# A handle on mass coincidence errors in *de novo* sequencing of antibodies by bottom-up proteomics

**DOI:** 10.1101/2024.02.20.581155

**Authors:** Douwe Schulte, Joost Snijder

## Abstract

Antibody sequences can be determined at 99% accuracy directly from the polypeptide product using bottom-up proteomics techniques. This circumvents the need to isolate the antibody-producing B-cell clone and enables reverse engineering of monoclonal antibodies from lost hybridoma cell lines, as well as the secreted protein in bodily fluid. Sequencing accuracy at the peptide level is limited by common mass coincidences of isobaric residues like leucine/isoleucine, but also by incomplete fragmentation spectra in which the order of two or more residues remains ambiguous due to lacking fragment ions for the intermediate positions. Likewise, different combinations of amino acids, of potentially different length, can also coincide to the same mass (*e*.*g*. GG=N, GA=Q *etc*.). Here we present several updates to Stitch (v1.5), which performs template-based assembly of *de novo* peptide reads to reconstruct antibody sequences. This version introduces a mass-based alignment algorithm that explicitly accounts for mass coincidence errors. In addition, it incorporates a postprocessing procedure to assign I/L residues based on secondary fragments (satellite ions, *i*.*e. w-*ions). Moreover, evidence for sequence assignments can now be directly evaluated with the addition of an integrated spectrum viewer. This version of Stitch also allows input data from a wider selection of *de novo* peptide sequencing algorithms, now including Casanovo, PEAKS, Novor.Cloud, pNovo, and MaxNovo, in addition to flat text and FASTA. Combined, these changes make Stitch compatible with a larger range of data processing pipelines and improve its tolerance to peptide-level sequencing errors.

## Introduction

Antibodies are essential components of the adaptive immune system that can recognize a vast array of antigens in the fight against pathogens or in autoimmune disease.^1–4^ The diversity of antigens recognized by antibodies is mirrored by the diversity in antibody sequences. This diversity is generated by somatic recombination and hypermutation of the paired heavy and light chains, taking place at singular B cells. Studying these unique antibody sequences therefore is crucial to understand the immune response in health and disease, and to develop diagnostics, therapeutics, and affinity reagents for life science research based on antibody-mediated recognition and binding of target antigens.

Established methods for antibody sequencing target the coding mRNA in single B cells.^5–12^ While these methods have laid the foundation for our current understanding of the antibody response, they do not directly access the functional secreted products found in bodily fluids in the same way as they are probed in common serological assays (to determine the binding and neutralization titres following natural infection or immunization). Additionally, B cells reside in spleen, bone marrow, and blood, of which only the latter population can be easily sampled in human subjects. In contrast, mass spectrometry-based methods can probe specific antibody sequences directly from the secreted polypeptide product, thereby circumventing the need to sample the antibody-producing B-cell clone and providing a direct glimpse into the so-called serum compartment of the immunoglobulin repertoire.^13–26^

Using a bottom-up proteomics approach, accurate peptide sequences can be determined and assembled into complete heavy and light chain sequences. We have recently developed the software tool Stitch to perform template-based assembly of peptide sequences against the coding gene segments of antibodies available in immunogenetics databases.^27^ Stitch has been shown to enable the accurate reconstruction of monoclonal antibody sequences, as well as the sequencing of isolated Fab fragments from patient serum, M-proteins in monoclonal gammopathies, antibody light chains from urine, and the profiling of whole IgG from COVID-19 patient sera.^28–31^ Sequence accuracies of ∼99% can be obtained, which is sufficient to reverse engineer functional antibody products.^13,22,23,32^ Remaining sequencing errors stem in large parts from common mass coincidences of isobaric residues like leucine/isoleucine, but also from incomplete fragmentation spectra in which the order of two or more residues remains ambiguous due to lacking fragment ions for the intermediate positions. Likewise, different combinations of amino acids, of potentially different length, can also coincide to the same mass (*e*.*g*. GG=N, GA=Q *etc*.). In addition to limiting accuracy of the input peptide sequences, these mass coincidence errors may also hamper proper assembly of the short peptides against the antibody template sequences.

Here we discuss several recent updates to Stitch (v1.1 up to and including v1.5) in the light of common MS-based *de novo* sequencing errors. These updates include a mass-based alignment algorithm (akin to Meta-SPS-contig ^33^), a post processing procedure to assign I/L residues based on secondary fragments, a built-in spectrum viewer, a graph based analysis which tracks the connectivity between identified sequence variants, and the ability to use a wider selection of *de novo* peptide sequencing programs as input, including Casanovo, MaxNovo, Novor.Cloud, PEAKS, and pNovo.^34–42^ Combined, these changes make Stitch compatible with a larger range of data processing pipelines and improve its tolerance to peptide-level sequencing errors.

## Results

Stitch assembles peptides from *de novo* sequencing programs for bottom-up LC-MS/MS data in the correct framework of the heavy and light chains of an antibody. An outline of the assembly and processing performed by Stitch, including the new features described here, is presented in Figure 1. The previously published version of Stitch was already able to handle PEAKS, FASTA, and plain text data, which is now extended to include the output from Casanovo, MaxNovo, Novor.Cloud, and pNovo. ^34–41^ These peptides are aligned to the germline sequences of the corresponding gene-segments (V/J/C) for the appropriate species and placed when the alignment score exceeds a user-defined cutoff. The consensus sequence of this alignment is weighed for the confidence and abundance of the individually placed reads as reported by the peptide *de novo* sequencing programs. This procedure robustly places peptide reads in the correct framework of the full antibody sequence, except for the hypervariable Complementarity Determining Region 3 of the heavy chain (CDRH3). CDRH3 is formed by the junction of three coding gene segments V, D and J, of which D is exceptionally short and variable, such that germline sequences do not provide a functional template to guide the assembly. Instead, Stitch extends the flanking V- and J-segments with a user-defined number of wildcard ‘X’ characters, such that overhanging reads from the V- and J-segments can be used to extend the consensus sequence into the CDRH3 region. The resulting overhanging V- and J-sequences are then aligned to find the overlap from both sides to reconstruct CDRH3. In contrast, CDR3 of the light chain is encoded only with V- and J-segments and can typically be reconstructed from the first order template matching step alone. These consensus sequences of the recombined V- and J-segments are then used for a second template matching step to reconstruct the final consensus sequence of the antibody. It should be noted that this procedure is only suitable for monoclonal antibodies (though it is tolerant to a moderate background of polyclonal sequences).^14–16^ The reconstruction of CDRH3 from disperse polyclonal mixtures currently still requires manual curation of candidate sequences. The template-based assembly (and recombination) results of Stitch are written out as an interactive report where all peptide alignments can be inspected and overview statistics like total score/area are collected for every template sequence. To aid in the analysis of disperse polyclonal mixtures, this new version of Stitch introduces the so-called ‘variant graph’, which shows how co-occurring sequence variations in reference to the templates are connected by peptide reads.

**Figure 1:**
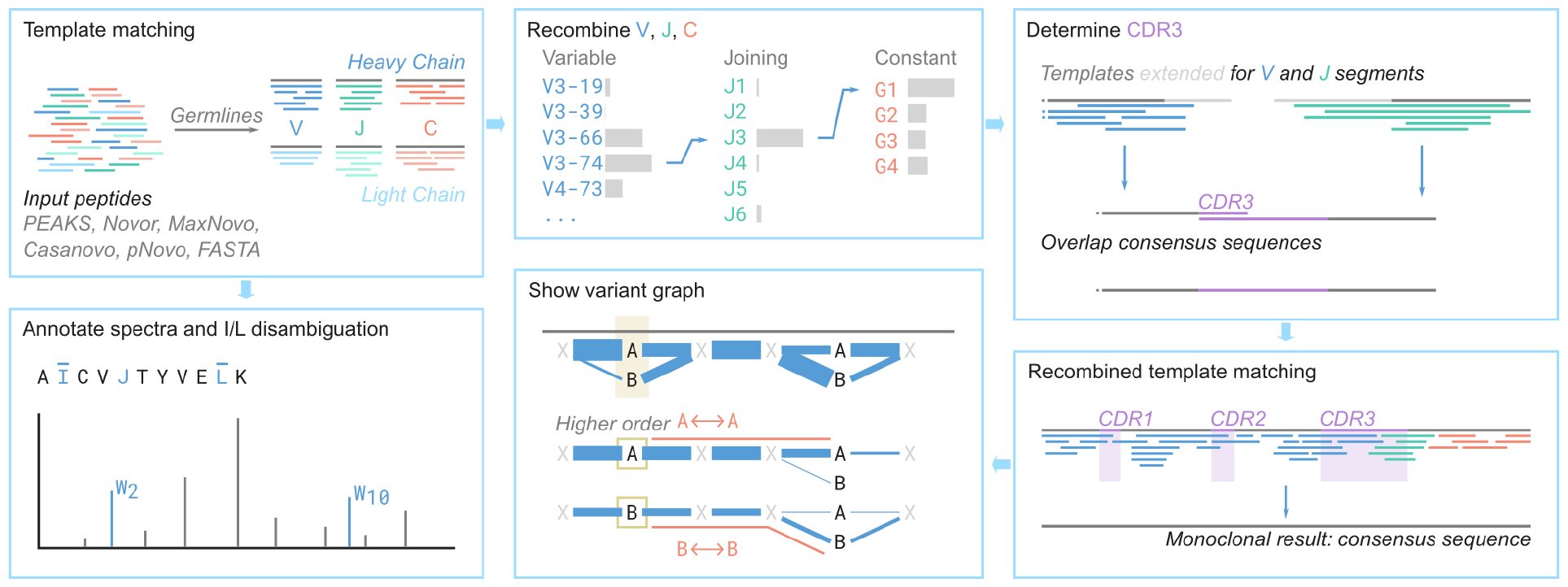
Overview of the program Stitch.

We also introduced a fundamentally different alignment algorithm for the placement of input reads against the template sequences. We previously used a modified version of the Smith-Waterman algorithm (SWA) using a matrix based on BLOSUM62.^43–45^ The SWA only accounts for mismatches between input reads and template sequence that stem from evolutionary processes like somatic hypermutation observed in mature antibody sequences compared to the germline precursors. In our experimental data, however, these mismatches are further convoluted with errors inherent to an MS-based approach to peptide sequencing.

To illustrate the sequencing errors that MS is prone to, Figure 2A plots the residue masses of the 20 common amino acids (with common modifications) on a scale of 50 to 200 Da, also including the amino acid combinations that fall within this range. Considering that mass analyzers in current proteomics setups typically operate within a precision range of 0.01-0.50 Da we can broadly distinguish between four categories of MS-based sequencing errors. The first category is the isobaric residues isoleucine and leucine, which simply have identical masses and cannot be distinguished without the presence of secondary fragment ions (as further discussed below). The second category of common MS-based errors includes larger isobaric *sets* of amino acids where different combinations of residues amount to identical masses and cannot be distinguished. This second category stems from missing fragment ions in the MS/MS spectra. There are three subcategories of errors in this second category: a) simple rotations within a set (*e*.*g*. AS=SA), b) isobaric sets (*e*.*g*. AS=GT), and c) isobaric sets of different lengths (*e*.*g*. N=GG, Q=GA). For sets of three residues or longer, combinations of these subcategories may occur. The third category of common MS-based errors includes (modified) residues that are not strictly identical in mass but are so similar that they cannot be distinguished given the experimental precision limits of the mass analyzer. For instance, the mass difference between K and Q is 0.036 Da, between M^ox^ and F 0.033 Da. This third category of MS-based errors can be eliminated with the use of a high precision mass analyzer (TOF, FT-MS), but poses a significant limitation to *de novo* sequencing at lower precision (ion trap). The fourth category of common MS-based errors do not necessarily stem from analytical limitations, but rather from artefacts of the sample processing pipeline in the form of deamidation of Q and especially N, converting these residues to E and D, respectively.

**Figure 2:**
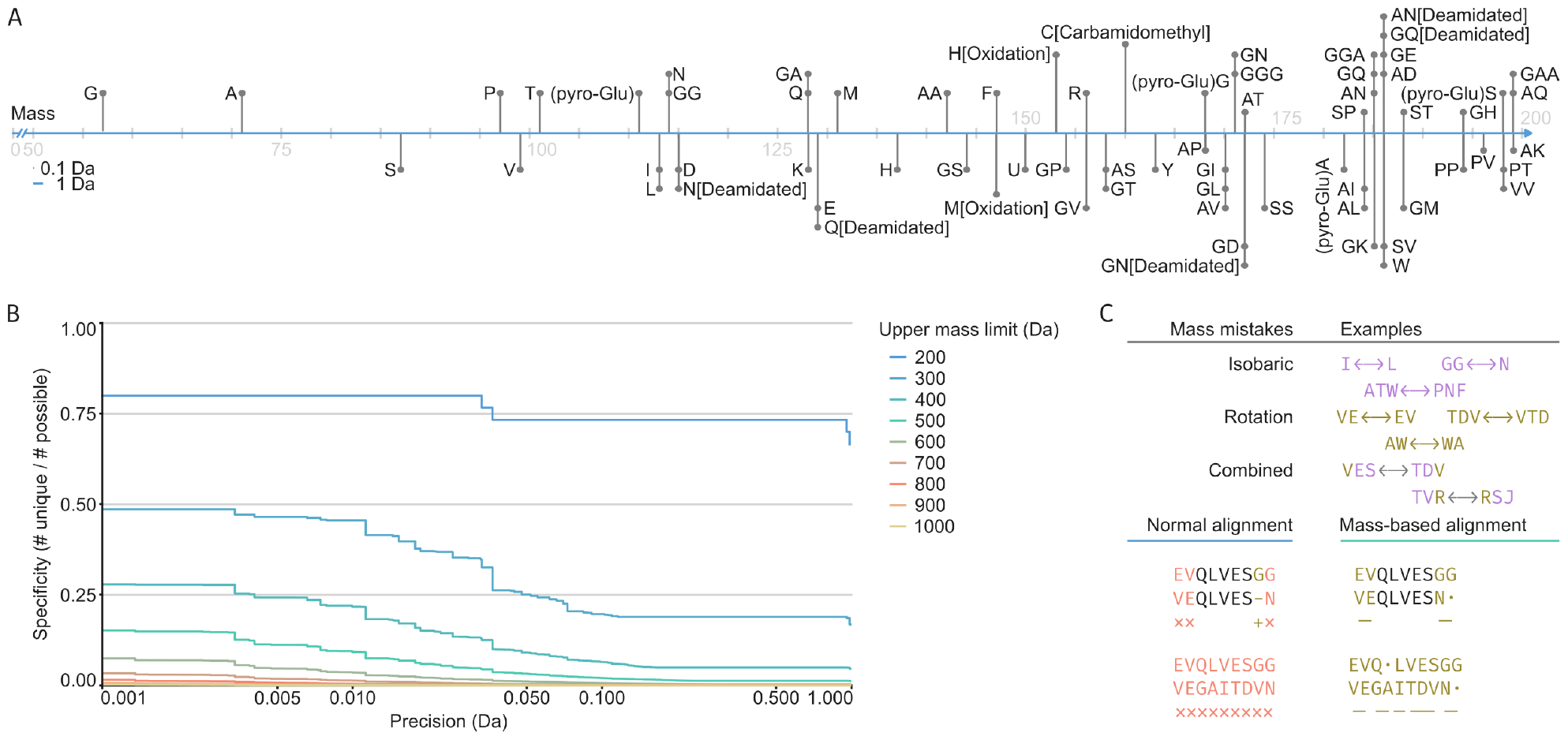
Mass coincidence errors in de novo peptide sequencing. A) All possible amino acid sequences up to 200 Da, including common modifications (carbamidomethyl fixed on C, pyro-Glu, deamidation on N and Q, oxidation on M and H). B) The fraction of sequences that can be told apart (specificity) based on the precision in Da and the upper mass limit (not counting rotations). C) Illustration of how the mass-based alignment provides a better handle on mass coincidence errors.

Figure 2B illustrates how these common MS-based errors accumulate as a function of precision for progressively larger mass intervals (*i*.*e*. a higher number of missing fragments). A few obvious, but fundamental requirements and limitations for *de novo* peptide sequencing can be observed from this ‘survival’ analysis, showing what fraction of possible unique residues (or residue combinations) can be distinguished based on their mass, not counting rotations. First, it is of the utmost importance to achieve complete fragmentation, as category 2 errors (isobaric subsets) readily accumulate over larger mass intervals. Second, category 3 errors (similar residue masses) can be eliminated with high precision mass analyzers. Important to note that the required precision to eliminate category 3 errors is readily achieved on modern mass analyzers for small peptide fragments with low charge states, but that this becomes a significant limitation for larger fragments with higher charge states in middle- and top-down approaches. Considering these common MS-based errors, we here propose a modification of the SWA to perform the template-based assembly for antibody sequencing.

One of the most important drawbacks in context of these MS-based errors is that in SWA a substitution can only be a single amino acid to another single amino acid, while the category 2 errors affect multiple consecutive residues. The new mass-based alignment we devised considers these errors by aligning not only single amino acids, but also larger sets, illustrated in Figure 2C. It does this by supplementing the alignment steps looked for in SWA (*i*.*e*. match/mismatch/insertion/deletion), with two additional steps: rotation and isobaric. These last two steps can be of length up to N steps on both template and peptide (*e*.*g*. match 2 amino acids on the template with 3 on the input peptide). With this addition the category 2 errors, up to N amino acids, can easily be found. The score for each aligned set of amino acids is looked up in a premade two-dimensional array with the same modified BLOSUM62 base matrix in the top left and all category 2 errors given a nonzero score in the array based on their mass identity. It is important for the behavior of the alignment that these rotations and isobaric matches score lower than any direct match. Hence, rotations score 3 per amino acid involved and isobaric matches score 2 times the maximal number of amino acids involved. The alignment algorithm looks for any nonzero score in the set of steps of up to N residues, alongside the normal cases for SWA, and the highest scoring of the possible steps is chosen. This approach is similar to Meta-SPS contig, with the main difference that alignment and assembly is performed on the level of the output peptide sequences, rather than the MS/MS data itself.^33^

Use of the MBA vs SWA improves the tolerance to mass coincidence errors during template-based assembly. This difference is illustrated in Figure 3 using PEAKS input data for the two monoclonal antibodies Herceptin and F59. PEAKS provides a peptide-level Average Local Confidence (ALC) score between 0 and 99, where peptides with lower ALC typically miss more fragment ions to support the given sequence. Figure 3A shows how the average length-normalized alignment score during the assembly develops as a function of ALC cutoff for MBA vs SWA. By design, the alignment scores are consistently higher using MBA vs SWA, the margin illustrating just how common mass coincidence errors are in the input data. Additionally, SWA cannot align any peptides below ALC 55, while these may still be placed using MBA. Peptides with higher ALC (and fewer errors) yield higher alignment scores, as expected, but to achieve optimal accuracy in the final consensus sequences there is an important trade off to be made between selection of the highest scoring input reads and achieving optimal coverage. As the coverage plummets with ALC cutoff scores >95, it follows that a substantial tolerance to mass coincidence errors is required to assemble the complete sequences (given the limitations of the experimental input data), by including lower ALC (<95) peptides (Figure 3B). The accuracy of the final consensus sequences of Herceptin and F59 are plotted as a function of ALC cutoff in Figure 3B, illustrating that MBA outperforms SWA across the sampled range. The accuracy increases from 0.95 (SWA) to 0.99 (MBA). While this may seem like a moderate increase at first, it represents a better than 100-fold improvement in the probability to determine the sequence of a 120 amino acid long variable domain with zero mistakes (*P=0*.*002* for accuracy 0.95 vs. *P=0*.*299* for accuracy 0.99). Of note, to achieve a probability of *P>0*.*99* to make zero mistakes in a sequence of this length, the accuracy needs to further improve to >0.9999 (see Figure 4).

**Figure 3:**
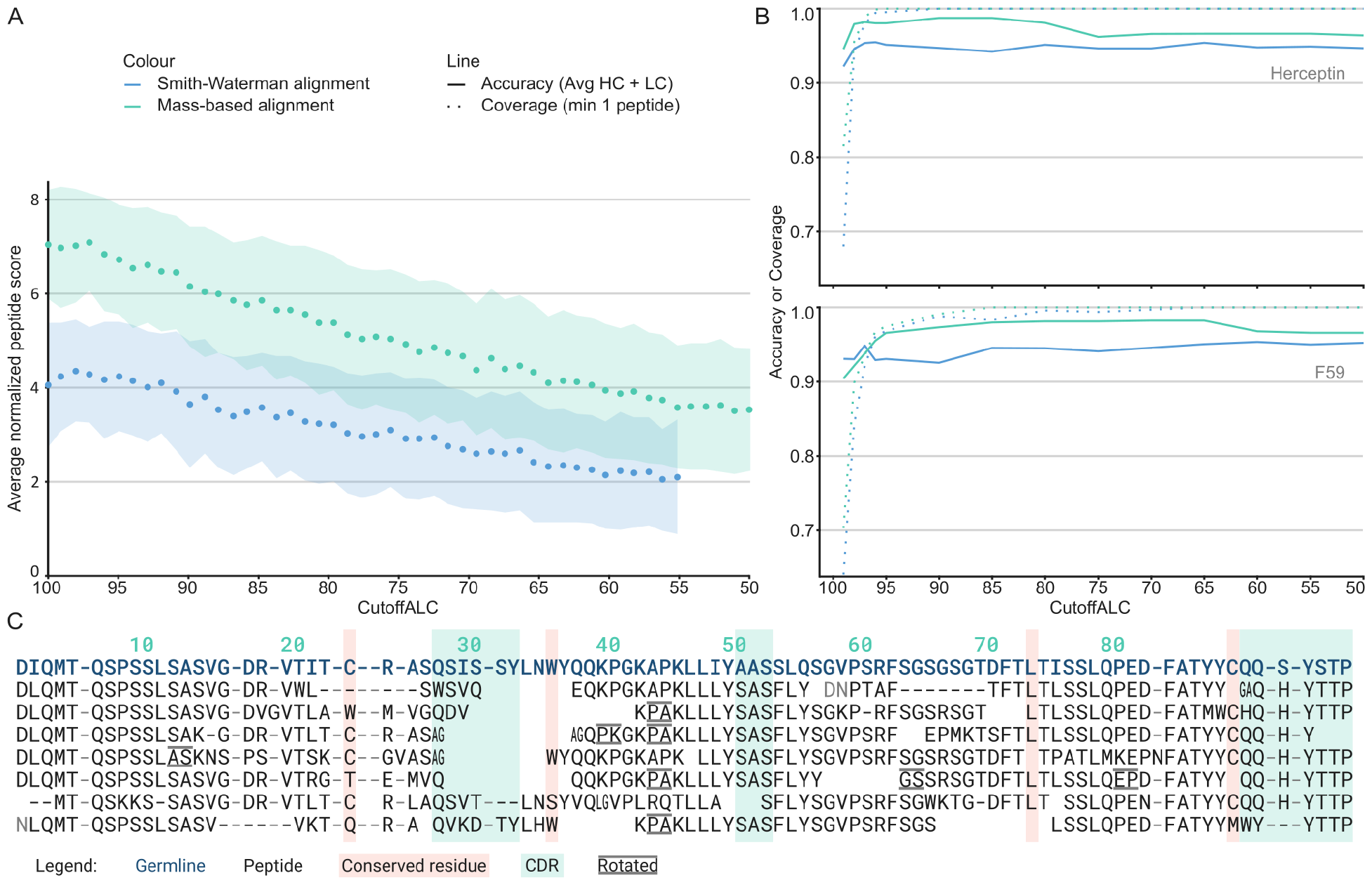
Mass based alignment results. A) The average normalized score (score / alignment length) as a function of PEAKS ALC. Note that the missing points for CutoffALC < 55 for Smith-Waterman alignment is because no peptides with these scores could be placed by this alignment algorithm. B) The result of running Stitch with the Herceptin and F59 sample data with different read cutoff scores presented as accuracy, defined here as identity against the true sequence, in a solid line, with the fraction of positions covered by at least one peptide drawn in a dotted line. Note that the results were generated running Stitch with a low cutoff score for template matching. C) An excerpt of the alignment of the highest scoring light chain variable gene from the Stitch report for Herceptin using mass-based alignment for cutoff ALC 85.

**Figure 4:**
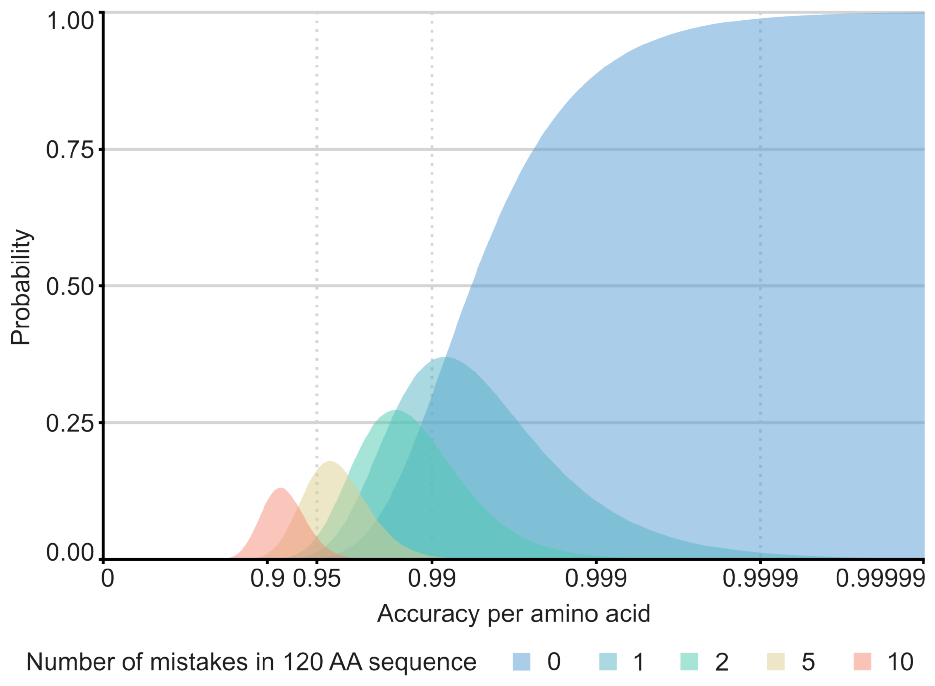
Probability of the indicated number of errors in a sequence of 120 amino acids given the sequencing accuracy per amino acid.

In addition to the new MBA, Stitch is also updated to read and display fragmentation spectra in an interactive spectrum viewer (see Figure 5A). This allows the user to manually inspect the underlying data for an assigned sequence. Moreover, accessing the fragmentation spectra in Stitch also allowed us to implement a new postprocessing procedure to differentiate between Leucine/Isoleucine based on secondary fragment ions that can be observed in some fragmentation modes, notably electron transfer high energy collision dissociation data.^42^ These diagnostic satellite ions (known as *w/d*-ions) are formed by secondary fragmentation of *z-* and *a-*ions, by transfer of the free electron to the side chain and breaking of a bond to the first C-atom in the side chain (Cβ). As the side chains are different between Isoleucine and Leucine the possible mass shifts of the satellite ions are different, being 15 and 29 Da for Isoleucine (loss of C_1_H_3_ and C_2_H_5_) and 43 Da for Leucine (loss of C_3_H_7_). Stitch gives the option to use these satellite ions to find the most likely candidate for each I/L position. This improves the fraction of correctly identified I/L positions from 0.81 (when defaulting to the germline template sequence, or to L in the case of newly introduced I/L mutations) to 0.94 when using the satellite ions as seen in Figure 5B.

**Figure 5:**
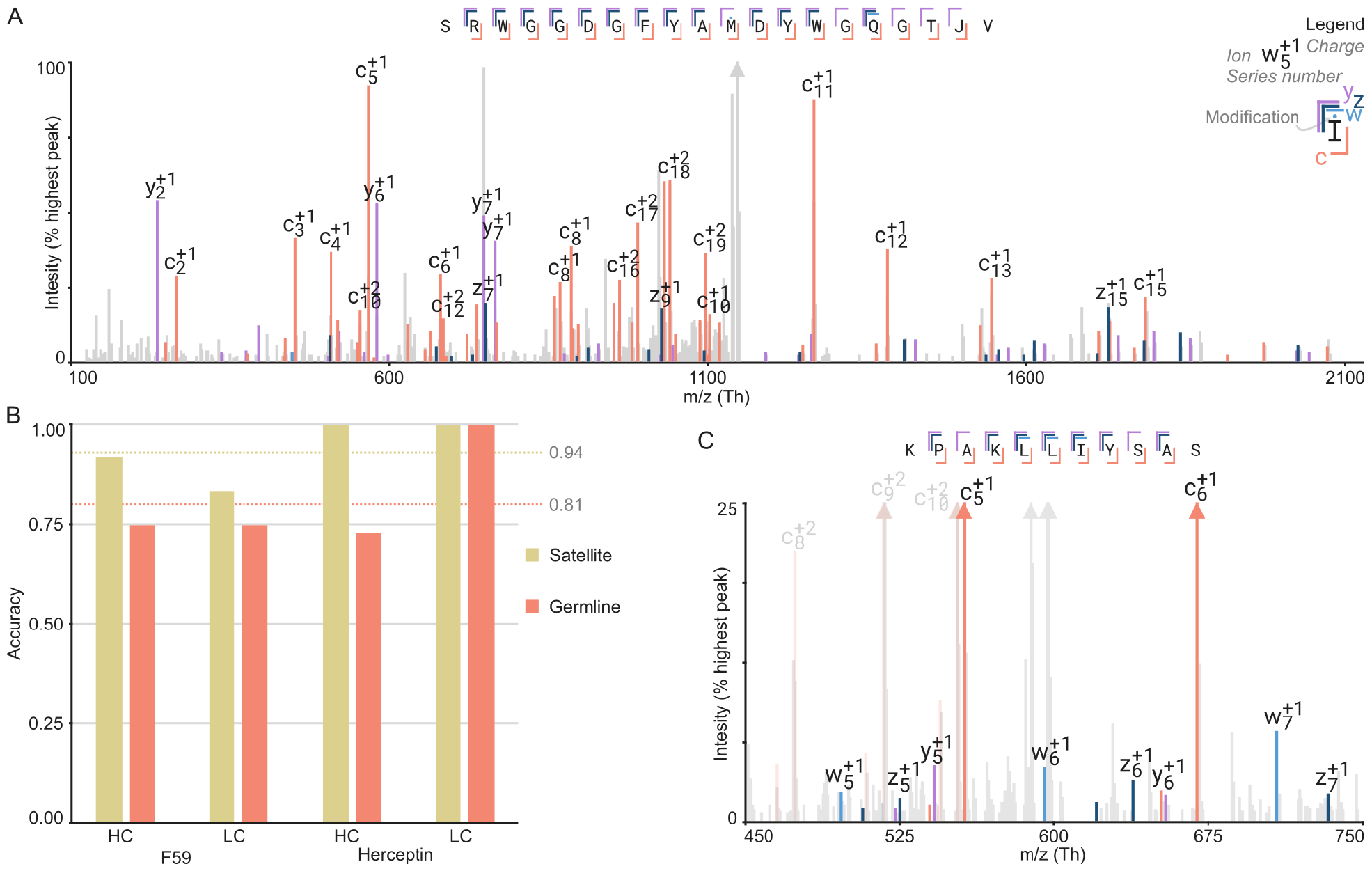
Satellite ions usage to identify I/L. A) Screen shot from the spectrum viewer of a CDRH3 peptide of Herceptin. B) The fraction of correct I/L identifications using just the germline (previous version of Stitch) or the new satellite ions, alignment identity is used as measure of accuracy. C) An example spectrum with satellite ions annotated as Stitch would display it.

## Discussion

These recent updates to Stitch provide a better handle on mass coincidence errors for *de novo* antibody sequencing by LC-MS/MS. These improvements extend to any MS-based *de novo* sequencing application beyond antibodies and indeed Stitch can perform the same tasks on an arbitrary set of template sequences.

Implementation of the mass-based alignment shed light on fundamental requirements and limitations for MS-based de novo sequencing and set out some important goals for the near future. To minimize peptide-level sequencing errors, it is crucially important to analyse at a precision of at least 0.02 Da, to achieve complete fragmentation, and obtain secondary fragment ions to differentiate I/L. The latter two points call for use of complementary fragmentation techniques beyond CID. Currently, accuracies of 0.99 can be achieved on the consensus sequence level, but to achieve a milestone 99% confidence in error-free sequences, this needs to improve to >0.9999 for the case of an antibody variable domain of 120 amino acids. Based on our own work, this requires further improvement of I/L assignments, but even more so elimination of deamidation errors (primarily N/D) from the sample processing workflows. Moreover, the 0.99 accuracy is currently based on consensus sequences from overlapping peptides and as we expand applications to complex polyclonal antibody mixtures, the depth of coverage will inevitably suffer, again calling for improved fragmentation to minimize sequencing errors at the peptide-level.

Stitch is now compatible with a wide range of current de novo sequencing algorithms, providing an improved strategy for sequence assembly with mass-based alignment, and the opportunity to browse spectrum-level evidence for the determined sequence to aid in manual curation. It is a versatile tool for MS-based antibody sequencing and beyond and may provide a springboard to dive into the serum compartment of the antibody repertoire with proteomics techniques.

## Methods

*Mass coincidences rate* – The mass coincidence rate over different precisions and mass ranges (Figure 2A and 2B) were calculated by generating all possible amino acid sequences from the canonical amino acids for each mass range (without rotations), including the following modifications: Carbamidomethyl (fixed on C), Oxidation (variable on WHM), Deamidated (variable on NQ), Glu->pyro-Glu (variable on N-term E), and Gln->pyro-Glu (variable on N-term Q). For each of these sets of possible sequences the number of unique masses was calculated by going through the possible sets sorted on mass and greedily combining any set with the previous if within the given precision.

*Smith-Waterman comparison to mass-based alignment* – The comparison between Smith-Waterman alignment (SWA) and mass-based alignment (MBA) (Figure 3A) was made by running Stitch with different PEAKS ALC peptide cutoff scores. The other parameters used were enforce unique 0.9, cutoff score template matching 5, cutoff score recombination 5, with the default templates for *Homo sapiens* heavy and light chain, and common contaminants. The identity was determined using a version of MBA by comparing the consensus sequence as given by these runs to the known sequence. The coverage was determined by counting the fractions of positions that had at least one peptide matching as given in the Stitch FASTA export.

The average scores for each ALC bin (Figure 2B) were calculated using the Herceptin data at PEAKS ALC cutoff 50 from Figure 2A. The score was normalized by dividing the absolute score by the length of each peptide it matched on the template. These scores were grouped by ALC of the peptide and the average and standard deviation calculated.

### Accuracy/error estimates

The likelihood of a given number of mistakes (Figure 4) was calculated following a simple binomial model, in which the probability *P*(*C, T*)of obtaining *C* correct amino acids in a sequence of length *T*, with a given accuracy *a*, equals *a*^*C*^ ⋅ (1 − *a*)^*T*−*C*^. Which can be simplified to *a*^*T*^ when *C* equals *T*, meaning no mistakes are made. Which can then be solved for *a* as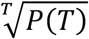.

### Accuracy of I/L assignments

The accuracy at I/L positions (Figure 5B) was determined by running Stitch with the same data as the SWA vs. MBA comparison, with PEAKS cutoff ALC 95, enforce unique 0.9, template matching and recombine cutoff score 10, mass-based alignment, the default templates for *Homo sapiens* heavy and light chain, and common contaminants, the raw data, and by turning on XleDisambiguation on the input file. For each I/L location the known true sequence, Stitch outcome and germline sequence were compared.

## Data and Code Availability

The source code of Stitch is available on GitHub: https://github.com/snijderlab/stitch. All Stitch HTML results related to this study are provided as Supporting Data. The raw data of the monoclonal antibodies Herceptin and F59 are available under identifier PXD023419, and doi 10.6084/m9.figshare.13194005, respectively.

## Acknowledgments

The authors thank Bastiaan de Graaf for the fruitful discussions regarding the design of Stitch and the mass-based alignment algorithm. We would also like to thank Lukas Käll for helpful input and feedback on the manuscript. This research was funded by the Dutch Research Council NWO Gravitation 2013 BOO, Institute for Chemical Immunology (ICI; 024.002.009), and the European Research Council Executive Agency HORIZON ERC-2022-STG (FLAVIR; 101077640).

## Notes

### Competing Interest Statement

The authors have declared no competing interest.

