## Supplementary material for "A handle on mass coincidence errors in *de novo* sequencing of antibodies by bottom-up proteomics": figure.r

```
shhh <- suppressPackageStartupMessages # It's a library, so shhh!
shhh(library(ggplot2))
shhh(library(ggpubr))
shhh(library(reshape2))
shhh(library(dplyr))
shhh(library("Cairo"))
shhh(library(extrafont))
shhh(library(tidyverse))
shhh(library("ggplotify"))

assembler_colours <- c("#569CD6", "#4EC9B0", "#F48771", "#DCD08F")


create_scale <- function(n) {
    # Simple cases when no extra colours need to be generated
    if (n < length(assembler_colours)) {
        return(assembler_colours[1:n])
    }
    if (n == length(assembler_colours)) {
        return(assembler_colours)
    }

    # Some extra colours need to be generated
    blocks <- length(assembler_colours) - 1
    steps <- floor((n - 1) / blocks)
    rem <- n - steps * blocks - 1
    last_colour <- assembler_colours[1]
    scale <- c(last_colour)
    for (colour in assembler_colours[2:length(assembler_colours)]) {
        actual_steps <- steps
        if (rem > 0) {
            actual_steps <- actual_steps + 1
            rem <- rem - 1
        }
        if (actual_steps == 1) {
            scale <- append(scale, colour)
        } else {
            extra_colours <- scales::seq_gradient_pal(last_colour, colour)(seq(0, 1, length.out = actual_steps + 1))

            scale <- append(
                scale,
                extra_colours[2:length(extra_colours)]
            )
        }
        last_colour <- colour
    }
    if (n != length(scale)) print(paste("Not the right number of colours, asked: ", n, " got: ", length(scale)))
    return(scale)
}

no_clipping <- function(plot) {
    pg <- ggplotGrob(plot)
    # print(pg)

    for (i in which(grepl("strip-t", pg$layout$name))) {
        pg$grobs[[i]]$layout$clip <- "off"
    }
    pg$layout$clip[pg$layout$name == "panel"] <- "off"

    # grid::grid.draw(pg)
    return(as.ggplot(pg))
}

open <- function(path) {
    data <- read.csv(path, header = TRUE, sep = ",")
    # data$Score <- data$Area / sum(data$Area)
    # print(data$Score[order(data$Point)])
    # frame <- data.frame(data$Point[order(data$Point)], data$Group[order(data$Point)], data$Score[order(data$Point)])
    # colnames(frame) <- c("Point", "Group", "Score")
    # print(frame)
    # data <- data.frame(data)
    return(data)
}


survival <- function(data) {
    return(no_clipping(
        # data_no_zero <- data %>% filter(Precision != 0)
        # data_zero <- data %>% filter(Precision == 0)
        ggplot(data = data %>% filter(Precision != 0), aes(x = Precision, y = Fraction, color = factor(Limit), label = Limit)) +
            geom_step() +
            geom_point(data = data %>% filter(Precision == 0)) +
            # geom_text(data = data %>% filter(Precision == 0), hjust = -0.4, vjust = 0.5) +
            # geom_smooth(method = lm, se = F) +
            # facet_wrap(~Group) +
            scale_fill_manual(
                values = create_scale(length(unique(data$Limit)))
            ) +
            scale_color_manual(
                values = create_scale(length(unique(data$Limit)))
            ) +
            scale_x_continuous(trans = "log2", breaks = c(0.001, 0.005, 0.01, 0.05, 0.1, 0.5, 1.0)) +
            # stat_cor(method = "pearson", cor.coef.name = "r", vjust = 1, size = 4, label.x = 4, label.y = 0.9) +
            ylim(0, 1.0) +
            theme_minimal() +
            labs(
                x = "Precision (Da) [log2]",
                y = "Fraction of unique masses (number unique masses / number of possible sequences)",
                color = "Upper mass limit (Da)"
            ) +
            theme(
                plot.background = element_rect(fill = "white", color = "white"),
                strip.background = element_blank(),
                # axis.text.x = element_text(angle = 90, vjust = 0.5, hjust = 1),
                panel.grid.major.x = element_blank(),
                panel.grid.major.y = element_line(linewidth = .1, color = "black"),
                panel.grid.minor = element_blank(),
                # legend.position = "none",
                text = element_text(size = 12, family = "roboto"),
                legend.position = c(0.9, .85)
            )
    ))
}

survival_y_log <- function(data) {
    return(no_clipping(
        # data_no_zero <- data %>% filter(Precision != 0)
        # data_zero <- data %>% filter(Precision == 0)
        ggplot(data = data %>% filter(Precision != 0), aes(x = Precision, y = Fraction, color = factor(Limit), label = Limit)) +
            geom_step() +
            geom_point(data = data %>% filter(Precision == 0)) +
            # geom_text(data = data %>% filter(Precision == 0), hjust = -0.4, vjust = 0.5) +
            # geom_smooth(method = lm, se = F) +
            # facet_wrap(~Group) +
            scale_fill_manual(
                values = create_scale(length(unique(data$Limit)))
            ) +
            scale_color_manual(
                values = create_scale(length(unique(data$Limit)))
            ) +
            scale_x_continuous(trans = "log2", breaks = c(0.0001, 0.0005, 0.001, 0.005, 0.01, 0.05, 0.1, 0.5, 1.0)) +
            scale_y_continuous(trans = "log2", breaks = c(0.001, 0.005, 0.01, 0.05, 0.1, 0.5, 1.0)) +
            # stat_cor(method = "pearson", cor.coef.name = "r", vjust = 1, size = 4, label.x = 4, label.y = 0.9) +
            # ylim(0, 1.0) +
            theme_minimal() +
            labs(
                x = "Precision (Da) [log2]",
                y = "Fraction of unique masses (number unique masses / number of possible sequences) [log2]",
                color = "Upper mass limit (Da)"
            ) +
            theme(
                plot.background = element_rect(fill = "white", color = "white"),
                strip.background = element_blank(),
                # axis.text.x = element_text(angle = 90, vjust = 0.5, hjust = 1),
                panel.grid.major.x = element_blank(),
                panel.grid.major.y = element_line(linewidth = .1, color = "black"),
                panel.grid.minor = element_blank(),
                # legend.position = "none",
                text = element_text(size = 12, family = "roboto"),
                legend.position = c(0.1, .15)
            )
    ))
}

data <- open("output.csv")
survival_plot <- survival(data)
ggsave("survival.png", plot = survival_plot, width = unit(x = 10, units = "cm"), height = unit(x = 5, units = "cm"))
ggsave("survival.svg", plot = survival_plot, width = unit(x = 10, units = "cm"), height = unit(x = 5, units = "cm"))
# survival_plot <- survival_y_log(data)
# ggsave("survival_y_log.png", plot = survival_plot, width = 10, height = 10)
```
