## Supplementary material for "A handle on mass coincidence errors in *de novo* sequencing of antibodies by bottom-up proteomics": main.rs

```
use std::{fmt::Display, io::BufWriter};

use rustyms::{
    ontologies::{unimod_ontology, OntologyLookup},
    placement_rule::{PlacementRule, Position},
    system::{dalton, Mass},
    AminoAcid, LinearPeptide, MassTolerance, Modification, SequenceElement,
};
use std::io::Write;

fn main() {
    let file = std::fs::File::create("output.csv").unwrap();
    let mut writer = BufWriter::new(file);
    writeln!(writer, "Precision,Limit,Surviving,Fraction").unwrap();
    for limit in (2..=10).map(|l| l * 100) {
        let state = find_isobarics(
            Mass::new::<dalton>(limit as f64),
            &[(
                unimod_ontology().find_name("Carbamidomethyl").unwrap(),
                Some(PlacementRule::AminoAcid(
                    vec![AminoAcid::C],
                    Position::Anywhere,
                )),
            )],
            &[
                (
                    unimod_ontology().find_name("Oxidation").unwrap(),
                    Some(PlacementRule::AminoAcid(
                        vec![AminoAcid::W, AminoAcid::H, AminoAcid::M],
                        Position::Anywhere,
                    )),
                ),
                (
                    unimod_ontology().find_name("Deamidated").unwrap(),
                    Some(PlacementRule::AminoAcid(
                        vec![AminoAcid::N, AminoAcid::Q],
                        Position::Anywhere,
                    )),
                ),
                (unimod_ontology().find_name("Glu->pyro-Glu").unwrap(), None),
                (unimod_ontology().find_name("Gln->pyro-Glu").unwrap(), None),
            ],
        );
        const SAMPLING_FACTOR: isize = 100;
        for precision in std::iter::once(0.0)
            .chain((-10 * SAMPLING_FACTOR..0).map(|p| (p as f64 / SAMPLING_FACTOR as f64).exp2()))
        {
            let isobarics = group_isobarics(
                &state,
                MassTolerance::Absolute(Mass::new::<dalton>(precision)),
            );
            writeln!(
                writer,
                "{},{},{},{}",
                precision,
                limit,
                isobarics.len(),
                isobarics.len() as f64 / state.len() as f64
            )
            .unwrap();
        }
    }
    // let isobarics = find_isobarics(
    //     Mass::new::<dalton>(300.0),
    //     MassTolerance::Ppm(10.0),
    //     &[(
    //         unimod_ontology().find_name("Carboxymethyl").unwrap(),
    //         Some(PlacementRule::AminoAcid(
    //             vec![AminoAcid::C],
    //             Position::Anywhere,
    //         )),
    //     )],
    //     &[
    //         (
    //             unimod_ontology().find_name("Oxidation").unwrap(),
    //             Some(PlacementRule::AminoAcid(
    //                 vec![AminoAcid::W, AminoAcid::H, AminoAcid::M],
    //                 Position::Anywhere,
    //             )),
    //         ),
    //         (
    //             unimod_ontology().find_name("Deamidated").unwrap(),
    //             Some(PlacementRule::AminoAcid(
    //                 vec![AminoAcid::N, AminoAcid::Q],
    //                 Position::Anywhere,
    //             )),
    //         ),
    //         (unimod_ontology().find_name("Glu->pyro-Glu").unwrap(), None),
    //     ],
    // );
    // for set in isobarics {
    //     println!("{set}");
    // }
}

struct Isobaric {
    pub mass: Mass,
    pub isobaric: usize,
}

// impl Display for Isobaric {
//     fn fmt(&self, f: &mut std::fmt::Formatter<'_>) -> std::fmt::Result {
//         write!(f, "{:.4};", self.mass.value)?;
//         let mut first = true;
//         for seq in &self.isobaric {
//             if !first {
//                 write!(f, ", ")?;
//             }
//             first = false;
//             write!(
//                 f,
//                 "{}",
//                 seq.to_string()
//                     .replace("U:", "")
//                     .replace("[Glu->pyro-Glu]-E", "(pyro-Glu)")
//             )?;
//         }
//         Ok(())
//     }
// }
#[derive(Clone)]
struct State {
    // n_term: Option<(SequenceElement, Modification)>,
    // c_term: Option<(SequenceElement, Modification)>,
    // middle: Vec<SequenceElement>,
    index: usize,
    mass: Mass,
}

impl State {
    fn with(&self, block: &(SequenceElement, Mass), index: usize) -> Self {
        // let mut new_seq = self.middle.clone();
        // new_seq.push(block.0.clone());
        Self {
            // middle: new_seq,
            index,
            mass: self.mass + block.1,
            ..self.clone()
        }
    }

    // fn peptide(&self) -> LinearPeptide {
    //     let n_term = self.n_term.as_ref().map(|n| n.1.clone());
    //     let c_term = self.c_term.as_ref().map(|c| c.1.clone());
    //     let mut seq = Vec::with_capacity(
    //         self.middle.len()
    //             + usize::from(self.n_term.is_some())
    //             + usize::from(self.c_term.is_some()),
    //     );
    //     if let Some((s, _)) = self.n_term.clone() {
    //         seq.push(s);
    //     }
    //     seq.append(&mut self.middle.clone());
    //     if let Some((s, _)) = self.c_term.clone() {
    //         seq.push(s);
    //     }

    //     LinearPeptide {
    //         global: Vec::new(),
    //         labile: Vec::new(),
    //         n_term,
    //         c_term,
    //         sequence: seq,
    //         ambiguous_modifications: Vec::new(),
    //         charge_carriers: None,
    //     }
    // }
}

fn find_isobarics(
    mass_limit: Mass,
    fixed: &[(Modification, Option<PlacementRule>)],
    variable: &[(Modification, Option<PlacementRule>)],
) -> Vec<State> {
    let (n_term, middle, c_term) =
        rustyms::building_blocks(AminoAcid::CANONICAL_AMINO_ACIDS, fixed, variable);
    let mut options = Vec::new();
    for element in n_term {
        options.push(State {
            // n_term: Some((element.0, element.1)),
            // c_term: None,
            // middle: Vec::new(),
            index: 0,
            mass: element.2,
        })
    }
    for element in c_term {
        options.push(State {
            // n_term: None,
            // c_term: Some((element.0, element.1)),
            // middle: Vec::new(),
            index: 0,
            mass: element.2,
        })
    }
    options.push(State {
        // n_term: None,
        // c_term: None,
        // middle: Vec::new(),
        index: 0,
        mass: Mass::new::<dalton>(0.0),
    });

    let mut next_layer;
    let mut finished = Vec::new();
    loop {
        if options[0].mass + middle[0].1 > mass_limit {
            break;
        }
        next_layer = Vec::with_capacity(options.len());
        for state in options {
            for (i, block) in middle.iter().enumerate().skip(state.index) {
                if state.mass + block.1 > mass_limit {
                    break; // skip this whole for loop as anything coming later is by definition heavier
                }
                next_layer.push(state.with(block, i));
            }
        }
        finished.extend_from_slice(&next_layer);
        options = next_layer;
    }

    finished.sort_unstable_by(|a, b| a.mass.value.total_cmp(&b.mass.value));
    finished
}

fn group_isobarics(finished: &[State], tolerance: MassTolerance) -> Vec<Isobaric> {
    let mut grouped: Vec<Isobaric> = Vec::new();

    for state in finished {
        let end = grouped.len() - 1;
        if !grouped.is_empty() && tolerance.within(grouped[end].mass, state.mass) {
            grouped[end].isobaric += 1;
        } else {
            grouped.push(Isobaric {
                mass: state.mass,
                isobaric: 1,
            });
        }
    }

    grouped
}
```
