## Supplementary material for "A handle on mass coincidence errors in *de novo* sequencing of antibodies by bottom-up proteomics": script.js

```
"use strict";
var assets_folder = undefined;

function sortTable(id, column_number, type) {
    var table, rows, switching, i, x, y, shouldSwitch, dir = 0;
    table = document.getElementById(id);
    switching = true;
    dir = "desc";
    let headers = table.getElementsByTagName("tr")[0].getElementsByTagName("th");

    var sorted = false;
    if (headers[column_number].getAttribute('data-sort-order') == "asc") {
        dir = "desc";
        sorted = true
    }
    if (headers[column_number].getAttribute('data-sort-order') == "desc") {
        dir = "asc";
        sorted = true
    }

    for (var j = 0; j < headers.length; j++) {
        headers[j].setAttribute('data-sort-order', "");
    }
    headers[column_number].setAttribute('data-sort-order', dir);

    rows = Array.from(table.getElementsByTagName("tr"));
    var values = [null];
    for (i = 1; i < rows.length; i++) {
        x = rows[i].getElementsByTagName("td")[column_number]
        switch (type) {
            case "string":
                values.push(x.innerHTML.toLowerCase())
                break
            case "id":
                values.push(x.firstChild.innerHTML.toLowerCase())
                break
            case "number":
                values.push(Number(x.innerHTML))
                break
        }
    }

    if (sorted) {
        rows = rows.reverse()
        rows.splice(0, 0, rows[rows.length - 1])
        rows.pop()
        switching = false
    } else {
        switching = true
    }

    while (switching) {
        switching = false;
        /* Loop through all table rows (except the
        first, which contains table headers): */
        for (i = 1; i < (rows.length - 1); i++) {
            shouldSwitch = false;
            if (typeof values[i] == "string") {
                if (values[i].localeCompare(values[i + 1], undefined, { numeric: true, sensitivity: "base" }) == 1) {
                    shouldSwitch = true;
                    break;
                }
            } else if (values[i] < values[i + 1]) {
                shouldSwitch = true;
                break;
            }
        }
        if (shouldSwitch) {
            var el = rows[i + 1]
            rows.splice(i + 1, 1)
            rows.splice(i, 0, el)
            el = values[i + 1]
            values.splice(i + 1, 1)
            values.splice(i, 0, el)
            //rows[i].parentNode.insertBefore(rows[i + 1], rows[i]);
            switching = true;
        }
    }
    // Remove old rows and append all new rows
    while (table.lastChild) {
        table.removeChild(table.lastChild);
    }
    var frag = document.createDocumentFragment();
    for (var i = 0; i < rows.length; ++i) {
        frag.appendChild(rows[i]);
    }
    table.appendChild(frag);
}

function Select(id) {
    window.location.href = assets_folder + "/contigs/" + id.replace(':', '-') + ".html";
}

var dragging = false;

function get(name) {
    if (name = (new RegExp('[?&]' + encodeURIComponent(name) + '=([^&]*)')).exec(location.search))
        return decodeURIComponent(name[1]);
}

var t;
function Setup() {
    if (assets_folder != undefined) {
        var back_button = document.getElementById("back-button");
        var ref = window.name.split('|').pop();
        if (back_button && ref) {
            back_button.href = ref;
            var id = ref.split('/').pop().split('.')[0].replace(/-/g, ':');
            back_button.innerText = id;
            if (id != assets_folder.replace(/-/g, ':')) back_button.style = "display: inline-block;"
        }

        window.name = window.name + "|" + window.location.href

        if (window.name.split('|').pop().split('/').pop().split('.')[0].replace(/-/g, ':') == assets_folder.replace(/-/g, ':')) window.name = window.location.href;
    }

    if (get('pos')) {
        var pos = parseInt(get('pos'));
        var text = document.getElementsByClassName('aside-seq')[0].innerText;
        document.getElementsByClassName('aside-seq')[0].innerHTML = text.substring(0, pos) + "<span class='highlight'>" + text.substring(pos, pos + 1) + "</span>" + text.substring(pos + 1);
    }

    var elements = document.getElementsByClassName("copy-data")
    for (var i = 0; i < elements.length; i++) {
        elements[i].addEventListener("click", CopyGraphData)
        var example = elements[i].getElementsByClassName("example")[0]
        if (example)
            example.innerHTML = GetGraphDataExample(elements[i]);
    }

    var elements = document.getElementsByClassName("user-help")
    for (var i = 0; i < elements.length; i++) {
        if (!elements[i].classList.contains("copy-data")) {
            elements[i].addEventListener("click", (event) => { event.stopPropagation() })
        }
        elements[i].addEventListener("focusin", openHelp)
        elements[i].addEventListener("focusout", closeHelp)
        elements[i].addEventListener("mouseenter", openHelp)
        elements[i].addEventListener("mouseleave", closeHelp)
    }

    // Highlight the correct nodes and table row on the homepage after clicking on the point graph
    var split = window.location.href.split('#')
    if (split.length == 2) {
        var parts = split[1].split('_');
        var target = parts[parts.length - 1];
        var elements = document.querySelectorAll("[id$=\"" + target + "\"]")
        for (var i = 0; i < elements.length; i++) {
            elements[i].classList.add("highlight");
        }
    }

    SpectrumSetUp()
}

function openHelp(event) {
    var content = event.target.children[1]
    content.classList.add("focus");
    var box = content.getBoundingClientRect()
    if (box.x < 0) {
        content.style.marginLeft = -box.x + "px";
    }
    if (box.right > window.innerWidth) {
        content.style.marginLeft = (window.innerWidth - box.right - 20) + "px";
    }
    if (box.bottom > window.innerHeight) {
        content.style.marginTop = (window.innerHeight - box.bottom) + "px";
    }
}

function closeHelp(event) {
    var content = event.target.children[1]
    content.classList.remove("focus")
}

function pauseEvent(e) {
    if (e.stopPropagation) e.stopPropagation();
    if (e.preventDefault) e.preventDefault();
    e.cancelBubble = true;
    e.returnValue = false;
    return false;
}

function GoBack() {
    var history = window.name.split('|');
    history.pop(); //remove current
    var url = history.pop();
    window.name = history.join('|');
    window.location.href = url;
}

function AlignmentDetails(number) {
    AlignmentDetailsClear();
    document.getElementById("alignment-details-" + number).className += " active";
}

function AlignmentDetailsClear() {
    document.getElementById("index-menus").childNodes.forEach(a => a.className = "alignment-details");
}

// Copy the graph data for the clicked on graph
function CopyGraphData(event) {
    var t = event.target;
    if (event.target.classList.contains("mark"))
        t = event.target.parentElement.parentElement;
    if (event.target.classList.contains("copy-data"))
        t = event.target.parentElement;

    var elements = t.getElementsByClassName("graph-data");
    for (var i = 0; i < elements.length; i++) {
        elements[i].select();
        elements[i].setSelectionRange(0, 99999);
        navigator.clipboard.writeText(elements[i].value);
        return;
    }
}

/** Get the first four lines of the data to work as an example of how the data looks.
 * @param {Element} element the .copy-data element to get the example for
*/
function GetGraphDataExample(element) {
    var children = element.children;
    if (element.classList.contains("mark"))
        children = element.parentElement.parentElement.children;
    if (element.classList.contains("copy-data"))
        children = element.parentElement.children;

    var example = undefined;
    for (var i = 0; i < children.length; i++) {
        if (children[i].classList.contains("graph-data")) {
            example = children[i].value.split("\n").slice(0, 4);
            break;
        }
    }

    if (example == undefined) {
        return "No data found";
    } else {
        var output = "";
        for (var line in example) {
            output += example[line].substring(0, Math.min(30, example[line].length)).replace(/ /gi, "<span class='sp'> </span>").replace(/\t/gi, "<span class='tab'>\t</span>")
            if (example[line].length <= 30) {
                output += "<span class='nl'></span>\n";
            } else {
                output += "...\n";
            }
        }
        return output;
    }
}

function ToggleCDRReads() {
    document.querySelector(".alignment-body").classList.toggle("cdr");
}

function ToggleAlignmentComic() {
    document.querySelector(".alignment-body").classList.toggle("comic");
}

/** Update the reads alignment to only show the read supporting the given position. Or if only this position is shown remove the highlight.
 * @param {String} position the class of the position eg for position 42 this would be "a42".
 * @param {Number} alignment_position the position of this node in the template alignment.
*/
function HighlightAmbiguous(position, alignment_position) {
    var body = document.querySelector(".alignment-body")
    if (body.classList.contains("highlight") && body.dataset.position == position) {
        body.style.setProperty("--highlight-pos", "-1ch");
        body.dataset.position = undefined;
        document.querySelectorAll(".highlight").forEach(e => e.classList.remove("highlight")); // Removes highlight from all reads, the .node, and .alignment-body
    } else {
        document.querySelectorAll(".highlight").forEach(e => e.classList.remove("highlight"));
        body.style.setProperty("--highlight-pos", String(alignment_position) + "ch");
        body.dataset.position = position;
        body.classList.add("highlight")
        document.querySelector(".higher-order-graphs").classList.add("highlight")
        document.querySelectorAll("." + position).forEach(e => e.classList.add("highlight")); // Highlights all reads and the .node
    }
}

/* Spectrum viewer code */
function SpectrumSetUp() {
    var elements = document.querySelectorAll(".spectrum .peptide span");
    for (let i = 0; i < elements.length; i++) {
        elements[i].addEventListener("mouseenter", e => { ToggleHighlight(e, false, true) });
        elements[i].addEventListener("mouseleave", e => { ToggleHighlight(e, false, false) });
        elements[i].addEventListener("focusin", e => { ToggleHighlight(e, false, true) });
        elements[i].addEventListener("focusout", e => { ToggleHighlight(e, false, false) });
        elements[i].addEventListener("mouseup", e => { ToggleHighlight(e, true) });
        elements[i].addEventListener("keyup", e => { if (e.key == "Enter") ToggleHighlight(e, true) });
    }
    var elements = document.querySelectorAll(".spectrum .legend .ion");
    for (let i = 0; i < elements.length; i++) {
        elements[i].addEventListener("mouseenter", e => { ToggleHighlight(e, false, true) });
        elements[i].addEventListener("mouseleave", e => { ToggleHighlight(e, false, false) });
        elements[i].addEventListener("focusin", e => { ToggleHighlight(e, false, true) });
        elements[i].addEventListener("focusout", e => { ToggleHighlight(e, false, false) });
        elements[i].addEventListener("mouseup", e => { ToggleHighlight(e, true) });
        elements[i].addEventListener("keyup", e => { if (e.key == "Enter") ToggleHighlight(e, true) });
    }
    var elements = document.querySelectorAll(".spectrum .legend .unassigned");
    for (let i = 0; i < elements.length; i++) {
        elements[i].addEventListener("keyup", e => { if (e.key == "Enter") e.target.click() });
    }
    var elements = document.querySelectorAll(".spectrum .legend input");
    for (let i = 0; i < elements.length; i++) {
        elements[i].addEventListener("change", SpectrumInputChange);
    }
    var elements = document.querySelectorAll(".spectrum>.manual-zoom input");
    for (let i = 0; i < elements.length; i++) {
        elements[i].addEventListener("change", ManualZoom);
    }
    var elements = document.querySelectorAll(".spectrum-graph .manual-zoom input");
    for (let i = 0; i < elements.length; i++) {
        elements[i].addEventListener("change", ManualZoomSpectrumGraph);
    }
    var elements = document.querySelectorAll(".spectrum-graph .plot");
    for (let i = 0; i < elements.length; i++) {
        elements[i].addEventListener("mousemove", SpectrumGraphMouseMove)
    }
    var elements = document.querySelectorAll(".spectrum .render-setup input");
    for (let i = 0; i < elements.length; i++) {
        elements[i].addEventListener("change", RenderSetup);
    }
    var elements = document.querySelectorAll(".spectrum .wrapper");
    for (let i = 0; i < elements.length; i++) {
        elements[i].addEventListener("mousedown", spectrumDragStart)
        elements[i].addEventListener("mousemove", spectrumDrag)
        elements[i].addEventListener("mouseup", spectrumDragEnd)
        elements[i].addEventListener("mouseout", spectrumDragOut)
    }
    var elements = document.querySelectorAll(".spectrum .canvas");
    for (let i = 0; i < elements.length; i++) {
        var d = elements[i].dataset;
        d.minMz = 0;
        d.maxMz = d.initialMaxMz;
        d.maxIntensity = d.initialMaxIntensity;
    }
    var elements = document.querySelectorAll(".spectrum .zoom-out");
    for (let i = 0; i < elements.length; i++) {
        elements[i].addEventListener("mouseup", spectrumZoomOut)
        elements[i].addEventListener("keyup", e => { if (e.key == "Enter") spectrumZoomOut(e) })
    }
}

var highlight;
function ToggleHighlight(event, permanent, start = null) {
    var t = event.target; // <span> with the sequence
    if (permanent) t.classList.toggle("permanent");
    if (t.classList.contains("permanent") && !permanent) return;

    highlight = t;
    var container = t.parentElement.parentElement;
    if (container.classList.contains("ion-series")) container = container.parentElement.parentElement;
    function toggle(element) {
        if (element.dataset.n == undefined) element.dataset.n = 0;
        if (element.dataset.n < 0) element.dataset.n = 0;
        if ((permanent && t.classList.contains("permanent"))) {
            element.dataset.n = Number(element.dataset.n) + 1;
            if (element.dataset.n == 1) element.classList.add("highlight");
        }
        else if (permanent) {
            element.dataset.n = Number(element.dataset.n) - 1;
            if (element.dataset.n <= 0) element.classList.remove("highlight");
        } else {
            if (element.dataset.n == undefined || element.dataset.n == 0) {
                if (start) {
                    element.classList.add("highlight");
                } else if (start == null) {
                    element.classList.toggle("highlight");
                } else {
                    element.classList.remove("highlight");
                }
            }
        }
    }
    container.querySelectorAll(".canvas").forEach(canvas => {
        if (t.classList.contains("ion")) {
            var cl = t.classList[1];
            var elements = canvas.querySelectorAll("." + cl);
            for (let i = 0; i < elements.length; i++) {
                toggle(elements[i]);
            }
        } else {
            var peaks = canvas.children;
            for (let i = 0; i < peaks.length; i++) {
                if (peaks[i].dataset.pos == t.dataset.pos) {
                    toggle(peaks[i]);
                }
            }
        }
    })

    if (start || container.querySelector(".permanent") != null) {
        container.querySelectorAll(".canvas").forEach(canvas => {
            canvas.classList.add("highlight");
        })
    } else {
        container.querySelectorAll(".canvas").forEach(canvas => {
            canvas.classList.remove("highlight");
        })
    }
}

/** Toggle features on or off in the spectrum (eg background peaks, peak labels)
 * @param {Event} event
*/
function SpectrumInputChange(event) {
    if (event.target.type == "checkbox") { // Background
        var wrapper = event.target.parentElement.parentElement;
        var canvas = wrapper.querySelector(".canvas");
        var update_axes = !wrapper.classList.contains("first") && !wrapper.classList.contains("second");
        if (event.target.checked) { // Will be adding all background peaks
            wrapper.classList.add(event.target.className);
            if (update_axes && canvas.dataset.maxIntensity == canvas.dataset.initialMaxIntensityAssigned) {
                canvas.dataset.maxIntensity = canvas.dataset.initialMaxIntensity;
                canvas.style.setProperty("--max-intensity", canvas.dataset.maxIntensity);
                UpdateSpectrumAxes(canvas)
            }
        } else { // Will be removing all background peaks
            wrapper.classList.remove(event.target.className);
            if (update_axes && canvas.dataset.maxIntensity == canvas.dataset.initialMaxIntensity) {
                canvas.dataset.maxIntensity = canvas.dataset.initialMaxIntensityAssigned;
                canvas.style.setProperty("--max-intensity", canvas.dataset.maxIntensity);
                UpdateSpectrumAxes(canvas)
            }
        }
    } else if (event.target.type == "range") { // Peak labels
        var ele = document.getElementById(event.target.id + "_value");
        ele.value = event.target.value;
        event.target.parentElement.parentElement.parentElement.querySelectorAll(".canvas").forEach(canvas => SpectrumUpdateLabels(Number(event.target.value), canvas))
    } else if (event.target.type == "number") { // Peak labels
        var ele = document.getElementById(event.target.id.substring(0, event.target.id.length - 6));
        ele.value = event.target.value;
        event.target.parentElement.parentElement.parentElement.querySelectorAll(".canvas").forEach(canvas => SpectrumUpdateLabels(Number(event.target.value), canvas))
    }
}

/** The mouse moved in the spectrum graph.
 * @param {MouseEvent} event
*/
function SpectrumGraphMouseMove(event) {
    var data = event.target;
    if (data.classList.contains("point")) data = data.parentElement;
    var spectrum = data.parentElement.parentElement;

    var baseline = data.getBoundingClientRect().top;
    var offset = event.clientY - baseline;

    var el = spectrum.querySelector(".ruler");
    el.style.top = offset + "px";
    var el = spectrum.querySelector("#ruler-value");

    var absolute = spectrum.querySelector("#absolute").checked;
    let y = absolute ? "abs" : "rel";

    let style = getComputedStyle(data)
    let min = Number(style.getPropertyValue(`--${y}-min`));
    let max = Number(style.getPropertyValue(`--${y}-max`));
    // TODO: on a point the offsetY is wrong (from itself instead of data)
    el.innerText = fancyRound(max, min, offset / data.clientHeight * - 1 * (max - min) + max);
}

/** Zoom the spectrum manually.
 * @param {Event} event
*/
function ManualZoom(event) {
    var spectrum = event.target.parentElement.parentElement;
    var min = Number(spectrum.querySelector(".mz-min").value);
    var max = Number(spectrum.querySelector(".mz-max").value);
    var maxI = Number(spectrum.querySelector(".intensity-max").value);

    spectrum.querySelectorAll(".canvas").forEach(canvas => Zoom(canvas, min, max, maxI));
}

/** Zoom the spectrum manually.
 * @param {Event} event
*/
function ManualZoomSpectrumGraph(event) {
    var spectrum = event.target.parentElement.parentElement;
    var min_x = Number(spectrum.querySelector(".x-min").value);
    var max_x = Number(spectrum.querySelector(".x-max").value);
    var min_y = Number(spectrum.querySelector(".y-min").value);
    var max_y = Number(spectrum.querySelector(".y-max").value);
    var absolute = spectrum.querySelector("#absolute").checked;
    var mz = spectrum.querySelector("#mz").checked;

    spectrum.querySelectorAll(".data").forEach(canvas => ZoomSpectrumGraph(canvas, min_x, max_x, min_y, max_y, absolute, mz));
}

/** Setup properties of the spectrum for publication
 * @param {Event} event
*/
function RenderSetup(event) {
    var cl = event.target.className;
    var spectrum = event.target.parentElement.parentElement;
    if (cl == "width") {
        spectrum.style.setProperty("--width", event.target.value);
    } else if (cl == "height") {
        spectrum.style.setProperty("--height", event.target.value);
    } else if (cl == "fs-peptide") {
        spectrum.style.setProperty("--fs-peptide", event.target.value);
    } else if (cl == "stroke-peptide") {
        spectrum.style.setProperty("--stroke-peptide", event.target.value);
    } else if (cl == "fs-spectrum") {
        spectrum.style.setProperty("--fs-spectrum", event.target.value);
    } else if (cl == "stroke-spectrum") {
        spectrum.style.setProperty("--stroke-spectrum", event.target.value);
    } else if (cl == "compact") {
        spectrum.classList.toggle("compact");
    }
}

/** Update the spectrum to only show the label for peaks within the given percentage group
 * @param {Number} value the percentage to include (0-100)
 * @param {Element} canvas the canvas to work on
*/
function SpectrumUpdateLabels(value, canvas) {
    var style = window.getComputedStyle(canvas);
    var max = Number(style.getPropertyValue("--max-intensity"));

    var peaks = canvas.children;
    for (let i = 0; i < peaks.length; i++) {
        var v = Number(window.getComputedStyle(peaks[i]).getPropertyValue("--intensity"));
        if (v > max * (100 - value) / 100) {
            peaks[i].classList.add("label");
        } else {
            peaks[i].classList.remove("label");
        }
    }
}

var startPoint;
var selection;
var linked_selection;
var last_selection;

function spectrumDragStart(event) {
    var wrapper = event.target;
    if (wrapper.classList.contains("peptide") || wrapper.classList.contains("canvas-wrapper")) wrapper = wrapper.parentElement;
    if (wrapper.classList.contains("canvas") || wrapper.classList.contains("y-axis") || wrapper.classList.contains("x-axis")) wrapper = wrapper.parentElement.parentElement;
    if (wrapper.classList.contains("wrapper")) {
        var canvas = wrapper.querySelector(".canvas");
        startPoint = event.pageX - wrapper.getBoundingClientRect().x;
        selection = canvas.querySelector(".selection");
        if (wrapper.classList.contains("first")) {
            linked_selection = wrapper.parentElement.querySelector(".selection.second")
        } else if (wrapper.classList.contains("second")) {
            linked_selection = wrapper.parentElement.querySelector(".selection.first")
        }
        wrapper.classList.add("dragging")
        selection.hidden = false;
        if (linked_selection != undefined) {
            linked_selection.hidden = false;
            linked_selection.classList.add("linked");
        }
        updateSelections(wrapper, canvas, startPoint, event.pageY - wrapper.getBoundingClientRect().y, startPoint);
    }
}

function updateSelections(wrapper, canvas, x, y, end_x) {
    var wrapper_box = wrapper.getBoundingClientRect();
    var canvas_box = canvas.getBoundingClientRect();
    var new_x = Math.min(Math.max(0, x - (canvas_box.left - wrapper_box.left)), canvas_box.width);
    var new_y = Math.min(Math.max(1, y - (canvas_box.top - wrapper_box.top)), canvas_box.height);
    var new_end_x = Math.min(Math.max(0, end_x - (canvas_box.left - wrapper_box.left)), canvas_box.width);
    var new_w = Math.min(Math.abs(new_end_x - new_x), canvas_box.width);
    var new_x = Math.min(new_x, new_end_x);
    if (selection != undefined) updateSelection(selection, new_x, new_y, new_w);
    if (linked_selection != undefined) updateSelection(linked_selection, new_x, new_y, new_w);
}


function updateSelection(select, offsetX, offsetY, width) {
    select.style.setProperty("left", offsetX + "px")
    select.style.setProperty("width", width + "px")
    select.style.setProperty("height", select.parentElement.getBoundingClientRect().height - offsetY + "px")

    if (select.classList.contains("first")) {
        select.style.setProperty("top", offsetY + "px")
    } else {
        select.style.setProperty("bottom", offsetY + "px")
    }
}

function spectrumDrag(event) {
    if (startPoint != undefined) {
        var canvas = event.target.querySelector(".canvas");
        var box = canvas.getBoundingClientRect();
        var offsetY = event.offsetY;
        if (selection.classList.contains("second")) offsetY = box.height - offsetY;
        updateSelections(event.target, canvas, startPoint, offsetY, event.offsetX)
    } else if (last_selection != undefined && event.target.classList.contains("canvas")) {
        if (last_selection.canvas == event.target) {
            selection = last_selection.selection;
            linked_selection = last_selection.linked_selection;
        } else {
            selection = last_selection.linked_selection;
            linked_selection = last_selection.selection;
        }
        startPoint = last_selection.startPoint;
        if (selection != undefined) selection.hidden = false;
        if (linked_selection != undefined) linked_selection.hidden = false;
        event.target.parentElement.parentElement.classList.add("dragging");
        last_selection = undefined;
    }
}

function spectrumDragOut(event) {
    if (event.target.classList.contains("wrapper")) {
        if (selection != undefined)
            last_selection = { selection: selection, linked_selection: linked_selection, startPoint: startPoint, canvas: event.target.querySelector(".canvas") };
        else last_selection = undefined;
        if (selection != undefined) selection.hidden = true;
        if (linked_selection != undefined) {
            linked_selection.hidden = true;
            linked_selection.classList.remove("linked");
        }
        selection = undefined
        linked_selection = undefined
        startPoint = undefined

        document.querySelectorAll(".wrapper.dragging").forEach(e => e.classList.remove("dragging"));
    }
}

function spectrumDragEnd(event) {
    var canvas = event.target.querySelector(".canvas");
    if (startPoint != undefined) {
        var d = canvas.dataset;
        var box = canvas.getBoundingClientRect();
        var wrapper_box = event.target.getBoundingClientRect();
        var width = box.width;
        var height = box.height;
        var boxY = box.y + window.scrollY;
        var boxX = box.x + window.scrollX;
        startPoint = startPoint + wrapper_box.x - boxX;
        var min = Math.max(0, Math.min(startPoint, event.pageX - boxX)) / width;
        var max = Math.min(width, Math.max(startPoint, event.pageX - boxX)) / width;
        var minMz = Number(d.minMz);
        var maxMz = Number(d.maxMz);
        var maxIntensity = Number(d.maxIntensity);
        var min = min * Math.max(1, maxMz - minMz) + minMz;
        var max = max * Math.max(1, maxMz - minMz) + minMz;
        var offsetY = Math.min(Math.max(0, event.pageY - boxY), height);
        if (selection.classList.contains("second")) offsetY = height - offsetY;
        var maxI = (1 - Math.max(0, offsetY / height)) * maxIntensity;

        Zoom(canvas, min, max, maxI);
        if (linked_selection != undefined) Zoom(linked_selection.parentElement, min, max, maxI);
        spectrumDragOut(event)
        last_selection = undefined;
    }
}

function Zoom(canvas, min, max, maxI) {
    canvas.classList.add("zoomed");
    canvas.dataset.minMz = min;
    canvas.dataset.maxMz = max;
    canvas.dataset.maxIntensity = maxI;
    canvas.style.setProperty("--min-mz", min);
    canvas.style.setProperty("--max-mz", max);
    canvas.style.setProperty("--max-intensity", canvas.dataset.maxIntensity);

    var spectrum = canvas.parentElement.parentElement.parentElement;
    spectrum.querySelector(".mz-min").value = fancyRound(max, min, min);
    spectrum.querySelector(".mz-max").value = fancyRound(max, min, max);
    spectrum.querySelector(".intensity-max").value = fancyRound(canvas.dataset.maxIntensity, 0, canvas.dataset.maxIntensity);

    UpdateSpectrumAxes(canvas)
}

function ZoomSpectrumGraph(canvas, min_x, max_x, min_y, max_y, absolute, mz) {
    canvas.classList.add("zoomed");
    // canvas.dataset.minMz = min;
    // canvas.dataset.maxMz = max;
    // canvas.dataset.maxIntensity = maxI;
    let x = mz ? "mz" : "mass";
    let y = absolute ? "abs" : "rel";
    canvas.style.setProperty(`--${x}-min`, min_x);
    canvas.style.setProperty(`--${x}-max`, max_x);
    canvas.style.setProperty(`--${y}-min`, min_y);
    canvas.style.setProperty(`--${y}-max`, max_y);

    var spectrum_graph = canvas.parentElement.parentElement;
    spectrum_graph.querySelector(".x-min").value = fancyRound(max_x, min_x, min_x);
    spectrum_graph.querySelector(".x-max").value = fancyRound(max_x, min_x, max_x);
    spectrum_graph.querySelector(".y-min").value = fancyRound(max_y, min_y, min_y);
    spectrum_graph.querySelector(".y-max").value = fancyRound(max_y, min_y, max_y);

    spectrum_graph.querySelector(`.min.${x}`).innerText = fancyRound(max_x, min_x, min_x);
    spectrum_graph.querySelector(`.max.${x}`).innerText = fancyRound(max_x, min_x, max_x);
    spectrum_graph.querySelector(`.min.${y}`).innerText = fancyRound(max_y, min_y, min_y);
    spectrum_graph.querySelector(`.max.${y}`).innerText = fancyRound(max_y, min_y, max_y);

    if (max_y > 0 && min_y > 0) spectrum_graph.querySelector(".x-axis").classList.add("hug-bottom");
    else spectrum_graph.querySelector(".x-axis").classList.remove("hug-bottom");
}

function spectrumZoomOut(event) {
    var spectrum = event.target.parentElement.parentElement.parentElement.parentElement;
    var min, max, maxI = 0;
    spectrum.querySelectorAll(".canvas").forEach(canvas => {
        var d = canvas.dataset;
        d.minMz = 0;
        d.maxMz = d.initialMaxMz;
        d.maxIntensity = d.initialMaxIntensity;
        canvas.style.setProperty("--min-mz", 0);
        canvas.style.setProperty("--max-mz", d.initialMaxMz);
        canvas.style.setProperty("--max-intensity", d.initialMaxIntensity);
        canvas.classList.remove("zoomed");

        UpdateSpectrumAxes(canvas)
        min = d.minMz;
        max = d.maxMz;
        maxI = d.maxIntensity;
    });
    spectrum.querySelector(".mz-min").value = fancyRound(max, min, min);
    spectrum.querySelector(".mz-max").value = fancyRound(max, min, max);
    spectrum.querySelector(".intensity-max").value = fancyRound(maxI, 0, maxI);
}

function fancyRound(max, min, value) {
    var factor = max - min < 5 ? 100 : max - min < 50 ? 10 : 1;
    return Math.round(value * factor) / factor;
}

// Give the canvas element
function UpdateSpectrumAxes(canvas) {
    // Update x-axis
    var axis = canvas.parentElement.getElementsByClassName("x-axis")[0];
    var ticks = axis.children;
    var min = Number(canvas.dataset.minMz);
    var max = Number(canvas.dataset.maxMz);
    for (let i = 0; i < ticks.length; i++) {
        ticks[i].innerText = fancyRound(max, min, min + i / 4 * (max - min))
    }

    // Update y-axis
    var axis = canvas.parentElement.getElementsByClassName("y-axis")[0];
    var ticks = axis.children;
    var max = Number(canvas.dataset.maxIntensity);
    for (let i = 0; i < ticks.length; i++) {
        if (i == 0)
            ticks[i].innerText = "0";
        else
            ticks[i].innerText = Math.round(i / 4 * (max)).toExponential(2);
    }

    var peaks = canvas.children;
    for (let i = 0; i < peaks.length; i++) {
        var v = Number(window.getComputedStyle(peaks[i]).getPropertyValue("--intensity"));
        if (v > max) {
            peaks[i].classList.add("cut");
        } else {
            peaks[i].classList.remove("cut");
        }
    }
}
```
