## Supplementary material for "A handle on mass coincidence errors in *de novo* sequencing of antibodies by bottom-up proteomics": Cargo.toml

```
[package]
name = "modelangelo-benchmark"
version = "0.1.0"
edition = "2021"

# See more keys and their definitions at https://doc.rust-lang.org/cargo/reference/manifest.html

[dependencies]
rustyms = { git = "https://github.com/snijderlab/rustyms", branch = "release" }
regex = "1"
```
