## Supplementary material for "A handle on mass coincidence errors in *de novo* sequencing of antibodies by bottom-up proteomics": main.rs

```
// #![warn(clippy::unwrap_used)]
use std::{
    fs::File,
    io::{BufWriter, Write},
    process::{Command, Stdio},
};

use rustyms::{align::Alignment, identifications::FastaData, ComplexPeptide, LinearPeptide};

fn main() {
    let actual_her: (LinearPeptide, LinearPeptide) = (
        ComplexPeptide::pro_forma("EVQLVESGGGLVQPGGSLRLSCAASGFNIKDTYIHWVRQAPGKGLEWVARIYPTNGYTRYADSVKGRFTISADTSKNTAYLQMNSLRAEDTAVYYCSRWGGDGFYAMDYWGQGTLVTVSSASTKGPSVFPLAPSSKSTSGGTAALGCLVKDYFPEPVTVSWNSGALTSGVHTFPAVLQSSGLYSLSSVVTVPSSSLGTQTYICNVNHKPSNTKVDKKVEPKSCDKTHTCPPCPAPELLGGPSVFLFPPKPKDTLMISRTPEVTCVVVDVSHEDPEVKFNWYVDGVEVHNAKTKPREEQYNSTYRVVSVLTVLHQDWLNGKEYKCKVSNKALPAPIEKTISKAKGQPREPQVYTLPPSREEMTKNQVSLTCLVKGFYPSDIAVEWESNGQPENNYKTTPPVLDSDGSFFLYSKLTVDKSRWQQGNVFSCSVMHEALHNHYTQKSLSLSPGK").unwrap().assume_linear(),
        ComplexPeptide::pro_forma("DIQMTQSPSSLSASVGDRVTITCRASQDVNTAVAWYQQKPGKAPKLLIYSASFLYSGVPSRFSGSRSGTDFTLTISSLQPEDFATYYCQQHYTTPPTFGQGTKVEIKRTVAAPSVFIFPPSDEQLKSGTASVVCLLNNFYPREAKVQWKVDNALQSGNSQESVTEQDSKDSTYSLSSTLTLSKADYEKHKVYACEVTHQGLSSPVTKSFNRGEC").unwrap().assume_linear(),
    );
    let actual_cr3022: (LinearPeptide, LinearPeptide) = (
        ComplexPeptide::pro_forma("EVQLVQSGTEVKKPGESLKISCKGSGYGFITYWIGWVRQMPGKGLEWMGIIYPGDSETRYSPSFQGQVTISADKSINTAYLQWSSLKASDTAIYYCAGGSGISTPMDVWGQGTTVTVSSASTKGPSVFPLAPSSKSTSGGTAALGCLVKDYFPEPVTVSWNSGALTSGVHTFPAVLQSSGLYSLSSVVTVPSSSLGTQTYICNVNHKPSNTKVDKKVEPKSCDK").unwrap().assume_linear(),
        ComplexPeptide::pro_forma("DIVMTQSPDSLAVSLGERATINCKSSQSVLYSSINKNYLAWYQQKPGQPPKLLIYWASTRESGVPDRFSGSGSGTDFTLTISSLQAEDVAVYYCQQYYSTPYTFGQGTKVEIKRTVAAPSVFIFPPSDEQLKSGTASVVCLLNNFYPREAKVQWKVDNALQSGNSQESVTEQDSKDSTYSLSSTLTLSKADYEKHKVYACEVTHQGLSSPVTKSFNRGEC").unwrap().assume_linear(),
    );
    let actual_f59: (LinearPeptide, LinearPeptide) = (
        ComplexPeptide::pro_forma("EPELVESGGGLAQPGTSLRLSCEASGFTFDDYAMHWVRQAPGRALEWVSGISWSSDNLAYSDSVEGRFTISRDNAKNSLYLQMNSLRLDDTAFYYCAKDVRPYYDFWAFDSWGRGTPVTVSSASTKGPSVFPLAPSSKSTSGGTAALGCLVKDYFPEPVTVSWNSGALTSGVHTFPAVLQSSGLYSLSSVVTVPSSSLGTQTYICNVNHKPSNTKVDKRVEPKSCDKTHTCPPCPAPELLGGPSVFLFPPKPKDTLMISRTPEVTCVVVDVSHEDPEVKFNWYVDGVEVHNAKTKPREEQYNSTYRVVSVLTVLHQDWLNGKEYKCKVSNKALPAPIEKTISKAKGQPREPQVYTLPPSREEMTKNQVSLTCLVKGFYPSDIAVEWESNGQPENNYKTTPPVLDSDGSFFLYSKLTVDKSRWQQGNVFSCSVMHEALHNHYTQKSLSLSPG").unwrap().assume_linear(),
        ComplexPeptide::pro_forma("QSALTQPASVSGSPGQSITISCTGTSSDVGGYNYVSWYQHHPGKAPKLLISEVSDRPSGVSSRFSGSKSGNTASLTISGLQAEDESMYFCSSYTDLTSFVVFGGGTKLTVLGQPKAAPSVTLFPPSSEELQANKATLVCLISDFYPGAVTVAWKADSSPVKAGVETTTPSKQSNNKYAASSYLSLTPEQWKSHRSYSCQVTHEGSTVEKTVAPTECS").unwrap().assume_linear(),
    );
    let mut file =
        BufWriter::new(File::create("stitch_scan_results.csv").expect("Open output CSV"));
    writeln!(file, "{},Resolution", Run::header(None)).unwrap();
    // The runs for final output based analysis
    for ab in [
        ("20210325_F59_3ug_DENOVO_12", actual_f59),
        ("CR3022_DENOVO_17", actual_cr3022),
        ("200305_HER_test_04_DENOVO", actual_her),
    ] {
        for alphabet in ["blosum62", "mass_alphabet"] {
            for cutoff_score in [99, 98, 97, 96, 95, 90, 85, 80, 75, 70, 65, 60, 55, 50] {
                let run = Run::new(
                    Settings::new(
                        format!("C{cutoff_score}-{alphabet}-{}", ab.0),
                        ab.0,
                        cutoff_score,
                        0.9,
                        5,
                        5,
                        alphabet,
                    ),
                    ab.1.clone(),
                );
                match run {
                    Ok(res) => writeln!(file, "{}", res.row()).unwrap(),
                    Err(e) => println!("{e}"),
                }
                file.flush().unwrap();
            }
        }
    }
    // The two runs for read based analysis
    // Run::new(
    //     Settings::new("N", 50, 1.0, 0, 0, "blosum62"),
    //     actual.clone(),
    // )
    // .unwrap();
    // Run::new(
    //     Settings::new("MB", 50, 1.0, 0, 0, "mass_alphabet"),
    //     actual.clone(),
    // )
    // .unwrap();
}

trait Save {
    fn header(prefix: Option<&str>) -> String;
    fn row(self) -> String;
}

#[derive(Debug)]
struct Run {
    hc: (Alignment, Vec<f64>),
    lc: (Alignment, Vec<f64>),
    settings: Settings,
}

impl Save for Run {
    fn header(prefix: Option<&str>) -> String {
        format!(
            "HC NScore,HC Score,HC Id,HC Sim,HC Gap,HC Length,HC Coverage,LC NScore,LC Score,LC Id,LC Sim,LC Gap,LC Length,LC Coverage,{}",
            Settings::header(prefix)
        )
    }
    fn row(self) -> String {
        let hc_stats = self.hc.0.stats();
        let lc_stats = self.lc.0.stats();
        format!(
            "{},{},{},{},{},{},{},{},{},{},{},{},{},{},{}",
            self.hc.0.normalised_score,
            self.hc.0.absolute_score,
            hc_stats.0,
            hc_stats.1,
            hc_stats.2,
            hc_stats.3,
            self.hc.1.iter().filter(|i| **i != 0.0).count(),
            self.lc.0.normalised_score,
            self.lc.0.absolute_score,
            lc_stats.0,
            lc_stats.1,
            lc_stats.2,
            lc_stats.3,
            self.lc.1.iter().filter(|i| **i != 0.0).count(),
            self.settings.row(),
        )
    }
}

impl Run {
    fn new(settings: Settings, actual: (LinearPeptide, LinearPeptide)) -> Result<Self, String> {
        let built = stitch_run(
            &settings.name,
            settings.data,
            settings.cutoff_alc,
            settings.enforce_unique,
            settings.cutoff_score_tm,
            settings.cutoff_score_rec,
            settings.alphabet,
        )?;

        let hc = built
            .iter()
            .map(|seq| {
                (
                    rustyms::align::align(
                        seq.peptide.clone(),
                        actual.0.clone(),
                        rustyms::align::BLOSUM62,
                        rustyms::MassTolerance::Ppm(10.0),
                        rustyms::align::Type::Global,
                    ),
                    seq.full_header
                        .split(' ')
                        .last()
                        .unwrap()
                        .split(':')
                        .last()
                        .unwrap()
                        .split(',')
                        .map(|s| s.parse::<f64>().unwrap())
                        .collect(),
                )
            })
            .max_by(|a, b| {
                a.0.normalised_score
                    .partial_cmp(&b.0.normalised_score)
                    .unwrap()
            })
            .unwrap();
        let lc = built
            .iter()
            .map(|seq| {
                (
                    rustyms::align::align(
                        seq.peptide.clone(),
                        actual.1.clone(),
                        rustyms::align::BLOSUM62,
                        rustyms::MassTolerance::Ppm(10.0),
                        rustyms::align::Type::Global,
                    ),
                    seq.full_header
                        .split(' ')
                        .last()
                        .unwrap()
                        .split(':')
                        .last()
                        .unwrap()
                        .split(',')
                        .map(|s| s.parse::<f64>().unwrap())
                        .collect(),
                )
            })
            .max_by(|a, b| {
                a.0.normalised_score
                    .partial_cmp(&b.0.normalised_score)
                    .unwrap()
            })
            .unwrap();

        Ok(Self { hc, lc, settings })
    }
}

#[derive(Debug)]
struct Settings {
    name: String,
    data: &'static str,
    cutoff_alc: usize,
    enforce_unique: f64,
    cutoff_score_tm: usize,
    cutoff_score_rec: usize,
    alphabet: &'static str,
}

impl Settings {
    fn new(
        name: impl Into<String>,
        data: &'static str,
        cutoff_alc: usize,
        enforce_unique: f64,
        cutoff_score_tm: usize,
        cutoff_score_rec: usize,
        alphabet: &'static str,
    ) -> Self {
        Settings {
            name: name.into(),
            data,
            cutoff_alc,
            enforce_unique,
            cutoff_score_tm,
            cutoff_score_rec,
            alphabet,
        }
    }
}

impl Save for Settings {
    fn header(_prefix: Option<&str>) -> String {
        String::from("Name,CutoffALC,EnforceUnique,CutoffScoreTM,CutoffScoreREC,Alphabet,Data")
    }
    fn row(self) -> String {
        format!(
            "{},{},{},{},{},{},{}",
            self.name,
            self.cutoff_alc,
            self.enforce_unique,
            self.cutoff_score_tm,
            self.cutoff_score_rec,
            self.alphabet,
            self.data,
        )
    }
}

fn stitch_run(
    name: &str,
    data: &str,
    cutoff_alc: usize,
    enforce_unique: f64,
    cutoff_score_tm: usize,
    cutoff_score_rec: usize,
    alphabet: &str,
) -> Result<Vec<FastaData>, String> {
    let template = include_str!("template.txt")
        .replace("{NAME}", name)
        .replace("{DATA}", data)
        .replace("{CUTOFFALC}", &cutoff_alc.to_string())
        .replace("{ENFORCEUNIQUE}", &enforce_unique.to_string())
        .replace("{CUTOFFSCORETM}", &cutoff_score_tm.to_string())
        .replace("{CUTOFFSCOREREC}", &cutoff_score_rec.to_string())
        .replace("{ALPHABET}", alphabet);
    let path = format!("results/{name}.txt");
    {
        let mut file =
            File::create(&path).map_err(|e| format!("Could not create stitch file {e}"))?;
        file.write_all(template.as_bytes())
            .map_err(|e| format!("Could not write to stitch file {e}"))?;
    }
    // Run stitch with the new batchfile
    let _status = Command::new("bin/stitch.exe")
        .args(["run", &path])
        .stdout(Stdio::null())
        .status()
        .map_err(|e| format!("Could not run stitch {e}"))?;
    println!("Stitch built status: {_status}");
    // Do something with the result, parse the files get the stuff
    let built_rec =
        FastaData::parse_file(&format!("results/{name}/data.fasta")).map_err(|e| e.to_string())?;
    Ok(built_rec)
}
```
