## Supplementary material for "A handle on mass coincidence errors in *de novo* sequencing of antibodies by bottom-up proteomics": readme.md

# Supplementary data to: A handle on mass coincidence errors in de novo sequencing of antibodies by bottom-up proteomics

Authors: Douwe Schulte, Joost Snijder


## Overview of the data

### isobarics-statistics
The script for the generation of the data and base figure for figure 2 in the manuscript.
### mass-based-benchmark
The script for the generation of the data and base figures for figure 3 in the manuscript. Note that the `bin` subfolder contains the stitch binaries used to run the analysis as well as all data necessary to run the whole test. Using `cargo run --release` in this folder (if Rust is set up) will rerun the same analysis.
### xln-disambiguation
The data, Stitch batchfiles, and Stitch results files for figure 5 in the manuscript. Note that the raw data necessary to run these batchfiles is not included dues to size limitations. To be able to rerun these files Stitch will have to be installed in a subfolder called `bin` (copy that folder over from the previous dataset) and the raw data will have to be put in a subfolder called `raw`. As specified in the manuscript the raw data can be found under identifier PXD023419, and doi 10.6084/m9.figshare.13194005, for Herceptin and F59 respectively.  
