## Supplementary material for "A handle on mass coincidence errors in *de novo* sequencing of antibodies by bottom-up proteomics": Combined.html

Details Combined | Stitch OverviewUndefined

### Read Combined

#### Sequence (length=10)

JSSPATJNSR

#### Spectrum 3548? Spectrum 3548 The raw spectrum of this peptide as annotated by Hecklib. The fragments are coloured according to ion type (see legend). Any peaks with a star '\*' as text can be hovered over to see the full details, first the ion type second the mass shift type. By hovering over the amino acids in the peptide or ions in the legend the corresponding peaks are highlighted. By toggling the 'Unassigned' label you can turn the background (unassigned) peaks on or off in the plot. By updating the slider in the Ion legend you can update the spectrum to only show the top X% of the peaks with labels. The top X% means any peak that is within X% of the highest intensity. By dragging in the spectrum you can zoom in to a specific part of the spectrum and use 'Zoom Out' to get back to the original zoom level. The annotation of the spectrum is based on the given sequence in the peptides file and is done with different software so inconsistencies are likely. The peaks are annotated based on the given sequence, with 20 ppm tolerance.

Copy Data

##### Spectrum 3548 (TSV)

###### Preview

```
Loading example...
```

*Click on the button to copy the data to your clipboard.*

Mz MinMz MaxIntensity Max

WidthHeightPeptide font sizePeptide stroke widthSpectrum font sizeSpectrum stroke widthCompact peptide

Ion legend

wxyz

abcd

OtherUnassignedIon^Charge^_Position_Show for top:%

JSSPATJNSR

01.59e+63.17e+64.76e+66.35e+6

Zoom Out

a^+1^_2_y^+1^_1_a^+1^_2_y^+1^_1_b^+1^_2_b^+2^_4_b^+1^_2_a^+1^_3_y^+1^_2_y^+1^_2_y^+2^_4_a^+1^_3_y^+1^_2_b^+1^_3_b^+2^_6_y^+2^_5_b^+1^_3_y^+2^_5_y^+2^_6_y^+2^_6_y^+1^_3_y^+1^_3_b^+1^_4_y^+2^_7_y^+1^_3_y^+2^_7_b^+1^_4_y^+2^_8_y^+2^_8_b^+1^_5_b^+1^_5_y^+2^_9_y^+2^_9_y^+1^_4_y^+1^_4_y^+1^_4_\*\*\*b^+1^_6_b^+1^_6_y^+1^_5_y^+1^_5_y^+1^_5_y^+1^_6_y^+1^_6_b^+1^_7_y^+1^_6_b^+1^_7_y^+1^_7_y^+1^_7_y^+1^_7_b^+1^_8_b^+1^_8_b^+1^_8_y^+1^_8_y^+1^_8_y^+1^_8_b^+1^_9_y^+1^_9_y^+1^_9_y^+1^_9_

0778155623343113

Fragment Matches Table

Show background peaks

| Position | Ion type | Intensity | mz Theoretical | mz Error (Th) | mz Error (ppm) | Charge | Series Number |
| --- | --- | --- | --- | --- | --- | --- | --- |
| - | - | 3148 | 123.1 | - | - | 0 | - |
| - | - | 2.908E+04 | 123.1 | - | - | 0 | - |
| - | - | 1.331E+05 | 125.1 | - | - | 0 | - |
| - | - | 1.016E+04 | 126.1 | - | - | 0 | - |
| - | - | 1.81E+04 | 127.1 | - | - | 0 | - |
| - | - | 6711 | 128.1 | - | - | 0 | - |
| - | - | 6.864E+04 | 128.1 | - | - | 0 | - |
| - | - | 5.3E+04 | 129.1 | - | - | 0 | - |
| - | - | 1.34E+04 | 129.1 | - | - | 0 | - |
| - | - | 3599 | 129.2 | - | - | 0 | - |
| - | - | 3474 | 130.1 | - | - | 0 | - |
| - | - | 4.563E+04 | 130.1 | - | - | 0 | - |
| - | - | 2.616E+04 | 130.1 | - | - | 0 | - |
| - | - | 3994 | 130.4 | - | - | 0 | - |
| - | - | 7651 | 131.1 | - | - | 0 | - |
| - | - | 3423 | 132 | - | - | 0 | - |
| - | - | 9635 | 133.1 | - | - | 0 | - |
| - | - | 1.309E+04 | 137.1 | - | - | 0 | - |
| - | - | 2.401E+04 | 138.1 | - | - | 0 | - |
| - | - | 1.114E+04 | 139.1 | - | - | 0 | - |
| - | - | 2.68E+05 | 139.1 | - | - | 0 | - |
| - | - | 6402 | 140.1 | - | - | 0 | - |
| - | - | 1.526E+04 | 140.1 | - | - | 0 | - |
| - | - | 3.473E+04 | 141.1 | - | - | 0 | - |
| - | - | 9.262E+05 | 141.1 | - | - | 0 | - |
| - | - | 6689 | 141.1 | - | - | 0 | - |
| - | - | 8477 | 142.1 | - | - | 0 | - |
| - | - | 5.955E+04 | 142.1 | - | - | 0 | - |
| - | - | 8157 | 142.1 | - | - | 0 | - |
| - | - | 2.245E+04 | 143.1 | - | - | 0 | - |
| - | - | 3814 | 145.1 | - | - | 0 | - |
| - | - | 6.154E+04 | 145.1 | - | - | 0 | - |
| - | - | 9.802E+04 | 147.1 | - | - | 0 | - |
| - | - | 5497 | 148.9 | - | - | 0 | - |
| - | - | 8176 | 150.1 | - | - | 0 | - |
| - | - | 9246 | 151.1 | - | - | 0 | - |
| - | - | 4404 | 152.1 | - | - | 0 | - |
| - | - | 1.715E+04 | 153.1 | - | - | 0 | - |
| - | - | 7413 | 153.1 | - | - | 0 | - |
| - | - | 4887 | 155 | - | - | 0 | - |
| - | - | 4.575E+04 | 155.1 | - | - | 0 | - |
| 2 | a | 2.906E+05 | 155.1 | 0.000305 | 1.967 | +1 | 2 |
| - | - | 1.059E+04 | 156.1 | - | - | 0 | - |
| - | - | 1.271E+04 | 156.1 | - | - | 0 | - |
| - | - | 1.828E+04 | 156.1 | - | - | 0 | - |
| - | - | 2.049E+05 | 157.1 | - | - | 0 | - |
| - | - | 8.81E+05 | 157.1 | - | - | 0 | - |
| - | - | 3.349E+04 | 157.1 | - | - | 0 | - |
| - | - | 1.001E+04 | 158.1 | - | - | 0 | - |
| 10 | y | 2.728E+05 | 158.1 | 0.0003551 | 2.246 | +1 | 1 |
| - | - | 6.997E+04 | 158.1 | - | - | 0 | - |
| - | - | 1.137E+04 | 159.1 | - | - | 0 | - |
| - | - | 1.752E+04 | 159.1 | - | - | 0 | - |
| - | - | 4280 | 160 | - | - | 0 | - |
| - | - | 4599 | 164.1 | - | - | 0 | - |
| - | - | 9550 | 165.1 | - | - | 0 | - |
| - | - | 1.187E+04 | 166.1 | - | - | 0 | - |
| - | - | 8535 | 167 | - | - | 0 | - |
| - | - | 2.964E+05 | 167.1 | - | - | 0 | - |
| - | - | 1.03E+04 | 167.1 | - | - | 0 | - |
| - | - | 6101 | 168.1 | - | - | 0 | - |
| - | - | 2.32E+04 | 168.1 | - | - | 0 | - |
| - | - | 2.65E+04 | 169.1 | - | - | 0 | - |
| - | - | 1.032E+06 | 169.1 | - | - | 0 | - |
| - | - | 4.56E+04 | 169.1 | - | - | 0 | - |
| - | - | 1.015E+04 | 170.1 | - | - | 0 | - |
| - | - | 8.311E+04 | 170.1 | - | - | 0 | - |
| - | - | 7869 | 171.1 | - | - | 0 | - |
| - | - | 4826 | 171.1 | - | - | 0 | - |
| - | - | 1.087E+04 | 171.1 | - | - | 0 | - |
| - | - | 1.562E+05 | 173.1 | - | - | 0 | - |
| 2 | a | 1.183E+06 | 173.1 | 0.0003452 | 1.994 | +1 | 2 |
| - | - | 1.487E+04 | 173.4 | - | - | 0 | - |
| - | - | 6.975E+04 | 174.1 | - | - | 0 | - |
| - | - | 1.029E+04 | 174.1 | - | - | 0 | - |
| - | - | 9.828E+04 | 174.1 | - | - | 0 | - |
| - | - | 8.377E+04 | 175.1 | - | - | 0 | - |
| 10 | y | 6.717E+05 | 175.1 | 0.0003258 | 1.86 | +1 | 1 |
| - | - | 4.464E+04 | 176.1 | - | - | 0 | - |
| - | - | 1.512E+04 | 179.1 | - | - | 0 | - |
| - | - | 1.382E+04 | 181.1 | - | - | 0 | - |
| - | - | 1.401E+04 | 182.1 | - | - | 0 | - |
| 2 | b | 1.847E+06 | 183.1 | 0.0003703 | 2.022 | +1 | 2 |
| - | - | 1.608E+04 | 183.1 | - | - | 0 | - |
| - | - | 2.319E+04 | 184.1 | - | - | 0 | - |
| 4 | b | 6189 | 184.1 | 0.002064 | 11.21 | +2 | 4 |
| - | - | 3.336E+04 | 184.1 | - | - | 0 | - |
| - | - | 1.817E+05 | 184.1 | - | - | 0 | - |
| - | - | 2.583E+04 | 185.1 | - | - | 0 | - |
| - | - | 7.829E+05 | 185.1 | - | - | 0 | - |
| - | - | 6114 | 186.1 | - | - | 0 | - |
| - | - | 6.009E+04 | 186.1 | - | - | 0 | - |
| - | - | 2.663E+04 | 186.1 | - | - | 0 | - |
| - | - | 8860 | 187.1 | - | - | 0 | - |
| - | - | 8796 | 187.1 | - | - | 0 | - |
| - | - | 3.234E+04 | 187.1 | - | - | 0 | - |
| - | - | 3.475E+05 | 187.1 | - | - | 0 | - |
| - | - | 2.92E+04 | 188.1 | - | - | 0 | - |
| - | - | 1.1E+04 | 192.1 | - | - | 0 | - |
| - | - | 4298 | 193.1 | - | - | 0 | - |
| - | - | 1.62E+04 | 193.1 | - | - | 0 | - |
| - | - | 9467 | 194.1 | - | - | 0 | - |
| - | - | 2.256E+04 | 195.1 | - | - | 0 | - |
| - | - | 4336 | 195.1 | - | - | 0 | - |
| - | - | 7678 | 196.1 | - | - | 0 | - |
| - | - | 8096 | 196.1 | - | - | 0 | - |
| - | - | 8017 | 197.1 | - | - | 0 | - |
| - | - | 8.557E+04 | 197.1 | - | - | 0 | - |
| - | - | 6778 | 198.1 | - | - | 0 | - |
| - | - | 6194 | 198.1 | - | - | 0 | - |
| - | - | 1.463E+04 | 199.1 | - | - | 0 | - |
| - | - | 1.984E+04 | 200.1 | - | - | 0 | - |
| 2 | b | 1.062E+06 | 201.1 | 0.0002731 | 1.358 | +1 | 2 |
| - | - | 1.662E+05 | 202.1 | - | - | 0 | - |
| - | - | 4.982E+04 | 202.1 | - | - | 0 | - |
| - | - | 1.011E+05 | 202.1 | - | - | 0 | - |
| - | - | 8398 | 203.1 | - | - | 0 | - |
| - | - | 1.411E+04 | 203.1 | - | - | 0 | - |
| - | - | 1.455E+04 | 203.1 | - | - | 0 | - |
| - | - | 6890 | 203.1 | - | - | 0 | - |
| - | - | 8439 | 207.1 | - | - | 0 | - |
| - | - | 7.687E+04 | 208.1 | - | - | 0 | - |
| - | - | 7398 | 208.1 | - | - | 0 | - |
| - | - | 1.688E+04 | 209.1 | - | - | 0 | - |
| - | - | 1.534E+04 | 209.1 | - | - | 0 | - |
| - | - | 6380 | 209.1 | - | - | 0 | - |
| - | - | 2.187E+04 | 210.1 | - | - | 0 | - |
| - | - | 1.419E+05 | 210.1 | - | - | 0 | - |
| - | - | 3.975E+04 | 211.1 | - | - | 0 | - |
| - | - | 1.473E+04 | 211.1 | - | - | 0 | - |
| - | - | 1.141E+04 | 211.1 | - | - | 0 | - |
| - | - | 1.546E+04 | 212.1 | - | - | 0 | - |
| - | - | 8025 | 213.1 | - | - | 0 | - |
| - | - | 5664 | 214.2 | - | - | 0 | - |
| - | - | 1.95E+05 | 215.1 | - | - | 0 | - |
| - | - | 2.83E+04 | 216.1 | - | - | 0 | - |
| - | - | 1.558E+04 | 219.1 | - | - | 0 | - |
| - | - | 6728 | 219.1 | - | - | 0 | - |
| - | - | 8040 | 220.1 | - | - | 0 | - |
| - | - | 1.936E+04 | 220.1 | - | - | 0 | - |
| - | - | 7.098E+04 | 221.1 | - | - | 0 | - |
| - | - | 3.403E+04 | 222.1 | - | - | 0 | - |
| - | - | 1.761E+04 | 223.1 | - | - | 0 | - |
| - | - | 7644 | 223.1 | - | - | 0 | - |
| - | - | 2.39E+04 | 224.1 | - | - | 0 | - |
| - | - | 8.901E+04 | 224.1 | - | - | 0 | - |
| - | - | 1.941E+04 | 225.1 | - | - | 0 | - |
| - | - | 2.85E+05 | 225.1 | - | - | 0 | - |
| - | - | 6362 | 225.2 | - | - | 0 | - |
| - | - | 1.058E+05 | 226.1 | - | - | 0 | - |
| - | - | 8254 | 226.2 | - | - | 0 | - |
| - | - | 6978 | 227.1 | - | - | 0 | - |
| - | - | 6.236E+04 | 227.1 | - | - | 0 | - |
| - | - | 2.079E+04 | 228.1 | - | - | 0 | - |
| - | - | 6181 | 228.1 | - | - | 0 | - |
| - | - | 1.342E+05 | 228.1 | - | - | 0 | - |
| - | - | 7692 | 229.1 | - | - | 0 | - |
| - | - | 1.763E+04 | 229.1 | - | - | 0 | - |
| - | - | 8629 | 232.1 | - | - | 0 | - |
| - | - | 4588 | 232.2 | - | - | 0 | - |
| - | - | 4.266E+04 | 236.1 | - | - | 0 | - |
| - | - | 1.31E+04 | 237.1 | - | - | 0 | - |
| - | - | 1.383E+04 | 237.1 | - | - | 0 | - |
| - | - | 5900 | 237.1 | - | - | 0 | - |
| - | - | 1.708E+06 | 238.1 | - | - | 0 | - |
| - | - | 1.966E+05 | 239.1 | - | - | 0 | - |
| - | - | 6854 | 240.1 | - | - | 0 | - |
| - | - | 1.137E+04 | 240.1 | - | - | 0 | - |
| - | - | 2.393E+04 | 240.1 | - | - | 0 | - |
| - | - | 2.264E+04 | 240.2 | - | - | 0 | - |
| - | - | 8090 | 241.2 | - | - | 0 | - |
| - | - | 2.087E+04 | 242.1 | - | - | 0 | - |
| 3 | a | 1.294E+05 | 242.1 | 0.0004269 | 1.763 | +1 | 3 |
| - | - | 1.536E+04 | 243.1 | - | - | 0 | - |
| - | - | 7991 | 243.2 | - | - | 0 | - |
| 9 | y | 4.45E+04 | 244.1 | 0.0002396 | 0.9815 | +1 | 2 |
| 9 | y | 2.951E+05 | 245.1 | 0.0004007 | 1.635 | +1 | 2 |
| 7 | y | 1.435E+04 | 245.1 | 0.0007327 | 2.989 | +2 | 4 |
| - | - | 8784 | 246.1 | - | - | 0 | - |
| - | - | 2.701E+04 | 246.1 | - | - | 0 | - |
| - | - | 4426 | 246.1 | - | - | 0 | - |
| - | - | 2.526E+04 | 250.1 | - | - | 0 | - |
| - | - | 3.606E+04 | 252.1 | - | - | 0 | - |
| - | - | 5.761E+05 | 252.1 | - | - | 0 | - |
| - | - | 2.723E+04 | 253.1 | - | - | 0 | - |
| - | - | 6.949E+04 | 253.1 | - | - | 0 | - |
| - | - | 8.504E+04 | 254.1 | - | - | 0 | - |
| - | - | 3.264E+04 | 255.1 | - | - | 0 | - |
| - | - | 1.197E+04 | 255.1 | - | - | 0 | - |
| - | - | 4.654E+05 | 256.1 | - | - | 0 | - |
| - | - | 4.675E+04 | 257.1 | - | - | 0 | - |
| 3 | a | 5494 | 260.2 | 0.0004671 | 1.795 | +1 | 3 |
| 9 | y | 4.041E+05 | 262.2 | 0.0002645 | 1.009 | +1 | 2 |
| - | - | 4.17E+04 | 263.2 | - | - | 0 | - |
| - | - | 4.84E+04 | 264.1 | - | - | 0 | - |
| - | - | 7224 | 266.1 | - | - | 0 | - |
| - | - | 7043 | 266.1 | - | - | 0 | - |
| - | - | 4.682E+04 | 268.1 | - | - | 0 | - |
| - | - | 1.502E+05 | 268.2 | - | - | 0 | - |
| - | - | 1.591E+04 | 269.2 | - | - | 0 | - |
| - | - | 6.767E+04 | 270.1 | - | - | 0 | - |
| 3 | b | 2.567E+06 | 270.1 | 0.0003396 | 1.257 | +1 | 3 |
| - | - | 3.501E+05 | 271.1 | - | - | 0 | - |
| - | - | 1.729E+04 | 272.1 | - | - | 0 | - |
| - | - | 1.411E+04 | 272.1 | - | - | 0 | - |
| - | - | 2.007E+04 | 272.2 | - | - | 0 | - |
| - | - | 8685 | 273.2 | - | - | 0 | - |
| - | - | 6712 | 276.1 | - | - | 0 | - |
| 6 | b | 7983 | 279.2 | 0.005248 | 18.8 | +2 | 6 |
| - | - | 1.559E+04 | 280.1 | - | - | 0 | - |
| - | - | 1.25E+04 | 280.2 | - | - | 0 | - |
| - | - | 2.198E+04 | 281.1 | - | - | 0 | - |
| - | - | 3.794E+04 | 282.1 | - | - | 0 | - |
| - | - | 6391 | 282.1 | - | - | 0 | - |
| - | - | 2.225E+04 | 282.2 | - | - | 0 | - |
| - | - | 4692 | 283.1 | - | - | 0 | - |
| - | - | 6797 | 283.1 | - | - | 0 | - |
| - | - | 1.652E+04 | 284.2 | - | - | 0 | - |
| - | - | 1.138E+04 | 285.2 | - | - | 0 | - |
| - | - | 1.606E+04 | 286.2 | - | - | 0 | - |
| 6 | y | 3.928E+04 | 286.7 | 0.0003179 | 1.109 | +2 | 5 |
| - | - | 6529 | 287.2 | - | - | 0 | - |
| 3 | b | 1.548E+05 | 288.2 | 0.0004255 | 1.477 | +1 | 3 |
| - | - | 1.588E+04 | 289.2 | - | - | 0 | - |
| - | - | 1.964E+04 | 294.1 | - | - | 0 | - |
| - | - | 1.499E+04 | 295.1 | - | - | 0 | - |
| 6 | y | 4.495E+04 | 295.7 | 0.0004677 | 1.582 | +2 | 5 |
| - | - | 6027 | 296.1 | - | - | 0 | - |
| - | - | 2.071E+04 | 296.2 | - | - | 0 | - |
| - | - | 8714 | 297.1 | - | - | 0 | - |
| - | - | 2.097E+04 | 297.2 | - | - | 0 | - |
| - | - | 8917 | 298.1 | - | - | 0 | - |
| - | - | 1.5E+04 | 298.2 | - | - | 0 | - |
| - | - | 8428 | 298.2 | - | - | 0 | - |
| - | - | 5.915E+04 | 299.1 | - | - | 0 | - |
| - | - | 2.243E+04 | 300.1 | - | - | 0 | - |
| - | - | 8505 | 300.1 | - | - | 0 | - |
| - | - | 1.129E+04 | 300.7 | - | - | 0 | - |
| - | - | 1.043E+04 | 301.2 | - | - | 0 | - |
| - | - | 1.041E+04 | 304.1 | - | - | 0 | - |
| - | - | 5102 | 304.2 | - | - | 0 | - |
| - | - | 1.212E+04 | 306.1 | - | - | 0 | - |
| - | - | 1.916E+04 | 307.1 | - | - | 0 | - |
| - | - | 2.253E+04 | 307.1 | - | - | 0 | - |
| - | - | 1.475E+04 | 308.1 | - | - | 0 | - |
| - | - | 6707 | 309.2 | - | - | 0 | - |
| - | - | 6258 | 310.2 | - | - | 0 | - |
| - | - | 8986 | 311.1 | - | - | 0 | - |
| - | - | 3.76E+04 | 311.2 | - | - | 0 | - |
| - | - | 3.546E+04 | 311.2 | - | - | 0 | - |
| - | - | 4.973E+04 | 312.2 | - | - | 0 | - |
| - | - | 2.694E+04 | 313.2 | - | - | 0 | - |
| - | - | 7157 | 313.7 | - | - | 0 | - |
| - | - | 2.462E+04 | 314.1 | - | - | 0 | - |
| - | - | 6250 | 315.1 | - | - | 0 | - |
| - | - | 6.074E+04 | 315.2 | - | - | 0 | - |
| - | - | 2.931E+04 | 316.2 | - | - | 0 | - |
| - | - | 1.713E+04 | 317.1 | - | - | 0 | - |
| - | - | 8714 | 317.2 | - | - | 0 | - |
| - | - | 7.201E+04 | 321.2 | - | - | 0 | - |
| - | - | 1.88E+04 | 322.1 | - | - | 0 | - |
| - | - | 1.051E+04 | 322.2 | - | - | 0 | - |
| 5 | y | 9.961E+04 | 322.2 | 0.0003462 | 1.075 | +2 | 6 |
| - | - | 2.452E+04 | 322.7 | - | - | 0 | - |
| - | - | 6028 | 323.1 | - | - | 0 | - |
| - | - | 6.706E+04 | 324.1 | - | - | 0 | - |
| - | - | 3.339E+04 | 325.1 | - | - | 0 | - |
| - | - | 4.801E+04 | 325.2 | - | - | 0 | - |
| - | - | 6404 | 326.1 | - | - | 0 | - |
| - | - | 7507 | 327.1 | - | - | 0 | - |
| - | - | 7838 | 327.2 | - | - | 0 | - |
| - | - | 1.233E+05 | 329.2 | - | - | 0 | - |
| - | - | 8802 | 329.2 | - | - | 0 | - |
| - | - | 2.236E+04 | 330.2 | - | - | 0 | - |
| 5 | y | 6.437E+04 | 331.2 | 0.0006181 | 1.866 | +2 | 6 |
| - | - | 2.105E+04 | 331.7 | - | - | 0 | - |
| - | - | 3.525E+04 | 334.2 | - | - | 0 | - |
| - | - | 8329 | 335.1 | - | - | 0 | - |
| - | - | 1.052E+04 | 335.2 | - | - | 0 | - |
| - | - | 1.308E+04 | 335.2 | - | - | 0 | - |
| - | - | 1.197E+04 | 336.2 | - | - | 0 | - |
| - | - | 1.707E+04 | 337.2 | - | - | 0 | - |
| - | - | 8.75E+04 | 337.2 | - | - | 0 | - |
| - | - | 1.531E+04 | 338.2 | - | - | 0 | - |
| - | - | 2.45E+05 | 339.2 | - | - | 0 | - |
| - | - | 3.417E+04 | 339.2 | - | - | 0 | - |
| - | - | 3.786E+04 | 340.2 | - | - | 0 | - |
| - | - | 1.354E+05 | 341.2 | - | - | 0 | - |
| - | - | 4834 | 341.2 | - | - | 0 | - |
| - | - | 2.702E+05 | 342.1 | - | - | 0 | - |
| - | - | 2.68E+04 | 343.1 | - | - | 0 | - |
| - | - | 4.991E+04 | 343.2 | - | - | 0 | - |
| - | - | 9890 | 344.2 | - | - | 0 | - |
| - | - | 9056 | 349.2 | - | - | 0 | - |
| - | - | 2.548E+04 | 349.2 | - | - | 0 | - |
| - | - | 6750 | 349.7 | - | - | 0 | - |
| - | - | 8523 | 351.2 | - | - | 0 | - |
| - | - | 6190 | 352.1 | - | - | 0 | - |
| - | - | 1.525E+04 | 353.1 | - | - | 0 | - |
| - | - | 9095 | 353.2 | - | - | 0 | - |
| - | - | 6194 | 353.2 | - | - | 0 | - |
| - | - | 6701 | 353.7 | - | - | 0 | - |
| - | - | 5.333E+04 | 355.2 | - | - | 0 | - |
| - | - | 7.757E+04 | 355.2 | - | - | 0 | - |
| - | - | 1.334E+04 | 356.2 | - | - | 0 | - |
| - | - | 1.303E+04 | 356.2 | - | - | 0 | - |
| - | - | 1.98E+05 | 357.2 | - | - | 0 | - |
| - | - | 8448 | 357.2 | - | - | 0 | - |
| 8 | y | 7.247E+04 | 358.2 | 0.001214 | 3.391 | +1 | 3 |
| 8 | y | 5.187E+05 | 359.2 | 0.0002436 | 0.6783 | +1 | 3 |
| - | - | 7.728E+04 | 360.2 | - | - | 0 | - |
| - | - | 6066 | 360.7 | - | - | 0 | - |
| - | - | 6868 | 361.7 | - | - | 0 | - |
| - | - | 1.743E+04 | 362.2 | - | - | 0 | - |
| - | - | 5578 | 362.5 | - | - | 0 | - |
| - | - | 6835 | 362.7 | - | - | 0 | - |
| - | - | 1.702E+04 | 363.2 | - | - | 0 | - |
| - | - | 5071 | 364.2 | - | - | 0 | - |
| - | - | 7078 | 365.1 | - | - | 0 | - |
| - | - | 1.22E+05 | 365.2 | - | - | 0 | - |
| - | - | 9572 | 366.2 | - | - | 0 | - |
| - | - | 1.797E+04 | 366.2 | - | - | 0 | - |
| - | - | 1.078E+04 | 367.2 | - | - | 0 | - |
| 4 | b | 2.534E+04 | 367.2 | 0.0003406 | 0.9275 | +1 | 4 |
| - | - | 4950 | 368.2 | - | - | 0 | - |
| - | - | 1.094E+05 | 369.7 | - | - | 0 | - |
| - | - | 4.897E+04 | 370.2 | - | - | 0 | - |
| 4 | y | 9.712E+05 | 370.7 | 0.0002704 | 0.7295 | +2 | 7 |
| - | - | 3.744E+05 | 371.2 | - | - | 0 | - |
| - | - | 6.489E+04 | 371.7 | - | - | 0 | - |
| - | - | 7536 | 372.2 | - | - | 0 | - |
| 8 | y | 1.121E+06 | 376.2 | 0.0003364 | 0.8941 | +1 | 3 |
| - | - | 1.895E+05 | 377.2 | - | - | 0 | - |
| - | - | 2.618E+04 | 378.2 | - | - | 0 | - |
| - | - | 2.083E+05 | 378.7 | - | - | 0 | - |
| - | - | 8.914E+04 | 379.2 | - | - | 0 | - |
| 4 | y | 5.408E+06 | 379.7 | 0.0002676 | 0.7048 | +2 | 7 |
| - | - | 2.136E+06 | 380.2 | - | - | 0 | - |
| - | - | 5.189E+05 | 380.7 | - | - | 0 | - |
| - | - | 1.296E+04 | 381.2 | - | - | 0 | - |
| - | - | 3.837E+04 | 381.2 | - | - | 0 | - |
| - | - | 1.175E+04 | 382.2 | - | - | 0 | - |
| - | - | 2.072E+04 | 382.2 | - | - | 0 | - |
| - | - | 1.021E+05 | 383.2 | - | - | 0 | - |
| - | - | 1.464E+04 | 384.2 | - | - | 0 | - |
| - | - | 1.842E+04 | 384.2 | - | - | 0 | - |
| - | - | 3.93E+04 | 384.7 | - | - | 0 | - |
| 4 | b | 5.798E+04 | 385.2 | 0.0009773 | 2.537 | +1 | 4 |
| - | - | 1.124E+04 | 386.2 | - | - | 0 | - |
| - | - | 2.755E+04 | 387.2 | - | - | 0 | - |
| - | - | 8015 | 388.2 | - | - | 0 | - |
| - | - | 6484 | 388.2 | - | - | 0 | - |
| - | - | 2.549E+04 | 390.2 | - | - | 0 | - |
| - | - | 3.45E+04 | 391.2 | - | - | 0 | - |
| - | - | 8683 | 391.7 | - | - | 0 | - |
| - | - | 5561 | 392.2 | - | - | 0 | - |
| - | - | 7892 | 393.2 | - | - | 0 | - |
| - | - | 1.223E+04 | 394.2 | - | - | 0 | - |
| - | - | 7757 | 396.2 | - | - | 0 | - |
| - | - | 3.498E+04 | 396.7 | - | - | 0 | - |
| - | - | 1.663E+04 | 397.2 | - | - | 0 | - |
| - | - | 2.625E+04 | 398.2 | - | - | 0 | - |
| - | - | 6078 | 399.2 | - | - | 0 | - |
| - | - | 1.467E+04 | 399.2 | - | - | 0 | - |
| - | - | 8270 | 400.2 | - | - | 0 | - |
| - | - | 3.807E+04 | 400.2 | - | - | 0 | - |
| - | - | 8110 | 401.2 | - | - | 0 | - |
| - | - | 2.791E+05 | 405.2 | - | - | 0 | - |
| - | - | 1.945E+05 | 405.7 | - | - | 0 | - |
| - | - | 7.232E+04 | 406.2 | - | - | 0 | - |
| - | - | 1.021E+04 | 406.2 | - | - | 0 | - |
| - | - | 1.901E+04 | 406.7 | - | - | 0 | - |
| - | - | 9478 | 407.2 | - | - | 0 | - |
| - | - | 5.833E+04 | 408.2 | - | - | 0 | - |
| - | - | 8901 | 408.2 | - | - | 0 | - |
| - | - | 1.536E+04 | 409.2 | - | - | 0 | - |
| - | - | 6457 | 410.2 | - | - | 0 | - |
| - | - | 1.134E+04 | 410.2 | - | - | 0 | - |
| - | - | 9338 | 410.2 | - | - | 0 | - |
| - | - | 9941 | 411.2 | - | - | 0 | - |
| - | - | 1.464E+04 | 412.2 | - | - | 0 | - |
| 3 | y | 3.767E+06 | 414.2 | 0.0006747 | 1.629 | +2 | 8 |
| - | - | 1.681E+06 | 414.7 | - | - | 0 | - |
| - | - | 4.26E+05 | 415.2 | - | - | 0 | - |
| - | - | 3.767E+04 | 415.7 | - | - | 0 | - |
| - | - | 4.432E+04 | 416.2 | - | - | 0 | - |
| - | - | 9462 | 417.2 | - | - | 0 | - |
| - | - | 8037 | 420.2 | - | - | 0 | - |
| - | - | 2.995E+04 | 420.2 | - | - | 0 | - |
| 3 | y | 3.297E+06 | 423.2 | 0.0005803 | 1.371 | +2 | 8 |
| - | - | 1.391E+06 | 423.7 | - | - | 0 | - |
| - | - | 3.031E+05 | 424.2 | - | - | 0 | - |
| - | - | 3.063E+04 | 424.7 | - | - | 0 | - |
| - | - | 5.662E+04 | 426.2 | - | - | 0 | - |
| - | - | 1.069E+04 | 427.2 | - | - | 0 | - |
| - | - | 1.051E+04 | 428.2 | - | - | 0 | - |
| - | - | 2.084E+04 | 428.3 | - | - | 0 | - |
| - | - | 8944 | 429.2 | - | - | 0 | - |
| - | - | 1.459E+04 | 430.2 | - | - | 0 | - |
| - | - | 7.165E+04 | 434.2 | - | - | 0 | - |
| - | - | 9604 | 435.2 | - | - | 0 | - |
| - | - | 2.07E+04 | 436.2 | - | - | 0 | - |
| - | - | 1.03E+04 | 436.2 | - | - | 0 | - |
| - | - | 3.102E+04 | 437.2 | - | - | 0 | - |
| - | - | 1.52E+04 | 438.2 | - | - | 0 | - |
| 5 | b | 5.127E+04 | 438.2 | 0.0006414 | 1.463 | +1 | 5 |
| - | - | 9158 | 440.2 | - | - | 0 | - |
| - | - | 7098 | 441.2 | - | - | 0 | - |
| - | - | 3.722E+04 | 442.3 | - | - | 0 | - |
| - | - | 9591 | 443.2 | - | - | 0 | - |
| - | - | 6947 | 443.3 | - | - | 0 | - |
| - | - | 1.89E+04 | 444.2 | - | - | 0 | - |
| - | - | 7326 | 445.2 | - | - | 0 | - |
| - | - | 1.251E+04 | 447.3 | - | - | 0 | - |
| - | - | 6.201E+04 | 448.7 | - | - | 0 | - |
| - | - | 3.193E+04 | 449.2 | - | - | 0 | - |
| - | - | 1.466E+04 | 449.7 | - | - | 0 | - |
| - | - | 8812 | 451.2 | - | - | 0 | - |
| - | - | 5824 | 452.2 | - | - | 0 | - |
| - | - | 9.765E+04 | 452.3 | - | - | 0 | - |
| - | - | 7549 | 453.2 | - | - | 0 | - |
| - | - | 3.515E+04 | 453.3 | - | - | 0 | - |
| - | - | 6943 | 453.7 | - | - | 0 | - |
| - | - | 1.251E+04 | 454.2 | - | - | 0 | - |
| - | - | 2.582E+04 | 454.2 | - | - | 0 | - |
| - | - | 7.483E+04 | 455.2 | - | - | 0 | - |
| 5 | b | 5.037E+04 | 456.2 | 0.0007681 | 1.683 | +1 | 5 |
| - | - | 1.549E+04 | 457.2 | - | - | 0 | - |
| 2 | y | 8.834E+04 | 457.7 | 0.0004991 | 1.09 | +2 | 9 |
| - | - | 3.58E+04 | 458.2 | - | - | 0 | - |
| - | - | 7586 | 458.7 | - | - | 0 | - |
| - | - | 1.382E+04 | 462.2 | - | - | 0 | - |
| - | - | 6.238E+04 | 462.7 | - | - | 0 | - |
| - | - | 3.558E+04 | 463.2 | - | - | 0 | - |
| - | - | 8753 | 463.7 | - | - | 0 | - |
| 2 | y | 5.744E+04 | 466.7 | 0.0009236 | 1.979 | +2 | 9 |
| - | - | 3.222E+04 | 467.2 | - | - | 0 | - |
| - | - | 6128 | 468.2 | - | - | 0 | - |
| - | - | 2.062E+04 | 469.2 | - | - | 0 | - |
| - | - | 1.081E+04 | 469.3 | - | - | 0 | - |
| - | - | 4.738E+04 | 470.3 | - | - | 0 | - |
| - | - | 6879 | 470.8 | - | - | 0 | - |
| 7 | y | 4.598E+04 | 471.3 | 0.0005006 | 1.062 | +1 | 4 |
| - | - | 7.492E+04 | 471.7 | - | - | 0 | - |
| 7 | y | 1.363E+05 | 472.3 | 0.000721 | 1.527 | +1 | 4 |
| - | - | 1.089E+04 | 472.7 | - | - | 0 | - |
| - | - | 2.704E+04 | 473.3 | - | - | 0 | - |
| - | - | 5897 | 477.3 | - | - | 0 | - |
| - | - | 2.437E+04 | 479.3 | - | - | 0 | - |
| - | - | 3.831E+04 | 480.2 | - | - | 0 | - |
| - | - | 1.047E+04 | 481.3 | - | - | 0 | - |
| - | - | 1.325E+04 | 483.2 | - | - | 0 | - |
| - | - | 8854 | 484.8 | - | - | 0 | - |
| - | - | 1.832E+04 | 487.3 | - | - | 0 | - |
| - | - | 8350 | 488.3 | - | - | 0 | - |
| 7 | y | 1.579E+06 | 489.3 | 0.0005619 | 1.148 | +1 | 4 |
| - | - | 3.89E+05 | 490.3 | - | - | 0 | - |
| - | - | 6.74E+04 | 491.3 | - | - | 0 | - |
| - | - | 1.201E+04 | 495.3 | - | - | 0 | - |
| - | - | 2.516E+04 | 496.8 | - | - | 0 | - |
| - | - | 7.267E+04 | 497.3 | - | - | 0 | - |
| - | - | 1.814E+04 | 498.3 | - | - | 0 | - |
| - | - | 1.711E+04 | 499.3 | - | - | 0 | - |
| - | - | 2.356E+04 | 503.3 | - | - | 0 | - |
| - | - | 1.147E+04 | 504.2 | - | - | 0 | - |
| - | - | 1.521E+05 | 505.3 | - | - | 0 | - |
| - | - | 9.572E+04 | 505.8 | - | - | 0 | - |
| - | - | 4.154E+04 | 506.3 | - | - | 0 | - |
| - | - | 1.373E+04 | 511.3 | - | - | 0 | - |
| - | - | 8857 | 512.2 | - | - | 0 | - |
| - | - | 1.254E+04 | 513.3 | - | - | 0 | - |
| 0 | Precursor | 2.649E+05 | 514.3 | 9.309E-05 | 0.181 | +2 | -1 |
| 0 | Precursor | 1.708E+05 | 514.8 | 0.009184 | 17.84 | +2 | -1 |
| - | - | 5.415E+04 | 515.3 | - | - | 0 | - |
| - | - | 5986 | 521.2 | - | - | 0 | - |
| - | - | 7.322E+04 | 521.3 | - | - | 0 | - |
| - | - | 2.468E+04 | 522.3 | - | - | 0 | - |
| 0 | Precursor | 5.034E+05 | 523.3 | 0.0003649 | 0.6974 | +2 | -1 |
| - | - | 2.734E+05 | 523.8 | - | - | 0 | - |
| - | - | 9.545E+04 | 524.3 | - | - | 0 | - |
| - | - | 1.404E+04 | 524.8 | - | - | 0 | - |
| - | - | 8367 | 528.3 | - | - | 0 | - |
| - | - | 1.524E+04 | 529.3 | - | - | 0 | - |
| - | - | 9922 | 530.3 | - | - | 0 | - |
| - | - | 7009 | 531.3 | - | - | 0 | - |
| - | - | 1.531E+04 | 537.3 | - | - | 0 | - |
| - | - | 1.797E+04 | 538.3 | - | - | 0 | - |
| 6 | b | 7.425E+04 | 539.3 | 0.0001011 | 0.1874 | +1 | 6 |
| - | - | 2.368E+04 | 540.3 | - | - | 0 | - |
| - | - | 6374 | 543.3 | - | - | 0 | - |
| - | - | 8099 | 545.8 | - | - | 0 | - |
| - | - | 1.468E+05 | 546.3 | - | - | 0 | - |
| - | - | 3.489E+04 | 547.3 | - | - | 0 | - |
| - | - | 1.732E+04 | 548.3 | - | - | 0 | - |
| - | - | 2.51E+04 | 549.3 | - | - | 0 | - |
| - | - | 9842 | 550.3 | - | - | 0 | - |
| - | - | 4.292E+04 | 555.3 | - | - | 0 | - |
| - | - | 4.767E+04 | 556.3 | - | - | 0 | - |
| 6 | b | 3.472E+04 | 557.3 | 0.002121 | 3.806 | +1 | 6 |
| - | - | 6518 | 558.3 | - | - | 0 | - |
| - | - | 4.533E+04 | 566.3 | - | - | 0 | - |
| - | - | 2.588E+04 | 567.3 | - | - | 0 | - |
| 6 | y | 1.866E+05 | 572.3 | 3.872E-05 | 0.06765 | +1 | 5 |
| 6 | y | 1.347E+05 | 573.3 | 0.003633 | 6.337 | +1 | 5 |
| - | - | 4.83E+04 | 574.3 | - | - | 0 | - |
| - | - | 3.208E+04 | 582.3 | - | - | 0 | - |
| - | - | 6.511E+04 | 584.3 | - | - | 0 | - |
| - | - | 1.701E+04 | 585.3 | - | - | 0 | - |
| 6 | y | 4.957E+06 | 590.3 | 0.0004604 | 0.7798 | +1 | 5 |
| - | - | 1.416E+06 | 591.3 | - | - | 0 | - |
| - | - | 2.701E+05 | 592.3 | - | - | 0 | - |
| - | - | 1.306E+04 | 593.3 | - | - | 0 | - |
| - | - | 7784 | 599.3 | - | - | 0 | - |
| - | - | 6.903E+04 | 600.3 | - | - | 0 | - |
| - | - | 3.045E+04 | 601.3 | - | - | 0 | - |
| - | - | 2.878E+04 | 602.3 | - | - | 0 | - |
| - | - | 8170 | 603.3 | - | - | 0 | - |
| - | - | 1.218E+04 | 609.3 | - | - | 0 | - |
| - | - | 5.885E+04 | 617.3 | - | - | 0 | - |
| - | - | 2.566E+04 | 618.3 | - | - | 0 | - |
| - | - | 1.201E+04 | 619.3 | - | - | 0 | - |
| - | - | 2.332E+04 | 625.3 | - | - | 0 | - |
| - | - | 3.599E+04 | 626.3 | - | - | 0 | - |
| - | - | 3.46E+04 | 627.3 | - | - | 0 | - |
| - | - | 9438 | 628.3 | - | - | 0 | - |
| - | - | 1.015E+04 | 631.3 | - | - | 0 | - |
| - | - | 1.682E+04 | 634.4 | - | - | 0 | - |
| - | - | 7734 | 635.3 | - | - | 0 | - |
| - | - | 6566 | 642.3 | - | - | 0 | - |
| 5 | y | 4.583E+05 | 643.4 | 0.0001564 | 0.2431 | +1 | 6 |
| 5 | y | 1.95E+05 | 644.3 | 0.0101 | 15.67 | +1 | 6 |
| - | - | 6.381E+04 | 645.3 | - | - | 0 | - |
| - | - | 7850 | 646.3 | - | - | 0 | - |
| 7 | b | 2.949E+04 | 652.4 | 0.0002466 | 0.3779 | +1 | 7 |
| - | - | 2.051E+04 | 653.3 | - | - | 0 | - |
| - | - | 1.123E+04 | 654.3 | - | - | 0 | - |
| 5 | y | 3.784E+06 | 661.4 | 0.0004559 | 0.6894 | +1 | 6 |
| - | - | 1.236E+06 | 662.4 | - | - | 0 | - |
| - | - | 2.251E+05 | 663.4 | - | - | 0 | - |
| - | - | 1.932E+04 | 664.4 | - | - | 0 | - |
| 7 | b | 1.31E+04 | 670.4 | 0.003665 | 5.468 | +1 | 7 |
| - | - | 1.592E+05 | 671.3 | - | - | 0 | - |
| - | - | 5.107E+04 | 672.3 | - | - | 0 | - |
| - | - | 1.227E+04 | 673.4 | - | - | 0 | - |
| - | - | 7700 | 680.4 | - | - | 0 | - |
| - | - | 1.506E+04 | 681.4 | - | - | 0 | - |
| - | - | 1.545E+04 | 689.4 | - | - | 0 | - |
| - | - | 1.424E+04 | 696.4 | - | - | 0 | - |
| - | - | 1.371E+04 | 697.4 | - | - | 0 | - |
| - | - | 3.636E+04 | 698.4 | - | - | 0 | - |
| - | - | 1.235E+04 | 699.4 | - | - | 0 | - |
| - | - | 9955 | 706.4 | - | - | 0 | - |
| - | - | 6484 | 713.4 | - | - | 0 | - |
| - | - | 7.179E+04 | 714.4 | - | - | 0 | - |
| - | - | 2.314E+04 | 715.4 | - | - | 0 | - |
| - | - | 1.945E+04 | 716.4 | - | - | 0 | - |
| - | - | 6973 | 722.4 | - | - | 0 | - |
| - | - | 2.038E+04 | 723.4 | - | - | 0 | - |
| - | - | 2.544E+04 | 724.4 | - | - | 0 | - |
| - | - | 6397 | 725.4 | - | - | 0 | - |
| - | - | 7259 | 728.4 | - | - | 0 | - |
| 4 | y | 1.63E+05 | 740.4 | 0.0008497 | 1.148 | +1 | 7 |
| 4 | y | 2.394E+05 | 741.4 | 0.003477 | 4.69 | +1 | 7 |
| - | - | 8.262E+04 | 742.4 | - | - | 0 | - |
| - | - | 2.467E+04 | 743.4 | - | - | 0 | - |
| - | - | 2.88E+04 | 756.4 | - | - | 0 | - |
| - | - | 1.304E+04 | 757.4 | - | - | 0 | - |
| 4 | y | 6.287E+06 | 758.4 | 0.0003044 | 0.4014 | +1 | 7 |
| - | - | 2.454E+06 | 759.4 | - | - | 0 | - |
| - | - | 5.454E+05 | 760.4 | - | - | 0 | - |
| - | - | 4.123E+04 | 761.4 | - | - | 0 | - |
| 8 | b | 7837 | 766.4 | 0.003645 | 4.756 | +1 | 8 |
| 8 | b | 7243 | 767.4 | 0.003455 | 4.502 | +1 | 8 |
| 8 | b | 8520 | 784.4 | 0.001793 | 2.286 | +1 | 8 |
| - | - | 6743 | 785.4 | - | - | 0 | - |
| - | - | 7736 | 786.4 | - | - | 0 | - |
| - | - | 8969 | 797.4 | - | - | 0 | - |
| - | - | 2.202E+04 | 801.4 | - | - | 0 | - |
| - | - | 1.114E+04 | 802.4 | - | - | 0 | - |
| - | - | 1.064E+04 | 803.4 | - | - | 0 | - |
| - | - | 9955 | 809.4 | - | - | 0 | - |
| - | - | 6909 | 810.4 | - | - | 0 | - |
| - | - | 9746 | 811.4 | - | - | 0 | - |
| - | - | 6.313E+04 | 815.4 | - | - | 0 | - |
| - | - | 2.864E+04 | 816.4 | - | - | 0 | - |
| - | - | 1.008E+04 | 817.4 | - | - | 0 | - |
| 3 | y | 1.553E+05 | 827.4 | 0.0003251 | 0.3928 | +1 | 8 |
| 3 | y | 9.63E+04 | 828.4 | 0.00752 | 9.078 | +1 | 8 |
| - | - | 4.15E+04 | 829.4 | - | - | 0 | - |
| 3 | y | 2.929E+06 | 845.4 | 4.677E-05 | 0.05531 | +1 | 8 |
| - | - | 1.291E+06 | 846.5 | - | - | 0 | - |
| - | - | 3.162E+05 | 847.5 | - | - | 0 | - |
| - | - | 3.266E+04 | 848.5 | - | - | 0 | - |
| - | - | 5.05E+04 | 855.4 | - | - | 0 | - |
| - | - | 2.376E+04 | 856.4 | - | - | 0 | - |
| 9 | b | 6665 | 871.5 | 0.0102 | 11.71 | +1 | 9 |
| - | - | 1.853E+04 | 888.5 | - | - | 0 | - |
| - | - | 9668 | 889.5 | - | - | 0 | - |
| - | - | 3.087E+04 | 896.5 | - | - | 0 | - |
| - | - | 1.791E+04 | 897.5 | - | - | 0 | - |
| - | - | 7935 | 898.5 | - | - | 0 | - |
| - | - | 3.173E+04 | 902.5 | - | - | 0 | - |
| - | - | 9907 | 903.5 | - | - | 0 | - |
| - | - | 8463 | 912.5 | - | - | 0 | - |
| - | - | 7690 | 913.5 | - | - | 0 | - |
| 2 | y | 2.593E+05 | 914.5 | 0.0003313 | 0.3623 | +1 | 9 |
| 2 | y | 1.365E+05 | 915.5 | 0.01474 | 16.1 | +1 | 9 |
| - | - | 4.061E+04 | 916.5 | - | - | 0 | - |
| - | - | 2.792E+04 | 924.5 | - | - | 0 | - |
| - | - | 1.735E+04 | 925.5 | - | - | 0 | - |
| 2 | y | 2.225E+06 | 932.5 | 0.0009472 | 1.016 | +1 | 9 |
| - | - | 1.048E+06 | 933.5 | - | - | 0 | - |
| - | - | 3.022E+05 | 934.5 | - | - | 0 | - |
| - | - | 3.37E+04 | 935.5 | - | - | 0 | - |
| - | - | 2.052E+05 | 942.5 | - | - | 0 | - |
| - | - | 1.016E+05 | 943.5 | - | - | 0 | - |
| - | - | 3.184E+04 | 944.5 | - | - | 0 | - |
| - | - | 5457 | 3081 | - | - | 0 | - |
| - | - | 6041 | 3082 | - | - | 0 | - |

m/z Charge Intensity FragmentType MassShift Position
123.08815002441406 0 3148.1763
123.09199523925781 0 29082.693
125.10768127441406 0 133115.42
126.11094665527344 0 10161.243
127.08684539794922 0 18100.299
128.0709991455078 0 6710.6514
128.10731506347656 0 68637.375
129.06617736816406 0 52998.58
129.11378479003906 0 13404.345
129.20755004882812 0 3598.749
130.05010986328125 0 3473.9805
130.0614013671875 0 45628.113
130.0977783203125 0 26161.281
130.41746520996094 0 3994.2017
131.11831665039062 0 7650.611
132.03350830078125 0 3422.6074
133.09747314453125 0 9635.222
137.07119750976562 0 13086.476
138.09176635742188 0 24014.516
139.0504913330078 0 11137.239
139.08694458007812 0 267967.22
140.08335876464844 0 6402.2524
140.09031677246094 0 15261.28
141.06619262695312 0 34731.934
141.1025848388672 0 926235.44
141.13902282714844 0 6688.7163
142.10011291503906 0 8477.386
142.10586547851562 0 59551.086
142.1228790283203 0 8157.173
143.11830139160156 0 22448.576
145.06117248535156 0 3813.7297
145.0974578857422 0 61539.727
147.07675170898438 0 98024.65
148.9469451904297 0 5496.507
150.05502319335938 0 8176.2144
151.0870361328125 0 9246.473
152.08221435546875 0 4403.561
153.06619262695312 0 17151.318
153.10281372070312 0 7412.692
155.04563903808594 0 4886.5654
155.08187866210938 0 45753.53
155.11819458007812 0 290561.06 a Water loss 1
156.07699584960938 0 10585.026
156.10220336914062 0 12708.507
156.12156677246094 0 18281.96
157.0611114501953 0 204903.12
157.0974884033203 0 880988.1
157.10911560058594 0 33491.793
158.0646514892578 0 10005.338
158.09275817871094 0 272818.16 y Ammonia loss 9
158.10076904296875 0 69973.875
159.07659912109375 0 11367.083
159.09608459472656 0 17524.94
159.9760284423828 0 4279.8745
164.0821990966797 0 4598.9126
165.1026153564453 0 9550.267
166.09764099121094 0 11866.778
167.04542541503906 0 8535.282
167.0818328857422 0 296408.47
167.11859130859375 0 10301.064
168.07826232910156 0 6101.2847
168.0853271484375 0 23201.03
169.0611114501953 0 26500.283
169.09751892089844 0 1032172.5
169.1338653564453 0 45603.184
170.09365844726562 0 10150.454
170.10089111328125 0 83106.766
171.0770263671875 0 7869.1846
171.10421752929688 0 4826.285
171.11331176757812 0 10869.063
173.0924072265625 0 156181.31
173.12879943847656 0 1182839.6 a 1
173.438720703125 0 14865.007
174.08766174316406 0 69745.945
174.0958709716797 0 10289.674
174.13217163085938 0 98284.51
175.07168579101562 0 83766.64
175.11927795410156 0 671749.8 y 9
176.12274169921875 0 44644.293
179.08209228515625 0 15118.982
181.09742736816406 0 13824.794
182.09266662597656 0 14013.604
183.11317443847656 0 1847264.5 b Water loss 1
183.1493377685547 0 16080.761
184.07192993164062 0 23188.682
184.10037231445312 0 6189.153 b Water loss 3
184.1086883544922 0 33360.07
184.11656188964844 0 181703.48
185.0559539794922 0 25834.41
185.09237670898438 0 782895.06
186.08851623535156 0 6113.922
186.09585571289062 0 60086.277
186.12413024902344 0 26628.492
187.0718536376953 0 8859.946
187.09886169433594 0 8796.142
187.1080322265625 0 32339.14
187.1444549560547 0 347484.56
188.14797973632812 0 29201.61
192.11309814453125 0 10995.735
193.06068420410156 0 4298.0356
193.0973358154297 0 16198.556
194.0927276611328 0 9466.764
195.0767822265625 0 22556.213
195.1499481201172 0 4335.613
196.10838317871094 0 7678.4585
196.14524841308594 0 8096.034
197.09263610839844 0 8017.293
197.12872314453125 0 85566.14
198.08700561523438 0 6777.9365
198.1331024169922 0 6193.8677
199.1192626953125 0 14627.029
200.13954162597656 0 19836.662
201.12364196777344 0 1062334 b 1
202.0824737548828 0 166212.19
202.1186065673828 0 49823.133
202.1271209716797 0 101070.9
203.0665740966797 0 8398.068
203.0857391357422 0 14110.493
203.10276794433594 0 14549.735
203.12939453125 0 6889.8257
207.1131134033203 0 8438.767
208.10829162597656 0 76870.99
208.14511108398438 0 7397.792
209.09239196777344 0 16884.69
209.103759765625 0 15341.617
209.12844848632812 0 6379.832
210.08738708496094 0 21873.877
210.1239471435547 0 141873.67
211.10791015625 0 39747.637
211.12750244140625 0 14732.102
211.14442443847656 0 11412.91
212.1033172607422 0 15459.908
213.08740234375 0 8024.6816
214.15509033203125 0 5664.3647
215.13934326171875 0 195016.72
216.14280700683594 0 28295.086
219.10934448242188 0 15584.63
219.13475036621094 0 6727.5923
220.10845947265625 0 8040.2163
220.12924194335938 0 19360.928
221.09239196777344 0 70975.09
222.1241455078125 0 34028.098
223.10792541503906 0 17608.594
223.14505004882812 0 7644.192
224.1033477783203 0 23901.555
224.13967895507812 0 89012.4
225.0874786376953 0 19413.031
225.12371826171875 0 284981
225.15899658203125 0 6362.4883
226.11891174316406 0 105790.75
226.15463256835938 0 8253.538
227.077392578125 0 6977.774
227.11419677734375 0 62358.367
228.09817504882812 0 20790.32
228.1210174560547 0 6181.1074
228.1346435546875 0 134186.77
229.1186065673828 0 7691.8794
229.13819885253906 0 17627.793
232.1404266357422 0 8629.135
232.16566467285156 0 4587.549
236.10336303710938 0 42662.652
237.0869140625 0 13095.108
237.0989990234375 0 13831.508
237.12557983398438 0 5900.121
238.11904907226562 0 1707888
239.12232971191406 0 196574.48
240.0976104736328 0 6854.2256
240.12286376953125 0 11372.299
240.13487243652344 0 23932.672
240.1709442138672 0 22641.215
241.15476989746094 0 8090.0464
242.11404418945312 0 20874.67
242.1503448486328 0 129373.26 a Water loss 2
243.13424682617188 0 15364.577
243.1540985107422 0 7991.021
244.14065551757812 0 44497.02 y Water loss 8
245.1248321533203 0 295093.62 y Ammonia loss 8
245.1418914794922 0 14346.25 y 6
246.08718872070312 0 8783.752
246.12791442871094 0 27009.484
246.1419219970703 0 4426.223
250.11895751953125 0 25257.172
252.09808349609375 0 36059.152
252.13465881347656 0 576093.3
253.11854553222656 0 27225.197
253.137939453125 0 69493.69
254.11383056640625 0 85043.445
255.1094970703125 0 32641.36
255.14552307128906 0 11965.283
256.1295166015625 0 465365.38
257.1328125 0 46754.496
260.16094970703125 0 5494.3774 a 2
262.1512451171875 0 404065.72 y 8
263.1542663574219 0 41704.51
264.0977783203125 0 48396.723
266.112548828125 0 7224.307
266.149658203125 0 7043.268
268.12939453125 0 46824.516
268.1658630371094 0 150185.19
269.1685791015625 0 15910.481
270.1095886230469 0 67667.055
270.1451721191406 0 2566994.2 b Water loss 2
271.1483154296875 0 350123.88
272.1239318847656 0 17288.314
272.1362609863281 0 14107.964
272.1510009765625 0 20074.059
273.15679931640625 0 8684.504
276.13470458984375 0 6712.3804
279.1448669433594 0 7983.338 b 5
280.1294250488281 0 15588.541
280.1662292480469 0 12502.157
281.1245422363281 0 21979.666
282.1090087890625 0 37940.754
282.1445007324219 0 6391.394
282.1822204589844 0 22247.56
283.1089782714844 0 4692.4424
283.1410827636719 0 6796.6636
284.1610107421875 0 16515.988
285.1556396484375 0 11383.904
286.1763610839844 0 16064.047
286.6614990234375 0 39281.85 y Water loss 5
287.17291259765625 0 6528.954
288.15582275390625 0 154795.33 b 2
289.1591796875 0 15882.106
294.1451110839844 0 19642.586
295.1416015625 0 14991.666
295.66693115234375 0 44953.27 y 5
296.1365966796875 0 6026.9126
296.16290283203125 0 20708.033
297.11932373046875 0 8713.544
297.1568603515625 0 20973.686
298.1374206542969 0 8916.693
298.15325927734375 0 15000.66
298.1774597167969 0 8428.434
299.1356201171875 0 59149.684
300.119873046875 0 22432.17
300.13824462890625 0 8504.625
300.65869140625 0 11285.16
301.1871337890625 0 10427.028
304.1292419433594 0 10413.926
304.1673889160156 0 5102.3423
306.1197814941406 0 12121.497
307.1044921875 0 19157.129
307.1404724121094 0 22528.19
308.1241455078125 0 14754.386
309.1557312011719 0 6707.4336
310.177734375 0 6257.8735
311.1473388671875 0 8985.862
311.17169189453125 0 37600.816
311.20843505859375 0 35464.7
312.15576171875 0 49731.293
313.1518859863281 0 26944.273
313.6653137207031 0 7156.5757
314.1468811035156 0 24620.965
315.1476745605469 0 6249.9297
315.16680908203125 0 60736.11
316.162841796875 0 29305.64
317.1452331542969 0 17125.682
317.1642761230469 0 8714.27
321.1562194824219 0 72008.45
322.1402587890625 0 18803.742
322.1597900390625 0 10506.176
322.1800842285156 0 99608.95 y Water loss 4
322.68060302734375 0 24523.729
323.14617919921875 0 6027.5957
324.1307678222656 0 67062.66
325.1149597167969 0 33386.26
325.151123046875 0 48013.457
326.1173095703125 0 6403.642
327.1305236816406 0 7506.6553
327.1667785644531 0 7838.235
329.1824035644531 0 123319.164
329.20257568359375 0 8802.377
330.1859130859375 0 22363.82
331.1856384277344 0 64368.86 y 4
331.686767578125 0 21050.232
334.17352294921875 0 35246.07
335.1354064941406 0 8329.376
335.1742858886719 0 10515.762
335.2086486816406 0 13076.332
336.1773376464844 0 11969.893
337.1870422363281 0 17074.86
337.2239990234375 0 87498.67
338.22686767578125 0 15313.167
339.1667785644531 0 245013.19
339.20361328125 0 34174.117
340.16925048828125 0 37864.742
341.1571044921875 0 135434.55
341.17938232421875 0 4833.9355
342.1412048339844 0 270158
343.1434020996094 0 26796.38
343.1618957519531 0 49914.02
344.1650695800781 0 9890.013
349.1506652832031 0 9055.622
349.18707275390625 0 25484.621
349.68695068359375 0 6749.796
351.20281982421875 0 8522.804
352.1242980957031 0 6189.8594
353.1459045410156 0 15247.73
353.18218994140625 0 9094.861
353.2185363769531 0 6194.014
353.6808166503906 0 6700.6675
355.19805908203125 0 53331.508
355.23443603515625 0 77566.74
356.1986083984375 0 13336.89
356.23638916015625 0 13031.489
357.177001953125 0 197981.94
357.2156066894531 0 8448.31
358.18212890625 0 72473.37 y Water loss 7
359.1676025390625 0 518698.88 y Ammonia loss 7
360.1705627441406 0 77280.29
360.6940002441406 0 6065.806
361.7012939453125 0 6867.5
362.1932373046875 0 17434.371
362.5487365722656 0 5578.1187
362.69366455078125 0 6835.1294
363.201416015625 0 17018.889
364.16162109375 0 5071.081
365.14581298828125 0 7077.59
365.2186584472656 0 122022.75
366.17987060546875 0 9571.506
366.2227478027344 0 17968.104
367.1601867675781 0 10775.353
367.19793701171875 0 25338.58 b Water loss 3
368.195068359375 0 4950.282
369.69842529296875 0 109367.39
370.20001220703125 0 48973.434
370.7063903808594 0 971193.6 y Water loss 3
371.2073669433594 0 374384.06
371.7082214355469 0 64889.65
372.20751953125 0 7535.7954
376.1942443847656 0 1120808.1 y 7
377.19677734375 0 189460.47
378.19921875 0 26177.852
378.70391845703125 0 208308.94
379.2047119140625 0 89139.43
379.711669921875 0 5408223.5 y 3
380.2130432128906 0 2136003.2
380.7143249511719 0 518900.78
381.1770324707031 0 12957.709
381.21441650390625 0 38374.77
382.1778869628906 0 11748.091
382.20941162109375 0 20720.338
383.2291259765625 0 102093.94
384.1885986328125 0 14636.24
384.23309326171875 0 18417.469
384.70391845703125 0 39302.48
385.2071838378906 0 57983.32 b 3
386.1795349121094 0 11235.126
387.1629638671875 0 27549.223
388.1917724609375 0 8015.392
388.21868896484375 0 6484.4365
390.1775207519531 0 25493.332
391.1617736816406 0 34504.633
391.7096862792969 0 8683.414
392.1636657714844 0 5561.1196
393.2127380371094 0 7891.7656
394.2096252441406 0 12232.641
396.2137145996094 0 7757.24
396.7042541503906 0 34976.938
397.2038879394531 0 16626.088
398.2036437988281 0 26252.64
399.1865234375 0 6078.1377
399.2173767089844 0 14669.609
400.1880798339844 0 8270.162
400.2194519042969 0 38072.688
401.22076416015625 0 8109.579
405.2172546386719 0 279057.62
405.71356201171875 0 194529.83
406.21270751953125 0 72320.016
406.2455139160156 0 10210.38
406.7117919921875 0 19012.469
407.2309875488281 0 9477.9375
408.18853759765625 0 58330.516
408.2222595214844 0 8901.471
409.1700134277344 0 15364.254
410.1739196777344 0 6456.9043
410.20379638671875 0 11340.232
410.2378234863281 0 9337.501
411.2336120605469 0 9940.588
412.21990966796875 0 14638.1455
414.2228088378906 0 3766517.8 y Water loss 2
414.7239074707031 0 1681395.2
415.22515869140625 0 425979.8
415.7254333496094 0 37665.64
416.21441650390625 0 44323.684
417.2163391113281 0 9462.05
420.1913757324219 0 8037.132
420.22503662109375 0 29954.87
423.2279968261719 0 3296633.2 y 2
423.7293701171875 0 1391306.2
424.2304382324219 0 303093.16
424.7315979003906 0 30630.574
426.1990051269531 0 56620.78
427.20257568359375 0 10694.305
428.21923828125 0 10514.212
428.2510070800781 0 20838.87
429.2472229003906 0 8943.813
430.2039794921875 0 14589.384
434.2401428222656 0 71645.83
435.2458801269531 0 9604.329
436.1827392578125 0 20699.236
436.2174377441406 0 10304.235
437.21502685546875 0 31022.373
438.200439453125 0 15202.644
438.2353515625 0 51268.695 b Water loss 4
440.2216796875 0 9157.977
441.24639892578125 0 7097.751
442.26641845703125 0 37221.99
443.23541259765625 0 9591.127
443.2699279785156 0 6946.903
444.2095947265625 0 18895.893
445.2126159667969 0 7325.582
447.2572021484375 0 12511.16
448.733642578125 0 62007.715
449.2333984375 0 31934.393
449.7351379394531 0 14657.666
451.229248046875 0 8812.202
452.21728515625 0 5823.779
452.25091552734375 0 97645.305
453.2184143066406 0 7549.4795
453.2526550292969 0 35150.36
453.723876953125 0 6943.272
454.1942138671875 0 12514.459
454.2414855957031 0 25818.096
455.22564697265625 0 74833.03
456.2445068359375 0 50371.773 b 4
457.24810791015625 0 15490.295
457.7386474609375 0 88344.914 y Water loss 1
458.23956298828125 0 35803.11
458.7421875 0 7585.903
462.2353820800781 0 13815.1
462.7310791015625 0 62375.88
463.2325439453125 0 35577.617
463.7345275878906 0 8752.812
466.7443542480469 0 57437.07 y 1
467.24591064453125 0 32217.975
468.24908447265625 0 6128.489
469.2404479980469 0 20624.645
469.2774658203125 0 10809.955
470.26129150390625 0 47378.043
470.7647705078125 0 6879.476
471.26690673828125 0 45975.742 y Water loss 6
471.7358703613281 0 74921.31
472.2507019042969 0 136325.2 y Ammonia loss 6
472.7376403808594 0 10885.611
473.2542419433594 0 27040.156
477.25006103515625 0 5896.969
479.2623596191406 0 24372.904
480.2463073730469 0 38313.918
481.2515869140625 0 10468.7705
483.2210998535156 0 13252.903
484.76153564453125 0 8853.776
487.2521057128906 0 18320.127
488.2587585449219 0 8350.312
489.2785339355469 0 1579362.2 y 6
490.2813415527344 0 388957.34
491.2838439941406 0 67403.99
495.25811767578125 0 12005.99
496.7632141113281 0 25157.906
497.2709045410156 0 72667.37
498.27374267578125 0 18144.66
499.26470947265625 0 17112.93
503.26123046875 0 23561.756
504.2456970214844 0 11469.158
505.2755432128906 0 152053.95
505.7751770019531 0 95715.57
506.2754821777344 0 41540.008
511.26165771484375 0 13725.212
512.2424926757812 0 8856.504
513.2669677734375 0 12539.333
514.2802734375 0 264860.7 Precursor Water loss
514.7813720703125 0 170814.33 Precursor Ammonia loss
515.2826538085938 0 54152.977
521.2321166992188 0 5986.349
521.2722778320312 0 73223.98
522.2723999023438 0 24679.26
523.2858276367188 0 503448.47 Precursor
523.7872314453125 0 273387.5
524.288330078125 0 95445.555
524.78955078125 0 14039.222
528.2905883789062 0 8366.905
529.2720336914062 0 15240.195
530.2928466796875 0 9921.744
531.2529907226562 0 7009.412
537.27734375 0 15312.761
538.2630615234375 0 17971.568
539.2822875976562 0 74249.13 b Water loss 5
540.2835693359375 0 23680.955
543.2871704101562 0 6373.5405
545.76611328125 0 8098.598
546.2998046875 0 146796.75
547.3031616210938 0 34886.766
548.3009643554688 0 17323.14
549.2673950195312 0 25104.8
550.2662353515625 0 9841.555
555.288818359375 0 42915.35
556.275390625 0 47665.68
557.2908325195312 0 34718.8 b 5
558.2928466796875 0 6517.876
566.2936401367188 0 45326.297
567.2799072265625 0 25880.715
572.3151245117188 0 186574.44 y Water loss 5
573.302734375 0 134718.97 y Ammonia loss 5
574.3026123046875 0 48296.37
582.3001098632812 0 32077.242
584.3048706054688 0 65111.42
585.305908203125 0 17013.367
590.3261108398438 0 4956763.5 y 5
591.3286743164062 0 1415905
592.3310546875 0 270132.62
593.3333740234375 0 13059.194
599.3240966796875 0 7784.033
600.31005859375 0 69029.13
601.3216552734375 0 30445.209
602.317626953125 0 28776.86
603.3223266601562 0 8169.6797
609.296630859375 0 12182.958
617.3370971679688 0 58848.152
618.3381958007812 0 25663.18
619.3414916992188 0 12013.154
625.3427734375 0 23320.475
626.3274536132812 0 35985.023
627.3158569335938 0 34603.66
628.313720703125 0 9437.866
631.34765625 0 10149.631
634.3565063476562 0 16817.984
635.3132934570312 0 7733.616
642.318359375 0 6566.0166
643.3523559570312 0 458256.97 y Water loss 4
644.3463134765625 0 195007.86 y Ammonia loss 4
645.3442993164062 0 63813.773
646.3493041992188 0 7849.9717
652.36669921875 0 29491.4 b Water loss 6
653.3287353515625 0 20506.527
654.3242797851562 0 11229.399
661.3632202148438 0 3783606.8 y 4
662.3656616210938 0 1235812.6
663.3677368164062 0 225056.58
664.3695678710938 0 19320.996
670.3733520507812 0 13095.569 b 6
671.3463745117188 0 159165.06
672.3482666015625 0 51070.027
673.3511962890625 0 12266.618
680.3704833984375 0 7699.99
681.3596801757812 0 15055.214
689.3590087890625 0 15451.97
696.376953125 0 14238.528
697.3721923828125 0 13711.471
698.383544921875 0 36362.973
699.3868408203125 0 12347.158
706.3515014648438 0 9955.369
713.4185791015625 0 6484.1235
714.3894653320312 0 71792.09
715.3931884765625 0 23138.613
716.392333984375 0 19446.203
722.399658203125 0 6973.3633
723.3822631835938 0 20381.06
724.3670043945312 0 25436.262
725.3741455078125 0 6397.236
728.406982421875 0 7258.649
740.4041137695312 0 162952.69 y Water loss 3
741.3924560546875 0 239362.19 y Ammonia loss 3
742.3942260742188 0 82621.53
743.3935546875 0 24671.482
756.4013061523438 0 28802.383
757.4049682617188 0 13037.113
758.4158325195312 0 6287084 y 3
759.4183959960938 0 2453811.8
760.4204711914062 0 545378.4
761.4227905273438 0 41225.742
766.4130249023438 0 7836.542 b Water loss 7
767.3968505859375 0 7243.0356 b Ammonia loss 7
784.4181518554688 0 8519.801 b 7
785.4259643554688 0 6742.53
786.4165649414062 0 7736.138
797.43017578125 0 8968.545
801.4197998046875 0 22020.139
802.424072265625 0 11137.692
803.4248046875 0 10641.875
809.425537109375 0 9955.084
810.4085693359375 0 6908.8613
811.4046020507812 0 9746.156
815.4368896484375 0 63133.387
816.4404296875 0 28638.463
817.4450073242188 0 10075.954
827.4373168945312 0 155308.08 y Water loss 2
828.4285278320312 0 96303.87 y Ammonia loss 2
829.4276733398438 0 41500.863
845.447509765625 0 2929080.8 y 2
846.4501342773438 0 1290599.8
847.4521484375 0 316162.66
848.4531860351562 0 32662.654
855.43115234375 0 50497.324
856.4342041015625 0 23757.34
871.4417724609375 0 6664.882 b 8
888.4506225585938 0 18532.91
889.4603271484375 0 9667.753
896.4602661132812 0 30866.684
897.4509887695312 0 17911.223
898.4552001953125 0 7934.6475
902.4668579101562 0 31727.021
903.4658203125 0 9906.563
912.4528198242188 0 8463.141
913.4564819335938 0 7690.369
914.4686889648438 0 259329.48 y Water loss 1
915.4677734375 0 136474.77 y Ammonia loss 1
916.4662475585938 0 40608.734
924.4517211914062 0 27918.541
925.4564819335938 0 17347.557
932.4786376953125 0 2225046.8 y 1
933.4815673828125 0 1048498.3
934.4837646484375 0 302239.72
935.4854125976562 0 33699.098
942.4627685546875 0 205249.02
943.4656982421875 0 101620.79
944.4683837890625 0 31844.916
3081.05322265625 0 5457.2397
3081.763427734375 0 6041.2637

Spectrum Details

|  |  |
| --- | --- |
| Matched peaks? Matched peaks The total absolute number of peaks matched. Additionally in brackets the total fraction of peaks matched and the total number of peaks is shown. | 62 (10.02% of 619) |
| FDR? FDR The false discovery rate estimated for this peptide. It is calculated by matching all theoretical fragments with a non-integer shift with the raw peaks for this spectrum. This is done with 40 different shifts. The resulting percentage is the average number of annotated peaks over the number of annotated peaks with the correct spectrum. | 0.00% |
| Satellite FDR? Satellite FDR See the FDR for details on its calculation. This satellite ion specific FDR only contains the satellite ions (d/w) for I/L/J positions. | - |
| PSM Score? PSM Score The PSM Score as given by Hecklib to this annotated spectrum. It is shown with three significant figures. | 857 |

#### Spectrum 3712? Spectrum 3712 The raw spectrum of this peptide as annotated by Hecklib. The fragments are coloured according to ion type (see legend). Any peaks with a star '\*' as text can be hovered over to see the full details, first the ion type second the mass shift type. By hovering over the amino acids in the peptide or ions in the legend the corresponding peaks are highlighted. By toggling the 'Unassigned' label you can turn the background (unassigned) peaks on or off in the plot. By updating the slider in the Ion legend you can update the spectrum to only show the top X% of the peaks with labels. The top X% means any peak that is within X% of the highest intensity. By dragging in the spectrum you can zoom in to a specific part of the spectrum and use 'Zoom Out' to get back to the original zoom level. The annotation of the spectrum is based on the given sequence in the peptides file and is done with different software so inconsistencies are likely. The peaks are annotated based on the given sequence, with 20 ppm tolerance.

Copy Data

##### Spectrum 3712 (TSV)

###### Preview

```
Loading example...
```

*Click on the button to copy the data to your clipboard.*

Mz MinMz MaxIntensity Max

WidthHeightPeptide font sizePeptide stroke widthSpectrum font sizeSpectrum stroke widthCompact peptide

Ion legend

wxyz

abcd

OtherUnassignedIon^Charge^_Position_Show for top:%

JSSPATJNSR

01.97e+43.94e+45.90e+47.87e+4

Zoom Out

a^+1^_2_y^+1^_1_a^+1^_2_y^+1^_1_b^+1^_2_b^+1^_2_a^+1^_3_y^+1^_2_y^+1^_2_y^+1^_2_b^+1^_3_b^+1^_3_y^+2^_6_y^+1^_3_y^+1^_3_b^+1^_4_y^+2^_7_y^+2^_7_y^+1^_3_y^+2^_7_y^+2^_8_y^+2^_8_y^+2^_8_b^+1^_5_b^+1^_5_y^+2^_9_y^+2^_9_y^+2^_9_y^+1^_4_y^+1^_4_\*\*\*b^+1^_6_y^+1^_5_y^+1^_5_y^+1^_5_y^+1^_6_y^+1^_6_y^+1^_6_y^+1^_7_y^+1^_7_y^+1^_7_y^+1^_8_y^+1^_8_y^+1^_8_y^+1^_9_y^+1^_9_y^+1^_9_

0807161424223229

Fragment Matches Table

Show background peaks

| Position | Ion type | Intensity | mz Theoretical | mz Error (Th) | mz Error (ppm) | Charge | Series Number |
| --- | --- | --- | --- | --- | --- | --- | --- |
| - | - | 515.8 | 123.1 | - | - | 0 | - |
| - | - | 3305 | 125.1 | - | - | 0 | - |
| - | - | 1007 | 128.1 | - | - | 0 | - |
| - | - | 1321 | 129.1 | - | - | 0 | - |
| - | - | 1501 | 129.1 | - | - | 0 | - |
| - | - | 400.4 | 129.1 | - | - | 0 | - |
| - | - | 366.6 | 129.8 | - | - | 0 | - |
| - | - | 1295 | 130.1 | - | - | 0 | - |
| - | - | 626.8 | 130.1 | - | - | 0 | - |
| - | - | 593.9 | 130.1 | - | - | 0 | - |
| - | - | 437 | 130.3 | - | - | 0 | - |
| - | - | 406.3 | 131.8 | - | - | 0 | - |
| - | - | 972.7 | 133.1 | - | - | 0 | - |
| - | - | 376.6 | 133.4 | - | - | 0 | - |
| - | - | 7560 | 136.1 | - | - | 0 | - |
| - | - | 361.3 | 136.7 | - | - | 0 | - |
| - | - | 869.3 | 137.1 | - | - | 0 | - |
| - | - | 445.4 | 138.1 | - | - | 0 | - |
| - | - | 370.1 | 138.7 | - | - | 0 | - |
| - | - | 6311 | 139.1 | - | - | 0 | - |
| - | - | 444.7 | 141.1 | - | - | 0 | - |
| - | - | 1.555E+04 | 141.1 | - | - | 0 | - |
| - | - | 1433 | 142.1 | - | - | 0 | - |
| - | - | 797.3 | 143.1 | - | - | 0 | - |
| - | - | 392.1 | 143.3 | - | - | 0 | - |
| - | - | 1325 | 145 | - | - | 0 | - |
| - | - | 876.4 | 145.1 | - | - | 0 | - |
| - | - | 526.4 | 145.7 | - | - | 0 | - |
| - | - | 1781 | 147.1 | - | - | 0 | - |
| - | - | 783 | 147.1 | - | - | 0 | - |
| - | - | 2555 | 149 | - | - | 0 | - |
| - | - | 824.8 | 150 | - | - | 0 | - |
| - | - | 1201 | 151 | - | - | 0 | - |
| - | - | 450.7 | 155.1 | - | - | 0 | - |
| 2 | a | 5059 | 155.1 | 0.0001067 | 0.6877 | +1 | 2 |
| - | - | 4232 | 157.1 | - | - | 0 | - |
| - | - | 1.502E+04 | 157.1 | - | - | 0 | - |
| - | - | 1187 | 157.1 | - | - | 0 | - |
| 10 | y | 6465 | 158.1 | 0.0002025 | 1.281 | +1 | 1 |
| - | - | 1473 | 158.1 | - | - | 0 | - |
| - | - | 415.1 | 161.5 | - | - | 0 | - |
| - | - | 501.8 | 166.1 | - | - | 0 | - |
| - | - | 566.9 | 167 | - | - | 0 | - |
| - | - | 2453 | 167.1 | - | - | 0 | - |
| - | - | 5333 | 167.1 | - | - | 0 | - |
| - | - | 465.3 | 167.2 | - | - | 0 | - |
| - | - | 698 | 168.1 | - | - | 0 | - |
| - | - | 471.6 | 168.1 | - | - | 0 | - |
| - | - | 1839 | 169.1 | - | - | 0 | - |
| - | - | 1.832E+04 | 169.1 | - | - | 0 | - |
| - | - | 1120 | 169.1 | - | - | 0 | - |
| - | - | 1158 | 170.1 | - | - | 0 | - |
| - | - | 2710 | 173.1 | - | - | 0 | - |
| 2 | a | 2.132E+04 | 173.1 | 0.0001469 | 0.8482 | +1 | 2 |
| - | - | 2282 | 173.5 | - | - | 0 | - |
| - | - | 654.3 | 174.1 | - | - | 0 | - |
| - | - | 2367 | 174.1 | - | - | 0 | - |
| - | - | 2095 | 175.1 | - | - | 0 | - |
| 10 | y | 1.181E+04 | 175.1 | 0.0001427 | 0.8148 | +1 | 1 |
| - | - | 738.9 | 177.1 | - | - | 0 | - |
| - | - | 615.6 | 183.1 | - | - | 0 | - |
| 2 | b | 3.341E+04 | 183.1 | 0.0001414 | 0.7722 | +1 | 2 |
| - | - | 813.4 | 184.1 | - | - | 0 | - |
| - | - | 2459 | 184.1 | - | - | 0 | - |
| - | - | 1.397E+04 | 185.1 | - | - | 0 | - |
| - | - | 1154 | 186.1 | - | - | 0 | - |
| - | - | 540.2 | 186.1 | - | - | 0 | - |
| - | - | 476.2 | 187.1 | - | - | 0 | - |
| - | - | 8867 | 187.1 | - | - | 0 | - |
| - | - | 1063 | 188.1 | - | - | 0 | - |
| - | - | 649 | 195.1 | - | - | 0 | - |
| - | - | 544.9 | 196.3 | - | - | 0 | - |
| - | - | 2111 | 197.1 | - | - | 0 | - |
| 2 | b | 1.95E+04 | 201.1 | 5.95E-05 | 0.2958 | +1 | 2 |
| - | - | 2701 | 202.1 | - | - | 0 | - |
| - | - | 1711 | 202.1 | - | - | 0 | - |
| - | - | 421.5 | 206 | - | - | 0 | - |
| - | - | 990.2 | 208.1 | - | - | 0 | - |
| - | - | 1088 | 209.1 | - | - | 0 | - |
| - | - | 2030 | 210.1 | - | - | 0 | - |
| - | - | 564.1 | 212.1 | - | - | 0 | - |
| - | - | 3551 | 215.1 | - | - | 0 | - |
| - | - | 1126 | 221.1 | - | - | 0 | - |
| - | - | 522.4 | 221.2 | - | - | 0 | - |
| - | - | 1110 | 223.1 | - | - | 0 | - |
| - | - | 1450 | 224.1 | - | - | 0 | - |
| - | - | 2509 | 225 | - | - | 0 | - |
| - | - | 1210 | 225 | - | - | 0 | - |
| - | - | 5604 | 225.1 | - | - | 0 | - |
| - | - | 637.4 | 226 | - | - | 0 | - |
| - | - | 672 | 226.1 | - | - | 0 | - |
| - | - | 1987 | 226.1 | - | - | 0 | - |
| - | - | 578 | 226.1 | - | - | 0 | - |
| - | - | 1214 | 227.1 | - | - | 0 | - |
| - | - | 549.1 | 228 | - | - | 0 | - |
| - | - | 2528 | 228.1 | - | - | 0 | - |
| - | - | 5398 | 235.1 | - | - | 0 | - |
| - | - | 677.8 | 236.1 | - | - | 0 | - |
| - | - | 585.2 | 236.1 | - | - | 0 | - |
| - | - | 630.3 | 237.1 | - | - | 0 | - |
| - | - | 3.003E+04 | 238.1 | - | - | 0 | - |
| - | - | 1200 | 239.1 | - | - | 0 | - |
| - | - | 3520 | 239.1 | - | - | 0 | - |
| - | - | 651.6 | 240.1 | - | - | 0 | - |
| - | - | 586.3 | 240.2 | - | - | 0 | - |
| - | - | 741.4 | 241.1 | - | - | 0 | - |
| 3 | a | 2304 | 242.1 | 0.0002743 | 1.133 | +1 | 3 |
| - | - | 511.5 | 243.2 | - | - | 0 | - |
| 9 | y | 738.7 | 244.1 | 0.0001418 | 0.581 | +1 | 2 |
| 9 | y | 6703 | 245.1 | 0.0001413 | 0.5764 | +1 | 2 |
| - | - | 709.5 | 247.1 | - | - | 0 | - |
| - | - | 561.6 | 247.6 | - | - | 0 | - |
| - | - | 629.6 | 248.2 | - | - | 0 | - |
| - | - | 549 | 250.1 | - | - | 0 | - |
| - | - | 648.8 | 252.1 | - | - | 0 | - |
| - | - | 1.012E+04 | 252.1 | - | - | 0 | - |
| - | - | 1169 | 253.1 | - | - | 0 | - |
| - | - | 1205 | 254.1 | - | - | 0 | - |
| - | - | 8718 | 256.1 | - | - | 0 | - |
| - | - | 754.4 | 257.1 | - | - | 0 | - |
| 9 | y | 4.718E+04 | 262.2 | 8.143E-05 | 0.3106 | +1 | 2 |
| - | - | 1380 | 263.1 | - | - | 0 | - |
| - | - | 3122 | 263.2 | - | - | 0 | - |
| - | - | 1160 | 264.1 | - | - | 0 | - |
| - | - | 757.1 | 268.1 | - | - | 0 | - |
| - | - | 2846 | 268.2 | - | - | 0 | - |
| - | - | 1197 | 270.1 | - | - | 0 | - |
| 3 | b | 4.398E+04 | 270.1 | 6.489E-05 | 0.2402 | +1 | 3 |
| - | - | 4874 | 271.1 | - | - | 0 | - |
| - | - | 603.9 | 281.1 | - | - | 0 | - |
| - | - | 718.8 | 282.1 | - | - | 0 | - |
| - | - | 715.1 | 283 | - | - | 0 | - |
| - | - | 588.9 | 283 | - | - | 0 | - |
| - | - | 771.4 | 284 | - | - | 0 | - |
| - | - | 660 | 284.2 | - | - | 0 | - |
| - | - | 815.9 | 288.1 | - | - | 0 | - |
| 3 | b | 3138 | 288.2 | 0.0003339 | 1.159 | +1 | 3 |
| - | - | 4252 | 299.1 | - | - | 0 | - |
| - | - | 5544 | 300.1 | - | - | 0 | - |
| - | - | 9121 | 301.1 | - | - | 0 | - |
| - | - | 6044 | 302.1 | - | - | 0 | - |
| - | - | 1440 | 303.1 | - | - | 0 | - |
| - | - | 572.4 | 306.8 | - | - | 0 | - |
| - | - | 564.8 | 307.1 | - | - | 0 | - |
| - | - | 585.5 | 311.2 | - | - | 0 | - |
| - | - | 824.3 | 312.2 | - | - | 0 | - |
| - | - | 1176 | 315.2 | - | - | 0 | - |
| - | - | 1574 | 321.2 | - | - | 0 | - |
| 5 | y | 1405 | 322.2 | 0.0001726 | 0.5357 | +2 | 6 |
| - | - | 1737 | 324.1 | - | - | 0 | - |
| - | - | 676.3 | 325.2 | - | - | 0 | - |
| - | - | 1057 | 327.2 | - | - | 0 | - |
| - | - | 2068 | 329.2 | - | - | 0 | - |
| - | - | 679.9 | 332.2 | - | - | 0 | - |
| - | - | 1074 | 337.2 | - | - | 0 | - |
| - | - | 2769 | 339.2 | - | - | 0 | - |
| - | - | 1623 | 341.2 | - | - | 0 | - |
| - | - | 4191 | 342.1 | - | - | 0 | - |
| - | - | 762.2 | 343.2 | - | - | 0 | - |
| - | - | 943.7 | 346 | - | - | 0 | - |
| - | - | 1125 | 346.2 | - | - | 0 | - |
| - | - | 2448 | 347 | - | - | 0 | - |
| - | - | 1794 | 348 | - | - | 0 | - |
| - | - | 1014 | 349 | - | - | 0 | - |
| - | - | 955.5 | 355.2 | - | - | 0 | - |
| - | - | 1186 | 355.2 | - | - | 0 | - |
| - | - | 3084 | 357.2 | - | - | 0 | - |
| 8 | y | 918.6 | 358.2 | 0.00213 | 5.947 | +1 | 3 |
| 8 | y | 7950 | 359.2 | 0.0001226 | 0.3413 | +1 | 3 |
| - | - | 1521 | 360.2 | - | - | 0 | - |
| - | - | 769.5 | 360.2 | - | - | 0 | - |
| - | - | 803 | 361 | - | - | 0 | - |
| - | - | 908.4 | 362 | - | - | 0 | - |
| - | - | 1071 | 363 | - | - | 0 | - |
| - | - | 1661 | 365.2 | - | - | 0 | - |
| 4 | b | 789.8 | 367.2 | 0.0003101 | 0.8444 | +1 | 4 |
| - | - | 1838 | 369.7 | - | - | 0 | - |
| 4 | y | 1.371E+04 | 370.7 | 0.0003399 | 0.9169 | +2 | 7 |
| 4 | y | 5031 | 371.2 | 0.004967 | 13.38 | +2 | 7 |
| - | - | 1463 | 371.7 | - | - | 0 | - |
| 8 | y | 1.523E+04 | 376.2 | 0.0001824 | 0.485 | +1 | 3 |
| - | - | 6046 | 377.2 | - | - | 0 | - |
| - | - | 1306 | 377.2 | - | - | 0 | - |
| - | - | 724.1 | 378.2 | - | - | 0 | - |
| - | - | 3546 | 378.7 | - | - | 0 | - |
| - | - | 926.6 | 379.2 | - | - | 0 | - |
| 4 | y | 7.453E+04 | 379.7 | 9.858E-05 | 0.2596 | +2 | 7 |
| - | - | 3.533E+04 | 380.2 | - | - | 0 | - |
| - | - | 9594 | 380.7 | - | - | 0 | - |
| - | - | 1149 | 381.2 | - | - | 0 | - |
| - | - | 2022 | 383.2 | - | - | 0 | - |
| - | - | 600.3 | 391.2 | - | - | 0 | - |
| - | - | 936.6 | 396.7 | - | - | 0 | - |
| - | - | 548.6 | 400.2 | - | - | 0 | - |
| - | - | 3293 | 405.2 | - | - | 0 | - |
| - | - | 4729 | 405.7 | - | - | 0 | - |
| - | - | 1496 | 406.2 | - | - | 0 | - |
| - | - | 686.7 | 408.2 | - | - | 0 | - |
| 3 | y | 5.381E+04 | 414.2 | 0.0002169 | 0.5237 | +2 | 8 |
| 3 | y | 2.806E+04 | 414.7 | 0.00647 | 15.6 | +2 | 8 |
| - | - | 9658 | 415.2 | - | - | 0 | - |
| - | - | 621.2 | 415.7 | - | - | 0 | - |
| - | - | 662.8 | 416.2 | - | - | 0 | - |
| - | - | 758.6 | 417 | - | - | 0 | - |
| - | - | 1688 | 418 | - | - | 0 | - |
| - | - | 1965 | 419 | - | - | 0 | - |
| - | - | 977.4 | 419 | - | - | 0 | - |
| - | - | 854.3 | 420 | - | - | 0 | - |
| - | - | 626.8 | 421 | - | - | 0 | - |
| 3 | y | 4.705E+04 | 423.2 | 0.0001836 | 0.4338 | +2 | 8 |
| - | - | 2.133E+04 | 423.7 | - | - | 0 | - |
| - | - | 7973 | 424.2 | - | - | 0 | - |
| - | - | 1043 | 434.2 | - | - | 0 | - |
| 5 | b | 593.4 | 438.2 | 0.0002437 | 0.556 | +1 | 5 |
| - | - | 792 | 442.3 | - | - | 0 | - |
| - | - | 880.8 | 448.7 | - | - | 0 | - |
| - | - | 765.3 | 449.2 | - | - | 0 | - |
| - | - | 1446 | 452.3 | - | - | 0 | - |
| - | - | 1363 | 455.2 | - | - | 0 | - |
| 5 | b | 727.1 | 456.2 | 0.001653 | 3.623 | +1 | 5 |
| 2 | y | 1934 | 457.7 | 0.0001634 | 0.357 | +2 | 9 |
| 2 | y | 852.2 | 458.2 | 0.008796 | 19.2 | +2 | 9 |
| 2 | y | 1513 | 466.7 | 0.000114 | 0.2443 | +2 | 9 |
| - | - | 816.1 | 470.3 | - | - | 0 | - |
| - | - | 1372 | 471.7 | - | - | 0 | - |
| 7 | y | 1486 | 472.3 | 0.0003166 | 0.6704 | +1 | 4 |
| - | - | 1194 | 480.2 | - | - | 0 | - |
| - | - | 566.5 | 482.1 | - | - | 0 | - |
| - | - | 1423 | 489.1 | - | - | 0 | - |
| 7 | y | 1.935E+04 | 489.3 | 0.0001957 | 0.4 | +1 | 4 |
| - | - | 3651 | 490.3 | - | - | 0 | - |
| - | - | 1124 | 497.3 | - | - | 0 | - |
| - | - | 618.4 | 504.8 | - | - | 0 | - |
| - | - | 1398 | 505.3 | - | - | 0 | - |
| - | - | 1814 | 505.8 | - | - | 0 | - |
| - | - | 2113 | 507.2 | - | - | 0 | - |
| - | - | 2108 | 513.8 | - | - | 0 | - |
| 0 | Precursor | 3633 | 514.3 | 0.002715 | 5.278 | +2 | -1 |
| 0 | Precursor | 2069 | 514.8 | 0.003996 | 7.763 | +2 | -1 |
| - | - | 1156 | 516.9 | - | - | 0 | - |
| - | - | 645 | 517.3 | - | - | 0 | - |
| - | - | 755.9 | 517.6 | - | - | 0 | - |
| - | - | 1799 | 521.3 | - | - | 0 | - |
| 0 | Precursor | 8214 | 523.3 | 0.0003675 | 0.7022 | +2 | -1 |
| - | - | 7457 | 523.8 | - | - | 0 | - |
| - | - | 1530 | 524.3 | - | - | 0 | - |
| 6 | b | 1462 | 539.3 | 2.099E-05 | 0.03892 | +1 | 6 |
| - | - | 651.9 | 540.3 | - | - | 0 | - |
| - | - | 7609 | 545.8 | - | - | 0 | - |
| - | - | 4547 | 546.3 | - | - | 0 | - |
| - | - | 996 | 546.3 | - | - | 0 | - |
| - | - | 1690 | 546.8 | - | - | 0 | - |
| - | - | 612.7 | 549.5 | - | - | 0 | - |
| - | - | 857.7 | 556.3 | - | - | 0 | - |
| 6 | y | 1891 | 572.3 | 0.0007101 | 1.241 | +1 | 5 |
| 6 | y | 1992 | 573.3 | 0.004182 | 7.295 | +1 | 5 |
| 6 | y | 6.111E+04 | 590.3 | 8.896E-05 | 0.1507 | +1 | 5 |
| - | - | 1.758E+04 | 591.3 | - | - | 0 | - |
| - | - | 4065 | 592.3 | - | - | 0 | - |
| - | - | 762.4 | 600.3 | - | - | 0 | - |
| - | - | 956.4 | 602.3 | - | - | 0 | - |
| - | - | 639.5 | 603.3 | - | - | 0 | - |
| - | - | 1206 | 617.3 | - | - | 0 | - |
| 5 | y | 5661 | 643.4 | 8.777E-05 | 0.1364 | +1 | 6 |
| 5 | y | 3202 | 644.3 | 0.008145 | 12.64 | +1 | 6 |
| - | - | 872.9 | 645.3 | - | - | 0 | - |
| - | - | 4379 | 652.8 | - | - | 0 | - |
| - | - | 2772 | 653.3 | - | - | 0 | - |
| - | - | 1714 | 653.8 | - | - | 0 | - |
| 5 | y | 4.835E+04 | 661.4 | 9.337E-05 | 0.1412 | +1 | 6 |
| - | - | 1.803E+04 | 662.4 | - | - | 0 | - |
| - | - | 4545 | 663.4 | - | - | 0 | - |
| - | - | 1807 | 671.3 | - | - | 0 | - |
| - | - | 1216 | 672.3 | - | - | 0 | - |
| - | - | 1023 | 734.4 | - | - | 0 | - |
| - | - | 1038 | 734.9 | - | - | 0 | - |
| 4 | y | 1901 | 740.4 | 0.0005445 | 0.7354 | +1 | 7 |
| 4 | y | 4471 | 741.4 | 0.001036 | 1.397 | +1 | 7 |
| - | - | 818.7 | 742.4 | - | - | 0 | - |
| 4 | y | 7.793E+04 | 758.4 | 0.000306 | 0.4034 | +1 | 7 |
| - | - | 4.065E+04 | 759.4 | - | - | 0 | - |
| - | - | 1.096E+04 | 760.4 | - | - | 0 | - |
| - | - | 820.4 | 761.4 | - | - | 0 | - |
| - | - | 944.1 | 815.4 | - | - | 0 | - |
| 3 | y | 1566 | 827.4 | 0.0007523 | 0.9092 | +1 | 8 |
| 3 | y | 1824 | 828.4 | 0.00398 | 4.805 | +1 | 8 |
| - | - | 858 | 829.4 | - | - | 0 | - |
| 3 | y | 3.948E+04 | 845.4 | 0.0002909 | 0.3441 | +1 | 8 |
| - | - | 2.096E+04 | 846.4 | - | - | 0 | - |
| - | - | 6860 | 847.4 | - | - | 0 | - |
| - | - | 857.7 | 848.5 | - | - | 0 | - |
| - | - | 663.9 | 902.5 | - | - | 0 | - |
| 2 | y | 3835 | 914.5 | 0.001796 | 1.964 | +1 | 9 |
| 2 | y | 1990 | 915.5 | 0.009611 | 10.5 | +1 | 9 |
| - | - | 778.2 | 916.5 | - | - | 0 | - |
| - | - | 1037 | 929.5 | - | - | 0 | - |
| 2 | y | 2.959E+04 | 932.5 | 0.001252 | 1.343 | +1 | 9 |
| - | - | 1.734E+04 | 933.5 | - | - | 0 | - |
| - | - | 5829 | 934.5 | - | - | 0 | - |
| - | - | 2759 | 942.5 | - | - | 0 | - |
| - | - | 2077 | 943.5 | - | - | 0 | - |
| - | - | 1381 | 993.5 | - | - | 0 | - |
| - | - | 1476 | 1027 | - | - | 0 | - |
| - | - | 2256 | 1091 | - | - | 0 | - |
| - | - | 806.8 | 1092 | - | - | 0 | - |
| - | - | 638.4 | 1583 | - | - | 0 | - |
| - | - | 644.9 | 2409 | - | - | 0 | - |
| - | - | 719.7 | 3080 | - | - | 0 | - |
| - | - | 1017 | 3080 | - | - | 0 | - |
| - | - | 663.3 | 3197 | - | - | 0 | - |

m/z Charge Intensity FragmentType MassShift Position
123.09166717529297 0 515.8336
125.10748291015625 0 3305.3335
128.1071319580078 0 1006.90625
129.06617736816406 0 1320.6681
129.1024627685547 0 1500.7758
129.11419677734375 0 400.35034
129.824462890625 0 366.57376
130.06114196777344 0 1295.074
130.0865020751953 0 626.81885
130.09768676757812 0 593.9173
130.33448791503906 0 436.9599
131.84657287597656 0 406.27493
133.08631896972656 0 972.709
133.36891174316406 0 376.55692
136.07586669921875 0 7560.374
136.69143676757812 0 361.3153
137.0792999267578 0 869.32947
138.09140014648438 0 445.4026
138.69886779785156 0 370.0939
139.08677673339844 0 6310.5444
141.06521606445312 0 444.74072
141.1024169921875 0 15550.852
142.10581970214844 0 1433.0519
143.1179656982422 0 797.2642
143.34942626953125 0 392.0608
145.0497283935547 0 1325.144
145.09718322753906 0 876.41675
145.70999145507812 0 526.43854
147.07650756835938 0 1781.1161
147.11314392089844 0 783.0464
149.0450439453125 0 2555.3633
150.04396057128906 0 824.84674
151.04220581054688 0 1201.1144
155.08140563964844 0 450.68262
155.1179962158203 0 5058.805 a Water loss 1
157.0609130859375 0 4231.9795
157.09730529785156 0 15019.972
157.1086883544922 0 1186.9613
158.0926055908203 0 6465.1533 y Ammonia loss 9
158.10084533691406 0 1472.743
161.5320587158203 0 415.08752
166.09808349609375 0 501.759
167.04566955566406 0 566.89764
167.05551147460938 0 2453.4995
167.0816650390625 0 5333.279
167.23001098632812 0 465.33093
168.05540466308594 0 698.00214
168.10301208496094 0 471.57217
169.05238342285156 0 1839.4985
169.09730529785156 0 18317.182
169.13345336914062 0 1119.8112
170.10073852539062 0 1157.748
173.09217834472656 0 2709.599
173.12860107421875 0 21324.96 a 1
173.450927734375 0 2281.9946
174.08724975585938 0 654.274
174.13217163085938 0 2366.643
175.07162475585938 0 2095.1284
175.1190948486328 0 11813.464 y 9
177.1126251220703 0 738.9374
183.07691955566406 0 615.56555
183.11294555664062 0 33406.754 b Water loss 1
184.10829162597656 0 813.3812
184.11656188964844 0 2459.0295
185.09219360351562 0 13967.114
186.09616088867188 0 1153.7178
186.12330627441406 0 540.19305
187.1074981689453 0 476.17035
187.14427185058594 0 8867.476
188.14752197265625 0 1062.7786
195.07693481445312 0 649.0248
196.26055908203125 0 544.89
197.1287841796875 0 2111.4163
201.12342834472656 0 19497.998 b 1
202.08229064941406 0 2700.5457
202.12680053710938 0 1711.2722
205.96542358398438 0 421.50006
208.1082763671875 0 990.2031
209.0922088623047 0 1087.6237
210.1238250732422 0 2029.6522
212.10267639160156 0 564.1325
215.13897705078125 0 3550.539
221.09225463867188 0 1126.1971
221.24929809570312 0 522.368
223.0640106201172 0 1110.2661
224.1394805908203 0 1450.4337
225.0062255859375 0 2509.4617
225.04327392578125 0 1210.2533
225.12356567382812 0 5603.5894
226.0435028076172 0 637.38293
226.08297729492188 0 672.0011
226.11842346191406 0 1986.5164
226.12911987304688 0 577.99634
227.11358642578125 0 1213.6648
227.97946166992188 0 549.06775
228.13421630859375 0 2527.5845
235.1441192626953 0 5397.6016
236.10296630859375 0 677.8222
236.14794921875 0 585.1889
237.1233367919922 0 630.2661
238.11880493164062 0 30033.316
239.09451293945312 0 1199.7289
239.1219482421875 0 3520.273
240.13479614257812 0 651.6154
240.17103576660156 0 586.3134
241.0918426513672 0 741.3969
242.1501922607422 0 2304.2598 a Water loss 2
243.1552276611328 0 511.4679
244.14027404785156 0 738.7404 y Water loss 8
245.12457275390625 0 6702.9995 y Ammonia loss 8
247.10694885253906 0 709.5097
247.6209716796875 0 561.5816
248.15989685058594 0 629.5736
250.11801147460938 0 549.0342
252.09786987304688 0 648.84143
252.13453674316406 0 10124.496
253.13853454589844 0 1169.0833
254.1132354736328 0 1204.8965
256.1293029785156 0 8717.638
257.13336181640625 0 754.398
262.15106201171875 0 47181.02 y 8
263.1383056640625 0 1380.3713
263.1546325683594 0 3121.9348
264.0978088378906 0 1160.2452
268.12847900390625 0 757.13666
268.1655578613281 0 2845.7007
270.1100769042969 0 1196.6962
270.1448974609375 0 43978.254 b Water loss 2
271.1481018066406 0 4873.756
281.0523681640625 0 603.94855
282.0515441894531 0 718.84625
283.0304870605469 0 715.1165
283.0493469238281 0 588.8817
284.0494689941406 0 771.39825
284.160888671875 0 659.97296
288.13360595703125 0 815.9215
288.1557312011719 0 3138.2683 b 2
299.0617370605469 0 4251.9883
300.06243896484375 0 5543.9663
301.05950927734375 0 9121.068
302.0594482421875 0 6044.042
303.0563049316406 0 1439.8351
306.7558898925781 0 572.4113
307.1045227050781 0 564.79144
311.1716003417969 0 585.48
312.156494140625 0 824.3445
315.1662902832031 0 1175.607
321.15618896484375 0 1574.1188
322.1795654296875 0 1404.6283 y Water loss 4
324.1309509277344 0 1736.9973
325.1510314941406 0 676.2832
327.20166015625 0 1056.8898
329.1821594238281 0 2067.7266
332.1965637207031 0 679.8529
337.22357177734375 0 1074.0515
339.16644287109375 0 2769.3186
341.157470703125 0 1623.0177
342.14093017578125 0 4191.0195
343.1619873046875 0 762.16565
345.9757385253906 0 943.7183
346.176025390625 0 1124.9863
346.97357177734375 0 2447.7197
347.9746398925781 0 1793.672
348.97259521484375 0 1014.0087
355.1968994140625 0 955.5054
355.2332763671875 0 1185.7603
357.17724609375 0 3084.073
358.18121337890625 0 918.6362 y Water loss 7
359.167236328125 0 7950.4507 y Ammonia loss 7
360.1517639160156 0 1521.3661
360.1724548339844 0 769.5237
361.0243835449219 0 802.95593
362.0240783691406 0 908.37885
363.0231018066406 0 1071.3964
365.2188415527344 0 1660.6765
367.1979064941406 0 789.80524 b Water loss 3
369.698486328125 0 1837.6918
370.7057800292969 0 13705.813 y Water loss 3
371.2030944824219 0 5031.201 y Ammonia loss 3
371.7055969238281 0 1463.2198
376.1937255859375 0 15231.812 y 7
377.177734375 0 6046.479
377.19915771484375 0 1305.7898
378.18011474609375 0 724.07367
378.7034606933594 0 3545.6406
379.1966247558594 0 926.62866
379.7113037109375 0 74527.016 y 3
380.2099914550781 0 35330.426
380.7098083496094 0 9594.149
381.2077941894531 0 1149.4639
383.2289123535156 0 2022.0958
391.1617126464844 0 600.26556
396.7031555175781 0 936.5945
400.21630859375 0 548.55426
405.2174377441406 0 3293.0723
405.7102966308594 0 4728.682
406.2123107910156 0 1496.4517
408.1886901855469 0 686.6912
414.22235107421875 0 53813.562 y Water loss 2
414.7206115722656 0 28061.184 y Ammonia loss 2
415.2206115722656 0 9657.891
415.7189636230469 0 621.24396
416.2131652832031 0 662.75464
417.03558349609375 0 758.61725
418.0354919433594 0 1688.2532
418.9952087402344 0 1965.3124
419.02972412109375 0 977.3591
419.99468994140625 0 854.32996
420.99053955078125 0 626.76324
423.22760009765625 0 47051.62 y 2
423.7258605957031 0 21327.312
424.2249755859375 0 7972.883
434.241455078125 0 1043.3661
438.2344665527344 0 593.35205 b Water loss 4
442.26483154296875 0 792.02954
448.73211669921875 0 880.81866
449.23260498046875 0 765.31476
452.25164794921875 0 1445.7985
455.2237243652344 0 1363.0946
456.2436218261719 0 727.09656 b 4
457.7383117675781 0 1934.423 y Water loss 1
458.23895263671875 0 852.18115 y Ammonia loss 1
466.7433166503906 0 1513.4688 y 1
470.2613830566406 0 816.1278
471.7359313964844 0 1371.5704
472.2517395019531 0 1486.0206 y Ammonia loss 6
480.24737548828125 0 1194.2051
482.14959716796875 0 566.5236
489.05670166015625 0 1423.2502
489.2781677246094 0 19348.459 y 6
490.2641906738281 0 3651.0693
497.2717590332031 0 1124.1847
504.7583923339844 0 618.43317
505.2745666503906 0 1397.954
505.77508544921875 0 1813.7949
507.1743469238281 0 2112.5776
513.7637329101562 0 2108.4976
514.2774658203125 0 3633.2144 Precursor Water loss
514.7761840820312 0 2068.7766 Precursor Ammonia loss
516.9281005859375 0 1156.0312
517.2593383789062 0 645.01575
517.5950317382812 0 755.9014
521.2722778320312 0 1799.3744
523.2850952148438 0 8214.253 Precursor
523.7842407226562 0 7456.7734
524.2799682617188 0 1529.579
539.2824096679688 0 1461.8591 b Water loss 5
540.2858276367188 0 651.8906
545.763916015625 0 7609.1357
546.264404296875 0 4546.641
546.3037719726562 0 996.0344
546.76513671875 0 1690.0491
549.53955078125 0 612.686
556.2740478515625 0 857.6894
572.3157958984375 0 1891.0563 y Water loss 5
573.3032836914062 0 1992.4933 y Ammonia loss 5
590.3255615234375 0 61108.457 y 5
591.3219604492188 0 17579.77
592.3248901367188 0 4064.5662
600.3095092773438 0 762.3979
602.3081665039062 0 956.39374
603.3076171875 0 639.5232
617.3364868164062 0 1205.6198
643.3521118164062 0 5660.686 y Water loss 4
644.3443603515625 0 3201.9788 y Ammonia loss 4
645.3446655273438 0 872.8832
652.8279418945312 0 4379.0566
653.3299560546875 0 2771.7422
653.8328247070312 0 1713.5139
661.3626708984375 0 48348 y 4
662.361328125 0 18033.508
663.3578491210938 0 4544.6343
671.3463745117188 0 1807.3905
672.345458984375 0 1216.0814
734.35888671875 0 1022.68787
734.8619995117188 0 1038.0271
740.4044189453125 0 1901.1356 y Water loss 3
741.3900146484375 0 4470.863 y Ammonia loss 3
742.39013671875 0 818.693
758.4152221679688 0 77934.05 y 3
759.412109375 0 40650.668
760.4122314453125 0 10959.133
761.4125366210938 0 820.40283
815.4371948242188 0 944.0614
827.437744140625 0 1565.5862 y Water loss 2
828.4249877929688 0 1823.5565 y Ammonia loss 2
829.4354248046875 0 858.03815
845.447265625 0 39480.555 y 2
846.4437255859375 0 20963.002
847.4432373046875 0 6859.8994
848.450927734375 0 857.7023
902.4645385742188 0 663.8953
914.4672241210938 0 3834.6226 y Water loss 1
915.462646484375 0 1990.4376 y Ammonia loss 1
916.4531860351562 0 778.22314
929.4680786132812 0 1036.604
932.4783325195312 0 29587.23 y 1
933.4755859375 0 17340.76
934.4765625 0 5829.3345
942.4636840820312 0 2759.207
943.4600830078125 0 2076.8857
993.4671630859375 0 1380.5302
1026.516357421875 0 1475.9287
1090.518310546875 0 2255.5708
1091.519287109375 0 806.7902
1583.4478759765625 0 638.3948
2408.6455078125 0 644.9292
3079.537353515625 0 719.69977
3080.34130859375 0 1016.7077
3196.740478515625 0 663.2527

Spectrum Details

|  |  |
| --- | --- |
| Matched peaks? Matched peaks The total absolute number of peaks matched. Additionally in brackets the total fraction of peaks matched and the total number of peaks is shown. | 49 (15.81% of 310) |
| FDR? FDR The false discovery rate estimated for this peptide. It is calculated by matching all theoretical fragments with a non-integer shift with the raw peaks for this spectrum. This is done with 40 different shifts. The resulting percentage is the average number of annotated peaks over the number of annotated peaks with the correct spectrum. | 0.53% |
| Satellite FDR? Satellite FDR See the FDR for details on its calculation. This satellite ion specific FDR only contains the satellite ions (d/w) for I/L/J positions. | - |
| PSM Score? PSM Score The PSM Score as given by Hecklib to this annotated spectrum. It is shown with three significant figures. | 596 |

#### Spectrum 3883? Spectrum 3883 The raw spectrum of this peptide as annotated by Hecklib. The fragments are coloured according to ion type (see legend). Any peaks with a star '\*' as text can be hovered over to see the full details, first the ion type second the mass shift type. By hovering over the amino acids in the peptide or ions in the legend the corresponding peaks are highlighted. By toggling the 'Unassigned' label you can turn the background (unassigned) peaks on or off in the plot. By updating the slider in the Ion legend you can update the spectrum to only show the top X% of the peaks with labels. The top X% means any peak that is within X% of the highest intensity. By dragging in the spectrum you can zoom in to a specific part of the spectrum and use 'Zoom Out' to get back to the original zoom level. The annotation of the spectrum is based on the given sequence in the peptides file and is done with different software so inconsistencies are likely. The peaks are annotated based on the given sequence, with 20 ppm tolerance.

Copy Data

##### Spectrum 3883 (TSV)

###### Preview

```
Loading example...
```

*Click on the button to copy the data to your clipboard.*

Mz MinMz MaxIntensity Max

WidthHeightPeptide font sizePeptide stroke widthSpectrum font sizeSpectrum stroke widthCompact peptide

Ion legend

wxyz

abcd

OtherUnassignedIon^Charge^_Position_Show for top:%

JSSPATJNSR

09.03e+31.81e+42.71e+43.61e+4

Zoom Out

a^+1^_2_y^+1^_1_a^+1^_2_y^+1^_1_b^+1^_2_b^+1^_2_a^+1^_3_y^+1^_2_y^+1^_2_y^+1^_2_b^+1^_3_b^+1^_3_y^+2^_6_y^+1^_3_y^+1^_3_y^+2^_7_y^+2^_7_y^+1^_3_y^+2^_7_y^+2^_8_y^+2^_8_y^+2^_8_y^+1^_4_\*\*y^+1^_5_y^+1^_5_y^+1^_5_y^+1^_6_y^+1^_6_y^+1^_6_y^+1^_7_y^+1^_7_y^+1^_7_y^+1^_8_y^+1^_8_y^+1^_9_y^+1^_9_

0666133219992665

Fragment Matches Table

Show background peaks

| Position | Ion type | Intensity | mz Theoretical | mz Error (Th) | mz Error (ppm) | Charge | Series Number |
| --- | --- | --- | --- | --- | --- | --- | --- |
| - | - | 400.9 | 121 | - | - | 0 | - |
| - | - | 371.9 | 123.8 | - | - | 0 | - |
| - | - | 1286 | 125.1 | - | - | 0 | - |
| - | - | 734.7 | 128.1 | - | - | 0 | - |
| - | - | 419.8 | 129.1 | - | - | 0 | - |
| - | - | 421.6 | 129.1 | - | - | 0 | - |
| - | - | 1362 | 130.1 | - | - | 0 | - |
| - | - | 420.5 | 134.1 | - | - | 0 | - |
| - | - | 4681 | 136.1 | - | - | 0 | - |
| - | - | 1973 | 139.1 | - | - | 0 | - |
| - | - | 8922 | 141.1 | - | - | 0 | - |
| - | - | 614.4 | 142.1 | - | - | 0 | - |
| - | - | 832.5 | 145.1 | - | - | 0 | - |
| - | - | 923 | 147.1 | - | - | 0 | - |
| - | - | 764.8 | 147.1 | - | - | 0 | - |
| - | - | 464 | 148.8 | - | - | 0 | - |
| - | - | 548.6 | 148.9 | - | - | 0 | - |
| - | - | 643.6 | 148.9 | - | - | 0 | - |
| - | - | 703.1 | 148.9 | - | - | 0 | - |
| - | - | 1211 | 148.9 | - | - | 0 | - |
| - | - | 1415 | 148.9 | - | - | 0 | - |
| - | - | 2687 | 148.9 | - | - | 0 | - |
| - | - | 4389 | 148.9 | - | - | 0 | - |
| - | - | 3941 | 149 | - | - | 0 | - |
| - | - | 2404 | 149 | - | - | 0 | - |
| - | - | 1129 | 149 | - | - | 0 | - |
| - | - | 1258 | 149 | - | - | 0 | - |
| - | - | 1134 | 149 | - | - | 0 | - |
| - | - | 736.9 | 149 | - | - | 0 | - |
| - | - | 443.8 | 149 | - | - | 0 | - |
| - | - | 503.1 | 149 | - | - | 0 | - |
| - | - | 1962 | 149 | - | - | 0 | - |
| - | - | 521.5 | 149.1 | - | - | 0 | - |
| - | - | 615.9 | 150 | - | - | 0 | - |
| - | - | 698.4 | 151 | - | - | 0 | - |
| - | - | 826.6 | 155.1 | - | - | 0 | - |
| 2 | a | 2471 | 155.1 | 7.616E-05 | 0.491 | +1 | 2 |
| - | - | 1731 | 157.1 | - | - | 0 | - |
| - | - | 6580 | 157.1 | - | - | 0 | - |
| - | - | 603.4 | 157.1 | - | - | 0 | - |
| 10 | y | 5015 | 158.1 | 8.045E-05 | 0.5089 | +1 | 1 |
| - | - | 694.7 | 158.1 | - | - | 0 | - |
| - | - | 2958 | 167.1 | - | - | 0 | - |
| - | - | 2337 | 167.1 | - | - | 0 | - |
| - | - | 733.9 | 168.1 | - | - | 0 | - |
| - | - | 1851 | 169.1 | - | - | 0 | - |
| - | - | 627.9 | 169.1 | - | - | 0 | - |
| - | - | 8568 | 169.1 | - | - | 0 | - |
| - | - | 2524 | 169.1 | - | - | 0 | - |
| - | - | 682.5 | 170.1 | - | - | 0 | - |
| - | - | 1745 | 173.1 | - | - | 0 | - |
| 2 | a | 9308 | 173.1 | 9.524E-06 | 0.05501 | +1 | 2 |
| - | - | 870.4 | 173.4 | - | - | 0 | - |
| - | - | 895.9 | 175.1 | - | - | 0 | - |
| 10 | y | 1.125E+04 | 175.1 | 9.69E-05 | 0.5534 | +1 | 1 |
| - | - | 559.8 | 176.1 | - | - | 0 | - |
| 2 | b | 1.542E+04 | 183.1 | 4.985E-05 | 0.2722 | +1 | 2 |
| - | - | 641.7 | 184.1 | - | - | 0 | - |
| - | - | 1572 | 184.1 | - | - | 0 | - |
| - | - | 6035 | 185.1 | - | - | 0 | - |
| - | - | 683.9 | 186.1 | - | - | 0 | - |
| - | - | 4572 | 187.1 | - | - | 0 | - |
| - | - | 573 | 192.1 | - | - | 0 | - |
| - | - | 618.2 | 195.1 | - | - | 0 | - |
| - | - | 488 | 197 | - | - | 0 | - |
| - | - | 1952 | 197.1 | - | - | 0 | - |
| 2 | b | 8591 | 201.1 | 1.679E-05 | 0.08349 | +1 | 2 |
| - | - | 945 | 202.1 | - | - | 0 | - |
| - | - | 1675 | 202.1 | - | - | 0 | - |
| - | - | 508.8 | 203 | - | - | 0 | - |
| - | - | 977.2 | 210.1 | - | - | 0 | - |
| - | - | 490.8 | 210.6 | - | - | 0 | - |
| - | - | 2208 | 215.1 | - | - | 0 | - |
| - | - | 462.3 | 220.1 | - | - | 0 | - |
| - | - | 1580 | 223.1 | - | - | 0 | - |
| - | - | 504.6 | 224.1 | - | - | 0 | - |
| - | - | 585.2 | 225 | - | - | 0 | - |
| - | - | 968.5 | 225.1 | - | - | 0 | - |
| - | - | 2505 | 225.1 | - | - | 0 | - |
| - | - | 491.1 | 226 | - | - | 0 | - |
| - | - | 663.3 | 226.1 | - | - | 0 | - |
| - | - | 512.6 | 227 | - | - | 0 | - |
| - | - | 685.6 | 227.1 | - | - | 0 | - |
| - | - | 696.1 | 228.1 | - | - | 0 | - |
| - | - | 1746 | 233.1 | - | - | 0 | - |
| - | - | 3430 | 235.1 | - | - | 0 | - |
| - | - | 540.2 | 236.1 | - | - | 0 | - |
| - | - | 655.7 | 236.1 | - | - | 0 | - |
| - | - | 1.332E+04 | 238.1 | - | - | 0 | - |
| - | - | 1343 | 239.1 | - | - | 0 | - |
| 3 | a | 1368 | 242.1 | 4.543E-05 | 0.1876 | +1 | 3 |
| 9 | y | 1090 | 244.1 | 0.0005996 | 2.456 | +1 | 2 |
| 9 | y | 6194 | 245.1 | 0.0001718 | 0.7009 | +1 | 2 |
| - | - | 645.1 | 246.1 | - | - | 0 | - |
| - | - | 3958 | 252.1 | - | - | 0 | - |
| - | - | 604.1 | 254.1 | - | - | 0 | - |
| - | - | 3597 | 256.1 | - | - | 0 | - |
| 9 | y | 2.139E+04 | 262.2 | 7.115E-05 | 0.2714 | +1 | 2 |
| - | - | 707.1 | 263.1 | - | - | 0 | - |
| - | - | 1819 | 263.2 | - | - | 0 | - |
| - | - | 626.4 | 268.2 | - | - | 0 | - |
| 3 | b | 1.873E+04 | 270.1 | 0.0001792 | 0.6635 | +1 | 3 |
| - | - | 2707 | 271.1 | - | - | 0 | - |
| - | - | 572.9 | 281.1 | - | - | 0 | - |
| - | - | 575.5 | 282.1 | - | - | 0 | - |
| - | - | 970 | 283 | - | - | 0 | - |
| - | - | 599.6 | 284.2 | - | - | 0 | - |
| 3 | b | 1094 | 288.2 | 0.0003339 | 1.159 | +1 | 3 |
| - | - | 3895 | 299.1 | - | - | 0 | - |
| - | - | 5170 | 300.1 | - | - | 0 | - |
| - | - | 7590 | 301.1 | - | - | 0 | - |
| - | - | 5522 | 302.1 | - | - | 0 | - |
| - | - | 923.5 | 303.1 | - | - | 0 | - |
| - | - | 634.4 | 312.2 | - | - | 0 | - |
| - | - | 595.5 | 321.2 | - | - | 0 | - |
| 5 | y | 701 | 322.2 | 0.0003252 | 1.009 | +2 | 6 |
| - | - | 1974 | 339.2 | - | - | 0 | - |
| - | - | 2091 | 342.1 | - | - | 0 | - |
| - | - | 697.5 | 346 | - | - | 0 | - |
| - | - | 2112 | 347 | - | - | 0 | - |
| - | - | 1742 | 348 | - | - | 0 | - |
| - | - | 561.6 | 349 | - | - | 0 | - |
| - | - | 660 | 355.2 | - | - | 0 | - |
| - | - | 1830 | 357.2 | - | - | 0 | - |
| 8 | y | 780.2 | 358.2 | 0.0005736 | 1.601 | +1 | 3 |
| 8 | y | 2666 | 359.2 | 0.001008 | 2.805 | +1 | 3 |
| - | - | 810.3 | 360.2 | - | - | 0 | - |
| - | - | 899.6 | 361 | - | - | 0 | - |
| - | - | 975 | 362 | - | - | 0 | - |
| - | - | 582.6 | 363 | - | - | 0 | - |
| - | - | 580.3 | 364.2 | - | - | 0 | - |
| - | - | 833.7 | 365.2 | - | - | 0 | - |
| 4 | y | 5008 | 370.7 | 0.0001873 | 0.5053 | +2 | 7 |
| 4 | y | 2720 | 371.2 | 0.006554 | 17.66 | +2 | 7 |
| - | - | 877.5 | 375.2 | - | - | 0 | - |
| 8 | y | 5567 | 376.2 | 0.0007623 | 2.026 | +1 | 3 |
| - | - | 2777 | 377.2 | - | - | 0 | - |
| - | - | 619.4 | 378.2 | - | - | 0 | - |
| 4 | y | 3.015E+04 | 379.7 | 0.0004953 | 1.304 | +2 | 7 |
| - | - | 1.5E+04 | 380.2 | - | - | 0 | - |
| - | - | 4763 | 380.7 | - | - | 0 | - |
| - | - | 747.9 | 383.2 | - | - | 0 | - |
| - | - | 609.1 | 391.9 | - | - | 0 | - |
| - | - | 711.3 | 395.7 | - | - | 0 | - |
| - | - | 2221 | 402.2 | - | - | 0 | - |
| - | - | 2070 | 405.2 | - | - | 0 | - |
| - | - | 2279 | 405.7 | - | - | 0 | - |
| - | - | 714.1 | 406.2 | - | - | 0 | - |
| 3 | y | 2.116E+04 | 414.2 | 0.0002103 | 0.5077 | +2 | 8 |
| 3 | y | 1.447E+04 | 414.7 | 0.004761 | 11.48 | +2 | 8 |
| - | - | 4649 | 415.2 | - | - | 0 | - |
| - | - | 949.3 | 415.7 | - | - | 0 | - |
| - | - | 1678 | 417 | - | - | 0 | - |
| - | - | 1647 | 418 | - | - | 0 | - |
| - | - | 1463 | 419 | - | - | 0 | - |
| - | - | 1293 | 419 | - | - | 0 | - |
| - | - | 936.8 | 420 | - | - | 0 | - |
| 3 | y | 1.907E+04 | 423.2 | 0.0001826 | 0.4315 | +2 | 8 |
| - | - | 1.22E+04 | 423.7 | - | - | 0 | - |
| - | - | 3932 | 424.2 | - | - | 0 | - |
| - | - | 4201 | 431.2 | - | - | 0 | - |
| - | - | 1054 | 432.3 | - | - | 0 | - |
| - | - | 999.2 | 440.3 | - | - | 0 | - |
| - | - | 1357 | 441.2 | - | - | 0 | - |
| - | - | 1371 | 452.2 | - | - | 0 | - |
| - | - | 2.844E+04 | 458.3 | - | - | 0 | - |
| - | - | 7432 | 459.3 | - | - | 0 | - |
| - | - | 1495 | 460.3 | - | - | 0 | - |
| - | - | 952.4 | 468.1 | - | - | 0 | - |
| - | - | 881.3 | 488.6 | - | - | 0 | - |
| - | - | 1183 | 489.1 | - | - | 0 | - |
| 7 | y | 7697 | 489.3 | 0.0005672 | 1.159 | +1 | 4 |
| - | - | 2197 | 490.3 | - | - | 0 | - |
| - | - | 860.9 | 497.2 | - | - | 0 | - |
| - | - | 1003 | 498.3 | - | - | 0 | - |
| - | - | 877.4 | 505.8 | - | - | 0 | - |
| - | - | 676.1 | 507.1 | - | - | 0 | - |
| - | - | 723.9 | 507.2 | - | - | 0 | - |
| - | - | 1548 | 513.8 | - | - | 0 | - |
| 0 | Precursor | 1577 | 514.3 | 0.004179 | 8.127 | +2 | -1 |
| 0 | Precursor | 703.7 | 514.8 | 0.001009 | 1.96 | +2 | -1 |
| - | - | 850.2 | 516.9 | - | - | 0 | - |
| - | - | 769.5 | 521.3 | - | - | 0 | - |
| - | - | 3.575E+04 | 522.3 | - | - | 0 | - |
| - | - | 1.333E+04 | 523.3 | - | - | 0 | - |
| - | - | 1882 | 523.8 | - | - | 0 | - |
| - | - | 3357 | 524.3 | - | - | 0 | - |
| - | - | 972.3 | 524.8 | - | - | 0 | - |
| - | - | 3308 | 545.8 | - | - | 0 | - |
| - | - | 1959 | 546.3 | - | - | 0 | - |
| - | - | 822.6 | 546.3 | - | - | 0 | - |
| - | - | 1350 | 546.8 | - | - | 0 | - |
| - | - | 947.5 | 567.3 | - | - | 0 | - |
| 6 | y | 686.6 | 572.3 | 0.001259 | 2.201 | +1 | 5 |
| 6 | y | 898.9 | 573.3 | 0.0001512 | 0.2637 | +1 | 5 |
| 6 | y | 2.314E+04 | 590.3 | 0.0004552 | 0.771 | +1 | 5 |
| - | - | 9748 | 591.3 | - | - | 0 | - |
| - | - | 1725 | 592.3 | - | - | 0 | - |
| - | - | 794.9 | 602.3 | - | - | 0 | - |
| 5 | y | 2587 | 643.4 | 0.0008202 | 1.275 | +1 | 6 |
| 5 | y | 1402 | 644.3 | 0.005765 | 8.947 | +1 | 6 |
| - | - | 2756 | 652.8 | - | - | 0 | - |
| - | - | 1810 | 653.3 | - | - | 0 | - |
| - | - | 1436 | 653.8 | - | - | 0 | - |
| 5 | y | 1.959E+04 | 661.4 | 0.0008258 | 1.249 | +1 | 6 |
| - | - | 9167 | 662.4 | - | - | 0 | - |
| - | - | 2313 | 663.4 | - | - | 0 | - |
| - | - | 768 | 725.4 | - | - | 0 | - |
| 4 | y | 798.7 | 740.4 | 0.0005445 | 0.7354 | +1 | 7 |
| 4 | y | 1862 | 741.4 | 0.00104 | 1.402 | +1 | 7 |
| - | - | 975.6 | 742.4 | - | - | 0 | - |
| 4 | y | 3.202E+04 | 758.4 | 0.0009773 | 1.289 | +1 | 7 |
| - | - | 2.031E+04 | 759.4 | - | - | 0 | - |
| - | - | 5758 | 760.4 | - | - | 0 | - |
| 3 | y | 1022 | 827.4 | 0.002522 | 3.048 | +1 | 8 |
| 3 | y | 1.46E+04 | 845.4 | 0.002 | 2.365 | +1 | 8 |
| - | - | 9649 | 846.4 | - | - | 0 | - |
| - | - | 3815 | 847.4 | - | - | 0 | - |
| 2 | y | 1458 | 914.5 | 0.002773 | 3.032 | +1 | 9 |
| - | - | 767 | 929.5 | - | - | 0 | - |
| 2 | y | 1.107E+04 | 932.5 | 0.002656 | 2.849 | +1 | 9 |
| - | - | 8623 | 933.5 | - | - | 0 | - |
| - | - | 2998 | 934.5 | - | - | 0 | - |
| - | - | 963.7 | 942.5 | - | - | 0 | - |
| - | - | 763.6 | 943.5 | - | - | 0 | - |
| - | - | 749.9 | 1027 | - | - | 0 | - |
| - | - | 701.4 | 2638 | - | - | 0 | - |

m/z Charge Intensity FragmentType MassShift Position
121.02857971191406 0 400.9017
123.81367492675781 0 371.89932
125.10758972167969 0 1286.1969
128.10687255859375 0 734.7195
129.06591796875 0 419.83444
129.1024932861328 0 421.58755
130.06130981445312 0 1362.0956
134.0599822998047 0 420.51767
136.07579040527344 0 4681.459
139.08668518066406 0 1973.4387
141.10235595703125 0 8922.452
142.10536193847656 0 614.389
145.096923828125 0 832.535
147.0766143798828 0 923.03015
147.11297607421875 0 764.81305
148.79269409179688 0 464.0265
148.89288330078125 0 548.62836
148.90756225585938 0 643.5523
148.91482543945312 0 703.104
148.92184448242188 0 1210.8618
148.9288787841797 0 1415.0619
148.93617248535156 0 2686.9507
148.94387817382812 0 4389.03
148.9603271484375 0 3941.3428
148.96804809570312 0 2403.6675
148.97499084472656 0 1129.3094
148.98243713378906 0 1257.7384
148.98936462402344 0 1134.2108
148.99644470214844 0 736.86145
149.00352478027344 0 443.79816
149.01202392578125 0 503.099
149.04531860351562 0 1961.9948
149.0690155029297 0 521.5201
150.04415893554688 0 615.8698
151.0419921875 0 698.4448
155.08139038085938 0 826.5616
155.1179656982422 0 2470.6648 a Water loss 1
157.060791015625 0 1730.5251
157.0972137451172 0 6580.349
157.10806274414062 0 603.40576
158.0924835205078 0 5014.8145 y Ammonia loss 9
158.09967041015625 0 694.6666
167.05543518066406 0 2957.622
167.08160400390625 0 2336.5264
168.0552520751953 0 733.863
169.05223083496094 0 1850.8934
169.05966186523438 0 627.877
169.0972137451172 0 8568.373
169.1336669921875 0 2524.0342
170.10012817382812 0 682.53314
173.09217834472656 0 1745.4244
173.1284637451172 0 9308.089 a 1
173.44146728515625 0 870.35834
175.1107940673828 0 895.86926
175.11904907226562 0 11247.136 y 9
176.1230926513672 0 559.77704
183.11285400390625 0 15419.365 b Water loss 1
184.10797119140625 0 641.6799
184.11619567871094 0 1572.1433
185.0920867919922 0 6034.7056
186.0953369140625 0 683.94696
187.14410400390625 0 4571.9326
192.10128784179688 0 572.9928
195.0773162841797 0 618.24365
197.01626586914062 0 487.97626
197.128662109375 0 1951.583
201.12335205078125 0 8590.541 b 1
202.08193969726562 0 945.0133
202.11875915527344 0 1674.9039
202.98678588867188 0 508.76855
210.12332153320312 0 977.22375
210.6256561279297 0 490.84717
215.13894653320312 0 2207.88
220.13021850585938 0 462.3023
223.0639190673828 0 1579.8007
224.10391235351562 0 504.56073
225.04347229003906 0 585.17883
225.0604248046875 0 968.4952
225.12335205078125 0 2505.3313
226.04525756835938 0 491.09277
226.1181182861328 0 663.2751
227.04026794433594 0 512.5601
227.11363220214844 0 685.5751
228.13400268554688 0 696.1149
233.13209533691406 0 1746.0372
235.14381408691406 0 3429.5688
236.10238647460938 0 540.17694
236.14724731445312 0 655.66473
238.11866760253906 0 13320.738
239.1221160888672 0 1342.6177
242.14996337890625 0 1367.7648 a Water loss 2
244.1398162841797 0 1089.9254 y Water loss 8
245.12460327148438 0 6193.7983 y Ammonia loss 8
246.13040161132812 0 645.1106
252.13436889648438 0 3958.4578
254.1137237548828 0 604.09845
256.1290283203125 0 3596.5525
262.1509094238281 0 21390.145 y 8
263.1385192871094 0 707.1144
263.1541748046875 0 1818.8223
268.1661071777344 0 626.3739
270.1446533203125 0 18728.92 b Water loss 2
271.1479187011719 0 2706.6821
281.12469482421875 0 572.9416
282.10626220703125 0 575.48914
283.0489501953125 0 969.9708
284.1600341796875 0 599.5938
288.1557312011719 0 1094.4797 b 2
299.06170654296875 0 3894.5251
300.0621337890625 0 5169.658
301.0595397949219 0 7590.3105
302.0597839355469 0 5522.1143
303.05572509765625 0 923.4563
312.1558532714844 0 634.41925
321.1570739746094 0 595.51154
322.1794128417969 0 701.0297 y Water loss 4
339.16656494140625 0 1973.6022
342.1402587890625 0 2091.3027
345.97552490234375 0 697.49536
346.974365234375 0 2111.955
347.97454833984375 0 1742.111
348.9722900390625 0 561.6207
355.2332458496094 0 659.9907
357.1767578125 0 1830.4851
358.1827697753906 0 780.18146 y Water loss 7
359.1663513183594 0 2665.73 y Ammonia loss 7
360.151611328125 0 810.2941
361.0265197753906 0 899.6436
362.0258483886719 0 975.01685
363.0261535644531 0 582.58136
364.1850280761719 0 580.3021
365.2189025878906 0 833.6836
370.7059326171875 0 5008.281 y Water loss 3
371.2046813964844 0 2720.4912 y Ammonia loss 3
375.2358703613281 0 877.5127
376.1931457519531 0 5566.661 y 7
377.1770935058594 0 2777.2773
378.1793518066406 0 619.4095
379.7109069824219 0 30154.932 y 3
380.2076416015625 0 14998.11
380.7079162597656 0 4762.6924
383.2276916503906 0 747.9479
391.8552551269531 0 609.10254
395.6831970214844 0 711.25256
402.208984375 0 2220.6265
405.21710205078125 0 2069.9563
405.7108459472656 0 2279.4932
406.2105712890625 0 714.11584
414.221923828125 0 21164.596 y Water loss 2
414.7189025878906 0 14474.307 y Ammonia loss 2
415.2187805175781 0 4648.6094
415.7193298339844 0 949.34125
417.0336608886719 0 1678.0701
418.034912109375 0 1646.6987
418.995361328125 0 1463.0552
419.0330505371094 0 1292.577
419.9976806640625 0 936.8158
423.22723388671875 0 19068.234 y 2
423.7232971191406 0 12201.896
424.2244873046875 0 3931.931
431.2486877441406 0 4200.7896
432.25115966796875 0 1054.3187
440.2626953125 0 999.1724
441.2436828613281 0 1357.0831
452.24908447265625 0 1371.4509
458.2721252441406 0 28439.125
459.2745361328125 0 7432.1035
460.2756652832031 0 1495.0605
468.1218566894531 0 952.43396
488.64501953125 0 881.3232
489.05426025390625 0 1183.1691
489.27740478515625 0 7697.238 y 6
490.2651672363281 0 2197.128
497.2358093261719 0 860.8798
498.2542419433594 0 1002.9855
505.7713317871094 0 877.3771
507.1025085449219 0 676.0888
507.2461853027344 0 723.89764
513.7659912109375 0 1548.2642
514.2760009765625 0 1577.0498 Precursor Water loss
514.7711791992188 0 703.7477 Precursor Ammonia loss
516.9295654296875 0 850.1678
521.2735595703125 0 769.5223
522.27001953125 0 35750.45
523.27392578125 0 13326.894
523.781982421875 0 1882.4589
524.2734375 0 3357.3784
524.7767333984375 0 972.3448
545.763427734375 0 3308.1406
546.263427734375 0 1958.5992
546.3042602539062 0 822.5567
546.765380859375 0 1350.1393
567.2754516601562 0 947.5155
572.3163452148438 0 686.63403 y Water loss 5
573.2989501953125 0 898.8681 y Ammonia loss 5
590.3251953125 0 23140.037 y 5
591.3172607421875 0 9747.765
592.3204345703125 0 1725.2554
602.3035278320312 0 794.94025
643.3513793945312 0 2587.2468 y Water loss 4
644.3419799804688 0 1401.9081 y Ammonia loss 4
652.8277587890625 0 2756.2505
653.3281860351562 0 1809.999
653.8309936523438 0 1435.7664
661.3619384765625 0 19594.031 y 4
662.3562622070312 0 9167.385
663.3543701171875 0 2313.031
725.3563232421875 0 767.97595
740.4044189453125 0 798.6967 y Water loss 3
741.387939453125 0 1861.9836 y Ammonia loss 3
742.39111328125 0 975.6214
758.41455078125 0 32020.77 y 3
759.4078979492188 0 20305.705
760.4074096679688 0 5758.4507
827.4395141601562 0 1022.18884 y Water loss 2
845.445556640625 0 14600.461 y 2
846.4401245117188 0 9649.308
847.4427490234375 0 3815.0374
914.4662475585938 0 1457.7323 y Water loss 1
929.473388671875 0 766.9899
932.4769287109375 0 11068.88 y 1
933.47314453125 0 8623.065
934.4708862304688 0 2998.4219
942.4594116210938 0 963.73016
943.4678955078125 0 763.564
1026.5140380859375 0 749.91705
2638.43994140625 0 701.43665

Spectrum Details

|  |  |
| --- | --- |
| Matched peaks? Matched peaks The total absolute number of peaks matched. Additionally in brackets the total fraction of peaks matched and the total number of peaks is shown. | 38 (16.74% of 227) |
| FDR? FDR The false discovery rate estimated for this peptide. It is calculated by matching all theoretical fragments with a non-integer shift with the raw peaks for this spectrum. This is done with 40 different shifts. The resulting percentage is the average number of annotated peaks over the number of annotated peaks with the correct spectrum. | 0.56% |
| Satellite FDR? Satellite FDR See the FDR for details on its calculation. This satellite ion specific FDR only contains the satellite ions (d/w) for I/L/J positions. | - |
| PSM Score? PSM Score The PSM Score as given by Hecklib to this annotated spectrum. It is shown with three significant figures. | 439 |

#### Spectrum 3404? Spectrum 3404 The raw spectrum of this peptide as annotated by Hecklib. The fragments are coloured according to ion type (see legend). Any peaks with a star '\*' as text can be hovered over to see the full details, first the ion type second the mass shift type. By hovering over the amino acids in the peptide or ions in the legend the corresponding peaks are highlighted. By toggling the 'Unassigned' label you can turn the background (unassigned) peaks on or off in the plot. By updating the slider in the Ion legend you can update the spectrum to only show the top X% of the peaks with labels. The top X% means any peak that is within X% of the highest intensity. By dragging in the spectrum you can zoom in to a specific part of the spectrum and use 'Zoom Out' to get back to the original zoom level. The annotation of the spectrum is based on the given sequence in the peptides file and is done with different software so inconsistencies are likely. The peaks are annotated based on the given sequence, with 20 ppm tolerance.

Copy Data

##### Spectrum 3404 (TSV)

###### Preview

```
Loading example...
```

*Click on the button to copy the data to your clipboard.*

Mz MinMz MaxIntensity Max

WidthHeightPeptide font sizePeptide stroke widthSpectrum font sizeSpectrum stroke widthCompact peptide

Ion legend

wxyz

abcd

OtherUnassignedIon^Charge^_Position_Show for top:%

JSSPATJNSR

04.10e+38.19e+31.23e+41.64e+4

Zoom Out

a^+1^_2_y^+1^_1_a^+1^_2_y^+1^_1_b^+1^_2_b^+1^_2_y^+1^_2_y^+1^_2_b^+1^_3_y^+1^_3_y^+1^_3_y^+2^_7_y^+1^_3_y^+2^_7_y^+2^_8_y^+2^_8_y^+1^_4_\*\*y^+1^_5_y^+1^_6_y^+1^_6_y^+1^_7_y^+1^_7_y^+1^_8_y^+1^_9_

0806161124173223

Fragment Matches Table

Show background peaks

| Position | Ion type | Intensity | mz Theoretical | mz Error (Th) | mz Error (ppm) | Charge | Series Number |
| --- | --- | --- | --- | --- | --- | --- | --- |
| - | - | 381.7 | 123.1 | - | - | 0 | - |
| - | - | 444.1 | 125.1 | - | - | 0 | - |
| - | - | 417.4 | 127.1 | - | - | 0 | - |
| - | - | 518.4 | 127.1 | - | - | 0 | - |
| - | - | 482.8 | 129.1 | - | - | 0 | - |
| - | - | 561.8 | 129.1 | - | - | 0 | - |
| - | - | 380.1 | 131.7 | - | - | 0 | - |
| - | - | 474.4 | 135 | - | - | 0 | - |
| - | - | 1178 | 136.1 | - | - | 0 | - |
| - | - | 573.5 | 139.1 | - | - | 0 | - |
| - | - | 3563 | 141.1 | - | - | 0 | - |
| - | - | 397.1 | 142.6 | - | - | 0 | - |
| - | - | 436.1 | 146.4 | - | - | 0 | - |
| - | - | 507.7 | 147.1 | - | - | 0 | - |
| - | - | 637.7 | 147.1 | - | - | 0 | - |
| - | - | 390 | 148.8 | - | - | 0 | - |
| - | - | 483.9 | 148.9 | - | - | 0 | - |
| - | - | 463.3 | 148.9 | - | - | 0 | - |
| - | - | 662.3 | 148.9 | - | - | 0 | - |
| - | - | 868.1 | 148.9 | - | - | 0 | - |
| - | - | 962.8 | 148.9 | - | - | 0 | - |
| - | - | 1188 | 148.9 | - | - | 0 | - |
| - | - | 1208 | 148.9 | - | - | 0 | - |
| - | - | 1618 | 148.9 | - | - | 0 | - |
| - | - | 3584 | 148.9 | - | - | 0 | - |
| - | - | 5709 | 149 | - | - | 0 | - |
| - | - | 3545 | 149 | - | - | 0 | - |
| - | - | 1509 | 149 | - | - | 0 | - |
| - | - | 994.9 | 149 | - | - | 0 | - |
| - | - | 837.4 | 149 | - | - | 0 | - |
| - | - | 1031 | 149 | - | - | 0 | - |
| - | - | 697.5 | 149 | - | - | 0 | - |
| - | - | 613.1 | 149 | - | - | 0 | - |
| - | - | 511.3 | 149 | - | - | 0 | - |
| - | - | 405.3 | 149 | - | - | 0 | - |
| - | - | 486.1 | 149 | - | - | 0 | - |
| - | - | 447.5 | 149 | - | - | 0 | - |
| - | - | 2092 | 149 | - | - | 0 | - |
| - | - | 453.9 | 149.2 | - | - | 0 | - |
| - | - | 1276 | 151 | - | - | 0 | - |
| - | - | 659 | 155.1 | - | - | 0 | - |
| 2 | a | 1552 | 155.1 | 1.305E-07 | 0.0008411 | +1 | 2 |
| - | - | 969.4 | 157.1 | - | - | 0 | - |
| - | - | 3111 | 157.1 | - | - | 0 | - |
| 10 | y | 697.7 | 158.1 | 9.571E-05 | 0.6054 | +1 | 1 |
| - | - | 3753 | 167.1 | - | - | 0 | - |
| - | - | 1109 | 167.1 | - | - | 0 | - |
| - | - | 900.1 | 168.1 | - | - | 0 | - |
| - | - | 1357 | 169.1 | - | - | 0 | - |
| - | - | 4172 | 169.1 | - | - | 0 | - |
| - | - | 634.8 | 170.1 | - | - | 0 | - |
| 2 | a | 5234 | 173.1 | 5.735E-06 | 0.03312 | +1 | 2 |
| 10 | y | 2207 | 175.1 | 0.0001472 | 0.8408 | +1 | 1 |
| 2 | b | 8530 | 183.1 | 1.119E-05 | 0.0611 | +1 | 2 |
| - | - | 2794 | 185.1 | - | - | 0 | - |
| - | - | 1731 | 187.1 | - | - | 0 | - |
| - | - | 771.5 | 197.1 | - | - | 0 | - |
| 2 | b | 4336 | 201.1 | 4.731E-05 | 0.2352 | +1 | 2 |
| - | - | 1014 | 202.1 | - | - | 0 | - |
| - | - | 493.2 | 206.1 | - | - | 0 | - |
| - | - | 1217 | 215.1 | - | - | 0 | - |
| - | - | 479.4 | 221.3 | - | - | 0 | - |
| - | - | 1665 | 223.1 | - | - | 0 | - |
| - | - | 627.4 | 224.1 | - | - | 0 | - |
| - | - | 1199 | 225 | - | - | 0 | - |
| - | - | 705.6 | 225.1 | - | - | 0 | - |
| - | - | 1024 | 225.1 | - | - | 0 | - |
| - | - | 920.8 | 226 | - | - | 0 | - |
| - | - | 940.9 | 228.1 | - | - | 0 | - |
| - | - | 5375 | 238.1 | - | - | 0 | - |
| - | - | 817.8 | 240.1 | - | - | 0 | - |
| 9 | y | 537.9 | 245.1 | 0.001103 | 4.498 | +1 | 2 |
| - | - | 2005 | 252.1 | - | - | 0 | - |
| - | - | 686.8 | 254.1 | - | - | 0 | - |
| - | - | 1222 | 256.1 | - | - | 0 | - |
| 9 | y | 1655 | 262.2 | 0.000112 | 0.4271 | +1 | 2 |
| 3 | b | 1.043E+04 | 270.1 | 0.0001487 | 0.5506 | +1 | 3 |
| - | - | 1437 | 271.1 | - | - | 0 | - |
| - | - | 887 | 281.1 | - | - | 0 | - |
| - | - | 847.9 | 282.1 | - | - | 0 | - |
| - | - | 687.1 | 282.1 | - | - | 0 | - |
| - | - | 563.9 | 282.2 | - | - | 0 | - |
| - | - | 1177 | 283 | - | - | 0 | - |
| - | - | 930.4 | 283 | - | - | 0 | - |
| - | - | 1083 | 285 | - | - | 0 | - |
| - | - | 6285 | 299.1 | - | - | 0 | - |
| - | - | 8085 | 300.1 | - | - | 0 | - |
| - | - | 1.043E+04 | 301.1 | - | - | 0 | - |
| - | - | 6863 | 302.1 | - | - | 0 | - |
| - | - | 1942 | 303.1 | - | - | 0 | - |
| - | - | 553.7 | 304.9 | - | - | 0 | - |
| - | - | 631.7 | 309.2 | - | - | 0 | - |
| - | - | 1127 | 339.2 | - | - | 0 | - |
| - | - | 1004 | 341.2 | - | - | 0 | - |
| - | - | 571.3 | 345 | - | - | 0 | - |
| - | - | 687.1 | 346 | - | - | 0 | - |
| - | - | 1870 | 347 | - | - | 0 | - |
| - | - | 1681 | 348 | - | - | 0 | - |
| - | - | 774.3 | 349 | - | - | 0 | - |
| 8 | y | 615 | 358.2 | 0.0006041 | 1.687 | +1 | 3 |
| 8 | y | 988.3 | 359.2 | 0.0005804 | 1.616 | +1 | 3 |
| - | - | 954.5 | 360 | - | - | 0 | - |
| - | - | 1008 | 361 | - | - | 0 | - |
| - | - | 1617 | 362 | - | - | 0 | - |
| - | - | 630 | 365.2 | - | - | 0 | - |
| - | - | 540.7 | 369.7 | - | - | 0 | - |
| - | - | 540.2 | 369.8 | - | - | 0 | - |
| - | - | 979.4 | 370.2 | - | - | 0 | - |
| 4 | y | 1742 | 370.7 | 0.0008808 | 2.376 | +2 | 7 |
| - | - | 829.7 | 371.2 | - | - | 0 | - |
| - | - | 1917 | 375.2 | - | - | 0 | - |
| 8 | y | 4332 | 376.2 | 0.0001519 | 0.4038 | +1 | 3 |
| - | - | 562.1 | 376.5 | - | - | 0 | - |
| - | - | 651.8 | 377.2 | - | - | 0 | - |
| - | - | 6943 | 379.2 | - | - | 0 | - |
| 4 | y | 1.402E+04 | 379.7 | 0.0003287 | 0.8656 | +2 | 7 |
| - | - | 5988 | 380.2 | - | - | 0 | - |
| - | - | 1144 | 380.7 | - | - | 0 | - |
| - | - | 606.4 | 393.9 | - | - | 0 | - |
| - | - | 1035 | 405.2 | - | - | 0 | - |
| - | - | 4155 | 413.7 | - | - | 0 | - |
| 3 | y | 1.045E+04 | 414.2 | 0.0007052 | 1.702 | +2 | 8 |
| - | - | 5784 | 414.7 | - | - | 0 | - |
| - | - | 872.2 | 415.2 | - | - | 0 | - |
| - | - | 597.7 | 415.9 | - | - | 0 | - |
| - | - | 1251 | 417 | - | - | 0 | - |
| - | - | 2539 | 418 | - | - | 0 | - |
| - | - | 2441 | 419 | - | - | 0 | - |
| - | - | 1434 | 419 | - | - | 0 | - |
| - | - | 1389 | 420 | - | - | 0 | - |
| - | - | 836.7 | 421 | - | - | 0 | - |
| - | - | 2850 | 422.7 | - | - | 0 | - |
| 3 | y | 9730 | 423.2 | 0.001008 | 2.381 | +2 | 8 |
| - | - | 3110 | 423.7 | - | - | 0 | - |
| - | - | 1075 | 424.2 | - | - | 0 | - |
| - | - | 2684 | 488.3 | - | - | 0 | - |
| - | - | 1295 | 489.1 | - | - | 0 | - |
| 7 | y | 3839 | 489.3 | 0.00014 | 0.2861 | +1 | 4 |
| - | - | 1437 | 490.3 | - | - | 0 | - |
| - | - | 815.6 | 491.1 | - | - | 0 | - |
| - | - | 726.4 | 505.3 | - | - | 0 | - |
| 0 | Precursor | 1197 | 514.3 | 9.309E-05 | 0.181 | +2 | -1 |
| - | - | 730.5 | 522.8 | - | - | 0 | - |
| 0 | Precursor | 2186 | 523.3 | 0.0006091 | 1.164 | +2 | -1 |
| - | - | 1123 | 523.8 | - | - | 0 | - |
| - | - | 602 | 560.9 | - | - | 0 | - |
| - | - | 5530 | 589.3 | - | - | 0 | - |
| 6 | y | 9510 | 590.3 | 0.001437 | 2.434 | +1 | 5 |
| - | - | 4203 | 591.3 | - | - | 0 | - |
| 5 | y | 1296 | 643.4 | 0.0008278 | 1.287 | +1 | 6 |
| - | - | 4073 | 660.4 | - | - | 0 | - |
| 5 | y | 9252 | 661.4 | 0.001555 | 2.351 | +1 | 6 |
| - | - | 2091 | 662.4 | - | - | 0 | - |
| - | - | 702.6 | 671.3 | - | - | 0 | - |
| 4 | y | 991.5 | 740.4 | 0.002803 | 3.785 | +1 | 7 |
| - | - | 5114 | 757.4 | - | - | 0 | - |
| 4 | y | 1.622E+04 | 758.4 | 0.001464 | 1.93 | +1 | 7 |
| - | - | 6103 | 759.4 | - | - | 0 | - |
| - | - | 1684 | 760.4 | - | - | 0 | - |
| - | - | 2202 | 844.5 | - | - | 0 | - |
| 3 | y | 7060 | 845.4 | 0.001662 | 1.966 | +1 | 8 |
| - | - | 3263 | 846.5 | - | - | 0 | - |
| - | - | 1527 | 847.5 | - | - | 0 | - |
| - | - | 1065 | 931.5 | - | - | 0 | - |
| 2 | y | 5958 | 932.5 | 0.0007007 | 0.7514 | +1 | 9 |
| - | - | 3343 | 933.5 | - | - | 0 | - |
| - | - | 1329 | 934.5 | - | - | 0 | - |
| - | - | 679.5 | 1057 | - | - | 0 | - |
| - | - | 717.6 | 1659 | - | - | 0 | - |
| - | - | 757.2 | 3191 | - | - | 0 | - |

m/z Charge Intensity FragmentType MassShift Position
123.0909423828125 0 381.7413
125.10746002197266 0 444.08646
127.07215118408203 0 417.40356
127.08634948730469 0 518.3917
129.0658416748047 0 482.78293
129.10223388671875 0 561.80396
131.693603515625 0 380.13983
135.04315185546875 0 474.44052
136.07566833496094 0 1178.1267
139.08676147460938 0 573.50653
141.102294921875 0 3562.7124
142.62734985351562 0 397.05298
146.3690185546875 0 436.1423
147.07638549804688 0 507.72693
147.11279296875 0 637.6849
148.8197021484375 0 390.0412
148.86924743652344 0 483.91583
148.89022827148438 0 463.26807
148.89736938476562 0 662.254
148.90467834472656 0 868.0748
148.91165161132812 0 962.82465
148.91888427734375 0 1187.502
148.92601013183594 0 1207.8196
148.93284606933594 0 1617.9974
148.9403533935547 0 3584.1973
148.9565887451172 0 5709.425
148.96420288085938 0 3544.8372
148.9717254638672 0 1508.531
148.978515625 0 994.93475
148.9857940673828 0 837.35913
148.99285888671875 0 1030.7571
148.999755859375 0 697.4791
149.00723266601562 0 613.1061
149.013671875 0 511.25714
149.02076721191406 0 405.31604
149.02813720703125 0 486.0738
149.03515625 0 447.53107
149.04473876953125 0 2092.3801
149.2476043701172 0 453.92365
151.04164123535156 0 1276.3807
155.08160400390625 0 658.9676
155.11788940429688 0 1552.2013 a Water loss 1
157.0606231689453 0 969.4223
157.09718322753906 0 3110.8381
158.09249877929688 0 697.68744 y Ammonia loss 9
167.05548095703125 0 3752.581
167.08177185058594 0 1108.9983
168.0547332763672 0 900.08453
169.05262756347656 0 1356.9111
169.09710693359375 0 4171.6826
170.05223083496094 0 634.8127
173.12844848632812 0 5233.9 a 1
175.11880493164062 0 2206.8457 y 9
183.11279296875 0 8530.07 b Water loss 1
185.09207153320312 0 2794.4382
187.14419555664062 0 1731.1536
197.12783813476562 0 771.5088
201.12332153320312 0 4336.04 b 1
202.0816650390625 0 1013.6451
206.1044921875 0 493.20557
215.1387176513672 0 1216.5446
221.2584991455078 0 479.40558
223.06349182128906 0 1665.0967
224.06370544433594 0 627.41473
225.04263305664062 0 1199.0787
225.0615997314453 0 705.64355
225.1233367919922 0 1023.92847
226.04312133789062 0 920.7528
228.13436889648438 0 940.89075
238.1186981201172 0 5375.462
240.09519958496094 0 817.8256
245.1255340576172 0 537.93427 y Ammonia loss 8
252.13426208496094 0 2005.4092
254.1129150390625 0 686.79224
256.1296081542969 0 1221.7229
262.1510925292969 0 1655.4921 y 8
270.1446838378906 0 10431.919 b Water loss 2
271.1479187011719 0 1437.4517
281.0508117675781 0 887.00555
282.0522766113281 0 847.8839
282.1082458496094 0 687.121
282.18145751953125 0 563.9381
283.0308837890625 0 1176.8708
283.047607421875 0 930.38776
285.0271911621094 0 1083.2472
299.0616455078125 0 6284.67
300.0620422363281 0 8085.4014
301.0593566894531 0 10426.507
302.06011962890625 0 6863.38
303.0569152832031 0 1941.6598
304.8558654785156 0 553.71436
309.2039794921875 0 631.69855
339.1671142578125 0 1126.842
341.1578674316406 0 1003.7752
344.97698974609375 0 571.25256
345.97601318359375 0 687.1293
346.9742126464844 0 1870.4563
347.9747009277344 0 1680.5352
348.97052001953125 0 774.28876
358.1827392578125 0 615.00085 y Water loss 7
359.1667785644531 0 988.3091 y Ammonia loss 7
360.0268859863281 0 954.5446
361.0257263183594 0 1007.6618
362.0267333984375 0 1617.3354
365.21710205078125 0 629.9827
369.6974182128906 0 540.66705
369.8280334472656 0 540.1629
370.2135009765625 0 979.3984
370.7070007324219 0 1742.3143 y Water loss 3
371.2086486816406 0 829.7115
375.2098083496094 0 1916.8889
376.1937561035156 0 4332.4927 y 7
376.5417785644531 0 562.1372
377.19647216796875 0 651.7842
379.2190856933594 0 6942.8296
379.71173095703125 0 14015.479 y 3
380.2126159667969 0 5987.775
380.7145690917969 0 1143.8777
393.8713073730469 0 606.3746
405.2171936035156 0 1034.9484
413.730224609375 0 4154.967
414.22283935546875 0 10447.132 y Water loss 2
414.7234191894531 0 5783.784
415.2234191894531 0 872.208
415.9082336425781 0 597.735
417.03607177734375 0 1251.1248
418.0356750488281 0 2539.3545
418.99493408203125 0 2441.206
419.032958984375 0 1433.9082
419.9963073730469 0 1388.8961
420.99395751953125 0 836.70074
422.7353820800781 0 2849.7202
423.2284240722656 0 9729.908 y 2
423.7294921875 0 3110.0862
424.2296142578125 0 1074.7482
488.2938537597656 0 2684.186
489.05352783203125 0 1294.748
489.27783203125 0 3839.487 y 6
490.2791442871094 0 1436.7351
491.0556640625 0 815.5754
505.2768249511719 0 726.43945
514.2802734375 0 1196.9009 Precursor Water loss
522.787109375 0 730.47125
523.2860717773438 0 2186.1177 Precursor
523.788330078125 0 1123.0876
560.8603515625 0 601.98975
589.3417358398438 0 5529.6587
590.3270874023438 0 9509.837 y 5
591.3287353515625 0 4203.1914
643.35302734375 0 1296.4929 y Water loss 4
660.3779907226562 0 4073.2126
661.3643188476562 0 9251.78 y 4
662.364990234375 0 2091.3506
671.34423828125 0 702.6133
740.4021606445312 0 991.4658 y Water loss 3
757.4306030273438 0 5114.2837
758.4169921875 0 16222.336 y 3
759.4187622070312 0 6102.6177
760.4189453125 0 1684.0646
844.4616088867188 0 2202.1748
845.44921875 0 7060.075 y 2
846.451416015625 0 3262.9944
847.455078125 0 1527.1923
931.4917602539062 0 1064.5918
932.4802856445312 0 5958.445 y 1
933.4832153320312 0 3343.2449
934.4777221679688 0 1328.5592
1057.306640625 0 679.5161
1658.852783203125 0 717.6458
3190.903076171875 0 757.1603

Spectrum Details

|  |  |
| --- | --- |
| Matched peaks? Matched peaks The total absolute number of peaks matched. Additionally in brackets the total fraction of peaks matched and the total number of peaks is shown. | 26 (15.29% of 170) |
| FDR? FDR The false discovery rate estimated for this peptide. It is calculated by matching all theoretical fragments with a non-integer shift with the raw peaks for this spectrum. This is done with 40 different shifts. The resulting percentage is the average number of annotated peaks over the number of annotated peaks with the correct spectrum. | 1.28% |
| Satellite FDR? Satellite FDR See the FDR for details on its calculation. This satellite ion specific FDR only contains the satellite ions (d/w) for I/L/J positions. | - |
| PSM Score? PSM Score The PSM Score as given by Hecklib to this annotated spectrum. It is shown with three significant figures. | 262 |

#### Spectrum 4042? Spectrum 4042 The raw spectrum of this peptide as annotated by Hecklib. The fragments are coloured according to ion type (see legend). Any peaks with a star '\*' as text can be hovered over to see the full details, first the ion type second the mass shift type. By hovering over the amino acids in the peptide or ions in the legend the corresponding peaks are highlighted. By toggling the 'Unassigned' label you can turn the background (unassigned) peaks on or off in the plot. By updating the slider in the Ion legend you can update the spectrum to only show the top X% of the peaks with labels. The top X% means any peak that is within X% of the highest intensity. By dragging in the spectrum you can zoom in to a specific part of the spectrum and use 'Zoom Out' to get back to the original zoom level. The annotation of the spectrum is based on the given sequence in the peptides file and is done with different software so inconsistencies are likely. The peaks are annotated based on the given sequence, with 20 ppm tolerance.

Copy Data

##### Spectrum 4042 (TSV)

###### Preview

```
Loading example...
```

*Click on the button to copy the data to your clipboard.*

Mz MinMz MaxIntensity Max

WidthHeightPeptide font sizePeptide stroke widthSpectrum font sizeSpectrum stroke widthCompact peptide

Ion legend

wxyz

abcd

OtherUnassignedIon^Charge^_Position_Show for top:%

JSSPATJNSR

04.34e+38.67e+31.30e+41.73e+4

Zoom Out

a^+1^_2_y^+1^_1_a^+1^_2_y^+1^_1_b^+1^_2_b^+1^_2_y^+1^_2_y^+1^_2_b^+1^_3_y^+1^_3_y^+2^_7_y^+1^_3_y^+2^_7_y^+2^_8_y^+2^_8_y^+2^_8_y^+2^_9_y^+1^_4_\*\*\*y^+1^_5_y^+1^_5_y^+1^_6_y^+1^_6_y^+1^_7_y^+1^_7_y^+1^_8_y^+1^_9_y^+1^_9_

0830166024903320

Fragment Matches Table

Show background peaks

| Position | Ion type | Intensity | mz Theoretical | mz Error (Th) | mz Error (ppm) | Charge | Series Number |
| --- | --- | --- | --- | --- | --- | --- | --- |
| - | - | 451.3 | 120.1 | - | - | 0 | - |
| - | - | 572.9 | 125.1 | - | - | 0 | - |
| - | - | 331.7 | 125.8 | - | - | 0 | - |
| - | - | 336.5 | 125.8 | - | - | 0 | - |
| - | - | 409.6 | 125.9 | - | - | 0 | - |
| - | - | 782.2 | 129.1 | - | - | 0 | - |
| - | - | 3072 | 136.1 | - | - | 0 | - |
| - | - | 376.6 | 138.6 | - | - | 0 | - |
| - | - | 1412 | 139.1 | - | - | 0 | - |
| - | - | 383.1 | 140.6 | - | - | 0 | - |
| - | - | 3155 | 141.1 | - | - | 0 | - |
| - | - | 569.6 | 142.1 | - | - | 0 | - |
| - | - | 398.1 | 147 | - | - | 0 | - |
| - | - | 676 | 147.1 | - | - | 0 | - |
| - | - | 536.8 | 148.8 | - | - | 0 | - |
| - | - | 570.4 | 148.9 | - | - | 0 | - |
| - | - | 378.9 | 148.9 | - | - | 0 | - |
| - | - | 436.3 | 148.9 | - | - | 0 | - |
| - | - | 637.4 | 148.9 | - | - | 0 | - |
| - | - | 774.7 | 148.9 | - | - | 0 | - |
| - | - | 1101 | 148.9 | - | - | 0 | - |
| - | - | 1294 | 148.9 | - | - | 0 | - |
| - | - | 1416 | 148.9 | - | - | 0 | - |
| - | - | 3534 | 148.9 | - | - | 0 | - |
| - | - | 6435 | 149 | - | - | 0 | - |
| - | - | 3933 | 149 | - | - | 0 | - |
| - | - | 1571 | 149 | - | - | 0 | - |
| - | - | 1192 | 149 | - | - | 0 | - |
| - | - | 1183 | 149 | - | - | 0 | - |
| - | - | 554.4 | 149 | - | - | 0 | - |
| - | - | 619.1 | 149 | - | - | 0 | - |
| - | - | 716.4 | 149 | - | - | 0 | - |
| - | - | 518.7 | 149 | - | - | 0 | - |
| - | - | 516.3 | 149 | - | - | 0 | - |
| - | - | 472.7 | 149 | - | - | 0 | - |
| - | - | 2313 | 149 | - | - | 0 | - |
| - | - | 416.7 | 149.2 | - | - | 0 | - |
| - | - | 1322 | 151 | - | - | 0 | - |
| - | - | 431.5 | 154.4 | - | - | 0 | - |
| 2 | a | 1197 | 155.1 | 9.142E-05 | 0.5894 | +1 | 2 |
| - | - | 846.1 | 157.1 | - | - | 0 | - |
| - | - | 3163 | 157.1 | - | - | 0 | - |
| 10 | y | 1549 | 158.1 | 9.571E-05 | 0.6054 | +1 | 1 |
| - | - | 4087 | 167.1 | - | - | 0 | - |
| - | - | 867.3 | 167.1 | - | - | 0 | - |
| - | - | 429.1 | 167.1 | - | - | 0 | - |
| - | - | 916.7 | 168.1 | - | - | 0 | - |
| - | - | 1998 | 169.1 | - | - | 0 | - |
| - | - | 4197 | 169.1 | - | - | 0 | - |
| - | - | 810.8 | 173.1 | - | - | 0 | - |
| 2 | a | 5364 | 173.1 | 2.099E-05 | 0.1213 | +1 | 2 |
| 10 | y | 3394 | 175.1 | 4.042E-05 | 0.2308 | +1 | 1 |
| 2 | b | 6825 | 183.1 | 4.171E-05 | 0.2278 | +1 | 2 |
| - | - | 680.1 | 183.1 | - | - | 0 | - |
| - | - | 2973 | 185.1 | - | - | 0 | - |
| - | - | 591.4 | 185.6 | - | - | 0 | - |
| - | - | 1476 | 187.1 | - | - | 0 | - |
| - | - | 622.5 | 192.1 | - | - | 0 | - |
| - | - | 879.3 | 197.1 | - | - | 0 | - |
| - | - | 456.9 | 197.3 | - | - | 0 | - |
| 2 | b | 4416 | 201.1 | 0.0001389 | 0.6904 | +1 | 2 |
| - | - | 608.5 | 202.1 | - | - | 0 | - |
| - | - | 897.8 | 215.1 | - | - | 0 | - |
| - | - | 482.8 | 217.5 | - | - | 0 | - |
| - | - | 587.6 | 222.1 | - | - | 0 | - |
| - | - | 1274 | 223.1 | - | - | 0 | - |
| - | - | 859.4 | 224.1 | - | - | 0 | - |
| - | - | 598.9 | 224.1 | - | - | 0 | - |
| - | - | 1176 | 225 | - | - | 0 | - |
| - | - | 646.6 | 225.1 | - | - | 0 | - |
| - | - | 869.5 | 225.1 | - | - | 0 | - |
| - | - | 892.5 | 226 | - | - | 0 | - |
| - | - | 642.4 | 226.1 | - | - | 0 | - |
| - | - | 802.2 | 227 | - | - | 0 | - |
| - | - | 1318 | 235.1 | - | - | 0 | - |
| - | - | 6552 | 238.1 | - | - | 0 | - |
| - | - | 850.9 | 240.1 | - | - | 0 | - |
| 9 | y | 1496 | 245.1 | 0.0003244 | 1.323 | +1 | 2 |
| - | - | 2244 | 252.1 | - | - | 0 | - |
| - | - | 1160 | 256.1 | - | - | 0 | - |
| 9 | y | 7500 | 262.2 | 0.0001322 | 0.5042 | +1 | 2 |
| - | - | 745.6 | 263.2 | - | - | 0 | - |
| 3 | b | 9101 | 270.1 | 0.0002403 | 0.8895 | +1 | 3 |
| - | - | 1107 | 271.1 | - | - | 0 | - |
| - | - | 1050 | 282.1 | - | - | 0 | - |
| - | - | 1412 | 283 | - | - | 0 | - |
| - | - | 609.9 | 284 | - | - | 0 | - |
| - | - | 945.8 | 284 | - | - | 0 | - |
| - | - | 5501 | 299.1 | - | - | 0 | - |
| - | - | 6792 | 300.1 | - | - | 0 | - |
| - | - | 1.193E+04 | 301.1 | - | - | 0 | - |
| - | - | 7393 | 302.1 | - | - | 0 | - |
| - | - | 2566 | 303.1 | - | - | 0 | - |
| - | - | 816.6 | 339.2 | - | - | 0 | - |
| - | - | 846.4 | 342.1 | - | - | 0 | - |
| - | - | 1596 | 346 | - | - | 0 | - |
| - | - | 1707 | 347 | - | - | 0 | - |
| - | - | 1946 | 348 | - | - | 0 | - |
| - | - | 938.2 | 349 | - | - | 0 | - |
| - | - | 726.7 | 355.2 | - | - | 0 | - |
| - | - | 826.6 | 357.2 | - | - | 0 | - |
| - | - | 568.1 | 357.5 | - | - | 0 | - |
| 8 | y | 962.2 | 359.2 | 0.0003362 | 0.9361 | +1 | 3 |
| - | - | 876.7 | 360 | - | - | 0 | - |
| - | - | 1604 | 361 | - | - | 0 | - |
| - | - | 1226 | 362 | - | - | 0 | - |
| 4 | y | 3073 | 370.7 | 0.0007977 | 2.152 | +2 | 7 |
| - | - | 901.3 | 371.2 | - | - | 0 | - |
| 8 | y | 3244 | 376.2 | 0.0001519 | 0.4038 | +1 | 3 |
| - | - | 1005 | 377.2 | - | - | 0 | - |
| 4 | y | 1.7E+04 | 379.7 | 0.0005869 | 1.546 | +2 | 7 |
| - | - | 6785 | 380.2 | - | - | 0 | - |
| - | - | 2285 | 380.7 | - | - | 0 | - |
| - | - | 1112 | 405.2 | - | - | 0 | - |
| - | - | 648.3 | 405.7 | - | - | 0 | - |
| 3 | y | 1.07E+04 | 414.2 | 0.0003019 | 0.7288 | +2 | 8 |
| 3 | y | 4639 | 414.7 | 0.006042 | 14.57 | +2 | 8 |
| - | - | 1730 | 415.2 | - | - | 0 | - |
| - | - | 1932 | 417 | - | - | 0 | - |
| - | - | 3493 | 418 | - | - | 0 | - |
| - | - | 864 | 418.2 | - | - | 0 | - |
| - | - | 1931 | 419 | - | - | 0 | - |
| - | - | 1766 | 419 | - | - | 0 | - |
| - | - | 716.8 | 420 | - | - | 0 | - |
| 3 | y | 1.148E+04 | 423.2 | 0.0001216 | 0.2872 | +2 | 8 |
| - | - | 4075 | 423.7 | - | - | 0 | - |
| - | - | 1067 | 424.2 | - | - | 0 | - |
| - | - | 1494 | 424.2 | - | - | 0 | - |
| - | - | 749.6 | 431.2 | - | - | 0 | - |
| - | - | 3020 | 458.3 | - | - | 0 | - |
| - | - | 696.5 | 459.3 | - | - | 0 | - |
| 2 | y | 652.7 | 466.7 | 0.0007099 | 1.521 | +2 | 9 |
| - | - | 1182 | 489.1 | - | - | 0 | - |
| 7 | y | 4860 | 489.3 | 0.00014 | 0.2861 | +1 | 4 |
| - | - | 886.5 | 490.3 | - | - | 0 | - |
| - | - | 638.3 | 506.1 | - | - | 0 | - |
| - | - | 599.9 | 506.3 | - | - | 0 | - |
| 0 | Precursor | 724.3 | 514.3 | 0.0005783 | 1.124 | +2 | -1 |
| 0 | Precursor | 841.6 | 514.8 | 0.004728 | 9.185 | +2 | -1 |
| - | - | 4805 | 522.3 | - | - | 0 | - |
| - | - | 852.3 | 522.8 | - | - | 0 | - |
| 0 | Precursor | 2340 | 523.3 | 0.005739 | 10.97 | +2 | -1 |
| - | - | 720.3 | 523.8 | - | - | 0 | - |
| - | - | 750.6 | 524.2 | - | - | 0 | - |
| - | - | 2295 | 545.8 | - | - | 0 | - |
| 6 | y | 593.6 | 572.3 | 0.0006937 | 1.212 | +1 | 5 |
| 6 | y | 1.221E+04 | 590.3 | 0.0008824 | 1.495 | +1 | 5 |
| - | - | 3915 | 591.3 | - | - | 0 | - |
| 5 | y | 1416 | 643.4 | 0.001614 | 2.508 | +1 | 6 |
| - | - | 1354 | 653.3 | - | - | 0 | - |
| 5 | y | 1.074E+04 | 661.4 | 0.0007037 | 1.064 | +1 | 6 |
| - | - | 4497 | 662.4 | - | - | 0 | - |
| - | - | 786 | 663.4 | - | - | 0 | - |
| - | - | 665.3 | 709.3 | - | - | 0 | - |
| - | - | 647 | 726.4 | - | - | 0 | - |
| 4 | y | 735.7 | 741.4 | 0.0002461 | 0.332 | +1 | 7 |
| 4 | y | 1.717E+04 | 758.4 | 0.00116 | 1.53 | +1 | 7 |
| - | - | 8224 | 759.4 | - | - | 0 | - |
| - | - | 2635 | 760.4 | - | - | 0 | - |
| 3 | y | 8330 | 845.4 | 0.001267 | 1.499 | +1 | 8 |
| - | - | 3826 | 846.4 | - | - | 0 | - |
| - | - | 1500 | 847.4 | - | - | 0 | - |
| 2 | y | 621.7 | 914.5 | 0.00272 | 2.975 | +1 | 9 |
| 2 | y | 5378 | 932.5 | 0.00229 | 2.456 | +1 | 9 |
| - | - | 2953 | 933.5 | - | - | 0 | - |
| - | - | 1835 | 934.5 | - | - | 0 | - |
| - | - | 615.5 | 1091 | - | - | 0 | - |
| - | - | 655.4 | 1131 | - | - | 0 | - |
| - | - | 588.2 | 2446 | - | - | 0 | - |
| - | - | 655.6 | 2963 | - | - | 0 | - |
| - | - | 688.5 | 3287 | - | - | 0 | - |

m/z Charge Intensity FragmentType MassShift Position
120.08075714111328 0 451.30807
125.1076889038086 0 572.9426
125.75416564941406 0 331.70627
125.83147430419922 0 336.51797
125.89170837402344 0 409.60095
129.1023406982422 0 782.1801
136.07571411132812 0 3071.7847
138.6365203857422 0 376.63043
139.08677673339844 0 1412.3193
140.6178436279297 0 383.1479
141.1023406982422 0 3154.7214
142.10609436035156 0 569.61694
147.0355987548828 0 398.06088
147.07635498046875 0 675.9764
148.84629821777344 0 536.80475
148.86778259277344 0 570.39185
148.88218688964844 0 378.90454
148.89671325683594 0 436.33395
148.90342712402344 0 637.423
148.91038513183594 0 774.7472
148.91781616210938 0 1100.9109
148.92491149902344 0 1293.9028
148.9319305419922 0 1415.571
148.93942260742188 0 3533.8757
148.95562744140625 0 6434.6025
148.9633331298828 0 3933.2979
148.9708709716797 0 1571.2218
148.97779846191406 0 1191.5981
148.9847869873047 0 1183.2611
148.99217224121094 0 554.3801
148.99940490722656 0 619.1156
149.00636291503906 0 716.41064
149.0131072998047 0 518.6513
149.0210418701172 0 516.26685
149.02822875976562 0 472.72873
149.0447998046875 0 2313.153
149.17039489746094 0 416.6775
151.04173278808594 0 1322.007
154.36434936523438 0 431.52576
155.11798095703125 0 1196.9475 a Water loss 1
157.0608367919922 0 846.0802
157.09725952148438 0 3163.045
158.09249877929688 0 1549 y Ammonia loss 9
167.05535888671875 0 4087.141
167.08132934570312 0 867.3063
167.08963012695312 0 429.13773
168.054931640625 0 916.6783
169.05210876464844 0 1998.2384
169.0971221923828 0 4197.4956
173.09185791015625 0 810.81464
173.12843322753906 0 5364.175 a 1
175.11891174316406 0 3393.6633 y 9
183.11276245117188 0 6824.6655 b Water loss 1
183.14895629882812 0 680.0579
185.09201049804688 0 2972.5085
185.63194274902344 0 591.38727
187.1441650390625 0 1476.2698
192.10348510742188 0 622.5442
197.12879943847656 0 879.3046
197.34768676757812 0 456.86783
201.12322998046875 0 4415.8506 b 1
202.08238220214844 0 608.522
215.1393280029297 0 897.767
217.4929962158203 0 482.8404
222.0842742919922 0 587.642
223.06362915039062 0 1273.703
224.06483459472656 0 859.4063
224.13877868652344 0 598.8792
225.04270935058594 0 1175.5969
225.06039428710938 0 646.61725
225.12286376953125 0 869.48816
226.04420471191406 0 892.46936
226.11907958984375 0 642.38367
227.03955078125 0 802.1855
235.14356994628906 0 1317.6952
238.1185302734375 0 6551.9087
240.09518432617188 0 850.8691
245.124755859375 0 1496.0892 y Ammonia loss 8
252.13436889648438 0 2244.0479
256.1283874511719 0 1159.8844
262.1508483886719 0 7500.4355 y 8
263.1539306640625 0 745.62714
270.14459228515625 0 9101.232 b Water loss 2
271.14874267578125 0 1107.1389
282.0511779785156 0 1049.5715
283.0487060546875 0 1412.3094
284.0298156738281 0 609.93414
284.04931640625 0 945.8107
299.0617370605469 0 5500.9805
300.0623474121094 0 6792.0024
301.05926513671875 0 11925.099
302.05963134765625 0 7392.6816
303.05682373046875 0 2565.7874
339.166748046875 0 816.5818
342.1402893066406 0 846.3541
345.9756164550781 0 1595.541
346.975341796875 0 1706.8873
347.9740905761719 0 1945.6661
348.971435546875 0 938.22034
355.2347412109375 0 726.6547
357.17681884765625 0 826.5757
357.4622802734375 0 568.0538
359.1670227050781 0 962.2374 y Ammonia loss 7
360.0283508300781 0 876.71094
361.02508544921875 0 1603.8735
362.02655029296875 0 1225.7911
370.705322265625 0 3072.8962 y Water loss 3
371.2066955566406 0 901.325
376.1937561035156 0 3243.9563 y 7
377.17626953125 0 1005.0933
379.7108154296875 0 17004.684 y 3
380.2094421386719 0 6785.4395
380.7087707519531 0 2285.1746
405.2174072265625 0 1111.6215
405.7183837890625 0 648.3498
414.2218322753906 0 10697.222 y Water loss 2
414.7201843261719 0 4639.1577 y Ammonia loss 2
415.2193298339844 0 1730.0929
417.0343017578125 0 1931.8573
418.0345764160156 0 3493.2705
418.1910095214844 0 863.9718
418.9952697753906 0 1931.3536
419.0332946777344 0 1765.8461
419.9954833984375 0 716.84686
423.227294921875 0 11477.609 y 2
423.72601318359375 0 4074.9019
424.19891357421875 0 1067.0972
424.2285461425781 0 1493.9409
431.248779296875 0 749.573
458.2714538574219 0 3020.292
459.2737121582031 0 696.4708
466.744140625 0 652.72156 y 1
489.053466796875 0 1181.8197
489.27783203125 0 4860.278 y 6
490.2796936035156 0 886.5085
506.106689453125 0 638.29517
506.2692565917969 0 599.89014
514.2796020507812 0 724.2556 Precursor Water loss
514.7769165039062 0 841.5532 Precursor Ammonia loss
522.27001953125 0 4805.3735
522.7814331054688 0 852.2977
523.2797241210938 0 2339.9182 Precursor
523.778076171875 0 720.2738
524.21875 0 750.6292
545.7625122070312 0 2294.6133
572.3143920898438 0 593.558 y Water loss 5
590.3247680664062 0 12207.421 y 5
591.3240356445312 0 3914.8857
643.3505859375 0 1416.4124 y Water loss 4
653.3284912109375 0 1353.9559
661.362060546875 0 10738.711 y 4
662.3615112304688 0 4497.332
663.3614501953125 0 786.0328
709.3253173828125 0 665.2794
726.3596801757812 0 647.03925
741.3887329101562 0 735.7296 y Ammonia loss 3
758.4143676757812 0 17169.502 y 3
759.4121704101562 0 8223.514
760.4140014648438 0 2635.2058
845.4462890625 0 8330.406 y 2
846.4425048828125 0 3825.998
847.4445190429688 0 1499.5725
914.4717407226562 0 621.67084 y Water loss 1
932.477294921875 0 5377.9346 y 1
933.4739990234375 0 2953.2458
934.4733276367188 0 1834.7332
1090.5343017578125 0 615.5425
1131.45361328125 0 655.4499
2446.1376953125 0 588.22186
2962.715576171875 0 655.57587
3286.819580078125 0 688.5048

Spectrum Details

|  |  |
| --- | --- |
| Matched peaks? Matched peaks The total absolute number of peaks matched. Additionally in brackets the total fraction of peaks matched and the total number of peaks is shown. | 30 (17.54% of 171) |
| FDR? FDR The false discovery rate estimated for this peptide. It is calculated by matching all theoretical fragments with a non-integer shift with the raw peaks for this spectrum. This is done with 40 different shifts. The resulting percentage is the average number of annotated peaks over the number of annotated peaks with the correct spectrum. | 1.03% |
| Satellite FDR? Satellite FDR See the FDR for details on its calculation. This satellite ion specific FDR only contains the satellite ions (d/w) for I/L/J positions. | - |
| PSM Score? PSM Score The PSM Score as given by Hecklib to this annotated spectrum. It is shown with three significant figures. | 281 |

#### Spectrum 4222? Spectrum 4222 The raw spectrum of this peptide as annotated by Hecklib. The fragments are coloured according to ion type (see legend). Any peaks with a star '\*' as text can be hovered over to see the full details, first the ion type second the mass shift type. By hovering over the amino acids in the peptide or ions in the legend the corresponding peaks are highlighted. By toggling the 'Unassigned' label you can turn the background (unassigned) peaks on or off in the plot. By updating the slider in the Ion legend you can update the spectrum to only show the top X% of the peaks with labels. The top X% means any peak that is within X% of the highest intensity. By dragging in the spectrum you can zoom in to a specific part of the spectrum and use 'Zoom Out' to get back to the original zoom level. The annotation of the spectrum is based on the given sequence in the peptides file and is done with different software so inconsistencies are likely. The peaks are annotated based on the given sequence, with 20 ppm tolerance.

Copy Data

##### Spectrum 4222 (TSV)

###### Preview

```
Loading example...
```

*Click on the button to copy the data to your clipboard.*

Mz MinMz MaxIntensity Max

WidthHeightPeptide font sizePeptide stroke widthSpectrum font sizeSpectrum stroke widthCompact peptide

Ion legend

wxyz

abcd

OtherUnassignedIon^Charge^_Position_Show for top:%

JSSPATJNSR

03.43e+36.85e+31.03e+41.37e+4

Zoom Out

a^+1^_2_y^+1^_1_a^+1^_2_y^+1^_1_b^+1^_2_b^+1^_2_a^+1^_3_y^+1^_2_y^+1^_2_b^+1^_3_y^+1^_3_y^+2^_7_y^+2^_7_y^+1^_3_y^+2^_7_y^+2^_8_y^+2^_8_y^+2^_8_y^+1^_4_\*y^+1^_5_y^+1^_6_y^+1^_6_y^+1^_7_y^+1^_8_y^+1^_9_

0774154823223096

Fragment Matches Table

Show background peaks

| Position | Ion type | Intensity | mz Theoretical | mz Error (Th) | mz Error (ppm) | Charge | Series Number |
| --- | --- | --- | --- | --- | --- | --- | --- |
| - | - | 566.8 | 120.1 | - | - | 0 | - |
| - | - | 347.4 | 125.8 | - | - | 0 | - |
| - | - | 632.7 | 129.1 | - | - | 0 | - |
| - | - | 1983 | 129.1 | - | - | 0 | - |
| - | - | 552.2 | 130.1 | - | - | 0 | - |
| - | - | 4358 | 133.1 | - | - | 0 | - |
| - | - | 464.1 | 134.1 | - | - | 0 | - |
| - | - | 4608 | 136.1 | - | - | 0 | - |
| - | - | 397 | 136.3 | - | - | 0 | - |
| - | - | 1030 | 139.1 | - | - | 0 | - |
| - | - | 3146 | 141.1 | - | - | 0 | - |
| - | - | 1641 | 143.1 | - | - | 0 | - |
| - | - | 372.6 | 145.8 | - | - | 0 | - |
| - | - | 807.1 | 147.1 | - | - | 0 | - |
| - | - | 1223 | 147.1 | - | - | 0 | - |
| - | - | 463.1 | 148.8 | - | - | 0 | - |
| - | - | 519.3 | 148.9 | - | - | 0 | - |
| - | - | 470.7 | 148.9 | - | - | 0 | - |
| - | - | 596.5 | 148.9 | - | - | 0 | - |
| - | - | 545.2 | 148.9 | - | - | 0 | - |
| - | - | 891 | 148.9 | - | - | 0 | - |
| - | - | 995.5 | 148.9 | - | - | 0 | - |
| - | - | 1537 | 148.9 | - | - | 0 | - |
| - | - | 3548 | 148.9 | - | - | 0 | - |
| - | - | 6049 | 149 | - | - | 0 | - |
| - | - | 3576 | 149 | - | - | 0 | - |
| - | - | 1665 | 149 | - | - | 0 | - |
| - | - | 1280 | 149 | - | - | 0 | - |
| - | - | 1049 | 149 | - | - | 0 | - |
| - | - | 617.1 | 149 | - | - | 0 | - |
| - | - | 849 | 149 | - | - | 0 | - |
| - | - | 749.7 | 149 | - | - | 0 | - |
| - | - | 451 | 149 | - | - | 0 | - |
| - | - | 732.2 | 149 | - | - | 0 | - |
| - | - | 660.2 | 151 | - | - | 0 | - |
| 2 | a | 822.3 | 155.1 | 0.0001372 | 0.8845 | +1 | 2 |
| - | - | 1644 | 157.1 | - | - | 0 | - |
| - | - | 2502 | 157.1 | - | - | 0 | - |
| 10 | y | 1355 | 158.1 | 9.571E-05 | 0.6054 | +1 | 1 |
| - | - | 470.1 | 158.1 | - | - | 0 | - |
| - | - | 535.7 | 160.2 | - | - | 0 | - |
| - | - | 2071 | 167.1 | - | - | 0 | - |
| - | - | 559.5 | 167.1 | - | - | 0 | - |
| - | - | 833.3 | 169.1 | - | - | 0 | - |
| - | - | 3847 | 169.1 | - | - | 0 | - |
| - | - | 534 | 169.1 | - | - | 0 | - |
| - | - | 520.7 | 171 | - | - | 0 | - |
| 2 | a | 4445 | 173.1 | 4.004E-05 | 0.2313 | +1 | 2 |
| - | - | 1446 | 174.1 | - | - | 0 | - |
| - | - | 482.1 | 174.1 | - | - | 0 | - |
| 10 | y | 4164 | 175.1 | 0.0001427 | 0.8148 | +1 | 1 |
| - | - | 427.2 | 177.6 | - | - | 0 | - |
| 2 | b | 6999 | 183.1 | 6.511E-05 | 0.3556 | +1 | 2 |
| - | - | 705.5 | 184.1 | - | - | 0 | - |
| - | - | 786.6 | 185.1 | - | - | 0 | - |
| - | - | 1447 | 185.1 | - | - | 0 | - |
| - | - | 907.1 | 185.2 | - | - | 0 | - |
| - | - | 679.4 | 187.1 | - | - | 0 | - |
| - | - | 1575 | 187.1 | - | - | 0 | - |
| - | - | 511 | 194.7 | - | - | 0 | - |
| 2 | b | 3746 | 201.1 | 3.205E-05 | 0.1594 | +1 | 2 |
| - | - | 1723 | 202.1 | - | - | 0 | - |
| - | - | 698.5 | 204.1 | - | - | 0 | - |
| - | - | 939.3 | 213.2 | - | - | 0 | - |
| - | - | 1164 | 215.1 | - | - | 0 | - |
| - | - | 528.3 | 219.5 | - | - | 0 | - |
| - | - | 1045 | 223.1 | - | - | 0 | - |
| - | - | 918.5 | 225 | - | - | 0 | - |
| - | - | 702.3 | 225.1 | - | - | 0 | - |
| - | - | 980.9 | 225.1 | - | - | 0 | - |
| - | - | 1237 | 227 | - | - | 0 | - |
| - | - | 636.2 | 227.1 | - | - | 0 | - |
| - | - | 496.8 | 228.4 | - | - | 0 | - |
| - | - | 672.4 | 230.1 | - | - | 0 | - |
| - | - | 4589 | 238.1 | - | - | 0 | - |
| - | - | 541.7 | 240.1 | - | - | 0 | - |
| 3 | a | 850.4 | 242.1 | 0.0004734 | 1.955 | +1 | 3 |
| - | - | 486.5 | 243.7 | - | - | 0 | - |
| - | - | 2023 | 244.1 | - | - | 0 | - |
| 9 | y | 1094 | 245.1 | 0.0001334 | 0.5441 | +1 | 2 |
| - | - | 1540 | 249.2 | - | - | 0 | - |
| - | - | 534.6 | 250.5 | - | - | 0 | - |
| - | - | 1385 | 252.1 | - | - | 0 | - |
| - | - | 1581 | 256.1 | - | - | 0 | - |
| 9 | y | 4777 | 262.2 | 5.092E-05 | 0.1942 | +1 | 2 |
| 3 | b | 7189 | 270.1 | 8.769E-05 | 0.3246 | +1 | 3 |
| - | - | 1442 | 271.1 | - | - | 0 | - |
| - | - | 2095 | 299.1 | - | - | 0 | - |
| - | - | 3442 | 300.1 | - | - | 0 | - |
| - | - | 7147 | 301.1 | - | - | 0 | - |
| - | - | 2698 | 302.1 | - | - | 0 | - |
| - | - | 503 | 302.5 | - | - | 0 | - |
| - | - | 1796 | 303.1 | - | - | 0 | - |
| - | - | 822.2 | 308.1 | - | - | 0 | - |
| - | - | 713.2 | 309.2 | - | - | 0 | - |
| - | - | 1089 | 339.2 | - | - | 0 | - |
| - | - | 929.9 | 342.1 | - | - | 0 | - |
| - | - | 1074 | 347 | - | - | 0 | - |
| - | - | 1291 | 349 | - | - | 0 | - |
| - | - | 601.1 | 357.2 | - | - | 0 | - |
| 8 | y | 1478 | 359.2 | 0.0005183 | 1.443 | +1 | 3 |
| - | - | 909.8 | 361 | - | - | 0 | - |
| 4 | y | 2148 | 370.7 | 0.0002704 | 0.7295 | +2 | 7 |
| 4 | y | 1414 | 371.2 | 0.006187 | 16.67 | +2 | 7 |
| 8 | y | 2525 | 376.2 | 3.118E-05 | 0.08289 | +1 | 3 |
| 4 | y | 1.16E+04 | 379.7 | 0.0001596 | 0.4203 | +2 | 7 |
| - | - | 4758 | 380.2 | - | - | 0 | - |
| - | - | 900.8 | 380.7 | - | - | 0 | - |
| - | - | 636.2 | 405.2 | - | - | 0 | - |
| 3 | y | 8778 | 414.2 | 3.382E-05 | 0.08165 | +2 | 8 |
| 3 | y | 4466 | 414.7 | 0.007202 | 17.37 | +2 | 8 |
| - | - | 1820 | 415.2 | - | - | 0 | - |
| - | - | 778.9 | 418 | - | - | 0 | - |
| - | - | 960.3 | 419 | - | - | 0 | - |
| - | - | 544 | 419 | - | - | 0 | - |
| - | - | 1310 | 420 | - | - | 0 | - |
| 3 | y | 8142 | 423.2 | 3.002E-05 | 0.07092 | +2 | 8 |
| - | - | 3282 | 423.7 | - | - | 0 | - |
| - | - | 1012 | 424.2 | - | - | 0 | - |
| - | - | 1172 | 458.3 | - | - | 0 | - |
| - | - | 922.5 | 482.7 | - | - | 0 | - |
| 7 | y | 4712 | 489.3 | 0.0003231 | 0.6603 | +1 | 4 |
| - | - | 840.5 | 490.3 | - | - | 0 | - |
| - | - | 1718 | 522.3 | - | - | 0 | - |
| 0 | Precursor | 2112 | 523.3 | 0.003541 | 6.767 | +2 | -1 |
| - | - | 777.7 | 523.8 | - | - | 0 | - |
| - | - | 1354 | 547.8 | - | - | 0 | - |
| 6 | y | 1.079E+04 | 590.3 | 2.792E-05 | 0.0473 | +1 | 5 |
| - | - | 2787 | 591.3 | - | - | 0 | - |
| - | - | 2766 | 629.3 | - | - | 0 | - |
| - | - | 2300 | 629.8 | - | - | 0 | - |
| 5 | y | 1210 | 643.4 | 0.001858 | 2.888 | +1 | 6 |
| 5 | y | 7953 | 661.4 | 0.0002154 | 0.3258 | +1 | 6 |
| - | - | 2984 | 662.4 | - | - | 0 | - |
| - | - | 1381 | 685.9 | - | - | 0 | - |
| - | - | 1466 | 686.3 | - | - | 0 | - |
| - | - | 696.1 | 686.8 | - | - | 0 | - |
| - | - | 667.4 | 729.4 | - | - | 0 | - |
| - | - | 1392 | 734.4 | - | - | 0 | - |
| - | - | 688.8 | 735.4 | - | - | 0 | - |
| 4 | y | 1.357E+04 | 758.4 | 0.000367 | 0.4839 | +1 | 7 |
| - | - | 6662 | 759.4 | - | - | 0 | - |
| - | - | 1761 | 760.4 | - | - | 0 | - |
| - | - | 669 | 817.4 | - | - | 0 | - |
| - | - | 561.5 | 836.1 | - | - | 0 | - |
| - | - | 1126 | 836.4 | - | - | 0 | - |
| 3 | y | 5934 | 845.4 | 0.001023 | 1.21 | +1 | 8 |
| - | - | 3425 | 846.4 | - | - | 0 | - |
| - | - | 1418 | 847.4 | - | - | 0 | - |
| 2 | y | 4373 | 932.5 | 0.000581 | 0.6231 | +1 | 9 |
| - | - | 2585 | 933.5 | - | - | 0 | - |
| - | - | 674.8 | 2280 | - | - | 0 | - |
| - | - | 655.1 | 2603 | - | - | 0 | - |
| - | - | 652.8 | 3065 | - | - | 0 | - |

m/z Charge Intensity FragmentType MassShift Position
120.08090209960938 0 566.83014
125.8163833618164 0 347.3847
129.0657196044922 0 632.6871
129.1022491455078 0 1983.0032
130.08660888671875 0 552.23627
133.06085205078125 0 4358.157
134.11001586914062 0 464.0715
136.0757598876953 0 4608.25
136.32261657714844 0 396.9911
139.0869140625 0 1030.4272
141.10232543945312 0 3145.8835
143.1177978515625 0 1641.4081
145.75291442871094 0 372.6094
147.07620239257812 0 807.0997
147.1132049560547 0 1222.7831
148.8472137451172 0 463.1285
148.87554931640625 0 519.32666
148.88255310058594 0 470.67834
148.89695739746094 0 596.5311
148.90350341796875 0 545.1501
148.91799926757812 0 891.0063
148.9254913330078 0 995.50934
148.93235778808594 0 1537.3824
148.93978881835938 0 3547.9932
148.9560089111328 0 6048.512
148.96368408203125 0 3576.0447
148.9711151123047 0 1664.6764
148.97817993164062 0 1279.566
148.98500061035156 0 1048.703
148.99220275878906 0 617.0819
148.99940490722656 0 849.00635
149.0064239501953 0 749.73627
149.03515625 0 451.0448
149.04440307617188 0 732.1724
151.04183959960938 0 660.1817
155.11802673339844 0 822.3089 a Water loss 1
157.0608367919922 0 1644.0146
157.09718322753906 0 2502.2266
158.09249877929688 0 1355.0399 y Ammonia loss 9
158.09934997558594 0 470.07953
160.2064208984375 0 535.6689
167.05545043945312 0 2070.9546
167.08169555664062 0 559.516
169.05230712890625 0 833.28864
169.09725952148438 0 3846.5999
169.13429260253906 0 533.9545
171.0492706298828 0 520.6901
173.1284942626953 0 4444.925 a 1
174.0874481201172 0 1445.7109
174.1321563720703 0 482.07498
175.1190948486328 0 4164.4165 y 9
177.61163330078125 0 427.18057
183.1128692626953 0 6998.6323 b Water loss 1
184.07151794433594 0 705.4618
185.05564880371094 0 786.6016
185.09214782714844 0 1447.3276
185.1646728515625 0 907.10547
187.071533203125 0 679.35547
187.14413452148438 0 1575.2407
194.6818084716797 0 510.97726
201.1233367919922 0 3745.7632 b 1
202.08245849609375 0 1722.759
204.098388671875 0 698.509
213.1598358154297 0 939.3439
215.13929748535156 0 1163.5479
219.53248596191406 0 528.2785
223.06362915039062 0 1044.5544
225.0430145263672 0 918.4882
225.06076049804688 0 702.25195
225.1235809326172 0 980.9219
227.0401153564453 0 1237.1302
227.1138153076172 0 636.2293
228.3655548095703 0 496.8245
230.07693481445312 0 672.371
238.1184844970703 0 4588.724
240.09609985351562 0 541.6862
242.14944458007812 0 850.43335 a Water loss 2
243.6772003173828 0 486.5083
244.12916564941406 0 2022.5767
245.12429809570312 0 1094.4774 y Ammonia loss 8
249.1598663330078 0 1539.833
250.49021911621094 0 534.5642
252.134033203125 0 1385.0713
256.1287536621094 0 1580.5012
262.1510314941406 0 4776.9023 y 8
270.1447448730469 0 7189.34 b Water loss 2
271.148193359375 0 1442.0236
299.06170654296875 0 2095.1409
300.0623474121094 0 3441.6555
301.05914306640625 0 7147.198
302.0599670410156 0 2698.3887
302.53656005859375 0 502.9669
303.0580749511719 0 1795.9036
308.1270751953125 0 822.2434
309.2032165527344 0 713.1769
339.1659240722656 0 1088.868
342.1402587890625 0 929.9424
346.9737243652344 0 1074.3671
348.9727783203125 0 1290.7311
357.17608642578125 0 601.0579
359.1678771972656 0 1477.9083 y Ammonia loss 7
361.0260925292969 0 909.8449
370.7063903808594 0 2147.6047 y Water loss 3
371.2043151855469 0 1413.8801 y Ammonia loss 3
376.1939392089844 0 2524.817 y 7
379.71124267578125 0 11598.819 y 3
380.2106018066406 0 4757.6665
380.7134094238281 0 900.7973
405.21807861328125 0 636.23145
414.22216796875 0 8778.414 y Water loss 2
414.7213439941406 0 4465.541 y Ammonia loss 2
415.22247314453125 0 1820.0469
418.03448486328125 0 778.8976
418.998046875 0 960.30896
419.0328674316406 0 543.97186
419.9967041015625 0 1309.5454
423.2273864746094 0 8142.45 y 2
423.72491455078125 0 3282.1123
424.22698974609375 0 1011.72253
458.2721862792969 0 1172.4088
482.7244873046875 0 922.51587
489.27764892578125 0 4712.1416 y 6
490.278076171875 0 840.45807
522.269287109375 0 1717.9291
523.2819213867188 0 2112.3066 Precursor
523.7847290039062 0 777.65356
547.7750854492188 0 1353.9045
590.3256225585938 0 10787.771 y 5
591.3245239257812 0 2786.7153
629.3057861328125 0 2765.8896
629.808349609375 0 2299.8013
643.350341796875 0 1210.344 y Water loss 4
661.362548828125 0 7953.192 y 4
662.3593139648438 0 2983.814
685.8504638671875 0 1380.7511
686.34765625 0 1465.593
686.8484497070312 0 696.1285
729.363037109375 0 667.3523
734.3585205078125 0 1391.9373
735.3640747070312 0 688.84656
758.4151611328125 0 13573.427 y 3
759.413818359375 0 6661.5728
760.4165649414062 0 1761.1082
817.3501586914062 0 669.0219
836.0673828125 0 561.5168
836.3942260742188 0 1125.5704
845.446533203125 0 5934.3237 y 2
846.446533203125 0 3424.8865
847.4446411132812 0 1418.1011
932.47900390625 0 4372.8555 y 1
933.4754638671875 0 2584.7168
2279.832275390625 0 674.81976
2602.807373046875 0 655.1256
3065.01025390625 0 652.8341

Spectrum Details

|  |  |
| --- | --- |
| Matched peaks? Matched peaks The total absolute number of peaks matched. Additionally in brackets the total fraction of peaks matched and the total number of peaks is shown. | 26 (16.88% of 154) |
| FDR? FDR The false discovery rate estimated for this peptide. It is calculated by matching all theoretical fragments with a non-integer shift with the raw peaks for this spectrum. This is done with 40 different shifts. The resulting percentage is the average number of annotated peaks over the number of annotated peaks with the correct spectrum. | 0.55% |
| Satellite FDR? Satellite FDR See the FDR for details on its calculation. This satellite ion specific FDR only contains the satellite ions (d/w) for I/L/J positions. | - |
| PSM Score? PSM Score The PSM Score as given by Hecklib to this annotated spectrum. It is shown with three significant figures. | 244 |

#### Spectrum 4272? Spectrum 4272 The raw spectrum of this peptide as annotated by Hecklib. The fragments are coloured according to ion type (see legend). Any peaks with a star '\*' as text can be hovered over to see the full details, first the ion type second the mass shift type. By hovering over the amino acids in the peptide or ions in the legend the corresponding peaks are highlighted. By toggling the 'Unassigned' label you can turn the background (unassigned) peaks on or off in the plot. By updating the slider in the Ion legend you can update the spectrum to only show the top X% of the peaks with labels. The top X% means any peak that is within X% of the highest intensity. By dragging in the spectrum you can zoom in to a specific part of the spectrum and use 'Zoom Out' to get back to the original zoom level. The annotation of the spectrum is based on the given sequence in the peptides file and is done with different software so inconsistencies are likely. The peaks are annotated based on the given sequence, with 20 ppm tolerance.

Copy Data

##### Spectrum 4272 (TSV)

###### Preview

```
Loading example...
```

*Click on the button to copy the data to your clipboard.*

Mz MinMz MaxIntensity Max

WidthHeightPeptide font sizePeptide stroke widthSpectrum font sizeSpectrum stroke widthCompact peptide

Ion legend

wxyz

abcd

OtherUnassignedIon^Charge^_Position_Show for top:%

JSSPATJNSR

03.18e+36.36e+39.54e+31.27e+4

Zoom Out

a^+1^_2_y^+1^_1_a^+1^_2_y^+1^_1_b^+1^_2_b^+1^_2_y^+1^_2_y^+1^_2_b^+1^_3_y^+1^_3_y^+2^_7_y^+1^_3_y^+2^_7_y^+2^_8_y^+2^_8_y^+2^_8_y^+1^_4_\*\*y^+1^_5_y^+1^_6_y^+1^_6_y^+1^_7_y^+1^_7_y^+1^_8_y^+1^_9_y^+1^_9_

0778155623353113

Fragment Matches Table

Show background peaks

| Position | Ion type | Intensity | mz Theoretical | mz Error (Th) | mz Error (ppm) | Charge | Series Number |
| --- | --- | --- | --- | --- | --- | --- | --- |
| - | - | 659.7 | 120.1 | - | - | 0 | - |
| - | - | 355.9 | 124.4 | - | - | 0 | - |
| - | - | 409.6 | 125.1 | - | - | 0 | - |
| - | - | 398.1 | 126.8 | - | - | 0 | - |
| - | - | 612.1 | 129.1 | - | - | 0 | - |
| - | - | 1402 | 129.1 | - | - | 0 | - |
| - | - | 459 | 134.2 | - | - | 0 | - |
| - | - | 3159 | 136.1 | - | - | 0 | - |
| - | - | 485.4 | 136.1 | - | - | 0 | - |
| - | - | 380.8 | 138.3 | - | - | 0 | - |
| - | - | 943.8 | 139.1 | - | - | 0 | - |
| - | - | 500.8 | 141.1 | - | - | 0 | - |
| - | - | 2042 | 141.1 | - | - | 0 | - |
| - | - | 471 | 143.8 | - | - | 0 | - |
| - | - | 457.9 | 145.3 | - | - | 0 | - |
| - | - | 686.9 | 147.1 | - | - | 0 | - |
| - | - | 529.5 | 147.1 | - | - | 0 | - |
| - | - | 406.8 | 148.3 | - | - | 0 | - |
| - | - | 538.2 | 148.9 | - | - | 0 | - |
| - | - | 544.7 | 148.9 | - | - | 0 | - |
| - | - | 606.7 | 148.9 | - | - | 0 | - |
| - | - | 545.6 | 148.9 | - | - | 0 | - |
| - | - | 724.6 | 148.9 | - | - | 0 | - |
| - | - | 896.2 | 148.9 | - | - | 0 | - |
| - | - | 1076 | 148.9 | - | - | 0 | - |
| - | - | 1284 | 148.9 | - | - | 0 | - |
| - | - | 2223 | 148.9 | - | - | 0 | - |
| - | - | 3939 | 148.9 | - | - | 0 | - |
| - | - | 4366 | 149 | - | - | 0 | - |
| - | - | 2898 | 149 | - | - | 0 | - |
| - | - | 1313 | 149 | - | - | 0 | - |
| - | - | 1060 | 149 | - | - | 0 | - |
| - | - | 896.3 | 149 | - | - | 0 | - |
| - | - | 846 | 149 | - | - | 0 | - |
| - | - | 700.1 | 149 | - | - | 0 | - |
| - | - | 472.1 | 149 | - | - | 0 | - |
| - | - | 498 | 149 | - | - | 0 | - |
| - | - | 603.3 | 149 | - | - | 0 | - |
| - | - | 670.5 | 149 | - | - | 0 | - |
| - | - | 2485 | 149 | - | - | 0 | - |
| - | - | 415.6 | 149.1 | - | - | 0 | - |
| - | - | 727.2 | 151 | - | - | 0 | - |
| - | - | 390.7 | 153.4 | - | - | 0 | - |
| 2 | a | 1050 | 155.1 | 0.0002135 | 1.376 | +1 | 2 |
| - | - | 525.1 | 155.6 | - | - | 0 | - |
| - | - | 843 | 157.1 | - | - | 0 | - |
| - | - | 2292 | 157.1 | - | - | 0 | - |
| 10 | y | 1034 | 158.1 | 0.0001789 | 1.132 | +1 | 1 |
| - | - | 431.2 | 159.2 | - | - | 0 | - |
| - | - | 4529 | 167.1 | - | - | 0 | - |
| - | - | 533.4 | 167.1 | - | - | 0 | - |
| - | - | 1920 | 168.1 | - | - | 0 | - |
| - | - | 1708 | 169.1 | - | - | 0 | - |
| - | - | 3212 | 169.1 | - | - | 0 | - |
| 2 | a | 3595 | 173.1 | 9.729E-05 | 0.5619 | +1 | 2 |
| - | - | 1161 | 174.1 | - | - | 0 | - |
| 10 | y | 2814 | 175.1 | 9.907E-06 | 0.05657 | +1 | 1 |
| - | - | 430.5 | 175.1 | - | - | 0 | - |
| 2 | b | 5464 | 183.1 | 8.748E-05 | 0.4777 | +1 | 2 |
| - | - | 1878 | 185.1 | - | - | 0 | - |
| - | - | 1327 | 187.1 | - | - | 0 | - |
| 2 | b | 3231 | 201.1 | 0.000322 | 1.601 | +1 | 2 |
| - | - | 632.6 | 202.1 | - | - | 0 | - |
| - | - | 707.3 | 215.1 | - | - | 0 | - |
| - | - | 639.9 | 216.1 | - | - | 0 | - |
| - | - | 669.1 | 217.1 | - | - | 0 | - |
| - | - | 1751 | 223.1 | - | - | 0 | - |
| - | - | 1208 | 224.1 | - | - | 0 | - |
| - | - | 1050 | 225 | - | - | 0 | - |
| - | - | 818 | 225.1 | - | - | 0 | - |
| - | - | 723 | 225.1 | - | - | 0 | - |
| - | - | 829.3 | 226 | - | - | 0 | - |
| - | - | 470.8 | 226.9 | - | - | 0 | - |
| - | - | 632.8 | 228.1 | - | - | 0 | - |
| - | - | 6157 | 238.1 | - | - | 0 | - |
| - | - | 1259 | 239.1 | - | - | 0 | - |
| - | - | 780.7 | 240.1 | - | - | 0 | - |
| - | - | 1018 | 241.1 | - | - | 0 | - |
| 9 | y | 1007 | 245.1 | 0.0003317 | 1.353 | +1 | 2 |
| - | - | 1245 | 251.1 | - | - | 0 | - |
| - | - | 1785 | 252.1 | - | - | 0 | - |
| - | - | 788.8 | 256.1 | - | - | 0 | - |
| 9 | y | 4301 | 262.2 | 0.0002848 | 1.086 | +1 | 2 |
| - | - | 531.9 | 262.9 | - | - | 0 | - |
| 3 | b | 6789 | 270.1 | 0.0004844 | 1.793 | +1 | 3 |
| - | - | 640.2 | 281.1 | - | - | 0 | - |
| - | - | 704.3 | 283 | - | - | 0 | - |
| - | - | 830.3 | 283 | - | - | 0 | - |
| - | - | 636.3 | 284 | - | - | 0 | - |
| - | - | 5586 | 299.1 | - | - | 0 | - |
| - | - | 8905 | 300.1 | - | - | 0 | - |
| - | - | 1.09E+04 | 301.1 | - | - | 0 | - |
| - | - | 6466 | 302.1 | - | - | 0 | - |
| - | - | 2618 | 303.1 | - | - | 0 | - |
| - | - | 707.9 | 339.2 | - | - | 0 | - |
| - | - | 780.7 | 346 | - | - | 0 | - |
| - | - | 1768 | 347 | - | - | 0 | - |
| - | - | 2180 | 348 | - | - | 0 | - |
| - | - | 736.2 | 357.2 | - | - | 0 | - |
| 8 | y | 944.9 | 359.2 | 0.000794 | 2.211 | +1 | 3 |
| - | - | 579.6 | 360 | - | - | 0 | - |
| - | - | 1338 | 361 | - | - | 0 | - |
| - | - | 1118 | 362 | - | - | 0 | - |
| 4 | y | 2368 | 370.7 | 0.0008282 | 2.234 | +2 | 7 |
| - | - | 986.4 | 371.2 | - | - | 0 | - |
| 8 | y | 3378 | 376.2 | 0.0005792 | 1.54 | +1 | 3 |
| - | - | 863.6 | 378.7 | - | - | 0 | - |
| 4 | y | 1.055E+04 | 379.7 | 0.0006784 | 1.787 | +2 | 7 |
| - | - | 3776 | 380.2 | - | - | 0 | - |
| - | - | 1455 | 380.7 | - | - | 0 | - |
| - | - | 689 | 405.7 | - | - | 0 | - |
| 3 | y | 9684 | 414.2 | 8.825E-05 | 0.213 | +2 | 8 |
| 3 | y | 3498 | 414.7 | 0.007294 | 17.59 | +2 | 8 |
| - | - | 821.9 | 415.2 | - | - | 0 | - |
| - | - | 1193 | 417 | - | - | 0 | - |
| - | - | 2636 | 418 | - | - | 0 | - |
| - | - | 2006 | 419 | - | - | 0 | - |
| - | - | 1638 | 419 | - | - | 0 | - |
| - | - | 1131 | 420 | - | - | 0 | - |
| 3 | y | 8179 | 423.2 | 0.0001521 | 0.3594 | +2 | 8 |
| - | - | 4010 | 423.7 | - | - | 0 | - |
| - | - | 833.6 | 424.2 | - | - | 0 | - |
| - | - | 1232 | 458.3 | - | - | 0 | - |
| - | - | 1320 | 489.1 | - | - | 0 | - |
| 7 | y | 3212 | 489.3 | 4.842E-05 | 0.09897 | +1 | 4 |
| - | - | 899.5 | 490.1 | - | - | 0 | - |
| 0 | Precursor | 704.4 | 514.3 | 0.0009476 | 1.843 | +2 | -1 |
| - | - | 1722 | 522.3 | - | - | 0 | - |
| 0 | Precursor | 1651 | 523.3 | 0.002076 | 3.968 | +2 | -1 |
| - | - | 769.1 | 523.8 | - | - | 0 | - |
| - | - | 672.4 | 545.8 | - | - | 0 | - |
| 6 | y | 1.042E+04 | 590.3 | 0.0003331 | 0.5643 | +1 | 5 |
| - | - | 2859 | 591.3 | - | - | 0 | - |
| 5 | y | 1201 | 643.4 | 0.002163 | 3.362 | +1 | 6 |
| 5 | y | 7157 | 661.4 | 0.0006427 | 0.9718 | +1 | 6 |
| - | - | 2066 | 662.4 | - | - | 0 | - |
| - | - | 683.2 | 730.9 | - | - | 0 | - |
| 4 | y | 928.6 | 741.4 | 0.00342 | 4.613 | +1 | 7 |
| 4 | y | 1.26E+04 | 758.4 | 0.001099 | 1.45 | +1 | 7 |
| - | - | 5621 | 759.4 | - | - | 0 | - |
| - | - | 1166 | 760.4 | - | - | 0 | - |
| 3 | y | 5679 | 845.4 | 0.000474 | 0.5607 | +1 | 8 |
| - | - | 3197 | 846.4 | - | - | 0 | - |
| 2 | y | 675.6 | 914.5 | 0.002537 | 2.775 | +1 | 9 |
| 2 | y | 4020 | 932.5 | 0.001741 | 1.867 | +1 | 9 |
| - | - | 2115 | 933.5 | - | - | 0 | - |
| - | - | 664.1 | 1033 | - | - | 0 | - |
| - | - | 651.5 | 1609 | - | - | 0 | - |
| - | - | 680.4 | 2861 | - | - | 0 | - |
| - | - | 831.1 | 3082 | - | - | 0 | - |

m/z Charge Intensity FragmentType MassShift Position
120.08082580566406 0 659.69434
124.41504669189453 0 355.88177
125.1072006225586 0 409.63724
126.81462097167969 0 398.14133
129.06602478027344 0 612.13654
129.10226440429688 0 1401.8649
134.19639587402344 0 458.97772
136.07569885253906 0 3158.533
136.08056640625 0 485.42685
138.3024139404297 0 380.80115
139.08651733398438 0 943.82025
141.06625366210938 0 500.8367
141.10232543945312 0 2041.8536
143.79795837402344 0 470.9748
145.33914184570312 0 457.9007
147.0764923095703 0 686.857
147.1122589111328 0 529.4934
148.30235290527344 0 406.78662
148.8712615966797 0 538.23804
148.8785858154297 0 544.71576
148.892578125 0 606.66095
148.9000701904297 0 545.61615
148.90713500976562 0 724.63763
148.9140625 0 896.2162
148.9210205078125 0 1075.9623
148.92823791503906 0 1283.6598
148.93521118164062 0 2223.3162
148.94288635253906 0 3939.0674
148.95913696289062 0 4366.003
148.96681213378906 0 2898.3035
148.97401428222656 0 1313.1533
148.98133850097656 0 1060.0121
148.9882354736328 0 896.32043
148.99505615234375 0 846.0135
149.00204467773438 0 700.0778
149.00914001464844 0 472.08734
149.01681518554688 0 497.99847
149.0233612060547 0 603.2666
149.0384063720703 0 670.46814
149.04486083984375 0 2485.2515
149.14353942871094 0 415.58914
151.04173278808594 0 727.1698
153.3665008544922 0 390.7216
155.11810302734375 0 1050.1086 a Water loss 1
155.5533447265625 0 525.1229
157.060546875 0 842.95215
157.09695434570312 0 2291.9644
158.09222412109375 0 1033.6615 y Ammonia loss 9
159.17552185058594 0 431.2393
167.0553741455078 0 4528.946
167.0820770263672 0 533.3894
168.05496215820312 0 1920.2906
169.05230712890625 0 1707.7524
169.09698486328125 0 3211.887
173.12835693359375 0 3594.6104 a 1
174.0875244140625 0 1160.5527
175.1189422607422 0 2813.9275 y 9
175.139404296875 0 430.48645
183.1127166748047 0 5463.6562 b Water loss 1
185.09194946289062 0 1878.1672
187.14422607421875 0 1327.4937
201.123046875 0 3231.4111 b 1
202.08181762695312 0 632.60474
215.13929748535156 0 707.3232
216.09878540039062 0 639.9288
217.08193969726562 0 669.0869
223.06344604492188 0 1751.1504
224.06324768066406 0 1208.0934
225.0422821044922 0 1049.5177
225.0595245361328 0 817.98126
225.12252807617188 0 723.0262
226.04383850097656 0 829.3274
226.8594970703125 0 470.83914
228.13345336914062 0 632.7593
238.1184539794922 0 6156.7993
239.0947723388672 0 1259.4744
240.09698486328125 0 780.6556
241.09205627441406 0 1017.60657
245.1240997314453 0 1006.9885 y Ammonia loss 8
251.1024627685547 0 1245.2683
252.1338348388672 0 1785.4668
256.1280517578125 0 788.7764
262.15069580078125 0 4301.1855 y 8
262.9435729980469 0 531.89905
270.14434814453125 0 6789.4 b Water loss 2
281.05108642578125 0 640.20245
283.03082275390625 0 704.33606
283.0482482910156 0 830.33093
284.047607421875 0 636.30066
299.0614318847656 0 5586.0854
300.06219482421875 0 8905.261
301.0590515136719 0 10895.299
302.0594787597656 0 6465.808
303.0559387207031 0 2618.0564
339.1667175292969 0 707.87665
345.9769287109375 0 780.67975
346.97357177734375 0 1768.1626
347.97393798828125 0 2180.4055
357.176025390625 0 736.19226
359.16656494140625 0 944.92413 y Ammonia loss 7
360.0257568359375 0 579.6367
361.02459716796875 0 1337.7107
362.02630615234375 0 1118.4637
370.7052917480469 0 2367.5222 y Water loss 3
371.2061767578125 0 986.3756
376.1933288574219 0 3378.377 y 7
378.7024230957031 0 863.61176
379.7107238769531 0 10554.331 y 3
380.2100830078125 0 3776.0706
380.7115478515625 0 1454.582
405.7095947265625 0 689.0037
414.2220458984375 0 9683.703 y Water loss 2
414.721435546875 0 3498.0706 y Ammonia loss 2
415.22198486328125 0 821.8642
417.03515625 0 1192.6736
418.03515625 0 2636.3215
418.9950866699219 0 2005.6592
419.0312194824219 0 1637.5131
419.9944152832031 0 1131.3549
423.2272644042969 0 8178.904 y 2
423.72711181640625 0 4009.993
424.2256164550781 0 833.60516
458.27093505859375 0 1231.8464
489.05419921875 0 1319.6498
489.2779235839844 0 3212.4126 y 6
490.05694580078125 0 899.4685
514.2811279296875 0 704.4487 Precursor Water loss
522.2706298828125 0 1721.8336
523.2833862304688 0 1651.0804 Precursor
523.784912109375 0 769.0579
545.7625122070312 0 672.3828
590.3253173828125 0 10422.824 y 5
591.3233032226562 0 2858.7795
643.3500366210938 0 1200.5667 y Water loss 4
661.3621215820312 0 7157.0024 y 4
662.3629150390625 0 2066.491
730.8989868164062 0 683.17975
741.3855590820312 0 928.64905 y Ammonia loss 3
758.4144287109375 0 12599.448 y 3
759.4130859375 0 5621.1016
760.4196166992188 0 1165.9569
845.4470825195312 0 5679.476 y 2
846.4411010742188 0 3196.9263
914.4715576171875 0 675.63605 y Water loss 1
932.4778442382812 0 4019.9656 y 1
933.4758911132812 0 2114.6877
1033.45068359375 0 664.06055
1608.56640625 0 651.46436
2860.924560546875 0 680.38226
3082.0166015625 0 831.0688

Spectrum Details

|  |  |
| --- | --- |
| Matched peaks? Matched peaks The total absolute number of peaks matched. Additionally in brackets the total fraction of peaks matched and the total number of peaks is shown. | 27 (18.00% of 150) |
| FDR? FDR The false discovery rate estimated for this peptide. It is calculated by matching all theoretical fragments with a non-integer shift with the raw peaks for this spectrum. This is done with 40 different shifts. The resulting percentage is the average number of annotated peaks over the number of annotated peaks with the correct spectrum. | 1.06% |
| Satellite FDR? Satellite FDR See the FDR for details on its calculation. This satellite ion specific FDR only contains the satellite ions (d/w) for I/L/J positions. | - |
| PSM Score? PSM Score The PSM Score as given by Hecklib to this annotated spectrum. It is shown with three significant figures. | 262 |

#### Spectrum 3097? Spectrum 3097 The raw spectrum of this peptide as annotated by Hecklib. The fragments are coloured according to ion type (see legend). Any peaks with a star '\*' as text can be hovered over to see the full details, first the ion type second the mass shift type. By hovering over the amino acids in the peptide or ions in the legend the corresponding peaks are highlighted. By toggling the 'Unassigned' label you can turn the background (unassigned) peaks on or off in the plot. By updating the slider in the Ion legend you can update the spectrum to only show the top X% of the peaks with labels. The top X% means any peak that is within X% of the highest intensity. By dragging in the spectrum you can zoom in to a specific part of the spectrum and use 'Zoom Out' to get back to the original zoom level. The annotation of the spectrum is based on the given sequence in the peptides file and is done with different software so inconsistencies are likely. The peaks are annotated based on the given sequence, with 20 ppm tolerance.

Copy Data

##### Spectrum 3097 (TSV)

###### Preview

```
Loading example...
```

*Click on the button to copy the data to your clipboard.*

Mz MinMz MaxIntensity Max

WidthHeightPeptide font sizePeptide stroke widthSpectrum font sizeSpectrum stroke widthCompact peptide

Ion legend

wxyz

abcd

OtherUnassignedIon^Charge^_Position_Show for top:%

JSSPATJNSR

05.69e+31.14e+41.71e+42.28e+4

Zoom Out

a^+1^_2_y^+1^_1_a^+1^_2_y^+1^_1_b^+1^_2_b^+1^_2_a^+1^_3_y^+1^_2_y^+1^_2_b^+1^_3_y^+1^_3_y^+2^_7_y^+1^_3_y^+2^_7_y^+2^_8_y^+2^_8_y^+1^_4_\*y^+1^_5_y^+1^_6_y^+1^_6_y^+1^_7_y^+1^_7_y^+1^_7_y^+1^_8_y^+1^_9_y^+1^_9_

0777155423323109

Fragment Matches Table

Show background peaks

| Position | Ion type | Intensity | mz Theoretical | mz Error (Th) | mz Error (ppm) | Charge | Series Number |
| --- | --- | --- | --- | --- | --- | --- | --- |
| - | - | 482.6 | 120.1 | - | - | 0 | - |
| - | - | 332.3 | 120.8 | - | - | 0 | - |
| - | - | 856.7 | 125.1 | - | - | 0 | - |
| - | - | 735.4 | 125.1 | - | - | 0 | - |
| - | - | 396.1 | 125.5 | - | - | 0 | - |
| - | - | 1780 | 129.1 | - | - | 0 | - |
| - | - | 899.5 | 130.1 | - | - | 0 | - |
| - | - | 483.8 | 130.7 | - | - | 0 | - |
| - | - | 365.5 | 131.7 | - | - | 0 | - |
| - | - | 433.1 | 135.3 | - | - | 0 | - |
| - | - | 526 | 136.1 | - | - | 0 | - |
| - | - | 987.7 | 139.1 | - | - | 0 | - |
| - | - | 3677 | 141.1 | - | - | 0 | - |
| - | - | 3834 | 141.1 | - | - | 0 | - |
| - | - | 741.3 | 147.1 | - | - | 0 | - |
| - | - | 1334 | 147.1 | - | - | 0 | - |
| - | - | 2437 | 149 | - | - | 0 | - |
| - | - | 1154 | 151 | - | - | 0 | - |
| - | - | 526.1 | 153.1 | - | - | 0 | - |
| - | - | 2283 | 153.1 | - | - | 0 | - |
| 2 | a | 1452 | 155.1 | 0.0001067 | 0.6877 | +1 | 2 |
| - | - | 1189 | 157.1 | - | - | 0 | - |
| - | - | 3191 | 157.1 | - | - | 0 | - |
| 10 | y | 1638 | 158.1 | 0.0001415 | 0.895 | +1 | 1 |
| - | - | 3712 | 167.1 | - | - | 0 | - |
| - | - | 909.9 | 167.1 | - | - | 0 | - |
| - | - | 973.5 | 168.1 | - | - | 0 | - |
| - | - | 1441 | 169.1 | - | - | 0 | - |
| - | - | 3803 | 169.1 | - | - | 0 | - |
| 2 | a | 4086 | 173.1 | 2.478E-05 | 0.1431 | +1 | 2 |
| - | - | 4301 | 173.5 | - | - | 0 | - |
| 10 | y | 1928 | 175.1 | 0.0001274 | 0.7276 | +1 | 1 |
| - | - | 1.63E+04 | 181.1 | - | - | 0 | - |
| - | - | 3896 | 181.1 | - | - | 0 | - |
| - | - | 628.3 | 181.1 | - | - | 0 | - |
| - | - | 1843 | 182.1 | - | - | 0 | - |
| 2 | b | 7777 | 183.1 | 9.562E-05 | 0.5222 | +1 | 2 |
| - | - | 3457 | 185.1 | - | - | 0 | - |
| - | - | 604 | 187.1 | - | - | 0 | - |
| - | - | 1309 | 187.1 | - | - | 0 | - |
| - | - | 451.5 | 188 | - | - | 0 | - |
| - | - | 470.3 | 192.1 | - | - | 0 | - |
| 2 | b | 4464 | 201.1 | 1.534E-06 | 0.007626 | +1 | 2 |
| - | - | 618.5 | 202.1 | - | - | 0 | - |
| - | - | 563.8 | 202.6 | - | - | 0 | - |
| - | - | 710.7 | 215.1 | - | - | 0 | - |
| - | - | 1655 | 223.1 | - | - | 0 | - |
| - | - | 988.3 | 224.1 | - | - | 0 | - |
| - | - | 1532 | 225 | - | - | 0 | - |
| - | - | 877.5 | 225.1 | - | - | 0 | - |
| - | - | 828 | 225.1 | - | - | 0 | - |
| - | - | 1056 | 226 | - | - | 0 | - |
| - | - | 663.4 | 227 | - | - | 0 | - |
| - | - | 473.1 | 228 | - | - | 0 | - |
| - | - | 942.9 | 228.1 | - | - | 0 | - |
| - | - | 2249 | 234.1 | - | - | 0 | - |
| - | - | 4064 | 238.1 | - | - | 0 | - |
| - | - | 1172 | 239.1 | - | - | 0 | - |
| - | - | 657 | 239.1 | - | - | 0 | - |
| - | - | 687.5 | 240.1 | - | - | 0 | - |
| 3 | a | 630.4 | 242.1 | 0.0003971 | 1.64 | +1 | 3 |
| 9 | y | 768.9 | 245.1 | 2.657E-05 | 0.1084 | +1 | 2 |
| - | - | 465.9 | 251.7 | - | - | 0 | - |
| - | - | 2453 | 252.1 | - | - | 0 | - |
| - | - | 870.8 | 256.1 | - | - | 0 | - |
| - | - | 546.2 | 260.8 | - | - | 0 | - |
| 9 | y | 1544 | 262.2 | 0.0002848 | 1.086 | +1 | 2 |
| - | - | 701.7 | 268.1 | - | - | 0 | - |
| 3 | b | 1.091E+04 | 270.1 | 0.0001487 | 0.5506 | +1 | 3 |
| - | - | 1251 | 271.1 | - | - | 0 | - |
| - | - | 823.1 | 273 | - | - | 0 | - |
| - | - | 821.2 | 280.2 | - | - | 0 | - |
| - | - | 785.2 | 281.1 | - | - | 0 | - |
| - | - | 1038 | 283 | - | - | 0 | - |
| - | - | 529.3 | 283.3 | - | - | 0 | - |
| - | - | 620.7 | 286.3 | - | - | 0 | - |
| - | - | 5897 | 299.1 | - | - | 0 | - |
| - | - | 6485 | 300.1 | - | - | 0 | - |
| - | - | 1.052E+04 | 301.1 | - | - | 0 | - |
| - | - | 6214 | 302.1 | - | - | 0 | - |
| - | - | 1821 | 303.1 | - | - | 0 | - |
| - | - | 607 | 309.2 | - | - | 0 | - |
| - | - | 579.7 | 329.2 | - | - | 0 | - |
| - | - | 635.5 | 330.2 | - | - | 0 | - |
| - | - | 5998 | 331.2 | - | - | 0 | - |
| - | - | 1563 | 332.2 | - | - | 0 | - |
| - | - | 866.1 | 339.2 | - | - | 0 | - |
| - | - | 835.3 | 346 | - | - | 0 | - |
| - | - | 2138 | 347 | - | - | 0 | - |
| - | - | 830.6 | 348 | - | - | 0 | - |
| - | - | 722.9 | 349 | - | - | 0 | - |
| 8 | y | 2119 | 359.2 | 0.0008245 | 2.296 | +1 | 3 |
| - | - | 641.4 | 361 | - | - | 0 | - |
| - | - | 592.5 | 363 | - | - | 0 | - |
| - | - | 641.1 | 365.2 | - | - | 0 | - |
| 4 | y | 4159 | 370.7 | 0.0006756 | 1.822 | +2 | 7 |
| - | - | 1133 | 371.2 | - | - | 0 | - |
| 8 | y | 3198 | 376.2 | 0.0001838 | 0.4885 | +1 | 3 |
| - | - | 854.2 | 378.7 | - | - | 0 | - |
| - | - | 687.1 | 379.4 | - | - | 0 | - |
| 4 | y | 2.253E+04 | 379.7 | 0.0005258 | 1.385 | +2 | 7 |
| - | - | 7242 | 380.2 | - | - | 0 | - |
| - | - | 2221 | 380.7 | - | - | 0 | - |
| - | - | 590.5 | 405.2 | - | - | 0 | - |
| 3 | y | 9403 | 414.2 | 3.382E-05 | 0.08165 | +2 | 8 |
| - | - | 3510 | 414.7 | - | - | 0 | - |
| - | - | 1767 | 415.2 | - | - | 0 | - |
| - | - | 1379 | 417 | - | - | 0 | - |
| - | - | 2769 | 418 | - | - | 0 | - |
| - | - | 2806 | 419 | - | - | 0 | - |
| - | - | 900.4 | 419 | - | - | 0 | - |
| - | - | 860.4 | 420 | - | - | 0 | - |
| 3 | y | 7766 | 423.2 | 0.0003352 | 0.792 | +2 | 8 |
| - | - | 4753 | 423.7 | - | - | 0 | - |
| - | - | 871.1 | 424.2 | - | - | 0 | - |
| - | - | 731 | 432.2 | - | - | 0 | - |
| - | - | 588.8 | 437.5 | - | - | 0 | - |
| - | - | 624.8 | 454.5 | - | - | 0 | - |
| 7 | y | 5057 | 489.3 | 0.0007808 | 1.596 | +1 | 4 |
| - | - | 1016 | 490.3 | - | - | 0 | - |
| - | - | 582.5 | 497.6 | - | - | 0 | - |
| - | - | 618.7 | 506.1 | - | - | 0 | - |
| - | - | 2306 | 521.8 | - | - | 0 | - |
| - | - | 1810 | 522.3 | - | - | 0 | - |
| - | - | 1706 | 522.8 | - | - | 0 | - |
| 0 | Precursor | 5418 | 523.3 | 0.0007337 | 1.402 | +2 | -1 |
| - | - | 2563 | 523.8 | - | - | 0 | - |
| 6 | y | 1.402E+04 | 590.3 | 0.0006383 | 1.081 | +1 | 5 |
| - | - | 3454 | 591.3 | - | - | 0 | - |
| - | - | 1922 | 634.3 | - | - | 0 | - |
| - | - | 958.3 | 635.3 | - | - | 0 | - |
| 5 | y | 620.9 | 643.4 | 0.005947 | 9.244 | +1 | 6 |
| 5 | y | 1.134E+04 | 661.4 | 0.001314 | 1.987 | +1 | 6 |
| - | - | 3135 | 662.4 | - | - | 0 | - |
| 4 | y | 694.8 | 740.4 | 0.0002393 | 0.3232 | +1 | 7 |
| 4 | y | 887.3 | 741.4 | 0.001894 | 2.555 | +1 | 7 |
| 4 | y | 2.028E+04 | 758.4 | 0.00171 | 2.254 | +1 | 7 |
| - | - | 8655 | 759.4 | - | - | 0 | - |
| - | - | 2005 | 760.4 | - | - | 0 | - |
| - | - | 1850 | 763.4 | - | - | 0 | - |
| - | - | 1118 | 764.4 | - | - | 0 | - |
| 3 | y | 8280 | 845.4 | 0.002549 | 3.015 | +1 | 8 |
| - | - | 4044 | 846.4 | - | - | 0 | - |
| - | - | 1118 | 847.5 | - | - | 0 | - |
| - | - | 5606 | 862.4 | - | - | 0 | - |
| - | - | 2845 | 863.5 | - | - | 0 | - |
| - | - | 1114 | 864.5 | - | - | 0 | - |
| - | - | 605.9 | 874.4 | - | - | 0 | - |
| 2 | y | 746.5 | 914.5 | 0.0006453 | 0.7056 | +1 | 9 |
| 2 | y | 6318 | 932.5 | 0.003083 | 3.307 | +1 | 9 |
| - | - | 2869 | 933.5 | - | - | 0 | - |
| - | - | 892 | 934.5 | - | - | 0 | - |
| - | - | 741.7 | 942.5 | - | - | 0 | - |
| - | - | 694.4 | 1212 | - | - | 0 | - |
| - | - | 800.5 | 3078 | - | - | 0 | - |

m/z Charge Intensity FragmentType MassShift Position
120.08080291748047 0 482.64328
120.8337173461914 0 332.3449
125.07128143310547 0 856.7159
125.10734558105469 0 735.44775
125.49581909179688 0 396.11847
129.10240173339844 0 1779.9973
130.08651733398438 0 899.5153
130.68971252441406 0 483.8341
131.70689392089844 0 365.53955
135.30923461914062 0 433.10913
136.0756072998047 0 526.046
139.08676147460938 0 987.6611
141.06594848632812 0 3677.4421
141.1023406982422 0 3834.403
147.07666015625 0 741.3049
147.1131134033203 0 1334.2069
149.0448760986328 0 2437.114
151.04183959960938 0 1154.0134
153.06558227539062 0 526.1365
153.1024169921875 0 2282.7844
155.1179962158203 0 1452.3513 a Water loss 1
157.06092834472656 0 1189.2438
157.09719848632812 0 3190.7852
158.09254455566406 0 1638.2502 y Ammonia loss 9
167.05560302734375 0 3711.9412
167.08203125 0 909.9014
168.0550994873047 0 973.52075
169.05227661132812 0 1440.6515
169.09730529785156 0 3803.2996
173.12847900390625 0 4086.3381 a 1
173.4508819580078 0 4300.842
175.11907958984375 0 1928.3354 y 9
181.0608367919922 0 16302.125
181.0972442626953 0 3895.7832
181.13323974609375 0 628.3232
182.06439208984375 0 1843.2898
183.11289978027344 0 7776.7563 b Water loss 1
185.09219360351562 0 3456.5042
187.0524444580078 0 603.95856
187.1441192626953 0 1308.6108
187.95858764648438 0 451.52493
192.06784057617188 0 470.2963
201.1233673095703 0 4464.18 b 1
202.0823211669922 0 618.5336
202.63433837890625 0 563.79706
215.13851928710938 0 710.6657
223.06353759765625 0 1654.641
224.06472778320312 0 988.31354
225.0428924560547 0 1532.0701
225.06103515625 0 877.50903
225.123291015625 0 828.0043
226.04415893554688 0 1056.3768
227.04025268554688 0 663.41223
228.03216552734375 0 473.1014
228.1337432861328 0 942.85596
234.14454650878906 0 2248.9983
238.11875915527344 0 4063.7788
239.09463500976562 0 1171.5924
239.1221160888672 0 656.9531
240.09567260742188 0 687.47003
242.14952087402344 0 630.41644 a Water loss 2
245.12440490722656 0 768.8675 y Ammonia loss 8
251.73765563964844 0 465.9219
252.13427734375 0 2452.9456
256.1294860839844 0 870.75146
260.78302001953125 0 546.2316
262.15069580078125 0 1544.039 y 8
268.1305236816406 0 701.7457
270.1446838378906 0 10913.186 b Water loss 2
271.14794921875 0 1250.8438
273.04730224609375 0 823.08356
280.16448974609375 0 821.2376
281.0503234863281 0 785.1989
283.04925537109375 0 1037.5164
283.3288269042969 0 529.32196
286.2860412597656 0 620.65393
299.0615234375 0 5897.043
300.0622253417969 0 6485.2295
301.05914306640625 0 10516.684
302.0599060058594 0 6214.3584
303.05670166015625 0 1820.5734
309.2023620605469 0 607.0321
329.17999267578125 0 579.6554
330.1665954589844 0 635.49567
331.197509765625 0 5998.4873
332.20111083984375 0 1562.711
339.16595458984375 0 866.12744
345.97808837890625 0 835.2816
346.974365234375 0 2138.2498
347.9732971191406 0 830.6417
348.9716491699219 0 722.9489
359.1665344238281 0 2119.2922 y Ammonia loss 7
361.02447509765625 0 641.40405
363.0240173339844 0 592.47797
365.2171325683594 0 641.12335
370.7054443359375 0 4158.5576 y Water loss 3
371.2075500488281 0 1132.6036
376.194091796875 0 3197.7952 y 7
378.7048645019531 0 854.2108
379.40362548828125 0 687.1108
379.71087646484375 0 22528.5 y 3
380.2121887207031 0 7242.294
380.713623046875 0 2221.243
405.2129821777344 0 590.47253
414.22216796875 0 9403.169 y Water loss 2
414.72393798828125 0 3510.1282
415.2240295410156 0 1766.837
417.0353088378906 0 1379.4277
418.0352478027344 0 2769.4478
418.99554443359375 0 2805.6702
419.02984619140625 0 900.375
419.9941711425781 0 860.3821
423.2270812988281 0 7766.1094 y 2
423.72900390625 0 4752.972
424.2292785644531 0 871.06445
432.2446594238281 0 730.99915
437.5302429199219 0 588.8272
454.4531555175781 0 624.7973
489.2771911621094 0 5056.8975 y 6
490.2814025878906 0 1015.5554
497.6377258300781 0 582.53455
506.10498046875 0 618.6583
521.7727661132812 0 2306.3435
522.2745361328125 0 1809.7
522.7778930664062 0 1706.1163
523.2847290039062 0 5418.0576 Precursor
523.7860107421875 0 2562.945
590.3250122070312 0 14018.27 y 5
591.3272094726562 0 3454.2615
634.3382568359375 0 1922.4425
635.3434448242188 0 958.29004
643.3462524414062 0 620.8992 y Water loss 4
661.3614501953125 0 11344.248 y 4
662.3652954101562 0 3134.6487
740.4047241210938 0 694.769 y Water loss 3
741.3870849609375 0 887.33685 y Ammonia loss 3
758.413818359375 0 20281.955 y 3
759.416259765625 0 8655.24
760.4190673828125 0 2005.2328
763.3816528320312 0 1849.9263
764.3878784179688 0 1117.87
845.4450073242188 0 8279.973 y 2
846.44921875 0 4044.1685
847.455078125 0 1117.6562
862.4490966796875 0 5605.8306
863.4528198242188 0 2844.8442
864.4578247070312 0 1114.33
874.44384765625 0 605.94855
914.4696655273438 0 746.4623 y Water loss 1
932.4765014648438 0 6317.7803 y 1
933.4788818359375 0 2869.2568
934.484375 0 891.9764
942.4523315429688 0 741.7492
1212.2799072265625 0 694.43335
3078.1201171875 0 800.4517

Spectrum Details

|  |  |
| --- | --- |
| Matched peaks? Matched peaks The total absolute number of peaks matched. Additionally in brackets the total fraction of peaks matched and the total number of peaks is shown. | 27 (17.42% of 155) |
| FDR? FDR The false discovery rate estimated for this peptide. It is calculated by matching all theoretical fragments with a non-integer shift with the raw peaks for this spectrum. This is done with 40 different shifts. The resulting percentage is the average number of annotated peaks over the number of annotated peaks with the correct spectrum. | 1.59% |
| Satellite FDR? Satellite FDR See the FDR for details on its calculation. This satellite ion specific FDR only contains the satellite ions (d/w) for I/L/J positions. | - |
| PSM Score? PSM Score The PSM Score as given by Hecklib to this annotated spectrum. It is shown with three significant figures. | 299 |

#### Spectrum 4326? Spectrum 4326 The raw spectrum of this peptide as annotated by Hecklib. The fragments are coloured according to ion type (see legend). Any peaks with a star '\*' as text can be hovered over to see the full details, first the ion type second the mass shift type. By hovering over the amino acids in the peptide or ions in the legend the corresponding peaks are highlighted. By toggling the 'Unassigned' label you can turn the background (unassigned) peaks on or off in the plot. By updating the slider in the Ion legend you can update the spectrum to only show the top X% of the peaks with labels. The top X% means any peak that is within X% of the highest intensity. By dragging in the spectrum you can zoom in to a specific part of the spectrum and use 'Zoom Out' to get back to the original zoom level. The annotation of the spectrum is based on the given sequence in the peptides file and is done with different software so inconsistencies are likely. The peaks are annotated based on the given sequence, with 20 ppm tolerance.

Copy Data

##### Spectrum 4326 (TSV)

###### Preview

```
Loading example...
```

*Click on the button to copy the data to your clipboard.*

Mz MinMz MaxIntensity Max

WidthHeightPeptide font sizePeptide stroke widthSpectrum font sizeSpectrum stroke widthCompact peptide

Ion legend

wxyz

abcd

OtherUnassignedIon^Charge^_Position_Show for top:%

JSSPATJNSR

03.10e+36.21e+39.31e+31.24e+4

Zoom Out

a^+1^_2_y^+1^_1_a^+1^_2_y^+1^_1_b^+1^_2_b^+1^_2_a^+1^_3_y^+1^_2_y^+1^_2_b^+1^_3_y^+1^_3_y^+2^_7_y^+2^_7_y^+1^_3_y^+2^_7_y^+2^_8_y^+2^_8_y^+2^_8_y^+1^_4_\*y^+1^_5_y^+1^_6_y^+1^_6_y^+1^_6_y^+1^_7_y^+1^_8_y^+1^_9_y^+1^_9_

0778155623333111

Fragment Matches Table

Show background peaks

| Position | Ion type | Intensity | mz Theoretical | mz Error (Th) | mz Error (ppm) | Charge | Series Number |
| --- | --- | --- | --- | --- | --- | --- | --- |
| - | - | 1247 | 120.1 | - | - | 0 | - |
| - | - | 477.2 | 121.1 | - | - | 0 | - |
| - | - | 337.9 | 121.7 | - | - | 0 | - |
| - | - | 355.5 | 127.3 | - | - | 0 | - |
| - | - | 442.7 | 127.4 | - | - | 0 | - |
| - | - | 604.2 | 129.1 | - | - | 0 | - |
| - | - | 1019 | 129.1 | - | - | 0 | - |
| - | - | 368.2 | 133.1 | - | - | 0 | - |
| - | - | 2842 | 136.1 | - | - | 0 | - |
| - | - | 506.2 | 139.1 | - | - | 0 | - |
| - | - | 2092 | 141.1 | - | - | 0 | - |
| - | - | 426.2 | 146.6 | - | - | 0 | - |
| - | - | 462 | 148.8 | - | - | 0 | - |
| - | - | 476.7 | 148.9 | - | - | 0 | - |
| - | - | 456.7 | 148.9 | - | - | 0 | - |
| - | - | 487.4 | 148.9 | - | - | 0 | - |
| - | - | 490.4 | 148.9 | - | - | 0 | - |
| - | - | 577.5 | 148.9 | - | - | 0 | - |
| - | - | 717.9 | 148.9 | - | - | 0 | - |
| - | - | 569.2 | 148.9 | - | - | 0 | - |
| - | - | 764.3 | 148.9 | - | - | 0 | - |
| - | - | 894.6 | 148.9 | - | - | 0 | - |
| - | - | 1191 | 148.9 | - | - | 0 | - |
| - | - | 1304 | 148.9 | - | - | 0 | - |
| - | - | 3475 | 148.9 | - | - | 0 | - |
| - | - | 5567 | 148.9 | - | - | 0 | - |
| - | - | 3968 | 149 | - | - | 0 | - |
| - | - | 1544 | 149 | - | - | 0 | - |
| - | - | 872.9 | 149 | - | - | 0 | - |
| - | - | 1038 | 149 | - | - | 0 | - |
| - | - | 922 | 149 | - | - | 0 | - |
| - | - | 665 | 149 | - | - | 0 | - |
| - | - | 450.5 | 149 | - | - | 0 | - |
| - | - | 454.5 | 149 | - | - | 0 | - |
| - | - | 1855 | 149 | - | - | 0 | - |
| - | - | 633 | 149.1 | - | - | 0 | - |
| - | - | 1755 | 151 | - | - | 0 | - |
| 2 | a | 1136 | 155.1 | 0.0002898 | 1.868 | +1 | 2 |
| - | - | 716.7 | 157.1 | - | - | 0 | - |
| - | - | 1976 | 157.1 | - | - | 0 | - |
| 10 | y | 700.6 | 158.1 | 0.0003399 | 2.15 | +1 | 1 |
| - | - | 465.4 | 161.1 | - | - | 0 | - |
| - | - | 417.1 | 161.6 | - | - | 0 | - |
| - | - | 2891 | 167.1 | - | - | 0 | - |
| - | - | 632.9 | 167.1 | - | - | 0 | - |
| - | - | 1189 | 168.1 | - | - | 0 | - |
| - | - | 1726 | 169.1 | - | - | 0 | - |
| - | - | 2609 | 169.1 | - | - | 0 | - |
| - | - | 413.2 | 170.6 | - | - | 0 | - |
| - | - | 917.4 | 173.1 | - | - | 0 | - |
| 2 | a | 3506 | 173.1 | 5.735E-06 | 0.03312 | +1 | 2 |
| - | - | 891.1 | 173.4 | - | - | 0 | - |
| - | - | 609.9 | 174.1 | - | - | 0 | - |
| 10 | y | 2228 | 175.1 | 5.568E-05 | 0.318 | +1 | 1 |
| - | - | 522.5 | 178.3 | - | - | 0 | - |
| 2 | b | 5386 | 183.1 | 1.933E-05 | 0.1056 | +1 | 2 |
| - | - | 1508 | 185.1 | - | - | 0 | - |
| - | - | 1083 | 187.1 | - | - | 0 | - |
| - | - | 685 | 189.1 | - | - | 0 | - |
| - | - | 499.2 | 193.7 | - | - | 0 | - |
| 2 | b | 3211 | 201.1 | 0.0001205 | 0.5993 | +1 | 2 |
| - | - | 948.9 | 202.1 | - | - | 0 | - |
| - | - | 645 | 216.1 | - | - | 0 | - |
| - | - | 2574 | 223.1 | - | - | 0 | - |
| - | - | 960.9 | 224.1 | - | - | 0 | - |
| - | - | 1409 | 225 | - | - | 0 | - |
| - | - | 986.1 | 226 | - | - | 0 | - |
| - | - | 661.8 | 226.1 | - | - | 0 | - |
| - | - | 810.8 | 227 | - | - | 0 | - |
| - | - | 4108 | 238.1 | - | - | 0 | - |
| - | - | 570.9 | 240.1 | - | - | 0 | - |
| - | - | 887.1 | 241.1 | - | - | 0 | - |
| 3 | a | 792 | 242.1 | 0.0006412 | 2.648 | +1 | 3 |
| 9 | y | 758.4 | 245.1 | 4.973E-05 | 0.2029 | +1 | 2 |
| - | - | 1089 | 252.1 | - | - | 0 | - |
| - | - | 1372 | 256.1 | - | - | 0 | - |
| 9 | y | 3643 | 262.2 | 0.0001322 | 0.5042 | +1 | 2 |
| - | - | 608.9 | 262.2 | - | - | 0 | - |
| 3 | b | 6461 | 270.1 | 8.769E-05 | 0.3246 | +1 | 3 |
| - | - | 693.9 | 271.2 | - | - | 0 | - |
| - | - | 659.1 | 281.1 | - | - | 0 | - |
| - | - | 921.5 | 282.1 | - | - | 0 | - |
| - | - | 1326 | 283 | - | - | 0 | - |
| - | - | 707 | 284 | - | - | 0 | - |
| - | - | 6226 | 299.1 | - | - | 0 | - |
| - | - | 6343 | 300.1 | - | - | 0 | - |
| - | - | 1.082E+04 | 301.1 | - | - | 0 | - |
| - | - | 6968 | 302.1 | - | - | 0 | - |
| - | - | 2151 | 303.1 | - | - | 0 | - |
| - | - | 756.9 | 339.2 | - | - | 0 | - |
| - | - | 717 | 342.1 | - | - | 0 | - |
| - | - | 1334 | 346 | - | - | 0 | - |
| - | - | 2384 | 347 | - | - | 0 | - |
| - | - | 1692 | 348 | - | - | 0 | - |
| - | - | 643.9 | 359 | - | - | 0 | - |
| 8 | y | 1122 | 359.2 | 0.0005804 | 1.616 | +1 | 3 |
| - | - | 1002 | 360 | - | - | 0 | - |
| - | - | 1447 | 361 | - | - | 0 | - |
| - | - | 1605 | 362 | - | - | 0 | - |
| - | - | 771.6 | 363 | - | - | 0 | - |
| 4 | y | 1918 | 370.7 | 0.0007977 | 2.152 | +2 | 7 |
| 4 | y | 783.3 | 371.2 | 0.005791 | 15.6 | +2 | 7 |
| - | - | 630.6 | 371.7 | - | - | 0 | - |
| 8 | y | 1622 | 376.2 | 0.0003974 | 1.056 | +1 | 3 |
| - | - | 696.4 | 378.7 | - | - | 0 | - |
| - | - | 520.8 | 379.2 | - | - | 0 | - |
| 4 | y | 1.03E+04 | 379.7 | 0.0001596 | 0.4203 | +2 | 7 |
| - | - | 3387 | 380.2 | - | - | 0 | - |
| - | - | 1115 | 380.7 | - | - | 0 | - |
| 3 | y | 6897 | 414.2 | 5.773E-05 | 0.1394 | +2 | 8 |
| 3 | y | 2508 | 414.7 | 0.004913 | 11.85 | +2 | 8 |
| - | - | 1329 | 415.2 | - | - | 0 | - |
| - | - | 1493 | 417 | - | - | 0 | - |
| - | - | 2928 | 418 | - | - | 0 | - |
| - | - | 2336 | 419 | - | - | 0 | - |
| - | - | 1434 | 419 | - | - | 0 | - |
| - | - | 673.6 | 420 | - | - | 0 | - |
| - | - | 937.5 | 421 | - | - | 0 | - |
| 3 | y | 7755 | 423.2 | 3.002E-05 | 0.07092 | +2 | 8 |
| - | - | 3657 | 423.7 | - | - | 0 | - |
| - | - | 1461 | 424.2 | - | - | 0 | - |
| - | - | 637.8 | 435.2 | - | - | 0 | - |
| - | - | 608 | 456.8 | - | - | 0 | - |
| - | - | 1320 | 458.3 | - | - | 0 | - |
| - | - | 679.3 | 471.7 | - | - | 0 | - |
| - | - | 704 | 489.1 | - | - | 0 | - |
| 7 | y | 2776 | 489.3 | 4.842E-05 | 0.09897 | +1 | 4 |
| - | - | 767.5 | 490.3 | - | - | 0 | - |
| - | - | 2409 | 522.3 | - | - | 0 | - |
| 0 | Precursor | 1333 | 523.3 | 0.006532 | 12.48 | +2 | -1 |
| - | - | 827.7 | 523.8 | - | - | 0 | - |
| - | - | 669.7 | 546.3 | - | - | 0 | - |
| 6 | y | 9533 | 590.3 | 0.0002773 | 0.4697 | +1 | 5 |
| - | - | 2194 | 591.3 | - | - | 0 | - |
| - | - | 716.1 | 592.3 | - | - | 0 | - |
| 5 | y | 851.5 | 643.4 | 8.777E-05 | 0.1364 | +1 | 6 |
| 5 | y | 629.9 | 644.3 | 0.009488 | 14.73 | +1 | 6 |
| 5 | y | 6810 | 661.4 | 0.0003375 | 0.5103 | +1 | 6 |
| - | - | 2755 | 662.4 | - | - | 0 | - |
| 4 | y | 1.229E+04 | 758.4 | 0.0001228 | 0.162 | +1 | 7 |
| - | - | 4934 | 759.4 | - | - | 0 | - |
| - | - | 1742 | 760.4 | - | - | 0 | - |
| 3 | y | 5135 | 845.4 | 0.0001363 | 0.1613 | +1 | 8 |
| - | - | 2741 | 846.4 | - | - | 0 | - |
| 2 | y | 726.9 | 914.5 | 0.001195 | 1.306 | +1 | 9 |
| 2 | y | 4306 | 932.5 | 0.0008862 | 0.9504 | +1 | 9 |
| - | - | 1995 | 933.5 | - | - | 0 | - |
| - | - | 708.3 | 1222 | - | - | 0 | - |
| - | - | 760.4 | 3081 | - | - | 0 | - |

m/z Charge Intensity FragmentType MassShift Position
120.08113098144531 0 1246.597
121.07630920410156 0 477.15265
121.66571044921875 0 337.85358
127.2706069946289 0 355.54428
127.43753051757812 0 442.73184
129.06594848632812 0 604.24304
129.1023712158203 0 1018.7833
133.14984130859375 0 368.22964
136.0758819580078 0 2842.405
139.08641052246094 0 506.1604
141.1023712158203 0 2092.4854
146.6175994873047 0 426.17957
148.82420349121094 0 462.00394
148.8598175048828 0 476.7282
148.86676025390625 0 456.66446
148.87408447265625 0 487.4339
148.8807830810547 0 490.4449
148.88790893554688 0 577.5354
148.89523315429688 0 717.9249
148.90240478515625 0 569.18494
148.90960693359375 0 764.2598
148.91709899902344 0 894.55554
148.92410278320312 0 1190.8013
148.9309844970703 0 1303.8145
148.93841552734375 0 3475.046
148.94606018066406 0 5566.566
148.9624481201172 0 3967.9346
148.969970703125 0 1543.611
148.97621154785156 0 872.9099
148.984130859375 0 1038.3265
148.99151611328125 0 921.96124
148.99839782714844 0 665.0255
149.0052947998047 0 450.54852
149.01307678222656 0 454.49557
149.04498291015625 0 1855.2693
149.06283569335938 0 632.9634
151.041748046875 0 1754.7667
155.11817932128906 0 1135.9261 a Water loss 1
157.06076049804688 0 716.6532
157.09725952148438 0 1976.2163
158.09274291992188 0 700.6036 y Ammonia loss 9
161.09266662597656 0 465.35547
161.62142944335938 0 417.1097
167.05538940429688 0 2891.3562
167.0814666748047 0 632.91
168.05484008789062 0 1189.3661
169.05258178710938 0 1726.48
169.09710693359375 0 2608.9084
170.5741729736328 0 413.21033
173.09231567382812 0 917.3515
173.12844848632812 0 3505.9714 a 1
173.44996643066406 0 891.0971
174.0872039794922 0 609.9299
175.118896484375 0 2227.8616 y 9
178.33657836914062 0 522.5014
183.11282348632812 0 5385.589 b Water loss 1
185.092041015625 0 1507.5604
187.14407348632812 0 1082.6951
189.0869903564453 0 685.001
193.6626434326172 0 499.1739
201.1234893798828 0 3210.8418 b 1
202.08251953125 0 948.9169
216.097900390625 0 644.9553
223.06375122070312 0 2573.7427
224.0637969970703 0 960.8511
225.04287719726562 0 1408.729
226.04351806640625 0 986.0876
226.11944580078125 0 661.80206
227.0399932861328 0 810.79224
238.11856079101562 0 4107.9243
240.09645080566406 0 570.9204
241.0921173095703 0 887.1112
242.14927673339844 0 792.01825 a Water loss 2
245.12448120117188 0 758.37286 y Ammonia loss 8
252.1347198486328 0 1089.0997
256.1290283203125 0 1371.7576
262.1508483886719 0 3642.9756 y 8
262.1647033691406 0 608.8757
270.1447448730469 0 6460.9297 b Water loss 2
271.15032958984375 0 693.9471
281.0517578125 0 659.0562
282.0522766113281 0 921.53174
283.0489501953125 0 1325.5944
284.0498352050781 0 707.00616
299.0616760253906 0 6225.5376
300.0622253417969 0 6342.9404
301.0595397949219 0 10823.018
302.0601501464844 0 6967.737
303.0563049316406 0 2150.7158
339.1653137207031 0 756.8669
342.13873291015625 0 717.04333
345.97601318359375 0 1333.7521
346.9738464355469 0 2384.0461
347.9732360839844 0 1692.1891
359.02862548828125 0 643.86694
359.1667785644531 0 1121.5245 y Ammonia loss 7
360.0281677246094 0 1001.91656
361.0249938964844 0 1447.4459
362.0257263183594 0 1605.2635
363.02410888671875 0 771.6177
370.705322265625 0 1917.9784 y Water loss 3
371.20391845703125 0 783.3358 y Ammonia loss 3
371.7076416015625 0 630.58435
376.1943054199219 0 1621.7412 y 7
378.7009582519531 0 696.3891
379.2071228027344 0 520.844
379.71124267578125 0 10301.702 y 3
380.21014404296875 0 3386.8005
380.7114562988281 0 1115.1547
414.2220764160156 0 6897.4746 y Water loss 2
414.71905517578125 0 2507.607 y Ammonia loss 2
415.2231140136719 0 1328.8688
417.0345458984375 0 1492.679
418.0348815917969 0 2928.3787
418.9955749511719 0 2335.5571
419.03204345703125 0 1433.8706
420.03155517578125 0 673.56384
420.9933776855469 0 937.50134
423.2273864746094 0 7754.7827 y 2
423.72698974609375 0 3657.4722
424.2261657714844 0 1461.447
435.22772216796875 0 637.83813
456.83575439453125 0 608.04834
458.272705078125 0 1320.1248
471.7342834472656 0 679.28705
489.055419921875 0 703.97815
489.2779235839844 0 2776.289 y 6
490.2850036621094 0 767.5083
522.2702026367188 0 2409.192
523.2789306640625 0 1332.5874 Precursor
523.7879028320312 0 827.6656
546.29052734375 0 669.6927
590.325927734375 0 9533.434 y 5
591.3225708007812 0 2193.9243
592.33056640625 0 716.1216
643.3521118164062 0 851.48486 y Water loss 4
644.345703125 0 629.88074 y Ammonia loss 4
661.3624267578125 0 6809.9556 y 4
662.3598022460938 0 2754.7495
758.4154052734375 0 12288.356 y 3
759.413818359375 0 4934.495
760.4155883789062 0 1741.5795
845.4476928710938 0 5134.865 y 2
846.4484252929688 0 2740.5789
914.47021484375 0 726.85223 y Water loss 1
932.4786987304688 0 4306.2285 y 1
933.4782104492188 0 1994.5754
1222.3505859375 0 708.34766
3080.51171875 0 760.3835

Spectrum Details

|  |  |
| --- | --- |
| Matched peaks? Matched peaks The total absolute number of peaks matched. Additionally in brackets the total fraction of peaks matched and the total number of peaks is shown. | 28 (18.79% of 149) |
| FDR? FDR The false discovery rate estimated for this peptide. It is calculated by matching all theoretical fragments with a non-integer shift with the raw peaks for this spectrum. This is done with 40 different shifts. The resulting percentage is the average number of annotated peaks over the number of annotated peaks with the correct spectrum. | 1.11% |
| Satellite FDR? Satellite FDR See the FDR for details on its calculation. This satellite ion specific FDR only contains the satellite ions (d/w) for I/L/J positions. | - |
| PSM Score? PSM Score The PSM Score as given by Hecklib to this annotated spectrum. It is shown with three significant figures. | 281 |

#### Spectrum 4517? Spectrum 4517 The raw spectrum of this peptide as annotated by Hecklib. The fragments are coloured according to ion type (see legend). Any peaks with a star '\*' as text can be hovered over to see the full details, first the ion type second the mass shift type. By hovering over the amino acids in the peptide or ions in the legend the corresponding peaks are highlighted. By toggling the 'Unassigned' label you can turn the background (unassigned) peaks on or off in the plot. By updating the slider in the Ion legend you can update the spectrum to only show the top X% of the peaks with labels. The top X% means any peak that is within X% of the highest intensity. By dragging in the spectrum you can zoom in to a specific part of the spectrum and use 'Zoom Out' to get back to the original zoom level. The annotation of the spectrum is based on the given sequence in the peptides file and is done with different software so inconsistencies are likely. The peaks are annotated based on the given sequence, with 20 ppm tolerance.

Copy Data

##### Spectrum 4517 (TSV)

###### Preview

```
Loading example...
```

*Click on the button to copy the data to your clipboard.*

Mz MinMz MaxIntensity Max

WidthHeightPeptide font sizePeptide stroke widthSpectrum font sizeSpectrum stroke widthCompact peptide

Ion legend

wxyz

abcd

OtherUnassignedIon^Charge^_Position_Show for top:%

JSSPATJNSR

02.78e+35.57e+38.35e+31.11e+4

Zoom Out

d^+1^_2_y^+1^_1_a^+1^_2_y^+1^_1_b^+1^_2_b^+1^_2_y^+1^_2_y^+1^_2_b^+1^_3_y^+1^_3_y^+2^_7_y^+1^_3_y^+2^_7_y^+2^_8_y^+2^_8_y^+2^_8_y^+1^_4_\*y^+1^_5_y^+1^_6_y^+1^_7_y^+1^_8_y^+1^_9_

0557111416712227

Fragment Matches Table

Show background peaks

| Position | Ion type | Intensity | mz Theoretical | mz Error (Th) | mz Error (ppm) | Charge | Series Number |
| --- | --- | --- | --- | --- | --- | --- | --- |
| - | - | 509.9 | 120.1 | - | - | 0 | - |
| - | - | 1068 | 129.1 | - | - | 0 | - |
| - | - | 415.4 | 131 | - | - | 0 | - |
| - | - | 376.6 | 134.5 | - | - | 0 | - |
| - | - | 353 | 136 | - | - | 0 | - |
| - | - | 989.1 | 136.1 | - | - | 0 | - |
| - | - | 377.4 | 137.4 | - | - | 0 | - |
| - | - | 435.5 | 139.1 | - | - | 0 | - |
| - | - | 451.5 | 140.6 | - | - | 0 | - |
| - | - | 1662 | 141.1 | - | - | 0 | - |
| - | - | 459.4 | 148.9 | - | - | 0 | - |
| - | - | 576.1 | 148.9 | - | - | 0 | - |
| - | - | 647 | 148.9 | - | - | 0 | - |
| - | - | 788.3 | 148.9 | - | - | 0 | - |
| - | - | 626 | 148.9 | - | - | 0 | - |
| - | - | 1188 | 148.9 | - | - | 0 | - |
| - | - | 1286 | 148.9 | - | - | 0 | - |
| - | - | 2261 | 148.9 | - | - | 0 | - |
| - | - | 3812 | 148.9 | - | - | 0 | - |
| - | - | 4436 | 149 | - | - | 0 | - |
| - | - | 2568 | 149 | - | - | 0 | - |
| - | - | 1336 | 149 | - | - | 0 | - |
| - | - | 1064 | 149 | - | - | 0 | - |
| - | - | 906.8 | 149 | - | - | 0 | - |
| - | - | 682 | 149 | - | - | 0 | - |
| - | - | 706.4 | 149 | - | - | 0 | - |
| - | - | 530.4 | 149 | - | - | 0 | - |
| - | - | 384.6 | 149 | - | - | 0 | - |
| - | - | 472.6 | 149 | - | - | 0 | - |
| - | - | 3489 | 149 | - | - | 0 | - |
| - | - | 957 | 150 | - | - | 0 | - |
| - | - | 958.6 | 151 | - | - | 0 | - |
| - | - | 566.9 | 157.1 | - | - | 0 | - |
| - | - | 1375 | 157.1 | - | - | 0 | - |
| 2 | d | 1567 | 157.1 | 0.0002189 | 1.393 | +1 | 2 |
| 10 | y | 672.1 | 158.1 | 0.0003399 | 2.15 | +1 | 1 |
| - | - | 413 | 158.9 | - | - | 0 | - |
| - | - | 514 | 162.2 | - | - | 0 | - |
| - | - | 440.2 | 163.5 | - | - | 0 | - |
| - | - | 3982 | 167.1 | - | - | 0 | - |
| - | - | 663.6 | 167.1 | - | - | 0 | - |
| - | - | 2018 | 169.1 | - | - | 0 | - |
| - | - | 2067 | 169.1 | - | - | 0 | - |
| 2 | a | 1900 | 173.1 | 2.099E-05 | 0.1213 | +1 | 2 |
| - | - | 616.5 | 173.5 | - | - | 0 | - |
| 10 | y | 1902 | 175.1 | 5.352E-06 | 0.03056 | +1 | 1 |
| 2 | b | 3913 | 183.1 | 4.985E-05 | 0.2722 | +1 | 2 |
| - | - | 1733 | 185.1 | - | - | 0 | - |
| - | - | 857.3 | 185.1 | - | - | 0 | - |
| - | - | 1511 | 187.1 | - | - | 0 | - |
| 2 | b | 2488 | 201.1 | 2.898E-05 | 0.1441 | +1 | 2 |
| - | - | 673.8 | 202.1 | - | - | 0 | - |
| - | - | 2299 | 223.1 | - | - | 0 | - |
| - | - | 868.1 | 225 | - | - | 0 | - |
| - | - | 815.9 | 226 | - | - | 0 | - |
| - | - | 775.8 | 227 | - | - | 0 | - |
| - | - | 2949 | 238.1 | - | - | 0 | - |
| - | - | 829.7 | 239.1 | - | - | 0 | - |
| - | - | 796.9 | 240.1 | - | - | 0 | - |
| - | - | 890.5 | 241.1 | - | - | 0 | - |
| 9 | y | 1103 | 245.1 | 2.657E-05 | 0.1084 | +1 | 2 |
| - | - | 1090 | 252.1 | - | - | 0 | - |
| 9 | y | 2650 | 262.2 | 0.0001425 | 0.5435 | +1 | 2 |
| - | - | 578 | 268.6 | - | - | 0 | - |
| 3 | b | 4432 | 270.1 | 0.0001259 | 0.4662 | +1 | 3 |
| - | - | 928.7 | 282.1 | - | - | 0 | - |
| - | - | 972.7 | 283 | - | - | 0 | - |
| - | - | 823.3 | 285 | - | - | 0 | - |
| - | - | 684.9 | 287.8 | - | - | 0 | - |
| - | - | 5801 | 299.1 | - | - | 0 | - |
| - | - | 764.8 | 299.1 | - | - | 0 | - |
| - | - | 7947 | 300.1 | - | - | 0 | - |
| - | - | 1.103E+04 | 301.1 | - | - | 0 | - |
| - | - | 6694 | 302.1 | - | - | 0 | - |
| - | - | 2069 | 303.1 | - | - | 0 | - |
| - | - | 780.2 | 309.2 | - | - | 0 | - |
| - | - | 1913 | 327.7 | - | - | 0 | - |
| - | - | 637.4 | 328.2 | - | - | 0 | - |
| - | - | 608.1 | 345 | - | - | 0 | - |
| - | - | 1297 | 346 | - | - | 0 | - |
| - | - | 538.7 | 346.9 | - | - | 0 | - |
| - | - | 2436 | 347 | - | - | 0 | - |
| - | - | 1871 | 348 | - | - | 0 | - |
| 8 | y | 1210 | 359.2 | 0.0007024 | 1.956 | +1 | 3 |
| - | - | 1436 | 362 | - | - | 0 | - |
| 4 | y | 1820 | 370.7 | 0.0006146 | 1.658 | +2 | 7 |
| - | - | 960.5 | 371.2 | - | - | 0 | - |
| 8 | y | 1650 | 376.2 | 9.222E-05 | 0.2451 | +1 | 3 |
| 4 | y | 7892 | 379.7 | 9.858E-05 | 0.2596 | +2 | 7 |
| - | - | 2729 | 380.2 | - | - | 0 | - |
| - | - | 732.6 | 380.7 | - | - | 0 | - |
| 3 | y | 5363 | 414.2 | 0.0002474 | 0.5974 | +2 | 8 |
| 3 | y | 2959 | 414.7 | 0.007477 | 18.03 | +2 | 8 |
| - | - | 1500 | 417 | - | - | 0 | - |
| - | - | 2702 | 418 | - | - | 0 | - |
| - | - | 2059 | 419 | - | - | 0 | - |
| - | - | 799.1 | 419 | - | - | 0 | - |
| - | - | 705.7 | 420 | - | - | 0 | - |
| 3 | y | 5121 | 423.2 | 0.0002141 | 0.5059 | +2 | 8 |
| - | - | 2943 | 423.7 | - | - | 0 | - |
| 7 | y | 2254 | 489.3 | 7.365E-05 | 0.1505 | +1 | 4 |
| - | - | 634.2 | 505.8 | - | - | 0 | - |
| - | - | 716.8 | 515.3 | - | - | 0 | - |
| - | - | 1784 | 522.3 | - | - | 0 | - |
| 0 | Precursor | 1526 | 523.3 | 0.00348 | 6.651 | +2 | -1 |
| - | - | 755.2 | 524.2 | - | - | 0 | - |
| 6 | y | 7161 | 590.3 | 2.792E-05 | 0.0473 | +1 | 5 |
| - | - | 2418 | 591.3 | - | - | 0 | - |
| - | - | 753.2 | 620 | - | - | 0 | - |
| 5 | y | 4283 | 661.4 | 0.0005206 | 0.7872 | +1 | 6 |
| - | - | 1732 | 662.4 | - | - | 0 | - |
| 4 | y | 8722 | 758.4 | 0.0001839 | 0.2425 | +1 | 7 |
| - | - | 4059 | 759.4 | - | - | 0 | - |
| - | - | 1024 | 760.4 | - | - | 0 | - |
| - | - | 646.1 | 774.2 | - | - | 0 | - |
| 3 | y | 3708 | 845.4 | 0.001329 | 1.571 | +1 | 8 |
| - | - | 2383 | 846.4 | - | - | 0 | - |
| 2 | y | 3555 | 932.5 | 0.001741 | 1.867 | +1 | 9 |
| - | - | 1288 | 933.5 | - | - | 0 | - |
| - | - | 781.1 | 934.5 | - | - | 0 | - |
| - | - | 716.2 | 1473 | - | - | 0 | - |
| - | - | 695.6 | 1501 | - | - | 0 | - |
| - | - | 632.4 | 1850 | - | - | 0 | - |
| - | - | 811.9 | 2037 | - | - | 0 | - |
| - | - | 708.6 | 2205 | - | - | 0 | - |

m/z Charge Intensity FragmentType MassShift Position
120.08042907714844 0 509.91602
129.10247802734375 0 1068.083
131.01199340820312 0 415.38565
134.4532470703125 0 376.61392
135.99319458007812 0 352.98602
136.0758056640625 0 989.07544
137.4030303955078 0 377.35474
139.08724975585938 0 435.53934
140.61610412597656 0 451.47842
141.10235595703125 0 1662.3785
148.88514709472656 0 459.38965
148.8924560546875 0 576.1003
148.89968872070312 0 647.005
148.9069061279297 0 788.31274
148.91419982910156 0 625.9975
148.92124938964844 0 1188.3652
148.92843627929688 0 1285.8235
148.93539428710938 0 2261.0994
148.94308471679688 0 3812.295
148.9595947265625 0 4436.338
148.96731567382812 0 2568.0427
148.97427368164062 0 1336.3151
148.98147583007812 0 1063.806
148.9889678955078 0 906.8429
148.99560546875 0 681.98364
149.00270080566406 0 706.3924
149.009521484375 0 530.4138
149.01670837402344 0 384.6021
149.0315704345703 0 472.58185
149.04507446289062 0 3489.2751
150.04469299316406 0 956.9989
151.04168701171875 0 958.60803
157.06065368652344 0 566.91766
157.09725952148438 0 1375.1685
157.13375854492188 0 1567.0642 d 1
158.09274291992188 0 672.0785 y Ammonia loss 9
158.85366821289062 0 413.04675
162.22161865234375 0 513.99884
163.5028839111328 0 440.24142
167.055419921875 0 3982.4214
167.08155822753906 0 663.6075
169.05233764648438 0 2017.6647
169.09719848632812 0 2066.954
173.12843322753906 0 1899.6997 a 1
173.4501495361328 0 616.53186
175.11895751953125 0 1901.9272 y 9
183.11285400390625 0 3912.5095 b Water loss 1
185.09214782714844 0 1733.4528
185.12867736816406 0 857.2792
187.10789489746094 0 1510.5607
201.12339782714844 0 2487.61 b 1
202.0822296142578 0 673.77966
223.0638885498047 0 2299.2236
225.04283142089844 0 868.14606
226.043701171875 0 815.9434
227.03871154785156 0 775.7596
238.11842346191406 0 2948.5925
239.09466552734375 0 829.66644
240.0952911376953 0 796.8885
241.09130859375 0 890.5082
245.12440490722656 0 1102.7386 y Ammonia loss 8
252.13421630859375 0 1089.6877
262.151123046875 0 2649.876 y 8
268.5558776855469 0 578.0005
270.14495849609375 0 4431.9604 b Water loss 2
282.0505676269531 0 928.6617
283.04876708984375 0 972.7405
285.0274963378906 0 823.3431
287.83355712890625 0 684.88416
299.0618591308594 0 5801.145
299.0802001953125 0 764.8328
300.0622253417969 0 7947.2437
301.0593566894531 0 11025.765
302.0599060058594 0 6693.535
303.0577697753906 0 2069.4746
309.2037353515625 0 780.155
327.69500732421875 0 1912.9836
328.1963195800781 0 637.37585
344.9752502441406 0 608.10425
345.97564697265625 0 1297.4775
346.91064453125 0 538.6618
346.9740905761719 0 2435.5984
347.9737854003906 0 1871.0928
359.1666564941406 0 1210.4702 y Ammonia loss 7
362.02484130859375 0 1435.9047
370.70550537109375 0 1820.3018 y Water loss 3
371.2060852050781 0 960.52435
376.1940002441406 0 1649.9512 y 7
379.7113037109375 0 7891.9497 y 3
380.2095642089844 0 2729.0542
380.7093200683594 0 732.60614
414.2223815917969 0 5363.26 y Water loss 2
414.72161865234375 0 2958.5044 y Ammonia loss 2
417.0355529785156 0 1499.8572
418.0350341796875 0 2701.714
418.994873046875 0 2059.154
419.0324401855469 0 799.1135
419.995849609375 0 705.727
423.2276306152344 0 5121.122 y 2
423.7271728515625 0 2942.911
489.2780456542969 0 2253.8855 y 6
505.78643798828125 0 634.2284
515.2814331054688 0 716.82983
522.271240234375 0 1783.8479
523.281982421875 0 1526.1921 Precursor
524.2103271484375 0 755.22217
590.3256225585938 0 7160.578 y 5
591.326416015625 0 2417.7002
619.982666015625 0 753.2222
661.3622436523438 0 4283.2197 y 4
662.3634033203125 0 1731.69
758.4153442382812 0 8721.963 y 3
759.4150390625 0 4059.081
760.409423828125 0 1023.99634
774.2234497070312 0 646.11444
845.4462280273438 0 3708.3286 y 2
846.44580078125 0 2382.9219
932.4778442382812 0 3555.462 y 1
933.4734497070312 0 1287.9823
934.4738159179688 0 781.09973
1473.4820556640625 0 716.2169
1501.36083984375 0 695.60266
1849.6075439453125 0 632.35614
2036.9124755859375 0 811.8527
2205.42041015625 0 708.6024

Spectrum Details

|  |  |
| --- | --- |
| Matched peaks? Matched peaks The total absolute number of peaks matched. Additionally in brackets the total fraction of peaks matched and the total number of peaks is shown. | 23 (18.40% of 125) |
| FDR? FDR The false discovery rate estimated for this peptide. It is calculated by matching all theoretical fragments with a non-integer shift with the raw peaks for this spectrum. This is done with 40 different shifts. The resulting percentage is the average number of annotated peaks over the number of annotated peaks with the correct spectrum. | 1.66% |
| Satellite FDR? Satellite FDR See the FDR for details on its calculation. This satellite ion specific FDR only contains the satellite ions (d/w) for I/L/J positions. | - |
| PSM Score? PSM Score The PSM Score as given by Hecklib to this annotated spectrum. It is shown with three significant figures. | 193 |

#### Spectrum 4461? Spectrum 4461 The raw spectrum of this peptide as annotated by Hecklib. The fragments are coloured according to ion type (see legend). Any peaks with a star '\*' as text can be hovered over to see the full details, first the ion type second the mass shift type. By hovering over the amino acids in the peptide or ions in the legend the corresponding peaks are highlighted. By toggling the 'Unassigned' label you can turn the background (unassigned) peaks on or off in the plot. By updating the slider in the Ion legend you can update the spectrum to only show the top X% of the peaks with labels. The top X% means any peak that is within X% of the highest intensity. By dragging in the spectrum you can zoom in to a specific part of the spectrum and use 'Zoom Out' to get back to the original zoom level. The annotation of the spectrum is based on the given sequence in the peptides file and is done with different software so inconsistencies are likely. The peaks are annotated based on the given sequence, with 20 ppm tolerance.

Copy Data

##### Spectrum 4461 (TSV)

###### Preview

```
Loading example...
```

*Click on the button to copy the data to your clipboard.*

Mz MinMz MaxIntensity Max

WidthHeightPeptide font sizePeptide stroke widthSpectrum font sizeSpectrum stroke widthCompact peptide

Ion legend

wxyz

abcd

OtherUnassignedIon^Charge^_Position_Show for top:%

JSSPATJNSR

02.89e+35.78e+38.67e+31.16e+4

Zoom Out

a^+1^_2_d^+1^_2_y^+1^_1_a^+1^_2_y^+1^_1_b^+1^_2_b^+1^_2_y^+1^_2_y^+1^_2_b^+1^_3_y^+1^_3_y^+2^_7_y^+1^_3_y^+2^_7_y^+2^_8_y^+2^_8_y^+2^_8_y^+1^_4_\*y^+1^_5_y^+1^_6_y^+1^_6_y^+1^_6_y^+1^_7_y^+1^_8_y^+1^_9_y^+1^_9_

0778155623333111

Fragment Matches Table

Show background peaks

| Position | Ion type | Intensity | mz Theoretical | mz Error (Th) | mz Error (ppm) | Charge | Series Number |
| --- | --- | --- | --- | --- | --- | --- | --- |
| - | - | 441.6 | 120.1 | - | - | 0 | - |
| - | - | 415.3 | 124.3 | - | - | 0 | - |
| - | - | 446 | 125.1 | - | - | 0 | - |
| - | - | 414.5 | 125.3 | - | - | 0 | - |
| - | - | 456.1 | 129.1 | - | - | 0 | - |
| - | - | 1861 | 129.1 | - | - | 0 | - |
| - | - | 566.3 | 130.1 | - | - | 0 | - |
| - | - | 1211 | 136.1 | - | - | 0 | - |
| - | - | 914.8 | 139.1 | - | - | 0 | - |
| - | - | 2965 | 141.1 | - | - | 0 | - |
| - | - | 440.5 | 144.6 | - | - | 0 | - |
| - | - | 795.3 | 147.1 | - | - | 0 | - |
| - | - | 505.8 | 148.9 | - | - | 0 | - |
| - | - | 460 | 148.9 | - | - | 0 | - |
| - | - | 816.9 | 148.9 | - | - | 0 | - |
| - | - | 776.7 | 148.9 | - | - | 0 | - |
| - | - | 956.1 | 148.9 | - | - | 0 | - |
| - | - | 1054 | 148.9 | - | - | 0 | - |
| - | - | 1082 | 148.9 | - | - | 0 | - |
| - | - | 1843 | 148.9 | - | - | 0 | - |
| - | - | 3595 | 148.9 | - | - | 0 | - |
| - | - | 5095 | 149 | - | - | 0 | - |
| - | - | 2980 | 149 | - | - | 0 | - |
| - | - | 1242 | 149 | - | - | 0 | - |
| - | - | 1278 | 149 | - | - | 0 | - |
| - | - | 967.3 | 149 | - | - | 0 | - |
| - | - | 727.2 | 149 | - | - | 0 | - |
| - | - | 703.8 | 149 | - | - | 0 | - |
| - | - | 541.8 | 149 | - | - | 0 | - |
| - | - | 480.9 | 149 | - | - | 0 | - |
| - | - | 2212 | 149 | - | - | 0 | - |
| - | - | 647.7 | 149.1 | - | - | 0 | - |
| - | - | 703.2 | 150 | - | - | 0 | - |
| - | - | 1080 | 151 | - | - | 0 | - |
| 2 | a | 730.8 | 155.1 | 0.0003053 | 1.968 | +1 | 2 |
| - | - | 568 | 157.1 | - | - | 0 | - |
| - | - | 1752 | 157.1 | - | - | 0 | - |
| 2 | d | 3228 | 157.1 | 0.0001884 | 1.199 | +1 | 2 |
| 10 | y | 831.8 | 158.1 | 0.0001262 | 0.7985 | +1 | 1 |
| - | - | 834.6 | 158.1 | - | - | 0 | - |
| - | - | 567.9 | 161.1 | - | - | 0 | - |
| - | - | 3901 | 167.1 | - | - | 0 | - |
| - | - | 486.2 | 167.1 | - | - | 0 | - |
| - | - | 554.7 | 168.1 | - | - | 0 | - |
| - | - | 2652 | 169.1 | - | - | 0 | - |
| - | - | 2999 | 169.1 | - | - | 0 | - |
| 2 | a | 2806 | 173.1 | 2.099E-05 | 0.1213 | +1 | 2 |
| - | - | 503.3 | 173.4 | - | - | 0 | - |
| 10 | y | 858.4 | 175.1 | 0.0001427 | 0.8148 | +1 | 1 |
| 2 | b | 4592 | 183.1 | 1.119E-05 | 0.0611 | +1 | 2 |
| - | - | 1050 | 183.1 | - | - | 0 | - |
| - | - | 1139 | 185.1 | - | - | 0 | - |
| - | - | 1379 | 185.1 | - | - | 0 | - |
| - | - | 479.6 | 191.2 | - | - | 0 | - |
| 2 | b | 2572 | 201.1 | 0.0002273 | 1.13 | +1 | 2 |
| - | - | 479.8 | 211.1 | - | - | 0 | - |
| - | - | 1823 | 223.1 | - | - | 0 | - |
| - | - | 738.4 | 224.1 | - | - | 0 | - |
| - | - | 1049 | 225 | - | - | 0 | - |
| - | - | 602.8 | 225.1 | - | - | 0 | - |
| - | - | 636.1 | 225.1 | - | - | 0 | - |
| - | - | 1068 | 227 | - | - | 0 | - |
| - | - | 4187 | 238.1 | - | - | 0 | - |
| - | - | 593.3 | 239.1 | - | - | 0 | - |
| - | - | 643.6 | 241.1 | - | - | 0 | - |
| 9 | y | 677.2 | 245.1 | 9.55E-05 | 0.3896 | +1 | 2 |
| - | - | 564 | 250.1 | - | - | 0 | - |
| - | - | 853.6 | 252.1 | - | - | 0 | - |
| - | - | 764.7 | 256.1 | - | - | 0 | - |
| - | - | 603.8 | 258.1 | - | - | 0 | - |
| - | - | 528.3 | 258.4 | - | - | 0 | - |
| 9 | y | 3432 | 262.2 | 0.0001932 | 0.7371 | +1 | 2 |
| - | - | 476.7 | 268.8 | - | - | 0 | - |
| 3 | b | 5704 | 270.1 | 0.0002403 | 0.8895 | +1 | 3 |
| - | - | 1021 | 271.1 | - | - | 0 | - |
| - | - | 804 | 276.2 | - | - | 0 | - |
| - | - | 495 | 278.2 | - | - | 0 | - |
| - | - | 710 | 281.1 | - | - | 0 | - |
| - | - | 553 | 282.1 | - | - | 0 | - |
| - | - | 523.2 | 283 | - | - | 0 | - |
| - | - | 1351 | 283 | - | - | 0 | - |
| - | - | 561.2 | 284.1 | - | - | 0 | - |
| - | - | 5730 | 299.1 | - | - | 0 | - |
| - | - | 5975 | 300.1 | - | - | 0 | - |
| - | - | 1.144E+04 | 301.1 | - | - | 0 | - |
| - | - | 6959 | 302.1 | - | - | 0 | - |
| - | - | 2437 | 303.1 | - | - | 0 | - |
| - | - | 1019 | 309.2 | - | - | 0 | - |
| - | - | 7584 | 327.7 | - | - | 0 | - |
| - | - | 2170 | 328.2 | - | - | 0 | - |
| - | - | 609.2 | 328.7 | - | - | 0 | - |
| - | - | 510.6 | 340.2 | - | - | 0 | - |
| - | - | 617.7 | 345 | - | - | 0 | - |
| - | - | 1001 | 346 | - | - | 0 | - |
| - | - | 2580 | 347 | - | - | 0 | - |
| - | - | 1727 | 348 | - | - | 0 | - |
| - | - | 962.9 | 349 | - | - | 0 | - |
| 8 | y | 633.3 | 359.2 | 0.001008 | 2.805 | +1 | 3 |
| - | - | 798.1 | 361 | - | - | 0 | - |
| - | - | 775 | 362 | - | - | 0 | - |
| 4 | y | 1596 | 370.7 | 0.0007061 | 1.905 | +2 | 7 |
| 8 | y | 1542 | 376.2 | 0.0004266 | 1.134 | +1 | 3 |
| 4 | y | 7824 | 379.7 | 0.0002371 | 0.6245 | +2 | 7 |
| - | - | 3806 | 380.2 | - | - | 0 | - |
| - | - | 690.7 | 401 | - | - | 0 | - |
| - | - | 828.5 | 405.2 | - | - | 0 | - |
| - | - | 776.5 | 405.7 | - | - | 0 | - |
| 3 | y | 6857 | 414.2 | 0.0004916 | 1.187 | +2 | 8 |
| 3 | y | 2694 | 414.7 | 0.008087 | 19.5 | +2 | 8 |
| - | - | 882 | 417 | - | - | 0 | - |
| - | - | 3088 | 418 | - | - | 0 | - |
| - | - | 1983 | 419 | - | - | 0 | - |
| - | - | 1428 | 419 | - | - | 0 | - |
| - | - | 752.2 | 420 | - | - | 0 | - |
| - | - | 729.1 | 421 | - | - | 0 | - |
| 3 | y | 4053 | 423.2 | 9.105E-05 | 0.2151 | +2 | 8 |
| - | - | 2540 | 423.7 | - | - | 0 | - |
| - | - | 556.8 | 424.2 | - | - | 0 | - |
| - | - | 633.9 | 430.7 | - | - | 0 | - |
| - | - | 1989 | 431.7 | - | - | 0 | - |
| - | - | 575 | 434.1 | - | - | 0 | - |
| - | - | 700.4 | 455.7 | - | - | 0 | - |
| - | - | 1985 | 486.3 | - | - | 0 | - |
| - | - | 910.8 | 489.1 | - | - | 0 | - |
| 7 | y | 2297 | 489.3 | 0.00014 | 0.2861 | +1 | 4 |
| - | - | 686.7 | 490.1 | - | - | 0 | - |
| - | - | 667.9 | 522.3 | - | - | 0 | - |
| 0 | Precursor | 742.1 | 523.3 | 0.0001844 | 0.3523 | +2 | -1 |
| - | - | 1116 | 523.8 | - | - | 0 | - |
| 6 | y | 6684 | 590.3 | 9.415E-05 | 0.1595 | +1 | 5 |
| - | - | 1908 | 591.3 | - | - | 0 | - |
| 5 | y | 821.7 | 643.4 | 0.003872 | 6.018 | +1 | 6 |
| 5 | y | 710.4 | 644.3 | 0.003329 | 5.167 | +1 | 6 |
| - | - | 4533 | 654.4 | - | - | 0 | - |
| - | - | 1544 | 655.4 | - | - | 0 | - |
| 5 | y | 5937 | 661.4 | 0.0001544 | 0.2335 | +1 | 6 |
| - | - | 2514 | 662.4 | - | - | 0 | - |
| 4 | y | 1.043E+04 | 758.4 | 7.769E-07 | 0.001024 | +1 | 7 |
| - | - | 3983 | 759.4 | - | - | 0 | - |
| - | - | 1035 | 760.4 | - | - | 0 | - |
| 3 | y | 4239 | 845.4 | 0.000413 | 0.4885 | +1 | 8 |
| - | - | 2183 | 846.4 | - | - | 0 | - |
| - | - | 567.5 | 879.1 | - | - | 0 | - |
| 2 | y | 634 | 915.5 | 0.006132 | 6.698 | +1 | 9 |
| 2 | y | 2941 | 932.5 | 0.0002148 | 0.2304 | +1 | 9 |
| - | - | 1366 | 933.5 | - | - | 0 | - |
| - | - | 593.2 | 986.8 | - | - | 0 | - |
| - | - | 736.3 | 1651 | - | - | 0 | - |
| - | - | 653.5 | 2076 | - | - | 0 | - |
| - | - | 752.7 | 3080 | - | - | 0 | - |

m/z Charge Intensity FragmentType MassShift Position
120.0809097290039 0 441.6314
124.3116226196289 0 415.2963
125.107666015625 0 446.04727
125.30028533935547 0 414.47043
129.06619262695312 0 456.0509
129.10231018066406 0 1861.0748
130.08619689941406 0 566.28766
136.07579040527344 0 1211.4948
139.0868377685547 0 914.8121
141.10231018066406 0 2965.1948
144.56788635253906 0 440.48032
147.1127471923828 0 795.3021
148.86927795410156 0 505.7981
148.88311767578125 0 459.97342
148.8979949951172 0 816.9075
148.90487670898438 0 776.73676
148.9120635986328 0 956.11035
148.9193878173828 0 1054.0048
148.92657470703125 0 1082.0264
148.93374633789062 0 1842.7616
148.9414825439453 0 3594.5881
148.95809936523438 0 5095.226
148.96591186523438 0 2979.9543
148.9733123779297 0 1241.6151
148.98045349121094 0 1277.564
148.98760986328125 0 967.3142
148.99497985839844 0 727.2094
149.00198364257812 0 703.755
149.00904846191406 0 541.84595
149.03880310058594 0 480.88748
149.04510498046875 0 2212.0117
149.05201721191406 0 647.713
150.04464721679688 0 703.24603
151.04188537597656 0 1080.2821
155.11758422851562 0 730.7952 a Water loss 1
157.06080627441406 0 567.98785
157.09715270996094 0 1751.8481
157.13372802734375 0 3227.6702 d 1
158.092529296875 0 831.7975 y Ammonia loss 9
158.13674926757812 0 834.6179
161.09246826171875 0 567.93286
167.0556640625 0 3901.1455
167.08140563964844 0 486.22595
168.0550537109375 0 554.6624
169.05250549316406 0 2652.3926
169.09718322753906 0 2998.603
173.12843322753906 0 2805.9812 a 1
173.44175720214844 0 503.25897
175.1190948486328 0 858.4251 y 9
183.11279296875 0 4591.7764 b Water loss 1
183.1494140625 0 1049.599
185.09237670898438 0 1139.1154
185.12890625 0 1378.7596
191.17706298828125 0 479.6105
201.12359619140625 0 2572.4392 b 1
211.14337158203125 0 479.8168
223.063720703125 0 1822.521
224.06503295898438 0 738.4257
225.04261779785156 0 1049.3384
225.06068420410156 0 602.789
225.12355041503906 0 636.1061
227.0396728515625 0 1068.1008
238.1186065673828 0 4187.067
239.0947265625 0 593.30133
241.09274291992188 0 643.5736
245.12452697753906 0 677.1904 y Ammonia loss 8
250.12351989746094 0 563.9748
252.13433837890625 0 853.55975
256.12969970703125 0 764.6594
258.14508056640625 0 603.84875
258.3761901855469 0 528.33
262.1507873535156 0 3432.0425 y 8
268.82562255859375 0 476.69122
270.14459228515625 0 5703.7754 b Water loss 2
271.1478271484375 0 1021.0441
276.15557861328125 0 803.9723
278.2366638183594 0 495.00555
281.05072021484375 0 709.9534
282.0521545410156 0 552.9538
283.0318908691406 0 523.2426
283.04864501953125 0 1351.4185
284.0509338378906 0 561.2419
299.0617980957031 0 5729.8457
300.0624084472656 0 5975.1504
301.05950927734375 0 11440.599
302.0601501464844 0 6958.523
303.05743408203125 0 2437.276
309.20404052734375 0 1018.659
327.6948547363281 0 7584.157
328.1964111328125 0 2170.215
328.69757080078125 0 609.2005
340.1869201660156 0 510.6004
344.97723388671875 0 617.6557
345.9755554199219 0 1000.7853
346.97406005859375 0 2580.3499
347.9740295410156 0 1726.8237
348.9709777832031 0 962.85266
359.1663513183594 0 633.2959 y Ammonia loss 7
361.0260009765625 0 798.13794
362.0251770019531 0 774.96686
370.7054138183594 0 1595.9364 y Water loss 3
376.1934814453125 0 1541.7772 y 7
379.7116394042969 0 7824.0483 y 3
380.2123718261719 0 3805.7114
400.98638916015625 0 690.73553
405.2158508300781 0 828.5171
405.7093811035156 0 776.49677
414.2226257324219 0 6856.5527 y Water loss 2
414.72222900390625 0 2693.6692 y Ammonia loss 2
417.0348815917969 0 882.0464
418.03521728515625 0 3088.0366
418.9950866699219 0 1983.2366
419.0315246582031 0 1427.8308
419.9962463378906 0 752.1738
420.99383544921875 0 729.05634
423.2273254394531 0 4053.1707 y 2
423.7270812988281 0 2539.6897
424.22857666015625 0 556.7812
430.7435302734375 0 633.89764
431.7348327636719 0 1989.221
434.1414794921875 0 574.9551
455.7052307128906 0 700.3593
486.2924499511719 0 1984.5491
489.0545654296875 0 910.7575
489.27783203125 0 2297.047 y 6
490.0557861328125 0 686.6873
522.2698364257812 0 667.9125
523.2852783203125 0 742.0527 Precursor
523.7886962890625 0 1115.5173
590.3257446289062 0 6684.4976 y 5
591.322265625 0 1908.1871
643.3483276367188 0 821.7169 y Water loss 4
644.3328857421875 0 710.4244 y Ammonia loss 4
654.3820190429688 0 4533.332
655.384765625 0 1543.9194
661.3626098632812 0 5936.701 y 4
662.3639526367188 0 2513.506
758.41552734375 0 10433.683 y 3
759.4135131835938 0 3982.9128
760.4161987304688 0 1034.7479
845.4471435546875 0 4239.3735 y 2
846.4444580078125 0 2183.1785
879.1298217773438 0 567.54974
915.4591674804688 0 633.9781 y Ammonia loss 1
932.4793701171875 0 2941.1238 y 1
933.4801635742188 0 1366.481
986.7864379882812 0 593.21155
1651.10205078125 0 736.2756
2075.723388671875 0 653.5347
3080.40771484375 0 752.6593

Spectrum Details

|  |  |
| --- | --- |
| Matched peaks? Matched peaks The total absolute number of peaks matched. Additionally in brackets the total fraction of peaks matched and the total number of peaks is shown. | 27 (18.00% of 150) |
| FDR? FDR The false discovery rate estimated for this peptide. It is calculated by matching all theoretical fragments with a non-integer shift with the raw peaks for this spectrum. This is done with 40 different shifts. The resulting percentage is the average number of annotated peaks over the number of annotated peaks with the correct spectrum. | 1.23% |
| Satellite FDR? Satellite FDR See the FDR for details on its calculation. This satellite ion specific FDR only contains the satellite ions (d/w) for I/L/J positions. | - |
| PSM Score? PSM Score The PSM Score as given by Hecklib to this annotated spectrum. It is shown with three significant figures. | 262 |

#### Spectrum 4573? Spectrum 4573 The raw spectrum of this peptide as annotated by Hecklib. The fragments are coloured according to ion type (see legend). Any peaks with a star '\*' as text can be hovered over to see the full details, first the ion type second the mass shift type. By hovering over the amino acids in the peptide or ions in the legend the corresponding peaks are highlighted. By toggling the 'Unassigned' label you can turn the background (unassigned) peaks on or off in the plot. By updating the slider in the Ion legend you can update the spectrum to only show the top X% of the peaks with labels. The top X% means any peak that is within X% of the highest intensity. By dragging in the spectrum you can zoom in to a specific part of the spectrum and use 'Zoom Out' to get back to the original zoom level. The annotation of the spectrum is based on the given sequence in the peptides file and is done with different software so inconsistencies are likely. The peaks are annotated based on the given sequence, with 20 ppm tolerance.

Copy Data

##### Spectrum 4573 (TSV)

###### Preview

```
Loading example...
```

*Click on the button to copy the data to your clipboard.*

Mz MinMz MaxIntensity Max

WidthHeightPeptide font sizePeptide stroke widthSpectrum font sizeSpectrum stroke widthCompact peptide

Ion legend

wxyz

abcd

OtherUnassignedIon^Charge^_Position_Show for top:%

JSSPATJNSR

03.17e+36.35e+39.52e+31.27e+4

Zoom Out

a^+1^_2_d^+1^_2_y^+1^_1_a^+1^_2_y^+1^_1_b^+1^_2_b^+1^_2_y^+1^_2_y^+1^_2_b^+1^_3_y^+1^_3_y^+2^_7_y^+1^_3_y^+2^_7_y^+2^_8_y^+2^_8_y^+2^_8_y^+1^_4_\*y^+1^_5_y^+1^_6_y^+1^_6_y^+1^_7_y^+1^_8_y^+1^_9_

0820164024603280

Fragment Matches Table

Show background peaks

| Position | Ion type | Intensity | mz Theoretical | mz Error (Th) | mz Error (ppm) | Charge | Series Number |
| --- | --- | --- | --- | --- | --- | --- | --- |
| - | - | 860.3 | 120.1 | - | - | 0 | - |
| - | - | 1089 | 129.1 | - | - | 0 | - |
| - | - | 407.4 | 133.3 | - | - | 0 | - |
| - | - | 1434 | 136.1 | - | - | 0 | - |
| - | - | 366.7 | 138.9 | - | - | 0 | - |
| - | - | 529.9 | 139.1 | - | - | 0 | - |
| - | - | 1663 | 141.1 | - | - | 0 | - |
| - | - | 445 | 141.9 | - | - | 0 | - |
| - | - | 434.6 | 147.1 | - | - | 0 | - |
| - | - | 482.9 | 147.4 | - | - | 0 | - |
| - | - | 2284 | 149 | - | - | 0 | - |
| - | - | 553 | 150 | - | - | 0 | - |
| - | - | 1690 | 151 | - | - | 0 | - |
| - | - | 506.4 | 151.1 | - | - | 0 | - |
| - | - | 438.1 | 152.1 | - | - | 0 | - |
| 2 | a | 582.8 | 155.1 | 0.0001525 | 0.9828 | +1 | 2 |
| - | - | 1010 | 157.1 | - | - | 0 | - |
| - | - | 1575 | 157.1 | - | - | 0 | - |
| 2 | d | 624 | 157.1 | 0.0001778 | 1.131 | +1 | 2 |
| 10 | y | 799.9 | 158.1 | 0.0005382 | 3.404 | +1 | 1 |
| - | - | 467.5 | 159.1 | - | - | 0 | - |
| - | - | 472.1 | 163.9 | - | - | 0 | - |
| - | - | 4549 | 167.1 | - | - | 0 | - |
| - | - | 680.7 | 167.1 | - | - | 0 | - |
| - | - | 1415 | 168.1 | - | - | 0 | - |
| - | - | 1900 | 169.1 | - | - | 0 | - |
| - | - | 1615 | 169.1 | - | - | 0 | - |
| 2 | a | 2605 | 173.1 | 3.625E-05 | 0.2094 | +1 | 2 |
| - | - | 3170 | 173.4 | - | - | 0 | - |
| 10 | y | 1605 | 175.1 | 0.0001015 | 0.5794 | +1 | 1 |
| 2 | b | 3405 | 183.1 | 1.933E-05 | 0.1056 | +1 | 2 |
| - | - | 1524 | 185.1 | - | - | 0 | - |
| 2 | b | 2298 | 201.1 | 4.424E-05 | 0.22 | +1 | 2 |
| - | - | 499 | 202 | - | - | 0 | - |
| - | - | 462.7 | 222.1 | - | - | 0 | - |
| - | - | 2766 | 223.1 | - | - | 0 | - |
| - | - | 1158 | 224.1 | - | - | 0 | - |
| - | - | 1515 | 225 | - | - | 0 | - |
| - | - | 949.4 | 225.1 | - | - | 0 | - |
| - | - | 741.6 | 225.1 | - | - | 0 | - |
| - | - | 775.5 | 226 | - | - | 0 | - |
| - | - | 966.8 | 227 | - | - | 0 | - |
| - | - | 539.8 | 228.1 | - | - | 0 | - |
| - | - | 2635 | 238.1 | - | - | 0 | - |
| - | - | 545.3 | 241.1 | - | - | 0 | - |
| 9 | y | 1017 | 245.1 | 0.00082 | 3.345 | +1 | 2 |
| - | - | 684.4 | 252.1 | - | - | 0 | - |
| - | - | 850.4 | 256.1 | - | - | 0 | - |
| 9 | y | 2176 | 262.2 | 0.0002848 | 1.086 | +1 | 2 |
| - | - | 658.7 | 264.1 | - | - | 0 | - |
| 3 | b | 4461 | 270.1 | 0.0002403 | 0.8895 | +1 | 3 |
| - | - | 608.1 | 281.1 | - | - | 0 | - |
| - | - | 880.6 | 282.1 | - | - | 0 | - |
| - | - | 702.5 | 283 | - | - | 0 | - |
| - | - | 1156 | 283 | - | - | 0 | - |
| - | - | 6436 | 299.1 | - | - | 0 | - |
| - | - | 7782 | 300.1 | - | - | 0 | - |
| - | - | 1.257E+04 | 301.1 | - | - | 0 | - |
| - | - | 7778 | 302.1 | - | - | 0 | - |
| - | - | 1965 | 303.1 | - | - | 0 | - |
| - | - | 654.4 | 309.2 | - | - | 0 | - |
| - | - | 567.3 | 327.5 | - | - | 0 | - |
| - | - | 1072 | 327.7 | - | - | 0 | - |
| - | - | 1604 | 346 | - | - | 0 | - |
| - | - | 2338 | 347 | - | - | 0 | - |
| - | - | 1896 | 348 | - | - | 0 | - |
| - | - | 863.2 | 349 | - | - | 0 | - |
| 8 | y | 561.8 | 359.2 | 0.001313 | 3.655 | +1 | 3 |
| - | - | 1004 | 360 | - | - | 0 | - |
| - | - | 1392 | 361 | - | - | 0 | - |
| - | - | 1051 | 362 | - | - | 0 | - |
| 4 | y | 1334 | 370.7 | 0.001378 | 3.716 | +2 | 7 |
| - | - | 661.6 | 371.2 | - | - | 0 | - |
| - | - | 538.3 | 371.4 | - | - | 0 | - |
| 8 | y | 766.4 | 376.2 | 0.001647 | 4.379 | +1 | 3 |
| 4 | y | 7195 | 379.7 | 7.023E-06 | 0.0185 | +2 | 7 |
| - | - | 2809 | 380.2 | - | - | 0 | - |
| - | - | 723.3 | 380.7 | - | - | 0 | - |
| - | - | 574.7 | 409.5 | - | - | 0 | - |
| 3 | y | 5712 | 414.2 | 0.0005526 | 1.334 | +2 | 8 |
| 3 | y | 1536 | 414.7 | 0.006958 | 16.78 | +2 | 8 |
| - | - | 733.4 | 416 | - | - | 0 | - |
| - | - | 867.2 | 417 | - | - | 0 | - |
| - | - | 3481 | 418 | - | - | 0 | - |
| - | - | 2354 | 419 | - | - | 0 | - |
| - | - | 1988 | 419 | - | - | 0 | - |
| - | - | 859.8 | 420 | - | - | 0 | - |
| 3 | y | 4569 | 423.2 | 0.0002436 | 0.5757 | +2 | 8 |
| - | - | 1903 | 423.7 | - | - | 0 | - |
| - | - | 1147 | 458.3 | - | - | 0 | - |
| - | - | 572 | 463.8 | - | - | 0 | - |
| - | - | 885.9 | 489.1 | - | - | 0 | - |
| 7 | y | 2549 | 489.3 | 0.0005672 | 1.159 | +1 | 4 |
| - | - | 906.2 | 490.1 | - | - | 0 | - |
| 0 | Precursor | 1111 | 523.3 | 0.007448 | 14.23 | +2 | -1 |
| - | - | 746 | 523.8 | - | - | 0 | - |
| - | - | 652.9 | 524.2 | - | - | 0 | - |
| 6 | y | 5406 | 590.3 | 2.792E-05 | 0.0473 | +1 | 5 |
| - | - | 2205 | 591.3 | - | - | 0 | - |
| - | - | 674.8 | 598.8 | - | - | 0 | - |
| 5 | y | 844.2 | 643.4 | 0.0008202 | 1.275 | +1 | 6 |
| 5 | y | 5695 | 661.4 | 0.0005817 | 0.8795 | +1 | 6 |
| - | - | 2426 | 662.4 | - | - | 0 | - |
| 4 | y | 8501 | 758.4 | 0.0001839 | 0.2425 | +1 | 7 |
| - | - | 4018 | 759.4 | - | - | 0 | - |
| 3 | y | 3174 | 845.4 | 0.0003805 | 0.45 | +1 | 8 |
| - | - | 2289 | 846.4 | - | - | 0 | - |
| - | - | 723.3 | 847.4 | - | - | 0 | - |
| - | - | 598.2 | 908 | - | - | 0 | - |
| 2 | y | 2471 | 932.5 | 0.0008252 | 0.8849 | +1 | 9 |
| - | - | 2036 | 933.5 | - | - | 0 | - |
| - | - | 581.1 | 1001 | - | - | 0 | - |
| - | - | 599.4 | 1048 | - | - | 0 | - |
| - | - | 625.7 | 1259 | - | - | 0 | - |
| - | - | 635.6 | 1568 | - | - | 0 | - |
| - | - | 734.2 | 2311 | - | - | 0 | - |
| - | - | 684.4 | 2948 | - | - | 0 | - |
| - | - | 906 | 3247 | - | - | 0 | - |

m/z Charge Intensity FragmentType MassShift Position
120.08101654052734 0 860.3399
129.10235595703125 0 1088.6665
133.32965087890625 0 407.35034
136.07579040527344 0 1434.0522
138.8619842529297 0 366.67175
139.0869598388672 0 529.86536
141.1023712158203 0 1662.831
141.92138671875 0 444.99698
147.1135711669922 0 434.62878
147.44020080566406 0 482.86716
149.0448760986328 0 2284.4863
150.04417419433594 0 552.99915
151.04176330566406 0 1689.9456
151.1481475830078 0 506.39337
152.05679321289062 0 438.0731
155.1180419921875 0 582.8396 a Water loss 1
157.06112670898438 0 1009.7205
157.09715270996094 0 1575.1754
157.13336181640625 0 624.03094 d 1
158.0929412841797 0 799.9189 y Ammonia loss 9
159.0847930908203 0 467.4875
163.9087677001953 0 472.1455
167.05548095703125 0 4549.463
167.0819854736328 0 680.68713
168.054931640625 0 1415.4452
169.0523223876953 0 1900.3147
169.09722900390625 0 1615.2665
173.12841796875 0 2605.0894 a 1
173.4398193359375 0 3170.2537
175.1188507080078 0 1605.3788 y 9
183.11282348632812 0 3404.6973 b Water loss 1
185.09205627441406 0 1523.925
201.1234130859375 0 2298.0195 b 1
202.0496063232422 0 499.04065
222.08563232421875 0 462.67905
223.06382751464844 0 2765.9844
224.0637969970703 0 1157.5442
225.04270935058594 0 1514.9397
225.06045532226562 0 949.4071
225.12281799316406 0 741.6307
226.0430908203125 0 775.484
227.0402374267578 0 966.77686
228.1346435546875 0 539.84247
238.11871337890625 0 2634.752
241.09239196777344 0 545.334
245.1236114501953 0 1016.74225 y Ammonia loss 8
252.13621520996094 0 684.39056
256.12884521484375 0 850.38837
262.15069580078125 0 2175.6736 y 8
264.0965576171875 0 658.6833
270.14459228515625 0 4461.051 b Water loss 2
281.050537109375 0 608.0792
282.05096435546875 0 880.6294
283.0312194824219 0 702.52277
283.0478515625 0 1155.9598
299.061767578125 0 6435.8135
300.0621032714844 0 7782.2495
301.059326171875 0 12570.912
302.0596008300781 0 7777.77
303.0570373535156 0 1964.9082
309.2055969238281 0 654.38947
327.4520263671875 0 567.2691
327.6950988769531 0 1072.3846
345.976806640625 0 1603.6725
346.9742126464844 0 2337.794
347.97406005859375 0 1895.7798
348.9723205566406 0 863.22614
359.1660461425781 0 561.8219 y Ammonia loss 7
360.02984619140625 0 1004.4304
361.0265197753906 0 1392.498
362.02593994140625 0 1050.9878
370.7047424316406 0 1333.8204 y Water loss 3
371.2059631347656 0 661.6143
371.3814697265625 0 538.29034
376.1922607421875 0 766.4157 y 7
379.7113952636719 0 7195.213 y 3
380.2106628417969 0 2809.4927
380.7126159667969 0 723.25507
409.4825439453125 0 574.6575
414.2226867675781 0 5712.1567 y Water loss 2
414.7210998535156 0 1536.3525 y Ammonia loss 2
416.0378112792969 0 733.38104
417.03692626953125 0 867.21155
418.035400390625 0 3481.3665
418.9959716796875 0 2353.847
419.03253173828125 0 1987.7142
419.99853515625 0 859.80194
423.2271728515625 0 4569.174 y 2
423.7281188964844 0 1903.0114
458.2717590332031 0 1147.2664
463.83721923828125 0 572.03357
489.05303955078125 0 885.9132
489.27740478515625 0 2548.883 y 6
490.0563659667969 0 906.1699
523.2780151367188 0 1110.8469 Precursor
523.7880859375 0 746.0183
524.2127685546875 0 652.9372
590.3256225585938 0 5405.6094 y 5
591.32470703125 0 2204.5437
598.8052368164062 0 674.7916
643.3513793945312 0 844.1783 y Water loss 4
661.3621826171875 0 5694.9556 y 4
662.3637084960938 0 2425.7695
758.4153442382812 0 8500.739 y 3
759.4137573242188 0 4018.4968
845.4479370117188 0 3173.9321 y 2
846.4469604492188 0 2288.8884
847.4468994140625 0 723.26874
907.9908447265625 0 598.2381
932.478759765625 0 2471.3484 y 1
933.4793701171875 0 2036.0989
1000.9503784179688 0 581.1318
1047.69775390625 0 599.38
1258.9556884765625 0 625.7455
1567.8355712890625 0 635.64453
2311.32666015625 0 734.1815
2948.484130859375 0 684.4172
3247.46484375 0 905.9985

Spectrum Details

|  |  |
| --- | --- |
| Matched peaks? Matched peaks The total absolute number of peaks matched. Additionally in brackets the total fraction of peaks matched and the total number of peaks is shown. | 25 (21.19% of 118) |
| FDR? FDR The false discovery rate estimated for this peptide. It is calculated by matching all theoretical fragments with a non-integer shift with the raw peaks for this spectrum. This is done with 40 different shifts. The resulting percentage is the average number of annotated peaks over the number of annotated peaks with the correct spectrum. | 1.33% |
| Satellite FDR? Satellite FDR See the FDR for details on its calculation. This satellite ion specific FDR only contains the satellite ions (d/w) for I/L/J positions. | - |
| PSM Score? PSM Score The PSM Score as given by Hecklib to this annotated spectrum. It is shown with three significant figures. | 227 |

#### Reverse Lookup? Reverse Lookup All places where this read could be placed.

| Group | Segment | Template | Template Part | Read Part | Score | Unique |
| --- | --- | --- | --- | --- | --- | --- |
| Decoy | Decoy | TRYP | [97..107] | [0..10] | 80 | True |

| Recombined | Template Part | Read Part | Score | Unique |
| --- | --- | --- | --- | --- |
| TRYP | [97..107] | [0..10] | 80 | True |

#### Meta Information from Multiple reads

##### Number of combined reads

12

##### Intensity

0.6538

##### TotalArea

1.091E+08

##### Changes to the peptide sequence

JSSPATJNSR

L→JNo support for either Leucine or Isoleucine based on side chain ions (Position: 7)

L→JNo support for either Leucine or Isoleucine based on side chain ions (Position: 1)

#### Positional Score

Copy Data

##### Positional Score (TSV)

###### Preview

```
Loading example...
```

*Click on the button to copy the data to your clipboard.*

100123456789

Label Value
"0" 0.803
"1" 0.822
"2" 0.829
"3" 0.827
"4" 0.822
"5" 0.814
"6" 0.796
"7" 0.778
"8" 0.802
"9" 0.82

#### Meta Information from PEAKS

##### Scan Identifier

F4:3548

##### Original sequence

L

S

S

P

A

T

L

N

S

R

##### Posttranslational Modifications

##### Source File

D:\separate\_stitch\_analyses\xle-disambiguation\raw\20210323\_F1\_UM1\_Peng0013\_SA\_F59\_ingel\_3ug\_tryp.raw

##### Fraction

4

##### Scan Feature

F4:3752

##### De Novo Score

99

##### ConfidenceScore

99

### m/z

523.2861

##### Mass

1044.5564

##### Charge

2

##### Retention Time

18.9

##### Predicted Retention Time

19.27

##### Area

9.802E+07

##### Parts Per Million

1.3

##### Fragmentation mode

HCD

##### Originating file

01 D:\separate\_stitch\_analyses\xle-disambiguation\20210325\_F59\_3ug\_DENOVO\_12.csv

#### Meta Information from PEAKS

##### Scan Identifier

F4:3712

##### Original sequence

L

S

S

P

A

T

L

N

S

R

##### Posttranslational Modifications

##### Source File

D:\separate\_stitch\_analyses\xle-disambiguation\raw\20210323\_F1\_UM1\_Peng0013\_SA\_F59\_ingel\_3ug\_tryp.raw

##### Fraction

4

##### Scan Feature

-

##### De Novo Score

99

##### ConfidenceScore

99

### m/z

523.2863

##### Mass

1044.5564

##### Charge

2

##### Retention Time

19.85

##### Predicted Retention Time

19.27

##### Area

0

##### Parts Per Million

1.6

##### Fragmentation mode

HCD

##### Originating file

01 D:\separate\_stitch\_analyses\xle-disambiguation\20210325\_F59\_3ug\_DENOVO\_12.csv

#### Meta Information from PEAKS

##### Scan Identifier

F4:3883

##### Original sequence

L

S

S

P

A

T

L

N

S

R

##### Posttranslational Modifications

##### Source File

D:\separate\_stitch\_analyses\xle-disambiguation\raw\20210323\_F1\_UM1\_Peng0013\_SA\_F59\_ingel\_3ug\_tryp.raw

##### Fraction

4

##### Scan Feature

F4:3750

##### De Novo Score

99

##### ConfidenceScore

98

### m/z

523.2853

##### Mass

1044.5564

##### Charge

2

##### Retention Time

20.88

##### Predicted Retention Time

19.27

##### Area

2.184E+06

##### Fragmentation mode

HCD

##### Originating file

01 D:\separate\_stitch\_analyses\xle-disambiguation\20210325\_F59\_3ug\_DENOVO\_12.csv

#### Meta Information from PEAKS

##### Scan Identifier

F4:3404

##### Original sequence

L

S

S

P

A

T

L

N

S

R

##### Posttranslational Modifications

##### Source File

D:\separate\_stitch\_analyses\xle-disambiguation\raw\20210323\_F1\_UM1\_Peng0013\_SA\_F59\_ingel\_3ug\_tryp.raw

##### Fraction

4

##### Scan Feature

F4:3753

##### De Novo Score

98

##### ConfidenceScore

97

### m/z

523.2883

##### Mass

1044.5564

##### Charge

2

##### Retention Time

18.06

##### Predicted Retention Time

19.27

##### Area

1.502E+05

##### Parts Per Million

5.4

##### Fragmentation mode

HCD

##### Originating file

01 D:\separate\_stitch\_analyses\xle-disambiguation\20210325\_F59\_3ug\_DENOVO\_12.csv

#### Meta Information from PEAKS

##### Scan Identifier

F4:4042

##### Original sequence

L

S

S

P

A

T

L

N

S

R

##### Posttranslational Modifications

##### Source File

D:\separate\_stitch\_analyses\xle-disambiguation\raw\20210323\_F1\_UM1\_Peng0013\_SA\_F59\_ingel\_3ug\_tryp.raw

##### Fraction

4

##### Scan Feature

F4:3750

##### De Novo Score

98

##### ConfidenceScore

97

### m/z

523.2853

##### Mass

1044.5564

##### Charge

2

##### Retention Time

20.88

##### Predicted Retention Time

19.27

##### Area

2.184E+06

##### Fragmentation mode

HCD

##### Originating file

01 D:\separate\_stitch\_analyses\xle-disambiguation\20210325\_F59\_3ug\_DENOVO\_12.csv

#### Meta Information from PEAKS

##### Scan Identifier

F4:4222

##### Original sequence

L

S

S

P

A

T

L

N

S

R

##### Posttranslational Modifications

##### Source File

D:\separate\_stitch\_analyses\xle-disambiguation\raw\20210323\_F1\_UM1\_Peng0013\_SA\_F59\_ingel\_3ug\_tryp.raw

##### Fraction

4

##### Scan Feature

F4:3750

##### De Novo Score

97

##### ConfidenceScore

98

### m/z

523.2853

##### Mass

1044.5564

##### Charge

2

##### Retention Time

20.88

##### Predicted Retention Time

19.27

##### Area

2.184E+06

##### Fragmentation mode

HCD

##### Originating file

01 D:\separate\_stitch\_analyses\xle-disambiguation\20210325\_F59\_3ug\_DENOVO\_12.csv

#### Meta Information from PEAKS

##### Scan Identifier

F4:4272

##### Original sequence

L

S

S

P

A

T

L

N

S

R

##### Posttranslational Modifications

##### Source File

D:\separate\_stitch\_analyses\xle-disambiguation\raw\20210323\_F1\_UM1\_Peng0013\_SA\_F59\_ingel\_3ug\_tryp.raw

##### Fraction

4

##### Scan Feature

F4:3750

##### De Novo Score

97

##### ConfidenceScore

98

### m/z

523.2853

##### Mass

1044.5564

##### Charge

2

##### Retention Time

20.88

##### Predicted Retention Time

19.27

##### Area

2.184E+06

##### Fragmentation mode

HCD

##### Originating file

01 D:\separate\_stitch\_analyses\xle-disambiguation\20210325\_F59\_3ug\_DENOVO\_12.csv

#### Meta Information from PEAKS

##### Scan Identifier

F4:3097

##### Original sequence

L

S

S

P

A

T

L

N

S

R

##### Posttranslational Modifications

##### Source File

D:\separate\_stitch\_analyses\xle-disambiguation\raw\20210323\_F1\_UM1\_Peng0013\_SA\_F59\_ingel\_3ug\_tryp.raw

##### Fraction

4

##### Scan Feature

F4:3751

##### De Novo Score

97

##### ConfidenceScore

97

### m/z

523.2859

##### Mass

1044.5564

##### Charge

2

##### Retention Time

16.45

##### Predicted Retention Time

19.27

##### Area

5.371E+04

##### Parts Per Million

0.9

##### Fragmentation mode

HCD

##### Originating file

01 D:\separate\_stitch\_analyses\xle-disambiguation\20210325\_F59\_3ug\_DENOVO\_12.csv

#### Meta Information from PEAKS

##### Scan Identifier

F4:4326

##### Original sequence

L

S

S

P

A

T

L

N

S

R

##### Posttranslational Modifications

##### Source File

D:\separate\_stitch\_analyses\xle-disambiguation\raw\20210323\_F1\_UM1\_Peng0013\_SA\_F59\_ingel\_3ug\_tryp.raw

##### Fraction

4

##### Scan Feature

F4:3750

##### De Novo Score

96

##### ConfidenceScore

96

### m/z

523.2853

##### Mass

1044.5564

##### Charge

2

##### Retention Time

20.88

##### Predicted Retention Time

19.27

##### Area

2.184E+06

##### Fragmentation mode

HCD

##### Originating file

01 D:\separate\_stitch\_analyses\xle-disambiguation\20210325\_F59\_3ug\_DENOVO\_12.csv

#### Meta Information from PEAKS

##### Scan Identifier

F4:4517

##### Original sequence

L

S

S

P

A

T

L

N

S

R

##### Posttranslational Modifications

##### Source File

D:\separate\_stitch\_analyses\xle-disambiguation\raw\20210323\_F1\_UM1\_Peng0013\_SA\_F59\_ingel\_3ug\_tryp.raw

##### Fraction

4

##### Scan Feature

-

##### De Novo Score

95

##### ConfidenceScore

97

### m/z

523.2858

##### Mass

1044.5564

##### Charge

2

##### Retention Time

24.19

##### Predicted Retention Time

19.27

##### Area

0

##### Parts Per Million

0.7

##### Fragmentation mode

HCD

##### Originating file

01 D:\separate\_stitch\_analyses\xle-disambiguation\20210325\_F59\_3ug\_DENOVO\_12.csv

#### Meta Information from PEAKS

##### Scan Identifier

F4:4461

##### Original sequence

L

S

S

P

A

T

L

N

S

R

##### Posttranslational Modifications

##### Source File

D:\separate\_stitch\_analyses\xle-disambiguation\raw\20210323\_F1\_UM1\_Peng0013\_SA\_F59\_ingel\_3ug\_tryp.raw

##### Fraction

4

##### Scan Feature

-

##### De Novo Score

94

##### ConfidenceScore

95

### m/z

523.2855

##### Mass

1044.5564

##### Charge

2

##### Retention Time

23.9

##### Predicted Retention Time

19.27

##### Area

0

##### Parts Per Million

0

##### Fragmentation mode

HCD

##### Originating file

01 D:\separate\_stitch\_analyses\xle-disambiguation\20210325\_F59\_3ug\_DENOVO\_12.csv

#### Meta Information from PEAKS

##### Scan Identifier

F4:4573

##### Original sequence

L

S

S

P

A

T

L

N

S

R

##### Posttranslational Modifications

##### Source File

D:\separate\_stitch\_analyses\xle-disambiguation\raw\20210323\_F1\_UM1\_Peng0013\_SA\_F59\_ingel\_3ug\_tryp.raw

##### Fraction

4

##### Scan Feature

-

##### De Novo Score

94

##### ConfidenceScore

96

### m/z

523.2866

##### Mass

1044.5564

##### Charge

2

##### Retention Time

24.49

##### Predicted Retention Time

19.27

##### Area

0

##### Parts Per Million

2.2

##### Fragmentation mode

HCD

##### Originating file

01 D:\separate\_stitch\_analyses\xle-disambiguation\20210325\_F59\_3ug\_DENOVO\_12.csv
