## Supplementary material for "A handle on mass coincidence errors in *de novo* sequencing of antibodies by bottom-up proteomics": Combined_002.html

Details Combined\_002 | Stitch OverviewUndefined

### Read Combined\_002

#### Sequence (length=11)

JFPPSSEEJQA

#### Spectrum 7848? Spectrum 7848 The raw spectrum of this peptide as annotated by Hecklib. The fragments are coloured according to ion type (see legend). Any peaks with a star '\*' as text can be hovered over to see the full details, first the ion type second the mass shift type. By hovering over the amino acids in the peptide or ions in the legend the corresponding peaks are highlighted. By toggling the 'Unassigned' label you can turn the background (unassigned) peaks on or off in the plot. By updating the slider in the Ion legend you can update the spectrum to only show the top X% of the peaks with labels. The top X% means any peak that is within X% of the highest intensity. By dragging in the spectrum you can zoom in to a specific part of the spectrum and use 'Zoom Out' to get back to the original zoom level. The annotation of the spectrum is based on the given sequence in the peptides file and is done with different software so inconsistencies are likely. The peaks are annotated based on the given sequence, with 20 ppm tolerance.

Copy Data

##### Spectrum 7848 (TSV)

###### Preview

```
Loading example...
```

*Click on the button to copy the data to your clipboard.*

Mz MinMz MaxIntensity Max

WidthHeightPeptide font sizePeptide stroke widthSpectrum font sizeSpectrum stroke widthCompact peptide

Ion legend

wxyz

abcd

OtherUnassignedIon^Charge^_Position_Show for top:%

JFPPSSEEJQA

06.00e+51.20e+61.80e+62.40e+6

Zoom Out

y^+1^_2_y^+1^_2_y^+1^_3_z^+1^_3_y^+1^_3_c^+2^_7_y^+1^_4_y^+1^_4_y^+1^_4_z^+2^_9_y^+2^_9_y^+2^_9_z^+2^_9_c^+1^_4_y^+2^_9_c^+2^_9_c^+1^_5_c^+2^_10_y^+1^_5_w^+1^_6_c^+1^_6_y^+1^_6_z^+1^_6_y^+1^_6_y^+1^_7_y^+1^_7_z^+1^_7_c^+1^_7_y^+1^_7_c^+1^_7_y^+1^_8_y^+1^_8_c^+1^_8_c^+1^_8_y^+1^_9_y^+1^_9_z^+1^_9_y^+1^_9_c^+1^_9_z^+1^_10_z^+1^_10_y^+1^_10_z^+1^_10_c^+1^_10_c^+1^_10_

0652130419562608

Fragment Matches Table

Show background peaks

| Position | Ion type | Intensity | mz Theoretical | mz Error (Th) | mz Error (ppm) | Charge | Series Number |
| --- | --- | --- | --- | --- | --- | --- | --- |
| - | - | 3159 | 120.1 | - | - | 0 | - |
| - | - | 3.204E+04 | 120.1 | - | - | 0 | - |
| - | - | 1614 | 120.7 | - | - | 0 | - |
| - | - | 3668 | 121.1 | - | - | 0 | - |
| - | - | 5.092E+04 | 129.1 | - | - | 0 | - |
| - | - | 2335 | 130.1 | - | - | 0 | - |
| - | - | 1.211E+04 | 131.1 | - | - | 0 | - |
| - | - | 1.03E+04 | 147.1 | - | - | 0 | - |
| - | - | 2214 | 148.9 | - | - | 0 | - |
| - | - | 2082 | 151.1 | - | - | 0 | - |
| - | - | 2433 | 155.1 | - | - | 0 | - |
| - | - | 1.001E+04 | 173.5 | - | - | 0 | - |
| - | - | 2519 | 173.9 | - | - | 0 | - |
| - | - | 2114 | 183.1 | - | - | 0 | - |
| - | - | 2035 | 186.4 | - | - | 0 | - |
| - | - | 3699 | 197.1 | - | - | 0 | - |
| - | - | 1.147E+05 | 200.1 | - | - | 0 | - |
| 10 | y | 2.099E+05 | 201.1 | 0.0003885 | 1.932 | +1 | 2 |
| - | - | 1.102E+04 | 201.1 | - | - | 0 | - |
| - | - | 1.611E+04 | 202.1 | - | - | 0 | - |
| - | - | 2902 | 212.1 | - | - | 0 | - |
| - | - | 2307 | 215.1 | - | - | 0 | - |
| 10 | y | 2.605E+05 | 218.1 | 0.0003897 | 1.787 | +1 | 2 |
| - | - | 2.149E+04 | 219.1 | - | - | 0 | - |
| - | - | 4599 | 221.1 | - | - | 0 | - |
| - | - | 1.305E+04 | 225.1 | - | - | 0 | - |
| - | - | 2.899E+05 | 233.2 | - | - | 0 | - |
| - | - | 4.177E+04 | 234.2 | - | - | 0 | - |
| - | - | 1.268E+05 | 242.2 | - | - | 0 | - |
| - | - | 5686 | 243.1 | - | - | 0 | - |
| - | - | 1.433E+04 | 243.2 | - | - | 0 | - |
| - | - | 4927 | 244.1 | - | - | 0 | - |
| - | - | 3501 | 245.1 | - | - | 0 | - |
| - | - | 5034 | 259.1 | - | - | 0 | - |
| - | - | 2948 | 260.2 | - | - | 0 | - |
| - | - | 2.816E+05 | 261.2 | - | - | 0 | - |
| - | - | 4.382E+04 | 262.2 | - | - | 0 | - |
| - | - | 2194 | 263.2 | - | - | 0 | - |
| - | - | 3424 | 264.1 | - | - | 0 | - |
| - | - | 3.462E+04 | 282.1 | - | - | 0 | - |
| - | - | 4998 | 283.1 | - | - | 0 | - |
| - | - | 2443 | 298 | - | - | 0 | - |
| - | - | 4022 | 299.1 | - | - | 0 | - |
| - | - | 9263 | 299.2 | - | - | 0 | - |
| - | - | 1.727E+04 | 314.1 | - | - | 0 | - |
| 9 | y | 4.738E+04 | 314.2 | 0.0006141 | 1.955 | +1 | 3 |
| - | - | 2488 | 314.2 | - | - | 0 | - |
| - | - | 3844 | 314.6 | - | - | 0 | - |
| 9 | z | 3710 | 315.2 | 0.004251 | 13.49 | +1 | 3 |
| - | - | 2345 | 318.4 | - | - | 0 | - |
| 9 | y | 4.467E+04 | 331.2 | 0.0005237 | 1.581 | +1 | 3 |
| - | - | 5757 | 332.2 | - | - | 0 | - |
| - | - | 2513 | 333.2 | - | - | 0 | - |
| - | - | 3541 | 338.7 | - | - | 0 | - |
| - | - | 4222 | 346.1 | - | - | 0 | - |
| - | - | 7514 | 347.7 | - | - | 0 | - |
| - | - | 3484 | 348.2 | - | - | 0 | - |
| - | - | 5596 | 351.2 | - | - | 0 | - |
| - | - | 1.63E+04 | 353.2 | - | - | 0 | - |
| - | - | 5.117E+04 | 355.1 | - | - | 0 | - |
| - | - | 7341 | 356.7 | - | - | 0 | - |
| - | - | 2.65E+04 | 358.2 | - | - | 0 | - |
| - | - | 6135 | 359.2 | - | - | 0 | - |
| - | - | 5724 | 361.7 | - | - | 0 | - |
| - | - | 4121 | 362.2 | - | - | 0 | - |
| - | - | 1.474E+04 | 369.2 | - | - | 0 | - |
| - | - | 2929 | 370.2 | - | - | 0 | - |
| - | - | 1.579E+05 | 370.7 | - | - | 0 | - |
| - | - | 5.222E+04 | 371.2 | - | - | 0 | - |
| - | - | 1.349E+04 | 371.7 | - | - | 0 | - |
| - | - | 3457 | 372.2 | - | - | 0 | - |
| 7 | c | 2.828E+04 | 379.2 | 0.007197 | 18.98 | +2 | 7 |
| - | - | 1.434E+04 | 379.7 | - | - | 0 | - |
| - | - | 7403 | 380.2 | - | - | 0 | - |
| - | - | 1.36E+04 | 386.2 | - | - | 0 | - |
| - | - | 8127 | 420.7 | - | - | 0 | - |
| - | - | 5440 | 421.2 | - | - | 0 | - |
| - | - | 2909 | 425.2 | - | - | 0 | - |
| - | - | 1.202E+04 | 425.7 | - | - | 0 | - |
| - | - | 1.111E+04 | 426.2 | - | - | 0 | - |
| - | - | 3679 | 426.7 | - | - | 0 | - |
| - | - | 5345 | 427.3 | - | - | 0 | - |
| - | - | 5581 | 428.3 | - | - | 0 | - |
| - | - | 5352 | 429.3 | - | - | 0 | - |
| - | - | 4727 | 430.2 | - | - | 0 | - |
| - | - | 4692 | 430.7 | - | - | 0 | - |
| - | - | 1.331E+05 | 434.7 | - | - | 0 | - |
| - | - | 5.838E+04 | 435.2 | - | - | 0 | - |
| - | - | 2.329E+04 | 435.7 | - | - | 0 | - |
| 8 | y | 3363 | 442.2 | 0.002461 | 5.566 | +1 | 4 |
| 8 | y | 4853 | 443.2 | 0.0016 | 3.61 | +1 | 4 |
| - | - | 5.137E+04 | 444.2 | - | - | 0 | - |
| - | - | 3.08E+04 | 444.7 | - | - | 0 | - |
| - | - | 5178 | 445.2 | - | - | 0 | - |
| - | - | 6091 | 447.7 | - | - | 0 | - |
| - | - | 3112 | 452.7 | - | - | 0 | - |
| - | - | 1.577E+04 | 455.3 | - | - | 0 | - |
| - | - | 3693 | 456.3 | - | - | 0 | - |
| - | - | 3596 | 459.2 | - | - | 0 | - |
| 8 | y | 9718 | 460.2 | 0.0006857 | 1.49 | +1 | 4 |
| - | - | 3520 | 461.2 | - | - | 0 | - |
| - | - | 5178 | 461.7 | - | - | 0 | - |
| 3 | z | 4941 | 462.2 | 0.0003072 | 0.6646 | +2 | 9 |
| 3 | y | 2.645E+04 | 470.2 | 0.001046 | 2.226 | +2 | 9 |
| 3 | y | 1.6E+04 | 470.7 | 0.007513 | 15.96 | +2 | 9 |
| 3 | z | 6787 | 471.2 | 0.004638 | 9.842 | +2 | 9 |
| - | - | 2.211E+04 | 471.3 | - | - | 0 | - |
| 4 | c | 2.742E+04 | 472.3 | 0.0002967 | 0.6283 | +1 | 4 |
| - | - | 6855 | 473.3 | - | - | 0 | - |
| - | - | 3130 | 477.8 | - | - | 0 | - |
| 3 | y | 2.77E+04 | 479.2 | 0.001166 | 2.433 | +2 | 9 |
| - | - | 1.233E+04 | 479.7 | - | - | 0 | - |
| - | - | 7789 | 480.2 | - | - | 0 | - |
| - | - | 2829 | 481.2 | - | - | 0 | - |
| - | - | 4258 | 482.2 | - | - | 0 | - |
| - | - | 3389 | 487.3 | - | - | 0 | - |
| - | - | 6.142E+04 | 498.2 | - | - | 0 | - |
| - | - | 3664 | 498.3 | - | - | 0 | - |
| - | - | 1.506E+04 | 499.2 | - | - | 0 | - |
| - | - | 7911 | 500.2 | - | - | 0 | - |
| - | - | 4.41E+04 | 500.8 | - | - | 0 | - |
| - | - | 2.271E+04 | 501.3 | - | - | 0 | - |
| - | - | 6053 | 501.8 | - | - | 0 | - |
| 9 | c | 4178 | 509.3 | 0.001345 | 2.642 | +2 | 9 |
| - | - | 2924 | 512.2 | - | - | 0 | - |
| - | - | 2.801E+04 | 515.2 | - | - | 0 | - |
| - | - | 9561 | 516.2 | - | - | 0 | - |
| - | - | 8205 | 516.3 | - | - | 0 | - |
| - | - | 8724 | 530.2 | - | - | 0 | - |
| - | - | 1.527E+04 | 542.3 | - | - | 0 | - |
| - | - | 5957 | 543.3 | - | - | 0 | - |
| - | - | 1.336E+04 | 544.3 | - | - | 0 | - |
| - | - | 5659 | 546.2 | - | - | 0 | - |
| - | - | 4128 | 546.8 | - | - | 0 | - |
| - | - | 3343 | 547.2 | - | - | 0 | - |
| - | - | 1.255E+05 | 558.3 | - | - | 0 | - |
| 5 | c | 1.411E+05 | 559.3 | 0.0002512 | 0.4491 | +1 | 5 |
| - | - | 4.161E+04 | 560.3 | - | - | 0 | - |
| - | - | 6236 | 561.3 | - | - | 0 | - |
| 10 | c | 4214 | 564.8 | 0.0006663 | 1.18 | +2 | 10 |
| - | - | 4211 | 565.3 | - | - | 0 | - |
| - | - | 3575 | 581.3 | - | - | 0 | - |
| - | - | 5923 | 587.3 | - | - | 0 | - |
| - | - | 3013 | 588.3 | - | - | 0 | - |
| 7 | y | 9047 | 589.3 | 0.0008478 | 1.439 | +1 | 5 |
| - | - | 5443 | 599.3 | - | - | 0 | - |
| - | - | 1.423E+04 | 603.4 | - | - | 0 | - |
| - | - | 6903 | 604.4 | - | - | 0 | - |
| - | - | 1.26E+04 | 609.3 | - | - | 0 | - |
| - | - | 1.217E+04 | 610.2 | - | - | 0 | - |
| - | - | 4844 | 610.3 | - | - | 0 | - |
| - | - | 4853 | 611.3 | - | - | 0 | - |
| - | - | 2911 | 612.3 | - | - | 0 | - |
| - | - | 4618 | 624.3 | - | - | 0 | - |
| - | - | 3.802E+05 | 627.3 | - | - | 0 | - |
| - | - | 1.209E+05 | 628.3 | - | - | 0 | - |
| - | - | 2.538E+04 | 629.3 | - | - | 0 | - |
| - | - | 1.867E+04 | 629.3 | - | - | 0 | - |
| - | - | 7582 | 630.3 | - | - | 0 | - |
| - | - | 3219 | 631.3 | - | - | 0 | - |
| 6 | w | 9700 | 643.3 | 0.0002318 | 0.3603 | +1 | 6 |
| - | - | 3.598E+04 | 644.3 | - | - | 0 | - |
| - | - | 3841 | 644.3 | - | - | 0 | - |
| - | - | 1.639E+04 | 645.3 | - | - | 0 | - |
| - | - | 1.034E+05 | 645.3 | - | - | 0 | - |
| 6 | c | 5.273E+05 | 646.4 | 0.0008015 | 1.24 | +1 | 6 |
| - | - | 1.9E+05 | 647.4 | - | - | 0 | - |
| - | - | 2774 | 647.4 | - | - | 0 | - |
| - | - | 2.917E+04 | 648.4 | - | - | 0 | - |
| - | - | 3184 | 649.4 | - | - | 0 | - |
| - | - | 2.717E+04 | 658.3 | - | - | 0 | - |
| 6 | y | 1.165E+04 | 659.3 | 0.0009084 | 1.378 | +1 | 6 |
| 6 | z | 6067 | 660.3 | 0.000727 | 1.101 | +1 | 6 |
| - | - | 4799 | 661.3 | - | - | 0 | - |
| - | - | 6011 | 672.4 | - | - | 0 | - |
| - | - | 6243 | 674.3 | - | - | 0 | - |
| - | - | 1.942E+04 | 675.3 | - | - | 0 | - |
| 6 | y | 3.083E+04 | 676.3 | 0.0001914 | 0.283 | +1 | 6 |
| - | - | 1.112E+04 | 677.3 | - | - | 0 | - |
| - | - | 4043 | 702.4 | - | - | 0 | - |
| - | - | 9651 | 704.3 | - | - | 0 | - |
| - | - | 3525 | 705.9 | - | - | 0 | - |
| - | - | 1.968E+04 | 712.4 | - | - | 0 | - |
| - | - | 6898 | 713.4 | - | - | 0 | - |
| - | - | 6391 | 714.4 | - | - | 0 | - |
| - | - | 4000 | 715.4 | - | - | 0 | - |
| - | - | 4695 | 719.4 | - | - | 0 | - |
| - | - | 5578 | 720.4 | - | - | 0 | - |
| - | - | 3.955E+04 | 722.3 | - | - | 0 | - |
| - | - | 1.816E+04 | 723.3 | - | - | 0 | - |
| - | - | 2914 | 730.4 | - | - | 0 | - |
| - | - | 2.151E+04 | 731.4 | - | - | 0 | - |
| - | - | 2.084E+04 | 732.4 | - | - | 0 | - |
| - | - | 5144 | 733.4 | - | - | 0 | - |
| - | - | 4628 | 739.3 | - | - | 0 | - |
| - | - | 4.367E+05 | 740.3 | - | - | 0 | - |
| - | - | 1.632E+05 | 741.4 | - | - | 0 | - |
| - | - | 3.135E+04 | 742.4 | - | - | 0 | - |
| - | - | 9858 | 744.3 | - | - | 0 | - |
| 5 | y | 7361 | 745.3 | 0.01169 | 15.68 | +1 | 7 |
| 5 | y | 3806 | 746.3 | 0.006675 | 8.944 | +1 | 7 |
| 5 | z | 1.859E+04 | 747.3 | 0.001291 | 1.728 | +1 | 7 |
| - | - | 1.644E+04 | 748.3 | - | - | 0 | - |
| - | - | 4491 | 749.3 | - | - | 0 | - |
| 7 | c | 6.286E+04 | 757.4 | 0.01389 | 18.34 | +1 | 7 |
| - | - | 9.779E+04 | 758.4 | - | - | 0 | - |
| - | - | 3.447E+04 | 759.4 | - | - | 0 | - |
| - | - | 7087 | 760.4 | - | - | 0 | - |
| - | - | 9116 | 762.3 | - | - | 0 | - |
| 5 | y | 6.728E+04 | 763.3 | 0.001244 | 1.63 | +1 | 7 |
| - | - | 2.301E+04 | 764.4 | - | - | 0 | - |
| - | - | 3617 | 765.4 | - | - | 0 | - |
| - | - | 2.103E+04 | 771.4 | - | - | 0 | - |
| - | - | 9842 | 772.4 | - | - | 0 | - |
| - | - | 4437 | 773.4 | - | - | 0 | - |
| - | - | 1.507E+05 | 774.4 | - | - | 0 | - |
| 7 | c | 8.981E+05 | 775.4 | 0.000872 | 1.125 | +1 | 7 |
| - | - | 3.877E+05 | 776.4 | - | - | 0 | - |
| - | - | 7.916E+04 | 777.4 | - | - | 0 | - |
| - | - | 8830 | 778.4 | - | - | 0 | - |
| - | - | 8719 | 801.4 | - | - | 0 | - |
| - | - | 5731 | 802.4 | - | - | 0 | - |
| - | - | 3538 | 831.4 | - | - | 0 | - |
| - | - | 4935 | 840.4 | - | - | 0 | - |
| - | - | 3673 | 841.4 | - | - | 0 | - |
| - | - | 9241 | 842.4 | - | - | 0 | - |
| 4 | y | 6780 | 843.4 | 0.006035 | 7.156 | +1 | 8 |
| - | - | 4179 | 844.4 | - | - | 0 | - |
| - | - | 3783 | 845.4 | - | - | 0 | - |
| - | - | 1.035E+04 | 850.4 | - | - | 0 | - |
| - | - | 1.173E+04 | 851.4 | - | - | 0 | - |
| - | - | 2.128E+04 | 853.4 | - | - | 0 | - |
| - | - | 1.317E+04 | 854.4 | - | - | 0 | - |
| - | - | 5075 | 858.4 | - | - | 0 | - |
| - | - | 1.472E+04 | 859.4 | - | - | 0 | - |
| 4 | y | 2.334E+05 | 860.4 | 0.003595 | 4.178 | +1 | 8 |
| - | - | 1.012E+05 | 861.4 | - | - | 0 | - |
| - | - | 2.912E+04 | 862.4 | - | - | 0 | - |
| - | - | 2745 | 863.4 | - | - | 0 | - |
| - | - | 1.338E+04 | 867.4 | - | - | 0 | - |
| - | - | 2.025E+05 | 868.4 | - | - | 0 | - |
| - | - | 1.069E+05 | 869.4 | - | - | 0 | - |
| - | - | 3.871E+04 | 870.4 | - | - | 0 | - |
| - | - | 2.265E+04 | 871.4 | - | - | 0 | - |
| - | - | 1.192E+04 | 872.4 | - | - | 0 | - |
| - | - | 5132 | 873.4 | - | - | 0 | - |
| - | - | 1.601E+04 | 884.4 | - | - | 0 | - |
| - | - | 3.249E+04 | 885.4 | - | - | 0 | - |
| 8 | c | 1.477E+04 | 886.4 | 0.01388 | 15.66 | +1 | 8 |
| - | - | 4.014E+05 | 887.4 | - | - | 0 | - |
| - | - | 1.948E+05 | 888.4 | - | - | 0 | - |
| - | - | 5.804E+04 | 889.4 | - | - | 0 | - |
| - | - | 6238 | 890.4 | - | - | 0 | - |
| - | - | 3.664E+04 | 901.5 | - | - | 0 | - |
| - | - | 1.85E+04 | 902.5 | - | - | 0 | - |
| - | - | 7.068E+04 | 903.4 | - | - | 0 | - |
| 8 | c | 9.062E+05 | 904.4 | 0.0009424 | 1.042 | +1 | 8 |
| - | - | 4.409E+05 | 905.4 | - | - | 0 | - |
| - | - | 1.251E+05 | 906.4 | - | - | 0 | - |
| - | - | 1.264E+04 | 907.4 | - | - | 0 | - |
| - | - | 3476 | 922.4 | - | - | 0 | - |
| - | - | 9649 | 930.5 | - | - | 0 | - |
| - | - | 4736 | 931.5 | - | - | 0 | - |
| - | - | 1.967E+04 | 938.4 | - | - | 0 | - |
| 3 | y | 2.596E+04 | 939.4 | 0.0007624 | 0.8116 | +1 | 9 |
| 3 | y | 3.808E+04 | 940.4 | 0.005029 | 5.348 | +1 | 9 |
| 3 | z | 2.077E+04 | 941.4 | 4.932E-05 | 0.05239 | +1 | 9 |
| - | - | 5902 | 942.4 | - | - | 0 | - |
| - | - | 8193 | 955.4 | - | - | 0 | - |
| - | - | 2.132E+05 | 956.4 | - | - | 0 | - |
| 3 | y | 1.097E+06 | 957.5 | 0.0006968 | 0.7278 | +1 | 9 |
| - | - | 5.063E+05 | 958.5 | - | - | 0 | - |
| - | - | 1.486E+05 | 959.5 | - | - | 0 | - |
| - | - | 1.895E+04 | 960.5 | - | - | 0 | - |
| - | - | 5851 | 964.5 | - | - | 0 | - |
| - | - | 4647 | 966.4 | - | - | 0 | - |
| - | - | 3.058E+04 | 972.5 | - | - | 0 | - |
| - | - | 1.585E+05 | 973.5 | - | - | 0 | - |
| - | - | 1.08E+05 | 974.5 | - | - | 0 | - |
| - | - | 3.729E+04 | 975.5 | - | - | 0 | - |
| - | - | 6734 | 976.5 | - | - | 0 | - |
| - | - | 2.287E+04 | 982.5 | - | - | 0 | - |
| - | - | 1.178E+04 | 983.5 | - | - | 0 | - |
| - | - | 1.632E+04 | 984.5 | - | - | 0 | - |
| - | - | 1.528E+04 | 985.5 | - | - | 0 | - |
| - | - | 7329 | 986.5 | - | - | 0 | - |
| - | - | 1.732E+04 | 999.5 | - | - | 0 | - |
| - | - | 2.651E+05 | 1000 | - | - | 0 | - |
| - | - | 1.488E+05 | 1002 | - | - | 0 | - |
| - | - | 5.061E+04 | 1003 | - | - | 0 | - |
| - | - | 5795 | 1004 | - | - | 0 | - |
| - | - | 7877 | 1016 | - | - | 0 | - |
| - | - | 1.477E+04 | 1016 | - | - | 0 | - |
| 9 | c | 6.727E+05 | 1018 | 0.001046 | 1.028 | +1 | 9 |
| - | - | 3.984E+05 | 1019 | - | - | 0 | - |
| - | - | 1.191E+05 | 1020 | - | - | 0 | - |
| - | - | 1.569E+04 | 1021 | - | - | 0 | - |
| - | - | 1.155E+05 | 1044 | - | - | 0 | - |
| - | - | 6.397E+04 | 1045 | - | - | 0 | - |
| - | - | 3.59E+04 | 1046 | - | - | 0 | - |
| - | - | 1.677E+04 | 1047 | - | - | 0 | - |
| - | - | 6098 | 1048 | - | - | 0 | - |
| - | - | 4741 | 1053 | - | - | 0 | - |
| 2 | z | 9660 | 1070 | 0.004479 | 4.184 | +1 | 10 |
| 2 | z | 7773 | 1071 | 0.01961 | 18.3 | +1 | 10 |
| 2 | y | 3963 | 1087 | 0.01212 | 11.14 | +1 | 10 |
| 2 | z | 2.423E+05 | 1089 | 0.001239 | 1.138 | +1 | 10 |
| - | - | 1.476E+05 | 1090 | - | - | 0 | - |
| - | - | 5.241E+04 | 1091 | - | - | 0 | - |
| - | - | 9257 | 1092 | - | - | 0 | - |
| - | - | 1.006E+05 | 1102 | - | - | 0 | - |
| - | - | 6.078E+04 | 1103 | - | - | 0 | - |
| - | - | 2.761E+04 | 1104 | - | - | 0 | - |
| - | - | 3838 | 1110 | - | - | 0 | - |
| - | - | 4558 | 1111 | - | - | 0 | - |
| - | - | 3.46E+04 | 1113 | - | - | 0 | - |
| - | - | 2.068E+04 | 1114 | - | - | 0 | - |
| - | - | 1.221E+04 | 1115 | - | - | 0 | - |
| - | - | 3.524E+04 | 1118 | - | - | 0 | - |
| - | - | 2.615E+04 | 1119 | - | - | 0 | - |
| - | - | 7794 | 1120 | - | - | 0 | - |
| 10 | c | 2.442E+04 | 1129 | 0.004113 | 3.644 | +1 | 10 |
| - | - | 3.801E+04 | 1130 | - | - | 0 | - |
| - | - | 7.192E+04 | 1131 | - | - | 0 | - |
| - | - | 3.965E+04 | 1132 | - | - | 0 | - |
| - | - | 2.048E+04 | 1133 | - | - | 0 | - |
| - | - | 1.067E+04 | 1134 | - | - | 0 | - |
| - | - | 3550 | 1135 | - | - | 0 | - |
| - | - | 1.445E+04 | 1144 | - | - | 0 | - |
| - | - | 1.44E+04 | 1145 | - | - | 0 | - |
| 10 | c | 2.378E+06 | 1146 | 0.0003416 | 0.2982 | +1 | 10 |
| - | - | 1.605E+06 | 1147 | - | - | 0 | - |
| - | - | 6.146E+05 | 1148 | - | - | 0 | - |
| - | - | 1.22E+05 | 1149 | - | - | 0 | - |
| - | - | 1.044E+04 | 1150 | - | - | 0 | - |
| - | - | 3873 | 1155 | - | - | 0 | - |
| - | - | 4685 | 1156 | - | - | 0 | - |
| - | - | 3.734E+04 | 1157 | - | - | 0 | - |
| - | - | 2.367E+04 | 1158 | - | - | 0 | - |
| - | - | 1.396E+04 | 1159 | - | - | 0 | - |
| - | - | 1.101E+04 | 1160 | - | - | 0 | - |
| - | - | 1.045E+04 | 1161 | - | - | 0 | - |
| - | - | 4307 | 1162 | - | - | 0 | - |
| - | - | 4.416E+04 | 1163 | - | - | 0 | - |
| - | - | 2.823E+04 | 1164 | - | - | 0 | - |
| - | - | 1.323E+04 | 1165 | - | - | 0 | - |
| - | - | 5988 | 1172 | - | - | 0 | - |
| - | - | 5.799E+04 | 1173 | - | - | 0 | - |
| - | - | 2.018E+05 | 1174 | - | - | 0 | - |
| - | - | 1.321E+05 | 1175 | - | - | 0 | - |
| - | - | 4.946E+04 | 1176 | - | - | 0 | - |
| - | - | 8612 | 1177 | - | - | 0 | - |
| - | - | 3.232E+04 | 1184 | - | - | 0 | - |
| - | - | 1.955E+04 | 1185 | - | - | 0 | - |
| - | - | 4370 | 1186 | - | - | 0 | - |
| - | - | 3779 | 1189 | - | - | 0 | - |
| - | - | 5214 | 1190 | - | - | 0 | - |
| - | - | 6.91E+04 | 1191 | - | - | 0 | - |
| - | - | 4.438E+04 | 1192 | - | - | 0 | - |
| - | - | 1.826E+04 | 1193 | - | - | 0 | - |
| - | - | 5113 | 1200 | - | - | 0 | - |
| - | - | 2.653E+04 | 1201 | - | - | 0 | - |
| - | - | 3.373E+05 | 1202 | - | - | 0 | - |
| - | - | 2.278E+05 | 1203 | - | - | 0 | - |
| - | - | 8.973E+04 | 1204 | - | - | 0 | - |
| - | - | 1.751E+04 | 1205 | - | - | 0 | - |
| - | - | 8675 | 1216 | - | - | 0 | - |
| - | - | 3.13E+04 | 1217 | - | - | 0 | - |
| - | - | 4.053E+05 | 1218 | - | - | 0 | - |
| - | - | 2.15E+06 | 1219 | - | - | 0 | - |
| - | - | 1.387E+06 | 1220 | - | - | 0 | - |
| - | - | 5.134E+05 | 1221 | - | - | 0 | - |
| - | - | 7.102E+04 | 1222 | - | - | 0 | - |
| - | - | 9448 | 1251 | - | - | 0 | - |
| - | - | 6646 | 1252 | - | - | 0 | - |
| - | - | 3181 | 1468 | - | - | 0 | - |
| - | - | 3498 | 1494 | - | - | 0 | - |
| - | - | 4287 | 1574 | - | - | 0 | - |
| - | - | 3994 | 1831 | - | - | 0 | - |
| - | - | 3398 | 2413 | - | - | 0 | - |
| - | - | 3158 | 2582 | - | - | 0 | - |

m/z Charge Intensity FragmentType MassShift Position
120.07682037353516 0 3158.9697
120.08111572265625 0 32037.303
120.72374725341797 0 1613.5679
121.08477020263672 0 3668.2395
129.066162109375 0 50916.25
130.05035400390625 0 2334.8757
131.1182098388672 0 12105.791
147.0767822265625 0 10302.744
148.9464569091797 0 2214.4248
151.06402587890625 0 2082.4492
155.08193969726562 0 2433.278
173.45138549804688 0 10013.423
173.9221649169922 0 2519.4204
183.07696533203125 0 2113.825
186.4086456298828 0 2035.2836
197.128662109375 0 3699.142
200.10328674316406 0 114661.27
201.08737182617188 0 209852.67 y Ammonia loss 9
201.1066131591797 0 11024.267
202.09056091308594 0 16107.058
212.14015197753906 0 2902.1106
215.13856506347656 0 2306.731
218.11392211914062 0 260519.17 y 9
219.1171875 0 21489.482
221.08468627929688 0 4599.497
225.12380981445312 0 13053.075
233.16531372070312 0 289885.4
234.1686248779297 0 41773.625
242.15037536621094 0 126763.59
243.13401794433594 0 5685.945
243.15357971191406 0 14332.637
244.1338348388672 0 4926.986
245.12881469726562 0 3500.6335
259.0927429199219 0 5034.3276
260.1616516113281 0 2947.672
261.16015625 0 281592.9
262.1634826660156 0 43816.18
263.1641540527344 0 2194.181
264.13507080078125 0 3424.015
282.1452941894531 0 34622.688
283.1490478515625 0 4997.846
297.97808837890625 0 2443.1213
299.0622253417969 0 4021.6384
299.17254638671875 0 9262.968
314.1355895996094 0 17273.05
314.1716613769531 0 47377.094 y Ammonia loss 8
314.19268798828125 0 2488.3545
314.63641357421875 0 3843.6653
315.17462158203125 0 3710.2424 z 8
318.3980407714844 0 2344.658
331.1981201171875 0 44671.188 y 8
332.20074462890625 0 5757.149
333.22528076171875 0 2512.5935
338.6693115234375 0 3540.6077
346.1258239746094 0 4221.7305
347.67388916015625 0 7513.6704
348.1769104003906 0 3484.4443
351.1666564941406 0 5596.238
353.1824645996094 0 16299.686
355.0706481933594 0 51166.62
356.67999267578125 0 7340.897
358.2129211425781 0 26503.238
359.2156982421875 0 6134.8228
361.673095703125 0 5723.9688
362.17431640625 0 4121.2173
369.1775817871094 0 14735.88
370.18487548828125 0 2929.1108
370.67718505859375 0 157881.2
371.1791076660156 0 52222.69
371.6794128417969 0 13490.189
372.19757080078125 0 3457.3328
379.1903991699219 0 28280.037 c Water loss 6
379.6910705566406 0 14340.003
380.1927185058594 0 7403.275
386.2044677734375 0 13595.775
420.7097473144531 0 8127.059
421.20904541015625 0 5440.0757
425.2029724121094 0 2909.2922
425.7017822265625 0 12021.022
426.1982421875 0 11105.085
426.70013427734375 0 3678.7332
427.2705078125 0 5344.523
428.2781677246094 0 5580.549
429.28643798828125 0 5351.532
430.2159729003906 0 4727.4185
430.7174072265625 0 4692.3394
434.70684814453125 0 133060.05
435.2082824707031 0 58376.902
435.708984375 0 23293.943
442.2320861816406 0 3363.4272 y Water loss 7
443.2152404785156 0 4852.511 y Ammonia loss 7
444.21173095703125 0 51373.688
444.7135009765625 0 30798.188
445.2138977050781 0 5178.0093
447.71551513671875 0 6090.9478
452.7065734863281 0 3112.1946
455.2668762207031 0 15772.771
456.26641845703125 0 3693.1477
459.2086181640625 0 3596.3047
460.2408752441406 0 9718.082 y 7
461.2436218261719 0 3519.6355
461.71002197265625 0 5178.4717
462.2154846191406 0 4940.708 z Water loss 2
470.2255859375 0 26446 y Water loss 2
470.72406005859375 0 16004.604 y Ammonia loss 2
471.22509765625 0 6786.9814 z 2
471.2854309082031 0 22110.74
472.2915344238281 0 27421.045 c 3
473.29681396484375 0 6855.3
477.7522277832031 0 3130.495
479.2309875488281 0 27695.639 y 2
479.7320556640625 0 12325.973
480.2093811035156 0 7789.0127
481.2108154296875 0 2828.5796
482.22369384765625 0 4257.5083
487.2585144042969 0 3389.141
498.2205505371094 0 61417.332
498.30859375 0 3664.0732
499.2237243652344 0 15055.502
500.23333740234375 0 7910.8945
500.7538146972656 0 44103.46
501.25494384765625 0 22707.25
501.7555236816406 0 6052.528
509.2648620605469 0 4178.266 c 8
512.2017211914062 0 2924.4954
515.2468872070312 0 28011.104
516.2479248046875 0 9560.559
516.3186645507812 0 8205.359
530.21044921875 0 8723.676
542.2979736328125 0 15265.132
543.3027954101562 0 5956.722
544.31396484375 0 13363.034
546.2398071289062 0 5658.8613
546.7734985351562 0 4127.7827
547.2437133789062 0 3343.023
558.3169555664062 0 125499.32
559.3236083984375 0 141054.2 c 4
560.326904296875 0 41606.754
561.3331298828125 0 6235.8506
564.7815551757812 0 4213.5396 c Ammonia loss 9
565.2835083007812 0 4211.017
581.2545166015625 0 3575.4497
587.2677001953125 0 5923.426
588.2764282226562 0 3012.7112
589.2836303710938 0 9046.869 y 6
599.2672729492188 0 5442.9775
603.3502807617188 0 14231.154
604.3535766601562 0 6902.8374
609.2514038085938 0 12599.2
610.1848754882812 0 12165.78
610.2537841796875 0 4843.819
611.3162231445312 0 4853.3477
612.319091796875 0 2910.574
624.339111328125 0 4617.6743
627.262939453125 0 380155.78
628.2660522460938 0 120942.34
629.2679443359375 0 25377.56
629.3308715820312 0 18673.889
630.3348999023438 0 7582.3813
631.3397216796875 0 3219.3857
643.2935791015625 0 9699.639 w 5
644.2904663085938 0 35983.832
644.3414306640625 0 3841.2761
645.291015625 0 16393.752
645.3493041992188 0 103376.2
646.356689453125 0 527317.75 c 5
647.3595581054688 0 189972.52
647.423583984375 0 2774.2385
648.3632202148438 0 29172.99
649.3630981445312 0 3184.3877
658.2822875976562 0 27167.613
659.287353515625 0 11651.017 y Ammonia loss 5
660.2968139648438 0 6067.3813 z 5
661.3040771484375 0 4799.3438
672.373046875 0 6011.1987
674.2974853515625 0 6243.0493
675.3082275390625 0 19418.826
676.3150024414062 0 30826.764 y 5
677.319580078125 0 11121.684
702.3666381835938 0 4043.238
704.3255004882812 0 9650.969
705.8951416015625 0 3525.3105
712.3518676757812 0 19683.08
713.3554077148438 0 6897.6416
714.3656005859375 0 6390.736
715.373291015625 0 4000.2097
719.3660888671875 0 4695.015
720.3668212890625 0 5578.056
722.33740234375 0 39553.176
723.34130859375 0 18163.123
730.3837890625 0 2914.2563
731.3848876953125 0 21511.127
732.3909301757812 0 20839.738
733.3927001953125 0 5143.998
739.3457641601562 0 4628.4487
740.347412109375 0 436684.12
741.3506469726562 0 163154.22
742.353515625 0 31346.45
744.32861328125 0 9858.423
745.3245849609375 0 7360.6147 y Water loss 4
746.3269653320312 0 3805.943 y Ammonia loss 4
747.3294067382812 0 18590.412 z 4
748.3353271484375 0 16437.24
749.3369140625 0 4490.516
757.3740234375 0 62855.16 c Water loss 6
758.3738403320312 0 97794.836
759.376708984375 0 34471.664
760.3790893554688 0 7087.3623
762.3396606445312 0 9116.262
763.3480834960938 0 67283.07 y 4
764.3511352539062 0 23010.137
765.3548583984375 0 3617.498
771.351806640625 0 21031.104
772.3555908203125 0 9842.122
773.3715209960938 0 4437.0396
774.3915405273438 0 150712.9
775.3993530273438 0 898114.8 c 6
776.4022827148438 0 387707.12
777.405029296875 0 79158.195
778.4072875976562 0 8830.459
801.4127197265625 0 8719.432
802.4196166992188 0 5731.0537
831.4072265625 0 3538.0444
840.4091186523438 0 4934.8164
841.4193115234375 0 3673.3816
842.417236328125 0 9240.897
843.3790893554688 0 6779.5327 y Ammonia loss 3
844.4280395507812 0 4179.186
845.4290161132812 0 3782.7095
850.3955078125 0 10347.0625
851.3886108398438 0 11728.248
853.4298095703125 0 21276.053
854.43408203125 0 13167.421
858.384033203125 0 5074.6943
859.4156494140625 0 14722.408
860.4031982421875 0 233426.06 y 3
861.4064331054688 0 101218.89
862.409912109375 0 29120.531
863.4169311523438 0 2744.5413
867.397705078125 0 13382.821
868.4058837890625 0 202490.66
869.4083862304688 0 106887.24
870.4105834960938 0 38705.434
871.3989868164062 0 22646.113
872.4063720703125 0 11915.617
873.404541015625 0 5132.3335
884.4235229492188 0 16014.71
885.415283203125 0 32487.455
886.4166259765625 0 14765.577 c Water loss 7
887.415771484375 0 401374.9
888.4187622070312 0 194824.8
889.4212036132812 0 58039.89
890.4246215820312 0 6238.4873
901.4915161132812 0 36639.39
902.4935302734375 0 18504.545
903.4349365234375 0 70681.14
904.4420166015625 0 906233 c 7
905.4447021484375 0 440929.28
906.4476318359375 0 125052.27
907.449951171875 0 12636.5625
922.4287719726562 0 3475.6575
930.4544677734375 0 9649.145
931.4583740234375 0 4736.136
938.4351806640625 0 19669.805
939.4410400390625 0 25956.69 y Water loss 2
940.4308471679688 0 38082.203 y Ammonia loss 2
941.43359375 0 20773.771 z 2
942.435791015625 0 5901.6514
955.4398193359375 0 8192.681
956.4456787109375 0 213200.48
957.4530639648438 0 1096549.8 y 2
958.4561157226562 0 506252.06
959.4592895507812 0 148579.83
960.4635009765625 0 18946.607
964.4783935546875 0 5850.983
966.4251708984375 0 4646.76
972.5050659179688 0 30578.834
973.5121459960938 0 158512.27
974.5169067382812 0 108002.25
975.52001953125 0 37285.746
976.5185546875 0 6734.2075
982.4896240234375 0 22865.77
983.4910278320312 0 11777.197
984.4754638671875 0 16322.859
985.4766845703125 0 15284.717
986.4744262695312 0 7329.443
999.458251953125 0 17320.475
1000.4989013671875 0 265108
1001.502685546875 0 148817.62
1002.5062866210938 0 50607.47
1003.5154418945312 0 5795.3896
1015.5040893554688 0 7876.7505
1016.49169921875 0 14773.543
1017.5261840820312 0 672738.25 c 8
1018.5293579101562 0 398424.66
1019.5316162109375 0 119093.91
1020.5357666015625 0 15691.389
1043.542236328125 0 115481.64
1044.5445556640625 0 63969.85
1045.525146484375 0 35901.086
1046.5185546875 0 16773.734
1047.513916015625 0 6098.0273
1052.5765380859375 0 4741.447
1070.4959716796875 0 9660.252 z Water loss 1
1071.4951171875 0 7773.002 z Ammonia loss 1
1087.50634765625 0 3962.6377 y Ammonia loss 1
1088.5032958984375 0 242304.88 z 1
1089.50634765625 0 147608.77
1090.5087890625 0 52410.29
1091.5155029296875 0 9256.974
1101.571533203125 0 100580.016
1102.573974609375 0 60776.78
1103.5771484375 0 27611.531
1109.5543212890625 0 3838.3538
1110.5439453125 0 4558.202
1112.5391845703125 0 34600.742
1113.5458984375 0 20684.2
1114.556640625 0 12206.786
1117.5313720703125 0 35235.64
1118.53369140625 0 26146.152
1119.54541015625 0 7793.5054
1128.561279296875 0 24422.584 c Ammonia loss 9
1129.5653076171875 0 38011.027
1130.5565185546875 0 71916.6
1131.560302734375 0 39650.715
1132.57421875 0 20484.97
1133.5841064453125 0 10666.124
1134.5863037109375 0 3549.9978
1143.5697021484375 0 14450.115
1144.5751953125 0 14398.278
1145.5833740234375 0 2378208.5 c 9
1146.5867919921875 0 1605390.9
1147.5889892578125 0 614582.1
1148.5906982421875 0 121973.54
1149.5838623046875 0 10436.046
1154.5919189453125 0 3872.6533
1155.5982666015625 0 4685.493
1156.6116943359375 0 37337.008
1157.6171875 0 23665.03
1158.5345458984375 0 13955.223
1159.608154296875 0 11012.194
1160.592041015625 0 10447.15
1161.587890625 0 4306.752
1162.5516357421875 0 44159.71
1163.5528564453125 0 28225.758
1164.5574951171875 0 13234.771
1171.5897216796875 0 5987.7236
1172.6082763671875 0 57987.582
1173.59521484375 0 201785.97
1174.59716796875 0 132057.22
1175.598388671875 0 49463.004
1176.5994873046875 0 8611.898
1183.576416015625 0 32315.42
1184.5782470703125 0 19552.367
1185.586181640625 0 4370.1436
1188.5826416015625 0 3778.7944
1189.6004638671875 0 5214.3438
1190.61865234375 0 69099.4
1191.6221923828125 0 44378.203
1192.6243896484375 0 18260.67
1199.56591796875 0 5112.871
1200.602294921875 0 26527.113
1201.5882568359375 0 337270.7
1202.59033203125 0 227811.56
1203.5921630859375 0 89730.64
1204.595458984375 0 17510.303
1215.6002197265625 0 8674.865
1216.59716796875 0 31304.217
1217.60498046875 0 405283
1218.613037109375 0 2149862
1219.615478515625 0 1386593.6
1220.61865234375 0 513423.06
1221.62109375 0 71024.59
1250.6043701171875 0 9447.998
1251.6082763671875 0 6645.879
1467.7164306640625 0 3181.2375
1493.77783203125 0 3498.3381
1573.8497314453125 0 4287.291
1831.0291748046875 0 3994.0378
2413.146728515625 0 3398.215
2581.8935546875 0 3158.178

Spectrum Details

|  |  |
| --- | --- |
| Matched peaks? Matched peaks The total absolute number of peaks matched. Additionally in brackets the total fraction of peaks matched and the total number of peaks is shown. | 45 (11.81% of 381) |
| FDR? FDR The false discovery rate estimated for this peptide. It is calculated by matching all theoretical fragments with a non-integer shift with the raw peaks for this spectrum. This is done with 40 different shifts. The resulting percentage is the average number of annotated peaks over the number of annotated peaks with the correct spectrum. | 0.79% |
| Satellite FDR? Satellite FDR See the FDR for details on its calculation. This satellite ion specific FDR only contains the satellite ions (d/w) for I/L/J positions. | - |
| PSM Score? PSM Score The PSM Score as given by Hecklib to this annotated spectrum. It is shown with three significant figures. | 527 |

#### Spectrum 7929? Spectrum 7929 The raw spectrum of this peptide as annotated by Hecklib. The fragments are coloured according to ion type (see legend). Any peaks with a star '\*' as text can be hovered over to see the full details, first the ion type second the mass shift type. By hovering over the amino acids in the peptide or ions in the legend the corresponding peaks are highlighted. By toggling the 'Unassigned' label you can turn the background (unassigned) peaks on or off in the plot. By updating the slider in the Ion legend you can update the spectrum to only show the top X% of the peaks with labels. The top X% means any peak that is within X% of the highest intensity. By dragging in the spectrum you can zoom in to a specific part of the spectrum and use 'Zoom Out' to get back to the original zoom level. The annotation of the spectrum is based on the given sequence in the peptides file and is done with different software so inconsistencies are likely. The peaks are annotated based on the given sequence, with 20 ppm tolerance.

Copy Data

##### Spectrum 7929 (TSV)

###### Preview

```
Loading example...
```

*Click on the button to copy the data to your clipboard.*

Mz MinMz MaxIntensity Max

WidthHeightPeptide font sizePeptide stroke widthSpectrum font sizeSpectrum stroke widthCompact peptide

Ion legend

wxyz

abcd

OtherUnassignedIon^Charge^_Position_Show for top:%

JFPPSSEEJQA

05.57e+51.11e+61.67e+62.23e+6

Zoom Out

y^+1^_2_y^+1^_2_y^+1^_3_z^+1^_3_y^+1^_3_c^+2^_7_y^+1^_4_y^+1^_4_y^+2^_9_y^+2^_9_z^+2^_9_c^+1^_4_y^+2^_9_c^+1^_5_c^+2^_10_y^+1^_5_w^+1^_6_c^+1^_6_y^+1^_6_z^+1^_6_y^+1^_6_y^+1^_7_y^+1^_7_z^+1^_7_c^+1^_7_y^+1^_7_c^+1^_7_y^+1^_8_c^+1^_8_c^+1^_8_y^+1^_9_y^+1^_9_z^+1^_9_y^+1^_9_c^+1^_9_z^+1^_10_z^+1^_10_c^+1^_10_c^+1^_10_

0583116717502334

Fragment Matches Table

Show background peaks

| Position | Ion type | Intensity | mz Theoretical | mz Error (Th) | mz Error (ppm) | Charge | Series Number |
| --- | --- | --- | --- | --- | --- | --- | --- |
| - | - | 2.889E+04 | 120.1 | - | - | 0 | - |
| - | - | 3380 | 121.1 | - | - | 0 | - |
| - | - | 1982 | 123 | - | - | 0 | - |
| - | - | 3.509E+04 | 129.1 | - | - | 0 | - |
| - | - | 1.201E+04 | 131.1 | - | - | 0 | - |
| - | - | 2165 | 134.5 | - | - | 0 | - |
| - | - | 6328 | 147.1 | - | - | 0 | - |
| - | - | 2877 | 148.9 | - | - | 0 | - |
| - | - | 2327 | 149 | - | - | 0 | - |
| - | - | 2333 | 152.5 | - | - | 0 | - |
| - | - | 1.428E+04 | 173.5 | - | - | 0 | - |
| - | - | 2085 | 176.7 | - | - | 0 | - |
| - | - | 6160 | 197.1 | - | - | 0 | - |
| - | - | 9.707E+04 | 200.1 | - | - | 0 | - |
| 10 | y | 1.565E+05 | 201.1 | 0.0002969 | 1.477 | +1 | 2 |
| - | - | 1.217E+04 | 202.1 | - | - | 0 | - |
| - | - | 2762 | 212.1 | - | - | 0 | - |
| - | - | 5791 | 215.1 | - | - | 0 | - |
| 10 | y | 2.054E+05 | 218.1 | 0.0003134 | 1.437 | +1 | 2 |
| - | - | 1.999E+04 | 219.1 | - | - | 0 | - |
| - | - | 3746 | 221.1 | - | - | 0 | - |
| - | - | 1.005E+04 | 225.1 | - | - | 0 | - |
| - | - | 4765 | 232.1 | - | - | 0 | - |
| - | - | 2.328E+05 | 233.2 | - | - | 0 | - |
| - | - | 2399 | 233.3 | - | - | 0 | - |
| - | - | 3.447E+04 | 234.2 | - | - | 0 | - |
| - | - | 9.182E+04 | 242.2 | - | - | 0 | - |
| - | - | 5873 | 243.1 | - | - | 0 | - |
| - | - | 1.185E+04 | 243.2 | - | - | 0 | - |
| - | - | 3902 | 256.1 | - | - | 0 | - |
| - | - | 2602 | 259.9 | - | - | 0 | - |
| - | - | 2.294E+05 | 261.2 | - | - | 0 | - |
| - | - | 3.721E+04 | 262.2 | - | - | 0 | - |
| - | - | 3065 | 264.1 | - | - | 0 | - |
| - | - | 2702 | 280.6 | - | - | 0 | - |
| - | - | 2.727E+04 | 282.1 | - | - | 0 | - |
| - | - | 5256 | 299.1 | - | - | 0 | - |
| - | - | 5666 | 299.2 | - | - | 0 | - |
| - | - | 1.356E+04 | 314.1 | - | - | 0 | - |
| 9 | y | 3.47E+04 | 314.2 | 0.0006446 | 2.052 | +1 | 3 |
| - | - | 4463 | 314.6 | - | - | 0 | - |
| 9 | z | 5513 | 315.2 | 0.004922 | 15.62 | +1 | 3 |
| 9 | y | 2.424E+04 | 331.2 | 0.0005847 | 1.766 | +1 | 3 |
| - | - | 3847 | 346.1 | - | - | 0 | - |
| - | - | 8675 | 347.7 | - | - | 0 | - |
| - | - | 5654 | 348.2 | - | - | 0 | - |
| - | - | 7606 | 351.2 | - | - | 0 | - |
| - | - | 8514 | 353.2 | - | - | 0 | - |
| - | - | 3.71E+04 | 355.1 | - | - | 0 | - |
| - | - | 3608 | 356.7 | - | - | 0 | - |
| - | - | 3338 | 357.2 | - | - | 0 | - |
| - | - | 2.398E+04 | 358.2 | - | - | 0 | - |
| - | - | 4253 | 359.2 | - | - | 0 | - |
| - | - | 2957 | 361.7 | - | - | 0 | - |
| - | - | 1.392E+04 | 369.2 | - | - | 0 | - |
| - | - | 3122 | 370.2 | - | - | 0 | - |
| - | - | 1.299E+05 | 370.7 | - | - | 0 | - |
| - | - | 4.538E+04 | 371.2 | - | - | 0 | - |
| - | - | 9904 | 371.7 | - | - | 0 | - |
| - | - | 3331 | 372.2 | - | - | 0 | - |
| 7 | c | 3.041E+04 | 379.2 | 0.007197 | 18.98 | +2 | 7 |
| - | - | 1.074E+04 | 379.7 | - | - | 0 | - |
| - | - | 1.476E+04 | 386.2 | - | - | 0 | - |
| - | - | 3247 | 418.2 | - | - | 0 | - |
| - | - | 2983 | 418.5 | - | - | 0 | - |
| - | - | 7119 | 420.7 | - | - | 0 | - |
| - | - | 5134 | 421.2 | - | - | 0 | - |
| - | - | 6262 | 425.7 | - | - | 0 | - |
| - | - | 5275 | 426.2 | - | - | 0 | - |
| - | - | 4772 | 427.3 | - | - | 0 | - |
| - | - | 3423 | 428.3 | - | - | 0 | - |
| - | - | 4198 | 429.3 | - | - | 0 | - |
| - | - | 3327 | 430.2 | - | - | 0 | - |
| - | - | 4205 | 430.7 | - | - | 0 | - |
| - | - | 1.18E+05 | 434.7 | - | - | 0 | - |
| - | - | 4.285E+04 | 435.2 | - | - | 0 | - |
| - | - | 1.597E+04 | 435.7 | - | - | 0 | - |
| 8 | y | 3440 | 442.2 | 0.001241 | 2.805 | +1 | 4 |
| - | - | 4.911E+04 | 444.2 | - | - | 0 | - |
| - | - | 2.589E+04 | 444.7 | - | - | 0 | - |
| - | - | 3075 | 445.2 | - | - | 0 | - |
| - | - | 1.365E+04 | 455.3 | - | - | 0 | - |
| - | - | 5394 | 456.3 | - | - | 0 | - |
| - | - | 3840 | 459.2 | - | - | 0 | - |
| 8 | y | 7998 | 460.2 | 0.0007486 | 1.627 | +1 | 4 |
| - | - | 3572 | 461.7 | - | - | 0 | - |
| - | - | 3924 | 464.7 | - | - | 0 | - |
| - | - | 2794 | 466.4 | - | - | 0 | - |
| 3 | y | 2.356E+04 | 470.2 | 0.0008939 | 1.901 | +2 | 9 |
| 3 | y | 1.217E+04 | 470.7 | 0.005926 | 12.59 | +2 | 9 |
| 3 | z | 5370 | 471.2 | 0.006377 | 13.53 | +2 | 9 |
| - | - | 2.263E+04 | 471.3 | - | - | 0 | - |
| 4 | c | 2.166E+04 | 472.3 | 0.0003578 | 0.7575 | +1 | 4 |
| - | - | 4798 | 473.3 | - | - | 0 | - |
| - | - | 3887 | 476.8 | - | - | 0 | - |
| - | - | 2910 | 478.2 | - | - | 0 | - |
| 3 | y | 1.354E+04 | 479.2 | 0.001196 | 2.496 | +2 | 9 |
| - | - | 8511 | 479.7 | - | - | 0 | - |
| - | - | 7707 | 480.2 | - | - | 0 | - |
| - | - | 2771 | 488.3 | - | - | 0 | - |
| - | - | 4.567E+04 | 498.2 | - | - | 0 | - |
| - | - | 8435 | 499.2 | - | - | 0 | - |
| - | - | 4174 | 500.2 | - | - | 0 | - |
| - | - | 3.5E+04 | 500.8 | - | - | 0 | - |
| - | - | 2.198E+04 | 501.3 | - | - | 0 | - |
| - | - | 3228 | 501.8 | - | - | 0 | - |
| - | - | 3420 | 505.2 | - | - | 0 | - |
| - | - | 2.442E+04 | 515.2 | - | - | 0 | - |
| - | - | 6756 | 516.2 | - | - | 0 | - |
| - | - | 7614 | 516.3 | - | - | 0 | - |
| - | - | 3125 | 520.3 | - | - | 0 | - |
| - | - | 8040 | 530.2 | - | - | 0 | - |
| - | - | 6221 | 535.8 | - | - | 0 | - |
| - | - | 9816 | 542.3 | - | - | 0 | - |
| - | - | 5724 | 543.3 | - | - | 0 | - |
| - | - | 1.469E+04 | 544.3 | - | - | 0 | - |
| - | - | 1.12E+05 | 558.3 | - | - | 0 | - |
| 5 | c | 1.398E+05 | 559.3 | 0.0006174 | 1.104 | +1 | 5 |
| - | - | 3.811E+04 | 560.3 | - | - | 0 | - |
| - | - | 7243 | 561.3 | - | - | 0 | - |
| 10 | c | 7295 | 564.8 | 0.0003001 | 0.5314 | +2 | 10 |
| - | - | 5570 | 587.3 | - | - | 0 | - |
| 7 | y | 6699 | 589.3 | 0.0008002 | 1.358 | +1 | 5 |
| - | - | 5163 | 591.2 | - | - | 0 | - |
| - | - | 3984 | 592.3 | - | - | 0 | - |
| - | - | 3840 | 593.3 | - | - | 0 | - |
| - | - | 5127 | 599.3 | - | - | 0 | - |
| - | - | 1.237E+04 | 603.4 | - | - | 0 | - |
| - | - | 3967 | 604.4 | - | - | 0 | - |
| - | - | 3570 | 606.3 | - | - | 0 | - |
| - | - | 1.284E+04 | 609.3 | - | - | 0 | - |
| - | - | 9624 | 610.2 | - | - | 0 | - |
| - | - | 5243 | 611.3 | - | - | 0 | - |
| - | - | 2.971E+05 | 627.3 | - | - | 0 | - |
| - | - | 9.47E+04 | 628.3 | - | - | 0 | - |
| - | - | 1.487E+04 | 629.3 | - | - | 0 | - |
| - | - | 1.949E+04 | 629.3 | - | - | 0 | - |
| - | - | 4727 | 630.3 | - | - | 0 | - |
| 6 | w | 1.433E+04 | 643.3 | 0.000537 | 0.8347 | +1 | 6 |
| - | - | 3.34E+04 | 644.3 | - | - | 0 | - |
| - | - | 4878 | 644.3 | - | - | 0 | - |
| - | - | 1.013E+04 | 645.3 | - | - | 0 | - |
| - | - | 1.099E+05 | 645.3 | - | - | 0 | - |
| 6 | c | 4.782E+05 | 646.4 | 0.0004963 | 0.7678 | +1 | 6 |
| - | - | 1.685E+05 | 647.4 | - | - | 0 | - |
| - | - | 3.962E+04 | 648.4 | - | - | 0 | - |
| - | - | 4039 | 657.3 | - | - | 0 | - |
| - | - | 2.355E+04 | 658.3 | - | - | 0 | - |
| 6 | y | 1.356E+04 | 659.3 | 0.001092 | 1.656 | +1 | 6 |
| 6 | z | 6195 | 660.3 | 0.002192 | 3.32 | +1 | 6 |
| - | - | 3993 | 661.3 | - | - | 0 | - |
| - | - | 8825 | 672.4 | - | - | 0 | - |
| - | - | 4733 | 674.3 | - | - | 0 | - |
| - | - | 2.083E+04 | 675.3 | - | - | 0 | - |
| 6 | y | 2.962E+04 | 676.3 | 0.000541 | 0.7999 | +1 | 6 |
| - | - | 8958 | 677.3 | - | - | 0 | - |
| - | - | 2890 | 679.4 | - | - | 0 | - |
| - | - | 3158 | 690.3 | - | - | 0 | - |
| - | - | 4002 | 697.4 | - | - | 0 | - |
| - | - | 3770 | 700.3 | - | - | 0 | - |
| - | - | 9444 | 704.3 | - | - | 0 | - |
| - | - | 3559 | 705.3 | - | - | 0 | - |
| - | - | 1.217E+04 | 712.4 | - | - | 0 | - |
| - | - | 7858 | 713.4 | - | - | 0 | - |
| - | - | 4834 | 714.4 | - | - | 0 | - |
| - | - | 3.262E+04 | 722.3 | - | - | 0 | - |
| - | - | 1.545E+04 | 723.3 | - | - | 0 | - |
| - | - | 3672 | 730.4 | - | - | 0 | - |
| - | - | 2.112E+04 | 731.4 | - | - | 0 | - |
| - | - | 1.601E+04 | 732.4 | - | - | 0 | - |
| - | - | 5281 | 733.4 | - | - | 0 | - |
| - | - | 3806 | 739.3 | - | - | 0 | - |
| - | - | 3.551E+05 | 740.3 | - | - | 0 | - |
| - | - | 1.37E+05 | 741.4 | - | - | 0 | - |
| - | - | 2.85E+04 | 742.4 | - | - | 0 | - |
| - | - | 9525 | 744.3 | - | - | 0 | - |
| 5 | y | 7196 | 745.3 | 0.01334 | 17.89 | +1 | 7 |
| 5 | y | 5139 | 746.3 | 0.007896 | 10.58 | +1 | 7 |
| 5 | z | 1.961E+04 | 747.3 | 0.0007838 | 1.049 | +1 | 7 |
| - | - | 1.53E+04 | 748.3 | - | - | 0 | - |
| 7 | c | 5.593E+04 | 757.4 | 0.01432 | 18.91 | +1 | 7 |
| - | - | 8.122E+04 | 758.4 | - | - | 0 | - |
| - | - | 2.772E+04 | 759.4 | - | - | 0 | - |
| - | - | 5860 | 760.4 | - | - | 0 | - |
| - | - | 7478 | 762.3 | - | - | 0 | - |
| 5 | y | 6.157E+04 | 763.3 | 0.0007558 | 0.9901 | +1 | 7 |
| - | - | 1.999E+04 | 764.4 | - | - | 0 | - |
| - | - | 4566 | 765.4 | - | - | 0 | - |
| - | - | 1.667E+04 | 771.4 | - | - | 0 | - |
| - | - | 1.004E+04 | 772.4 | - | - | 0 | - |
| - | - | 5009 | 773.4 | - | - | 0 | - |
| - | - | 1.194E+05 | 774.4 | - | - | 0 | - |
| 7 | c | 8.444E+05 | 775.4 | 0.0006278 | 0.8097 | +1 | 7 |
| - | - | 3.589E+05 | 776.4 | - | - | 0 | - |
| - | - | 8.652E+04 | 777.4 | - | - | 0 | - |
| - | - | 5825 | 778.4 | - | - | 0 | - |
| - | - | 4883 | 786.3 | - | - | 0 | - |
| - | - | 4152 | 787.4 | - | - | 0 | - |
| - | - | 1.428E+04 | 801.4 | - | - | 0 | - |
| - | - | 4026 | 802.4 | - | - | 0 | - |
| - | - | 7250 | 826.4 | - | - | 0 | - |
| - | - | 3094 | 827.4 | - | - | 0 | - |
| - | - | 7986 | 842.4 | - | - | 0 | - |
| - | - | 3437 | 844.4 | - | - | 0 | - |
| - | - | 1.04E+04 | 850.4 | - | - | 0 | - |
| - | - | 9158 | 851.4 | - | - | 0 | - |
| - | - | 1.332E+04 | 853.4 | - | - | 0 | - |
| - | - | 6918 | 854.4 | - | - | 0 | - |
| - | - | 4058 | 855.4 | - | - | 0 | - |
| - | - | 1.014E+04 | 859.4 | - | - | 0 | - |
| 4 | y | 2.177E+05 | 860.4 | 0.003412 | 3.965 | +1 | 8 |
| - | - | 1.003E+05 | 861.4 | - | - | 0 | - |
| - | - | 2.778E+04 | 862.4 | - | - | 0 | - |
| - | - | 1.329E+04 | 867.4 | - | - | 0 | - |
| - | - | 1.669E+05 | 868.4 | - | - | 0 | - |
| - | - | 8.472E+04 | 869.4 | - | - | 0 | - |
| - | - | 2.717E+04 | 870.4 | - | - | 0 | - |
| - | - | 2.154E+04 | 871.4 | - | - | 0 | - |
| - | - | 1.406E+04 | 872.4 | - | - | 0 | - |
| - | - | 5026 | 873.4 | - | - | 0 | - |
| - | - | 1.343E+04 | 884.4 | - | - | 0 | - |
| - | - | 3.135E+04 | 885.4 | - | - | 0 | - |
| 8 | c | 1.195E+04 | 886.4 | 0.01321 | 14.9 | +1 | 8 |
| - | - | 3.261E+05 | 887.4 | - | - | 0 | - |
| - | - | 1.483E+05 | 888.4 | - | - | 0 | - |
| - | - | 4.331E+04 | 889.4 | - | - | 0 | - |
| - | - | 4840 | 890.4 | - | - | 0 | - |
| - | - | 4380 | 894.4 | - | - | 0 | - |
| - | - | 3456 | 895.4 | - | - | 0 | - |
| - | - | 3.483E+04 | 901.5 | - | - | 0 | - |
| - | - | 1.353E+04 | 902.5 | - | - | 0 | - |
| - | - | 6.786E+04 | 903.4 | - | - | 0 | - |
| 8 | c | 8.259E+05 | 904.4 | 0.0007593 | 0.8396 | +1 | 8 |
| - | - | 4.15E+05 | 905.4 | - | - | 0 | - |
| - | - | 6457 | 905.6 | - | - | 0 | - |
| - | - | 1.184E+05 | 906.4 | - | - | 0 | - |
| - | - | 1.146E+04 | 907.5 | - | - | 0 | - |
| - | - | 4530 | 910.5 | - | - | 0 | - |
| - | - | 8440 | 913.5 | - | - | 0 | - |
| - | - | 7330 | 914.5 | - | - | 0 | - |
| - | - | 3673 | 916.5 | - | - | 0 | - |
| - | - | 8855 | 930.5 | - | - | 0 | - |
| - | - | 7877 | 931.5 | - | - | 0 | - |
| - | - | 2.108E+04 | 938.4 | - | - | 0 | - |
| 3 | y | 2.106E+04 | 939.4 | 0.002349 | 2.501 | +1 | 9 |
| 3 | y | 3.132E+04 | 940.4 | 0.00625 | 6.646 | +1 | 9 |
| 3 | z | 1.422E+04 | 941.4 | 0.0006221 | 0.6608 | +1 | 9 |
| - | - | 4153 | 942.4 | - | - | 0 | - |
| - | - | 1.063E+04 | 955.4 | - | - | 0 | - |
| - | - | 3670 | 956.3 | - | - | 0 | - |
| - | - | 1.887E+05 | 956.4 | - | - | 0 | - |
| 3 | y | 9.966E+05 | 957.5 | 0.0003917 | 0.4091 | +1 | 9 |
| - | - | 4.794E+05 | 958.5 | - | - | 0 | - |
| - | - | 1.438E+05 | 959.5 | - | - | 0 | - |
| - | - | 2.005E+04 | 960.5 | - | - | 0 | - |
| - | - | 3568 | 965.5 | - | - | 0 | - |
| - | - | 3734 | 969.5 | - | - | 0 | - |
| - | - | 2.336E+04 | 972.5 | - | - | 0 | - |
| - | - | 1.587E+05 | 973.5 | - | - | 0 | - |
| - | - | 9.134E+04 | 974.5 | - | - | 0 | - |
| - | - | 2.931E+04 | 975.5 | - | - | 0 | - |
| - | - | 3988 | 976.5 | - | - | 0 | - |
| - | - | 1.344E+04 | 982.5 | - | - | 0 | - |
| - | - | 1.126E+04 | 983.5 | - | - | 0 | - |
| - | - | 1.747E+04 | 984.5 | - | - | 0 | - |
| - | - | 1.518E+04 | 985.5 | - | - | 0 | - |
| - | - | 2.059E+04 | 999.5 | - | - | 0 | - |
| - | - | 2.046E+05 | 1000 | - | - | 0 | - |
| - | - | 1.063E+05 | 1002 | - | - | 0 | - |
| - | - | 4.089E+04 | 1003 | - | - | 0 | - |
| - | - | 3518 | 1003 | - | - | 0 | - |
| - | - | 6117 | 1004 | - | - | 0 | - |
| - | - | 4308 | 1013 | - | - | 0 | - |
| - | - | 3967 | 1016 | - | - | 0 | - |
| - | - | 1.568E+04 | 1016 | - | - | 0 | - |
| 9 | c | 6.489E+05 | 1018 | 0.0006187 | 0.608 | +1 | 9 |
| - | - | 3.684E+05 | 1019 | - | - | 0 | - |
| - | - | 1.169E+05 | 1020 | - | - | 0 | - |
| - | - | 1.48E+04 | 1021 | - | - | 0 | - |
| - | - | 4977 | 1025 | - | - | 0 | - |
| - | - | 8606 | 1032 | - | - | 0 | - |
| - | - | 4777 | 1033 | - | - | 0 | - |
| - | - | 7336 | 1043 | - | - | 0 | - |
| - | - | 1.14E+05 | 1044 | - | - | 0 | - |
| - | - | 6.224E+04 | 1045 | - | - | 0 | - |
| - | - | 3.733E+04 | 1046 | - | - | 0 | - |
| - | - | 1.447E+04 | 1047 | - | - | 0 | - |
| - | - | 5620 | 1048 | - | - | 0 | - |
| - | - | 3674 | 1060 | - | - | 0 | - |
| - | - | 7035 | 1067 | - | - | 0 | - |
| 2 | z | 1.89E+04 | 1070 | 0.01229 | 11.48 | +1 | 10 |
| - | - | 1.028E+04 | 1072 | - | - | 0 | - |
| - | - | 4209 | 1073 | - | - | 0 | - |
| - | - | 6971 | 1081 | - | - | 0 | - |
| - | - | 7599 | 1082 | - | - | 0 | - |
| - | - | 7018 | 1085 | - | - | 0 | - |
| 2 | z | 2.218E+05 | 1089 | 0.001361 | 1.25 | +1 | 10 |
| - | - | 1.357E+05 | 1090 | - | - | 0 | - |
| - | - | 5.117E+04 | 1091 | - | - | 0 | - |
| - | - | 5756 | 1092 | - | - | 0 | - |
| - | - | 9665 | 1099 | - | - | 0 | - |
| - | - | 6777 | 1100 | - | - | 0 | - |
| - | - | 6591 | 1101 | - | - | 0 | - |
| - | - | 9.882E+04 | 1102 | - | - | 0 | - |
| - | - | 5.792E+04 | 1103 | - | - | 0 | - |
| - | - | 2.89E+04 | 1104 | - | - | 0 | - |
| - | - | 7256 | 1105 | - | - | 0 | - |
| - | - | 7212 | 1110 | - | - | 0 | - |
| - | - | 5192 | 1111 | - | - | 0 | - |
| - | - | 2.597E+04 | 1113 | - | - | 0 | - |
| - | - | 2.276E+04 | 1114 | - | - | 0 | - |
| - | - | 1.139E+04 | 1115 | - | - | 0 | - |
| - | - | 4919 | 1116 | - | - | 0 | - |
| - | - | 3.809E+04 | 1118 | - | - | 0 | - |
| - | - | 2.863E+04 | 1119 | - | - | 0 | - |
| - | - | 7252 | 1120 | - | - | 0 | - |
| - | - | 3798 | 1125 | - | - | 0 | - |
| 10 | c | 2.586E+04 | 1129 | 0.00216 | 1.914 | +1 | 10 |
| - | - | 4.192E+04 | 1130 | - | - | 0 | - |
| - | - | 6.306E+04 | 1131 | - | - | 0 | - |
| - | - | 3.809E+04 | 1132 | - | - | 0 | - |
| - | - | 2.198E+04 | 1133 | - | - | 0 | - |
| - | - | 7401 | 1134 | - | - | 0 | - |
| - | - | 3378 | 1141 | - | - | 0 | - |
| - | - | 1.69E+04 | 1144 | - | - | 0 | - |
| - | - | 1.53E+04 | 1145 | - | - | 0 | - |
| 10 | c | 2.207E+06 | 1146 | 0.0005858 | 0.5113 | +1 | 10 |
| - | - | 1.553E+06 | 1147 | - | - | 0 | - |
| - | - | 6.027E+05 | 1148 | - | - | 0 | - |
| - | - | 1.149E+05 | 1149 | - | - | 0 | - |
| - | - | 1.463E+04 | 1150 | - | - | 0 | - |
| - | - | 3920 | 1151 | - | - | 0 | - |
| - | - | 1.066E+04 | 1155 | - | - | 0 | - |
| - | - | 9256 | 1156 | - | - | 0 | - |
| - | - | 3.343E+04 | 1157 | - | - | 0 | - |
| - | - | 2.663E+04 | 1158 | - | - | 0 | - |
| - | - | 9068 | 1159 | - | - | 0 | - |
| - | - | 1.763E+04 | 1160 | - | - | 0 | - |
| - | - | 1.008E+04 | 1161 | - | - | 0 | - |
| - | - | 4.663E+04 | 1163 | - | - | 0 | - |
| - | - | 2.374E+04 | 1164 | - | - | 0 | - |
| - | - | 1.144E+04 | 1165 | - | - | 0 | - |
| - | - | 1.096E+04 | 1167 | - | - | 0 | - |
| - | - | 2.842E+04 | 1168 | - | - | 0 | - |
| - | - | 1.727E+04 | 1169 | - | - | 0 | - |
| - | - | 4.589E+04 | 1173 | - | - | 0 | - |
| - | - | 1.929E+05 | 1174 | - | - | 0 | - |
| - | - | 1.379E+05 | 1175 | - | - | 0 | - |
| - | - | 4.599E+04 | 1176 | - | - | 0 | - |
| - | - | 8386 | 1177 | - | - | 0 | - |
| - | - | 2.452E+04 | 1184 | - | - | 0 | - |
| - | - | 1.498E+05 | 1185 | - | - | 0 | - |
| - | - | 1.047E+05 | 1186 | - | - | 0 | - |
| - | - | 2.508E+04 | 1187 | - | - | 0 | - |
| - | - | 6646 | 1188 | - | - | 0 | - |
| - | - | 4615 | 1190 | - | - | 0 | - |
| - | - | 6.058E+04 | 1191 | - | - | 0 | - |
| - | - | 4.269E+04 | 1192 | - | - | 0 | - |
| - | - | 1.233E+04 | 1193 | - | - | 0 | - |
| - | - | 5158 | 1199 | - | - | 0 | - |
| - | - | 9197 | 1200 | - | - | 0 | - |
| - | - | 3.803E+04 | 1201 | - | - | 0 | - |
| - | - | 3.734E+05 | 1202 | - | - | 0 | - |
| - | - | 3.918E+05 | 1203 | - | - | 0 | - |
| - | - | 1.817E+05 | 1204 | - | - | 0 | - |
| - | - | 3.335E+04 | 1205 | - | - | 0 | - |
| - | - | 8878 | 1216 | - | - | 0 | - |
| - | - | 3.818E+04 | 1217 | - | - | 0 | - |
| - | - | 3.728E+05 | 1218 | - | - | 0 | - |
| - | - | 2.025E+06 | 1219 | - | - | 0 | - |
| - | - | 1.303E+06 | 1220 | - | - | 0 | - |
| - | - | 4.732E+05 | 1221 | - | - | 0 | - |
| - | - | 7.896E+04 | 1222 | - | - | 0 | - |
| - | - | 8680 | 1251 | - | - | 0 | - |
| - | - | 8243 | 1252 | - | - | 0 | - |
| - | - | 8792 | 1830 | - | - | 0 | - |
| - | - | 3552 | 2035 | - | - | 0 | - |
| - | - | 3256 | 2311 | - | - | 0 | - |

m/z Charge Intensity FragmentType MassShift Position
120.08109283447266 0 28885.633
121.08468627929688 0 3379.8743
122.98612213134766 0 1981.6328
129.0661163330078 0 35093.77
131.1180877685547 0 12010.179
134.54293823242188 0 2164.799
147.07701110839844 0 6328.4194
148.94778442382812 0 2876.8079
149.0452423095703 0 2327.1
152.51307678222656 0 2333.3198
173.4508819580078 0 14278.984
176.66856384277344 0 2085.2144
197.12875366210938 0 6159.848
200.1031951904297 0 97069.414
201.0872802734375 0 156471.8 y Ammonia loss 9
202.0906524658203 0 12171.906
212.13967895507812 0 2761.5935
215.1392364501953 0 5790.9946
218.1138458251953 0 205372.2 y 9
219.11721801757812 0 19987.443
221.0841522216797 0 3746.1897
225.12356567382812 0 10045.783
232.09339904785156 0 4765.2026
233.16525268554688 0 232772.81
233.25416564941406 0 2399.0862
234.16864013671875 0 34470.51
242.15028381347656 0 91816.13
243.13339233398438 0 5872.6924
243.1537628173828 0 11851.79
256.1290283203125 0 3902.2722
259.9483947753906 0 2601.605
261.1600646972656 0 229390.77
262.1634216308594 0 37211.4
264.13525390625 0 3065.2283
280.6141052246094 0 2702.232
282.14544677734375 0 27274.654
299.0627136230469 0 5255.511
299.172119140625 0 5666.0396
314.13531494140625 0 13559.75
314.17169189453125 0 34703.06 y Ammonia loss 8
314.6364440917969 0 4462.8853
315.1739501953125 0 5513.4106 z 8
331.19818115234375 0 24242.805 y 8
346.1241455078125 0 3847.3767
347.67474365234375 0 8675.226
348.17626953125 0 5653.7134
351.1669006347656 0 7606.414
353.18206787109375 0 8513.855
355.0702209472656 0 37098.004
356.6800537109375 0 3608.2605
357.1806335449219 0 3338.0347
358.212890625 0 23975.564
359.2151184082031 0 4252.598
361.674560546875 0 2957.416
369.17742919921875 0 13917.933
370.1791687011719 0 3122.3384
370.6769714355469 0 129931.24
371.1785888671875 0 45375.277
371.6791076660156 0 9903.757
372.19818115234375 0 3331.387
379.1903991699219 0 30408.22 c Water loss 6
379.6916198730469 0 10741.976
386.2048034667969 0 14755.943
418.1939697265625 0 3246.9922
418.5021057128906 0 2982.8318
420.7091064453125 0 7118.9087
421.2108459472656 0 5133.9404
425.6999206542969 0 6261.6665
426.1996154785156 0 5275.0264
427.2718200683594 0 4771.6353
428.2764892578125 0 3422.6396
429.2861022949219 0 4198.311
430.21575927734375 0 3326.6653
430.71490478515625 0 4205.4395
434.70660400390625 0 117974.57
435.208251953125 0 42849.64
435.71026611328125 0 15969.232
442.2308654785156 0 3440.4622 y Water loss 7
444.2117004394531 0 49112.465
444.712646484375 0 25892.361
445.2121276855469 0 3075.2083
455.2658996582031 0 13652.236
456.2698059082031 0 5393.8354
459.2102966308594 0 3840.403
460.23944091796875 0 7997.6177 y 7
461.70947265625 0 3571.684
464.7420349121094 0 3924.4727
466.3830261230469 0 2793.841
470.2254333496094 0 23555.508 y Water loss 2
470.72247314453125 0 12172.023 y Ammonia loss 2
471.2268371582031 0 5370.473 z 2
471.285400390625 0 22634.846
472.2914733886719 0 21657.016 c 3
473.2980041503906 0 4797.655
476.7581787109375 0 3886.5547
478.2168884277344 0 2910.2554
479.23101806640625 0 13540.135 y 2
479.7326965332031 0 8510.753
480.20855712890625 0 7706.5854
488.2731018066406 0 2770.8633
498.2202453613281 0 45669.82
499.22369384765625 0 8434.522
500.2352600097656 0 4174.491
500.7539978027344 0 34998.94
501.2550964355469 0 21983.832
501.75927734375 0 3227.8804
505.2254943847656 0 3420.2263
515.2464599609375 0 24420.785
516.2473754882812 0 6755.8003
516.3189086914062 0 7614.418
520.2787475585938 0 3125.3516
530.2090454101562 0 8040.288
535.7617797851562 0 6221.2344
542.298828125 0 9816.069
543.3054809570312 0 5724.2734
544.3128662109375 0 14687.915
558.3169555664062 0 112006.82
559.3232421875 0 139848.22 c 4
560.3272705078125 0 38105.633
561.333740234375 0 7243.3574
564.7819213867188 0 7295.4634 c Ammonia loss 9
587.2703247070312 0 5570.4053
589.281982421875 0 6699.3306 y 6
591.2400512695312 0 5162.6567
592.3025512695312 0 3983.928
593.30126953125 0 3840.1562
599.268310546875 0 5127.239
603.3505859375 0 12367.384
604.3551635742188 0 3967.016
606.2761840820312 0 3570.3496
609.2527465820312 0 12839.0625
610.1849365234375 0 9623.827
611.3176879882812 0 5242.5845
627.2628173828125 0 297108.47
628.26611328125 0 94695.51
629.2672729492188 0 14866.12
629.3303833007812 0 19492.127
630.3342895507812 0 4727.199
643.2938842773438 0 14329.293 w 5
644.2894897460938 0 33401.117
644.3434448242188 0 4878.031
645.2913208007812 0 10130.1045
645.3487548828125 0 109909.34
646.3563842773438 0 478157.6 c 5
647.3594360351562 0 168473.53
648.3623657226562 0 39621.605
657.2886352539062 0 4039.3655
658.2821655273438 0 23554.918
659.2871704101562 0 13556.602 y Ammonia loss 5
660.2982788085938 0 6195.2266 z 5
661.304931640625 0 3993.4312
672.3734130859375 0 8825.102
674.2980346679688 0 4732.9756
675.308349609375 0 20827.818
676.3142700195312 0 29624.416 y 5
677.3184204101562 0 8957.876
679.3749389648438 0 2890.2935
690.3242797851562 0 3157.836
697.3846435546875 0 4002.298
700.3321533203125 0 3770.0322
704.3238525390625 0 9444.256
705.328125 0 3559.0598
712.3502197265625 0 12172.661
713.3555908203125 0 7857.9487
714.363037109375 0 4834.141
722.3377075195312 0 32618.744
723.3419799804688 0 15448.633
730.3818969726562 0 3672.3115
731.3849487304688 0 21124.818
732.3916625976562 0 16006.844
733.3931884765625 0 5281.089
739.3464965820312 0 3806.0198
740.3473510742188 0 355091.38
741.35009765625 0 136982.66
742.3524780273438 0 28498.719
744.3298950195312 0 9524.914
745.3229370117188 0 7195.7456 y Water loss 4
746.3281860351562 0 5139.091 y Ammonia loss 4
747.3273315429688 0 19606.732 z 4
748.335205078125 0 15301.155
757.3735961914062 0 55926.465 c Water loss 6
758.3737182617188 0 81219.16
759.3764038085938 0 27720.629
760.3773803710938 0 5860.236
762.3408203125 0 7478.302
763.3475952148438 0 61571.277 y 4
764.3502807617188 0 19988.033
765.3523559570312 0 4565.8853
771.3527221679688 0 16669.172
772.3563842773438 0 10042.947
773.3786010742188 0 5008.82
774.3916015625 0 119408.34
775.3991088867188 0 844435.9 c 6
776.4019165039062 0 358873.62
777.4048461914062 0 86520.74
778.4039306640625 0 5824.698
786.349853515625 0 4883.1455
787.3604736328125 0 4151.626
801.4144897460938 0 14276.66
802.4237060546875 0 4026.0854
826.4293212890625 0 7250.387
827.4375610351562 0 3094.3103
842.41943359375 0 7985.968
844.415283203125 0 3436.7007
850.3965454101562 0 10400.112
851.3922119140625 0 9157.978
853.4329833984375 0 13321.396
854.4349975585938 0 6917.6455
855.4315185546875 0 4058.1677
859.4140014648438 0 10143.804
860.4030151367188 0 217731.44 y 3
861.4071655273438 0 100287.96
862.4096069335938 0 27775.861
867.398681640625 0 13290.859
868.4054565429688 0 166914.95
869.4076538085938 0 84718.63
870.4086303710938 0 27165.693
871.4010620117188 0 21536.574
872.4010009765625 0 14057.501
873.4032592773438 0 5025.8184
884.4254760742188 0 13430.091
885.4149169921875 0 31349.994
886.4172973632812 0 11947.803 c Water loss 7
887.4154052734375 0 326122.7
888.4183349609375 0 148311.69
889.4209594726562 0 43308.098
890.4209594726562 0 4840.422
894.447021484375 0 4379.954
895.4493408203125 0 3455.6108
901.4921264648438 0 34834.594
902.4943237304688 0 13531.401
903.43505859375 0 67864.016
904.4418334960938 0 825906.94 c 7
905.444580078125 0 415016.28
905.5504150390625 0 6457
906.4474487304688 0 118395.03
907.452392578125 0 11456.232
910.4647827148438 0 4530.3115
913.4600219726562 0 8440.366
914.4675903320312 0 7330.076
916.5109252929688 0 3673.2341
930.4615478515625 0 8855.017
931.4656372070312 0 7877.1846
938.4358520507812 0 21081.777
939.439453125 0 21059.604 y Water loss 2
940.4320678710938 0 31317.068 y Ammonia loss 2
941.4342651367188 0 14224.915 z 2
942.4381103515625 0 4153.299
955.4409790039062 0 10633.295
956.2787475585938 0 3670.3643
956.4451904296875 0 188729.17
957.4527587890625 0 996629.44 y 2
958.4557495117188 0 479363.25
959.4592895507812 0 143803.73
960.463623046875 0 20047.572
965.4703369140625 0 3567.5427
969.4664306640625 0 3733.954
972.5046997070312 0 23358.928
973.511962890625 0 158697.36
974.51513671875 0 91343.85
975.5211181640625 0 29307.273
976.5203857421875 0 3987.9294
982.4881591796875 0 13439.765
983.4844360351562 0 11258.014
984.4764404296875 0 17473.621
985.4784545898438 0 15184.161
999.4561157226562 0 20587.88
1000.4983520507812 0 204606.19
1001.5015869140625 0 106270.96
1002.5047607421875 0 40890.535
1002.7713623046875 0 3518.3286
1003.522216796875 0 6116.7446
1012.533203125 0 4308.1934
1015.500732421875 0 3966.7678
1016.49658203125 0 15681.429
1017.5257568359375 0 648871.9 c 8
1018.5287475585938 0 368363.1
1019.5314331054688 0 116916.18
1020.5343017578125 0 14797.486
1024.535888671875 0 4976.9404
1031.510009765625 0 8606.479
1032.5308837890625 0 4776.8125
1042.5389404296875 0 7335.8496
1043.5416259765625 0 113995.36
1044.54443359375 0 62237.156
1045.5250244140625 0 37327.13
1046.5189208984375 0 14471.924
1047.5172119140625 0 5619.7544
1059.5606689453125 0 3674.349
1066.514404296875 0 7035.432
1070.5037841796875 0 18896.568 z Water loss 1
1071.5113525390625 0 10276.568
1072.5008544921875 0 4209.0054
1080.544189453125 0 6971.269
1081.5494384765625 0 7599.447
1084.5313720703125 0 7017.5654
1088.50341796875 0 221830.16 z 1
1089.50634765625 0 135671.34
1090.508544921875 0 51168.1
1091.513427734375 0 5755.7114
1098.5528564453125 0 9665.248
1099.5574951171875 0 6776.759
1100.5693359375 0 6591.4136
1101.5709228515625 0 98819.8
1102.5736083984375 0 57921.97
1103.576904296875 0 28903.695
1104.589599609375 0 7255.555
1109.5457763671875 0 7212.0117
1110.5419921875 0 5191.7354
1112.539306640625 0 25967.986
1113.547119140625 0 22759.428
1114.5501708984375 0 11392.735
1115.5555419921875 0 4919.433
1117.53125 0 38090.6
1118.5323486328125 0 28632.121
1119.540283203125 0 7252.2695
1124.5584716796875 0 3797.97
1128.559326171875 0 25863.83 c Ammonia loss 9
1129.5640869140625 0 41920.953
1130.5576171875 0 63064.777
1131.5611572265625 0 38092.688
1132.57666015625 0 21979.738
1133.581787109375 0 7401.2983
1140.5841064453125 0 3378.2205
1143.566650390625 0 16899.889
1144.5697021484375 0 15302.13
1145.5831298828125 0 2206629.2 c 9
1146.5863037109375 0 1552608
1147.58837890625 0 602707.94
1148.5899658203125 0 114853.42
1149.5838623046875 0 14630.331
1150.5732421875 0 3919.9316
1154.5916748046875 0 10657.111
1155.5950927734375 0 9256.4375
1156.6104736328125 0 33429.062
1157.6134033203125 0 26627.238
1158.530029296875 0 9067.893
1159.591796875 0 17630.78
1160.591064453125 0 10075.849
1162.551025390625 0 46633.22
1163.552978515625 0 23744.398
1164.5518798828125 0 11435.785
1166.594482421875 0 10958.036
1167.5814208984375 0 28419.066
1168.5806884765625 0 17267.102
1172.6075439453125 0 45889.66
1173.5947265625 0 192938.44
1174.596923828125 0 137921.39
1175.59912109375 0 45989.08
1176.6002197265625 0 8386.113
1183.5751953125 0 24516.236
1184.6015625 0 149786.39
1185.6004638671875 0 104674.83
1186.5982666015625 0 25078.945
1187.5947265625 0 6646.2104
1189.597900390625 0 4615.2324
1190.6171875 0 60576.83
1191.621337890625 0 42689.1
1192.6220703125 0 12333.695
1198.5771484375 0 5158.403
1199.577392578125 0 9196.738
1200.59912109375 0 38031.59
1201.5899658203125 0 373392.75
1202.600830078125 0 391825.1
1203.60546875 0 181748.27
1204.6083984375 0 33345.652
1215.58740234375 0 8877.706
1216.59521484375 0 38182.707
1217.6043701171875 0 372789.62
1218.6123046875 0 2025071.2
1219.614990234375 0 1302694.6
1220.6182861328125 0 473225.47
1221.6190185546875 0 78963.97
1250.6019287109375 0 8679.762
1251.60400390625 0 8243.232
1830.033447265625 0 8791.875
2034.5167236328125 0 3552.0107
2310.708251953125 0 3256.3765

Spectrum Details

|  |  |
| --- | --- |
| Matched peaks? Matched peaks The total absolute number of peaks matched. Additionally in brackets the total fraction of peaks matched and the total number of peaks is shown. | 39 (10.32% of 378) |
| FDR? FDR The false discovery rate estimated for this peptide. It is calculated by matching all theoretical fragments with a non-integer shift with the raw peaks for this spectrum. This is done with 40 different shifts. The resulting percentage is the average number of annotated peaks over the number of annotated peaks with the correct spectrum. | 0.98% |
| Satellite FDR? Satellite FDR See the FDR for details on its calculation. This satellite ion specific FDR only contains the satellite ions (d/w) for I/L/J positions. | - |
| PSM Score? PSM Score The PSM Score as given by Hecklib to this annotated spectrum. It is shown with three significant figures. | 447 |

#### Spectrum 8007? Spectrum 8007 The raw spectrum of this peptide as annotated by Hecklib. The fragments are coloured according to ion type (see legend). Any peaks with a star '\*' as text can be hovered over to see the full details, first the ion type second the mass shift type. By hovering over the amino acids in the peptide or ions in the legend the corresponding peaks are highlighted. By toggling the 'Unassigned' label you can turn the background (unassigned) peaks on or off in the plot. By updating the slider in the Ion legend you can update the spectrum to only show the top X% of the peaks with labels. The top X% means any peak that is within X% of the highest intensity. By dragging in the spectrum you can zoom in to a specific part of the spectrum and use 'Zoom Out' to get back to the original zoom level. The annotation of the spectrum is based on the given sequence in the peptides file and is done with different software so inconsistencies are likely. The peaks are annotated based on the given sequence, with 20 ppm tolerance.

Copy Data

##### Spectrum 8007 (TSV)

###### Preview

```
Loading example...
```

*Click on the button to copy the data to your clipboard.*

Mz MinMz MaxIntensity Max

WidthHeightPeptide font sizePeptide stroke widthSpectrum font sizeSpectrum stroke widthCompact peptide

Ion legend

wxyz

abcd

OtherUnassignedIon^Charge^_Position_Show for top:%

JFPPSSEEJQA

03.55e+57.11e+51.07e+61.42e+6

Zoom Out

y^+1^_2_z^+1^_2_y^+1^_2_y^+1^_3_z^+1^_3_y^+1^_3_c^+1^_3_c^+2^_7_y^+1^_4_y^+1^_4_y^+1^_4_y^+2^_9_y^+2^_9_z^+2^_9_c^+1^_4_y^+2^_9_c^+1^_5_y^+1^_5_w^+1^_6_c^+1^_6_y^+1^_6_z^+1^_6_y^+1^_6_z^+1^_7_c^+1^_7_y^+1^_7_c^+1^_7_y^+1^_8_y^+1^_8_c^+1^_8_c^+1^_8_y^+1^_9_y^+1^_9_z^+1^_9_y^+1^_9_c^+1^_9_z^+1^_10_z^+1^_10_c^+1^_10_c^+1^_10_

0783156623503133

Fragment Matches Table

Show background peaks

| Position | Ion type | Intensity | mz Theoretical | mz Error (Th) | mz Error (ppm) | Charge | Series Number |
| --- | --- | --- | --- | --- | --- | --- | --- |
| - | - | 1342 | 120.1 | - | - | 0 | - |
| - | - | 1.61E+04 | 120.1 | - | - | 0 | - |
| - | - | 1.86E+04 | 129.1 | - | - | 0 | - |
| - | - | 4955 | 131.1 | - | - | 0 | - |
| - | - | 1709 | 142.3 | - | - | 0 | - |
| - | - | 4644 | 147.1 | - | - | 0 | - |
| - | - | 1642 | 147.5 | - | - | 0 | - |
| - | - | 1695 | 148.5 | - | - | 0 | - |
| - | - | 6547 | 173.5 | - | - | 0 | - |
| - | - | 1979 | 186.3 | - | - | 0 | - |
| - | - | 5.607E+04 | 200.1 | - | - | 0 | - |
| 10 | y | 8.642E+04 | 201.1 | 0.0002969 | 1.477 | +1 | 2 |
| - | - | 4214 | 201.1 | - | - | 0 | - |
| 10 | z | 7738 | 202.1 | 0.003927 | 19.43 | +1 | 2 |
| - | - | 2690 | 209 | - | - | 0 | - |
| 10 | y | 1.188E+05 | 218.1 | 0.0003439 | 1.577 | +1 | 2 |
| - | - | 9733 | 219.1 | - | - | 0 | - |
| - | - | 4563 | 221.1 | - | - | 0 | - |
| - | - | 7232 | 225.1 | - | - | 0 | - |
| - | - | 1.334E+05 | 233.2 | - | - | 0 | - |
| - | - | 1.957E+04 | 234.2 | - | - | 0 | - |
| - | - | 6753 | 239.1 | - | - | 0 | - |
| - | - | 5.591E+04 | 242.2 | - | - | 0 | - |
| - | - | 2443 | 243.1 | - | - | 0 | - |
| - | - | 6707 | 243.2 | - | - | 0 | - |
| - | - | 1.272E+05 | 261.2 | - | - | 0 | - |
| - | - | 1918 | 261.6 | - | - | 0 | - |
| - | - | 1.689E+04 | 262.2 | - | - | 0 | - |
| - | - | 2664 | 264.1 | - | - | 0 | - |
| - | - | 2133 | 281.1 | - | - | 0 | - |
| - | - | 2400 | 281.4 | - | - | 0 | - |
| - | - | 1.588E+04 | 282.1 | - | - | 0 | - |
| - | - | 2903 | 283.1 | - | - | 0 | - |
| - | - | 6858 | 299.1 | - | - | 0 | - |
| - | - | 4545 | 299.2 | - | - | 0 | - |
| - | - | 7238 | 314.1 | - | - | 0 | - |
| 9 | y | 1.557E+04 | 314.2 | 0.000431 | 1.372 | +1 | 3 |
| - | - | 3204 | 314.6 | - | - | 0 | - |
| 9 | z | 4294 | 315.2 | 0.0048 | 15.23 | +1 | 3 |
| - | - | 2340 | 323.9 | - | - | 0 | - |
| 9 | y | 1.517E+04 | 331.2 | 0.0002796 | 0.8441 | +1 | 3 |
| - | - | 2119 | 347.7 | - | - | 0 | - |
| - | - | 2673 | 351.2 | - | - | 0 | - |
| - | - | 5139 | 353.2 | - | - | 0 | - |
| - | - | 4.916E+04 | 355.1 | - | - | 0 | - |
| - | - | 3326 | 356.7 | - | - | 0 | - |
| - | - | 2456 | 357.2 | - | - | 0 | - |
| - | - | 1.316E+04 | 358.2 | - | - | 0 | - |
| - | - | 3770 | 361.7 | - | - | 0 | - |
| - | - | 8873 | 369.2 | - | - | 0 | - |
| - | - | 7.769E+04 | 370.7 | - | - | 0 | - |
| - | - | 2675 | 371.1 | - | - | 0 | - |
| - | - | 3.235E+04 | 371.2 | - | - | 0 | - |
| - | - | 7546 | 371.7 | - | - | 0 | - |
| 3 | c | 2340 | 375.2 | 0.004402 | 11.73 | +1 | 3 |
| 7 | c | 1.996E+04 | 379.2 | 0.007197 | 18.98 | +2 | 7 |
| - | - | 4778 | 379.7 | - | - | 0 | - |
| - | - | 2306 | 380.2 | - | - | 0 | - |
| - | - | 9302 | 386.2 | - | - | 0 | - |
| - | - | 3406 | 415 | - | - | 0 | - |
| - | - | 3964 | 421.2 | - | - | 0 | - |
| - | - | 4179 | 425.7 | - | - | 0 | - |
| - | - | 3833 | 426.2 | - | - | 0 | - |
| - | - | 2302 | 427.3 | - | - | 0 | - |
| - | - | 4696 | 430.2 | - | - | 0 | - |
| - | - | 5.592E+04 | 434.7 | - | - | 0 | - |
| - | - | 2.966E+04 | 435.2 | - | - | 0 | - |
| - | - | 5710 | 435.7 | - | - | 0 | - |
| - | - | 2285 | 436.2 | - | - | 0 | - |
| 8 | y | 2697 | 442.2 | 0.00121 | 2.736 | +1 | 4 |
| 8 | y | 3015 | 443.2 | 0.0009287 | 2.095 | +1 | 4 |
| - | - | 3.112E+04 | 444.2 | - | - | 0 | - |
| - | - | 1.086E+04 | 444.7 | - | - | 0 | - |
| - | - | 5459 | 445.2 | - | - | 0 | - |
| - | - | 2284 | 447.7 | - | - | 0 | - |
| - | - | 7826 | 455.3 | - | - | 0 | - |
| - | - | 3145 | 459.2 | - | - | 0 | - |
| 8 | y | 4205 | 460.2 | 0.0002298 | 0.4993 | +1 | 4 |
| 3 | y | 1.57E+04 | 470.2 | 0.0005887 | 1.252 | +2 | 9 |
| 3 | y | 1.024E+04 | 470.7 | 0.007238 | 15.38 | +2 | 9 |
| 3 | z | 3135 | 471.2 | 0.005462 | 11.59 | +2 | 9 |
| - | - | 1.326E+04 | 471.3 | - | - | 0 | - |
| 4 | c | 1.53E+04 | 472.3 | 0.0006019 | 1.274 | +1 | 4 |
| - | - | 4595 | 473.3 | - | - | 0 | - |
| 3 | y | 1.107E+04 | 479.2 | 0.0008606 | 1.796 | +2 | 9 |
| - | - | 5060 | 479.7 | - | - | 0 | - |
| - | - | 3184 | 480.2 | - | - | 0 | - |
| - | - | 3048 | 489.1 | - | - | 0 | - |
| - | - | 2.847E+04 | 498.2 | - | - | 0 | - |
| - | - | 6033 | 499.2 | - | - | 0 | - |
| - | - | 1.853E+04 | 500.8 | - | - | 0 | - |
| - | - | 6363 | 501.3 | - | - | 0 | - |
| - | - | 7921 | 501.8 | - | - | 0 | - |
| - | - | 2.116E+04 | 515.2 | - | - | 0 | - |
| - | - | 2793 | 516.2 | - | - | 0 | - |
| - | - | 5540 | 516.3 | - | - | 0 | - |
| - | - | 5247 | 542.3 | - | - | 0 | - |
| - | - | 8588 | 544.3 | - | - | 0 | - |
| - | - | 2816 | 545.3 | - | - | 0 | - |
| - | - | 7.676E+04 | 558.3 | - | - | 0 | - |
| 5 | c | 8.712E+04 | 559.3 | 0.0008005 | 1.431 | +1 | 5 |
| - | - | 2.347E+04 | 560.3 | - | - | 0 | - |
| - | - | 2982 | 561.3 | - | - | 0 | - |
| 7 | y | 2761 | 589.3 | 0.002692 | 4.569 | +1 | 5 |
| - | - | 2819 | 590.3 | - | - | 0 | - |
| - | - | 3522 | 591.2 | - | - | 0 | - |
| - | - | 6785 | 603.4 | - | - | 0 | - |
| - | - | 3793 | 604.4 | - | - | 0 | - |
| - | - | 2.559E+04 | 608.3 | - | - | 0 | - |
| - | - | 6632 | 609.3 | - | - | 0 | - |
| - | - | 3757 | 609.3 | - | - | 0 | - |
| - | - | 1.441E+04 | 610.2 | - | - | 0 | - |
| - | - | 2818 | 610.3 | - | - | 0 | - |
| - | - | 1.662E+05 | 627.3 | - | - | 0 | - |
| - | - | 4.865E+04 | 628.3 | - | - | 0 | - |
| - | - | 7233 | 629.3 | - | - | 0 | - |
| - | - | 8407 | 629.3 | - | - | 0 | - |
| - | - | 3377 | 630.3 | - | - | 0 | - |
| 6 | w | 6242 | 643.3 | 0.0005617 | 0.8731 | +1 | 6 |
| - | - | 1.966E+04 | 644.3 | - | - | 0 | - |
| - | - | 6403 | 645.3 | - | - | 0 | - |
| - | - | 6.733E+04 | 645.3 | - | - | 0 | - |
| 6 | c | 3.192E+05 | 646.4 | 0.0001911 | 0.2957 | +1 | 6 |
| - | - | 1.142E+05 | 647.4 | - | - | 0 | - |
| - | - | 2.002E+04 | 648.4 | - | - | 0 | - |
| - | - | 3608 | 657.3 | - | - | 0 | - |
| - | - | 1.511E+04 | 658.3 | - | - | 0 | - |
| 6 | y | 6471 | 659.3 | 0.00335 | 5.081 | +1 | 6 |
| - | - | 2376 | 659.5 | - | - | 0 | - |
| 6 | z | 3122 | 660.3 | 0.004278 | 6.479 | +1 | 6 |
| - | - | 2608 | 661.3 | - | - | 0 | - |
| - | - | 3801 | 672.4 | - | - | 0 | - |
| - | - | 2921 | 673.4 | - | - | 0 | - |
| - | - | 2864 | 674.3 | - | - | 0 | - |
| - | - | 1.283E+04 | 675.3 | - | - | 0 | - |
| 6 | y | 1.503E+04 | 676.3 | 0.001212 | 1.793 | +1 | 6 |
| - | - | 3139 | 677.3 | - | - | 0 | - |
| - | - | 2788 | 704.3 | - | - | 0 | - |
| - | - | 8589 | 712.4 | - | - | 0 | - |
| - | - | 2941 | 713.4 | - | - | 0 | - |
| - | - | 3540 | 714.4 | - | - | 0 | - |
| - | - | 2.295E+04 | 722.3 | - | - | 0 | - |
| - | - | 6467 | 723.3 | - | - | 0 | - |
| - | - | 1.318E+04 | 731.4 | - | - | 0 | - |
| - | - | 1.211E+04 | 732.4 | - | - | 0 | - |
| - | - | 1.876E+05 | 740.3 | - | - | 0 | - |
| - | - | 7.082E+04 | 741.3 | - | - | 0 | - |
| - | - | 1.717E+04 | 742.4 | - | - | 0 | - |
| - | - | 4804 | 744.3 | - | - | 0 | - |
| - | - | 2743 | 745.3 | - | - | 0 | - |
| 5 | z | 8582 | 747.3 | 0.001047 | 1.401 | +1 | 7 |
| - | - | 1.314E+04 | 748.3 | - | - | 0 | - |
| 7 | c | 2.784E+04 | 757.4 | 0.01487 | 19.63 | +1 | 7 |
| - | - | 5.138E+04 | 758.4 | - | - | 0 | - |
| - | - | 2.026E+04 | 759.4 | - | - | 0 | - |
| - | - | 6281 | 762.3 | - | - | 0 | - |
| 5 | y | 4.09E+04 | 763.3 | 0.0005116 | 0.6703 | +1 | 7 |
| - | - | 1.555E+04 | 764.4 | - | - | 0 | - |
| - | - | 2804 | 765.4 | - | - | 0 | - |
| - | - | 8367 | 771.4 | - | - | 0 | - |
| - | - | 4407 | 772.4 | - | - | 0 | - |
| - | - | 3355 | 773.4 | - | - | 0 | - |
| - | - | 8.466E+04 | 774.4 | - | - | 0 | - |
| 7 | c | 5.341E+05 | 775.4 | 0.0001046 | 0.1349 | +1 | 7 |
| - | - | 2.223E+05 | 776.4 | - | - | 0 | - |
| - | - | 4.936E+04 | 777.4 | - | - | 0 | - |
| - | - | 4155 | 778.4 | - | - | 0 | - |
| - | - | 2484 | 796.8 | - | - | 0 | - |
| - | - | 4311 | 801.4 | - | - | 0 | - |
| - | - | 2356 | 802.4 | - | - | 0 | - |
| - | - | 4671 | 840.4 | - | - | 0 | - |
| - | - | 4001 | 842.4 | - | - | 0 | - |
| 4 | y | 5382 | 843.4 | 0.002068 | 2.452 | +1 | 8 |
| - | - | 6300 | 850.4 | - | - | 0 | - |
| - | - | 5125 | 851.4 | - | - | 0 | - |
| - | - | 3497 | 852.4 | - | - | 0 | - |
| - | - | 9911 | 853.4 | - | - | 0 | - |
| - | - | 5209 | 854.4 | - | - | 0 | - |
| - | - | 3080 | 855.4 | - | - | 0 | - |
| - | - | 7291 | 859.4 | - | - | 0 | - |
| 4 | y | 1.336E+05 | 860.4 | 0.002679 | 3.114 | +1 | 8 |
| - | - | 5.334E+04 | 861.4 | - | - | 0 | - |
| - | - | 1.531E+04 | 862.4 | - | - | 0 | - |
| - | - | 7857 | 867.4 | - | - | 0 | - |
| - | - | 9.193E+04 | 868.4 | - | - | 0 | - |
| - | - | 4.587E+04 | 869.4 | - | - | 0 | - |
| - | - | 1.513E+04 | 870.4 | - | - | 0 | - |
| - | - | 1.561E+04 | 871.4 | - | - | 0 | - |
| - | - | 6507 | 872.4 | - | - | 0 | - |
| - | - | 1.121E+04 | 884.4 | - | - | 0 | - |
| - | - | 1.916E+04 | 885.4 | - | - | 0 | - |
| 8 | c | 6345 | 886.4 | 0.01498 | 16.9 | +1 | 8 |
| - | - | 1.754E+05 | 887.4 | - | - | 0 | - |
| - | - | 8.617E+04 | 888.4 | - | - | 0 | - |
| - | - | 2.65E+04 | 889.4 | - | - | 0 | - |
| - | - | 1.791E+04 | 901.5 | - | - | 0 | - |
| - | - | 9932 | 902.5 | - | - | 0 | - |
| - | - | 4.201E+04 | 903.4 | - | - | 0 | - |
| 8 | c | 5.26E+05 | 904.4 | 2.69E-05 | 0.02975 | +1 | 8 |
| - | - | 2.719E+05 | 905.4 | - | - | 0 | - |
| - | - | 7.921E+04 | 906.4 | - | - | 0 | - |
| - | - | 7596 | 907.5 | - | - | 0 | - |
| - | - | 6384 | 930.5 | - | - | 0 | - |
| - | - | 4314 | 931.5 | - | - | 0 | - |
| - | - | 1.453E+04 | 938.4 | - | - | 0 | - |
| 3 | y | 1.41E+04 | 939.4 | 0.0009455 | 1.006 | +1 | 9 |
| 3 | y | 2.05E+04 | 940.4 | 0.005701 | 6.062 | +1 | 9 |
| 3 | z | 9777 | 941.4 | 0.003711 | 3.942 | +1 | 9 |
| - | - | 3415 | 955.4 | - | - | 0 | - |
| - | - | 1.212E+05 | 956.4 | - | - | 0 | - |
| 3 | y | 6.122E+05 | 957.5 | 0.0004018 | 0.4197 | +1 | 9 |
| - | - | 2.858E+05 | 958.5 | - | - | 0 | - |
| - | - | 7.695E+04 | 959.5 | - | - | 0 | - |
| - | - | 8241 | 960.5 | - | - | 0 | - |
| - | - | 1.253E+04 | 972.5 | - | - | 0 | - |
| - | - | 9.469E+04 | 973.5 | - | - | 0 | - |
| - | - | 5.917E+04 | 974.5 | - | - | 0 | - |
| - | - | 2.366E+04 | 975.5 | - | - | 0 | - |
| - | - | 3107 | 976.5 | - | - | 0 | - |
| - | - | 8758 | 982.5 | - | - | 0 | - |
| - | - | 8904 | 984.5 | - | - | 0 | - |
| - | - | 1.039E+04 | 985.5 | - | - | 0 | - |
| - | - | 4228 | 986.5 | - | - | 0 | - |
| - | - | 8148 | 999.5 | - | - | 0 | - |
| - | - | 1.143E+05 | 1000 | - | - | 0 | - |
| - | - | 6.971E+04 | 1002 | - | - | 0 | - |
| - | - | 2.05E+04 | 1003 | - | - | 0 | - |
| - | - | 2752 | 1004 | - | - | 0 | - |
| - | - | 5528 | 1016 | - | - | 0 | - |
| - | - | 1.097E+04 | 1016 | - | - | 0 | - |
| 9 | c | 4E+05 | 1018 | 0.0003579 | 0.3517 | +1 | 9 |
| - | - | 2.359E+05 | 1019 | - | - | 0 | - |
| - | - | 4686 | 1019 | - | - | 0 | - |
| - | - | 7.208E+04 | 1020 | - | - | 0 | - |
| - | - | 8647 | 1021 | - | - | 0 | - |
| - | - | 6.154E+04 | 1044 | - | - | 0 | - |
| - | - | 3.772E+04 | 1045 | - | - | 0 | - |
| - | - | 2.322E+04 | 1046 | - | - | 0 | - |
| - | - | 9804 | 1047 | - | - | 0 | - |
| - | - | 3230 | 1069 | - | - | 0 | - |
| 2 | z | 4832 | 1070 | 0.004432 | 4.14 | +1 | 10 |
| - | - | 3585 | 1071 | - | - | 0 | - |
| 2 | z | 1.496E+05 | 1089 | 0.0001039 | 0.09541 | +1 | 10 |
| - | - | 9.292E+04 | 1090 | - | - | 0 | - |
| - | - | 2.788E+04 | 1091 | - | - | 0 | - |
| - | - | 3517 | 1092 | - | - | 0 | - |
| - | - | 3809 | 1101 | - | - | 0 | - |
| - | - | 6.721E+04 | 1102 | - | - | 0 | - |
| - | - | 3.996E+04 | 1103 | - | - | 0 | - |
| - | - | 1.831E+04 | 1104 | - | - | 0 | - |
| - | - | 2890 | 1105 | - | - | 0 | - |
| - | - | 4395 | 1110 | - | - | 0 | - |
| - | - | 3783 | 1112 | - | - | 0 | - |
| - | - | 1.627E+04 | 1113 | - | - | 0 | - |
| - | - | 8683 | 1114 | - | - | 0 | - |
| - | - | 5814 | 1115 | - | - | 0 | - |
| - | - | 2.473E+04 | 1118 | - | - | 0 | - |
| - | - | 1.661E+04 | 1119 | - | - | 0 | - |
| - | - | 5610 | 1120 | - | - | 0 | - |
| 10 | c | 1.289E+04 | 1129 | 0.003136 | 2.779 | +1 | 10 |
| - | - | 2.626E+04 | 1130 | - | - | 0 | - |
| - | - | 4.267E+04 | 1131 | - | - | 0 | - |
| - | - | 2.364E+04 | 1132 | - | - | 0 | - |
| - | - | 1.312E+04 | 1133 | - | - | 0 | - |
| - | - | 8798 | 1134 | - | - | 0 | - |
| - | - | 7849 | 1144 | - | - | 0 | - |
| - | - | 9553 | 1145 | - | - | 0 | - |
| 10 | c | 1.408E+06 | 1146 | 0.002051 | 1.79 | +1 | 10 |
| - | - | 9.494E+05 | 1147 | - | - | 0 | - |
| - | - | 3.712E+05 | 1148 | - | - | 0 | - |
| - | - | 7.028E+04 | 1149 | - | - | 0 | - |
| - | - | 4172 | 1150 | - | - | 0 | - |
| - | - | 2.22E+04 | 1157 | - | - | 0 | - |
| - | - | 1.112E+04 | 1158 | - | - | 0 | - |
| - | - | 7979 | 1159 | - | - | 0 | - |
| - | - | 1.134E+04 | 1160 | - | - | 0 | - |
| - | - | 6434 | 1161 | - | - | 0 | - |
| - | - | 2.256E+04 | 1163 | - | - | 0 | - |
| - | - | 1.66E+04 | 1164 | - | - | 0 | - |
| - | - | 8191 | 1165 | - | - | 0 | - |
| - | - | 3.145E+04 | 1173 | - | - | 0 | - |
| - | - | 1.187E+05 | 1174 | - | - | 0 | - |
| - | - | 8.042E+04 | 1175 | - | - | 0 | - |
| - | - | 2.765E+04 | 1176 | - | - | 0 | - |
| - | - | 5713 | 1177 | - | - | 0 | - |
| - | - | 1.595E+04 | 1184 | - | - | 0 | - |
| - | - | 1.413E+04 | 1185 | - | - | 0 | - |
| - | - | 6117 | 1186 | - | - | 0 | - |
| - | - | 3445 | 1190 | - | - | 0 | - |
| - | - | 3.25E+04 | 1191 | - | - | 0 | - |
| - | - | 2.745E+04 | 1192 | - | - | 0 | - |
| - | - | 1.114E+04 | 1193 | - | - | 0 | - |
| - | - | 2.419E+04 | 1201 | - | - | 0 | - |
| - | - | 1.87E+05 | 1202 | - | - | 0 | - |
| - | - | 1.366E+05 | 1203 | - | - | 0 | - |
| - | - | 5.644E+04 | 1204 | - | - | 0 | - |
| - | - | 8235 | 1205 | - | - | 0 | - |
| - | - | 3793 | 1216 | - | - | 0 | - |
| - | - | 2.002E+04 | 1217 | - | - | 0 | - |
| - | - | 2.254E+05 | 1218 | - | - | 0 | - |
| - | - | 1.254E+06 | 1219 | - | - | 0 | - |
| - | - | 8.235E+05 | 1220 | - | - | 0 | - |
| - | - | 2.947E+05 | 1221 | - | - | 0 | - |
| - | - | 4.595E+04 | 1222 | - | - | 0 | - |
| - | - | 3107 | 1251 | - | - | 0 | - |
| - | - | 4400 | 1830 | - | - | 0 | - |
| - | - | 3154 | 1831 | - | - | 0 | - |
| - | - | 2708 | 2831 | - | - | 0 | - |
| - | - | 3041 | 3002 | - | - | 0 | - |
| - | - | 2869 | 3040 | - | - | 0 | - |
| - | - | 2448 | 3102 | - | - | 0 | - |

m/z Charge Intensity FragmentType MassShift Position
120.0775375366211 0 1341.9307
120.08109283447266 0 16095.596
129.06613159179688 0 18595.287
131.11810302734375 0 4954.549
142.2928009033203 0 1709.2384
147.07679748535156 0 4643.8477
147.49334716796875 0 1642.4292
148.5498046875 0 1695.0135
173.4516143798828 0 6547.1343
186.28689575195312 0 1979.0419
200.10321044921875 0 56069.582
201.0872802734375 0 86418.14 y Ammonia loss 9
201.10513305664062 0 4214.2476
202.09088134765625 0 7737.904 z 9
208.9923095703125 0 2690.0547
218.11387634277344 0 118821.06 y 9
219.1174774169922 0 9733.355
221.08445739746094 0 4562.594
225.12355041503906 0 7231.6587
233.1652069091797 0 133406.44
234.1685333251953 0 19572.957
239.09559631347656 0 6752.6885
242.15025329589844 0 55910.043
243.1338653564453 0 2442.5103
243.15379333496094 0 6707.404
261.15997314453125 0 127247.12
261.5692443847656 0 1918.342
262.1629333496094 0 16894.172
264.1347961425781 0 2664.0266
281.05084228515625 0 2133.1494
281.3722839355469 0 2399.7124
282.145263671875 0 15881.068
283.14715576171875 0 2902.6267
299.0627746582031 0 6858.3687
299.1715087890625 0 4544.678
314.1357116699219 0 7237.7075
314.1714782714844 0 15574.868 y Ammonia loss 8
314.6372375488281 0 3204.41
315.174072265625 0 4293.769 z 8
323.9090881347656 0 2339.753
331.1978759765625 0 15170.778 y 8
347.674560546875 0 2118.884
351.1663818359375 0 2672.657
353.1818542480469 0 5139.246
355.0703125 0 49162.33
356.6795349121094 0 3326.0408
357.18182373046875 0 2456.2515
358.2130126953125 0 13155.475
361.6720275878906 0 3769.8984
369.1773376464844 0 8872.687
370.67694091796875 0 77685.664
371.1026306152344 0 2675.2803
371.1786804199219 0 32347.826
371.6795959472656 0 7546.1787
375.24346923828125 0 2339.8179 c 2
379.1903991699219 0 19960.428 c Water loss 6
379.6910095214844 0 4777.802
380.1917724609375 0 2306.2017
386.2039489746094 0 9302.306
415.0375671386719 0 3405.5425
421.2108154296875 0 3963.6091
425.7003479003906 0 4179.4365
426.2021484375 0 3833.1758
427.2704162597656 0 2301.5254
430.21380615234375 0 4695.5337
434.70648193359375 0 55924.145
435.2077941894531 0 29660.416
435.70849609375 0 5709.955
436.2121276855469 0 2284.54
442.2308349609375 0 2697.182 y Water loss 7
443.2145690917969 0 3015.083 y Ammonia loss 7
444.2115478515625 0 31115.654
444.7125244140625 0 10856.446
445.2149658203125 0 5459.0767
447.7146911621094 0 2283.9902
455.266845703125 0 7826.0723
459.20831298828125 0 3145.2087
460.2399597167969 0 4205.153 y 7
470.2251281738281 0 15700.594 y Water loss 2
470.7237854003906 0 10235.612 y Ammonia loss 2
471.2259216308594 0 3134.998 z 2
471.2846984863281 0 13256.441
472.2912292480469 0 15299.996 c 3
473.29669189453125 0 4595.4204
479.2306823730469 0 11067.533 y 2
479.7314147949219 0 5060.1226
480.20745849609375 0 3184.5
489.0562744140625 0 3048.436
498.2196044921875 0 28466.504
499.2243347167969 0 6033.417
500.75390625 0 18531.738
501.25494384765625 0 6362.69
501.75604248046875 0 7920.581
515.24658203125 0 21159.693
516.2496948242188 0 2793.3037
516.317138671875 0 5539.6274
542.299560546875 0 5247.3276
544.3138427734375 0 8588.248
545.3107299804688 0 2816.2532
558.3165283203125 0 76756.555
559.3230590820312 0 87117.7 c 4
560.3259887695312 0 23468.236
561.3265380859375 0 2982.099
589.2800903320312 0 2760.6206 y 6
590.2918701171875 0 2818.6226
591.2430419921875 0 3521.6611
603.350830078125 0 6784.551
604.3526611328125 0 3792.6807
608.3079223632812 0 25592.34
609.2523193359375 0 6632.4644
609.3082275390625 0 3757.1106
610.1849975585938 0 14412.576
610.2536010742188 0 2818.0479
627.2625122070312 0 166191.08
628.265380859375 0 48646.082
629.2678833007812 0 7232.5654
629.3301391601562 0 8407.246
630.336669921875 0 3377.0916
643.2927856445312 0 6241.7827 w 5
644.2894287109375 0 19659.182
645.2925415039062 0 6402.8984
645.348388671875 0 67334.086
646.3560791015625 0 319173.72 c 5
647.3590087890625 0 114231.96
648.3612670898438 0 20016.955
657.2969970703125 0 3608.027
658.2814331054688 0 15111.911
659.284912109375 0 6470.6934 y Ammonia loss 5
659.534423828125 0 2375.7922
660.2918090820312 0 3121.5894 z 5
661.30908203125 0 2607.6033
672.3742065429688 0 3801.499
673.3652954101562 0 2921.1406
674.2993774414062 0 2863.5137
675.3076171875 0 12827.117
676.3135986328125 0 15033.011 y 5
677.3170166015625 0 3139.264
704.3300170898438 0 2787.9312
712.3527221679688 0 8589.2295
713.3568115234375 0 2941.2632
714.3644409179688 0 3539.829
722.3364868164062 0 22954.213
723.3402709960938 0 6466.7144
731.3839111328125 0 13177.556
732.3914184570312 0 12111.851
740.3465576171875 0 187632.17
741.349853515625 0 70818.36
742.3522338867188 0 17168.512
744.3296508789062 0 4803.936
745.3211059570312 0 2743.3071
747.3291625976562 0 8582.425 z 4
748.3350830078125 0 13139.847
757.373046875 0 27838.87 c Water loss 6
758.373779296875 0 51381.855
759.3762817382812 0 20257.787
762.339111328125 0 6280.517
763.3473510742188 0 40898.83 y 4
764.3501586914062 0 15548.099
765.35302734375 0 2804.3547
771.3540649414062 0 8366.732
772.3560180664062 0 4406.726
773.3817138671875 0 3355.0037
774.39111328125 0 84663.97
775.3983764648438 0 534105.6 c 6
776.401611328125 0 222303.14
777.404296875 0 49355.062
778.4051513671875 0 4154.863
796.7598876953125 0 2483.6694
801.414306640625 0 4311.416
802.4175415039062 0 2356.1372
840.4073486328125 0 4671.2056
842.423583984375 0 4000.995
843.3751220703125 0 5382.045 y Ammonia loss 3
850.3932495117188 0 6299.8535
851.3866577148438 0 5124.6885
852.3916625976562 0 3497.444
853.4284057617188 0 9910.855
854.4335327148438 0 5209.298
855.4371337890625 0 3080.1104
859.409423828125 0 7290.673
860.4022827148438 0 133595.27 y 3
861.4055786132812 0 53342.785
862.4097900390625 0 15308.315
867.3950805664062 0 7856.6445
868.4050903320312 0 91932.01
869.4071655273438 0 45872.406
870.408935546875 0 15134.12
871.400146484375 0 15611.319
872.4027099609375 0 6506.693
884.4239501953125 0 11212.38
885.4127197265625 0 19164.068
886.41552734375 0 6345.3564 c Water loss 7
887.4143676757812 0 175409.03
888.417724609375 0 86165.26
889.4201049804688 0 26502.512
901.4883422851562 0 17910.848
902.4938354492188 0 9931.69
903.4332885742188 0 42005.207
904.4411010742188 0 525984 c 7
905.4437255859375 0 271926.97
906.4468994140625 0 79211.42
907.4501342773438 0 7595.96
930.457763671875 0 6383.5444
931.4622802734375 0 4313.834
938.43212890625 0 14529.259
939.4408569335938 0 14095.825 y Water loss 2
940.4315185546875 0 20495.83 y Ammonia loss 2
941.429931640625 0 9776.778 z 2
955.4343872070312 0 3414.8215
956.4442749023438 0 121159.55
957.4519653320312 0 612197.5 y 2
958.4551391601562 0 285816.16
959.4584350585938 0 76954.125
960.45849609375 0 8240.695
972.502685546875 0 12525.674
973.5108032226562 0 94689.375
974.5155029296875 0 59170.973
975.517822265625 0 23658.973
976.5150146484375 0 3106.8154
982.4903564453125 0 8757.582
984.4744873046875 0 8903.518
985.4775390625 0 10389.778
986.4744262695312 0 4228.0347
999.4600830078125 0 8148.3076
1000.497802734375 0 114293.26
1001.5012817382812 0 69712.16
1002.5050659179688 0 20498.078
1003.5269775390625 0 2752.0037
1015.5069580078125 0 5528.395
1016.4925537109375 0 10969.828
1017.5247802734375 0 400032.2 c 8
1018.52783203125 0 235857.22
1018.6547241210938 0 4686.2144
1019.531005859375 0 72082.93
1020.5365600585938 0 8647.452
1043.5406494140625 0 61544.902
1044.54296875 0 37720.715
1045.524658203125 0 23223.729
1046.5130615234375 0 9803.866
1069.4776611328125 0 3230.1736
1070.487060546875 0 4831.8076 z Water loss 1
1071.4993896484375 0 3584.6638
1088.501953125 0 149564.06 z 1
1089.5050048828125 0 92918.8
1090.5079345703125 0 27882.05
1091.51611328125 0 3516.8057
1100.5745849609375 0 3808.8967
1101.5697021484375 0 67205.4
1102.57275390625 0 39957.734
1103.5782470703125 0 18312.217
1104.58935546875 0 2890.182
1109.549072265625 0 4394.9316
1111.55419921875 0 3783.1643
1112.537109375 0 16266.515
1113.5400390625 0 8682.653
1114.5606689453125 0 5813.7764
1117.5289306640625 0 24728.127
1118.5338134765625 0 16613.857
1119.541259765625 0 5609.7476
1128.560302734375 0 12885.98 c Ammonia loss 9
1129.5634765625 0 26257.445
1130.556884765625 0 42665.46
1131.561767578125 0 23636.096
1132.5684814453125 0 13119.706
1133.5821533203125 0 8798.486
1143.5692138671875 0 7849.046
1144.568359375 0 9552.874
1145.5816650390625 0 1407616.5 c 9
1146.585205078125 0 949365.2
1147.58740234375 0 371175.34
1148.5888671875 0 70283.234
1149.583984375 0 4171.5693
1156.6099853515625 0 22198.066
1157.61181640625 0 11124.59
1158.52978515625 0 7979.2637
1159.5872802734375 0 11338.807
1160.5963134765625 0 6434.24
1162.549560546875 0 22557.568
1163.5521240234375 0 16604.285
1164.5523681640625 0 8191.3643
1172.6055908203125 0 31451.832
1173.5933837890625 0 118688.94
1174.5955810546875 0 80416.85
1175.5965576171875 0 27653.367
1176.5997314453125 0 5713.3037
1183.5738525390625 0 15947.51
1184.581787109375 0 14127.345
1185.5836181640625 0 6117.4873
1189.605224609375 0 3444.5679
1190.615966796875 0 32499.824
1191.6201171875 0 27451.516
1192.622314453125 0 11142.984
1200.59521484375 0 24194.482
1201.5865478515625 0 187007.28
1202.58984375 0 136553.6
1203.5914306640625 0 56442.754
1204.597412109375 0 8235.059
1215.579345703125 0 3792.973
1216.5936279296875 0 20019.268
1217.603271484375 0 225384.42
1218.6107177734375 0 1253684.4
1219.6138916015625 0 823527.3
1220.6170654296875 0 294704.72
1221.6187744140625 0 45947.12
1250.6024169921875 0 3106.6672
1830.032470703125 0 4399.7993
1831.0390625 0 3154.1277
2830.7509765625 0 2707.5244
3001.578125 0 3041.052
3039.888916015625 0 2868.8503
3101.775146484375 0 2448.001

Spectrum Details

|  |  |
| --- | --- |
| Matched peaks? Matched peaks The total absolute number of peaks matched. Additionally in brackets the total fraction of peaks matched and the total number of peaks is shown. | 40 (12.86% of 311) |
| FDR? FDR The false discovery rate estimated for this peptide. It is calculated by matching all theoretical fragments with a non-integer shift with the raw peaks for this spectrum. This is done with 40 different shifts. The resulting percentage is the average number of annotated peaks over the number of annotated peaks with the correct spectrum. | 0.71% |
| Satellite FDR? Satellite FDR See the FDR for details on its calculation. This satellite ion specific FDR only contains the satellite ions (d/w) for I/L/J positions. | - |
| PSM Score? PSM Score The PSM Score as given by Hecklib to this annotated spectrum. It is shown with three significant figures. | 487 |

#### Spectrum 7659? Spectrum 7659 The raw spectrum of this peptide as annotated by Hecklib. The fragments are coloured according to ion type (see legend). Any peaks with a star '\*' as text can be hovered over to see the full details, first the ion type second the mass shift type. By hovering over the amino acids in the peptide or ions in the legend the corresponding peaks are highlighted. By toggling the 'Unassigned' label you can turn the background (unassigned) peaks on or off in the plot. By updating the slider in the Ion legend you can update the spectrum to only show the top X% of the peaks with labels. The top X% means any peak that is within X% of the highest intensity. By dragging in the spectrum you can zoom in to a specific part of the spectrum and use 'Zoom Out' to get back to the original zoom level. The annotation of the spectrum is based on the given sequence in the peptides file and is done with different software so inconsistencies are likely. The peaks are annotated based on the given sequence, with 20 ppm tolerance.

Copy Data

##### Spectrum 7659 (TSV)

###### Preview

```
Loading example...
```

*Click on the button to copy the data to your clipboard.*

Mz MinMz MaxIntensity Max

WidthHeightPeptide font sizePeptide stroke widthSpectrum font sizeSpectrum stroke widthCompact peptide

Ion legend

wxyz

abcd

OtherUnassignedIon^Charge^_Position_Show for top:%

JFPPSSEEJQA

08.89e+41.78e+52.67e+53.55e+5

Zoom Out

y^+1^_2_z^+1^_2_y^+1^_2_y^+1^_3_z^+1^_3_y^+1^_3_y^+1^_4_y^+1^_4_y^+2^_9_y^+2^_9_z^+2^_9_c^+1^_4_y^+2^_9_c^+1^_5_c^+2^_10_y^+1^_5_w^+1^_6_c^+1^_6_y^+1^_6_z^+1^_6_y^+1^_6_y^+1^_7_z^+1^_7_c^+1^_7_y^+1^_7_c^+1^_7_y^+1^_8_c^+1^_8_y^+1^_9_y^+1^_9_z^+1^_9_y^+1^_9_c^+1^_9_z^+1^_10_z^+1^_10_y^+1^_10_z^+1^_10_c^+1^_10_c^+1^_10_

0770154123113082

Fragment Matches Table

Show background peaks

| Position | Ion type | Intensity | mz Theoretical | mz Error (Th) | mz Error (ppm) | Charge | Series Number |
| --- | --- | --- | --- | --- | --- | --- | --- |
| - | - | 4488 | 120.1 | - | - | 0 | - |
| - | - | 380.8 | 127.8 | - | - | 0 | - |
| - | - | 6229 | 129.1 | - | - | 0 | - |
| - | - | 500.1 | 129.1 | - | - | 0 | - |
| - | - | 1158 | 131.1 | - | - | 0 | - |
| - | - | 841.1 | 133.1 | - | - | 0 | - |
| - | - | 418.1 | 148.9 | - | - | 0 | - |
| - | - | 402.7 | 148.9 | - | - | 0 | - |
| - | - | 680.2 | 148.9 | - | - | 0 | - |
| - | - | 845.3 | 148.9 | - | - | 0 | - |
| - | - | 778.6 | 148.9 | - | - | 0 | - |
| - | - | 1049 | 148.9 | - | - | 0 | - |
| - | - | 1304 | 148.9 | - | - | 0 | - |
| - | - | 3098 | 148.9 | - | - | 0 | - |
| - | - | 6329 | 149 | - | - | 0 | - |
| - | - | 3295 | 149 | - | - | 0 | - |
| - | - | 1441 | 149 | - | - | 0 | - |
| - | - | 1199 | 149 | - | - | 0 | - |
| - | - | 1120 | 149 | - | - | 0 | - |
| - | - | 853 | 149 | - | - | 0 | - |
| - | - | 501.8 | 149 | - | - | 0 | - |
| - | - | 1167 | 149 | - | - | 0 | - |
| - | - | 493 | 149.1 | - | - | 0 | - |
| - | - | 471.5 | 149.1 | - | - | 0 | - |
| - | - | 419.4 | 158.9 | - | - | 0 | - |
| - | - | 1162 | 173.4 | - | - | 0 | - |
| - | - | 461 | 192.5 | - | - | 0 | - |
| - | - | 519.7 | 195.5 | - | - | 0 | - |
| - | - | 715.7 | 197.1 | - | - | 0 | - |
| - | - | 501.2 | 197.3 | - | - | 0 | - |
| - | - | 1.465E+04 | 200.1 | - | - | 0 | - |
| 10 | y | 2.438E+04 | 201.1 | 0.0002664 | 1.325 | +1 | 2 |
| - | - | 851.1 | 201.1 | - | - | 0 | - |
| 10 | z | 1894 | 202.1 | 0.003835 | 18.98 | +1 | 2 |
| - | - | 483.7 | 203 | - | - | 0 | - |
| - | - | 968.1 | 215.1 | - | - | 0 | - |
| 10 | y | 3.189E+04 | 218.1 | 0.0002676 | 1.227 | +1 | 2 |
| - | - | 2252 | 219.1 | - | - | 0 | - |
| - | - | 5137 | 221.1 | - | - | 0 | - |
| - | - | 1151 | 225 | - | - | 0 | - |
| - | - | 1066 | 225.1 | - | - | 0 | - |
| - | - | 3.369E+04 | 233.2 | - | - | 0 | - |
| - | - | 4627 | 234.2 | - | - | 0 | - |
| - | - | 473.6 | 237.4 | - | - | 0 | - |
| - | - | 5778 | 239.1 | - | - | 0 | - |
| - | - | 1264 | 240.1 | - | - | 0 | - |
| - | - | 559.4 | 241.1 | - | - | 0 | - |
| - | - | 1.434E+04 | 242.2 | - | - | 0 | - |
| - | - | 1650 | 243.2 | - | - | 0 | - |
| - | - | 3.384E+04 | 261.2 | - | - | 0 | - |
| - | - | 5808 | 262.2 | - | - | 0 | - |
| - | - | 1216 | 281.1 | - | - | 0 | - |
| - | - | 5073 | 282.1 | - | - | 0 | - |
| - | - | 680.6 | 283.1 | - | - | 0 | - |
| - | - | 1416 | 295.1 | - | - | 0 | - |
| - | - | 1414 | 296.1 | - | - | 0 | - |
| - | - | 613.7 | 297.1 | - | - | 0 | - |
| - | - | 5510 | 299.1 | - | - | 0 | - |
| - | - | 988.8 | 299.2 | - | - | 0 | - |
| - | - | 691.4 | 300.1 | - | - | 0 | - |
| - | - | 1526 | 300.1 | - | - | 0 | - |
| - | - | 2387 | 314.1 | - | - | 0 | - |
| 9 | y | 6291 | 314.2 | 0.000431 | 1.372 | +1 | 3 |
| - | - | 759.9 | 314.6 | - | - | 0 | - |
| 9 | z | 1149 | 315.2 | 0.00364 | 11.55 | +1 | 3 |
| 9 | y | 5001 | 331.2 | 0.0003711 | 1.121 | +1 | 3 |
| - | - | 1073 | 332.2 | - | - | 0 | - |
| - | - | 954.9 | 346.1 | - | - | 0 | - |
| - | - | 669.6 | 347.7 | - | - | 0 | - |
| - | - | 727.6 | 351.2 | - | - | 0 | - |
| - | - | 2290 | 353.2 | - | - | 0 | - |
| - | - | 4.022E+04 | 355.1 | - | - | 0 | - |
| - | - | 608.6 | 356.7 | - | - | 0 | - |
| - | - | 3850 | 358.2 | - | - | 0 | - |
| - | - | 879.4 | 359.2 | - | - | 0 | - |
| - | - | 1106 | 360.2 | - | - | 0 | - |
| - | - | 2985 | 369.2 | - | - | 0 | - |
| - | - | 815.4 | 370.2 | - | - | 0 | - |
| - | - | 1.935E+04 | 370.7 | - | - | 0 | - |
| - | - | 1022 | 371.1 | - | - | 0 | - |
| - | - | 5207 | 371.2 | - | - | 0 | - |
| - | - | 1814 | 371.7 | - | - | 0 | - |
| - | - | 903.9 | 373.1 | - | - | 0 | - |
| - | - | 2272 | 379.2 | - | - | 0 | - |
| - | - | 2070 | 379.7 | - | - | 0 | - |
| - | - | 887.1 | 380.2 | - | - | 0 | - |
| - | - | 537.8 | 385.5 | - | - | 0 | - |
| - | - | 2522 | 386.2 | - | - | 0 | - |
| - | - | 558 | 411.5 | - | - | 0 | - |
| - | - | 2194 | 415 | - | - | 0 | - |
| - | - | 669.9 | 419 | - | - | 0 | - |
| - | - | 820.3 | 420.7 | - | - | 0 | - |
| - | - | 954.7 | 425.2 | - | - | 0 | - |
| - | - | 1198 | 425.7 | - | - | 0 | - |
| - | - | 1208 | 426.2 | - | - | 0 | - |
| - | - | 924 | 428.3 | - | - | 0 | - |
| - | - | 964.3 | 429.3 | - | - | 0 | - |
| - | - | 892.6 | 430.2 | - | - | 0 | - |
| - | - | 1.75E+04 | 434.7 | - | - | 0 | - |
| - | - | 8330 | 435.2 | - | - | 0 | - |
| - | - | 640.7 | 435.2 | - | - | 0 | - |
| - | - | 1947 | 435.7 | - | - | 0 | - |
| 8 | y | 705.6 | 442.2 | 0.0005998 | 1.356 | +1 | 4 |
| - | - | 8147 | 444.2 | - | - | 0 | - |
| - | - | 2918 | 444.7 | - | - | 0 | - |
| - | - | 1041 | 445.2 | - | - | 0 | - |
| - | - | 882.7 | 447.7 | - | - | 0 | - |
| - | - | 2071 | 455.3 | - | - | 0 | - |
| - | - | 672.5 | 459.2 | - | - | 0 | - |
| 8 | y | 709.6 | 460.2 | 0.001052 | 2.286 | +1 | 4 |
| - | - | 601.3 | 461.7 | - | - | 0 | - |
| 3 | y | 2768 | 470.2 | 0.0009244 | 1.966 | +2 | 9 |
| 3 | y | 1767 | 470.7 | 0.007391 | 15.7 | +2 | 9 |
| 3 | z | 790.4 | 471.2 | 0.0009468 | 2.009 | +2 | 9 |
| - | - | 3437 | 471.3 | - | - | 0 | - |
| 4 | c | 3895 | 472.3 | 0.0005409 | 1.145 | +1 | 4 |
| - | - | 622.9 | 474.3 | - | - | 0 | - |
| 3 | y | 2902 | 479.2 | 0.0009216 | 1.923 | +2 | 9 |
| - | - | 1236 | 479.7 | - | - | 0 | - |
| - | - | 1589 | 480.2 | - | - | 0 | - |
| - | - | 730.7 | 487.3 | - | - | 0 | - |
| - | - | 1894 | 488.3 | - | - | 0 | - |
| - | - | 2084 | 489.1 | - | - | 0 | - |
| - | - | 746 | 491.7 | - | - | 0 | - |
| - | - | 7189 | 498.2 | - | - | 0 | - |
| - | - | 1557 | 499.2 | - | - | 0 | - |
| - | - | 921.1 | 500.2 | - | - | 0 | - |
| - | - | 4793 | 500.8 | - | - | 0 | - |
| - | - | 3029 | 501.3 | - | - | 0 | - |
| - | - | 675.2 | 501.8 | - | - | 0 | - |
| - | - | 609.8 | 512.2 | - | - | 0 | - |
| - | - | 4259 | 515.2 | - | - | 0 | - |
| - | - | 602.5 | 516.2 | - | - | 0 | - |
| - | - | 971 | 516.3 | - | - | 0 | - |
| - | - | 653.4 | 530.2 | - | - | 0 | - |
| - | - | 685 | 536.3 | - | - | 0 | - |
| - | - | 2181 | 542.3 | - | - | 0 | - |
| - | - | 800.3 | 543.3 | - | - | 0 | - |
| - | - | 1704 | 544.3 | - | - | 0 | - |
| - | - | 1202 | 545.3 | - | - | 0 | - |
| - | - | 1078 | 546.2 | - | - | 0 | - |
| - | - | 1.707E+04 | 558.3 | - | - | 0 | - |
| 5 | c | 2.182E+04 | 559.3 | 0.0008615 | 1.54 | +1 | 5 |
| - | - | 6908 | 560.3 | - | - | 0 | - |
| - | - | 1061 | 561.3 | - | - | 0 | - |
| 10 | c | 558.5 | 564.8 | 0.0007375 | 1.306 | +2 | 10 |
| - | - | 1112 | 577.1 | - | - | 0 | - |
| - | - | 985.8 | 587.3 | - | - | 0 | - |
| 7 | y | 1079 | 589.3 | 0.001105 | 1.876 | +1 | 5 |
| - | - | 1082 | 591.4 | - | - | 0 | - |
| - | - | 804.9 | 599.3 | - | - | 0 | - |
| - | - | 2472 | 603.4 | - | - | 0 | - |
| - | - | 2013 | 609.3 | - | - | 0 | - |
| - | - | 8794 | 610.2 | - | - | 0 | - |
| - | - | 854.6 | 610.2 | - | - | 0 | - |
| - | - | 824.7 | 611.3 | - | - | 0 | - |
| - | - | 611.4 | 611.9 | - | - | 0 | - |
| - | - | 2271 | 616.3 | - | - | 0 | - |
| - | - | 2226 | 624.3 | - | - | 0 | - |
| - | - | 4.627E+04 | 627.3 | - | - | 0 | - |
| - | - | 1.35E+04 | 628.3 | - | - | 0 | - |
| - | - | 2589 | 629.3 | - | - | 0 | - |
| - | - | 2677 | 629.3 | - | - | 0 | - |
| - | - | 615.5 | 640.3 | - | - | 0 | - |
| 6 | w | 1595 | 643.3 | 0.0007811 | 1.214 | +1 | 6 |
| - | - | 4527 | 644.3 | - | - | 0 | - |
| - | - | 2227 | 645.3 | - | - | 0 | - |
| - | - | 1.608E+04 | 645.3 | - | - | 0 | - |
| 6 | c | 8.059E+04 | 646.4 | 0.0002522 | 0.3901 | +1 | 6 |
| - | - | 2.68E+04 | 647.4 | - | - | 0 | - |
| - | - | 6225 | 648.4 | - | - | 0 | - |
| - | - | 3815 | 658.3 | - | - | 0 | - |
| 6 | y | 2502 | 659.3 | 0.002068 | 3.137 | +1 | 6 |
| 6 | z | 1098 | 660.3 | 0.002863 | 4.336 | +1 | 6 |
| - | - | 867.5 | 661.3 | - | - | 0 | - |
| - | - | 1580 | 664.4 | - | - | 0 | - |
| - | - | 903.4 | 672.4 | - | - | 0 | - |
| - | - | 5517 | 673.4 | - | - | 0 | - |
| - | - | 1591 | 674.4 | - | - | 0 | - |
| - | - | 2953 | 675.3 | - | - | 0 | - |
| 6 | y | 4852 | 676.3 | 0.0006187 | 0.9148 | +1 | 6 |
| - | - | 1524 | 677.3 | - | - | 0 | - |
| - | - | 650.9 | 682.4 | - | - | 0 | - |
| - | - | 728 | 691.3 | - | - | 0 | - |
| - | - | 1125 | 704.3 | - | - | 0 | - |
| - | - | 672.6 | 705.3 | - | - | 0 | - |
| - | - | 2460 | 712.4 | - | - | 0 | - |
| - | - | 810.6 | 713.4 | - | - | 0 | - |
| - | - | 1103 | 714.4 | - | - | 0 | - |
| - | - | 658.7 | 717.9 | - | - | 0 | - |
| - | - | 5609 | 722.3 | - | - | 0 | - |
| - | - | 1793 | 723.3 | - | - | 0 | - |
| - | - | 941.7 | 727.4 | - | - | 0 | - |
| - | - | 3719 | 731.4 | - | - | 0 | - |
| - | - | 2441 | 732.4 | - | - | 0 | - |
| - | - | 763 | 739.4 | - | - | 0 | - |
| - | - | 5.155E+04 | 740.3 | - | - | 0 | - |
| - | - | 2.107E+04 | 741.3 | - | - | 0 | - |
| - | - | 5697 | 742.4 | - | - | 0 | - |
| - | - | 689 | 743.4 | - | - | 0 | - |
| - | - | 1036 | 744.3 | - | - | 0 | - |
| 5 | y | 842.4 | 745.3 | 0.01456 | 19.53 | +1 | 7 |
| 5 | z | 3474 | 747.3 | 0.0002538 | 0.3396 | +1 | 7 |
| - | - | 2756 | 748.3 | - | - | 0 | - |
| - | - | 1064 | 749.3 | - | - | 0 | - |
| - | - | 862.9 | 750.4 | - | - | 0 | - |
| - | - | 2499 | 750.9 | - | - | 0 | - |
| - | - | 755.9 | 751.4 | - | - | 0 | - |
| 7 | c | 8844 | 757.4 | 0.01463 | 19.31 | +1 | 7 |
| - | - | 1.157E+04 | 758.4 | - | - | 0 | - |
| - | - | 4395 | 759.4 | - | - | 0 | - |
| - | - | 1146 | 760.4 | - | - | 0 | - |
| - | - | 1199 | 762.3 | - | - | 0 | - |
| 5 | y | 9958 | 763.3 | 3.767E-05 | 0.04935 | +1 | 7 |
| - | - | 3253 | 764.4 | - | - | 0 | - |
| - | - | 1233 | 765.4 | - | - | 0 | - |
| - | - | 2838 | 771.4 | - | - | 0 | - |
| - | - | 1013 | 772.4 | - | - | 0 | - |
| - | - | 2.101E+04 | 774.4 | - | - | 0 | - |
| 7 | c | 1.354E+05 | 775.4 | 0.0001395 | 0.1799 | +1 | 7 |
| - | - | 5.801E+04 | 776.4 | - | - | 0 | - |
| - | - | 1.379E+04 | 777.4 | - | - | 0 | - |
| - | - | 1123 | 778.4 | - | - | 0 | - |
| - | - | 1202 | 783.9 | - | - | 0 | - |
| - | - | 1048 | 784.4 | - | - | 0 | - |
| - | - | 1398 | 786.5 | - | - | 0 | - |
| - | - | 1507 | 787 | - | - | 0 | - |
| - | - | 914.8 | 791.5 | - | - | 0 | - |
| - | - | 900.2 | 801.4 | - | - | 0 | - |
| - | - | 980.1 | 802.4 | - | - | 0 | - |
| - | - | 1192 | 809.4 | - | - | 0 | - |
| - | - | 1120 | 840.4 | - | - | 0 | - |
| - | - | 1186 | 842.4 | - | - | 0 | - |
| - | - | 1545 | 850.4 | - | - | 0 | - |
| - | - | 2291 | 851.4 | - | - | 0 | - |
| - | - | 786.1 | 852.4 | - | - | 0 | - |
| - | - | 1825 | 853.4 | - | - | 0 | - |
| - | - | 1432 | 854.4 | - | - | 0 | - |
| - | - | 1782 | 859.4 | - | - | 0 | - |
| 4 | y | 3.329E+04 | 860.4 | 0.003107 | 3.611 | +1 | 8 |
| - | - | 1.615E+04 | 861.4 | - | - | 0 | - |
| - | - | 5584 | 862.4 | - | - | 0 | - |
| - | - | 951.5 | 863.4 | - | - | 0 | - |
| - | - | 2483 | 867.4 | - | - | 0 | - |
| - | - | 2.559E+04 | 868.4 | - | - | 0 | - |
| - | - | 1.344E+04 | 869.4 | - | - | 0 | - |
| - | - | 4879 | 870.4 | - | - | 0 | - |
| - | - | 719.8 | 870.5 | - | - | 0 | - |
| - | - | 2411 | 871.4 | - | - | 0 | - |
| - | - | 1666 | 872.4 | - | - | 0 | - |
| - | - | 761.4 | 873.4 | - | - | 0 | - |
| - | - | 1846 | 878.5 | - | - | 0 | - |
| - | - | 1073 | 879.5 | - | - | 0 | - |
| - | - | 2866 | 884.4 | - | - | 0 | - |
| - | - | 5031 | 885.4 | - | - | 0 | - |
| - | - | 1812 | 886.4 | - | - | 0 | - |
| - | - | 787.1 | 886.5 | - | - | 0 | - |
| - | - | 4.768E+04 | 887.4 | - | - | 0 | - |
| - | - | 2.428E+04 | 888.4 | - | - | 0 | - |
| - | - | 7291 | 889.4 | - | - | 0 | - |
| - | - | 812.5 | 901.4 | - | - | 0 | - |
| - | - | 5031 | 901.5 | - | - | 0 | - |
| - | - | 2083 | 902.5 | - | - | 0 | - |
| - | - | 1.216E+04 | 903.4 | - | - | 0 | - |
| 8 | c | 1.302E+05 | 904.4 | 0.000271 | 0.2997 | +1 | 8 |
| - | - | 6.584E+04 | 905.4 | - | - | 0 | - |
| - | - | 1.633E+04 | 906.4 | - | - | 0 | - |
| - | - | 837.3 | 907 | - | - | 0 | - |
| - | - | 2521 | 907.4 | - | - | 0 | - |
| - | - | 704.8 | 912.5 | - | - | 0 | - |
| - | - | 1851 | 914.5 | - | - | 0 | - |
| - | - | 3794 | 915 | - | - | 0 | - |
| - | - | 2501 | 915.5 | - | - | 0 | - |
| - | - | 868 | 930.5 | - | - | 0 | - |
| - | - | 693.5 | 931.5 | - | - | 0 | - |
| - | - | 2594 | 938.4 | - | - | 0 | - |
| 3 | y | 2920 | 939.4 | 0.0008845 | 0.9415 | +1 | 9 |
| 3 | y | 3946 | 940.4 | 0.006006 | 6.386 | +1 | 9 |
| 3 | z | 2124 | 941.4 | 0.001171 | 1.244 | +1 | 9 |
| - | - | 947.9 | 942.4 | - | - | 0 | - |
| - | - | 968.2 | 955.4 | - | - | 0 | - |
| - | - | 3.274E+04 | 956.4 | - | - | 0 | - |
| 3 | y | 1.512E+05 | 957.5 | 3.559E-05 | 0.03718 | +1 | 9 |
| - | - | 7.658E+04 | 958.5 | - | - | 0 | - |
| - | - | 2.23E+04 | 959.5 | - | - | 0 | - |
| - | - | 2346 | 960.5 | - | - | 0 | - |
| - | - | 894.3 | 966.4 | - | - | 0 | - |
| - | - | 3674 | 972.5 | - | - | 0 | - |
| - | - | 2.147E+04 | 973.5 | - | - | 0 | - |
| - | - | 1.54E+04 | 974.5 | - | - | 0 | - |
| - | - | 6224 | 975.5 | - | - | 0 | - |
| - | - | 851.5 | 976.5 | - | - | 0 | - |
| - | - | 1344 | 977.6 | - | - | 0 | - |
| - | - | 619.2 | 978.6 | - | - | 0 | - |
| - | - | 3217 | 982.5 | - | - | 0 | - |
| - | - | 1036 | 983.5 | - | - | 0 | - |
| - | - | 2799 | 984.5 | - | - | 0 | - |
| - | - | 1806 | 985.5 | - | - | 0 | - |
| - | - | 1454 | 986.5 | - | - | 0 | - |
| - | - | 1364 | 996.8 | - | - | 0 | - |
| - | - | 781.8 | 997.1 | - | - | 0 | - |
| - | - | 2457 | 999.5 | - | - | 0 | - |
| - | - | 3.194E+04 | 1000 | - | - | 0 | - |
| - | - | 1.911E+04 | 1002 | - | - | 0 | - |
| - | - | 6879 | 1003 | - | - | 0 | - |
| - | - | 902.9 | 1004 | - | - | 0 | - |
| - | - | 809.9 | 1006 | - | - | 0 | - |
| - | - | 834.2 | 1015 | - | - | 0 | - |
| - | - | 2885 | 1016 | - | - | 0 | - |
| - | - | 736.7 | 1017 | - | - | 0 | - |
| 9 | c | 1.006E+05 | 1018 | 0.0001137 | 0.1118 | +1 | 9 |
| - | - | 5.672E+04 | 1019 | - | - | 0 | - |
| - | - | 1.803E+04 | 1020 | - | - | 0 | - |
| - | - | 2619 | 1021 | - | - | 0 | - |
| - | - | 1.534E+04 | 1044 | - | - | 0 | - |
| - | - | 1.021E+04 | 1045 | - | - | 0 | - |
| - | - | 5208 | 1046 | - | - | 0 | - |
| - | - | 2694 | 1047 | - | - | 0 | - |
| - | - | 982.1 | 1054 | - | - | 0 | - |
| 2 | z | 785.4 | 1070 | 0.001183 | 1.106 | +1 | 10 |
| 2 | z | 1349 | 1071 | 0.01473 | 13.74 | +1 | 10 |
| - | - | 1084 | 1079 | - | - | 0 | - |
| 2 | y | 941 | 1087 | 0.008942 | 8.222 | +1 | 10 |
| 2 | z | 3.614E+04 | 1089 | 1.821E-05 | 0.01673 | +1 | 10 |
| - | - | 2.336E+04 | 1090 | - | - | 0 | - |
| - | - | 7047 | 1091 | - | - | 0 | - |
| - | - | 1281 | 1092 | - | - | 0 | - |
| - | - | 1.702E+04 | 1102 | - | - | 0 | - |
| - | - | 1.054E+04 | 1103 | - | - | 0 | - |
| - | - | 3822 | 1104 | - | - | 0 | - |
| - | - | 742.8 | 1106 | - | - | 0 | - |
| - | - | 846.4 | 1110 | - | - | 0 | - |
| - | - | 1146 | 1112 | - | - | 0 | - |
| - | - | 4775 | 1113 | - | - | 0 | - |
| - | - | 4495 | 1114 | - | - | 0 | - |
| - | - | 1226 | 1115 | - | - | 0 | - |
| - | - | 6177 | 1118 | - | - | 0 | - |
| - | - | 3754 | 1119 | - | - | 0 | - |
| - | - | 1140 | 1120 | - | - | 0 | - |
| 10 | c | 4105 | 1129 | 0.002282 | 2.022 | +1 | 10 |
| - | - | 6541 | 1130 | - | - | 0 | - |
| - | - | 1.011E+04 | 1131 | - | - | 0 | - |
| - | - | 5013 | 1132 | - | - | 0 | - |
| - | - | 3130 | 1133 | - | - | 0 | - |
| - | - | 2893 | 1144 | - | - | 0 | - |
| - | - | 2676 | 1145 | - | - | 0 | - |
| 10 | c | 3.519E+05 | 1146 | 0.00144 | 1.257 | +1 | 10 |
| - | - | 2.416E+05 | 1147 | - | - | 0 | - |
| - | - | 9.478E+04 | 1148 | - | - | 0 | - |
| - | - | 1.658E+04 | 1149 | - | - | 0 | - |
| - | - | 1126 | 1150 | - | - | 0 | - |
| - | - | 4703 | 1157 | - | - | 0 | - |
| - | - | 4139 | 1158 | - | - | 0 | - |
| - | - | 1804 | 1159 | - | - | 0 | - |
| - | - | 1578 | 1160 | - | - | 0 | - |
| - | - | 2294 | 1161 | - | - | 0 | - |
| - | - | 789.8 | 1162 | - | - | 0 | - |
| - | - | 6695 | 1163 | - | - | 0 | - |
| - | - | 4004 | 1164 | - | - | 0 | - |
| - | - | 1380 | 1165 | - | - | 0 | - |
| - | - | 1582 | 1166 | - | - | 0 | - |
| - | - | 8955 | 1173 | - | - | 0 | - |
| - | - | 2.8E+04 | 1174 | - | - | 0 | - |
| - | - | 2.139E+04 | 1175 | - | - | 0 | - |
| - | - | 7667 | 1176 | - | - | 0 | - |
| - | - | 1031 | 1177 | - | - | 0 | - |
| - | - | 4146 | 1184 | - | - | 0 | - |
| - | - | 2572 | 1185 | - | - | 0 | - |
| - | - | 1533 | 1186 | - | - | 0 | - |
| - | - | 1338 | 1188 | - | - | 0 | - |
| - | - | 1099 | 1189 | - | - | 0 | - |
| - | - | 1187 | 1189 | - | - | 0 | - |
| - | - | 9320 | 1191 | - | - | 0 | - |
| - | - | 5657 | 1192 | - | - | 0 | - |
| - | - | 2230 | 1193 | - | - | 0 | - |
| - | - | 909 | 1200 | - | - | 0 | - |
| - | - | 3827 | 1201 | - | - | 0 | - |
| - | - | 5.071E+04 | 1202 | - | - | 0 | - |
| - | - | 3.575E+04 | 1203 | - | - | 0 | - |
| - | - | 718.6 | 1203 | - | - | 0 | - |
| - | - | 1.308E+04 | 1204 | - | - | 0 | - |
| - | - | 2658 | 1205 | - | - | 0 | - |
| - | - | 1519 | 1216 | - | - | 0 | - |
| - | - | 4289 | 1217 | - | - | 0 | - |
| - | - | 6.246E+04 | 1218 | - | - | 0 | - |
| - | - | 3.138E+05 | 1219 | - | - | 0 | - |
| - | - | 2.124E+05 | 1220 | - | - | 0 | - |
| - | - | 7.835E+04 | 1221 | - | - | 0 | - |
| - | - | 1.158E+04 | 1222 | - | - | 0 | - |
| - | - | 1608 | 1251 | - | - | 0 | - |
| - | - | 1165 | 1252 | - | - | 0 | - |
| - | - | 1635 | 1282 | - | - | 0 | - |
| - | - | 1834 | 1294 | - | - | 0 | - |
| - | - | 1293 | 1295 | - | - | 0 | - |
| - | - | 1052 | 1296 | - | - | 0 | - |
| - | - | 730.5 | 1310 | - | - | 0 | - |
| - | - | 1775 | 1313 | - | - | 0 | - |
| - | - | 1227 | 1314 | - | - | 0 | - |
| - | - | 1093 | 1373 | - | - | 0 | - |
| - | - | 1054 | 1374 | - | - | 0 | - |
| - | - | 2063 | 1396 | - | - | 0 | - |
| - | - | 1102 | 1397 | - | - | 0 | - |
| - | - | 814.8 | 1417 | - | - | 0 | - |
| - | - | 2133 | 1453 | - | - | 0 | - |
| - | - | 1569 | 1454 | - | - | 0 | - |
| - | - | 1030 | 1455 | - | - | 0 | - |
| - | - | 1176 | 1494 | - | - | 0 | - |
| - | - | 1974 | 1495 | - | - | 0 | - |
| - | - | 2055 | 1495 | - | - | 0 | - |
| - | - | 1061 | 1496 | - | - | 0 | - |
| - | - | 2212 | 1496 | - | - | 0 | - |
| - | - | 1643 | 1501 | - | - | 0 | - |
| - | - | 949.1 | 1502 | - | - | 0 | - |
| - | - | 1815 | 1503 | - | - | 0 | - |
| - | - | 1571 | 1503 | - | - | 0 | - |
| - | - | 1099 | 1504 | - | - | 0 | - |
| - | - | 1643 | 1510 | - | - | 0 | - |
| - | - | 1345 | 1516 | - | - | 0 | - |
| - | - | 1739 | 1517 | - | - | 0 | - |
| - | - | 1834 | 1517 | - | - | 0 | - |
| - | - | 1017 | 1518 | - | - | 0 | - |
| - | - | 1184 | 1524 | - | - | 0 | - |
| - | - | 1923 | 1525 | - | - | 0 | - |
| - | - | 2559 | 1525 | - | - | 0 | - |
| - | - | 2257 | 1526 | - | - | 0 | - |
| - | - | 1352 | 1526 | - | - | 0 | - |
| - | - | 2288 | 1567 | - | - | 0 | - |
| - | - | 1879 | 1568 | - | - | 0 | - |
| - | - | 956.9 | 1573 | - | - | 0 | - |
| - | - | 1453 | 1714 | - | - | 0 | - |
| - | - | 837.2 | 1737 | - | - | 0 | - |
| - | - | 1365 | 1813 | - | - | 0 | - |
| - | - | 1567 | 1814 | - | - | 0 | - |
| - | - | 927 | 1825 | - | - | 0 | - |
| - | - | 2665 | 1829 | - | - | 0 | - |
| - | - | 4730 | 1830 | - | - | 0 | - |
| - | - | 2668 | 1831 | - | - | 0 | - |
| - | - | 1180 | 1832 | - | - | 0 | - |
| - | - | 1005 | 2377 | - | - | 0 | - |
| - | - | 790.7 | 2378 | - | - | 0 | - |
| - | - | 1137 | 3051 | - | - | 0 | - |

m/z Charge Intensity FragmentType MassShift Position
120.0810775756836 0 4488.041
127.79950714111328 0 380.76196
129.0660858154297 0 6229.185
129.10235595703125 0 500.12814
131.11827087402344 0 1157.6143
133.08615112304688 0 841.08527
148.88084411621094 0 418.08563
148.89488220214844 0 402.65192
148.90225219726562 0 680.1735
148.90924072265625 0 845.3442
148.9168243408203 0 778.55273
148.92401123046875 0 1048.5894
148.93113708496094 0 1304.2834
148.93850708007812 0 3097.7952
148.9550018310547 0 6328.5767
148.96278381347656 0 3294.62
148.97064208984375 0 1441.4904
148.97775268554688 0 1199.4695
148.9849090576172 0 1120.4723
148.99252319335938 0 852.9651
148.99964904785156 0 501.83554
149.04481506347656 0 1166.9989
149.0718994140625 0 493.00903
149.10772705078125 0 471.51266
158.87066650390625 0 419.39218
173.44970703125 0 1161.6528
192.46746826171875 0 460.97327
195.49261474609375 0 519.66614
197.1281280517578 0 715.6879
197.3432159423828 0 501.19196
200.1031494140625 0 14645.938
201.08724975585938 0 24377.486 y Ammonia loss 9
201.10516357421875 0 851.11066
202.09097290039062 0 1893.9601 z 9
202.97479248046875 0 483.686
215.13893127441406 0 968.1074
218.11380004882812 0 31887.213 y 9
219.11732482910156 0 2252.101
221.0846710205078 0 5137.4253
225.0428924560547 0 1150.7052
225.1236572265625 0 1065.7823
233.16517639160156 0 33688.95
234.16845703125 0 4627.44
237.4429168701172 0 473.59473
239.09532165527344 0 5777.6987
240.0955810546875 0 1264.0908
241.0924072265625 0 559.3921
242.1502227783203 0 14342.394
243.1534881591797 0 1650.0521
261.1600036621094 0 33836.152
262.1632385253906 0 5808.224
281.05120849609375 0 1216.06
282.1449890136719 0 5073.0874
283.1473388671875 0 680.602
295.10302734375 0 1415.852
296.103271484375 0 1414.0153
297.1024169921875 0 613.72626
299.0620422363281 0 5509.9883
299.1701354980469 0 988.7991
300.06280517578125 0 691.4175
300.11956787109375 0 1526.4374
314.1352233886719 0 2386.52
314.1714782714844 0 6290.844 y Ammonia loss 8
314.6376037597656 0 759.90607
315.17523193359375 0 1149.2269 z 8
331.1979675292969 0 5000.7817 y 8
332.20184326171875 0 1072.523
346.1253967285156 0 954.89734
347.674072265625 0 669.5842
351.16741943359375 0 727.62823
353.1827697753906 0 2289.8691
355.0702209472656 0 40220.188
356.6799621582031 0 608.62335
358.2129821777344 0 3850.1616
359.21575927734375 0 879.43005
360.21484375 0 1105.6927
369.17694091796875 0 2984.6938
370.1789245605469 0 815.446
370.6769104003906 0 19351.68
371.10113525390625 0 1022.1954
371.1784362792969 0 5207.1074
371.6790771484375 0 1814.2034
373.07958984375 0 903.8833
379.18994140625 0 2271.5298
379.69000244140625 0 2070.2373
380.1929626464844 0 887.1261
385.4922790527344 0 537.83545
386.203857421875 0 2522.1313
411.549560546875 0 557.9805
415.0372314453125 0 2194.4868
418.9920349121094 0 669.8957
420.7084655761719 0 820.3383
425.2052001953125 0 954.69116
425.7019348144531 0 1197.66
426.1985168457031 0 1208.4666
428.2783508300781 0 924.03455
429.2843933105469 0 964.3192
430.2139587402344 0 892.5884
434.70654296875 0 17499.92
435.2079772949219 0 8329.712
435.2386474609375 0 640.6864
435.7104797363281 0 1946.7234
442.230224609375 0 705.649 y Water loss 7
444.2115478515625 0 8146.8315
444.7129211425781 0 2917.5786
445.2143249511719 0 1041.306
447.7171630859375 0 882.70416
455.2652893066406 0 2071.0217
459.208251953125 0 672.49243
460.2412414550781 0 709.61127 y 7
461.7098693847656 0 601.3291
470.2254638671875 0 2768.296 y Water loss 2
470.72393798828125 0 1767.311 y Ammonia loss 2
471.2195129394531 0 790.4436 z 2
471.2845153808594 0 3437.4634
472.2912902832031 0 3895.0378 c 3
474.3086853027344 0 622.9094
479.2307434082031 0 2902.0066 y 2
479.7320556640625 0 1235.7358
480.2095031738281 0 1589.3923
487.2572021484375 0 730.71924
488.2724609375 0 1894.1578
489.056396484375 0 2084.2722
491.74993896484375 0 745.9679
498.2198791503906 0 7188.775
499.22344970703125 0 1556.871
500.2369384765625 0 921.1204
500.75311279296875 0 4792.8306
501.25421142578125 0 3028.5864
501.7557067871094 0 675.22736
512.1986694335938 0 609.8352
515.2466430664062 0 4259.44
516.2465209960938 0 602.4743
516.3167724609375 0 970.9955
530.20654296875 0 653.4051
536.2828979492188 0 685.0158
542.2958984375 0 2181.4185
543.3101196289062 0 800.2507
544.3147583007812 0 1704.4303
545.3151245117188 0 1201.6781
546.2407836914062 0 1077.6122
558.3165283203125 0 17073.893
559.322998046875 0 21816.697 c 4
560.3267822265625 0 6907.559
561.334228515625 0 1061.1543
564.782958984375 0 558.4879 c Ammonia loss 9
577.1259765625 0 1111.9534
587.2694091796875 0 985.82385
589.2816772460938 0 1078.52 y 6
591.36083984375 0 1081.5374
599.2664184570312 0 804.89215
603.3500366210938 0 2471.71
609.2507934570312 0 2013.1041
610.1845703125 0 8793.88
610.2446899414062 0 854.57025
611.313720703125 0 824.6853
611.909423828125 0 611.41833
616.3324584960938 0 2270.812
624.3363037109375 0 2225.9656
627.2626342773438 0 46267.848
628.2655639648438 0 13504.9
629.2681274414062 0 2588.6643
629.3311157226562 0 2676.9587
640.294677734375 0 615.541
643.2941284179688 0 1595.1573 w 5
644.28955078125 0 4527.194
645.2907104492188 0 2227.4634
645.3486938476562 0 16078.273
646.3561401367188 0 80592.23 c 5
647.3592529296875 0 26804.104
648.3624267578125 0 6224.6045
658.28173828125 0 3814.867
659.2861938476562 0 2501.726 y Ammonia loss 5
660.2989501953125 0 1098.212 z 5
661.3080444335938 0 867.53955
664.3777465820312 0 1580.3341
672.3682250976562 0 903.39624
673.3546752929688 0 5516.996
674.35791015625 0 1591.3225
675.3069458007812 0 2953.4094
676.3154296875 0 4851.751 y 5
677.3178100585938 0 1524.1376
682.3682861328125 0 650.8803
691.2667846679688 0 727.9714
704.3245849609375 0 1124.9125
705.3276977539062 0 672.5658
712.3504638671875 0 2460.4468
713.3576049804688 0 810.55316
714.36669921875 0 1103.1134
717.894287109375 0 658.7047
722.3363037109375 0 5608.9575
723.3390502929688 0 1792.8319
727.3984985351562 0 941.69055
731.3842163085938 0 3718.9597
732.3893432617188 0 2440.9211
739.356201171875 0 762.9976
740.3466796875 0 51551.938
741.3499755859375 0 21070.115
742.3530883789062 0 5696.526
743.3527221679688 0 689.0469
744.3282470703125 0 1035.8506
745.3217163085938 0 842.3908 y Water loss 4
747.328369140625 0 3473.846 z 4
748.3348999023438 0 2755.8994
749.3319091796875 0 1064.406
750.4409790039062 0 862.9431
750.9483032226562 0 2499.0867
751.4494018554688 0 755.90137
757.373291015625 0 8844.073 c Water loss 6
758.3734741210938 0 11574.924
759.3758544921875 0 4394.728
760.3798217773438 0 1146.2794
762.3411865234375 0 1199.2366
763.3468017578125 0 9957.931 y 4
764.351318359375 0 3253.3093
765.3511352539062 0 1233.102
771.3511352539062 0 2838.0278
772.3594970703125 0 1012.5197
774.3911743164062 0 21006.574
775.3986206054688 0 135437.03 c 6
776.4016723632812 0 58011.02
777.4044189453125 0 13789.202
778.4022216796875 0 1123.0354
783.9456787109375 0 1202.235
784.4490966796875 0 1047.828
786.4694213867188 0 1397.9482
786.9727783203125 0 1507.0507
791.4532470703125 0 914.84357
801.4138793945312 0 900.21655
802.4268798828125 0 980.1317
809.4121704101562 0 1192.2432
840.4034423828125 0 1119.7535
842.4212646484375 0 1185.6665
850.3981323242188 0 1545.3646
851.3920288085938 0 2291.3296
852.3942260742188 0 786.14435
853.4308471679688 0 1825.2949
854.427978515625 0 1431.5659
859.4119873046875 0 1781.9308
860.4027099609375 0 33292.26 y 3
861.406494140625 0 16148.905
862.4082641601562 0 5584.148
863.3985595703125 0 951.4861
867.3983154296875 0 2482.5967
868.4049072265625 0 25588.984
869.4080810546875 0 13437.434
870.4097900390625 0 4878.774
870.5089721679688 0 719.8113
871.4004516601562 0 2410.5906
872.4002685546875 0 1666.045
873.4060668945312 0 761.36035
878.5060424804688 0 1846.3923
879.5145263671875 0 1072.689
884.4246215820312 0 2865.8367
885.4151000976562 0 5030.7236
886.408935546875 0 1812.1183
886.4679565429688 0 787.0664
887.4148559570312 0 47682.93
888.4181518554688 0 24281.986
889.4205322265625 0 7291.3843
901.3985595703125 0 812.5483
901.489990234375 0 5031.2627
902.4906616210938 0 2083.4348
903.43359375 0 12162.642
904.4413452148438 0 130224.39 c 7
905.444091796875 0 65836.836
906.4471435546875 0 16327.908
907.0149536132812 0 837.2504
907.4484252929688 0 2521.2527
912.4616088867188 0 704.797
914.5096435546875 0 1850.5448
915.022705078125 0 3794.202
915.5245361328125 0 2500.8682
930.4537353515625 0 868.0054
931.4641723632812 0 693.47644
938.435302734375 0 2594.287
939.44091796875 0 2920.3584 y Water loss 2
940.4318237304688 0 3946.4487 y Ammonia loss 2
941.434814453125 0 2124.3835 z 2
942.433837890625 0 947.91064
955.4376220703125 0 968.16003
956.4444580078125 0 32741.152
957.4523315429688 0 151239.52 y 2
958.455322265625 0 76582.29
959.4589233398438 0 22303.729
960.461669921875 0 2345.5703
966.4284057617188 0 894.3455
972.5044555664062 0 3674.193
973.5110473632812 0 21471.164
974.515625 0 15404.696
975.5185546875 0 6224.4854
976.5170288085938 0 851.4848
977.5802612304688 0 1343.8087
978.583740234375 0 619.21246
982.48779296875 0 3216.5725
983.4942016601562 0 1036.0713
984.4814453125 0 2798.6787
985.4849243164062 0 1806.1564
986.4779663085938 0 1453.9683
996.7920532226562 0 1364.2377
997.1334838867188 0 781.8151
999.457763671875 0 2457.2385
1000.4976806640625 0 31938.201
1001.501220703125 0 19107.773
1002.5048217773438 0 6878.991
1003.5132446289062 0 902.912
1005.5040893554688 0 809.9139
1015.4908447265625 0 834.24994
1016.4883422851562 0 2885.12
1016.7969970703125 0 736.6533
1017.5250244140625 0 100584.24 c 8
1018.5281372070312 0 56720.977
1019.5310668945312 0 18031.908
1020.53369140625 0 2618.9631
1043.5408935546875 0 15339.773
1044.54443359375 0 10209.189
1045.52197265625 0 5207.803
1046.5177001953125 0 2693.8506
1053.59228515625 0 982.11383
1070.49267578125 0 785.3556 z Water loss 1
1071.490234375 0 1349.3715 z Ammonia loss 1
1079.4981689453125 0 1084.3649
1087.503173828125 0 941.04254 y Ammonia loss 1
1088.5020751953125 0 36135.24 z 1
1089.505126953125 0 23358.475
1090.5084228515625 0 7046.5493
1091.5107421875 0 1281.4601
1101.5701904296875 0 17023.533
1102.573486328125 0 10536.832
1103.5736083984375 0 3822.109
1105.5511474609375 0 742.808
1109.5579833984375 0 846.4292
1111.5528564453125 0 1145.8561
1112.5389404296875 0 4774.805
1113.5430908203125 0 4494.8984
1114.5506591796875 0 1225.9338
1117.5289306640625 0 6176.539
1118.53076171875 0 3753.6597
1119.5345458984375 0 1140.2504
1128.5594482421875 0 4105.494 c Ammonia loss 9
1129.5655517578125 0 6541.2905
1130.5556640625 0 10112.589
1131.561767578125 0 5013.0947
1132.5697021484375 0 3130.0686
1143.562744140625 0 2892.7893
1144.568115234375 0 2676.2153
1145.582275390625 0 351939.53 c 9
1146.585693359375 0 241620.34
1147.5877685546875 0 94780.33
1148.5889892578125 0 16581.154
1149.5880126953125 0 1125.7139
1156.611328125 0 4702.5654
1157.6116943359375 0 4138.6772
1158.525634765625 0 1803.7959
1159.60498046875 0 1577.5815
1160.594482421875 0 2293.8022
1161.60498046875 0 789.77094
1162.5511474609375 0 6694.6987
1163.5528564453125 0 4004.133
1164.5560302734375 0 1379.9819
1165.671630859375 0 1581.8973
1172.6064453125 0 8954.544
1173.5938720703125 0 27997.088
1174.5966796875 0 21387.582
1175.597412109375 0 7667.1953
1176.59619140625 0 1030.8657
1183.578125 0 4146.415
1184.5797119140625 0 2572.2275
1185.5753173828125 0 1532.5657
1188.029052734375 0 1337.7837
1188.5330810546875 0 1098.9078
1189.0205078125 0 1187.2842
1190.6180419921875 0 9319.958
1191.6201171875 0 5656.7715
1192.6256103515625 0 2229.7385
1199.6162109375 0 908.95795
1200.5985107421875 0 3827.2466
1201.58642578125 0 50705.207
1202.588623046875 0 35752.867
1203.441162109375 0 718.62506
1203.5902099609375 0 13080.823
1204.586669921875 0 2658.1458
1215.594970703125 0 1519.0709
1216.589111328125 0 4289.369
1217.603271484375 0 62462.67
1218.611083984375 0 313815.8
1219.6141357421875 0 212394.88
1220.61767578125 0 78351.58
1221.6199951171875 0 11584.831
1250.595947265625 0 1608.0043
1251.599853515625 0 1164.6088
1281.5753173828125 0 1635.3159
1293.76416015625 0 1834.0964
1294.7718505859375 0 1293.3563
1295.7869873046875 0 1051.6287
1309.7796630859375 0 730.52924
1312.6212158203125 0 1774.904
1313.631591796875 0 1227.1174
1372.7935791015625 0 1093.2296
1373.7978515625 0 1053.9147
1395.8162841796875 0 2062.876
1396.818115234375 0 1102.4972
1416.6505126953125 0 814.8273
1452.84228515625 0 2133.4692
1453.8408203125 0 1569.0426
1454.8363037109375 0 1029.898
1494.1650390625 0 1176.3971
1494.693115234375 0 1974.0162
1495.187255859375 0 2054.869
1495.6832275390625 0 1061.4834
1496.2041015625 0 2212.3438
1500.882568359375 0 1642.8829
1501.896240234375 0 949.0672
1502.693359375 0 1814.504
1503.1953125 0 1571.3417
1503.6982421875 0 1099.0173
1509.8621826171875 0 1643.4294
1516.183837890625 0 1344.7971
1516.6973876953125 0 1738.6189
1517.1925048828125 0 1833.9457
1517.6904296875 0 1017.1924
1524.1949462890625 0 1184.464
1524.698486328125 0 1922.7172
1525.209716796875 0 2559.096
1525.7069091796875 0 2256.9763
1526.201904296875 0 1352.1654
1566.88330078125 0 2288.3577
1567.88037109375 0 1878.6643
1572.923583984375 0 956.9228
1713.9600830078125 0 1452.5132
1736.78076171875 0 837.2162
1813.0252685546875 0 1364.5651
1814.0274658203125 0 1566.9877
1824.877197265625 0 927.02185
1829.03125 0 2665.0261
1830.0377197265625 0 4729.8574
1831.049560546875 0 2667.808
1832.0570068359375 0 1180.3173
2377.05859375 0 1005.148
2378.056884765625 0 790.67365
3051.408447265625 0 1136.5754

Spectrum Details

|  |  |
| --- | --- |
| Matched peaks? Matched peaks The total absolute number of peaks matched. Additionally in brackets the total fraction of peaks matched and the total number of peaks is shown. | 39 (8.84% of 441) |
| FDR? FDR The false discovery rate estimated for this peptide. It is calculated by matching all theoretical fragments with a non-integer shift with the raw peaks for this spectrum. This is done with 40 different shifts. The resulting percentage is the average number of annotated peaks over the number of annotated peaks with the correct spectrum. | 1.89% |
| Satellite FDR? Satellite FDR See the FDR for details on its calculation. This satellite ion specific FDR only contains the satellite ions (d/w) for I/L/J positions. | - |
| PSM Score? PSM Score The PSM Score as given by Hecklib to this annotated spectrum. It is shown with three significant figures. | 467 |

#### Spectrum 8210? Spectrum 8210 The raw spectrum of this peptide as annotated by Hecklib. The fragments are coloured according to ion type (see legend). Any peaks with a star '\*' as text can be hovered over to see the full details, first the ion type second the mass shift type. By hovering over the amino acids in the peptide or ions in the legend the corresponding peaks are highlighted. By toggling the 'Unassigned' label you can turn the background (unassigned) peaks on or off in the plot. By updating the slider in the Ion legend you can update the spectrum to only show the top X% of the peaks with labels. The top X% means any peak that is within X% of the highest intensity. By dragging in the spectrum you can zoom in to a specific part of the spectrum and use 'Zoom Out' to get back to the original zoom level. The annotation of the spectrum is based on the given sequence in the peptides file and is done with different software so inconsistencies are likely. The peaks are annotated based on the given sequence, with 20 ppm tolerance.

Copy Data

##### Spectrum 8210 (TSV)

###### Preview

```
Loading example...
```

*Click on the button to copy the data to your clipboard.*

Mz MinMz MaxIntensity Max

WidthHeightPeptide font sizePeptide stroke widthSpectrum font sizeSpectrum stroke widthCompact peptide

Ion legend

wxyz

abcd

OtherUnassignedIon^Charge^_Position_Show for top:%

JFPPSSEEJQA

04.40e+48.81e+41.32e+51.76e+5

Zoom Out

y^+1^_2_z^+1^_2_y^+1^_2_y^+1^_3_y^+1^_3_c^+2^_7_y^+1^_4_y^+2^_9_c^+1^_4_y^+2^_9_c^+1^_5_w^+1^_6_c^+1^_6_y^+1^_6_y^+1^_6_z^+1^_7_c^+1^_7_y^+1^_7_c^+1^_7_y^+1^_8_c^+1^_8_c^+1^_8_y^+1^_9_y^+1^_9_z^+1^_9_y^+1^_9_c^+1^_9_y^+1^_10_z^+1^_10_c^+1^_10_

0766153322993065

Fragment Matches Table

Show background peaks

| Position | Ion type | Intensity | mz Theoretical | mz Error (Th) | mz Error (ppm) | Charge | Series Number |
| --- | --- | --- | --- | --- | --- | --- | --- |
| - | - | 2259 | 120.1 | - | - | 0 | - |
| - | - | 352.3 | 124.3 | - | - | 0 | - |
| - | - | 2665 | 129.1 | - | - | 0 | - |
| - | - | 2200 | 129.1 | - | - | 0 | - |
| - | - | 766.3 | 131.1 | - | - | 0 | - |
| - | - | 451.7 | 140.1 | - | - | 0 | - |
| - | - | 408.2 | 146.6 | - | - | 0 | - |
| - | - | 972.9 | 149 | - | - | 0 | - |
| - | - | 552.7 | 157.1 | - | - | 0 | - |
| - | - | 438.9 | 160.7 | - | - | 0 | - |
| - | - | 470 | 162.8 | - | - | 0 | - |
| - | - | 641.6 | 167.1 | - | - | 0 | - |
| - | - | 730.9 | 169.1 | - | - | 0 | - |
| - | - | 958.3 | 173.4 | - | - | 0 | - |
| - | - | 446 | 176.9 | - | - | 0 | - |
| - | - | 586 | 177.1 | - | - | 0 | - |
| - | - | 433.9 | 179.5 | - | - | 0 | - |
| - | - | 910.2 | 183.1 | - | - | 0 | - |
| - | - | 428.5 | 186.2 | - | - | 0 | - |
| - | - | 6703 | 200.1 | - | - | 0 | - |
| 10 | y | 1.317E+04 | 201.1 | 0.0002206 | 1.097 | +1 | 2 |
| 10 | z | 980.1 | 202.1 | 0.003866 | 19.13 | +1 | 2 |
| - | - | 4.38E+04 | 203.1 | - | - | 0 | - |
| - | - | 3873 | 204.1 | - | - | 0 | - |
| - | - | 572.5 | 211.1 | - | - | 0 | - |
| - | - | 600.5 | 212.1 | - | - | 0 | - |
| - | - | 1121 | 217.1 | - | - | 0 | - |
| 10 | y | 1.325E+04 | 218.1 | 0.0002829 | 1.297 | +1 | 2 |
| - | - | 1035 | 219.1 | - | - | 0 | - |
| - | - | 1411 | 219.1 | - | - | 0 | - |
| - | - | 4825 | 221.1 | - | - | 0 | - |
| - | - | 1041 | 225 | - | - | 0 | - |
| - | - | 546.1 | 225.1 | - | - | 0 | - |
| - | - | 1328 | 230.2 | - | - | 0 | - |
| - | - | 1.63E+04 | 233.2 | - | - | 0 | - |
| - | - | 2042 | 234.2 | - | - | 0 | - |
| - | - | 525.7 | 234.6 | - | - | 0 | - |
| - | - | 6025 | 239.1 | - | - | 0 | - |
| - | - | 975.7 | 240.1 | - | - | 0 | - |
| - | - | 7528 | 242.2 | - | - | 0 | - |
| - | - | 738.1 | 243.1 | - | - | 0 | - |
| - | - | 1.378E+04 | 261.2 | - | - | 0 | - |
| - | - | 2033 | 262.2 | - | - | 0 | - |
| - | - | 4961 | 274.1 | - | - | 0 | - |
| - | - | 709.1 | 275.1 | - | - | 0 | - |
| - | - | 1442 | 281.1 | - | - | 0 | - |
| - | - | 2166 | 282.1 | - | - | 0 | - |
| - | - | 688.2 | 283.1 | - | - | 0 | - |
| - | - | 1161 | 287.2 | - | - | 0 | - |
| - | - | 1965 | 288.1 | - | - | 0 | - |
| - | - | 1329 | 295.1 | - | - | 0 | - |
| - | - | 579.9 | 296.1 | - | - | 0 | - |
| - | - | 612.4 | 297.1 | - | - | 0 | - |
| - | - | 6917 | 299.1 | - | - | 0 | - |
| - | - | 724 | 299.2 | - | - | 0 | - |
| - | - | 629.4 | 300.1 | - | - | 0 | - |
| - | - | 684.8 | 314.1 | - | - | 0 | - |
| 9 | y | 1807 | 314.2 | 0.0007666 | 2.44 | +1 | 3 |
| 9 | y | 1667 | 331.2 | 5.614E-05 | 0.1695 | +1 | 3 |
| - | - | 658.2 | 353.2 | - | - | 0 | - |
| - | - | 4.02E+04 | 355.1 | - | - | 0 | - |
| - | - | 1235 | 358.2 | - | - | 0 | - |
| - | - | 972 | 369.1 | - | - | 0 | - |
| - | - | 708.4 | 370.1 | - | - | 0 | - |
| - | - | 9143 | 370.7 | - | - | 0 | - |
| - | - | 1179 | 371.1 | - | - | 0 | - |
| - | - | 2915 | 371.2 | - | - | 0 | - |
| - | - | 892.8 | 373.1 | - | - | 0 | - |
| 7 | c | 1045 | 379.2 | 0.006282 | 16.57 | +2 | 7 |
| - | - | 630.4 | 383.2 | - | - | 0 | - |
| - | - | 1187 | 386.2 | - | - | 0 | - |
| - | - | 3065 | 400.2 | - | - | 0 | - |
| - | - | 2466 | 401.3 | - | - | 0 | - |
| - | - | 2811 | 415 | - | - | 0 | - |
| - | - | 529.7 | 415.2 | - | - | 0 | - |
| - | - | 705.7 | 429.2 | - | - | 0 | - |
| - | - | 6112 | 434.7 | - | - | 0 | - |
| - | - | 3638 | 435.2 | - | - | 0 | - |
| - | - | 1029 | 435.7 | - | - | 0 | - |
| 8 | y | 566.1 | 443.2 | 0.003313 | 7.476 | +1 | 4 |
| - | - | 2327 | 444.2 | - | - | 0 | - |
| - | - | 1087 | 444.7 | - | - | 0 | - |
| - | - | 1875 | 445.3 | - | - | 0 | - |
| - | - | 643.2 | 450.8 | - | - | 0 | - |
| - | - | 863.9 | 455.3 | - | - | 0 | - |
| 3 | y | 1347 | 470.2 | 0.00109 | 2.317 | +2 | 9 |
| - | - | 2016 | 471.3 | - | - | 0 | - |
| 4 | c | 1623 | 472.3 | 0.00225 | 4.764 | +1 | 4 |
| 3 | y | 811.2 | 479.2 | 0.002112 | 4.407 | +2 | 9 |
| - | - | 812.9 | 482.3 | - | - | 0 | - |
| - | - | 2672 | 484.3 | - | - | 0 | - |
| - | - | 669.9 | 485.3 | - | - | 0 | - |
| - | - | 1413 | 486.3 | - | - | 0 | - |
| - | - | 1474 | 487.3 | - | - | 0 | - |
| - | - | 5955 | 488.3 | - | - | 0 | - |
| - | - | 1895 | 489.1 | - | - | 0 | - |
| - | - | 1540 | 489.3 | - | - | 0 | - |
| - | - | 667.6 | 492.2 | - | - | 0 | - |
| - | - | 2559 | 498.2 | - | - | 0 | - |
| - | - | 1725 | 498.3 | - | - | 0 | - |
| - | - | 2243 | 500.8 | - | - | 0 | - |
| - | - | 1898 | 501.3 | - | - | 0 | - |
| - | - | 1183 | 502.3 | - | - | 0 | - |
| - | - | 1244 | 507.8 | - | - | 0 | - |
| - | - | 2002 | 515.2 | - | - | 0 | - |
| - | - | 737.4 | 516.2 | - | - | 0 | - |
| - | - | 659.2 | 542.3 | - | - | 0 | - |
| - | - | 682.8 | 543.3 | - | - | 0 | - |
| - | - | 1190 | 544.3 | - | - | 0 | - |
| - | - | 1.157E+04 | 558.3 | - | - | 0 | - |
| 5 | c | 1.108E+04 | 559.3 | 0.0004343 | 0.7764 | +1 | 5 |
| - | - | 3384 | 560.3 | - | - | 0 | - |
| - | - | 696.4 | 561.3 | - | - | 0 | - |
| - | - | 1183 | 568.3 | - | - | 0 | - |
| - | - | 3191 | 570.3 | - | - | 0 | - |
| - | - | 1324 | 571.3 | - | - | 0 | - |
| - | - | 1938 | 577.1 | - | - | 0 | - |
| - | - | 825.2 | 584.3 | - | - | 0 | - |
| - | - | 7473 | 586.3 | - | - | 0 | - |
| - | - | 757.9 | 587.3 | - | - | 0 | - |
| - | - | 1.7E+04 | 587.4 | - | - | 0 | - |
| - | - | 5302 | 588.4 | - | - | 0 | - |
| - | - | 1478 | 589.4 | - | - | 0 | - |
| - | - | 1001 | 591.4 | - | - | 0 | - |
| - | - | 1162 | 599.8 | - | - | 0 | - |
| - | - | 3903 | 600.3 | - | - | 0 | - |
| - | - | 3178 | 601.3 | - | - | 0 | - |
| - | - | 3235 | 602.3 | - | - | 0 | - |
| - | - | 3412 | 603.3 | - | - | 0 | - |
| - | - | 999.3 | 604.4 | - | - | 0 | - |
| - | - | 1310 | 607.8 | - | - | 0 | - |
| - | - | 1414 | 608.3 | - | - | 0 | - |
| - | - | 1934 | 608.8 | - | - | 0 | - |
| - | - | 1157 | 609.2 | - | - | 0 | - |
| - | - | 8596 | 610.2 | - | - | 0 | - |
| - | - | 663.8 | 623.5 | - | - | 0 | - |
| - | - | 826.3 | 624.3 | - | - | 0 | - |
| - | - | 2.06E+04 | 627.3 | - | - | 0 | - |
| - | - | 6005 | 628.3 | - | - | 0 | - |
| - | - | 1251 | 629.3 | - | - | 0 | - |
| - | - | 1409 | 629.3 | - | - | 0 | - |
| - | - | 772.6 | 631.3 | - | - | 0 | - |
| 6 | w | 733.6 | 643.3 | 0.0004149 | 0.645 | +1 | 6 |
| - | - | 2447 | 644.3 | - | - | 0 | - |
| - | - | 823.5 | 645.3 | - | - | 0 | - |
| - | - | 8947 | 645.3 | - | - | 0 | - |
| 6 | c | 3.971E+04 | 646.4 | 0.0001751 | 0.2709 | +1 | 6 |
| - | - | 1.424E+04 | 647.4 | - | - | 0 | - |
| - | - | 2900 | 648.4 | - | - | 0 | - |
| - | - | 1693 | 658.3 | - | - | 0 | - |
| 6 | y | 1439 | 659.3 | 0.002739 | 4.155 | +1 | 6 |
| - | - | 559.2 | 661.3 | - | - | 0 | - |
| - | - | 1080 | 664.4 | - | - | 0 | - |
| - | - | 823.3 | 672.4 | - | - | 0 | - |
| - | - | 1998 | 675.3 | - | - | 0 | - |
| 6 | y | 2041 | 676.3 | 0.00048 | 0.7097 | +1 | 6 |
| - | - | 887.7 | 690.4 | - | - | 0 | - |
| - | - | 829.2 | 691.4 | - | - | 0 | - |
| - | - | 1134 | 712.4 | - | - | 0 | - |
| - | - | 656.7 | 713.4 | - | - | 0 | - |
| - | - | 645.6 | 714.4 | - | - | 0 | - |
| - | - | 638.5 | 715.4 | - | - | 0 | - |
| - | - | 2454 | 717.4 | - | - | 0 | - |
| - | - | 2060 | 718.4 | - | - | 0 | - |
| - | - | 1799 | 722.3 | - | - | 0 | - |
| - | - | 828.2 | 723.3 | - | - | 0 | - |
| - | - | 1719 | 731.4 | - | - | 0 | - |
| - | - | 1405 | 732.4 | - | - | 0 | - |
| - | - | 1113 | 733.4 | - | - | 0 | - |
| - | - | 1116 | 734.4 | - | - | 0 | - |
| - | - | 2.16E+04 | 740.3 | - | - | 0 | - |
| - | - | 9568 | 741.3 | - | - | 0 | - |
| - | - | 2073 | 742.4 | - | - | 0 | - |
| 5 | z | 1145 | 747.3 | 0.0006007 | 0.8038 | +1 | 7 |
| - | - | 1635 | 748.3 | - | - | 0 | - |
| - | - | 1364 | 750.9 | - | - | 0 | - |
| 7 | c | 3591 | 757.4 | 0.01463 | 19.31 | +1 | 7 |
| - | - | 5766 | 758.4 | - | - | 0 | - |
| - | - | 1628 | 759.4 | - | - | 0 | - |
| 5 | y | 4652 | 763.3 | 0.001136 | 1.489 | +1 | 7 |
| - | - | 1768 | 764.3 | - | - | 0 | - |
| - | - | 836.6 | 771.3 | - | - | 0 | - |
| - | - | 801.7 | 772.3 | - | - | 0 | - |
| - | - | 823.1 | 774.3 | - | - | 0 | - |
| - | - | 1.054E+04 | 774.4 | - | - | 0 | - |
| 7 | c | 6.494E+04 | 775.4 | 0.000715 | 0.9221 | +1 | 7 |
| - | - | 2.99E+04 | 776.4 | - | - | 0 | - |
| - | - | 6209 | 777.4 | - | - | 0 | - |
| - | - | 897.2 | 778.4 | - | - | 0 | - |
| - | - | 773.2 | 786.5 | - | - | 0 | - |
| - | - | 658 | 787 | - | - | 0 | - |
| - | - | 842.6 | 809.4 | - | - | 0 | - |
| - | - | 1572 | 814.4 | - | - | 0 | - |
| - | - | 653 | 815.4 | - | - | 0 | - |
| - | - | 851.8 | 831.5 | - | - | 0 | - |
| - | - | 562.5 | 832.4 | - | - | 0 | - |
| - | - | 899.4 | 832.5 | - | - | 0 | - |
| - | - | 1999 | 833.4 | - | - | 0 | - |
| - | - | 1135 | 834.4 | - | - | 0 | - |
| - | - | 2210 | 845.4 | - | - | 0 | - |
| - | - | 2093 | 846.5 | - | - | 0 | - |
| - | - | 4297 | 847.5 | - | - | 0 | - |
| - | - | 1037 | 848.5 | - | - | 0 | - |
| - | - | 1114 | 850.4 | - | - | 0 | - |
| - | - | 981.2 | 853.4 | - | - | 0 | - |
| - | - | 616 | 859.4 | - | - | 0 | - |
| 4 | y | 1.567E+04 | 860.4 | 0.002008 | 2.334 | +1 | 8 |
| - | - | 5990 | 861.4 | - | - | 0 | - |
| - | - | 2433 | 862.4 | - | - | 0 | - |
| - | - | 926.1 | 867.4 | - | - | 0 | - |
| - | - | 1.112E+04 | 868.4 | - | - | 0 | - |
| - | - | 5864 | 869.4 | - | - | 0 | - |
| - | - | 1606 | 870.4 | - | - | 0 | - |
| - | - | 696.3 | 870.5 | - | - | 0 | - |
| - | - | 752.5 | 871 | - | - | 0 | - |
| - | - | 1541 | 871.4 | - | - | 0 | - |
| - | - | 685.7 | 871.5 | - | - | 0 | - |
| - | - | 812.2 | 872.4 | - | - | 0 | - |
| - | - | 641.1 | 873.4 | - | - | 0 | - |
| - | - | 678.1 | 878.5 | - | - | 0 | - |
| - | - | 1033 | 879.5 | - | - | 0 | - |
| - | - | 1085 | 884.4 | - | - | 0 | - |
| - | - | 1612 | 885.4 | - | - | 0 | - |
| 8 | c | 714.4 | 886.4 | 0.01547 | 17.45 | +1 | 8 |
| - | - | 2.028E+04 | 887.4 | - | - | 0 | - |
| - | - | 1.083E+04 | 888.4 | - | - | 0 | - |
| - | - | 3374 | 889.4 | - | - | 0 | - |
| - | - | 1115 | 891.4 | - | - | 0 | - |
| - | - | 2601 | 901.5 | - | - | 0 | - |
| - | - | 1155 | 902.5 | - | - | 0 | - |
| - | - | 4658 | 903.4 | - | - | 0 | - |
| 8 | c | 6.598E+04 | 904.4 | 0.0007055 | 0.7801 | +1 | 8 |
| - | - | 3.27E+04 | 905.4 | - | - | 0 | - |
| - | - | 7705 | 906.4 | - | - | 0 | - |
| - | - | 1166 | 907 | - | - | 0 | - |
| - | - | 2213 | 914.5 | - | - | 0 | - |
| - | - | 2158 | 915 | - | - | 0 | - |
| - | - | 2095 | 915.5 | - | - | 0 | - |
| - | - | 641.8 | 916.5 | - | - | 0 | - |
| - | - | 2331 | 927.5 | - | - | 0 | - |
| - | - | 888.6 | 928.5 | - | - | 0 | - |
| - | - | 723.7 | 930.5 | - | - | 0 | - |
| - | - | 1569 | 938.4 | - | - | 0 | - |
| 3 | y | 1129 | 939.4 | 0.003692 | 3.93 | +1 | 9 |
| 3 | y | 1858 | 940.4 | 0.005517 | 5.867 | +1 | 9 |
| 3 | z | 804.1 | 941.4 | 0.01628 | 17.3 | +1 | 9 |
| - | - | 1135 | 941.5 | - | - | 0 | - |
| - | - | 1122 | 942.5 | - | - | 0 | - |
| - | - | 4506 | 943.5 | - | - | 0 | - |
| - | - | 6738 | 944.5 | - | - | 0 | - |
| - | - | 3026 | 945.6 | - | - | 0 | - |
| - | - | 690.6 | 946.6 | - | - | 0 | - |
| - | - | 1.423E+04 | 956.4 | - | - | 0 | - |
| 3 | y | 7.661E+04 | 957.5 | 0.001195 | 1.248 | +1 | 9 |
| - | - | 3.424E+04 | 958.5 | - | - | 0 | - |
| - | - | 3221 | 958.5 | - | - | 0 | - |
| - | - | 8463 | 959.5 | - | - | 0 | - |
| - | - | 2426 | 959.5 | - | - | 0 | - |
| - | - | 1473 | 960.5 | - | - | 0 | - |
| - | - | 1.406E+04 | 960.6 | - | - | 0 | - |
| - | - | 7672 | 961.6 | - | - | 0 | - |
| - | - | 2504 | 962.6 | - | - | 0 | - |
| - | - | 1051 | 969.5 | - | - | 0 | - |
| - | - | 1442 | 970.5 | - | - | 0 | - |
| - | - | 1852 | 971.5 | - | - | 0 | - |
| - | - | 2763 | 972.5 | - | - | 0 | - |
| - | - | 1.065E+04 | 973.5 | - | - | 0 | - |
| - | - | 7607 | 974.5 | - | - | 0 | - |
| - | - | 3115 | 975.5 | - | - | 0 | - |
| - | - | 807.2 | 977.6 | - | - | 0 | - |
| - | - | 1287 | 982.5 | - | - | 0 | - |
| - | - | 1422 | 984.5 | - | - | 0 | - |
| - | - | 1892 | 985.5 | - | - | 0 | - |
| - | - | 662.3 | 986.5 | - | - | 0 | - |
| - | - | 775.6 | 987.5 | - | - | 0 | - |
| - | - | 2276 | 994.5 | - | - | 0 | - |
| - | - | 1343 | 995.5 | - | - | 0 | - |
| - | - | 3201 | 996.5 | - | - | 0 | - |
| - | - | 1419 | 997.5 | - | - | 0 | - |
| - | - | 3982 | 998.6 | - | - | 0 | - |
| - | - | 1490 | 999.5 | - | - | 0 | - |
| - | - | 1677 | 999.6 | - | - | 0 | - |
| - | - | 1.393E+04 | 1000 | - | - | 0 | - |
| - | - | 8293 | 1002 | - | - | 0 | - |
| - | - | 2961 | 1003 | - | - | 0 | - |
| - | - | 8982 | 1013 | - | - | 0 | - |
| - | - | 5885 | 1014 | - | - | 0 | - |
| - | - | 1.565E+04 | 1015 | - | - | 0 | - |
| - | - | 8983 | 1016 | - | - | 0 | - |
| - | - | 3280 | 1017 | - | - | 0 | - |
| 9 | c | 5.057E+04 | 1018 | 0.001212 | 1.191 | +1 | 9 |
| - | - | 2.848E+04 | 1019 | - | - | 0 | - |
| - | - | 485.8 | 1019 | - | - | 0 | - |
| - | - | 855.5 | 1019 | - | - | 0 | - |
| - | - | 9208 | 1020 | - | - | 0 | - |
| - | - | 1082 | 1021 | - | - | 0 | - |
| - | - | 8677 | 1044 | - | - | 0 | - |
| - | - | 5453 | 1045 | - | - | 0 | - |
| - | - | 2231 | 1046 | - | - | 0 | - |
| - | - | 1386 | 1047 | - | - | 0 | - |
| - | - | 846.8 | 1053 | - | - | 0 | - |
| - | - | 663.1 | 1071 | - | - | 0 | - |
| - | - | 1217 | 1082 | - | - | 0 | - |
| - | - | 645.7 | 1085 | - | - | 0 | - |
| 2 | y | 765.7 | 1087 | 0.002655 | 2.441 | +1 | 10 |
| 2 | z | 1.711E+04 | 1089 | 0.000348 | 0.3197 | +1 | 10 |
| - | - | 9962 | 1090 | - | - | 0 | - |
| - | - | 3446 | 1091 | - | - | 0 | - |
| - | - | 1004 | 1092 | - | - | 0 | - |
| - | - | 3910 | 1098 | - | - | 0 | - |
| - | - | 4401 | 1099 | - | - | 0 | - |
| - | - | 8132 | 1100 | - | - | 0 | - |
| - | - | 6903 | 1101 | - | - | 0 | - |
| - | - | 9799 | 1102 | - | - | 0 | - |
| - | - | 4841 | 1103 | - | - | 0 | - |
| - | - | 2628 | 1104 | - | - | 0 | - |
| - | - | 716.9 | 1110 | - | - | 0 | - |
| - | - | 2221 | 1113 | - | - | 0 | - |
| - | - | 1716 | 1114 | - | - | 0 | - |
| - | - | 864.8 | 1116 | - | - | 0 | - |
| - | - | 737.6 | 1117 | - | - | 0 | - |
| - | - | 2723 | 1118 | - | - | 0 | - |
| - | - | 1122 | 1119 | - | - | 0 | - |
| - | - | 868.3 | 1126 | - | - | 0 | - |
| - | - | 1.034E+04 | 1127 | - | - | 0 | - |
| - | - | 1.006E+04 | 1128 | - | - | 0 | - |
| - | - | 1.541E+04 | 1129 | - | - | 0 | - |
| - | - | 8840 | 1130 | - | - | 0 | - |
| - | - | 3868 | 1131 | - | - | 0 | - |
| - | - | 2494 | 1131 | - | - | 0 | - |
| - | - | 2848 | 1132 | - | - | 0 | - |
| - | - | 2782 | 1133 | - | - | 0 | - |
| - | - | 893.3 | 1143 | - | - | 0 | - |
| - | - | 2073 | 1144 | - | - | 0 | - |
| - | - | 2355 | 1145 | - | - | 0 | - |
| 10 | c | 1.744E+05 | 1146 | 0.003149 | 2.749 | +1 | 10 |
| - | - | 1.242E+05 | 1147 | - | - | 0 | - |
| - | - | 5E+04 | 1148 | - | - | 0 | - |
| - | - | 9162 | 1149 | - | - | 0 | - |
| - | - | 947.5 | 1150 | - | - | 0 | - |
| - | - | 728.8 | 1155 | - | - | 0 | - |
| - | - | 1142 | 1156 | - | - | 0 | - |
| - | - | 3560 | 1157 | - | - | 0 | - |
| - | - | 3125 | 1158 | - | - | 0 | - |
| - | - | 1835 | 1159 | - | - | 0 | - |
| - | - | 2770 | 1160 | - | - | 0 | - |
| - | - | 2179 | 1161 | - | - | 0 | - |
| - | - | 1668 | 1162 | - | - | 0 | - |
| - | - | 4005 | 1163 | - | - | 0 | - |
| - | - | 3598 | 1164 | - | - | 0 | - |
| - | - | 1706 | 1165 | - | - | 0 | - |
| - | - | 2269 | 1166 | - | - | 0 | - |
| - | - | 887.6 | 1170 | - | - | 0 | - |
| - | - | 1856 | 1171 | - | - | 0 | - |
| - | - | 2644 | 1172 | - | - | 0 | - |
| - | - | 5411 | 1173 | - | - | 0 | - |
| - | - | 1.341E+04 | 1174 | - | - | 0 | - |
| - | - | 9281 | 1175 | - | - | 0 | - |
| - | - | 4225 | 1176 | - | - | 0 | - |
| - | - | 1053 | 1177 | - | - | 0 | - |
| - | - | 1128 | 1182 | - | - | 0 | - |
| - | - | 1934 | 1183 | - | - | 0 | - |
| - | - | 2136 | 1184 | - | - | 0 | - |
| - | - | 3311 | 1185 | - | - | 0 | - |
| - | - | 2712 | 1186 | - | - | 0 | - |
| - | - | 1482 | 1187 | - | - | 0 | - |
| - | - | 2651 | 1188 | - | - | 0 | - |
| - | - | 2864 | 1189 | - | - | 0 | - |
| - | - | 3799 | 1190 | - | - | 0 | - |
| - | - | 5423 | 1191 | - | - | 0 | - |
| - | - | 3284 | 1192 | - | - | 0 | - |
| - | - | 904.7 | 1193 | - | - | 0 | - |
| - | - | 8169 | 1198 | - | - | 0 | - |
| - | - | 2.271E+04 | 1199 | - | - | 0 | - |
| - | - | 2.606E+04 | 1200 | - | - | 0 | - |
| - | - | 3.987E+04 | 1201 | - | - | 0 | - |
| - | - | 3.091E+04 | 1202 | - | - | 0 | - |
| - | - | 2.122E+04 | 1203 | - | - | 0 | - |
| - | - | 9035 | 1204 | - | - | 0 | - |
| - | - | 4720 | 1205 | - | - | 0 | - |
| - | - | 9073 | 1215 | - | - | 0 | - |
| - | - | 5.215E+04 | 1216 | - | - | 0 | - |
| - | - | 6.39E+04 | 1217 | - | - | 0 | - |
| - | - | 8.153E+04 | 1218 | - | - | 0 | - |
| - | - | 1.633E+05 | 1219 | - | - | 0 | - |
| - | - | 1.037E+05 | 1220 | - | - | 0 | - |
| - | - | 4.274E+04 | 1221 | - | - | 0 | - |
| - | - | 8079 | 1222 | - | - | 0 | - |
| - | - | 964.1 | 1294 | - | - | 0 | - |
| - | - | 1037 | 1295 | - | - | 0 | - |
| - | - | 748.3 | 1310 | - | - | 0 | - |
| - | - | 1115 | 1373 | - | - | 0 | - |
| - | - | 1340 | 1396 | - | - | 0 | - |
| - | - | 1345 | 1397 | - | - | 0 | - |
| - | - | 1409 | 1501 | - | - | 0 | - |
| - | - | 1070 | 1502 | - | - | 0 | - |
| - | - | 831.7 | 1510 | - | - | 0 | - |
| - | - | 906.1 | 1567 | - | - | 0 | - |
| - | - | 871.7 | 1568 | - | - | 0 | - |
| - | - | 1203 | 1572 | - | - | 0 | - |
| - | - | 1247 | 1573 | - | - | 0 | - |
| - | - | 1109 | 1694 | - | - | 0 | - |
| - | - | 991 | 1695 | - | - | 0 | - |
| - | - | 1193 | 1714 | - | - | 0 | - |
| - | - | 694.1 | 1715 | - | - | 0 | - |
| - | - | 1067 | 1813 | - | - | 0 | - |
| - | - | 1664 | 1814 | - | - | 0 | - |
| - | - | 3557 | 1830 | - | - | 0 | - |
| - | - | 3305 | 1831 | - | - | 0 | - |
| - | - | 621.6 | 3034 | - | - | 0 | - |
| - | - | 853.4 | 3035 | - | - | 0 | - |

m/z Charge Intensity FragmentType MassShift Position
120.08116149902344 0 2258.836
124.25291442871094 0 352.30115
129.0662078857422 0 2665.061
129.1024627685547 0 2199.7666
131.11825561523438 0 766.2684
140.08189392089844 0 451.71832
146.5814208984375 0 408.17056
149.04501342773438 0 972.926
157.09754943847656 0 552.6667
160.6846466064453 0 438.88446
162.76234436035156 0 469.9555
167.05555725097656 0 641.61786
169.0974884033203 0 730.86847
173.43861389160156 0 958.31055
176.9459686279297 0 445.99338
177.1124725341797 0 586.0139
179.54730224609375 0 433.9226
183.0771484375 0 910.20483
186.19985961914062 0 428.51016
200.1031951904297 0 6702.7637
201.0872039794922 0 13172.518 y Ammonia loss 9
202.0909423828125 0 980.1125 z 9
203.10287475585938 0 43804.53
204.10623168945312 0 3872.6086
211.1442413330078 0 572.4866
212.13980102539062 0 600.50494
217.08255004882812 0 1120.7651
218.1138153076172 0 13245.646 y 9
219.09739685058594 0 1035.0773
219.11770629882812 0 1410.8528
221.08462524414062 0 4824.7666
225.04327392578125 0 1041.1045
225.12281799316406 0 546.1005
230.1502227783203 0 1328.1681
233.16517639160156 0 16297.453
234.168212890625 0 2041.7429
234.55601501464844 0 525.69904
239.0951385498047 0 6025.149
240.0960235595703 0 975.69684
242.15036010742188 0 7528.357
243.1344757080078 0 738.0609
261.1598815917969 0 13777.797
262.1636657714844 0 2033.1392
274.1399230957031 0 4960.704
275.1431884765625 0 709.0727
281.05157470703125 0 1441.7705
282.14459228515625 0 2166.0906
283.14422607421875 0 688.153
287.171875 0 1160.5411
288.1192932128906 0 1964.9374
295.1038818359375 0 1328.7588
296.1043701171875 0 579.85425
297.101318359375 0 612.3745
299.0621032714844 0 6916.7065
299.1722412109375 0 723.9891
300.0605773925781 0 629.4211
314.1354064941406 0 684.8164
314.17181396484375 0 1807.0857 y Ammonia loss 8
331.1975402832031 0 1667.1754 y 8
353.1817321777344 0 658.2129
355.0701599121094 0 40200.79
358.2131042480469 0 1234.966
369.1214904785156 0 972.0188
370.12310791015625 0 708.3512
370.6767272949219 0 9143.259
371.10113525390625 0 1178.7578
371.1788024902344 0 2914.9207
373.08087158203125 0 892.82855
379.1913146972656 0 1044.5597 c Water loss 6
383.1936340332031 0 630.3789
386.2034606933594 0 1186.5348
400.24334716796875 0 3065.373
401.2503967285156 0 2466.3
415.0367431640625 0 2811.3984
415.22021484375 0 529.67816
429.248046875 0 705.66895
434.70660400390625 0 6112.3296
435.2078857421875 0 3637.6406
435.7083740234375 0 1029.3553
443.2103271484375 0 566.11884 y Ammonia loss 7
444.21124267578125 0 2326.9233
444.71331787109375 0 1086.9183
445.2781066894531 0 1875.4065
450.77313232421875 0 643.18866
455.2668151855469 0 863.94916
470.22344970703125 0 1346.6207 y Water loss 2
471.2850036621094 0 2015.7821
472.2895812988281 0 1622.7388 c 3
479.23193359375 0 811.2148 y 2
482.2600402832031 0 812.8846
484.2773132324219 0 2671.7534
485.278564453125 0 669.86975
486.3040771484375 0 1412.5272
487.2745361328125 0 1474.116
488.28265380859375 0 5955.4893
489.0553894042969 0 1894.7972
489.2834167480469 0 1539.821
492.2454833984375 0 667.63934
498.2200012207031 0 2558.9038
498.2579650878906 0 1725.2413
500.7518005371094 0 2243.4956
501.25384521484375 0 1897.8549
502.25994873046875 0 1183.2987
507.78619384765625 0 1243.754
515.2459716796875 0 2002.414
516.2444458007812 0 737.4324
542.2958984375 0 659.17523
543.3359375 0 682.78644
544.3132934570312 0 1189.7676
558.3162841796875 0 11572.736
559.3234252929688 0 11084.833 c 4
560.3268432617188 0 3384.2798
561.3294067382812 0 696.43994
568.3330688476562 0 1182.5797
570.3240966796875 0 3190.543
571.3270263671875 0 1324.0516
577.124755859375 0 1938.1592
584.3280639648438 0 825.2223
586.3436279296875 0 7472.8813
587.2685546875 0 757.85876
587.3505859375 0 16999.406
588.3538208007812 0 5301.54
589.3562622070312 0 1478.2155
591.3597412109375 0 1000.9969
599.82666015625 0 1161.6028
600.3233642578125 0 3902.9717
601.3287963867188 0 3178.022
602.336181640625 0 3234.6846
603.34716796875 0 3411.8691
604.3502197265625 0 999.27875
607.8248901367188 0 1309.5645
608.326416015625 0 1413.9789
608.8325805664062 0 1934.1426
609.2496948242188 0 1156.8687
610.1839599609375 0 8596.25
623.4679565429688 0 663.75415
624.3350219726562 0 826.32434
627.2620849609375 0 20597.613
628.265625 0 6005.4023
629.2681884765625 0 1250.9722
629.3286743164062 0 1409.1489
631.3414306640625 0 772.579
643.2937622070312 0 733.5608 w 5
644.2898559570312 0 2447.0725
645.29296875 0 823.45557
645.3471069335938 0 8947.289
646.355712890625 0 39709.09 c 5
647.3584594726562 0 14244.01
648.3612060546875 0 2899.6934
658.2811279296875 0 1692.8247
659.2855224609375 0 1438.8928 y Ammonia loss 5
661.3097534179688 0 559.1708
664.37548828125 0 1080.438
672.375732421875 0 823.26056
675.305419921875 0 1998.3003
676.3143310546875 0 2041.3435 y 5
690.4061279296875 0 887.7316
691.407470703125 0 829.241
712.3519897460938 0 1134.4855
713.3590698242188 0 656.74384
714.3693237304688 0 645.567
715.369140625 0 638.5483
717.3919677734375 0 2453.6836
718.3944091796875 0 2060.0571
722.3358154296875 0 1798.529
723.3384399414062 0 828.19794
731.3803100585938 0 1718.9746
732.3902587890625 0 1404.6802
733.392822265625 0 1113.4447
734.3915405273438 0 1115.9712
740.345947265625 0 21596.477
741.3489990234375 0 9568.041
742.3531494140625 0 2073.499
747.3275146484375 0 1144.5916 z 4
748.3330078125 0 1635.4355
750.9476318359375 0 1363.92
757.373291015625 0 3591.046 c Water loss 6
758.373046875 0 5765.7886
759.3761596679688 0 1628.0778
763.345703125 0 4651.902 y 4
764.349609375 0 1768.1641
771.3487548828125 0 836.60065
772.3482055664062 0 801.6792
774.3204956054688 0 823.0514
774.390380859375 0 10544.087
775.3977661132812 0 64940.684 c 6
776.4009399414062 0 29899.383
777.4034423828125 0 6208.6465
778.4095458984375 0 897.1969
786.4699096679688 0 773.15204
786.9713745117188 0 658.0255
809.4096069335938 0 842.59766
814.44482421875 0 1571.8097
815.4340209960938 0 653.00726
831.4751586914062 0 851.81177
832.4263305664062 0 562.50354
832.4790649414062 0 899.3706
833.4389038085938 0 1998.6246
834.4422607421875 0 1135.4349
845.4475708007812 0 2209.7217
846.4544067382812 0 2093.428
847.4617309570312 0 4297.111
848.4679565429688 0 1036.7823
850.3867797851562 0 1114.3774
853.4351806640625 0 981.249
859.3984375 0 616.0449
860.401611328125 0 15669.591 y 3
861.4039916992188 0 5989.8623
862.40966796875 0 2432.703
867.3977661132812 0 926.1052
868.4046630859375 0 11115.42
869.4041137695312 0 5864.184
870.408203125 0 1606.4716
870.5077514648438 0 696.27997
871.0073852539062 0 752.5056
871.3992309570312 0 1540.558
871.5106811523438 0 685.708
872.3960571289062 0 812.18805
873.4077758789062 0 641.0596
878.5133666992188 0 678.06146
879.5088500976562 0 1033.3768
884.4210815429688 0 1085.398
885.4139404296875 0 1611.6715
886.4150390625 0 714.42615 c Water loss 7
887.4134521484375 0 20279.832
888.4177856445312 0 10825.74
889.4216918945312 0 3374.011
891.4078369140625 0 1114.864
901.4896240234375 0 2600.6716
902.49365234375 0 1154.9663
903.4326171875 0 4658.171
904.4403686523438 0 65979.9 c 7
905.443115234375 0 32696.918
906.4457397460938 0 7705.2
907.008056640625 0 1165.7955
914.5139770507812 0 2212.6746
915.0195922851562 0 2158.4014
915.513916015625 0 2094.9973
916.5270385742188 0 641.79614
927.5286865234375 0 2330.6763
928.5438842773438 0 888.61035
930.4541625976562 0 723.7446
938.4384155273438 0 1569.0879
939.4381103515625 0 1128.628 y Water loss 2
940.4313354492188 0 1858.091 y Ammonia loss 2
941.4173583984375 0 804.076 z 2
941.51171875 0 1134.7655
942.52099609375 0 1121.88
943.5241088867188 0 4505.8945
944.5496215820312 0 6737.529
945.555419921875 0 3025.5688
946.5609741210938 0 690.60706
956.443359375 0 14232.529
957.451171875 0 76609.29 y 2
958.4536743164062 0 34240.758
958.5297241210938 0 3220.701
959.4570922851562 0 8463.263
959.5410766601562 0 2425.9265
960.4567260742188 0 1472.9186
960.5502319335938 0 14060.576
961.5513916015625 0 7671.9175
962.557373046875 0 2504.225
969.46728515625 0 1051.4248
970.4871215820312 0 1442.3333
971.495849609375 0 1852.3167
972.4986572265625 0 2762.7512
973.50927734375 0 10650.535
974.51318359375 0 7607.1504
975.5166625976562 0 3115.177
977.5767822265625 0 807.1732
982.49267578125 0 1287.2202
984.4801025390625 0 1421.5061
985.4915771484375 0 1892.1371
986.4873046875 0 662.2578
987.5123291015625 0 775.62164
994.5330200195312 0 2276.1414
995.534912109375 0 1342.667
996.546875 0 3201.2886
997.5457763671875 0 1418.896
998.5628662109375 0 3982.146
999.4531860351562 0 1490.487
999.5709228515625 0 1676.6223
1000.496337890625 0 13931.95
1001.5003662109375 0 8292.816
1002.50341796875 0 2961.4998
1012.5468139648438 0 8982.104
1013.5471801757812 0 5885.059
1014.558837890625 0 15653.827
1015.5634765625 0 8982.715
1016.564208984375 0 3280.1274
1017.52392578125 0 50569 c 8
1018.5263061523438 0 28479.232
1018.623046875 0 485.75873
1018.65283203125 0 855.496
1019.5294189453125 0 9208.357
1020.5267944335938 0 1081.6309
1043.5390625 0 8677.087
1044.5400390625 0 5453.454
1045.5281982421875 0 2231.2131
1046.50927734375 0 1386.2117
1052.578369140625 0 846.7565
1070.6058349609375 0 663.08594
1081.5792236328125 0 1217.1151
1084.60205078125 0 645.7125
1087.4915771484375 0 765.7064 y Ammonia loss 1
1088.501708984375 0 17111.504 z 1
1089.5042724609375 0 9962.305
1090.509521484375 0 3446.4412
1091.507568359375 0 1003.7027
1097.5743408203125 0 3909.8416
1098.574462890625 0 4401.1255
1099.5897216796875 0 8132.0728
1100.591796875 0 6902.7227
1101.572509765625 0 9798.671
1102.5716552734375 0 4841.3916
1103.57568359375 0 2627.6477
1109.5550537109375 0 716.8947
1112.54248046875 0 2220.6833
1113.542724609375 0 1715.5195
1115.6099853515625 0 864.805
1116.6060791015625 0 737.64343
1117.5321044921875 0 2723.4583
1118.5361328125 0 1121.7825
1125.57080078125 0 868.2715
1126.6231689453125 0 10339.151
1127.62744140625 0 10063.791
1128.6356201171875 0 15406.256
1129.638427734375 0 8840.498
1130.5570068359375 0 3867.5476
1130.65673828125 0 2494.3953
1131.566162109375 0 2848.0837
1132.5814208984375 0 2782.1125
1142.5679931640625 0 893.262
1143.5711669921875 0 2073.4832
1144.5736083984375 0 2355.2522
1145.58056640625 0 174408.45 c 9
1146.5836181640625 0 124227.8
1147.586181640625 0 50003.316
1148.5880126953125 0 9161.534
1149.5872802734375 0 947.54755
1154.6270751953125 0 728.80237
1155.64892578125 0 1142.183
1156.61376953125 0 3560.2693
1157.6070556640625 0 3125.3257
1158.6134033203125 0 1834.657
1159.602783203125 0 2769.9749
1160.598876953125 0 2179.4453
1161.5963134765625 0 1667.7593
1162.560791015625 0 4004.7383
1163.5693359375 0 3597.8384
1164.5810546875 0 1705.8574
1165.60791015625 0 2269.0396
1169.6124267578125 0 887.642
1170.6329345703125 0 1855.6
1171.635986328125 0 2644.3303
1172.6153564453125 0 5411.2563
1173.5943603515625 0 13407.803
1174.5948486328125 0 9281.223
1175.5985107421875 0 4225.257
1176.605712890625 0 1053.3141
1181.5994873046875 0 1127.7886
1182.6102294921875 0 1934.0487
1183.59130859375 0 2135.6829
1184.5999755859375 0 3311.0642
1185.5908203125 0 2711.822
1186.597900390625 0 1481.7795
1187.63818359375 0 2650.798
1188.6466064453125 0 2864.4397
1189.661376953125 0 3799.49
1190.62744140625 0 5422.872
1191.6253662109375 0 3283.78
1192.6207275390625 0 904.72125
1197.6378173828125 0 8169.194
1198.6259765625 0 22709.934
1199.634765625 0 26061.883
1200.63671875 0 39868.004
1201.6058349609375 0 30911.904
1202.5986328125 0 21215.076
1203.5977783203125 0 9035.426
1204.6068115234375 0 4720.182
1214.6385498046875 0 9072.768
1215.64501953125 0 52146.363
1216.6495361328125 0 63897.95
1217.6488037109375 0 81534.59
1218.615966796875 0 163295.94
1219.615966796875 0 103745.74
1220.617919921875 0 42744.62
1221.6180419921875 0 8079.3394
1293.7696533203125 0 964.1141
1294.775146484375 0 1037.1224
1309.795654296875 0 748.3442
1372.78759765625 0 1115.1964
1395.8209228515625 0 1339.5989
1396.8232421875 0 1345.2263
1500.8800048828125 0 1409.1855
1501.8858642578125 0 1070.2555
1509.8546142578125 0 831.66345
1566.8643798828125 0 906.0982
1567.889892578125 0 871.6909
1571.91796875 0 1203.4446
1572.932861328125 0 1246.8909
1693.8624267578125 0 1109.1448
1694.8663330078125 0 991.0106
1713.9482421875 0 1192.7473
1714.984130859375 0 694.1306
1813.0164794921875 0 1067.1422
1813.9935302734375 0 1663.6235
1830.0369873046875 0 3556.8052
1831.0269775390625 0 3304.6594
3034.077880859375 0 621.58716
3034.861083984375 0 853.4279

Spectrum Details

|  |  |
| --- | --- |
| Matched peaks? Matched peaks The total absolute number of peaks matched. Additionally in brackets the total fraction of peaks matched and the total number of peaks is shown. | 30 (7.30% of 411) |
| FDR? FDR The false discovery rate estimated for this peptide. It is calculated by matching all theoretical fragments with a non-integer shift with the raw peaks for this spectrum. This is done with 40 different shifts. The resulting percentage is the average number of annotated peaks over the number of annotated peaks with the correct spectrum. | 5.95% |
| Satellite FDR? Satellite FDR See the FDR for details on its calculation. This satellite ion specific FDR only contains the satellite ions (d/w) for I/L/J positions. | - |
| PSM Score? PSM Score The PSM Score as given by Hecklib to this annotated spectrum. It is shown with three significant figures. | 335 |

#### Reverse Lookup? Reverse Lookup All places where this read could be placed.

| Group | Segment | Template | Template Part | Read Part | Score | Unique |
| --- | --- | --- | --- | --- | --- | --- |
| Homo sapiens Light Chain | IGLC | IGLC2 | [10..21] | [0..11] | 88 | False |
| Homo sapiens Light Chain | IGLC | IGLC3 | [8..19] | [0..11] | 88 | False |
| Homo sapiens Light Chain | IGLC | IGLC6 | [10..21] | [0..11] | 88 | False |
| Homo sapiens Light Chain | IGLC | IGLC7 | [10..21] | [0..11] | 88 | False |

| Recombined | Template Part | Read Part | Score | Unique |
| --- | --- | --- | --- | --- |
| REC-0-1\_002 | [121..132] | [0..11] | 88 | True |

#### Meta Information from Multiple reads

##### Number of combined reads

5

##### Intensity

0.8627

##### TotalArea

2.379E+09

##### Changes to the peptide sequence

JFPPSSEEJQA

L→JNo support for either Leucine or Isoleucine based on side chain ions (Position: 9)

L→JNo support for either Leucine or Isoleucine based on side chain ions (Position: 1)

#### Positional Score

Copy Data

##### Positional Score (TSV)

###### Preview

```
Loading example...
```

*Click on the button to copy the data to your clipboard.*

10012345678910

Label Value
"0" 0.6
"1" 0.596
"2" 0.598
"3" 0.598
"4" 0.598
"5" 0.594
"6" 0.598
"7" 0.6
"8" 0.6
"9" 0.598
"10" 0.598

#### Meta Information from PEAKS

##### Scan Identifier

F1:7848

##### Original sequence

L

F

P

P

S

S

E

E

L

Q

A

##### Posttranslational Modifications

##### Source File

D:\separate\_stitch\_analyses\xle-disambiguation\raw\20210323\_F1\_UM1\_Peng0013\_SA\_F59\_ingel\_3ug\_ELA.raw

##### Fraction

1

##### Scan Feature

F1:8758

##### De Novo Score

99

##### ConfidenceScore

99

### m/z

609.3065

##### Mass

1216.5974

##### Charge

2

##### Retention Time

42.74

##### Predicted Retention Time

-

##### Area

4.758E+08

##### Parts Per Million

0.9

##### Fragmentation mode

ETHCD

##### Originating file

01 D:\separate\_stitch\_analyses\xle-disambiguation\20210325\_F59\_3ug\_DENOVO\_12.csv

#### Meta Information from PEAKS

##### Scan Identifier

F1:7929

##### Original sequence

L

F

P

P

S

S

E

E

L

Q

A

##### Posttranslational Modifications

##### Source File

D:\separate\_stitch\_analyses\xle-disambiguation\raw\20210323\_F1\_UM1\_Peng0013\_SA\_F59\_ingel\_3ug\_ELA.raw

##### Fraction

1

##### Scan Feature

F1:8758

##### De Novo Score

99

##### ConfidenceScore

99

### m/z

609.3065

##### Mass

1216.5974

##### Charge

2

##### Retention Time

42.74

##### Predicted Retention Time

-

##### Area

4.758E+08

##### Parts Per Million

0.9

##### Fragmentation mode

ETHCD

##### Originating file

01 D:\separate\_stitch\_analyses\xle-disambiguation\20210325\_F59\_3ug\_DENOVO\_12.csv

#### Meta Information from PEAKS

##### Scan Identifier

F1:8007

##### Original sequence

L

F

P

P

S

S

E

E

L

Q

A

##### Posttranslational Modifications

##### Source File

D:\separate\_stitch\_analyses\xle-disambiguation\raw\20210323\_F1\_UM1\_Peng0013\_SA\_F59\_ingel\_3ug\_ELA.raw

##### Fraction

1

##### Scan Feature

F1:8758

##### De Novo Score

99

##### ConfidenceScore

99

### m/z

609.3065

##### Mass

1216.5974

##### Charge

2

##### Retention Time

42.74

##### Predicted Retention Time

-

##### Area

4.758E+08

##### Parts Per Million

0.9

##### Fragmentation mode

ETHCD

##### Originating file

01 D:\separate\_stitch\_analyses\xle-disambiguation\20210325\_F59\_3ug\_DENOVO\_12.csv

#### Meta Information from PEAKS

##### Scan Identifier

F1:7659

##### Original sequence

L

F

P

P

S

S

E

E

L

Q

A

##### Posttranslational Modifications

##### Source File

D:\separate\_stitch\_analyses\xle-disambiguation\raw\20210323\_F1\_UM1\_Peng0013\_SA\_F59\_ingel\_3ug\_ELA.raw

##### Fraction

1

##### Scan Feature

F1:8758

##### De Novo Score

99

##### ConfidenceScore

99

### m/z

609.3065

##### Mass

1216.5974

##### Charge

2

##### Retention Time

42.74

##### Predicted Retention Time

-

##### Area

4.758E+08

##### Parts Per Million

0.9

##### Fragmentation mode

ETHCD

##### Originating file

01 D:\separate\_stitch\_analyses\xle-disambiguation\20210325\_F59\_3ug\_DENOVO\_12.csv

#### Meta Information from PEAKS

##### Scan Identifier

F1:8210

##### Original sequence

L

F

P

P

S

S

E

E

L

Q

A

##### Posttranslational Modifications

##### Source File

D:\separate\_stitch\_analyses\xle-disambiguation\raw\20210323\_F1\_UM1\_Peng0013\_SA\_F59\_ingel\_3ug\_ELA.raw

##### Fraction

1

##### Scan Feature

F1:8758

##### De Novo Score

99

##### ConfidenceScore

99

### m/z

609.3065

##### Mass

1216.5974

##### Charge

2

##### Retention Time

42.74

##### Predicted Retention Time

-

##### Area

4.758E+08

##### Parts Per Million

0.9

##### Fragmentation mode

ETHCD

##### Originating file

01 D:\separate\_stitch\_analyses\xle-disambiguation\20210325\_F59\_3ug\_DENOVO\_12.csv
