## Supplementary material for "A handle on mass coincidence errors in *de novo* sequencing of antibodies by bottom-up proteomics": Combined_003.html

Details Combined\_003 | Stitch OverviewUndefined

### Read Combined\_003

#### Sequence (length=10)

FTFDDYAMHW

#### Spectrum 9743? Spectrum 9743 The raw spectrum of this peptide as annotated by Hecklib. The fragments are coloured according to ion type (see legend). Any peaks with a star '\*' as text can be hovered over to see the full details, first the ion type second the mass shift type. By hovering over the amino acids in the peptide or ions in the legend the corresponding peaks are highlighted. By toggling the 'Unassigned' label you can turn the background (unassigned) peaks on or off in the plot. By updating the slider in the Ion legend you can update the spectrum to only show the top X% of the peaks with labels. The top X% means any peak that is within X% of the highest intensity. By dragging in the spectrum you can zoom in to a specific part of the spectrum and use 'Zoom Out' to get back to the original zoom level. The annotation of the spectrum is based on the given sequence in the peptides file and is done with different software so inconsistencies are likely. The peaks are annotated based on the given sequence, with 20 ppm tolerance.

Copy Data

##### Spectrum 9743 (TSV)

###### Preview

```
Loading example...
```

*Click on the button to copy the data to your clipboard.*

Mz MinMz MaxIntensity Max

WidthHeightPeptide font sizePeptide stroke widthSpectrum font sizeSpectrum stroke widthCompact peptide

Ion legend

wxyz

abcd

OtherUnassignedIon^Charge^_Position_Show for top:%

FTFDDYAMHW

02.56e+55.11e+57.67e+51.02e+6

Zoom Out

b^+2^_3_a^+1^_2_a^+1^_2_y^+1^_1_d^+1^_2_d^+1^_2_a^+1^_2_b^+1^_2_y^+2^_3_b^+2^_4_b^+1^_2_y^+1^_2_a^+1^_3_a^+1^_3_b^+1^_3_b^+2^_6_b^+1^_3_y^+2^_7_y^+1^_3_b^+1^_4_b^+2^_8_b^+1^_4_y^+2^_8_y^+2^_8_y^+1^_4_b^+2^_9_b^+2^_9_b^+2^_9_y^+2^_9_y^+2^_9_b^+1^_5_b^+1^_5_\*\*y^+1^_5_b^+1^_6_b^+1^_6_y^+1^_6_y^+1^_6_b^+1^_7_b^+1^_7_b^+1^_7_y^+1^_7_y^+1^_7_b^+1^_8_b^+1^_8_y^+1^_8_y^+1^_8_b^+1^_9_b^+1^_9_y^+1^_9_y^+1^_9_

0739147922182958

Fragment Matches Table

Show background peaks

| Position | Ion type | Intensity | mz Theoretical | mz Error (Th) | mz Error (ppm) | Charge | Series Number |
| --- | --- | --- | --- | --- | --- | --- | --- |
| - | - | 1.012E+06 | 120.1 | - | - | 0 | - |
| - | - | 800.6 | 121 | - | - | 0 | - |
| - | - | 3112 | 121.1 | - | - | 0 | - |
| - | - | 8.606E+04 | 121.1 | - | - | 0 | - |
| - | - | 1335 | 122.1 | - | - | 0 | - |
| - | - | 1790 | 122.1 | - | - | 0 | - |
| - | - | 1527 | 129.1 | - | - | 0 | - |
| - | - | 2264 | 130.1 | - | - | 0 | - |
| - | - | 1.007E+04 | 130.1 | - | - | 0 | - |
| - | - | 2320 | 131 | - | - | 0 | - |
| - | - | 4163 | 132.1 | - | - | 0 | - |
| - | - | 7.768E+04 | 136.1 | - | - | 0 | - |
| - | - | 2218 | 136.1 | - | - | 0 | - |
| - | - | 6060 | 137.1 | - | - | 0 | - |
| - | - | 7448 | 138.1 | - | - | 0 | - |
| - | - | 1.141E+04 | 138.1 | - | - | 0 | - |
| - | - | 3123 | 143.1 | - | - | 0 | - |
| - | - | 904.2 | 143.9 | - | - | 0 | - |
| - | - | 1442 | 144.1 | - | - | 0 | - |
| - | - | 3673 | 146.1 | - | - | 0 | - |
| - | - | 3090 | 146.1 | - | - | 0 | - |
| - | - | 1014 | 147.1 | - | - | 0 | - |
| - | - | 3707 | 148.1 | - | - | 0 | - |
| - | - | 1943 | 148.9 | - | - | 0 | - |
| - | - | 2268 | 155.1 | - | - | 0 | - |
| - | - | 4526 | 155.1 | - | - | 0 | - |
| - | - | 2.766E+04 | 156.1 | - | - | 0 | - |
| - | - | 1857 | 157.1 | - | - | 0 | - |
| - | - | 2.297E+04 | 158.1 | - | - | 0 | - |
| - | - | 4.943E+04 | 159.1 | - | - | 0 | - |
| - | - | 3008 | 159.1 | - | - | 0 | - |
| - | - | 2048 | 160.1 | - | - | 0 | - |
| - | - | 4749 | 160.1 | - | - | 0 | - |
| - | - | 2419 | 160.1 | - | - | 0 | - |
| - | - | 2886 | 161.1 | - | - | 0 | - |
| - | - | 1172 | 162.1 | - | - | 0 | - |
| - | - | 1403 | 165.1 | - | - | 0 | - |
| - | - | 5825 | 165.1 | - | - | 0 | - |
| - | - | 1276 | 165.1 | - | - | 0 | - |
| - | - | 2120 | 166.1 | - | - | 0 | - |
| - | - | 4420 | 166.1 | - | - | 0 | - |
| - | - | 2.287E+04 | 166.1 | - | - | 0 | - |
| - | - | 5867 | 167.1 | - | - | 0 | - |
| - | - | 2293 | 169.1 | - | - | 0 | - |
| - | - | 1207 | 169.1 | - | - | 0 | - |
| - | - | 2544 | 171.1 | - | - | 0 | - |
| - | - | 1071 | 171.1 | - | - | 0 | - |
| - | - | 1278 | 172.1 | - | - | 0 | - |
| - | - | 999.7 | 173.1 | - | - | 0 | - |
| - | - | 1083 | 173.4 | - | - | 0 | - |
| - | - | 3029 | 174.1 | - | - | 0 | - |
| - | - | 1975 | 175.1 | - | - | 0 | - |
| - | - | 2576 | 175.1 | - | - | 0 | - |
| - | - | 8393 | 176.1 | - | - | 0 | - |
| - | - | 4.638E+05 | 176.1 | - | - | 0 | - |
| - | - | 7889 | 177.1 | - | - | 0 | - |
| - | - | 5.059E+04 | 177.1 | - | - | 0 | - |
| - | - | 1788 | 178.1 | - | - | 0 | - |
| - | - | 1181 | 178.1 | - | - | 0 | - |
| - | - | 3648 | 180.1 | - | - | 0 | - |
| - | - | 3788 | 185.1 | - | - | 0 | - |
| - | - | 6.113E+04 | 186.1 | - | - | 0 | - |
| - | - | 1886 | 187.1 | - | - | 0 | - |
| - | - | 7304 | 187.1 | - | - | 0 | - |
| - | - | 3.457E+04 | 188.1 | - | - | 0 | - |
| - | - | 3293 | 189.1 | - | - | 0 | - |
| - | - | 5075 | 189.1 | - | - | 0 | - |
| - | - | 3701 | 189.1 | - | - | 0 | - |
| 3 | b | 1554 | 190.1 | 0.0003996 | 2.102 | +2 | 3 |
| - | - | 1422 | 191.1 | - | - | 0 | - |
| - | - | 3183 | 191.1 | - | - | 0 | - |
| - | - | 7472 | 191.1 | - | - | 0 | - |
| - | - | 1354 | 192.1 | - | - | 0 | - |
| - | - | 1771 | 193.1 | - | - | 0 | - |
| - | - | 2.936E+04 | 193.1 | - | - | 0 | - |
| - | - | 2446 | 194.1 | - | - | 0 | - |
| - | - | 1384 | 194.1 | - | - | 0 | - |
| - | - | 6867 | 195.1 | - | - | 0 | - |
| - | - | 2572 | 197.1 | - | - | 0 | - |
| - | - | 1091 | 199.1 | - | - | 0 | - |
| - | - | 2171 | 201.1 | - | - | 0 | - |
| - | - | 1929 | 202.1 | - | - | 0 | - |
| - | - | 1108 | 202.2 | - | - | 0 | - |
| - | - | 4267 | 203.1 | - | - | 0 | - |
| - | - | 2348 | 203.1 | - | - | 0 | - |
| 2 | a | 3.367E+04 | 203.1 | 0.0001677 | 0.8257 | +1 | 2 |
| - | - | 1.486E+04 | 204.1 | - | - | 0 | - |
| 2 | a | 1072 | 204.1 | 6.926E-06 | 0.03393 | +1 | 2 |
| - | - | 4181 | 204.1 | - | - | 0 | - |
| - | - | 1746 | 205.1 | - | - | 0 | - |
| 10 | y | 2.388E+04 | 205.1 | 0.0001665 | 0.8116 | +1 | 1 |
| - | - | 1108 | 205.1 | - | - | 0 | - |
| 2 | d | 1343 | 205.1 | 0.0002083 | 1.015 | +1 | 2 |
| - | - | 2710 | 206.1 | - | - | 0 | - |
| - | - | 893.3 | 206.5 | - | - | 0 | - |
| 2 | d | 9453 | 207.1 | 0.0001872 | 0.9037 | +1 | 2 |
| - | - | 2414 | 208.1 | - | - | 0 | - |
| - | - | 909.9 | 209.1 | - | - | 0 | - |
| - | - | 7512 | 212.1 | - | - | 0 | - |
| - | - | 1216 | 213.1 | - | - | 0 | - |
| - | - | 1122 | 213.1 | - | - | 0 | - |
| - | - | 3768 | 214.1 | - | - | 0 | - |
| - | - | 4734 | 215.1 | - | - | 0 | - |
| - | - | 1304 | 215.1 | - | - | 0 | - |
| - | - | 3.237E+04 | 217.1 | - | - | 0 | - |
| - | - | 4778 | 218.1 | - | - | 0 | - |
| - | - | 4595 | 219.1 | - | - | 0 | - |
| - | - | 3411 | 219.1 | - | - | 0 | - |
| - | - | 7.879E+04 | 221.1 | - | - | 0 | - |
| 2 | a | 5.843E+05 | 221.1 | 0.0002994 | 1.354 | +1 | 2 |
| - | - | 7470 | 222.1 | - | - | 0 | - |
| - | - | 7.269E+04 | 222.1 | - | - | 0 | - |
| - | - | 1638 | 223.1 | - | - | 0 | - |
| - | - | 1635 | 223.1 | - | - | 0 | - |
| - | - | 4548 | 223.1 | - | - | 0 | - |
| - | - | 1339 | 226.1 | - | - | 0 | - |
| - | - | 1003 | 227.1 | - | - | 0 | - |
| - | - | 945 | 227.3 | - | - | 0 | - |
| - | - | 3156 | 228.1 | - | - | 0 | - |
| - | - | 2167 | 229.1 | - | - | 0 | - |
| - | - | 2265 | 229.1 | - | - | 0 | - |
| - | - | 1527 | 229.1 | - | - | 0 | - |
| - | - | 9876 | 231.1 | - | - | 0 | - |
| 2 | b | 1.456E+05 | 231.1 | 0.0002482 | 1.074 | +1 | 2 |
| - | - | 1.842E+04 | 232.1 | - | - | 0 | - |
| - | - | 1134 | 233.1 | - | - | 0 | - |
| - | - | 5.068E+04 | 235.1 | - | - | 0 | - |
| - | - | 6910 | 236.1 | - | - | 0 | - |
| - | - | 2172 | 238.1 | - | - | 0 | - |
| - | - | 8815 | 239.1 | - | - | 0 | - |
| - | - | 1769 | 240.1 | - | - | 0 | - |
| 8 | y | 2764 | 245.1 | 0.004125 | 16.83 | +2 | 3 |
| - | - | 1708 | 246.1 | - | - | 0 | - |
| 4 | b | 1460 | 247.1 | 0.0009378 | 3.795 | +2 | 4 |
| - | - | 1210 | 247.1 | - | - | 0 | - |
| - | - | 2063 | 247.1 | - | - | 0 | - |
| - | - | 3013 | 248.2 | - | - | 0 | - |
| 2 | b | 3.025E+05 | 249.1 | 0.0002884 | 1.158 | +1 | 2 |
| - | - | 4.019E+04 | 250.1 | - | - | 0 | - |
| - | - | 2.606E+04 | 251.1 | - | - | 0 | - |
| - | - | 4841 | 251.1 | - | - | 0 | - |
| - | - | 2915 | 252.1 | - | - | 0 | - |
| - | - | 1.682E+04 | 253.1 | - | - | 0 | - |
| - | - | 5646 | 254.1 | - | - | 0 | - |
| - | - | 1.39E+04 | 257.1 | - | - | 0 | - |
| - | - | 4620 | 259.1 | - | - | 0 | - |
| - | - | 3499 | 261.1 | - | - | 0 | - |
| - | - | 3003 | 261.1 | - | - | 0 | - |
| - | - | 1217 | 262.1 | - | - | 0 | - |
| - | - | 6.134E+04 | 263.1 | - | - | 0 | - |
| - | - | 3911 | 263.1 | - | - | 0 | - |
| - | - | 8068 | 264.1 | - | - | 0 | - |
| - | - | 3715 | 264.1 | - | - | 0 | - |
| - | - | 2808 | 267.1 | - | - | 0 | - |
| - | - | 2399 | 268.1 | - | - | 0 | - |
| - | - | 1977 | 268.1 | - | - | 0 | - |
| - | - | 1194 | 269.1 | - | - | 0 | - |
| - | - | 2057 | 271.1 | - | - | 0 | - |
| - | - | 1082 | 272.1 | - | - | 0 | - |
| - | - | 1727 | 273.1 | - | - | 0 | - |
| - | - | 1.442E+04 | 274.1 | - | - | 0 | - |
| - | - | 1.967E+04 | 279.1 | - | - | 0 | - |
| - | - | 1728 | 279.1 | - | - | 0 | - |
| - | - | 2678 | 280.1 | - | - | 0 | - |
| - | - | 1421 | 280.1 | - | - | 0 | - |
| - | - | 1363 | 283.1 | - | - | 0 | - |
| - | - | 1362 | 284.2 | - | - | 0 | - |
| - | - | 2.832E+04 | 285.1 | - | - | 0 | - |
| - | - | 2641 | 286.1 | - | - | 0 | - |
| - | - | 1780 | 287.1 | - | - | 0 | - |
| - | - | 1516 | 289.1 | - | - | 0 | - |
| - | - | 1723 | 289.1 | - | - | 0 | - |
| - | - | 3751 | 289.1 | - | - | 0 | - |
| - | - | 1.832E+04 | 292.1 | - | - | 0 | - |
| - | - | 1178 | 292.2 | - | - | 0 | - |
| - | - | 1068 | 292.2 | - | - | 0 | - |
| - | - | 2753 | 293.1 | - | - | 0 | - |
| - | - | 2.28E+04 | 296.2 | - | - | 0 | - |
| - | - | 3060 | 297.2 | - | - | 0 | - |
| - | - | 2623 | 300.1 | - | - | 0 | - |
| - | - | 1150 | 302.2 | - | - | 0 | - |
| - | - | 4609 | 303.1 | - | - | 0 | - |
| - | - | 2184 | 304.1 | - | - | 0 | - |
| - | - | 1973 | 306.1 | - | - | 0 | - |
| - | - | 1523 | 307.1 | - | - | 0 | - |
| - | - | 1604 | 307.1 | - | - | 0 | - |
| - | - | 3281 | 310.2 | - | - | 0 | - |
| - | - | 1511 | 312.2 | - | - | 0 | - |
| - | - | 1472 | 314.1 | - | - | 0 | - |
| - | - | 1928 | 315.1 | - | - | 0 | - |
| - | - | 2058 | 316.2 | - | - | 0 | - |
| - | - | 4250 | 317.1 | - | - | 0 | - |
| - | - | 1.006E+04 | 318.1 | - | - | 0 | - |
| - | - | 1480 | 319.1 | - | - | 0 | - |
| - | - | 2161 | 320.1 | - | - | 0 | - |
| - | - | 3900 | 323.1 | - | - | 0 | - |
| - | - | 1625 | 323.2 | - | - | 0 | - |
| - | - | 5.042E+04 | 324.1 | - | - | 0 | - |
| - | - | 8166 | 325.1 | - | - | 0 | - |
| - | - | 1657 | 327.1 | - | - | 0 | - |
| - | - | 1931 | 328.1 | - | - | 0 | - |
| - | - | 1064 | 329.1 | - | - | 0 | - |
| - | - | 2402 | 330.1 | - | - | 0 | - |
| - | - | 2778 | 330.2 | - | - | 0 | - |
| - | - | 3729 | 332.1 | - | - | 0 | - |
| - | - | 2767 | 333.1 | - | - | 0 | - |
| - | - | 9399 | 333.2 | - | - | 0 | - |
| - | - | 1006 | 334.1 | - | - | 0 | - |
| - | - | 2142 | 334.1 | - | - | 0 | - |
| - | - | 1295 | 334.2 | - | - | 0 | - |
| - | - | 2538 | 338.1 | - | - | 0 | - |
| - | - | 4844 | 339.2 | - | - | 0 | - |
| 9 | y | 2.596E+05 | 342.2 | 0.0004282 | 1.251 | +1 | 2 |
| - | - | 4.964E+04 | 343.2 | - | - | 0 | - |
| - | - | 1850 | 344.1 | - | - | 0 | - |
| - | - | 5936 | 344.2 | - | - | 0 | - |
| - | - | 8099 | 346.1 | - | - | 0 | - |
| - | - | 1096 | 347.1 | - | - | 0 | - |
| - | - | 4176 | 348.1 | - | - | 0 | - |
| - | - | 1437 | 349.1 | - | - | 0 | - |
| - | - | 1523 | 349.2 | - | - | 0 | - |
| - | - | 1.798E+04 | 350.1 | - | - | 0 | - |
| 3 | a | 3681 | 350.2 | 0.0007083 | 2.023 | +1 | 3 |
| - | - | 3363 | 351.1 | - | - | 0 | - |
| 3 | a | 2427 | 351.2 | 0.0003361 | 0.9572 | +1 | 3 |
| - | - | 6300 | 352.1 | - | - | 0 | - |
| - | - | 1140 | 352.2 | - | - | 0 | - |
| - | - | 1136 | 354.2 | - | - | 0 | - |
| - | - | 2726 | 355.1 | - | - | 0 | - |
| - | - | 1.24E+04 | 356.1 | - | - | 0 | - |
| - | - | 2200 | 357.1 | - | - | 0 | - |
| - | - | 2336 | 358.1 | - | - | 0 | - |
| - | - | 5122 | 360.1 | - | - | 0 | - |
| - | - | 990.9 | 360.1 | - | - | 0 | - |
| - | - | 1930 | 362.1 | - | - | 0 | - |
| - | - | 2.239E+04 | 364.2 | - | - | 0 | - |
| - | - | 4775 | 365.2 | - | - | 0 | - |
| - | - | 4277 | 366.1 | - | - | 0 | - |
| - | - | 2658 | 367.2 | - | - | 0 | - |
| - | - | 1380 | 371.1 | - | - | 0 | - |
| - | - | 4301 | 374.1 | - | - | 0 | - |
| - | - | 7546 | 376.1 | - | - | 0 | - |
| - | - | 7.549E+04 | 378.1 | - | - | 0 | - |
| 3 | b | 5.679E+04 | 378.2 | 0.0003005 | 0.7945 | +1 | 3 |
| - | - | 1.568E+04 | 379.1 | - | - | 0 | - |
| - | - | 1.486E+04 | 379.2 | - | - | 0 | - |
| - | - | 4128 | 380.1 | - | - | 0 | - |
| - | - | 1708 | 381.1 | - | - | 0 | - |
| - | - | 6536 | 382.1 | - | - | 0 | - |
| - | - | 978.8 | 382.2 | - | - | 0 | - |
| - | - | 4217 | 390.2 | - | - | 0 | - |
| - | - | 1579 | 391.2 | - | - | 0 | - |
| - | - | 3.015E+04 | 394.1 | - | - | 0 | - |
| - | - | 3836 | 395.1 | - | - | 0 | - |
| 6 | b | 1816 | 395.2 | 0.00381 | 9.64 | +2 | 6 |
| 3 | b | 3591 | 396.2 | 0.0009663 | 2.439 | +1 | 3 |
| - | - | 1520 | 397.2 | - | - | 0 | - |
| - | - | 1680 | 398.2 | - | - | 0 | - |
| - | - | 2799 | 399.2 | - | - | 0 | - |
| - | - | 2346 | 401.2 | - | - | 0 | - |
| - | - | 1558 | 404.1 | - | - | 0 | - |
| - | - | 5597 | 404.1 | - | - | 0 | - |
| - | - | 1351 | 405.1 | - | - | 0 | - |
| - | - | 1799 | 406.2 | - | - | 0 | - |
| - | - | 5281 | 407.2 | - | - | 0 | - |
| - | - | 9255 | 408.2 | - | - | 0 | - |
| - | - | 2859 | 409.2 | - | - | 0 | - |
| - | - | 2473 | 410.1 | - | - | 0 | - |
| - | - | 1501 | 413.2 | - | - | 0 | - |
| - | - | 1600 | 415.2 | - | - | 0 | - |
| - | - | 2302 | 416.1 | - | - | 0 | - |
| - | - | 2709 | 419.2 | - | - | 0 | - |
| - | - | 4.78E+04 | 425.2 | - | - | 0 | - |
| - | - | 9474 | 426.2 | - | - | 0 | - |
| - | - | 7659 | 427.2 | - | - | 0 | - |
| - | - | 4259 | 433.2 | - | - | 0 | - |
| - | - | 1314 | 434.2 | - | - | 0 | - |
| - | - | 1617 | 442.2 | - | - | 0 | - |
| - | - | 2721 | 443.2 | - | - | 0 | - |
| - | - | 5042 | 447.2 | - | - | 0 | - |
| - | - | 3962 | 449.2 | - | - | 0 | - |
| - | - | 2353 | 454.2 | - | - | 0 | - |
| - | - | 6783 | 455.2 | - | - | 0 | - |
| - | - | 8435 | 461.2 | - | - | 0 | - |
| - | - | 3195 | 461.2 | - | - | 0 | - |
| - | - | 1846 | 462.2 | - | - | 0 | - |
| - | - | 2199 | 462.2 | - | - | 0 | - |
| - | - | 2.073E+04 | 465.2 | - | - | 0 | - |
| - | - | 4103 | 466.2 | - | - | 0 | - |
| - | - | 4318 | 470.2 | - | - | 0 | - |
| - | - | 4034 | 471.2 | - | - | 0 | - |
| - | - | 1317 | 473.2 | - | - | 0 | - |
| - | - | 2835 | 473.2 | - | - | 0 | - |
| 4 | y | 3740 | 477.2 | 0.0006768 | 1.418 | +2 | 7 |
| - | - | 1.304E+04 | 479.2 | - | - | 0 | - |
| - | - | 2788 | 480.2 | - | - | 0 | - |
| - | - | 1739 | 481.2 | - | - | 0 | - |
| - | - | 3256 | 486.2 | - | - | 0 | - |
| - | - | 3152 | 489.2 | - | - | 0 | - |
| 8 | y | 6.557E+04 | 489.2 | 0.005619 | 11.49 | +1 | 3 |
| - | - | 1.871E+04 | 490.2 | - | - | 0 | - |
| - | - | 7293 | 491.2 | - | - | 0 | - |
| - | - | 1830 | 492.2 | - | - | 0 | - |
| 4 | b | 6354 | 493.2 | 0.0007012 | 1.422 | +1 | 4 |
| - | - | 3741 | 494.2 | - | - | 0 | - |
| 8 | b | 7585 | 495.2 | 0.001396 | 2.818 | +2 | 8 |
| - | - | 2569 | 496.2 | - | - | 0 | - |
| - | - | 3.223E+04 | 496.2 | - | - | 0 | - |
| - | - | 4597 | 497.2 | - | - | 0 | - |
| - | - | 7339 | 497.2 | - | - | 0 | - |
| - | - | 1521 | 498.2 | - | - | 0 | - |
| - | - | 2310 | 509.7 | - | - | 0 | - |
| 4 | b | 2.934E+04 | 511.2 | 0.0001158 | 0.2264 | +1 | 4 |
| - | - | 6964 | 512.2 | - | - | 0 | - |
| - | - | 1.259E+04 | 513.2 | - | - | 0 | - |
| - | - | 5053 | 514.2 | - | - | 0 | - |
| - | - | 1236 | 518.5 | - | - | 0 | - |
| - | - | 4638 | 518.7 | - | - | 0 | - |
| - | - | 9793 | 519.2 | - | - | 0 | - |
| - | - | 4752 | 520.2 | - | - | 0 | - |
| - | - | 1849 | 521.2 | - | - | 0 | - |
| - | - | 2638 | 523.2 | - | - | 0 | - |
| - | - | 2135 | 527.7 | - | - | 0 | - |
| - | - | 1936 | 530.2 | - | - | 0 | - |
| - | - | 1331 | 532.2 | - | - | 0 | - |
| - | - | 1968 | 533.2 | - | - | 0 | - |
| - | - | 2723 | 537.2 | - | - | 0 | - |
| - | - | 1457 | 539.2 | - | - | 0 | - |
| - | - | 8584 | 541.2 | - | - | 0 | - |
| 3 | y | 7554 | 541.7 | 0.003113 | 5.747 | +2 | 8 |
| - | - | 5936 | 542.2 | - | - | 0 | - |
| - | - | 4878 | 542.2 | - | - | 0 | - |
| - | - | 3463 | 542.7 | - | - | 0 | - |
| - | - | 6949 | 548.2 | - | - | 0 | - |
| - | - | 3124 | 549.2 | - | - | 0 | - |
| - | - | 1427 | 550.2 | - | - | 0 | - |
| 3 | y | 7.684E+04 | 550.7 | 0.002592 | 4.706 | +2 | 8 |
| - | - | 5.193E+04 | 551.2 | - | - | 0 | - |
| - | - | 1.868E+04 | 551.7 | - | - | 0 | - |
| - | - | 3954 | 552.2 | - | - | 0 | - |
| - | - | 1461 | 555.2 | - | - | 0 | - |
| - | - | 2329 | 557.2 | - | - | 0 | - |
| - | - | 1229 | 558.2 | - | - | 0 | - |
| 7 | y | 9.125E+04 | 560.2 | 0.004973 | 8.877 | +1 | 4 |
| - | - | 2.96E+04 | 561.2 | - | - | 0 | - |
| - | - | 1.008E+04 | 562.2 | - | - | 0 | - |
| 9 | b | 2458 | 563.7 | 0.003983 | 7.066 | +2 | 9 |
| 9 | b | 2680 | 564.2 | 0.00874 | 15.49 | +2 | 9 |
| - | - | 2827 | 567.2 | - | - | 0 | - |
| - | - | 1375 | 569.2 | - | - | 0 | - |
| - | - | 1469 | 569.7 | - | - | 0 | - |
| - | - | 7278 | 570.2 | - | - | 0 | - |
| - | - | 1798 | 571.2 | - | - | 0 | - |
| 9 | b | 6155 | 572.7 | 0.001875 | 3.273 | +2 | 9 |
| - | - | 3816 | 573.2 | - | - | 0 | - |
| - | - | 1738 | 579.2 | - | - | 0 | - |
| - | - | 4098 | 580.2 | - | - | 0 | - |
| - | - | 1179 | 581.2 | - | - | 0 | - |
| - | - | 2010 | 581.7 | - | - | 0 | - |
| - | - | 2190 | 582.2 | - | - | 0 | - |
| - | - | 3793 | 584.2 | - | - | 0 | - |
| - | - | 1843 | 584.7 | - | - | 0 | - |
| - | - | 3696 | 590.2 | - | - | 0 | - |
| 2 | y | 6189 | 592.2 | 0.002468 | 4.167 | +2 | 9 |
| - | - | 3161 | 592.7 | - | - | 0 | - |
| - | - | 2154 | 594.2 | - | - | 0 | - |
| - | - | 2785 | 596.2 | - | - | 0 | - |
| - | - | 2220 | 597.2 | - | - | 0 | - |
| 2 | y | 3.168E+04 | 601.2 | 0.002678 | 4.455 | +2 | 9 |
| - | - | 1.795E+04 | 601.7 | - | - | 0 | - |
| - | - | 1.032E+04 | 602.2 | - | - | 0 | - |
| - | - | 1744 | 605.2 | - | - | 0 | - |
| - | - | 1.164E+04 | 606.2 | - | - | 0 | - |
| - | - | 3279 | 607.2 | - | - | 0 | - |
| 5 | b | 1.249E+04 | 608.2 | 0.0007357 | 1.21 | +1 | 5 |
| - | - | 3854 | 609.2 | - | - | 0 | - |
| - | - | 6890 | 612.2 | - | - | 0 | - |
| - | - | 6448 | 612.2 | - | - | 0 | - |
| - | - | 1638 | 613.3 | - | - | 0 | - |
| - | - | 3123 | 614.2 | - | - | 0 | - |
| - | - | 1495 | 615.2 | - | - | 0 | - |
| - | - | 1613 | 622.2 | - | - | 0 | - |
| - | - | 1472 | 623.2 | - | - | 0 | - |
| - | - | 8759 | 624.2 | - | - | 0 | - |
| - | - | 2064 | 625.2 | - | - | 0 | - |
| 5 | b | 2.112E+04 | 626.2 | 0.0005516 | 0.8809 | +1 | 5 |
| - | - | 7350 | 627.3 | - | - | 0 | - |
| - | - | 1518 | 628.3 | - | - | 0 | - |
| - | - | 3030 | 631.2 | - | - | 0 | - |
| - | - | 1974 | 633.2 | - | - | 0 | - |
| - | - | 5607 | 633.8 | - | - | 0 | - |
| - | - | 8278 | 634.2 | - | - | 0 | - |
| - | - | 3851 | 634.3 | - | - | 0 | - |
| - | - | 2905 | 634.8 | - | - | 0 | - |
| - | - | 5242 | 635.2 | - | - | 0 | - |
| - | - | 1380 | 639.3 | - | - | 0 | - |
| - | - | 1902 | 641.3 | - | - | 0 | - |
| - | - | 5153 | 642.2 | - | - | 0 | - |
| - | - | 1.07E+04 | 642.8 | - | - | 0 | - |
| - | - | 8286 | 643.3 | - | - | 0 | - |
| - | - | 4392 | 643.8 | - | - | 0 | - |
| - | - | 1252 | 644.2 | - | - | 0 | - |
| - | - | 2099 | 650.2 | - | - | 0 | - |
| - | - | 6819 | 651.8 | - | - | 0 | - |
| - | - | 3677 | 652.2 | - | - | 0 | - |
| - | - | 3659 | 652.3 | - | - | 0 | - |
| - | - | 2150 | 652.8 | - | - | 0 | - |
| - | - | 1279 | 653.2 | - | - | 0 | - |
| - | - | 4450 | 656.8 | - | - | 0 | - |
| - | - | 1.495E+04 | 657.3 | - | - | 0 | - |
| - | - | 1443 | 657.8 | - | - | 0 | - |
| - | - | 2975 | 658.3 | - | - | 0 | - |
| - | - | 3.463E+04 | 659.3 | - | - | 0 | - |
| - | - | 1.187E+04 | 660.3 | - | - | 0 | - |
| - | - | 1525 | 661.3 | - | - | 0 | - |
| 0 | Precursor | 5.026E+04 | 665.8 | 0.002379 | 3.574 | +2 | -1 |
| - | - | 3.969E+04 | 666.3 | - | - | 0 | - |
| - | - | 1.541E+04 | 666.8 | - | - | 0 | - |
| - | - | 7074 | 667.3 | - | - | 0 | - |
| - | - | 1167 | 669.3 | - | - | 0 | - |
| 0 | Precursor | 9.167E+04 | 674.8 | 0.002468 | 3.658 | +2 | -1 |
| - | - | 7.628E+04 | 675.3 | - | - | 0 | - |
| - | - | 3.435E+04 | 675.8 | - | - | 0 | - |
| - | - | 8002 | 676.3 | - | - | 0 | - |
| - | - | 2708 | 677.3 | - | - | 0 | - |
| - | - | 1.361E+04 | 685.3 | - | - | 0 | - |
| - | - | 6575 | 686.3 | - | - | 0 | - |
| - | - | 1524 | 687.3 | - | - | 0 | - |
| - | - | 1.461E+04 | 695.3 | - | - | 0 | - |
| - | - | 5838 | 696.3 | - | - | 0 | - |
| - | - | 3471 | 703.3 | - | - | 0 | - |
| - | - | 6437 | 704.3 | - | - | 0 | - |
| - | - | 4330 | 705.3 | - | - | 0 | - |
| - | - | 2155 | 706.3 | - | - | 0 | - |
| - | - | 3301 | 713.3 | - | - | 0 | - |
| - | - | 1646 | 714.3 | - | - | 0 | - |
| - | - | 1.175E+04 | 721.3 | - | - | 0 | - |
| - | - | 5235 | 722.3 | - | - | 0 | - |
| 6 | y | 9.616E+04 | 723.3 | 0.004694 | 6.49 | +1 | 5 |
| - | - | 3.655E+04 | 724.3 | - | - | 0 | - |
| - | - | 1.101E+04 | 725.3 | - | - | 0 | - |
| - | - | 1760 | 726.3 | - | - | 0 | - |
| - | - | 1402 | 728.3 | - | - | 0 | - |
| - | - | 1531 | 731.3 | - | - | 0 | - |
| - | - | 1584 | 733.3 | - | - | 0 | - |
| - | - | 2225 | 741.3 | - | - | 0 | - |
| - | - | 1187 | 742.3 | - | - | 0 | - |
| - | - | 3208 | 743.3 | - | - | 0 | - |
| - | - | 3737 | 744.3 | - | - | 0 | - |
| - | - | 1598 | 745.3 | - | - | 0 | - |
| - | - | 1.834E+04 | 749.3 | - | - | 0 | - |
| - | - | 5564 | 750.3 | - | - | 0 | - |
| - | - | 1357 | 751.3 | - | - | 0 | - |
| - | - | 1561 | 753.3 | - | - | 0 | - |
| - | - | 3115 | 756.3 | - | - | 0 | - |
| - | - | 8390 | 759.3 | - | - | 0 | - |
| - | - | 3965 | 760.3 | - | - | 0 | - |
| - | - | 4295 | 761.3 | - | - | 0 | - |
| - | - | 1830 | 762.3 | - | - | 0 | - |
| - | - | 2976 | 767.3 | - | - | 0 | - |
| - | - | 1489 | 768.3 | - | - | 0 | - |
| - | - | 1602 | 770.3 | - | - | 0 | - |
| 6 | b | 1.014E+04 | 771.3 | 9.027E-05 | 0.117 | +1 | 6 |
| - | - | 3820 | 772.3 | - | - | 0 | - |
| - | - | 4.382E+04 | 774.3 | - | - | 0 | - |
| - | - | 1.594E+04 | 775.3 | - | - | 0 | - |
| - | - | 4596 | 776.3 | - | - | 0 | - |
| - | - | 3473 | 778.3 | - | - | 0 | - |
| - | - | 1781 | 779.3 | - | - | 0 | - |
| - | - | 2725 | 787.3 | - | - | 0 | - |
| 6 | b | 6106 | 789.3 | 0.0004646 | 0.5887 | +1 | 6 |
| - | - | 3089 | 790.3 | - | - | 0 | - |
| - | - | 2500 | 792.3 | - | - | 0 | - |
| - | - | 2927 | 796.3 | - | - | 0 | - |
| - | - | 1345 | 803.3 | - | - | 0 | - |
| - | - | 1.382E+04 | 804.3 | - | - | 0 | - |
| - | - | 6697 | 805.3 | - | - | 0 | - |
| - | - | 1254 | 806.3 | - | - | 0 | - |
| - | - | 2242 | 814.3 | - | - | 0 | - |
| - | - | 2221 | 815.3 | - | - | 0 | - |
| 5 | y | 6511 | 820.3 | 0.004185 | 5.102 | +1 | 6 |
| - | - | 3377 | 821.3 | - | - | 0 | - |
| - | - | 1303 | 822.3 | - | - | 0 | - |
| - | - | 1733 | 824.3 | - | - | 0 | - |
| - | - | 1368 | 825.9 | - | - | 0 | - |
| - | - | 1423 | 827.3 | - | - | 0 | - |
| - | - | 2.023E+04 | 832.3 | - | - | 0 | - |
| - | - | 1.035E+04 | 833.3 | - | - | 0 | - |
| - | - | 2163 | 834.3 | - | - | 0 | - |
| 5 | y | 1.293E+05 | 838.3 | 0.004912 | 5.859 | +1 | 6 |
| - | - | 5.551E+04 | 839.3 | - | - | 0 | - |
| - | - | 1.844E+04 | 840.3 | - | - | 0 | - |
| - | - | 1294 | 841.3 | - | - | 0 | - |
| 7 | b | 5089 | 842.3 | 0.008398 | 9.97 | +1 | 7 |
| 7 | b | 4031 | 843.3 | 0.01558 | 18.48 | +1 | 7 |
| - | - | 2335 | 844.3 | - | - | 0 | - |
| - | - | 6648 | 850.3 | - | - | 0 | - |
| - | - | 3616 | 851.3 | - | - | 0 | - |
| - | - | 1987 | 854.3 | - | - | 0 | - |
| 7 | b | 4153 | 860.3 | 0.009197 | 10.69 | +1 | 7 |
| - | - | 1783 | 861.4 | - | - | 0 | - |
| - | - | 2292 | 862.3 | - | - | 0 | - |
| - | - | 1526 | 863.3 | - | - | 0 | - |
| - | - | 1.478E+04 | 868.3 | - | - | 0 | - |
| - | - | 7915 | 869.3 | - | - | 0 | - |
| - | - | 3385 | 870.3 | - | - | 0 | - |
| - | - | 7412 | 871.3 | - | - | 0 | - |
| - | - | 5435 | 872.3 | - | - | 0 | - |
| - | - | 1822 | 873.3 | - | - | 0 | - |
| - | - | 2229 | 878.3 | - | - | 0 | - |
| - | - | 1580 | 886.4 | - | - | 0 | - |
| - | - | 1343 | 888.3 | - | - | 0 | - |
| - | - | 7.599E+04 | 889.3 | - | - | 0 | - |
| - | - | 3.505E+04 | 890.4 | - | - | 0 | - |
| - | - | 8711 | 891.4 | - | - | 0 | - |
| - | - | 1668 | 893.3 | - | - | 0 | - |
| - | - | 2.767E+04 | 896.3 | - | - | 0 | - |
| - | - | 1.49E+04 | 897.3 | - | - | 0 | - |
| - | - | 4862 | 898.3 | - | - | 0 | - |
| - | - | 2130 | 899.3 | - | - | 0 | - |
| - | - | 2500 | 905.4 | - | - | 0 | - |
| - | - | 1693 | 906.4 | - | - | 0 | - |
| - | - | 3575 | 907.3 | - | - | 0 | - |
| - | - | 1981 | 908.3 | - | - | 0 | - |
| - | - | 7055 | 914.3 | - | - | 0 | - |
| - | - | 2580 | 915.3 | - | - | 0 | - |
| - | - | 1873 | 918.3 | - | - | 0 | - |
| - | - | 4269 | 933.4 | - | - | 0 | - |
| - | - | 1431 | 934.4 | - | - | 0 | - |
| 4 | y | 1.487E+04 | 935.3 | 0.00544 | 5.816 | +1 | 7 |
| - | - | 8770 | 936.3 | - | - | 0 | - |
| - | - | 1557 | 937.3 | - | - | 0 | - |
| - | - | 2555 | 938.3 | - | - | 0 | - |
| - | - | 1448 | 943.4 | - | - | 0 | - |
| - | - | 1847 | 951.4 | - | - | 0 | - |
| - | - | 1212 | 952.4 | - | - | 0 | - |
| 4 | y | 2.266E+05 | 953.3 | 0.004702 | 4.932 | +1 | 7 |
| - | - | 1.098E+05 | 954.3 | - | - | 0 | - |
| - | - | 1201 | 954.5 | - | - | 0 | - |
| - | - | 4.188E+04 | 955.3 | - | - | 0 | - |
| - | - | 5424 | 956.3 | - | - | 0 | - |
| - | - | 4056 | 963.3 | - | - | 0 | - |
| - | - | 1631 | 964.3 | - | - | 0 | - |
| - | - | 1342 | 967.4 | - | - | 0 | - |
| - | - | 3206 | 969.4 | - | - | 0 | - |
| - | - | 1503 | 970.4 | - | - | 0 | - |
| - | - | 1940 | 979.4 | - | - | 0 | - |
| 8 | b | 2691 | 989.4 | 0.002133 | 2.156 | +1 | 8 |
| - | - | 3841 | 990.4 | - | - | 0 | - |
| - | - | 2084 | 991.4 | - | - | 0 | - |
| - | - | 4895 | 997.4 | - | - | 0 | - |
| - | - | 3335 | 998.4 | - | - | 0 | - |
| 8 | b | 2522 | 1007 | 0.004648 | 4.613 | +1 | 8 |
| - | - | 2937 | 1008 | - | - | 0 | - |
| - | - | 3394 | 1009 | - | - | 0 | - |
| - | - | 8730 | 1018 | - | - | 0 | - |
| - | - | 4838 | 1019 | - | - | 0 | - |
| - | - | 1675 | 1021 | - | - | 0 | - |
| - | - | 1.013E+05 | 1036 | - | - | 0 | - |
| - | - | 6.593E+04 | 1037 | - | - | 0 | - |
| - | - | 1.741E+04 | 1038 | - | - | 0 | - |
| - | - | 2143 | 1039 | - | - | 0 | - |
| - | - | 3896 | 1054 | - | - | 0 | - |
| - | - | 3426 | 1055 | - | - | 0 | - |
| - | - | 1417 | 1065 | - | - | 0 | - |
| - | - | 1411 | 1080 | - | - | 0 | - |
| 3 | y | 1.759E+04 | 1082 | 0.004714 | 4.355 | +1 | 8 |
| - | - | 1.243E+04 | 1083 | - | - | 0 | - |
| - | - | 6189 | 1084 | - | - | 0 | - |
| - | - | 3208 | 1085 | - | - | 0 | - |
| 3 | y | 3.692E+05 | 1100 | 0.004403 | 4.002 | +1 | 8 |
| - | - | 2.277E+05 | 1101 | - | - | 0 | - |
| - | - | 1586 | 1102 | - | - | 0 | - |
| - | - | 9.06E+04 | 1102 | - | - | 0 | - |
| - | - | 1.205E+04 | 1103 | - | - | 0 | - |
| - | - | 1822 | 1110 | - | - | 0 | - |
| - | - | 2078 | 1116 | - | - | 0 | - |
| - | - | 4808 | 1119 | - | - | 0 | - |
| - | - | 2699 | 1120 | - | - | 0 | - |
| 9 | b | 1488 | 1126 | 0.003068 | 2.724 | +1 | 9 |
| - | - | 1931 | 1137 | - | - | 0 | - |
| - | - | 2.453E+04 | 1137 | - | - | 0 | - |
| - | - | 1.488E+04 | 1138 | - | - | 0 | - |
| - | - | 4817 | 1139 | - | - | 0 | - |
| 9 | b | 5239 | 1144 | 0.005777 | 5.048 | +1 | 9 |
| - | - | 4774 | 1145 | - | - | 0 | - |
| - | - | 2213 | 1146 | - | - | 0 | - |
| - | - | 4867 | 1147 | - | - | 0 | - |
| - | - | 1947 | 1148 | - | - | 0 | - |
| - | - | 1792 | 1157 | - | - | 0 | - |
| - | - | 2628 | 1162 | - | - | 0 | - |
| 2 | y | 1.136E+04 | 1183 | 0.004155 | 3.511 | +1 | 9 |
| - | - | 7110 | 1184 | - | - | 0 | - |
| - | - | 3107 | 1185 | - | - | 0 | - |
| - | - | 2018 | 1193 | - | - | 0 | - |
| 2 | y | 6.781E+04 | 1201 | 0.003478 | 2.895 | +1 | 9 |
| - | - | 4.592E+04 | 1202 | - | - | 0 | - |
| - | - | 1.758E+04 | 1203 | - | - | 0 | - |
| - | - | 3262 | 1204 | - | - | 0 | - |
| - | - | 9710 | 1211 | - | - | 0 | - |
| - | - | 7068 | 1212 | - | - | 0 | - |
| - | - | 3357 | 1213 | - | - | 0 | - |
| - | - | 1145 | 1741 | - | - | 0 | - |
| - | - | 1342 | 1955 | - | - | 0 | - |
| - | - | 1373 | 2253 | - | - | 0 | - |
| - | - | 1569 | 2341 | - | - | 0 | - |
| - | - | 1445 | 2928 | - | - | 0 | - |

m/z Charge Intensity FragmentType MassShift Position
120.08111572265625 0 1011909.3
121.04015350341797 0 800.5969
121.07847595214844 0 3111.836
121.0843505859375 0 86058.16
122.07144165039062 0 1334.6449
122.08763122558594 0 1789.998
129.10243225097656 0 1527.3761
130.0500030517578 0 2264.3198
130.06539916992188 0 10071.656
131.04957580566406 0 2319.5654
132.0810089111328 0 4163.401
136.07594299316406 0 77675.37
136.08624267578125 0 2218.1504
137.07923889160156 0 6059.643
138.0550994873047 0 7448.1904
138.06639099121094 0 11405.172
143.07313537597656 0 3122.631
143.89749145507812 0 904.2271
144.0811767578125 0 1441.7068
146.06033325195312 0 3672.8848
146.09661865234375 0 3090.1357
147.0760498046875 0 1014.0512
148.08702087402344 0 3706.6384
148.94715881347656 0 1943.1345
155.08177185058594 0 2267.883
155.09286499023438 0 4525.6343
156.0769500732422 0 27662.594
157.08026123046875 0 1857.26
158.0966339111328 0 22969.906
159.0918731689453 0 49434.75
159.09945678710938 0 3008.4702
160.07611083984375 0 2047.6398
160.09535217285156 0 4748.68
160.1119842529297 0 2418.7422
161.0924835205078 0 2886.0369
162.0923614501953 0 1171.6154
165.07015991210938 0 1402.9543
165.0773468017578 0 5824.9575
165.1026153564453 0 1275.8132
166.05401611328125 0 2119.6396
166.0614013671875 0 4420.4663
166.08645629882812 0 22874.387
167.091552734375 0 5866.5747
169.07611083984375 0 2292.9077
169.13360595703125 0 1206.5851
171.07664489746094 0 2544.2769
171.091796875 0 1070.7112
172.07598876953125 0 1277.8202
173.09225463867188 0 999.7487
173.43756103515625 0 1083.4485
174.06674194335938 0 3028.8042
175.08706665039062 0 1974.8721
175.12318420410156 0 2575.5715
176.0823974609375 0 8392.727
176.1072998046875 0 463810.78
177.1024932861328 0 7889.4414
177.11073303222656 0 50590.254
178.0615234375 0 1787.951
178.11383056640625 0 1180.5133
180.0770263671875 0 3647.5593
185.05604553222656 0 3788.0728
186.09153747558594 0 61128.785
187.06272888183594 0 1886.1523
187.09495544433594 0 7304.1094
188.07078552246094 0 34568.367
189.07456970214844 0 3292.828
189.0870361328125 0 5074.5854
189.10240173339844 0 3700.616
190.08665466308594 0 1554.0424 b Ammonia loss 2
191.08285522460938 0 1421.8253
191.0931396484375 0 3182.9194
191.1181640625 0 7471.6
192.10189819335938 0 1353.8031
193.0723114013672 0 1770.9476
193.1085662841797 0 29358.666
194.080078125 0 2446.1543
194.11236572265625 0 1384.0205
195.087646484375 0 6867.3184
197.1284942626953 0 2572.4377
199.07171630859375 0 1090.7557
201.12368774414062 0 2171.0127
202.05300903320312 0 1928.629
202.23037719726562 0 1108.1859
203.06643676757812 0 4266.759
203.092529296875 0 2348.3242
203.11805725097656 0 33672.508 a Water loss 1
204.07691955566406 0 14862.97
204.10189819335938 0 1071.9939 a Ammonia loss 1
204.12152099609375 0 4181.041
205.08096313476562 0 1745.9664
205.09732055664062 0 23875.668 y 9
205.10772705078125 0 1108.2175
205.13333129882812 0 1343.1473 d 1
206.10096740722656 0 2710.004
206.49945068359375 0 893.2968
207.1129913330078 0 9452.942 d 1
208.0717315673828 0 2413.7214
209.1029052734375 0 909.8917
212.11817932128906 0 7512.0957
213.1006317138672 0 1215.9869
213.12088012695312 0 1121.8536
214.086181640625 0 3767.8987
215.0818328857422 0 4733.891
215.13876342773438 0 1303.7788
217.09738159179688 0 32367.45
218.10107421875 0 4778.456
219.079833984375 0 4595.3413
219.11329650878906 0 3411.202
221.10353088378906 0 78787.484
221.12875366210938 0 584273.4 a 1
222.10707092285156 0 7470.027
222.1320037841797 0 72690.7
223.10818481445312 0 1638.0381
223.11988830566406 0 1635.1724
223.13458251953125 0 4548.057
226.0825958251953 0 1338.6433
227.06686401367188 0 1003.2709
227.2652587890625 0 945.00385
228.1131591796875 0 3156.0747
229.097412109375 0 2167.0232
229.10784912109375 0 2264.9438
229.11962890625 0 1526.9471
231.0615234375 0 9875.885
231.11305236816406 0 145634.61 b Water loss 1
232.11630249023438 0 18417.9
233.09197998046875 0 1134.4064
235.10800170898438 0 50680.21
236.11109924316406 0 6910.168
238.1217803955078 0 2172.4456
239.11410522460938 0 8814.659
240.11416625976562 0 1769.2845
245.09278869628906 0 2764.2915 y 7
246.13519287109375 0 1707.9756
247.10678100585938 0 1459.58 b Water loss 3
247.11976623535156 0 1210.3922
247.144775390625 0 2062.9934
248.1507110595703 0 3012.8696
249.1236572265625 0 302547.06 b 1
250.12693786621094 0 40188.453
251.1028289794922 0 26062.19
251.129638671875 0 4840.9707
252.10577392578125 0 2914.6155
253.11849975585938 0 16815.572
254.12203979492188 0 5646.337
257.10626220703125 0 13895.886
259.1075439453125 0 4619.8413
261.08697509765625 0 3498.6375
261.123046875 0 3002.7517
262.07196044921875 0 1217.0194
263.1027526855469 0 61339.266
263.1287536621094 0 3910.7136
264.106201171875 0 8068.2246
264.14581298828125 0 3715.118
267.0913391113281 0 2808.0896
268.07489013671875 0 2398.5015
268.1401062011719 0 1977.2731
269.0766296386719 0 1194.4468
271.1083679199219 0 2057.4922
272.1139221191406 0 1081.7043
273.12408447265625 0 1726.5566
274.1300048828125 0 14422.486
279.0976867675781 0 19669.842
279.1241455078125 0 1728.0452
280.1007385253906 0 2678.1067
280.1295471191406 0 1420.6328
283.14398193359375 0 1362.6985
284.16143798828125 0 1361.6399
285.101806640625 0 28318.1
286.104736328125 0 2640.6438
287.1017761230469 0 1779.9695
289.08209228515625 0 1516.4568
289.1177062988281 0 1722.9829
289.13446044921875 0 3751.361
292.1406555175781 0 18320.814
292.1623840332031 0 1178.1476
292.16571044921875 0 1068.4259
293.14385986328125 0 2753.1143
296.15087890625 0 22804.068
297.1539306640625 0 3060.4788
300.1349792480469 0 2623.4727
302.1510314941406 0 1149.8749
303.1125793457031 0 4608.58
304.1303405761719 0 2183.539
306.1442565917969 0 1972.633
307.11981201171875 0 1523.084
307.1436462402344 0 1603.6017
310.1508483886719 0 3280.843
312.1553955078125 0 1510.8204
314.0974426269531 0 1472.2732
315.0982360839844 0 1928.3743
316.1865539550781 0 2058.2102
317.12896728515625 0 4250.4014
318.1452331542969 0 10056.784
319.1475830078125 0 1479.9913
320.1359558105469 0 2160.54
323.0987548828125 0 3899.574
323.1762390136719 0 1625.4161
324.1458435058594 0 50415.32
325.1488342285156 0 8165.8467
327.1459655761719 0 1657.3414
328.14569091796875 0 1930.5935
329.1131591796875 0 1064.4269
330.10833740234375 0 2401.6433
330.16650390625 0 2778.0254
332.1247253417969 0 3729.0417
333.1079406738281 0 2766.544
333.16021728515625 0 9399.371
334.1087951660156 0 1006.1539
334.1379699707031 0 2141.5151
334.1629333496094 0 1295.2153
338.12841796875 0 2538.3271
339.1709289550781 0 4844.05
342.156494140625 0 259618.92 y 8
343.1594543457031 0 49638.457
344.12408447265625 0 1850.398
344.1618347167969 0 5936.143
346.1396789550781 0 8099.3223
347.1440734863281 0 1096.0353
348.1193542480469 0 4176.1787
349.1239929199219 0 1436.9291
349.15411376953125 0 1523.2056
350.1346130371094 0 17983.268
350.18701171875 0 3681.3718 a Water loss 2
351.1382141113281 0 3363.0757
351.16998291015625 0 2427.239 a Ammonia loss 2
352.140869140625 0 6299.59
352.1651306152344 0 1140.0436
354.1606140136719 0 1135.5544
355.1405029296875 0 2726.3188
356.1388854980469 0 12402.051
357.1423034667969 0 2199.5857
358.1024475097656 0 2336.0737
360.11920166015625 0 5121.857
360.1436767578125 0 990.87683
362.13507080078125 0 1929.8513
364.1505432128906 0 22386.754
365.1535949707031 0 4775.286
366.1293640136719 0 4276.972
367.165771484375 0 2657.6213
371.1356506347656 0 1380.0981
374.134521484375 0 4300.7974
376.11370849609375 0 7545.7695
378.1298828125 0 75491.39
378.1815185546875 0 56785.004 b Water loss 2
379.1329650878906 0 15676.851
379.184814453125 0 14856.163
380.13665771484375 0 4127.94
381.14556884765625 0 1707.8005
382.14361572265625 0 6536.4224
382.1730041503906 0 978.7632
390.1556396484375 0 4217.282
391.15863037109375 0 1579.3746
394.1248779296875 0 30146.426
395.12713623046875 0 3836.0776
395.1543273925781 0 1816.1732 b 5
396.1927490234375 0 3591.4033 b 2
397.16314697265625 0 1520.0771
398.170166015625 0 1679.8599
399.17498779296875 0 2798.9907
401.1922607421875 0 2346.3987
404.11248779296875 0 1558.1223
404.14556884765625 0 5596.59
405.14337158203125 0 1351.1459
406.15106201171875 0 1798.8575
407.1836853027344 0 5281.0703
408.1667175292969 0 9254.553
409.1702575683594 0 2858.6855
410.1381530761719 0 2472.6887
413.2025451660156 0 1500.9423
415.15936279296875 0 1600.4534
416.145263671875 0 2302.4475
419.1559143066406 0 2709.157
425.19354248046875 0 47801.05
426.1963806152344 0 9473.534
427.2082214355469 0 7658.832
433.17181396484375 0 4258.7686
434.17694091796875 0 1314.2539
442.17242431640625 0 1617.1357
443.1561584472656 0 2721.1982
447.1519470214844 0 5041.608
449.1664733886719 0 3962.173
454.1952209472656 0 2352.8835
455.20361328125 0 6782.8857
461.16668701171875 0 8434.556
461.19476318359375 0 3194.5178
462.16754150390625 0 1845.8201
462.2001953125 0 2198.545
465.162353515625 0 20730.63
466.1650085449219 0 4102.659
470.1669921875 0 4317.5474
471.179931640625 0 4034.2122
473.1728515625 0 1316.6116
473.2162780761719 0 2834.812
477.17340087890625 0 3739.8008 y 3
479.1776428222656 0 13037.996
480.18182373046875 0 2788.0205
481.17791748046875 0 1739.4581
486.1612854003906 0 3256.2173
489.162353515625 0 3151.9495
489.1921691894531 0 65567.9 y 7
490.1946105957031 0 18705.043
491.20452880859375 0 7293.1597
492.2125244140625 0 1829.8861
493.2088623046875 0 6354.449 b Water loss 3
494.2125244140625 0 3740.6687
495.18804931640625 0 7585.3228 b Water loss 7
496.1893005371094 0 2569.0232
496.2306823730469 0 32226.79
497.17138671875 0 4597.26
497.2335510253906 0 7338.9424
498.17388916015625 0 1521.4022
509.705078125 0 2309.632
511.2188415527344 0 29337.143 b 3
512.2218627929688 0 6963.589
513.19775390625 0 12587.367
514.201904296875 0 5053.4683
518.5222778320312 0 1235.7482
518.7113647460938 0 4637.702
519.204345703125 0 9792.609
520.2027587890625 0 4752.306
521.2022094726562 0 1848.5247
523.1806640625 0 2637.946
527.70751953125 0 2134.5867
530.1912231445312 0 1936.056
532.20556640625 0 1330.6027
533.2030029296875 0 1968.1058
537.2119140625 0 2723.2383
539.2142944335938 0 1457.2548
541.192626953125 0 8584.178
541.7061157226562 0 7553.9395 y Water loss 2
542.2029418945312 0 5935.983
542.2374267578125 0 4878.1006
542.70751953125 0 3463.0051
548.1984252929688 0 6948.85
549.2005615234375 0 3124.2095
550.2171630859375 0 1427.0991
550.7108764648438 0 76837.75 y 2
551.2122802734375 0 51926.89
551.712646484375 0 18676.469
552.213134765625 0 3954.1804
555.2153930664062 0 1460.5164
557.2009887695312 0 2329.2615
558.20458984375 0 1229.089
560.2286376953125 0 91254.21 y 6
561.2320556640625 0 29598.719
562.2302856445312 0 10081.466
563.7200927734375 0 2458.24 b Water loss 8
564.2168579101562 0 2679.5679 b Ammonia loss 8
567.20947265625 0 2826.9617
569.2263793945312 0 1375.1475
569.7394409179688 0 1469.134
570.2285766601562 0 7278.361
571.2276611328125 0 1797.9573
572.7232666015625 0 6155.388 b 8
573.2260131835938 0 3816.2827
579.208984375 0 1737.8315
580.2405395507812 0 4098.06
581.2286376953125 0 1179.0918
581.7274169921875 0 2010.1123
582.2296142578125 0 2189.6682
584.2347412109375 0 3792.7336
584.71484375 0 1843.1073
590.222900390625 0 3695.8855
592.2293090820312 0 6188.746 y Water loss 1
592.72998046875 0 3160.641
594.2235107421875 0 2154.0005
596.233154296875 0 2785.0283
597.2269287109375 0 2219.6274
601.2348022460938 0 31681.883 y 1
601.7361450195312 0 17948.092
602.23681640625 0 10317.033
605.2362060546875 0 1744.2206
606.2320556640625 0 11644.801
607.2352294921875 0 3278.9185
608.23583984375 0 12493.492 b Water loss 4
609.238037109375 0 3853.6895
612.1947021484375 0 6890.1216
612.2352905273438 0 6448.068
613.28955078125 0 1637.514
614.2439575195312 0 3123.1782
615.2421875 0 1495.0331
622.219482421875 0 1613.3408
623.2275390625 0 1471.8623
624.230712890625 0 8758.737
625.2269287109375 0 2063.8826
626.2451171875 0 21123.4 b 4
627.2501831054688 0 7349.901
628.2544555664062 0 1517.634
631.247802734375 0 3030.128
633.2286987304688 0 1973.7141
633.7637329101562 0 5606.8696
634.228759765625 0 8278.241
634.2716674804688 0 3850.5496
634.7642211914062 0 2904.7925
635.2338256835938 0 5241.753
639.2526245117188 0 1379.9788
641.28515625 0 1901.7325
642.2404174804688 0 5152.5093
642.7677001953125 0 10702.853
643.2664184570312 0 8286.099
643.7682495117188 0 4391.908
644.2369995117188 0 1251.6827
650.2455444335938 0 2099.201
651.7660522460938 0 6819.257
652.2432250976562 0 3676.9858
652.2546997070312 0 3659.2227
652.7606201171875 0 2149.9553
653.2421875 0 1278.5884
656.7579345703125 0 4449.797
657.2613525390625 0 14945.061
657.7548217773438 0 1443.01
658.2643432617188 0 2975.3552
659.2939453125 0 34633.49
660.297119140625 0 11868.331
661.29736328125 0 1524.5696
665.763427734375 0 50255.76 Precursor Water loss
666.2651977539062 0 39688.67
666.7659301757812 0 15407.552
667.2630004882812 0 7074.2993
669.2695922851562 0 1166.6119
674.768798828125 0 91665.09 Precursor
675.2702026367188 0 76275.4
675.7708129882812 0 34350.9
676.2713623046875 0 8002.0664
677.2610473632812 0 2707.5103
685.2576904296875 0 13612.544
686.2600708007812 0 6574.992
687.2664184570312 0 1524.3936
695.2667846679688 0 14606.373
696.270263671875 0 5838.4785
703.2682495117188 0 3471.4734
704.268798828125 0 6437.382
705.2778930664062 0 4330.4453
706.2827758789062 0 2155.4028
713.271484375 0 3300.9182
714.274658203125 0 1645.9061
721.2604370117188 0 11752.3955
722.2628173828125 0 5235.004
723.2916870117188 0 96157.42 y 5
724.29443359375 0 36554.895
725.2951049804688 0 11012.473
726.2921142578125 0 1759.9298
728.3182373046875 0 1402.487
731.265869140625 0 1530.6193
733.2780151367188 0 1583.9219
741.2525024414062 0 2224.6824
742.2559814453125 0 1187.1967
743.3035278320312 0 3207.8318
744.2985229492188 0 3737.3462
745.3091430664062 0 1597.7057
749.2554931640625 0 18335.824
750.259033203125 0 5564.0215
751.2512817382812 0 1356.6627
753.285400390625 0 1561.4672
756.3104248046875 0 3115.2446
759.2651977539062 0 8390.413
760.27001953125 0 3964.7178
761.3106079101562 0 4294.8296
762.3157348632812 0 1829.8458
767.2696533203125 0 2975.6113
768.2698364257812 0 1489.3083
770.2962036132812 0 1601.7247
771.2985229492188 0 10143.387 b Water loss 5
772.3012084960938 0 3820.1614
774.3203125 0 43824.984
775.3236083984375 0 15943.15
776.3274536132812 0 4595.824
778.30712890625 0 3473.1375
779.3143310546875 0 1781.2866
787.307861328125 0 2725.4817
789.3085327148438 0 6106.228 b 5
790.3095092773438 0 3088.8118
792.3113403320312 0 2499.8853
796.3130493164062 0 2927.3752
803.2737426757812 0 1344.7913
804.3309326171875 0 13820.785
805.3331298828125 0 6696.5513
806.3203125 0 1253.5145
814.31689453125 0 2241.6455
815.323974609375 0 2221.1335
820.3075561523438 0 6511.3804 y Water loss 4
821.3052368164062 0 3376.5906
822.303955078125 0 1302.7467
824.3128051757812 0 1733.1147
825.9221801757812 0 1368.0431
827.3177490234375 0 1422.8633
832.3265380859375 0 20231.48
833.3290405273438 0 10352.051
834.3252563476562 0 2162.6833
838.31884765625 0 129342.305 y 4
839.3218383789062 0 55512.08
840.3203125 0 18440.295
841.3242797851562 0 1294.4133
842.3271484375 0 5088.573 b Water loss 6
843.3351440429688 0 4031.2283 b Ammonia loss 6
844.3392333984375 0 2335.0178
850.333984375 0 6647.9746
851.3308715820312 0 3616.4856
854.3064575195312 0 1987.4333
860.3369140625 0 4153.2715 b 6
861.3533935546875 0 1782.7552
862.3224487304688 0 2291.5222
863.3361206054688 0 1525.9
868.3286743164062 0 14779.404
869.3321533203125 0 7914.9756
870.3271484375 0 3385.3735
871.3348999023438 0 7412.2144
872.3307495117188 0 5435.046
873.3357543945312 0 1822.4014
878.3104858398438 0 2228.654
886.412841796875 0 1579.902
888.3423461914062 0 1342.9092
889.346923828125 0 75988.17
890.3500366210938 0 35045.91
891.3538818359375 0 8710.58
893.3452758789062 0 1668.2771
896.32470703125 0 27668.121
897.327392578125 0 14897.307
898.3257446289062 0 4861.775
899.331298828125 0 2129.928
905.3786010742188 0 2500.3296
906.3892211914062 0 1692.8339
907.34765625 0 3574.6348
908.3414916992188 0 1981.0651
914.3359375 0 7055.0747
915.3478393554688 0 2579.5042
918.3128051757812 0 1872.9193
933.3726196289062 0 4268.5234
934.3703002929688 0 1431.3212
935.3357543945312 0 14870.629 y Water loss 3
936.3325805664062 0 8769.823
937.3373413085938 0 1557.2317
938.3219604492188 0 2554.9714
943.3702392578125 0 1447.6642
951.3748168945312 0 1847.2275
952.3677978515625 0 1212.0383
953.3455810546875 0 226585.97 y 3
954.3482055664062 0 109781.484
954.4660034179688 0 1200.7639
955.347900390625 0 41882
956.34912109375 0 5424.2007
963.3289794921875 0 4055.87
964.3311157226562 0 1631.1459
967.4237670898438 0 1341.6404
969.3748168945312 0 3205.7717
970.3775024414062 0 1502.9973
979.3627319335938 0 1939.5435
989.3681640625 0 2690.582 b Water loss 7
990.4011840820312 0 3841.2017
991.4099731445312 0 2084.1907
997.3717651367188 0 4894.839
998.3746948242188 0 3335.4468
1007.3719482421875 0 2521.7095 b 7
1008.3839721679688 0 2936.5652
1009.3878784179688 0 3394.365
1018.40380859375 0 8729.947
1019.4107666015625 0 4837.9146
1021.3958740234375 0 1675.109
1036.4150390625 0 101338.836
1037.4180908203125 0 65925.96
1038.419921875 0 17411.82
1039.4222412109375 0 2143.391
1054.41015625 0 3895.9841
1055.413818359375 0 3426.165
1065.3851318359375 0 1417.346
1080.4375 0 1411.3514
1082.4034423828125 0 17594.51 y Water loss 2
1083.4072265625 0 12427.685
1084.40625 0 6189.1494
1085.399169921875 0 3207.5977
1100.4136962890625 0 369219 y 2
1101.4166259765625 0 227662.14
1101.698486328125 0 1585.8381
1102.4169921875 0 90604.4
1103.416748046875 0 12046.641
1110.395263671875 0 1821.7186
1116.4444580078125 0 2078.0393
1119.447265625 0 4808.459
1120.4530029296875 0 2698.7742
1126.421875 0 1487.616 b Water loss 8
1137.3271484375 0 1930.519
1137.462890625 0 24527.98
1138.46484375 0 14879.782
1139.4696044921875 0 4816.531
1144.4412841796875 0 5239.329 b 8
1145.438232421875 0 4773.5796
1146.4476318359375 0 2212.5154
1147.443603515625 0 4867.0474
1148.4443359375 0 1946.7723
1157.439697265625 0 1791.6317
1162.45263671875 0 2628.1912
1183.4505615234375 0 11363.926 y Water loss 1
1184.45166015625 0 7110.4233
1185.4615478515625 0 3106.9888
1193.43359375 0 2017.5292
1201.46044921875 0 67807.18 y 1
1202.4638671875 0 45922.81
1203.4630126953125 0 17579.121
1204.464111328125 0 3262.4653
1211.4473876953125 0 9710.437
1212.4476318359375 0 7068.264
1213.453857421875 0 3357.322
1740.5755615234375 0 1145.3129
1954.7794189453125 0 1342.3052
2253.40673828125 0 1372.8938
2340.8828125 0 1569.187
2928.376220703125 0 1445.1279

Spectrum Details

|  |  |
| --- | --- |
| Matched peaks? Matched peaks The total absolute number of peaks matched. Additionally in brackets the total fraction of peaks matched and the total number of peaks is shown. | 52 (8.57% of 607) |
| FDR? FDR The false discovery rate estimated for this peptide. It is calculated by matching all theoretical fragments with a non-integer shift with the raw peaks for this spectrum. This is done with 40 different shifts. The resulting percentage is the average number of annotated peaks over the number of annotated peaks with the correct spectrum. | 0.05% |
| Satellite FDR? Satellite FDR See the FDR for details on its calculation. This satellite ion specific FDR only contains the satellite ions (d/w) for I/L/J positions. | - |
| PSM Score? PSM Score The PSM Score as given by Hecklib to this annotated spectrum. It is shown with three significant figures. | 605 |

#### Spectrum 9689? Spectrum 9689 The raw spectrum of this peptide as annotated by Hecklib. The fragments are coloured according to ion type (see legend). Any peaks with a star '\*' as text can be hovered over to see the full details, first the ion type second the mass shift type. By hovering over the amino acids in the peptide or ions in the legend the corresponding peaks are highlighted. By toggling the 'Unassigned' label you can turn the background (unassigned) peaks on or off in the plot. By updating the slider in the Ion legend you can update the spectrum to only show the top X% of the peaks with labels. The top X% means any peak that is within X% of the highest intensity. By dragging in the spectrum you can zoom in to a specific part of the spectrum and use 'Zoom Out' to get back to the original zoom level. The annotation of the spectrum is based on the given sequence in the peptides file and is done with different software so inconsistencies are likely. The peaks are annotated based on the given sequence, with 20 ppm tolerance.

Copy Data

##### Spectrum 9689 (TSV)

###### Preview

```
Loading example...
```

*Click on the button to copy the data to your clipboard.*

Mz MinMz MaxIntensity Max

WidthHeightPeptide font sizePeptide stroke widthSpectrum font sizeSpectrum stroke widthCompact peptide

Ion legend

wxyz

abcd

OtherUnassignedIon^Charge^_Position_Show for top:%

FTFDDYAMHW

06.28e+41.26e+51.89e+52.51e+5

Zoom Out

b^+2^_3_a^+1^_2_a^+1^_2_y^+1^_1_d^+1^_2_a^+1^_2_b^+1^_2_b^+1^_2_y^+1^_2_a^+1^_3_a^+1^_3_b^+1^_3_b^+1^_3_b^+1^_3_y^+1^_3_b^+1^_4_b^+2^_8_b^+1^_4_y^+2^_8_y^+2^_8_y^+1^_4_b^+2^_9_b^+2^_9_y^+2^_9_y^+2^_9_b^+1^_5_b^+1^_5_\*\*y^+1^_5_b^+1^_6_b^+1^_6_y^+1^_6_y^+1^_6_b^+1^_7_y^+1^_7_y^+1^_7_b^+1^_8_y^+1^_8_y^+1^_8_b^+1^_9_y^+1^_9_y^+1^_9_

0803160624093212

Fragment Matches Table

Show background peaks

| Position | Ion type | Intensity | mz Theoretical | mz Error (Th) | mz Error (ppm) | Charge | Series Number |
| --- | --- | --- | --- | --- | --- | --- | --- |
| - | - | 2.489E+05 | 120.1 | - | - | 0 | - |
| - | - | 367.2 | 120.8 | - | - | 0 | - |
| - | - | 431.6 | 121 | - | - | 0 | - |
| - | - | 2.09E+04 | 121.1 | - | - | 0 | - |
| - | - | 328 | 122 | - | - | 0 | - |
| - | - | 415.9 | 122.1 | - | - | 0 | - |
| - | - | 524.8 | 127.1 | - | - | 0 | - |
| - | - | 488 | 128.1 | - | - | 0 | - |
| - | - | 2111 | 129.1 | - | - | 0 | - |
| - | - | 1510 | 130.1 | - | - | 0 | - |
| - | - | 2601 | 130.1 | - | - | 0 | - |
| - | - | 1171 | 132.1 | - | - | 0 | - |
| - | - | 452 | 133.1 | - | - | 0 | - |
| - | - | 742.8 | 133.1 | - | - | 0 | - |
| - | - | 2.18E+04 | 136.1 | - | - | 0 | - |
| - | - | 742.9 | 136.1 | - | - | 0 | - |
| - | - | 1599 | 137.1 | - | - | 0 | - |
| - | - | 2458 | 138.1 | - | - | 0 | - |
| - | - | 3113 | 138.1 | - | - | 0 | - |
| - | - | 500.7 | 142.3 | - | - | 0 | - |
| - | - | 723.5 | 143.1 | - | - | 0 | - |
| - | - | 465.5 | 145.2 | - | - | 0 | - |
| - | - | 1393 | 146.1 | - | - | 0 | - |
| - | - | 1445 | 146.1 | - | - | 0 | - |
| - | - | 2145 | 147.1 | - | - | 0 | - |
| - | - | 732.4 | 148.1 | - | - | 0 | - |
| - | - | 891.3 | 149 | - | - | 0 | - |
| - | - | 603.6 | 152.1 | - | - | 0 | - |
| - | - | 501 | 153.1 | - | - | 0 | - |
| - | - | 1033 | 155.1 | - | - | 0 | - |
| - | - | 7847 | 156.1 | - | - | 0 | - |
| - | - | 5732 | 158.1 | - | - | 0 | - |
| - | - | 1.185E+04 | 159.1 | - | - | 0 | - |
| - | - | 787 | 159.1 | - | - | 0 | - |
| - | - | 1091 | 160.1 | - | - | 0 | - |
| - | - | 612.4 | 160.1 | - | - | 0 | - |
| - | - | 751.9 | 165.1 | - | - | 0 | - |
| - | - | 681.6 | 166.1 | - | - | 0 | - |
| - | - | 4343 | 166.1 | - | - | 0 | - |
| - | - | 499.9 | 166.1 | - | - | 0 | - |
| - | - | 507.4 | 166.1 | - | - | 0 | - |
| - | - | 947.8 | 167.1 | - | - | 0 | - |
| - | - | 439 | 169.1 | - | - | 0 | - |
| - | - | 2194 | 169.1 | - | - | 0 | - |
| - | - | 642.4 | 171.1 | - | - | 0 | - |
| - | - | 697 | 173.1 | - | - | 0 | - |
| - | - | 967.4 | 175.1 | - | - | 0 | - |
| - | - | 524.9 | 175.1 | - | - | 0 | - |
| - | - | 2751 | 176.1 | - | - | 0 | - |
| - | - | 1.156E+05 | 176.1 | - | - | 0 | - |
| - | - | 2474 | 177.1 | - | - | 0 | - |
| - | - | 1.262E+04 | 177.1 | - | - | 0 | - |
| - | - | 1101 | 180.1 | - | - | 0 | - |
| - | - | 917 | 183.1 | - | - | 0 | - |
| - | - | 1.486E+04 | 186.1 | - | - | 0 | - |
| - | - | 1714 | 187.1 | - | - | 0 | - |
| - | - | 7514 | 188.1 | - | - | 0 | - |
| - | - | 1198 | 189.1 | - | - | 0 | - |
| - | - | 1224 | 189.1 | - | - | 0 | - |
| 3 | b | 531.3 | 190.1 | 0.0008574 | 4.51 | +2 | 3 |
| - | - | 824.7 | 191.1 | - | - | 0 | - |
| - | - | 2719 | 191.1 | - | - | 0 | - |
| - | - | 7920 | 193.1 | - | - | 0 | - |
| - | - | 1434 | 194.1 | - | - | 0 | - |
| - | - | 2849 | 195.1 | - | - | 0 | - |
| - | - | 591.7 | 197.1 | - | - | 0 | - |
| - | - | 1872 | 197.1 | - | - | 0 | - |
| - | - | 1035 | 201.1 | - | - | 0 | - |
| - | - | 721.6 | 203.1 | - | - | 0 | - |
| - | - | 579.4 | 203.1 | - | - | 0 | - |
| 2 | a | 8736 | 203.1 | 0.000244 | 1.201 | +1 | 2 |
| - | - | 4145 | 204.1 | - | - | 0 | - |
| 2 | a | 523.3 | 204.1 | 2.359E-05 | 0.1156 | +1 | 2 |
| - | - | 729.7 | 204.1 | - | - | 0 | - |
| - | - | 579.5 | 205.1 | - | - | 0 | - |
| 10 | y | 5907 | 205.1 | 0.0001817 | 0.886 | +1 | 1 |
| - | - | 940.1 | 205.1 | - | - | 0 | - |
| - | - | 660.2 | 206.1 | - | - | 0 | - |
| 2 | d | 2247 | 207.1 | 7.222E-05 | 0.3487 | +1 | 2 |
| - | - | 905.6 | 208.1 | - | - | 0 | - |
| - | - | 730.1 | 208.1 | - | - | 0 | - |
| - | - | 932.2 | 209.2 | - | - | 0 | - |
| - | - | 2341 | 212.1 | - | - | 0 | - |
| - | - | 951.4 | 217.1 | - | - | 0 | - |
| - | - | 7726 | 217.1 | - | - | 0 | - |
| - | - | 1088 | 218.1 | - | - | 0 | - |
| - | - | 1141 | 219.1 | - | - | 0 | - |
| - | - | 475.7 | 220.4 | - | - | 0 | - |
| - | - | 1.691E+04 | 221.1 | - | - | 0 | - |
| 2 | a | 1.433E+05 | 221.1 | 0.00033 | 1.492 | +1 | 2 |
| - | - | 1912 | 222.1 | - | - | 0 | - |
| - | - | 1.701E+04 | 222.1 | - | - | 0 | - |
| - | - | 1250 | 223.1 | - | - | 0 | - |
| - | - | 1536 | 227.1 | - | - | 0 | - |
| - | - | 578.2 | 227.2 | - | - | 0 | - |
| - | - | 1005 | 228.1 | - | - | 0 | - |
| - | - | 762.9 | 229.1 | - | - | 0 | - |
| - | - | 601.4 | 229.1 | - | - | 0 | - |
| - | - | 2496 | 231.1 | - | - | 0 | - |
| 2 | b | 3.444E+04 | 231.1 | 0.0003398 | 1.47 | +1 | 2 |
| - | - | 4146 | 232.1 | - | - | 0 | - |
| - | - | 1.221E+04 | 235.1 | - | - | 0 | - |
| - | - | 1640 | 236.1 | - | - | 0 | - |
| - | - | 1674 | 237.2 | - | - | 0 | - |
| - | - | 918.3 | 238.1 | - | - | 0 | - |
| - | - | 2298 | 239.1 | - | - | 0 | - |
| - | - | 514.5 | 244.1 | - | - | 0 | - |
| 2 | b | 7.535E+04 | 249.1 | 0.0003494 | 1.403 | +1 | 2 |
| - | - | 1.105E+04 | 250.1 | - | - | 0 | - |
| - | - | 7731 | 251.1 | - | - | 0 | - |
| - | - | 815 | 251.1 | - | - | 0 | - |
| - | - | 3672 | 253.1 | - | - | 0 | - |
| - | - | 1358 | 254.1 | - | - | 0 | - |
| - | - | 3886 | 257.1 | - | - | 0 | - |
| - | - | 1532 | 259.1 | - | - | 0 | - |
| - | - | 729.7 | 261.1 | - | - | 0 | - |
| - | - | 1.323E+04 | 263.1 | - | - | 0 | - |
| - | - | 1027 | 263.1 | - | - | 0 | - |
| - | - | 2103 | 264.1 | - | - | 0 | - |
| - | - | 1098 | 264.1 | - | - | 0 | - |
| - | - | 827.3 | 267.1 | - | - | 0 | - |
| - | - | 3337 | 274.1 | - | - | 0 | - |
| - | - | 6298 | 279.1 | - | - | 0 | - |
| - | - | 640 | 279.1 | - | - | 0 | - |
| - | - | 639.6 | 280.1 | - | - | 0 | - |
| - | - | 5297 | 285.1 | - | - | 0 | - |
| - | - | 801.1 | 286.1 | - | - | 0 | - |
| - | - | 708.3 | 289.1 | - | - | 0 | - |
| - | - | 4153 | 292.1 | - | - | 0 | - |
| - | - | 6810 | 296.2 | - | - | 0 | - |
| - | - | 1250 | 296.2 | - | - | 0 | - |
| - | - | 717.8 | 297.2 | - | - | 0 | - |
| - | - | 651.1 | 302.1 | - | - | 0 | - |
| - | - | 675.7 | 303.1 | - | - | 0 | - |
| - | - | 642.8 | 309.2 | - | - | 0 | - |
| - | - | 1087 | 310.2 | - | - | 0 | - |
| - | - | 2594 | 316.2 | - | - | 0 | - |
| - | - | 919.9 | 317.1 | - | - | 0 | - |
| - | - | 2001 | 318.1 | - | - | 0 | - |
| - | - | 1262 | 323.1 | - | - | 0 | - |
| - | - | 1.222E+04 | 324.1 | - | - | 0 | - |
| - | - | 2417 | 325.1 | - | - | 0 | - |
| - | - | 1219 | 333.1 | - | - | 0 | - |
| - | - | 2037 | 333.2 | - | - | 0 | - |
| - | - | 1352 | 338.1 | - | - | 0 | - |
| - | - | 855 | 339.2 | - | - | 0 | - |
| 9 | y | 6.294E+04 | 342.2 | 0.0005197 | 1.519 | +1 | 2 |
| - | - | 1.223E+04 | 343.2 | - | - | 0 | - |
| - | - | 863.2 | 344.2 | - | - | 0 | - |
| - | - | 1470 | 346.1 | - | - | 0 | - |
| - | - | 1323 | 348.1 | - | - | 0 | - |
| - | - | 4944 | 350.1 | - | - | 0 | - |
| 3 | a | 1444 | 350.2 | 0.0002988 | 0.8533 | +1 | 3 |
| - | - | 796.6 | 351.1 | - | - | 0 | - |
| 3 | a | 1151 | 351.2 | 0.0005794 | 1.65 | +1 | 3 |
| - | - | 3502 | 356.1 | - | - | 0 | - |
| - | - | 1030 | 360.1 | - | - | 0 | - |
| - | - | 4949 | 364.2 | - | - | 0 | - |
| - | - | 931.5 | 365.2 | - | - | 0 | - |
| - | - | 730.8 | 366.1 | - | - | 0 | - |
| - | - | 1079 | 374.1 | - | - | 0 | - |
| - | - | 1453 | 376.1 | - | - | 0 | - |
| - | - | 1.724E+04 | 378.1 | - | - | 0 | - |
| 3 | b | 1.47E+04 | 378.2 | 0.0003615 | 0.9559 | +1 | 3 |
| - | - | 3473 | 379.1 | - | - | 0 | - |
| 3 | b | 764.2 | 379.2 | 0.003613 | 9.528 | +1 | 3 |
| - | - | 3919 | 379.2 | - | - | 0 | - |
| - | - | 2227 | 382.1 | - | - | 0 | - |
| - | - | 799.4 | 388.1 | - | - | 0 | - |
| - | - | 964.8 | 390.2 | - | - | 0 | - |
| - | - | 6617 | 394.1 | - | - | 0 | - |
| - | - | 802.5 | 395.1 | - | - | 0 | - |
| 3 | b | 755.5 | 396.2 | 8.125E-05 | 0.2051 | +1 | 3 |
| - | - | 826 | 399.2 | - | - | 0 | - |
| - | - | 1072 | 404.1 | - | - | 0 | - |
| - | - | 823.9 | 407.2 | - | - | 0 | - |
| - | - | 2752 | 408.2 | - | - | 0 | - |
| - | - | 644.4 | 409.2 | - | - | 0 | - |
| - | - | 822.7 | 410.1 | - | - | 0 | - |
| - | - | 617.5 | 416.2 | - | - | 0 | - |
| - | - | 1.081E+04 | 425.2 | - | - | 0 | - |
| - | - | 2661 | 426.2 | - | - | 0 | - |
| - | - | 1209 | 427.2 | - | - | 0 | - |
| - | - | 734.8 | 428.2 | - | - | 0 | - |
| - | - | 957.8 | 433.2 | - | - | 0 | - |
| - | - | 588.6 | 435.2 | - | - | 0 | - |
| - | - | 776.8 | 443.2 | - | - | 0 | - |
| - | - | 1154 | 447.2 | - | - | 0 | - |
| - | - | 1189 | 449.2 | - | - | 0 | - |
| - | - | 608.4 | 452.2 | - | - | 0 | - |
| - | - | 622.1 | 454.2 | - | - | 0 | - |
| - | - | 1961 | 455.2 | - | - | 0 | - |
| - | - | 2585 | 461.2 | - | - | 0 | - |
| - | - | 5390 | 465.2 | - | - | 0 | - |
| - | - | 951.7 | 470.2 | - | - | 0 | - |
| - | - | 1749 | 471.2 | - | - | 0 | - |
| - | - | 656.5 | 473.4 | - | - | 0 | - |
| - | - | 685 | 477.2 | - | - | 0 | - |
| - | - | 3750 | 479.2 | - | - | 0 | - |
| - | - | 974.6 | 486.2 | - | - | 0 | - |
| 8 | y | 1.546E+04 | 489.2 | 0.005649 | 11.55 | +1 | 3 |
| - | - | 4366 | 490.2 | - | - | 0 | - |
| - | - | 1593 | 491.2 | - | - | 0 | - |
| 4 | b | 1783 | 493.2 | 0.0006416 | 1.301 | +1 | 4 |
| 8 | b | 2722 | 495.2 | 0.001365 | 2.757 | +2 | 8 |
| - | - | 6283 | 496.2 | - | - | 0 | - |
| - | - | 1893 | 497.2 | - | - | 0 | - |
| - | - | 1950 | 497.2 | - | - | 0 | - |
| 4 | b | 6160 | 511.2 | 0.0008482 | 1.659 | +1 | 4 |
| - | - | 2045 | 512.2 | - | - | 0 | - |
| - | - | 2029 | 513.2 | - | - | 0 | - |
| - | - | 1531 | 518.7 | - | - | 0 | - |
| - | - | 1895 | 519.2 | - | - | 0 | - |
| - | - | 743 | 520.2 | - | - | 0 | - |
| - | - | 836.6 | 532.2 | - | - | 0 | - |
| - | - | 777.7 | 541.2 | - | - | 0 | - |
| 3 | y | 2870 | 541.7 | 0.002625 | 4.846 | +2 | 8 |
| - | - | 1584 | 542.2 | - | - | 0 | - |
| - | - | 895.3 | 542.3 | - | - | 0 | - |
| - | - | 660.5 | 542.7 | - | - | 0 | - |
| - | - | 1470 | 548.2 | - | - | 0 | - |
| 3 | y | 1.498E+04 | 550.7 | 0.003019 | 5.482 | +2 | 8 |
| - | - | 1.255E+04 | 551.2 | - | - | 0 | - |
| - | - | 5124 | 551.7 | - | - | 0 | - |
| - | - | 905.7 | 552.2 | - | - | 0 | - |
| - | - | 1078 | 557.2 | - | - | 0 | - |
| 7 | y | 2.246E+04 | 560.2 | 0.005095 | 9.095 | +1 | 4 |
| - | - | 7520 | 561.2 | - | - | 0 | - |
| - | - | 2371 | 562.2 | - | - | 0 | - |
| 9 | b | 706.6 | 564.2 | 0.006604 | 11.71 | +2 | 9 |
| - | - | 711 | 567.2 | - | - | 0 | - |
| - | - | 1642 | 570.2 | - | - | 0 | - |
| 9 | b | 885 | 572.7 | 0.003339 | 5.831 | +2 | 9 |
| - | - | 997.4 | 573.2 | - | - | 0 | - |
| 2 | y | 1208 | 592.2 | 0.0008284 | 1.399 | +2 | 9 |
| - | - | 1107 | 592.7 | - | - | 0 | - |
| - | - | 1046 | 594.2 | - | - | 0 | - |
| 2 | y | 6117 | 601.2 | 0.002861 | 4.759 | +2 | 9 |
| - | - | 4526 | 601.7 | - | - | 0 | - |
| - | - | 1483 | 602.2 | - | - | 0 | - |
| - | - | 902.5 | 602.7 | - | - | 0 | - |
| - | - | 2278 | 606.2 | - | - | 0 | - |
| 5 | b | 2794 | 608.2 | 3.284E-06 | 0.005399 | +1 | 5 |
| - | - | 936.9 | 609.2 | - | - | 0 | - |
| - | - | 642.9 | 610.4 | - | - | 0 | - |
| - | - | 1813 | 612.2 | - | - | 0 | - |
| - | - | 1142 | 612.2 | - | - | 0 | - |
| - | - | 1032 | 614.2 | - | - | 0 | - |
| - | - | 2071 | 624.2 | - | - | 0 | - |
| - | - | 1057 | 625.2 | - | - | 0 | - |
| 5 | b | 4285 | 626.2 | 0.0003075 | 0.491 | +1 | 5 |
| - | - | 1224 | 627.3 | - | - | 0 | - |
| - | - | 1624 | 633.8 | - | - | 0 | - |
| - | - | 2190 | 634.2 | - | - | 0 | - |
| - | - | 1412 | 635.2 | - | - | 0 | - |
| - | - | 768.1 | 642.2 | - | - | 0 | - |
| - | - | 3086 | 642.8 | - | - | 0 | - |
| - | - | 1091 | 643.3 | - | - | 0 | - |
| - | - | 798.5 | 643.8 | - | - | 0 | - |
| - | - | 1095 | 651.8 | - | - | 0 | - |
| - | - | 747.2 | 656.8 | - | - | 0 | - |
| - | - | 2398 | 657.3 | - | - | 0 | - |
| - | - | 877.1 | 658.3 | - | - | 0 | - |
| - | - | 6968 | 659.3 | - | - | 0 | - |
| - | - | 2704 | 660.3 | - | - | 0 | - |
| - | - | 970.3 | 661.3 | - | - | 0 | - |
| 0 | Precursor | 1.102E+04 | 665.8 | 0.002806 | 4.215 | +2 | -1 |
| - | - | 9433 | 666.3 | - | - | 0 | - |
| - | - | 3499 | 666.8 | - | - | 0 | - |
| - | - | 1622 | 667.3 | - | - | 0 | - |
| 0 | Precursor | 2.025E+04 | 674.8 | 0.002712 | 4.019 | +2 | -1 |
| - | - | 1.943E+04 | 675.3 | - | - | 0 | - |
| - | - | 8441 | 675.8 | - | - | 0 | - |
| - | - | 2169 | 676.3 | - | - | 0 | - |
| - | - | 1503 | 685.3 | - | - | 0 | - |
| - | - | 1174 | 686.3 | - | - | 0 | - |
| - | - | 2400 | 695.3 | - | - | 0 | - |
| - | - | 801.8 | 696.3 | - | - | 0 | - |
| - | - | 748.6 | 703.3 | - | - | 0 | - |
| - | - | 823.7 | 704.3 | - | - | 0 | - |
| - | - | 871.2 | 705.3 | - | - | 0 | - |
| - | - | 643.4 | 708.3 | - | - | 0 | - |
| - | - | 1195 | 713.3 | - | - | 0 | - |
| - | - | 2447 | 721.3 | - | - | 0 | - |
| - | - | 968.8 | 722.3 | - | - | 0 | - |
| 6 | y | 1.968E+04 | 723.3 | 0.004816 | 6.659 | +1 | 5 |
| - | - | 8128 | 724.3 | - | - | 0 | - |
| - | - | 3127 | 725.3 | - | - | 0 | - |
| - | - | 869.2 | 726.3 | - | - | 0 | - |
| - | - | 690.2 | 743.3 | - | - | 0 | - |
| - | - | 743.4 | 744.3 | - | - | 0 | - |
| - | - | 3577 | 749.3 | - | - | 0 | - |
| - | - | 1512 | 750.3 | - | - | 0 | - |
| - | - | 917.6 | 756.3 | - | - | 0 | - |
| - | - | 1560 | 759.3 | - | - | 0 | - |
| - | - | 939.6 | 761.3 | - | - | 0 | - |
| - | - | 689.6 | 767.3 | - | - | 0 | - |
| 6 | b | 2451 | 771.3 | 0.0005786 | 0.7501 | +1 | 6 |
| - | - | 954.1 | 772.3 | - | - | 0 | - |
| - | - | 9098 | 774.3 | - | - | 0 | - |
| - | - | 3870 | 775.3 | - | - | 0 | - |
| - | - | 744.4 | 776.3 | - | - | 0 | - |
| - | - | 760.5 | 778.3 | - | - | 0 | - |
| - | - | 968.5 | 780.4 | - | - | 0 | - |
| 6 | b | 1167 | 789.3 | 0.003028 | 3.836 | +1 | 6 |
| - | - | 3436 | 804.3 | - | - | 0 | - |
| - | - | 2030 | 805.3 | - | - | 0 | - |
| - | - | 789.9 | 814.3 | - | - | 0 | - |
| 5 | y | 1360 | 820.3 | 0.006626 | 8.078 | +1 | 6 |
| - | - | 863.2 | 821.3 | - | - | 0 | - |
| - | - | 708.9 | 828.4 | - | - | 0 | - |
| - | - | 3971 | 832.3 | - | - | 0 | - |
| - | - | 1505 | 833.3 | - | - | 0 | - |
| 5 | y | 2.918E+04 | 838.3 | 0.004973 | 5.932 | +1 | 6 |
| - | - | 1.321E+04 | 839.3 | - | - | 0 | - |
| - | - | 3593 | 840.3 | - | - | 0 | - |
| - | - | 842 | 841.3 | - | - | 0 | - |
| - | - | 1197 | 843.3 | - | - | 0 | - |
| - | - | 1648 | 850.3 | - | - | 0 | - |
| 7 | b | 1277 | 860.3 | 0.002155 | 2.505 | +1 | 7 |
| - | - | 3260 | 868.3 | - | - | 0 | - |
| - | - | 1170 | 869.3 | - | - | 0 | - |
| - | - | 2127 | 871.3 | - | - | 0 | - |
| - | - | 2284 | 886.4 | - | - | 0 | - |
| - | - | 987.8 | 887.4 | - | - | 0 | - |
| - | - | 1.433E+04 | 889.3 | - | - | 0 | - |
| - | - | 7322 | 890.4 | - | - | 0 | - |
| - | - | 1799 | 891.4 | - | - | 0 | - |
| - | - | 5212 | 896.3 | - | - | 0 | - |
| - | - | 2788 | 897.3 | - | - | 0 | - |
| - | - | 883.3 | 898.3 | - | - | 0 | - |
| - | - | 654 | 905.4 | - | - | 0 | - |
| - | - | 1224 | 914.3 | - | - | 0 | - |
| - | - | 1126 | 915.3 | - | - | 0 | - |
| 4 | y | 2727 | 935.3 | 0.005379 | 5.751 | +1 | 7 |
| - | - | 1334 | 936.3 | - | - | 0 | - |
| 4 | y | 4.796E+04 | 953.3 | 0.004824 | 5.06 | +1 | 7 |
| - | - | 2.607E+04 | 954.3 | - | - | 0 | - |
| - | - | 8823 | 955.3 | - | - | 0 | - |
| - | - | 1149 | 956.3 | - | - | 0 | - |
| - | - | 779.9 | 963.3 | - | - | 0 | - |
| - | - | 909.7 | 969.4 | - | - | 0 | - |
| 8 | b | 1429 | 989.4 | 0.001834 | 1.854 | +1 | 8 |
| - | - | 1430 | 997.4 | - | - | 0 | - |
| - | - | 2134 | 1018 | - | - | 0 | - |
| - | - | 1024 | 1019 | - | - | 0 | - |
| - | - | 720.3 | 1036 | - | - | 0 | - |
| - | - | 2.246E+04 | 1036 | - | - | 0 | - |
| - | - | 1.575E+04 | 1037 | - | - | 0 | - |
| - | - | 4695 | 1038 | - | - | 0 | - |
| - | - | 1121 | 1054 | - | - | 0 | - |
| 3 | y | 3912 | 1082 | 0.005813 | 5.37 | +1 | 8 |
| - | - | 2935 | 1083 | - | - | 0 | - |
| - | - | 1379 | 1084 | - | - | 0 | - |
| - | - | 633.2 | 1085 | - | - | 0 | - |
| 3 | y | 8.051E+04 | 1100 | 0.00477 | 4.334 | +1 | 8 |
| - | - | 5.675E+04 | 1101 | - | - | 0 | - |
| - | - | 2.487E+04 | 1102 | - | - | 0 | - |
| - | - | 4076 | 1103 | - | - | 0 | - |
| - | - | 983.8 | 1116 | - | - | 0 | - |
| - | - | 1097 | 1119 | - | - | 0 | - |
| - | - | 851.6 | 1120 | - | - | 0 | - |
| - | - | 5362 | 1137 | - | - | 0 | - |
| - | - | 3252 | 1138 | - | - | 0 | - |
| - | - | 981.2 | 1139 | - | - | 0 | - |
| 9 | b | 1129 | 1144 | 0.0002834 | 0.2476 | +1 | 9 |
| - | - | 1365 | 1145 | - | - | 0 | - |
| - | - | 1185 | 1147 | - | - | 0 | - |
| - | - | 990.3 | 1148 | - | - | 0 | - |
| 2 | y | 1970 | 1183 | 0.001958 | 1.654 | +1 | 9 |
| - | - | 1527 | 1184 | - | - | 0 | - |
| 2 | y | 1.541E+04 | 1201 | 0.003722 | 3.098 | +1 | 9 |
| - | - | 1.149E+04 | 1202 | - | - | 0 | - |
| - | - | 4419 | 1203 | - | - | 0 | - |
| - | - | 924.9 | 1204 | - | - | 0 | - |
| - | - | 2066 | 1211 | - | - | 0 | - |
| - | - | 2540 | 1212 | - | - | 0 | - |
| - | - | 770.3 | 1213 | - | - | 0 | - |
| - | - | 768 | 3180 | - | - | 0 | - |

m/z Charge Intensity FragmentType MassShift Position
120.08113861083984 0 248892.98
120.75008392333984 0 367.16858
121.03997802734375 0 431.58145
121.08440399169922 0 20899.408
122.04613494873047 0 327.9599
122.07164001464844 0 415.9098
127.08705139160156 0 524.75525
128.10694885253906 0 487.96606
129.10252380371094 0 2111.0771
130.05014038085938 0 1509.9312
130.0654296875 0 2601.1208
132.08116149902344 0 1170.98
133.06114196777344 0 452.03635
133.08607482910156 0 742.8266
136.0759735107422 0 21804.857
136.0867919921875 0 742.8641
137.07943725585938 0 1599.2986
138.05517578125 0 2458.1787
138.0664520263672 0 3112.7979
142.2949676513672 0 500.6951
143.0733642578125 0 723.5117
145.22573852539062 0 465.4715
146.06036376953125 0 1393.167
146.0968017578125 0 1444.6377
147.07676696777344 0 2145.197
148.087158203125 0 732.38464
148.95472717285156 0 891.2722
152.0709686279297 0 603.61804
153.0652618408203 0 501.0165
155.0818328857422 0 1032.5635
156.07693481445312 0 7847.0337
158.09674072265625 0 5732.387
159.09190368652344 0 11850.297
159.09938049316406 0 786.9789
160.0953826904297 0 1090.6243
160.11224365234375 0 612.39465
165.07699584960938 0 751.9162
166.0615692138672 0 681.6274
166.08645629882812 0 4342.867
166.0941162109375 0 499.86118
166.09716796875 0 507.41418
167.0916748046875 0 947.7647
169.07278442382812 0 439.04614
169.13385009765625 0 2193.8286
171.07693481445312 0 642.41833
173.12823486328125 0 696.9753
175.0871124267578 0 967.4299
175.0980987548828 0 524.9136
176.0822296142578 0 2751.239
176.10731506347656 0 115590.47
177.1024169921875 0 2474.3855
177.1107940673828 0 12623.534
180.0771484375 0 1101.4371
183.1134490966797 0 916.97534
186.09158325195312 0 14860.768
187.09518432617188 0 1714.2839
188.07083129882812 0 7513.697
189.07432556152344 0 1198.1849
189.08737182617188 0 1224.2247
190.0871124267578 0 531.26416 b Ammonia loss 2
191.09307861328125 0 824.7227
191.11801147460938 0 2718.586
193.10862731933594 0 7920.361
194.11178588867188 0 1434.0781
195.0878143310547 0 2849.0312
197.10394287109375 0 591.74396
197.12847900390625 0 1871.8606
201.12356567382812 0 1035.2778
203.06613159179688 0 721.58966
203.09295654296875 0 579.4196
203.11813354492188 0 8735.99 a Water loss 1
204.0770721435547 0 4144.642
204.1019287109375 0 523.317 a Ammonia loss 1
204.1212921142578 0 729.7081
205.0809326171875 0 579.5054
205.0973358154297 0 5906.5625 y 9
205.10769653320312 0 940.1223
206.1011505126953 0 660.1891
207.11273193359375 0 2247.3723 d 1
208.07205200195312 0 905.5927
208.1163330078125 0 730.10156
209.16500854492188 0 932.1808
212.11839294433594 0 2341.0063
217.0845489501953 0 951.36597
217.09739685058594 0 7726.1865
218.10140991210938 0 1087.9297
219.07957458496094 0 1140.5972
220.4041290283203 0 475.73422
221.10357666015625 0 16907.34
221.1287841796875 0 143259.47 a 1
222.10728454589844 0 1912.2261
222.13209533691406 0 17008.258
223.1342315673828 0 1249.97
227.1025848388672 0 1535.9536
227.1751708984375 0 578.15936
228.1129913330078 0 1005.2138
229.097412109375 0 762.8733
229.10826110839844 0 601.36957
231.06158447265625 0 2496.1638
231.11314392089844 0 34443.098 b Water loss 1
232.1166229248047 0 4146.1963
235.1080322265625 0 12211.9
236.1116943359375 0 1639.7385
237.1598358154297 0 1673.9874
238.12246704101562 0 918.2526
239.11415100097656 0 2298.3157
244.12881469726562 0 514.47894
249.12371826171875 0 75347.55 b 1
250.12716674804688 0 11045.716
251.10293579101562 0 7731.174
251.12950134277344 0 815.02576
253.1184844970703 0 3672.008
254.1216583251953 0 1358.4319
257.1062927246094 0 3885.7693
259.1066589355469 0 1531.5962
261.0879211425781 0 729.7267
263.1028137207031 0 13225.301
263.12750244140625 0 1027.3922
264.10638427734375 0 2102.6082
264.145263671875 0 1098.1705
267.0909423828125 0 827.28723
274.12969970703125 0 3336.5542
279.09796142578125 0 6298.251
279.1236572265625 0 639.97327
280.1007385253906 0 639.55396
285.10186767578125 0 5297.139
286.1048278808594 0 801.07776
289.1350402832031 0 708.2708
292.1408996582031 0 4153.1636
296.15093994140625 0 6809.6226
296.19677734375 0 1249.5243
297.15399169921875 0 717.75354
302.0994873046875 0 651.0586
303.113037109375 0 675.7037
309.1600036621094 0 642.83417
310.151123046875 0 1086.7367
316.1869201660156 0 2593.9722
317.1278991699219 0 919.8704
318.1446228027344 0 2000.5795
323.0985107421875 0 1261.6306
324.1461181640625 0 12218.643
325.1484680175781 0 2416.6414
333.1081848144531 0 1219.2368
333.1596374511719 0 2036.6683
338.12945556640625 0 1352.3851
339.1708068847656 0 855.0022
342.1565856933594 0 62938.082 y 8
343.1595153808594 0 12231.441
344.16387939453125 0 863.18854
346.14056396484375 0 1469.533
348.1194152832031 0 1322.5725
350.13482666015625 0 4943.698
350.1860046386719 0 1443.8229 a Water loss 2
351.1382751464844 0 796.59546
351.1708984375 0 1150.9159 a Ammonia loss 2
356.1391296386719 0 3502.4258
360.119384765625 0 1029.8633
364.15032958984375 0 4949.428
365.15411376953125 0 931.47406
366.128662109375 0 730.7545
374.13311767578125 0 1079.4011
376.1141357421875 0 1452.9406
378.1298828125 0 17239.105
378.18157958984375 0 14699.1875 b Water loss 2
379.1331481933594 0 3473.4028
379.16162109375 0 764.2263 b Ammonia loss 2
379.1857604980469 0 3919.4177
382.14324951171875 0 2226.9531
388.11553955078125 0 799.35785
390.1584777832031 0 964.8394
394.125 0 6616.7754
395.1291809082031 0 802.46674
396.1918640136719 0 755.5065 b 2
399.17877197265625 0 825.9618
404.14556884765625 0 1071.6555
407.1846618652344 0 823.8633
408.16717529296875 0 2751.7139
409.17169189453125 0 644.37537
410.1383056640625 0 822.658
416.1509704589844 0 617.5426
425.1937561035156 0 10812.561
426.19659423828125 0 2660.5479
427.20849609375 0 1208.7314
428.2114562988281 0 734.78076
433.1725158691406 0 957.8221
435.17803955078125 0 588.63214
443.1569519042969 0 776.7764
447.1515808105469 0 1154.2305
449.16778564453125 0 1189.2062
452.18902587890625 0 608.40717
454.19720458984375 0 622.0808
455.20428466796875 0 1961.0171
461.166748046875 0 2584.6462
465.1622314453125 0 5390.409
470.1668701171875 0 951.72516
471.18023681640625 0 1748.8911
473.43304443359375 0 656.47656
477.1640625 0 684.9541
479.17755126953125 0 3750.2644
486.1606140136719 0 974.6441
489.19219970703125 0 15459.46 y 7
490.1949157714844 0 4366.496
491.20635986328125 0 1593.3159
493.20751953125 0 1782.8773 b Water loss 3
495.1880187988281 0 2722.2231 b Water loss 7
496.2306823730469 0 6282.911
497.1724548339844 0 1893.2784
497.2343444824219 0 1949.5571
511.2195739746094 0 6159.513 b 3
512.2233276367188 0 2045.2802
513.197265625 0 2028.5502
518.7135009765625 0 1530.5579
519.2046508789062 0 1895.4955
520.2019653320312 0 743.02216
532.2169189453125 0 836.61975
541.1944580078125 0 777.704
541.7056274414062 0 2870.4639 y Water loss 2
542.2059936523438 0 1584.1584
542.2808837890625 0 895.30475
542.7086791992188 0 660.51636
548.19970703125 0 1470.4785
550.7113037109375 0 14979.002 y 2
551.2125854492188 0 12545.563
551.7130737304688 0 5123.8467
552.2164306640625 0 905.6857
557.198486328125 0 1078.2277
560.228759765625 0 22460.725 y 6
561.2320556640625 0 7519.87
562.2293090820312 0 2370.7783
564.2147216796875 0 706.59656 b Ammonia loss 8
567.2074584960938 0 710.9679
570.2274169921875 0 1641.8981
572.7247314453125 0 885.0061 b 8
573.2254028320312 0 997.3953
592.2260131835938 0 1207.807 y Water loss 1
592.7294921875 0 1106.6696
594.2229614257812 0 1046.4064
601.2349853515625 0 6116.93 y 1
601.7362670898438 0 4525.759
602.2393798828125 0 1482.8645
602.73828125 0 902.53033
606.2338256835938 0 2277.7515
608.235107421875 0 2794.2437 b Water loss 4
609.2409057617188 0 936.8573
610.4353637695312 0 642.9495
612.1947021484375 0 1812.7759
612.240234375 0 1141.8761
614.2462158203125 0 1032.17
624.232421875 0 2071.2505
625.2383422851562 0 1056.9056
626.245361328125 0 4284.6226 b 4
627.2527465820312 0 1224.4039
633.7643432617188 0 1624.293
634.22900390625 0 2190.0962
635.2355346679688 0 1412.4994
642.2413940429688 0 768.1457
642.7686157226562 0 3086.3499
643.2711181640625 0 1090.9521
643.7708740234375 0 798.4808
651.7697143554688 0 1094.8462
656.755126953125 0 747.1766
657.2614135742188 0 2397.8123
658.2619018554688 0 877.0962
659.2941284179688 0 6967.563
660.2958984375 0 2704.492
661.3021240234375 0 970.2542
665.7638549804688 0 11021.254 Precursor Water loss
666.2650756835938 0 9432.502
666.7659301757812 0 3498.8022
667.26904296875 0 1622.3896
674.76904296875 0 20251.508 Precursor
675.2703247070312 0 19426.387
675.7703247070312 0 8441.1045
676.2713012695312 0 2169.2249
685.2567749023438 0 1502.9885
686.25927734375 0 1174.357
695.2670288085938 0 2400.0654
696.269775390625 0 801.7801
703.26806640625 0 748.6378
704.2685546875 0 823.66534
705.281494140625 0 871.2435
708.2642211914062 0 643.4278
713.2758178710938 0 1195.291
721.2616577148438 0 2446.6223
722.2622680664062 0 968.826
723.2918090820312 0 19678.73 y 5
724.2947387695312 0 8128.4663
725.2935180664062 0 3126.6013
726.2932739257812 0 869.2195
743.307861328125 0 690.2029
744.2978515625 0 743.39557
749.2554931640625 0 3577.0596
750.2586059570312 0 1511.927
756.30712890625 0 917.5965
759.2626953125 0 1560.1877
761.3171997070312 0 939.6126
767.263427734375 0 689.5621
771.2990112304688 0 2450.9304 b Water loss 5
772.3016357421875 0 954.1051
774.3203735351562 0 9097.788
775.3244018554688 0 3870.2966
776.327392578125 0 744.42316
778.3108520507812 0 760.5322
780.3536987304688 0 968.45276
789.3059692382812 0 1166.5386 b 5
804.3298950195312 0 3435.773
805.33056640625 0 2029.8463
814.3158569335938 0 789.90936
820.3099975585938 0 1359.6544 y Water loss 4
821.30517578125 0 863.15533
828.4295043945312 0 708.9233
832.3260498046875 0 3971.4946
833.328857421875 0 1505.1414
838.3189086914062 0 29180.104 y 4
839.32177734375 0 13209.714
840.3207397460938 0 3593.2034
841.32421875 0 842.0475
843.3369140625 0 1196.6814
850.331787109375 0 1647.7385
860.3482666015625 0 1276.9323 b 6
868.3286743164062 0 3260.174
869.3309936523438 0 1169.9133
871.336669921875 0 2127.4602
886.4157104492188 0 2284.2683
887.4151000976562 0 987.79974
889.3472290039062 0 14333.008
890.35009765625 0 7321.8203
891.3529663085938 0 1799.2576
896.322998046875 0 5211.633
897.3274536132812 0 2787.8901
898.322265625 0 883.3078
905.3799438476562 0 654.04474
914.335693359375 0 1224.3138
915.3391723632812 0 1126.1094
935.335693359375 0 2726.9673 y Water loss 3
936.335205078125 0 1333.7272
953.345703125 0 47963.234 y 3
954.3482666015625 0 26072.693
955.348876953125 0 8823.21
956.3458862304688 0 1148.8939
963.3314208984375 0 779.8688
969.3763427734375 0 909.6979
989.3641967773438 0 1428.9841 b Water loss 7
997.3726806640625 0 1429.5715
1018.4063720703125 0 2133.6992
1019.412353515625 0 1024.148
1035.517822265625 0 720.2693
1036.41552734375 0 22458.865
1037.41845703125 0 15753.941
1038.42041015625 0 4694.6904
1054.40673828125 0 1121.2407
1082.404541015625 0 3912.0881 y Water loss 2
1083.4072265625 0 2935.3052
1084.4039306640625 0 1379.1327
1085.3916015625 0 633.19977
1100.4140625 0 80514.414 y 2
1101.4168701171875 0 56749.547
1102.4176025390625 0 24873.482
1103.4178466796875 0 4075.5273
1116.452392578125 0 983.834
1119.4417724609375 0 1096.594
1120.4542236328125 0 851.5525
1137.462646484375 0 5361.6245
1138.4669189453125 0 3251.5852
1139.472412109375 0 981.2207
1144.435791015625 0 1128.6748 b 8
1145.4434814453125 0 1365.4208
1147.44482421875 0 1185.37
1148.449462890625 0 990.26154
1183.4483642578125 0 1969.8328 y Water loss 1
1184.45263671875 0 1527.2808
1201.460693359375 0 15405.752 y 1
1202.464111328125 0 11489.48
1203.4638671875 0 4419.3057
1204.467041015625 0 924.9039
1211.446533203125 0 2066.3813
1212.4456787109375 0 2539.6675
1213.4459228515625 0 770.3379
3180.157958984375 0 767.9836

Spectrum Details

|  |  |
| --- | --- |
| Matched peaks? Matched peaks The total absolute number of peaks matched. Additionally in brackets the total fraction of peaks matched and the total number of peaks is shown. | 43 (11.35% of 379) |
| FDR? FDR The false discovery rate estimated for this peptide. It is calculated by matching all theoretical fragments with a non-integer shift with the raw peaks for this spectrum. This is done with 40 different shifts. The resulting percentage is the average number of annotated peaks over the number of annotated peaks with the correct spectrum. | 0.00% |
| Satellite FDR? Satellite FDR See the FDR for details on its calculation. This satellite ion specific FDR only contains the satellite ions (d/w) for I/L/J positions. | - |
| PSM Score? PSM Score The PSM Score as given by Hecklib to this annotated spectrum. It is shown with three significant figures. | 532 |

#### Spectrum 8424? Spectrum 8424 The raw spectrum of this peptide as annotated by Hecklib. The fragments are coloured according to ion type (see legend). Any peaks with a star '\*' as text can be hovered over to see the full details, first the ion type second the mass shift type. By hovering over the amino acids in the peptide or ions in the legend the corresponding peaks are highlighted. By toggling the 'Unassigned' label you can turn the background (unassigned) peaks on or off in the plot. By updating the slider in the Ion legend you can update the spectrum to only show the top X% of the peaks with labels. The top X% means any peak that is within X% of the highest intensity. By dragging in the spectrum you can zoom in to a specific part of the spectrum and use 'Zoom Out' to get back to the original zoom level. The annotation of the spectrum is based on the given sequence in the peptides file and is done with different software so inconsistencies are likely. The peaks are annotated based on the given sequence, with 20 ppm tolerance.

Copy Data

##### Spectrum 8424 (TSV)

###### Preview

```
Loading example...
```

*Click on the button to copy the data to your clipboard.*

Mz MinMz MaxIntensity Max

WidthHeightPeptide font sizePeptide stroke widthSpectrum font sizeSpectrum stroke widthCompact peptide

Ion legend

wxyz

abcd

OtherUnassignedIon^Charge^_Position_Show for top:%

FTFDDYAMHW

03.51e+47.02e+41.05e+51.40e+5

Zoom Out

a^+1^_2_d^+1^_2_a^+1^_2_b^+1^_2_b^+1^_2_a^+1^_3_a^+1^_3_y^+1^_2_b^+1^_3_b^+1^_3_b^+2^_8_y^+1^_3_b^+1^_4_y^+2^_8_y^+2^_8_b^+2^_9_y^+1^_4_y^+2^_9_b^+1^_5_y^+2^_9_b^+1^_5_\*\*y^+1^_5_b^+1^_6_b^+1^_6_y^+1^_6_y^+1^_7_y^+1^_7_y^+1^_8_y^+1^_8_b^+1^_9_y^+1^_9_y^+1^_9_

0535106916042138

Fragment Matches Table

Show background peaks

| Position | Ion type | Intensity | mz Theoretical | mz Error (Th) | mz Error (ppm) | Charge | Series Number |
| --- | --- | --- | --- | --- | --- | --- | --- |
| - | - | 1.39E+05 | 120.1 | - | - | 0 | - |
| - | - | 1.174E+04 | 121.1 | - | - | 0 | - |
| - | - | 395.1 | 128.1 | - | - | 0 | - |
| - | - | 2483 | 129.1 | - | - | 0 | - |
| - | - | 1557 | 130.1 | - | - | 0 | - |
| - | - | 452.8 | 133.1 | - | - | 0 | - |
| - | - | 1.116E+04 | 136.1 | - | - | 0 | - |
| - | - | 971 | 137.1 | - | - | 0 | - |
| - | - | 493.5 | 138.1 | - | - | 0 | - |
| - | - | 1007 | 138.1 | - | - | 0 | - |
| - | - | 1022 | 143.1 | - | - | 0 | - |
| - | - | 696.3 | 146.1 | - | - | 0 | - |
| - | - | 522.1 | 147.1 | - | - | 0 | - |
| - | - | 584.8 | 148.1 | - | - | 0 | - |
| - | - | 1825 | 149 | - | - | 0 | - |
| - | - | 619.5 | 155.1 | - | - | 0 | - |
| - | - | 511.1 | 155.1 | - | - | 0 | - |
| - | - | 4847 | 156.1 | - | - | 0 | - |
| - | - | 426.1 | 157.8 | - | - | 0 | - |
| - | - | 741.3 | 158.1 | - | - | 0 | - |
| - | - | 3444 | 158.1 | - | - | 0 | - |
| - | - | 3837 | 159.1 | - | - | 0 | - |
| - | - | 800.2 | 166.1 | - | - | 0 | - |
| - | - | 1060 | 166.1 | - | - | 0 | - |
| - | - | 3053 | 167.1 | - | - | 0 | - |
| - | - | 674.5 | 167.1 | - | - | 0 | - |
| - | - | 937 | 169.1 | - | - | 0 | - |
| - | - | 1143 | 171.1 | - | - | 0 | - |
| - | - | 8842 | 175.1 | - | - | 0 | - |
| - | - | 1634 | 176.1 | - | - | 0 | - |
| - | - | 6.726E+04 | 176.1 | - | - | 0 | - |
| - | - | 1415 | 177.1 | - | - | 0 | - |
| - | - | 6170 | 177.1 | - | - | 0 | - |
| - | - | 7621 | 186.1 | - | - | 0 | - |
| - | - | 763.1 | 187.1 | - | - | 0 | - |
| - | - | 563.7 | 187.1 | - | - | 0 | - |
| - | - | 536.5 | 191.1 | - | - | 0 | - |
| - | - | 1290 | 191.1 | - | - | 0 | - |
| - | - | 4218 | 193.1 | - | - | 0 | - |
| - | - | 555.5 | 193.8 | - | - | 0 | - |
| - | - | 589.4 | 195.1 | - | - | 0 | - |
| - | - | 542.4 | 197.1 | - | - | 0 | - |
| - | - | 475.7 | 198.7 | - | - | 0 | - |
| - | - | 733.8 | 201.1 | - | - | 0 | - |
| - | - | 749.6 | 203.1 | - | - | 0 | - |
| - | - | 3525 | 203.1 | - | - | 0 | - |
| - | - | 973.1 | 203.1 | - | - | 0 | - |
| 2 | a | 4572 | 203.1 | 0.0002135 | 1.051 | +1 | 2 |
| - | - | 540.5 | 204.1 | - | - | 0 | - |
| - | - | 1321 | 204.1 | - | - | 0 | - |
| - | - | 955.2 | 205.1 | - | - | 0 | - |
| 2 | d | 613.8 | 207.1 | 0.0004313 | 2.083 | +1 | 2 |
| - | - | 934.2 | 212.1 | - | - | 0 | - |
| - | - | 554.9 | 215.1 | - | - | 0 | - |
| - | - | 482 | 215.1 | - | - | 0 | - |
| - | - | 4271 | 217.1 | - | - | 0 | - |
| - | - | 743.6 | 218.1 | - | - | 0 | - |
| - | - | 710.7 | 219.1 | - | - | 0 | - |
| - | - | 540.2 | 219.1 | - | - | 0 | - |
| - | - | 1.713E+04 | 221.1 | - | - | 0 | - |
| - | - | 9926 | 221.1 | - | - | 0 | - |
| 2 | a | 8.388E+04 | 221.1 | 0.0002689 | 1.216 | +1 | 2 |
| - | - | 2104 | 222.1 | - | - | 0 | - |
| - | - | 561.1 | 222.1 | - | - | 0 | - |
| - | - | 9644 | 222.1 | - | - | 0 | - |
| - | - | 6317 | 225 | - | - | 0 | - |
| - | - | 574.1 | 226 | - | - | 0 | - |
| - | - | 583.3 | 227 | - | - | 0 | - |
| - | - | 484.7 | 227.7 | - | - | 0 | - |
| - | - | 620.9 | 228.1 | - | - | 0 | - |
| - | - | 1935 | 231.1 | - | - | 0 | - |
| 2 | b | 2.15E+04 | 231.1 | 0.0003245 | 1.404 | +1 | 2 |
| - | - | 2902 | 232.1 | - | - | 0 | - |
| - | - | 6465 | 235.1 | - | - | 0 | - |
| - | - | 755.1 | 236.1 | - | - | 0 | - |
| - | - | 6914 | 239.1 | - | - | 0 | - |
| - | - | 1066 | 239.1 | - | - | 0 | - |
| - | - | 1041 | 240.1 | - | - | 0 | - |
| 2 | b | 4.375E+04 | 249.1 | 0.0003036 | 1.219 | +1 | 2 |
| - | - | 5548 | 250.1 | - | - | 0 | - |
| - | - | 3961 | 251.1 | - | - | 0 | - |
| - | - | 850.3 | 252.1 | - | - | 0 | - |
| - | - | 1851 | 257.1 | - | - | 0 | - |
| - | - | 1.052E+04 | 263.1 | - | - | 0 | - |
| - | - | 707.2 | 264.1 | - | - | 0 | - |
| - | - | 1893 | 274.1 | - | - | 0 | - |
| - | - | 493.7 | 275.3 | - | - | 0 | - |
| - | - | 2216 | 279.1 | - | - | 0 | - |
| - | - | 803.2 | 281.1 | - | - | 0 | - |
| - | - | 742.6 | 285 | - | - | 0 | - |
| - | - | 557.2 | 285.1 | - | - | 0 | - |
| - | - | 3321 | 285.1 | - | - | 0 | - |
| - | - | 2439 | 292.1 | - | - | 0 | - |
| - | - | 1658 | 295.1 | - | - | 0 | - |
| - | - | 4630 | 299.1 | - | - | 0 | - |
| - | - | 1859 | 312.1 | - | - | 0 | - |
| - | - | 632.5 | 313.1 | - | - | 0 | - |
| - | - | 593.5 | 315.1 | - | - | 0 | - |
| - | - | 883.1 | 318.1 | - | - | 0 | - |
| - | - | 669.7 | 322.1 | - | - | 0 | - |
| - | - | 733.2 | 323.1 | - | - | 0 | - |
| - | - | 606.8 | 323.1 | - | - | 0 | - |
| - | - | 1452 | 325.1 | - | - | 0 | - |
| - | - | 775.2 | 327.1 | - | - | 0 | - |
| - | - | 513.4 | 332.1 | - | - | 0 | - |
| - | - | 762 | 333.1 | - | - | 0 | - |
| - | - | 1157 | 333.2 | - | - | 0 | - |
| - | - | 691.8 | 339.2 | - | - | 0 | - |
| - | - | 7845 | 340.1 | - | - | 0 | - |
| - | - | 972.3 | 341 | - | - | 0 | - |
| - | - | 965.3 | 341.1 | - | - | 0 | - |
| - | - | 622.4 | 346.1 | - | - | 0 | - |
| - | - | 769.5 | 348.1 | - | - | 0 | - |
| - | - | 2605 | 350.1 | - | - | 0 | - |
| 3 | a | 987.1 | 350.2 | 0.0004819 | 1.376 | +1 | 3 |
| 3 | a | 711 | 351.2 | 0.0005803 | 1.652 | +1 | 3 |
| - | - | 4078 | 355.1 | - | - | 0 | - |
| - | - | 2435 | 356.1 | - | - | 0 | - |
| 9 | y | 4.003E+04 | 358.1 | 0.005149 | 14.38 | +1 | 2 |
| - | - | 7432 | 359 | - | - | 0 | - |
| - | - | 7678 | 359.2 | - | - | 0 | - |
| - | - | 955 | 360.1 | - | - | 0 | - |
| - | - | 588.7 | 360.2 | - | - | 0 | - |
| - | - | 3525 | 364.2 | - | - | 0 | - |
| - | - | 699.9 | 365.2 | - | - | 0 | - |
| - | - | 1119 | 368.1 | - | - | 0 | - |
| - | - | 781.8 | 369.1 | - | - | 0 | - |
| - | - | 586.3 | 370.1 | - | - | 0 | - |
| - | - | 596 | 370.2 | - | - | 0 | - |
| - | - | 675 | 376.1 | - | - | 0 | - |
| - | - | 1.041E+04 | 378.1 | - | - | 0 | - |
| 3 | b | 7275 | 378.2 | 0.0001479 | 0.3911 | +1 | 3 |
| - | - | 2181 | 379.1 | - | - | 0 | - |
| - | - | 1700 | 379.2 | - | - | 0 | - |
| - | - | 982.9 | 382.1 | - | - | 0 | - |
| - | - | 4575 | 394.1 | - | - | 0 | - |
| 3 | b | 828.7 | 396.2 | 0.002116 | 5.341 | +1 | 3 |
| - | - | 1612 | 406.2 | - | - | 0 | - |
| - | - | 564.4 | 416.8 | - | - | 0 | - |
| - | - | 1186 | 419 | - | - | 0 | - |
| - | - | 1058 | 423.2 | - | - | 0 | - |
| - | - | 976.9 | 424.2 | - | - | 0 | - |
| - | - | 803.6 | 427.2 | - | - | 0 | - |
| - | - | 7415 | 429.1 | - | - | 0 | - |
| - | - | 785.9 | 433.2 | - | - | 0 | - |
| - | - | 638.1 | 439.2 | - | - | 0 | - |
| - | - | 8294 | 441.2 | - | - | 0 | - |
| - | - | 1308 | 442.2 | - | - | 0 | - |
| - | - | 1025 | 449.2 | - | - | 0 | - |
| - | - | 582.4 | 454.2 | - | - | 0 | - |
| - | - | 661.7 | 455.2 | - | - | 0 | - |
| - | - | 601.4 | 459.8 | - | - | 0 | - |
| - | - | 1241 | 461.2 | - | - | 0 | - |
| - | - | 2914 | 465.2 | - | - | 0 | - |
| - | - | 585.9 | 477.2 | - | - | 0 | - |
| - | - | 1957 | 479.2 | - | - | 0 | - |
| - | - | 633.8 | 487.2 | - | - | 0 | - |
| - | - | 1013 | 491.2 | - | - | 0 | - |
| 8 | b | 1011 | 495.2 | 0.00048 | 0.9694 | +2 | 8 |
| - | - | 1230 | 497.2 | - | - | 0 | - |
| 8 | y | 9393 | 505.2 | 0.009851 | 19.5 | +1 | 3 |
| - | - | 2628 | 506.2 | - | - | 0 | - |
| - | - | 1200 | 507.2 | - | - | 0 | - |
| 4 | b | 4699 | 511.2 | 0.0002683 | 0.5249 | +1 | 4 |
| - | - | 6794 | 512.2 | - | - | 0 | - |
| - | - | 1314 | 513.2 | - | - | 0 | - |
| - | - | 1802 | 513.2 | - | - | 0 | - |
| - | - | 1889 | 519.2 | - | - | 0 | - |
| - | - | 832.6 | 520.2 | - | - | 0 | - |
| - | - | 1323 | 526.7 | - | - | 0 | - |
| - | - | 1101 | 527.2 | - | - | 0 | - |
| - | - | 588.9 | 530.2 | - | - | 0 | - |
| - | - | 1602 | 541.2 | - | - | 0 | - |
| - | - | 698.1 | 548.2 | - | - | 0 | - |
| 3 | y | 1026 | 549.7 | 0.005855 | 10.65 | +2 | 8 |
| - | - | 1047 | 550.2 | - | - | 0 | - |
| 3 | y | 1.124E+04 | 558.7 | 0.005028 | 9 | +2 | 8 |
| - | - | 7011 | 559.2 | - | - | 0 | - |
| - | - | 2404 | 559.7 | - | - | 0 | - |
| - | - | 715.9 | 560.2 | - | - | 0 | - |
| - | - | 661.2 | 563.2 | - | - | 0 | - |
| - | - | 1079 | 570.2 | - | - | 0 | - |
| 9 | b | 698.9 | 572.7 | 0.003339 | 5.831 | +2 | 9 |
| 7 | y | 1.595E+04 | 576.2 | 0.009846 | 17.09 | +1 | 4 |
| - | - | 4463 | 577.2 | - | - | 0 | - |
| - | - | 875.1 | 578.2 | - | - | 0 | - |
| - | - | 670.1 | 590.2 | - | - | 0 | - |
| 2 | y | 673.7 | 600.2 | 0.006552 | 10.92 | +2 | 9 |
| - | - | 1390 | 606.2 | - | - | 0 | - |
| 5 | b | 1683 | 608.2 | 0.0007291 | 1.199 | +1 | 5 |
| 2 | y | 6279 | 609.2 | 0.005359 | 8.796 | +2 | 9 |
| - | - | 3284 | 609.7 | - | - | 0 | - |
| - | - | 1434 | 610.2 | - | - | 0 | - |
| - | - | 869.4 | 612.2 | - | - | 0 | - |
| - | - | 1197 | 624.2 | - | - | 0 | - |
| 5 | b | 2317 | 626.2 | 0.0006737 | 1.076 | +1 | 5 |
| - | - | 1305 | 627.2 | - | - | 0 | - |
| - | - | 692.6 | 633.2 | - | - | 0 | - |
| - | - | 1232 | 634.2 | - | - | 0 | - |
| - | - | 932.1 | 641.8 | - | - | 0 | - |
| - | - | 2264 | 650.8 | - | - | 0 | - |
| - | - | 1832 | 651.3 | - | - | 0 | - |
| - | - | 1205 | 657.3 | - | - | 0 | - |
| 0 | Precursor | 7540 | 673.8 | 0.004877 | 7.238 | +2 | -1 |
| - | - | 5205 | 674.3 | - | - | 0 | - |
| - | - | 2709 | 674.8 | - | - | 0 | - |
| - | - | 5965 | 675.3 | - | - | 0 | - |
| - | - | 1875 | 676.3 | - | - | 0 | - |
| - | - | 870.3 | 681.8 | - | - | 0 | - |
| - | - | 1579 | 682.3 | - | - | 0 | - |
| 0 | Precursor | 1.673E+04 | 682.8 | 0.004721 | 6.915 | +2 | -1 |
| - | - | 778.1 | 682.9 | - | - | 0 | - |
| - | - | 1.157E+04 | 683.3 | - | - | 0 | - |
| - | - | 2231 | 683.4 | - | - | 0 | - |
| - | - | 6091 | 683.8 | - | - | 0 | - |
| - | - | 2045 | 684.2 | - | - | 0 | - |
| - | - | 1058 | 684.3 | - | - | 0 | - |
| - | - | 1778 | 685.3 | - | - | 0 | - |
| - | - | 1023 | 686.3 | - | - | 0 | - |
| - | - | 1726 | 695.3 | - | - | 0 | - |
| - | - | 1739 | 721.3 | - | - | 0 | - |
| - | - | 900.6 | 722.3 | - | - | 0 | - |
| 6 | y | 1.397E+04 | 739.3 | 0.009872 | 13.35 | +1 | 5 |
| - | - | 5507 | 740.3 | - | - | 0 | - |
| - | - | 1208 | 741.3 | - | - | 0 | - |
| - | - | 892.1 | 744.3 | - | - | 0 | - |
| - | - | 1629 | 749.3 | - | - | 0 | - |
| - | - | 1154 | 750.3 | - | - | 0 | - |
| - | - | 861.1 | 759.3 | - | - | 0 | - |
| - | - | 911 | 761.3 | - | - | 0 | - |
| 6 | b | 1879 | 771.3 | 0.001741 | 2.257 | +1 | 6 |
| - | - | 1295 | 772.3 | - | - | 0 | - |
| 6 | b | 1332 | 789.3 | 0.001929 | 2.445 | +1 | 6 |
| - | - | 6868 | 790.3 | - | - | 0 | - |
| - | - | 3127 | 791.3 | - | - | 0 | - |
| - | - | 1742 | 804.3 | - | - | 0 | - |
| - | - | 948.6 | 805.3 | - | - | 0 | - |
| - | - | 2599 | 832.3 | - | - | 0 | - |
| - | - | 1304 | 833.3 | - | - | 0 | - |
| - | - | 757 | 842.3 | - | - | 0 | - |
| 5 | y | 1.945E+04 | 854.3 | 0.009419 | 11.02 | +1 | 6 |
| - | - | 8447 | 855.3 | - | - | 0 | - |
| - | - | 2668 | 856.3 | - | - | 0 | - |
| - | - | 732.6 | 857.2 | - | - | 0 | - |
| - | - | 1554 | 868.3 | - | - | 0 | - |
| - | - | 909.8 | 887.3 | - | - | 0 | - |
| - | - | 4308 | 896.3 | - | - | 0 | - |
| - | - | 1470 | 897.3 | - | - | 0 | - |
| - | - | 733.5 | 898.3 | - | - | 0 | - |
| - | - | 9567 | 905.3 | - | - | 0 | - |
| - | - | 4598 | 906.3 | - | - | 0 | - |
| - | - | 1367 | 907.3 | - | - | 0 | - |
| 4 | y | 1983 | 951.3 | 0.0119 | 12.51 | +1 | 7 |
| - | - | 1347 | 952.3 | - | - | 0 | - |
| 4 | y | 2.973E+04 | 969.3 | 0.009148 | 9.437 | +1 | 7 |
| - | - | 1.564E+04 | 970.3 | - | - | 0 | - |
| - | - | 5734 | 971.3 | - | - | 0 | - |
| - | - | 709.4 | 972.4 | - | - | 0 | - |
| - | - | 1290 | 1034 | - | - | 0 | - |
| - | - | 1.509E+04 | 1052 | - | - | 0 | - |
| - | - | 8928 | 1053 | - | - | 0 | - |
| - | - | 2537 | 1054 | - | - | 0 | - |
| 3 | y | 2510 | 1098 | 0.009343 | 8.506 | +1 | 8 |
| - | - | 1968 | 1099 | - | - | 0 | - |
| 3 | y | 4.496E+04 | 1116 | 0.008422 | 7.544 | +1 | 8 |
| - | - | 2.923E+04 | 1117 | - | - | 0 | - |
| - | - | 1.212E+04 | 1118 | - | - | 0 | - |
| - | - | 1115 | 1119 | - | - | 0 | - |
| 9 | b | 1848 | 1144 | 0.003335 | 2.914 | +1 | 9 |
| - | - | 1395 | 1145 | - | - | 0 | - |
| - | - | 871.6 | 1146 | - | - | 0 | - |
| - | - | 2457 | 1153 | - | - | 0 | - |
| - | - | 1526 | 1154 | - | - | 0 | - |
| - | - | 649.2 | 1163 | - | - | 0 | - |
| 2 | y | 723.3 | 1199 | 0.01562 | 13.02 | +1 | 9 |
| 2 | y | 7398 | 1217 | 0.008107 | 6.659 | +1 | 9 |
| - | - | 6411 | 1218 | - | - | 0 | - |
| - | - | 2439 | 1219 | - | - | 0 | - |
| - | - | 1473 | 1227 | - | - | 0 | - |
| - | - | 1169 | 1228 | - | - | 0 | - |
| - | - | 610.9 | 2081 | - | - | 0 | - |
| - | - | 662.7 | 2117 | - | - | 0 | - |

m/z Charge Intensity FragmentType MassShift Position
120.08110809326172 0 138995.48
121.08438873291016 0 11744.153
128.10739135742188 0 395.07816
129.1024932861328 0 2483.278
130.06541442871094 0 1556.509
133.06085205078125 0 452.77115
136.07598876953125 0 11158.84
137.07945251464844 0 971.00696
138.05511474609375 0 493.53772
138.06646728515625 0 1006.5518
143.11790466308594 0 1022.49835
146.09706115722656 0 696.3162
147.1132354736328 0 522.12915
148.08718872070312 0 584.7726
149.04510498046875 0 1824.6205
155.0819091796875 0 619.4562
155.09327697753906 0 511.06876
156.07704162597656 0 4846.9707
157.7808837890625 0 426.10504
158.0606689453125 0 741.3021
158.09661865234375 0 3444.0938
159.09188842773438 0 3837.437
166.0535888671875 0 800.1837
166.061279296875 0 1060.0679
167.0557098388672 0 3052.6272
167.093505859375 0 674.4774
169.13372802734375 0 936.96967
171.11312866210938 0 1143.0991
175.08682250976562 0 8841.979
176.0825653076172 0 1634.2152
176.10728454589844 0 67263.58
177.1024169921875 0 1414.8386
177.1107940673828 0 6170.2944
186.09156799316406 0 7620.6587
187.09478759765625 0 763.08826
187.10800170898438 0 563.72797
191.09262084960938 0 536.53326
191.11801147460938 0 1290.2468
193.10862731933594 0 4218.1577
193.79933166503906 0 555.4704
195.08782958984375 0 589.3756
197.12918090820312 0 542.4266
198.7136688232422 0 475.66855
201.1240997314453 0 733.8159
203.06617736816406 0 749.60767
203.08189392089844 0 3525.2104
203.09280395507812 0 973.1492
203.11810302734375 0 4571.9775 a Water loss 1
204.0673828125 0 540.54565
204.0770721435547 0 1321.2258
205.0972900390625 0 955.233
207.1132354736328 0 613.8425 d 1
212.1189727783203 0 934.2318
215.08169555664062 0 554.87445
215.13919067382812 0 482.00253
217.0974578857422 0 4271.0127
218.10044860839844 0 743.6192
219.07943725585938 0 710.7438
219.11297607421875 0 540.2292
221.09226989746094 0 17130.803
221.10369873046875 0 9925.986
221.12872314453125 0 83882.65 a 1
222.09576416015625 0 2103.7769
222.1070098876953 0 561.13354
222.13211059570312 0 9643.975
225.04322814941406 0 6317.1934
226.0440673828125 0 574.0844
227.02174377441406 0 583.29834
227.70724487304688 0 484.71802
228.1136932373047 0 620.9154
231.0614471435547 0 1934.6271
231.11312866210938 0 21504.934 b Water loss 1
232.11622619628906 0 2902.2583
235.10797119140625 0 6464.6836
236.11138916015625 0 755.10223
239.09523010253906 0 6913.813
239.1144256591797 0 1066.1571
240.09666442871094 0 1041.2966
249.12367248535156 0 43753.883 b 1
250.1270751953125 0 5547.9883
251.10321044921875 0 3961.2754
252.10740661621094 0 850.279
257.1062927246094 0 1850.7162
263.102783203125 0 10521.767
264.107177734375 0 707.2401
274.1299133300781 0 1893.124
275.2738037109375 0 493.73575
279.09765625 0 2215.5654
281.0513000488281 0 803.24243
285.0097961425781 0 742.6101
285.08660888671875 0 557.2283
285.1020202636719 0 3321.412
292.14068603515625 0 2438.8843
295.1043395996094 0 1657.5454
299.0619812011719 0 4629.6045
312.1461486816406 0 1858.9408
313.1498718261719 0 632.4819
315.0998229980469 0 593.54504
318.1448059082031 0 883.14923
322.130126953125 0 669.7479
323.11517333984375 0 733.1845
323.1347351074219 0 606.84064
325.138671875 0 1452.4901
327.1449890136719 0 775.2008
332.1275634765625 0 513.40247
333.1087646484375 0 762.0121
333.1601257324219 0 1157.3477
339.1718444824219 0 691.761
340.1409606933594 0 7845.21
341.0175476074219 0 972.3119
341.14385986328125 0 965.27155
346.1410217285156 0 622.4163
348.11865234375 0 769.4576
350.1344909667969 0 2605.4988
350.1858215332031 0 987.14197 a Water loss 2
351.16973876953125 0 710.9925 a Ammonia loss 2
355.06988525390625 0 4077.9014
356.1387939453125 0 2434.7856
358.1512145996094 0 40030.89 y 8
359.0286560058594 0 7431.7705
359.1542663574219 0 7677.5454
360.11871337890625 0 954.98663
360.15802001953125 0 588.68787
364.15069580078125 0 3525.3833
365.1529846191406 0 699.8594
368.134765625 0 1118.9027
369.1230163574219 0 781.7607
370.1250915527344 0 586.329
370.1503601074219 0 596.0205
376.1148376464844 0 675.0032
378.1299133300781 0 10405.71
378.1813659667969 0 7274.9297 b Water loss 2
379.1328125 0 2181.4824
379.1847839355469 0 1699.5153
382.1449279785156 0 982.9234
394.12451171875 0 4574.732
396.1896667480469 0 828.7167 b 2
406.15203857421875 0 1612.2762
416.8260803222656 0 564.3691
418.9941101074219 0 1186.3217
423.1773986816406 0 1057.5964
424.1591491699219 0 976.88934
427.20947265625 0 803.56915
429.08917236328125 0 7415.4097
433.1719970703125 0 785.92847
439.1708679199219 0 638.06354
441.188720703125 0 8293.743
442.19305419921875 0 1308.291
449.1680908203125 0 1024.8219
454.238037109375 0 582.4394
455.2023620605469 0 661.7244
459.79705810546875 0 601.44446
461.16827392578125 0 1241.0939
465.16229248046875 0 2913.9097
477.1859130859375 0 585.89496
479.1766662597656 0 1957.0002
487.172119140625 0 633.80035
491.2068786621094 0 1012.58545
495.1871337890625 0 1011.15814 b Water loss 7
497.1706237792969 0 1230.2498
505.1864013671875 0 9392.742 y 7
506.1889953613281 0 2627.6948
507.1865539550781 0 1200.0745
511.218994140625 0 4698.6157 b 3
512.2247924804688 0 6794.2817
513.1954956054688 0 1313.6154
513.22900390625 0 1802.2504
519.2019653320312 0 1889.189
520.2055053710938 0 832.61285
526.70849609375 0 1322.9546
527.2096557617188 0 1100.557
530.1920166015625 0 588.8664
541.1934814453125 0 1602.167
548.197021484375 0 698.0814
549.703857421875 0 1025.5112 y Water loss 2
550.2068481445312 0 1047.0812
558.7083129882812 0 11239.324 y 2
559.2099609375 0 7010.602
559.7117309570312 0 2403.7708
560.208251953125 0 715.87726
563.2218627929688 0 661.23816
570.231689453125 0 1078.968
572.7247314453125 0 698.8734 b 8
576.2235107421875 0 15952.301 y 6
577.2264404296875 0 4463.0884
578.2254638671875 0 875.1353
590.2241821289062 0 670.05176
600.2283935546875 0 673.70306 y Water loss 1
606.2337646484375 0 1390.2821
608.234375 0 1683.4926 b Water loss 4
609.2324829101562 0 6279.1763 y 1
609.7339477539062 0 3284.1394
610.2357788085938 0 1434.0142
612.1914672851562 0 869.35297
624.2271728515625 0 1197.3124
626.2449951171875 0 2317.0347 b 4
627.2496948242188 0 1304.6938
633.2271728515625 0 692.6056
634.2284545898438 0 1231.5521
641.7594604492188 0 932.11237
650.7675170898438 0 2264.4414
651.2684326171875 0 1832.2561
657.263916015625 0 1205.4672
673.7609252929688 0 7540.0176 Precursor Water loss
674.2627563476562 0 5205.037
674.761962890625 0 2709.4246
675.2880859375 0 5965.3276
676.2913208007812 0 1874.6139
681.8353271484375 0 870.2706
682.3377075195312 0 1578.9573
682.7660522460938 0 16726.393 Precursor
682.8942260742188 0 778.0586
683.2673950195312 0 11567.606
683.3741455078125 0 2230.8284
683.7683715820312 0 6091.2637
684.2028198242188 0 2044.5941
684.2680053710938 0 1058.104
685.2603149414062 0 1778.249
686.2582397460938 0 1022.6566
695.2662963867188 0 1726.2944
721.2632446289062 0 1739.4622
722.261962890625 0 900.5877
739.286865234375 0 13966.846 y 5
740.2898559570312 0 5506.6475
741.289794921875 0 1207.9124
744.2955932617188 0 892.0534
749.2548828125 0 1629.2665
750.2574462890625 0 1153.8529
759.2621459960938 0 861.0667
761.313720703125 0 910.98676
771.2966918945312 0 1879.2701 b Water loss 5
772.3043212890625 0 1295.1118
789.3070678710938 0 1332.2654 b 5
790.3148803710938 0 6867.5586
791.318603515625 0 3127.3018
804.3331298828125 0 1741.6066
805.33349609375 0 948.57806
832.3270263671875 0 2599.2773
833.3273315429688 0 1303.6073
842.3167724609375 0 756.9872
854.3133544921875 0 19452.123 y 4
855.3158569335938 0 8447.194
856.31396484375 0 2667.8342
857.1583862304688 0 732.6043
868.3307495117188 0 1553.5786
887.3341674804688 0 909.831
896.3239135742188 0 4307.8237
897.3275146484375 0 1470.021
898.33544921875 0 733.52277
905.3419189453125 0 9566.718
906.3446655273438 0 4597.683
907.3479614257812 0 1367.2365
951.3322143554688 0 1982.5576 y Water loss 3
952.329345703125 0 1347.0225
969.3400268554688 0 29734.207 y 3
970.3431396484375 0 15644.431
971.3425903320312 0 5733.883
972.3508911132812 0 709.3838
1034.3980712890625 0 1290.4622
1052.4105224609375 0 15094.298
1053.412841796875 0 8927.843
1054.41259765625 0 2536.6912
1098.3980712890625 0 2510.1821 y Water loss 2
1099.4012451171875 0 1968.4015
1116.40771484375 0 44961.727 y 2
1117.411376953125 0 29228.396
1118.4111328125 0 12124.328
1119.4088134765625 0 1114.774
1144.4388427734375 0 1847.8821 b 8
1145.4456787109375 0 1394.9476
1146.444580078125 0 871.6083
1153.4578857421875 0 2457.1465
1154.460693359375 0 1526.4569
1163.4473876953125 0 649.17535
1199.4520263671875 0 723.34955 y Water loss 1
1217.455078125 0 7398.27 y 1
1218.457275390625 0 6411.0513
1219.461669921875 0 2439.038
1227.438232421875 0 1472.7057
1228.45068359375 0 1168.8516
2081.0205078125 0 610.94476
2116.94482421875 0 662.6688

Spectrum Details

|  |  |
| --- | --- |
| Matched peaks? Matched peaks The total absolute number of peaks matched. Additionally in brackets the total fraction of peaks matched and the total number of peaks is shown. | 34 (12.06% of 282) |
| FDR? FDR The false discovery rate estimated for this peptide. It is calculated by matching all theoretical fragments with a non-integer shift with the raw peaks for this spectrum. This is done with 40 different shifts. The resulting percentage is the average number of annotated peaks over the number of annotated peaks with the correct spectrum. | 0.14% |
| Satellite FDR? Satellite FDR See the FDR for details on its calculation. This satellite ion specific FDR only contains the satellite ions (d/w) for I/L/J positions. | - |
| PSM Score? PSM Score The PSM Score as given by Hecklib to this annotated spectrum. It is shown with three significant figures. | 376 |

#### Spectrum 8805? Spectrum 8805 The raw spectrum of this peptide as annotated by Hecklib. The fragments are coloured according to ion type (see legend). Any peaks with a star '\*' as text can be hovered over to see the full details, first the ion type second the mass shift type. By hovering over the amino acids in the peptide or ions in the legend the corresponding peaks are highlighted. By toggling the 'Unassigned' label you can turn the background (unassigned) peaks on or off in the plot. By updating the slider in the Ion legend you can update the spectrum to only show the top X% of the peaks with labels. The top X% means any peak that is within X% of the highest intensity. By dragging in the spectrum you can zoom in to a specific part of the spectrum and use 'Zoom Out' to get back to the original zoom level. The annotation of the spectrum is based on the given sequence in the peptides file and is done with different software so inconsistencies are likely. The peaks are annotated based on the given sequence, with 20 ppm tolerance.

Copy Data

##### Spectrum 8805 (TSV)

###### Preview

```
Loading example...
```

*Click on the button to copy the data to your clipboard.*

Mz MinMz MaxIntensity Max

WidthHeightPeptide font sizePeptide stroke widthSpectrum font sizeSpectrum stroke widthCompact peptide

Ion legend

wxyz

abcd

OtherUnassignedIon^Charge^_Position_Show for top:%

FTFDDYAMHW

05.07e+41.01e+51.52e+52.03e+5

Zoom Out

a^+1^_2_d^+1^_2_a^+1^_2_b^+1^_2_b^+1^_2_a^+1^_3_y^+1^_2_b^+1^_3_b^+1^_3_b^+1^_4_b^+2^_8_y^+1^_3_b^+1^_4_y^+2^_8_y^+2^_8_y^+1^_4_y^+2^_9_b^+1^_5_y^+2^_9_b^+1^_5_\*\*y^+1^_5_b^+1^_6_b^+1^_6_y^+1^_6_b^+1^_7_y^+1^_6_b^+1^_7_y^+1^_7_y^+1^_7_b^+1^_8_y^+1^_8_y^+1^_8_b^+1^_9_y^+1^_9_y^+1^_9_

0776155323293105

Fragment Matches Table

Show background peaks

| Position | Ion type | Intensity | mz Theoretical | mz Error (Th) | mz Error (ppm) | Charge | Series Number |
| --- | --- | --- | --- | --- | --- | --- | --- |
| - | - | 2.006E+05 | 120.1 | - | - | 0 | - |
| - | - | 784.2 | 121.1 | - | - | 0 | - |
| - | - | 1.489E+04 | 121.1 | - | - | 0 | - |
| - | - | 398 | 122.1 | - | - | 0 | - |
| - | - | 578.9 | 122.1 | - | - | 0 | - |
| - | - | 482.8 | 127.1 | - | - | 0 | - |
| - | - | 1966 | 129.1 | - | - | 0 | - |
| - | - | 1211 | 130.1 | - | - | 0 | - |
| - | - | 465.4 | 131 | - | - | 0 | - |
| - | - | 382.7 | 132.4 | - | - | 0 | - |
| - | - | 575.6 | 133.1 | - | - | 0 | - |
| - | - | 1.465E+04 | 136.1 | - | - | 0 | - |
| - | - | 545.9 | 136.1 | - | - | 0 | - |
| - | - | 1377 | 137.1 | - | - | 0 | - |
| - | - | 1683 | 138.1 | - | - | 0 | - |
| - | - | 1112 | 138.1 | - | - | 0 | - |
| - | - | 510.4 | 138.1 | - | - | 0 | - |
| - | - | 614 | 146.1 | - | - | 0 | - |
| - | - | 434.7 | 146.1 | - | - | 0 | - |
| - | - | 1185 | 149 | - | - | 0 | - |
| - | - | 1737 | 149 | - | - | 0 | - |
| - | - | 414.8 | 151.9 | - | - | 0 | - |
| - | - | 5930 | 156.1 | - | - | 0 | - |
| - | - | 1488 | 158.1 | - | - | 0 | - |
| - | - | 4693 | 158.1 | - | - | 0 | - |
| - | - | 6569 | 159.1 | - | - | 0 | - |
| - | - | 959.8 | 159.1 | - | - | 0 | - |
| - | - | 834.7 | 160.1 | - | - | 0 | - |
| - | - | 713.8 | 160.1 | - | - | 0 | - |
| - | - | 1348 | 166.1 | - | - | 0 | - |
| - | - | 780.7 | 166.1 | - | - | 0 | - |
| - | - | 2264 | 167.1 | - | - | 0 | - |
| - | - | 1531 | 167.1 | - | - | 0 | - |
| - | - | 601.7 | 169.1 | - | - | 0 | - |
| - | - | 558.7 | 171.1 | - | - | 0 | - |
| - | - | 447.8 | 174.1 | - | - | 0 | - |
| - | - | 1.082E+04 | 175.1 | - | - | 0 | - |
| - | - | 703.9 | 175.1 | - | - | 0 | - |
| - | - | 2221 | 176.1 | - | - | 0 | - |
| - | - | 9.401E+04 | 176.1 | - | - | 0 | - |
| - | - | 1897 | 177.1 | - | - | 0 | - |
| - | - | 9447 | 177.1 | - | - | 0 | - |
| - | - | 489.2 | 180.1 | - | - | 0 | - |
| - | - | 618 | 185.2 | - | - | 0 | - |
| - | - | 1.223E+04 | 186.1 | - | - | 0 | - |
| - | - | 1664 | 187.1 | - | - | 0 | - |
| - | - | 870.9 | 191.1 | - | - | 0 | - |
| - | - | 1461 | 191.1 | - | - | 0 | - |
| - | - | 5778 | 193.1 | - | - | 0 | - |
| - | - | 598.3 | 194.1 | - | - | 0 | - |
| - | - | 1033 | 195.1 | - | - | 0 | - |
| - | - | 493.5 | 196.5 | - | - | 0 | - |
| - | - | 523.9 | 197.1 | - | - | 0 | - |
| - | - | 861.3 | 203.1 | - | - | 0 | - |
| - | - | 1631 | 203.1 | - | - | 0 | - |
| 2 | a | 6418 | 203.1 | 0.0001525 | 0.7506 | +1 | 2 |
| - | - | 2421 | 204.1 | - | - | 0 | - |
| - | - | 932 | 204.1 | - | - | 0 | - |
| - | - | 1258 | 205.1 | - | - | 0 | - |
| 2 | d | 1422 | 207.1 | 0.0002787 | 1.346 | +1 | 2 |
| - | - | 990 | 208.1 | - | - | 0 | - |
| - | - | 1481 | 212.1 | - | - | 0 | - |
| - | - | 525.8 | 213.1 | - | - | 0 | - |
| - | - | 691 | 215.1 | - | - | 0 | - |
| - | - | 507.1 | 215.4 | - | - | 0 | - |
| - | - | 6459 | 217.1 | - | - | 0 | - |
| - | - | 957.7 | 218.1 | - | - | 0 | - |
| - | - | 1143 | 219.1 | - | - | 0 | - |
| - | - | 551.1 | 219.1 | - | - | 0 | - |
| - | - | 2.757E+04 | 221.1 | - | - | 0 | - |
| - | - | 1.526E+04 | 221.1 | - | - | 0 | - |
| 2 | a | 1.146E+05 | 221.1 | 0.0002689 | 1.216 | +1 | 2 |
| - | - | 2129 | 222.1 | - | - | 0 | - |
| - | - | 1475 | 222.1 | - | - | 0 | - |
| - | - | 1.3E+04 | 222.1 | - | - | 0 | - |
| - | - | 586.2 | 223.1 | - | - | 0 | - |
| - | - | 4149 | 225 | - | - | 0 | - |
| - | - | 709.5 | 228.1 | - | - | 0 | - |
| - | - | 843.4 | 229.1 | - | - | 0 | - |
| - | - | 801.9 | 229.1 | - | - | 0 | - |
| - | - | 2553 | 231.1 | - | - | 0 | - |
| 2 | b | 2.993E+04 | 231.1 | 0.0002635 | 1.14 | +1 | 2 |
| - | - | 3854 | 232.1 | - | - | 0 | - |
| - | - | 8976 | 235.1 | - | - | 0 | - |
| - | - | 966.9 | 236.1 | - | - | 0 | - |
| - | - | 5806 | 239.1 | - | - | 0 | - |
| - | - | 1512 | 239.1 | - | - | 0 | - |
| - | - | 607.2 | 241.1 | - | - | 0 | - |
| - | - | 641.8 | 245.1 | - | - | 0 | - |
| - | - | 585.8 | 247.1 | - | - | 0 | - |
| - | - | 908.6 | 248.2 | - | - | 0 | - |
| 2 | b | 6.079E+04 | 249.1 | 0.0002579 | 1.035 | +1 | 2 |
| - | - | 8375 | 250.1 | - | - | 0 | - |
| - | - | 4952 | 251.1 | - | - | 0 | - |
| - | - | 855.8 | 251.1 | - | - | 0 | - |
| - | - | 603.9 | 252.1 | - | - | 0 | - |
| - | - | 2577 | 257.1 | - | - | 0 | - |
| - | - | 554.7 | 258.1 | - | - | 0 | - |
| - | - | 836.4 | 259.1 | - | - | 0 | - |
| - | - | 1.287E+04 | 263.1 | - | - | 0 | - |
| - | - | 1973 | 264.1 | - | - | 0 | - |
| - | - | 823.5 | 264.1 | - | - | 0 | - |
| - | - | 2686 | 274.1 | - | - | 0 | - |
| - | - | 742.9 | 275.1 | - | - | 0 | - |
| - | - | 4510 | 279.1 | - | - | 0 | - |
| - | - | 601.3 | 281.1 | - | - | 0 | - |
| - | - | 1439 | 285 | - | - | 0 | - |
| - | - | 5625 | 285.1 | - | - | 0 | - |
| - | - | 805.6 | 287.1 | - | - | 0 | - |
| - | - | 701.1 | 289.1 | - | - | 0 | - |
| - | - | 4141 | 292.1 | - | - | 0 | - |
| - | - | 631.4 | 294.1 | - | - | 0 | - |
| - | - | 1125 | 295.1 | - | - | 0 | - |
| - | - | 3833 | 299.1 | - | - | 0 | - |
| - | - | 2506 | 312.1 | - | - | 0 | - |
| - | - | 872.1 | 313.1 | - | - | 0 | - |
| - | - | 614.9 | 314.1 | - | - | 0 | - |
| - | - | 1500 | 318.1 | - | - | 0 | - |
| - | - | 574.9 | 319.1 | - | - | 0 | - |
| - | - | 793 | 320.3 | - | - | 0 | - |
| - | - | 1445 | 322.1 | - | - | 0 | - |
| - | - | 857.8 | 323.1 | - | - | 0 | - |
| - | - | 1649 | 325.1 | - | - | 0 | - |
| - | - | 704.6 | 332.1 | - | - | 0 | - |
| - | - | 1925 | 333.2 | - | - | 0 | - |
| - | - | 882.9 | 338.1 | - | - | 0 | - |
| - | - | 1164 | 339.2 | - | - | 0 | - |
| - | - | 1.251E+04 | 340.1 | - | - | 0 | - |
| - | - | 2244 | 341.1 | - | - | 0 | - |
| - | - | 1205 | 346.1 | - | - | 0 | - |
| - | - | 1092 | 348.1 | - | - | 0 | - |
| - | - | 2988 | 350.1 | - | - | 0 | - |
| 3 | a | 841.9 | 350.2 | 0.0008609 | 2.458 | +1 | 3 |
| - | - | 3524 | 355.1 | - | - | 0 | - |
| - | - | 750.2 | 355.1 | - | - | 0 | - |
| - | - | 3138 | 356.1 | - | - | 0 | - |
| - | - | 962.2 | 357.1 | - | - | 0 | - |
| 9 | y | 6.294E+04 | 358.1 | 0.005027 | 14.03 | +1 | 2 |
| - | - | 5231 | 359 | - | - | 0 | - |
| - | - | 1.055E+04 | 359.2 | - | - | 0 | - |
| - | - | 1047 | 360.1 | - | - | 0 | - |
| - | - | 4798 | 364.2 | - | - | 0 | - |
| - | - | 1030 | 365.2 | - | - | 0 | - |
| - | - | 677 | 367.2 | - | - | 0 | - |
| - | - | 1596 | 368.1 | - | - | 0 | - |
| - | - | 1291 | 374.1 | - | - | 0 | - |
| - | - | 1303 | 376.1 | - | - | 0 | - |
| - | - | 664.4 | 377.1 | - | - | 0 | - |
| - | - | 730.8 | 378.1 | - | - | 0 | - |
| - | - | 1.521E+04 | 378.1 | - | - | 0 | - |
| 3 | b | 1.102E+04 | 378.2 | 0.00027 | 0.7138 | +1 | 3 |
| - | - | 1899 | 379.1 | - | - | 0 | - |
| - | - | 3197 | 379.2 | - | - | 0 | - |
| - | - | 1321 | 382.1 | - | - | 0 | - |
| - | - | 634 | 388.1 | - | - | 0 | - |
| - | - | 802.5 | 388.1 | - | - | 0 | - |
| - | - | 5333 | 394.1 | - | - | 0 | - |
| 3 | b | 1246 | 396.2 | 5.074E-05 | 0.1281 | +1 | 3 |
| - | - | 834.7 | 404.1 | - | - | 0 | - |
| - | - | 1798 | 406.2 | - | - | 0 | - |
| - | - | 1057 | 416.1 | - | - | 0 | - |
| - | - | 810.6 | 422.1 | - | - | 0 | - |
| - | - | 1097 | 423.2 | - | - | 0 | - |
| - | - | 2593 | 424.2 | - | - | 0 | - |
| - | - | 1226 | 427.2 | - | - | 0 | - |
| - | - | 5756 | 429.1 | - | - | 0 | - |
| - | - | 1370 | 433.2 | - | - | 0 | - |
| - | - | 1.093E+04 | 441.2 | - | - | 0 | - |
| - | - | 2607 | 442.2 | - | - | 0 | - |
| - | - | 1112 | 443.2 | - | - | 0 | - |
| - | - | 1324 | 447.2 | - | - | 0 | - |
| - | - | 891.7 | 449.2 | - | - | 0 | - |
| - | - | 967.9 | 455.2 | - | - | 0 | - |
| - | - | 1974 | 461.2 | - | - | 0 | - |
| - | - | 5009 | 465.2 | - | - | 0 | - |
| - | - | 865.9 | 466.2 | - | - | 0 | - |
| - | - | 878.6 | 470.2 | - | - | 0 | - |
| - | - | 654.6 | 477.2 | - | - | 0 | - |
| - | - | 2709 | 479.2 | - | - | 0 | - |
| - | - | 726.1 | 480.2 | - | - | 0 | - |
| - | - | 1346 | 487.2 | - | - | 0 | - |
| - | - | 1135 | 489.2 | - | - | 0 | - |
| - | - | 1480 | 491.2 | - | - | 0 | - |
| 4 | b | 1826 | 493.2 | 0.0008857 | 1.796 | +1 | 4 |
| 8 | b | 1976 | 495.2 | 0.001396 | 2.818 | +2 | 8 |
| - | - | 1472 | 497.2 | - | - | 0 | - |
| 8 | y | 1.55E+04 | 505.2 | 0.01003 | 19.86 | +1 | 3 |
| - | - | 4581 | 506.2 | - | - | 0 | - |
| - | - | 1008 | 507.2 | - | - | 0 | - |
| 4 | b | 8157 | 511.2 | 0.0002073 | 0.4055 | +1 | 4 |
| - | - | 9116 | 512.2 | - | - | 0 | - |
| - | - | 1591 | 513.2 | - | - | 0 | - |
| - | - | 1162 | 513.2 | - | - | 0 | - |
| - | - | 1941 | 519.2 | - | - | 0 | - |
| - | - | 623 | 523.2 | - | - | 0 | - |
| - | - | 1042 | 526.7 | - | - | 0 | - |
| - | - | 745.5 | 533.8 | - | - | 0 | - |
| - | - | 2098 | 541.2 | - | - | 0 | - |
| - | - | 1235 | 542.2 | - | - | 0 | - |
| - | - | 1355 | 548.2 | - | - | 0 | - |
| 3 | y | 2121 | 549.7 | 0.006771 | 12.32 | +2 | 8 |
| - | - | 1216 | 550.2 | - | - | 0 | - |
| - | - | 694.2 | 557.2 | - | - | 0 | - |
| - | - | 1193 | 558.2 | - | - | 0 | - |
| - | - | 917.7 | 558.7 | - | - | 0 | - |
| 3 | y | 1.858E+04 | 558.7 | 0.005028 | 9 | +2 | 8 |
| - | - | 1.118E+04 | 559.2 | - | - | 0 | - |
| - | - | 4822 | 559.7 | - | - | 0 | - |
| - | - | 658.9 | 560.2 | - | - | 0 | - |
| - | - | 1537 | 570.2 | - | - | 0 | - |
| - | - | 709 | 571.2 | - | - | 0 | - |
| - | - | 674.8 | 573.2 | - | - | 0 | - |
| 7 | y | 2.401E+04 | 576.2 | 0.009846 | 17.09 | +1 | 4 |
| - | - | 6855 | 577.2 | - | - | 0 | - |
| - | - | 1943 | 578.2 | - | - | 0 | - |
| - | - | 1017 | 584.2 | - | - | 0 | - |
| - | - | 725 | 590.2 | - | - | 0 | - |
| - | - | 887.7 | 592.2 | - | - | 0 | - |
| 2 | y | 865.6 | 600.2 | 0.005942 | 9.899 | +2 | 9 |
| - | - | 1429 | 600.7 | - | - | 0 | - |
| - | - | 632.6 | 601.2 | - | - | 0 | - |
| - | - | 2355 | 606.2 | - | - | 0 | - |
| 5 | b | 2536 | 608.2 | 0.001773 | 2.915 | +1 | 5 |
| 2 | y | 5834 | 609.2 | 0.005847 | 9.598 | +2 | 9 |
| - | - | 4744 | 609.7 | - | - | 0 | - |
| - | - | 2363 | 610.2 | - | - | 0 | - |
| - | - | 1958 | 612.2 | - | - | 0 | - |
| - | - | 1753 | 624.2 | - | - | 0 | - |
| 5 | b | 4572 | 626.2 | 0.0001244 | 0.1986 | +1 | 5 |
| - | - | 1625 | 627.3 | - | - | 0 | - |
| - | - | 915.4 | 631.2 | - | - | 0 | - |
| - | - | 2045 | 634.2 | - | - | 0 | - |
| - | - | 933.5 | 635.2 | - | - | 0 | - |
| - | - | 810.6 | 641.8 | - | - | 0 | - |
| - | - | 2129 | 642.3 | - | - | 0 | - |
| - | - | 2806 | 650.8 | - | - | 0 | - |
| - | - | 2093 | 651.3 | - | - | 0 | - |
| - | - | 916.5 | 652.2 | - | - | 0 | - |
| - | - | 2339 | 657.3 | - | - | 0 | - |
| - | - | 1347 | 665.3 | - | - | 0 | - |
| 0 | Precursor | 1.034E+04 | 673.8 | 0.004816 | 7.148 | +2 | -1 |
| - | - | 6269 | 674.3 | - | - | 0 | - |
| - | - | 4478 | 674.8 | - | - | 0 | - |
| - | - | 6763 | 675.3 | - | - | 0 | - |
| - | - | 3317 | 676.3 | - | - | 0 | - |
| - | - | 621.2 | 681.8 | - | - | 0 | - |
| 0 | Precursor | 2.128E+04 | 682.8 | 0.004843 | 7.094 | +2 | -1 |
| - | - | 1012 | 682.8 | - | - | 0 | - |
| - | - | 1.504E+04 | 683.3 | - | - | 0 | - |
| - | - | 650.5 | 683.3 | - | - | 0 | - |
| - | - | 8825 | 683.8 | - | - | 0 | - |
| - | - | 1280 | 684.2 | - | - | 0 | - |
| - | - | 1580 | 684.3 | - | - | 0 | - |
| - | - | 2656 | 685.3 | - | - | 0 | - |
| - | - | 1125 | 686.3 | - | - | 0 | - |
| - | - | 1729 | 695.3 | - | - | 0 | - |
| - | - | 1120 | 696.3 | - | - | 0 | - |
| - | - | 704.3 | 703.3 | - | - | 0 | - |
| - | - | 1190 | 713.3 | - | - | 0 | - |
| - | - | 2072 | 721.3 | - | - | 0 | - |
| - | - | 1121 | 722.3 | - | - | 0 | - |
| 6 | y | 2.056E+04 | 739.3 | 0.00975 | 13.19 | +1 | 5 |
| - | - | 7982 | 740.3 | - | - | 0 | - |
| - | - | 2494 | 741.3 | - | - | 0 | - |
| - | - | 1662 | 743.3 | - | - | 0 | - |
| - | - | 740.7 | 744.3 | - | - | 0 | - |
| - | - | 3553 | 749.3 | - | - | 0 | - |
| - | - | 1898 | 750.3 | - | - | 0 | - |
| - | - | 1747 | 759.3 | - | - | 0 | - |
| - | - | 874.8 | 760.3 | - | - | 0 | - |
| - | - | 1162 | 761.3 | - | - | 0 | - |
| 6 | b | 2252 | 771.3 | 0.0005811 | 0.7534 | +1 | 6 |
| - | - | 1514 | 772.3 | - | - | 0 | - |
| - | - | 898.5 | 773.3 | - | - | 0 | - |
| - | - | 754 | 778.3 | - | - | 0 | - |
| 6 | b | 1950 | 789.3 | 0.00155 | 1.963 | +1 | 6 |
| - | - | 1.105E+04 | 790.3 | - | - | 0 | - |
| - | - | 4095 | 791.3 | - | - | 0 | - |
| - | - | 699.1 | 792.3 | - | - | 0 | - |
| - | - | 2685 | 804.3 | - | - | 0 | - |
| - | - | 633.8 | 824.3 | - | - | 0 | - |
| - | - | 3685 | 832.3 | - | - | 0 | - |
| - | - | 1951 | 833.3 | - | - | 0 | - |
| 5 | y | 1101 | 836.3 | 0.01064 | 12.73 | +1 | 6 |
| 7 | b | 749.2 | 842.3 | 0.005224 | 6.202 | +1 | 7 |
| 5 | y | 2.623E+04 | 854.3 | 0.009358 | 10.95 | +1 | 6 |
| - | - | 1.207E+04 | 855.3 | - | - | 0 | - |
| - | - | 3963 | 856.3 | - | - | 0 | - |
| 7 | b | 723.3 | 860.3 | 0.001446 | 1.68 | +1 | 7 |
| - | - | 2172 | 868.3 | - | - | 0 | - |
| - | - | 1506 | 869.3 | - | - | 0 | - |
| - | - | 1869 | 887.3 | - | - | 0 | - |
| - | - | 720.7 | 888.3 | - | - | 0 | - |
| - | - | 4540 | 896.3 | - | - | 0 | - |
| - | - | 2188 | 897.3 | - | - | 0 | - |
| - | - | 1080 | 898.3 | - | - | 0 | - |
| - | - | 1.467E+04 | 905.3 | - | - | 0 | - |
| - | - | 7602 | 906.3 | - | - | 0 | - |
| - | - | 1343 | 907.4 | - | - | 0 | - |
| 4 | y | 2638 | 951.3 | 0.008909 | 9.365 | +1 | 7 |
| - | - | 1672 | 952.3 | - | - | 0 | - |
| - | - | 957.6 | 953.3 | - | - | 0 | - |
| 4 | y | 4.009E+04 | 969.3 | 0.009026 | 9.311 | +1 | 7 |
| - | - | 2.29E+04 | 970.3 | - | - | 0 | - |
| - | - | 7939 | 971.3 | - | - | 0 | - |
| - | - | 1073 | 972.3 | - | - | 0 | - |
| - | - | 1134 | 979.3 | - | - | 0 | - |
| 8 | b | 1436 | 989.4 | 0.005368 | 5.426 | +1 | 8 |
| - | - | 951.2 | 997.4 | - | - | 0 | - |
| - | - | 2507 | 1034 | - | - | 0 | - |
| - | - | 1074 | 1035 | - | - | 0 | - |
| - | - | 1.866E+04 | 1052 | - | - | 0 | - |
| - | - | 1.134E+04 | 1053 | - | - | 0 | - |
| - | - | 3153 | 1054 | - | - | 0 | - |
| 3 | y | 3567 | 1098 | 0.008122 | 7.395 | +1 | 8 |
| - | - | 2660 | 1099 | - | - | 0 | - |
| - | - | 1070 | 1100 | - | - | 0 | - |
| 3 | y | 6.491E+04 | 1116 | 0.008544 | 7.653 | +1 | 8 |
| - | - | 3.779E+04 | 1117 | - | - | 0 | - |
| - | - | 1.629E+04 | 1118 | - | - | 0 | - |
| - | - | 2221 | 1119 | - | - | 0 | - |
| - | - | 1029 | 1135 | - | - | 0 | - |
| 9 | b | 2104 | 1144 | 0.001382 | 1.208 | +1 | 9 |
| - | - | 1653 | 1145 | - | - | 0 | - |
| - | - | 4674 | 1153 | - | - | 0 | - |
| - | - | 2431 | 1154 | - | - | 0 | - |
| 2 | y | 1592 | 1199 | 0.008418 | 7.018 | +1 | 9 |
| - | - | 1115 | 1200 | - | - | 0 | - |
| 2 | y | 1.228E+04 | 1217 | 0.007252 | 5.957 | +1 | 9 |
| - | - | 7808 | 1218 | - | - | 0 | - |
| - | - | 3404 | 1219 | - | - | 0 | - |
| - | - | 805.8 | 1220 | - | - | 0 | - |
| - | - | 1901 | 1227 | - | - | 0 | - |
| - | - | 1016 | 1228 | - | - | 0 | - |
| - | - | 705.1 | 1361 | - | - | 0 | - |
| - | - | 864.4 | 3050 | - | - | 0 | - |
| - | - | 834.7 | 3074 | - | - | 0 | - |

m/z Charge Intensity FragmentType MassShift Position
120.08110809326172 0 200604.19
121.07903289794922 0 784.15967
121.08436584472656 0 14893.53
122.0713882446289 0 397.98703
122.08777618408203 0 578.93896
127.08642578125 0 482.79053
129.10263061523438 0 1965.911
130.06549072265625 0 1210.5714
131.04969787597656 0 465.41046
132.37399291992188 0 382.71176
133.08555603027344 0 575.63947
136.075927734375 0 14652.343
136.0879364013672 0 545.8669
137.07948303222656 0 1377.0669
138.05517578125 0 1683.1967
138.06617736816406 0 1111.6888
138.09165954589844 0 510.40085
146.06080627441406 0 613.99274
146.0966033935547 0 434.67157
148.95458984375 0 1184.8302
149.04505920410156 0 1736.8518
151.85348510742188 0 414.77213
156.07699584960938 0 5930.2803
158.06016540527344 0 1487.9436
158.0966339111328 0 4692.5435
159.0919189453125 0 6568.8447
159.09910583496094 0 959.8297
160.0950469970703 0 834.7335
160.1126708984375 0 713.84784
166.06106567382812 0 1347.9343
166.08627319335938 0 780.72833
167.05567932128906 0 2263.8733
167.09304809570312 0 1531.451
169.13421630859375 0 601.7121
171.11289978027344 0 558.6995
174.06654357910156 0 447.83463
175.0868377685547 0 10818.709
175.12298583984375 0 703.88196
176.08221435546875 0 2220.671
176.10726928710938 0 94008.93
177.1023712158203 0 1897.2079
177.11082458496094 0 9446.61
180.07693481445312 0 489.24664
185.16482543945312 0 617.97205
186.09149169921875 0 12232.562
187.0950469970703 0 1663.9761
191.09283447265625 0 870.9442
191.1183319091797 0 1460.5762
193.10858154296875 0 5777.67
194.0797882080078 0 598.3421
195.0878143310547 0 1032.9332
196.4595489501953 0 493.45007
197.12869262695312 0 523.9233
203.06634521484375 0 861.2557
203.08164978027344 0 1631.157
203.1180419921875 0 6417.6934 a Water loss 1
204.076904296875 0 2421.2168
204.12149047851562 0 932.0397
205.09765625 0 1258.0544
207.1130828857422 0 1421.8516 d 1
208.0714569091797 0 990.002
212.11865234375 0 1480.5825
213.09869384765625 0 525.81287
215.08204650878906 0 690.96234
215.38841247558594 0 507.09735
217.0973663330078 0 6459.0146
218.1011962890625 0 957.68585
219.0795440673828 0 1143.3423
219.1134490966797 0 551.09296
221.0922393798828 0 27572.477
221.1036376953125 0 15260.323
221.12872314453125 0 114635.88 a 1
222.09564208984375 0 2128.9631
222.10702514648438 0 1475.4955
222.13197326660156 0 12999.017
223.1334991455078 0 586.2243
225.0431671142578 0 4149.498
228.1132049560547 0 709.48303
229.09715270996094 0 843.44
229.10882568359375 0 801.8907
231.06163024902344 0 2553.1863
231.11306762695312 0 29930.004 b Water loss 1
232.11651611328125 0 3853.772
235.1080322265625 0 8976.194
236.1114501953125 0 966.881
239.09518432617188 0 5805.6772
239.11361694335938 0 1511.911
241.09439086914062 0 607.2349
245.09141540527344 0 641.7968
247.14529418945312 0 585.8021
248.151123046875 0 908.55505
249.12362670898438 0 60793.793 b 1
250.12686157226562 0 8374.779
251.10293579101562 0 4952.4307
251.1282958984375 0 855.8128
252.10765075683594 0 603.88293
257.1067810058594 0 2577.2244
258.1088562011719 0 554.69696
259.1064453125 0 836.37134
263.10272216796875 0 12866.63
264.1062316894531 0 1972.9019
264.14508056640625 0 823.4875
274.13018798828125 0 2686.0215
275.1150207519531 0 742.93146
279.0973815917969 0 4510.0215
281.05126953125 0 601.3201
285.0093688964844 0 1438.8752
285.10186767578125 0 5625.246
287.10137939453125 0 805.6084
289.1346130371094 0 701.1492
292.14080810546875 0 4140.8916
294.1357116699219 0 631.4173
295.1036376953125 0 1125.1177
299.06207275390625 0 3832.5798
312.1463623046875 0 2505.8616
313.114013671875 0 872.12225
314.098388671875 0 614.85504
318.1455078125 0 1500.1439
319.1494445800781 0 574.89954
320.2980041503906 0 793.02277
322.1308288574219 0 1444.984
323.11541748046875 0 857.77606
325.1380920410156 0 1648.5159
332.1248779296875 0 704.57135
333.15985107421875 0 1925.1704
338.1297607421875 0 882.94214
339.1700439453125 0 1163.7786
340.14080810546875 0 12508.539
341.1439208984375 0 2244.3862
346.1389465332031 0 1204.7695
348.1192932128906 0 1092.388
350.13427734375 0 2987.9016
350.1871643066406 0 841.9487 a Water loss 2
355.07000732421875 0 3523.586
355.1384582519531 0 750.2492
356.1385192871094 0 3138.3904
357.1426086425781 0 962.16754
358.1510925292969 0 62938.7 y 8
359.0281677246094 0 5231.1616
359.154052734375 0 10554.767
360.1193542480469 0 1047.4725
364.1503601074219 0 4797.9043
365.1534118652344 0 1030.024
367.1631164550781 0 677.0129
368.1355285644531 0 1595.6935
374.1325988769531 0 1290.8545
376.1138916015625 0 1303.3452
377.0859069824219 0 664.44885
378.10101318359375 0 730.7771
378.1297302246094 0 15207.54
378.1814880371094 0 11019.967 b Water loss 2
379.1328430175781 0 1898.8644
379.18487548828125 0 3196.969
382.1435241699219 0 1321.0399
388.114013671875 0 634.04193
388.14007568359375 0 802.4787
394.1248779296875 0 5332.558
396.19183349609375 0 1246.4519 b 2
404.1459655761719 0 834.7338
406.15093994140625 0 1798.2107
416.1445617675781 0 1056.9912
422.146484375 0 810.5874
423.1781005859375 0 1097.2682
424.1621398925781 0 2592.873
427.2093811035156 0 1225.6141
429.08905029296875 0 5755.5654
433.174072265625 0 1369.8237
441.1884765625 0 10932.8125
442.19134521484375 0 2606.5425
443.1559753417969 0 1112.4933
447.1514587402344 0 1324.2332
449.1676330566406 0 891.69073
455.2030334472656 0 967.94495
461.167236328125 0 1974.4651
465.1624450683594 0 5009.3853
466.16455078125 0 865.8972
470.16632080078125 0 878.5909
477.18756103515625 0 654.5945
479.1767272949219 0 2709.3696
480.1795959472656 0 726.1309
487.177001953125 0 1346.1656
489.161376953125 0 1135.2628
491.2089538574219 0 1479.9739
493.207275390625 0 1825.8745 b Water loss 3
495.18804931640625 0 1975.7949 b Water loss 7
497.16888427734375 0 1472.0038
505.18658447265625 0 15504.97 y 7
506.1897277832031 0 4581.112
507.1900939941406 0 1008.31226
511.21893310546875 0 8156.7397 b 3
512.2245483398438 0 9116.43
513.1964721679688 0 1591.3843
513.2328491210938 0 1161.7091
519.2020263671875 0 1940.7252
523.1826171875 0 622.9958
526.7083129882812 0 1042.3066
533.7797241210938 0 745.46747
541.1941528320312 0 2098.2075
542.236572265625 0 1234.6841
548.198486328125 0 1355.2473
549.7047729492188 0 2120.9487 y Water loss 2
550.2035522460938 0 1216.4078
557.1986694335938 0 694.21686
558.208740234375 0 1192.9537
558.6588745117188 0 917.7186
558.7083129882812 0 18577.135 y 2
559.2101440429688 0 11182.87
559.71044921875 0 4822.495
560.21044921875 0 658.8676
570.2317504882812 0 1537.1188
571.230712890625 0 709.0115
573.2269287109375 0 674.81665
576.2235107421875 0 24007.201 y 6
577.2261962890625 0 6855.255
578.225341796875 0 1943.0475
584.2362670898438 0 1017.4817
590.224365234375 0 725.0269
592.2130737304688 0 887.7189
600.227783203125 0 865.5582 y Water loss 1
600.728271484375 0 1428.5837
601.2286987304688 0 632.5512
606.2328491210938 0 2355.192
608.2368774414062 0 2535.8618 b Water loss 4
609.2329711914062 0 5833.87 y 1
609.7338256835938 0 4743.6147
610.2357788085938 0 2362.7964
612.2324829101562 0 1958.0645
624.2299194335938 0 1752.6523
626.2455444335938 0 4572.428 b 4
627.2505493164062 0 1624.7168
631.2477416992188 0 915.3577
634.2315063476562 0 2045.2688
635.2265014648438 0 933.4542
641.7617797851562 0 810.598
642.2568969726562 0 2129.255
650.7660522460938 0 2806.1797
651.2658081054688 0 2092.6445
652.2269897460938 0 916.5394
657.2664794921875 0 2339.0332
665.2587890625 0 1346.9199
673.7608642578125 0 10344.551 Precursor Water loss
674.262451171875 0 6268.602
674.7623901367188 0 4478.245
675.2875366210938 0 6762.683
676.2904663085938 0 3316.9834
681.8367919921875 0 621.23505
682.7661743164062 0 21278.885 Precursor
682.8358764648438 0 1012.3927
683.2677001953125 0 15038.369
683.3373413085938 0 650.52673
683.7680053710938 0 8825.277
684.2042236328125 0 1279.5344
684.2703247070312 0 1580.2388
685.25927734375 0 2655.6394
686.2619018554688 0 1124.8577
695.2658081054688 0 1728.8379
696.271484375 0 1120.2307
703.2682495117188 0 704.3132
713.2752075195312 0 1190.3158
721.266357421875 0 2071.936
722.2703247070312 0 1120.7336
739.2867431640625 0 20561.6 y 5
740.2900390625 0 7981.967
741.287353515625 0 2493.688
743.303466796875 0 1662.166
744.2962646484375 0 740.7148
749.2554321289062 0 3552.8438
750.2579956054688 0 1897.9019
759.2664794921875 0 1747.3774
760.2672119140625 0 874.79877
761.3137817382812 0 1161.5089
771.2978515625 0 2251.7095 b Water loss 5
772.3007202148438 0 1513.6469
773.3037719726562 0 898.5117
778.311767578125 0 754.0154
789.310546875 0 1949.7322 b 5
790.3150024414062 0 11050.704
791.3172607421875 0 4095.0679
792.3255004882812 0 699.0854
804.3286743164062 0 2684.7368
824.3219604492188 0 633.7746
832.3262939453125 0 3684.9995
833.3276977539062 0 1950.7272
836.3040161132812 0 1101.006 y Water loss 4
842.330322265625 0 749.2403 b Water loss 6
854.3132934570312 0 26230.326 y 4
855.3162231445312 0 12070.824
856.3155517578125 0 3963.1768
860.3446655273438 0 723.3195 b 6
868.3276977539062 0 2172.175
869.3309326171875 0 1506.4985
887.3305053710938 0 1869.0844
888.3173828125 0 720.668
896.3241577148438 0 4540.379
897.3239135742188 0 2187.6514
898.3297729492188 0 1080.0209
905.34130859375 0 14673.772
906.3450317382812 0 7601.7173
907.3502807617188 0 1343.0319
951.3292236328125 0 2637.5132 y Water loss 3
952.326171875 0 1672.4617
953.33203125 0 957.5794
969.3399047851562 0 40089.844 y 3
970.3429565429688 0 22898.816
971.3432006835938 0 7939.1226
972.344482421875 0 1072.9114
979.3302001953125 0 1134.1124
989.3713989257812 0 1436.423 b Water loss 7
997.3720703125 0 951.23395
1034.3990478515625 0 2506.636
1035.4066162109375 0 1073.9829
1052.4100341796875 0 18664.928
1053.41259765625 0 11344.824
1054.4173583984375 0 3153.2402
1098.3968505859375 0 3566.7563 y Water loss 2
1099.4013671875 0 2659.516
1100.3978271484375 0 1069.6736
1116.4078369140625 0 64906.547 y 2
1117.410888671875 0 37785.37
1118.4112548828125 0 16288.928
1119.413818359375 0 2220.5964
1135.4461669921875 0 1029.1523
1144.4368896484375 0 2103.8782 b 8
1145.433837890625 0 1653.3862
1153.456787109375 0 4673.9146
1154.4609375 0 2430.8088
1199.44482421875 0 1592.4595 y Water loss 1
1200.4447021484375 0 1114.9575
1217.4542236328125 0 12283.522 y 1
1218.45751953125 0 7808.3496
1219.4559326171875 0 3404.3755
1220.4656982421875 0 805.75323
1227.437255859375 0 1900.6038
1228.4398193359375 0 1015.91
1361.068603515625 0 705.0881
3050.26708984375 0 864.3698
3074.473388671875 0 834.66327

Spectrum Details

|  |  |
| --- | --- |
| Matched peaks? Matched peaks The total absolute number of peaks matched. Additionally in brackets the total fraction of peaks matched and the total number of peaks is shown. | 37 (10.98% of 337) |
| FDR? FDR The false discovery rate estimated for this peptide. It is calculated by matching all theoretical fragments with a non-integer shift with the raw peaks for this spectrum. This is done with 40 different shifts. The resulting percentage is the average number of annotated peaks over the number of annotated peaks with the correct spectrum. | 0.19% |
| Satellite FDR? Satellite FDR See the FDR for details on its calculation. This satellite ion specific FDR only contains the satellite ions (d/w) for I/L/J positions. | - |
| PSM Score? PSM Score The PSM Score as given by Hecklib to this annotated spectrum. It is shown with three significant figures. | 463 |

#### Spectrum 9795? Spectrum 9795 The raw spectrum of this peptide as annotated by Hecklib. The fragments are coloured according to ion type (see legend). Any peaks with a star '\*' as text can be hovered over to see the full details, first the ion type second the mass shift type. By hovering over the amino acids in the peptide or ions in the legend the corresponding peaks are highlighted. By toggling the 'Unassigned' label you can turn the background (unassigned) peaks on or off in the plot. By updating the slider in the Ion legend you can update the spectrum to only show the top X% of the peaks with labels. The top X% means any peak that is within X% of the highest intensity. By dragging in the spectrum you can zoom in to a specific part of the spectrum and use 'Zoom Out' to get back to the original zoom level. The annotation of the spectrum is based on the given sequence in the peptides file and is done with different software so inconsistencies are likely. The peaks are annotated based on the given sequence, with 20 ppm tolerance.

Copy Data

##### Spectrum 9795 (TSV)

###### Preview

```
Loading example...
```

*Click on the button to copy the data to your clipboard.*

Mz MinMz MaxIntensity Max

WidthHeightPeptide font sizePeptide stroke widthSpectrum font sizeSpectrum stroke widthCompact peptide

Ion legend

wxyz

abcd

OtherUnassignedIon^Charge^_Position_Show for top:%

FTFDDYAMHW

03.57e+47.13e+41.07e+51.43e+5

Zoom Out

y^+1^_1_c^+1^_2_y^+1^_2_z^+1^_3_y^+1^_3_c^+1^_4_z^+1^_4_y^+2^_8_y^+1^_4_y^+2^_9_c^+1^_5_z^+1^_5_y^+1^_5_c^+1^_6_z^+1^_6_y^+1^_6_c^+1^_7_z^+1^_7_y^+1^_7_c^+1^_8_z^+1^_8_y^+1^_8_c^+1^_9_y^+1^_9_z^+1^_9_y^+1^_9_

0836167225093345

Fragment Matches Table

Show background peaks

| Position | Ion type | Intensity | mz Theoretical | mz Error (Th) | mz Error (ppm) | Charge | Series Number |
| --- | --- | --- | --- | --- | --- | --- | --- |
| - | - | 837.4 | 120.1 | - | - | 0 | - |
| - | - | 1.439E+04 | 120.1 | - | - | 0 | - |
| - | - | 1036 | 121.1 | - | - | 0 | - |
| - | - | 431.6 | 135.2 | - | - | 0 | - |
| - | - | 542.7 | 148.9 | - | - | 0 | - |
| - | - | 935.9 | 148.9 | - | - | 0 | - |
| - | - | 582.1 | 148.9 | - | - | 0 | - |
| - | - | 1014 | 148.9 | - | - | 0 | - |
| - | - | 2013 | 148.9 | - | - | 0 | - |
| - | - | 1989 | 148.9 | - | - | 0 | - |
| - | - | 5845 | 149 | - | - | 0 | - |
| - | - | 2540 | 149 | - | - | 0 | - |
| - | - | 1853 | 149 | - | - | 0 | - |
| - | - | 1086 | 149 | - | - | 0 | - |
| - | - | 1091 | 149 | - | - | 0 | - |
| - | - | 780.9 | 159.1 | - | - | 0 | - |
| - | - | 931.5 | 166.1 | - | - | 0 | - |
| - | - | 4803 | 176.1 | - | - | 0 | - |
| - | - | 442.5 | 184.7 | - | - | 0 | - |
| 10 | y | 1008 | 205.1 | 0.0001512 | 0.7372 | +1 | 1 |
| - | - | 501.6 | 205.4 | - | - | 0 | - |
| - | - | 469.9 | 216.2 | - | - | 0 | - |
| - | - | 1.928E+04 | 221.1 | - | - | 0 | - |
| - | - | 2750 | 222.1 | - | - | 0 | - |
| - | - | 2697 | 231.1 | - | - | 0 | - |
| - | - | 653.7 | 232.1 | - | - | 0 | - |
| 2 | c | 1.76E+04 | 249.1 | 0.0002579 | 1.035 | +1 | 2 |
| - | - | 1645 | 250.1 | - | - | 0 | - |
| - | - | 518.2 | 252.9 | - | - | 0 | - |
| - | - | 1250 | 253.1 | - | - | 0 | - |
| - | - | 1591 | 263.1 | - | - | 0 | - |
| - | - | 486.9 | 299.1 | - | - | 0 | - |
| - | - | 498.7 | 316.2 | - | - | 0 | - |
| - | - | 807.7 | 327.1 | - | - | 0 | - |
| - | - | 541.2 | 331.9 | - | - | 0 | - |
| - | - | 536.8 | 342.1 | - | - | 0 | - |
| 9 | y | 3018 | 342.2 | 0.0004282 | 1.251 | +1 | 2 |
| - | - | 1091 | 355.1 | - | - | 0 | - |
| - | - | 687.2 | 364.2 | - | - | 0 | - |
| - | - | 577.3 | 365.2 | - | - | 0 | - |
| - | - | 976.8 | 378.1 | - | - | 0 | - |
| - | - | 4023 | 378.2 | - | - | 0 | - |
| - | - | 599.9 | 379.2 | - | - | 0 | - |
| 8 | z | 1826 | 473.2 | 0.004903 | 10.36 | +1 | 3 |
| - | - | 5662 | 474.2 | - | - | 0 | - |
| - | - | 1641 | 475.2 | - | - | 0 | - |
| 8 | y | 2215 | 489.2 | 0.005374 | 10.99 | +1 | 3 |
| - | - | 599.9 | 498.8 | - | - | 0 | - |
| 4 | c | 1775 | 511.2 | 0.0001894 | 0.3705 | +1 | 4 |
| 7 | z | 5566 | 544.2 | 0.005082 | 9.338 | +1 | 4 |
| - | - | 2943 | 545.2 | - | - | 0 | - |
| - | - | 765.8 | 546.2 | - | - | 0 | - |
| 3 | y | 6464 | 550.7 | 0.002897 | 5.26 | +2 | 8 |
| - | - | 3270 | 551.2 | - | - | 0 | - |
| - | - | 1377 | 551.7 | - | - | 0 | - |
| 7 | y | 5025 | 560.2 | 0.004912 | 8.768 | +1 | 4 |
| - | - | 1023 | 561.2 | - | - | 0 | - |
| 2 | y | 1892 | 601.2 | 0.002373 | 3.947 | +2 | 9 |
| - | - | 1919 | 601.7 | - | - | 0 | - |
| - | - | 1030 | 608.2 | - | - | 0 | - |
| 5 | c | 1426 | 626.2 | 0.002378 | 3.797 | +1 | 5 |
| - | - | 590.2 | 633.8 | - | - | 0 | - |
| - | - | 707.6 | 659.3 | - | - | 0 | - |
| - | - | 947 | 663.3 | - | - | 0 | - |
| - | - | 3554 | 665.8 | - | - | 0 | - |
| - | - | 3048 | 666.3 | - | - | 0 | - |
| - | - | 1107 | 666.8 | - | - | 0 | - |
| - | - | 730.7 | 674.4 | - | - | 0 | - |
| - | - | 7194 | 674.8 | - | - | 0 | - |
| - | - | 4699 | 675.3 | - | - | 0 | - |
| - | - | 2930 | 675.8 | - | - | 0 | - |
| - | - | 745.8 | 676.4 | - | - | 0 | - |
| 6 | z | 1.611E+04 | 707.3 | 0.004558 | 6.445 | +1 | 5 |
| - | - | 7791 | 708.3 | - | - | 0 | - |
| - | - | 2653 | 709.3 | - | - | 0 | - |
| 6 | y | 5775 | 723.3 | 0.00506 | 6.996 | +1 | 5 |
| - | - | 2143 | 724.3 | - | - | 0 | - |
| - | - | 2458 | 750.3 | - | - | 0 | - |
| - | - | 1070 | 751.3 | - | - | 0 | - |
| - | - | 833.8 | 774.3 | - | - | 0 | - |
| - | - | 7330 | 778.3 | - | - | 0 | - |
| - | - | 4169 | 779.3 | - | - | 0 | - |
| - | - | 925.2 | 780.3 | - | - | 0 | - |
| 6 | c | 1111 | 789.3 | 0.001258 | 1.594 | +1 | 6 |
| 5 | z | 2.013E+04 | 822.3 | 0.004471 | 5.437 | +1 | 6 |
| - | - | 2.205E+04 | 823.3 | - | - | 0 | - |
| - | - | 9224 | 824.3 | - | - | 0 | - |
| - | - | 2723 | 825.3 | - | - | 0 | - |
| 5 | y | 8894 | 838.3 | 0.004301 | 5.131 | +1 | 6 |
| - | - | 3917 | 839.3 | - | - | 0 | - |
| - | - | 1338 | 840.3 | - | - | 0 | - |
| - | - | 1137 | 849.4 | - | - | 0 | - |
| - | - | 984.9 | 876.4 | - | - | 0 | - |
| 7 | c | 2905 | 877.4 | 0.0005697 | 0.6494 | +1 | 7 |
| - | - | 1310 | 878.4 | - | - | 0 | - |
| - | - | 1194 | 889.3 | - | - | 0 | - |
| - | - | 7303 | 893.3 | - | - | 0 | - |
| - | - | 3105 | 894.3 | - | - | 0 | - |
| - | - | 1775 | 896.3 | - | - | 0 | - |
| - | - | 722.2 | 918.4 | - | - | 0 | - |
| 4 | z | 8532 | 937.3 | 0.00359 | 3.83 | +1 | 7 |
| - | - | 1.605E+04 | 938.3 | - | - | 0 | - |
| - | - | 7414 | 939.3 | - | - | 0 | - |
| - | - | 2561 | 940.3 | - | - | 0 | - |
| 4 | y | 1.199E+04 | 953.3 | 0.00458 | 4.804 | +1 | 7 |
| - | - | 7172 | 954.3 | - | - | 0 | - |
| - | - | 1915 | 955.3 | - | - | 0 | - |
| - | - | 6552 | 980.4 | - | - | 0 | - |
| - | - | 3606 | 981.4 | - | - | 0 | - |
| - | - | 1303 | 982.4 | - | - | 0 | - |
| - | - | 734.2 | 983.4 | - | - | 0 | - |
| - | - | 646.3 | 997.4 | - | - | 0 | - |
| - | - | 762.7 | 1021 | - | - | 0 | - |
| - | - | 381.5 | 1023 | - | - | 0 | - |
| - | - | 1.298E+04 | 1023 | - | - | 0 | - |
| 8 | c | 2.327E+04 | 1024 | 0.002251 | 2.197 | +1 | 8 |
| - | - | 1.063E+04 | 1025 | - | - | 0 | - |
| - | - | 3137 | 1026 | - | - | 0 | - |
| - | - | 1991 | 1036 | - | - | 0 | - |
| - | - | 2372 | 1037 | - | - | 0 | - |
| - | - | 1162 | 1040 | - | - | 0 | - |
| - | - | 741.1 | 1041 | - | - | 0 | - |
| - | - | 694.4 | 1083 | - | - | 0 | - |
| 3 | z | 2799 | 1084 | 0.004329 | 3.992 | +1 | 8 |
| - | - | 2009 | 1085 | - | - | 0 | - |
| - | - | 987.3 | 1086 | - | - | 0 | - |
| 3 | y | 2.35E+04 | 1100 | 0.003915 | 3.558 | +1 | 8 |
| - | - | 1.591E+04 | 1101 | - | - | 0 | - |
| - | - | 5346 | 1102 | - | - | 0 | - |
| - | - | 685.8 | 1110 | - | - | 0 | - |
| - | - | 6004 | 1117 | - | - | 0 | - |
| - | - | 3365 | 1118 | - | - | 0 | - |
| - | - | 1374 | 1119 | - | - | 0 | - |
| - | - | 718.7 | 1137 | - | - | 0 | - |
| - | - | 3984 | 1139 | - | - | 0 | - |
| - | - | 2487 | 1140 | - | - | 0 | - |
| - | - | 2307 | 1141 | - | - | 0 | - |
| - | - | 990.7 | 1142 | - | - | 0 | - |
| - | - | 851.6 | 1145 | - | - | 0 | - |
| - | - | 1102 | 1146 | - | - | 0 | - |
| - | - | 974 | 1147 | - | - | 0 | - |
| 9 | c | 4.099E+04 | 1161 | 0.003642 | 3.135 | +1 | 9 |
| - | - | 2.658E+04 | 1162 | - | - | 0 | - |
| - | - | 1.027E+04 | 1163 | - | - | 0 | - |
| - | - | 951.6 | 1164 | - | - | 0 | - |
| 2 | y | 1326 | 1183 | 0.01196 | 10.1 | +1 | 9 |
| - | - | 6076 | 1184 | - | - | 0 | - |
| 2 | z | 1.128E+04 | 1185 | 0.002182 | 1.841 | +1 | 9 |
| - | - | 7790 | 1186 | - | - | 0 | - |
| - | - | 2050 | 1187 | - | - | 0 | - |
| - | - | 933.4 | 1188 | - | - | 0 | - |
| - | - | 851.6 | 1197 | - | - | 0 | - |
| - | - | 639.6 | 1198 | - | - | 0 | - |
| 2 | y | 4934 | 1201 | 0.003844 | 3.2 | +1 | 9 |
| - | - | 3668 | 1202 | - | - | 0 | - |
| - | - | 1818 | 1203 | - | - | 0 | - |
| - | - | 1030 | 1211 | - | - | 0 | - |
| - | - | 849 | 1219 | - | - | 0 | - |
| - | - | 963.6 | 1240 | - | - | 0 | - |
| - | - | 1026 | 1244 | - | - | 0 | - |
| - | - | 1080 | 1250 | - | - | 0 | - |
| - | - | 775.1 | 1252 | - | - | 0 | - |
| - | - | 1163 | 1257 | - | - | 0 | - |
| - | - | 987.4 | 1258 | - | - | 0 | - |
| - | - | 1368 | 1270 | - | - | 0 | - |
| - | - | 2038 | 1271 | - | - | 0 | - |
| - | - | 2215 | 1272 | - | - | 0 | - |
| - | - | 1106 | 1273 | - | - | 0 | - |
| - | - | 1346 | 1286 | - | - | 0 | - |
| - | - | 2575 | 1287 | - | - | 0 | - |
| - | - | 1.231E+04 | 1288 | - | - | 0 | - |
| - | - | 2.093E+04 | 1289 | - | - | 0 | - |
| - | - | 3.047E+04 | 1290 | - | - | 0 | - |
| - | - | 1.688E+04 | 1291 | - | - | 0 | - |
| - | - | 6831 | 1292 | - | - | 0 | - |
| - | - | 791 | 1293 | - | - | 0 | - |
| - | - | 675.2 | 1300 | - | - | 0 | - |
| - | - | 1069 | 1302 | - | - | 0 | - |
| - | - | 2438 | 1304 | - | - | 0 | - |
| - | - | 2359 | 1305 | - | - | 0 | - |
| - | - | 2317 | 1306 | - | - | 0 | - |
| - | - | 1194 | 1307 | - | - | 0 | - |
| - | - | 928.5 | 1315 | - | - | 0 | - |
| - | - | 1140 | 1315 | - | - | 0 | - |
| - | - | 913.7 | 1317 | - | - | 0 | - |
| - | - | 1313 | 1317 | - | - | 0 | - |
| - | - | 863.2 | 1319 | - | - | 0 | - |
| - | - | 689.5 | 1322 | - | - | 0 | - |
| - | - | 1430 | 1330 | - | - | 0 | - |
| - | - | 2085 | 1331 | - | - | 0 | - |
| - | - | 7522 | 1332 | - | - | 0 | - |
| - | - | 7.925E+04 | 1333 | - | - | 0 | - |
| - | - | 6.538E+04 | 1334 | - | - | 0 | - |
| - | - | 3.142E+04 | 1335 | - | - | 0 | - |
| - | - | 6201 | 1336 | - | - | 0 | - |
| - | - | 2509 | 1347 | - | - | 0 | - |
| - | - | 2292 | 1348 | - | - | 0 | - |
| - | - | 3.782E+04 | 1349 | - | - | 0 | - |
| - | - | 1.413E+05 | 1350 | - | - | 0 | - |
| - | - | 1.085E+05 | 1351 | - | - | 0 | - |
| - | - | 2844 | 1351 | - | - | 0 | - |
| - | - | 4.563E+04 | 1352 | - | - | 0 | - |
| - | - | 8816 | 1353 | - | - | 0 | - |
| - | - | 677.6 | 2500 | - | - | 0 | - |
| - | - | 681.7 | 3018 | - | - | 0 | - |
| - | - | 707 | 3312 | - | - | 0 | - |

m/z Charge Intensity FragmentType MassShift Position
120.07628631591797 0 837.3833
120.08102416992188 0 14386.282
121.08438110351562 0 1036.3032
135.1670379638672 0 431.57104
148.89755249023438 0 542.65466
148.91983032226562 0 935.92395
148.92562866210938 0 582.068
148.93064880371094 0 1014.14825
148.93624877929688 0 2012.9735
148.94248962402344 0 1988.9193
148.9550323486328 0 5845.0225
148.9607391357422 0 2540.351
148.96697998046875 0 1852.5148
148.97250366210938 0 1085.9614
148.9833526611328 0 1091.3356
159.0919189453125 0 780.88654
166.08627319335938 0 931.5091
176.10733032226562 0 4803.4917
184.6937255859375 0 442.4594
205.09730529785156 0 1008.3886 y 9
205.3511962890625 0 501.60532
216.1676788330078 0 469.899
221.12863159179688 0 19279.412
222.1318817138672 0 2749.9895
231.11318969726562 0 2697.3506
232.11663818359375 0 653.6853
249.12362670898438 0 17598.85 c Ammonia loss 1
250.12677001953125 0 1644.7759
252.8597869873047 0 518.17255
253.117919921875 0 1249.9733
263.1029357910156 0 1590.6958
299.10302734375 0 486.9394
316.1827392578125 0 498.66995
327.14569091796875 0 807.748
331.8740234375 0 541.2416
342.1363525390625 0 536.8054
342.156494140625 0 3018.4834 y 8
355.0715026855469 0 1091.1666
364.151611328125 0 687.1629
365.15679931640625 0 577.26483
378.1294860839844 0 976.79126
378.1810302734375 0 4023.4937
379.1838073730469 0 599.86444
473.1727294921875 0 1826.0265 z 7
474.1806335449219 0 5661.6353
475.1837463378906 0 1640.8497
489.1919250488281 0 2215.0476 y 7
498.76171875 0 599.9195
511.2185363769531 0 1774.6522 c Ammonia loss 3
544.2100219726562 0 5566.24 z 6
545.2142333984375 0 2943.1885
546.2199096679688 0 765.84784
550.711181640625 0 6464.0757 y 2
551.21240234375 0 3269.8928
551.71337890625 0 1376.759
560.2285766601562 0 5025.27 y 6
561.2322998046875 0 1022.9434
601.2344970703125 0 1891.678 y 1
601.737548828125 0 1919.2504
608.2349853515625 0 1030.1953
626.248046875 0 1426.4132 c Ammonia loss 4
633.7656860351562 0 590.24725
659.291259765625 0 707.6257
663.28466796875 0 946.9908
665.762939453125 0 3554.4993
666.2638549804688 0 3047.5293
666.7667236328125 0 1106.6726
674.4367065429688 0 730.6566
674.7681884765625 0 7194.4487
675.2706298828125 0 4699.4453
675.770263671875 0 2929.6846
676.3677368164062 0 745.8178
707.2728271484375 0 16106.751 z 5
708.2783203125 0 7790.8267
709.2783813476562 0 2653.4717
723.2920532226562 0 5774.9536 y 5
724.2958374023438 0 2142.6785
750.2774658203125 0 2458.1677
751.2874755859375 0 1070.0548
774.3225708007812 0 833.76117
778.3095703125 0 7330.482
779.31494140625 0 4168.9062
780.3123168945312 0 925.1785
789.3077392578125 0 1110.9089 c Ammonia loss 5
822.2996826171875 0 20131.898 z 4
823.3056030273438 0 22045.254
824.3084106445312 0 9224.278
825.309326171875 0 2723.204
838.3182373046875 0 8893.634 y 4
839.3199462890625 0 3917.4436
840.320068359375 0 1337.8364
849.3509521484375 0 1136.949
876.3662109375 0 984.85406
877.3732299804688 0 2904.8403 c 6
878.3763427734375 0 1310.2013
889.3477783203125 0 1193.6725
893.3373413085938 0 7303.2944
894.3372802734375 0 3104.805
896.3239135742188 0 1774.8253
918.4083251953125 0 722.2391
937.3257446289062 0 8531.906 z 3
938.3323974609375 0 16047.976
939.3345336914062 0 7413.506
940.3365478515625 0 2561.4453
953.345458984375 0 11987.78 y 3
954.3485717773438 0 7172.062
955.3489990234375 0 1915.2195
980.3921508789062 0 6551.5854
981.3965454101562 0 3605.8145
982.4035034179688 0 1303.2882
983.4063720703125 0 734.18475
997.3739013671875 0 646.25287
1021.3790283203125 0 762.69305
1023.30615234375 0 381.46448
1023.3999633789062 0 12976.466
1024.4053955078125 0 23273.87 c 7
1025.4091796875 0 10634.917
1026.410888671875 0 3136.8938
1036.4156494140625 0 1991.0352
1037.4156494140625 0 2372.033
1040.397705078125 0 1161.9908
1041.407958984375 0 741.05096
1083.4010009765625 0 694.4209
1084.3948974609375 0 2798.7986 z 2
1085.39208984375 0 2008.6943
1086.3917236328125 0 987.3338
1100.4132080078125 0 23495.771 y 2
1101.41650390625 0 15913.54
1102.41796875 0 5346.411
1110.4466552734375 0 685.7509
1117.4508056640625 0 6003.8
1118.454345703125 0 3364.6838
1119.4554443359375 0 1373.8867
1137.4512939453125 0 718.7262
1139.43603515625 0 3984.4934
1140.44091796875 0 2487.4968
1141.4488525390625 0 2307.1282
1142.455322265625 0 990.68134
1145.4530029296875 0 851.5912
1146.457763671875 0 1102.3342
1147.4554443359375 0 973.9533
1161.4656982421875 0 40993.895 c 8
1162.4676513671875 0 26575.424
1163.46826171875 0 10273.32
1164.45166015625 0 951.58575
1183.4344482421875 0 1325.9713 y Water loss 1
1184.4342041015625 0 6075.9062
1185.4404296875 0 11281.371 z 1
1186.441162109375 0 7790.221
1187.4473876953125 0 2049.9934
1188.4632568359375 0 933.41656
1197.4713134765625 0 851.58453
1198.468505859375 0 639.56805
1201.4608154296875 0 4933.6826 y 1
1202.46240234375 0 3668.1377
1203.4605712890625 0 1817.779
1211.43896484375 0 1030.4083
1219.47509765625 0 848.9558
1239.5582275390625 0 963.6395
1243.559814453125 0 1025.5032
1250.4775390625 0 1080.1034
1251.50146484375 0 775.0656
1257.486572265625 0 1162.7333
1258.48486328125 0 987.3779
1269.5208740234375 0 1367.5417
1270.518310546875 0 2037.6383
1271.508544921875 0 2214.942
1272.5179443359375 0 1105.8668
1285.530517578125 0 1345.8862
1286.5316162109375 0 2574.7537
1287.5338134765625 0 12306.574
1288.5264892578125 0 20929.145
1289.5196533203125 0 30471.82
1290.5186767578125 0 16877.58
1291.518310546875 0 6831.27
1292.5157470703125 0 790.99536
1299.5281982421875 0 675.2027
1301.5296630859375 0 1069.3928
1303.5325927734375 0 2438.3457
1304.5341796875 0 2358.6938
1305.540771484375 0 2316.786
1306.5438232421875 0 1193.7549
1314.5101318359375 0 928.4737
1315.4991455078125 0 1140.2722
1316.51318359375 0 913.66516
1317.4986572265625 0 1313.2748
1318.515625 0 863.188
1321.541015625 0 689.49835
1329.5291748046875 0 1429.6959
1330.516845703125 0 2084.9211
1331.52294921875 0 7522.178
1332.51171875 0 79253.8
1333.513916015625 0 65383.957
1334.515869140625 0 31424.195
1335.516845703125 0 6200.5356
1346.5379638671875 0 2508.8188
1347.53955078125 0 2291.5476
1348.527099609375 0 37821.383
1349.5341796875 0 141263.05
1350.5362548828125 0 108514.266
1350.682373046875 0 2844.305
1351.53857421875 0 45625.945
1352.540771484375 0 8815.875
2500.344970703125 0 677.6129
3018.380859375 0 681.6789
3311.613037109375 0 706.98627

Spectrum Details

|  |  |
| --- | --- |
| Matched peaks? Matched peaks The total absolute number of peaks matched. Additionally in brackets the total fraction of peaks matched and the total number of peaks is shown. | 26 (12.62% of 206) |
| FDR? FDR The false discovery rate estimated for this peptide. It is calculated by matching all theoretical fragments with a non-integer shift with the raw peaks for this spectrum. This is done with 40 different shifts. The resulting percentage is the average number of annotated peaks over the number of annotated peaks with the correct spectrum. | 0.09% |
| Satellite FDR? Satellite FDR See the FDR for details on its calculation. This satellite ion specific FDR only contains the satellite ions (d/w) for I/L/J positions. | - |
| PSM Score? PSM Score The PSM Score as given by Hecklib to this annotated spectrum. It is shown with three significant figures. | 340 |

#### Spectrum 8586? Spectrum 8586 The raw spectrum of this peptide as annotated by Hecklib. The fragments are coloured according to ion type (see legend). Any peaks with a star '\*' as text can be hovered over to see the full details, first the ion type second the mass shift type. By hovering over the amino acids in the peptide or ions in the legend the corresponding peaks are highlighted. By toggling the 'Unassigned' label you can turn the background (unassigned) peaks on or off in the plot. By updating the slider in the Ion legend you can update the spectrum to only show the top X% of the peaks with labels. The top X% means any peak that is within X% of the highest intensity. By dragging in the spectrum you can zoom in to a specific part of the spectrum and use 'Zoom Out' to get back to the original zoom level. The annotation of the spectrum is based on the given sequence in the peptides file and is done with different software so inconsistencies are likely. The peaks are annotated based on the given sequence, with 20 ppm tolerance.

Copy Data

##### Spectrum 8586 (TSV)

###### Preview

```
Loading example...
```

*Click on the button to copy the data to your clipboard.*

Mz MinMz MaxIntensity Max

WidthHeightPeptide font sizePeptide stroke widthSpectrum font sizeSpectrum stroke widthCompact peptide

Ion legend

wxyz

abcd

OtherUnassignedIon^Charge^_Position_Show for top:%

FTFDDYAMHW

04.80e+49.59e+41.44e+51.92e+5

Zoom Out

a^+1^_2_d^+1^_2_a^+1^_2_b^+1^_2_b^+1^_2_a^+1^_3_y^+1^_2_b^+1^_3_b^+1^_4_b^+2^_8_y^+1^_3_b^+1^_4_y^+2^_8_y^+2^_8_b^+2^_9_y^+1^_4_y^+2^_9_b^+1^_5_y^+2^_9_b^+1^_5_\*\*y^+1^_5_b^+1^_6_b^+1^_6_y^+1^_6_b^+1^_7_b^+1^_7_y^+1^_6_b^+1^_7_y^+1^_7_y^+1^_7_b^+1^_8_y^+1^_8_y^+1^_8_b^+1^_9_y^+1^_9_y^+1^_9_

0829165924883317

Fragment Matches Table

Show background peaks

| Position | Ion type | Intensity | mz Theoretical | mz Error (Th) | mz Error (ppm) | Charge | Series Number |
| --- | --- | --- | --- | --- | --- | --- | --- |
| - | - | 1.899E+05 | 120.1 | - | - | 0 | - |
| - | - | 764.4 | 121.1 | - | - | 0 | - |
| - | - | 1.446E+04 | 121.1 | - | - | 0 | - |
| - | - | 2473 | 129.1 | - | - | 0 | - |
| - | - | 1726 | 130.1 | - | - | 0 | - |
| - | - | 777 | 132.1 | - | - | 0 | - |
| - | - | 431.5 | 133.1 | - | - | 0 | - |
| - | - | 1917 | 133.1 | - | - | 0 | - |
| - | - | 1.523E+04 | 136.1 | - | - | 0 | - |
| - | - | 374.1 | 136.9 | - | - | 0 | - |
| - | - | 1137 | 137.1 | - | - | 0 | - |
| - | - | 836.2 | 138.1 | - | - | 0 | - |
| - | - | 1511 | 138.1 | - | - | 0 | - |
| - | - | 908.2 | 146.1 | - | - | 0 | - |
| - | - | 762.7 | 146.1 | - | - | 0 | - |
| - | - | 586.8 | 148.1 | - | - | 0 | - |
| - | - | 452.1 | 148.9 | - | - | 0 | - |
| - | - | 509 | 148.9 | - | - | 0 | - |
| - | - | 758.1 | 148.9 | - | - | 0 | - |
| - | - | 661 | 148.9 | - | - | 0 | - |
| - | - | 474.3 | 148.9 | - | - | 0 | - |
| - | - | 1193 | 148.9 | - | - | 0 | - |
| - | - | 1443 | 148.9 | - | - | 0 | - |
| - | - | 2630 | 148.9 | - | - | 0 | - |
| - | - | 4406 | 148.9 | - | - | 0 | - |
| - | - | 4369 | 149 | - | - | 0 | - |
| - | - | 2468 | 149 | - | - | 0 | - |
| - | - | 1303 | 149 | - | - | 0 | - |
| - | - | 1250 | 149 | - | - | 0 | - |
| - | - | 817.6 | 149 | - | - | 0 | - |
| - | - | 652.8 | 149 | - | - | 0 | - |
| - | - | 606.9 | 149 | - | - | 0 | - |
| - | - | 464.9 | 149 | - | - | 0 | - |
| - | - | 451.6 | 149 | - | - | 0 | - |
| - | - | 537.1 | 149 | - | - | 0 | - |
| - | - | 1906 | 149 | - | - | 0 | - |
| - | - | 6439 | 156.1 | - | - | 0 | - |
| - | - | 1140 | 158.1 | - | - | 0 | - |
| - | - | 2934 | 158.1 | - | - | 0 | - |
| - | - | 5897 | 159.1 | - | - | 0 | - |
| - | - | 841.9 | 160.1 | - | - | 0 | - |
| - | - | 949.5 | 160.1 | - | - | 0 | - |
| - | - | 501.8 | 165.1 | - | - | 0 | - |
| - | - | 727 | 166.1 | - | - | 0 | - |
| - | - | 1451 | 166.1 | - | - | 0 | - |
| - | - | 534.3 | 166.1 | - | - | 0 | - |
| - | - | 2490 | 167.1 | - | - | 0 | - |
| - | - | 1115 | 167.1 | - | - | 0 | - |
| - | - | 1124 | 169.1 | - | - | 0 | - |
| - | - | 480.6 | 172.1 | - | - | 0 | - |
| - | - | 539.5 | 174.1 | - | - | 0 | - |
| - | - | 463 | 174.9 | - | - | 0 | - |
| - | - | 1.236E+04 | 175.1 | - | - | 0 | - |
| - | - | 497 | 175.1 | - | - | 0 | - |
| - | - | 2009 | 176.1 | - | - | 0 | - |
| - | - | 8.904E+04 | 176.1 | - | - | 0 | - |
| - | - | 1657 | 177.1 | - | - | 0 | - |
| - | - | 8547 | 177.1 | - | - | 0 | - |
| - | - | 526.6 | 180.1 | - | - | 0 | - |
| - | - | 1.12E+04 | 186.1 | - | - | 0 | - |
| - | - | 1270 | 187.1 | - | - | 0 | - |
| - | - | 1707 | 191.1 | - | - | 0 | - |
| - | - | 6336 | 193.1 | - | - | 0 | - |
| - | - | 611.5 | 194.1 | - | - | 0 | - |
| - | - | 678.1 | 195.1 | - | - | 0 | - |
| - | - | 566.6 | 197.1 | - | - | 0 | - |
| - | - | 435.5 | 200.4 | - | - | 0 | - |
| - | - | 667.5 | 201.1 | - | - | 0 | - |
| - | - | 1114 | 203.1 | - | - | 0 | - |
| - | - | 2735 | 203.1 | - | - | 0 | - |
| 2 | a | 6820 | 203.1 | 0.0001677 | 0.8257 | +1 | 2 |
| - | - | 3235 | 204.1 | - | - | 0 | - |
| - | - | 756.6 | 204.1 | - | - | 0 | - |
| - | - | 895.1 | 205.1 | - | - | 0 | - |
| - | - | 653.9 | 205.1 | - | - | 0 | - |
| 2 | d | 1056 | 207.1 | 0.0001414 | 0.6827 | +1 | 2 |
| - | - | 874.3 | 208.1 | - | - | 0 | - |
| - | - | 566.2 | 209.2 | - | - | 0 | - |
| - | - | 522.7 | 211.1 | - | - | 0 | - |
| - | - | 1681 | 212.1 | - | - | 0 | - |
| - | - | 5275 | 217.1 | - | - | 0 | - |
| - | - | 868.9 | 218.1 | - | - | 0 | - |
| - | - | 723.1 | 219.1 | - | - | 0 | - |
| - | - | 560.4 | 219.1 | - | - | 0 | - |
| - | - | 2.398E+04 | 221.1 | - | - | 0 | - |
| - | - | 1.302E+04 | 221.1 | - | - | 0 | - |
| 2 | a | 1.14E+05 | 221.1 | 0.0002231 | 1.009 | +1 | 2 |
| - | - | 2424 | 222.1 | - | - | 0 | - |
| - | - | 1559 | 222.1 | - | - | 0 | - |
| - | - | 1.328E+04 | 222.1 | - | - | 0 | - |
| - | - | 710.7 | 223.1 | - | - | 0 | - |
| - | - | 5031 | 225 | - | - | 0 | - |
| - | - | 494.1 | 226 | - | - | 0 | - |
| - | - | 636.2 | 226.2 | - | - | 0 | - |
| - | - | 566.2 | 227 | - | - | 0 | - |
| - | - | 537.2 | 229.1 | - | - | 0 | - |
| - | - | 545.8 | 229.1 | - | - | 0 | - |
| - | - | 1949 | 231.1 | - | - | 0 | - |
| 2 | b | 2.788E+04 | 231.1 | 0.0002787 | 1.206 | +1 | 2 |
| - | - | 3992 | 232.1 | - | - | 0 | - |
| - | - | 8405 | 235.1 | - | - | 0 | - |
| - | - | 619.8 | 236.1 | - | - | 0 | - |
| - | - | 5902 | 239.1 | - | - | 0 | - |
| - | - | 1358 | 239.1 | - | - | 0 | - |
| - | - | 1033 | 240.1 | - | - | 0 | - |
| - | - | 621.1 | 240.1 | - | - | 0 | - |
| - | - | 534.5 | 246.1 | - | - | 0 | - |
| - | - | 625.6 | 248.2 | - | - | 0 | - |
| 2 | b | 5.774E+04 | 249.1 | 0.0002426 | 0.9738 | +1 | 2 |
| - | - | 7779 | 250.1 | - | - | 0 | - |
| - | - | 6073 | 251.1 | - | - | 0 | - |
| - | - | 474.4 | 251.1 | - | - | 0 | - |
| - | - | 835.2 | 252.1 | - | - | 0 | - |
| - | - | 2528 | 257.1 | - | - | 0 | - |
| - | - | 965.2 | 259.1 | - | - | 0 | - |
| - | - | 1.174E+04 | 263.1 | - | - | 0 | - |
| - | - | 1488 | 264.1 | - | - | 0 | - |
| - | - | 652.5 | 267.1 | - | - | 0 | - |
| - | - | 566.2 | 269.1 | - | - | 0 | - |
| - | - | 1812 | 274.1 | - | - | 0 | - |
| - | - | 4860 | 279.1 | - | - | 0 | - |
| - | - | 694.7 | 280.1 | - | - | 0 | - |
| - | - | 914.1 | 285 | - | - | 0 | - |
| - | - | 5825 | 285.1 | - | - | 0 | - |
| - | - | 639.3 | 289.1 | - | - | 0 | - |
| - | - | 3815 | 292.1 | - | - | 0 | - |
| - | - | 521.2 | 294.1 | - | - | 0 | - |
| - | - | 533.8 | 295.1 | - | - | 0 | - |
| - | - | 1557 | 295.1 | - | - | 0 | - |
| - | - | 4172 | 299.1 | - | - | 0 | - |
| - | - | 556.9 | 300.1 | - | - | 0 | - |
| - | - | 616.1 | 303.1 | - | - | 0 | - |
| - | - | 732.3 | 306.1 | - | - | 0 | - |
| - | - | 2981 | 312.1 | - | - | 0 | - |
| - | - | 1415 | 318.1 | - | - | 0 | - |
| - | - | 896.3 | 322.1 | - | - | 0 | - |
| - | - | 1255 | 323.1 | - | - | 0 | - |
| - | - | 1298 | 325.1 | - | - | 0 | - |
| - | - | 677.3 | 330.1 | - | - | 0 | - |
| - | - | 1134 | 332.1 | - | - | 0 | - |
| - | - | 1374 | 333.2 | - | - | 0 | - |
| - | - | 577.7 | 339.2 | - | - | 0 | - |
| - | - | 1.079E+04 | 340.1 | - | - | 0 | - |
| - | - | 665.8 | 341 | - | - | 0 | - |
| - | - | 2234 | 341.1 | - | - | 0 | - |
| - | - | 1175 | 346.1 | - | - | 0 | - |
| - | - | 1231 | 348.1 | - | - | 0 | - |
| - | - | 2979 | 350.1 | - | - | 0 | - |
| 3 | a | 1226 | 350.2 | 0.0001462 | 0.4176 | +1 | 3 |
| - | - | 3358 | 355.1 | - | - | 0 | - |
| - | - | 2751 | 356.1 | - | - | 0 | - |
| 9 | y | 5.522E+04 | 358.1 | 0.005057 | 14.12 | +1 | 2 |
| - | - | 5428 | 359 | - | - | 0 | - |
| - | - | 9864 | 359.2 | - | - | 0 | - |
| - | - | 919.2 | 360.1 | - | - | 0 | - |
| - | - | 1393 | 360.2 | - | - | 0 | - |
| - | - | 4555 | 364.2 | - | - | 0 | - |
| - | - | 996.9 | 365.2 | - | - | 0 | - |
| - | - | 773.5 | 366.1 | - | - | 0 | - |
| - | - | 1235 | 368.1 | - | - | 0 | - |
| - | - | 553.6 | 369.1 | - | - | 0 | - |
| - | - | 889.3 | 376.1 | - | - | 0 | - |
| - | - | 1.368E+04 | 378.1 | - | - | 0 | - |
| 3 | b | 1.377E+04 | 378.2 | 8.686E-05 | 0.2297 | +1 | 3 |
| - | - | 3191 | 379.1 | - | - | 0 | - |
| - | - | 2664 | 379.2 | - | - | 0 | - |
| - | - | 1100 | 382.1 | - | - | 0 | - |
| - | - | 4709 | 394.1 | - | - | 0 | - |
| - | - | 980.4 | 395.1 | - | - | 0 | - |
| - | - | 650.6 | 404.1 | - | - | 0 | - |
| - | - | 742.3 | 404.1 | - | - | 0 | - |
| - | - | 1161 | 406.2 | - | - | 0 | - |
| - | - | 1358 | 423.2 | - | - | 0 | - |
| - | - | 2166 | 424.2 | - | - | 0 | - |
| - | - | 1510 | 427.2 | - | - | 0 | - |
| - | - | 7385 | 429.1 | - | - | 0 | - |
| - | - | 683.6 | 433.2 | - | - | 0 | - |
| - | - | 8965 | 441.2 | - | - | 0 | - |
| - | - | 2199 | 442.2 | - | - | 0 | - |
| - | - | 678.5 | 447.2 | - | - | 0 | - |
| - | - | 659.8 | 449.2 | - | - | 0 | - |
| - | - | 644.1 | 451.2 | - | - | 0 | - |
| - | - | 1588 | 455.2 | - | - | 0 | - |
| - | - | 2063 | 461.2 | - | - | 0 | - |
| - | - | 3189 | 465.2 | - | - | 0 | - |
| - | - | 1493 | 477.2 | - | - | 0 | - |
| - | - | 2472 | 479.2 | - | - | 0 | - |
| - | - | 817.6 | 480.2 | - | - | 0 | - |
| - | - | 1232 | 487.2 | - | - | 0 | - |
| - | - | 1534 | 489.2 | - | - | 0 | - |
| - | - | 1740 | 491.2 | - | - | 0 | - |
| 4 | b | 1147 | 493.2 | 0.001067 | 2.164 | +1 | 4 |
| 8 | b | 1436 | 495.2 | 0.001365 | 2.757 | +2 | 8 |
| - | - | 1169 | 497.2 | - | - | 0 | - |
| 8 | y | 1.549E+04 | 505.2 | 0.01006 | 19.92 | +1 | 3 |
| - | - | 4746 | 506.2 | - | - | 0 | - |
| - | - | 1415 | 507.2 | - | - | 0 | - |
| 4 | b | 5638 | 511.2 | 6.734E-05 | 0.1317 | +1 | 4 |
| - | - | 8524 | 512.2 | - | - | 0 | - |
| - | - | 1914 | 513.2 | - | - | 0 | - |
| - | - | 2077 | 513.2 | - | - | 0 | - |
| - | - | 1488 | 519.2 | - | - | 0 | - |
| - | - | 1365 | 526.7 | - | - | 0 | - |
| - | - | 880.1 | 527.2 | - | - | 0 | - |
| - | - | 1967 | 541.2 | - | - | 0 | - |
| - | - | 955.4 | 542.2 | - | - | 0 | - |
| - | - | 1130 | 548.2 | - | - | 0 | - |
| 3 | y | 1220 | 549.7 | 0.002925 | 5.322 | +2 | 8 |
| - | - | 926.8 | 550.2 | - | - | 0 | - |
| - | - | 897.8 | 558.2 | - | - | 0 | - |
| 3 | y | 1.582E+04 | 558.7 | 0.004845 | 8.672 | +2 | 8 |
| - | - | 1E+04 | 559.2 | - | - | 0 | - |
| - | - | 3304 | 559.7 | - | - | 0 | - |
| - | - | 968.2 | 560.2 | - | - | 0 | - |
| - | - | 678.8 | 562.7 | - | - | 0 | - |
| - | - | 604.8 | 567.2 | - | - | 0 | - |
| - | - | 1730 | 570 | - | - | 0 | - |
| - | - | 1050 | 570.2 | - | - | 0 | - |
| 9 | b | 870.2 | 572.7 | 0.0006279 | 1.096 | +2 | 9 |
| 7 | y | 2.25E+04 | 576.2 | 0.009907 | 17.19 | +1 | 4 |
| - | - | 6894 | 577.2 | - | - | 0 | - |
| - | - | 1518 | 578.2 | - | - | 0 | - |
| - | - | 829 | 580.2 | - | - | 0 | - |
| - | - | 801.3 | 590.2 | - | - | 0 | - |
| - | - | 1183 | 595 | - | - | 0 | - |
| 2 | y | 1211 | 600.2 | 0.005148 | 8.577 | +2 | 9 |
| - | - | 615.9 | 600.7 | - | - | 0 | - |
| - | - | 1582 | 606.2 | - | - | 0 | - |
| 5 | b | 1997 | 608.2 | 0.0001798 | 0.2956 | +1 | 5 |
| 2 | y | 7371 | 609.2 | 0.005847 | 9.598 | +2 | 9 |
| - | - | 3629 | 609.7 | - | - | 0 | - |
| - | - | 1919 | 610.2 | - | - | 0 | - |
| - | - | 795.1 | 612.2 | - | - | 0 | - |
| - | - | 959.4 | 614.2 | - | - | 0 | - |
| - | - | 714.6 | 625.2 | - | - | 0 | - |
| 5 | b | 3559 | 626.2 | 0.0006737 | 1.076 | +1 | 5 |
| - | - | 1568 | 627.3 | - | - | 0 | - |
| - | - | 1242 | 634.2 | - | - | 0 | - |
| - | - | 693 | 640.2 | - | - | 0 | - |
| - | - | 1182 | 641.8 | - | - | 0 | - |
| - | - | 902 | 642.3 | - | - | 0 | - |
| - | - | 2589 | 650.8 | - | - | 0 | - |
| - | - | 1402 | 651.3 | - | - | 0 | - |
| - | - | 1509 | 657.3 | - | - | 0 | - |
| - | - | 713.2 | 659.8 | - | - | 0 | - |
| - | - | 854.6 | 660.3 | - | - | 0 | - |
| - | - | 1451 | 664.8 | - | - | 0 | - |
| - | - | 670.5 | 665.3 | - | - | 0 | - |
| 0 | Precursor | 9532 | 673.8 | 0.004755 | 7.057 | +2 | -1 |
| - | - | 7370 | 674.3 | - | - | 0 | - |
| - | - | 3664 | 674.8 | - | - | 0 | - |
| - | - | 7516 | 675.3 | - | - | 0 | - |
| - | - | 2669 | 676.3 | - | - | 0 | - |
| - | - | 827.9 | 681.4 | - | - | 0 | - |
| - | - | 1284 | 682.3 | - | - | 0 | - |
| 0 | Precursor | 1.811E+04 | 682.8 | 0.00466 | 6.826 | +2 | -1 |
| - | - | 1.416E+04 | 683.3 | - | - | 0 | - |
| - | - | 678.7 | 683.3 | - | - | 0 | - |
| - | - | 600.7 | 683.4 | - | - | 0 | - |
| - | - | 6919 | 683.8 | - | - | 0 | - |
| - | - | 1496 | 684.2 | - | - | 0 | - |
| - | - | 1837 | 684.3 | - | - | 0 | - |
| - | - | 709.1 | 685.2 | - | - | 0 | - |
| - | - | 3010 | 685.3 | - | - | 0 | - |
| - | - | 1176 | 686.3 | - | - | 0 | - |
| - | - | 2202 | 695.3 | - | - | 0 | - |
| - | - | 810.2 | 696.3 | - | - | 0 | - |
| - | - | 709 | 704.3 | - | - | 0 | - |
| - | - | 919.7 | 713.3 | - | - | 0 | - |
| - | - | 2405 | 721.3 | - | - | 0 | - |
| - | - | 741.2 | 722.3 | - | - | 0 | - |
| - | - | 651.1 | 723.3 | - | - | 0 | - |
| 6 | y | 1.908E+04 | 739.3 | 0.009689 | 13.11 | +1 | 5 |
| - | - | 7814 | 740.3 | - | - | 0 | - |
| - | - | 2018 | 741.3 | - | - | 0 | - |
| - | - | 723.9 | 743.3 | - | - | 0 | - |
| - | - | 2732 | 749.3 | - | - | 0 | - |
| - | - | 760.9 | 750.3 | - | - | 0 | - |
| - | - | 1121 | 759.3 | - | - | 0 | - |
| - | - | 821.7 | 761.3 | - | - | 0 | - |
| - | - | 1125 | 767.3 | - | - | 0 | - |
| 6 | b | 1348 | 771.3 | 0.003511 | 4.552 | +1 | 6 |
| - | - | 1135 | 772.3 | - | - | 0 | - |
| - | - | 942.1 | 778.3 | - | - | 0 | - |
| 6 | b | 1537 | 789.3 | 0.001977 | 2.504 | +1 | 6 |
| - | - | 8833 | 790.3 | - | - | 0 | - |
| - | - | 3933 | 791.3 | - | - | 0 | - |
| - | - | 773 | 792.3 | - | - | 0 | - |
| - | - | 1277 | 804.3 | - | - | 0 | - |
| - | - | 898.6 | 805.3 | - | - | 0 | - |
| - | - | 4195 | 832.3 | - | - | 0 | - |
| - | - | 1138 | 833.3 | - | - | 0 | - |
| 5 | y | 1307 | 836.3 | 0.008387 | 10.03 | +1 | 6 |
| 7 | b | 956.7 | 842.3 | 0.005224 | 6.202 | +1 | 7 |
| 7 | b | 811.1 | 843.3 | 0.009662 | 11.46 | +1 | 7 |
| 5 | y | 2.476E+04 | 854.3 | 0.009419 | 11.02 | +1 | 6 |
| - | - | 1.089E+04 | 855.3 | - | - | 0 | - |
| - | - | 3928 | 856.3 | - | - | 0 | - |
| 7 | b | 818.3 | 860.3 | 0.009624 | 11.19 | +1 | 7 |
| - | - | 914.5 | 861.3 | - | - | 0 | - |
| - | - | 2477 | 868.3 | - | - | 0 | - |
| - | - | 1712 | 869.3 | - | - | 0 | - |
| - | - | 1502 | 887.3 | - | - | 0 | - |
| - | - | 1138 | 888.3 | - | - | 0 | - |
| - | - | 3831 | 896.3 | - | - | 0 | - |
| - | - | 1931 | 897.3 | - | - | 0 | - |
| - | - | 1.254E+04 | 905.3 | - | - | 0 | - |
| - | - | 7145 | 906.3 | - | - | 0 | - |
| - | - | 1693 | 907.3 | - | - | 0 | - |
| - | - | 1143 | 914.3 | - | - | 0 | - |
| - | - | 937.7 | 915.3 | - | - | 0 | - |
| 4 | y | 2549 | 951.3 | 0.01068 | 11.23 | +1 | 7 |
| - | - | 2187 | 952.3 | - | - | 0 | - |
| 4 | y | 4.358E+04 | 969.3 | 0.008904 | 9.185 | +1 | 7 |
| - | - | 1.998E+04 | 970.3 | - | - | 0 | - |
| - | - | 5619 | 971.3 | - | - | 0 | - |
| - | - | 746.4 | 979.3 | - | - | 0 | - |
| 8 | b | 866.5 | 989.4 | 0.006039 | 6.104 | +1 | 8 |
| - | - | 816.8 | 1008 | - | - | 0 | - |
| - | - | 2262 | 1034 | - | - | 0 | - |
| - | - | 758.5 | 1051 | - | - | 0 | - |
| - | - | 1.833E+04 | 1052 | - | - | 0 | - |
| - | - | 1.108E+04 | 1053 | - | - | 0 | - |
| - | - | 3271 | 1054 | - | - | 0 | - |
| 3 | y | 4111 | 1098 | 0.01008 | 9.173 | +1 | 8 |
| - | - | 1894 | 1099 | - | - | 0 | - |
| - | - | 875.2 | 1100 | - | - | 0 | - |
| 3 | y | 6.254E+04 | 1116 | 0.008544 | 7.653 | +1 | 8 |
| - | - | 4.167E+04 | 1117 | - | - | 0 | - |
| - | - | 1.475E+04 | 1118 | - | - | 0 | - |
| - | - | 2887 | 1119 | - | - | 0 | - |
| 9 | b | 2088 | 1144 | 0.002725 | 2.381 | +1 | 9 |
| - | - | 1653 | 1145 | - | - | 0 | - |
| - | - | 3988 | 1153 | - | - | 0 | - |
| - | - | 3059 | 1154 | - | - | 0 | - |
| - | - | 966.9 | 1155 | - | - | 0 | - |
| 2 | y | 1660 | 1199 | 0.01403 | 11.7 | +1 | 9 |
| - | - | 940.1 | 1200 | - | - | 0 | - |
| 2 | y | 1.148E+04 | 1217 | 0.007374 | 6.057 | +1 | 9 |
| - | - | 8011 | 1218 | - | - | 0 | - |
| - | - | 2903 | 1219 | - | - | 0 | - |
| - | - | 1841 | 1227 | - | - | 0 | - |
| - | - | 2115 | 1228 | - | - | 0 | - |
| - | - | 696.6 | 1374 | - | - | 0 | - |
| - | - | 725.4 | 1997 | - | - | 0 | - |
| - | - | 641.4 | 2460 | - | - | 0 | - |
| - | - | 692.4 | 3074 | - | - | 0 | - |
| - | - | 789.1 | 3075 | - | - | 0 | - |
| - | - | 828.9 | 3075 | - | - | 0 | - |
| - | - | 627 | 3284 | - | - | 0 | - |

m/z Charge Intensity FragmentType MassShift Position
120.08108520507812 0 189929.39
121.07908630371094 0 764.36884
121.08434295654297 0 14463.035
129.1023712158203 0 2472.6008
130.0653533935547 0 1725.7583
132.08082580566406 0 777.04126
133.06138610839844 0 431.46774
133.08615112304688 0 1917.4691
136.075927734375 0 15225.343
136.8873291015625 0 374.06464
137.07911682128906 0 1137.0541
138.05540466308594 0 836.2472
138.0664520263672 0 1510.6232
146.06007385253906 0 908.2333
146.09642028808594 0 762.6639
148.08709716796875 0 586.78625
148.87159729003906 0 452.08188
148.89315795898438 0 508.9674
148.9007568359375 0 758.0541
148.90745544433594 0 660.964
148.9150848388672 0 474.30328
148.92236328125 0 1193.4846
148.92942810058594 0 1442.9143
148.93646240234375 0 2630.4402
148.94424438476562 0 4406.1055
148.96072387695312 0 4368.794
148.96852111816406 0 2468.2102
148.97555541992188 0 1302.7933
148.98275756835938 0 1249.8679
148.98983764648438 0 817.58563
148.99696350097656 0 652.8102
149.00425720214844 0 606.93353
149.01124572753906 0 464.90826
149.01885986328125 0 451.55737
149.02603149414062 0 537.0591
149.0453643798828 0 1906.2155
156.0769500732422 0 6439.205
158.0603485107422 0 1139.5164
158.09661865234375 0 2934.3635
159.09182739257812 0 5897.149
160.09527587890625 0 841.85364
160.1126251220703 0 949.53595
165.0770721435547 0 501.835
166.05416870117188 0 727.0016
166.061279296875 0 1451.4761
166.09768676757812 0 534.31067
167.0556640625 0 2490.1458
167.0930633544922 0 1114.9739
169.1342010498047 0 1124.1714
172.1083984375 0 480.64767
174.0660858154297 0 539.4965
174.91201782226562 0 463.04907
175.08677673339844 0 12358.522
175.1231689453125 0 496.96313
176.08233642578125 0 2008.7223
176.1072540283203 0 89043.4
177.10232543945312 0 1656.6062
177.11080932617188 0 8547.15
180.07662963867188 0 526.6416
186.09149169921875 0 11204.611
187.09523010253906 0 1270.2465
191.11813354492188 0 1707.4011
193.1085968017578 0 6336.3174
194.079345703125 0 611.5491
195.08792114257812 0 678.05774
197.0795135498047 0 566.6196
200.35543823242188 0 435.4723
201.1229705810547 0 667.54205
203.06643676757812 0 1113.9365
203.08177185058594 0 2734.659
203.11805725097656 0 6819.776 a Water loss 1
204.0767364501953 0 3234.6057
204.12132263183594 0 756.6355
205.09703063964844 0 895.1474
205.10716247558594 0 653.92883
207.11294555664062 0 1055.5825 d 1
208.07261657714844 0 874.25006
209.1649627685547 0 566.17926
211.0825958251953 0 522.69495
212.11830139160156 0 1681.4113
217.0973358154297 0 5274.5234
218.10055541992188 0 868.88763
219.07960510253906 0 723.1022
219.11297607421875 0 560.4377
221.0922088623047 0 23980.47
221.1036376953125 0 13024.197
221.12867736816406 0 113989.44 a 1
222.09561157226562 0 2424.3918
222.1073455810547 0 1559.2363
222.1319580078125 0 13280.209
223.13475036621094 0 710.73566
225.0430908203125 0 5030.945
226.04385375976562 0 494.1073
226.1555938720703 0 636.19385
227.02284240722656 0 566.23615
229.09719848632812 0 537.16504
229.13211059570312 0 545.7918
231.06173706054688 0 1949.2233
231.1130828857422 0 27879.326 b Water loss 1
232.11624145507812 0 3991.8772
235.10789489746094 0 8405.057
236.1122589111328 0 619.7987
239.09506225585938 0 5902.417
239.1141357421875 0 1357.5039
240.096923828125 0 1033.1194
240.1130828857422 0 621.11676
246.13397216796875 0 534.51105
248.15098571777344 0 625.62775
249.1236114501953 0 57740.7 b 1
250.12680053710938 0 7778.643
251.10279846191406 0 6073.45
251.1303253173828 0 474.3829
252.10537719726562 0 835.23065
257.106689453125 0 2527.674
259.1083984375 0 965.1885
263.102783203125 0 11744.112
264.1065368652344 0 1487.5183
267.1254577636719 0 652.5242
269.0762634277344 0 566.1589
274.1294860839844 0 1812.136
279.0973815917969 0 4859.528
280.1015319824219 0 694.703
285.0094909667969 0 914.146
285.1018981933594 0 5824.562
289.1346740722656 0 639.3216
292.1407470703125 0 3814.7344
294.13616943359375 0 521.16376
295.0861511230469 0 533.84937
295.1043395996094 0 1556.7303
299.06195068359375 0 4171.6743
300.0633239746094 0 556.8669
303.1114196777344 0 616.10565
306.1440734863281 0 732.2523
312.1458740234375 0 2981.2905
318.1451416015625 0 1415.0687
322.13140869140625 0 896.2703
323.0985107421875 0 1254.9323
325.1377258300781 0 1298.3983
330.10845947265625 0 677.26013
332.12493896484375 0 1133.616
333.1611328125 0 1374.379
339.16925048828125 0 577.655
340.14068603515625 0 10786.8
341.0167236328125 0 665.8139
341.14410400390625 0 2233.7158
346.1400451660156 0 1174.9418
348.1184997558594 0 1230.6683
350.13470458984375 0 2979.0269
350.1861572265625 0 1226.4373 a Water loss 2
355.0702819824219 0 3358.1785
356.1387939453125 0 2750.8718
358.151123046875 0 55217.74 y 8
359.02850341796875 0 5428.492
359.15411376953125 0 9864.327
360.1206970214844 0 919.23615
360.1546936035156 0 1392.7224
364.1504211425781 0 4554.9976
365.1532287597656 0 996.93536
366.1285400390625 0 773.52673
368.1334228515625 0 1234.6108
369.12200927734375 0 553.62756
376.1151123046875 0 889.3484
378.1297607421875 0 13684.948
378.1813049316406 0 13765.477 b Water loss 2
379.1327819824219 0 3191.1262
379.18408203125 0 2663.8462
382.1445007324219 0 1099.5175
394.12432861328125 0 4708.903
395.12774658203125 0 980.4124
404.10845947265625 0 650.55536
404.1434631347656 0 742.2989
406.15118408203125 0 1161.343
423.1778869628906 0 1358.2225
424.1614074707031 0 2166.3352
427.2085876464844 0 1510.48
429.0890808105469 0 7384.951
433.17291259765625 0 683.6145
441.18853759765625 0 8965.006
442.191162109375 0 2199.1174
447.1507568359375 0 678.54974
449.16595458984375 0 659.8324
451.1697998046875 0 644.1318
455.20343017578125 0 1587.8273
461.16845703125 0 2063.2446
465.161865234375 0 3189.3584
477.18707275390625 0 1493.473
479.17755126953125 0 2472.1729
480.1814270019531 0 817.6428
487.17694091796875 0 1231.7733
489.1618347167969 0 1533.7042
491.2066955566406 0 1740.1561
493.209228515625 0 1146.5105 b Water loss 3
495.1880187988281 0 1435.5714 b Water loss 7
497.16766357421875 0 1168.5365
505.1866149902344 0 15490.085 y 7
506.18896484375 0 4746.3247
507.18603515625 0 1415.3152
511.2186584472656 0 5637.84 b 3
512.224853515625 0 8524.302
513.1970825195312 0 1914.3748
513.22802734375 0 2076.8364
519.2028198242188 0 1487.9248
526.7068481445312 0 1364.5243
527.207275390625 0 880.06226
541.1944580078125 0 1967.0607
542.2371215820312 0 955.41345
548.1986083984375 0 1130.3947
549.700927734375 0 1219.8315 y Water loss 2
550.20703125 0 926.765
558.2135620117188 0 897.785
558.7081298828125 0 15820.688 y 2
559.2095947265625 0 10004.782
559.7102661132812 0 3303.8333
560.2072143554688 0 968.157
562.7041625976562 0 678.832
567.209228515625 0 604.8385
570.02490234375 0 1730.2124
570.2302856445312 0 1050.3102
572.7207641601562 0 870.2144 b 8
576.2235717773438 0 22497.273 y 6
577.2269287109375 0 6894.426
578.2233276367188 0 1517.5043
580.2402954101562 0 829.04224
590.2221069335938 0 801.3186
594.955078125 0 1182.9623
600.2269897460938 0 1211.4976 y Water loss 1
600.723876953125 0 615.94354
606.2323608398438 0 1582.1267
608.2349243164062 0 1997.2621 b Water loss 4
609.2329711914062 0 7371.0864 y 1
609.73291015625 0 3628.5132
610.2328491210938 0 1918.5242
612.1941528320312 0 795.1281
614.2408447265625 0 959.4076
625.2344360351562 0 714.63574
626.2449951171875 0 3559.4011 b 4
627.251708984375 0 1567.7013
634.2302856445312 0 1242.4083
640.248291015625 0 692.996
641.7593383789062 0 1182.0402
642.2601318359375 0 901.96655
650.7652587890625 0 2589.2056
651.267822265625 0 1402.1677
657.2682495117188 0 1509.0294
659.7596435546875 0 713.2263
660.2589721679688 0 854.6153
664.756591796875 0 1450.7078
665.2596435546875 0 670.5441
673.7608032226562 0 9531.887 Precursor Water loss
674.2633666992188 0 7370.001
674.7630615234375 0 3663.9968
675.2875366210938 0 7516.4062
676.2902221679688 0 2669.0076
681.4077758789062 0 827.91125
682.340576171875 0 1284.2943
682.7659912109375 0 18109.506 Precursor
683.267578125 0 14157.365
683.3343505859375 0 678.6974
683.3956909179688 0 600.68207
683.76806640625 0 6919.382
684.2026977539062 0 1496.3231
684.269775390625 0 1836.6255
685.1943359375 0 709.13184
685.259765625 0 3009.7356
686.264892578125 0 1175.9724
695.2669067382812 0 2202.1582
696.2723388671875 0 810.16254
704.2593994140625 0 709.0314
713.2760009765625 0 919.73224
721.26708984375 0 2405.42
722.2711181640625 0 741.1549
723.263916015625 0 651.0637
739.2866821289062 0 19075.736 y 5
740.289306640625 0 7814.4526
741.2864379882812 0 2018.1504
743.3049926757812 0 723.88165
749.2567138671875 0 2732.43
750.25927734375 0 760.9043
759.2653198242188 0 1121.1788
761.312744140625 0 821.74585
767.2664794921875 0 1125.2343
771.294921875 0 1348.0819 b Water loss 5
772.306884765625 0 1135.0847
778.3090209960938 0 942.11475
789.3109741210938 0 1536.9829 b 5
790.3148803710938 0 8833.321
791.3189086914062 0 3933.4902
792.321533203125 0 773.03094
804.3312377929688 0 1276.9314
805.3320922851562 0 898.5746
832.3265991210938 0 4194.876
833.3280639648438 0 1138.214
836.3017578125 0 1307.064 y Water loss 4
842.330322265625 0 956.72736 b Water loss 6
843.3292236328125 0 811.0663 b Ammonia loss 6
854.3133544921875 0 24760.543 y 4
855.3160400390625 0 10891.396
856.3162231445312 0 3928.4812
860.3364868164062 0 818.279 b 6
861.3373413085938 0 914.5366
868.3271484375 0 2476.7053
869.3310546875 0 1711.9031
887.3290405273438 0 1501.6049
888.3270874023438 0 1138.163
896.3226318359375 0 3831.406
897.3264770507812 0 1930.5327
905.3418579101562 0 12543.188
906.34423828125 0 7144.5693
907.3480224609375 0 1693.2877
914.3389892578125 0 1143.4176
915.3350219726562 0 937.68726
951.3309936523438 0 2549.4429 y Water loss 3
952.3267822265625 0 2186.9856
969.3397827148438 0 43581.176 y 3
970.3430786132812 0 19978.16
971.3426513671875 0 5619.2095
979.3300170898438 0 746.39923
989.3720703125 0 866.4693 b Water loss 7
1008.3899536132812 0 816.84564
1034.397705078125 0 2262.3718
1050.9588623046875 0 758.509
1052.41064453125 0 18327.74
1053.4132080078125 0 11076.044
1054.4188232421875 0 3270.864
1098.3988037109375 0 4110.989 y Water loss 2
1099.39892578125 0 1894.4642
1100.4073486328125 0 875.17053
1116.4078369140625 0 62543.445 y 2
1117.4112548828125 0 41671.715
1118.4111328125 0 14753.156
1119.409912109375 0 2886.8823
1144.438232421875 0 2088.4214 b 8
1145.44287109375 0 1652.9642
1153.45654296875 0 3988.483
1154.457275390625 0 3058.5903
1155.463134765625 0 966.85895
1199.450439453125 0 1659.7478 y Water loss 1
1200.4495849609375 0 940.12024
1217.454345703125 0 11482.513 y 1
1218.4576416015625 0 8010.744
1219.4566650390625 0 2902.754
1227.43603515625 0 1840.6432
1228.4410400390625 0 2114.5513
1373.5811767578125 0 696.6446
1997.0435791015625 0 725.3534
2459.696533203125 0 641.4488
3074.0458984375 0 692.4073
3074.757568359375 0 789.10284
3075.41064453125 0 828.9067
3284.385498046875 0 626.9627

Spectrum Details

|  |  |
| --- | --- |
| Matched peaks? Matched peaks The total absolute number of peaks matched. Additionally in brackets the total fraction of peaks matched and the total number of peaks is shown. | 38 (10.86% of 350) |
| FDR? FDR The false discovery rate estimated for this peptide. It is calculated by matching all theoretical fragments with a non-integer shift with the raw peaks for this spectrum. This is done with 40 different shifts. The resulting percentage is the average number of annotated peaks over the number of annotated peaks with the correct spectrum. | 0.19% |
| Satellite FDR? Satellite FDR See the FDR for details on its calculation. This satellite ion specific FDR only contains the satellite ions (d/w) for I/L/J positions. | - |
| PSM Score? PSM Score The PSM Score as given by Hecklib to this annotated spectrum. It is shown with three significant figures. | 463 |

#### Spectrum 9848? Spectrum 9848 The raw spectrum of this peptide as annotated by Hecklib. The fragments are coloured according to ion type (see legend). Any peaks with a star '\*' as text can be hovered over to see the full details, first the ion type second the mass shift type. By hovering over the amino acids in the peptide or ions in the legend the corresponding peaks are highlighted. By toggling the 'Unassigned' label you can turn the background (unassigned) peaks on or off in the plot. By updating the slider in the Ion legend you can update the spectrum to only show the top X% of the peaks with labels. The top X% means any peak that is within X% of the highest intensity. By dragging in the spectrum you can zoom in to a specific part of the spectrum and use 'Zoom Out' to get back to the original zoom level. The annotation of the spectrum is based on the given sequence in the peptides file and is done with different software so inconsistencies are likely. The peaks are annotated based on the given sequence, with 20 ppm tolerance.

Copy Data

##### Spectrum 9848 (TSV)

###### Preview

```
Loading example...
```

*Click on the button to copy the data to your clipboard.*

Mz MinMz MaxIntensity Max

WidthHeightPeptide font sizePeptide stroke widthSpectrum font sizeSpectrum stroke widthCompact peptide

Ion legend

wxyz

abcd

OtherUnassignedIon^Charge^_Position_Show for top:%

FTFDDYAMHW

04.81e+49.62e+41.44e+51.92e+5

Zoom Out

a^+1^_2_a^+1^_2_y^+1^_1_d^+1^_2_a^+1^_2_b^+1^_2_b^+2^_4_b^+1^_2_y^+1^_2_b^+1^_3_y^+1^_3_b^+1^_4_b^+2^_8_b^+1^_4_y^+2^_8_y^+1^_4_b^+2^_9_b^+2^_9_y^+2^_9_b^+1^_5_b^+1^_5_\*\*y^+1^_5_b^+1^_6_b^+1^_6_y^+1^_6_b^+1^_7_y^+1^_7_y^+1^_7_y^+1^_8_y^+1^_8_b^+1^_9_y^+1^_9_y^+1^_9_

0849169825473397

Fragment Matches Table

Show background peaks

| Position | Ion type | Intensity | mz Theoretical | mz Error (Th) | mz Error (ppm) | Charge | Series Number |
| --- | --- | --- | --- | --- | --- | --- | --- |
| - | - | 1.905E+05 | 120.1 | - | - | 0 | - |
| - | - | 956.9 | 121.1 | - | - | 0 | - |
| - | - | 1.587E+04 | 121.1 | - | - | 0 | - |
| - | - | 342.7 | 121.8 | - | - | 0 | - |
| - | - | 394.7 | 122.1 | - | - | 0 | - |
| - | - | 544.6 | 129.1 | - | - | 0 | - |
| - | - | 1624 | 129.1 | - | - | 0 | - |
| - | - | 700.9 | 130.1 | - | - | 0 | - |
| - | - | 1818 | 130.1 | - | - | 0 | - |
| - | - | 793.4 | 131 | - | - | 0 | - |
| - | - | 421.1 | 131.1 | - | - | 0 | - |
| - | - | 607.4 | 132.1 | - | - | 0 | - |
| - | - | 780.5 | 133.1 | - | - | 0 | - |
| - | - | 1463 | 134 | - | - | 0 | - |
| - | - | 2.589E+04 | 136.1 | - | - | 0 | - |
| - | - | 2176 | 137.1 | - | - | 0 | - |
| - | - | 1163 | 138.1 | - | - | 0 | - |
| - | - | 1951 | 138.1 | - | - | 0 | - |
| - | - | 896.1 | 142.1 | - | - | 0 | - |
| - | - | 824.4 | 143.1 | - | - | 0 | - |
| - | - | 854.1 | 143.1 | - | - | 0 | - |
| - | - | 1001 | 146.1 | - | - | 0 | - |
| - | - | 892 | 147.1 | - | - | 0 | - |
| - | - | 965.7 | 148.1 | - | - | 0 | - |
| - | - | 547.1 | 148.9 | - | - | 0 | - |
| - | - | 521.7 | 152.1 | - | - | 0 | - |
| - | - | 926.1 | 155.1 | - | - | 0 | - |
| - | - | 4450 | 156.1 | - | - | 0 | - |
| - | - | 800.2 | 157.1 | - | - | 0 | - |
| - | - | 3106 | 158.1 | - | - | 0 | - |
| - | - | 8957 | 159.1 | - | - | 0 | - |
| - | - | 809.2 | 160.1 | - | - | 0 | - |
| - | - | 643.1 | 160.1 | - | - | 0 | - |
| - | - | 2245 | 161.1 | - | - | 0 | - |
| - | - | 578.6 | 162.1 | - | - | 0 | - |
| - | - | 1252 | 165.1 | - | - | 0 | - |
| - | - | 1081 | 166.1 | - | - | 0 | - |
| - | - | 2.057E+04 | 166.1 | - | - | 0 | - |
| - | - | 2559 | 167.1 | - | - | 0 | - |
| - | - | 510.9 | 169.1 | - | - | 0 | - |
| - | - | 1255 | 169.1 | - | - | 0 | - |
| - | - | 2178 | 171.1 | - | - | 0 | - |
| - | - | 682.7 | 173.1 | - | - | 0 | - |
| - | - | 882.7 | 175.1 | - | - | 0 | - |
| - | - | 1636 | 176.1 | - | - | 0 | - |
| - | - | 7.895E+04 | 176.1 | - | - | 0 | - |
| - | - | 1439 | 177.1 | - | - | 0 | - |
| - | - | 9384 | 177.1 | - | - | 0 | - |
| - | - | 793.8 | 180.1 | - | - | 0 | - |
| - | - | 538.2 | 181.1 | - | - | 0 | - |
| - | - | 941 | 181.2 | - | - | 0 | - |
| - | - | 981.2 | 182.1 | - | - | 0 | - |
| - | - | 1022 | 185.1 | - | - | 0 | - |
| - | - | 9323 | 186.1 | - | - | 0 | - |
| - | - | 1215 | 187.1 | - | - | 0 | - |
| - | - | 1757 | 187.1 | - | - | 0 | - |
| - | - | 5945 | 188.1 | - | - | 0 | - |
| - | - | 4249 | 189.1 | - | - | 0 | - |
| - | - | 1308 | 189.1 | - | - | 0 | - |
| - | - | 679.1 | 191.1 | - | - | 0 | - |
| - | - | 1413 | 191.1 | - | - | 0 | - |
| - | - | 4593 | 193.1 | - | - | 0 | - |
| - | - | 1937 | 195.1 | - | - | 0 | - |
| - | - | 557.3 | 195.1 | - | - | 0 | - |
| - | - | 1399 | 197.1 | - | - | 0 | - |
| - | - | 1914 | 201.1 | - | - | 0 | - |
| - | - | 737.4 | 202.1 | - | - | 0 | - |
| - | - | 640.5 | 203.1 | - | - | 0 | - |
| 2 | a | 5453 | 203.1 | 0.000305 | 1.502 | +1 | 2 |
| - | - | 2735 | 204.1 | - | - | 0 | - |
| 2 | a | 641 | 204.1 | 0.0004966 | 2.433 | +1 | 2 |
| - | - | 865.2 | 204.1 | - | - | 0 | - |
| 10 | y | 4902 | 205.1 | 0.0003953 | 1.928 | +1 | 1 |
| - | - | 834.7 | 205.1 | - | - | 0 | - |
| 2 | d | 5079 | 207.1 | 0.0004771 | 2.304 | +1 | 2 |
| - | - | 879.7 | 208.1 | - | - | 0 | - |
| - | - | 743.8 | 209.2 | - | - | 0 | - |
| - | - | 1054 | 212.1 | - | - | 0 | - |
| - | - | 555.9 | 214.1 | - | - | 0 | - |
| - | - | 499.8 | 215.1 | - | - | 0 | - |
| - | - | 1777 | 215.1 | - | - | 0 | - |
| - | - | 1622 | 217.1 | - | - | 0 | - |
| - | - | 6052 | 217.1 | - | - | 0 | - |
| - | - | 646.9 | 218.1 | - | - | 0 | - |
| - | - | 893.5 | 218.1 | - | - | 0 | - |
| - | - | 645.4 | 219.1 | - | - | 0 | - |
| - | - | 1.281E+04 | 221.1 | - | - | 0 | - |
| 2 | a | 1.04E+05 | 221.1 | 0.0004825 | 2.182 | +1 | 2 |
| - | - | 1063 | 222.1 | - | - | 0 | - |
| - | - | 1.277E+04 | 222.1 | - | - | 0 | - |
| - | - | 2072 | 223.1 | - | - | 0 | - |
| - | - | 826.3 | 223.1 | - | - | 0 | - |
| - | - | 605.7 | 226.1 | - | - | 0 | - |
| - | - | 835.1 | 227.1 | - | - | 0 | - |
| - | - | 844.2 | 228.1 | - | - | 0 | - |
| - | - | 1322 | 229.1 | - | - | 0 | - |
| - | - | 712 | 231 | - | - | 0 | - |
| - | - | 2115 | 231.1 | - | - | 0 | - |
| 2 | b | 2.459E+04 | 231.1 | 0.0004771 | 2.064 | +1 | 2 |
| - | - | 2743 | 232.1 | - | - | 0 | - |
| - | - | 526.2 | 235.1 | - | - | 0 | - |
| - | - | 1.108E+04 | 235.1 | - | - | 0 | - |
| - | - | 1149 | 236.1 | - | - | 0 | - |
| - | - | 760.5 | 236.1 | - | - | 0 | - |
| - | - | 973.7 | 237.2 | - | - | 0 | - |
| - | - | 1428 | 239.1 | - | - | 0 | - |
| - | - | 549.9 | 240.1 | - | - | 0 | - |
| - | - | 647.7 | 245.1 | - | - | 0 | - |
| 4 | b | 901.5 | 247.1 | 6.802E-05 | 0.2753 | +2 | 4 |
| - | - | 517.7 | 247.1 | - | - | 0 | - |
| 2 | b | 5.25E+04 | 249.1 | 0.0004562 | 1.831 | +1 | 2 |
| - | - | 6518 | 250.1 | - | - | 0 | - |
| - | - | 5902 | 251.1 | - | - | 0 | - |
| - | - | 647.5 | 251.1 | - | - | 0 | - |
| - | - | 1.921E+04 | 253.1 | - | - | 0 | - |
| - | - | 3392 | 254.1 | - | - | 0 | - |
| - | - | 2305 | 257.1 | - | - | 0 | - |
| - | - | 591.1 | 257.1 | - | - | 0 | - |
| - | - | 1088 | 259.1 | - | - | 0 | - |
| - | - | 1.051E+04 | 263.1 | - | - | 0 | - |
| - | - | 713.4 | 263.1 | - | - | 0 | - |
| - | - | 1317 | 264.1 | - | - | 0 | - |
| - | - | 1636 | 265.1 | - | - | 0 | - |
| - | - | 1951 | 269.1 | - | - | 0 | - |
| - | - | 2064 | 274.1 | - | - | 0 | - |
| - | - | 3701 | 279.1 | - | - | 0 | - |
| - | - | 896.1 | 280.1 | - | - | 0 | - |
| - | - | 8154 | 281.1 | - | - | 0 | - |
| - | - | 1190 | 282.1 | - | - | 0 | - |
| - | - | 842.4 | 283.1 | - | - | 0 | - |
| - | - | 1026 | 284.2 | - | - | 0 | - |
| - | - | 510.2 | 285 | - | - | 0 | - |
| - | - | 5183 | 285.1 | - | - | 0 | - |
| - | - | 638.6 | 286.1 | - | - | 0 | - |
| - | - | 2384 | 292.1 | - | - | 0 | - |
| - | - | 557.4 | 295.1 | - | - | 0 | - |
| - | - | 4654 | 296.2 | - | - | 0 | - |
| - | - | 1122 | 297.2 | - | - | 0 | - |
| - | - | 1503 | 304.1 | - | - | 0 | - |
| - | - | 4423 | 309.1 | - | - | 0 | - |
| - | - | 1828 | 312.2 | - | - | 0 | - |
| - | - | 1727 | 316.2 | - | - | 0 | - |
| - | - | 1547 | 318.1 | - | - | 0 | - |
| - | - | 2343 | 318.1 | - | - | 0 | - |
| - | - | 927.5 | 320.1 | - | - | 0 | - |
| - | - | 8666 | 324.1 | - | - | 0 | - |
| - | - | 1003 | 325.1 | - | - | 0 | - |
| - | - | 548 | 326.4 | - | - | 0 | - |
| - | - | 2017 | 330.2 | - | - | 0 | - |
| - | - | 1498 | 333.2 | - | - | 0 | - |
| - | - | 1125 | 334.1 | - | - | 0 | - |
| - | - | 980.6 | 336.2 | - | - | 0 | - |
| - | - | 671.2 | 339.2 | - | - | 0 | - |
| 9 | y | 4.545E+04 | 342.2 | 0.0007028 | 2.054 | +1 | 2 |
| - | - | 7279 | 343.2 | - | - | 0 | - |
| - | - | 998.5 | 344.2 | - | - | 0 | - |
| - | - | 881.7 | 346.1 | - | - | 0 | - |
| - | - | 1523 | 348.1 | - | - | 0 | - |
| - | - | 3282 | 350.1 | - | - | 0 | - |
| - | - | 535.4 | 352.1 | - | - | 0 | - |
| - | - | 2234 | 354.2 | - | - | 0 | - |
| - | - | 783 | 355.1 | - | - | 0 | - |
| - | - | 2137 | 356.1 | - | - | 0 | - |
| - | - | 827.9 | 360.1 | - | - | 0 | - |
| - | - | 1198 | 362.1 | - | - | 0 | - |
| - | - | 4052 | 364.2 | - | - | 0 | - |
| - | - | 788.7 | 365.2 | - | - | 0 | - |
| - | - | 711.4 | 366.1 | - | - | 0 | - |
| - | - | 1347 | 376.1 | - | - | 0 | - |
| - | - | 1.329E+04 | 378.1 | - | - | 0 | - |
| 3 | b | 9651 | 378.2 | 0.0004836 | 1.279 | +1 | 3 |
| - | - | 2093 | 379.1 | - | - | 0 | - |
| - | - | 2417 | 379.2 | - | - | 0 | - |
| - | - | 1133 | 380.1 | - | - | 0 | - |
| - | - | 740.3 | 382.1 | - | - | 0 | - |
| - | - | 1109 | 390.2 | - | - | 0 | - |
| - | - | 5063 | 394.1 | - | - | 0 | - |
| - | - | 992.5 | 395.1 | - | - | 0 | - |
| - | - | 559.7 | 399.2 | - | - | 0 | - |
| - | - | 555.4 | 400.1 | - | - | 0 | - |
| - | - | 875.4 | 404.1 | - | - | 0 | - |
| - | - | 1666 | 408.2 | - | - | 0 | - |
| - | - | 1089 | 421.2 | - | - | 0 | - |
| - | - | 6972 | 425.2 | - | - | 0 | - |
| - | - | 1798 | 426.2 | - | - | 0 | - |
| - | - | 547.3 | 427.1 | - | - | 0 | - |
| - | - | 1039 | 427.2 | - | - | 0 | - |
| - | - | 735.9 | 431.2 | - | - | 0 | - |
| - | - | 736.4 | 433.2 | - | - | 0 | - |
| - | - | 733.7 | 439.2 | - | - | 0 | - |
| - | - | 657.4 | 443.2 | - | - | 0 | - |
| - | - | 675.9 | 445.1 | - | - | 0 | - |
| - | - | 836 | 449.2 | - | - | 0 | - |
| - | - | 929.7 | 455.2 | - | - | 0 | - |
| - | - | 1294 | 461.2 | - | - | 0 | - |
| - | - | 848 | 461.2 | - | - | 0 | - |
| - | - | 2672 | 465.2 | - | - | 0 | - |
| - | - | 1211 | 471.2 | - | - | 0 | - |
| - | - | 2924 | 479.2 | - | - | 0 | - |
| - | - | 728.2 | 489.2 | - | - | 0 | - |
| 8 | y | 9301 | 489.2 | 0.006168 | 12.61 | +1 | 3 |
| - | - | 862.4 | 489.2 | - | - | 0 | - |
| - | - | 2245 | 490.2 | - | - | 0 | - |
| - | - | 1049 | 491.2 | - | - | 0 | - |
| 4 | b | 909.5 | 493.2 | 0.0005486 | 1.112 | +1 | 4 |
| 8 | b | 982.9 | 495.2 | 0.002036 | 4.112 | +2 | 8 |
| - | - | 4876 | 496.2 | - | - | 0 | - |
| - | - | 1890 | 497.2 | - | - | 0 | - |
| 4 | b | 5656 | 511.2 | 0.0006346 | 1.241 | +1 | 4 |
| - | - | 1328 | 512.2 | - | - | 0 | - |
| - | - | 1646 | 513.2 | - | - | 0 | - |
| - | - | 1410 | 518.7 | - | - | 0 | - |
| - | - | 1192 | 519.2 | - | - | 0 | - |
| - | - | 763.2 | 520.2 | - | - | 0 | - |
| - | - | 815.2 | 529.7 | - | - | 0 | - |
| - | - | 730.1 | 536.2 | - | - | 0 | - |
| - | - | 789 | 537.2 | - | - | 0 | - |
| - | - | 652.9 | 538.7 | - | - | 0 | - |
| - | - | 1163 | 541.2 | - | - | 0 | - |
| - | - | 758.6 | 542.2 | - | - | 0 | - |
| 3 | y | 1.267E+04 | 550.7 | 0.003141 | 5.704 | +2 | 8 |
| - | - | 7946 | 551.2 | - | - | 0 | - |
| - | - | 2093 | 551.7 | - | - | 0 | - |
| - | - | 1593 | 552.2 | - | - | 0 | - |
| 7 | y | 1.702E+04 | 560.2 | 0.005462 | 9.749 | +1 | 4 |
| - | - | 4531 | 561.2 | - | - | 0 | - |
| - | - | 1415 | 562.2 | - | - | 0 | - |
| 9 | b | 775.8 | 563.7 | 0.002335 | 4.142 | +2 | 9 |
| - | - | 1075 | 570.2 | - | - | 0 | - |
| 9 | b | 731.6 | 572.7 | 0.002851 | 4.978 | +2 | 9 |
| - | - | 1589 | 573.2 | - | - | 0 | - |
| - | - | 1340 | 573.7 | - | - | 0 | - |
| - | - | 916.2 | 580.2 | - | - | 0 | - |
| - | - | 758.1 | 592.7 | - | - | 0 | - |
| - | - | 672.2 | 597.2 | - | - | 0 | - |
| 2 | y | 5113 | 601.2 | 0.003472 | 5.774 | +2 | 9 |
| - | - | 2754 | 601.7 | - | - | 0 | - |
| - | - | 960.6 | 602.2 | - | - | 0 | - |
| - | - | 1761 | 606.2 | - | - | 0 | - |
| 5 | b | 1387 | 608.2 | 0.0005526 | 0.9085 | +1 | 5 |
| - | - | 682.4 | 609.2 | - | - | 0 | - |
| - | - | 1543 | 612.2 | - | - | 0 | - |
| - | - | 901 | 613.2 | - | - | 0 | - |
| - | - | 848.3 | 614.2 | - | - | 0 | - |
| - | - | 1189 | 624.2 | - | - | 0 | - |
| 5 | b | 1687 | 626.2 | 0.0006127 | 0.9783 | +1 | 5 |
| - | - | 727.1 | 627.2 | - | - | 0 | - |
| - | - | 1401 | 633.8 | - | - | 0 | - |
| - | - | 1101 | 634.2 | - | - | 0 | - |
| - | - | 810.8 | 642.2 | - | - | 0 | - |
| - | - | 1614 | 642.8 | - | - | 0 | - |
| - | - | 1587 | 643.3 | - | - | 0 | - |
| - | - | 1002 | 651.8 | - | - | 0 | - |
| - | - | 807.7 | 652.8 | - | - | 0 | - |
| - | - | 759.9 | 655.8 | - | - | 0 | - |
| - | - | 701.9 | 656.8 | - | - | 0 | - |
| - | - | 2440 | 657.3 | - | - | 0 | - |
| - | - | 4809 | 659.3 | - | - | 0 | - |
| - | - | 2924 | 660.3 | - | - | 0 | - |
| 0 | Precursor | 7729 | 665.8 | 0.002929 | 4.399 | +2 | -1 |
| - | - | 5271 | 666.3 | - | - | 0 | - |
| - | - | 4111 | 666.8 | - | - | 0 | - |
| - | - | 1031 | 667.3 | - | - | 0 | - |
| 0 | Precursor | 1.356E+04 | 674.8 | 0.002834 | 4.2 | +2 | -1 |
| - | - | 1.163E+04 | 675.3 | - | - | 0 | - |
| - | - | 6031 | 675.8 | - | - | 0 | - |
| - | - | 1382 | 676.3 | - | - | 0 | - |
| - | - | 807.2 | 683.3 | - | - | 0 | - |
| - | - | 1654 | 685.3 | - | - | 0 | - |
| - | - | 680.8 | 686.3 | - | - | 0 | - |
| - | - | 2223 | 695.3 | - | - | 0 | - |
| - | - | 928.5 | 696.3 | - | - | 0 | - |
| - | - | 774.4 | 704.3 | - | - | 0 | - |
| - | - | 2421 | 721.3 | - | - | 0 | - |
| 6 | y | 1.506E+04 | 723.3 | 0.00506 | 6.996 | +1 | 5 |
| - | - | 5848 | 724.3 | - | - | 0 | - |
| - | - | 1993 | 725.3 | - | - | 0 | - |
| - | - | 1181 | 743.3 | - | - | 0 | - |
| - | - | 3116 | 749.3 | - | - | 0 | - |
| - | - | 1210 | 750.3 | - | - | 0 | - |
| - | - | 740.6 | 759.3 | - | - | 0 | - |
| 6 | b | 1014 | 771.3 | 0.003572 | 4.631 | +1 | 6 |
| - | - | 6502 | 774.3 | - | - | 0 | - |
| - | - | 2225 | 775.3 | - | - | 0 | - |
| - | - | 1104 | 787.3 | - | - | 0 | - |
| 6 | b | 1115 | 789.3 | 0.000695 | 0.8805 | +1 | 6 |
| - | - | 2490 | 804.3 | - | - | 0 | - |
| - | - | 952.2 | 805.3 | - | - | 0 | - |
| - | - | 969.7 | 821.3 | - | - | 0 | - |
| - | - | 2840 | 832.3 | - | - | 0 | - |
| - | - | 1888 | 833.3 | - | - | 0 | - |
| 5 | y | 1.889E+04 | 838.3 | 0.005827 | 6.951 | +1 | 6 |
| - | - | 9409 | 839.3 | - | - | 0 | - |
| - | - | 3033 | 840.3 | - | - | 0 | - |
| 7 | b | 1094 | 842.3 | 0.00675 | 8.014 | +1 | 7 |
| - | - | 908.3 | 843.3 | - | - | 0 | - |
| - | - | 835.6 | 850.3 | - | - | 0 | - |
| - | - | 718.9 | 861.4 | - | - | 0 | - |
| - | - | 2344 | 862.3 | - | - | 0 | - |
| - | - | 1305 | 863.3 | - | - | 0 | - |
| - | - | 2584 | 868.3 | - | - | 0 | - |
| - | - | 985.6 | 869.3 | - | - | 0 | - |
| - | - | 835 | 871.3 | - | - | 0 | - |
| - | - | 1101 | 877.4 | - | - | 0 | - |
| - | - | 1.029E+04 | 889.3 | - | - | 0 | - |
| - | - | 5375 | 890.4 | - | - | 0 | - |
| - | - | 1568 | 891.4 | - | - | 0 | - |
| - | - | 4301 | 896.3 | - | - | 0 | - |
| - | - | 2229 | 897.3 | - | - | 0 | - |
| - | - | 786.2 | 898.3 | - | - | 0 | - |
| - | - | 822 | 907.3 | - | - | 0 | - |
| - | - | 1021 | 914.3 | - | - | 0 | - |
| - | - | 672.5 | 915.4 | - | - | 0 | - |
| - | - | 912.7 | 933.4 | - | - | 0 | - |
| 4 | y | 2072 | 935.3 | 0.005806 | 6.208 | +1 | 7 |
| - | - | 1441 | 936.3 | - | - | 0 | - |
| 4 | y | 3.297E+04 | 953.3 | 0.005373 | 5.636 | +1 | 7 |
| - | - | 1.735E+04 | 954.3 | - | - | 0 | - |
| - | - | 5633 | 955.3 | - | - | 0 | - |
| - | - | 927 | 963.3 | - | - | 0 | - |
| - | - | 998.7 | 975.4 | - | - | 0 | - |
| - | - | 1178 | 997.4 | - | - | 0 | - |
| - | - | 1200 | 1018 | - | - | 0 | - |
| - | - | 1.619E+04 | 1036 | - | - | 0 | - |
| - | - | 9535 | 1037 | - | - | 0 | - |
| - | - | 3694 | 1038 | - | - | 0 | - |
| - | - | 1739 | 1042 | - | - | 0 | - |
| - | - | 836.9 | 1049 | - | - | 0 | - |
| - | - | 859.3 | 1054 | - | - | 0 | - |
| - | - | 1152 | 1058 | - | - | 0 | - |
| 3 | y | 2905 | 1082 | 0.007644 | 7.062 | +1 | 8 |
| - | - | 1451 | 1083 | - | - | 0 | - |
| 3 | y | 5.406E+04 | 1100 | 0.005624 | 5.111 | +1 | 8 |
| - | - | 3.441E+04 | 1101 | - | - | 0 | - |
| - | - | 1.373E+04 | 1102 | - | - | 0 | - |
| - | - | 2890 | 1103 | - | - | 0 | - |
| - | - | 728.4 | 1120 | - | - | 0 | - |
| - | - | 3942 | 1137 | - | - | 0 | - |
| - | - | 3408 | 1138 | - | - | 0 | - |
| 9 | b | 1617 | 1144 | 0.003946 | 3.448 | +1 | 9 |
| - | - | 1464 | 1145 | - | - | 0 | - |
| - | - | 1513 | 1147 | - | - | 0 | - |
| 2 | y | 1655 | 1183 | 0.003789 | 3.201 | +1 | 9 |
| - | - | 1267 | 1185 | - | - | 0 | - |
| 2 | y | 1.079E+04 | 1201 | 0.003844 | 3.2 | +1 | 9 |
| - | - | 6834 | 1202 | - | - | 0 | - |
| - | - | 3685 | 1203 | - | - | 0 | - |
| - | - | 1757 | 1211 | - | - | 0 | - |
| - | - | 1613 | 1212 | - | - | 0 | - |
| - | - | 706.2 | 2727 | - | - | 0 | - |
| - | - | 662.1 | 2815 | - | - | 0 | - |
| - | - | 764.7 | 3168 | - | - | 0 | - |
| - | - | 785.3 | 3363 | - | - | 0 | - |

m/z Charge Intensity FragmentType MassShift Position
120.08121490478516 0 190549.73
121.0790786743164 0 956.8506
121.0844955444336 0 15867.234
121.77378845214844 0 342.67807
122.08782196044922 0 394.6738
129.06640625 0 544.56177
129.1025390625 0 1624.3337
130.05014038085938 0 700.8511
130.06556701660156 0 1817.5863
131.04965209960938 0 793.4211
131.0690460205078 0 421.05264
132.08111572265625 0 607.37976
133.08596801757812 0 780.5003
134.02725219726562 0 1462.7249
136.07608032226562 0 25886.424
137.0794677734375 0 2176.2913
138.0552520751953 0 1162.7905
138.06655883789062 0 1950.5762
142.12298583984375 0 896.0754
143.07337951660156 0 824.3962
143.08193969726562 0 854.0723
146.06057739257812 0 1001.3824
147.0767059326172 0 892.029
148.08750915527344 0 965.74493
148.94683837890625 0 547.06415
152.07106018066406 0 521.7439
155.09312438964844 0 926.1016
156.07713317871094 0 4449.5776
157.06121826171875 0 800.24725
158.09683227539062 0 3106.362
159.09207153320312 0 8957.383
160.09530639648438 0 809.168
160.1124267578125 0 643.08185
161.09237670898438 0 2245.4038
162.09124755859375 0 578.59625
165.0773468017578 0 1252.0573
166.0615234375 0 1081.1693
166.08660888671875 0 20567.37
167.09034729003906 0 2559.232
169.0975799560547 0 510.85645
169.1339111328125 0 1254.7102
171.0768280029297 0 2178.4844
173.12887573242188 0 682.7081
175.0869140625 0 882.7157
176.0825653076172 0 1635.8091
176.10743713378906 0 78949.445
177.10264587402344 0 1439.4166
177.11093139648438 0 9383.984
180.07733154296875 0 793.7762
181.06126403808594 0 538.2008
181.1704559326172 0 940.9968
182.08152770996094 0 981.1797
185.0559539794922 0 1021.5326
186.0917205810547 0 9323.419
187.09483337402344 0 1214.6667
187.1445770263672 0 1756.707
188.0709991455078 0 5944.951
189.08737182617188 0 4249.324
189.10287475585938 0 1307.6404
191.09327697753906 0 679.0542
191.1182403564453 0 1412.7289
193.10879516601562 0 4592.947
195.0880889892578 0 1936.971
195.1124725341797 0 557.26227
197.12855529785156 0 1398.6624
201.12393188476562 0 1913.7493
202.0536651611328 0 737.39417
203.0922088623047 0 640.4776
203.11819458007812 0 5452.772 a Water loss 1
204.0770721435547 0 2734.9546
204.10240173339844 0 640.9605 a Ammonia loss 1
204.12185668945312 0 865.2163
205.09754943847656 0 4901.7925 y 9
205.1077423095703 0 834.69196
207.11328125 0 5078.579 d 1
208.11627197265625 0 879.6503
209.16482543945312 0 743.84924
212.11903381347656 0 1053.8706
214.0847930908203 0 555.90186
215.08139038085938 0 499.81772
215.13951110839844 0 1777.0424
217.08250427246094 0 1622.0752
217.09750366210938 0 6051.699
218.0634002685547 0 646.9279
218.1020050048828 0 893.50055
219.0796661376953 0 645.3872
221.10369873046875 0 12810.934
221.12893676757812 0 103973.89 a 1
222.10682678222656 0 1063.393
222.13223266601562 0 12767.774
223.1082000732422 0 2072.0828
223.1345672607422 0 826.3111
226.1185302734375 0 605.6955
227.10377502441406 0 835.1287
228.11331176757812 0 844.20087
229.11862182617188 0 1322.4606
231.04461669921875 0 712.00824
231.06155395507812 0 2114.9705
231.11328125 0 24594.014 b Water loss 1
232.1168212890625 0 2743.4617
235.0871124267578 0 526.2268
235.1082305908203 0 11080.266
236.07452392578125 0 1149.484
236.11245727539062 0 760.50714
237.159912109375 0 973.65607
239.11436462402344 0 1427.9673
240.13511657714844 0 549.9317
245.0918426513672 0 647.6817
247.10765075683594 0 901.46576 b Water loss 3
247.1447296142578 0 517.6533
249.1238250732422 0 52500.973 b 1
250.1272735595703 0 6518.4023
251.10305786132812 0 5901.504
251.12905883789062 0 647.4672
253.1187286376953 0 19211.186
254.12203979492188 0 3391.972
257.1072692871094 0 2305.4866
257.12774658203125 0 591.1461
259.108154296875 0 1087.7922
263.1029357910156 0 10507.041
263.127685546875 0 713.4136
264.1069030761719 0 1316.8207
265.1183776855469 0 1635.8386
269.1138610839844 0 1950.6289
274.13018798828125 0 2064.206
279.09796142578125 0 3701.0293
280.1010437011719 0 896.1358
281.0959167480469 0 8154.043
282.0988464355469 0 1189.6881
283.0908203125 0 842.4135
284.1614074707031 0 1026.3625
285.0447692871094 0 510.16135
285.1021728515625 0 5183.0933
286.1202697753906 0 638.5902
292.1412353515625 0 2383.5618
295.1109619140625 0 557.36237
296.15118408203125 0 4654.4663
297.15496826171875 0 1122.3656
304.1302490234375 0 1503.322
309.09088134765625 0 4422.941
312.1561279296875 0 1827.6934
316.1878356933594 0 1726.729
318.0760192871094 0 1547.05
318.1452331542969 0 2343.4082
320.1247863769531 0 927.5273
324.14617919921875 0 8665.959
325.1485290527344 0 1003.25323
326.3993835449219 0 548.0374
330.1666259765625 0 2017.1099
333.160888671875 0 1497.8436
334.13958740234375 0 1125.4612
336.1562805175781 0 980.5551
339.1696472167969 0 671.1583
342.1567687988281 0 45453.84 y 8
343.15985107421875 0 7279.0083
344.1619873046875 0 998.4918
346.1413269042969 0 881.717
348.12042236328125 0 1523.4093
350.135498046875 0 3281.8022
352.141845703125 0 535.4097
354.16619873046875 0 2233.5325
355.0698547363281 0 783.03296
356.1391296386719 0 2136.7168
360.1197509765625 0 827.85974
362.1351623535156 0 1198.0084
364.1507263183594 0 4052.3545
365.1528625488281 0 788.7356
366.1322937011719 0 711.3906
376.1145935058594 0 1346.591
378.1300048828125 0 13294.92
378.18170166015625 0 9651.042 b Water loss 2
379.13323974609375 0 2093.4412
379.18536376953125 0 2416.8708
380.145263671875 0 1133.2192
382.145751953125 0 740.31445
390.15679931640625 0 1108.9407
394.1252136230469 0 5062.9297
395.1287536621094 0 992.5262
399.1824035644531 0 559.6658
400.1425476074219 0 555.4378
404.14544677734375 0 875.42523
408.1663818359375 0 1666.1764
421.1729736328125 0 1089.2096
425.1938781738281 0 6972.4546
426.1979675292969 0 1797.5365
427.1325988769531 0 547.28674
427.2087707519531 0 1038.5496
431.2135009765625 0 735.9394
433.1701354980469 0 736.4315
439.1837158203125 0 733.73773
443.15838623046875 0 657.4476
445.1370849609375 0 675.8808
449.1675109863281 0 836.02826
455.20416259765625 0 929.69934
461.16552734375 0 1294.1545
461.1971740722656 0 847.952
465.1617126464844 0 2672.2932
471.18475341796875 0 1210.8241
479.1775207519531 0 2924.3506
489.15740966796875 0 728.2445
489.1927185058594 0 9301.478 y 7
489.23236083984375 0 862.42615
490.1955871582031 0 2244.5798
491.2063903808594 0 1049.1671
493.2087097167969 0 909.49805 b Water loss 3
495.1886901855469 0 982.9235 b Water loss 7
496.2315979003906 0 4875.7075
497.2341613769531 0 1890.1758
511.2193603515625 0 5655.8545 b 3
512.2203979492188 0 1328.2068
513.1987915039062 0 1645.6768
518.7113037109375 0 1410.4216
519.2010498046875 0 1192.3098
520.2030639648438 0 763.2371
529.708740234375 0 815.1733
536.2002563476562 0 730.1067
537.2116088867188 0 788.9801
538.7120361328125 0 652.85547
541.191650390625 0 1163.462
542.2396850585938 0 758.5716
550.71142578125 0 12674.664 y 2
551.2127685546875 0 7945.5
551.714599609375 0 2093.1877
552.2167358398438 0 1592.7877
560.2291259765625 0 17024.523 y 6
561.2322387695312 0 4530.7
562.2344970703125 0 1414.612
563.7184448242188 0 775.79333 b Water loss 8
570.22705078125 0 1074.9684
572.7242431640625 0 731.6167 b 8
573.2266235351562 0 1588.5603
573.7267456054688 0 1340.4231
580.2479858398438 0 916.1895
592.7302856445312 0 758.14685
597.2144165039062 0 672.24066
601.235595703125 0 5112.6235 y 1
601.7366943359375 0 2753.698
602.236328125 0 960.64374
606.2323608398438 0 1760.9548
608.2356567382812 0 1386.5039 b Water loss 4
609.2346801757812 0 682.4009
612.1943969726562 0 1542.6388
613.2402954101562 0 901.0383
614.2464599609375 0 848.308
624.2299194335938 0 1188.5005
626.2450561523438 0 1687.0068 b 4
627.2478637695312 0 727.1098
633.7652587890625 0 1401.4421
634.2271728515625 0 1100.6798
642.2351684570312 0 810.7595
642.7684936523438 0 1614.4924
643.2683715820312 0 1586.8589
651.7651977539062 0 1001.78503
652.7689208984375 0 807.72473
655.760009765625 0 759.85406
656.7588500976562 0 701.8618
657.2626342773438 0 2440.3008
659.294677734375 0 4808.91
660.2963256835938 0 2923.6821
665.7639770507812 0 7728.727 Precursor Water loss
666.2654418945312 0 5270.824
666.7651977539062 0 4110.846
667.2711791992188 0 1030.8817
674.7691650390625 0 13559.38 Precursor
675.27099609375 0 11626.52
675.7700805664062 0 6031.0107
676.2664794921875 0 1381.7372
683.3278198242188 0 807.17377
685.2583618164062 0 1653.5967
686.2591552734375 0 680.7854
695.2666625976562 0 2222.5364
696.2737426757812 0 928.48505
704.2738647460938 0 774.40137
721.2615356445312 0 2421.0627
723.2920532226562 0 15063.749 y 5
724.2953491210938 0 5848.107
725.2952270507812 0 1992.6564
743.3004760742188 0 1180.5514
749.256591796875 0 3115.94
750.260498046875 0 1210.4955
759.26318359375 0 740.5737
771.2948608398438 0 1013.6597 b Water loss 5
774.3209838867188 0 6502.269
775.3240356445312 0 2224.889
787.3117065429688 0 1103.9562
789.3096923828125 0 1115.345 b 5
804.3311767578125 0 2490.2808
805.3316040039062 0 952.2044
821.3016357421875 0 969.68384
832.3278198242188 0 2839.7544
833.3294067382812 0 1887.8722
838.3197631835938 0 18894.268 y 4
839.3230590820312 0 9408.775
840.32275390625 0 3032.8267
842.3287963867188 0 1094.2424 b Water loss 6
843.3392333984375 0 908.3137
850.3291625976562 0 835.6215
861.3519287109375 0 718.90546
862.292724609375 0 2343.8774
863.292236328125 0 1304.9141
868.33056640625 0 2583.5312
869.332763671875 0 985.63586
871.3372192382812 0 834.97766
877.4158935546875 0 1100.7714
889.3475952148438 0 10285.072
890.3507080078125 0 5374.641
891.3545532226562 0 1567.5532
896.3250122070312 0 4300.9194
897.3280639648438 0 2228.7708
898.32275390625 0 786.21967
907.3433837890625 0 821.9825
914.3436279296875 0 1021.2774
915.3619384765625 0 672.51965
933.370361328125 0 912.6687
935.3361206054688 0 2072.2446 y Water loss 3
936.336669921875 0 1441.1936
953.3462524414062 0 32968.195 y 3
954.3489379882812 0 17345.156
955.3496704101562 0 5632.7354
963.3421630859375 0 926.97833
975.384765625 0 998.70605
997.3753051757812 0 1177.9918
1018.4052734375 0 1199.7533
1036.4163818359375 0 16191.339
1037.4183349609375 0 9534.674
1038.42041015625 0 3694.1765
1042.4326171875 0 1739.486
1049.4254150390625 0 836.866
1054.4017333984375 0 859.3022
1058.41748046875 0 1152.2897
1082.4063720703125 0 2904.6543 y Water loss 2
1083.4091796875 0 1450.5292
1100.4149169921875 0 54061.24 y 2
1101.417236328125 0 34413.64
1102.4176025390625 0 13727.125
1103.4185791015625 0 2889.6362
1120.4583740234375 0 728.40155
1137.464111328125 0 3941.8965
1138.466552734375 0 3408.204
1144.439453125 0 1616.807 b 8
1145.44970703125 0 1463.7361
1147.44775390625 0 1512.8575
1183.4501953125 0 1655.2899 y Water loss 1
1185.4390869140625 0 1266.9038
1201.4608154296875 0 10792.179 y 1
1202.46435546875 0 6833.7764
1203.4617919921875 0 3685.09
1211.447509765625 0 1757.173
1212.449951171875 0 1613.2766
2727.266845703125 0 706.2046
2815.0615234375 0 662.05316
3168.0732421875 0 764.73267
3363.032958984375 0 785.3424

Spectrum Details

|  |  |
| --- | --- |
| Matched peaks? Matched peaks The total absolute number of peaks matched. Additionally in brackets the total fraction of peaks matched and the total number of peaks is shown. | 35 (9.92% of 353) |
| FDR? FDR The false discovery rate estimated for this peptide. It is calculated by matching all theoretical fragments with a non-integer shift with the raw peaks for this spectrum. This is done with 40 different shifts. The resulting percentage is the average number of annotated peaks over the number of annotated peaks with the correct spectrum. | 0.00% |
| Satellite FDR? Satellite FDR See the FDR for details on its calculation. This satellite ion specific FDR only contains the satellite ions (d/w) for I/L/J positions. | - |
| PSM Score? PSM Score The PSM Score as given by Hecklib to this annotated spectrum. It is shown with three significant figures. | 397 |

#### Spectrum 9898? Spectrum 9898 The raw spectrum of this peptide as annotated by Hecklib. The fragments are coloured according to ion type (see legend). Any peaks with a star '\*' as text can be hovered over to see the full details, first the ion type second the mass shift type. By hovering over the amino acids in the peptide or ions in the legend the corresponding peaks are highlighted. By toggling the 'Unassigned' label you can turn the background (unassigned) peaks on or off in the plot. By updating the slider in the Ion legend you can update the spectrum to only show the top X% of the peaks with labels. The top X% means any peak that is within X% of the highest intensity. By dragging in the spectrum you can zoom in to a specific part of the spectrum and use 'Zoom Out' to get back to the original zoom level. The annotation of the spectrum is based on the given sequence in the peptides file and is done with different software so inconsistencies are likely. The peaks are annotated based on the given sequence, with 20 ppm tolerance.

Copy Data

##### Spectrum 9898 (TSV)

###### Preview

```
Loading example...
```

*Click on the button to copy the data to your clipboard.*

Mz MinMz MaxIntensity Max

WidthHeightPeptide font sizePeptide stroke widthSpectrum font sizeSpectrum stroke widthCompact peptide

Ion legend

wxyz

abcd

OtherUnassignedIon^Charge^_Position_Show for top:%

FTFDDYAMHW

01.92e+43.83e+45.75e+47.67e+4

Zoom Out

c^+1^_2_y^+1^_2_z^+1^_3_y^+1^_3_c^+1^_4_z^+1^_4_y^+2^_8_y^+1^_4_y^+2^_9_c^+1^_5_z^+1^_5_y^+1^_5_z^+1^_6_y^+1^_6_c^+1^_7_z^+1^_7_y^+1^_7_c^+1^_8_y^+1^_8_z^+1^_8_y^+1^_8_c^+1^_9_z^+1^_9_y^+1^_9_

0860172025803440

Fragment Matches Table

Show background peaks

| Position | Ion type | Intensity | mz Theoretical | mz Error (Th) | mz Error (ppm) | Charge | Series Number |
| --- | --- | --- | --- | --- | --- | --- | --- |
| - | - | 7170 | 120.1 | - | - | 0 | - |
| - | - | 655.5 | 148.9 | - | - | 0 | - |
| - | - | 524.4 | 148.9 | - | - | 0 | - |
| - | - | 823.7 | 148.9 | - | - | 0 | - |
| - | - | 549.4 | 148.9 | - | - | 0 | - |
| - | - | 840.4 | 148.9 | - | - | 0 | - |
| - | - | 1840 | 148.9 | - | - | 0 | - |
| - | - | 2351 | 148.9 | - | - | 0 | - |
| - | - | 5501 | 149 | - | - | 0 | - |
| - | - | 2625 | 149 | - | - | 0 | - |
| - | - | 1616 | 149 | - | - | 0 | - |
| - | - | 938.5 | 149 | - | - | 0 | - |
| - | - | 720.8 | 149 | - | - | 0 | - |
| - | - | 670.8 | 149 | - | - | 0 | - |
| - | - | 445.8 | 149 | - | - | 0 | - |
| - | - | 584 | 149 | - | - | 0 | - |
| - | - | 563 | 149 | - | - | 0 | - |
| - | - | 638.7 | 149 | - | - | 0 | - |
| - | - | 417.1 | 151.6 | - | - | 0 | - |
| - | - | 2121 | 176.1 | - | - | 0 | - |
| - | - | 492 | 198.7 | - | - | 0 | - |
| - | - | 1.009E+04 | 221.1 | - | - | 0 | - |
| - | - | 841.2 | 222.1 | - | - | 0 | - |
| - | - | 1956 | 231.1 | - | - | 0 | - |
| - | - | 550.3 | 232.7 | - | - | 0 | - |
| - | - | 724.5 | 235.1 | - | - | 0 | - |
| 2 | c | 9595 | 249.1 | 0.0001053 | 0.4226 | +1 | 2 |
| - | - | 1064 | 250.1 | - | - | 0 | - |
| - | - | 1696 | 253.1 | - | - | 0 | - |
| - | - | 1081 | 263.1 | - | - | 0 | - |
| 9 | y | 1723 | 342.2 | 9.061E-05 | 0.2648 | +1 | 2 |
| - | - | 557.3 | 365 | - | - | 0 | - |
| - | - | 1458 | 378.2 | - | - | 0 | - |
| 8 | z | 1128 | 473.2 | 0.005971 | 12.62 | +1 | 3 |
| - | - | 2817 | 474.2 | - | - | 0 | - |
| 8 | y | 1118 | 489.2 | 0.006229 | 12.73 | +1 | 3 |
| 4 | c | 686.8 | 511.2 | 0.002722 | 5.325 | +1 | 4 |
| - | - | 603.4 | 533.2 | - | - | 0 | - |
| 7 | z | 4154 | 544.2 | 0.005143 | 9.45 | +1 | 4 |
| - | - | 2170 | 545.2 | - | - | 0 | - |
| - | - | 621.3 | 546.2 | - | - | 0 | - |
| 3 | y | 3462 | 550.7 | 0.002287 | 4.152 | +2 | 8 |
| - | - | 1018 | 551.2 | - | - | 0 | - |
| - | - | 999.6 | 551.7 | - | - | 0 | - |
| 7 | y | 2282 | 560.2 | 0.003997 | 7.134 | +1 | 4 |
| - | - | 627.1 | 561.2 | - | - | 0 | - |
| 2 | y | 1089 | 601.2 | 0.001519 | 2.526 | +2 | 9 |
| 5 | c | 1160 | 626.2 | 0.0001808 | 0.2887 | +1 | 5 |
| - | - | 632.4 | 633.8 | - | - | 0 | - |
| - | - | 1933 | 665.8 | - | - | 0 | - |
| - | - | 915.5 | 666.3 | - | - | 0 | - |
| - | - | 2915 | 674.8 | - | - | 0 | - |
| - | - | 3571 | 675.3 | - | - | 0 | - |
| - | - | 1660 | 675.8 | - | - | 0 | - |
| 6 | z | 8175 | 707.3 | 0.004497 | 6.359 | +1 | 5 |
| - | - | 5585 | 708.3 | - | - | 0 | - |
| - | - | 1461 | 709.3 | - | - | 0 | - |
| 6 | y | 3087 | 723.3 | 0.004816 | 6.659 | +1 | 5 |
| - | - | 1056 | 724.3 | - | - | 0 | - |
| - | - | 817.1 | 750.3 | - | - | 0 | - |
| - | - | 703.3 | 751.3 | - | - | 0 | - |
| - | - | 613.6 | 761.3 | - | - | 0 | - |
| - | - | 599.7 | 761.8 | - | - | 0 | - |
| - | - | 856.5 | 763.3 | - | - | 0 | - |
| - | - | 1374 | 764.3 | - | - | 0 | - |
| - | - | 999.6 | 765.3 | - | - | 0 | - |
| - | - | 4273 | 778.3 | - | - | 0 | - |
| - | - | 2096 | 779.3 | - | - | 0 | - |
| - | - | 576 | 782.4 | - | - | 0 | - |
| 5 | z | 1.119E+04 | 822.3 | 0.004837 | 5.882 | +1 | 6 |
| - | - | 1.197E+04 | 823.3 | - | - | 0 | - |
| - | - | 5158 | 824.3 | - | - | 0 | - |
| - | - | 1292 | 825.3 | - | - | 0 | - |
| - | - | 599.3 | 830.3 | - | - | 0 | - |
| 5 | y | 4543 | 838.3 | 0.005095 | 6.077 | +1 | 6 |
| - | - | 1662 | 839.3 | - | - | 0 | - |
| - | - | 901 | 840.3 | - | - | 0 | - |
| - | - | 1006 | 849.3 | - | - | 0 | - |
| 7 | c | 1787 | 877.4 | 0.000773 | 0.8811 | +1 | 7 |
| - | - | 1066 | 889.3 | - | - | 0 | - |
| - | - | 4744 | 893.3 | - | - | 0 | - |
| - | - | 2041 | 894.3 | - | - | 0 | - |
| - | - | 626.8 | 896.3 | - | - | 0 | - |
| - | - | 701.4 | 918.4 | - | - | 0 | - |
| 4 | z | 5581 | 937.3 | 0.004994 | 5.327 | +1 | 7 |
| - | - | 9419 | 938.3 | - | - | 0 | - |
| - | - | 4351 | 939.3 | - | - | 0 | - |
| - | - | 2086 | 940.3 | - | - | 0 | - |
| 4 | y | 6189 | 953.3 | 0.00458 | 4.804 | +1 | 7 |
| - | - | 3817 | 954.3 | - | - | 0 | - |
| - | - | 728.8 | 955.3 | - | - | 0 | - |
| - | - | 3262 | 980.4 | - | - | 0 | - |
| - | - | 2091 | 981.4 | - | - | 0 | - |
| - | - | 1097 | 982.4 | - | - | 0 | - |
| - | - | 623 | 1012 | - | - | 0 | - |
| - | - | 7707 | 1023 | - | - | 0 | - |
| 8 | c | 1.185E+04 | 1024 | 0.002861 | 2.793 | +1 | 8 |
| - | - | 5290 | 1025 | - | - | 0 | - |
| - | - | 1959 | 1026 | - | - | 0 | - |
| - | - | 2197 | 1036 | - | - | 0 | - |
| - | - | 1037 | 1037 | - | - | 0 | - |
| - | - | 902.1 | 1040 | - | - | 0 | - |
| 3 | y | 807.2 | 1082 | 0.003098 | 2.862 | +1 | 8 |
| 3 | z | 934.1 | 1084 | 0.003108 | 2.866 | +1 | 8 |
| - | - | 916.4 | 1085 | - | - | 0 | - |
| - | - | 1929 | 1093 | - | - | 0 | - |
| - | - | 1317 | 1094 | - | - | 0 | - |
| 3 | y | 1.3E+04 | 1100 | 0.004648 | 4.223 | +1 | 8 |
| - | - | 8048 | 1101 | - | - | 0 | - |
| - | - | 2989 | 1102 | - | - | 0 | - |
| - | - | 2887 | 1117 | - | - | 0 | - |
| - | - | 2235 | 1118 | - | - | 0 | - |
| - | - | 1332 | 1119 | - | - | 0 | - |
| - | - | 2136 | 1139 | - | - | 0 | - |
| - | - | 1641 | 1140 | - | - | 0 | - |
| - | - | 1207 | 1141 | - | - | 0 | - |
| - | - | 778.2 | 1146 | - | - | 0 | - |
| 9 | c | 1.998E+04 | 1161 | 0.00413 | 3.556 | +1 | 9 |
| - | - | 1.351E+04 | 1162 | - | - | 0 | - |
| - | - | 7222 | 1163 | - | - | 0 | - |
| - | - | 1918 | 1180 | - | - | 0 | - |
| - | - | 1845 | 1181 | - | - | 0 | - |
| - | - | 780.6 | 1182 | - | - | 0 | - |
| - | - | 3161 | 1184 | - | - | 0 | - |
| 2 | z | 6383 | 1185 | 0.001816 | 1.532 | +1 | 9 |
| - | - | 3419 | 1186 | - | - | 0 | - |
| - | - | 1497 | 1187 | - | - | 0 | - |
| 2 | y | 2323 | 1201 | 0.001159 | 0.9643 | +1 | 9 |
| - | - | 1639 | 1202 | - | - | 0 | - |
| - | - | 1035 | 1203 | - | - | 0 | - |
| - | - | 657.9 | 1232 | - | - | 0 | - |
| - | - | 872.5 | 1270 | - | - | 0 | - |
| - | - | 2156 | 1271 | - | - | 0 | - |
| - | - | 1627 | 1272 | - | - | 0 | - |
| - | - | 944.2 | 1273 | - | - | 0 | - |
| - | - | 1115 | 1287 | - | - | 0 | - |
| - | - | 7579 | 1288 | - | - | 0 | - |
| - | - | 1.307E+04 | 1289 | - | - | 0 | - |
| - | - | 1.674E+04 | 1290 | - | - | 0 | - |
| - | - | 1.002E+04 | 1291 | - | - | 0 | - |
| - | - | 2718 | 1292 | - | - | 0 | - |
| - | - | 1375 | 1305 | - | - | 0 | - |
| - | - | 1294 | 1306 | - | - | 0 | - |
| - | - | 841.1 | 1313 | - | - | 0 | - |
| - | - | 1142 | 1314 | - | - | 0 | - |
| - | - | 1218 | 1314 | - | - | 0 | - |
| - | - | 924.4 | 1318 | - | - | 0 | - |
| - | - | 1036 | 1329 | - | - | 0 | - |
| - | - | 3217 | 1330 | - | - | 0 | - |
| - | - | 2226 | 1331 | - | - | 0 | - |
| - | - | 3438 | 1332 | - | - | 0 | - |
| - | - | 4.427E+04 | 1333 | - | - | 0 | - |
| - | - | 3.289E+04 | 1334 | - | - | 0 | - |
| - | - | 1.652E+04 | 1335 | - | - | 0 | - |
| - | - | 3520 | 1336 | - | - | 0 | - |
| - | - | 1139 | 1347 | - | - | 0 | - |
| - | - | 1270 | 1348 | - | - | 0 | - |
| - | - | 2.086E+04 | 1349 | - | - | 0 | - |
| - | - | 1196 | 1349 | - | - | 0 | - |
| - | - | 7.594E+04 | 1350 | - | - | 0 | - |
| - | - | 5.423E+04 | 1351 | - | - | 0 | - |
| - | - | 1941 | 1351 | - | - | 0 | - |
| - | - | 2.566E+04 | 1352 | - | - | 0 | - |
| - | - | 1112 | 1352 | - | - | 0 | - |
| - | - | 4768 | 1353 | - | - | 0 | - |
| - | - | 749 | 1368 | - | - | 0 | - |
| - | - | 709 | 3406 | - | - | 0 | - |

m/z Charge Intensity FragmentType MassShift Position
120.08091735839844 0 7169.822
148.90188598632812 0 655.4995
148.90765380859375 0 524.3969
148.91903686523438 0 823.7428
148.92495727539062 0 549.3832
148.93038940429688 0 840.38586
148.93600463867188 0 1839.7638
148.94235229492188 0 2350.7473
148.95526123046875 0 5500.8687
148.96102905273438 0 2625.3086
148.967529296875 0 1616.1512
148.97299194335938 0 938.53687
148.97869873046875 0 720.7964
148.9841766357422 0 670.8211
148.99534606933594 0 445.75507
149.0013427734375 0 583.9905
149.00711059570312 0 563.03906
149.0188446044922 0 638.72375
151.60964965820312 0 417.0529
176.1071014404297 0 2121.1802
198.73123168945312 0 491.9855
221.12843322753906 0 10094.701
222.13217163085938 0 841.18866
231.11260986328125 0 1955.8555
232.67901611328125 0 550.337
235.10755920410156 0 724.4744
249.12347412109375 0 9595.219 c Ammonia loss 1
250.1261749267578 0 1063.7766
253.11825561523438 0 1695.9036
263.10272216796875 0 1080.652
342.1559753417969 0 1723.4357 y 8
365.01513671875 0 557.3126
378.1822204589844 0 1458.4767
473.1737976074219 0 1128.0433 z 7
474.1804504394531 0 2816.696
489.1927795410156 0 1117.8184 y 7
511.21600341796875 0 686.80597 c Ammonia loss 3
533.1700439453125 0 603.4394
544.2100830078125 0 4154.1963 z 6
545.2142333984375 0 2169.8447
546.21240234375 0 621.27686
550.7105712890625 0 3461.9456 y 2
551.2116088867188 0 1018.0018
551.712158203125 0 999.63574
560.2276611328125 0 2281.7712 y 6
561.2313842773438 0 627.0563
601.233642578125 0 1089.4623 y 1
626.245849609375 0 1159.9957 c Ammonia loss 4
633.763671875 0 632.35925
665.7630615234375 0 1932.7561
666.2688598632812 0 915.5425
674.7689208984375 0 2915.149
675.269287109375 0 3571.0786
675.7702026367188 0 1660.4614
707.2727661132812 0 8174.7227 z 5
708.2777709960938 0 5585.2964
709.2781372070312 0 1461.1276
723.2918090820312 0 3087.2812 y 5
724.2955932617188 0 1056.2794
750.2796630859375 0 817.1055
751.2767944335938 0 703.3434
761.3206176757812 0 613.6385
761.7706298828125 0 599.69104
763.2813110351562 0 856.4926
764.2879028320312 0 1374.1069
765.2948608398438 0 999.64484
778.3101196289062 0 4273.4824
779.3125610351562 0 2096.2014
782.419921875 0 575.9595
822.300048828125 0 11186.674 z 4
823.3057861328125 0 11972.698
824.3076782226562 0 5158.1562
825.3096923828125 0 1291.8729
830.3358764648438 0 599.2611
838.3190307617188 0 4543.4014 y 4
839.32177734375 0 1661.7367
840.3247680664062 0 900.9827
849.3474731445312 0 1005.9958
877.3718872070312 0 1787.2163 c 6
889.3450317382812 0 1066.4725
893.336181640625 0 4744.399
894.3394775390625 0 2041.4102
896.3184814453125 0 626.822
918.4082641601562 0 701.35956
937.3271484375 0 5581.3228 z 3
938.3323364257812 0 9419.37
939.3363647460938 0 4350.9795
940.3392333984375 0 2085.847
953.345458984375 0 6188.6426 y 3
954.3475952148438 0 3816.5168
955.348388671875 0 728.81793
980.392333984375 0 3262.1873
981.400634765625 0 2090.6318
982.3984985351562 0 1096.8824
1012.3809814453125 0 623.0157
1023.3990478515625 0 7707.4614
1024.406005859375 0 11848.692 c 7
1025.408447265625 0 5289.8257
1026.411376953125 0 1958.8276
1036.41015625 0 2196.8267
1037.414794921875 0 1037.4922
1040.415771484375 0 902.1478
1082.3956298828125 0 807.19763 y Water loss 2
1084.3936767578125 0 934.0948 z 2
1085.4017333984375 0 916.36615
1093.4493408203125 0 1929.0411
1094.453125 0 1317.035
1100.4139404296875 0 13001.612 y 2
1101.4176025390625 0 8048.0244
1102.4173583984375 0 2988.6272
1117.451416015625 0 2887.0068
1118.4559326171875 0 2235.1619
1119.4658203125 0 1331.6571
1139.4383544921875 0 2135.504
1140.4417724609375 0 1640.8201
1141.4498291015625 0 1207.4609
1146.4581298828125 0 778.19946
1161.4661865234375 0 19980.896 c 8
1162.4686279296875 0 13507.742
1163.469970703125 0 7221.8696
1180.4815673828125 0 1917.8693
1181.48681640625 0 1844.9965
1182.487548828125 0 780.6369
1184.4334716796875 0 3161.2078
1185.4400634765625 0 6382.971 z 1
1186.44287109375 0 3419.485
1187.44775390625 0 1497.1001
1201.4581298828125 0 2323.0051 y 1
1202.460693359375 0 1638.7882
1203.4647216796875 0 1034.9747
1232.296142578125 0 657.9404
1269.5123291015625 0 872.51794
1270.521240234375 0 2156.3027
1271.515869140625 0 1626.8629
1272.5125732421875 0 944.2186
1286.530517578125 0 1114.9486
1287.5361328125 0 7579.4893
1288.5279541015625 0 13069.361
1289.5201416015625 0 16742.428
1290.5185546875 0 10020.826
1291.520751953125 0 2717.6587
1304.5343017578125 0 1375.4225
1305.534423828125 0 1293.7804
1312.5076904296875 0 841.07007
1313.508544921875 0 1141.7405
1314.48779296875 0 1218.4896
1317.5015869140625 0 924.39746
1328.534423828125 0 1035.8478
1329.537841796875 0 3217.042
1330.533203125 0 2226.3887
1331.5264892578125 0 3438.4624
1332.513427734375 0 44273.99
1333.514892578125 0 32887.95
1334.51611328125 0 16523.537
1335.5179443359375 0 3520.3862
1346.5267333984375 0 1139.1019
1347.52197265625 0 1269.9956
1348.5279541015625 0 20857.064
1348.7119140625 0 1195.8138
1349.5345458984375 0 75939.71
1350.5369873046875 0 54230.336
1350.68017578125 0 1940.5687
1351.5391845703125 0 25659.838
1351.701904296875 0 1112.0074
1352.5400390625 0 4767.618
1368.32177734375 0 748.995
3405.544921875 0 708.99207

Spectrum Details

|  |  |
| --- | --- |
| Matched peaks? Matched peaks The total absolute number of peaks matched. Additionally in brackets the total fraction of peaks matched and the total number of peaks is shown. | 24 (14.37% of 167) |
| FDR? FDR The false discovery rate estimated for this peptide. It is calculated by matching all theoretical fragments with a non-integer shift with the raw peaks for this spectrum. This is done with 40 different shifts. The resulting percentage is the average number of annotated peaks over the number of annotated peaks with the correct spectrum. | 0.00% |
| Satellite FDR? Satellite FDR See the FDR for details on its calculation. This satellite ion specific FDR only contains the satellite ions (d/w) for I/L/J positions. | - |
| PSM Score? PSM Score The PSM Score as given by Hecklib to this annotated spectrum. It is shown with three significant figures. | 302 |

#### Spectrum 8533? Spectrum 8533 The raw spectrum of this peptide as annotated by Hecklib. The fragments are coloured according to ion type (see legend). Any peaks with a star '\*' as text can be hovered over to see the full details, first the ion type second the mass shift type. By hovering over the amino acids in the peptide or ions in the legend the corresponding peaks are highlighted. By toggling the 'Unassigned' label you can turn the background (unassigned) peaks on or off in the plot. By updating the slider in the Ion legend you can update the spectrum to only show the top X% of the peaks with labels. The top X% means any peak that is within X% of the highest intensity. By dragging in the spectrum you can zoom in to a specific part of the spectrum and use 'Zoom Out' to get back to the original zoom level. The annotation of the spectrum is based on the given sequence in the peptides file and is done with different software so inconsistencies are likely. The peaks are annotated based on the given sequence, with 20 ppm tolerance.

Copy Data

##### Spectrum 8533 (TSV)

###### Preview

```
Loading example...
```

*Click on the button to copy the data to your clipboard.*

Mz MinMz MaxIntensity Max

WidthHeightPeptide font sizePeptide stroke widthSpectrum font sizeSpectrum stroke widthCompact peptide

Ion legend

wxyz

abcd

OtherUnassignedIon^Charge^_Position_Show for top:%

FTFDDYAMHW

04.63e+49.26e+41.39e+51.85e+5

Zoom Out

a^+1^_2_d^+1^_2_a^+1^_2_b^+1^_2_b^+1^_2_a^+1^_3_a^+1^_3_y^+1^_2_y^+2^_5_b^+1^_3_b^+1^_4_b^+2^_8_y^+1^_3_b^+1^_4_y^+2^_8_y^+2^_8_b^+2^_9_y^+1^_4_y^+2^_9_b^+1^_5_y^+2^_9_b^+1^_5_\*\*y^+1^_5_b^+1^_6_b^+1^_6_y^+1^_6_b^+1^_7_y^+1^_6_b^+1^_7_y^+1^_7_y^+1^_7_b^+1^_8_y^+1^_8_y^+1^_8_b^+1^_9_y^+1^_9_y^+1^_9_

0776155323293105

Fragment Matches Table

Show background peaks

| Position | Ion type | Intensity | mz Theoretical | mz Error (Th) | mz Error (ppm) | Charge | Series Number |
| --- | --- | --- | --- | --- | --- | --- | --- |
| - | - | 1.834E+05 | 120.1 | - | - | 0 | - |
| - | - | 1.46E+04 | 121.1 | - | - | 0 | - |
| - | - | 399.8 | 121.7 | - | - | 0 | - |
| - | - | 563.2 | 122.1 | - | - | 0 | - |
| - | - | 2053 | 129.1 | - | - | 0 | - |
| - | - | 1626 | 130.1 | - | - | 0 | - |
| - | - | 1202 | 131.1 | - | - | 0 | - |
| - | - | 410.3 | 132.1 | - | - | 0 | - |
| - | - | 564.8 | 132.1 | - | - | 0 | - |
| - | - | 1.741E+04 | 133.1 | - | - | 0 | - |
| - | - | 3814 | 134.1 | - | - | 0 | - |
| - | - | 1.336E+04 | 136.1 | - | - | 0 | - |
| - | - | 601.5 | 136.1 | - | - | 0 | - |
| - | - | 1153 | 137.1 | - | - | 0 | - |
| - | - | 1141 | 138.1 | - | - | 0 | - |
| - | - | 1686 | 138.1 | - | - | 0 | - |
| - | - | 482.2 | 143.1 | - | - | 0 | - |
| - | - | 450 | 146.1 | - | - | 0 | - |
| - | - | 495.8 | 148.1 | - | - | 0 | - |
| - | - | 596.9 | 149 | - | - | 0 | - |
| - | - | 2276 | 149 | - | - | 0 | - |
| - | - | 423.3 | 150.9 | - | - | 0 | - |
| - | - | 433.3 | 152.6 | - | - | 0 | - |
| - | - | 5683 | 156.1 | - | - | 0 | - |
| - | - | 523.7 | 157.1 | - | - | 0 | - |
| - | - | 1027 | 158.1 | - | - | 0 | - |
| - | - | 4250 | 158.1 | - | - | 0 | - |
| - | - | 4892 | 159.1 | - | - | 0 | - |
| - | - | 860.8 | 159.1 | - | - | 0 | - |
| - | - | 512.3 | 160.1 | - | - | 0 | - |
| - | - | 1075 | 161.1 | - | - | 0 | - |
| - | - | 517.9 | 164.1 | - | - | 0 | - |
| - | - | 1605 | 166.1 | - | - | 0 | - |
| - | - | 694.5 | 166.1 | - | - | 0 | - |
| - | - | 3160 | 167.1 | - | - | 0 | - |
| - | - | 1523 | 167.1 | - | - | 0 | - |
| - | - | 1153 | 169.1 | - | - | 0 | - |
| - | - | 575.5 | 171.1 | - | - | 0 | - |
| - | - | 2620 | 173.4 | - | - | 0 | - |
| - | - | 689.4 | 174.1 | - | - | 0 | - |
| - | - | 1.049E+04 | 175.1 | - | - | 0 | - |
| - | - | 1378 | 175.1 | - | - | 0 | - |
| - | - | 628.2 | 175.1 | - | - | 0 | - |
| - | - | 2156 | 176.1 | - | - | 0 | - |
| - | - | 8.443E+04 | 176.1 | - | - | 0 | - |
| - | - | 1691 | 177.1 | - | - | 0 | - |
| - | - | 1.267E+04 | 177.1 | - | - | 0 | - |
| - | - | 2912 | 178.1 | - | - | 0 | - |
| - | - | 532 | 180.1 | - | - | 0 | - |
| - | - | 553.2 | 181.6 | - | - | 0 | - |
| - | - | 1.045E+04 | 186.1 | - | - | 0 | - |
| - | - | 1738 | 187.1 | - | - | 0 | - |
| - | - | 446 | 187.1 | - | - | 0 | - |
| - | - | 1905 | 191.1 | - | - | 0 | - |
| - | - | 6131 | 193.1 | - | - | 0 | - |
| - | - | 541.7 | 195 | - | - | 0 | - |
| - | - | 1374 | 197.1 | - | - | 0 | - |
| - | - | 502.6 | 199.9 | - | - | 0 | - |
| - | - | 1201 | 203.1 | - | - | 0 | - |
| - | - | 2481 | 203.1 | - | - | 0 | - |
| 2 | a | 6718 | 203.1 | 0.0002135 | 1.051 | +1 | 2 |
| - | - | 3528 | 204.1 | - | - | 0 | - |
| - | - | 1011 | 204.1 | - | - | 0 | - |
| - | - | 538.8 | 205.1 | - | - | 0 | - |
| - | - | 6379 | 205.1 | - | - | 0 | - |
| - | - | 2739 | 206.1 | - | - | 0 | - |
| 2 | d | 1649 | 207.1 | 0.0002553 | 1.233 | +1 | 2 |
| - | - | 1494 | 212.1 | - | - | 0 | - |
| - | - | 781.3 | 214.1 | - | - | 0 | - |
| - | - | 5351 | 217.1 | - | - | 0 | - |
| - | - | 1338 | 218.1 | - | - | 0 | - |
| - | - | 1079 | 219.1 | - | - | 0 | - |
| - | - | 526.3 | 219.1 | - | - | 0 | - |
| - | - | 2.36E+04 | 221.1 | - | - | 0 | - |
| - | - | 1.348E+04 | 221.1 | - | - | 0 | - |
| 2 | a | 1.088E+05 | 221.1 | 0.0001926 | 0.8711 | +1 | 2 |
| - | - | 632.6 | 222.1 | - | - | 0 | - |
| - | - | 2108 | 222.1 | - | - | 0 | - |
| - | - | 682.6 | 222.1 | - | - | 0 | - |
| - | - | 1.326E+04 | 222.1 | - | - | 0 | - |
| - | - | 1115 | 222.1 | - | - | 0 | - |
| - | - | 4973 | 225 | - | - | 0 | - |
| - | - | 626.2 | 226 | - | - | 0 | - |
| - | - | 906.4 | 228.1 | - | - | 0 | - |
| - | - | 2559 | 231.1 | - | - | 0 | - |
| 2 | b | 2.87E+04 | 231.1 | 0.0002177 | 0.9419 | +1 | 2 |
| - | - | 3746 | 232.1 | - | - | 0 | - |
| - | - | 7893 | 235.1 | - | - | 0 | - |
| - | - | 912.4 | 236.1 | - | - | 0 | - |
| - | - | 7033 | 239.1 | - | - | 0 | - |
| - | - | 682.8 | 239.1 | - | - | 0 | - |
| - | - | 1214 | 240.1 | - | - | 0 | - |
| - | - | 540.2 | 248.2 | - | - | 0 | - |
| 2 | b | 5.715E+04 | 249.1 | 0.0002426 | 0.9738 | +1 | 2 |
| - | - | 6390 | 250.1 | - | - | 0 | - |
| - | - | 547.8 | 250.2 | - | - | 0 | - |
| - | - | 5386 | 251.1 | - | - | 0 | - |
| - | - | 793.6 | 251.1 | - | - | 0 | - |
| - | - | 568.1 | 253.1 | - | - | 0 | - |
| - | - | 1815 | 257.1 | - | - | 0 | - |
| - | - | 1231 | 259.1 | - | - | 0 | - |
| - | - | 1.195E+04 | 263.1 | - | - | 0 | - |
| - | - | 1469 | 264.1 | - | - | 0 | - |
| - | - | 701.7 | 265.2 | - | - | 0 | - |
| - | - | 2467 | 274.1 | - | - | 0 | - |
| - | - | 559 | 277.5 | - | - | 0 | - |
| - | - | 5075 | 279.1 | - | - | 0 | - |
| - | - | 1275 | 285 | - | - | 0 | - |
| - | - | 5718 | 285.1 | - | - | 0 | - |
| - | - | 731.8 | 287.1 | - | - | 0 | - |
| - | - | 736.4 | 289.1 | - | - | 0 | - |
| - | - | 3307 | 292.1 | - | - | 0 | - |
| - | - | 699.1 | 293.1 | - | - | 0 | - |
| - | - | 1589 | 295.1 | - | - | 0 | - |
| - | - | 3918 | 299.1 | - | - | 0 | - |
| - | - | 774.7 | 300.1 | - | - | 0 | - |
| - | - | 699.9 | 301.1 | - | - | 0 | - |
| - | - | 605.7 | 309.2 | - | - | 0 | - |
| - | - | 641.5 | 310.2 | - | - | 0 | - |
| - | - | 2320 | 312.1 | - | - | 0 | - |
| - | - | 990.4 | 316.2 | - | - | 0 | - |
| - | - | 984.6 | 318.1 | - | - | 0 | - |
| - | - | 773.9 | 322.1 | - | - | 0 | - |
| - | - | 1033 | 323.1 | - | - | 0 | - |
| - | - | 1758 | 325.1 | - | - | 0 | - |
| - | - | 712.1 | 332.1 | - | - | 0 | - |
| - | - | 693.6 | 333.1 | - | - | 0 | - |
| - | - | 1533 | 333.2 | - | - | 0 | - |
| - | - | 1.121E+04 | 340.1 | - | - | 0 | - |
| - | - | 610.1 | 341 | - | - | 0 | - |
| - | - | 1517 | 341.1 | - | - | 0 | - |
| - | - | 629.3 | 343.1 | - | - | 0 | - |
| - | - | 1425 | 346.1 | - | - | 0 | - |
| - | - | 948.8 | 348.1 | - | - | 0 | - |
| - | - | 619.9 | 349.2 | - | - | 0 | - |
| - | - | 4440 | 350.1 | - | - | 0 | - |
| 3 | a | 803.1 | 350.2 | 0.0007083 | 2.023 | +1 | 3 |
| 3 | a | 643.6 | 351.2 | 0.0001835 | 0.5226 | +1 | 3 |
| - | - | 680.5 | 354.2 | - | - | 0 | - |
| - | - | 3600 | 355.1 | - | - | 0 | - |
| - | - | 1877 | 356.1 | - | - | 0 | - |
| 9 | y | 5.369E+04 | 358.1 | 0.005027 | 14.03 | +1 | 2 |
| - | - | 5960 | 359 | - | - | 0 | - |
| - | - | 9973 | 359.2 | - | - | 0 | - |
| - | - | 1190 | 360.1 | - | - | 0 | - |
| - | - | 799.2 | 360.2 | - | - | 0 | - |
| - | - | 4252 | 364.2 | - | - | 0 | - |
| - | - | 624.2 | 367.2 | - | - | 0 | - |
| - | - | 1379 | 368.1 | - | - | 0 | - |
| - | - | 947.6 | 369.1 | - | - | 0 | - |
| 6 | y | 747.4 | 370.1 | 0.003312 | 8.948 | +2 | 5 |
| - | - | 1649 | 374.1 | - | - | 0 | - |
| - | - | 1177 | 376.1 | - | - | 0 | - |
| - | - | 1.397E+04 | 378.1 | - | - | 0 | - |
| 3 | b | 1.22E+04 | 378.2 | 8.686E-05 | 0.2297 | +1 | 3 |
| - | - | 2262 | 379.1 | - | - | 0 | - |
| - | - | 2319 | 379.2 | - | - | 0 | - |
| - | - | 602.1 | 382.1 | - | - | 0 | - |
| - | - | 4906 | 394.1 | - | - | 0 | - |
| - | - | 1062 | 395.1 | - | - | 0 | - |
| - | - | 938 | 400.2 | - | - | 0 | - |
| - | - | 953.5 | 404.1 | - | - | 0 | - |
| - | - | 1764 | 406.2 | - | - | 0 | - |
| - | - | 819.6 | 419 | - | - | 0 | - |
| - | - | 798.4 | 419.2 | - | - | 0 | - |
| - | - | 967 | 423.2 | - | - | 0 | - |
| - | - | 1747 | 424.2 | - | - | 0 | - |
| - | - | 1206 | 427.2 | - | - | 0 | - |
| - | - | 7185 | 429.1 | - | - | 0 | - |
| - | - | 713.7 | 439.2 | - | - | 0 | - |
| - | - | 1.039E+04 | 441.2 | - | - | 0 | - |
| - | - | 2428 | 442.2 | - | - | 0 | - |
| - | - | 851.3 | 449.2 | - | - | 0 | - |
| - | - | 993.4 | 455.2 | - | - | 0 | - |
| - | - | 2087 | 461.2 | - | - | 0 | - |
| - | - | 3497 | 465.2 | - | - | 0 | - |
| - | - | 867.5 | 477.2 | - | - | 0 | - |
| - | - | 2793 | 479.2 | - | - | 0 | - |
| - | - | 1081 | 480.2 | - | - | 0 | - |
| - | - | 811.4 | 483.3 | - | - | 0 | - |
| - | - | 805 | 489.2 | - | - | 0 | - |
| - | - | 1499 | 491.2 | - | - | 0 | - |
| 4 | b | 1058 | 493.2 | 0.0003059 | 0.6202 | +1 | 4 |
| - | - | 803.6 | 494.2 | - | - | 0 | - |
| 8 | b | 727.6 | 495.2 | 0.002175 | 4.392 | +2 | 8 |
| - | - | 628.6 | 496.2 | - | - | 0 | - |
| - | - | 744.7 | 497.2 | - | - | 0 | - |
| 8 | y | 1.456E+04 | 505.2 | 0.00979 | 19.38 | +1 | 3 |
| - | - | 3994 | 506.2 | - | - | 0 | - |
| - | - | 1096 | 507.2 | - | - | 0 | - |
| 4 | b | 6999 | 511.2 | 0.0001768 | 0.3458 | +1 | 4 |
| - | - | 8811 | 512.2 | - | - | 0 | - |
| - | - | 2123 | 513.2 | - | - | 0 | - |
| - | - | 1682 | 513.2 | - | - | 0 | - |
| - | - | 781.2 | 514.2 | - | - | 0 | - |
| - | - | 1279 | 519.2 | - | - | 0 | - |
| - | - | 830.1 | 520.2 | - | - | 0 | - |
| - | - | 784.2 | 526.7 | - | - | 0 | - |
| - | - | 1146 | 527.2 | - | - | 0 | - |
| - | - | 2137 | 541.2 | - | - | 0 | - |
| - | - | 948.2 | 542.2 | - | - | 0 | - |
| - | - | 1683 | 548.2 | - | - | 0 | - |
| 3 | y | 1866 | 549.7 | 0.00555 | 10.1 | +2 | 8 |
| - | - | 1496 | 558.2 | - | - | 0 | - |
| 3 | y | 1.552E+04 | 558.7 | 0.004723 | 8.454 | +2 | 8 |
| - | - | 1.011E+04 | 559.2 | - | - | 0 | - |
| - | - | 5518 | 559.7 | - | - | 0 | - |
| - | - | 1340 | 560.2 | - | - | 0 | - |
| - | - | 1032 | 570.2 | - | - | 0 | - |
| 9 | b | 1128 | 572.7 | 0.003034 | 5.298 | +2 | 9 |
| - | - | 740.3 | 573.2 | - | - | 0 | - |
| 7 | y | 2.118E+04 | 576.2 | 0.009846 | 17.09 | +1 | 4 |
| - | - | 7344 | 577.2 | - | - | 0 | - |
| - | - | 1033 | 578.2 | - | - | 0 | - |
| - | - | 1182 | 580.2 | - | - | 0 | - |
| - | - | 777.9 | 590.2 | - | - | 0 | - |
| - | - | 772.1 | 592.2 | - | - | 0 | - |
| 2 | y | 978.5 | 600.2 | 0.005575 | 9.289 | +2 | 9 |
| - | - | 1144 | 600.7 | - | - | 0 | - |
| - | - | 1848 | 606.2 | - | - | 0 | - |
| 5 | b | 2147 | 608.2 | 0.0001254 | 0.2061 | +1 | 5 |
| 2 | y | 7095 | 609.2 | 0.005542 | 9.097 | +2 | 9 |
| - | - | 4164 | 609.7 | - | - | 0 | - |
| - | - | 1824 | 610.2 | - | - | 0 | - |
| - | - | 1718 | 612.2 | - | - | 0 | - |
| - | - | 630.6 | 614.2 | - | - | 0 | - |
| - | - | 1377 | 624.2 | - | - | 0 | - |
| - | - | 812.5 | 625.2 | - | - | 0 | - |
| 5 | b | 4200 | 626.2 | 0.0009178 | 1.466 | +1 | 5 |
| - | - | 1142 | 627.3 | - | - | 0 | - |
| - | - | 788.2 | 633.2 | - | - | 0 | - |
| - | - | 1216 | 634.2 | - | - | 0 | - |
| - | - | 2894 | 636.4 | - | - | 0 | - |
| - | - | 1349 | 637.4 | - | - | 0 | - |
| - | - | 789.6 | 640.2 | - | - | 0 | - |
| - | - | 616 | 641.3 | - | - | 0 | - |
| - | - | 1753 | 641.8 | - | - | 0 | - |
| - | - | 847.1 | 642.2 | - | - | 0 | - |
| - | - | 2070 | 650.8 | - | - | 0 | - |
| - | - | 1851 | 651.3 | - | - | 0 | - |
| - | - | 744.1 | 651.8 | - | - | 0 | - |
| - | - | 668.1 | 652.2 | - | - | 0 | - |
| - | - | 1488 | 657.3 | - | - | 0 | - |
| - | - | 6934 | 664.4 | - | - | 0 | - |
| - | - | 894.6 | 664.8 | - | - | 0 | - |
| - | - | 1030 | 665.3 | - | - | 0 | - |
| - | - | 2429 | 665.4 | - | - | 0 | - |
| - | - | 1052 | 666.4 | - | - | 0 | - |
| - | - | 689.8 | 673.3 | - | - | 0 | - |
| 0 | Precursor | 9739 | 673.8 | 0.004877 | 7.238 | +2 | -1 |
| - | - | 6111 | 674.3 | - | - | 0 | - |
| - | - | 4061 | 674.8 | - | - | 0 | - |
| - | - | 7509 | 675.3 | - | - | 0 | - |
| - | - | 2118 | 676.3 | - | - | 0 | - |
| - | - | 1.261E+04 | 681.4 | - | - | 0 | - |
| - | - | 1297 | 681.5 | - | - | 0 | - |
| - | - | 817.3 | 681.8 | - | - | 0 | - |
| - | - | 1550 | 682.3 | - | - | 0 | - |
| - | - | 4848 | 682.4 | - | - | 0 | - |
| 0 | Precursor | 1.921E+04 | 682.8 | 0.004416 | 6.468 | +2 | -1 |
| - | - | 689.2 | 682.8 | - | - | 0 | - |
| - | - | 1.561E+04 | 683.3 | - | - | 0 | - |
| - | - | 792.7 | 683.3 | - | - | 0 | - |
| - | - | 1660 | 683.4 | - | - | 0 | - |
| - | - | 6824 | 683.8 | - | - | 0 | - |
| - | - | 1048 | 684.2 | - | - | 0 | - |
| - | - | 1597 | 684.3 | - | - | 0 | - |
| - | - | 1953 | 685.3 | - | - | 0 | - |
| - | - | 666.3 | 686.3 | - | - | 0 | - |
| - | - | 1989 | 695.3 | - | - | 0 | - |
| - | - | 899.9 | 704.3 | - | - | 0 | - |
| - | - | 712.9 | 713.3 | - | - | 0 | - |
| - | - | 3264 | 721.3 | - | - | 0 | - |
| 6 | y | 1.892E+04 | 739.3 | 0.009628 | 13.02 | +1 | 5 |
| - | - | 7548 | 740.3 | - | - | 0 | - |
| - | - | 2184 | 741.3 | - | - | 0 | - |
| - | - | 1416 | 743.3 | - | - | 0 | - |
| - | - | 3334 | 749.3 | - | - | 0 | - |
| - | - | 790.7 | 750.3 | - | - | 0 | - |
| - | - | 1512 | 759.3 | - | - | 0 | - |
| - | - | 771.4 | 760.3 | - | - | 0 | - |
| - | - | 880.2 | 761.3 | - | - | 0 | - |
| 6 | b | 1941 | 771.3 | 0.000337 | 0.4369 | +1 | 6 |
| - | - | 1790 | 772.3 | - | - | 0 | - |
| - | - | 794.4 | 782.2 | - | - | 0 | - |
| 6 | b | 1516 | 789.3 | 0.001075 | 1.362 | +1 | 6 |
| - | - | 9110 | 790.3 | - | - | 0 | - |
| - | - | 2996 | 791.3 | - | - | 0 | - |
| - | - | 819.5 | 792.3 | - | - | 0 | - |
| - | - | 2011 | 804.3 | - | - | 0 | - |
| - | - | 950.2 | 805.3 | - | - | 0 | - |
| - | - | 2966 | 832.3 | - | - | 0 | - |
| 5 | y | 901.1 | 836.3 | 0.0101 | 12.07 | +1 | 6 |
| - | - | 820 | 837.3 | - | - | 0 | - |
| 7 | b | 645.8 | 842.3 | 0.01316 | 15.62 | +1 | 7 |
| 5 | y | 2.231E+04 | 854.3 | 0.009174 | 10.74 | +1 | 6 |
| - | - | 1.084E+04 | 855.3 | - | - | 0 | - |
| - | - | 2162 | 856.3 | - | - | 0 | - |
| 7 | b | 705.9 | 860.3 | 0.01426 | 16.58 | +1 | 7 |
| - | - | 2217 | 868.3 | - | - | 0 | - |
| - | - | 1317 | 869.3 | - | - | 0 | - |
| - | - | 1816 | 887.3 | - | - | 0 | - |
| - | - | 814.5 | 888.3 | - | - | 0 | - |
| - | - | 3644 | 896.3 | - | - | 0 | - |
| - | - | 2304 | 897.3 | - | - | 0 | - |
| - | - | 708.2 | 898.3 | - | - | 0 | - |
| - | - | 1.255E+04 | 905.3 | - | - | 0 | - |
| - | - | 6012 | 906.3 | - | - | 0 | - |
| - | - | 1731 | 907.3 | - | - | 0 | - |
| - | - | 734.4 | 914.3 | - | - | 0 | - |
| - | - | 1035 | 915.3 | - | - | 0 | - |
| - | - | 723.5 | 933.4 | - | - | 0 | - |
| 4 | y | 2622 | 951.3 | 0.007933 | 8.339 | +1 | 7 |
| - | - | 907.8 | 952.3 | - | - | 0 | - |
| 4 | y | 3.912E+04 | 969.3 | 0.008904 | 9.185 | +1 | 7 |
| - | - | 1.927E+04 | 970.3 | - | - | 0 | - |
| - | - | 7170 | 971.3 | - | - | 0 | - |
| - | - | 1210 | 979.3 | - | - | 0 | - |
| 8 | b | 841 | 989.4 | 0.002743 | 2.773 | +1 | 8 |
| - | - | 1496 | 997.4 | - | - | 0 | - |
| - | - | 1973 | 1034 | - | - | 0 | - |
| - | - | 979.1 | 1035 | - | - | 0 | - |
| - | - | 1.926E+04 | 1052 | - | - | 0 | - |
| - | - | 1.181E+04 | 1053 | - | - | 0 | - |
| - | - | 3761 | 1054 | - | - | 0 | - |
| 3 | y | 3319 | 1098 | 0.0102 | 9.284 | +1 | 8 |
| - | - | 2249 | 1099 | - | - | 0 | - |
| - | - | 756.6 | 1100 | - | - | 0 | - |
| 3 | y | 5.955E+04 | 1116 | 0.008544 | 7.653 | +1 | 8 |
| - | - | 3.897E+04 | 1117 | - | - | 0 | - |
| - | - | 1.593E+04 | 1118 | - | - | 0 | - |
| - | - | 2237 | 1119 | - | - | 0 | - |
| - | - | 745.8 | 1135 | - | - | 0 | - |
| 9 | b | 1931 | 1144 | 0.0008152 | 0.7123 | +1 | 9 |
| - | - | 1573 | 1145 | - | - | 0 | - |
| - | - | 3553 | 1153 | - | - | 0 | - |
| - | - | 2625 | 1154 | - | - | 0 | - |
| - | - | 1038 | 1155 | - | - | 0 | - |
| - | - | 1418 | 1163 | - | - | 0 | - |
| 2 | y | 1686 | 1199 | 0.00915 | 7.629 | +1 | 9 |
| - | - | 847 | 1200 | - | - | 0 | - |
| 2 | y | 1.188E+04 | 1217 | 0.006764 | 5.556 | +1 | 9 |
| - | - | 7240 | 1218 | - | - | 0 | - |
| - | - | 3149 | 1219 | - | - | 0 | - |
| - | - | 1491 | 1227 | - | - | 0 | - |
| - | - | 1693 | 1228 | - | - | 0 | - |
| - | - | 660 | 1278 | - | - | 0 | - |
| - | - | 751.3 | 1602 | - | - | 0 | - |
| - | - | 750.6 | 2401 | - | - | 0 | - |
| - | - | 828.8 | 3075 | - | - | 0 | - |

m/z Charge Intensity FragmentType MassShift Position
120.08106994628906 0 183373.34
121.08434295654297 0 14598.768
121.73993682861328 0 399.808
122.07140350341797 0 563.2446
129.1024932861328 0 2053.0488
130.06541442871094 0 1625.6017
131.0703887939453 0 1202.1243
132.07334899902344 0 410.2781
132.08091735839844 0 564.7664
133.08615112304688 0 17405.475
134.0895233154297 0 3813.5735
136.07594299316406 0 13361.296
136.0868377685547 0 601.48145
137.0791015625 0 1153.1569
138.05519104003906 0 1140.5746
138.06639099121094 0 1685.621
143.11793518066406 0 482.23605
146.06053161621094 0 449.9725
148.08685302734375 0 495.8498
148.95433044433594 0 596.8578
149.04502868652344 0 2276.0378
150.8808135986328 0 423.3473
152.60389709472656 0 433.2806
156.07687377929688 0 5682.7217
157.0760955810547 0 523.7197
158.0603485107422 0 1026.6503
158.0965576171875 0 4249.838
159.09181213378906 0 4891.534
159.09910583496094 0 860.78455
160.09506225585938 0 512.2858
161.08094787597656 0 1075.301
164.08200073242188 0 517.9231
166.0610809326172 0 1604.5621
166.08616638183594 0 694.46155
167.05567932128906 0 3160.4758
167.09303283691406 0 1523.221
169.13385009765625 0 1152.9813
171.0770721435547 0 575.5041
173.4392852783203 0 2620.215
174.0665283203125 0 689.3666
175.0867919921875 0 10488.275
175.0966339111328 0 1378.4423
175.12289428710938 0 628.15485
176.0821075439453 0 2156.0676
176.1072235107422 0 84426.47
177.10238647460938 0 1690.8782
177.11134338378906 0 12673.054
178.11546325683594 0 2911.5627
180.11279296875 0 532.0165
181.58705139160156 0 553.189
186.09149169921875 0 10446.753
187.09523010253906 0 1738.1709
187.10780334472656 0 446.03516
191.11801147460938 0 1904.8442
193.1085205078125 0 6131.417
195.0343475341797 0 541.7045
197.1287384033203 0 1374.474
199.861083984375 0 502.62674
203.06607055664062 0 1201.3837
203.08180236816406 0 2480.6733
203.11810302734375 0 6717.927 a Water loss 1
204.0771026611328 0 3528.0706
204.1212158203125 0 1011.36597
205.0977325439453 0 538.77356
205.10726928710938 0 6378.918
206.11070251464844 0 2739.474
207.112548828125 0 1648.6603 d 1
212.11865234375 0 1494.2439
214.08689880371094 0 781.32495
217.0973358154297 0 5350.8535
218.10035705566406 0 1338.0868
219.0797576904297 0 1078.5964
219.11151123046875 0 526.2583
221.09217834472656 0 23598.975
221.10360717773438 0 13481.107
221.12864685058594 0 108792.89 a 1
222.08456420898438 0 632.58136
222.0955047607422 0 2108.1628
222.1068572998047 0 682.5836
222.13197326660156 0 13258.076
222.14393615722656 0 1114.8171
225.0430908203125 0 4972.994
226.04405212402344 0 626.15466
228.1127471923828 0 906.44415
231.06155395507812 0 2559.1172
231.11302185058594 0 28700.246 b Water loss 1
232.11643981933594 0 3745.839
235.10792541503906 0 7892.855
236.11099243164062 0 912.40466
239.0951385498047 0 7032.6978
239.114013671875 0 682.8347
240.09642028808594 0 1213.5425
248.15338134765625 0 540.17175
249.1236114501953 0 57154.195 b 1
250.1269073486328 0 6389.666
250.15513610839844 0 547.78143
251.1028289794922 0 5385.6357
251.1280517578125 0 793.61224
253.1175994873047 0 568.0811
257.10675048828125 0 1815.1027
259.107177734375 0 1231.0809
263.10272216796875 0 11953.254
264.1059875488281 0 1468.5665
265.164306640625 0 701.7225
274.1297912597656 0 2467.0032
277.4535217285156 0 558.98303
279.09747314453125 0 5074.645
285.0086669921875 0 1275.1903
285.101806640625 0 5718.2993
287.10089111328125 0 731.7821
289.1331787109375 0 736.3614
292.140625 0 3307.2747
293.1453552246094 0 699.11957
295.10357666015625 0 1589.259
299.0615539550781 0 3918.3499
300.0643310546875 0 774.7446
301.0581359863281 0 699.94293
309.19049072265625 0 605.6777
310.19354248046875 0 641.5192
312.14581298828125 0 2319.5308
316.1864929199219 0 990.3695
318.14508056640625 0 984.5987
322.12982177734375 0 773.9028
323.1156311035156 0 1033.2917
325.1388854980469 0 1758.4974
332.1239929199219 0 712.1064
333.1089172363281 0 693.5687
333.1602478027344 0 1533.0322
340.1407470703125 0 11206.813
341.0172119140625 0 610.1476
341.14447021484375 0 1516.7769
343.1407470703125 0 629.344
346.14044189453125 0 1425.2643
348.1197814941406 0 948.7537
349.1554870605469 0 619.91046
350.1346435546875 0 4439.709
350.18701171875 0 803.1147 a Water loss 2
351.1701354980469 0 643.60645 a Ammonia loss 2
354.2196044921875 0 680.4764
355.06964111328125 0 3600.326
356.138671875 0 1877.2211
358.1510925292969 0 53692.28 y 8
359.02838134765625 0 5960.1313
359.1540222167969 0 9973.186
360.1190490722656 0 1189.6486
360.1566467285156 0 799.1681
364.15045166015625 0 4252.093
367.1652526855469 0 624.20605
368.1361083984375 0 1379.3772
369.12066650390625 0 947.5935
370.14544677734375 0 747.3916 y 5
374.1354675292969 0 1649.4504
376.1139221191406 0 1176.6454
378.1296081542969 0 13972.594
378.1813049316406 0 12203.383 b Water loss 2
379.1324768066406 0 2261.606
379.1841735839844 0 2318.8718
382.1445007324219 0 602.06244
394.124755859375 0 4905.8228
395.1271667480469 0 1061.9484
400.1570739746094 0 937.96155
404.1447448730469 0 953.53406
406.15106201171875 0 1764.1019
418.9978942871094 0 819.60065
419.1567687988281 0 798.355
423.17840576171875 0 967.03076
424.1613464355469 0 1746.7837
427.20965576171875 0 1206.154
429.0887451171875 0 7184.849
439.1728210449219 0 713.6789
441.1886291503906 0 10390.033
442.19110107421875 0 2427.809
449.16668701171875 0 851.2509
455.2043151855469 0 993.3919
461.1670837402344 0 2087.3765
465.1622009277344 0 3496.8716
477.1911926269531 0 867.4619
479.1773986816406 0 2793.0989
480.1805114746094 0 1081.0618
483.2620544433594 0 811.3632
489.1613464355469 0 805.03784
491.20721435546875 0 1498.6846
493.2078552246094 0 1058.4784 b Water loss 3
494.21466064453125 0 803.6038
495.1844787597656 0 727.6201 b Water loss 7
496.17205810546875 0 628.60767
497.1689453125 0 744.7032
505.18634033203125 0 14562.172 y 7
506.1898498535156 0 3994.0837
507.1864013671875 0 1095.6603
511.2189025878906 0 6998.7666 b 3
512.224609375 0 8811.177
513.1968383789062 0 2123.0186
513.2294921875 0 1681.686
514.196044921875 0 781.21094
519.201171875 0 1279.0033
520.2037353515625 0 830.14386
526.7066650390625 0 784.15485
527.2116088867188 0 1145.5256
541.1939697265625 0 2137.3765
542.2369995117188 0 948.2181
548.1981201171875 0 1682.7307
549.7035522460938 0 1866.2039 y Water loss 2
558.2125854492188 0 1495.5311
558.7080078125 0 15519.139 y 2
559.2096557617188 0 10109.512
559.7103881835938 0 5517.5083
560.2120971679688 0 1340.2765
570.2289428710938 0 1031.7306
572.7244262695312 0 1127.8456 b 8
573.2210693359375 0 740.30975
576.2235107421875 0 21176.787 y 6
577.2269287109375 0 7343.8994
578.2265625 0 1033.1285
580.240478515625 0 1181.5402
590.2218627929688 0 777.8547
592.215576171875 0 772.08514
600.2274169921875 0 978.4616 y Water loss 1
600.7291259765625 0 1144.341
606.2301635742188 0 1847.8513
608.2352294921875 0 2146.7354 b Water loss 4
609.232666015625 0 7094.6943 y 1
609.7333984375 0 4163.6255
610.2355346679688 0 1824.4734
612.1940307617188 0 1717.6959
614.236083984375 0 630.641
624.23388671875 0 1376.797
625.2378540039062 0 812.47864
626.2447509765625 0 4200.1846 b 4
627.2500610351562 0 1141.5385
633.2332153320312 0 788.1818
634.2315673828125 0 1216.3534
636.3889770507812 0 2894.228
637.390869140625 0 1348.764
640.2424926757812 0 789.60126
641.2547607421875 0 616.01154
641.7619018554688 0 1753.346
642.2489624023438 0 847.0925
650.7686767578125 0 2069.501
651.2654418945312 0 1851.3954
651.7694091796875 0 744.103
652.230712890625 0 668.1453
657.2640380859375 0 1488.3441
664.382568359375 0 6933.994
664.7538452148438 0 894.5671
665.2522583007812 0 1030.1122
665.3848876953125 0 2428.623
666.3896484375 0 1051.5189
673.2752685546875 0 689.79517
673.7609252929688 0 9739.042 Precursor Water loss
674.2628173828125 0 6110.6465
674.7634887695312 0 4061.301
675.2877807617188 0 7508.831
676.2926025390625 0 2117.6492
681.4098510742188 0 12607.118
681.5170288085938 0 1297.0753
681.8369140625 0 817.25385
682.3422241210938 0 1549.6277
682.4126586914062 0 4848.3203
682.7657470703125 0 19208.387 Precursor
682.8322143554688 0 689.1836
683.2671508789062 0 15608.515
683.3396606445312 0 792.6625
683.4122314453125 0 1659.6104
683.7684936523438 0 6823.9424
684.2098999023438 0 1048.4604
684.2688598632812 0 1596.5647
685.2581176757812 0 1952.5154
686.2605590820312 0 666.2954
695.2666015625 0 1989.056
704.264404296875 0 899.93353
713.2791137695312 0 712.9315
721.2644653320312 0 3264.3123
739.28662109375 0 18918.205 y 5
740.2893676757812 0 7547.7
741.2884521484375 0 2183.6682
743.3023681640625 0 1416.195
749.2572631835938 0 3333.53
750.255859375 0 790.6717
759.2642822265625 0 1512.0848
760.2711181640625 0 771.3857
761.3126220703125 0 880.19946
771.298095703125 0 1941.0687 b Water loss 5
772.3017578125 0 1790.2346
782.2124633789062 0 794.38354
789.3079223632812 0 1515.7168 b 5
790.3148803710938 0 9110.07
791.3178100585938 0 2995.6792
792.3196411132812 0 819.5328
804.3309326171875 0 2010.9451
805.3322143554688 0 950.15063
832.3270874023438 0 2965.9087
836.303466796875 0 901.1326 y Water loss 4
837.3010864257812 0 819.96313
842.3223876953125 0 645.84314 b Water loss 6
854.3131103515625 0 22311.371 y 4
855.3162841796875 0 10838.582
856.31591796875 0 2162.2212
860.3318481445312 0 705.8903 b 6
868.3280639648438 0 2216.655
869.3340454101562 0 1317.4032
887.3306884765625 0 1815.5348
888.3333740234375 0 814.51404
896.3255004882812 0 3643.7646
897.3258666992188 0 2304.0789
898.320068359375 0 708.2451
905.3416748046875 0 12545.163
906.3447265625 0 6011.952
907.3474731445312 0 1731.4652
914.33544921875 0 734.37384
915.3385620117188 0 1034.5825
933.3680419921875 0 723.46954
951.3282470703125 0 2621.7412 y Water loss 3
952.3335571289062 0 907.83405
969.3397827148438 0 39116.03 y 3
970.3428344726562 0 19270.086
971.3432006835938 0 7169.6963
979.3231811523438 0 1210.4824
989.3687744140625 0 840.9968 b Water loss 7
997.3776245117188 0 1496.1147
1034.3973388671875 0 1972.9694
1035.3975830078125 0 979.09863
1052.40966796875 0 19264.453
1053.41259765625 0 11809.764
1054.415771484375 0 3761.4607
1098.39892578125 0 3318.9407 y Water loss 2
1099.4012451171875 0 2248.6694
1100.399169921875 0 756.5604
1116.4078369140625 0 59552.37 y 2
1117.4110107421875 0 38972.742
1118.411865234375 0 15925.104
1119.4144287109375 0 2236.6753
1135.45751953125 0 745.7962
1144.4346923828125 0 1930.5347 b 8
1145.4423828125 0 1573.379
1153.4573974609375 0 3552.9036
1154.46044921875 0 2624.6902
1155.4696044921875 0 1038.0386
1163.440185546875 0 1417.9633
1199.445556640625 0 1685.5173 y Water loss 1
1200.4517822265625 0 846.9735
1217.4537353515625 0 11882.021 y 1
1218.456298828125 0 7240.2153
1219.459228515625 0 3149.2517
1227.43603515625 0 1490.9922
1228.446533203125 0 1693.3721
1277.6546630859375 0 660.03625
1602.3121337890625 0 751.26996
2401.22509765625 0 750.62537
3074.66064453125 0 828.77454

Spectrum Details

|  |  |
| --- | --- |
| Matched peaks? Matched peaks The total absolute number of peaks matched. Additionally in brackets the total fraction of peaks matched and the total number of peaks is shown. | 39 (11.14% of 350) |
| FDR? FDR The false discovery rate estimated for this peptide. It is calculated by matching all theoretical fragments with a non-integer shift with the raw peaks for this spectrum. This is done with 40 different shifts. The resulting percentage is the average number of annotated peaks over the number of annotated peaks with the correct spectrum. | 0.18% |
| Satellite FDR? Satellite FDR See the FDR for details on its calculation. This satellite ion specific FDR only contains the satellite ions (d/w) for I/L/J positions. | - |
| PSM Score? PSM Score The PSM Score as given by Hecklib to this annotated spectrum. It is shown with three significant figures. | 463 |

#### Spectrum 8477? Spectrum 8477 The raw spectrum of this peptide as annotated by Hecklib. The fragments are coloured according to ion type (see legend). Any peaks with a star '\*' as text can be hovered over to see the full details, first the ion type second the mass shift type. By hovering over the amino acids in the peptide or ions in the legend the corresponding peaks are highlighted. By toggling the 'Unassigned' label you can turn the background (unassigned) peaks on or off in the plot. By updating the slider in the Ion legend you can update the spectrum to only show the top X% of the peaks with labels. The top X% means any peak that is within X% of the highest intensity. By dragging in the spectrum you can zoom in to a specific part of the spectrum and use 'Zoom Out' to get back to the original zoom level. The annotation of the spectrum is based on the given sequence in the peptides file and is done with different software so inconsistencies are likely. The peaks are annotated based on the given sequence, with 20 ppm tolerance.

Copy Data

##### Spectrum 8477 (TSV)

###### Preview

```
Loading example...
```

*Click on the button to copy the data to your clipboard.*

Mz MinMz MaxIntensity Max

WidthHeightPeptide font sizePeptide stroke widthSpectrum font sizeSpectrum stroke widthCompact peptide

Ion legend

wxyz

abcd

OtherUnassignedIon^Charge^_Position_Show for top:%

FTFDDYAMHW

04.49e+48.97e+41.35e+51.79e+5

Zoom Out

a^+1^_2_d^+1^_2_a^+1^_2_b^+1^_2_b^+1^_2_a^+1^_3_y^+1^_2_b^+1^_3_b^+1^_3_b^+1^_4_b^+2^_8_b^+1^_4_y^+2^_8_y^+2^_8_y^+1^_4_y^+2^_9_b^+1^_5_y^+2^_9_b^+1^_5_\*\*y^+1^_5_b^+1^_6_b^+1^_6_y^+1^_6_y^+1^_6_y^+1^_7_y^+1^_7_y^+1^_8_y^+1^_8_b^+1^_9_y^+1^_9_y^+1^_9_

0776155323293105

Fragment Matches Table

Show background peaks

| Position | Ion type | Intensity | mz Theoretical | mz Error (Th) | mz Error (ppm) | Charge | Series Number |
| --- | --- | --- | --- | --- | --- | --- | --- |
| - | - | 1.777E+05 | 120.1 | - | - | 0 | - |
| - | - | 1.374E+04 | 121.1 | - | - | 0 | - |
| - | - | 421.1 | 121.9 | - | - | 0 | - |
| - | - | 2607 | 129.1 | - | - | 0 | - |
| - | - | 1707 | 130.1 | - | - | 0 | - |
| - | - | 557.5 | 130.1 | - | - | 0 | - |
| - | - | 1520 | 133.1 | - | - | 0 | - |
| - | - | 1.266E+04 | 136.1 | - | - | 0 | - |
| - | - | 1267 | 137.1 | - | - | 0 | - |
| - | - | 1259 | 138.1 | - | - | 0 | - |
| - | - | 1355 | 138.1 | - | - | 0 | - |
| - | - | 412.7 | 140 | - | - | 0 | - |
| - | - | 2979 | 143.1 | - | - | 0 | - |
| - | - | 580.9 | 146.1 | - | - | 0 | - |
| - | - | 837.8 | 146.1 | - | - | 0 | - |
| - | - | 685.8 | 147.1 | - | - | 0 | - |
| - | - | 559.2 | 147.1 | - | - | 0 | - |
| - | - | 742.7 | 148.1 | - | - | 0 | - |
| - | - | 1355 | 149 | - | - | 0 | - |
| - | - | 645.4 | 153.1 | - | - | 0 | - |
| - | - | 764.6 | 155.1 | - | - | 0 | - |
| - | - | 596.3 | 155.1 | - | - | 0 | - |
| - | - | 5329 | 156.1 | - | - | 0 | - |
| - | - | 1108 | 158.1 | - | - | 0 | - |
| - | - | 3925 | 158.1 | - | - | 0 | - |
| - | - | 4468 | 159.1 | - | - | 0 | - |
| - | - | 812.8 | 159.1 | - | - | 0 | - |
| - | - | 645.8 | 160.1 | - | - | 0 | - |
| - | - | 567.5 | 160.1 | - | - | 0 | - |
| - | - | 662.5 | 166.1 | - | - | 0 | - |
| - | - | 1647 | 166.1 | - | - | 0 | - |
| - | - | 574.9 | 166.1 | - | - | 0 | - |
| - | - | 2702 | 167.1 | - | - | 0 | - |
| - | - | 1212 | 167.1 | - | - | 0 | - |
| - | - | 1672 | 169.1 | - | - | 0 | - |
| - | - | 1567 | 171.1 | - | - | 0 | - |
| - | - | 4473 | 173.5 | - | - | 0 | - |
| - | - | 563.8 | 174.1 | - | - | 0 | - |
| - | - | 1.084E+04 | 175.1 | - | - | 0 | - |
| - | - | 1095 | 175.1 | - | - | 0 | - |
| - | - | 1220 | 176.1 | - | - | 0 | - |
| - | - | 7.981E+04 | 176.1 | - | - | 0 | - |
| - | - | 534.3 | 176.4 | - | - | 0 | - |
| - | - | 831.4 | 177.1 | - | - | 0 | - |
| - | - | 9289 | 177.1 | - | - | 0 | - |
| - | - | 536.7 | 180.1 | - | - | 0 | - |
| - | - | 775.1 | 185.1 | - | - | 0 | - |
| - | - | 1.237E+04 | 186.1 | - | - | 0 | - |
| - | - | 1689 | 187.1 | - | - | 0 | - |
| - | - | 1676 | 191.1 | - | - | 0 | - |
| - | - | 4931 | 193.1 | - | - | 0 | - |
| - | - | 820.8 | 194.1 | - | - | 0 | - |
| - | - | 1191 | 197.1 | - | - | 0 | - |
| - | - | 700 | 202.1 | - | - | 0 | - |
| - | - | 864.7 | 203.1 | - | - | 0 | - |
| - | - | 2756 | 203.1 | - | - | 0 | - |
| - | - | 586.6 | 203.1 | - | - | 0 | - |
| 2 | a | 5865 | 203.1 | 0.0002593 | 1.276 | +1 | 2 |
| - | - | 2621 | 204.1 | - | - | 0 | - |
| - | - | 736.4 | 205.1 | - | - | 0 | - |
| 2 | d | 1266 | 207.1 | 0.0005076 | 2.451 | +1 | 2 |
| - | - | 570.4 | 211.1 | - | - | 0 | - |
| - | - | 1767 | 212.1 | - | - | 0 | - |
| - | - | 979 | 214.1 | - | - | 0 | - |
| - | - | 4949 | 217.1 | - | - | 0 | - |
| - | - | 889.9 | 219.1 | - | - | 0 | - |
| - | - | 610.7 | 219.1 | - | - | 0 | - |
| - | - | 2.261E+04 | 221.1 | - | - | 0 | - |
| - | - | 1.192E+04 | 221.1 | - | - | 0 | - |
| 2 | a | 1.039E+05 | 221.1 | 0.0003147 | 1.423 | +1 | 2 |
| - | - | 2078 | 222.1 | - | - | 0 | - |
| - | - | 1314 | 222.1 | - | - | 0 | - |
| - | - | 1.225E+04 | 222.1 | - | - | 0 | - |
| - | - | 596.2 | 223.1 | - | - | 0 | - |
| - | - | 557 | 223.1 | - | - | 0 | - |
| - | - | 4713 | 225 | - | - | 0 | - |
| - | - | 821.3 | 226 | - | - | 0 | - |
| - | - | 614 | 228.1 | - | - | 0 | - |
| - | - | 1336 | 231.1 | - | - | 0 | - |
| 2 | b | 2.644E+04 | 231.1 | 0.0003398 | 1.47 | +1 | 2 |
| - | - | 3198 | 232.1 | - | - | 0 | - |
| - | - | 8047 | 235.1 | - | - | 0 | - |
| - | - | 870.6 | 236.1 | - | - | 0 | - |
| - | - | 7224 | 239.1 | - | - | 0 | - |
| - | - | 1063 | 239.1 | - | - | 0 | - |
| - | - | 1120 | 240.1 | - | - | 0 | - |
| - | - | 812.5 | 240.1 | - | - | 0 | - |
| - | - | 562.6 | 247.1 | - | - | 0 | - |
| - | - | 550.6 | 248.2 | - | - | 0 | - |
| 2 | b | 5.61E+04 | 249.1 | 0.0003189 | 1.28 | +1 | 2 |
| - | - | 7594 | 250.1 | - | - | 0 | - |
| - | - | 6073 | 251.1 | - | - | 0 | - |
| - | - | 586.9 | 251.1 | - | - | 0 | - |
| - | - | 2717 | 257.1 | - | - | 0 | - |
| - | - | 948.2 | 258.1 | - | - | 0 | - |
| - | - | 833.3 | 259.1 | - | - | 0 | - |
| - | - | 529.5 | 259.5 | - | - | 0 | - |
| - | - | 1.125E+04 | 263.1 | - | - | 0 | - |
| - | - | 1487 | 264.1 | - | - | 0 | - |
| - | - | 537.4 | 267 | - | - | 0 | - |
| - | - | 641.2 | 267.1 | - | - | 0 | - |
| - | - | 2976 | 274.1 | - | - | 0 | - |
| - | - | 935.1 | 276.2 | - | - | 0 | - |
| - | - | 3762 | 279.1 | - | - | 0 | - |
| - | - | 553 | 281.5 | - | - | 0 | - |
| - | - | 1348 | 285 | - | - | 0 | - |
| - | - | 5375 | 285.1 | - | - | 0 | - |
| - | - | 559.7 | 287.1 | - | - | 0 | - |
| - | - | 589.4 | 289.1 | - | - | 0 | - |
| - | - | 3760 | 292.1 | - | - | 0 | - |
| - | - | 1458 | 295.1 | - | - | 0 | - |
| - | - | 544.9 | 296.1 | - | - | 0 | - |
| - | - | 4154 | 299.1 | - | - | 0 | - |
| - | - | 680.6 | 300.1 | - | - | 0 | - |
| - | - | 835.3 | 306.1 | - | - | 0 | - |
| - | - | 2769 | 312.1 | - | - | 0 | - |
| - | - | 1337 | 316.2 | - | - | 0 | - |
| - | - | 1221 | 318.1 | - | - | 0 | - |
| - | - | 1214 | 322.1 | - | - | 0 | - |
| - | - | 1476 | 325.1 | - | - | 0 | - |
| - | - | 1553 | 333.2 | - | - | 0 | - |
| - | - | 957.9 | 338.1 | - | - | 0 | - |
| - | - | 979.7 | 339.2 | - | - | 0 | - |
| - | - | 1.14E+04 | 340.1 | - | - | 0 | - |
| - | - | 1049 | 341 | - | - | 0 | - |
| - | - | 1718 | 341.1 | - | - | 0 | - |
| - | - | 1137 | 346.1 | - | - | 0 | - |
| - | - | 800.4 | 348.1 | - | - | 0 | - |
| - | - | 3220 | 350.1 | - | - | 0 | - |
| 3 | a | 1218 | 350.2 | 0.001136 | 3.243 | +1 | 3 |
| - | - | 3774 | 355.1 | - | - | 0 | - |
| - | - | 2881 | 356.1 | - | - | 0 | - |
| 9 | y | 5.065E+04 | 358.1 | 0.00521 | 14.55 | +1 | 2 |
| - | - | 6318 | 359 | - | - | 0 | - |
| - | - | 9506 | 359.2 | - | - | 0 | - |
| - | - | 876.7 | 360.2 | - | - | 0 | - |
| - | - | 4665 | 364.2 | - | - | 0 | - |
| - | - | 722.1 | 365.2 | - | - | 0 | - |
| - | - | 1157 | 368.1 | - | - | 0 | - |
| - | - | 921.6 | 369.1 | - | - | 0 | - |
| - | - | 631.8 | 373.1 | - | - | 0 | - |
| - | - | 1062 | 376.1 | - | - | 0 | - |
| - | - | 1.469E+04 | 378.1 | - | - | 0 | - |
| 3 | b | 1.078E+04 | 378.2 | 0.0002089 | 0.5525 | +1 | 3 |
| - | - | 2353 | 379.1 | - | - | 0 | - |
| - | - | 2752 | 379.2 | - | - | 0 | - |
| - | - | 566.6 | 381.1 | - | - | 0 | - |
| - | - | 1209 | 382.1 | - | - | 0 | - |
| - | - | 4748 | 394.1 | - | - | 0 | - |
| 3 | b | 719 | 396.2 | 0.001048 | 2.645 | +1 | 3 |
| - | - | 693.2 | 406.1 | - | - | 0 | - |
| - | - | 716.5 | 412.7 | - | - | 0 | - |
| - | - | 688 | 413.2 | - | - | 0 | - |
| - | - | 792.6 | 415.2 | - | - | 0 | - |
| - | - | 571.7 | 417.2 | - | - | 0 | - |
| - | - | 864.1 | 419 | - | - | 0 | - |
| - | - | 1435 | 423.2 | - | - | 0 | - |
| - | - | 1611 | 424.2 | - | - | 0 | - |
| - | - | 611.1 | 425.2 | - | - | 0 | - |
| - | - | 916 | 426.2 | - | - | 0 | - |
| - | - | 1142 | 427.2 | - | - | 0 | - |
| - | - | 6985 | 429.1 | - | - | 0 | - |
| - | - | 647.1 | 433.2 | - | - | 0 | - |
| - | - | 625.5 | 439.2 | - | - | 0 | - |
| - | - | 8832 | 441.2 | - | - | 0 | - |
| - | - | 1891 | 442.2 | - | - | 0 | - |
| - | - | 774.6 | 447.2 | - | - | 0 | - |
| - | - | 546.9 | 447.2 | - | - | 0 | - |
| - | - | 1059 | 455.2 | - | - | 0 | - |
| - | - | 1800 | 461.2 | - | - | 0 | - |
| - | - | 3291 | 465.2 | - | - | 0 | - |
| - | - | 537.1 | 466.1 | - | - | 0 | - |
| - | - | 1202 | 466.2 | - | - | 0 | - |
| - | - | 1596 | 477.2 | - | - | 0 | - |
| - | - | 2717 | 479.2 | - | - | 0 | - |
| - | - | 787.1 | 489.2 | - | - | 0 | - |
| - | - | 1291 | 491.2 | - | - | 0 | - |
| 4 | b | 1516 | 493.2 | 0.0004571 | 0.9267 | +1 | 4 |
| - | - | 952.3 | 494.2 | - | - | 0 | - |
| 8 | b | 1175 | 495.2 | 0.003288 | 6.639 | +2 | 8 |
| - | - | 824.5 | 497.2 | - | - | 0 | - |
| - | - | 1.401E+04 | 505.2 | - | - | 0 | - |
| - | - | 4263 | 506.2 | - | - | 0 | - |
| 4 | b | 5492 | 511.2 | 5.473E-05 | 0.107 | +1 | 4 |
| - | - | 7481 | 512.2 | - | - | 0 | - |
| - | - | 1760 | 513.2 | - | - | 0 | - |
| - | - | 1716 | 513.2 | - | - | 0 | - |
| - | - | 1058 | 514.2 | - | - | 0 | - |
| - | - | 1391 | 519.2 | - | - | 0 | - |
| - | - | 1168 | 526.7 | - | - | 0 | - |
| - | - | 1072 | 527.2 | - | - | 0 | - |
| - | - | 1972 | 541.2 | - | - | 0 | - |
| - | - | 1147 | 542.2 | - | - | 0 | - |
| - | - | 915.5 | 548.2 | - | - | 0 | - |
| 3 | y | 1753 | 549.7 | 0.004634 | 8.431 | +2 | 8 |
| - | - | 812.2 | 550.2 | - | - | 0 | - |
| - | - | 1335 | 558.2 | - | - | 0 | - |
| 3 | y | 1.695E+04 | 558.7 | 0.005028 | 9 | +2 | 8 |
| - | - | 1.004E+04 | 559.2 | - | - | 0 | - |
| - | - | 3926 | 559.7 | - | - | 0 | - |
| - | - | 760.4 | 560.2 | - | - | 0 | - |
| - | - | 1219 | 570.2 | - | - | 0 | - |
| 7 | y | 1.886E+04 | 576.2 | 0.01009 | 17.51 | +1 | 4 |
| - | - | 6683 | 577.2 | - | - | 0 | - |
| - | - | 1580 | 578.2 | - | - | 0 | - |
| - | - | 678.2 | 584.2 | - | - | 0 | - |
| - | - | 1004 | 596.8 | - | - | 0 | - |
| 2 | y | 1718 | 600.2 | 0.005759 | 9.594 | +2 | 9 |
| - | - | 1384 | 600.7 | - | - | 0 | - |
| - | - | 623 | 601.2 | - | - | 0 | - |
| - | - | 1418 | 606.2 | - | - | 0 | - |
| - | - | 666.9 | 607.2 | - | - | 0 | - |
| 5 | b | 2381 | 608.2 | 0.00159 | 2.614 | +1 | 5 |
| 2 | y | 7727 | 609.2 | 0.005542 | 9.097 | +2 | 9 |
| - | - | 3157 | 609.7 | - | - | 0 | - |
| - | - | 942.5 | 610.2 | - | - | 0 | - |
| - | - | 1078 | 612.2 | - | - | 0 | - |
| - | - | 909.4 | 612.2 | - | - | 0 | - |
| - | - | 1034 | 623.9 | - | - | 0 | - |
| - | - | 1033 | 624.2 | - | - | 0 | - |
| 5 | b | 4419 | 626.2 | 0.0001854 | 0.2961 | +1 | 5 |
| - | - | 926.2 | 627.3 | - | - | 0 | - |
| - | - | 1650 | 634.2 | - | - | 0 | - |
| - | - | 669.3 | 643.2 | - | - | 0 | - |
| - | - | 2322 | 650.8 | - | - | 0 | - |
| - | - | 1436 | 651.3 | - | - | 0 | - |
| - | - | 1036 | 651.8 | - | - | 0 | - |
| - | - | 1018 | 657.3 | - | - | 0 | - |
| - | - | 713.8 | 659.8 | - | - | 0 | - |
| - | - | 656.3 | 667.2 | - | - | 0 | - |
| 0 | Precursor | 8589 | 673.8 | 0.005121 | 7.601 | +2 | -1 |
| - | - | 9205 | 674.3 | - | - | 0 | - |
| - | - | 3330 | 674.8 | - | - | 0 | - |
| - | - | 6389 | 675.3 | - | - | 0 | - |
| - | - | 2212 | 676.3 | - | - | 0 | - |
| - | - | 1393 | 681.8 | - | - | 0 | - |
| - | - | 1773 | 682.3 | - | - | 0 | - |
| 0 | Precursor | 1.902E+04 | 682.8 | 0.004599 | 6.736 | +2 | -1 |
| - | - | 1.443E+04 | 683.3 | - | - | 0 | - |
| - | - | 8655 | 683.8 | - | - | 0 | - |
| - | - | 1789 | 684.2 | - | - | 0 | - |
| - | - | 1196 | 684.3 | - | - | 0 | - |
| - | - | 2438 | 685.3 | - | - | 0 | - |
| - | - | 1772 | 695.3 | - | - | 0 | - |
| - | - | 769.2 | 703.8 | - | - | 0 | - |
| - | - | 2564 | 721.3 | - | - | 0 | - |
| - | - | 617.6 | 722.3 | - | - | 0 | - |
| - | - | 821.6 | 723.3 | - | - | 0 | - |
| - | - | 713.8 | 725.3 | - | - | 0 | - |
| 6 | y | 1.778E+04 | 739.3 | 0.01006 | 13.6 | +1 | 5 |
| - | - | 7835 | 740.3 | - | - | 0 | - |
| - | - | 1792 | 741.3 | - | - | 0 | - |
| - | - | 2640 | 749.3 | - | - | 0 | - |
| - | - | 837.1 | 750.3 | - | - | 0 | - |
| - | - | 1587 | 759.3 | - | - | 0 | - |
| 6 | b | 1629 | 771.3 | 0.000398 | 0.516 | +1 | 6 |
| - | - | 1322 | 772.3 | - | - | 0 | - |
| 6 | b | 1267 | 789.3 | 0.004981 | 6.311 | +1 | 6 |
| - | - | 7402 | 790.3 | - | - | 0 | - |
| - | - | 3276 | 791.3 | - | - | 0 | - |
| - | - | 649.2 | 792.3 | - | - | 0 | - |
| - | - | 2525 | 804.3 | - | - | 0 | - |
| - | - | 1279 | 805.3 | - | - | 0 | - |
| - | - | 3380 | 832.3 | - | - | 0 | - |
| - | - | 1040 | 833.3 | - | - | 0 | - |
| 5 | y | 1405 | 836.3 | 0.0137 | 16.38 | +1 | 6 |
| - | - | 805.6 | 837.3 | - | - | 0 | - |
| - | - | 798.9 | 851.3 | - | - | 0 | - |
| 5 | y | 2.257E+04 | 854.3 | 0.00948 | 11.1 | +1 | 6 |
| - | - | 9884 | 855.3 | - | - | 0 | - |
| - | - | 2487 | 856.3 | - | - | 0 | - |
| - | - | 1776 | 868.3 | - | - | 0 | - |
| - | - | 759.5 | 869.3 | - | - | 0 | - |
| - | - | 2037 | 887.3 | - | - | 0 | - |
| - | - | 848.6 | 888.3 | - | - | 0 | - |
| - | - | 4152 | 896.3 | - | - | 0 | - |
| - | - | 2472 | 897.3 | - | - | 0 | - |
| - | - | 1.304E+04 | 905.3 | - | - | 0 | - |
| - | - | 6820 | 906.3 | - | - | 0 | - |
| - | - | 1669 | 907.3 | - | - | 0 | - |
| - | - | 739 | 914.3 | - | - | 0 | - |
| - | - | 726.3 | 933.4 | - | - | 0 | - |
| 4 | y | 2428 | 951.3 | 0.01184 | 12.44 | +1 | 7 |
| - | - | 1611 | 952.3 | - | - | 0 | - |
| 4 | y | 3.834E+04 | 969.3 | 0.009331 | 9.626 | +1 | 7 |
| - | - | 1.89E+04 | 970.3 | - | - | 0 | - |
| - | - | 6719 | 971.3 | - | - | 0 | - |
| - | - | 706 | 997.4 | - | - | 0 | - |
| - | - | 1700 | 1034 | - | - | 0 | - |
| - | - | 762 | 1035 | - | - | 0 | - |
| - | - | 1141 | 1050 | - | - | 0 | - |
| - | - | 1.909E+04 | 1052 | - | - | 0 | - |
| - | - | 1047 | 1053 | - | - | 0 | - |
| - | - | 1.142E+04 | 1053 | - | - | 0 | - |
| - | - | 3542 | 1054 | - | - | 0 | - |
| 3 | y | 3588 | 1098 | 0.01093 | 9.951 | +1 | 8 |
| - | - | 2314 | 1099 | - | - | 0 | - |
| - | - | 860 | 1100 | - | - | 0 | - |
| 3 | y | 6.017E+04 | 1116 | 0.009032 | 8.091 | +1 | 8 |
| - | - | 3.638E+04 | 1117 | - | - | 0 | - |
| - | - | 1.201E+04 | 1118 | - | - | 0 | - |
| - | - | 1952 | 1119 | - | - | 0 | - |
| 9 | b | 2186 | 1144 | 0.004434 | 3.874 | +1 | 9 |
| - | - | 1187 | 1145 | - | - | 0 | - |
| - | - | 832.6 | 1146 | - | - | 0 | - |
| - | - | 3984 | 1153 | - | - | 0 | - |
| - | - | 2479 | 1154 | - | - | 0 | - |
| 2 | y | 1953 | 1199 | 5.291E-06 | 0.004411 | +1 | 9 |
| 2 | y | 1.105E+04 | 1217 | 0.007985 | 6.559 | +1 | 9 |
| - | - | 7513 | 1218 | - | - | 0 | - |
| - | - | 2933 | 1219 | - | - | 0 | - |
| - | - | 2098 | 1227 | - | - | 0 | - |
| - | - | 1677 | 1228 | - | - | 0 | - |
| - | - | 661 | 2251 | - | - | 0 | - |
| - | - | 796.1 | 3074 | - | - | 0 | - |

m/z Charge Intensity FragmentType MassShift Position
120.08113098144531 0 177664.86
121.08438873291016 0 13735.702
121.86904907226562 0 421.1499
129.10238647460938 0 2607.2388
130.06546020507812 0 1707.4832
130.08663940429688 0 557.5014
133.08627319335938 0 1520.4159
136.07598876953125 0 12656.978
137.079345703125 0 1267.138
138.05514526367188 0 1258.8282
138.06649780273438 0 1354.9158
140.02520751953125 0 412.72394
143.1180419921875 0 2979.2156
146.06040954589844 0 580.89185
146.09669494628906 0 837.7841
147.10174560546875 0 685.763
147.11373901367188 0 559.173
148.08721923828125 0 742.6661
149.04513549804688 0 1354.6783
153.06625366210938 0 645.4091
155.08184814453125 0 764.6464
155.09310913085938 0 596.3332
156.07705688476562 0 5329.1846
158.06094360351562 0 1107.5771
158.09674072265625 0 3925.162
159.09194946289062 0 4467.876
159.0990447998047 0 812.8482
160.09478759765625 0 645.833
160.1122589111328 0 567.47687
166.05360412597656 0 662.4822
166.06134033203125 0 1646.569
166.08705139160156 0 574.8697
167.05567932128906 0 2702.4392
167.09317016601562 0 1211.935
169.13381958007812 0 1672.086
171.113037109375 0 1567.2434
173.4502716064453 0 4473.4263
174.06642150878906 0 563.82465
175.08689880371094 0 10843.246
175.09564208984375 0 1094.8517
176.08270263671875 0 1219.7618
176.1072998046875 0 79811.5
176.44451904296875 0 534.3179
177.1024932861328 0 831.4197
177.11080932617188 0 9288.647
180.07720947265625 0 536.6552
185.0564727783203 0 775.09924
186.0916290283203 0 12372.394
187.0950469970703 0 1688.6888
191.11837768554688 0 1675.5886
193.10861206054688 0 4931.373
194.1117401123047 0 820.8132
197.12857055664062 0 1191.1787
202.05300903320312 0 700.0217
203.06610107421875 0 864.73737
203.081787109375 0 2756.2476
203.09181213378906 0 586.56537
203.11814880371094 0 5864.9795 a Water loss 1
204.07708740234375 0 2621.1387
205.09716796875 0 736.4126
207.11331176757812 0 1265.7327 d 1
211.0826416015625 0 570.42535
212.1183319091797 0 1767.2554
214.08668518066406 0 978.9646
217.097412109375 0 4948.992
219.07980346679688 0 889.9487
219.11233520507812 0 610.6722
221.09230041503906 0 22612.895
221.10369873046875 0 11917.685
221.12876892089844 0 103941.77 a 1
222.09571838378906 0 2077.6086
222.1068572998047 0 1313.533
222.1320037841797 0 12245.945
223.0838165283203 0 596.21747
223.13433837890625 0 556.96106
225.04310607910156 0 4713.2134
226.0439453125 0 821.2726
228.11325073242188 0 614.0186
231.0617218017578 0 1336.0159
231.11314392089844 0 26435.236 b Water loss 1
232.1162567138672 0 3198.1418
235.10806274414062 0 8047.4546
236.1112823486328 0 870.63074
239.09519958496094 0 7223.9746
239.11415100097656 0 1062.7773
240.09625244140625 0 1119.9689
240.11288452148438 0 812.51575
247.1209716796875 0 562.5623
248.1503143310547 0 550.61554
249.12368774414062 0 56101.09 b 1
250.126953125 0 7594.0103
251.10289001464844 0 6072.7637
251.1273193359375 0 586.9216
257.1064453125 0 2717.2786
258.14501953125 0 948.1863
259.10736083984375 0 833.3194
259.48150634765625 0 529.4908
263.1028137207031 0 11250.178
264.1061096191406 0 1486.9275
267.02197265625 0 537.4048
267.0914306640625 0 641.16254
274.12994384765625 0 2976.4397
276.1552429199219 0 935.14795
279.09771728515625 0 3762.211
281.4518127441406 0 553.0112
285.00946044921875 0 1347.9098
285.1019592285156 0 5374.852
287.1040954589844 0 559.6604
289.13568115234375 0 589.42596
292.14080810546875 0 3759.8342
295.1033630371094 0 1458.2574
296.1038513183594 0 544.9168
299.0621643066406 0 4154.23
300.0621032714844 0 680.61395
306.1449890136719 0 835.302
312.14581298828125 0 2769.3857
316.1869201660156 0 1336.9342
318.1461486816406 0 1220.754
322.13006591796875 0 1214.1141
325.1385192871094 0 1475.5312
333.1603698730469 0 1553.1863
338.12744140625 0 957.93524
339.17010498046875 0 979.6855
340.1409606933594 0 11402.613
341.0181579589844 0 1049.3816
341.1441345214844 0 1717.9016
346.1415710449219 0 1137.267
348.1190185546875 0 800.40594
350.13482666015625 0 3219.5168
350.18743896484375 0 1218.4893 a Water loss 2
355.0703430175781 0 3773.799
356.1385498046875 0 2880.6333
358.1512756347656 0 50652.11 y 8
359.02862548828125 0 6318.104
359.1543273925781 0 9505.637
360.1545715332031 0 876.68976
364.15069580078125 0 4664.968
365.15325927734375 0 722.10425
368.1368408203125 0 1157.468
369.1208801269531 0 921.5819
373.08135986328125 0 631.7514
376.11370849609375 0 1061.8114
378.1298522949219 0 14694.216
378.1814270019531 0 10778.79 b Water loss 2
379.1332092285156 0 2352.9465
379.1846923828125 0 2751.5425
381.144775390625 0 566.58795
382.1441650390625 0 1209.3127
394.12493896484375 0 4747.7837
396.19073486328125 0 718.99274 b 2
406.1498718261719 0 693.1605
412.7087707519531 0 716.5209
413.1875 0 688.0195
415.1722106933594 0 792.6319
417.1733093261719 0 571.6803
418.9972839355469 0 864.068
423.1779479980469 0 1435.2592
424.16253662109375 0 1611.3483
425.1673889160156 0 611.1136
426.166748046875 0 916.02167
427.21044921875 0 1142.0446
429.0893249511719 0 6984.558
433.17083740234375 0 647.0918
439.17041015625 0 625.5272
441.18896484375 0 8832.042
442.19158935546875 0 1891.4872
447.1515808105469 0 774.63934
447.1959533691406 0 546.89996
455.2037658691406 0 1058.7396
461.166015625 0 1799.5879
465.1627197265625 0 3291.3235
466.124267578125 0 537.1229
466.1656799316406 0 1202.2698
477.1888427734375 0 1596.0781
479.17864990234375 0 2716.5337
489.16217041015625 0 787.1335
491.2098083496094 0 1290.8505
493.2086181640625 0 1516.2322 b Water loss 3
494.21514892578125 0 952.3412
495.18994140625 0 1174.988 b Water loss 7
497.1703186035156 0 824.51685
505.1868896484375 0 14012.899
506.1899719238281 0 4263.275
511.2187805175781 0 5491.8823 b 3
512.22509765625 0 7480.628
513.1969604492188 0 1759.541
513.2295532226562 0 1715.9846
514.1974487304688 0 1057.8068
519.2044067382812 0 1391.2551
526.7080078125 0 1168.0242
527.2080688476562 0 1072.3868
541.1923828125 0 1972.253
542.2357177734375 0 1146.6132
548.1968383789062 0 915.4745
549.70263671875 0 1753.34 y Water loss 2
550.2031860351562 0 812.1914
558.212158203125 0 1334.9851
558.7083129882812 0 16945.25 y 2
559.2102661132812 0 10037.436
559.7106323242188 0 3925.5981
560.2107543945312 0 760.4213
570.2320556640625 0 1218.9624
576.2237548828125 0 18861.309 y 6
577.22705078125 0 6683.4756
578.2286376953125 0 1579.648
584.2340698242188 0 678.22076
596.765869140625 0 1003.8844
600.2276000976562 0 1717.5947 y Water loss 1
600.730224609375 0 1383.5244
601.2239990234375 0 622.9559
606.234619140625 0 1418.355
607.2383422851562 0 666.8569
608.2366943359375 0 2381.056 b Water loss 4
609.232666015625 0 7727.3213 y 1
609.73486328125 0 3156.8977
610.2332763671875 0 942.53107
612.1904907226562 0 1078.237
612.2400512695312 0 909.3543
623.929931640625 0 1033.9615
624.2299194335938 0 1033.1447
626.2454833984375 0 4419.486 b 4
627.2525634765625 0 926.1912
634.23046875 0 1650.4448
643.2438354492188 0 669.3355
650.7670288085938 0 2322.4963
651.2680053710938 0 1436.1885
651.76904296875 0 1036.3726
657.26318359375 0 1018.2985
659.7651977539062 0 713.77844
667.1773071289062 0 656.271
673.7611694335938 0 8588.835 Precursor Water loss
674.2628784179688 0 9204.581
674.7631225585938 0 3329.5645
675.2880859375 0 6389.411
676.2924194335938 0 2212.4988
681.8344116210938 0 1392.6533
682.3367919921875 0 1773.2295
682.7659301757812 0 19015.457 Precursor
683.2680053710938 0 14429.48
683.7688598632812 0 8655.426
684.2023315429688 0 1789.3442
684.2642211914062 0 1195.9229
685.2587890625 0 2437.9885
695.2695922851562 0 1771.9546
703.7692260742188 0 769.2413
721.2637329101562 0 2564.455
722.2666625976562 0 617.55725
723.2561645507812 0 821.59454
725.288330078125 0 713.76935
739.2870483398438 0 17778.26 y 5
740.2901611328125 0 7835.348
741.2879638671875 0 1791.5288
749.2562866210938 0 2640
750.2598266601562 0 837.0805
759.2678833007812 0 1587.1161
771.2980346679688 0 1629.4297 b Water loss 5
772.3043823242188 0 1321.9995
789.3040161132812 0 1266.546 b 5
790.3143920898438 0 7402.166
791.3193969726562 0 3275.9639
792.3204345703125 0 649.21857
804.3330078125 0 2524.8293
805.33056640625 0 1278.8815
832.3270874023438 0 3380.3542
833.3302001953125 0 1039.6903
836.3070678710938 0 1404.872 y Water loss 4
837.304443359375 0 805.61774
851.3344116210938 0 798.9441
854.3134155273438 0 22574.246 y 4
855.31689453125 0 9884.377
856.3187255859375 0 2486.7078
868.329345703125 0 1775.6964
869.32373046875 0 759.4731
887.3333740234375 0 2036.935
888.3358764648438 0 848.5506
896.3241577148438 0 4151.7354
897.3242797851562 0 2472.3267
905.3417358398438 0 13039.412
906.3453979492188 0 6819.6963
907.3462524414062 0 1669.2754
914.3291625976562 0 739.0493
933.3732299804688 0 726.29065
951.3321533203125 0 2427.515 y Water loss 3
952.3246459960938 0 1610.9525
969.3402099609375 0 38342.133 y 3
970.3434448242188 0 18904.574
971.3433837890625 0 6719.1655
997.3721313476562 0 706.0288
1034.3992919921875 0 1699.6619
1035.399658203125 0 761.9728
1049.5377197265625 0 1140.879
1052.4100341796875 0 19087.596
1053.2947998046875 0 1046.611
1053.413330078125 0 11421.611
1054.4171142578125 0 3541.5928
1098.399658203125 0 3588.0315 y Water loss 2
1099.3997802734375 0 2313.871
1100.4095458984375 0 859.97015
1116.4083251953125 0 60168.324 y 2
1117.411376953125 0 36382.98
1118.4117431640625 0 12013.907
1119.4117431640625 0 1951.9192
1144.43994140625 0 2186.2903 b 8
1145.4365234375 0 1186.6302
1146.44091796875 0 832.55554
1153.4566650390625 0 3984.276
1154.460693359375 0 2479.1125
1199.4364013671875 0 1952.5161 y Water loss 1
1217.4549560546875 0 11045.182 y 1
1218.457275390625 0 7513.0947
1219.456787109375 0 2932.7554
1227.4364013671875 0 2097.5754
1228.4385986328125 0 1676.9897
2251.30517578125 0 660.9842
3074.271728515625 0 796.1485

Spectrum Details

|  |  |
| --- | --- |
| Matched peaks? Matched peaks The total absolute number of peaks matched. Additionally in brackets the total fraction of peaks matched and the total number of peaks is shown. | 33 (10.48% of 315) |
| FDR? FDR The false discovery rate estimated for this peptide. It is calculated by matching all theoretical fragments with a non-integer shift with the raw peaks for this spectrum. This is done with 40 different shifts. The resulting percentage is the average number of annotated peaks over the number of annotated peaks with the correct spectrum. | 0.29% |
| Satellite FDR? Satellite FDR See the FDR for details on its calculation. This satellite ion specific FDR only contains the satellite ions (d/w) for I/L/J positions. | - |
| PSM Score? PSM Score The PSM Score as given by Hecklib to this annotated spectrum. It is shown with three significant figures. | 376 |

#### Spectrum 8649? Spectrum 8649 The raw spectrum of this peptide as annotated by Hecklib. The fragments are coloured according to ion type (see legend). Any peaks with a star '\*' as text can be hovered over to see the full details, first the ion type second the mass shift type. By hovering over the amino acids in the peptide or ions in the legend the corresponding peaks are highlighted. By toggling the 'Unassigned' label you can turn the background (unassigned) peaks on or off in the plot. By updating the slider in the Ion legend you can update the spectrum to only show the top X% of the peaks with labels. The top X% means any peak that is within X% of the highest intensity. By dragging in the spectrum you can zoom in to a specific part of the spectrum and use 'Zoom Out' to get back to the original zoom level. The annotation of the spectrum is based on the given sequence in the peptides file and is done with different software so inconsistencies are likely. The peaks are annotated based on the given sequence, with 20 ppm tolerance.

Copy Data

##### Spectrum 8649 (TSV)

###### Preview

```
Loading example...
```

*Click on the button to copy the data to your clipboard.*

Mz MinMz MaxIntensity Max

WidthHeightPeptide font sizePeptide stroke widthSpectrum font sizeSpectrum stroke widthCompact peptide

Ion legend

wxyz

abcd

OtherUnassignedIon^Charge^_Position_Show for top:%

FTFDDYAMHW

03.15e+46.30e+49.46e+41.26e+5

Zoom Out

y^+1^_1_c^+1^_2_y^+1^_2_y^+1^_3_c^+1^_4_y^+2^_8_y^+2^_8_z^+1^_4_y^+1^_4_y^+2^_9_c^+1^_5_z^+1^_5_y^+1^_5_c^+1^_6_z^+1^_6_y^+1^_6_c^+1^_7_z^+1^_7_y^+1^_7_c^+1^_8_y^+1^_8_z^+1^_8_y^+1^_8_c^+1^_9_c^+1^_9_y^+1^_9_z^+1^_9_y^+1^_9_

0709141921282837

Fragment Matches Table

Show background peaks

| Position | Ion type | Intensity | mz Theoretical | mz Error (Th) | mz Error (ppm) | Charge | Series Number |
| --- | --- | --- | --- | --- | --- | --- | --- |
| - | - | 1.202E+04 | 120.1 | - | - | 0 | - |
| - | - | 529.4 | 121.1 | - | - | 0 | - |
| - | - | 369.4 | 123.9 | - | - | 0 | - |
| - | - | 377.5 | 125.2 | - | - | 0 | - |
| - | - | 416.6 | 126.7 | - | - | 0 | - |
| - | - | 452.6 | 129.3 | - | - | 0 | - |
| - | - | 394.5 | 130.6 | - | - | 0 | - |
| - | - | 503.8 | 131.1 | - | - | 0 | - |
| - | - | 3089 | 133.1 | - | - | 0 | - |
| - | - | 805.8 | 149 | - | - | 0 | - |
| - | - | 410.2 | 149.2 | - | - | 0 | - |
| - | - | 3250 | 173.4 | - | - | 0 | - |
| - | - | 3665 | 176.1 | - | - | 0 | - |
| - | - | 1630 | 177.1 | - | - | 0 | - |
| - | - | 607 | 205.1 | - | - | 0 | - |
| 10 | y | 1769 | 221.1 | 0.003048 | 13.78 | +1 | 1 |
| - | - | 1630 | 221.1 | - | - | 0 | - |
| - | - | 1.522E+04 | 221.1 | - | - | 0 | - |
| - | - | 761 | 221.1 | - | - | 0 | - |
| - | - | 1753 | 222.1 | - | - | 0 | - |
| - | - | 1310 | 225 | - | - | 0 | - |
| - | - | 594.4 | 229.7 | - | - | 0 | - |
| - | - | 2510 | 231.1 | - | - | 0 | - |
| - | - | 568.9 | 235.1 | - | - | 0 | - |
| - | - | 519.6 | 239 | - | - | 0 | - |
| - | - | 2764 | 239.1 | - | - | 0 | - |
| 2 | c | 1.552E+04 | 249.1 | 2.898E-05 | 0.1163 | +1 | 2 |
| - | - | 1575 | 250.1 | - | - | 0 | - |
| - | - | 1813 | 263.1 | - | - | 0 | - |
| - | - | 582.5 | 272.3 | - | - | 0 | - |
| - | - | 1885 | 295.1 | - | - | 0 | - |
| - | - | 1729 | 299.1 | - | - | 0 | - |
| - | - | 611.1 | 313.1 | - | - | 0 | - |
| - | - | 2917 | 355.1 | - | - | 0 | - |
| 9 | y | 4037 | 358.1 | 0.00466 | 13.01 | +1 | 2 |
| - | - | 1675 | 359 | - | - | 0 | - |
| - | - | 708.3 | 364.2 | - | - | 0 | - |
| - | - | 846.3 | 369.1 | - | - | 0 | - |
| - | - | 940.9 | 378.1 | - | - | 0 | - |
| - | - | 2857 | 378.2 | - | - | 0 | - |
| - | - | 1006 | 379.2 | - | - | 0 | - |
| - | - | 628 | 410.1 | - | - | 0 | - |
| - | - | 1113 | 415.2 | - | - | 0 | - |
| - | - | 8101 | 429.1 | - | - | 0 | - |
| - | - | 571.9 | 479.2 | - | - | 0 | - |
| - | - | 756.9 | 480.3 | - | - | 0 | - |
| - | - | 1493 | 489.2 | - | - | 0 | - |
| - | - | 3239 | 490.2 | - | - | 0 | - |
| - | - | 780.5 | 491.2 | - | - | 0 | - |
| - | - | 1687 | 504.2 | - | - | 0 | - |
| 8 | y | 2733 | 505.2 | 0.009301 | 18.41 | +1 | 3 |
| - | - | 616.9 | 506.2 | - | - | 0 | - |
| 4 | c | 1554 | 511.2 | 0.001459 | 2.853 | +1 | 4 |
| - | - | 1163 | 512.2 | - | - | 0 | - |
| 3 | y | 877.1 | 549.7 | 0.00616 | 11.21 | +2 | 8 |
| 3 | y | 7414 | 558.7 | 0.004784 | 8.563 | +2 | 8 |
| - | - | 3931 | 559.2 | - | - | 0 | - |
| - | - | 1874 | 559.7 | - | - | 0 | - |
| 7 | z | 4291 | 560.2 | 0.01093 | 19.51 | +1 | 4 |
| - | - | 1460 | 561.2 | - | - | 0 | - |
| 7 | y | 5401 | 576.2 | 0.009785 | 16.98 | +1 | 4 |
| - | - | 1520 | 577.2 | - | - | 0 | - |
| - | - | 828.6 | 578.2 | - | - | 0 | - |
| 2 | y | 1509 | 609.2 | 0.003711 | 6.091 | +2 | 9 |
| - | - | 1215 | 609.7 | - | - | 0 | - |
| 5 | c | 1410 | 626.2 | 0.0001854 | 0.2961 | +1 | 5 |
| - | - | 1250 | 635.4 | - | - | 0 | - |
| - | - | 1086 | 636.4 | - | - | 0 | - |
| - | - | 860.4 | 649.3 | - | - | 0 | - |
| - | - | 685.2 | 650.3 | - | - | 0 | - |
| - | - | 921.2 | 650.8 | - | - | 0 | - |
| - | - | 1108 | 651.3 | - | - | 0 | - |
| - | - | 3121 | 673.8 | - | - | 0 | - |
| - | - | 2119 | 674.3 | - | - | 0 | - |
| - | - | 1070 | 674.8 | - | - | 0 | - |
| - | - | 2151 | 681.4 | - | - | 0 | - |
| - | - | 1157 | 682.3 | - | - | 0 | - |
| - | - | 1286 | 682.4 | - | - | 0 | - |
| - | - | 4591 | 682.8 | - | - | 0 | - |
| - | - | 5746 | 683.3 | - | - | 0 | - |
| - | - | 2310 | 683.8 | - | - | 0 | - |
| - | - | 1996 | 684.2 | - | - | 0 | - |
| - | - | 1481 | 693.2 | - | - | 0 | - |
| - | - | 1263 | 694.2 | - | - | 0 | - |
| 6 | z | 1.394E+04 | 723.3 | 0.009676 | 13.38 | +1 | 5 |
| - | - | 9898 | 724.3 | - | - | 0 | - |
| - | - | 2748 | 725.3 | - | - | 0 | - |
| - | - | 942.4 | 738.3 | - | - | 0 | - |
| 6 | y | 6009 | 739.3 | 0.009689 | 13.11 | +1 | 5 |
| - | - | 2203 | 740.3 | - | - | 0 | - |
| - | - | 804.4 | 741.3 | - | - | 0 | - |
| - | - | 1162 | 764.3 | - | - | 0 | - |
| - | - | 1128 | 766.3 | - | - | 0 | - |
| 6 | c | 743 | 789.3 | 0.001136 | 1.439 | +1 | 6 |
| - | - | 686.2 | 790.3 | - | - | 0 | - |
| - | - | 5302 | 794.3 | - | - | 0 | - |
| - | - | 1944 | 795.3 | - | - | 0 | - |
| - | - | 728.1 | 796.3 | - | - | 0 | - |
| - | - | 1188 | 808.3 | - | - | 0 | - |
| 5 | z | 1.76E+04 | 838.3 | 0.009771 | 11.66 | +1 | 6 |
| - | - | 2.659E+04 | 839.3 | - | - | 0 | - |
| - | - | 9841 | 840.3 | - | - | 0 | - |
| - | - | 2299 | 841.3 | - | - | 0 | - |
| - | - | 560 | 847.3 | - | - | 0 | - |
| 5 | y | 6906 | 854.3 | 0.009113 | 10.67 | +1 | 6 |
| - | - | 3147 | 855.3 | - | - | 0 | - |
| - | - | 780.3 | 856.3 | - | - | 0 | - |
| 7 | c | 1327 | 877.4 | 0.005392 | 6.145 | +1 | 7 |
| - | - | 1294 | 896.3 | - | - | 0 | - |
| - | - | 1319 | 905.3 | - | - | 0 | - |
| - | - | 2683 | 909.3 | - | - | 0 | - |
| - | - | 1213 | 910.3 | - | - | 0 | - |
| - | - | 597.8 | 952.3 | - | - | 0 | - |
| 4 | z | 3797 | 953.3 | 0.01023 | 10.73 | +1 | 7 |
| - | - | 1.466E+04 | 954.3 | - | - | 0 | - |
| - | - | 5920 | 955.3 | - | - | 0 | - |
| - | - | 2768 | 956.3 | - | - | 0 | - |
| 4 | y | 1.221E+04 | 969.3 | 0.008782 | 9.06 | +1 | 7 |
| - | - | 5915 | 970.3 | - | - | 0 | - |
| - | - | 1767 | 971.3 | - | - | 0 | - |
| - | - | 1074 | 980.4 | - | - | 0 | - |
| - | - | 1498 | 981.4 | - | - | 0 | - |
| - | - | 1595 | 982.4 | - | - | 0 | - |
| - | - | 8903 | 1023 | - | - | 0 | - |
| 8 | c | 1.159E+04 | 1024 | 0.003471 | 3.389 | +1 | 8 |
| - | - | 6111 | 1025 | - | - | 0 | - |
| - | - | 2793 | 1026 | - | - | 0 | - |
| - | - | 3235 | 1052 | - | - | 0 | - |
| - | - | 1672 | 1053 | - | - | 0 | - |
| - | - | 3711 | 1056 | - | - | 0 | - |
| - | - | 1871 | 1057 | - | - | 0 | - |
| - | - | 993.4 | 1058 | - | - | 0 | - |
| 3 | y | 943.3 | 1098 | 0.00739 | 6.728 | +1 | 8 |
| 3 | z | 1999 | 1100 | 0.007737 | 7.031 | +1 | 8 |
| - | - | 807.5 | 1101 | - | - | 0 | - |
| 3 | y | 2.149E+04 | 1116 | 0.009765 | 8.747 | +1 | 8 |
| - | - | 1.123E+04 | 1117 | - | - | 0 | - |
| - | - | 5046 | 1118 | - | - | 0 | - |
| - | - | 795.1 | 1119 | - | - | 0 | - |
| - | - | 1903 | 1128 | - | - | 0 | - |
| - | - | 1056 | 1129 | - | - | 0 | - |
| 9 | c | 1595 | 1144 | 0.003946 | 3.448 | +1 | 9 |
| - | - | 8905 | 1146 | - | - | 0 | - |
| - | - | 6067 | 1147 | - | - | 0 | - |
| - | - | 2367 | 1148 | - | - | 0 | - |
| - | - | 1061 | 1158 | - | - | 0 | - |
| 9 | c | 3.891E+04 | 1161 | 0.004252 | 3.661 | +1 | 9 |
| - | - | 2.537E+04 | 1162 | - | - | 0 | - |
| - | - | 9774 | 1163 | - | - | 0 | - |
| - | - | 1520 | 1164 | - | - | 0 | - |
| - | - | 788.7 | 1172 | - | - | 0 | - |
| - | - | 4808 | 1187 | - | - | 0 | - |
| - | - | 3006 | 1188 | - | - | 0 | - |
| - | - | 1045 | 1189 | - | - | 0 | - |
| 2 | y | 2426 | 1199 | 0.01111 | 9.266 | +1 | 9 |
| - | - | 4567 | 1200 | - | - | 0 | - |
| 2 | z | 8386 | 1201 | 0.006445 | 5.365 | +1 | 9 |
| - | - | 9728 | 1202 | - | - | 0 | - |
| - | - | 1.697E+04 | 1203 | - | - | 0 | - |
| - | - | 9664 | 1204 | - | - | 0 | - |
| - | - | 3164 | 1205 | - | - | 0 | - |
| - | - | 2162 | 1216 | - | - | 0 | - |
| 2 | y | 3981 | 1217 | 0.01262 | 10.37 | +1 | 9 |
| - | - | 7277 | 1218 | - | - | 0 | - |
| - | - | 3227 | 1219 | - | - | 0 | - |
| - | - | 1668 | 1220 | - | - | 0 | - |
| - | - | 1229 | 1227 | - | - | 0 | - |
| - | - | 2362 | 1287 | - | - | 0 | - |
| - | - | 835.2 | 1288 | - | - | 0 | - |
| - | - | 710.1 | 1289 | - | - | 0 | - |
| - | - | 981.8 | 1302 | - | - | 0 | - |
| - | - | 755.9 | 1303 | - | - | 0 | - |
| - | - | 1.361E+04 | 1304 | - | - | 0 | - |
| - | - | 1.059E+04 | 1305 | - | - | 0 | - |
| - | - | 2.596E+04 | 1306 | - | - | 0 | - |
| - | - | 1.901E+04 | 1307 | - | - | 0 | - |
| - | - | 8899 | 1308 | - | - | 0 | - |
| - | - | 1672 | 1309 | - | - | 0 | - |
| - | - | 782.1 | 1332 | - | - | 0 | - |
| - | - | 836.9 | 1333 | - | - | 0 | - |
| - | - | 827 | 1339 | - | - | 0 | - |
| - | - | 1.71E+04 | 1348 | - | - | 0 | - |
| - | - | 7.46E+04 | 1349 | - | - | 0 | - |
| - | - | 5.506E+04 | 1350 | - | - | 0 | - |
| - | - | 2.763E+04 | 1351 | - | - | 0 | - |
| - | - | 4770 | 1352 | - | - | 0 | - |
| - | - | 762.9 | 1353 | - | - | 0 | - |
| - | - | 3.514E+04 | 1365 | - | - | 0 | - |
| - | - | 1.248E+05 | 1366 | - | - | 0 | - |
| - | - | 8.784E+04 | 1367 | - | - | 0 | - |
| - | - | 4.123E+04 | 1368 | - | - | 0 | - |
| - | - | 7268 | 1369 | - | - | 0 | - |
| - | - | 640.5 | 2809 | - | - | 0 | - |

m/z Charge Intensity FragmentType MassShift Position
120.08088684082031 0 12021.711
121.08404541015625 0 529.41174
123.86109924316406 0 369.40692
125.2017822265625 0 377.5494
126.74169921875 0 416.6041
129.30226135253906 0 452.63226
130.56298828125 0 394.51617
131.07077026367188 0 503.78384
133.0859832763672 0 3089.2861
149.04493713378906 0 805.7879
149.22193908691406 0 410.1668
173.43934631347656 0 3249.85
176.10707092285156 0 3665.28
177.11146545410156 0 1630.3119
205.1072998046875 0 606.9627
221.0841064453125 0 1769.1487 y 9
221.0930633544922 0 1630.4526
221.12844848632812 0 15215.456
221.13894653320312 0 761.0217
222.13168334960938 0 1752.7465
225.0426025390625 0 1309.6172
229.68214416503906 0 594.3839
231.11273193359375 0 2510.0134
235.1079864501953 0 568.9251
239.0400390625 0 519.6232
239.09481811523438 0 2763.7515
249.12339782714844 0 15523.334 c Ammonia loss 1
250.1263885498047 0 1575.3688
263.10223388671875 0 1812.7078
272.2875671386719 0 582.5207
295.1033020019531 0 1884.6997
299.0614318847656 0 1729.0972
313.114013671875 0 611.08594
355.0692138671875 0 2916.8176
358.1507263183594 0 4037.0168 y 8
359.0282287597656 0 1674.8541
364.1514892578125 0 708.27625
369.1211853027344 0 846.34174
378.1292419433594 0 940.92926
378.1816101074219 0 2856.5078
379.1861267089844 0 1006.18823
410.06195068359375 0 628.0212
415.1518249511719 0 1112.5564
429.08905029296875 0 8100.737
479.1809387207031 0 571.9181
480.2669982910156 0 756.9048
489.1682434082031 0 1493.0361
490.1750793457031 0 3239.1821
491.1757507324219 0 780.4856
504.1784362792969 0 1686.8948
505.18585205078125 0 2733.0344 y 7
506.19024658203125 0 616.9407
511.2201843261719 0 1554.2261 c Ammonia loss 3
512.2244262695312 0 1163.3428
549.7041625976562 0 877.1337 y Water loss 2
558.7080688476562 0 7414.2144 y 2
559.2099609375 0 3931.0571
559.70849609375 0 1874.2365
560.2058715820312 0 4290.674 z 6
561.2080078125 0 1459.8167
576.2234497070312 0 5400.5654 y 6
577.225341796875 0 1519.5773
578.2202758789062 0 828.59216
609.2308349609375 0 1508.9623 y 1
609.7347412109375 0 1214.9727
626.2454833984375 0 1410.2991 c Ammonia loss 4
635.3849487304688 0 1250.1519
636.3886108398438 0 1086.4075
649.256591796875 0 860.4106
650.2557983398438 0 685.1964
650.7679443359375 0 921.2255
651.267578125 0 1107.5913
673.7608642578125 0 3120.9993
674.260009765625 0 2118.5164
674.7627563476562 0 1070.3857
681.4102172851562 0 2151.3208
682.3407592773438 0 1156.5887
682.4185180664062 0 1286.4014
682.7661743164062 0 4591.03
683.2681274414062 0 5745.5723
683.7676391601562 0 2310.443
684.20068359375 0 1995.637
693.2425537109375 0 1481.0027
694.247802734375 0 1262.6249
723.2679443359375 0 13938.076 z 5
724.2723999023438 0 9898.184
725.2764892578125 0 2747.7996
738.2810668945312 0 942.3914
739.2866821289062 0 6008.667 y 5
740.2896118164062 0 2202.6116
741.2923583984375 0 804.4345
764.27734375 0 1162.2814
766.2743530273438 0 1127.5282
789.307861328125 0 742.9903 c Ammonia loss 5
790.3174438476562 0 686.247
794.3043823242188 0 5301.7627
795.3055419921875 0 1944.4495
796.3136596679688 0 728.0983
808.2637939453125 0 1188.1206
838.2949829101562 0 17601.646 z 4
839.3010864257812 0 26587.057
840.3040771484375 0 9840.712
841.3035278320312 0 2298.869
847.343017578125 0 560.0279
854.3130493164062 0 6905.68 y 4
855.3164672851562 0 3146.758
856.3153686523438 0 780.2747
877.3780517578125 0 1327.3923 c 6
896.3245849609375 0 1293.5017
905.3436279296875 0 1318.9965
909.3322143554688 0 2683.2598
910.3291625976562 0 1213.4365
952.3196411132812 0 597.8384
953.3223876953125 0 3797.4717 z 3
954.3290405273438 0 14662.502
955.3338012695312 0 5920.3467
956.335693359375 0 2768.2327
969.3396606445312 0 12209.138 y 3
970.3443603515625 0 5914.974
971.3418579101562 0 1766.895
980.4002685546875 0 1074.0227
981.4022216796875 0 1497.8986
982.4031372070312 0 1594.6176
1023.3995971679688 0 8903.316
1024.4066162109375 0 11590.699 c 7
1025.40869140625 0 6111.0483
1026.41015625 0 2792.712
1052.4124755859375 0 3234.7085
1053.415771484375 0 1672.4271
1056.3873291015625 0 3710.519
1057.3905029296875 0 1870.5807
1058.3837890625 0 993.39453
1098.3961181640625 0 943.26154 y Water loss 2
1100.3883056640625 0 1999.1187 z 2
1101.391845703125 0 807.4785
1116.4090576171875 0 21490.916 y 2
1117.41259765625 0 11225.018
1118.4139404296875 0 5045.6245
1119.4122314453125 0 795.1266
1128.444580078125 0 1902.7499
1129.4512939453125 0 1055.7566
1144.439453125 0 1594.9176 c Ammonia loss 8
1146.4554443359375 0 8904.602
1147.45849609375 0 6067.293
1148.4593505859375 0 2367.2551
1158.46728515625 0 1061.0665
1161.46630859375 0 38914.297 c 8
1162.4686279296875 0 25369.668
1163.468994140625 0 9773.822
1164.470703125 0 1519.5199
1172.483642578125 0 788.6897
1187.482666015625 0 4808.2334
1188.4810791015625 0 3005.6838
1189.4732666015625 0 1044.7635
1199.42529296875 0 2425.6472 y Water loss 1
1200.4283447265625 0 4567.0513
1201.4346923828125 0 8385.619 z 1
1202.4527587890625 0 9727.867
1203.455810546875 0 16972.994
1204.4586181640625 0 9663.582
1205.45849609375 0 3164.096
1216.460205078125 0 2162.221
1217.4595947265625 0 3980.7732 y 1
1218.461181640625 0 7277.3257
1219.46533203125 0 3227.0735
1220.468505859375 0 1668.1235
1227.44384765625 0 1228.9421
1287.49951171875 0 2362.4194
1288.494384765625 0 835.21
1289.4906005859375 0 710.0925
1301.5335693359375 0 981.8247
1302.5335693359375 0 755.9324
1303.53125 0 13614.072
1304.53076171875 0 10593.242
1305.5157470703125 0 25962.385
1306.5137939453125 0 19010.963
1307.5147705078125 0 8898.927
1308.5159912109375 0 1671.5344
1331.5013427734375 0 782.1297
1333.4920654296875 0 836.87854
1338.528564453125 0 827.0369
1347.5208740234375 0 17099.762
1348.5106201171875 0 74603.82
1349.511474609375 0 55057.586
1350.51123046875 0 27633.57
1351.514404296875 0 4769.88
1352.5096435546875 0 762.8938
1364.523681640625 0 35144.44
1365.5294189453125 0 124836.16
1366.532470703125 0 87835.33
1367.5345458984375 0 41230.89
1368.53564453125 0 7268.4585
2809.06005859375 0 640.54553

Spectrum Details

|  |  |
| --- | --- |
| Matched peaks? Matched peaks The total absolute number of peaks matched. Additionally in brackets the total fraction of peaks matched and the total number of peaks is shown. | 28 (14.51% of 193) |
| FDR? FDR The false discovery rate estimated for this peptide. It is calculated by matching all theoretical fragments with a non-integer shift with the raw peaks for this spectrum. This is done with 40 different shifts. The resulting percentage is the average number of annotated peaks over the number of annotated peaks with the correct spectrum. | 0.26% |
| Satellite FDR? Satellite FDR See the FDR for details on its calculation. This satellite ion specific FDR only contains the satellite ions (d/w) for I/L/J positions. | - |
| PSM Score? PSM Score The PSM Score as given by Hecklib to this annotated spectrum. It is shown with three significant figures. | 360 |

#### Spectrum 9949? Spectrum 9949 The raw spectrum of this peptide as annotated by Hecklib. The fragments are coloured according to ion type (see legend). Any peaks with a star '\*' as text can be hovered over to see the full details, first the ion type second the mass shift type. By hovering over the amino acids in the peptide or ions in the legend the corresponding peaks are highlighted. By toggling the 'Unassigned' label you can turn the background (unassigned) peaks on or off in the plot. By updating the slider in the Ion legend you can update the spectrum to only show the top X% of the peaks with labels. The top X% means any peak that is within X% of the highest intensity. By dragging in the spectrum you can zoom in to a specific part of the spectrum and use 'Zoom Out' to get back to the original zoom level. The annotation of the spectrum is based on the given sequence in the peptides file and is done with different software so inconsistencies are likely. The peaks are annotated based on the given sequence, with 20 ppm tolerance.

Copy Data

##### Spectrum 9949 (TSV)

###### Preview

```
Loading example...
```

*Click on the button to copy the data to your clipboard.*

Mz MinMz MaxIntensity Max

WidthHeightPeptide font sizePeptide stroke widthSpectrum font sizeSpectrum stroke widthCompact peptide

Ion legend

wxyz

abcd

OtherUnassignedIon^Charge^_Position_Show for top:%

FTFDDYAMHW

01.66e+43.33e+44.99e+46.65e+4

Zoom Out

c^+1^_2_y^+1^_2_z^+1^_3_y^+1^_3_c^+1^_4_z^+1^_4_y^+2^_8_y^+1^_4_y^+2^_9_c^+1^_5_z^+1^_5_y^+1^_5_z^+1^_6_y^+1^_6_c^+1^_7_z^+1^_7_y^+1^_7_c^+1^_8_z^+1^_8_y^+1^_8_c^+1^_9_z^+1^_9_y^+1^_9_

048196214431924

Fragment Matches Table

Show background peaks

| Position | Ion type | Intensity | mz Theoretical | mz Error (Th) | mz Error (ppm) | Charge | Series Number |
| --- | --- | --- | --- | --- | --- | --- | --- |
| - | - | 7284 | 120.1 | - | - | 0 | - |
| - | - | 339.3 | 120.2 | - | - | 0 | - |
| - | - | 476.8 | 135.3 | - | - | 0 | - |
| - | - | 616.7 | 148.9 | - | - | 0 | - |
| - | - | 557 | 148.9 | - | - | 0 | - |
| - | - | 887.7 | 148.9 | - | - | 0 | - |
| - | - | 949.8 | 148.9 | - | - | 0 | - |
| - | - | 843.5 | 148.9 | - | - | 0 | - |
| - | - | 1556 | 148.9 | - | - | 0 | - |
| - | - | 2721 | 148.9 | - | - | 0 | - |
| - | - | 5241 | 149 | - | - | 0 | - |
| - | - | 2949 | 149 | - | - | 0 | - |
| - | - | 1654 | 149 | - | - | 0 | - |
| - | - | 894.3 | 149 | - | - | 0 | - |
| - | - | 776.7 | 149 | - | - | 0 | - |
| - | - | 746.1 | 149 | - | - | 0 | - |
| - | - | 533 | 149 | - | - | 0 | - |
| - | - | 489.4 | 149 | - | - | 0 | - |
| - | - | 459.6 | 149 | - | - | 0 | - |
| - | - | 453.5 | 152.3 | - | - | 0 | - |
| - | - | 591.8 | 166.1 | - | - | 0 | - |
| - | - | 473.5 | 173.4 | - | - | 0 | - |
| - | - | 1876 | 176.1 | - | - | 0 | - |
| - | - | 522.8 | 194.8 | - | - | 0 | - |
| - | - | 8809 | 221.1 | - | - | 0 | - |
| - | - | 715.5 | 222.1 | - | - | 0 | - |
| - | - | 1971 | 231.1 | - | - | 0 | - |
| 2 | c | 8569 | 249.1 | 0.0004593 | 1.844 | +1 | 2 |
| - | - | 881 | 250.1 | - | - | 0 | - |
| - | - | 1092 | 253.1 | - | - | 0 | - |
| - | - | 953.1 | 263.1 | - | - | 0 | - |
| - | - | 627.1 | 264.1 | - | - | 0 | - |
| - | - | 619.6 | 303.4 | - | - | 0 | - |
| - | - | 585.3 | 319.5 | - | - | 0 | - |
| 9 | y | 1952 | 342.2 | 0.0003958 | 1.157 | +1 | 2 |
| - | - | 891 | 355.1 | - | - | 0 | - |
| - | - | 1852 | 378.2 | - | - | 0 | - |
| - | - | 562.4 | 401.5 | - | - | 0 | - |
| - | - | 540.9 | 447.6 | - | - | 0 | - |
| 8 | z | 1497 | 473.2 | 0.004598 | 9.717 | +1 | 3 |
| - | - | 2946 | 474.2 | - | - | 0 | - |
| - | - | 680.2 | 475.2 | - | - | 0 | - |
| 8 | y | 1168 | 489.2 | 0.004581 | 9.365 | +1 | 3 |
| 4 | c | 1171 | 511.2 | 0.0005556 | 1.087 | +1 | 4 |
| - | - | 514.2 | 518.7 | - | - | 0 | - |
| 7 | z | 3562 | 544.2 | 0.004288 | 7.88 | +1 | 4 |
| - | - | 1172 | 545.2 | - | - | 0 | - |
| - | - | 555 | 546.2 | - | - | 0 | - |
| 3 | y | 2147 | 550.7 | 0.001371 | 2.49 | +2 | 8 |
| - | - | 1449 | 551.2 | - | - | 0 | - |
| 7 | y | 2371 | 560.2 | 0.003325 | 5.936 | +1 | 4 |
| - | - | 629.4 | 573.2 | - | - | 0 | - |
| - | - | 620.7 | 587.4 | - | - | 0 | - |
| 2 | y | 1078 | 601.2 | 0.001763 | 2.932 | +2 | 9 |
| - | - | 766.1 | 601.7 | - | - | 0 | - |
| 5 | c | 606.7 | 626.2 | 0.002627 | 4.195 | +1 | 5 |
| - | - | 623 | 633.8 | - | - | 0 | - |
| - | - | 574 | 657.2 | - | - | 0 | - |
| - | - | 1350 | 665.8 | - | - | 0 | - |
| - | - | 1462 | 666.3 | - | - | 0 | - |
| - | - | 745 | 666.8 | - | - | 0 | - |
| - | - | 2148 | 674.8 | - | - | 0 | - |
| - | - | 1863 | 675.3 | - | - | 0 | - |
| - | - | 759.7 | 675.8 | - | - | 0 | - |
| - | - | 805.4 | 676.4 | - | - | 0 | - |
| - | - | 592.7 | 706.3 | - | - | 0 | - |
| 6 | z | 6476 | 707.3 | 0.003582 | 5.064 | +1 | 5 |
| - | - | 4833 | 708.3 | - | - | 0 | - |
| - | - | 956.9 | 709.3 | - | - | 0 | - |
| 6 | y | 2712 | 723.3 | 0.004816 | 6.659 | +1 | 5 |
| - | - | 905.7 | 724.3 | - | - | 0 | - |
| - | - | 888.9 | 750.3 | - | - | 0 | - |
| - | - | 698.8 | 764.3 | - | - | 0 | - |
| - | - | 638.1 | 775.3 | - | - | 0 | - |
| - | - | 3605 | 778.3 | - | - | 0 | - |
| - | - | 1043 | 779.3 | - | - | 0 | - |
| - | - | 1091 | 780.3 | - | - | 0 | - |
| 5 | z | 8429 | 822.3 | 0.003799 | 4.62 | +1 | 6 |
| - | - | 9311 | 823.3 | - | - | 0 | - |
| - | - | 4475 | 824.3 | - | - | 0 | - |
| 5 | y | 4501 | 838.3 | 0.002409 | 2.874 | +1 | 6 |
| - | - | 1191 | 839.3 | - | - | 0 | - |
| - | - | 646.5 | 849.3 | - | - | 0 | - |
| 7 | c | 867.5 | 877.4 | 0.005595 | 6.377 | +1 | 7 |
| - | - | 773 | 878.4 | - | - | 0 | - |
| - | - | 3138 | 893.3 | - | - | 0 | - |
| - | - | 1502 | 894.3 | - | - | 0 | - |
| - | - | 651.8 | 917.4 | - | - | 0 | - |
| 4 | z | 3453 | 937.3 | 0.002918 | 3.114 | +1 | 7 |
| - | - | 7297 | 938.3 | - | - | 0 | - |
| - | - | 3420 | 939.3 | - | - | 0 | - |
| - | - | 786.6 | 940.3 | - | - | 0 | - |
| 4 | y | 5616 | 953.3 | 0.003176 | 3.332 | +1 | 7 |
| - | - | 3439 | 954.3 | - | - | 0 | - |
| - | - | 3277 | 980.4 | - | - | 0 | - |
| - | - | 1986 | 981.4 | - | - | 0 | - |
| - | - | 1280 | 982.4 | - | - | 0 | - |
| - | - | 5685 | 1023 | - | - | 0 | - |
| 8 | c | 7961 | 1024 | 0.001884 | 1.84 | +1 | 8 |
| - | - | 4735 | 1025 | - | - | 0 | - |
| - | - | 1282 | 1026 | - | - | 0 | - |
| - | - | 665.1 | 1027 | - | - | 0 | - |
| - | - | 2050 | 1036 | - | - | 0 | - |
| 3 | z | 1099 | 1084 | 0.00385 | 3.55 | +1 | 8 |
| - | - | 1002 | 1094 | - | - | 0 | - |
| - | - | 656.8 | 1095 | - | - | 0 | - |
| 3 | y | 1.151E+04 | 1100 | 0.003061 | 2.781 | +1 | 8 |
| - | - | 6693 | 1101 | - | - | 0 | - |
| - | - | 2373 | 1102 | - | - | 0 | - |
| - | - | 2349 | 1117 | - | - | 0 | - |
| - | - | 2155 | 1118 | - | - | 0 | - |
| - | - | 1690 | 1139 | - | - | 0 | - |
| - | - | 928.7 | 1140 | - | - | 0 | - |
| - | - | 1009 | 1141 | - | - | 0 | - |
| - | - | 931.2 | 1146 | - | - | 0 | - |
| - | - | 1095 | 1147 | - | - | 0 | - |
| 9 | c | 1.739E+04 | 1161 | 0.002177 | 1.874 | +1 | 9 |
| - | - | 1.093E+04 | 1162 | - | - | 0 | - |
| - | - | 3822 | 1163 | - | - | 0 | - |
| - | - | 2658 | 1184 | - | - | 0 | - |
| 2 | z | 5116 | 1185 | 0.001358 | 1.145 | +1 | 9 |
| - | - | 2951 | 1186 | - | - | 0 | - |
| - | - | 1552 | 1187 | - | - | 0 | - |
| 2 | y | 1628 | 1201 | 0.001525 | 1.269 | +1 | 9 |
| - | - | 1603 | 1202 | - | - | 0 | - |
| - | - | 720.5 | 1203 | - | - | 0 | - |
| - | - | 1401 | 1271 | - | - | 0 | - |
| - | - | 1506 | 1272 | - | - | 0 | - |
| - | - | 1035 | 1273 | - | - | 0 | - |
| - | - | 854.2 | 1286 | - | - | 0 | - |
| - | - | 6489 | 1288 | - | - | 0 | - |
| - | - | 8862 | 1289 | - | - | 0 | - |
| - | - | 1.142E+04 | 1290 | - | - | 0 | - |
| - | - | 7854 | 1291 | - | - | 0 | - |
| - | - | 3482 | 1292 | - | - | 0 | - |
| - | - | 871.3 | 1304 | - | - | 0 | - |
| - | - | 844 | 1305 | - | - | 0 | - |
| - | - | 1160 | 1306 | - | - | 0 | - |
| - | - | 677.8 | 1322 | - | - | 0 | - |
| - | - | 1157 | 1329 | - | - | 0 | - |
| - | - | 2184 | 1330 | - | - | 0 | - |
| - | - | 1632 | 1331 | - | - | 0 | - |
| - | - | 4279 | 1332 | - | - | 0 | - |
| - | - | 943.3 | 1332 | - | - | 0 | - |
| - | - | 3.661E+04 | 1333 | - | - | 0 | - |
| - | - | 2.678E+04 | 1334 | - | - | 0 | - |
| - | - | 1.056E+04 | 1335 | - | - | 0 | - |
| - | - | 3636 | 1336 | - | - | 0 | - |
| - | - | 963 | 1348 | - | - | 0 | - |
| - | - | 1.699E+04 | 1349 | - | - | 0 | - |
| - | - | 674.6 | 1349 | - | - | 0 | - |
| - | - | 6.585E+04 | 1350 | - | - | 0 | - |
| - | - | 4.572E+04 | 1351 | - | - | 0 | - |
| - | - | 2025 | 1351 | - | - | 0 | - |
| - | - | 2.04E+04 | 1352 | - | - | 0 | - |
| - | - | 1027 | 1352 | - | - | 0 | - |
| - | - | 4179 | 1353 | - | - | 0 | - |
| - | - | 607.2 | 1905 | - | - | 0 | - |

m/z Charge Intensity FragmentType MassShift Position
120.08064270019531 0 7283.8477
120.206298828125 0 339.29984
135.337890625 0 476.81305
148.90200805664062 0 616.66833
148.90802001953125 0 557.025
148.91896057128906 0 887.7462
148.92466735839844 0 949.82007
148.93020629882812 0 843.4889
148.9356689453125 0 1556.4847
148.94204711914062 0 2721.084
148.95472717285156 0 5240.613
148.9604949951172 0 2949.0776
148.96678161621094 0 1653.7944
148.97225952148438 0 894.3114
148.9779052734375 0 776.6939
148.98336791992188 0 746.10156
148.98910522460938 0 532.9985
149.00030517578125 0 489.3698
149.02337646484375 0 459.57092
152.25344848632812 0 453.46017
166.0861358642578 0 591.78845
173.43023681640625 0 473.5119
176.10665893554688 0 1876.2184
194.7610321044922 0 522.7801
221.1280517578125 0 8808.771
222.13186645507812 0 715.53033
231.1123504638672 0 1970.6685
249.12290954589844 0 8568.744 c Ammonia loss 1
250.12664794921875 0 881.02075
253.1177520751953 0 1092.193
263.10198974609375 0 953.12445
264.13677978515625 0 627.10114
303.42181396484375 0 619.5557
319.5334167480469 0 585.29694
342.1556701660156 0 1952.2305 y 8
355.06878662109375 0 891.0065
378.17999267578125 0 1851.5312
401.4523620605469 0 562.42554
447.5718078613281 0 540.87494
473.17242431640625 0 1497.1604 z 7
474.1794738769531 0 2946.121
475.1830749511719 0 680.2112
489.1911315917969 0 1167.6115 y 7
511.2181701660156 0 1170.939 c Ammonia loss 3
518.7078857421875 0 514.1993
544.209228515625 0 3561.7344 z 6
545.2152099609375 0 1171.6105
546.2201538085938 0 555.0443
550.7096557617188 0 2146.8765 y 2
551.2120971679688 0 1449.3284
560.2269897460938 0 2370.8203 y 6
573.2245483398438 0 629.4382
587.438720703125 0 620.65454
601.23388671875 0 1077.6982 y 1
601.7259521484375 0 766.06354
626.2430419921875 0 606.6501 c Ammonia loss 4
633.76220703125 0 623.03674
657.2105102539062 0 573.9938
665.7657470703125 0 1349.6692
666.2675170898438 0 1461.6586
666.7666625976562 0 744.952
674.7681884765625 0 2147.5586
675.2677612304688 0 1862.8748
675.765380859375 0 759.7049
676.359619140625 0 805.35596
706.2728881835938 0 592.70746
707.2718505859375 0 6475.5522 z 5
708.2772216796875 0 4833.411
709.2823486328125 0 956.8886
723.2918090820312 0 2711.8923 y 5
724.2919921875 0 905.66626
750.2796020507812 0 888.9175
764.2874145507812 0 698.7538
775.3319091796875 0 638.1117
778.30859375 0 3605.2888
779.3123168945312 0 1043.0387
780.3143310546875 0 1090.7235
822.2990112304688 0 8429.071 z 4
823.3047485351562 0 9310.594
824.3072509765625 0 4474.615
838.3163452148438 0 4500.7114 y 4
839.3242797851562 0 1191.3431
849.3478393554688 0 646.4963
877.3670654296875 0 867.4752 c 6
878.375244140625 0 773.01465
893.3355102539062 0 3137.8318
894.3408813476562 0 1501.731
917.3965454101562 0 651.8448
937.3250732421875 0 3452.9292 z 3
938.331787109375 0 7296.7466
939.3368530273438 0 3420.4226
940.332275390625 0 786.58746
953.3440551757812 0 5616.269 y 3
954.3475952148438 0 3439.0378
980.3941040039062 0 3276.77
981.3984375 0 1986.0483
982.3966674804688 0 1280.2626
1023.3980102539062 0 5684.693
1024.405029296875 0 7960.9844 c 7
1025.408203125 0 4735.4487
1026.41015625 0 1282.2657
1027.4122314453125 0 665.1363
1036.4168701171875 0 2050.3816
1084.38671875 0 1099.4788 z 2
1094.4591064453125 0 1001.7003
1095.4453125 0 656.8246
1100.412353515625 0 11505.015 y 2
1101.415283203125 0 6692.767
1102.4139404296875 0 2373.3499
1117.452392578125 0 2349.2644
1118.456298828125 0 2154.5806
1139.436767578125 0 1690.237
1140.442138671875 0 928.721
1141.4439697265625 0 1008.8171
1146.447509765625 0 931.19867
1147.450927734375 0 1094.7805
1161.4642333984375 0 17390.775 c 8
1162.466796875 0 10933.094
1163.468017578125 0 3821.5544
1184.4332275390625 0 2657.5247
1185.4368896484375 0 5116.277 z 1
1186.4415283203125 0 2951.1326
1187.448486328125 0 1552.263
1201.45849609375 0 1627.5537 y 1
1202.4627685546875 0 1602.514
1203.457763671875 0 720.5151
1270.5179443359375 0 1400.5529
1271.5128173828125 0 1505.7025
1272.520751953125 0 1034.5735
1285.5147705078125 0 854.17413
1287.5355224609375 0 6489.2793
1288.522216796875 0 8861.842
1289.5186767578125 0 11419.349
1290.5177001953125 0 7854.029
1291.51513671875 0 3481.5618
1303.52880859375 0 871.3057
1304.535400390625 0 844.0185
1305.5408935546875 0 1160.1189
1321.51708984375 0 677.7938
1328.52490234375 0 1156.7582
1329.5361328125 0 2183.658
1330.533203125 0 1632.4165
1331.5267333984375 0 4278.5166
1331.69482421875 0 943.2865
1332.51220703125 0 36606.387
1333.5142822265625 0 26783.607
1334.51611328125 0 10558.049
1335.5140380859375 0 3635.5254
1347.5267333984375 0 962.9924
1348.5264892578125 0 16985.25
1348.6844482421875 0 674.6116
1349.5338134765625 0 65853.87
1350.535888671875 0 45716.137
1350.6893310546875 0 2024.9912
1351.537353515625 0 20399.602
1351.703125 0 1027.0746
1352.5391845703125 0 4179.2344
1904.897216796875 0 607.2244

Spectrum Details

|  |  |
| --- | --- |
| Matched peaks? Matched peaks The total absolute number of peaks matched. Additionally in brackets the total fraction of peaks matched and the total number of peaks is shown. | 23 (14.56% of 158) |
| FDR? FDR The false discovery rate estimated for this peptide. It is calculated by matching all theoretical fragments with a non-integer shift with the raw peaks for this spectrum. This is done with 40 different shifts. The resulting percentage is the average number of annotated peaks over the number of annotated peaks with the correct spectrum. | 0.00% |
| Satellite FDR? Satellite FDR See the FDR for details on its calculation. This satellite ion specific FDR only contains the satellite ions (d/w) for I/L/J positions. | - |
| PSM Score? PSM Score The PSM Score as given by Hecklib to this annotated spectrum. It is shown with three significant figures. | 283 |

#### Spectrum 8969? Spectrum 8969 The raw spectrum of this peptide as annotated by Hecklib. The fragments are coloured according to ion type (see legend). Any peaks with a star '\*' as text can be hovered over to see the full details, first the ion type second the mass shift type. By hovering over the amino acids in the peptide or ions in the legend the corresponding peaks are highlighted. By toggling the 'Unassigned' label you can turn the background (unassigned) peaks on or off in the plot. By updating the slider in the Ion legend you can update the spectrum to only show the top X% of the peaks with labels. The top X% means any peak that is within X% of the highest intensity. By dragging in the spectrum you can zoom in to a specific part of the spectrum and use 'Zoom Out' to get back to the original zoom level. The annotation of the spectrum is based on the given sequence in the peptides file and is done with different software so inconsistencies are likely. The peaks are annotated based on the given sequence, with 20 ppm tolerance.

Copy Data

##### Spectrum 8969 (TSV)

###### Preview

```
Loading example...
```

*Click on the button to copy the data to your clipboard.*

Mz MinMz MaxIntensity Max

WidthHeightPeptide font sizePeptide stroke widthSpectrum font sizeSpectrum stroke widthCompact peptide

Ion legend

wxyz

abcd

OtherUnassignedIon^Charge^_Position_Show for top:%

FTFDDYAMHW

01.14e+42.27e+43.41e+44.54e+4

Zoom Out

a^+1^_2_y^+1^_1_a^+1^_2_b^+1^_2_b^+1^_2_y^+1^_2_b^+1^_3_b^+1^_4_y^+2^_8_y^+1^_4_b^+1^_5_\*\*y^+1^_5_b^+1^_6_y^+1^_6_y^+1^_7_y^+1^_8_y^+1^_8_b^+1^_9_y^+1^_9_

0694138720812774

Fragment Matches Table

Show background peaks

| Position | Ion type | Intensity | mz Theoretical | mz Error (Th) | mz Error (ppm) | Charge | Series Number |
| --- | --- | --- | --- | --- | --- | --- | --- |
| - | - | 4.496E+04 | 120.1 | - | - | 0 | - |
| - | - | 3048 | 121.1 | - | - | 0 | - |
| - | - | 352.1 | 125.4 | - | - | 0 | - |
| - | - | 2306 | 129.1 | - | - | 0 | - |
| - | - | 446.9 | 130 | - | - | 0 | - |
| - | - | 1420 | 130.1 | - | - | 0 | - |
| - | - | 2760 | 133.1 | - | - | 0 | - |
| - | - | 6628 | 136.1 | - | - | 0 | - |
| - | - | 538.6 | 146.1 | - | - | 0 | - |
| - | - | 639 | 147 | - | - | 0 | - |
| - | - | 707.3 | 148.9 | - | - | 0 | - |
| - | - | 2676 | 149 | - | - | 0 | - |
| - | - | 447.9 | 152.1 | - | - | 0 | - |
| - | - | 523.3 | 152.7 | - | - | 0 | - |
| - | - | 489.5 | 153.1 | - | - | 0 | - |
| - | - | 1491 | 156.1 | - | - | 0 | - |
| - | - | 482.5 | 157.1 | - | - | 0 | - |
| - | - | 1066 | 158.1 | - | - | 0 | - |
| - | - | 726.5 | 158.1 | - | - | 0 | - |
| - | - | 1840 | 159.1 | - | - | 0 | - |
| - | - | 401.9 | 161.4 | - | - | 0 | - |
| - | - | 450.8 | 162.1 | - | - | 0 | - |
| - | - | 565.5 | 166.1 | - | - | 0 | - |
| - | - | 5366 | 167.1 | - | - | 0 | - |
| - | - | 1393 | 169.1 | - | - | 0 | - |
| - | - | 829.3 | 173.1 | - | - | 0 | - |
| - | - | 3564 | 175.1 | - | - | 0 | - |
| - | - | 539.5 | 175.1 | - | - | 0 | - |
| - | - | 1.786E+04 | 176.1 | - | - | 0 | - |
| - | - | 856.3 | 177.1 | - | - | 0 | - |
| - | - | 2457 | 177.1 | - | - | 0 | - |
| - | - | 523.6 | 181.1 | - | - | 0 | - |
| - | - | 2450 | 181.2 | - | - | 0 | - |
| - | - | 578.1 | 185.1 | - | - | 0 | - |
| - | - | 1145 | 185.1 | - | - | 0 | - |
| - | - | 2163 | 186.1 | - | - | 0 | - |
| - | - | 482.6 | 186.1 | - | - | 0 | - |
| - | - | 808.7 | 191.1 | - | - | 0 | - |
| - | - | 459.3 | 191.1 | - | - | 0 | - |
| - | - | 1647 | 193.1 | - | - | 0 | - |
| - | - | 749.3 | 197.1 | - | - | 0 | - |
| - | - | 1426 | 203.1 | - | - | 0 | - |
| 2 | a | 1141 | 203.1 | 0.0002443 | 1.203 | +1 | 2 |
| - | - | 2364 | 209.2 | - | - | 0 | - |
| - | - | 1333 | 217.1 | - | - | 0 | - |
| 10 | y | 1833 | 221.1 | 0.004039 | 18.27 | +1 | 1 |
| - | - | 4120 | 221.1 | - | - | 0 | - |
| - | - | 2387 | 221.1 | - | - | 0 | - |
| 2 | a | 2.459E+04 | 221.1 | 3.625E-05 | 0.1639 | +1 | 2 |
| - | - | 2932 | 222.1 | - | - | 0 | - |
| - | - | 8516 | 225 | - | - | 0 | - |
| - | - | 1046 | 226 | - | - | 0 | - |
| - | - | 628.7 | 227 | - | - | 0 | - |
| - | - | 746.3 | 227.1 | - | - | 0 | - |
| - | - | 687.1 | 231.1 | - | - | 0 | - |
| 2 | b | 5641 | 231.1 | 5.696E-05 | 0.2465 | +1 | 2 |
| - | - | 2493 | 235.1 | - | - | 0 | - |
| - | - | 1703 | 237.2 | - | - | 0 | - |
| - | - | 9183 | 239.1 | - | - | 0 | - |
| - | - | 3574 | 239.2 | - | - | 0 | - |
| - | - | 1863 | 240.1 | - | - | 0 | - |
| 2 | b | 1.178E+04 | 249.1 | 3.205E-05 | 0.1287 | +1 | 2 |
| - | - | 1747 | 250.1 | - | - | 0 | - |
| - | - | 1122 | 251.1 | - | - | 0 | - |
| - | - | 1028 | 256.2 | - | - | 0 | - |
| - | - | 1796 | 263.1 | - | - | 0 | - |
| - | - | 634.5 | 272.9 | - | - | 0 | - |
| - | - | 1080 | 279.1 | - | - | 0 | - |
| - | - | 562.9 | 281.1 | - | - | 0 | - |
| - | - | 1586 | 285 | - | - | 0 | - |
| - | - | 726.1 | 285.1 | - | - | 0 | - |
| - | - | 673.5 | 292.1 | - | - | 0 | - |
| - | - | 1868 | 295.1 | - | - | 0 | - |
| - | - | 637.5 | 296.1 | - | - | 0 | - |
| - | - | 5877 | 299.1 | - | - | 0 | - |
| - | - | 883 | 300.1 | - | - | 0 | - |
| - | - | 696.3 | 301.1 | - | - | 0 | - |
| - | - | 1128 | 302.1 | - | - | 0 | - |
| - | - | 622.5 | 313.1 | - | - | 0 | - |
| - | - | 1114 | 316.2 | - | - | 0 | - |
| - | - | 578.9 | 334.1 | - | - | 0 | - |
| - | - | 2866 | 340.1 | - | - | 0 | - |
| - | - | 879.4 | 341 | - | - | 0 | - |
| - | - | 3057 | 355.1 | - | - | 0 | - |
| - | - | 564.3 | 356.1 | - | - | 0 | - |
| 9 | y | 1.116E+04 | 358.1 | 0.004782 | 13.35 | +1 | 2 |
| - | - | 6257 | 359 | - | - | 0 | - |
| - | - | 2025 | 359.2 | - | - | 0 | - |
| - | - | 841.7 | 364.1 | - | - | 0 | - |
| - | - | 1052 | 369.1 | - | - | 0 | - |
| - | - | 2093 | 378.1 | - | - | 0 | - |
| 3 | b | 2108 | 378.2 | 0.0004531 | 1.198 | +1 | 3 |
| - | - | 648.1 | 379.1 | - | - | 0 | - |
| - | - | 1223 | 394.1 | - | - | 0 | - |
| - | - | 566.7 | 406 | - | - | 0 | - |
| - | - | 593 | 417.4 | - | - | 0 | - |
| - | - | 1220 | 419 | - | - | 0 | - |
| - | - | 7732 | 429.1 | - | - | 0 | - |
| - | - | 2095 | 441.2 | - | - | 0 | - |
| - | - | 576.3 | 456.8 | - | - | 0 | - |
| - | - | 1986 | 505.2 | - | - | 0 | - |
| - | - | 696.7 | 506.2 | - | - | 0 | - |
| 4 | b | 824.6 | 511.2 | 0.001868 | 3.654 | +1 | 4 |
| - | - | 1312 | 512.2 | - | - | 0 | - |
| - | - | 764.6 | 513.2 | - | - | 0 | - |
| 3 | y | 3889 | 558.7 | 0.004479 | 8.017 | +2 | 8 |
| - | - | 2224 | 559.2 | - | - | 0 | - |
| - | - | 1032 | 564.4 | - | - | 0 | - |
| - | - | 623.6 | 574.1 | - | - | 0 | - |
| 7 | y | 3988 | 576.2 | 0.008992 | 15.61 | +1 | 4 |
| - | - | 1717 | 577.2 | - | - | 0 | - |
| - | - | 753 | 583.4 | - | - | 0 | - |
| - | - | 736.3 | 609.7 | - | - | 0 | - |
| 5 | b | 982.7 | 626.2 | 0.005373 | 8.58 | +1 | 5 |
| - | - | 643.5 | 634.2 | - | - | 0 | - |
| - | - | 636.5 | 643.8 | - | - | 0 | - |
| - | - | 679.4 | 645.9 | - | - | 0 | - |
| - | - | 579.3 | 649.4 | - | - | 0 | - |
| - | - | 936.1 | 649.5 | - | - | 0 | - |
| - | - | 819 | 649.5 | - | - | 0 | - |
| - | - | 608.1 | 651.3 | - | - | 0 | - |
| - | - | 678.4 | 658.3 | - | - | 0 | - |
| - | - | 667.2 | 662.5 | - | - | 0 | - |
| - | - | 708.8 | 662.7 | - | - | 0 | - |
| - | - | 667.5 | 665.3 | - | - | 0 | - |
| - | - | 659 | 672.7 | - | - | 0 | - |
| 0 | Precursor | 1377 | 673.8 | 0.005731 | 8.506 | +2 | -1 |
| - | - | 955.3 | 675.3 | - | - | 0 | - |
| - | - | 859.9 | 676.3 | - | - | 0 | - |
| - | - | 1040 | 681.8 | - | - | 0 | - |
| - | - | 1303 | 682.3 | - | - | 0 | - |
| 0 | Precursor | 3523 | 682.8 | 0.004172 | 6.111 | +2 | -1 |
| - | - | 2189 | 683.3 | - | - | 0 | - |
| - | - | 594.1 | 683.4 | - | - | 0 | - |
| - | - | 1292 | 683.8 | - | - | 0 | - |
| - | - | 991.2 | 684.2 | - | - | 0 | - |
| - | - | 642 | 684.3 | - | - | 0 | - |
| 6 | y | 3232 | 739.3 | 0.01036 | 14.01 | +1 | 5 |
| - | - | 1608 | 740.3 | - | - | 0 | - |
| - | - | 1189 | 742.1 | - | - | 0 | - |
| - | - | 668.9 | 742.1 | - | - | 0 | - |
| 6 | b | 1015 | 771.3 | 0.004241 | 5.498 | +1 | 6 |
| - | - | 1901 | 790.3 | - | - | 0 | - |
| - | - | 677.9 | 810.4 | - | - | 0 | - |
| 5 | y | 3922 | 854.3 | 0.008137 | 9.525 | +1 | 6 |
| - | - | 2279 | 855.3 | - | - | 0 | - |
| - | - | 805.3 | 856.3 | - | - | 0 | - |
| - | - | 2138 | 905.3 | - | - | 0 | - |
| - | - | 733.2 | 906.3 | - | - | 0 | - |
| 4 | y | 7714 | 969.3 | 0.007927 | 8.178 | +1 | 7 |
| - | - | 3295 | 970.3 | - | - | 0 | - |
| - | - | 940.9 | 971.3 | - | - | 0 | - |
| - | - | 3990 | 1052 | - | - | 0 | - |
| - | - | 2606 | 1053 | - | - | 0 | - |
| - | - | 926.6 | 1054 | - | - | 0 | - |
| - | - | 605.9 | 1079 | - | - | 0 | - |
| 3 | y | 798.2 | 1098 | 0.006413 | 5.839 | +1 | 8 |
| - | - | 1154 | 1099 | - | - | 0 | - |
| 3 | y | 1.331E+04 | 1116 | 0.008666 | 7.763 | +1 | 8 |
| - | - | 7164 | 1117 | - | - | 0 | - |
| - | - | 2124 | 1118 | - | - | 0 | - |
| 9 | b | 1035 | 1144 | 0.006265 | 5.474 | +1 | 9 |
| - | - | 627.4 | 1200 | - | - | 0 | - |
| 2 | y | 2014 | 1217 | 0.007252 | 5.957 | +1 | 9 |
| - | - | 1280 | 1218 | - | - | 0 | - |
| - | - | 645.6 | 2747 | - | - | 0 | - |

m/z Charge Intensity FragmentType MassShift Position
120.08089447021484 0 44963.668
121.08419799804688 0 3047.5066
125.44956970214844 0 352.13882
129.10227966308594 0 2306.474
130.0499725341797 0 446.88513
130.06521606445312 0 1420.1821
133.08604431152344 0 2760.1377
136.0758056640625 0 6628.2446
146.06039428710938 0 538.5836
147.0446014404297 0 639.0487
148.9475555419922 0 707.3417
149.04490661621094 0 2676.2358
152.08206176757812 0 447.86957
152.72671508789062 0 523.308
153.10227966308594 0 489.49454
156.07684326171875 0 1490.7457
157.07582092285156 0 482.4811
158.06002807617188 0 1066.275
158.09646606445312 0 726.49896
159.09188842773438 0 1840.4021
161.4298095703125 0 401.92026
162.05389404296875 0 450.83115
166.0859832763672 0 565.46655
167.055419921875 0 5365.598
169.1334228515625 0 1392.942
173.07054138183594 0 829.34283
175.08677673339844 0 3564.2441
175.093994140625 0 539.4536
176.10696411132812 0 17862.59
177.1025390625 0 856.31555
177.1110382080078 0 2456.5552
181.09703063964844 0 523.5728
181.16995239257812 0 2449.916
185.05621337890625 0 578.0714
185.0712890625 0 1145.3477
186.09104919433594 0 2162.5159
186.146484375 0 482.58832
191.08135986328125 0 808.7086
191.09194946289062 0 459.26697
193.10821533203125 0 1646.9155
197.1279296875 0 749.3011
203.0811004638672 0 1426.4275
203.11764526367188 0 1141.0575 a Water loss 1
209.1649169921875 0 2364.2173
217.09707641601562 0 1332.6726
221.08311462402344 0 1832.7421 y 9
221.09207153320312 0 4119.555
221.10342407226562 0 2386.824
221.12841796875 0 24586.03 a 1
222.1317901611328 0 2931.515
225.0428466796875 0 8516.35
226.04371643066406 0 1046.0225
227.02188110351562 0 628.7046
227.1025848388672 0 746.33844
231.06112670898438 0 687.1404
231.1127471923828 0 5640.595 b Water loss 1
235.1077117919922 0 2492.6199
237.16004943847656 0 1703.2847
239.09503173828125 0 9182.647
239.15016174316406 0 3574.1965
240.09567260742188 0 1862.7362
249.1233367919922 0 11775.643 b 1
250.126953125 0 1746.9998
251.1029052734375 0 1122.149
256.17816162109375 0 1028.1622
263.1025695800781 0 1796.2538
272.8559875488281 0 634.5375
279.0978088378906 0 1080.326
281.05145263671875 0 562.87115
285.0096740722656 0 1585.94
285.1012268066406 0 726.0525
292.1405334472656 0 673.534
295.10296630859375 0 1868.1948
296.10345458984375 0 637.4613
299.0617370605469 0 5877.0293
300.0621032714844 0 883.0395
301.0600891113281 0 696.3024
302.11309814453125 0 1127.6627
313.11431884765625 0 622.51984
316.1875305175781 0 1114.2877
334.055908203125 0 578.94336
340.14105224609375 0 2866.045
341.01806640625 0 879.3941
355.0693054199219 0 3056.6123
356.1380920410156 0 564.3256
358.1508483886719 0 11158.21 y 8
359.0282897949219 0 6257.327
359.15374755859375 0 2025.3329
364.1482849121094 0 841.7236
369.12060546875 0 1052.2588
378.1289367675781 0 2093.216
378.1816711425781 0 2107.8198 b Water loss 2
379.13323974609375 0 648.06036
394.1246032714844 0 1222.7395
405.9954528808594 0 566.72156
417.38958740234375 0 592.9836
418.99615478515625 0 1220.3986
429.0882263183594 0 7732.3574
441.18798828125 0 2095.095
456.821533203125 0 576.29987
505.1876220703125 0 1985.6768
506.1895751953125 0 696.66504
511.21685791015625 0 824.6479 b 3
512.2230834960938 0 1312.3389
513.1953735351562 0 764.6311
558.707763671875 0 3888.6729 y 2
559.210205078125 0 2224.3953
564.3622436523438 0 1032.1743
574.0592041015625 0 623.5562
576.22265625 0 3987.6184 y 6
577.2272338867188 0 1716.6307
583.3697509765625 0 752.9941
609.7337646484375 0 736.25165
626.2402954101562 0 982.67224 b 4
634.2352905273438 0 643.48474
643.7575073242188 0 636.5292
645.8506469726562 0 679.4426
649.38623046875 0 579.339
649.4641723632812 0 936.06274
649.539794921875 0 818.99194
651.264404296875 0 608.0835
658.2686767578125 0 678.44617
662.4913940429688 0 667.2301
662.6567993164062 0 708.84094
665.2556762695312 0 667.5145
672.7333984375 0 658.96124
673.7617797851562 0 1376.5175 Precursor Water loss
675.2868041992188 0 955.3397
676.2858276367188 0 859.9378
681.8397827148438 0 1040.4745
682.3361206054688 0 1303.153
682.7655029296875 0 3522.9695 Precursor
683.266357421875 0 2189.0303
683.3500366210938 0 594.0795
683.7693481445312 0 1292.1036
684.1983642578125 0 991.24994
684.260009765625 0 641.9781
739.287353515625 0 3232.3794 y 5
740.2896728515625 0 1608.2012
742.0551147460938 0 1188.8029
742.1236572265625 0 668.90137
771.3026733398438 0 1014.5518 b Water loss 5
790.313720703125 0 1900.6749
810.3809204101562 0 677.9365
854.3120727539062 0 3922.4858 y 4
855.3165893554688 0 2278.8096
856.3199462890625 0 805.2582
905.339111328125 0 2138.4578
906.3485107421875 0 733.20355
969.3388061523438 0 7713.852 y 3
970.3438110351562 0 3294.8657
971.3430786132812 0 940.92554
1052.4110107421875 0 3989.549
1053.412841796875 0 2606.1406
1054.419189453125 0 926.63293
1078.9130859375 0 605.9067
1098.3951416015625 0 798.22125 y Water loss 2
1099.404296875 0 1153.9176
1116.407958984375 0 13313.278 y 2
1117.4095458984375 0 7163.733
1118.41015625 0 2124.3718
1144.4417724609375 0 1035.2078 b 8
1200.4345703125 0 627.37067
1217.4542236328125 0 2014.4285 y 1
1218.456787109375 0 1280.2501
2746.812744140625 0 645.59937

Spectrum Details

|  |  |
| --- | --- |
| Matched peaks? Matched peaks The total absolute number of peaks matched. Additionally in brackets the total fraction of peaks matched and the total number of peaks is shown. | 21 (12.65% of 166) |
| FDR? FDR The false discovery rate estimated for this peptide. It is calculated by matching all theoretical fragments with a non-integer shift with the raw peaks for this spectrum. This is done with 40 different shifts. The resulting percentage is the average number of annotated peaks over the number of annotated peaks with the correct spectrum. | 0.45% |
| Satellite FDR? Satellite FDR See the FDR for details on its calculation. This satellite ion specific FDR only contains the satellite ions (d/w) for I/L/J positions. | - |
| PSM Score? PSM Score The PSM Score as given by Hecklib to this annotated spectrum. It is shown with three significant figures. | 239 |

#### Spectrum 10048? Spectrum 10048 The raw spectrum of this peptide as annotated by Hecklib. The fragments are coloured according to ion type (see legend). Any peaks with a star '\*' as text can be hovered over to see the full details, first the ion type second the mass shift type. By hovering over the amino acids in the peptide or ions in the legend the corresponding peaks are highlighted. By toggling the 'Unassigned' label you can turn the background (unassigned) peaks on or off in the plot. By updating the slider in the Ion legend you can update the spectrum to only show the top X% of the peaks with labels. The top X% means any peak that is within X% of the highest intensity. By dragging in the spectrum you can zoom in to a specific part of the spectrum and use 'Zoom Out' to get back to the original zoom level. The annotation of the spectrum is based on the given sequence in the peptides file and is done with different software so inconsistencies are likely. The peaks are annotated based on the given sequence, with 20 ppm tolerance.

Copy Data

##### Spectrum 10048 (TSV)

###### Preview

```
Loading example...
```

*Click on the button to copy the data to your clipboard.*

Mz MinMz MaxIntensity Max

WidthHeightPeptide font sizePeptide stroke widthSpectrum font sizeSpectrum stroke widthCompact peptide

Ion legend

wxyz

abcd

OtherUnassignedIon^Charge^_Position_Show for top:%

FTFDDYAMHW

02.01e+44.02e+46.02e+48.03e+4

Zoom Out

a^+1^_2_y^+1^_1_d^+1^_2_a^+1^_2_b^+1^_2_b^+1^_2_y^+1^_2_b^+1^_3_y^+2^_7_y^+1^_3_b^+1^_4_y^+2^_8_y^+2^_8_y^+1^_4_y^+2^_9_b^+1^_5_\*\*y^+1^_5_y^+1^_6_y^+1^_6_y^+1^_7_y^+1^_7_y^+1^_8_y^+1^_8_y^+1^_9_

0639127819172556

Fragment Matches Table

Show background peaks

| Position | Ion type | Intensity | mz Theoretical | mz Error (Th) | mz Error (ppm) | Charge | Series Number |
| --- | --- | --- | --- | --- | --- | --- | --- |
| - | - | 7.953E+04 | 120.1 | - | - | 0 | - |
| - | - | 356.2 | 120.8 | - | - | 0 | - |
| - | - | 7882 | 121.1 | - | - | 0 | - |
| - | - | 421.7 | 126.1 | - | - | 0 | - |
| - | - | 674 | 127.1 | - | - | 0 | - |
| - | - | 485.1 | 128.1 | - | - | 0 | - |
| - | - | 401.7 | 128.3 | - | - | 0 | - |
| - | - | 546.2 | 129.1 | - | - | 0 | - |
| - | - | 1743 | 129.1 | - | - | 0 | - |
| - | - | 992 | 130.1 | - | - | 0 | - |
| - | - | 574.2 | 131 | - | - | 0 | - |
| - | - | 1503 | 133.1 | - | - | 0 | - |
| - | - | 7481 | 136.1 | - | - | 0 | - |
| - | - | 547.5 | 137.1 | - | - | 0 | - |
| - | - | 693.5 | 138.1 | - | - | 0 | - |
| - | - | 422 | 141.1 | - | - | 0 | - |
| - | - | 430.8 | 142.9 | - | - | 0 | - |
| - | - | 405 | 144.2 | - | - | 0 | - |
| - | - | 544 | 145.1 | - | - | 0 | - |
| - | - | 460.5 | 146.7 | - | - | 0 | - |
| - | - | 502.5 | 148.9 | - | - | 0 | - |
| - | - | 509.3 | 148.9 | - | - | 0 | - |
| - | - | 745 | 148.9 | - | - | 0 | - |
| - | - | 594.7 | 148.9 | - | - | 0 | - |
| - | - | 855.5 | 148.9 | - | - | 0 | - |
| - | - | 681.2 | 148.9 | - | - | 0 | - |
| - | - | 1058 | 148.9 | - | - | 0 | - |
| - | - | 1012 | 148.9 | - | - | 0 | - |
| - | - | 2368 | 148.9 | - | - | 0 | - |
| - | - | 4197 | 148.9 | - | - | 0 | - |
| - | - | 4257 | 149 | - | - | 0 | - |
| - | - | 2435 | 149 | - | - | 0 | - |
| - | - | 1468 | 149 | - | - | 0 | - |
| - | - | 1293 | 149 | - | - | 0 | - |
| - | - | 1078 | 149 | - | - | 0 | - |
| - | - | 626.8 | 149 | - | - | 0 | - |
| - | - | 697.1 | 149 | - | - | 0 | - |
| - | - | 699.4 | 149 | - | - | 0 | - |
| - | - | 450.9 | 149 | - | - | 0 | - |
| - | - | 451.1 | 149 | - | - | 0 | - |
| - | - | 469.9 | 149.1 | - | - | 0 | - |
| - | - | 446.4 | 149.2 | - | - | 0 | - |
| - | - | 619.8 | 155.1 | - | - | 0 | - |
| - | - | 2284 | 156.1 | - | - | 0 | - |
| - | - | 1207 | 158.1 | - | - | 0 | - |
| - | - | 4026 | 159.1 | - | - | 0 | - |
| - | - | 663.1 | 160.1 | - | - | 0 | - |
| - | - | 2773 | 166.1 | - | - | 0 | - |
| - | - | 454.3 | 167.7 | - | - | 0 | - |
| - | - | 549.4 | 169.1 | - | - | 0 | - |
| - | - | 1093 | 169.1 | - | - | 0 | - |
| - | - | 748.9 | 171.1 | - | - | 0 | - |
| - | - | 830 | 171.1 | - | - | 0 | - |
| - | - | 555.2 | 172.1 | - | - | 0 | - |
| - | - | 712.1 | 175.1 | - | - | 0 | - |
| - | - | 3.526E+04 | 176.1 | - | - | 0 | - |
| - | - | 792.5 | 177.1 | - | - | 0 | - |
| - | - | 3775 | 177.1 | - | - | 0 | - |
| - | - | 440.5 | 178.2 | - | - | 0 | - |
| - | - | 536.6 | 181.2 | - | - | 0 | - |
| - | - | 5005 | 186.1 | - | - | 0 | - |
| - | - | 680.1 | 187.1 | - | - | 0 | - |
| - | - | 2533 | 188.1 | - | - | 0 | - |
| - | - | 2188 | 193.1 | - | - | 0 | - |
| - | - | 872.1 | 195.1 | - | - | 0 | - |
| - | - | 1144 | 197.1 | - | - | 0 | - |
| - | - | 751.3 | 201.1 | - | - | 0 | - |
| - | - | 681.6 | 203.1 | - | - | 0 | - |
| 2 | a | 3290 | 203.1 | 0.0002745 | 1.352 | +1 | 2 |
| - | - | 543.4 | 204.1 | - | - | 0 | - |
| 10 | y | 2624 | 205.1 | 7.491E-05 | 0.3652 | +1 | 1 |
| 2 | d | 838.1 | 207.1 | 2.645E-05 | 0.1277 | +1 | 2 |
| - | - | 908.1 | 209.2 | - | - | 0 | - |
| - | - | 519.5 | 212.1 | - | - | 0 | - |
| - | - | 537.6 | 214.1 | - | - | 0 | - |
| - | - | 2152 | 217.1 | - | - | 0 | - |
| - | - | 656.6 | 219.1 | - | - | 0 | - |
| - | - | 7055 | 221.1 | - | - | 0 | - |
| 2 | a | 4.355E+04 | 221.1 | 0.0002689 | 1.216 | +1 | 2 |
| - | - | 5103 | 222.1 | - | - | 0 | - |
| - | - | 649.6 | 226.1 | - | - | 0 | - |
| - | - | 553.4 | 227.1 | - | - | 0 | - |
| - | - | 645.7 | 231.1 | - | - | 0 | - |
| 2 | b | 1.003E+04 | 231.1 | 0.0003245 | 1.404 | +1 | 2 |
| - | - | 1105 | 232.1 | - | - | 0 | - |
| - | - | 4474 | 235.1 | - | - | 0 | - |
| - | - | 482.4 | 235.1 | - | - | 0 | - |
| - | - | 1208 | 237.2 | - | - | 0 | - |
| - | - | 901.9 | 239.1 | - | - | 0 | - |
| - | - | 563.7 | 249.1 | - | - | 0 | - |
| 2 | b | 2.469E+04 | 249.1 | 0.0002884 | 1.158 | +1 | 2 |
| - | - | 2926 | 250.1 | - | - | 0 | - |
| - | - | 2448 | 251.1 | - | - | 0 | - |
| - | - | 2319 | 253.1 | - | - | 0 | - |
| - | - | 1489 | 257.1 | - | - | 0 | - |
| - | - | 4223 | 263.1 | - | - | 0 | - |
| - | - | 767.6 | 264.1 | - | - | 0 | - |
| - | - | 697.5 | 267.1 | - | - | 0 | - |
| - | - | 633.1 | 270.1 | - | - | 0 | - |
| - | - | 845.1 | 274.1 | - | - | 0 | - |
| - | - | 2059 | 279.1 | - | - | 0 | - |
| - | - | 731 | 281.1 | - | - | 0 | - |
| - | - | 2093 | 285.1 | - | - | 0 | - |
| - | - | 1436 | 292.1 | - | - | 0 | - |
| - | - | 1705 | 296.2 | - | - | 0 | - |
| - | - | 660.4 | 300.1 | - | - | 0 | - |
| - | - | 779.8 | 303.1 | - | - | 0 | - |
| - | - | 1286 | 316.2 | - | - | 0 | - |
| - | - | 703.6 | 318.1 | - | - | 0 | - |
| - | - | 3213 | 324.1 | - | - | 0 | - |
| - | - | 722.1 | 333.2 | - | - | 0 | - |
| 9 | y | 1.849E+04 | 342.2 | 0.0004892 | 1.43 | +1 | 2 |
| - | - | 1016 | 342.2 | - | - | 0 | - |
| - | - | 4230 | 343.2 | - | - | 0 | - |
| - | - | 558.2 | 344.4 | - | - | 0 | - |
| - | - | 476.5 | 344.7 | - | - | 0 | - |
| - | - | 961.9 | 346.1 | - | - | 0 | - |
| - | - | 960.8 | 350.1 | - | - | 0 | - |
| - | - | 791.9 | 356.1 | - | - | 0 | - |
| - | - | 1325 | 364.2 | - | - | 0 | - |
| - | - | 5189 | 378.1 | - | - | 0 | - |
| 3 | b | 3190 | 378.2 | 0.0006057 | 1.601 | +1 | 3 |
| - | - | 1088 | 379.1 | - | - | 0 | - |
| - | - | 1245 | 379.2 | - | - | 0 | - |
| - | - | 609.4 | 387.3 | - | - | 0 | - |
| - | - | 2325 | 394.1 | - | - | 0 | - |
| - | - | 716.8 | 395.1 | - | - | 0 | - |
| - | - | 793.7 | 395.2 | - | - | 0 | - |
| - | - | 644.1 | 407.2 | - | - | 0 | - |
| - | - | 631.7 | 408.2 | - | - | 0 | - |
| - | - | 3186 | 425.2 | - | - | 0 | - |
| - | - | 585.9 | 449.2 | - | - | 0 | - |
| - | - | 666.3 | 455.2 | - | - | 0 | - |
| - | - | 1375 | 465.2 | - | - | 0 | - |
| - | - | 603.5 | 471.2 | - | - | 0 | - |
| 4 | y | 784.9 | 477.2 | 0.0001472 | 0.3084 | +2 | 7 |
| - | - | 843.2 | 479.2 | - | - | 0 | - |
| 8 | y | 4214 | 489.2 | 0.005802 | 11.86 | +1 | 3 |
| - | - | 1136 | 490.2 | - | - | 0 | - |
| - | - | 650.5 | 490.3 | - | - | 0 | - |
| - | - | 2277 | 496.2 | - | - | 0 | - |
| 4 | b | 1829 | 511.2 | 0.0001158 | 0.2264 | +1 | 4 |
| - | - | 948.2 | 512.2 | - | - | 0 | - |
| - | - | 965.9 | 513.2 | - | - | 0 | - |
| - | - | 686.8 | 519.2 | - | - | 0 | - |
| 3 | y | 886 | 541.7 | 0.00232 | 4.283 | +2 | 8 |
| - | - | 602.3 | 542.2 | - | - | 0 | - |
| - | - | 921.6 | 548.2 | - | - | 0 | - |
| 3 | y | 5425 | 550.7 | 0.002836 | 5.15 | +2 | 8 |
| - | - | 3295 | 551.2 | - | - | 0 | - |
| - | - | 1459 | 551.7 | - | - | 0 | - |
| - | - | 561.9 | 559.2 | - | - | 0 | - |
| 7 | y | 6629 | 560.2 | 0.00479 | 8.551 | +1 | 4 |
| - | - | 1700 | 561.2 | - | - | 0 | - |
| - | - | 800.9 | 570.2 | - | - | 0 | - |
| 2 | y | 1537 | 601.2 | 0.004631 | 7.703 | +2 | 9 |
| - | - | 1183 | 601.7 | - | - | 0 | - |
| - | - | 703.5 | 606.2 | - | - | 0 | - |
| 5 | b | 726.4 | 626.2 | 0.002688 | 4.292 | +1 | 5 |
| - | - | 833.7 | 634.2 | - | - | 0 | - |
| - | - | 631 | 642.8 | - | - | 0 | - |
| - | - | 906.3 | 643.3 | - | - | 0 | - |
| - | - | 984.3 | 657.3 | - | - | 0 | - |
| - | - | 2172 | 659.3 | - | - | 0 | - |
| - | - | 1225 | 660.3 | - | - | 0 | - |
| 0 | Precursor | 3703 | 665.8 | 0.002562 | 3.849 | +2 | -1 |
| - | - | 1823 | 666.3 | - | - | 0 | - |
| - | - | 1252 | 666.8 | - | - | 0 | - |
| 0 | Precursor | 4409 | 674.8 | 0.001919 | 2.843 | +2 | -1 |
| - | - | 5069 | 675.3 | - | - | 0 | - |
| - | - | 2154 | 675.8 | - | - | 0 | - |
| - | - | 1032 | 676.3 | - | - | 0 | - |
| - | - | 1178 | 685.3 | - | - | 0 | - |
| - | - | 829.1 | 695.3 | - | - | 0 | - |
| 6 | y | 6548 | 723.3 | 0.005427 | 7.503 | +1 | 5 |
| - | - | 2952 | 724.3 | - | - | 0 | - |
| - | - | 671 | 725.3 | - | - | 0 | - |
| - | - | 847.8 | 749.3 | - | - | 0 | - |
| - | - | 589.9 | 759.3 | - | - | 0 | - |
| - | - | 3070 | 774.3 | - | - | 0 | - |
| - | - | 992.3 | 775.3 | - | - | 0 | - |
| - | - | 947 | 804.3 | - | - | 0 | - |
| 5 | y | 579.5 | 820.3 | 0.01456 | 17.75 | +1 | 6 |
| - | - | 1847 | 832.3 | - | - | 0 | - |
| - | - | 763.2 | 833.3 | - | - | 0 | - |
| 5 | y | 8985 | 838.3 | 0.005644 | 6.733 | +1 | 6 |
| - | - | 3989 | 839.3 | - | - | 0 | - |
| - | - | 1203 | 840.3 | - | - | 0 | - |
| - | - | 587.1 | 856.5 | - | - | 0 | - |
| - | - | 1116 | 868.3 | - | - | 0 | - |
| - | - | 5111 | 889.3 | - | - | 0 | - |
| - | - | 2157 | 890.4 | - | - | 0 | - |
| - | - | 1959 | 896.3 | - | - | 0 | - |
| - | - | 802.4 | 897.3 | - | - | 0 | - |
| 4 | y | 1209 | 935.3 | 0.005379 | 5.751 | +1 | 7 |
| 4 | y | 1.534E+04 | 953.3 | 0.005984 | 6.277 | +1 | 7 |
| - | - | 8185 | 954.3 | - | - | 0 | - |
| - | - | 2922 | 955.3 | - | - | 0 | - |
| - | - | 972.4 | 1018 | - | - | 0 | - |
| - | - | 7250 | 1036 | - | - | 0 | - |
| - | - | 4300 | 1037 | - | - | 0 | - |
| - | - | 1349 | 1038 | - | - | 0 | - |
| 3 | y | 1573 | 1082 | 0.004836 | 4.468 | +1 | 8 |
| - | - | 1209 | 1083 | - | - | 0 | - |
| 3 | y | 2.582E+04 | 1100 | 0.00477 | 4.334 | +1 | 8 |
| - | - | 1.616E+04 | 1101 | - | - | 0 | - |
| - | - | 6386 | 1102 | - | - | 0 | - |
| - | - | 629.1 | 1103 | - | - | 0 | - |
| - | - | 1954 | 1137 | - | - | 0 | - |
| - | - | 672.6 | 1138 | - | - | 0 | - |
| 2 | y | 4974 | 1201 | 0.005187 | 4.317 | +1 | 9 |
| - | - | 3212 | 1202 | - | - | 0 | - |
| - | - | 733.5 | 1203 | - | - | 0 | - |
| - | - | 910.8 | 1211 | - | - | 0 | - |
| - | - | 1032 | 1212 | - | - | 0 | - |
| - | - | 620.1 | 1496 | - | - | 0 | - |
| - | - | 773 | 1839 | - | - | 0 | - |
| - | - | 652.7 | 2530 | - | - | 0 | - |

m/z Charge Intensity FragmentType MassShift Position
120.0810546875 0 79532.81
120.78719329833984 0 356.23557
121.0843734741211 0 7881.94
126.0666275024414 0 421.69238
127.05052947998047 0 673.97174
128.10736083984375 0 485.11795
128.32928466796875 0 401.67886
129.06607055664062 0 546.2016
129.10252380371094 0 1743.1355
130.06549072265625 0 991.95404
131.04515075683594 0 574.18964
133.0861358642578 0 1503.3955
136.07598876953125 0 7481.3037
137.0792694091797 0 547.4742
138.0659637451172 0 693.53906
141.06674194335938 0 421.99207
142.91677856445312 0 430.75806
144.1814727783203 0 404.99307
145.0612030029297 0 544.04156
146.65634155273438 0 460.473
148.87811279296875 0 502.4908
148.8853759765625 0 509.28134
148.8924560546875 0 744.96936
148.90025329589844 0 594.71027
148.90701293945312 0 855.5494
148.9136962890625 0 681.15985
148.9209442138672 0 1057.9574
148.92880249023438 0 1012.3705
148.9355010986328 0 2368.2012
148.94314575195312 0 4197.086
148.959716796875 0 4256.6997
148.96746826171875 0 2434.7317
148.97451782226562 0 1468.1775
148.98194885253906 0 1293.1481
148.98924255371094 0 1078.3262
148.99606323242188 0 626.76587
149.00340270996094 0 697.081
149.01031494140625 0 699.4401
149.02561950683594 0 450.90524
149.03245544433594 0 451.07324
149.08229064941406 0 469.93802
149.18943786621094 0 446.37277
155.09317016601562 0 619.79474
156.07711791992188 0 2284.2417
158.09664916992188 0 1207.1537
159.0919647216797 0 4025.6877
160.0955047607422 0 663.1037
166.08641052246094 0 2773.2097
167.69125366210938 0 454.34695
169.0975799560547 0 549.44556
169.13380432128906 0 1092.5184
171.07717895507812 0 748.92126
171.11328125 0 829.98
172.07211303710938 0 555.2388
175.08633422851562 0 712.09265
176.1072540283203 0 35259.99
177.10289001464844 0 792.50037
177.1110076904297 0 3774.9836
178.2306671142578 0 440.46826
181.17013549804688 0 536.5552
186.09149169921875 0 5005.0234
187.09523010253906 0 680.05853
188.07093811035156 0 2532.8066
193.10862731933594 0 2187.9536
195.0878448486328 0 872.1124
197.12844848632812 0 1143.6227
201.12359619140625 0 751.28973
203.06663513183594 0 681.551
203.1181640625 0 3289.74 a Water loss 1
204.07723999023438 0 543.3986
205.09722900390625 0 2624.259 y 9
207.11277770996094 0 838.10736 d 1
209.1654815673828 0 908.1469
212.1175537109375 0 519.4551
214.0860137939453 0 537.59186
217.0973358154297 0 2152.379
219.0792694091797 0 656.5776
221.10348510742188 0 7055.0728
221.12872314453125 0 43546.58 a 1
222.1319122314453 0 5102.9497
226.11837768554688 0 649.6326
227.10305786132812 0 553.41766
231.0618438720703 0 645.7122
231.11312866210938 0 10030.285 b Water loss 1
232.11648559570312 0 1104.9623
235.10784912109375 0 4473.9004
235.1435546875 0 482.4307
237.16055297851562 0 1208.3722
239.1136932373047 0 901.86975
249.0714569091797 0 563.69574
249.1236572265625 0 24694.273 b 1
250.12672424316406 0 2925.695
251.10276794433594 0 2448.4893
253.11880493164062 0 2318.9192
257.1062316894531 0 1488.5911
263.1026611328125 0 4222.703
264.1061096191406 0 767.56616
267.11053466796875 0 697.54425
270.1446228027344 0 633.1231
274.1300048828125 0 845.07697
279.0975036621094 0 2059.1792
281.0954895019531 0 730.9593
285.1015319824219 0 2092.5408
292.14044189453125 0 1435.8606
296.1515197753906 0 1704.7084
300.1351623535156 0 660.4168
303.1124267578125 0 779.8351
316.1871337890625 0 1286.4315
318.14544677734375 0 703.62085
324.1462707519531 0 3213.3657
333.1597595214844 0 722.1183
342.15655517578125 0 18486.707 y 8
342.17926025390625 0 1015.8741
343.1593933105469 0 4230.3896
344.43048095703125 0 558.1826
344.7193908691406 0 476.53058
346.1401672363281 0 961.8715
350.13543701171875 0 960.83234
356.1396484375 0 791.85425
364.1507873535156 0 1325.3943
378.1300048828125 0 5189.1943
378.18182373046875 0 3190.1624 b Water loss 2
379.13189697265625 0 1088.207
379.1846008300781 0 1245.2859
387.3259582519531 0 609.44183
394.125 0 2325.0676
395.12677001953125 0 716.7905
395.22991943359375 0 793.6734
407.1835021972656 0 644.09564
408.1675109863281 0 631.66187
425.1939392089844 0 3185.7568
449.1676940917969 0 585.93384
455.2032775878906 0 666.34406
465.16217041015625 0 1375.221
471.1833801269531 0 603.50146
477.1742248535156 0 784.87463 y 3
479.1755065917969 0 843.1952
489.1923522949219 0 4214.286 y 7
490.1958312988281 0 1135.7686
490.2880859375 0 650.45844
496.2302551269531 0 2276.5535
511.2188415527344 0 1828.9918 b 3
512.2200927734375 0 948.23206
513.1959228515625 0 965.93787
519.21142578125 0 686.8
541.705322265625 0 886.0161 y Water loss 2
542.21435546875 0 602.2795
548.1956176757812 0 921.6471
550.7111206054688 0 5425.0327 y 2
551.2127685546875 0 3294.9094
551.7130126953125 0 1458.6188
559.23046875 0 561.8879
560.2284545898438 0 6629.082 y 6
561.2316284179688 0 1700.3273
570.2301025390625 0 800.87555
601.2367553710938 0 1537.1731 y 1
601.7345581054688 0 1183.0638
606.22900390625 0 703.54565
626.2429809570312 0 726.3847 b 4
634.2281494140625 0 833.7131
642.7670288085938 0 630.98444
643.2685546875 0 906.2976
657.2601928710938 0 984.3079
659.2926025390625 0 2172.4712
660.295166015625 0 1224.8981
665.7636108398438 0 3703.2617 Precursor Water loss
666.2645874023438 0 1823.3706
666.7637939453125 0 1251.5641
674.7682495117188 0 4409.4 Precursor
675.2695922851562 0 5068.58
675.7726440429688 0 2154.0461
676.2735595703125 0 1031.6387
685.2578125 0 1177.6943
695.2633056640625 0 829.0537
723.2924194335938 0 6547.848 y 5
724.2947998046875 0 2952.1726
725.2979736328125 0 670.9852
749.2532348632812 0 847.816
759.260498046875 0 589.853
774.3201904296875 0 3070.3086
775.3226928710938 0 992.301
804.3316040039062 0 947.03986
820.3179321289062 0 579.4923 y Water loss 4
832.3285522460938 0 1847.3755
833.329345703125 0 763.19116
838.319580078125 0 8984.968 y 4
839.3220825195312 0 3988.7888
840.3231201171875 0 1203.0354
856.5242309570312 0 587.09955
868.3345947265625 0 1116.2286
889.34716796875 0 5110.5166
890.3529663085938 0 2157.3726
896.3264770507812 0 1958.7903
897.3302001953125 0 802.38306
935.335693359375 0 1209.2666 y Water loss 3
953.3468627929688 0 15338.392 y 3
954.3486328125 0 8184.677
955.3496704101562 0 2922.035
1018.4050903320312 0 972.39014
1036.4150390625 0 7249.7847
1037.418212890625 0 4299.8247
1038.421630859375 0 1349.4812
1082.403564453125 0 1572.833 y Water loss 2
1083.4033203125 0 1209.4597
1100.4140625 0 25823.562 y 2
1101.4168701171875 0 16161.724
1102.41748046875 0 6385.924
1103.4229736328125 0 629.14355
1137.4576416015625 0 1953.9504
1138.4638671875 0 672.63525
1201.462158203125 0 4973.519 y 1
1202.46484375 0 3212.0818
1203.4720458984375 0 733.45557
1211.4366455078125 0 910.76935
1212.4581298828125 0 1032.4825
1496.19482421875 0 620.06866
1838.6353759765625 0 773.0488
2530.335693359375 0 652.6857

Spectrum Details

|  |  |
| --- | --- |
| Matched peaks? Matched peaks The total absolute number of peaks matched. Additionally in brackets the total fraction of peaks matched and the total number of peaks is shown. | 26 (11.93% of 218) |
| FDR? FDR The false discovery rate estimated for this peptide. It is calculated by matching all theoretical fragments with a non-integer shift with the raw peaks for this spectrum. This is done with 40 different shifts. The resulting percentage is the average number of annotated peaks over the number of annotated peaks with the correct spectrum. | 0.00% |
| Satellite FDR? Satellite FDR See the FDR for details on its calculation. This satellite ion specific FDR only contains the satellite ions (d/w) for I/L/J positions. | - |
| PSM Score? PSM Score The PSM Score as given by Hecklib to this annotated spectrum. It is shown with three significant figures. | 257 |

#### Spectrum 8861? Spectrum 8861 The raw spectrum of this peptide as annotated by Hecklib. The fragments are coloured according to ion type (see legend). Any peaks with a star '\*' as text can be hovered over to see the full details, first the ion type second the mass shift type. By hovering over the amino acids in the peptide or ions in the legend the corresponding peaks are highlighted. By toggling the 'Unassigned' label you can turn the background (unassigned) peaks on or off in the plot. By updating the slider in the Ion legend you can update the spectrum to only show the top X% of the peaks with labels. The top X% means any peak that is within X% of the highest intensity. By dragging in the spectrum you can zoom in to a specific part of the spectrum and use 'Zoom Out' to get back to the original zoom level. The annotation of the spectrum is based on the given sequence in the peptides file and is done with different software so inconsistencies are likely. The peaks are annotated based on the given sequence, with 20 ppm tolerance.

Copy Data

##### Spectrum 8861 (TSV)

###### Preview

```
Loading example...
```

*Click on the button to copy the data to your clipboard.*

Mz MinMz MaxIntensity Max

WidthHeightPeptide font sizePeptide stroke widthSpectrum font sizeSpectrum stroke widthCompact peptide

Ion legend

wxyz

abcd

OtherUnassignedIon^Charge^_Position_Show for top:%

FTFDDYAMHW

03.28e+46.56e+49.84e+41.31e+5

Zoom Out

y^+1^_1_c^+1^_2_y^+1^_2_z^+1^_3_y^+1^_3_c^+1^_4_y^+2^_8_y^+2^_8_z^+1^_4_y^+1^_4_y^+2^_9_c^+1^_5_z^+1^_5_y^+1^_5_z^+1^_6_y^+1^_6_c^+1^_7_z^+1^_7_y^+1^_7_c^+1^_8_y^+1^_8_z^+1^_8_y^+1^_8_c^+1^_9_w^+1^_9_y^+1^_9_z^+1^_9_y^+1^_9_

0785157023553140

Fragment Matches Table

Show background peaks

| Position | Ion type | Intensity | mz Theoretical | mz Error (Th) | mz Error (ppm) | Charge | Series Number |
| --- | --- | --- | --- | --- | --- | --- | --- |
| - | - | 1.132E+04 | 120.1 | - | - | 0 | - |
| - | - | 1376 | 121.1 | - | - | 0 | - |
| - | - | 380.4 | 122.7 | - | - | 0 | - |
| - | - | 375.7 | 122.9 | - | - | 0 | - |
| - | - | 576.4 | 122.9 | - | - | 0 | - |
| - | - | 376.4 | 125.3 | - | - | 0 | - |
| - | - | 343.8 | 128.9 | - | - | 0 | - |
| - | - | 444.5 | 140.2 | - | - | 0 | - |
| - | - | 407.8 | 145.5 | - | - | 0 | - |
| - | - | 405.8 | 147 | - | - | 0 | - |
| - | - | 612.4 | 148.9 | - | - | 0 | - |
| - | - | 1890 | 148.9 | - | - | 0 | - |
| - | - | 1902 | 148.9 | - | - | 0 | - |
| - | - | 3648 | 149 | - | - | 0 | - |
| - | - | 2303 | 149 | - | - | 0 | - |
| - | - | 1564 | 149 | - | - | 0 | - |
| - | - | 581.6 | 149 | - | - | 0 | - |
| - | - | 1354 | 149 | - | - | 0 | - |
| - | - | 472.5 | 160.3 | - | - | 0 | - |
| - | - | 441.1 | 162.9 | - | - | 0 | - |
| - | - | 494 | 173.4 | - | - | 0 | - |
| - | - | 653.9 | 173.5 | - | - | 0 | - |
| - | - | 468.8 | 175.7 | - | - | 0 | - |
| - | - | 3051 | 176.1 | - | - | 0 | - |
| - | - | 518.1 | 177.1 | - | - | 0 | - |
| - | - | 556.7 | 180.9 | - | - | 0 | - |
| - | - | 470.3 | 194.3 | - | - | 0 | - |
| 10 | y | 1017 | 221.1 | 0.003856 | 17.44 | +1 | 1 |
| - | - | 848.5 | 221.1 | - | - | 0 | - |
| - | - | 1.74E+04 | 221.1 | - | - | 0 | - |
| - | - | 1986 | 222.1 | - | - | 0 | - |
| - | - | 1624 | 225 | - | - | 0 | - |
| - | - | 3284 | 231.1 | - | - | 0 | - |
| - | - | 524.3 | 232.1 | - | - | 0 | - |
| - | - | 503.3 | 233.9 | - | - | 0 | - |
| - | - | 980.9 | 235.1 | - | - | 0 | - |
| - | - | 2792 | 239.1 | - | - | 0 | - |
| - | - | 729.2 | 240.1 | - | - | 0 | - |
| 2 | c | 1.605E+04 | 249.1 | 0.0001511 | 0.6063 | +1 | 2 |
| - | - | 2578 | 250.1 | - | - | 0 | - |
| - | - | 1880 | 263.1 | - | - | 0 | - |
| - | - | 2259 | 295.1 | - | - | 0 | - |
| - | - | 931.5 | 296.1 | - | - | 0 | - |
| - | - | 560.3 | 296.2 | - | - | 0 | - |
| - | - | 1702 | 299.1 | - | - | 0 | - |
| - | - | 708 | 313.1 | - | - | 0 | - |
| - | - | 607.3 | 333.2 | - | - | 0 | - |
| - | - | 3438 | 355.1 | - | - | 0 | - |
| 9 | y | 3516 | 358.1 | 0.004538 | 12.67 | +1 | 2 |
| - | - | 1298 | 359 | - | - | 0 | - |
| - | - | 754.7 | 369.1 | - | - | 0 | - |
| - | - | 760 | 370.1 | - | - | 0 | - |
| - | - | 1224 | 378.1 | - | - | 0 | - |
| - | - | 3681 | 378.2 | - | - | 0 | - |
| - | - | 641.9 | 415.2 | - | - | 0 | - |
| - | - | 9204 | 429.1 | - | - | 0 | - |
| - | - | 656.2 | 479.2 | - | - | 0 | - |
| 8 | z | 864.7 | 489.2 | 0.00941 | 19.24 | +1 | 3 |
| - | - | 3262 | 490.2 | - | - | 0 | - |
| - | - | 784.5 | 491.2 | - | - | 0 | - |
| - | - | 1348 | 504.2 | - | - | 0 | - |
| 8 | y | 2696 | 505.2 | 0.009363 | 18.53 | +1 | 3 |
| - | - | 1104 | 506.2 | - | - | 0 | - |
| 4 | c | 1207 | 511.2 | 0.001123 | 2.196 | +1 | 4 |
| 3 | y | 819.2 | 549.7 | 0.004085 | 7.432 | +2 | 8 |
| 3 | y | 6681 | 558.7 | 0.005272 | 9.437 | +2 | 8 |
| - | - | 5090 | 559.2 | - | - | 0 | - |
| - | - | 1332 | 559.7 | - | - | 0 | - |
| 7 | z | 3029 | 560.2 | 0.01002 | 17.88 | +1 | 4 |
| - | - | 521.9 | 561.1 | - | - | 0 | - |
| - | - | 1920 | 561.2 | - | - | 0 | - |
| 7 | y | 6237 | 576.2 | 0.009846 | 17.09 | +1 | 4 |
| - | - | 1810 | 577.2 | - | - | 0 | - |
| - | - | 1352 | 578.2 | - | - | 0 | - |
| 2 | y | 2586 | 609.2 | 0.005054 | 8.295 | +2 | 9 |
| - | - | 1429 | 609.7 | - | - | 0 | - |
| - | - | 897.4 | 610.2 | - | - | 0 | - |
| 5 | c | 1694 | 626.2 | 0.0008522 | 1.361 | +1 | 5 |
| - | - | 621.5 | 641.8 | - | - | 0 | - |
| - | - | 1134 | 649.3 | - | - | 0 | - |
| - | - | 776.7 | 650.8 | - | - | 0 | - |
| - | - | 2761 | 673.8 | - | - | 0 | - |
| - | - | 2894 | 674.3 | - | - | 0 | - |
| - | - | 835.6 | 682.3 | - | - | 0 | - |
| - | - | 7252 | 682.8 | - | - | 0 | - |
| - | - | 7415 | 683.3 | - | - | 0 | - |
| - | - | 3042 | 683.8 | - | - | 0 | - |
| - | - | 1168 | 684.2 | - | - | 0 | - |
| - | - | 1097 | 693.2 | - | - | 0 | - |
| - | - | 643.2 | 703.7 | - | - | 0 | - |
| - | - | 641.2 | 704.3 | - | - | 0 | - |
| 6 | z | 1.524E+04 | 723.3 | 0.009981 | 13.8 | +1 | 5 |
| - | - | 8143 | 724.3 | - | - | 0 | - |
| - | - | 2910 | 725.3 | - | - | 0 | - |
| 6 | y | 6366 | 739.3 | 0.009262 | 12.53 | +1 | 5 |
| - | - | 2201 | 740.3 | - | - | 0 | - |
| - | - | 1358 | 764.3 | - | - | 0 | - |
| - | - | 930.1 | 766.3 | - | - | 0 | - |
| - | - | 577.2 | 772.3 | - | - | 0 | - |
| - | - | 561.6 | 790.3 | - | - | 0 | - |
| - | - | 6605 | 794.3 | - | - | 0 | - |
| - | - | 2649 | 795.3 | - | - | 0 | - |
| - | - | 678.3 | 796.3 | - | - | 0 | - |
| - | - | 1352 | 808.3 | - | - | 0 | - |
| 5 | z | 2.152E+04 | 838.3 | 0.009771 | 11.66 | +1 | 6 |
| - | - | 2.617E+04 | 839.3 | - | - | 0 | - |
| - | - | 1.068E+04 | 840.3 | - | - | 0 | - |
| - | - | 2652 | 841.3 | - | - | 0 | - |
| 5 | y | 6137 | 854.3 | 0.009358 | 10.95 | +1 | 6 |
| - | - | 3283 | 855.3 | - | - | 0 | - |
| - | - | 1410 | 856.3 | - | - | 0 | - |
| 7 | c | 937.5 | 877.4 | 0.00236 | 2.69 | +1 | 7 |
| - | - | 1410 | 896.3 | - | - | 0 | - |
| - | - | 1459 | 905.3 | - | - | 0 | - |
| - | - | 850.3 | 906.4 | - | - | 0 | - |
| - | - | 3184 | 909.3 | - | - | 0 | - |
| - | - | 1845 | 910.3 | - | - | 0 | - |
| - | - | 656.5 | 920.3 | - | - | 0 | - |
| 4 | z | 4014 | 953.3 | 0.0095 | 9.966 | +1 | 7 |
| - | - | 1.599E+04 | 954.3 | - | - | 0 | - |
| - | - | 8104 | 955.3 | - | - | 0 | - |
| - | - | 2955 | 956.3 | - | - | 0 | - |
| 4 | y | 1.099E+04 | 969.3 | 0.009453 | 9.752 | +1 | 7 |
| - | - | 6717 | 970.3 | - | - | 0 | - |
| - | - | 1750 | 971.3 | - | - | 0 | - |
| - | - | 3301 | 981.4 | - | - | 0 | - |
| - | - | 1719 | 982.4 | - | - | 0 | - |
| - | - | 1.033E+04 | 1023 | - | - | 0 | - |
| 8 | c | 1.299E+04 | 1024 | 0.002739 | 2.674 | +1 | 8 |
| - | - | 6631 | 1025 | - | - | 0 | - |
| - | - | 2189 | 1026 | - | - | 0 | - |
| - | - | 2892 | 1052 | - | - | 0 | - |
| - | - | 1304 | 1053 | - | - | 0 | - |
| - | - | 937.6 | 1054 | - | - | 0 | - |
| - | - | 6076 | 1056 | - | - | 0 | - |
| - | - | 3245 | 1057 | - | - | 0 | - |
| - | - | 962 | 1058 | - | - | 0 | - |
| 3 | y | 737.3 | 1098 | 0.01349 | 12.28 | +1 | 8 |
| 3 | z | 2654 | 1100 | 0.01018 | 9.25 | +1 | 8 |
| - | - | 1463 | 1101 | - | - | 0 | - |
| 3 | y | 1.981E+04 | 1116 | 0.009399 | 8.419 | +1 | 8 |
| - | - | 1.25E+04 | 1117 | - | - | 0 | - |
| - | - | 4763 | 1118 | - | - | 0 | - |
| - | - | 657.9 | 1119 | - | - | 0 | - |
| - | - | 2853 | 1128 | - | - | 0 | - |
| - | - | 2147 | 1129 | - | - | 0 | - |
| - | - | 695.6 | 1130 | - | - | 0 | - |
| - | - | 680.9 | 1145 | - | - | 0 | - |
| - | - | 1.144E+04 | 1146 | - | - | 0 | - |
| - | - | 1.037E+04 | 1147 | - | - | 0 | - |
| - | - | 2811 | 1148 | - | - | 0 | - |
| - | - | 976.2 | 1158 | - | - | 0 | - |
| 9 | c | 4.071E+04 | 1161 | 0.004374 | 3.766 | +1 | 9 |
| - | - | 2.451E+04 | 1162 | - | - | 0 | - |
| - | - | 9908 | 1163 | - | - | 0 | - |
| - | - | 1884 | 1164 | - | - | 0 | - |
| - | - | 736.9 | 1170 | - | - | 0 | - |
| 2 | w | 604.6 | 1184 | 0.02249 | 18.99 | +1 | 9 |
| - | - | 877.9 | 1186 | - | - | 0 | - |
| - | - | 2162 | 1187 | - | - | 0 | - |
| - | - | 1779 | 1188 | - | - | 0 | - |
| 2 | y | 1725 | 1199 | 0.0105 | 8.757 | +1 | 9 |
| - | - | 3588 | 1200 | - | - | 0 | - |
| 2 | z | 8568 | 1201 | 0.00852 | 7.092 | +1 | 9 |
| - | - | 1.27E+04 | 1202 | - | - | 0 | - |
| - | - | 1.984E+04 | 1203 | - | - | 0 | - |
| - | - | 1.561E+04 | 1204 | - | - | 0 | - |
| - | - | 4975 | 1205 | - | - | 0 | - |
| - | - | 950.2 | 1206 | - | - | 0 | - |
| - | - | 1226 | 1216 | - | - | 0 | - |
| 2 | y | 3912 | 1217 | 0.009205 | 7.561 | +1 | 9 |
| - | - | 5272 | 1218 | - | - | 0 | - |
| - | - | 2520 | 1219 | - | - | 0 | - |
| - | - | 1043 | 1220 | - | - | 0 | - |
| - | - | 791.5 | 1221 | - | - | 0 | - |
| - | - | 869.6 | 1228 | - | - | 0 | - |
| - | - | 1569 | 1273 | - | - | 0 | - |
| - | - | 876.7 | 1286 | - | - | 0 | - |
| - | - | 1634 | 1287 | - | - | 0 | - |
| - | - | 2260 | 1287 | - | - | 0 | - |
| - | - | 1075 | 1288 | - | - | 0 | - |
| - | - | 824.2 | 1302 | - | - | 0 | - |
| - | - | 1247 | 1303 | - | - | 0 | - |
| - | - | 8546 | 1304 | - | - | 0 | - |
| - | - | 7506 | 1305 | - | - | 0 | - |
| - | - | 2.338E+04 | 1306 | - | - | 0 | - |
| - | - | 1.617E+04 | 1307 | - | - | 0 | - |
| - | - | 7357 | 1308 | - | - | 0 | - |
| - | - | 937.7 | 1309 | - | - | 0 | - |
| - | - | 1200 | 1320 | - | - | 0 | - |
| - | - | 1480 | 1321 | - | - | 0 | - |
| - | - | 845.3 | 1322 | - | - | 0 | - |
| - | - | 719.2 | 1323 | - | - | 0 | - |
| - | - | 848.7 | 1331 | - | - | 0 | - |
| - | - | 1501 | 1332 | - | - | 0 | - |
| - | - | 1936 | 1332 | - | - | 0 | - |
| - | - | 2161 | 1334 | - | - | 0 | - |
| - | - | 1900 | 1335 | - | - | 0 | - |
| - | - | 967.3 | 1336 | - | - | 0 | - |
| - | - | 1004 | 1338 | - | - | 0 | - |
| - | - | 1026 | 1339 | - | - | 0 | - |
| - | - | 1199 | 1347 | - | - | 0 | - |
| - | - | 2.306E+04 | 1348 | - | - | 0 | - |
| - | - | 7.928E+04 | 1349 | - | - | 0 | - |
| - | - | 6.599E+04 | 1350 | - | - | 0 | - |
| - | - | 3.507E+04 | 1351 | - | - | 0 | - |
| - | - | 6758 | 1352 | - | - | 0 | - |
| - | - | 1083 | 1353 | - | - | 0 | - |
| - | - | 1126 | 1364 | - | - | 0 | - |
| - | - | 3.547E+04 | 1365 | - | - | 0 | - |
| - | - | 1.299E+05 | 1366 | - | - | 0 | - |
| - | - | 9.779E+04 | 1367 | - | - | 0 | - |
| - | - | 5.227E+04 | 1368 | - | - | 0 | - |
| - | - | 9188 | 1369 | - | - | 0 | - |
| - | - | 822.7 | 3108 | - | - | 0 | - |

m/z Charge Intensity FragmentType MassShift Position
120.08096313476562 0 11323.759
121.08431243896484 0 1375.548
122.68122863769531 0 380.3672
122.88201904296875 0 375.67245
122.89640808105469 0 576.42584
125.26056671142578 0 376.39352
128.8741912841797 0 343.8069
140.15280151367188 0 444.5083
145.4541473388672 0 407.8112
147.0406036376953 0 405.78635
148.92832946777344 0 612.3549
148.93853759765625 0 1890.2891
148.94383239746094 0 1901.639
148.95545959472656 0 3647.818
148.96041870117188 0 2303.177
148.966064453125 0 1563.8511
148.9807586669922 0 581.6311
149.04498291015625 0 1353.6157
160.2872314453125 0 472.50067
162.85183715820312 0 441.14783
173.4234619140625 0 493.98364
173.4573974609375 0 653.9161
175.6533660888672 0 468.7709
176.10716247558594 0 3051.233
177.11097717285156 0 518.13135
180.94918823242188 0 556.6773
194.25167846679688 0 470.27493
221.0832977294922 0 1016.57764 y 9
221.09376525878906 0 848.4802
221.12857055664062 0 17396.686
222.1317138671875 0 1985.9379
225.0432891845703 0 1624.3757
231.1130828857422 0 3284.3872
232.05654907226562 0 524.3281
233.9435272216797 0 503.2686
235.10757446289062 0 980.90985
239.0949249267578 0 2791.79
240.09500122070312 0 729.20215
249.12351989746094 0 16050.343 c Ammonia loss 1
250.12692260742188 0 2577.5962
263.1025695800781 0 1880.0945
295.1028747558594 0 2258.8613
296.1033020019531 0 931.4962
296.1906433105469 0 560.28925
299.0614013671875 0 1701.8641
313.1134948730469 0 707.9802
333.1600036621094 0 607.2524
355.0697937011719 0 3438.4114
358.1506042480469 0 3515.8677 y 8
359.02874755859375 0 1298.2903
369.12078857421875 0 754.74677
370.1227722167969 0 759.9604
378.1302490234375 0 1223.7579
378.1811828613281 0 3680.7185
415.1535949707031 0 641.86774
429.0892333984375 0 9204.047
479.1785583496094 0 656.1879
489.167236328125 0 864.6955 z 7
490.17547607421875 0 3262.3643
491.1759948730469 0 784.53046
504.17926025390625 0 1348.2126
505.1859130859375 0 2696.1326 y 7
506.1885681152344 0 1103.9188
511.2198486328125 0 1206.7081 c Ammonia loss 3
549.7020874023438 0 819.17615 y Water loss 2
558.7085571289062 0 6681.43 y 2
559.2099609375 0 5089.9106
559.7106323242188 0 1332.2614
560.2049560546875 0 3028.9204 z 6
561.1243896484375 0 521.87103
561.2092895507812 0 1920.3383
576.2235107421875 0 6237.278 y 6
577.2257690429688 0 1810.2871
578.2157592773438 0 1352.2762
609.232177734375 0 2586.149 y 1
609.7322998046875 0 1429.068
610.234375 0 897.39294
626.2465209960938 0 1694.0615 c Ammonia loss 4
641.7607421875 0 621.5151
649.2523803710938 0 1134.4393
650.7674560546875 0 776.6987
673.760986328125 0 2760.9197
674.2620239257812 0 2894.197
682.3352661132812 0 835.5631
682.766357421875 0 7251.6953
683.2672729492188 0 7414.785
683.7674560546875 0 3042.4783
684.2012329101562 0 1168.223
693.2364501953125 0 1097.3844
703.713134765625 0 643.2487
704.3027954101562 0 641.1834
723.2682495117188 0 15236.031 z 5
724.2730102539062 0 8142.8096
725.2761840820312 0 2909.514
739.2862548828125 0 6365.571 y 5
740.2879638671875 0 2200.6196
764.2788696289062 0 1358.2843
766.2783203125 0 930.0604
772.2980346679688 0 577.23773
790.314453125 0 561.6433
794.30517578125 0 6604.719
795.3082275390625 0 2649.1465
796.3029174804688 0 678.3202
808.2672119140625 0 1351.8673
838.2949829101562 0 21515.125 z 4
839.300537109375 0 26169.11
840.3031005859375 0 10684.686
841.3053588867188 0 2652.097
854.3132934570312 0 6137.3267 y 4
855.3154907226562 0 3282.581
856.3162231445312 0 1409.9882
877.3703002929688 0 937.4729 c 6
896.3248291015625 0 1409.9683
905.3414916992188 0 1458.8486
906.3531494140625 0 850.3342
909.3312377929688 0 3184.4097
910.3322143554688 0 1845.4565
920.3078002929688 0 656.4702
953.3216552734375 0 4014.114 z 3
954.3287353515625 0 15991.871
955.3320922851562 0 8103.86
956.3336181640625 0 2955.1057
969.34033203125 0 10985.199 y 3
970.343994140625 0 6717.113
971.34716796875 0 1749.542
981.4004516601562 0 3300.515
982.4077758789062 0 1719.0461
1023.400146484375 0 10333.402
1024.4058837890625 0 12987.48 c 7
1025.409423828125 0 6630.5415
1026.4068603515625 0 2189.2344
1052.4093017578125 0 2892.024
1053.411376953125 0 1303.5187
1054.4283447265625 0 937.61945
1056.3868408203125 0 6075.542
1057.390869140625 0 3244.628
1058.3807373046875 0 962.0478
1098.4022216796875 0 737.25287 y Water loss 2
1100.3907470703125 0 2654.021 z 2
1101.39453125 0 1462.9402
1116.40869140625 0 19814.04 y 2
1117.41357421875 0 12500.92
1118.4146728515625 0 4763.0396
1119.423095703125 0 657.8587
1128.443115234375 0 2853.2473
1129.4498291015625 0 2146.5737
1130.4525146484375 0 695.6119
1145.4454345703125 0 680.935
1146.4544677734375 0 11437.681
1147.4578857421875 0 10367.014
1148.4605712890625 0 2811.3433
1158.4822998046875 0 976.18506
1161.4664306640625 0 40706.105 c 8
1162.4686279296875 0 24505.844
1163.4693603515625 0 9908.465
1164.4658203125 0 1883.7758
1170.458251953125 0 736.8836
1184.447998046875 0 604.57684 w 1
1186.4376220703125 0 877.918
1187.4761962890625 0 2162.092
1188.480224609375 0 1778.5437
1199.4259033203125 0 1725.2632 y Water loss 1
1200.429931640625 0 3588.427
1201.436767578125 0 8567.552 z 1
1202.4552001953125 0 12698.437
1203.457275390625 0 19842.426
1204.4593505859375 0 15606.36
1205.4578857421875 0 4974.7173
1206.4566650390625 0 950.1692
1216.4559326171875 0 1225.8468
1217.4561767578125 0 3912.4338 y 1
1218.4599609375 0 5272.0215
1219.4608154296875 0 2520.312
1220.46630859375 0 1043.186
1221.4732666015625 0 791.5302
1228.4522705078125 0 869.5641
1273.489990234375 0 1569.0573
1285.5269775390625 0 876.6512
1286.5057373046875 0 1633.9248
1287.4979248046875 0 2259.5774
1288.49560546875 0 1074.5786
1301.5399169921875 0 824.20197
1302.53271484375 0 1247.0564
1303.5303955078125 0 8545.585
1304.5316162109375 0 7505.738
1305.513427734375 0 23379.898
1306.513916015625 0 16165.041
1307.5130615234375 0 7356.999
1308.5164794921875 0 937.7493
1319.512451171875 0 1199.7007
1320.51416015625 0 1480.171
1321.516357421875 0 845.27844
1322.524169921875 0 719.15393
1330.51318359375 0 848.7415
1331.507080078125 0 1500.6527
1332.498046875 0 1935.7668
1333.5064697265625 0 2161.235
1334.5244140625 0 1900.2134
1335.541259765625 0 967.2661
1337.5267333984375 0 1003.8314
1338.53857421875 0 1025.9642
1346.505615234375 0 1199.0272
1347.5189208984375 0 23063.395
1348.5103759765625 0 79280.85
1349.5123291015625 0 65990.195
1350.51220703125 0 35065.465
1351.51171875 0 6758.2417
1352.5157470703125 0 1083.3347
1363.512939453125 0 1125.8231
1364.522705078125 0 35469.363
1365.529296875 0 129916.805
1366.5316162109375 0 97791.53
1367.532470703125 0 52266.86
1368.534423828125 0 9188.073
3108.419921875 0 822.7287

Spectrum Details

|  |  |
| --- | --- |
| Matched peaks? Matched peaks The total absolute number of peaks matched. Additionally in brackets the total fraction of peaks matched and the total number of peaks is shown. | 28 (13.02% of 215) |
| FDR? FDR The false discovery rate estimated for this peptide. It is calculated by matching all theoretical fragments with a non-integer shift with the raw peaks for this spectrum. This is done with 40 different shifts. The resulting percentage is the average number of annotated peaks over the number of annotated peaks with the correct spectrum. | 0.26% |
| Satellite FDR? Satellite FDR See the FDR for details on its calculation. This satellite ion specific FDR only contains the satellite ions (d/w) for I/L/J positions. | - |
| PSM Score? PSM Score The PSM Score as given by Hecklib to this annotated spectrum. It is shown with three significant figures. | 340 |

#### Spectrum 8914? Spectrum 8914 The raw spectrum of this peptide as annotated by Hecklib. The fragments are coloured according to ion type (see legend). Any peaks with a star '\*' as text can be hovered over to see the full details, first the ion type second the mass shift type. By hovering over the amino acids in the peptide or ions in the legend the corresponding peaks are highlighted. By toggling the 'Unassigned' label you can turn the background (unassigned) peaks on or off in the plot. By updating the slider in the Ion legend you can update the spectrum to only show the top X% of the peaks with labels. The top X% means any peak that is within X% of the highest intensity. By dragging in the spectrum you can zoom in to a specific part of the spectrum and use 'Zoom Out' to get back to the original zoom level. The annotation of the spectrum is based on the given sequence in the peptides file and is done with different software so inconsistencies are likely. The peaks are annotated based on the given sequence, with 20 ppm tolerance.

Copy Data

##### Spectrum 8914 (TSV)

###### Preview

```
Loading example...
```

*Click on the button to copy the data to your clipboard.*

Mz MinMz MaxIntensity Max

WidthHeightPeptide font sizePeptide stroke widthSpectrum font sizeSpectrum stroke widthCompact peptide

Ion legend

wxyz

abcd

OtherUnassignedIon^Charge^_Position_Show for top:%

FTFDDYAMHW

02.03e+44.06e+46.09e+48.12e+4

Zoom Out

a^+1^_2_d^+1^_2_a^+1^_2_b^+1^_2_b^+1^_2_y^+1^_2_b^+1^_3_b^+2^_8_b^+1^_4_y^+2^_8_y^+1^_4_b^+1^_5_y^+2^_9_b^+1^_5_\*\*y^+1^_5_y^+1^_6_y^+1^_7_y^+1^_7_b^+1^_8_y^+1^_8_y^+1^_8_y^+1^_9_

0830166124913321

Fragment Matches Table

Show background peaks

| Position | Ion type | Intensity | mz Theoretical | mz Error (Th) | mz Error (ppm) | Charge | Series Number |
| --- | --- | --- | --- | --- | --- | --- | --- |
| - | - | 8.04E+04 | 120.1 | - | - | 0 | - |
| - | - | 793 | 121.1 | - | - | 0 | - |
| - | - | 7051 | 121.1 | - | - | 0 | - |
| - | - | 359 | 127.1 | - | - | 0 | - |
| - | - | 1904 | 129.1 | - | - | 0 | - |
| - | - | 518.3 | 130.1 | - | - | 0 | - |
| - | - | 946.6 | 130.1 | - | - | 0 | - |
| - | - | 403.9 | 130.4 | - | - | 0 | - |
| - | - | 419.3 | 131.1 | - | - | 0 | - |
| - | - | 433.6 | 132.8 | - | - | 0 | - |
| - | - | 699.8 | 133.1 | - | - | 0 | - |
| - | - | 477 | 134.1 | - | - | 0 | - |
| - | - | 496.9 | 134.1 | - | - | 0 | - |
| - | - | 7879 | 136.1 | - | - | 0 | - |
| - | - | 474.9 | 136.8 | - | - | 0 | - |
| - | - | 623.9 | 138.1 | - | - | 0 | - |
| - | - | 395.9 | 144.4 | - | - | 0 | - |
| - | - | 747.6 | 146.1 | - | - | 0 | - |
| - | - | 624.4 | 148.9 | - | - | 0 | - |
| - | - | 599.9 | 148.9 | - | - | 0 | - |
| - | - | 667.9 | 148.9 | - | - | 0 | - |
| - | - | 592.3 | 148.9 | - | - | 0 | - |
| - | - | 1209 | 148.9 | - | - | 0 | - |
| - | - | 1290 | 148.9 | - | - | 0 | - |
| - | - | 3070 | 148.9 | - | - | 0 | - |
| - | - | 4971 | 148.9 | - | - | 0 | - |
| - | - | 3789 | 149 | - | - | 0 | - |
| - | - | 1735 | 149 | - | - | 0 | - |
| - | - | 1235 | 149 | - | - | 0 | - |
| - | - | 1027 | 149 | - | - | 0 | - |
| - | - | 1087 | 149 | - | - | 0 | - |
| - | - | 1034 | 149 | - | - | 0 | - |
| - | - | 617.9 | 149 | - | - | 0 | - |
| - | - | 412.9 | 149 | - | - | 0 | - |
| - | - | 1756 | 149 | - | - | 0 | - |
| - | - | 2387 | 156.1 | - | - | 0 | - |
| - | - | 1444 | 158.1 | - | - | 0 | - |
| - | - | 482.3 | 159.1 | - | - | 0 | - |
| - | - | 2866 | 159.1 | - | - | 0 | - |
| - | - | 551.6 | 160.1 | - | - | 0 | - |
| - | - | 732.4 | 166.1 | - | - | 0 | - |
| - | - | 3369 | 167.1 | - | - | 0 | - |
| - | - | 638.5 | 167.1 | - | - | 0 | - |
| - | - | 451.3 | 173.3 | - | - | 0 | - |
| - | - | 4892 | 175.1 | - | - | 0 | - |
| - | - | 3.889E+04 | 176.1 | - | - | 0 | - |
| - | - | 427.5 | 176.4 | - | - | 0 | - |
| - | - | 819.6 | 177.1 | - | - | 0 | - |
| - | - | 2819 | 177.1 | - | - | 0 | - |
| - | - | 3941 | 186.1 | - | - | 0 | - |
| - | - | 547.7 | 186.7 | - | - | 0 | - |
| - | - | 971.3 | 191.1 | - | - | 0 | - |
| - | - | 2506 | 193.1 | - | - | 0 | - |
| - | - | 529.9 | 198.8 | - | - | 0 | - |
| - | - | 1474 | 203.1 | - | - | 0 | - |
| 2 | a | 2242 | 203.1 | 0.0001677 | 0.8257 | +1 | 2 |
| - | - | 852.2 | 204.1 | - | - | 0 | - |
| 2 | d | 544.7 | 207.1 | 0.0006297 | 3.04 | +1 | 2 |
| - | - | 640.6 | 209.2 | - | - | 0 | - |
| - | - | 537.6 | 214.1 | - | - | 0 | - |
| - | - | 2057 | 217.1 | - | - | 0 | - |
| - | - | 490.6 | 217.1 | - | - | 0 | - |
| - | - | 764.4 | 221.1 | - | - | 0 | - |
| - | - | 8601 | 221.1 | - | - | 0 | - |
| - | - | 6133 | 221.1 | - | - | 0 | - |
| 2 | a | 4.376E+04 | 221.1 | 0.00033 | 1.492 | +1 | 2 |
| - | - | 1381 | 222.1 | - | - | 0 | - |
| - | - | 614.7 | 222.1 | - | - | 0 | - |
| - | - | 5482 | 222.1 | - | - | 0 | - |
| - | - | 5913 | 225 | - | - | 0 | - |
| - | - | 962.5 | 226 | - | - | 0 | - |
| 2 | b | 1.09E+04 | 231.1 | 0.0004161 | 1.8 | +1 | 2 |
| - | - | 1151 | 232.1 | - | - | 0 | - |
| - | - | 4699 | 235.1 | - | - | 0 | - |
| - | - | 7256 | 239.1 | - | - | 0 | - |
| - | - | 1325 | 240.1 | - | - | 0 | - |
| 2 | b | 2.292E+04 | 249.1 | 0.0003494 | 1.403 | +1 | 2 |
| - | - | 2719 | 250.1 | - | - | 0 | - |
| - | - | 1590 | 251.1 | - | - | 0 | - |
| - | - | 997.5 | 257.1 | - | - | 0 | - |
| - | - | 570.3 | 259.1 | - | - | 0 | - |
| - | - | 3530 | 263.1 | - | - | 0 | - |
| - | - | 522.2 | 272.5 | - | - | 0 | - |
| - | - | 1211 | 274.1 | - | - | 0 | - |
| - | - | 1970 | 279.1 | - | - | 0 | - |
| - | - | 757.6 | 281.1 | - | - | 0 | - |
| - | - | 485.8 | 282.8 | - | - | 0 | - |
| - | - | 1499 | 285 | - | - | 0 | - |
| - | - | 1670 | 285.1 | - | - | 0 | - |
| - | - | 1767 | 292.1 | - | - | 0 | - |
| - | - | 1797 | 295.1 | - | - | 0 | - |
| - | - | 536.1 | 297.2 | - | - | 0 | - |
| - | - | 5029 | 299.1 | - | - | 0 | - |
| - | - | 775.3 | 300.1 | - | - | 0 | - |
| - | - | 685.1 | 301.1 | - | - | 0 | - |
| - | - | 870.7 | 313.1 | - | - | 0 | - |
| - | - | 701.5 | 325.1 | - | - | 0 | - |
| - | - | 641.3 | 327.8 | - | - | 0 | - |
| - | - | 609 | 333.2 | - | - | 0 | - |
| - | - | 4022 | 340.1 | - | - | 0 | - |
| - | - | 814.4 | 341 | - | - | 0 | - |
| - | - | 720.7 | 341.1 | - | - | 0 | - |
| - | - | 749.9 | 346.1 | - | - | 0 | - |
| - | - | 1587 | 350.1 | - | - | 0 | - |
| - | - | 4782 | 355.1 | - | - | 0 | - |
| - | - | 1296 | 356.1 | - | - | 0 | - |
| 9 | y | 2.175E+04 | 358.1 | 0.005271 | 14.72 | +1 | 2 |
| - | - | 7951 | 359 | - | - | 0 | - |
| - | - | 4085 | 359.2 | - | - | 0 | - |
| - | - | 1781 | 364.2 | - | - | 0 | - |
| - | - | 828.9 | 368.1 | - | - | 0 | - |
| - | - | 994.9 | 369.1 | - | - | 0 | - |
| - | - | 680.2 | 374.1 | - | - | 0 | - |
| - | - | 5408 | 378.1 | - | - | 0 | - |
| 3 | b | 4085 | 378.2 | 0.0005446 | 1.44 | +1 | 3 |
| - | - | 1393 | 379.1 | - | - | 0 | - |
| - | - | 737.7 | 379.2 | - | - | 0 | - |
| - | - | 737.1 | 382.1 | - | - | 0 | - |
| - | - | 1555 | 394.1 | - | - | 0 | - |
| - | - | 608.3 | 404.1 | - | - | 0 | - |
| - | - | 1028 | 419 | - | - | 0 | - |
| - | - | 882.7 | 423.2 | - | - | 0 | - |
| - | - | 708.8 | 424.2 | - | - | 0 | - |
| - | - | 7699 | 429.1 | - | - | 0 | - |
| - | - | 549.7 | 431.5 | - | - | 0 | - |
| - | - | 3918 | 441.2 | - | - | 0 | - |
| - | - | 1073 | 442.2 | - | - | 0 | - |
| - | - | 680.4 | 461.2 | - | - | 0 | - |
| - | - | 1098 | 465.2 | - | - | 0 | - |
| - | - | 1322 | 479.2 | - | - | 0 | - |
| - | - | 643.1 | 482.2 | - | - | 0 | - |
| 8 | b | 746.9 | 495.2 | 0.002372 | 4.79 | +2 | 8 |
| - | - | 593.5 | 497.2 | - | - | 0 | - |
| - | - | 5260 | 505.2 | - | - | 0 | - |
| - | - | 1364 | 506.2 | - | - | 0 | - |
| 4 | b | 2214 | 511.2 | 0.001275 | 2.495 | +1 | 4 |
| - | - | 3338 | 512.2 | - | - | 0 | - |
| - | - | 610.4 | 513.8 | - | - | 0 | - |
| - | - | 646.3 | 519.2 | - | - | 0 | - |
| 3 | y | 7525 | 558.7 | 0.005456 | 9.765 | +2 | 8 |
| - | - | 4137 | 559.2 | - | - | 0 | - |
| - | - | 869 | 559.7 | - | - | 0 | - |
| 7 | y | 9575 | 576.2 | 0.01033 | 17.94 | +1 | 4 |
| - | - | 2313 | 577.2 | - | - | 0 | - |
| - | - | 714 | 587.2 | - | - | 0 | - |
| - | - | 951.3 | 606.2 | - | - | 0 | - |
| 5 | b | 1306 | 608.2 | 0.0002409 | 0.396 | +1 | 5 |
| 2 | y | 2223 | 609.2 | 0.005054 | 8.295 | +2 | 9 |
| - | - | 1079 | 609.7 | - | - | 0 | - |
| - | - | 786.7 | 610.2 | - | - | 0 | - |
| 5 | b | 1436 | 626.2 | 0.0009132 | 1.458 | +1 | 5 |
| - | - | 871.9 | 627.2 | - | - | 0 | - |
| - | - | 738.6 | 642.3 | - | - | 0 | - |
| - | - | 1243 | 650.8 | - | - | 0 | - |
| - | - | 739.6 | 657.3 | - | - | 0 | - |
| 0 | Precursor | 3596 | 673.8 | 0.005426 | 8.054 | +2 | -1 |
| - | - | 2548 | 674.3 | - | - | 0 | - |
| - | - | 1573 | 674.8 | - | - | 0 | - |
| - | - | 2663 | 675.3 | - | - | 0 | - |
| - | - | 1045 | 676.3 | - | - | 0 | - |
| - | - | 1119 | 681.8 | - | - | 0 | - |
| - | - | 1162 | 682.3 | - | - | 0 | - |
| 0 | Precursor | 8091 | 682.8 | 0.005332 | 7.809 | +2 | -1 |
| - | - | 694.9 | 682.8 | - | - | 0 | - |
| - | - | 4563 | 683.3 | - | - | 0 | - |
| - | - | 646.8 | 683.3 | - | - | 0 | - |
| - | - | 2941 | 683.8 | - | - | 0 | - |
| - | - | 1845 | 684.2 | - | - | 0 | - |
| - | - | 611 | 686.3 | - | - | 0 | - |
| - | - | 923.9 | 695.3 | - | - | 0 | - |
| - | - | 725.6 | 721.3 | - | - | 0 | - |
| - | - | 935.1 | 724.1 | - | - | 0 | - |
| 6 | y | 6371 | 739.3 | 0.01115 | 15.09 | +1 | 5 |
| - | - | 2296 | 740.3 | - | - | 0 | - |
| - | - | 697.3 | 741.3 | - | - | 0 | - |
| - | - | 963.2 | 749.3 | - | - | 0 | - |
| - | - | 720.8 | 772.3 | - | - | 0 | - |
| - | - | 3061 | 790.3 | - | - | 0 | - |
| - | - | 1177 | 791.3 | - | - | 0 | - |
| - | - | 948.1 | 832.3 | - | - | 0 | - |
| - | - | 748.8 | 833.3 | - | - | 0 | - |
| 5 | y | 9517 | 854.3 | 0.01058 | 12.38 | +1 | 6 |
| - | - | 4127 | 855.3 | - | - | 0 | - |
| - | - | 1155 | 856.3 | - | - | 0 | - |
| - | - | 1492 | 896.3 | - | - | 0 | - |
| - | - | 5729 | 905.3 | - | - | 0 | - |
| - | - | 2533 | 906.3 | - | - | 0 | - |
| 4 | y | 823.9 | 951.3 | 0.01239 | 13.02 | +1 | 7 |
| - | - | 626.2 | 952.3 | - | - | 0 | - |
| 4 | y | 1.587E+04 | 969.3 | 0.01025 | 10.57 | +1 | 7 |
| - | - | 7358 | 970.3 | - | - | 0 | - |
| - | - | 3167 | 971.3 | - | - | 0 | - |
| 8 | b | 691.2 | 1007 | 0.0001309 | 0.13 | +1 | 8 |
| - | - | 1139 | 1034 | - | - | 0 | - |
| - | - | 8027 | 1052 | - | - | 0 | - |
| - | - | 4306 | 1053 | - | - | 0 | - |
| - | - | 1682 | 1054 | - | - | 0 | - |
| 3 | y | 1223 | 1098 | 0.0152 | 13.84 | +1 | 8 |
| 3 | y | 2.174E+04 | 1116 | 0.01086 | 9.731 | +1 | 8 |
| - | - | 1.596E+04 | 1117 | - | - | 0 | - |
| - | - | 5598 | 1118 | - | - | 0 | - |
| - | - | 787.4 | 1119 | - | - | 0 | - |
| - | - | 1373 | 1153 | - | - | 0 | - |
| - | - | 901 | 1154 | - | - | 0 | - |
| - | - | 755.1 | 1155 | - | - | 0 | - |
| 2 | y | 3432 | 1217 | 0.01067 | 8.764 | +1 | 9 |
| - | - | 3019 | 1218 | - | - | 0 | - |
| - | - | 937.6 | 1219 | - | - | 0 | - |
| - | - | 778.2 | 1227 | - | - | 0 | - |
| - | - | 689.2 | 1362 | - | - | 0 | - |
| - | - | 843.6 | 1731 | - | - | 0 | - |
| - | - | 662.3 | 2159 | - | - | 0 | - |
| - | - | 746.7 | 3289 | - | - | 0 | - |

m/z Charge Intensity FragmentType MassShift Position
120.08106994628906 0 80397.29
121.07962036132812 0 792.99634
121.08438873291016 0 7051.3647
127.11666107177734 0 358.9826
129.1025390625 0 1904.435
130.06076049804688 0 518.32904
130.0655975341797 0 946.6451
130.40126037597656 0 403.89548
131.1040802001953 0 419.3202
132.84246826171875 0 433.6273
133.0860137939453 0 699.7983
134.05967712402344 0 477.03128
134.0650177001953 0 496.88632
136.0759735107422 0 7879.057
136.80311584472656 0 474.94785
138.0664520263672 0 623.88135
144.41590881347656 0 395.8831
146.060546875 0 747.639
148.89404296875 0 624.3707
148.90118408203125 0 599.89764
148.90850830078125 0 667.8915
148.91610717773438 0 592.34106
148.92330932617188 0 1208.6283
148.93032836914062 0 1289.8187
148.93768310546875 0 3069.653
148.94544982910156 0 4970.6484
148.96212768554688 0 3789.188
148.9697723388672 0 1734.8021
148.97662353515625 0 1234.9153
148.98428344726562 0 1027.3058
148.99159240722656 0 1086.8898
148.99884033203125 0 1033.6265
149.0059356689453 0 617.9499
149.01292419433594 0 412.88126
149.04498291015625 0 1755.9185
156.0770721435547 0 2386.7444
158.0965576171875 0 1444.2434
159.07666015625 0 482.3164
159.09182739257812 0 2866.3052
160.09556579589844 0 551.5642
166.06101989746094 0 732.4405
167.05557250976562 0 3369.4768
167.09317016601562 0 638.5218
173.29673767089844 0 451.28815
175.0869903564453 0 4891.569
176.1072998046875 0 38889.773
176.3779296875 0 427.453
177.1026611328125 0 819.5836
177.11083984375 0 2819.0752
186.09164428710938 0 3940.8987
186.6786346435547 0 547.6824
191.11793518066406 0 971.2768
193.10885620117188 0 2505.834
198.7702178955078 0 529.8576
203.08192443847656 0 1473.9735
203.11805725097656 0 2242.3447 a Water loss 1
204.07699584960938 0 852.16705
207.11343383789062 0 544.6607 d 1
209.16531372070312 0 640.6498
214.08566284179688 0 537.59015
217.0974884033203 0 2056.5269
217.1334228515625 0 490.64435
221.08238220214844 0 764.4491
221.09230041503906 0 8600.611
221.10377502441406 0 6132.879
221.1287841796875 0 43758.586 a 1
222.09617614746094 0 1380.896
222.10789489746094 0 614.7338
222.1322784423828 0 5481.9766
225.04319763183594 0 5912.799
226.04415893554688 0 962.5197
231.11322021484375 0 10895.629 b Water loss 1
232.11607360839844 0 1151.2704
235.10806274414062 0 4698.865
239.0953369140625 0 7255.7524
240.0970916748047 0 1324.6191
249.12371826171875 0 22918.533 b 1
250.1269989013672 0 2718.766
251.10357666015625 0 1589.7917
257.1072692871094 0 997.4699
259.107421875 0 570.2745
263.1029052734375 0 3530.4683
272.48675537109375 0 522.1609
274.1302795410156 0 1210.7362
279.0977783203125 0 1969.8792
281.0506286621094 0 757.6278
282.8125305175781 0 485.7737
285.01019287109375 0 1498.5364
285.1016845703125 0 1669.9393
292.1407775878906 0 1766.5004
295.1036376953125 0 1796.927
297.15777587890625 0 536.1258
299.0621337890625 0 5028.5234
300.0613098144531 0 775.3204
301.0603942871094 0 685.1337
313.11566162109375 0 870.7372
325.139892578125 0 701.456
327.8438720703125 0 641.3453
333.1605529785156 0 608.97797
340.1407470703125 0 4021.8228
341.02044677734375 0 814.4092
341.1414489746094 0 720.7197
346.1401062011719 0 749.9228
350.1353759765625 0 1587.2434
355.070556640625 0 4782.3247
356.1388244628906 0 1295.5537
358.1513366699219 0 21750.238 y 8
359.0288391113281 0 7950.8433
359.1542663574219 0 4084.8384
364.1503601074219 0 1780.9906
368.13580322265625 0 828.9041
369.1230163574219 0 994.8513
374.1478271484375 0 680.1834
378.13006591796875 0 5408.171
378.1817626953125 0 4084.602 b Water loss 2
379.1330261230469 0 1392.8281
379.18377685546875 0 737.6579
382.1455383300781 0 737.121
394.1255187988281 0 1555.1179
404.1435852050781 0 608.2808
418.9975891113281 0 1028.1064
423.1793518066406 0 882.6664
424.1632080078125 0 708.76575
429.0897216796875 0 7699.1885
431.5069885253906 0 549.70514
441.18896484375 0 3917.6558
442.1910095214844 0 1073.2869
461.1666259765625 0 680.4013
465.1622619628906 0 1097.9515
479.1773681640625 0 1321.601
482.19122314453125 0 643.1399
495.18902587890625 0 746.90015 b Water loss 7
497.1737365722656 0 593.45593
505.1871643066406 0 5259.6016
506.1899719238281 0 1363.6147
511.2200012207031 0 2213.9033 b 3
512.2255249023438 0 3337.8567
513.8385620117188 0 610.41144
519.2024536132812 0 646.34406
558.708740234375 0 7525.3174 y 2
559.2108764648438 0 4136.972
559.7119750976562 0 869.00885
576.2239990234375 0 9575.156 y 6
577.2265014648438 0 2312.69
587.2177124023438 0 714.0493
606.2329711914062 0 951.27625
608.23486328125 0 1306.4141 b Water loss 4
609.232177734375 0 2223.3906 y 1
609.73193359375 0 1078.8252
610.237548828125 0 786.7326
626.24658203125 0 1435.5319 b 4
627.2471923828125 0 871.904
642.262939453125 0 738.57245
650.765625 0 1242.9922
657.264892578125 0 739.6268
673.761474609375 0 3595.6194 Precursor Water loss
674.2630615234375 0 2547.8604
674.764404296875 0 1572.9912
675.289794921875 0 2662.6404
676.2947387695312 0 1044.7833
681.847900390625 0 1119.1095
682.3423461914062 0 1161.771
682.7666625976562 0 8090.5713 Precursor
682.8295288085938 0 694.947
683.2684936523438 0 4563.1143
683.3240966796875 0 646.7913
683.7686157226562 0 2940.773
684.2051391601562 0 1844.8915
686.2606201171875 0 611.0408
695.2706298828125 0 923.9156
721.268798828125 0 725.5542
724.1284790039062 0 935.1232
739.2881469726562 0 6370.9746 y 5
740.2898559570312 0 2295.8494
741.2839965820312 0 697.26965
749.2535400390625 0 963.2047
772.307861328125 0 720.8259
790.3162841796875 0 3060.6506
791.3189697265625 0 1177.0288
832.330078125 0 948.0554
833.3276977539062 0 748.78735
854.3145141601562 0 9517.296 y 4
855.3178100585938 0 4126.8467
856.3130493164062 0 1154.8711
896.324462890625 0 1492.1943
905.3433227539062 0 5728.8994
906.3463745117188 0 2533.2043
951.3327026367188 0 823.88837 y Water loss 3
952.3242797851562 0 626.21063
969.3411254882812 0 15873.836 y 3
970.3436889648438 0 7357.8843
971.3445434570312 0 3167.3108
1007.37646484375 0 691.20276 b 7
1034.4022216796875 0 1139.4973
1052.4114990234375 0 8026.964
1053.4156494140625 0 4306.4893
1054.41748046875 0 1681.7493
1098.4039306640625 0 1223.2307 y Water loss 2
1116.41015625 0 21738.17 y 2
1117.4129638671875 0 15958.048
1118.4150390625 0 5598.482
1119.413818359375 0 787.35785
1153.455810546875 0 1373.3936
1154.458984375 0 900.98987
1155.4613037109375 0 755.13495
1217.4576416015625 0 3431.566 y 1
1218.4615478515625 0 3018.6145
1219.4453125 0 937.5534
1227.440673828125 0 778.2477
1362.1031494140625 0 689.22577
1731.263916015625 0 843.5835
2158.8271484375 0 662.30975
3288.5546875 0 746.6815

Spectrum Details

|  |  |
| --- | --- |
| Matched peaks? Matched peaks The total absolute number of peaks matched. Additionally in brackets the total fraction of peaks matched and the total number of peaks is shown. | 24 (11.27% of 213) |
| FDR? FDR The false discovery rate estimated for this peptide. It is calculated by matching all theoretical fragments with a non-integer shift with the raw peaks for this spectrum. This is done with 40 different shifts. The resulting percentage is the average number of annotated peaks over the number of annotated peaks with the correct spectrum. | 0.40% |
| Satellite FDR? Satellite FDR See the FDR for details on its calculation. This satellite ion specific FDR only contains the satellite ions (d/w) for I/L/J positions. | - |
| PSM Score? PSM Score The PSM Score as given by Hecklib to this annotated spectrum. It is shown with three significant figures. | 239 |

#### Spectrum 9998? Spectrum 9998 The raw spectrum of this peptide as annotated by Hecklib. The fragments are coloured according to ion type (see legend). Any peaks with a star '\*' as text can be hovered over to see the full details, first the ion type second the mass shift type. By hovering over the amino acids in the peptide or ions in the legend the corresponding peaks are highlighted. By toggling the 'Unassigned' label you can turn the background (unassigned) peaks on or off in the plot. By updating the slider in the Ion legend you can update the spectrum to only show the top X% of the peaks with labels. The top X% means any peak that is within X% of the highest intensity. By dragging in the spectrum you can zoom in to a specific part of the spectrum and use 'Zoom Out' to get back to the original zoom level. The annotation of the spectrum is based on the given sequence in the peptides file and is done with different software so inconsistencies are likely. The peaks are annotated based on the given sequence, with 20 ppm tolerance.

Copy Data

##### Spectrum 9998 (TSV)

###### Preview

```
Loading example...
```

*Click on the button to copy the data to your clipboard.*

Mz MinMz MaxIntensity Max

WidthHeightPeptide font sizePeptide stroke widthSpectrum font sizeSpectrum stroke widthCompact peptide

Ion legend

wxyz

abcd

OtherUnassignedIon^Charge^_Position_Show for top:%

FTFDDYAMHW

01.36e+42.72e+44.09e+45.45e+4

Zoom Out

c^+1^_2_y^+1^_2_y^+1^_3_z^+1^_4_y^+2^_8_y^+1^_4_y^+2^_9_c^+1^_5_z^+1^_5_y^+1^_5_z^+1^_6_y^+1^_6_c^+1^_7_z^+1^_7_y^+1^_7_c^+1^_8_z^+1^_8_y^+1^_8_c^+1^_9_z^+1^_9_y^+1^_9_

0874174726213495

Fragment Matches Table

Show background peaks

| Position | Ion type | Intensity | mz Theoretical | mz Error (Th) | mz Error (ppm) | Charge | Series Number |
| --- | --- | --- | --- | --- | --- | --- | --- |
| - | - | 378.9 | 120.1 | - | - | 0 | - |
| - | - | 5031 | 120.1 | - | - | 0 | - |
| - | - | 418.4 | 121.1 | - | - | 0 | - |
| - | - | 734.6 | 148.9 | - | - | 0 | - |
| - | - | 676.2 | 148.9 | - | - | 0 | - |
| - | - | 805.7 | 148.9 | - | - | 0 | - |
| - | - | 940.1 | 148.9 | - | - | 0 | - |
| - | - | 1080 | 148.9 | - | - | 0 | - |
| - | - | 2549 | 148.9 | - | - | 0 | - |
| - | - | 2983 | 148.9 | - | - | 0 | - |
| - | - | 3718 | 149 | - | - | 0 | - |
| - | - | 3147 | 149 | - | - | 0 | - |
| - | - | 1069 | 149 | - | - | 0 | - |
| - | - | 1120 | 149 | - | - | 0 | - |
| - | - | 727.6 | 149 | - | - | 0 | - |
| - | - | 625.1 | 149 | - | - | 0 | - |
| - | - | 737.1 | 149 | - | - | 0 | - |
| - | - | 506.1 | 149 | - | - | 0 | - |
| - | - | 432.5 | 149 | - | - | 0 | - |
| - | - | 512.6 | 149 | - | - | 0 | - |
| - | - | 586.8 | 166.1 | - | - | 0 | - |
| - | - | 1057 | 176.1 | - | - | 0 | - |
| - | - | 449.6 | 185.6 | - | - | 0 | - |
| - | - | 7104 | 221.1 | - | - | 0 | - |
| - | - | 1690 | 231.1 | - | - | 0 | - |
| 2 | c | 6249 | 249.1 | 4.731E-05 | 0.1899 | +1 | 2 |
| - | - | 617.6 | 250.1 | - | - | 0 | - |
| - | - | 715.5 | 253.1 | - | - | 0 | - |
| - | - | 473.6 | 261.1 | - | - | 0 | - |
| - | - | 525.2 | 282.4 | - | - | 0 | - |
| - | - | 583.4 | 316.2 | - | - | 0 | - |
| 9 | y | 1132 | 342.2 | 0.0006094 | 1.781 | +1 | 2 |
| - | - | 1212 | 378.2 | - | - | 0 | - |
| - | - | 2662 | 474.2 | - | - | 0 | - |
| 8 | y | 1095 | 489.2 | 0.004947 | 10.11 | +1 | 3 |
| 7 | z | 2791 | 544.2 | 0.004838 | 8.889 | +1 | 4 |
| - | - | 853.1 | 545.2 | - | - | 0 | - |
| 3 | y | 2085 | 550.7 | 0.002042 | 3.709 | +2 | 8 |
| - | - | 850.9 | 551.2 | - | - | 0 | - |
| - | - | 684.2 | 551.7 | - | - | 0 | - |
| 7 | y | 1733 | 560.2 | 0.00357 | 6.372 | +1 | 4 |
| 2 | y | 813.6 | 601.2 | 0.0028 | 4.658 | +2 | 9 |
| 5 | c | 648.7 | 626.2 | 0.004214 | 6.729 | +1 | 5 |
| - | - | 1396 | 665.8 | - | - | 0 | - |
| - | - | 1511 | 666.3 | - | - | 0 | - |
| - | - | 705.5 | 666.8 | - | - | 0 | - |
| - | - | 2728 | 674.8 | - | - | 0 | - |
| - | - | 2824 | 675.3 | - | - | 0 | - |
| - | - | 1302 | 675.8 | - | - | 0 | - |
| 6 | z | 5046 | 707.3 | 0.00468 | 6.618 | +1 | 5 |
| - | - | 3381 | 708.3 | - | - | 0 | - |
| - | - | 1010 | 709.3 | - | - | 0 | - |
| 6 | y | 1685 | 723.3 | 0.001154 | 1.596 | +1 | 5 |
| - | - | 865.9 | 724.3 | - | - | 0 | - |
| - | - | 619.9 | 727 | - | - | 0 | - |
| - | - | 734.7 | 761.5 | - | - | 0 | - |
| - | - | 2700 | 778.3 | - | - | 0 | - |
| - | - | 1349 | 779.3 | - | - | 0 | - |
| 5 | z | 8355 | 822.3 | 0.004654 | 5.66 | +1 | 6 |
| - | - | 8553 | 823.3 | - | - | 0 | - |
| - | - | 3160 | 824.3 | - | - | 0 | - |
| - | - | 1352 | 825.3 | - | - | 0 | - |
| - | - | 581.4 | 832.4 | - | - | 0 | - |
| 5 | y | 2892 | 838.3 | 0.005034 | 6.005 | +1 | 6 |
| - | - | 1486 | 839.3 | - | - | 0 | - |
| 7 | c | 1172 | 877.4 | 0.00297 | 3.385 | +1 | 7 |
| - | - | 2570 | 893.3 | - | - | 0 | - |
| - | - | 996.2 | 894.3 | - | - | 0 | - |
| - | - | 1007 | 896.3 | - | - | 0 | - |
| - | - | 752.6 | 923.5 | - | - | 0 | - |
| 4 | z | 3049 | 937.3 | 0.004322 | 4.611 | +1 | 7 |
| - | - | 6587 | 938.3 | - | - | 0 | - |
| - | - | 2509 | 939.3 | - | - | 0 | - |
| - | - | 888.1 | 940.3 | - | - | 0 | - |
| 4 | y | 4101 | 953.3 | 0.003848 | 4.036 | +1 | 7 |
| - | - | 2321 | 954.3 | - | - | 0 | - |
| - | - | 2364 | 980.4 | - | - | 0 | - |
| - | - | 1326 | 981.4 | - | - | 0 | - |
| - | - | 967.5 | 982.4 | - | - | 0 | - |
| - | - | 3976 | 1023 | - | - | 0 | - |
| 8 | c | 8013 | 1024 | 0.00164 | 1.601 | +1 | 8 |
| - | - | 3139 | 1025 | - | - | 0 | - |
| - | - | 1253 | 1026 | - | - | 0 | - |
| - | - | 1336 | 1036 | - | - | 0 | - |
| - | - | 1116 | 1037 | - | - | 0 | - |
| 3 | z | 749.7 | 1084 | 0.004573 | 4.217 | +1 | 8 |
| 3 | y | 8233 | 1100 | 0.004281 | 3.891 | +1 | 8 |
| - | - | 5083 | 1101 | - | - | 0 | - |
| - | - | 2252 | 1102 | - | - | 0 | - |
| - | - | 2632 | 1117 | - | - | 0 | - |
| - | - | 1171 | 1118 | - | - | 0 | - |
| - | - | 1308 | 1119 | - | - | 0 | - |
| - | - | 937 | 1146 | - | - | 0 | - |
| 9 | c | 1.287E+04 | 1161 | 0.003886 | 3.345 | +1 | 9 |
| - | - | 9683 | 1162 | - | - | 0 | - |
| - | - | 3630 | 1163 | - | - | 0 | - |
| - | - | 959 | 1164 | - | - | 0 | - |
| - | - | 763.5 | 1180 | - | - | 0 | - |
| - | - | 2077 | 1184 | - | - | 0 | - |
| 2 | z | 4687 | 1185 | 0.003037 | 2.562 | +1 | 9 |
| - | - | 2706 | 1186 | - | - | 0 | - |
| - | - | 890.5 | 1187 | - | - | 0 | - |
| 2 | y | 1126 | 1201 | 0.005433 | 4.522 | +1 | 9 |
| - | - | 1315 | 1202 | - | - | 0 | - |
| - | - | 670.5 | 1218 | - | - | 0 | - |
| - | - | 1042 | 1272 | - | - | 0 | - |
| - | - | 614 | 1287 | - | - | 0 | - |
| - | - | 3596 | 1288 | - | - | 0 | - |
| - | - | 7748 | 1289 | - | - | 0 | - |
| - | - | 1.203E+04 | 1290 | - | - | 0 | - |
| - | - | 7051 | 1291 | - | - | 0 | - |
| - | - | 2593 | 1292 | - | - | 0 | - |
| - | - | 1111 | 1304 | - | - | 0 | - |
| - | - | 712.7 | 1306 | - | - | 0 | - |
| - | - | 902.3 | 1316 | - | - | 0 | - |
| - | - | 666.7 | 1328 | - | - | 0 | - |
| - | - | 536.2 | 1330 | - | - | 0 | - |
| - | - | 1031 | 1331 | - | - | 0 | - |
| - | - | 2722 | 1332 | - | - | 0 | - |
| - | - | 3.076E+04 | 1333 | - | - | 0 | - |
| - | - | 2.298E+04 | 1334 | - | - | 0 | - |
| - | - | 1.197E+04 | 1335 | - | - | 0 | - |
| - | - | 2376 | 1336 | - | - | 0 | - |
| - | - | 1.191E+04 | 1349 | - | - | 0 | - |
| - | - | 5.395E+04 | 1350 | - | - | 0 | - |
| - | - | 4.059E+04 | 1351 | - | - | 0 | - |
| - | - | 2081 | 1351 | - | - | 0 | - |
| - | - | 1.615E+04 | 1352 | - | - | 0 | - |
| - | - | 1024 | 1352 | - | - | 0 | - |
| - | - | 2821 | 1353 | - | - | 0 | - |
| - | - | 712.7 | 1445 | - | - | 0 | - |
| - | - | 606.2 | 1929 | - | - | 0 | - |
| - | - | 820 | 3460 | - | - | 0 | - |

m/z Charge Intensity FragmentType MassShift Position
120.07707977294922 0 378.92206
120.08084869384766 0 5030.953
121.08422088623047 0 418.41397
148.90377807617188 0 734.56055
148.90972900390625 0 676.19476
148.9210205078125 0 805.69617
148.92674255371094 0 940.0813
148.932373046875 0 1080.4052
148.93817138671875 0 2549.0251
148.9439697265625 0 2982.6797
148.9573211669922 0 3717.9421
148.96298217773438 0 3147.308
148.9691925048828 0 1068.5469
148.9746551513672 0 1120.4406
148.98048400878906 0 727.64856
148.98577880859375 0 625.1075
148.99159240722656 0 737.12866
149.00294494628906 0 506.11304
149.0198211669922 0 432.47183
149.04258728027344 0 512.64404
166.08668518066406 0 586.83093
176.1074676513672 0 1056.7035
185.55613708496094 0 449.57947
221.1283416748047 0 7104.1934
231.1129608154297 0 1689.8163
249.12332153320312 0 6248.6367 c Ammonia loss 1
250.1269989013672 0 617.6118
253.11764526367188 0 715.4681
261.070556640625 0 473.55124
282.4227600097656 0 525.22614
316.1888122558594 0 583.356
342.15545654296875 0 1132.2123 y 8
378.1805114746094 0 1212.2715
474.18072509765625 0 2661.6743
489.1914978027344 0 1094.8577 y 7
544.2097778320312 0 2790.6216 z 6
545.216552734375 0 853.0639
550.7103271484375 0 2084.6978 y 2
551.212646484375 0 850.89984
551.7125854492188 0 684.1961
560.2272338867188 0 1732.6218 y 6
601.2349243164062 0 813.5953 y 1
626.241455078125 0 648.6645 c Ammonia loss 4
665.7649536132812 0 1395.6992
666.2628784179688 0 1511.3198
666.7686767578125 0 705.5469
674.7677001953125 0 2727.6516
675.27001953125 0 2823.588
675.7698364257812 0 1302.1877
707.27294921875 0 5046.1494 z 5
708.2782592773438 0 3380.7617
709.27685546875 0 1010.4565
723.2881469726562 0 1685.4938 y 5
724.2860717773438 0 865.9332
727.0345458984375 0 619.88354
761.5482788085938 0 734.6878
778.309814453125 0 2700.237
779.3130493164062 0 1349.2573
822.2998657226562 0 8355.434 z 4
823.3054809570312 0 8553.08
824.306640625 0 3160.2327
825.309814453125 0 1352.363
832.422119140625 0 581.371
838.3189697265625 0 2892.1267 y 4
839.3228149414062 0 1486.0837
877.3696899414062 0 1172.2378 c 6
893.33740234375 0 2569.7402
894.3334350585938 0 996.17444
896.3289184570312 0 1007.4154
923.501220703125 0 752.6004
937.3264770507812 0 3049.1594 z 3
938.3330078125 0 6586.8726
939.3351440429688 0 2508.9023
940.3314208984375 0 888.0632
953.3447265625 0 4101.28 y 3
954.34814453125 0 2321.2798
980.3949584960938 0 2364.3533
981.3990478515625 0 1326.2927
982.4077758789062 0 967.4563
1023.3978881835938 0 3975.7234
1024.40478515625 0 8013.2793 c 7
1025.4073486328125 0 3138.6907
1026.4132080078125 0 1252.9863
1036.4146728515625 0 1335.5074
1037.425537109375 0 1115.7528
1084.3951416015625 0 749.6807 z 2
1100.41357421875 0 8232.673 y 2
1101.41650390625 0 5083.434
1102.4190673828125 0 2252.4185
1117.451171875 0 2632.2954
1118.4547119140625 0 1170.6692
1119.4620361328125 0 1307.5267
1146.4517822265625 0 937.04083
1161.4659423828125 0 12869.514 c 8
1162.4678955078125 0 9682.929
1163.470703125 0 3630.449
1164.4798583984375 0 958.95746
1180.4803466796875 0 763.47614
1184.4398193359375 0 2076.7004
1185.4412841796875 0 4687.464 z 1
1186.44384765625 0 2705.97
1187.453369140625 0 890.4866
1201.4515380859375 0 1126.4124 y 1
1202.4659423828125 0 1314.8798
1218.4765625 0 670.4537
1271.5125732421875 0 1041.8383
1286.54736328125 0 613.987
1287.5343017578125 0 3596.4258
1288.52685546875 0 7748.318
1289.519775390625 0 12027.041
1290.520263671875 0 7051.411
1291.52001953125 0 2593.4375
1303.5277099609375 0 1110.5538
1305.5416259765625 0 712.7372
1315.505859375 0 902.3496
1327.795654296875 0 666.7337
1329.575439453125 0 536.23224
1330.5328369140625 0 1031.4811
1331.5283203125 0 2721.6125
1332.5123291015625 0 30758.965
1333.516357421875 0 22975.342
1334.5157470703125 0 11970.451
1335.51806640625 0 2376.2788
1348.5286865234375 0 11911.283
1349.534912109375 0 53947.85
1350.5367431640625 0 40593.88
1350.692626953125 0 2081.0227
1351.5386962890625 0 16153.295
1351.6995849609375 0 1024.3706
1352.541748046875 0 2821.0605
1444.5950927734375 0 712.6526
1928.552001953125 0 606.22833
3460.127685546875 0 819.97266

Spectrum Details

|  |  |
| --- | --- |
| Matched peaks? Matched peaks The total absolute number of peaks matched. Additionally in brackets the total fraction of peaks matched and the total number of peaks is shown. | 21 (15.79% of 133) |
| FDR? FDR The false discovery rate estimated for this peptide. It is calculated by matching all theoretical fragments with a non-integer shift with the raw peaks for this spectrum. This is done with 40 different shifts. The resulting percentage is the average number of annotated peaks over the number of annotated peaks with the correct spectrum. | 0.11% |
| Satellite FDR? Satellite FDR See the FDR for details on its calculation. This satellite ion specific FDR only contains the satellite ions (d/w) for I/L/J positions. | - |
| PSM Score? PSM Score The PSM Score as given by Hecklib to this annotated spectrum. It is shown with three significant figures. | 246 |

#### Spectrum 9024? Spectrum 9024 The raw spectrum of this peptide as annotated by Hecklib. The fragments are coloured according to ion type (see legend). Any peaks with a star '\*' as text can be hovered over to see the full details, first the ion type second the mass shift type. By hovering over the amino acids in the peptide or ions in the legend the corresponding peaks are highlighted. By toggling the 'Unassigned' label you can turn the background (unassigned) peaks on or off in the plot. By updating the slider in the Ion legend you can update the spectrum to only show the top X% of the peaks with labels. The top X% means any peak that is within X% of the highest intensity. By dragging in the spectrum you can zoom in to a specific part of the spectrum and use 'Zoom Out' to get back to the original zoom level. The annotation of the spectrum is based on the given sequence in the peptides file and is done with different software so inconsistencies are likely. The peaks are annotated based on the given sequence, with 20 ppm tolerance.

Copy Data

##### Spectrum 9024 (TSV)

###### Preview

```
Loading example...
```

*Click on the button to copy the data to your clipboard.*

Mz MinMz MaxIntensity Max

WidthHeightPeptide font sizePeptide stroke widthSpectrum font sizeSpectrum stroke widthCompact peptide

Ion legend

wxyz

abcd

OtherUnassignedIon^Charge^_Position_Show for top:%

FTFDDYAMHW

08.70e+31.74e+42.61e+43.48e+4

Zoom Out

a^+1^_2_y^+1^_1_a^+1^_2_b^+1^_2_b^+1^_2_y^+1^_2_b^+1^_3_y^+1^_3_b^+1^_4_y^+2^_8_y^+1^_4_y^+2^_9_b^+1^_5_\*\*y^+1^_5_y^+1^_6_y^+1^_7_y^+1^_8_b^+1^_9_y^+1^_9_

0803160624093211

Fragment Matches Table

Show background peaks

| Position | Ion type | Intensity | mz Theoretical | mz Error (Th) | mz Error (ppm) | Charge | Series Number |
| --- | --- | --- | --- | --- | --- | --- | --- |
| - | - | 3.446E+04 | 120.1 | - | - | 0 | - |
| - | - | 2593 | 121.1 | - | - | 0 | - |
| - | - | 653.3 | 129.1 | - | - | 0 | - |
| - | - | 1882 | 129.1 | - | - | 0 | - |
| - | - | 860.6 | 130.1 | - | - | 0 | - |
| - | - | 438.1 | 130.1 | - | - | 0 | - |
| - | - | 1111 | 133.1 | - | - | 0 | - |
| - | - | 481.8 | 135 | - | - | 0 | - |
| - | - | 5205 | 136.1 | - | - | 0 | - |
| - | - | 518.9 | 138.1 | - | - | 0 | - |
| - | - | 581.3 | 138.1 | - | - | 0 | - |
| - | - | 760.4 | 141.1 | - | - | 0 | - |
| - | - | 1898 | 149 | - | - | 0 | - |
| - | - | 456.9 | 152.1 | - | - | 0 | - |
| - | - | 477.4 | 154.6 | - | - | 0 | - |
| - | - | 1413 | 156.1 | - | - | 0 | - |
| - | - | 631.2 | 158.1 | - | - | 0 | - |
| - | - | 987.8 | 158.1 | - | - | 0 | - |
| - | - | 1304 | 159.1 | - | - | 0 | - |
| - | - | 468.5 | 166.1 | - | - | 0 | - |
| - | - | 3143 | 167.1 | - | - | 0 | - |
| - | - | 1411 | 169.1 | - | - | 0 | - |
| - | - | 443.1 | 170 | - | - | 0 | - |
| - | - | 449.5 | 171.8 | - | - | 0 | - |
| - | - | 716.6 | 173.1 | - | - | 0 | - |
| - | - | 1546 | 173.4 | - | - | 0 | - |
| - | - | 463.8 | 174.1 | - | - | 0 | - |
| - | - | 2539 | 175.1 | - | - | 0 | - |
| - | - | 1.633E+04 | 176.1 | - | - | 0 | - |
| - | - | 806.3 | 176.1 | - | - | 0 | - |
| - | - | 480.5 | 177.1 | - | - | 0 | - |
| - | - | 1967 | 177.1 | - | - | 0 | - |
| - | - | 1373 | 186.1 | - | - | 0 | - |
| - | - | 737.2 | 193.1 | - | - | 0 | - |
| - | - | 518 | 195.5 | - | - | 0 | - |
| - | - | 606.6 | 197.1 | - | - | 0 | - |
| - | - | 759.3 | 203.1 | - | - | 0 | - |
| 2 | a | 835.8 | 203.1 | 0.0002443 | 1.203 | +1 | 2 |
| - | - | 960.8 | 204.1 | - | - | 0 | - |
| - | - | 552.5 | 209.2 | - | - | 0 | - |
| - | - | 1222 | 217.1 | - | - | 0 | - |
| 10 | y | 2075 | 221.1 | 0.003795 | 17.17 | +1 | 1 |
| - | - | 3210 | 221.1 | - | - | 0 | - |
| - | - | 2618 | 221.1 | - | - | 0 | - |
| 2 | a | 1.858E+04 | 221.1 | 0.0001316 | 0.5951 | +1 | 2 |
| - | - | 729.7 | 222.1 | - | - | 0 | - |
| - | - | 2480 | 222.1 | - | - | 0 | - |
| - | - | 503.7 | 224.1 | - | - | 0 | - |
| - | - | 5916 | 225 | - | - | 0 | - |
| - | - | 1430 | 226 | - | - | 0 | - |
| - | - | 634.1 | 227 | - | - | 0 | - |
| - | - | 774.4 | 227.1 | - | - | 0 | - |
| - | - | 506.1 | 228.3 | - | - | 0 | - |
| 2 | b | 5266 | 231.1 | 0.000233 | 1.008 | +1 | 2 |
| - | - | 1773 | 235.1 | - | - | 0 | - |
| - | - | 8906 | 239.1 | - | - | 0 | - |
| - | - | 778.4 | 239.1 | - | - | 0 | - |
| - | - | 1017 | 240.1 | - | - | 0 | - |
| - | - | 508.2 | 241.1 | - | - | 0 | - |
| 2 | b | 9640 | 249.1 | 0.0001511 | 0.6063 | +1 | 2 |
| - | - | 512.9 | 249.9 | - | - | 0 | - |
| - | - | 1187 | 250.1 | - | - | 0 | - |
| - | - | 980.1 | 251.1 | - | - | 0 | - |
| - | - | 2402 | 263.1 | - | - | 0 | - |
| - | - | 681.3 | 279.1 | - | - | 0 | - |
| - | - | 1840 | 285 | - | - | 0 | - |
| - | - | 741.3 | 285.1 | - | - | 0 | - |
| - | - | 1642 | 295.1 | - | - | 0 | - |
| - | - | 669.5 | 296.1 | - | - | 0 | - |
| - | - | 4799 | 299.1 | - | - | 0 | - |
| - | - | 587.2 | 300.1 | - | - | 0 | - |
| - | - | 818.3 | 316.2 | - | - | 0 | - |
| - | - | 663.5 | 325.1 | - | - | 0 | - |
| - | - | 1309 | 340.1 | - | - | 0 | - |
| - | - | 528.4 | 348.2 | - | - | 0 | - |
| - | - | 732.1 | 350.1 | - | - | 0 | - |
| - | - | 4000 | 355.1 | - | - | 0 | - |
| 9 | y | 8271 | 358.1 | 0.004996 | 13.95 | +1 | 2 |
| - | - | 8208 | 359 | - | - | 0 | - |
| - | - | 1303 | 359.2 | - | - | 0 | - |
| - | - | 965.8 | 369.1 | - | - | 0 | - |
| - | - | 519 | 370.1 | - | - | 0 | - |
| - | - | 2493 | 378.1 | - | - | 0 | - |
| 3 | b | 1766 | 378.2 | 0.000392 | 1.037 | +1 | 3 |
| - | - | 1001 | 394.1 | - | - | 0 | - |
| - | - | 916.4 | 419 | - | - | 0 | - |
| - | - | 6867 | 429.1 | - | - | 0 | - |
| - | - | 1572 | 441.2 | - | - | 0 | - |
| - | - | 785.5 | 465.2 | - | - | 0 | - |
| - | - | 660.3 | 482 | - | - | 0 | - |
| - | - | 605.8 | 487.2 | - | - | 0 | - |
| 8 | y | 1860 | 505.2 | 0.009851 | 19.5 | +1 | 3 |
| 4 | b | 1112 | 511.2 | 0.0002199 | 0.4302 | +1 | 4 |
| - | - | 1675 | 512.2 | - | - | 0 | - |
| 3 | y | 3308 | 558.7 | 0.005639 | 10.09 | +2 | 8 |
| - | - | 1592 | 559.2 | - | - | 0 | - |
| 7 | y | 3253 | 576.2 | 0.01015 | 17.62 | +1 | 4 |
| - | - | 919.6 | 577.2 | - | - | 0 | - |
| 2 | y | 1504 | 609.2 | 0.005603 | 9.197 | +2 | 9 |
| - | - | 728 | 609.7 | - | - | 0 | - |
| 5 | b | 829.1 | 626.2 | 0.002993 | 4.779 | +1 | 5 |
| - | - | 1034 | 651.3 | - | - | 0 | - |
| 0 | Precursor | 1537 | 673.8 | 0.005792 | 8.597 | +2 | -1 |
| - | - | 1062 | 674.3 | - | - | 0 | - |
| - | - | 707.6 | 674.8 | - | - | 0 | - |
| - | - | 2019 | 675.3 | - | - | 0 | - |
| - | - | 1233 | 681.8 | - | - | 0 | - |
| - | - | 1482 | 682.3 | - | - | 0 | - |
| 0 | Precursor | 2715 | 682.8 | 0.005698 | 8.345 | +2 | -1 |
| - | - | 767.1 | 682.8 | - | - | 0 | - |
| - | - | 2279 | 683.3 | - | - | 0 | - |
| - | - | 958.6 | 683.3 | - | - | 0 | - |
| - | - | 1439 | 683.8 | - | - | 0 | - |
| - | - | 1970 | 684.2 | - | - | 0 | - |
| 6 | y | 2815 | 739.3 | 0.007553 | 10.22 | +1 | 5 |
| - | - | 1705 | 740.3 | - | - | 0 | - |
| - | - | 933 | 772.3 | - | - | 0 | - |
| - | - | 999.6 | 790.3 | - | - | 0 | - |
| 5 | y | 4132 | 854.3 | 0.0118 | 13.81 | +1 | 6 |
| - | - | 1879 | 855.3 | - | - | 0 | - |
| - | - | 939.3 | 856.3 | - | - | 0 | - |
| - | - | 622.4 | 877.9 | - | - | 0 | - |
| - | - | 835.7 | 891.9 | - | - | 0 | - |
| - | - | 2217 | 905.3 | - | - | 0 | - |
| - | - | 737.8 | 906.3 | - | - | 0 | - |
| 4 | y | 5830 | 969.3 | 0.009514 | 9.815 | +1 | 7 |
| - | - | 3049 | 970.3 | - | - | 0 | - |
| - | - | 1423 | 971.3 | - | - | 0 | - |
| - | - | 2675 | 1052 | - | - | 0 | - |
| - | - | 1784 | 1053 | - | - | 0 | - |
| 3 | y | 9292 | 1116 | 0.0105 | 9.403 | +1 | 8 |
| - | - | 6748 | 1117 | - | - | 0 | - |
| - | - | 2411 | 1118 | - | - | 0 | - |
| 9 | b | 729.8 | 1144 | 0.01742 | 15.22 | +1 | 9 |
| 2 | y | 1476 | 1217 | 0.01116 | 9.166 | +1 | 9 |
| - | - | 1430 | 1218 | - | - | 0 | - |
| - | - | 711.4 | 3180 | - | - | 0 | - |

m/z Charge Intensity FragmentType MassShift Position
120.08098602294922 0 34464.727
121.08427429199219 0 2593.366
129.0658416748047 0 653.28546
129.1024169921875 0 1882.0499
130.0653533935547 0 860.5786
130.10546875 0 438.13214
133.08628845214844 0 1110.9102
134.99221801757812 0 481.79523
136.0758819580078 0 5205.035
138.05494689941406 0 518.8743
138.06625366210938 0 581.2921
141.06588745117188 0 760.40576
149.0449676513672 0 1898.2389
152.0706787109375 0 456.91116
154.6448516845703 0 477.44327
156.07693481445312 0 1412.9838
158.05996704101562 0 631.182
158.09689331054688 0 987.8413
159.09176635742188 0 1303.5619
166.06170654296875 0 468.4866
167.0556182861328 0 3142.9744
169.13397216796875 0 1411.218
170.02899169921875 0 443.0605
171.82150268554688 0 449.4629
173.12789916992188 0 716.6471
173.43878173828125 0 1545.9465
174.06712341308594 0 463.80768
175.08665466308594 0 2539.0635
176.10719299316406 0 16327.66
176.11605834960938 0 806.3447
177.10275268554688 0 480.49243
177.11114501953125 0 1967.2069
186.0919189453125 0 1373.2462
193.10855102539062 0 737.2422
195.52572631835938 0 517.9547
197.1292266845703 0 606.5952
203.08169555664062 0 759.26294
203.11764526367188 0 835.79724 a Water loss 1
204.0762939453125 0 960.8173
209.16513061523438 0 552.5062
217.09678649902344 0 1221.6497
221.08335876464844 0 2075.2158 y 9
221.09228515625 0 3209.7646
221.1035614013672 0 2617.933
221.1285858154297 0 18578.459 a 1
222.08505249023438 0 729.74506
222.1318359375 0 2479.6863
224.10255432128906 0 503.6921
225.0431671142578 0 5916.3774
226.0440673828125 0 1429.5746
227.03955078125 0 634.1438
227.10263061523438 0 774.439
228.34658813476562 0 506.11362
231.113037109375 0 5266.2114 b Water loss 1
235.1079559326172 0 1772.6321
239.0951690673828 0 8905.754
239.14979553222656 0 778.3663
240.0956268310547 0 1017.36066
241.0921630859375 0 508.2206
249.12351989746094 0 9640.219 b 1
249.9182891845703 0 512.89417
250.12741088867188 0 1187.3124
251.10324096679688 0 980.1307
263.10260009765625 0 2401.9612
279.09661865234375 0 681.32794
285.0093994140625 0 1839.8647
285.1017150878906 0 741.3168
295.1034851074219 0 1641.6034
296.10211181640625 0 669.5458
299.06195068359375 0 4799.004
300.06378173828125 0 587.15717
316.1847229003906 0 818.3185
325.1380615234375 0 663.46533
340.1397705078125 0 1309.2804
348.1558532714844 0 528.418
350.1332092285156 0 732.0659
355.070068359375 0 4000.2932
358.15106201171875 0 8270.869 y 8
359.0284118652344 0 8208.152
359.1539306640625 0 1303.2686
369.12042236328125 0 965.81396
370.1242980957031 0 518.96674
378.1298828125 0 2492.6748
378.1816101074219 0 1766.4402 b Water loss 2
394.1243896484375 0 1001.3164
418.99700927734375 0 916.39844
429.0893859863281 0 6867.021
441.1869812011719 0 1572.3164
465.16400146484375 0 785.472
482.0011901855469 0 660.3042
487.17333984375 0 605.7516
505.1864013671875 0 1860.1576 y 7
511.218505859375 0 1112.3983 b 3
512.2246704101562 0 1674.7471
558.7089233398438 0 3308.384 y 2
559.2081298828125 0 1592.0828
576.2238159179688 0 3253.4553 y 6
577.2261352539062 0 919.60394
609.2327270507812 0 1504.4742 y 1
609.7339477539062 0 727.9789
626.24267578125 0 829.1134 b 4
651.2733764648438 0 1034.1315
673.7618408203125 0 1536.7191 Precursor Water loss
674.2610473632812 0 1062.0898
674.7620849609375 0 707.6163
675.2880249023438 0 2019.3613
681.8356323242188 0 1233.4606
682.341796875 0 1482.2952
682.7670288085938 0 2714.6077 Precursor
682.8366088867188 0 767.11536
683.2686157226562 0 2278.9043
683.347900390625 0 958.5645
683.7697143554688 0 1439.048
684.2031860351562 0 1969.802
739.2845458984375 0 2814.705 y 5
740.2902221679688 0 1704.9358
772.3047485351562 0 933.0061
790.3128051757812 0 999.59296
854.3157348632812 0 4132.482 y 4
855.3168334960938 0 1879.2257
856.3153686523438 0 939.2837
877.898193359375 0 622.3926
891.9222412109375 0 835.6774
905.34033203125 0 2216.671
906.3433837890625 0 737.84985
969.3403930664062 0 5830.4463 y 3
970.3433837890625 0 3049.2314
971.3442993164062 0 1422.5637
1052.40966796875 0 2675.1272
1053.4130859375 0 1783.5452
1116.4097900390625 0 9292.279 y 2
1117.412353515625 0 6748.232
1118.4122314453125 0 2410.6313
1144.4180908203125 0 729.7653 b 8
1217.4581298828125 0 1476.1661 y 1
1218.4613037109375 0 1429.8595
3179.54638671875 0 711.3722

Spectrum Details

|  |  |
| --- | --- |
| Matched peaks? Matched peaks The total absolute number of peaks matched. Additionally in brackets the total fraction of peaks matched and the total number of peaks is shown. | 21 (15.33% of 137) |
| FDR? FDR The false discovery rate estimated for this peptide. It is calculated by matching all theoretical fragments with a non-integer shift with the raw peaks for this spectrum. This is done with 40 different shifts. The resulting percentage is the average number of annotated peaks over the number of annotated peaks with the correct spectrum. | 0.23% |
| Satellite FDR? Satellite FDR See the FDR for details on its calculation. This satellite ion specific FDR only contains the satellite ions (d/w) for I/L/J positions. | - |
| PSM Score? PSM Score The PSM Score as given by Hecklib to this annotated spectrum. It is shown with three significant figures. | 221 |

#### Spectrum 10100? Spectrum 10100 The raw spectrum of this peptide as annotated by Hecklib. The fragments are coloured according to ion type (see legend). Any peaks with a star '\*' as text can be hovered over to see the full details, first the ion type second the mass shift type. By hovering over the amino acids in the peptide or ions in the legend the corresponding peaks are highlighted. By toggling the 'Unassigned' label you can turn the background (unassigned) peaks on or off in the plot. By updating the slider in the Ion legend you can update the spectrum to only show the top X% of the peaks with labels. The top X% means any peak that is within X% of the highest intensity. By dragging in the spectrum you can zoom in to a specific part of the spectrum and use 'Zoom Out' to get back to the original zoom level. The annotation of the spectrum is based on the given sequence in the peptides file and is done with different software so inconsistencies are likely. The peaks are annotated based on the given sequence, with 20 ppm tolerance.

Copy Data

##### Spectrum 10100 (TSV)

###### Preview

```
Loading example...
```

*Click on the button to copy the data to your clipboard.*

Mz MinMz MaxIntensity Max

WidthHeightPeptide font sizePeptide stroke widthSpectrum font sizeSpectrum stroke widthCompact peptide

Ion legend

wxyz

abcd

OtherUnassignedIon^Charge^_Position_Show for top:%

FTFDDYAMHW

08.55e+31.71e+42.57e+43.42e+4

Zoom Out

c^+1^_2_z^+1^_3_y^+1^_3_z^+1^_4_y^+2^_8_y^+1^_4_z^+1^_5_y^+1^_5_z^+1^_6_y^+1^_6_c^+1^_7_z^+1^_7_y^+1^_7_c^+1^_8_y^+1^_8_c^+1^_9_y^+1^_9_z^+1^_9_y^+1^_9_

0775154923243098

Fragment Matches Table

Show background peaks

| Position | Ion type | Intensity | mz Theoretical | mz Error (Th) | mz Error (ppm) | Charge | Series Number |
| --- | --- | --- | --- | --- | --- | --- | --- |
| - | - | 3770 | 120.1 | - | - | 0 | - |
| - | - | 356.6 | 121 | - | - | 0 | - |
| - | - | 457.3 | 123.4 | - | - | 0 | - |
| - | - | 373.3 | 132.4 | - | - | 0 | - |
| - | - | 423.6 | 142.8 | - | - | 0 | - |
| - | - | 386.1 | 147.3 | - | - | 0 | - |
| - | - | 418 | 149.9 | - | - | 0 | - |
| - | - | 431.3 | 159.3 | - | - | 0 | - |
| - | - | 466.4 | 161.1 | - | - | 0 | - |
| - | - | 1201 | 173.4 | - | - | 0 | - |
| - | - | 1088 | 176.1 | - | - | 0 | - |
| - | - | 472.3 | 189.1 | - | - | 0 | - |
| - | - | 440.6 | 197.4 | - | - | 0 | - |
| - | - | 4868 | 221.1 | - | - | 0 | - |
| - | - | 566.3 | 231.1 | - | - | 0 | - |
| - | - | 516.4 | 238.7 | - | - | 0 | - |
| - | - | 440.4 | 240 | - | - | 0 | - |
| 2 | c | 4093 | 249.1 | 6.257E-05 | 0.2512 | +1 | 2 |
| - | - | 469 | 300.5 | - | - | 0 | - |
| - | - | 592.2 | 305.1 | - | - | 0 | - |
| - | - | 1087 | 355.1 | - | - | 0 | - |
| - | - | 1752 | 378.2 | - | - | 0 | - |
| - | - | 588.4 | 405.6 | - | - | 0 | - |
| 8 | z | 685.4 | 473.2 | 0.004384 | 9.266 | +1 | 3 |
| - | - | 857.6 | 474.2 | - | - | 0 | - |
| 8 | y | 565.2 | 489.2 | 0.008243 | 16.85 | +1 | 3 |
| 7 | z | 955.3 | 544.2 | 0.004227 | 7.768 | +1 | 4 |
| - | - | 864.9 | 545.2 | - | - | 0 | - |
| 3 | y | 1816 | 550.7 | 0.002165 | 3.93 | +2 | 8 |
| - | - | 1068 | 551.2 | - | - | 0 | - |
| 7 | y | 1463 | 560.2 | 0.004485 | 8.006 | +1 | 4 |
| - | - | 535.7 | 567.7 | - | - | 0 | - |
| - | - | 774.9 | 601.7 | - | - | 0 | - |
| - | - | 665.3 | 665.8 | - | - | 0 | - |
| - | - | 1941 | 674.8 | - | - | 0 | - |
| - | - | 1673 | 675.3 | - | - | 0 | - |
| 6 | z | 3000 | 707.3 | 0.004925 | 6.963 | +1 | 5 |
| - | - | 2146 | 708.3 | - | - | 0 | - |
| - | - | 745 | 709.3 | - | - | 0 | - |
| 6 | y | 1464 | 723.3 | 0.001459 | 2.018 | +1 | 5 |
| - | - | 768.7 | 724.3 | - | - | 0 | - |
| - | - | 1975 | 778.3 | - | - | 0 | - |
| - | - | 678.7 | 779.3 | - | - | 0 | - |
| 5 | z | 4229 | 822.3 | 0.004166 | 5.066 | +1 | 6 |
| - | - | 4884 | 823.3 | - | - | 0 | - |
| - | - | 2174 | 824.3 | - | - | 0 | - |
| 5 | y | 1363 | 838.3 | 0.003386 | 4.039 | +1 | 6 |
| 7 | c | 802 | 877.4 | 0.003642 | 4.151 | +1 | 7 |
| - | - | 2088 | 893.3 | - | - | 0 | - |
| - | - | 711.7 | 894.3 | - | - | 0 | - |
| - | - | 604.2 | 917.4 | - | - | 0 | - |
| 4 | z | 2149 | 937.3 | 0.004566 | 4.872 | +1 | 7 |
| - | - | 3792 | 938.3 | - | - | 0 | - |
| - | - | 2558 | 939.3 | - | - | 0 | - |
| 4 | y | 2816 | 953.3 | 0.002444 | 2.563 | +1 | 7 |
| - | - | 1326 | 954.3 | - | - | 0 | - |
| - | - | 1035 | 980.4 | - | - | 0 | - |
| - | - | 3262 | 1023 | - | - | 0 | - |
| 8 | c | 5153 | 1024 | 0.002983 | 2.912 | +1 | 8 |
| - | - | 2287 | 1025 | - | - | 0 | - |
| - | - | 1107 | 1026 | - | - | 0 | - |
| - | - | 705.8 | 1036 | - | - | 0 | - |
| 3 | y | 5742 | 1100 | 0.004892 | 4.445 | +1 | 8 |
| - | - | 3636 | 1101 | - | - | 0 | - |
| - | - | 1290 | 1102 | - | - | 0 | - |
| - | - | 1488 | 1117 | - | - | 0 | - |
| - | - | 1070 | 1139 | - | - | 0 | - |
| 9 | c | 7877 | 1161 | 0.00413 | 3.556 | +1 | 9 |
| - | - | 6369 | 1162 | - | - | 0 | - |
| - | - | 2846 | 1163 | - | - | 0 | - |
| 2 | y | 923.6 | 1183 | 0.01318 | 11.14 | +1 | 9 |
| - | - | 1012 | 1184 | - | - | 0 | - |
| 2 | z | 1961 | 1185 | 0.002182 | 1.841 | +1 | 9 |
| - | - | 2144 | 1186 | - | - | 0 | - |
| - | - | 654.9 | 1187 | - | - | 0 | - |
| 2 | y | 1435 | 1201 | 0.00775 | 6.451 | +1 | 9 |
| - | - | 837.1 | 1202 | - | - | 0 | - |
| - | - | 2489 | 1288 | - | - | 0 | - |
| - | - | 4988 | 1289 | - | - | 0 | - |
| - | - | 7002 | 1290 | - | - | 0 | - |
| - | - | 4477 | 1291 | - | - | 0 | - |
| - | - | 2204 | 1292 | - | - | 0 | - |
| - | - | 606 | 1305 | - | - | 0 | - |
| - | - | 879.8 | 1306 | - | - | 0 | - |
| - | - | 1214 | 1313 | - | - | 0 | - |
| - | - | 1141 | 1314 | - | - | 0 | - |
| - | - | 656.8 | 1323 | - | - | 0 | - |
| - | - | 2517 | 1329 | - | - | 0 | - |
| - | - | 711.8 | 1330 | - | - | 0 | - |
| - | - | 1140 | 1330 | - | - | 0 | - |
| - | - | 1727 | 1332 | - | - | 0 | - |
| - | - | 949.3 | 1332 | - | - | 0 | - |
| - | - | 2.062E+04 | 1333 | - | - | 0 | - |
| - | - | 1075 | 1333 | - | - | 0 | - |
| - | - | 1.579E+04 | 1334 | - | - | 0 | - |
| - | - | 513.1 | 1334 | - | - | 0 | - |
| - | - | 7694 | 1335 | - | - | 0 | - |
| - | - | 1473 | 1336 | - | - | 0 | - |
| - | - | 716 | 1348 | - | - | 0 | - |
| - | - | 8539 | 1349 | - | - | 0 | - |
| - | - | 679.4 | 1349 | - | - | 0 | - |
| - | - | 3.387E+04 | 1350 | - | - | 0 | - |
| - | - | 2.58E+04 | 1351 | - | - | 0 | - |
| - | - | 1713 | 1351 | - | - | 0 | - |
| - | - | 1.136E+04 | 1352 | - | - | 0 | - |
| - | - | 930.1 | 1352 | - | - | 0 | - |
| - | - | 1736 | 1353 | - | - | 0 | - |
| - | - | 662.4 | 2551 | - | - | 0 | - |
| - | - | 673.4 | 2719 | - | - | 0 | - |
| - | - | 829.2 | 3068 | - | - | 0 | - |

m/z Charge Intensity FragmentType MassShift Position
120.08079528808594 0 3770.459
120.96550750732422 0 356.63354
123.36460876464844 0 457.27823
132.43289184570312 0 373.2801
142.80384826660156 0 423.63077
147.26104736328125 0 386.084
149.90927124023438 0 417.99454
159.25875854492188 0 431.29068
161.08973693847656 0 466.44003
173.43763732910156 0 1200.8573
176.1072998046875 0 1088.0884
189.08694458007812 0 472.3215
197.44735717773438 0 440.63092
221.12835693359375 0 4868.198
231.11228942871094 0 566.2515
238.74798583984375 0 516.4328
240.01661682128906 0 440.42294
249.12330627441406 0 4093.354 c Ammonia loss 1
300.5356750488281 0 469.0388
305.13775634765625 0 592.24646
355.07061767578125 0 1086.8444
378.1807556152344 0 1751.9882
405.60333251953125 0 588.3864
473.1722106933594 0 685.40674 z 7
474.1807556152344 0 857.55347
489.1947937011719 0 565.1607 y 7
544.2091674804688 0 955.33325 z 6
545.2138671875 0 864.85095
550.71044921875 0 1815.7986 y 2
551.2130126953125 0 1067.6877
560.2281494140625 0 1462.726 y 6
567.6968383789062 0 535.7445
601.7411499023438 0 774.85034
665.7596435546875 0 665.2931
674.7681884765625 0 1941.4578
675.2709350585938 0 1672.7427
707.273193359375 0 3000.1147 z 5
708.2769775390625 0 2146.1492
709.2805786132812 0 744.9958
723.2884521484375 0 1464.3582 y 5
724.2910766601562 0 768.69806
778.3082885742188 0 1975.0382
779.3173217773438 0 678.6743
822.2993774414062 0 4229.037 z 4
823.3057861328125 0 4884.4663
824.3074951171875 0 2173.9026
838.3173217773438 0 1363.296 y 4
877.3690185546875 0 802.02325 c 6
893.3367309570312 0 2087.6306
894.3410034179688 0 711.6932
917.4061279296875 0 604.2439
937.3267211914062 0 2149.463 z 3
938.3320922851562 0 3792.158
939.3360595703125 0 2557.7368
953.3433227539062 0 2815.9517 y 3
954.34912109375 0 1326.0764
980.3945922851562 0 1035.2498
1023.3997192382812 0 3261.505
1024.4061279296875 0 5152.621 c 7
1025.408203125 0 2287.1555
1026.4102783203125 0 1106.9198
1036.4183349609375 0 705.79333
1100.4141845703125 0 5742.1084 y 2
1101.416259765625 0 3635.7324
1102.414794921875 0 1289.5809
1117.447998046875 0 1488.4426
1139.441650390625 0 1070.1145
1161.4661865234375 0 7876.5923 c 8
1162.470703125 0 6369.4424
1163.4681396484375 0 2846.3855
1183.4332275390625 0 923.61957 y Water loss 1
1184.4425048828125 0 1011.89087
1185.4404296875 0 1960.7264 z 1
1186.44140625 0 2144.3855
1187.4573974609375 0 654.8959
1201.4647216796875 0 1435.1233 y 1
1202.4547119140625 0 837.1077
1287.5362548828125 0 2488.5542
1288.529296875 0 4987.5356
1289.5208740234375 0 7001.5894
1290.51904296875 0 4476.993
1291.5167236328125 0 2204.0186
1304.525146484375 0 605.966
1305.5361328125 0 879.8473
1312.7640380859375 0 1214.1415
1313.76025390625 0 1141.2611
1322.5185546875 0 656.76355
1328.785888671875 0 2517.3237
1329.5374755859375 0 711.7644
1329.7900390625 0 1140.292
1331.5328369140625 0 1726.7876
1331.7388916015625 0 949.272
1332.512939453125 0 20615.68
1332.6895751953125 0 1075.1412
1333.5137939453125 0 15791.636
1333.6956787109375 0 513.1227
1334.515380859375 0 7694.011
1335.5235595703125 0 1472.7645
1347.70703125 0 716.0097
1348.5284423828125 0 8538.942
1348.6845703125 0 679.4022
1349.5345458984375 0 33868.71
1350.53662109375 0 25798.79
1350.6978759765625 0 1713.3638
1351.5379638671875 0 11361.636
1351.6937255859375 0 930.0521
1352.5447998046875 0 1736.2809
2551.15478515625 0 662.4375
2719.111328125 0 673.4417
3067.515625 0 829.15076

Spectrum Details

|  |  |
| --- | --- |
| Matched peaks? Matched peaks The total absolute number of peaks matched. Additionally in brackets the total fraction of peaks matched and the total number of peaks is shown. | 19 (17.27% of 110) |
| FDR? FDR The false discovery rate estimated for this peptide. It is calculated by matching all theoretical fragments with a non-integer shift with the raw peaks for this spectrum. This is done with 40 different shifts. The resulting percentage is the average number of annotated peaks over the number of annotated peaks with the correct spectrum. | 0.13% |
| Satellite FDR? Satellite FDR See the FDR for details on its calculation. This satellite ion specific FDR only contains the satellite ions (d/w) for I/L/J positions. | - |
| PSM Score? PSM Score The PSM Score as given by Hecklib to this annotated spectrum. It is shown with three significant figures. | 229 |

#### Spectrum 8754? Spectrum 8754 The raw spectrum of this peptide as annotated by Hecklib. The fragments are coloured according to ion type (see legend). Any peaks with a star '\*' as text can be hovered over to see the full details, first the ion type second the mass shift type. By hovering over the amino acids in the peptide or ions in the legend the corresponding peaks are highlighted. By toggling the 'Unassigned' label you can turn the background (unassigned) peaks on or off in the plot. By updating the slider in the Ion legend you can update the spectrum to only show the top X% of the peaks with labels. The top X% means any peak that is within X% of the highest intensity. By dragging in the spectrum you can zoom in to a specific part of the spectrum and use 'Zoom Out' to get back to the original zoom level. The annotation of the spectrum is based on the given sequence in the peptides file and is done with different software so inconsistencies are likely. The peaks are annotated based on the given sequence, with 20 ppm tolerance.

Copy Data

##### Spectrum 8754 (TSV)

###### Preview

```
Loading example...
```

*Click on the button to copy the data to your clipboard.*

Mz MinMz MaxIntensity Max

WidthHeightPeptide font sizePeptide stroke widthSpectrum font sizeSpectrum stroke widthCompact peptide

Ion legend

wxyz

abcd

OtherUnassignedIon^Charge^_Position_Show for top:%

FTFDDYAMHW

02.85e+45.71e+48.56e+41.14e+5

Zoom Out

y^+1^_1_c^+1^_2_y^+1^_2_c^+1^_4_y^+2^_8_y^+2^_8_z^+1^_4_y^+1^_4_y^+2^_9_c^+1^_5_z^+1^_5_y^+1^_5_z^+1^_6_y^+1^_6_z^+1^_7_y^+1^_7_c^+1^_8_y^+1^_8_z^+1^_8_y^+1^_8_c^+1^_9_c^+1^_9_y^+1^_9_z^+1^_9_y^+1^_9_

034669110371382

Fragment Matches Table

Show background peaks

| Position | Ion type | Intensity | mz Theoretical | mz Error (Th) | mz Error (ppm) | Charge | Series Number |
| --- | --- | --- | --- | --- | --- | --- | --- |
| - | - | 1.043E+04 | 120.1 | - | - | 0 | - |
| - | - | 774.1 | 121.1 | - | - | 0 | - |
| - | - | 418.8 | 136.1 | - | - | 0 | - |
| - | - | 455.1 | 143.4 | - | - | 0 | - |
| - | - | 411.7 | 144.4 | - | - | 0 | - |
| - | - | 427.9 | 149 | - | - | 0 | - |
| - | - | 431.9 | 156.4 | - | - | 0 | - |
| - | - | 425.8 | 157 | - | - | 0 | - |
| - | - | 558.9 | 166.2 | - | - | 0 | - |
| - | - | 453.5 | 171.9 | - | - | 0 | - |
| - | - | 462.8 | 172.5 | - | - | 0 | - |
| - | - | 1314 | 173.5 | - | - | 0 | - |
| - | - | 3124 | 176.1 | - | - | 0 | - |
| - | - | 438.8 | 192.4 | - | - | 0 | - |
| - | - | 488.5 | 206.4 | - | - | 0 | - |
| 10 | y | 1976 | 221.1 | 0.003002 | 13.58 | +1 | 1 |
| - | - | 1144 | 221.1 | - | - | 0 | - |
| - | - | 1.426E+04 | 221.1 | - | - | 0 | - |
| - | - | 1933 | 222.1 | - | - | 0 | - |
| - | - | 2331 | 225 | - | - | 0 | - |
| - | - | 486.9 | 228 | - | - | 0 | - |
| - | - | 535.4 | 230.1 | - | - | 0 | - |
| - | - | 3417 | 231.1 | - | - | 0 | - |
| - | - | 3212 | 239.1 | - | - | 0 | - |
| - | - | 773.3 | 240.1 | - | - | 0 | - |
| 2 | c | 1.418E+04 | 249.1 | 0.0001053 | 0.4226 | +1 | 2 |
| - | - | 2192 | 250.1 | - | - | 0 | - |
| - | - | 1457 | 263.1 | - | - | 0 | - |
| - | - | 1670 | 295.1 | - | - | 0 | - |
| - | - | 2058 | 299.1 | - | - | 0 | - |
| - | - | 4462 | 355.1 | - | - | 0 | - |
| 9 | y | 2774 | 358.1 | 0.004691 | 13.1 | +1 | 2 |
| - | - | 1636 | 359 | - | - | 0 | - |
| - | - | 726.1 | 359.2 | - | - | 0 | - |
| - | - | 1048 | 369.1 | - | - | 0 | - |
| - | - | 1565 | 378.1 | - | - | 0 | - |
| - | - | 4088 | 378.2 | - | - | 0 | - |
| - | - | 784 | 379.2 | - | - | 0 | - |
| - | - | 720.6 | 415.2 | - | - | 0 | - |
| - | - | 9253 | 429.1 | - | - | 0 | - |
| - | - | 538.1 | 448.7 | - | - | 0 | - |
| - | - | 672 | 485.8 | - | - | 0 | - |
| - | - | 1697 | 489.2 | - | - | 0 | - |
| - | - | 3860 | 490.2 | - | - | 0 | - |
| - | - | 1176 | 504.2 | - | - | 0 | - |
| - | - | 2682 | 505.2 | - | - | 0 | - |
| - | - | 1410 | 506.2 | - | - | 0 | - |
| 4 | c | 1316 | 511.2 | 0.0009397 | 1.838 | +1 | 4 |
| - | - | 907.5 | 512.2 | - | - | 0 | - |
| - | - | 674.8 | 513.2 | - | - | 0 | - |
| 3 | y | 780.4 | 549.7 | 0.009334 | 16.98 | +2 | 8 |
| 3 | y | 6560 | 558.7 | 0.005272 | 9.437 | +2 | 8 |
| - | - | 3913 | 559.2 | - | - | 0 | - |
| - | - | 1652 | 559.7 | - | - | 0 | - |
| 7 | z | 3490 | 560.2 | 0.01032 | 18.42 | +1 | 4 |
| - | - | 1048 | 561.2 | - | - | 0 | - |
| 7 | y | 4716 | 576.2 | 0.00948 | 16.45 | +1 | 4 |
| - | - | 1676 | 577.2 | - | - | 0 | - |
| - | - | 795.4 | 578.2 | - | - | 0 | - |
| 2 | y | 1449 | 609.2 | 0.006824 | 11.2 | +2 | 9 |
| - | - | 1010 | 609.7 | - | - | 0 | - |
| - | - | 998.7 | 610.2 | - | - | 0 | - |
| 5 | c | 1477 | 626.2 | 0.001463 | 2.335 | +1 | 5 |
| - | - | 735.2 | 627.2 | - | - | 0 | - |
| - | - | 620.7 | 649.3 | - | - | 0 | - |
| - | - | 1021 | 650.8 | - | - | 0 | - |
| - | - | 689.2 | 664.3 | - | - | 0 | - |
| - | - | 3300 | 673.8 | - | - | 0 | - |
| - | - | 1601 | 674.3 | - | - | 0 | - |
| - | - | 1176 | 674.8 | - | - | 0 | - |
| - | - | 1379 | 682.3 | - | - | 0 | - |
| - | - | 5865 | 682.8 | - | - | 0 | - |
| - | - | 5646 | 683.3 | - | - | 0 | - |
| - | - | 901.6 | 683.3 | - | - | 0 | - |
| - | - | 2601 | 683.8 | - | - | 0 | - |
| - | - | 2524 | 684.2 | - | - | 0 | - |
| - | - | 1153 | 693.2 | - | - | 0 | - |
| 6 | z | 1.335E+04 | 723.3 | 0.009737 | 13.46 | +1 | 5 |
| - | - | 7702 | 724.3 | - | - | 0 | - |
| - | - | 2707 | 725.3 | - | - | 0 | - |
| - | - | 810.6 | 738.3 | - | - | 0 | - |
| 6 | y | 5775 | 739.3 | 0.009994 | 13.52 | +1 | 5 |
| - | - | 2204 | 740.3 | - | - | 0 | - |
| - | - | 952.2 | 764.3 | - | - | 0 | - |
| - | - | 1364 | 766.3 | - | - | 0 | - |
| - | - | 793.2 | 767.3 | - | - | 0 | - |
| - | - | 1103 | 790.3 | - | - | 0 | - |
| - | - | 6232 | 794.3 | - | - | 0 | - |
| - | - | 2087 | 795.3 | - | - | 0 | - |
| - | - | 1182 | 808.3 | - | - | 0 | - |
| 5 | z | 1.777E+04 | 838.3 | 0.00971 | 11.58 | +1 | 6 |
| - | - | 2.296E+04 | 839.3 | - | - | 0 | - |
| - | - | 8950 | 840.3 | - | - | 0 | - |
| - | - | 2352 | 841.3 | - | - | 0 | - |
| - | - | 711.2 | 842.3 | - | - | 0 | - |
| 5 | y | 5366 | 854.3 | 0.009541 | 11.17 | +1 | 6 |
| - | - | 2418 | 855.3 | - | - | 0 | - |
| - | - | 829.9 | 856.3 | - | - | 0 | - |
| - | - | 1178 | 896.3 | - | - | 0 | - |
| - | - | 1240 | 905.3 | - | - | 0 | - |
| - | - | 2374 | 909.3 | - | - | 0 | - |
| - | - | 1441 | 910.3 | - | - | 0 | - |
| 4 | z | 3241 | 953.3 | 0.009622 | 10.09 | +1 | 7 |
| - | - | 1.117E+04 | 954.3 | - | - | 0 | - |
| - | - | 5655 | 955.3 | - | - | 0 | - |
| - | - | 1938 | 956.3 | - | - | 0 | - |
| 4 | y | 9022 | 969.3 | 0.008965 | 9.248 | +1 | 7 |
| - | - | 6473 | 970.3 | - | - | 0 | - |
| - | - | 2154 | 971.3 | - | - | 0 | - |
| - | - | 921.4 | 980.4 | - | - | 0 | - |
| - | - | 2228 | 981.4 | - | - | 0 | - |
| - | - | 850.4 | 982.4 | - | - | 0 | - |
| - | - | 651.6 | 991 | - | - | 0 | - |
| - | - | 9474 | 1023 | - | - | 0 | - |
| 8 | c | 1.128E+04 | 1024 | 0.002495 | 2.435 | +1 | 8 |
| - | - | 5824 | 1025 | - | - | 0 | - |
| - | - | 1346 | 1026 | - | - | 0 | - |
| - | - | 2933 | 1052 | - | - | 0 | - |
| - | - | 1122 | 1053 | - | - | 0 | - |
| - | - | 3332 | 1056 | - | - | 0 | - |
| - | - | 2068 | 1057 | - | - | 0 | - |
| - | - | 928.9 | 1058 | - | - | 0 | - |
| 3 | y | 1067 | 1098 | 0.01398 | 12.73 | +1 | 8 |
| 3 | z | 1883 | 1100 | 0.01213 | 11.02 | +1 | 8 |
| 3 | y | 1.763E+04 | 1116 | 0.009887 | 8.856 | +1 | 8 |
| - | - | 1.22E+04 | 1117 | - | - | 0 | - |
| - | - | 4459 | 1118 | - | - | 0 | - |
| - | - | 2908 | 1128 | - | - | 0 | - |
| - | - | 1433 | 1129 | - | - | 0 | - |
| 9 | c | 1030 | 1144 | 0.001181 | 1.032 | +1 | 9 |
| - | - | 1.166E+04 | 1146 | - | - | 0 | - |
| - | - | 6911 | 1147 | - | - | 0 | - |
| - | - | 2767 | 1148 | - | - | 0 | - |
| - | - | 863 | 1158 | - | - | 0 | - |
| 9 | c | 3.594E+04 | 1161 | 0.003886 | 3.345 | +1 | 9 |
| - | - | 2.324E+04 | 1162 | - | - | 0 | - |
| - | - | 9512 | 1163 | - | - | 0 | - |
| - | - | 1530 | 1164 | - | - | 0 | - |
| - | - | 3532 | 1187 | - | - | 0 | - |
| - | - | 2408 | 1188 | - | - | 0 | - |
| - | - | 940.9 | 1189 | - | - | 0 | - |
| 2 | y | 2400 | 1199 | 0.01526 | 12.73 | +1 | 9 |
| - | - | 4259 | 1200 | - | - | 0 | - |
| 2 | z | 7989 | 1201 | 0.007666 | 6.381 | +1 | 9 |
| - | - | 1.1E+04 | 1202 | - | - | 0 | - |
| - | - | 1.737E+04 | 1203 | - | - | 0 | - |
| - | - | 8685 | 1204 | - | - | 0 | - |
| - | - | 2793 | 1205 | - | - | 0 | - |
| - | - | 1703 | 1216 | - | - | 0 | - |
| 2 | y | 4081 | 1217 | 0.01152 | 9.466 | +1 | 9 |
| - | - | 5058 | 1218 | - | - | 0 | - |
| - | - | 3208 | 1219 | - | - | 0 | - |
| - | - | 1039 | 1220 | - | - | 0 | - |
| - | - | 777.6 | 1221 | - | - | 0 | - |
| - | - | 1123 | 1227 | - | - | 0 | - |
| - | - | 717.8 | 1228 | - | - | 0 | - |
| - | - | 874.7 | 1274 | - | - | 0 | - |
| - | - | 1546 | 1287 | - | - | 0 | - |
| - | - | 1487 | 1287 | - | - | 0 | - |
| - | - | 1971 | 1288 | - | - | 0 | - |
| - | - | 1599 | 1302 | - | - | 0 | - |
| - | - | 1058 | 1303 | - | - | 0 | - |
| - | - | 1.003E+04 | 1304 | - | - | 0 | - |
| - | - | 8812 | 1305 | - | - | 0 | - |
| - | - | 2.284E+04 | 1306 | - | - | 0 | - |
| - | - | 1.637E+04 | 1307 | - | - | 0 | - |
| - | - | 6532 | 1308 | - | - | 0 | - |
| - | - | 1143 | 1330 | - | - | 0 | - |
| - | - | 678.4 | 1331 | - | - | 0 | - |
| - | - | 982.2 | 1332 | - | - | 0 | - |
| - | - | 920.1 | 1335 | - | - | 0 | - |
| - | - | 1099 | 1338 | - | - | 0 | - |
| - | - | 742.5 | 1346 | - | - | 0 | - |
| - | - | 1.637E+04 | 1348 | - | - | 0 | - |
| - | - | 7.224E+04 | 1349 | - | - | 0 | - |
| - | - | 5.608E+04 | 1350 | - | - | 0 | - |
| - | - | 2.616E+04 | 1351 | - | - | 0 | - |
| - | - | 5276 | 1352 | - | - | 0 | - |
| - | - | 3.292E+04 | 1365 | - | - | 0 | - |
| - | - | 1.13E+05 | 1366 | - | - | 0 | - |
| - | - | 8.223E+04 | 1367 | - | - | 0 | - |
| - | - | 3.941E+04 | 1368 | - | - | 0 | - |
| - | - | 5586 | 1369 | - | - | 0 | - |

m/z Charge Intensity FragmentType MassShift Position
120.0809555053711 0 10428.348
121.08450317382812 0 774.0529
136.14727783203125 0 418.76535
143.35044860839844 0 455.11804
144.4131622314453 0 411.74542
149.0450439453125 0 427.903
156.35133361816406 0 431.87997
156.9976043701172 0 425.82156
166.1777801513672 0 558.8754
171.92909240722656 0 453.48724
172.4510498046875 0 462.8392
173.4519805908203 0 1313.855
176.10716247558594 0 3123.632
192.4090576171875 0 438.83624
206.35792541503906 0 488.5164
221.0841522216797 0 1976.3893 y 9
221.09397888183594 0 1143.7288
221.1285400390625 0 14260.388
222.13165283203125 0 1932.9283
225.04302978515625 0 2330.8528
227.97483825683594 0 486.92596
230.1152801513672 0 535.4005
231.11277770996094 0 3417.096
239.09490966796875 0 3212.339
240.0958709716797 0 773.31604
249.12347412109375 0 14182.698 c Ammonia loss 1
250.1267547607422 0 2191.891
263.102294921875 0 1456.6228
295.1028137207031 0 1669.8617
299.0614929199219 0 2058.3718
355.0698547363281 0 4461.9434
358.1507568359375 0 2774.2593 y 8
359.0283508300781 0 1635.7174
359.1541442871094 0 726.1446
369.1222229003906 0 1048.21
378.1307678222656 0 1564.9542
378.1810607910156 0 4087.8428
379.1830749511719 0 783.95624
415.1513671875 0 720.6101
429.0889587402344 0 9253.276
448.68157958984375 0 538.1415
485.8171081542969 0 671.9556
489.1678771972656 0 1696.7778
490.1752014160156 0 3860.1843
504.1779479980469 0 1176.0701
505.186767578125 0 2682.3184
506.1884460449219 0 1409.722
511.21966552734375 0 1315.9962 c Ammonia loss 3
512.223876953125 0 907.5029
513.1934814453125 0 674.77124
549.7073364257812 0 780.3923 y Water loss 2
558.7085571289062 0 6559.6035 y 2
559.2089233398438 0 3912.8262
559.7100830078125 0 1652.2476
560.2052612304688 0 3489.5635 z 6
561.2113037109375 0 1048.1127
576.22314453125 0 4715.7637 y 6
577.228271484375 0 1675.6471
578.21630859375 0 795.39166
609.2339477539062 0 1449.2252 y 1
609.731201171875 0 1009.8445
610.2321166992188 0 998.6887
626.2471313476562 0 1477.24 c Ammonia loss 4
627.246337890625 0 735.1556
649.2529296875 0 620.68365
650.7675170898438 0 1021.31836
664.3350219726562 0 689.16644
673.76025390625 0 3299.908
674.2628784179688 0 1600.9612
674.7658081054688 0 1176.141
682.3365478515625 0 1378.8092
682.7658081054688 0 5865.377
683.26708984375 0 5645.9106
683.3359375 0 901.5954
683.768798828125 0 2600.6277
684.2020263671875 0 2524.4812
693.2420654296875 0 1152.7352
723.2680053710938 0 13349.812 z 5
724.2724609375 0 7702.2607
725.27490234375 0 2707.346
738.2777099609375 0 810.6417
739.2869873046875 0 5775.3374 y 5
740.289794921875 0 2203.9197
764.2821655273438 0 952.1618
766.2774658203125 0 1363.5796
767.2725219726562 0 793.2174
790.3139038085938 0 1102.9127
794.3043823242188 0 6232.218
795.3088989257812 0 2087.2698
808.267822265625 0 1181.5092
838.294921875 0 17772.842 z 4
839.30078125 0 22961.906
840.304443359375 0 8950.391
841.3011474609375 0 2351.6785
842.3216552734375 0 711.1909
854.3134765625 0 5365.966 y 4
855.317626953125 0 2417.9958
856.3164672851562 0 829.92926
896.3260498046875 0 1178.3258
905.3386840820312 0 1239.8209
909.331298828125 0 2374.3796
910.3364868164062 0 1441.3634
953.32177734375 0 3241.1206 z 3
954.3281860351562 0 11174.645
955.3333129882812 0 5654.9707
956.3338623046875 0 1937.7499
969.33984375 0 9022.026 y 3
970.3430786132812 0 6473.113
971.341796875 0 2154.04
980.3973999023438 0 921.407
981.4005126953125 0 2227.882
982.4025268554688 0 850.41187
991.0432739257812 0 651.63855
1023.400634765625 0 9473.634
1024.4056396484375 0 11279.549 c 7
1025.408447265625 0 5823.798
1026.4136962890625 0 1346.4866
1052.4132080078125 0 2933.0547
1053.4129638671875 0 1121.7393
1056.3857421875 0 3332.1016
1057.3853759765625 0 2068.4746
1058.384521484375 0 928.94257
1098.4027099609375 0 1066.5508 y Water loss 2
1100.3927001953125 0 1882.9159 z 2
1116.4091796875 0 17630.672 y 2
1117.412353515625 0 12197.88
1118.4122314453125 0 4459.165
1128.43994140625 0 2908.419
1129.4498291015625 0 1433.1055
1144.434326171875 0 1029.5996 c Ammonia loss 8
1146.45458984375 0 11657.313
1147.4576416015625 0 6910.532
1148.4561767578125 0 2766.9683
1158.47314453125 0 863.03284
1161.4659423828125 0 35943.094 c 8
1162.468505859375 0 23237.148
1163.46923828125 0 9511.761
1164.4715576171875 0 1529.9258
1187.4796142578125 0 3531.8499
1188.4835205078125 0 2407.7903
1189.4947509765625 0 940.8588
1199.421142578125 0 2399.9966 y Water loss 1
1200.4266357421875 0 4259.492
1201.4359130859375 0 7988.587 z 1
1202.4547119140625 0 11000.683
1203.4547119140625 0 17367.809
1204.4583740234375 0 8685.304
1205.457275390625 0 2792.8945
1216.4622802734375 0 1702.8516
1217.45849609375 0 4081.4983 y 1
1218.4608154296875 0 5057.8193
1219.4603271484375 0 3208.367
1220.4674072265625 0 1038.8163
1221.4683837890625 0 777.623
1227.438232421875 0 1122.853
1228.449462890625 0 717.81024
1274.4910888671875 0 874.7156
1286.5169677734375 0 1546.1421
1287.4901123046875 0 1486.7385
1288.49072265625 0 1971.2188
1301.5233154296875 0 1599.3152
1302.5220947265625 0 1057.9574
1303.531494140625 0 10034.108
1304.532470703125 0 8811.745
1305.5120849609375 0 22841.514
1306.5135498046875 0 16367.618
1307.5113525390625 0 6532.4375
1330.4971923828125 0 1142.8707
1330.68701171875 0 678.4489
1331.5072021484375 0 982.19183
1335.4920654296875 0 920.097
1337.525390625 0 1099.4353
1346.491455078125 0 742.53595
1347.519775390625 0 16366.255
1348.5089111328125 0 72238.195
1349.5108642578125 0 56079.07
1350.510986328125 0 26155.203
1351.5108642578125 0 5275.938
1364.5225830078125 0 32917.594
1365.5289306640625 0 112988.22
1366.53173828125 0 82225.734
1367.533203125 0 39408.52
1368.5340576171875 0 5585.5405

Spectrum Details

|  |  |
| --- | --- |
| Matched peaks? Matched peaks The total absolute number of peaks matched. Additionally in brackets the total fraction of peaks matched and the total number of peaks is shown. | 25 (13.66% of 183) |
| FDR? FDR The false discovery rate estimated for this peptide. It is calculated by matching all theoretical fragments with a non-integer shift with the raw peaks for this spectrum. This is done with 40 different shifts. The resulting percentage is the average number of annotated peaks over the number of annotated peaks with the correct spectrum. | 0.19% |
| Satellite FDR? Satellite FDR See the FDR for details on its calculation. This satellite ion specific FDR only contains the satellite ions (d/w) for I/L/J positions. | - |
| PSM Score? PSM Score The PSM Score as given by Hecklib to this annotated spectrum. It is shown with three significant figures. | 302 |

#### Spectrum 10360? Spectrum 10360 The raw spectrum of this peptide as annotated by Hecklib. The fragments are coloured according to ion type (see legend). Any peaks with a star '\*' as text can be hovered over to see the full details, first the ion type second the mass shift type. By hovering over the amino acids in the peptide or ions in the legend the corresponding peaks are highlighted. By toggling the 'Unassigned' label you can turn the background (unassigned) peaks on or off in the plot. By updating the slider in the Ion legend you can update the spectrum to only show the top X% of the peaks with labels. The top X% means any peak that is within X% of the highest intensity. By dragging in the spectrum you can zoom in to a specific part of the spectrum and use 'Zoom Out' to get back to the original zoom level. The annotation of the spectrum is based on the given sequence in the peptides file and is done with different software so inconsistencies are likely. The peaks are annotated based on the given sequence, with 20 ppm tolerance.

Copy Data

##### Spectrum 10360 (TSV)

###### Preview

```
Loading example...
```

*Click on the button to copy the data to your clipboard.*

Mz MinMz MaxIntensity Max

WidthHeightPeptide font sizePeptide stroke widthSpectrum font sizeSpectrum stroke widthCompact peptide

Ion legend

wxyz

abcd

OtherUnassignedIon^Charge^_Position_Show for top:%

FTFDDYAMHW

09.16e+31.83e+42.75e+43.66e+4

Zoom Out

a^+1^_2_y^+1^_1_a^+1^_2_b^+1^_2_b^+1^_2_y^+1^_2_b^+1^_3_y^+1^_3_b^+1^_4_y^+2^_8_y^+1^_4_y^+2^_9_b^+1^_5_\*\*y^+1^_5_y^+1^_6_y^+1^_7_y^+1^_8_y^+1^_8_y^+1^_9_

0777155423313109

Fragment Matches Table

Show background peaks

| Position | Ion type | Intensity | mz Theoretical | mz Error (Th) | mz Error (ppm) | Charge | Series Number |
| --- | --- | --- | --- | --- | --- | --- | --- |
| - | - | 3.627E+04 | 120.1 | - | - | 0 | - |
| - | - | 2860 | 121.1 | - | - | 0 | - |
| - | - | 409.1 | 124.4 | - | - | 0 | - |
| - | - | 427.4 | 126.1 | - | - | 0 | - |
| - | - | 412.6 | 126.6 | - | - | 0 | - |
| - | - | 470.9 | 127.1 | - | - | 0 | - |
| - | - | 499.9 | 129.1 | - | - | 0 | - |
| - | - | 1970 | 129.1 | - | - | 0 | - |
| - | - | 528.2 | 130.1 | - | - | 0 | - |
| - | - | 554.2 | 130.1 | - | - | 0 | - |
| - | - | 915.7 | 133.1 | - | - | 0 | - |
| - | - | 4342 | 136.1 | - | - | 0 | - |
| - | - | 399.6 | 137.9 | - | - | 0 | - |
| - | - | 798.1 | 138.1 | - | - | 0 | - |
| - | - | 504.7 | 140.6 | - | - | 0 | - |
| - | - | 477 | 145.9 | - | - | 0 | - |
| - | - | 551.3 | 148.9 | - | - | 0 | - |
| - | - | 505.4 | 148.9 | - | - | 0 | - |
| - | - | 614.5 | 148.9 | - | - | 0 | - |
| - | - | 797.5 | 148.9 | - | - | 0 | - |
| - | - | 820.5 | 148.9 | - | - | 0 | - |
| - | - | 1208 | 148.9 | - | - | 0 | - |
| - | - | 1251 | 148.9 | - | - | 0 | - |
| - | - | 3243 | 148.9 | - | - | 0 | - |
| - | - | 5451 | 148.9 | - | - | 0 | - |
| - | - | 4081 | 149 | - | - | 0 | - |
| - | - | 1613 | 149 | - | - | 0 | - |
| - | - | 1161 | 149 | - | - | 0 | - |
| - | - | 1135 | 149 | - | - | 0 | - |
| - | - | 620.2 | 149 | - | - | 0 | - |
| - | - | 679.3 | 149 | - | - | 0 | - |
| - | - | 566.1 | 149 | - | - | 0 | - |
| - | - | 679.4 | 149 | - | - | 0 | - |
| - | - | 496.2 | 149 | - | - | 0 | - |
| - | - | 561 | 149 | - | - | 0 | - |
| - | - | 613.9 | 149.1 | - | - | 0 | - |
| - | - | 505.4 | 149.1 | - | - | 0 | - |
| - | - | 473.3 | 149.1 | - | - | 0 | - |
| - | - | 400.8 | 149.2 | - | - | 0 | - |
| - | - | 1397 | 156.1 | - | - | 0 | - |
| - | - | 2532 | 159.1 | - | - | 0 | - |
| - | - | 1153 | 166.1 | - | - | 0 | - |
| - | - | 547.7 | 169.1 | - | - | 0 | - |
| - | - | 788.7 | 169.1 | - | - | 0 | - |
| - | - | 1.511E+04 | 176.1 | - | - | 0 | - |
| - | - | 1381 | 177.1 | - | - | 0 | - |
| - | - | 459.5 | 179.1 | - | - | 0 | - |
| - | - | 744.1 | 181.2 | - | - | 0 | - |
| - | - | 439.7 | 182.8 | - | - | 0 | - |
| - | - | 1651 | 186.1 | - | - | 0 | - |
| - | - | 502 | 188.1 | - | - | 0 | - |
| - | - | 557.5 | 191.1 | - | - | 0 | - |
| - | - | 767.1 | 193.1 | - | - | 0 | - |
| - | - | 665.8 | 197.1 | - | - | 0 | - |
| - | - | 817.6 | 197.1 | - | - | 0 | - |
| 2 | a | 1247 | 203.1 | 0.000229 | 1.127 | +1 | 2 |
| - | - | 565.6 | 204.1 | - | - | 0 | - |
| 10 | y | 644.7 | 205.1 | 0.0004439 | 2.164 | +1 | 1 |
| - | - | 937.2 | 209.2 | - | - | 0 | - |
| - | - | 887.3 | 217.1 | - | - | 0 | - |
| - | - | 2249 | 221.1 | - | - | 0 | - |
| 2 | a | 1.889E+04 | 221.1 | 0.0001736 | 0.785 | +1 | 2 |
| - | - | 2680 | 222.1 | - | - | 0 | - |
| - | - | 514.4 | 226 | - | - | 0 | - |
| - | - | 923.1 | 227.1 | - | - | 0 | - |
| - | - | 506.9 | 231.1 | - | - | 0 | - |
| 2 | b | 4556 | 231.1 | 0.0002401 | 1.039 | +1 | 2 |
| - | - | 939.5 | 232.1 | - | - | 0 | - |
| - | - | 1796 | 235.1 | - | - | 0 | - |
| - | - | 997.9 | 237.2 | - | - | 0 | - |
| 2 | b | 9927 | 249.1 | 0.0001694 | 0.6799 | +1 | 2 |
| - | - | 1176 | 250.1 | - | - | 0 | - |
| - | - | 972.4 | 251.1 | - | - | 0 | - |
| - | - | 1215 | 253.1 | - | - | 0 | - |
| - | - | 2507 | 263.1 | - | - | 0 | - |
| - | - | 812.7 | 279.1 | - | - | 0 | - |
| - | - | 611.4 | 281.1 | - | - | 0 | - |
| - | - | 774.8 | 285.1 | - | - | 0 | - |
| - | - | 595.1 | 296.2 | - | - | 0 | - |
| - | - | 1642 | 316.2 | - | - | 0 | - |
| - | - | 1606 | 324.1 | - | - | 0 | - |
| 9 | y | 7981 | 342.2 | 6.01E-05 | 0.1756 | +1 | 2 |
| - | - | 667.7 | 343.2 | - | - | 0 | - |
| - | - | 672.9 | 350.1 | - | - | 0 | - |
| - | - | 563.5 | 355.1 | - | - | 0 | - |
| - | - | 3181 | 378.1 | - | - | 0 | - |
| 3 | b | 2526 | 378.2 | 6.573E-05 | 0.1738 | +1 | 3 |
| - | - | 596.2 | 379.2 | - | - | 0 | - |
| - | - | 1127 | 394.1 | - | - | 0 | - |
| - | - | 536 | 417.2 | - | - | 0 | - |
| - | - | 1399 | 425.2 | - | - | 0 | - |
| 8 | y | 2561 | 489.2 | 0.004917 | 10.05 | +1 | 3 |
| - | - | 1161 | 496.2 | - | - | 0 | - |
| - | - | 620.7 | 505.8 | - | - | 0 | - |
| 4 | b | 674.3 | 511.2 | 0.002478 | 4.848 | +1 | 4 |
| - | - | 711.5 | 513.2 | - | - | 0 | - |
| - | - | 679.3 | 529.6 | - | - | 0 | - |
| 3 | y | 2701 | 550.7 | 0.002409 | 4.374 | +2 | 8 |
| - | - | 736.6 | 551.2 | - | - | 0 | - |
| - | - | 1419 | 551.7 | - | - | 0 | - |
| 7 | y | 4099 | 560.2 | 0.00479 | 8.551 | +1 | 4 |
| - | - | 851.9 | 561.2 | - | - | 0 | - |
| - | - | 598.1 | 564.2 | - | - | 0 | - |
| - | - | 591 | 598.9 | - | - | 0 | - |
| 2 | y | 941.1 | 601.2 | 0.0007252 | 1.206 | +2 | 9 |
| - | - | 747.1 | 601.7 | - | - | 0 | - |
| 5 | b | 763.2 | 626.2 | 0.00342 | 5.462 | +1 | 5 |
| - | - | 683.1 | 642.8 | - | - | 0 | - |
| - | - | 865.5 | 659.3 | - | - | 0 | - |
| 0 | Precursor | 1464 | 665.8 | 0.003783 | 5.682 | +2 | -1 |
| - | - | 923.7 | 666.3 | - | - | 0 | - |
| 0 | Precursor | 2722 | 674.8 | 0.002102 | 3.115 | +2 | -1 |
| - | - | 2609 | 675.3 | - | - | 0 | - |
| - | - | 1014 | 675.8 | - | - | 0 | - |
| 6 | y | 2226 | 723.3 | 0.005243 | 7.249 | +1 | 5 |
| - | - | 913.4 | 724.3 | - | - | 0 | - |
| - | - | 687.4 | 725.3 | - | - | 0 | - |
| - | - | 1187 | 774.3 | - | - | 0 | - |
| - | - | 644.6 | 808.4 | - | - | 0 | - |
| 5 | y | 3326 | 838.3 | 0.006804 | 8.116 | +1 | 6 |
| - | - | 1291 | 839.3 | - | - | 0 | - |
| - | - | 528.7 | 867.7 | - | - | 0 | - |
| - | - | 2552 | 889.3 | - | - | 0 | - |
| - | - | 1119 | 890.3 | - | - | 0 | - |
| - | - | 903 | 896.3 | - | - | 0 | - |
| - | - | 679.6 | 913.5 | - | - | 0 | - |
| - | - | 665.5 | 930.9 | - | - | 0 | - |
| 4 | y | 7320 | 953.3 | 0.005251 | 5.508 | +1 | 7 |
| - | - | 3387 | 954.4 | - | - | 0 | - |
| - | - | 1335 | 955.4 | - | - | 0 | - |
| - | - | 3484 | 1036 | - | - | 0 | - |
| - | - | 1464 | 1037 | - | - | 0 | - |
| - | - | 724.3 | 1038 | - | - | 0 | - |
| 3 | y | 907.7 | 1082 | 0.003005 | 2.776 | +1 | 8 |
| - | - | 724.8 | 1096 | - | - | 0 | - |
| 3 | y | 1.083E+04 | 1100 | 0.004403 | 4.002 | +1 | 8 |
| - | - | 6645 | 1101 | - | - | 0 | - |
| - | - | 3325 | 1102 | - | - | 0 | - |
| 2 | y | 1952 | 1201 | 0.007506 | 6.248 | +1 | 9 |
| - | - | 1007 | 1202 | - | - | 0 | - |
| - | - | 629.1 | 1211 | - | - | 0 | - |
| - | - | 800.8 | 2418 | - | - | 0 | - |
| - | - | 681.5 | 2926 | - | - | 0 | - |
| - | - | 819.2 | 3069 | - | - | 0 | - |
| - | - | 702.4 | 3078 | - | - | 0 | - |

m/z Charge Intensity FragmentType MassShift Position
120.0807876586914 0 36270.78
121.08415222167969 0 2859.89
124.41207885742188 0 409.06583
126.05467224121094 0 427.3874
126.56034851074219 0 412.5876
127.08750915527344 0 470.90356
129.06590270996094 0 499.8869
129.1021728515625 0 1969.9377
130.06057739257812 0 528.1881
130.0654296875 0 554.19727
133.08590698242188 0 915.6954
136.07571411132812 0 4342.1885
137.94769287109375 0 399.5619
138.06617736816406 0 798.0565
140.6222686767578 0 504.6991
145.9359893798828 0 477.02563
148.88035583496094 0 551.32404
148.88760375976562 0 505.4096
148.90219116210938 0 614.5229
148.90908813476562 0 797.5001
148.91677856445312 0 820.451
148.9237518310547 0 1208.3027
148.9308624267578 0 1250.6151
148.9381561279297 0 3243.2654
148.94593811035156 0 5451.1025
148.96243286132812 0 4081.2915
148.97018432617188 0 1612.602
148.97718811035156 0 1160.8948
148.98422241210938 0 1135.3948
148.99136352539062 0 620.2149
148.99905395507812 0 679.32635
149.00621032714844 0 566.13495
149.01316833496094 0 679.439
149.04251098632812 0 496.20825
149.04962158203125 0 560.99084
149.05581665039062 0 613.9342
149.07070922851562 0 505.3642
149.10690307617188 0 473.25146
149.1504669189453 0 400.75177
156.07666015625 0 1396.5674
159.09164428710938 0 2532.4668
166.08615112304688 0 1152.8068
169.09689331054688 0 547.6605
169.13368225097656 0 788.70074
176.10682678222656 0 15114.413
177.11094665527344 0 1380.6337
179.09222412109375 0 459.5218
181.16995239257812 0 744.11505
182.78871154785156 0 439.6639
186.0912628173828 0 1651.134
188.07041931152344 0 502.01447
191.11769104003906 0 557.4714
193.1079864501953 0 767.1354
197.10305786132812 0 665.82
197.1283416748047 0 817.6297
203.11766052246094 0 1246.6526 a Water loss 1
204.0772705078125 0 565.57587
205.09671020507812 0 644.68176 y 9
209.1641387939453 0 937.164
217.09666442871094 0 887.27374
221.10316467285156 0 2248.83
221.12828063964844 0 18889.328 a 1
222.1315155029297 0 2680.459
225.99026489257812 0 514.4298
227.10240173339844 0 923.096
231.0606689453125 0 506.94467
231.11256408691406 0 4556.448 b Water loss 1
232.11598205566406 0 939.51996
235.10777282714844 0 1795.6812
237.15948486328125 0 997.9387
249.12319946289062 0 9926.795 b 1
250.126708984375 0 1175.6486
251.102783203125 0 972.3915
253.1182098388672 0 1215.4653
263.1024475097656 0 2507.0608
279.0956115722656 0 812.6607
281.0959777832031 0 611.42804
285.1024475097656 0 774.8033
296.1501159667969 0 595.13513
316.18707275390625 0 1642.0242
324.1462707519531 0 1606.2509
342.156005859375 0 7980.571 y 8
343.1583557128906 0 667.73
350.1333923339844 0 672.9018
355.0677795410156 0 563.5325
378.12933349609375 0 3180.6565
378.18115234375 0 2526.1377 b Water loss 2
379.18414306640625 0 596.2467
394.12445068359375 0 1127.3389
417.17852783203125 0 536.0064
425.1936340332031 0 1398.6989
489.19146728515625 0 2560.9988 y 7
496.2268981933594 0 1160.8655
505.8134460449219 0 620.6963
511.21624755859375 0 674.34265 b 3
513.1968383789062 0 711.4693
529.640380859375 0 679.3469
550.710693359375 0 2701.13 y 2
551.2107543945312 0 736.61536
551.712890625 0 1418.7227
560.2284545898438 0 4098.747 y 6
561.2301025390625 0 851.864
564.2200927734375 0 598.11707
598.9016723632812 0 591.03394
601.2328491210938 0 941.1463 y 1
601.7354125976562 0 747.07837
626.2422485351562 0 763.2186 b 4
642.77294921875 0 683.11316
659.29736328125 0 865.4832
665.7648315429688 0 1463.5293 Precursor Water loss
666.260498046875 0 923.7445
674.7684326171875 0 2721.6357 Precursor
675.2703247070312 0 2609.1614
675.7691040039062 0 1013.9804
723.292236328125 0 2226.066 y 5
724.2933349609375 0 913.40643
725.2899780273438 0 687.42444
774.321533203125 0 1186.6527
808.4212036132812 0 644.55884
838.3207397460938 0 3325.825 y 4
839.3236083984375 0 1291.3876
867.6823120117188 0 528.7478
889.3469848632812 0 2552.2583
890.3478393554688 0 1118.9535
896.3272094726562 0 903.0087
913.4502563476562 0 679.59827
930.94677734375 0 665.4525
953.3461303710938 0 7319.569 y 3
954.35009765625 0 3386.8428
955.3516235351562 0 1335.0529
1036.4151611328125 0 3484.3748
1037.42041015625 0 1463.9514
1038.4259033203125 0 724.32904
1082.4017333984375 0 907.65063 y Water loss 2
1095.527099609375 0 724.75037
1100.4136962890625 0 10829.371 y 2
1101.416015625 0 6645.1416
1102.417236328125 0 3324.652
1201.4644775390625 0 1951.6285 y 1
1202.4580078125 0 1007.3106
1211.4342041015625 0 629.13416
2418.269287109375 0 800.81256
2926.07666015625 0 681.4842
3068.763671875 0 819.22534
3077.793212890625 0 702.3507

Spectrum Details

|  |  |
| --- | --- |
| Matched peaks? Matched peaks The total absolute number of peaks matched. Additionally in brackets the total fraction of peaks matched and the total number of peaks is shown. | 21 (14.48% of 145) |
| FDR? FDR The false discovery rate estimated for this peptide. It is calculated by matching all theoretical fragments with a non-integer shift with the raw peaks for this spectrum. This is done with 40 different shifts. The resulting percentage is the average number of annotated peaks over the number of annotated peaks with the correct spectrum. | 0.45% |
| Satellite FDR? Satellite FDR See the FDR for details on its calculation. This satellite ion specific FDR only contains the satellite ions (d/w) for I/L/J positions. | - |
| PSM Score? PSM Score The PSM Score as given by Hecklib to this annotated spectrum. It is shown with three significant figures. | 221 |

#### Spectrum 8704? Spectrum 8704 The raw spectrum of this peptide as annotated by Hecklib. The fragments are coloured according to ion type (see legend). Any peaks with a star '\*' as text can be hovered over to see the full details, first the ion type second the mass shift type. By hovering over the amino acids in the peptide or ions in the legend the corresponding peaks are highlighted. By toggling the 'Unassigned' label you can turn the background (unassigned) peaks on or off in the plot. By updating the slider in the Ion legend you can update the spectrum to only show the top X% of the peaks with labels. The top X% means any peak that is within X% of the highest intensity. By dragging in the spectrum you can zoom in to a specific part of the spectrum and use 'Zoom Out' to get back to the original zoom level. The annotation of the spectrum is based on the given sequence in the peptides file and is done with different software so inconsistencies are likely. The peaks are annotated based on the given sequence, with 20 ppm tolerance.

Copy Data

##### Spectrum 8704 (TSV)

###### Preview

```
Loading example...
```

*Click on the button to copy the data to your clipboard.*

Mz MinMz MaxIntensity Max

WidthHeightPeptide font sizePeptide stroke widthSpectrum font sizeSpectrum stroke widthCompact peptide

Ion legend

wxyz

abcd

OtherUnassignedIon^Charge^_Position_Show for top:%

FTFDDYAMHW

03.06e+46.12e+49.17e+41.22e+5

Zoom Out

y^+1^_1_c^+1^_2_y^+1^_2_z^+1^_3_y^+1^_3_c^+1^_4_y^+2^_8_z^+1^_4_y^+1^_4_y^+2^_9_c^+1^_5_z^+1^_5_y^+1^_5_z^+1^_6_y^+1^_6_c^+1^_7_z^+1^_7_y^+1^_7_c^+1^_8_y^+1^_8_z^+1^_8_y^+1^_8_c^+1^_9_c^+1^_9_y^+1^_9_z^+1^_9_y^+1^_9_

0834166825023336

Fragment Matches Table

Show background peaks

| Position | Ion type | Intensity | mz Theoretical | mz Error (Th) | mz Error (ppm) | Charge | Series Number |
| --- | --- | --- | --- | --- | --- | --- | --- |
| - | - | 1.006E+04 | 120.1 | - | - | 0 | - |
| - | - | 638.9 | 121.1 | - | - | 0 | - |
| - | - | 393.5 | 124.6 | - | - | 0 | - |
| - | - | 398.8 | 125.4 | - | - | 0 | - |
| - | - | 679.7 | 133.1 | - | - | 0 | - |
| - | - | 1132 | 149 | - | - | 0 | - |
| - | - | 732.2 | 149 | - | - | 0 | - |
| - | - | 487 | 169.8 | - | - | 0 | - |
| - | - | 2818 | 173.4 | - | - | 0 | - |
| - | - | 3247 | 176.1 | - | - | 0 | - |
| - | - | 405.4 | 182.4 | - | - | 0 | - |
| - | - | 496.7 | 203.1 | - | - | 0 | - |
| 10 | y | 1670 | 221.1 | 0.002926 | 13.23 | +1 | 1 |
| - | - | 638.8 | 221.1 | - | - | 0 | - |
| - | - | 1.375E+04 | 221.1 | - | - | 0 | - |
| - | - | 2103 | 222.1 | - | - | 0 | - |
| - | - | 1797 | 225 | - | - | 0 | - |
| - | - | 3368 | 231.1 | - | - | 0 | - |
| - | - | 585.9 | 235.1 | - | - | 0 | - |
| - | - | 3361 | 239.1 | - | - | 0 | - |
| - | - | 534.9 | 241.1 | - | - | 0 | - |
| 2 | c | 1.427E+04 | 249.1 | 0.0001694 | 0.6799 | +1 | 2 |
| - | - | 1960 | 250.1 | - | - | 0 | - |
| - | - | 2109 | 263.1 | - | - | 0 | - |
| - | - | 532.2 | 269 | - | - | 0 | - |
| - | - | 746.1 | 281.1 | - | - | 0 | - |
| - | - | 2153 | 295.1 | - | - | 0 | - |
| - | - | 1640 | 299.1 | - | - | 0 | - |
| - | - | 604.2 | 313.1 | - | - | 0 | - |
| - | - | 528.3 | 317.7 | - | - | 0 | - |
| - | - | 3551 | 355.1 | - | - | 0 | - |
| 9 | y | 4947 | 358.1 | 0.004599 | 12.84 | +1 | 2 |
| - | - | 1808 | 359 | - | - | 0 | - |
| - | - | 818.3 | 359.2 | - | - | 0 | - |
| - | - | 955 | 364.2 | - | - | 0 | - |
| - | - | 1310 | 369.1 | - | - | 0 | - |
| - | - | 1825 | 378.1 | - | - | 0 | - |
| - | - | 2940 | 378.2 | - | - | 0 | - |
| - | - | 788.9 | 379.2 | - | - | 0 | - |
| - | - | 709.2 | 415.2 | - | - | 0 | - |
| - | - | 8548 | 429.1 | - | - | 0 | - |
| - | - | 649.2 | 489.1 | - | - | 0 | - |
| 8 | z | 2111 | 489.2 | 0.009532 | 19.49 | +1 | 3 |
| - | - | 4559 | 490.2 | - | - | 0 | - |
| - | - | 1885 | 504.2 | - | - | 0 | - |
| 8 | y | 2704 | 505.2 | 0.008172 | 16.18 | +1 | 3 |
| 4 | c | 1623 | 511.2 | 0.0007692 | 1.505 | +1 | 4 |
| - | - | 959.1 | 512.2 | - | - | 0 | - |
| - | - | 652.4 | 548.7 | - | - | 0 | - |
| 3 | y | 6621 | 558.7 | 0.004479 | 8.017 | +2 | 8 |
| - | - | 4645 | 559.2 | - | - | 0 | - |
| - | - | 2010 | 559.7 | - | - | 0 | - |
| 7 | z | 3747 | 560.2 | 0.008856 | 15.81 | +1 | 4 |
| - | - | 1823 | 561.2 | - | - | 0 | - |
| 7 | y | 4757 | 576.2 | 0.00887 | 15.39 | +1 | 4 |
| - | - | 1593 | 577.2 | - | - | 0 | - |
| - | - | 893.5 | 578.2 | - | - | 0 | - |
| - | - | 719.1 | 608.2 | - | - | 0 | - |
| 2 | y | 2663 | 609.2 | 0.004871 | 7.995 | +2 | 9 |
| - | - | 2406 | 609.7 | - | - | 0 | - |
| 5 | c | 839.6 | 626.2 | 0.002383 | 3.805 | +1 | 5 |
| - | - | 695.5 | 634.2 | - | - | 0 | - |
| - | - | 834.9 | 649.3 | - | - | 0 | - |
| - | - | 768.3 | 651.3 | - | - | 0 | - |
| - | - | 615.6 | 664.3 | - | - | 0 | - |
| - | - | 2671 | 673.8 | - | - | 0 | - |
| - | - | 2567 | 674.3 | - | - | 0 | - |
| - | - | 586.3 | 675.3 | - | - | 0 | - |
| - | - | 1659 | 682.3 | - | - | 0 | - |
| - | - | 6300 | 682.8 | - | - | 0 | - |
| - | - | 5555 | 683.3 | - | - | 0 | - |
| - | - | 2529 | 683.8 | - | - | 0 | - |
| - | - | 1968 | 684.2 | - | - | 0 | - |
| - | - | 1174 | 684.3 | - | - | 0 | - |
| - | - | 1007 | 693.2 | - | - | 0 | - |
| - | - | 624.4 | 694.3 | - | - | 0 | - |
| 6 | z | 1.235E+04 | 723.3 | 0.009126 | 12.62 | +1 | 5 |
| - | - | 8753 | 724.3 | - | - | 0 | - |
| - | - | 2756 | 725.3 | - | - | 0 | - |
| - | - | 672.6 | 726.3 | - | - | 0 | - |
| - | - | 1002 | 738.3 | - | - | 0 | - |
| 6 | y | 5201 | 739.3 | 0.009262 | 12.53 | +1 | 5 |
| - | - | 2058 | 740.3 | - | - | 0 | - |
| - | - | 829.1 | 764.3 | - | - | 0 | - |
| - | - | 856.5 | 765.3 | - | - | 0 | - |
| - | - | 1130 | 766.3 | - | - | 0 | - |
| - | - | 1097 | 790.3 | - | - | 0 | - |
| - | - | 5770 | 794.3 | - | - | 0 | - |
| - | - | 3600 | 795.3 | - | - | 0 | - |
| - | - | 1224 | 808.3 | - | - | 0 | - |
| 5 | z | 1.816E+04 | 838.3 | 0.008978 | 10.71 | +1 | 6 |
| - | - | 2.457E+04 | 839.3 | - | - | 0 | - |
| - | - | 1.039E+04 | 840.3 | - | - | 0 | - |
| - | - | 2753 | 841.3 | - | - | 0 | - |
| - | - | 815 | 853.3 | - | - | 0 | - |
| 5 | y | 5867 | 854.3 | 0.007771 | 9.096 | +1 | 6 |
| - | - | 3280 | 855.3 | - | - | 0 | - |
| - | - | 1365 | 856.3 | - | - | 0 | - |
| - | - | 829 | 876.4 | - | - | 0 | - |
| 7 | c | 1048 | 877.4 | 0.0002848 | 0.3246 | +1 | 7 |
| - | - | 913.6 | 896.3 | - | - | 0 | - |
| - | - | 1170 | 905.3 | - | - | 0 | - |
| - | - | 850.1 | 906.3 | - | - | 0 | - |
| - | - | 2756 | 909.3 | - | - | 0 | - |
| - | - | 1432 | 910.3 | - | - | 0 | - |
| 4 | z | 3821 | 953.3 | 0.009378 | 9.838 | +1 | 7 |
| - | - | 1.275E+04 | 954.3 | - | - | 0 | - |
| - | - | 7546 | 955.3 | - | - | 0 | - |
| - | - | 2188 | 956.3 | - | - | 0 | - |
| 4 | y | 1.046E+04 | 969.3 | 0.00811 | 8.367 | +1 | 7 |
| - | - | 6500 | 970.3 | - | - | 0 | - |
| - | - | 2083 | 971.3 | - | - | 0 | - |
| - | - | 726.4 | 980.4 | - | - | 0 | - |
| - | - | 3534 | 981.4 | - | - | 0 | - |
| - | - | 1008 | 982.4 | - | - | 0 | - |
| - | - | 8341 | 1023 | - | - | 0 | - |
| 8 | c | 1.508E+04 | 1024 | 0.00164 | 1.601 | +1 | 8 |
| - | - | 5732 | 1025 | - | - | 0 | - |
| - | - | 1720 | 1026 | - | - | 0 | - |
| - | - | 3140 | 1052 | - | - | 0 | - |
| - | - | 1452 | 1053 | - | - | 0 | - |
| - | - | 3413 | 1056 | - | - | 0 | - |
| - | - | 2485 | 1057 | - | - | 0 | - |
| - | - | 727.5 | 1058 | - | - | 0 | - |
| 3 | y | 730.4 | 1098 | 0.01996 | 18.18 | +1 | 8 |
| 3 | z | 2257 | 1100 | 0.0136 | 12.36 | +1 | 8 |
| - | - | 1614 | 1101 | - | - | 0 | - |
| - | - | 646.7 | 1112 | - | - | 0 | - |
| 3 | y | 1.921E+04 | 1116 | 0.008788 | 7.872 | +1 | 8 |
| - | - | 1.173E+04 | 1117 | - | - | 0 | - |
| - | - | 5308 | 1118 | - | - | 0 | - |
| - | - | 2231 | 1128 | - | - | 0 | - |
| - | - | 1622 | 1129 | - | - | 0 | - |
| - | - | 963.7 | 1130 | - | - | 0 | - |
| 9 | c | 1051 | 1144 | 0.0008152 | 0.7123 | +1 | 9 |
| - | - | 1.034E+04 | 1146 | - | - | 0 | - |
| - | - | 7381 | 1147 | - | - | 0 | - |
| - | - | 2682 | 1148 | - | - | 0 | - |
| - | - | 872.8 | 1158 | - | - | 0 | - |
| - | - | 1207 | 1160 | - | - | 0 | - |
| 9 | c | 3.776E+04 | 1161 | 0.003153 | 2.715 | +1 | 9 |
| - | - | 2.439E+04 | 1162 | - | - | 0 | - |
| - | - | 498.4 | 1163 | - | - | 0 | - |
| - | - | 9227 | 1163 | - | - | 0 | - |
| - | - | 1240 | 1164 | - | - | 0 | - |
| - | - | 670.3 | 1186 | - | - | 0 | - |
| - | - | 4531 | 1187 | - | - | 0 | - |
| - | - | 2975 | 1188 | - | - | 0 | - |
| - | - | 1308 | 1189 | - | - | 0 | - |
| 2 | y | 2491 | 1199 | 0.01356 | 11.3 | +1 | 9 |
| - | - | 4305 | 1200 | - | - | 0 | - |
| 2 | z | 8573 | 1201 | 0.006567 | 5.466 | +1 | 9 |
| - | - | 9140 | 1202 | - | - | 0 | - |
| - | - | 1.784E+04 | 1203 | - | - | 0 | - |
| - | - | 8152 | 1204 | - | - | 0 | - |
| - | - | 3451 | 1205 | - | - | 0 | - |
| - | - | 783.6 | 1215 | - | - | 0 | - |
| - | - | 942.5 | 1216 | - | - | 0 | - |
| 2 | y | 4520 | 1217 | 0.01214 | 9.968 | +1 | 9 |
| - | - | 7053 | 1218 | - | - | 0 | - |
| - | - | 3280 | 1219 | - | - | 0 | - |
| - | - | 1570 | 1220 | - | - | 0 | - |
| - | - | 1026 | 1286 | - | - | 0 | - |
| - | - | 2084 | 1287 | - | - | 0 | - |
| - | - | 1109 | 1288 | - | - | 0 | - |
| - | - | 1459 | 1302 | - | - | 0 | - |
| - | - | 1368 | 1303 | - | - | 0 | - |
| - | - | 1.238E+04 | 1304 | - | - | 0 | - |
| - | - | 1.143E+04 | 1305 | - | - | 0 | - |
| - | - | 2.713E+04 | 1306 | - | - | 0 | - |
| - | - | 1.808E+04 | 1307 | - | - | 0 | - |
| - | - | 7475 | 1308 | - | - | 0 | - |
| - | - | 1029 | 1309 | - | - | 0 | - |
| - | - | 635.9 | 1310 | - | - | 0 | - |
| - | - | 925.2 | 1320 | - | - | 0 | - |
| - | - | 977.6 | 1330 | - | - | 0 | - |
| - | - | 815.4 | 1332 | - | - | 0 | - |
| - | - | 1123 | 1338 | - | - | 0 | - |
| - | - | 1.608E+04 | 1348 | - | - | 0 | - |
| - | - | 7.178E+04 | 1349 | - | - | 0 | - |
| - | - | 5.758E+04 | 1350 | - | - | 0 | - |
| - | - | 2.828E+04 | 1351 | - | - | 0 | - |
| - | - | 5184 | 1352 | - | - | 0 | - |
| - | - | 639.6 | 1364 | - | - | 0 | - |
| - | - | 3.291E+04 | 1365 | - | - | 0 | - |
| - | - | 1.211E+05 | 1366 | - | - | 0 | - |
| - | - | 9.037E+04 | 1367 | - | - | 0 | - |
| - | - | 3.897E+04 | 1368 | - | - | 0 | - |
| - | - | 996.2 | 1368 | - | - | 0 | - |
| - | - | 5434 | 1369 | - | - | 0 | - |
| - | - | 664.7 | 3303 | - | - | 0 | - |

m/z Charge Intensity FragmentType MassShift Position
120.080810546875 0 10058.83
121.08416748046875 0 638.938
124.63272094726562 0 393.4626
125.40113067626953 0 398.79974
133.0859832763672 0 679.74316
148.95436096191406 0 1131.8529
149.0452117919922 0 732.1832
169.77474975585938 0 486.98862
173.4395751953125 0 2817.7622
176.10691833496094 0 3246.5981
182.3540496826172 0 405.35284
203.1177215576172 0 496.6808
221.084228515625 0 1670.2213 y 9
221.0940399169922 0 638.8324
221.12828063964844 0 13750.107
222.13165283203125 0 2103.389
225.04278564453125 0 1797.156
231.1125946044922 0 3368.0403
235.10818481445312 0 585.8616
239.09490966796875 0 3361.0774
241.10577392578125 0 534.91754
249.12319946289062 0 14267.748 c Ammonia loss 1
250.12655639648438 0 1960.3815
263.1024475097656 0 2108.5393
268.98687744140625 0 532.2488
281.0510559082031 0 746.0607
295.1031188964844 0 2152.7373
299.0614318847656 0 1639.6337
313.1137390136719 0 604.1779
317.73455810546875 0 528.31964
355.0695495605469 0 3550.8875
358.1506652832031 0 4946.8 y 8
359.0278015136719 0 1807.7765
359.1545104980469 0 818.34174
364.1503601074219 0 954.9839
369.1205749511719 0 1309.7589
378.1280517578125 0 1824.6616
378.18109130859375 0 2939.998
379.1846008300781 0 788.921
415.1534423828125 0 709.2289
429.08843994140625 0 8547.585
489.0535888671875 0 649.17993
489.1673583984375 0 2110.7834 z 7
490.1746520996094 0 4558.9604
504.176513671875 0 1884.5933
505.1847229003906 0 2704.3538 y 7
511.21795654296875 0 1622.9978 c Ammonia loss 3
512.2217407226562 0 959.0855
548.6607666015625 0 652.37805
558.707763671875 0 6620.6196 y 2
559.2091674804688 0 4644.712
559.709228515625 0 2009.8225
560.2037963867188 0 3747.1382 z 6
561.2096557617188 0 1823.1953
576.2225341796875 0 4756.859 y 6
577.22705078125 0 1592.9104
578.2202758789062 0 893.46704
608.2339477539062 0 719.1257
609.2319946289062 0 2663.2056 y 1
609.733154296875 0 2405.599
626.2432861328125 0 839.62067 c Ammonia loss 4
634.226318359375 0 695.54736
649.2505493164062 0 834.91907
651.2686767578125 0 768.2514
664.3318481445312 0 615.5534
673.7605590820312 0 2670.7693
674.261474609375 0 2567.2607
675.2893676757812 0 586.3489
682.3410034179688 0 1658.7465
682.7653198242188 0 6300.279
683.2669067382812 0 5554.9624
683.7675170898438 0 2528.8506
684.2019653320312 0 1968.401
684.2680053710938 0 1174.1307
693.242919921875 0 1006.6377
694.2506713867188 0 624.36835
723.2673950195312 0 12350.814 z 5
724.2718505859375 0 8753.022
725.2757568359375 0 2755.5625
726.2783813476562 0 672.63556
738.2787475585938 0 1001.7274
739.2862548828125 0 5200.993 y 5
740.2874755859375 0 2058.0752
764.2760009765625 0 829.1469
765.2828369140625 0 856.4522
766.275390625 0 1129.621
790.31298828125 0 1096.6965
794.3045654296875 0 5769.659
795.3090209960938 0 3599.778
808.26806640625 0 1224.1324
838.294189453125 0 18160.172 z 4
839.3001708984375 0 24572.059
840.3027954101562 0 10386.74
841.3012084960938 0 2753.1545
853.3086547851562 0 814.9764
854.3117065429688 0 5866.7666 y 4
855.3163452148438 0 3280.2437
856.3140869140625 0 1365.4916
876.3668212890625 0 829.048
877.3723754882812 0 1048.0052 c 6
896.3192749023438 0 913.634
905.3401489257812 0 1170.2035
906.34130859375 0 850.14197
909.3287353515625 0 2755.9666
910.3343505859375 0 1431.6583
953.321533203125 0 3821.2114 z 3
954.3277587890625 0 12750.449
955.3306884765625 0 7546.1714
956.3305053710938 0 2188.486
969.3389892578125 0 10461.167 y 3
970.3431396484375 0 6499.9077
971.34521484375 0 2083.238
980.3936157226562 0 726.42737
981.4015502929688 0 3534.1707
982.3972778320312 0 1007.5236
1023.3989868164062 0 8340.928
1024.40478515625 0 15079.318 c 7
1025.4071044921875 0 5732.266
1026.4080810546875 0 1720.3752
1052.406494140625 0 3140.472
1053.409423828125 0 1451.9612
1056.38818359375 0 3413.0208
1057.3875732421875 0 2484.6802
1058.398193359375 0 727.5235
1098.40869140625 0 730.4205 y Water loss 2
1100.3941650390625 0 2256.6492 z 2
1101.385986328125 0 1614.2743
1111.8961181640625 0 646.69354
1116.4080810546875 0 19208.018 y 2
1117.411376953125 0 11726.836
1118.4127197265625 0 5307.5312
1128.4454345703125 0 2231.303
1129.444091796875 0 1621.6013
1130.4384765625 0 963.68976
1144.4346923828125 0 1051.4456 c Ammonia loss 8
1146.4541015625 0 10340.639
1147.457763671875 0 7380.546
1148.4599609375 0 2682.28
1158.47705078125 0 872.84393
1160.4632568359375 0 1207.2648
1161.4652099609375 0 37760.574 c 8
1162.4676513671875 0 24385.79
1162.5908203125 0 498.41504
1163.468994140625 0 9226.541
1164.4659423828125 0 1240.4623
1186.4427490234375 0 670.2994
1187.47998046875 0 4530.6226
1188.485595703125 0 2975.4746
1189.4864501953125 0 1307.5227
1199.4228515625 0 2490.956 y Water loss 1
1200.4293212890625 0 4304.855
1201.434814453125 0 8573.232 z 1
1202.454345703125 0 9139.77
1203.4537353515625 0 17836.031
1204.4586181640625 0 8152.027
1205.4576416015625 0 3450.9194
1215.20703125 0 783.63684
1216.4571533203125 0 942.52264
1217.4591064453125 0 4520.4834 y 1
1218.4598388671875 0 7053.187
1219.4617919921875 0 3280.0093
1220.4688720703125 0 1569.9928
1285.530517578125 0 1025.9027
1287.4901123046875 0 2083.9258
1288.48388671875 0 1109.3413
1301.5299072265625 0 1458.8468
1302.54052734375 0 1368.3354
1303.5313720703125 0 12384.88
1304.530029296875 0 11432.617
1305.512451171875 0 27133.484
1306.5115966796875 0 18075.744
1307.5125732421875 0 7474.9395
1308.5155029296875 0 1028.6875
1309.626953125 0 635.89795
1319.52197265625 0 925.15137
1330.48681640625 0 977.62866
1331.51171875 0 815.4399
1337.5313720703125 0 1122.916
1347.5184326171875 0 16082.624
1348.5084228515625 0 71781.195
1349.5108642578125 0 57583.5
1350.510009765625 0 28277.645
1351.5126953125 0 5183.7827
1363.6806640625 0 639.56305
1364.5216064453125 0 32911.535
1365.5284423828125 0 121092.25
1366.5313720703125 0 90369.29
1367.5328369140625 0 38972.566
1367.6837158203125 0 996.17017
1368.5328369140625 0 5434.297
3302.86279296875 0 664.6888

Spectrum Details

|  |  |
| --- | --- |
| Matched peaks? Matched peaks The total absolute number of peaks matched. Additionally in brackets the total fraction of peaks matched and the total number of peaks is shown. | 27 (14.14% of 191) |
| FDR? FDR The false discovery rate estimated for this peptide. It is calculated by matching all theoretical fragments with a non-integer shift with the raw peaks for this spectrum. This is done with 40 different shifts. The resulting percentage is the average number of annotated peaks over the number of annotated peaks with the correct spectrum. | 0.09% |
| Satellite FDR? Satellite FDR See the FDR for details on its calculation. This satellite ion specific FDR only contains the satellite ions (d/w) for I/L/J positions. | - |
| PSM Score? PSM Score The PSM Score as given by Hecklib to this annotated spectrum. It is shown with three significant figures. | 360 |

#### Spectrum 10560? Spectrum 10560 The raw spectrum of this peptide as annotated by Hecklib. The fragments are coloured according to ion type (see legend). Any peaks with a star '\*' as text can be hovered over to see the full details, first the ion type second the mass shift type. By hovering over the amino acids in the peptide or ions in the legend the corresponding peaks are highlighted. By toggling the 'Unassigned' label you can turn the background (unassigned) peaks on or off in the plot. By updating the slider in the Ion legend you can update the spectrum to only show the top X% of the peaks with labels. The top X% means any peak that is within X% of the highest intensity. By dragging in the spectrum you can zoom in to a specific part of the spectrum and use 'Zoom Out' to get back to the original zoom level. The annotation of the spectrum is based on the given sequence in the peptides file and is done with different software so inconsistencies are likely. The peaks are annotated based on the given sequence, with 20 ppm tolerance.

Copy Data

##### Spectrum 10560 (TSV)

###### Preview

```
Loading example...
```

*Click on the button to copy the data to your clipboard.*

Mz MinMz MaxIntensity Max

WidthHeightPeptide font sizePeptide stroke widthSpectrum font sizeSpectrum stroke widthCompact peptide

Ion legend

wxyz

abcd

OtherUnassignedIon^Charge^_Position_Show for top:%

FTFDDYAMHW

06.19e+31.24e+41.86e+42.48e+4

Zoom Out

a^+1^_2_y^+1^_1_a^+1^_2_b^+1^_2_b^+1^_2_y^+1^_2_b^+1^_3_y^+1^_3_y^+1^_4_\*\*y^+1^_5_y^+1^_6_y^+1^_7_y^+1^_8_y^+1^_9_

048496914531938

Fragment Matches Table

Show background peaks

| Position | Ion type | Intensity | mz Theoretical | mz Error (Th) | mz Error (ppm) | Charge | Series Number |
| --- | --- | --- | --- | --- | --- | --- | --- |
| - | - | 2.452E+04 | 120.1 | - | - | 0 | - |
| - | - | 1658 | 121.1 | - | - | 0 | - |
| - | - | 351.1 | 122.7 | - | - | 0 | - |
| - | - | 397.8 | 129 | - | - | 0 | - |
| - | - | 1981 | 129.1 | - | - | 0 | - |
| - | - | 446.9 | 132.1 | - | - | 0 | - |
| - | - | 544.7 | 133.1 | - | - | 0 | - |
| - | - | 2597 | 133.1 | - | - | 0 | - |
| - | - | 398.1 | 134 | - | - | 0 | - |
| - | - | 4201 | 136.1 | - | - | 0 | - |
| - | - | 654.9 | 138.1 | - | - | 0 | - |
| - | - | 531.6 | 145.1 | - | - | 0 | - |
| - | - | 519.3 | 147.4 | - | - | 0 | - |
| - | - | 495.6 | 148.9 | - | - | 0 | - |
| - | - | 600.9 | 148.9 | - | - | 0 | - |
| - | - | 592.4 | 148.9 | - | - | 0 | - |
| - | - | 565 | 148.9 | - | - | 0 | - |
| - | - | 648 | 148.9 | - | - | 0 | - |
| - | - | 980.9 | 148.9 | - | - | 0 | - |
| - | - | 1248 | 148.9 | - | - | 0 | - |
| - | - | 1397 | 148.9 | - | - | 0 | - |
| - | - | 3260 | 148.9 | - | - | 0 | - |
| - | - | 5022 | 148.9 | - | - | 0 | - |
| - | - | 3927 | 149 | - | - | 0 | - |
| - | - | 1621 | 149 | - | - | 0 | - |
| - | - | 1226 | 149 | - | - | 0 | - |
| - | - | 996 | 149 | - | - | 0 | - |
| - | - | 801.4 | 149 | - | - | 0 | - |
| - | - | 666.7 | 149 | - | - | 0 | - |
| - | - | 717 | 149 | - | - | 0 | - |
| - | - | 522 | 149 | - | - | 0 | - |
| - | - | 450.6 | 149 | - | - | 0 | - |
| - | - | 570.1 | 149.1 | - | - | 0 | - |
| - | - | 1256 | 156.1 | - | - | 0 | - |
| - | - | 496.6 | 158.1 | - | - | 0 | - |
| - | - | 1257 | 159.1 | - | - | 0 | - |
| - | - | 2484 | 166.1 | - | - | 0 | - |
| - | - | 1057 | 169.1 | - | - | 0 | - |
| - | - | 566.2 | 172.1 | - | - | 0 | - |
| - | - | 491.8 | 172.8 | - | - | 0 | - |
| - | - | 523.2 | 174.1 | - | - | 0 | - |
| - | - | 558.5 | 175.1 | - | - | 0 | - |
| - | - | 1.04E+04 | 176.1 | - | - | 0 | - |
| - | - | 1719 | 177.1 | - | - | 0 | - |
| - | - | 457.2 | 181.2 | - | - | 0 | - |
| - | - | 1015 | 186.1 | - | - | 0 | - |
| - | - | 589.1 | 188.1 | - | - | 0 | - |
| - | - | 594.6 | 191.1 | - | - | 0 | - |
| - | - | 677.5 | 193.1 | - | - | 0 | - |
| - | - | 1051 | 197.1 | - | - | 0 | - |
| - | - | 761 | 201.1 | - | - | 0 | - |
| 2 | a | 665.7 | 203.1 | 0.0005342 | 2.63 | +1 | 2 |
| 10 | y | 614.8 | 205.1 | 0.0001359 | 0.6628 | +1 | 1 |
| - | - | 545.6 | 209.2 | - | - | 0 | - |
| - | - | 1288 | 221.1 | - | - | 0 | - |
| 2 | a | 1.398E+04 | 221.1 | 3.625E-05 | 0.1639 | +1 | 2 |
| - | - | 1691 | 222.1 | - | - | 0 | - |
| 2 | b | 3063 | 231.1 | 0.0001638 | 0.7086 | +1 | 2 |
| - | - | 1517 | 235.1 | - | - | 0 | - |
| 2 | b | 6813 | 249.1 | 3.205E-05 | 0.1287 | +1 | 2 |
| - | - | 638.7 | 251.1 | - | - | 0 | - |
| - | - | 596.3 | 251.1 | - | - | 0 | - |
| - | - | 2153 | 253.1 | - | - | 0 | - |
| - | - | 553 | 257.1 | - | - | 0 | - |
| - | - | 1623 | 263.1 | - | - | 0 | - |
| - | - | 682 | 279.1 | - | - | 0 | - |
| - | - | 557.3 | 297.8 | - | - | 0 | - |
| - | - | 1323 | 316.2 | - | - | 0 | - |
| - | - | 772.4 | 324.1 | - | - | 0 | - |
| 9 | y | 5572 | 342.2 | 0.000184 | 0.5379 | +1 | 2 |
| - | - | 1227 | 343.2 | - | - | 0 | - |
| - | - | 690 | 350.1 | - | - | 0 | - |
| - | - | 1454 | 355.1 | - | - | 0 | - |
| - | - | 1572 | 378.1 | - | - | 0 | - |
| 3 | b | 1365 | 378.2 | 0.0002089 | 0.5525 | +1 | 3 |
| - | - | 761 | 379.1 | - | - | 0 | - |
| - | - | 590 | 394.1 | - | - | 0 | - |
| - | - | 985.7 | 425.2 | - | - | 0 | - |
| - | - | 622.9 | 442.1 | - | - | 0 | - |
| 8 | y | 1124 | 489.2 | 0.003971 | 8.117 | +1 | 3 |
| - | - | 756.1 | 551.2 | - | - | 0 | - |
| 7 | y | 1681 | 560.2 | 0.002959 | 5.282 | +1 | 4 |
| - | - | 870.7 | 656.3 | - | - | 0 | - |
| - | - | 681.5 | 657.3 | - | - | 0 | - |
| - | - | 751.9 | 659.3 | - | - | 0 | - |
| - | - | 659.3 | 660.4 | - | - | 0 | - |
| - | - | 690.4 | 661.6 | - | - | 0 | - |
| 0 | Precursor | 920.8 | 665.8 | 0.00183 | 2.749 | +2 | -1 |
| - | - | 755.5 | 666.3 | - | - | 0 | - |
| 0 | Precursor | 1759 | 674.8 | 0.002102 | 3.115 | +2 | -1 |
| - | - | 1853 | 675.3 | - | - | 0 | - |
| - | - | 659.5 | 675.4 | - | - | 0 | - |
| - | - | 785.5 | 675.8 | - | - | 0 | - |
| - | - | 678.2 | 712.1 | - | - | 0 | - |
| 6 | y | 1190 | 723.3 | 0.002497 | 3.452 | +1 | 5 |
| - | - | 897.3 | 774.3 | - | - | 0 | - |
| 5 | y | 2028 | 838.3 | 0.007292 | 8.699 | +1 | 6 |
| - | - | 1520 | 839.3 | - | - | 0 | - |
| - | - | 708.3 | 868.3 | - | - | 0 | - |
| - | - | 1044 | 889.3 | - | - | 0 | - |
| - | - | 810.8 | 890.3 | - | - | 0 | - |
| - | - | 745.9 | 896.3 | - | - | 0 | - |
| 4 | y | 3610 | 953.3 | 0.004397 | 4.612 | +1 | 7 |
| - | - | 2267 | 954.3 | - | - | 0 | - |
| - | - | 3135 | 1036 | - | - | 0 | - |
| - | - | 1631 | 1037 | - | - | 0 | - |
| - | - | 609.7 | 1038 | - | - | 0 | - |
| 3 | y | 7662 | 1100 | 0.003061 | 2.781 | +1 | 8 |
| - | - | 4393 | 1101 | - | - | 0 | - |
| - | - | 1849 | 1102 | - | - | 0 | - |
| - | - | 986 | 1137 | - | - | 0 | - |
| 2 | y | 1560 | 1201 | 0.004699 | 3.911 | +1 | 9 |
| - | - | 622.8 | 1596 | - | - | 0 | - |
| - | - | 628.5 | 1918 | - | - | 0 | - |

m/z Charge Intensity FragmentType MassShift Position
120.08086395263672 0 24520.174
121.08421325683594 0 1658.114
122.74864959716797 0 351.1169
129.02374267578125 0 397.78647
129.10227966308594 0 1981.1135
132.0887908935547 0 446.87384
133.06109619140625 0 544.6946
133.08595275878906 0 2597.2173
134.0453338623047 0 398.13623
136.0757598876953 0 4201.2876
138.0658721923828 0 654.86884
145.06080627441406 0 531.5999
147.44761657714844 0 519.30963
148.87265014648438 0 495.6081
148.87930297851562 0 600.893
148.88632202148438 0 592.39685
148.9013671875 0 564.98517
148.9086456298828 0 647.99445
148.91574096679688 0 980.8867
148.9228515625 0 1247.6748
148.93008422851562 0 1396.8364
148.93748474121094 0 3260.2834
148.94522094726562 0 5022.0117
148.96185302734375 0 3927.249
148.96954345703125 0 1620.9489
148.97671508789062 0 1225.9164
148.9840545654297 0 996.0103
148.9913330078125 0 801.41925
148.99868774414062 0 666.7495
149.0059051513672 0 717.04785
149.0133056640625 0 522.02405
149.03472900390625 0 450.62408
149.0557098388672 0 570.062
156.07647705078125 0 1255.6562
158.09169006347656 0 496.55734
159.0914764404297 0 1257.4915
166.0863494873047 0 2484.4385
169.13356018066406 0 1057.1464
172.07151794433594 0 566.18555
172.77235412597656 0 491.79645
174.08729553222656 0 523.2261
175.096435546875 0 558.50226
176.10694885253906 0 10402.988
177.11122131347656 0 1719.2096
181.16993713378906 0 457.23105
186.0919647216797 0 1014.9224
188.07073974609375 0 589.05505
191.11788940429688 0 594.64923
193.10797119140625 0 677.54315
197.12832641601562 0 1050.6666
201.12319946289062 0 761.0331
203.1173553466797 0 665.6775 a Water loss 1
205.0972900390625 0 614.75586 y 9
209.1650390625 0 545.6415
221.10313415527344 0 1288.0706
221.12841796875 0 13984.599 a 1
222.13153076171875 0 1691.0055
231.11264038085938 0 3062.7075 b Water loss 1
235.10733032226562 0 1516.8988
249.1233367919922 0 6812.598 b 1
251.0843505859375 0 638.67163
251.10214233398438 0 596.2853
253.11849975585938 0 2153.0225
257.1061706542969 0 552.99994
263.10223388671875 0 1622.6995
279.0975646972656 0 682.0352
297.79583740234375 0 557.2651
316.1866149902344 0 1322.64
324.1446533203125 0 772.4297
342.15625 0 5571.538 y 8
343.16070556640625 0 1227.3676
350.1338195800781 0 690.0487
355.0693054199219 0 1453.6951
378.1292419433594 0 1571.6239
378.1814270019531 0 1364.5186 b Water loss 2
379.1354675292969 0 761.04913
394.1234436035156 0 590.03906
425.1942138671875 0 985.66504
442.1024169921875 0 622.9258
489.1905212402344 0 1124.3392 y 7
551.2116088867188 0 756.1027
560.2266235351562 0 1681.3046 y 6
656.3316650390625 0 870.7354
657.332763671875 0 681.4817
659.2933959960938 0 751.85443
660.3538818359375 0 659.2963
661.626708984375 0 690.4202
665.7628784179688 0 920.7638 Precursor Water loss
666.2628784179688 0 755.5104
674.7684326171875 0 1759.0767 Precursor
675.2705688476562 0 1852.5017
675.3531494140625 0 659.5373
675.7656860351562 0 785.5448
712.1300048828125 0 678.1913
723.2894897460938 0 1189.6947 y 5
774.3201293945312 0 897.30273
838.3212280273438 0 2028.4097 y 4
839.3240966796875 0 1520.0997
868.3237915039062 0 708.3409
889.3411254882812 0 1043.7146
890.3425903320312 0 810.8494
896.3193359375 0 745.8602
953.3452758789062 0 3610.1772 y 3
954.3458862304688 0 2266.8616
1036.4150390625 0 3134.8638
1037.4215087890625 0 1631.474
1038.4061279296875 0 609.67004
1100.412353515625 0 7662.2173 y 2
1101.4169921875 0 4393.1646
1102.418701171875 0 1848.8232
1137.4654541015625 0 986.01587
1201.461669921875 0 1559.9542 y 1
1596.1915283203125 0 622.84827
1918.3560791015625 0 628.48505

Spectrum Details

|  |  |
| --- | --- |
| Matched peaks? Matched peaks The total absolute number of peaks matched. Additionally in brackets the total fraction of peaks matched and the total number of peaks is shown. | 16 (14.04% of 114) |
| FDR? FDR The false discovery rate estimated for this peptide. It is calculated by matching all theoretical fragments with a non-integer shift with the raw peaks for this spectrum. This is done with 40 different shifts. The resulting percentage is the average number of annotated peaks over the number of annotated peaks with the correct spectrum. | 0.00% |
| Satellite FDR? Satellite FDR See the FDR for details on its calculation. This satellite ion specific FDR only contains the satellite ions (d/w) for I/L/J positions. | - |
| PSM Score? PSM Score The PSM Score as given by Hecklib to this annotated spectrum. It is shown with three significant figures. | 169 |

#### Spectrum 9715? Spectrum 9715 The raw spectrum of this peptide as annotated by Hecklib. The fragments are coloured according to ion type (see legend). Any peaks with a star '\*' as text can be hovered over to see the full details, first the ion type second the mass shift type. By hovering over the amino acids in the peptide or ions in the legend the corresponding peaks are highlighted. By toggling the 'Unassigned' label you can turn the background (unassigned) peaks on or off in the plot. By updating the slider in the Ion legend you can update the spectrum to only show the top X% of the peaks with labels. The top X% means any peak that is within X% of the highest intensity. By dragging in the spectrum you can zoom in to a specific part of the spectrum and use 'Zoom Out' to get back to the original zoom level. The annotation of the spectrum is based on the given sequence in the peptides file and is done with different software so inconsistencies are likely. The peaks are annotated based on the given sequence, with 20 ppm tolerance.

Copy Data

##### Spectrum 9715 (TSV)

###### Preview

```
Loading example...
```

*Click on the button to copy the data to your clipboard.*

Mz MinMz MaxIntensity Max

WidthHeightPeptide font sizePeptide stroke widthSpectrum font sizeSpectrum stroke widthCompact peptide

Ion legend

wxyz

abcd

OtherUnassignedIon^Charge^_Position_Show for top:%

FTFDDYAMHW

02.03e+34.06e+36.09e+38.12e+3

Zoom Out

c^+1^_2_y^+1^_3_c^+1^_4_c^+1^_4_y^+1^_4_c^+2^_9_c^+1^_5_c^+1^_5_z^+1^_5_c^+1^_6_z^+1^_6_c^+1^_7_z^+1^_7_c^+1^_8_z^+1^_8_c^+1^_9_c^+1^_9_z^+1^_9_

034669210371383

Fragment Matches Table

Show background peaks

| Position | Ion type | Intensity | mz Theoretical | mz Error (Th) | mz Error (ppm) | Charge | Series Number |
| --- | --- | --- | --- | --- | --- | --- | --- |
| - | - | 5136 | 120.1 | - | - | 0 | - |
| - | - | 534.5 | 124.1 | - | - | 0 | - |
| - | - | 401 | 124.7 | - | - | 0 | - |
| - | - | 441.5 | 125.1 | - | - | 0 | - |
| - | - | 1576 | 126.1 | - | - | 0 | - |
| - | - | 361.7 | 127.1 | - | - | 0 | - |
| - | - | 529.2 | 127.1 | - | - | 0 | - |
| - | - | 616.6 | 128.1 | - | - | 0 | - |
| - | - | 444.2 | 128.1 | - | - | 0 | - |
| - | - | 581.1 | 129.1 | - | - | 0 | - |
| - | - | 708.7 | 130.1 | - | - | 0 | - |
| - | - | 682.1 | 130.1 | - | - | 0 | - |
| - | - | 691.9 | 131 | - | - | 0 | - |
| - | - | 625.8 | 133.1 | - | - | 0 | - |
| - | - | 1617 | 133.1 | - | - | 0 | - |
| - | - | 2335 | 136.1 | - | - | 0 | - |
| - | - | 425.7 | 137.1 | - | - | 0 | - |
| - | - | 537.7 | 137.1 | - | - | 0 | - |
| - | - | 394.6 | 137.1 | - | - | 0 | - |
| - | - | 669.7 | 138.1 | - | - | 0 | - |
| - | - | 442.7 | 139.7 | - | - | 0 | - |
| - | - | 508.3 | 140 | - | - | 0 | - |
| - | - | 1390 | 140.1 | - | - | 0 | - |
| - | - | 709.4 | 142.1 | - | - | 0 | - |
| - | - | 511.9 | 142.1 | - | - | 0 | - |
| - | - | 1418 | 143.1 | - | - | 0 | - |
| - | - | 566.5 | 144.1 | - | - | 0 | - |
| - | - | 1142 | 146.1 | - | - | 0 | - |
| - | - | 522 | 153.1 | - | - | 0 | - |
| - | - | 500.7 | 153.1 | - | - | 0 | - |
| - | - | 479.1 | 153.5 | - | - | 0 | - |
| - | - | 770.8 | 156.1 | - | - | 0 | - |
| - | - | 590 | 158.1 | - | - | 0 | - |
| - | - | 415.8 | 163.4 | - | - | 0 | - |
| - | - | 1652 | 165.1 | - | - | 0 | - |
| - | - | 1590 | 169.1 | - | - | 0 | - |
| - | - | 440.8 | 169.9 | - | - | 0 | - |
| - | - | 731.9 | 170.1 | - | - | 0 | - |
| - | - | 963.7 | 170.1 | - | - | 0 | - |
| - | - | 548.3 | 172.1 | - | - | 0 | - |
| - | - | 1438 | 173.4 | - | - | 0 | - |
| - | - | 781.2 | 177.1 | - | - | 0 | - |
| - | - | 498.4 | 180.8 | - | - | 0 | - |
| - | - | 1195 | 202.1 | - | - | 0 | - |
| - | - | 2167 | 203.1 | - | - | 0 | - |
| - | - | 2507 | 221.1 | - | - | 0 | - |
| - | - | 817.3 | 231.1 | - | - | 0 | - |
| - | - | 617.6 | 239.1 | - | - | 0 | - |
| - | - | 485.1 | 241.2 | - | - | 0 | - |
| 2 | c | 4249 | 249.1 | 4.424E-05 | 0.1776 | +1 | 2 |
| - | - | 976.7 | 280.1 | - | - | 0 | - |
| - | - | 668.1 | 282.1 | - | - | 0 | - |
| - | - | 976.1 | 338.2 | - | - | 0 | - |
| - | - | 755.8 | 366.1 | - | - | 0 | - |
| - | - | 1817 | 378.2 | - | - | 0 | - |
| - | - | 527.2 | 379.8 | - | - | 0 | - |
| - | - | 560.9 | 385.2 | - | - | 0 | - |
| - | - | 697.5 | 403 | - | - | 0 | - |
| - | - | 560.8 | 415.1 | - | - | 0 | - |
| - | - | 1428 | 455.2 | - | - | 0 | - |
| - | - | 2580 | 457.2 | - | - | 0 | - |
| - | - | 658.5 | 475.2 | - | - | 0 | - |
| 8 | y | 1212 | 505.2 | 0.008996 | 17.81 | +1 | 3 |
| 4 | c | 2009 | 511.2 | 0.0006956 | 1.361 | +1 | 4 |
| - | - | 1023 | 512.2 | - | - | 0 | - |
| 4 | c | 1288 | 528.2 | 0.0009512 | 1.801 | +1 | 4 |
| - | - | 563.3 | 565.7 | - | - | 0 | - |
| 7 | y | 909.7 | 576.2 | 0.007832 | 13.59 | +1 | 4 |
| - | - | 609.6 | 577.2 | - | - | 0 | - |
| 9 | c | 1494 | 581.2 | 0.001295 | 2.229 | +2 | 9 |
| - | - | 842.1 | 581.7 | - | - | 0 | - |
| - | - | 1065 | 594.2 | - | - | 0 | - |
| - | - | 788.5 | 595.2 | - | - | 0 | - |
| - | - | 932.2 | 601.7 | - | - | 0 | - |
| - | - | 1996 | 609.7 | - | - | 0 | - |
| - | - | 1189 | 610.2 | - | - | 0 | - |
| - | - | 971.5 | 610.7 | - | - | 0 | - |
| - | - | 664.4 | 617.7 | - | - | 0 | - |
| 5 | c | 1477 | 626.2 | 0.0007347 | 1.173 | +1 | 5 |
| - | - | 570.6 | 643.2 | - | - | 0 | - |
| 5 | c | 2408 | 643.3 | 0.000243 | 0.3778 | +1 | 5 |
| - | - | 949.8 | 644.3 | - | - | 0 | - |
| - | - | 749.9 | 651.3 | - | - | 0 | - |
| - | - | 968.6 | 652.3 | - | - | 0 | - |
| - | - | 1071 | 652.8 | - | - | 0 | - |
| - | - | 785.8 | 653.3 | - | - | 0 | - |
| - | - | 630.9 | 656.3 | - | - | 0 | - |
| - | - | 1201 | 660.8 | - | - | 0 | - |
| - | - | 1093 | 661.3 | - | - | 0 | - |
| - | - | 825.4 | 666.4 | - | - | 0 | - |
| - | - | 644.9 | 667.3 | - | - | 0 | - |
| - | - | 1141 | 673.8 | - | - | 0 | - |
| - | - | 2210 | 674.3 | - | - | 0 | - |
| - | - | 3620 | 674.8 | - | - | 0 | - |
| - | - | 3047 | 675.3 | - | - | 0 | - |
| - | - | 848.1 | 675.8 | - | - | 0 | - |
| - | - | 739.9 | 681.8 | - | - | 0 | - |
| - | - | 2890 | 682.3 | - | - | 0 | - |
| - | - | 5624 | 682.8 | - | - | 0 | - |
| - | - | 7584 | 683.3 | - | - | 0 | - |
| - | - | 3880 | 683.8 | - | - | 0 | - |
| - | - | 2465 | 684.3 | - | - | 0 | - |
| - | - | 794.6 | 684.8 | - | - | 0 | - |
| - | - | 646.3 | 688.8 | - | - | 0 | - |
| 6 | z | 1329 | 723.3 | 0.0076 | 10.51 | +1 | 5 |
| - | - | 940.4 | 794.3 | - | - | 0 | - |
| - | - | 2275 | 795.5 | - | - | 0 | - |
| - | - | 904.7 | 796.5 | - | - | 0 | - |
| 6 | c | 1696 | 806.3 | 0.001196 | 1.483 | +1 | 6 |
| - | - | 637.9 | 807.3 | - | - | 0 | - |
| - | - | 4716 | 812.5 | - | - | 0 | - |
| - | - | 1321 | 813.5 | - | - | 0 | - |
| 5 | z | 1182 | 838.3 | 0.006597 | 7.87 | +1 | 6 |
| - | - | 1263 | 839.3 | - | - | 0 | - |
| 7 | c | 1294 | 877.4 | 0.004679 | 5.333 | +1 | 7 |
| - | - | 898.1 | 878.4 | - | - | 0 | - |
| - | - | 740.9 | 895.5 | - | - | 0 | - |
| - | - | 1008 | 909.3 | - | - | 0 | - |
| - | - | 1915 | 913.5 | - | - | 0 | - |
| - | - | 771.2 | 930.1 | - | - | 0 | - |
| 4 | z | 1128 | 953.3 | 0.0178 | 18.67 | +1 | 7 |
| - | - | 928.4 | 954.3 | - | - | 0 | - |
| - | - | 709.9 | 955.3 | - | - | 0 | - |
| 8 | c | 858.7 | 1024 | 0.007988 | 7.798 | +1 | 8 |
| - | - | 752.3 | 1025 | - | - | 0 | - |
| - | - | 602.9 | 1068 | - | - | 0 | - |
| 3 | z | 787.2 | 1100 | 0.0136 | 12.36 | +1 | 8 |
| - | - | 735.9 | 1118 | - | - | 0 | - |
| - | - | 650.5 | 1128 | - | - | 0 | - |
| - | - | 1112 | 1129 | - | - | 0 | - |
| 9 | c | 705.6 | 1144 | 0.01314 | 11.49 | +1 | 9 |
| - | - | 717.4 | 1145 | - | - | 0 | - |
| - | - | 1105 | 1146 | - | - | 0 | - |
| 9 | c | 2077 | 1161 | 0.005228 | 4.502 | +1 | 9 |
| - | - | 1500 | 1162 | - | - | 0 | - |
| - | - | 830.9 | 1163 | - | - | 0 | - |
| - | - | 2878 | 1187 | - | - | 0 | - |
| - | - | 1980 | 1188 | - | - | 0 | - |
| - | - | 1386 | 1189 | - | - | 0 | - |
| 2 | z | 1286 | 1201 | 0.004126 | 3.434 | +1 | 9 |
| - | - | 1314 | 1202 | - | - | 0 | - |
| - | - | 1075 | 1219 | - | - | 0 | - |
| - | - | 753.8 | 1220 | - | - | 0 | - |
| - | - | 805.8 | 1287 | - | - | 0 | - |
| - | - | 909 | 1288 | - | - | 0 | - |
| - | - | 978 | 1289 | - | - | 0 | - |
| - | - | 1021 | 1290 | - | - | 0 | - |
| - | - | 621.8 | 1303 | - | - | 0 | - |
| - | - | 1175 | 1304 | - | - | 0 | - |
| - | - | 3595 | 1305 | - | - | 0 | - |
| - | - | 1919 | 1306 | - | - | 0 | - |
| - | - | 1962 | 1307 | - | - | 0 | - |
| - | - | 1342 | 1308 | - | - | 0 | - |
| - | - | 865.2 | 1321 | - | - | 0 | - |
| - | - | 844.8 | 1332 | - | - | 0 | - |
| - | - | 1447 | 1333 | - | - | 0 | - |
| - | - | 943 | 1334 | - | - | 0 | - |
| - | - | 2601 | 1348 | - | - | 0 | - |
| - | - | 6685 | 1349 | - | - | 0 | - |
| - | - | 8041 | 1350 | - | - | 0 | - |
| - | - | 6061 | 1351 | - | - | 0 | - |
| - | - | 2926 | 1352 | - | - | 0 | - |
| - | - | 1216 | 1353 | - | - | 0 | - |
| - | - | 666.1 | 1354 | - | - | 0 | - |
| - | - | 755.9 | 1359 | - | - | 0 | - |
| - | - | 2588 | 1365 | - | - | 0 | - |
| - | - | 3683 | 1366 | - | - | 0 | - |
| - | - | 4364 | 1367 | - | - | 0 | - |
| - | - | 3252 | 1368 | - | - | 0 | - |
| - | - | 1463 | 1369 | - | - | 0 | - |
| - | - | 718.5 | 1370 | - | - | 0 | - |

m/z Charge Intensity FragmentType MassShift Position
120.08087158203125 0 5136.1084
124.08695220947266 0 534.4858
124.74072265625 0 401.0134
125.06051635742188 0 441.5252
126.10265350341797 0 1575.853
127.05052185058594 0 361.71704
127.07528686523438 0 529.15485
128.0820770263672 0 616.5918
128.11819458007812 0 444.1603
129.05455017089844 0 581.0974
130.06097412109375 0 708.68475
130.0655975341797 0 682.1473
131.0452880859375 0 691.87396
133.060791015625 0 625.8175
133.0859375 0 1616.7283
136.0758056640625 0 2335.3203
137.05923461914062 0 425.72922
137.08253479003906 0 537.73035
137.1481475830078 0 394.60745
138.06600952148438 0 669.66394
139.66102600097656 0 442.65594
140.0455780029297 0 508.33847
140.08189392089844 0 1389.7521
142.06121826171875 0 709.43054
142.097900390625 0 511.9251
143.07286071777344 0 1417.9076
144.07666015625 0 566.4803
146.06027221679688 0 1142.4163
153.0548553466797 0 521.9537
153.07725524902344 0 500.71255
153.5349578857422 0 479.06384
156.0769805908203 0 770.7727
158.08396911621094 0 589.98535
163.3751678466797 0 415.7695
165.10226440429688 0 1652.4369
169.07205200195312 0 1589.7188
169.9373321533203 0 440.78195
170.05612182617188 0 731.851
170.0925750732422 0 963.6644
172.0721435546875 0 548.2595
173.4389190673828 0 1437.9532
177.11175537109375 0 781.2231
180.7921905517578 0 498.42502
202.07374572753906 0 1194.6343
203.0814666748047 0 2167.4368
221.1285400390625 0 2506.7747
231.11285400390625 0 817.3013
239.0888671875 0 617.57166
241.24110412597656 0 485.14835
249.1234130859375 0 4249.357 c Ammonia loss 1
280.12982177734375 0 976.65356
282.13177490234375 0 668.1359
338.15069580078125 0 976.09607
366.1465759277344 0 755.7676
378.1817932128906 0 1816.5522
379.81793212890625 0 527.1752
385.19873046875 0 560.90125
402.96575927734375 0 697.4768
415.14862060546875 0 560.76416
455.1994323730469 0 1427.7528
457.1590270996094 0 2579.5728
475.1558532714844 0 658.45856
505.185546875 0 1212.4108 y 7
511.21942138671875 0 2008.8269 c Ammonia loss 3
512.2239379882812 0 1022.7719
528.2443237304688 0 1288.041 c 3
565.656494140625 0 563.33295
576.2214965820312 0 909.655 y 6
577.2311401367188 0 609.562
581.2359619140625 0 1493.8226 c 8
581.7388916015625 0 842.05774
594.244873046875 0 1064.6816
595.2460327148438 0 788.5392
601.7344360351562 0 932.17523
609.7369995117188 0 1995.8422
610.2356567382812 0 1188.7598
610.7395629882812 0 971.54834
617.7288818359375 0 664.4376
626.2449340820312 0 1476.9099 c Ammonia loss 4
643.22021484375 0 570.5716
643.2724609375 0 2407.9001 c 4
644.2706909179688 0 949.8426
651.2617797851562 0 749.85956
652.27099609375 0 968.6066
652.766845703125 0 1071.4268
653.2700805664062 0 785.7674
656.2569580078125 0 630.9368
660.7704467773438 0 1200.9263
661.2693481445312 0 1093.3761
666.363525390625 0 825.41907
667.2571411132812 0 644.8924
673.7597045898438 0 1141.0242
674.2557983398438 0 2210.416
674.7625732421875 0 3619.5427
675.2647705078125 0 3047.3672
675.76953125 0 848.12524
681.7540283203125 0 739.8982
682.2622680664062 0 2889.981
682.7654418945312 0 5623.6445
683.2674560546875 0 7583.5947
683.7678833007812 0 3879.621
684.2664184570312 0 2464.6765
684.7664184570312 0 794.62866
688.80615234375 0 646.28125
723.265869140625 0 1329.3921 z 5
794.3080444335938 0 940.4204
795.4566650390625 0 2275.4226
796.4671630859375 0 904.6927
806.3343505859375 0 1696.1946 c 5
807.3330078125 0 637.88696
812.4841918945312 0 4716.497
813.489013671875 0 1321.2336
838.2918090820312 0 1182.4623 z 4
839.2954711914062 0 1262.5376
877.3679809570312 0 1293.5986 c 6
878.376953125 0 898.08685
895.478759765625 0 740.8981
909.3309936523438 0 1008.0216
913.5183715820312 0 1915.3468
930.1487426757812 0 771.18
953.3299560546875 0 1127.9408 z 3
954.325927734375 0 928.40436
955.3270263671875 0 709.9013
1024.4111328125 0 858.6884 c 7
1025.4154052734375 0 752.2509
1068.09423828125 0 602.9124
1100.3941650390625 0 787.2077 z 2
1118.44189453125 0 735.9276
1128.45556640625 0 650.54254
1129.4256591796875 0 1111.549
1144.42236328125 0 705.607 c Ammonia loss 8
1145.44873046875 0 717.41394
1146.458251953125 0 1104.6918
1161.46728515625 0 2076.6086 c 8
1162.4713134765625 0 1500.2886
1163.48095703125 0 830.9429
1187.4833984375 0 2878.393
1188.481689453125 0 1980.1954
1189.488525390625 0 1386.1427
1201.432373046875 0 1286.0879 z 1
1202.4444580078125 0 1314.4159
1219.4715576171875 0 1074.5509
1220.4686279296875 0 753.8471
1286.510986328125 0 805.8356
1287.5245361328125 0 909.04956
1288.5196533203125 0 977.97266
1289.5025634765625 0 1020.9895
1302.5361328125 0 621.77795
1303.5394287109375 0 1174.5286
1304.5316162109375 0 3595.3147
1305.52734375 0 1918.608
1306.5235595703125 0 1961.8339
1307.5296630859375 0 1342.0054
1320.534423828125 0 865.1737
1331.50732421875 0 844.84454
1332.5069580078125 0 1447.2092
1333.5220947265625 0 943.0007
1347.505126953125 0 2601.441
1348.51318359375 0 6685.041
1349.5159912109375 0 8040.6514
1350.519287109375 0 6061.0874
1351.5179443359375 0 2926.4036
1352.53271484375 0 1215.658
1353.5269775390625 0 666.0628
1359.2750244140625 0 755.8666
1364.52490234375 0 2587.7966
1365.5262451171875 0 3682.6697
1366.5338134765625 0 4363.991
1367.533935546875 0 3251.52
1368.5294189453125 0 1463.3976
1369.5338134765625 0 718.5166

Spectrum Details

|  |  |
| --- | --- |
| Matched peaks? Matched peaks The total absolute number of peaks matched. Additionally in brackets the total fraction of peaks matched and the total number of peaks is shown. | 18 (10.53% of 171) |
| FDR? FDR The false discovery rate estimated for this peptide. It is calculated by matching all theoretical fragments with a non-integer shift with the raw peaks for this spectrum. This is done with 40 different shifts. The resulting percentage is the average number of annotated peaks over the number of annotated peaks with the correct spectrum. | 0.26% |
| Satellite FDR? Satellite FDR See the FDR for details on its calculation. This satellite ion specific FDR only contains the satellite ions (d/w) for I/L/J positions. | - |
| PSM Score? PSM Score The PSM Score as given by Hecklib to this annotated spectrum. It is shown with three significant figures. | 211 |

#### Spectrum 8264? Spectrum 8264 The raw spectrum of this peptide as annotated by Hecklib. The fragments are coloured according to ion type (see legend). Any peaks with a star '\*' as text can be hovered over to see the full details, first the ion type second the mass shift type. By hovering over the amino acids in the peptide or ions in the legend the corresponding peaks are highlighted. By toggling the 'Unassigned' label you can turn the background (unassigned) peaks on or off in the plot. By updating the slider in the Ion legend you can update the spectrum to only show the top X% of the peaks with labels. The top X% means any peak that is within X% of the highest intensity. By dragging in the spectrum you can zoom in to a specific part of the spectrum and use 'Zoom Out' to get back to the original zoom level. The annotation of the spectrum is based on the given sequence in the peptides file and is done with different software so inconsistencies are likely. The peaks are annotated based on the given sequence, with 20 ppm tolerance.

Copy Data

##### Spectrum 8264 (TSV)

###### Preview

```
Loading example...
```

*Click on the button to copy the data to your clipboard.*

Mz MinMz MaxIntensity Max

WidthHeightPeptide font sizePeptide stroke widthSpectrum font sizeSpectrum stroke widthCompact peptide

Ion legend

wxyz

abcd

OtherUnassignedIon^Charge^_Position_Show for top:%

FTFDDYAMHW

08.22e+31.64e+42.47e+43.29e+4

Zoom Out

a^+1^_2_y^+1^_1_a^+1^_2_b^+1^_2_b^+1^_2_y^+1^_2_b^+1^_3_b^+1^_4_y^+2^_8_y^+1^_4_\*y^+1^_5_y^+1^_6_y^+1^_7_y^+1^_8_y^+1^_8_b^+1^_9_y^+1^_9_

0646129119372583

Fragment Matches Table

Show background peaks

| Position | Ion type | Intensity | mz Theoretical | mz Error (Th) | mz Error (ppm) | Charge | Series Number |
| --- | --- | --- | --- | --- | --- | --- | --- |
| - | - | 3.257E+04 | 120.1 | - | - | 0 | - |
| - | - | 1706 | 121.1 | - | - | 0 | - |
| - | - | 433.8 | 128.6 | - | - | 0 | - |
| - | - | 699.5 | 129.1 | - | - | 0 | - |
| - | - | 3291 | 129.1 | - | - | 0 | - |
| - | - | 483.6 | 130.1 | - | - | 0 | - |
| - | - | 675.4 | 130.1 | - | - | 0 | - |
| - | - | 1670 | 133.1 | - | - | 0 | - |
| - | - | 5261 | 136.1 | - | - | 0 | - |
| - | - | 457.3 | 143.2 | - | - | 0 | - |
| - | - | 482 | 145.1 | - | - | 0 | - |
| - | - | 817.3 | 147 | - | - | 0 | - |
| - | - | 405.4 | 147.1 | - | - | 0 | - |
| - | - | 976.4 | 148.9 | - | - | 0 | - |
| - | - | 2767 | 149 | - | - | 0 | - |
| - | - | 1139 | 156.1 | - | - | 0 | - |
| - | - | 592.1 | 158.1 | - | - | 0 | - |
| - | - | 1762 | 159.1 | - | - | 0 | - |
| - | - | 3888 | 167.1 | - | - | 0 | - |
| - | - | 428.2 | 167.1 | - | - | 0 | - |
| - | - | 810.4 | 169.1 | - | - | 0 | - |
| - | - | 616.6 | 173.1 | - | - | 0 | - |
| - | - | 1789 | 173.4 | - | - | 0 | - |
| - | - | 2010 | 175.1 | - | - | 0 | - |
| - | - | 1.191E+04 | 176.1 | - | - | 0 | - |
| - | - | 1465 | 177.1 | - | - | 0 | - |
| - | - | 488.3 | 181.1 | - | - | 0 | - |
| - | - | 716.9 | 182.1 | - | - | 0 | - |
| - | - | 654.4 | 183.1 | - | - | 0 | - |
| - | - | 486 | 185.2 | - | - | 0 | - |
| - | - | 2160 | 186.1 | - | - | 0 | - |
| - | - | 498.4 | 187.1 | - | - | 0 | - |
| - | - | 622.2 | 187.1 | - | - | 0 | - |
| - | - | 593.5 | 195.1 | - | - | 0 | - |
| - | - | 1550 | 197.1 | - | - | 0 | - |
| - | - | 674.6 | 201.1 | - | - | 0 | - |
| - | - | 4823 | 203.1 | - | - | 0 | - |
| 2 | a | 930.1 | 203.1 | 0.0002135 | 1.051 | +1 | 2 |
| - | - | 559.5 | 204.1 | - | - | 0 | - |
| - | - | 587.4 | 213.1 | - | - | 0 | - |
| 10 | y | 1743 | 221.1 | 0.002895 | 13.09 | +1 | 1 |
| - | - | 1495 | 221.1 | - | - | 0 | - |
| 2 | a | 1.386E+04 | 221.1 | 8.582E-05 | 0.3881 | +1 | 2 |
| - | - | 986.5 | 221.1 | - | - | 0 | - |
| - | - | 1885 | 222.1 | - | - | 0 | - |
| - | - | 6632 | 225 | - | - | 0 | - |
| - | - | 1022 | 226 | - | - | 0 | - |
| - | - | 784.5 | 227 | - | - | 0 | - |
| - | - | 710.6 | 229.2 | - | - | 0 | - |
| 2 | b | 2612 | 231.1 | 0.0004924 | 2.13 | +1 | 2 |
| - | - | 1115 | 235.1 | - | - | 0 | - |
| - | - | 8212 | 239.1 | - | - | 0 | - |
| - | - | 1367 | 240.1 | - | - | 0 | - |
| - | - | 758.8 | 241.1 | - | - | 0 | - |
| 2 | b | 8289 | 249.1 | 0.0001358 | 0.5451 | +1 | 2 |
| - | - | 938.9 | 250.1 | - | - | 0 | - |
| - | - | 730.7 | 251.1 | - | - | 0 | - |
| - | - | 502.7 | 256.2 | - | - | 0 | - |
| - | - | 1714 | 263.1 | - | - | 0 | - |
| - | - | 575.9 | 270.3 | - | - | 0 | - |
| - | - | 686.3 | 279.1 | - | - | 0 | - |
| - | - | 2046 | 285 | - | - | 0 | - |
| - | - | 900.5 | 285.1 | - | - | 0 | - |
| - | - | 2435 | 295.1 | - | - | 0 | - |
| - | - | 5394 | 299.1 | - | - | 0 | - |
| - | - | 902.5 | 300.1 | - | - | 0 | - |
| - | - | 560.3 | 313.1 | - | - | 0 | - |
| - | - | 1007 | 316.2 | - | - | 0 | - |
| - | - | 578.3 | 322.1 | - | - | 0 | - |
| - | - | 1097 | 340.1 | - | - | 0 | - |
| - | - | 801.5 | 341 | - | - | 0 | - |
| - | - | 694.6 | 341.1 | - | - | 0 | - |
| - | - | 576.7 | 341.2 | - | - | 0 | - |
| - | - | 5572 | 355.1 | - | - | 0 | - |
| 9 | y | 5797 | 358.1 | 0.004843 | 13.52 | +1 | 2 |
| - | - | 1.074E+04 | 359 | - | - | 0 | - |
| - | - | 908.7 | 359.2 | - | - | 0 | - |
| - | - | 787.4 | 364.1 | - | - | 0 | - |
| - | - | 696.6 | 369.1 | - | - | 0 | - |
| - | - | 2599 | 378.1 | - | - | 0 | - |
| 3 | b | 1141 | 378.2 | 0.0004319 | 1.142 | +1 | 3 |
| - | - | 753.7 | 382.1 | - | - | 0 | - |
| - | - | 953.4 | 394.1 | - | - | 0 | - |
| - | - | 553.7 | 415 | - | - | 0 | - |
| - | - | 710.2 | 419 | - | - | 0 | - |
| - | - | 1.169E+04 | 429.1 | - | - | 0 | - |
| - | - | 929.6 | 441.2 | - | - | 0 | - |
| - | - | 673.6 | 501.3 | - | - | 0 | - |
| - | - | 1809 | 505.2 | - | - | 0 | - |
| 4 | b | 875.1 | 511.2 | 0.001123 | 2.196 | +1 | 4 |
| - | - | 1022 | 512.2 | - | - | 0 | - |
| 3 | y | 1220 | 558.7 | 0.003014 | 5.395 | +2 | 8 |
| - | - | 577.9 | 568.7 | - | - | 0 | - |
| 7 | y | 2717 | 576.2 | 0.01003 | 17.41 | +1 | 4 |
| - | - | 931.7 | 577.2 | - | - | 0 | - |
| - | - | 656.2 | 610.2 | - | - | 0 | - |
| - | - | 618.9 | 674.3 | - | - | 0 | - |
| - | - | 893.9 | 675.3 | - | - | 0 | - |
| - | - | 973.4 | 681.8 | - | - | 0 | - |
| - | - | 1982 | 682.3 | - | - | 0 | - |
| 0 | Precursor | 1396 | 682.8 | 0.005332 | 7.809 | +2 | -1 |
| - | - | 1439 | 683.3 | - | - | 0 | - |
| - | - | 1405 | 683.3 | - | - | 0 | - |
| - | - | 2442 | 684.2 | - | - | 0 | - |
| 6 | y | 1869 | 739.3 | 0.009262 | 12.53 | +1 | 5 |
| - | - | 981.5 | 790.3 | - | - | 0 | - |
| - | - | 803.6 | 791.3 | - | - | 0 | - |
| 5 | y | 2686 | 854.3 | 0.01052 | 12.31 | +1 | 6 |
| - | - | 739.6 | 855.3 | - | - | 0 | - |
| - | - | 1125 | 896.3 | - | - | 0 | - |
| - | - | 908.2 | 905.3 | - | - | 0 | - |
| 4 | y | 3813 | 969.3 | 0.009575 | 9.878 | +1 | 7 |
| - | - | 1849 | 970.3 | - | - | 0 | - |
| - | - | 1944 | 1052 | - | - | 0 | - |
| - | - | 1042 | 1053 | - | - | 0 | - |
| 3 | y | 629.1 | 1098 | 0.01288 | 11.73 | +1 | 8 |
| 3 | y | 6023 | 1116 | 0.01062 | 9.512 | +1 | 8 |
| - | - | 3586 | 1117 | - | - | 0 | - |
| - | - | 1688 | 1118 | - | - | 0 | - |
| 9 | b | 1190 | 1144 | 0.003213 | 2.808 | +1 | 9 |
| 2 | y | 1680 | 1217 | 0.007496 | 6.157 | +1 | 9 |
| - | - | 1024 | 1218 | - | - | 0 | - |
| - | - | 575.6 | 2020 | - | - | 0 | - |
| - | - | 642.7 | 2557 | - | - | 0 | - |

m/z Charge Intensity FragmentType MassShift Position
120.0809555053711 0 32572.223
121.084228515625 0 1706.4974
128.6337127685547 0 433.78345
129.066162109375 0 699.4852
129.10243225097656 0 3290.768
130.05014038085938 0 483.59308
130.06558227539062 0 675.38745
133.08624267578125 0 1669.5471
136.0758056640625 0 5261.387
143.22470092773438 0 457.3153
145.0607452392578 0 482.01117
147.04769897460938 0 817.3107
147.07589721679688 0 405.36642
148.94720458984375 0 976.38916
149.0450897216797 0 2766.689
156.07717895507812 0 1139.2273
158.0605010986328 0 592.10504
159.09170532226562 0 1762.0939
167.0556182861328 0 3887.856
167.09246826171875 0 428.15964
169.13394165039062 0 810.37274
173.12933349609375 0 616.60425
173.43882751464844 0 1789.2738
175.08677673339844 0 2010.4358
176.10711669921875 0 11914.939
177.11083984375 0 1465.3374
181.1333465576172 0 488.31918
182.08123779296875 0 716.89014
183.1497344970703 0 654.3738
185.1651153564453 0 486.03198
186.09124755859375 0 2160.4766
187.0704345703125 0 498.4292
187.10752868652344 0 622.1627
195.11277770996094 0 593.48047
197.12860107421875 0 1550.2386
201.1229705810547 0 674.6008
203.08164978027344 0 4822.8965
203.11810302734375 0 930.0934 a Water loss 1
204.08494567871094 0 559.53424
213.12269592285156 0 587.3852
221.08425903320312 0 1743.0303 y 9
221.10350036621094 0 1494.5142
221.1285400390625 0 13856.77 a 1
221.1399383544922 0 986.52423
222.13235473632812 0 1884.8324
225.04307556152344 0 6632.2715
226.04380798339844 0 1021.8239
227.0399932861328 0 784.521
229.1549530029297 0 710.62085
231.11329650878906 0 2611.5579 b Water loss 1
235.10791015625 0 1114.5098
239.09507751464844 0 8211.521
240.09664916992188 0 1367.3301
241.0924835205078 0 758.82
249.12350463867188 0 8289.337 b 1
250.12725830078125 0 938.93866
251.10255432128906 0 730.70123
256.16143798828125 0 502.69263
263.10272216796875 0 1713.8364
270.33392333984375 0 575.89905
279.0972900390625 0 686.2567
285.010009765625 0 2045.7122
285.10247802734375 0 900.45416
295.1036071777344 0 2435.4744
299.0617370605469 0 5394.104
300.0618896484375 0 902.5247
313.1142578125 0 560.2639
316.188232421875 0 1006.8323
322.1305236816406 0 578.30475
340.1410217285156 0 1097.3958
341.01788330078125 0 801.4748
341.14630126953125 0 694.6156
341.17254638671875 0 576.74603
355.0699768066406 0 5571.948
358.1509094238281 0 5796.8096 y 8
359.02838134765625 0 10738.27
359.15472412109375 0 908.72815
364.1493835449219 0 787.3828
369.12164306640625 0 696.58496
378.1290588378906 0 2598.7551
378.1807861328125 0 1141.3794 b Water loss 2
382.1431579589844 0 753.7012
394.124755859375 0 953.4344
415.0320129394531 0 553.7417
418.9953918457031 0 710.19635
429.08929443359375 0 11690.137
441.18756103515625 0 929.55963
501.26654052734375 0 673.59607
505.1867980957031 0 1808.9493
511.2198486328125 0 875.0625 b 3
512.2236938476562 0 1022.08655
558.706298828125 0 1219.9727 y 2
568.7232666015625 0 577.9256
576.2236938476562 0 2716.579 y 6
577.2298583984375 0 931.69366
610.22412109375 0 656.1871
674.2606811523438 0 618.9084
675.2869873046875 0 893.8526
681.8368530273438 0 973.4234
682.3380126953125 0 1981.6672
682.7666625976562 0 1395.6003 Precursor
683.2678833007812 0 1438.5801
683.3455810546875 0 1404.7742
684.2029418945312 0 2442.0842
739.2862548828125 0 1869.1604 y 5
790.313720703125 0 981.5041
791.3224487304688 0 803.556
854.314453125 0 2686.1904 y 4
855.3193969726562 0 739.6008
896.327392578125 0 1125.2245
905.3411254882812 0 908.2392
969.3404541015625 0 3812.892 y 3
970.3436279296875 0 1849.182
1052.412109375 0 1944.2972
1053.40673828125 0 1041.6477
1098.401611328125 0 629.12476 y Water loss 2
1116.409912109375 0 6022.779 y 2
1117.4146728515625 0 3585.7224
1118.4151611328125 0 1687.6315
1144.438720703125 0 1189.8947 b 8
1217.4544677734375 0 1680.4688 y 1
1218.4515380859375 0 1023.5142
2019.715576171875 0 575.603
2556.944091796875 0 642.67444

Spectrum Details

|  |  |
| --- | --- |
| Matched peaks? Matched peaks The total absolute number of peaks matched. Additionally in brackets the total fraction of peaks matched and the total number of peaks is shown. | 18 (14.52% of 124) |
| FDR? FDR The false discovery rate estimated for this peptide. It is calculated by matching all theoretical fragments with a non-integer shift with the raw peaks for this spectrum. This is done with 40 different shifts. The resulting percentage is the average number of annotated peaks over the number of annotated peaks with the correct spectrum. | 0.40% |
| Satellite FDR? Satellite FDR See the FDR for details on its calculation. This satellite ion specific FDR only contains the satellite ions (d/w) for I/L/J positions. | - |
| PSM Score? PSM Score The PSM Score as given by Hecklib to this annotated spectrum. It is shown with three significant figures. | 203 |

#### Spectrum 9159? Spectrum 9159 The raw spectrum of this peptide as annotated by Hecklib. The fragments are coloured according to ion type (see legend). Any peaks with a star '\*' as text can be hovered over to see the full details, first the ion type second the mass shift type. By hovering over the amino acids in the peptide or ions in the legend the corresponding peaks are highlighted. By toggling the 'Unassigned' label you can turn the background (unassigned) peaks on or off in the plot. By updating the slider in the Ion legend you can update the spectrum to only show the top X% of the peaks with labels. The top X% means any peak that is within X% of the highest intensity. By dragging in the spectrum you can zoom in to a specific part of the spectrum and use 'Zoom Out' to get back to the original zoom level. The annotation of the spectrum is based on the given sequence in the peptides file and is done with different software so inconsistencies are likely. The peaks are annotated based on the given sequence, with 20 ppm tolerance.

Copy Data

##### Spectrum 9159 (TSV)

###### Preview

```
Loading example...
```

*Click on the button to copy the data to your clipboard.*

Mz MinMz MaxIntensity Max

WidthHeightPeptide font sizePeptide stroke widthSpectrum font sizeSpectrum stroke widthCompact peptide

Ion legend

wxyz

abcd

OtherUnassignedIon^Charge^_Position_Show for top:%

FTFDDYAMHW

08.68e+31.74e+42.60e+43.47e+4

Zoom Out

a^+1^_2_y^+1^_1_a^+1^_2_b^+1^_2_b^+1^_2_y^+1^_2_a^+1^_3_b^+1^_3_b^+1^_4_y^+2^_8_y^+1^_4_b^+1^_5_\*y^+1^_5_y^+1^_6_y^+1^_7_y^+1^_8_y^+1^_9_

0775155123263102

Fragment Matches Table

Show background peaks

| Position | Ion type | Intensity | mz Theoretical | mz Error (Th) | mz Error (ppm) | Charge | Series Number |
| --- | --- | --- | --- | --- | --- | --- | --- |
| - | - | 3.438E+04 | 120.1 | - | - | 0 | - |
| - | - | 298 | 121.1 | - | - | 0 | - |
| - | - | 2651 | 121.1 | - | - | 0 | - |
| - | - | 2706 | 129.1 | - | - | 0 | - |
| - | - | 1779 | 130.1 | - | - | 0 | - |
| - | - | 352.9 | 131.2 | - | - | 0 | - |
| - | - | 963 | 133.1 | - | - | 0 | - |
| - | - | 446.4 | 134.1 | - | - | 0 | - |
| - | - | 424 | 134.8 | - | - | 0 | - |
| - | - | 1.006E+04 | 136.1 | - | - | 0 | - |
| - | - | 771.4 | 137.1 | - | - | 0 | - |
| - | - | 455.3 | 138.1 | - | - | 0 | - |
| - | - | 412.1 | 143.7 | - | - | 0 | - |
| - | - | 736.2 | 146.1 | - | - | 0 | - |
| - | - | 4688 | 147.1 | - | - | 0 | - |
| - | - | 476.8 | 148.8 | - | - | 0 | - |
| - | - | 442.1 | 148.8 | - | - | 0 | - |
| - | - | 453.4 | 148.9 | - | - | 0 | - |
| - | - | 621.2 | 148.9 | - | - | 0 | - |
| - | - | 833.7 | 148.9 | - | - | 0 | - |
| - | - | 894.5 | 148.9 | - | - | 0 | - |
| - | - | 1326 | 148.9 | - | - | 0 | - |
| - | - | 1605 | 148.9 | - | - | 0 | - |
| - | - | 3571 | 148.9 | - | - | 0 | - |
| - | - | 5340 | 149 | - | - | 0 | - |
| - | - | 3207 | 149 | - | - | 0 | - |
| - | - | 1529 | 149 | - | - | 0 | - |
| - | - | 1197 | 149 | - | - | 0 | - |
| - | - | 719.6 | 149 | - | - | 0 | - |
| - | - | 890.7 | 149 | - | - | 0 | - |
| - | - | 604.2 | 149 | - | - | 0 | - |
| - | - | 577.4 | 149 | - | - | 0 | - |
| - | - | 408.2 | 149 | - | - | 0 | - |
| - | - | 416 | 149 | - | - | 0 | - |
| - | - | 2505 | 149 | - | - | 0 | - |
| - | - | 1342 | 152.1 | - | - | 0 | - |
| - | - | 834.6 | 158.1 | - | - | 0 | - |
| - | - | 1098 | 158.1 | - | - | 0 | - |
| - | - | 1347 | 159.1 | - | - | 0 | - |
| - | - | 659.8 | 164.1 | - | - | 0 | - |
| - | - | 3563 | 167.1 | - | - | 0 | - |
| - | - | 462.7 | 173.1 | - | - | 0 | - |
| - | - | 639.2 | 174.1 | - | - | 0 | - |
| - | - | 1759 | 175.1 | - | - | 0 | - |
| - | - | 458.6 | 175.1 | - | - | 0 | - |
| - | - | 630 | 176.1 | - | - | 0 | - |
| - | - | 1.294E+04 | 176.1 | - | - | 0 | - |
| - | - | 534.3 | 177.1 | - | - | 0 | - |
| - | - | 1853 | 177.1 | - | - | 0 | - |
| - | - | 469.2 | 179.4 | - | - | 0 | - |
| - | - | 1883 | 182.1 | - | - | 0 | - |
| - | - | 983.3 | 183.1 | - | - | 0 | - |
| - | - | 1554 | 186.1 | - | - | 0 | - |
| - | - | 1204 | 193.1 | - | - | 0 | - |
| - | - | 530.3 | 195.9 | - | - | 0 | - |
| - | - | 662.8 | 201.1 | - | - | 0 | - |
| 2 | a | 824.6 | 203.1 | 7.642E-05 | 0.3763 | +1 | 2 |
| - | - | 603.8 | 209.1 | - | - | 0 | - |
| - | - | 502.1 | 216.1 | - | - | 0 | - |
| - | - | 593.2 | 217.1 | - | - | 0 | - |
| - | - | 705.1 | 217.1 | - | - | 0 | - |
| - | - | 475.5 | 220 | - | - | 0 | - |
| 10 | y | 1903 | 221.1 | 0.002987 | 13.51 | +1 | 1 |
| - | - | 1880 | 221.1 | - | - | 0 | - |
| - | - | 2357 | 221.1 | - | - | 0 | - |
| 2 | a | 1.813E+04 | 221.1 | 0.0001163 | 0.5261 | +1 | 2 |
| - | - | 2408 | 222.1 | - | - | 0 | - |
| - | - | 6318 | 225 | - | - | 0 | - |
| - | - | 931.3 | 226 | - | - | 0 | - |
| - | - | 560.7 | 226.2 | - | - | 0 | - |
| 2 | b | 4008 | 231.1 | 3.459E-05 | 0.1497 | +1 | 2 |
| - | - | 1663 | 235.1 | - | - | 0 | - |
| - | - | 556 | 237.3 | - | - | 0 | - |
| - | - | 8696 | 239.1 | - | - | 0 | - |
| - | - | 639.1 | 239.2 | - | - | 0 | - |
| - | - | 1614 | 240.1 | - | - | 0 | - |
| 2 | b | 9354 | 249.1 | 0.0001511 | 0.6063 | +1 | 2 |
| - | - | 602.7 | 250.1 | - | - | 0 | - |
| - | - | 766.4 | 251.1 | - | - | 0 | - |
| - | - | 509.5 | 252.9 | - | - | 0 | - |
| - | - | 670.1 | 253.2 | - | - | 0 | - |
| - | - | 815.5 | 260.2 | - | - | 0 | - |
| - | - | 2636 | 263.1 | - | - | 0 | - |
| - | - | 647.3 | 279.1 | - | - | 0 | - |
| - | - | 638.2 | 283.6 | - | - | 0 | - |
| - | - | 1759 | 285 | - | - | 0 | - |
| - | - | 917.2 | 285.1 | - | - | 0 | - |
| - | - | 700.5 | 292.1 | - | - | 0 | - |
| - | - | 2074 | 295.1 | - | - | 0 | - |
| - | - | 747.6 | 296.1 | - | - | 0 | - |
| - | - | 2896 | 296.2 | - | - | 0 | - |
| - | - | 493.2 | 296.3 | - | - | 0 | - |
| - | - | 601.2 | 297.1 | - | - | 0 | - |
| - | - | 4393 | 299.1 | - | - | 0 | - |
| - | - | 864.7 | 300.1 | - | - | 0 | - |
| - | - | 960 | 314.2 | - | - | 0 | - |
| - | - | 598.9 | 323.1 | - | - | 0 | - |
| - | - | 875.4 | 340.1 | - | - | 0 | - |
| - | - | 882.5 | 341 | - | - | 0 | - |
| - | - | 584.7 | 350.1 | - | - | 0 | - |
| - | - | 3957 | 355.1 | - | - | 0 | - |
| - | - | 2927 | 356.1 | - | - | 0 | - |
| - | - | 548.5 | 357.1 | - | - | 0 | - |
| 9 | y | 5124 | 358.1 | 0.004752 | 13.27 | +1 | 2 |
| - | - | 7320 | 359 | - | - | 0 | - |
| - | - | 775.2 | 359.1 | - | - | 0 | - |
| - | - | 1468 | 359.2 | - | - | 0 | - |
| - | - | 572.9 | 364.1 | - | - | 0 | - |
| 3 | a | 536.8 | 368.2 | 0.002441 | 6.629 | +1 | 3 |
| - | - | 687.9 | 369.1 | - | - | 0 | - |
| - | - | 2332 | 378.1 | - | - | 0 | - |
| 3 | b | 2195 | 378.2 | 0.0003709 | 0.9808 | +1 | 3 |
| - | - | 1004 | 394.1 | - | - | 0 | - |
| - | - | 1011 | 419 | - | - | 0 | - |
| - | - | 6908 | 429.1 | - | - | 0 | - |
| - | - | 697.3 | 439.2 | - | - | 0 | - |
| - | - | 1389 | 441.2 | - | - | 0 | - |
| - | - | 672.1 | 459 | - | - | 0 | - |
| - | - | 818.2 | 465.2 | - | - | 0 | - |
| - | - | 775.2 | 475 | - | - | 0 | - |
| - | - | 644.6 | 477.2 | - | - | 0 | - |
| - | - | 726.5 | 479.2 | - | - | 0 | - |
| - | - | 675 | 503.2 | - | - | 0 | - |
| - | - | 748.6 | 505.2 | - | - | 0 | - |
| 4 | b | 761 | 511.2 | 0.0008482 | 1.659 | +1 | 4 |
| - | - | 553.4 | 512.2 | - | - | 0 | - |
| - | - | 935.7 | 557.7 | - | - | 0 | - |
| 3 | y | 895.1 | 558.7 | 0.005395 | 9.655 | +2 | 8 |
| - | - | 1393 | 559.2 | - | - | 0 | - |
| - | - | 898 | 574.2 | - | - | 0 | - |
| 7 | y | 1853 | 576.2 | 0.01088 | 18.89 | +1 | 4 |
| - | - | 679.7 | 577.2 | - | - | 0 | - |
| - | - | 779.5 | 613.3 | - | - | 0 | - |
| 5 | b | 859.1 | 626.2 | 0.000608 | 0.9709 | +1 | 5 |
| - | - | 910.1 | 627.3 | - | - | 0 | - |
| - | - | 588.6 | 634.2 | - | - | 0 | - |
| - | - | 717.1 | 674.3 | - | - | 0 | - |
| - | - | 776.2 | 675.3 | - | - | 0 | - |
| - | - | 555.6 | 676.3 | - | - | 0 | - |
| - | - | 703.6 | 681.8 | - | - | 0 | - |
| - | - | 1144 | 681.8 | - | - | 0 | - |
| - | - | 637 | 682.3 | - | - | 0 | - |
| - | - | 1462 | 682.3 | - | - | 0 | - |
| 0 | Precursor | 2101 | 682.8 | 0.003318 | 4.859 | +2 | -1 |
| - | - | 688.5 | 682.8 | - | - | 0 | - |
| - | - | 1526 | 683.3 | - | - | 0 | - |
| - | - | 1833 | 684.2 | - | - | 0 | - |
| - | - | 648.6 | 714.2 | - | - | 0 | - |
| - | - | 1305 | 737.3 | - | - | 0 | - |
| 6 | y | 1485 | 739.3 | 0.01018 | 13.77 | +1 | 5 |
| - | - | 998.7 | 740.3 | - | - | 0 | - |
| - | - | 748.8 | 740.4 | - | - | 0 | - |
| - | - | 590.4 | 757.4 | - | - | 0 | - |
| - | - | 544.6 | 780.5 | - | - | 0 | - |
| - | - | 1634 | 785.4 | - | - | 0 | - |
| - | - | 782.4 | 786.4 | - | - | 0 | - |
| - | - | 1161 | 790.3 | - | - | 0 | - |
| - | - | 1012 | 804.3 | - | - | 0 | - |
| - | - | 646.8 | 809.4 | - | - | 0 | - |
| - | - | 922 | 852.3 | - | - | 0 | - |
| 5 | y | 1923 | 854.3 | 0.01088 | 12.74 | +1 | 6 |
| - | - | 1012 | 855.3 | - | - | 0 | - |
| - | - | 4242 | 886.4 | - | - | 0 | - |
| - | - | 2531 | 887.4 | - | - | 0 | - |
| - | - | 912.3 | 896.3 | - | - | 0 | - |
| - | - | 993.5 | 905.3 | - | - | 0 | - |
| - | - | 926.3 | 905.5 | - | - | 0 | - |
| - | - | 680.4 | 906.3 | - | - | 0 | - |
| - | - | 1783 | 932.5 | - | - | 0 | - |
| - | - | 1121 | 933.5 | - | - | 0 | - |
| - | - | 817 | 953.3 | - | - | 0 | - |
| - | - | 1338 | 967.3 | - | - | 0 | - |
| - | - | 545.3 | 968.3 | - | - | 0 | - |
| 4 | y | 4048 | 969.3 | 0.01177 | 12.14 | +1 | 7 |
| - | - | 1855 | 970.3 | - | - | 0 | - |
| - | - | 803.8 | 971.3 | - | - | 0 | - |
| - | - | 685.6 | 984.4 | - | - | 0 | - |
| - | - | 586.7 | 996.3 | - | - | 0 | - |
| - | - | 865.6 | 1050 | - | - | 0 | - |
| - | - | 934.7 | 1051 | - | - | 0 | - |
| - | - | 1681 | 1052 | - | - | 0 | - |
| - | - | 1078 | 1053 | - | - | 0 | - |
| - | - | 2443 | 1114 | - | - | 0 | - |
| - | - | 712.3 | 1115 | - | - | 0 | - |
| 3 | y | 5430 | 1116 | 0.01038 | 9.293 | +1 | 8 |
| - | - | 3377 | 1117 | - | - | 0 | - |
| - | - | 1537 | 1118 | - | - | 0 | - |
| 2 | y | 1123 | 1217 | 0.005299 | 4.353 | +1 | 9 |
| - | - | 677.7 | 2655 | - | - | 0 | - |
| - | - | 730.1 | 3071 | - | - | 0 | - |

m/z Charge Intensity FragmentType MassShift Position
120.08092498779297 0 34384.844
121.080810546875 0 297.98007
121.08428192138672 0 2650.8198
129.10232543945312 0 2705.7249
130.050048828125 0 1779.4387
131.1768341064453 0 352.9232
133.08602905273438 0 963.01416
134.0889129638672 0 446.43472
134.77650451660156 0 423.95142
136.0758056640625 0 10055.87
137.0791778564453 0 771.41034
138.06594848632812 0 455.34717
143.69309997558594 0 412.09302
146.0604705810547 0 736.21643
147.07652282714844 0 4688.494
148.82598876953125 0 476.81116
148.84722900390625 0 442.12064
148.89759826660156 0 453.42218
148.90452575683594 0 621.2324
148.91183471679688 0 833.74896
148.91908264160156 0 894.52386
148.9261474609375 0 1326.4087
148.93325805664062 0 1605.177
148.94078063964844 0 3570.561
148.95718383789062 0 5340.396
148.9649200439453 0 3206.5237
148.97232055664062 0 1529.3917
148.97938537597656 0 1196.7695
148.9866485595703 0 719.574
148.9940643310547 0 890.66174
149.00070190429688 0 604.17126
149.00794982910156 0 577.3614
149.02273559570312 0 408.17755
149.02947998046875 0 415.9507
149.04470825195312 0 2504.811
152.070556640625 0 1342.0956
158.0597686767578 0 834.6377
158.09677124023438 0 1098.271
159.09197998046875 0 1347.3691
164.0706329345703 0 659.77655
167.0555877685547 0 3563.0325
173.1286163330078 0 462.74057
174.054931640625 0 639.15753
175.0868377685547 0 1758.7722
175.09555053710938 0 458.59363
176.08224487304688 0 629.9743
176.10711669921875 0 12941.268
177.1027374267578 0 534.3399
177.11077880859375 0 1852.55
179.427978515625 0 469.17093
182.08140563964844 0 1882.759
183.1131134033203 0 983.2702
186.09144592285156 0 1553.5916
193.10841369628906 0 1203.5691
195.9257049560547 0 530.33484
201.12368774414062 0 662.81384
203.11781311035156 0 824.64624 a Water loss 1
209.09112548828125 0 603.79877
216.09765625 0 502.07535
217.08126831054688 0 593.1617
217.0970916748047 0 705.0641
220.00413513183594 0 475.4615
221.08416748046875 0 1903.3567 y 9
221.09295654296875 0 1880.3066
221.10350036621094 0 2356.665
221.12857055664062 0 18128.285 a 1
222.13192749023438 0 2407.976
225.0430450439453 0 6318.0664
226.0426788330078 0 931.33887
226.15504455566406 0 560.6993
231.1128387451172 0 4007.5283 b Water loss 1
235.1078338623047 0 1663.0569
237.3192901611328 0 556.0011
239.09507751464844 0 8695.948
239.15052795410156 0 639.1109
240.09649658203125 0 1613.9194
249.12351989746094 0 9354.252 b 1
250.12652587890625 0 602.6837
251.1026153564453 0 766.4166
252.8696746826172 0 509.50385
253.1659393310547 0 670.0763
260.1603698730469 0 815.4882
263.1026916503906 0 2636.4473
279.09674072265625 0 647.34467
283.599609375 0 638.2231
285.010009765625 0 1759.0302
285.1019287109375 0 917.19147
292.1409912109375 0 700.49414
295.1035461425781 0 2073.5444
296.1045837402344 0 747.56146
296.1969909667969 0 2895.949
296.3175048828125 0 493.15628
297.1001892089844 0 601.1903
299.0619201660156 0 4393.0786
300.0626525878906 0 864.7217
314.2078552246094 0 960.0131
323.09844970703125 0 598.89954
340.1412048339844 0 875.4324
341.0187072753906 0 882.534
350.1343078613281 0 584.7211
355.070068359375 0 3956.7217
356.13555908203125 0 2926.6492
357.1382751464844 0 548.54553
358.15081787109375 0 5123.6826 y 8
359.0284729003906 0 7320.1597
359.05181884765625 0 775.246
359.1545715332031 0 1468.231
364.14990234375 0 572.9283
368.1944274902344 0 536.79047 a 2
369.12310791015625 0 687.9415
378.1301574707031 0 2332.4817
378.18084716796875 0 2194.77 b Water loss 2
394.12432861328125 0 1003.9632
418.99554443359375 0 1010.9499
429.08941650390625 0 6907.9146
439.1727600097656 0 697.2968
441.18939208984375 0 1388.9788
458.9822998046875 0 672.0785
465.1640930175781 0 818.2167
475.001953125 0 775.23083
477.189453125 0 644.6455
479.1781005859375 0 726.46045
503.1747741699219 0 675.03546
505.18817138671875 0 748.5665
511.2195739746094 0 760.99603 b 3
512.2246704101562 0 553.3818
557.7015380859375 0 935.6539
558.7086791992188 0 895.0949 y 2
559.2120971679688 0 1392.532
574.208984375 0 897.9716
576.2245483398438 0 1853.1083 y 6
577.2286987304688 0 679.7073
613.33349609375 0 779.4926
626.2462768554688 0 859.06354 b 4
627.2593383789062 0 910.0638
634.2376098632812 0 588.5671
674.2640991210938 0 717.14136
675.2901611328125 0 776.2093
676.2949829101562 0 555.6476
681.7567138671875 0 703.6401
681.84228515625 0 1144.414
682.2639770507812 0 637.0489
682.3388671875 0 1462.477
682.7646484375 0 2101.2756 Precursor
682.8287963867188 0 688.5465
683.2665405273438 0 1525.9185
684.2035522460938 0 1832.823
714.2254028320312 0 648.63306
737.2745971679688 0 1304.5715
739.2871704101562 0 1485.4683 y 5
740.2833251953125 0 998.6802
740.351806640625 0 748.83997
757.4130859375 0 590.38
780.4602661132812 0 544.5876
785.4174194335938 0 1633.6963
786.4161376953125 0 782.43176
790.3140869140625 0 1161.1578
804.3392944335938 0 1012.36945
809.3800659179688 0 646.8485
852.2940673828125 0 921.9766
854.3148193359375 0 1923.1425 y 4
855.3182983398438 0 1011.8
886.4169311523438 0 4242.4473
887.4169921875 0 2530.91
896.3229370117188 0 912.2949
905.3484497070312 0 993.45996
905.4891357421875 0 926.2652
906.342041015625 0 680.4006
932.4863891601562 0 1782.9276
933.4901733398438 0 1120.6327
953.301025390625 0 816.99695
967.3262329101562 0 1338.4003
968.330078125 0 545.304
969.3426513671875 0 4047.5554 y 3
970.3442993164062 0 1855.0674
971.3307495117188 0 803.8013
984.4232177734375 0 685.62317
996.2566528320312 0 586.6883
1050.3931884765625 0 865.557
1051.3956298828125 0 934.68506
1052.4088134765625 0 1681.13
1053.423583984375 0 1077.6139
1114.3931884765625 0 2443.0615
1115.39453125 0 712.2921
1116.40966796875 0 5429.926 y 2
1117.409912109375 0 3376.8657
1118.4039306640625 0 1536.6171
1217.4522705078125 0 1122.899 y 1
2655.254150390625 0 677.7012
3070.893310546875 0 730.05164

Spectrum Details

|  |  |
| --- | --- |
| Matched peaks? Matched peaks The total absolute number of peaks matched. Additionally in brackets the total fraction of peaks matched and the total number of peaks is shown. | 18 (9.47% of 190) |
| FDR? FDR The false discovery rate estimated for this peptide. It is calculated by matching all theoretical fragments with a non-integer shift with the raw peaks for this spectrum. This is done with 40 different shifts. The resulting percentage is the average number of annotated peaks over the number of annotated peaks with the correct spectrum. | 1.06% |
| Satellite FDR? Satellite FDR See the FDR for details on its calculation. This satellite ion specific FDR only contains the satellite ions (d/w) for I/L/J positions. | - |
| PSM Score? PSM Score The PSM Score as given by Hecklib to this annotated spectrum. It is shown with three significant figures. | 203 |

#### Spectrum 8970? Spectrum 8970 The raw spectrum of this peptide as annotated by Hecklib. The fragments are coloured according to ion type (see legend). Any peaks with a star '\*' as text can be hovered over to see the full details, first the ion type second the mass shift type. By hovering over the amino acids in the peptide or ions in the legend the corresponding peaks are highlighted. By toggling the 'Unassigned' label you can turn the background (unassigned) peaks on or off in the plot. By updating the slider in the Ion legend you can update the spectrum to only show the top X% of the peaks with labels. The top X% means any peak that is within X% of the highest intensity. By dragging in the spectrum you can zoom in to a specific part of the spectrum and use 'Zoom Out' to get back to the original zoom level. The annotation of the spectrum is based on the given sequence in the peptides file and is done with different software so inconsistencies are likely. The peaks are annotated based on the given sequence, with 20 ppm tolerance.

Copy Data

##### Spectrum 8970 (TSV)

###### Preview

```
Loading example...
```

*Click on the button to copy the data to your clipboard.*

Mz MinMz MaxIntensity Max

WidthHeightPeptide font sizePeptide stroke widthSpectrum font sizeSpectrum stroke widthCompact peptide

Ion legend

wxyz

abcd

OtherUnassignedIon^Charge^_Position_Show for top:%

FTFDDYAMHW

02.59e+45.18e+47.77e+41.04e+5

Zoom Out

a^+1^_2_a^+1^_2_b^+1^_2_b^+1^_2_b^+1^_3_y^+2^_6_b^+1^_4_y^+2^_8_y^+2^_8_b^+1^_5_y^+2^_9_b^+1^_5_\*\*y^+1^_5_b^+1^_6_b^+1^_7_y^+1^_6_y^+1^_7_y^+1^_8_y^+1^_8_y^+1^_9_y^+1^_9_

0731146221932924

Fragment Matches Table

Show background peaks

| Position | Ion type | Intensity | mz Theoretical | mz Error (Th) | mz Error (ppm) | Charge | Series Number |
| --- | --- | --- | --- | --- | --- | --- | --- |
| - | - | 1.025E+05 | 120.1 | - | - | 0 | - |
| - | - | 6638 | 121.1 | - | - | 0 | - |
| - | - | 363.9 | 121.9 | - | - | 0 | - |
| - | - | 444.5 | 123.7 | - | - | 0 | - |
| - | - | 2053 | 129.1 | - | - | 0 | - |
| - | - | 601.8 | 130.1 | - | - | 0 | - |
| - | - | 424.7 | 131 | - | - | 0 | - |
| - | - | 601.4 | 132.1 | - | - | 0 | - |
| - | - | 388.7 | 133.1 | - | - | 0 | - |
| - | - | 679.2 | 133.1 | - | - | 0 | - |
| - | - | 501.9 | 136.1 | - | - | 0 | - |
| - | - | 9063 | 136.1 | - | - | 0 | - |
| - | - | 486.7 | 136.1 | - | - | 0 | - |
| - | - | 858.7 | 137.1 | - | - | 0 | - |
| - | - | 629.4 | 138.1 | - | - | 0 | - |
| - | - | 816.4 | 138.1 | - | - | 0 | - |
| - | - | 1247 | 146.1 | - | - | 0 | - |
| - | - | 502.1 | 148.9 | - | - | 0 | - |
| - | - | 524 | 148.9 | - | - | 0 | - |
| - | - | 697.6 | 148.9 | - | - | 0 | - |
| - | - | 666.5 | 148.9 | - | - | 0 | - |
| - | - | 974.7 | 148.9 | - | - | 0 | - |
| - | - | 1016 | 148.9 | - | - | 0 | - |
| - | - | 1966 | 148.9 | - | - | 0 | - |
| - | - | 3842 | 148.9 | - | - | 0 | - |
| - | - | 5030 | 149 | - | - | 0 | - |
| - | - | 3292 | 149 | - | - | 0 | - |
| - | - | 1291 | 149 | - | - | 0 | - |
| - | - | 1377 | 149 | - | - | 0 | - |
| - | - | 1120 | 149 | - | - | 0 | - |
| - | - | 913.9 | 149 | - | - | 0 | - |
| - | - | 770.5 | 149 | - | - | 0 | - |
| - | - | 511 | 149 | - | - | 0 | - |
| - | - | 466.2 | 149 | - | - | 0 | - |
| - | - | 471.1 | 149 | - | - | 0 | - |
| - | - | 426.6 | 149 | - | - | 0 | - |
| - | - | 578 | 152.1 | - | - | 0 | - |
| - | - | 394.6 | 152.3 | - | - | 0 | - |
| - | - | 452.6 | 154.2 | - | - | 0 | - |
| - | - | 1786 | 155.1 | - | - | 0 | - |
| - | - | 3211 | 156.1 | - | - | 0 | - |
| - | - | 593.1 | 158.1 | - | - | 0 | - |
| - | - | 1970 | 158.1 | - | - | 0 | - |
| - | - | 7054 | 159.1 | - | - | 0 | - |
| - | - | 756.7 | 160.1 | - | - | 0 | - |
| - | - | 589.5 | 160.1 | - | - | 0 | - |
| - | - | 439.9 | 165.1 | - | - | 0 | - |
| - | - | 1080 | 166.1 | - | - | 0 | - |
| - | - | 717.4 | 167.1 | - | - | 0 | - |
| - | - | 1119 | 169.1 | - | - | 0 | - |
| - | - | 1943 | 171.1 | - | - | 0 | - |
| - | - | 1662 | 174.1 | - | - | 0 | - |
| - | - | 477.4 | 174.1 | - | - | 0 | - |
| - | - | 1922 | 175.1 | - | - | 0 | - |
| - | - | 1424 | 176.1 | - | - | 0 | - |
| - | - | 4.526E+04 | 176.1 | - | - | 0 | - |
| - | - | 1681 | 177.1 | - | - | 0 | - |
| - | - | 4355 | 177.1 | - | - | 0 | - |
| - | - | 449.1 | 178.4 | - | - | 0 | - |
| - | - | 908 | 181.2 | - | - | 0 | - |
| - | - | 480.8 | 181.3 | - | - | 0 | - |
| - | - | 832.1 | 185.1 | - | - | 0 | - |
| - | - | 456.4 | 185.3 | - | - | 0 | - |
| - | - | 5486 | 186.1 | - | - | 0 | - |
| - | - | 601.4 | 187.1 | - | - | 0 | - |
| - | - | 1041 | 191.1 | - | - | 0 | - |
| - | - | 743.3 | 191.1 | - | - | 0 | - |
| - | - | 3346 | 193.1 | - | - | 0 | - |
| - | - | 804.7 | 195.1 | - | - | 0 | - |
| - | - | 799.8 | 197.1 | - | - | 0 | - |
| - | - | 1224 | 199.1 | - | - | 0 | - |
| - | - | 481.8 | 200.9 | - | - | 0 | - |
| - | - | 525.7 | 201.1 | - | - | 0 | - |
| - | - | 713.6 | 202.1 | - | - | 0 | - |
| - | - | 773.1 | 203.1 | - | - | 0 | - |
| 2 | a | 2725 | 203.1 | 6.09E-05 | 0.2998 | +1 | 2 |
| - | - | 1296 | 204.1 | - | - | 0 | - |
| - | - | 955.8 | 209.2 | - | - | 0 | - |
| - | - | 569.6 | 213.2 | - | - | 0 | - |
| - | - | 1024 | 215.1 | - | - | 0 | - |
| - | - | 2439 | 217.1 | - | - | 0 | - |
| - | - | 1497 | 220.1 | - | - | 0 | - |
| - | - | 8391 | 221.1 | - | - | 0 | - |
| 2 | a | 5.719E+04 | 221.1 | 2.099E-05 | 0.09494 | +1 | 2 |
| - | - | 718.4 | 222.1 | - | - | 0 | - |
| - | - | 6770 | 222.1 | - | - | 0 | - |
| - | - | 564.5 | 225.1 | - | - | 0 | - |
| - | - | 1254 | 227.1 | - | - | 0 | - |
| - | - | 968.8 | 231.1 | - | - | 0 | - |
| 2 | b | 1.341E+04 | 231.1 | 3.459E-05 | 0.1497 | +1 | 2 |
| - | - | 1169 | 232.1 | - | - | 0 | - |
| - | - | 5264 | 235.1 | - | - | 0 | - |
| - | - | 664 | 236.1 | - | - | 0 | - |
| - | - | 2792 | 237.1 | - | - | 0 | - |
| - | - | 692.6 | 237.2 | - | - | 0 | - |
| - | - | 828.7 | 238.1 | - | - | 0 | - |
| - | - | 822.4 | 239.1 | - | - | 0 | - |
| - | - | 2112 | 239.2 | - | - | 0 | - |
| - | - | 1347 | 243.1 | - | - | 0 | - |
| 2 | b | 2.939E+04 | 249.1 | 1.534E-06 | 0.006156 | +1 | 2 |
| - | - | 3590 | 250.1 | - | - | 0 | - |
| - | - | 3161 | 251.1 | - | - | 0 | - |
| - | - | 1016 | 256.2 | - | - | 0 | - |
| - | - | 1008 | 257.1 | - | - | 0 | - |
| - | - | 1323 | 258.2 | - | - | 0 | - |
| - | - | 6508 | 263.1 | - | - | 0 | - |
| - | - | 916.4 | 264.1 | - | - | 0 | - |
| - | - | 583.8 | 268.1 | - | - | 0 | - |
| - | - | 1002 | 274.1 | - | - | 0 | - |
| - | - | 2603 | 279.1 | - | - | 0 | - |
| - | - | 2210 | 285.1 | - | - | 0 | - |
| - | - | 674 | 286.2 | - | - | 0 | - |
| - | - | 638.3 | 292.1 | - | - | 0 | - |
| - | - | 787.3 | 302.1 | - | - | 0 | - |
| - | - | 713.3 | 303.1 | - | - | 0 | - |
| - | - | 734.2 | 306.1 | - | - | 0 | - |
| - | - | 789.8 | 310.2 | - | - | 0 | - |
| - | - | 874.2 | 316.2 | - | - | 0 | - |
| - | - | 571.7 | 325 | - | - | 0 | - |
| - | - | 1594 | 328.1 | - | - | 0 | - |
| - | - | 1160 | 333.2 | - | - | 0 | - |
| - | - | 614.9 | 339.2 | - | - | 0 | - |
| - | - | 749.9 | 346.1 | - | - | 0 | - |
| - | - | 668.6 | 348.1 | - | - | 0 | - |
| - | - | 1006 | 350.1 | - | - | 0 | - |
| - | - | 3428 | 356.1 | - | - | 0 | - |
| - | - | 911.7 | 357.1 | - | - | 0 | - |
| - | - | 1141 | 358.2 | - | - | 0 | - |
| - | - | 2648 | 364.2 | - | - | 0 | - |
| - | - | 2.877E+04 | 374.1 | - | - | 0 | - |
| - | - | 5001 | 375.1 | - | - | 0 | - |
| - | - | 863.6 | 376.2 | - | - | 0 | - |
| - | - | 6334 | 378.1 | - | - | 0 | - |
| 3 | b | 5548 | 378.2 | 0.0001268 | 0.3352 | +1 | 3 |
| - | - | 1888 | 379.1 | - | - | 0 | - |
| - | - | 1033 | 379.2 | - | - | 0 | - |
| - | - | 777.6 | 382.1 | - | - | 0 | - |
| - | - | 2482 | 394.1 | - | - | 0 | - |
| - | - | 2193 | 429.2 | - | - | 0 | - |
| - | - | 598.7 | 433.2 | - | - | 0 | - |
| 5 | y | 721.1 | 435.7 | 0.006773 | 15.55 | +2 | 6 |
| - | - | 1322 | 440.2 | - | - | 0 | - |
| - | - | 9009 | 457.2 | - | - | 0 | - |
| - | - | 1321 | 458.2 | - | - | 0 | - |
| - | - | 1681 | 465.2 | - | - | 0 | - |
| - | - | 1666 | 479.2 | - | - | 0 | - |
| - | - | 696.9 | 480.2 | - | - | 0 | - |
| - | - | 770.8 | 493.2 | - | - | 0 | - |
| - | - | 729 | 503.2 | - | - | 0 | - |
| 4 | b | 4082 | 511.2 | 0.0002505 | 0.4899 | +1 | 4 |
| - | - | 927.4 | 512.2 | - | - | 0 | - |
| - | - | 1125 | 513.2 | - | - | 0 | - |
| - | - | 824.3 | 514.2 | - | - | 0 | - |
| - | - | 7968 | 521.2 | - | - | 0 | - |
| - | - | 2504 | 522.2 | - | - | 0 | - |
| - | - | 880.2 | 523.2 | - | - | 0 | - |
| - | - | 4918 | 528.2 | - | - | 0 | - |
| - | - | 815.9 | 528.3 | - | - | 0 | - |
| - | - | 1374 | 529.2 | - | - | 0 | - |
| - | - | 654.1 | 534.7 | - | - | 0 | - |
| - | - | 783.7 | 535.2 | - | - | 0 | - |
| - | - | 911.9 | 541.2 | - | - | 0 | - |
| - | - | 884.8 | 542.2 | - | - | 0 | - |
| 3 | y | 1299 | 557.7 | 0.00701 | 12.57 | +2 | 8 |
| - | - | 576.8 | 561 | - | - | 0 | - |
| 3 | y | 9578 | 566.7 | 0.006793 | 11.99 | +2 | 8 |
| - | - | 6501 | 567.2 | - | - | 0 | - |
| - | - | 2840 | 567.7 | - | - | 0 | - |
| - | - | 1.041E+04 | 592.2 | - | - | 0 | - |
| - | - | 3093 | 593.2 | - | - | 0 | - |
| - | - | 720.4 | 606.2 | - | - | 0 | - |
| 5 | b | 1411 | 608.2 | 0.00195 | 3.206 | +1 | 5 |
| - | - | 873.2 | 615.3 | - | - | 0 | - |
| 2 | y | 1956 | 617.2 | 0.008223 | 13.32 | +2 | 9 |
| - | - | 1855 | 617.7 | - | - | 0 | - |
| - | - | 1000 | 624.2 | - | - | 0 | - |
| 5 | b | 1431 | 626.2 | 0.001223 | 1.953 | +1 | 5 |
| - | - | 704.6 | 634.2 | - | - | 0 | - |
| - | - | 718.8 | 649.8 | - | - | 0 | - |
| - | - | 649.3 | 650.3 | - | - | 0 | - |
| - | - | 670.3 | 658.8 | - | - | 0 | - |
| - | - | 882.5 | 659.8 | - | - | 0 | - |
| - | - | 658.4 | 673.3 | - | - | 0 | - |
| 0 | Precursor | 3692 | 681.8 | 0.007191 | 10.55 | +2 | -1 |
| - | - | 2431 | 682.3 | - | - | 0 | - |
| - | - | 596.9 | 682.3 | - | - | 0 | - |
| - | - | 1338 | 682.8 | - | - | 0 | - |
| - | - | 1261 | 690.3 | - | - | 0 | - |
| 0 | Precursor | 1.047E+04 | 690.8 | 0.006364 | 9.214 | +2 | -1 |
| - | - | 8397 | 691.3 | - | - | 0 | - |
| - | - | 3719 | 691.8 | - | - | 0 | - |
| - | - | 735.5 | 691.9 | - | - | 0 | - |
| - | - | 1913 | 692.3 | - | - | 0 | - |
| - | - | 1104 | 695.3 | - | - | 0 | - |
| 6 | y | 9736 | 755.3 | 0.01377 | 18.23 | +1 | 5 |
| - | - | 3694 | 756.3 | - | - | 0 | - |
| - | - | 791.3 | 761.3 | - | - | 0 | - |
| 6 | b | 914.2 | 789.3 | 0.002357 | 2.986 | +1 | 6 |
| - | - | 809.4 | 804.3 | - | - | 0 | - |
| - | - | 4732 | 806.3 | - | - | 0 | - |
| - | - | 2004 | 807.3 | - | - | 0 | - |
| - | - | 1355 | 832.3 | - | - | 0 | - |
| 7 | b | 661.4 | 860.3 | 0.0007516 | 0.8737 | +1 | 7 |
| 5 | y | 1.066E+04 | 870.3 | 0.01344 | 15.44 | +1 | 6 |
| - | - | 6166 | 871.3 | - | - | 0 | - |
| - | - | 1879 | 872.3 | - | - | 0 | - |
| - | - | 1330 | 896.3 | - | - | 0 | - |
| - | - | 1440 | 903.3 | - | - | 0 | - |
| - | - | 6781 | 921.3 | - | - | 0 | - |
| - | - | 2894 | 922.3 | - | - | 0 | - |
| - | - | 836.5 | 967.3 | - | - | 0 | - |
| - | - | 897 | 968.3 | - | - | 0 | - |
| 4 | y | 1.98E+04 | 985.3 | 0.01298 | 13.18 | +1 | 7 |
| - | - | 1.158E+04 | 986.3 | - | - | 0 | - |
| - | - | 3347 | 987.3 | - | - | 0 | - |
| - | - | 888.3 | 988.3 | - | - | 0 | - |
| - | - | 665.3 | 995.3 | - | - | 0 | - |
| - | - | 685.2 | 1050 | - | - | 0 | - |
| - | - | 1.071E+04 | 1068 | - | - | 0 | - |
| - | - | 5715 | 1069 | - | - | 0 | - |
| - | - | 2291 | 1070 | - | - | 0 | - |
| 3 | y | 1110 | 1114 | 0.01422 | 12.76 | +1 | 8 |
| - | - | 1119 | 1115 | - | - | 0 | - |
| 3 | y | 3.192E+04 | 1132 | 0.01207 | 10.66 | +1 | 8 |
| - | - | 1.831E+04 | 1133 | - | - | 0 | - |
| - | - | 7365 | 1134 | - | - | 0 | - |
| - | - | 1091 | 1135 | - | - | 0 | - |
| - | - | 656.4 | 1151 | - | - | 0 | - |
| - | - | 1500 | 1169 | - | - | 0 | - |
| 2 | y | 805.9 | 1215 | 0.01659 | 13.65 | +1 | 9 |
| - | - | 841.5 | 1217 | - | - | 0 | - |
| 2 | y | 5171 | 1233 | 0.008219 | 6.664 | +1 | 9 |
| - | - | 4446 | 1234 | - | - | 0 | - |
| - | - | 1838 | 1235 | - | - | 0 | - |
| - | - | 956.1 | 1243 | - | - | 0 | - |
| - | - | 727.4 | 1509 | - | - | 0 | - |
| - | - | 696.6 | 1594 | - | - | 0 | - |
| - | - | 685.3 | 2402 | - | - | 0 | - |
| - | - | 747.9 | 2895 | - | - | 0 | - |

m/z Charge Intensity FragmentType MassShift Position
120.0809555053711 0 102535.95
121.08425903320312 0 6638.0537
121.8948974609375 0 363.9369
123.65139770507812 0 444.5333
129.10232543945312 0 2053.3794
130.0652313232422 0 601.8009
131.04876708984375 0 424.65308
132.08087158203125 0 601.4357
133.0608673095703 0 388.7
133.085693359375 0 679.2144
136.0701904296875 0 501.93433
136.0758056640625 0 9062.819
136.08692932128906 0 486.72693
137.07916259765625 0 858.6554
138.05517578125 0 629.4075
138.0663299560547 0 816.4424
146.0604705810547 0 1247.3495
148.876220703125 0 502.08463
148.89779663085938 0 524.02637
148.9046630859375 0 697.62994
148.91226196289062 0 666.47705
148.9196319580078 0 974.74493
148.92681884765625 0 1016.09973
148.9336700439453 0 1965.8398
148.9413299560547 0 3842.0088
148.95770263671875 0 5030.22
148.9655303955078 0 3292.4924
148.9728240966797 0 1290.5573
148.97998046875 0 1377.4458
148.98724365234375 0 1120.0294
148.99452209472656 0 913.8631
149.00152587890625 0 770.454
149.0088348388672 0 510.9633
149.01588439941406 0 466.1674
149.02325439453125 0 471.06766
149.04539489746094 0 426.60867
152.07102966308594 0 578.04443
152.26141357421875 0 394.58438
154.2385711669922 0 452.64627
155.09275817871094 0 1785.7303
156.07687377929688 0 3211.3792
158.05987548828125 0 593.11926
158.09645080566406 0 1970.3854
159.09176635742188 0 7053.6113
160.0951690673828 0 756.66266
160.11302185058594 0 589.45233
165.07664489746094 0 439.90012
166.06105041503906 0 1080.0184
167.09249877929688 0 717.4102
169.13343811035156 0 1119.2034
171.1493377685547 0 1942.9369
174.05511474609375 0 1662.2151
174.06764221191406 0 477.43954
175.08656311035156 0 1922.3613
176.08204650878906 0 1423.7781
176.10704040527344 0 45263.56
177.1024169921875 0 1680.885
177.11070251464844 0 4355.064
178.42376708984375 0 449.13272
181.16998291015625 0 908.0295
181.34791564941406 0 480.78564
185.07080078125 0 832.10345
185.27444458007812 0 456.35468
186.09136962890625 0 5485.582
187.09469604492188 0 601.4343
191.08200073242188 0 1040.8474
191.1181182861328 0 743.31726
193.1083526611328 0 3346.4062
195.08790588378906 0 804.71106
197.1284637451172 0 799.82544
199.14454650878906 0 1223.8547
200.899658203125 0 481.838
201.1229248046875 0 525.7065
202.05018615722656 0 713.59393
203.10220336914062 0 773.1374
203.11795043945312 0 2725.4685 a Water loss 1
204.07656860351562 0 1296.1466
209.1650390625 0 955.8422
213.1601104736328 0 569.59753
215.13922119140625 0 1024.3273
217.09695434570312 0 2438.822
220.06072998046875 0 1497.24
221.10330200195312 0 8391.109
221.12843322753906 0 57190.777 a 1
222.10699462890625 0 718.39636
222.13182067871094 0 6769.726
225.09933471679688 0 564.50934
227.10272216796875 0 1253.6083
231.06198120117188 0 968.795
231.1128387451172 0 13409.679 b Water loss 1
232.11651611328125 0 1169.0337
235.10784912109375 0 5263.6353
236.11175537109375 0 663.9948
237.08694458007812 0 2792.3247
237.1595916748047 0 692.6227
238.0912322998047 0 828.7028
239.11447143554688 0 822.41815
239.1500701904297 0 2112.434
243.133544921875 0 1346.6132
249.1233673095703 0 29393.527 b 1
250.12649536132812 0 3589.847
251.10250854492188 0 3161.1113
256.1764221191406 0 1015.5614
257.1061096191406 0 1007.93176
258.16015625 0 1323.0839
263.1023254394531 0 6508.1978
264.10748291015625 0 916.3927
268.129150390625 0 583.7804
274.13079833984375 0 1002.0167
279.0971374511719 0 2602.683
285.101806640625 0 2209.6438
286.155029296875 0 674.02637
292.1416931152344 0 638.2533
302.113525390625 0 787.29736
303.1128234863281 0 713.3179
306.1457824707031 0 734.1577
310.1527099609375 0 789.7658
316.1867980957031 0 874.2425
324.9699401855469 0 571.7421
328.1404113769531 0 1593.7037
333.15850830078125 0 1159.5945
339.1690368652344 0 614.8878
346.1404113769531 0 749.8525
348.11871337890625 0 668.56134
350.134521484375 0 1005.97095
356.1351623535156 0 3428.138
357.1379699707031 0 911.74365
358.1507568359375 0 1140.9945
364.15020751953125 0 2648.3567
374.14556884765625 0 28770.113
375.1485290527344 0 5000.9043
376.1525573730469 0 863.5784
378.1294250488281 0 6333.585
378.18109130859375 0 5548.319 b Water loss 2
379.13299560546875 0 1887.6298
379.1824035644531 0 1033.3706
382.14227294921875 0 777.55896
394.1246032714844 0 2481.7974
429.21307373046875 0 2192.9277
433.174072265625 0 598.6668
435.6573791503906 0 721.0555 y 4
440.1557922363281 0 1321.5841
457.18255615234375 0 9008.85
458.1873474121094 0 1320.5516
465.16131591796875 0 1681.1653
479.17626953125 0 1665.7454
480.1820983886719 0 696.8904
493.1817626953125 0 770.77167
503.1720275878906 0 729.01294
511.2184753417969 0 4081.6323 b 3
512.2208251953125 0 927.354
513.197998046875 0 1124.9146
514.1995849609375 0 824.2861
521.1809692382812 0 7967.823
522.1830444335938 0 2503.6309
523.1829833984375 0 880.2454
528.2191162109375 0 4917.7866
528.280029296875 0 815.93024
529.22265625 0 1373.7549
534.7128295898438 0 654.08954
535.2107543945312 0 783.6865
541.1940307617188 0 911.9209
542.23828125 0 884.75134
557.7000122070312 0 1298.6212 y Water loss 2
560.989013671875 0 576.77356
566.705078125 0 9577.772 y 2
567.2066040039062 0 6500.832
567.7062377929688 0 2839.964
592.2178344726562 0 10406.599
593.22119140625 0 3092.7073
606.2261352539062 0 720.41046
608.233154296875 0 1411.1448 b Water loss 4
615.3101196289062 0 873.1601
617.2303466796875 0 1956.1814 y 1
617.7302856445312 0 1854.574
624.2294921875 0 1000.1458
626.2444458007812 0 1431.2593 b 4
634.2280883789062 0 704.58826
649.76123046875 0 718.84204
650.2549438476562 0 649.3398
658.76416015625 0 670.3335
659.7662963867188 0 882.4654
673.255859375 0 658.38873
681.7582397460938 0 3692.3887 Precursor Water loss
682.2584228515625 0 2430.6343
682.322265625 0 596.88513
682.7576293945312 0 1337.6603
690.34326171875 0 1261.4874
690.7626953125 0 10474.128 Precursor
691.2711181640625 0 8396.973
691.7645874023438 0 3719.1677
691.8577270507812 0 735.4662
692.2826538085938 0 1913.1234
695.26513671875 0 1103.8042
755.28076171875 0 9735.514 y 5
756.283935546875 0 3693.9536
761.3087768554688 0 791.2906
789.306640625 0 914.1554 b 5
804.3364868164062 0 809.391
806.30859375 0 4732.148
807.3118896484375 0 2003.6753
832.3274536132812 0 1354.7883
860.3468627929688 0 661.4178 b 6
870.307373046875 0 10657.206 y 4
871.3099975585938 0 6165.9453
872.3103637695312 0 1878.8992
896.3202514648438 0 1329.9324
903.3265991210938 0 1440.4988
921.3365478515625 0 6781.3413
922.33642578125 0 2893.7456
967.3331298828125 0 836.5202
968.3211059570312 0 897.0093
985.3338623046875 0 19797.969 y 3
986.3372802734375 0 11583.264
987.3372802734375 0 3347.4644
988.3326416015625 0 888.26526
995.3157958984375 0 665.2912
1050.3798828125 0 685.20355
1068.4039306640625 0 10711.425
1069.405517578125 0 5715.1377
1070.4073486328125 0 2291.122
1114.3929443359375 0 1109.9924 y Water loss 2
1115.3948974609375 0 1118.533
1132.4013671875 0 31924.773 y 2
1133.40478515625 0 18311.014
1134.405029296875 0 7365.187
1135.3953857421875 0 1090.6259
1151.436767578125 0 656.4193
1169.44970703125 0 1500.3003
1215.4429931640625 0 805.91956 y Water loss 1
1217.4378662109375 0 841.4542
1233.4451904296875 0 5170.84 y 1
1234.4501953125 0 4446.167
1235.452392578125 0 1837.715
1243.4334716796875 0 956.1158
1509.2047119140625 0 727.3564
1593.90625 0 696.5536
2402.06005859375 0 685.31757
2894.621337890625 0 747.8861

Spectrum Details

|  |  |
| --- | --- |
| Matched peaks? Matched peaks The total absolute number of peaks matched. Additionally in brackets the total fraction of peaks matched and the total number of peaks is shown. | 23 (9.62% of 239) |
| FDR? FDR The false discovery rate estimated for this peptide. It is calculated by matching all theoretical fragments with a non-integer shift with the raw peaks for this spectrum. This is done with 40 different shifts. The resulting percentage is the average number of annotated peaks over the number of annotated peaks with the correct spectrum. | 0.41% |
| Satellite FDR? Satellite FDR See the FDR for details on its calculation. This satellite ion specific FDR only contains the satellite ions (d/w) for I/L/J positions. | - |
| PSM Score? PSM Score The PSM Score as given by Hecklib to this annotated spectrum. It is shown with three significant figures. | 221 |

#### Spectrum 9262? Spectrum 9262 The raw spectrum of this peptide as annotated by Hecklib. The fragments are coloured according to ion type (see legend). Any peaks with a star '\*' as text can be hovered over to see the full details, first the ion type second the mass shift type. By hovering over the amino acids in the peptide or ions in the legend the corresponding peaks are highlighted. By toggling the 'Unassigned' label you can turn the background (unassigned) peaks on or off in the plot. By updating the slider in the Ion legend you can update the spectrum to only show the top X% of the peaks with labels. The top X% means any peak that is within X% of the highest intensity. By dragging in the spectrum you can zoom in to a specific part of the spectrum and use 'Zoom Out' to get back to the original zoom level. The annotation of the spectrum is based on the given sequence in the peptides file and is done with different software so inconsistencies are likely. The peaks are annotated based on the given sequence, with 20 ppm tolerance.

Copy Data

##### Spectrum 9262 (TSV)

###### Preview

```
Loading example...
```

*Click on the button to copy the data to your clipboard.*

Mz MinMz MaxIntensity Max

WidthHeightPeptide font sizePeptide stroke widthSpectrum font sizeSpectrum stroke widthCompact peptide

Ion legend

wxyz

abcd

OtherUnassignedIon^Charge^_Position_Show for top:%

FTFDDYAMHW

08.47e+31.69e+42.54e+43.39e+4

Zoom Out

y^+1^_1_a^+1^_2_b^+1^_2_b^+1^_2_y^+1^_2_b^+1^_3_b^+2^_8_y^+1^_3_b^+1^_4_y^+2^_8_b^+1^_5_\*\*y^+1^_5_y^+1^_6_y^+1^_7_y^+1^_8_b^+1^_9_y^+1^_9_

0830166024903320

Fragment Matches Table

Show background peaks

| Position | Ion type | Intensity | mz Theoretical | mz Error (Th) | mz Error (ppm) | Charge | Series Number |
| --- | --- | --- | --- | --- | --- | --- | --- |
| - | - | 1979 | 120.1 | - | - | 0 | - |
| - | - | 3.356E+04 | 120.1 | - | - | 0 | - |
| - | - | 2795 | 121.1 | - | - | 0 | - |
| - | - | 412.9 | 126.5 | - | - | 0 | - |
| - | - | 2071 | 129.1 | - | - | 0 | - |
| - | - | 3322 | 130.1 | - | - | 0 | - |
| - | - | 771.8 | 130.1 | - | - | 0 | - |
| - | - | 1426 | 133.1 | - | - | 0 | - |
| - | - | 7869 | 136.1 | - | - | 0 | - |
| - | - | 913.2 | 137.1 | - | - | 0 | - |
| - | - | 580.7 | 138.1 | - | - | 0 | - |
| - | - | 553.4 | 138.1 | - | - | 0 | - |
| - | - | 453.4 | 140.1 | - | - | 0 | - |
| - | - | 461 | 142.6 | - | - | 0 | - |
| - | - | 869.3 | 146.1 | - | - | 0 | - |
| - | - | 504.6 | 147 | - | - | 0 | - |
| - | - | 6140 | 147.1 | - | - | 0 | - |
| - | - | 498.4 | 148.9 | - | - | 0 | - |
| - | - | 2020 | 149 | - | - | 0 | - |
| - | - | 424.7 | 149.6 | - | - | 0 | - |
| - | - | 498.3 | 150.1 | - | - | 0 | - |
| - | - | 1190 | 152.1 | - | - | 0 | - |
| - | - | 668 | 153.1 | - | - | 0 | - |
| - | - | 449.3 | 153.4 | - | - | 0 | - |
| - | - | 1610 | 155.1 | - | - | 0 | - |
| - | - | 1059 | 156.1 | - | - | 0 | - |
| - | - | 594.3 | 157.1 | - | - | 0 | - |
| - | - | 988.7 | 158.1 | - | - | 0 | - |
| - | - | 586.1 | 159.1 | - | - | 0 | - |
| - | - | 1727 | 159.1 | - | - | 0 | - |
| - | - | 938.1 | 162.1 | - | - | 0 | - |
| - | - | 3850 | 167.1 | - | - | 0 | - |
| - | - | 727.1 | 169.1 | - | - | 0 | - |
| - | - | 517.1 | 173.1 | - | - | 0 | - |
| - | - | 4039 | 173.4 | - | - | 0 | - |
| - | - | 4323 | 175.1 | - | - | 0 | - |
| - | - | 1.178E+04 | 176.1 | - | - | 0 | - |
| - | - | 582.1 | 177.1 | - | - | 0 | - |
| - | - | 1950 | 177.1 | - | - | 0 | - |
| - | - | 670.1 | 181.1 | - | - | 0 | - |
| - | - | 819.3 | 182.1 | - | - | 0 | - |
| - | - | 486.2 | 183.1 | - | - | 0 | - |
| - | - | 1465 | 183.1 | - | - | 0 | - |
| - | - | 540.7 | 185.1 | - | - | 0 | - |
| - | - | 1352 | 186.1 | - | - | 0 | - |
| - | - | 516.2 | 187.1 | - | - | 0 | - |
| - | - | 1628 | 191.1 | - | - | 0 | - |
| - | - | 514.2 | 193.1 | - | - | 0 | - |
| - | - | 1026 | 193.1 | - | - | 0 | - |
| - | - | 469.4 | 195.1 | - | - | 0 | - |
| - | - | 913.5 | 197.1 | - | - | 0 | - |
| - | - | 2951 | 199.2 | - | - | 0 | - |
| - | - | 1153 | 201.1 | - | - | 0 | - |
| - | - | 857.9 | 203.1 | - | - | 0 | - |
| - | - | 1176 | 203.1 | - | - | 0 | - |
| - | - | 535.6 | 207.1 | - | - | 0 | - |
| - | - | 593.9 | 215.1 | - | - | 0 | - |
| - | - | 759.5 | 217.1 | - | - | 0 | - |
| - | - | 1282 | 219.1 | - | - | 0 | - |
| 10 | y | 1753 | 221.1 | 0.002834 | 12.82 | +1 | 1 |
| - | - | 1008 | 221.1 | - | - | 0 | - |
| - | - | 1757 | 221.1 | - | - | 0 | - |
| 2 | a | 1.597E+04 | 221.1 | 0.0002079 | 0.9401 | +1 | 2 |
| - | - | 499.9 | 222.1 | - | - | 0 | - |
| - | - | 550.3 | 222.1 | - | - | 0 | - |
| - | - | 1853 | 222.1 | - | - | 0 | - |
| - | - | 6506 | 225 | - | - | 0 | - |
| - | - | 1071 | 226 | - | - | 0 | - |
| - | - | 3473 | 226.1 | - | - | 0 | - |
| - | - | 561.2 | 227 | - | - | 0 | - |
| - | - | 959.7 | 227.2 | - | - | 0 | - |
| - | - | 471.1 | 228.1 | - | - | 0 | - |
| - | - | 559.9 | 230.3 | - | - | 0 | - |
| 2 | b | 4239 | 231.1 | 0.0001567 | 0.6778 | +1 | 2 |
| - | - | 587.6 | 233.1 | - | - | 0 | - |
| - | - | 680.6 | 233.2 | - | - | 0 | - |
| - | - | 1622 | 235.1 | - | - | 0 | - |
| - | - | 8556 | 239.1 | - | - | 0 | - |
| - | - | 3914 | 239.2 | - | - | 0 | - |
| - | - | 506.3 | 239.6 | - | - | 0 | - |
| - | - | 1153 | 240.1 | - | - | 0 | - |
| - | - | 454.7 | 240.6 | - | - | 0 | - |
| 2 | b | 7389 | 249.1 | 0.0003036 | 1.219 | +1 | 2 |
| - | - | 1954 | 250.1 | - | - | 0 | - |
| - | - | 1016 | 251.1 | - | - | 0 | - |
| - | - | 1287 | 256.2 | - | - | 0 | - |
| - | - | 538.7 | 259.1 | - | - | 0 | - |
| - | - | 1340 | 260.2 | - | - | 0 | - |
| - | - | 558.5 | 261.2 | - | - | 0 | - |
| - | - | 2415 | 263.1 | - | - | 0 | - |
| - | - | 706.4 | 279.1 | - | - | 0 | - |
| - | - | 1850 | 285 | - | - | 0 | - |
| - | - | 711.6 | 290.1 | - | - | 0 | - |
| - | - | 2072 | 295.1 | - | - | 0 | - |
| - | - | 3145 | 296.2 | - | - | 0 | - |
| - | - | 663 | 297.1 | - | - | 0 | - |
| - | - | 846.2 | 297.2 | - | - | 0 | - |
| - | - | 4455 | 299.1 | - | - | 0 | - |
| - | - | 940.9 | 300.1 | - | - | 0 | - |
| - | - | 950.3 | 314.2 | - | - | 0 | - |
| - | - | 611 | 316.2 | - | - | 0 | - |
| - | - | 697.3 | 325.2 | - | - | 0 | - |
| - | - | 586.4 | 340.1 | - | - | 0 | - |
| - | - | 814 | 345.1 | - | - | 0 | - |
| - | - | 1326 | 350.1 | - | - | 0 | - |
| - | - | 3861 | 355.1 | - | - | 0 | - |
| - | - | 4160 | 356.1 | - | - | 0 | - |
| - | - | 859 | 357.1 | - | - | 0 | - |
| 9 | y | 4295 | 358.1 | 0.005301 | 14.8 | +1 | 2 |
| - | - | 5872 | 359 | - | - | 0 | - |
| - | - | 864 | 359.2 | - | - | 0 | - |
| - | - | 2332 | 378.1 | - | - | 0 | - |
| 3 | b | 1499 | 378.2 | 0.0001573 | 0.4159 | +1 | 3 |
| - | - | 7279 | 429.1 | - | - | 0 | - |
| - | - | 1128 | 433.3 | - | - | 0 | - |
| - | - | 1038 | 441.2 | - | - | 0 | - |
| - | - | 2782 | 444.2 | - | - | 0 | - |
| - | - | 554 | 457.2 | - | - | 0 | - |
| - | - | 702.1 | 461.3 | - | - | 0 | - |
| - | - | 574.5 | 463.2 | - | - | 0 | - |
| 8 | b | 573.3 | 495.2 | 0.0004965 | 1.003 | +2 | 8 |
| - | - | 706.3 | 498.2 | - | - | 0 | - |
| 8 | y | 790.7 | 505.2 | 0.009698 | 19.2 | +1 | 3 |
| 4 | b | 1291 | 511.2 | 0.00199 | 3.893 | +1 | 4 |
| - | - | 926.7 | 512.2 | - | - | 0 | - |
| - | - | 722.5 | 558.2 | - | - | 0 | - |
| 3 | y | 1566 | 558.7 | 0.005944 | 10.64 | +2 | 8 |
| - | - | 655.4 | 559.2 | - | - | 0 | - |
| - | - | 1185 | 574.2 | - | - | 0 | - |
| - | - | 1634 | 576.2 | - | - | 0 | - |
| - | - | 684.7 | 599.3 | - | - | 0 | - |
| 5 | b | 1016 | 609.2 | 0.01196 | 19.63 | +1 | 5 |
| - | - | 1358 | 627.3 | - | - | 0 | - |
| - | - | 635.6 | 664.6 | - | - | 0 | - |
| - | - | 631.2 | 672.6 | - | - | 0 | - |
| 0 | Precursor | 927.4 | 673.8 | 0.002435 | 3.615 | +2 | -1 |
| - | - | 912.6 | 674.3 | - | - | 0 | - |
| - | - | 1055 | 675.3 | - | - | 0 | - |
| - | - | 796.8 | 677.2 | - | - | 0 | - |
| - | - | 1226 | 681.8 | - | - | 0 | - |
| - | - | 647.1 | 682.3 | - | - | 0 | - |
| - | - | 819.2 | 682.3 | - | - | 0 | - |
| 0 | Precursor | 2137 | 682.8 | 0.003745 | 5.485 | +2 | -1 |
| - | - | 1273 | 683.3 | - | - | 0 | - |
| - | - | 604.6 | 683.3 | - | - | 0 | - |
| - | - | 1702 | 684.2 | - | - | 0 | - |
| - | - | 958.2 | 712.4 | - | - | 0 | - |
| - | - | 827.5 | 714.4 | - | - | 0 | - |
| - | - | 612.8 | 737.3 | - | - | 0 | - |
| 6 | y | 1238 | 739.3 | 0.01012 | 13.68 | +1 | 5 |
| - | - | 1203 | 740.3 | - | - | 0 | - |
| - | - | 825 | 785.4 | - | - | 0 | - |
| - | - | 1046 | 794.4 | - | - | 0 | - |
| - | - | 1330 | 852.3 | - | - | 0 | - |
| 5 | y | 1927 | 854.3 | 0.009541 | 11.17 | +1 | 6 |
| - | - | 935.4 | 855.3 | - | - | 0 | - |
| - | - | 669 | 872.4 | - | - | 0 | - |
| - | - | 4772 | 886.4 | - | - | 0 | - |
| - | - | 2460 | 887.4 | - | - | 0 | - |
| - | - | 994.1 | 888.4 | - | - | 0 | - |
| - | - | 612.7 | 904.3 | - | - | 0 | - |
| - | - | 945.6 | 905.3 | - | - | 0 | - |
| - | - | 702.9 | 906.3 | - | - | 0 | - |
| - | - | 1669 | 967.3 | - | - | 0 | - |
| - | - | 873.8 | 968.3 | - | - | 0 | - |
| 4 | y | 3176 | 969.3 | 0.01092 | 11.26 | +1 | 7 |
| - | - | 1839 | 970.3 | - | - | 0 | - |
| - | - | 821 | 1020 | - | - | 0 | - |
| - | - | 642.9 | 1025 | - | - | 0 | - |
| - | - | 827.5 | 1050 | - | - | 0 | - |
| - | - | 1763 | 1052 | - | - | 0 | - |
| - | - | 2421 | 1114 | - | - | 0 | - |
| - | - | 1611 | 1115 | - | - | 0 | - |
| 3 | y | 5624 | 1116 | 0.01099 | 9.84 | +1 | 8 |
| - | - | 3075 | 1117 | - | - | 0 | - |
| - | - | 2124 | 1118 | - | - | 0 | - |
| 9 | b | 775.9 | 1144 | 0.0005275 | 0.461 | +1 | 9 |
| - | - | 773.4 | 1202 | - | - | 0 | - |
| - | - | 718.2 | 1211 | - | - | 0 | - |
| - | - | 618.1 | 1215 | - | - | 0 | - |
| 2 | y | 1105 | 1217 | 0.01079 | 8.865 | +1 | 9 |
| - | - | 649.3 | 1218 | - | - | 0 | - |
| - | - | 624.5 | 1240 | - | - | 0 | - |
| - | - | 693.3 | 3287 | - | - | 0 | - |

m/z Charge Intensity FragmentType MassShift Position
120.06563568115234 0 1978.7825
120.08096313476562 0 33560.926
121.08426666259766 0 2794.9111
126.46712493896484 0 412.9001
129.10247802734375 0 2071.428
130.0500030517578 0 3321.6128
130.06541442871094 0 771.77155
133.0861358642578 0 1426.3638
136.07583618164062 0 7868.518
137.07901000976562 0 913.18494
138.06689453125 0 580.70416
138.0916748046875 0 553.42126
140.0816650390625 0 453.42337
142.63027954101562 0 461.0089
146.060302734375 0 869.2529
147.04367065429688 0 504.58972
147.0766143798828 0 6139.7153
148.94737243652344 0 498.39212
149.04490661621094 0 2019.9987
149.61819458007812 0 424.7475
150.05532836914062 0 498.3429
152.07077026367188 0 1190.2181
153.06661987304688 0 668.04126
153.37509155273438 0 449.2566
155.08177185058594 0 1609.714
156.0769500732422 0 1059.4653
157.09727478027344 0 594.30585
158.06016540527344 0 988.6577
159.07662963867188 0 586.1281
159.09165954589844 0 1727.2388
162.05502319335938 0 938.10657
167.0555877685547 0 3850.4607
169.13348388671875 0 727.05536
173.12893676757812 0 517.11163
173.44041442871094 0 4039.1594
175.08682250976562 0 4323.269
176.10719299316406 0 11779.579
177.1025390625 0 582.1052
177.11111450195312 0 1950.1766
181.10841369628906 0 670.14594
182.08160400390625 0 819.30835
183.08639526367188 0 486.18906
183.11325073242188 0 1465.0814
185.10260009765625 0 540.6986
186.09158325195312 0 1351.6814
187.14447021484375 0 516.1754
191.08177185058594 0 1628.3276
193.0974884033203 0 514.17456
193.1083984375 0 1025.5316
195.11373901367188 0 469.42026
197.12875366210938 0 913.47705
199.1806640625 0 2951.0156
201.12388610839844 0 1152.6753
203.0660858154297 0 857.89716
203.08163452148438 0 1175.5222
207.07749938964844 0 535.63794
215.13888549804688 0 593.8883
217.0974578857422 0 759.53467
219.1342010498047 0 1282.1488
221.08432006835938 0 1753.4658 y 9
221.09375 0 1008.1986
221.1039276123047 0 1756.7487
221.128662109375 0 15969.452 a 1
222.0842742919922 0 499.90692
222.10726928710938 0 550.28467
222.13201904296875 0 1852.9222
225.0431671142578 0 6506.0977
226.043212890625 0 1071.4861
226.1187286376953 0 3473.2012
227.03981018066406 0 561.19574
227.17538452148438 0 959.6579
228.1343231201172 0 471.14337
230.28594970703125 0 559.87354
231.1129608154297 0 4239.1772 b Water loss 1
233.12887573242188 0 587.619
233.1658172607422 0 680.6106
235.10818481445312 0 1621.6093
239.09523010253906 0 8555.529
239.1505126953125 0 3914.0151
239.5657958984375 0 506.27933
240.09548950195312 0 1153.1395
240.564697265625 0 454.71335
249.12367248535156 0 7389.1274 b 1
250.12716674804688 0 1953.927
251.1029815673828 0 1016.39453
256.1767272949219 0 1286.6881
259.09222412109375 0 538.65314
260.1605529785156 0 1339.656
261.1587219238281 0 558.46014
263.1028747558594 0 2415.4653
279.0980224609375 0 706.3632
285.0103454589844 0 1849.7822
290.1152648925781 0 711.64154
295.1033630371094 0 2071.6396
296.1971130371094 0 3144.5166
297.100830078125 0 662.95856
297.19512939453125 0 846.1736
299.06201171875 0 4455.032
300.0628356933594 0 940.90204
314.2080078125 0 950.2924
316.15185546875 0 610.98706
325.1874694824219 0 697.2614
340.1410217285156 0 586.4432
345.1462097167969 0 814.0453
350.1341247558594 0 1326.19
355.070068359375 0 3860.5093
356.135986328125 0 4159.925
357.1401062011719 0 859.01587
358.1513671875 0 4295.332 y 8
359.02850341796875 0 5871.7266
359.1533203125 0 863.9688
378.1297607421875 0 2331.696
378.1810607910156 0 1498.6825 b Water loss 2
429.08929443359375 0 7279.3247
433.2789306640625 0 1128.163
441.18756103515625 0 1038.3053
444.24578857421875 0 2782.1335
457.1689453125 0 554.0413
461.2762756347656 0 702.0937
463.2190246582031 0 574.47284
495.1861572265625 0 573.33527 b Water loss 7
498.219970703125 0 706.2771
505.1862487792969 0 790.6541 y 7
511.21673583984375 0 1290.9177 b 3
512.2251586914062 0 926.7004
558.2057495117188 0 722.4931
558.709228515625 0 1566.0594 y 2
559.2119140625 0 655.3543
574.2071533203125 0 1184.903
576.2266845703125 0 1633.6439
599.2647705078125 0 684.70135
609.2310791015625 0 1016.1267 b Ammonia loss 4
627.263916015625 0 1357.8749
664.5741577148438 0 635.5903
672.6499633789062 0 631.2253
673.7584838867188 0 927.40875 Precursor Water loss
674.2661743164062 0 912.63416
675.2892456054688 0 1055.0901
677.1676635742188 0 796.7903
681.7556762695312 0 1226.2845
682.259521484375 0 647.1187
682.337646484375 0 819.2411
682.7650756835938 0 2137.4011 Precursor
683.2673950195312 0 1273.289
683.333740234375 0 604.62366
684.2045288085938 0 1702.2938
712.36376953125 0 958.1733
714.3793334960938 0 827.51166
737.2769775390625 0 612.76764
739.287109375 0 1238.3766 y 5
740.3492431640625 0 1203.3265
785.4193725585938 0 825.03265
794.3831787109375 0 1045.654
852.3055419921875 0 1329.9073
854.3134765625 0 1926.6084 y 4
855.3158569335938 0 935.35956
872.4326171875 0 668.9853
886.4174194335938 0 4771.895
887.4197387695312 0 2460.2634
888.4192504882812 0 994.11584
904.3300170898438 0 612.74677
905.3445434570312 0 945.61926
906.3482666015625 0 702.91345
967.328125 0 1669.4783
968.3295288085938 0 873.8294
969.341796875 0 3176.3235 y 3
970.3433837890625 0 1838.5143
1019.5123291015625 0 821.022
1024.8892822265625 0 642.8901
1050.3961181640625 0 827.5114
1052.416015625 0 1763.2151
1114.39697265625 0 2420.8232
1115.39453125 0 1610.9227
1116.4102783203125 0 5624.094 y 2
1117.4130859375 0 3074.7422
1118.4150390625 0 2123.5107
1144.43603515625 0 775.90466 b 8
1201.6204833984375 0 773.4287
1210.5782470703125 0 718.1679
1215.4739990234375 0 618.1409
1217.457763671875 0 1104.6016 y 1
1218.4501953125 0 649.2591
1240.3828125 0 624.4926
3286.69921875 0 693.3336

Spectrum Details

|  |  |
| --- | --- |
| Matched peaks? Matched peaks The total absolute number of peaks matched. Additionally in brackets the total fraction of peaks matched and the total number of peaks is shown. | 19 (10.33% of 184) |
| FDR? FDR The false discovery rate estimated for this peptide. It is calculated by matching all theoretical fragments with a non-integer shift with the raw peaks for this spectrum. This is done with 40 different shifts. The resulting percentage is the average number of annotated peaks over the number of annotated peaks with the correct spectrum. | 0.50% |
| Satellite FDR? Satellite FDR See the FDR for details on its calculation. This satellite ion specific FDR only contains the satellite ions (d/w) for I/L/J positions. | - |
| PSM Score? PSM Score The PSM Score as given by Hecklib to this annotated spectrum. It is shown with three significant figures. | 186 |

#### Reverse Lookup? Reverse Lookup All places where this read could be placed.

| Group | Segment | Template | Template Part | Read Part | Score | Unique |
| --- | --- | --- | --- | --- | --- | --- |
| Homo sapiens Heavy Chain | IGHV | IGHV3-9 | [26..36] | [0..10] | 80 | True |

| Recombined | Template Part | Read Part | Score | Unique |
| --- | --- | --- | --- | --- |
| REC-0-1 | [26..36] | [0..10] | 80 | True |

#### Meta Information from Multiple reads

##### Number of combined reads

28

##### Intensity

0.7938

##### TotalArea

3.906E+08

#### Positional Score

Copy Data

##### Positional Score (TSV)

###### Preview

```
Loading example...
```

*Click on the button to copy the data to your clipboard.*

100123456789

Label Value
"0" 0.927
"1" 0.927
"2" 0.926
"3" 0.927
"4" 0.926
"5" 0.913
"6" 0.903
"7" 0.904
"8" 0.874
"9" 0.886

#### Meta Information from PEAKS

##### Scan Identifier

F2:9743

##### Original sequence

F

T

F

D

D

Y

A

M

+15.99

H

W

##### Posttranslational Modifications

Oxidation (M)

##### Source File

D:\separate\_stitch\_analyses\xle-disambiguation\raw\20210323\_F1\_UM1\_Peng0013\_SA\_F59\_ingel\_3ug\_TL.raw

##### Fraction

2

##### Scan Feature

F2:11435

##### De Novo Score

99

##### ConfidenceScore

99

### m/z

674.7697

##### Mass

1347.5229

##### Charge

2

##### Retention Time

54.69

##### Predicted Retention Time

-

##### Area

5.238E+07

##### Parts Per Million

1.3

##### Fragmentation mode

HCD

##### Originating file

01 D:\separate\_stitch\_analyses\xle-disambiguation\20210325\_F59\_3ug\_DENOVO\_12.csv

#### Meta Information from PEAKS

##### Scan Identifier

F2:9689

##### Original sequence

F

T

F

D

D

Y

A

M

+15.99

H

W

##### Posttranslational Modifications

Oxidation (M)

##### Source File

D:\separate\_stitch\_analyses\xle-disambiguation\raw\20210323\_F1\_UM1\_Peng0013\_SA\_F59\_ingel\_3ug\_TL.raw

##### Fraction

2

##### Scan Feature

F2:11435

##### De Novo Score

99

##### ConfidenceScore

99

### m/z

674.7697

##### Mass

1347.5229

##### Charge

2

##### Retention Time

54.69

##### Predicted Retention Time

-

##### Area

5.238E+07

##### Parts Per Million

1.3

##### Fragmentation mode

HCD

##### Originating file

01 D:\separate\_stitch\_analyses\xle-disambiguation\20210325\_F59\_3ug\_DENOVO\_12.csv

#### Meta Information from PEAKS

##### Scan Identifier

F2:8424

##### Original sequence

F

T

F

D

D

Y

A

M

+15.99

H

W

+15.99

##### Posttranslational Modifications

Oxidation (M); Oxidation (HW)

##### Source File

D:\separate\_stitch\_analyses\xle-disambiguation\raw\20210323\_F1\_UM1\_Peng0013\_SA\_F59\_ingel\_3ug\_TL.raw

##### Fraction

2

##### Scan Feature

-

##### De Novo Score

99

##### ConfidenceScore

99

### m/z

682.7671

##### Mass

1363.5178

##### Charge

2

##### Retention Time

47.46

##### Predicted Retention Time

-

##### Area

0

##### Parts Per Million

1.3

##### Fragmentation mode

HCD

##### Originating file

01 D:\separate\_stitch\_analyses\xle-disambiguation\20210325\_F59\_3ug\_DENOVO\_12.csv

#### Meta Information from PEAKS

##### Scan Identifier

F2:8805

##### Original sequence

F

T

F

D

D

Y

A

M

+15.99

H

W

+15.99

##### Posttranslational Modifications

Oxidation (M); Oxidation (HW)

##### Source File

D:\separate\_stitch\_analyses\xle-disambiguation\raw\20210323\_F1\_UM1\_Peng0013\_SA\_F59\_ingel\_3ug\_TL.raw

##### Fraction

2

##### Scan Feature

F2:11788

##### De Novo Score

99

##### ConfidenceScore

99

### m/z

682.7664

##### Mass

1363.5178

##### Charge

2

##### Retention Time

49.24

##### Predicted Retention Time

-

##### Area

9.51E+06

##### Parts Per Million

0.3

##### Fragmentation mode

HCD

##### Originating file

01 D:\separate\_stitch\_analyses\xle-disambiguation\20210325\_F59\_3ug\_DENOVO\_12.csv

#### Meta Information from PEAKS

##### Scan Identifier

F2:9795

##### Original sequence

F

T

F

D

D

Y

A

M

+15.99

H

W

##### Posttranslational Modifications

Oxidation (M)

##### Source File

D:\separate\_stitch\_analyses\xle-disambiguation\raw\20210323\_F1\_UM1\_Peng0013\_SA\_F59\_ingel\_3ug\_TL.raw

##### Fraction

2

##### Scan Feature

F2:11435

##### De Novo Score

99

##### ConfidenceScore

99

### m/z

674.7697

##### Mass

1347.5229

##### Charge

2

##### Retention Time

54.69

##### Predicted Retention Time

-

##### Area

5.238E+07

##### Parts Per Million

1.3

##### Fragmentation mode

ETHCD

##### Originating file

01 D:\separate\_stitch\_analyses\xle-disambiguation\20210325\_F59\_3ug\_DENOVO\_12.csv

#### Meta Information from PEAKS

##### Scan Identifier

F2:8586

##### Original sequence

F

T

F

D

D

Y

A

M

+15.99

H

W

+15.99

##### Posttranslational Modifications

Oxidation (M); Oxidation (HW)

##### Source File

D:\separate\_stitch\_analyses\xle-disambiguation\raw\20210323\_F1\_UM1\_Peng0013\_SA\_F59\_ingel\_3ug\_TL.raw

##### Fraction

2

##### Scan Feature

-

##### De Novo Score

99

##### ConfidenceScore

99

### m/z

682.7668

##### Mass

1363.5178

##### Charge

2

##### Retention Time

48.35

##### Predicted Retention Time

-

##### Area

0

##### Parts Per Million

1

##### Fragmentation mode

HCD

##### Originating file

01 D:\separate\_stitch\_analyses\xle-disambiguation\20210325\_F59\_3ug\_DENOVO\_12.csv

#### Meta Information from PEAKS

##### Scan Identifier

F2:9848

##### Original sequence

F

T

F

D

D

Y

A

M

+15.99

H

W

##### Posttranslational Modifications

Oxidation (M)

##### Source File

D:\separate\_stitch\_analyses\xle-disambiguation\raw\20210323\_F1\_UM1\_Peng0013\_SA\_F59\_ingel\_3ug\_TL.raw

##### Fraction

2

##### Scan Feature

F2:11435

##### De Novo Score

99

##### ConfidenceScore

99

### m/z

674.7697

##### Mass

1347.5229

##### Charge

2

##### Retention Time

54.69

##### Predicted Retention Time

-

##### Area

5.238E+07

##### Parts Per Million

1.3

##### Fragmentation mode

HCD

##### Originating file

01 D:\separate\_stitch\_analyses\xle-disambiguation\20210325\_F59\_3ug\_DENOVO\_12.csv

#### Meta Information from PEAKS

##### Scan Identifier

F2:9898

##### Original sequence

F

T

F

D

D

Y

A

M

+15.99

H

W

##### Posttranslational Modifications

Oxidation (M)

##### Source File

D:\separate\_stitch\_analyses\xle-disambiguation\raw\20210323\_F1\_UM1\_Peng0013\_SA\_F59\_ingel\_3ug\_TL.raw

##### Fraction

2

##### Scan Feature

F2:11435

##### De Novo Score

99

##### ConfidenceScore

99

### m/z

674.7697

##### Mass

1347.5229

##### Charge

2

##### Retention Time

54.69

##### Predicted Retention Time

-

##### Area

5.238E+07

##### Parts Per Million

1.3

##### Fragmentation mode

ETHCD

##### Originating file

01 D:\separate\_stitch\_analyses\xle-disambiguation\20210325\_F59\_3ug\_DENOVO\_12.csv

#### Meta Information from PEAKS

##### Scan Identifier

F2:8533

##### Original sequence

F

T

F

D

D

Y

A

M

+15.99

H

W

+15.99

##### Posttranslational Modifications

Oxidation (M); Oxidation (HW)

##### Source File

D:\separate\_stitch\_analyses\xle-disambiguation\raw\20210323\_F1\_UM1\_Peng0013\_SA\_F59\_ingel\_3ug\_TL.raw

##### Fraction

2

##### Scan Feature

-

##### De Novo Score

99

##### ConfidenceScore

99

### m/z

682.767

##### Mass

1363.5178

##### Charge

2

##### Retention Time

48.05

##### Predicted Retention Time

-

##### Area

0

##### Parts Per Million

1.1

##### Fragmentation mode

HCD

##### Originating file

01 D:\separate\_stitch\_analyses\xle-disambiguation\20210325\_F59\_3ug\_DENOVO\_12.csv

#### Meta Information from PEAKS

##### Scan Identifier

F2:8477

##### Original sequence

F

T

F

D

D

Y

A

M

+15.99

H

W

+15.99

##### Posttranslational Modifications

Oxidation (M); Oxidation (HW)

##### Source File

D:\separate\_stitch\_analyses\xle-disambiguation\raw\20210323\_F1\_UM1\_Peng0013\_SA\_F59\_ingel\_3ug\_TL.raw

##### Fraction

2

##### Scan Feature

-

##### De Novo Score

99

##### ConfidenceScore

99

### m/z

682.767

##### Mass

1363.5178

##### Charge

2

##### Retention Time

47.75

##### Predicted Retention Time

-

##### Area

0

##### Parts Per Million

1.1

##### Fragmentation mode

HCD

##### Originating file

01 D:\separate\_stitch\_analyses\xle-disambiguation\20210325\_F59\_3ug\_DENOVO\_12.csv

#### Meta Information from PEAKS

##### Scan Identifier

F2:8649

##### Original sequence

F

T

F

D

D

Y

A

M

+15.99

H

W

+15.99

##### Posttranslational Modifications

Oxidation (M); Oxidation (HW)

##### Source File

D:\separate\_stitch\_analyses\xle-disambiguation\raw\20210323\_F1\_UM1\_Peng0013\_SA\_F59\_ingel\_3ug\_TL.raw

##### Fraction

2

##### Scan Feature

-

##### De Novo Score

99

##### ConfidenceScore

99

### m/z

682.7672

##### Mass

1363.5178

##### Charge

2

##### Retention Time

48.69

##### Predicted Retention Time

-

##### Area

0

##### Parts Per Million

1.4

##### Fragmentation mode

ETHCD

##### Originating file

01 D:\separate\_stitch\_analyses\xle-disambiguation\20210325\_F59\_3ug\_DENOVO\_12.csv

#### Meta Information from PEAKS

##### Scan Identifier

F2:9949

##### Original sequence

F

T

F

D

D

Y

A

M

+15.99

H

W

##### Posttranslational Modifications

Oxidation (M)

##### Source File

D:\separate\_stitch\_analyses\xle-disambiguation\raw\20210323\_F1\_UM1\_Peng0013\_SA\_F59\_ingel\_3ug\_TL.raw

##### Fraction

2

##### Scan Feature

F2:11435

##### De Novo Score

99

##### ConfidenceScore

99

### m/z

674.7697

##### Mass

1347.5229

##### Charge

2

##### Retention Time

54.69

##### Predicted Retention Time

-

##### Area

5.238E+07

##### Parts Per Million

1.3

##### Fragmentation mode

ETHCD

##### Originating file

01 D:\separate\_stitch\_analyses\xle-disambiguation\20210325\_F59\_3ug\_DENOVO\_12.csv

#### Meta Information from PEAKS

##### Scan Identifier

F2:8969

##### Original sequence

F

T

F

D

D

Y

A

M

+15.99

H

W

+15.99

##### Posttranslational Modifications

Oxidation (M); Oxidation (HW)

##### Source File

D:\separate\_stitch\_analyses\xle-disambiguation\raw\20210323\_F1\_UM1\_Peng0013\_SA\_F59\_ingel\_3ug\_TL.raw

##### Fraction

2

##### Scan Feature

F2:11788

##### De Novo Score

98

##### ConfidenceScore

98

### m/z

682.7664

##### Mass

1363.5178

##### Charge

2

##### Retention Time

49.24

##### Predicted Retention Time

-

##### Area

9.51E+06

##### Parts Per Million

0.3

##### Fragmentation mode

HCD

##### Originating file

01 D:\separate\_stitch\_analyses\xle-disambiguation\20210325\_F59\_3ug\_DENOVO\_12.csv

#### Meta Information from PEAKS

##### Scan Identifier

F2:10048

##### Original sequence

F

T

F

D

D

Y

A

M

+15.99

H

W

##### Posttranslational Modifications

Oxidation (M)

##### Source File

D:\separate\_stitch\_analyses\xle-disambiguation\raw\20210323\_F1\_UM1\_Peng0013\_SA\_F59\_ingel\_3ug\_TL.raw

##### Fraction

2

##### Scan Feature

-

##### De Novo Score

98

##### ConfidenceScore

98

### m/z

674.77

##### Mass

1347.5229

##### Charge

2

##### Retention Time

56.51

##### Predicted Retention Time

-

##### Area

0

##### Parts Per Million

1.8

##### Fragmentation mode

HCD

##### Originating file

01 D:\separate\_stitch\_analyses\xle-disambiguation\20210325\_F59\_3ug\_DENOVO\_12.csv

#### Meta Information from PEAKS

##### Scan Identifier

F2:8861

##### Original sequence

F

T

F

D

D

Y

A

M

+15.99

H

W

+15.99

##### Posttranslational Modifications

Oxidation (M); Oxidation (HW)

##### Source File

D:\separate\_stitch\_analyses\xle-disambiguation\raw\20210323\_F1\_UM1\_Peng0013\_SA\_F59\_ingel\_3ug\_TL.raw

##### Fraction

2

##### Scan Feature

F2:11788

##### De Novo Score

98

##### ConfidenceScore

98

### m/z

682.7664

##### Mass

1363.5178

##### Charge

2

##### Retention Time

49.24

##### Predicted Retention Time

-

##### Area

9.51E+06

##### Parts Per Million

0.3

##### Fragmentation mode

ETHCD

##### Originating file

01 D:\separate\_stitch\_analyses\xle-disambiguation\20210325\_F59\_3ug\_DENOVO\_12.csv

#### Meta Information from PEAKS

##### Scan Identifier

F2:8914

##### Original sequence

F

T

F

D

D

Y

A

M

+15.99

H

W

+15.99

##### Posttranslational Modifications

Oxidation (M); Oxidation (HW)

##### Source File

D:\separate\_stitch\_analyses\xle-disambiguation\raw\20210323\_F1\_UM1\_Peng0013\_SA\_F59\_ingel\_3ug\_TL.raw

##### Fraction

2

##### Scan Feature

F2:11788

##### De Novo Score

98

##### ConfidenceScore

98

### m/z

682.7664

##### Mass

1363.5178

##### Charge

2

##### Retention Time

49.24

##### Predicted Retention Time

-

##### Area

9.51E+06

##### Parts Per Million

0.3

##### Fragmentation mode

HCD

##### Originating file

01 D:\separate\_stitch\_analyses\xle-disambiguation\20210325\_F59\_3ug\_DENOVO\_12.csv

#### Meta Information from PEAKS

##### Scan Identifier

F2:9998

##### Original sequence

F

T

F

D

D

Y

A

M

+15.99

H

W

##### Posttranslational Modifications

Oxidation (M)

##### Source File

D:\separate\_stitch\_analyses\xle-disambiguation\raw\20210323\_F1\_UM1\_Peng0013\_SA\_F59\_ingel\_3ug\_TL.raw

##### Fraction

2

##### Scan Feature

-

##### De Novo Score

98

##### ConfidenceScore

98

### m/z

674.7701

##### Mass

1347.5229

##### Charge

2

##### Retention Time

56.22

##### Predicted Retention Time

-

##### Area

0

##### Parts Per Million

2.1

##### Fragmentation mode

ETHCD

##### Originating file

01 D:\separate\_stitch\_analyses\xle-disambiguation\20210325\_F59\_3ug\_DENOVO\_12.csv

#### Meta Information from PEAKS

##### Scan Identifier

F2:9024

##### Original sequence

F

T

F

D

D

Y

A

M

+15.99

H

W

+15.99

##### Posttranslational Modifications

Oxidation (M); Oxidation (HW)

##### Source File

D:\separate\_stitch\_analyses\xle-disambiguation\raw\20210323\_F1\_UM1\_Peng0013\_SA\_F59\_ingel\_3ug\_TL.raw

##### Fraction

2

##### Scan Feature

-

##### De Novo Score

98

##### ConfidenceScore

98

### m/z

682.7673

##### Mass

1363.5178

##### Charge

2

##### Retention Time

50.8

##### Predicted Retention Time

-

##### Area

0

##### Parts Per Million

1.6

##### Fragmentation mode

HCD

##### Originating file

01 D:\separate\_stitch\_analyses\xle-disambiguation\20210325\_F59\_3ug\_DENOVO\_12.csv

#### Meta Information from PEAKS

##### Scan Identifier

F2:10100

##### Original sequence

F

T

F

D

D

Y

A

M

+15.99

H

W

##### Posttranslational Modifications

Oxidation (M)

##### Source File

D:\separate\_stitch\_analyses\xle-disambiguation\raw\20210323\_F1\_UM1\_Peng0013\_SA\_F59\_ingel\_3ug\_TL.raw

##### Fraction

2

##### Scan Feature

-

##### De Novo Score

98

##### ConfidenceScore

98

### m/z

674.7689

##### Mass

1347.5229

##### Charge

2

##### Retention Time

56.79

##### Predicted Retention Time

-

##### Area

0

##### Parts Per Million

0.3

##### Fragmentation mode

ETHCD

##### Originating file

01 D:\separate\_stitch\_analyses\xle-disambiguation\20210325\_F59\_3ug\_DENOVO\_12.csv

#### Meta Information from PEAKS

##### Scan Identifier

F2:8754

##### Original sequence

F

T

F

D

D

Y

A

M

+15.99

H

W

+15.99

##### Posttranslational Modifications

Oxidation (M); Oxidation (HW)

##### Source File

D:\separate\_stitch\_analyses\xle-disambiguation\raw\20210323\_F1\_UM1\_Peng0013\_SA\_F59\_ingel\_3ug\_TL.raw

##### Fraction

2

##### Scan Feature

F2:11788

##### De Novo Score

97

##### ConfidenceScore

97

### m/z

682.7664

##### Mass

1363.5178

##### Charge

2

##### Retention Time

49.24

##### Predicted Retention Time

-

##### Area

9.51E+06

##### Parts Per Million

0.3

##### Fragmentation mode

ETHCD

##### Originating file

01 D:\separate\_stitch\_analyses\xle-disambiguation\20210325\_F59\_3ug\_DENOVO\_12.csv

#### Meta Information from PEAKS

##### Scan Identifier

F2:10360

##### Original sequence

F

T

F

D

D

Y

A

M

+15.99

H

W

##### Posttranslational Modifications

Oxidation (M)

##### Source File

D:\separate\_stitch\_analyses\xle-disambiguation\raw\20210323\_F1\_UM1\_Peng0013\_SA\_F59\_ingel\_3ug\_TL.raw

##### Fraction

2

##### Scan Feature

-

##### De Novo Score

97

##### ConfidenceScore

97

### m/z

674.7698

##### Mass

1347.5229

##### Charge

2

##### Retention Time

58.22

##### Predicted Retention Time

-

##### Area

0

##### Parts Per Million

1.6

##### Fragmentation mode

HCD

##### Originating file

01 D:\separate\_stitch\_analyses\xle-disambiguation\20210325\_F59\_3ug\_DENOVO\_12.csv

#### Meta Information from PEAKS

##### Scan Identifier

F2:8704

##### Original sequence

F

T

F

D

D

Y

A

M

+15.99

H

W

+15.99

##### Posttranslational Modifications

Oxidation (M); Oxidation (HW)

##### Source File

D:\separate\_stitch\_analyses\xle-disambiguation\raw\20210323\_F1\_UM1\_Peng0013\_SA\_F59\_ingel\_3ug\_TL.raw

##### Fraction

2

##### Scan Feature

F2:11788

##### De Novo Score

96

##### ConfidenceScore

96

### m/z

682.7664

##### Mass

1363.5178

##### Charge

2

##### Retention Time

49.24

##### Predicted Retention Time

-

##### Area

9.51E+06

##### Parts Per Million

0.3

##### Fragmentation mode

ETHCD

##### Originating file

01 D:\separate\_stitch\_analyses\xle-disambiguation\20210325\_F59\_3ug\_DENOVO\_12.csv

#### Meta Information from PEAKS

##### Scan Identifier

F2:10560

##### Original sequence

F

T

F

D

D

Y

A

M

+15.99

H

W

##### Posttranslational Modifications

Oxidation (M)

##### Source File

D:\separate\_stitch\_analyses\xle-disambiguation\raw\20210323\_F1\_UM1\_Peng0013\_SA\_F59\_ingel\_3ug\_TL.raw

##### Fraction

2

##### Scan Feature

-

##### De Novo Score

96

##### ConfidenceScore

96

### m/z

674.7698

##### Mass

1347.5229

##### Charge

2

##### Retention Time

59.31

##### Predicted Retention Time

-

##### Area

0

##### Parts Per Million

1.5

##### Fragmentation mode

HCD

##### Originating file

01 D:\separate\_stitch\_analyses\xle-disambiguation\20210325\_F59\_3ug\_DENOVO\_12.csv

#### Meta Information from PEAKS

##### Scan Identifier

F2:9715

##### Original sequence

F

T

F

D

D

Y

A

M

+15.99

H

W

+15.99

##### Posttranslational Modifications

Oxidation (M); Oxidation (HW)

##### Source File

D:\separate\_stitch\_analyses\xle-disambiguation\raw\20210323\_F1\_UM1\_Peng0013\_SA\_F59\_ingel\_3ug\_TL.raw

##### Fraction

2

##### Scan Feature

F2:2457

##### De Novo Score

96

##### ConfidenceScore

96

### m/z

455.5132

##### Mass

1363.5178

##### Charge

3

##### Retention Time

54.69

##### Predicted Retention Time

-

##### Area

1.139E+05

##### Parts Per Million

0.1

##### Fragmentation mode

ETHCD

##### Originating file

01 D:\separate\_stitch\_analyses\xle-disambiguation\20210325\_F59\_3ug\_DENOVO\_12.csv

#### Meta Information from PEAKS

##### Scan Identifier

F2:8264

##### Original sequence

F

T

F

D

D

Y

A

M

+15.99

H

W

+15.99

##### Posttranslational Modifications

Oxidation (M); Oxidation (HW)

##### Source File

D:\separate\_stitch\_analyses\xle-disambiguation\raw\20210323\_F1\_UM1\_Peng0013\_SA\_F59\_ingel\_3ug\_TL.raw

##### Fraction

2

##### Scan Feature

F2:11797

##### De Novo Score

95

##### ConfidenceScore

95

### m/z

682.7676

##### Mass

1363.5178

##### Charge

2

##### Retention Time

46.65

##### Predicted Retention Time

-

##### Area

3.557E+05

##### Parts Per Million

2.1

##### Fragmentation mode

HCD

##### Originating file

01 D:\separate\_stitch\_analyses\xle-disambiguation\20210325\_F59\_3ug\_DENOVO\_12.csv

#### Meta Information from PEAKS

##### Scan Identifier

F2:9159

##### Original sequence

F

T

F

D

D

Y

A

M

+15.99

H

W

+15.99

##### Posttranslational Modifications

Oxidation (M); Oxidation (HW)

##### Source File

D:\separate\_stitch\_analyses\xle-disambiguation\raw\20210323\_F1\_UM1\_Peng0013\_SA\_F59\_ingel\_3ug\_TL.raw

##### Fraction

2

##### Scan Feature

-

##### De Novo Score

95

##### ConfidenceScore

95

### m/z

682.7665

##### Mass

1363.5178

##### Charge

2

##### Retention Time

51.57

##### Predicted Retention Time

-

##### Area

0

##### Parts Per Million

0.5

##### Fragmentation mode

HCD

##### Originating file

01 D:\separate\_stitch\_analyses\xle-disambiguation\20210325\_F59\_3ug\_DENOVO\_12.csv

#### Meta Information from PEAKS

##### Scan Identifier

F2:8970

##### Original sequence

F

T

F

D

D

Y

A

M

+15.99

H

+15.99

W

+15.99

##### Posttranslational Modifications

Oxidation (M); Oxidation (HW)

##### Source File

D:\separate\_stitch\_analyses\xle-disambiguation\raw\20210323\_F1\_UM1\_Peng0013\_SA\_F59\_ingel\_3ug\_TL.raw

##### Fraction

2

##### Scan Feature

F2:12180

##### De Novo Score

95

##### ConfidenceScore

95

### m/z

690.7643

##### Mass

1379.5129

##### Charge

2

##### Retention Time

49.9

##### Predicted Retention Time

-

##### Area

1.879E+07

##### Parts Per Million

0.8

##### Fragmentation mode

HCD

##### Originating file

01 D:\separate\_stitch\_analyses\xle-disambiguation\20210325\_F59\_3ug\_DENOVO\_12.csv

#### Meta Information from PEAKS

##### Scan Identifier

F2:9262

##### Original sequence

F

T

F

D

D

Y

A

M

+15.99

H

W

+15.99

##### Posttranslational Modifications

Oxidation (M); Oxidation (HW)

##### Source File

D:\separate\_stitch\_analyses\xle-disambiguation\raw\20210323\_F1\_UM1\_Peng0013\_SA\_F59\_ingel\_3ug\_TL.raw

##### Fraction

2

##### Scan Feature

-

##### De Novo Score

95

##### ConfidenceScore

95

### m/z

682.7656

##### Mass

1363.5178

##### Charge

2

##### Retention Time

52.15

##### Predicted Retention Time

-

##### Area

0

##### Fragmentation mode

HCD

##### Originating file

01 D:\separate\_stitch\_analyses\xle-disambiguation\20210325\_F59\_3ug\_DENOVO\_12.csv
