## Supplementary material for "A handle on mass coincidence errors in *de novo* sequencing of antibodies by bottom-up proteomics": Combined_004.html

Details Combined\_004 | Stitch OverviewUndefined

### Read Combined\_004

#### Sequence (length=12)

AVMDDFAAFVEK

#### Spectrum 8408? Spectrum 8408 The raw spectrum of this peptide as annotated by Hecklib. The fragments are coloured according to ion type (see legend). Any peaks with a star '\*' as text can be hovered over to see the full details, first the ion type second the mass shift type. By hovering over the amino acids in the peptide or ions in the legend the corresponding peaks are highlighted. By toggling the 'Unassigned' label you can turn the background (unassigned) peaks on or off in the plot. By updating the slider in the Ion legend you can update the spectrum to only show the top X% of the peaks with labels. The top X% means any peak that is within X% of the highest intensity. By dragging in the spectrum you can zoom in to a specific part of the spectrum and use 'Zoom Out' to get back to the original zoom level. The annotation of the spectrum is based on the given sequence in the peptides file and is done with different software so inconsistencies are likely. The peaks are annotated based on the given sequence, with 20 ppm tolerance.

Copy Data

##### Spectrum 8408 (TSV)

###### Preview

```
Loading example...
```

*Click on the button to copy the data to your clipboard.*

Mz MinMz MaxIntensity Max

WidthHeightPeptide font sizePeptide stroke widthSpectrum font sizeSpectrum stroke widthCompact peptide

Ion legend

wxyz

abcd

OtherUnassignedIon^Charge^_Position_Show for top:%

AVMDDFAAFVEK

01.33e+42.66e+43.99e+45.32e+4

Zoom Out

d^+1^_2_y^+1^_1_a^+1^_2_y^+1^_1_b^+1^_2_y^+1^_2_y^+1^_2_a^+1^_3_b^+1^_3_y^+1^_3_b^+1^_4_b^+1^_4_y^+1^_4_b^+1^_5_b^+1^_5_y^+2^_10_y^+2^_10_y^+1^_5_y^+2^_10_y^+2^_11_y^+1^_6_\*b^+1^_6_\*b^+1^_6_b^+1^_7_b^+1^_7_y^+1^_7_b^+1^_8_y^+1^_8_y^+1^_8_b^+1^_9_y^+1^_9_y^+1^_9_y^+1^_9_y^+1^_10_y^+1^_10_

0577115417312308

Fragment Matches Table

Show background peaks

| Position | Ion type | Intensity | mz Theoretical | mz Error (Th) | mz Error (ppm) | Charge | Series Number |
| --- | --- | --- | --- | --- | --- | --- | --- |
| - | - | 593.4 | 120 | - | - | 0 | - |
| - | - | 638.7 | 120.1 | - | - | 0 | - |
| - | - | 2.483E+04 | 120.1 | - | - | 0 | - |
| - | - | 2187 | 121.1 | - | - | 0 | - |
| - | - | 391.8 | 122.1 | - | - | 0 | - |
| - | - | 412.1 | 122.7 | - | - | 0 | - |
| - | - | 1537 | 127.1 | - | - | 0 | - |
| 2 | d | 1.324E+04 | 129.1 | 0.0001775 | 1.375 | +1 | 2 |
| 12 | y | 5919 | 130.1 | 0.0001707 | 1.312 | +1 | 1 |
| - | - | 1291 | 130.1 | - | - | 0 | - |
| - | - | 441.9 | 131.1 | - | - | 0 | - |
| - | - | 490.7 | 133.1 | - | - | 0 | - |
| - | - | 393.7 | 133.1 | - | - | 0 | - |
| - | - | 2590 | 136.1 | - | - | 0 | - |
| - | - | 806.1 | 141.1 | - | - | 0 | - |
| - | - | 560 | 143 | - | - | 0 | - |
| - | - | 1128 | 143.1 | - | - | 0 | - |
| 2 | a | 5.263E+04 | 143.1 | 0.0001525 | 1.065 | +1 | 2 |
| - | - | 3074 | 144.1 | - | - | 0 | - |
| - | - | 415.9 | 144.4 | - | - | 0 | - |
| 12 | y | 1.246E+04 | 147.1 | 0.0001567 | 1.065 | +1 | 1 |
| - | - | 485.8 | 148.9 | - | - | 0 | - |
| - | - | 604.1 | 148.9 | - | - | 0 | - |
| - | - | 834.2 | 148.9 | - | - | 0 | - |
| - | - | 908.2 | 148.9 | - | - | 0 | - |
| - | - | 1050 | 148.9 | - | - | 0 | - |
| - | - | 1462 | 148.9 | - | - | 0 | - |
| - | - | 3482 | 148.9 | - | - | 0 | - |
| - | - | 5821 | 149 | - | - | 0 | - |
| - | - | 3530 | 149 | - | - | 0 | - |
| - | - | 1600 | 149 | - | - | 0 | - |
| - | - | 1065 | 149 | - | - | 0 | - |
| - | - | 1028 | 149 | - | - | 0 | - |
| - | - | 622.8 | 149 | - | - | 0 | - |
| - | - | 934.6 | 149 | - | - | 0 | - |
| - | - | 895.2 | 149 | - | - | 0 | - |
| - | - | 636.7 | 149 | - | - | 0 | - |
| - | - | 477.7 | 149 | - | - | 0 | - |
| - | - | 491.2 | 149 | - | - | 0 | - |
| - | - | 860.3 | 149 | - | - | 0 | - |
| - | - | 438.2 | 149.1 | - | - | 0 | - |
| - | - | 484 | 149.1 | - | - | 0 | - |
| - | - | 1233 | 153.1 | - | - | 0 | - |
| - | - | 710.9 | 155.1 | - | - | 0 | - |
| - | - | 1619 | 155.1 | - | - | 0 | - |
| - | - | 472.2 | 162.4 | - | - | 0 | - |
| - | - | 623 | 165.1 | - | - | 0 | - |
| - | - | 577.9 | 166.1 | - | - | 0 | - |
| - | - | 8150 | 166.1 | - | - | 0 | - |
| - | - | 663.8 | 167.1 | - | - | 0 | - |
| - | - | 634.1 | 167.1 | - | - | 0 | - |
| - | - | 467.1 | 167.1 | - | - | 0 | - |
| - | - | 4646 | 169.1 | - | - | 0 | - |
| - | - | 518 | 170.1 | - | - | 0 | - |
| - | - | 664.1 | 171 | - | - | 0 | - |
| 2 | b | 2.85E+04 | 171.1 | 9.562E-05 | 0.5588 | +1 | 2 |
| - | - | 2106 | 172.1 | - | - | 0 | - |
| - | - | 1078 | 181.1 | - | - | 0 | - |
| - | - | 1694 | 183.1 | - | - | 0 | - |
| - | - | 517.9 | 183.7 | - | - | 0 | - |
| - | - | 1855 | 185.2 | - | - | 0 | - |
| - | - | 850.1 | 191.1 | - | - | 0 | - |
| - | - | 4577 | 191.1 | - | - | 0 | - |
| - | - | 558.7 | 192.1 | - | - | 0 | - |
| - | - | 523.6 | 193.1 | - | - | 0 | - |
| - | - | 791 | 195.1 | - | - | 0 | - |
| - | - | 4302 | 197.1 | - | - | 0 | - |
| - | - | 4983 | 199.1 | - | - | 0 | - |
| - | - | 814.8 | 199.1 | - | - | 0 | - |
| - | - | 2188 | 201.1 | - | - | 0 | - |
| - | - | 868.6 | 209.1 | - | - | 0 | - |
| - | - | 2164 | 211.1 | - | - | 0 | - |
| - | - | 1035 | 213.2 | - | - | 0 | - |
| - | - | 2881 | 217.1 | - | - | 0 | - |
| - | - | 720.4 | 217.1 | - | - | 0 | - |
| - | - | 8421 | 219.1 | - | - | 0 | - |
| - | - | 4984 | 219.1 | - | - | 0 | - |
| - | - | 582.4 | 220.1 | - | - | 0 | - |
| - | - | 2665 | 225 | - | - | 0 | - |
| - | - | 565.8 | 226.1 | - | - | 0 | - |
| - | - | 1554 | 226.1 | - | - | 0 | - |
| - | - | 1678 | 227.1 | - | - | 0 | - |
| - | - | 2234 | 229.1 | - | - | 0 | - |
| - | - | 843.3 | 230.1 | - | - | 0 | - |
| - | - | 1793 | 231.1 | - | - | 0 | - |
| - | - | 632.6 | 231.1 | - | - | 0 | - |
| - | - | 738.2 | 235.1 | - | - | 0 | - |
| - | - | 3270 | 235.1 | - | - | 0 | - |
| - | - | 2698 | 239.1 | - | - | 0 | - |
| - | - | 3945 | 240.1 | - | - | 0 | - |
| - | - | 664.6 | 241.1 | - | - | 0 | - |
| - | - | 756.3 | 244.1 | - | - | 0 | - |
| - | - | 1091 | 245.1 | - | - | 0 | - |
| - | - | 2442 | 245.1 | - | - | 0 | - |
| - | - | 770.8 | 246 | - | - | 0 | - |
| - | - | 2214 | 247.1 | - | - | 0 | - |
| - | - | 2925 | 247.1 | - | - | 0 | - |
| - | - | 501.6 | 252.1 | - | - | 0 | - |
| - | - | 4008 | 254.1 | - | - | 0 | - |
| 11 | y | 1.423E+04 | 258.1 | 3.859E-06 | 0.01495 | +1 | 2 |
| - | - | 1187 | 259.1 | - | - | 0 | - |
| - | - | 1.637E+04 | 263.1 | - | - | 0 | - |
| - | - | 5005 | 263.1 | - | - | 0 | - |
| - | - | 1375 | 264.1 | - | - | 0 | - |
| - | - | 578.6 | 268.9 | - | - | 0 | - |
| - | - | 6463 | 275.1 | - | - | 0 | - |
| - | - | 1004 | 276.1 | - | - | 0 | - |
| 11 | y | 1.752E+04 | 276.2 | 3.226E-05 | 0.1168 | +1 | 2 |
| - | - | 2206 | 277.2 | - | - | 0 | - |
| - | - | 512.3 | 283.1 | - | - | 0 | - |
| - | - | 1772 | 286.1 | - | - | 0 | - |
| - | - | 685.4 | 286.1 | - | - | 0 | - |
| 3 | a | 7792 | 290.1 | 0.001528 | 5.267 | +1 | 3 |
| - | - | 887.9 | 291.2 | - | - | 0 | - |
| - | - | 1070 | 295.1 | - | - | 0 | - |
| - | - | 1979 | 296.1 | - | - | 0 | - |
| - | - | 985.6 | 298.1 | - | - | 0 | - |
| - | - | 884.7 | 298.2 | - | - | 0 | - |
| - | - | 1114 | 299.1 | - | - | 0 | - |
| - | - | 782.1 | 300.1 | - | - | 0 | - |
| - | - | 709 | 313.1 | - | - | 0 | - |
| - | - | 5931 | 314.1 | - | - | 0 | - |
| - | - | 863.4 | 315.1 | - | - | 0 | - |
| - | - | 930.5 | 316.1 | - | - | 0 | - |
| - | - | 4684 | 316.2 | - | - | 0 | - |
| - | - | 764 | 317.2 | - | - | 0 | - |
| 3 | b | 1.532E+04 | 318.1 | 0.004996 | 15.7 | +1 | 3 |
| - | - | 1569 | 318.2 | - | - | 0 | - |
| - | - | 2455 | 319.2 | - | - | 0 | - |
| - | - | 804.8 | 324.2 | - | - | 0 | - |
| - | - | 617.1 | 332.1 | - | - | 0 | - |
| - | - | 6069 | 334.1 | - | - | 0 | - |
| - | - | 1152 | 335.1 | - | - | 0 | - |
| - | - | 533.3 | 335.8 | - | - | 0 | - |
| - | - | 726.1 | 339.2 | - | - | 0 | - |
| - | - | 741.1 | 343.2 | - | - | 0 | - |
| - | - | 927.6 | 344.1 | - | - | 0 | - |
| - | - | 682.5 | 344.2 | - | - | 0 | - |
| - | - | 789.4 | 350.1 | - | - | 0 | - |
| - | - | 1667 | 360.1 | - | - | 0 | - |
| - | - | 1724 | 360.1 | - | - | 0 | - |
| - | - | 958.6 | 361.2 | - | - | 0 | - |
| - | - | 1404 | 362.1 | - | - | 0 | - |
| - | - | 1127 | 369.1 | - | - | 0 | - |
| - | - | 742.4 | 369.2 | - | - | 0 | - |
| - | - | 802.7 | 370.1 | - | - | 0 | - |
| - | - | 581.5 | 373.1 | - | - | 0 | - |
| 10 | y | 1.647E+04 | 375.2 | 8.68E-05 | 0.2313 | +1 | 3 |
| - | - | 1200 | 376.2 | - | - | 0 | - |
| - | - | 2712 | 376.2 | - | - | 0 | - |
| - | - | 1.41E+04 | 378.1 | - | - | 0 | - |
| - | - | 5538 | 378.1 | - | - | 0 | - |
| - | - | 1122 | 379.1 | - | - | 0 | - |
| - | - | 816 | 379.1 | - | - | 0 | - |
| - | - | 608.1 | 387.1 | - | - | 0 | - |
| - | - | 2548 | 388.1 | - | - | 0 | - |
| - | - | 627.2 | 389.1 | - | - | 0 | - |
| - | - | 1448 | 389.2 | - | - | 0 | - |
| - | - | 591.1 | 390.2 | - | - | 0 | - |
| - | - | 606.3 | 392.2 | - | - | 0 | - |
| - | - | 647.8 | 403.2 | - | - | 0 | - |
| - | - | 683.3 | 404.1 | - | - | 0 | - |
| - | - | 3554 | 405.2 | - | - | 0 | - |
| - | - | 860.9 | 409.2 | - | - | 0 | - |
| - | - | 1109 | 414.1 | - | - | 0 | - |
| 4 | b | 1185 | 415.2 | 0.002137 | 5.148 | +1 | 4 |
| - | - | 641.9 | 421.2 | - | - | 0 | - |
| - | - | 1623 | 431.2 | - | - | 0 | - |
| - | - | 668.8 | 433.1 | - | - | 0 | - |
| 4 | b | 3130 | 433.2 | 0.00381 | 8.796 | +1 | 4 |
| - | - | 664.3 | 434.2 | - | - | 0 | - |
| - | - | 1203 | 437.2 | - | - | 0 | - |
| - | - | 931.4 | 443.2 | - | - | 0 | - |
| - | - | 1020 | 447.2 | - | - | 0 | - |
| - | - | 915.7 | 449.1 | - | - | 0 | - |
| - | - | 5430 | 449.2 | - | - | 0 | - |
| - | - | 1087 | 450.2 | - | - | 0 | - |
| - | - | 1163 | 456.3 | - | - | 0 | - |
| - | - | 2122 | 459.2 | - | - | 0 | - |
| - | - | 916.5 | 461.2 | - | - | 0 | - |
| - | - | 1006 | 466.2 | - | - | 0 | - |
| - | - | 708 | 475.2 | - | - | 0 | - |
| - | - | 1467 | 477.2 | - | - | 0 | - |
| - | - | 1013 | 479.2 | - | - | 0 | - |
| - | - | 1429 | 484.2 | - | - | 0 | - |
| - | - | 610.9 | 486.2 | - | - | 0 | - |
| - | - | 648 | 492.2 | - | - | 0 | - |
| - | - | 1558 | 497.2 | - | - | 0 | - |
| - | - | 665.9 | 498.2 | - | - | 0 | - |
| - | - | 1182 | 502.2 | - | - | 0 | - |
| - | - | 644.4 | 508.1 | - | - | 0 | - |
| - | - | 692.9 | 508.3 | - | - | 0 | - |
| - | - | 771 | 518.3 | - | - | 0 | - |
| - | - | 5464 | 520.2 | - | - | 0 | - |
| - | - | 1420 | 521.2 | - | - | 0 | - |
| 9 | y | 1.507E+04 | 522.3 | 0.0002329 | 0.4459 | +1 | 4 |
| - | - | 3371 | 523.3 | - | - | 0 | - |
| - | - | 855.3 | 524.3 | - | - | 0 | - |
| - | - | 3261 | 525.2 | - | - | 0 | - |
| - | - | 698.9 | 526.2 | - | - | 0 | - |
| - | - | 720.3 | 526.2 | - | - | 0 | - |
| 5 | b | 2668 | 530.2 | 0.002843 | 5.362 | +1 | 5 |
| - | - | 642.2 | 531.2 | - | - | 0 | - |
| - | - | 1535 | 532.2 | - | - | 0 | - |
| - | - | 990.6 | 542.3 | - | - | 0 | - |
| 5 | b | 3749 | 548.2 | 0.004058 | 7.403 | +1 | 5 |
| - | - | 977.3 | 549.2 | - | - | 0 | - |
| - | - | 1401 | 552.2 | - | - | 0 | - |
| - | - | 805.4 | 553.2 | - | - | 0 | - |
| - | - | 813.6 | 553.8 | - | - | 0 | - |
| - | - | 644.4 | 557.6 | - | - | 0 | - |
| - | - | 8037 | 562.8 | - | - | 0 | - |
| - | - | 6510 | 563.3 | - | - | 0 | - |
| - | - | 1426 | 563.8 | - | - | 0 | - |
| - | - | 564 | 567.3 | - | - | 0 | - |
| - | - | 712 | 585.2 | - | - | 0 | - |
| 3 | y | 1870 | 585.8 | 0.00173 | 2.954 | +2 | 10 |
| 3 | y | 2355 | 586.2 | 0.007891 | 13.46 | +2 | 10 |
| - | - | 911.3 | 586.8 | - | - | 0 | - |
| 8 | y | 1.608E+04 | 593.3 | 0.0002373 | 0.4 | +1 | 5 |
| - | - | 4637 | 594.3 | - | - | 0 | - |
| 3 | y | 3.032E+04 | 594.8 | 0.002124 | 3.571 | +2 | 10 |
| - | - | 2.001E+04 | 595.3 | - | - | 0 | - |
| - | - | 622.3 | 595.3 | - | - | 0 | - |
| - | - | 9172 | 595.8 | - | - | 0 | - |
| - | - | 2509 | 596.2 | - | - | 0 | - |
| - | - | 1065 | 596.3 | - | - | 0 | - |
| - | - | 745.1 | 597.2 | - | - | 0 | - |
| - | - | 1709 | 603.2 | - | - | 0 | - |
| - | - | 837 | 604.2 | - | - | 0 | - |
| - | - | 1474 | 612.3 | - | - | 0 | - |
| - | - | 641.6 | 622.2 | - | - | 0 | - |
| - | - | 1134 | 631.3 | - | - | 0 | - |
| - | - | 653.2 | 635.8 | - | - | 0 | - |
| - | - | 729.9 | 638.8 | - | - | 0 | - |
| 2 | y | 4174 | 644.3 | 0.001853 | 2.875 | +2 | 11 |
| - | - | 3274 | 644.8 | - | - | 0 | - |
| - | - | 1074 | 645.3 | - | - | 0 | - |
| - | - | 6440 | 647.8 | - | - | 0 | - |
| - | - | 4776 | 648.3 | - | - | 0 | - |
| - | - | 1462 | 648.8 | - | - | 0 | - |
| - | - | 1458 | 649.3 | - | - | 0 | - |
| - | - | 734.4 | 650.2 | - | - | 0 | - |
| - | - | 791.9 | 651.3 | - | - | 0 | - |
| - | - | 622.3 | 663.3 | - | - | 0 | - |
| 7 | y | 1.976E+04 | 664.4 | 0.0006079 | 0.9151 | +1 | 6 |
| - | - | 7493 | 665.4 | - | - | 0 | - |
| - | - | 1211 | 666.4 | - | - | 0 | - |
| - | - | 922.8 | 667.2 | - | - | 0 | - |
| - | - | 3115 | 667.3 | - | - | 0 | - |
| - | - | 884 | 668.3 | - | - | 0 | - |
| 0 | Precursor | 849.3 | 671.3 | 0.01177 | 17.53 | +2 | -1 |
| 6 | b | 1773 | 677.3 | 0.004497 | 6.641 | +1 | 6 |
| 0 | Precursor | 7032 | 679.8 | 0.001972 | 2.901 | +2 | -1 |
| - | - | 795.8 | 680 | - | - | 0 | - |
| - | - | 5861 | 680.3 | - | - | 0 | - |
| - | - | 3479 | 680.8 | - | - | 0 | - |
| - | - | 743.4 | 689.3 | - | - | 0 | - |
| 6 | b | 2227 | 695.3 | 0.005285 | 7.602 | +1 | 6 |
| - | - | 773.3 | 696.3 | - | - | 0 | - |
| - | - | 935.1 | 702.3 | - | - | 0 | - |
| - | - | 677.4 | 732.3 | - | - | 0 | - |
| 7 | b | 1020 | 748.3 | 0.002845 | 3.802 | +1 | 7 |
| - | - | 1333 | 750.3 | - | - | 0 | - |
| 7 | b | 1425 | 766.3 | 0.01236 | 16.13 | +1 | 7 |
| - | - | 798.3 | 767.3 | - | - | 0 | - |
| - | - | 1033 | 773.3 | - | - | 0 | - |
| - | - | 960.4 | 782.4 | - | - | 0 | - |
| 6 | y | 1.699E+04 | 811.4 | 0.0008456 | 1.042 | +1 | 7 |
| - | - | 6903 | 812.4 | - | - | 0 | - |
| - | - | 1720 | 813.4 | - | - | 0 | - |
| - | - | 1426 | 814.3 | - | - | 0 | - |
| - | - | 947 | 815.3 | - | - | 0 | - |
| - | - | 636.9 | 820.4 | - | - | 0 | - |
| 8 | b | 2031 | 837.3 | 0.003995 | 4.771 | +1 | 8 |
| - | - | 1209 | 838.3 | - | - | 0 | - |
| - | - | 1324 | 849.4 | - | - | 0 | - |
| - | - | 866.7 | 895.4 | - | - | 0 | - |
| 5 | y | 912.7 | 908.5 | 0.002796 | 3.077 | +1 | 8 |
| - | - | 1369 | 913.4 | - | - | 0 | - |
| - | - | 929.3 | 920.4 | - | - | 0 | - |
| 5 | y | 1.32E+04 | 926.5 | 0.00136 | 1.468 | +1 | 8 |
| - | - | 6777 | 927.5 | - | - | 0 | - |
| - | - | 1363 | 928.5 | - | - | 0 | - |
| 9 | b | 2422 | 984.4 | 0.004917 | 4.995 | +1 | 9 |
| - | - | 1319 | 985.4 | - | - | 0 | - |
| - | - | 1000 | 1008 | - | - | 0 | - |
| 4 | y | 1015 | 1023 | 0.007851 | 7.671 | +1 | 9 |
| 4 | y | 1498 | 1024 | 0.01534 | 14.97 | +1 | 9 |
| 4 | y | 3.642E+04 | 1041 | 0.001814 | 1.742 | +1 | 9 |
| - | - | 2.156E+04 | 1042 | - | - | 0 | - |
| - | - | 5812 | 1043 | - | - | 0 | - |
| - | - | 706.6 | 1045 | - | - | 0 | - |
| - | - | 1942 | 1051 | - | - | 0 | - |
| - | - | 1741 | 1052 | - | - | 0 | - |
| - | - | 631.6 | 1072 | - | - | 0 | - |
| - | - | 690.6 | 1097 | - | - | 0 | - |
| - | - | 6518 | 1125 | - | - | 0 | - |
| - | - | 4396 | 1126 | - | - | 0 | - |
| - | - | 802.7 | 1127 | - | - | 0 | - |
| 3 | y | 1187 | 1171 | 0.0001165 | 0.09953 | +1 | 10 |
| 3 | y | 2.112E+04 | 1189 | 0.002247 | 1.891 | +1 | 10 |
| - | - | 1.46E+04 | 1190 | - | - | 0 | - |
| - | - | 6203 | 1191 | - | - | 0 | - |
| - | - | 964.3 | 1192 | - | - | 0 | - |
| - | - | 638.6 | 2016 | - | - | 0 | - |
| - | - | 683.6 | 2031 | - | - | 0 | - |
| - | - | 758.5 | 2285 | - | - | 0 | - |

m/z Charge Intensity FragmentType MassShift Position
120.04817962646484 0 593.4333
120.06566619873047 0 638.7104
120.08096313476562 0 24833.426
121.08425903320312 0 2186.8367
122.09637451171875 0 391.78998
122.73540496826172 0 412.14896
127.08683776855469 0 1536.9686
129.1024169921875 0 13241.167 d 1
130.08642578125 0 5919.291 y Ammonia loss 11
130.10572814941406 0 1290.8955
131.0900421142578 0 441.9107
133.06097412109375 0 490.7395
133.0863800048828 0 393.65082
136.07582092285156 0 2590.1235
141.06578063964844 0 806.0754
143.04518127441406 0 560.0077
143.08152770996094 0 1128.287
143.1180419921875 0 52627.445 a 1
144.12144470214844 0 3074.2068
144.36865234375 0 415.93137
147.1129608154297 0 12457.742 y 11
148.8824005126953 0 485.7608
148.89706420898438 0 604.1313
148.91087341308594 0 834.21216
148.91847229003906 0 908.22327
148.92556762695312 0 1049.9054
148.93272399902344 0 1462.3685
148.94007873535156 0 3482.466
148.95651245117188 0 5821.455
148.96437072753906 0 3530.4421
148.9720458984375 0 1599.6747
148.9789581298828 0 1065.2538
148.98573303222656 0 1028.0046
148.99319458007812 0 622.82745
149.00042724609375 0 934.6188
149.0076446533203 0 895.1524
149.0148162841797 0 636.73535
149.02154541015625 0 477.67206
149.0282745361328 0 491.19583
149.0446014404297 0 860.3105
149.0653839111328 0 438.22092
149.09385681152344 0 483.98462
153.0663604736328 0 1233.3417
155.08145141601562 0 710.88495
155.11807250976562 0 1618.5306
162.3587646484375 0 472.22784
165.1024932861328 0 623.0199
166.05389404296875 0 577.88477
166.08645629882812 0 8150.4893
167.0556640625 0 663.79004
167.0893096923828 0 634.1222
167.1177520751953 0 467.103
169.13368225097656 0 4645.901
170.12908935546875 0 517.9781
171.0401153564453 0 664.0778
171.11289978027344 0 28496.488 b 1
172.11636352539062 0 2106.0137
181.0607147216797 0 1078.2173
183.11297607421875 0 1693.6887
183.740478515625 0 517.9231
185.16497802734375 0 1854.926
191.0845947265625 0 850.1263
191.1180419921875 0 4577.454
192.1211700439453 0 558.7259
193.09674072265625 0 523.6203
195.11277770996094 0 790.96625
197.12847900390625 0 4302.079
199.0714569091797 0 4982.9385
199.10818481445312 0 814.8288
201.1234130859375 0 2187.8003
209.1292724609375 0 868.61346
211.10748291015625 0 2163.5623
213.15975952148438 0 1034.7926
217.0641632080078 0 2880.8225
217.1337890625 0 720.38226
219.11280822753906 0 8420.787
219.14930725097656 0 4984.092
220.11647033691406 0 582.3659
225.04283142089844 0 2664.7524
226.0948028564453 0 565.82904
226.11886596679688 0 1553.815
227.10250854492188 0 1677.6294
229.11839294433594 0 2233.927
230.0843963623047 0 843.3296
231.061279296875 0 1792.907
231.1127166748047 0 632.5706
235.07516479492188 0 738.1812
235.1077880859375 0 3270.4287
239.09487915039062 0 2698.4873
240.13429260253906 0 3944.6912
241.11880493164062 0 664.60223
244.12966918945312 0 756.3432
245.0589599609375 0 1090.8322
245.12869262695312 0 2442.0454
246.04222106933594 0 770.8287
247.1111297607422 0 2214.1313
247.14413452148438 0 2924.9133
252.135009765625 0 501.58093
254.14993286132812 0 4008.175
258.14483642578125 0 14228.701 y Water loss 10
259.14837646484375 0 1186.8397
263.069580078125 0 16368.472
263.1026611328125 0 5005.1416
264.07330322265625 0 1375.3973
268.93304443359375 0 578.61664
275.1058349609375 0 6462.728
276.1097106933594 0 1003.72375
276.1553649902344 0 17522.715 y 10
277.1582946777344 0 2206.0447
283.1396179199219 0 512.2515
286.0668029785156 0 1772.1091
286.1033020019531 0 685.3662
290.14990234375 0 7791.9873 a 2
291.1526794433594 0 887.9333
295.1036682128906 0 1069.8665
296.0875244140625 0 1978.9114
298.1395568847656 0 985.56305
298.17626953125 0 884.65894
299.0614318847656 0 1113.8123
300.1368103027344 0 782.0651
313.11328125 0 708.99164
314.0985107421875 0 5930.862
315.1007995605469 0 863.4206
316.130615234375 0 930.4828
316.1868591308594 0 4683.983
317.1893310546875 0 763.9768
318.1482849121094 0 15318.202 b 2
318.18194580078125 0 1568.5847
319.1509094238281 0 2455.0173
324.15521240234375 0 804.7864
332.1238098144531 0 617.0676
334.14007568359375 0 6069.0186
335.1434631347656 0 1152.258
335.83343505859375 0 533.3038
339.2042541503906 0 726.1361
343.2026672363281 0 741.1333
344.1225891113281 0 927.63257
344.20025634765625 0 682.5391
350.134765625 0 789.3751
360.0867004394531 0 1667.123
360.11871337890625 0 1724.1747
361.2234802246094 0 958.61914
362.1357727050781 0 1404.1302
369.1202392578125 0 1127.2264
369.175048828125 0 742.373
370.102783203125 0 802.7383
373.1356201171875 0 581.5047
375.2237243652344 0 16466.49 y 9
376.1862487792969 0 1199.5515
376.22662353515625 0 2712.0454
378.0965576171875 0 14099.946
378.1292724609375 0 5537.9014
379.0980529785156 0 1122.489
379.1340026855469 0 816.0146
387.1357727050781 0 608.13116
388.11419677734375 0 2548.358
389.1170349121094 0 627.1683
389.21783447265625 0 1447.733
390.220703125 0 591.1489
392.1964416503906 0 606.3057
403.1639099121094 0 647.816
404.1112976074219 0 683.273
405.1768493652344 0 3553.8757
409.224365234375 0 860.94104
414.1298828125 0 1109.0795
415.16180419921875 0 1185.2926 b Water loss 3
421.17071533203125 0 641.8767
431.1551818847656 0 1622.8193
433.1408996582031 0 668.7624
433.1740417480469 0 3129.7666 b 3
434.1780700683594 0 664.2565
437.21734619140625 0 1203.3009
443.1575012207031 0 931.4097
447.2225036621094 0 1020.4037
449.1333312988281 0 915.65985
449.16705322265625 0 5430.0693
450.1695556640625 0 1087.1407
456.2803039550781 0 1162.9857
459.1514587402344 0 2121.553
461.16693115234375 0 916.5307
466.1935729980469 0 1005.9336
475.18304443359375 0 707.9869
477.1650695800781 0 1467.3667
479.1600341796875 0 1012.7419
484.2041015625 0 1428.638
486.1990051269531 0 610.9235
492.2103576660156 0 648.04956
497.16961669921875 0 1558.2361
498.17279052734375 0 665.897
502.19403076171875 0 1182.482
508.1496276855469 0 644.3885
508.2925109863281 0 692.88965
518.2601318359375 0 770.9672
520.2039184570312 0 5464.1064
521.206787109375 0 1419.6102
522.2919921875 0 15072.242 y 8
523.2950439453125 0 3370.6514
524.2969360351562 0 855.3288
525.1647338867188 0 3260.639
526.1690673828125 0 698.9079
526.2413940429688 0 720.2765
530.189453125 0 2667.9714 b Water loss 4
531.1920776367188 0 642.1888
532.2018432617188 0 1534.9343
542.2833251953125 0 990.5944
548.2012329101562 0 3748.9023 b 4
549.2042846679688 0 977.32544
552.2447509765625 0 1400.5986
553.24658203125 0 805.44104
553.763427734375 0 813.569
557.6250610351562 0 644.4088
562.7666015625 0 8036.502
563.2677612304688 0 6510.21
563.76953125 0 1425.9658
567.312744140625 0 564.0261
585.2293090820312 0 712.01385
585.7597045898438 0 1870.0828 y Water loss 2
586.2578735351562 0 2355.2764 y Ammonia loss 2
586.7559204101562 0 911.33905
593.3291015625 0 16081.989 y 7
594.33203125 0 4637.4067
594.765380859375 0 30317.701 y 2
595.2667236328125 0 20014.125
595.3253173828125 0 622.3462
595.7672729492188 0 9172.426
596.2024536132812 0 2508.6448
596.2716064453125 0 1065.3105
597.2073974609375 0 745.0805
603.2409057617188 0 1709.4307
604.2399291992188 0 837.0066
612.30029296875 0 1474.4702
622.245849609375 0 641.61615
631.267578125 0 1133.7341
635.7635498046875 0 653.1837
638.8145141601562 0 729.91034
644.29931640625 0 4173.527 y 1
644.80078125 0 3273.7869
645.3043212890625 0 1074.1271
647.8187866210938 0 6440.33
648.31982421875 0 4776.315
648.82080078125 0 1461.7363
649.2669677734375 0 1457.8599
650.2468872070312 0 734.3729
651.3117065429688 0 791.90186
663.26171875 0 622.3388
664.3658447265625 0 19761.744 y 6
665.3687744140625 0 7493.4736
666.3723754882812 0 1211.2847
667.2317504882812 0 922.7943
667.2728271484375 0 3114.9268
668.275390625 0 883.9554
671.3145141601562 0 849.25323 Precursor Ammonia loss
677.259521484375 0 1773.1987 b Water loss 5
679.8179931640625 0 7031.847 Precursor
680.0125732421875 0 795.8344
680.31884765625 0 5861.2607
680.8223266601562 0 3478.63
689.3436889648438 0 743.40753
695.2708740234375 0 2227.3567 b 5
696.2694702148438 0 773.3183
702.3096923828125 0 935.09656
732.2943725585938 0 677.3957
748.2949829101562 0 1020.2015 b Water loss 6
750.3079833984375 0 1332.6099
766.3150634765625 0 1425.1399 b 6
767.3125 0 798.30084
773.3458862304688 0 1032.6791
782.4066772460938 0 960.43726
811.4340209960938 0 16985.074 y 5
812.4375 0 6902.813
813.4403076171875 0 1720.2976
814.3084716796875 0 1426.3715
815.31298828125 0 947.0009
820.3524780273438 0 636.8831
837.3438110351562 0 2031.0116 b 7
838.346923828125 0 1208.8212
849.3748168945312 0 1323.7306
895.3619384765625 0 866.6689
908.4540405273438 0 912.7357 y Water loss 4
913.37451171875 0 1369.0221
920.41455078125 0 929.3182
926.46044921875 0 13196.195 y 4
927.46337890625 0 6777.473
928.4628295898438 0 1362.5742
984.4131469726562 0 2422.312 b 8
985.4142456054688 0 1318.811
1008.4966430664062 0 1000.42145
1023.4703369140625 0 1014.93365 y Water loss 3
1024.4775390625 0 1497.7955 y Ammonia loss 3
1041.4869384765625 0 36424.914 y 3
1042.490234375 0 21563.947
1043.4927978515625 0 5812.39
1044.5003662109375 0 706.6292
1051.4715576171875 0 1941.9719
1052.476806640625 0 1741.1365
1072.48681640625 0 631.581
1097.491455078125 0 690.56006
1124.5225830078125 0 6518.161
1125.526123046875 0 4395.901
1126.52880859375 0 802.67523
1170.5087890625 0 1187.2467 y Water loss 2
1188.521484375 0 21124.146 y 2
1189.525146484375 0 14596.2295
1190.5230712890625 0 6203.3237
1191.5272216796875 0 964.31177
2015.8380126953125 0 638.6099
2031.2301025390625 0 683.62714
2284.666259765625 0 758.4738

Spectrum Details

|  |  |
| --- | --- |
| Matched peaks? Matched peaks The total absolute number of peaks matched. Additionally in brackets the total fraction of peaks matched and the total number of peaks is shown. | 37 (12.01% of 308) |
| FDR? FDR The false discovery rate estimated for this peptide. It is calculated by matching all theoretical fragments with a non-integer shift with the raw peaks for this spectrum. This is done with 40 different shifts. The resulting percentage is the average number of annotated peaks over the number of annotated peaks with the correct spectrum. | 1.61% |
| Satellite FDR? Satellite FDR See the FDR for details on its calculation. This satellite ion specific FDR only contains the satellite ions (d/w) for I/L/J positions. | - |
| PSM Score? PSM Score The PSM Score as given by Hecklib to this annotated spectrum. It is shown with three significant figures. | 446 |

#### Spectrum 8461? Spectrum 8461 The raw spectrum of this peptide as annotated by Hecklib. The fragments are coloured according to ion type (see legend). Any peaks with a star '\*' as text can be hovered over to see the full details, first the ion type second the mass shift type. By hovering over the amino acids in the peptide or ions in the legend the corresponding peaks are highlighted. By toggling the 'Unassigned' label you can turn the background (unassigned) peaks on or off in the plot. By updating the slider in the Ion legend you can update the spectrum to only show the top X% of the peaks with labels. The top X% means any peak that is within X% of the highest intensity. By dragging in the spectrum you can zoom in to a specific part of the spectrum and use 'Zoom Out' to get back to the original zoom level. The annotation of the spectrum is based on the given sequence in the peptides file and is done with different software so inconsistencies are likely. The peaks are annotated based on the given sequence, with 20 ppm tolerance.

Copy Data

##### Spectrum 8461 (TSV)

###### Preview

```
Loading example...
```

*Click on the button to copy the data to your clipboard.*

Mz MinMz MaxIntensity Max

WidthHeightPeptide font sizePeptide stroke widthSpectrum font sizeSpectrum stroke widthCompact peptide

Ion legend

wxyz

abcd

OtherUnassignedIon^Charge^_Position_Show for top:%

AVMDDFAAFVEK

07.68e+31.54e+42.30e+43.07e+4

Zoom Out

d^+1^_2_y^+1^_1_a^+1^_2_y^+1^_1_b^+1^_2_y^+1^_2_y^+1^_2_a^+1^_3_b^+1^_3_y^+1^_3_b^+1^_4_b^+1^_4_y^+1^_4_b^+1^_5_b^+1^_5_y^+2^_10_y^+2^_10_y^+1^_5_y^+2^_10_y^+2^_11_y^+1^_6_\*b^+1^_6_\*b^+1^_6_b^+1^_7_y^+1^_7_b^+1^_8_y^+1^_8_b^+1^_9_y^+1^_9_b^+1^_10_y^+1^_10_

0776155223283105

Fragment Matches Table

Show background peaks

| Position | Ion type | Intensity | mz Theoretical | mz Error (Th) | mz Error (ppm) | Charge | Series Number |
| --- | --- | --- | --- | --- | --- | --- | --- |
| - | - | 926.6 | 120.1 | - | - | 0 | - |
| - | - | 1.498E+04 | 120.1 | - | - | 0 | - |
| - | - | 1023 | 121.1 | - | - | 0 | - |
| - | - | 421 | 126.9 | - | - | 0 | - |
| - | - | 861.6 | 127.1 | - | - | 0 | - |
| 2 | d | 8053 | 129.1 | 0.0001165 | 0.9023 | +1 | 2 |
| - | - | 375.5 | 129.1 | - | - | 0 | - |
| 12 | y | 3543 | 130.1 | 0.0001249 | 0.9605 | +1 | 1 |
| - | - | 2424 | 133.1 | - | - | 0 | - |
| - | - | 2351 | 136.1 | - | - | 0 | - |
| - | - | 412.7 | 140.1 | - | - | 0 | - |
| 2 | a | 3.04E+04 | 143.1 | 9.142E-05 | 0.6388 | +1 | 2 |
| - | - | 1842 | 144.1 | - | - | 0 | - |
| - | - | 413.8 | 144.4 | - | - | 0 | - |
| - | - | 545.5 | 147.1 | - | - | 0 | - |
| 12 | y | 6948 | 147.1 | 0.0001261 | 0.8574 | +1 | 1 |
| - | - | 453.2 | 147.8 | - | - | 0 | - |
| - | - | 454.1 | 148.9 | - | - | 0 | - |
| - | - | 650.9 | 149 | - | - | 0 | - |
| - | - | 607.8 | 152.1 | - | - | 0 | - |
| - | - | 733.9 | 153.1 | - | - | 0 | - |
| - | - | 930.4 | 155.1 | - | - | 0 | - |
| - | - | 3290 | 166.1 | - | - | 0 | - |
| - | - | 1147 | 167.1 | - | - | 0 | - |
| - | - | 491.2 | 167.1 | - | - | 0 | - |
| - | - | 3714 | 169.1 | - | - | 0 | - |
| - | - | 570.2 | 170.1 | - | - | 0 | - |
| - | - | 1157 | 171.1 | - | - | 0 | - |
| 2 | b | 1.331E+04 | 171.1 | 3.459E-05 | 0.2021 | +1 | 2 |
| - | - | 1154 | 172.1 | - | - | 0 | - |
| - | - | 1579 | 173.4 | - | - | 0 | - |
| - | - | 872.5 | 177.1 | - | - | 0 | - |
| - | - | 491.2 | 181.1 | - | - | 0 | - |
| - | - | 547 | 182.1 | - | - | 0 | - |
| - | - | 1003 | 183.1 | - | - | 0 | - |
| - | - | 1446 | 185.2 | - | - | 0 | - |
| - | - | 562.5 | 191.1 | - | - | 0 | - |
| - | - | 2632 | 191.1 | - | - | 0 | - |
| - | - | 3167 | 197.1 | - | - | 0 | - |
| - | - | 2770 | 199.1 | - | - | 0 | - |
| - | - | 607 | 199.1 | - | - | 0 | - |
| - | - | 1735 | 201.1 | - | - | 0 | - |
| - | - | 480.3 | 207.3 | - | - | 0 | - |
| - | - | 613.1 | 209.1 | - | - | 0 | - |
| - | - | 1106 | 211.1 | - | - | 0 | - |
| - | - | 504 | 213.1 | - | - | 0 | - |
| - | - | 673.5 | 213.2 | - | - | 0 | - |
| - | - | 1731 | 217.1 | - | - | 0 | - |
| - | - | 5296 | 219.1 | - | - | 0 | - |
| - | - | 529.7 | 219.1 | - | - | 0 | - |
| - | - | 2624 | 219.1 | - | - | 0 | - |
| - | - | 1312 | 221.1 | - | - | 0 | - |
| - | - | 602 | 222.1 | - | - | 0 | - |
| - | - | 2907 | 225 | - | - | 0 | - |
| - | - | 1013 | 226.1 | - | - | 0 | - |
| - | - | 1223 | 227.1 | - | - | 0 | - |
| - | - | 1363 | 229.1 | - | - | 0 | - |
| - | - | 1029 | 231.1 | - | - | 0 | - |
| - | - | 696.2 | 235.1 | - | - | 0 | - |
| - | - | 1695 | 235.1 | - | - | 0 | - |
| - | - | 3834 | 239.1 | - | - | 0 | - |
| - | - | 2815 | 240.1 | - | - | 0 | - |
| - | - | 842.5 | 244.1 | - | - | 0 | - |
| - | - | 869.5 | 245.1 | - | - | 0 | - |
| - | - | 1463 | 245.1 | - | - | 0 | - |
| - | - | 1028 | 247.1 | - | - | 0 | - |
| - | - | 854 | 247.1 | - | - | 0 | - |
| - | - | 1482 | 254.2 | - | - | 0 | - |
| 11 | y | 9521 | 258.1 | 2.666E-05 | 0.1033 | +1 | 2 |
| - | - | 7014 | 263.1 | - | - | 0 | - |
| - | - | 2189 | 263.1 | - | - | 0 | - |
| - | - | 554 | 273.2 | - | - | 0 | - |
| - | - | 3089 | 275.1 | - | - | 0 | - |
| 11 | y | 9622 | 276.2 | 3.226E-05 | 0.1168 | +1 | 2 |
| 3 | a | 4615 | 290.1 | 0.001681 | 5.793 | +1 | 3 |
| - | - | 1047 | 295.1 | - | - | 0 | - |
| - | - | 1384 | 296.1 | - | - | 0 | - |
| - | - | 574 | 298.8 | - | - | 0 | - |
| - | - | 2060 | 299.1 | - | - | 0 | - |
| - | - | 568.1 | 313.1 | - | - | 0 | - |
| - | - | 3490 | 314.1 | - | - | 0 | - |
| - | - | 4240 | 316.2 | - | - | 0 | - |
| 3 | b | 6224 | 318.1 | 0.004935 | 15.51 | +1 | 3 |
| - | - | 904.1 | 318.2 | - | - | 0 | - |
| - | - | 883.8 | 319.2 | - | - | 0 | - |
| - | - | 2754 | 334.1 | - | - | 0 | - |
| - | - | 790.7 | 335.1 | - | - | 0 | - |
| - | - | 731.9 | 341.2 | - | - | 0 | - |
| - | - | 514.2 | 349.7 | - | - | 0 | - |
| - | - | 525.9 | 350.1 | - | - | 0 | - |
| - | - | 577.4 | 350.7 | - | - | 0 | - |
| - | - | 563.1 | 355.1 | - | - | 0 | - |
| - | - | 600.2 | 355.1 | - | - | 0 | - |
| - | - | 1454 | 360.1 | - | - | 0 | - |
| - | - | 620.1 | 360.1 | - | - | 0 | - |
| - | - | 739.9 | 362.1 | - | - | 0 | - |
| - | - | 1125 | 369.1 | - | - | 0 | - |
| - | - | 574.4 | 374.2 | - | - | 0 | - |
| 10 | y | 7521 | 375.2 | 0.0002394 | 0.638 | +1 | 3 |
| - | - | 606.7 | 376.2 | - | - | 0 | - |
| - | - | 1072 | 376.2 | - | - | 0 | - |
| - | - | 6845 | 378.1 | - | - | 0 | - |
| - | - | 2877 | 378.1 | - | - | 0 | - |
| - | - | 690.3 | 379.1 | - | - | 0 | - |
| - | - | 580.4 | 379.1 | - | - | 0 | - |
| - | - | 1371 | 388.1 | - | - | 0 | - |
| - | - | 642 | 388.2 | - | - | 0 | - |
| - | - | 906.8 | 389.2 | - | - | 0 | - |
| - | - | 743.6 | 403.2 | - | - | 0 | - |
| - | - | 790 | 404.1 | - | - | 0 | - |
| - | - | 2352 | 405.2 | - | - | 0 | - |
| - | - | 629.8 | 409.2 | - | - | 0 | - |
| 4 | b | 675.6 | 415.2 | 0.002442 | 5.883 | +1 | 4 |
| - | - | 614.7 | 421.2 | - | - | 0 | - |
| - | - | 1046 | 431.2 | - | - | 0 | - |
| 4 | b | 2146 | 433.2 | 0.004695 | 10.84 | +1 | 4 |
| - | - | 656.6 | 437.2 | - | - | 0 | - |
| - | - | 716.1 | 447.2 | - | - | 0 | - |
| - | - | 2002 | 449.2 | - | - | 0 | - |
| - | - | 630.3 | 456.3 | - | - | 0 | - |
| - | - | 967.5 | 459.2 | - | - | 0 | - |
| - | - | 701.8 | 461.2 | - | - | 0 | - |
| - | - | 1102 | 477.2 | - | - | 0 | - |
| - | - | 603.7 | 487.1 | - | - | 0 | - |
| - | - | 1354 | 497.2 | - | - | 0 | - |
| - | - | 658.8 | 507.3 | - | - | 0 | - |
| - | - | 2865 | 520.2 | - | - | 0 | - |
| - | - | 627.5 | 521.2 | - | - | 0 | - |
| 9 | y | 7521 | 522.3 | 0.000355 | 0.6796 | +1 | 4 |
| - | - | 1548 | 523.3 | - | - | 0 | - |
| - | - | 1435 | 525.2 | - | - | 0 | - |
| 5 | b | 1525 | 530.2 | 0.004308 | 8.125 | +1 | 5 |
| - | - | 933.2 | 532.2 | - | - | 0 | - |
| - | - | 1284 | 541.6 | - | - | 0 | - |
| 5 | b | 1772 | 548.2 | 0.006194 | 11.3 | +1 | 5 |
| - | - | 852.3 | 549.2 | - | - | 0 | - |
| - | - | 690.5 | 550.2 | - | - | 0 | - |
| - | - | 1115 | 552.2 | - | - | 0 | - |
| - | - | 722.7 | 553.8 | - | - | 0 | - |
| - | - | 3511 | 562.8 | - | - | 0 | - |
| - | - | 2251 | 563.3 | - | - | 0 | - |
| - | - | 1471 | 563.8 | - | - | 0 | - |
| - | - | 773.7 | 585.2 | - | - | 0 | - |
| 3 | y | 957.4 | 585.8 | 0.002585 | 4.412 | +2 | 10 |
| 3 | y | 2086 | 586.2 | 0.005938 | 10.13 | +2 | 10 |
| 8 | y | 7994 | 593.3 | 0.0002983 | 0.5028 | +1 | 5 |
| - | - | 2507 | 594.3 | - | - | 0 | - |
| 3 | y | 1.547E+04 | 594.8 | 0.002307 | 3.879 | +2 | 10 |
| - | - | 1.139E+04 | 595.3 | - | - | 0 | - |
| - | - | 4320 | 595.8 | - | - | 0 | - |
| - | - | 1576 | 596.2 | - | - | 0 | - |
| - | - | 1193 | 603.2 | - | - | 0 | - |
| - | - | 642.1 | 604.2 | - | - | 0 | - |
| - | - | 910.3 | 612.8 | - | - | 0 | - |
| - | - | 679.1 | 631.3 | - | - | 0 | - |
| - | - | 753.6 | 638 | - | - | 0 | - |
| 2 | y | 1724 | 644.3 | 0.003012 | 4.675 | +2 | 11 |
| - | - | 1710 | 644.8 | - | - | 0 | - |
| - | - | 783 | 645.3 | - | - | 0 | - |
| - | - | 4398 | 647.8 | - | - | 0 | - |
| - | - | 1649 | 648.3 | - | - | 0 | - |
| - | - | 678.4 | 663.4 | - | - | 0 | - |
| 7 | y | 9751 | 664.4 | 0.0001855 | 0.2792 | +1 | 6 |
| - | - | 3250 | 665.4 | - | - | 0 | - |
| - | - | 800.1 | 667.2 | - | - | 0 | - |
| - | - | 527.9 | 667.3 | - | - | 0 | - |
| 0 | Precursor | 1023 | 670.8 | 0.001945 | 2.899 | +2 | -1 |
| 6 | b | 973.4 | 677.3 | 0.004192 | 6.19 | +1 | 6 |
| - | - | 791.4 | 679.5 | - | - | 0 | - |
| 0 | Precursor | 4250 | 679.8 | 0.001789 | 2.632 | +2 | -1 |
| - | - | 4682 | 680.3 | - | - | 0 | - |
| - | - | 907.5 | 680.4 | - | - | 0 | - |
| - | - | 2529 | 680.8 | - | - | 0 | - |
| - | - | 623.7 | 684.3 | - | - | 0 | - |
| 6 | b | 1499 | 695.3 | 0.005651 | 8.129 | +1 | 6 |
| - | - | 730.6 | 704.3 | - | - | 0 | - |
| 7 | b | 773.7 | 748.3 | 0.001672 | 2.234 | +1 | 7 |
| - | - | 1162 | 750.3 | - | - | 0 | - |
| - | - | 676.9 | 766.1 | - | - | 0 | - |
| - | - | 704.5 | 782.4 | - | - | 0 | - |
| 6 | y | 9716 | 811.4 | 6.994E-05 | 0.0862 | +1 | 7 |
| - | - | 2719 | 812.4 | - | - | 0 | - |
| - | - | 1204 | 813.4 | - | - | 0 | - |
| 8 | b | 808.1 | 837.3 | 0.01199 | 14.32 | +1 | 8 |
| - | - | 737.1 | 849.4 | - | - | 0 | - |
| - | - | 1024 | 913.4 | - | - | 0 | - |
| 5 | y | 7743 | 926.5 | 0.001055 | 1.139 | +1 | 8 |
| - | - | 2922 | 927.5 | - | - | 0 | - |
| - | - | 1069 | 928.5 | - | - | 0 | - |
| 9 | b | 862.5 | 984.4 | 0.004734 | 4.809 | +1 | 9 |
| - | - | 864.9 | 1007 | - | - | 0 | - |
| 4 | y | 1.447E+04 | 1041 | 0.001448 | 1.39 | +1 | 9 |
| - | - | 1087 | 1042 | - | - | 0 | - |
| - | - | 1.016E+04 | 1042 | - | - | 0 | - |
| - | - | 3228 | 1043 | - | - | 0 | - |
| - | - | 709 | 1045 | - | - | 0 | - |
| - | - | 1604 | 1051 | - | - | 0 | - |
| - | - | 933 | 1052 | - | - | 0 | - |
| 10 | b | 782.6 | 1083 | 0.001261 | 1.164 | +1 | 10 |
| - | - | 799.2 | 1098 | - | - | 0 | - |
| - | - | 4395 | 1125 | - | - | 0 | - |
| - | - | 2875 | 1126 | - | - | 0 | - |
| - | - | 974.9 | 1127 | - | - | 0 | - |
| 3 | y | 1.266E+04 | 1189 | 0.003346 | 2.815 | +1 | 10 |
| - | - | 9655 | 1190 | - | - | 0 | - |
| - | - | 3612 | 1191 | - | - | 0 | - |
| - | - | 682.4 | 1505 | - | - | 0 | - |
| - | - | 780.4 | 3074 | - | - | 0 | - |

m/z Charge Intensity FragmentType MassShift Position
120.06566619873047 0 926.63403
120.08090209960938 0 14977.192
121.08431243896484 0 1022.9051
126.91268920898438 0 421.02933
127.08673858642578 0 861.6387
129.10235595703125 0 8052.851 d 1
129.1102752685547 0 375.54385
130.0863800048828 0 3542.9592 y Ammonia loss 11
133.08602905273438 0 2423.8052
136.0758056640625 0 2350.7222
140.082275390625 0 412.7395
143.11798095703125 0 30403.822 a 1
144.12118530273438 0 1841.6993
144.4171905517578 0 413.7928
147.10208129882812 0 545.4872
147.11293029785156 0 6948.3906 y 11
147.78240966796875 0 453.22925
148.94644165039062 0 454.09906
149.04469299316406 0 650.8536
152.07080078125 0 607.7802
153.06546020507812 0 733.8666
155.11778259277344 0 930.4109
166.08627319335938 0 3289.6506
167.05589294433594 0 1146.6266
167.11782836914062 0 491.24176
169.13375854492188 0 3714.4712
170.1366729736328 0 570.23706
171.07669067382812 0 1157.0963
171.1128387451172 0 13308.112 b 1
172.1162872314453 0 1154.4602
173.43862915039062 0 1579.3839
177.11227416992188 0 872.5007
181.13360595703125 0 491.24466
182.09226989746094 0 547.02075
183.11302185058594 0 1003.4329
185.1649932861328 0 1446.2252
191.08438110351562 0 562.5177
191.11807250976562 0 2631.5044
197.1285400390625 0 3167.2346
199.07138061523438 0 2770.341
199.10760498046875 0 606.9776
201.1232452392578 0 1734.62
207.27215576171875 0 480.2678
209.1290740966797 0 613.0679
211.107421875 0 1106.4276
213.12400817871094 0 504.00235
213.1598358154297 0 673.5152
217.064208984375 0 1730.976
219.11279296875 0 5296.1304
219.13475036621094 0 529.67456
219.14947509765625 0 2623.5984
221.08436584472656 0 1311.5454
222.12435913085938 0 601.9627
225.04283142089844 0 2907.333
226.11892700195312 0 1012.81976
227.10302734375 0 1222.6167
229.11839294433594 0 1362.913
231.0611572265625 0 1028.7006
235.07395935058594 0 696.2055
235.1074676513672 0 1694.5638
239.09490966796875 0 3834.3599
240.1345977783203 0 2814.5789
244.12864685058594 0 842.4534
245.05824279785156 0 869.47455
245.1283416748047 0 1462.8315
247.11062622070312 0 1027.7142
247.14401245117188 0 854.0405
254.15049743652344 0 1481.7554
258.1448059082031 0 9521.45 y Water loss 10
263.0696716308594 0 7013.8994
263.1027526855469 0 2188.9873
273.15826416015625 0 554.033
275.1062927246094 0 3089.1287
276.1553649902344 0 9622.04 y 10
290.1500549316406 0 4615.222 a 2
295.1031494140625 0 1047.1425
296.08734130859375 0 1383.706
298.82568359375 0 573.97675
299.06170654296875 0 2060.3513
313.11395263671875 0 568.0829
314.0979919433594 0 3489.5762
316.1866149902344 0 4240.066
318.1482238769531 0 6224.452 b 2
318.17938232421875 0 904.0939
319.1524963378906 0 883.82733
334.1394958496094 0 2754.2363
335.1449890136719 0 790.6884
341.18121337890625 0 731.90924
349.6614685058594 0 514.2327
350.1351318359375 0 525.85004
350.6555480957031 0 577.4242
355.0682373046875 0 563.0668
355.1224060058594 0 600.1808
360.0849304199219 0 1454.3632
360.1175537109375 0 620.1261
362.1367492675781 0 739.855
369.1209716796875 0 1124.9457
374.1720886230469 0 574.4045
375.22357177734375 0 7521.309 y 9
376.18414306640625 0 606.65326
376.2279052734375 0 1072.3425
378.0963439941406 0 6845.2104
378.1291809082031 0 2877.3604
379.10040283203125 0 690.32764
379.1350402832031 0 580.435
388.1136474609375 0 1371.1392
388.1824645996094 0 642.0012
389.21856689453125 0 906.8493
403.1625671386719 0 743.5918
404.1119079589844 0 790.02673
405.1770935058594 0 2352.2397
409.22509765625 0 629.7772
415.162109375 0 675.5813 b Water loss 3
421.1720275878906 0 614.7378
431.1568908691406 0 1046.3087
433.1749267578125 0 2146.4036 b 3
437.21807861328125 0 656.58734
447.22393798828125 0 716.0714
449.16717529296875 0 2002.0219
456.28326416015625 0 630.31885
459.15216064453125 0 967.5251
461.16802978515625 0 701.8115
477.1656799316406 0 1101.9303
487.1452331542969 0 603.652
497.17059326171875 0 1354.0797
507.2576599121094 0 658.7678
520.2042236328125 0 2864.8271
521.2027587890625 0 627.48016
522.2918701171875 0 7521.0225 y 8
523.29638671875 0 1548.0793
525.16552734375 0 1435.4314
530.19091796875 0 1524.5514 b Water loss 4
532.2028198242188 0 933.1961
541.6131591796875 0 1283.7006
548.203369140625 0 1771.7384 b 4
549.2053833007812 0 852.3241
550.1959228515625 0 690.5485
552.2456665039062 0 1114.5327
553.7603149414062 0 722.7316
562.7666625976562 0 3510.616
563.2695922851562 0 2251.4287
563.7702026367188 0 1471.2588
585.2326049804688 0 773.70874
585.7605590820312 0 957.4076 y Water loss 2
586.2559204101562 0 2086.4685 y Ammonia loss 2
593.3290405273438 0 7994.303 y 7
594.333740234375 0 2506.9165
594.7655639648438 0 15467.966 y 2
595.2672119140625 0 11390.46
595.767578125 0 4319.595
596.2030639648438 0 1575.6492
603.2388916015625 0 1192.6155
604.2431640625 0 642.09546
612.80224609375 0 910.25934
631.2709350585938 0 679.1182
637.975341796875 0 753.5544
644.3004760742188 0 1723.6932 y 1
644.8001708984375 0 1710.2289
645.298828125 0 782.9748
647.8186645507812 0 4397.9004
648.31982421875 0 1649.4265
663.3766479492188 0 678.42487
664.3666381835938 0 9750.894 y 6
665.3692626953125 0 3250.176
667.2344360351562 0 800.0865
667.275146484375 0 527.8656
670.8126831054688 0 1023.0568 Precursor Water loss
677.2592163085938 0 973.4453 b Water loss 5
679.5140380859375 0 791.41907
679.8178100585938 0 4249.9814 Precursor
680.3206787109375 0 4682.3906
680.399658203125 0 907.5473
680.8212280273438 0 2529.4758
684.305419921875 0 623.6753
695.271240234375 0 1498.7177 b 5
704.32373046875 0 730.6127
748.2904663085938 0 773.7483 b Water loss 6
750.3056640625 0 1161.7788
766.0935668945312 0 676.8847
782.4051513671875 0 704.4952
811.4349365234375 0 9716.316 y 5
812.4351806640625 0 2719.3484
813.4403076171875 0 1203.7686
837.351806640625 0 808.076 b 7
849.3750610351562 0 737.0828
913.3740234375 0 1024.3287
926.4607543945312 0 7742.7344 y 4
927.4646606445312 0 2921.7097
928.4639892578125 0 1069.1652
984.4129638671875 0 862.5053 b 8
1007.4874267578125 0 864.9476
1041.4873046875 0 14465.924 y 3
1041.6063232421875 0 1087.3654
1042.490966796875 0 10159.208
1043.494140625 0 3228.1667
1044.50634765625 0 709.01074
1051.4722900390625 0 1604.2053
1052.474853515625 0 933.0307
1083.4779052734375 0 782.5872 b 9
1097.5052490234375 0 799.1784
1124.5250244140625 0 4394.7695
1125.529541015625 0 2874.5142
1126.5281982421875 0 974.93604
1188.5225830078125 0 12664.245 y 2
1189.526123046875 0 9654.982
1190.525634765625 0 3611.5415
1504.9208984375 0 682.4498
3073.8984375 0 780.41284

Spectrum Details

|  |  |
| --- | --- |
| Matched peaks? Matched peaks The total absolute number of peaks matched. Additionally in brackets the total fraction of peaks matched and the total number of peaks is shown. | 33 (15.87% of 208) |
| FDR? FDR The false discovery rate estimated for this peptide. It is calculated by matching all theoretical fragments with a non-integer shift with the raw peaks for this spectrum. This is done with 40 different shifts. The resulting percentage is the average number of annotated peaks over the number of annotated peaks with the correct spectrum. | 1.01% |
| Satellite FDR? Satellite FDR See the FDR for details on its calculation. This satellite ion specific FDR only contains the satellite ions (d/w) for I/L/J positions. | - |
| PSM Score? PSM Score The PSM Score as given by Hecklib to this annotated spectrum. It is shown with three significant figures. | 366 |

#### Spectrum 8638? Spectrum 8638 The raw spectrum of this peptide as annotated by Hecklib. The fragments are coloured according to ion type (see legend). Any peaks with a star '\*' as text can be hovered over to see the full details, first the ion type second the mass shift type. By hovering over the amino acids in the peptide or ions in the legend the corresponding peaks are highlighted. By toggling the 'Unassigned' label you can turn the background (unassigned) peaks on or off in the plot. By updating the slider in the Ion legend you can update the spectrum to only show the top X% of the peaks with labels. The top X% means any peak that is within X% of the highest intensity. By dragging in the spectrum you can zoom in to a specific part of the spectrum and use 'Zoom Out' to get back to the original zoom level. The annotation of the spectrum is based on the given sequence in the peptides file and is done with different software so inconsistencies are likely. The peaks are annotated based on the given sequence, with 20 ppm tolerance.

Copy Data

##### Spectrum 8638 (TSV)

###### Preview

```
Loading example...
```

*Click on the button to copy the data to your clipboard.*

Mz MinMz MaxIntensity Max

WidthHeightPeptide font sizePeptide stroke widthSpectrum font sizeSpectrum stroke widthCompact peptide

Ion legend

wxyz

abcd

OtherUnassignedIon^Charge^_Position_Show for top:%

AVMDDFAAFVEK

05.87e+31.17e+41.76e+42.35e+4

Zoom Out

y^+1^_1_y^+1^_2_z^+1^_3_y^+1^_3_z^+1^_4_y^+1^_4_z^+1^_5_y^+1^_5_y^+2^_10_y^+2^_11_z^+1^_6_y^+1^_6_z^+1^_7_y^+1^_7_c^+1^_8_z^+1^_8_y^+1^_8_c^+1^_9_z^+1^_9_y^+1^_9_c^+1^_10_z^+1^_10_y^+1^_10_c^+1^_11_z^+1^_11_

0688137620642753

Fragment Matches Table

Show background peaks

| Position | Ion type | Intensity | mz Theoretical | mz Error (Th) | mz Error (ppm) | Charge | Series Number |
| --- | --- | --- | --- | --- | --- | --- | --- |
| - | - | 395.2 | 132.9 | - | - | 0 | - |
| - | - | 592.5 | 133.1 | - | - | 0 | - |
| - | - | 426.3 | 135.8 | - | - | 0 | - |
| - | - | 400.8 | 136.2 | - | - | 0 | - |
| - | - | 360.3 | 136.5 | - | - | 0 | - |
| - | - | 413.8 | 138.6 | - | - | 0 | - |
| - | - | 3053 | 143.1 | - | - | 0 | - |
| - | - | 380.6 | 144 | - | - | 0 | - |
| 12 | y | 603.7 | 147.1 | 0.0004924 | 3.347 | +1 | 1 |
| - | - | 417.4 | 148.5 | - | - | 0 | - |
| - | - | 547.5 | 148.9 | - | - | 0 | - |
| - | - | 504.4 | 148.9 | - | - | 0 | - |
| - | - | 399.9 | 148.9 | - | - | 0 | - |
| - | - | 659.6 | 148.9 | - | - | 0 | - |
| - | - | 810.4 | 148.9 | - | - | 0 | - |
| - | - | 1051 | 148.9 | - | - | 0 | - |
| - | - | 1338 | 148.9 | - | - | 0 | - |
| - | - | 2580 | 148.9 | - | - | 0 | - |
| - | - | 4052 | 148.9 | - | - | 0 | - |
| - | - | 3700 | 149 | - | - | 0 | - |
| - | - | 1688 | 149 | - | - | 0 | - |
| - | - | 1198 | 149 | - | - | 0 | - |
| - | - | 999.5 | 149 | - | - | 0 | - |
| - | - | 763.8 | 149 | - | - | 0 | - |
| - | - | 481.8 | 149 | - | - | 0 | - |
| - | - | 483.8 | 149 | - | - | 0 | - |
| - | - | 485 | 149 | - | - | 0 | - |
| - | - | 387.1 | 149.1 | - | - | 0 | - |
| - | - | 429.5 | 151.6 | - | - | 0 | - |
| - | - | 413.5 | 153.6 | - | - | 0 | - |
| - | - | 2670 | 171.1 | - | - | 0 | - |
| - | - | 746.7 | 203.1 | - | - | 0 | - |
| - | - | 740.9 | 221.1 | - | - | 0 | - |
| - | - | 605.7 | 224.1 | - | - | 0 | - |
| - | - | 1014 | 225 | - | - | 0 | - |
| - | - | 522.6 | 232.3 | - | - | 0 | - |
| - | - | 528.4 | 233.7 | - | - | 0 | - |
| - | - | 1320 | 239.1 | - | - | 0 | - |
| - | - | 887.2 | 263.1 | - | - | 0 | - |
| - | - | 583.8 | 275.1 | - | - | 0 | - |
| 11 | y | 1510 | 276.2 | 0.0007002 | 2.535 | +1 | 2 |
| - | - | 1432 | 295.1 | - | - | 0 | - |
| - | - | 1073 | 299.1 | - | - | 0 | - |
| - | - | 2322 | 318.1 | - | - | 0 | - |
| - | - | 683.9 | 319.2 | - | - | 0 | - |
| - | - | 643.2 | 350.1 | - | - | 0 | - |
| 10 | z | 587.5 | 359.2 | 0.001425 | 3.968 | +1 | 3 |
| - | - | 2000 | 369.1 | - | - | 0 | - |
| 10 | y | 1919 | 375.2 | 8.68E-05 | 0.2313 | +1 | 3 |
| - | - | 1441 | 378.1 | - | - | 0 | - |
| - | - | 718.8 | 404.2 | - | - | 0 | - |
| - | - | 533.8 | 445.2 | - | - | 0 | - |
| 9 | z | 871.7 | 506.3 | 0.00192 | 3.793 | +1 | 4 |
| - | - | 3268 | 507.3 | - | - | 0 | - |
| - | - | 972 | 521.3 | - | - | 0 | - |
| 9 | y | 2844 | 522.3 | 0.000355 | 0.6796 | +1 | 4 |
| - | - | 1102 | 523.3 | - | - | 0 | - |
| - | - | 764.7 | 548.2 | - | - | 0 | - |
| - | - | 1348 | 562.8 | - | - | 0 | - |
| - | - | 644.2 | 563.3 | - | - | 0 | - |
| 8 | z | 1796 | 577.3 | 0.0006035 | 1.045 | +1 | 5 |
| - | - | 1339 | 578.3 | - | - | 0 | - |
| 8 | y | 1958 | 593.3 | 0.0004951 | 0.8345 | +1 | 5 |
| - | - | 808.3 | 594.3 | - | - | 0 | - |
| 3 | y | 6383 | 594.8 | 0.002551 | 4.29 | +2 | 10 |
| - | - | 3794 | 595.3 | - | - | 0 | - |
| - | - | 1784 | 595.8 | - | - | 0 | - |
| 2 | y | 1048 | 644.3 | 0.001303 | 2.023 | +2 | 11 |
| - | - | 564 | 644.8 | - | - | 0 | - |
| - | - | 1649 | 647.8 | - | - | 0 | - |
| 7 | z | 8370 | 648.3 | 0.001232 | 1.9 | +1 | 6 |
| - | - | 3774 | 649.4 | - | - | 0 | - |
| - | - | 1315 | 650.4 | - | - | 0 | - |
| 7 | y | 3762 | 664.4 | 0.0001807 | 0.272 | +1 | 6 |
| - | - | 774.4 | 665.4 | - | - | 0 | - |
| - | - | 2248 | 679.8 | - | - | 0 | - |
| - | - | 2176 | 680.3 | - | - | 0 | - |
| 6 | z | 6291 | 795.4 | 0.0005445 | 0.6845 | +1 | 7 |
| - | - | 3576 | 796.4 | - | - | 0 | - |
| - | - | 993.9 | 797.4 | - | - | 0 | - |
| 6 | y | 2123 | 811.4 | 0.0008456 | 1.042 | +1 | 7 |
| - | - | 1072 | 812.4 | - | - | 0 | - |
| - | - | 685.7 | 822.3 | - | - | 0 | - |
| - | - | 1314 | 853.4 | - | - | 0 | - |
| 8 | c | 1551 | 854.4 | 0.002836 | 3.32 | +1 | 8 |
| - | - | 1187 | 866.5 | - | - | 0 | - |
| 5 | z | 4519 | 910.4 | 0.0003349 | 0.3678 | +1 | 8 |
| - | - | 1.975E+04 | 911.4 | - | - | 0 | - |
| - | - | 9696 | 912.5 | - | - | 0 | - |
| - | - | 2245 | 913.5 | - | - | 0 | - |
| - | - | 2013 | 925.5 | - | - | 0 | - |
| 5 | y | 3372 | 926.5 | 0.002093 | 2.259 | +1 | 8 |
| - | - | 1561 | 927.5 | - | - | 0 | - |
| - | - | 3006 | 957.4 | - | - | 0 | - |
| - | - | 2199 | 958.4 | - | - | 0 | - |
| - | - | 660.3 | 959.4 | - | - | 0 | - |
| - | - | 769.5 | 981.5 | - | - | 0 | - |
| - | - | 797 | 1000 | - | - | 0 | - |
| 9 | c | 3175 | 1001 | 0.005467 | 5.459 | +1 | 9 |
| - | - | 1623 | 1002 | - | - | 0 | - |
| - | - | 1073 | 1003 | - | - | 0 | - |
| 4 | z | 827 | 1025 | 0.002011 | 1.961 | +1 | 9 |
| - | - | 8155 | 1026 | - | - | 0 | - |
| - | - | 3320 | 1027 | - | - | 0 | - |
| - | - | 1389 | 1028 | - | - | 0 | - |
| - | - | 1032 | 1032 | - | - | 0 | - |
| 4 | y | 6374 | 1041 | 0.002058 | 1.976 | +1 | 9 |
| - | - | 3626 | 1042 | - | - | 0 | - |
| - | - | 928.8 | 1044 | - | - | 0 | - |
| - | - | 3478 | 1056 | - | - | 0 | - |
| - | - | 4557 | 1057 | - | - | 0 | - |
| - | - | 1506 | 1058 | - | - | 0 | - |
| - | - | 635.8 | 1100 | - | - | 0 | - |
| 10 | c | 2965 | 1101 | 0.006328 | 5.75 | +1 | 10 |
| - | - | 1612 | 1102 | - | - | 0 | - |
| - | - | 709.3 | 1108 | - | - | 0 | - |
| - | - | 1804 | 1110 | - | - | 0 | - |
| - | - | 4190 | 1111 | - | - | 0 | - |
| - | - | 1803 | 1112 | - | - | 0 | - |
| - | - | 1082 | 1113 | - | - | 0 | - |
| - | - | 735.9 | 1125 | - | - | 0 | - |
| - | - | 847.2 | 1130 | - | - | 0 | - |
| 3 | z | 2056 | 1173 | 0.003271 | 2.79 | +1 | 10 |
| - | - | 1329 | 1174 | - | - | 0 | - |
| 3 | y | 4834 | 1189 | 0.002369 | 1.993 | +1 | 10 |
| - | - | 3373 | 1190 | - | - | 0 | - |
| - | - | 878.4 | 1191 | - | - | 0 | - |
| - | - | 722.5 | 1213 | - | - | 0 | - |
| - | - | 930.1 | 1218 | - | - | 0 | - |
| 11 | c | 1.63E+04 | 1230 | 0.002309 | 1.878 | +1 | 11 |
| - | - | 890.7 | 1230 | - | - | 0 | - |
| - | - | 1.27E+04 | 1231 | - | - | 0 | - |
| - | - | 5737 | 1232 | - | - | 0 | - |
| - | - | 882.5 | 1233 | - | - | 0 | - |
| - | - | 926.2 | 1254 | - | - | 0 | - |
| 2 | z | 2226 | 1272 | 0.003705 | 2.914 | +1 | 11 |
| - | - | 2013 | 1273 | - | - | 0 | - |
| - | - | 784.6 | 1274 | - | - | 0 | - |
| - | - | 4824 | 1300 | - | - | 0 | - |
| - | - | 3708 | 1301 | - | - | 0 | - |
| - | - | 2045 | 1302 | - | - | 0 | - |
| - | - | 657.5 | 1325 | - | - | 0 | - |
| - | - | 1394 | 1327 | - | - | 0 | - |
| - | - | 2687 | 1332 | - | - | 0 | - |
| - | - | 2161 | 1333 | - | - | 0 | - |
| - | - | 1419 | 1334 | - | - | 0 | - |
| - | - | 724 | 1341 | - | - | 0 | - |
| - | - | 1853 | 1342 | - | - | 0 | - |
| - | - | 2.326E+04 | 1343 | - | - | 0 | - |
| - | - | 1.925E+04 | 1344 | - | - | 0 | - |
| - | - | 9425 | 1345 | - | - | 0 | - |
| - | - | 2094 | 1346 | - | - | 0 | - |
| - | - | 1156 | 1354 | - | - | 0 | - |
| - | - | 855.9 | 1357 | - | - | 0 | - |
| - | - | 903.2 | 1358 | - | - | 0 | - |
| - | - | 1.404E+04 | 1359 | - | - | 0 | - |
| - | - | 2.013E+04 | 1360 | - | - | 0 | - |
| - | - | 1.459E+04 | 1361 | - | - | 0 | - |
| - | - | 1030 | 1361 | - | - | 0 | - |
| - | - | 4791 | 1362 | - | - | 0 | - |
| - | - | 2142 | 1363 | - | - | 0 | - |
| - | - | 831.1 | 1804 | - | - | 0 | - |
| - | - | 899.7 | 2025 | - | - | 0 | - |
| - | - | 744.6 | 2032 | - | - | 0 | - |
| - | - | 1355 | 2039 | - | - | 0 | - |
| - | - | 926.3 | 2040 | - | - | 0 | - |
| - | - | 890.6 | 2043 | - | - | 0 | - |
| - | - | 891.7 | 2064 | - | - | 0 | - |
| - | - | 704.9 | 2725 | - | - | 0 | - |

m/z Charge Intensity FragmentType MassShift Position
132.9029083251953 0 395.16333
133.08645629882812 0 592.52045
135.7625732421875 0 426.25848
136.1800537109375 0 400.7769
136.47718811035156 0 360.33832
138.57070922851562 0 413.81076
143.11834716796875 0 3052.9084
144.02914428710938 0 380.59525
147.11329650878906 0 603.68604 y 11
148.54501342773438 0 417.44476
148.8792724609375 0 547.5419
148.89358520507812 0 504.3737
148.90139770507812 0 399.87613
148.90866088867188 0 659.6498
148.91552734375 0 810.4067
148.92263793945312 0 1050.947
148.92974853515625 0 1338.2369
148.93716430664062 0 2579.9006
148.94482421875 0 4051.898
148.9613800048828 0 3700.14
148.96923828125 0 1688.1057
148.9761505126953 0 1198.0244
148.9833984375 0 999.4881
148.99073791503906 0 763.82086
148.99765014648438 0 481.79553
149.0049285888672 0 483.77795
149.01937866210938 0 484.99313
149.06365966796875 0 387.06393
151.57469177246094 0 429.45175
153.61647033691406 0 413.53036
171.11302185058594 0 2670.4998
203.10256958007812 0 746.6645
221.08480834960938 0 740.93713
224.08001708984375 0 605.6812
225.0428466796875 0 1013.5889
232.25299072265625 0 522.59454
233.65463256835938 0 528.3713
239.09461975097656 0 1320.4501
263.06976318359375 0 887.23676
275.1063537597656 0 583.7721
276.1560974121094 0 1510.086 y 10
295.1038818359375 0 1432.0499
299.0620422363281 0 1072.6993
318.1486511230469 0 2321.9805
319.150634765625 0 683.91895
350.13714599609375 0 643.1889
359.2065124511719 0 587.5467 z 9
369.12188720703125 0 2000.2544
375.2237243652344 0 1919.0029 y 9
378.097412109375 0 1440.958
404.21868896484375 0 718.8484
445.21307373046875 0 533.7931
506.2754211425781 0 871.6913 z 8
507.2822265625 0 3268.2214
521.2855834960938 0 971.9958
522.2918701171875 0 2843.8904 y 8
523.2947998046875 0 1101.7338
548.2000122070312 0 764.69037
562.7699584960938 0 1347.5524
563.2682495117188 0 644.22107
577.3112182617188 0 1795.9446 z 7
578.3143310546875 0 1339.3474
593.329833984375 0 1957.6572 y 7
594.3321533203125 0 808.2867
594.7658081054688 0 6382.7524 y 2
595.268310546875 0 3794.5
595.7681274414062 0 1783.9706
644.2987670898438 0 1048.2115 y 1
644.8009033203125 0 564.0276
647.8193359375 0 1648.9213
648.3464965820312 0 8369.678 z 6
649.3514404296875 0 3773.7964
650.3573608398438 0 1314.7917
664.3662719726562 0 3762.1387 y 6
665.3684692382812 0 774.44946
679.8187866210938 0 2247.8003
680.3186645507812 0 2175.5786
795.4166870117188 0 6290.669 z 5
796.4202270507812 0 3576.1943
797.4208984375 0 993.8505
811.4340209960938 0 2122.7305 y 5
812.4367065429688 0 1071.5851
822.3117065429688 0 685.6594
853.3623046875 0 1314.0571
854.3692016601562 0 1551.1646 c 7
866.4501342773438 0 1186.8324
910.4434204101562 0 4519.164 z 4
911.44970703125 0 19752.107
912.4539184570312 0 9695.6045
913.4567260742188 0 2245.293
925.4542236328125 0 2012.7028
926.459716796875 0 3372.4082 y 4
927.4642944335938 0 1561.3315
957.424560546875 0 3006.3665
958.4284057617188 0 2199.0972
959.4292602539062 0 660.31024
981.4840698242188 0 769.4668
1000.428466796875 0 796.974
1001.4402465820312 0 3175.3472 c 8
1002.44287109375 0 1622.6003
1003.4407958984375 0 1073.3462
1025.468017578125 0 827.03937 z 3
1026.47607421875 0 8154.5596
1027.4788818359375 0 3320.0955
1028.4830322265625 0 1389.0425
1032.4976806640625 0 1031.7053
1041.4866943359375 0 6373.94 y 3
1042.491943359375 0 3625.614
1043.5025634765625 0 928.78357
1056.4915771484375 0 3477.6404
1057.497314453125 0 4557.1387
1058.49951171875 0 1505.9211
1099.5013427734375 0 635.7508
1100.509521484375 0 2964.8271 c 9
1101.5107421875 0 1612.4556
1108.492431640625 0 709.2641
1109.5142822265625 0 1803.8386
1110.519775390625 0 4190.481
1111.5250244140625 0 1802.72
1112.5272216796875 0 1082.4038
1124.520751953125 0 735.8592
1129.5140380859375 0 847.1584
1172.5037841796875 0 2056.1602 z 2
1173.5087890625 0 1328.6301
1188.5216064453125 0 4833.7637 y 2
1189.5260009765625 0 3373.4133
1190.525390625 0 878.43976
1212.5322265625 0 722.49225
1217.6024169921875 0 930.0591
1229.548095703125 0 16303.071 c 10
1229.7086181640625 0 890.72565
1230.5509033203125 0 12695.233
1231.5511474609375 0 5736.9814
1232.551513671875 0 882.5201
1253.643798828125 0 926.20746
1271.5726318359375 0 2226.2166 z 1
1272.5740966796875 0 2013.2955
1273.578857421875 0 784.606
1299.613525390625 0 4824.031
1300.61474609375 0 3707.7893
1301.6134033203125 0 2044.6603
1324.591552734375 0 657.543
1326.62255859375 0 1393.5344
1331.638916015625 0 2686.8994
1332.640380859375 0 2160.8362
1333.64892578125 0 1418.6553
1340.667236328125 0 723.975
1341.617919921875 0 1852.668
1342.60986328125 0 23262.662
1343.613037109375 0 19254.227
1344.615966796875 0 9425.461
1345.623291015625 0 2093.5913
1354.174072265625 0 1155.7893
1356.7171630859375 0 855.9401
1357.6666259765625 0 903.1544
1358.626708984375 0 14037.277
1359.63232421875 0 20129.67
1360.635498046875 0 14586.953
1361.1341552734375 0 1030.046
1361.639892578125 0 4790.596
1362.6611328125 0 2141.5857
1803.9151611328125 0 831.1067
2024.889404296875 0 899.6514
2032.4281005859375 0 744.6301
2038.9447021484375 0 1354.5388
2039.943359375 0 926.27277
2042.955078125 0 890.633
2063.972900390625 0 891.69855
2725.36767578125 0 704.877

Spectrum Details

|  |  |
| --- | --- |
| Matched peaks? Matched peaks The total absolute number of peaks matched. Additionally in brackets the total fraction of peaks matched and the total number of peaks is shown. | 25 (14.79% of 169) |
| FDR? FDR The false discovery rate estimated for this peptide. It is calculated by matching all theoretical fragments with a non-integer shift with the raw peaks for this spectrum. This is done with 40 different shifts. The resulting percentage is the average number of annotated peaks over the number of annotated peaks with the correct spectrum. | 0.86% |
| Satellite FDR? Satellite FDR See the FDR for details on its calculation. This satellite ion specific FDR only contains the satellite ions (d/w) for I/L/J positions. | - |
| PSM Score? PSM Score The PSM Score as given by Hecklib to this annotated spectrum. It is shown with three significant figures. | 261 |

#### Reverse Lookup? Reverse Lookup All places where this read could be placed.

| Group | Segment | Template | Template Part | Read Part | Score | Unique |
| --- | --- | --- | --- | --- | --- | --- |
| Decoy | Decoy | PRLA | [247..259] | [0..12] | 46 | True |

| Recombined | Template Part | Read Part | Score | Unique |
| --- | --- | --- | --- | --- |
| PRLA | [247..259] | [0..12] | 46 | True |

#### Meta Information from Multiple reads

##### Number of combined reads

3

##### Intensity

0.3957

##### TotalArea

1.368E+06

#### Positional Score

Copy Data

##### Positional Score (TSV)

###### Preview

```
Loading example...
```

*Click on the button to copy the data to your clipboard.*

1001234567891011

Label Value
"0" 0.32
"1" 0.32
"2" 0.33
"3" 0.33
"4" 0.333
"5" 0.33
"6" 0.333
"7" 0.333
"8" 0.333
"9" 0.33
"10" 0.33
"11" 0.327

#### Meta Information from PEAKS

##### Scan Identifier

F2:8408

##### Original sequence

A

V

M

+15.99

D

D

F

A

A

F

V

E

K

##### Posttranslational Modifications

Oxidation (M)

##### Source File

D:\separate\_stitch\_analyses\xle-disambiguation\raw\20210323\_F1\_UM1\_Peng0013\_SA\_F59\_ingel\_3ug\_TL.raw

##### Fraction

2

##### Scan Feature

F2:11636

##### De Novo Score

99

##### ConfidenceScore

99

### m/z

679.8196

##### Mass

1357.6223

##### Charge

2

##### Retention Time

47.4

##### Predicted Retention Time

-

##### Area

4.518E+05

##### Parts Per Million

1.7

##### Fragmentation mode

HCD

##### Originating file

01 D:\separate\_stitch\_analyses\xle-disambiguation\20210325\_F59\_3ug\_DENOVO\_12.csv

#### Meta Information from PEAKS

##### Scan Identifier

F2:8461

##### Original sequence

A

V

M

+15.99

D

D

F

A

A

F

V

E

K

##### Posttranslational Modifications

Oxidation (M)

##### Source File

D:\separate\_stitch\_analyses\xle-disambiguation\raw\20210323\_F1\_UM1\_Peng0013\_SA\_F59\_ingel\_3ug\_TL.raw

##### Fraction

2

##### Scan Feature

F2:11635

##### De Novo Score

99

##### ConfidenceScore

99

### m/z

679.8192

##### Mass

1357.6223

##### Charge

2

##### Retention Time

47.74

##### Predicted Retention Time

-

##### Area

5.806E+05

##### Parts Per Million

1.2

##### Fragmentation mode

HCD

##### Originating file

01 D:\separate\_stitch\_analyses\xle-disambiguation\20210325\_F59\_3ug\_DENOVO\_12.csv

#### Meta Information from PEAKS

##### Scan Identifier

F1:8638

##### Original sequence

A

V

M

+15.99

D

D

F

A

A

F

V

E

K

##### Posttranslational Modifications

Oxidation (M)

##### Source File

D:\separate\_stitch\_analyses\xle-disambiguation\raw\20210323\_F1\_UM1\_Peng0013\_SA\_F59\_ingel\_3ug\_ELA.raw

##### Fraction

1

##### Scan Feature

F1:11828

##### De Novo Score

98

##### ConfidenceScore

98

### m/z

679.8201

##### Mass

1357.6223

##### Charge

2

##### Retention Time

47.4

##### Predicted Retention Time

-

##### Area

3.358E+05

##### Parts Per Million

2.4

##### Fragmentation mode

ETHCD

##### Originating file

01 D:\separate\_stitch\_analyses\xle-disambiguation\20210325\_F59\_3ug\_DENOVO\_12.csv
