## Supplementary material for "A handle on mass coincidence errors in *de novo* sequencing of antibodies by bottom-up proteomics": Combined_005.html

Details Combined\_005 | Stitch OverviewUndefined

### Read Combined\_005

#### Sequence (length=9)

TFDDYAMHW

#### Spectrum 7317? Spectrum 7317 The raw spectrum of this peptide as annotated by Hecklib. The fragments are coloured according to ion type (see legend). Any peaks with a star '\*' as text can be hovered over to see the full details, first the ion type second the mass shift type. By hovering over the amino acids in the peptide or ions in the legend the corresponding peaks are highlighted. By toggling the 'Unassigned' label you can turn the background (unassigned) peaks on or off in the plot. By updating the slider in the Ion legend you can update the spectrum to only show the top X% of the peaks with labels. The top X% means any peak that is within X% of the highest intensity. By dragging in the spectrum you can zoom in to a specific part of the spectrum and use 'Zoom Out' to get back to the original zoom level. The annotation of the spectrum is based on the given sequence in the peptides file and is done with different software so inconsistencies are likely. The peaks are annotated based on the given sequence, with 20 ppm tolerance.

Copy Data

##### Spectrum 7317 (TSV)

###### Preview

```
Loading example...
```

*Click on the button to copy the data to your clipboard.*

Mz MinMz MaxIntensity Max

WidthHeightPeptide font sizePeptide stroke widthSpectrum font sizeSpectrum stroke widthCompact peptide

Ion legend

wxyz

abcd

OtherUnassignedIon^Charge^_Position_Show for top:%

TFDDYAMHW

07.44e+61.49e+72.23e+72.98e+7

Zoom Out

b^+2^_3_a^+1^_2_a^+1^_2_y^+1^_1_a^+1^_2_b^+2^_4_b^+1^_2_y^+2^_3_b^+1^_2_d^+1^_3_a^+1^_3_a^+1^_3_y^+1^_2_b^+1^_3_b^+2^_6_y^+2^_5_b^+1^_3_b^+1^_4_y^+2^_7_y^+2^_7_b^+1^_4_y^+1^_3_b^+2^_8_y^+2^_8_y^+2^_8_y^+1^_4_\*\*b^+1^_5_b^+1^_5_b^+1^_6_b^+1^_6_y^+1^_5_y^+1^_6_y^+1^_6_b^+1^_7_b^+1^_7_y^+1^_7_y^+1^_7_b^+1^_8_b^+1^_8_b^+1^_8_y^+1^_8_y^+1^_8_

0616123318492466

Fragment Matches Table

Show background peaks

| Position | Ion type | Intensity | mz Theoretical | mz Error (Th) | mz Error (ppm) | Charge | Series Number |
| --- | --- | --- | --- | --- | --- | --- | --- |
| - | - | 7.092E+06 | 120.1 | - | - | 0 | - |
| - | - | 2.212E+04 | 121 | - | - | 0 | - |
| - | - | 2.139E+04 | 121.1 | - | - | 0 | - |
| - | - | 5.55E+05 | 121.1 | - | - | 0 | - |
| - | - | 3.172E+04 | 122.1 | - | - | 0 | - |
| - | - | 2.007E+04 | 122.1 | - | - | 0 | - |
| - | - | 1.502E+04 | 125.1 | - | - | 0 | - |
| - | - | 3.206E+04 | 127.1 | - | - | 0 | - |
| - | - | 1.358E+04 | 129.1 | - | - | 0 | - |
| - | - | 1.994E+05 | 130.1 | - | - | 0 | - |
| - | - | 1.681E+04 | 131.1 | - | - | 0 | - |
| - | - | 3.675E+05 | 132.1 | - | - | 0 | - |
| - | - | 4.066E+04 | 133.1 | - | - | 0 | - |
| - | - | 4.033E+04 | 134.1 | - | - | 0 | - |
| - | - | 1.395E+04 | 135.1 | - | - | 0 | - |
| - | - | 2.585E+06 | 136.1 | - | - | 0 | - |
| - | - | 7.759E+04 | 136.1 | - | - | 0 | - |
| - | - | 1.827E+05 | 137.1 | - | - | 0 | - |
| - | - | 1.259E+04 | 137.1 | - | - | 0 | - |
| - | - | 1.942E+05 | 138.1 | - | - | 0 | - |
| - | - | 3.076E+05 | 138.1 | - | - | 0 | - |
| - | - | 1.509E+04 | 139.1 | - | - | 0 | - |
| - | - | 1.679E+04 | 139.1 | - | - | 0 | - |
| - | - | 1.01E+04 | 141.6 | - | - | 0 | - |
| - | - | 4.82E+04 | 143.1 | - | - | 0 | - |
| - | - | 1.208E+04 | 143.1 | - | - | 0 | - |
| - | - | 4.587E+04 | 143.1 | - | - | 0 | - |
| - | - | 5.166E+04 | 144.1 | - | - | 0 | - |
| - | - | 9.894E+04 | 146.1 | - | - | 0 | - |
| - | - | 5.26E+04 | 146.1 | - | - | 0 | - |
| - | - | 1.122E+04 | 147 | - | - | 0 | - |
| - | - | 1.846E+04 | 147.1 | - | - | 0 | - |
| - | - | 3.351E+04 | 148.1 | - | - | 0 | - |
| - | - | 9.316E+04 | 148.1 | - | - | 0 | - |
| - | - | 1.315E+04 | 148.1 | - | - | 0 | - |
| - | - | 1.219E+04 | 150.1 | - | - | 0 | - |
| - | - | 2.26E+04 | 153.1 | - | - | 0 | - |
| - | - | 4.355E+04 | 155.1 | - | - | 0 | - |
| - | - | 6.302E+04 | 155.1 | - | - | 0 | - |
| - | - | 6.427E+05 | 156.1 | - | - | 0 | - |
| - | - | 4.064E+04 | 157.1 | - | - | 0 | - |
| - | - | 1.079E+06 | 158.1 | - | - | 0 | - |
| - | - | 1.471E+04 | 159.1 | - | - | 0 | - |
| - | - | 5.712E+05 | 159.1 | - | - | 0 | - |
| - | - | 1.182E+05 | 159.1 | - | - | 0 | - |
| - | - | 2.066E+05 | 160.1 | - | - | 0 | - |
| - | - | 2.173E+04 | 160.1 | - | - | 0 | - |
| - | - | 4.241E+04 | 160.1 | - | - | 0 | - |
| - | - | 4.865E+05 | 160.1 | - | - | 0 | - |
| - | - | 5.282E+04 | 161.1 | - | - | 0 | - |
| - | - | 1.968E+04 | 162.1 | - | - | 0 | - |
| - | - | 1.821E+04 | 164.1 | - | - | 0 | - |
| - | - | 2.865E+04 | 165.1 | - | - | 0 | - |
| - | - | 1.31E+05 | 165.1 | - | - | 0 | - |
| - | - | 1.376E+04 | 165.1 | - | - | 0 | - |
| - | - | 4.225E+04 | 166.1 | - | - | 0 | - |
| - | - | 9.407E+04 | 166.1 | - | - | 0 | - |
| - | - | 1.359E+04 | 166.1 | - | - | 0 | - |
| - | - | 1.393E+04 | 167 | - | - | 0 | - |
| - | - | 1.165E+05 | 167.1 | - | - | 0 | - |
| - | - | 3.248E+04 | 169.1 | - | - | 0 | - |
| - | - | 1.672E+04 | 170.1 | - | - | 0 | - |
| - | - | 8.393E+04 | 171.1 | - | - | 0 | - |
| - | - | 1.76E+04 | 172.1 | - | - | 0 | - |
| - | - | 1.727E+04 | 172.1 | - | - | 0 | - |
| - | - | 1.398E+04 | 173.1 | - | - | 0 | - |
| - | - | 2.036E+04 | 173.4 | - | - | 0 | - |
| 3 | b | 7.996E+04 | 174.1 | 0.001177 | 6.759 | +2 | 3 |
| - | - | 1.345E+04 | 174.1 | - | - | 0 | - |
| - | - | 1.164E+04 | 175.1 | - | - | 0 | - |
| - | - | 3.813E+04 | 175.1 | - | - | 0 | - |
| - | - | 1.432E+04 | 175.1 | - | - | 0 | - |
| - | - | 2.83E+04 | 175.1 | - | - | 0 | - |
| - | - | 1.78E+05 | 176.1 | - | - | 0 | - |
| - | - | 2.176E+06 | 176.1 | - | - | 0 | - |
| - | - | 1.492E+06 | 177.1 | - | - | 0 | - |
| - | - | 2.127E+05 | 177.1 | - | - | 0 | - |
| - | - | 2.102E+04 | 178.1 | - | - | 0 | - |
| - | - | 1.242E+04 | 178.1 | - | - | 0 | - |
| - | - | 1.605E+05 | 178.1 | - | - | 0 | - |
| - | - | 1.121E+05 | 180.1 | - | - | 0 | - |
| - | - | 1.805E+05 | 185.1 | - | - | 0 | - |
| - | - | 1.852E+05 | 186.1 | - | - | 0 | - |
| - | - | 2.875E+04 | 187.1 | - | - | 0 | - |
| - | - | 7.801E+05 | 188.1 | - | - | 0 | - |
| - | - | 1.059E+05 | 189.1 | - | - | 0 | - |
| - | - | 1.18E+04 | 189.1 | - | - | 0 | - |
| - | - | 2.943E+04 | 190.1 | - | - | 0 | - |
| - | - | 7.766E+04 | 191.1 | - | - | 0 | - |
| - | - | 1.339E+05 | 191.1 | - | - | 0 | - |
| - | - | 2.19E+04 | 192.1 | - | - | 0 | - |
| - | - | 3.934E+04 | 193.1 | - | - | 0 | - |
| - | - | 6.274E+05 | 193.1 | - | - | 0 | - |
| - | - | 2.437E+04 | 194.1 | - | - | 0 | - |
| - | - | 1.129E+04 | 194.1 | - | - | 0 | - |
| - | - | 5.268E+04 | 194.1 | - | - | 0 | - |
| - | - | 1.686E+05 | 195.1 | - | - | 0 | - |
| - | - | 1.523E+04 | 197.1 | - | - | 0 | - |
| - | - | 7.945E+04 | 202.1 | - | - | 0 | - |
| - | - | 1.805E+05 | 203.1 | - | - | 0 | - |
| - | - | 6.063E+04 | 203.1 | - | - | 0 | - |
| 2 | a | 7.366E+05 | 203.1 | 0.0003813 | 1.877 | +1 | 2 |
| - | - | 3.337E+05 | 204.1 | - | - | 0 | - |
| 2 | a | 5.33E+04 | 204.1 | 0.0001762 | 0.8632 | +1 | 2 |
| - | - | 1.045E+05 | 204.1 | - | - | 0 | - |
| - | - | 2.748E+04 | 205.1 | - | - | 0 | - |
| 9 | y | 5.802E+05 | 205.1 | 0.0004716 | 2.3 | +1 | 1 |
| - | - | 4.376E+04 | 205.1 | - | - | 0 | - |
| - | - | 5.878E+04 | 206.1 | - | - | 0 | - |
| - | - | 1.516E+05 | 207.1 | - | - | 0 | - |
| - | - | 9.289E+04 | 208.1 | - | - | 0 | - |
| - | - | 2.695E+04 | 208.1 | - | - | 0 | - |
| - | - | 3.891E+04 | 212.1 | - | - | 0 | - |
| - | - | 1.594E+04 | 213.1 | - | - | 0 | - |
| - | - | 1.425E+04 | 213.1 | - | - | 0 | - |
| - | - | 1.264E+04 | 214 | - | - | 0 | - |
| - | - | 3.747E+04 | 215.1 | - | - | 0 | - |
| - | - | 4.05E+05 | 217.1 | - | - | 0 | - |
| - | - | 2.258E+04 | 218.1 | - | - | 0 | - |
| - | - | 5.618E+04 | 218.1 | - | - | 0 | - |
| - | - | 1.216E+05 | 219.1 | - | - | 0 | - |
| - | - | 1.8E+04 | 219.1 | - | - | 0 | - |
| - | - | 1.889E+04 | 220.1 | - | - | 0 | - |
| - | - | 1.702E+06 | 221.1 | - | - | 0 | - |
| 2 | a | 2.947E+07 | 221.1 | 0.0006199 | 2.803 | +1 | 2 |
| - | - | 1.618E+05 | 222.1 | - | - | 0 | - |
| - | - | 3.583E+06 | 222.1 | - | - | 0 | - |
| - | - | 4.136E+04 | 223.1 | - | - | 0 | - |
| - | - | 2.203E+05 | 223.1 | - | - | 0 | - |
| - | - | 2.111E+04 | 224.1 | - | - | 0 | - |
| - | - | 1.647E+04 | 225.1 | - | - | 0 | - |
| - | - | 8.469E+04 | 228.1 | - | - | 0 | - |
| - | - | 6.141E+04 | 229.1 | - | - | 0 | - |
| - | - | 4.534E+05 | 231.1 | - | - | 0 | - |
| 4 | b | 1.493E+04 | 231.1 | 0.001319 | 5.709 | +2 | 4 |
| 2 | b | 3.008E+05 | 231.1 | 0.0004771 | 2.064 | +1 | 2 |
| - | - | 1.981E+04 | 231.1 | - | - | 0 | - |
| - | - | 3.57E+04 | 232.1 | - | - | 0 | - |
| - | - | 3.004E+04 | 232.1 | - | - | 0 | - |
| - | - | 3.643E+04 | 233.1 | - | - | 0 | - |
| - | - | 8.076E+05 | 235.1 | - | - | 0 | - |
| - | - | 9.08E+04 | 236.1 | - | - | 0 | - |
| - | - | 2.61E+04 | 237.1 | - | - | 0 | - |
| - | - | 2.192E+04 | 237.1 | - | - | 0 | - |
| - | - | 2.848E+04 | 238.1 | - | - | 0 | - |
| - | - | 1.664E+05 | 239.1 | - | - | 0 | - |
| - | - | 1.806E+04 | 240.1 | - | - | 0 | - |
| - | - | 1.779E+04 | 240.1 | - | - | 0 | - |
| 7 | y | 3.482E+04 | 245.1 | 0.004094 | 16.7 | +2 | 3 |
| - | - | 2.855E+04 | 245.1 | - | - | 0 | - |
| - | - | 1.542E+04 | 246.1 | - | - | 0 | - |
| - | - | 5.131E+04 | 246.1 | - | - | 0 | - |
| - | - | 3.4E+04 | 247.1 | - | - | 0 | - |
| - | - | 3.905E+04 | 247.1 | - | - | 0 | - |
| - | - | 1.851E+04 | 247.1 | - | - | 0 | - |
| - | - | 2.053E+04 | 248.1 | - | - | 0 | - |
| - | - | 7.327E+04 | 248.2 | - | - | 0 | - |
| 2 | b | 5.132E+06 | 249.1 | 0.0005936 | 2.383 | +1 | 2 |
| - | - | 6.919E+05 | 250.1 | - | - | 0 | - |
| - | - | 5.155E+05 | 251.1 | - | - | 0 | - |
| - | - | 8.308E+04 | 251.1 | - | - | 0 | - |
| - | - | 5.55E+04 | 252.1 | - | - | 0 | - |
| - | - | 2.388E+04 | 253.1 | - | - | 0 | - |
| - | - | 1.495E+04 | 255.1 | - | - | 0 | - |
| - | - | 3.327E+04 | 256.1 | - | - | 0 | - |
| - | - | 2.774E+04 | 257.1 | - | - | 0 | - |
| - | - | 3.441E+05 | 257.1 | - | - | 0 | - |
| - | - | 3.34E+04 | 258.1 | - | - | 0 | - |
| - | - | 1.99E+04 | 258.1 | - | - | 0 | - |
| - | - | 9.03E+04 | 261.1 | - | - | 0 | - |
| - | - | 4.817E+04 | 263.1 | - | - | 0 | - |
| - | - | 1.039E+06 | 263.1 | - | - | 0 | - |
| - | - | 9.945E+04 | 263.1 | - | - | 0 | - |
| - | - | 1.675E+05 | 264.1 | - | - | 0 | - |
| - | - | 2.169E+04 | 264.1 | - | - | 0 | - |
| - | - | 1.052E+05 | 264.1 | - | - | 0 | - |
| - | - | 4.273E+04 | 265.1 | - | - | 0 | - |
| - | - | 2.31E+04 | 266.1 | - | - | 0 | - |
| - | - | 6.202E+04 | 267.1 | - | - | 0 | - |
| - | - | 5.77E+04 | 268.1 | - | - | 0 | - |
| - | - | 1.899E+04 | 268.1 | - | - | 0 | - |
| - | - | 3.667E+04 | 268.1 | - | - | 0 | - |
| - | - | 4.996E+04 | 269.1 | - | - | 0 | - |
| - | - | 1.399E+04 | 270.1 | - | - | 0 | - |
| - | - | 2.534E+04 | 270.1 | - | - | 0 | - |
| - | - | 4.612E+04 | 271.1 | - | - | 0 | - |
| - | - | 2.447E+04 | 272.1 | - | - | 0 | - |
| - | - | 1.651E+04 | 273.1 | - | - | 0 | - |
| - | - | 2.277E+04 | 273.1 | - | - | 0 | - |
| - | - | 2.828E+05 | 274.1 | - | - | 0 | - |
| - | - | 2.51E+04 | 274.2 | - | - | 0 | - |
| - | - | 4.325E+04 | 275.1 | - | - | 0 | - |
| - | - | 3.553E+04 | 275.1 | - | - | 0 | - |
| - | - | 1.738E+04 | 277.6 | - | - | 0 | - |
| - | - | 1.428E+04 | 277.6 | - | - | 0 | - |
| - | - | 4.893E+05 | 279.1 | - | - | 0 | - |
| - | - | 4.284E+04 | 279.1 | - | - | 0 | - |
| - | - | 4.353E+04 | 280.1 | - | - | 0 | - |
| - | - | 1.881E+04 | 283.1 | - | - | 0 | - |
| - | - | 2E+04 | 283.2 | - | - | 0 | - |
| - | - | 6.77E+05 | 285.1 | - | - | 0 | - |
| - | - | 7.442E+04 | 286.1 | - | - | 0 | - |
| - | - | 1.439E+04 | 286.1 | - | - | 0 | - |
| - | - | 3.454E+04 | 287.1 | - | - | 0 | - |
| - | - | 3.383E+04 | 289.1 | - | - | 0 | - |
| - | - | 1.591E+04 | 289.1 | - | - | 0 | - |
| - | - | 2.72E+04 | 290.1 | - | - | 0 | - |
| - | - | 4.082E+05 | 292.1 | - | - | 0 | - |
| 3 | d | 9.963E+04 | 292.2 | 0.0006612 | 2.263 | +1 | 3 |
| - | - | 4.242E+04 | 293.1 | - | - | 0 | - |
| - | - | 2.002E+04 | 294.2 | - | - | 0 | - |
| - | - | 1.352E+04 | 295.1 | - | - | 0 | - |
| - | - | 5.391E+05 | 296.2 | - | - | 0 | - |
| - | - | 2.216E+04 | 297.1 | - | - | 0 | - |
| - | - | 8.926E+04 | 297.2 | - | - | 0 | - |
| - | - | 3.438E+04 | 299.2 | - | - | 0 | - |
| - | - | 6.348E+04 | 300.1 | - | - | 0 | - |
| - | - | 3.336E+04 | 301.1 | - | - | 0 | - |
| - | - | 4.723E+04 | 302.1 | - | - | 0 | - |
| - | - | 9.16E+04 | 303.1 | - | - | 0 | - |
| - | - | 1.723E+04 | 304.1 | - | - | 0 | - |
| - | - | 1.57E+04 | 305.5 | - | - | 0 | - |
| - | - | 2.102E+04 | 306.1 | - | - | 0 | - |
| - | - | 1.584E+04 | 306.1 | - | - | 0 | - |
| - | - | 7.32E+04 | 310.2 | - | - | 0 | - |
| - | - | 4.283E+04 | 313.1 | - | - | 0 | - |
| - | - | 7.011E+04 | 314.1 | - | - | 0 | - |
| - | - | 2.502E+04 | 315.1 | - | - | 0 | - |
| 3 | a | 1.661E+05 | 318.1 | 0.0006752 | 2.122 | +1 | 3 |
| - | - | 3.497E+04 | 319.1 | - | - | 0 | - |
| - | - | 2.64E+04 | 320.1 | - | - | 0 | - |
| - | - | 1.977E+05 | 323.1 | - | - | 0 | - |
| - | - | 1.485E+04 | 323.1 | - | - | 0 | - |
| - | - | 1.181E+06 | 324.1 | - | - | 0 | - |
| - | - | 2.016E+05 | 325.1 | - | - | 0 | - |
| - | - | 1.857E+04 | 326.2 | - | - | 0 | - |
| - | - | 5.518E+04 | 327.1 | - | - | 0 | - |
| - | - | 1.14E+05 | 328.1 | - | - | 0 | - |
| - | - | 4.926E+04 | 328.1 | - | - | 0 | - |
| - | - | 1.846E+04 | 329.1 | - | - | 0 | - |
| - | - | 6.51E+04 | 330.1 | - | - | 0 | - |
| - | - | 2.631E+04 | 331.1 | - | - | 0 | - |
| - | - | 1.334E+04 | 331.1 | - | - | 0 | - |
| - | - | 8.633E+04 | 332.1 | - | - | 0 | - |
| - | - | 3.808E+04 | 333.1 | - | - | 0 | - |
| - | - | 2.211E+04 | 334.1 | - | - | 0 | - |
| - | - | 2.245E+04 | 334.1 | - | - | 0 | - |
| - | - | 2.385E+04 | 335.2 | - | - | 0 | - |
| 3 | a | 2.595E+04 | 336.2 | 0.0007002 | 2.083 | +1 | 3 |
| - | - | 7.542E+04 | 338.1 | - | - | 0 | - |
| - | - | 1.095E+05 | 339.2 | - | - | 0 | - |
| - | - | 1.705E+04 | 340.2 | - | - | 0 | - |
| 8 | y | 6.117E+06 | 342.2 | 0.0007334 | 2.143 | +1 | 2 |
| - | - | 1.121E+06 | 343.2 | - | - | 0 | - |
| - | - | 8.662E+04 | 344.2 | - | - | 0 | - |
| 3 | b | 3.654E+05 | 346.1 | 0.0006032 | 1.743 | +1 | 3 |
| - | - | 6.295E+04 | 347.1 | - | - | 0 | - |
| - | - | 1.655E+05 | 348.1 | - | - | 0 | - |
| - | - | 1.797E+04 | 349.1 | - | - | 0 | - |
| - | - | 1.668E+04 | 349.1 | - | - | 0 | - |
| - | - | 3.538E+05 | 350.1 | - | - | 0 | - |
| - | - | 5.513E+04 | 351.1 | - | - | 0 | - |
| - | - | 1.085E+05 | 352.1 | - | - | 0 | - |
| - | - | 2.38E+04 | 353.1 | - | - | 0 | - |
| - | - | 1.513E+04 | 354.2 | - | - | 0 | - |
| - | - | 3.383E+05 | 356.1 | - | - | 0 | - |
| 6 | b | 4.834E+04 | 357.1 | 0.0002749 | 0.7698 | +2 | 6 |
| - | - | 2.037E+04 | 358.1 | - | - | 0 | - |
| - | - | 2.094E+04 | 359.1 | - | - | 0 | - |
| - | - | 9.46E+04 | 360.1 | - | - | 0 | - |
| - | - | 2.147E+04 | 360.1 | - | - | 0 | - |
| - | - | 1.94E+04 | 361.1 | - | - | 0 | - |
| 5 | y | 5.52E+04 | 362.1 | 0.002981 | 8.232 | +2 | 5 |
| 3 | b | 1.533E+06 | 364.2 | 0.0005365 | 1.473 | +1 | 3 |
| - | - | 2.6E+05 | 365.2 | - | - | 0 | - |
| - | - | 1.356E+05 | 366.1 | - | - | 0 | - |
| - | - | 4.048E+04 | 366.2 | - | - | 0 | - |
| - | - | 3.769E+04 | 367.1 | - | - | 0 | - |
| - | - | 7.089E+04 | 367.2 | - | - | 0 | - |
| - | - | 1.789E+04 | 368.1 | - | - | 0 | - |
| - | - | 4.528E+04 | 371.1 | - | - | 0 | - |
| - | - | 6.732E+04 | 374.1 | - | - | 0 | - |
| - | - | 1.589E+05 | 376.1 | - | - | 0 | - |
| - | - | 1.499E+04 | 377.1 | - | - | 0 | - |
| - | - | 2.22E+04 | 377.1 | - | - | 0 | - |
| - | - | 5.557E+04 | 378.1 | - | - | 0 | - |
| - | - | 8.416E+05 | 378.1 | - | - | 0 | - |
| - | - | 1.674E+05 | 379.1 | - | - | 0 | - |
| - | - | 5.396E+04 | 380.1 | - | - | 0 | - |
| - | - | 1.678E+05 | 382.1 | - | - | 0 | - |
| - | - | 1.961E+04 | 383.1 | - | - | 0 | - |
| - | - | 4.301E+04 | 384.1 | - | - | 0 | - |
| - | - | 5.008E+04 | 385.1 | - | - | 0 | - |
| - | - | 1.765E+04 | 386.1 | - | - | 0 | - |
| - | - | 2.527E+04 | 388.1 | - | - | 0 | - |
| - | - | 3.338E+04 | 388.1 | - | - | 0 | - |
| - | - | 1.235E+05 | 390.2 | - | - | 0 | - |
| - | - | 2.105E+04 | 391.2 | - | - | 0 | - |
| - | - | 1.093E+06 | 394.1 | - | - | 0 | - |
| - | - | 2.052E+05 | 395.1 | - | - | 0 | - |
| - | - | 2.468E+04 | 395.2 | - | - | 0 | - |
| - | - | 2.619E+04 | 396.1 | - | - | 0 | - |
| - | - | 4.089E+04 | 397.2 | - | - | 0 | - |
| - | - | 2.386E+04 | 398.2 | - | - | 0 | - |
| - | - | 6.006E+04 | 399.2 | - | - | 0 | - |
| - | - | 1.8E+04 | 401.2 | - | - | 0 | - |
| - | - | 3.083E+04 | 404.1 | - | - | 0 | - |
| - | - | 2.902E+04 | 404.1 | - | - | 0 | - |
| - | - | 1.586E+04 | 405.1 | - | - | 0 | - |
| - | - | 4.157E+04 | 406.2 | - | - | 0 | - |
| - | - | 1.139E+05 | 407.2 | - | - | 0 | - |
| - | - | 2.417E+05 | 408.2 | - | - | 0 | - |
| - | - | 5.339E+04 | 409.2 | - | - | 0 | - |
| - | - | 2.086E+04 | 410.1 | - | - | 0 | - |
| - | - | 1.758E+04 | 410.2 | - | - | 0 | - |
| - | - | 7.861E+04 | 415.2 | - | - | 0 | - |
| - | - | 1.625E+04 | 416.1 | - | - | 0 | - |
| - | - | 1.587E+04 | 417.1 | - | - | 0 | - |
| - | - | 8.144E+04 | 419.2 | - | - | 0 | - |
| - | - | 1.63E+04 | 423.2 | - | - | 0 | - |
| - | - | 9.406E+05 | 425.2 | - | - | 0 | - |
| - | - | 1.993E+05 | 426.2 | - | - | 0 | - |
| - | - | 1.834E+05 | 427.2 | - | - | 0 | - |
| - | - | 3.113E+04 | 428.2 | - | - | 0 | - |
| - | - | 1.86E+04 | 429.1 | - | - | 0 | - |
| - | - | 1.451E+04 | 429.2 | - | - | 0 | - |
| - | - | 2.208E+04 | 430.1 | - | - | 0 | - |
| - | - | 1.919E+04 | 431.2 | - | - | 0 | - |
| - | - | 4.005E+04 | 432.1 | - | - | 0 | - |
| - | - | 1.276E+05 | 433.2 | - | - | 0 | - |
| - | - | 3.345E+04 | 434.2 | - | - | 0 | - |
| - | - | 5.625E+04 | 435.2 | - | - | 0 | - |
| - | - | 1.809E+04 | 436.1 | - | - | 0 | - |
| - | - | 2.252E+04 | 436.2 | - | - | 0 | - |
| - | - | 3.364E+04 | 437.2 | - | - | 0 | - |
| - | - | 2.536E+04 | 438.2 | - | - | 0 | - |
| - | - | 4.036E+04 | 442.2 | - | - | 0 | - |
| - | - | 4.566E+04 | 443.2 | - | - | 0 | - |
| - | - | 2.614E+04 | 443.2 | - | - | 0 | - |
| - | - | 1.217E+05 | 447.2 | - | - | 0 | - |
| - | - | 4.753E+04 | 449.1 | - | - | 0 | - |
| - | - | 3.203E+04 | 449.2 | - | - | 0 | - |
| - | - | 1.774E+04 | 452.7 | - | - | 0 | - |
| - | - | 7.414E+04 | 454.2 | - | - | 0 | - |
| - | - | 1.883E+05 | 455.2 | - | - | 0 | - |
| - | - | 3.899E+04 | 456.2 | - | - | 0 | - |
| - | - | 4.331E+04 | 458.2 | - | - | 0 | - |
| - | - | 3.582E+04 | 459.2 | - | - | 0 | - |
| 4 | b | 2.671E+05 | 461.2 | 0.0003935 | 0.8533 | +1 | 4 |
| - | - | 9.997E+04 | 461.2 | - | - | 0 | - |
| - | - | 5.612E+04 | 462.2 | - | - | 0 | - |
| - | - | 3.915E+04 | 462.2 | - | - | 0 | - |
| - | - | 8.267E+05 | 465.2 | - | - | 0 | - |
| - | - | 1.699E+05 | 466.2 | - | - | 0 | - |
| - | - | 2.192E+04 | 467.2 | - | - | 0 | - |
| 3 | y | 3.754E+04 | 468.2 | 0.00485 | 10.36 | +2 | 7 |
| - | - | 2.796E+04 | 469.2 | - | - | 0 | - |
| - | - | 1.894E+04 | 470.2 | - | - | 0 | - |
| - | - | 1.397E+04 | 470.2 | - | - | 0 | - |
| - | - | 1.095E+05 | 471.2 | - | - | 0 | - |
| - | - | 3.591E+04 | 472.2 | - | - | 0 | - |
| - | - | 6.613E+04 | 473.2 | - | - | 0 | - |
| - | - | 1.996E+04 | 475.1 | - | - | 0 | - |
| 3 | y | 1.476E+05 | 477.2 | 0.004095 | 8.581 | +2 | 7 |
| - | - | 4.074E+04 | 477.7 | - | - | 0 | - |
| - | - | 3.939E+04 | 478.2 | - | - | 0 | - |
| - | - | 5.092E+04 | 478.2 | - | - | 0 | - |
| 4 | b | 8.281E+05 | 479.2 | 0.0008762 | 1.829 | +1 | 4 |
| - | - | 1.143E+04 | 480.1 | - | - | 0 | - |
| - | - | 2.054E+05 | 480.2 | - | - | 0 | - |
| - | - | 4.133E+04 | 481.2 | - | - | 0 | - |
| - | - | 5.482E+04 | 482.2 | - | - | 0 | - |
| - | - | 2.416E+04 | 483.2 | - | - | 0 | - |
| - | - | 1.135E+05 | 486.2 | - | - | 0 | - |
| - | - | 3.548E+04 | 487.2 | - | - | 0 | - |
| 7 | y | 1.403E+06 | 489.2 | 0.005893 | 12.05 | +1 | 3 |
| - | - | 3.981E+05 | 490.2 | - | - | 0 | - |
| - | - | 2.377E+04 | 490.7 | - | - | 0 | - |
| - | - | 1.94E+05 | 491.2 | - | - | 0 | - |
| - | - | 4.192E+04 | 492.2 | - | - | 0 | - |
| - | - | 1.792E+04 | 493.2 | - | - | 0 | - |
| - | - | 1.137E+05 | 495.2 | - | - | 0 | - |
| - | - | 1.481E+04 | 495.7 | - | - | 0 | - |
| - | - | 3.726E+04 | 496.2 | - | - | 0 | - |
| - | - | 3.955E+04 | 496.2 | - | - | 0 | - |
| - | - | 7.567E+05 | 496.2 | - | - | 0 | - |
| - | - | 1.322E+05 | 497.2 | - | - | 0 | - |
| - | - | 1.926E+05 | 497.2 | - | - | 0 | - |
| - | - | 2.299E+04 | 498.2 | - | - | 0 | - |
| - | - | 3.492E+04 | 498.2 | - | - | 0 | - |
| 8 | b | 6.528E+04 | 499.2 | 0.0002344 | 0.4696 | +2 | 8 |
| - | - | 2.089E+04 | 500.2 | - | - | 0 | - |
| - | - | 2.664E+04 | 503.2 | - | - | 0 | - |
| - | - | 2.94E+04 | 504.2 | - | - | 0 | - |
| - | - | 2.858E+04 | 505.2 | - | - | 0 | - |
| - | - | 2.154E+04 | 506.2 | - | - | 0 | - |
| - | - | 2.922E+04 | 508.2 | - | - | 0 | - |
| - | - | 2.442E+04 | 511.2 | - | - | 0 | - |
| - | - | 1.603E+04 | 513.2 | - | - | 0 | - |
| - | - | 1.015E+05 | 513.2 | - | - | 0 | - |
| - | - | 2.234E+04 | 514.2 | - | - | 0 | - |
| - | - | 2.131E+04 | 514.2 | - | - | 0 | - |
| - | - | 3.847E+04 | 516.2 | - | - | 0 | - |
| - | - | 1.665E+04 | 517.2 | - | - | 0 | - |
| - | - | 4.493E+04 | 518.7 | - | - | 0 | - |
| - | - | 2.14E+05 | 519.2 | - | - | 0 | - |
| - | - | 1.191E+05 | 520.2 | - | - | 0 | - |
| - | - | 3.548E+04 | 521.2 | - | - | 0 | - |
| - | - | 2.967E+04 | 523.2 | - | - | 0 | - |
| - | - | 2.831E+04 | 524.2 | - | - | 0 | - |
| - | - | 2.161E+04 | 525.2 | - | - | 0 | - |
| - | - | 5.251E+04 | 530.2 | - | - | 0 | - |
| - | - | 2.855E+04 | 531.2 | - | - | 0 | - |
| - | - | 2.324E+04 | 532.7 | - | - | 0 | - |
| - | - | 6.267E+04 | 537.2 | - | - | 0 | - |
| - | - | 2.535E+04 | 538.2 | - | - | 0 | - |
| - | - | 5.843E+04 | 541.2 | - | - | 0 | - |
| 2 | y | 6.441E+04 | 541.7 | 0.003235 | 5.973 | +2 | 8 |
| - | - | 1.114E+05 | 542.2 | - | - | 0 | - |
| - | - | 2.786E+04 | 542.7 | - | - | 0 | - |
| - | - | 2.599E+04 | 543.2 | - | - | 0 | - |
| - | - | 2.082E+04 | 544.2 | - | - | 0 | - |
| - | - | 2.358E+04 | 547.2 | - | - | 0 | - |
| - | - | 2.501E+05 | 548.2 | - | - | 0 | - |
| - | - | 6.937E+04 | 549.2 | - | - | 0 | - |
| - | - | 1.694E+04 | 550.2 | - | - | 0 | - |
| 2 | y | 7.237E+05 | 550.7 | 0.003019 | 5.482 | +2 | 8 |
| - | - | 5.213E+05 | 551.2 | - | - | 0 | - |
| - | - | 2.326E+05 | 551.7 | - | - | 0 | - |
| - | - | 5.052E+04 | 552.2 | - | - | 0 | - |
| - | - | 2.155E+04 | 553.2 | - | - | 0 | - |
| - | - | 1.925E+04 | 556.2 | - | - | 0 | - |
| - | - | 1.255E+05 | 557.2 | - | - | 0 | - |
| - | - | 4.191E+04 | 558.2 | - | - | 0 | - |
| 6 | y | 2.332E+06 | 560.2 | 0.00534 | 9.531 | +1 | 4 |
| - | - | 7.284E+04 | 560.7 | - | - | 0 | - |
| - | - | 7.474E+05 | 561.2 | - | - | 0 | - |
| - | - | 1.769E+05 | 562.2 | - | - | 0 | - |
| - | - | 1.506E+05 | 569.2 | - | - | 0 | - |
| - | - | 1.024E+05 | 569.7 | - | - | 0 | - |
| - | - | 2.426E+05 | 570.2 | - | - | 0 | - |
| - | - | 3.708E+04 | 570.7 | - | - | 0 | - |
| - | - | 5.908E+04 | 571.2 | - | - | 0 | - |
| - | - | 1.813E+04 | 574.7 | - | - | 0 | - |
| - | - | 8.928E+04 | 578.2 | - | - | 0 | - |
| - | - | 4.611E+04 | 578.7 | - | - | 0 | - |
| - | - | 1.877E+04 | 579.2 | - | - | 0 | - |
| - | - | 7.152E+04 | 583.2 | - | - | 0 | - |
| - | - | 6.132E+04 | 583.7 | - | - | 0 | - |
| - | - | 3.915E+04 | 584.2 | - | - | 0 | - |
| - | - | 3.852E+04 | 588.2 | - | - | 0 | - |
| 0 | Precursor | 4.085E+05 | 592.2 | 0.003139 | 5.3 | +2 | -1 |
| - | - | 2.967E+05 | 592.7 | - | - | 0 | - |
| - | - | 1.13E+05 | 593.2 | - | - | 0 | - |
| - | - | 2.883E+04 | 593.7 | - | - | 0 | - |
| - | - | 7.999E+04 | 594.2 | - | - | 0 | - |
| - | - | 3.373E+04 | 595.2 | - | - | 0 | - |
| - | - | 1.357E+05 | 596.2 | - | - | 0 | - |
| - | - | 1.044E+05 | 597.2 | - | - | 0 | - |
| - | - | 5.224E+04 | 598.2 | - | - | 0 | - |
| 0 | Precursor | 1.071E+06 | 601.2 | 0.003411 | 5.673 | +2 | -1 |
| - | - | 7.931E+05 | 601.7 | - | - | 0 | - |
| - | - | 3.461E+05 | 602.2 | - | - | 0 | - |
| - | - | 4.7E+04 | 602.7 | - | - | 0 | - |
| - | - | 2.309E+05 | 606.2 | - | - | 0 | - |
| - | - | 6.022E+04 | 607.2 | - | - | 0 | - |
| - | - | 2.704E+04 | 608.2 | - | - | 0 | - |
| - | - | 3.648E+05 | 612.2 | - | - | 0 | - |
| - | - | 1.057E+05 | 613.2 | - | - | 0 | - |
| - | - | 2.277E+04 | 613.3 | - | - | 0 | - |
| - | - | 1.613E+04 | 614.2 | - | - | 0 | - |
| - | - | 2.876E+05 | 614.2 | - | - | 0 | - |
| - | - | 1.093E+05 | 615.2 | - | - | 0 | - |
| - | - | 1.92E+04 | 622.2 | - | - | 0 | - |
| 5 | b | 2.023E+05 | 624.2 | 0.00106 | 1.699 | +1 | 5 |
| - | - | 5.032E+04 | 625.2 | - | - | 0 | - |
| - | - | 1.696E+04 | 626.2 | - | - | 0 | - |
| - | - | 1.03E+05 | 631.2 | - | - | 0 | - |
| - | - | 4.347E+04 | 632.3 | - | - | 0 | - |
| - | - | 2.651E+05 | 634.2 | - | - | 0 | - |
| - | - | 5.802E+04 | 635.2 | - | - | 0 | - |
| - | - | 1.855E+04 | 636.2 | - | - | 0 | - |
| - | - | 6.004E+04 | 639.3 | - | - | 0 | - |
| - | - | 1.167E+05 | 640.2 | - | - | 0 | - |
| - | - | 4.97E+04 | 641.3 | - | - | 0 | - |
| 5 | b | 1.716E+05 | 642.2 | 0.0006885 | 1.072 | +1 | 5 |
| - | - | 6.447E+04 | 643.2 | - | - | 0 | - |
| - | - | 1.97E+04 | 644.2 | - | - | 0 | - |
| - | - | 1.906E+04 | 650.2 | - | - | 0 | - |
| - | - | 6.632E+04 | 652.2 | - | - | 0 | - |
| - | - | 1.582E+04 | 653.2 | - | - | 0 | - |
| - | - | 5.362E+05 | 657.3 | - | - | 0 | - |
| - | - | 1.618E+05 | 658.3 | - | - | 0 | - |
| - | - | 8.746E+05 | 659.3 | - | - | 0 | - |
| - | - | 3.251E+05 | 660.3 | - | - | 0 | - |
| - | - | 5.664E+04 | 661.3 | - | - | 0 | - |
| - | - | 9.632E+04 | 667.3 | - | - | 0 | - |
| - | - | 4.595E+04 | 668.3 | - | - | 0 | - |
| - | - | 1.328E+04 | 669.3 | - | - | 0 | - |
| - | - | 1.422E+05 | 676.3 | - | - | 0 | - |
| - | - | 7.123E+04 | 677.3 | - | - | 0 | - |
| - | - | 2.169E+04 | 678.3 | - | - | 0 | - |
| - | - | 7.51E+05 | 685.3 | - | - | 0 | - |
| - | - | 2.57E+05 | 686.3 | - | - | 0 | - |
| - | - | 4.771E+04 | 687.3 | - | - | 0 | - |
| 6 | b | 1.385E+05 | 695.3 | 0.0008362 | 1.203 | +1 | 6 |
| - | - | 4.502E+04 | 696.3 | - | - | 0 | - |
| - | - | 2.257E+04 | 702.3 | - | - | 0 | - |
| - | - | 2.12E+05 | 703.3 | - | - | 0 | - |
| - | - | 4.014E+04 | 704.2 | - | - | 0 | - |
| - | - | 4.193E+04 | 704.3 | - | - | 0 | - |
| - | - | 1.002E+05 | 705.3 | - | - | 0 | - |
| - | - | 6.066E+04 | 706.3 | - | - | 0 | - |
| - | - | 1.47E+04 | 708.3 | - | - | 0 | - |
| 6 | b | 9.379E+04 | 713.3 | 0.0001958 | 0.2746 | +1 | 6 |
| - | - | 2.682E+04 | 714.3 | - | - | 0 | - |
| - | - | 7.167E+05 | 721.3 | - | - | 0 | - |
| - | - | 2.844E+05 | 722.3 | - | - | 0 | - |
| 5 | y | 2.251E+06 | 723.3 | 0.005182 | 7.165 | +1 | 5 |
| - | - | 8.954E+05 | 724.3 | - | - | 0 | - |
| - | - | 3.052E+05 | 725.3 | - | - | 0 | - |
| - | - | 3.125E+04 | 728.3 | - | - | 0 | - |
| - | - | 8.181E+04 | 731.2 | - | - | 0 | - |
| - | - | 3.635E+04 | 732.2 | - | - | 0 | - |
| - | - | 2.874E+04 | 733.3 | - | - | 0 | - |
| - | - | 2.123E+04 | 747.3 | - | - | 0 | - |
| - | - | 1.72E+04 | 748.3 | - | - | 0 | - |
| - | - | 1.032E+06 | 749.3 | - | - | 0 | - |
| - | - | 4.035E+05 | 750.3 | - | - | 0 | - |
| - | - | 1.261E+05 | 751.3 | - | - | 0 | - |
| - | - | 2.281E+04 | 752.3 | - | - | 0 | - |
| - | - | 7.121E+04 | 756.3 | - | - | 0 | - |
| - | - | 4.273E+04 | 757.3 | - | - | 0 | - |
| - | - | 2.544E+04 | 759.3 | - | - | 0 | - |
| - | - | 2.618E+05 | 767.3 | - | - | 0 | - |
| - | - | 9.797E+04 | 768.3 | - | - | 0 | - |
| - | - | 4.099E+04 | 769.3 | - | - | 0 | - |
| - | - | 8.773E+05 | 774.3 | - | - | 0 | - |
| - | - | 3.821E+05 | 775.3 | - | - | 0 | - |
| - | - | 1.017E+05 | 776.3 | - | - | 0 | - |
| - | - | 3.291E+04 | 778.3 | - | - | 0 | - |
| - | - | 2.295E+04 | 779.3 | - | - | 0 | - |
| - | - | 1.664E+04 | 787.3 | - | - | 0 | - |
| - | - | 5.595E+04 | 792.3 | - | - | 0 | - |
| - | - | 1.981E+04 | 793.3 | - | - | 0 | - |
| - | - | 2.736E+04 | 796.3 | - | - | 0 | - |
| - | - | 2.257E+04 | 803.3 | - | - | 0 | - |
| - | - | 4.01E+04 | 804.3 | - | - | 0 | - |
| - | - | 1.68E+04 | 805.3 | - | - | 0 | - |
| - | - | 2.656E+04 | 814.3 | - | - | 0 | - |
| - | - | 2.153E+04 | 817.3 | - | - | 0 | - |
| - | - | 2.32E+04 | 818.3 | - | - | 0 | - |
| - | - | 1.75E+04 | 819.3 | - | - | 0 | - |
| 4 | y | 1.423E+05 | 820.3 | 0.006077 | 7.408 | +1 | 6 |
| - | - | 7.973E+04 | 821.3 | - | - | 0 | - |
| - | - | 2.994E+04 | 822.3 | - | - | 0 | - |
| - | - | 2.172E+04 | 823.3 | - | - | 0 | - |
| - | - | 1.675E+04 | 825.3 | - | - | 0 | - |
| - | - | 1.848E+04 | 827.3 | - | - | 0 | - |
| - | - | 8.053E+04 | 832.3 | - | - | 0 | - |
| - | - | 3.477E+04 | 833.3 | - | - | 0 | - |
| 4 | y | 2.833E+06 | 838.3 | 0.005278 | 6.296 | +1 | 6 |
| - | - | 1.291E+06 | 839.3 | - | - | 0 | - |
| - | - | 4.27E+05 | 840.3 | - | - | 0 | - |
| - | - | 4.093E+04 | 841.3 | - | - | 0 | - |
| 7 | b | 6.276E+04 | 842.3 | 0.00408 | 4.843 | +1 | 7 |
| - | - | 1.1E+05 | 843.3 | - | - | 0 | - |
| - | - | 5.416E+04 | 844.3 | - | - | 0 | - |
| - | - | 1.746E+04 | 845.4 | - | - | 0 | - |
| - | - | 1.85E+04 | 846.4 | - | - | 0 | - |
| - | - | 2.48E+04 | 848.3 | - | - | 0 | - |
| - | - | 4.082E+04 | 850.2 | - | - | 0 | - |
| - | - | 2.694E+04 | 850.3 | - | - | 0 | - |
| - | - | 2.214E+04 | 853.3 | - | - | 0 | - |
| - | - | 2.714E+04 | 854.3 | - | - | 0 | - |
| 7 | b | 6.683E+04 | 860.3 | 0.009323 | 10.84 | +1 | 7 |
| - | - | 3.282E+04 | 861.3 | - | - | 0 | - |
| - | - | 8.008E+04 | 862.3 | - | - | 0 | - |
| - | - | 4.335E+04 | 863.3 | - | - | 0 | - |
| - | - | 6.28E+04 | 868.3 | - | - | 0 | - |
| - | - | 4.239E+05 | 871.3 | - | - | 0 | - |
| - | - | 1.931E+05 | 872.3 | - | - | 0 | - |
| - | - | 7.782E+04 | 873.3 | - | - | 0 | - |
| - | - | 3.969E+06 | 889.3 | - | - | 0 | - |
| - | - | 1.996E+06 | 890.4 | - | - | 0 | - |
| - | - | 3.105E+04 | 890.4 | - | - | 0 | - |
| - | - | 5.132E+05 | 891.4 | - | - | 0 | - |
| - | - | 4.751E+04 | 892.3 | - | - | 0 | - |
| - | - | 8.08E+04 | 896.3 | - | - | 0 | - |
| - | - | 3.911E+04 | 897.3 | - | - | 0 | - |
| - | - | 2.225E+04 | 898.3 | - | - | 0 | - |
| - | - | 2.341E+04 | 899.3 | - | - | 0 | - |
| - | - | 1.49E+05 | 907.3 | - | - | 0 | - |
| - | - | 8.263E+04 | 908.3 | - | - | 0 | - |
| - | - | 2.834E+04 | 909.3 | - | - | 0 | - |
| - | - | 1.98E+04 | 914.3 | - | - | 0 | - |
| - | - | 2.392E+04 | 917.3 | - | - | 0 | - |
| - | - | 3.898E+04 | 918.3 | - | - | 0 | - |
| - | - | 2.767E+04 | 919.3 | - | - | 0 | - |
| - | - | 3.676E+04 | 933.4 | - | - | 0 | - |
| 3 | y | 7.442E+05 | 935.3 | 0.005196 | 5.555 | +1 | 7 |
| - | - | 4.882E+05 | 936.3 | - | - | 0 | - |
| - | - | 1.706E+05 | 937.3 | - | - | 0 | - |
| - | - | 1.222E+05 | 938.3 | - | - | 0 | - |
| - | - | 3.497E+04 | 939.3 | - | - | 0 | - |
| - | - | 2.076E+04 | 951.4 | - | - | 0 | - |
| 3 | y | 1.21E+07 | 953.3 | 0.00519 | 5.444 | +1 | 7 |
| - | - | 6.461E+06 | 954.3 | - | - | 0 | - |
| - | - | 2.286E+06 | 955.3 | - | - | 0 | - |
| - | - | 2.312E+05 | 956.3 | - | - | 0 | - |
| - | - | 9.231E+04 | 963.3 | - | - | 0 | - |
| - | - | 3.356E+04 | 964.3 | - | - | 0 | - |
| - | - | 5.118E+04 | 969.4 | - | - | 0 | - |
| - | - | 3.517E+04 | 970.4 | - | - | 0 | - |
| 8 | b | 2.374E+04 | 979.4 | 0.001991 | 2.033 | +1 | 8 |
| 8 | b | 1.897E+04 | 980.3 | 0.0195 | 19.89 | +1 | 8 |
| 8 | b | 1.296E+05 | 997.4 | 0.004366 | 4.378 | +1 | 8 |
| - | - | 7.063E+04 | 998.4 | - | - | 0 | - |
| - | - | 2.095E+04 | 999.4 | - | - | 0 | - |
| - | - | 4.72E+04 | 1015 | - | - | 0 | - |
| - | - | 2.586E+04 | 1016 | - | - | 0 | - |
| - | - | 6.461E+04 | 1018 | - | - | 0 | - |
| - | - | 5.421E+04 | 1019 | - | - | 0 | - |
| - | - | 2.625E+05 | 1036 | - | - | 0 | - |
| - | - | 1.689E+05 | 1037 | - | - | 0 | - |
| - | - | 3.802E+04 | 1038 | - | - | 0 | - |
| - | - | 1.787E+04 | 1046 | - | - | 0 | - |
| - | - | 1.761E+04 | 1056 | - | - | 0 | - |
| 2 | y | 9.476E+04 | 1082 | 0.005447 | 5.032 | +1 | 8 |
| - | - | 5.339E+04 | 1083 | - | - | 0 | - |
| 2 | y | 7.1E+05 | 1100 | 0.004648 | 4.223 | +1 | 8 |
| - | - | 4.524E+05 | 1101 | - | - | 0 | - |
| - | - | 1.852E+05 | 1102 | - | - | 0 | - |
| - | - | 2.792E+04 | 1103 | - | - | 0 | - |
| - | - | 1.209E+05 | 1110 | - | - | 0 | - |
| - | - | 8.447E+04 | 1111 | - | - | 0 | - |
| - | - | 4.446E+04 | 1112 | - | - | 0 | - |
| - | - | 1.49E+04 | 2401 | - | - | 0 | - |
| - | - | 1.719E+04 | 2441 | - | - | 0 | - |

m/z Charge Intensity FragmentType MassShift Position
120.08125305175781 0 7091824
121.04000091552734 0 22120.4
121.07906341552734 0 21391.3
121.08455657958984 0 554972.5
122.07174682617188 0 31723.475
122.0879898071289 0 20072.482
125.07132720947266 0 15021.966
127.0870361328125 0 32059.105
129.07052612304688 0 13581.879
130.06556701660156 0 199376.14
131.0684814453125 0 16810.377
132.08120727539062 0 367484.75
133.0846405029297 0 40655.637
134.09693908691406 0 40326.64
135.0999755859375 0 13953.187
136.07614135742188 0 2584963.5
136.08746337890625 0 77592.38
137.0794677734375 0 182685.56
137.09120178222656 0 12589.119
138.0553741455078 0 194238.75
138.06661987304688 0 307647.88
139.0592498779297 0 15089.203
139.0699462890625 0 16790.79
141.5588836669922 0 10099.392
143.07350158691406 0 48200.76
143.080078125 0 12080.138
143.08595275878906 0 45874.727
144.08120727539062 0 51663.48
146.06051635742188 0 98938.21
146.0968475341797 0 52597.51
147.04437255859375 0 11223.638
147.09190368652344 0 18455.066
148.0763702392578 0 33506.887
148.0873565673828 0 93157.78
148.1126708984375 0 13146.571
150.06591796875 0 12190.28
153.06643676757812 0 22596.104
155.08197021484375 0 43545.14
155.09312438964844 0 63021.57
156.0771942138672 0 642724.06
157.0805206298828 0 40636.016
158.0968475341797 0 1079209.1
159.05596923828125 0 14712.73
159.09213256835938 0 571159.9
159.10009765625 0 118211.93
160.07614135742188 0 206598.5
160.08779907226562 0 21734.955
160.0957489013672 0 42408.652
160.11248779296875 0 486542.5
161.11590576171875 0 52815.125
162.09225463867188 0 19684.922
164.0823516845703 0 18208.635
165.070068359375 0 28652.992
165.07757568359375 0 131011.414
165.10256958007812 0 13757.527
166.05368041992188 0 42249.56
166.06158447265625 0 94070.18
166.0863037109375 0 13592.713
167.045166015625 0 13930.623
167.0932159423828 0 116456.34
169.0765380859375 0 32475.537
170.0606689453125 0 16724.562
171.07687377929688 0 83934.62
172.07650756835938 0 17595.84
172.10855102539062 0 17268.973
173.0921173095703 0 13975.182
173.43838500976562 0 20356.822
174.0666961669922 0 79959.51 b Ammonia loss 2
174.09226989746094 0 13449.933
175.0697479248047 0 11640.266
175.0870361328125 0 38126.297
175.0979766845703 0 14319.765
175.12355041503906 0 28299.873
176.0824432373047 0 177990.14
176.10748291015625 0 2176386.8
177.10272216796875 0 1491951.9
177.11134338378906 0 212685.4
178.0616455078125 0 21021.857
178.09829711914062 0 12423.441
178.10606384277344 0 160506.55
180.0772705078125 0 112085.43
185.05618286132812 0 180524.08
186.0919189453125 0 185211.06
187.0631561279297 0 28753.375
188.07106018066406 0 780071.6
189.07443237304688 0 105914.8
189.10354614257812 0 11797.865
190.08688354492188 0 29427.6
191.0931854248047 0 77657.49
191.1183624267578 0 133873.02
192.1022186279297 0 21900.92
193.0727996826172 0 39338.87
193.10877990722656 0 627374.44
194.08038330078125 0 24371.896
194.0914306640625 0 11294.174
194.11231994628906 0 52683.793
195.08807373046875 0 168648.25
197.10464477539062 0 15232.787
202.05380249023438 0 79446.984
203.0665740966797 0 180502.05
203.0934295654297 0 60632.39
203.11827087402344 0 736588.94 a Water loss 1
204.0771484375 0 333707.66
204.10208129882812 0 53299.23 a Ammonia loss 1
204.1217498779297 0 104456.35
205.08135986328125 0 27482.154
205.09762573242188 0 580222.1 y 8
205.10818481445312 0 43762.242
206.10128784179688 0 58784.086
207.11334228515625 0 151594.2
208.07217407226562 0 92894.48
208.1168212890625 0 26954.988
212.11720275878906 0 38909.61
213.0502166748047 0 15942.259
213.09934997558594 0 14254.291
214.03570556640625 0 12637.427
215.08201599121094 0 37469.164
217.09765625 0 405020.78
218.08216857910156 0 22576.346
218.10110473632812 0 56179.3
219.08041381835938 0 121560.33
219.11318969726562 0 18001.064
220.12094116210938 0 18886.168
221.1041259765625 0 1701863
221.1290740966797 0 29472186 a 1
222.1071014404297 0 161774.6
222.1322479248047 0 3583316.8
223.11935424804688 0 41356.246
223.1347198486328 0 220344.48
224.11778259277344 0 21107.822
225.0990753173828 0 16474.602
228.11337280273438 0 84686.37
229.10865783691406 0 61412.816
231.0616912841797 0 453406.94
231.0883026123047 0 14929.592 b Water loss 3
231.11328125 0 300844.5 b Water loss 1
231.12620544433594 0 19814.086
232.06497192382812 0 35698.066
232.11688232421875 0 30037.113
233.0927734375 0 36433.324
235.10826110839844 0 807556.9
236.1109619140625 0 90795.734
237.08973693847656 0 26099.092
237.1241912841797 0 21918.414
238.1313018798828 0 28482.307
239.11448669433594 0 166438.62
240.08131408691406 0 18056.799
240.11700439453125 0 17790.857
245.0928192138672 0 34820.273 y 6
245.1290740966797 0 28546.729
246.07447814941406 0 15415.251
246.13531494140625 0 51312.098
247.10736083984375 0 33999.992
247.1199493408203 0 39053.004
247.14503479003906 0 18505.752
248.11526489257812 0 20531.105
248.15106201171875 0 73272.016
249.12396240234375 0 5132272.5 b 1
250.12730407714844 0 691860.06
251.1031494140625 0 515539.6
251.1292724609375 0 83075.69
252.10678100585938 0 55501.97
253.09353637695312 0 23877.201
255.11231994628906 0 14946.366
256.1084289550781 0 33272.297
257.0752868652344 0 27740.34
257.1068115234375 0 344114.3
258.11016845703125 0 33404.67
258.12432861328125 0 19896.186
261.0875244140625 0 90300.875
263.07012939453125 0 48171.45
263.10302734375 0 1039388.9
263.12933349609375 0 99452.945
264.1066589355469 0 167543.64
264.13177490234375 0 21689.162
264.1461181640625 0 105177.13
265.1173095703125 0 42726.055
266.1235046386719 0 23095.809
267.0914306640625 0 62016.17
268.0751953125 0 57698.758
268.09197998046875 0 18991.299
268.14013671875 0 36671.152
269.0773010253906 0 49964.836
270.0788269042969 0 13992.194
270.109375 0 25339.457
271.1087341308594 0 46116.293
272.1133728027344 0 24471.035
273.0876159667969 0 16507.945
273.1239013671875 0 22773.203
274.13018798828125 0 282823.7
274.1558837890625 0 25103.445
275.11444091796875 0 43252.918
275.1336364746094 0 35526.137
277.59674072265625 0 17379.633
277.6205749511719 0 14277.675
279.09808349609375 0 489292.44
279.125244140625 0 42838.312
280.10137939453125 0 43526.074
283.1064453125 0 18806.129
283.15252685546875 0 19998.6
285.1021423339844 0 676977.2
286.1048889160156 0 74415.36
286.12017822265625 0 14392.861
287.100830078125 0 34535.527
289.0824890136719 0 33828.72
289.1190185546875 0 15912.827
290.1260986328125 0 27203.725
292.1409912109375 0 408179.62
292.1662292480469 0 99628.516 d 2
293.1435852050781 0 42419.7
294.1546936035156 0 20023.74
295.14385986328125 0 13518.481
296.1512145996094 0 539114.7
297.13525390625 0 22160.12
297.15478515625 0 89264.84
299.1508483886719 0 34379.918
300.13482666015625 0 63476.777
301.1190490722656 0 33364.152
302.114501953125 0 47228.824
303.1136474609375 0 91601.23
304.1147155761719 0 17232.934
305.4649353027344 0 15703.436
306.1333923339844 0 21017.219
306.1468200683594 0 15839.155
310.15167236328125 0 73200.15
313.0967712402344 0 42826.98
314.0987854003906 0 70105.1
315.0997009277344 0 25023.266
318.1455078125 0 166064.39 a Water loss 2
319.1489562988281 0 34968.24
320.1355285644531 0 26396.791
323.09942626953125 0 197730.75
323.14013671875 0 14853.643
324.1462707519531 0 1181274.5
325.14910888671875 0 201630.5
326.154296875 0 18567.598
327.14593505859375 0 55179.363
328.1294860839844 0 113971.17
328.1466064453125 0 49255.79
329.13189697265625 0 18456.887
330.1100158691406 0 65099.27
331.0929260253906 0 26305.56
331.1131286621094 0 13343.931
332.12457275390625 0 86330.23
333.10882568359375 0 38082.9
334.1083984375 0 22110.426
334.1302490234375 0 22453.457
335.17132568359375 0 23854.68
336.1560974121094 0 25949.467 a 2
338.12884521484375 0 75424.28
339.171142578125 0 109545.414
340.1752014160156 0 17046.537
342.15679931640625 0 6117012 y 7
343.1598815917969 0 1121017.2
344.162353515625 0 86624.555
346.1403503417969 0 365387.1 b Water loss 2
347.14337158203125 0 62950.97
348.1194152832031 0 165547.98
349.10198974609375 0 17972.873
349.1243591308594 0 16675.88
350.13525390625 0 353847.25
351.1380310058594 0 55128.45
352.1407470703125 0 108494.516
353.1465759277344 0 23797.516
354.1554870605469 0 15125.087
356.1390686035156 0 338278.34
357.1422119140625 0 48344.164 b 5
358.1046447753906 0 20365.488
359.0865173339844 0 20942.352
360.1195373535156 0 94603.32
360.14306640625 0 21470.03
361.123779296875 0 19404.912
362.1501159667969 0 55202.16 y 4
364.1508483886719 0 1533393.8 b 2
365.15411376953125 0 259976.33
366.1294860839844 0 135582.11
366.15631103515625 0 40478.703
367.13250732421875 0 37693.54
367.16571044921875 0 70892.51
368.1190490722656 0 17890.582
371.1353759765625 0 45279.965
374.14923095703125 0 67324.73
376.11456298828125 0 158904.62
377.0954895019531 0 14987.46
377.11981201171875 0 22201.352
378.09979248046875 0 55572.652
378.1301574707031 0 841624.2
379.1331787109375 0 167445.56
380.1393127441406 0 53956.055
382.14373779296875 0 167816.05
383.14617919921875 0 19612.463
384.13519287109375 0 43006.934
385.1361389160156 0 50082.742
386.13726806640625 0 17651.154
388.11602783203125 0 25273.578
388.149658203125 0 33380.54
390.1570129394531 0 123475.86
391.15966796875 0 21049.473
394.12518310546875 0 1092798.9
395.12823486328125 0 205218.47
395.1567687988281 0 24677.285
396.13177490234375 0 26186.62
397.16314697265625 0 40892.2
398.1690368652344 0 23864.39
399.17681884765625 0 60064.41
401.1961975097656 0 18002.166
404.1130065917969 0 30831.734
404.145263671875 0 29022.496
405.1146545410156 0 15861.225
406.1518249511719 0 41573.016
407.1836242675781 0 113943.336
408.1673278808594 0 241731
409.1696472167969 0 53388.66
410.1404113769531 0 20862.219
410.18212890625 0 17581.076
415.1618347167969 0 78611.77
416.14508056640625 0 16252.024
417.144775390625 0 15870.451
419.1568603515625 0 81439.83
423.17840576171875 0 16295.685
425.1940612792969 0 940629
426.1971130371094 0 199303.53
427.2091064453125 0 183421.38
428.2106628417969 0 31130.363
429.143310546875 0 18602.459
429.19097900390625 0 14505.265
430.1235656738281 0 22080.316
431.1575927734375 0 19186.416
432.1413269042969 0 40046.9
433.1729736328125 0 127627.29
434.1763916015625 0 33446.184
435.1800842285156 0 56252.543
436.14752197265625 0 18093.51
436.181640625 0 22522.129
437.1673889160156 0 33639.03
438.1706848144531 0 25360.713
442.17303466796875 0 40355.234
443.1556701660156 0 45661.68
443.1866455078125 0 26143.566
447.15142822265625 0 121650.49
449.1344909667969 0 47526.46
449.1679382324219 0 32030.584
452.6690368652344 0 17740.193
454.1968688964844 0 74139.195
455.2041931152344 0 188332.23
456.2077331542969 0 38989.684
458.1688232421875 0 43310.41
459.15362548828125 0 35817.305
461.1670837402344 0 267069.72 b Water loss 3
461.1960144042969 0 99971.234
462.1698303222656 0 56123.58
462.20025634765625 0 39146.586
465.1625061035156 0 826656.2
466.16571044921875 0 169914.06
467.16748046875 0 21922.553
468.17364501953125 0 37536.184 y Water loss 2
469.20965576171875 0 27963.074
470.1679992675781 0 18938.62
470.21240234375 0 13965.2705
471.1808776855469 0 109504.43
472.1824645996094 0 35905.223
473.2148742675781 0 66127.01
475.1444091796875 0 19962.004
477.16998291015625 0 147606.14 y 2
477.6781005859375 0 40736.805
478.16595458984375 0 39389.117
478.22113037109375 0 50917.164
479.1781311035156 0 828099.2 b 3
480.1445617675781 0 11430.541
480.1812438964844 0 205365.92
481.18206787109375 0 41329.89
482.19342041015625 0 54823.266
483.19647216796875 0 24159.973
486.1631164550781 0 113471.32
487.16668701171875 0 35479.88
489.19244384765625 0 1402828.8 y 6
490.19464111328125 0 398135.12
490.68585205078125 0 23773.553
491.2055358886719 0 193974.7
492.20977783203125 0 41917.445
493.20306396484375 0 17922.95
495.1868896484375 0 113702.586
495.69329833984375 0 14810.388
496.1729431152344 0 37258.367
496.1885070800781 0 39546.875
496.2311096191406 0 756706.56
497.1711730957031 0 132197.81
497.2343444824219 0 192576.52
498.1737976074219 0 22990.46
498.2389221191406 0 34921.066
499.18695068359375 0 65282.605 b 7
500.18804931640625 0 20894.54
503.1781311035156 0 26642.234
504.1814880371094 0 29397.338
505.19317626953125 0 28582.809
506.21563720703125 0 21542.047
508.19476318359375 0 29215.78
511.19354248046875 0 24422.688
513.1630249023438 0 16025.253
513.1991577148438 0 101534.78
514.1925659179688 0 22339.537
514.22900390625 0 21307.936
516.2232055664062 0 38473.617
517.2260131835938 0 16646.162
518.7092895507812 0 44931.723
519.2041015625 0 214008.55
520.2052612304688 0 119136.49
521.20556640625 0 35481.543
523.1859741210938 0 29667.137
524.223388671875 0 28305.275
525.1992797851562 0 21614.746
530.1896362304688 0 52507.496
531.18994140625 0 28549.568
532.70068359375 0 23238.268
537.2135620117188 0 62672.75
538.2155151367188 0 25351.807
541.1942138671875 0 58426.08
541.7062377929688 0 64413.434 y Water loss 1
542.234375 0 111427.586
542.710205078125 0 27860.082
543.2239990234375 0 25986.758
544.2119750976562 0 20820.752
547.195068359375 0 23580.62
548.1995849609375 0 250130.2
549.2020874023438 0 69374.15
550.2078247070312 0 16941.42
550.7113037109375 0 723724.7 y 1
551.2130737304688 0 521344.88
551.7138671875 0 232558.61
552.215576171875 0 50518.207
553.2003173828125 0 21550.092
556.210693359375 0 19246.875
557.19970703125 0 125481.59
558.2023315429688 0 41912
560.22900390625 0 2331863.5 y 5
560.7299194335938 0 72842.92
561.2321166992188 0 747350.2
562.2308959960938 0 176872.72
569.2340698242188 0 150616.28
569.7352294921875 0 102422.25
570.2291870117188 0 242603.75
570.720458984375 0 37080.816
571.2305297851562 0 59076.535
574.7166137695312 0 18129.379
578.2305297851562 0 89280.84
578.7325439453125 0 46113.383
579.2156982421875 0 18765.475
583.2258911132812 0 71517.17
583.7255249023438 0 61316.992
584.2031860351562 0 39154.95
588.240234375 0 38517.82
592.22998046875 0 408459.25 Precursor Water loss
592.7318115234375 0 296696.03
593.2322387695312 0 113009.51
593.7312622070312 0 28834.582
594.186279296875 0 79985.91
595.1869506835938 0 33729.34
596.2355346679688 0 135728.6
597.2254638671875 0 104437.01
598.2254638671875 0 52235.19
601.2355346679688 0 1071055 Precursor
601.7366943359375 0 793070.94
602.2374267578125 0 346144.7
602.737060546875 0 47001.742
606.233154296875 0 230944.55
607.23681640625 0 60216.496
608.2337646484375 0 27038.58
612.1978149414062 0 364796
613.2012329101562 0 105690.48
613.29052734375 0 22766.865
614.2008666992188 0 16126.004
614.246337890625 0 287555
615.2483520507812 0 109298.76
622.2219848632812 0 19202.357
624.2310791015625 0 202283.47 b Water loss 4
625.2332153320312 0 50317.55
626.2349243164062 0 16957.771
631.2467041015625 0 103007.23
632.2583618164062 0 43473.555
634.2295532226562 0 265102.2
635.2333374023438 0 58021.3
636.2343139648438 0 18546.445
639.2531127929688 0 60044.01
640.2382202148438 0 116724.33
641.2804565429688 0 49695.445
642.2412719726562 0 171637.66 b 4
643.2451171875 0 64466.04
644.24658203125 0 19699.273
650.2454833984375 0 19056.543
652.2403564453125 0 66319.29
653.2449340820312 0 15817.383
657.263427734375 0 536225.7
658.2657470703125 0 161762.14
659.294189453125 0 874613.5
660.297119140625 0 325069
661.29931640625 0 56643.938
667.2515258789062 0 96323.93
668.251220703125 0 45951.023
669.27783203125 0 13282.995
676.2696533203125 0 142213.83
677.2758178710938 0 71227.414
678.2951049804688 0 21691.428
685.2583618164062 0 750989.3
686.2612915039062 0 257029.88
687.2650146484375 0 47711.516
695.2662963867188 0 138460.03 b Water loss 5
696.2725830078125 0 45015.1
702.28662109375 0 22572.508
703.2653198242188 0 212025.97
704.2440185546875 0 40143.445
704.2664794921875 0 41927.61
705.2815551757812 0 100177.26
706.2854614257812 0 60657.54
708.2720947265625 0 14695.937
713.2778930664062 0 93788.875 b 5
714.2777099609375 0 26820.889
721.2614135742188 0 716726.25
722.2645874023438 0 284427.78
723.2921752929688 0 2251368.8 y 4
724.2952880859375 0 895409.75
725.295166015625 0 305183.9
728.3198852539062 0 31251.404
731.2474975585938 0 81812.29
732.2432250976562 0 36352.227
733.2720336914062 0 28742.07
747.2918090820312 0 21234.957
748.2979125976562 0 17199.387
749.2561645507812 0 1032349.4
750.2588500976562 0 403495.75
751.259521484375 0 126069.5
752.25341796875 0 22811.729
756.310546875 0 71207.36
757.30517578125 0 42727.7
759.259521484375 0 25444.688
767.2667236328125 0 261791.95
768.2706298828125 0 97966.02
769.2672729492188 0 40989.246
774.320556640625 0 877278.3
775.323486328125 0 382132.34
776.32666015625 0 101700.03
778.298095703125 0 32909.516
779.31103515625 0 22947.367
787.3148803710938 0 16640.75
792.3156127929688 0 55948.45
793.3134765625 0 19811.488
796.3153686523438 0 27357.713
803.284912109375 0 22574.84
804.32763671875 0 40102.793
805.3328247070312 0 16797.027
814.3160400390625 0 26555.697
817.3228759765625 0 21534.867
818.33447265625 0 23195.697
819.3413696289062 0 17500.926
820.3094482421875 0 142270 y Water loss 3
821.3070068359375 0 79731.14
822.299560546875 0 29935.748
823.2855224609375 0 21721.725
825.3106079101562 0 16753.254
827.3470458984375 0 18483.537
832.3264770507812 0 80530.99
833.3284301757812 0 34774.934
838.3192138671875 0 2832887.5 y 3
839.3218383789062 0 1290682.8
840.321533203125 0 427000.9
841.3250732421875 0 40934.477
842.3016967773438 0 62756.58 b Water loss 6
843.3391723632812 0 109953.484
844.34228515625 0 54159.844
845.3539428710938 0 17456.76
846.3591918945312 0 18496.676
848.30224609375 0 24801.084
850.1714477539062 0 40816.23
850.3236083984375 0 26940.258
853.3203735351562 0 22138.746
854.3191528320312 0 27143.225
860.3175048828125 0 66830.13 b 6
861.314697265625 0 32816.91
862.3238525390625 0 80082.89
863.3256225585938 0 43352.4
868.3311767578125 0 62796.49
871.3368530273438 0 423891.62
872.3364868164062 0 193145.03
873.3384399414062 0 77824.11
889.3473510742188 0 3969402.8
890.3502197265625 0 1995913.8
890.4302978515625 0 31046.857
891.35302734375 0 513234.4
892.3488159179688 0 47507.44
896.3245239257812 0 80802.3
897.3297729492188 0 39106.176
898.3217163085938 0 22253.932
899.3314819335938 0 23411.113
907.3394775390625 0 149010.17
908.342041015625 0 82629.92
909.3403930664062 0 28336.707
914.3306274414062 0 19799.23
917.3245849609375 0 23919.973
918.3199462890625 0 38980.496
919.318603515625 0 27668.904
933.3786010742188 0 36758.055
935.3355102539062 0 744199.2 y Water loss 2
936.3350219726562 0 488167.2
937.3347778320312 0 170649.77
938.3265991210938 0 122225.1
939.3259887695312 0 34972.613
951.3657836914062 0 20760.234
953.3460693359375 0 12102720 y 2
954.3486328125 0 6460993.5
955.348876953125 0 2285785.2
956.3493041992188 0 231213.11
963.3313598632812 0 92309.11
964.3329467773438 0 33563.17
969.376953125 0 51178.1
970.3757934570312 0 35174.973
979.3585205078125 0 23744.436 b Water loss 7
980.3600463867188 0 18966.85 b Ammonia loss 7
997.3714599609375 0 129582.09 b 7
998.375732421875 0 70625.555
999.3756713867188 0 20945.385
1015.3811645507812 0 47200.61
1016.39111328125 0 25858.322
1018.3936157226562 0 64606.22
1019.4008178710938 0 54206.047
1036.413818359375 0 262479.56
1037.419189453125 0 168900.1
1038.4210205078125 0 38022.637
1046.3929443359375 0 17869.68
1055.9176025390625 0 17607.543
1082.4041748046875 0 94762.484 y Water loss 1
1083.4022216796875 0 53388.46
1100.4139404296875 0 709954.5 y 1
1101.4166259765625 0 452356.28
1102.417236328125 0 185164.64
1103.4202880859375 0 27918.594
1110.3992919921875 0 120917.45
1111.4007568359375 0 84474.38
1112.4019775390625 0 44457.62
2401.114013671875 0 14902.679
2441.31640625 0 17188.86

Spectrum Details

|  |  |
| --- | --- |
| Matched peaks? Matched peaks The total absolute number of peaks matched. Additionally in brackets the total fraction of peaks matched and the total number of peaks is shown. | 44 (6.89% of 639) |
| FDR? FDR The false discovery rate estimated for this peptide. It is calculated by matching all theoretical fragments with a non-integer shift with the raw peaks for this spectrum. This is done with 40 different shifts. The resulting percentage is the average number of annotated peaks over the number of annotated peaks with the correct spectrum. | 0.00% |
| Satellite FDR? Satellite FDR See the FDR for details on its calculation. This satellite ion specific FDR only contains the satellite ions (d/w) for I/L/J positions. | - |
| PSM Score? PSM Score The PSM Score as given by Hecklib to this annotated spectrum. It is shown with three significant figures. | 536 |

#### Spectrum 7379? Spectrum 7379 The raw spectrum of this peptide as annotated by Hecklib. The fragments are coloured according to ion type (see legend). Any peaks with a star '\*' as text can be hovered over to see the full details, first the ion type second the mass shift type. By hovering over the amino acids in the peptide or ions in the legend the corresponding peaks are highlighted. By toggling the 'Unassigned' label you can turn the background (unassigned) peaks on or off in the plot. By updating the slider in the Ion legend you can update the spectrum to only show the top X% of the peaks with labels. The top X% means any peak that is within X% of the highest intensity. By dragging in the spectrum you can zoom in to a specific part of the spectrum and use 'Zoom Out' to get back to the original zoom level. The annotation of the spectrum is based on the given sequence in the peptides file and is done with different software so inconsistencies are likely. The peaks are annotated based on the given sequence, with 20 ppm tolerance.

Copy Data

##### Spectrum 7379 (TSV)

###### Preview

```
Loading example...
```

*Click on the button to copy the data to your clipboard.*

Mz MinMz MaxIntensity Max

WidthHeightPeptide font sizePeptide stroke widthSpectrum font sizeSpectrum stroke widthCompact peptide

Ion legend

wxyz

abcd

OtherUnassignedIon^Charge^_Position_Show for top:%

TFDDYAMHW

02.23e+64.47e+66.70e+68.94e+6

Zoom Out

b^+2^_3_a^+1^_2_a^+1^_2_y^+1^_1_a^+1^_2_b^+2^_4_b^+1^_2_y^+2^_3_b^+1^_2_d^+1^_3_a^+1^_3_y^+1^_2_b^+1^_3_b^+2^_6_b^+2^_6_y^+2^_5_b^+1^_3_b^+1^_4_y^+2^_7_y^+2^_7_b^+1^_4_y^+1^_3_b^+2^_8_y^+2^_8_y^+2^_8_y^+1^_4_\*\*b^+1^_5_b^+1^_5_b^+1^_6_b^+1^_6_y^+1^_5_y^+1^_6_y^+1^_6_b^+1^_7_b^+1^_7_y^+1^_7_y^+1^_7_b^+1^_8_b^+1^_8_y^+1^_8_y^+1^_8_

0818163624543272

Fragment Matches Table

Show background peaks

| Position | Ion type | Intensity | mz Theoretical | mz Error (Th) | mz Error (ppm) | Charge | Series Number |
| --- | --- | --- | --- | --- | --- | --- | --- |
| - | - | 2.182E+06 | 120.1 | - | - | 0 | - |
| - | - | 5911 | 121 | - | - | 0 | - |
| - | - | 1.627E+05 | 121.1 | - | - | 0 | - |
| - | - | 1.277E+04 | 122.1 | - | - | 0 | - |
| - | - | 6464 | 122.1 | - | - | 0 | - |
| - | - | 6640 | 127.1 | - | - | 0 | - |
| - | - | 9324 | 129.1 | - | - | 0 | - |
| - | - | 3154 | 129.2 | - | - | 0 | - |
| - | - | 4.657E+04 | 130.1 | - | - | 0 | - |
| - | - | 5443 | 131.1 | - | - | 0 | - |
| - | - | 1.12E+05 | 132.1 | - | - | 0 | - |
| - | - | 6168 | 133.1 | - | - | 0 | - |
| - | - | 8821 | 133.1 | - | - | 0 | - |
| - | - | 9515 | 134.1 | - | - | 0 | - |
| - | - | 7.861E+05 | 136.1 | - | - | 0 | - |
| - | - | 2.627E+04 | 136.1 | - | - | 0 | - |
| - | - | 5548 | 137.1 | - | - | 0 | - |
| - | - | 6.285E+04 | 137.1 | - | - | 0 | - |
| - | - | 5369 | 137.1 | - | - | 0 | - |
| - | - | 5.913E+04 | 138.1 | - | - | 0 | - |
| - | - | 8.726E+04 | 138.1 | - | - | 0 | - |
| - | - | 4916 | 139.1 | - | - | 0 | - |
| - | - | 6577 | 141.1 | - | - | 0 | - |
| - | - | 5459 | 143 | - | - | 0 | - |
| - | - | 1.395E+04 | 143.1 | - | - | 0 | - |
| - | - | 1.259E+04 | 143.1 | - | - | 0 | - |
| - | - | 3006 | 144.1 | - | - | 0 | - |
| - | - | 1.091E+04 | 144.1 | - | - | 0 | - |
| - | - | 3.001E+04 | 146.1 | - | - | 0 | - |
| - | - | 4785 | 146.1 | - | - | 0 | - |
| - | - | 1.386E+04 | 146.1 | - | - | 0 | - |
| - | - | 7203 | 148.1 | - | - | 0 | - |
| - | - | 2.491E+04 | 148.1 | - | - | 0 | - |
| - | - | 6613 | 150.1 | - | - | 0 | - |
| - | - | 1.391E+04 | 155.1 | - | - | 0 | - |
| - | - | 2.172E+04 | 155.1 | - | - | 0 | - |
| - | - | 2.044E+05 | 156.1 | - | - | 0 | - |
| - | - | 1.233E+04 | 157.1 | - | - | 0 | - |
| - | - | 3.26E+05 | 158.1 | - | - | 0 | - |
| - | - | 3697 | 159.1 | - | - | 0 | - |
| - | - | 1.649E+05 | 159.1 | - | - | 0 | - |
| - | - | 3.406E+04 | 159.1 | - | - | 0 | - |
| - | - | 5.868E+04 | 160.1 | - | - | 0 | - |
| - | - | 6060 | 160.1 | - | - | 0 | - |
| - | - | 1.54E+04 | 160.1 | - | - | 0 | - |
| - | - | 1.486E+05 | 160.1 | - | - | 0 | - |
| - | - | 3859 | 161.1 | - | - | 0 | - |
| - | - | 1.65E+04 | 161.1 | - | - | 0 | - |
| - | - | 5179 | 162.1 | - | - | 0 | - |
| - | - | 4468 | 164.1 | - | - | 0 | - |
| - | - | 9347 | 165.1 | - | - | 0 | - |
| - | - | 3.29E+04 | 165.1 | - | - | 0 | - |
| - | - | 1.337E+04 | 166.1 | - | - | 0 | - |
| - | - | 2.806E+04 | 166.1 | - | - | 0 | - |
| - | - | 5245 | 166.1 | - | - | 0 | - |
| - | - | 2.515E+04 | 167.1 | - | - | 0 | - |
| - | - | 1.202E+04 | 169.1 | - | - | 0 | - |
| - | - | 2.719E+04 | 171.1 | - | - | 0 | - |
| - | - | 5293 | 171.1 | - | - | 0 | - |
| - | - | 5993 | 172.1 | - | - | 0 | - |
| - | - | 5835 | 172.1 | - | - | 0 | - |
| - | - | 8810 | 173.1 | - | - | 0 | - |
| 3 | b | 2.021E+04 | 174.1 | 0.001222 | 7.022 | +2 | 3 |
| - | - | 4125 | 174.1 | - | - | 0 | - |
| - | - | 1.27E+04 | 175.1 | - | - | 0 | - |
| - | - | 5952 | 175.1 | - | - | 0 | - |
| - | - | 8749 | 175.1 | - | - | 0 | - |
| - | - | 5.372E+04 | 176.1 | - | - | 0 | - |
| - | - | 6.578E+05 | 176.1 | - | - | 0 | - |
| - | - | 4.675E+05 | 177.1 | - | - | 0 | - |
| - | - | 7.02E+04 | 177.1 | - | - | 0 | - |
| - | - | 4.455E+04 | 178.1 | - | - | 0 | - |
| - | - | 4998 | 178.1 | - | - | 0 | - |
| - | - | 2.927E+04 | 180.1 | - | - | 0 | - |
| - | - | 5605 | 182 | - | - | 0 | - |
| - | - | 5.017E+04 | 185.1 | - | - | 0 | - |
| - | - | 5.44E+04 | 186.1 | - | - | 0 | - |
| - | - | 1.201E+04 | 187.1 | - | - | 0 | - |
| - | - | 2.233E+05 | 188.1 | - | - | 0 | - |
| - | - | 2.768E+04 | 189.1 | - | - | 0 | - |
| - | - | 8221 | 190.1 | - | - | 0 | - |
| - | - | 5370 | 191.1 | - | - | 0 | - |
| - | - | 2.356E+04 | 191.1 | - | - | 0 | - |
| - | - | 4.673E+04 | 191.1 | - | - | 0 | - |
| - | - | 1.12E+04 | 192.1 | - | - | 0 | - |
| - | - | 4599 | 192.1 | - | - | 0 | - |
| - | - | 1.346E+04 | 193.1 | - | - | 0 | - |
| - | - | 1.859E+05 | 193.1 | - | - | 0 | - |
| - | - | 9673 | 194.1 | - | - | 0 | - |
| - | - | 1.411E+04 | 194.1 | - | - | 0 | - |
| - | - | 4155 | 194.3 | - | - | 0 | - |
| - | - | 4.846E+04 | 195.1 | - | - | 0 | - |
| - | - | 3839 | 196.7 | - | - | 0 | - |
| - | - | 1.592E+04 | 202.1 | - | - | 0 | - |
| - | - | 5.248E+04 | 203.1 | - | - | 0 | - |
| - | - | 1.184E+04 | 203.1 | - | - | 0 | - |
| 2 | a | 2.237E+05 | 203.1 | 0.0004729 | 2.328 | +1 | 2 |
| - | - | 1.023E+05 | 204.1 | - | - | 0 | - |
| 2 | a | 1.362E+04 | 204.1 | 0.0003898 | 1.91 | +1 | 2 |
| - | - | 3.078E+04 | 204.1 | - | - | 0 | - |
| - | - | 8510 | 205.1 | - | - | 0 | - |
| 9 | y | 1.731E+05 | 205.1 | 0.0005327 | 2.597 | +1 | 1 |
| - | - | 1.346E+04 | 205.1 | - | - | 0 | - |
| - | - | 2.532E+04 | 206.1 | - | - | 0 | - |
| - | - | 4.775E+04 | 207.1 | - | - | 0 | - |
| - | - | 2.809E+04 | 208.1 | - | - | 0 | - |
| - | - | 4858 | 209.1 | - | - | 0 | - |
| - | - | 5479 | 211.1 | - | - | 0 | - |
| - | - | 1.209E+04 | 212.1 | - | - | 0 | - |
| - | - | 1.131E+04 | 215.1 | - | - | 0 | - |
| - | - | 7610 | 216.1 | - | - | 0 | - |
| - | - | 1.171E+05 | 217.1 | - | - | 0 | - |
| - | - | 1.368E+04 | 218.1 | - | - | 0 | - |
| - | - | 3.737E+04 | 219.1 | - | - | 0 | - |
| - | - | 7069 | 219.1 | - | - | 0 | - |
| - | - | 4828 | 220.1 | - | - | 0 | - |
| - | - | 5.394E+05 | 221.1 | - | - | 0 | - |
| 2 | a | 8.848E+06 | 221.1 | 0.0007267 | 3.286 | +1 | 2 |
| - | - | 5.126E+04 | 222.1 | - | - | 0 | - |
| - | - | 1.075E+06 | 222.1 | - | - | 0 | - |
| - | - | 8257 | 223.1 | - | - | 0 | - |
| - | - | 6.325E+04 | 223.1 | - | - | 0 | - |
| - | - | 8979 | 224.1 | - | - | 0 | - |
| - | - | 6291 | 225.1 | - | - | 0 | - |
| - | - | 2.429E+04 | 228.1 | - | - | 0 | - |
| - | - | 1.875E+04 | 229.1 | - | - | 0 | - |
| - | - | 5973 | 230.1 | - | - | 0 | - |
| - | - | 1.493E+05 | 231.1 | - | - | 0 | - |
| 4 | b | 5860 | 231.1 | 0.001899 | 8.218 | +2 | 4 |
| 2 | b | 8.454E+04 | 231.1 | 0.0005839 | 2.526 | +1 | 2 |
| - | - | 4046 | 231.1 | - | - | 0 | - |
| - | - | 1.373E+04 | 232.1 | - | - | 0 | - |
| - | - | 1.832E+04 | 232.1 | - | - | 0 | - |
| - | - | 4682 | 233.1 | - | - | 0 | - |
| - | - | 5954 | 234.1 | - | - | 0 | - |
| - | - | 2.085E+05 | 235.1 | - | - | 0 | - |
| - | - | 2.329E+04 | 236.1 | - | - | 0 | - |
| - | - | 8882 | 237.1 | - | - | 0 | - |
| - | - | 6521 | 237.1 | - | - | 0 | - |
| - | - | 1.373E+04 | 238.1 | - | - | 0 | - |
| - | - | 5.744E+04 | 239.1 | - | - | 0 | - |
| - | - | 8397 | 240.1 | - | - | 0 | - |
| - | - | 4632 | 241.1 | - | - | 0 | - |
| - | - | 5882 | 241.1 | - | - | 0 | - |
| 7 | y | 8183 | 245.1 | 0.003514 | 14.34 | +2 | 3 |
| - | - | 8516 | 245.1 | - | - | 0 | - |
| - | - | 6426 | 246.1 | - | - | 0 | - |
| - | - | 1.646E+04 | 246.1 | - | - | 0 | - |
| - | - | 7315 | 247.1 | - | - | 0 | - |
| - | - | 7701 | 247.1 | - | - | 0 | - |
| - | - | 5871 | 248.1 | - | - | 0 | - |
| - | - | 1.699E+04 | 248.2 | - | - | 0 | - |
| 2 | b | 1.626E+06 | 249.1 | 0.0007156 | 2.873 | +1 | 2 |
| - | - | 2.247E+05 | 250.1 | - | - | 0 | - |
| - | - | 1.538E+05 | 251.1 | - | - | 0 | - |
| - | - | 2.944E+04 | 251.1 | - | - | 0 | - |
| - | - | 2.557E+04 | 252.1 | - | - | 0 | - |
| - | - | 1.233E+04 | 253.1 | - | - | 0 | - |
| - | - | 5021 | 255.1 | - | - | 0 | - |
| - | - | 7279 | 256.1 | - | - | 0 | - |
| - | - | 4329 | 257.1 | - | - | 0 | - |
| - | - | 8.671E+04 | 257.1 | - | - | 0 | - |
| - | - | 1.677E+04 | 261.1 | - | - | 0 | - |
| - | - | 1.117E+04 | 263.1 | - | - | 0 | - |
| - | - | 3.091E+05 | 263.1 | - | - | 0 | - |
| - | - | 3.163E+04 | 263.1 | - | - | 0 | - |
| - | - | 4.37E+04 | 264.1 | - | - | 0 | - |
| - | - | 6830 | 264.1 | - | - | 0 | - |
| - | - | 2.738E+04 | 264.1 | - | - | 0 | - |
| - | - | 1.321E+04 | 265.1 | - | - | 0 | - |
| - | - | 4818 | 265.1 | - | - | 0 | - |
| - | - | 2.037E+04 | 267.1 | - | - | 0 | - |
| - | - | 1.207E+04 | 268.1 | - | - | 0 | - |
| - | - | 4823 | 268.1 | - | - | 0 | - |
| - | - | 9172 | 268.1 | - | - | 0 | - |
| - | - | 1.544E+04 | 269.1 | - | - | 0 | - |
| - | - | 6127 | 270.1 | - | - | 0 | - |
| - | - | 5975 | 270.1 | - | - | 0 | - |
| - | - | 7538 | 271.1 | - | - | 0 | - |
| - | - | 6759 | 273.1 | - | - | 0 | - |
| - | - | 7786 | 273.1 | - | - | 0 | - |
| - | - | 7.739E+04 | 274.1 | - | - | 0 | - |
| - | - | 5396 | 277.7 | - | - | 0 | - |
| - | - | 1.318E+05 | 279.1 | - | - | 0 | - |
| - | - | 1.455E+04 | 279.1 | - | - | 0 | - |
| - | - | 1.184E+04 | 280.1 | - | - | 0 | - |
| - | - | 6148 | 283.1 | - | - | 0 | - |
| - | - | 6417 | 283.2 | - | - | 0 | - |
| - | - | 1.92E+05 | 285.1 | - | - | 0 | - |
| - | - | 2.465E+04 | 286.1 | - | - | 0 | - |
| - | - | 6761 | 287.1 | - | - | 0 | - |
| - | - | 1.731E+04 | 289.1 | - | - | 0 | - |
| - | - | 5360 | 289.1 | - | - | 0 | - |
| - | - | 1.001E+04 | 290.1 | - | - | 0 | - |
| - | - | 1.382E+05 | 292.1 | - | - | 0 | - |
| 3 | d | 3.028E+04 | 292.2 | 0.000295 | 1.01 | +1 | 3 |
| - | - | 1.682E+04 | 293.1 | - | - | 0 | - |
| - | - | 1.567E+05 | 296.2 | - | - | 0 | - |
| - | - | 2.645E+04 | 297.2 | - | - | 0 | - |
| - | - | 6630 | 299.2 | - | - | 0 | - |
| - | - | 1.264E+04 | 300.1 | - | - | 0 | - |
| - | - | 7678 | 301.1 | - | - | 0 | - |
| - | - | 8491 | 302.1 | - | - | 0 | - |
| - | - | 1.486E+04 | 302.1 | - | - | 0 | - |
| - | - | 4904 | 303.1 | - | - | 0 | - |
| - | - | 2.627E+04 | 303.1 | - | - | 0 | - |
| - | - | 1.773E+04 | 306.1 | - | - | 0 | - |
| - | - | 7465 | 307.1 | - | - | 0 | - |
| - | - | 2.044E+04 | 310.2 | - | - | 0 | - |
| - | - | 5165 | 311.1 | - | - | 0 | - |
| - | - | 3980 | 311.2 | - | - | 0 | - |
| - | - | 8922 | 313.1 | - | - | 0 | - |
| - | - | 1.893E+04 | 314.1 | - | - | 0 | - |
| - | - | 1.066E+04 | 315.1 | - | - | 0 | - |
| - | - | 4818 | 316.1 | - | - | 0 | - |
| 3 | a | 5.111E+04 | 318.1 | 0.0008889 | 2.794 | +1 | 3 |
| - | - | 1.03E+04 | 319.1 | - | - | 0 | - |
| - | - | 9409 | 320.1 | - | - | 0 | - |
| - | - | 6.266E+04 | 323.1 | - | - | 0 | - |
| - | - | 6259 | 323.1 | - | - | 0 | - |
| - | - | 5492 | 323.1 | - | - | 0 | - |
| - | - | 3.813E+05 | 324.1 | - | - | 0 | - |
| - | - | 5.56E+04 | 325.1 | - | - | 0 | - |
| - | - | 1.271E+04 | 327.1 | - | - | 0 | - |
| - | - | 2.977E+04 | 328.1 | - | - | 0 | - |
| - | - | 1.415E+04 | 328.1 | - | - | 0 | - |
| - | - | 1.755E+04 | 330.1 | - | - | 0 | - |
| - | - | 2.406E+04 | 332.1 | - | - | 0 | - |
| - | - | 1.385E+04 | 333.1 | - | - | 0 | - |
| - | - | 4755 | 333.2 | - | - | 0 | - |
| - | - | 1.603E+04 | 334.1 | - | - | 0 | - |
| - | - | 8096 | 334.1 | - | - | 0 | - |
| - | - | 7293 | 335.2 | - | - | 0 | - |
| - | - | 2.355E+04 | 338.1 | - | - | 0 | - |
| - | - | 7153 | 339.1 | - | - | 0 | - |
| - | - | 2.704E+04 | 339.2 | - | - | 0 | - |
| 8 | y | 1.827E+06 | 342.2 | 0.000947 | 2.768 | +1 | 2 |
| - | - | 3.588E+05 | 343.2 | - | - | 0 | - |
| - | - | 2.856E+04 | 344.2 | - | - | 0 | - |
| 3 | b | 9.36E+04 | 346.1 | 0.0008778 | 2.536 | +1 | 3 |
| - | - | 7926 | 346.2 | - | - | 0 | - |
| - | - | 1.607E+04 | 347.1 | - | - | 0 | - |
| - | - | 5.845E+04 | 348.1 | - | - | 0 | - |
| 6 | b | 4545 | 348.1 | 0.006747 | 19.38 | +2 | 6 |
| - | - | 1.006E+05 | 350.1 | - | - | 0 | - |
| - | - | 2.365E+04 | 351.1 | - | - | 0 | - |
| - | - | 4.29E+04 | 352.1 | - | - | 0 | - |
| - | - | 8560 | 353.1 | - | - | 0 | - |
| - | - | 9569 | 354.2 | - | - | 0 | - |
| - | - | 9.969E+04 | 356.1 | - | - | 0 | - |
| 6 | b | 1.456E+04 | 357.1 | 9.128E-05 | 0.2556 | +2 | 6 |
| - | - | 5464 | 359.1 | - | - | 0 | - |
| - | - | 2.419E+04 | 360.1 | - | - | 0 | - |
| - | - | 6571 | 361.1 | - | - | 0 | - |
| 5 | y | 1.5E+04 | 362.1 | 0.004049 | 11.18 | +2 | 5 |
| - | - | 5483 | 362.7 | - | - | 0 | - |
| 3 | b | 4.671E+05 | 364.2 | 0.0007807 | 2.144 | +1 | 3 |
| - | - | 7.811E+04 | 365.2 | - | - | 0 | - |
| - | - | 4.784E+04 | 366.1 | - | - | 0 | - |
| - | - | 1.13E+04 | 366.2 | - | - | 0 | - |
| - | - | 7633 | 367.1 | - | - | 0 | - |
| - | - | 1.805E+04 | 367.2 | - | - | 0 | - |
| - | - | 5990 | 368.2 | - | - | 0 | - |
| - | - | 1.536E+04 | 371.1 | - | - | 0 | - |
| - | - | 2.138E+04 | 374.1 | - | - | 0 | - |
| - | - | 5521 | 375.2 | - | - | 0 | - |
| - | - | 4.046E+04 | 376.1 | - | - | 0 | - |
| - | - | 1.396E+04 | 378.1 | - | - | 0 | - |
| - | - | 2.521E+05 | 378.1 | - | - | 0 | - |
| - | - | 4.303E+04 | 379.1 | - | - | 0 | - |
| - | - | 1.891E+04 | 380.1 | - | - | 0 | - |
| - | - | 5.375E+04 | 382.1 | - | - | 0 | - |
| - | - | 1.33E+04 | 383.1 | - | - | 0 | - |
| - | - | 1.519E+04 | 384.1 | - | - | 0 | - |
| - | - | 1.582E+04 | 385.1 | - | - | 0 | - |
| - | - | 8243 | 388.1 | - | - | 0 | - |
| - | - | 9694 | 388.2 | - | - | 0 | - |
| - | - | 3.445E+04 | 390.2 | - | - | 0 | - |
| - | - | 3.366E+05 | 394.1 | - | - | 0 | - |
| - | - | 5.749E+04 | 395.1 | - | - | 0 | - |
| - | - | 7100 | 396.1 | - | - | 0 | - |
| - | - | 8875 | 397.2 | - | - | 0 | - |
| - | - | 5902 | 398.2 | - | - | 0 | - |
| - | - | 1.251E+04 | 399.2 | - | - | 0 | - |
| - | - | 1.013E+04 | 401.2 | - | - | 0 | - |
| - | - | 6794 | 404.1 | - | - | 0 | - |
| - | - | 7389 | 404.1 | - | - | 0 | - |
| - | - | 1.022E+04 | 406.1 | - | - | 0 | - |
| - | - | 1.396E+04 | 406.2 | - | - | 0 | - |
| - | - | 2.945E+04 | 407.2 | - | - | 0 | - |
| - | - | 8.365E+04 | 408.2 | - | - | 0 | - |
| - | - | 2.411E+04 | 409.2 | - | - | 0 | - |
| - | - | 6922 | 409.2 | - | - | 0 | - |
| - | - | 9429 | 410.1 | - | - | 0 | - |
| - | - | 5756 | 410.2 | - | - | 0 | - |
| - | - | 4947 | 414.7 | - | - | 0 | - |
| - | - | 1.565E+04 | 415.2 | - | - | 0 | - |
| - | - | 1.953E+04 | 419.2 | - | - | 0 | - |
| - | - | 7481 | 423.2 | - | - | 0 | - |
| - | - | 2.871E+05 | 425.2 | - | - | 0 | - |
| - | - | 6.62E+04 | 426.2 | - | - | 0 | - |
| - | - | 5.123E+04 | 427.2 | - | - | 0 | - |
| - | - | 9265 | 428.2 | - | - | 0 | - |
| - | - | 6007 | 429.2 | - | - | 0 | - |
| - | - | 4599 | 431.2 | - | - | 0 | - |
| - | - | 6746 | 432.1 | - | - | 0 | - |
| - | - | 4.036E+04 | 433.2 | - | - | 0 | - |
| - | - | 1.175E+04 | 434.2 | - | - | 0 | - |
| - | - | 1.11E+04 | 435.2 | - | - | 0 | - |
| - | - | 8973 | 437.2 | - | - | 0 | - |
| - | - | 1.14E+04 | 442.2 | - | - | 0 | - |
| - | - | 9807 | 443.2 | - | - | 0 | - |
| - | - | 5243 | 443.2 | - | - | 0 | - |
| - | - | 7072 | 446.2 | - | - | 0 | - |
| - | - | 4.13E+04 | 447.2 | - | - | 0 | - |
| - | - | 1.449E+04 | 449.1 | - | - | 0 | - |
| - | - | 1.572E+04 | 449.2 | - | - | 0 | - |
| - | - | 8409 | 450.2 | - | - | 0 | - |
| - | - | 5890 | 451.2 | - | - | 0 | - |
| - | - | 6856 | 452.7 | - | - | 0 | - |
| - | - | 5417 | 453.2 | - | - | 0 | - |
| - | - | 8726 | 454.2 | - | - | 0 | - |
| - | - | 2.236E+04 | 454.2 | - | - | 0 | - |
| - | - | 6.239E+04 | 455.2 | - | - | 0 | - |
| - | - | 1.536E+04 | 456.2 | - | - | 0 | - |
| - | - | 7958 | 458.2 | - | - | 0 | - |
| - | - | 9358 | 459.1 | - | - | 0 | - |
| 4 | b | 6.972E+04 | 461.2 | 0.0004546 | 0.9857 | +1 | 4 |
| - | - | 3.024E+04 | 461.2 | - | - | 0 | - |
| - | - | 2.187E+04 | 462.2 | - | - | 0 | - |
| - | - | 5442 | 464.2 | - | - | 0 | - |
| - | - | 2.29E+05 | 465.2 | - | - | 0 | - |
| - | - | 5.377E+04 | 466.2 | - | - | 0 | - |
| - | - | 9939 | 467.2 | - | - | 0 | - |
| 3 | y | 8171 | 468.2 | 0.004178 | 8.925 | +2 | 7 |
| - | - | 7272 | 469.2 | - | - | 0 | - |
| - | - | 1.082E+04 | 469.2 | - | - | 0 | - |
| - | - | 8448 | 470.2 | - | - | 0 | - |
| - | - | 9669 | 470.2 | - | - | 0 | - |
| - | - | 3.734E+04 | 471.2 | - | - | 0 | - |
| - | - | 8905 | 472.2 | - | - | 0 | - |
| - | - | 1.911E+04 | 473.2 | - | - | 0 | - |
| - | - | 4548 | 473.6 | - | - | 0 | - |
| - | - | 7060 | 475.1 | - | - | 0 | - |
| 3 | y | 4.41E+04 | 477.2 | 0.006231 | 13.06 | +2 | 7 |
| - | - | 1.808E+04 | 477.7 | - | - | 0 | - |
| - | - | 1.651E+04 | 478.2 | - | - | 0 | - |
| - | - | 1.431E+04 | 478.2 | - | - | 0 | - |
| 4 | b | 2.495E+05 | 479.2 | 0.001181 | 2.465 | +1 | 4 |
| - | - | 4.857E+04 | 480.2 | - | - | 0 | - |
| - | - | 9983 | 481.2 | - | - | 0 | - |
| - | - | 1.439E+04 | 482.2 | - | - | 0 | - |
| - | - | 8179 | 483.2 | - | - | 0 | - |
| - | - | 3.468E+04 | 486.2 | - | - | 0 | - |
| - | - | 8830 | 487.2 | - | - | 0 | - |
| 7 | y | 4.556E+05 | 489.2 | 0.006259 | 12.8 | +1 | 3 |
| - | - | 1.115E+05 | 490.2 | - | - | 0 | - |
| - | - | 5.633E+04 | 491.2 | - | - | 0 | - |
| - | - | 1.757E+04 | 492.2 | - | - | 0 | - |
| - | - | 3.459E+04 | 495.2 | - | - | 0 | - |
| - | - | 8806 | 496.2 | - | - | 0 | - |
| - | - | 1.222E+04 | 496.2 | - | - | 0 | - |
| - | - | 2.324E+05 | 496.2 | - | - | 0 | - |
| - | - | 4.232E+04 | 497.2 | - | - | 0 | - |
| - | - | 6.379E+04 | 497.2 | - | - | 0 | - |
| - | - | 8224 | 498.2 | - | - | 0 | - |
| 8 | b | 1.873E+04 | 499.2 | 0.0007727 | 1.548 | +2 | 8 |
| - | - | 1.241E+04 | 499.7 | - | - | 0 | - |
| - | - | 8652 | 502.2 | - | - | 0 | - |
| - | - | 5070 | 504.2 | - | - | 0 | - |
| - | - | 5862 | 507.2 | - | - | 0 | - |
| - | - | 8343 | 508.2 | - | - | 0 | - |
| - | - | 7708 | 511.2 | - | - | 0 | - |
| - | - | 2.85E+04 | 513.2 | - | - | 0 | - |
| - | - | 1.276E+04 | 514.2 | - | - | 0 | - |
| - | - | 9907 | 516.2 | - | - | 0 | - |
| - | - | 1.845E+04 | 518.7 | - | - | 0 | - |
| - | - | 6.22E+04 | 519.2 | - | - | 0 | - |
| - | - | 3.636E+04 | 520.2 | - | - | 0 | - |
| - | - | 1.461E+04 | 521.2 | - | - | 0 | - |
| - | - | 1.015E+04 | 523.2 | - | - | 0 | - |
| - | - | 5838 | 525.2 | - | - | 0 | - |
| - | - | 6058 | 525.2 | - | - | 0 | - |
| - | - | 5501 | 528.2 | - | - | 0 | - |
| - | - | 2.24E+04 | 530.2 | - | - | 0 | - |
| - | - | 6908 | 531.2 | - | - | 0 | - |
| - | - | 8045 | 532.2 | - | - | 0 | - |
| - | - | 8174 | 533.2 | - | - | 0 | - |
| - | - | 2.26E+04 | 537.2 | - | - | 0 | - |
| - | - | 7732 | 538.2 | - | - | 0 | - |
| - | - | 1.476E+04 | 541.2 | - | - | 0 | - |
| 2 | y | 3.855E+04 | 541.7 | 0.003174 | 5.86 | +2 | 8 |
| - | - | 3.343E+04 | 542.2 | - | - | 0 | - |
| - | - | 1.31E+04 | 542.7 | - | - | 0 | - |
| - | - | 9713 | 543.2 | - | - | 0 | - |
| - | - | 5650 | 547.2 | - | - | 0 | - |
| - | - | 7.836E+04 | 548.2 | - | - | 0 | - |
| - | - | 1.983E+04 | 549.2 | - | - | 0 | - |
| - | - | 9859 | 550.2 | - | - | 0 | - |
| 2 | y | 2.159E+05 | 550.7 | 0.003629 | 6.59 | +2 | 8 |
| - | - | 1.633E+05 | 551.2 | - | - | 0 | - |
| - | - | 6E+04 | 551.7 | - | - | 0 | - |
| - | - | 1.543E+04 | 552.2 | - | - | 0 | - |
| - | - | 7353 | 555.7 | - | - | 0 | - |
| - | - | 4.859E+04 | 557.2 | - | - | 0 | - |
| - | - | 1.196E+04 | 558.2 | - | - | 0 | - |
| 6 | y | 7.084E+05 | 560.2 | 0.005889 | 10.51 | +1 | 4 |
| - | - | 1.665E+04 | 560.7 | - | - | 0 | - |
| - | - | 2.326E+05 | 561.2 | - | - | 0 | - |
| - | - | 5.797E+04 | 562.2 | - | - | 0 | - |
| - | - | 8525 | 566.2 | - | - | 0 | - |
| - | - | 4.321E+04 | 569.2 | - | - | 0 | - |
| - | - | 3.177E+04 | 569.7 | - | - | 0 | - |
| - | - | 7.277E+04 | 570.2 | - | - | 0 | - |
| - | - | 1.432E+04 | 571.2 | - | - | 0 | - |
| - | - | 1.206E+04 | 574.7 | - | - | 0 | - |
| - | - | 2.953E+04 | 578.2 | - | - | 0 | - |
| - | - | 1.57E+04 | 578.7 | - | - | 0 | - |
| - | - | 7582 | 579.2 | - | - | 0 | - |
| - | - | 5964 | 580.2 | - | - | 0 | - |
| - | - | 3.034E+04 | 583.2 | - | - | 0 | - |
| - | - | 1.553E+04 | 583.7 | - | - | 0 | - |
| - | - | 8031 | 584.2 | - | - | 0 | - |
| - | - | 1.723E+04 | 588.2 | - | - | 0 | - |
| - | - | 6184 | 589.2 | - | - | 0 | - |
| 0 | Precursor | 1.125E+05 | 592.2 | 0.003505 | 5.919 | +2 | -1 |
| - | - | 1.005E+05 | 592.7 | - | - | 0 | - |
| - | - | 3.886E+04 | 593.2 | - | - | 0 | - |
| - | - | 7093 | 593.7 | - | - | 0 | - |
| - | - | 2.049E+04 | 594.2 | - | - | 0 | - |
| - | - | 9600 | 595.2 | - | - | 0 | - |
| - | - | 5.067E+04 | 596.2 | - | - | 0 | - |
| - | - | 3.03E+04 | 597.2 | - | - | 0 | - |
| - | - | 1.494E+04 | 598.2 | - | - | 0 | - |
| 0 | Precursor | 3.377E+05 | 601.2 | 0.003838 | 6.384 | +2 | -1 |
| - | - | 2.666E+05 | 601.7 | - | - | 0 | - |
| - | - | 1.009E+05 | 602.2 | - | - | 0 | - |
| - | - | 1.212E+04 | 602.7 | - | - | 0 | - |
| - | - | 7.198E+04 | 606.2 | - | - | 0 | - |
| - | - | 1.832E+04 | 607.2 | - | - | 0 | - |
| - | - | 8878 | 608.2 | - | - | 0 | - |
| - | - | 1.147E+05 | 612.2 | - | - | 0 | - |
| - | - | 3.383E+04 | 613.2 | - | - | 0 | - |
| - | - | 9533 | 613.3 | - | - | 0 | - |
| - | - | 7.626E+04 | 614.2 | - | - | 0 | - |
| - | - | 2.18E+04 | 615.2 | - | - | 0 | - |
| - | - | 6415 | 616.3 | - | - | 0 | - |
| - | - | 5554 | 622.2 | - | - | 0 | - |
| 5 | b | 5.808E+04 | 624.2 | 0.001854 | 2.97 | +1 | 5 |
| - | - | 1.992E+04 | 625.2 | - | - | 0 | - |
| - | - | 2.318E+04 | 631.2 | - | - | 0 | - |
| - | - | 1.197E+04 | 632.3 | - | - | 0 | - |
| - | - | 7.041E+04 | 634.2 | - | - | 0 | - |
| - | - | 1.974E+04 | 635.2 | - | - | 0 | - |
| - | - | 1.676E+04 | 639.3 | - | - | 0 | - |
| - | - | 3.276E+04 | 640.2 | - | - | 0 | - |
| - | - | 1.458E+04 | 641.3 | - | - | 0 | - |
| 5 | b | 5.193E+04 | 642.2 | 0.001848 | 2.878 | +1 | 5 |
| - | - | 2.062E+04 | 643.2 | - | - | 0 | - |
| - | - | 2.488E+04 | 652.2 | - | - | 0 | - |
| - | - | 7899 | 653.2 | - | - | 0 | - |
| - | - | 1.667E+05 | 657.3 | - | - | 0 | - |
| - | - | 5.107E+04 | 658.3 | - | - | 0 | - |
| - | - | 2.62E+05 | 659.3 | - | - | 0 | - |
| - | - | 6420 | 660.2 | - | - | 0 | - |
| - | - | 1.051E+05 | 660.3 | - | - | 0 | - |
| - | - | 1.467E+04 | 661.3 | - | - | 0 | - |
| - | - | 1.992E+04 | 667.3 | - | - | 0 | - |
| - | - | 1.518E+04 | 668.2 | - | - | 0 | - |
| - | - | 7123 | 669.3 | - | - | 0 | - |
| - | - | 3.371E+04 | 676.3 | - | - | 0 | - |
| - | - | 2.143E+04 | 677.3 | - | - | 0 | - |
| - | - | 6506 | 678.3 | - | - | 0 | - |
| - | - | 2.119E+05 | 685.3 | - | - | 0 | - |
| - | - | 7.631E+04 | 686.3 | - | - | 0 | - |
| - | - | 1.852E+04 | 687.3 | - | - | 0 | - |
| 6 | b | 4.35E+04 | 695.3 | 0.0001037 | 0.1492 | +1 | 6 |
| - | - | 1.183E+04 | 696.3 | - | - | 0 | - |
| - | - | 6.582E+04 | 703.3 | - | - | 0 | - |
| - | - | 1.884E+04 | 704.3 | - | - | 0 | - |
| - | - | 2.502E+04 | 705.3 | - | - | 0 | - |
| - | - | 9097 | 706.3 | - | - | 0 | - |
| 6 | b | 2.368E+04 | 713.3 | 0.002088 | 2.927 | +1 | 6 |
| - | - | 5567 | 714.3 | - | - | 0 | - |
| - | - | 9375 | 721.2 | - | - | 0 | - |
| - | - | 2.119E+05 | 721.3 | - | - | 0 | - |
| - | - | 6.895E+04 | 722.3 | - | - | 0 | - |
| 5 | y | 6.873E+05 | 723.3 | 0.005854 | 8.093 | +1 | 5 |
| - | - | 2.711E+05 | 724.3 | - | - | 0 | - |
| - | - | 7.824E+04 | 725.3 | - | - | 0 | - |
| - | - | 9781 | 726.3 | - | - | 0 | - |
| - | - | 9625 | 728.3 | - | - | 0 | - |
| - | - | 2.115E+04 | 731.2 | - | - | 0 | - |
| - | - | 1.312E+04 | 732.2 | - | - | 0 | - |
| - | - | 1.262E+04 | 733.3 | - | - | 0 | - |
| - | - | 6316 | 734.3 | - | - | 0 | - |
| - | - | 6185 | 740.3 | - | - | 0 | - |
| - | - | 1.183E+04 | 746.3 | - | - | 0 | - |
| - | - | 5763 | 747.3 | - | - | 0 | - |
| - | - | 7229 | 748.3 | - | - | 0 | - |
| - | - | 2.899E+05 | 749.3 | - | - | 0 | - |
| - | - | 1.166E+05 | 750.3 | - | - | 0 | - |
| - | - | 3.809E+04 | 751.3 | - | - | 0 | - |
| - | - | 1.859E+04 | 756.3 | - | - | 0 | - |
| - | - | 1.425E+04 | 757.3 | - | - | 0 | - |
| - | - | 1.223E+04 | 759.3 | - | - | 0 | - |
| - | - | 7.267E+04 | 767.3 | - | - | 0 | - |
| - | - | 2.929E+04 | 768.3 | - | - | 0 | - |
| - | - | 7544 | 769.3 | - | - | 0 | - |
| - | - | 2.825E+05 | 774.3 | - | - | 0 | - |
| - | - | 1.158E+05 | 775.3 | - | - | 0 | - |
| - | - | 2.259E+04 | 776.3 | - | - | 0 | - |
| - | - | 1.433E+04 | 778.3 | - | - | 0 | - |
| - | - | 6974 | 779.3 | - | - | 0 | - |
| - | - | 7479 | 784.3 | - | - | 0 | - |
| - | - | 9612 | 792.3 | - | - | 0 | - |
| - | - | 5774 | 793.3 | - | - | 0 | - |
| - | - | 7951 | 796.3 | - | - | 0 | - |
| - | - | 8156 | 803.3 | - | - | 0 | - |
| - | - | 9148 | 804.3 | - | - | 0 | - |
| - | - | 8710 | 805.3 | - | - | 0 | - |
| - | - | 8866 | 818.3 | - | - | 0 | - |
| 4 | y | 4.236E+04 | 820.3 | 0.005894 | 7.185 | +1 | 6 |
| - | - | 2.216E+04 | 821.3 | - | - | 0 | - |
| - | - | 1.089E+04 | 822.3 | - | - | 0 | - |
| - | - | 6426 | 823.3 | - | - | 0 | - |
| - | - | 7256 | 824.3 | - | - | 0 | - |
| - | - | 9211 | 828.4 | - | - | 0 | - |
| - | - | 2.446E+04 | 832.3 | - | - | 0 | - |
| - | - | 1.677E+04 | 833.3 | - | - | 0 | - |
| 4 | y | 8.721E+05 | 838.3 | 0.006193 | 7.388 | +1 | 6 |
| - | - | 3.874E+05 | 839.3 | - | - | 0 | - |
| - | - | 1.309E+05 | 840.3 | - | - | 0 | - |
| - | - | 9753 | 841.3 | - | - | 0 | - |
| 7 | b | 1.44E+04 | 842.3 | 0.01049 | 12.45 | +1 | 7 |
| - | - | 3.66E+04 | 843.3 | - | - | 0 | - |
| - | - | 1.578E+04 | 844.3 | - | - | 0 | - |
| - | - | 1.336E+04 | 848.3 | - | - | 0 | - |
| - | - | 6242 | 850.3 | - | - | 0 | - |
| - | - | 5755 | 851.3 | - | - | 0 | - |
| - | - | 5538 | 853.3 | - | - | 0 | - |
| - | - | 1.026E+04 | 854.3 | - | - | 0 | - |
| 7 | b | 1.436E+04 | 860.3 | 0.003891 | 4.523 | +1 | 7 |
| - | - | 9270 | 861.3 | - | - | 0 | - |
| - | - | 2.743E+04 | 862.3 | - | - | 0 | - |
| - | - | 1.377E+04 | 863.3 | - | - | 0 | - |
| - | - | 1.58E+04 | 868.3 | - | - | 0 | - |
| - | - | 9692 | 869.3 | - | - | 0 | - |
| - | - | 1.168E+05 | 871.3 | - | - | 0 | - |
| - | - | 6.629E+04 | 872.3 | - | - | 0 | - |
| - | - | 2.3E+04 | 873.3 | - | - | 0 | - |
| - | - | 1.203E+06 | 889.3 | - | - | 0 | - |
| - | - | 5.57E+05 | 890.4 | - | - | 0 | - |
| - | - | 9090 | 890.5 | - | - | 0 | - |
| - | - | 1.548E+05 | 891.4 | - | - | 0 | - |
| - | - | 1.574E+04 | 892.4 | - | - | 0 | - |
| - | - | 2.32E+04 | 896.3 | - | - | 0 | - |
| - | - | 1.297E+04 | 897.3 | - | - | 0 | - |
| - | - | 7818 | 899.3 | - | - | 0 | - |
| - | - | 4.579E+04 | 907.3 | - | - | 0 | - |
| - | - | 2.637E+04 | 908.3 | - | - | 0 | - |
| - | - | 1.115E+04 | 909.3 | - | - | 0 | - |
| - | - | 7221 | 914.3 | - | - | 0 | - |
| - | - | 1.26E+04 | 917.3 | - | - | 0 | - |
| - | - | 1.291E+04 | 918.3 | - | - | 0 | - |
| - | - | 1.241E+04 | 933.4 | - | - | 0 | - |
| 3 | y | 2.38E+05 | 935.3 | 0.005989 | 6.404 | +1 | 7 |
| - | - | 1.365E+05 | 936.3 | - | - | 0 | - |
| - | - | 5.475E+04 | 937.3 | - | - | 0 | - |
| - | - | 2.544E+04 | 938.3 | - | - | 0 | - |
| - | - | 8975 | 939.3 | - | - | 0 | - |
| 3 | y | 3.722E+06 | 953.3 | 0.006289 | 6.597 | +1 | 7 |
| - | - | 1.945E+06 | 954.3 | - | - | 0 | - |
| - | - | 6.993E+05 | 955.4 | - | - | 0 | - |
| - | - | 6.848E+04 | 956.4 | - | - | 0 | - |
| - | - | 2.345E+04 | 963.3 | - | - | 0 | - |
| - | - | 1.18E+04 | 964.3 | - | - | 0 | - |
| - | - | 1.423E+04 | 969.4 | - | - | 0 | - |
| - | - | 5420 | 970.4 | - | - | 0 | - |
| 8 | b | 1.028E+04 | 979.4 | 0.003639 | 3.716 | +1 | 8 |
| - | - | 5809 | 990.1 | - | - | 0 | - |
| 8 | b | 4.149E+04 | 997.4 | 0.005587 | 5.602 | +1 | 8 |
| - | - | 2.141E+04 | 998.4 | - | - | 0 | - |
| - | - | 1.088E+04 | 999.4 | - | - | 0 | - |
| - | - | 1.248E+04 | 1015 | - | - | 0 | - |
| - | - | 5830 | 1016 | - | - | 0 | - |
| - | - | 1.879E+04 | 1018 | - | - | 0 | - |
| - | - | 9061 | 1019 | - | - | 0 | - |
| - | - | 5689 | 1020 | - | - | 0 | - |
| - | - | 8.097E+04 | 1036 | - | - | 0 | - |
| - | - | 5.15E+04 | 1037 | - | - | 0 | - |
| - | - | 1.791E+04 | 1038 | - | - | 0 | - |
| - | - | 6202 | 1046 | - | - | 0 | - |
| 2 | y | 2.351E+04 | 1082 | 0.006545 | 6.047 | +1 | 8 |
| - | - | 1.369E+04 | 1083 | - | - | 0 | - |
| 2 | y | 2.073E+05 | 1100 | 0.006479 | 5.887 | +1 | 8 |
| - | - | 1.385E+05 | 1101 | - | - | 0 | - |
| - | - | 5.836E+04 | 1102 | - | - | 0 | - |
| - | - | 9629 | 1103 | - | - | 0 | - |
| - | - | 3.557E+04 | 1110 | - | - | 0 | - |
| - | - | 2.573E+04 | 1111 | - | - | 0 | - |
| - | - | 9870 | 1112 | - | - | 0 | - |
| - | - | 5642 | 1956 | - | - | 0 | - |
| - | - | 6472 | 3240 | - | - | 0 | - |

m/z Charge Intensity FragmentType MassShift Position
120.08128356933594 0 2181933.5
121.04022216796875 0 5911.368
121.08458709716797 0 162669.42
122.0720443725586 0 12771.19
122.08824920654297 0 6464.0454
127.08721923828125 0 6639.904
129.07052612304688 0 9323.835
129.16705322265625 0 3154.443
130.0655517578125 0 46569.223
131.0685272216797 0 5443.433
132.08123779296875 0 111954.1
133.06103515625 0 6168.472
133.0843963623047 0 8820.505
134.09689331054688 0 9514.965
136.076171875 0 786052.4
136.08749389648438 0 26270.564
137.07366943359375 0 5548.151
137.0795440673828 0 62848.367
137.0844268798828 0 5368.931
138.05543518066406 0 59126.77
138.06666564941406 0 87262.48
139.07008361816406 0 4916.3735
141.0663299560547 0 6576.707
143.0456085205078 0 5458.708
143.073486328125 0 13951.242
143.08607482910156 0 12589.368
144.0749969482422 0 3006.4016
144.08119201660156 0 10914.035
146.06056213378906 0 30014.475
146.0717315673828 0 4784.646
146.09686279296875 0 13855.525
148.07640075683594 0 7202.824
148.08741760253906 0 24912.656
150.06700134277344 0 6612.655
155.0819854736328 0 13912.131
155.0931854248047 0 21720.398
156.0772247314453 0 204384.56
157.08058166503906 0 12332.426
158.09690856933594 0 325995.34
159.054931640625 0 3697.449
159.0922088623047 0 164924.53
159.10018920898438 0 34061.094
160.07615661621094 0 58682.53
160.08753967285156 0 6059.764
160.09539794921875 0 15395.32
160.112548828125 0 148562.6
161.0790557861328 0 3859.1362
161.1159210205078 0 16495.67
162.09239196777344 0 5179.444
164.08201599121094 0 4468.3647
165.0701141357422 0 9346.841
165.07760620117188 0 32898.152
166.05369567871094 0 13367.219
166.0616455078125 0 28061.467
166.0860137939453 0 5245.241
167.09320068359375 0 25148.924
169.07664489746094 0 12021.771
171.076904296875 0 27188.268
171.0919647216797 0 5292.718
172.07723999023438 0 5993.219
172.10867309570312 0 5835.1836
173.09262084960938 0 8810.467
174.06674194335938 0 20212.145 b Ammonia loss 2
174.09173583984375 0 4125.1265
175.0872802734375 0 12704.858
175.09779357910156 0 5951.9893
175.12351989746094 0 8748.858
176.0824432373047 0 53717.37
176.1075439453125 0 657830.3
177.102783203125 0 467468.94
177.11135864257812 0 70200.25
178.10614013671875 0 44548.84
178.11419677734375 0 4997.9736
180.0771942138672 0 29272.6
182.0456085205078 0 5604.774
185.05624389648438 0 50171.668
186.09185791015625 0 54400.4
187.06356811523438 0 12011.196
188.0711212158203 0 223348.25
189.0745849609375 0 27677.74
190.0868682861328 0 8221.209
191.08358764648438 0 5370.167
191.0935821533203 0 23561.326
191.11842346191406 0 46733.445
192.1027069091797 0 11195.881
192.1204376220703 0 4598.6733
193.07254028320312 0 13456.464
193.10888671875 0 185870
194.0802459716797 0 9672.893
194.11248779296875 0 14105.567
194.33055114746094 0 4155.0615
195.0881805419922 0 48464.77
196.6621856689453 0 3839.406
202.05381774902344 0 15919.494
203.0667266845703 0 52484.848
203.0931854248047 0 11838.531
203.1183624267578 0 223670.89 a Water loss 1
204.07725524902344 0 102332.87
204.102294921875 0 13616.174 a Ammonia loss 1
204.12156677246094 0 30775.748
205.08135986328125 0 8510.439
205.09768676757812 0 173131.36 y 8
205.10829162597656 0 13461.29
206.1012420654297 0 25323.28
207.11334228515625 0 47748.52
208.0721893310547 0 28086.008
209.07562255859375 0 4857.9556
211.08340454101562 0 5478.818
212.11679077148438 0 12089.758
215.0823211669922 0 11311.131
216.0655517578125 0 7609.9453
217.0977325439453 0 117130.99
218.10150146484375 0 13684.508
219.08035278320312 0 37374.92
219.09072875976562 0 7069.462
220.12049865722656 0 4827.6895
221.10406494140625 0 539384.9
221.12918090820312 0 8847761 a 1
222.10714721679688 0 51256.75
222.13238525390625 0 1074879.2
223.1199951171875 0 8257.199
223.1348419189453 0 63254.65
224.118408203125 0 8979.486
225.09967041015625 0 6291.1206
228.11351013183594 0 24287.598
229.1089324951172 0 18746.22
230.12936401367188 0 5973.041
231.06179809570312 0 149280.97
231.08888244628906 0 5860.252 b Water loss 3
231.11338806152344 0 84543.94 b Water loss 1
231.1240997314453 0 4046.3032
232.0647735595703 0 13725.806
232.1166534423828 0 18324.592
233.0930938720703 0 4681.994
234.12408447265625 0 5954.1973
235.10838317871094 0 208485.81
236.11109924316406 0 23285.402
237.09042358398438 0 8882.131
237.1244354248047 0 6521.125
238.1304473876953 0 13730.005
239.11444091796875 0 57436.402
240.1169891357422 0 8396.95
241.08460998535156 0 4632.0483
241.09864807128906 0 5881.937
245.09339904785156 0 8182.7495 y 6
245.1292724609375 0 8516.108
246.09893798828125 0 6426.07
246.1354522705078 0 16460.455
247.10707092285156 0 7315.119
247.12034606933594 0 7700.661
248.08914184570312 0 5871.0977
248.15103149414062 0 16991.596
249.12408447265625 0 1625605.9 b 1
250.1273956298828 0 224715.83
251.10324096679688 0 153772.47
251.12974548339844 0 29440.498
252.10691833496094 0 25574.51
253.09397888183594 0 12325.319
255.11279296875 0 5021.4565
256.1087951660156 0 7279.006
257.07879638671875 0 4329.331
257.1069641113281 0 86709.24
261.0870361328125 0 16772.875
263.0703430175781 0 11170.293
263.10321044921875 0 309139.16
263.1295471191406 0 31627.879
264.106689453125 0 43696.414
264.1312561035156 0 6829.831
264.14605712890625 0 27382.709
265.1171875 0 13212.423
265.14349365234375 0 4817.633
267.0916748046875 0 20366.047
268.0757751464844 0 12071.028
268.0950622558594 0 4822.7007
268.140380859375 0 9172.347
269.077392578125 0 15436.0205
270.07244873046875 0 6126.639
270.1083679199219 0 5975.0366
271.1079406738281 0 7537.863
273.088134765625 0 6759.008
273.12298583984375 0 7786.3438
274.1304626464844 0 77390.11
277.6948547363281 0 5395.6963
279.0982666015625 0 131750.97
279.12542724609375 0 14545.06
280.1016845703125 0 11844.262
283.10784912109375 0 6148.2227
283.15167236328125 0 6416.617
285.1023254394531 0 192045.72
286.1045227050781 0 24645.846
287.100830078125 0 6760.728
289.08270263671875 0 17313.047
289.1191101074219 0 5359.882
290.1257019042969 0 10007.213
292.1412353515625 0 138245.48
292.1658630371094 0 30276.068 d 2
293.144287109375 0 16823.74
296.1513671875 0 156707.73
297.154052734375 0 26454.1
299.1521301269531 0 6630.267
300.1346130371094 0 12635.563
301.11859130859375 0 7677.919
302.09783935546875 0 8491.267
302.1150207519531 0 14855.417
303.09619140625 0 4903.748
303.113525390625 0 26272.299
306.14520263671875 0 17727.746
307.12115478515625 0 7464.934
310.15185546875 0 20438.406
311.11895751953125 0 5164.9688
311.155029296875 0 3979.7737
313.0979309082031 0 8922.309
314.0994567871094 0 18933.344
315.09881591796875 0 10661.101
316.1296081542969 0 4818.3384
318.1457214355469 0 51108.703 a Water loss 2
319.1494445800781 0 10298.334
320.1371154785156 0 9408.925
323.0995178222656 0 62656.844
323.11871337890625 0 6258.8984
323.1402587890625 0 5492.2476
324.1464538574219 0 381259.78
325.1490478515625 0 55595.285
327.14642333984375 0 12713.654
328.1296081542969 0 29765.594
328.1465148925781 0 14154.25
330.10955810546875 0 17545.176
332.12530517578125 0 24060.355
333.1093444824219 0 13852.653
333.21673583984375 0 4754.676
334.1075439453125 0 16033.018
334.1295471191406 0 8095.9546
335.1717224121094 0 7292.821
338.1288757324219 0 23553.598
339.133056640625 0 7153.0366
339.17108154296875 0 27036.098
342.1570129394531 0 1826543.5 y 7
343.1600646972656 0 358795.5
344.1622314453125 0 28558.496
346.140625 0 93602.09 b Water loss 2
346.1625061035156 0 7926.03
347.14337158203125 0 16065.131
348.119873046875 0 58449.57
348.1439514160156 0 4544.755 b Water loss 5
350.1353759765625 0 100575.73
351.137939453125 0 23653.082
352.1408996582031 0 42897.49
353.1481628417969 0 8559.761
354.1556396484375 0 9569.318
356.1396484375 0 99694.03
357.142578125 0 14563.707 b 5
359.0858154296875 0 5464.26
360.1199951171875 0 24191.043
361.1219787597656 0 6570.986
362.15118408203125 0 15000.353 y 4
362.6501770019531 0 5483.0293
364.1510925292969 0 467080.4 b 2
365.1545104980469 0 78110.125
366.1298522949219 0 47838.87
366.1559753417969 0 11298.902
367.1334533691406 0 7633.1934
367.1657409667969 0 18054.309
368.16650390625 0 5989.5073
371.13525390625 0 15358.471
374.1499938964844 0 21378.459
375.1528625488281 0 5520.721
376.11468505859375 0 40462.17
378.1008605957031 0 13960.353
378.13043212890625 0 252102.14
379.13348388671875 0 43030.25
380.1390075683594 0 18908.71
382.1435546875 0 53745.918
383.1467590332031 0 13303.781
384.1355895996094 0 15185.886
385.1370849609375 0 15824.381
388.11572265625 0 8243.153
388.15032958984375 0 9694.376
390.15667724609375 0 34450.492
394.12554931640625 0 336600
395.1290588378906 0 57494.51
396.1305236816406 0 7100.391
397.1635437011719 0 8875.369
398.1702880859375 0 5901.903
399.1772155761719 0 12514.386
401.193603515625 0 10126.145
404.11346435546875 0 6794.243
404.1451110839844 0 7388.6323
406.1238708496094 0 10217.56
406.1522216796875 0 13962.809
407.1835021972656 0 29445.678
408.16754150390625 0 83649.59
409.1707458496094 0 24114.117
409.1991271972656 0 6921.582
410.1392822265625 0 9429.155
410.1814270019531 0 5755.977
414.72283935546875 0 4947.156
415.1619873046875 0 15654.642
419.1566162109375 0 19527.266
423.1805114746094 0 7480.9185
425.1943664550781 0 287123.34
426.1971740722656 0 66202.86
427.2097473144531 0 51234.81
428.2128601074219 0 9265.369
429.1883239746094 0 6006.9043
431.1598205566406 0 4598.7935
432.1419982910156 0 6745.7275
433.1728820800781 0 40360.484
434.1766357421875 0 11753.126
435.1807861328125 0 11100.952
437.1676025390625 0 8973.459
442.1737976074219 0 11400.148
443.1556091308594 0 9806.575
443.18743896484375 0 5242.531
446.21630859375 0 7072.4116
447.1521301269531 0 41297.05
449.1341552734375 0 14489.806
449.1668395996094 0 15722.004
450.22686767578125 0 8408.951
451.1778564453125 0 5889.5913
452.6684875488281 0 6855.554
453.1895751953125 0 5417.3354
454.158935546875 0 8725.863
454.1971740722656 0 22361.383
455.2043151855469 0 62394.336
456.2062072753906 0 15362.8125
458.1673583984375 0 7957.529
459.1479797363281 0 9357.519
461.1671447753906 0 69722.35 b Water loss 3
461.1961975097656 0 30241.57
462.1709899902344 0 21872.447
464.1863098144531 0 5441.8887
465.162841796875 0 228991.92
466.1658630371094 0 53768.285
467.1670837402344 0 9938.92
468.1729736328125 0 8171.347 y Water loss 2
469.1731872558594 0 7272.144
469.2090759277344 0 10817.193
470.16693115234375 0 8447.771
470.2126770019531 0 9669.2295
471.18182373046875 0 37344.355
472.1853332519531 0 8904.687
473.2153625488281 0 19110.312
473.5928039550781 0 4547.584
475.14483642578125 0 7059.5293
477.1678466796875 0 44098.54 y 2
477.6787109375 0 18076.326
478.1681213378906 0 16512.145
478.22021484375 0 14312.334
479.1784362792969 0 249481.45 b 3
480.1817932128906 0 48570.465
481.1827087402344 0 9982.827
482.19232177734375 0 14385.575
483.1963806152344 0 8179.4663
486.1636657714844 0 34680.895
487.1650695800781 0 8830.274
489.19281005859375 0 455575.84 y 6
490.19512939453125 0 111502.94
491.205078125 0 56330.855
492.20989990234375 0 17569.576
495.1868896484375 0 34591.03
496.1698913574219 0 8806.172
496.1889343261719 0 12222.2
496.2315979003906 0 232371.8
497.1713562011719 0 42315.387
497.23455810546875 0 63789.766
498.23785400390625 0 8224.265
499.1879577636719 0 18729.596 b 7
499.6936950683594 0 12409.62
502.1936340332031 0 8651.736
504.1790771484375 0 5069.562
507.1520080566406 0 5861.7856
508.196533203125 0 8342.51
511.1936950683594 0 7707.979
513.1993408203125 0 28504.137
514.224365234375 0 12756.905
516.2210083007812 0 9907.169
518.7117919921875 0 18454.428
519.20458984375 0 62202.71
520.2054443359375 0 36355.47
521.20654296875 0 14613.535
523.189697265625 0 10149.605
525.1622314453125 0 5837.811
525.2067260742188 0 6057.9106
528.2100830078125 0 5501.368
530.189697265625 0 22400.621
531.1885375976562 0 6907.521
532.2052001953125 0 8044.824
533.2040405273438 0 8174.3096
537.212646484375 0 22604.74
538.21337890625 0 7731.9727
541.1947021484375 0 14759.629
541.7061767578125 0 38553.758 y Water loss 1
542.2341918945312 0 33432.74
542.7105102539062 0 13098.348
543.2423095703125 0 9712.734
547.19677734375 0 5650.0557
548.199951171875 0 78361.055
549.2035522460938 0 19827.977
550.2059936523438 0 9859.31
550.7119140625 0 215872.4 y 1
551.2137451171875 0 163313.36
551.714111328125 0 59995.45
552.2168579101562 0 15428.408
555.7022705078125 0 7352.665
557.2001953125 0 48587.24
558.2012329101562 0 11958.626
560.2295532226562 0 708430.7 y 5
560.730712890625 0 16649.041
561.2324829101562 0 232633.45
562.230712890625 0 57973.145
566.1953125 0 8524.865
569.2343139648438 0 43209.883
569.7352905273438 0 31766.398
570.229736328125 0 72771.39
571.2306518554688 0 14321.262
574.7137451171875 0 12060.506
578.2310791015625 0 29527.422
578.7354125976562 0 15696.626
579.2308349609375 0 7581.79
580.2115478515625 0 5964.396
583.2246704101562 0 30344.54
583.72607421875 0 15527.608
584.1993408203125 0 8031.3174
588.241943359375 0 17231.535
589.2054443359375 0 6183.9473
592.2303466796875 0 112470.54 Precursor Water loss
592.73193359375 0 100496.05
593.2328491210938 0 38861.484
593.728515625 0 7092.8706
594.1886596679688 0 20487.16
595.1889038085938 0 9600.0625
596.236328125 0 50667.094
597.2257080078125 0 30298.082
598.2246704101562 0 14941.212
601.2359619140625 0 337733.66 Precursor
601.7373657226562 0 266579.88
602.2379150390625 0 100882.13
602.7376098632812 0 12118.864
606.2332153320312 0 71978.78
607.2353515625 0 18319.09
608.2390747070312 0 8877.875
612.1982421875 0 114740.48
613.200927734375 0 33832.63
613.2879028320312 0 9532.743
614.2467651367188 0 76263.96
615.2488403320312 0 21799.9
616.2564086914062 0 6414.5103
622.2164916992188 0 5553.948
624.2318725585938 0 58077.707 b Water loss 4
625.2364501953125 0 19916.186
631.2484130859375 0 23184.887
632.2551879882812 0 11965.216
634.23046875 0 70413.61
635.2330322265625 0 19742.078
639.2538452148438 0 16762.365
640.2393798828125 0 32756.66
641.2820434570312 0 14575.529
642.242431640625 0 51928.777 b 4
643.244140625 0 20617.04
652.2413330078125 0 24881.35
653.245361328125 0 7898.9634
657.2639770507812 0 166738.8
658.2658081054688 0 51069.844
659.2947998046875 0 262025.52
660.2374877929688 0 6420.137
660.2978515625 0 105090.125
661.3015747070312 0 14666.085
667.2503662109375 0 19922.95
668.2452392578125 0 15183.845
669.2774658203125 0 7122.9614
676.2698364257812 0 33711.523
677.27685546875 0 21433.71
678.27880859375 0 6506.113
685.2589721679688 0 211878.38
686.2616577148438 0 76306.805
687.2659301757812 0 18517.84
695.2670288085938 0 43497.242 b Water loss 5
696.2720336914062 0 11833.97
703.2664794921875 0 65822.734
704.2676391601562 0 18837.438
705.2794189453125 0 25022.406
706.2910766601562 0 9097.447
713.27978515625 0 23675.873 b 5
714.280029296875 0 5566.757
721.1887817382812 0 9374.528
721.2622680664062 0 211906.77
722.2652587890625 0 68950.97
723.2928466796875 0 687285.06 y 4
724.2958984375 0 271128.66
725.2962036132812 0 78237.64
726.2960815429688 0 9780.581
728.3192749023438 0 9625.308
731.2472534179688 0 21148.012
732.2402954101562 0 13118.889
733.2714233398438 0 12615.959
734.2821044921875 0 6316.319
740.297607421875 0 6184.7153
746.2752685546875 0 11829.718
747.295166015625 0 5763.4155
748.30712890625 0 7229.3857
749.2572021484375 0 289933.2
750.259765625 0 116633.945
751.2608032226562 0 38087.63
756.310302734375 0 18586.658
757.3065185546875 0 14253.49
759.2623291015625 0 12226.231
767.2681884765625 0 72669.625
768.2711181640625 0 29294.533
769.2700805664062 0 7544.035
774.3216552734375 0 282454.1
775.3247680664062 0 115798.516
776.3257446289062 0 22587.059
778.3037719726562 0 14330.035
779.3119506835938 0 6973.727
784.3017578125 0 7478.7085
792.322021484375 0 9612.122
793.31640625 0 5773.977
796.3175048828125 0 7951.3936
803.2791748046875 0 8155.8433
804.3356323242188 0 9148.472
805.333740234375 0 8709.848
818.3333740234375 0 8866.148
820.3092651367188 0 42355.02 y Water loss 3
821.3036499023438 0 22160.145
822.305908203125 0 10893.616
823.2994384765625 0 6426.346
824.3014526367188 0 7256.0527
828.4381103515625 0 9210.656
832.32666015625 0 24464.412
833.32470703125 0 16772.865
838.3201293945312 0 872098.5 y 3
839.3228759765625 0 387434.5
840.323974609375 0 130885.41
841.3272094726562 0 9753.028
842.30810546875 0 14399.812 b Water loss 6
843.3414916992188 0 36602.555
844.34228515625 0 15778.613
848.3036499023438 0 13359.339
850.3196411132812 0 6242.312
851.3267822265625 0 5755.324
853.3162841796875 0 5538.022
854.3164672851562 0 10263.047
860.3120727539062 0 14361.596 b 6
861.3201293945312 0 9269.537
862.3258056640625 0 27431.871
863.3323974609375 0 13768.885
868.3306884765625 0 15801.212
869.3309936523438 0 9691.689
871.3385620117188 0 116843.28
872.3367919921875 0 66285.31
873.33837890625 0 23002.104
889.3484497070312 0 1202962
890.3511352539062 0 556980.1
890.4560546875 0 9089.946
891.3541259765625 0 154758.25
892.3532104492188 0 15740.692
896.3237915039062 0 23196.402
897.3292236328125 0 12974.866
899.3331298828125 0 7817.9595
907.3412475585938 0 45789.594
908.3453369140625 0 26367.908
909.3482055664062 0 11150.911
914.3378295898438 0 7221.38
917.3282470703125 0 12598.105
918.3123779296875 0 12909.952
933.375 0 12414.952
935.3363037109375 0 237980.16 y Water loss 2
936.3361206054688 0 136542.23
937.33544921875 0 54746.68
938.3240356445312 0 25444.453
939.3244018554688 0 8975.313
953.34716796875 0 3721935.5 y 2
954.3497314453125 0 1945402.8
955.3500366210938 0 699255.94
956.3515014648438 0 68475.93
963.3317260742188 0 23447.814
964.3347778320312 0 11802.063
969.37841796875 0 14226.93
970.3892211914062 0 5419.886
979.3601684570312 0 10277.703 b Water loss 7
990.0547485351562 0 5808.9736
997.3726806640625 0 41487.027 b 7
998.37353515625 0 21413.25
999.3815307617188 0 10884.492
1015.3798217773438 0 12477.065
1016.3836059570312 0 5829.796
1018.3960571289062 0 18790.969
1019.4061279296875 0 9060.508
1020.3939208984375 0 5689.033
1036.4158935546875 0 80972.93
1037.418212890625 0 51501.32
1038.42333984375 0 17914.045
1046.398681640625 0 6202.001
1082.4052734375 0 23512.863 y Water loss 1
1083.40771484375 0 13691.651
1100.415771484375 0 207322.73 y 1
1101.41796875 0 138519.31
1102.4185791015625 0 58362.56
1103.4189453125 0 9629.02
1110.399658203125 0 35571.395
1111.4024658203125 0 25730.951
1112.4095458984375 0 9870.391
1955.6422119140625 0 5642.3633
3239.639404296875 0 6471.89

Spectrum Details

|  |  |
| --- | --- |
| Matched peaks? Matched peaks The total absolute number of peaks matched. Additionally in brackets the total fraction of peaks matched and the total number of peaks is shown. | 43 (7.12% of 604) |
| FDR? FDR The false discovery rate estimated for this peptide. It is calculated by matching all theoretical fragments with a non-integer shift with the raw peaks for this spectrum. This is done with 40 different shifts. The resulting percentage is the average number of annotated peaks over the number of annotated peaks with the correct spectrum. | 0.06% |
| Satellite FDR? Satellite FDR See the FDR for details on its calculation. This satellite ion specific FDR only contains the satellite ions (d/w) for I/L/J positions. | - |
| PSM Score? PSM Score The PSM Score as given by Hecklib to this annotated spectrum. It is shown with three significant figures. | 487 |

#### Spectrum 7537? Spectrum 7537 The raw spectrum of this peptide as annotated by Hecklib. The fragments are coloured according to ion type (see legend). Any peaks with a star '\*' as text can be hovered over to see the full details, first the ion type second the mass shift type. By hovering over the amino acids in the peptide or ions in the legend the corresponding peaks are highlighted. By toggling the 'Unassigned' label you can turn the background (unassigned) peaks on or off in the plot. By updating the slider in the Ion legend you can update the spectrum to only show the top X% of the peaks with labels. The top X% means any peak that is within X% of the highest intensity. By dragging in the spectrum you can zoom in to a specific part of the spectrum and use 'Zoom Out' to get back to the original zoom level. The annotation of the spectrum is based on the given sequence in the peptides file and is done with different software so inconsistencies are likely. The peaks are annotated based on the given sequence, with 20 ppm tolerance.

Copy Data

##### Spectrum 7537 (TSV)

###### Preview

```
Loading example...
```

*Click on the button to copy the data to your clipboard.*

Mz MinMz MaxIntensity Max

WidthHeightPeptide font sizePeptide stroke widthSpectrum font sizeSpectrum stroke widthCompact peptide

Ion legend

wxyz

abcd

OtherUnassignedIon^Charge^_Position_Show for top:%

TFDDYAMHW

06.64e+51.33e+61.99e+62.65e+6

Zoom Out

b^+2^_3_a^+1^_2_a^+1^_2_y^+1^_1_a^+1^_2_b^+1^_2_y^+2^_3_b^+1^_2_d^+1^_3_a^+1^_3_a^+1^_3_y^+1^_2_b^+1^_3_b^+2^_6_y^+2^_5_b^+1^_3_y^+2^_6_b^+1^_4_y^+2^_7_y^+2^_7_b^+1^_4_y^+1^_3_b^+2^_8_y^+2^_8_y^+2^_8_y^+1^_4_\*\*b^+1^_5_b^+1^_5_b^+1^_6_b^+1^_6_y^+1^_5_y^+1^_6_y^+1^_6_b^+1^_7_b^+1^_7_y^+1^_7_y^+1^_7_b^+1^_8_b^+1^_8_y^+1^_8_y^+1^_8_

02815628431124

Fragment Matches Table

Show background peaks

| Position | Ion type | Intensity | mz Theoretical | mz Error (Th) | mz Error (ppm) | Charge | Series Number |
| --- | --- | --- | --- | --- | --- | --- | --- |
| - | - | 6.55E+05 | 120.1 | - | - | 0 | - |
| - | - | 1944 | 121 | - | - | 0 | - |
| - | - | 4.978E+04 | 121.1 | - | - | 0 | - |
| - | - | 2414 | 122.1 | - | - | 0 | - |
| - | - | 910 | 122.1 | - | - | 0 | - |
| - | - | 1979 | 127.1 | - | - | 0 | - |
| - | - | 1676 | 129.1 | - | - | 0 | - |
| - | - | 3201 | 129.1 | - | - | 0 | - |
| - | - | 894.4 | 130.1 | - | - | 0 | - |
| - | - | 1196 | 130.1 | - | - | 0 | - |
| - | - | 2.041E+04 | 130.1 | - | - | 0 | - |
| - | - | 920.2 | 131 | - | - | 0 | - |
| - | - | 1375 | 131.1 | - | - | 0 | - |
| - | - | 883.7 | 131.1 | - | - | 0 | - |
| - | - | 3.287E+04 | 132.1 | - | - | 0 | - |
| - | - | 3510 | 133.1 | - | - | 0 | - |
| - | - | 2417 | 134.1 | - | - | 0 | - |
| - | - | 2.231E+05 | 136.1 | - | - | 0 | - |
| - | - | 7133 | 136.1 | - | - | 0 | - |
| - | - | 1.57E+04 | 137.1 | - | - | 0 | - |
| - | - | 1.636E+04 | 138.1 | - | - | 0 | - |
| - | - | 2.571E+04 | 138.1 | - | - | 0 | - |
| - | - | 863.8 | 138.4 | - | - | 0 | - |
| - | - | 1418 | 139.1 | - | - | 0 | - |
| - | - | 1374 | 141.1 | - | - | 0 | - |
| - | - | 1988 | 142.1 | - | - | 0 | - |
| - | - | 1429 | 143 | - | - | 0 | - |
| - | - | 4598 | 143.1 | - | - | 0 | - |
| - | - | 4643 | 143.1 | - | - | 0 | - |
| - | - | 3032 | 144.1 | - | - | 0 | - |
| - | - | 8304 | 146.1 | - | - | 0 | - |
| - | - | 1313 | 146.1 | - | - | 0 | - |
| - | - | 3289 | 146.1 | - | - | 0 | - |
| - | - | 1020 | 147 | - | - | 0 | - |
| - | - | 1023 | 147.1 | - | - | 0 | - |
| - | - | 1112 | 147.1 | - | - | 0 | - |
| - | - | 1775 | 148.1 | - | - | 0 | - |
| - | - | 4720 | 148.1 | - | - | 0 | - |
| - | - | 1512 | 150.1 | - | - | 0 | - |
| - | - | 4895 | 155.1 | - | - | 0 | - |
| - | - | 4759 | 155.1 | - | - | 0 | - |
| - | - | 5.819E+04 | 156.1 | - | - | 0 | - |
| - | - | 3457 | 157.1 | - | - | 0 | - |
| - | - | 2822 | 157.1 | - | - | 0 | - |
| - | - | 9.55E+04 | 158.1 | - | - | 0 | - |
| - | - | 1132 | 159.1 | - | - | 0 | - |
| - | - | 4.987E+04 | 159.1 | - | - | 0 | - |
| - | - | 8698 | 159.1 | - | - | 0 | - |
| - | - | 1.871E+04 | 160.1 | - | - | 0 | - |
| - | - | 2453 | 160.1 | - | - | 0 | - |
| - | - | 4462 | 160.1 | - | - | 0 | - |
| - | - | 4.322E+04 | 160.1 | - | - | 0 | - |
| - | - | 5381 | 161.1 | - | - | 0 | - |
| - | - | 2025 | 165.1 | - | - | 0 | - |
| - | - | 1.285E+04 | 165.1 | - | - | 0 | - |
| - | - | 1260 | 165.1 | - | - | 0 | - |
| - | - | 3096 | 166.1 | - | - | 0 | - |
| - | - | 8882 | 166.1 | - | - | 0 | - |
| - | - | 2558 | 166.1 | - | - | 0 | - |
| - | - | 1416 | 167.1 | - | - | 0 | - |
| - | - | 8704 | 167.1 | - | - | 0 | - |
| - | - | 4054 | 169.1 | - | - | 0 | - |
| - | - | 1259 | 169.1 | - | - | 0 | - |
| - | - | 8600 | 171.1 | - | - | 0 | - |
| - | - | 1567 | 171.1 | - | - | 0 | - |
| - | - | 2313 | 172.1 | - | - | 0 | - |
| - | - | 1199 | 172.1 | - | - | 0 | - |
| - | - | 2844 | 173.1 | - | - | 0 | - |
| - | - | 1137 | 173.1 | - | - | 0 | - |
| - | - | 2225 | 173.4 | - | - | 0 | - |
| 3 | b | 4974 | 174.1 | 0.0009782 | 5.62 | +2 | 3 |
| - | - | 1138 | 174.1 | - | - | 0 | - |
| - | - | 3101 | 175.1 | - | - | 0 | - |
| - | - | 2131 | 175.1 | - | - | 0 | - |
| - | - | 2967 | 175.1 | - | - | 0 | - |
| - | - | 1.713E+04 | 176.1 | - | - | 0 | - |
| - | - | 1.873E+05 | 176.1 | - | - | 0 | - |
| - | - | 1.357E+05 | 177.1 | - | - | 0 | - |
| - | - | 1.94E+04 | 177.1 | - | - | 0 | - |
| - | - | 1224 | 178.1 | - | - | 0 | - |
| - | - | 1734 | 178.1 | - | - | 0 | - |
| - | - | 1.352E+04 | 178.1 | - | - | 0 | - |
| - | - | 9863 | 180.1 | - | - | 0 | - |
| - | - | 1.794E+04 | 185.1 | - | - | 0 | - |
| - | - | 1.432E+04 | 186.1 | - | - | 0 | - |
| - | - | 2544 | 187.1 | - | - | 0 | - |
| - | - | 1055 | 187.1 | - | - | 0 | - |
| - | - | 6.426E+04 | 188.1 | - | - | 0 | - |
| - | - | 7657 | 189.1 | - | - | 0 | - |
| - | - | 1353 | 189.1 | - | - | 0 | - |
| - | - | 1613 | 190.1 | - | - | 0 | - |
| - | - | 2040 | 191.1 | - | - | 0 | - |
| - | - | 6178 | 191.1 | - | - | 0 | - |
| - | - | 1.389E+04 | 191.1 | - | - | 0 | - |
| - | - | 2709 | 192.1 | - | - | 0 | - |
| - | - | 1277 | 192.1 | - | - | 0 | - |
| - | - | 3328 | 193.1 | - | - | 0 | - |
| - | - | 6.075E+04 | 193.1 | - | - | 0 | - |
| - | - | 3312 | 194.1 | - | - | 0 | - |
| - | - | 1100 | 194.1 | - | - | 0 | - |
| - | - | 3393 | 194.1 | - | - | 0 | - |
| - | - | 1.37E+04 | 195.1 | - | - | 0 | - |
| - | - | 1497 | 196.1 | - | - | 0 | - |
| - | - | 1208 | 199.1 | - | - | 0 | - |
| - | - | 1120 | 201.1 | - | - | 0 | - |
| - | - | 5986 | 202.1 | - | - | 0 | - |
| - | - | 1.536E+04 | 203.1 | - | - | 0 | - |
| - | - | 4591 | 203.1 | - | - | 0 | - |
| 2 | a | 6.406E+04 | 203.1 | 0.0003661 | 1.802 | +1 | 2 |
| - | - | 1992 | 204.1 | - | - | 0 | - |
| - | - | 2.468E+04 | 204.1 | - | - | 0 | - |
| 2 | a | 4105 | 204.1 | 0.0001304 | 0.6389 | +1 | 2 |
| - | - | 9034 | 204.1 | - | - | 0 | - |
| - | - | 2442 | 205.1 | - | - | 0 | - |
| 9 | y | 4.798E+04 | 205.1 | 0.0003648 | 1.779 | +1 | 1 |
| - | - | 2655 | 205.1 | - | - | 0 | - |
| - | - | 6278 | 206.1 | - | - | 0 | - |
| - | - | 1.366E+04 | 207.1 | - | - | 0 | - |
| - | - | 8575 | 208.1 | - | - | 0 | - |
| - | - | 1527 | 208.1 | - | - | 0 | - |
| - | - | 1175 | 208.9 | - | - | 0 | - |
| - | - | 1326 | 211.1 | - | - | 0 | - |
| - | - | 4132 | 212.1 | - | - | 0 | - |
| - | - | 1362 | 213.1 | - | - | 0 | - |
| - | - | 2010 | 213.1 | - | - | 0 | - |
| - | - | 3970 | 215.1 | - | - | 0 | - |
| - | - | 1325 | 216.1 | - | - | 0 | - |
| - | - | 3.176E+04 | 217.1 | - | - | 0 | - |
| - | - | 1642 | 218.1 | - | - | 0 | - |
| - | - | 3419 | 218.1 | - | - | 0 | - |
| - | - | 7784 | 219.1 | - | - | 0 | - |
| - | - | 1840 | 219.1 | - | - | 0 | - |
| - | - | 1091 | 219.1 | - | - | 0 | - |
| - | - | 1997 | 220.1 | - | - | 0 | - |
| - | - | 1.588E+05 | 221.1 | - | - | 0 | - |
| 2 | a | 2.628E+06 | 221.1 | 0.0005588 | 2.527 | +1 | 2 |
| - | - | 1.327E+04 | 222.1 | - | - | 0 | - |
| - | - | 3.172E+05 | 222.1 | - | - | 0 | - |
| - | - | 1230 | 223.1 | - | - | 0 | - |
| - | - | 2452 | 223.1 | - | - | 0 | - |
| - | - | 1.788E+04 | 223.1 | - | - | 0 | - |
| - | - | 1015 | 223.2 | - | - | 0 | - |
| - | - | 2042 | 224.1 | - | - | 0 | - |
| - | - | 8330 | 228.1 | - | - | 0 | - |
| - | - | 5112 | 229.1 | - | - | 0 | - |
| - | - | 1122 | 230.1 | - | - | 0 | - |
| - | - | 4.22E+04 | 231.1 | - | - | 0 | - |
| 2 | b | 2.794E+04 | 231.1 | 0.0004618 | 1.998 | +1 | 2 |
| - | - | 3191 | 232.1 | - | - | 0 | - |
| - | - | 3905 | 232.1 | - | - | 0 | - |
| - | - | 2660 | 233.1 | - | - | 0 | - |
| - | - | 6.408E+04 | 235.1 | - | - | 0 | - |
| - | - | 8763 | 236.1 | - | - | 0 | - |
| - | - | 4179 | 238.1 | - | - | 0 | - |
| - | - | 1062 | 239.1 | - | - | 0 | - |
| - | - | 1.41E+04 | 239.1 | - | - | 0 | - |
| - | - | 1342 | 240.1 | - | - | 0 | - |
| - | - | 1346 | 240.1 | - | - | 0 | - |
| - | - | 1396 | 241.1 | - | - | 0 | - |
| 7 | y | 3247 | 245.1 | 0.004842 | 19.76 | +2 | 3 |
| - | - | 1397 | 245.1 | - | - | 0 | - |
| - | - | 1395 | 246.1 | - | - | 0 | - |
| - | - | 4958 | 246.1 | - | - | 0 | - |
| - | - | 1236 | 247.1 | - | - | 0 | - |
| - | - | 1867 | 247.1 | - | - | 0 | - |
| - | - | 2580 | 248.1 | - | - | 0 | - |
| - | - | 4227 | 248.2 | - | - | 0 | - |
| 2 | b | 4.648E+05 | 249.1 | 0.0005478 | 2.199 | +1 | 2 |
| - | - | 2467 | 250.1 | - | - | 0 | - |
| - | - | 5.837E+04 | 250.1 | - | - | 0 | - |
| - | - | 4.963E+04 | 251.1 | - | - | 0 | - |
| - | - | 7028 | 251.1 | - | - | 0 | - |
| - | - | 5055 | 252.1 | - | - | 0 | - |
| - | - | 2347 | 253.1 | - | - | 0 | - |
| - | - | 2342 | 255.1 | - | - | 0 | - |
| - | - | 3702 | 256.1 | - | - | 0 | - |
| - | - | 2.781E+04 | 257.1 | - | - | 0 | - |
| - | - | 2964 | 258.1 | - | - | 0 | - |
| - | - | 1381 | 260.7 | - | - | 0 | - |
| - | - | 7077 | 261.1 | - | - | 0 | - |
| - | - | 1348 | 262.1 | - | - | 0 | - |
| - | - | 1675 | 262.1 | - | - | 0 | - |
| - | - | 2826 | 263.1 | - | - | 0 | - |
| - | - | 9.19E+04 | 263.1 | - | - | 0 | - |
| - | - | 9810 | 263.1 | - | - | 0 | - |
| - | - | 1.171E+04 | 264.1 | - | - | 0 | - |
| - | - | 9057 | 264.1 | - | - | 0 | - |
| - | - | 3167 | 265.1 | - | - | 0 | - |
| - | - | 6861 | 267.1 | - | - | 0 | - |
| - | - | 3701 | 268.1 | - | - | 0 | - |
| - | - | 2343 | 268.1 | - | - | 0 | - |
| - | - | 3139 | 268.1 | - | - | 0 | - |
| - | - | 2029 | 269.1 | - | - | 0 | - |
| - | - | 1399 | 270.1 | - | - | 0 | - |
| - | - | 1797 | 270.1 | - | - | 0 | - |
| - | - | 4815 | 271.1 | - | - | 0 | - |
| - | - | 1551 | 272.1 | - | - | 0 | - |
| - | - | 2254 | 273.1 | - | - | 0 | - |
| - | - | 2.325E+04 | 274.1 | - | - | 0 | - |
| - | - | 1695 | 274.2 | - | - | 0 | - |
| - | - | 4435 | 275.1 | - | - | 0 | - |
| - | - | 3682 | 275.1 | - | - | 0 | - |
| - | - | 1598 | 277.7 | - | - | 0 | - |
| - | - | 1128 | 277.8 | - | - | 0 | - |
| - | - | 1012 | 277.8 | - | - | 0 | - |
| - | - | 1482 | 278.1 | - | - | 0 | - |
| - | - | 3.822E+04 | 279.1 | - | - | 0 | - |
| - | - | 3821 | 279.1 | - | - | 0 | - |
| - | - | 6055 | 280.1 | - | - | 0 | - |
| - | - | 1645 | 282.1 | - | - | 0 | - |
| - | - | 1355 | 283.2 | - | - | 0 | - |
| - | - | 5.931E+04 | 285.1 | - | - | 0 | - |
| - | - | 8055 | 286.1 | - | - | 0 | - |
| - | - | 2215 | 287.1 | - | - | 0 | - |
| - | - | 1569 | 288.1 | - | - | 0 | - |
| - | - | 4265 | 289.1 | - | - | 0 | - |
| - | - | 1639 | 290.1 | - | - | 0 | - |
| - | - | 3.869E+04 | 292.1 | - | - | 0 | - |
| 3 | d | 9110 | 292.2 | 0.0005392 | 1.845 | +1 | 3 |
| - | - | 4635 | 293.1 | - | - | 0 | - |
| - | - | 1863 | 296.1 | - | - | 0 | - |
| - | - | 4.329E+04 | 296.2 | - | - | 0 | - |
| - | - | 1574 | 297.1 | - | - | 0 | - |
| - | - | 7805 | 297.2 | - | - | 0 | - |
| - | - | 4326 | 300.1 | - | - | 0 | - |
| - | - | 3466 | 301.1 | - | - | 0 | - |
| - | - | 4370 | 302.1 | - | - | 0 | - |
| - | - | 6029 | 303.1 | - | - | 0 | - |
| - | - | 1698 | 306.1 | - | - | 0 | - |
| - | - | 1134 | 306.1 | - | - | 0 | - |
| - | - | 1879 | 307.1 | - | - | 0 | - |
| - | - | 6705 | 310.2 | - | - | 0 | - |
| - | - | 2776 | 313.1 | - | - | 0 | - |
| - | - | 4091 | 314.1 | - | - | 0 | - |
| - | - | 3427 | 315.1 | - | - | 0 | - |
| - | - | 1762 | 316.1 | - | - | 0 | - |
| 3 | a | 1.516E+04 | 318.1 | 0.0006447 | 2.027 | +1 | 3 |
| - | - | 1537 | 319.1 | - | - | 0 | - |
| - | - | 3902 | 320.1 | - | - | 0 | - |
| - | - | 1.761E+04 | 323.1 | - | - | 0 | - |
| - | - | 1.031E+05 | 324.1 | - | - | 0 | - |
| - | - | 1.56E+04 | 325.1 | - | - | 0 | - |
| - | - | 1233 | 327.1 | - | - | 0 | - |
| - | - | 2513 | 327.1 | - | - | 0 | - |
| - | - | 7216 | 328.1 | - | - | 0 | - |
| - | - | 4932 | 328.1 | - | - | 0 | - |
| - | - | 5824 | 330.1 | - | - | 0 | - |
| - | - | 2568 | 331.1 | - | - | 0 | - |
| - | - | 5319 | 332.1 | - | - | 0 | - |
| - | - | 3791 | 333.1 | - | - | 0 | - |
| - | - | 1675 | 333.1 | - | - | 0 | - |
| - | - | 2844 | 334.1 | - | - | 0 | - |
| - | - | 3270 | 334.1 | - | - | 0 | - |
| - | - | 2638 | 335.2 | - | - | 0 | - |
| 3 | a | 1325 | 336.2 | 0.0001814 | 0.5395 | +1 | 3 |
| - | - | 7243 | 338.1 | - | - | 0 | - |
| - | - | 9064 | 339.2 | - | - | 0 | - |
| - | - | 1649 | 340.2 | - | - | 0 | - |
| 8 | y | 5.377E+05 | 342.2 | 0.0007944 | 2.322 | +1 | 2 |
| - | - | 9.701E+04 | 343.2 | - | - | 0 | - |
| - | - | 9883 | 344.2 | - | - | 0 | - |
| 3 | b | 3.078E+04 | 346.1 | 0.0004811 | 1.39 | +1 | 3 |
| - | - | 4527 | 347.1 | - | - | 0 | - |
| - | - | 1.408E+04 | 348.1 | - | - | 0 | - |
| - | - | 1574 | 349.1 | - | - | 0 | - |
| - | - | 3093 | 349.1 | - | - | 0 | - |
| - | - | 3.146E+04 | 350.1 | - | - | 0 | - |
| - | - | 4398 | 351.1 | - | - | 0 | - |
| - | - | 9164 | 352.1 | - | - | 0 | - |
| - | - | 2013 | 353.1 | - | - | 0 | - |
| - | - | 2629 | 354.2 | - | - | 0 | - |
| - | - | 3.236E+04 | 356.1 | - | - | 0 | - |
| 6 | b | 4701 | 357.1 | 0.000458 | 1.282 | +2 | 6 |
| - | - | 2559 | 358.1 | - | - | 0 | - |
| - | - | 2605 | 359.1 | - | - | 0 | - |
| - | - | 6722 | 360.1 | - | - | 0 | - |
| 5 | y | 4639 | 362.1 | 0.003531 | 9.749 | +2 | 5 |
| - | - | 3286 | 362.7 | - | - | 0 | - |
| 3 | b | 1.298E+05 | 364.2 | 0.0006281 | 1.725 | +1 | 3 |
| - | - | 2.64E+04 | 365.2 | - | - | 0 | - |
| - | - | 1.422E+04 | 366.1 | - | - | 0 | - |
| - | - | 2124 | 366.2 | - | - | 0 | - |
| - | - | 2159 | 367.1 | - | - | 0 | - |
| - | - | 6502 | 367.2 | - | - | 0 | - |
| - | - | 1870 | 368.2 | - | - | 0 | - |
| - | - | 3382 | 371.1 | - | - | 0 | - |
| - | - | 6205 | 374.1 | - | - | 0 | - |
| - | - | 1.36E+04 | 376.1 | - | - | 0 | - |
| - | - | 2851 | 377.1 | - | - | 0 | - |
| - | - | 3634 | 378.1 | - | - | 0 | - |
| - | - | 7.319E+04 | 378.1 | - | - | 0 | - |
| - | - | 1.244E+04 | 379.1 | - | - | 0 | - |
| - | - | 5090 | 380.1 | - | - | 0 | - |
| - | - | 2640 | 381.1 | - | - | 0 | - |
| - | - | 1.564E+04 | 382.1 | - | - | 0 | - |
| - | - | 3144 | 383.1 | - | - | 0 | - |
| - | - | 4377 | 384.1 | - | - | 0 | - |
| - | - | 3858 | 385.1 | - | - | 0 | - |
| - | - | 1882 | 386.1 | - | - | 0 | - |
| - | - | 1893 | 388.1 | - | - | 0 | - |
| - | - | 1988 | 388.1 | - | - | 0 | - |
| - | - | 1720 | 389.1 | - | - | 0 | - |
| - | - | 9620 | 390.2 | - | - | 0 | - |
| - | - | 1813 | 391.2 | - | - | 0 | - |
| - | - | 9.138E+04 | 394.1 | - | - | 0 | - |
| - | - | 1.792E+04 | 395.1 | - | - | 0 | - |
| - | - | 1789 | 395.2 | - | - | 0 | - |
| - | - | 1773 | 396.1 | - | - | 0 | - |
| - | - | 1707 | 397.2 | - | - | 0 | - |
| - | - | 2599 | 398.2 | - | - | 0 | - |
| - | - | 4252 | 399.2 | - | - | 0 | - |
| - | - | 2578 | 404.1 | - | - | 0 | - |
| - | - | 2373 | 404.1 | - | - | 0 | - |
| - | - | 2146 | 406.1 | - | - | 0 | - |
| - | - | 2772 | 406.2 | - | - | 0 | - |
| - | - | 9792 | 407.2 | - | - | 0 | - |
| - | - | 2.123E+04 | 408.2 | - | - | 0 | - |
| - | - | 6368 | 409.2 | - | - | 0 | - |
| - | - | 2926 | 410.1 | - | - | 0 | - |
| - | - | 2098 | 410.2 | - | - | 0 | - |
| - | - | 4603 | 415.2 | - | - | 0 | - |
| - | - | 2253 | 416.1 | - | - | 0 | - |
| - | - | 1530 | 417.2 | - | - | 0 | - |
| - | - | 3928 | 419.2 | - | - | 0 | - |
| 4 | y | 1427 | 419.7 | 0.00129 | 3.073 | +2 | 6 |
| - | - | 1495 | 420.2 | - | - | 0 | - |
| - | - | 2198 | 423.2 | - | - | 0 | - |
| - | - | 7.709E+04 | 425.2 | - | - | 0 | - |
| - | - | 1.676E+04 | 426.2 | - | - | 0 | - |
| - | - | 1.604E+04 | 427.2 | - | - | 0 | - |
| - | - | 3659 | 428.2 | - | - | 0 | - |
| - | - | 2145 | 430.1 | - | - | 0 | - |
| - | - | 1654 | 431.1 | - | - | 0 | - |
| - | - | 1529 | 431.2 | - | - | 0 | - |
| - | - | 2240 | 432.1 | - | - | 0 | - |
| - | - | 1567 | 433.1 | - | - | 0 | - |
| - | - | 1.47E+04 | 433.2 | - | - | 0 | - |
| - | - | 2187 | 434.2 | - | - | 0 | - |
| - | - | 4260 | 435.2 | - | - | 0 | - |
| - | - | 3920 | 437.2 | - | - | 0 | - |
| - | - | 1942 | 438.2 | - | - | 0 | - |
| - | - | 2966 | 442.2 | - | - | 0 | - |
| - | - | 3066 | 443.2 | - | - | 0 | - |
| - | - | 1423 | 445.2 | - | - | 0 | - |
| - | - | 1449 | 446.2 | - | - | 0 | - |
| - | - | 1.155E+04 | 447.2 | - | - | 0 | - |
| - | - | 2343 | 448.2 | - | - | 0 | - |
| - | - | 2895 | 449.1 | - | - | 0 | - |
| - | - | 3223 | 449.2 | - | - | 0 | - |
| - | - | 2548 | 450.2 | - | - | 0 | - |
| - | - | 1671 | 451.2 | - | - | 0 | - |
| - | - | 2608 | 454.2 | - | - | 0 | - |
| - | - | 5796 | 454.2 | - | - | 0 | - |
| - | - | 1.448E+04 | 455.2 | - | - | 0 | - |
| - | - | 3441 | 456.2 | - | - | 0 | - |
| - | - | 3602 | 458.2 | - | - | 0 | - |
| - | - | 3522 | 459.2 | - | - | 0 | - |
| 4 | b | 2.036E+04 | 461.2 | 0.0006377 | 1.383 | +1 | 4 |
| - | - | 7611 | 461.2 | - | - | 0 | - |
| - | - | 6412 | 462.2 | - | - | 0 | - |
| - | - | 2194 | 462.2 | - | - | 0 | - |
| - | - | 7.285E+04 | 465.2 | - | - | 0 | - |
| - | - | 1.681E+04 | 466.2 | - | - | 0 | - |
| - | - | 1570 | 467.2 | - | - | 0 | - |
| 3 | y | 3130 | 468.2 | 0.002805 | 5.991 | +2 | 7 |
| - | - | 3204 | 469.2 | - | - | 0 | - |
| - | - | 1593 | 470.2 | - | - | 0 | - |
| - | - | 1485 | 470.2 | - | - | 0 | - |
| - | - | 1.176E+04 | 471.2 | - | - | 0 | - |
| - | - | 3264 | 472.2 | - | - | 0 | - |
| - | - | 1270 | 472.2 | - | - | 0 | - |
| - | - | 6696 | 473.2 | - | - | 0 | - |
| - | - | 1544 | 475.1 | - | - | 0 | - |
| - | - | 1516 | 476.2 | - | - | 0 | - |
| 3 | y | 1.204E+04 | 477.2 | 0.006933 | 14.53 | +2 | 7 |
| - | - | 3512 | 477.7 | - | - | 0 | - |
| - | - | 3535 | 478.2 | - | - | 0 | - |
| - | - | 2294 | 478.2 | - | - | 0 | - |
| 4 | b | 6.895E+04 | 479.2 | 0.0008762 | 1.829 | +1 | 4 |
| - | - | 1.559E+04 | 480.2 | - | - | 0 | - |
| - | - | 1777 | 481.2 | - | - | 0 | - |
| - | - | 3813 | 482.2 | - | - | 0 | - |
| - | - | 1704 | 483.2 | - | - | 0 | - |
| - | - | 1513 | 485.7 | - | - | 0 | - |
| - | - | 1.199E+04 | 486.2 | - | - | 0 | - |
| - | - | 2175 | 487.2 | - | - | 0 | - |
| 7 | y | 1.283E+05 | 489.2 | 0.005954 | 12.17 | +1 | 3 |
| - | - | 3.609E+04 | 490.2 | - | - | 0 | - |
| - | - | 2378 | 490.7 | - | - | 0 | - |
| - | - | 1.684E+04 | 491.2 | - | - | 0 | - |
| - | - | 4277 | 492.2 | - | - | 0 | - |
| - | - | 1284 | 493.2 | - | - | 0 | - |
| - | - | 9251 | 495.2 | - | - | 0 | - |
| - | - | 2297 | 496.2 | - | - | 0 | - |
| - | - | 3675 | 496.2 | - | - | 0 | - |
| - | - | 6.561E+04 | 496.2 | - | - | 0 | - |
| - | - | 1.201E+04 | 497.2 | - | - | 0 | - |
| - | - | 1.965E+04 | 497.2 | - | - | 0 | - |
| - | - | 2571 | 498.2 | - | - | 0 | - |
| - | - | 2010 | 498.2 | - | - | 0 | - |
| 8 | b | 5624 | 499.2 | 0.001719 | 3.443 | +2 | 8 |
| - | - | 1745 | 502.2 | - | - | 0 | - |
| - | - | 3198 | 503.2 | - | - | 0 | - |
| - | - | 1653 | 506.2 | - | - | 0 | - |
| - | - | 1687 | 509.7 | - | - | 0 | - |
| - | - | 1611 | 511.2 | - | - | 0 | - |
| - | - | 1.103E+04 | 513.2 | - | - | 0 | - |
| - | - | 2730 | 514.2 | - | - | 0 | - |
| - | - | 3745 | 516.2 | - | - | 0 | - |
| - | - | 4520 | 518.7 | - | - | 0 | - |
| - | - | 1.938E+04 | 519.2 | - | - | 0 | - |
| - | - | 1483 | 519.7 | - | - | 0 | - |
| - | - | 8478 | 520.2 | - | - | 0 | - |
| - | - | 2053 | 521.2 | - | - | 0 | - |
| - | - | 3271 | 523.2 | - | - | 0 | - |
| - | - | 1393 | 525.2 | - | - | 0 | - |
| - | - | 4007 | 525.2 | - | - | 0 | - |
| - | - | 1613 | 526.2 | - | - | 0 | - |
| - | - | 1604 | 527.2 | - | - | 0 | - |
| - | - | 2068 | 527.7 | - | - | 0 | - |
| - | - | 3920 | 530.2 | - | - | 0 | - |
| - | - | 1533 | 532.2 | - | - | 0 | - |
| - | - | 1710 | 533.2 | - | - | 0 | - |
| - | - | 6359 | 537.2 | - | - | 0 | - |
| - | - | 5089 | 541.2 | - | - | 0 | - |
| 2 | y | 9249 | 541.7 | 0.002686 | 4.959 | +2 | 8 |
| - | - | 1.191E+04 | 542.2 | - | - | 0 | - |
| - | - | 5138 | 542.7 | - | - | 0 | - |
| - | - | 3207 | 543.2 | - | - | 0 | - |
| - | - | 2025 | 545.2 | - | - | 0 | - |
| - | - | 2.392E+04 | 548.2 | - | - | 0 | - |
| - | - | 8286 | 549.2 | - | - | 0 | - |
| - | - | 2009 | 550.2 | - | - | 0 | - |
| 2 | y | 6.756E+04 | 550.7 | 0.003141 | 5.704 | +2 | 8 |
| - | - | 4.973E+04 | 551.2 | - | - | 0 | - |
| - | - | 1.997E+04 | 551.7 | - | - | 0 | - |
| - | - | 4202 | 552.2 | - | - | 0 | - |
| - | - | 2317 | 553.2 | - | - | 0 | - |
| - | - | 3081 | 555.7 | - | - | 0 | - |
| - | - | 1.17E+04 | 557.2 | - | - | 0 | - |
| - | - | 1615 | 558.2 | - | - | 0 | - |
| 6 | y | 2.042E+05 | 560.2 | 0.005462 | 9.749 | +1 | 4 |
| - | - | 5372 | 560.7 | - | - | 0 | - |
| - | - | 6.736E+04 | 561.2 | - | - | 0 | - |
| - | - | 1.413E+04 | 562.2 | - | - | 0 | - |
| - | - | 1558 | 563.2 | - | - | 0 | - |
| - | - | 1.388E+04 | 569.2 | - | - | 0 | - |
| - | - | 1.131E+04 | 569.7 | - | - | 0 | - |
| - | - | 2.025E+04 | 570.2 | - | - | 0 | - |
| - | - | 1356 | 570.7 | - | - | 0 | - |
| - | - | 3570 | 571.2 | - | - | 0 | - |
| - | - | 2606 | 574.7 | - | - | 0 | - |
| - | - | 1.048E+04 | 578.2 | - | - | 0 | - |
| - | - | 5561 | 578.7 | - | - | 0 | - |
| - | - | 1760 | 579.2 | - | - | 0 | - |
| - | - | 6215 | 583.2 | - | - | 0 | - |
| - | - | 3611 | 583.7 | - | - | 0 | - |
| - | - | 4942 | 588.2 | - | - | 0 | - |
| 0 | Precursor | 4.421E+04 | 592.2 | 0.003444 | 5.815 | +2 | -1 |
| - | - | 2.84E+04 | 592.7 | - | - | 0 | - |
| - | - | 1.416E+04 | 593.2 | - | - | 0 | - |
| - | - | 2856 | 593.7 | - | - | 0 | - |
| - | - | 5815 | 594.2 | - | - | 0 | - |
| - | - | 1.41E+04 | 596.2 | - | - | 0 | - |
| - | - | 9118 | 597.2 | - | - | 0 | - |
| - | - | 3522 | 598.2 | - | - | 0 | - |
| 0 | Precursor | 1.054E+05 | 601.2 | 0.003411 | 5.673 | +2 | -1 |
| - | - | 7.35E+04 | 601.7 | - | - | 0 | - |
| - | - | 1814 | 601.8 | - | - | 0 | - |
| - | - | 3.26E+04 | 602.2 | - | - | 0 | - |
| - | - | 5006 | 602.7 | - | - | 0 | - |
| - | - | 2.057E+04 | 606.2 | - | - | 0 | - |
| - | - | 6713 | 607.2 | - | - | 0 | - |
| - | - | 2552 | 608.2 | - | - | 0 | - |
| - | - | 3.572E+04 | 612.2 | - | - | 0 | - |
| - | - | 1.12E+04 | 613.2 | - | - | 0 | - |
| - | - | 2301 | 613.3 | - | - | 0 | - |
| - | - | 2079 | 614.2 | - | - | 0 | - |
| - | - | 2.479E+04 | 614.2 | - | - | 0 | - |
| - | - | 1319 | 614.3 | - | - | 0 | - |
| - | - | 6558 | 615.2 | - | - | 0 | - |
| - | - | 1661 | 622.2 | - | - | 0 | - |
| 5 | b | 1.603E+04 | 624.2 | 0.001243 | 1.992 | +1 | 5 |
| - | - | 5388 | 625.2 | - | - | 0 | - |
| - | - | 9085 | 631.2 | - | - | 0 | - |
| - | - | 4627 | 632.3 | - | - | 0 | - |
| - | - | 1370 | 633.3 | - | - | 0 | - |
| - | - | 1.864E+04 | 634.2 | - | - | 0 | - |
| - | - | 7531 | 635.2 | - | - | 0 | - |
| - | - | 1572 | 636.2 | - | - | 0 | - |
| - | - | 6264 | 639.3 | - | - | 0 | - |
| - | - | 8924 | 640.2 | - | - | 0 | - |
| - | - | 2170 | 641.2 | - | - | 0 | - |
| - | - | 2126 | 641.3 | - | - | 0 | - |
| 5 | b | 1.523E+04 | 642.2 | 0.001299 | 2.022 | +1 | 5 |
| - | - | 3374 | 643.2 | - | - | 0 | - |
| - | - | 5391 | 652.2 | - | - | 0 | - |
| - | - | 1383 | 653.2 | - | - | 0 | - |
| - | - | 4.736E+04 | 657.3 | - | - | 0 | - |
| - | - | 1.353E+04 | 658.3 | - | - | 0 | - |
| - | - | 7.355E+04 | 659.3 | - | - | 0 | - |
| - | - | 2.692E+04 | 660.3 | - | - | 0 | - |
| - | - | 6523 | 661.3 | - | - | 0 | - |
| - | - | 5791 | 667.3 | - | - | 0 | - |
| - | - | 4322 | 668.3 | - | - | 0 | - |
| - | - | 9196 | 676.3 | - | - | 0 | - |
| - | - | 5837 | 677.3 | - | - | 0 | - |
| - | - | 2333 | 678.3 | - | - | 0 | - |
| - | - | 6.358E+04 | 685.3 | - | - | 0 | - |
| - | - | 2.12E+04 | 686.3 | - | - | 0 | - |
| - | - | 5234 | 687.3 | - | - | 0 | - |
| 6 | b | 1.241E+04 | 695.3 | 0.0002258 | 0.3248 | +1 | 6 |
| - | - | 4211 | 696.3 | - | - | 0 | - |
| - | - | 2680 | 702.3 | - | - | 0 | - |
| - | - | 1.678E+04 | 703.3 | - | - | 0 | - |
| - | - | 3969 | 704.2 | - | - | 0 | - |
| - | - | 4331 | 704.3 | - | - | 0 | - |
| - | - | 8059 | 705.3 | - | - | 0 | - |
| - | - | 3007 | 706.3 | - | - | 0 | - |
| - | - | 1516 | 708.3 | - | - | 0 | - |
| 6 | b | 6364 | 713.3 | 0.001818 | 2.549 | +1 | 6 |
| - | - | 1825 | 714.3 | - | - | 0 | - |
| - | - | 5.745E+04 | 721.3 | - | - | 0 | - |
| - | - | 2.343E+04 | 722.3 | - | - | 0 | - |
| 5 | y | 1.891E+05 | 723.3 | 0.005366 | 7.418 | +1 | 5 |
| - | - | 7.931E+04 | 724.3 | - | - | 0 | - |
| - | - | 2.496E+04 | 725.3 | - | - | 0 | - |
| - | - | 1610 | 728.3 | - | - | 0 | - |
| - | - | 6410 | 731.2 | - | - | 0 | - |
| - | - | 3536 | 732.2 | - | - | 0 | - |
| - | - | 4372 | 733.3 | - | - | 0 | - |
| - | - | 1845 | 739.3 | - | - | 0 | - |
| - | - | 3377 | 746.3 | - | - | 0 | - |
| - | - | 1816 | 747.3 | - | - | 0 | - |
| - | - | 2006 | 748.3 | - | - | 0 | - |
| - | - | 8.518E+04 | 749.3 | - | - | 0 | - |
| - | - | 2.974E+04 | 750.3 | - | - | 0 | - |
| - | - | 8775 | 751.3 | - | - | 0 | - |
| - | - | 5940 | 756.3 | - | - | 0 | - |
| - | - | 3725 | 757.3 | - | - | 0 | - |
| - | - | 2607 | 759.3 | - | - | 0 | - |
| - | - | 2.077E+04 | 767.3 | - | - | 0 | - |
| - | - | 8136 | 768.3 | - | - | 0 | - |
| - | - | 2013 | 769.3 | - | - | 0 | - |
| - | - | 7.819E+04 | 774.3 | - | - | 0 | - |
| - | - | 3.359E+04 | 775.3 | - | - | 0 | - |
| - | - | 8560 | 776.3 | - | - | 0 | - |
| - | - | 3662 | 778.3 | - | - | 0 | - |
| - | - | 4651 | 792.3 | - | - | 0 | - |
| - | - | 3281 | 793.3 | - | - | 0 | - |
| - | - | 2225 | 796.3 | - | - | 0 | - |
| - | - | 2442 | 803.3 | - | - | 0 | - |
| - | - | 1930 | 804.3 | - | - | 0 | - |
| - | - | 2114 | 814.3 | - | - | 0 | - |
| - | - | 1564 | 817.3 | - | - | 0 | - |
| - | - | 2959 | 818.3 | - | - | 0 | - |
| - | - | 1825 | 819.3 | - | - | 0 | - |
| 4 | y | 1.077E+04 | 820.3 | 0.005711 | 6.962 | +1 | 6 |
| - | - | 7344 | 821.3 | - | - | 0 | - |
| - | - | 2508 | 822.3 | - | - | 0 | - |
| - | - | 1475 | 823.3 | - | - | 0 | - |
| - | - | 5917 | 832.3 | - | - | 0 | - |
| - | - | 2501 | 833.3 | - | - | 0 | - |
| - | - | 1791 | 834.3 | - | - | 0 | - |
| 4 | y | 2.356E+05 | 838.3 | 0.005522 | 6.587 | +1 | 6 |
| - | - | 1.093E+05 | 839.3 | - | - | 0 | - |
| - | - | 3.664E+04 | 840.3 | - | - | 0 | - |
| - | - | 4770 | 841.3 | - | - | 0 | - |
| 7 | b | 6596 | 842.3 | 0.006582 | 7.814 | +1 | 7 |
| - | - | 7051 | 843.3 | - | - | 0 | - |
| - | - | 4077 | 844.3 | - | - | 0 | - |
| - | - | 2320 | 845.4 | - | - | 0 | - |
| - | - | 2442 | 848.3 | - | - | 0 | - |
| - | - | 2490 | 850.6 | - | - | 0 | - |
| - | - | 2675 | 853.3 | - | - | 0 | - |
| - | - | 3938 | 854.3 | - | - | 0 | - |
| 7 | b | 4325 | 860.3 | 0.005356 | 6.225 | +1 | 7 |
| - | - | 5395 | 862.3 | - | - | 0 | - |
| - | - | 2968 | 863.3 | - | - | 0 | - |
| - | - | 4006 | 868.3 | - | - | 0 | - |
| - | - | 1676 | 869.3 | - | - | 0 | - |
| - | - | 3.236E+04 | 871.3 | - | - | 0 | - |
| - | - | 1.933E+04 | 872.3 | - | - | 0 | - |
| - | - | 7252 | 873.3 | - | - | 0 | - |
| - | - | 2044 | 874.3 | - | - | 0 | - |
| - | - | 1901 | 888.3 | - | - | 0 | - |
| - | - | 3.297E+05 | 889.3 | - | - | 0 | - |
| - | - | 1.685E+05 | 890.4 | - | - | 0 | - |
| - | - | 2206 | 890.5 | - | - | 0 | - |
| - | - | 4.351E+04 | 891.4 | - | - | 0 | - |
| - | - | 4450 | 892.4 | - | - | 0 | - |
| - | - | 7324 | 896.3 | - | - | 0 | - |
| - | - | 3911 | 897.3 | - | - | 0 | - |
| - | - | 1600 | 899.3 | - | - | 0 | - |
| - | - | 1.535E+04 | 907.3 | - | - | 0 | - |
| - | - | 9560 | 908.3 | - | - | 0 | - |
| - | - | 3097 | 909.3 | - | - | 0 | - |
| - | - | 2909 | 915.4 | - | - | 0 | - |
| - | - | 2028 | 917.3 | - | - | 0 | - |
| - | - | 3593 | 918.3 | - | - | 0 | - |
| - | - | 3012 | 933.4 | - | - | 0 | - |
| 3 | y | 5.99E+04 | 935.3 | 0.005562 | 5.947 | +1 | 7 |
| - | - | 3.975E+04 | 936.3 | - | - | 0 | - |
| - | - | 1.533E+04 | 937.3 | - | - | 0 | - |
| - | - | 8958 | 938.3 | - | - | 0 | - |
| - | - | 3917 | 939.3 | - | - | 0 | - |
| 3 | y | 1.085E+06 | 953.3 | 0.005679 | 5.957 | +1 | 7 |
| - | - | 5.653E+05 | 954.3 | - | - | 0 | - |
| - | - | 1.926E+05 | 955.3 | - | - | 0 | - |
| - | - | 2.591E+04 | 956.4 | - | - | 0 | - |
| - | - | 8431 | 963.3 | - | - | 0 | - |
| - | - | 3868 | 964.3 | - | - | 0 | - |
| - | - | 4569 | 969.4 | - | - | 0 | - |
| - | - | 2930 | 970.4 | - | - | 0 | - |
| 8 | b | 3178 | 979.4 | 0.001686 | 1.722 | +1 | 8 |
| - | - | 1576 | 980.4 | - | - | 0 | - |
| - | - | 1594 | 981.4 | - | - | 0 | - |
| 8 | b | 1.297E+04 | 997.4 | 0.005587 | 5.602 | +1 | 8 |
| - | - | 6537 | 998.4 | - | - | 0 | - |
| - | - | 2532 | 999.4 | - | - | 0 | - |
| - | - | 4391 | 1015 | - | - | 0 | - |
| - | - | 5215 | 1018 | - | - | 0 | - |
| - | - | 1805 | 1019 | - | - | 0 | - |
| - | - | 2.415E+04 | 1036 | - | - | 0 | - |
| - | - | 1.513E+04 | 1037 | - | - | 0 | - |
| - | - | 4800 | 1038 | - | - | 0 | - |
| - | - | 1720 | 1046 | - | - | 0 | - |
| 2 | y | 7427 | 1082 | 0.005202 | 4.806 | +1 | 8 |
| - | - | 4194 | 1083 | - | - | 0 | - |
| 2 | y | 6.038E+04 | 1100 | 0.005136 | 4.667 | +1 | 8 |
| - | - | 3.974E+04 | 1101 | - | - | 0 | - |
| - | - | 1.643E+04 | 1102 | - | - | 0 | - |
| - | - | 1836 | 1103 | - | - | 0 | - |
| - | - | 9855 | 1110 | - | - | 0 | - |
| - | - | 4929 | 1111 | - | - | 0 | - |
| - | - | 2976 | 1112 | - | - | 0 | - |

m/z Charge Intensity FragmentType MassShift Position
120.08120727539062 0 655006.7
121.03990173339844 0 1943.9277
121.08448791503906 0 49775.227
122.07177734375 0 2414.2556
122.08861541748047 0 910.02686
127.08715057373047 0 1979.1365
129.07061767578125 0 1675.9445
129.10269165039062 0 3201.2234
130.050537109375 0 894.4042
130.05996704101562 0 1196.4757
130.06552124023438 0 20409.764
131.0491180419922 0 920.2438
131.0688934326172 0 1374.8514
131.08584594726562 0 883.71545
132.0811309814453 0 32865.227
133.084716796875 0 3510.1946
134.09674072265625 0 2416.5183
136.0760955810547 0 223087.67
136.08729553222656 0 7132.913
137.0794677734375 0 15697.696
138.0552978515625 0 16358.1455
138.0666046142578 0 25712.2
138.36717224121094 0 863.8476
139.06997680664062 0 1417.796
141.10238647460938 0 1373.5
142.06533813476562 0 1988.4291
143.0454864501953 0 1429.1755
143.0734100341797 0 4597.922
143.08592224121094 0 4643.113
144.0811767578125 0 3032.2705
146.06044006347656 0 8304.412
146.0718994140625 0 1312.8082
146.09693908691406 0 3289.0571
147.04412841796875 0 1020.25903
147.0635223388672 0 1023.02454
147.09231567382812 0 1111.8215
148.07606506347656 0 1775.0371
148.08721923828125 0 4720.4697
150.06663513183594 0 1512.2742
155.08193969726562 0 4894.689
155.09315490722656 0 4758.6963
156.07713317871094 0 58194.367
157.08053588867188 0 3457.0464
157.1342315673828 0 2821.9756
158.09678649902344 0 95504.77
159.05563354492188 0 1131.5476
159.09207153320312 0 49865.984
159.0999755859375 0 8698.0205
160.07606506347656 0 18714.943
160.08749389648438 0 2453.4377
160.09555053710938 0 4462.1016
160.11241149902344 0 43224.086
161.11570739746094 0 5381.053
165.070068359375 0 2024.6824
165.07742309570312 0 12845.015
165.10313415527344 0 1259.9515
166.0536346435547 0 3096.393
166.06153869628906 0 8881.914
166.08627319335938 0 2557.6475
167.0567626953125 0 1416.1543
167.0932159423828 0 8704.479
169.07643127441406 0 4053.958
169.09738159179688 0 1258.6539
171.07691955566406 0 8599.688
171.09185791015625 0 1566.6725
172.0762481689453 0 2312.754
172.10833740234375 0 1199.0736
173.0924072265625 0 2843.531
173.12904357910156 0 1137.3196
173.4388427734375 0 2224.8162
174.06649780273438 0 4973.751 b Ammonia loss 2
174.09214782714844 0 1137.8798
175.08743286132812 0 3100.5913
175.09837341308594 0 2131.1687
175.1234893798828 0 2967.1675
176.08236694335938 0 17134.12
176.10743713378906 0 187296.58
177.10264587402344 0 135737.72
177.11109924316406 0 19404.389
178.06134033203125 0 1224.4556
178.09814453125 0 1733.5239
178.10610961914062 0 13517.441
180.0772247314453 0 9862.59
185.05615234375 0 17935.03
186.09173583984375 0 14324.095
187.0633087158203 0 2544.0771
187.0952911376953 0 1055.0028
188.0709991455078 0 64259.207
189.07432556152344 0 7656.75
189.10198974609375 0 1352.9362
190.08663940429688 0 1613.4431
191.08285522460938 0 2040.1997
191.09329223632812 0 6177.99
191.11837768554688 0 13888.793
192.10272216796875 0 2709.0974
192.12205505371094 0 1276.7059
193.07276916503906 0 3328.0964
193.10877990722656 0 60748.617
194.080078125 0 3311.878
194.0930938720703 0 1099.8557
194.1122589111328 0 3392.9294
195.0880584716797 0 13703.3955
196.09194946289062 0 1496.7706
199.07142639160156 0 1207.5057
201.06582641601562 0 1119.8401
202.05355834960938 0 5986.0903
203.06666564941406 0 15357.032
203.0931396484375 0 4591.217
203.11825561523438 0 64058.535 a Water loss 1
204.06809997558594 0 1991.7843
204.0771484375 0 24677.11
204.10203552246094 0 4104.7876 a Ammonia loss 1
204.12164306640625 0 9033.745
205.08132934570312 0 2441.938
205.09751892089844 0 47983.45 y 8
205.10781860351562 0 2655.1733
206.10101318359375 0 6278.4917
207.11331176757812 0 13659.186
208.0720672607422 0 8574.618
208.11724853515625 0 1527.1578
208.926513671875 0 1174.64
211.05856323242188 0 1326.2236
212.11610412597656 0 4131.636
213.05136108398438 0 1362.4452
213.10006713867188 0 2010.371
215.08216857910156 0 3970.1853
216.06680297851562 0 1324.7069
217.09759521484375 0 31759.615
218.08206176757812 0 1641.8201
218.10177612304688 0 3419.3108
219.0801239013672 0 7784.0356
219.09005737304688 0 1839.91
219.11367797851562 0 1090.6678
220.12022399902344 0 1996.6882
221.10406494140625 0 158804.08
221.12901306152344 0 2628171.2 a 1
222.10694885253906 0 13270.981
222.13217163085938 0 317154.7
223.10699462890625 0 1230.0045
223.1195068359375 0 2451.7454
223.13462829589844 0 17880.385
223.18272399902344 0 1015.42053
224.11795043945312 0 2042.1597
228.1134033203125 0 8330.455
229.10882568359375 0 5111.579
230.1139678955078 0 1121.8392
231.0615997314453 0 42202.7
231.11326599121094 0 27938.076 b Water loss 1
232.0651092529297 0 3190.6362
232.11660766601562 0 3904.7039
233.0914764404297 0 2659.8142
235.10821533203125 0 64083.48
236.1111602783203 0 8762.927
238.13038635253906 0 4178.581
239.0952911376953 0 1062.4667
239.11428833007812 0 14097.047
240.08151245117188 0 1342.1515
240.117431640625 0 1346.0073
241.08407592773438 0 1395.9797
245.09207153320312 0 3246.8574 y 6
245.1301727294922 0 1396.6715
246.09873962402344 0 1395.065
246.13519287109375 0 4957.724
247.10675048828125 0 1236.2273
247.12025451660156 0 1866.5511
248.11569213867188 0 2579.8066
248.1510009765625 0 4227.208
249.12391662597656 0 464768.44 b 1
250.10537719726562 0 2466.8389
250.12721252441406 0 58368.453
251.10311889648438 0 49630.31
251.1297149658203 0 7027.679
252.10684204101562 0 5055.468
253.09371948242188 0 2347.303
255.11199951171875 0 2342.2397
256.1084289550781 0 3701.948
257.1068420410156 0 27806.127
258.1099548339844 0 2963.686
260.7489013671875 0 1380.918
261.087646484375 0 7077.4253
262.070068359375 0 1347.5609
262.1191101074219 0 1674.8418
263.0707092285156 0 2825.8135
263.10302734375 0 91900.73
263.1294250488281 0 9809.604
264.10662841796875 0 11706.672
264.14593505859375 0 9056.768
265.11785888671875 0 3167.3584
267.0914611816406 0 6860.9126
268.07568359375 0 3701.3262
268.0929870605469 0 2342.748
268.141845703125 0 3139.3254
269.07708740234375 0 2028.8745
270.07342529296875 0 1399.3889
270.10821533203125 0 1797.0527
271.108642578125 0 4815.0146
272.1141662597656 0 1551.4655
273.1228332519531 0 2254.2485
274.13031005859375 0 23247.102
274.1563415527344 0 1694.586
275.1146240234375 0 4435.3286
275.1341857910156 0 3681.8403
277.7330017089844 0 1598.2203
277.7564697265625 0 1128.4482
277.7767639160156 0 1011.8174
278.1391296386719 0 1482.3912
279.0981140136719 0 38224.727
279.125 0 3820.945
280.1014099121094 0 6055.007
282.1217346191406 0 1645.3188
283.1540222167969 0 1354.6404
285.1021423339844 0 59312.684
286.1051330566406 0 8055.236
287.0997009277344 0 2214.7285
288.13421630859375 0 1568.8612
289.0825500488281 0 4265.169
290.1258239746094 0 1639.1357
292.1409912109375 0 38686.08
292.1661071777344 0 9109.908 d 2
293.1432800292969 0 4634.535
296.0887145996094 0 1862.8988
296.151123046875 0 43289.906
297.1373291015625 0 1573.6985
297.15496826171875 0 7804.9487
300.13525390625 0 4325.6377
301.1189270019531 0 3466.4255
302.1147155761719 0 4370.415
303.1133728027344 0 6028.7393
306.1334533691406 0 1697.8877
306.1471862792969 0 1133.5023
307.1199645996094 0 1878.5795
310.151611328125 0 6705.1694
313.09783935546875 0 2775.9673
314.0992736816406 0 4090.8848
315.0984191894531 0 3427.1433
316.1307373046875 0 1762.3309
318.1454772949219 0 15163.056 a Water loss 2
319.14837646484375 0 1536.9923
320.13507080078125 0 3902.01
323.09942626953125 0 17606.857
324.1462707519531 0 103113.125
325.14898681640625 0 15601.766
327.09722900390625 0 1232.9373
327.145751953125 0 2513.2034
328.12921142578125 0 7216.108
328.1454772949219 0 4931.6323
330.110107421875 0 5823.602
331.09295654296875 0 2568.4587
332.12506103515625 0 5318.584
333.10882568359375 0 3791.35
333.129638671875 0 1675.2139
334.1072692871094 0 2843.7715
334.132568359375 0 3269.737
335.1714782714844 0 2637.89
336.15557861328125 0 1324.6097 a 2
338.1288757324219 0 7243.4263
339.1712646484375 0 9064.027
340.1754455566406 0 1648.5146
342.1568603515625 0 537688.75 y 7
343.1598815917969 0 97014.96
344.16192626953125 0 9882.805
346.1402282714844 0 30784.844 b Water loss 2
347.1429138183594 0 4526.8184
348.1197814941406 0 14077.768
349.10113525390625 0 1573.5166
349.12322998046875 0 3092.6528
350.1351318359375 0 31457.81
351.1382751464844 0 4397.6357
352.1407470703125 0 9164.232
353.1473083496094 0 2012.7504
354.1562194824219 0 2628.785
356.13916015625 0 32357.812
357.14202880859375 0 4701.4863 b 5
358.1041259765625 0 2558.8076
359.08807373046875 0 2605.2234
360.11981201171875 0 6722.1733
362.1506652832031 0 4639.373 y 4
362.6512451171875 0 3285.5906
364.15093994140625 0 129821.72 b 2
365.1541442871094 0 26402.525
366.12969970703125 0 14222.144
366.1544189453125 0 2124.0303
367.13250732421875 0 2159.3225
367.165771484375 0 6501.922
368.1688232421875 0 1869.5585
371.1365966796875 0 3382.0254
374.1493835449219 0 6204.742
376.1145935058594 0 13599.233
377.12005615234375 0 2850.7893
378.1006164550781 0 3634.4292
378.13018798828125 0 73193.71
379.1331787109375 0 12444.761
380.13916015625 0 5090.3335
381.14508056640625 0 2640.296
382.1438903808594 0 15643.522
383.14715576171875 0 3144.0332
384.1354064941406 0 4376.6367
385.1358642578125 0 3857.9119
386.1371765136719 0 1881.9977
388.1155700683594 0 1893.4998
388.1489562988281 0 1987.8489
389.1481628417969 0 1720.214
390.1567687988281 0 9620.155
391.15997314453125 0 1812.7428
394.1252746582031 0 91379.18
395.128173828125 0 17918.506
395.1558837890625 0 1788.8707
396.12908935546875 0 1772.8109
397.1629943847656 0 1706.7963
398.170166015625 0 2599.4084
399.1752624511719 0 4252.1816
404.110595703125 0 2577.737
404.1455078125 0 2372.5444
406.12567138671875 0 2146.3435
406.15289306640625 0 2771.889
407.18359375 0 9791.642
408.1675109863281 0 21227.877
409.1713562011719 0 6368.0522
410.1392822265625 0 2926.4688
410.18267822265625 0 2097.652
415.1620178222656 0 4602.6855
416.1457214355469 0 2252.651
417.1748352050781 0 1530.1332
419.1566162109375 0 3927.7764
419.6618957519531 0 1426.5105 y 3
420.1636047363281 0 1494.8773
423.17791748046875 0 2198.1235
425.194091796875 0 77093.33
426.196533203125 0 16760.396
427.20880126953125 0 16044.458
428.214599609375 0 3659.0024
430.12548828125 0 2144.6853
431.1253356933594 0 1653.9756
431.1571044921875 0 1529.4131
432.13909912109375 0 2239.7896
433.1419677734375 0 1566.7491
433.17266845703125 0 14703.074
434.1787414550781 0 2187.4734
435.17864990234375 0 4260.065
437.16748046875 0 3920.4036
438.1724548339844 0 1942.2416
442.17413330078125 0 2966.106
443.156982421875 0 3066.0146
445.1757507324219 0 1422.5725
446.2176818847656 0 1448.9598
447.15167236328125 0 11545.06
448.1573791503906 0 2342.854
449.1327209472656 0 2894.6155
449.1666259765625 0 3223.3372
450.2287902832031 0 2548.3936
451.1802673339844 0 1671.1417
454.1561584472656 0 2607.8362
454.1972961425781 0 5796.191
455.2039489746094 0 14480.469
456.2068786621094 0 3440.8374
458.16729736328125 0 3602.0552
459.1531066894531 0 3522.2688
461.1673278808594 0 20361.676 b Water loss 3
461.19586181640625 0 7610.704
462.1693420410156 0 6412.4155
462.2013244628906 0 2194.3726
465.16253662109375 0 72848.61
466.1656188964844 0 16808.992
467.1675109863281 0 1569.6323
468.1716003417969 0 3129.6274 y Water loss 2
469.2063903808594 0 3203.7458
470.1700744628906 0 1592.5618
470.208984375 0 1484.776
471.181884765625 0 11756.759
472.18560791015625 0 3263.8557
472.2305603027344 0 1269.6356
473.2149963378906 0 6696.3965
475.1459045410156 0 1543.9972
476.1869201660156 0 1516.2251
477.1671447753906 0 12035.732 y 2
477.678955078125 0 3512.0713
478.169677734375 0 3534.6606
478.2201843261719 0 2293.587
479.1781311035156 0 68951.01 b 3
480.1817321777344 0 15586.481
481.18243408203125 0 1776.5442
482.1914978027344 0 3813.0486
483.1986389160156 0 1704.2844
485.6944274902344 0 1513.2848
486.1632080078125 0 11992.325
487.1629943847656 0 2175.3762
489.1925048828125 0 128345.06 y 6
490.19482421875 0 36088.45
490.68499755859375 0 2377.7703
491.2050476074219 0 16837.896
492.20904541015625 0 4277.1616
493.156494140625 0 1283.8553
495.18499755859375 0 9251.376
496.1690368652344 0 2296.5547
496.1887512207031 0 3675.4946
496.231201171875 0 65613.85
497.1714782714844 0 12005.437
497.2343444824219 0 19647.162
498.1733093261719 0 2570.7952
498.2394104003906 0 2009.8298
499.18890380859375 0 5624.365 b 7
502.19696044921875 0 1745.2897
503.1819763183594 0 3198.4514
506.214111328125 0 1652.6471
509.7073059082031 0 1687.0619
511.1950378417969 0 1610.9193
513.198486328125 0 11031.513
514.196533203125 0 2730.1594
516.2207641601562 0 3745.3845
518.7103881835938 0 4519.645
519.2041015625 0 19376.031
519.7161254882812 0 1483.3477
520.2056884765625 0 8478.418
521.2060546875 0 2053.3203
523.1860961914062 0 3271.4307
525.1605834960938 0 1393.0721
525.2042846679688 0 4007.4792
526.206298828125 0 1613.4353
527.2122802734375 0 1604.4269
527.7095947265625 0 2067.975
530.1912231445312 0 3919.9192
532.203369140625 0 1532.8994
533.2030029296875 0 1710.099
537.2139892578125 0 6359.2275
541.1947631835938 0 5089.152
541.7056884765625 0 9249.052 y Water loss 1
542.2332153320312 0 11913.791
542.7101440429688 0 5137.8726
543.2404174804688 0 3206.689
545.209716796875 0 2024.921
548.1997680664062 0 23922.46
549.2026977539062 0 8285.62
550.2078247070312 0 2009.0723
550.71142578125 0 67558.28 y 1
551.2130737304688 0 49729.35
551.7137451171875 0 19965.602
552.2146606445312 0 4201.944
553.1980590820312 0 2317.0164
555.7045288085938 0 3080.674
557.19970703125 0 11698.4
558.2073364257812 0 1615.367
560.2291259765625 0 204233.61 y 5
560.73095703125 0 5372.221
561.2323608398438 0 67360.266
562.230712890625 0 14126.073
563.2342529296875 0 1558.4706
569.2338256835938 0 13877.834
569.7357788085938 0 11305.058
570.22998046875 0 20245.309
570.7149658203125 0 1355.5648
571.233154296875 0 3569.8162
574.7113037109375 0 2606.3413
578.2305908203125 0 10482.999
578.7332153320312 0 5560.869
579.2340698242188 0 1760.1838
583.2241821289062 0 6214.603
583.72607421875 0 3611.2769
588.24072265625 0 4941.6636
592.2302856445312 0 44207.453 Precursor Water loss
592.7319946289062 0 28396.787
593.232421875 0 14160.639
593.7296142578125 0 2856.4438
594.1875610351562 0 5814.7773
596.2363891601562 0 14097.564
597.2255249023438 0 9117.759
598.2293701171875 0 3521.769
601.2355346679688 0 105427.33 Precursor
601.7369384765625 0 73497.78
601.8181762695312 0 1813.9718
602.2374267578125 0 32600.924
602.73876953125 0 5006.261
606.233642578125 0 20569.645
607.235595703125 0 6713.252
608.2372436523438 0 2551.7012
612.1978759765625 0 35722.22
613.2008056640625 0 11200.814
613.2879028320312 0 2301.4966
614.2005004882812 0 2078.974
614.2465209960938 0 24793.73
614.29296875 0 1318.7683
615.24951171875 0 6558.428
622.2181396484375 0 1661.1803
624.2312622070312 0 16030.322 b Water loss 4
625.235595703125 0 5388.152
631.2476196289062 0 9084.823
632.2593383789062 0 4627.26
633.2838134765625 0 1370.0878
634.2299194335938 0 18639.95
635.2338256835938 0 7530.553
636.2266845703125 0 1571.9669
639.251220703125 0 6264.0513
640.2380981445312 0 8924.194
641.2344970703125 0 2169.5774
641.2880859375 0 2126.2847
642.2418823242188 0 15233.086 b 4
643.2450561523438 0 3374.1445
652.2405395507812 0 5390.76
653.2381591796875 0 1382.7831
657.263427734375 0 47360.008
658.265380859375 0 13532.226
659.294189453125 0 73548.19
660.2971801757812 0 26923.234
661.2997436523438 0 6523.4214
667.2503051757812 0 5790.733
668.2513427734375 0 4321.5464
676.2689208984375 0 9195.509
677.2723999023438 0 5836.85
678.2920532226562 0 2333.2332
685.2586059570312 0 63575.582
686.2617797851562 0 21203.24
687.2622680664062 0 5233.762
695.2669067382812 0 12405.139 b Water loss 5
696.27099609375 0 4210.7373
702.2882080078125 0 2679.7637
703.2667846679688 0 16775.934
704.2403564453125 0 3968.647
704.2681884765625 0 4330.847
705.27783203125 0 8059.436
706.2854614257812 0 3006.7356
708.270263671875 0 1516.3378
713.27587890625 0 6363.864 b 5
714.2827758789062 0 1824.9425
721.2615356445312 0 57452.547
722.2646484375 0 23430.174
723.2923583984375 0 189148.81 y 4
724.2955932617188 0 79311.45
725.2946166992188 0 24959.102
728.3179321289062 0 1610.1587
731.2471313476562 0 6410.461
732.238037109375 0 3535.897
733.2744750976562 0 4372.475
739.2919921875 0 1845.1597
746.2737426757812 0 3377.276
747.2976684570312 0 1815.8197
748.302978515625 0 2006.1334
749.2564086914062 0 85176.51
750.2590942382812 0 29740.576
751.2589111328125 0 8775.1
756.3106689453125 0 5939.6123
757.3074340820312 0 3725.1243
759.2630004882812 0 2606.8542
767.2672729492188 0 20765.27
768.2706909179688 0 8136.4985
769.2611083984375 0 2012.7195
774.3212280273438 0 78194.26
775.3237915039062 0 33592.58
776.326416015625 0 8559.693
778.302001953125 0 3661.6836
792.315673828125 0 4650.9434
793.3177490234375 0 3280.5852
796.3170776367188 0 2224.9224
803.2808837890625 0 2441.9912
804.3418579101562 0 1929.7942
814.3135986328125 0 2113.7737
817.32080078125 0 1563.6152
818.3331298828125 0 2958.5852
819.3389282226562 0 1825.1858
820.30908203125 0 10771.703 y Water loss 3
821.30224609375 0 7343.6265
822.3143920898438 0 2508.3843
823.2891845703125 0 1474.9518
832.3289794921875 0 5916.859
833.3307495117188 0 2500.9763
834.3251953125 0 1790.6102
838.3194580078125 0 235587.19 y 3
839.3220825195312 0 109252.94
840.3231201171875 0 36638.305
841.32275390625 0 4769.812
842.30419921875 0 6595.5825 b Water loss 6
843.3377685546875 0 7051.218
844.3375854492188 0 4077.3523
845.3507080078125 0 2319.513
848.3008422851562 0 2441.8115
850.5880126953125 0 2489.7927
853.3255615234375 0 2675.1775
854.321533203125 0 3937.7664
860.3135375976562 0 4325.3833 b 6
862.3243408203125 0 5395.102
863.3316650390625 0 2967.7612
868.3290405273438 0 4005.861
869.3340454101562 0 1675.8552
871.337158203125 0 32358.209
872.3369750976562 0 19332.559
873.3384399414062 0 7252.373
874.3203125 0 2044.0474
888.3489990234375 0 1901.258
889.3477783203125 0 329666.22
890.3504638671875 0 168475.12
890.4552612304688 0 2205.647
891.353759765625 0 43508.2
892.3524780273438 0 4449.9253
896.3252563476562 0 7323.5894
897.3292846679688 0 3911.262
899.3383178710938 0 1599.8584
907.3411865234375 0 15351.666
908.3438110351562 0 9560.227
909.3446044921875 0 3097.1619
915.354248046875 0 2909.054
917.3283081054688 0 2027.724
918.3145751953125 0 3592.6091
933.3751831054688 0 3012.0293
935.3358764648438 0 59900.863 y Water loss 2
936.3358764648438 0 39751.543
937.3339233398438 0 15325.871
938.3248291015625 0 8957.561
939.3258666992188 0 3916.6414
953.3465576171875 0 1085217.2 y 2
954.3491821289062 0 565254.56
955.3494873046875 0 192562.45
956.3505249023438 0 25907.246
963.3322143554688 0 8430.684
964.33056640625 0 3868.374
969.381103515625 0 4569.3354
970.381591796875 0 2930.327
979.3582153320312 0 3177.5261 b Water loss 7
980.3811645507812 0 1576.1697
981.3692016601562 0 1594.4335
997.3726806640625 0 12968.312 b 7
998.3759765625 0 6536.64
999.3818969726562 0 2531.641
1015.3814086914062 0 4390.8877
1018.3994140625 0 5214.942
1019.4024658203125 0 1804.873
1036.4156494140625 0 24147.96
1037.418212890625 0 15128.36
1038.4237060546875 0 4800.4204
1046.39501953125 0 1720.4977
1082.4039306640625 0 7427.4375 y Water loss 1
1083.4061279296875 0 4194.392
1100.4144287109375 0 60379.797 y 1
1101.41748046875 0 39741.734
1102.417724609375 0 16431.43
1103.4149169921875 0 1836.0184
1110.3994140625 0 9854.918
1111.40380859375 0 4928.8857
1112.4013671875 0 2975.5261

Spectrum Details

|  |  |
| --- | --- |
| Matched peaks? Matched peaks The total absolute number of peaks matched. Additionally in brackets the total fraction of peaks matched and the total number of peaks is shown. | 43 (6.76% of 636) |
| FDR? FDR The false discovery rate estimated for this peptide. It is calculated by matching all theoretical fragments with a non-integer shift with the raw peaks for this spectrum. This is done with 40 different shifts. The resulting percentage is the average number of annotated peaks over the number of annotated peaks with the correct spectrum. | 0.00% |
| Satellite FDR? Satellite FDR See the FDR for details on its calculation. This satellite ion specific FDR only contains the satellite ions (d/w) for I/L/J positions. | - |
| PSM Score? PSM Score The PSM Score as given by Hecklib to this annotated spectrum. It is shown with three significant figures. | 511 |

#### Spectrum 7636? Spectrum 7636 The raw spectrum of this peptide as annotated by Hecklib. The fragments are coloured according to ion type (see legend). Any peaks with a star '\*' as text can be hovered over to see the full details, first the ion type second the mass shift type. By hovering over the amino acids in the peptide or ions in the legend the corresponding peaks are highlighted. By toggling the 'Unassigned' label you can turn the background (unassigned) peaks on or off in the plot. By updating the slider in the Ion legend you can update the spectrum to only show the top X% of the peaks with labels. The top X% means any peak that is within X% of the highest intensity. By dragging in the spectrum you can zoom in to a specific part of the spectrum and use 'Zoom Out' to get back to the original zoom level. The annotation of the spectrum is based on the given sequence in the peptides file and is done with different software so inconsistencies are likely. The peaks are annotated based on the given sequence, with 20 ppm tolerance.

Copy Data

##### Spectrum 7636 (TSV)

###### Preview

```
Loading example...
```

*Click on the button to copy the data to your clipboard.*

Mz MinMz MaxIntensity Max

WidthHeightPeptide font sizePeptide stroke widthSpectrum font sizeSpectrum stroke widthCompact peptide

Ion legend

wxyz

abcd

OtherUnassignedIon^Charge^_Position_Show for top:%

TFDDYAMHW

03.87e+57.73e+51.16e+61.55e+6

Zoom Out

b^+2^_3_a^+1^_2_a^+1^_2_y^+1^_1_a^+1^_2_b^+2^_4_b^+1^_2_y^+2^_3_b^+1^_2_d^+1^_3_a^+1^_3_a^+1^_3_y^+1^_2_b^+1^_3_b^+2^_6_y^+2^_5_b^+1^_3_b^+1^_4_y^+2^_7_b^+1^_4_y^+1^_3_b^+2^_8_y^+2^_8_y^+2^_8_y^+1^_4_\*\*b^+1^_5_b^+1^_5_b^+1^_6_b^+1^_6_y^+1^_5_y^+1^_6_y^+1^_6_b^+1^_7_b^+1^_7_y^+1^_7_y^+1^_7_b^+1^_8_y^+1^_8_y^+1^_8_

0830166024903320

Fragment Matches Table

Show background peaks

| Position | Ion type | Intensity | mz Theoretical | mz Error (Th) | mz Error (ppm) | Charge | Series Number |
| --- | --- | --- | --- | --- | --- | --- | --- |
| - | - | 3.695E+05 | 120.1 | - | - | 0 | - |
| - | - | 1208 | 121 | - | - | 0 | - |
| - | - | 1307 | 121.1 | - | - | 0 | - |
| - | - | 2.755E+04 | 121.1 | - | - | 0 | - |
| - | - | 489.4 | 121.6 | - | - | 0 | - |
| - | - | 1817 | 122.1 | - | - | 0 | - |
| - | - | 1008 | 122.1 | - | - | 0 | - |
| - | - | 511 | 124.8 | - | - | 0 | - |
| - | - | 1513 | 127.1 | - | - | 0 | - |
| - | - | 1445 | 129.1 | - | - | 0 | - |
| - | - | 3941 | 129.1 | - | - | 0 | - |
| - | - | 1.162E+04 | 130.1 | - | - | 0 | - |
| - | - | 789.1 | 131.1 | - | - | 0 | - |
| - | - | 2.035E+04 | 132.1 | - | - | 0 | - |
| - | - | 539.5 | 132.1 | - | - | 0 | - |
| - | - | 645.7 | 133.1 | - | - | 0 | - |
| - | - | 1658 | 133.1 | - | - | 0 | - |
| - | - | 1629 | 134.1 | - | - | 0 | - |
| - | - | 1.339E+05 | 136.1 | - | - | 0 | - |
| - | - | 4454 | 136.1 | - | - | 0 | - |
| - | - | 9722 | 137.1 | - | - | 0 | - |
| - | - | 667.1 | 137.1 | - | - | 0 | - |
| - | - | 1.066E+04 | 138.1 | - | - | 0 | - |
| - | - | 1.381E+04 | 138.1 | - | - | 0 | - |
| - | - | 736.1 | 139.1 | - | - | 0 | - |
| - | - | 1008 | 140.1 | - | - | 0 | - |
| - | - | 699.9 | 141.1 | - | - | 0 | - |
| - | - | 883.4 | 141.1 | - | - | 0 | - |
| - | - | 660.9 | 143 | - | - | 0 | - |
| - | - | 2011 | 143.1 | - | - | 0 | - |
| - | - | 2261 | 143.1 | - | - | 0 | - |
| - | - | 2276 | 144.1 | - | - | 0 | - |
| - | - | 4361 | 146.1 | - | - | 0 | - |
| - | - | 2561 | 146.1 | - | - | 0 | - |
| - | - | 701.7 | 147.1 | - | - | 0 | - |
| - | - | 1456 | 148.1 | - | - | 0 | - |
| - | - | 4070 | 148.1 | - | - | 0 | - |
| - | - | 552.5 | 148.1 | - | - | 0 | - |
| - | - | 693.3 | 148.3 | - | - | 0 | - |
| - | - | 694.8 | 152.1 | - | - | 0 | - |
| - | - | 1965 | 155.1 | - | - | 0 | - |
| - | - | 3944 | 155.1 | - | - | 0 | - |
| - | - | 3.29E+04 | 156.1 | - | - | 0 | - |
| - | - | 2000 | 157.1 | - | - | 0 | - |
| - | - | 824.7 | 157.1 | - | - | 0 | - |
| - | - | 5.631E+04 | 158.1 | - | - | 0 | - |
| - | - | 1150 | 159.1 | - | - | 0 | - |
| - | - | 2.757E+04 | 159.1 | - | - | 0 | - |
| - | - | 7101 | 159.1 | - | - | 0 | - |
| - | - | 9129 | 160.1 | - | - | 0 | - |
| - | - | 1156 | 160.1 | - | - | 0 | - |
| - | - | 2227 | 160.1 | - | - | 0 | - |
| - | - | 2.67E+04 | 160.1 | - | - | 0 | - |
| - | - | 1487 | 161.1 | - | - | 0 | - |
| - | - | 2637 | 161.1 | - | - | 0 | - |
| - | - | 606 | 161.7 | - | - | 0 | - |
| - | - | 1040 | 162.1 | - | - | 0 | - |
| - | - | 1580 | 165.1 | - | - | 0 | - |
| - | - | 6783 | 165.1 | - | - | 0 | - |
| - | - | 2634 | 166.1 | - | - | 0 | - |
| - | - | 6618 | 166.1 | - | - | 0 | - |
| - | - | 2793 | 166.1 | - | - | 0 | - |
| - | - | 776 | 167 | - | - | 0 | - |
| - | - | 663.8 | 167.1 | - | - | 0 | - |
| - | - | 4959 | 167.1 | - | - | 0 | - |
| - | - | 1900 | 169.1 | - | - | 0 | - |
| - | - | 1172 | 170.1 | - | - | 0 | - |
| - | - | 4927 | 171.1 | - | - | 0 | - |
| - | - | 862.4 | 172.1 | - | - | 0 | - |
| - | - | 1531 | 173.1 | - | - | 0 | - |
| - | - | 885.4 | 173.1 | - | - | 0 | - |
| 3 | b | 4072 | 174.1 | 0.001283 | 7.373 | +2 | 3 |
| - | - | 884.5 | 174.1 | - | - | 0 | - |
| - | - | 1880 | 175.1 | - | - | 0 | - |
| - | - | 1640 | 175.1 | - | - | 0 | - |
| - | - | 1808 | 175.1 | - | - | 0 | - |
| - | - | 9822 | 176.1 | - | - | 0 | - |
| - | - | 1.107E+05 | 176.1 | - | - | 0 | - |
| - | - | 7.463E+04 | 177.1 | - | - | 0 | - |
| - | - | 1.32E+04 | 177.1 | - | - | 0 | - |
| - | - | 1443 | 178.1 | - | - | 0 | - |
| - | - | 1218 | 178.1 | - | - | 0 | - |
| - | - | 6980 | 178.1 | - | - | 0 | - |
| - | - | 6032 | 180.1 | - | - | 0 | - |
| - | - | 1883 | 183.1 | - | - | 0 | - |
| - | - | 9426 | 185.1 | - | - | 0 | - |
| - | - | 887.4 | 185.1 | - | - | 0 | - |
| - | - | 9670 | 186.1 | - | - | 0 | - |
| - | - | 1405 | 187.1 | - | - | 0 | - |
| - | - | 3.707E+04 | 188.1 | - | - | 0 | - |
| - | - | 4291 | 189.1 | - | - | 0 | - |
| - | - | 999.5 | 189.1 | - | - | 0 | - |
| - | - | 1639 | 190.1 | - | - | 0 | - |
| - | - | 1085 | 191.1 | - | - | 0 | - |
| - | - | 3587 | 191.1 | - | - | 0 | - |
| - | - | 6999 | 191.1 | - | - | 0 | - |
| - | - | 2095 | 192.1 | - | - | 0 | - |
| - | - | 621 | 192.1 | - | - | 0 | - |
| - | - | 1324 | 193.1 | - | - | 0 | - |
| - | - | 3.298E+04 | 193.1 | - | - | 0 | - |
| - | - | 2916 | 194.1 | - | - | 0 | - |
| - | - | 2691 | 194.1 | - | - | 0 | - |
| - | - | 8696 | 195.1 | - | - | 0 | - |
| - | - | 958.8 | 195.1 | - | - | 0 | - |
| - | - | 710.7 | 198.1 | - | - | 0 | - |
| - | - | 772.8 | 199.1 | - | - | 0 | - |
| - | - | 1023 | 201.1 | - | - | 0 | - |
| - | - | 3500 | 202.1 | - | - | 0 | - |
| - | - | 8499 | 203.1 | - | - | 0 | - |
| - | - | 3310 | 203.1 | - | - | 0 | - |
| 2 | a | 3.726E+04 | 203.1 | 0.0004424 | 2.178 | +1 | 2 |
| - | - | 1.856E+04 | 204.1 | - | - | 0 | - |
| 2 | a | 2109 | 204.1 | 0.0001137 | 0.5573 | +1 | 2 |
| - | - | 4949 | 204.1 | - | - | 0 | - |
| - | - | 660.6 | 204.5 | - | - | 0 | - |
| - | - | 1687 | 205.1 | - | - | 0 | - |
| 9 | y | 3.026E+04 | 205.1 | 0.0005327 | 2.597 | +1 | 1 |
| - | - | 1157 | 205.1 | - | - | 0 | - |
| - | - | 3787 | 206.1 | - | - | 0 | - |
| - | - | 8167 | 207.1 | - | - | 0 | - |
| - | - | 4753 | 208.1 | - | - | 0 | - |
| - | - | 1505 | 212.1 | - | - | 0 | - |
| - | - | 811.7 | 213.1 | - | - | 0 | - |
| - | - | 758.6 | 213.1 | - | - | 0 | - |
| - | - | 1433 | 213.1 | - | - | 0 | - |
| - | - | 2272 | 215.1 | - | - | 0 | - |
| - | - | 887.9 | 216.1 | - | - | 0 | - |
| - | - | 983.3 | 216.1 | - | - | 0 | - |
| - | - | 1.938E+04 | 217.1 | - | - | 0 | - |
| - | - | 852.3 | 218.1 | - | - | 0 | - |
| - | - | 2325 | 218.1 | - | - | 0 | - |
| - | - | 5949 | 219.1 | - | - | 0 | - |
| - | - | 1131 | 219.1 | - | - | 0 | - |
| - | - | 675.5 | 220.1 | - | - | 0 | - |
| - | - | 1883 | 220.1 | - | - | 0 | - |
| - | - | 8.811E+04 | 221.1 | - | - | 0 | - |
| 2 | a | 1.531E+06 | 221.1 | 0.0006962 | 3.148 | +1 | 2 |
| - | - | 7442 | 222.1 | - | - | 0 | - |
| - | - | 1.858E+05 | 222.1 | - | - | 0 | - |
| - | - | 9969 | 223.1 | - | - | 0 | - |
| - | - | 1163 | 224.1 | - | - | 0 | - |
| - | - | 1031 | 225.1 | - | - | 0 | - |
| - | - | 920.2 | 226.2 | - | - | 0 | - |
| - | - | 6017 | 228.1 | - | - | 0 | - |
| - | - | 3038 | 229.1 | - | - | 0 | - |
| - | - | 2.067E+04 | 229.2 | - | - | 0 | - |
| - | - | 1276 | 229.2 | - | - | 0 | - |
| - | - | 3329 | 230.2 | - | - | 0 | - |
| - | - | 2.336E+04 | 231.1 | - | - | 0 | - |
| 4 | b | 806.5 | 231.1 | 0.0004495 | 1.945 | +2 | 4 |
| 2 | b | 1.482E+04 | 231.1 | 0.0005686 | 2.46 | +1 | 2 |
| - | - | 1953 | 232.1 | - | - | 0 | - |
| - | - | 1685 | 232.1 | - | - | 0 | - |
| - | - | 1782 | 233.1 | - | - | 0 | - |
| - | - | 837 | 233.1 | - | - | 0 | - |
| - | - | 4.054E+04 | 235.1 | - | - | 0 | - |
| - | - | 5356 | 236.1 | - | - | 0 | - |
| - | - | 713.1 | 237.1 | - | - | 0 | - |
| - | - | 1746 | 238.1 | - | - | 0 | - |
| - | - | 1115 | 239.1 | - | - | 0 | - |
| - | - | 1.031E+04 | 239.1 | - | - | 0 | - |
| - | - | 786.7 | 240.1 | - | - | 0 | - |
| - | - | 1277 | 240.1 | - | - | 0 | - |
| 7 | y | 1302 | 245.1 | 0.004537 | 18.51 | +2 | 3 |
| - | - | 1931 | 245.1 | - | - | 0 | - |
| - | - | 2953 | 246.1 | - | - | 0 | - |
| - | - | 1637 | 247.1 | - | - | 0 | - |
| - | - | 949.3 | 247.1 | - | - | 0 | - |
| - | - | 1016 | 247.1 | - | - | 0 | - |
| - | - | 959 | 248.1 | - | - | 0 | - |
| - | - | 3423 | 248.2 | - | - | 0 | - |
| 2 | b | 2.682E+05 | 249.1 | 0.0006699 | 2.689 | +1 | 2 |
| - | - | 3.373E+04 | 250.1 | - | - | 0 | - |
| - | - | 2.895E+04 | 251.1 | - | - | 0 | - |
| - | - | 4737 | 251.1 | - | - | 0 | - |
| - | - | 4439 | 252.1 | - | - | 0 | - |
| - | - | 874.8 | 252.1 | - | - | 0 | - |
| - | - | 1236 | 253.1 | - | - | 0 | - |
| - | - | 1093 | 253.1 | - | - | 0 | - |
| - | - | 1805 | 256.1 | - | - | 0 | - |
| - | - | 1.666E+04 | 257.1 | - | - | 0 | - |
| - | - | 2214 | 258.1 | - | - | 0 | - |
| - | - | 1053 | 258.1 | - | - | 0 | - |
| - | - | 3149 | 261.1 | - | - | 0 | - |
| - | - | 1021 | 261.1 | - | - | 0 | - |
| - | - | 1436 | 262.1 | - | - | 0 | - |
| - | - | 5.39E+04 | 263.1 | - | - | 0 | - |
| - | - | 5990 | 263.1 | - | - | 0 | - |
| - | - | 5898 | 264.1 | - | - | 0 | - |
| - | - | 4897 | 264.1 | - | - | 0 | - |
| - | - | 2700 | 265.1 | - | - | 0 | - |
| - | - | 765.8 | 265.1 | - | - | 0 | - |
| - | - | 1123 | 266.1 | - | - | 0 | - |
| - | - | 923 | 266.2 | - | - | 0 | - |
| - | - | 3883 | 267.1 | - | - | 0 | - |
| - | - | 1445 | 268.1 | - | - | 0 | - |
| - | - | 915.7 | 268.1 | - | - | 0 | - |
| - | - | 2450 | 268.1 | - | - | 0 | - |
| - | - | 1599 | 269.1 | - | - | 0 | - |
| - | - | 1150 | 271.1 | - | - | 0 | - |
| - | - | 875.2 | 273.1 | - | - | 0 | - |
| - | - | 1552 | 273.1 | - | - | 0 | - |
| - | - | 1.466E+04 | 274.1 | - | - | 0 | - |
| - | - | 1259 | 274.2 | - | - | 0 | - |
| - | - | 1807 | 275.1 | - | - | 0 | - |
| - | - | 1067 | 278 | - | - | 0 | - |
| - | - | 1064 | 278 | - | - | 0 | - |
| - | - | 2.131E+04 | 279.1 | - | - | 0 | - |
| - | - | 1952 | 279.1 | - | - | 0 | - |
| - | - | 3030 | 280.1 | - | - | 0 | - |
| - | - | 870.9 | 283.1 | - | - | 0 | - |
| - | - | 1223 | 283.2 | - | - | 0 | - |
| - | - | 1696 | 285.1 | - | - | 0 | - |
| - | - | 3.478E+04 | 285.1 | - | - | 0 | - |
| - | - | 4216 | 286.1 | - | - | 0 | - |
| - | - | 1156 | 287.1 | - | - | 0 | - |
| - | - | 2037 | 289.1 | - | - | 0 | - |
| - | - | 1133 | 290.1 | - | - | 0 | - |
| - | - | 2.086E+04 | 292.1 | - | - | 0 | - |
| 3 | d | 5014 | 292.2 | 0.0008749 | 2.994 | +1 | 3 |
| - | - | 2410 | 293.1 | - | - | 0 | - |
| - | - | 947.9 | 293.2 | - | - | 0 | - |
| - | - | 876.1 | 296.1 | - | - | 0 | - |
| - | - | 2.767E+04 | 296.2 | - | - | 0 | - |
| - | - | 891.3 | 297.1 | - | - | 0 | - |
| - | - | 4534 | 297.2 | - | - | 0 | - |
| - | - | 940.8 | 299.2 | - | - | 0 | - |
| - | - | 2094 | 300.1 | - | - | 0 | - |
| - | - | 1478 | 301.1 | - | - | 0 | - |
| - | - | 1134 | 302.1 | - | - | 0 | - |
| - | - | 2541 | 302.1 | - | - | 0 | - |
| - | - | 837.1 | 302.2 | - | - | 0 | - |
| - | - | 5272 | 303.1 | - | - | 0 | - |
| - | - | 914.5 | 306.1 | - | - | 0 | - |
| - | - | 719.9 | 307.1 | - | - | 0 | - |
| - | - | 1513 | 307.1 | - | - | 0 | - |
| - | - | 3125 | 310.2 | - | - | 0 | - |
| - | - | 2129 | 313.1 | - | - | 0 | - |
| - | - | 2669 | 314.1 | - | - | 0 | - |
| - | - | 1982 | 315.1 | - | - | 0 | - |
| 3 | a | 7854 | 318.1 | 0.0008278 | 2.602 | +1 | 3 |
| - | - | 1689 | 319.1 | - | - | 0 | - |
| - | - | 3065 | 320.1 | - | - | 0 | - |
| - | - | 1.102E+04 | 323.1 | - | - | 0 | - |
| - | - | 2066 | 324.1 | - | - | 0 | - |
| - | - | 6.16E+04 | 324.1 | - | - | 0 | - |
| - | - | 9130 | 325.1 | - | - | 0 | - |
| - | - | 2402 | 327.1 | - | - | 0 | - |
| - | - | 4849 | 328.1 | - | - | 0 | - |
| - | - | 1983 | 328.1 | - | - | 0 | - |
| - | - | 1611 | 329.1 | - | - | 0 | - |
| - | - | 2553 | 330.1 | - | - | 0 | - |
| - | - | 1373 | 331.1 | - | - | 0 | - |
| - | - | 4138 | 332.1 | - | - | 0 | - |
| - | - | 2002 | 333.1 | - | - | 0 | - |
| - | - | 1879 | 334.1 | - | - | 0 | - |
| - | - | 2220 | 334.1 | - | - | 0 | - |
| - | - | 1477 | 335.2 | - | - | 0 | - |
| 3 | a | 1255 | 336.2 | 0.0005781 | 1.72 | +1 | 3 |
| - | - | 4200 | 338.1 | - | - | 0 | - |
| - | - | 1347 | 339.1 | - | - | 0 | - |
| - | - | 6747 | 339.2 | - | - | 0 | - |
| 8 | y | 3.164E+05 | 342.2 | 0.0009165 | 2.679 | +1 | 2 |
| - | - | 5.928E+04 | 343.2 | - | - | 0 | - |
| - | - | 5625 | 344.2 | - | - | 0 | - |
| 3 | b | 1.616E+04 | 346.1 | 0.0008168 | 2.36 | +1 | 3 |
| - | - | 2660 | 347.1 | - | - | 0 | - |
| - | - | 7438 | 348.1 | - | - | 0 | - |
| - | - | 1.786E+04 | 350.1 | - | - | 0 | - |
| - | - | 4759 | 351.1 | - | - | 0 | - |
| - | - | 6267 | 352.1 | - | - | 0 | - |
| - | - | 1214 | 353.1 | - | - | 0 | - |
| - | - | 1154 | 354.2 | - | - | 0 | - |
| - | - | 1.687E+04 | 356.1 | - | - | 0 | - |
| 6 | b | 2611 | 357.1 | 0.0002134 | 0.5974 | +2 | 6 |
| - | - | 1446 | 358.1 | - | - | 0 | - |
| - | - | 1615 | 359.1 | - | - | 0 | - |
| - | - | 5005 | 360.1 | - | - | 0 | - |
| - | - | 914 | 360.1 | - | - | 0 | - |
| - | - | 962.3 | 361.1 | - | - | 0 | - |
| 5 | y | 2732 | 362.1 | 0.004721 | 13.04 | +2 | 5 |
| - | - | 1283 | 362.7 | - | - | 0 | - |
| 3 | b | 7.65E+04 | 364.2 | 0.0008112 | 2.228 | +1 | 3 |
| - | - | 1.455E+04 | 365.2 | - | - | 0 | - |
| - | - | 8593 | 366.1 | - | - | 0 | - |
| - | - | 1083 | 366.2 | - | - | 0 | - |
| - | - | 1028 | 367.1 | - | - | 0 | - |
| - | - | 3859 | 367.2 | - | - | 0 | - |
| - | - | 2587 | 371.1 | - | - | 0 | - |
| - | - | 5006 | 374.2 | - | - | 0 | - |
| - | - | 9233 | 376.1 | - | - | 0 | - |
| - | - | 2403 | 378.1 | - | - | 0 | - |
| - | - | 4.191E+04 | 378.1 | - | - | 0 | - |
| - | - | 7957 | 379.1 | - | - | 0 | - |
| - | - | 2785 | 380.1 | - | - | 0 | - |
| - | - | 947.1 | 381.1 | - | - | 0 | - |
| - | - | 7597 | 382.1 | - | - | 0 | - |
| - | - | 2036 | 383.1 | - | - | 0 | - |
| - | - | 1897 | 384.1 | - | - | 0 | - |
| - | - | 2403 | 385.1 | - | - | 0 | - |
| - | - | 1589 | 388.2 | - | - | 0 | - |
| - | - | 759.3 | 389.1 | - | - | 0 | - |
| - | - | 5691 | 390.2 | - | - | 0 | - |
| - | - | 1352 | 391.2 | - | - | 0 | - |
| - | - | 1049 | 392.2 | - | - | 0 | - |
| - | - | 5.327E+04 | 394.1 | - | - | 0 | - |
| - | - | 1.098E+04 | 395.1 | - | - | 0 | - |
| - | - | 1376 | 396.1 | - | - | 0 | - |
| - | - | 2721 | 397.2 | - | - | 0 | - |
| - | - | 897.9 | 398.1 | - | - | 0 | - |
| - | - | 1187 | 398.2 | - | - | 0 | - |
| - | - | 1929 | 399.2 | - | - | 0 | - |
| - | - | 1212 | 401.2 | - | - | 0 | - |
| - | - | 1502 | 404.1 | - | - | 0 | - |
| - | - | 1829 | 404.1 | - | - | 0 | - |
| - | - | 1570 | 406.1 | - | - | 0 | - |
| - | - | 2104 | 406.2 | - | - | 0 | - |
| - | - | 4915 | 407.2 | - | - | 0 | - |
| - | - | 1.302E+04 | 408.2 | - | - | 0 | - |
| - | - | 3536 | 409.2 | - | - | 0 | - |
| - | - | 838 | 409.2 | - | - | 0 | - |
| - | - | 1813 | 410.1 | - | - | 0 | - |
| - | - | 1315 | 410.2 | - | - | 0 | - |
| - | - | 2637 | 415.2 | - | - | 0 | - |
| - | - | 853 | 417.2 | - | - | 0 | - |
| - | - | 2158 | 419.2 | - | - | 0 | - |
| - | - | 1282 | 420.2 | - | - | 0 | - |
| - | - | 5.024E+04 | 425.2 | - | - | 0 | - |
| - | - | 1.101E+04 | 426.2 | - | - | 0 | - |
| - | - | 8383 | 427.2 | - | - | 0 | - |
| - | - | 1774 | 428.2 | - | - | 0 | - |
| - | - | 1068 | 429.2 | - | - | 0 | - |
| - | - | 1167 | 430.1 | - | - | 0 | - |
| - | - | 938 | 431.1 | - | - | 0 | - |
| - | - | 931.3 | 431.2 | - | - | 0 | - |
| - | - | 1809 | 432.1 | - | - | 0 | - |
| - | - | 8106 | 433.2 | - | - | 0 | - |
| - | - | 973.7 | 434.2 | - | - | 0 | - |
| - | - | 1504 | 435.2 | - | - | 0 | - |
| - | - | 2363 | 437.2 | - | - | 0 | - |
| - | - | 910.1 | 438.2 | - | - | 0 | - |
| - | - | 2481 | 442.2 | - | - | 0 | - |
| - | - | 1843 | 443.2 | - | - | 0 | - |
| - | - | 1765 | 443.2 | - | - | 0 | - |
| - | - | 871.6 | 446.2 | - | - | 0 | - |
| - | - | 6363 | 447.2 | - | - | 0 | - |
| - | - | 2050 | 449.1 | - | - | 0 | - |
| - | - | 2741 | 449.2 | - | - | 0 | - |
| - | - | 1152 | 450.2 | - | - | 0 | - |
| - | - | 3278 | 454.2 | - | - | 0 | - |
| - | - | 1.081E+04 | 455.2 | - | - | 0 | - |
| - | - | 1770 | 456.2 | - | - | 0 | - |
| - | - | 1881 | 458.2 | - | - | 0 | - |
| - | - | 1130 | 459.1 | - | - | 0 | - |
| 4 | b | 1.197E+04 | 461.2 | 0.0006682 | 1.449 | +1 | 4 |
| - | - | 4093 | 461.2 | - | - | 0 | - |
| - | - | 3766 | 462.2 | - | - | 0 | - |
| - | - | 1639 | 462.2 | - | - | 0 | - |
| - | - | 4.128E+04 | 465.2 | - | - | 0 | - |
| - | - | 1482 | 466.1 | - | - | 0 | - |
| - | - | 9517 | 466.2 | - | - | 0 | - |
| - | - | 946.5 | 467.2 | - | - | 0 | - |
| - | - | 1776 | 469.2 | - | - | 0 | - |
| - | - | 888.5 | 470.2 | - | - | 0 | - |
| - | - | 6042 | 471.2 | - | - | 0 | - |
| - | - | 1233 | 472.2 | - | - | 0 | - |
| - | - | 3577 | 473.2 | - | - | 0 | - |
| - | - | 1237 | 475.1 | - | - | 0 | - |
| - | - | 1102 | 476.2 | - | - | 0 | - |
| 3 | y | 7626 | 477.2 | 0.006689 | 14.02 | +2 | 7 |
| - | - | 2134 | 477.7 | - | - | 0 | - |
| - | - | 2217 | 478.2 | - | - | 0 | - |
| - | - | 2039 | 478.2 | - | - | 0 | - |
| 4 | b | 4.211E+04 | 479.2 | 0.001273 | 2.656 | +1 | 4 |
| - | - | 1.008E+04 | 480.2 | - | - | 0 | - |
| - | - | 1179 | 481.2 | - | - | 0 | - |
| - | - | 2739 | 482.2 | - | - | 0 | - |
| - | - | 958.2 | 483.2 | - | - | 0 | - |
| - | - | 6749 | 486.2 | - | - | 0 | - |
| - | - | 1199 | 487.2 | - | - | 0 | - |
| 7 | y | 7.481E+04 | 489.2 | 0.006259 | 12.8 | +1 | 3 |
| - | - | 2.084E+04 | 490.2 | - | - | 0 | - |
| - | - | 1.053E+04 | 491.2 | - | - | 0 | - |
| - | - | 2938 | 492.2 | - | - | 0 | - |
| - | - | 971.8 | 493.2 | - | - | 0 | - |
| - | - | 5530 | 495.2 | - | - | 0 | - |
| - | - | 2309 | 496.2 | - | - | 0 | - |
| - | - | 3.618E+04 | 496.2 | - | - | 0 | - |
| - | - | 6676 | 497.2 | - | - | 0 | - |
| - | - | 9366 | 497.2 | - | - | 0 | - |
| - | - | 2003 | 498.2 | - | - | 0 | - |
| - | - | 1214 | 498.2 | - | - | 0 | - |
| 8 | b | 3376 | 499.2 | 0.002726 | 5.46 | +2 | 8 |
| - | - | 1022 | 499.7 | - | - | 0 | - |
| - | - | 1286 | 502.2 | - | - | 0 | - |
| - | - | 1899 | 506.2 | - | - | 0 | - |
| - | - | 1628 | 511.2 | - | - | 0 | - |
| - | - | 6676 | 513.2 | - | - | 0 | - |
| - | - | 1931 | 514.2 | - | - | 0 | - |
| - | - | 1446 | 516.2 | - | - | 0 | - |
| - | - | 2632 | 518.7 | - | - | 0 | - |
| - | - | 1.16E+04 | 519.2 | - | - | 0 | - |
| - | - | 6822 | 520.2 | - | - | 0 | - |
| - | - | 2406 | 521.2 | - | - | 0 | - |
| - | - | 1424 | 522.2 | - | - | 0 | - |
| - | - | 1971 | 523.2 | - | - | 0 | - |
| - | - | 865.6 | 525.2 | - | - | 0 | - |
| - | - | 1176 | 525.2 | - | - | 0 | - |
| - | - | 1556 | 529.2 | - | - | 0 | - |
| - | - | 3209 | 530.2 | - | - | 0 | - |
| - | - | 1358 | 531.2 | - | - | 0 | - |
| - | - | 854.8 | 532.2 | - | - | 0 | - |
| - | - | 994.9 | 533.2 | - | - | 0 | - |
| - | - | 1014 | 535.3 | - | - | 0 | - |
| - | - | 1016 | 536.2 | - | - | 0 | - |
| - | - | 3250 | 537.2 | - | - | 0 | - |
| - | - | 2329 | 538.2 | - | - | 0 | - |
| - | - | 2384 | 541.2 | - | - | 0 | - |
| 2 | y | 5630 | 541.7 | 0.003174 | 5.86 | +2 | 8 |
| - | - | 6189 | 542.2 | - | - | 0 | - |
| - | - | 2030 | 542.7 | - | - | 0 | - |
| - | - | 2125 | 543.2 | - | - | 0 | - |
| - | - | 1073 | 544.2 | - | - | 0 | - |
| - | - | 1.249E+04 | 548.2 | - | - | 0 | - |
| - | - | 4172 | 549.2 | - | - | 0 | - |
| 2 | y | 3.93E+04 | 550.7 | 0.003568 | 6.48 | +2 | 8 |
| - | - | 2.756E+04 | 551.2 | - | - | 0 | - |
| - | - | 1.052E+04 | 551.7 | - | - | 0 | - |
| - | - | 3398 | 552.2 | - | - | 0 | - |
| - | - | 824.2 | 553.2 | - | - | 0 | - |
| - | - | 1193 | 555.7 | - | - | 0 | - |
| - | - | 6603 | 557.2 | - | - | 0 | - |
| - | - | 2580 | 558.2 | - | - | 0 | - |
| 6 | y | 1.287E+05 | 560.2 | 0.00595 | 10.62 | +1 | 4 |
| - | - | 2456 | 560.7 | - | - | 0 | - |
| - | - | 3.694E+04 | 561.2 | - | - | 0 | - |
| - | - | 8376 | 562.2 | - | - | 0 | - |
| - | - | 811 | 563.2 | - | - | 0 | - |
| - | - | 7590 | 569.2 | - | - | 0 | - |
| - | - | 7508 | 569.7 | - | - | 0 | - |
| - | - | 1.076E+04 | 570.2 | - | - | 0 | - |
| - | - | 1073 | 570.7 | - | - | 0 | - |
| - | - | 3311 | 571.2 | - | - | 0 | - |
| - | - | 982.2 | 574.7 | - | - | 0 | - |
| - | - | 6543 | 578.2 | - | - | 0 | - |
| - | - | 2676 | 578.7 | - | - | 0 | - |
| - | - | 1816 | 579.2 | - | - | 0 | - |
| - | - | 3450 | 583.2 | - | - | 0 | - |
| - | - | 2543 | 583.7 | - | - | 0 | - |
| - | - | 1991 | 584.2 | - | - | 0 | - |
| - | - | 2155 | 588.2 | - | - | 0 | - |
| 0 | Precursor | 2.049E+04 | 592.2 | 0.003688 | 6.228 | +2 | -1 |
| - | - | 1.566E+04 | 592.7 | - | - | 0 | - |
| - | - | 7889 | 593.2 | - | - | 0 | - |
| - | - | 2025 | 593.7 | - | - | 0 | - |
| - | - | 3013 | 594.2 | - | - | 0 | - |
| - | - | 1233 | 596.2 | - | - | 0 | - |
| - | - | 7803 | 596.2 | - | - | 0 | - |
| - | - | 3462 | 597.2 | - | - | 0 | - |
| - | - | 1864 | 598.2 | - | - | 0 | - |
| 0 | Precursor | 5.926E+04 | 601.2 | 0.003777 | 6.282 | +2 | -1 |
| - | - | 3.875E+04 | 601.7 | - | - | 0 | - |
| - | - | 1.883E+04 | 602.2 | - | - | 0 | - |
| - | - | 1237 | 602.3 | - | - | 0 | - |
| - | - | 2594 | 602.7 | - | - | 0 | - |
| - | - | 1106 | 605.2 | - | - | 0 | - |
| - | - | 1.045E+04 | 606.2 | - | - | 0 | - |
| - | - | 3691 | 607.2 | - | - | 0 | - |
| - | - | 2.14E+04 | 612.2 | - | - | 0 | - |
| - | - | 959.5 | 612.4 | - | - | 0 | - |
| - | - | 6548 | 613.2 | - | - | 0 | - |
| - | - | 1800 | 613.3 | - | - | 0 | - |
| - | - | 1.194E+04 | 614.2 | - | - | 0 | - |
| - | - | 4013 | 615.3 | - | - | 0 | - |
| - | - | 956.8 | 623.2 | - | - | 0 | - |
| 5 | b | 8819 | 624.2 | 0.002037 | 3.263 | +1 | 5 |
| - | - | 3843 | 625.2 | - | - | 0 | - |
| - | - | 5014 | 631.2 | - | - | 0 | - |
| - | - | 1244 | 632.3 | - | - | 0 | - |
| - | - | 1.509E+04 | 634.2 | - | - | 0 | - |
| - | - | 3841 | 635.2 | - | - | 0 | - |
| - | - | 3210 | 639.3 | - | - | 0 | - |
| - | - | 5461 | 640.2 | - | - | 0 | - |
| - | - | 2172 | 641.2 | - | - | 0 | - |
| - | - | 1543 | 641.3 | - | - | 0 | - |
| 5 | b | 1.035E+04 | 642.2 | 0.001421 | 2.212 | +1 | 5 |
| - | - | 2494 | 643.2 | - | - | 0 | - |
| - | - | 1023 | 649.3 | - | - | 0 | - |
| - | - | 3368 | 652.2 | - | - | 0 | - |
| - | - | 951.3 | 653.2 | - | - | 0 | - |
| - | - | 903.8 | 654.7 | - | - | 0 | - |
| - | - | 2.903E+04 | 657.3 | - | - | 0 | - |
| - | - | 8206 | 658.3 | - | - | 0 | - |
| - | - | 4.596E+04 | 659.3 | - | - | 0 | - |
| - | - | 1.683E+04 | 660.3 | - | - | 0 | - |
| - | - | 4315 | 661.3 | - | - | 0 | - |
| - | - | 888.7 | 662.2 | - | - | 0 | - |
| - | - | 1060 | 663.3 | - | - | 0 | - |
| - | - | 960.9 | 663.8 | - | - | 0 | - |
| - | - | 4282 | 667.3 | - | - | 0 | - |
| - | - | 2574 | 668.3 | - | - | 0 | - |
| - | - | 1939 | 669.3 | - | - | 0 | - |
| - | - | 6093 | 676.3 | - | - | 0 | - |
| - | - | 3853 | 677.3 | - | - | 0 | - |
| - | - | 1182 | 678.3 | - | - | 0 | - |
| - | - | 3.768E+04 | 685.3 | - | - | 0 | - |
| - | - | 1.427E+04 | 686.3 | - | - | 0 | - |
| - | - | 1643 | 687.3 | - | - | 0 | - |
| 6 | b | 9091 | 695.3 | 0.001239 | 1.782 | +1 | 6 |
| - | - | 3661 | 696.3 | - | - | 0 | - |
| - | - | 1052 | 700.8 | - | - | 0 | - |
| - | - | 1522 | 702.3 | - | - | 0 | - |
| - | - | 1.133E+04 | 703.3 | - | - | 0 | - |
| - | - | 3365 | 704.3 | - | - | 0 | - |
| - | - | 4484 | 705.3 | - | - | 0 | - |
| - | - | 1597 | 706.3 | - | - | 0 | - |
| - | - | 1449 | 712.3 | - | - | 0 | - |
| 6 | b | 4822 | 713.3 | 0.00105 | 1.473 | +1 | 6 |
| - | - | 1680 | 714.3 | - | - | 0 | - |
| - | - | 1678 | 719.4 | - | - | 0 | - |
| - | - | 3.465E+04 | 721.3 | - | - | 0 | - |
| - | - | 809.6 | 721.8 | - | - | 0 | - |
| - | - | 1.286E+04 | 722.3 | - | - | 0 | - |
| 5 | y | 1.179E+05 | 723.3 | 0.005854 | 8.093 | +1 | 5 |
| - | - | 4.906E+04 | 724.3 | - | - | 0 | - |
| - | - | 1.507E+04 | 725.3 | - | - | 0 | - |
| - | - | 1464 | 725.8 | - | - | 0 | - |
| - | - | 3752 | 726.3 | - | - | 0 | - |
| - | - | 848.2 | 727.8 | - | - | 0 | - |
| - | - | 1455 | 728.3 | - | - | 0 | - |
| - | - | 877.6 | 729.3 | - | - | 0 | - |
| - | - | 3899 | 731.2 | - | - | 0 | - |
| - | - | 2286 | 732.2 | - | - | 0 | - |
| - | - | 1924 | 733.3 | - | - | 0 | - |
| - | - | 924.2 | 734.8 | - | - | 0 | - |
| - | - | 1052 | 735.8 | - | - | 0 | - |
| - | - | 1610 | 739.3 | - | - | 0 | - |
| - | - | 888.4 | 742.3 | - | - | 0 | - |
| - | - | 1731 | 746.3 | - | - | 0 | - |
| - | - | 1285 | 747.3 | - | - | 0 | - |
| - | - | 1086 | 747.8 | - | - | 0 | - |
| - | - | 2272 | 748.3 | - | - | 0 | - |
| - | - | 5.429E+04 | 749.3 | - | - | 0 | - |
| - | - | 1.972E+04 | 750.3 | - | - | 0 | - |
| - | - | 5816 | 751.3 | - | - | 0 | - |
| - | - | 5139 | 756.3 | - | - | 0 | - |
| - | - | 2259 | 756.8 | - | - | 0 | - |
| - | - | 4747 | 757.3 | - | - | 0 | - |
| - | - | 1033 | 758.3 | - | - | 0 | - |
| - | - | 1294 | 759.3 | - | - | 0 | - |
| - | - | 1028 | 760.3 | - | - | 0 | - |
| - | - | 1.28E+04 | 767.3 | - | - | 0 | - |
| - | - | 4197 | 768.3 | - | - | 0 | - |
| - | - | 1379 | 769.3 | - | - | 0 | - |
| - | - | 4.646E+04 | 774.3 | - | - | 0 | - |
| - | - | 2.013E+04 | 775.3 | - | - | 0 | - |
| - | - | 5502 | 776.3 | - | - | 0 | - |
| - | - | 959.7 | 777.3 | - | - | 0 | - |
| - | - | 1868 | 778.3 | - | - | 0 | - |
| - | - | 2529 | 792.3 | - | - | 0 | - |
| - | - | 1524 | 796.3 | - | - | 0 | - |
| - | - | 1398 | 803.3 | - | - | 0 | - |
| - | - | 2173 | 804.3 | - | - | 0 | - |
| - | - | 1312 | 804.4 | - | - | 0 | - |
| - | - | 1492 | 805.3 | - | - | 0 | - |
| - | - | 1628 | 810.3 | - | - | 0 | - |
| - | - | 1164 | 814.3 | - | - | 0 | - |
| - | - | 2633 | 818.3 | - | - | 0 | - |
| 4 | y | 6215 | 820.3 | 0.007054 | 8.599 | +1 | 6 |
| - | - | 3480 | 821.3 | - | - | 0 | - |
| - | - | 986.8 | 822.3 | - | - | 0 | - |
| - | - | 1081 | 827.3 | - | - | 0 | - |
| - | - | 3755 | 832.3 | - | - | 0 | - |
| - | - | 2085 | 833.3 | - | - | 0 | - |
| 4 | y | 1.4E+05 | 838.3 | 0.006193 | 7.388 | +1 | 6 |
| - | - | 6.677E+04 | 839.3 | - | - | 0 | - |
| - | - | 2.288E+04 | 840.3 | - | - | 0 | - |
| - | - | 2323 | 841.3 | - | - | 0 | - |
| 7 | b | 2692 | 842.3 | 0.004446 | 5.278 | +1 | 7 |
| - | - | 4919 | 843.3 | - | - | 0 | - |
| - | - | 2719 | 844.3 | - | - | 0 | - |
| - | - | 1596 | 850.3 | - | - | 0 | - |
| - | - | 959.4 | 851.4 | - | - | 0 | - |
| - | - | 1912 | 853.3 | - | - | 0 | - |
| - | - | 1118 | 854.3 | - | - | 0 | - |
| 7 | b | 2917 | 860.3 | 0.005478 | 6.367 | +1 | 7 |
| - | - | 3511 | 862.3 | - | - | 0 | - |
| - | - | 1037 | 863.3 | - | - | 0 | - |
| - | - | 3153 | 868.3 | - | - | 0 | - |
| - | - | 978.6 | 869.3 | - | - | 0 | - |
| - | - | 2.249E+04 | 871.3 | - | - | 0 | - |
| - | - | 1.093E+04 | 872.3 | - | - | 0 | - |
| - | - | 3102 | 873.3 | - | - | 0 | - |
| - | - | 2.004E+05 | 889.3 | - | - | 0 | - |
| - | - | 9.568E+04 | 890.4 | - | - | 0 | - |
| - | - | 2.642E+04 | 891.4 | - | - | 0 | - |
| - | - | 2954 | 892.4 | - | - | 0 | - |
| - | - | 4769 | 896.3 | - | - | 0 | - |
| - | - | 1576 | 897.3 | - | - | 0 | - |
| - | - | 1309 | 898.3 | - | - | 0 | - |
| - | - | 1331 | 899.3 | - | - | 0 | - |
| - | - | 9516 | 907.3 | - | - | 0 | - |
| - | - | 4059 | 908.3 | - | - | 0 | - |
| - | - | 2404 | 909.3 | - | - | 0 | - |
| - | - | 1375 | 914.3 | - | - | 0 | - |
| - | - | 1273 | 915.3 | - | - | 0 | - |
| - | - | 1323 | 917.3 | - | - | 0 | - |
| - | - | 3350 | 918.3 | - | - | 0 | - |
| - | - | 1301 | 919.3 | - | - | 0 | - |
| - | - | 822.2 | 924.3 | - | - | 0 | - |
| - | - | 1764 | 933.4 | - | - | 0 | - |
| 3 | y | 3.812E+04 | 935.3 | 0.006234 | 6.665 | +1 | 7 |
| - | - | 2.54E+04 | 936.3 | - | - | 0 | - |
| - | - | 8626 | 937.3 | - | - | 0 | - |
| - | - | 5909 | 938.3 | - | - | 0 | - |
| - | - | 1301 | 939.3 | - | - | 0 | - |
| - | - | 1120 | 940.3 | - | - | 0 | - |
| 3 | y | 6.363E+05 | 953.3 | 0.006289 | 6.597 | +1 | 7 |
| - | - | 3.334E+05 | 954.3 | - | - | 0 | - |
| - | - | 1.212E+05 | 955.4 | - | - | 0 | - |
| - | - | 1.425E+04 | 956.4 | - | - | 0 | - |
| - | - | 4602 | 963.3 | - | - | 0 | - |
| - | - | 2018 | 964.3 | - | - | 0 | - |
| - | - | 1806 | 969.4 | - | - | 0 | - |
| - | - | 1773 | 970.4 | - | - | 0 | - |
| - | - | 1752 | 980.4 | - | - | 0 | - |
| 8 | b | 7970 | 997.4 | 0.005587 | 5.602 | +1 | 8 |
| - | - | 4168 | 998.4 | - | - | 0 | - |
| - | - | 1109 | 999.4 | - | - | 0 | - |
| - | - | 948.7 | 1009 | - | - | 0 | - |
| - | - | 2494 | 1015 | - | - | 0 | - |
| - | - | 1341 | 1016 | - | - | 0 | - |
| - | - | 3240 | 1018 | - | - | 0 | - |
| - | - | 1405 | 1019 | - | - | 0 | - |
| - | - | 1.468E+04 | 1036 | - | - | 0 | - |
| - | - | 8789 | 1037 | - | - | 0 | - |
| - | - | 2054 | 1038 | - | - | 0 | - |
| - | - | 1252 | 1046 | - | - | 0 | - |
| - | - | 903 | 1047 | - | - | 0 | - |
| 2 | y | 3918 | 1082 | 0.003249 | 3.002 | +1 | 8 |
| - | - | 2442 | 1083 | - | - | 0 | - |
| 2 | y | 3.58E+04 | 1100 | 0.005868 | 5.333 | +1 | 8 |
| - | - | 2.288E+04 | 1101 | - | - | 0 | - |
| - | - | 9240 | 1102 | - | - | 0 | - |
| - | - | 1295 | 1103 | - | - | 0 | - |
| - | - | 4962 | 1110 | - | - | 0 | - |
| - | - | 2754 | 1111 | - | - | 0 | - |
| - | - | 1391 | 1112 | - | - | 0 | - |
| - | - | 779.4 | 1241 | - | - | 0 | - |
| - | - | 975.3 | 2080 | - | - | 0 | - |
| - | - | 1035 | 3081 | - | - | 0 | - |
| - | - | 1093 | 3287 | - | - | 0 | - |

m/z Charge Intensity FragmentType MassShift Position
120.08126831054688 0 369533.38
121.03996276855469 0 1207.5842
121.07914733886719 0 1307.13
121.0845718383789 0 27553.94
121.58333587646484 0 489.36572
122.07172393798828 0 1816.6228
122.0877456665039 0 1008.4808
124.76318359375 0 511.0499
127.08699798583984 0 1512.5953
129.0704803466797 0 1445.2957
129.10267639160156 0 3941.0527
130.0655517578125 0 11624.403
131.08583068847656 0 789.0686
132.08123779296875 0 20348.256
132.10214233398438 0 539.47314
133.0612335205078 0 645.7221
133.0846405029297 0 1657.7229
134.096923828125 0 1628.8036
136.07615661621094 0 133914.56
136.08743286132812 0 4453.6353
137.0795135498047 0 9721.661
137.09075927734375 0 667.1475
138.055419921875 0 10661.192
138.06663513183594 0 13808.415
139.05889892578125 0 736.14294
140.08250427246094 0 1007.7129
141.0662384033203 0 699.9486
141.10267639160156 0 883.359
143.0457305908203 0 660.8744
143.07350158691406 0 2010.8842
143.0859832763672 0 2261.086
144.08126831054688 0 2275.7314
146.0605010986328 0 4360.7217
146.096923828125 0 2561.2817
147.09213256835938 0 701.6676
148.07635498046875 0 1456.346
148.08741760253906 0 4069.7698
148.09335327148438 0 552.5485
148.30393981933594 0 693.29504
152.08248901367188 0 694.8452
155.08175659179688 0 1964.6946
155.09326171875 0 3943.6726
156.07723999023438 0 32895.633
157.08053588867188 0 1999.7927
157.1342010498047 0 824.65845
158.09689331054688 0 56312.305
159.0561981201172 0 1150.2598
159.09217834472656 0 27573.992
159.1001739501953 0 7101.0522
160.07620239257812 0 9128.996
160.08778381347656 0 1156.4255
160.0957489013672 0 2227.4683
160.11253356933594 0 26700.236
161.07945251464844 0 1486.6498
161.1157684326172 0 2637.4126
161.71182250976562 0 605.98486
162.0917205810547 0 1040.3629
165.07012939453125 0 1580.3054
165.07762145996094 0 6783.0425
166.0537109375 0 2633.8076
166.06155395507812 0 6618.1636
166.0865020751953 0 2792.7449
167.0464324951172 0 775.9618
167.08160400390625 0 663.8125
167.09304809570312 0 4959.469
169.07640075683594 0 1899.6855
170.0605926513672 0 1172.2335
171.07687377929688 0 4927.483
172.10845947265625 0 862.3955
173.09259033203125 0 1531.2952
173.12953186035156 0 885.43854
174.06680297851562 0 4071.7725 b Ammonia loss 2
174.09210205078125 0 884.48395
175.0873565673828 0 1880.0366
175.09823608398438 0 1640.1676
175.1236114501953 0 1808.3518
176.08238220214844 0 9822.318
176.10751342773438 0 110690.33
177.1027374267578 0 74629.82
177.11129760742188 0 13197.494
178.06163024902344 0 1443.2832
178.09793090820312 0 1218.2572
178.10609436035156 0 6980.274
180.07725524902344 0 6031.501
183.14955139160156 0 1883.2098
185.0562744140625 0 9425.522
185.0924835205078 0 887.3963
186.09185791015625 0 9669.72
187.06353759765625 0 1404.5299
188.07115173339844 0 37074.05
189.07435607910156 0 4290.7837
189.0865478515625 0 999.5287
190.08741760253906 0 1639.286
191.0828857421875 0 1084.9984
191.09323120117188 0 3586.5825
191.118408203125 0 6999.119
192.10276794433594 0 2095.4119
192.1211700439453 0 620.9766
193.07276916503906 0 1324.1307
193.10882568359375 0 32978.082
194.0802001953125 0 2916.1338
194.1125030517578 0 2691.2256
195.08815002441406 0 8695.608
195.1239776611328 0 958.7778
198.08831787109375 0 710.66974
199.0720977783203 0 772.81177
201.12393188476562 0 1023.177
202.05340576171875 0 3499.8098
203.0668182373047 0 8499.061
203.09336853027344 0 3310.113
203.1183319091797 0 37262.375 a Water loss 1
204.07725524902344 0 18559.23
204.10179138183594 0 2109.0461 a Ammonia loss 1
204.12158203125 0 4948.6943
204.48049926757812 0 660.60724
205.08106994628906 0 1687.0348
205.09768676757812 0 30262.715 y 8
205.10733032226562 0 1156.7955
206.10122680664062 0 3787.3303
207.1133270263672 0 8167.493
208.07223510742188 0 4752.7104
212.1177520751953 0 1504.5682
213.05177307128906 0 811.6828
213.0876922607422 0 758.60657
213.09881591796875 0 1433.1163
215.08187866210938 0 2271.8845
216.06610107421875 0 887.8647
216.09754943847656 0 983.276
217.09768676757812 0 19376.842
218.0824432373047 0 852.27936
218.1015625 0 2325.0254
219.08033752441406 0 5949.175
219.1140594482422 0 1131.0269
220.095458984375 0 675.50903
220.1198272705078 0 1882.5734
221.10421752929688 0 88107.99
221.129150390625 0 1530951.4 a 1
222.10691833496094 0 7442.231
222.13235473632812 0 185797.19
223.1346893310547 0 9968.911
224.11740112304688 0 1163.0834
225.09957885742188 0 1030.9445
226.1554718017578 0 920.17944
228.1136932373047 0 6017.3306
229.10902404785156 0 3038.138
229.15524291992188 0 20674.908
229.16775512695312 0 1276.4375
230.1586456298828 0 3329.2817
231.06179809570312 0 23359.87
231.08743286132812 0 806.5371 b Water loss 3
231.11337280273438 0 14821.493 b Water loss 1
232.0648651123047 0 1952.6062
232.11656188964844 0 1685.351
233.0924530029297 0 1781.8008
233.10464477539062 0 836.97797
235.1083526611328 0 40535.35
236.1116943359375 0 5356.4795
237.0895233154297 0 713.07947
238.13145446777344 0 1745.8398
239.10044860839844 0 1115.4666
239.11441040039062 0 10313.442
240.0814666748047 0 786.6572
240.11676025390625 0 1277.3099
245.09237670898438 0 1302.2407 y 6
245.129638671875 0 1930.8486
246.13571166992188 0 2952.579
247.10647583007812 0 1636.8016
247.1199493408203 0 949.3331
247.1453094482422 0 1016.357
248.08912658691406 0 959.03503
248.15127563476562 0 3423.3188
249.12403869628906 0 268155.62 b 1
250.1273193359375 0 33727.574
251.1032257080078 0 28947.514
251.12966918945312 0 4736.941
252.10678100585938 0 4438.872
252.13409423828125 0 874.75214
253.0929718017578 0 1236.4319
253.1184844970703 0 1092.7721
256.10797119140625 0 1804.8723
257.1069641113281 0 16655.578
258.1100158691406 0 2213.5793
258.1242370605469 0 1052.5659
261.0874328613281 0 3149.185
261.112060546875 0 1021.34625
262.11883544921875 0 1435.9072
263.1031494140625 0 53903.137
263.1294250488281 0 5990.3394
264.1065979003906 0 5898.2134
264.1459045410156 0 4897.021
265.1182556152344 0 2700.2832
265.14788818359375 0 765.82745
266.1229248046875 0 1123.4323
266.15118408203125 0 923.0448
267.0918273925781 0 3882.6865
268.0762634277344 0 1444.7598
268.09332275390625 0 915.6614
268.1407165527344 0 2450.2798
269.07757568359375 0 1598.798
271.1065368652344 0 1149.6643
273.0872497558594 0 875.21796
273.123779296875 0 1552.0111
274.13037109375 0 14663.498
274.156005859375 0 1258.5438
275.1146240234375 0 1806.707
278.0209045410156 0 1067.2247
278.0382995605469 0 1064.3228
279.0982666015625 0 21312.654
279.125244140625 0 1951.6755
280.1018371582031 0 3029.918
283.1063537597656 0 870.8762
283.1531066894531 0 1223.4875
285.0841064453125 0 1696.1274
285.102294921875 0 34776.36
286.1051330566406 0 4216.1016
287.0995788574219 0 1155.8438
289.082275390625 0 2037.4564
290.1264343261719 0 1133.0947
292.14129638671875 0 20863.588
292.16644287109375 0 5013.9966 d 2
293.14404296875 0 2409.54
293.1700744628906 0 947.91785
296.0880126953125 0 876.11975
296.15142822265625 0 27672.64
297.1358337402344 0 891.31415
297.1543884277344 0 4533.5146
299.15228271484375 0 940.8059
300.1352844238281 0 2093.5005
301.11822509765625 0 1477.5521
302.09783935546875 0 1133.7694
302.1151123046875 0 2540.864
302.1516418457031 0 837.1236
303.1136474609375 0 5271.7695
306.14306640625 0 914.52106
307.0963439941406 0 719.8869
307.1200256347656 0 1513.0942
310.1520080566406 0 3124.6387
313.09759521484375 0 2129.0994
314.09954833984375 0 2668.8567
315.09832763671875 0 1981.8035
318.1456604003906 0 7854.228 a Water loss 2
319.1490783691406 0 1688.895
320.13543701171875 0 3065.0176
323.099609375 0 11018.823
324.1020812988281 0 2065.5024
324.1464538574219 0 61604.68
325.1490173339844 0 9129.726
327.1454772949219 0 2402.0051
328.12969970703125 0 4849.2134
328.1479797363281 0 1982.6252
329.13348388671875 0 1610.8302
330.1103820800781 0 2553.0107
331.0937194824219 0 1373.278
332.12518310546875 0 4137.6934
333.10894775390625 0 2001.7806
334.1077575683594 0 1878.8301
334.1322021484375 0 2219.5613
335.1708679199219 0 1477.1292
336.1559753417969 0 1254.5404 a 2
338.1290283203125 0 4199.815
339.1321716308594 0 1346.6335
339.1714782714844 0 6746.824
342.156982421875 0 316370.22 y 7
343.1600646972656 0 59281.637
344.1623229980469 0 5625.2837
346.14056396484375 0 16157.04 b Water loss 2
347.1429748535156 0 2660.439
348.11968994140625 0 7437.813
350.13543701171875 0 17856.102
351.1390075683594 0 4758.9644
352.1410827636719 0 6267.3076
353.1467590332031 0 1213.8318
354.1556091308594 0 1153.541
356.13946533203125 0 16870.473
357.1427001953125 0 2610.7542 b 5
358.10333251953125 0 1445.6514
359.089111328125 0 1615.2916
360.11968994140625 0 5005.278
360.1441345214844 0 913.9657
361.1220397949219 0 962.3293
362.15185546875 0 2731.8896 y 4
362.65325927734375 0 1283.485
364.151123046875 0 76503.54 b 2
365.1541748046875 0 14549.737
366.1302795410156 0 8593.221
366.1549987792969 0 1083.0033
367.1327209472656 0 1028.1401
367.1661376953125 0 3858.5066
371.13690185546875 0 2586.782
374.15008544921875 0 5006.3804
376.1148681640625 0 9233.423
378.1004638671875 0 2402.787
378.13043212890625 0 41906.812
379.1329040527344 0 7956.971
380.1398010253906 0 2784.6562
381.14642333984375 0 947.1171
382.14398193359375 0 7596.799
383.1474304199219 0 2035.9601
384.13580322265625 0 1897.1499
385.13751220703125 0 2402.5125
388.1507263183594 0 1589.0054
389.11767578125 0 759.3144
390.1572265625 0 5690.772
391.15838623046875 0 1352.0978
392.17352294921875 0 1049.3707
394.1255187988281 0 53269.188
395.1285400390625 0 10979.697
396.13116455078125 0 1375.9882
397.1633605957031 0 2721.1553
398.1398620605469 0 897.91064
398.17071533203125 0 1186.6538
399.1749267578125 0 1929.0332
401.1942443847656 0 1211.9545
404.112060546875 0 1501.6896
404.14599609375 0 1828.7499
406.1242980957031 0 1569.6888
406.1534729003906 0 2104.084
407.1837463378906 0 4915.261
408.16766357421875 0 13019.779
409.1710510253906 0 3535.6943
409.1977844238281 0 837.9531
410.1404113769531 0 1812.8235
410.1813659667969 0 1315.3694
415.1625061035156 0 2636.865
417.1740417480469 0 852.9834
419.15643310546875 0 2157.5671
420.16363525390625 0 1282.3754
425.1943054199219 0 50237.66
426.1969299316406 0 11010.228
427.208984375 0 8382.899
428.213623046875 0 1774.469
429.1896667480469 0 1068.0969
430.1252136230469 0 1166.7334
431.1240234375 0 937.98706
431.15643310546875 0 931.3225
432.1409606933594 0 1808.9692
433.1726379394531 0 8106.2856
434.17864990234375 0 973.6549
435.1795349121094 0 1504.2036
437.16851806640625 0 2362.804
438.1750793457031 0 910.119
442.1737365722656 0 2481.106
443.15625 0 1843.137
443.1873779296875 0 1764.5148
446.2175598144531 0 871.6047
447.15216064453125 0 6363.314
449.13446044921875 0 2050.3755
449.16802978515625 0 2741.459
450.2285461425781 0 1152.1323
454.1971740722656 0 3278.291
455.2046203613281 0 10811.344
456.2062683105469 0 1769.8401
458.168212890625 0 1880.6782
459.14984130859375 0 1129.992
461.1673583984375 0 11967.915 b Water loss 3
461.1963195800781 0 4093.338
462.169921875 0 3765.7307
462.20166015625 0 1639.2656
465.1628112792969 0 41278.234
466.1289367675781 0 1481.5679
466.16607666015625 0 9517.224
467.1696472167969 0 946.54297
469.20831298828125 0 1775.8745
470.21221923828125 0 888.49384
471.18231201171875 0 6041.95
472.1827087402344 0 1233.26
473.2152404785156 0 3576.5654
475.1448974609375 0 1236.7594
476.18682861328125 0 1102.4692
477.1673889160156 0 7625.798 y 2
477.679443359375 0 2134.4534
478.1668701171875 0 2216.8718
478.221435546875 0 2039.3588
479.17852783203125 0 42109.875 b 3
480.1812744140625 0 10079.487
481.1817321777344 0 1178.7336
482.1903076171875 0 2738.9888
483.19970703125 0 958.2207
486.1637878417969 0 6749.3257
487.16259765625 0 1199.3423
489.19281005859375 0 74809.32 y 6
490.1950378417969 0 20842.783
491.2054443359375 0 10532.051
492.2101135253906 0 2937.9282
493.2091979980469 0 971.8331
495.1867980957031 0 5530.059
496.1894226074219 0 2309.069
496.2315673828125 0 36176.816
497.17144775390625 0 6676.275
497.2343444824219 0 9366.469
498.17352294921875 0 2003.0073
498.23870849609375 0 1213.9717
499.1899108886719 0 3376.3655 b 7
499.6943664550781 0 1022.3287
502.19677734375 0 1286.1273
506.21728515625 0 1898.6476
511.19232177734375 0 1628.1487
513.19873046875 0 6675.7793
514.2283935546875 0 1930.519
516.2210693359375 0 1446.2681
518.7106323242188 0 2632.191
519.2039794921875 0 11599.8
520.2058715820312 0 6822.062
521.205322265625 0 2405.828
522.1897583007812 0 1424.2976
523.1834716796875 0 1971.0592
525.1611328125 0 865.58105
525.2023315429688 0 1176.2136
529.2060546875 0 1556.417
530.1907348632812 0 3208.763
531.19091796875 0 1357.9058
532.2051391601562 0 854.78754
533.2044677734375 0 994.91003
535.3050537109375 0 1014.20654
536.2176513671875 0 1015.9198
537.214111328125 0 3250.3013
538.2188110351562 0 2329.1519
541.1937255859375 0 2383.6445
541.7061767578125 0 5630.32 y Water loss 1
542.23388671875 0 6189.2827
542.70849609375 0 2030.2551
543.236328125 0 2124.8123
544.2117309570312 0 1072.816
548.2002563476562 0 12491.029
549.2032470703125 0 4172.1953
550.7118530273438 0 39302.68 y 1
551.2133178710938 0 27559.225
551.713623046875 0 10524.197
552.2169799804688 0 3398.0234
553.202392578125 0 824.1542
555.7022094726562 0 1192.6975
557.200927734375 0 6603.143
558.2044067382812 0 2580.2576
560.2296142578125 0 128711.914 y 5
560.7275390625 0 2455.7017
561.2327880859375 0 36939.15
562.2308959960938 0 8375.878
563.2348022460938 0 811.01294
569.2343139648438 0 7590.2856
569.736328125 0 7507.6626
570.2300415039062 0 10755.997
570.7168579101562 0 1073.3827
571.231689453125 0 3310.9805
574.7139892578125 0 982.2282
578.2308959960938 0 6543.3623
578.7346801757812 0 2675.6045
579.2105102539062 0 1816.3082
583.2252807617188 0 3449.6099
583.7277221679688 0 2543.187
584.237060546875 0 1991.4823
588.2396850585938 0 2154.5925
592.2305297851562 0 20486.133 Precursor Water loss
592.7318725585938 0 15655.008
593.23291015625 0 7888.6533
593.7310180664062 0 2025.2734
594.1871337890625 0 3012.7734
596.1884765625 0 1233.18
596.236572265625 0 7803.232
597.231201171875 0 3462.2043
598.2265625 0 1864.394
601.2359008789062 0 59257.78 Precursor
601.7373046875 0 38745.81
602.2379760742188 0 18832.604
602.3209228515625 0 1236.6936
602.7395629882812 0 2593.6545
605.244384765625 0 1105.5488
606.2343139648438 0 10445.888
607.2372436523438 0 3691.0815
612.1981201171875 0 21397.86
612.3595581054688 0 959.4559
613.2005004882812 0 6548.3833
613.289794921875 0 1800.4471
614.2462158203125 0 11935.501
615.2507934570312 0 4013.2712
623.2496337890625 0 956.814
624.2320556640625 0 8819.381 b Water loss 4
625.2352294921875 0 3843.3455
631.2471923828125 0 5013.7607
632.2630004882812 0 1244.1031
634.2301635742188 0 15089.239
635.2333374023438 0 3840.8965
639.2542724609375 0 3209.8357
640.2406005859375 0 5461.028
641.2369384765625 0 2172.3188
641.289306640625 0 1542.7241
642.2420043945312 0 10354.506 b 4
643.245361328125 0 2493.9797
649.2626342773438 0 1022.5062
652.2424926757812 0 3368.285
653.2479248046875 0 951.3083
654.7432250976562 0 903.814
657.2640991210938 0 29034.69
658.2659301757812 0 8205.683
659.2947387695312 0 45962.508
660.2977294921875 0 16825.96
661.3001098632812 0 4314.89
662.2227783203125 0 888.6825
663.2703247070312 0 1059.9816
663.7672729492188 0 960.94336
667.2503662109375 0 4281.8354
668.2520141601562 0 2573.588
669.2732543945312 0 1939.2961
676.27001953125 0 6093.258
677.2738647460938 0 3852.7227
678.2713012695312 0 1181.7362
685.259033203125 0 37680.383
686.2620239257812 0 14270.177
687.2640991210938 0 1643.0481
695.2683715820312 0 9090.67 b Water loss 5
696.2720336914062 0 3661.4866
700.7830200195312 0 1051.925
702.2838745117188 0 1521.8458
703.2669677734375 0 11329.895
704.2689208984375 0 3364.8455
705.279296875 0 4483.6533
706.292236328125 0 1597.1151
712.305419921875 0 1448.8716
713.2787475585938 0 4821.6553 b 5
714.2803955078125 0 1680.313
719.3951416015625 0 1678.0222
721.2621459960938 0 34652.168
721.8115844726562 0 809.61865
722.2652587890625 0 12858.927
723.2928466796875 0 117859.73 y 4
724.2960815429688 0 49055.336
725.2963256835938 0 15066.804
725.8079223632812 0 1464.296
726.2982788085938 0 3751.536
727.8041381835938 0 848.18317
728.3137817382812 0 1454.5986
729.3131103515625 0 877.56964
731.2482299804688 0 3899.429
732.238525390625 0 2286.066
733.2740478515625 0 1924.1624
734.8175048828125 0 924.1877
735.816650390625 0 1051.7399
739.2874755859375 0 1609.9445
742.3193969726562 0 888.37103
746.2747802734375 0 1731.3643
747.292724609375 0 1285.4813
747.8209838867188 0 1085.9565
748.3140258789062 0 2271.9734
749.257080078125 0 54289.83
750.2601928710938 0 19720.3
751.2606201171875 0 5816.048
756.3131103515625 0 5138.903
756.825927734375 0 2258.8167
757.31640625 0 4747.445
758.3151245117188 0 1032.8044
759.2647705078125 0 1294.3291
760.2976684570312 0 1028.1599
767.267578125 0 12796.494
768.2699584960938 0 4196.7026
769.2705078125 0 1379.4155
774.3215942382812 0 46459.81
775.324462890625 0 20125.848
776.3280029296875 0 5501.7573
777.32421875 0 959.6713
778.3043823242188 0 1867.6146
792.3145141601562 0 2529.065
796.3184204101562 0 1523.8451
803.2844848632812 0 1397.7631
804.3298950195312 0 2173.0798
804.4264526367188 0 1312.0624
805.3302001953125 0 1492.4342
810.3225708007812 0 1627.7957
814.3148803710938 0 1163.7596
818.3355712890625 0 2633.1123
820.3104248046875 0 6215.3037 y Water loss 3
821.3055419921875 0 3480.0525
822.303466796875 0 986.82733
827.3479614257812 0 1080.79
832.3258666992188 0 3755.109
833.3271484375 0 2085.02
838.3201293945312 0 139990.25 y 3
839.3231201171875 0 66769.11
840.3230590820312 0 22884.54
841.3257446289062 0 2322.8323
842.3020629882812 0 2691.6553 b Water loss 6
843.33984375 0 4918.6504
844.3394775390625 0 2719.1736
850.3287353515625 0 1595.6416
851.4258422851562 0 959.3688
853.3267211914062 0 1911.5938
854.31201171875 0 1117.5721
860.3136596679688 0 2917.1064 b 6
862.3233032226562 0 3511.4058
863.3312377929688 0 1036.9823
868.3313598632812 0 3152.7627
869.328857421875 0 978.5715
871.3381958007812 0 22486.387
872.3373413085938 0 10930.817
873.3395385742188 0 3102.1072
889.3484497070312 0 200363.47
890.3511352539062 0 95679.19
891.3543090820312 0 26418.492
892.354248046875 0 2953.562
896.3260498046875 0 4768.5537
897.3248901367188 0 1575.6599
898.3250122070312 0 1308.507
899.3304443359375 0 1330.6941
907.3412475585938 0 9515.653
908.3432006835938 0 4059.1858
909.3422241210938 0 2404.067
914.3370971679688 0 1374.9932
915.3482055664062 0 1273.2108
917.3256225585938 0 1322.5947
918.3145751953125 0 3349.876
919.3212890625 0 1300.5671
924.3329467773438 0 822.2163
933.3740844726562 0 1764.0918
935.3365478515625 0 38116.48 y Water loss 2
936.3357543945312 0 25396.135
937.3359985351562 0 8625.703
938.325439453125 0 5909.4966
939.3244018554688 0 1300.7046
940.3269653320312 0 1120.2905
953.34716796875 0 636301.56 y 2
954.3497314453125 0 333384.22
955.350341796875 0 121194.43
956.3521118164062 0 14254.829
963.3311767578125 0 4601.5195
964.3370361328125 0 2017.5824
969.3761596679688 0 1806.0498
970.3794555664062 0 1773.4933
980.3678588867188 0 1752.0121
997.3726806640625 0 7969.669 b 7
998.3761596679688 0 4168.1143
999.3746948242188 0 1108.7039
1009.3821411132812 0 948.6528
1015.384521484375 0 2494.2285
1016.3828125 0 1340.649
1018.3985595703125 0 3239.5188
1019.400146484375 0 1405.0002
1036.4154052734375 0 14678.127
1037.41845703125 0 8788.928
1038.4276123046875 0 2054.348
1046.4007568359375 0 1251.6556
1047.40625 0 903.0405
1082.4019775390625 0 3918.0427 y Water loss 1
1083.403564453125 0 2441.8914
1100.4151611328125 0 35804.97 y 1
1101.4185791015625 0 22878.395
1102.4185791015625 0 9240.098
1103.4200439453125 0 1295.085
1110.3988037109375 0 4962.005
1111.403076171875 0 2753.9102
1112.40283203125 0 1391.1752
1240.9527587890625 0 779.44104
2079.87109375 0 975.27826
3080.990966796875 0 1035.0864
3287.3818359375 0 1093.2383

Spectrum Details

|  |  |
| --- | --- |
| Matched peaks? Matched peaks The total absolute number of peaks matched. Additionally in brackets the total fraction of peaks matched and the total number of peaks is shown. | 41 (6.29% of 652) |
| FDR? FDR The false discovery rate estimated for this peptide. It is calculated by matching all theoretical fragments with a non-integer shift with the raw peaks for this spectrum. This is done with 40 different shifts. The resulting percentage is the average number of annotated peaks over the number of annotated peaks with the correct spectrum. | 0.17% |
| Satellite FDR? Satellite FDR See the FDR for details on its calculation. This satellite ion specific FDR only contains the satellite ions (d/w) for I/L/J positions. | - |
| PSM Score? PSM Score The PSM Score as given by Hecklib to this annotated spectrum. It is shown with three significant figures. | 487 |

#### Spectrum 9198? Spectrum 9198 The raw spectrum of this peptide as annotated by Hecklib. The fragments are coloured according to ion type (see legend). Any peaks with a star '\*' as text can be hovered over to see the full details, first the ion type second the mass shift type. By hovering over the amino acids in the peptide or ions in the legend the corresponding peaks are highlighted. By toggling the 'Unassigned' label you can turn the background (unassigned) peaks on or off in the plot. By updating the slider in the Ion legend you can update the spectrum to only show the top X% of the peaks with labels. The top X% means any peak that is within X% of the highest intensity. By dragging in the spectrum you can zoom in to a specific part of the spectrum and use 'Zoom Out' to get back to the original zoom level. The annotation of the spectrum is based on the given sequence in the peptides file and is done with different software so inconsistencies are likely. The peaks are annotated based on the given sequence, with 20 ppm tolerance.

Copy Data

##### Spectrum 9198 (TSV)

###### Preview

```
Loading example...
```

*Click on the button to copy the data to your clipboard.*

Mz MinMz MaxIntensity Max

WidthHeightPeptide font sizePeptide stroke widthSpectrum font sizeSpectrum stroke widthCompact peptide

Ion legend

wxyz

abcd

OtherUnassignedIon^Charge^_Position_Show for top:%

TFDDYAMHW

05.48e+41.10e+51.64e+52.19e+5

Zoom Out

a^+1^_2_y^+1^_1_a^+1^_2_b^+1^_2_b^+2^_4_b^+1^_2_a^+1^_3_y^+1^_2_b^+1^_3_b^+1^_3_b^+1^_4_y^+1^_3_b^+1^_4_b^+2^_8_b^+2^_8_y^+2^_8_y^+1^_4_\*\*b^+1^_5_b^+1^_5_b^+1^_6_y^+1^_5_b^+1^_6_y^+1^_6_y^+1^_6_y^+1^_7_y^+1^_7_b^+1^_8_y^+1^_8_

0675135020252700

Fragment Matches Table

Show background peaks

| Position | Ion type | Intensity | mz Theoretical | mz Error (Th) | mz Error (ppm) | Charge | Series Number |
| --- | --- | --- | --- | --- | --- | --- | --- |
| - | - | 6.346E+04 | 120.1 | - | - | 0 | - |
| - | - | 4943 | 121.1 | - | - | 0 | - |
| - | - | 608.7 | 127.1 | - | - | 0 | - |
| - | - | 352 | 129 | - | - | 0 | - |
| - | - | 2873 | 129.1 | - | - | 0 | - |
| - | - | 2027 | 130.1 | - | - | 0 | - |
| - | - | 2811 | 132.1 | - | - | 0 | - |
| - | - | 1692 | 132.1 | - | - | 0 | - |
| - | - | 3877 | 133.1 | - | - | 0 | - |
| - | - | 539.5 | 134 | - | - | 0 | - |
| - | - | 418.4 | 134.1 | - | - | 0 | - |
| - | - | 411.7 | 134.2 | - | - | 0 | - |
| - | - | 2.992E+04 | 136.1 | - | - | 0 | - |
| - | - | 2423 | 137.1 | - | - | 0 | - |
| - | - | 2507 | 138.1 | - | - | 0 | - |
| - | - | 427.1 | 139.1 | - | - | 0 | - |
| - | - | 465.2 | 141.1 | - | - | 0 | - |
| - | - | 553.2 | 142.4 | - | - | 0 | - |
| - | - | 916.5 | 146.1 | - | - | 0 | - |
| - | - | 775.1 | 146.1 | - | - | 0 | - |
| - | - | 482.1 | 149 | - | - | 0 | - |
| - | - | 3175 | 149 | - | - | 0 | - |
| - | - | 6671 | 156.1 | - | - | 0 | - |
| - | - | 8258 | 158.1 | - | - | 0 | - |
| - | - | 4077 | 159.1 | - | - | 0 | - |
| - | - | 905.7 | 159.1 | - | - | 0 | - |
| - | - | 2023 | 160.1 | - | - | 0 | - |
| - | - | 3481 | 160.1 | - | - | 0 | - |
| - | - | 417.8 | 161.1 | - | - | 0 | - |
| - | - | 565.5 | 165.1 | - | - | 0 | - |
| - | - | 698.5 | 165.1 | - | - | 0 | - |
| - | - | 2724 | 166.1 | - | - | 0 | - |
| - | - | 3949 | 166.1 | - | - | 0 | - |
| - | - | 4600 | 167.1 | - | - | 0 | - |
| - | - | 573.4 | 167.1 | - | - | 0 | - |
| - | - | 1389 | 167.1 | - | - | 0 | - |
| - | - | 1093 | 168.1 | - | - | 0 | - |
| - | - | 620.5 | 169.1 | - | - | 0 | - |
| - | - | 904.7 | 171.1 | - | - | 0 | - |
| - | - | 437 | 171.1 | - | - | 0 | - |
| - | - | 1032 | 173.1 | - | - | 0 | - |
| - | - | 540 | 173.4 | - | - | 0 | - |
| - | - | 616 | 175.1 | - | - | 0 | - |
| - | - | 579.9 | 175.1 | - | - | 0 | - |
| - | - | 1.616E+04 | 176.1 | - | - | 0 | - |
| - | - | 1.234E+04 | 177.1 | - | - | 0 | - |
| - | - | 2977 | 177.1 | - | - | 0 | - |
| - | - | 905.8 | 178.1 | - | - | 0 | - |
| - | - | 834.3 | 180.1 | - | - | 0 | - |
| - | - | 496.6 | 181.1 | - | - | 0 | - |
| - | - | 874.6 | 182.1 | - | - | 0 | - |
| - | - | 452.4 | 182.5 | - | - | 0 | - |
| - | - | 462.1 | 183.1 | - | - | 0 | - |
| - | - | 1656 | 185.1 | - | - | 0 | - |
| - | - | 1129 | 186.1 | - | - | 0 | - |
| - | - | 528.7 | 187.1 | - | - | 0 | - |
| - | - | 421.2 | 187.1 | - | - | 0 | - |
| - | - | 777.1 | 188.1 | - | - | 0 | - |
| - | - | 7627 | 188.1 | - | - | 0 | - |
| - | - | 1096 | 189.1 | - | - | 0 | - |
| - | - | 736.5 | 189.1 | - | - | 0 | - |
| - | - | 1160 | 191.1 | - | - | 0 | - |
| - | - | 464.9 | 193.1 | - | - | 0 | - |
| - | - | 1188 | 195.1 | - | - | 0 | - |
| - | - | 1217 | 196.1 | - | - | 0 | - |
| - | - | 583.6 | 197.1 | - | - | 0 | - |
| - | - | 651.3 | 199.2 | - | - | 0 | - |
| - | - | 1004 | 201.1 | - | - | 0 | - |
| - | - | 1421 | 203.1 | - | - | 0 | - |
| 2 | a | 4906 | 203.1 | 0.0002135 | 1.051 | +1 | 2 |
| - | - | 1161 | 204.1 | - | - | 0 | - |
| 9 | y | 4672 | 205.1 | 0.0001665 | 0.8116 | +1 | 1 |
| - | - | 1929 | 207.1 | - | - | 0 | - |
| - | - | 651.5 | 208.1 | - | - | 0 | - |
| - | - | 477.2 | 211.5 | - | - | 0 | - |
| - | - | 617.8 | 212.1 | - | - | 0 | - |
| - | - | 698 | 215.1 | - | - | 0 | - |
| - | - | 2526 | 217.1 | - | - | 0 | - |
| - | - | 4749 | 221.1 | - | - | 0 | - |
| 2 | a | 2.168E+05 | 221.1 | 0.00033 | 1.492 | +1 | 2 |
| - | - | 538.8 | 221.6 | - | - | 0 | - |
| - | - | 940.7 | 222.1 | - | - | 0 | - |
| - | - | 2.81E+04 | 222.1 | - | - | 0 | - |
| - | - | 839.3 | 223.1 | - | - | 0 | - |
| - | - | 1644 | 223.1 | - | - | 0 | - |
| - | - | 688 | 224.1 | - | - | 0 | - |
| - | - | 1830 | 225 | - | - | 0 | - |
| - | - | 1202 | 226.2 | - | - | 0 | - |
| - | - | 3768 | 231.1 | - | - | 0 | - |
| 2 | b | 2111 | 231.1 | 0.0004771 | 2.064 | +1 | 2 |
| - | - | 576.1 | 231.2 | - | - | 0 | - |
| - | - | 1077 | 231.2 | - | - | 0 | - |
| - | - | 648.2 | 233.1 | - | - | 0 | - |
| - | - | 1522 | 233.2 | - | - | 0 | - |
| - | - | 6465 | 235.1 | - | - | 0 | - |
| - | - | 608.2 | 236.1 | - | - | 0 | - |
| - | - | 582.8 | 238.1 | - | - | 0 | - |
| - | - | 1.155E+04 | 239.1 | - | - | 0 | - |
| 4 | b | 2229 | 240.1 | 0.003102 | 12.92 | +2 | 4 |
| - | - | 4150 | 241.1 | - | - | 0 | - |
| - | - | 625.8 | 243.1 | - | - | 0 | - |
| - | - | 529 | 244.1 | - | - | 0 | - |
| 2 | b | 3.617E+04 | 249.1 | 0.0002731 | 1.096 | +1 | 2 |
| - | - | 3992 | 250.1 | - | - | 0 | - |
| - | - | 5188 | 251.1 | - | - | 0 | - |
| - | - | 3314 | 252.1 | - | - | 0 | - |
| - | - | 1623 | 253.1 | - | - | 0 | - |
| - | - | 791.3 | 261.1 | - | - | 0 | - |
| - | - | 7855 | 263.1 | - | - | 0 | - |
| - | - | 700.7 | 263.1 | - | - | 0 | - |
| - | - | 559.5 | 265.1 | - | - | 0 | - |
| - | - | 513.1 | 265.3 | - | - | 0 | - |
| - | - | 732.8 | 267 | - | - | 0 | - |
| - | - | 1590 | 269 | - | - | 0 | - |
| - | - | 1.629E+04 | 269.1 | - | - | 0 | - |
| - | - | 1872 | 270.1 | - | - | 0 | - |
| - | - | 3523 | 279.1 | - | - | 0 | - |
| - | - | 1667 | 281.1 | - | - | 0 | - |
| - | - | 949.9 | 283 | - | - | 0 | - |
| - | - | 2.275E+04 | 285 | - | - | 0 | - |
| - | - | 3602 | 286 | - | - | 0 | - |
| - | - | 1567 | 287.1 | - | - | 0 | - |
| - | - | 4763 | 296.2 | - | - | 0 | - |
| - | - | 641.2 | 297.2 | - | - | 0 | - |
| - | - | 1.483E+04 | 299.1 | - | - | 0 | - |
| - | - | 641.9 | 299.2 | - | - | 0 | - |
| - | - | 2556 | 300.1 | - | - | 0 | - |
| - | - | 706.9 | 301.1 | - | - | 0 | - |
| - | - | 2906 | 303 | - | - | 0 | - |
| - | - | 673.9 | 304 | - | - | 0 | - |
| - | - | 1004 | 315.2 | - | - | 0 | - |
| 3 | a | 708.1 | 318.1 | 0.0006065 | 1.906 | +1 | 3 |
| - | - | 1645 | 322.1 | - | - | 0 | - |
| - | - | 1335 | 323.1 | - | - | 0 | - |
| - | - | 632.4 | 323.2 | - | - | 0 | - |
| - | - | 1.135E+04 | 324.1 | - | - | 0 | - |
| - | - | 2259 | 325.1 | - | - | 0 | - |
| - | - | 726.1 | 328.1 | - | - | 0 | - |
| - | - | 642.3 | 332.1 | - | - | 0 | - |
| - | - | 710 | 339.2 | - | - | 0 | - |
| - | - | 5241 | 340.1 | - | - | 0 | - |
| - | - | 1492 | 341.1 | - | - | 0 | - |
| 8 | y | 5.677E+04 | 342.2 | 0.0003977 | 1.162 | +1 | 2 |
| - | - | 1.151E+04 | 343.2 | - | - | 0 | - |
| - | - | 619.7 | 344.2 | - | - | 0 | - |
| - | - | 2814 | 345 | - | - | 0 | - |
| - | - | 927.2 | 346 | - | - | 0 | - |
| 3 | b | 2065 | 346.1 | 0.0002675 | 0.7727 | +1 | 3 |
| - | - | 1774 | 348.1 | - | - | 0 | - |
| - | - | 4231 | 350.1 | - | - | 0 | - |
| - | - | 1201 | 352.1 | - | - | 0 | - |
| - | - | 5.824E+04 | 355.1 | - | - | 0 | - |
| - | - | 1.514E+04 | 356.1 | - | - | 0 | - |
| - | - | 589.1 | 357.6 | - | - | 0 | - |
| - | - | 770.1 | 358.2 | - | - | 0 | - |
| 3 | b | 8930 | 364.2 | 0.0003839 | 1.054 | +1 | 3 |
| - | - | 1581 | 365.2 | - | - | 0 | - |
| - | - | 1063 | 366.1 | - | - | 0 | - |
| - | - | 1403 | 371.1 | - | - | 0 | - |
| - | - | 5452 | 373.1 | - | - | 0 | - |
| - | - | 2008 | 374.1 | - | - | 0 | - |
| - | - | 755.9 | 376.1 | - | - | 0 | - |
| - | - | 6225 | 378.1 | - | - | 0 | - |
| - | - | 1.205E+04 | 394.1 | - | - | 0 | - |
| - | - | 2471 | 395.1 | - | - | 0 | - |
| - | - | 1636 | 401 | - | - | 0 | - |
| - | - | 2899 | 415 | - | - | 0 | - |
| - | - | 582.3 | 415.2 | - | - | 0 | - |
| - | - | 783.9 | 416 | - | - | 0 | - |
| - | - | 4754 | 419 | - | - | 0 | - |
| - | - | 1667 | 420 | - | - | 0 | - |
| - | - | 968.3 | 432.1 | - | - | 0 | - |
| - | - | 989.3 | 433.2 | - | - | 0 | - |
| - | - | 825.8 | 438.2 | - | - | 0 | - |
| - | - | 537.1 | 449.2 | - | - | 0 | - |
| - | - | 1156 | 455.2 | - | - | 0 | - |
| 4 | b | 1014 | 461.2 | 0.001065 | 2.309 | +1 | 4 |
| - | - | 3163 | 462.3 | - | - | 0 | - |
| - | - | 1267 | 462.8 | - | - | 0 | - |
| - | - | 959.4 | 463.3 | - | - | 0 | - |
| - | - | 1.189E+04 | 465.2 | - | - | 0 | - |
| - | - | 2821 | 466.2 | - | - | 0 | - |
| 7 | y | 1.229E+04 | 473.2 | 0.0001352 | 0.2858 | +1 | 3 |
| - | - | 3181 | 474.2 | - | - | 0 | - |
| - | - | 1681 | 475.2 | - | - | 0 | - |
| - | - | 783.7 | 477.2 | - | - | 0 | - |
| - | - | 706.8 | 478.8 | - | - | 0 | - |
| 4 | b | 5486 | 479.2 | 0.0006626 | 1.383 | +1 | 4 |
| - | - | 1006 | 480.2 | - | - | 0 | - |
| 8 | b | 772.5 | 482.2 | 0.002062 | 4.277 | +2 | 8 |
| - | - | 695.5 | 483.2 | - | - | 0 | - |
| - | - | 1740 | 486.2 | - | - | 0 | - |
| 8 | b | 787.7 | 491.2 | 0.001327 | 2.701 | +2 | 8 |
| - | - | 2371 | 503.2 | - | - | 0 | - |
| - | - | 888.4 | 513.2 | - | - | 0 | - |
| - | - | 789.3 | 521.2 | - | - | 0 | - |
| - | - | 1029 | 540.2 | - | - | 0 | - |
| 2 | y | 4115 | 542.7 | 0.0006436 | 1.186 | +2 | 8 |
| - | - | 3237 | 543.2 | - | - | 0 | - |
| - | - | 1046 | 543.7 | - | - | 0 | - |
| 6 | y | 1.334E+04 | 544.2 | 3.926E-05 | 0.07214 | +1 | 4 |
| - | - | 3484 | 545.2 | - | - | 0 | - |
| - | - | 938.2 | 546.2 | - | - | 0 | - |
| - | - | 741.9 | 548.2 | - | - | 0 | - |
| - | - | 2166 | 557.2 | - | - | 0 | - |
| - | - | 2208 | 575.4 | - | - | 0 | - |
| 0 | Precursor | 2923 | 584.2 | 0.0005803 | 0.9933 | +2 | -1 |
| - | - | 1381 | 584.7 | - | - | 0 | - |
| - | - | 855.4 | 585.2 | - | - | 0 | - |
| - | - | 1595 | 590.2 | - | - | 0 | - |
| - | - | 1052 | 591.2 | - | - | 0 | - |
| - | - | 771.8 | 592.3 | - | - | 0 | - |
| - | - | 910.6 | 592.3 | - | - | 0 | - |
| - | - | 796.2 | 593.1 | - | - | 0 | - |
| 0 | Precursor | 2693 | 593.2 | 0.0003685 | 0.6212 | +2 | -1 |
| - | - | 826 | 593.3 | - | - | 0 | - |
| - | - | 1856 | 593.7 | - | - | 0 | - |
| - | - | 1433 | 594.2 | - | - | 0 | - |
| - | - | 2064 | 596.2 | - | - | 0 | - |
| - | - | 1549 | 614.2 | - | - | 0 | - |
| - | - | 3635 | 618.2 | - | - | 0 | - |
| - | - | 803.6 | 619.2 | - | - | 0 | - |
| 5 | b | 1582 | 624.2 | 0.001686 | 2.701 | +1 | 5 |
| - | - | 660.7 | 636.2 | - | - | 0 | - |
| 5 | b | 824.6 | 642.2 | 0.0009594 | 1.494 | +1 | 5 |
| - | - | 667.1 | 649.3 | - | - | 0 | - |
| - | - | 917.3 | 656.3 | - | - | 0 | - |
| - | - | 1309 | 688.2 | - | - | 0 | - |
| - | - | 747.6 | 689.2 | - | - | 0 | - |
| 6 | b | 1224 | 695.3 | 0.003033 | 4.363 | +1 | 6 |
| - | - | 953.6 | 699.4 | - | - | 0 | - |
| - | - | 7835 | 705.3 | - | - | 0 | - |
| - | - | 2238 | 706.3 | - | - | 0 | - |
| 5 | y | 1.451E+04 | 707.3 | 0.0007282 | 1.03 | +1 | 5 |
| - | - | 6918 | 708.3 | - | - | 0 | - |
| - | - | 1083 | 709.3 | - | - | 0 | - |
| 6 | b | 838.1 | 713.3 | 0.001086 | 1.522 | +1 | 6 |
| - | - | 1.426E+04 | 733.3 | - | - | 0 | - |
| - | - | 5974 | 734.3 | - | - | 0 | - |
| - | - | 943 | 735.3 | - | - | 0 | - |
| - | - | 2819 | 751.3 | - | - | 0 | - |
| - | - | 1071 | 752.3 | - | - | 0 | - |
| - | - | 1252 | 788.4 | - | - | 0 | - |
| 4 | y | 888.5 | 804.3 | 0.002458 | 3.056 | +1 | 6 |
| 4 | y | 1.758E+04 | 822.3 | 8.338E-05 | 0.1014 | +1 | 6 |
| - | - | 8637 | 823.3 | - | - | 0 | - |
| - | - | 2328 | 824.3 | - | - | 0 | - |
| - | - | 1315 | 880.3 | - | - | 0 | - |
| - | - | 963.1 | 891.3 | - | - | 0 | - |
| 3 | y | 5696 | 919.3 | 0.001996 | 2.172 | +1 | 7 |
| - | - | 3422 | 920.3 | - | - | 0 | - |
| - | - | 1411 | 921.3 | - | - | 0 | - |
| - | - | 1019 | 923.5 | - | - | 0 | - |
| - | - | 735.8 | 924.5 | - | - | 0 | - |
| 3 | y | 9.121E+04 | 937.4 | 0.0007203 | 0.7684 | +1 | 7 |
| - | - | 4.925E+04 | 938.4 | - | - | 0 | - |
| - | - | 1.478E+04 | 939.4 | - | - | 0 | - |
| - | - | 1972 | 940.4 | - | - | 0 | - |
| - | - | 668.9 | 942 | - | - | 0 | - |
| - | - | 3484 | 956.4 | - | - | 0 | - |
| - | - | 1228 | 957.5 | - | - | 0 | - |
| 8 | b | 1359 | 981.4 | 0.001789 | 1.822 | +1 | 8 |
| - | - | 793.1 | 1056 | - | - | 0 | - |
| 2 | y | 6851 | 1084 | 0.00169 | 1.559 | +1 | 8 |
| - | - | 4589 | 1085 | - | - | 0 | - |
| - | - | 1529 | 1086 | - | - | 0 | - |
| - | - | 683.7 | 1394 | - | - | 0 | - |
| - | - | 743.1 | 2033 | - | - | 0 | - |
| - | - | 601.6 | 2304 | - | - | 0 | - |
| - | - | 651.1 | 2674 | - | - | 0 | - |

m/z Charge Intensity FragmentType MassShift Position
120.08106994628906 0 63463.07
121.0844497680664 0 4943.2837
127.08684539794922 0 608.72955
129.0480499267578 0 351.9599
129.10250854492188 0 2872.5771
130.06546020507812 0 2027.4053
132.08103942871094 0 2811.0896
132.1022491455078 0 1691.9514
133.08612060546875 0 3876.8486
134.02732849121094 0 539.45844
134.08946228027344 0 418.37253
134.2315673828125 0 411.71304
136.0759735107422 0 29924.674
137.07931518554688 0 2422.7576
138.06637573242188 0 2506.8855
139.11697387695312 0 427.0782
141.10264587402344 0 465.15707
142.43756103515625 0 553.2468
146.060302734375 0 916.5293
146.09640502929688 0 775.13605
148.95509338378906 0 482.08994
149.0452117919922 0 3174.7915
156.07701110839844 0 6670.889
158.09664916992188 0 8257.678
159.09201049804688 0 4076.647
159.09933471679688 0 905.6695
160.07601928710938 0 2023.3156
160.11236572265625 0 3481.4277
161.09228515625 0 417.78287
165.05467224121094 0 565.4593
165.07720947265625 0 698.54944
166.06134033203125 0 2723.8357
166.0865020751953 0 3948.7222
167.05569458007812 0 4600.2383
167.08206176757812 0 573.3935
167.08985900878906 0 1388.6356
168.07701110839844 0 1092.6461
169.09764099121094 0 620.5149
171.0771026611328 0 904.69434
171.11512756347656 0 436.99844
173.1285400390625 0 1032.042
173.43765258789062 0 540.00024
175.08937072753906 0 616.0417
175.0977020263672 0 579.88074
176.10726928710938 0 16156.024
177.10243225097656 0 12342.484
177.11134338378906 0 2976.642
178.1061553955078 0 905.77905
180.0771942138672 0 834.323
181.09780883789062 0 496.6479
182.08145141601562 0 874.6306
182.45204162597656 0 452.39743
183.06336975097656 0 462.0876
185.0559844970703 0 1655.6514
186.0914306640625 0 1129.24
187.13427734375 0 528.706
187.14480590820312 0 421.1652
188.06227111816406 0 777.0703
188.07086181640625 0 7626.7744
189.07437133789062 0 1096.1381
189.0865936279297 0 736.50525
191.1183319091797 0 1160.4044
193.1083984375 0 464.90628
195.0879669189453 0 1188.4327
196.09059143066406 0 1217.4354
197.1280975341797 0 583.5654
199.17019653320312 0 651.28864
201.1236572265625 0 1003.7478
203.0663299560547 0 1420.6113
203.11810302734375 0 4905.576 a Water loss 1
204.1216583251953 0 1161.4464
205.09732055664062 0 4672.1895 y 8
207.11289978027344 0 1928.6068
208.07208251953125 0 651.4979
211.45167541503906 0 477.1709
212.1179656982422 0 617.8308
215.13818359375 0 697.9901
217.09759521484375 0 2525.92
221.08480834960938 0 4748.5376
221.1287841796875 0 216849.97 a 1
221.55271911621094 0 538.78064
222.08445739746094 0 940.6876
222.1320037841797 0 28102.133
223.1081085205078 0 839.2829
223.1342315673828 0 1643.8467
224.10289001464844 0 688.0253
225.0430908203125 0 1830.4349
226.1554412841797 0 1202.3064
231.0614471435547 0 3767.5603
231.11328125 0 2110.6055 b Water loss 1
231.15846252441406 0 576.0734
231.17051696777344 0 1076.9641
233.09072875976562 0 648.17334
233.16531372070312 0 1522.2173
235.1080780029297 0 6464.576
236.1117401123047 0 608.1571
238.1208953857422 0 582.79474
239.09530639648438 0 11554.261
240.09536743164062 0 2229.1902 b 3
241.11212158203125 0 4149.667
243.1343231201172 0 625.81744
244.0926513671875 0 529.0495
249.12364196777344 0 36172.62 b 1
250.126953125 0 3992.4045
251.10276794433594 0 5188.27
252.0802764892578 0 3313.9465
253.1182403564453 0 1623.4304
261.0870056152344 0 791.3241
263.1028747558594 0 7854.6074
263.12744140625 0 700.6518
265.11883544921875 0 559.52765
265.3362731933594 0 513.10034
266.9991760253906 0 732.81335
268.9784851074219 0 1589.8026
269.1069030761719 0 16290.364
270.1093444824219 0 1872.3152
279.0976257324219 0 3522.6038
281.05169677734375 0 1666.7754
283.0304870605469 0 949.8768
285.0100402832031 0 22752.56
286.0101013183594 0 3601.6416
287.11712646484375 0 1566.9789
296.15106201171875 0 4763.4004
297.15411376953125 0 641.2104
299.0619201660156 0 14827.998
299.15118408203125 0 641.9195
300.06317138671875 0 2555.8398
301.13873291015625 0 706.9432
303.0205078125 0 2905.7004
304.0209045410156 0 673.85565
315.1669921875 0 1004.1676
318.14422607421875 0 708.101 a Water loss 2
322.1342468261719 0 1645.1953
323.0985107421875 0 1334.8047
323.17138671875 0 632.4083
324.14581298828125 0 11349.111
325.14898681640625 0 2258.9412
328.13006591796875 0 726.12476
332.123779296875 0 642.3066
339.17034912109375 0 710.04065
340.14410400390625 0 5241.2554
341.14544677734375 0 1491.9161
342.1564636230469 0 56771.38 y 7
343.15960693359375 0 11513.827
344.1634521484375 0 619.72595
344.9764709472656 0 2813.6372
345.9755554199219 0 927.2224
346.1400146484375 0 2064.596 b Water loss 2
348.1185607910156 0 1773.586
350.13494873046875 0 4230.8325
352.1404113769531 0 1201.3813
355.07000732421875 0 58242.668
356.0707092285156 0 15135.749
357.6465759277344 0 589.0754
358.155517578125 0 770.14795
364.15069580078125 0 8930.311 b 2
365.15277099609375 0 1580.6095
366.13037109375 0 1063.0286
371.1005554199219 0 1403.0896
373.0807189941406 0 5451.9146
374.0818176269531 0 2008.4823
376.1155700683594 0 755.89496
378.1297607421875 0 6225.06
394.1248779296875 0 12047.196
395.12841796875 0 2471.0498
400.98565673828125 0 1635.9397
415.0373840332031 0 2899.445
415.15325927734375 0 582.2796
416.0365295410156 0 783.8743
418.99566650390625 0 4754.246
419.99530029296875 0 1667.429
432.1413269042969 0 968.265
433.1722412109375 0 989.3294
438.1603698730469 0 825.7883
449.16668701171875 0 537.0791
455.1861572265625 0 1155.6952
461.1677551269531 0 1014.16907 b Water loss 3
462.26416015625 0 3163.4797
462.7658996582031 0 1266.7484
463.2648010253906 0 959.4008
465.1621398925781 0 11885.224
466.1655578613281 0 2821.2695
473.1966857910156 0 12285.362 y 6
474.1999206542969 0 3180.8892
475.2105712890625 0 1680.7576
477.1611328125 0 783.6565
478.8359375 0 706.81885
479.17791748046875 0 5485.563 b 3
480.1810302734375 0 1006.1308
482.18896484375 0 772.545 b Water loss 7
483.1791687011719 0 695.54004
486.1610412597656 0 1740.234
491.1935119628906 0 787.653 b 7
503.20782470703125 0 2370.6223
513.19775390625 0 888.4142
521.2176513671875 0 789.2529
540.24267578125 0 1029.2953
542.7139282226562 0 4115.433 y 1
543.2135620117188 0 3236.8538
543.7171020507812 0 1045.558
544.2337036132812 0 13343.203 y 5
545.2364501953125 0 3483.6943
546.232666015625 0 938.16583
548.1981811523438 0 741.8588
557.1986083984375 0 2165.679
575.363525390625 0 2207.8738
584.232421875 0 2922.7405 Precursor Water loss
584.7340698242188 0 1381.0369
585.2349243164062 0 855.44965
590.2384033203125 0 1594.6599
591.23974609375 0 1051.869
592.2561645507812 0 771.786
592.33984375 0 910.61237
593.1439208984375 0 796.1597
593.2367553710938 0 2692.8782 Precursor
593.33984375 0 825.98883
593.7389526367188 0 1856.157
594.24072265625 0 1432.7947
596.20263671875 0 2064.4514
614.2450561523438 0 1549.4419
618.2338256835938 0 3634.9995
619.2378540039062 0 803.6471
624.2283325195312 0 1581.6228 b Water loss 4
636.2454833984375 0 660.6835
642.2396240234375 0 824.6229 b 4
649.3399047851562 0 667.14014
656.3020629882812 0 917.26965
688.2399291992188 0 1308.6904
689.2371215820312 0 747.64136
695.2640991210938 0 1223.6624 b Water loss 5
699.4032592773438 0 953.61896
705.2656860351562 0 7834.577
706.2687377929688 0 2237.7646
707.2962646484375 0 14508.633 y 4
708.2996826171875 0 6917.6025
709.297119140625 0 1083.1698
713.276611328125 0 838.12177 b 5
733.2605590820312 0 14260.914
734.2639770507812 0 5973.7974
735.264404296875 0 942.97534
751.27099609375 0 2818.8066
752.2725830078125 0 1071.257
788.3558959960938 0 1252.1725
804.3109130859375 0 888.4528 y Water loss 3
822.3238525390625 0 17582.615 y 3
823.326171875 0 8637.4375
824.3276977539062 0 2327.5647
880.330078125 0 1314.8188
891.3427124023438 0 963.0831
919.3383178710938 0 5695.8975 y Water loss 2
920.3377685546875 0 3422.2842
921.3384399414062 0 1410.7673
923.5155639648438 0 1019.2341
924.5242919921875 0 735.79034
937.3501586914062 0 91207.125 y 2
938.3529663085938 0 49246.152
939.3532104492188 0 14775.649
940.3558959960938 0 1971.8398
941.9912109375 0 668.9095
956.4464111328125 0 3484.233
957.4514770507812 0 1228.4435
981.3753051757812 0 1358.687 b 7
1055.515625 0 793.11145
1084.4176025390625 0 6850.86 y 1
1085.4207763671875 0 4588.953
1086.425537109375 0 1528.6871
1394.168212890625 0 683.69366
2033.496826171875 0 743.0636
2303.85595703125 0 601.57996
2673.571044921875 0 651.0887

Spectrum Details

|  |  |
| --- | --- |
| Matched peaks? Matched peaks The total absolute number of peaks matched. Additionally in brackets the total fraction of peaks matched and the total number of peaks is shown. | 30 (11.11% of 270) |
| FDR? FDR The false discovery rate estimated for this peptide. It is calculated by matching all theoretical fragments with a non-integer shift with the raw peaks for this spectrum. This is done with 40 different shifts. The resulting percentage is the average number of annotated peaks over the number of annotated peaks with the correct spectrum. | 0.71% |
| Satellite FDR? Satellite FDR See the FDR for details on its calculation. This satellite ion specific FDR only contains the satellite ions (d/w) for I/L/J positions. | - |
| PSM Score? PSM Score The PSM Score as given by Hecklib to this annotated spectrum. It is shown with three significant figures. | 373 |

#### Spectrum 8296? Spectrum 8296 The raw spectrum of this peptide as annotated by Hecklib. The fragments are coloured according to ion type (see legend). Any peaks with a star '\*' as text can be hovered over to see the full details, first the ion type second the mass shift type. By hovering over the amino acids in the peptide or ions in the legend the corresponding peaks are highlighted. By toggling the 'Unassigned' label you can turn the background (unassigned) peaks on or off in the plot. By updating the slider in the Ion legend you can update the spectrum to only show the top X% of the peaks with labels. The top X% means any peak that is within X% of the highest intensity. By dragging in the spectrum you can zoom in to a specific part of the spectrum and use 'Zoom Out' to get back to the original zoom level. The annotation of the spectrum is based on the given sequence in the peptides file and is done with different software so inconsistencies are likely. The peaks are annotated based on the given sequence, with 20 ppm tolerance.

Copy Data

##### Spectrum 8296 (TSV)

###### Preview

```
Loading example...
```

*Click on the button to copy the data to your clipboard.*

Mz MinMz MaxIntensity Max

WidthHeightPeptide font sizePeptide stroke widthSpectrum font sizeSpectrum stroke widthCompact peptide

Ion legend

wxyz

abcd

OtherUnassignedIon^Charge^_Position_Show for top:%

TFDDYAMHW

09.90e+41.98e+52.97e+53.96e+5

Zoom Out

b^+2^_3_a^+1^_2_a^+1^_2_y^+1^_1_a^+1^_2_b^+1^_2_b^+1^_2_d^+1^_3_a^+1^_3_a^+1^_3_y^+1^_2_b^+1^_3_b^+1^_3_b^+1^_4_y^+2^_7_b^+1^_4_y^+1^_3_y^+2^_8_y^+2^_8_y^+1^_4_\*\*b^+1^_5_b^+1^_5_b^+1^_6_b^+1^_6_y^+1^_5_y^+1^_6_y^+1^_6_b^+1^_7_y^+1^_7_y^+1^_7_b^+1^_8_y^+1^_8_

0840167925193358

Fragment Matches Table

Show background peaks

| Position | Ion type | Intensity | mz Theoretical | mz Error (Th) | mz Error (ppm) | Charge | Series Number |
| --- | --- | --- | --- | --- | --- | --- | --- |
| - | - | 1.068E+05 | 120.1 | - | - | 0 | - |
| - | - | 7907 | 121.1 | - | - | 0 | - |
| - | - | 798.4 | 122.1 | - | - | 0 | - |
| - | - | 1048 | 127.1 | - | - | 0 | - |
| - | - | 388.6 | 128.1 | - | - | 0 | - |
| - | - | 1343 | 128.1 | - | - | 0 | - |
| - | - | 567.5 | 129.1 | - | - | 0 | - |
| - | - | 5974 | 129.1 | - | - | 0 | - |
| - | - | 584.6 | 129.1 | - | - | 0 | - |
| - | - | 4779 | 130.1 | - | - | 0 | - |
| - | - | 696.4 | 131.1 | - | - | 0 | - |
| - | - | 438 | 131.1 | - | - | 0 | - |
| - | - | 6124 | 132.1 | - | - | 0 | - |
| - | - | 673.5 | 133.1 | - | - | 0 | - |
| - | - | 3.762E+04 | 136.1 | - | - | 0 | - |
| - | - | 651.5 | 137 | - | - | 0 | - |
| - | - | 3156 | 137.1 | - | - | 0 | - |
| - | - | 3073 | 138.1 | - | - | 0 | - |
| - | - | 3554 | 138.1 | - | - | 0 | - |
| - | - | 475.1 | 139.1 | - | - | 0 | - |
| - | - | 382.3 | 139.1 | - | - | 0 | - |
| - | - | 344.7 | 140.6 | - | - | 0 | - |
| - | - | 444.3 | 141.1 | - | - | 0 | - |
| - | - | 1036 | 141.1 | - | - | 0 | - |
| - | - | 428.3 | 142.3 | - | - | 0 | - |
| - | - | 873.4 | 143.1 | - | - | 0 | - |
| - | - | 501.7 | 143.1 | - | - | 0 | - |
| - | - | 1683 | 143.1 | - | - | 0 | - |
| - | - | 547.1 | 144.1 | - | - | 0 | - |
| - | - | 448.1 | 145.1 | - | - | 0 | - |
| - | - | 2461 | 146.1 | - | - | 0 | - |
| - | - | 432.4 | 146.5 | - | - | 0 | - |
| - | - | 1151 | 148.1 | - | - | 0 | - |
| - | - | 384.4 | 150.1 | - | - | 0 | - |
| - | - | 397.9 | 154.2 | - | - | 0 | - |
| - | - | 1622 | 155.1 | - | - | 0 | - |
| - | - | 651.8 | 155.1 | - | - | 0 | - |
| - | - | 482.2 | 155.1 | - | - | 0 | - |
| - | - | 1.142E+04 | 156.1 | - | - | 0 | - |
| - | - | 587.1 | 157.1 | - | - | 0 | - |
| - | - | 2275 | 157.1 | - | - | 0 | - |
| - | - | 1.356E+04 | 158.1 | - | - | 0 | - |
| - | - | 543 | 159.1 | - | - | 0 | - |
| - | - | 9096 | 159.1 | - | - | 0 | - |
| - | - | 784 | 159.1 | - | - | 0 | - |
| - | - | 2230 | 160.1 | - | - | 0 | - |
| - | - | 758.2 | 160.1 | - | - | 0 | - |
| - | - | 1089 | 160.1 | - | - | 0 | - |
| - | - | 5823 | 160.1 | - | - | 0 | - |
| - | - | 410.1 | 160.4 | - | - | 0 | - |
| - | - | 1232 | 161.1 | - | - | 0 | - |
| - | - | 468.4 | 165.1 | - | - | 0 | - |
| - | - | 1144 | 165.1 | - | - | 0 | - |
| - | - | 795.8 | 165.1 | - | - | 0 | - |
| - | - | 793 | 166.1 | - | - | 0 | - |
| - | - | 1629 | 166.1 | - | - | 0 | - |
| - | - | 3554 | 166.1 | - | - | 0 | - |
| - | - | 832.9 | 167.1 | - | - | 0 | - |
| - | - | 2501 | 167.1 | - | - | 0 | - |
| - | - | 885.2 | 167.1 | - | - | 0 | - |
| - | - | 869.3 | 169.1 | - | - | 0 | - |
| - | - | 616.4 | 169.1 | - | - | 0 | - |
| - | - | 1001 | 169.1 | - | - | 0 | - |
| - | - | 936.6 | 169.1 | - | - | 0 | - |
| - | - | 1449 | 171.1 | - | - | 0 | - |
| - | - | 938.3 | 171.1 | - | - | 0 | - |
| - | - | 428.5 | 172.1 | - | - | 0 | - |
| - | - | 3725 | 173.1 | - | - | 0 | - |
| - | - | 699.8 | 173.4 | - | - | 0 | - |
| 3 | b | 1337 | 174.1 | 9.318E-05 | 0.5353 | +2 | 3 |
| - | - | 1358 | 175.1 | - | - | 0 | - |
| - | - | 2273 | 176.1 | - | - | 0 | - |
| - | - | 2.683E+04 | 176.1 | - | - | 0 | - |
| - | - | 2.204E+04 | 177.1 | - | - | 0 | - |
| - | - | 3302 | 177.1 | - | - | 0 | - |
| - | - | 2199 | 178.1 | - | - | 0 | - |
| - | - | 1583 | 180.1 | - | - | 0 | - |
| - | - | 1254 | 182.1 | - | - | 0 | - |
| - | - | 535.3 | 183.1 | - | - | 0 | - |
| - | - | 2666 | 185.1 | - | - | 0 | - |
| - | - | 992.1 | 185.1 | - | - | 0 | - |
| - | - | 1049 | 186.1 | - | - | 0 | - |
| - | - | 699 | 187.1 | - | - | 0 | - |
| - | - | 6590 | 187.1 | - | - | 0 | - |
| - | - | 1.532E+04 | 188.1 | - | - | 0 | - |
| - | - | 1582 | 189.1 | - | - | 0 | - |
| - | - | 1711 | 191.1 | - | - | 0 | - |
| - | - | 755.7 | 192.1 | - | - | 0 | - |
| - | - | 472.5 | 193 | - | - | 0 | - |
| - | - | 523.3 | 193.1 | - | - | 0 | - |
| - | - | 8679 | 193.1 | - | - | 0 | - |
| - | - | 1472 | 195.1 | - | - | 0 | - |
| - | - | 557.8 | 197.1 | - | - | 0 | - |
| - | - | 1998 | 199.1 | - | - | 0 | - |
| - | - | 512.6 | 201.1 | - | - | 0 | - |
| - | - | 1662 | 201.1 | - | - | 0 | - |
| - | - | 1085 | 202.1 | - | - | 0 | - |
| - | - | 2878 | 203.1 | - | - | 0 | - |
| - | - | 621.8 | 203.1 | - | - | 0 | - |
| - | - | 981.2 | 203.1 | - | - | 0 | - |
| - | - | 1616 | 203.1 | - | - | 0 | - |
| 2 | a | 1.085E+04 | 203.1 | 9.168E-05 | 0.4514 | +1 | 2 |
| - | - | 485.6 | 204.1 | - | - | 0 | - |
| - | - | 3508 | 204.1 | - | - | 0 | - |
| 2 | a | 647.1 | 204.1 | 0.000222 | 1.087 | +1 | 2 |
| - | - | 1046 | 204.1 | - | - | 0 | - |
| - | - | 559.6 | 205.1 | - | - | 0 | - |
| 9 | y | 8562 | 205.1 | 0.0001082 | 0.5275 | +1 | 1 |
| - | - | 848.5 | 205.1 | - | - | 0 | - |
| - | - | 1033 | 206.1 | - | - | 0 | - |
| - | - | 2722 | 207.1 | - | - | 0 | - |
| - | - | 1068 | 208.1 | - | - | 0 | - |
| - | - | 603.6 | 211.1 | - | - | 0 | - |
| - | - | 638.9 | 212.1 | - | - | 0 | - |
| - | - | 646.4 | 213.2 | - | - | 0 | - |
| - | - | 831.3 | 215.1 | - | - | 0 | - |
| - | - | 7291 | 215.1 | - | - | 0 | - |
| - | - | 585.9 | 217.1 | - | - | 0 | - |
| - | - | 5322 | 217.1 | - | - | 0 | - |
| - | - | 2222 | 219.1 | - | - | 0 | - |
| - | - | 519.8 | 220.1 | - | - | 0 | - |
| - | - | 2.162E+04 | 221.1 | - | - | 0 | - |
| 2 | a | 3.922E+05 | 221.1 | 0.0001011 | 0.4571 | +1 | 2 |
| - | - | 2049 | 222.1 | - | - | 0 | - |
| - | - | 4.885E+04 | 222.1 | - | - | 0 | - |
| - | - | 603.6 | 223.1 | - | - | 0 | - |
| - | - | 528.4 | 223.1 | - | - | 0 | - |
| - | - | 2929 | 223.1 | - | - | 0 | - |
| - | - | 3085 | 226.2 | - | - | 0 | - |
| - | - | 3344 | 227.1 | - | - | 0 | - |
| - | - | 728.9 | 227.2 | - | - | 0 | - |
| - | - | 1529 | 228.1 | - | - | 0 | - |
| - | - | 769.7 | 229.1 | - | - | 0 | - |
| - | - | 2228 | 229.2 | - | - | 0 | - |
| - | - | 5983 | 231.1 | - | - | 0 | - |
| 2 | b | 3246 | 231.1 | 0.0002024 | 0.8759 | +1 | 2 |
| - | - | 1316 | 231.2 | - | - | 0 | - |
| - | - | 880.6 | 232.1 | - | - | 0 | - |
| - | - | 1453 | 233.2 | - | - | 0 | - |
| - | - | 8716 | 235.1 | - | - | 0 | - |
| - | - | 577.3 | 235.1 | - | - | 0 | - |
| - | - | 1351 | 236.1 | - | - | 0 | - |
| - | - | 584.2 | 238.1 | - | - | 0 | - |
| - | - | 904.2 | 239.1 | - | - | 0 | - |
| - | - | 2745 | 239.1 | - | - | 0 | - |
| - | - | 571.6 | 240.1 | - | - | 0 | - |
| - | - | 567 | 242.2 | - | - | 0 | - |
| - | - | 716.5 | 244.1 | - | - | 0 | - |
| - | - | 563.4 | 245.1 | - | - | 0 | - |
| - | - | 649.4 | 246.1 | - | - | 0 | - |
| - | - | 514.7 | 247.1 | - | - | 0 | - |
| - | - | 2562 | 248.1 | - | - | 0 | - |
| - | - | 871.2 | 248.2 | - | - | 0 | - |
| 2 | b | 6.796E+04 | 249.1 | 5.95E-05 | 0.2388 | +1 | 2 |
| - | - | 9360 | 250.1 | - | - | 0 | - |
| - | - | 6503 | 251.1 | - | - | 0 | - |
| - | - | 911 | 251.1 | - | - | 0 | - |
| - | - | 862.1 | 254.1 | - | - | 0 | - |
| - | - | 949.4 | 254.2 | - | - | 0 | - |
| - | - | 843.7 | 255.1 | - | - | 0 | - |
| - | - | 926.8 | 256.1 | - | - | 0 | - |
| - | - | 4609 | 257.1 | - | - | 0 | - |
| - | - | 519.3 | 257.2 | - | - | 0 | - |
| - | - | 957.4 | 261.1 | - | - | 0 | - |
| - | - | 700.1 | 261.1 | - | - | 0 | - |
| - | - | 1.406E+04 | 263.1 | - | - | 0 | - |
| - | - | 1719 | 263.1 | - | - | 0 | - |
| - | - | 1666 | 264.1 | - | - | 0 | - |
| - | - | 760.7 | 264.1 | - | - | 0 | - |
| - | - | 871.3 | 264.1 | - | - | 0 | - |
| - | - | 638.2 | 265.2 | - | - | 0 | - |
| - | - | 776.5 | 266.1 | - | - | 0 | - |
| - | - | 984.5 | 267.1 | - | - | 0 | - |
| - | - | 1312 | 270.2 | - | - | 0 | - |
| - | - | 548.5 | 271.1 | - | - | 0 | - |
| - | - | 2494 | 273.1 | - | - | 0 | - |
| - | - | 3525 | 274.1 | - | - | 0 | - |
| - | - | 678.5 | 275.2 | - | - | 0 | - |
| - | - | 6598 | 279.1 | - | - | 0 | - |
| - | - | 689.8 | 280.1 | - | - | 0 | - |
| - | - | 641.2 | 284.2 | - | - | 0 | - |
| - | - | 8046 | 285.1 | - | - | 0 | - |
| - | - | 914.6 | 286.1 | - | - | 0 | - |
| - | - | 6190 | 292.1 | - | - | 0 | - |
| 3 | d | 978.8 | 292.2 | 0.0001424 | 0.4875 | +1 | 3 |
| - | - | 749.9 | 293.1 | - | - | 0 | - |
| - | - | 775.8 | 293.1 | - | - | 0 | - |
| - | - | 6940 | 296.2 | - | - | 0 | - |
| - | - | 506.1 | 297 | - | - | 0 | - |
| - | - | 1101 | 297.2 | - | - | 0 | - |
| - | - | 719.9 | 302.1 | - | - | 0 | - |
| - | - | 6181 | 303.1 | - | - | 0 | - |
| - | - | 1115 | 310.2 | - | - | 0 | - |
| - | - | 1043 | 314.1 | - | - | 0 | - |
| - | - | 2706 | 314.2 | - | - | 0 | - |
| - | - | 928.3 | 315.1 | - | - | 0 | - |
| - | - | 1242 | 316.1 | - | - | 0 | - |
| 3 | a | 2412 | 318.1 | 0.0003013 | 0.9471 | +1 | 3 |
| - | - | 756.4 | 321.1 | - | - | 0 | - |
| - | - | 2695 | 323.1 | - | - | 0 | - |
| - | - | 596.5 | 323.2 | - | - | 0 | - |
| - | - | 996.3 | 323.2 | - | - | 0 | - |
| - | - | 1.611E+04 | 324.1 | - | - | 0 | - |
| - | - | 1818 | 325.1 | - | - | 0 | - |
| - | - | 882.2 | 328.1 | - | - | 0 | - |
| - | - | 1066 | 328.1 | - | - | 0 | - |
| - | - | 750.4 | 330.1 | - | - | 0 | - |
| - | - | 806.3 | 332.1 | - | - | 0 | - |
| - | - | 772.8 | 333.2 | - | - | 0 | - |
| 3 | a | 783.1 | 336.2 | 0.0001849 | 0.5499 | +1 | 3 |
| - | - | 1365 | 339.2 | - | - | 0 | - |
| - | - | 658.8 | 339.3 | - | - | 0 | - |
| 8 | y | 8.034E+04 | 342.2 | 6.01E-05 | 0.1756 | +1 | 2 |
| - | - | 1.387E+04 | 343.2 | - | - | 0 | - |
| - | - | 1029 | 344.2 | - | - | 0 | - |
| 3 | b | 4850 | 346.1 | 7.199E-06 | 0.0208 | +1 | 3 |
| - | - | 644.1 | 347.1 | - | - | 0 | - |
| - | - | 1798 | 348.1 | - | - | 0 | - |
| - | - | 4687 | 350.1 | - | - | 0 | - |
| - | - | 902.3 | 351.1 | - | - | 0 | - |
| - | - | 1885 | 352.1 | - | - | 0 | - |
| - | - | 1306 | 353.2 | - | - | 0 | - |
| - | - | 1706 | 355.2 | - | - | 0 | - |
| - | - | 3847 | 356.1 | - | - | 0 | - |
| - | - | 986.9 | 358.2 | - | - | 0 | - |
| - | - | 688.9 | 359.1 | - | - | 0 | - |
| - | - | 1375 | 360.1 | - | - | 0 | - |
| 3 | b | 2.006E+04 | 364.2 | 0.0002264 | 0.6218 | +1 | 3 |
| - | - | 3561 | 365.2 | - | - | 0 | - |
| - | - | 2199 | 366.1 | - | - | 0 | - |
| - | - | 987 | 367.2 | - | - | 0 | - |
| - | - | 744.5 | 367.2 | - | - | 0 | - |
| - | - | 849.1 | 368.2 | - | - | 0 | - |
| - | - | 3462 | 371.2 | - | - | 0 | - |
| - | - | 616.2 | 374.1 | - | - | 0 | - |
| - | - | 2164 | 376.1 | - | - | 0 | - |
| - | - | 8227 | 378.1 | - | - | 0 | - |
| - | - | 1874 | 379.1 | - | - | 0 | - |
| - | - | 2256 | 382.1 | - | - | 0 | - |
| - | - | 984.7 | 385.2 | - | - | 0 | - |
| - | - | 1660 | 387.3 | - | - | 0 | - |
| - | - | 1394 | 390.2 | - | - | 0 | - |
| - | - | 1.435E+04 | 394.1 | - | - | 0 | - |
| - | - | 2090 | 395.1 | - | - | 0 | - |
| - | - | 4270 | 402.2 | - | - | 0 | - |
| - | - | 1172 | 407.2 | - | - | 0 | - |
| - | - | 3249 | 408.2 | - | - | 0 | - |
| - | - | 855.2 | 409.2 | - | - | 0 | - |
| - | - | 566.1 | 413.8 | - | - | 0 | - |
| - | - | 1190 | 420.2 | - | - | 0 | - |
| - | - | 636.2 | 421.2 | - | - | 0 | - |
| - | - | 1.167E+04 | 425.2 | - | - | 0 | - |
| - | - | 2159 | 426.2 | - | - | 0 | - |
| - | - | 2253 | 427.2 | - | - | 0 | - |
| - | - | 771.2 | 428.2 | - | - | 0 | - |
| - | - | 2331 | 433.2 | - | - | 0 | - |
| - | - | 1962 | 447.2 | - | - | 0 | - |
| - | - | 1352 | 448.8 | - | - | 0 | - |
| - | - | 873.9 | 449.1 | - | - | 0 | - |
| - | - | 652.8 | 449.3 | - | - | 0 | - |
| - | - | 1043 | 454.2 | - | - | 0 | - |
| - | - | 2166 | 455.2 | - | - | 0 | - |
| - | - | 2344 | 456.3 | - | - | 0 | - |
| - | - | 864.8 | 457.3 | - | - | 0 | - |
| - | - | 728 | 460.8 | - | - | 0 | - |
| 4 | b | 3413 | 461.2 | 0.0003694 | 0.8011 | +1 | 4 |
| - | - | 1043 | 461.2 | - | - | 0 | - |
| - | - | 887.2 | 462.2 | - | - | 0 | - |
| - | - | 703.7 | 462.3 | - | - | 0 | - |
| - | - | 1.006E+04 | 465.2 | - | - | 0 | - |
| - | - | 1989 | 466.2 | - | - | 0 | - |
| - | - | 1601 | 466.3 | - | - | 0 | - |
| - | - | 1470 | 471.2 | - | - | 0 | - |
| 3 | y | 1648 | 477.2 | 0.006719 | 14.08 | +2 | 7 |
| - | - | 680 | 477.7 | - | - | 0 | - |
| 4 | b | 1.122E+04 | 479.2 | 0.0001309 | 0.2731 | +1 | 4 |
| - | - | 2685 | 480.2 | - | - | 0 | - |
| - | - | 966.9 | 482.2 | - | - | 0 | - |
| - | - | 5594 | 484.3 | - | - | 0 | - |
| - | - | 1461 | 485.3 | - | - | 0 | - |
| - | - | 2051 | 486.2 | - | - | 0 | - |
| 7 | y | 2.061E+04 | 489.2 | 0.004825 | 9.864 | +1 | 3 |
| - | - | 5550 | 490.2 | - | - | 0 | - |
| - | - | 2079 | 491.2 | - | - | 0 | - |
| - | - | 1699 | 495.2 | - | - | 0 | - |
| - | - | 8064 | 496.2 | - | - | 0 | - |
| - | - | 1605 | 497.2 | - | - | 0 | - |
| - | - | 2838 | 497.2 | - | - | 0 | - |
| - | - | 1816 | 513.2 | - | - | 0 | - |
| - | - | 912.9 | 513.3 | - | - | 0 | - |
| - | - | 630.7 | 518.7 | - | - | 0 | - |
| - | - | 3569 | 519.2 | - | - | 0 | - |
| - | - | 1981 | 520.2 | - | - | 0 | - |
| - | - | 638.3 | 527.7 | - | - | 0 | - |
| - | - | 978.9 | 534.3 | - | - | 0 | - |
| - | - | 934.9 | 537.2 | - | - | 0 | - |
| 2 | y | 1095 | 541.7 | 0.004212 | 7.776 | +2 | 8 |
| - | - | 2043 | 542.2 | - | - | 0 | - |
| - | - | 3426 | 548.2 | - | - | 0 | - |
| - | - | 722.2 | 549.2 | - | - | 0 | - |
| 2 | y | 1.129E+04 | 550.7 | 0.002042 | 3.709 | +2 | 8 |
| - | - | 6500 | 551.2 | - | - | 0 | - |
| - | - | 2273 | 551.7 | - | - | 0 | - |
| - | - | 1250 | 553.3 | - | - | 0 | - |
| - | - | 1421 | 557.2 | - | - | 0 | - |
| - | - | 1590 | 559.3 | - | - | 0 | - |
| 6 | y | 3.067E+04 | 560.2 | 0.004485 | 8.006 | +1 | 4 |
| - | - | 9028 | 561.2 | - | - | 0 | - |
| - | - | 2067 | 562.2 | - | - | 0 | - |
| - | - | 2410 | 569.2 | - | - | 0 | - |
| - | - | 1471 | 569.7 | - | - | 0 | - |
| - | - | 3464 | 570.2 | - | - | 0 | - |
| - | - | 1107 | 574.3 | - | - | 0 | - |
| - | - | 1348 | 578.2 | - | - | 0 | - |
| - | - | 1103 | 579.2 | - | - | 0 | - |
| - | - | 1055 | 590.8 | - | - | 0 | - |
| - | - | 764.2 | 591.3 | - | - | 0 | - |
| 0 | Precursor | 5903 | 592.2 | 0.002468 | 4.167 | +2 | -1 |
| - | - | 4704 | 592.7 | - | - | 0 | - |
| - | - | 1860 | 593.2 | - | - | 0 | - |
| - | - | 1693 | 596.2 | - | - | 0 | - |
| - | - | 795.1 | 599.8 | - | - | 0 | - |
| - | - | 754.3 | 600.3 | - | - | 0 | - |
| - | - | 744.9 | 601 | - | - | 0 | - |
| 0 | Precursor | 1.496E+04 | 601.2 | 0.001946 | 3.237 | +2 | -1 |
| - | - | 854.2 | 601.3 | - | - | 0 | - |
| - | - | 3141 | 601.4 | - | - | 0 | - |
| - | - | 9649 | 601.7 | - | - | 0 | - |
| - | - | 4875 | 602.2 | - | - | 0 | - |
| - | - | 3507 | 606.2 | - | - | 0 | - |
| - | - | 5060 | 612.2 | - | - | 0 | - |
| - | - | 1385 | 613.2 | - | - | 0 | - |
| - | - | 2866 | 614.2 | - | - | 0 | - |
| - | - | 1118 | 615.2 | - | - | 0 | - |
| 5 | b | 2535 | 624.2 | 0.001793 | 2.872 | +1 | 5 |
| - | - | 801.2 | 631.2 | - | - | 0 | - |
| - | - | 3021 | 634.2 | - | - | 0 | - |
| - | - | 789 | 635.2 | - | - | 0 | - |
| - | - | 1589 | 640.2 | - | - | 0 | - |
| - | - | 1106 | 641.3 | - | - | 0 | - |
| 5 | b | 2091 | 642.2 | 0.0008984 | 1.399 | +1 | 5 |
| - | - | 780.9 | 655.3 | - | - | 0 | - |
| - | - | 5450 | 657.3 | - | - | 0 | - |
| - | - | 1829 | 658.3 | - | - | 0 | - |
| - | - | 790.4 | 659.2 | - | - | 0 | - |
| - | - | 9749 | 659.3 | - | - | 0 | - |
| - | - | 3940 | 660.3 | - | - | 0 | - |
| - | - | 645.5 | 672.3 | - | - | 0 | - |
| - | - | 1206 | 676.3 | - | - | 0 | - |
| - | - | 685 | 677.3 | - | - | 0 | - |
| - | - | 8208 | 685.3 | - | - | 0 | - |
| - | - | 2918 | 686.3 | - | - | 0 | - |
| - | - | 931.5 | 691.3 | - | - | 0 | - |
| 6 | b | 1310 | 695.3 | 0.0008972 | 1.29 | +1 | 6 |
| - | - | 982 | 696.3 | - | - | 0 | - |
| - | - | 673.6 | 696.8 | - | - | 0 | - |
| - | - | 2945 | 703.3 | - | - | 0 | - |
| - | - | 591.8 | 704.2 | - | - | 0 | - |
| - | - | 1872 | 705.3 | - | - | 0 | - |
| 6 | b | 1371 | 713.3 | 0.002062 | 2.892 | +1 | 6 |
| - | - | 9393 | 721.3 | - | - | 0 | - |
| - | - | 2801 | 722.3 | - | - | 0 | - |
| 5 | y | 2.955E+04 | 723.3 | 0.003718 | 5.14 | +1 | 5 |
| - | - | 1.119E+04 | 724.3 | - | - | 0 | - |
| - | - | 3052 | 725.3 | - | - | 0 | - |
| - | - | 1337 | 731.2 | - | - | 0 | - |
| - | - | 803.7 | 733.3 | - | - | 0 | - |
| - | - | 1.254E+04 | 749.3 | - | - | 0 | - |
| - | - | 3753 | 750.3 | - | - | 0 | - |
| - | - | 1394 | 751.3 | - | - | 0 | - |
| - | - | 709.5 | 754.4 | - | - | 0 | - |
| - | - | 1042 | 756.3 | - | - | 0 | - |
| - | - | 3433 | 767.3 | - | - | 0 | - |
| - | - | 1.092E+04 | 774.3 | - | - | 0 | - |
| - | - | 5146 | 775.3 | - | - | 0 | - |
| - | - | 900.8 | 776.3 | - | - | 0 | - |
| - | - | 649.4 | 796.3 | - | - | 0 | - |
| - | - | 768.2 | 804.3 | - | - | 0 | - |
| - | - | 1646 | 810 | - | - | 0 | - |
| 4 | y | 1649 | 820.3 | 0.004856 | 5.92 | +1 | 6 |
| - | - | 1133 | 821.3 | - | - | 0 | - |
| - | - | 994.5 | 832.3 | - | - | 0 | - |
| 4 | y | 3.342E+04 | 838.3 | 0.003752 | 4.476 | +1 | 6 |
| - | - | 1.755E+04 | 839.3 | - | - | 0 | - |
| - | - | 5476 | 840.3 | - | - | 0 | - |
| 7 | b | 869.2 | 842.3 | 0.007498 | 8.901 | +1 | 7 |
| - | - | 1041 | 862.3 | - | - | 0 | - |
| - | - | 727 | 865.7 | - | - | 0 | - |
| - | - | 4133 | 871.3 | - | - | 0 | - |
| - | - | 3015 | 872.3 | - | - | 0 | - |
| - | - | 902.2 | 872.4 | - | - | 0 | - |
| - | - | 5.008E+04 | 889.3 | - | - | 0 | - |
| - | - | 2.257E+04 | 890.3 | - | - | 0 | - |
| - | - | 5787 | 891.4 | - | - | 0 | - |
| - | - | 776.2 | 892.3 | - | - | 0 | - |
| - | - | 1577 | 895.5 | - | - | 0 | - |
| - | - | 2957 | 896.5 | - | - | 0 | - |
| - | - | 1042 | 897.5 | - | - | 0 | - |
| - | - | 1784 | 899.4 | - | - | 0 | - |
| - | - | 1466 | 900.4 | - | - | 0 | - |
| - | - | 2043 | 907.3 | - | - | 0 | - |
| - | - | 1210 | 908.3 | - | - | 0 | - |
| - | - | 856.1 | 918.3 | - | - | 0 | - |
| - | - | 750.2 | 933.4 | - | - | 0 | - |
| 3 | y | 9800 | 935.3 | 0.003121 | 3.337 | +1 | 7 |
| - | - | 7265 | 936.3 | - | - | 0 | - |
| - | - | 1944 | 937.3 | - | - | 0 | - |
| - | - | 1233 | 938.3 | - | - | 0 | - |
| 3 | y | 1.558E+05 | 953.3 | 0.003726 | 3.908 | +1 | 7 |
| - | - | 8.248E+04 | 954.3 | - | - | 0 | - |
| - | - | 3.161E+04 | 955.3 | - | - | 0 | - |
| - | - | 724.5 | 955.5 | - | - | 0 | - |
| - | - | 3411 | 956.3 | - | - | 0 | - |
| - | - | 1051 | 969.4 | - | - | 0 | - |
| 8 | b | 2419 | 997.4 | 0.00162 | 1.624 | +1 | 8 |
| - | - | 1184 | 998.4 | - | - | 0 | - |
| - | - | 979.5 | 998.6 | - | - | 0 | - |
| - | - | 765.3 | 1018 | - | - | 0 | - |
| - | - | 2713 | 1036 | - | - | 0 | - |
| - | - | 2455 | 1037 | - | - | 0 | - |
| 2 | y | 8248 | 1100 | 0.00184 | 1.672 | +1 | 8 |
| - | - | 5654 | 1101 | - | - | 0 | - |
| - | - | 2818 | 1102 | - | - | 0 | - |
| - | - | 820.9 | 1110 | - | - | 0 | - |
| - | - | 1181 | 1111 | - | - | 0 | - |
| - | - | 600.6 | 2147 | - | - | 0 | - |
| - | - | 670.2 | 2426 | - | - | 0 | - |
| - | - | 903.8 | 3080 | - | - | 0 | - |
| - | - | 1005 | 3080 | - | - | 0 | - |
| - | - | 667.2 | 3325 | - | - | 0 | - |

m/z Charge Intensity FragmentType MassShift Position
120.0809555053711 0 106798.125
121.08427429199219 0 7907.037
122.07159423828125 0 798.3794
127.08661651611328 0 1047.9135
128.0900115966797 0 388.5552
128.1071319580078 0 1343.0938
129.06581115722656 0 567.5327
129.10238647460938 0 5974.3735
129.1105194091797 0 584.62994
130.0652313232422 0 4779.4883
131.0684356689453 0 696.4083
131.11795043945312 0 437.97498
132.08094787597656 0 6124.4326
133.0845489501953 0 673.5101
136.0758056640625 0 37618.215
137.03475952148438 0 651.54333
137.0791473388672 0 3156.461
138.0549774169922 0 3072.54
138.0662841796875 0 3553.585
139.06988525390625 0 475.0821
139.08644104003906 0 382.2755
140.5623321533203 0 344.67453
141.0657958984375 0 444.293
141.10215759277344 0 1035.8916
142.28428649902344 0 428.31143
143.07275390625 0 873.36145
143.08580017089844 0 501.7298
143.11805725097656 0 1682.9429
144.08096313476562 0 547.1384
145.06141662597656 0 448.12396
146.0601348876953 0 2460.9429
146.47555541992188 0 432.37518
148.0869598388672 0 1151.4911
150.0662384033203 0 384.39108
154.20001220703125 0 397.94513
155.08160400390625 0 1621.7831
155.09317016601562 0 651.79205
155.11773681640625 0 482.2
156.0768280029297 0 11422.032
157.08035278320312 0 587.1345
157.09716796875 0 2274.8584
158.09645080566406 0 13560.104
159.07635498046875 0 543.0043
159.0917205810547 0 9095.888
159.09921264648438 0 784.0109
160.0758056640625 0 2229.6206
160.08726501464844 0 758.1922
160.09506225585938 0 1088.7548
160.11207580566406 0 5822.62
160.3658905029297 0 410.06024
161.1158447265625 0 1232.2871
165.05516052246094 0 468.4261
165.0771484375 0 1143.668
165.10238647460938 0 795.8435
166.05360412597656 0 792.97516
166.06117248535156 0 1629.354
166.08631896972656 0 3553.8992
167.0552520751953 0 832.88916
167.0814666748047 0 2501.1953
167.0926055908203 0 885.1744
169.06063842773438 0 869.3293
169.07638549804688 0 616.42957
169.09718322753906 0 1000.86365
169.13348388671875 0 936.63367
171.07667541503906 0 1449.1265
171.11276245117188 0 938.30396
172.10821533203125 0 428.49704
173.12850952148438 0 3725.2449
173.4374237060547 0 699.81055
174.06561279296875 0 1336.8674 b Ammonia loss 2
175.0865936279297 0 1358.4573
176.0819854736328 0 2273.4126
176.1070556640625 0 26827.625
177.10223388671875 0 22042.564
177.11099243164062 0 3301.6367
178.10572814941406 0 2198.8784
180.0768585205078 0 1582.5162
182.0806884765625 0 1254.3962
183.1133575439453 0 535.2904
185.05581665039062 0 2665.7734
185.09207153320312 0 992.0648
186.0902099609375 0 1049.0244
187.0629119873047 0 698.99536
187.14413452148438 0 6589.6265
188.0706787109375 0 15317.38
189.0739288330078 0 1582.2743
191.11810302734375 0 1711.3572
192.10218811035156 0 755.7131
193.02513122558594 0 472.50452
193.07249450683594 0 523.2602
193.10848999023438 0 8679.317
195.08731079101562 0 1471.7247
197.12850952148438 0 557.83417
199.1079559326172 0 1997.9836
201.0657958984375 0 512.6197
201.12353515625 0 1662.1328
202.053466796875 0 1084.7106
203.0662384033203 0 2878.0747
203.08241271972656 0 621.8125
203.09239196777344 0 981.2484
203.10226440429688 0 1616.1309
203.1177978515625 0 10850.271 a Water loss 1
204.0677032470703 0 485.64728
204.07675170898438 0 3507.5642
204.1021270751953 0 647.1122 a Ammonia loss 1
204.12156677246094 0 1046.3539
205.0806884765625 0 559.5864
205.0970458984375 0 8561.772 y 8
205.10775756835938 0 848.46814
206.10113525390625 0 1032.6315
207.1127471923828 0 2721.6196
208.0716552734375 0 1067.526
211.1444091796875 0 603.5995
212.1160888671875 0 638.90857
213.1599578857422 0 646.4364
215.08143615722656 0 831.2855
215.1389923095703 0 7291.468
217.08456420898438 0 585.9422
217.09707641601562 0 5321.521
219.07972717285156 0 2221.5332
220.1200408935547 0 519.7583
221.10333251953125 0 21615.977
221.12855529785156 0 392150.8 a 1
222.10650634765625 0 2048.5159
222.13182067871094 0 48848.6
223.0623779296875 0 603.6405
223.1194305419922 0 528.42053
223.13409423828125 0 2928.8975
226.15501403808594 0 3085.1448
227.1022491455078 0 3344.0151
227.15826416015625 0 728.9442
228.11285400390625 0 1529.2017
229.10751342773438 0 769.68585
229.15484619140625 0 2227.9058
231.06121826171875 0 5982.9087
231.11300659179688 0 3245.5784 b Water loss 1
231.169921875 0 1315.6846
232.1165771484375 0 880.556
233.1654815673828 0 1453.3765
235.1077423095703 0 8715.595
235.14303588867188 0 577.31976
236.11155700683594 0 1350.6312
238.13026428222656 0 584.2482
239.0955047607422 0 904.23486
239.1136932373047 0 2745.357
240.1328582763672 0 571.56854
242.1865997314453 0 566.96594
244.12994384765625 0 716.529
245.1290283203125 0 563.4058
246.13482666015625 0 649.3727
247.1070098876953 0 514.70844
248.10714721679688 0 2561.645
248.15003967285156 0 871.19025
249.12342834472656 0 67958.22 b 1
250.12660217285156 0 9360.445
251.10260009765625 0 6503.3867
251.12738037109375 0 910.97894
254.11370849609375 0 862.10565
254.15025329589844 0 949.3522
255.14501953125 0 843.706
256.10870361328125 0 926.75116
257.106201171875 0 4609.4844
257.1631164550781 0 519.31537
261.08697509765625 0 957.37494
261.1255187988281 0 700.1145
263.1025390625 0 14064.565
263.1288146972656 0 1718.8212
264.10595703125 0 1666.0881
264.1312561035156 0 760.66266
264.14556884765625 0 871.2748
265.1538391113281 0 638.2119
266.1490478515625 0 776.4583
267.0907287597656 0 984.4569
270.1815185546875 0 1311.5237
271.10760498046875 0 548.5063
273.134521484375 0 2493.6023
274.1297302246094 0 3524.9482
275.1746826171875 0 678.54364
279.0975036621094 0 6598.422
280.12677001953125 0 689.822
284.1605224609375 0 641.23816
285.1016540527344 0 8045.669
286.1040954589844 0 914.59656
292.1403503417969 0 6190.029
292.16571044921875 0 978.76764 d 2
293.12347412109375 0 749.89935
293.1430358886719 0 775.7666
296.1506042480469 0 6939.9614
297.0419006347656 0 506.06216
297.1539001464844 0 1101.3086
302.1139221191406 0 719.91766
303.10986328125 0 6181.2495
310.1507873535156 0 1114.6819
314.0989990234375 0 1043.0653
314.2077941894531 0 2706.1172
315.1462707519531 0 928.2921
316.12890625 0 1241.8998
318.14453125 0 2412.3315 a Water loss 2
321.11883544921875 0 756.3512
323.098388671875 0 2695.265
323.1731262207031 0 596.5278
323.208251953125 0 996.32904
324.1455078125 0 16107.247
325.1485900878906 0 1818.3453
328.12646484375 0 882.2406
328.14703369140625 0 1065.941
330.11004638671875 0 750.4062
332.124267578125 0 806.2669
333.1574401855469 0 772.82074
336.15521240234375 0 783.0653 a 2
339.1710205078125 0 1364.535
339.26507568359375 0 658.8195
342.156005859375 0 80335.18 y 7
343.1591796875 0 13873.534
344.1614685058594 0 1029.3921
346.1397399902344 0 4850.243 b Water loss 2
347.1414794921875 0 644.0719
348.1189880371094 0 1797.7812
350.1344909667969 0 4687.2666
351.1360778808594 0 902.3134
352.1401062011719 0 1884.7158
353.2185974121094 0 1305.8281
355.1610412597656 0 1705.9282
356.13848876953125 0 3846.6428
358.15032958984375 0 986.9089
359.08892822265625 0 688.89484
360.1199645996094 0 1375.3713
364.15008544921875 0 20064.127 b 2
365.1529235839844 0 3560.6316
366.1287536621094 0 2199.0137
367.1640319824219 0 987.0173
367.23480224609375 0 744.51746
368.22955322265625 0 849.05817
371.22882080078125 0 3461.5208
374.1497802734375 0 616.20447
376.11395263671875 0 2163.6155
378.12945556640625 0 8227.12
379.13226318359375 0 1873.7908
382.1429748535156 0 2256.0588
385.2430114746094 0 984.66187
387.2594299316406 0 1660.2007
390.15545654296875 0 1394.3318
394.1242980957031 0 14349.946
395.1276550292969 0 2089.5925
402.1767883300781 0 4270.1533
407.1838073730469 0 1172.2126
408.16680908203125 0 3248.9326
409.168701171875 0 855.24585
413.84857177734375 0 566.1343
420.1870422363281 0 1189.8822
421.2234191894531 0 636.2462
425.1929931640625 0 11668.6045
426.1958923339844 0 2159.391
427.20806884765625 0 2252.5232
428.21051025390625 0 771.1779
433.1723327636719 0 2330.671
447.1510314941406 0 1961.799
448.7696533203125 0 1352.0507
449.1335144042969 0 873.888
449.2691650390625 0 652.8371
454.1957702636719 0 1043.0098
455.2050476074219 0 2166.4033
456.3185119628906 0 2343.8958
457.32025146484375 0 864.7553
460.76593017578125 0 727.99756
461.16632080078125 0 3413.1406 b Water loss 3
461.1976318359375 0 1042.962
462.1693420410156 0 887.22205
462.2654724121094 0 703.66705
465.1617736816406 0 10061.072
466.16363525390625 0 1988.8601
466.30316162109375 0 1601.2472
471.1806640625 0 1470.0734
477.1673583984375 0 1647.5569 y 2
477.6753845214844 0 679.99023
479.1771240234375 0 11218.919 b 3
480.1800537109375 0 2684.5432
482.19146728515625 0 966.91626
484.3128662109375 0 5594.161
485.3162536621094 0 1461.3495
486.16107177734375 0 2050.5159
489.1913757324219 0 20612.762 y 6
490.1935729980469 0 5549.6357
491.20208740234375 0 2079.196
495.18499755859375 0 1698.5656
496.2303466796875 0 8063.629
497.1705017089844 0 1604.7181
497.2333068847656 0 2838.2986
513.2000122070312 0 1815.6592
513.3134155273438 0 912.8737
518.7142333984375 0 630.6696
519.2032470703125 0 3569.3074
520.2039794921875 0 1980.5526
527.7089233398438 0 638.34595
534.3006591796875 0 978.894
537.2144165039062 0 934.8719
541.7072143554688 0 1095.1273 y Water loss 1
542.22900390625 0 2042.6029
548.1991577148438 0 3426.424
549.1998901367188 0 722.24286
550.7103271484375 0 11285.784 y 1
551.2119140625 0 6499.652
551.7120971679688 0 2273.2437
553.3102416992188 0 1249.7638
557.1978149414062 0 1420.8402
559.2501831054688 0 1590.0543
560.2281494140625 0 30672.615 y 5
561.23095703125 0 9028.359
562.2313842773438 0 2067.2012
569.2329711914062 0 2410.043
569.7352294921875 0 1470.5496
570.2276611328125 0 3464.2346
574.3278198242188 0 1106.6741
578.2301025390625 0 1348.0715
579.232666015625 0 1102.7257
590.8449096679688 0 1054.7443
591.3427734375 0 764.169
592.2293090820312 0 5903.066 Precursor Water loss
592.7311401367188 0 4704.3877
593.2296752929688 0 1860.423
596.2340087890625 0 1693.0798
599.8474731445312 0 795.06085
600.347412109375 0 754.2893
601.046630859375 0 744.9211
601.2340698242188 0 14955.613 Precursor
601.316162109375 0 854.217
601.3919067382812 0 3140.6465
601.7355346679688 0 9649.133
602.2360229492188 0 4875.017
606.2330322265625 0 3507.1619
612.1964111328125 0 5060.417
613.1980590820312 0 1385.3534
614.245361328125 0 2866.2852
615.248046875 0 1117.5612
624.2318115234375 0 2535.0344 b Water loss 4
631.2437133789062 0 801.222
634.2276000976562 0 3021.0513
635.2360229492188 0 788.9862
640.2359008789062 0 1588.5123
641.2886962890625 0 1105.9518
642.2396850585938 0 2090.7974 b 4
655.3031005859375 0 780.91486
657.2623901367188 0 5450.128
658.2647094726562 0 1829.0197
659.2359008789062 0 790.396
659.2926025390625 0 9748.505
660.2957763671875 0 3940.1294
672.3223876953125 0 645.4686
676.2640991210938 0 1205.8364
677.2772216796875 0 684.98926
685.25732421875 0 8207.969
686.2610473632812 0 2917.8591
691.33837890625 0 931.4651
695.2662353515625 0 1310.0208 b Water loss 5
696.2726440429688 0 982.00073
696.7733764648438 0 673.5925
703.2628173828125 0 2944.9424
704.2459716796875 0 591.837
705.2770385742188 0 1871.7349
713.275634765625 0 1370.9537 b 5
721.2601318359375 0 9392.79
722.2640380859375 0 2800.8389
723.2907104492188 0 29548.973 y 4
724.2935180664062 0 11186.157
725.293212890625 0 3052.0513
731.2449340820312 0 1337.0775
733.2622680664062 0 803.749
749.2546997070312 0 12535.747
750.2576293945312 0 3753.1423
751.25732421875 0 1394.0764
754.3675537109375 0 709.5104
756.3111572265625 0 1041.9985
767.2667236328125 0 3433.0637
774.3196411132812 0 10922.18
775.3225708007812 0 5145.764
776.3246459960938 0 900.76044
796.305419921875 0 649.3784
804.3320922851562 0 768.22723
810.0420532226562 0 1645.8474
820.3082275390625 0 1649.4116 y Water loss 3
821.3095703125 0 1132.5508
832.3189697265625 0 994.5229
838.3176879882812 0 33416.992 y 3
839.3207397460938 0 17552.451
840.3198852539062 0 5476.0605
842.3051147460938 0 869.23975 b Water loss 6
862.3184204101562 0 1041.0304
865.7373046875 0 727.017
871.3367309570312 0 4133.103
872.3362426757812 0 3014.8445
872.4375610351562 0 902.2493
889.3460693359375 0 50080.74
890.3488159179688 0 22569.238
891.35107421875 0 5787.34
892.3444213867188 0 776.23254
895.5223999023438 0 1576.5991
896.52392578125 0 2956.6208
897.5274047851562 0 1041.9243
899.4246826171875 0 1783.887
900.4262084960938 0 1465.7043
907.3358154296875 0 2042.5051
908.340087890625 0 1210.252
918.31201171875 0 856.0934
933.3724975585938 0 750.22675
935.3334350585938 0 9799.988 y Water loss 2
936.3350830078125 0 7265.265
937.32763671875 0 1944.0225
938.3173828125 0 1232.7847
953.3446044921875 0 155779.64 y 2
954.3472290039062 0 82478.914
955.3474731445312 0 31608.64
955.4636840820312 0 724.50244
956.3473510742188 0 3410.9895
969.3678588867188 0 1050.834
997.3687133789062 0 2419.1653 b 7
998.3734741210938 0 1183.9127
998.5703735351562 0 979.48804
1018.4050903320312 0 765.2556
1036.413330078125 0 2712.9668
1037.4161376953125 0 2454.9006
1100.4111328125 0 8247.889 y 1
1101.416259765625 0 5654.3867
1102.4173583984375 0 2817.632
1110.3946533203125 0 820.9077
1111.3974609375 0 1181.3713
2146.81005859375 0 600.5825
2425.674560546875 0 670.19977
3079.611328125 0 903.78436
3080.35498046875 0 1004.76794
3325.0634765625 0 667.1938

Spectrum Details

|  |  |
| --- | --- |
| Matched peaks? Matched peaks The total absolute number of peaks matched. Additionally in brackets the total fraction of peaks matched and the total number of peaks is shown. | 34 (7.91% of 430) |
| FDR? FDR The false discovery rate estimated for this peptide. It is calculated by matching all theoretical fragments with a non-integer shift with the raw peaks for this spectrum. This is done with 40 different shifts. The resulting percentage is the average number of annotated peaks over the number of annotated peaks with the correct spectrum. | 0.77% |
| Satellite FDR? Satellite FDR See the FDR for details on its calculation. This satellite ion specific FDR only contains the satellite ions (d/w) for I/L/J positions. | - |
| PSM Score? PSM Score The PSM Score as given by Hecklib to this annotated spectrum. It is shown with three significant figures. | 440 |

#### Spectrum 10292? Spectrum 10292 The raw spectrum of this peptide as annotated by Hecklib. The fragments are coloured according to ion type (see legend). Any peaks with a star '\*' as text can be hovered over to see the full details, first the ion type second the mass shift type. By hovering over the amino acids in the peptide or ions in the legend the corresponding peaks are highlighted. By toggling the 'Unassigned' label you can turn the background (unassigned) peaks on or off in the plot. By updating the slider in the Ion legend you can update the spectrum to only show the top X% of the peaks with labels. The top X% means any peak that is within X% of the highest intensity. By dragging in the spectrum you can zoom in to a specific part of the spectrum and use 'Zoom Out' to get back to the original zoom level. The annotation of the spectrum is based on the given sequence in the peptides file and is done with different software so inconsistencies are likely. The peaks are annotated based on the given sequence, with 20 ppm tolerance.

Copy Data

##### Spectrum 10292 (TSV)

###### Preview

```
Loading example...
```

*Click on the button to copy the data to your clipboard.*

Mz MinMz MaxIntensity Max

WidthHeightPeptide font sizePeptide stroke widthSpectrum font sizeSpectrum stroke widthCompact peptide

Ion legend

wxyz

abcd

OtherUnassignedIon^Charge^_Position_Show for top:%

TFDDYAMHW

02.11e+44.23e+46.34e+48.45e+4

Zoom Out

a^+1^_2_y^+1^_1_a^+1^_2_b^+1^_2_b^+1^_2_a^+1^_3_y^+1^_2_b^+1^_3_b^+1^_3_b^+1^_4_y^+1^_3_y^+2^_8_y^+1^_4_\*\*b^+1^_5_y^+1^_5_y^+1^_6_y^+1^_7_y^+1^_7_y^+1^_8_

0779155723363115

Fragment Matches Table

Show background peaks

| Position | Ion type | Intensity | mz Theoretical | mz Error (Th) | mz Error (ppm) | Charge | Series Number |
| --- | --- | --- | --- | --- | --- | --- | --- |
| - | - | 7.483E+04 | 120.1 | - | - | 0 | - |
| - | - | 760.6 | 121.1 | - | - | 0 | - |
| - | - | 6623 | 121.1 | - | - | 0 | - |
| - | - | 524.7 | 122.1 | - | - | 0 | - |
| - | - | 355.3 | 122.5 | - | - | 0 | - |
| - | - | 937.8 | 123 | - | - | 0 | - |
| - | - | 464.9 | 127.1 | - | - | 0 | - |
| - | - | 689.6 | 129.1 | - | - | 0 | - |
| - | - | 7613 | 129.1 | - | - | 0 | - |
| - | - | 2188 | 130.1 | - | - | 0 | - |
| - | - | 592 | 130.1 | - | - | 0 | - |
| - | - | 2820 | 132.1 | - | - | 0 | - |
| - | - | 867.5 | 132.1 | - | - | 0 | - |
| - | - | 1193 | 133.1 | - | - | 0 | - |
| - | - | 3.091E+04 | 136.1 | - | - | 0 | - |
| - | - | 2776 | 137.1 | - | - | 0 | - |
| - | - | 685.1 | 138.1 | - | - | 0 | - |
| - | - | 1118 | 138.1 | - | - | 0 | - |
| - | - | 474.7 | 141.9 | - | - | 0 | - |
| - | - | 448.5 | 143 | - | - | 0 | - |
| - | - | 876.8 | 144.1 | - | - | 0 | - |
| - | - | 5361 | 146.1 | - | - | 0 | - |
| - | - | 1138 | 147 | - | - | 0 | - |
| - | - | 1042 | 147.1 | - | - | 0 | - |
| - | - | 645.8 | 148.9 | - | - | 0 | - |
| - | - | 491.9 | 148.9 | - | - | 0 | - |
| - | - | 643.4 | 148.9 | - | - | 0 | - |
| - | - | 611.9 | 148.9 | - | - | 0 | - |
| - | - | 963.5 | 148.9 | - | - | 0 | - |
| - | - | 1191 | 148.9 | - | - | 0 | - |
| - | - | 1329 | 148.9 | - | - | 0 | - |
| - | - | 3216 | 148.9 | - | - | 0 | - |
| - | - | 5391 | 148.9 | - | - | 0 | - |
| - | - | 3706 | 149 | - | - | 0 | - |
| - | - | 1854 | 149 | - | - | 0 | - |
| - | - | 1121 | 149 | - | - | 0 | - |
| - | - | 979.6 | 149 | - | - | 0 | - |
| - | - | 870.9 | 149 | - | - | 0 | - |
| - | - | 660.8 | 149 | - | - | 0 | - |
| - | - | 554 | 149 | - | - | 0 | - |
| - | - | 511.5 | 149 | - | - | 0 | - |
| - | - | 529.6 | 149 | - | - | 0 | - |
| - | - | 425.3 | 149 | - | - | 0 | - |
| - | - | 1159 | 149 | - | - | 0 | - |
| - | - | 427.8 | 155.1 | - | - | 0 | - |
| - | - | 2041 | 156.1 | - | - | 0 | - |
| - | - | 1340 | 157.1 | - | - | 0 | - |
| - | - | 2966 | 158.1 | - | - | 0 | - |
| - | - | 8511 | 159.1 | - | - | 0 | - |
| - | - | 775.6 | 160.1 | - | - | 0 | - |
| - | - | 1163 | 160.1 | - | - | 0 | - |
| - | - | 1506 | 160.1 | - | - | 0 | - |
| - | - | 8466 | 165.1 | - | - | 0 | - |
| - | - | 665.4 | 165.1 | - | - | 0 | - |
| - | - | 530.5 | 165.1 | - | - | 0 | - |
| - | - | 1577 | 166.1 | - | - | 0 | - |
| - | - | 1767 | 167.1 | - | - | 0 | - |
| - | - | 703.3 | 169.1 | - | - | 0 | - |
| - | - | 768.3 | 170.1 | - | - | 0 | - |
| - | - | 3968 | 173.1 | - | - | 0 | - |
| - | - | 735.8 | 174.1 | - | - | 0 | - |
| - | - | 694.3 | 175.1 | - | - | 0 | - |
| - | - | 593.4 | 176.1 | - | - | 0 | - |
| - | - | 7048 | 176.1 | - | - | 0 | - |
| - | - | 4356 | 177.1 | - | - | 0 | - |
| - | - | 916.5 | 177.1 | - | - | 0 | - |
| - | - | 844.3 | 178.1 | - | - | 0 | - |
| - | - | 569.3 | 181.1 | - | - | 0 | - |
| - | - | 1.642E+04 | 182.1 | - | - | 0 | - |
| - | - | 1306 | 183.1 | - | - | 0 | - |
| - | - | 1004 | 185.1 | - | - | 0 | - |
| - | - | 1715 | 185.1 | - | - | 0 | - |
| - | - | 697.6 | 186.1 | - | - | 0 | - |
| - | - | 652.4 | 187.1 | - | - | 0 | - |
| - | - | 763.9 | 187.1 | - | - | 0 | - |
| - | - | 2.84E+04 | 188.1 | - | - | 0 | - |
| - | - | 3589 | 189.1 | - | - | 0 | - |
| - | - | 480.3 | 189.1 | - | - | 0 | - |
| - | - | 460 | 189.1 | - | - | 0 | - |
| - | - | 1692 | 191.1 | - | - | 0 | - |
| - | - | 1671 | 193.1 | - | - | 0 | - |
| - | - | 772.5 | 195.1 | - | - | 0 | - |
| - | - | 497.2 | 199.1 | - | - | 0 | - |
| - | - | 941.8 | 200.1 | - | - | 0 | - |
| - | - | 2961 | 201.1 | - | - | 0 | - |
| - | - | 1168 | 203.1 | - | - | 0 | - |
| 2 | a | 3032 | 203.1 | 1.539E-05 | 0.07577 | +1 | 2 |
| - | - | 1183 | 204.1 | - | - | 0 | - |
| 9 | y | 1.078E+04 | 205.1 | 0.0001665 | 0.8116 | +1 | 1 |
| - | - | 1691 | 206.1 | - | - | 0 | - |
| - | - | 749 | 214.1 | - | - | 0 | - |
| - | - | 1441 | 217.1 | - | - | 0 | - |
| - | - | 774.9 | 219.1 | - | - | 0 | - |
| - | - | 5434 | 221.1 | - | - | 0 | - |
| 2 | a | 8.37E+04 | 221.1 | 0.0002079 | 0.9401 | +1 | 2 |
| - | - | 941 | 222.1 | - | - | 0 | - |
| - | - | 751.1 | 222.1 | - | - | 0 | - |
| - | - | 9170 | 222.1 | - | - | 0 | - |
| - | - | 1494 | 223.1 | - | - | 0 | - |
| - | - | 681.7 | 223.1 | - | - | 0 | - |
| - | - | 503.7 | 224.6 | - | - | 0 | - |
| - | - | 478 | 225.9 | - | - | 0 | - |
| - | - | 765.9 | 226.2 | - | - | 0 | - |
| - | - | 712.9 | 229.1 | - | - | 0 | - |
| - | - | 500.6 | 229.2 | - | - | 0 | - |
| - | - | 1941 | 231.1 | - | - | 0 | - |
| - | - | 637.3 | 231.1 | - | - | 0 | - |
| 2 | b | 1469 | 231.1 | 3.459E-05 | 0.1497 | +1 | 2 |
| - | - | 2241 | 233.1 | - | - | 0 | - |
| - | - | 1553 | 233.1 | - | - | 0 | - |
| - | - | 827.3 | 233.2 | - | - | 0 | - |
| - | - | 538.9 | 233.2 | - | - | 0 | - |
| - | - | 2433 | 235.1 | - | - | 0 | - |
| - | - | 640.9 | 237.1 | - | - | 0 | - |
| - | - | 927.1 | 239.1 | - | - | 0 | - |
| - | - | 815.7 | 239.1 | - | - | 0 | - |
| - | - | 1284 | 239.2 | - | - | 0 | - |
| - | - | 981.1 | 241.1 | - | - | 0 | - |
| 2 | b | 1.368E+04 | 249.1 | 0.0002579 | 1.035 | +1 | 2 |
| - | - | 1981 | 250.1 | - | - | 0 | - |
| - | - | 2138 | 251.1 | - | - | 0 | - |
| - | - | 898.6 | 251.2 | - | - | 0 | - |
| - | - | 771.7 | 253.1 | - | - | 0 | - |
| - | - | 1058 | 253.2 | - | - | 0 | - |
| - | - | 942 | 257.1 | - | - | 0 | - |
| - | - | 559.3 | 257.2 | - | - | 0 | - |
| - | - | 1399 | 261.1 | - | - | 0 | - |
| - | - | 2791 | 263.1 | - | - | 0 | - |
| - | - | 632.8 | 274.1 | - | - | 0 | - |
| - | - | 2063 | 279.1 | - | - | 0 | - |
| - | - | 581.4 | 282.1 | - | - | 0 | - |
| - | - | 598.9 | 284.2 | - | - | 0 | - |
| - | - | 958.7 | 285 | - | - | 0 | - |
| - | - | 1604 | 285.1 | - | - | 0 | - |
| - | - | 1051 | 292.1 | - | - | 0 | - |
| - | - | 1590 | 296.2 | - | - | 0 | - |
| 3 | a | 738.4 | 318.1 | 0.0008278 | 2.602 | +1 | 3 |
| - | - | 1203 | 319.1 | - | - | 0 | - |
| - | - | 682 | 321.2 | - | - | 0 | - |
| - | - | 3610 | 324.1 | - | - | 0 | - |
| - | - | 717.2 | 325.1 | - | - | 0 | - |
| - | - | 1.07E+04 | 329.1 | - | - | 0 | - |
| - | - | 2242 | 330.2 | - | - | 0 | - |
| - | - | 1813 | 340.2 | - | - | 0 | - |
| 8 | y | 1.815E+04 | 342.2 | 0.0002451 | 0.7163 | +1 | 2 |
| - | - | 1131 | 342.2 | - | - | 0 | - |
| - | - | 3447 | 343.2 | - | - | 0 | - |
| 3 | b | 1294 | 346.1 | 0.00135 | 3.9 | +1 | 3 |
| - | - | 554.2 | 349.4 | - | - | 0 | - |
| - | - | 1092 | 350.1 | - | - | 0 | - |
| - | - | 986.7 | 352.2 | - | - | 0 | - |
| - | - | 1284 | 355.1 | - | - | 0 | - |
| - | - | 673.1 | 356.1 | - | - | 0 | - |
| - | - | 1163 | 356.1 | - | - | 0 | - |
| - | - | 2458 | 357.2 | - | - | 0 | - |
| 3 | b | 3299 | 364.2 | 7.384E-05 | 0.2028 | +1 | 3 |
| - | - | 527.7 | 365.2 | - | - | 0 | - |
| - | - | 630.1 | 366.1 | - | - | 0 | - |
| - | - | 608.3 | 366.2 | - | - | 0 | - |
| - | - | 842.7 | 368.2 | - | - | 0 | - |
| - | - | 2795 | 378.1 | - | - | 0 | - |
| - | - | 2905 | 394.1 | - | - | 0 | - |
| - | - | 1012 | 398.2 | - | - | 0 | - |
| - | - | 728.5 | 401.7 | - | - | 0 | - |
| - | - | 709.7 | 408.2 | - | - | 0 | - |
| - | - | 1102 | 413.3 | - | - | 0 | - |
| - | - | 661.5 | 414.3 | - | - | 0 | - |
| - | - | 794.8 | 422.7 | - | - | 0 | - |
| - | - | 612.6 | 423.2 | - | - | 0 | - |
| - | - | 2175 | 425.2 | - | - | 0 | - |
| - | - | 1050 | 427.7 | - | - | 0 | - |
| - | - | 1082 | 433.3 | - | - | 0 | - |
| - | - | 629 | 436.7 | - | - | 0 | - |
| - | - | 954.4 | 437.2 | - | - | 0 | - |
| - | - | 820.6 | 455.2 | - | - | 0 | - |
| - | - | 2181 | 465.2 | - | - | 0 | - |
| 4 | b | 2218 | 479.2 | 0.0007846 | 1.637 | +1 | 4 |
| - | - | 923.7 | 480.2 | - | - | 0 | - |
| - | - | 592.7 | 480.8 | - | - | 0 | - |
| - | - | 729.1 | 483.2 | - | - | 0 | - |
| 7 | y | 5124 | 489.2 | 0.005313 | 10.86 | +1 | 3 |
| - | - | 1371 | 490.2 | - | - | 0 | - |
| - | - | 1136 | 496.2 | - | - | 0 | - |
| - | - | 693.8 | 497.2 | - | - | 0 | - |
| - | - | 625.3 | 499.7 | - | - | 0 | - |
| - | - | 692.4 | 501.3 | - | - | 0 | - |
| - | - | 742.6 | 513.4 | - | - | 0 | - |
| - | - | 624.6 | 540.2 | - | - | 0 | - |
| - | - | 640 | 547.1 | - | - | 0 | - |
| - | - | 1236 | 547.3 | - | - | 0 | - |
| 2 | y | 1748 | 550.7 | 0.003141 | 5.704 | +2 | 8 |
| - | - | 1694 | 551.2 | - | - | 0 | - |
| - | - | 1020 | 551.7 | - | - | 0 | - |
| - | - | 655.1 | 557.2 | - | - | 0 | - |
| 6 | y | 6552 | 560.2 | 0.004607 | 8.224 | +1 | 4 |
| - | - | 2376 | 561.2 | - | - | 0 | - |
| - | - | 735 | 562.2 | - | - | 0 | - |
| - | - | 691 | 569.2 | - | - | 0 | - |
| - | - | 577.2 | 570.2 | - | - | 0 | - |
| - | - | 4210 | 585.3 | - | - | 0 | - |
| - | - | 1443 | 586.3 | - | - | 0 | - |
| 0 | Precursor | 1025 | 592.2 | 0.001674 | 2.827 | +2 | -1 |
| - | - | 707.3 | 592.7 | - | - | 0 | - |
| 0 | Precursor | 2982 | 601.2 | 0.002984 | 4.962 | +2 | -1 |
| - | - | 629.1 | 601.3 | - | - | 0 | - |
| - | - | 1892 | 601.7 | - | - | 0 | - |
| - | - | 732.3 | 601.8 | - | - | 0 | - |
| - | - | 1388 | 612.2 | - | - | 0 | - |
| - | - | 771.6 | 614.2 | - | - | 0 | - |
| - | - | 628.3 | 614.8 | - | - | 0 | - |
| 5 | b | 1149 | 624.2 | 0.0004045 | 0.648 | +1 | 5 |
| - | - | 1021 | 628.4 | - | - | 0 | - |
| - | - | 605.7 | 640.2 | - | - | 0 | - |
| - | - | 1074 | 657.3 | - | - | 0 | - |
| - | - | 644 | 658.3 | - | - | 0 | - |
| - | - | 2634 | 659.3 | - | - | 0 | - |
| - | - | 5850 | 670.8 | - | - | 0 | - |
| - | - | 6108 | 671.3 | - | - | 0 | - |
| - | - | 3015 | 671.8 | - | - | 0 | - |
| - | - | 663.5 | 672.3 | - | - | 0 | - |
| - | - | 656.4 | 682.3 | - | - | 0 | - |
| - | - | 1524 | 685.3 | - | - | 0 | - |
| - | - | 1390 | 693.8 | - | - | 0 | - |
| - | - | 1201 | 694.3 | - | - | 0 | - |
| - | - | 683.8 | 695.3 | - | - | 0 | - |
| - | - | 9676 | 700.4 | - | - | 0 | - |
| - | - | 4312 | 701.4 | - | - | 0 | - |
| - | - | 1419 | 721.3 | - | - | 0 | - |
| 5 | y | 5580 | 723.3 | 0.003351 | 4.633 | +1 | 5 |
| - | - | 1889 | 724.3 | - | - | 0 | - |
| - | - | 826.8 | 725.3 | - | - | 0 | - |
| - | - | 1509 | 730.4 | - | - | 0 | - |
| - | - | 1832 | 730.9 | - | - | 0 | - |
| - | - | 3412 | 744.4 | - | - | 0 | - |
| - | - | 3664 | 744.9 | - | - | 0 | - |
| - | - | 1371 | 745.4 | - | - | 0 | - |
| - | - | 2395 | 749.3 | - | - | 0 | - |
| - | - | 883.2 | 750.3 | - | - | 0 | - |
| - | - | 1016 | 757.4 | - | - | 0 | - |
| - | - | 638.9 | 767.3 | - | - | 0 | - |
| - | - | 2449 | 774.3 | - | - | 0 | - |
| - | - | 1270 | 775.3 | - | - | 0 | - |
| - | - | 2833 | 783.4 | - | - | 0 | - |
| - | - | 1387 | 784.4 | - | - | 0 | - |
| - | - | 2816 | 794.4 | - | - | 0 | - |
| - | - | 1345 | 795.4 | - | - | 0 | - |
| - | - | 1387 | 800.4 | - | - | 0 | - |
| - | - | 4901 | 801.4 | - | - | 0 | - |
| - | - | 2529 | 802.4 | - | - | 0 | - |
| - | - | 2811 | 828.4 | - | - | 0 | - |
| - | - | 792.2 | 829.4 | - | - | 0 | - |
| 4 | y | 7332 | 838.3 | 0.003752 | 4.476 | +1 | 6 |
| - | - | 3518 | 839.3 | - | - | 0 | - |
| - | - | 1201 | 840.3 | - | - | 0 | - |
| - | - | 6471 | 844.5 | - | - | 0 | - |
| - | - | 3356 | 845.5 | - | - | 0 | - |
| - | - | 664.6 | 846.5 | - | - | 0 | - |
| - | - | 926.9 | 854.4 | - | - | 0 | - |
| - | - | 714.7 | 855.4 | - | - | 0 | - |
| - | - | 616.5 | 862.3 | - | - | 0 | - |
| - | - | 837.2 | 871.3 | - | - | 0 | - |
| - | - | 983.5 | 871.4 | - | - | 0 | - |
| - | - | 2.642E+04 | 872.4 | - | - | 0 | - |
| - | - | 1.363E+04 | 873.4 | - | - | 0 | - |
| - | - | 3138 | 874.5 | - | - | 0 | - |
| - | - | 8885 | 889.3 | - | - | 0 | - |
| - | - | 6132 | 890.3 | - | - | 0 | - |
| - | - | 1067 | 891.4 | - | - | 0 | - |
| - | - | 814.2 | 896.3 | - | - | 0 | - |
| - | - | 2245 | 909.4 | - | - | 0 | - |
| - | - | 1001 | 910.4 | - | - | 0 | - |
| 3 | y | 1996 | 935.3 | 0.005013 | 5.359 | +1 | 7 |
| - | - | 1452 | 936.3 | - | - | 0 | - |
| 3 | y | 3.244E+04 | 953.3 | 0.003481 | 3.652 | +1 | 7 |
| - | - | 1.776E+04 | 954.3 | - | - | 0 | - |
| - | - | 6996 | 955.3 | - | - | 0 | - |
| - | - | 5203 | 991.5 | - | - | 0 | - |
| - | - | 1936 | 992.5 | - | - | 0 | - |
| - | - | 1.172E+04 | 1020 | - | - | 0 | - |
| - | - | 7033 | 1021 | - | - | 0 | - |
| - | - | 2884 | 1022 | - | - | 0 | - |
| - | - | 750.1 | 1036 | - | - | 0 | - |
| - | - | 809.2 | 1038 | - | - | 0 | - |
| - | - | 655.8 | 1039 | - | - | 0 | - |
| - | - | 768.1 | 1056 | - | - | 0 | - |
| 2 | y | 1785 | 1100 | 0.001718 | 1.561 | +1 | 8 |
| - | - | 1644 | 1101 | - | - | 0 | - |
| - | - | 805 | 1178 | - | - | 0 | - |
| - | - | 640 | 1220 | - | - | 0 | - |
| - | - | 739.4 | 1325 | - | - | 0 | - |
| - | - | 1351 | 1341 | - | - | 0 | - |
| - | - | 727.2 | 1342 | - | - | 0 | - |
| - | - | 667.2 | 1343 | - | - | 0 | - |
| - | - | 793.5 | 1387 | - | - | 0 | - |
| - | - | 640.8 | 2239 | - | - | 0 | - |
| - | - | 621.3 | 3007 | - | - | 0 | - |
| - | - | 706.5 | 3083 | - | - | 0 | - |
| - | - | 736.6 | 3084 | - | - | 0 | - |

m/z Charge Intensity FragmentType MassShift Position
120.0810775756836 0 74825.69
121.07963562011719 0 760.59607
121.0843734741211 0 6622.862
122.08763885498047 0 524.7402
122.54342651367188 0 355.2934
123.04446411132812 0 937.8296
127.08678436279297 0 464.88043
129.06622314453125 0 689.5675
129.1024932861328 0 7612.9683
130.06544494628906 0 2187.7705
130.1060791015625 0 591.957
132.0810089111328 0 2819.6099
132.10191345214844 0 867.5407
133.0860595703125 0 1193.0459
136.0759735107422 0 30910.54
137.07937622070312 0 2776.1365
138.05543518066406 0 685.0808
138.06640625 0 1118.3257
141.92776489257812 0 474.72717
143.04627990722656 0 448.52484
144.08108520507812 0 876.7502
146.06028747558594 0 5361.3223
147.0443115234375 0 1137.6934
147.06393432617188 0 1042.0813
148.88705444335938 0 645.76886
148.89427185058594 0 491.92996
148.90174865722656 0 643.44
148.90902709960938 0 611.87714
148.91615295410156 0 963.5283
148.92298889160156 0 1190.9082
148.93060302734375 0 1329.0435
148.9380645751953 0 3216.1963
148.94590759277344 0 5391.2007
148.9626007080078 0 3706.0193
148.97032165527344 0 1854.2363
148.97735595703125 0 1121.4265
148.98477172851562 0 979.57355
148.9920196533203 0 870.8611
148.99908447265625 0 660.7506
149.00631713867188 0 554.0174
149.0138397216797 0 511.4777
149.02175903320312 0 529.6324
149.03555297851562 0 425.28384
149.04469299316406 0 1159.2622
155.1179962158203 0 427.8363
156.07696533203125 0 2040.6235
157.1342010498047 0 1340.2266
158.09669494628906 0 2965.566
159.0919189453125 0 8510.533
160.07569885253906 0 775.5916
160.09539794921875 0 1163.3433
160.1122589111328 0 1505.7382
165.05484008789062 0 8465.788
165.07716369628906 0 665.42255
165.1027374267578 0 530.53723
166.08636474609375 0 1577.4423
167.05567932128906 0 1767.2898
169.09759521484375 0 703.3233
170.0601806640625 0 768.2877
173.12869262695312 0 3968.0977
174.0552520751953 0 735.7778
175.08738708496094 0 694.34564
176.08270263671875 0 593.35864
176.1072235107422 0 7047.6416
177.10256958007812 0 4356.3384
177.1110076904297 0 916.468
178.1062774658203 0 844.27704
181.09707641601562 0 569.3438
182.0813751220703 0 16423.047
183.0848388671875 0 1306.2047
185.05593872070312 0 1003.5048
185.128662109375 0 1715.4496
186.09144592285156 0 697.56824
187.08726501464844 0 652.3694
187.14486694335938 0 763.931
188.07081604003906 0 28400.797
189.07432556152344 0 3588.8406
189.083251953125 0 480.27866
189.08689880371094 0 459.99008
191.1179962158203 0 1691.8612
193.10899353027344 0 1670.6501
195.0878448486328 0 772.4851
199.14369201660156 0 497.22894
200.13986206054688 0 941.7689
201.12347412109375 0 2961.0789
203.0663299560547 0 1168.0022
203.1178741455078 0 3031.9849 a Water loss 1
204.07701110839844 0 1183.024
205.09732055664062 0 10779.662 y 8
206.10086059570312 0 1691.1487
214.1192169189453 0 749.0025
217.0974578857422 0 1440.784
219.113037109375 0 774.94556
221.10348510742188 0 5433.93
221.128662109375 0 83697.1 a 1
222.1064453125 0 941.0348
222.12191772460938 0 751.05615
222.13201904296875 0 9169.681
223.06394958496094 0 1493.6649
223.13414001464844 0 681.68536
224.63760375976562 0 503.66226
225.85354614257812 0 477.96643
226.1544952392578 0 765.889
229.11915588378906 0 712.93756
229.1718292236328 0 500.56186
231.06143188476562 0 1941.256
231.09791564941406 0 637.337
231.1128387451172 0 1469.3253 b Water loss 1
233.09207153320312 0 2241.3352
233.12884521484375 0 1552.7098
233.15052795410156 0 827.29083
233.22940063476562 0 538.9385
235.1080780029297 0 2433.3772
237.13531494140625 0 640.8737
239.09564208984375 0 927.1436
239.1146697998047 0 815.6695
239.1500244140625 0 1283.7881
241.0742950439453 0 981.1117
249.12362670898438 0 13681.495 b 1
250.12677001953125 0 1980.6306
251.1028594970703 0 2137.5713
251.15045166015625 0 898.6455
253.11892700195312 0 771.69116
253.16542053222656 0 1057.6295
257.1064147949219 0 941.9957
257.1636047363281 0 559.2632
261.12408447265625 0 1398.5833
263.1026916503906 0 2791.0024
274.12921142578125 0 632.78656
279.097900390625 0 2062.9368
282.0733642578125 0 581.42426
284.19696044921875 0 598.9029
285.0096435546875 0 958.7087
285.102294921875 0 1603.6631
292.14013671875 0 1050.6718
296.15057373046875 0 1589.5886
318.1456604003906 0 738.38727 a Water loss 2
319.1406555175781 0 1202.5692
321.15643310546875 0 682.01855
324.1455078125 0 3609.6624
325.1466979980469 0 717.17334
329.1498718261719 0 10697.475
330.1529846191406 0 2242.4946
340.1976013183594 0 1812.782
342.15631103515625 0 18146.129 y 7
342.17938232421875 0 1130.6898
343.1593933105469 0 3446.9165
346.1383972167969 0 1294.2783 b Water loss 2
349.4438781738281 0 554.20886
350.1356506347656 0 1092.267
352.16522216796875 0 986.74945
355.0696105957031 0 1284.3989
356.0703430175781 0 673.05475
356.13836669921875 0 1163.2494
357.22430419921875 0 2457.7888
364.1502380371094 0 3298.979 b 2
365.1522521972656 0 527.68
366.1318664550781 0 630.059
366.1788635253906 0 608.3415
368.19207763671875 0 842.69257
378.1294860839844 0 2795.4949
394.12469482421875 0 2904.7056
398.2403869628906 0 1012.112
401.7113952636719 0 728.5161
408.1669921875 0 709.70416
413.2662353515625 0 1101.8892
414.2691650390625 0 661.485
422.7315979003906 0 794.81476
423.2313232421875 0 612.64325
425.19342041015625 0 2174.502
427.7229309082031 0 1050.3813
433.2784423828125 0 1082.2023
436.72760009765625 0 628.99384
437.2298889160156 0 954.4045
455.1998596191406 0 820.5557
465.16162109375 0 2181.1929
479.17803955078125 0 2218.1455 b 3
480.1819763183594 0 923.6663
480.7956848144531 0 592.6813
483.2198791503906 0 729.08966
489.1918640136719 0 5124.308 y 6
490.1963806152344 0 1371.2264
496.231201171875 0 1136.0952
497.2349853515625 0 693.8341
499.67803955078125 0 625.3261
501.2513122558594 0 692.42896
513.3526611328125 0 742.5673
540.2152099609375 0 624.597
547.10205078125 0 639.99725
547.2503051757812 0 1236.1854
550.71142578125 0 1748.3896 y 1
551.21142578125 0 1693.643
551.7146606445312 0 1020.2867
557.1973876953125 0 655.0757
560.228271484375 0 6551.6577 y 5
561.2307739257812 0 2376.007
562.2269287109375 0 735.0182
569.2363891601562 0 691.0455
570.224365234375 0 577.17126
585.3350830078125 0 4209.502
586.3395385742188 0 1442.9493
592.228515625 0 1025.3196 Precursor Water loss
592.7317504882812 0 707.33093
601.235107421875 0 2982.3735 Precursor
601.3348999023438 0 629.0687
601.7377319335938 0 1891.8937
601.8200073242188 0 732.2514
612.1963500976562 0 1387.7468
614.24267578125 0 771.609
614.7567138671875 0 628.27606
624.2296142578125 0 1148.6498 b Water loss 4
628.3759765625 0 1020.86426
640.24072265625 0 605.66595
657.2596435546875 0 1074.2959
658.263671875 0 644.01294
659.29248046875 0 2634.085
670.8169555664062 0 5849.8486
671.318359375 0 6107.933
671.818359375 0 3015.292
672.320068359375 0 663.53125
682.3495483398438 0 656.44965
685.2571411132812 0 1523.9951
693.8124389648438 0 1390.0758
694.3053588867188 0 1200.5438
695.316162109375 0 683.7854
700.3618774414062 0 9676.276
701.3641967773438 0 4311.9946
721.2625732421875 0 1418.9855
723.2903442382812 0 5580.415 y 4
724.2950439453125 0 1888.9052
725.3018798828125 0 826.75714
730.3539428710938 0 1509.2874
730.8550415039062 0 1831.6211
744.350830078125 0 3412.2964
744.852294921875 0 3664.3735
745.3519897460938 0 1371.0907
749.2554931640625 0 2395.3867
750.2579956054688 0 883.1646
757.3824462890625 0 1015.9397
767.2647705078125 0 638.89655
774.3203735351562 0 2448.7825
775.3240356445312 0 1269.5634
783.3988037109375 0 2833.4917
784.4013061523438 0 1386.6251
794.3824462890625 0 2815.9001
795.385986328125 0 1344.6469
800.4276733398438 0 1387.0732
801.4111938476562 0 4901.196
802.4136352539062 0 2528.6985
828.4185791015625 0 2811.206
829.419921875 0 792.17456
838.3176879882812 0 7332.0864 y 3
839.3213500976562 0 3518.31
840.3238525390625 0 1201.4911
844.4503784179688 0 6470.7227
845.4526977539062 0 3356.4448
846.4612426757812 0 664.6456
854.4331665039062 0 926.9094
855.4400024414062 0 714.722
862.3141479492188 0 616.46576
871.3304443359375 0 837.1832
871.414306640625 0 983.4592
872.4462280273438 0 26415.834
873.44873046875 0 13626.743
874.451416015625 0 3138.4548
889.3466796875 0 8884.795
890.349609375 0 6131.971
891.3570556640625 0 1067.3441
896.32763671875 0 814.19745
909.4090576171875 0 2245.028
910.4092407226562 0 1000.9867
935.3353271484375 0 1995.5905 y Water loss 2
936.3323974609375 0 1451.911
953.3443603515625 0 32441.254 y 2
954.3469848632812 0 17762.115
955.347412109375 0 6996.055
991.5183715820312 0 5202.6807
992.519775390625 0 1935.9857
1019.5133056640625 0 11717.781
1020.515869140625 0 7032.5596
1021.5178833007812 0 2884.462
1036.4190673828125 0 750.0618
1037.5079345703125 0 809.18756
1038.513427734375 0 655.7966
1056.4842529296875 0 768.11694
1100.4110107421875 0 1784.5399 y 1
1101.41650390625 0 1643.7509
1177.568359375 0 805.03235
1220.1669921875 0 639.9993
1324.6290283203125 0 739.3899
1340.6229248046875 0 1350.9779
1341.6287841796875 0 727.211
1342.593017578125 0 667.2108
1386.580810546875 0 793.5208
2238.658447265625 0 640.8371
3007.208740234375 0 621.2717
3083.005615234375 0 706.52844
3083.75537109375 0 736.63104

Spectrum Details

|  |  |
| --- | --- |
| Matched peaks? Matched peaks The total absolute number of peaks matched. Additionally in brackets the total fraction of peaks matched and the total number of peaks is shown. | 21 (7.05% of 298) |
| FDR? FDR The false discovery rate estimated for this peptide. It is calculated by matching all theoretical fragments with a non-integer shift with the raw peaks for this spectrum. This is done with 40 different shifts. The resulting percentage is the average number of annotated peaks over the number of annotated peaks with the correct spectrum. | 1.47% |
| Satellite FDR? Satellite FDR See the FDR for details on its calculation. This satellite ion specific FDR only contains the satellite ions (d/w) for I/L/J positions. | - |
| PSM Score? PSM Score The PSM Score as given by Hecklib to this annotated spectrum. It is shown with three significant figures. | 250 |

#### Spectrum 10402? Spectrum 10402 The raw spectrum of this peptide as annotated by Hecklib. The fragments are coloured according to ion type (see legend). Any peaks with a star '\*' as text can be hovered over to see the full details, first the ion type second the mass shift type. By hovering over the amino acids in the peptide or ions in the legend the corresponding peaks are highlighted. By toggling the 'Unassigned' label you can turn the background (unassigned) peaks on or off in the plot. By updating the slider in the Ion legend you can update the spectrum to only show the top X% of the peaks with labels. The top X% means any peak that is within X% of the highest intensity. By dragging in the spectrum you can zoom in to a specific part of the spectrum and use 'Zoom Out' to get back to the original zoom level. The annotation of the spectrum is based on the given sequence in the peptides file and is done with different software so inconsistencies are likely. The peaks are annotated based on the given sequence, with 20 ppm tolerance.

Copy Data

##### Spectrum 10402 (TSV)

###### Preview

```
Loading example...
```

*Click on the button to copy the data to your clipboard.*

Mz MinMz MaxIntensity Max

WidthHeightPeptide font sizePeptide stroke widthSpectrum font sizeSpectrum stroke widthCompact peptide

Ion legend

wxyz

abcd

OtherUnassignedIon^Charge^_Position_Show for top:%

TFDDYAMHW

02.10e+44.20e+46.30e+48.40e+4

Zoom Out

a^+1^_2_y^+1^_1_a^+1^_2_b^+1^_2_b^+1^_2_a^+1^_3_y^+1^_2_b^+1^_3_b^+1^_3_b^+1^_4_b^+1^_4_y^+1^_3_y^+2^_8_y^+1^_4_\*\*y^+1^_5_y^+1^_6_y^+1^_7_y^+1^_7_y^+1^_8_

0698139520932790

Fragment Matches Table

Show background peaks

| Position | Ion type | Intensity | mz Theoretical | mz Error (Th) | mz Error (ppm) | Charge | Series Number |
| --- | --- | --- | --- | --- | --- | --- | --- |
| - | - | 528.4 | 120.1 | - | - | 0 | - |
| - | - | 7.228E+04 | 120.1 | - | - | 0 | - |
| - | - | 5420 | 121.1 | - | - | 0 | - |
| - | - | 694.8 | 123 | - | - | 0 | - |
| - | - | 8080 | 129.1 | - | - | 0 | - |
| - | - | 2503 | 130.1 | - | - | 0 | - |
| - | - | 397.7 | 130.1 | - | - | 0 | - |
| - | - | 396.9 | 130.1 | - | - | 0 | - |
| - | - | 2595 | 132.1 | - | - | 0 | - |
| - | - | 1083 | 132.1 | - | - | 0 | - |
| - | - | 1009 | 133.1 | - | - | 0 | - |
| - | - | 2.942E+04 | 136.1 | - | - | 0 | - |
| - | - | 2448 | 137.1 | - | - | 0 | - |
| - | - | 662.6 | 138.1 | - | - | 0 | - |
| - | - | 950.2 | 138.1 | - | - | 0 | - |
| - | - | 394.4 | 140.1 | - | - | 0 | - |
| - | - | 478.1 | 141.6 | - | - | 0 | - |
| - | - | 388.6 | 141.7 | - | - | 0 | - |
| - | - | 557.2 | 142.1 | - | - | 0 | - |
| - | - | 485.3 | 144.1 | - | - | 0 | - |
| - | - | 6072 | 146.1 | - | - | 0 | - |
| - | - | 1249 | 147 | - | - | 0 | - |
| - | - | 988.7 | 148.1 | - | - | 0 | - |
| - | - | 832.8 | 148.9 | - | - | 0 | - |
| - | - | 1362 | 149 | - | - | 0 | - |
| - | - | 397.9 | 149.7 | - | - | 0 | - |
| - | - | 451 | 154.2 | - | - | 0 | - |
| - | - | 496.2 | 155.1 | - | - | 0 | - |
| - | - | 1303 | 156.1 | - | - | 0 | - |
| - | - | 1435 | 157.1 | - | - | 0 | - |
| - | - | 2427 | 158.1 | - | - | 0 | - |
| - | - | 8936 | 159.1 | - | - | 0 | - |
| - | - | 1024 | 159.1 | - | - | 0 | - |
| - | - | 657.9 | 160.1 | - | - | 0 | - |
| - | - | 1177 | 160.1 | - | - | 0 | - |
| - | - | 1272 | 160.1 | - | - | 0 | - |
| - | - | 8156 | 165.1 | - | - | 0 | - |
| - | - | 442.3 | 165.1 | - | - | 0 | - |
| - | - | 839.1 | 166.1 | - | - | 0 | - |
| - | - | 1727 | 166.1 | - | - | 0 | - |
| - | - | 1360 | 167.1 | - | - | 0 | - |
| - | - | 539.9 | 167.1 | - | - | 0 | - |
| - | - | 550.9 | 169.1 | - | - | 0 | - |
| - | - | 649.4 | 170.1 | - | - | 0 | - |
| - | - | 520.5 | 173.1 | - | - | 0 | - |
| - | - | 5419 | 173.1 | - | - | 0 | - |
| - | - | 912.1 | 173.5 | - | - | 0 | - |
| - | - | 1421 | 175.1 | - | - | 0 | - |
| - | - | 744.7 | 175.1 | - | - | 0 | - |
| - | - | 501.8 | 176.1 | - | - | 0 | - |
| - | - | 5825 | 176.1 | - | - | 0 | - |
| - | - | 4576 | 177.1 | - | - | 0 | - |
| - | - | 791 | 177.1 | - | - | 0 | - |
| - | - | 1.552E+04 | 182.1 | - | - | 0 | - |
| - | - | 813.3 | 182.1 | - | - | 0 | - |
| - | - | 1101 | 183.1 | - | - | 0 | - |
| - | - | 533.2 | 185.1 | - | - | 0 | - |
| - | - | 1758 | 185.1 | - | - | 0 | - |
| - | - | 432.1 | 186.1 | - | - | 0 | - |
| - | - | 2.779E+04 | 188.1 | - | - | 0 | - |
| - | - | 3451 | 189.1 | - | - | 0 | - |
| - | - | 484.9 | 189.8 | - | - | 0 | - |
| - | - | 1519 | 191.1 | - | - | 0 | - |
| - | - | 1492 | 193.1 | - | - | 0 | - |
| - | - | 539.3 | 195.1 | - | - | 0 | - |
| - | - | 557 | 195.1 | - | - | 0 | - |
| - | - | 907.7 | 200.1 | - | - | 0 | - |
| - | - | 2484 | 201.1 | - | - | 0 | - |
| - | - | 595.1 | 202.1 | - | - | 0 | - |
| - | - | 493.5 | 203.1 | - | - | 0 | - |
| 2 | a | 2394 | 203.1 | 0.0002745 | 1.352 | +1 | 2 |
| - | - | 943.6 | 204.1 | - | - | 0 | - |
| 9 | y | 1.143E+04 | 205.1 | 0.0003648 | 1.779 | +1 | 1 |
| - | - | 1210 | 206.1 | - | - | 0 | - |
| - | - | 911.3 | 207.1 | - | - | 0 | - |
| - | - | 669.8 | 211.1 | - | - | 0 | - |
| - | - | 730.5 | 214.1 | - | - | 0 | - |
| - | - | 531.9 | 214.9 | - | - | 0 | - |
| - | - | 696.8 | 215.1 | - | - | 0 | - |
| - | - | 828.3 | 216.1 | - | - | 0 | - |
| - | - | 1165 | 217.1 | - | - | 0 | - |
| - | - | 500.7 | 218.7 | - | - | 0 | - |
| - | - | 1029 | 219.1 | - | - | 0 | - |
| - | - | 4260 | 221.1 | - | - | 0 | - |
| 2 | a | 8.316E+04 | 221.1 | 0.0003605 | 1.63 | +1 | 2 |
| - | - | 595.1 | 222.1 | - | - | 0 | - |
| - | - | 8928 | 222.1 | - | - | 0 | - |
| - | - | 1513 | 223.1 | - | - | 0 | - |
| - | - | 539.1 | 223.1 | - | - | 0 | - |
| - | - | 485.2 | 224.1 | - | - | 0 | - |
| - | - | 604.7 | 225.1 | - | - | 0 | - |
| - | - | 1174 | 225.2 | - | - | 0 | - |
| - | - | 558.9 | 226.1 | - | - | 0 | - |
| - | - | 796.7 | 227.1 | - | - | 0 | - |
| - | - | 545 | 228.1 | - | - | 0 | - |
| - | - | 1321 | 231.1 | - | - | 0 | - |
| 2 | b | 1103 | 231.1 | 0.0002706 | 1.171 | +1 | 2 |
| - | - | 1092 | 233.1 | - | - | 0 | - |
| - | - | 1583 | 233.1 | - | - | 0 | - |
| - | - | 605.4 | 233.1 | - | - | 0 | - |
| - | - | 3151 | 235.1 | - | - | 0 | - |
| - | - | 545.6 | 239.1 | - | - | 0 | - |
| - | - | 1825 | 239.2 | - | - | 0 | - |
| - | - | 641.7 | 241.6 | - | - | 0 | - |
| - | - | 484 | 243.1 | - | - | 0 | - |
| - | - | 817.7 | 249.1 | - | - | 0 | - |
| 2 | b | 1.525E+04 | 249.1 | 0.0004105 | 1.648 | +1 | 2 |
| - | - | 2221 | 250.1 | - | - | 0 | - |
| - | - | 1442 | 251.1 | - | - | 0 | - |
| - | - | 497 | 251.1 | - | - | 0 | - |
| - | - | 743.6 | 252.1 | - | - | 0 | - |
| - | - | 671.8 | 253.1 | - | - | 0 | - |
| - | - | 1515 | 253.2 | - | - | 0 | - |
| - | - | 800.2 | 256.2 | - | - | 0 | - |
| - | - | 1040 | 257.1 | - | - | 0 | - |
| - | - | 1458 | 261.1 | - | - | 0 | - |
| - | - | 3714 | 263.1 | - | - | 0 | - |
| - | - | 505.4 | 268 | - | - | 0 | - |
| - | - | 2262 | 279.1 | - | - | 0 | - |
| - | - | 514.9 | 280.2 | - | - | 0 | - |
| - | - | 1362 | 285 | - | - | 0 | - |
| - | - | 594.4 | 285.1 | - | - | 0 | - |
| - | - | 1404 | 285.1 | - | - | 0 | - |
| - | - | 805.3 | 292.1 | - | - | 0 | - |
| - | - | 515.3 | 294.9 | - | - | 0 | - |
| - | - | 1884 | 296.2 | - | - | 0 | - |
| - | - | 809.4 | 302.1 | - | - | 0 | - |
| - | - | 1229 | 304.1 | - | - | 0 | - |
| - | - | 628 | 315.2 | - | - | 0 | - |
| 3 | a | 652.8 | 318.1 | 0.001072 | 3.369 | +1 | 3 |
| - | - | 1300 | 319.1 | - | - | 0 | - |
| - | - | 731.4 | 323.1 | - | - | 0 | - |
| - | - | 2614 | 324.1 | - | - | 0 | - |
| - | - | 1019 | 325.1 | - | - | 0 | - |
| - | - | 9764 | 329.2 | - | - | 0 | - |
| - | - | 2147 | 330.2 | - | - | 0 | - |
| - | - | 758.2 | 335.2 | - | - | 0 | - |
| - | - | 636.5 | 336.2 | - | - | 0 | - |
| - | - | 689 | 337.2 | - | - | 0 | - |
| - | - | 2320 | 340.2 | - | - | 0 | - |
| 8 | y | 1.642E+04 | 342.2 | 0.0005808 | 1.697 | +1 | 2 |
| - | - | 1107 | 342.2 | - | - | 0 | - |
| - | - | 2873 | 343.2 | - | - | 0 | - |
| 3 | b | 685.8 | 346.1 | 0.0009694 | 2.8 | +1 | 3 |
| - | - | 905.1 | 350.1 | - | - | 0 | - |
| - | - | 1030 | 352.2 | - | - | 0 | - |
| - | - | 1295 | 355.1 | - | - | 0 | - |
| - | - | 663.4 | 356.1 | - | - | 0 | - |
| - | - | 1949 | 357.2 | - | - | 0 | - |
| 3 | b | 3490 | 364.2 | 0.0006281 | 1.725 | +1 | 3 |
| - | - | 839.3 | 365.2 | - | - | 0 | - |
| - | - | 1082 | 368.2 | - | - | 0 | - |
| - | - | 625.7 | 374.2 | - | - | 0 | - |
| - | - | 2142 | 378.1 | - | - | 0 | - |
| - | - | 627.3 | 390.2 | - | - | 0 | - |
| - | - | 1110 | 392.2 | - | - | 0 | - |
| - | - | 2469 | 394.1 | - | - | 0 | - |
| - | - | 554.5 | 401.2 | - | - | 0 | - |
| - | - | 680.4 | 401.7 | - | - | 0 | - |
| - | - | 626.7 | 410.8 | - | - | 0 | - |
| - | - | 1197 | 413.3 | - | - | 0 | - |
| - | - | 2511 | 425.2 | - | - | 0 | - |
| - | - | 967.5 | 427.7 | - | - | 0 | - |
| - | - | 634.2 | 428.2 | - | - | 0 | - |
| - | - | 549 | 436.7 | - | - | 0 | - |
| - | - | 729 | 437.2 | - | - | 0 | - |
| 4 | b | 759.8 | 461.2 | 0.0007597 | 1.647 | +1 | 4 |
| - | - | 1764 | 465.2 | - | - | 0 | - |
| 4 | b | 2160 | 479.2 | 0.0008152 | 1.701 | +1 | 4 |
| - | - | 826.5 | 480.2 | - | - | 0 | - |
| - | - | 839 | 483.2 | - | - | 0 | - |
| - | - | 636 | 487.1 | - | - | 0 | - |
| - | - | 681.6 | 487.3 | - | - | 0 | - |
| 7 | y | 3419 | 489.2 | 0.006076 | 12.42 | +1 | 3 |
| - | - | 1060 | 490.2 | - | - | 0 | - |
| - | - | 1754 | 496.2 | - | - | 0 | - |
| - | - | 704.5 | 510.3 | - | - | 0 | - |
| - | - | 881.6 | 519.2 | - | - | 0 | - |
| - | - | 1153 | 539.2 | - | - | 0 | - |
| - | - | 902.7 | 547.3 | - | - | 0 | - |
| - | - | 963.9 | 548.2 | - | - | 0 | - |
| 2 | y | 2251 | 550.7 | 0.00369 | 6.701 | +2 | 8 |
| - | - | 1651 | 551.2 | - | - | 0 | - |
| - | - | 665.3 | 559.3 | - | - | 0 | - |
| 6 | y | 6158 | 560.2 | 0.00534 | 9.531 | +1 | 4 |
| - | - | 2002 | 561.2 | - | - | 0 | - |
| - | - | 576.7 | 579.8 | - | - | 0 | - |
| - | - | 2376 | 585.3 | - | - | 0 | - |
| - | - | 1070 | 586.3 | - | - | 0 | - |
| - | - | 741.3 | 589.3 | - | - | 0 | - |
| 0 | Precursor | 1350 | 592.2 | 0.003627 | 6.125 | +2 | -1 |
| 0 | Precursor | 3811 | 601.2 | 0.003472 | 5.774 | +2 | -1 |
| - | - | 798.4 | 601.3 | - | - | 0 | - |
| - | - | 1646 | 601.7 | - | - | 0 | - |
| - | - | 1301 | 612.2 | - | - | 0 | - |
| - | - | 675.4 | 614.2 | - | - | 0 | - |
| - | - | 947.4 | 628.4 | - | - | 0 | - |
| - | - | 864.1 | 651.4 | - | - | 0 | - |
| - | - | 1461 | 657.3 | - | - | 0 | - |
| - | - | 1961 | 659.3 | - | - | 0 | - |
| - | - | 1073 | 660.3 | - | - | 0 | - |
| - | - | 7254 | 670.8 | - | - | 0 | - |
| - | - | 3637 | 671.3 | - | - | 0 | - |
| - | - | 702.7 | 671.4 | - | - | 0 | - |
| - | - | 2367 | 671.8 | - | - | 0 | - |
| - | - | 897.1 | 672.3 | - | - | 0 | - |
| - | - | 2344 | 685.3 | - | - | 0 | - |
| - | - | 1148 | 686.3 | - | - | 0 | - |
| - | - | 1553 | 693.8 | - | - | 0 | - |
| - | - | 1249 | 694.3 | - | - | 0 | - |
| - | - | 646.8 | 696.3 | - | - | 0 | - |
| - | - | 9650 | 700.4 | - | - | 0 | - |
| - | - | 3366 | 701.4 | - | - | 0 | - |
| - | - | 1202 | 702.4 | - | - | 0 | - |
| - | - | 1524 | 721.3 | - | - | 0 | - |
| - | - | 1012 | 722.4 | - | - | 0 | - |
| 5 | y | 6350 | 723.3 | 0.004755 | 6.574 | +1 | 5 |
| - | - | 2091 | 724.3 | - | - | 0 | - |
| - | - | 1654 | 730.4 | - | - | 0 | - |
| - | - | 1476 | 730.9 | - | - | 0 | - |
| - | - | 1167 | 731.4 | - | - | 0 | - |
| - | - | 4143 | 744.4 | - | - | 0 | - |
| - | - | 2368 | 744.9 | - | - | 0 | - |
| - | - | 1599 | 745.4 | - | - | 0 | - |
| - | - | 2821 | 749.3 | - | - | 0 | - |
| - | - | 705.7 | 754.9 | - | - | 0 | - |
| - | - | 846.8 | 757.4 | - | - | 0 | - |
| - | - | 706.6 | 768.3 | - | - | 0 | - |
| - | - | 2300 | 774.3 | - | - | 0 | - |
| - | - | 1063 | 775.3 | - | - | 0 | - |
| - | - | 3082 | 783.4 | - | - | 0 | - |
| - | - | 1322 | 784.4 | - | - | 0 | - |
| - | - | 3670 | 794.4 | - | - | 0 | - |
| - | - | 1748 | 795.4 | - | - | 0 | - |
| - | - | 1822 | 800.4 | - | - | 0 | - |
| - | - | 5687 | 801.4 | - | - | 0 | - |
| - | - | 2161 | 802.4 | - | - | 0 | - |
| - | - | 1814 | 828.4 | - | - | 0 | - |
| - | - | 889.4 | 829.4 | - | - | 0 | - |
| 4 | y | 7480 | 838.3 | 0.004973 | 5.932 | +1 | 6 |
| - | - | 3211 | 839.3 | - | - | 0 | - |
| - | - | 1151 | 840.3 | - | - | 0 | - |
| - | - | 8452 | 844.5 | - | - | 0 | - |
| - | - | 2167 | 845.5 | - | - | 0 | - |
| - | - | 834.6 | 846.5 | - | - | 0 | - |
| - | - | 1187 | 854.4 | - | - | 0 | - |
| - | - | 824.9 | 855.4 | - | - | 0 | - |
| - | - | 1446 | 871.3 | - | - | 0 | - |
| - | - | 2.648E+04 | 872.4 | - | - | 0 | - |
| - | - | 1.258E+04 | 873.5 | - | - | 0 | - |
| - | - | 4349 | 874.5 | - | - | 0 | - |
| - | - | 9290 | 889.3 | - | - | 0 | - |
| - | - | 4976 | 890.4 | - | - | 0 | - |
| - | - | 1106 | 891.4 | - | - | 0 | - |
| - | - | 788.6 | 896.5 | - | - | 0 | - |
| - | - | 2530 | 909.4 | - | - | 0 | - |
| - | - | 1626 | 910.4 | - | - | 0 | - |
| 3 | y | 1289 | 935.3 | 0.003487 | 3.728 | +1 | 7 |
| - | - | 723.3 | 936.3 | - | - | 0 | - |
| 3 | y | 3.01E+04 | 953.3 | 0.004885 | 5.124 | +1 | 7 |
| - | - | 1.588E+04 | 954.3 | - | - | 0 | - |
| - | - | 5985 | 955.3 | - | - | 0 | - |
| - | - | 4260 | 991.5 | - | - | 0 | - |
| - | - | 2378 | 992.5 | - | - | 0 | - |
| - | - | 644.1 | 1006 | - | - | 0 | - |
| - | - | 1.172E+04 | 1020 | - | - | 0 | - |
| - | - | 6955 | 1021 | - | - | 0 | - |
| - | - | 2403 | 1022 | - | - | 0 | - |
| - | - | 1016 | 1038 | - | - | 0 | - |
| 2 | y | 2195 | 1100 | 0.005014 | 4.556 | +1 | 8 |
| - | - | 789.2 | 1101 | - | - | 0 | - |
| - | - | 691.3 | 1178 | - | - | 0 | - |
| - | - | 891 | 1179 | - | - | 0 | - |
| - | - | 796 | 1325 | - | - | 0 | - |
| - | - | 935.3 | 1326 | - | - | 0 | - |
| - | - | 977.5 | 1341 | - | - | 0 | - |
| - | - | 761.7 | 1388 | - | - | 0 | - |
| - | - | 725 | 1691 | - | - | 0 | - |
| - | - | 599.4 | 1743 | - | - | 0 | - |
| - | - | 663.6 | 2454 | - | - | 0 | - |
| - | - | 668.1 | 2763 | - | - | 0 | - |

m/z Charge Intensity FragmentType MassShift Position
120.05274963378906 0 528.3845
120.08113098144531 0 72283.88
121.08444213867188 0 5420.418
123.04449462890625 0 694.8314
129.10255432128906 0 8079.9556
130.06546020507812 0 2502.8594
130.08653259277344 0 397.6913
130.10586547851562 0 396.9001
132.0811004638672 0 2595.2246
132.10231018066406 0 1082.6255
133.0862274169922 0 1008.84467
136.07601928710938 0 29422.805
137.07936096191406 0 2447.597
138.05517578125 0 662.6209
138.06666564941406 0 950.1739
140.08758544921875 0 394.40488
141.6322479248047 0 478.147
141.73451232910156 0 388.62628
142.0651092529297 0 557.1789
144.08099365234375 0 485.2975
146.06036376953125 0 6072.176
147.04452514648438 0 1248.9517
148.07606506347656 0 988.68085
148.947998046875 0 832.76105
149.04525756835938 0 1362.4324
149.69105529785156 0 397.88406
154.24655151367188 0 450.9836
155.0819091796875 0 496.1757
156.07687377929688 0 1302.9392
157.1336212158203 0 1435.4164
158.09664916992188 0 2427.0198
159.09202575683594 0 8936.187
159.09890747070312 0 1024.0952
160.0762481689453 0 657.9106
160.0948944091797 0 1177.3562
160.1123046875 0 1271.7322
165.05491638183594 0 8156.28
165.10301208496094 0 442.3448
166.0581512451172 0 839.14166
166.08644104003906 0 1726.8243
167.05564880371094 0 1360.4402
167.09288024902344 0 539.87634
169.07603454589844 0 550.8842
170.0604705810547 0 649.37164
173.09291076660156 0 520.5413
173.1288299560547 0 5418.854
173.4529571533203 0 912.0618
175.08685302734375 0 1421.4373
175.1190948486328 0 744.69147
176.08245849609375 0 501.8116
176.10736083984375 0 5824.5522
177.10247802734375 0 4575.506
177.1109619140625 0 791.033
182.08157348632812 0 15517.532
182.0906982421875 0 813.3195
183.08505249023438 0 1101.259
185.05615234375 0 533.2248
185.12864685058594 0 1758.3844
186.09120178222656 0 432.06287
188.0709228515625 0 27792.283
189.07440185546875 0 3450.7537
189.8454132080078 0 484.87366
191.1185760498047 0 1518.7468
193.10879516601562 0 1492.2798
195.0874481201172 0 539.3215
195.11257934570312 0 556.99
200.13954162597656 0 907.6705
201.1234130859375 0 2484.2866
202.05197143554688 0 595.08496
203.06784057617188 0 493.50003
203.1181640625 0 2394.265 a Water loss 1
204.07745361328125 0 943.59393
205.09751892089844 0 11434.23 y 8
206.10116577148438 0 1210.4684
207.1129913330078 0 911.27277
211.06040954589844 0 669.77386
214.11880493164062 0 730.528
214.88795471191406 0 531.8751
215.13987731933594 0 696.8246
216.06533813476562 0 828.3242
217.0972900390625 0 1164.5037
218.7314910888672 0 500.73312
219.1132354736328 0 1029.3645
221.10365295410156 0 4259.7983
221.12881469726562 0 83159.5 a 1
222.1060791015625 0 595.061
222.13217163085938 0 8927.736
223.06358337402344 0 1513.0472
223.13572692871094 0 539.11383
224.13980102539062 0 485.23343
225.1227264404297 0 604.712
225.1717529296875 0 1173.8484
226.11996459960938 0 558.90546
227.1028289794922 0 796.657
228.1125030517578 0 545.018
231.0616455078125 0 1320.6053
231.11253356933594 0 1103.0365 b Water loss 1
233.0924072265625 0 1091.7612
233.129150390625 0 1583.0413
233.14979553222656 0 605.41144
235.1080322265625 0 3151.181
239.0836639404297 0 545.58124
239.15052795410156 0 1825.4329
241.58250427246094 0 641.678
243.11087036132812 0 483.9956
249.10897827148438 0 817.7079
249.123779296875 0 15247.668 b 1
250.1268310546875 0 2221.3718
251.10305786132812 0 1442.2742
251.14939880371094 0 496.98138
252.13380432128906 0 743.5798
253.11874389648438 0 671.7779
253.1660614013672 0 1514.731
256.17706298828125 0 800.20776
257.1069030761719 0 1040.4934
261.1233825683594 0 1458.0654
263.1028747558594 0 3713.7124
267.95928955078125 0 505.4244
279.09771728515625 0 2261.883
280.16400146484375 0 514.8922
285.0101013183594 0 1361.8192
285.08544921875 0 594.3824
285.1020812988281 0 1403.9703
292.1411437988281 0 805.3313
294.921142578125 0 515.2848
296.1507873535156 0 1884.0399
302.11407470703125 0 809.4395
304.1287841796875 0 1229.244
315.167236328125 0 628.04663
318.1459045410156 0 652.75806 a Water loss 2
319.1400451660156 0 1300.4761
323.09832763671875 0 731.44885
324.1459045410156 0 2614.4216
325.1480712890625 0 1019.0516
329.1501159667969 0 9764.449
330.1533508300781 0 2147.1729
335.1808776855469 0 758.1664
336.18145751953125 0 636.5052
337.1866149902344 0 689.022
340.1982421875 0 2319.8186
342.1566467285156 0 16416.9 y 7
342.17962646484375 0 1107.1902
343.1593322753906 0 2872.6091
346.1407165527344 0 685.75616 b Water loss 2
350.13433837890625 0 905.10504
352.1661071777344 0 1030.4681
355.07025146484375 0 1294.9414
356.1387023925781 0 663.3715
357.2247619628906 0 1949.4021
364.15093994140625 0 3490.025 b 2
365.1542663574219 0 839.34576
368.1927795410156 0 1082.0171
374.15081787109375 0 625.6587
378.1298828125 0 2142.2017
390.15673828125 0 627.2715
392.20428466796875 0 1109.9601
394.12457275390625 0 2468.5225
401.20745849609375 0 554.49805
401.71282958984375 0 680.402
410.8466491699219 0 626.7346
413.2677307128906 0 1196.8252
425.1930236816406 0 2511.1375
427.7223815917969 0 967.4936
428.223876953125 0 634.2296
436.727294921875 0 549.02264
437.2280578613281 0 729.0428
461.1674499511719 0 759.8383 b Water loss 3
465.162841796875 0 1763.8226
479.1780700683594 0 2160.253 b 3
480.1837463378906 0 826.4528
483.2218017578125 0 838.9502
487.1086120605469 0 636.0113
487.33331298828125 0 681.57
489.192626953125 0 3418.6392 y 6
490.1949157714844 0 1059.9303
496.2297668457031 0 1754.1301
510.2618713378906 0 704.46735
519.2019653320312 0 881.6235
539.217041015625 0 1152.7689
547.25244140625 0 902.6691
548.1970825195312 0 963.91846
550.7119750976562 0 2251.4963 y 1
551.21240234375 0 1651.2352
559.2646484375 0 665.2907
560.22900390625 0 6157.582 y 5
561.2323608398438 0 2002.11
579.794921875 0 576.72424
585.3348999023438 0 2376.4978
586.338623046875 0 1070.0841
589.2849731445312 0 741.25653
592.23046875 0 1349.8389 Precursor Water loss
601.235595703125 0 3811.3972 Precursor
601.3343505859375 0 798.39105
601.738037109375 0 1645.5261
612.1961669921875 0 1300.6646
614.244384765625 0 675.41174
628.3781127929688 0 947.407
651.3648681640625 0 864.133
657.2647094726562 0 1460.7903
659.2937622070312 0 1960.5596
660.2958374023438 0 1072.5854
670.8173217773438 0 7253.5312
671.3195190429688 0 3637.4927
671.4031982421875 0 702.6588
671.819580078125 0 2366.8293
672.319580078125 0 897.11774
685.2590942382812 0 2343.6028
686.2600708007812 0 1147.5306
693.8112182617188 0 1552.8441
694.315185546875 0 1249.0684
696.3206787109375 0 646.77325
700.3623657226562 0 9649.598
701.3651123046875 0 3366.1868
702.3687744140625 0 1202.4656
721.26513671875 0 1524.0164
722.40087890625 0 1012.3358
723.291748046875 0 6350.138 y 4
724.2950439453125 0 2091.479
730.353759765625 0 1654.1277
730.8547973632812 0 1476.0222
731.3551635742188 0 1166.9066
744.351318359375 0 4142.8105
744.8533325195312 0 2368.1338
745.3521728515625 0 1599.4075
749.2551879882812 0 2821.3032
754.8512573242188 0 705.72156
757.3843994140625 0 846.8036
768.336669921875 0 706.55444
774.31982421875 0 2299.9895
775.3220825195312 0 1063.4358
783.399658203125 0 3082.0486
784.3994750976562 0 1321.6467
794.3839111328125 0 3670.3308
795.3856811523438 0 1747.9928
800.4281616210938 0 1822.284
801.41259765625 0 5686.5625
802.4133911132812 0 2161.1755
828.4200439453125 0 1813.7668
829.4193725585938 0 889.39844
838.3189086914062 0 7480.1587 y 3
839.3217163085938 0 3210.8767
840.3141479492188 0 1150.6272
844.45263671875 0 8451.682
845.4541625976562 0 2166.8442
846.4559326171875 0 834.59564
854.4364013671875 0 1186.7605
855.4330444335938 0 824.8736
871.3399047851562 0 1445.5566
872.4476318359375 0 26481.18
873.4501342773438 0 12581.717
874.45361328125 0 4349.2827
889.3473510742188 0 9289.597
890.3502197265625 0 4976.3765
891.3535766601562 0 1106.0988
896.52490234375 0 788.5963
909.4109497070312 0 2530.1116
910.4146728515625 0 1625.7242
935.3338012695312 0 1288.8478 y Water loss 2
936.3380126953125 0 723.33575
953.3457641601562 0 30104.13 y 2
954.3482666015625 0 15883.617
955.3494873046875 0 5984.744
991.5211791992188 0 4260.4575
992.5239868164062 0 2377.7476
1006.4817504882812 0 644.06494
1019.5150146484375 0 11715.347
1020.5181274414062 0 6954.882
1021.5191040039062 0 2403.1746
1037.50927734375 0 1016.1239
1100.414306640625 0 2195.4097 y 1
1101.4200439453125 0 789.1866
1177.5618896484375 0 691.29736
1178.5701904296875 0 891.00305
1324.6229248046875 0 796.02985
1325.6304931640625 0 935.32733
1340.61962890625 0 977.45764
1387.629638671875 0 761.7111
1691.113037109375 0 725.02747
1743.4324951171875 0 599.4033
2454.062744140625 0 663.6124
2762.576416015625 0 668.1058

Spectrum Details

|  |  |
| --- | --- |
| Matched peaks? Matched peaks The total absolute number of peaks matched. Additionally in brackets the total fraction of peaks matched and the total number of peaks is shown. | 21 (7.47% of 281) |
| FDR? FDR The false discovery rate estimated for this peptide. It is calculated by matching all theoretical fragments with a non-integer shift with the raw peaks for this spectrum. This is done with 40 different shifts. The resulting percentage is the average number of annotated peaks over the number of annotated peaks with the correct spectrum. | 1.36% |
| Satellite FDR? Satellite FDR See the FDR for details on its calculation. This satellite ion specific FDR only contains the satellite ions (d/w) for I/L/J positions. | - |
| PSM Score? PSM Score The PSM Score as given by Hecklib to this annotated spectrum. It is shown with three significant figures. | 250 |

#### Spectrum 10187? Spectrum 10187 The raw spectrum of this peptide as annotated by Hecklib. The fragments are coloured according to ion type (see legend). Any peaks with a star '\*' as text can be hovered over to see the full details, first the ion type second the mass shift type. By hovering over the amino acids in the peptide or ions in the legend the corresponding peaks are highlighted. By toggling the 'Unassigned' label you can turn the background (unassigned) peaks on or off in the plot. By updating the slider in the Ion legend you can update the spectrum to only show the top X% of the peaks with labels. The top X% means any peak that is within X% of the highest intensity. By dragging in the spectrum you can zoom in to a specific part of the spectrum and use 'Zoom Out' to get back to the original zoom level. The annotation of the spectrum is based on the given sequence in the peptides file and is done with different software so inconsistencies are likely. The peaks are annotated based on the given sequence, with 20 ppm tolerance.

Copy Data

##### Spectrum 10187 (TSV)

###### Preview

```
Loading example...
```

*Click on the button to copy the data to your clipboard.*

Mz MinMz MaxIntensity Max

WidthHeightPeptide font sizePeptide stroke widthSpectrum font sizeSpectrum stroke widthCompact peptide

Ion legend

wxyz

abcd

OtherUnassignedIon^Charge^_Position_Show for top:%

TFDDYAMHW

02.20e+44.40e+46.60e+48.80e+4

Zoom Out

a^+1^_2_y^+1^_1_a^+1^_2_b^+1^_2_b^+1^_2_y^+1^_2_b^+1^_3_b^+1^_4_y^+1^_3_y^+2^_8_y^+1^_4_\*\*y^+1^_5_y^+1^_6_y^+1^_7_y^+1^_7_y^+1^_8_

0837167325103347

Fragment Matches Table

Show background peaks

| Position | Ion type | Intensity | mz Theoretical | mz Error (Th) | mz Error (ppm) | Charge | Series Number |
| --- | --- | --- | --- | --- | --- | --- | --- |
| - | - | 7.869E+04 | 120.1 | - | - | 0 | - |
| - | - | 570.1 | 121.1 | - | - | 0 | - |
| - | - | 5937 | 121.1 | - | - | 0 | - |
| - | - | 341.5 | 121.2 | - | - | 0 | - |
| - | - | 1364 | 123 | - | - | 0 | - |
| - | - | 430.3 | 127.1 | - | - | 0 | - |
| - | - | 606.7 | 129.1 | - | - | 0 | - |
| - | - | 5627 | 129.1 | - | - | 0 | - |
| - | - | 1614 | 130.1 | - | - | 0 | - |
| - | - | 1819 | 132.1 | - | - | 0 | - |
| - | - | 1171 | 132.1 | - | - | 0 | - |
| - | - | 1127 | 133.1 | - | - | 0 | - |
| - | - | 2.689E+04 | 136.1 | - | - | 0 | - |
| - | - | 2874 | 137.1 | - | - | 0 | - |
| - | - | 567.7 | 138.1 | - | - | 0 | - |
| - | - | 1109 | 138.1 | - | - | 0 | - |
| - | - | 602.7 | 141.1 | - | - | 0 | - |
| - | - | 898.4 | 143.1 | - | - | 0 | - |
| - | - | 2294 | 146.1 | - | - | 0 | - |
| - | - | 813.2 | 147 | - | - | 0 | - |
| - | - | 401 | 148.3 | - | - | 0 | - |
| - | - | 415.6 | 148.9 | - | - | 0 | - |
| - | - | 751.2 | 149 | - | - | 0 | - |
| - | - | 462.9 | 155.1 | - | - | 0 | - |
| - | - | 2291 | 156.1 | - | - | 0 | - |
| - | - | 2024 | 157.1 | - | - | 0 | - |
| - | - | 3361 | 158.1 | - | - | 0 | - |
| - | - | 5102 | 159.1 | - | - | 0 | - |
| - | - | 812.8 | 159.1 | - | - | 0 | - |
| - | - | 640.6 | 160.1 | - | - | 0 | - |
| - | - | 1048 | 160.1 | - | - | 0 | - |
| - | - | 1614 | 160.1 | - | - | 0 | - |
| - | - | 531.9 | 161 | - | - | 0 | - |
| - | - | 1.024E+04 | 165.1 | - | - | 0 | - |
| - | - | 420.8 | 165.3 | - | - | 0 | - |
| - | - | 848.1 | 166.1 | - | - | 0 | - |
| - | - | 1292 | 166.1 | - | - | 0 | - |
| - | - | 1787 | 167.1 | - | - | 0 | - |
| - | - | 520.7 | 169.1 | - | - | 0 | - |
| - | - | 698.6 | 169.1 | - | - | 0 | - |
| - | - | 657.4 | 170.1 | - | - | 0 | - |
| - | - | 583.9 | 171.1 | - | - | 0 | - |
| - | - | 669.6 | 171.1 | - | - | 0 | - |
| - | - | 4873 | 173.1 | - | - | 0 | - |
| - | - | 2286 | 173.5 | - | - | 0 | - |
| - | - | 811.8 | 174.1 | - | - | 0 | - |
| - | - | 588.6 | 174.1 | - | - | 0 | - |
| - | - | 755.9 | 175.1 | - | - | 0 | - |
| - | - | 6179 | 176.1 | - | - | 0 | - |
| - | - | 5512 | 177.1 | - | - | 0 | - |
| - | - | 1345 | 177.1 | - | - | 0 | - |
| - | - | 536.5 | 181.1 | - | - | 0 | - |
| - | - | 1.754E+04 | 182.1 | - | - | 0 | - |
| - | - | 433.2 | 182.1 | - | - | 0 | - |
| - | - | 2337 | 183.1 | - | - | 0 | - |
| - | - | 500.4 | 183.2 | - | - | 0 | - |
| - | - | 638.1 | 184.1 | - | - | 0 | - |
| - | - | 556.7 | 185.1 | - | - | 0 | - |
| - | - | 540.4 | 185.1 | - | - | 0 | - |
| - | - | 2152 | 185.1 | - | - | 0 | - |
| - | - | 818.3 | 186.1 | - | - | 0 | - |
| - | - | 1.199E+04 | 188.1 | - | - | 0 | - |
| - | - | 1414 | 189.1 | - | - | 0 | - |
| - | - | 679.5 | 191.1 | - | - | 0 | - |
| - | - | 1677 | 191.1 | - | - | 0 | - |
| - | - | 1406 | 192.1 | - | - | 0 | - |
| - | - | 2029 | 193.1 | - | - | 0 | - |
| - | - | 723.5 | 197.1 | - | - | 0 | - |
| - | - | 709.9 | 199.1 | - | - | 0 | - |
| - | - | 3472 | 201.1 | - | - | 0 | - |
| - | - | 532.3 | 202.1 | - | - | 0 | - |
| - | - | 804.7 | 203.1 | - | - | 0 | - |
| 2 | a | 3111 | 203.1 | 7.616E-05 | 0.375 | +1 | 2 |
| - | - | 1061 | 204.1 | - | - | 0 | - |
| 9 | y | 5882 | 205.1 | 0.0001054 | 0.514 | +1 | 1 |
| - | - | 951.2 | 205.1 | - | - | 0 | - |
| - | - | 603.8 | 206.1 | - | - | 0 | - |
| - | - | 1051 | 207.1 | - | - | 0 | - |
| - | - | 2290 | 209.1 | - | - | 0 | - |
| - | - | 712.6 | 209.1 | - | - | 0 | - |
| - | - | 839.1 | 214.1 | - | - | 0 | - |
| - | - | 701.5 | 215.1 | - | - | 0 | - |
| - | - | 571.8 | 217.1 | - | - | 0 | - |
| - | - | 1555 | 217.1 | - | - | 0 | - |
| - | - | 1605 | 219.1 | - | - | 0 | - |
| - | - | 5238 | 221.1 | - | - | 0 | - |
| 2 | a | 8.717E+04 | 221.1 | 0.0002842 | 1.285 | +1 | 2 |
| - | - | 838.6 | 222.1 | - | - | 0 | - |
| - | - | 1.129E+04 | 222.1 | - | - | 0 | - |
| - | - | 1093 | 223.1 | - | - | 0 | - |
| - | - | 671.9 | 224.1 | - | - | 0 | - |
| - | - | 720.3 | 225.1 | - | - | 0 | - |
| - | - | 604.1 | 225.2 | - | - | 0 | - |
| - | - | 1213 | 229.1 | - | - | 0 | - |
| - | - | 1311 | 231.1 | - | - | 0 | - |
| 2 | b | 850.2 | 231.1 | 5.696E-05 | 0.2465 | +1 | 2 |
| - | - | 1197 | 233.1 | - | - | 0 | - |
| - | - | 3096 | 235.1 | - | - | 0 | - |
| - | - | 501.6 | 239.1 | - | - | 0 | - |
| - | - | 792.9 | 239.1 | - | - | 0 | - |
| - | - | 1160 | 239.2 | - | - | 0 | - |
| - | - | 913.9 | 241.1 | - | - | 0 | - |
| - | - | 468.3 | 242.1 | - | - | 0 | - |
| 2 | b | 1.57E+04 | 249.1 | 0.0003189 | 1.28 | +1 | 2 |
| - | - | 2079 | 250.1 | - | - | 0 | - |
| - | - | 2123 | 251.1 | - | - | 0 | - |
| - | - | 1314 | 253.2 | - | - | 0 | - |
| - | - | 542.9 | 257.1 | - | - | 0 | - |
| - | - | 2920 | 263.1 | - | - | 0 | - |
| - | - | 694 | 270.1 | - | - | 0 | - |
| - | - | 1824 | 279.1 | - | - | 0 | - |
| - | - | 718.1 | 284.2 | - | - | 0 | - |
| - | - | 2132 | 285.1 | - | - | 0 | - |
| - | - | 534.9 | 285.2 | - | - | 0 | - |
| - | - | 612.1 | 287.1 | - | - | 0 | - |
| - | - | 1080 | 292.1 | - | - | 0 | - |
| - | - | 1409 | 296.2 | - | - | 0 | - |
| - | - | 881.5 | 300.1 | - | - | 0 | - |
| - | - | 656.9 | 300.2 | - | - | 0 | - |
| - | - | 995.2 | 302.1 | - | - | 0 | - |
| - | - | 614.6 | 309.2 | - | - | 0 | - |
| - | - | 1307 | 319.1 | - | - | 0 | - |
| - | - | 865.4 | 320.1 | - | - | 0 | - |
| - | - | 4120 | 324.1 | - | - | 0 | - |
| - | - | 722.5 | 325.1 | - | - | 0 | - |
| - | - | 665.4 | 329.1 | - | - | 0 | - |
| - | - | 1.262E+04 | 329.2 | - | - | 0 | - |
| - | - | 2248 | 330.2 | - | - | 0 | - |
| - | - | 663.2 | 333.1 | - | - | 0 | - |
| - | - | 520.7 | 333.2 | - | - | 0 | - |
| - | - | 724.3 | 338.1 | - | - | 0 | - |
| - | - | 554.7 | 339.2 | - | - | 0 | - |
| - | - | 2425 | 340.2 | - | - | 0 | - |
| 8 | y | 1.684E+04 | 342.2 | 0.0004892 | 1.43 | +1 | 2 |
| - | - | 3021 | 343.2 | - | - | 0 | - |
| - | - | 793.4 | 350.1 | - | - | 0 | - |
| - | - | 708.7 | 355.1 | - | - | 0 | - |
| - | - | 1094 | 356.1 | - | - | 0 | - |
| - | - | 3504 | 357.2 | - | - | 0 | - |
| 3 | b | 4451 | 364.2 | 4.823E-05 | 0.1325 | +1 | 3 |
| - | - | 912.4 | 365.2 | - | - | 0 | - |
| - | - | 759.7 | 366.1 | - | - | 0 | - |
| - | - | 695.8 | 368.2 | - | - | 0 | - |
| - | - | 828.1 | 376.1 | - | - | 0 | - |
| - | - | 2377 | 378.1 | - | - | 0 | - |
| - | - | 637.5 | 392.2 | - | - | 0 | - |
| - | - | 2542 | 394.1 | - | - | 0 | - |
| - | - | 589.6 | 409.2 | - | - | 0 | - |
| - | - | 681.8 | 418.7 | - | - | 0 | - |
| - | - | 694.6 | 422.7 | - | - | 0 | - |
| - | - | 2565 | 425.2 | - | - | 0 | - |
| - | - | 1763 | 427.7 | - | - | 0 | - |
| - | - | 1106 | 428.2 | - | - | 0 | - |
| - | - | 980.6 | 432.2 | - | - | 0 | - |
| - | - | 1548 | 436.7 | - | - | 0 | - |
| - | - | 1193 | 437.2 | - | - | 0 | - |
| - | - | 710.5 | 441.3 | - | - | 0 | - |
| - | - | 572.3 | 447.2 | - | - | 0 | - |
| - | - | 2146 | 465.2 | - | - | 0 | - |
| 4 | b | 2229 | 479.2 | 0.0006321 | 1.319 | +1 | 4 |
| - | - | 1028 | 483.2 | - | - | 0 | - |
| 7 | y | 3884 | 489.2 | 0.00571 | 11.67 | +1 | 3 |
| - | - | 1563 | 490.2 | - | - | 0 | - |
| - | - | 1713 | 496.2 | - | - | 0 | - |
| - | - | 680.7 | 500.2 | - | - | 0 | - |
| - | - | 669.4 | 501.3 | - | - | 0 | - |
| - | - | 1339 | 519.2 | - | - | 0 | - |
| - | - | 902 | 524.3 | - | - | 0 | - |
| - | - | 971.7 | 539.2 | - | - | 0 | - |
| - | - | 615.1 | 547.3 | - | - | 0 | - |
| - | - | 743.3 | 548.2 | - | - | 0 | - |
| 2 | y | 2301 | 550.7 | 0.002897 | 5.26 | +2 | 8 |
| - | - | 1198 | 551.2 | - | - | 0 | - |
| - | - | 706.3 | 551.7 | - | - | 0 | - |
| 6 | y | 6772 | 560.2 | 0.005523 | 9.858 | +1 | 4 |
| - | - | 1690 | 561.2 | - | - | 0 | - |
| - | - | 4166 | 585.3 | - | - | 0 | - |
| - | - | 1133 | 586.3 | - | - | 0 | - |
| 0 | Precursor | 824.9 | 592.2 | 0.003078 | 5.197 | +2 | -1 |
| - | - | 1089 | 592.7 | - | - | 0 | - |
| - | - | 866.9 | 595.3 | - | - | 0 | - |
| - | - | 781.8 | 600.3 | - | - | 0 | - |
| - | - | 895.5 | 601 | - | - | 0 | - |
| 0 | Precursor | 2598 | 601.2 | 0.001946 | 3.237 | +2 | -1 |
| - | - | 892.7 | 601.3 | - | - | 0 | - |
| - | - | 1597 | 601.7 | - | - | 0 | - |
| - | - | 748.9 | 602.2 | - | - | 0 | - |
| - | - | 808.5 | 606.9 | - | - | 0 | - |
| - | - | 1143 | 612.2 | - | - | 0 | - |
| - | - | 813 | 614.2 | - | - | 0 | - |
| - | - | 853.3 | 628.4 | - | - | 0 | - |
| - | - | 1539 | 657.3 | - | - | 0 | - |
| - | - | 1614 | 659.3 | - | - | 0 | - |
| - | - | 1422 | 660.3 | - | - | 0 | - |
| - | - | 2579 | 670.8 | - | - | 0 | - |
| - | - | 875.4 | 671.3 | - | - | 0 | - |
| - | - | 1014 | 671.8 | - | - | 0 | - |
| - | - | 642.7 | 672.4 | - | - | 0 | - |
| - | - | 717.7 | 682.4 | - | - | 0 | - |
| - | - | 2131 | 685.3 | - | - | 0 | - |
| - | - | 819 | 686.3 | - | - | 0 | - |
| - | - | 1.144E+04 | 700.4 | - | - | 0 | - |
| - | - | 3866 | 701.4 | - | - | 0 | - |
| - | - | 906.8 | 702.4 | - | - | 0 | - |
| - | - | 1715 | 721.3 | - | - | 0 | - |
| - | - | 950.2 | 722.3 | - | - | 0 | - |
| - | - | 634.1 | 722.4 | - | - | 0 | - |
| 5 | y | 4872 | 723.3 | 0.005366 | 7.418 | +1 | 5 |
| - | - | 2594 | 724.3 | - | - | 0 | - |
| - | - | 713.1 | 725.3 | - | - | 0 | - |
| - | - | 1463 | 744.3 | - | - | 0 | - |
| - | - | 2670 | 749.3 | - | - | 0 | - |
| - | - | 1542 | 750.3 | - | - | 0 | - |
| - | - | 928.2 | 757.4 | - | - | 0 | - |
| - | - | 753.9 | 773.4 | - | - | 0 | - |
| - | - | 3437 | 774.3 | - | - | 0 | - |
| - | - | 2734 | 783.4 | - | - | 0 | - |
| - | - | 1186 | 784.4 | - | - | 0 | - |
| - | - | 1254 | 794.4 | - | - | 0 | - |
| - | - | 1367 | 800.4 | - | - | 0 | - |
| - | - | 6096 | 801.4 | - | - | 0 | - |
| - | - | 2937 | 802.4 | - | - | 0 | - |
| - | - | 656.7 | 803.2 | - | - | 0 | - |
| - | - | 807.5 | 803.4 | - | - | 0 | - |
| - | - | 2357 | 828.4 | - | - | 0 | - |
| - | - | 901.7 | 829.4 | - | - | 0 | - |
| 4 | y | 6770 | 838.3 | 0.005827 | 6.951 | +1 | 6 |
| - | - | 3053 | 839.3 | - | - | 0 | - |
| - | - | 707.7 | 840.3 | - | - | 0 | - |
| - | - | 8928 | 844.5 | - | - | 0 | - |
| - | - | 2823 | 845.5 | - | - | 0 | - |
| - | - | 1206 | 846.5 | - | - | 0 | - |
| - | - | 1783 | 854.4 | - | - | 0 | - |
| - | - | 3.28E+04 | 872.4 | - | - | 0 | - |
| - | - | 1.368E+04 | 873.4 | - | - | 0 | - |
| - | - | 3211 | 874.5 | - | - | 0 | - |
| - | - | 9799 | 889.3 | - | - | 0 | - |
| - | - | 4965 | 890.4 | - | - | 0 | - |
| - | - | 1468 | 891.4 | - | - | 0 | - |
| - | - | 658.8 | 907.3 | - | - | 0 | - |
| - | - | 843.9 | 909.4 | - | - | 0 | - |
| 3 | y | 2271 | 935.3 | 0.003487 | 3.728 | +1 | 7 |
| - | - | 1176 | 936.3 | - | - | 0 | - |
| 3 | y | 3.337E+04 | 953.3 | 0.004763 | 4.996 | +1 | 7 |
| - | - | 1.865E+04 | 954.3 | - | - | 0 | - |
| - | - | 521 | 954.4 | - | - | 0 | - |
| - | - | 6277 | 955.3 | - | - | 0 | - |
| - | - | 906.7 | 956.3 | - | - | 0 | - |
| - | - | 5555 | 991.5 | - | - | 0 | - |
| - | - | 2919 | 992.5 | - | - | 0 | - |
| - | - | 1096 | 993.5 | - | - | 0 | - |
| - | - | 924.9 | 1003 | - | - | 0 | - |
| - | - | 723.2 | 1004 | - | - | 0 | - |
| - | - | 1.181E+04 | 1020 | - | - | 0 | - |
| - | - | 8219 | 1021 | - | - | 0 | - |
| - | - | 3371 | 1022 | - | - | 0 | - |
| 2 | y | 1870 | 1100 | 0.006357 | 5.777 | +1 | 8 |
| - | - | 1003 | 1101 | - | - | 0 | - |
| - | - | 646.6 | 1810 | - | - | 0 | - |
| - | - | 684.2 | 2603 | - | - | 0 | - |
| - | - | 759.5 | 2744 | - | - | 0 | - |
| - | - | 726.4 | 3314 | - | - | 0 | - |

m/z Charge Intensity FragmentType MassShift Position
120.0810775756836 0 78690.08
121.0796127319336 0 570.1216
121.08438110351562 0 5936.549
121.24666595458984 0 341.5262
123.04437255859375 0 1364.3822
127.0870132446289 0 430.3237
129.06597900390625 0 606.6612
129.10247802734375 0 5627.443
130.0653076171875 0 1613.845
132.08106994628906 0 1818.9316
132.10206604003906 0 1171.1364
133.08627319335938 0 1126.7461
136.0759735107422 0 26892.732
137.07919311523438 0 2874.2207
138.05477905273438 0 567.6762
138.06639099121094 0 1108.9232
141.10243225097656 0 602.7204
143.1180419921875 0 898.37695
146.0603485107422 0 2294.2375
147.04412841796875 0 813.2065
148.3268585205078 0 401.04553
148.89614868164062 0 415.6424
149.04530334472656 0 751.2287
155.09286499023438 0 462.9214
156.07708740234375 0 2291.2764
157.13385009765625 0 2023.5339
158.0967559814453 0 3360.6655
159.0919952392578 0 5102.24
159.0991973876953 0 812.7911
160.07601928710938 0 640.6468
160.09532165527344 0 1047.9307
160.1121368408203 0 1614.3533
161.04547119140625 0 531.866
165.0548858642578 0 10237.037
165.3275604248047 0 420.7727
166.058837890625 0 848.0728
166.08644104003906 0 1292.0577
167.0557861328125 0 1787.2887
169.09725952148438 0 520.6956
169.1336212158203 0 698.6039
170.05984497070312 0 657.4058
171.07598876953125 0 583.92944
171.1129608154297 0 669.58997
173.12867736816406 0 4873.456
173.4512176513672 0 2285.898
174.05511474609375 0 811.8099
174.13253784179688 0 588.62305
175.08656311035156 0 755.9475
176.1072540283203 0 6179.049
177.10250854492188 0 5512.4385
177.111572265625 0 1345.1095
181.0974578857422 0 536.55
182.0814666748047 0 17540.992
182.1283721923828 0 433.1541
183.084716796875 0 2337.0542
183.15020751953125 0 500.42737
184.07203674316406 0 638.1284
185.05528259277344 0 556.7222
185.0916290283203 0 540.4078
185.128662109375 0 2152.0977
186.09158325195312 0 818.329
188.07086181640625 0 11993.623
189.07421875 0 1414.0117
191.08175659179688 0 679.5059
191.1178741455078 0 1676.8235
192.06570434570312 0 1406.2291
193.10861206054688 0 2029.1111
197.12828063964844 0 723.4927
199.07119750976562 0 709.94507
201.1236114501953 0 3472.134
202.05113220214844 0 532.3108
203.066162109375 0 804.69666
203.1179656982422 0 3110.9902 a Water loss 1
204.07708740234375 0 1060.64
205.09725952148438 0 5882.139 y 8
205.10763549804688 0 951.2351
206.1006622314453 0 603.7553
207.11279296875 0 1050.553
209.09217834472656 0 2290.264
209.1028594970703 0 712.59283
214.11915588378906 0 839.08887
215.13909912109375 0 701.522
217.08303833007812 0 571.7717
217.09751892089844 0 1555.207
219.11309814453125 0 1605.1743
221.103515625 0 5237.607
221.1287384033203 0 87165.14 a 1
222.10638427734375 0 838.64185
222.13211059570312 0 11292.048
223.06382751464844 0 1092.5588
224.14012145996094 0 671.9308
225.10963439941406 0 720.29333
225.1722869873047 0 604.1222
229.11862182617188 0 1212.7778
231.06182861328125 0 1311.1582
231.1127471923828 0 850.16327 b Water loss 1
233.12872314453125 0 1197.2301
235.1080322265625 0 3095.9243
239.0952606201172 0 501.5524
239.11410522460938 0 792.9403
239.15048217773438 0 1160.3221
241.07444763183594 0 913.933
242.11427307128906 0 468.3336
249.12368774414062 0 15697.872 b 1
250.12696838378906 0 2079.0918
251.1028594970703 0 2122.9773
253.16636657714844 0 1313.5834
257.105224609375 0 542.94446
263.10272216796875 0 2920.3162
270.1068115234375 0 693.9526
279.09814453125 0 1823.9956
284.19696044921875 0 718.1012
285.1020812988281 0 2132.3987
285.1540222167969 0 534.8652
287.1007995605469 0 612.1033
292.14068603515625 0 1080.0632
296.15142822265625 0 1408.5326
300.11767578125 0 881.51904
300.15631103515625 0 656.89484
302.11395263671875 0 995.2231
309.203369140625 0 614.6229
319.1399230957031 0 1306.7936
320.1241760253906 0 865.42737
324.1459045410156 0 4120.1733
325.1491394042969 0 722.54987
329.127685546875 0 665.43866
329.15008544921875 0 12620.763
330.1534423828125 0 2247.9902
333.12054443359375 0 663.2038
333.19293212890625 0 520.6717
338.1355895996094 0 724.34717
339.1690979003906 0 554.66095
340.19854736328125 0 2424.7886
342.15655517578125 0 16843.191 y 7
343.15960693359375 0 3020.8516
350.1343688964844 0 793.3789
355.0683898925781 0 708.6547
356.1380615234375 0 1093.8615
357.2247009277344 0 3504.449
364.1503601074219 0 4451.24 b 2
365.1554260253906 0 912.44214
366.13006591796875 0 759.6565
368.19134521484375 0 695.8249
376.114990234375 0 828.0991
378.1288757324219 0 2376.748
392.20635986328125 0 637.5134
394.12481689453125 0 2541.795
409.1697692871094 0 589.5602
418.7174377441406 0 681.78894
422.7294006347656 0 694.5545
425.19378662109375 0 2565.2214
427.7220764160156 0 1762.855
428.22369384765625 0 1105.5438
432.1868591308594 0 980.6327
436.7281188964844 0 1548.3325
437.22845458984375 0 1192.6357
441.2966003417969 0 710.5404
447.1556091308594 0 572.2836
465.16253662109375 0 2145.6528
479.1778869628906 0 2229.3816 b 3
483.21923828125 0 1027.8107
489.1922607421875 0 3883.888 y 6
490.1950378417969 0 1563.1538
496.229736328125 0 1712.7854
500.2446594238281 0 680.7254
501.2556457519531 0 669.3983
519.2006225585938 0 1338.5464
524.251220703125 0 901.9906
539.2145385742188 0 971.72424
547.252685546875 0 615.08765
548.1990966796875 0 743.30066
550.711181640625 0 2301.1035 y 1
551.21337890625 0 1198.2449
551.713623046875 0 706.2917
560.2291870117188 0 6772.2935 y 5
561.230712890625 0 1690.0574
585.3355712890625 0 4165.8843
586.340576171875 0 1133.0725
592.2299194335938 0 824.87085 Precursor Water loss
592.7260131835938 0 1088.9312
595.3255004882812 0 866.89215
600.3356323242188 0 781.809
600.9881591796875 0 895.5088
601.2340698242188 0 2597.7944 Precursor
601.3220825195312 0 892.7103
601.7366333007812 0 1597.3378
602.236328125 0 748.9384
606.9269409179688 0 808.49603
612.197509765625 0 1143.132
614.2463989257812 0 812.9945
628.3775024414062 0 853.2888
657.26123046875 0 1538.764
659.2939453125 0 1613.9319
660.2996826171875 0 1421.8481
670.8168334960938 0 2578.762
671.3197631835938 0 875.3535
671.8202514648438 0 1013.8709
672.392333984375 0 642.7142
682.3529663085938 0 717.7321
685.2567138671875 0 2130.5425
686.2603759765625 0 818.9505
700.3623657226562 0 11441.81
701.36572265625 0 3865.6538
702.3703002929688 0 906.77594
721.2598266601562 0 1715.3807
722.263427734375 0 950.2233
722.4005126953125 0 634.1129
723.2923583984375 0 4871.7 y 4
724.293212890625 0 2593.922
725.2947998046875 0 713.13336
744.346435546875 0 1462.5355
749.2566528320312 0 2669.5156
750.2601318359375 0 1541.9106
757.3831787109375 0 928.20575
773.4103393554688 0 753.9234
774.318359375 0 3436.627
783.3984375 0 2733.8423
784.4073486328125 0 1186.3274
794.3832397460938 0 1253.7246
800.4257202148438 0 1367.3721
801.4131469726562 0 6095.712
802.413330078125 0 2937.092
803.1714477539062 0 656.65173
803.41650390625 0 807.532
828.419189453125 0 2356.8916
829.427001953125 0 901.7207
838.3197631835938 0 6769.509 y 3
839.32177734375 0 3052.6948
840.3307495117188 0 707.6905
844.4519653320312 0 8927.713
845.4529418945312 0 2823.262
846.4580078125 0 1206.1481
854.4329223632812 0 1783.331
872.4470825195312 0 32802.242
873.4498291015625 0 13678.922
874.454345703125 0 3210.5642
889.3472290039062 0 9798.565
890.3507690429688 0 4965.0034
891.3523559570312 0 1468.2529
907.3327026367188 0 658.83746
909.408447265625 0 843.8872
935.3338012695312 0 2271.268 y Water loss 2
936.3345336914062 0 1175.8663
953.3456420898438 0 33368.1 y 2
954.3486328125 0 18651.074
954.4338989257812 0 520.9684
955.3496704101562 0 6276.839
956.3479614257812 0 906.70667
991.5186157226562 0 5555.4976
992.523681640625 0 2919.2688
993.5260620117188 0 1095.9982
1003.3854370117188 0 924.9484
1004.3820190429688 0 723.1774
1019.5140991210938 0 11814.348
1020.5172729492188 0 8218.838
1021.5198364257812 0 3371.0964
1100.4156494140625 0 1870.2688 y 1
1101.4136962890625 0 1002.5679
1810.0751953125 0 646.63135
2603.2861328125 0 684.22723
2744.458740234375 0 759.4819
3313.851806640625 0 726.4017

Spectrum Details

|  |  |
| --- | --- |
| Matched peaks? Matched peaks The total absolute number of peaks matched. Additionally in brackets the total fraction of peaks matched and the total number of peaks is shown. | 18 (6.87% of 262) |
| FDR? FDR The false discovery rate estimated for this peptide. It is calculated by matching all theoretical fragments with a non-integer shift with the raw peaks for this spectrum. This is done with 40 different shifts. The resulting percentage is the average number of annotated peaks over the number of annotated peaks with the correct spectrum. | 1.46% |
| Satellite FDR? Satellite FDR See the FDR for details on its calculation. This satellite ion specific FDR only contains the satellite ions (d/w) for I/L/J positions. | - |
| PSM Score? PSM Score The PSM Score as given by Hecklib to this annotated spectrum. It is shown with three significant figures. | 195 |

#### Spectrum 10126? Spectrum 10126 The raw spectrum of this peptide as annotated by Hecklib. The fragments are coloured according to ion type (see legend). Any peaks with a star '\*' as text can be hovered over to see the full details, first the ion type second the mass shift type. By hovering over the amino acids in the peptide or ions in the legend the corresponding peaks are highlighted. By toggling the 'Unassigned' label you can turn the background (unassigned) peaks on or off in the plot. By updating the slider in the Ion legend you can update the spectrum to only show the top X% of the peaks with labels. The top X% means any peak that is within X% of the highest intensity. By dragging in the spectrum you can zoom in to a specific part of the spectrum and use 'Zoom Out' to get back to the original zoom level. The annotation of the spectrum is based on the given sequence in the peptides file and is done with different software so inconsistencies are likely. The peaks are annotated based on the given sequence, with 20 ppm tolerance.

Copy Data

##### Spectrum 10126 (TSV)

###### Preview

```
Loading example...
```

*Click on the button to copy the data to your clipboard.*

Mz MinMz MaxIntensity Max

WidthHeightPeptide font sizePeptide stroke widthSpectrum font sizeSpectrum stroke widthCompact peptide

Ion legend

wxyz

abcd

OtherUnassignedIon^Charge^_Position_Show for top:%

TFDDYAMHW

02.32e+44.64e+46.96e+49.29e+4

Zoom Out

a^+1^_2_y^+1^_1_a^+1^_2_b^+1^_2_b^+1^_2_a^+1^_3_y^+1^_2_b^+1^_3_b^+1^_3_y^+2^_7_b^+1^_4_y^+1^_3_y^+2^_8_y^+1^_4_\*\*b^+1^_5_y^+1^_5_y^+1^_6_y^+1^_7_y^+1^_7_y^+1^_8_

0835167025043339

Fragment Matches Table

Show background peaks

| Position | Ion type | Intensity | mz Theoretical | mz Error (Th) | mz Error (ppm) | Charge | Series Number |
| --- | --- | --- | --- | --- | --- | --- | --- |
| - | - | 7.531E+04 | 120.1 | - | - | 0 | - |
| - | - | 837.2 | 121.1 | - | - | 0 | - |
| - | - | 7596 | 121.1 | - | - | 0 | - |
| - | - | 1149 | 123 | - | - | 0 | - |
| - | - | 577.5 | 127.1 | - | - | 0 | - |
| - | - | 4718 | 129.1 | - | - | 0 | - |
| - | - | 426.8 | 129.3 | - | - | 0 | - |
| - | - | 368.1 | 130.1 | - | - | 0 | - |
| - | - | 1740 | 130.1 | - | - | 0 | - |
| - | - | 929.9 | 132.1 | - | - | 0 | - |
| - | - | 864.9 | 132.1 | - | - | 0 | - |
| - | - | 824.1 | 133.1 | - | - | 0 | - |
| - | - | 490.3 | 133.4 | - | - | 0 | - |
| - | - | 385.1 | 134.5 | - | - | 0 | - |
| - | - | 2.549E+04 | 136.1 | - | - | 0 | - |
| - | - | 2440 | 137.1 | - | - | 0 | - |
| - | - | 893.2 | 138.1 | - | - | 0 | - |
| - | - | 1182 | 138.1 | - | - | 0 | - |
| - | - | 424.5 | 141.1 | - | - | 0 | - |
| - | - | 448 | 141.1 | - | - | 0 | - |
| - | - | 1358 | 146.1 | - | - | 0 | - |
| - | - | 1232 | 147 | - | - | 0 | - |
| - | - | 577.6 | 147.1 | - | - | 0 | - |
| - | - | 467.5 | 149 | - | - | 0 | - |
| - | - | 441 | 149 | - | - | 0 | - |
| - | - | 1036 | 155.1 | - | - | 0 | - |
| - | - | 2821 | 156.1 | - | - | 0 | - |
| - | - | 2073 | 157.1 | - | - | 0 | - |
| - | - | 656.4 | 158.1 | - | - | 0 | - |
| - | - | 3860 | 158.1 | - | - | 0 | - |
| - | - | 3280 | 159.1 | - | - | 0 | - |
| - | - | 456.2 | 159.1 | - | - | 0 | - |
| - | - | 1027 | 160.1 | - | - | 0 | - |
| - | - | 1346 | 160.1 | - | - | 0 | - |
| - | - | 407.4 | 163.5 | - | - | 0 | - |
| - | - | 1.194E+04 | 165.1 | - | - | 0 | - |
| - | - | 662.5 | 166.1 | - | - | 0 | - |
| - | - | 3014 | 166.1 | - | - | 0 | - |
| - | - | 518.4 | 167 | - | - | 0 | - |
| - | - | 1472 | 167.1 | - | - | 0 | - |
| - | - | 513.8 | 167.1 | - | - | 0 | - |
| - | - | 863.6 | 171.1 | - | - | 0 | - |
| - | - | 547.6 | 173.1 | - | - | 0 | - |
| - | - | 5826 | 173.1 | - | - | 0 | - |
| - | - | 2628 | 173.5 | - | - | 0 | - |
| - | - | 1423 | 175.1 | - | - | 0 | - |
| - | - | 459.5 | 175.6 | - | - | 0 | - |
| - | - | 653.3 | 176.1 | - | - | 0 | - |
| - | - | 7376 | 176.1 | - | - | 0 | - |
| - | - | 3778 | 177.1 | - | - | 0 | - |
| - | - | 637 | 178.1 | - | - | 0 | - |
| - | - | 438.2 | 181.1 | - | - | 0 | - |
| - | - | 1.992E+04 | 182.1 | - | - | 0 | - |
| - | - | 1364 | 183.1 | - | - | 0 | - |
| - | - | 736.4 | 183.1 | - | - | 0 | - |
| - | - | 677.6 | 183.1 | - | - | 0 | - |
| - | - | 1839 | 185.1 | - | - | 0 | - |
| - | - | 604.6 | 186.1 | - | - | 0 | - |
| - | - | 519.7 | 187.1 | - | - | 0 | - |
| - | - | 4678 | 188.1 | - | - | 0 | - |
| - | - | 655.1 | 189.1 | - | - | 0 | - |
| - | - | 2568 | 191.1 | - | - | 0 | - |
| - | - | 1929 | 193.1 | - | - | 0 | - |
| - | - | 773.3 | 195.1 | - | - | 0 | - |
| - | - | 648.4 | 199.1 | - | - | 0 | - |
| - | - | 456.4 | 200.6 | - | - | 0 | - |
| - | - | 3609 | 201.1 | - | - | 0 | - |
| - | - | 895.6 | 203.1 | - | - | 0 | - |
| 2 | a | 2430 | 203.1 | 0.0002135 | 1.051 | +1 | 2 |
| - | - | 724.2 | 204.1 | - | - | 0 | - |
| 9 | y | 3383 | 205.1 | 0.0004411 | 2.151 | +1 | 1 |
| - | - | 564.2 | 207.1 | - | - | 0 | - |
| - | - | 1049 | 207.1 | - | - | 0 | - |
| - | - | 1103 | 211.1 | - | - | 0 | - |
| - | - | 909 | 214.1 | - | - | 0 | - |
| - | - | 634 | 215.1 | - | - | 0 | - |
| - | - | 1108 | 217.1 | - | - | 0 | - |
| - | - | 1364 | 219.1 | - | - | 0 | - |
| - | - | 5088 | 221.1 | - | - | 0 | - |
| 2 | a | 9.194E+04 | 221.1 | 0.00033 | 1.492 | +1 | 2 |
| - | - | 528.4 | 222.1 | - | - | 0 | - |
| - | - | 645.6 | 222.1 | - | - | 0 | - |
| - | - | 1.184E+04 | 222.1 | - | - | 0 | - |
| - | - | 1401 | 223.1 | - | - | 0 | - |
| - | - | 688.5 | 225.2 | - | - | 0 | - |
| - | - | 546.3 | 226.2 | - | - | 0 | - |
| - | - | 816.6 | 229.2 | - | - | 0 | - |
| - | - | 1574 | 231.1 | - | - | 0 | - |
| 2 | b | 804.8 | 231.1 | 0.001026 | 4.441 | +1 | 2 |
| - | - | 1198 | 233.1 | - | - | 0 | - |
| - | - | 515.2 | 234.8 | - | - | 0 | - |
| - | - | 2090 | 235.1 | - | - | 0 | - |
| - | - | 656 | 238.1 | - | - | 0 | - |
| - | - | 919.8 | 239.1 | - | - | 0 | - |
| - | - | 878.8 | 239.1 | - | - | 0 | - |
| - | - | 967.4 | 241.1 | - | - | 0 | - |
| - | - | 562.1 | 241.5 | - | - | 0 | - |
| - | - | 594.9 | 246.9 | - | - | 0 | - |
| 2 | b | 1.583E+04 | 249.1 | 0.0003036 | 1.219 | +1 | 2 |
| - | - | 2425 | 250.1 | - | - | 0 | - |
| - | - | 2035 | 251.1 | - | - | 0 | - |
| - | - | 1397 | 253.1 | - | - | 0 | - |
| - | - | 660.4 | 253.2 | - | - | 0 | - |
| - | - | 880 | 257.1 | - | - | 0 | - |
| - | - | 3294 | 263.1 | - | - | 0 | - |
| - | - | 690.4 | 264.1 | - | - | 0 | - |
| - | - | 1465 | 279.1 | - | - | 0 | - |
| - | - | 689.2 | 283.1 | - | - | 0 | - |
| - | - | 1238 | 285 | - | - | 0 | - |
| - | - | 2285 | 285.1 | - | - | 0 | - |
| - | - | 612.2 | 286.1 | - | - | 0 | - |
| - | - | 1545 | 292.1 | - | - | 0 | - |
| - | - | 1354 | 296.2 | - | - | 0 | - |
| - | - | 732.3 | 302.1 | - | - | 0 | - |
| - | - | 823.1 | 314.1 | - | - | 0 | - |
| 3 | a | 903.6 | 318.1 | 0.0005532 | 1.739 | +1 | 3 |
| - | - | 1066 | 319.1 | - | - | 0 | - |
| - | - | 767.2 | 323.1 | - | - | 0 | - |
| - | - | 2928 | 324.1 | - | - | 0 | - |
| - | - | 788.2 | 325.1 | - | - | 0 | - |
| - | - | 1.294E+04 | 329.1 | - | - | 0 | - |
| - | - | 2529 | 330.2 | - | - | 0 | - |
| - | - | 798.1 | 333.1 | - | - | 0 | - |
| - | - | 2857 | 340.2 | - | - | 0 | - |
| - | - | 1069 | 342.1 | - | - | 0 | - |
| 8 | y | 1.986E+04 | 342.2 | 0.0005197 | 1.519 | +1 | 2 |
| - | - | 3349 | 343.2 | - | - | 0 | - |
| 3 | b | 1450 | 346.1 | 2.332E-05 | 0.06737 | +1 | 3 |
| - | - | 994.5 | 350.1 | - | - | 0 | - |
| - | - | 928.2 | 355.1 | - | - | 0 | - |
| - | - | 957.4 | 356.1 | - | - | 0 | - |
| - | - | 2731 | 357.2 | - | - | 0 | - |
| 3 | b | 4849 | 364.2 | 0.0002569 | 0.7056 | +1 | 3 |
| - | - | 825.4 | 368.2 | - | - | 0 | - |
| - | - | 2045 | 378.1 | - | - | 0 | - |
| - | - | 760.9 | 379.1 | - | - | 0 | - |
| - | - | 3939 | 394.1 | - | - | 0 | - |
| - | - | 602.5 | 395.1 | - | - | 0 | - |
| - | - | 889.2 | 401.2 | - | - | 0 | - |
| - | - | 679.9 | 401.7 | - | - | 0 | - |
| - | - | 560.4 | 402.2 | - | - | 0 | - |
| - | - | 619.3 | 408.2 | - | - | 0 | - |
| - | - | 705.3 | 413.3 | - | - | 0 | - |
| - | - | 2708 | 425.2 | - | - | 0 | - |
| - | - | 832.7 | 426.2 | - | - | 0 | - |
| - | - | 878.6 | 427.7 | - | - | 0 | - |
| - | - | 1042 | 428.2 | - | - | 0 | - |
| - | - | 991.7 | 432.2 | - | - | 0 | - |
| - | - | 2198 | 436.7 | - | - | 0 | - |
| - | - | 786.8 | 455.2 | - | - | 0 | - |
| - | - | 2628 | 465.2 | - | - | 0 | - |
| - | - | 692.4 | 470.3 | - | - | 0 | - |
| - | - | 804.8 | 471.3 | - | - | 0 | - |
| 3 | y | 646.1 | 477.2 | 0.004553 | 9.541 | +2 | 7 |
| - | - | 624.9 | 479 | - | - | 0 | - |
| 4 | b | 1665 | 479.2 | 0.002066 | 4.312 | +1 | 4 |
| - | - | 1253 | 483.2 | - | - | 0 | - |
| - | - | 734.7 | 487.3 | - | - | 0 | - |
| 7 | y | 4047 | 489.2 | 0.005893 | 12.05 | +1 | 3 |
| - | - | 1326 | 490.2 | - | - | 0 | - |
| - | - | 2057 | 496.2 | - | - | 0 | - |
| - | - | 721.4 | 497.2 | - | - | 0 | - |
| - | - | 805.3 | 497.2 | - | - | 0 | - |
| - | - | 838.4 | 500.2 | - | - | 0 | - |
| - | - | 939.5 | 501.3 | - | - | 0 | - |
| - | - | 885 | 510.3 | - | - | 0 | - |
| - | - | 703.5 | 513.4 | - | - | 0 | - |
| - | - | 671.1 | 519.2 | - | - | 0 | - |
| - | - | 568.1 | 525.5 | - | - | 0 | - |
| - | - | 675 | 527.7 | - | - | 0 | - |
| - | - | 773 | 533.2 | - | - | 0 | - |
| - | - | 938.5 | 548.2 | - | - | 0 | - |
| 2 | y | 2511 | 550.7 | 0.002531 | 4.595 | +2 | 8 |
| - | - | 1513 | 551.2 | - | - | 0 | - |
| 6 | y | 5597 | 560.2 | 0.005279 | 9.422 | +1 | 4 |
| - | - | 1502 | 561.2 | - | - | 0 | - |
| - | - | 835.5 | 570.2 | - | - | 0 | - |
| - | - | 658.6 | 580.3 | - | - | 0 | - |
| - | - | 736.1 | 583.3 | - | - | 0 | - |
| - | - | 4948 | 585.3 | - | - | 0 | - |
| - | - | 1583 | 586.3 | - | - | 0 | - |
| 0 | Precursor | 1343 | 592.2 | 0.0009416 | 1.59 | +2 | -1 |
| - | - | 1009 | 593.3 | - | - | 0 | - |
| - | - | 718.2 | 600.3 | - | - | 0 | - |
| 0 | Precursor | 1626 | 601.2 | 0.001702 | 2.83 | +2 | -1 |
| - | - | 737 | 601.7 | - | - | 0 | - |
| - | - | 1851 | 601.7 | - | - | 0 | - |
| - | - | 1073 | 612.2 | - | - | 0 | - |
| - | - | 885.3 | 613.2 | - | - | 0 | - |
| - | - | 1107 | 614.2 | - | - | 0 | - |
| - | - | 938.5 | 616.3 | - | - | 0 | - |
| 5 | b | 845.1 | 624.2 | 0.002647 | 4.241 | +1 | 5 |
| - | - | 1522 | 628.4 | - | - | 0 | - |
| - | - | 1333 | 657.3 | - | - | 0 | - |
| - | - | 2433 | 659.3 | - | - | 0 | - |
| - | - | 902.6 | 682.3 | - | - | 0 | - |
| - | - | 1934 | 685.3 | - | - | 0 | - |
| - | - | 776.4 | 699.4 | - | - | 0 | - |
| - | - | 1.372E+04 | 700.4 | - | - | 0 | - |
| - | - | 3564 | 701.4 | - | - | 0 | - |
| - | - | 828.1 | 702.4 | - | - | 0 | - |
| - | - | 823.4 | 703.3 | - | - | 0 | - |
| - | - | 668.6 | 715.4 | - | - | 0 | - |
| - | - | 1796 | 721.3 | - | - | 0 | - |
| 5 | y | 6159 | 723.3 | 0.004755 | 6.574 | +1 | 5 |
| - | - | 2921 | 724.3 | - | - | 0 | - |
| - | - | 2097 | 749.3 | - | - | 0 | - |
| - | - | 997.7 | 750.3 | - | - | 0 | - |
| - | - | 1638 | 757.4 | - | - | 0 | - |
| - | - | 2827 | 774.3 | - | - | 0 | - |
| - | - | 1367 | 775.3 | - | - | 0 | - |
| - | - | 2691 | 783.4 | - | - | 0 | - |
| - | - | 1407 | 784.4 | - | - | 0 | - |
| - | - | 1822 | 800.4 | - | - | 0 | - |
| - | - | 6413 | 801.4 | - | - | 0 | - |
| - | - | 2057 | 802.4 | - | - | 0 | - |
| - | - | 878.1 | 803.4 | - | - | 0 | - |
| - | - | 2795 | 828.4 | - | - | 0 | - |
| 4 | y | 7659 | 838.3 | 0.004973 | 5.932 | +1 | 6 |
| - | - | 3405 | 839.3 | - | - | 0 | - |
| - | - | 875.9 | 840.3 | - | - | 0 | - |
| - | - | 8475 | 844.5 | - | - | 0 | - |
| - | - | 4523 | 845.5 | - | - | 0 | - |
| - | - | 1433 | 846.5 | - | - | 0 | - |
| - | - | 1226 | 854.4 | - | - | 0 | - |
| - | - | 911.9 | 871.3 | - | - | 0 | - |
| - | - | 3.383E+04 | 872.4 | - | - | 0 | - |
| - | - | 1.489E+04 | 873.4 | - | - | 0 | - |
| - | - | 5080 | 874.5 | - | - | 0 | - |
| - | - | 862.3 | 875.5 | - | - | 0 | - |
| - | - | 1.058E+04 | 889.3 | - | - | 0 | - |
| - | - | 5781 | 890.3 | - | - | 0 | - |
| - | - | 1202 | 891.4 | - | - | 0 | - |
| 3 | y | 1998 | 935.3 | 0.003914 | 4.185 | +1 | 7 |
| - | - | 1346 | 936.3 | - | - | 0 | - |
| 3 | y | 3.642E+04 | 953.3 | 0.004336 | 4.548 | +1 | 7 |
| - | - | 1.762E+04 | 954.3 | - | - | 0 | - |
| - | - | 5814 | 955.3 | - | - | 0 | - |
| - | - | 981.3 | 956.3 | - | - | 0 | - |
| - | - | 5348 | 991.5 | - | - | 0 | - |
| - | - | 3000 | 992.5 | - | - | 0 | - |
| - | - | 1094 | 993.5 | - | - | 0 | - |
| - | - | 942.7 | 1002 | - | - | 0 | - |
| - | - | 1.604E+04 | 1020 | - | - | 0 | - |
| - | - | 8426 | 1021 | - | - | 0 | - |
| - | - | 3202 | 1022 | - | - | 0 | - |
| - | - | 655.9 | 1023 | - | - | 0 | - |
| 2 | y | 2080 | 1100 | 0.004037 | 3.669 | +1 | 8 |
| - | - | 1685 | 1101 | - | - | 0 | - |
| - | - | 725.1 | 1981 | - | - | 0 | - |
| - | - | 647 | 3306 | - | - | 0 | - |

m/z Charge Intensity FragmentType MassShift Position
120.08108520507812 0 75313.11
121.07976531982422 0 837.21857
121.08442687988281 0 7595.673
123.04444885253906 0 1149.4572
127.0868911743164 0 577.4934
129.10252380371094 0 4718.2197
129.33848571777344 0 426.78607
130.05010986328125 0 368.10193
130.06539916992188 0 1740.4673
132.08108520507812 0 929.94666
132.10220336914062 0 864.90906
133.08599853515625 0 824.11664
133.42918395996094 0 490.32425
134.47633361816406 0 385.08597
136.0760040283203 0 25494.3
137.0794219970703 0 2440.355
138.05502319335938 0 893.19275
138.06643676757812 0 1182.4369
141.06655883789062 0 424.4572
141.10264587402344 0 448.0341
146.0603485107422 0 1357.7173
147.0443115234375 0 1231.6442
147.11294555664062 0 577.6242
149.0234832763672 0 467.47702
149.04510498046875 0 441.03702
155.08145141601562 0 1036.4138
156.0769805908203 0 2821.449
157.1337432861328 0 2073.303
158.06048583984375 0 656.42865
158.0966796875 0 3859.863
159.09190368652344 0 3280.147
159.09854125976562 0 456.1568
160.07579040527344 0 1027.09
160.1123504638672 0 1345.5756
163.53880310058594 0 407.44986
165.05490112304688 0 11935.008
166.0605926513672 0 662.48395
166.08663940429688 0 3014.46
167.04444885253906 0 518.38873
167.05569458007812 0 1472.06
167.08197021484375 0 513.77594
171.07691955566406 0 863.616
173.09217834472656 0 547.5696
173.12879943847656 0 5825.8994
173.45138549804688 0 2628.3096
175.0868377685547 0 1422.6965
175.55300903320312 0 459.52643
176.0820770263672 0 653.2757
176.1072540283203 0 7376.197
177.1026153564453 0 3777.718
178.10610961914062 0 637.04504
181.0970001220703 0 438.2238
182.0814666748047 0 19922.693
183.0849151611328 0 1363.787
183.11300659179688 0 736.3883
183.1492919921875 0 677.59973
185.12881469726562 0 1839.3284
186.09201049804688 0 604.5733
187.14466857910156 0 519.7314
188.07093811035156 0 4678.211
189.08729553222656 0 655.10175
191.11834716796875 0 2567.9182
193.10858154296875 0 1928.8748
195.08773803710938 0 773.31915
199.07147216796875 0 648.40216
200.64329528808594 0 456.39062
201.12355041503906 0 3609.0654
203.06631469726562 0 895.57587
203.11810302734375 0 2430.1292 a Water loss 1
204.07730102539062 0 724.2379
205.09759521484375 0 3383.2754 y 8
207.07684326171875 0 564.1747
207.1131591796875 0 1048.6593
211.14385986328125 0 1102.5006
214.11924743652344 0 908.9845
215.1389617919922 0 633.9618
217.0970001220703 0 1107.5686
219.11302185058594 0 1363.551
221.10354614257812 0 5088.451
221.1287841796875 0 91939.45 a 1
222.0859375 0 528.3535
222.10691833496094 0 645.6466
222.13214111328125 0 11838.674
223.06398010253906 0 1400.7198
225.17141723632812 0 688.5134
226.15469360351562 0 546.2718
229.1710205078125 0 816.6411
231.06175231933594 0 1574.2847
231.11383056640625 0 804.81604 b Water loss 1
233.14993286132812 0 1197.5289
234.7512664794922 0 515.1508
235.10784912109375 0 2090.2585
238.11915588378906 0 656.041
239.09535217285156 0 919.7835
239.11416625976562 0 878.8365
241.0741729736328 0 967.41943
241.4734344482422 0 562.09717
246.91885375976562 0 594.88544
249.12367248535156 0 15827.041 b 1
250.12725830078125 0 2425.1958
251.102783203125 0 2034.9456
253.11866760253906 0 1397.4177
253.1654815673828 0 660.4244
257.1065673828125 0 880.0471
263.10272216796875 0 3294.147
264.14666748046875 0 690.3915
279.09808349609375 0 1465.3009
283.1440734863281 0 689.2331
285.0103759765625 0 1238.1849
285.10205078125 0 2285.3196
286.10321044921875 0 612.21
292.1413269042969 0 1545.1833
296.15032958984375 0 1354.423
302.1145935058594 0 732.3015
314.0978088378906 0 823.12897
318.1453857421875 0 903.63586 a Water loss 2
319.14129638671875 0 1065.7781
323.0990905761719 0 767.1947
324.1459655761719 0 2928.2283
325.14990234375 0 788.2388
329.1499938964844 0 12936.524
330.15350341796875 0 2529.2366
333.120849609375 0 798.1457
340.19830322265625 0 2857.0867
342.13311767578125 0 1069.4796
342.1565856933594 0 19856.361 y 7
343.159423828125 0 3349.1792
346.1397705078125 0 1449.9137 b Water loss 2
350.1351013183594 0 994.5183
355.0696105957031 0 928.22644
356.1395263671875 0 957.3887
357.22442626953125 0 2731.333
364.1500549316406 0 4849.104 b 2
368.1927795410156 0 825.38855
378.12939453125 0 2045.2319
379.1338806152344 0 760.92883
394.125 0 3938.9583
395.128173828125 0 602.48145
401.20928955078125 0 889.2479
401.7116394042969 0 679.928
402.215087890625 0 560.35284
408.1673583984375 0 619.32697
413.2671813964844 0 705.27216
425.19384765625 0 2707.5747
426.1971130371094 0 832.7487
427.72308349609375 0 878.6435
428.22314453125 0 1041.8668
432.1887512207031 0 991.674
436.7279357910156 0 2198.057
455.20330810546875 0 786.75214
465.1629333496094 0 2627.5413
470.3082580566406 0 692.4232
471.2555847167969 0 804.8368
477.1695251464844 0 646.06146 y 2
479.0285949707031 0 624.91486
479.1793212890625 0 1664.7766 b 3
483.21990966796875 0 1253.4845
487.3341064453125 0 734.6992
489.19244384765625 0 4047.4382 y 6
490.1951904296875 0 1326.0809
496.2319641113281 0 2057.4912
497.1672668457031 0 721.4251
497.23199462890625 0 805.26874
500.24615478515625 0 838.44543
501.2557373046875 0 939.5417
510.26220703125 0 885.02026
513.3521118164062 0 703.49426
519.207763671875 0 671.14197
525.5235595703125 0 568.09985
527.66943359375 0 674.9945
533.2044067382812 0 772.9859
548.1991577148438 0 938.51874
550.7108154296875 0 2511.055 y 1
551.2138061523438 0 1512.8279
560.2289428710938 0 5597.2495 y 5
561.2322998046875 0 1501.6659
570.2266845703125 0 835.51587
580.2970581054688 0 658.5911
583.28173828125 0 736.0546
585.3357543945312 0 4948.1997
586.3386840820312 0 1582.5524
592.227783203125 0 1343.161 Precursor Water loss
593.3148193359375 0 1008.6783
600.2871704101562 0 718.23486
601.2338256835938 0 1625.7046 Precursor
601.6558227539062 0 736.9764
601.7380981445312 0 1850.7911
612.198486328125 0 1073.254
613.1975708007812 0 885.3042
614.2483520507812 0 1106.5028
616.3095092773438 0 938.51245
624.232666015625 0 845.07227 b Water loss 4
628.3784790039062 0 1521.8993
657.2609252929688 0 1332.6609
659.2958374023438 0 2432.685
682.3488159179688 0 902.55676
685.2589721679688 0 1934.3044
699.4033203125 0 776.4241
700.3629760742188 0 13715.5625
701.364990234375 0 3563.889
702.3705444335938 0 828.13367
703.2666015625 0 823.4165
715.3754272460938 0 668.56683
721.2595825195312 0 1795.7366
723.291748046875 0 6158.6953 y 4
724.2938842773438 0 2921.4302
749.2564086914062 0 2096.7004
750.2608642578125 0 997.7303
757.3872680664062 0 1637.9763
774.3208618164062 0 2826.5117
775.3209228515625 0 1367.2075
783.3997802734375 0 2691.378
784.39990234375 0 1407.4503
800.4234008789062 0 1821.8845
801.4130859375 0 6413.291
802.4126586914062 0 2056.5269
803.4159545898438 0 878.058
828.4219360351562 0 2794.8774
838.3189086914062 0 7658.6465 y 3
839.321533203125 0 3405.1143
840.3233642578125 0 875.9065
844.451904296875 0 8474.893
845.4556884765625 0 4522.677
846.4552612304688 0 1432.6406
854.4349365234375 0 1225.779
871.3381958007812 0 911.9389
872.4473876953125 0 33826.96
873.4498291015625 0 14887.167
874.451171875 0 5079.5034
875.4544677734375 0 862.2736
889.3475341796875 0 10578.498
890.3499755859375 0 5780.8477
891.3543090820312 0 1201.7155
935.334228515625 0 1997.8395 y Water loss 2
936.3299560546875 0 1346.3616
953.34521484375 0 36415.29 y 2
954.347900390625 0 17624.684
955.3485717773438 0 5814.4946
956.3483276367188 0 981.2815
991.5196533203125 0 5347.761
992.5234985351562 0 3000.0815
993.5233154296875 0 1094.038
1001.5028076171875 0 942.6851
1019.5145874023438 0 16041.866
1020.5187377929688 0 8426.085
1021.5185546875 0 3201.665
1022.5311889648438 0 655.93243
1100.413330078125 0 2079.8105 y 1
1101.4190673828125 0 1684.9229
1980.8382568359375 0 725.11957
3306.2109375 0 646.9964

Spectrum Details

|  |  |
| --- | --- |
| Matched peaks? Matched peaks The total absolute number of peaks matched. Additionally in brackets the total fraction of peaks matched and the total number of peaks is shown. | 22 (8.76% of 251) |
| FDR? FDR The false discovery rate estimated for this peptide. It is calculated by matching all theoretical fragments with a non-integer shift with the raw peaks for this spectrum. This is done with 40 different shifts. The resulting percentage is the average number of annotated peaks over the number of annotated peaks with the correct spectrum. | 1.08% |
| Satellite FDR? Satellite FDR See the FDR for details on its calculation. This satellite ion specific FDR only contains the satellite ions (d/w) for I/L/J positions. | - |
| PSM Score? PSM Score The PSM Score as given by Hecklib to this annotated spectrum. It is shown with three significant figures. | 250 |

#### Spectrum 7474? Spectrum 7474 The raw spectrum of this peptide as annotated by Hecklib. The fragments are coloured according to ion type (see legend). Any peaks with a star '\*' as text can be hovered over to see the full details, first the ion type second the mass shift type. By hovering over the amino acids in the peptide or ions in the legend the corresponding peaks are highlighted. By toggling the 'Unassigned' label you can turn the background (unassigned) peaks on or off in the plot. By updating the slider in the Ion legend you can update the spectrum to only show the top X% of the peaks with labels. The top X% means any peak that is within X% of the highest intensity. By dragging in the spectrum you can zoom in to a specific part of the spectrum and use 'Zoom Out' to get back to the original zoom level. The annotation of the spectrum is based on the given sequence in the peptides file and is done with different software so inconsistencies are likely. The peaks are annotated based on the given sequence, with 20 ppm tolerance.

Copy Data

##### Spectrum 7474 (TSV)

###### Preview

```
Loading example...
```

*Click on the button to copy the data to your clipboard.*

Mz MinMz MaxIntensity Max

WidthHeightPeptide font sizePeptide stroke widthSpectrum font sizeSpectrum stroke widthCompact peptide

Ion legend

wxyz

abcd

OtherUnassignedIon^Charge^_Position_Show for top:%

TFDDYAMHW

04.03e+58.05e+51.21e+61.61e+6

Zoom Out

y^+1^_1_c^+1^_2_y^+1^_2_c^+1^_3_z^+1^_3_y^+2^_7_c^+1^_4_y^+1^_3_c^+2^_8_y^+2^_8_z^+2^_8_z^+1^_4_y^+2^_8_y^+1^_4_c^+1^_5_z^+1^_5_c^+1^_6_y^+1^_5_c^+1^_6_w^+1^_6_y^+1^_6_z^+1^_6_y^+1^_6_c^+1^_7_c^+1^_7_c^+1^_7_w^+1^_7_y^+1^_7_z^+1^_7_y^+1^_7_c^+1^_8_c^+1^_8_c^+1^_8_z^+1^_8_y^+1^_8_

0801160224043205

Fragment Matches Table

Show background peaks

| Position | Ion type | Intensity | mz Theoretical | mz Error (Th) | mz Error (ppm) | Charge | Series Number |
| --- | --- | --- | --- | --- | --- | --- | --- |
| - | - | 1597 | 120.1 | - | - | 0 | - |
| - | - | 1.762E+04 | 120.1 | - | - | 0 | - |
| - | - | 1405 | 121.1 | - | - | 0 | - |
| - | - | 953.6 | 124.6 | - | - | 0 | - |
| - | - | 944.9 | 126.9 | - | - | 0 | - |
| - | - | 3660 | 136.1 | - | - | 0 | - |
| - | - | 1169 | 143.1 | - | - | 0 | - |
| - | - | 2183 | 148.9 | - | - | 0 | - |
| - | - | 1167 | 152.5 | - | - | 0 | - |
| - | - | 6063 | 159.1 | - | - | 0 | - |
| - | - | 1414 | 160.1 | - | - | 0 | - |
| - | - | 1387 | 171.1 | - | - | 0 | - |
| - | - | 1107 | 171.3 | - | - | 0 | - |
| - | - | 1360 | 173.4 | - | - | 0 | - |
| - | - | 1399 | 173.4 | - | - | 0 | - |
| - | - | 1965 | 173.5 | - | - | 0 | - |
| - | - | 3460 | 176.1 | - | - | 0 | - |
| - | - | 6950 | 177.1 | - | - | 0 | - |
| - | - | 4618 | 188.1 | - | - | 0 | - |
| - | - | 4979 | 203.1 | - | - | 0 | - |
| 9 | y | 1.111E+04 | 205.1 | 0.0002122 | 1.035 | +1 | 1 |
| - | - | 5.573E+05 | 221.1 | - | - | 0 | - |
| - | - | 6.481E+04 | 222.1 | - | - | 0 | - |
| - | - | 3097 | 223.1 | - | - | 0 | - |
| - | - | 3890 | 231.1 | - | - | 0 | - |
| - | - | 6074 | 231.1 | - | - | 0 | - |
| - | - | 3334 | 235.1 | - | - | 0 | - |
| 2 | c | 2.167E+05 | 249.1 | 0.0002731 | 1.096 | +1 | 2 |
| - | - | 2.516E+04 | 250.1 | - | - | 0 | - |
| - | - | 1825 | 251.1 | - | - | 0 | - |
| - | - | 2254 | 251.1 | - | - | 0 | - |
| - | - | 1726 | 262 | - | - | 0 | - |
| - | - | 1.364E+04 | 263.1 | - | - | 0 | - |
| - | - | 2251 | 264.1 | - | - | 0 | - |
| - | - | 3.208E+04 | 264.1 | - | - | 0 | - |
| - | - | 7258 | 265.1 | - | - | 0 | - |
| - | - | 1.197E+04 | 282.1 | - | - | 0 | - |
| - | - | 1.622E+04 | 283.2 | - | - | 0 | - |
| - | - | 1330 | 283.7 | - | - | 0 | - |
| - | - | 2085 | 284.2 | - | - | 0 | - |
| - | - | 1397 | 287 | - | - | 0 | - |
| - | - | 2177 | 292.2 | - | - | 0 | - |
| - | - | 2110 | 324.1 | - | - | 0 | - |
| - | - | 1.343E+04 | 327.1 | - | - | 0 | - |
| - | - | 2367 | 328.1 | - | - | 0 | - |
| 8 | y | 5.771E+04 | 342.2 | 0.0003977 | 1.162 | +1 | 2 |
| - | - | 1.147E+04 | 343.2 | - | - | 0 | - |
| - | - | 5073 | 346.1 | - | - | 0 | - |
| - | - | 2814 | 356.1 | - | - | 0 | - |
| - | - | 4179 | 358.2 | - | - | 0 | - |
| 3 | c | 5.133E+04 | 364.2 | 0.0001093 | 0.3001 | +1 | 3 |
| - | - | 8093 | 365.2 | - | - | 0 | - |
| - | - | 8951 | 378.1 | - | - | 0 | - |
| - | - | 2240 | 379.1 | - | - | 0 | - |
| - | - | 5356 | 411.2 | - | - | 0 | - |
| - | - | 2545 | 412.2 | - | - | 0 | - |
| - | - | 5178 | 415.2 | - | - | 0 | - |
| - | - | 4173 | 425.2 | - | - | 0 | - |
| - | - | 9013 | 429.2 | - | - | 0 | - |
| - | - | 3927 | 430.2 | - | - | 0 | - |
| - | - | 1760 | 433.2 | - | - | 0 | - |
| - | - | 1676 | 454.2 | - | - | 0 | - |
| - | - | 1631 | 454.5 | - | - | 0 | - |
| - | - | 1781 | 455.2 | - | - | 0 | - |
| - | - | 7121 | 461.2 | - | - | 0 | - |
| - | - | 2215 | 465.2 | - | - | 0 | - |
| - | - | 1784 | 467.2 | - | - | 0 | - |
| - | - | 2884 | 471.2 | - | - | 0 | - |
| 7 | z | 1.965E+04 | 473.2 | 0.005513 | 11.65 | +1 | 3 |
| - | - | 1.037E+05 | 474.2 | - | - | 0 | - |
| - | - | 2.744E+04 | 475.2 | - | - | 0 | - |
| - | - | 4031 | 476.2 | - | - | 0 | - |
| 3 | y | 4593 | 477.2 | 0.001074 | 2.25 | +2 | 7 |
| - | - | 2175 | 477.7 | - | - | 0 | - |
| 4 | c | 3.069E+04 | 479.2 | 0.00051 | 1.064 | +1 | 4 |
| - | - | 6058 | 480.2 | - | - | 0 | - |
| - | - | 4385 | 481.2 | - | - | 0 | - |
| - | - | 4569 | 488.2 | - | - | 0 | - |
| 7 | y | 5.572E+04 | 489.2 | 0.005222 | 10.67 | +1 | 3 |
| - | - | 1.579E+04 | 490.2 | - | - | 0 | - |
| - | - | 2778 | 491.2 | - | - | 0 | - |
| - | - | 7223 | 496.2 | - | - | 0 | - |
| 8 | c | 3442 | 499.2 | 0.003275 | 6.561 | +2 | 8 |
| - | - | 2219 | 499.7 | - | - | 0 | - |
| - | - | 1.427E+04 | 500.2 | - | - | 0 | - |
| - | - | 5421 | 501.2 | - | - | 0 | - |
| - | - | 3280 | 513.2 | - | - | 0 | - |
| - | - | 4277 | 516.2 | - | - | 0 | - |
| - | - | 5666 | 519.2 | - | - | 0 | - |
| - | - | 1974 | 520.2 | - | - | 0 | - |
| - | - | 1.203E+04 | 521.2 | - | - | 0 | - |
| - | - | 3198 | 522.2 | - | - | 0 | - |
| 2 | y | 4614 | 541.7 | 0.002747 | 5.071 | +2 | 8 |
| - | - | 4490 | 542.2 | - | - | 0 | - |
| 2 | z | 2422 | 542.7 | 0.007193 | 13.25 | +2 | 8 |
| 6 | z | 3.448E+05 | 544.2 | 0.005326 | 9.786 | +1 | 4 |
| - | - | 1.638E+05 | 545.2 | - | - | 0 | - |
| - | - | 3.637E+04 | 546.2 | - | - | 0 | - |
| - | - | 6422 | 547.2 | - | - | 0 | - |
| 2 | y | 3.82E+04 | 550.7 | 0.002775 | 5.039 | +2 | 8 |
| - | - | 2.355E+04 | 551.2 | - | - | 0 | - |
| - | - | 8282 | 551.7 | - | - | 0 | - |
| - | - | 1537 | 552.2 | - | - | 0 | - |
| - | - | 1693 | 559.2 | - | - | 0 | - |
| 6 | y | 9.274E+04 | 560.2 | 0.004973 | 8.877 | +1 | 4 |
| - | - | 2.497E+04 | 561.2 | - | - | 0 | - |
| - | - | 6961 | 562.2 | - | - | 0 | - |
| - | - | 8226 | 569.2 | - | - | 0 | - |
| - | - | 3827 | 569.7 | - | - | 0 | - |
| - | - | 7979 | 578.2 | - | - | 0 | - |
| - | - | 2840 | 578.7 | - | - | 0 | - |
| - | - | 2542 | 579.2 | - | - | 0 | - |
| - | - | 2936 | 583.2 | - | - | 0 | - |
| - | - | 1.298E+04 | 587.2 | - | - | 0 | - |
| - | - | 4284 | 588.2 | - | - | 0 | - |
| - | - | 2117 | 591.2 | - | - | 0 | - |
| - | - | 2.335E+04 | 592.2 | - | - | 0 | - |
| - | - | 1.459E+04 | 592.7 | - | - | 0 | - |
| - | - | 7148 | 593.2 | - | - | 0 | - |
| - | - | 4062 | 596.2 | - | - | 0 | - |
| - | - | 1577 | 597.2 | - | - | 0 | - |
| - | - | 7.403E+04 | 601.2 | - | - | 0 | - |
| - | - | 3071 | 601.3 | - | - | 0 | - |
| - | - | 4.208E+04 | 601.7 | - | - | 0 | - |
| - | - | 1.692E+04 | 602.2 | - | - | 0 | - |
| - | - | 4590 | 602.7 | - | - | 0 | - |
| - | - | 1579 | 606.2 | - | - | 0 | - |
| - | - | 2458 | 612.2 | - | - | 0 | - |
| - | - | 1.03E+04 | 614.2 | - | - | 0 | - |
| - | - | 4855 | 615.2 | - | - | 0 | - |
| - | - | 2121 | 618.3 | - | - | 0 | - |
| - | - | 1560 | 619.3 | - | - | 0 | - |
| - | - | 5975 | 624.2 | - | - | 0 | - |
| - | - | 4205 | 634.2 | - | - | 0 | - |
| - | - | 1.325E+04 | 636.2 | - | - | 0 | - |
| - | - | 4040 | 637.2 | - | - | 0 | - |
| 5 | c | 8901 | 642.2 | 0.001482 | 2.308 | +1 | 5 |
| - | - | 1825 | 643.2 | - | - | 0 | - |
| - | - | 3285 | 649.3 | - | - | 0 | - |
| - | - | 9779 | 659.3 | - | - | 0 | - |
| - | - | 2685 | 660.3 | - | - | 0 | - |
| - | - | 4681 | 662.3 | - | - | 0 | - |
| - | - | 1.364E+04 | 663.3 | - | - | 0 | - |
| - | - | 3716 | 664.3 | - | - | 0 | - |
| - | - | 2174 | 665.3 | - | - | 0 | - |
| - | - | 4529 | 685.3 | - | - | 0 | - |
| - | - | 3983 | 688.3 | - | - | 0 | - |
| - | - | 3580 | 693.2 | - | - | 0 | - |
| - | - | 2106 | 695.3 | - | - | 0 | - |
| - | - | 2370 | 696.3 | - | - | 0 | - |
| - | - | 5141 | 706.3 | - | - | 0 | - |
| 5 | z | 1.916E+05 | 707.3 | 0.005352 | 7.567 | +1 | 5 |
| - | - | 9.103E+04 | 708.3 | - | - | 0 | - |
| - | - | 2204 | 708.4 | - | - | 0 | - |
| - | - | 2.721E+04 | 709.3 | - | - | 0 | - |
| - | - | 4263 | 710.3 | - | - | 0 | - |
| 6 | c | 3791 | 713.3 | 0.003187 | 4.467 | +1 | 6 |
| - | - | 1763 | 714.3 | - | - | 0 | - |
| - | - | 1.1E+04 | 715.3 | - | - | 0 | - |
| - | - | 6051 | 716.3 | - | - | 0 | - |
| - | - | 8310 | 721.3 | - | - | 0 | - |
| - | - | 7641 | 722.3 | - | - | 0 | - |
| 5 | y | 9.911E+04 | 723.3 | 0.004755 | 6.574 | +1 | 5 |
| - | - | 3.806E+04 | 724.3 | - | - | 0 | - |
| - | - | 1.136E+04 | 725.3 | - | - | 0 | - |
| - | - | 3844 | 729.3 | - | - | 0 | - |
| 6 | c | 5403 | 730.3 | 0.001512 | 2.07 | +1 | 6 |
| - | - | 2197 | 733.3 | - | - | 0 | - |
| - | - | 5378 | 734.3 | - | - | 0 | - |
| - | - | 3483 | 735.3 | - | - | 0 | - |
| - | - | 2.694E+04 | 749.3 | - | - | 0 | - |
| - | - | 2.203E+04 | 750.3 | - | - | 0 | - |
| - | - | 1.505E+04 | 751.3 | - | - | 0 | - |
| - | - | 3104 | 752.3 | - | - | 0 | - |
| - | - | 7859 | 759.3 | - | - | 0 | - |
| - | - | 5807 | 760.3 | - | - | 0 | - |
| - | - | 1841 | 761.3 | - | - | 0 | - |
| - | - | 3247 | 763.3 | - | - | 0 | - |
| - | - | 5416 | 764.3 | - | - | 0 | - |
| - | - | 1878 | 765.3 | - | - | 0 | - |
| - | - | 3834 | 767.3 | - | - | 0 | - |
| - | - | 2642 | 771.3 | - | - | 0 | - |
| - | - | 2318 | 772.3 | - | - | 0 | - |
| - | - | 1.007E+04 | 774.3 | - | - | 0 | - |
| - | - | 5489 | 775.3 | - | - | 0 | - |
| 4 | w | 4865 | 777.3 | 0.0006215 | 0.7996 | +1 | 6 |
| - | - | 1.067E+05 | 778.3 | - | - | 0 | - |
| - | - | 4.7E+04 | 779.3 | - | - | 0 | - |
| - | - | 1.317E+04 | 780.3 | - | - | 0 | - |
| - | - | 2247 | 781.3 | - | - | 0 | - |
| - | - | 3611 | 787.4 | - | - | 0 | - |
| - | - | 1723 | 793.3 | - | - | 0 | - |
| - | - | 2405 | 805.4 | - | - | 0 | - |
| - | - | 4762 | 808.3 | - | - | 0 | - |
| - | - | 2502 | 815.3 | - | - | 0 | - |
| - | - | 2997 | 816.3 | - | - | 0 | - |
| 4 | y | 3693 | 820.3 | 0.004368 | 5.325 | +1 | 6 |
| - | - | 2202 | 821.3 | - | - | 0 | - |
| 4 | z | 1.32E+05 | 822.3 | 0.004715 | 5.734 | +1 | 6 |
| - | - | 1.328E+05 | 823.3 | - | - | 0 | - |
| - | - | 5.717E+04 | 824.3 | - | - | 0 | - |
| - | - | 1.237E+04 | 825.3 | - | - | 0 | - |
| - | - | 2035 | 829.3 | - | - | 0 | - |
| - | - | 2.479E+04 | 830.3 | - | - | 0 | - |
| - | - | 1.543E+04 | 831.3 | - | - | 0 | - |
| - | - | 4591 | 832.4 | - | - | 0 | - |
| - | - | 5.641E+04 | 833.3 | - | - | 0 | - |
| - | - | 3.89E+04 | 834.3 | - | - | 0 | - |
| - | - | 1.423E+04 | 835.3 | - | - | 0 | - |
| - | - | 2525 | 836.3 | - | - | 0 | - |
| 4 | y | 1.044E+05 | 838.3 | 0.004729 | 5.641 | +1 | 6 |
| - | - | 4.823E+04 | 839.3 | - | - | 0 | - |
| - | - | 1.567E+04 | 840.3 | - | - | 0 | - |
| - | - | 7.329E+04 | 849.3 | - | - | 0 | - |
| - | - | 3.518E+04 | 850.3 | - | - | 0 | - |
| - | - | 1.1E+04 | 851.4 | - | - | 0 | - |
| - | - | 9916 | 858.3 | - | - | 0 | - |
| 7 | c | 7109 | 859.3 | 0.007028 | 8.178 | +1 | 7 |
| 7 | c | 8271 | 860.3 | 0.006699 | 7.786 | +1 | 7 |
| - | - | 3147 | 861.3 | - | - | 0 | - |
| - | - | 2073 | 862.3 | - | - | 0 | - |
| - | - | 4907 | 866.3 | - | - | 0 | - |
| - | - | 4190 | 871.3 | - | - | 0 | - |
| - | - | 2136 | 872.3 | - | - | 0 | - |
| - | - | 1.021E+04 | 874.3 | - | - | 0 | - |
| - | - | 7585 | 875.3 | - | - | 0 | - |
| - | - | 6.708E+04 | 876.3 | - | - | 0 | - |
| 7 | c | 2.877E+05 | 877.3 | 0.004136 | 4.715 | +1 | 7 |
| - | - | 1.305E+05 | 878.3 | - | - | 0 | - |
| - | - | 4.212E+04 | 879.3 | - | - | 0 | - |
| - | - | 4919 | 880.3 | - | - | 0 | - |
| - | - | 6.529E+04 | 889.3 | - | - | 0 | - |
| - | - | 3.484E+04 | 890.3 | - | - | 0 | - |
| - | - | 9392 | 891.4 | - | - | 0 | - |
| 3 | w | 4743 | 892.3 | 0.01034 | 11.59 | +1 | 7 |
| - | - | 1.66E+05 | 893.3 | - | - | 0 | - |
| - | - | 8.978E+04 | 894.3 | - | - | 0 | - |
| - | - | 2.695E+04 | 895.3 | - | - | 0 | - |
| - | - | 5284 | 896.3 | - | - | 0 | - |
| - | - | 4520 | 898.3 | - | - | 0 | - |
| - | - | 1990 | 907.3 | - | - | 0 | - |
| - | - | 2628 | 907.4 | - | - | 0 | - |
| - | - | 2261 | 908.4 | - | - | 0 | - |
| - | - | 2209 | 925.3 | - | - | 0 | - |
| - | - | 6387 | 926.4 | - | - | 0 | - |
| - | - | 3728 | 927.4 | - | - | 0 | - |
| 3 | y | 2.135E+04 | 935.3 | 0.004158 | 4.446 | +1 | 7 |
| - | - | 1.717E+04 | 936.3 | - | - | 0 | - |
| 3 | z | 3.81E+04 | 937.3 | 0.005604 | 5.979 | +1 | 7 |
| - | - | 2.348E+04 | 938.3 | - | - | 0 | - |
| - | - | 8793 | 939.3 | - | - | 0 | - |
| - | - | 2136 | 940.3 | - | - | 0 | - |
| - | - | 1.113E+04 | 952.4 | - | - | 0 | - |
| 3 | y | 5.05E+05 | 953.3 | 0.005129 | 5.38 | +1 | 7 |
| - | - | 2.567E+05 | 954.3 | - | - | 0 | - |
| - | - | 1.01E+05 | 955.3 | - | - | 0 | - |
| - | - | 1.156E+04 | 956.3 | - | - | 0 | - |
| - | - | 2287 | 963.3 | - | - | 0 | - |
| - | - | 6.007E+04 | 970.4 | - | - | 0 | - |
| - | - | 3.168E+04 | 971.4 | - | - | 0 | - |
| - | - | 1.174E+04 | 972.4 | - | - | 0 | - |
| - | - | 1658 | 981.4 | - | - | 0 | - |
| 8 | c | 2085 | 996.4 | 0.01866 | 18.72 | +1 | 8 |
| 8 | c | 8350 | 997.4 | 0.01151 | 11.54 | +1 | 8 |
| - | - | 5648 | 998.4 | - | - | 0 | - |
| - | - | 9167 | 999.4 | - | - | 0 | - |
| - | - | 3494 | 1000 | - | - | 0 | - |
| 8 | c | 5.235E+05 | 1014 | 0.004612 | 4.546 | +1 | 8 |
| - | - | 2.997E+05 | 1015 | - | - | 0 | - |
| - | - | 1.007E+05 | 1016 | - | - | 0 | - |
| - | - | 1.455E+04 | 1017 | - | - | 0 | - |
| - | - | 2015 | 1026 | - | - | 0 | - |
| - | - | 4976 | 1036 | - | - | 0 | - |
| - | - | 4221 | 1037 | - | - | 0 | - |
| - | - | 1831 | 1038 | - | - | 0 | - |
| - | - | 2.401E+04 | 1040 | - | - | 0 | - |
| - | - | 1.546E+04 | 1041 | - | - | 0 | - |
| - | - | 7035 | 1042 | - | - | 0 | - |
| - | - | 2400 | 1056 | - | - | 0 | - |
| - | - | 3140 | 1069 | - | - | 0 | - |
| - | - | 6563 | 1071 | - | - | 0 | - |
| - | - | 3622 | 1072 | - | - | 0 | - |
| - | - | 2045 | 1079 | - | - | 0 | - |
| - | - | 2749 | 1080 | - | - | 0 | - |
| 2 | z | 5.528E+04 | 1084 | 0.00384 | 3.542 | +1 | 8 |
| - | - | 3.55E+04 | 1085 | - | - | 0 | - |
| - | - | 1.367E+04 | 1086 | - | - | 0 | - |
| - | - | 7629 | 1094 | - | - | 0 | - |
| - | - | 3807 | 1095 | - | - | 0 | - |
| - | - | 3408 | 1096 | - | - | 0 | - |
| - | - | 4148 | 1097 | - | - | 0 | - |
| - | - | 2840 | 1098 | - | - | 0 | - |
| 2 | y | 2.606E+04 | 1100 | 0.004648 | 4.223 | +1 | 8 |
| - | - | 1.532E+04 | 1101 | - | - | 0 | - |
| - | - | 5510 | 1102 | - | - | 0 | - |
| - | - | 1.995E+04 | 1103 | - | - | 0 | - |
| - | - | 1.245E+04 | 1104 | - | - | 0 | - |
| - | - | 4096 | 1105 | - | - | 0 | - |
| - | - | 9414 | 1110 | - | - | 0 | - |
| - | - | 8167 | 1111 | - | - | 0 | - |
| - | - | 1910 | 1112 | - | - | 0 | - |
| - | - | 1971 | 1113 | - | - | 0 | - |
| - | - | 1.202E+04 | 1122 | - | - | 0 | - |
| - | - | 1.083E+04 | 1123 | - | - | 0 | - |
| - | - | 8797 | 1124 | - | - | 0 | - |
| - | - | 2217 | 1128 | - | - | 0 | - |
| - | - | 1.513E+04 | 1138 | - | - | 0 | - |
| - | - | 9183 | 1139 | - | - | 0 | - |
| - | - | 1.093E+05 | 1140 | - | - | 0 | - |
| - | - | 1.699E+05 | 1141 | - | - | 0 | - |
| - | - | 1.928E+05 | 1142 | - | - | 0 | - |
| - | - | 1.045E+05 | 1143 | - | - | 0 | - |
| - | - | 3.782E+04 | 1144 | - | - | 0 | - |
| - | - | 6187 | 1145 | - | - | 0 | - |
| - | - | 7394 | 1156 | - | - | 0 | - |
| - | - | 4223 | 1157 | - | - | 0 | - |
| - | - | 1.168E+04 | 1158 | - | - | 0 | - |
| - | - | 5509 | 1159 | - | - | 0 | - |
| - | - | 2931 | 1160 | - | - | 0 | - |
| - | - | 3928 | 1166 | - | - | 0 | - |
| - | - | 2950 | 1167 | - | - | 0 | - |
| - | - | 3118 | 1168 | - | - | 0 | - |
| - | - | 2414 | 1169 | - | - | 0 | - |
| - | - | 6672 | 1170 | - | - | 0 | - |
| - | - | 5212 | 1171 | - | - | 0 | - |
| - | - | 2076 | 1172 | - | - | 0 | - |
| - | - | 1.418E+04 | 1174 | - | - | 0 | - |
| - | - | 9386 | 1175 | - | - | 0 | - |
| - | - | 3883 | 1176 | - | - | 0 | - |
| - | - | 8.446E+04 | 1184 | - | - | 0 | - |
| - | - | 7.281E+05 | 1185 | - | - | 0 | - |
| - | - | 5.004E+05 | 1186 | - | - | 0 | - |
| - | - | 2.239E+05 | 1187 | - | - | 0 | - |
| - | - | 4.364E+04 | 1188 | - | - | 0 | - |
| - | - | 4706 | 1189 | - | - | 0 | - |
| - | - | 4.917E+05 | 1201 | - | - | 0 | - |
| - | - | 1.595E+06 | 1202 | - | - | 0 | - |
| - | - | 1.028E+06 | 1203 | - | - | 0 | - |
| - | - | 4.076E+05 | 1204 | - | - | 0 | - |
| - | - | 5.561E+04 | 1205 | - | - | 0 | - |
| - | - | 2235 | 3083 | - | - | 0 | - |
| - | - | 1883 | 3173 | - | - | 0 | - |

m/z Charge Intensity FragmentType MassShift Position
120.0763931274414 0 1597.0543
120.0810317993164 0 17615.98
121.08451843261719 0 1404.7292
124.58456420898438 0 953.57666
126.87700653076172 0 944.8678
136.07603454589844 0 3659.5237
143.0876922607422 0 1168.69
148.9464569091797 0 2182.7212
152.51382446289062 0 1167.3524
159.091796875 0 6062.777
160.11241149902344 0 1413.7346
171.0773162841797 0 1387.3049
171.34498596191406 0 1107.3647
173.4191131591797 0 1359.5571
173.4320831298828 0 1398.6136
173.45870971679688 0 1964.608
176.1072540283203 0 3460.2454
177.10243225097656 0 6949.8306
188.07095336914062 0 4618.0757
203.11810302734375 0 4978.7285
205.0973663330078 0 11108.559 y 8
221.1287384033203 0 557328.44
222.1320343017578 0 64813.535
223.13406372070312 0 3097.472
231.06126403808594 0 3890.468
231.11306762695312 0 6074.2295
235.1077117919922 0 3334.0747
249.12364196777344 0 216681.56 c Ammonia loss 1
250.12680053710938 0 25158.428
251.1029510498047 0 1824.9471
251.12948608398438 0 2254.347
262.03521728515625 0 1725.7743
263.10272216796875 0 13639.7
264.1064453125 0 2251.4421
264.1369934082031 0 32078.24
265.1413879394531 0 7257.9087
282.1478576660156 0 11965.072
283.1552734375 0 16222.265
283.7259216308594 0 1330.1963
284.1591491699219 0 2085.0156
287.0005187988281 0 1397.1726
292.16595458984375 0 2177.206
324.14654541015625 0 2110.3877
327.14556884765625 0 13426.602
328.14837646484375 0 2366.9255
342.1564636230469 0 57711.605 y 7
343.1595153808594 0 11465.715
346.1398010253906 0 5073.472
356.138916015625 0 2814.2498
358.1552734375 0 4178.832
364.1504211425781 0 51327.16 c Ammonia loss 2
365.15313720703125 0 8092.882
378.12969970703125 0 8951.372
379.1343078613281 0 2239.6248
411.1907043457031 0 5356.2207
412.1964111328125 0 2545.2217
415.1522216796875 0 5177.705
425.19317626953125 0 4172.9526
429.1831970214844 0 9013.047
430.18829345703125 0 3927.4434
433.1720886230469 0 1759.7737
454.197021484375 0 1676.2046
454.48876953125 0 1630.9086
455.20367431640625 0 1780.7893
461.16632080078125 0 7121.136
465.1632385253906 0 2214.968
467.2041015625 0 1784.1581
471.1825256347656 0 2884.402
473.17333984375 0 19650.596 z 6
474.1807556152344 0 103713.55
475.18359375 0 27435.912
476.18438720703125 0 4031.4663
477.1730041503906 0 4592.758 y 2
477.6753845214844 0 2175.0872
479.1777648925781 0 30694.164 c Ammonia loss 3
480.1806335449219 0 6058.0024
481.2210388183594 0 4384.7837
488.1833190917969 0 4568.978
489.1917724609375 0 55717.38 y 6
490.19384765625 0 15793.053
491.1948547363281 0 2778.3582
496.23040771484375 0 7222.529
499.1904602050781 0 3442.2793 c Ammonia loss 7
499.68914794921875 0 2218.8774
500.2200927734375 0 14270.626
501.2238464355469 0 5421.379
513.199462890625 0 3279.5068
516.178466796875 0 4276.696
519.2054443359375 0 5665.834
520.2051391601562 0 1973.9126
521.217529296875 0 12025.353
522.2206420898438 0 3198.1738
541.7057495117188 0 4613.9746 y Water loss 1
542.2055053710938 0 4490.375
542.7061157226562 0 2421.9854 z 1
544.2102661132812 0 344751.7 z 5
545.21484375 0 163803.11
546.21630859375 0 36367.37
547.224609375 0 6421.6987
550.7110595703125 0 38201.516 y 1
551.212646484375 0 23546.27
551.7138061523438 0 8281.598
552.2130737304688 0 1536.5061
559.21875 0 1692.6699
560.2286376953125 0 92735.39 y 5
561.23095703125 0 24971.309
562.2316284179688 0 6961.0674
569.2324829101562 0 8225.671
569.7335205078125 0 3826.651
578.22216796875 0 7978.8623
578.7315063476562 0 2840.4521
579.224853515625 0 2542.0461
583.2222290039062 0 2936.0356
587.216064453125 0 12983.528
588.222412109375 0 4283.953
591.2242431640625 0 2117.0596
592.2308349609375 0 23350.852
592.731201171875 0 14591.516
593.233154296875 0 7148.121
596.2354125976562 0 4061.8787
597.2260131835938 0 1577.225
601.2352905273438 0 74027.125
601.2835693359375 0 3070.556
601.736572265625 0 42082.266
602.2372436523438 0 16918.95
602.7361450195312 0 4589.638
606.2230834960938 0 1579.2554
612.196533203125 0 2458.268
614.2454223632812 0 10297.064
615.2467651367188 0 4855.448
618.271240234375 0 2120.9849
619.2765502929688 0 1559.7692
624.229736328125 0 5974.616
634.229248046875 0 4205.4146
636.2446899414062 0 13250.695
637.2457275390625 0 4040.0508
642.2420654296875 0 8901.32 c Ammonia loss 4
643.2413330078125 0 1824.7704
649.251220703125 0 3284.5652
659.2946166992188 0 9779.032
660.2965698242188 0 2685.4949
662.2590942382812 0 4680.7495
663.281494140625 0 13639.992
664.2841186523438 0 3715.7314
665.28857421875 0 2174.4072
685.2592163085938 0 4528.993
688.2975463867188 0 3982.9556
693.2442626953125 0 3579.5244
695.2677001953125 0 2105.8315
696.2699584960938 0 2370.4941
706.2610473632812 0 5141.3374
707.2736206054688 0 191584.39 z 4
708.2772827148438 0 91031.37
708.381103515625 0 2204.0024
709.2769165039062 0 27212.291
710.2749633789062 0 4262.874
713.2808837890625 0 3790.659 c Ammonia loss 5
714.3076171875 0 1762.5967
715.3191528320312 0 11000.93
716.3233642578125 0 6050.7954
721.2646484375 0 8309.877
722.2774047851562 0 7641.135
723.291748046875 0 99109.336 y 4
724.2948608398438 0 38055.344
725.294921875 0 11356.094
729.2942504882812 0 3843.6746
730.302734375 0 5402.939 c 5
733.29345703125 0 2196.6577
734.3177490234375 0 5378.452
735.323486328125 0 3483.4216
749.2562255859375 0 26937.627
750.2725219726562 0 22028.885
751.2739868164062 0 15046.661
752.2736206054688 0 3103.999
759.3095092773438 0 7858.878
760.313720703125 0 5807.1367
761.3201293945312 0 1840.9949
763.2886962890625 0 3247.2373
764.2849731445312 0 5416.136
765.2915649414062 0 1878.4666
767.2691650390625 0 3834.0593
771.3499145507812 0 2642.4888
772.347900390625 0 2317.8865
774.3197631835938 0 10073.877
775.3224487304688 0 5489.1045
777.2969360351562 0 4864.698 w 3
778.3099365234375 0 106737.71
779.3131103515625 0 46995.07
780.3137817382812 0 13169.536
781.3087158203125 0 2247.3118
787.3616943359375 0 3611.454
793.3167724609375 0 1723.1306
805.3583984375 0 2404.6553
808.2723388671875 0 4761.777
815.308349609375 0 2502.107
816.3087768554688 0 2997.4678
820.3077392578125 0 3693.0762 y Water loss 3
821.2907104492188 0 2201.553
822.2999267578125 0 131984.4 z 3
823.3057250976562 0 132803.73
824.3082275390625 0 57170.227
825.310791015625 0 12372.384
829.33740234375 0 2035.3494
830.3468627929688 0 24786.312
831.349853515625 0 15432.03
832.3513793945312 0 4591.1353
833.3257446289062 0 56410.293
834.3298950195312 0 38902.76
835.3318481445312 0 14227.51
836.3388671875 0 2524.6023
838.3186645507812 0 104423.09 y 3
839.3214111328125 0 48232.805
840.3222045898438 0 15674.408
849.34716796875 0 73294.58
850.3485717773438 0 35180.69
851.3502807617188 0 11001.07
858.3218383789062 0 9915.586
859.317138671875 0 7108.556 c Water loss 6
860.3148803710938 0 8270.832 c Ammonia loss 6
861.320068359375 0 3146.5435
862.3247680664062 0 2072.6572
866.313720703125 0 4907.376
871.3350219726562 0 4190.0537
872.3369750976562 0 2136.0176
874.3389892578125 0 10209.176
875.3345336914062 0 7584.87
876.3321533203125 0 67084.586
877.3388671875 0 287736.38 c 6
878.3416748046875 0 130465.79
879.3411865234375 0 42122.504
880.33935546875 0 4919.1855
889.346923828125 0 65286.207
890.3499755859375 0 34838.31
891.352783203125 0 9392.203
892.3348388671875 0 4743.4097 w 2
893.3369140625 0 165971.12
894.3400268554688 0 89777.7
895.3400268554688 0 26948.412
896.3323974609375 0 5284.4775
898.3367919921875 0 4519.5977
907.3273315429688 0 1989.6703
907.4049072265625 0 2627.7842
908.4090576171875 0 2261.0588
925.322998046875 0 2209.2256
926.396484375 0 6386.915
927.39599609375 0 3727.9612
935.33447265625 0 21353.395 y Water loss 2
936.3356323242188 0 17169.137
937.3277587890625 0 38096.42 z 2
938.329345703125 0 23477.918
939.3325805664062 0 8792.837
940.3386840820312 0 2135.9404
952.3734741210938 0 11131.198
953.3460083007812 0 504993 y 2
954.348876953125 0 256701.5
955.3491821289062 0 100988.016
956.3491821289062 0 11558.796
963.3301391601562 0 2286.812
970.38427734375 0 60065.094
971.38671875 0 31679.46
972.3889770507812 0 11738.292
981.3685302734375 0 1657.6431
996.4017333984375 0 2085.241 c Water loss 7
997.3786010742188 0 8350.193 c Ammonia loss 7
998.3869018554688 0 5647.6187
999.3867797851562 0 9167.1045
1000.393310546875 0 3493.549
1014.3982543945312 0 523461.03 c 7
1015.4010009765625 0 299738.38
1016.4011840820312 0 100694.23
1017.4039916992188 0 14552.26
1026.404541015625 0 2014.9081
1036.414306640625 0 4975.741
1037.4326171875 0 4221.003
1038.437255859375 0 1831.331
1040.405029296875 0 24005.008
1041.408203125 0 15463.059
1042.411376953125 0 7035.342
1056.3870849609375 0 2400.2166
1069.39013671875 0 3139.6824
1071.3927001953125 0 6562.8604
1072.400634765625 0 3621.5208
1079.466796875 0 2044.6904
1080.4573974609375 0 2749.2424
1084.3944091796875 0 55282.957 z 1
1085.398681640625 0 35500.156
1086.3988037109375 0 13671.765
1094.4775390625 0 7629.344
1095.4859619140625 0 3807.4016
1096.482177734375 0 3408.2117
1097.46142578125 0 4148.365
1098.4625244140625 0 2839.8127
1100.4139404296875 0 26058.34 y 1
1101.415771484375 0 15318.007
1102.4195556640625 0 5509.8823
1103.413330078125 0 19950.416
1104.4156494140625 0 12445.658
1105.420166015625 0 4096.07
1110.42236328125 0 9413.883
1111.42919921875 0 8166.6343
1112.4608154296875 0 1910.3658
1113.4951171875 0 1971.0558
1122.453857421875 0 12021.51
1123.4521484375 0 10832.114
1124.4522705078125 0 8796.666
1128.435302734375 0 2217.3965
1138.47265625 0 15125.29
1139.46826171875 0 9183.48
1140.467529296875 0 109341.3
1141.4598388671875 0 169922.12
1142.453369140625 0 192751.39
1143.452392578125 0 104523.12
1144.45263671875 0 37817.246
1145.453125 0 6187.409
1156.4620361328125 0 7393.529
1157.4688720703125 0 4222.756
1158.4853515625 0 11676.415
1159.49365234375 0 5508.6426
1160.4832763671875 0 2930.585
1166.44287109375 0 3928.3643
1167.452880859375 0 2949.732
1168.4361572265625 0 3118.3965
1169.4443359375 0 2413.972
1170.426025390625 0 6671.845
1171.426513671875 0 5212.219
1172.42822265625 0 2076.184
1174.4736328125 0 14184.857
1175.4749755859375 0 9385.6455
1176.4739990234375 0 3883.0718
1184.457763671875 0 84460.04
1185.446533203125 0 728107.06
1186.449462890625 0 500403.78
1187.4503173828125 0 223928.34
1188.45068359375 0 43638.516
1189.448974609375 0 4705.818
1201.4608154296875 0 491710.34
1202.4677734375 0 1594982.4
1203.47119140625 0 1028292.7
1204.4720458984375 0 407550.5
1205.473876953125 0 55607.14
3083.405517578125 0 2234.9524
3173.217041015625 0 1883.3218

Spectrum Details

|  |  |
| --- | --- |
| Matched peaks? Matched peaks The total absolute number of peaks matched. Additionally in brackets the total fraction of peaks matched and the total number of peaks is shown. | 35 (10.23% of 342) |
| FDR? FDR The false discovery rate estimated for this peptide. It is calculated by matching all theoretical fragments with a non-integer shift with the raw peaks for this spectrum. This is done with 40 different shifts. The resulting percentage is the average number of annotated peaks over the number of annotated peaks with the correct spectrum. | 0.48% |
| Satellite FDR? Satellite FDR See the FDR for details on its calculation. This satellite ion specific FDR only contains the satellite ions (d/w) for I/L/J positions. | - |
| PSM Score? PSM Score The PSM Score as given by Hecklib to this annotated spectrum. It is shown with three significant figures. | 449 |

#### Spectrum 10919? Spectrum 10919 The raw spectrum of this peptide as annotated by Hecklib. The fragments are coloured according to ion type (see legend). Any peaks with a star '\*' as text can be hovered over to see the full details, first the ion type second the mass shift type. By hovering over the amino acids in the peptide or ions in the legend the corresponding peaks are highlighted. By toggling the 'Unassigned' label you can turn the background (unassigned) peaks on or off in the plot. By updating the slider in the Ion legend you can update the spectrum to only show the top X% of the peaks with labels. The top X% means any peak that is within X% of the highest intensity. By dragging in the spectrum you can zoom in to a specific part of the spectrum and use 'Zoom Out' to get back to the original zoom level. The annotation of the spectrum is based on the given sequence in the peptides file and is done with different software so inconsistencies are likely. The peaks are annotated based on the given sequence, with 20 ppm tolerance.

Copy Data

##### Spectrum 10919 (TSV)

###### Preview

```
Loading example...
```

*Click on the button to copy the data to your clipboard.*

Mz MinMz MaxIntensity Max

WidthHeightPeptide font sizePeptide stroke widthSpectrum font sizeSpectrum stroke widthCompact peptide

Ion legend

wxyz

abcd

OtherUnassignedIon^Charge^_Position_Show for top:%

TFDDYAMHW

01.63e+43.26e+44.89e+46.52e+4

Zoom Out

a^+1^_2_y^+1^_1_a^+1^_2_b^+1^_2_b^+1^_2_a^+1^_3_y^+1^_2_b^+1^_3_b^+1^_4_y^+1^_3_y^+2^_8_y^+1^_4_\*y^+1^_5_y^+1^_6_y^+1^_7_y^+1^_7_y^+1^_8_

0778155623343112

Fragment Matches Table

Show background peaks

| Position | Ion type | Intensity | mz Theoretical | mz Error (Th) | mz Error (ppm) | Charge | Series Number |
| --- | --- | --- | --- | --- | --- | --- | --- |
| - | - | 5.054E+04 | 120.1 | - | - | 0 | - |
| - | - | 435.3 | 120.7 | - | - | 0 | - |
| - | - | 345 | 120.9 | - | - | 0 | - |
| - | - | 4087 | 121.1 | - | - | 0 | - |
| - | - | 418.9 | 122.1 | - | - | 0 | - |
| - | - | 776.2 | 123 | - | - | 0 | - |
| - | - | 414.9 | 127.1 | - | - | 0 | - |
| - | - | 560.8 | 129.1 | - | - | 0 | - |
| - | - | 7488 | 129.1 | - | - | 0 | - |
| - | - | 1568 | 130.1 | - | - | 0 | - |
| - | - | 717.6 | 130.1 | - | - | 0 | - |
| - | - | 816.6 | 132.1 | - | - | 0 | - |
| - | - | 466 | 133.1 | - | - | 0 | - |
| - | - | 1534 | 133.1 | - | - | 0 | - |
| - | - | 1.635E+04 | 136.1 | - | - | 0 | - |
| - | - | 1556 | 137.1 | - | - | 0 | - |
| - | - | 765.6 | 138.1 | - | - | 0 | - |
| - | - | 916.4 | 138.1 | - | - | 0 | - |
| - | - | 499.3 | 146.1 | - | - | 0 | - |
| - | - | 564.5 | 147 | - | - | 0 | - |
| - | - | 943.5 | 147.1 | - | - | 0 | - |
| - | - | 858.2 | 149 | - | - | 0 | - |
| - | - | 569.9 | 155.1 | - | - | 0 | - |
| - | - | 1853 | 156.1 | - | - | 0 | - |
| - | - | 1331 | 157.1 | - | - | 0 | - |
| - | - | 2479 | 158.1 | - | - | 0 | - |
| - | - | 1873 | 159.1 | - | - | 0 | - |
| - | - | 504.3 | 160.1 | - | - | 0 | - |
| - | - | 1002 | 160.1 | - | - | 0 | - |
| - | - | 654.1 | 162.1 | - | - | 0 | - |
| - | - | 5031 | 165.1 | - | - | 0 | - |
| - | - | 494.9 | 165.1 | - | - | 0 | - |
| - | - | 1485 | 166.1 | - | - | 0 | - |
| - | - | 1169 | 167.1 | - | - | 0 | - |
| - | - | 531.8 | 171.1 | - | - | 0 | - |
| - | - | 2984 | 173.1 | - | - | 0 | - |
| - | - | 551 | 173.5 | - | - | 0 | - |
| - | - | 5543 | 176.1 | - | - | 0 | - |
| - | - | 3586 | 177.1 | - | - | 0 | - |
| - | - | 1483 | 177.1 | - | - | 0 | - |
| - | - | 1.169E+04 | 182.1 | - | - | 0 | - |
| - | - | 994.4 | 183.1 | - | - | 0 | - |
| - | - | 3745 | 183.1 | - | - | 0 | - |
| - | - | 1006 | 185.1 | - | - | 0 | - |
| - | - | 3567 | 188.1 | - | - | 0 | - |
| - | - | 927.1 | 191.1 | - | - | 0 | - |
| - | - | 1745 | 193.1 | - | - | 0 | - |
| - | - | 797.5 | 197.1 | - | - | 0 | - |
| - | - | 890.5 | 200.1 | - | - | 0 | - |
| - | - | 3674 | 201.1 | - | - | 0 | - |
| - | - | 1060 | 203.1 | - | - | 0 | - |
| 2 | a | 923.4 | 203.1 | 0.0003203 | 1.577 | +1 | 2 |
| - | - | 606.7 | 204.1 | - | - | 0 | - |
| 9 | y | 2349 | 205.1 | 0.000197 | 0.9604 | +1 | 1 |
| - | - | 898.4 | 211.1 | - | - | 0 | - |
| - | - | 866.4 | 211.1 | - | - | 0 | - |
| - | - | 680.3 | 213.1 | - | - | 0 | - |
| - | - | 688.2 | 213.1 | - | - | 0 | - |
| - | - | 617.3 | 213.1 | - | - | 0 | - |
| - | - | 4348 | 221.1 | - | - | 0 | - |
| 2 | a | 6.459E+04 | 221.1 | 0.0001926 | 0.8711 | +1 | 2 |
| - | - | 8727 | 222.1 | - | - | 0 | - |
| - | - | 620.2 | 222.1 | - | - | 0 | - |
| - | - | 1207 | 223.1 | - | - | 0 | - |
| - | - | 541.1 | 223.1 | - | - | 0 | - |
| - | - | 606.7 | 227.1 | - | - | 0 | - |
| - | - | 917.2 | 231.1 | - | - | 0 | - |
| - | - | 548.2 | 231.1 | - | - | 0 | - |
| 2 | b | 683.2 | 231.1 | 0.0002024 | 0.8759 | +1 | 2 |
| - | - | 881.6 | 233.1 | - | - | 0 | - |
| - | - | 1523 | 235.1 | - | - | 0 | - |
| - | - | 1116 | 239.1 | - | - | 0 | - |
| - | - | 626.1 | 239.1 | - | - | 0 | - |
| - | - | 816 | 241.1 | - | - | 0 | - |
| 2 | b | 1.061E+04 | 249.1 | 0.0001968 | 0.7901 | +1 | 2 |
| - | - | 1408 | 250.1 | - | - | 0 | - |
| - | - | 1112 | 251.1 | - | - | 0 | - |
| - | - | 949.4 | 253.1 | - | - | 0 | - |
| - | - | 828.7 | 253.2 | - | - | 0 | - |
| - | - | 600.3 | 257.2 | - | - | 0 | - |
| - | - | 2300 | 263.1 | - | - | 0 | - |
| - | - | 1204 | 279.1 | - | - | 0 | - |
| - | - | 1441 | 285.1 | - | - | 0 | - |
| - | - | 894 | 296.2 | - | - | 0 | - |
| - | - | 701.9 | 302.1 | - | - | 0 | - |
| 3 | a | 601 | 318.1 | 0.002537 | 7.974 | +1 | 3 |
| - | - | 2313 | 324.1 | - | - | 0 | - |
| - | - | 5988 | 329.1 | - | - | 0 | - |
| - | - | 1514 | 330.2 | - | - | 0 | - |
| - | - | 1481 | 340.2 | - | - | 0 | - |
| 8 | y | 1.196E+04 | 342.2 | 0.0003672 | 1.073 | +1 | 2 |
| - | - | 2525 | 343.2 | - | - | 0 | - |
| - | - | 705.8 | 350.1 | - | - | 0 | - |
| - | - | 1526 | 355.1 | - | - | 0 | - |
| - | - | 711.6 | 356.1 | - | - | 0 | - |
| - | - | 1464 | 357.2 | - | - | 0 | - |
| 3 | b | 2932 | 364.2 | 0.000379 | 1.041 | +1 | 3 |
| - | - | 872.8 | 368.2 | - | - | 0 | - |
| - | - | 1271 | 378.1 | - | - | 0 | - |
| - | - | 670.4 | 392.2 | - | - | 0 | - |
| - | - | 2161 | 394.1 | - | - | 0 | - |
| - | - | 746.3 | 413.3 | - | - | 0 | - |
| - | - | 632.7 | 422.7 | - | - | 0 | - |
| - | - | 2310 | 425.2 | - | - | 0 | - |
| - | - | 1053 | 427.7 | - | - | 0 | - |
| - | - | 688.3 | 436.7 | - | - | 0 | - |
| - | - | 638.2 | 458.8 | - | - | 0 | - |
| - | - | 1546 | 465.2 | - | - | 0 | - |
| - | - | 677.5 | 472 | - | - | 0 | - |
| 4 | b | 2197 | 479.2 | 0.000571 | 1.192 | +1 | 4 |
| - | - | 757.8 | 483.2 | - | - | 0 | - |
| - | - | 780.6 | 487.3 | - | - | 0 | - |
| 7 | y | 2717 | 489.2 | 0.005374 | 10.99 | +1 | 3 |
| - | - | 754.8 | 490.2 | - | - | 0 | - |
| - | - | 1399 | 496.2 | - | - | 0 | - |
| - | - | 788.5 | 511.8 | - | - | 0 | - |
| 2 | y | 1499 | 550.7 | 0.002836 | 5.15 | +2 | 8 |
| - | - | 705.8 | 551.2 | - | - | 0 | - |
| 6 | y | 5290 | 560.2 | 0.004607 | 8.224 | +1 | 4 |
| - | - | 1398 | 561.2 | - | - | 0 | - |
| - | - | 2264 | 585.3 | - | - | 0 | - |
| - | - | 880.1 | 586.3 | - | - | 0 | - |
| - | - | 616.7 | 588 | - | - | 0 | - |
| - | - | 695.2 | 590.9 | - | - | 0 | - |
| - | - | 733.9 | 600.3 | - | - | 0 | - |
| 0 | Precursor | 2033 | 601.2 | 0.004143 | 6.891 | +2 | -1 |
| - | - | 636 | 602.2 | - | - | 0 | - |
| - | - | 1411 | 612.2 | - | - | 0 | - |
| - | - | 705 | 639.7 | - | - | 0 | - |
| - | - | 1068 | 643 | - | - | 0 | - |
| - | - | 641.3 | 649.3 | - | - | 0 | - |
| - | - | 967.3 | 657.3 | - | - | 0 | - |
| - | - | 972.2 | 659.3 | - | - | 0 | - |
| - | - | 727.3 | 660.3 | - | - | 0 | - |
| - | - | 1028 | 663.4 | - | - | 0 | - |
| - | - | 906.9 | 682.3 | - | - | 0 | - |
| - | - | 1913 | 685.3 | - | - | 0 | - |
| - | - | 5982 | 700.4 | - | - | 0 | - |
| - | - | 3165 | 701.4 | - | - | 0 | - |
| - | - | 966.6 | 702.4 | - | - | 0 | - |
| - | - | 776 | 705.1 | - | - | 0 | - |
| - | - | 1309 | 721.3 | - | - | 0 | - |
| 5 | y | 3837 | 723.3 | 0.004572 | 6.321 | +1 | 5 |
| - | - | 1744 | 724.3 | - | - | 0 | - |
| - | - | 2198 | 749.3 | - | - | 0 | - |
| - | - | 763 | 750.4 | - | - | 0 | - |
| - | - | 706.3 | 753.5 | - | - | 0 | - |
| - | - | 822.8 | 757.4 | - | - | 0 | - |
| - | - | 1930 | 774.3 | - | - | 0 | - |
| - | - | 1916 | 783.4 | - | - | 0 | - |
| - | - | 1006 | 785.4 | - | - | 0 | - |
| - | - | 3273 | 801.4 | - | - | 0 | - |
| - | - | 1084 | 802.4 | - | - | 0 | - |
| - | - | 1327 | 828.4 | - | - | 0 | - |
| 4 | y | 5596 | 838.3 | 0.003569 | 4.257 | +1 | 6 |
| - | - | 3052 | 839.3 | - | - | 0 | - |
| - | - | 1126 | 840.3 | - | - | 0 | - |
| - | - | 4929 | 844.5 | - | - | 0 | - |
| - | - | 3103 | 845.5 | - | - | 0 | - |
| - | - | 1070 | 854.4 | - | - | 0 | - |
| - | - | 952.8 | 871.3 | - | - | 0 | - |
| - | - | 1.802E+04 | 872.4 | - | - | 0 | - |
| - | - | 8073 | 873.4 | - | - | 0 | - |
| - | - | 1793 | 874.5 | - | - | 0 | - |
| - | - | 8877 | 889.3 | - | - | 0 | - |
| - | - | 3383 | 890.3 | - | - | 0 | - |
| - | - | 791.4 | 891.3 | - | - | 0 | - |
| 3 | y | 845.7 | 935.3 | 0.007515 | 8.035 | +1 | 7 |
| - | - | 639.2 | 936.3 | - | - | 0 | - |
| 3 | y | 2.446E+04 | 953.3 | 0.004336 | 4.548 | +1 | 7 |
| - | - | 1.321E+04 | 954.3 | - | - | 0 | - |
| - | - | 3556 | 955.3 | - | - | 0 | - |
| - | - | 2852 | 991.5 | - | - | 0 | - |
| - | - | 1760 | 992.5 | - | - | 0 | - |
| - | - | 680 | 995.6 | - | - | 0 | - |
| - | - | 7993 | 1020 | - | - | 0 | - |
| - | - | 4054 | 1021 | - | - | 0 | - |
| - | - | 1657 | 1022 | - | - | 0 | - |
| 2 | y | 1808 | 1100 | 0.005258 | 4.778 | +1 | 8 |
| - | - | 817.3 | 1101 | - | - | 0 | - |
| - | - | 963.6 | 3081 | - | - | 0 | - |

m/z Charge Intensity FragmentType MassShift Position
120.08102416992188 0 50540.28
120.68478393554688 0 435.31186
120.9048843383789 0 344.9729
121.08440399169922 0 4086.7212
122.0712890625 0 418.92365
123.04439544677734 0 776.1738
127.0868148803711 0 414.93036
129.06593322753906 0 560.80524
129.1024627685547 0 7487.6235
130.06521606445312 0 1568.4825
130.08657836914062 0 717.6031
132.0808563232422 0 816.61993
133.06092834472656 0 465.99066
133.08627319335938 0 1534.1483
136.07591247558594 0 16348.97
137.079345703125 0 1555.5374
138.06651306152344 0 765.636
138.0917510986328 0 916.4373
146.0603790283203 0 499.3232
147.04443359375 0 564.51135
147.11317443847656 0 943.5045
149.04519653320312 0 858.1831
155.08126831054688 0 569.8621
156.0769500732422 0 1852.9752
157.13365173339844 0 1330.8273
158.09664916992188 0 2478.6638
159.09194946289062 0 1873.1482
160.07586669921875 0 504.28635
160.11224365234375 0 1001.827
162.09173583984375 0 654.1201
165.05484008789062 0 5031.024
165.07684326171875 0 494.90668
166.08645629882812 0 1484.9368
167.05604553222656 0 1169.0874
171.0763397216797 0 531.7902
173.12876892089844 0 2984.0554
173.45396423339844 0 550.9648
176.1072235107422 0 5542.8228
177.10232543945312 0 3586.0159
177.1114959716797 0 1483.0769
182.08139038085938 0 11688.185
183.0849151611328 0 994.3702
183.1129608154297 0 3744.7197
185.12875366210938 0 1005.69464
188.07061767578125 0 3566.7585
191.11766052246094 0 927.11273
193.108642578125 0 1744.8597
197.1287078857422 0 797.5241
200.13914489746094 0 890.5057
201.12342834472656 0 3673.7085
203.06642150878906 0 1059.9158
203.1182098388672 0 923.4112 a Water loss 1
204.0770263671875 0 606.6942
205.09735107421875 0 2348.9526 y 8
211.10748291015625 0 898.4252
211.14480590820312 0 866.39746
213.08782958984375 0 680.2529
213.09959411621094 0 688.192
213.11138916015625 0 617.2801
221.1034698486328 0 4347.877
221.12864685058594 0 64589.9 a 1
222.1320037841797 0 8726.68
222.14422607421875 0 620.2389
223.0635223388672 0 1206.6619
223.13502502441406 0 541.0863
227.10328674316406 0 606.73755
231.06109619140625 0 917.21
231.0987091064453 0 548.16156
231.11300659179688 0 683.15454 b Water loss 1
233.09197998046875 0 881.58136
235.1082305908203 0 1523.2767
239.0950469970703 0 1116.334
239.1151123046875 0 626.0793
241.07382202148438 0 816.0028
249.12356567382812 0 10614.167 b 1
250.12684631347656 0 1408.0583
251.10317993164062 0 1112.2915
253.1184844970703 0 949.4325
253.16648864746094 0 828.71375
257.16357421875 0 600.26776
263.1028747558594 0 2299.5806
279.0977478027344 0 1203.876
285.1022644042969 0 1440.8046
296.150634765625 0 893.9806
302.1145935058594 0 701.87756
318.1473693847656 0 600.9904 a Water loss 2
324.1459045410156 0 2313.028
329.14971923828125 0 5988.3125
330.153076171875 0 1513.7043
340.1980895996094 0 1480.9598
342.15643310546875 0 11959.498 y 7
343.1598205566406 0 2524.564
350.1357421875 0 705.7987
355.070068359375 0 1525.6775
356.13916015625 0 711.6282
357.2242431640625 0 1463.7773
364.1499328613281 0 2931.907 b 2
368.19281005859375 0 872.80664
378.12939453125 0 1271.241
392.2042236328125 0 670.36066
394.1248779296875 0 2161.1377
413.266845703125 0 746.309
422.7298889160156 0 632.6772
425.1930236816406 0 2309.826
427.7208251953125 0 1053.0953
436.7270202636719 0 688.2705
458.7972717285156 0 638.15704
465.1615905761719 0 1545.6748
471.9722900390625 0 677.4635
479.1778259277344 0 2196.9753 b 3
483.2227783203125 0 757.8341
487.3342590332031 0 780.60284
489.1919250488281 0 2716.6753 y 6
490.19342041015625 0 754.81433
496.2298278808594 0 1399.2815
511.76568603515625 0 788.506
550.7111206054688 0 1498.575 y 1
551.2097778320312 0 705.8169
560.228271484375 0 5289.619 y 5
561.2304077148438 0 1398.2249
585.3350830078125 0 2264.1042
586.338134765625 0 880.1071
587.9902954101562 0 616.7396
590.85400390625 0 695.2135
600.310791015625 0 733.9289
601.2362670898438 0 2033.1282 Precursor
602.2351684570312 0 635.98236
612.1967163085938 0 1411.1196
639.6577758789062 0 705.03424
642.9663696289062 0 1068.4215
649.3262939453125 0 641.2659
657.2603759765625 0 967.28754
659.2932739257812 0 972.2236
660.2963256835938 0 727.3058
663.36376953125 0 1027.7887
682.3455200195312 0 906.87213
685.2564697265625 0 1912.824
700.3617553710938 0 5981.5635
701.3648681640625 0 3164.6738
702.3656005859375 0 966.6085
705.091796875 0 776.02405
721.2635498046875 0 1308.644
723.2915649414062 0 3837.2505 y 4
724.29296875 0 1744.2095
749.255615234375 0 2197.6213
750.4104614257812 0 763.0373
753.5126342773438 0 706.3173
757.3824462890625 0 822.7959
774.3187255859375 0 1930.1649
783.3973999023438 0 1916.4873
785.4105834960938 0 1005.7189
801.40869140625 0 3272.9
802.4091796875 0 1084.0396
828.4214477539062 0 1326.6582
838.3175048828125 0 5596.2104 y 3
839.3206787109375 0 3052.0068
840.3231811523438 0 1126.2335
844.4522094726562 0 4929.4014
845.4537353515625 0 3103.1523
854.4343872070312 0 1069.6958
871.3311157226562 0 952.7786
872.447021484375 0 18021.78
873.4497680664062 0 8073.206
874.452880859375 0 1793.049
889.345947265625 0 8877.343
890.34912109375 0 3383.4092
891.3487548828125 0 791.39417
935.3378295898438 0 845.6565 y Water loss 2
936.3406372070312 0 639.19055
953.34521484375 0 24456.56 y 2
954.34814453125 0 13208.013
955.3465576171875 0 3555.9307
991.5183715820312 0 2852.0442
992.5261840820312 0 1759.953
995.56689453125 0 679.9838
1019.5144653320312 0 7992.6353
1020.518310546875 0 4054.4326
1021.5200805664062 0 1656.7563
1100.41455078125 0 1807.6875 y 1
1101.4144287109375 0 817.34894
3081.4970703125 0 963.6072

Spectrum Details

|  |  |
| --- | --- |
| Matched peaks? Matched peaks The total absolute number of peaks matched. Additionally in brackets the total fraction of peaks matched and the total number of peaks is shown. | 18 (9.94% of 181) |
| FDR? FDR The false discovery rate estimated for this peptide. It is calculated by matching all theoretical fragments with a non-integer shift with the raw peaks for this spectrum. This is done with 40 different shifts. The resulting percentage is the average number of annotated peaks over the number of annotated peaks with the correct spectrum. | 1.46% |
| Satellite FDR? Satellite FDR See the FDR for details on its calculation. This satellite ion specific FDR only contains the satellite ions (d/w) for I/L/J positions. | - |
| PSM Score? PSM Score The PSM Score as given by Hecklib to this annotated spectrum. It is shown with three significant figures. | 213 |

#### Spectrum 9794? Spectrum 9794 The raw spectrum of this peptide as annotated by Hecklib. The fragments are coloured according to ion type (see legend). Any peaks with a star '\*' as text can be hovered over to see the full details, first the ion type second the mass shift type. By hovering over the amino acids in the peptide or ions in the legend the corresponding peaks are highlighted. By toggling the 'Unassigned' label you can turn the background (unassigned) peaks on or off in the plot. By updating the slider in the Ion legend you can update the spectrum to only show the top X% of the peaks with labels. The top X% means any peak that is within X% of the highest intensity. By dragging in the spectrum you can zoom in to a specific part of the spectrum and use 'Zoom Out' to get back to the original zoom level. The annotation of the spectrum is based on the given sequence in the peptides file and is done with different software so inconsistencies are likely. The peaks are annotated based on the given sequence, with 20 ppm tolerance.

Copy Data

##### Spectrum 9794 (TSV)

###### Preview

```
Loading example...
```

*Click on the button to copy the data to your clipboard.*

Mz MinMz MaxIntensity Max

WidthHeightPeptide font sizePeptide stroke widthSpectrum font sizeSpectrum stroke widthCompact peptide

Ion legend

wxyz

abcd

OtherUnassignedIon^Charge^_Position_Show for top:%

TFDDYAMHW

02.83e+45.67e+48.50e+41.13e+5

Zoom Out

a^+1^_2_y^+1^_1_a^+1^_2_b^+1^_2_b^+1^_2_y^+1^_2_b^+1^_3_b^+1^_3_b^+1^_4_b^+1^_4_y^+1^_3_y^+2^_8_y^+1^_4_\*\*y^+1^_5_y^+1^_6_y^+1^_7_y^+1^_7_b^+1^_8_y^+1^_8_

036272310851447

Fragment Matches Table

Show background peaks

| Position | Ion type | Intensity | mz Theoretical | mz Error (Th) | mz Error (ppm) | Charge | Series Number |
| --- | --- | --- | --- | --- | --- | --- | --- |
| - | - | 1.041E+05 | 120.1 | - | - | 0 | - |
| - | - | 9603 | 121.1 | - | - | 0 | - |
| - | - | 1334 | 123 | - | - | 0 | - |
| - | - | 464 | 128.6 | - | - | 0 | - |
| - | - | 510.7 | 129.1 | - | - | 0 | - |
| - | - | 5479 | 129.1 | - | - | 0 | - |
| - | - | 649.7 | 130.1 | - | - | 0 | - |
| - | - | 2924 | 130.1 | - | - | 0 | - |
| - | - | 461.2 | 132 | - | - | 0 | - |
| - | - | 1651 | 132.1 | - | - | 0 | - |
| - | - | 1062 | 132.1 | - | - | 0 | - |
| - | - | 592.7 | 133.1 | - | - | 0 | - |
| - | - | 3.281E+04 | 136.1 | - | - | 0 | - |
| - | - | 3393 | 137.1 | - | - | 0 | - |
| - | - | 1290 | 138.1 | - | - | 0 | - |
| - | - | 422.1 | 139.9 | - | - | 0 | - |
| - | - | 448.1 | 144 | - | - | 0 | - |
| - | - | 2132 | 146.1 | - | - | 0 | - |
| - | - | 939.8 | 147 | - | - | 0 | - |
| - | - | 622.1 | 148.1 | - | - | 0 | - |
| - | - | 1989 | 149 | - | - | 0 | - |
| - | - | 456 | 151.1 | - | - | 0 | - |
| - | - | 401.2 | 151.5 | - | - | 0 | - |
| - | - | 522.2 | 153.1 | - | - | 0 | - |
| - | - | 511 | 155.1 | - | - | 0 | - |
| - | - | 552 | 155.1 | - | - | 0 | - |
| - | - | 2853 | 156.1 | - | - | 0 | - |
| - | - | 403.4 | 156.1 | - | - | 0 | - |
| - | - | 567.1 | 157.1 | - | - | 0 | - |
| - | - | 669 | 157.1 | - | - | 0 | - |
| - | - | 1669 | 157.1 | - | - | 0 | - |
| - | - | 664.9 | 158.1 | - | - | 0 | - |
| - | - | 3979 | 158.1 | - | - | 0 | - |
| - | - | 5139 | 159.1 | - | - | 0 | - |
| - | - | 1124 | 160.1 | - | - | 0 | - |
| - | - | 651.4 | 160.1 | - | - | 0 | - |
| - | - | 1898 | 160.1 | - | - | 0 | - |
| - | - | 1.3E+04 | 165.1 | - | - | 0 | - |
| - | - | 1061 | 166.1 | - | - | 0 | - |
| - | - | 3231 | 166.1 | - | - | 0 | - |
| - | - | 2046 | 167.1 | - | - | 0 | - |
| - | - | 641.6 | 167.1 | - | - | 0 | - |
| - | - | 467.6 | 169.1 | - | - | 0 | - |
| - | - | 1035 | 169.1 | - | - | 0 | - |
| - | - | 759.4 | 171.1 | - | - | 0 | - |
| - | - | 567.8 | 171.1 | - | - | 0 | - |
| - | - | 639.3 | 173.1 | - | - | 0 | - |
| - | - | 7253 | 173.1 | - | - | 0 | - |
| - | - | 2364 | 175.1 | - | - | 0 | - |
| - | - | 766.9 | 176.1 | - | - | 0 | - |
| - | - | 8346 | 176.1 | - | - | 0 | - |
| - | - | 4917 | 177.1 | - | - | 0 | - |
| - | - | 809.5 | 177.1 | - | - | 0 | - |
| - | - | 504.5 | 178.1 | - | - | 0 | - |
| - | - | 599.4 | 181.1 | - | - | 0 | - |
| - | - | 2.494E+04 | 182.1 | - | - | 0 | - |
| - | - | 2932 | 183.1 | - | - | 0 | - |
| - | - | 539.4 | 183.1 | - | - | 0 | - |
| - | - | 1949 | 185.1 | - | - | 0 | - |
| - | - | 1108 | 186.1 | - | - | 0 | - |
| - | - | 7506 | 188.1 | - | - | 0 | - |
| - | - | 753.7 | 189.1 | - | - | 0 | - |
| - | - | 3242 | 191.1 | - | - | 0 | - |
| - | - | 2108 | 193.1 | - | - | 0 | - |
| - | - | 743.2 | 195.1 | - | - | 0 | - |
| - | - | 486.5 | 195.1 | - | - | 0 | - |
| - | - | 960.7 | 197.1 | - | - | 0 | - |
| - | - | 518.6 | 199.1 | - | - | 0 | - |
| - | - | 535.4 | 199.1 | - | - | 0 | - |
| - | - | 917.1 | 200.1 | - | - | 0 | - |
| - | - | 6189 | 201.1 | - | - | 0 | - |
| - | - | 585.7 | 203.1 | - | - | 0 | - |
| - | - | 1200 | 203.1 | - | - | 0 | - |
| 2 | a | 2170 | 203.1 | 1.513E-05 | 0.07448 | +1 | 2 |
| - | - | 1277 | 204.1 | - | - | 0 | - |
| 9 | y | 4310 | 205.1 | 9.017E-05 | 0.4396 | +1 | 1 |
| - | - | 1491 | 207.1 | - | - | 0 | - |
| - | - | 768 | 209.1 | - | - | 0 | - |
| - | - | 973.4 | 214.1 | - | - | 0 | - |
| - | - | 821.8 | 215.1 | - | - | 0 | - |
| - | - | 565.7 | 217.1 | - | - | 0 | - |
| - | - | 1667 | 217.1 | - | - | 0 | - |
| - | - | 1485 | 219.1 | - | - | 0 | - |
| - | - | 517.7 | 219.7 | - | - | 0 | - |
| - | - | 2345 | 221.1 | - | - | 0 | - |
| - | - | 7170 | 221.1 | - | - | 0 | - |
| 2 | a | 1.123E+05 | 221.1 | 0.0002231 | 1.009 | +1 | 2 |
| - | - | 1.388E+04 | 222.1 | - | - | 0 | - |
| - | - | 1131 | 223.1 | - | - | 0 | - |
| - | - | 940.9 | 223.1 | - | - | 0 | - |
| - | - | 1003 | 225.2 | - | - | 0 | - |
| - | - | 821 | 226.1 | - | - | 0 | - |
| - | - | 771.3 | 226.2 | - | - | 0 | - |
| - | - | 565.6 | 227.1 | - | - | 0 | - |
| - | - | 1092 | 229.1 | - | - | 0 | - |
| - | - | 736.4 | 229.2 | - | - | 0 | - |
| - | - | 533.7 | 229.2 | - | - | 0 | - |
| - | - | 1828 | 231.1 | - | - | 0 | - |
| - | - | 556.6 | 231.1 | - | - | 0 | - |
| 2 | b | 1403 | 231.1 | 0.000355 | 1.536 | +1 | 2 |
| - | - | 915.5 | 233.1 | - | - | 0 | - |
| - | - | 938 | 233.1 | - | - | 0 | - |
| - | - | 627.1 | 233.1 | - | - | 0 | - |
| - | - | 3325 | 235.1 | - | - | 0 | - |
| - | - | 1229 | 239.1 | - | - | 0 | - |
| - | - | 726.7 | 239.1 | - | - | 0 | - |
| - | - | 845.8 | 239.2 | - | - | 0 | - |
| - | - | 717.5 | 240.1 | - | - | 0 | - |
| - | - | 1485 | 241.1 | - | - | 0 | - |
| - | - | 647.4 | 248.1 | - | - | 0 | - |
| 2 | b | 1.777E+04 | 249.1 | 0.0002579 | 1.035 | +1 | 2 |
| - | - | 2485 | 250.1 | - | - | 0 | - |
| - | - | 2040 | 251.1 | - | - | 0 | - |
| - | - | 1859 | 253.1 | - | - | 0 | - |
| - | - | 2294 | 253.2 | - | - | 0 | - |
| - | - | 1221 | 257.1 | - | - | 0 | - |
| - | - | 789.2 | 257.2 | - | - | 0 | - |
| - | - | 3952 | 263.1 | - | - | 0 | - |
| - | - | 510.3 | 263.6 | - | - | 0 | - |
| - | - | 668.6 | 274.1 | - | - | 0 | - |
| - | - | 1406 | 279.1 | - | - | 0 | - |
| - | - | 687.6 | 283 | - | - | 0 | - |
| - | - | 974.3 | 283.1 | - | - | 0 | - |
| - | - | 1456 | 285 | - | - | 0 | - |
| - | - | 2904 | 285.1 | - | - | 0 | - |
| - | - | 1449 | 292.1 | - | - | 0 | - |
| - | - | 2365 | 296.2 | - | - | 0 | - |
| - | - | 746.6 | 297.1 | - | - | 0 | - |
| - | - | 1331 | 302.1 | - | - | 0 | - |
| - | - | 1288 | 312.2 | - | - | 0 | - |
| - | - | 1100 | 314.1 | - | - | 0 | - |
| - | - | 2033 | 319.1 | - | - | 0 | - |
| - | - | 4277 | 324.1 | - | - | 0 | - |
| - | - | 1051 | 325.1 | - | - | 0 | - |
| - | - | 934.9 | 325.1 | - | - | 0 | - |
| - | - | 1.805E+04 | 329.1 | - | - | 0 | - |
| - | - | 3933 | 330.2 | - | - | 0 | - |
| - | - | 839 | 331.2 | - | - | 0 | - |
| - | - | 3365 | 340.2 | - | - | 0 | - |
| 8 | y | 2.259E+04 | 342.2 | 0.0003061 | 0.8947 | +1 | 2 |
| - | - | 3802 | 343.2 | - | - | 0 | - |
| 3 | b | 1361 | 346.1 | 0.0009388 | 2.712 | +1 | 3 |
| - | - | 1565 | 350.1 | - | - | 0 | - |
| - | - | 1709 | 355.1 | - | - | 0 | - |
| - | - | 674.9 | 356.1 | - | - | 0 | - |
| - | - | 1313 | 356.1 | - | - | 0 | - |
| - | - | 4092 | 357.2 | - | - | 0 | - |
| - | - | 881 | 358.2 | - | - | 0 | - |
| - | - | 857.8 | 358.2 | - | - | 0 | - |
| 3 | b | 4417 | 364.2 | 0.0001044 | 0.2866 | +1 | 3 |
| - | - | 1136 | 365.2 | - | - | 0 | - |
| - | - | 620.5 | 366.1 | - | - | 0 | - |
| - | - | 1412 | 368.2 | - | - | 0 | - |
| - | - | 3098 | 378.1 | - | - | 0 | - |
| - | - | 584.6 | 379.1 | - | - | 0 | - |
| - | - | 725.2 | 383.2 | - | - | 0 | - |
| - | - | 868.2 | 392.2 | - | - | 0 | - |
| - | - | 4320 | 394.1 | - | - | 0 | - |
| - | - | 1238 | 395.1 | - | - | 0 | - |
| - | - | 615.9 | 397.2 | - | - | 0 | - |
| - | - | 676.7 | 401.2 | - | - | 0 | - |
| - | - | 659.3 | 413.7 | - | - | 0 | - |
| - | - | 2412 | 425.2 | - | - | 0 | - |
| - | - | 1046 | 425.2 | - | - | 0 | - |
| - | - | 2179 | 427.7 | - | - | 0 | - |
| - | - | 1497 | 428.2 | - | - | 0 | - |
| - | - | 2760 | 436.7 | - | - | 0 | - |
| - | - | 689.7 | 453.3 | - | - | 0 | - |
| 4 | b | 655.1 | 461.2 | 0.0001253 | 0.2717 | +1 | 4 |
| - | - | 2962 | 465.2 | - | - | 0 | - |
| - | - | 751.9 | 466.2 | - | - | 0 | - |
| - | - | 713.6 | 470.3 | - | - | 0 | - |
| 4 | b | 3203 | 479.2 | 0.0006015 | 1.255 | +1 | 4 |
| - | - | 952.7 | 480.2 | - | - | 0 | - |
| - | - | 1191 | 483.2 | - | - | 0 | - |
| - | - | 712 | 484.2 | - | - | 0 | - |
| - | - | 798.1 | 487.3 | - | - | 0 | - |
| - | - | 801.7 | 488.3 | - | - | 0 | - |
| 7 | y | 4448 | 489.2 | 0.005436 | 11.11 | +1 | 3 |
| - | - | 968.7 | 490.2 | - | - | 0 | - |
| - | - | 729.7 | 491.2 | - | - | 0 | - |
| - | - | 2595 | 496.2 | - | - | 0 | - |
| - | - | 676.3 | 497.3 | - | - | 0 | - |
| - | - | 1126 | 510.3 | - | - | 0 | - |
| - | - | 741.4 | 510.8 | - | - | 0 | - |
| - | - | 1587 | 513.4 | - | - | 0 | - |
| - | - | 1309 | 519.2 | - | - | 0 | - |
| - | - | 558 | 519.3 | - | - | 0 | - |
| 2 | y | 3392 | 550.7 | 0.001676 | 3.044 | +2 | 8 |
| - | - | 1809 | 551.2 | - | - | 0 | - |
| 6 | y | 9633 | 560.2 | 0.004851 | 8.66 | +1 | 4 |
| - | - | 691.1 | 561 | - | - | 0 | - |
| - | - | 2485 | 561.2 | - | - | 0 | - |
| - | - | 792.6 | 569.2 | - | - | 0 | - |
| - | - | 7161 | 585.3 | - | - | 0 | - |
| - | - | 2445 | 586.3 | - | - | 0 | - |
| - | - | 689.3 | 587.3 | - | - | 0 | - |
| - | - | 823.7 | 590.3 | - | - | 0 | - |
| 0 | Precursor | 1733 | 592.2 | 0.001369 | 2.311 | +2 | -1 |
| - | - | 1251 | 592.7 | - | - | 0 | - |
| 0 | Precursor | 4378 | 601.2 | 0.001336 | 2.221 | +2 | -1 |
| - | - | 2041 | 601.7 | - | - | 0 | - |
| - | - | 694.6 | 602.3 | - | - | 0 | - |
| - | - | 814.9 | 606.2 | - | - | 0 | - |
| - | - | 721.5 | 612.2 | - | - | 0 | - |
| - | - | 1170 | 614.2 | - | - | 0 | - |
| - | - | 1036 | 628.4 | - | - | 0 | - |
| - | - | 768 | 639.2 | - | - | 0 | - |
| - | - | 2418 | 657.3 | - | - | 0 | - |
| - | - | 3238 | 659.3 | - | - | 0 | - |
| - | - | 1509 | 660.3 | - | - | 0 | - |
| - | - | 731.1 | 665.3 | - | - | 0 | - |
| - | - | 781.2 | 671.3 | - | - | 0 | - |
| - | - | 1050 | 682.3 | - | - | 0 | - |
| - | - | 1666 | 685.3 | - | - | 0 | - |
| - | - | 1004 | 686.3 | - | - | 0 | - |
| - | - | 1.831E+04 | 700.4 | - | - | 0 | - |
| - | - | 6119 | 701.4 | - | - | 0 | - |
| - | - | 2104 | 702.4 | - | - | 0 | - |
| - | - | 1039 | 703.3 | - | - | 0 | - |
| - | - | 1441 | 716.8 | - | - | 0 | - |
| - | - | 1165 | 717.3 | - | - | 0 | - |
| - | - | 1905 | 721.3 | - | - | 0 | - |
| 5 | y | 6525 | 723.3 | 0.004633 | 6.406 | +1 | 5 |
| - | - | 2135 | 724.3 | - | - | 0 | - |
| - | - | 1371 | 725.3 | - | - | 0 | - |
| - | - | 1048 | 725.8 | - | - | 0 | - |
| - | - | 1080 | 744.3 | - | - | 0 | - |
| - | - | 3269 | 749.3 | - | - | 0 | - |
| - | - | 1740 | 750.3 | - | - | 0 | - |
| - | - | 1679 | 757.4 | - | - | 0 | - |
| - | - | 807.9 | 767.3 | - | - | 0 | - |
| - | - | 1376 | 773.4 | - | - | 0 | - |
| - | - | 3217 | 774.3 | - | - | 0 | - |
| - | - | 1335 | 775.3 | - | - | 0 | - |
| - | - | 5165 | 783.4 | - | - | 0 | - |
| - | - | 1815 | 784.4 | - | - | 0 | - |
| - | - | 888.7 | 785.4 | - | - | 0 | - |
| - | - | 980.9 | 794.4 | - | - | 0 | - |
| - | - | 816.1 | 798.8 | - | - | 0 | - |
| - | - | 631.8 | 799.6 | - | - | 0 | - |
| - | - | 3064 | 800.4 | - | - | 0 | - |
| - | - | 9321 | 801.4 | - | - | 0 | - |
| - | - | 3872 | 802.4 | - | - | 0 | - |
| - | - | 1368 | 803.4 | - | - | 0 | - |
| - | - | 3435 | 828.4 | - | - | 0 | - |
| - | - | 1770 | 829.4 | - | - | 0 | - |
| 4 | y | 9893 | 838.3 | 0.004607 | 5.495 | +1 | 6 |
| - | - | 3513 | 839.3 | - | - | 0 | - |
| - | - | 968 | 840.3 | - | - | 0 | - |
| - | - | 1.28E+04 | 844.5 | - | - | 0 | - |
| - | - | 5833 | 845.5 | - | - | 0 | - |
| - | - | 1492 | 846.5 | - | - | 0 | - |
| - | - | 851.7 | 847.4 | - | - | 0 | - |
| - | - | 1721 | 854.4 | - | - | 0 | - |
| - | - | 986.6 | 871.3 | - | - | 0 | - |
| - | - | 5.074E+04 | 872.4 | - | - | 0 | - |
| - | - | 2.289E+04 | 873.4 | - | - | 0 | - |
| - | - | 7186 | 874.5 | - | - | 0 | - |
| - | - | 1537 | 875.5 | - | - | 0 | - |
| - | - | 1.388E+04 | 889.3 | - | - | 0 | - |
| - | - | 6885 | 890.3 | - | - | 0 | - |
| - | - | 883.9 | 891.4 | - | - | 0 | - |
| - | - | 652 | 896.5 | - | - | 0 | - |
| - | - | 955.2 | 907.3 | - | - | 0 | - |
| - | - | 765.8 | 909.4 | - | - | 0 | - |
| 3 | y | 2604 | 935.3 | 0.002633 | 2.815 | +1 | 7 |
| - | - | 1232 | 936.3 | - | - | 0 | - |
| 3 | y | 3.986E+04 | 953.3 | 0.004214 | 4.42 | +1 | 7 |
| - | - | 1.961E+04 | 954.3 | - | - | 0 | - |
| - | - | 7123 | 955.3 | - | - | 0 | - |
| - | - | 1073 | 956.4 | - | - | 0 | - |
| - | - | 7556 | 991.5 | - | - | 0 | - |
| - | - | 4609 | 992.5 | - | - | 0 | - |
| 8 | b | 770.1 | 997.4 | 0.006991 | 7.009 | +1 | 8 |
| - | - | 1.919E+04 | 1020 | - | - | 0 | - |
| - | - | 1.033E+04 | 1021 | - | - | 0 | - |
| - | - | 3536 | 1022 | - | - | 0 | - |
| 2 | y | 2369 | 1100 | 0.003061 | 2.781 | +1 | 8 |
| - | - | 1281 | 1101 | - | - | 0 | - |
| - | - | 683.5 | 1237 | - | - | 0 | - |
| - | - | 848.9 | 1433 | - | - | 0 | - |

m/z Charge Intensity FragmentType MassShift Position
120.0810546875 0 104114.72
121.08436584472656 0 9602.626
123.04429626464844 0 1333.5529
128.6042938232422 0 464.0482
129.06637573242188 0 510.7285
129.10244750976562 0 5478.9043
130.05029296875 0 649.68353
130.06527709960938 0 2923.775
132.04466247558594 0 461.2053
132.08096313476562 0 1650.6034
132.1018829345703 0 1061.5425
133.0858612060547 0 592.66327
136.075927734375 0 32806.004
137.0792999267578 0 3392.7036
138.06622314453125 0 1290.0785
139.92318725585938 0 422.05066
143.9642333984375 0 448.13434
146.06036376953125 0 2131.811
147.04385375976562 0 939.7581
148.07553100585938 0 622.10126
149.04525756835938 0 1989.1748
151.0503692626953 0 455.98053
151.49871826171875 0 401.18
153.10220336914062 0 522.18176
155.08184814453125 0 510.98477
155.11781311035156 0 551.99445
156.07696533203125 0 2853.1716
156.0828857421875 0 403.41824
157.0761260986328 0 567.09485
157.10903930664062 0 668.9983
157.13385009765625 0 1668.9965
158.0599822998047 0 664.9104
158.09664916992188 0 3978.874
159.09190368652344 0 5138.637
160.07554626464844 0 1123.645
160.09521484375 0 651.3601
160.11228942871094 0 1897.9475
165.05482482910156 0 12999.201
166.05894470214844 0 1061.1721
166.0864715576172 0 3230.7153
167.05557250976562 0 2046.271
167.09222412109375 0 641.6091
169.0763397216797 0 467.5583
169.097412109375 0 1035.3615
171.07618713378906 0 759.35565
171.11306762695312 0 567.83417
173.09225463867188 0 639.2519
173.128662109375 0 7253.3496
175.08688354492188 0 2364.1821
176.08187866210938 0 766.89984
176.10723876953125 0 8345.631
177.1023712158203 0 4917.2085
177.11061096191406 0 809.46924
178.1055145263672 0 504.5026
181.09799194335938 0 599.40546
182.0813751220703 0 24936.914
183.08493041992188 0 2932.212
183.14938354492188 0 539.3877
185.12852478027344 0 1948.8322
186.09181213378906 0 1108.1785
188.0707550048828 0 7505.779
189.07418823242188 0 753.71436
191.1180877685547 0 3242.4868
193.1085968017578 0 2108.012
195.08787536621094 0 743.1674
195.1121368408203 0 486.45206
197.1286163330078 0 960.736
199.07191467285156 0 518.6438
199.10813903808594 0 535.357
200.1395721435547 0 917.14514
201.1236114501953 0 6188.9453
203.06639099121094 0 585.72943
203.0821533203125 0 1199.5996
203.11790466308594 0 2169.7 a Water loss 1
204.07681274414062 0 1277.0883
205.0972442626953 0 4309.8154 y 8
207.11273193359375 0 1490.8567
209.09231567382812 0 768.0165
214.11851501464844 0 973.4099
215.1387939453125 0 821.84424
217.08282470703125 0 565.7449
217.0973663330078 0 1667.1122
219.11306762695312 0 1485.3649
219.6790771484375 0 517.6883
221.0924530029297 0 2345.469
221.10354614257812 0 7170.143
221.12867736816406 0 112272.2 a 1
222.1320343017578 0 13878.053
223.06324768066406 0 1130.9833
223.13482666015625 0 940.874
225.1708984375 0 1002.8386
226.11883544921875 0 821.0474
226.1553955078125 0 771.33356
227.10296630859375 0 565.63025
229.11924743652344 0 1092.4957
229.1553192138672 0 736.3564
229.16964721679688 0 533.72186
231.06126403808594 0 1827.6443
231.09835815429688 0 556.62177
231.1131591796875 0 1403.186 b Water loss 1
233.09262084960938 0 915.4711
233.12966918945312 0 938.0166
233.14984130859375 0 627.0654
235.10797119140625 0 3325.4536
239.09466552734375 0 1229.0442
239.11390686035156 0 726.6672
239.15048217773438 0 845.8354
240.0978546142578 0 717.5034
241.0746612548828 0 1485.4344
248.1074676513672 0 647.4402
249.12362670898438 0 17773.9 b 1
250.12713623046875 0 2484.5454
251.10301208496094 0 2040.2366
253.11846923828125 0 1858.8403
253.16622924804688 0 2294.4753
257.106689453125 0 1220.559
257.1644287109375 0 789.1604
263.1029968261719 0 3952.49
263.61456298828125 0 510.28317
274.1302795410156 0 668.6045
279.0976257324219 0 1406.2706
283.0304260253906 0 687.5699
283.1434631347656 0 974.3229
285.00897216796875 0 1456.2021
285.1020812988281 0 2903.5728
292.1404724121094 0 1448.8242
296.15081787109375 0 2364.993
297.0902404785156 0 746.57916
302.1127624511719 0 1330.6241
312.15594482421875 0 1287.5862
314.0980224609375 0 1100.4788
319.1410217285156 0 2032.7936
324.14581298828125 0 4277.036
325.0856628417969 0 1050.9757
325.1478271484375 0 934.9462
329.1497802734375 0 18051.193
330.1534118652344 0 3932.5337
331.1547546386719 0 839.04926
340.1982727050781 0 3365.0635
342.1563720703125 0 22591.855 y 7
343.15972900390625 0 3802.3008
346.14068603515625 0 1360.5894 b Water loss 2
350.13446044921875 0 1565.3601
355.0701599121094 0 1708.5886
356.06903076171875 0 674.9286
356.1379089355469 0 1312.5367
357.22454833984375 0 4091.9202
358.1608581542969 0 880.9584
358.227294921875 0 857.7548
364.15020751953125 0 4417.3774 b 2
365.1531066894531 0 1136.3918
366.1287536621094 0 620.4656
368.19287109375 0 1412.249
378.1296691894531 0 3098.4004
379.1315002441406 0 584.6254
383.1966247558594 0 725.1707
392.2014465332031 0 868.2216
394.1249084472656 0 4319.9463
395.12841796875 0 1238.4144
397.1687316894531 0 615.9254
401.2079772949219 0 676.71716
413.7244873046875 0 659.28125
425.1927490234375 0 2411.7893
425.2409362792969 0 1045.8763
427.72314453125 0 2179.3745
428.22369384765625 0 1497.1334
436.7278747558594 0 2760.1409
453.2826843261719 0 689.7163
461.16656494140625 0 655.14185 b Water loss 3
465.1619873046875 0 2962.1362
466.1633605957031 0 751.9426
470.30694580078125 0 713.617
479.1778564453125 0 3202.7036 b 3
480.18096923828125 0 952.67
483.2207336425781 0 1190.8092
484.22515869140625 0 711.96356
487.3345642089844 0 798.1324
488.3362121582031 0 801.7082
489.1919860839844 0 4448.1006 y 6
490.19390869140625 0 968.70245
491.2014465332031 0 729.6943
496.2311706542969 0 2594.7847
497.2679443359375 0 676.3051
510.2645263671875 0 1126.1779
510.76226806640625 0 741.42505
513.3516845703125 0 1587.1082
519.205078125 0 1308.7316
519.2548828125 0 557.9591
550.7099609375 0 3391.7073 y 1
551.2139892578125 0 1809.1945
560.228515625 0 9632.941 y 5
561.01171875 0 691.09973
561.2311401367188 0 2485.1714
569.235107421875 0 792.59155
585.3353881835938 0 7160.7197
586.3377685546875 0 2445.0142
587.3384399414062 0 689.2904
590.3425903320312 0 823.7181
592.2282104492188 0 1732.9236 Precursor Water loss
592.73193359375 0 1250.5747
601.2334594726562 0 4377.7847 Precursor
601.7389526367188 0 2040.7903
602.3233032226562 0 694.6114
606.2306518554688 0 814.8682
612.1948852539062 0 721.5003
614.24609375 0 1170.2267
628.3785400390625 0 1035.8755
639.2438354492188 0 767.9876
657.2625122070312 0 2418.2876
659.2936401367188 0 3237.743
660.2982788085938 0 1508.8562
665.343017578125 0 731.0884
671.3223876953125 0 781.17786
682.349609375 0 1049.6357
685.2593994140625 0 1666.1006
686.259033203125 0 1004.0304
700.3626708984375 0 18310.375
701.364990234375 0 6118.587
702.3690185546875 0 2104.4248
703.2655029296875 0 1039.0724
716.8154296875 0 1440.5618
717.3123168945312 0 1165.4515
721.2612915039062 0 1905.4148
723.2916259765625 0 6525.2974 y 4
724.294677734375 0 2135.2126
725.29833984375 0 1371.2102
725.8156127929688 0 1048.0778
744.3456420898438 0 1080.4241
749.25537109375 0 3269.318
750.2588500976562 0 1740.2332
757.3818359375 0 1679.0292
767.2642822265625 0 807.91364
773.41015625 0 1375.6421
774.3198852539062 0 3217.3818
775.3218383789062 0 1334.6079
783.3971557617188 0 5165.286
784.4011840820312 0 1814.5382
785.4063110351562 0 888.67883
794.3782958984375 0 980.9057
798.8494873046875 0 816.1034
799.6142578125 0 631.75183
800.427490234375 0 3064.0918
801.4119873046875 0 9320.763
802.41259765625 0 3871.7695
803.4159545898438 0 1367.563
828.4201049804688 0 3435.4766
829.4217529296875 0 1769.932
838.3185424804688 0 9893.263 y 3
839.3207397460938 0 3513.4536
840.3226318359375 0 968.0123
844.4522705078125 0 12801.054
845.45556640625 0 5832.556
846.4613037109375 0 1492.2693
847.4415893554688 0 851.66534
854.4362182617188 0 1721.185
871.3370361328125 0 986.5899
872.447021484375 0 50738.56
873.4494018554688 0 22893.896
874.4523315429688 0 7186.118
875.4573364257812 0 1537.339
889.3463745117188 0 13881.966
890.3487548828125 0 6885.1084
891.3561401367188 0 883.9037
896.5275268554688 0 652.0187
907.3391723632812 0 955.2487
909.4083862304688 0 765.7693
935.3329467773438 0 2603.9807 y Water loss 2
936.3309936523438 0 1232.0898
953.3450927734375 0 39860.695 y 2
954.3478393554688 0 19606.355
955.3488159179688 0 7123.09
956.3530883789062 0 1072.9542
991.52001953125 0 7556.1294
992.5220336914062 0 4608.8193
997.3740844726562 0 770.1421 b 7
1019.5145263671875 0 19189.91
1020.5177612304688 0 10331.13
1021.5189819335938 0 3536.1677
1100.412353515625 0 2368.9531 y 1
1101.421630859375 0 1281.1235
1236.966064453125 0 683.53204
1432.6041259765625 0 848.8859

Spectrum Details

|  |  |
| --- | --- |
| Matched peaks? Matched peaks The total absolute number of peaks matched. Additionally in brackets the total fraction of peaks matched and the total number of peaks is shown. | 21 (7.45% of 282) |
| FDR? FDR The false discovery rate estimated for this peptide. It is calculated by matching all theoretical fragments with a non-integer shift with the raw peaks for this spectrum. This is done with 40 different shifts. The resulting percentage is the average number of annotated peaks over the number of annotated peaks with the correct spectrum. | 1.47% |
| Satellite FDR? Satellite FDR See the FDR for details on its calculation. This satellite ion specific FDR only contains the satellite ions (d/w) for I/L/J positions. | - |
| PSM Score? PSM Score The PSM Score as given by Hecklib to this annotated spectrum. It is shown with three significant figures. | 250 |

#### Spectrum 10004? Spectrum 10004 The raw spectrum of this peptide as annotated by Hecklib. The fragments are coloured according to ion type (see legend). Any peaks with a star '\*' as text can be hovered over to see the full details, first the ion type second the mass shift type. By hovering over the amino acids in the peptide or ions in the legend the corresponding peaks are highlighted. By toggling the 'Unassigned' label you can turn the background (unassigned) peaks on or off in the plot. By updating the slider in the Ion legend you can update the spectrum to only show the top X% of the peaks with labels. The top X% means any peak that is within X% of the highest intensity. By dragging in the spectrum you can zoom in to a specific part of the spectrum and use 'Zoom Out' to get back to the original zoom level. The annotation of the spectrum is based on the given sequence in the peptides file and is done with different software so inconsistencies are likely. The peaks are annotated based on the given sequence, with 20 ppm tolerance.

Copy Data

##### Spectrum 10004 (TSV)

###### Preview

```
Loading example...
```

*Click on the button to copy the data to your clipboard.*

Mz MinMz MaxIntensity Max

WidthHeightPeptide font sizePeptide stroke widthSpectrum font sizeSpectrum stroke widthCompact peptide

Ion legend

wxyz

abcd

OtherUnassignedIon^Charge^_Position_Show for top:%

TFDDYAMHW

02.36e+44.73e+47.09e+49.46e+4

Zoom Out

a^+1^_2_y^+1^_1_a^+1^_2_b^+1^_2_b^+1^_2_a^+1^_3_y^+1^_2_b^+1^_3_b^+1^_3_b^+1^_4_b^+1^_4_y^+1^_3_y^+2^_8_y^+2^_8_y^+1^_4_\*\*y^+1^_5_y^+1^_6_y^+1^_7_y^+1^_7_y^+1^_8_

0778155723353113

Fragment Matches Table

Show background peaks

| Position | Ion type | Intensity | mz Theoretical | mz Error (Th) | mz Error (ppm) | Charge | Series Number |
| --- | --- | --- | --- | --- | --- | --- | --- |
| - | - | 8.584E+04 | 120.1 | - | - | 0 | - |
| - | - | 6813 | 121.1 | - | - | 0 | - |
| - | - | 347 | 121.7 | - | - | 0 | - |
| - | - | 428.8 | 122.1 | - | - | 0 | - |
| - | - | 412 | 122.2 | - | - | 0 | - |
| - | - | 1147 | 123 | - | - | 0 | - |
| - | - | 353.6 | 128.1 | - | - | 0 | - |
| - | - | 552.1 | 128.1 | - | - | 0 | - |
| - | - | 759.8 | 129.1 | - | - | 0 | - |
| - | - | 3785 | 129.1 | - | - | 0 | - |
| - | - | 597.3 | 130.1 | - | - | 0 | - |
| - | - | 1955 | 130.1 | - | - | 0 | - |
| - | - | 475 | 130.1 | - | - | 0 | - |
| - | - | 1536 | 132.1 | - | - | 0 | - |
| - | - | 2311 | 132.1 | - | - | 0 | - |
| - | - | 484.6 | 133.1 | - | - | 0 | - |
| - | - | 563.9 | 133.3 | - | - | 0 | - |
| - | - | 2.592E+04 | 136.1 | - | - | 0 | - |
| - | - | 1662 | 137.1 | - | - | 0 | - |
| - | - | 797.4 | 138.1 | - | - | 0 | - |
| - | - | 1196 | 138.1 | - | - | 0 | - |
| - | - | 476.7 | 141.1 | - | - | 0 | - |
| - | - | 1164 | 146.1 | - | - | 0 | - |
| - | - | 430.7 | 146.4 | - | - | 0 | - |
| - | - | 1556 | 147 | - | - | 0 | - |
| - | - | 1151 | 149 | - | - | 0 | - |
| - | - | 409.4 | 151 | - | - | 0 | - |
| - | - | 434.2 | 155.1 | - | - | 0 | - |
| - | - | 2253 | 156.1 | - | - | 0 | - |
| - | - | 4982 | 157.1 | - | - | 0 | - |
| - | - | 733.5 | 158.1 | - | - | 0 | - |
| - | - | 4060 | 158.1 | - | - | 0 | - |
| - | - | 969.6 | 159.1 | - | - | 0 | - |
| - | - | 529.7 | 159.1 | - | - | 0 | - |
| - | - | 3231 | 159.1 | - | - | 0 | - |
| - | - | 671 | 160.1 | - | - | 0 | - |
| - | - | 1457 | 160.1 | - | - | 0 | - |
| - | - | 435.1 | 162.2 | - | - | 0 | - |
| - | - | 1.159E+04 | 165.1 | - | - | 0 | - |
| - | - | 1046 | 166.1 | - | - | 0 | - |
| - | - | 1689 | 166.1 | - | - | 0 | - |
| - | - | 1933 | 167.1 | - | - | 0 | - |
| - | - | 481.1 | 167.1 | - | - | 0 | - |
| - | - | 713.3 | 169.1 | - | - | 0 | - |
| - | - | 900.5 | 171.1 | - | - | 0 | - |
| - | - | 549.3 | 171.1 | - | - | 0 | - |
| - | - | 6143 | 173.1 | - | - | 0 | - |
| - | - | 3820 | 173.5 | - | - | 0 | - |
| - | - | 2041 | 174.1 | - | - | 0 | - |
| - | - | 1621 | 175.1 | - | - | 0 | - |
| - | - | 751.5 | 176.1 | - | - | 0 | - |
| - | - | 7310 | 176.1 | - | - | 0 | - |
| - | - | 4449 | 177.1 | - | - | 0 | - |
| - | - | 915.9 | 177.1 | - | - | 0 | - |
| - | - | 468.7 | 179.1 | - | - | 0 | - |
| - | - | 487 | 181 | - | - | 0 | - |
| - | - | 2.553E+04 | 182.1 | - | - | 0 | - |
| - | - | 552.4 | 183.1 | - | - | 0 | - |
| - | - | 2252 | 183.1 | - | - | 0 | - |
| - | - | 551 | 183.1 | - | - | 0 | - |
| - | - | 991.9 | 185.1 | - | - | 0 | - |
| - | - | 944.7 | 185.1 | - | - | 0 | - |
| - | - | 6167 | 185.1 | - | - | 0 | - |
| - | - | 457.6 | 185.2 | - | - | 0 | - |
| - | - | 512.3 | 186.1 | - | - | 0 | - |
| - | - | 4001 | 188.1 | - | - | 0 | - |
| - | - | 776.1 | 189.1 | - | - | 0 | - |
| - | - | 1885 | 191.1 | - | - | 0 | - |
| - | - | 1666 | 193.1 | - | - | 0 | - |
| - | - | 475.2 | 197.1 | - | - | 0 | - |
| - | - | 422 | 197.1 | - | - | 0 | - |
| - | - | 593 | 197.1 | - | - | 0 | - |
| - | - | 522.4 | 199.1 | - | - | 0 | - |
| - | - | 618.9 | 200.1 | - | - | 0 | - |
| - | - | 3093 | 201.1 | - | - | 0 | - |
| - | - | 786.3 | 202.1 | - | - | 0 | - |
| - | - | 810.7 | 203.1 | - | - | 0 | - |
| 2 | a | 2150 | 203.1 | 0.0004424 | 2.178 | +1 | 2 |
| - | - | 799.2 | 204.1 | - | - | 0 | - |
| 9 | y | 2645 | 205.1 | 0.0004716 | 2.3 | +1 | 1 |
| - | - | 694.4 | 207.1 | - | - | 0 | - |
| - | - | 1114 | 208.1 | - | - | 0 | - |
| - | - | 673.6 | 212.1 | - | - | 0 | - |
| - | - | 1034 | 213.1 | - | - | 0 | - |
| - | - | 1226 | 214.1 | - | - | 0 | - |
| - | - | 1445 | 215.1 | - | - | 0 | - |
| - | - | 784.6 | 217.1 | - | - | 0 | - |
| - | - | 1147 | 217.1 | - | - | 0 | - |
| - | - | 922.7 | 219.1 | - | - | 0 | - |
| - | - | 6240 | 221.1 | - | - | 0 | - |
| 2 | a | 9.362E+04 | 221.1 | 0.0004978 | 2.251 | +1 | 2 |
| - | - | 1.152E+04 | 222.1 | - | - | 0 | - |
| - | - | 1707 | 223.1 | - | - | 0 | - |
| - | - | 1490 | 225.2 | - | - | 0 | - |
| - | - | 1078 | 229.1 | - | - | 0 | - |
| - | - | 646.5 | 229.2 | - | - | 0 | - |
| - | - | 570.7 | 229.2 | - | - | 0 | - |
| - | - | 955.9 | 230.1 | - | - | 0 | - |
| - | - | 2355 | 231.1 | - | - | 0 | - |
| 2 | b | 902.5 | 231.1 | 0.000355 | 1.536 | +1 | 2 |
| - | - | 3980 | 233.2 | - | - | 0 | - |
| - | - | 2418 | 235.1 | - | - | 0 | - |
| - | - | 1418 | 239.1 | - | - | 0 | - |
| - | - | 683.3 | 239.1 | - | - | 0 | - |
| - | - | 992.6 | 239.2 | - | - | 0 | - |
| - | - | 827.9 | 240.1 | - | - | 0 | - |
| - | - | 1207 | 241.1 | - | - | 0 | - |
| - | - | 658.8 | 243.1 | - | - | 0 | - |
| - | - | 550.5 | 248.1 | - | - | 0 | - |
| 2 | b | 1.648E+04 | 249.1 | 0.0005478 | 2.199 | +1 | 2 |
| - | - | 1597 | 250.1 | - | - | 0 | - |
| - | - | 2044 | 251.1 | - | - | 0 | - |
| - | - | 818.7 | 253.1 | - | - | 0 | - |
| - | - | 877.1 | 253.2 | - | - | 0 | - |
| - | - | 1172 | 257.1 | - | - | 0 | - |
| - | - | 800.5 | 257.1 | - | - | 0 | - |
| - | - | 617.8 | 257.2 | - | - | 0 | - |
| - | - | 586.3 | 260.1 | - | - | 0 | - |
| - | - | 3762 | 263.1 | - | - | 0 | - |
| - | - | 587.9 | 270.1 | - | - | 0 | - |
| - | - | 586.5 | 274.1 | - | - | 0 | - |
| - | - | 1767 | 279.1 | - | - | 0 | - |
| - | - | 859.4 | 283.1 | - | - | 0 | - |
| - | - | 1651 | 284.2 | - | - | 0 | - |
| - | - | 791.1 | 285 | - | - | 0 | - |
| - | - | 2088 | 285.1 | - | - | 0 | - |
| - | - | 910 | 287.1 | - | - | 0 | - |
| - | - | 709 | 290.1 | - | - | 0 | - |
| - | - | 1090 | 292.1 | - | - | 0 | - |
| - | - | 1659 | 296.2 | - | - | 0 | - |
| - | - | 577.2 | 297.1 | - | - | 0 | - |
| - | - | 656.1 | 301 | - | - | 0 | - |
| - | - | 640.3 | 311.1 | - | - | 0 | - |
| 3 | a | 742.9 | 318.1 | 0.001255 | 3.945 | +1 | 3 |
| - | - | 1397 | 319.1 | - | - | 0 | - |
| - | - | 674.2 | 323.1 | - | - | 0 | - |
| - | - | 4445 | 324.1 | - | - | 0 | - |
| - | - | 631.6 | 325.1 | - | - | 0 | - |
| - | - | 1.516E+04 | 329.2 | - | - | 0 | - |
| - | - | 693.1 | 329.2 | - | - | 0 | - |
| - | - | 2920 | 330.2 | - | - | 0 | - |
| - | - | 617.7 | 332.1 | - | - | 0 | - |
| - | - | 1962 | 333.1 | - | - | 0 | - |
| - | - | 981.2 | 334.1 | - | - | 0 | - |
| - | - | 623 | 337.2 | - | - | 0 | - |
| - | - | 2431 | 340.2 | - | - | 0 | - |
| - | - | 639.5 | 341.2 | - | - | 0 | - |
| 8 | y | 1.669E+04 | 342.2 | 0.0006418 | 1.876 | +1 | 2 |
| - | - | 3500 | 343.2 | - | - | 0 | - |
| - | - | 1370 | 343.2 | - | - | 0 | - |
| 3 | b | 846.4 | 346.1 | 0.001702 | 4.916 | +1 | 3 |
| - | - | 854.4 | 348.1 | - | - | 0 | - |
| - | - | 1306 | 350.1 | - | - | 0 | - |
| - | - | 2360 | 355.1 | - | - | 0 | - |
| - | - | 1179 | 355.2 | - | - | 0 | - |
| - | - | 578.6 | 355.2 | - | - | 0 | - |
| - | - | 899.4 | 356.1 | - | - | 0 | - |
| - | - | 697.7 | 356.1 | - | - | 0 | - |
| - | - | 2502 | 357.2 | - | - | 0 | - |
| - | - | 756.2 | 359.2 | - | - | 0 | - |
| 3 | b | 4879 | 364.2 | 0.0001398 | 0.3839 | +1 | 3 |
| - | - | 1144 | 365.2 | - | - | 0 | - |
| - | - | 1777 | 368.2 | - | - | 0 | - |
| - | - | 2340 | 378.1 | - | - | 0 | - |
| - | - | 675.4 | 385.2 | - | - | 0 | - |
| - | - | 858.3 | 392.2 | - | - | 0 | - |
| - | - | 3794 | 394.1 | - | - | 0 | - |
| - | - | 842.1 | 399.2 | - | - | 0 | - |
| - | - | 749.8 | 400.2 | - | - | 0 | - |
| - | - | 771.2 | 412.2 | - | - | 0 | - |
| - | - | 633.7 | 413.3 | - | - | 0 | - |
| - | - | 706 | 417.1 | - | - | 0 | - |
| - | - | 2384 | 425.2 | - | - | 0 | - |
| - | - | 672.8 | 426.2 | - | - | 0 | - |
| - | - | 2261 | 427.7 | - | - | 0 | - |
| - | - | 1503 | 428.2 | - | - | 0 | - |
| - | - | 2261 | 432.2 | - | - | 0 | - |
| - | - | 1128 | 433.2 | - | - | 0 | - |
| - | - | 1288 | 436.7 | - | - | 0 | - |
| - | - | 1092 | 454.2 | - | - | 0 | - |
| 4 | b | 967.7 | 461.2 | 0.001987 | 4.308 | +1 | 4 |
| - | - | 745.3 | 461.2 | - | - | 0 | - |
| - | - | 2906 | 465.2 | - | - | 0 | - |
| - | - | 795.6 | 466.2 | - | - | 0 | - |
| - | - | 897.5 | 471.3 | - | - | 0 | - |
| 4 | b | 3047 | 479.2 | 0.00167 | 3.484 | +1 | 4 |
| - | - | 1020 | 480.2 | - | - | 0 | - |
| - | - | 983.4 | 483.2 | - | - | 0 | - |
| - | - | 707.6 | 487.3 | - | - | 0 | - |
| 7 | y | 5074 | 489.2 | 0.005985 | 12.23 | +1 | 3 |
| - | - | 1205 | 490.2 | - | - | 0 | - |
| - | - | 1994 | 496.2 | - | - | 0 | - |
| - | - | 991.4 | 497.2 | - | - | 0 | - |
| - | - | 832.6 | 500.2 | - | - | 0 | - |
| - | - | 894.6 | 501.3 | - | - | 0 | - |
| - | - | 785.9 | 501.8 | - | - | 0 | - |
| - | - | 1376 | 510.3 | - | - | 0 | - |
| - | - | 891.1 | 513.3 | - | - | 0 | - |
| 2 | y | 791.6 | 541.7 | 0.007142 | 13.18 | +2 | 8 |
| - | - | 907.2 | 548.2 | - | - | 0 | - |
| - | - | 623.7 | 549.2 | - | - | 0 | - |
| 2 | y | 2590 | 550.7 | 0.00308 | 5.593 | +2 | 8 |
| - | - | 1457 | 551.2 | - | - | 0 | - |
| - | - | 1102 | 551.7 | - | - | 0 | - |
| 6 | y | 9276 | 560.2 | 0.005645 | 10.08 | +1 | 4 |
| - | - | 2100 | 561.2 | - | - | 0 | - |
| - | - | 618.6 | 573.8 | - | - | 0 | - |
| - | - | 1059 | 584.8 | - | - | 0 | - |
| - | - | 4553 | 585.3 | - | - | 0 | - |
| - | - | 1437 | 586.3 | - | - | 0 | - |
| 0 | Precursor | 791.4 | 592.2 | 0.002468 | 4.167 | +2 | -1 |
| - | - | 858.4 | 593.2 | - | - | 0 | - |
| - | - | 750.4 | 601.2 | - | - | 0 | - |
| 0 | Precursor | 3211 | 601.2 | 0.002434 | 4.049 | +2 | -1 |
| - | - | 687.7 | 601.3 | - | - | 0 | - |
| - | - | 684.8 | 601.6 | - | - | 0 | - |
| - | - | 1451 | 601.7 | - | - | 0 | - |
| - | - | 1161 | 612.2 | - | - | 0 | - |
| - | - | 670.2 | 614.3 | - | - | 0 | - |
| - | - | 669.6 | 616.6 | - | - | 0 | - |
| - | - | 1529 | 628.4 | - | - | 0 | - |
| - | - | 1305 | 657.3 | - | - | 0 | - |
| - | - | 2794 | 659.3 | - | - | 0 | - |
| - | - | 1368 | 660.3 | - | - | 0 | - |
| - | - | 749.3 | 682.4 | - | - | 0 | - |
| - | - | 2323 | 685.3 | - | - | 0 | - |
| - | - | 1452 | 691.4 | - | - | 0 | - |
| - | - | 1.329E+04 | 700.4 | - | - | 0 | - |
| - | - | 5053 | 701.4 | - | - | 0 | - |
| - | - | 1126 | 702.4 | - | - | 0 | - |
| - | - | 809.1 | 717.3 | - | - | 0 | - |
| - | - | 2089 | 721.3 | - | - | 0 | - |
| - | - | 1086 | 722.3 | - | - | 0 | - |
| 5 | y | 6382 | 723.3 | 0.005366 | 7.418 | +1 | 5 |
| - | - | 1932 | 724.3 | - | - | 0 | - |
| - | - | 716 | 725.3 | - | - | 0 | - |
| - | - | 2668 | 749.3 | - | - | 0 | - |
| - | - | 778.8 | 750.3 | - | - | 0 | - |
| - | - | 1513 | 757.4 | - | - | 0 | - |
| - | - | 889.3 | 758.4 | - | - | 0 | - |
| - | - | 930.6 | 767.3 | - | - | 0 | - |
| - | - | 1097 | 773.4 | - | - | 0 | - |
| - | - | 2879 | 774.3 | - | - | 0 | - |
| - | - | 1758 | 775.3 | - | - | 0 | - |
| - | - | 4207 | 783.4 | - | - | 0 | - |
| - | - | 1694 | 784.4 | - | - | 0 | - |
| - | - | 834.4 | 785.4 | - | - | 0 | - |
| - | - | 2650 | 800.4 | - | - | 0 | - |
| - | - | 5654 | 801.4 | - | - | 0 | - |
| - | - | 2122 | 802.4 | - | - | 0 | - |
| - | - | 2167 | 804.4 | - | - | 0 | - |
| - | - | 749.2 | 826.4 | - | - | 0 | - |
| - | - | 2530 | 828.4 | - | - | 0 | - |
| - | - | 1100 | 829.4 | - | - | 0 | - |
| 4 | y | 6982 | 838.3 | 0.006316 | 7.534 | +1 | 6 |
| - | - | 4139 | 839.3 | - | - | 0 | - |
| - | - | 1171 | 840.3 | - | - | 0 | - |
| - | - | 8386 | 844.5 | - | - | 0 | - |
| - | - | 4941 | 845.5 | - | - | 0 | - |
| - | - | 1852 | 854.4 | - | - | 0 | - |
| - | - | 1532 | 871.3 | - | - | 0 | - |
| - | - | 3.806E+04 | 872.4 | - | - | 0 | - |
| - | - | 1.763E+04 | 873.5 | - | - | 0 | - |
| - | - | 5595 | 874.5 | - | - | 0 | - |
| - | - | 1021 | 875.5 | - | - | 0 | - |
| - | - | 1.049E+04 | 889.3 | - | - | 0 | - |
| - | - | 4777 | 890.3 | - | - | 0 | - |
| - | - | 1110 | 891.3 | - | - | 0 | - |
| - | - | 934.9 | 902.4 | - | - | 0 | - |
| 3 | y | 2023 | 935.3 | 0.004891 | 5.229 | +1 | 7 |
| 3 | y | 3.398E+04 | 953.3 | 0.005496 | 5.765 | +1 | 7 |
| - | - | 1.838E+04 | 954.3 | - | - | 0 | - |
| - | - | 431.4 | 954.4 | - | - | 0 | - |
| - | - | 6591 | 955.3 | - | - | 0 | - |
| - | - | 701 | 971.5 | - | - | 0 | - |
| - | - | 6436 | 991.5 | - | - | 0 | - |
| - | - | 3930 | 992.5 | - | - | 0 | - |
| - | - | 1157 | 993.5 | - | - | 0 | - |
| - | - | 791.6 | 1003 | - | - | 0 | - |
| - | - | 1.657E+04 | 1020 | - | - | 0 | - |
| - | - | 1.039E+04 | 1021 | - | - | 0 | - |
| - | - | 3970 | 1022 | - | - | 0 | - |
| 2 | y | 2301 | 1100 | 0.007577 | 6.886 | +1 | 8 |
| - | - | 1120 | 1101 | - | - | 0 | - |
| - | - | 609.5 | 1834 | - | - | 0 | - |
| - | - | 668.3 | 2612 | - | - | 0 | - |
| - | - | 646.1 | 3082 | - | - | 0 | - |

m/z Charge Intensity FragmentType MassShift Position
120.08118438720703 0 85839.805
121.08451843261719 0 6812.8096
121.74286651611328 0 346.99515
122.07133483886719 0 428.82504
122.24674987792969 0 411.95993
123.04441833496094 0 1147.1357
128.08160400390625 0 353.59607
128.10752868652344 0 552.1188
129.0662078857422 0 759.83875
129.1025848388672 0 3785.0134
130.05030822753906 0 597.3354
130.0655059814453 0 1955.3904
130.0865020751953 0 474.97247
132.08114624023438 0 1536.4974
132.10227966308594 0 2311.1262
133.08639526367188 0 484.5675
133.25631713867188 0 563.9269
136.07608032226562 0 25916.523
137.0794219970703 0 1662.1161
138.055419921875 0 797.4038
138.06655883789062 0 1195.6337
141.10263061523438 0 476.6812
146.0602569580078 0 1163.545
146.42510986328125 0 430.74628
147.0443115234375 0 1555.5366
149.04537963867188 0 1150.924
150.9563446044922 0 409.36896
155.0817108154297 0 434.15076
156.07717895507812 0 2252.7554
157.13392639160156 0 4982.331
158.06039428710938 0 733.51154
158.0968475341797 0 4060.36
159.0767059326172 0 969.5763
159.0855712890625 0 529.72174
159.09210205078125 0 3230.6228
160.07586669921875 0 671.0355
160.1123046875 0 1457.3146
162.181884765625 0 435.11148
165.0549774169922 0 11586.828
166.05812072753906 0 1046.24
166.08660888671875 0 1689.2865
167.0558319091797 0 1932.5485
167.08206176757812 0 481.13663
169.09732055664062 0 713.32764
171.0770721435547 0 900.4572
171.113037109375 0 549.3036
173.1288299560547 0 6142.5513
173.4504852294922 0 3820.0278
174.05540466308594 0 2040.9199
175.08692932128906 0 1621.3452
176.0821075439453 0 751.5149
176.10739135742188 0 7310.207
177.10267639160156 0 4448.87
177.1106414794922 0 915.87115
179.10595703125 0 468.69864
181.02777099609375 0 487.0405
182.0815887451172 0 25525.676
183.0767059326172 0 552.44574
183.08505249023438 0 2251.9521
183.14956665039062 0 550.9743
185.05606079101562 0 991.8833
185.09271240234375 0 944.6916
185.12890625 0 6167.1553
185.16482543945312 0 457.566
186.0918426513672 0 512.3195
188.0709228515625 0 4000.8608
189.087158203125 0 776.09827
191.11827087402344 0 1885.0813
193.10911560058594 0 1666.4451
197.0928955078125 0 475.2347
197.1051788330078 0 422.01736
197.1290283203125 0 593.04816
199.0716552734375 0 522.3861
200.1399383544922 0 618.9031
201.12362670898438 0 3092.9722
202.05087280273438 0 786.2647
203.0663604736328 0 810.70306
203.1183319091797 0 2149.6133 a Water loss 1
204.07708740234375 0 799.15546
205.09762573242188 0 2644.7524 y 8
207.11215209960938 0 694.36505
208.0723114013672 0 1114.2543
212.10357666015625 0 673.58905
213.0880126953125 0 1033.5632
214.11907958984375 0 1225.5486
215.13938903808594 0 1444.8365
217.08230590820312 0 784.6386
217.0974578857422 0 1147.3284
219.11305236816406 0 922.6844
221.10365295410156 0 6239.5522
221.1289520263672 0 93619.79 a 1
222.13229370117188 0 11516.987
223.06410217285156 0 1707.3032
225.17156982421875 0 1489.5896
229.1190948486328 0 1078.4211
229.15516662597656 0 646.46576
229.17152404785156 0 570.73987
230.0771942138672 0 955.8952
231.06126403808594 0 2355.15
231.1131591796875 0 902.51215 b Water loss 1
233.15008544921875 0 3979.8635
235.10816955566406 0 2417.7607
239.0945587158203 0 1418.3085
239.11474609375 0 683.3108
239.1754913330078 0 992.564
240.09735107421875 0 827.8871
241.07510375976562 0 1206.6327
243.13392639160156 0 658.83923
248.1076202392578 0 550.5056
249.12391662597656 0 16481.455 b 1
250.12738037109375 0 1596.53
251.10321044921875 0 2043.843
253.1190185546875 0 818.673
253.16690063476562 0 877.129
257.1068420410156 0 1172.3582
257.1246643066406 0 800.4855
257.1649475097656 0 617.84644
260.1394348144531 0 586.3343
263.10302734375 0 3761.6152
270.1080322265625 0 587.9484
274.13037109375 0 586.5064
279.0980529785156 0 1766.9169
283.14495849609375 0 859.3714
284.1972351074219 0 1650.5845
285.0082702636719 0 791.1004
285.102294921875 0 2088.2341
287.0983581542969 0 910.04205
290.149658203125 0 709.01
292.14093017578125 0 1089.522
296.1513977050781 0 1658.8558
297.0910949707031 0 577.22894
301.0414123535156 0 656.13367
311.12921142578125 0 640.3397
318.1460876464844 0 742.89514 a Water loss 2
319.1416320800781 0 1396.7081
323.0986022949219 0 674.16156
324.14599609375 0 4445.1484
325.1460266113281 0 631.6447
329.1502990722656 0 15156.766
329.22955322265625 0 693.0879
330.1534118652344 0 2920.486
332.1241455078125 0 617.6618
333.1207275390625 0 1961.7118
334.1037292480469 0 981.1511
337.15069580078125 0 623.0071
340.19891357421875 0 2431.277
341.2022399902344 0 639.5304
342.1567077636719 0 16693.031 y 7
343.1601867675781 0 3499.8877
343.1983337402344 0 1370.4675
346.1414489746094 0 846.37384 b Water loss 2
348.119140625 0 854.4266
350.1344299316406 0 1305.9447
355.07037353515625 0 2360.1934
355.1613464355469 0 1178.7428
355.1982727050781 0 578.62665
356.07073974609375 0 899.39154
356.14019775390625 0 697.66907
357.2254943847656 0 2501.7078
359.1943054199219 0 756.17267
364.15045166015625 0 4878.786 b 2
365.1556396484375 0 1144.4711
368.19287109375 0 1776.5784
378.1305847167969 0 2339.961
385.2208251953125 0 675.39954
392.2035827636719 0 858.25134
394.125244140625 0 3793.7515
399.2264404296875 0 842.0595
400.1852722167969 0 749.80096
412.2204284667969 0 771.1614
413.2655944824219 0 633.74445
417.08526611328125 0 705.97723
425.1944885253906 0 2383.731
426.1967468261719 0 672.754
427.7226257324219 0 2261.4573
428.22357177734375 0 1502.7781
432.1889343261719 0 2260.6584
433.1734313964844 0 1128.2085
436.7277526855469 0 1287.9979
454.2301330566406 0 1091.7192
461.1647033691406 0 967.6505 b Water loss 3
461.19696044921875 0 745.3065
465.1627502441406 0 2906.201
466.1669006347656 0 795.6269
471.25726318359375 0 897.4743
479.1789245605469 0 3047.3416 b 3
480.1822204589844 0 1019.75287
483.22198486328125 0 983.3808
487.33734130859375 0 707.57056
489.1925354003906 0 5074.2285 y 6
490.193603515625 0 1204.6403
496.2315368652344 0 1993.6086
497.2328186035156 0 991.3597
500.2470703125 0 832.6429
501.25457763671875 0 894.61096
501.75244140625 0 785.90125
510.2627258300781 0 1376.2928
513.3464965820312 0 891.1221
541.7101440429688 0 791.62585 y Water loss 1
548.2003784179688 0 907.2349
549.1998291015625 0 623.7431
550.7113647460938 0 2590.3992 y 1
551.2127685546875 0 1457.3712
551.7130126953125 0 1101.5997
560.2293090820312 0 9276.174 y 5
561.2310791015625 0 2099.8906
573.7811279296875 0 618.5556
584.7970581054688 0 1059.2611
585.3367919921875 0 4552.581
586.3375854492188 0 1436.7334
592.2293090820312 0 791.4353 Precursor Water loss
593.22998046875 0 858.39166
601.1536865234375 0 750.4379
601.2345581054688 0 3211.4106 Precursor
601.3271484375 0 687.67004
601.6194458007812 0 684.8356
601.73828125 0 1450.9275
612.1967163085938 0 1160.907
614.252197265625 0 670.2404
616.627685546875 0 669.56024
628.3782958984375 0 1529.4943
657.2642822265625 0 1305.3965
659.293701171875 0 2794.4836
660.2972412109375 0 1368.3606
682.3526000976562 0 749.29425
685.2589721679688 0 2322.8152
691.3540649414062 0 1452.0438
700.3628540039062 0 13286.747
701.3663940429688 0 5052.968
702.3703002929688 0 1125.6857
717.3176879882812 0 809.0937
721.2618408203125 0 2088.7856
722.2605590820312 0 1086.251
723.2923583984375 0 6382.404 y 4
724.2952270507812 0 1931.5231
725.297607421875 0 716.02997
749.255126953125 0 2668.2
750.2628173828125 0 778.8193
757.3876342773438 0 1513.0511
758.386474609375 0 889.3033
767.2659301757812 0 930.5928
773.4063110351562 0 1096.7649
774.3208618164062 0 2879.49
775.3252563476562 0 1758.1553
783.3987426757812 0 4206.655
784.3997802734375 0 1693.7538
785.4099731445312 0 834.4093
800.4262084960938 0 2650.0623
801.4127197265625 0 5654.45
802.414794921875 0 2121.9497
804.4415893554688 0 2167.1719
826.4202270507812 0 749.2192
828.4229736328125 0 2530.1094
829.421875 0 1099.795
838.3202514648438 0 6981.929 y 3
839.3236083984375 0 4139.365
840.3235473632812 0 1171.4037
844.4526977539062 0 8385.553
845.4556274414062 0 4940.997
854.436279296875 0 1851.7703
871.34033203125 0 1532.4243
872.44775390625 0 38060.133
873.4503784179688 0 17630.604
874.4523315429688 0 5595.307
875.454833984375 0 1020.63684
889.3474731445312 0 10486.484
890.3499145507812 0 4776.525
891.3478393554688 0 1110.4387
902.43896484375 0 934.8608
935.335205078125 0 2023.4967 y Water loss 2
953.3463745117188 0 33975.035 y 2
954.3490600585938 0 18378.441
954.4338989257812 0 431.37915
955.3493041992188 0 6590.56
971.4940185546875 0 700.957
991.5216674804688 0 6435.8203
992.5239868164062 0 3929.7793
993.5242309570312 0 1156.6764
1003.3740844726562 0 791.57544
1019.5159912109375 0 16574.805
1020.5178833007812 0 10392.166
1021.5194702148438 0 3970.2637
1100.4168701171875 0 2300.6638 y 1
1101.4119873046875 0 1119.861
1834.2681884765625 0 609.5157
2611.850341796875 0 668.3366
3082.417724609375 0 646.10895

Spectrum Details

|  |  |
| --- | --- |
| Matched peaks? Matched peaks The total absolute number of peaks matched. Additionally in brackets the total fraction of peaks matched and the total number of peaks is shown. | 22 (7.67% of 287) |
| FDR? FDR The false discovery rate estimated for this peptide. It is calculated by matching all theoretical fragments with a non-integer shift with the raw peaks for this spectrum. This is done with 40 different shifts. The resulting percentage is the average number of annotated peaks over the number of annotated peaks with the correct spectrum. | 1.52% |
| Satellite FDR? Satellite FDR See the FDR for details on its calculation. This satellite ion specific FDR only contains the satellite ions (d/w) for I/L/J positions. | - |
| PSM Score? PSM Score The PSM Score as given by Hecklib to this annotated spectrum. It is shown with three significant figures. | 250 |

#### Spectrum 9585? Spectrum 9585 The raw spectrum of this peptide as annotated by Hecklib. The fragments are coloured according to ion type (see legend). Any peaks with a star '\*' as text can be hovered over to see the full details, first the ion type second the mass shift type. By hovering over the amino acids in the peptide or ions in the legend the corresponding peaks are highlighted. By toggling the 'Unassigned' label you can turn the background (unassigned) peaks on or off in the plot. By updating the slider in the Ion legend you can update the spectrum to only show the top X% of the peaks with labels. The top X% means any peak that is within X% of the highest intensity. By dragging in the spectrum you can zoom in to a specific part of the spectrum and use 'Zoom Out' to get back to the original zoom level. The annotation of the spectrum is based on the given sequence in the peptides file and is done with different software so inconsistencies are likely. The peaks are annotated based on the given sequence, with 20 ppm tolerance.

Copy Data

##### Spectrum 9585 (TSV)

###### Preview

```
Loading example...
```

*Click on the button to copy the data to your clipboard.*

Mz MinMz MaxIntensity Max

WidthHeightPeptide font sizePeptide stroke widthSpectrum font sizeSpectrum stroke widthCompact peptide

Ion legend

wxyz

abcd

OtherUnassignedIon^Charge^_Position_Show for top:%

TFDDYAMHW

03.48e+46.95e+41.04e+51.39e+5

Zoom Out

a^+1^_2_y^+1^_1_a^+1^_2_b^+1^_2_b^+1^_2_a^+1^_3_y^+1^_2_b^+1^_3_b^+1^_3_b^+1^_4_b^+1^_4_y^+1^_3_y^+2^_8_y^+1^_4_\*\*b^+1^_5_b^+1^_5_y^+1^_5_y^+1^_6_y^+1^_6_y^+1^_7_y^+1^_7_y^+1^_8_

0758151622743032

Fragment Matches Table

Show background peaks

| Position | Ion type | Intensity | mz Theoretical | mz Error (Th) | mz Error (ppm) | Charge | Series Number |
| --- | --- | --- | --- | --- | --- | --- | --- |
| - | - | 661.6 | 120 | - | - | 0 | - |
| - | - | 1.376E+05 | 120.1 | - | - | 0 | - |
| - | - | 569.6 | 121.1 | - | - | 0 | - |
| - | - | 1.079E+04 | 121.1 | - | - | 0 | - |
| - | - | 2141 | 123 | - | - | 0 | - |
| - | - | 339.9 | 129.1 | - | - | 0 | - |
| - | - | 8128 | 129.1 | - | - | 0 | - |
| - | - | 4271 | 130.1 | - | - | 0 | - |
| - | - | 413 | 130.1 | - | - | 0 | - |
| - | - | 601.3 | 131.1 | - | - | 0 | - |
| - | - | 2199 | 132.1 | - | - | 0 | - |
| - | - | 1111 | 132.1 | - | - | 0 | - |
| - | - | 531 | 133.1 | - | - | 0 | - |
| - | - | 463.1 | 135.5 | - | - | 0 | - |
| - | - | 5.007E+04 | 136.1 | - | - | 0 | - |
| - | - | 3957 | 137.1 | - | - | 0 | - |
| - | - | 1100 | 138.1 | - | - | 0 | - |
| - | - | 1211 | 138.1 | - | - | 0 | - |
| - | - | 425.4 | 140.6 | - | - | 0 | - |
| - | - | 574.7 | 141.1 | - | - | 0 | - |
| - | - | 410 | 144.1 | - | - | 0 | - |
| - | - | 611.1 | 145.1 | - | - | 0 | - |
| - | - | 5065 | 146.1 | - | - | 0 | - |
| - | - | 1743 | 147 | - | - | 0 | - |
| - | - | 806.2 | 149 | - | - | 0 | - |
| - | - | 533.6 | 149 | - | - | 0 | - |
| - | - | 452.4 | 150.1 | - | - | 0 | - |
| - | - | 856.9 | 155.1 | - | - | 0 | - |
| - | - | 472.1 | 155.1 | - | - | 0 | - |
| - | - | 5050 | 156.1 | - | - | 0 | - |
| - | - | 573.8 | 157.1 | - | - | 0 | - |
| - | - | 800.7 | 157.1 | - | - | 0 | - |
| - | - | 1248 | 157.1 | - | - | 0 | - |
| - | - | 802.4 | 158.1 | - | - | 0 | - |
| - | - | 4239 | 158.1 | - | - | 0 | - |
| - | - | 6864 | 159.1 | - | - | 0 | - |
| - | - | 885.5 | 160.1 | - | - | 0 | - |
| - | - | 590.9 | 160.1 | - | - | 0 | - |
| - | - | 2341 | 160.1 | - | - | 0 | - |
| - | - | 803.8 | 164.1 | - | - | 0 | - |
| - | - | 1.824E+04 | 165.1 | - | - | 0 | - |
| - | - | 690.8 | 165.1 | - | - | 0 | - |
| - | - | 972.1 | 166.1 | - | - | 0 | - |
| - | - | 3218 | 166.1 | - | - | 0 | - |
| - | - | 1350 | 167.1 | - | - | 0 | - |
| - | - | 526.1 | 167.1 | - | - | 0 | - |
| - | - | 607.5 | 169.1 | - | - | 0 | - |
| - | - | 690.4 | 169.1 | - | - | 0 | - |
| - | - | 710.8 | 169.1 | - | - | 0 | - |
| - | - | 684.6 | 171.1 | - | - | 0 | - |
| - | - | 1037 | 173.1 | - | - | 0 | - |
| - | - | 1.057E+04 | 173.1 | - | - | 0 | - |
| - | - | 1946 | 174.1 | - | - | 0 | - |
| - | - | 548.6 | 174.1 | - | - | 0 | - |
| - | - | 4363 | 175.1 | - | - | 0 | - |
| - | - | 697.5 | 176.1 | - | - | 0 | - |
| - | - | 9327 | 176.1 | - | - | 0 | - |
| - | - | 4971 | 177.1 | - | - | 0 | - |
| - | - | 1269 | 177.1 | - | - | 0 | - |
| - | - | 878.8 | 178.1 | - | - | 0 | - |
| - | - | 490.5 | 180.1 | - | - | 0 | - |
| - | - | 709.2 | 181.1 | - | - | 0 | - |
| - | - | 3.163E+04 | 182.1 | - | - | 0 | - |
| - | - | 2851 | 183.1 | - | - | 0 | - |
| - | - | 850.4 | 183.1 | - | - | 0 | - |
| - | - | 513.3 | 184.6 | - | - | 0 | - |
| - | - | 1027 | 185.1 | - | - | 0 | - |
| - | - | 2081 | 185.1 | - | - | 0 | - |
| - | - | 544.6 | 186.1 | - | - | 0 | - |
| - | - | 6627 | 188.1 | - | - | 0 | - |
| - | - | 728.1 | 189.1 | - | - | 0 | - |
| - | - | 1121 | 189.1 | - | - | 0 | - |
| - | - | 669 | 191.1 | - | - | 0 | - |
| - | - | 2922 | 191.1 | - | - | 0 | - |
| - | - | 4096 | 192.1 | - | - | 0 | - |
| - | - | 800.4 | 192.1 | - | - | 0 | - |
| - | - | 508.2 | 193.1 | - | - | 0 | - |
| - | - | 2342 | 193.1 | - | - | 0 | - |
| - | - | 937.9 | 195.1 | - | - | 0 | - |
| - | - | 485.4 | 196.2 | - | - | 0 | - |
| - | - | 524.2 | 197.1 | - | - | 0 | - |
| - | - | 764.8 | 199.1 | - | - | 0 | - |
| - | - | 911.9 | 199.1 | - | - | 0 | - |
| - | - | 714.4 | 199.1 | - | - | 0 | - |
| - | - | 1218 | 200.1 | - | - | 0 | - |
| - | - | 6744 | 201.1 | - | - | 0 | - |
| - | - | 535.8 | 202.1 | - | - | 0 | - |
| - | - | 1184 | 203.1 | - | - | 0 | - |
| - | - | 1604 | 203.1 | - | - | 0 | - |
| 2 | a | 3008 | 203.1 | 0.0002593 | 1.276 | +1 | 2 |
| - | - | 1517 | 204.1 | - | - | 0 | - |
| 9 | y | 5028 | 205.1 | 0.0002733 | 1.332 | +1 | 1 |
| - | - | 1080 | 206.1 | - | - | 0 | - |
| - | - | 1489 | 207.1 | - | - | 0 | - |
| - | - | 527.7 | 208.1 | - | - | 0 | - |
| - | - | 2994 | 209.1 | - | - | 0 | - |
| - | - | 532.2 | 210.9 | - | - | 0 | - |
| - | - | 581.4 | 212.1 | - | - | 0 | - |
| - | - | 644.8 | 214.1 | - | - | 0 | - |
| - | - | 1015 | 215.1 | - | - | 0 | - |
| - | - | 654.2 | 217.1 | - | - | 0 | - |
| - | - | 2481 | 217.1 | - | - | 0 | - |
| - | - | 533.5 | 218.6 | - | - | 0 | - |
| - | - | 1544 | 219.1 | - | - | 0 | - |
| - | - | 2791 | 221.1 | - | - | 0 | - |
| - | - | 6374 | 221.1 | - | - | 0 | - |
| 2 | a | 1.196E+05 | 221.1 | 0.000391 | 1.768 | +1 | 2 |
| - | - | 478.8 | 221.5 | - | - | 0 | - |
| - | - | 1.495E+04 | 222.1 | - | - | 0 | - |
| - | - | 953.4 | 223.1 | - | - | 0 | - |
| - | - | 673.3 | 223.1 | - | - | 0 | - |
| - | - | 1053 | 225.2 | - | - | 0 | - |
| - | - | 562.8 | 226.1 | - | - | 0 | - |
| - | - | 1459 | 226.2 | - | - | 0 | - |
| - | - | 987.2 | 227.1 | - | - | 0 | - |
| - | - | 757.6 | 229.1 | - | - | 0 | - |
| - | - | 715.6 | 229.2 | - | - | 0 | - |
| - | - | 2893 | 231.1 | - | - | 0 | - |
| - | - | 622.4 | 231.1 | - | - | 0 | - |
| 2 | b | 1929 | 231.1 | 0.0005381 | 2.328 | +1 | 2 |
| - | - | 1214 | 233.1 | - | - | 0 | - |
| - | - | 1808 | 233.1 | - | - | 0 | - |
| - | - | 3498 | 235.1 | - | - | 0 | - |
| - | - | 533.5 | 236.8 | - | - | 0 | - |
| - | - | 1087 | 239.1 | - | - | 0 | - |
| - | - | 1240 | 239.1 | - | - | 0 | - |
| - | - | 1364 | 239.2 | - | - | 0 | - |
| - | - | 695.9 | 247.1 | - | - | 0 | - |
| 2 | b | 2.09E+04 | 249.1 | 0.0003647 | 1.464 | +1 | 2 |
| - | - | 2550 | 250.1 | - | - | 0 | - |
| - | - | 3111 | 251.1 | - | - | 0 | - |
| - | - | 1482 | 253.1 | - | - | 0 | - |
| - | - | 2471 | 253.2 | - | - | 0 | - |
| - | - | 790.6 | 256.2 | - | - | 0 | - |
| - | - | 1365 | 257.1 | - | - | 0 | - |
| - | - | 587.7 | 258.1 | - | - | 0 | - |
| - | - | 883.4 | 261.1 | - | - | 0 | - |
| - | - | 4781 | 263.1 | - | - | 0 | - |
| - | - | 670.8 | 264.1 | - | - | 0 | - |
| - | - | 639.5 | 270.1 | - | - | 0 | - |
| - | - | 1257 | 273.1 | - | - | 0 | - |
| - | - | 828.4 | 274.1 | - | - | 0 | - |
| - | - | 3245 | 279.1 | - | - | 0 | - |
| - | - | 1322 | 283.1 | - | - | 0 | - |
| - | - | 643.7 | 284.1 | - | - | 0 | - |
| - | - | 631 | 284.2 | - | - | 0 | - |
| - | - | 795.5 | 285.1 | - | - | 0 | - |
| - | - | 2625 | 285.1 | - | - | 0 | - |
| - | - | 511.7 | 286.1 | - | - | 0 | - |
| - | - | 1558 | 292.1 | - | - | 0 | - |
| - | - | 2753 | 296.2 | - | - | 0 | - |
| - | - | 1316 | 297.1 | - | - | 0 | - |
| - | - | 1053 | 302.1 | - | - | 0 | - |
| - | - | 589.8 | 312.2 | - | - | 0 | - |
| - | - | 886.1 | 314.1 | - | - | 0 | - |
| - | - | 984.4 | 316.1 | - | - | 0 | - |
| 3 | a | 1445 | 318.1 | 0.001133 | 3.561 | +1 | 3 |
| - | - | 2173 | 319.1 | - | - | 0 | - |
| - | - | 3652 | 324.1 | - | - | 0 | - |
| - | - | 1370 | 325.1 | - | - | 0 | - |
| - | - | 624.6 | 327.2 | - | - | 0 | - |
| - | - | 2.488E+04 | 329.2 | - | - | 0 | - |
| - | - | 4877 | 330.2 | - | - | 0 | - |
| - | - | 4291 | 340.2 | - | - | 0 | - |
| 8 | y | 2.278E+04 | 342.2 | 0.0006113 | 1.787 | +1 | 2 |
| - | - | 3769 | 343.2 | - | - | 0 | - |
| 3 | b | 1396 | 346.1 | 0.001061 | 3.065 | +1 | 3 |
| - | - | 660.9 | 348.1 | - | - | 0 | - |
| - | - | 576.1 | 350.1 | - | - | 0 | - |
| - | - | 1317 | 355.1 | - | - | 0 | - |
| - | - | 806.7 | 355.2 | - | - | 0 | - |
| - | - | 752.3 | 356.1 | - | - | 0 | - |
| - | - | 1780 | 356.1 | - | - | 0 | - |
| - | - | 5220 | 357.2 | - | - | 0 | - |
| - | - | 561.9 | 361.8 | - | - | 0 | - |
| 3 | b | 6336 | 364.2 | 0.0001398 | 0.3839 | +1 | 3 |
| - | - | 1361 | 365.2 | - | - | 0 | - |
| - | - | 1140 | 366.1 | - | - | 0 | - |
| - | - | 641 | 366.2 | - | - | 0 | - |
| - | - | 1458 | 368.2 | - | - | 0 | - |
| - | - | 536.6 | 371.1 | - | - | 0 | - |
| - | - | 3087 | 378.1 | - | - | 0 | - |
| - | - | 886.3 | 382.1 | - | - | 0 | - |
| - | - | 846.5 | 384.2 | - | - | 0 | - |
| - | - | 808.2 | 385.2 | - | - | 0 | - |
| - | - | 1602 | 392.2 | - | - | 0 | - |
| - | - | 5161 | 394.1 | - | - | 0 | - |
| - | - | 1567 | 395.1 | - | - | 0 | - |
| - | - | 973.7 | 398.2 | - | - | 0 | - |
| - | - | 1239 | 401.2 | - | - | 0 | - |
| - | - | 562.9 | 401.7 | - | - | 0 | - |
| - | - | 622.1 | 413.7 | - | - | 0 | - |
| - | - | 613.2 | 414.2 | - | - | 0 | - |
| - | - | 943.3 | 415.3 | - | - | 0 | - |
| - | - | 822.2 | 418.2 | - | - | 0 | - |
| - | - | 1132 | 422.7 | - | - | 0 | - |
| - | - | 790.8 | 423.2 | - | - | 0 | - |
| - | - | 3023 | 425.2 | - | - | 0 | - |
| - | - | 678.7 | 426.2 | - | - | 0 | - |
| - | - | 996.4 | 427.2 | - | - | 0 | - |
| - | - | 2720 | 427.7 | - | - | 0 | - |
| - | - | 983 | 428.2 | - | - | 0 | - |
| - | - | 2576 | 436.7 | - | - | 0 | - |
| - | - | 575 | 437.2 | - | - | 0 | - |
| - | - | 1746 | 437.2 | - | - | 0 | - |
| - | - | 681.9 | 437.7 | - | - | 0 | - |
| - | - | 632.7 | 448.3 | - | - | 0 | - |
| - | - | 793.8 | 448.8 | - | - | 0 | - |
| - | - | 1023 | 453.3 | - | - | 0 | - |
| 4 | b | 715.2 | 461.2 | 0.00137 | 2.971 | +1 | 4 |
| - | - | 3184 | 465.2 | - | - | 0 | - |
| - | - | 628.4 | 466.2 | - | - | 0 | - |
| - | - | 704.9 | 470.3 | - | - | 0 | - |
| 4 | b | 3250 | 479.2 | 0.0006321 | 1.319 | +1 | 4 |
| - | - | 2307 | 483.2 | - | - | 0 | - |
| - | - | 1094 | 486.2 | - | - | 0 | - |
| - | - | 949.2 | 487.3 | - | - | 0 | - |
| - | - | 666.5 | 487.8 | - | - | 0 | - |
| 7 | y | 5923 | 489.2 | 0.006076 | 12.42 | +1 | 3 |
| - | - | 1423 | 490.2 | - | - | 0 | - |
| - | - | 3315 | 496.2 | - | - | 0 | - |
| - | - | 1129 | 496.8 | - | - | 0 | - |
| - | - | 716.3 | 497.2 | - | - | 0 | - |
| - | - | 986.6 | 497.2 | - | - | 0 | - |
| - | - | 657.6 | 499.1 | - | - | 0 | - |
| - | - | 930.6 | 500.2 | - | - | 0 | - |
| - | - | 960 | 501.8 | - | - | 0 | - |
| - | - | 633.4 | 506.7 | - | - | 0 | - |
| - | - | 958.7 | 510.8 | - | - | 0 | - |
| - | - | 1868 | 513.4 | - | - | 0 | - |
| - | - | 993.4 | 514.2 | - | - | 0 | - |
| - | - | 761.1 | 519.2 | - | - | 0 | - |
| - | - | 734.9 | 525.3 | - | - | 0 | - |
| - | - | 661.3 | 532.3 | - | - | 0 | - |
| - | - | 651.9 | 536.2 | - | - | 0 | - |
| - | - | 795.8 | 545.2 | - | - | 0 | - |
| - | - | 1044 | 548.2 | - | - | 0 | - |
| 2 | y | 2401 | 550.7 | 0.002775 | 5.039 | +2 | 8 |
| - | - | 2024 | 551.2 | - | - | 0 | - |
| - | - | 1484 | 559.3 | - | - | 0 | - |
| 6 | y | 8662 | 560.2 | 0.005279 | 9.422 | +1 | 4 |
| - | - | 2436 | 561.2 | - | - | 0 | - |
| - | - | 1093 | 562.2 | - | - | 0 | - |
| - | - | 899.6 | 569.7 | - | - | 0 | - |
| - | - | 788 | 570.2 | - | - | 0 | - |
| - | - | 896.2 | 583.3 | - | - | 0 | - |
| - | - | 8821 | 585.3 | - | - | 0 | - |
| - | - | 2821 | 586.3 | - | - | 0 | - |
| - | - | 644.4 | 587.3 | - | - | 0 | - |
| - | - | 838.2 | 589.8 | - | - | 0 | - |
| 0 | Precursor | 2009 | 592.2 | 0.002406 | 4.063 | +2 | -1 |
| - | - | 1214 | 592.7 | - | - | 0 | - |
| 0 | Precursor | 4670 | 601.2 | 0.00396 | 6.587 | +2 | -1 |
| - | - | 784.7 | 601.3 | - | - | 0 | - |
| - | - | 1008 | 601.3 | - | - | 0 | - |
| - | - | 3267 | 601.7 | - | - | 0 | - |
| - | - | 1039 | 602.3 | - | - | 0 | - |
| - | - | 853.4 | 606.2 | - | - | 0 | - |
| - | - | 692.1 | 612.2 | - | - | 0 | - |
| - | - | 973.8 | 614.2 | - | - | 0 | - |
| 5 | b | 1103 | 624.2 | 0.001671 | 2.676 | +1 | 5 |
| - | - | 2730 | 628.4 | - | - | 0 | - |
| - | - | 1050 | 629.4 | - | - | 0 | - |
| 5 | b | 619.3 | 642.2 | 0.0006275 | 0.977 | +1 | 5 |
| - | - | 1233 | 655.3 | - | - | 0 | - |
| - | - | 1906 | 657.3 | - | - | 0 | - |
| - | - | 621.7 | 657.8 | - | - | 0 | - |
| - | - | 695.9 | 658.3 | - | - | 0 | - |
| - | - | 3459 | 659.3 | - | - | 0 | - |
| - | - | 779.5 | 660.3 | - | - | 0 | - |
| - | - | 632.6 | 664.8 | - | - | 0 | - |
| - | - | 604.9 | 665.3 | - | - | 0 | - |
| - | - | 6539 | 670.8 | - | - | 0 | - |
| - | - | 4861 | 671.3 | - | - | 0 | - |
| - | - | 1779 | 671.8 | - | - | 0 | - |
| - | - | 756.7 | 682.3 | - | - | 0 | - |
| - | - | 735.5 | 683.3 | - | - | 0 | - |
| - | - | 2755 | 685.3 | - | - | 0 | - |
| - | - | 714.6 | 686.3 | - | - | 0 | - |
| - | - | 922.7 | 693.8 | - | - | 0 | - |
| - | - | 843.2 | 694.3 | - | - | 0 | - |
| - | - | 691.1 | 696.4 | - | - | 0 | - |
| - | - | 2.187E+04 | 700.4 | - | - | 0 | - |
| - | - | 7400 | 701.4 | - | - | 0 | - |
| - | - | 2110 | 702.4 | - | - | 0 | - |
| - | - | 2394 | 716.8 | - | - | 0 | - |
| - | - | 1218 | 717.3 | - | - | 0 | - |
| - | - | 2232 | 721.3 | - | - | 0 | - |
| - | - | 1302 | 722.3 | - | - | 0 | - |
| 5 | y | 6979 | 723.3 | 0.005121 | 7.081 | +1 | 5 |
| - | - | 3386 | 724.3 | - | - | 0 | - |
| - | - | 1460 | 725.3 | - | - | 0 | - |
| - | - | 943.5 | 730.3 | - | - | 0 | - |
| - | - | 721.5 | 730.9 | - | - | 0 | - |
| - | - | 1447 | 744.4 | - | - | 0 | - |
| - | - | 1679 | 744.9 | - | - | 0 | - |
| - | - | 711.1 | 745.4 | - | - | 0 | - |
| - | - | 4487 | 749.3 | - | - | 0 | - |
| - | - | 1143 | 750.3 | - | - | 0 | - |
| - | - | 2295 | 752.9 | - | - | 0 | - |
| - | - | 725.4 | 754.4 | - | - | 0 | - |
| - | - | 2469 | 757.4 | - | - | 0 | - |
| - | - | 869.6 | 767.3 | - | - | 0 | - |
| - | - | 813.9 | 767.4 | - | - | 0 | - |
| - | - | 808.5 | 773.4 | - | - | 0 | - |
| - | - | 2522 | 774.3 | - | - | 0 | - |
| - | - | 621.1 | 774.4 | - | - | 0 | - |
| - | - | 1734 | 775.3 | - | - | 0 | - |
| - | - | 922.3 | 775.4 | - | - | 0 | - |
| - | - | 6176 | 783.4 | - | - | 0 | - |
| - | - | 2704 | 784.4 | - | - | 0 | - |
| - | - | 678.5 | 785.4 | - | - | 0 | - |
| - | - | 734.4 | 793.4 | - | - | 0 | - |
| - | - | 2855 | 794.4 | - | - | 0 | - |
| - | - | 1071 | 795.4 | - | - | 0 | - |
| - | - | 3788 | 800.4 | - | - | 0 | - |
| - | - | 1.129E+04 | 801.4 | - | - | 0 | - |
| - | - | 689.2 | 801.9 | - | - | 0 | - |
| - | - | 4965 | 802.4 | - | - | 0 | - |
| - | - | 1393 | 803.4 | - | - | 0 | - |
| 4 | y | 660.7 | 820.3 | 0.000828 | 1.009 | +1 | 6 |
| - | - | 4561 | 828.4 | - | - | 0 | - |
| - | - | 1663 | 829.4 | - | - | 0 | - |
| 4 | y | 1.007E+04 | 838.3 | 0.005278 | 6.296 | +1 | 6 |
| - | - | 4657 | 839.3 | - | - | 0 | - |
| - | - | 1376 | 840.3 | - | - | 0 | - |
| - | - | 1.694E+04 | 844.5 | - | - | 0 | - |
| - | - | 7310 | 845.5 | - | - | 0 | - |
| - | - | 2109 | 846.5 | - | - | 0 | - |
| - | - | 3447 | 854.4 | - | - | 0 | - |
| - | - | 1961 | 855.4 | - | - | 0 | - |
| - | - | 1151 | 871.3 | - | - | 0 | - |
| - | - | 6.67E+04 | 872.4 | - | - | 0 | - |
| - | - | 2.913E+04 | 873.4 | - | - | 0 | - |
| - | - | 8550 | 874.5 | - | - | 0 | - |
| - | - | 1772 | 875.5 | - | - | 0 | - |
| - | - | 1.239E+04 | 889.3 | - | - | 0 | - |
| - | - | 7051 | 890.4 | - | - | 0 | - |
| - | - | 1364 | 891.4 | - | - | 0 | - |
| - | - | 1512 | 895.5 | - | - | 0 | - |
| - | - | 1547 | 896.5 | - | - | 0 | - |
| - | - | 1012 | 897.5 | - | - | 0 | - |
| - | - | 809.7 | 899.4 | - | - | 0 | - |
| - | - | 2050 | 909.4 | - | - | 0 | - |
| - | - | 611.9 | 910.4 | - | - | 0 | - |
| - | - | 595 | 914.7 | - | - | 0 | - |
| 3 | y | 2263 | 935.3 | 0.003975 | 4.25 | +1 | 7 |
| - | - | 863.8 | 936.3 | - | - | 0 | - |
| - | - | 907.4 | 937.3 | - | - | 0 | - |
| 3 | y | 4.823E+04 | 953.3 | 0.004946 | 5.188 | +1 | 7 |
| - | - | 2.385E+04 | 954.3 | - | - | 0 | - |
| - | - | 9281 | 955.3 | - | - | 0 | - |
| - | - | 1222 | 956.4 | - | - | 0 | - |
| - | - | 1.14E+04 | 991.5 | - | - | 0 | - |
| - | - | 5854 | 992.5 | - | - | 0 | - |
| - | - | 2500 | 993.5 | - | - | 0 | - |
| - | - | 841.9 | 994.5 | - | - | 0 | - |
| - | - | 2.55E+04 | 1020 | - | - | 0 | - |
| - | - | 1.565E+04 | 1021 | - | - | 0 | - |
| - | - | 4964 | 1022 | - | - | 0 | - |
| - | - | 821.8 | 1023 | - | - | 0 | - |
| - | - | 1072 | 1036 | - | - | 0 | - |
| - | - | 776.2 | 1038 | - | - | 0 | - |
| - | - | 649.6 | 1039 | - | - | 0 | - |
| 2 | y | 2895 | 1100 | 0.002939 | 2.67 | +1 | 8 |
| - | - | 2063 | 1101 | - | - | 0 | - |
| - | - | 730.8 | 2430 | - | - | 0 | - |
| - | - | 674.8 | 2501 | - | - | 0 | - |
| - | - | 821.1 | 3002 | - | - | 0 | - |

m/z Charge Intensity FragmentType MassShift Position
120.04457092285156 0 661.6229
120.08114624023438 0 137649.89
121.07923126220703 0 569.6064
121.08443450927734 0 10785.149
123.0443344116211 0 2141.1104
129.06622314453125 0 339.93027
129.10255432128906 0 8127.604
130.0654296875 0 4271.161
130.10609436035156 0 412.98795
131.06842041015625 0 601.25214
132.08099365234375 0 2198.746
132.1022491455078 0 1111.037
133.0858612060547 0 531.02045
135.4722442626953 0 463.0609
136.07603454589844 0 50071.312
137.07936096191406 0 3957.0908
138.05531311035156 0 1100.4327
138.06610107421875 0 1210.9207
140.6341552734375 0 425.41666
141.1026611328125 0 574.728
144.08114624023438 0 410.03998
145.09716796875 0 611.0722
146.06033325195312 0 5065.0967
147.04449462890625 0 1743.4374
148.95480346679688 0 806.1863
149.0450439453125 0 533.5748
150.0550079345703 0 452.42178
155.0819549560547 0 856.9233
155.11813354492188 0 472.07425
156.0770721435547 0 5050.2188
157.07627868652344 0 573.8227
157.10848999023438 0 800.6563
157.13418579101562 0 1248.4407
158.0603790283203 0 802.3817
158.09671020507812 0 4238.9854
159.0919647216797 0 6864.2314
160.0757293701172 0 885.4887
160.09518432617188 0 590.93115
160.11233520507812 0 2340.7444
164.07147216796875 0 803.82
165.054931640625 0 18236.865
165.0771026611328 0 690.7981
166.05868530273438 0 972.1273
166.08651733398438 0 3218.3777
167.05555725097656 0 1349.9312
167.08145141601562 0 526.1335
169.0763702392578 0 607.4896
169.0979766845703 0 690.4245
169.13421630859375 0 710.7999
171.0767822265625 0 684.6184
173.09242248535156 0 1036.7484
173.1287841796875 0 10567.647
174.05517578125 0 1945.803
174.1319580078125 0 548.6225
175.08688354492188 0 4363.4985
176.08238220214844 0 697.4512
176.1073455810547 0 9327.464
177.10252380371094 0 4971.0938
177.11109924316406 0 1269.1586
178.1059112548828 0 878.75305
180.07708740234375 0 490.50198
181.0972137451172 0 709.1828
182.08152770996094 0 31633.557
183.0848388671875 0 2851.024
183.1133575439453 0 850.4226
184.6470489501953 0 513.3064
185.05580139160156 0 1027.1183
185.12872314453125 0 2080.9443
186.09152221679688 0 544.6465
188.07098388671875 0 6626.857
189.0745391845703 0 728.1292
189.08714294433594 0 1121.1013
191.08229064941406 0 669.0111
191.11822509765625 0 2921.58
192.0658416748047 0 4096.486
192.12164306640625 0 800.3629
193.07003784179688 0 508.2169
193.1086883544922 0 2341.9058
195.08767700195312 0 937.87524
196.24781799316406 0 485.4491
197.12799072265625 0 524.24396
199.07154846191406 0 764.801
199.10865783691406 0 911.9174
199.1439208984375 0 714.44775
200.13963317871094 0 1218.1553
201.1236572265625 0 6743.774
202.0530548095703 0 535.8267
203.06642150878906 0 1183.7019
203.08187866210938 0 1604.3491
203.11814880371094 0 3008.0203 a Water loss 1
204.07705688476562 0 1516.614
205.09742736816406 0 5028.264 y 8
206.1016845703125 0 1079.5566
207.1128692626953 0 1489.1945
208.0724334716797 0 527.6559
209.09237670898438 0 2993.5154
210.92779541015625 0 532.19
212.1392822265625 0 581.42816
214.11859130859375 0 644.84937
215.1398468017578 0 1014.68896
217.08251953125 0 654.16705
217.09744262695312 0 2480.955
218.55201721191406 0 533.53925
219.11300659179688 0 1543.5637
221.0923614501953 0 2790.808
221.10366821289062 0 6374.1543
221.12884521484375 0 119612.41 a 1
221.49929809570312 0 478.84125
222.13214111328125 0 14948.961
223.06381225585938 0 953.39246
223.13406372070312 0 673.30646
225.1709747314453 0 1053.3568
226.11871337890625 0 562.84607
226.1556854248047 0 1459.4584
227.1031951904297 0 987.2141
229.11903381347656 0 757.6268
229.15602111816406 0 715.6015
231.06166076660156 0 2893.1348
231.10006713867188 0 622.43085
231.11334228515625 0 1928.5557 b Water loss 1
233.0923614501953 0 1214.4122
233.1293182373047 0 1808.1595
235.1081085205078 0 3497.8684
236.79969787597656 0 533.5078
239.09532165527344 0 1087.2561
239.11422729492188 0 1239.692
239.15078735351562 0 1364.023
247.0933837890625 0 695.9479
249.1237335205078 0 20903.744 b 1
250.1272735595703 0 2550.0278
251.1029815673828 0 3110.5635
253.11863708496094 0 1481.5757
253.16627502441406 0 2470.6484
256.177978515625 0 790.587
257.1064758300781 0 1365.467
258.10882568359375 0 587.6873
261.1231994628906 0 883.3968
263.1029357910156 0 4781.258
264.1059875488281 0 670.7535
270.1087951660156 0 639.45386
273.1346740722656 0 1257.1823
274.1298828125 0 828.367
279.0977478027344 0 3245.4885
283.1441345214844 0 1322.1229
284.1036682128906 0 643.6616
284.1605529785156 0 631.04816
285.08636474609375 0 795.47095
285.1019287109375 0 2624.5078
286.10400390625 0 511.6667
292.1405334472656 0 1558.0714
296.15087890625 0 2752.6694
297.0904235839844 0 1315.876
302.1139831542969 0 1052.9974
312.2049560546875 0 589.78674
314.09918212890625 0 886.07294
316.1305236816406 0 984.42175
318.1459655761719 0 1444.5635 a Water loss 2
319.14044189453125 0 2172.7686
324.1460876464844 0 3651.8691
325.1485595703125 0 1370.3885
327.20379638671875 0 624.57477
329.15008544921875 0 24883.822
330.1533203125 0 4877.0703
340.19879150390625 0 4290.7515
342.15667724609375 0 22779.736 y 7
343.15997314453125 0 3769.1802
346.14080810546875 0 1395.9987 b Water loss 2
348.1177673339844 0 660.9459
350.1354675292969 0 576.1475
355.07000732421875 0 1316.503
355.16168212890625 0 806.743
356.07061767578125 0 752.2525
356.1392822265625 0 1780.3773
357.22491455078125 0 5220.095
361.7706604003906 0 561.9315
364.15045166015625 0 6336.173 b 2
365.1519775390625 0 1361.4443
366.1297302246094 0 1139.985
366.17755126953125 0 640.95123
368.193115234375 0 1457.8594
371.1325988769531 0 536.64606
378.1295166015625 0 3087.084
382.142333984375 0 886.2872
384.1881103515625 0 846.5083
385.220703125 0 808.21985
392.2034912109375 0 1601.9252
394.12493896484375 0 5161.4473
395.126708984375 0 1566.845
398.2408752441406 0 973.74554
401.2106018066406 0 1239.3842
401.71234130859375 0 562.93445
413.7236633300781 0 622.13995
414.2193603515625 0 613.24493
415.2558898925781 0 943.2878
418.20977783203125 0 822.1575
422.7311706542969 0 1132.0641
423.2320251464844 0 790.7895
425.1934814453125 0 3023.3677
426.19659423828125 0 678.7032
427.2103271484375 0 996.38226
427.7222900390625 0 2720.1482
428.223388671875 0 982.9934
436.7275390625 0 2576.4238
437.1940612792969 0 575.02264
437.22918701171875 0 1745.9578
437.73138427734375 0 681.9348
448.26800537109375 0 632.704
448.76959228515625 0 793.82623
453.2820129394531 0 1023.3474
461.1680603027344 0 715.1523 b Water loss 3
465.1618347167969 0 3183.5706
466.16607666015625 0 628.3581
470.31048583984375 0 704.88727
479.1778869628906 0 3249.6008 b 3
483.22113037109375 0 2307.1335
486.1631164550781 0 1093.8129
487.3345031738281 0 949.1724
487.7565002441406 0 666.45276
489.192626953125 0 5923.374 y 6
490.193603515625 0 1423.3392
496.23101806640625 0 3314.504
496.7649230957031 0 1129.439
497.17626953125 0 716.3331
497.23321533203125 0 986.5983
499.14892578125 0 657.59033
500.2440490722656 0 930.57367
501.7563781738281 0 959.9598
506.73077392578125 0 633.3992
510.7648620605469 0 958.7489
513.3509521484375 0 1867.9851
514.1560668945312 0 993.3717
519.2034912109375 0 761.0856
525.2930297851562 0 734.9017
532.2747802734375 0 661.3326
536.1661987304688 0 651.8627
545.1754150390625 0 795.78015
548.1995239257812 0 1043.7968
550.7110595703125 0 2401.0222 y 1
551.2131958007812 0 2024.0813
559.25146484375 0 1483.769
560.2289428710938 0 8661.528 y 5
561.2315673828125 0 2436.0095
562.22705078125 0 1093.4355
569.7314453125 0 899.5739
570.2286376953125 0 787.9711
583.283935546875 0 896.2136
585.3356323242188 0 8820.762
586.3386840820312 0 2821.274
587.3423461914062 0 644.3905
589.78759765625 0 838.15027
592.229248046875 0 2008.8528 Precursor Water loss
592.7274780273438 0 1213.6056
601.236083984375 0 4669.511 Precursor
601.28955078125 0 784.7041
601.3072509765625 0 1007.5106
601.7349853515625 0 3266.7095
602.3231811523438 0 1038.5708
606.23291015625 0 853.36786
612.1946411132812 0 692.1247
614.2467651367188 0 973.8323
624.231689453125 0 1102.88 b Water loss 4
628.3783569335938 0 2730.0332
629.3779907226562 0 1050.2528
642.2412109375 0 619.30176 b 4
655.3031005859375 0 1232.6089
657.2621459960938 0 1905.7721
657.765625 0 621.69824
658.2691040039062 0 695.8675
659.2928466796875 0 3459.2441
660.298583984375 0 779.5456
664.8027954101562 0 632.58246
665.3209228515625 0 604.8815
670.817626953125 0 6538.9004
671.3186645507812 0 4861.2466
671.8207397460938 0 1778.8993
682.3485107421875 0 756.6774
683.339599609375 0 735.4985
685.2564086914062 0 2754.5215
686.2615356445312 0 714.6072
693.8168334960938 0 922.6867
694.3120727539062 0 843.1863
696.3810424804688 0 691.0575
700.3628540039062 0 21867.318
701.3651733398438 0 7400.172
702.365234375 0 2109.897
716.8150634765625 0 2394.372
717.3174438476562 0 1218.3055
721.2606201171875 0 2231.9727
722.26220703125 0 1302.4849
723.2921142578125 0 6979.2983 y 4
724.2937622070312 0 3386.197
725.296142578125 0 1460.4198
730.3499145507812 0 943.4622
730.8523559570312 0 721.5077
744.3523559570312 0 1447.3025
744.8535766601562 0 1678.5851
745.3562622070312 0 711.1262
749.2557983398438 0 4486.6924
750.2620849609375 0 1143.4628
752.8634033203125 0 2294.594
754.365966796875 0 725.42395
757.385986328125 0 2469.4893
767.2681274414062 0 869.62177
767.398193359375 0 813.9153
773.4146118164062 0 808.48444
774.3203125 0 2521.6235
774.4061889648438 0 621.1004
775.3272705078125 0 1734.0453
775.405029296875 0 922.25745
783.3991088867188 0 6175.9214
784.401123046875 0 2704.2231
785.4097290039062 0 678.50433
793.3661499023438 0 734.43585
794.3817138671875 0 2855.1045
795.384521484375 0 1071.4248
800.427490234375 0 3788.1084
801.4119262695312 0 11291.094
801.8660888671875 0 689.1625
802.4136962890625 0 4964.5273
803.4151000976562 0 1393.286
820.30419921875 0 660.66534 y Water loss 3
828.4221801757812 0 4560.659
829.4281616210938 0 1662.8535
838.3192138671875 0 10072.07 y 3
839.3226318359375 0 4656.8696
840.3189697265625 0 1376.3572
844.45263671875 0 16938.453
845.4550170898438 0 7309.722
846.4609375 0 2108.526
854.4373168945312 0 3446.5698
855.4385986328125 0 1961.1355
871.340087890625 0 1151.3381
872.447509765625 0 66700.88
873.4498291015625 0 29129.984
874.4534912109375 0 8550.391
875.4576416015625 0 1771.5906
889.347412109375 0 12386.65
890.3505249023438 0 7051.189
891.3515014648438 0 1364.4226
895.5235595703125 0 1511.7782
896.5267944335938 0 1547.4403
897.528076171875 0 1012.49603
899.4263916015625 0 809.6631
909.4069213867188 0 2049.7317
910.4173583984375 0 611.87726
914.6800537109375 0 595.01874
935.3342895507812 0 2262.7212 y Water loss 2
936.3352661132812 0 863.78265
937.3380126953125 0 907.418
953.3458251953125 0 48233.387 y 2
954.3485717773438 0 23850.426
955.3489990234375 0 9281.183
956.3529663085938 0 1221.7411
991.5208740234375 0 11399.057
992.5225830078125 0 5854.214
993.5228271484375 0 2499.864
994.515869140625 0 841.8812
1019.5155029296875 0 25503.482
1020.5182495117188 0 15652.101
1021.5205078125 0 4964.3887
1022.5220947265625 0 821.7518
1036.4176025390625 0 1071.7389
1037.511474609375 0 776.15125
1038.5140380859375 0 649.5872
1100.4122314453125 0 2894.9707 y 1
1101.41845703125 0 2063.1133
2429.617431640625 0 730.7893
2501.01416015625 0 674.81177
3001.750732421875 0 821.1299

Spectrum Details

|  |  |
| --- | --- |
| Matched peaks? Matched peaks The total absolute number of peaks matched. Additionally in brackets the total fraction of peaks matched and the total number of peaks is shown. | 24 (6.50% of 369) |
| FDR? FDR The false discovery rate estimated for this peptide. It is calculated by matching all theoretical fragments with a non-integer shift with the raw peaks for this spectrum. This is done with 40 different shifts. The resulting percentage is the average number of annotated peaks over the number of annotated peaks with the correct spectrum. | 1.09% |
| Satellite FDR? Satellite FDR See the FDR for details on its calculation. This satellite ion specific FDR only contains the satellite ions (d/w) for I/L/J positions. | - |
| PSM Score? PSM Score The PSM Score as given by Hecklib to this annotated spectrum. It is shown with three significant figures. | 310 |

#### Spectrum 9669? Spectrum 9669 The raw spectrum of this peptide as annotated by Hecklib. The fragments are coloured according to ion type (see legend). Any peaks with a star '\*' as text can be hovered over to see the full details, first the ion type second the mass shift type. By hovering over the amino acids in the peptide or ions in the legend the corresponding peaks are highlighted. By toggling the 'Unassigned' label you can turn the background (unassigned) peaks on or off in the plot. By updating the slider in the Ion legend you can update the spectrum to only show the top X% of the peaks with labels. The top X% means any peak that is within X% of the highest intensity. By dragging in the spectrum you can zoom in to a specific part of the spectrum and use 'Zoom Out' to get back to the original zoom level. The annotation of the spectrum is based on the given sequence in the peptides file and is done with different software so inconsistencies are likely. The peaks are annotated based on the given sequence, with 20 ppm tolerance.

Copy Data

##### Spectrum 9669 (TSV)

###### Preview

```
Loading example...
```

*Click on the button to copy the data to your clipboard.*

Mz MinMz MaxIntensity Max

WidthHeightPeptide font sizePeptide stroke widthSpectrum font sizeSpectrum stroke widthCompact peptide

Ion legend

wxyz

abcd

OtherUnassignedIon^Charge^_Position_Show for top:%

TFDDYAMHW

03.08e+46.16e+49.23e+41.23e+5

Zoom Out

a^+1^_2_y^+1^_1_a^+1^_2_b^+1^_2_b^+1^_2_a^+1^_3_y^+1^_2_b^+1^_3_b^+1^_3_b^+1^_4_b^+1^_4_y^+1^_3_y^+2^_8_y^+1^_4_\*\*y^+1^_5_y^+1^_6_y^+1^_7_y^+1^_7_y^+1^_8_

02785568341112

Fragment Matches Table

Show background peaks

| Position | Ion type | Intensity | mz Theoretical | mz Error (Th) | mz Error (ppm) | Charge | Series Number |
| --- | --- | --- | --- | --- | --- | --- | --- |
| - | - | 1.219E+05 | 120.1 | - | - | 0 | - |
| - | - | 1.005E+04 | 121.1 | - | - | 0 | - |
| - | - | 404.4 | 121.3 | - | - | 0 | - |
| - | - | 1956 | 123 | - | - | 0 | - |
| - | - | 390.2 | 126.2 | - | - | 0 | - |
| - | - | 658.4 | 127.1 | - | - | 0 | - |
| - | - | 446.5 | 127.8 | - | - | 0 | - |
| - | - | 505.6 | 128.1 | - | - | 0 | - |
| - | - | 6695 | 129.1 | - | - | 0 | - |
| - | - | 458.8 | 130.1 | - | - | 0 | - |
| - | - | 2978 | 130.1 | - | - | 0 | - |
| - | - | 1964 | 132.1 | - | - | 0 | - |
| - | - | 1113 | 132.1 | - | - | 0 | - |
| - | - | 1848 | 133.1 | - | - | 0 | - |
| - | - | 555.9 | 134 | - | - | 0 | - |
| - | - | 517 | 135 | - | - | 0 | - |
| - | - | 3.853E+04 | 136.1 | - | - | 0 | - |
| - | - | 398 | 137.1 | - | - | 0 | - |
| - | - | 2486 | 137.1 | - | - | 0 | - |
| - | - | 605.8 | 138.1 | - | - | 0 | - |
| - | - | 1486 | 138.1 | - | - | 0 | - |
| - | - | 567.7 | 145.1 | - | - | 0 | - |
| - | - | 1622 | 146.1 | - | - | 0 | - |
| - | - | 1337 | 147 | - | - | 0 | - |
| - | - | 461.4 | 149 | - | - | 0 | - |
| - | - | 1268 | 149 | - | - | 0 | - |
| - | - | 557.6 | 150.9 | - | - | 0 | - |
| - | - | 643.1 | 155.1 | - | - | 0 | - |
| - | - | 500.3 | 155.1 | - | - | 0 | - |
| - | - | 4022 | 156.1 | - | - | 0 | - |
| - | - | 749.5 | 157.1 | - | - | 0 | - |
| - | - | 488.6 | 157.1 | - | - | 0 | - |
| - | - | 1489 | 157.1 | - | - | 0 | - |
| - | - | 1414 | 158.1 | - | - | 0 | - |
| - | - | 4338 | 158.1 | - | - | 0 | - |
| - | - | 4955 | 159.1 | - | - | 0 | - |
| - | - | 615.1 | 160.1 | - | - | 0 | - |
| - | - | 1734 | 160.1 | - | - | 0 | - |
| - | - | 410.7 | 163.1 | - | - | 0 | - |
| - | - | 1.587E+04 | 165.1 | - | - | 0 | - |
| - | - | 694.1 | 165.1 | - | - | 0 | - |
| - | - | 1643 | 166.1 | - | - | 0 | - |
| - | - | 2242 | 166.1 | - | - | 0 | - |
| - | - | 1727 | 167.1 | - | - | 0 | - |
| - | - | 534.6 | 167.1 | - | - | 0 | - |
| - | - | 706 | 169.1 | - | - | 0 | - |
| - | - | 650 | 171.1 | - | - | 0 | - |
| - | - | 506.3 | 171.1 | - | - | 0 | - |
| - | - | 458.9 | 173.1 | - | - | 0 | - |
| - | - | 7993 | 173.1 | - | - | 0 | - |
| - | - | 3697 | 173.4 | - | - | 0 | - |
| - | - | 637.7 | 174.1 | - | - | 0 | - |
| - | - | 551.9 | 174.1 | - | - | 0 | - |
| - | - | 3507 | 175.1 | - | - | 0 | - |
| - | - | 768.3 | 175.1 | - | - | 0 | - |
| - | - | 965.7 | 176.1 | - | - | 0 | - |
| - | - | 8766 | 176.1 | - | - | 0 | - |
| - | - | 5616 | 177.1 | - | - | 0 | - |
| - | - | 945.3 | 177.1 | - | - | 0 | - |
| - | - | 676.1 | 178.1 | - | - | 0 | - |
| - | - | 3.204E+04 | 182.1 | - | - | 0 | - |
| - | - | 3617 | 183.1 | - | - | 0 | - |
| - | - | 800.6 | 183.1 | - | - | 0 | - |
| - | - | 431 | 185.1 | - | - | 0 | - |
| - | - | 501.4 | 185.1 | - | - | 0 | - |
| - | - | 1517 | 185.1 | - | - | 0 | - |
| - | - | 502.8 | 185.2 | - | - | 0 | - |
| - | - | 649.7 | 186.1 | - | - | 0 | - |
| - | - | 461.5 | 187.6 | - | - | 0 | - |
| - | - | 6345 | 188.1 | - | - | 0 | - |
| - | - | 950.9 | 189.1 | - | - | 0 | - |
| - | - | 549.9 | 191.1 | - | - | 0 | - |
| - | - | 2679 | 191.1 | - | - | 0 | - |
| - | - | 1118 | 192.1 | - | - | 0 | - |
| - | - | 2186 | 193.1 | - | - | 0 | - |
| - | - | 1116 | 195.1 | - | - | 0 | - |
| - | - | 513.4 | 195.1 | - | - | 0 | - |
| - | - | 722.1 | 200.1 | - | - | 0 | - |
| - | - | 4824 | 201.1 | - | - | 0 | - |
| - | - | 911.8 | 203.1 | - | - | 0 | - |
| - | - | 1968 | 203.1 | - | - | 0 | - |
| - | - | 486.5 | 203.1 | - | - | 0 | - |
| 2 | a | 2714 | 203.1 | 0.0002288 | 1.126 | +1 | 2 |
| - | - | 1021 | 204.1 | - | - | 0 | - |
| 9 | y | 3856 | 205.1 | 0.0002428 | 1.184 | +1 | 1 |
| - | - | 796.7 | 207.1 | - | - | 0 | - |
| - | - | 757 | 209.1 | - | - | 0 | - |
| - | - | 878.2 | 214.1 | - | - | 0 | - |
| - | - | 752.7 | 215.1 | - | - | 0 | - |
| - | - | 563.4 | 217.1 | - | - | 0 | - |
| - | - | 1857 | 217.1 | - | - | 0 | - |
| - | - | 1799 | 219.1 | - | - | 0 | - |
| - | - | 3275 | 221.1 | - | - | 0 | - |
| - | - | 7220 | 221.1 | - | - | 0 | - |
| 2 | a | 1.173E+05 | 221.1 | 0.0003147 | 1.423 | +1 | 2 |
| - | - | 1.472E+04 | 222.1 | - | - | 0 | - |
| - | - | 2023 | 223.1 | - | - | 0 | - |
| - | - | 791.8 | 223.1 | - | - | 0 | - |
| - | - | 808.5 | 223.1 | - | - | 0 | - |
| - | - | 1664 | 225.2 | - | - | 0 | - |
| - | - | 563.6 | 226.1 | - | - | 0 | - |
| - | - | 907.2 | 226.2 | - | - | 0 | - |
| - | - | 859.8 | 227.1 | - | - | 0 | - |
| - | - | 845.1 | 229.1 | - | - | 0 | - |
| - | - | 1056 | 229.2 | - | - | 0 | - |
| - | - | 1563 | 231.1 | - | - | 0 | - |
| 2 | b | 1576 | 231.1 | 0.0001109 | 0.4798 | +1 | 2 |
| - | - | 497.9 | 232.9 | - | - | 0 | - |
| - | - | 881.1 | 233.1 | - | - | 0 | - |
| - | - | 1071 | 233.1 | - | - | 0 | - |
| - | - | 3337 | 235.1 | - | - | 0 | - |
| - | - | 541.7 | 236.1 | - | - | 0 | - |
| - | - | 1840 | 239.1 | - | - | 0 | - |
| - | - | 659.3 | 239.1 | - | - | 0 | - |
| - | - | 1140 | 239.2 | - | - | 0 | - |
| - | - | 872.5 | 241.1 | - | - | 0 | - |
| 2 | b | 2.049E+04 | 249.1 | 0.0003342 | 1.341 | +1 | 2 |
| - | - | 2936 | 250.1 | - | - | 0 | - |
| - | - | 2011 | 251.1 | - | - | 0 | - |
| - | - | 614.7 | 251.2 | - | - | 0 | - |
| - | - | 1617 | 253.1 | - | - | 0 | - |
| - | - | 2104 | 253.2 | - | - | 0 | - |
| - | - | 1360 | 257.1 | - | - | 0 | - |
| - | - | 987.1 | 261.1 | - | - | 0 | - |
| - | - | 4816 | 263.1 | - | - | 0 | - |
| - | - | 1097 | 274.1 | - | - | 0 | - |
| - | - | 2329 | 279.1 | - | - | 0 | - |
| - | - | 619.5 | 280.9 | - | - | 0 | - |
| - | - | 1101 | 283.1 | - | - | 0 | - |
| - | - | 1153 | 285 | - | - | 0 | - |
| - | - | 2097 | 285.1 | - | - | 0 | - |
| - | - | 749.3 | 286.1 | - | - | 0 | - |
| - | - | 714.1 | 288.1 | - | - | 0 | - |
| - | - | 1710 | 292.1 | - | - | 0 | - |
| - | - | 2225 | 296.2 | - | - | 0 | - |
| - | - | 1507 | 297.1 | - | - | 0 | - |
| - | - | 754.2 | 299.1 | - | - | 0 | - |
| - | - | 858.3 | 302.1 | - | - | 0 | - |
| - | - | 791.3 | 304.1 | - | - | 0 | - |
| - | - | 542.6 | 305.5 | - | - | 0 | - |
| 3 | a | 968.2 | 318.1 | 0.0007973 | 2.506 | +1 | 3 |
| - | - | 1636 | 319.1 | - | - | 0 | - |
| - | - | 616.4 | 323.1 | - | - | 0 | - |
| - | - | 3968 | 324.1 | - | - | 0 | - |
| - | - | 825.9 | 325.1 | - | - | 0 | - |
| - | - | 528.4 | 325.1 | - | - | 0 | - |
| - | - | 2.211E+04 | 329.1 | - | - | 0 | - |
| - | - | 830.6 | 329.2 | - | - | 0 | - |
| - | - | 4403 | 330.2 | - | - | 0 | - |
| - | - | 960 | 338.1 | - | - | 0 | - |
| - | - | 704.5 | 339.2 | - | - | 0 | - |
| - | - | 3202 | 340.2 | - | - | 0 | - |
| 8 | y | 2.174E+04 | 342.2 | 0.0005503 | 1.608 | +1 | 2 |
| - | - | 4122 | 343.2 | - | - | 0 | - |
| 3 | b | 1831 | 346.1 | 6.823E-05 | 0.1971 | +1 | 3 |
| - | - | 911.1 | 348.1 | - | - | 0 | - |
| - | - | 1292 | 350.1 | - | - | 0 | - |
| - | - | 543.1 | 352.1 | - | - | 0 | - |
| - | - | 1122 | 355.1 | - | - | 0 | - |
| - | - | 1506 | 356.1 | - | - | 0 | - |
| - | - | 621.6 | 357.1 | - | - | 0 | - |
| - | - | 4699 | 357.2 | - | - | 0 | - |
| - | - | 706.4 | 358.2 | - | - | 0 | - |
| - | - | 582.7 | 360 | - | - | 0 | - |
| - | - | 671.5 | 362.1 | - | - | 0 | - |
| 3 | b | 5162 | 364.2 | 0.0001044 | 0.2866 | +1 | 3 |
| - | - | 587.2 | 365.2 | - | - | 0 | - |
| - | - | 629.3 | 367.2 | - | - | 0 | - |
| - | - | 1210 | 368.2 | - | - | 0 | - |
| - | - | 3468 | 378.1 | - | - | 0 | - |
| - | - | 870.9 | 392.2 | - | - | 0 | - |
| - | - | 4504 | 394.1 | - | - | 0 | - |
| - | - | 722.9 | 395.1 | - | - | 0 | - |
| - | - | 1423 | 401.2 | - | - | 0 | - |
| - | - | 665.1 | 401.7 | - | - | 0 | - |
| - | - | 1167 | 408.2 | - | - | 0 | - |
| - | - | 706.6 | 413.3 | - | - | 0 | - |
| - | - | 856 | 413.7 | - | - | 0 | - |
| - | - | 685.3 | 418.2 | - | - | 0 | - |
| - | - | 674.9 | 423.2 | - | - | 0 | - |
| - | - | 3550 | 425.2 | - | - | 0 | - |
| - | - | 1013 | 426.2 | - | - | 0 | - |
| - | - | 2228 | 427.7 | - | - | 0 | - |
| - | - | 1135 | 428.2 | - | - | 0 | - |
| - | - | 2948 | 436.7 | - | - | 0 | - |
| - | - | 1027 | 437.2 | - | - | 0 | - |
| - | - | 829.3 | 448.8 | - | - | 0 | - |
| - | - | 766.8 | 455.2 | - | - | 0 | - |
| 4 | b | 1259 | 461.2 | 0.0007051 | 1.529 | +1 | 4 |
| - | - | 2967 | 465.2 | - | - | 0 | - |
| - | - | 790.2 | 466.2 | - | - | 0 | - |
| - | - | 903 | 470.3 | - | - | 0 | - |
| 4 | b | 4242 | 479.2 | 0.0009983 | 2.083 | +1 | 4 |
| - | - | 688.3 | 480.2 | - | - | 0 | - |
| - | - | 2338 | 483.2 | - | - | 0 | - |
| - | - | 618.8 | 487.3 | - | - | 0 | - |
| - | - | 1587 | 487.3 | - | - | 0 | - |
| - | - | 657.5 | 487.8 | - | - | 0 | - |
| 7 | y | 5795 | 489.2 | 0.005466 | 11.17 | +1 | 3 |
| - | - | 1816 | 490.2 | - | - | 0 | - |
| - | - | 741.3 | 491.2 | - | - | 0 | - |
| - | - | 602.2 | 495.2 | - | - | 0 | - |
| - | - | 2030 | 496.2 | - | - | 0 | - |
| - | - | 1001 | 501.3 | - | - | 0 | - |
| - | - | 784.4 | 501.8 | - | - | 0 | - |
| - | - | 1453 | 510.3 | - | - | 0 | - |
| - | - | 832.2 | 510.8 | - | - | 0 | - |
| - | - | 807.8 | 511.3 | - | - | 0 | - |
| - | - | 587.7 | 513.2 | - | - | 0 | - |
| - | - | 1833 | 513.4 | - | - | 0 | - |
| - | - | 640.9 | 548.2 | - | - | 0 | - |
| - | - | 780.9 | 549.2 | - | - | 0 | - |
| 2 | y | 2558 | 550.7 | 0.002714 | 4.928 | +2 | 8 |
| - | - | 1976 | 551.2 | - | - | 0 | - |
| - | - | 1562 | 551.7 | - | - | 0 | - |
| 6 | y | 9353 | 560.2 | 0.004851 | 8.66 | +1 | 4 |
| - | - | 2623 | 561.2 | - | - | 0 | - |
| - | - | 746.9 | 562.2 | - | - | 0 | - |
| - | - | 1081 | 568.3 | - | - | 0 | - |
| - | - | 787 | 570.2 | - | - | 0 | - |
| - | - | 685.9 | 583.2 | - | - | 0 | - |
| - | - | 8283 | 585.3 | - | - | 0 | - |
| - | - | 2056 | 586.3 | - | - | 0 | - |
| 0 | Precursor | 1799 | 592.2 | 0.003383 | 5.712 | +2 | -1 |
| - | - | 1134 | 596.2 | - | - | 0 | - |
| 0 | Precursor | 5133 | 601.2 | 0.003655 | 6.079 | +2 | -1 |
| - | - | 658.2 | 601.3 | - | - | 0 | - |
| - | - | 1465 | 601.3 | - | - | 0 | - |
| - | - | 2812 | 601.7 | - | - | 0 | - |
| - | - | 932.3 | 602.4 | - | - | 0 | - |
| - | - | 1029 | 606.2 | - | - | 0 | - |
| - | - | 1312 | 612.2 | - | - | 0 | - |
| - | - | 743.5 | 613.2 | - | - | 0 | - |
| - | - | 1067 | 614.2 | - | - | 0 | - |
| - | - | 2774 | 628.4 | - | - | 0 | - |
| - | - | 828.8 | 629.4 | - | - | 0 | - |
| - | - | 1045 | 634.2 | - | - | 0 | - |
| - | - | 1697 | 657.3 | - | - | 0 | - |
| - | - | 1126 | 658.3 | - | - | 0 | - |
| - | - | 3525 | 659.3 | - | - | 0 | - |
| - | - | 1293 | 660.3 | - | - | 0 | - |
| - | - | 1209 | 670.8 | - | - | 0 | - |
| - | - | 1185 | 682.4 | - | - | 0 | - |
| - | - | 3252 | 685.3 | - | - | 0 | - |
| - | - | 753.1 | 686.3 | - | - | 0 | - |
| - | - | 1.947E+04 | 700.4 | - | - | 0 | - |
| - | - | 7268 | 701.4 | - | - | 0 | - |
| - | - | 1850 | 702.4 | - | - | 0 | - |
| - | - | 2434 | 716.8 | - | - | 0 | - |
| - | - | 1667 | 717.3 | - | - | 0 | - |
| - | - | 3119 | 721.3 | - | - | 0 | - |
| - | - | 861.5 | 722.3 | - | - | 0 | - |
| 5 | y | 7251 | 723.3 | 0.004267 | 5.899 | +1 | 5 |
| - | - | 2621 | 724.3 | - | - | 0 | - |
| - | - | 1177 | 725.3 | - | - | 0 | - |
| - | - | 2961 | 749.3 | - | - | 0 | - |
| - | - | 1688 | 750.3 | - | - | 0 | - |
| - | - | 1731 | 757.4 | - | - | 0 | - |
| - | - | 837.6 | 758.4 | - | - | 0 | - |
| - | - | 1231 | 767.3 | - | - | 0 | - |
| - | - | 1317 | 773.4 | - | - | 0 | - |
| - | - | 2663 | 774.3 | - | - | 0 | - |
| - | - | 1248 | 775.3 | - | - | 0 | - |
| - | - | 4454 | 783.4 | - | - | 0 | - |
| - | - | 2627 | 784.4 | - | - | 0 | - |
| - | - | 1035 | 794.4 | - | - | 0 | - |
| - | - | 3157 | 800.4 | - | - | 0 | - |
| - | - | 1.052E+04 | 801.4 | - | - | 0 | - |
| - | - | 3594 | 802.4 | - | - | 0 | - |
| - | - | 1567 | 803.4 | - | - | 0 | - |
| - | - | 4385 | 828.4 | - | - | 0 | - |
| - | - | 1962 | 829.4 | - | - | 0 | - |
| 4 | y | 1.046E+04 | 838.3 | 0.004057 | 4.84 | +1 | 6 |
| - | - | 5753 | 839.3 | - | - | 0 | - |
| - | - | 1175 | 840.3 | - | - | 0 | - |
| - | - | 1.315E+04 | 844.5 | - | - | 0 | - |
| - | - | 7471 | 845.5 | - | - | 0 | - |
| - | - | 2123 | 846.5 | - | - | 0 | - |
| - | - | 918.9 | 847.4 | - | - | 0 | - |
| - | - | 1882 | 854.4 | - | - | 0 | - |
| - | - | 1367 | 855.4 | - | - | 0 | - |
| - | - | 781.8 | 856.4 | - | - | 0 | - |
| - | - | 1977 | 871.3 | - | - | 0 | - |
| - | - | 5.676E+04 | 872.4 | - | - | 0 | - |
| - | - | 2.979E+04 | 873.4 | - | - | 0 | - |
| - | - | 8356 | 874.5 | - | - | 0 | - |
| - | - | 1416 | 875.5 | - | - | 0 | - |
| - | - | 556.7 | 877.2 | - | - | 0 | - |
| - | - | 1.545E+04 | 889.3 | - | - | 0 | - |
| - | - | 6392 | 890.4 | - | - | 0 | - |
| - | - | 1576 | 891.4 | - | - | 0 | - |
| - | - | 650.1 | 895.5 | - | - | 0 | - |
| - | - | 1405 | 896.5 | - | - | 0 | - |
| - | - | 689.8 | 897.4 | - | - | 0 | - |
| - | - | 698.6 | 908.3 | - | - | 0 | - |
| - | - | 832.6 | 909.4 | - | - | 0 | - |
| - | - | 749 | 910.4 | - | - | 0 | - |
| 3 | y | 2737 | 935.3 | 0.002938 | 3.141 | +1 | 7 |
| - | - | 1595 | 936.3 | - | - | 0 | - |
| - | - | 1129 | 937.3 | - | - | 0 | - |
| 3 | y | 4.441E+04 | 953.3 | 0.00458 | 4.804 | +1 | 7 |
| - | - | 2.297E+04 | 954.3 | - | - | 0 | - |
| - | - | 7585 | 955.3 | - | - | 0 | - |
| - | - | 885.5 | 973.5 | - | - | 0 | - |
| - | - | 1.005E+04 | 991.5 | - | - | 0 | - |
| - | - | 5970 | 992.5 | - | - | 0 | - |
| - | - | 1686 | 993.5 | - | - | 0 | - |
| - | - | 2.635E+04 | 1020 | - | - | 0 | - |
| - | - | 1.477E+04 | 1021 | - | - | 0 | - |
| - | - | 4186 | 1022 | - | - | 0 | - |
| - | - | 713.9 | 1023 | - | - | 0 | - |
| - | - | 900.1 | 1036 | - | - | 0 | - |
| 2 | y | 2545 | 1100 | 0.00538 | 4.889 | +1 | 8 |
| - | - | 1168 | 1101 | - | - | 0 | - |

m/z Charge Intensity FragmentType MassShift Position
120.08111572265625 0 121887.27
121.08439636230469 0 10054.572
121.2587661743164 0 404.4023
123.04425048828125 0 1955.8926
126.17923736572266 0 390.19485
127.08714294433594 0 658.3833
127.75723266601562 0 446.50916
128.10702514648438 0 505.57123
129.1024932861328 0 6695.101
130.05003356933594 0 458.83096
130.0654296875 0 2978.0955
132.0810089111328 0 1964.2384
132.10211181640625 0 1112.8014
133.08615112304688 0 1847.6632
134.02818298339844 0 555.9104
134.99301147460938 0 517.01135
136.0760040283203 0 38528.848
137.0741424560547 0 397.9769
137.07937622070312 0 2485.7937
138.05555725097656 0 605.81177
138.06642150878906 0 1485.5358
145.09732055664062 0 567.6686
146.0603790283203 0 1622.3722
147.0443115234375 0 1337.2943
148.95538330078125 0 461.39044
149.0451202392578 0 1267.5463
150.9181671142578 0 557.622
155.0818328857422 0 643.1004
155.11814880371094 0 500.32156
156.076904296875 0 4021.9731
157.07630920410156 0 749.50433
157.1087188720703 0 488.6455
157.13375854492188 0 1489.2396
158.06048583984375 0 1414.317
158.0966796875 0 4338.2275
159.09194946289062 0 4955.1455
160.07577514648438 0 615.0734
160.11231994628906 0 1734.2671
163.11077880859375 0 410.677
165.05491638183594 0 15873.42
165.10264587402344 0 694.103
166.05882263183594 0 1642.7715
166.0864715576172 0 2242.3118
167.05552673339844 0 1727.4424
167.11679077148438 0 534.60876
169.09732055664062 0 705.97186
171.07688903808594 0 650.0448
171.1490020751953 0 506.25513
173.0921173095703 0 458.89294
173.12875366210938 0 7993.224
173.44049072265625 0 3697.1626
174.05511474609375 0 637.7462
174.13201904296875 0 551.9094
175.0868377685547 0 3507.3809
175.11949157714844 0 768.2899
176.08216857910156 0 965.6722
176.10733032226562 0 8765.997
177.10255432128906 0 5616.462
177.11082458496094 0 945.32245
178.10641479492188 0 676.0969
182.0814666748047 0 32041.518
183.084716796875 0 3617.2473
183.1128692626953 0 800.57495
185.05555725097656 0 430.9749
185.07015991210938 0 501.39838
185.12879943847656 0 1517.3835
185.16526794433594 0 502.82993
186.08937072753906 0 649.7408
187.58668518066406 0 461.4753
188.07086181640625 0 6345.129
189.08692932128906 0 950.94324
191.0818328857422 0 549.8979
191.1182861328125 0 2678.9973
192.06629943847656 0 1118.4951
193.10874938964844 0 2186.4324
195.0880584716797 0 1116.1078
195.11163330078125 0 513.4315
200.13958740234375 0 722.08374
201.1236114501953 0 4824.4487
203.06619262695312 0 911.82214
203.08180236816406 0 1968.3801
203.0915985107422 0 486.5087
203.1181182861328 0 2714.1038 a Water loss 1
204.07662963867188 0 1021.1066
205.09739685058594 0 3856.449 y 8
207.1133575439453 0 796.6626
209.09246826171875 0 756.9649
214.11949157714844 0 878.19666
215.13917541503906 0 752.65533
217.0812530517578 0 563.37335
217.09771728515625 0 1857.0608
219.11329650878906 0 1798.7251
221.0921630859375 0 3274.6133
221.1035919189453 0 7219.986
221.12876892089844 0 117302.67 a 1
222.1321563720703 0 14715.757
223.0638885498047 0 2022.7076
223.1077880859375 0 791.78375
223.1336212158203 0 808.5227
225.1708526611328 0 1664.2534
226.1187744140625 0 563.5997
226.15542602539062 0 907.21967
227.10287475585938 0 859.8071
229.11886596679688 0 845.066
229.1698760986328 0 1056.2397
231.06141662597656 0 1563.2274
231.1129150390625 0 1576.0642 b Water loss 1
232.8965606689453 0 497.9194
233.0922088623047 0 881.14874
233.12954711914062 0 1071.0764
235.1081085205078 0 3337.374
236.11048889160156 0 541.743
239.09559631347656 0 1840.1276
239.1143035888672 0 659.3012
239.1507568359375 0 1139.6376
241.0745849609375 0 872.4855
249.1237030029297 0 20493.375 b 1
250.127197265625 0 2935.5928
251.1031036376953 0 2010.5745
251.15135192871094 0 614.68994
253.11862182617188 0 1617.2493
253.16639709472656 0 2103.6306
257.1064453125 0 1360.1116
261.1238098144531 0 987.14197
263.10296630859375 0 4816.3306
274.1295166015625 0 1097.4636
279.0978698730469 0 2329.15
280.91131591796875 0 619.52136
283.1437072753906 0 1101.3308
285.0097961425781 0 1152.524
285.1019592285156 0 2096.8423
286.1044006347656 0 749.34564
288.1194763183594 0 714.1045
292.1405334472656 0 1710.306
296.15093994140625 0 2225.4846
297.09063720703125 0 1506.9406
299.06280517578125 0 754.24536
302.11358642578125 0 858.3175
304.12933349609375 0 791.3073
305.4905700683594 0 542.5705
318.1456298828125 0 968.1814 a Water loss 2
319.1407775878906 0 1636.4976
323.09893798828125 0 616.3786
324.1457824707031 0 3968.1099
325.08563232421875 0 825.8908
325.1495056152344 0 528.3875
329.1499938964844 0 22111.615
329.22930908203125 0 830.55865
330.1534729003906 0 4402.5474
338.1316833496094 0 960.01086
339.1695861816406 0 704.5417
340.19805908203125 0 3201.7861
342.1566162109375 0 21744.055 y 7
343.1596984863281 0 4121.67
346.1396789550781 0 1831.3445 b Water loss 2
348.119384765625 0 911.1399
350.13543701171875 0 1291.7378
352.1410827636719 0 543.06146
355.0702209472656 0 1122.3525
356.1403503417969 0 1506.0518
357.0666198730469 0 621.6409
357.224609375 0 4698.6597
358.2258605957031 0 706.44073
359.95782470703125 0 582.65405
362.1358337402344 0 671.5392
364.15020751953125 0 5162.1865 b 2
365.1530456542969 0 587.22
367.20013427734375 0 629.2969
368.1922302246094 0 1210.0248
378.12994384765625 0 3467.8447
392.2030029296875 0 870.87134
394.124755859375 0 4504.461
395.1279602050781 0 722.8593
401.20806884765625 0 1422.5055
401.7121276855469 0 665.0927
408.1658020019531 0 1166.7736
413.26556396484375 0 706.5738
413.72625732421875 0 855.9946
418.2081604003906 0 685.2645
423.2315673828125 0 674.8592
425.19342041015625 0 3549.7605
426.1979675292969 0 1012.69
427.7222900390625 0 2228.2803
428.22314453125 0 1134.826
436.728271484375 0 2948.3982
437.23046875 0 1026.7559
448.7655334472656 0 829.31433
455.19976806640625 0 766.78394
461.1659851074219 0 1259.491 b Water loss 3
465.16217041015625 0 2967.2473
466.1626892089844 0 790.21466
470.3086853027344 0 902.99365
479.1782531738281 0 4242.42 b 3
480.17816162109375 0 688.25226
483.2206115722656 0 2337.6216
487.2594909667969 0 618.81433
487.3352355957031 0 1587.1567
487.7518310546875 0 657.51984
489.1920166015625 0 5794.7256 y 6
490.1940002441406 0 1815.8387
491.20599365234375 0 741.2828
495.1899719238281 0 602.16284
496.23028564453125 0 2030.4104
501.2530517578125 0 1001.32697
501.7591857910156 0 784.3947
510.26239013671875 0 1452.5851
510.7657470703125 0 832.2032
511.2589111328125 0 807.822
513.1941528320312 0 587.7425
513.3502807617188 0 1832.9777
548.2015380859375 0 640.88745
549.203369140625 0 780.90265
550.7109985351562 0 2558.1777 y 1
551.2135009765625 0 1976.0677
551.7150268554688 0 1561.9172
560.228515625 0 9353.469 y 5
561.2327270507812 0 2623.2175
562.2305297851562 0 746.928
568.310302734375 0 1081.3126
570.226318359375 0 786.9652
583.2274780273438 0 685.91583
585.3360595703125 0 8283.192
586.3378295898438 0 2055.638
592.230224609375 0 1799.4857 Precursor Water loss
596.237060546875 0 1134.2428
601.2357788085938 0 5132.755 Precursor
601.2890014648438 0 658.1632
601.3204956054688 0 1464.6062
601.738525390625 0 2811.5608
602.3660278320312 0 932.2571
606.2314453125 0 1028.61
612.195068359375 0 1311.8768
613.1983032226562 0 743.48706
614.2457885742188 0 1066.7909
628.3773193359375 0 2773.6013
629.3809204101562 0 828.8175
634.2291259765625 0 1045.4178
657.2625122070312 0 1697.2483
658.2626342773438 0 1126.4714
659.2931518554688 0 3525.129
660.2973022460938 0 1293.4464
670.8152465820312 0 1209.206
682.3519897460938 0 1185.0898
685.257568359375 0 3251.668
686.2625732421875 0 753.0821
700.3624877929688 0 19469.965
701.3646240234375 0 7267.8755
702.3667602539062 0 1849.7744
716.81298828125 0 2433.9062
717.3137817382812 0 1666.577
721.2612915039062 0 3118.85
722.2617797851562 0 861.47705
723.291259765625 0 7250.825 y 4
724.2938842773438 0 2621.354
725.3016967773438 0 1176.6315
749.2550048828125 0 2961.1497
750.2605590820312 0 1688.0883
757.3867797851562 0 1730.753
758.3869018554688 0 837.59875
767.2659912109375 0 1230.5167
773.413330078125 0 1316.7279
774.3222045898438 0 2662.959
775.3250122070312 0 1248.151
783.399658203125 0 4453.527
784.4015502929688 0 2627.1409
794.3843383789062 0 1035.1555
800.4276733398438 0 3157.2043
801.4124755859375 0 10521.164
802.4142456054688 0 3594.2383
803.418212890625 0 1566.6342
828.4218139648438 0 4385.038
829.4244384765625 0 1961.5642
838.3179931640625 0 10458.867 y 3
839.3221435546875 0 5752.7373
840.3238525390625 0 1175.3826
844.4519653320312 0 13151.015
845.4542846679688 0 7470.787
846.45849609375 0 2122.5925
847.4395141601562 0 918.92224
854.4354248046875 0 1881.6819
855.435546875 0 1366.732
856.4445190429688 0 781.76324
871.3387451171875 0 1976.9554
872.4470825195312 0 56756.91
873.4498901367188 0 29787.154
874.4517822265625 0 8355.528
875.4501342773438 0 1415.8612
877.2146606445312 0 556.653
889.3472900390625 0 15447.883
890.3502197265625 0 6391.559
891.3514404296875 0 1576.1038
895.525390625 0 650.1257
896.5274047851562 0 1405.4756
897.4140014648438 0 689.84467
908.3406982421875 0 698.6313
909.4054565429688 0 832.6187
910.416259765625 0 749.0204
935.333251953125 0 2737.4033 y Water loss 2
936.330810546875 0 1595.284
937.3338623046875 0 1128.959
953.345458984375 0 44407.867 y 2
954.3483276367188 0 22969.553
955.3477783203125 0 7585.238
973.5131225585938 0 885.4863
991.5194702148438 0 10051.601
992.5232543945312 0 5969.643
993.5241088867188 0 1686.4863
1019.514892578125 0 26346.396
1020.5184326171875 0 14766.084
1021.5228881835938 0 4185.6445
1022.5260620117188 0 713.90344
1036.407470703125 0 900.13525
1100.4146728515625 0 2545.1333 y 1
1101.4141845703125 0 1167.7505

Spectrum Details

|  |  |
| --- | --- |
| Matched peaks? Matched peaks The total absolute number of peaks matched. Additionally in brackets the total fraction of peaks matched and the total number of peaks is shown. | 21 (6.69% of 314) |
| FDR? FDR The false discovery rate estimated for this peptide. It is calculated by matching all theoretical fragments with a non-integer shift with the raw peaks for this spectrum. This is done with 40 different shifts. The resulting percentage is the average number of annotated peaks over the number of annotated peaks with the correct spectrum. | 1.47% |
| Satellite FDR? Satellite FDR See the FDR for details on its calculation. This satellite ion specific FDR only contains the satellite ions (d/w) for I/L/J positions. | - |
| PSM Score? PSM Score The PSM Score as given by Hecklib to this annotated spectrum. It is shown with three significant figures. | 250 |

#### Spectrum 6163? Spectrum 6163 The raw spectrum of this peptide as annotated by Hecklib. The fragments are coloured according to ion type (see legend). Any peaks with a star '\*' as text can be hovered over to see the full details, first the ion type second the mass shift type. By hovering over the amino acids in the peptide or ions in the legend the corresponding peaks are highlighted. By toggling the 'Unassigned' label you can turn the background (unassigned) peaks on or off in the plot. By updating the slider in the Ion legend you can update the spectrum to only show the top X% of the peaks with labels. The top X% means any peak that is within X% of the highest intensity. By dragging in the spectrum you can zoom in to a specific part of the spectrum and use 'Zoom Out' to get back to the original zoom level. The annotation of the spectrum is based on the given sequence in the peptides file and is done with different software so inconsistencies are likely. The peaks are annotated based on the given sequence, with 20 ppm tolerance.

Copy Data

##### Spectrum 6163 (TSV)

###### Preview

```
Loading example...
```

*Click on the button to copy the data to your clipboard.*

Mz MinMz MaxIntensity Max

WidthHeightPeptide font sizePeptide stroke widthSpectrum font sizeSpectrum stroke widthCompact peptide

Ion legend

wxyz

abcd

OtherUnassignedIon^Charge^_Position_Show for top:%

TFDDYAMHW

06.70e+41.34e+52.01e+52.68e+5

Zoom Out

y^+1^_1_c^+2^_4_c^+1^_2_z^+1^_2_y^+1^_2_c^+1^_3_c^+1^_4_y^+2^_7_y^+1^_3_y^+2^_8_z^+1^_4_y^+1^_4_c^+1^_5_z^+1^_5_y^+1^_5_z^+1^_6_y^+1^_6_c^+1^_7_c^+1^_7_c^+1^_7_y^+1^_7_z^+1^_7_y^+1^_7_c^+1^_8_c^+1^_8_z^+1^_8_y^+1^_8_

0778155623343112

Fragment Matches Table

Show background peaks

| Position | Ion type | Intensity | mz Theoretical | mz Error (Th) | mz Error (ppm) | Charge | Series Number |
| --- | --- | --- | --- | --- | --- | --- | --- |
| - | - | 3087 | 120.1 | - | - | 0 | - |
| - | - | 400.5 | 130.2 | - | - | 0 | - |
| - | - | 1054 | 133.1 | - | - | 0 | - |
| - | - | 387.1 | 134.9 | - | - | 0 | - |
| - | - | 585.6 | 136.1 | - | - | 0 | - |
| - | - | 513.6 | 148.9 | - | - | 0 | - |
| - | - | 455.7 | 148.9 | - | - | 0 | - |
| - | - | 461.9 | 148.9 | - | - | 0 | - |
| - | - | 861.2 | 148.9 | - | - | 0 | - |
| - | - | 1261 | 148.9 | - | - | 0 | - |
| - | - | 1165 | 148.9 | - | - | 0 | - |
| - | - | 2136 | 148.9 | - | - | 0 | - |
| - | - | 3686 | 148.9 | - | - | 0 | - |
| - | - | 4632 | 149 | - | - | 0 | - |
| - | - | 2873 | 149 | - | - | 0 | - |
| - | - | 1272 | 149 | - | - | 0 | - |
| - | - | 1282 | 149 | - | - | 0 | - |
| - | - | 748.1 | 149 | - | - | 0 | - |
| - | - | 838.9 | 149 | - | - | 0 | - |
| - | - | 526.4 | 149 | - | - | 0 | - |
| - | - | 650.6 | 149 | - | - | 0 | - |
| - | - | 551.3 | 149 | - | - | 0 | - |
| - | - | 524 | 149 | - | - | 0 | - |
| - | - | 2102 | 149 | - | - | 0 | - |
| - | - | 448.6 | 149.9 | - | - | 0 | - |
| - | - | 499.2 | 155.6 | - | - | 0 | - |
| - | - | 495.2 | 156.5 | - | - | 0 | - |
| - | - | 617 | 167.1 | - | - | 0 | - |
| - | - | 937.6 | 173.5 | - | - | 0 | - |
| - | - | 788.8 | 175.1 | - | - | 0 | - |
| - | - | 873.5 | 177.1 | - | - | 0 | - |
| - | - | 528.4 | 177.1 | - | - | 0 | - |
| - | - | 1372 | 203.1 | - | - | 0 | - |
| - | - | 798.2 | 203.1 | - | - | 0 | - |
| 9 | y | 5912 | 221.1 | 0.002697 | 12.2 | +1 | 1 |
| - | - | 1676 | 221.1 | - | - | 0 | - |
| - | - | 8.054E+04 | 221.1 | - | - | 0 | - |
| - | - | 861.2 | 222.1 | - | - | 0 | - |
| - | - | 1.049E+04 | 222.1 | - | - | 0 | - |
| - | - | 506.3 | 222.9 | - | - | 0 | - |
| - | - | 611.9 | 223.1 | - | - | 0 | - |
| - | - | 2275 | 225 | - | - | 0 | - |
| - | - | 1152 | 226 | - | - | 0 | - |
| - | - | 1016 | 231.1 | - | - | 0 | - |
| - | - | 756.3 | 235.1 | - | - | 0 | - |
| - | - | 9520 | 239.1 | - | - | 0 | - |
| 4 | c | 1702 | 240.1 | 0.003575 | 14.89 | +2 | 4 |
| - | - | 1040 | 241.1 | - | - | 0 | - |
| 2 | c | 3.28E+04 | 249.1 | 0.0002884 | 1.158 | +1 | 2 |
| - | - | 3443 | 250.1 | - | - | 0 | - |
| - | - | 520.7 | 254.8 | - | - | 0 | - |
| - | - | 607.4 | 257.1 | - | - | 0 | - |
| - | - | 1723 | 263.1 | - | - | 0 | - |
| - | - | 1448 | 281.1 | - | - | 0 | - |
| - | - | 1011 | 285.1 | - | - | 0 | - |
| - | - | 3358 | 295.1 | - | - | 0 | - |
| - | - | 1413 | 296.1 | - | - | 0 | - |
| - | - | 1222 | 297.1 | - | - | 0 | - |
| - | - | 6057 | 299.1 | - | - | 0 | - |
| - | - | 526.2 | 309.1 | - | - | 0 | - |
| - | - | 555.6 | 313.1 | - | - | 0 | - |
| - | - | 572.4 | 314.1 | - | - | 0 | - |
| 8 | z | 2496 | 342.1 | 0.004952 | 14.47 | +1 | 2 |
| - | - | 6275 | 343.1 | - | - | 0 | - |
| - | - | 538.5 | 344.1 | - | - | 0 | - |
| - | - | 1033 | 346.1 | - | - | 0 | - |
| - | - | 4.466E+04 | 355.1 | - | - | 0 | - |
| 8 | y | 9493 | 358.1 | 0.00524 | 14.63 | +1 | 2 |
| - | - | 652.8 | 359 | - | - | 0 | - |
| - | - | 1820 | 359.2 | - | - | 0 | - |
| - | - | 479.3 | 362.6 | - | - | 0 | - |
| 3 | c | 7017 | 364.2 | 0.0001398 | 0.3839 | +1 | 3 |
| - | - | 1478 | 365.2 | - | - | 0 | - |
| - | - | 1354 | 369.1 | - | - | 0 | - |
| - | - | 949.6 | 370.1 | - | - | 0 | - |
| - | - | 2184 | 371.1 | - | - | 0 | - |
| - | - | 870.8 | 371.1 | - | - | 0 | - |
| - | - | 1687 | 378.1 | - | - | 0 | - |
| - | - | 1011 | 394.1 | - | - | 0 | - |
| - | - | 2310 | 415 | - | - | 0 | - |
| - | - | 6111 | 415.2 | - | - | 0 | - |
| - | - | 1414 | 416.2 | - | - | 0 | - |
| - | - | 973.7 | 419 | - | - | 0 | - |
| - | - | 594 | 433.2 | - | - | 0 | - |
| - | - | 984.1 | 441.2 | - | - | 0 | - |
| - | - | 1490 | 458.2 | - | - | 0 | - |
| - | - | 662.4 | 461.2 | - | - | 0 | - |
| 4 | c | 4452 | 479.2 | 0.00109 | 2.274 | +1 | 4 |
| - | - | 733.6 | 480.2 | - | - | 0 | - |
| 3 | y | 1028 | 485.2 | 0.006673 | 13.75 | +2 | 7 |
| - | - | 1832 | 489.1 | - | - | 0 | - |
| - | - | 2680 | 489.2 | - | - | 0 | - |
| - | - | 1.311E+04 | 490.2 | - | - | 0 | - |
| - | - | 3364 | 491.2 | - | - | 0 | - |
| - | - | 4836 | 504.2 | - | - | 0 | - |
| 7 | y | 6457 | 505.2 | 0.01006 | 19.92 | +1 | 3 |
| - | - | 1132 | 506.2 | - | - | 0 | - |
| - | - | 1246 | 512.2 | - | - | 0 | - |
| - | - | 704.1 | 532.2 | - | - | 0 | - |
| 2 | y | 5783 | 558.7 | 0.005456 | 9.765 | +2 | 8 |
| - | - | 3783 | 559.2 | - | - | 0 | - |
| - | - | 1401 | 559.7 | - | - | 0 | - |
| 6 | z | 2.363E+04 | 560.2 | 0.0102 | 18.21 | +1 | 4 |
| - | - | 1.044E+04 | 561.2 | - | - | 0 | - |
| - | - | 2166 | 562.2 | - | - | 0 | - |
| - | - | 628.1 | 565.2 | - | - | 0 | - |
| - | - | 1163 | 575.2 | - | - | 0 | - |
| 6 | y | 1.309E+04 | 576.2 | 0.009846 | 17.09 | +1 | 4 |
| - | - | 1467 | 577.1 | - | - | 0 | - |
| - | - | 5372 | 577.2 | - | - | 0 | - |
| - | - | 6599 | 578.2 | - | - | 0 | - |
| - | - | 2143 | 579.2 | - | - | 0 | - |
| - | - | 1170 | 591.2 | - | - | 0 | - |
| - | - | 1932 | 591.4 | - | - | 0 | - |
| - | - | 700.9 | 596.2 | - | - | 0 | - |
| - | - | 673.6 | 598.8 | - | - | 0 | - |
| - | - | 2666 | 600.2 | - | - | 0 | - |
| - | - | 2623 | 600.7 | - | - | 0 | - |
| - | - | 1498 | 601.2 | - | - | 0 | - |
| - | - | 758.6 | 603.2 | - | - | 0 | - |
| - | - | 1033 | 608.4 | - | - | 0 | - |
| - | - | 9193 | 609.2 | - | - | 0 | - |
| - | - | 4986 | 609.7 | - | - | 0 | - |
| - | - | 9262 | 610.2 | - | - | 0 | - |
| - | - | 2464 | 610.2 | - | - | 0 | - |
| - | - | 2107 | 614.2 | - | - | 0 | - |
| - | - | 696.2 | 615.2 | - | - | 0 | - |
| - | - | 1085 | 624.2 | - | - | 0 | - |
| 5 | c | 1664 | 642.2 | 0.0002613 | 0.4068 | +1 | 5 |
| - | - | 1.041E+04 | 649.3 | - | - | 0 | - |
| - | - | 3689 | 650.3 | - | - | 0 | - |
| - | - | 714.1 | 651.3 | - | - | 0 | - |
| - | - | 1159 | 662.3 | - | - | 0 | - |
| - | - | 807.3 | 663.3 | - | - | 0 | - |
| - | - | 1081 | 675.3 | - | - | 0 | - |
| - | - | 1.019E+04 | 693.2 | - | - | 0 | - |
| - | - | 3445 | 694.2 | - | - | 0 | - |
| - | - | 1213 | 695.2 | - | - | 0 | - |
| - | - | 735.1 | 707.3 | - | - | 0 | - |
| - | - | 707.5 | 708.3 | - | - | 0 | - |
| - | - | 1402 | 721.3 | - | - | 0 | - |
| 5 | z | 3.233E+04 | 723.3 | 0.01035 | 14.31 | +1 | 5 |
| - | - | 1.367E+04 | 724.3 | - | - | 0 | - |
| - | - | 3988 | 725.3 | - | - | 0 | - |
| - | - | 1653 | 731.3 | - | - | 0 | - |
| - | - | 672.6 | 731.4 | - | - | 0 | - |
| - | - | 669.6 | 733.3 | - | - | 0 | - |
| - | - | 3238 | 738.3 | - | - | 0 | - |
| 5 | y | 1.268E+04 | 739.3 | 0.009689 | 13.11 | +1 | 5 |
| - | - | 5156 | 740.3 | - | - | 0 | - |
| - | - | 1726 | 741.3 | - | - | 0 | - |
| - | - | 3461 | 749.3 | - | - | 0 | - |
| - | - | 7843 | 764.3 | - | - | 0 | - |
| - | - | 3713 | 765.3 | - | - | 0 | - |
| - | - | 1869 | 766.3 | - | - | 0 | - |
| - | - | 735.9 | 767.3 | - | - | 0 | - |
| - | - | 1941 | 775.3 | - | - | 0 | - |
| - | - | 1677 | 776.3 | - | - | 0 | - |
| - | - | 1828 | 777.3 | - | - | 0 | - |
| - | - | 789 | 779.3 | - | - | 0 | - |
| - | - | 1639 | 790.3 | - | - | 0 | - |
| - | - | 643.6 | 791.3 | - | - | 0 | - |
| - | - | 1.427E+04 | 794.3 | - | - | 0 | - |
| - | - | 6454 | 795.3 | - | - | 0 | - |
| - | - | 2384 | 796.3 | - | - | 0 | - |
| - | - | 4610 | 808.3 | - | - | 0 | - |
| - | - | 2152 | 809.3 | - | - | 0 | - |
| - | - | 871.5 | 810.3 | - | - | 0 | - |
| - | - | 2651 | 833.3 | - | - | 0 | - |
| - | - | 9719 | 834.3 | - | - | 0 | - |
| - | - | 3714 | 835.3 | - | - | 0 | - |
| - | - | 764.5 | 836.3 | - | - | 0 | - |
| - | - | 1131 | 836.4 | - | - | 0 | - |
| - | - | 650.6 | 838.2 | - | - | 0 | - |
| 4 | z | 3.504E+04 | 838.3 | 0.01038 | 12.38 | +1 | 6 |
| - | - | 3.683E+04 | 839.3 | - | - | 0 | - |
| - | - | 1.576E+04 | 840.3 | - | - | 0 | - |
| - | - | 3575 | 841.3 | - | - | 0 | - |
| - | - | 1489 | 846.3 | - | - | 0 | - |
| - | - | 1256 | 847.3 | - | - | 0 | - |
| - | - | 1108 | 853.3 | - | - | 0 | - |
| 4 | y | 1.354E+04 | 854.3 | 0.01003 | 11.74 | +1 | 6 |
| - | - | 6388 | 855.3 | - | - | 0 | - |
| - | - | 1962 | 856.3 | - | - | 0 | - |
| - | - | 2289 | 858.3 | - | - | 0 | - |
| 7 | c | 782.7 | 859.3 | 0.005013 | 5.834 | +1 | 7 |
| 7 | c | 792.6 | 860.3 | 0.009994 | 11.62 | +1 | 7 |
| - | - | 4143 | 865.3 | - | - | 0 | - |
| - | - | 2086 | 866.3 | - | - | 0 | - |
| - | - | 6001 | 876.3 | - | - | 0 | - |
| 7 | c | 2.162E+04 | 877.3 | 0.004441 | 5.062 | +1 | 7 |
| - | - | 873.4 | 877.5 | - | - | 0 | - |
| - | - | 1.142E+04 | 878.3 | - | - | 0 | - |
| - | - | 3677 | 879.3 | - | - | 0 | - |
| - | - | 886.3 | 890.3 | - | - | 0 | - |
| - | - | 1272 | 891.3 | - | - | 0 | - |
| - | - | 9264 | 905.3 | - | - | 0 | - |
| - | - | 4671 | 906.3 | - | - | 0 | - |
| - | - | 1666 | 907.4 | - | - | 0 | - |
| - | - | 1.511E+04 | 909.3 | - | - | 0 | - |
| - | - | 7277 | 910.3 | - | - | 0 | - |
| - | - | 2557 | 911.3 | - | - | 0 | - |
| 3 | y | 3671 | 951.3 | 0.00836 | 8.788 | +1 | 7 |
| - | - | 1506 | 952.3 | - | - | 0 | - |
| 3 | z | 4800 | 953.3 | 0.01035 | 10.86 | +1 | 7 |
| - | - | 4611 | 954.3 | - | - | 0 | - |
| - | - | 2011 | 955.3 | - | - | 0 | - |
| - | - | 667.1 | 956.3 | - | - | 0 | - |
| 3 | y | 6.731E+04 | 969.3 | 0.01019 | 10.51 | +1 | 7 |
| - | - | 3.725E+04 | 970.3 | - | - | 0 | - |
| - | - | 1.071E+04 | 971.3 | - | - | 0 | - |
| - | - | 1636 | 972.3 | - | - | 0 | - |
| - | - | 3151 | 981.4 | - | - | 0 | - |
| - | - | 2480 | 982.4 | - | - | 0 | - |
| - | - | 1002 | 983.4 | - | - | 0 | - |
| 8 | c | 2021 | 997.4 | 0.008028 | 8.05 | +1 | 8 |
| - | - | 786.4 | 998.4 | - | - | 0 | - |
| - | - | 1.14E+04 | 999.4 | - | - | 0 | - |
| - | - | 4545 | 1000 | - | - | 0 | - |
| - | - | 1086 | 1000 | - | - | 0 | - |
| - | - | 1669 | 1001 | - | - | 0 | - |
| - | - | 978.2 | 1001 | - | - | 0 | - |
| - | - | 915.2 | 1011 | - | - | 0 | - |
| - | - | 1256 | 1013 | - | - | 0 | - |
| 8 | c | 7.76E+04 | 1014 | 0.005466 | 5.389 | +1 | 8 |
| - | - | 4.287E+04 | 1015 | - | - | 0 | - |
| - | - | 1.528E+04 | 1016 | - | - | 0 | - |
| - | - | 1846 | 1017 | - | - | 0 | - |
| - | - | 8969 | 1040 | - | - | 0 | - |
| - | - | 5061 | 1041 | - | - | 0 | - |
| - | - | 2254 | 1042 | - | - | 0 | - |
| - | - | 817.9 | 1055 | - | - | 0 | - |
| - | - | 9891 | 1055 | - | - | 0 | - |
| - | - | 2.733E+04 | 1056 | - | - | 0 | - |
| - | - | 1.284E+04 | 1057 | - | - | 0 | - |
| - | - | 3758 | 1058 | - | - | 0 | - |
| - | - | 2960 | 1069 | - | - | 0 | - |
| - | - | 1560 | 1070 | - | - | 0 | - |
| - | - | 6467 | 1071 | - | - | 0 | - |
| - | - | 2402 | 1072 | - | - | 0 | - |
| - | - | 1521 | 1073 | - | - | 0 | - |
| - | - | 846 | 1084 | - | - | 0 | - |
| - | - | 714.9 | 1085 | - | - | 0 | - |
| - | - | 635.5 | 1089 | - | - | 0 | - |
| 2 | z | 2619 | 1100 | 0.01115 | 10.14 | +1 | 8 |
| - | - | 1845 | 1101 | - | - | 0 | - |
| - | - | 1534 | 1102 | - | - | 0 | - |
| - | - | 844.9 | 1110 | - | - | 0 | - |
| - | - | 815.8 | 1112 | - | - | 0 | - |
| 2 | y | 3460 | 1116 | 0.01001 | 8.965 | +1 | 8 |
| - | - | 1557 | 1117 | - | - | 0 | - |
| - | - | 1329 | 1126 | - | - | 0 | - |
| - | - | 843.6 | 1127 | - | - | 0 | - |
| - | - | 2497 | 1138 | - | - | 0 | - |
| - | - | 1913 | 1139 | - | - | 0 | - |
| - | - | 1955 | 1140 | - | - | 0 | - |
| - | - | 2344 | 1154 | - | - | 0 | - |
| - | - | 2232 | 1155 | - | - | 0 | - |
| - | - | 1.788E+04 | 1156 | - | - | 0 | - |
| - | - | 1.583E+04 | 1157 | - | - | 0 | - |
| - | - | 3.169E+04 | 1158 | - | - | 0 | - |
| - | - | 1.916E+04 | 1159 | - | - | 0 | - |
| - | - | 773.7 | 1160 | - | - | 0 | - |
| - | - | 6730 | 1160 | - | - | 0 | - |
| - | - | 887.9 | 1161 | - | - | 0 | - |
| - | - | 1087 | 1174 | - | - | 0 | - |
| - | - | 1448 | 1182 | - | - | 0 | - |
| - | - | 2066 | 1183 | - | - | 0 | - |
| - | - | 1022 | 1184 | - | - | 0 | - |
| - | - | 1442 | 1186 | - | - | 0 | - |
| - | - | 2453 | 1190 | - | - | 0 | - |
| - | - | 2248 | 1191 | - | - | 0 | - |
| - | - | 715.4 | 1199 | - | - | 0 | - |
| - | - | 2.773E+04 | 1200 | - | - | 0 | - |
| - | - | 1.283E+05 | 1201 | - | - | 0 | - |
| - | - | 8.404E+04 | 1202 | - | - | 0 | - |
| - | - | 3.623E+04 | 1203 | - | - | 0 | - |
| - | - | 5963 | 1204 | - | - | 0 | - |
| - | - | 996.3 | 1216 | - | - | 0 | - |
| - | - | 1529 | 1217 | - | - | 0 | - |
| - | - | 7.742E+04 | 1217 | - | - | 0 | - |
| - | - | 2.652E+05 | 1218 | - | - | 0 | - |
| - | - | 1.738E+05 | 1219 | - | - | 0 | - |
| - | - | 6.833E+04 | 1220 | - | - | 0 | - |
| - | - | 988.4 | 1221 | - | - | 0 | - |
| - | - | 8267 | 1221 | - | - | 0 | - |
| - | - | 730.9 | 1521 | - | - | 0 | - |
| - | - | 771.9 | 1787 | - | - | 0 | - |
| - | - | 684.3 | 1807 | - | - | 0 | - |
| - | - | 1939 | 1808 | - | - | 0 | - |
| - | - | 830.9 | 1809 | - | - | 0 | - |
| - | - | 784.2 | 1824 | - | - | 0 | - |
| - | - | 1521 | 1825 | - | - | 0 | - |
| - | - | 2105 | 1826 | - | - | 0 | - |
| - | - | 829.3 | 1830 | - | - | 0 | - |
| - | - | 1718 | 1832 | - | - | 0 | - |
| - | - | 788.7 | 3082 | - | - | 0 | - |

m/z Charge Intensity FragmentType MassShift Position
120.08102416992188 0 3087.1738
130.21246337890625 0 400.45792
133.08596801757812 0 1054.4983
134.9217071533203 0 387.05798
136.0757293701172 0 585.5661
148.87608337402344 0 513.5768
148.88314819335938 0 455.6857
148.8981475830078 0 461.863
148.9123077392578 0 861.21313
148.91986083984375 0 1261.0875
148.92710876464844 0 1165.0702
148.93411254882812 0 2136.0747
148.94195556640625 0 3685.8528
148.95864868164062 0 4632.292
148.9665985107422 0 2872.7727
148.9737548828125 0 1272.1937
148.9812774658203 0 1281.7722
148.98841857910156 0 748.0733
148.99571228027344 0 838.8642
149.00282287597656 0 526.41724
149.01719665527344 0 650.5548
149.0252227783203 0 551.3353
149.03213500976562 0 523.99207
149.04537963867188 0 2102.0527
149.90536499023438 0 448.5664
155.6481475830078 0 499.16226
156.50576782226562 0 495.17877
167.05526733398438 0 617.00385
173.4500274658203 0 937.6385
175.086669921875 0 788.79694
177.10243225097656 0 873.4961
177.11160278320312 0 528.4455
203.08184814453125 0 1372.243
203.117431640625 0 798.1928
221.08445739746094 0 5911.852 y 8
221.0944366455078 0 1676.328
221.1287078857422 0 80537.08
222.08523559570312 0 861.1775
222.13206481933594 0 10487.22
222.93612670898438 0 506.32114
223.08209228515625 0 611.8914
225.0431671142578 0 2274.9612
226.04278564453125 0 1151.741
231.11288452148438 0 1015.5178
235.10765075683594 0 756.3197
239.0952606201172 0 9520.307
240.09584045410156 0 1701.6954 c Ammonia loss 3
241.09140014648438 0 1039.652
249.1236572265625 0 32797.25 c Ammonia loss 1
250.12698364257812 0 3442.787
254.76686096191406 0 520.7064
257.1073303222656 0 607.41705
263.1033020019531 0 1723.167
281.0517272949219 0 1448.0713
285.1017761230469 0 1011.1363
295.1034851074219 0 3358.225
296.1043701171875 0 1413.0085
297.10125732421875 0 1222.238
299.06195068359375 0 6056.9155
309.05950927734375 0 526.17444
313.1148681640625 0 555.6436
314.11297607421875 0 572.409
342.1322937011719 0 2496.2532 z 7
343.1401672363281 0 6275.1963
344.142333984375 0 538.5323
346.1400451660156 0 1032.5917
355.07012939453125 0 44660.383
358.15130615234375 0 9493.223 y 7
359.0285949707031 0 652.7579
359.154541015625 0 1820.1432
362.5925598144531 0 479.2654
364.15045166015625 0 7016.5635 c Ammonia loss 2
365.15374755859375 0 1477.6677
369.12249755859375 0 1354.4175
370.1229248046875 0 949.5735
371.1010437011719 0 2184.126
371.1236267089844 0 870.8105
378.12933349609375 0 1686.885
394.1224670410156 0 1011.0138
415.0367736816406 0 2309.6787
415.15264892578125 0 6110.6157
416.1546325683594 0 1414.2573
418.99713134765625 0 973.67694
433.1729431152344 0 594.03375
441.1882019042969 0 984.1087
458.1576843261719 0 1490.4219
461.1676025390625 0 662.4313
479.1783447265625 0 4451.7295 c Ammonia loss 3
480.1811218261719 0 733.56494
485.1757507324219 0 1028.4863 y 2
489.05712890625 0 1832.229
489.1680908203125 0 2680.12
490.17596435546875 0 13109.705
491.1783447265625 0 3364.22
504.1792297363281 0 4835.9053
505.1866149902344 0 6456.94 y 6
506.18682861328125 0 1131.8932
512.223388671875 0 1245.8838
532.1727905273438 0 704.1077
558.708740234375 0 5782.856 y 1
559.210205078125 0 3782.611
559.7119140625 0 1401.2374
560.2051391601562 0 23629.957 z 5
561.2096557617188 0 10435.787
562.2102661132812 0 2166.3794
565.2301025390625 0 628.058
575.2183227539062 0 1163.3955
576.2235107421875 0 13089.122 y 5
577.1270141601562 0 1467.3136
577.2276611328125 0 5371.54
578.2166137695312 0 6599.3647
579.2183227539062 0 2142.6572
591.2235717773438 0 1170.0247
591.3590087890625 0 1932.4402
596.23388671875 0 700.8879
598.7767944335938 0 673.6429
600.2274169921875 0 2666.448
600.7284545898438 0 2622.791
601.22900390625 0 1497.7584
603.210205078125 0 758.56305
608.3872680664062 0 1032.9331
609.2326049804688 0 9192.704
609.733642578125 0 4986.037
610.1842041015625 0 9261.783
610.2355346679688 0 2463.76
614.245361328125 0 2106.9912
615.2490844726562 0 696.17664
624.2296142578125 0 1084.9076
642.2408447265625 0 1664.0586 c Ammonia loss 4
649.2531127929688 0 10408.197
650.25634765625 0 3689.0464
651.2557373046875 0 714.1246
662.2593994140625 0 1159.3331
663.2680053710938 0 807.3454
675.288330078125 0 1081.3668
693.2429809570312 0 10189.591
694.24609375 0 3445.1113
695.2494506835938 0 1212.8442
707.251708984375 0 735.1492
708.2534790039062 0 707.5194
721.264404296875 0 1401.6631
723.2686157226562 0 32334.268 z 4
724.2719116210938 0 13673.56
725.27294921875 0 3987.6797
731.3138427734375 0 1653.2057
731.3859252929688 0 672.57837
733.3148803710938 0 669.58636
738.2785034179688 0 3237.7417
739.2866821289062 0 12683.578 y 4
740.2898559570312 0 5156.171
741.2864379882812 0 1725.6375
749.25634765625 0 3460.578
764.2802124023438 0 7843.0264
765.28271484375 0 3713.2537
766.2755737304688 0 1869.1105
767.2708740234375 0 735.919
775.3058471679688 0 1941.4021
776.3077392578125 0 1677.4408
777.289794921875 0 1828.2991
779.2830810546875 0 788.9699
790.3167114257812 0 1638.6868
791.313720703125 0 643.6096
794.3055419921875 0 14270.44
795.3087158203125 0 6453.9053
796.306396484375 0 2383.8901
808.2689208984375 0 4610.302
809.2734375 0 2152.2993
810.271240234375 0 871.5294
833.328125 0 2651.4841
834.3338623046875 0 9719.484
835.3366088867188 0 3714.4358
836.3323974609375 0 764.4805
836.4207763671875 0 1130.683
838.2189331054688 0 650.5994
838.2955932617188 0 35042.406 z 3
839.3010864257812 0 36831.062
840.3040161132812 0 15762.229
841.3040161132812 0 3575.495
846.3419799804688 0 1488.581
847.3456420898438 0 1255.8619
853.3095092773438 0 1108.2987
854.31396484375 0 13542.444 y 3
855.3173217773438 0 6388.026
856.3163452148438 0 1961.9329
858.3218383789062 0 2289.2104
859.3191528320312 0 782.67255 c Water loss 6
860.3181762695312 0 792.6124 c Ammonia loss 6
865.3407592773438 0 4142.795
866.3475952148438 0 2085.744
876.3322143554688 0 6000.99
877.3391723632812 0 21621.266 c 6
877.501220703125 0 873.4075
878.342041015625 0 11416.562
879.3435668945312 0 3677.1829
890.3287353515625 0 886.3213
891.3344116210938 0 1271.6808
905.3423461914062 0 9263.682
906.3455810546875 0 4671.4585
907.3523559570312 0 1666.2224
909.3322143554688 0 15109.669
910.3344116210938 0 7276.578
911.33349609375 0 2557.1594
951.3286743164062 0 3670.8865 y Water loss 2
952.338623046875 0 1506.2355
953.322509765625 0 4800.2563 z 2
954.3250122070312 0 4610.852
955.3304443359375 0 2011.0184
956.3442993164062 0 667.0937
969.341064453125 0 67311.16 y 2
970.3442993164062 0 37254.645
971.3450317382812 0 10705.941
972.344970703125 0 1636.1586
981.379150390625 0 3151.1572
982.3656616210938 0 2480.4836
983.3773803710938 0 1002.1014
997.3751220703125 0 2021.2144 c Ammonia loss 7
998.3732299804688 0 786.39215
999.3880615234375 0 11395.223
1000.3901977539062 0 4544.7495
1000.4921264648438 0 1086.4712
1001.3880004882812 0 1668.7856
1001.4832153320312 0 978.20715
1011.4093017578125 0 915.1638
1013.3929443359375 0 1256.1831
1014.3991088867188 0 77603.266 c 7
1015.4015502929688 0 42871.504
1016.4022827148438 0 15278.631
1017.4065551757812 0 1846.3152
1040.413818359375 0 8968.828
1041.41650390625 0 5061.0796
1042.4122314453125 0 2253.6057
1054.5140380859375 0 817.8969
1055.4013671875 0 9891.164
1056.39013671875 0 27326.115
1057.391357421875 0 12839.128
1058.3897705078125 0 3757.6492
1069.393310546875 0 2959.9204
1070.4010009765625 0 1560.181
1071.395751953125 0 6467.074
1072.4017333984375 0 2402.2454
1073.4044189453125 0 1521.0597
1084.4039306640625 0 845.9669
1085.3909912109375 0 714.85046
1089.4923095703125 0 635.5154
1100.3917236328125 0 2618.5342 z 1
1101.3944091796875 0 1844.686
1102.3924560546875 0 1533.8954
1110.4764404296875 0 844.90546
1112.468505859375 0 815.77686
1116.4093017578125 0 3460.2266 y 1
1117.412841796875 0 1557.0408
1126.43212890625 0 1329.0889
1127.434814453125 0 843.5994
1138.4498291015625 0 2496.6777
1139.447998046875 0 1913.4404
1140.44677734375 0 1954.8716
1154.463134765625 0 2344.2993
1155.4625244140625 0 2231.948
1156.46337890625 0 17881.389
1157.462890625 0 15829.107
1158.4476318359375 0 31686.678
1159.44677734375 0 19161.125
1159.56982421875 0 773.73425
1160.448974609375 0 6729.862
1161.44482421875 0 887.8979
1174.4658203125 0 1086.6763
1182.44580078125 0 1448.361
1183.4329833984375 0 2065.5342
1184.4417724609375 0 1022.2715
1186.4161376953125 0 1442.3724
1190.4688720703125 0 2452.764
1191.469970703125 0 2248.1614
1198.694580078125 0 715.39606
1200.4534912109375 0 27731.352
1201.4437255859375 0 128261.45
1202.4454345703125 0 84044.586
1203.44580078125 0 36226.723
1204.447021484375 0 5962.5776
1215.543212890625 0 996.3202
1216.5723876953125 0 1529.2294
1217.45654296875 0 77417.74
1218.4625244140625 0 265171.75
1219.4654541015625 0 173808.98
1220.4674072265625 0 68332.37
1221.1962890625 0 988.4116
1221.4696044921875 0 8266.6
1521.33447265625 0 730.9146
1786.9515380859375 0 771.93396
1806.935791015625 0 684.34375
1807.9267578125 0 1938.5234
1808.936279296875 0 830.934
1823.94140625 0 784.2481
1824.9456787109375 0 1521.0908
1825.95068359375 0 2104.7468
1829.9046630859375 0 829.30334
1831.947021484375 0 1718.1456
3081.650634765625 0 788.681

Spectrum Details

|  |  |
| --- | --- |
| Matched peaks? Matched peaks The total absolute number of peaks matched. Additionally in brackets the total fraction of peaks matched and the total number of peaks is shown. | 27 (9.09% of 297) |
| FDR? FDR The false discovery rate estimated for this peptide. It is calculated by matching all theoretical fragments with a non-integer shift with the raw peaks for this spectrum. This is done with 40 different shifts. The resulting percentage is the average number of annotated peaks over the number of annotated peaks with the correct spectrum. | 0.88% |
| Satellite FDR? Satellite FDR See the FDR for details on its calculation. This satellite ion specific FDR only contains the satellite ions (d/w) for I/L/J positions. | - |
| PSM Score? PSM Score The PSM Score as given by Hecklib to this annotated spectrum. It is shown with three significant figures. | 362 |

#### Spectrum 11784? Spectrum 11784 The raw spectrum of this peptide as annotated by Hecklib. The fragments are coloured according to ion type (see legend). Any peaks with a star '\*' as text can be hovered over to see the full details, first the ion type second the mass shift type. By hovering over the amino acids in the peptide or ions in the legend the corresponding peaks are highlighted. By toggling the 'Unassigned' label you can turn the background (unassigned) peaks on or off in the plot. By updating the slider in the Ion legend you can update the spectrum to only show the top X% of the peaks with labels. The top X% means any peak that is within X% of the highest intensity. By dragging in the spectrum you can zoom in to a specific part of the spectrum and use 'Zoom Out' to get back to the original zoom level. The annotation of the spectrum is based on the given sequence in the peptides file and is done with different software so inconsistencies are likely. The peaks are annotated based on the given sequence, with 20 ppm tolerance.

Copy Data

##### Spectrum 11784 (TSV)

###### Preview

```
Loading example...
```

*Click on the button to copy the data to your clipboard.*

Mz MinMz MaxIntensity Max

WidthHeightPeptide font sizePeptide stroke widthSpectrum font sizeSpectrum stroke widthCompact peptide

Ion legend

wxyz

abcd

OtherUnassignedIon^Charge^_Position_Show for top:%

TFDDYAMHW

01.24e+42.48e+43.72e+44.96e+4

Zoom Out

a^+1^_2_y^+1^_1_a^+1^_2_b^+1^_2_y^+1^_2_b^+1^_3_b^+1^_4_y^+1^_3_y^+2^_8_y^+1^_4_\*y^+1^_5_y^+1^_6_y^+1^_7_y^+1^_7_y^+1^_8_

0778155523333111

Fragment Matches Table

Show background peaks

| Position | Ion type | Intensity | mz Theoretical | mz Error (Th) | mz Error (ppm) | Charge | Series Number |
| --- | --- | --- | --- | --- | --- | --- | --- |
| - | - | 3.213E+04 | 120.1 | - | - | 0 | - |
| - | - | 3438 | 121.1 | - | - | 0 | - |
| - | - | 616.9 | 123 | - | - | 0 | - |
| - | - | 377.7 | 129 | - | - | 0 | - |
| - | - | 3198 | 129.1 | - | - | 0 | - |
| - | - | 862.4 | 130.1 | - | - | 0 | - |
| - | - | 424.3 | 132.1 | - | - | 0 | - |
| - | - | 410.1 | 132.3 | - | - | 0 | - |
| - | - | 1079 | 133.1 | - | - | 0 | - |
| - | - | 474 | 134.1 | - | - | 0 | - |
| - | - | 1.191E+04 | 136.1 | - | - | 0 | - |
| - | - | 644.6 | 137.1 | - | - | 0 | - |
| - | - | 741.1 | 138.1 | - | - | 0 | - |
| - | - | 574 | 145.1 | - | - | 0 | - |
| - | - | 1012 | 149 | - | - | 0 | - |
| - | - | 470.2 | 150.2 | - | - | 0 | - |
| - | - | 480.5 | 153 | - | - | 0 | - |
| - | - | 1369 | 156.1 | - | - | 0 | - |
| - | - | 438.4 | 157.1 | - | - | 0 | - |
| - | - | 1652 | 158.1 | - | - | 0 | - |
| - | - | 427.8 | 158.8 | - | - | 0 | - |
| - | - | 1515 | 159.1 | - | - | 0 | - |
| - | - | 448.6 | 159.6 | - | - | 0 | - |
| - | - | 732.7 | 160.1 | - | - | 0 | - |
| - | - | 775.4 | 160.1 | - | - | 0 | - |
| - | - | 538.6 | 162.1 | - | - | 0 | - |
| - | - | 519.4 | 164.4 | - | - | 0 | - |
| - | - | 4029 | 165.1 | - | - | 0 | - |
| - | - | 549.2 | 166.1 | - | - | 0 | - |
| - | - | 1282 | 166.1 | - | - | 0 | - |
| - | - | 1783 | 167.1 | - | - | 0 | - |
| - | - | 610.5 | 167.1 | - | - | 0 | - |
| - | - | 848.6 | 169.1 | - | - | 0 | - |
| - | - | 419.7 | 170 | - | - | 0 | - |
| - | - | 3003 | 173.1 | - | - | 0 | - |
| - | - | 531 | 176.1 | - | - | 0 | - |
| - | - | 3377 | 176.1 | - | - | 0 | - |
| - | - | 2490 | 177.1 | - | - | 0 | - |
| - | - | 628.1 | 177.1 | - | - | 0 | - |
| - | - | 6871 | 182.1 | - | - | 0 | - |
| - | - | 699.6 | 183.1 | - | - | 0 | - |
| - | - | 684.4 | 183.1 | - | - | 0 | - |
| - | - | 736.3 | 185.1 | - | - | 0 | - |
| - | - | 1858 | 188.1 | - | - | 0 | - |
| - | - | 677.7 | 191.1 | - | - | 0 | - |
| - | - | 1050 | 193.1 | - | - | 0 | - |
| - | - | 620.2 | 197.1 | - | - | 0 | - |
| - | - | 627.8 | 199.1 | - | - | 0 | - |
| - | - | 1417 | 201.1 | - | - | 0 | - |
| - | - | 691.3 | 203.1 | - | - | 0 | - |
| - | - | 493 | 203.1 | - | - | 0 | - |
| 2 | a | 1691 | 203.1 | 0.0005339 | 2.629 | +1 | 2 |
| 9 | y | 1390 | 205.1 | 0.0003191 | 1.556 | +1 | 1 |
| - | - | 529.6 | 207.9 | - | - | 0 | - |
| - | - | 729.9 | 219.1 | - | - | 0 | - |
| - | - | 2942 | 221.1 | - | - | 0 | - |
| 2 | a | 4.914E+04 | 221.1 | 0.0001621 | 0.7331 | +1 | 2 |
| - | - | 6902 | 222.1 | - | - | 0 | - |
| - | - | 1302 | 223.1 | - | - | 0 | - |
| - | - | 495.2 | 223.1 | - | - | 0 | - |
| - | - | 486.5 | 224.3 | - | - | 0 | - |
| - | - | 1349 | 227.1 | - | - | 0 | - |
| - | - | 520.5 | 229.2 | - | - | 0 | - |
| - | - | 584.5 | 229.2 | - | - | 0 | - |
| - | - | 1319 | 235.1 | - | - | 0 | - |
| - | - | 1323 | 239.1 | - | - | 0 | - |
| - | - | 820.2 | 241.1 | - | - | 0 | - |
| 2 | b | 7889 | 249.1 | 0.0002273 | 0.9126 | +1 | 2 |
| - | - | 1091 | 250.1 | - | - | 0 | - |
| - | - | 1084 | 251.1 | - | - | 0 | - |
| - | - | 1648 | 263.1 | - | - | 0 | - |
| - | - | 993.1 | 279.1 | - | - | 0 | - |
| - | - | 874.6 | 281.1 | - | - | 0 | - |
| - | - | 539.8 | 282.9 | - | - | 0 | - |
| - | - | 1507 | 285 | - | - | 0 | - |
| - | - | 1431 | 285.1 | - | - | 0 | - |
| - | - | 663.5 | 285.2 | - | - | 0 | - |
| - | - | 750.2 | 292.1 | - | - | 0 | - |
| - | - | 830.1 | 309.2 | - | - | 0 | - |
| - | - | 1416 | 324.1 | - | - | 0 | - |
| - | - | 784.9 | 325.1 | - | - | 0 | - |
| - | - | 4591 | 329.1 | - | - | 0 | - |
| - | - | 966.1 | 330.2 | - | - | 0 | - |
| - | - | 882.7 | 340.2 | - | - | 0 | - |
| 8 | y | 9786 | 342.2 | 0.0002756 | 0.8055 | +1 | 2 |
| - | - | 1953 | 343.2 | - | - | 0 | - |
| - | - | 2563 | 355.1 | - | - | 0 | - |
| - | - | 704.9 | 356.1 | - | - | 0 | - |
| - | - | 1075 | 357.2 | - | - | 0 | - |
| 3 | b | 2213 | 364.2 | 0.0001959 | 0.538 | +1 | 3 |
| - | - | 1067 | 378.1 | - | - | 0 | - |
| - | - | 573.6 | 390.2 | - | - | 0 | - |
| - | - | 1151 | 394.1 | - | - | 0 | - |
| - | - | 754.6 | 395.2 | - | - | 0 | - |
| - | - | 833.4 | 398.2 | - | - | 0 | - |
| - | - | 969.6 | 413.3 | - | - | 0 | - |
| - | - | 704.2 | 417.1 | - | - | 0 | - |
| - | - | 1103 | 425.2 | - | - | 0 | - |
| - | - | 532.2 | 425.3 | - | - | 0 | - |
| - | - | 686.3 | 427.7 | - | - | 0 | - |
| - | - | 891.3 | 453.3 | - | - | 0 | - |
| - | - | 776.8 | 465.2 | - | - | 0 | - |
| - | - | 593.4 | 466.2 | - | - | 0 | - |
| 4 | b | 1480 | 479.2 | 0.001578 | 3.293 | +1 | 4 |
| 7 | y | 2509 | 489.2 | 0.004947 | 10.11 | +1 | 3 |
| - | - | 686.3 | 490.2 | - | - | 0 | - |
| - | - | 648.7 | 497.1 | - | - | 0 | - |
| - | - | 623.8 | 506.3 | - | - | 0 | - |
| - | - | 1056 | 508.3 | - | - | 0 | - |
| - | - | 607.3 | 533.4 | - | - | 0 | - |
| 2 | y | 1670 | 550.7 | 0.003935 | 7.145 | +2 | 8 |
| - | - | 1189 | 551.2 | - | - | 0 | - |
| 6 | y | 3744 | 560.2 | 0.004302 | 7.679 | +1 | 4 |
| - | - | 964.8 | 561.2 | - | - | 0 | - |
| - | - | 2175 | 585.3 | - | - | 0 | - |
| - | - | 743.7 | 592.7 | - | - | 0 | - |
| 0 | Precursor | 2016 | 601.2 | 0.0028 | 4.658 | +2 | -1 |
| - | - | 872.2 | 601.3 | - | - | 0 | - |
| - | - | 1317 | 601.7 | - | - | 0 | - |
| - | - | 863 | 612.2 | - | - | 0 | - |
| - | - | 920 | 621.4 | - | - | 0 | - |
| - | - | 642.5 | 622.4 | - | - | 0 | - |
| - | - | 738.2 | 634.2 | - | - | 0 | - |
| - | - | 591.2 | 657.3 | - | - | 0 | - |
| - | - | 1344 | 659.3 | - | - | 0 | - |
| - | - | 691.8 | 676.4 | - | - | 0 | - |
| - | - | 984.7 | 685.3 | - | - | 0 | - |
| - | - | 3833 | 700.4 | - | - | 0 | - |
| - | - | 1574 | 701.4 | - | - | 0 | - |
| - | - | 896.8 | 703.3 | - | - | 0 | - |
| - | - | 1174 | 721.3 | - | - | 0 | - |
| 5 | y | 3962 | 723.3 | 0.003779 | 5.224 | +1 | 5 |
| - | - | 1927 | 724.3 | - | - | 0 | - |
| - | - | 1112 | 749.3 | - | - | 0 | - |
| - | - | 1115 | 767.3 | - | - | 0 | - |
| - | - | 1281 | 774.3 | - | - | 0 | - |
| - | - | 804.8 | 783.4 | - | - | 0 | - |
| - | - | 565.7 | 794.5 | - | - | 0 | - |
| - | - | 864 | 800.4 | - | - | 0 | - |
| - | - | 2931 | 801.4 | - | - | 0 | - |
| - | - | 741.2 | 802.4 | - | - | 0 | - |
| - | - | 1186 | 828.4 | - | - | 0 | - |
| - | - | 711.3 | 834.4 | - | - | 0 | - |
| 4 | y | 3970 | 838.3 | 0.003935 | 4.694 | +1 | 6 |
| - | - | 1757 | 839.3 | - | - | 0 | - |
| - | - | 838.7 | 840.3 | - | - | 0 | - |
| - | - | 2531 | 844.5 | - | - | 0 | - |
| - | - | 1189 | 845.5 | - | - | 0 | - |
| - | - | 1.247E+04 | 872.4 | - | - | 0 | - |
| - | - | 5707 | 873.4 | - | - | 0 | - |
| - | - | 1404 | 874.5 | - | - | 0 | - |
| - | - | 611.3 | 877.6 | - | - | 0 | - |
| - | - | 6983 | 889.3 | - | - | 0 | - |
| - | - | 2388 | 890.3 | - | - | 0 | - |
| - | - | 735.8 | 901.5 | - | - | 0 | - |
| 3 | y | 1384 | 935.3 | 0.006661 | 7.121 | +1 | 7 |
| 3 | y | 1.786E+04 | 953.3 | 0.004214 | 4.42 | +1 | 7 |
| - | - | 9577 | 954.3 | - | - | 0 | - |
| - | - | 2510 | 955.3 | - | - | 0 | - |
| - | - | 2241 | 991.5 | - | - | 0 | - |
| - | - | 875.9 | 992.5 | - | - | 0 | - |
| - | - | 5715 | 1020 | - | - | 0 | - |
| - | - | 3278 | 1021 | - | - | 0 | - |
| - | - | 1100 | 1022 | - | - | 0 | - |
| 2 | y | 695.9 | 1100 | 0.001334 | 1.212 | +1 | 8 |
| - | - | 747.4 | 1101 | - | - | 0 | - |
| - | - | 712.3 | 1323 | - | - | 0 | - |
| - | - | 642.6 | 1328 | - | - | 0 | - |
| - | - | 652.6 | 1471 | - | - | 0 | - |
| - | - | 576.2 | 1801 | - | - | 0 | - |
| - | - | 675.3 | 2913 | - | - | 0 | - |
| - | - | 813.7 | 3080 | - | - | 0 | - |

m/z Charge Intensity FragmentType MassShift Position
120.08100128173828 0 32133.264
121.0843276977539 0 3438.3682
123.04447937011719 0 616.8564
129.04312133789062 0 377.69168
129.1024627685547 0 3198.1868
130.06529235839844 0 862.4178
132.08135986328125 0 424.31342
132.30181884765625 0 410.14633
133.08612060546875 0 1079.4764
134.0964813232422 0 473.99277
136.07591247558594 0 11910.074
137.0791473388672 0 644.64844
138.06634521484375 0 741.0541
145.06112670898438 0 574.0074
149.0447235107422 0 1012.2521
150.23724365234375 0 470.1606
152.95843505859375 0 480.49597
156.07675170898438 0 1368.9883
157.13380432128906 0 438.35767
158.0965576171875 0 1652.3613
158.7579803466797 0 427.78952
159.09190368652344 0 1514.6674
159.62619018554688 0 448.55258
160.07595825195312 0 732.7113
160.11231994628906 0 775.3976
162.09194946289062 0 538.5587
164.38345336914062 0 519.4012
165.0546875 0 4029.447
166.0600128173828 0 549.1867
166.0863494873047 0 1281.6539
167.05555725097656 0 1783.281
167.08168029785156 0 610.511
169.09718322753906 0 848.6355
170.04591369628906 0 419.66956
173.128662109375 0 3003.3513
176.0823516845703 0 531.04694
176.1072235107422 0 3376.7444
177.1022491455078 0 2489.9187
177.1106719970703 0 628.05536
182.0813446044922 0 6870.8223
183.0848388671875 0 699.60034
183.11282348632812 0 684.37286
185.05523681640625 0 736.32214
188.0707550048828 0 1857.7139
191.1179962158203 0 677.7445
193.10877990722656 0 1050.2161
197.12973022460938 0 620.1883
199.10768127441406 0 627.7664
201.12368774414062 0 1416.7063
203.06675720214844 0 691.3123
203.10870361328125 0 493.04007
203.11842346191406 0 1691.2695 a Water loss 1
205.09747314453125 0 1389.8384 y 8
207.86160278320312 0 529.62787
219.11280822753906 0 729.90704
221.10333251953125 0 2942.369
221.1286163330078 0 49144.246 a 1
222.1320343017578 0 6901.56
223.06338500976562 0 1302.2968
223.1331024169922 0 495.1831
224.25148010253906 0 486.47318
227.10324096679688 0 1349.028
229.15646362304688 0 520.5354
229.1693572998047 0 584.53235
235.1082305908203 0 1318.6643
239.0948028564453 0 1323.0598
241.07455444335938 0 820.24207
249.12359619140625 0 7889.448 b 1
250.1266326904297 0 1090.7823
251.1029510498047 0 1084.2341
263.1024475097656 0 1648.3793
279.0976867675781 0 993.0923
281.12384033203125 0 874.6175
282.8840637207031 0 539.7677
285.0096130371094 0 1506.9927
285.1018981933594 0 1431.2144
285.1605224609375 0 663.4894
292.1403503417969 0 750.2134
309.2038879394531 0 830.14777
324.14630126953125 0 1415.961
325.1470031738281 0 784.8627
329.1498107910156 0 4591.371
330.154296875 0 966.1454
340.1851501464844 0 882.7275
342.1563415527344 0 9786.219 y 7
343.1602478027344 0 1953.1995
355.0698547363281 0 2563.1272
356.07012939453125 0 704.87354
357.2247619628906 0 1075.0089
364.1501159667969 0 2213.4016 b 2
378.12908935546875 0 1066.7913
390.1591796875 0 573.62366
394.12353515625 0 1150.8707
395.19268798828125 0 754.62085
398.2038269042969 0 833.3741
413.2662048339844 0 969.63934
417.08636474609375 0 704.159
425.1935729980469 0 1103.16
425.2804870605469 0 532.2071
427.7217102050781 0 686.303
453.2732849121094 0 891.3401
465.1622314453125 0 776.84033
466.1628723144531 0 593.38556
479.1788330078125 0 1480.369 b 3
489.1914978027344 0 2508.7366 y 6
490.19866943359375 0 686.288
497.12384033203125 0 648.70105
506.2651672363281 0 623.79865
508.2781066894531 0 1055.5771
533.4295654296875 0 607.3395
550.7122192382812 0 1669.8796 y 1
551.2134399414062 0 1189.0593
560.2279663085938 0 3743.5083 y 5
561.23095703125 0 964.79315
585.3367919921875 0 2175.107
592.7293701171875 0 743.68176
601.2349243164062 0 2015.7694 Precursor
601.322021484375 0 872.16425
601.73583984375 0 1317.103
612.19677734375 0 863.035
621.3601684570312 0 920.00525
622.3590698242188 0 642.45294
634.2255859375 0 738.1815
657.2584838867188 0 591.2122
659.294677734375 0 1343.8323
676.3715209960938 0 691.83
685.26171875 0 984.73456
700.363525390625 0 3833.0532
701.3638305664062 0 1574.2661
703.2584838867188 0 896.847
721.2622680664062 0 1174.2646
723.290771484375 0 3961.9644 y 4
724.2938232421875 0 1927.1637
749.2564697265625 0 1111.9677
767.267578125 0 1115.2473
774.32080078125 0 1281.3992
783.3973388671875 0 804.7826
794.5311889648438 0 565.70013
800.4251708984375 0 863.951
801.4111938476562 0 2930.5037
802.4146118164062 0 741.23596
828.4244995117188 0 1186.4989
834.4466552734375 0 711.3373
838.31787109375 0 3969.8594 y 3
839.3228759765625 0 1757.3335
840.3159790039062 0 838.68616
844.451904296875 0 2530.8015
845.4529418945312 0 1189.3423
872.4469604492188 0 12469.7
873.449951171875 0 5706.9424
874.4564819335938 0 1404.1067
877.6029052734375 0 611.2739
889.346435546875 0 6982.6284
890.3480224609375 0 2388.1829
901.4808959960938 0 735.7531
935.3369750976562 0 1384.3452 y Water loss 2
953.3450927734375 0 17863.797 y 2
954.348388671875 0 9576.546
955.3485717773438 0 2510.456
991.5202026367188 0 2240.504
992.523193359375 0 875.9365
1019.5158081054688 0 5714.878
1020.5198974609375 0 3277.57
1021.5248413085938 0 1099.5148
1100.407958984375 0 695.8538 y 1
1101.416015625 0 747.38354
1323.264892578125 0 712.2598
1328.188232421875 0 642.5519
1470.92529296875 0 652.624
1800.772705078125 0 576.16406
2913.122314453125 0 675.32385
3080.150146484375 0 813.711

Spectrum Details

|  |  |
| --- | --- |
| Matched peaks? Matched peaks The total absolute number of peaks matched. Additionally in brackets the total fraction of peaks matched and the total number of peaks is shown. | 16 (9.30% of 172) |
| FDR? FDR The false discovery rate estimated for this peptide. It is calculated by matching all theoretical fragments with a non-integer shift with the raw peaks for this spectrum. This is done with 40 different shifts. The resulting percentage is the average number of annotated peaks over the number of annotated peaks with the correct spectrum. | 1.19% |
| Satellite FDR? Satellite FDR See the FDR for details on its calculation. This satellite ion specific FDR only contains the satellite ions (d/w) for I/L/J positions. | - |
| PSM Score? PSM Score The PSM Score as given by Hecklib to this annotated spectrum. It is shown with three significant figures. | 177 |

#### Spectrum 9943? Spectrum 9943 The raw spectrum of this peptide as annotated by Hecklib. The fragments are coloured according to ion type (see legend). Any peaks with a star '\*' as text can be hovered over to see the full details, first the ion type second the mass shift type. By hovering over the amino acids in the peptide or ions in the legend the corresponding peaks are highlighted. By toggling the 'Unassigned' label you can turn the background (unassigned) peaks on or off in the plot. By updating the slider in the Ion legend you can update the spectrum to only show the top X% of the peaks with labels. The top X% means any peak that is within X% of the highest intensity. By dragging in the spectrum you can zoom in to a specific part of the spectrum and use 'Zoom Out' to get back to the original zoom level. The annotation of the spectrum is based on the given sequence in the peptides file and is done with different software so inconsistencies are likely. The peaks are annotated based on the given sequence, with 20 ppm tolerance.

Copy Data

##### Spectrum 9943 (TSV)

###### Preview

```
Loading example...
```

*Click on the button to copy the data to your clipboard.*

Mz MinMz MaxIntensity Max

WidthHeightPeptide font sizePeptide stroke widthSpectrum font sizeSpectrum stroke widthCompact peptide

Ion legend

wxyz

abcd

OtherUnassignedIon^Charge^_Position_Show for top:%

TFDDYAMHW

02.38e+44.77e+47.15e+49.54e+4

Zoom Out

a^+1^_2_y^+1^_1_a^+1^_2_b^+1^_2_b^+1^_2_a^+1^_3_y^+1^_2_b^+1^_3_b^+1^_3_b^+1^_4_b^+1^_4_y^+1^_3_y^+2^_8_y^+1^_4_\*y^+1^_5_y^+1^_6_y^+1^_7_y^+1^_7_y^+1^_8_

0800159923993199

Fragment Matches Table

Show background peaks

| Position | Ion type | Intensity | mz Theoretical | mz Error (Th) | mz Error (ppm) | Charge | Series Number |
| --- | --- | --- | --- | --- | --- | --- | --- |
| - | - | 8.438E+04 | 120.1 | - | - | 0 | - |
| - | - | 6748 | 121.1 | - | - | 0 | - |
| - | - | 1931 | 123 | - | - | 0 | - |
| - | - | 419.4 | 125.1 | - | - | 0 | - |
| - | - | 454.3 | 128.1 | - | - | 0 | - |
| - | - | 1154 | 129.1 | - | - | 0 | - |
| - | - | 4728 | 129.1 | - | - | 0 | - |
| - | - | 2163 | 130.1 | - | - | 0 | - |
| - | - | 591.8 | 130.1 | - | - | 0 | - |
| - | - | 719 | 132.1 | - | - | 0 | - |
| - | - | 1187 | 132.1 | - | - | 0 | - |
| - | - | 3376 | 132.1 | - | - | 0 | - |
| - | - | 743.3 | 133.1 | - | - | 0 | - |
| - | - | 899 | 133.1 | - | - | 0 | - |
| - | - | 3.036E+04 | 136.1 | - | - | 0 | - |
| - | - | 1825 | 137.1 | - | - | 0 | - |
| - | - | 628.8 | 138.1 | - | - | 0 | - |
| - | - | 874 | 138.1 | - | - | 0 | - |
| - | - | 1588 | 141.1 | - | - | 0 | - |
| - | - | 492.5 | 143.1 | - | - | 0 | - |
| - | - | 948.9 | 143.1 | - | - | 0 | - |
| - | - | 2090 | 146.1 | - | - | 0 | - |
| - | - | 469.6 | 146.4 | - | - | 0 | - |
| - | - | 1544 | 147 | - | - | 0 | - |
| - | - | 1021 | 149 | - | - | 0 | - |
| - | - | 1040 | 149 | - | - | 0 | - |
| - | - | 503.5 | 154.9 | - | - | 0 | - |
| - | - | 533.7 | 155.1 | - | - | 0 | - |
| - | - | 598.6 | 155.1 | - | - | 0 | - |
| - | - | 1954 | 156.1 | - | - | 0 | - |
| - | - | 1.004E+04 | 157.1 | - | - | 0 | - |
| - | - | 1049 | 158.1 | - | - | 0 | - |
| - | - | 3615 | 158.1 | - | - | 0 | - |
| - | - | 2710 | 159.1 | - | - | 0 | - |
| - | - | 549.1 | 159.1 | - | - | 0 | - |
| - | - | 1839 | 159.1 | - | - | 0 | - |
| - | - | 424.5 | 159.4 | - | - | 0 | - |
| - | - | 1238 | 160.1 | - | - | 0 | - |
| - | - | 1938 | 161.1 | - | - | 0 | - |
| - | - | 433.4 | 164.6 | - | - | 0 | - |
| - | - | 1.4E+04 | 165.1 | - | - | 0 | - |
| - | - | 628.4 | 165.1 | - | - | 0 | - |
| - | - | 885.1 | 166.1 | - | - | 0 | - |
| - | - | 2062 | 166.1 | - | - | 0 | - |
| - | - | 2121 | 167.1 | - | - | 0 | - |
| - | - | 427.4 | 169.1 | - | - | 0 | - |
| - | - | 2369 | 169.1 | - | - | 0 | - |
| - | - | 1379 | 171.1 | - | - | 0 | - |
| - | - | 865.2 | 171.1 | - | - | 0 | - |
| - | - | 6449 | 173.1 | - | - | 0 | - |
| - | - | 2467 | 174.1 | - | - | 0 | - |
| - | - | 1493 | 175.1 | - | - | 0 | - |
| - | - | 518.7 | 176.1 | - | - | 0 | - |
| - | - | 6665 | 176.1 | - | - | 0 | - |
| - | - | 588.9 | 177.1 | - | - | 0 | - |
| - | - | 5612 | 177.1 | - | - | 0 | - |
| - | - | 892.7 | 177.1 | - | - | 0 | - |
| - | - | 576.7 | 181.1 | - | - | 0 | - |
| - | - | 2.298E+04 | 182.1 | - | - | 0 | - |
| - | - | 801.3 | 183.1 | - | - | 0 | - |
| - | - | 2074 | 183.1 | - | - | 0 | - |
| - | - | 550.8 | 183.1 | - | - | 0 | - |
| - | - | 1032 | 183.1 | - | - | 0 | - |
| - | - | 1372 | 185.1 | - | - | 0 | - |
| - | - | 7335 | 185.1 | - | - | 0 | - |
| - | - | 478.1 | 185.2 | - | - | 0 | - |
| - | - | 515.5 | 186.1 | - | - | 0 | - |
| - | - | 866.7 | 186.1 | - | - | 0 | - |
| - | - | 2467 | 187.1 | - | - | 0 | - |
| - | - | 871.4 | 187.1 | - | - | 0 | - |
| - | - | 1227 | 187.1 | - | - | 0 | - |
| - | - | 3835 | 188.1 | - | - | 0 | - |
| - | - | 541.7 | 188.9 | - | - | 0 | - |
| - | - | 2063 | 189.1 | - | - | 0 | - |
| - | - | 529.4 | 191.1 | - | - | 0 | - |
| - | - | 2268 | 191.1 | - | - | 0 | - |
| - | - | 486.5 | 191.9 | - | - | 0 | - |
| - | - | 2805 | 193.1 | - | - | 0 | - |
| - | - | 1254 | 195.1 | - | - | 0 | - |
| - | - | 700.9 | 197.1 | - | - | 0 | - |
| - | - | 637.9 | 197.1 | - | - | 0 | - |
| - | - | 794.2 | 199.1 | - | - | 0 | - |
| - | - | 3243 | 201.1 | - | - | 0 | - |
| - | - | 1243 | 202.1 | - | - | 0 | - |
| - | - | 729.7 | 203.1 | - | - | 0 | - |
| - | - | 624.8 | 203.1 | - | - | 0 | - |
| 2 | a | 1805 | 203.1 | 0.000244 | 1.201 | +1 | 2 |
| - | - | 835.3 | 204.1 | - | - | 0 | - |
| 9 | y | 2950 | 205.1 | 0.0003191 | 1.556 | +1 | 1 |
| - | - | 616.9 | 207.1 | - | - | 0 | - |
| - | - | 537.2 | 208.1 | - | - | 0 | - |
| - | - | 523 | 209.1 | - | - | 0 | - |
| - | - | 920.9 | 211.1 | - | - | 0 | - |
| - | - | 1366 | 212.1 | - | - | 0 | - |
| - | - | 471.1 | 213.1 | - | - | 0 | - |
| - | - | 1764 | 214.1 | - | - | 0 | - |
| - | - | 1411 | 215.1 | - | - | 0 | - |
| - | - | 509.9 | 217.1 | - | - | 0 | - |
| - | - | 1073 | 217.1 | - | - | 0 | - |
| - | - | 502.4 | 219.1 | - | - | 0 | - |
| - | - | 861.2 | 219.1 | - | - | 0 | - |
| - | - | 6009 | 221.1 | - | - | 0 | - |
| 2 | a | 9.442E+04 | 221.1 | 0.0002994 | 1.354 | +1 | 2 |
| - | - | 650.5 | 222.1 | - | - | 0 | - |
| - | - | 1.145E+04 | 222.1 | - | - | 0 | - |
| - | - | 1445 | 223.1 | - | - | 0 | - |
| - | - | 807.8 | 223.1 | - | - | 0 | - |
| - | - | 1194 | 225.2 | - | - | 0 | - |
| - | - | 1823 | 228.1 | - | - | 0 | - |
| - | - | 878.6 | 228.1 | - | - | 0 | - |
| - | - | 961.5 | 229.1 | - | - | 0 | - |
| - | - | 660.1 | 229.2 | - | - | 0 | - |
| - | - | 908.4 | 229.2 | - | - | 0 | - |
| - | - | 1567 | 231.1 | - | - | 0 | - |
| 2 | b | 1119 | 231.1 | 0.0007589 | 3.284 | +1 | 2 |
| - | - | 877.9 | 231.1 | - | - | 0 | - |
| - | - | 551.6 | 233.1 | - | - | 0 | - |
| - | - | 6708 | 233.1 | - | - | 0 | - |
| - | - | 3234 | 235.1 | - | - | 0 | - |
| - | - | 791.3 | 239.1 | - | - | 0 | - |
| - | - | 1664 | 239.1 | - | - | 0 | - |
| - | - | 714.4 | 239.1 | - | - | 0 | - |
| - | - | 950.1 | 239.2 | - | - | 0 | - |
| - | - | 1333 | 240.1 | - | - | 0 | - |
| - | - | 931.8 | 240.1 | - | - | 0 | - |
| - | - | 782.6 | 241.1 | - | - | 0 | - |
| - | - | 677.9 | 246.1 | - | - | 0 | - |
| 2 | b | 1.64E+04 | 249.1 | 0.0003189 | 1.28 | +1 | 2 |
| - | - | 2213 | 250.1 | - | - | 0 | - |
| - | - | 1578 | 251.1 | - | - | 0 | - |
| - | - | 824.1 | 253.1 | - | - | 0 | - |
| - | - | 1040 | 253.2 | - | - | 0 | - |
| - | - | 763.7 | 255.1 | - | - | 0 | - |
| - | - | 594.2 | 255.1 | - | - | 0 | - |
| - | - | 497.3 | 257.1 | - | - | 0 | - |
| - | - | 712.5 | 257.2 | - | - | 0 | - |
| - | - | 716.1 | 258.1 | - | - | 0 | - |
| - | - | 531.3 | 260.1 | - | - | 0 | - |
| - | - | 3746 | 263.1 | - | - | 0 | - |
| - | - | 659.5 | 274.1 | - | - | 0 | - |
| - | - | 853.3 | 279.1 | - | - | 0 | - |
| - | - | 644.1 | 282.2 | - | - | 0 | - |
| - | - | 803.7 | 283 | - | - | 0 | - |
| - | - | 1043 | 283.1 | - | - | 0 | - |
| - | - | 2976 | 284.2 | - | - | 0 | - |
| - | - | 701.9 | 285 | - | - | 0 | - |
| - | - | 2364 | 285.1 | - | - | 0 | - |
| - | - | 539.4 | 287.5 | - | - | 0 | - |
| - | - | 523.9 | 290.1 | - | - | 0 | - |
| - | - | 1523 | 292.1 | - | - | 0 | - |
| - | - | 1911 | 296.2 | - | - | 0 | - |
| - | - | 846.4 | 297.1 | - | - | 0 | - |
| - | - | 805.7 | 302.1 | - | - | 0 | - |
| - | - | 768.1 | 305.1 | - | - | 0 | - |
| - | - | 645.8 | 317.2 | - | - | 0 | - |
| 3 | a | 1613 | 318.1 | 0.000248 | 0.7795 | +1 | 3 |
| - | - | 1590 | 319.1 | - | - | 0 | - |
| - | - | 658.7 | 322.2 | - | - | 0 | - |
| - | - | 596.2 | 323.1 | - | - | 0 | - |
| - | - | 4301 | 324.1 | - | - | 0 | - |
| - | - | 836.2 | 325.1 | - | - | 0 | - |
| - | - | 658.8 | 325.2 | - | - | 0 | - |
| - | - | 1.596E+04 | 329.1 | - | - | 0 | - |
| - | - | 2883 | 330.2 | - | - | 0 | - |
| - | - | 3345 | 333.1 | - | - | 0 | - |
| - | - | 556.7 | 338.1 | - | - | 0 | - |
| - | - | 683.5 | 340.2 | - | - | 0 | - |
| - | - | 2641 | 340.2 | - | - | 0 | - |
| - | - | 828.7 | 341.1 | - | - | 0 | - |
| 8 | y | 1.722E+04 | 342.2 | 0.0004892 | 1.43 | +1 | 2 |
| - | - | 2967 | 343.2 | - | - | 0 | - |
| - | - | 2034 | 343.2 | - | - | 0 | - |
| 3 | b | 1185 | 346.1 | 0.0008168 | 2.36 | +1 | 3 |
| - | - | 1095 | 350.1 | - | - | 0 | - |
| - | - | 1518 | 355.1 | - | - | 0 | - |
| - | - | 732.5 | 355.2 | - | - | 0 | - |
| - | - | 1919 | 356.1 | - | - | 0 | - |
| - | - | 633.8 | 357.1 | - | - | 0 | - |
| - | - | 3707 | 357.2 | - | - | 0 | - |
| - | - | 703 | 361.2 | - | - | 0 | - |
| 3 | b | 4325 | 364.2 | 4.823E-05 | 0.1325 | +1 | 3 |
| - | - | 851.4 | 365.2 | - | - | 0 | - |
| - | - | 700.4 | 366.1 | - | - | 0 | - |
| - | - | 1216 | 368.2 | - | - | 0 | - |
| - | - | 3141 | 378.1 | - | - | 0 | - |
| - | - | 591.9 | 382.1 | - | - | 0 | - |
| - | - | 3819 | 394.1 | - | - | 0 | - |
| - | - | 1499 | 398.2 | - | - | 0 | - |
| - | - | 691.1 | 419.2 | - | - | 0 | - |
| - | - | 761.7 | 422.2 | - | - | 0 | - |
| - | - | 878.4 | 422.7 | - | - | 0 | - |
| - | - | 1985 | 424.3 | - | - | 0 | - |
| - | - | 652.6 | 424.8 | - | - | 0 | - |
| - | - | 1989 | 425.2 | - | - | 0 | - |
| - | - | 628.5 | 426.2 | - | - | 0 | - |
| - | - | 1014 | 427.2 | - | - | 0 | - |
| - | - | 1880 | 427.7 | - | - | 0 | - |
| - | - | 1188 | 428.2 | - | - | 0 | - |
| - | - | 5535 | 432.2 | - | - | 0 | - |
| - | - | 1561 | 436.7 | - | - | 0 | - |
| - | - | 742.8 | 444.7 | - | - | 0 | - |
| - | - | 556.4 | 451.9 | - | - | 0 | - |
| - | - | 1040 | 454.2 | - | - | 0 | - |
| 4 | b | 681.8 | 461.2 | 0.0002714 | 0.5886 | +1 | 4 |
| - | - | 2747 | 465.2 | - | - | 0 | - |
| - | - | 733 | 470.3 | - | - | 0 | - |
| - | - | 1819 | 471.3 | - | - | 0 | - |
| 4 | b | 2540 | 479.2 | 0.0004489 | 0.9369 | +1 | 4 |
| - | - | 628.2 | 482.2 | - | - | 0 | - |
| - | - | 881.7 | 483.2 | - | - | 0 | - |
| - | - | 946.9 | 487.3 | - | - | 0 | - |
| - | - | 712.8 | 487.7 | - | - | 0 | - |
| 7 | y | 3844 | 489.2 | 0.004764 | 9.739 | +1 | 3 |
| - | - | 1085 | 490.2 | - | - | 0 | - |
| - | - | 698.6 | 495.2 | - | - | 0 | - |
| - | - | 1939 | 496.2 | - | - | 0 | - |
| - | - | 873.9 | 496.3 | - | - | 0 | - |
| - | - | 662 | 497.2 | - | - | 0 | - |
| - | - | 795.9 | 500.2 | - | - | 0 | - |
| - | - | 1863 | 501.3 | - | - | 0 | - |
| - | - | 3026 | 508.3 | - | - | 0 | - |
| - | - | 1381 | 508.8 | - | - | 0 | - |
| - | - | 1126 | 510.3 | - | - | 0 | - |
| - | - | 757.3 | 510.8 | - | - | 0 | - |
| - | - | 1124 | 531.3 | - | - | 0 | - |
| - | - | 912.1 | 536.8 | - | - | 0 | - |
| - | - | 913 | 538.6 | - | - | 0 | - |
| - | - | 737.9 | 539.3 | - | - | 0 | - |
| - | - | 725.6 | 547.6 | - | - | 0 | - |
| 2 | y | 2163 | 550.7 | 0.003507 | 6.369 | +2 | 8 |
| - | - | 2071 | 551.2 | - | - | 0 | - |
| 6 | y | 6473 | 560.2 | 0.007171 | 12.8 | +1 | 4 |
| - | - | 1606 | 561.2 | - | - | 0 | - |
| - | - | 660.2 | 562.2 | - | - | 0 | - |
| - | - | 1309 | 567.6 | - | - | 0 | - |
| - | - | 1468 | 567.9 | - | - | 0 | - |
| - | - | 749.8 | 583.3 | - | - | 0 | - |
| - | - | 920.3 | 584.3 | - | - | 0 | - |
| - | - | 4920 | 585.3 | - | - | 0 | - |
| - | - | 1851 | 586.3 | - | - | 0 | - |
| - | - | 620.2 | 588.3 | - | - | 0 | - |
| - | - | 725.6 | 589.3 | - | - | 0 | - |
| - | - | 1234 | 592.7 | - | - | 0 | - |
| - | - | 921.4 | 593.3 | - | - | 0 | - |
| - | - | 1724 | 595.3 | - | - | 0 | - |
| - | - | 842.3 | 595.6 | - | - | 0 | - |
| - | - | 724.8 | 596.2 | - | - | 0 | - |
| 0 | Precursor | 2847 | 601.2 | 0.002312 | 3.846 | +2 | -1 |
| - | - | 6711 | 601.3 | - | - | 0 | - |
| - | - | 1.087E+04 | 601.6 | - | - | 0 | - |
| - | - | 1948 | 601.7 | - | - | 0 | - |
| - | - | 3536 | 602 | - | - | 0 | - |
| - | - | 886.7 | 602.2 | - | - | 0 | - |
| - | - | 1066 | 602.3 | - | - | 0 | - |
| - | - | 809.2 | 602.4 | - | - | 0 | - |
| - | - | 1206 | 602.6 | - | - | 0 | - |
| - | - | 2171 | 612.2 | - | - | 0 | - |
| - | - | 4424 | 616.3 | - | - | 0 | - |
| - | - | 846.4 | 628.4 | - | - | 0 | - |
| - | - | 774.5 | 656.3 | - | - | 0 | - |
| - | - | 1129 | 657.3 | - | - | 0 | - |
| - | - | 2302 | 659.3 | - | - | 0 | - |
| - | - | 900.6 | 660.3 | - | - | 0 | - |
| - | - | 701.6 | 660.8 | - | - | 0 | - |
| - | - | 1106 | 661.3 | - | - | 0 | - |
| - | - | 908.4 | 670.4 | - | - | 0 | - |
| - | - | 1923 | 685.3 | - | - | 0 | - |
| - | - | 1392 | 687.4 | - | - | 0 | - |
| - | - | 782.5 | 696.8 | - | - | 0 | - |
| - | - | 1.392E+04 | 700.4 | - | - | 0 | - |
| - | - | 3699 | 701.4 | - | - | 0 | - |
| - | - | 988.2 | 702.4 | - | - | 0 | - |
| - | - | 2666 | 715.4 | - | - | 0 | - |
| - | - | 1081 | 716.8 | - | - | 0 | - |
| - | - | 1408 | 721.3 | - | - | 0 | - |
| - | - | 843.7 | 722.3 | - | - | 0 | - |
| 5 | y | 6579 | 723.3 | 0.004389 | 6.068 | +1 | 5 |
| - | - | 2452 | 724.3 | - | - | 0 | - |
| - | - | 2978 | 729.3 | - | - | 0 | - |
| - | - | 2999 | 729.8 | - | - | 0 | - |
| - | - | 1631 | 730.3 | - | - | 0 | - |
| - | - | 957.5 | 731.3 | - | - | 0 | - |
| - | - | 861.4 | 741.4 | - | - | 0 | - |
| - | - | 3247 | 749.3 | - | - | 0 | - |
| - | - | 1354 | 750.3 | - | - | 0 | - |
| - | - | 1933 | 754.4 | - | - | 0 | - |
| - | - | 1500 | 754.9 | - | - | 0 | - |
| - | - | 657 | 755.3 | - | - | 0 | - |
| - | - | 691.1 | 756.4 | - | - | 0 | - |
| - | - | 1486 | 757.4 | - | - | 0 | - |
| - | - | 753 | 767.3 | - | - | 0 | - |
| - | - | 947.7 | 773.4 | - | - | 0 | - |
| - | - | 3588 | 774.3 | - | - | 0 | - |
| - | - | 658.8 | 774.4 | - | - | 0 | - |
| - | - | 676.9 | 774.9 | - | - | 0 | - |
| - | - | 1473 | 775.3 | - | - | 0 | - |
| - | - | 702.3 | 776.3 | - | - | 0 | - |
| - | - | 3443 | 783.4 | - | - | 0 | - |
| - | - | 1386 | 784.4 | - | - | 0 | - |
| - | - | 832.7 | 789.4 | - | - | 0 | - |
| - | - | 1816 | 796.9 | - | - | 0 | - |
| - | - | 1468 | 797.4 | - | - | 0 | - |
| - | - | 1181 | 800.4 | - | - | 0 | - |
| - | - | 8154 | 801.4 | - | - | 0 | - |
| - | - | 3946 | 802.4 | - | - | 0 | - |
| - | - | 2033 | 807.4 | - | - | 0 | - |
| - | - | 2431 | 807.9 | - | - | 0 | - |
| - | - | 1778 | 808.4 | - | - | 0 | - |
| - | - | 1033 | 808.9 | - | - | 0 | - |
| - | - | 833.7 | 810.9 | - | - | 0 | - |
| - | - | 1533 | 811.4 | - | - | 0 | - |
| - | - | 956.8 | 811.9 | - | - | 0 | - |
| - | - | 3886 | 828.4 | - | - | 0 | - |
| - | - | 1129 | 829.4 | - | - | 0 | - |
| 4 | y | 7140 | 838.3 | 0.004362 | 5.204 | +1 | 6 |
| - | - | 3921 | 839.3 | - | - | 0 | - |
| - | - | 1072 | 840.3 | - | - | 0 | - |
| - | - | 2177 | 843.4 | - | - | 0 | - |
| - | - | 1.165E+04 | 844.5 | - | - | 0 | - |
| - | - | 4464 | 845.5 | - | - | 0 | - |
| - | - | 1064 | 846.5 | - | - | 0 | - |
| - | - | 1190 | 847.5 | - | - | 0 | - |
| - | - | 1026 | 848.5 | - | - | 0 | - |
| - | - | 1060 | 850.9 | - | - | 0 | - |
| - | - | 1193 | 851.4 | - | - | 0 | - |
| - | - | 680.7 | 854.4 | - | - | 0 | - |
| - | - | 949 | 855.4 | - | - | 0 | - |
| - | - | 1028 | 871.3 | - | - | 0 | - |
| - | - | 4.109E+04 | 872.4 | - | - | 0 | - |
| - | - | 1.791E+04 | 873.4 | - | - | 0 | - |
| - | - | 5411 | 874.5 | - | - | 0 | - |
| - | - | 820.8 | 875.4 | - | - | 0 | - |
| - | - | 834.6 | 888.4 | - | - | 0 | - |
| - | - | 1.115E+04 | 889.3 | - | - | 0 | - |
| - | - | 4951 | 890.3 | - | - | 0 | - |
| - | - | 1801 | 891.4 | - | - | 0 | - |
| - | - | 718.9 | 899.4 | - | - | 0 | - |
| - | - | 726.8 | 900.4 | - | - | 0 | - |
| - | - | 1446 | 902.4 | - | - | 0 | - |
| 3 | y | 1973 | 935.3 | 0.003792 | 4.054 | +1 | 7 |
| - | - | 1219 | 936.3 | - | - | 0 | - |
| - | - | 987.7 | 943.5 | - | - | 0 | - |
| 3 | y | 3.624E+04 | 953.3 | 0.004397 | 4.612 | +1 | 7 |
| - | - | 1.882E+04 | 954.3 | - | - | 0 | - |
| - | - | 592.5 | 954.5 | - | - | 0 | - |
| - | - | 6996 | 955.3 | - | - | 0 | - |
| - | - | 1007 | 956.4 | - | - | 0 | - |
| - | - | 1787 | 971.5 | - | - | 0 | - |
| - | - | 7653 | 991.5 | - | - | 0 | - |
| - | - | 3604 | 992.5 | - | - | 0 | - |
| - | - | 1469 | 993.5 | - | - | 0 | - |
| - | - | 767.9 | 997.5 | - | - | 0 | - |
| - | - | 1642 | 1002 | - | - | 0 | - |
| - | - | 1.731E+04 | 1020 | - | - | 0 | - |
| - | - | 9238 | 1021 | - | - | 0 | - |
| - | - | 3756 | 1022 | - | - | 0 | - |
| - | - | 722.9 | 1036 | - | - | 0 | - |
| 2 | y | 2287 | 1100 | 0.005014 | 4.556 | +1 | 8 |
| - | - | 1237 | 1101 | - | - | 0 | - |
| - | - | 724.5 | 1103 | - | - | 0 | - |
| - | - | 607.4 | 1441 | - | - | 0 | - |
| - | - | 668.8 | 3094 | - | - | 0 | - |
| - | - | 692.7 | 3167 | - | - | 0 | - |

m/z Charge Intensity FragmentType MassShift Position
120.08109283447266 0 84377.16
121.08441162109375 0 6747.8174
123.0443115234375 0 1931.0756
125.0598373413086 0 419.38785
128.10731506347656 0 454.33154
129.06605529785156 0 1153.8373
129.10252380371094 0 4727.716
130.06539916992188 0 2162.5554
130.0857696533203 0 591.77057
132.07659912109375 0 718.9911
132.08148193359375 0 1186.8032
132.10220336914062 0 3375.9617
133.06103515625 0 743.2634
133.08621215820312 0 898.97864
136.07598876953125 0 30361.514
137.0792694091797 0 1825.1897
138.05548095703125 0 628.75714
138.0662841796875 0 874.0405
141.10255432128906 0 1587.572
143.07138061523438 0 492.4985
143.08154296875 0 948.92944
146.0602264404297 0 2090.0579
146.3544158935547 0 469.60477
147.04454040527344 0 1543.5112
148.9545440673828 0 1021.33234
149.04534912109375 0 1040.2001
154.9463348388672 0 503.52194
155.08177185058594 0 533.7113
155.11795043945312 0 598.6034
156.07687377929688 0 1954.2659
157.1338348388672 0 10040.945
158.05979919433594 0 1049.3506
158.09664916992188 0 3615.0884
159.09193420410156 0 2710.2295
159.0987548828125 0 549.07135
159.11302185058594 0 1839.1873
159.44754028320312 0 424.5315
160.11219787597656 0 1238.274
161.0921630859375 0 1938.3281
164.55088806152344 0 433.40002
165.05487060546875 0 14003.825
165.07716369628906 0 628.3702
166.05897521972656 0 885.1397
166.08645629882812 0 2062.1917
167.05567932128906 0 2121.0574
169.08670043945312 0 427.3777
169.09735107421875 0 2368.94
171.07666015625 0 1379.111
171.11302185058594 0 865.22174
173.12875366210938 0 6449.236
174.05519104003906 0 2466.681
175.0868377685547 0 1493.164
176.0826873779297 0 518.6818
176.10723876953125 0 6665.126
177.06788635253906 0 588.9233
177.1024932861328 0 5611.5396
177.1107940673828 0 892.6892
181.0973663330078 0 576.6925
182.08145141601562 0 22978.404
183.0763702392578 0 801.3069
183.0847930908203 0 2073.753
183.11268615722656 0 550.7589
183.1495361328125 0 1031.614
185.092529296875 0 1371.661
185.1287841796875 0 7335.392
185.1650390625 0 478.13217
186.09188842773438 0 515.4683
186.12416076660156 0 866.65936
187.0966339111328 0 2467.4116
187.1069793701172 0 871.37646
187.1446990966797 0 1226.6127
188.07086181640625 0 3835.187
188.93125915527344 0 541.6681
189.08712768554688 0 2063.395
191.08226013183594 0 529.3681
191.11810302734375 0 2268.1646
191.8586883544922 0 486.49857
193.108642578125 0 2804.869
195.0875244140625 0 1254.3289
197.0916748046875 0 700.8789
197.12875366210938 0 637.91254
199.07208251953125 0 794.19995
201.12359619140625 0 3242.5762
202.05035400390625 0 1243.0736
203.06605529785156 0 729.74414
203.08187866210938 0 624.81824
203.11813354492188 0 1804.7166 a Water loss 1
204.0771026611328 0 835.30884
205.09747314453125 0 2950.4045 y 8
207.11248779296875 0 616.94885
208.07237243652344 0 537.20874
209.09249877929688 0 523.03107
211.11940002441406 0 920.8854
212.1033172607422 0 1366.0754
213.08726501464844 0 471.08795
214.11875915527344 0 1764.1282
215.13949584960938 0 1410.8147
217.08152770996094 0 509.92474
217.0972442626953 0 1073.2524
219.07713317871094 0 502.38968
219.113525390625 0 861.19275
221.1035614013672 0 6009.008
221.12875366210938 0 94420.19 a 1
222.10684204101562 0 650.4859
222.1320037841797 0 11450.509
223.06349182128906 0 1445.2437
223.13465881347656 0 807.76074
225.17135620117188 0 1193.6768
228.0982666015625 0 1823.0839
228.1338653564453 0 878.6179
229.11846923828125 0 961.5241
229.1558837890625 0 660.1139
229.17050170898438 0 908.352
231.0612030029297 0 1566.9028
231.11204528808594 0 1118.6587 b Water loss 1
231.1236114501953 0 877.85004
233.09194946289062 0 551.60016
233.14988708496094 0 6707.854
235.1080322265625 0 3233.776
239.0946502685547 0 791.27277
239.1138916015625 0 1664.2424
239.14991760253906 0 714.36426
239.1759033203125 0 950.0868
240.09796142578125 0 1333.1414
240.13473510742188 0 931.8392
241.0740509033203 0 782.60504
246.10874938964844 0 677.88696
249.12368774414062 0 16400.084 b 1
250.12705993652344 0 2212.6511
251.10255432128906 0 1578.3958
253.1184844970703 0 824.0683
253.16549682617188 0 1039.6638
255.1121826171875 0 763.68524
255.14674377441406 0 594.18195
257.1058349609375 0 497.34167
257.1646423339844 0 712.5102
258.1453552246094 0 716.0636
260.1018371582031 0 531.28406
263.10284423828125 0 3745.713
274.130126953125 0 659.4628
279.0973815917969 0 853.3164
282.1800842285156 0 644.1456
283.03094482421875 0 803.6657
283.14373779296875 0 1043.3499
284.1972351074219 0 2976.1519
285.0096130371094 0 701.86816
285.10186767578125 0 2364.425
287.5415344238281 0 539.395
290.1499938964844 0 523.9167
292.1405334472656 0 1523.1375
296.1511535644531 0 1911.4194
297.0906677246094 0 846.4114
302.11395263671875 0 805.6766
305.124267578125 0 768.13477
317.181884765625 0 645.8233
318.14508056640625 0 1613.4965 a Water loss 2
319.1407470703125 0 1589.9908
322.1855773925781 0 658.69476
323.1390380859375 0 596.1574
324.1459045410156 0 4300.8535
325.1484069824219 0 836.16736
325.1861877441406 0 658.82745
329.14996337890625 0 15955.288
330.1534118652344 0 2882.5154
333.11981201171875 0 3344.6677
338.0787658691406 0 556.6986
340.1605529785156 0 683.45245
340.1985778808594 0 2641.0146
341.1455993652344 0 828.6534
342.15655517578125 0 17215.613 y 7
343.1593933105469 0 2967.0837
343.19793701171875 0 2034.3693
346.14056396484375 0 1185.1045 b Water loss 2
350.13507080078125 0 1095.048
355.0697021484375 0 1517.6881
355.19696044921875 0 732.5089
356.1383056640625 0 1918.8451
357.0667724609375 0 633.8139
357.2252197265625 0 3707.0051
361.2096862792969 0 702.9805
364.1503601074219 0 4324.7544 b 2
365.15411376953125 0 851.3808
366.1285095214844 0 700.3528
368.1923828125 0 1215.759
378.1300048828125 0 3140.834
382.1404724121094 0 591.8818
394.12506103515625 0 3818.6648
398.2415771484375 0 1499.28
419.219482421875 0 691.12164
422.22283935546875 0 761.6752
422.7330322265625 0 878.4039
424.2506103515625 0 1984.7704
424.75262451171875 0 652.6123
425.1927185058594 0 1988.8322
426.1969299316406 0 628.4675
427.2082214355469 0 1014.0812
427.7223815917969 0 1879.9829
428.2242126464844 0 1188.4994
432.1881408691406 0 5534.641
436.72784423828125 0 1560.5308
444.71173095703125 0 742.79193
451.9441223144531 0 556.42816
454.2301025390625 0 1040.4685
461.1669616699219 0 681.8122 b Water loss 3
465.1609191894531 0 2747.3403
470.30859375 0 733.0479
471.25665283203125 0 1819.2699
479.1777038574219 0 2540.0674 b 3
482.2362365722656 0 628.2439
483.220947265625 0 881.67755
487.33294677734375 0 946.87897
487.74969482421875 0 712.79565
489.1913146972656 0 3844.1516 y 6
490.1937561035156 0 1085.4465
495.1818542480469 0 698.63806
496.2292785644531 0 1938.5321
496.2665710449219 0 873.8642
497.2332763671875 0 661.9873
500.2471008300781 0 795.94885
501.2558898925781 0 1863.4197
508.2958679199219 0 3025.562
508.7975769042969 0 1380.5709
510.2642517089844 0 1126.3998
510.7622375488281 0 757.2567
531.254150390625 0 1124.2029
536.7721557617188 0 912.1361
538.5938720703125 0 912.9607
539.2625122070312 0 737.9281
547.599365234375 0 725.5518
550.7117919921875 0 2163.1145 y 1
551.2119140625 0 2070.5464
560.2308349609375 0 6473.346 y 5
561.2335815429688 0 1605.6011
562.233154296875 0 660.1617
567.6046752929688 0 1309.2253
567.939208984375 0 1467.5251
583.286376953125 0 749.8414
584.2794189453125 0 920.30475
585.3359375 0 4919.976
586.340087890625 0 1850.5298
588.3207397460938 0 620.1567
589.2804565429688 0 725.62744
592.7311401367188 0 1233.9275
593.313720703125 0 921.3515
595.2836303710938 0 1723.8335
595.6198120117188 0 842.2567
596.2384643554688 0 724.7706
601.2344360351562 0 2847.0454 Precursor
601.2879028320312 0 6711.077
601.6199340820312 0 10870.2
601.7377319335938 0 1948.4177
601.9552612304688 0 3536.1672
602.2344360351562 0 886.7306
602.286376953125 0 1066.4647
602.369873046875 0 809.24316
602.6236572265625 0 1205.8203
612.1978149414062 0 2170.528
616.3091430664062 0 4423.6885
628.3799438476562 0 846.4058
656.3054809570312 0 774.54266
657.2611694335938 0 1128.7593
659.2958374023438 0 2301.7917
660.2931518554688 0 900.63776
660.8154907226562 0 701.59344
661.3113403320312 0 1105.5068
670.3563842773438 0 908.4461
685.2583618164062 0 1923.4572
687.3829345703125 0 1391.6028
696.8287353515625 0 782.5057
700.3622436523438 0 13923.117
701.3657836914062 0 3699.1086
702.3670043945312 0 988.20135
715.3764038085938 0 2665.8079
716.8175659179688 0 1080.6475
721.2598876953125 0 1407.5626
722.26318359375 0 843.71936
723.2913818359375 0 6578.554 y 4
724.2948608398438 0 2451.9028
729.3405151367188 0 2977.5906
729.8412475585938 0 2999.1184
730.3417358398438 0 1630.89
731.3422241210938 0 957.5192
741.3506469726562 0 861.42474
749.25537109375 0 3247.455
750.261474609375 0 1353.8934
754.3501586914062 0 1932.6776
754.852294921875 0 1500.3517
755.3450927734375 0 656.9958
756.4061279296875 0 691.09814
757.384765625 0 1485.5845
767.2718505859375 0 753.0154
773.41357421875 0 947.7406
774.3206787109375 0 3587.887
774.3977661132812 0 658.77765
774.8839111328125 0 676.93066
775.323974609375 0 1473.1042
776.342041015625 0 702.3085
783.3992309570312 0 3442.5632
784.4022827148438 0 1386.3687
789.35693359375 0 832.6951
796.8870849609375 0 1816.4386
797.3888549804688 0 1468.4756
800.4261474609375 0 1180.5745
801.4113159179688 0 8153.5737
802.4136962890625 0 3946.2966
807.38671875 0 2033.4304
807.8876953125 0 2430.5146
808.3899536132812 0 1778.3451
808.892333984375 0 1033.3091
810.8821411132812 0 833.652
811.388671875 0 1533.1132
811.8907470703125 0 956.78064
828.419921875 0 3886.059
829.4197998046875 0 1128.6284
838.3182983398438 0 7139.5444 y 3
839.3215942382812 0 3920.858
840.315185546875 0 1071.9734
843.4374389648438 0 2177.203
844.4520874023438 0 11646.342
845.4551391601562 0 4463.901
846.4580688476562 0 1064.1664
847.4970092773438 0 1189.5065
848.4903564453125 0 1025.9958
850.9002075195312 0 1059.9884
851.4080200195312 0 1192.5928
854.4429321289062 0 680.66077
855.4400634765625 0 948.98456
871.3380737304688 0 1027.7102
872.4468994140625 0 41089.832
873.4498291015625 0 17906.633
874.4535522460938 0 5411.318
875.4498291015625 0 820.7597
888.4214477539062 0 834.64935
889.3464965820312 0 11148.031
890.3499145507812 0 4950.717
891.3511352539062 0 1800.693
899.4354248046875 0 718.86414
900.4380493164062 0 726.79193
902.4385375976562 0 1445.7292
935.3341064453125 0 1973.313 y Water loss 2
936.337158203125 0 1218.9966
943.490234375 0 987.6995
953.3452758789062 0 36240 y 2
954.348388671875 0 18823.635
954.4562377929688 0 592.4538
955.3485107421875 0 6996.0225
956.355712890625 0 1007.4072
971.49560546875 0 1786.7012
991.5191040039062 0 7653.19
992.5230712890625 0 3604.4492
993.5254516601562 0 1468.6499
997.5113525390625 0 767.9066
1001.50439453125 0 1642.493
1019.5149536132812 0 17312.955
1020.5178833007812 0 9237.941
1021.520263671875 0 3756.111
1036.41845703125 0 722.8847
1100.414306640625 0 2286.7246 y 1
1101.4185791015625 0 1237.1233
1103.42822265625 0 724.53796
1440.650634765625 0 607.43
3094.454833984375 0 668.7796
3166.9931640625 0 692.6549

Spectrum Details

|  |  |
| --- | --- |
| Matched peaks? Matched peaks The total absolute number of peaks matched. Additionally in brackets the total fraction of peaks matched and the total number of peaks is shown. | 20 (5.51% of 363) |
| FDR? FDR The false discovery rate estimated for this peptide. It is calculated by matching all theoretical fragments with a non-integer shift with the raw peaks for this spectrum. This is done with 40 different shifts. The resulting percentage is the average number of annotated peaks over the number of annotated peaks with the correct spectrum. | 2.14% |
| Satellite FDR? Satellite FDR See the FDR for details on its calculation. This satellite ion specific FDR only contains the satellite ions (d/w) for I/L/J positions. | - |
| PSM Score? PSM Score The PSM Score as given by Hecklib to this annotated spectrum. It is shown with three significant figures. | 250 |

#### Spectrum 10780? Spectrum 10780 The raw spectrum of this peptide as annotated by Hecklib. The fragments are coloured according to ion type (see legend). Any peaks with a star '\*' as text can be hovered over to see the full details, first the ion type second the mass shift type. By hovering over the amino acids in the peptide or ions in the legend the corresponding peaks are highlighted. By toggling the 'Unassigned' label you can turn the background (unassigned) peaks on or off in the plot. By updating the slider in the Ion legend you can update the spectrum to only show the top X% of the peaks with labels. The top X% means any peak that is within X% of the highest intensity. By dragging in the spectrum you can zoom in to a specific part of the spectrum and use 'Zoom Out' to get back to the original zoom level. The annotation of the spectrum is based on the given sequence in the peptides file and is done with different software so inconsistencies are likely. The peaks are annotated based on the given sequence, with 20 ppm tolerance.

Copy Data

##### Spectrum 10780 (TSV)

###### Preview

```
Loading example...
```

*Click on the button to copy the data to your clipboard.*

Mz MinMz MaxIntensity Max

WidthHeightPeptide font sizePeptide stroke widthSpectrum font sizeSpectrum stroke widthCompact peptide

Ion legend

wxyz

abcd

OtherUnassignedIon^Charge^_Position_Show for top:%

TFDDYAMHW

01.70e+43.40e+45.09e+46.79e+4

Zoom Out

a^+1^_2_y^+1^_1_a^+1^_2_b^+1^_2_b^+1^_2_y^+1^_2_b^+1^_3_b^+1^_3_b^+1^_4_y^+1^_3_y^+2^_8_y^+1^_4_\*\*y^+1^_5_y^+1^_6_y^+1^_7_y^+1^_7_y^+1^_8_

0778155723353114

Fragment Matches Table

Show background peaks

| Position | Ion type | Intensity | mz Theoretical | mz Error (Th) | mz Error (ppm) | Charge | Series Number |
| --- | --- | --- | --- | --- | --- | --- | --- |
| - | - | 5.535E+04 | 120.1 | - | - | 0 | - |
| - | - | 4737 | 121.1 | - | - | 0 | - |
| - | - | 750.4 | 123 | - | - | 0 | - |
| - | - | 665.2 | 126 | - | - | 0 | - |
| - | - | 406.4 | 126.7 | - | - | 0 | - |
| - | - | 417.5 | 129.1 | - | - | 0 | - |
| - | - | 420.7 | 129.1 | - | - | 0 | - |
| - | - | 4395 | 129.1 | - | - | 0 | - |
| - | - | 359.1 | 129.4 | - | - | 0 | - |
| - | - | 1081 | 130.1 | - | - | 0 | - |
| - | - | 437.6 | 130.1 | - | - | 0 | - |
| - | - | 399.4 | 130.1 | - | - | 0 | - |
| - | - | 957.5 | 132.1 | - | - | 0 | - |
| - | - | 614.1 | 132.1 | - | - | 0 | - |
| - | - | 643.4 | 133.1 | - | - | 0 | - |
| - | - | 2.438E+04 | 136.1 | - | - | 0 | - |
| - | - | 525.5 | 137.1 | - | - | 0 | - |
| - | - | 1891 | 137.1 | - | - | 0 | - |
| - | - | 481.5 | 138.1 | - | - | 0 | - |
| - | - | 783.4 | 138.1 | - | - | 0 | - |
| - | - | 786.1 | 146.1 | - | - | 0 | - |
| - | - | 738.1 | 147.1 | - | - | 0 | - |
| - | - | 935.7 | 149 | - | - | 0 | - |
| - | - | 1571 | 156.1 | - | - | 0 | - |
| - | - | 694.5 | 157.1 | - | - | 0 | - |
| - | - | 2314 | 158.1 | - | - | 0 | - |
| - | - | 2235 | 159.1 | - | - | 0 | - |
| - | - | 508.7 | 159.1 | - | - | 0 | - |
| - | - | 1157 | 160.1 | - | - | 0 | - |
| - | - | 642.8 | 162.1 | - | - | 0 | - |
| - | - | 6162 | 165.1 | - | - | 0 | - |
| - | - | 7815 | 166.1 | - | - | 0 | - |
| - | - | 1560 | 167.1 | - | - | 0 | - |
| - | - | 1299 | 171.1 | - | - | 0 | - |
| - | - | 506.9 | 171.1 | - | - | 0 | - |
| - | - | 577.3 | 173.1 | - | - | 0 | - |
| - | - | 3143 | 173.1 | - | - | 0 | - |
| - | - | 1469 | 173.4 | - | - | 0 | - |
| - | - | 608.1 | 175.1 | - | - | 0 | - |
| - | - | 5648 | 176.1 | - | - | 0 | - |
| - | - | 3655 | 177.1 | - | - | 0 | - |
| - | - | 807.6 | 177.1 | - | - | 0 | - |
| - | - | 659.5 | 181.1 | - | - | 0 | - |
| - | - | 1.106E+04 | 182.1 | - | - | 0 | - |
| - | - | 980.1 | 183.1 | - | - | 0 | - |
| - | - | 690.5 | 183.1 | - | - | 0 | - |
| - | - | 652.8 | 185.1 | - | - | 0 | - |
| - | - | 655 | 185.1 | - | - | 0 | - |
| - | - | 1244 | 185.1 | - | - | 0 | - |
| - | - | 1290 | 187.1 | - | - | 0 | - |
| - | - | 3001 | 188.1 | - | - | 0 | - |
| - | - | 493.4 | 189.1 | - | - | 0 | - |
| - | - | 1505 | 189.1 | - | - | 0 | - |
| - | - | 642.3 | 189.1 | - | - | 0 | - |
| - | - | 1103 | 191.1 | - | - | 0 | - |
| - | - | 1662 | 193.1 | - | - | 0 | - |
| - | - | 3539 | 201.1 | - | - | 0 | - |
| 2 | a | 1731 | 203.1 | 6.09E-05 | 0.2998 | +1 | 2 |
| - | - | 896.7 | 204.1 | - | - | 0 | - |
| 9 | y | 2825 | 205.1 | 0.0001359 | 0.6628 | +1 | 1 |
| - | - | 1367 | 207.1 | - | - | 0 | - |
| - | - | 651.2 | 211.1 | - | - | 0 | - |
| - | - | 3290 | 213.1 | - | - | 0 | - |
| - | - | 649.9 | 214.1 | - | - | 0 | - |
| - | - | 1165 | 215.1 | - | - | 0 | - |
| - | - | 1198 | 217.1 | - | - | 0 | - |
| - | - | 1190 | 217.1 | - | - | 0 | - |
| - | - | 3997 | 221.1 | - | - | 0 | - |
| 2 | a | 6.724E+04 | 221.1 | 0.0001316 | 0.5951 | +1 | 2 |
| - | - | 7926 | 222.1 | - | - | 0 | - |
| - | - | 1397 | 223.1 | - | - | 0 | - |
| - | - | 770.6 | 223.1 | - | - | 0 | - |
| - | - | 631.9 | 225.2 | - | - | 0 | - |
| - | - | 827.1 | 229.2 | - | - | 0 | - |
| - | - | 1866 | 231.1 | - | - | 0 | - |
| 2 | b | 786.4 | 231.1 | 0.0001638 | 0.7086 | +1 | 2 |
| - | - | 2826 | 235.1 | - | - | 0 | - |
| - | - | 552.3 | 237.1 | - | - | 0 | - |
| - | - | 1098 | 239.1 | - | - | 0 | - |
| - | - | 530.5 | 239.2 | - | - | 0 | - |
| - | - | 2534 | 241.1 | - | - | 0 | - |
| 2 | b | 1.253E+04 | 249.1 | 0.0001816 | 0.7288 | +1 | 2 |
| - | - | 1186 | 250.1 | - | - | 0 | - |
| - | - | 1149 | 251.1 | - | - | 0 | - |
| - | - | 645 | 251.2 | - | - | 0 | - |
| - | - | 5585 | 253.1 | - | - | 0 | - |
| - | - | 1085 | 257.1 | - | - | 0 | - |
| - | - | 2682 | 263.1 | - | - | 0 | - |
| - | - | 2013 | 269.1 | - | - | 0 | - |
| - | - | 573.1 | 270.1 | - | - | 0 | - |
| - | - | 549.3 | 276.2 | - | - | 0 | - |
| - | - | 1509 | 279.1 | - | - | 0 | - |
| - | - | 756.5 | 279.1 | - | - | 0 | - |
| - | - | 1378 | 285 | - | - | 0 | - |
| - | - | 1411 | 285.1 | - | - | 0 | - |
| - | - | 761.1 | 292.1 | - | - | 0 | - |
| - | - | 1516 | 296.2 | - | - | 0 | - |
| - | - | 541.9 | 301.5 | - | - | 0 | - |
| - | - | 1837 | 310.1 | - | - | 0 | - |
| - | - | 822.2 | 319.1 | - | - | 0 | - |
| - | - | 746.6 | 320.1 | - | - | 0 | - |
| - | - | 534.3 | 322.2 | - | - | 0 | - |
| - | - | 2765 | 324.1 | - | - | 0 | - |
| - | - | 516.8 | 326.1 | - | - | 0 | - |
| - | - | 1181 | 328.1 | - | - | 0 | - |
| - | - | 8800 | 329.1 | - | - | 0 | - |
| - | - | 993.7 | 330.2 | - | - | 0 | - |
| - | - | 576.6 | 334.5 | - | - | 0 | - |
| - | - | 1163 | 338.1 | - | - | 0 | - |
| - | - | 1158 | 340.2 | - | - | 0 | - |
| 8 | y | 1.394E+04 | 342.2 | 6.197E-05 | 0.1811 | +1 | 2 |
| - | - | 2281 | 343.2 | - | - | 0 | - |
| 3 | b | 925.4 | 346.1 | 0.00135 | 3.9 | +1 | 3 |
| - | - | 660.6 | 348.1 | - | - | 0 | - |
| - | - | 1550 | 355.1 | - | - | 0 | - |
| - | - | 844.4 | 356.1 | - | - | 0 | - |
| - | - | 1705 | 356.1 | - | - | 0 | - |
| - | - | 987.6 | 356.1 | - | - | 0 | - |
| - | - | 1931 | 357.2 | - | - | 0 | - |
| - | - | 575.4 | 363.9 | - | - | 0 | - |
| 3 | b | 2865 | 364.2 | 0.0006586 | 1.809 | +1 | 3 |
| - | - | 804.5 | 365.2 | - | - | 0 | - |
| - | - | 2258 | 378.1 | - | - | 0 | - |
| - | - | 1486 | 380.1 | - | - | 0 | - |
| - | - | 1680 | 394.1 | - | - | 0 | - |
| - | - | 696.5 | 413.3 | - | - | 0 | - |
| - | - | 2076 | 425.2 | - | - | 0 | - |
| - | - | 1052 | 428.2 | - | - | 0 | - |
| - | - | 853 | 436.7 | - | - | 0 | - |
| - | - | 626.7 | 448.8 | - | - | 0 | - |
| - | - | 658.5 | 455.2 | - | - | 0 | - |
| - | - | 743.5 | 456.3 | - | - | 0 | - |
| - | - | 1716 | 465.2 | - | - | 0 | - |
| 4 | b | 1326 | 479.2 | 0.0007541 | 1.574 | +1 | 4 |
| 7 | y | 3360 | 489.2 | 0.005161 | 10.55 | +1 | 3 |
| - | - | 930.8 | 490.2 | - | - | 0 | - |
| - | - | 628.9 | 491.2 | - | - | 0 | - |
| - | - | 1311 | 496.2 | - | - | 0 | - |
| - | - | 903.3 | 500.2 | - | - | 0 | - |
| - | - | 869.5 | 501.1 | - | - | 0 | - |
| - | - | 1011 | 519.2 | - | - | 0 | - |
| - | - | 747.6 | 519.2 | - | - | 0 | - |
| 2 | y | 2219 | 550.7 | 0.002775 | 5.039 | +2 | 8 |
| 6 | y | 4678 | 560.2 | 0.004668 | 8.333 | +1 | 4 |
| - | - | 2104 | 561.2 | - | - | 0 | - |
| - | - | 768.8 | 565.7 | - | - | 0 | - |
| - | - | 699.7 | 569.2 | - | - | 0 | - |
| - | - | 814.1 | 570.2 | - | - | 0 | - |
| - | - | 2598 | 585.3 | - | - | 0 | - |
| 0 | Precursor | 1038 | 592.2 | 0.001979 | 3.342 | +2 | -1 |
| 0 | Precursor | 2146 | 601.2 | 0.001336 | 2.221 | +2 | -1 |
| - | - | 989.2 | 601.4 | - | - | 0 | - |
| - | - | 1165 | 601.7 | - | - | 0 | - |
| - | - | 1028 | 602.4 | - | - | 0 | - |
| - | - | 924.6 | 612.2 | - | - | 0 | - |
| - | - | 799.7 | 620.2 | - | - | 0 | - |
| - | - | 857 | 647.3 | - | - | 0 | - |
| - | - | 1286 | 657.3 | - | - | 0 | - |
| - | - | 1155 | 659.3 | - | - | 0 | - |
| - | - | 1209 | 660.3 | - | - | 0 | - |
| - | - | 693.5 | 682.3 | - | - | 0 | - |
| - | - | 1312 | 685.3 | - | - | 0 | - |
| - | - | 6202 | 700.4 | - | - | 0 | - |
| - | - | 2119 | 701.4 | - | - | 0 | - |
| - | - | 1004 | 702.4 | - | - | 0 | - |
| - | - | 1009 | 703.3 | - | - | 0 | - |
| - | - | 670.8 | 713.8 | - | - | 0 | - |
| - | - | 585.9 | 717.2 | - | - | 0 | - |
| - | - | 1694 | 721.3 | - | - | 0 | - |
| 5 | y | 4777 | 723.3 | 0.004267 | 5.899 | +1 | 5 |
| - | - | 2715 | 724.3 | - | - | 0 | - |
| - | - | 2256 | 749.3 | - | - | 0 | - |
| - | - | 1451 | 774.3 | - | - | 0 | - |
| - | - | 857.9 | 775.3 | - | - | 0 | - |
| - | - | 2092 | 783.4 | - | - | 0 | - |
| - | - | 928 | 784.4 | - | - | 0 | - |
| - | - | 5120 | 801.4 | - | - | 0 | - |
| - | - | 1617 | 802.4 | - | - | 0 | - |
| - | - | 1640 | 828.4 | - | - | 0 | - |
| 4 | y | 5663 | 838.3 | 0.003813 | 4.549 | +1 | 6 |
| - | - | 2723 | 839.3 | - | - | 0 | - |
| - | - | 5962 | 844.5 | - | - | 0 | - |
| - | - | 2701 | 845.5 | - | - | 0 | - |
| - | - | 786.7 | 871.3 | - | - | 0 | - |
| - | - | 2.179E+04 | 872.4 | - | - | 0 | - |
| - | - | 7459 | 873.4 | - | - | 0 | - |
| - | - | 2130 | 874.5 | - | - | 0 | - |
| - | - | 8895 | 889.3 | - | - | 0 | - |
| - | - | 3708 | 890.3 | - | - | 0 | - |
| - | - | 1440 | 891.4 | - | - | 0 | - |
| - | - | 693.9 | 897.4 | - | - | 0 | - |
| 3 | y | 1448 | 935.3 | 0.006295 | 6.73 | +1 | 7 |
| - | - | 1540 | 936.3 | - | - | 0 | - |
| 3 | y | 2.362E+04 | 953.3 | 0.003787 | 3.972 | +1 | 7 |
| - | - | 1.356E+04 | 954.3 | - | - | 0 | - |
| - | - | 4542 | 955.3 | - | - | 0 | - |
| - | - | 1120 | 956.4 | - | - | 0 | - |
| - | - | 3311 | 991.5 | - | - | 0 | - |
| - | - | 1596 | 992.5 | - | - | 0 | - |
| - | - | 1179 | 993.5 | - | - | 0 | - |
| - | - | 7859 | 1020 | - | - | 0 | - |
| - | - | 5281 | 1021 | - | - | 0 | - |
| - | - | 1638 | 1022 | - | - | 0 | - |
| - | - | 685.9 | 1036 | - | - | 0 | - |
| - | - | 659.6 | 1090 | - | - | 0 | - |
| 2 | y | 1473 | 1100 | 0.0004972 | 0.4518 | +1 | 8 |
| - | - | 1361 | 1101 | - | - | 0 | - |
| - | - | 844.3 | 1711 | - | - | 0 | - |
| - | - | 628.1 | 2030 | - | - | 0 | - |
| - | - | 656.1 | 2223 | - | - | 0 | - |
| - | - | 1032 | 3082 | - | - | 0 | - |
| - | - | 779 | 3083 | - | - | 0 | - |

m/z Charge Intensity FragmentType MassShift Position
120.08100891113281 0 55348.418
121.08432006835938 0 4737.293
123.04415130615234 0 750.3804
126.03753662109375 0 665.18146
126.7004623413086 0 406.43396
129.0660400390625 0 417.54147
129.09823608398438 0 420.74203
129.10243225097656 0 4395.2803
129.42835998535156 0 359.143
130.0653839111328 0 1080.891
130.08673095703125 0 437.55728
130.1056365966797 0 399.40527
132.08103942871094 0 957.5484
132.10223388671875 0 614.1433
133.08595275878906 0 643.39355
136.07589721679688 0 24378.566
137.06044006347656 0 525.4578
137.079345703125 0 1891.4116
138.05545043945312 0 481.54575
138.0662384033203 0 783.39667
146.06011962890625 0 786.089
147.1126251220703 0 738.08765
149.0452423095703 0 935.6795
156.07701110839844 0 1571.3937
157.1336212158203 0 694.4893
158.0965118408203 0 2313.8647
159.09181213378906 0 2234.6455
159.09938049316406 0 508.6633
160.11248779296875 0 1156.5562
162.09129333496094 0 642.81824
165.05484008789062 0 6162.4795
166.08644104003906 0 7815.4854
167.05575561523438 0 1559.8611
171.07635498046875 0 1298.924
171.1493682861328 0 506.94852
173.09226989746094 0 577.2958
173.1285400390625 0 3142.5356
173.43881225585938 0 1469.0361
175.0867919921875 0 608.0791
176.10720825195312 0 5648.2944
177.10247802734375 0 3654.8303
177.11053466796875 0 807.55145
181.09742736816406 0 659.4537
182.081298828125 0 11064.308
183.08485412597656 0 980.0641
183.11322021484375 0 690.464
185.05517578125 0 652.8125
185.07412719726562 0 654.9974
185.12847900390625 0 1243.6558
187.14453125 0 1290.0228
188.07081604003906 0 3001.0254
189.0751953125 0 493.375
189.0871124267578 0 1505.2312
189.10276794433594 0 642.293
191.11830139160156 0 1103.1022
193.1083984375 0 1661.7771
201.1234893798828 0 3539.4836
203.11795043945312 0 1730.6232 a Water loss 1
204.07667541503906 0 896.7066
205.0972900390625 0 2824.9285 y 8
207.11302185058594 0 1367.4039
211.1439208984375 0 651.2001
213.06932067871094 0 3290.0554
214.1192169189453 0 649.92645
215.13900756835938 0 1164.5674
217.08218383789062 0 1197.5541
217.09727478027344 0 1189.9087
221.1034393310547 0 3997.4492
221.1285858154297 0 67243.74 a 1
222.1319580078125 0 7926.3457
223.05348205566406 0 1396.8864
223.0638885498047 0 770.6015
225.171142578125 0 631.9215
229.1693115234375 0 827.1462
231.06112670898438 0 1865.8577
231.11264038085938 0 786.3929 b Water loss 1
235.10780334472656 0 2825.766
237.12265014648438 0 552.2613
239.09425354003906 0 1098.0068
239.1504669189453 0 530.52954
241.0641632080078 0 2533.6133
249.12355041503906 0 12525.423 b 1
250.12721252441406 0 1186.2683
251.1026611328125 0 1149.1211
251.15078735351562 0 645.0466
253.11842346191406 0 5584.839
257.1070251464844 0 1085.1414
263.10247802734375 0 2681.8413
269.05877685546875 0 2013.0912
270.1085205078125 0 573.0739
276.1778259277344 0 549.33624
279.09674072265625 0 1508.8812
279.13482666015625 0 756.451
285.0095520019531 0 1377.5403
285.1016845703125 0 1411.1115
292.07415771484375 0 761.0689
296.15057373046875 0 1515.5236
301.4803466796875 0 541.88995
310.0857849121094 0 1837.3013
319.1399841308594 0 822.21375
320.0714416503906 0 746.58276
322.1863098144531 0 534.3383
324.14599609375 0 2765.4836
326.08905029296875 0 516.7643
328.0963439941406 0 1181.1448
329.149658203125 0 8800.169
330.1515197753906 0 993.7031
334.5445556640625 0 576.58295
338.0804748535156 0 1163.333
340.19757080078125 0 1158.2465
342.1561279296875 0 13936.82 y 7
343.1582946777344 0 2280.5693
346.1383972167969 0 925.40826 b Water loss 2
348.1175842285156 0 660.64404
355.069580078125 0 1550.0377
356.06878662109375 0 844.4258
356.09130859375 0 1704.588
356.13897705078125 0 987.62537
357.2242126464844 0 1930.583
363.8935546875 0 575.42334
364.1509704589844 0 2865.4258 b 2
365.15087890625 0 804.538
378.1300354003906 0 2258.3013
380.1449890136719 0 1485.5496
394.1236877441406 0 1679.7853
413.2630310058594 0 696.526
425.1932373046875 0 2075.7258
428.22247314453125 0 1051.7614
436.7289733886719 0 852.9833
448.7664794921875 0 626.7224
455.2243347167969 0 658.4938
456.318115234375 0 743.4759
465.1618957519531 0 1715.9888
479.1780090332031 0 1326.1548 b 3
489.19171142578125 0 3359.9775 y 6
490.1950378417969 0 930.7648
491.209716796875 0 628.9371
496.2299499511719 0 1311.3413
500.2459411621094 0 903.2539
501.1449890136719 0 869.4831
519.1543579101562 0 1010.8598
519.1986083984375 0 747.61365
550.7110595703125 0 2219.3218 y 1
560.2283325195312 0 4677.7812 y 5
561.2313232421875 0 2103.6594
565.70361328125 0 768.7512
569.2349243164062 0 699.71387
570.2298583984375 0 814.07385
585.3355712890625 0 2598.4353
592.2288208007812 0 1037.805 Precursor Water loss
601.2334594726562 0 2145.9822 Precursor
601.3612670898438 0 989.1799
601.7342529296875 0 1165.2142
602.361083984375 0 1027.6754
612.1982421875 0 924.56635
620.204833984375 0 799.71735
647.2987670898438 0 856.9604
657.2630615234375 0 1285.671
659.2940063476562 0 1155.4598
660.295654296875 0 1209.0492
682.343017578125 0 693.4723
685.257080078125 0 1312.0282
700.3618774414062 0 6202.087
701.3653564453125 0 2119.1035
702.3668823242188 0 1003.5395
703.2682495117188 0 1009.3299
713.834228515625 0 670.76465
717.220458984375 0 585.9472
721.259521484375 0 1693.6569
723.291259765625 0 4777.062 y 4
724.2958374023438 0 2715.0024
749.2550048828125 0 2256.0884
774.3194580078125 0 1451.4473
775.318603515625 0 857.8717
783.4002685546875 0 2092.4092
784.4019165039062 0 927.97217
801.4105834960938 0 5119.644
802.4143676757812 0 1616.7427
828.4169921875 0 1640.0674
838.3177490234375 0 5663.2754 y 3
839.3201904296875 0 2722.901
844.4508666992188 0 5961.5537
845.45556640625 0 2701.0618
871.3345947265625 0 786.7062
872.4464111328125 0 21793.824
873.44921875 0 7459.122
874.4513549804688 0 2130.3423
889.3468017578125 0 8895.186
890.3494262695312 0 3707.9495
891.351806640625 0 1439.6359
897.403564453125 0 693.88873
935.3366088867188 0 1447.8857 y Water loss 2
936.3355102539062 0 1539.521
953.3446655273438 0 23619.758 y 2
954.3477172851562 0 13555.669
955.3475952148438 0 4541.5845
956.3511962890625 0 1119.6127
991.5204467773438 0 3310.582
992.5210571289062 0 1596.4961
993.5294189453125 0 1178.657
1019.5142822265625 0 7859.2686
1020.515869140625 0 5281.0503
1021.5206298828125 0 1637.7672
1036.41650390625 0 685.8518
1090.182861328125 0 659.62036
1100.4097900390625 0 1472.9374 y 1
1101.412109375 0 1361.2173
1710.8590087890625 0 844.3092
2029.798583984375 0 628.11487
2222.6708984375 0 656.11395
3082.13330078125 0 1032.3737
3082.702880859375 0 778.971

Spectrum Details

|  |  |
| --- | --- |
| Matched peaks? Matched peaks The total absolute number of peaks matched. Additionally in brackets the total fraction of peaks matched and the total number of peaks is shown. | 19 (8.96% of 212) |
| FDR? FDR The false discovery rate estimated for this peptide. It is calculated by matching all theoretical fragments with a non-integer shift with the raw peaks for this spectrum. This is done with 40 different shifts. The resulting percentage is the average number of annotated peaks over the number of annotated peaks with the correct spectrum. | 1.25% |
| Satellite FDR? Satellite FDR See the FDR for details on its calculation. This satellite ion specific FDR only contains the satellite ions (d/w) for I/L/J positions. | - |
| PSM Score? PSM Score The PSM Score as given by Hecklib to this annotated spectrum. It is shown with three significant figures. | 213 |

#### Spectrum 9135? Spectrum 9135 The raw spectrum of this peptide as annotated by Hecklib. The fragments are coloured according to ion type (see legend). Any peaks with a star '\*' as text can be hovered over to see the full details, first the ion type second the mass shift type. By hovering over the amino acids in the peptide or ions in the legend the corresponding peaks are highlighted. By toggling the 'Unassigned' label you can turn the background (unassigned) peaks on or off in the plot. By updating the slider in the Ion legend you can update the spectrum to only show the top X% of the peaks with labels. The top X% means any peak that is within X% of the highest intensity. By dragging in the spectrum you can zoom in to a specific part of the spectrum and use 'Zoom Out' to get back to the original zoom level. The annotation of the spectrum is based on the given sequence in the peptides file and is done with different software so inconsistencies are likely. The peaks are annotated based on the given sequence, with 20 ppm tolerance.

Copy Data

##### Spectrum 9135 (TSV)

###### Preview

```
Loading example...
```

*Click on the button to copy the data to your clipboard.*

Mz MinMz MaxIntensity Max

WidthHeightPeptide font sizePeptide stroke widthSpectrum font sizeSpectrum stroke widthCompact peptide

Ion legend

wxyz

abcd

OtherUnassignedIon^Charge^_Position_Show for top:%

TFDDYAMHW

01.29e+52.58e+53.86e+55.15e+5

Zoom Out

y^+1^_1_c^+1^_2_z^+1^_2_y^+1^_2_z^+2^_5_c^+1^_3_w^+1^_3_z^+1^_3_z^+2^_7_y^+1^_3_c^+1^_4_z^+1^_4_y^+2^_8_y^+2^_8_y^+1^_4_c^+1^_5_z^+1^_5_y^+1^_5_c^+1^_6_c^+1^_6_z^+1^_6_y^+1^_6_z^+1^_6_y^+1^_6_c^+1^_7_c^+1^_7_c^+1^_7_w^+1^_7_y^+1^_7_z^+1^_7_y^+1^_7_c^+1^_8_c^+1^_8_y^+1^_8_z^+1^_8_y^+1^_8_

0778155623343112

Fragment Matches Table

Show background peaks

| Position | Ion type | Intensity | mz Theoretical | mz Error (Th) | mz Error (ppm) | Charge | Series Number |
| --- | --- | --- | --- | --- | --- | --- | --- |
| - | - | 6660 | 120.1 | - | - | 0 | - |
| - | - | 2802 | 133.1 | - | - | 0 | - |
| - | - | 1133 | 136.1 | - | - | 0 | - |
| - | - | 415.3 | 141.4 | - | - | 0 | - |
| - | - | 679 | 143.1 | - | - | 0 | - |
| - | - | 450.9 | 146.8 | - | - | 0 | - |
| - | - | 1297 | 149 | - | - | 0 | - |
| - | - | 426.7 | 158.1 | - | - | 0 | - |
| - | - | 1222 | 159.1 | - | - | 0 | - |
| - | - | 422.7 | 162.4 | - | - | 0 | - |
| - | - | 439 | 163.3 | - | - | 0 | - |
| - | - | 790.7 | 166.1 | - | - | 0 | - |
| - | - | 1441 | 176.1 | - | - | 0 | - |
| - | - | 2119 | 177.1 | - | - | 0 | - |
| - | - | 2471 | 177.1 | - | - | 0 | - |
| - | - | 492.5 | 183.1 | - | - | 0 | - |
| - | - | 2285 | 187.1 | - | - | 0 | - |
| - | - | 1176 | 188.1 | - | - | 0 | - |
| - | - | 1692 | 203.1 | - | - | 0 | - |
| 9 | y | 4657 | 205.1 | 0.0001817 | 0.886 | +1 | 1 |
| - | - | 2440 | 221.1 | - | - | 0 | - |
| - | - | 1.765E+05 | 221.1 | - | - | 0 | - |
| - | - | 607.3 | 222.1 | - | - | 0 | - |
| - | - | 2.223E+04 | 222.1 | - | - | 0 | - |
| - | - | 1419 | 223.1 | - | - | 0 | - |
| - | - | 1152 | 231.1 | - | - | 0 | - |
| - | - | 2189 | 231.1 | - | - | 0 | - |
| - | - | 971.3 | 231.2 | - | - | 0 | - |
| - | - | 1690 | 235.1 | - | - | 0 | - |
| - | - | 4209 | 239.1 | - | - | 0 | - |
| 2 | c | 5.814E+04 | 249.1 | 0.0001816 | 0.7288 | +1 | 2 |
| - | - | 8265 | 250.1 | - | - | 0 | - |
| - | - | 786.5 | 251.1 | - | - | 0 | - |
| - | - | 6450 | 263.1 | - | - | 0 | - |
| - | - | 654.8 | 264.1 | - | - | 0 | - |
| - | - | 1.864E+04 | 264.1 | - | - | 0 | - |
| - | - | 2080 | 265.1 | - | - | 0 | - |
| - | - | 2562 | 269.1 | - | - | 0 | - |
| - | - | 1252 | 271.1 | - | - | 0 | - |
| - | - | 1383 | 275.2 | - | - | 0 | - |
| - | - | 604.3 | 280.1 | - | - | 0 | - |
| - | - | 1270 | 281.1 | - | - | 0 | - |
| - | - | 5534 | 282.1 | - | - | 0 | - |
| - | - | 4498 | 283.2 | - | - | 0 | - |
| - | - | 2334 | 285 | - | - | 0 | - |
| - | - | 625.2 | 292.2 | - | - | 0 | - |
| - | - | 5400 | 299.1 | - | - | 0 | - |
| - | - | 689.6 | 300.1 | - | - | 0 | - |
| - | - | 1505 | 324.1 | - | - | 0 | - |
| 8 | z | 1229 | 326.1 | 0.0003534 | 1.084 | +1 | 2 |
| - | - | 7340 | 327.1 | - | - | 0 | - |
| - | - | 1027 | 328.1 | - | - | 0 | - |
| - | - | 1606 | 337.2 | - | - | 0 | - |
| - | - | 1813 | 340.1 | - | - | 0 | - |
| 8 | y | 3.411E+04 | 342.2 | 0.0003672 | 1.073 | +1 | 2 |
| - | - | 6435 | 343.2 | - | - | 0 | - |
| - | - | 768.1 | 344.2 | - | - | 0 | - |
| 5 | z | 1480 | 346.1 | 0.002819 | 8.144 | +2 | 5 |
| - | - | 4.996E+04 | 355.1 | - | - | 0 | - |
| - | - | 1.291E+04 | 356.1 | - | - | 0 | - |
| 3 | c | 1.29E+04 | 364.2 | 0.0001349 | 0.3704 | +1 | 3 |
| - | - | 2308 | 365.2 | - | - | 0 | - |
| - | - | 646.3 | 366.2 | - | - | 0 | - |
| - | - | 533.9 | 367.2 | - | - | 0 | - |
| - | - | 2156 | 371.1 | - | - | 0 | - |
| - | - | 3253 | 378.1 | - | - | 0 | - |
| - | - | 644 | 379.1 | - | - | 0 | - |
| - | - | 1033 | 381.2 | - | - | 0 | - |
| - | - | 539.3 | 382.3 | - | - | 0 | - |
| - | - | 1274 | 394.1 | - | - | 0 | - |
| 7 | w | 957.5 | 396.2 | 0.0005235 | 1.321 | +1 | 3 |
| - | - | 1172 | 399.2 | - | - | 0 | - |
| - | - | 7961 | 413.2 | - | - | 0 | - |
| - | - | 5108 | 414.2 | - | - | 0 | - |
| - | - | 2474 | 415 | - | - | 0 | - |
| - | - | 1510 | 415.2 | - | - | 0 | - |
| - | - | 758.2 | 416 | - | - | 0 | - |
| - | - | 702.1 | 419 | - | - | 0 | - |
| - | - | 777.5 | 425.3 | - | - | 0 | - |
| - | - | 847 | 433.2 | - | - | 0 | - |
| 7 | z | 3062 | 457.2 | 0.0006414 | 1.403 | +1 | 3 |
| - | - | 1.346E+04 | 458.2 | - | - | 0 | - |
| - | - | 3430 | 459.2 | - | - | 0 | - |
| - | - | 884.8 | 460.2 | - | - | 0 | - |
| 3 | z | 1414 | 461.2 | 0.00431 | 9.347 | +2 | 7 |
| - | - | 2227 | 465.2 | - | - | 0 | - |
| - | - | 2438 | 467.2 | - | - | 0 | - |
| - | - | 798.9 | 469.3 | - | - | 0 | - |
| - | - | 850 | 472.2 | - | - | 0 | - |
| 7 | y | 1.732E+04 | 473.2 | 0.0001352 | 0.2858 | +1 | 3 |
| - | - | 4820 | 474.2 | - | - | 0 | - |
| - | - | 617.7 | 475.2 | - | - | 0 | - |
| - | - | 613.4 | 476.2 | - | - | 0 | - |
| - | - | 725.8 | 477.2 | - | - | 0 | - |
| 4 | c | 7104 | 479.2 | 0.0004795 | 1.001 | +1 | 4 |
| - | - | 1290 | 480.2 | - | - | 0 | - |
| - | - | 589.6 | 481.2 | - | - | 0 | - |
| - | - | 4557 | 484.2 | - | - | 0 | - |
| - | - | 2065 | 485.2 | - | - | 0 | - |
| - | - | 679 | 486.2 | - | - | 0 | - |
| - | - | 630.4 | 487.3 | - | - | 0 | - |
| - | - | 860.7 | 491.7 | - | - | 0 | - |
| - | - | 946.7 | 500.2 | - | - | 0 | - |
| - | - | 1845 | 503.2 | - | - | 0 | - |
| - | - | 816.9 | 504.2 | - | - | 0 | - |
| - | - | 2184 | 505.2 | - | - | 0 | - |
| - | - | 997.8 | 506.2 | - | - | 0 | - |
| 6 | z | 1.198E+05 | 528.2 | 8.657E-05 | 0.1639 | +1 | 4 |
| - | - | 5.096E+04 | 529.2 | - | - | 0 | - |
| - | - | 1.203E+04 | 530.2 | - | - | 0 | - |
| - | - | 1721 | 531.2 | - | - | 0 | - |
| 2 | y | 1206 | 533.7 | 0.0002386 | 0.4471 | +2 | 8 |
| - | - | 595.2 | 537.1 | - | - | 0 | - |
| - | - | 586.9 | 542.2 | - | - | 0 | - |
| 2 | y | 7263 | 542.7 | 0.0002774 | 0.511 | +2 | 8 |
| - | - | 4819 | 543.2 | - | - | 0 | - |
| - | - | 1620 | 543.7 | - | - | 0 | - |
| 6 | y | 2.222E+04 | 544.2 | 0.0001613 | 0.2964 | +1 | 4 |
| - | - | 6934 | 545.2 | - | - | 0 | - |
| - | - | 2009 | 546.2 | - | - | 0 | - |
| - | - | 700.8 | 558.2 | - | - | 0 | - |
| - | - | 1016 | 560.2 | - | - | 0 | - |
| - | - | 1556 | 562.2 | - | - | 0 | - |
| - | - | 1605 | 571.2 | - | - | 0 | - |
| - | - | 669.5 | 572.2 | - | - | 0 | - |
| - | - | 1509 | 575.2 | - | - | 0 | - |
| - | - | 1.289E+04 | 575.4 | - | - | 0 | - |
| - | - | 2527 | 576.3 | - | - | 0 | - |
| - | - | 5519 | 576.4 | - | - | 0 | - |
| - | - | 746.9 | 577.3 | - | - | 0 | - |
| - | - | 737.6 | 577.4 | - | - | 0 | - |
| - | - | 4324 | 584.2 | - | - | 0 | - |
| - | - | 3911 | 584.7 | - | - | 0 | - |
| - | - | 1965 | 585.2 | - | - | 0 | - |
| - | - | 1079 | 590.2 | - | - | 0 | - |
| - | - | 1233 | 592.4 | - | - | 0 | - |
| - | - | 1.022E+04 | 593.2 | - | - | 0 | - |
| - | - | 8735 | 593.7 | - | - | 0 | - |
| - | - | 2859 | 594.2 | - | - | 0 | - |
| - | - | 1076 | 596.2 | - | - | 0 | - |
| - | - | 663.1 | 596.2 | - | - | 0 | - |
| - | - | 850.2 | 599.2 | - | - | 0 | - |
| - | - | 1991 | 614.2 | - | - | 0 | - |
| - | - | 810.4 | 617.3 | - | - | 0 | - |
| - | - | 2432 | 618.2 | - | - | 0 | - |
| - | - | 4789 | 620.2 | - | - | 0 | - |
| - | - | 2005 | 621.3 | - | - | 0 | - |
| - | - | 1630 | 624.2 | - | - | 0 | - |
| - | - | 779.2 | 625.2 | - | - | 0 | - |
| - | - | 3152 | 630.3 | - | - | 0 | - |
| - | - | 907.2 | 631.3 | - | - | 0 | - |
| - | - | 1235 | 633.3 | - | - | 0 | - |
| 5 | c | 1958 | 642.2 | 0.001081 | 1.684 | +1 | 5 |
| - | - | 2273 | 646.3 | - | - | 0 | - |
| - | - | 3673 | 647.3 | - | - | 0 | - |
| - | - | 1853 | 648.3 | - | - | 0 | - |
| - | - | 1827 | 677.2 | - | - | 0 | - |
| - | - | 945.4 | 686.3 | - | - | 0 | - |
| - | - | 6838 | 688.3 | - | - | 0 | - |
| - | - | 3221 | 689.3 | - | - | 0 | - |
| - | - | 1054 | 690.3 | - | - | 0 | - |
| 5 | z | 6.431E+04 | 691.3 | 7.058E-05 | 0.1021 | +1 | 5 |
| - | - | 3.502E+04 | 692.3 | - | - | 0 | - |
| - | - | 1.054E+04 | 693.3 | - | - | 0 | - |
| - | - | 1708 | 694.3 | - | - | 0 | - |
| - | - | 2310 | 695.3 | - | - | 0 | - |
| - | - | 8373 | 701.3 | - | - | 0 | - |
| - | - | 2558 | 702.3 | - | - | 0 | - |
| - | - | 1014 | 703.3 | - | - | 0 | - |
| - | - | 4265 | 705.3 | - | - | 0 | - |
| - | - | 2520 | 706.3 | - | - | 0 | - |
| 5 | y | 3.524E+04 | 707.3 | 0.0007282 | 1.03 | +1 | 5 |
| - | - | 1.386E+04 | 708.3 | - | - | 0 | - |
| - | - | 3925 | 709.3 | - | - | 0 | - |
| 6 | c | 1704 | 713.3 | 0.0006231 | 0.8735 | +1 | 6 |
| - | - | 946.9 | 714.3 | - | - | 0 | - |
| - | - | 1306 | 715.3 | - | - | 0 | - |
| - | - | 685.1 | 716.3 | - | - | 0 | - |
| - | - | 827.2 | 718.3 | - | - | 0 | - |
| - | - | 5132 | 729.3 | - | - | 0 | - |
| 6 | c | 1.918E+04 | 730.3 | 0.0008406 | 1.151 | +1 | 6 |
| - | - | 6843 | 731.3 | - | - | 0 | - |
| - | - | 8709 | 732.3 | - | - | 0 | - |
| - | - | 1.626E+04 | 733.3 | - | - | 0 | - |
| - | - | 8248 | 734.3 | - | - | 0 | - |
| - | - | 7216 | 735.3 | - | - | 0 | - |
| - | - | 2470 | 736.3 | - | - | 0 | - |
| - | - | 785 | 737.3 | - | - | 0 | - |
| - | - | 1.531E+04 | 745.3 | - | - | 0 | - |
| - | - | 6071 | 746.3 | - | - | 0 | - |
| - | - | 1867 | 747.3 | - | - | 0 | - |
| - | - | 978 | 748.3 | - | - | 0 | - |
| - | - | 1775 | 751.3 | - | - | 0 | - |
| - | - | 1654 | 759.3 | - | - | 0 | - |
| - | - | 1075 | 760.3 | - | - | 0 | - |
| - | - | 1013 | 761.3 | - | - | 0 | - |
| - | - | 2.294E+04 | 762.3 | - | - | 0 | - |
| - | - | 1.049E+04 | 763.3 | - | - | 0 | - |
| - | - | 3875 | 764.3 | - | - | 0 | - |
| - | - | 708.7 | 776.3 | - | - | 0 | - |
| - | - | 1077 | 786.3 | - | - | 0 | - |
| 4 | z | 897.6 | 788.3 | 0.001984 | 2.516 | +1 | 6 |
| - | - | 1189 | 792.3 | - | - | 0 | - |
| - | - | 8757 | 803.3 | - | - | 0 | - |
| 4 | y | 3594 | 804.3 | 0.00981 | 12.2 | +1 | 6 |
| - | - | 1428 | 805.3 | - | - | 0 | - |
| 4 | z | 4.185E+04 | 806.3 | 0.0004023 | 0.4989 | +1 | 6 |
| - | - | 4.392E+04 | 807.3 | - | - | 0 | - |
| - | - | 1.753E+04 | 808.3 | - | - | 0 | - |
| - | - | 4185 | 809.3 | - | - | 0 | - |
| - | - | 6570 | 816.3 | - | - | 0 | - |
| - | - | 8379 | 817.3 | - | - | 0 | - |
| - | - | 6127 | 818.3 | - | - | 0 | - |
| - | - | 2136 | 819.3 | - | - | 0 | - |
| 4 | y | 3.942E+04 | 822.3 | 0.0006327 | 0.7694 | +1 | 6 |
| - | - | 1.763E+04 | 823.3 | - | - | 0 | - |
| - | - | 5862 | 824.3 | - | - | 0 | - |
| - | - | 686.3 | 825.3 | - | - | 0 | - |
| - | - | 4714 | 830.3 | - | - | 0 | - |
| - | - | 2616 | 831.4 | - | - | 0 | - |
| - | - | 7177 | 833.4 | - | - | 0 | - |
| - | - | 3751 | 834.4 | - | - | 0 | - |
| - | - | 642.8 | 835.4 | - | - | 0 | - |
| - | - | 820.6 | 842.3 | - | - | 0 | - |
| 7 | c | 717.3 | 843.3 | 0.003295 | 3.907 | +1 | 7 |
| 7 | c | 2384 | 844.3 | 0.003046 | 3.608 | +1 | 7 |
| - | - | 1323 | 845.3 | - | - | 0 | - |
| - | - | 6085 | 847.3 | - | - | 0 | - |
| - | - | 2705 | 848.3 | - | - | 0 | - |
| - | - | 2029 | 850.3 | - | - | 0 | - |
| - | - | 1090 | 851.3 | - | - | 0 | - |
| - | - | 2.57E+04 | 860.3 | - | - | 0 | - |
| 7 | c | 7.24E+04 | 861.3 | 0.002446 | 2.839 | +1 | 7 |
| - | - | 3.119E+04 | 862.3 | - | - | 0 | - |
| - | - | 1.01E+04 | 863.3 | - | - | 0 | - |
| - | - | 1106 | 864.3 | - | - | 0 | - |
| - | - | 3846 | 874.3 | - | - | 0 | - |
| - | - | 2640 | 875.3 | - | - | 0 | - |
| 3 | w | 1214 | 876.3 | 0.004183 | 4.774 | +1 | 7 |
| - | - | 2.647E+04 | 877.3 | - | - | 0 | - |
| - | - | 1.384E+04 | 878.3 | - | - | 0 | - |
| - | - | 4223 | 879.3 | - | - | 0 | - |
| - | - | 1658 | 880.3 | - | - | 0 | - |
| - | - | 1423 | 881.4 | - | - | 0 | - |
| - | - | 1445 | 882.3 | - | - | 0 | - |
| - | - | 964.2 | 883.4 | - | - | 0 | - |
| - | - | 1219 | 891.3 | - | - | 0 | - |
| - | - | 2523 | 893.4 | - | - | 0 | - |
| - | - | 1490 | 894.4 | - | - | 0 | - |
| - | - | 3498 | 910.4 | - | - | 0 | - |
| - | - | 1872 | 911.4 | - | - | 0 | - |
| 3 | y | 7302 | 919.3 | 0.0009588 | 1.043 | +1 | 7 |
| - | - | 4495 | 920.3 | - | - | 0 | - |
| 3 | z | 1.492E+04 | 921.3 | 0.0002457 | 0.2667 | +1 | 7 |
| - | - | 8374 | 922.3 | - | - | 0 | - |
| - | - | 2828 | 923.3 | - | - | 0 | - |
| - | - | 6273 | 936.4 | - | - | 0 | - |
| 3 | y | 1.617E+05 | 937.4 | 0.0001099 | 0.1173 | +1 | 7 |
| - | - | 8.665E+04 | 938.4 | - | - | 0 | - |
| - | - | 3.029E+04 | 939.4 | - | - | 0 | - |
| - | - | 3226 | 940.4 | - | - | 0 | - |
| - | - | 1477 | 953.4 | - | - | 0 | - |
| - | - | 2.162E+04 | 954.4 | - | - | 0 | - |
| - | - | 1.048E+04 | 955.4 | - | - | 0 | - |
| - | - | 2471 | 956.4 | - | - | 0 | - |
| 8 | c | 3125 | 981.4 | 0.002118 | 2.158 | +1 | 8 |
| - | - | 1645 | 982.4 | - | - | 0 | - |
| - | - | 5275 | 983.4 | - | - | 0 | - |
| - | - | 2526 | 984.4 | - | - | 0 | - |
| - | - | 1114 | 985.4 | - | - | 0 | - |
| - | - | 3760 | 994.4 | - | - | 0 | - |
| - | - | 2779 | 995.4 | - | - | 0 | - |
| - | - | 805.7 | 996.4 | - | - | 0 | - |
| 8 | c | 2.065E+05 | 998.4 | 0.0006887 | 0.6898 | +1 | 8 |
| - | - | 1.148E+05 | 999.4 | - | - | 0 | - |
| - | - | 4.232E+04 | 1000 | - | - | 0 | - |
| - | - | 4913 | 1001 | - | - | 0 | - |
| - | - | 7227 | 1024 | - | - | 0 | - |
| - | - | 4991 | 1025 | - | - | 0 | - |
| - | - | 1922 | 1026 | - | - | 0 | - |
| - | - | 1085 | 1040 | - | - | 0 | - |
| - | - | 1169 | 1041 | - | - | 0 | - |
| - | - | 733.5 | 1052 | - | - | 0 | - |
| - | - | 3810 | 1055 | - | - | 0 | - |
| - | - | 2263 | 1056 | - | - | 0 | - |
| - | - | 661.9 | 1057 | - | - | 0 | - |
| - | - | 784.2 | 1065 | - | - | 0 | - |
| 2 | y | 1389 | 1066 | 0.01144 | 10.73 | +1 | 8 |
| - | - | 1085 | 1067 | - | - | 0 | - |
| 2 | z | 2.709E+04 | 1068 | 0.001643 | 1.538 | +1 | 8 |
| - | - | 1.603E+04 | 1069 | - | - | 0 | - |
| - | - | 6245 | 1070 | - | - | 0 | - |
| - | - | 1126 | 1079 | - | - | 0 | - |
| - | - | 1059 | 1081 | - | - | 0 | - |
| - | - | 1427 | 1082 | - | - | 0 | - |
| 2 | y | 1.038E+04 | 1084 | 0.00108 | 0.9959 | +1 | 8 |
| - | - | 7124 | 1085 | - | - | 0 | - |
| - | - | 2884 | 1086 | - | - | 0 | - |
| - | - | 2562 | 1087 | - | - | 0 | - |
| - | - | 1749 | 1088 | - | - | 0 | - |
| - | - | 1103 | 1093 | - | - | 0 | - |
| - | - | 1.866E+04 | 1095 | - | - | 0 | - |
| - | - | 1.068E+04 | 1096 | - | - | 0 | - |
| - | - | 3834 | 1097 | - | - | 0 | - |
| - | - | 979.6 | 1105 | - | - | 0 | - |
| - | - | 1142 | 1106 | - | - | 0 | - |
| - | - | 769.8 | 1107 | - | - | 0 | - |
| - | - | 2711 | 1108 | - | - | 0 | - |
| - | - | 2043 | 1109 | - | - | 0 | - |
| - | - | 778.2 | 1110 | - | - | 0 | - |
| - | - | 1097 | 1112 | - | - | 0 | - |
| - | - | 912.2 | 1115 | - | - | 0 | - |
| - | - | 1650 | 1122 | - | - | 0 | - |
| - | - | 2082 | 1123 | - | - | 0 | - |
| - | - | 1.631E+04 | 1124 | - | - | 0 | - |
| - | - | 5.875E+04 | 1125 | - | - | 0 | - |
| - | - | 7.352E+04 | 1126 | - | - | 0 | - |
| - | - | 3.748E+04 | 1127 | - | - | 0 | - |
| - | - | 1.329E+04 | 1128 | - | - | 0 | - |
| - | - | 2255 | 1129 | - | - | 0 | - |
| - | - | 1470 | 1129 | - | - | 0 | - |
| - | - | 1192 | 1130 | - | - | 0 | - |
| - | - | 1579 | 1140 | - | - | 0 | - |
| - | - | 1879 | 1141 | - | - | 0 | - |
| - | - | 1.007E+04 | 1142 | - | - | 0 | - |
| - | - | 6163 | 1143 | - | - | 0 | - |
| - | - | 1054 | 1144 | - | - | 0 | - |
| - | - | 3075 | 1144 | - | - | 0 | - |
| - | - | 851.3 | 1145 | - | - | 0 | - |
| - | - | 1906 | 1152 | - | - | 0 | - |
| - | - | 1253 | 1153 | - | - | 0 | - |
| - | - | 2948 | 1154 | - | - | 0 | - |
| - | - | 1833 | 1155 | - | - | 0 | - |
| - | - | 1557 | 1156 | - | - | 0 | - |
| - | - | 4982 | 1158 | - | - | 0 | - |
| - | - | 3590 | 1159 | - | - | 0 | - |
| - | - | 793.5 | 1168 | - | - | 0 | - |
| - | - | 2.52E+04 | 1168 | - | - | 0 | - |
| - | - | 1.968E+05 | 1169 | - | - | 0 | - |
| - | - | 1.333E+05 | 1170 | - | - | 0 | - |
| - | - | 5.909E+04 | 1171 | - | - | 0 | - |
| - | - | 937.8 | 1172 | - | - | 0 | - |
| - | - | 8705 | 1172 | - | - | 0 | - |
| - | - | 966.9 | 1185 | - | - | 0 | - |
| - | - | 1.709E+05 | 1185 | - | - | 0 | - |
| - | - | 5.101E+05 | 1186 | - | - | 0 | - |
| - | - | 3.285E+05 | 1187 | - | - | 0 | - |
| - | - | 1.251E+05 | 1188 | - | - | 0 | - |
| - | - | 2866 | 1189 | - | - | 0 | - |
| - | - | 1.571E+04 | 1189 | - | - | 0 | - |
| - | - | 663 | 1413 | - | - | 0 | - |
| - | - | 750.2 | 1777 | - | - | 0 | - |
| - | - | 904.6 | 1783 | - | - | 0 | - |
| - | - | 649.2 | 2342 | - | - | 0 | - |
| - | - | 786.2 | 3081 | - | - | 0 | - |

m/z Charge Intensity FragmentType MassShift Position
120.08103942871094 0 6659.774
133.086181640625 0 2801.9902
136.07595825195312 0 1133.3768
141.3522491455078 0 415.34302
143.10684204101562 0 678.9949
146.8499755859375 0 450.8965
149.04498291015625 0 1297.2683
158.09652709960938 0 426.738
159.09202575683594 0 1222.0673
162.39537048339844 0 422.67834
163.2989501953125 0 438.97314
166.09747314453125 0 790.65283
176.1070556640625 0 1440.5273
177.102294921875 0 2119.444
177.11199951171875 0 2470.5986
183.08364868164062 0 492.52573
187.1331024169922 0 2284.6267
188.07080078125 0 1176.1382
203.11795043945312 0 1692.3893
205.0973358154297 0 4657.362 y 8
221.0846710205078 0 2439.8206
221.1286163330078 0 176476.36
222.08554077148438 0 607.335
222.1319122314453 0 22227.982
223.13485717773438 0 1418.9805
231.0608673095703 0 1151.5762
231.11328125 0 2189.245
231.15847778320312 0 971.2586
235.10751342773438 0 1689.8506
239.09519958496094 0 4209.38
249.12355041503906 0 58139.285 c Ammonia loss 1
250.12681579589844 0 8264.973
251.1029815673828 0 786.46
263.10272216796875 0 6450.4077
264.10577392578125 0 654.78265
264.1369323730469 0 18641.504
265.1412353515625 0 2080.108
269.10675048828125 0 2562.259
271.1225280761719 0 1252.3293
275.1849060058594 0 1382.7357
280.13238525390625 0 604.27844
281.0515441894531 0 1270.367
282.1474304199219 0 5533.6787
283.1547546386719 0 4498.2173
285.0103454589844 0 2334.326
292.16461181640625 0 625.17737
299.0619201660156 0 5399.9463
300.0626525878906 0 689.6296
324.1456604003906 0 1504.6199
326.1376953125 0 1228.9313 z 7
327.1453552246094 0 7340.014
328.14813232421875 0 1026.8137
337.2220764160156 0 1605.9913
340.14459228515625 0 1813.1571
342.15643310546875 0 34109.527 y 7
343.1595458984375 0 6434.9326
344.1607971191406 0 768.14575
346.13995361328125 0 1480.2222 z 4
355.06982421875 0 49955.586
356.070556640625 0 12907.128
364.1501770019531 0 12900.305 c Ammonia loss 2
365.1541748046875 0 2307.9517
366.15380859375 0 646.34015
367.1606140136719 0 533.86176
371.1009826660156 0 2155.6492
378.1294860839844 0 3253.3682
379.13092041015625 0 643.9668
381.2489318847656 0 1033.2477
382.2539367675781 0 539.30237
394.1256103515625 0 1273.9993
396.1661071777344 0 957.48334 w 6
399.15753173828125 0 1171.8903
413.1883544921875 0 7960.572
414.1944580078125 0 5107.6494
415.03704833984375 0 2474.1472
415.1981506347656 0 1510.1547
416.0384826660156 0 758.17163
418.9931640625 0 702.0776
425.27642822265625 0 777.49866
433.1722412109375 0 847.02954
457.17718505859375 0 3062.3262 z 6
458.185791015625 0 13460.28
459.1889953613281 0 3429.6042
460.18585205078125 0 884.834
461.1654052734375 0 1413.8352 z 2
465.16217041015625 0 2226.5125
467.2042541503906 0 2438.4653
469.3009033203125 0 798.93225
472.188232421875 0 849.99854
473.1966857910156 0 17318.49 y 6
474.199951171875 0 4820.495
475.20587158203125 0 617.6866
476.2150573730469 0 613.3503
477.1583251953125 0 725.84576
479.177734375 0 7104.0127 c Ammonia loss 3
480.1797790527344 0 1290.3967
481.2182922363281 0 589.6441
484.22509765625 0 4557.058
485.2300109863281 0 2064.6191
486.2328796386719 0 678.97687
487.31011962890625 0 630.3688
491.6974182128906 0 860.71265
500.183837890625 0 946.7122
503.2087097167969 0 1845.36
504.2139892578125 0 816.86945
505.22320556640625 0 2183.5789
506.226318359375 0 997.81976
528.2150268554688 0 119765.3 z 5
529.2196655273438 0 50964.11
530.2196044921875 0 12031.179
531.2224731445312 0 1721.1365
533.707763671875 0 1206.1831 y Water loss 1
537.1450805664062 0 595.23267
542.2200927734375 0 586.90454
542.7135620117188 0 7262.773 y 1
543.2161254882812 0 4818.882
543.7151489257812 0 1620.0867
544.2338256835938 0 22222.281 y 5
545.2361450195312 0 6934.452
546.233642578125 0 2008.722
558.2250366210938 0 700.7791
560.2030029296875 0 1016.1636
562.221923828125 0 1556.402
571.22216796875 0 1604.758
572.2236328125 0 669.5216
575.2265014648438 0 1509.1085
575.3634643554688 0 12885.845
576.2589111328125 0 2526.7158
576.3670043945312 0 5519.365
577.2658081054688 0 746.9343
577.3671875 0 737.6488
584.2313232421875 0 4324.263
584.7327880859375 0 3910.7156
585.2325439453125 0 1965.339
590.2376098632812 0 1079.2628
592.390625 0 1233.054
593.237060546875 0 10223.868
593.7383422851562 0 8735.416
594.239501953125 0 2858.5981
596.196044921875 0 1075.5327
596.2444458007812 0 663.0559
599.2362670898438 0 850.24585
614.24462890625 0 1990.5436
617.263671875 0 810.4148
618.2350463867188 0 2432.2444
620.2492065429688 0 4788.9023
621.25244140625 0 2004.9025
624.2301635742188 0 1629.8458
625.233154296875 0 779.22833
630.2673950195312 0 3151.703
631.2678833007812 0 907.1932
633.2548217773438 0 1235.275
642.239501953125 0 1957.7759 c Ammonia loss 4
646.2647094726562 0 2273.3086
647.2867431640625 0 3672.8564
648.291259765625 0 1852.6542
677.2478637695312 0 1826.9482
686.28759765625 0 945.43945
688.2960205078125 0 6837.575
689.2974243164062 0 3220.965
690.2584838867188 0 1053.5739
691.2781982421875 0 64310.27 z 4
692.282470703125 0 35016.324
693.2832641601562 0 10543.728
694.2842407226562 0 1708.3572
695.2680053710938 0 2310.0994
701.3036499023438 0 8373.269
702.3071899414062 0 2558.3853
703.3095703125 0 1014.1348
705.2681884765625 0 4265.418
706.2794799804688 0 2519.8555
707.2962646484375 0 35238.348 y 4
708.29931640625 0 13857.154
709.29833984375 0 3924.9888
713.2783203125 0 1704.0115 c Ammonia loss 5
714.2765502929688 0 946.943
715.3200073242188 0 1305.8391
716.3309326171875 0 685.107
718.3275756835938 0 827.249
729.29541015625 0 5132.111
730.3034057617188 0 19184.78 c 5
731.3071899414062 0 6842.8555
732.2877197265625 0 8709.342
733.2625732421875 0 16262.508
734.2713623046875 0 8248.147
735.2760620117188 0 7216.211
736.280029296875 0 2470.305
737.2865600585938 0 784.9769
745.2940063476562 0 15306.227
746.2969360351562 0 6071.287
747.2949829101562 0 1867.0677
748.2880249023438 0 978.01385
751.2733154296875 0 1775.3273
759.3179321289062 0 1653.5938
760.3140869140625 0 1075.0055
761.2886352539062 0 1012.53174
762.3146362304688 0 22944.484
763.3179321289062 0 10485.392
764.3193969726562 0 3875.0515
776.3189086914062 0 708.6912
786.3190307617188 0 1077.0234
788.296630859375 0 897.63324 z Water loss 3
792.2803955078125 0 1189.3109
803.3223876953125 0 8756.536
804.3231811523438 0 3594.4866 y Water loss 3
805.3179321289062 0 1428.4471
806.3048095703125 0 41851.164 z 3
807.3104858398438 0 43919.27
808.3124389648438 0 17531.205
809.314453125 0 4185.0146
816.3307495117188 0 6570.494
817.3314208984375 0 8378.91
818.3361206054688 0 6126.68
819.3411254882812 0 2135.6213
822.3233032226562 0 39423.93 y 3
823.32666015625 0 17631.164
824.3276977539062 0 5861.6294
825.3242797851562 0 686.33606
830.3453369140625 0 4713.6807
831.3502197265625 0 2615.722
833.3516845703125 0 7176.8804
834.3555908203125 0 3751.3813
835.3530883789062 0 642.8049
842.3280029296875 0 820.6215
843.3308715820312 0 717.3324 c Water loss 6
844.3212280273438 0 2384.3103 c Ammonia loss 6
845.3208618164062 0 1323.104
847.3129272460938 0 6085.203
848.3169555664062 0 2705.3452
850.3192749023438 0 2029.3829
851.323486328125 0 1090.0448
860.3289794921875 0 25696.957
861.34228515625 0 72400.625 c 6
862.345458984375 0 31193.89
863.3453979492188 0 10101.661
864.3495483398438 0 1106.1189
874.3361206054688 0 3845.8372
875.3409423828125 0 2639.6873
876.3386840820312 0 1213.9961 w 2
877.3416137695312 0 26465.838
878.344482421875 0 13838.534
879.3450927734375 0 4222.7817
880.3431396484375 0 1657.6942
881.3524780273438 0 1422.7986
882.3464965820312 0 1444.5603
883.3501586914062 0 964.2001
891.3428344726562 0 1218.7081
893.3742065429688 0 2522.6326
894.384033203125 0 1490.2432
910.3972778320312 0 3497.5752
911.400146484375 0 1871.5603
919.33935546875 0 7301.562 y Water loss 2
920.3389282226562 0 4495.0195
921.3319091796875 0 14921.061 z 2
922.33447265625 0 8374.451
923.3363647460938 0 2828.2153
936.3775024414062 0 6273.1416
937.3507690429688 0 161739.62 y 2
938.3534545898438 0 86646.266
939.3535766601562 0 30293.445
940.3522338867188 0 3226.1104
953.376953125 0 1477.0554
954.3893432617188 0 21619.156
955.3917236328125 0 10475.497
956.3934936523438 0 2471.4187
981.3792114257812 0 3124.7688 c Ammonia loss 7
982.38037109375 0 1644.8734
983.390869140625 0 5275.095
984.3926391601562 0 2526.2402
985.3845825195312 0 1113.9823
994.3810424804688 0 3760.203
995.388916015625 0 2778.9937
996.3912353515625 0 805.73364
998.4029541015625 0 206509.62 c 7
999.4053955078125 0 114818.984
1000.4061889648438 0 42319.4
1001.4066162109375 0 4912.5664
1024.41162109375 0 7226.9004
1025.4127197265625 0 4991.197
1026.4161376953125 0 1922.1693
1040.3861083984375 0 1085.3694
1041.4000244140625 0 1168.7288
1052.4339599609375 0 733.5446
1055.3980712890625 0 3810.12
1056.4027099609375 0 2262.9473
1057.399658203125 0 661.8781
1065.4532470703125 0 784.19543
1066.420166015625 0 1389.1499 y Water loss 1
1067.4110107421875 0 1084.9817
1068.39892578125 0 27089.47 z 1
1069.40234375 0 16034.819
1070.40380859375 0 6245.1123
1079.4635009765625 0 1125.7516
1081.47119140625 0 1059.4993
1082.4639892578125 0 1426.8035
1084.418212890625 0 10378.832 y 1
1085.4212646484375 0 7123.958
1086.4208984375 0 2884.389
1087.416259765625 0 2561.5283
1088.4195556640625 0 1749.1475
1093.4774169921875 0 1103.1774
1095.427490234375 0 18663.473
1096.4312744140625 0 10682.128
1097.4375 0 3834.113
1105.424072265625 0 979.63525
1106.444580078125 0 1141.9613
1107.4541015625 0 769.7992
1108.4393310546875 0 2710.5305
1109.4447021484375 0 2042.5267
1110.452880859375 0 778.21564
1112.4453125 0 1097.0826
1115.4754638671875 0 912.18353
1122.4512939453125 0 1650.0123
1123.4532470703125 0 2082.4553
1124.4710693359375 0 16311.995
1125.460205078125 0 58748.566
1126.456787109375 0 73515.125
1127.4578857421875 0 37476.707
1128.4580078125 0 13288.557
1128.60595703125 0 2254.7126
1129.4637451171875 0 1469.755
1129.5987548828125 0 1192.2141
1140.47314453125 0 1578.8595
1141.4708251953125 0 1879.3945
1142.4825439453125 0 10073.145
1143.486572265625 0 6162.604
1143.62158203125 0 1053.9562
1144.489990234375 0 3074.7642
1144.6279296875 0 851.3057
1152.4444580078125 0 1906.2732
1153.44775390625 0 1252.8942
1154.4254150390625 0 2948.1833
1155.425537109375 0 1832.5106
1156.4217529296875 0 1556.6702
1158.4765625 0 4982.4956
1159.4791259765625 0 3589.6633
1167.646484375 0 793.50116
1168.4625244140625 0 25202.506
1169.4483642578125 0 196846.3
1170.4501953125 0 133298.39
1171.4505615234375 0 59094.523
1171.5802001953125 0 937.8105
1172.4500732421875 0 8705.146
1184.6856689453125 0 966.9116
1185.465087890625 0 170946.62
1186.4722900390625 0 510113.66
1187.47509765625 0 328509.8
1188.476318359375 0 125112.7
1188.58984375 0 2865.933
1189.478759765625 0 15711.053
1412.7396240234375 0 662.974
1776.8309326171875 0 750.2061
1782.93603515625 0 904.5846
2341.721435546875 0 649.21
3080.768798828125 0 786.2408

Spectrum Details

|  |  |
| --- | --- |
| Matched peaks? Matched peaks The total absolute number of peaks matched. Additionally in brackets the total fraction of peaks matched and the total number of peaks is shown. | 36 (10.14% of 355) |
| FDR? FDR The false discovery rate estimated for this peptide. It is calculated by matching all theoretical fragments with a non-integer shift with the raw peaks for this spectrum. This is done with 40 different shifts. The resulting percentage is the average number of annotated peaks over the number of annotated peaks with the correct spectrum. | 0.40% |
| Satellite FDR? Satellite FDR See the FDR for details on its calculation. This satellite ion specific FDR only contains the satellite ions (d/w) for I/L/J positions. | - |
| PSM Score? PSM Score The PSM Score as given by Hecklib to this annotated spectrum. It is shown with three significant figures. | 495 |

#### Spectrum 7880? Spectrum 7880 The raw spectrum of this peptide as annotated by Hecklib. The fragments are coloured according to ion type (see legend). Any peaks with a star '\*' as text can be hovered over to see the full details, first the ion type second the mass shift type. By hovering over the amino acids in the peptide or ions in the legend the corresponding peaks are highlighted. By toggling the 'Unassigned' label you can turn the background (unassigned) peaks on or off in the plot. By updating the slider in the Ion legend you can update the spectrum to only show the top X% of the peaks with labels. The top X% means any peak that is within X% of the highest intensity. By dragging in the spectrum you can zoom in to a specific part of the spectrum and use 'Zoom Out' to get back to the original zoom level. The annotation of the spectrum is based on the given sequence in the peptides file and is done with different software so inconsistencies are likely. The peaks are annotated based on the given sequence, with 20 ppm tolerance.

Copy Data

##### Spectrum 7880 (TSV)

###### Preview

```
Loading example...
```

*Click on the button to copy the data to your clipboard.*

Mz MinMz MaxIntensity Max

WidthHeightPeptide font sizePeptide stroke widthSpectrum font sizeSpectrum stroke widthCompact peptide

Ion legend

wxyz

abcd

OtherUnassignedIon^Charge^_Position_Show for top:%

TFDDYAMHW

09.65e+41.93e+52.90e+53.86e+5

Zoom Out

y^+1^_1_c^+1^_2_y^+1^_2_c^+1^_3_z^+1^_3_y^+2^_7_c^+1^_4_y^+1^_3_c^+2^_8_z^+1^_4_y^+2^_8_y^+1^_4_c^+1^_5_z^+1^_5_c^+1^_6_y^+1^_5_c^+1^_6_w^+1^_6_z^+1^_6_y^+1^_6_c^+1^_7_c^+1^_7_c^+1^_7_w^+1^_7_y^+1^_7_z^+1^_7_y^+1^_7_c^+1^_8_c^+1^_8_z^+1^_8_y^+1^_8_

0631126218922523

Fragment Matches Table

Show background peaks

| Position | Ion type | Intensity | mz Theoretical | mz Error (Th) | mz Error (ppm) | Charge | Series Number |
| --- | --- | --- | --- | --- | --- | --- | --- |
| - | - | 3061 | 120.1 | - | - | 0 | - |
| - | - | 348.3 | 124.1 | - | - | 0 | - |
| - | - | 473 | 134.9 | - | - | 0 | - |
| - | - | 804.3 | 136.1 | - | - | 0 | - |
| - | - | 405.1 | 137 | - | - | 0 | - |
| - | - | 399.7 | 138.5 | - | - | 0 | - |
| - | - | 367.8 | 139.5 | - | - | 0 | - |
| - | - | 388.3 | 140.7 | - | - | 0 | - |
| - | - | 403.8 | 145.7 | - | - | 0 | - |
| - | - | 744.8 | 149 | - | - | 0 | - |
| - | - | 788.8 | 159.1 | - | - | 0 | - |
| - | - | 690.6 | 166.1 | - | - | 0 | - |
| - | - | 441.3 | 166.5 | - | - | 0 | - |
| - | - | 436.5 | 172.6 | - | - | 0 | - |
| - | - | 599.3 | 176.1 | - | - | 0 | - |
| - | - | 797.7 | 177.1 | - | - | 0 | - |
| - | - | 706.2 | 188.1 | - | - | 0 | - |
| - | - | 1680 | 203.1 | - | - | 0 | - |
| 9 | y | 2282 | 205.1 | 0.000258 | 1.258 | +1 | 1 |
| - | - | 1.095E+05 | 221.1 | - | - | 0 | - |
| - | - | 1.37E+04 | 222.1 | - | - | 0 | - |
| - | - | 728.8 | 223.1 | - | - | 0 | - |
| - | - | 1165 | 223.1 | - | - | 0 | - |
| - | - | 598.2 | 229.2 | - | - | 0 | - |
| - | - | 713.5 | 231.1 | - | - | 0 | - |
| - | - | 770.5 | 235.1 | - | - | 0 | - |
| - | - | 536.7 | 236.5 | - | - | 0 | - |
| - | - | 536.3 | 243.7 | - | - | 0 | - |
| - | - | 503.5 | 245.8 | - | - | 0 | - |
| 2 | c | 3.915E+04 | 249.1 | 0.0002273 | 0.9126 | +1 | 2 |
| - | - | 5581 | 250.1 | - | - | 0 | - |
| - | - | 2129 | 263.1 | - | - | 0 | - |
| - | - | 674.2 | 263.1 | - | - | 0 | - |
| - | - | 7863 | 264.1 | - | - | 0 | - |
| - | - | 1078 | 265.1 | - | - | 0 | - |
| - | - | 521.3 | 278 | - | - | 0 | - |
| - | - | 2979 | 282.1 | - | - | 0 | - |
| - | - | 3568 | 283.2 | - | - | 0 | - |
| - | - | 931.1 | 284.2 | - | - | 0 | - |
| - | - | 1086 | 292.2 | - | - | 0 | - |
| - | - | 3443 | 327.1 | - | - | 0 | - |
| 8 | y | 1.122E+04 | 342.2 | 0.0003366 | 0.9839 | +1 | 2 |
| - | - | 2519 | 343.2 | - | - | 0 | - |
| - | - | 896.9 | 346.1 | - | - | 0 | - |
| - | - | 772.2 | 355.1 | - | - | 0 | - |
| - | - | 906.4 | 356.1 | - | - | 0 | - |
| - | - | 1166 | 358.2 | - | - | 0 | - |
| 3 | c | 1.028E+04 | 364.2 | 4.823E-05 | 0.1325 | +1 | 3 |
| - | - | 2824 | 365.2 | - | - | 0 | - |
| - | - | 1806 | 378.1 | - | - | 0 | - |
| - | - | 901.9 | 394.1 | - | - | 0 | - |
| - | - | 959.4 | 411.2 | - | - | 0 | - |
| - | - | 675.2 | 412.2 | - | - | 0 | - |
| - | - | 3733 | 413.3 | - | - | 0 | - |
| - | - | 758.9 | 414.3 | - | - | 0 | - |
| - | - | 880 | 415.2 | - | - | 0 | - |
| - | - | 1942 | 429.2 | - | - | 0 | - |
| - | - | 942.9 | 430.2 | - | - | 0 | - |
| - | - | 546.9 | 432 | - | - | 0 | - |
| - | - | 713.4 | 433.2 | - | - | 0 | - |
| - | - | 644.1 | 454.8 | - | - | 0 | - |
| - | - | 865.5 | 461.2 | - | - | 0 | - |
| 7 | z | 5135 | 473.2 | 0.005086 | 10.75 | +1 | 3 |
| - | - | 2.324E+04 | 474.2 | - | - | 0 | - |
| - | - | 6194 | 475.2 | - | - | 0 | - |
| - | - | 1020 | 476.2 | - | - | 0 | - |
| 3 | y | 908.8 | 477.2 | 0.0005744 | 1.204 | +2 | 7 |
| - | - | 788.3 | 477.7 | - | - | 0 | - |
| 4 | c | 6758 | 479.2 | 0.0006015 | 1.255 | +1 | 4 |
| - | - | 1035 | 480.2 | - | - | 0 | - |
| - | - | 1065 | 488.2 | - | - | 0 | - |
| 7 | y | 1.095E+04 | 489.2 | 0.005161 | 10.55 | +1 | 3 |
| - | - | 3023 | 490.2 | - | - | 0 | - |
| - | - | 2448 | 496.2 | - | - | 0 | - |
| 8 | c | 1086 | 499.2 | 0.0002844 | 0.5697 | +2 | 8 |
| - | - | 3661 | 500.2 | - | - | 0 | - |
| - | - | 1899 | 501.2 | - | - | 0 | - |
| - | - | 879.5 | 516.2 | - | - | 0 | - |
| - | - | 703.6 | 518.7 | - | - | 0 | - |
| - | - | 596.7 | 520.2 | - | - | 0 | - |
| - | - | 2511 | 521.2 | - | - | 0 | - |
| - | - | 808.2 | 522.2 | - | - | 0 | - |
| - | - | 662.2 | 542.2 | - | - | 0 | - |
| 6 | z | 7.481E+04 | 544.2 | 0.005021 | 9.226 | +1 | 4 |
| - | - | 3.399E+04 | 545.2 | - | - | 0 | - |
| - | - | 9352 | 546.2 | - | - | 0 | - |
| - | - | 1505 | 547.2 | - | - | 0 | - |
| 2 | y | 6199 | 550.7 | 0.002042 | 3.709 | +2 | 8 |
| - | - | 3498 | 551.2 | - | - | 0 | - |
| - | - | 1371 | 551.7 | - | - | 0 | - |
| 6 | y | 1.695E+04 | 560.2 | 0.00479 | 8.551 | +1 | 4 |
| - | - | 5584 | 561.2 | - | - | 0 | - |
| - | - | 1320 | 562.2 | - | - | 0 | - |
| - | - | 2311 | 569.2 | - | - | 0 | - |
| - | - | 1153 | 569.7 | - | - | 0 | - |
| - | - | 640.7 | 578.2 | - | - | 0 | - |
| - | - | 832.1 | 583.2 | - | - | 0 | - |
| - | - | 2432 | 587.2 | - | - | 0 | - |
| - | - | 3979 | 592.2 | - | - | 0 | - |
| - | - | 2435 | 592.7 | - | - | 0 | - |
| - | - | 978.6 | 593.3 | - | - | 0 | - |
| - | - | 849.9 | 596.2 | - | - | 0 | - |
| - | - | 1.472E+04 | 601.2 | - | - | 0 | - |
| - | - | 6694 | 601.7 | - | - | 0 | - |
| - | - | 4062 | 602.2 | - | - | 0 | - |
| - | - | 1731 | 614.2 | - | - | 0 | - |
| - | - | 1136 | 624.2 | - | - | 0 | - |
| - | - | 3190 | 636.2 | - | - | 0 | - |
| 5 | c | 1369 | 642.2 | 0.0007153 | 1.114 | +1 | 5 |
| - | - | 1173 | 649.3 | - | - | 0 | - |
| - | - | 805.5 | 650.3 | - | - | 0 | - |
| - | - | 1876 | 659.3 | - | - | 0 | - |
| - | - | 1403 | 662.3 | - | - | 0 | - |
| - | - | 2290 | 663.3 | - | - | 0 | - |
| - | - | 854.1 | 664.3 | - | - | 0 | - |
| - | - | 632.7 | 687.9 | - | - | 0 | - |
| - | - | 1331 | 688.3 | - | - | 0 | - |
| - | - | 795.8 | 693.2 | - | - | 0 | - |
| - | - | 734.8 | 706.3 | - | - | 0 | - |
| 5 | z | 4.168E+04 | 707.3 | 0.005169 | 7.308 | +1 | 5 |
| - | - | 1.878E+04 | 708.3 | - | - | 0 | - |
| - | - | 6746 | 709.3 | - | - | 0 | - |
| - | - | 640.6 | 710.3 | - | - | 0 | - |
| 6 | c | 679.3 | 713.3 | 0.002673 | 3.747 | +1 | 6 |
| - | - | 2243 | 715.3 | - | - | 0 | - |
| - | - | 1572 | 716.3 | - | - | 0 | - |
| - | - | 779.4 | 721.3 | - | - | 0 | - |
| - | - | 1810 | 722.3 | - | - | 0 | - |
| 5 | y | 1.964E+04 | 723.3 | 0.00445 | 6.152 | +1 | 5 |
| - | - | 8011 | 724.3 | - | - | 0 | - |
| - | - | 2906 | 725.3 | - | - | 0 | - |
| 6 | c | 1401 | 730.3 | 0.004259 | 5.831 | +1 | 6 |
| - | - | 697.1 | 732.3 | - | - | 0 | - |
| - | - | 743.9 | 734.3 | - | - | 0 | - |
| - | - | 5372 | 749.3 | - | - | 0 | - |
| - | - | 3614 | 750.3 | - | - | 0 | - |
| - | - | 2694 | 751.3 | - | - | 0 | - |
| - | - | 1155 | 752.3 | - | - | 0 | - |
| - | - | 1708 | 759.3 | - | - | 0 | - |
| - | - | 1067 | 760.3 | - | - | 0 | - |
| - | - | 2563 | 764.3 | - | - | 0 | - |
| - | - | 1101 | 767.3 | - | - | 0 | - |
| - | - | 1081 | 771.3 | - | - | 0 | - |
| - | - | 817.7 | 772.3 | - | - | 0 | - |
| - | - | 2399 | 774.3 | - | - | 0 | - |
| - | - | 1338 | 775.3 | - | - | 0 | - |
| 4 | w | 1140 | 777.3 | 0.004406 | 5.668 | +1 | 6 |
| - | - | 2.451E+04 | 778.3 | - | - | 0 | - |
| - | - | 1.164E+04 | 779.3 | - | - | 0 | - |
| - | - | 3067 | 780.3 | - | - | 0 | - |
| 4 | z | 3.327E+04 | 822.3 | 0.004593 | 5.585 | +1 | 6 |
| - | - | 3.355E+04 | 823.3 | - | - | 0 | - |
| - | - | 1.276E+04 | 824.3 | - | - | 0 | - |
| - | - | 3839 | 825.3 | - | - | 0 | - |
| - | - | 4604 | 830.3 | - | - | 0 | - |
| - | - | 3329 | 831.3 | - | - | 0 | - |
| - | - | 1185 | 832.4 | - | - | 0 | - |
| - | - | 1.226E+04 | 833.3 | - | - | 0 | - |
| - | - | 9320 | 834.3 | - | - | 0 | - |
| - | - | 2612 | 835.3 | - | - | 0 | - |
| - | - | 617.5 | 836.3 | - | - | 0 | - |
| - | - | 836.7 | 837.3 | - | - | 0 | - |
| 4 | y | 2.285E+04 | 838.3 | 0.004179 | 4.985 | +1 | 6 |
| - | - | 9257 | 839.3 | - | - | 0 | - |
| - | - | 3635 | 840.3 | - | - | 0 | - |
| - | - | 859.4 | 842.3 | - | - | 0 | - |
| - | - | 667.4 | 848.3 | - | - | 0 | - |
| - | - | 1.426E+04 | 849.3 | - | - | 0 | - |
| - | - | 8424 | 850.3 | - | - | 0 | - |
| - | - | 2404 | 851.3 | - | - | 0 | - |
| - | - | 2561 | 858.3 | - | - | 0 | - |
| 7 | c | 1820 | 859.3 | 0.01118 | 13.01 | +1 | 7 |
| 7 | c | 2319 | 860.3 | 0.003036 | 3.529 | +1 | 7 |
| - | - | 745 | 866.3 | - | - | 0 | - |
| - | - | 784.3 | 871.3 | - | - | 0 | - |
| - | - | 2572 | 874.3 | - | - | 0 | - |
| - | - | 1197 | 875.3 | - | - | 0 | - |
| - | - | 1.548E+04 | 876.3 | - | - | 0 | - |
| 7 | c | 6.715E+04 | 877.3 | 0.003892 | 4.436 | +1 | 7 |
| - | - | 3.087E+04 | 878.3 | - | - | 0 | - |
| - | - | 1.07E+04 | 879.3 | - | - | 0 | - |
| - | - | 1.387E+04 | 889.3 | - | - | 0 | - |
| - | - | 6019 | 890.3 | - | - | 0 | - |
| - | - | 1911 | 891.4 | - | - | 0 | - |
| 3 | w | 969.9 | 892.3 | 0.005211 | 5.84 | +1 | 7 |
| - | - | 3.93E+04 | 893.3 | - | - | 0 | - |
| - | - | 1.902E+04 | 894.3 | - | - | 0 | - |
| - | - | 6944 | 895.3 | - | - | 0 | - |
| - | - | 889.3 | 896.3 | - | - | 0 | - |
| - | - | 1209 | 898.3 | - | - | 0 | - |
| - | - | 1535 | 899.9 | - | - | 0 | - |
| - | - | 1966 | 900.4 | - | - | 0 | - |
| - | - | 862.2 | 900.9 | - | - | 0 | - |
| - | - | 1464 | 901.4 | - | - | 0 | - |
| - | - | 1412 | 902.9 | - | - | 0 | - |
| - | - | 1207 | 903.4 | - | - | 0 | - |
| - | - | 1113 | 926.4 | - | - | 0 | - |
| - | - | 810.1 | 927.4 | - | - | 0 | - |
| 3 | y | 4592 | 935.3 | 0.003487 | 3.728 | +1 | 7 |
| - | - | 2701 | 936.3 | - | - | 0 | - |
| 3 | z | 9513 | 937.3 | 0.004994 | 5.327 | +1 | 7 |
| - | - | 4355 | 938.3 | - | - | 0 | - |
| - | - | 2398 | 939.3 | - | - | 0 | - |
| - | - | 2342 | 952.4 | - | - | 0 | - |
| 3 | y | 9.691E+04 | 953.3 | 0.004641 | 4.868 | +1 | 7 |
| - | - | 4.746E+04 | 954.3 | - | - | 0 | - |
| - | - | 1.82E+04 | 955.3 | - | - | 0 | - |
| - | - | 2625 | 956.4 | - | - | 0 | - |
| - | - | 1.387E+04 | 970.4 | - | - | 0 | - |
| - | - | 833.1 | 970.5 | - | - | 0 | - |
| - | - | 6951 | 971.4 | - | - | 0 | - |
| - | - | 1882 | 972.4 | - | - | 0 | - |
| 8 | c | 1707 | 997.4 | 0.005343 | 5.357 | +1 | 8 |
| - | - | 2154 | 998.4 | - | - | 0 | - |
| - | - | 1777 | 999.4 | - | - | 0 | - |
| 8 | c | 1.243E+05 | 1014 | 0.003879 | 3.824 | +1 | 8 |
| - | - | 6.761E+04 | 1015 | - | - | 0 | - |
| - | - | 2.586E+04 | 1016 | - | - | 0 | - |
| - | - | 2509 | 1017 | - | - | 0 | - |
| - | - | 944 | 1036 | - | - | 0 | - |
| - | - | 6930 | 1040 | - | - | 0 | - |
| - | - | 3764 | 1041 | - | - | 0 | - |
| - | - | 1707 | 1042 | - | - | 0 | - |
| - | - | 1531 | 1069 | - | - | 0 | - |
| - | - | 1751 | 1071 | - | - | 0 | - |
| - | - | 807.9 | 1072 | - | - | 0 | - |
| 2 | z | 1.284E+04 | 1084 | 0.003718 | 3.429 | +1 | 8 |
| - | - | 7079 | 1085 | - | - | 0 | - |
| - | - | 2548 | 1086 | - | - | 0 | - |
| - | - | 1848 | 1094 | - | - | 0 | - |
| - | - | 1380 | 1095 | - | - | 0 | - |
| - | - | 737 | 1096 | - | - | 0 | - |
| - | - | 1056 | 1097 | - | - | 0 | - |
| 2 | y | 5744 | 1100 | 0.003671 | 3.336 | +1 | 8 |
| - | - | 2917 | 1101 | - | - | 0 | - |
| - | - | 1546 | 1102 | - | - | 0 | - |
| - | - | 3787 | 1103 | - | - | 0 | - |
| - | - | 2473 | 1104 | - | - | 0 | - |
| - | - | 730.6 | 1105 | - | - | 0 | - |
| - | - | 2199 | 1110 | - | - | 0 | - |
| - | - | 886.1 | 1111 | - | - | 0 | - |
| - | - | 3355 | 1122 | - | - | 0 | - |
| - | - | 3017 | 1123 | - | - | 0 | - |
| - | - | 1116 | 1124 | - | - | 0 | - |
| - | - | 757 | 1128 | - | - | 0 | - |
| - | - | 3738 | 1138 | - | - | 0 | - |
| - | - | 2627 | 1139 | - | - | 0 | - |
| - | - | 2.476E+04 | 1140 | - | - | 0 | - |
| - | - | 3.628E+04 | 1141 | - | - | 0 | - |
| - | - | 4.229E+04 | 1142 | - | - | 0 | - |
| - | - | 2.279E+04 | 1143 | - | - | 0 | - |
| - | - | 8341 | 1144 | - | - | 0 | - |
| - | - | 527.8 | 1145 | - | - | 0 | - |
| - | - | 1182 | 1145 | - | - | 0 | - |
| - | - | 1061 | 1156 | - | - | 0 | - |
| - | - | 2114 | 1158 | - | - | 0 | - |
| - | - | 1525 | 1159 | - | - | 0 | - |
| - | - | 758.2 | 1166 | - | - | 0 | - |
| - | - | 2995 | 1170 | - | - | 0 | - |
| - | - | 1088 | 1171 | - | - | 0 | - |
| - | - | 2672 | 1174 | - | - | 0 | - |
| - | - | 2139 | 1175 | - | - | 0 | - |
| - | - | 1.997E+04 | 1184 | - | - | 0 | - |
| - | - | 1.733E+05 | 1185 | - | - | 0 | - |
| - | - | 1.181E+05 | 1186 | - | - | 0 | - |
| - | - | 5.055E+04 | 1187 | - | - | 0 | - |
| - | - | 8971 | 1188 | - | - | 0 | - |
| - | - | 779 | 1189 | - | - | 0 | - |
| - | - | 1.134E+05 | 1201 | - | - | 0 | - |
| - | - | 3.823E+05 | 1202 | - | - | 0 | - |
| - | - | 2.34E+05 | 1203 | - | - | 0 | - |
| - | - | 2895 | 1204 | - | - | 0 | - |
| - | - | 9.147E+04 | 1204 | - | - | 0 | - |
| - | - | 1.438E+04 | 1205 | - | - | 0 | - |
| - | - | 877.4 | 1782 | - | - | 0 | - |
| - | - | 783.2 | 1783 | - | - | 0 | - |
| - | - | 1009 | 1784 | - | - | 0 | - |
| - | - | 715.2 | 1785 | - | - | 0 | - |
| - | - | 1664 | 1788 | - | - | 0 | - |
| - | - | 2682 | 1789 | - | - | 0 | - |
| - | - | 786.2 | 1790 | - | - | 0 | - |
| - | - | 2001 | 1800 | - | - | 0 | - |
| - | - | 1862 | 1801 | - | - | 0 | - |
| - | - | 2123 | 1802 | - | - | 0 | - |
| - | - | 1168 | 1803 | - | - | 0 | - |
| - | - | 881.6 | 1805 | - | - | 0 | - |
| - | - | 1151 | 1806 | - | - | 0 | - |
| - | - | 704.5 | 1969 | - | - | 0 | - |
| - | - | 684.2 | 2132 | - | - | 0 | - |
| - | - | 777.4 | 2142 | - | - | 0 | - |
| - | - | 1481 | 2406 | - | - | 0 | - |
| - | - | 1999 | 2407 | - | - | 0 | - |
| - | - | 862.5 | 2408 | - | - | 0 | - |
| - | - | 696.8 | 2498 | - | - | 0 | - |

m/z Charge Intensity FragmentType MassShift Position
120.08098602294922 0 3060.619
124.13399505615234 0 348.33688
134.90725708007812 0 473.0005
136.07611083984375 0 804.2848
136.98207092285156 0 405.14786
138.53561401367188 0 399.74307
139.50379943847656 0 367.7749
140.7045440673828 0 388.33395
145.66432189941406 0 403.75238
148.9550323486328 0 744.8471
159.09182739257812 0 788.8148
166.09835815429688 0 690.583
166.46791076660156 0 441.34067
172.63453674316406 0 436.50793
176.10723876953125 0 599.33655
177.10267639160156 0 797.74384
188.07081604003906 0 706.19
203.117919921875 0 1680.4261
205.097412109375 0 2281.942 y 8
221.12863159179688 0 109451.67
222.132080078125 0 13698.19
223.06399536132812 0 728.793
223.13478088378906 0 1164.5566
229.15499877929688 0 598.16547
231.11338806152344 0 713.4639
235.10777282714844 0 770.46173
236.47164916992188 0 536.7272
243.7183837890625 0 536.3003
245.82318115234375 0 503.51514
249.12359619140625 0 39151.977 c Ammonia loss 1
250.126953125 0 5580.9
263.1025085449219 0 2128.6423
263.12933349609375 0 674.1803
264.1369934082031 0 7863.314
265.1427001953125 0 1078.3494
277.9958801269531 0 521.2621
282.1479187011719 0 2979.2715
283.1551208496094 0 3567.6555
284.1589660644531 0 931.1158
292.166015625 0 1085.8943
327.1454162597656 0 3443.3003
342.1564025878906 0 11222.507 y 7
343.15899658203125 0 2519.3245
346.138916015625 0 896.86237
355.0700378417969 0 772.15076
356.1385498046875 0 906.3528
358.1551818847656 0 1165.984
364.1503601074219 0 10278.171 c Ammonia loss 2
365.1535339355469 0 2823.5461
378.1302185058594 0 1805.7346
394.124267578125 0 901.87994
411.1891784667969 0 959.39954
412.197021484375 0 675.2073
413.2666015625 0 3733.035
414.2679138183594 0 758.942
415.1535339355469 0 879.99835
429.1824951171875 0 1941.754
430.1888122558594 0 942.93066
431.9551086425781 0 546.93286
433.1736755371094 0 713.3917
454.768798828125 0 644.084
461.1658630371094 0 865.5308
473.17291259765625 0 5134.668 z 6
474.1806335449219 0 23240.94
475.1833190917969 0 6194.1953
476.1863708496094 0 1020.21436
477.1746520996094 0 908.81274 y 2
477.6778564453125 0 788.289
479.1778564453125 0 6758.465 c Ammonia loss 3
480.17974853515625 0 1034.586
488.18359375 0 1064.9645
489.19171142578125 0 10946.383 y 6
490.1945495605469 0 3023.177
496.23101806640625 0 2448.3047
499.1874694824219 0 1086.313 c Ammonia loss 7
500.2204284667969 0 3660.7166
501.22393798828125 0 1899.2701
516.1791381835938 0 879.49695
518.7103881835938 0 703.617
520.2035522460938 0 596.7316
521.2174682617188 0 2510.8809
522.2218627929688 0 808.1527
542.212646484375 0 662.24243
544.2099609375 0 74806.88 z 5
545.2144775390625 0 33992.47
546.2155151367188 0 9351.676
547.2203369140625 0 1504.6086
550.7103271484375 0 6198.539 y 1
551.2127075195312 0 3497.588
551.714599609375 0 1370.671
560.2284545898438 0 16950.57 y 5
561.2319946289062 0 5584.1875
562.2291259765625 0 1319.8857
569.2348022460938 0 2311.2817
569.7340087890625 0 1152.7816
578.2210083007812 0 640.69684
583.2195434570312 0 832.0686
587.2156982421875 0 2432.4028
592.230712890625 0 3978.7422
592.7313842773438 0 2435.293
593.2577514648438 0 978.58435
596.233642578125 0 849.8672
601.2341918945312 0 14717.502
601.7356567382812 0 6693.524
602.236572265625 0 4061.984
614.245361328125 0 1731.4336
624.22998046875 0 1135.7877
636.244140625 0 3190.1353
642.2398681640625 0 1368.7621 c Ammonia loss 4
649.2518920898438 0 1172.7698
650.256591796875 0 805.4578
659.2933349609375 0 1876.3617
662.2632446289062 0 1402.717
663.2808227539062 0 2289.773
664.2907104492188 0 854.1404
687.9082641601562 0 632.7289
688.2968139648438 0 1330.7673
693.2398681640625 0 795.75586
706.2517700195312 0 734.771
707.2734375 0 41678.652 z 4
708.2770385742188 0 18780.164
709.276611328125 0 6745.66
710.2809448242188 0 640.6423
713.2750244140625 0 679.29175 c Ammonia loss 5
715.3191528320312 0 2242.8174
716.3231201171875 0 1572.0696
721.2662353515625 0 779.3528
722.2767333984375 0 1809.6259
723.2914428710938 0 19637.256 y 4
724.293701171875 0 8010.729
725.2938232421875 0 2905.5923
730.2999877929688 0 1400.9496 c 5
732.289306640625 0 697.0602
734.322021484375 0 743.89575
749.2554321289062 0 5372.1104
750.2734985351562 0 3613.553
751.272705078125 0 2693.6533
752.2772216796875 0 1154.7963
759.3082885742188 0 1707.8639
760.3128662109375 0 1067.2362
764.283203125 0 2562.6992
767.2627563476562 0 1101.1882
771.3394165039062 0 1080.7834
772.3485107421875 0 817.6634
774.3196411132812 0 2398.5437
775.3201293945312 0 1338.3865
777.2931518554688 0 1140.0103 w 3
778.3096923828125 0 24513.86
779.3125 0 11640.497
780.3121337890625 0 3067.4253
822.2998046875 0 33270.312 z 3
823.3052978515625 0 33554.86
824.3075561523438 0 12755.859
825.30859375 0 3838.8801
830.3480834960938 0 4604.05
831.3489379882812 0 3328.5188
832.3529663085938 0 1185.3363
833.3257446289062 0 12261.084
834.3292846679688 0 9320.294
835.3326416015625 0 2611.925
836.3417358398438 0 617.4656
837.313720703125 0 836.7346
838.318115234375 0 22852.367 y 3
839.3216552734375 0 9257.424
840.3202514648438 0 3635.368
842.3056030273438 0 859.3575
848.3391723632812 0 667.4254
849.3463745117188 0 14258.464
850.349609375 0 8423.987
851.349853515625 0 2404.068
858.3196411132812 0 2560.7717
859.31298828125 0 1820.379 c Water loss 6
860.3112182617188 0 2318.699 c Ammonia loss 6
866.3165283203125 0 744.9964
871.340576171875 0 784.28394
874.3343505859375 0 2572.0564
875.3389892578125 0 1196.6439
876.3316650390625 0 15480.641
877.338623046875 0 67151.6 c 6
878.3412475585938 0 30868.67
879.3408813476562 0 10699.24
889.3464965820312 0 13867.771
890.349365234375 0 6018.59
891.3515625 0 1911.021
892.3297119140625 0 969.8977 w 2
893.3364868164062 0 39303.047
894.3392333984375 0 19015.545
895.3399047851562 0 6943.6084
896.3345336914062 0 889.3094
898.3339233398438 0 1209.453
899.9144897460938 0 1534.6107
900.4140625 0 1966.4198
900.9137573242188 0 862.2364
901.4083862304688 0 1464.0323
902.9102172851562 0 1411.7269
903.412353515625 0 1206.5609
926.3944702148438 0 1112.862
927.3994750976562 0 810.0764
935.3338012695312 0 4592.353 y Water loss 2
936.3336181640625 0 2700.8042
937.3271484375 0 9512.559 z 2
938.3294067382812 0 4354.9204
939.33203125 0 2398.1123
952.3742065429688 0 2342.4736
953.3455200195312 0 96908.945 y 2
954.3482055664062 0 47461.03
955.3486328125 0 18196.023
956.3512573242188 0 2625.2153
970.3837890625 0 13866.136
970.4956665039062 0 833.0862
971.3865966796875 0 6950.7373
972.3898315429688 0 1882.2947
997.3724365234375 0 1707.4928 c Ammonia loss 7
998.3950805664062 0 2153.784
999.395751953125 0 1777.4672
1014.3975219726562 0 124261.5 c 7
1015.400390625 0 67611.164
1016.4004516601562 0 25859.275
1017.4012451171875 0 2509.3064
1036.41796875 0 943.95026
1040.4044189453125 0 6930.0996
1041.4088134765625 0 3764.101
1042.4071044921875 0 1706.7328
1069.38330078125 0 1530.9783
1071.3985595703125 0 1750.768
1072.4088134765625 0 807.94104
1084.394287109375 0 12837.123 z 1
1085.39697265625 0 7078.9893
1086.3997802734375 0 2547.8025
1094.4793701171875 0 1847.7076
1095.482666015625 0 1379.9641
1096.482666015625 0 736.95276
1097.4556884765625 0 1056.0956
1100.4129638671875 0 5744.0005 y 1
1101.417236328125 0 2917.1365
1102.4190673828125 0 1545.8807
1103.4100341796875 0 3786.7437
1104.4132080078125 0 2473.4316
1105.420654296875 0 730.6393
1110.4183349609375 0 2199.2957
1111.428466796875 0 886.0517
1122.45458984375 0 3355.2195
1123.4552001953125 0 3017.319
1124.459228515625 0 1115.8353
1128.441162109375 0 756.97705
1138.465576171875 0 3738.05
1139.469970703125 0 2626.6558
1140.46728515625 0 24756.096
1141.458740234375 0 36277.94
1142.45263671875 0 42286.336
1143.4515380859375 0 22794.545
1144.45166015625 0 8341.495
1144.588134765625 0 527.787
1145.4573974609375 0 1182.2568
1156.4527587890625 0 1061.0323
1158.47412109375 0 2113.5322
1159.484375 0 1524.5957
1166.4495849609375 0 758.17725
1170.4158935546875 0 2995.261
1171.43115234375 0 1087.7437
1174.4713134765625 0 2672.141
1175.4769287109375 0 2138.5984
1184.45703125 0 19968.52
1185.4456787109375 0 173318.92
1186.4490966796875 0 118125.42
1187.4493408203125 0 50547.31
1188.448974609375 0 8970.595
1189.4549560546875 0 778.96625
1201.4600830078125 0 113425.89
1202.4669189453125 0 382345
1203.4703369140625 0 233998.22
1204.1387939453125 0 2894.683
1204.4713134765625 0 91466.87
1205.4727783203125 0 14379.555
1781.82275390625 0 877.3538
1782.8197021484375 0 783.19885
1783.7786865234375 0 1008.5559
1784.802001953125 0 715.2238
1787.8284912109375 0 1663.9252
1788.81884765625 0 2682.0396
1789.810546875 0 786.15137
1799.8262939453125 0 2000.5494
1800.835205078125 0 1862.0087
1801.822509765625 0 2123.4922
1802.835205078125 0 1167.7369
1804.8223876953125 0 881.64795
1805.845703125 0 1150.7537
1969.26904296875 0 704.48627
2132.02490234375 0 684.1966
2142.384521484375 0 777.39056
2406.24609375 0 1481.3699
2407.26416015625 0 1998.8923
2408.261474609375 0 862.51294
2498.257568359375 0 696.75543

Spectrum Details

|  |  |
| --- | --- |
| Matched peaks? Matched peaks The total absolute number of peaks matched. Additionally in brackets the total fraction of peaks matched and the total number of peaks is shown. | 31 (10.54% of 294) |
| FDR? FDR The false discovery rate estimated for this peptide. It is calculated by matching all theoretical fragments with a non-integer shift with the raw peaks for this spectrum. This is done with 40 different shifts. The resulting percentage is the average number of annotated peaks over the number of annotated peaks with the correct spectrum. | 0.77% |
| Satellite FDR? Satellite FDR See the FDR for details on its calculation. This satellite ion specific FDR only contains the satellite ions (d/w) for I/L/J positions. | - |
| PSM Score? PSM Score The PSM Score as given by Hecklib to this annotated spectrum. It is shown with three significant figures. | 404 |

#### Spectrum 9729? Spectrum 9729 The raw spectrum of this peptide as annotated by Hecklib. The fragments are coloured according to ion type (see legend). Any peaks with a star '\*' as text can be hovered over to see the full details, first the ion type second the mass shift type. By hovering over the amino acids in the peptide or ions in the legend the corresponding peaks are highlighted. By toggling the 'Unassigned' label you can turn the background (unassigned) peaks on or off in the plot. By updating the slider in the Ion legend you can update the spectrum to only show the top X% of the peaks with labels. The top X% means any peak that is within X% of the highest intensity. By dragging in the spectrum you can zoom in to a specific part of the spectrum and use 'Zoom Out' to get back to the original zoom level. The annotation of the spectrum is based on the given sequence in the peptides file and is done with different software so inconsistencies are likely. The peaks are annotated based on the given sequence, with 20 ppm tolerance.

Copy Data

##### Spectrum 9729 (TSV)

###### Preview

```
Loading example...
```

*Click on the button to copy the data to your clipboard.*

Mz MinMz MaxIntensity Max

WidthHeightPeptide font sizePeptide stroke widthSpectrum font sizeSpectrum stroke widthCompact peptide

Ion legend

wxyz

abcd

OtherUnassignedIon^Charge^_Position_Show for top:%

TFDDYAMHW

03.08e+46.15e+49.23e+41.23e+5

Zoom Out

a^+1^_2_y^+1^_1_a^+1^_2_b^+1^_2_b^+1^_2_d^+1^_3_y^+1^_2_b^+1^_3_b^+1^_3_b^+1^_4_b^+1^_4_y^+1^_3_y^+2^_8_y^+1^_4_\*\*b^+1^_5_y^+1^_5_y^+1^_6_y^+1^_6_y^+1^_7_y^+1^_7_y^+1^_8_

0506101115172023

Fragment Matches Table

Show background peaks

| Position | Ion type | Intensity | mz Theoretical | mz Error (Th) | mz Error (ppm) | Charge | Series Number |
| --- | --- | --- | --- | --- | --- | --- | --- |
| - | - | 1154 | 120 | - | - | 0 | - |
| - | - | 1.219E+05 | 120.1 | - | - | 0 | - |
| - | - | 9315 | 121.1 | - | - | 0 | - |
| - | - | 1561 | 123 | - | - | 0 | - |
| - | - | 811 | 127.1 | - | - | 0 | - |
| - | - | 440.3 | 129.1 | - | - | 0 | - |
| - | - | 6178 | 129.1 | - | - | 0 | - |
| - | - | 2713 | 130.1 | - | - | 0 | - |
| - | - | 691.4 | 132 | - | - | 0 | - |
| - | - | 1650 | 132.1 | - | - | 0 | - |
| - | - | 828.6 | 132.1 | - | - | 0 | - |
| - | - | 1347 | 133.1 | - | - | 0 | - |
| - | - | 4.546E+04 | 136.1 | - | - | 0 | - |
| - | - | 3719 | 137.1 | - | - | 0 | - |
| - | - | 1393 | 138.1 | - | - | 0 | - |
| - | - | 759.9 | 141.1 | - | - | 0 | - |
| - | - | 449.8 | 143.1 | - | - | 0 | - |
| - | - | 7421 | 146.1 | - | - | 0 | - |
| - | - | 450.9 | 147 | - | - | 0 | - |
| - | - | 1323 | 147 | - | - | 0 | - |
| - | - | 977.9 | 149 | - | - | 0 | - |
| - | - | 808.6 | 149 | - | - | 0 | - |
| - | - | 541.1 | 150.1 | - | - | 0 | - |
| - | - | 404.8 | 151.8 | - | - | 0 | - |
| - | - | 1187 | 155.1 | - | - | 0 | - |
| - | - | 2722 | 156.1 | - | - | 0 | - |
| - | - | 445.1 | 157.1 | - | - | 0 | - |
| - | - | 715.3 | 157.1 | - | - | 0 | - |
| - | - | 517.2 | 157.1 | - | - | 0 | - |
| - | - | 1382 | 157.1 | - | - | 0 | - |
| - | - | 1325 | 158.1 | - | - | 0 | - |
| - | - | 4070 | 158.1 | - | - | 0 | - |
| - | - | 3988 | 159.1 | - | - | 0 | - |
| - | - | 822.9 | 159.1 | - | - | 0 | - |
| - | - | 1154 | 160.1 | - | - | 0 | - |
| - | - | 513.8 | 160.1 | - | - | 0 | - |
| - | - | 398.6 | 160.1 | - | - | 0 | - |
| - | - | 1674 | 160.1 | - | - | 0 | - |
| - | - | 451.4 | 162.1 | - | - | 0 | - |
| - | - | 1208 | 163.1 | - | - | 0 | - |
| - | - | 760.9 | 164.1 | - | - | 0 | - |
| - | - | 1.363E+04 | 165.1 | - | - | 0 | - |
| - | - | 696 | 165.1 | - | - | 0 | - |
| - | - | 1139 | 166.1 | - | - | 0 | - |
| - | - | 2156 | 166.1 | - | - | 0 | - |
| - | - | 1339 | 167.1 | - | - | 0 | - |
| - | - | 677.4 | 167.1 | - | - | 0 | - |
| - | - | 1293 | 169.1 | - | - | 0 | - |
| - | - | 569.9 | 169.1 | - | - | 0 | - |
| - | - | 935.8 | 171.1 | - | - | 0 | - |
| - | - | 1168 | 173.1 | - | - | 0 | - |
| - | - | 8442 | 173.1 | - | - | 0 | - |
| - | - | 2066 | 173.5 | - | - | 0 | - |
| - | - | 2885 | 174.1 | - | - | 0 | - |
| - | - | 697.3 | 174.1 | - | - | 0 | - |
| - | - | 472.1 | 175.1 | - | - | 0 | - |
| - | - | 2784 | 175.1 | - | - | 0 | - |
| - | - | 552.2 | 176.1 | - | - | 0 | - |
| - | - | 7863 | 176.1 | - | - | 0 | - |
| - | - | 585.3 | 177.1 | - | - | 0 | - |
| - | - | 6611 | 177.1 | - | - | 0 | - |
| - | - | 1189 | 177.1 | - | - | 0 | - |
| - | - | 703 | 178.1 | - | - | 0 | - |
| - | - | 826.7 | 180.1 | - | - | 0 | - |
| - | - | 3.102E+04 | 182.1 | - | - | 0 | - |
| - | - | 2679 | 183.1 | - | - | 0 | - |
| - | - | 1029 | 185.1 | - | - | 0 | - |
| - | - | 1181 | 185.1 | - | - | 0 | - |
| - | - | 751.1 | 186.1 | - | - | 0 | - |
| - | - | 668.5 | 187.1 | - | - | 0 | - |
| - | - | 7529 | 188.1 | - | - | 0 | - |
| - | - | 1188 | 189.1 | - | - | 0 | - |
| - | - | 706.6 | 189.1 | - | - | 0 | - |
| - | - | 485.5 | 190.6 | - | - | 0 | - |
| - | - | 1556 | 191.1 | - | - | 0 | - |
| - | - | 2018 | 191.1 | - | - | 0 | - |
| - | - | 6773 | 192.1 | - | - | 0 | - |
| - | - | 690.3 | 193.1 | - | - | 0 | - |
| - | - | 2195 | 193.1 | - | - | 0 | - |
| - | - | 855.6 | 195.1 | - | - | 0 | - |
| - | - | 616.1 | 195.1 | - | - | 0 | - |
| - | - | 646 | 197.1 | - | - | 0 | - |
| - | - | 661.4 | 197.1 | - | - | 0 | - |
| - | - | 1233 | 199.2 | - | - | 0 | - |
| - | - | 1231 | 200.1 | - | - | 0 | - |
| - | - | 800.3 | 200.2 | - | - | 0 | - |
| - | - | 622.3 | 201.1 | - | - | 0 | - |
| - | - | 4280 | 201.1 | - | - | 0 | - |
| - | - | 480.1 | 201.3 | - | - | 0 | - |
| - | - | 645.7 | 202.1 | - | - | 0 | - |
| - | - | 1072 | 203.1 | - | - | 0 | - |
| - | - | 1430 | 203.1 | - | - | 0 | - |
| 2 | a | 2506 | 203.1 | 0.0001982 | 0.976 | +1 | 2 |
| - | - | 764.2 | 204.1 | - | - | 0 | - |
| 9 | y | 4849 | 205.1 | 0.0001207 | 0.5884 | +1 | 1 |
| - | - | 996.7 | 207.1 | - | - | 0 | - |
| - | - | 4730 | 209.1 | - | - | 0 | - |
| - | - | 506.5 | 210.1 | - | - | 0 | - |
| - | - | 1052 | 214.1 | - | - | 0 | - |
| - | - | 638.8 | 216.1 | - | - | 0 | - |
| - | - | 718.5 | 217.1 | - | - | 0 | - |
| - | - | 1552 | 217.1 | - | - | 0 | - |
| - | - | 491.7 | 217.3 | - | - | 0 | - |
| - | - | 1772 | 219.1 | - | - | 0 | - |
| - | - | 1687 | 221.1 | - | - | 0 | - |
| - | - | 4873 | 221.1 | - | - | 0 | - |
| 2 | a | 1.126E+05 | 221.1 | 0.0001469 | 0.6641 | +1 | 2 |
| - | - | 1.273E+04 | 222.1 | - | - | 0 | - |
| - | - | 776.1 | 223.1 | - | - | 0 | - |
| - | - | 1045 | 223.1 | - | - | 0 | - |
| - | - | 578.1 | 223.1 | - | - | 0 | - |
| - | - | 1114 | 225.2 | - | - | 0 | - |
| - | - | 538.6 | 226.1 | - | - | 0 | - |
| - | - | 1424 | 226.2 | - | - | 0 | - |
| - | - | 928.9 | 229.1 | - | - | 0 | - |
| - | - | 675.9 | 229.2 | - | - | 0 | - |
| - | - | 844.1 | 229.2 | - | - | 0 | - |
| - | - | 2240 | 231.1 | - | - | 0 | - |
| 2 | b | 1225 | 231.1 | 0.0002482 | 1.074 | +1 | 2 |
| - | - | 1646 | 233.1 | - | - | 0 | - |
| - | - | 2143 | 233.1 | - | - | 0 | - |
| - | - | 3686 | 235.1 | - | - | 0 | - |
| - | - | 832.2 | 239.1 | - | - | 0 | - |
| - | - | 826.2 | 239.1 | - | - | 0 | - |
| - | - | 1119 | 239.1 | - | - | 0 | - |
| - | - | 2701 | 239.2 | - | - | 0 | - |
| - | - | 660.3 | 240.1 | - | - | 0 | - |
| - | - | 701.1 | 240.2 | - | - | 0 | - |
| - | - | 502.1 | 241.1 | - | - | 0 | - |
| - | - | 612.5 | 242.1 | - | - | 0 | - |
| 2 | b | 1.958E+04 | 249.1 | 0.0001358 | 0.5451 | +1 | 2 |
| - | - | 2687 | 250.1 | - | - | 0 | - |
| - | - | 2029 | 251.1 | - | - | 0 | - |
| - | - | 1470 | 253.1 | - | - | 0 | - |
| - | - | 971 | 253.2 | - | - | 0 | - |
| - | - | 873 | 256.2 | - | - | 0 | - |
| - | - | 1440 | 257.1 | - | - | 0 | - |
| - | - | 1724 | 261.1 | - | - | 0 | - |
| - | - | 4771 | 263.1 | - | - | 0 | - |
| - | - | 828.2 | 264.1 | - | - | 0 | - |
| - | - | 824.7 | 270.1 | - | - | 0 | - |
| - | - | 1376 | 274.1 | - | - | 0 | - |
| - | - | 622.4 | 279.1 | - | - | 0 | - |
| - | - | 2463 | 279.1 | - | - | 0 | - |
| - | - | 639.8 | 282.2 | - | - | 0 | - |
| - | - | 664.4 | 283.1 | - | - | 0 | - |
| - | - | 649.5 | 284.2 | - | - | 0 | - |
| - | - | 1027 | 285 | - | - | 0 | - |
| - | - | 2531 | 285.1 | - | - | 0 | - |
| - | - | 775.3 | 286.1 | - | - | 0 | - |
| - | - | 1146 | 292.1 | - | - | 0 | - |
| 3 | d | 561.4 | 292.2 | 0.001444 | 4.944 | +1 | 3 |
| - | - | 1852 | 296.2 | - | - | 0 | - |
| - | - | 977.6 | 296.2 | - | - | 0 | - |
| - | - | 1182 | 297.1 | - | - | 0 | - |
| - | - | 805.4 | 302.1 | - | - | 0 | - |
| - | - | 605.3 | 304.1 | - | - | 0 | - |
| - | - | 2900 | 312.3 | - | - | 0 | - |
| - | - | 2514 | 313.3 | - | - | 0 | - |
| - | - | 1709 | 319.1 | - | - | 0 | - |
| - | - | 695.9 | 323.1 | - | - | 0 | - |
| - | - | 4235 | 324.1 | - | - | 0 | - |
| - | - | 983.9 | 329.1 | - | - | 0 | - |
| - | - | 2.002E+04 | 329.1 | - | - | 0 | - |
| - | - | 3064 | 330.2 | - | - | 0 | - |
| - | - | 542 | 333.1 | - | - | 0 | - |
| - | - | 521.6 | 338.1 | - | - | 0 | - |
| - | - | 3385 | 340.2 | - | - | 0 | - |
| 8 | y | 2.145E+04 | 342.2 | 0.000123 | 0.3595 | +1 | 2 |
| - | - | 3788 | 343.2 | - | - | 0 | - |
| 3 | b | 856.4 | 346.1 | 0.001838 | 5.311 | +1 | 3 |
| - | - | 1194 | 350.1 | - | - | 0 | - |
| - | - | 533.2 | 351.1 | - | - | 0 | - |
| - | - | 1304 | 355.1 | - | - | 0 | - |
| - | - | 1504 | 356.1 | - | - | 0 | - |
| - | - | 4266 | 357.2 | - | - | 0 | - |
| 3 | b | 5617 | 364.2 | 4.823E-05 | 0.1325 | +1 | 3 |
| - | - | 553.2 | 366.1 | - | - | 0 | - |
| - | - | 1353 | 368.2 | - | - | 0 | - |
| - | - | 566.5 | 376.1 | - | - | 0 | - |
| - | - | 3258 | 378.1 | - | - | 0 | - |
| - | - | 667.9 | 384.2 | - | - | 0 | - |
| - | - | 826.6 | 385.2 | - | - | 0 | - |
| - | - | 607.2 | 390 | - | - | 0 | - |
| - | - | 703.8 | 392.2 | - | - | 0 | - |
| - | - | 3012 | 394.1 | - | - | 0 | - |
| - | - | 828.7 | 395.1 | - | - | 0 | - |
| - | - | 1070 | 401.2 | - | - | 0 | - |
| - | - | 727.7 | 401.7 | - | - | 0 | - |
| - | - | 757.1 | 407.2 | - | - | 0 | - |
| - | - | 1278 | 408.2 | - | - | 0 | - |
| - | - | 580.4 | 410.4 | - | - | 0 | - |
| - | - | 549 | 414.7 | - | - | 0 | - |
| - | - | 612.7 | 417.1 | - | - | 0 | - |
| - | - | 667.6 | 418.7 | - | - | 0 | - |
| - | - | 694.5 | 423.2 | - | - | 0 | - |
| - | - | 3471 | 425.2 | - | - | 0 | - |
| - | - | 657.4 | 426.2 | - | - | 0 | - |
| - | - | 1963 | 427.7 | - | - | 0 | - |
| - | - | 726.8 | 428.2 | - | - | 0 | - |
| - | - | 847.8 | 428.3 | - | - | 0 | - |
| - | - | 2591 | 436.7 | - | - | 0 | - |
| - | - | 1390 | 437.2 | - | - | 0 | - |
| - | - | 824.8 | 442.3 | - | - | 0 | - |
| - | - | 666.1 | 453.3 | - | - | 0 | - |
| - | - | 677.3 | 458.2 | - | - | 0 | - |
| - | - | 596.4 | 460.8 | - | - | 0 | - |
| 4 | b | 1031 | 461.2 | 0.0001799 | 0.3901 | +1 | 4 |
| - | - | 3547 | 465.2 | - | - | 0 | - |
| - | - | 715.1 | 466.2 | - | - | 0 | - |
| - | - | 655.8 | 473.2 | - | - | 0 | - |
| 4 | b | 3031 | 479.2 | 6.985E-05 | 0.1458 | +1 | 4 |
| - | - | 1393 | 483.2 | - | - | 0 | - |
| - | - | 921.6 | 484.2 | - | - | 0 | - |
| - | - | 1028 | 487.3 | - | - | 0 | - |
| - | - | 794.7 | 487.8 | - | - | 0 | - |
| - | - | 576.7 | 488.3 | - | - | 0 | - |
| 7 | y | 6009 | 489.2 | 0.005008 | 10.24 | +1 | 3 |
| - | - | 1468 | 490.2 | - | - | 0 | - |
| - | - | 3045 | 496.2 | - | - | 0 | - |
| - | - | 811.8 | 497.2 | - | - | 0 | - |
| - | - | 790.9 | 501.3 | - | - | 0 | - |
| - | - | 1351 | 510.3 | - | - | 0 | - |
| - | - | 1167 | 513.4 | - | - | 0 | - |
| - | - | 707.1 | 516.3 | - | - | 0 | - |
| - | - | 895.8 | 519.2 | - | - | 0 | - |
| - | - | 1152 | 531.8 | - | - | 0 | - |
| - | - | 740.1 | 539.2 | - | - | 0 | - |
| - | - | 755 | 542.2 | - | - | 0 | - |
| - | - | 1115 | 547.3 | - | - | 0 | - |
| - | - | 896.8 | 548.2 | - | - | 0 | - |
| 2 | y | 2884 | 550.7 | 0.00192 | 3.487 | +2 | 8 |
| - | - | 2092 | 551.2 | - | - | 0 | - |
| 6 | y | 8596 | 560.2 | 0.004973 | 8.877 | +1 | 4 |
| - | - | 2503 | 561.2 | - | - | 0 | - |
| - | - | 765.6 | 566.3 | - | - | 0 | - |
| - | - | 583.9 | 570.2 | - | - | 0 | - |
| - | - | 701.7 | 583.3 | - | - | 0 | - |
| - | - | 6750 | 585.3 | - | - | 0 | - |
| - | - | 1744 | 586.3 | - | - | 0 | - |
| - | - | 1016 | 589.3 | - | - | 0 | - |
| 0 | Precursor | 1943 | 592.2 | 0.001308 | 2.208 | +2 | -1 |
| - | - | 1019 | 599.4 | - | - | 0 | - |
| - | - | 668.7 | 600.3 | - | - | 0 | - |
| - | - | 8838 | 600.4 | - | - | 0 | - |
| 0 | Precursor | 3280 | 601.2 | 0.002251 | 3.744 | +2 | -1 |
| - | - | 600.4 | 601.3 | - | - | 0 | - |
| - | - | 1971 | 601.4 | - | - | 0 | - |
| - | - | 2492 | 601.7 | - | - | 0 | - |
| - | - | 1015 | 601.8 | - | - | 0 | - |
| - | - | 1179 | 602.2 | - | - | 0 | - |
| - | - | 612.8 | 603.3 | - | - | 0 | - |
| - | - | 1440 | 606.2 | - | - | 0 | - |
| - | - | 916.3 | 612.2 | - | - | 0 | - |
| - | - | 768.6 | 614.2 | - | - | 0 | - |
| - | - | 963 | 615.2 | - | - | 0 | - |
| - | - | 1976 | 628.4 | - | - | 0 | - |
| 5 | b | 688 | 642.2 | 0.001997 | 3.109 | +1 | 5 |
| - | - | 1772 | 657.3 | - | - | 0 | - |
| - | - | 1055 | 658.3 | - | - | 0 | - |
| - | - | 2935 | 659.3 | - | - | 0 | - |
| - | - | 1623 | 660.3 | - | - | 0 | - |
| - | - | 1.012E+04 | 670.8 | - | - | 0 | - |
| - | - | 4924 | 671.3 | - | - | 0 | - |
| - | - | 1475 | 671.8 | - | - | 0 | - |
| - | - | 805.3 | 682.4 | - | - | 0 | - |
| - | - | 656.5 | 684.8 | - | - | 0 | - |
| - | - | 2508 | 685.3 | - | - | 0 | - |
| - | - | 997.5 | 686.3 | - | - | 0 | - |
| - | - | 718.1 | 688.3 | - | - | 0 | - |
| - | - | 672.7 | 689.4 | - | - | 0 | - |
| - | - | 1677 | 693.8 | - | - | 0 | - |
| - | - | 1402 | 694.3 | - | - | 0 | - |
| - | - | 1063 | 694.8 | - | - | 0 | - |
| - | - | 1.953E+04 | 700.4 | - | - | 0 | - |
| - | - | 6967 | 701.4 | - | - | 0 | - |
| - | - | 1354 | 702.4 | - | - | 0 | - |
| - | - | 1048 | 703.3 | - | - | 0 | - |
| - | - | 947.4 | 716.8 | - | - | 0 | - |
| - | - | 1826 | 717.3 | - | - | 0 | - |
| - | - | 776.6 | 717.8 | - | - | 0 | - |
| - | - | 2390 | 721.3 | - | - | 0 | - |
| - | - | 787.4 | 722.4 | - | - | 0 | - |
| 5 | y | 6725 | 723.3 | 0.003046 | 4.212 | +1 | 5 |
| - | - | 707.3 | 723.4 | - | - | 0 | - |
| - | - | 2827 | 724.3 | - | - | 0 | - |
| - | - | 1858 | 725.3 | - | - | 0 | - |
| - | - | 1632 | 730.4 | - | - | 0 | - |
| - | - | 770.2 | 730.9 | - | - | 0 | - |
| - | - | 2357 | 744.4 | - | - | 0 | - |
| - | - | 1652 | 744.9 | - | - | 0 | - |
| - | - | 3204 | 749.3 | - | - | 0 | - |
| - | - | 1219 | 750.3 | - | - | 0 | - |
| - | - | 2084 | 752.9 | - | - | 0 | - |
| - | - | 1656 | 753.4 | - | - | 0 | - |
| - | - | 1813 | 757.4 | - | - | 0 | - |
| - | - | 847.3 | 767.3 | - | - | 0 | - |
| - | - | 762.9 | 771.4 | - | - | 0 | - |
| - | - | 1019 | 773.4 | - | - | 0 | - |
| - | - | 3048 | 774.3 | - | - | 0 | - |
| - | - | 942.7 | 775.3 | - | - | 0 | - |
| - | - | 4867 | 783.4 | - | - | 0 | - |
| - | - | 1625 | 784.4 | - | - | 0 | - |
| - | - | 3171 | 794.4 | - | - | 0 | - |
| - | - | 1099 | 795.4 | - | - | 0 | - |
| - | - | 2431 | 800.4 | - | - | 0 | - |
| - | - | 1.002E+04 | 801.4 | - | - | 0 | - |
| - | - | 1246 | 801.9 | - | - | 0 | - |
| - | - | 4210 | 802.4 | - | - | 0 | - |
| - | - | 1085 | 803.4 | - | - | 0 | - |
| - | - | 660.8 | 814.4 | - | - | 0 | - |
| 4 | y | 974.6 | 820.3 | 0.0112 | 13.66 | +1 | 6 |
| - | - | 4162 | 828.4 | - | - | 0 | - |
| - | - | 2091 | 829.4 | - | - | 0 | - |
| - | - | 1076 | 830.4 | - | - | 0 | - |
| 4 | y | 9085 | 838.3 | 0.004179 | 4.985 | +1 | 6 |
| - | - | 4031 | 839.3 | - | - | 0 | - |
| - | - | 934.2 | 840.3 | - | - | 0 | - |
| - | - | 1.395E+04 | 844.5 | - | - | 0 | - |
| - | - | 5709 | 845.5 | - | - | 0 | - |
| - | - | 1258 | 846.5 | - | - | 0 | - |
| - | - | 2489 | 854.4 | - | - | 0 | - |
| - | - | 1003 | 855.4 | - | - | 0 | - |
| - | - | 1348 | 871.3 | - | - | 0 | - |
| - | - | 5.069E+04 | 872.4 | - | - | 0 | - |
| - | - | 2.646E+04 | 873.4 | - | - | 0 | - |
| - | - | 7697 | 874.5 | - | - | 0 | - |
| - | - | 1325 | 875.5 | - | - | 0 | - |
| - | - | 1.331E+04 | 889.3 | - | - | 0 | - |
| - | - | 7346 | 890.3 | - | - | 0 | - |
| - | - | 1960 | 891.4 | - | - | 0 | - |
| - | - | 1083 | 896.5 | - | - | 0 | - |
| - | - | 3680 | 909.4 | - | - | 0 | - |
| - | - | 1184 | 910.4 | - | - | 0 | - |
| - | - | 765.9 | 916.7 | - | - | 0 | - |
| 3 | y | 2270 | 935.3 | 0.001335 | 1.427 | +1 | 7 |
| - | - | 1992 | 936.3 | - | - | 0 | - |
| 3 | y | 4.254E+04 | 953.3 | 0.004031 | 4.228 | +1 | 7 |
| - | - | 2.269E+04 | 954.3 | - | - | 0 | - |
| - | - | 6925 | 955.3 | - | - | 0 | - |
| - | - | 1183 | 956.4 | - | - | 0 | - |
| - | - | 8082 | 991.5 | - | - | 0 | - |
| - | - | 4900 | 992.5 | - | - | 0 | - |
| - | - | 1500 | 993.5 | - | - | 0 | - |
| - | - | 2.51E+04 | 1020 | - | - | 0 | - |
| - | - | 1.254E+04 | 1021 | - | - | 0 | - |
| - | - | 4965 | 1022 | - | - | 0 | - |
| - | - | 740.2 | 1023 | - | - | 0 | - |
| - | - | 948.2 | 1038 | - | - | 0 | - |
| - | - | 963.7 | 1056 | - | - | 0 | - |
| - | - | 625.7 | 1075 | - | - | 0 | - |
| 2 | y | 2588 | 1100 | 0.0008634 | 0.7846 | +1 | 8 |
| - | - | 1081 | 1101 | - | - | 0 | - |
| - | - | 1110 | 1109 | - | - | 0 | - |
| - | - | 859.8 | 1178 | - | - | 0 | - |
| - | - | 630.2 | 2003 | - | - | 0 | - |

m/z Charge Intensity FragmentType MassShift Position
120.04467010498047 0 1154.1261
120.0810317993164 0 121864.29
121.08431243896484 0 9314.98
123.04441833496094 0 1561.2311
127.08679962158203 0 810.9835
129.0662841796875 0 440.34708
129.1024932861328 0 6177.801
130.06546020507812 0 2712.7444
132.04483032226562 0 691.4035
132.08091735839844 0 1650.3647
132.10186767578125 0 828.55524
133.08604431152344 0 1347.1455
136.07589721679688 0 45460.125
137.07923889160156 0 3718.598
138.0663604736328 0 1392.9513
141.10235595703125 0 759.9148
143.08175659179688 0 449.81757
146.06024169921875 0 7421.4717
147.03004455566406 0 450.94925
147.0445556640625 0 1322.8544
148.95472717285156 0 977.9432
149.04525756835938 0 808.64905
150.05526733398438 0 541.1233
151.76986694335938 0 404.8321
155.0818328857422 0 1186.5656
156.07696533203125 0 2721.6772
157.0609588623047 0 445.09418
157.07620239257812 0 715.25543
157.10850524902344 0 517.21533
157.13388061523438 0 1382.3165
158.05999755859375 0 1325.2739
158.09658813476562 0 4070.4128
159.09182739257812 0 3988.1448
159.09918212890625 0 822.8643
160.0757293701172 0 1154.2247
160.09539794921875 0 513.7941
160.1044464111328 0 398.58945
160.11236572265625 0 1673.8939
162.0550079345703 0 451.41928
163.08673095703125 0 1208.1371
164.0704803466797 0 760.88214
165.05477905273438 0 13634.9795
165.1023406982422 0 696.039
166.05825805664062 0 1138.599
166.0865020751953 0 2155.9895
167.0556640625 0 1339.1879
167.09323120117188 0 677.39075
169.09719848632812 0 1292.929
169.13352966308594 0 569.8722
171.07632446289062 0 935.7673
173.09226989746094 0 1168.3796
173.12864685058594 0 8442.468
173.4516143798828 0 2065.9565
174.0551300048828 0 2884.6768
174.13243103027344 0 697.32605
175.05857849121094 0 472.07382
175.08670043945312 0 2783.9395
176.0812530517578 0 552.1885
176.10711669921875 0 7862.9917
177.0675811767578 0 585.30524
177.10240173339844 0 6611.29
177.11105346679688 0 1189.4454
178.10585021972656 0 703.03735
180.07728576660156 0 826.66437
182.08131408691406 0 31021.05
183.08453369140625 0 2678.593
185.05599975585938 0 1028.9065
185.1283416748047 0 1180.6901
186.09120178222656 0 751.0918
187.10791015625 0 668.51056
188.07077026367188 0 7528.5503
189.07427978515625 0 1187.9083
189.08702087402344 0 706.602
190.58804321289062 0 485.47867
191.08154296875 0 1556.0159
191.11805725097656 0 2017.6628
192.06564331054688 0 6772.7505
193.06895446777344 0 690.2574
193.10867309570312 0 2195.2827
195.08767700195312 0 855.57623
195.12222290039062 0 616.087
197.0919952392578 0 645.9617
197.12835693359375 0 661.3907
199.18040466308594 0 1232.9802
200.13938903808594 0 1231.075
200.18399047851562 0 800.30975
201.11390686035156 0 622.2871
201.12335205078125 0 4279.8354
201.31942749023438 0 480.12277
202.12709045410156 0 645.66547
203.0662078857422 0 1072.4333
203.0817108154297 0 1429.5634
203.1180877685547 0 2506.0605 a Water loss 1
204.07684326171875 0 764.17267
205.09727478027344 0 4848.92 y 8
207.1131591796875 0 996.73425
209.09207153320312 0 4729.6743
210.09654235839844 0 506.479
214.1186981201172 0 1052.4788
216.06561279296875 0 638.7596
217.08160400390625 0 718.53375
217.09706115722656 0 1551.7742
217.34991455078125 0 491.74945
219.11285400390625 0 1772.2006
221.0923614501953 0 1687.3627
221.10342407226562 0 4873.362
221.12860107421875 0 112644.64 a 1
222.13201904296875 0 12732.092
223.0637969970703 0 776.0843
223.10784912109375 0 1045.4583
223.134765625 0 578.086
225.17111206054688 0 1113.729
226.11880493164062 0 538.63696
226.1551513671875 0 1423.7063
229.11886596679688 0 928.885
229.1561737060547 0 675.87463
229.1700439453125 0 844.07745
231.0613555908203 0 2240.4082
231.11305236816406 0 1225.4856 b Water loss 1
233.0924072265625 0 1645.9286
233.1288604736328 0 2143.4102
235.10780334472656 0 3685.9197
239.08351135253906 0 832.15875
239.09593200683594 0 826.1822
239.1143035888672 0 1118.831
239.15017700195312 0 2700.6165
240.13592529296875 0 660.29865
240.1531982421875 0 701.0974
241.0739288330078 0 502.14233
242.11309814453125 0 612.4513
249.12350463867188 0 19583.05 b 1
250.1269989013672 0 2687.0247
251.10305786132812 0 2029.2051
253.11862182617188 0 1470.3081
253.16647338867188 0 971.0309
256.1769714355469 0 872.9766
257.1065979003906 0 1440.2571
261.1234436035156 0 1724.1831
263.10260009765625 0 4770.9004
264.10638427734375 0 828.22217
270.1087646484375 0 824.67444
274.13079833984375 0 1375.971
279.08160400390625 0 622.4281
279.097900390625 0 2462.6494
282.1827087402344 0 639.79474
283.1446838378906 0 664.39844
284.1968078613281 0 649.5197
285.00994873046875 0 1026.5453
285.101806640625 0 2530.962
286.10540771484375 0 775.2827
292.140869140625 0 1145.5793
292.16412353515625 0 561.41473 d 2
296.1504821777344 0 1851.7324
296.2091369628906 0 977.6313
297.0903015136719 0 1182.2081
302.1137390136719 0 805.40894
304.128662109375 0 605.3471
312.2648620605469 0 2900.4788
313.2679138183594 0 2514.3623
319.1407470703125 0 1709.3151
323.09893798828125 0 695.94446
324.14581298828125 0 4235.416
329.1275939941406 0 983.89886
329.1498107910156 0 20020.115
330.15338134765625 0 3063.5908
333.10821533203125 0 541.9513
338.13201904296875 0 521.6171
340.1983947753906 0 3385.3013
342.15618896484375 0 21449.508 y 7
343.1593933105469 0 3788.1448
346.1379089355469 0 856.4088 b Water loss 2
350.1348571777344 0 1193.6249
351.1359558105469 0 533.1829
355.06976318359375 0 1304.387
356.1386413574219 0 1504.1062
357.22393798828125 0 4266.152
364.1503601074219 0 5616.8516 b 2
366.1285705566406 0 553.18286
368.1924743652344 0 1353.2322
376.1134033203125 0 566.53815
378.1290588378906 0 3258.1616
384.18939208984375 0 667.922
385.2178649902344 0 826.5528
389.9964599609375 0 607.16956
392.20574951171875 0 703.8239
394.1249084472656 0 3011.7031
395.12786865234375 0 828.7412
401.2088928222656 0 1070.3636
401.7098083496094 0 727.707
407.18438720703125 0 757.1445
408.16729736328125 0 1277.5071
410.44158935546875 0 580.3916
414.7137451171875 0 549.0367
417.0843505859375 0 612.7196
418.7189025878906 0 667.5667
423.2313537597656 0 694.49994
425.1933288574219 0 3471.3337
426.1964416503906 0 657.4098
427.7221374511719 0 1963.0463
428.2223205566406 0 726.8062
428.29205322265625 0 847.8334
436.7268981933594 0 2591.2954
437.2273864746094 0 1390.3243
442.3133850097656 0 824.7824
453.28302001953125 0 666.12836
458.2305603027344 0 677.29
460.767333984375 0 596.4405
461.1668701171875 0 1030.8502 b Water loss 3
465.1612854003906 0 3547.305
466.1650695800781 0 715.13214
473.2139892578125 0 655.7787
479.17718505859375 0 3031.4626 b 3
483.22076416015625 0 1392.9822
484.22393798828125 0 921.5589
487.3355712890625 0 1028.1573
487.7531433105469 0 794.6619
488.2970275878906 0 576.71735
489.1915588378906 0 6008.6523 y 6
490.1954040527344 0 1467.832
496.23052978515625 0 3045.2175
497.2310791015625 0 811.8435
501.2536315917969 0 790.8938
510.2643127441406 0 1351.1797
513.3513793945312 0 1166.7585
516.2924194335938 0 707.14734
519.2061157226562 0 895.7549
531.7725219726562 0 1152.1882
539.2152099609375 0 740.0835
542.2310180664062 0 755.0015
547.2510986328125 0 1115.2919
548.1998291015625 0 896.7548
550.710205078125 0 2884.404 y 1
551.21240234375 0 2091.896
560.2286376953125 0 8595.667 y 5
561.2305908203125 0 2503.279
566.25244140625 0 765.643
570.2257080078125 0 583.8575
583.281005859375 0 701.66187
585.3355102539062 0 6749.7983
586.3391723632812 0 1743.6416
589.2847900390625 0 1015.99506
592.2281494140625 0 1943.3306 Precursor Water loss
599.3773803710938 0 1019.13574
600.2879028320312 0 668.7285
600.3787231445312 0 8838.312
601.234375 0 3280.1638 Precursor
601.3260498046875 0 600.35596
601.3822021484375 0 1971.291
601.7379150390625 0 2492.3135
601.8153076171875 0 1014.51495
602.2340087890625 0 1179.3593
603.3482666015625 0 612.7733
606.2297973632812 0 1440.0319
612.1952514648438 0 916.30536
614.2449951171875 0 768.5572
615.2464599609375 0 963.01483
628.3765258789062 0 1975.6699
642.2385864257812 0 687.9753 b 4
657.2640380859375 0 1771.9828
658.2610473632812 0 1055.3075
659.29296875 0 2935.3394
660.2964477539062 0 1623.4218
670.81689453125 0 10124.056
671.3187255859375 0 4923.564
671.8197021484375 0 1475.1506
682.353515625 0 805.2923
684.8060302734375 0 656.53534
685.257080078125 0 2507.8066
686.26220703125 0 997.4932
688.33447265625 0 718.0517
689.4219360351562 0 672.73706
693.8106689453125 0 1676.9847
694.3128051757812 0 1401.8228
694.8155517578125 0 1062.7755
700.362060546875 0 19527.596
701.3640747070312 0 6967.233
702.3671264648438 0 1354.0428
703.263671875 0 1047.5476
716.8134155273438 0 947.422
717.3148803710938 0 1826.2037
717.8121337890625 0 776.6346
721.2615356445312 0 2389.9143
722.4022827148438 0 787.38635
723.2900390625 0 6724.547 y 4
723.4029541015625 0 707.3454
724.2943115234375 0 2826.7153
725.305908203125 0 1858.4235
730.353271484375 0 1631.554
730.8569946289062 0 770.218
744.3507080078125 0 2357.2678
744.8532104492188 0 1652.4982
749.2557983398438 0 3203.9622
750.2554321289062 0 1218.5497
752.8637084960938 0 2083.7869
753.3627319335938 0 1655.9366
757.3849487304688 0 1812.7067
767.2684936523438 0 847.3203
771.3935546875 0 762.8951
773.4144897460938 0 1019.3051
774.320068359375 0 3047.5842
775.322998046875 0 942.68463
783.3970947265625 0 4866.8267
784.3995971679688 0 1624.8722
794.3834838867188 0 3171.3098
795.3831787109375 0 1099.4171
800.4263916015625 0 2431.1047
801.4111328125 0 10019.446
801.8678588867188 0 1245.8688
802.4150390625 0 4210.1807
803.4205932617188 0 1085.1718
814.4442749023438 0 660.769
820.3145751953125 0 974.6116 y Water loss 3
828.4216918945312 0 4161.8735
829.4231567382812 0 2091.3267
830.4279174804688 0 1075.6652
838.318115234375 0 9084.676 y 3
839.320068359375 0 4030.6387
840.3185424804688 0 934.2393
844.4515380859375 0 13948.725
845.454345703125 0 5709.4033
846.4572143554688 0 1258.2479
854.4342651367188 0 2489.0952
855.4385375976562 0 1002.7858
871.3386840820312 0 1348.0544
872.446533203125 0 50694.3
873.4493408203125 0 26457.63
874.4511108398438 0 7697.4277
875.4521484375 0 1324.822
889.346435546875 0 13306.489
890.3489990234375 0 7346.3335
891.3502197265625 0 1959.8682
896.5291748046875 0 1082.6182
909.4083862304688 0 3679.6619
910.4127197265625 0 1184.172
916.6693725585938 0 765.90045
935.3289794921875 0 2270.3472 y Water loss 2
936.3351440429688 0 1991.6887
953.3449096679688 0 42541.94 y 2
954.3475341796875 0 22690.822
955.349365234375 0 6924.7017
956.3507080078125 0 1183.4653
991.5196533203125 0 8081.9136
992.522216796875 0 4900.4536
993.5245971679688 0 1500.2554
1019.5139770507812 0 25102.902
1020.5169677734375 0 12543.835
1021.5191040039062 0 4965.4297
1022.5285034179688 0 740.24506
1037.514404296875 0 948.2111
1056.4810791015625 0 963.6801
1075.211181640625 0 625.6905
1100.41015625 0 2587.8271 y 1
1101.4171142578125 0 1081.1195
1108.52490234375 0 1109.7988
1177.5611572265625 0 859.78796
2002.52734375 0 630.2146

Spectrum Details

|  |  |
| --- | --- |
| Matched peaks? Matched peaks The total absolute number of peaks matched. Additionally in brackets the total fraction of peaks matched and the total number of peaks is shown. | 23 (6.46% of 356) |
| FDR? FDR The false discovery rate estimated for this peptide. It is calculated by matching all theoretical fragments with a non-integer shift with the raw peaks for this spectrum. This is done with 40 different shifts. The resulting percentage is the average number of annotated peaks over the number of annotated peaks with the correct spectrum. | 1.55% |
| Satellite FDR? Satellite FDR See the FDR for details on its calculation. This satellite ion specific FDR only contains the satellite ions (d/w) for I/L/J positions. | - |
| PSM Score? PSM Score The PSM Score as given by Hecklib to this annotated spectrum. It is shown with three significant figures. | 270 |

#### Spectrum 11731? Spectrum 11731 The raw spectrum of this peptide as annotated by Hecklib. The fragments are coloured according to ion type (see legend). Any peaks with a star '\*' as text can be hovered over to see the full details, first the ion type second the mass shift type. By hovering over the amino acids in the peptide or ions in the legend the corresponding peaks are highlighted. By toggling the 'Unassigned' label you can turn the background (unassigned) peaks on or off in the plot. By updating the slider in the Ion legend you can update the spectrum to only show the top X% of the peaks with labels. The top X% means any peak that is within X% of the highest intensity. By dragging in the spectrum you can zoom in to a specific part of the spectrum and use 'Zoom Out' to get back to the original zoom level. The annotation of the spectrum is based on the given sequence in the peptides file and is done with different software so inconsistencies are likely. The peaks are annotated based on the given sequence, with 20 ppm tolerance.

Copy Data

##### Spectrum 11731 (TSV)

###### Preview

```
Loading example...
```

*Click on the button to copy the data to your clipboard.*

Mz MinMz MaxIntensity Max

WidthHeightPeptide font sizePeptide stroke widthSpectrum font sizeSpectrum stroke widthCompact peptide

Ion legend

wxyz

abcd

OtherUnassignedIon^Charge^_Position_Show for top:%

TFDDYAMHW

01.25e+42.51e+43.76e+45.01e+4

Zoom Out

a^+1^_2_y^+1^_1_a^+1^_2_b^+1^_2_b^+1^_2_y^+1^_2_b^+1^_3_b^+1^_4_y^+1^_3_b^+2^_8_y^+2^_8_y^+1^_4_\*y^+1^_5_y^+1^_6_y^+1^_7_y^+1^_7_y^+1^_8_

0778155523333111

Fragment Matches Table

Show background peaks

| Position | Ion type | Intensity | mz Theoretical | mz Error (Th) | mz Error (ppm) | Charge | Series Number |
| --- | --- | --- | --- | --- | --- | --- | --- |
| - | - | 3.453E+04 | 120.1 | - | - | 0 | - |
| - | - | 2929 | 121.1 | - | - | 0 | - |
| - | - | 549.7 | 123 | - | - | 0 | - |
| - | - | 436.8 | 127.1 | - | - | 0 | - |
| - | - | 3128 | 129.1 | - | - | 0 | - |
| - | - | 549.7 | 130.1 | - | - | 0 | - |
| - | - | 537.8 | 132.1 | - | - | 0 | - |
| - | - | 984.8 | 133.1 | - | - | 0 | - |
| - | - | 414.7 | 134.2 | - | - | 0 | - |
| - | - | 1.08E+04 | 136.1 | - | - | 0 | - |
| - | - | 782.7 | 137.1 | - | - | 0 | - |
| - | - | 545.5 | 141.1 | - | - | 0 | - |
| - | - | 505.9 | 146.1 | - | - | 0 | - |
| - | - | 1252 | 149 | - | - | 0 | - |
| - | - | 459.7 | 152.1 | - | - | 0 | - |
| - | - | 406.6 | 152.4 | - | - | 0 | - |
| - | - | 478.2 | 155.1 | - | - | 0 | - |
| - | - | 846.8 | 156.1 | - | - | 0 | - |
| - | - | 430.5 | 156.5 | - | - | 0 | - |
| - | - | 987.3 | 157.1 | - | - | 0 | - |
| - | - | 2087 | 158.1 | - | - | 0 | - |
| - | - | 1930 | 159.1 | - | - | 0 | - |
| - | - | 977.8 | 160.1 | - | - | 0 | - |
| - | - | 3712 | 165.1 | - | - | 0 | - |
| - | - | 2356 | 166.1 | - | - | 0 | - |
| - | - | 2327 | 167.1 | - | - | 0 | - |
| - | - | 868.1 | 167.1 | - | - | 0 | - |
| - | - | 477.2 | 167.1 | - | - | 0 | - |
| - | - | 575 | 169.1 | - | - | 0 | - |
| - | - | 449.5 | 171.1 | - | - | 0 | - |
| - | - | 2596 | 173.1 | - | - | 0 | - |
| - | - | 2758 | 173.4 | - | - | 0 | - |
| - | - | 646.1 | 176.1 | - | - | 0 | - |
| - | - | 3668 | 176.1 | - | - | 0 | - |
| - | - | 2705 | 177.1 | - | - | 0 | - |
| - | - | 6452 | 182.1 | - | - | 0 | - |
| - | - | 471.7 | 183.1 | - | - | 0 | - |
| - | - | 690.8 | 183.1 | - | - | 0 | - |
| - | - | 534.1 | 183.1 | - | - | 0 | - |
| - | - | 475.8 | 185.1 | - | - | 0 | - |
| - | - | 571.8 | 185.1 | - | - | 0 | - |
| - | - | 1893 | 188.1 | - | - | 0 | - |
| - | - | 634.5 | 191.1 | - | - | 0 | - |
| - | - | 922.3 | 193.1 | - | - | 0 | - |
| - | - | 622.7 | 199.1 | - | - | 0 | - |
| - | - | 483.2 | 200.1 | - | - | 0 | - |
| - | - | 1827 | 201.1 | - | - | 0 | - |
| 2 | a | 1149 | 203.1 | 0.0004271 | 2.103 | +1 | 2 |
| - | - | 648.2 | 204.1 | - | - | 0 | - |
| 9 | y | 1793 | 205.1 | 0.0003801 | 1.853 | +1 | 1 |
| - | - | 1844 | 212.2 | - | - | 0 | - |
| - | - | 530.8 | 215.1 | - | - | 0 | - |
| - | - | 660.9 | 217.1 | - | - | 0 | - |
| - | - | 2941 | 221.1 | - | - | 0 | - |
| 2 | a | 4.965E+04 | 221.1 | 0.0001469 | 0.6641 | +1 | 2 |
| - | - | 6101 | 222.1 | - | - | 0 | - |
| - | - | 1649 | 223.1 | - | - | 0 | - |
| - | - | 589.3 | 223.1 | - | - | 0 | - |
| - | - | 499.5 | 223.4 | - | - | 0 | - |
| - | - | 506.6 | 224.1 | - | - | 0 | - |
| - | - | 2171 | 227.1 | - | - | 0 | - |
| - | - | 659.8 | 229.1 | - | - | 0 | - |
| - | - | 856.1 | 231.1 | - | - | 0 | - |
| 2 | b | 722.8 | 231.1 | 0.0003011 | 1.303 | +1 | 2 |
| - | - | 950.9 | 235.1 | - | - | 0 | - |
| - | - | 1617 | 239.1 | - | - | 0 | - |
| - | - | 1182 | 241.1 | - | - | 0 | - |
| - | - | 620.7 | 241.7 | - | - | 0 | - |
| 2 | b | 8078 | 249.1 | 0.0001816 | 0.7288 | +1 | 2 |
| - | - | 944.2 | 250.1 | - | - | 0 | - |
| - | - | 940.7 | 251.1 | - | - | 0 | - |
| - | - | 547 | 253.1 | - | - | 0 | - |
| - | - | 1396 | 263.1 | - | - | 0 | - |
| - | - | 697.1 | 265.2 | - | - | 0 | - |
| - | - | 944.1 | 279.1 | - | - | 0 | - |
| - | - | 879.2 | 281.1 | - | - | 0 | - |
| - | - | 650.9 | 283 | - | - | 0 | - |
| - | - | 683.2 | 283.1 | - | - | 0 | - |
| - | - | 506.4 | 284.1 | - | - | 0 | - |
| - | - | 2174 | 285 | - | - | 0 | - |
| - | - | 1210 | 285.1 | - | - | 0 | - |
| - | - | 781.8 | 292.1 | - | - | 0 | - |
| - | - | 649.3 | 299.1 | - | - | 0 | - |
| - | - | 735.4 | 319.1 | - | - | 0 | - |
| - | - | 1754 | 324.1 | - | - | 0 | - |
| - | - | 4359 | 329.1 | - | - | 0 | - |
| - | - | 973.9 | 330.2 | - | - | 0 | - |
| - | - | 545.2 | 336 | - | - | 0 | - |
| - | - | 1948 | 340.2 | - | - | 0 | - |
| 8 | y | 1.079E+04 | 342.2 | 3.146E-05 | 0.09194 | +1 | 2 |
| - | - | 1776 | 343.2 | - | - | 0 | - |
| - | - | 782.8 | 350.1 | - | - | 0 | - |
| - | - | 2426 | 355.1 | - | - | 0 | - |
| - | - | 1156 | 356.1 | - | - | 0 | - |
| - | - | 905.3 | 356.1 | - | - | 0 | - |
| - | - | 1100 | 357.2 | - | - | 0 | - |
| 3 | b | 2201 | 364.2 | 0.000445 | 1.222 | +1 | 3 |
| - | - | 920.5 | 378.1 | - | - | 0 | - |
| - | - | 558.8 | 380.2 | - | - | 0 | - |
| - | - | 480.9 | 387.8 | - | - | 0 | - |
| - | - | 948.1 | 393.2 | - | - | 0 | - |
| - | - | 2377 | 394.1 | - | - | 0 | - |
| - | - | 889.7 | 395.2 | - | - | 0 | - |
| - | - | 778.7 | 398.2 | - | - | 0 | - |
| - | - | 713.4 | 403.2 | - | - | 0 | - |
| - | - | 493 | 411.2 | - | - | 0 | - |
| - | - | 1120 | 413.3 | - | - | 0 | - |
| - | - | 576.8 | 415.3 | - | - | 0 | - |
| - | - | 1904 | 425.2 | - | - | 0 | - |
| - | - | 743.6 | 427.7 | - | - | 0 | - |
| - | - | 712.7 | 453.3 | - | - | 0 | - |
| - | - | 594.3 | 458.4 | - | - | 0 | - |
| - | - | 1539 | 465.2 | - | - | 0 | - |
| 4 | b | 1531 | 479.2 | 0.0001743 | 0.3637 | +1 | 4 |
| 7 | y | 2880 | 489.2 | 0.004764 | 9.739 | +1 | 3 |
| 8 | b | 635.3 | 490.2 | 0.006971 | 14.22 | +2 | 8 |
| - | - | 901.4 | 496.2 | - | - | 0 | - |
| - | - | 551.2 | 501.5 | - | - | 0 | - |
| - | - | 1082 | 508.3 | - | - | 0 | - |
| - | - | 548.7 | 519.3 | - | - | 0 | - |
| 2 | y | 883.4 | 550.7 | 0.001249 | 2.268 | +2 | 8 |
| 6 | y | 3364 | 560.2 | 0.005156 | 9.204 | +1 | 4 |
| - | - | 655.8 | 561.2 | - | - | 0 | - |
| - | - | 758.7 | 563.3 | - | - | 0 | - |
| - | - | 593.3 | 568.3 | - | - | 0 | - |
| - | - | 860.7 | 569.2 | - | - | 0 | - |
| - | - | 2002 | 585.3 | - | - | 0 | - |
| 0 | Precursor | 1976 | 601.2 | 0.001885 | 3.135 | +2 | -1 |
| - | - | 983.5 | 601.7 | - | - | 0 | - |
| - | - | 729.7 | 601.8 | - | - | 0 | - |
| - | - | 774.1 | 602.3 | - | - | 0 | - |
| - | - | 1157 | 602.4 | - | - | 0 | - |
| - | - | 1388 | 621.4 | - | - | 0 | - |
| - | - | 744.6 | 628.4 | - | - | 0 | - |
| - | - | 730.6 | 640.3 | - | - | 0 | - |
| - | - | 734.5 | 657 | - | - | 0 | - |
| - | - | 1449 | 657.3 | - | - | 0 | - |
| - | - | 1141 | 659.3 | - | - | 0 | - |
| - | - | 1129 | 685.3 | - | - | 0 | - |
| - | - | 4874 | 700.4 | - | - | 0 | - |
| - | - | 1355 | 701.4 | - | - | 0 | - |
| - | - | 670.5 | 721.3 | - | - | 0 | - |
| 5 | y | 2896 | 723.3 | 0.004633 | 6.406 | +1 | 5 |
| - | - | 1470 | 724.3 | - | - | 0 | - |
| - | - | 735.7 | 735.4 | - | - | 0 | - |
| - | - | 1996 | 749.3 | - | - | 0 | - |
| - | - | 743.1 | 750.3 | - | - | 0 | - |
| - | - | 867.4 | 774.3 | - | - | 0 | - |
| - | - | 890.8 | 783.4 | - | - | 0 | - |
| - | - | 1648 | 801.4 | - | - | 0 | - |
| - | - | 1011 | 802.4 | - | - | 0 | - |
| 4 | y | 3542 | 838.3 | 0.003081 | 3.675 | +1 | 6 |
| - | - | 1248 | 839.3 | - | - | 0 | - |
| - | - | 3010 | 844.5 | - | - | 0 | - |
| - | - | 1420 | 845.5 | - | - | 0 | - |
| - | - | 848.4 | 854.4 | - | - | 0 | - |
| - | - | 1.128E+04 | 872.4 | - | - | 0 | - |
| - | - | 5380 | 873.4 | - | - | 0 | - |
| - | - | 1138 | 874.5 | - | - | 0 | - |
| - | - | 6301 | 889.3 | - | - | 0 | - |
| - | - | 2601 | 890.3 | - | - | 0 | - |
| - | - | 1169 | 901.5 | - | - | 0 | - |
| - | - | 1185 | 902.5 | - | - | 0 | - |
| - | - | 645.9 | 908.3 | - | - | 0 | - |
| 3 | y | 1068 | 935.3 | 0.003487 | 3.728 | +1 | 7 |
| 3 | y | 1.814E+04 | 953.3 | 0.003848 | 4.036 | +1 | 7 |
| - | - | 9386 | 954.3 | - | - | 0 | - |
| - | - | 4546 | 955.3 | - | - | 0 | - |
| - | - | 766.8 | 956.4 | - | - | 0 | - |
| - | - | 2172 | 991.5 | - | - | 0 | - |
| - | - | 883.5 | 992.5 | - | - | 0 | - |
| - | - | 604.6 | 1006 | - | - | 0 | - |
| - | - | 5579 | 1020 | - | - | 0 | - |
| - | - | 2499 | 1021 | - | - | 0 | - |
| - | - | 949.3 | 1022 | - | - | 0 | - |
| 2 | y | 1293 | 1100 | 0.00245 | 2.227 | +1 | 8 |
| - | - | 608.5 | 1206 | - | - | 0 | - |
| - | - | 714.8 | 3079 | - | - | 0 | - |
| - | - | 808 | 3080 | - | - | 0 | - |

m/z Charge Intensity FragmentType MassShift Position
120.08100891113281 0 34526.418
121.0843276977539 0 2928.877
123.04418182373047 0 549.6511
127.08744812011719 0 436.80527
129.1024627685547 0 3128.1716
130.0654296875 0 549.6973
132.0812225341797 0 537.8353
133.08595275878906 0 984.82446
134.2151336669922 0 414.70932
136.07589721679688 0 10797.627
137.0792236328125 0 782.66016
141.1025848388672 0 545.5393
146.06024169921875 0 505.9305
149.04513549804688 0 1252.1249
152.09445190429688 0 459.70633
152.40768432617188 0 406.604
155.0817413330078 0 478.15674
156.07672119140625 0 846.77136
156.5098876953125 0 430.50806
157.1336669921875 0 987.3196
158.0966033935547 0 2086.8828
159.09170532226562 0 1929.5443
160.1123046875 0 977.8325
165.0548095703125 0 3712.1484
166.08642578125 0 2356.26
167.05572509765625 0 2326.9055
167.0814971923828 0 868.1317
167.09304809570312 0 477.21057
169.0972442626953 0 575.00867
171.0765380859375 0 449.52432
173.12860107421875 0 2596.487
173.439208984375 0 2757.587
176.0823974609375 0 646.05133
176.1070556640625 0 3667.7222
177.10255432128906 0 2704.9424
182.081298828125 0 6452.4204
183.08474731445312 0 471.72647
183.1132049560547 0 690.7806
183.1493682861328 0 534.1099
185.05572509765625 0 475.7769
185.12892150878906 0 571.7761
188.0706787109375 0 1892.6799
191.11753845214844 0 634.51825
193.10870361328125 0 922.317
199.10888671875 0 622.7083
200.138916015625 0 483.20413
201.12327575683594 0 1826.846
203.11831665039062 0 1148.7513 a Water loss 1
204.07740783691406 0 648.15497
205.0975341796875 0 1792.7393 y 8
212.1646270751953 0 1844.3536
215.1280059814453 0 530.7755
217.09689331054688 0 660.91626
221.1034698486328 0 2940.8232
221.12860107421875 0 49646.27 a 1
222.13194274902344 0 6100.809
223.06350708007812 0 1649.465
223.1083984375 0 589.33136
223.393798828125 0 499.50882
224.10342407226562 0 506.61737
227.10292053222656 0 2171.148
229.11813354492188 0 659.7682
231.06161499023438 0 856.0901
231.1125030517578 0 722.79456 b Water loss 1
235.10791015625 0 950.9411
239.0948486328125 0 1616.9264
241.0735626220703 0 1182.3079
241.6948699951172 0 620.6982
249.12355041503906 0 8077.7134 b 1
250.1276397705078 0 944.2147
251.10304260253906 0 940.74335
253.09507751464844 0 547.0397
263.10211181640625 0 1395.899
265.15478515625 0 697.0653
279.0971374511719 0 944.0685
281.1240539550781 0 879.1753
283.0300598144531 0 650.8549
283.14337158203125 0 683.22003
284.1197814941406 0 506.44244
285.0098571777344 0 2173.9507
285.1009521484375 0 1209.8026
292.1417541503906 0 781.80664
299.06097412109375 0 649.3262
319.13958740234375 0 735.3962
324.1453857421875 0 1753.9926
329.1496276855469 0 4359.335
330.15093994140625 0 973.93256
336.0159606933594 0 545.22327
340.1875915527344 0 1948.2274
342.1560974121094 0 10790.124 y 7
343.1585998535156 0 1775.5026
350.1337890625 0 782.8423
355.070068359375 0 2425.9177
356.070068359375 0 1156.023
356.1392517089844 0 905.2645
357.2234191894531 0 1100.3286
364.1507568359375 0 2200.6255 b 2
378.12860107421875 0 920.5304
380.19287109375 0 558.7776
387.8145751953125 0 480.88275
393.177490234375 0 948.146
394.1243591308594 0 2377.0312
395.19293212890625 0 889.6825
398.2027893066406 0 778.723
403.2129821777344 0 713.35455
411.22637939453125 0 493.04715
413.2662658691406 0 1119.9772
415.2510070800781 0 576.84625
425.1923828125 0 1904.344
427.72332763671875 0 743.6061
453.2705993652344 0 712.70984
458.447021484375 0 594.3085
465.16204833984375 0 1538.8003
479.17742919921875 0 1531.2368 b 3
489.1913146972656 0 2880.402 y 6
490.1888732910156 0 635.2602 b Water loss 7
496.22869873046875 0 901.40326
501.4621276855469 0 551.2067
508.27703857421875 0 1082.2108
519.3052978515625 0 548.65015
550.7095336914062 0 883.4402 y 1
560.2288208007812 0 3363.5723 y 5
561.231201171875 0 655.8128
563.2845458984375 0 758.7407
568.3045043945312 0 593.2674
569.2314453125 0 860.7286
585.3436889648438 0 2001.6266
601.2340087890625 0 1975.7867 Precursor
601.7387084960938 0 983.4542
601.8202514648438 0 729.70264
602.3211059570312 0 774.07855
602.373046875 0 1157.2571
621.3596801757812 0 1388.2021
628.37744140625 0 744.5526
640.3369750976562 0 730.5516
656.9848022460938 0 734.53674
657.262939453125 0 1448.9437
659.2944946289062 0 1140.5695
685.2551879882812 0 1129.481
700.3612670898438 0 4873.957
701.3599853515625 0 1354.888
721.2635498046875 0 670.52893
723.2916259765625 0 2895.7136 y 4
724.2930908203125 0 1469.6562
735.4017944335938 0 735.66327
749.2553100585938 0 1995.5106
750.2520751953125 0 743.0628
774.3224487304688 0 867.3942
783.39453125 0 890.8427
801.4149780273438 0 1647.6123
802.4148559570312 0 1011.30316
838.3170166015625 0 3542.1704 y 3
839.3207397460938 0 1248.0114
844.4525756835938 0 3010.3267
845.4552001953125 0 1420.0806
854.428466796875 0 848.42163
872.4471435546875 0 11282.513
873.4490966796875 0 5380.2383
874.4519653320312 0 1138.0265
889.3458862304688 0 6300.8804
890.3485107421875 0 2600.6458
901.477783203125 0 1168.6802
902.47900390625 0 1184.9323
908.3421020507812 0 645.94745
935.3338012695312 0 1067.9196 y Water loss 2
953.3447265625 0 18143.316 y 2
954.3472290039062 0 9385.505
955.3499145507812 0 4546.257
956.3502807617188 0 766.77496
991.5173950195312 0 2172.1455
992.5176391601562 0 883.4621
1005.7100830078125 0 604.5746
1019.51513671875 0 5578.8403
1020.5202026367188 0 2499.4663
1021.5153198242188 0 949.2647
1100.4117431640625 0 1293.3981 y 1
1206.4422607421875 0 608.47174
3079.388671875 0 714.76355
3080.038818359375 0 807.9606

Spectrum Details

|  |  |
| --- | --- |
| Matched peaks? Matched peaks The total absolute number of peaks matched. Additionally in brackets the total fraction of peaks matched and the total number of peaks is shown. | 18 (10.06% of 179) |
| FDR? FDR The false discovery rate estimated for this peptide. It is calculated by matching all theoretical fragments with a non-integer shift with the raw peaks for this spectrum. This is done with 40 different shifts. The resulting percentage is the average number of annotated peaks over the number of annotated peaks with the correct spectrum. | 1.32% |
| Satellite FDR? Satellite FDR See the FDR for details on its calculation. This satellite ion specific FDR only contains the satellite ions (d/w) for I/L/J positions. | - |
| PSM Score? PSM Score The PSM Score as given by Hecklib to this annotated spectrum. It is shown with three significant figures. | 195 |

#### Spectrum 10660? Spectrum 10660 The raw spectrum of this peptide as annotated by Hecklib. The fragments are coloured according to ion type (see legend). Any peaks with a star '\*' as text can be hovered over to see the full details, first the ion type second the mass shift type. By hovering over the amino acids in the peptide or ions in the legend the corresponding peaks are highlighted. By toggling the 'Unassigned' label you can turn the background (unassigned) peaks on or off in the plot. By updating the slider in the Ion legend you can update the spectrum to only show the top X% of the peaks with labels. The top X% means any peak that is within X% of the highest intensity. By dragging in the spectrum you can zoom in to a specific part of the spectrum and use 'Zoom Out' to get back to the original zoom level. The annotation of the spectrum is based on the given sequence in the peptides file and is done with different software so inconsistencies are likely. The peaks are annotated based on the given sequence, with 20 ppm tolerance.

Copy Data

##### Spectrum 10660 (TSV)

###### Preview

```
Loading example...
```

*Click on the button to copy the data to your clipboard.*

Mz MinMz MaxIntensity Max

WidthHeightPeptide font sizePeptide stroke widthSpectrum font sizeSpectrum stroke widthCompact peptide

Ion legend

wxyz

abcd

OtherUnassignedIon^Charge^_Position_Show for top:%

TFDDYAMHW

01.96e+43.93e+45.89e+47.85e+4

Zoom Out

a^+1^_2_y^+1^_1_a^+1^_2_b^+1^_2_b^+1^_2_d^+1^_3_y^+1^_2_b^+1^_3_b^+1^_3_b^+1^_4_y^+1^_3_y^+2^_8_y^+1^_4_\*\*y^+1^_5_y^+1^_6_y^+1^_7_y^+1^_7_y^+1^_8_

0740148022202959

Fragment Matches Table

Show background peaks

| Position | Ion type | Intensity | mz Theoretical | mz Error (Th) | mz Error (ppm) | Charge | Series Number |
| --- | --- | --- | --- | --- | --- | --- | --- |
| - | - | 5.536E+04 | 120.1 | - | - | 0 | - |
| - | - | 4014 | 121.1 | - | - | 0 | - |
| - | - | 1327 | 123 | - | - | 0 | - |
| - | - | 384.4 | 128.1 | - | - | 0 | - |
| - | - | 3470 | 129.1 | - | - | 0 | - |
| - | - | 518.1 | 130.1 | - | - | 0 | - |
| - | - | 1044 | 130.1 | - | - | 0 | - |
| - | - | 547.1 | 130.1 | - | - | 0 | - |
| - | - | 955.3 | 132.1 | - | - | 0 | - |
| - | - | 1210 | 132.1 | - | - | 0 | - |
| - | - | 842.7 | 133.1 | - | - | 0 | - |
| - | - | 416.5 | 135.9 | - | - | 0 | - |
| - | - | 1.819E+04 | 136.1 | - | - | 0 | - |
| - | - | 1260 | 137.1 | - | - | 0 | - |
| - | - | 664 | 138.1 | - | - | 0 | - |
| - | - | 1040 | 138.1 | - | - | 0 | - |
| - | - | 387.3 | 143.1 | - | - | 0 | - |
| - | - | 908.5 | 146.1 | - | - | 0 | - |
| - | - | 886.7 | 147 | - | - | 0 | - |
| - | - | 649.3 | 148.9 | - | - | 0 | - |
| - | - | 1122 | 149 | - | - | 0 | - |
| - | - | 1990 | 156.1 | - | - | 0 | - |
| - | - | 621.3 | 157.1 | - | - | 0 | - |
| - | - | 2698 | 158.1 | - | - | 0 | - |
| - | - | 467.7 | 158.1 | - | - | 0 | - |
| - | - | 2855 | 159.1 | - | - | 0 | - |
| - | - | 468.6 | 159.1 | - | - | 0 | - |
| - | - | 747.1 | 160.1 | - | - | 0 | - |
| - | - | 1043 | 160.1 | - | - | 0 | - |
| - | - | 6543 | 165.1 | - | - | 0 | - |
| - | - | 447.1 | 165.1 | - | - | 0 | - |
| - | - | 781.7 | 166.1 | - | - | 0 | - |
| - | - | 1996 | 167.1 | - | - | 0 | - |
| - | - | 689.7 | 169.1 | - | - | 0 | - |
| - | - | 3438 | 173.1 | - | - | 0 | - |
| - | - | 542.5 | 175.1 | - | - | 0 | - |
| - | - | 4303 | 176.1 | - | - | 0 | - |
| - | - | 4407 | 177.1 | - | - | 0 | - |
| - | - | 1033 | 177.1 | - | - | 0 | - |
| - | - | 484.9 | 177.3 | - | - | 0 | - |
| - | - | 1.345E+04 | 182.1 | - | - | 0 | - |
| - | - | 929.4 | 183.1 | - | - | 0 | - |
| - | - | 654.1 | 183.1 | - | - | 0 | - |
| - | - | 819.5 | 183.1 | - | - | 0 | - |
| - | - | 455.6 | 183.8 | - | - | 0 | - |
| - | - | 873.1 | 185.1 | - | - | 0 | - |
| - | - | 4939 | 188.1 | - | - | 0 | - |
| - | - | 1216 | 191.1 | - | - | 0 | - |
| - | - | 1859 | 193.1 | - | - | 0 | - |
| - | - | 613.5 | 197.1 | - | - | 0 | - |
| - | - | 589 | 200.1 | - | - | 0 | - |
| - | - | 1382 | 201.1 | - | - | 0 | - |
| - | - | 760.9 | 203.1 | - | - | 0 | - |
| - | - | 565.7 | 203.1 | - | - | 0 | - |
| 2 | a | 1402 | 203.1 | 0.000244 | 1.201 | +1 | 2 |
| - | - | 695.2 | 204.1 | - | - | 0 | - |
| 9 | y | 2407 | 205.1 | 0.0005937 | 2.895 | +1 | 1 |
| - | - | 809.9 | 207.1 | - | - | 0 | - |
| - | - | 517.1 | 215.1 | - | - | 0 | - |
| - | - | 1153 | 217.1 | - | - | 0 | - |
| - | - | 4556 | 221.1 | - | - | 0 | - |
| 2 | a | 7.773E+04 | 221.1 | 0.0001163 | 0.5261 | +1 | 2 |
| - | - | 8505 | 222.1 | - | - | 0 | - |
| - | - | 838.6 | 223.1 | - | - | 0 | - |
| - | - | 764 | 225.2 | - | - | 0 | - |
| - | - | 746.3 | 226.1 | - | - | 0 | - |
| - | - | 613.4 | 226.2 | - | - | 0 | - |
| - | - | 721.5 | 228.1 | - | - | 0 | - |
| - | - | 548.1 | 229.2 | - | - | 0 | - |
| - | - | 723 | 229.2 | - | - | 0 | - |
| 2 | b | 1012 | 231.1 | 0.0005381 | 2.328 | +1 | 2 |
| - | - | 1071 | 233.1 | - | - | 0 | - |
| - | - | 540.7 | 233.1 | - | - | 0 | - |
| - | - | 1861 | 235.1 | - | - | 0 | - |
| - | - | 720.5 | 239.1 | - | - | 0 | - |
| - | - | 749.7 | 240.1 | - | - | 0 | - |
| - | - | 496.5 | 241.1 | - | - | 0 | - |
| 2 | b | 1.208E+04 | 249.1 | 0.0002121 | 0.8513 | +1 | 2 |
| - | - | 2018 | 250.1 | - | - | 0 | - |
| - | - | 1608 | 251.1 | - | - | 0 | - |
| - | - | 774.4 | 253.2 | - | - | 0 | - |
| - | - | 764.4 | 257.1 | - | - | 0 | - |
| - | - | 1891 | 263.1 | - | - | 0 | - |
| - | - | 665.4 | 263.1 | - | - | 0 | - |
| - | - | 540.9 | 268.1 | - | - | 0 | - |
| - | - | 664.6 | 274.1 | - | - | 0 | - |
| - | - | 1234 | 279.1 | - | - | 0 | - |
| - | - | 1113 | 285 | - | - | 0 | - |
| - | - | 1759 | 285.1 | - | - | 0 | - |
| - | - | 943.6 | 292.1 | - | - | 0 | - |
| 3 | d | 593.7 | 292.2 | 0.001058 | 3.621 | +1 | 3 |
| - | - | 949.7 | 296.1 | - | - | 0 | - |
| - | - | 1183 | 302.1 | - | - | 0 | - |
| - | - | 570.6 | 319 | - | - | 0 | - |
| - | - | 788.5 | 319.1 | - | - | 0 | - |
| - | - | 3357 | 324.1 | - | - | 0 | - |
| - | - | 627.1 | 325.1 | - | - | 0 | - |
| - | - | 8366 | 329.1 | - | - | 0 | - |
| - | - | 2547 | 330.2 | - | - | 0 | - |
| - | - | 1311 | 340.2 | - | - | 0 | - |
| 8 | y | 1.419E+04 | 342.2 | 0.0003061 | 0.8947 | +1 | 2 |
| - | - | 1881 | 343.2 | - | - | 0 | - |
| 3 | b | 859.7 | 346.1 | 0.0004506 | 1.302 | +1 | 3 |
| - | - | 679.6 | 348 | - | - | 0 | - |
| - | - | 545.1 | 354.9 | - | - | 0 | - |
| - | - | 899.4 | 355.1 | - | - | 0 | - |
| - | - | 651.7 | 356.1 | - | - | 0 | - |
| - | - | 2123 | 357.2 | - | - | 0 | - |
| 3 | b | 3124 | 364.2 | 0.0001349 | 0.3704 | +1 | 3 |
| - | - | 560.9 | 367.2 | - | - | 0 | - |
| - | - | 965.1 | 368.2 | - | - | 0 | - |
| - | - | 2037 | 378.1 | - | - | 0 | - |
| - | - | 2331 | 394.1 | - | - | 0 | - |
| - | - | 753.8 | 395.1 | - | - | 0 | - |
| - | - | 881.9 | 401.2 | - | - | 0 | - |
| - | - | 919 | 413.3 | - | - | 0 | - |
| - | - | 2300 | 425.2 | - | - | 0 | - |
| - | - | 621.1 | 427.7 | - | - | 0 | - |
| - | - | 1210 | 428.2 | - | - | 0 | - |
| - | - | 578.2 | 434.9 | - | - | 0 | - |
| - | - | 1092 | 436.7 | - | - | 0 | - |
| - | - | 2229 | 465.2 | - | - | 0 | - |
| 4 | b | 1572 | 479.2 | 0.0002048 | 0.4274 | +1 | 4 |
| - | - | 1104 | 487.3 | - | - | 0 | - |
| 7 | y | 4158 | 489.2 | 0.005283 | 10.8 | +1 | 3 |
| - | - | 799 | 490.2 | - | - | 0 | - |
| - | - | 1179 | 496.2 | - | - | 0 | - |
| - | - | 825.4 | 510.3 | - | - | 0 | - |
| 2 | y | 1405 | 550.7 | 0.001432 | 2.601 | +2 | 8 |
| - | - | 632.8 | 551.7 | - | - | 0 | - |
| 6 | y | 4631 | 560.2 | 0.005095 | 9.095 | +1 | 4 |
| - | - | 1788 | 561.2 | - | - | 0 | - |
| - | - | 577.9 | 570.2 | - | - | 0 | - |
| - | - | 726.9 | 578.3 | - | - | 0 | - |
| - | - | 636.8 | 578.7 | - | - | 0 | - |
| - | - | 2996 | 585.3 | - | - | 0 | - |
| - | - | 940.1 | 586.3 | - | - | 0 | - |
| 0 | Precursor | 808.1 | 592.2 | 0.004238 | 7.155 | +2 | -1 |
| 0 | Precursor | 2090 | 601.2 | 0.00457 | 7.602 | +2 | -1 |
| - | - | 788.4 | 601.7 | - | - | 0 | - |
| - | - | 701.5 | 611.2 | - | - | 0 | - |
| - | - | 1023 | 612.2 | - | - | 0 | - |
| - | - | 643.1 | 628.4 | - | - | 0 | - |
| - | - | 1140 | 657.3 | - | - | 0 | - |
| - | - | 1740 | 659.3 | - | - | 0 | - |
| - | - | 1563 | 685.3 | - | - | 0 | - |
| - | - | 8526 | 700.4 | - | - | 0 | - |
| - | - | 2995 | 701.4 | - | - | 0 | - |
| - | - | 804.4 | 702.4 | - | - | 0 | - |
| - | - | 853.8 | 721.3 | - | - | 0 | - |
| - | - | 901.4 | 722.3 | - | - | 0 | - |
| 5 | y | 4050 | 723.3 | 0.004267 | 5.899 | +1 | 5 |
| - | - | 2369 | 724.3 | - | - | 0 | - |
| - | - | 2540 | 749.3 | - | - | 0 | - |
| - | - | 1272 | 757.4 | - | - | 0 | - |
| - | - | 1802 | 774.3 | - | - | 0 | - |
| - | - | 1014 | 775.3 | - | - | 0 | - |
| - | - | 2436 | 783.4 | - | - | 0 | - |
| - | - | 964.7 | 784.4 | - | - | 0 | - |
| - | - | 814.9 | 800.4 | - | - | 0 | - |
| - | - | 3736 | 801.4 | - | - | 0 | - |
| - | - | 1486 | 802.4 | - | - | 0 | - |
| - | - | 991.9 | 803.4 | - | - | 0 | - |
| - | - | 1500 | 828.4 | - | - | 0 | - |
| - | - | 581.2 | 834.4 | - | - | 0 | - |
| 4 | y | 5558 | 838.3 | 0.003996 | 4.767 | +1 | 6 |
| - | - | 2789 | 839.3 | - | - | 0 | - |
| - | - | 1574 | 840.3 | - | - | 0 | - |
| - | - | 7318 | 844.5 | - | - | 0 | - |
| - | - | 3079 | 845.5 | - | - | 0 | - |
| - | - | 987.2 | 846.5 | - | - | 0 | - |
| - | - | 806 | 854.4 | - | - | 0 | - |
| - | - | 1144 | 871.3 | - | - | 0 | - |
| - | - | 2.121E+04 | 872.4 | - | - | 0 | - |
| - | - | 1.062E+04 | 873.4 | - | - | 0 | - |
| - | - | 3542 | 874.5 | - | - | 0 | - |
| - | - | 9310 | 889.3 | - | - | 0 | - |
| - | - | 3995 | 890.3 | - | - | 0 | - |
| - | - | 1683 | 891.4 | - | - | 0 | - |
| - | - | 769.9 | 904.7 | - | - | 0 | - |
| 3 | y | 1446 | 935.3 | 0.003487 | 3.728 | +1 | 7 |
| - | - | 1596 | 936.3 | - | - | 0 | - |
| 3 | y | 2.529E+04 | 953.3 | 0.00397 | 4.164 | +1 | 7 |
| - | - | 1.323E+04 | 954.3 | - | - | 0 | - |
| - | - | 4377 | 955.3 | - | - | 0 | - |
| - | - | 884.2 | 956.3 | - | - | 0 | - |
| - | - | 3818 | 991.5 | - | - | 0 | - |
| - | - | 2669 | 992.5 | - | - | 0 | - |
| - | - | 666.6 | 993.5 | - | - | 0 | - |
| - | - | 9075 | 1020 | - | - | 0 | - |
| - | - | 5417 | 1021 | - | - | 0 | - |
| - | - | 1597 | 1022 | - | - | 0 | - |
| - | - | 662.1 | 1023 | - | - | 0 | - |
| 2 | y | 1535 | 1100 | 0.004159 | 3.78 | +1 | 8 |
| - | - | 1198 | 1101 | - | - | 0 | - |
| - | - | 695.8 | 1358 | - | - | 0 | - |
| - | - | 623.5 | 2341 | - | - | 0 | - |
| - | - | 671.2 | 2930 | - | - | 0 | - |

m/z Charge Intensity FragmentType MassShift Position
120.08100128173828 0 55358.11
121.08433532714844 0 4013.8186
123.04421997070312 0 1327.4475
128.1073760986328 0 384.41592
129.1024169921875 0 3469.8083
130.0503692626953 0 518.1412
130.06544494628906 0 1043.8112
130.08656311035156 0 547.05927
132.08103942871094 0 955.29877
132.1019287109375 0 1209.8638
133.08599853515625 0 842.70953
135.89303588867188 0 416.45276
136.0758819580078 0 18192.678
137.07923889160156 0 1259.7833
138.05523681640625 0 663.9752
138.06654357910156 0 1040.1874
143.07144165039062 0 387.2707
146.0601348876953 0 908.5303
147.0443878173828 0 886.69904
148.94786071777344 0 649.3486
149.0451202392578 0 1122.17
156.07679748535156 0 1990.2806
157.133544921875 0 621.2803
158.0966033935547 0 2697.6174
158.10292053222656 0 467.7121
159.09181213378906 0 2855.0598
159.0984344482422 0 468.58136
160.07589721679688 0 747.1265
160.11160278320312 0 1043.3076
165.0548095703125 0 6542.9
165.0666046142578 0 447.0938
166.0867462158203 0 781.6681
167.05577087402344 0 1995.732
169.09735107421875 0 689.6592
173.12875366210938 0 3438.2944
175.08680725097656 0 542.5421
176.10723876953125 0 4302.8965
177.10235595703125 0 4407.096
177.11105346679688 0 1033.3115
177.27061462402344 0 484.94687
182.08132934570312 0 13453.891
183.0851287841797 0 929.3851
183.11370849609375 0 654.1043
183.1493682861328 0 819.5283
183.80487060546875 0 455.6246
185.128662109375 0 873.0729
188.0707550048828 0 4939.365
191.11790466308594 0 1215.7402
193.1083984375 0 1859.1858
197.128662109375 0 613.5291
200.13978576660156 0 588.972
201.12295532226562 0 1382.4906
203.0667724609375 0 760.87854
203.08285522460938 0 565.6705
203.11813354492188 0 1402.0347 a Water loss 1
204.0771026611328 0 695.15735
205.09774780273438 0 2406.5745 y 8
207.11288452148438 0 809.88293
215.13929748535156 0 517.0746
217.09744262695312 0 1152.9966
221.10350036621094 0 4556.221
221.12857055664062 0 77733.24 a 1
222.13197326660156 0 8504.87
223.06375122070312 0 838.56366
225.17160034179688 0 764.0479
226.11878967285156 0 746.2913
226.15521240234375 0 613.4174
228.1126708984375 0 721.46265
229.15524291992188 0 548.1438
229.1702880859375 0 723.0132
231.11334228515625 0 1012.13525 b Water loss 1
233.0911102294922 0 1070.945
233.11422729492188 0 540.70856
235.1078338623047 0 1860.8854
239.094482421875 0 720.4805
240.13412475585938 0 749.69495
241.07443237304688 0 496.50092
249.1235809326172 0 12079.993 b 1
250.12672424316406 0 2017.625
251.1034698486328 0 1608.3091
253.16632080078125 0 774.3533
257.1073913574219 0 764.41235
263.10260009765625 0 1891.3813
263.12713623046875 0 665.3817
268.0752258300781 0 540.86316
274.12921142578125 0 664.5646
279.0972595214844 0 1234.282
285.0101013183594 0 1112.5219
285.1016845703125 0 1759.0165
292.14044189453125 0 943.61865
292.1666259765625 0 593.65924 d 2
296.1497802734375 0 949.68396
302.11407470703125 0 1182.815
319.02496337890625 0 570.6195
319.13958740234375 0 788.5455
324.14556884765625 0 3356.803
325.1468505859375 0 627.13916
329.149658203125 0 8366.2705
330.1534118652344 0 2547.4175
340.197509765625 0 1310.8853
342.1563720703125 0 14192.737 y 7
343.1595153808594 0 1880.6971
346.14019775390625 0 859.73865 b Water loss 2
347.96923828125 0 679.6111
354.8597412109375 0 545.148
355.06976318359375 0 899.41266
356.1382141113281 0 651.72064
357.2245178222656 0 2123.1267
364.1501770019531 0 3123.9631 b 2
367.1649475097656 0 560.9306
368.192626953125 0 965.14636
378.1286315917969 0 2037.111
394.1242370605469 0 2331.1433
395.1277770996094 0 753.7538
401.20654296875 0 881.8512
413.2653503417969 0 918.96747
425.1929626464844 0 2300.126
427.7213134765625 0 621.1069
428.2231750488281 0 1210.4131
434.8704528808594 0 578.15625
436.7275390625 0 1091.8871
465.1613464355469 0 2229.2136
479.1774597167969 0 1571.8738 b 3
487.3370361328125 0 1104.3529
489.19183349609375 0 4157.873 y 6
490.1940002441406 0 798.9831
496.2279357910156 0 1179.3616
510.2624206542969 0 825.37573
550.709716796875 0 1405.4618 y 1
551.7132568359375 0 632.7574
560.228759765625 0 4630.9272 y 5
561.231689453125 0 1787.5046
570.2254028320312 0 577.94763
578.2530517578125 0 726.87775
578.7491455078125 0 636.8097
585.3347778320312 0 2996.304
586.3383178710938 0 940.0564
592.2310791015625 0 808.05286 Precursor Water loss
601.2366943359375 0 2089.9282 Precursor
601.7318725585938 0 788.3748
611.2240600585938 0 701.4636
612.1954345703125 0 1022.845
628.376708984375 0 643.1095
657.2611694335938 0 1139.9674
659.292724609375 0 1739.9832
685.2549438476562 0 1563.4119
700.36181640625 0 8526.036
701.36376953125 0 2995.2515
702.3643188476562 0 804.4427
721.2589111328125 0 853.84235
722.2630004882812 0 901.37787
723.291259765625 0 4050.23 y 4
724.292724609375 0 2368.7068
749.255615234375 0 2539.6616
757.3825073242188 0 1272.1235
774.3204345703125 0 1802.0188
775.316162109375 0 1013.5439
783.398681640625 0 2435.6711
784.3945922851562 0 964.69086
800.419921875 0 814.8762
801.4132690429688 0 3735.6052
802.4110717773438 0 1486.3142
803.415771484375 0 991.93713
828.4189453125 0 1500.0935
834.3959350585938 0 581.2356
838.3179321289062 0 5558.165 y 3
839.3214721679688 0 2789.487
840.3226318359375 0 1574.2026
844.4517822265625 0 7317.9688
845.4539794921875 0 3079.1426
846.4537963867188 0 987.1529
854.435546875 0 806.0112
871.33642578125 0 1143.8691
872.4464721679688 0 21210.537
873.4490966796875 0 10623.343
874.4535522460938 0 3542.4692
889.346435546875 0 9309.899
890.349609375 0 3995.4102
891.3544311523438 0 1683.2528
904.6875610351562 0 769.91223
935.3338012695312 0 1446.3477 y Water loss 2
936.3353881835938 0 1595.9635
953.3448486328125 0 25291.182 y 2
954.3478393554688 0 13227.09
955.3482055664062 0 4376.819
956.3494873046875 0 884.2259
991.5200805664062 0 3818.396
992.5233764648438 0 2669.2266
993.5277709960938 0 666.626
1019.5140380859375 0 9074.676
1020.516845703125 0 5416.798
1021.5244750976562 0 1596.9979
1022.513427734375 0 662.07654
1100.4134521484375 0 1535.2041 y 1
1101.41748046875 0 1197.7913
1358.2684326171875 0 695.8492
2340.927490234375 0 623.4804
2930.16650390625 0 671.20355

Spectrum Details

|  |  |
| --- | --- |
| Matched peaks? Matched peaks The total absolute number of peaks matched. Additionally in brackets the total fraction of peaks matched and the total number of peaks is shown. | 20 (10.10% of 198) |
| FDR? FDR The false discovery rate estimated for this peptide. It is calculated by matching all theoretical fragments with a non-integer shift with the raw peaks for this spectrum. This is done with 40 different shifts. The resulting percentage is the average number of annotated peaks over the number of annotated peaks with the correct spectrum. | 1.07% |
| Satellite FDR? Satellite FDR See the FDR for details on its calculation. This satellite ion specific FDR only contains the satellite ions (d/w) for I/L/J positions. | - |
| PSM Score? PSM Score The PSM Score as given by Hecklib to this annotated spectrum. It is shown with three significant figures. | 213 |

#### Spectrum 11253? Spectrum 11253 The raw spectrum of this peptide as annotated by Hecklib. The fragments are coloured according to ion type (see legend). Any peaks with a star '\*' as text can be hovered over to see the full details, first the ion type second the mass shift type. By hovering over the amino acids in the peptide or ions in the legend the corresponding peaks are highlighted. By toggling the 'Unassigned' label you can turn the background (unassigned) peaks on or off in the plot. By updating the slider in the Ion legend you can update the spectrum to only show the top X% of the peaks with labels. The top X% means any peak that is within X% of the highest intensity. By dragging in the spectrum you can zoom in to a specific part of the spectrum and use 'Zoom Out' to get back to the original zoom level. The annotation of the spectrum is based on the given sequence in the peptides file and is done with different software so inconsistencies are likely. The peaks are annotated based on the given sequence, with 20 ppm tolerance.

Copy Data

##### Spectrum 11253 (TSV)

###### Preview

```
Loading example...
```

*Click on the button to copy the data to your clipboard.*

Mz MinMz MaxIntensity Max

WidthHeightPeptide font sizePeptide stroke widthSpectrum font sizeSpectrum stroke widthCompact peptide

Ion legend

wxyz

abcd

OtherUnassignedIon^Charge^_Position_Show for top:%

TFDDYAMHW

01.48e+42.96e+44.44e+45.92e+4

Zoom Out

a^+1^_2_y^+1^_1_a^+1^_2_b^+1^_2_b^+1^_2_a^+1^_3_y^+1^_2_b^+1^_3_b^+1^_3_b^+1^_4_y^+1^_3_y^+2^_8_y^+1^_4_\*\*y^+1^_5_y^+1^_6_y^+1^_7_y^+1^_7_y^+1^_8_

0778155723353114

Fragment Matches Table

Show background peaks

| Position | Ion type | Intensity | mz Theoretical | mz Error (Th) | mz Error (ppm) | Charge | Series Number |
| --- | --- | --- | --- | --- | --- | --- | --- |
| - | - | 4.977E+04 | 120.1 | - | - | 0 | - |
| - | - | 4411 | 121.1 | - | - | 0 | - |
| - | - | 814.9 | 127.1 | - | - | 0 | - |
| - | - | 3569 | 129.1 | - | - | 0 | - |
| - | - | 1277 | 130.1 | - | - | 0 | - |
| - | - | 410.8 | 130.1 | - | - | 0 | - |
| - | - | 1042 | 132.1 | - | - | 0 | - |
| - | - | 457.5 | 133.1 | - | - | 0 | - |
| - | - | 884.6 | 136.1 | - | - | 0 | - |
| - | - | 1.652E+04 | 136.1 | - | - | 0 | - |
| - | - | 1343 | 137.1 | - | - | 0 | - |
| - | - | 483.4 | 138.1 | - | - | 0 | - |
| - | - | 601.5 | 138.1 | - | - | 0 | - |
| - | - | 384 | 140.7 | - | - | 0 | - |
| - | - | 1419 | 141.1 | - | - | 0 | - |
| - | - | 690.1 | 147 | - | - | 0 | - |
| - | - | 530.4 | 147.1 | - | - | 0 | - |
| - | - | 1032 | 149 | - | - | 0 | - |
| - | - | 505.9 | 155.1 | - | - | 0 | - |
| - | - | 786.1 | 155.1 | - | - | 0 | - |
| - | - | 1694 | 156.1 | - | - | 0 | - |
| - | - | 864 | 157.1 | - | - | 0 | - |
| - | - | 698 | 157.1 | - | - | 0 | - |
| - | - | 2654 | 158.1 | - | - | 0 | - |
| - | - | 457.1 | 158.9 | - | - | 0 | - |
| - | - | 1510 | 159.1 | - | - | 0 | - |
| - | - | 473.1 | 160.1 | - | - | 0 | - |
| - | - | 739.1 | 160.1 | - | - | 0 | - |
| - | - | 554.9 | 162.1 | - | - | 0 | - |
| - | - | 5190 | 165.1 | - | - | 0 | - |
| - | - | 487.5 | 165.1 | - | - | 0 | - |
| - | - | 704.5 | 166.1 | - | - | 0 | - |
| - | - | 8633 | 166.1 | - | - | 0 | - |
| - | - | 2362 | 167.1 | - | - | 0 | - |
| - | - | 3175 | 167.1 | - | - | 0 | - |
| - | - | 1078 | 167.1 | - | - | 0 | - |
| - | - | 2609 | 169.1 | - | - | 0 | - |
| - | - | 608.1 | 171.1 | - | - | 0 | - |
| - | - | 2589 | 173.1 | - | - | 0 | - |
| - | - | 2606 | 173.5 | - | - | 0 | - |
| - | - | 505.7 | 175.1 | - | - | 0 | - |
| - | - | 672.4 | 176.1 | - | - | 0 | - |
| - | - | 4367 | 176.1 | - | - | 0 | - |
| - | - | 496.4 | 176.4 | - | - | 0 | - |
| - | - | 2499 | 177.1 | - | - | 0 | - |
| - | - | 678 | 177.1 | - | - | 0 | - |
| - | - | 430.3 | 181.2 | - | - | 0 | - |
| - | - | 8962 | 182.1 | - | - | 0 | - |
| - | - | 723.5 | 183.1 | - | - | 0 | - |
| - | - | 1091 | 183.1 | - | - | 0 | - |
| - | - | 924.9 | 185.1 | - | - | 0 | - |
| - | - | 514.5 | 185.1 | - | - | 0 | - |
| - | - | 543.3 | 187.1 | - | - | 0 | - |
| - | - | 616.8 | 187.1 | - | - | 0 | - |
| - | - | 2224 | 188.1 | - | - | 0 | - |
| - | - | 677.9 | 189.1 | - | - | 0 | - |
| - | - | 1046 | 191.1 | - | - | 0 | - |
| - | - | 498.1 | 192.3 | - | - | 0 | - |
| - | - | 1167 | 193.1 | - | - | 0 | - |
| - | - | 541.9 | 195.8 | - | - | 0 | - |
| - | - | 1567 | 199.1 | - | - | 0 | - |
| - | - | 1832 | 201.1 | - | - | 0 | - |
| - | - | 826.4 | 203.1 | - | - | 0 | - |
| 2 | a | 1434 | 203.1 | 0.0005187 | 2.554 | +1 | 2 |
| - | - | 718.3 | 204.1 | - | - | 0 | - |
| - | - | 495.9 | 204.4 | - | - | 0 | - |
| 9 | y | 1451 | 205.1 | 0.0001054 | 0.514 | +1 | 1 |
| - | - | 1247 | 207.1 | - | - | 0 | - |
| - | - | 514.3 | 209 | - | - | 0 | - |
| - | - | 519.7 | 211.1 | - | - | 0 | - |
| - | - | 480.2 | 212.1 | - | - | 0 | - |
| - | - | 935.9 | 215.1 | - | - | 0 | - |
| - | - | 780.3 | 215.1 | - | - | 0 | - |
| - | - | 716.4 | 217.1 | - | - | 0 | - |
| - | - | 2766 | 221.1 | - | - | 0 | - |
| 2 | a | 5.861E+04 | 221.1 | 0.0001163 | 0.5261 | +1 | 2 |
| - | - | 7510 | 222.1 | - | - | 0 | - |
| - | - | 1548 | 223.1 | - | - | 0 | - |
| - | - | 2169 | 224.1 | - | - | 0 | - |
| - | - | 776.6 | 226.2 | - | - | 0 | - |
| - | - | 6264 | 227.1 | - | - | 0 | - |
| - | - | 589.6 | 229.2 | - | - | 0 | - |
| - | - | 1450 | 231.1 | - | - | 0 | - |
| 2 | b | 1011 | 231.1 | 0.0004924 | 2.13 | +1 | 2 |
| - | - | 778.1 | 233.1 | - | - | 0 | - |
| - | - | 1924 | 235.1 | - | - | 0 | - |
| - | - | 1500 | 239.1 | - | - | 0 | - |
| - | - | 664.9 | 241.1 | - | - | 0 | - |
| - | - | 1009 | 242.1 | - | - | 0 | - |
| - | - | 729.5 | 243.1 | - | - | 0 | - |
| - | - | 628.4 | 243.1 | - | - | 0 | - |
| - | - | 715 | 244.1 | - | - | 0 | - |
| 2 | b | 1.052E+04 | 249.1 | 0.0001205 | 0.4838 | +1 | 2 |
| - | - | 1621 | 250.1 | - | - | 0 | - |
| - | - | 2132 | 251.1 | - | - | 0 | - |
| - | - | 1711 | 253.1 | - | - | 0 | - |
| - | - | 612.6 | 253.2 | - | - | 0 | - |
| - | - | 603.2 | 261.2 | - | - | 0 | - |
| - | - | 1729 | 263.1 | - | - | 0 | - |
| - | - | 1543 | 265.2 | - | - | 0 | - |
| - | - | 1119 | 266.1 | - | - | 0 | - |
| - | - | 504.8 | 279.1 | - | - | 0 | - |
| - | - | 2415 | 279.1 | - | - | 0 | - |
| - | - | 3910 | 281.1 | - | - | 0 | - |
| - | - | 898.4 | 285 | - | - | 0 | - |
| - | - | 1054 | 285.1 | - | - | 0 | - |
| - | - | 575.5 | 286 | - | - | 0 | - |
| - | - | 919.6 | 292.1 | - | - | 0 | - |
| - | - | 762.3 | 295.2 | - | - | 0 | - |
| - | - | 588.8 | 296.1 | - | - | 0 | - |
| - | - | 942.4 | 299.1 | - | - | 0 | - |
| - | - | 809 | 302.1 | - | - | 0 | - |
| - | - | 573.4 | 309.2 | - | - | 0 | - |
| 3 | a | 589.3 | 318.1 | 0.000248 | 0.7795 | +1 | 3 |
| - | - | 1023 | 319.1 | - | - | 0 | - |
| - | - | 645 | 321.1 | - | - | 0 | - |
| - | - | 602.2 | 322.1 | - | - | 0 | - |
| - | - | 1142 | 322.2 | - | - | 0 | - |
| - | - | 669.6 | 323.1 | - | - | 0 | - |
| - | - | 506.9 | 323.2 | - | - | 0 | - |
| - | - | 1925 | 324.1 | - | - | 0 | - |
| - | - | 613 | 325.2 | - | - | 0 | - |
| - | - | 5468 | 329.1 | - | - | 0 | - |
| - | - | 1297 | 330.2 | - | - | 0 | - |
| - | - | 5700 | 340.2 | - | - | 0 | - |
| - | - | 1621 | 341.2 | - | - | 0 | - |
| - | - | 767.2 | 342.1 | - | - | 0 | - |
| 8 | y | 1.132E+04 | 342.2 | 0.0002756 | 0.8055 | +1 | 2 |
| - | - | 2054 | 343.2 | - | - | 0 | - |
| 3 | b | 1079 | 346.1 | 0.001152 | 3.329 | +1 | 3 |
| - | - | 1081 | 352.2 | - | - | 0 | - |
| - | - | 1272 | 355.1 | - | - | 0 | - |
| - | - | 919.6 | 356.1 | - | - | 0 | - |
| - | - | 1101 | 356.1 | - | - | 0 | - |
| - | - | 1060 | 357.2 | - | - | 0 | - |
| 3 | b | 3407 | 364.2 | 4.332E-05 | 0.119 | +1 | 3 |
| - | - | 851.4 | 365.2 | - | - | 0 | - |
| - | - | 983.3 | 370.2 | - | - | 0 | - |
| - | - | 782.1 | 376.1 | - | - | 0 | - |
| - | - | 1703 | 378.1 | - | - | 0 | - |
| - | - | 4535 | 380.2 | - | - | 0 | - |
| - | - | 872.3 | 381.2 | - | - | 0 | - |
| - | - | 622.6 | 383.2 | - | - | 0 | - |
| - | - | 743.5 | 392.2 | - | - | 0 | - |
| - | - | 2197 | 394.1 | - | - | 0 | - |
| - | - | 758.6 | 394.2 | - | - | 0 | - |
| - | - | 2342 | 395.2 | - | - | 0 | - |
| - | - | 1114 | 396.2 | - | - | 0 | - |
| - | - | 3485 | 398.2 | - | - | 0 | - |
| - | - | 533.6 | 398.2 | - | - | 0 | - |
| - | - | 688.2 | 399.2 | - | - | 0 | - |
| - | - | 703.5 | 407.2 | - | - | 0 | - |
| - | - | 600.8 | 413.2 | - | - | 0 | - |
| - | - | 792.9 | 413.3 | - | - | 0 | - |
| - | - | 995.3 | 425.2 | - | - | 0 | - |
| - | - | 863.2 | 425.3 | - | - | 0 | - |
| - | - | 803.6 | 427.7 | - | - | 0 | - |
| - | - | 618.7 | 431.2 | - | - | 0 | - |
| - | - | 907.1 | 449.2 | - | - | 0 | - |
| - | - | 2151 | 453.3 | - | - | 0 | - |
| - | - | 696.4 | 454.3 | - | - | 0 | - |
| - | - | 1231 | 465.2 | - | - | 0 | - |
| - | - | 634.2 | 472.4 | - | - | 0 | - |
| - | - | 576.9 | 474.2 | - | - | 0 | - |
| 4 | b | 1708 | 479.2 | 0.0001919 | 0.4005 | +1 | 4 |
| - | - | 651 | 479.3 | - | - | 0 | - |
| - | - | 854.3 | 480.3 | - | - | 0 | - |
| 7 | y | 2858 | 489.2 | 0.005222 | 10.67 | +1 | 3 |
| - | - | 706.2 | 491.2 | - | - | 0 | - |
| - | - | 865.1 | 493.3 | - | - | 0 | - |
| - | - | 1645 | 496.2 | - | - | 0 | - |
| - | - | 2019 | 500.8 | - | - | 0 | - |
| - | - | 808.7 | 501.3 | - | - | 0 | - |
| - | - | 645.6 | 503.2 | - | - | 0 | - |
| - | - | 803.3 | 506.3 | - | - | 0 | - |
| - | - | 5543 | 508.3 | - | - | 0 | - |
| - | - | 1680 | 509.3 | - | - | 0 | - |
| - | - | 864.4 | 511.3 | - | - | 0 | - |
| 2 | y | 1747 | 550.7 | 0.002226 | 4.041 | +2 | 8 |
| - | - | 708 | 551.2 | - | - | 0 | - |
| 6 | y | 4711 | 560.2 | 0.004912 | 8.768 | +1 | 4 |
| - | - | 1188 | 561.2 | - | - | 0 | - |
| - | - | 2810 | 563.3 | - | - | 0 | - |
| - | - | 749.7 | 569.2 | - | - | 0 | - |
| - | - | 2359 | 585.3 | - | - | 0 | - |
| - | - | 1438 | 586.3 | - | - | 0 | - |
| 0 | Precursor | 1199 | 592.2 | 0.003139 | 5.3 | +2 | -1 |
| - | - | 1746 | 593.4 | - | - | 0 | - |
| 0 | Precursor | 2175 | 601.2 | 0.001152 | 1.917 | +2 | -1 |
| - | - | 735.2 | 601.6 | - | - | 0 | - |
| - | - | 1043 | 601.7 | - | - | 0 | - |
| - | - | 578 | 609.3 | - | - | 0 | - |
| - | - | 1072 | 612.2 | - | - | 0 | - |
| - | - | 9040 | 621.4 | - | - | 0 | - |
| - | - | 2787 | 622.4 | - | - | 0 | - |
| - | - | 615.2 | 634.2 | - | - | 0 | - |
| - | - | 1140 | 657.3 | - | - | 0 | - |
| - | - | 1472 | 659.3 | - | - | 0 | - |
| - | - | 1157 | 676.4 | - | - | 0 | - |
| - | - | 1593 | 678.4 | - | - | 0 | - |
| - | - | 810.8 | 679.4 | - | - | 0 | - |
| - | - | 1385 | 685.3 | - | - | 0 | - |
| - | - | 5227 | 700.4 | - | - | 0 | - |
| - | - | 1187 | 701.4 | - | - | 0 | - |
| - | - | 626.5 | 701.6 | - | - | 0 | - |
| - | - | 653.6 | 702.4 | - | - | 0 | - |
| - | - | 881 | 721.3 | - | - | 0 | - |
| 5 | y | 3841 | 723.3 | 0.004206 | 5.815 | +1 | 5 |
| - | - | 1793 | 724.3 | - | - | 0 | - |
| - | - | 701.2 | 733.4 | - | - | 0 | - |
| - | - | 3335 | 735.4 | - | - | 0 | - |
| - | - | 1070 | 736.4 | - | - | 0 | - |
| - | - | 1626 | 749.3 | - | - | 0 | - |
| - | - | 1437 | 774.3 | - | - | 0 | - |
| - | - | 1343 | 783.4 | - | - | 0 | - |
| - | - | 925.8 | 800.4 | - | - | 0 | - |
| - | - | 2744 | 801.4 | - | - | 0 | - |
| - | - | 1196 | 802.4 | - | - | 0 | - |
| - | - | 728.6 | 826.4 | - | - | 0 | - |
| - | - | 811.4 | 828.4 | - | - | 0 | - |
| 4 | y | 5974 | 838.3 | 0.003935 | 4.694 | +1 | 6 |
| - | - | 1989 | 839.3 | - | - | 0 | - |
| - | - | 755.7 | 840.3 | - | - | 0 | - |
| - | - | 3556 | 844.5 | - | - | 0 | - |
| - | - | 2200 | 845.5 | - | - | 0 | - |
| - | - | 1037 | 871.3 | - | - | 0 | - |
| - | - | 1.47E+04 | 872.4 | - | - | 0 | - |
| - | - | 6619 | 873.4 | - | - | 0 | - |
| - | - | 1984 | 874.5 | - | - | 0 | - |
| - | - | 7570 | 889.3 | - | - | 0 | - |
| - | - | 4354 | 890.3 | - | - | 0 | - |
| - | - | 4840 | 901.5 | - | - | 0 | - |
| - | - | 2742 | 902.5 | - | - | 0 | - |
| - | - | 622.4 | 903.5 | - | - | 0 | - |
| - | - | 1813 | 919.5 | - | - | 0 | - |
| 3 | y | 1325 | 935.3 | 0.0006794 | 0.7264 | +1 | 7 |
| - | - | 1116 | 936.3 | - | - | 0 | - |
| - | - | 789.4 | 951.4 | - | - | 0 | - |
| - | - | 781.5 | 951.5 | - | - | 0 | - |
| 3 | y | 2.3E+04 | 953.3 | 0.003665 | 3.844 | +1 | 7 |
| - | - | 1.124E+04 | 954.3 | - | - | 0 | - |
| - | - | 3727 | 955.3 | - | - | 0 | - |
| - | - | 895.2 | 956.3 | - | - | 0 | - |
| - | - | 780.3 | 964.3 | - | - | 0 | - |
| - | - | 1968 | 991.5 | - | - | 0 | - |
| - | - | 1438 | 992.5 | - | - | 0 | - |
| - | - | 1561 | 1001 | - | - | 0 | - |
| - | - | 1169 | 1002 | - | - | 0 | - |
| - | - | 1836 | 1019 | - | - | 0 | - |
| - | - | 5548 | 1020 | - | - | 0 | - |
| - | - | 3918 | 1021 | - | - | 0 | - |
| - | - | 945.7 | 1022 | - | - | 0 | - |
| 2 | y | 1776 | 1100 | 0.005624 | 5.111 | +1 | 8 |
| - | - | 962.9 | 1101 | - | - | 0 | - |
| - | - | 977.9 | 3083 | - | - | 0 | - |

m/z Charge Intensity FragmentType MassShift Position
120.08099365234375 0 49770.938
121.08433532714844 0 4410.8467
127.08676147460938 0 814.8809
129.10243225097656 0 3568.6604
130.06521606445312 0 1276.5181
130.10598754882812 0 410.79123
132.08148193359375 0 1042.3502
133.06057739257812 0 457.46136
136.07009887695312 0 884.61005
136.07586669921875 0 16516.234
137.07920837402344 0 1342.8933
138.0549774169922 0 483.37405
138.06639099121094 0 601.5213
140.72178649902344 0 384.01614
141.1024169921875 0 1419.0087
147.04429626464844 0 690.13684
147.11300659179688 0 530.38715
149.04507446289062 0 1032.0059
155.07058715820312 0 505.94635
155.0813446044922 0 786.0606
156.07681274414062 0 1693.5486
157.09698486328125 0 863.9947
157.13375854492188 0 698.0244
158.0967254638672 0 2653.8467
158.9319610595703 0 457.0841
159.091796875 0 1509.5134
160.0951385498047 0 473.10614
160.11231994628906 0 739.10223
162.09140014648438 0 554.9042
165.05479431152344 0 5189.837
165.10281372070312 0 487.54996
166.05905151367188 0 704.49347
166.08642578125 0 8633.337
167.05548095703125 0 2362.0625
167.0816650390625 0 3174.9966
167.0892791748047 0 1077.8535
169.09722900390625 0 2609.4058
171.07595825195312 0 608.1487
173.12875366210938 0 2588.5754
173.45115661621094 0 2606.0825
175.08692932128906 0 505.68427
176.08164978027344 0 672.4475
176.10708618164062 0 4367.2495
176.4478302001953 0 496.37772
177.1023712158203 0 2499.2651
177.11048889160156 0 677.9751
181.23684692382812 0 430.26373
182.08128356933594 0 8962.105
183.08509826660156 0 723.45526
183.11279296875 0 1091.053
185.09173583984375 0 924.85767
185.1290740966797 0 514.53827
187.1078643798828 0 543.26086
187.1439666748047 0 616.8182
188.07081604003906 0 2224.2036
189.0866241455078 0 677.8816
191.1179656982422 0 1046.0172
192.32188415527344 0 498.08105
193.1087646484375 0 1166.9823
195.7607879638672 0 541.94073
199.1080322265625 0 1567.3047
201.1236572265625 0 1831.6672
203.06602478027344 0 826.4077
203.118408203125 0 1433.6636 a Water loss 1
204.07725524902344 0 718.30884
204.3808135986328 0 495.8887
205.09725952148438 0 1451.4539 y 8
207.11322021484375 0 1247.2562
209.01022338867188 0 514.3235
211.10739135742188 0 519.7323
212.10232543945312 0 480.16525
215.08534240722656 0 935.86426
215.1398162841797 0 780.297
217.0962677001953 0 716.35315
221.10333251953125 0 2766.031
221.12857055664062 0 58612.68 a 1
222.1318359375 0 7510.4624
223.0637969970703 0 1548.0015
224.10321044921875 0 2169.133
226.15403747558594 0 776.581
227.10275268554688 0 6264.3955
229.17019653320312 0 589.61346
231.06138610839844 0 1450.2933
231.11329650878906 0 1011.00226 b Water loss 1
233.09190368652344 0 778.08856
235.10752868652344 0 1923.5873
239.09494018554688 0 1499.8658
241.0738983154297 0 664.93884
242.11239624023438 0 1008.8603
243.07984924316406 0 729.47955
243.1332550048828 0 628.4409
244.12905883789062 0 714.9816
249.1234893798828 0 10520.829 b 1
250.1268310546875 0 1620.5331
251.10284423828125 0 2131.7742
253.11830139160156 0 1710.7174
253.1664581298828 0 612.59076
261.15972900390625 0 603.17126
263.1023254394531 0 1728.8644
265.1552429199219 0 1543.2235
266.1495361328125 0 1119.0488
279.0815734863281 0 504.8185
279.0975646972656 0 2415.3054
281.1243896484375 0 3910.425
285.009521484375 0 898.4252
285.1018981933594 0 1053.7756
286.0072937011719 0 575.50323
292.1409912109375 0 919.6481
295.1768798828125 0 762.31464
296.1499328613281 0 588.8121
299.1348571777344 0 942.3772
302.1141662597656 0 808.9673
309.2057800292969 0 573.3508
318.14508056640625 0 589.2689 a Water loss 2
319.1409912109375 0 1022.7687
321.1222839355469 0 645.00525
322.08477783203125 0 602.2111
322.1557312011719 0 1142.332
323.09808349609375 0 669.60333
323.1690368652344 0 506.85007
324.1458740234375 0 1924.7164
325.1876525878906 0 612.9706
329.1497802734375 0 5468.1196
330.1534118652344 0 1297.0173
340.1873779296875 0 5699.6416
341.1879577636719 0 1621.2108
342.1365051269531 0 767.1825
342.1563415527344 0 11317.211 y 7
343.1595764160156 0 2054.2275
346.1408996582031 0 1079.3461 b Water loss 2
352.1973571777344 0 1080.6222
355.0696716308594 0 1272.1056
356.0700378417969 0 919.5574
356.138671875 0 1100.5103
357.22412109375 0 1060.244
364.1502685546875 0 3407.3054 b 2
365.1536865234375 0 851.41754
370.206298828125 0 983.25934
376.1119689941406 0 782.0822
378.12921142578125 0 1703.3561
380.1925964355469 0 4535.0806
381.1957702636719 0 872.31586
383.2310791015625 0 622.63525
392.20458984375 0 743.5087
394.1238708496094 0 2197.2686
394.20989990234375 0 758.5554
395.192626953125 0 2341.692
396.1943664550781 0 1114.3818
398.20330810546875 0 3485.2803
398.2314453125 0 533.61115
399.2042236328125 0 688.2354
407.1844177246094 0 703.5226
413.1590576171875 0 600.8344
413.2674560546875 0 792.8647
425.1934814453125 0 995.3243
425.2752990722656 0 863.2126
427.7231140136719 0 803.6369
431.21221923828125 0 618.71106
449.2399597167969 0 907.07996
453.2713317871094 0 2151.2632
454.27423095703125 0 696.3866
465.1607666015625 0 1231.4545
472.3750915527344 0 634.1583
474.19677734375 0 576.8579
479.17706298828125 0 1707.7489 b 3
479.2884521484375 0 650.96893
480.2801513671875 0 854.29626
489.1917724609375 0 2858.1133 y 6
491.2061767578125 0 706.19495
493.27716064453125 0 865.1153
496.2301025390625 0 1645.4246
500.7768249511719 0 2018.5848
501.2802734375 0 808.7252
503.15948486328125 0 645.63934
506.26007080078125 0 803.31244
508.2763366699219 0 5543.15
509.2786560058594 0 1679.9028
511.28997802734375 0 864.35376
550.7105102539062 0 1746.9296 y 1
551.2106323242188 0 707.9779
560.2285766601562 0 4711.2207 y 5
561.2325439453125 0 1188.0961
563.2817993164062 0 2810.0442
569.22900390625 0 749.7366
585.3341064453125 0 2358.5186
586.3397216796875 0 1438.0802
592.22998046875 0 1199.2206 Precursor Water loss
593.3641967773438 0 1746.3015
601.2332763671875 0 2174.882 Precursor
601.6322631835938 0 735.2027
601.737548828125 0 1043.2943
609.3295288085938 0 577.98895
612.1942749023438 0 1072.4302
621.3599243164062 0 9040.391
622.36474609375 0 2787.3274
634.2235717773438 0 615.17114
657.2620849609375 0 1139.6378
659.2911987304688 0 1471.6871
676.3682250976562 0 1156.8523
678.3807983398438 0 1593.458
679.3840942382812 0 810.7748
685.2593994140625 0 1384.648
700.3618774414062 0 5227.3535
701.3671875 0 1187.4849
701.5818481445312 0 626.54083
702.3712768554688 0 653.5643
721.264404296875 0 881.03754
723.2911987304688 0 3841.0637 y 4
724.2927856445312 0 1792.9911
733.3931884765625 0 701.1517
735.40234375 0 3335.3723
736.4143676757812 0 1069.8638
749.2579956054688 0 1626.0835
774.3184204101562 0 1436.8116
783.3977661132812 0 1342.7653
800.4246215820312 0 925.75543
801.4111938476562 0 2743.643
802.41357421875 0 1196.0962
826.4347534179688 0 728.5768
828.4188842773438 0 811.4318
838.31787109375 0 5973.574 y 3
839.32275390625 0 1988.5441
840.3211059570312 0 755.69885
844.4519653320312 0 3556.1133
845.453857421875 0 2200.0483
871.3375244140625 0 1036.8954
872.4464721679688 0 14702.253
873.4489135742188 0 6618.7
874.4523315429688 0 1984.3494
889.346923828125 0 7570.36
890.3486938476562 0 4354.448
901.4776000976562 0 4839.761
902.47802734375 0 2742.1714
903.4839477539062 0 622.3651
919.484619140625 0 1812.7076
935.3309936523438 0 1324.9855 y Water loss 2
936.3321533203125 0 1115.9438
951.3766479492188 0 789.37006
951.49365234375 0 781.4739
953.3445434570312 0 22999.936 y 2
954.3479614257812 0 11244.149
955.3480834960938 0 3726.6238
956.3492431640625 0 895.1881
964.3429565429688 0 780.29956
991.5177612304688 0 1968.2817
992.5230102539062 0 1437.5122
1000.5443115234375 0 1561.4486
1001.5528564453125 0 1169.1714
1018.5521850585938 0 1835.8196
1019.5183715820312 0 5548.145
1020.5189819335938 0 3917.703
1021.5226440429688 0 945.66034
1100.4149169921875 0 1775.5901 y 1
1101.416748046875 0 962.8632
3082.76318359375 0 977.8784

Spectrum Details

|  |  |
| --- | --- |
| Matched peaks? Matched peaks The total absolute number of peaks matched. Additionally in brackets the total fraction of peaks matched and the total number of peaks is shown. | 20 (7.84% of 255) |
| FDR? FDR The false discovery rate estimated for this peptide. It is calculated by matching all theoretical fragments with a non-integer shift with the raw peaks for this spectrum. This is done with 40 different shifts. The resulting percentage is the average number of annotated peaks over the number of annotated peaks with the correct spectrum. | 3.21% |
| Satellite FDR? Satellite FDR See the FDR for details on its calculation. This satellite ion specific FDR only contains the satellite ions (d/w) for I/L/J positions. | - |
| PSM Score? PSM Score The PSM Score as given by Hecklib to this annotated spectrum. It is shown with three significant figures. | 231 |

#### Spectrum 11490? Spectrum 11490 The raw spectrum of this peptide as annotated by Hecklib. The fragments are coloured according to ion type (see legend). Any peaks with a star '\*' as text can be hovered over to see the full details, first the ion type second the mass shift type. By hovering over the amino acids in the peptide or ions in the legend the corresponding peaks are highlighted. By toggling the 'Unassigned' label you can turn the background (unassigned) peaks on or off in the plot. By updating the slider in the Ion legend you can update the spectrum to only show the top X% of the peaks with labels. The top X% means any peak that is within X% of the highest intensity. By dragging in the spectrum you can zoom in to a specific part of the spectrum and use 'Zoom Out' to get back to the original zoom level. The annotation of the spectrum is based on the given sequence in the peptides file and is done with different software so inconsistencies are likely. The peaks are annotated based on the given sequence, with 20 ppm tolerance.

Copy Data

##### Spectrum 11490 (TSV)

###### Preview

```
Loading example...
```

*Click on the button to copy the data to your clipboard.*

Mz MinMz MaxIntensity Max

WidthHeightPeptide font sizePeptide stroke widthSpectrum font sizeSpectrum stroke widthCompact peptide

Ion legend

wxyz

abcd

OtherUnassignedIon^Charge^_Position_Show for top:%

TFDDYAMHW

01.37e+42.73e+44.10e+45.47e+4

Zoom Out

a^+1^_2_y^+1^_1_a^+1^_2_b^+1^_2_b^+1^_2_a^+1^_3_y^+1^_2_b^+1^_3_b^+1^_4_y^+1^_3_y^+2^_8_y^+1^_4_\*y^+1^_5_y^+1^_6_y^+1^_7_y^+1^_7_y^+1^_8_

0675134920242699

Fragment Matches Table

Show background peaks

| Position | Ion type | Intensity | mz Theoretical | mz Error (Th) | mz Error (ppm) | Charge | Series Number |
| --- | --- | --- | --- | --- | --- | --- | --- |
| - | - | 3.959E+04 | 120.1 | - | - | 0 | - |
| - | - | 3203 | 121.1 | - | - | 0 | - |
| - | - | 816.3 | 123 | - | - | 0 | - |
| - | - | 390.3 | 124.5 | - | - | 0 | - |
| - | - | 715.9 | 127.1 | - | - | 0 | - |
| - | - | 3203 | 129.1 | - | - | 0 | - |
| - | - | 1398 | 130.1 | - | - | 0 | - |
| - | - | 405.2 | 131.7 | - | - | 0 | - |
| - | - | 704.7 | 132.1 | - | - | 0 | - |
| - | - | 1095 | 133.1 | - | - | 0 | - |
| - | - | 1.19E+04 | 136.1 | - | - | 0 | - |
| - | - | 1003 | 137.1 | - | - | 0 | - |
| - | - | 622.1 | 138.1 | - | - | 0 | - |
| - | - | 2488 | 139.1 | - | - | 0 | - |
| - | - | 787 | 141.1 | - | - | 0 | - |
| - | - | 503.3 | 142.1 | - | - | 0 | - |
| - | - | 420.3 | 142.3 | - | - | 0 | - |
| - | - | 563.8 | 143.1 | - | - | 0 | - |
| - | - | 389.6 | 144.9 | - | - | 0 | - |
| - | - | 799.9 | 146.1 | - | - | 0 | - |
| - | - | 585.1 | 147 | - | - | 0 | - |
| - | - | 1423 | 149 | - | - | 0 | - |
| - | - | 510.4 | 155.1 | - | - | 0 | - |
| - | - | 516.6 | 155.1 | - | - | 0 | - |
| - | - | 1209 | 156.1 | - | - | 0 | - |
| - | - | 790.8 | 157.1 | - | - | 0 | - |
| - | - | 1842 | 158.1 | - | - | 0 | - |
| - | - | 2599 | 159.1 | - | - | 0 | - |
| - | - | 446.9 | 159.1 | - | - | 0 | - |
| - | - | 625.6 | 159.1 | - | - | 0 | - |
| - | - | 479 | 160.1 | - | - | 0 | - |
| - | - | 975 | 160.1 | - | - | 0 | - |
| - | - | 696.8 | 162.1 | - | - | 0 | - |
| - | - | 3865 | 165.1 | - | - | 0 | - |
| - | - | 7846 | 166.1 | - | - | 0 | - |
| - | - | 2455 | 167.1 | - | - | 0 | - |
| - | - | 7281 | 167.1 | - | - | 0 | - |
| - | - | 1173 | 167.1 | - | - | 0 | - |
| - | - | 739.6 | 168.1 | - | - | 0 | - |
| - | - | 1377 | 169.1 | - | - | 0 | - |
| - | - | 465.6 | 169.1 | - | - | 0 | - |
| - | - | 618.6 | 173.1 | - | - | 0 | - |
| - | - | 2560 | 173.1 | - | - | 0 | - |
| - | - | 2195 | 173.4 | - | - | 0 | - |
| - | - | 721.6 | 174.1 | - | - | 0 | - |
| - | - | 589.7 | 175.1 | - | - | 0 | - |
| - | - | 4099 | 176.1 | - | - | 0 | - |
| - | - | 453 | 176.4 | - | - | 0 | - |
| - | - | 2566 | 177.1 | - | - | 0 | - |
| - | - | 813.2 | 177.1 | - | - | 0 | - |
| - | - | 456.4 | 180.1 | - | - | 0 | - |
| - | - | 476.4 | 180.3 | - | - | 0 | - |
| - | - | 8871 | 182.1 | - | - | 0 | - |
| - | - | 562.7 | 183.1 | - | - | 0 | - |
| - | - | 506.7 | 185.1 | - | - | 0 | - |
| - | - | 879.1 | 185.1 | - | - | 0 | - |
| - | - | 1387 | 185.1 | - | - | 0 | - |
| - | - | 571.7 | 187.1 | - | - | 0 | - |
| - | - | 1834 | 188.1 | - | - | 0 | - |
| - | - | 698.1 | 191.1 | - | - | 0 | - |
| - | - | 1300 | 193.1 | - | - | 0 | - |
| - | - | 1726 | 199.1 | - | - | 0 | - |
| - | - | 1643 | 201.1 | - | - | 0 | - |
| - | - | 514.8 | 203.1 | - | - | 0 | - |
| 2 | a | 1614 | 203.1 | 0.0002898 | 1.427 | +1 | 2 |
| - | - | 646.1 | 204.1 | - | - | 0 | - |
| 9 | y | 1671 | 205.1 | 0.0002733 | 1.332 | +1 | 1 |
| - | - | 919.5 | 214.1 | - | - | 0 | - |
| - | - | 953.8 | 215.1 | - | - | 0 | - |
| - | - | 922.1 | 217.1 | - | - | 0 | - |
| - | - | 2794 | 221.1 | - | - | 0 | - |
| 2 | a | 5.415E+04 | 221.1 | 0.0001163 | 0.5261 | +1 | 2 |
| - | - | 6639 | 222.1 | - | - | 0 | - |
| - | - | 1102 | 223.1 | - | - | 0 | - |
| - | - | 1763 | 224.1 | - | - | 0 | - |
| - | - | 7058 | 227.1 | - | - | 0 | - |
| - | - | 1180 | 228.1 | - | - | 0 | - |
| - | - | 753.8 | 229.2 | - | - | 0 | - |
| - | - | 987.6 | 231.1 | - | - | 0 | - |
| 2 | b | 964.4 | 231.1 | 0.00053 | 2.293 | +1 | 2 |
| - | - | 1073 | 235.1 | - | - | 0 | - |
| - | - | 1087 | 239.1 | - | - | 0 | - |
| - | - | 1097 | 241.1 | - | - | 0 | - |
| - | - | 762.7 | 243.1 | - | - | 0 | - |
| - | - | 525 | 248.7 | - | - | 0 | - |
| 2 | b | 7826 | 249.1 | 0.0002579 | 1.035 | +1 | 2 |
| - | - | 779.4 | 250.1 | - | - | 0 | - |
| - | - | 915.5 | 251.1 | - | - | 0 | - |
| - | - | 1991 | 263.1 | - | - | 0 | - |
| - | - | 1886 | 264.1 | - | - | 0 | - |
| - | - | 1190 | 265.2 | - | - | 0 | - |
| - | - | 667.6 | 266.1 | - | - | 0 | - |
| - | - | 627.5 | 267.2 | - | - | 0 | - |
| - | - | 635.1 | 270.1 | - | - | 0 | - |
| - | - | 593.2 | 273.1 | - | - | 0 | - |
| - | - | 533.2 | 274.1 | - | - | 0 | - |
| - | - | 1551 | 279.1 | - | - | 0 | - |
| - | - | 2968 | 281.1 | - | - | 0 | - |
| - | - | 1162 | 285 | - | - | 0 | - |
| - | - | 1364 | 285.1 | - | - | 0 | - |
| - | - | 654.6 | 286 | - | - | 0 | - |
| - | - | 750.4 | 296.2 | - | - | 0 | - |
| - | - | 630.5 | 299.1 | - | - | 0 | - |
| - | - | 568.1 | 302.1 | - | - | 0 | - |
| 3 | a | 641.4 | 318.1 | 0.000187 | 0.5877 | +1 | 3 |
| - | - | 1100 | 319.1 | - | - | 0 | - |
| - | - | 629.6 | 322.2 | - | - | 0 | - |
| - | - | 617.1 | 323.2 | - | - | 0 | - |
| - | - | 1692 | 324.1 | - | - | 0 | - |
| - | - | 5517 | 329.1 | - | - | 0 | - |
| - | - | 997 | 330.2 | - | - | 0 | - |
| - | - | 5940 | 340.2 | - | - | 0 | - |
| - | - | 1365 | 341.2 | - | - | 0 | - |
| 8 | y | 1.034E+04 | 342.2 | 0.0002756 | 0.8055 | +1 | 2 |
| - | - | 530.7 | 342.7 | - | - | 0 | - |
| - | - | 1725 | 343.2 | - | - | 0 | - |
| - | - | 1820 | 355.1 | - | - | 0 | - |
| - | - | 597.5 | 355.3 | - | - | 0 | - |
| - | - | 671.2 | 357.2 | - | - | 0 | - |
| 3 | b | 3150 | 364.2 | 0.0007501 | 2.06 | +1 | 3 |
| - | - | 1040 | 365.2 | - | - | 0 | - |
| - | - | 691 | 366.2 | - | - | 0 | - |
| - | - | 756.3 | 367.2 | - | - | 0 | - |
| - | - | 879.1 | 370.2 | - | - | 0 | - |
| - | - | 1192 | 378.1 | - | - | 0 | - |
| - | - | 3561 | 380.2 | - | - | 0 | - |
| - | - | 670.6 | 392.7 | - | - | 0 | - |
| - | - | 6936 | 393.2 | - | - | 0 | - |
| - | - | 1500 | 394.1 | - | - | 0 | - |
| - | - | 1453 | 394.2 | - | - | 0 | - |
| - | - | 599.5 | 395.1 | - | - | 0 | - |
| - | - | 677.1 | 395.2 | - | - | 0 | - |
| - | - | 2405 | 398.2 | - | - | 0 | - |
| - | - | 773.6 | 404.1 | - | - | 0 | - |
| - | - | 748.1 | 412.2 | - | - | 0 | - |
| - | - | 1104 | 413.3 | - | - | 0 | - |
| - | - | 1009 | 425.2 | - | - | 0 | - |
| - | - | 1324 | 427.7 | - | - | 0 | - |
| - | - | 923.6 | 428.2 | - | - | 0 | - |
| - | - | 785.6 | 434.7 | - | - | 0 | - |
| - | - | 897.2 | 436.7 | - | - | 0 | - |
| - | - | 678.1 | 441.3 | - | - | 0 | - |
| - | - | 886.3 | 449.2 | - | - | 0 | - |
| - | - | 1066 | 450.2 | - | - | 0 | - |
| - | - | 628.8 | 451.2 | - | - | 0 | - |
| - | - | 733.7 | 453.3 | - | - | 0 | - |
| - | - | 648.7 | 454.3 | - | - | 0 | - |
| - | - | 748.5 | 457.2 | - | - | 0 | - |
| - | - | 922.3 | 465.2 | - | - | 0 | - |
| - | - | 867.8 | 465.7 | - | - | 0 | - |
| - | - | 668.2 | 471.3 | - | - | 0 | - |
| - | - | 854.7 | 478.3 | - | - | 0 | - |
| 4 | b | 1425 | 479.2 | 0.0006015 | 1.255 | +1 | 4 |
| 7 | y | 2573 | 489.2 | 0.005161 | 10.55 | +1 | 3 |
| - | - | 1070 | 490.2 | - | - | 0 | - |
| - | - | 597.4 | 493.3 | - | - | 0 | - |
| - | - | 994.1 | 496.2 | - | - | 0 | - |
| - | - | 820.1 | 496.3 | - | - | 0 | - |
| - | - | 7026 | 506.3 | - | - | 0 | - |
| - | - | 2025 | 507.3 | - | - | 0 | - |
| - | - | 1318 | 508.3 | - | - | 0 | - |
| 2 | y | 717.5 | 550.7 | 0.0002158 | 0.3919 | +2 | 8 |
| - | - | 762.5 | 551.2 | - | - | 0 | - |
| 6 | y | 3913 | 560.2 | 0.005767 | 10.29 | +1 | 4 |
| - | - | 1100 | 561.2 | - | - | 0 | - |
| - | - | 1654 | 563.3 | - | - | 0 | - |
| - | - | 801.7 | 564.3 | - | - | 0 | - |
| - | - | 910.7 | 567.3 | - | - | 0 | - |
| - | - | 2671 | 585.3 | - | - | 0 | - |
| - | - | 848.8 | 586.3 | - | - | 0 | - |
| - | - | 981.6 | 593.4 | - | - | 0 | - |
| 0 | Precursor | 1917 | 601.2 | 0.001397 | 2.323 | +2 | -1 |
| - | - | 942.4 | 601.3 | - | - | 0 | - |
| - | - | 703.7 | 601.5 | - | - | 0 | - |
| - | - | 822.5 | 601.7 | - | - | 0 | - |
| - | - | 872 | 602.2 | - | - | 0 | - |
| - | - | 700.7 | 602.3 | - | - | 0 | - |
| - | - | 984.6 | 612.2 | - | - | 0 | - |
| - | - | 2701 | 619.3 | - | - | 0 | - |
| - | - | 1143 | 620.4 | - | - | 0 | - |
| - | - | 3159 | 621.4 | - | - | 0 | - |
| - | - | 635.9 | 622.2 | - | - | 0 | - |
| - | - | 1164 | 622.4 | - | - | 0 | - |
| - | - | 1625 | 659.3 | - | - | 0 | - |
| - | - | 563.8 | 665.6 | - | - | 0 | - |
| - | - | 2811 | 676.4 | - | - | 0 | - |
| - | - | 738.7 | 677.4 | - | - | 0 | - |
| - | - | 822.4 | 678.4 | - | - | 0 | - |
| - | - | 812.4 | 685.3 | - | - | 0 | - |
| - | - | 4878 | 700.4 | - | - | 0 | - |
| - | - | 1592 | 701.4 | - | - | 0 | - |
| - | - | 1092 | 721.3 | - | - | 0 | - |
| 5 | y | 3243 | 723.3 | 0.003473 | 4.802 | +1 | 5 |
| - | - | 1900 | 724.3 | - | - | 0 | - |
| - | - | 2944 | 733.4 | - | - | 0 | - |
| - | - | 706.8 | 734.4 | - | - | 0 | - |
| - | - | 1372 | 735.4 | - | - | 0 | - |
| - | - | 1377 | 749.3 | - | - | 0 | - |
| - | - | 744.2 | 751.3 | - | - | 0 | - |
| - | - | 1161 | 774.3 | - | - | 0 | - |
| - | - | 1438 | 783.4 | - | - | 0 | - |
| - | - | 1780 | 801.4 | - | - | 0 | - |
| - | - | 647.1 | 802.4 | - | - | 0 | - |
| - | - | 595.1 | 817.5 | - | - | 0 | - |
| - | - | 712.6 | 829.4 | - | - | 0 | - |
| - | - | 1095 | 832.5 | - | - | 0 | - |
| 4 | y | 4030 | 838.3 | 0.003813 | 4.549 | +1 | 6 |
| - | - | 1593 | 839.3 | - | - | 0 | - |
| - | - | 4926 | 844.5 | - | - | 0 | - |
| - | - | 1753 | 845.5 | - | - | 0 | - |
| - | - | 751.4 | 846.5 | - | - | 0 | - |
| - | - | 495.3 | 872.3 | - | - | 0 | - |
| - | - | 1.3E+04 | 872.4 | - | - | 0 | - |
| - | - | 5945 | 873.4 | - | - | 0 | - |
| - | - | 2159 | 874.5 | - | - | 0 | - |
| - | - | 6005 | 889.3 | - | - | 0 | - |
| - | - | 3728 | 890.3 | - | - | 0 | - |
| - | - | 665 | 896.5 | - | - | 0 | - |
| - | - | 783 | 899.5 | - | - | 0 | - |
| - | - | 1298 | 901.5 | - | - | 0 | - |
| - | - | 1822 | 902.5 | - | - | 0 | - |
| - | - | 921.3 | 919.5 | - | - | 0 | - |
| - | - | 630.1 | 920.5 | - | - | 0 | - |
| 3 | y | 1258 | 935.3 | 0.001351 | 1.444 | +1 | 7 |
| - | - | 1070 | 936.3 | - | - | 0 | - |
| 3 | y | 1.969E+04 | 953.3 | 0.004214 | 4.42 | +1 | 7 |
| - | - | 1.123E+04 | 954.3 | - | - | 0 | - |
| - | - | 3415 | 955.3 | - | - | 0 | - |
| - | - | 631.5 | 980 | - | - | 0 | - |
| - | - | 2664 | 991.5 | - | - | 0 | - |
| - | - | 1134 | 992.5 | - | - | 0 | - |
| - | - | 5814 | 1020 | - | - | 0 | - |
| - | - | 4067 | 1021 | - | - | 0 | - |
| - | - | 1200 | 1022 | - | - | 0 | - |
| 2 | y | 1787 | 1100 | 0.00477 | 4.334 | +1 | 8 |
| - | - | 1167 | 1101 | - | - | 0 | - |
| - | - | 704.7 | 1971 | - | - | 0 | - |
| - | - | 631.8 | 2309 | - | - | 0 | - |
| - | - | 735.8 | 2672 | - | - | 0 | - |

m/z Charge Intensity FragmentType MassShift Position
120.08099365234375 0 39589.477
121.08430480957031 0 3202.7346
123.04419708251953 0 816.2878
124.45535278320312 0 390.26236
127.0868911743164 0 715.931
129.1023712158203 0 3203.4773
130.0652618408203 0 1398.081
131.69361877441406 0 405.16287
132.08108520507812 0 704.6677
133.0862274169922 0 1094.819
136.0758819580078 0 11898.046
137.0793914794922 0 1002.56506
138.06639099121094 0 622.1362
139.08682250976562 0 2488.2642
141.10260009765625 0 786.9843
142.1233367919922 0 503.26117
142.29066467285156 0 420.3191
143.11741638183594 0 563.7991
144.8822021484375 0 389.61166
146.05999755859375 0 799.87067
147.04408264160156 0 585.1457
149.04490661621094 0 1423.4816
155.08152770996094 0 510.37717
155.1181182861328 0 516.60974
156.0767059326172 0 1208.6315
157.1331787109375 0 790.77515
158.09661865234375 0 1842.3561
159.09173583984375 0 2598.5835
159.09844970703125 0 446.8792
159.1131134033203 0 625.64594
160.07603454589844 0 479.01163
160.11190795898438 0 975.02594
162.09182739257812 0 696.8139
165.05479431152344 0 3864.8987
166.08644104003906 0 7845.7397
167.05551147460938 0 2455.471
167.0816192626953 0 7280.705
167.08961486816406 0 1173.2755
168.08494567871094 0 739.6353
169.09722900390625 0 1377.3477
169.13392639160156 0 465.6422
173.09226989746094 0 618.5837
173.12864685058594 0 2560.4573
173.43858337402344 0 2195.1643
174.13233947753906 0 721.63074
175.1194610595703 0 589.6789
176.10711669921875 0 4099.433
176.36404418945312 0 453.03525
177.10235595703125 0 2566.1973
177.1105499267578 0 813.1626
180.0765380859375 0 456.40427
180.25820922851562 0 476.41113
182.081298828125 0 8871.452
183.0845489501953 0 562.65216
185.05517578125 0 506.66864
185.09228515625 0 879.0775
185.12869262695312 0 1387.1687
187.1073455810547 0 571.686
188.07095336914062 0 1833.8121
191.1175537109375 0 698.14636
193.10885620117188 0 1299.6876
199.10784912109375 0 1726.092
201.12353515625 0 1643.3772
203.08184814453125 0 514.7691
203.11817932128906 0 1613.6102 a Water loss 1
204.07676696777344 0 646.0908
205.09742736816406 0 1670.8291 y 8
214.1185302734375 0 919.4986
215.13946533203125 0 953.8323
217.09674072265625 0 922.0948
221.1033477783203 0 2794.1626
221.12857055664062 0 54150.105 a 1
222.1319580078125 0 6639.005
223.06387329101562 0 1102.1139
224.10362243652344 0 1762.9958
227.1027069091797 0 7057.8926
228.10589599609375 0 1179.6036
229.17010498046875 0 753.80865
231.06155395507812 0 987.5988
231.11227416992188 0 964.3767 b Water loss 1
235.10757446289062 0 1072.663
239.09524536132812 0 1086.5033
241.07411193847656 0 1096.5658
243.1342010498047 0 762.6852
248.74118041992188 0 524.9756
249.12362670898438 0 7825.642 b 1
250.12713623046875 0 779.41174
251.10316467285156 0 915.48724
263.102294921875 0 1990.6279
264.1338195800781 0 1886.4805
265.1543273925781 0 1190.3647
266.1485595703125 0 667.61786
267.2196960449219 0 627.51575
270.1097717285156 0 635.1034
273.1391296386719 0 593.169
274.1300048828125 0 533.1794
279.09765625 0 1550.5383
281.1242980957031 0 2967.621
285.0090026855469 0 1161.5426
285.1007995605469 0 1363.8175
286.01123046875 0 654.6323
296.15020751953125 0 750.4041
299.1330871582031 0 630.5307
302.1129455566406 0 568.1005
318.14501953125 0 641.4072 a Water loss 2
319.14068603515625 0 1099.8816
322.175048828125 0 629.55365
323.17010498046875 0 617.1353
324.1452331542969 0 1692.126
329.14996337890625 0 5516.765
330.15423583984375 0 997.0099
340.18701171875 0 5940.4844
341.190673828125 0 1365.0691
342.1563415527344 0 10343.025 y 7
342.6596984863281 0 530.7416
343.1595458984375 0 1724.9358
355.0698547363281 0 1820.3755
355.2723388671875 0 597.5262
357.2249450683594 0 671.2243
364.15106201171875 0 3150.1675 b 2
365.1815185546875 0 1040.164
366.2146301269531 0 691.01556
367.1986999511719 0 756.29895
370.20855712890625 0 879.08167
378.1308288574219 0 1191.5648
380.1927795410156 0 3560.8127
392.7001037597656 0 670.5632
393.1767578125 0 6935.697
394.1242980957031 0 1500.3292
394.1788024902344 0 1453.4689
395.1283874511719 0 599.46747
395.19287109375 0 677.1284
398.2041015625 0 2405.3093
404.1076354980469 0 773.56213
412.2177734375 0 748.05634
413.26611328125 0 1103.6189
425.1938781738281 0 1008.51904
427.7217102050781 0 1323.893
428.22235107421875 0 923.6157
434.71514892578125 0 785.5709
436.7267761230469 0 897.1806
441.2962341308594 0 678.1292
449.2397155761719 0 886.2741
450.2372741699219 0 1065.7576
451.2422790527344 0 628.839
453.2715759277344 0 733.7141
454.2697448730469 0 648.73517
457.22991943359375 0 748.5418
465.16058349609375 0 922.3005
465.7156066894531 0 867.81555
471.33428955078125 0 668.1708
478.26617431640625 0 854.6894
479.1778564453125 0 1425.0989 b 3
489.19171142578125 0 2572.9766 y 6
490.19244384765625 0 1069.586
493.2807922363281 0 597.36414
496.2298278808594 0 994.1055
496.26629638671875 0 820.14215
506.2607421875 0 7025.984
507.26397705078125 0 2025.0355
508.2771911621094 0 1317.5977
550.7080688476562 0 717.50745 y 1
551.21484375 0 762.5018
560.2294311523438 0 3912.5076 y 5
561.2343139648438 0 1100.4569
563.281982421875 0 1654.2224
564.2879028320312 0 801.7476
567.3101806640625 0 910.6804
585.3365478515625 0 2670.5862
586.3375854492188 0 848.81714
593.3628540039062 0 981.6458
601.2335205078125 0 1917.4198 Precursor
601.3309326171875 0 942.38745
601.476806640625 0 703.65027
601.7352905273438 0 822.5366
602.2427368164062 0 871.9734
602.3199462890625 0 700.71857
612.1988525390625 0 984.597
619.344970703125 0 2701.3145
620.3507080078125 0 1142.5719
621.3595581054688 0 3158.6309
622.2160034179688 0 635.89764
622.3644409179688 0 1163.6992
659.2940063476562 0 1624.9667
665.5662841796875 0 563.8264
676.3655395507812 0 2810.7168
677.3687744140625 0 738.6949
678.3768920898438 0 822.37555
685.2586059570312 0 812.4054
700.3613891601562 0 4878.281
701.3650512695312 0 1591.7015
721.2575073242188 0 1091.8413
723.2904663085938 0 3243.482 y 4
724.295166015625 0 1900.018
733.3882446289062 0 2944.1936
734.3911743164062 0 706.7733
735.3998413085938 0 1372.3008
749.2554321289062 0 1376.6039
751.2611083984375 0 744.19556
774.3178100585938 0 1161.1964
783.3969116210938 0 1437.7307
801.412353515625 0 1779.5931
802.4114379882812 0 647.143
817.517578125 0 595.05597
829.427490234375 0 712.57745
832.453369140625 0 1094.8059
838.3177490234375 0 4029.5466 y 3
839.3212890625 0 1593.3901
844.4517822265625 0 4925.826
845.4570922851562 0 1752.6122
846.457763671875 0 751.36993
872.3494873046875 0 495.26202
872.4465942382812 0 13003.826
873.44873046875 0 5944.5737
874.4501953125 0 2158.682
889.3477783203125 0 6005.263
890.3490600585938 0 3727.834
896.5249633789062 0 664.965
899.4600219726562 0 782.9886
901.47509765625 0 1298.2982
902.4791259765625 0 1821.6532
919.489501953125 0 921.3224
920.4972534179688 0 630.064
935.3316650390625 0 1258.0115 y Water loss 2
936.3350219726562 0 1069.52
953.3450927734375 0 19687.908 y 2
954.3472290039062 0 11231.653
955.34765625 0 3414.6765
979.9837646484375 0 631.47894
991.5183715820312 0 2664.1655
992.5225219726562 0 1133.6542
1019.5150756835938 0 5814.341
1020.5194091796875 0 4066.913
1021.5269165039062 0 1199.9702
1100.4140625 0 1787.4666 y 1
1101.413818359375 0 1167.4663
1971.1463623046875 0 704.6773
2308.557861328125 0 631.8063
2672.154052734375 0 735.8439

Spectrum Details

|  |  |
| --- | --- |
| Matched peaks? Matched peaks The total absolute number of peaks matched. Additionally in brackets the total fraction of peaks matched and the total number of peaks is shown. | 18 (7.53% of 239) |
| FDR? FDR The false discovery rate estimated for this peptide. It is calculated by matching all theoretical fragments with a non-integer shift with the raw peaks for this spectrum. This is done with 40 different shifts. The resulting percentage is the average number of annotated peaks over the number of annotated peaks with the correct spectrum. | 2.51% |
| Satellite FDR? Satellite FDR See the FDR for details on its calculation. This satellite ion specific FDR only contains the satellite ions (d/w) for I/L/J positions. | - |
| PSM Score? PSM Score The PSM Score as given by Hecklib to this annotated spectrum. It is shown with three significant figures. | 213 |

#### Spectrum 9421? Spectrum 9421 The raw spectrum of this peptide as annotated by Hecklib. The fragments are coloured according to ion type (see legend). Any peaks with a star '\*' as text can be hovered over to see the full details, first the ion type second the mass shift type. By hovering over the amino acids in the peptide or ions in the legend the corresponding peaks are highlighted. By toggling the 'Unassigned' label you can turn the background (unassigned) peaks on or off in the plot. By updating the slider in the Ion legend you can update the spectrum to only show the top X% of the peaks with labels. The top X% means any peak that is within X% of the highest intensity. By dragging in the spectrum you can zoom in to a specific part of the spectrum and use 'Zoom Out' to get back to the original zoom level. The annotation of the spectrum is based on the given sequence in the peptides file and is done with different software so inconsistencies are likely. The peaks are annotated based on the given sequence, with 20 ppm tolerance.

Copy Data

##### Spectrum 9421 (TSV)

###### Preview

```
Loading example...
```

*Click on the button to copy the data to your clipboard.*

Mz MinMz MaxIntensity Max

WidthHeightPeptide font sizePeptide stroke widthSpectrum font sizeSpectrum stroke widthCompact peptide

Ion legend

wxyz

abcd

OtherUnassignedIon^Charge^_Position_Show for top:%

TFDDYAMHW

01.44e+42.88e+44.31e+45.75e+4

Zoom Out

a^+1^_2_y^+1^_1_a^+1^_2_b^+1^_2_b^+2^_4_b^+1^_2_y^+1^_2_b^+1^_3_b^+1^_4_y^+1^_3_b^+1^_4_y^+2^_8_y^+1^_4_\*y^+1^_5_y^+1^_6_y^+1^_7_y^+1^_7_y^+1^_8_

0778155523333111

Fragment Matches Table

Show background peaks

| Position | Ion type | Intensity | mz Theoretical | mz Error (Th) | mz Error (ppm) | Charge | Series Number |
| --- | --- | --- | --- | --- | --- | --- | --- |
| - | - | 1068 | 120.1 | - | - | 0 | - |
| - | - | 2.047E+04 | 120.1 | - | - | 0 | - |
| - | - | 1845 | 121.1 | - | - | 0 | - |
| - | - | 546.3 | 126.1 | - | - | 0 | - |
| - | - | 719.8 | 127.1 | - | - | 0 | - |
| - | - | 545 | 129.1 | - | - | 0 | - |
| - | - | 3283 | 129.1 | - | - | 0 | - |
| - | - | 585.6 | 130.1 | - | - | 0 | - |
| - | - | 663.1 | 132.1 | - | - | 0 | - |
| - | - | 2288 | 132.1 | - | - | 0 | - |
| - | - | 505.1 | 133.1 | - | - | 0 | - |
| - | - | 1574 | 133.1 | - | - | 0 | - |
| - | - | 403.9 | 134.3 | - | - | 0 | - |
| - | - | 628.7 | 136.1 | - | - | 0 | - |
| - | - | 1.112E+04 | 136.1 | - | - | 0 | - |
| - | - | 776.7 | 137.1 | - | - | 0 | - |
| - | - | 544.4 | 138.1 | - | - | 0 | - |
| - | - | 605 | 141.1 | - | - | 0 | - |
| - | - | 795.1 | 146.1 | - | - | 0 | - |
| - | - | 631.1 | 146.1 | - | - | 0 | - |
| - | - | 512.4 | 147.1 | - | - | 0 | - |
| - | - | 3306 | 149 | - | - | 0 | - |
| - | - | 698 | 155.1 | - | - | 0 | - |
| - | - | 1812 | 156.1 | - | - | 0 | - |
| - | - | 1622 | 157.1 | - | - | 0 | - |
| - | - | 2331 | 158.1 | - | - | 0 | - |
| - | - | 1655 | 159.1 | - | - | 0 | - |
| - | - | 1012 | 160.1 | - | - | 0 | - |
| - | - | 400.5 | 160.3 | - | - | 0 | - |
| - | - | 570.1 | 165.1 | - | - | 0 | - |
| - | - | 720.6 | 166.1 | - | - | 0 | - |
| - | - | 3857 | 166.1 | - | - | 0 | - |
| - | - | 5008 | 167.1 | - | - | 0 | - |
| - | - | 1102 | 169.1 | - | - | 0 | - |
| - | - | 1010 | 169.1 | - | - | 0 | - |
| - | - | 982.4 | 171.1 | - | - | 0 | - |
| - | - | 1203 | 173.1 | - | - | 0 | - |
| - | - | 3715 | 173.4 | - | - | 0 | - |
| - | - | 4002 | 176.1 | - | - | 0 | - |
| - | - | 3160 | 177.1 | - | - | 0 | - |
| - | - | 738.8 | 177.1 | - | - | 0 | - |
| - | - | 598.4 | 181.1 | - | - | 0 | - |
| - | - | 1278 | 182.1 | - | - | 0 | - |
| - | - | 3112 | 183.1 | - | - | 0 | - |
| - | - | 610.2 | 183.1 | - | - | 0 | - |
| - | - | 499.9 | 185.1 | - | - | 0 | - |
| - | - | 1674 | 185.1 | - | - | 0 | - |
| - | - | 528.2 | 185.2 | - | - | 0 | - |
| - | - | 653.7 | 187.1 | - | - | 0 | - |
| - | - | 3078 | 188.1 | - | - | 0 | - |
| - | - | 611.5 | 191.1 | - | - | 0 | - |
| - | - | 785.1 | 195.1 | - | - | 0 | - |
| - | - | 667.4 | 195.1 | - | - | 0 | - |
| - | - | 992.3 | 197.1 | - | - | 0 | - |
| - | - | 523.7 | 200.1 | - | - | 0 | - |
| - | - | 1185 | 201.1 | - | - | 0 | - |
| - | - | 979.4 | 203.1 | - | - | 0 | - |
| 2 | a | 1119 | 203.1 | 0.0003203 | 1.577 | +1 | 2 |
| 9 | y | 1700 | 205.1 | 0.0003191 | 1.556 | +1 | 1 |
| - | - | 1188 | 207.1 | - | - | 0 | - |
| - | - | 822.4 | 209.1 | - | - | 0 | - |
| - | - | 757.6 | 214.1 | - | - | 0 | - |
| - | - | 611.8 | 217.1 | - | - | 0 | - |
| - | - | 1120 | 219.1 | - | - | 0 | - |
| - | - | 3732 | 221.1 | - | - | 0 | - |
| 2 | a | 5.128E+04 | 221.1 | 0.0001469 | 0.6641 | +1 | 2 |
| - | - | 4931 | 222.1 | - | - | 0 | - |
| - | - | 625.9 | 223.1 | - | - | 0 | - |
| - | - | 547.9 | 224.8 | - | - | 0 | - |
| - | - | 1745 | 225 | - | - | 0 | - |
| - | - | 707.5 | 226.1 | - | - | 0 | - |
| - | - | 1210 | 226.2 | - | - | 0 | - |
| - | - | 612.3 | 227.1 | - | - | 0 | - |
| - | - | 550.4 | 229.1 | - | - | 0 | - |
| - | - | 1141 | 231.1 | - | - | 0 | - |
| 2 | b | 610.7 | 231.1 | 0.000828 | 3.583 | +1 | 2 |
| - | - | 1342 | 233.2 | - | - | 0 | - |
| - | - | 689.6 | 233.2 | - | - | 0 | - |
| - | - | 2523 | 235.1 | - | - | 0 | - |
| - | - | 1.172E+04 | 239.1 | - | - | 0 | - |
| 4 | b | 2038 | 240.1 | 0.004094 | 17.05 | +2 | 4 |
| - | - | 613.9 | 241.1 | - | - | 0 | - |
| 2 | b | 8685 | 249.1 | 0.0002579 | 1.035 | +1 | 2 |
| - | - | 1030 | 250.1 | - | - | 0 | - |
| - | - | 1354 | 251.1 | - | - | 0 | - |
| - | - | 1165 | 252.1 | - | - | 0 | - |
| - | - | 1883 | 253.1 | - | - | 0 | - |
| - | - | 2135 | 263.1 | - | - | 0 | - |
| - | - | 973.4 | 267 | - | - | 0 | - |
| - | - | 2045 | 269 | - | - | 0 | - |
| - | - | 3876 | 269.1 | - | - | 0 | - |
| - | - | 1026 | 279.1 | - | - | 0 | - |
| - | - | 1674 | 281.1 | - | - | 0 | - |
| - | - | 2.223E+04 | 285 | - | - | 0 | - |
| - | - | 3219 | 286 | - | - | 0 | - |
| - | - | 581.6 | 296.2 | - | - | 0 | - |
| - | - | 1.439E+04 | 299.1 | - | - | 0 | - |
| - | - | 2687 | 300.1 | - | - | 0 | - |
| - | - | 4234 | 303 | - | - | 0 | - |
| - | - | 653.2 | 304 | - | - | 0 | - |
| - | - | 674 | 322.1 | - | - | 0 | - |
| - | - | 2607 | 324.1 | - | - | 0 | - |
| - | - | 1326 | 340.1 | - | - | 0 | - |
| 8 | y | 1.335E+04 | 342.2 | 0.0003061 | 0.8947 | +1 | 2 |
| - | - | 1956 | 343.2 | - | - | 0 | - |
| - | - | 624.3 | 344.2 | - | - | 0 | - |
| - | - | 3013 | 345 | - | - | 0 | - |
| - | - | 622.7 | 346 | - | - | 0 | - |
| - | - | 1372 | 350.1 | - | - | 0 | - |
| - | - | 5.695E+04 | 355.1 | - | - | 0 | - |
| - | - | 1.657E+04 | 356.1 | - | - | 0 | - |
| 3 | b | 1670 | 364.2 | 0.0007501 | 2.06 | +1 | 3 |
| - | - | 987.1 | 366.1 | - | - | 0 | - |
| - | - | 1337 | 371.1 | - | - | 0 | - |
| - | - | 5591 | 373.1 | - | - | 0 | - |
| - | - | 723.5 | 374.1 | - | - | 0 | - |
| - | - | 1421 | 378.1 | - | - | 0 | - |
| - | - | 832.1 | 389.2 | - | - | 0 | - |
| - | - | 571.8 | 392.2 | - | - | 0 | - |
| - | - | 2772 | 394.1 | - | - | 0 | - |
| - | - | 1013 | 401 | - | - | 0 | - |
| - | - | 1897 | 415 | - | - | 0 | - |
| - | - | 715.7 | 415.2 | - | - | 0 | - |
| - | - | 1049 | 416 | - | - | 0 | - |
| - | - | 603.8 | 418.2 | - | - | 0 | - |
| - | - | 5294 | 419 | - | - | 0 | - |
| - | - | 1625 | 420 | - | - | 0 | - |
| - | - | 948 | 433.2 | - | - | 0 | - |
| - | - | 836.2 | 448.8 | - | - | 0 | - |
| 4 | b | 1028 | 461.2 | 0.008084 | 17.53 | +1 | 4 |
| - | - | 2700 | 462.3 | - | - | 0 | - |
| - | - | 1431 | 462.8 | - | - | 0 | - |
| - | - | 2107 | 465.2 | - | - | 0 | - |
| - | - | 613.2 | 465.7 | - | - | 0 | - |
| - | - | 759.4 | 466.2 | - | - | 0 | - |
| 7 | y | 2693 | 473.2 | 0.0001352 | 0.2858 | +1 | 3 |
| 4 | b | 942.9 | 479.2 | 0.002066 | 4.312 | +1 | 4 |
| - | - | 705.8 | 480.2 | - | - | 0 | - |
| - | - | 845.5 | 534.3 | - | - | 0 | - |
| 2 | y | 766 | 542.7 | 0.002103 | 3.875 | +2 | 8 |
| - | - | 814.1 | 543.2 | - | - | 0 | - |
| 6 | y | 2911 | 544.2 | 0.0002834 | 0.5207 | +1 | 4 |
| - | - | 763.4 | 545.2 | - | - | 0 | - |
| - | - | 559.2 | 575.4 | - | - | 0 | - |
| - | - | 553.4 | 582.4 | - | - | 0 | - |
| - | - | 594.1 | 584.3 | - | - | 0 | - |
| 0 | Precursor | 871.3 | 593.2 | 0.0008568 | 1.444 | +2 | -1 |
| - | - | 960.4 | 593.3 | - | - | 0 | - |
| - | - | 977.6 | 593.5 | - | - | 0 | - |
| - | - | 836.2 | 597.3 | - | - | 0 | - |
| - | - | 956.4 | 618.2 | - | - | 0 | - |
| - | - | 884.9 | 699.4 | - | - | 0 | - |
| - | - | 1337 | 705.3 | - | - | 0 | - |
| 5 | y | 3874 | 707.3 | 0.0008503 | 1.202 | +1 | 5 |
| - | - | 1104 | 708.3 | - | - | 0 | - |
| - | - | 2787 | 733.3 | - | - | 0 | - |
| - | - | 1449 | 734.3 | - | - | 0 | - |
| - | - | 914.7 | 770.4 | - | - | 0 | - |
| - | - | 605.4 | 811.9 | - | - | 0 | - |
| 4 | y | 5173 | 822.3 | 0.0006937 | 0.8436 | +1 | 6 |
| - | - | 2398 | 823.3 | - | - | 0 | - |
| - | - | 832.4 | 824.3 | - | - | 0 | - |
| - | - | 840.1 | 854.4 | - | - | 0 | - |
| 3 | y | 1358 | 919.3 | 0.002241 | 2.437 | +1 | 7 |
| 3 | y | 2.095E+04 | 937.4 | 0.0009034 | 0.9637 | +1 | 7 |
| - | - | 1.062E+04 | 938.4 | - | - | 0 | - |
| - | - | 4647 | 939.4 | - | - | 0 | - |
| - | - | 2081 | 956.4 | - | - | 0 | - |
| - | - | 831.8 | 957.4 | - | - | 0 | - |
| 2 | y | 1508 | 1084 | 0.003559 | 3.282 | +1 | 8 |
| - | - | 688.3 | 1085 | - | - | 0 | - |
| - | - | 663 | 1501 | - | - | 0 | - |
| - | - | 667.7 | 2426 | - | - | 0 | - |
| - | - | 874.6 | 3080 | - | - | 0 | - |

m/z Charge Intensity FragmentType MassShift Position
120.07634735107422 0 1068.0015
120.08101654052734 0 20467.873
121.08434295654297 0 1844.778
126.05583190917969 0 546.34
127.08704376220703 0 719.7735
129.0661163330078 0 544.9546
129.10243225097656 0 3283.252
130.0654754638672 0 585.5993
132.0811309814453 0 663.09344
132.10211181640625 0 2288.365
133.06134033203125 0 505.1261
133.0861358642578 0 1573.5586
134.2555694580078 0 403.90717
136.07015991210938 0 628.6605
136.07591247558594 0 11120.9795
137.07913208007812 0 776.73224
138.06622314453125 0 544.4114
141.1027069091797 0 604.979
146.0604248046875 0 795.0725
146.096923828125 0 631.1043
147.11326599121094 0 512.3726
149.04505920410156 0 3306.1
155.08189392089844 0 697.9976
156.07708740234375 0 1811.8634
157.13381958007812 0 1622.1971
158.0966033935547 0 2331.078
159.0917205810547 0 1654.9299
160.11248779296875 0 1012.0032
160.32533264160156 0 400.4806
165.05535888671875 0 570.0867
166.06150817871094 0 720.56616
166.08642578125 0 3857.3853
167.05569458007812 0 5007.97
169.0975799560547 0 1101.7135
169.13368225097656 0 1010.40045
171.07652282714844 0 982.42053
173.1288299560547 0 1202.7262
173.44044494628906 0 3714.7488
176.10716247558594 0 4002.342
177.1026153564453 0 3160.2668
177.11106872558594 0 738.75433
181.09713745117188 0 598.40515
182.08131408691406 0 1278.3772
183.0554656982422 0 3112.1587
183.1128387451172 0 610.16003
185.09202575683594 0 499.92056
185.12867736816406 0 1673.7968
185.1654510498047 0 528.181
187.144287109375 0 653.67755
188.07077026367188 0 3078.1033
191.11764526367188 0 611.4872
195.1130828857422 0 785.1091
195.12295532226562 0 667.3828
197.12875366210938 0 992.2731
200.09703063964844 0 523.74396
201.12364196777344 0 1185.3856
203.06649780273438 0 979.4408
203.1182098388672 0 1118.799 a Water loss 1
205.09747314453125 0 1699.5743 y 8
207.1125946044922 0 1187.9127
209.09242248535156 0 822.3676
214.11868286132812 0 757.5881
217.0975341796875 0 611.807
219.13430786132812 0 1119.9531
221.08448791503906 0 3732.1997
221.12860107421875 0 51282.69 a 1
222.13189697265625 0 4930.705
223.10826110839844 0 625.8684
224.82452392578125 0 547.9247
225.04327392578125 0 1745.1577
226.11917114257812 0 707.506
226.1551055908203 0 1210.381
227.10382080078125 0 612.30334
229.11805725097656 0 550.4112
231.0614471435547 0 1141.4889
231.11363220214844 0 610.7066 b Water loss 1
233.15008544921875 0 1341.6008
233.16456604003906 0 689.6431
235.1079559326172 0 2522.6572
239.0950469970703 0 11722.72
240.0963592529297 0 2038.0388 b 3
241.1117706298828 0 613.901
249.12362670898438 0 8684.952 b 1
250.12637329101562 0 1030.2687
251.10267639160156 0 1354.1565
252.080322265625 0 1164.8662
253.118408203125 0 1882.6492
263.1028137207031 0 2134.532
266.99932861328125 0 973.3929
268.9787292480469 0 2044.5508
269.10693359375 0 3876.0276
279.0981140136719 0 1026.127
281.0509338378906 0 1674.0122
285.0098571777344 0 22232.549
286.01007080078125 0 3218.858
296.1504211425781 0 581.60614
299.0618896484375 0 14385.087
300.06268310546875 0 2686.6372
303.02069091796875 0 4233.6426
304.02093505859375 0 653.2386
322.1339416503906 0 673.9737
324.1454772949219 0 2607.4573
340.1432800292969 0 1326.0243
342.1563720703125 0 13345.458 y 7
343.1606140136719 0 1956.338
344.1593322753906 0 624.321
344.97601318359375 0 3013.4631
345.9772644042969 0 622.7317
350.13446044921875 0 1372.2391
355.0699157714844 0 56947.36
356.0706787109375 0 16565.832
364.15106201171875 0 1669.8506 b 2
366.1290588378906 0 987.11304
371.1003723144531 0 1337.0273
373.08001708984375 0 5590.7793
374.07989501953125 0 723.5139
378.1297302246094 0 1421.2505
389.1585693359375 0 832.1231
392.2299499511719 0 571.8183
394.1242980957031 0 2772.1309
400.9844970703125 0 1013.0893
415.036865234375 0 1897.0011
415.1727600097656 0 715.7344
416.0367126464844 0 1048.7952
418.2315368652344 0 603.8492
418.99542236328125 0 5294.0347
419.9951171875 0 1625.4302
433.1821594238281 0 947.95764
448.7678527832031 0 836.1553
461.1747741699219 0 1028.2577 b Water loss 3
462.2633361816406 0 2700.2124
462.7642517089844 0 1431.2263
465.16253662109375 0 2107.4568
465.670654296875 0 613.2005
466.1605529785156 0 759.38086
473.1966857910156 0 2692.8118 y 6
479.1793212890625 0 942.89075 b 3
480.1824035644531 0 705.8425
534.3054809570312 0 845.5418
542.711181640625 0 766.018 y 1
543.214111328125 0 814.10596
544.2339477539062 0 2911.2222 y 5
545.23583984375 0 763.3729
575.3623657226562 0 559.1807
582.3525390625 0 553.37585
584.32763671875 0 594.0703
593.2362670898438 0 871.26874 Precursor
593.341552734375 0 960.35144
593.5416870117188 0 977.56305
597.3163452148438 0 836.15436
618.2362670898438 0 956.3776
699.4015502929688 0 884.94574
705.2681884765625 0 1337.1074
707.296142578125 0 3873.846 y 4
708.298828125 0 1103.9762
733.2610473632812 0 2786.7786
734.2619018554688 0 1449.1316
770.3516235351562 0 914.69617
811.8689575195312 0 605.3644
822.3232421875 0 5172.754 y 3
823.3270263671875 0 2397.596
824.3250122070312 0 832.4321
854.4390258789062 0 840.05365
919.3380737304688 0 1357.8976 y Water loss 2
937.3499755859375 0 20954.123 y 2
938.3532104492188 0 10617.96
939.3546142578125 0 4647.3003
956.4424438476562 0 2080.5388
957.4497680664062 0 831.77985
1084.4228515625 0 1508.398 y 1
1085.423095703125 0 688.3241
1501.28125 0 663.01996
2426.256103515625 0 667.696
3079.867919921875 0 874.6193

Spectrum Details

|  |  |
| --- | --- |
| Matched peaks? Matched peaks The total absolute number of peaks matched. Additionally in brackets the total fraction of peaks matched and the total number of peaks is shown. | 19 (10.92% of 174) |
| FDR? FDR The false discovery rate estimated for this peptide. It is calculated by matching all theoretical fragments with a non-integer shift with the raw peaks for this spectrum. This is done with 40 different shifts. The resulting percentage is the average number of annotated peaks over the number of annotated peaks with the correct spectrum. | 0.75% |
| Satellite FDR? Satellite FDR See the FDR for details on its calculation. This satellite ion specific FDR only contains the satellite ions (d/w) for I/L/J positions. | - |
| PSM Score? PSM Score The PSM Score as given by Hecklib to this annotated spectrum. It is shown with three significant figures. | 213 |

#### Spectrum 9404? Spectrum 9404 The raw spectrum of this peptide as annotated by Hecklib. The fragments are coloured according to ion type (see legend). Any peaks with a star '\*' as text can be hovered over to see the full details, first the ion type second the mass shift type. By hovering over the amino acids in the peptide or ions in the legend the corresponding peaks are highlighted. By toggling the 'Unassigned' label you can turn the background (unassigned) peaks on or off in the plot. By updating the slider in the Ion legend you can update the spectrum to only show the top X% of the peaks with labels. The top X% means any peak that is within X% of the highest intensity. By dragging in the spectrum you can zoom in to a specific part of the spectrum and use 'Zoom Out' to get back to the original zoom level. The annotation of the spectrum is based on the given sequence in the peptides file and is done with different software so inconsistencies are likely. The peaks are annotated based on the given sequence, with 20 ppm tolerance.

Copy Data

##### Spectrum 9404 (TSV)

###### Preview

```
Loading example...
```

*Click on the button to copy the data to your clipboard.*

Mz MinMz MaxIntensity Max

WidthHeightPeptide font sizePeptide stroke widthSpectrum font sizeSpectrum stroke widthCompact peptide

Ion legend

wxyz

abcd

OtherUnassignedIon^Charge^_Position_Show for top:%

TFDDYAMHW

03.78e+47.56e+41.13e+51.51e+5

Zoom Out

a^+1^_2_y^+1^_1_a^+1^_2_b^+1^_2_b^+1^_2_a^+1^_3_y^+1^_2_b^+1^_3_b^+1^_3_b^+1^_4_b^+1^_4_y^+1^_3_y^+2^_8_y^+1^_4_\*\*b^+1^_5_y^+1^_5_y^+1^_6_y^+1^_7_y^+1^_7_y^+1^_8_

035170210531404

Fragment Matches Table

Show background peaks

| Position | Ion type | Intensity | mz Theoretical | mz Error (Th) | mz Error (ppm) | Charge | Series Number |
| --- | --- | --- | --- | --- | --- | --- | --- |
| - | - | 1.497E+05 | 120.1 | - | - | 0 | - |
| - | - | 991.9 | 121.1 | - | - | 0 | - |
| - | - | 1.513E+04 | 121.1 | - | - | 0 | - |
| - | - | 546 | 122.1 | - | - | 0 | - |
| - | - | 2580 | 123 | - | - | 0 | - |
| - | - | 371.6 | 125.5 | - | - | 0 | - |
| - | - | 710.8 | 127.1 | - | - | 0 | - |
| - | - | 827.7 | 129.1 | - | - | 0 | - |
| - | - | 8568 | 129.1 | - | - | 0 | - |
| - | - | 498.3 | 130.1 | - | - | 0 | - |
| - | - | 2634 | 130.1 | - | - | 0 | - |
| - | - | 974.3 | 130.1 | - | - | 0 | - |
| - | - | 1824 | 132.1 | - | - | 0 | - |
| - | - | 1558 | 132.1 | - | - | 0 | - |
| - | - | 1694 | 133.1 | - | - | 0 | - |
| - | - | 487.4 | 134 | - | - | 0 | - |
| - | - | 4.733E+04 | 136.1 | - | - | 0 | - |
| - | - | 4405 | 137.1 | - | - | 0 | - |
| - | - | 952 | 138.1 | - | - | 0 | - |
| - | - | 1438 | 138.1 | - | - | 0 | - |
| - | - | 1512 | 141.1 | - | - | 0 | - |
| - | - | 8203 | 143.1 | - | - | 0 | - |
| - | - | 627.6 | 144.1 | - | - | 0 | - |
| - | - | 455.9 | 145.1 | - | - | 0 | - |
| - | - | 1469 | 146.1 | - | - | 0 | - |
| - | - | 1771 | 147 | - | - | 0 | - |
| - | - | 460.6 | 147.8 | - | - | 0 | - |
| - | - | 1019 | 148.9 | - | - | 0 | - |
| - | - | 600.6 | 149 | - | - | 0 | - |
| - | - | 934.1 | 155.1 | - | - | 0 | - |
| - | - | 5872 | 156.1 | - | - | 0 | - |
| - | - | 555 | 157.1 | - | - | 0 | - |
| - | - | 2497 | 157.1 | - | - | 0 | - |
| - | - | 1064 | 158.1 | - | - | 0 | - |
| - | - | 5171 | 158.1 | - | - | 0 | - |
| - | - | 6037 | 159.1 | - | - | 0 | - |
| - | - | 1236 | 160.1 | - | - | 0 | - |
| - | - | 646.8 | 160.1 | - | - | 0 | - |
| - | - | 1732 | 160.1 | - | - | 0 | - |
| - | - | 583.4 | 162.1 | - | - | 0 | - |
| - | - | 2.118E+04 | 165.1 | - | - | 0 | - |
| - | - | 1644 | 166.1 | - | - | 0 | - |
| - | - | 3952 | 166.1 | - | - | 0 | - |
| - | - | 1045 | 167.1 | - | - | 0 | - |
| - | - | 668.7 | 167.1 | - | - | 0 | - |
| - | - | 438.4 | 168.5 | - | - | 0 | - |
| - | - | 416.3 | 168.8 | - | - | 0 | - |
| - | - | 863.5 | 169.1 | - | - | 0 | - |
| - | - | 1737 | 169.1 | - | - | 0 | - |
| - | - | 418.6 | 170.9 | - | - | 0 | - |
| - | - | 1202 | 171.1 | - | - | 0 | - |
| - | - | 6202 | 171.1 | - | - | 0 | - |
| - | - | 1168 | 173.1 | - | - | 0 | - |
| - | - | 650.6 | 173.1 | - | - | 0 | - |
| - | - | 1.051E+04 | 173.1 | - | - | 0 | - |
| - | - | 1336 | 174.1 | - | - | 0 | - |
| - | - | 3732 | 175.1 | - | - | 0 | - |
| - | - | 972.7 | 176.1 | - | - | 0 | - |
| - | - | 7975 | 176.1 | - | - | 0 | - |
| - | - | 6707 | 177.1 | - | - | 0 | - |
| - | - | 1350 | 177.1 | - | - | 0 | - |
| - | - | 532 | 180.1 | - | - | 0 | - |
| - | - | 546.6 | 181.1 | - | - | 0 | - |
| - | - | 4.19E+04 | 182.1 | - | - | 0 | - |
| - | - | 4102 | 183.1 | - | - | 0 | - |
| - | - | 925.8 | 183.1 | - | - | 0 | - |
| - | - | 607 | 183.1 | - | - | 0 | - |
| - | - | 1712 | 185.1 | - | - | 0 | - |
| - | - | 1878 | 185.1 | - | - | 0 | - |
| - | - | 879.2 | 186.1 | - | - | 0 | - |
| - | - | 2445 | 186.1 | - | - | 0 | - |
| - | - | 593.9 | 187.1 | - | - | 0 | - |
| - | - | 598.1 | 187.1 | - | - | 0 | - |
| - | - | 6572 | 188.1 | - | - | 0 | - |
| - | - | 1077 | 189.1 | - | - | 0 | - |
| - | - | 3540 | 191.1 | - | - | 0 | - |
| - | - | 3586 | 193.1 | - | - | 0 | - |
| - | - | 468.6 | 194.3 | - | - | 0 | - |
| - | - | 493.5 | 195.1 | - | - | 0 | - |
| - | - | 516.8 | 195.1 | - | - | 0 | - |
| - | - | 1224 | 197.1 | - | - | 0 | - |
| - | - | 1245 | 199.1 | - | - | 0 | - |
| - | - | 6323 | 199.1 | - | - | 0 | - |
| - | - | 940.1 | 199.1 | - | - | 0 | - |
| - | - | 1048 | 200.1 | - | - | 0 | - |
| - | - | 562.9 | 200.3 | - | - | 0 | - |
| - | - | 6910 | 201.1 | - | - | 0 | - |
| - | - | 1075 | 202.1 | - | - | 0 | - |
| - | - | 1737 | 203.1 | - | - | 0 | - |
| 2 | a | 3108 | 203.1 | 0.0004271 | 2.103 | +1 | 2 |
| - | - | 1506 | 204.1 | - | - | 0 | - |
| 9 | y | 4894 | 205.1 | 0.000197 | 0.9604 | +1 | 1 |
| - | - | 467.2 | 207 | - | - | 0 | - |
| - | - | 1204 | 207.1 | - | - | 0 | - |
| - | - | 690.7 | 210.2 | - | - | 0 | - |
| - | - | 1540 | 214.1 | - | - | 0 | - |
| - | - | 663.8 | 215.1 | - | - | 0 | - |
| - | - | 648.3 | 216.1 | - | - | 0 | - |
| - | - | 1489 | 217.1 | - | - | 0 | - |
| - | - | 1096 | 217.1 | - | - | 0 | - |
| - | - | 825 | 217.1 | - | - | 0 | - |
| - | - | 1.532E+04 | 217.1 | - | - | 0 | - |
| - | - | 548.3 | 219.1 | - | - | 0 | - |
| - | - | 1972 | 219.1 | - | - | 0 | - |
| - | - | 2401 | 221.1 | - | - | 0 | - |
| - | - | 7393 | 221.1 | - | - | 0 | - |
| 2 | a | 1.343E+05 | 221.1 | 0.000391 | 1.768 | +1 | 2 |
| - | - | 1.696E+04 | 222.1 | - | - | 0 | - |
| - | - | 1379 | 223.1 | - | - | 0 | - |
| - | - | 761.4 | 223.1 | - | - | 0 | - |
| - | - | 1048 | 223.1 | - | - | 0 | - |
| - | - | 2609 | 225.2 | - | - | 0 | - |
| - | - | 1571 | 226.1 | - | - | 0 | - |
| - | - | 2780 | 226.2 | - | - | 0 | - |
| - | - | 705 | 227.1 | - | - | 0 | - |
| - | - | 806 | 227.1 | - | - | 0 | - |
| - | - | 1.044E+04 | 227.1 | - | - | 0 | - |
| - | - | 652.6 | 227.2 | - | - | 0 | - |
| - | - | 721.1 | 228.1 | - | - | 0 | - |
| - | - | 2498 | 229.1 | - | - | 0 | - |
| - | - | 1106 | 229.2 | - | - | 0 | - |
| - | - | 741.4 | 229.2 | - | - | 0 | - |
| - | - | 3659 | 231.1 | - | - | 0 | - |
| - | - | 582.7 | 231.1 | - | - | 0 | - |
| 2 | b | 1019 | 231.1 | 4.985E-05 | 0.2157 | +1 | 2 |
| - | - | 515.9 | 232.1 | - | - | 0 | - |
| - | - | 637.2 | 233.1 | - | - | 0 | - |
| - | - | 801.1 | 233.1 | - | - | 0 | - |
| - | - | 545.4 | 233.2 | - | - | 0 | - |
| - | - | 713 | 233.2 | - | - | 0 | - |
| - | - | 4746 | 235.1 | - | - | 0 | - |
| - | - | 589.5 | 235.1 | - | - | 0 | - |
| - | - | 675.9 | 239.1 | - | - | 0 | - |
| - | - | 905.3 | 239.1 | - | - | 0 | - |
| - | - | 1267 | 239.1 | - | - | 0 | - |
| - | - | 1297 | 240.1 | - | - | 0 | - |
| - | - | 1487 | 243.1 | - | - | 0 | - |
| - | - | 1330 | 244.1 | - | - | 0 | - |
| - | - | 7207 | 245.1 | - | - | 0 | - |
| - | - | 919.1 | 246.1 | - | - | 0 | - |
| - | - | 852.9 | 247.1 | - | - | 0 | - |
| - | - | 1017 | 248.1 | - | - | 0 | - |
| - | - | 1141 | 249.1 | - | - | 0 | - |
| 2 | b | 2.35E+04 | 249.1 | 0.0003952 | 1.586 | +1 | 2 |
| - | - | 3127 | 250.1 | - | - | 0 | - |
| - | - | 2496 | 251.1 | - | - | 0 | - |
| - | - | 2638 | 253.1 | - | - | 0 | - |
| - | - | 3348 | 253.2 | - | - | 0 | - |
| - | - | 778.7 | 254.1 | - | - | 0 | - |
| - | - | 556.6 | 254.6 | - | - | 0 | - |
| - | - | 1018 | 256.1 | - | - | 0 | - |
| - | - | 888.7 | 257.1 | - | - | 0 | - |
| - | - | 513.7 | 258.1 | - | - | 0 | - |
| - | - | 4829 | 263.1 | - | - | 0 | - |
| - | - | 899.1 | 264.1 | - | - | 0 | - |
| - | - | 792.2 | 271.1 | - | - | 0 | - |
| - | - | 4376 | 273.1 | - | - | 0 | - |
| - | - | 1156 | 274.1 | - | - | 0 | - |
| - | - | 2225 | 279.1 | - | - | 0 | - |
| - | - | 708.1 | 280.1 | - | - | 0 | - |
| - | - | 1471 | 283.1 | - | - | 0 | - |
| - | - | 875.4 | 283.2 | - | - | 0 | - |
| - | - | 673.9 | 284.1 | - | - | 0 | - |
| - | - | 656.3 | 285 | - | - | 0 | - |
| - | - | 2250 | 285.1 | - | - | 0 | - |
| - | - | 543.1 | 288.1 | - | - | 0 | - |
| - | - | 977.9 | 292.1 | - | - | 0 | - |
| - | - | 2466 | 296.2 | - | - | 0 | - |
| - | - | 651 | 296.2 | - | - | 0 | - |
| - | - | 833.6 | 297.1 | - | - | 0 | - |
| - | - | 890.6 | 297.2 | - | - | 0 | - |
| - | - | 2054 | 301.1 | - | - | 0 | - |
| - | - | 1674 | 302.1 | - | - | 0 | - |
| - | - | 606.1 | 303.1 | - | - | 0 | - |
| - | - | 963.9 | 304.1 | - | - | 0 | - |
| - | - | 566.4 | 315.2 | - | - | 0 | - |
| 3 | a | 1060 | 318.1 | 0.0006752 | 2.122 | +1 | 3 |
| - | - | 2633 | 319.1 | - | - | 0 | - |
| - | - | 1363 | 323.1 | - | - | 0 | - |
| - | - | 1589 | 323.2 | - | - | 0 | - |
| - | - | 4801 | 324.1 | - | - | 0 | - |
| - | - | 891.4 | 325.1 | - | - | 0 | - |
| - | - | 833.5 | 325.2 | - | - | 0 | - |
| - | - | 4366 | 326.2 | - | - | 0 | - |
| - | - | 1094 | 327.2 | - | - | 0 | - |
| - | - | 867.8 | 327.2 | - | - | 0 | - |
| - | - | 2.828E+04 | 329.2 | - | - | 0 | - |
| - | - | 636.4 | 329.2 | - | - | 0 | - |
| - | - | 5332 | 330.2 | - | - | 0 | - |
| - | - | 689.8 | 339.2 | - | - | 0 | - |
| - | - | 5243 | 340.2 | - | - | 0 | - |
| - | - | 819.9 | 341.2 | - | - | 0 | - |
| 8 | y | 2.724E+04 | 342.2 | 0.0005808 | 1.697 | +1 | 2 |
| - | - | 5183 | 343.2 | - | - | 0 | - |
| - | - | 1205 | 344.1 | - | - | 0 | - |
| 3 | b | 1550 | 346.1 | 0.0003895 | 1.125 | +1 | 3 |
| - | - | 1050 | 348.1 | - | - | 0 | - |
| - | - | 1640 | 350.1 | - | - | 0 | - |
| - | - | 1146 | 355.1 | - | - | 0 | - |
| - | - | 619.9 | 355.2 | - | - | 0 | - |
| - | - | 1066 | 356.1 | - | - | 0 | - |
| - | - | 7169 | 357.2 | - | - | 0 | - |
| - | - | 800.6 | 358.2 | - | - | 0 | - |
| 3 | b | 5811 | 364.2 | 0.0002619 | 0.7191 | +1 | 3 |
| - | - | 1033 | 365.2 | - | - | 0 | - |
| - | - | 1049 | 366.1 | - | - | 0 | - |
| - | - | 2143 | 368.2 | - | - | 0 | - |
| - | - | 613.2 | 369.2 | - | - | 0 | - |
| - | - | 668.1 | 371.7 | - | - | 0 | - |
| - | - | 694.5 | 374.2 | - | - | 0 | - |
| - | - | 747.4 | 375.2 | - | - | 0 | - |
| - | - | 575 | 375.2 | - | - | 0 | - |
| - | - | 752.5 | 376.1 | - | - | 0 | - |
| - | - | 3427 | 378.1 | - | - | 0 | - |
| - | - | 937 | 382.1 | - | - | 0 | - |
| - | - | 714.1 | 383.2 | - | - | 0 | - |
| - | - | 1050 | 385.2 | - | - | 0 | - |
| - | - | 1486 | 392.2 | - | - | 0 | - |
| - | - | 7239 | 394.1 | - | - | 0 | - |
| - | - | 923 | 395.1 | - | - | 0 | - |
| - | - | 759.8 | 398.2 | - | - | 0 | - |
| - | - | 939 | 400.2 | - | - | 0 | - |
| - | - | 1929 | 401.2 | - | - | 0 | - |
| - | - | 1142 | 401.7 | - | - | 0 | - |
| - | - | 730.3 | 405.7 | - | - | 0 | - |
| - | - | 564.5 | 414.7 | - | - | 0 | - |
| - | - | 864.6 | 415 | - | - | 0 | - |
| - | - | 783.5 | 418.2 | - | - | 0 | - |
| - | - | 987.1 | 422.7 | - | - | 0 | - |
| - | - | 3522 | 425.2 | - | - | 0 | - |
| - | - | 3807 | 427.7 | - | - | 0 | - |
| - | - | 2608 | 428.2 | - | - | 0 | - |
| - | - | 836.6 | 428.7 | - | - | 0 | - |
| - | - | 3206 | 436.7 | - | - | 0 | - |
| - | - | 1659 | 437.2 | - | - | 0 | - |
| - | - | 877 | 442.3 | - | - | 0 | - |
| - | - | 1460 | 443.2 | - | - | 0 | - |
| - | - | 760.1 | 444.2 | - | - | 0 | - |
| - | - | 704.3 | 445.2 | - | - | 0 | - |
| - | - | 1012 | 445.2 | - | - | 0 | - |
| - | - | 1469 | 448.3 | - | - | 0 | - |
| - | - | 1190 | 448.8 | - | - | 0 | - |
| - | - | 1325 | 453.3 | - | - | 0 | - |
| 4 | b | 900.8 | 461.2 | 0.00101 | 2.191 | +1 | 4 |
| - | - | 712.9 | 461.2 | - | - | 0 | - |
| - | - | 760.3 | 462.2 | - | - | 0 | - |
| - | - | 3124 | 465.2 | - | - | 0 | - |
| - | - | 990.2 | 466.2 | - | - | 0 | - |
| - | - | 1305 | 470.3 | - | - | 0 | - |
| - | - | 875.7 | 473.2 | - | - | 0 | - |
| 4 | b | 3337 | 479.2 | 0.0005405 | 1.128 | +1 | 4 |
| - | - | 803.3 | 480.2 | - | - | 0 | - |
| - | - | 706.8 | 482.2 | - | - | 0 | - |
| - | - | 2707 | 483.2 | - | - | 0 | - |
| - | - | 1838 | 487.3 | - | - | 0 | - |
| 7 | y | 6126 | 489.2 | 0.005466 | 11.17 | +1 | 3 |
| - | - | 1469 | 490.2 | - | - | 0 | - |
| - | - | 760.1 | 491.2 | - | - | 0 | - |
| - | - | 2399 | 496.2 | - | - | 0 | - |
| - | - | 997.5 | 496.3 | - | - | 0 | - |
| - | - | 788.1 | 496.8 | - | - | 0 | - |
| - | - | 675.6 | 498.8 | - | - | 0 | - |
| - | - | 1152 | 500.2 | - | - | 0 | - |
| - | - | 1911 | 501.3 | - | - | 0 | - |
| - | - | 1403 | 501.8 | - | - | 0 | - |
| - | - | 939.8 | 507.2 | - | - | 0 | - |
| - | - | 1537 | 510.3 | - | - | 0 | - |
| - | - | 1303 | 510.8 | - | - | 0 | - |
| - | - | 2053 | 513.4 | - | - | 0 | - |
| - | - | 869.5 | 539.3 | - | - | 0 | - |
| - | - | 1014 | 539.6 | - | - | 0 | - |
| - | - | 1.515E+04 | 545 | - | - | 0 | - |
| - | - | 1.611E+04 | 545.3 | - | - | 0 | - |
| - | - | 4471 | 545.6 | - | - | 0 | - |
| - | - | 2710 | 546 | - | - | 0 | - |
| - | - | 638.9 | 548.2 | - | - | 0 | - |
| 2 | y | 3168 | 550.7 | 0.003874 | 7.034 | +2 | 8 |
| - | - | 1928 | 551.2 | - | - | 0 | - |
| - | - | 846.7 | 551.7 | - | - | 0 | - |
| - | - | 686.2 | 557.2 | - | - | 0 | - |
| 6 | y | 9053 | 560.2 | 0.005217 | 9.313 | +1 | 4 |
| - | - | 2707 | 561.2 | - | - | 0 | - |
| - | - | 875.1 | 561.3 | - | - | 0 | - |
| - | - | 884.4 | 564.8 | - | - | 0 | - |
| - | - | 1489 | 568.3 | - | - | 0 | - |
| - | - | 791.5 | 568.6 | - | - | 0 | - |
| - | - | 727.6 | 569.2 | - | - | 0 | - |
| - | - | 808.1 | 569.3 | - | - | 0 | - |
| - | - | 1332 | 573.3 | - | - | 0 | - |
| - | - | 1504 | 574.3 | - | - | 0 | - |
| - | - | 698.3 | 580.3 | - | - | 0 | - |
| - | - | 652.8 | 582.3 | - | - | 0 | - |
| - | - | 859.3 | 583.3 | - | - | 0 | - |
| - | - | 793.6 | 584.8 | - | - | 0 | - |
| - | - | 1.113E+04 | 585.3 | - | - | 0 | - |
| - | - | 3864 | 586.3 | - | - | 0 | - |
| 0 | Precursor | 1623 | 592.2 | 0.002529 | 4.27 | +2 | -1 |
| - | - | 1505 | 592.7 | - | - | 0 | - |
| - | - | 695.8 | 596 | - | - | 0 | - |
| - | - | 968.8 | 596.2 | - | - | 0 | - |
| - | - | 876.9 | 600.3 | - | - | 0 | - |
| 0 | Precursor | 4708 | 601.2 | 0.004082 | 6.79 | +2 | -1 |
| - | - | 1078 | 601.3 | - | - | 0 | - |
| - | - | 2616 | 601.7 | - | - | 0 | - |
| - | - | 738.1 | 602.2 | - | - | 0 | - |
| - | - | 1706 | 602.4 | - | - | 0 | - |
| - | - | 1072 | 603.4 | - | - | 0 | - |
| - | - | 768.1 | 606.2 | - | - | 0 | - |
| - | - | 2261 | 612.2 | - | - | 0 | - |
| - | - | 979.3 | 614.2 | - | - | 0 | - |
| - | - | 2499 | 628.4 | - | - | 0 | - |
| - | - | 1750 | 629.3 | - | - | 0 | - |
| - | - | 833.1 | 629.4 | - | - | 0 | - |
| - | - | 834.3 | 631.2 | - | - | 0 | - |
| - | - | 794 | 634.2 | - | - | 0 | - |
| - | - | 748.9 | 637.3 | - | - | 0 | - |
| - | - | 658.2 | 640.2 | - | - | 0 | - |
| 5 | b | 797.4 | 642.2 | 0.0005932 | 0.9237 | +1 | 5 |
| - | - | 777.2 | 654.3 | - | - | 0 | - |
| - | - | 1700 | 657.3 | - | - | 0 | - |
| - | - | 779.4 | 658.3 | - | - | 0 | - |
| - | - | 2737 | 659.3 | - | - | 0 | - |
| - | - | 1132 | 660.3 | - | - | 0 | - |
| - | - | 1205 | 682.4 | - | - | 0 | - |
| - | - | 883.4 | 683.3 | - | - | 0 | - |
| - | - | 2943 | 685.3 | - | - | 0 | - |
| - | - | 687 | 685.9 | - | - | 0 | - |
| - | - | 822.8 | 686.4 | - | - | 0 | - |
| - | - | 1222 | 686.9 | - | - | 0 | - |
| - | - | 1025 | 689.4 | - | - | 0 | - |
| - | - | 1.632E+04 | 694.9 | - | - | 0 | - |
| - | - | 1.474E+04 | 695.4 | - | - | 0 | - |
| - | - | 4078 | 695.9 | - | - | 0 | - |
| - | - | 1962 | 696.4 | - | - | 0 | - |
| - | - | 2.792E+04 | 700.4 | - | - | 0 | - |
| - | - | 9674 | 701.4 | - | - | 0 | - |
| - | - | 2346 | 702.4 | - | - | 0 | - |
| - | - | 1582 | 716.8 | - | - | 0 | - |
| - | - | 740.4 | 717.3 | - | - | 0 | - |
| - | - | 807.5 | 717.8 | - | - | 0 | - |
| - | - | 3013 | 721.3 | - | - | 0 | - |
| - | - | 793.1 | 722.3 | - | - | 0 | - |
| 5 | y | 7765 | 723.3 | 0.005427 | 7.503 | +1 | 5 |
| - | - | 3533 | 724.3 | - | - | 0 | - |
| - | - | 1011 | 725.3 | - | - | 0 | - |
| - | - | 803.7 | 726.4 | - | - | 0 | - |
| - | - | 1179 | 736.4 | - | - | 0 | - |
| - | - | 3762 | 749.3 | - | - | 0 | - |
| - | - | 848.6 | 750.3 | - | - | 0 | - |
| - | - | 2913 | 757.4 | - | - | 0 | - |
| - | - | 1499 | 758.4 | - | - | 0 | - |
| - | - | 1101 | 759.4 | - | - | 0 | - |
| - | - | 1100 | 767.3 | - | - | 0 | - |
| - | - | 3627 | 768.4 | - | - | 0 | - |
| - | - | 3703 | 768.9 | - | - | 0 | - |
| - | - | 979.9 | 769.4 | - | - | 0 | - |
| - | - | 2203 | 773.4 | - | - | 0 | - |
| - | - | 2743 | 774.3 | - | - | 0 | - |
| - | - | 801.6 | 774.9 | - | - | 0 | - |
| - | - | 2048 | 775.3 | - | - | 0 | - |
| - | - | 2207 | 775.4 | - | - | 0 | - |
| - | - | 1941 | 775.9 | - | - | 0 | - |
| - | - | 705.2 | 776.4 | - | - | 0 | - |
| - | - | 6892 | 783.4 | - | - | 0 | - |
| - | - | 3757 | 784.4 | - | - | 0 | - |
| - | - | 1331 | 785.4 | - | - | 0 | - |
| - | - | 907.6 | 787.4 | - | - | 0 | - |
| - | - | 1029 | 798.5 | - | - | 0 | - |
| - | - | 930 | 799.5 | - | - | 0 | - |
| - | - | 4928 | 800.4 | - | - | 0 | - |
| - | - | 1.374E+04 | 801.4 | - | - | 0 | - |
| - | - | 6616 | 802.4 | - | - | 0 | - |
| - | - | 1881 | 803.4 | - | - | 0 | - |
| - | - | 7816 | 816.9 | - | - | 0 | - |
| - | - | 8861 | 817.4 | - | - | 0 | - |
| - | - | 4583 | 817.9 | - | - | 0 | - |
| - | - | 1304 | 818.4 | - | - | 0 | - |
| - | - | 903.8 | 827.4 | - | - | 0 | - |
| - | - | 6783 | 828.4 | - | - | 0 | - |
| - | - | 2788 | 829.4 | - | - | 0 | - |
| - | - | 1031 | 830.4 | - | - | 0 | - |
| 4 | y | 1.014E+04 | 838.3 | 0.005095 | 6.077 | +1 | 6 |
| - | - | 5094 | 839.3 | - | - | 0 | - |
| - | - | 1317 | 840.3 | - | - | 0 | - |
| - | - | 2.313E+04 | 844.5 | - | - | 0 | - |
| - | - | 9903 | 845.5 | - | - | 0 | - |
| - | - | 2223 | 846.5 | - | - | 0 | - |
| - | - | 958.5 | 847.4 | - | - | 0 | - |
| - | - | 854.3 | 852.4 | - | - | 0 | - |
| - | - | 2894 | 854.4 | - | - | 0 | - |
| - | - | 2392 | 855.4 | - | - | 0 | - |
| - | - | 1668 | 871.3 | - | - | 0 | - |
| - | - | 8.099E+04 | 872.4 | - | - | 0 | - |
| - | - | 3.434E+04 | 873.5 | - | - | 0 | - |
| - | - | 9306 | 874.5 | - | - | 0 | - |
| - | - | 1779 | 875.5 | - | - | 0 | - |
| - | - | 1.432E+04 | 889.3 | - | - | 0 | - |
| - | - | 871.1 | 889.4 | - | - | 0 | - |
| - | - | 6992 | 890.4 | - | - | 0 | - |
| - | - | 1916 | 891.4 | - | - | 0 | - |
| - | - | 3987 | 895.5 | - | - | 0 | - |
| - | - | 4384 | 896.5 | - | - | 0 | - |
| - | - | 1545 | 897.5 | - | - | 0 | - |
| - | - | 2752 | 899.4 | - | - | 0 | - |
| - | - | 2552 | 900.4 | - | - | 0 | - |
| - | - | 1644 | 916.5 | - | - | 0 | - |
| - | - | 1053 | 917.5 | - | - | 0 | - |
| - | - | 811.8 | 932.5 | - | - | 0 | - |
| 3 | y | 3089 | 935.3 | 0.00251 | 2.684 | +1 | 7 |
| - | - | 1313 | 936.3 | - | - | 0 | - |
| 3 | y | 5.066E+04 | 953.3 | 0.004885 | 5.124 | +1 | 7 |
| - | - | 2.545E+04 | 954.3 | - | - | 0 | - |
| - | - | 9453 | 955.3 | - | - | 0 | - |
| - | - | 1251 | 956.4 | - | - | 0 | - |
| - | - | 1.386E+04 | 991.5 | - | - | 0 | - |
| - | - | 7329 | 992.5 | - | - | 0 | - |
| - | - | 3287 | 993.5 | - | - | 0 | - |
| - | - | 877.3 | 1000 | - | - | 0 | - |
| - | - | 838.7 | 1001 | - | - | 0 | - |
| - | - | 3.468E+04 | 1020 | - | - | 0 | - |
| - | - | 2.144E+04 | 1021 | - | - | 0 | - |
| - | - | 7274 | 1022 | - | - | 0 | - |
| - | - | 934.7 | 1023 | - | - | 0 | - |
| - | - | 1413 | 1036 | - | - | 0 | - |
| - | - | 806.6 | 1043 | - | - | 0 | - |
| - | - | 1022 | 1044 | - | - | 0 | - |
| - | - | 2860 | 1064 | - | - | 0 | - |
| - | - | 1279 | 1065 | - | - | 0 | - |
| 2 | y | 2718 | 1100 | 0.007699 | 6.997 | +1 | 8 |
| - | - | 1402 | 1101 | - | - | 0 | - |
| - | - | 1201 | 1163 | - | - | 0 | - |
| - | - | 760.9 | 1164 | - | - | 0 | - |
| - | - | 614 | 1202 | - | - | 0 | - |
| - | - | 1394 | 1389 | - | - | 0 | - |
| - | - | 1449 | 1390 | - | - | 0 | - |

m/z Charge Intensity FragmentType MassShift Position
120.08114624023438 0 149663.05
121.07901763916016 0 991.8849
121.08442687988281 0 15129.369
122.08782958984375 0 546.0405
123.04434967041016 0 2579.8386
125.51982116699219 0 371.58914
127.08719635009766 0 710.8262
129.0660400390625 0 827.7058
129.1025390625 0 8568.234
130.05018615722656 0 498.28323
130.06539916992188 0 2633.8154
130.105712890625 0 974.30865
132.0811767578125 0 1824.0991
132.1021728515625 0 1558.3164
133.0861358642578 0 1694.1172
134.0272674560547 0 487.42233
136.07601928710938 0 47333.254
137.07936096191406 0 4404.7188
138.0553436279297 0 952.0391
138.06649780273438 0 1438.0417
141.10260009765625 0 1512.2529
143.11817932128906 0 8202.734
144.12168884277344 0 627.6058
145.06143188476562 0 455.93338
146.06033325195312 0 1468.7363
147.04452514648438 0 1771.1996
147.7678680419922 0 460.63812
148.94723510742188 0 1018.99036
149.0451202392578 0 600.57715
155.08172607421875 0 934.0727
156.0770263671875 0 5871.543
157.10850524902344 0 555.02734
157.1338653564453 0 2496.592
158.06039428710938 0 1064.386
158.0967559814453 0 5171.06
159.09207153320312 0 6037.383
160.075927734375 0 1235.8894
160.0959014892578 0 646.7725
160.11245727539062 0 1731.9382
162.05540466308594 0 583.3778
165.054931640625 0 21182.412
166.0587615966797 0 1643.9478
166.0866241455078 0 3952.0525
167.05564880371094 0 1045.324
167.0926971435547 0 668.7429
168.5148468017578 0 438.41525
168.77005004882812 0 416.26303
169.0762176513672 0 863.4757
169.09716796875 0 1737.261
170.93405151367188 0 418.58633
171.0765380859375 0 1202.0021
171.1131134033203 0 6202.475
173.09217834472656 0 1168.31
173.12049865722656 0 650.5661
173.12881469726562 0 10513.731
174.13232421875 0 1336.4951
175.0869140625 0 3732.2349
176.08253479003906 0 972.7127
176.1073455810547 0 7975.153
177.10250854492188 0 6707.1367
177.11138916015625 0 1349.6046
180.07723999023438 0 531.99365
181.0967254638672 0 546.6219
182.08151245117188 0 41904.863
183.08499145507812 0 4102.009
183.1132354736328 0 925.7697
183.14930725097656 0 606.9738
185.0559539794922 0 1712.2972
185.12879943847656 0 1878.0171
186.09152221679688 0 879.2297
186.12399291992188 0 2444.771
187.08645629882812 0 593.9142
187.1446075439453 0 598.06946
188.07086181640625 0 6571.7314
189.0875244140625 0 1077.1289
191.1183319091797 0 3540.0269
193.108642578125 0 3585.7588
194.30099487304688 0 468.6123
195.08755493164062 0 493.46667
195.1116943359375 0 516.77466
197.1287384033203 0 1223.9677
199.0719757080078 0 1245.48
199.10800170898438 0 6322.7524
199.14450073242188 0 940.09357
200.13990783691406 0 1047.9127
200.26329040527344 0 562.87964
201.1237030029297 0 6909.936
202.12750244140625 0 1074.56
203.0665740966797 0 1736.9078
203.11831665039062 0 3108.2244 a Water loss 1
204.07676696777344 0 1505.7253
205.09735107421875 0 4893.832 y 8
206.96041870117188 0 467.18964
207.11288452148438 0 1204.1549
210.15997314453125 0 690.67377
214.1190185546875 0 1540.0995
215.1398162841797 0 663.7653
216.0982666015625 0 648.2848
217.08250427246094 0 1488.9391
217.09713745117188 0 1096.3171
217.12167358398438 0 824.9615
217.1338653564453 0 15322.339
219.07879638671875 0 548.2874
219.113037109375 0 1971.9102
221.09231567382812 0 2401.0227
221.10360717773438 0 7392.6895
221.12884521484375 0 134294.06 a 1
222.13211059570312 0 16955.146
223.0640869140625 0 1378.5669
223.10829162597656 0 761.3948
223.1343994140625 0 1047.6608
225.17144775390625 0 2609.2004
226.1190948486328 0 1571.1904
226.15550231933594 0 2780.0173
227.06655883789062 0 704.98224
227.09109497070312 0 806.03894
227.10299682617188 0 10441.607
227.1586456298828 0 652.56134
228.1070556640625 0 721.08026
229.11856079101562 0 2498.286
229.15487670898438 0 1105.6583
229.1692352294922 0 741.3694
231.06153869628906 0 3658.8037
231.0730743408203 0 582.69965
231.11285400390625 0 1019.00745 b Water loss 1
232.06541442871094 0 515.949
233.09228515625 0 637.2267
233.12887573242188 0 801.09656
233.15103149414062 0 545.3992
233.1648712158203 0 713.03534
235.1079864501953 0 4745.574
235.1437530517578 0 589.53894
239.08229064941406 0 675.943
239.11471557617188 0 905.3437
239.1496124267578 0 1266.9406
240.13491821289062 0 1297.431
243.13421630859375 0 1486.7493
244.0931854248047 0 1330.1117
245.1287841796875 0 7206.5464
246.13352966308594 0 919.0581
247.14427185058594 0 852.8948
248.1072235107422 0 1016.6225
249.10903930664062 0 1141.1416
249.12376403808594 0 23504.785 b 1
250.12716674804688 0 3126.847
251.10308837890625 0 2496.0938
253.118896484375 0 2637.7605
253.16619873046875 0 3347.6448
254.1136932373047 0 778.66516
254.6184844970703 0 556.6331
256.1087341308594 0 1018.13116
257.106689453125 0 888.74927
258.10906982421875 0 513.6639
263.10302734375 0 4829.341
264.105712890625 0 899.0761
271.10986328125 0 792.1504
273.13494873046875 0 4376.1206
274.1307678222656 0 1156.4741
279.0974426269531 0 2225.2546
280.129638671875 0 708.0651
283.1435241699219 0 1471.3218
283.1758728027344 0 875.43396
284.10382080078125 0 673.85803
285.01068115234375 0 656.2598
285.1026916503906 0 2250.2476
288.11773681640625 0 543.0632
292.1412048339844 0 977.8965
296.15106201171875 0 2465.8833
296.2091979980469 0 650.99713
297.0906066894531 0 833.63556
297.1557312011719 0 890.6209
301.1293640136719 0 2053.666
302.1142578125 0 1674.24
303.1136169433594 0 606.1153
304.1318359375 0 963.89215
315.1654357910156 0 566.3647
318.1455078125 0 1060.3121 a Water loss 2
319.14068603515625 0 2633.1294
323.098388671875 0 1363.4072
323.20806884765625 0 1589.2379
324.1458740234375 0 4800.699
325.1481018066406 0 891.41675
325.1878967285156 0 833.4858
326.1716613769531 0 4366.0693
327.17584228515625 0 1093.5905
327.2008972167969 0 867.81366
329.1501159667969 0 28275.426
329.23309326171875 0 636.43726
330.1532897949219 0 5332.341
339.2052001953125 0 689.8233
340.19842529296875 0 5243.214
341.202880859375 0 819.92957
342.1566467285156 0 27240.764 y 7
343.159912109375 0 5182.8193
344.1441345214844 0 1205.477
346.14013671875 0 1549.9154 b Water loss 2
348.1201477050781 0 1049.6881
350.1345520019531 0 1640.4741
355.06903076171875 0 1146.1003
355.1574401855469 0 619.93896
356.13958740234375 0 1066.4408
357.2247314453125 0 7169.1973
358.22918701171875 0 800.62933
364.15057373046875 0 5811.2183 b 2
365.1541748046875 0 1032.723
366.1300354003906 0 1048.8958
368.1923522949219 0 2142.5793
369.1971740722656 0 613.2145
371.68438720703125 0 668.1069
374.17376708984375 0 694.51117
375.185791015625 0 747.35223
375.2353515625 0 574.9757
376.115234375 0 752.4506
378.1295471191406 0 3427.4365
382.1430358886719 0 937.0292
383.197509765625 0 714.05054
385.2179870605469 0 1049.7496
392.2034912109375 0 1486.1094
394.1249694824219 0 7239.484
395.12774658203125 0 923.0274
398.24139404296875 0 759.8294
400.1876220703125 0 939.04736
401.20989990234375 0 1928.747
401.71075439453125 0 1142.0894
405.7116394042969 0 730.3046
414.72393798828125 0 564.4876
414.962646484375 0 864.56555
418.2095642089844 0 783.54626
422.7307434082031 0 987.1025
425.1943054199219 0 3521.655
427.72259521484375 0 3806.9153
428.2228698730469 0 2608.0623
428.7229919433594 0 836.6057
436.7278137207031 0 3206.1414
437.22833251953125 0 1658.6787
442.3141174316406 0 876.9662
443.2291564941406 0 1459.723
444.2240295410156 0 760.0627
445.19140625 0 704.2906
445.24603271484375 0 1011.603
448.26611328125 0 1469.0385
448.7684631347656 0 1189.9268
453.28143310546875 0 1325.0299
461.1656799316406 0 900.81775 b Water loss 3
461.1972961425781 0 712.929
462.2357482910156 0 760.2568
465.16265869140625 0 3124.3262
466.16436767578125 0 990.1536
470.3088073730469 0 1304.8204
473.24029541015625 0 875.6594
479.17779541015625 0 3336.9172 b 3
480.1807556152344 0 803.3407
482.2320556640625 0 706.81757
483.2200927734375 0 2706.6475
487.3359069824219 0 1838.2694
489.1920166015625 0 6125.5273 y 6
490.1942443847656 0 1468.6243
491.20367431640625 0 760.09937
496.230712890625 0 2398.7998
496.2657775878906 0 997.50995
496.7674255371094 0 788.0736
498.7798156738281 0 675.5663
500.2472229003906 0 1151.7893
501.255126953125 0 1911.4004
501.7568054199219 0 1403.0167
507.2093811035156 0 939.77014
510.26177978515625 0 1536.5474
510.7641296386719 0 1303.076
513.35205078125 0 2052.5317
539.2919311523438 0 869.4545
539.6224975585938 0 1013.7053
544.959716796875 0 15153.922
545.293701171875 0 16106.975
545.6289672851562 0 4471.393
545.9625244140625 0 2710.109
548.2018432617188 0 638.92773
550.712158203125 0 3168.2764 y 1
551.2119140625 0 1928.061
551.7160034179688 0 846.66534
557.1970825195312 0 686.18976
560.2288818359375 0 9053.472 y 5
561.2305297851562 0 2706.9182
561.2984619140625 0 875.05194
564.8046264648438 0 884.4315
568.3046875 0 1488.5621
568.6372680664062 0 791.51874
569.236083984375 0 727.6364
569.3054809570312 0 808.08105
573.3247680664062 0 1331.6268
574.3271484375 0 1503.9572
580.2604370117188 0 698.30255
582.3284301757812 0 652.80975
583.2872314453125 0 859.3126
584.7937622070312 0 793.623
585.3357543945312 0 11133.355
586.3391723632812 0 3864.459
592.2293701171875 0 1623.2488 Precursor Water loss
592.7311401367188 0 1505.0225
595.9959716796875 0 695.79095
596.2313232421875 0 968.8486
600.3078002929688 0 876.94543
601.2362060546875 0 4708.1777 Precursor
601.2943115234375 0 1078.3268
601.7377319335938 0 2616.1768
602.2347412109375 0 738.0883
602.354736328125 0 1706.1956
603.3511962890625 0 1071.7787
606.2313232421875 0 768.0947
612.1978149414062 0 2260.9297
614.2453002929688 0 979.3366
628.3779296875 0 2499.0403
629.3265991210938 0 1750.2765
629.3817749023438 0 833.14844
631.2463989257812 0 834.2711
634.2285766601562 0 794.0115
637.2964477539062 0 748.9032
640.2377319335938 0 658.2312
642.239990234375 0 797.4021 b 4
654.2767333984375 0 777.2018
657.2637329101562 0 1700.2249
658.2620239257812 0 779.44977
659.2918090820312 0 2737.1743
660.3010864257812 0 1132.3639
682.35107421875 0 1204.5024
683.3433227539062 0 883.3656
685.2576293945312 0 2943.0986
685.8657836914062 0 687.0266
686.3644409179688 0 822.8434
686.8655395507812 0 1222.1906
689.397705078125 0 1024.5723
694.874755859375 0 16319.706
695.3767700195312 0 14738.387
695.8768310546875 0 4077.8752
696.3753051757812 0 1961.7993
700.3627319335938 0 27921.434
701.36572265625 0 9674.128
702.3680419921875 0 2346.302
716.814453125 0 1582.1445
717.311279296875 0 740.3596
717.8165283203125 0 807.52203
721.26123046875 0 3012.971
722.2625122070312 0 793.0554
723.2924194335938 0 7764.5083 y 4
724.295654296875 0 3532.5254
725.2981567382812 0 1011.1939
726.3773803710938 0 803.70734
736.3602905273438 0 1179.2863
749.2564697265625 0 3762.2131
750.2589111328125 0 848.62964
757.3887329101562 0 2913.1648
758.390625 0 1498.5771
759.3674926757812 0 1101.0476
767.2645874023438 0 1099.6444
768.4091796875 0 3627.1372
768.9110107421875 0 3703.2153
769.4049072265625 0 979.9136
773.4129638671875 0 2202.7947
774.3194580078125 0 2743.2393
774.9010009765625 0 801.58386
775.3244018554688 0 2047.675
775.4049072265625 0 2207.273
775.9033203125 0 1941.4617
776.4103393554688 0 705.1766
783.3995971679688 0 6892.2847
784.401123046875 0 3756.7214
785.4150390625 0 1330.7518
787.4070434570312 0 907.61914
798.470947265625 0 1029.2422
799.4810791015625 0 929.9805
800.4268188476562 0 4927.8096
801.411376953125 0 13740.485
802.4136352539062 0 6616.4346
803.4221801757812 0 1880.9297
816.9354248046875 0 7815.7305
817.4363403320312 0 8861.462
817.9371948242188 0 4582.748
818.4417114257812 0 1303.7477
827.4385986328125 0 903.82715
828.422119140625 0 6783.3755
829.4232177734375 0 2788.399
830.42626953125 0 1030.9961
838.3190307617188 0 10144.812 y 3
839.3218383789062 0 5094.111
840.3213500976562 0 1316.769
844.4525146484375 0 23129.225
845.4552001953125 0 9902.993
846.45458984375 0 2223.3235
847.445556640625 0 958.493
852.4486083984375 0 854.3237
854.4368286132812 0 2894.1204
855.4363403320312 0 2392.045
871.3367309570312 0 1668.2611
872.4473266601562 0 80986.516
873.4500122070312 0 34340.902
874.4525756835938 0 9306.078
875.4577026367188 0 1779.422
889.3475952148438 0 14318.22
889.4436645507812 0 871.0878
890.3500366210938 0 6992.1733
891.3550415039062 0 1916.3699
895.5241088867188 0 3986.705
896.5274047851562 0 4383.916
897.5276489257812 0 1544.7432
899.4271240234375 0 2751.937
900.4293212890625 0 2551.7615
916.508544921875 0 1643.81
917.5164184570312 0 1053.4856
932.4830322265625 0 811.78375
935.3328247070312 0 3089.4087 y Water loss 2
936.3329467773438 0 1312.5332
953.3457641601562 0 50657.43 y 2
954.3486938476562 0 25451.215
955.3485107421875 0 9452.804
956.3505249023438 0 1250.6759
991.5202026367188 0 13862.166
992.5240478515625 0 7328.771
993.5272827148438 0 3286.7915
1000.4750366210938 0 877.34753
1001.48828125 0 838.7152
1019.51513671875 0 34675.645
1020.5181884765625 0 21437.04
1021.5218505859375 0 7274.246
1022.51953125 0 934.72034
1036.4122314453125 0 1412.8232
1042.587890625 0 806.5558
1043.601806640625 0 1022.4746
1063.5765380859375 0 2860.2837
1064.58056640625 0 1279.3564
1100.4169921875 0 2717.6392 y 1
1101.41650390625 0 1402.2301
1162.641845703125 0 1200.8357
1163.652099609375 0 760.86505
1201.751953125 0 614
1388.73486328125 0 1393.8767
1389.7381591796875 0 1448.7837

Spectrum Details

|  |  |
| --- | --- |
| Matched peaks? Matched peaks The total absolute number of peaks matched. Additionally in brackets the total fraction of peaks matched and the total number of peaks is shown. | 22 (5.06% of 435) |
| FDR? FDR The false discovery rate estimated for this peptide. It is calculated by matching all theoretical fragments with a non-integer shift with the raw peaks for this spectrum. This is done with 40 different shifts. The resulting percentage is the average number of annotated peaks over the number of annotated peaks with the correct spectrum. | 2.38% |
| Satellite FDR? Satellite FDR See the FDR for details on its calculation. This satellite ion specific FDR only contains the satellite ions (d/w) for I/L/J positions. | - |
| PSM Score? PSM Score The PSM Score as given by Hecklib to this annotated spectrum. It is shown with three significant figures. | 270 |

#### Spectrum 11192? Spectrum 11192 The raw spectrum of this peptide as annotated by Hecklib. The fragments are coloured according to ion type (see legend). Any peaks with a star '\*' as text can be hovered over to see the full details, first the ion type second the mass shift type. By hovering over the amino acids in the peptide or ions in the legend the corresponding peaks are highlighted. By toggling the 'Unassigned' label you can turn the background (unassigned) peaks on or off in the plot. By updating the slider in the Ion legend you can update the spectrum to only show the top X% of the peaks with labels. The top X% means any peak that is within X% of the highest intensity. By dragging in the spectrum you can zoom in to a specific part of the spectrum and use 'Zoom Out' to get back to the original zoom level. The annotation of the spectrum is based on the given sequence in the peptides file and is done with different software so inconsistencies are likely. The peaks are annotated based on the given sequence, with 20 ppm tolerance.

Copy Data

##### Spectrum 11192 (TSV)

###### Preview

```
Loading example...
```

*Click on the button to copy the data to your clipboard.*

Mz MinMz MaxIntensity Max

WidthHeightPeptide font sizePeptide stroke widthSpectrum font sizeSpectrum stroke widthCompact peptide

Ion legend

wxyz

abcd

OtherUnassignedIon^Charge^_Position_Show for top:%

TFDDYAMHW

01.50e+42.99e+44.49e+45.99e+4

Zoom Out

a^+1^_2_y^+1^_1_a^+1^_2_b^+1^_2_y^+1^_2_b^+1^_3_b^+1^_4_y^+1^_3_b^+2^_8_y^+2^_8_y^+1^_4_\*\*b^+1^_5_y^+1^_5_y^+1^_6_y^+1^_7_y^+1^_7_y^+1^_8_

0836167225083344

Fragment Matches Table

Show background peaks

| Position | Ion type | Intensity | mz Theoretical | mz Error (Th) | mz Error (ppm) | Charge | Series Number |
| --- | --- | --- | --- | --- | --- | --- | --- |
| - | - | 5.002E+04 | 120.1 | - | - | 0 | - |
| - | - | 4417 | 121.1 | - | - | 0 | - |
| - | - | 693.2 | 123 | - | - | 0 | - |
| - | - | 347.8 | 125 | - | - | 0 | - |
| - | - | 1125 | 127.1 | - | - | 0 | - |
| - | - | 4334 | 129.1 | - | - | 0 | - |
| - | - | 1533 | 130.1 | - | - | 0 | - |
| - | - | 735.6 | 132.1 | - | - | 0 | - |
| - | - | 1221 | 133.1 | - | - | 0 | - |
| - | - | 1.362E+04 | 136.1 | - | - | 0 | - |
| - | - | 936.1 | 137.1 | - | - | 0 | - |
| - | - | 447.9 | 138.1 | - | - | 0 | - |
| - | - | 795.9 | 138.1 | - | - | 0 | - |
| - | - | 438.1 | 138.4 | - | - | 0 | - |
| - | - | 546.7 | 139.1 | - | - | 0 | - |
| - | - | 2592 | 141.1 | - | - | 0 | - |
| - | - | 545.6 | 146.1 | - | - | 0 | - |
| - | - | 704.8 | 147 | - | - | 0 | - |
| - | - | 515.8 | 147.1 | - | - | 0 | - |
| - | - | 860 | 149 | - | - | 0 | - |
| - | - | 1219 | 155.1 | - | - | 0 | - |
| - | - | 1008 | 156.1 | - | - | 0 | - |
| - | - | 1971 | 157.1 | - | - | 0 | - |
| - | - | 1153 | 157.1 | - | - | 0 | - |
| - | - | 2599 | 158.1 | - | - | 0 | - |
| - | - | 2073 | 159.1 | - | - | 0 | - |
| - | - | 760.8 | 159.1 | - | - | 0 | - |
| - | - | 1147 | 160.1 | - | - | 0 | - |
| - | - | 437 | 162 | - | - | 0 | - |
| - | - | 659.5 | 162.1 | - | - | 0 | - |
| - | - | 5223 | 165.1 | - | - | 0 | - |
| - | - | 434.7 | 165.1 | - | - | 0 | - |
| - | - | 1.207E+04 | 166.1 | - | - | 0 | - |
| - | - | 2301 | 167.1 | - | - | 0 | - |
| - | - | 5020 | 167.1 | - | - | 0 | - |
| - | - | 1130 | 167.1 | - | - | 0 | - |
| - | - | 3662 | 169.1 | - | - | 0 | - |
| - | - | 427.7 | 170.1 | - | - | 0 | - |
| - | - | 639.5 | 171.1 | - | - | 0 | - |
| - | - | 2215 | 173.1 | - | - | 0 | - |
| - | - | 3002 | 173.4 | - | - | 0 | - |
| - | - | 708.7 | 175.1 | - | - | 0 | - |
| - | - | 3914 | 176.1 | - | - | 0 | - |
| - | - | 2821 | 177.1 | - | - | 0 | - |
| - | - | 763.6 | 177.1 | - | - | 0 | - |
| - | - | 8275 | 182.1 | - | - | 0 | - |
| - | - | 648.6 | 183.1 | - | - | 0 | - |
| - | - | 1369 | 185.1 | - | - | 0 | - |
| - | - | 711.8 | 185.1 | - | - | 0 | - |
| - | - | 2083 | 188.1 | - | - | 0 | - |
| - | - | 839.7 | 191.1 | - | - | 0 | - |
| - | - | 644.6 | 193.1 | - | - | 0 | - |
| - | - | 812 | 197.1 | - | - | 0 | - |
| - | - | 2576 | 199.1 | - | - | 0 | - |
| - | - | 510.3 | 200.1 | - | - | 0 | - |
| - | - | 2984 | 201.1 | - | - | 0 | - |
| - | - | 2343 | 201.1 | - | - | 0 | - |
| 2 | a | 1234 | 203.1 | 0.0004424 | 2.178 | +1 | 2 |
| 9 | y | 1536 | 205.1 | 1.383E-06 | 0.006741 | +1 | 1 |
| - | - | 726.1 | 207.1 | - | - | 0 | - |
| - | - | 735.2 | 209.2 | - | - | 0 | - |
| - | - | 628.1 | 211.1 | - | - | 0 | - |
| - | - | 741.6 | 212.1 | - | - | 0 | - |
| - | - | 1123 | 215.1 | - | - | 0 | - |
| - | - | 1112 | 217.1 | - | - | 0 | - |
| - | - | 3132 | 221.1 | - | - | 0 | - |
| 2 | a | 5.927E+04 | 221.1 | 0.0002079 | 0.9401 | +1 | 2 |
| - | - | 6447 | 222.1 | - | - | 0 | - |
| - | - | 1344 | 223.1 | - | - | 0 | - |
| - | - | 2337 | 224.1 | - | - | 0 | - |
| - | - | 623 | 225.2 | - | - | 0 | - |
| - | - | 9864 | 227.1 | - | - | 0 | - |
| - | - | 710.2 | 228.1 | - | - | 0 | - |
| - | - | 502.3 | 228.1 | - | - | 0 | - |
| - | - | 1073 | 231.1 | - | - | 0 | - |
| - | - | 1879 | 235.1 | - | - | 0 | - |
| - | - | 540 | 238.2 | - | - | 0 | - |
| - | - | 1045 | 239.1 | - | - | 0 | - |
| - | - | 1150 | 241.1 | - | - | 0 | - |
| - | - | 509.6 | 242.1 | - | - | 0 | - |
| - | - | 891.6 | 243.1 | - | - | 0 | - |
| 2 | b | 1.022E+04 | 249.1 | 0.0003189 | 1.28 | +1 | 2 |
| - | - | 1450 | 250.1 | - | - | 0 | - |
| - | - | 1023 | 251.1 | - | - | 0 | - |
| - | - | 682.8 | 254.1 | - | - | 0 | - |
| - | - | 539.3 | 258.1 | - | - | 0 | - |
| - | - | 2751 | 263.1 | - | - | 0 | - |
| - | - | 552.9 | 264.1 | - | - | 0 | - |
| - | - | 2783 | 265.2 | - | - | 0 | - |
| - | - | 1029 | 266.1 | - | - | 0 | - |
| - | - | 1215 | 279.1 | - | - | 0 | - |
| - | - | 6161 | 281.1 | - | - | 0 | - |
| - | - | 1049 | 282.1 | - | - | 0 | - |
| - | - | 795.2 | 283.1 | - | - | 0 | - |
| - | - | 523.1 | 284.1 | - | - | 0 | - |
| - | - | 1050 | 284.2 | - | - | 0 | - |
| - | - | 664.9 | 285 | - | - | 0 | - |
| - | - | 1160 | 285.1 | - | - | 0 | - |
| - | - | 564.5 | 286.1 | - | - | 0 | - |
| - | - | 846 | 292.1 | - | - | 0 | - |
| - | - | 781.9 | 294.2 | - | - | 0 | - |
| - | - | 543.8 | 295.2 | - | - | 0 | - |
| - | - | 818.8 | 296.2 | - | - | 0 | - |
| - | - | 738.7 | 298.1 | - | - | 0 | - |
| - | - | 1513 | 299.1 | - | - | 0 | - |
| - | - | 628.4 | 302.1 | - | - | 0 | - |
| - | - | 630.4 | 309.2 | - | - | 0 | - |
| - | - | 1197 | 312.2 | - | - | 0 | - |
| - | - | 871.3 | 319.1 | - | - | 0 | - |
| - | - | 763.6 | 322.2 | - | - | 0 | - |
| - | - | 824.7 | 323.2 | - | - | 0 | - |
| - | - | 1814 | 324.1 | - | - | 0 | - |
| - | - | 593.3 | 325.1 | - | - | 0 | - |
| - | - | 6121 | 329.1 | - | - | 0 | - |
| - | - | 1143 | 330.2 | - | - | 0 | - |
| - | - | 999 | 339.7 | - | - | 0 | - |
| - | - | 1.079E+04 | 340.2 | - | - | 0 | - |
| - | - | 1894 | 341.2 | - | - | 0 | - |
| 8 | y | 1.207E+04 | 342.2 | 0.0004282 | 1.251 | +1 | 2 |
| - | - | 1770 | 343.2 | - | - | 0 | - |
| - | - | 649.2 | 344.2 | - | - | 0 | - |
| - | - | 528.1 | 349.2 | - | - | 0 | - |
| - | - | 505 | 351.7 | - | - | 0 | - |
| - | - | 2050 | 352.2 | - | - | 0 | - |
| - | - | 1842 | 355.1 | - | - | 0 | - |
| - | - | 908.7 | 356.1 | - | - | 0 | - |
| - | - | 783.8 | 356.1 | - | - | 0 | - |
| - | - | 1021 | 357.2 | - | - | 0 | - |
| 3 | b | 2707 | 364.2 | 0.0002924 | 0.8029 | +1 | 3 |
| - | - | 1177 | 367.2 | - | - | 0 | - |
| - | - | 1103 | 370.2 | - | - | 0 | - |
| - | - | 1045 | 378.1 | - | - | 0 | - |
| - | - | 6425 | 380.2 | - | - | 0 | - |
| - | - | 878.9 | 381.2 | - | - | 0 | - |
| - | - | 636.8 | 393.3 | - | - | 0 | - |
| - | - | 549.6 | 393.3 | - | - | 0 | - |
| - | - | 2115 | 394.1 | - | - | 0 | - |
| - | - | 732.4 | 394.2 | - | - | 0 | - |
| - | - | 5242 | 395.2 | - | - | 0 | - |
| - | - | 998.4 | 396.2 | - | - | 0 | - |
| - | - | 5458 | 398.2 | - | - | 0 | - |
| - | - | 1033 | 399.2 | - | - | 0 | - |
| - | - | 823.9 | 412.2 | - | - | 0 | - |
| - | - | 1173 | 413.3 | - | - | 0 | - |
| - | - | 669.4 | 414.2 | - | - | 0 | - |
| - | - | 640.7 | 416.7 | - | - | 0 | - |
| - | - | 728.9 | 417.2 | - | - | 0 | - |
| - | - | 1268 | 425.2 | - | - | 0 | - |
| - | - | 1136 | 425.3 | - | - | 0 | - |
| - | - | 610.9 | 427.7 | - | - | 0 | - |
| - | - | 836.6 | 433.2 | - | - | 0 | - |
| - | - | 920 | 436.7 | - | - | 0 | - |
| - | - | 622.9 | 446.3 | - | - | 0 | - |
| - | - | 2276 | 449.2 | - | - | 0 | - |
| - | - | 1512 | 451.2 | - | - | 0 | - |
| - | - | 1243 | 451.7 | - | - | 0 | - |
| - | - | 691 | 452.2 | - | - | 0 | - |
| - | - | 3076 | 453.3 | - | - | 0 | - |
| - | - | 1007 | 454.3 | - | - | 0 | - |
| - | - | 1200 | 465.2 | - | - | 0 | - |
| 4 | b | 1262 | 479.2 | 0.0006321 | 1.319 | +1 | 4 |
| - | - | 1747 | 480.3 | - | - | 0 | - |
| 7 | y | 3078 | 489.2 | 0.005497 | 11.24 | +1 | 3 |
| 8 | b | 609.2 | 490.2 | 0.005231 | 10.67 | +2 | 8 |
| - | - | 1153 | 493.3 | - | - | 0 | - |
| - | - | 1368 | 496.2 | - | - | 0 | - |
| - | - | 3398 | 500.8 | - | - | 0 | - |
| - | - | 1316 | 501.3 | - | - | 0 | - |
| - | - | 1572 | 506.3 | - | - | 0 | - |
| - | - | 9211 | 508.3 | - | - | 0 | - |
| - | - | 2086 | 509.3 | - | - | 0 | - |
| - | - | 630 | 516.4 | - | - | 0 | - |
| 2 | y | 945.6 | 550.7 | 0.004179 | 7.588 | +2 | 8 |
| - | - | 1143 | 551.2 | - | - | 0 | - |
| 6 | y | 4924 | 560.2 | 0.005034 | 8.986 | +1 | 4 |
| - | - | 1651 | 561.2 | - | - | 0 | - |
| - | - | 4710 | 563.3 | - | - | 0 | - |
| - | - | 1387 | 564.3 | - | - | 0 | - |
| - | - | 2392 | 585.3 | - | - | 0 | - |
| 0 | Precursor | 673.3 | 592.2 | 0.005885 | 9.938 | +2 | -1 |
| - | - | 3020 | 593.4 | - | - | 0 | - |
| - | - | 1377 | 594.4 | - | - | 0 | - |
| 0 | Precursor | 1479 | 601.2 | 0.002129 | 3.541 | +2 | -1 |
| - | - | 913.3 | 601.3 | - | - | 0 | - |
| - | - | 738.3 | 601.6 | - | - | 0 | - |
| - | - | 871.8 | 601.7 | - | - | 0 | - |
| - | - | 678 | 601.8 | - | - | 0 | - |
| - | - | 552.4 | 602.2 | - | - | 0 | - |
| - | - | 1195 | 602.3 | - | - | 0 | - |
| - | - | 1.546E+04 | 621.4 | - | - | 0 | - |
| - | - | 4584 | 622.4 | - | - | 0 | - |
| - | - | 1540 | 623.4 | - | - | 0 | - |
| 5 | b | 612 | 624.2 | 0.002236 | 3.581 | +1 | 5 |
| - | - | 829 | 628.4 | - | - | 0 | - |
| - | - | 622.2 | 650.4 | - | - | 0 | - |
| - | - | 1210 | 657.3 | - | - | 0 | - |
| - | - | 999.9 | 659.3 | - | - | 0 | - |
| - | - | 669.3 | 660.3 | - | - | 0 | - |
| - | - | 769.6 | 660.4 | - | - | 0 | - |
| - | - | 2665 | 676.4 | - | - | 0 | - |
| - | - | 737 | 677.4 | - | - | 0 | - |
| - | - | 2802 | 678.4 | - | - | 0 | - |
| - | - | 1059 | 679.4 | - | - | 0 | - |
| - | - | 796.8 | 680.4 | - | - | 0 | - |
| - | - | 932.6 | 685.3 | - | - | 0 | - |
| - | - | 5778 | 700.4 | - | - | 0 | - |
| - | - | 2415 | 701.4 | - | - | 0 | - |
| - | - | 663.2 | 702.4 | - | - | 0 | - |
| - | - | 1086 | 717.4 | - | - | 0 | - |
| - | - | 1520 | 721.3 | - | - | 0 | - |
| 5 | y | 3507 | 723.3 | 0.004755 | 6.574 | +1 | 5 |
| - | - | 1112 | 724.3 | - | - | 0 | - |
| - | - | 721.7 | 725.3 | - | - | 0 | - |
| - | - | 5431 | 735.4 | - | - | 0 | - |
| - | - | 2312 | 736.4 | - | - | 0 | - |
| - | - | 1949 | 749.3 | - | - | 0 | - |
| - | - | 641.9 | 757.4 | - | - | 0 | - |
| - | - | 1811 | 774.3 | - | - | 0 | - |
| - | - | 683.7 | 775.3 | - | - | 0 | - |
| - | - | 1559 | 783.4 | - | - | 0 | - |
| - | - | 677.2 | 784.4 | - | - | 0 | - |
| - | - | 705.7 | 800.4 | - | - | 0 | - |
| - | - | 2649 | 801.4 | - | - | 0 | - |
| - | - | 851.9 | 802.4 | - | - | 0 | - |
| - | - | 662.9 | 817.7 | - | - | 0 | - |
| - | - | 934.9 | 828.4 | - | - | 0 | - |
| - | - | 894.4 | 829.4 | - | - | 0 | - |
| - | - | 686.1 | 832.5 | - | - | 0 | - |
| 4 | y | 4827 | 838.3 | 0.003874 | 4.621 | +1 | 6 |
| - | - | 2174 | 839.3 | - | - | 0 | - |
| - | - | 3531 | 844.5 | - | - | 0 | - |
| - | - | 1473 | 845.5 | - | - | 0 | - |
| - | - | 618.2 | 871.3 | - | - | 0 | - |
| - | - | 1.686E+04 | 872.4 | - | - | 0 | - |
| - | - | 7699 | 873.5 | - | - | 0 | - |
| - | - | 2271 | 874.5 | - | - | 0 | - |
| - | - | 7007 | 889.3 | - | - | 0 | - |
| - | - | 798 | 889.4 | - | - | 0 | - |
| - | - | 3554 | 890.4 | - | - | 0 | - |
| - | - | 1114 | 891.4 | - | - | 0 | - |
| - | - | 8565 | 901.5 | - | - | 0 | - |
| - | - | 5053 | 902.5 | - | - | 0 | - |
| - | - | 1891 | 903.5 | - | - | 0 | - |
| - | - | 4299 | 919.5 | - | - | 0 | - |
| - | - | 2345 | 920.5 | - | - | 0 | - |
| 3 | y | 1484 | 935.3 | 0.003914 | 4.185 | +1 | 7 |
| 3 | y | 2.08E+04 | 953.3 | 0.004519 | 4.74 | +1 | 7 |
| - | - | 1.21E+04 | 954.3 | - | - | 0 | - |
| - | - | 3160 | 955.3 | - | - | 0 | - |
| - | - | 3078 | 991.5 | - | - | 0 | - |
| - | - | 1474 | 992.5 | - | - | 0 | - |
| - | - | 2013 | 1001 | - | - | 0 | - |
| - | - | 1592 | 1002 | - | - | 0 | - |
| - | - | 725.7 | 1003 | - | - | 0 | - |
| - | - | 3234 | 1019 | - | - | 0 | - |
| - | - | 7009 | 1020 | - | - | 0 | - |
| - | - | 3708 | 1021 | - | - | 0 | - |
| - | - | 1001 | 1022 | - | - | 0 | - |
| 2 | y | 1852 | 1100 | 0.005502 | 5 | +1 | 8 |
| - | - | 1290 | 1101 | - | - | 0 | - |
| - | - | 599.2 | 1313 | - | - | 0 | - |
| - | - | 739 | 1727 | - | - | 0 | - |
| - | - | 731.6 | 1755 | - | - | 0 | - |
| - | - | 638.5 | 2206 | - | - | 0 | - |
| - | - | 800.9 | 2351 | - | - | 0 | - |
| - | - | 789.7 | 2371 | - | - | 0 | - |
| - | - | 1015 | 2989 | - | - | 0 | - |
| - | - | 665.4 | 3311 | - | - | 0 | - |

m/z Charge Intensity FragmentType MassShift Position
120.08103942871094 0 50018.832
121.08438873291016 0 4417.297
123.04415130615234 0 693.2209
125.02149200439453 0 347.8258
127.08683013916016 0 1125.0844
129.10247802734375 0 4334.2188
130.06524658203125 0 1533.1052
132.10223388671875 0 735.5654
133.0862274169922 0 1221.3807
136.075927734375 0 13622.635
137.07949829101562 0 936.0939
138.05552673339844 0 447.85242
138.06642150878906 0 795.94086
138.36703491210938 0 438.14938
139.0864715576172 0 546.68274
141.10238647460938 0 2592.1216
146.05995178222656 0 545.5873
147.0446319580078 0 704.8207
147.1122589111328 0 515.83746
149.0453643798828 0 860.00146
155.0819091796875 0 1219.3114
156.0768585205078 0 1008.40765
157.09742736816406 0 1971.398
157.13388061523438 0 1153.321
158.09657287597656 0 2598.959
159.09201049804688 0 2073.065
159.1128692626953 0 760.8124
160.1122589111328 0 1146.8184
161.9861297607422 0 436.9563
162.09117126464844 0 659.5316
165.0548095703125 0 5222.911
165.1019287109375 0 434.73578
166.08648681640625 0 12071.828
167.0557098388672 0 2301.3792
167.0817413330078 0 5019.8223
167.0896759033203 0 1130.1444
169.0973663330078 0 3662.2378
170.1009063720703 0 427.72403
171.07667541503906 0 639.5091
173.12879943847656 0 2214.6445
173.44003295898438 0 3002.3516
175.11923217773438 0 708.6946
176.10720825195312 0 3914.408
177.10243225097656 0 2820.5312
177.1109619140625 0 763.6052
182.08139038085938 0 8275.2705
183.11297607421875 0 648.60925
185.09251403808594 0 1369.2208
185.12901306152344 0 711.8298
188.07093811035156 0 2082.526
191.11810302734375 0 839.70575
193.10809326171875 0 644.60565
197.1289520263672 0 811.98413
199.1078338623047 0 2575.5676
200.11160278320312 0 510.32156
201.11239624023438 0 2983.5776
201.12368774414062 0 2343.1255
203.1183319091797 0 1233.9171 a Water loss 1
205.09715270996094 0 1535.986 y 8
207.11294555664062 0 726.1236
209.1650848388672 0 735.2323
211.1188201904297 0 628.1347
212.1029815673828 0 741.6439
215.13961791992188 0 1122.6307
217.09735107421875 0 1111.763
221.10353088378906 0 3131.9636
221.128662109375 0 59274.32 a 1
222.13204956054688 0 6447.4556
223.06419372558594 0 1344.0664
224.10289001464844 0 2336.8066
225.17169189453125 0 623.01227
227.10281372070312 0 9863.549
228.10617065429688 0 710.176
228.13462829589844 0 502.33447
231.0615234375 0 1072.6152
235.1075897216797 0 1878.82
238.1554718017578 0 540.0397
239.0942840576172 0 1044.5416
241.0740203857422 0 1149.5482
242.11444091796875 0 509.61374
243.13406372070312 0 891.55786
249.12368774414062 0 10217.39 b 1
250.12722778320312 0 1449.646
251.10296630859375 0 1023.4751
254.1138916015625 0 682.7538
258.1097106933594 0 539.3269
263.1029357910156 0 2750.8887
264.1459655761719 0 552.89636
265.15472412109375 0 2783.0227
266.149169921875 0 1028.9128
279.09765625 0 1215.4276
281.12469482421875 0 6160.5225
282.1277160644531 0 1048.7933
283.14202880859375 0 795.167
284.1246032714844 0 523.08496
284.1605224609375 0 1049.6659
285.0102844238281 0 664.9298
285.1017761230469 0 1160.2306
286.1065368652344 0 564.4653
292.14093017578125 0 846.04736
294.18218994140625 0 781.88367
295.17877197265625 0 543.773
296.15069580078125 0 818.7617
298.1401062011719 0 738.6936
299.13519287109375 0 1512.5693
302.11279296875 0 628.40393
309.2032775878906 0 630.4253
312.1920166015625 0 1196.679
319.1396179199219 0 871.2589
322.17620849609375 0 763.5879
323.1711120605469 0 824.69995
324.1462097167969 0 1813.886
325.14923095703125 0 593.2576
329.14984130859375 0 6121.3164
330.1532897949219 0 1142.9524
339.69415283203125 0 998.95654
340.1874694824219 0 10794.052
341.1893005371094 0 1894.3347
342.156494140625 0 12072.279 y 7
343.1595458984375 0 1770.4698
344.1659851074219 0 649.1988
349.1869812011719 0 528.0827
351.7264099121094 0 504.98935
352.197998046875 0 2050.317
355.0698547363281 0 1841.9479
356.0702819824219 0 908.69147
356.139892578125 0 783.75757
357.22467041015625 0 1020.6808
364.1506042480469 0 2707.434 b 2
367.1980285644531 0 1177.1957
370.20745849609375 0 1103.0052
378.1300048828125 0 1044.7675
380.19287109375 0 6425.2607
381.1953125 0 878.91644
393.2856750488281 0 636.76056
393.3422546386719 0 549.57587
394.1243896484375 0 2115.003
394.20806884765625 0 732.37854
395.19287109375 0 5241.653
396.1948547363281 0 998.41174
398.20379638671875 0 5458.294
399.2057189941406 0 1032.635
412.21929931640625 0 823.91174
413.2659912109375 0 1172.9518
414.2258605957031 0 669.4383
416.7328186035156 0 640.702
417.232421875 0 728.87616
425.19329833984375 0 1267.8779
425.27545166015625 0 1135.9204
427.723876953125 0 610.90906
433.1734924316406 0 836.6246
436.72705078125 0 920.0314
446.2521057128906 0 622.92
449.2397766113281 0 2275.618
451.2441711425781 0 1511.7454
451.7442321777344 0 1243.3306
452.2469177246094 0 691.03674
453.27166748046875 0 3075.8953
454.27337646484375 0 1006.7861
465.1603088378906 0 1199.8451
479.1778869628906 0 1262.4995 b 3
480.2821044921875 0 1747.1089
489.1920471191406 0 3077.9238 y 6
490.1871337890625 0 609.1996 b Water loss 7
493.2782287597656 0 1153.0187
496.23040771484375 0 1368.0881
500.77685546875 0 3398.259
501.279296875 0 1316.4641
506.26129150390625 0 1571.7214
508.2768859863281 0 9211.144
509.281494140625 0 2085.7117
516.4049682617188 0 630.0066
550.7124633789062 0 945.5784 y 1
551.2138671875 0 1142.8191
560.2286987304688 0 4924.0234 y 5
561.231689453125 0 1651.0519
563.2823486328125 0 4709.7974
564.2849731445312 0 1386.7627
585.3350219726562 0 2391.548
592.2327270507812 0 673.2659 Precursor Water loss
593.3665771484375 0 3019.9072
594.3682861328125 0 1377.2318
601.2342529296875 0 1479.1523 Precursor
601.3138427734375 0 913.3395
601.6483154296875 0 738.33673
601.7388916015625 0 871.8157
601.8137817382812 0 678.0253
602.248779296875 0 552.40234
602.3342895507812 0 1194.8263
621.36083984375 0 15463.133
622.364501953125 0 4583.962
623.3662109375 0 1540.2174
624.227783203125 0 611.99164 b Water loss 4
628.3780517578125 0 829.00024
650.382080078125 0 622.1925
657.2620239257812 0 1210.4839
659.2949829101562 0 999.94507
660.3004150390625 0 669.2798
660.3646240234375 0 769.56714
676.3671264648438 0 2665.4705
677.3697509765625 0 736.9765
678.3812866210938 0 2801.65
679.3818969726562 0 1058.5688
680.3812255859375 0 796.799
685.2568359375 0 932.57214
700.3622436523438 0 5777.9727
701.36474609375 0 2415.0283
702.3673706054688 0 663.159
717.3923950195312 0 1085.6128
721.2611694335938 0 1519.6173
723.291748046875 0 3507.3699 y 4
724.2958984375 0 1111.6301
725.3002319335938 0 721.72034
735.404052734375 0 5431.3647
736.4056396484375 0 2312.4905
749.2533569335938 0 1948.6641
757.3892822265625 0 641.8764
774.3203125 0 1810.5135
775.3251342773438 0 683.74365
783.397705078125 0 1558.6146
784.39453125 0 677.1531
800.4262084960938 0 705.6725
801.4139404296875 0 2648.8933
802.41015625 0 851.85834
817.7122192382812 0 662.8512
828.4297485351562 0 934.8567
829.421142578125 0 894.38763
832.4624633789062 0 686.1186
838.3178100585938 0 4826.788 y 3
839.32177734375 0 2173.8774
844.4515380859375 0 3530.6255
845.45263671875 0 1473.0266
871.3350219726562 0 618.2356
872.4470825195312 0 16863.49
873.4500732421875 0 7699.2515
874.4514770507812 0 2270.8591
889.3466796875 0 7007.306
889.43408203125 0 798.02466
890.3505249023438 0 3554.4854
891.3507080078125 0 1113.5444
901.47802734375 0 8565.271
902.47900390625 0 5053.091
903.4803466796875 0 1891.225
919.4860229492188 0 4298.9653
920.4901733398438 0 2344.5728
935.334228515625 0 1483.5879 y Water loss 2
953.3453979492188 0 20802.166 y 2
954.3483276367188 0 12100.933
955.3477172851562 0 3159.5088
991.5206909179688 0 3078.2002
992.5234985351562 0 1473.7756
1000.5452880859375 0 2012.5142
1001.5458984375 0 1591.509
1003.3688354492188 0 725.68164
1018.5554809570312 0 3233.8564
1019.5189819335938 0 7008.718
1020.5218505859375 0 3708.407
1021.5166015625 0 1001.294
1100.414794921875 0 1852.4517 y 1
1101.413330078125 0 1289.9756
1313.3682861328125 0 599.2122
1726.6494140625 0 738.991
1754.9508056640625 0 731.57623
2205.6162109375 0 638.5398
2350.88037109375 0 800.9068
2370.9677734375 0 789.7363
2989.365478515625 0 1014.9787
3311.008544921875 0 665.35266

Spectrum Details

|  |  |
| --- | --- |
| Matched peaks? Matched peaks The total absolute number of peaks matched. Additionally in brackets the total fraction of peaks matched and the total number of peaks is shown. | 19 (7.09% of 268) |
| FDR? FDR The false discovery rate estimated for this peptide. It is calculated by matching all theoretical fragments with a non-integer shift with the raw peaks for this spectrum. This is done with 40 different shifts. The resulting percentage is the average number of annotated peaks over the number of annotated peaks with the correct spectrum. | 3.01% |
| Satellite FDR? Satellite FDR See the FDR for details on its calculation. This satellite ion specific FDR only contains the satellite ions (d/w) for I/L/J positions. | - |
| PSM Score? PSM Score The PSM Score as given by Hecklib to this annotated spectrum. It is shown with three significant figures. | 195 |

#### Spectrum 11306? Spectrum 11306 The raw spectrum of this peptide as annotated by Hecklib. The fragments are coloured according to ion type (see legend). Any peaks with a star '\*' as text can be hovered over to see the full details, first the ion type second the mass shift type. By hovering over the amino acids in the peptide or ions in the legend the corresponding peaks are highlighted. By toggling the 'Unassigned' label you can turn the background (unassigned) peaks on or off in the plot. By updating the slider in the Ion legend you can update the spectrum to only show the top X% of the peaks with labels. The top X% means any peak that is within X% of the highest intensity. By dragging in the spectrum you can zoom in to a specific part of the spectrum and use 'Zoom Out' to get back to the original zoom level. The annotation of the spectrum is based on the given sequence in the peptides file and is done with different software so inconsistencies are likely. The peaks are annotated based on the given sequence, with 20 ppm tolerance.

Copy Data

##### Spectrum 11306 (TSV)

###### Preview

```
Loading example...
```

*Click on the button to copy the data to your clipboard.*

Mz MinMz MaxIntensity Max

WidthHeightPeptide font sizePeptide stroke widthSpectrum font sizeSpectrum stroke widthCompact peptide

Ion legend

wxyz

abcd

OtherUnassignedIon^Charge^_Position_Show for top:%

TFDDYAMHW

01.43e+42.87e+44.30e+45.74e+4

Zoom Out

a^+1^_2_y^+1^_1_a^+1^_2_b^+1^_2_y^+1^_2_b^+1^_3_b^+1^_4_y^+1^_3_y^+2^_8_y^+1^_4_\*b^+1^_5_y^+1^_5_y^+1^_6_y^+1^_7_y^+1^_7_y^+1^_8_

0882176526473529

Fragment Matches Table

Show background peaks

| Position | Ion type | Intensity | mz Theoretical | mz Error (Th) | mz Error (ppm) | Charge | Series Number |
| --- | --- | --- | --- | --- | --- | --- | --- |
| - | - | 4.304E+04 | 120.1 | - | - | 0 | - |
| - | - | 3378 | 121.1 | - | - | 0 | - |
| - | - | 676.4 | 123 | - | - | 0 | - |
| - | - | 821.4 | 127.1 | - | - | 0 | - |
| - | - | 3758 | 129.1 | - | - | 0 | - |
| - | - | 1006 | 130.1 | - | - | 0 | - |
| - | - | 1054 | 132.1 | - | - | 0 | - |
| - | - | 718.7 | 132.1 | - | - | 0 | - |
| - | - | 1.357E+04 | 136.1 | - | - | 0 | - |
| - | - | 808.1 | 137.1 | - | - | 0 | - |
| - | - | 792.6 | 138.1 | - | - | 0 | - |
| - | - | 754.2 | 138.1 | - | - | 0 | - |
| - | - | 359.5 | 141.1 | - | - | 0 | - |
| - | - | 1350 | 141.1 | - | - | 0 | - |
| - | - | 461.6 | 141.9 | - | - | 0 | - |
| - | - | 481.6 | 143.1 | - | - | 0 | - |
| - | - | 1173 | 147 | - | - | 0 | - |
| - | - | 430 | 147.1 | - | - | 0 | - |
| - | - | 384 | 147.9 | - | - | 0 | - |
| - | - | 441.3 | 148.5 | - | - | 0 | - |
| - | - | 1021 | 149 | - | - | 0 | - |
| - | - | 1092 | 149 | - | - | 0 | - |
| - | - | 724.2 | 155.1 | - | - | 0 | - |
| - | - | 1101 | 156.1 | - | - | 0 | - |
| - | - | 923.3 | 157.1 | - | - | 0 | - |
| - | - | 900 | 157.1 | - | - | 0 | - |
| - | - | 1758 | 158.1 | - | - | 0 | - |
| - | - | 1545 | 159.1 | - | - | 0 | - |
| - | - | 477.2 | 159.1 | - | - | 0 | - |
| - | - | 1220 | 160.1 | - | - | 0 | - |
| - | - | 777.4 | 162.1 | - | - | 0 | - |
| - | - | 491.4 | 163.1 | - | - | 0 | - |
| - | - | 5067 | 165.1 | - | - | 0 | - |
| - | - | 6136 | 166.1 | - | - | 0 | - |
| - | - | 1526 | 167.1 | - | - | 0 | - |
| - | - | 1946 | 167.1 | - | - | 0 | - |
| - | - | 970.2 | 167.1 | - | - | 0 | - |
| - | - | 1836 | 169.1 | - | - | 0 | - |
| - | - | 520.6 | 171.2 | - | - | 0 | - |
| - | - | 527.1 | 173 | - | - | 0 | - |
| - | - | 2812 | 173.1 | - | - | 0 | - |
| - | - | 2721 | 173.4 | - | - | 0 | - |
| - | - | 506.4 | 174.1 | - | - | 0 | - |
| - | - | 549.6 | 175.1 | - | - | 0 | - |
| - | - | 3753 | 176.1 | - | - | 0 | - |
| - | - | 3301 | 177.1 | - | - | 0 | - |
| - | - | 776.8 | 177.1 | - | - | 0 | - |
| - | - | 467.2 | 178.1 | - | - | 0 | - |
| - | - | 8977 | 182.1 | - | - | 0 | - |
| - | - | 1145 | 183.1 | - | - | 0 | - |
| - | - | 448.5 | 183.1 | - | - | 0 | - |
| - | - | 659 | 183.1 | - | - | 0 | - |
| - | - | 490.5 | 183.2 | - | - | 0 | - |
| - | - | 1245 | 185.1 | - | - | 0 | - |
| - | - | 1308 | 185.1 | - | - | 0 | - |
| - | - | 432.5 | 185.2 | - | - | 0 | - |
| - | - | 2757 | 188.1 | - | - | 0 | - |
| - | - | 715.5 | 191.1 | - | - | 0 | - |
| - | - | 1552 | 193.1 | - | - | 0 | - |
| - | - | 1363 | 199.1 | - | - | 0 | - |
| - | - | 1557 | 201.1 | - | - | 0 | - |
| 2 | a | 1325 | 203.1 | 6.117E-05 | 0.3011 | +1 | 2 |
| 9 | y | 1870 | 205.1 | 3.19E-05 | 0.1555 | +1 | 1 |
| - | - | 729 | 207.1 | - | - | 0 | - |
| - | - | 614.2 | 211.1 | - | - | 0 | - |
| - | - | 686.8 | 212.1 | - | - | 0 | - |
| - | - | 834.3 | 214.1 | - | - | 0 | - |
| - | - | 3103 | 221.1 | - | - | 0 | - |
| 2 | a | 5.68E+04 | 221.1 | 0.0002994 | 1.354 | +1 | 2 |
| - | - | 5350 | 222.1 | - | - | 0 | - |
| - | - | 1101 | 223.1 | - | - | 0 | - |
| - | - | 474.1 | 223.6 | - | - | 0 | - |
| - | - | 1828 | 224.1 | - | - | 0 | - |
| - | - | 4604 | 227.1 | - | - | 0 | - |
| - | - | 592.5 | 229.1 | - | - | 0 | - |
| - | - | 552.1 | 229.2 | - | - | 0 | - |
| - | - | 895.5 | 231.1 | - | - | 0 | - |
| - | - | 1793 | 235.1 | - | - | 0 | - |
| - | - | 767.3 | 239.1 | - | - | 0 | - |
| - | - | 1471 | 241.1 | - | - | 0 | - |
| - | - | 663 | 242.1 | - | - | 0 | - |
| 2 | b | 9951 | 249.1 | 0.0003799 | 1.525 | +1 | 2 |
| - | - | 841.6 | 250.1 | - | - | 0 | - |
| - | - | 1868 | 251.1 | - | - | 0 | - |
| - | - | 773.3 | 253.2 | - | - | 0 | - |
| - | - | 643.2 | 255.6 | - | - | 0 | - |
| - | - | 527.6 | 257.2 | - | - | 0 | - |
| - | - | 2019 | 263.1 | - | - | 0 | - |
| - | - | 1191 | 265.2 | - | - | 0 | - |
| - | - | 575.1 | 275.2 | - | - | 0 | - |
| - | - | 1238 | 279.1 | - | - | 0 | - |
| - | - | 2536 | 281.1 | - | - | 0 | - |
| - | - | 522.2 | 283 | - | - | 0 | - |
| - | - | 1172 | 283.1 | - | - | 0 | - |
| - | - | 555.1 | 284.2 | - | - | 0 | - |
| - | - | 1167 | 285 | - | - | 0 | - |
| - | - | 1166 | 285.1 | - | - | 0 | - |
| - | - | 704.9 | 286 | - | - | 0 | - |
| - | - | 871.1 | 292.1 | - | - | 0 | - |
| - | - | 1121 | 296.2 | - | - | 0 | - |
| - | - | 519.3 | 303.1 | - | - | 0 | - |
| - | - | 786.6 | 319.1 | - | - | 0 | - |
| - | - | 1537 | 324.1 | - | - | 0 | - |
| - | - | 530.4 | 324.2 | - | - | 0 | - |
| - | - | 5322 | 329.1 | - | - | 0 | - |
| - | - | 1505 | 330.2 | - | - | 0 | - |
| - | - | 3661 | 340.2 | - | - | 0 | - |
| 8 | y | 1.269E+04 | 342.2 | 0.0004892 | 1.43 | +1 | 2 |
| - | - | 2100 | 343.2 | - | - | 0 | - |
| - | - | 679.9 | 350.1 | - | - | 0 | - |
| - | - | 716.1 | 352.2 | - | - | 0 | - |
| - | - | 1863 | 355.1 | - | - | 0 | - |
| - | - | 1076 | 357.2 | - | - | 0 | - |
| - | - | 550.1 | 359.2 | - | - | 0 | - |
| - | - | 587.9 | 363.6 | - | - | 0 | - |
| 3 | b | 1826 | 364.2 | 1.28E-05 | 0.03516 | +1 | 3 |
| - | - | 665.7 | 378.1 | - | - | 0 | - |
| - | - | 3458 | 380.2 | - | - | 0 | - |
| - | - | 1782 | 394.1 | - | - | 0 | - |
| - | - | 2685 | 395.2 | - | - | 0 | - |
| - | - | 2326 | 398.2 | - | - | 0 | - |
| - | - | 775 | 398.2 | - | - | 0 | - |
| - | - | 712 | 413.3 | - | - | 0 | - |
| - | - | 1481 | 425.2 | - | - | 0 | - |
| - | - | 572.2 | 425.3 | - | - | 0 | - |
| - | - | 717.5 | 436.7 | - | - | 0 | - |
| - | - | 979.1 | 449.2 | - | - | 0 | - |
| - | - | 858.7 | 451.2 | - | - | 0 | - |
| - | - | 1559 | 453.3 | - | - | 0 | - |
| - | - | 2103 | 465.2 | - | - | 0 | - |
| 4 | b | 1724 | 479.2 | 0.0001133 | 0.2364 | +1 | 4 |
| - | - | 817.2 | 480.3 | - | - | 0 | - |
| 7 | y | 1674 | 489.2 | 0.00513 | 10.49 | +1 | 3 |
| - | - | 835 | 490.2 | - | - | 0 | - |
| - | - | 940.3 | 500.8 | - | - | 0 | - |
| - | - | 803.4 | 501.3 | - | - | 0 | - |
| - | - | 2970 | 508.3 | - | - | 0 | - |
| - | - | 682.4 | 509.3 | - | - | 0 | - |
| - | - | 782.3 | 510.8 | - | - | 0 | - |
| 2 | y | 1062 | 550.7 | 0.005399 | 9.805 | +2 | 8 |
| - | - | 938.1 | 551.2 | - | - | 0 | - |
| 6 | y | 4904 | 560.2 | 0.004912 | 8.768 | +1 | 4 |
| - | - | 1200 | 561.2 | - | - | 0 | - |
| - | - | 1078 | 563.3 | - | - | 0 | - |
| - | - | 722.3 | 564.3 | - | - | 0 | - |
| - | - | 743.8 | 567.3 | - | - | 0 | - |
| - | - | 1771 | 585.3 | - | - | 0 | - |
| - | - | 905.9 | 586.3 | - | - | 0 | - |
| - | - | 1535 | 593.4 | - | - | 0 | - |
| - | - | 765.5 | 594.4 | - | - | 0 | - |
| - | - | 644.3 | 596.2 | - | - | 0 | - |
| - | - | 635.6 | 600.2 | - | - | 0 | - |
| 0 | Precursor | 2839 | 601.2 | 0.0028 | 4.658 | +2 | -1 |
| - | - | 1047 | 601.3 | - | - | 0 | - |
| - | - | 1415 | 601.7 | - | - | 0 | - |
| - | - | 841.3 | 603.4 | - | - | 0 | - |
| - | - | 1091 | 612.2 | - | - | 0 | - |
| - | - | 805.5 | 613.2 | - | - | 0 | - |
| - | - | 5798 | 621.4 | - | - | 0 | - |
| - | - | 626 | 621.8 | - | - | 0 | - |
| - | - | 1546 | 622.4 | - | - | 0 | - |
| 5 | b | 734.3 | 642.2 | 0.0002002 | 0.3118 | +1 | 5 |
| - | - | 1639 | 659.3 | - | - | 0 | - |
| - | - | 591.3 | 675.8 | - | - | 0 | - |
| - | - | 1052 | 678.4 | - | - | 0 | - |
| - | - | 672 | 678.9 | - | - | 0 | - |
| - | - | 1236 | 685.3 | - | - | 0 | - |
| - | - | 6063 | 700.4 | - | - | 0 | - |
| - | - | 1899 | 701.4 | - | - | 0 | - |
| - | - | 1271 | 721.3 | - | - | 0 | - |
| 5 | y | 4004 | 723.3 | 0.003657 | 5.055 | +1 | 5 |
| - | - | 1313 | 724.3 | - | - | 0 | - |
| - | - | 2214 | 735.4 | - | - | 0 | - |
| - | - | 877.8 | 736.4 | - | - | 0 | - |
| - | - | 1995 | 749.3 | - | - | 0 | - |
| - | - | 857.6 | 750.3 | - | - | 0 | - |
| - | - | 645 | 751.4 | - | - | 0 | - |
| - | - | 1095 | 774.3 | - | - | 0 | - |
| - | - | 634.9 | 775.3 | - | - | 0 | - |
| - | - | 639.3 | 781.9 | - | - | 0 | - |
| - | - | 1833 | 783.4 | - | - | 0 | - |
| - | - | 687.3 | 784.4 | - | - | 0 | - |
| - | - | 699.6 | 796.9 | - | - | 0 | - |
| - | - | 2442 | 801.4 | - | - | 0 | - |
| - | - | 633.1 | 809.6 | - | - | 0 | - |
| - | - | 1018 | 828.4 | - | - | 0 | - |
| 4 | y | 4131 | 838.3 | 0.004729 | 5.641 | +1 | 6 |
| - | - | 2781 | 839.3 | - | - | 0 | - |
| - | - | 4322 | 844.5 | - | - | 0 | - |
| - | - | 1852 | 845.5 | - | - | 0 | - |
| - | - | 713 | 872.3 | - | - | 0 | - |
| - | - | 1.552E+04 | 872.4 | - | - | 0 | - |
| - | - | 7551 | 873.4 | - | - | 0 | - |
| - | - | 1653 | 874.5 | - | - | 0 | - |
| - | - | 6800 | 889.3 | - | - | 0 | - |
| - | - | 2932 | 890.4 | - | - | 0 | - |
| - | - | 3153 | 901.5 | - | - | 0 | - |
| - | - | 1782 | 902.5 | - | - | 0 | - |
| - | - | 759.7 | 903.5 | - | - | 0 | - |
| - | - | 678.9 | 911.2 | - | - | 0 | - |
| - | - | 1204 | 919.5 | - | - | 0 | - |
| - | - | 832.9 | 920.5 | - | - | 0 | - |
| 3 | y | 727.2 | 935.3 | 0.006295 | 6.73 | +1 | 7 |
| 3 | y | 2.03E+04 | 953.3 | 0.004458 | 4.676 | +1 | 7 |
| - | - | 1.132E+04 | 954.3 | - | - | 0 | - |
| - | - | 3584 | 955.4 | - | - | 0 | - |
| - | - | 3527 | 991.5 | - | - | 0 | - |
| - | - | 1368 | 992.5 | - | - | 0 | - |
| - | - | 848.5 | 1019 | - | - | 0 | - |
| - | - | 6128 | 1020 | - | - | 0 | - |
| - | - | 3542 | 1021 | - | - | 0 | - |
| - | - | 1393 | 1022 | - | - | 0 | - |
| 2 | y | 1791 | 1100 | 0.001474 | 1.339 | +1 | 8 |
| - | - | 878.5 | 1101 | - | - | 0 | - |
| - | - | 929.2 | 1830 | - | - | 0 | - |
| - | - | 739.4 | 3398 | - | - | 0 | - |
| - | - | 707.2 | 3494 | - | - | 0 | - |

m/z Charge Intensity FragmentType MassShift Position
120.0810775756836 0 43042.64
121.0844497680664 0 3378.4368
123.04462432861328 0 676.3553
127.08680725097656 0 821.3827
129.10252380371094 0 3757.8982
130.06536865234375 0 1005.9776
132.08123779296875 0 1054.3945
132.10227966308594 0 718.7392
136.0760040283203 0 13572.615
137.0792236328125 0 808.13385
138.0553741455078 0 792.55206
138.0665740966797 0 754.2046
141.092529296875 0 359.53107
141.10243225097656 0 1350.0834
141.93482971191406 0 461.55997
143.0808868408203 0 481.62943
147.0442352294922 0 1173.047
147.1134033203125 0 430.0218
147.92141723632812 0 383.9926
148.53054809570312 0 441.28802
148.95465087890625 0 1021.41595
149.0452117919922 0 1091.772
155.08163452148438 0 724.153
156.0772247314453 0 1101.4363
157.0972442626953 0 923.2548
157.1337432861328 0 899.9891
158.09674072265625 0 1757.5563
159.0919647216797 0 1544.7117
159.11279296875 0 477.16904
160.11239624023438 0 1220.178
162.09164428710938 0 777.3988
163.13516235351562 0 491.41116
165.0548858642578 0 5066.8896
166.0865478515625 0 6136.2974
167.05575561523438 0 1525.6028
167.0817413330078 0 1945.9934
167.08966064453125 0 970.2409
169.09756469726562 0 1836.3223
171.15028381347656 0 520.62714
173.04335021972656 0 527.13293
173.12867736816406 0 2812.3223
173.44003295898438 0 2721.3533
174.05552673339844 0 506.3529
175.11917114257812 0 549.6468
176.10726928710938 0 3753.439
177.10256958007812 0 3300.8665
177.11048889160156 0 776.79395
178.0876922607422 0 467.23624
182.08155822753906 0 8976.668
183.08465576171875 0 1144.8331
183.11279296875 0 448.512
183.14906311035156 0 658.97687
183.18556213378906 0 490.53357
185.0924072265625 0 1244.7562
185.1288604736328 0 1307.692
185.16542053222656 0 432.54346
188.0708770751953 0 2757.0525
191.1183319091797 0 715.49536
193.10861206054688 0 1552.3868
199.1082305908203 0 1363.2733
201.12339782714844 0 1557.0271
203.11782836914062 0 1324.715 a Water loss 1
205.0971221923828 0 1869.5428 y 8
207.11331176757812 0 729.03937
211.14358520507812 0 614.19476
212.1031036376953 0 686.7825
214.11965942382812 0 834.2846
221.1035919189453 0 3103.125
221.12875366210938 0 56799.03 a 1
222.13197326660156 0 5349.8906
223.06369018554688 0 1101.4656
223.59352111816406 0 474.10043
224.1033935546875 0 1828.4014
227.10289001464844 0 4604.373
229.1182403564453 0 592.4732
229.1549835205078 0 552.1072
231.09716796875 0 895.45636
235.10792541503906 0 1792.5587
239.09561157226562 0 767.3331
241.07460021972656 0 1471.0104
242.11376953125 0 662.97723
249.12374877929688 0 9951.432 b 1
250.12705993652344 0 841.6053
251.1027374267578 0 1868.352
253.16644287109375 0 773.281
255.61032104492188 0 643.2252
257.16583251953125 0 527.6108
263.1027526855469 0 2018.568
265.15460205078125 0 1191.1691
275.17510986328125 0 575.10266
279.0971984863281 0 1237.6072
281.1247253417969 0 2535.804
283.0293273925781 0 522.20074
283.14019775390625 0 1171.8977
284.1607360839844 0 555.1034
285.01025390625 0 1167.4509
285.1019287109375 0 1166.2583
286.01031494140625 0 704.9175
292.1408996582031 0 871.1184
296.1521911621094 0 1120.8159
303.1190185546875 0 519.32153
319.13934326171875 0 786.57587
324.1456604003906 0 1537.3174
324.2262878417969 0 530.3649
329.1499938964844 0 5321.9985
330.1530456542969 0 1504.8086
340.1875305175781 0 3661.1584
342.15655517578125 0 12690.885 y 7
343.15948486328125 0 2100.0935
350.13494873046875 0 679.86426
352.1990661621094 0 716.13007
355.07037353515625 0 1862.8787
357.223388671875 0 1075.6425
359.1916198730469 0 550.0626
363.59490966796875 0 587.92804
364.1502990722656 0 1825.7084 b 2
378.1292419433594 0 665.67017
380.1929931640625 0 3457.544
394.1252746582031 0 1782.0779
395.19281005859375 0 2684.8506
398.2030944824219 0 2325.6838
398.2399597167969 0 774.99023
413.267822265625 0 711.9718
425.1947937011719 0 1481.3146
425.2771911621094 0 572.22314
436.7300109863281 0 717.4797
449.2399597167969 0 979.06396
451.2447204589844 0 858.68317
453.27191162109375 0 1559.3563
465.162353515625 0 2102.7805
479.1773681640625 0 1723.6266 b 3
480.2834777832031 0 817.21545
489.1916809082031 0 1674.3925 y 6
490.1962890625 0 834.9932
500.77734375 0 940.2617
501.2785339355469 0 803.364
508.275634765625 0 2969.7393
509.2808532714844 0 682.4384
510.7610778808594 0 782.2601
550.7136840820312 0 1062.2429 y 1
551.2103271484375 0 938.06165
560.2285766601562 0 4904.493 y 5
561.2317504882812 0 1199.9495
563.2833862304688 0 1078.4718
564.28271484375 0 722.2513
567.3170776367188 0 743.75006
585.3340454101562 0 1770.9487
586.3377685546875 0 905.86865
593.36328125 0 1535.2708
594.3695068359375 0 765.50397
596.2373046875 0 644.33624
600.2323608398438 0 635.5913
601.2349243164062 0 2839.257 Precursor
601.322509765625 0 1046.7278
601.7376098632812 0 1414.6404
603.3533325195312 0 841.3258
612.1981811523438 0 1091.3011
613.203369140625 0 805.47363
621.3605346679688 0 5797.914
621.8054809570312 0 626.03467
622.3661499023438 0 1545.7168
642.2407836914062 0 734.27466 b 4
659.2943115234375 0 1638.5974
675.7855834960938 0 591.25476
678.3824462890625 0 1052.3496
678.9306030273438 0 672.0483
685.25927734375 0 1235.7192
700.363037109375 0 6062.589
701.364501953125 0 1898.7035
721.2608032226562 0 1271.0306
723.2906494140625 0 4003.9436 y 4
724.2962646484375 0 1312.899
735.4031372070312 0 2214.2725
736.4114990234375 0 877.8113
749.25537109375 0 1994.6376
750.257568359375 0 857.61224
751.3839721679688 0 644.98627
774.3231811523438 0 1095.2909
775.3240356445312 0 634.9364
781.9374389648438 0 639.31396
783.399169921875 0 1832.584
784.3970336914062 0 687.2592
796.9010620117188 0 699.5844
801.412841796875 0 2442.0889
809.64990234375 0 633.09485
828.420654296875 0 1017.9921
838.3186645507812 0 4131.468 y 3
839.321533203125 0 2781.0479
844.4532470703125 0 4322.282
845.454833984375 0 1852.3275
872.3486938476562 0 712.9964
872.4473266601562 0 15524.353
873.44921875 0 7550.85
874.4555053710938 0 1653.1934
889.3470458984375 0 6800.19
890.3511352539062 0 2932.1248
901.4772338867188 0 3152.5374
902.4796752929688 0 1782.3823
903.4878540039062 0 759.72296
911.1678466796875 0 678.8664
919.4871826171875 0 1203.9365
920.4923095703125 0 832.8996
935.3366088867188 0 727.2058 y Water loss 2
953.3453369140625 0 20300.72 y 2
954.3477783203125 0 11323.076
955.3505859375 0 3584.1726
991.5206298828125 0 3527.2385
992.5198364257812 0 1367.7676
1018.5595703125 0 848.456
1019.517578125 0 6127.5537
1020.51953125 0 3541.6882
1021.5197143554688 0 1392.5596
1100.4107666015625 0 1790.7354 y 1
1101.4229736328125 0 878.5456
1829.8433837890625 0 929.2252
3397.75927734375 0 739.41907
3494.4677734375 0 707.20605

Spectrum Details

|  |  |
| --- | --- |
| Matched peaks? Matched peaks The total absolute number of peaks matched. Additionally in brackets the total fraction of peaks matched and the total number of peaks is shown. | 17 (7.83% of 217) |
| FDR? FDR The false discovery rate estimated for this peptide. It is calculated by matching all theoretical fragments with a non-integer shift with the raw peaks for this spectrum. This is done with 40 different shifts. The resulting percentage is the average number of annotated peaks over the number of annotated peaks with the correct spectrum. | 2.10% |
| Satellite FDR? Satellite FDR See the FDR for details on its calculation. This satellite ion specific FDR only contains the satellite ions (d/w) for I/L/J positions. | - |
| PSM Score? PSM Score The PSM Score as given by Hecklib to this annotated spectrum. It is shown with three significant figures. | 195 |

#### Spectrum 9335? Spectrum 9335 The raw spectrum of this peptide as annotated by Hecklib. The fragments are coloured according to ion type (see legend). Any peaks with a star '\*' as text can be hovered over to see the full details, first the ion type second the mass shift type. By hovering over the amino acids in the peptide or ions in the legend the corresponding peaks are highlighted. By toggling the 'Unassigned' label you can turn the background (unassigned) peaks on or off in the plot. By updating the slider in the Ion legend you can update the spectrum to only show the top X% of the peaks with labels. The top X% means any peak that is within X% of the highest intensity. By dragging in the spectrum you can zoom in to a specific part of the spectrum and use 'Zoom Out' to get back to the original zoom level. The annotation of the spectrum is based on the given sequence in the peptides file and is done with different software so inconsistencies are likely. The peaks are annotated based on the given sequence, with 20 ppm tolerance.

Copy Data

##### Spectrum 9335 (TSV)

###### Preview

```
Loading example...
```

*Click on the button to copy the data to your clipboard.*

Mz MinMz MaxIntensity Max

WidthHeightPeptide font sizePeptide stroke widthSpectrum font sizeSpectrum stroke widthCompact peptide

Ion legend

wxyz

abcd

OtherUnassignedIon^Charge^_Position_Show for top:%

TFDDYAMHW

01.72e+43.43e+45.15e+46.87e+4

Zoom Out

a^+1^_2_y^+1^_1_a^+1^_2_b^+1^_2_b^+2^_4_b^+1^_2_a^+1^_3_y^+1^_2_b^+1^_3_y^+1^_3_b^+1^_4_y^+2^_8_y^+1^_4_\*\*y^+1^_5_y^+1^_6_y^+1^_7_y^+1^_7_y^+1^_8_

0794158723813174

Fragment Matches Table

Show background peaks

| Position | Ion type | Intensity | mz Theoretical | mz Error (Th) | mz Error (ppm) | Charge | Series Number |
| --- | --- | --- | --- | --- | --- | --- | --- |
| - | - | 4.185E+04 | 120.1 | - | - | 0 | - |
| - | - | 2916 | 121.1 | - | - | 0 | - |
| - | - | 340.5 | 121.2 | - | - | 0 | - |
| - | - | 600.8 | 127.1 | - | - | 0 | - |
| - | - | 525.7 | 128.1 | - | - | 0 | - |
| - | - | 588.7 | 129.1 | - | - | 0 | - |
| - | - | 3415 | 129.1 | - | - | 0 | - |
| - | - | 1334 | 130.1 | - | - | 0 | - |
| - | - | 940.7 | 132.1 | - | - | 0 | - |
| - | - | 736.5 | 132.1 | - | - | 0 | - |
| - | - | 1514 | 133.1 | - | - | 0 | - |
| - | - | 3.164E+04 | 136.1 | - | - | 0 | - |
| - | - | 1915 | 137.1 | - | - | 0 | - |
| - | - | 668.1 | 138.1 | - | - | 0 | - |
| - | - | 1557 | 141.1 | - | - | 0 | - |
| - | - | 947.9 | 146.1 | - | - | 0 | - |
| - | - | 928.6 | 146.1 | - | - | 0 | - |
| - | - | 513.9 | 148.9 | - | - | 0 | - |
| - | - | 3084 | 149 | - | - | 0 | - |
| - | - | 415 | 149.2 | - | - | 0 | - |
| - | - | 474.7 | 151.5 | - | - | 0 | - |
| - | - | 5012 | 155.1 | - | - | 0 | - |
| - | - | 659.3 | 155.1 | - | - | 0 | - |
| - | - | 1795 | 156.1 | - | - | 0 | - |
| - | - | 690 | 157.1 | - | - | 0 | - |
| - | - | 2958 | 157.1 | - | - | 0 | - |
| - | - | 2632 | 158.1 | - | - | 0 | - |
| - | - | 503.1 | 158.4 | - | - | 0 | - |
| - | - | 2721 | 159.1 | - | - | 0 | - |
| - | - | 492 | 160.1 | - | - | 0 | - |
| - | - | 1351 | 160.1 | - | - | 0 | - |
| - | - | 1695 | 161.1 | - | - | 0 | - |
| - | - | 470.9 | 162.1 | - | - | 0 | - |
| - | - | 420.6 | 164.2 | - | - | 0 | - |
| - | - | 1.861E+04 | 166.1 | - | - | 0 | - |
| - | - | 460.9 | 166.9 | - | - | 0 | - |
| - | - | 4323 | 167.1 | - | - | 0 | - |
| - | - | 1023 | 167.1 | - | - | 0 | - |
| - | - | 2693 | 169.1 | - | - | 0 | - |
| - | - | 522.9 | 169.1 | - | - | 0 | - |
| - | - | 932.9 | 171.1 | - | - | 0 | - |
| - | - | 617.7 | 171.1 | - | - | 0 | - |
| - | - | 726.4 | 172.1 | - | - | 0 | - |
| - | - | 7423 | 173.1 | - | - | 0 | - |
| - | - | 1140 | 174.1 | - | - | 0 | - |
| - | - | 538.4 | 175.1 | - | - | 0 | - |
| - | - | 5796 | 176.1 | - | - | 0 | - |
| - | - | 4382 | 177.1 | - | - | 0 | - |
| - | - | 753 | 177.1 | - | - | 0 | - |
| - | - | 647.9 | 181.1 | - | - | 0 | - |
| - | - | 758.5 | 182 | - | - | 0 | - |
| - | - | 855.5 | 182.1 | - | - | 0 | - |
| - | - | 960.7 | 183.1 | - | - | 0 | - |
| - | - | 1465 | 183.1 | - | - | 0 | - |
| - | - | 547.8 | 184 | - | - | 0 | - |
| - | - | 567.2 | 185.1 | - | - | 0 | - |
| - | - | 923.8 | 185.1 | - | - | 0 | - |
| - | - | 612.7 | 185.2 | - | - | 0 | - |
| - | - | 5374 | 188.1 | - | - | 0 | - |
| - | - | 549.2 | 188.1 | - | - | 0 | - |
| - | - | 2919 | 189.1 | - | - | 0 | - |
| - | - | 447.6 | 190.9 | - | - | 0 | - |
| - | - | 546.8 | 191.1 | - | - | 0 | - |
| - | - | 732.1 | 192.1 | - | - | 0 | - |
| - | - | 590.1 | 195.1 | - | - | 0 | - |
| - | - | 1168 | 197.1 | - | - | 0 | - |
| - | - | 909.7 | 198.1 | - | - | 0 | - |
| - | - | 764.2 | 199.1 | - | - | 0 | - |
| - | - | 1.042E+04 | 201.1 | - | - | 0 | - |
| - | - | 1057 | 202.1 | - | - | 0 | - |
| - | - | 1599 | 203.1 | - | - | 0 | - |
| 2 | a | 1378 | 203.1 | 0.0001069 | 0.5265 | +1 | 2 |
| - | - | 641.4 | 204.1 | - | - | 0 | - |
| 9 | y | 3521 | 205.1 | 0.0001207 | 0.5884 | +1 | 1 |
| - | - | 546.6 | 205.3 | - | - | 0 | - |
| - | - | 3394 | 207.1 | - | - | 0 | - |
| - | - | 886.6 | 209.1 | - | - | 0 | - |
| - | - | 511.1 | 209.6 | - | - | 0 | - |
| - | - | 1240 | 211.1 | - | - | 0 | - |
| - | - | 797.8 | 212.1 | - | - | 0 | - |
| - | - | 2157 | 215.1 | - | - | 0 | - |
| - | - | 1879 | 217.1 | - | - | 0 | - |
| - | - | 1179 | 217.1 | - | - | 0 | - |
| - | - | 2.156E+04 | 219.1 | - | - | 0 | - |
| - | - | 2183 | 220.1 | - | - | 0 | - |
| - | - | 4238 | 221.1 | - | - | 0 | - |
| 2 | a | 6.802E+04 | 221.1 | 0.0001163 | 0.5261 | +1 | 2 |
| - | - | 835.9 | 222.1 | - | - | 0 | - |
| - | - | 8217 | 222.1 | - | - | 0 | - |
| - | - | 2792 | 223.1 | - | - | 0 | - |
| - | - | 658.8 | 223.1 | - | - | 0 | - |
| - | - | 1380 | 225 | - | - | 0 | - |
| - | - | 1.16E+04 | 226.1 | - | - | 0 | - |
| - | - | 617.7 | 226.2 | - | - | 0 | - |
| - | - | 3440 | 227 | - | - | 0 | - |
| - | - | 1029 | 227.1 | - | - | 0 | - |
| - | - | 1009 | 227.1 | - | - | 0 | - |
| - | - | 1126 | 229.1 | - | - | 0 | - |
| - | - | 781.6 | 229.2 | - | - | 0 | - |
| - | - | 1181 | 231.1 | - | - | 0 | - |
| 2 | b | 870 | 231.1 | 4.171E-05 | 0.1805 | +1 | 2 |
| - | - | 526.9 | 233.1 | - | - | 0 | - |
| - | - | 1064 | 233.2 | - | - | 0 | - |
| - | - | 5518 | 235.1 | - | - | 0 | - |
| - | - | 877.1 | 236.1 | - | - | 0 | - |
| - | - | 649.1 | 237 | - | - | 0 | - |
| - | - | 586 | 237.1 | - | - | 0 | - |
| - | - | 1.25E+04 | 239.1 | - | - | 0 | - |
| 4 | b | 2050 | 240.1 | 0.003193 | 13.3 | +2 | 4 |
| - | - | 663.4 | 240.1 | - | - | 0 | - |
| - | - | 1442 | 241.1 | - | - | 0 | - |
| - | - | 668 | 243.1 | - | - | 0 | - |
| - | - | 515.5 | 243.1 | - | - | 0 | - |
| - | - | 992 | 244.1 | - | - | 0 | - |
| - | - | 1068 | 247.1 | - | - | 0 | - |
| 2 | b | 1.188E+04 | 249.1 | 0.0001663 | 0.6676 | +1 | 2 |
| - | - | 1135 | 250.1 | - | - | 0 | - |
| - | - | 2594 | 251.1 | - | - | 0 | - |
| - | - | 747.1 | 252.1 | - | - | 0 | - |
| - | - | 1.214E+04 | 253.1 | - | - | 0 | - |
| - | - | 771.8 | 254.1 | - | - | 0 | - |
| - | - | 1861 | 255 | - | - | 0 | - |
| - | - | 607.5 | 256.1 | - | - | 0 | - |
| - | - | 659.2 | 260.1 | - | - | 0 | - |
| - | - | 3010 | 263.1 | - | - | 0 | - |
| - | - | 887.7 | 265.1 | - | - | 0 | - |
| - | - | 737.8 | 266.1 | - | - | 0 | - |
| - | - | 1440 | 269 | - | - | 0 | - |
| - | - | 4415 | 269.1 | - | - | 0 | - |
| - | - | 658 | 270 | - | - | 0 | - |
| - | - | 816.3 | 278 | - | - | 0 | - |
| - | - | 654.7 | 279 | - | - | 0 | - |
| - | - | 794.3 | 279.1 | - | - | 0 | - |
| - | - | 1557 | 281.1 | - | - | 0 | - |
| - | - | 1098 | 282.2 | - | - | 0 | - |
| - | - | 1220 | 283 | - | - | 0 | - |
| - | - | 2135 | 283.1 | - | - | 0 | - |
| - | - | 1086 | 284.2 | - | - | 0 | - |
| - | - | 2.321E+04 | 285 | - | - | 0 | - |
| - | - | 3976 | 286 | - | - | 0 | - |
| - | - | 727.8 | 291.2 | - | - | 0 | - |
| - | - | 1977 | 296 | - | - | 0 | - |
| - | - | 1576 | 296.2 | - | - | 0 | - |
| - | - | 5862 | 297.2 | - | - | 0 | - |
| - | - | 2377 | 298.1 | - | - | 0 | - |
| - | - | 1019 | 298.2 | - | - | 0 | - |
| - | - | 1.432E+04 | 299.1 | - | - | 0 | - |
| - | - | 1122 | 299.1 | - | - | 0 | - |
| - | - | 3245 | 300.1 | - | - | 0 | - |
| - | - | 822.2 | 300.2 | - | - | 0 | - |
| - | - | 2686 | 303 | - | - | 0 | - |
| - | - | 2255 | 306 | - | - | 0 | - |
| - | - | 1304 | 311.1 | - | - | 0 | - |
| - | - | 691.6 | 311.2 | - | - | 0 | - |
| - | - | 803.8 | 312.2 | - | - | 0 | - |
| - | - | 982 | 316.1 | - | - | 0 | - |
| - | - | 3449 | 316.2 | - | - | 0 | - |
| 3 | a | 1559 | 318.1 | 0.0005532 | 1.739 | +1 | 3 |
| - | - | 1067 | 318.2 | - | - | 0 | - |
| - | - | 4446 | 324 | - | - | 0 | - |
| - | - | 3232 | 324.1 | - | - | 0 | - |
| - | - | 7029 | 325.2 | - | - | 0 | - |
| - | - | 889.3 | 326.2 | - | - | 0 | - |
| - | - | 1687 | 330.2 | - | - | 0 | - |
| - | - | 574.3 | 333.2 | - | - | 0 | - |
| - | - | 749.6 | 334.1 | - | - | 0 | - |
| - | - | 1529 | 338.1 | - | - | 0 | - |
| - | - | 569.4 | 339 | - | - | 0 | - |
| - | - | 622.6 | 339.2 | - | - | 0 | - |
| - | - | 1537 | 340.1 | - | - | 0 | - |
| - | - | 864.6 | 340.2 | - | - | 0 | - |
| - | - | 638.8 | 341.1 | - | - | 0 | - |
| - | - | 2122 | 342 | - | - | 0 | - |
| 8 | y | 1.666E+04 | 342.2 | 9.249E-05 | 0.2703 | +1 | 2 |
| - | - | 3564 | 343.2 | - | - | 0 | - |
| - | - | 2248 | 345 | - | - | 0 | - |
| - | - | 685.7 | 346 | - | - | 0 | - |
| - | - | 6037 | 350.1 | - | - | 0 | - |
| - | - | 898 | 351.1 | - | - | 0 | - |
| - | - | 603.1 | 351.2 | - | - | 0 | - |
| - | - | 1660 | 354.2 | - | - | 0 | - |
| - | - | 5.509E+04 | 355.1 | - | - | 0 | - |
| - | - | 1080 | 355.2 | - | - | 0 | - |
| - | - | 1.639E+04 | 356.1 | - | - | 0 | - |
| - | - | 1312 | 362.1 | - | - | 0 | - |
| 3 | b | 2473 | 364.2 | 0.0001959 | 0.538 | +1 | 3 |
| - | - | 865.7 | 366.1 | - | - | 0 | - |
| - | - | 1042 | 371.1 | - | - | 0 | - |
| - | - | 4412 | 373.1 | - | - | 0 | - |
| - | - | 1516 | 374.1 | - | - | 0 | - |
| - | - | 2275 | 378.1 | - | - | 0 | - |
| - | - | 946.6 | 380.1 | - | - | 0 | - |
| - | - | 573.3 | 381.1 | - | - | 0 | - |
| - | - | 612 | 382.2 | - | - | 0 | - |
| - | - | 853.1 | 389.2 | - | - | 0 | - |
| - | - | 4347 | 394.1 | - | - | 0 | - |
| - | - | 989.7 | 395.1 | - | - | 0 | - |
| - | - | 714 | 398.2 | - | - | 0 | - |
| - | - | 966.9 | 401 | - | - | 0 | - |
| - | - | 697.8 | 405.2 | - | - | 0 | - |
| - | - | 2095 | 408.2 | - | - | 0 | - |
| - | - | 779.3 | 412.2 | - | - | 0 | - |
| - | - | 1654 | 413.2 | - | - | 0 | - |
| - | - | 1773 | 415 | - | - | 0 | - |
| - | - | 6569 | 419 | - | - | 0 | - |
| - | - | 1820 | 420 | - | - | 0 | - |
| - | - | 979.1 | 423.7 | - | - | 0 | - |
| - | - | 1491 | 424.7 | - | - | 0 | - |
| - | - | 844.9 | 426.2 | - | - | 0 | - |
| - | - | 1069 | 426.2 | - | - | 0 | - |
| - | - | 2693 | 426.2 | - | - | 0 | - |
| - | - | 709.6 | 427.2 | - | - | 0 | - |
| - | - | 1214 | 431.2 | - | - | 0 | - |
| - | - | 987 | 434.2 | - | - | 0 | - |
| - | - | 1646 | 435.2 | - | - | 0 | - |
| - | - | 965.6 | 438.3 | - | - | 0 | - |
| - | - | 600.4 | 447.2 | - | - | 0 | - |
| - | - | 705.6 | 449.2 | - | - | 0 | - |
| - | - | 787.7 | 459.1 | - | - | 0 | - |
| - | - | 754.3 | 460.3 | - | - | 0 | - |
| - | - | 2237 | 462.3 | - | - | 0 | - |
| - | - | 1152 | 462.8 | - | - | 0 | - |
| - | - | 1826 | 463.2 | - | - | 0 | - |
| - | - | 604.3 | 463.3 | - | - | 0 | - |
| - | - | 4247 | 465.2 | - | - | 0 | - |
| 7 | y | 3459 | 473.2 | 0.0001394 | 0.2947 | +1 | 3 |
| - | - | 1226 | 474.2 | - | - | 0 | - |
| - | - | 650.7 | 478.8 | - | - | 0 | - |
| 4 | b | 1254 | 479.2 | 8.274E-05 | 0.1727 | +1 | 4 |
| - | - | 1593 | 485.2 | - | - | 0 | - |
| - | - | 1824 | 487.1 | - | - | 0 | - |
| - | - | 700.5 | 489.1 | - | - | 0 | - |
| - | - | 766.4 | 495.2 | - | - | 0 | - |
| - | - | 853.3 | 500.2 | - | - | 0 | - |
| - | - | 729.7 | 504.2 | - | - | 0 | - |
| - | - | 3961 | 513.2 | - | - | 0 | - |
| - | - | 1696 | 514.2 | - | - | 0 | - |
| - | - | 639.6 | 522.8 | - | - | 0 | - |
| - | - | 2588 | 525.3 | - | - | 0 | - |
| - | - | 849.5 | 526.3 | - | - | 0 | - |
| 2 | y | 1475 | 542.7 | 0.0007602 | 1.401 | +2 | 8 |
| - | - | 598 | 542.8 | - | - | 0 | - |
| - | - | 1.457E+04 | 543.3 | - | - | 0 | - |
| 6 | y | 4391 | 544.2 | 0.0002049 | 0.3765 | +1 | 4 |
| - | - | 4098 | 544.3 | - | - | 0 | - |
| - | - | 1799 | 545.2 | - | - | 0 | - |
| - | - | 790.3 | 553.3 | - | - | 0 | - |
| - | - | 676.2 | 553.8 | - | - | 0 | - |
| - | - | 935.3 | 559.3 | - | - | 0 | - |
| - | - | 839.9 | 562.3 | - | - | 0 | - |
| - | - | 1140 | 571.3 | - | - | 0 | - |
| - | - | 813.8 | 575.1 | - | - | 0 | - |
| - | - | 660.5 | 581.3 | - | - | 0 | - |
| 0 | Precursor | 1349 | 584.2 | 0.001922 | 3.29 | +2 | -1 |
| - | - | 834.6 | 590.2 | - | - | 0 | - |
| - | - | 885.1 | 592.9 | - | - | 0 | - |
| - | - | 836.6 | 593.1 | - | - | 0 | - |
| 0 | Precursor | 972.7 | 593.2 | 0.001772 | 2.988 | +2 | -1 |
| - | - | 1186 | 593.3 | - | - | 0 | - |
| - | - | 693.2 | 593.7 | - | - | 0 | - |
| - | - | 690.9 | 596.2 | - | - | 0 | - |
| - | - | 1936 | 598.3 | - | - | 0 | - |
| - | - | 950.3 | 599.3 | - | - | 0 | - |
| - | - | 1011 | 606.2 | - | - | 0 | - |
| - | - | 842.1 | 608.3 | - | - | 0 | - |
| - | - | 3842 | 626.3 | - | - | 0 | - |
| - | - | 1136 | 627.3 | - | - | 0 | - |
| - | - | 655.2 | 679.4 | - | - | 0 | - |
| - | - | 816.3 | 697.4 | - | - | 0 | - |
| - | - | 1584 | 705.3 | - | - | 0 | - |
| - | - | 1016 | 706.4 | - | - | 0 | - |
| 5 | y | 4936 | 707.3 | 0.001827 | 2.583 | +1 | 5 |
| - | - | 765.8 | 707.4 | - | - | 0 | - |
| - | - | 2224 | 708.3 | - | - | 0 | - |
| - | - | 3951 | 733.3 | - | - | 0 | - |
| - | - | 786.3 | 734.3 | - | - | 0 | - |
| - | - | 930 | 738.3 | - | - | 0 | - |
| - | - | 779.7 | 749.4 | - | - | 0 | - |
| - | - | 935.5 | 751.3 | - | - | 0 | - |
| - | - | 779.9 | 780.4 | - | - | 0 | - |
| - | - | 663 | 798.9 | - | - | 0 | - |
| 4 | y | 5220 | 822.3 | 0.001426 | 1.734 | +1 | 6 |
| - | - | 2084 | 823.3 | - | - | 0 | - |
| - | - | 1009 | 823.4 | - | - | 0 | - |
| - | - | 895.9 | 824.4 | - | - | 0 | - |
| - | - | 1973 | 846.5 | - | - | 0 | - |
| - | - | 1039 | 848.5 | - | - | 0 | - |
| - | - | 5318 | 851.4 | - | - | 0 | - |
| - | - | 2174 | 852.4 | - | - | 0 | - |
| - | - | 815.8 | 872.4 | - | - | 0 | - |
| - | - | 929.8 | 873.5 | - | - | 0 | - |
| 3 | y | 1694 | 919.3 | 0.003156 | 3.433 | +1 | 7 |
| - | - | 837.7 | 920.3 | - | - | 0 | - |
| - | - | 719.5 | 922.5 | - | - | 0 | - |
| - | - | 875 | 923.5 | - | - | 0 | - |
| - | - | 850.2 | 932.5 | - | - | 0 | - |
| 3 | y | 2.82E+04 | 937.4 | 0.002124 | 2.266 | +1 | 7 |
| - | - | 1.565E+04 | 938.4 | - | - | 0 | - |
| - | - | 5033 | 939.4 | - | - | 0 | - |
| - | - | 833.4 | 940.4 | - | - | 0 | - |
| - | - | 4634 | 950.5 | - | - | 0 | - |
| - | - | 2982 | 951.5 | - | - | 0 | - |
| - | - | 753.2 | 954.4 | - | - | 0 | - |
| - | - | 867.2 | 1020 | - | - | 0 | - |
| - | - | 1663 | 1021 | - | - | 0 | - |
| - | - | 1064 | 1033 | - | - | 0 | - |
| - | - | 866.7 | 1053 | - | - | 0 | - |
| 2 | y | 2137 | 1084 | 0.004986 | 4.598 | +1 | 8 |
| - | - | 1522 | 1085 | - | - | 0 | - |
| - | - | 831.8 | 1249 | - | - | 0 | - |
| - | - | 742.3 | 1894 | - | - | 0 | - |
| - | - | 612.9 | 2514 | - | - | 0 | - |
| - | - | 669.5 | 2834 | - | - | 0 | - |
| - | - | 729.2 | 3143 | - | - | 0 | - |

m/z Charge Intensity FragmentType MassShift Position
120.0810317993164 0 41851.85
121.08446502685547 0 2915.6294
121.1670913696289 0 340.46545
127.08702087402344 0 600.7828
128.1072235107422 0 525.6824
129.06605529785156 0 588.7019
129.10247802734375 0 3415.4639
130.06541442871094 0 1334.0564
132.0809326171875 0 940.6658
132.10191345214844 0 736.4537
133.08596801757812 0 1513.9929
136.075927734375 0 31644.715
137.0792236328125 0 1914.9309
138.0666046142578 0 668.1016
141.10235595703125 0 1556.5034
146.06040954589844 0 947.9097
146.09666442871094 0 928.58624
148.9451446533203 0 513.919
149.0451202392578 0 3084.4365
149.18589782714844 0 414.98123
151.5478515625 0 474.74118
155.08172607421875 0 5012.2505
155.11790466308594 0 659.3069
156.07684326171875 0 1795.4706
157.06082153320312 0 690.02167
157.13369750976562 0 2957.5388
158.09674072265625 0 2632.4988
158.41229248046875 0 503.0994
159.09178161621094 0 2721.0662
160.07615661621094 0 491.9587
160.1123504638672 0 1351.1947
161.09222412109375 0 1695.1129
162.0914306640625 0 470.9252
164.188720703125 0 420.56152
166.0864715576172 0 18607.78
166.88368225097656 0 460.93253
167.05563354492188 0 4323.136
167.08985900878906 0 1022.77563
169.0972900390625 0 2693.061
169.13363647460938 0 522.8982
171.0768585205078 0 932.8598
171.11288452148438 0 617.68665
172.10873413085938 0 726.38806
173.12857055664062 0 7423.105
174.13204956054688 0 1140.1797
175.0886688232422 0 538.427
176.10728454589844 0 5796.313
177.10226440429688 0 4381.5728
177.1108856201172 0 752.9554
181.09751892089844 0 647.9227
181.9940185546875 0 758.49915
182.08160400390625 0 855.4803
183.11318969726562 0 960.70276
183.1492462158203 0 1465.4064
183.96005249023438 0 547.8101
185.05531311035156 0 567.16925
185.12863159179688 0 923.84674
185.1647491455078 0 612.7156
188.0708465576172 0 5373.8853
188.14308166503906 0 549.1689
189.08705139160156 0 2919.136
190.91184997558594 0 447.63135
191.1180877685547 0 546.7622
192.1019744873047 0 732.13226
195.1123809814453 0 590.11
197.12879943847656 0 1168.3528
198.1238250732422 0 909.74274
199.1077880859375 0 764.22064
201.12355041503906 0 10416.042
202.1261444091797 0 1056.6327
203.0664825439453 0 1598.9076
203.11778259277344 0 1378.389 a Water loss 1
204.1387939453125 0 641.4115
205.09727478027344 0 3521.0718 y 8
205.30162048339844 0 546.56354
207.1128387451172 0 3393.6624
209.0924530029297 0 886.64453
209.5644073486328 0 511.06967
211.14427185058594 0 1239.7157
212.10231018066406 0 797.8156
215.139404296875 0 2156.5356
217.0821533203125 0 1878.5139
217.0973358154297 0 1178.5854
219.1340789794922 0 21560.889
220.13743591308594 0 2182.7207
221.08450317382812 0 4237.9854
221.12857055664062 0 68015.04 a 1
222.086181640625 0 835.8859
222.13192749023438 0 8217.242
223.1075439453125 0 2792.4995
223.13482666015625 0 658.7553
225.0430145263672 0 1379.7649
226.11871337890625 0 11600.423
226.1545867919922 0 617.7049
227.01565551757812 0 3440.3787
227.1023406982422 0 1029.2463
227.12261962890625 0 1009.4418
229.11807250976562 0 1126.0216
229.15579223632812 0 781.61945
231.06085205078125 0 1181.1805
231.11276245117188 0 870.02563 b Water loss 1
233.09202575683594 0 526.9367
233.16500854492188 0 1063.9003
235.10775756835938 0 5518.1953
236.112060546875 0 877.0815
236.9998779296875 0 649.1156
237.12322998046875 0 585.97437
239.09507751464844 0 12501.774
240.095458984375 0 2050.2905 b 3
240.13385009765625 0 663.37604
241.1116180419922 0 1441.9863
243.13368225097656 0 667.9726
243.14712524414062 0 515.5365
244.12969970703125 0 992.0008
247.10800170898438 0 1067.805
249.12353515625 0 11875.557 b 1
250.1266632080078 0 1135.117
251.10269165039062 0 2593.8093
252.0799102783203 0 747.1022
253.1184539794922 0 12140.877
254.1220703125 0 771.75494
255.01077270507812 0 1861.4392
256.1097717285156 0 607.5442
260.1064758300781 0 659.193
263.10260009765625 0 3010.0686
265.1180419921875 0 887.6769
266.1490478515625 0 737.7815
268.9788513183594 0 1440.2206
269.1064147949219 0 4415.183
269.97882080078125 0 658.0077
278.0260925292969 0 816.335
279.0109558105469 0 654.6856
279.09698486328125 0 794.32526
281.05157470703125 0 1556.8628
282.1814880371094 0 1098.1486
283.03021240234375 0 1220.3774
283.14385986328125 0 2135.3945
284.16180419921875 0 1085.7849
285.009765625 0 23209.012
286.00994873046875 0 3975.8264
291.1707763671875 0 727.83356
296.036865234375 0 1976.8925
296.15106201171875 0 1576.2833
297.1923522949219 0 5861.789
298.1397705078125 0 2376.7246
298.1960754394531 0 1018.57837
299.061767578125 0 14321.801
299.0802001953125 0 1122.1888
300.06243896484375 0 3245.4475
300.19122314453125 0 822.1746
303.0206298828125 0 2686.1985
306.0212707519531 0 2254.8984
311.13873291015625 0 1303.9299
311.20819091796875 0 691.5997
312.1545715332031 0 803.7881
316.13104248046875 0 982.0309
316.1503601074219 0 3449.22
318.1453857421875 0 1558.6398 a Water loss 2
318.2021484375 0 1067.3463
324.03216552734375 0 4446.008
324.1452331542969 0 3232.1677
325.1871643066406 0 7028.9097
326.19091796875 0 889.27527
330.1659240722656 0 1687.1609
333.155029296875 0 574.26154
334.13983154296875 0 749.5773
338.1349792480469 0 1529.3505
339.0411071777344 0 569.3946
339.2021484375 0 622.587
340.1434631347656 0 1536.7389
340.1863708496094 0 864.59875
341.14678955078125 0 638.829
342.0424499511719 0 2122.2979
342.1561584472656 0 16655.535 y 7
343.1587219238281 0 3564.37
344.97637939453125 0 2248.3855
345.97454833984375 0 685.66876
350.1343994140625 0 6037.1743
351.13848876953125 0 897.96027
351.16583251953125 0 603.12573
354.1661071777344 0 1660.2795
355.0697326660156 0 55087.727
355.1622619628906 0 1080.3489
356.0704040527344 0 16390.826
362.1343994140625 0 1312.2552
364.1501159667969 0 2472.731 b 2
366.1296081542969 0 865.6573
371.1000061035156 0 1041.7875
373.0802001953125 0 4411.7915
374.08111572265625 0 1515.5991
378.12884521484375 0 2275.0193
380.1448669433594 0 946.57074
381.1493835449219 0 573.3334
382.243408203125 0 611.9975
389.18206787109375 0 853.096
394.12432861328125 0 4346.6377
395.1292724609375 0 989.7488
398.2406921386719 0 714.01263
400.98468017578125 0 966.8907
405.2147216796875 0 697.794
408.2238464355469 0 2094.5386
412.21820068359375 0 779.3411
413.2036437988281 0 1653.7448
415.0356750488281 0 1772.6318
418.9949951171875 0 6568.8315
419.9952697753906 0 1820.4233
423.7364501953125 0 979.08105
424.72918701171875 0 1490.7377
426.1654357910156 0 844.8861
426.200927734375 0 1068.8931
426.2347717285156 0 2692.9062
427.2372131347656 0 709.56635
431.21514892578125 0 1214.2592
434.19525146484375 0 986.97723
435.2237854003906 0 1646.2795
438.271728515625 0 965.6009
447.1549072265625 0 600.3899
449.16705322265625 0 705.6204
459.1007080078125 0 787.73773
460.2564697265625 0 754.25775
462.264404296875 0 2237.3804
462.7630920410156 0 1151.9861
463.2188720703125 0 1826.2717
463.2665710449219 0 604.28046
465.1619567871094 0 4247.4365
473.1964111328125 0 3458.9978 y 6
474.20001220703125 0 1225.6761
478.83160400390625 0 650.66345
479.1773376464844 0 1254.2832 b 3
485.202392578125 0 1593.0109
487.0954284667969 0 1824.0098
489.05438232421875 0 700.4885
495.18621826171875 0 766.4224
500.2352600097656 0 853.3397
504.21295166015625 0 729.6921
513.197265625 0 3961.0662
514.201171875 0 1695.6384
522.7973022460938 0 639.60724
525.3032836914062 0 2587.7388
526.2604370117188 0 849.52466
542.7125244140625 0 1474.815 y 1
542.7623901367188 0 598.045
543.3131103515625 0 14568.361
544.2334594726562 0 4390.9243 y 5
544.31591796875 0 4097.52
545.235107421875 0 1798.8235
553.2719116210938 0 790.34595
553.7818603515625 0 676.16473
559.2505493164062 0 935.30334
562.2508544921875 0 839.9008
571.28466796875 0 1139.7705
575.1188354492188 0 813.7692
581.2603149414062 0 660.5291
584.2299194335938 0 1348.608 Precursor Water loss
590.2426147460938 0 834.6342
592.9386596679688 0 885.07104
593.137939453125 0 836.6082
593.2353515625 0 972.65247 Precursor
593.3395385742188 0 1186.181
593.7373046875 0 693.17316
596.1978149414062 0 690.8974
598.2850952148438 0 1936.4019
599.2910766601562 0 950.2952
606.1541137695312 0 1010.804
608.2737426757812 0 842.10345
626.2814331054688 0 3841.6917
627.281494140625 0 1135.5562
679.3740234375 0 655.1914
697.3732299804688 0 816.33
705.2631225585938 0 1583.7554
706.3734741210938 0 1016.2264
707.295166015625 0 4935.6357 y 4
707.3658447265625 0 765.8029
708.2982788085938 0 2223.652
733.2593383789062 0 3951.443
734.2633056640625 0 786.3308
738.308837890625 0 930.0021
749.4140014648438 0 779.70496
751.2750854492188 0 935.453
780.3554077148438 0 779.9322
798.9137573242188 0 662.96265
822.322509765625 0 5219.5503 y 3
823.3263549804688 0 2083.7534
823.4002685546875 0 1009.2438
824.40869140625 0 895.8928
846.4682006835938 0 1972.9744
848.4514770507812 0 1039.3643
851.3919677734375 0 5317.595
852.3927612304688 0 2173.781
872.4454345703125 0 815.83636
873.4536743164062 0 929.8445
919.337158203125 0 1694.062 y Water loss 2
920.333251953125 0 837.729
922.46142578125 0 719.5238
923.5283203125 0 874.95984
932.4539794921875 0 850.17346
937.3487548828125 0 28199.57 y 2
938.3514404296875 0 15652.63
939.351318359375 0 5032.7983
940.3512573242188 0 833.42755
950.4601440429688 0 4633.833
951.460205078125 0 2981.8413
954.4361572265625 0 753.21
1019.50927734375 0 867.2297
1020.5157470703125 0 1662.9773
1033.498291015625 0 1063.5386
1052.513427734375 0 866.66705
1084.414306640625 0 2136.8047 y 1
1085.4150390625 0 1521.736
1248.5721435546875 0 831.7762
1894.146240234375 0 742.34045
2514.22509765625 0 612.9059
2834.08349609375 0 669.49963
3142.83544921875 0 729.2367

Spectrum Details

|  |  |
| --- | --- |
| Matched peaks? Matched peaks The total absolute number of peaks matched. Additionally in brackets the total fraction of peaks matched and the total number of peaks is shown. | 20 (6.37% of 314) |
| FDR? FDR The false discovery rate estimated for this peptide. It is calculated by matching all theoretical fragments with a non-integer shift with the raw peaks for this spectrum. This is done with 40 different shifts. The resulting percentage is the average number of annotated peaks over the number of annotated peaks with the correct spectrum. | 3.33% |
| Satellite FDR? Satellite FDR See the FDR for details on its calculation. This satellite ion specific FDR only contains the satellite ions (d/w) for I/L/J positions. | - |
| PSM Score? PSM Score The PSM Score as given by Hecklib to this annotated spectrum. It is shown with three significant figures. | 213 |

#### Spectrum 12075? Spectrum 12075 The raw spectrum of this peptide as annotated by Hecklib. The fragments are coloured according to ion type (see legend). Any peaks with a star '\*' as text can be hovered over to see the full details, first the ion type second the mass shift type. By hovering over the amino acids in the peptide or ions in the legend the corresponding peaks are highlighted. By toggling the 'Unassigned' label you can turn the background (unassigned) peaks on or off in the plot. By updating the slider in the Ion legend you can update the spectrum to only show the top X% of the peaks with labels. The top X% means any peak that is within X% of the highest intensity. By dragging in the spectrum you can zoom in to a specific part of the spectrum and use 'Zoom Out' to get back to the original zoom level. The annotation of the spectrum is based on the given sequence in the peptides file and is done with different software so inconsistencies are likely. The peaks are annotated based on the given sequence, with 20 ppm tolerance.

Copy Data

##### Spectrum 12075 (TSV)

###### Preview

```
Loading example...
```

*Click on the button to copy the data to your clipboard.*

Mz MinMz MaxIntensity Max

WidthHeightPeptide font sizePeptide stroke widthSpectrum font sizeSpectrum stroke widthCompact peptide

Ion legend

wxyz

abcd

OtherUnassignedIon^Charge^_Position_Show for top:%

TFDDYAMHW

01.19e+42.38e+43.57e+44.76e+4

Zoom Out

a^+1^_2_y^+1^_1_a^+1^_2_b^+1^_2_y^+1^_2_b^+1^_3_b^+1^_4_y^+1^_3_y^+2^_8_y^+1^_4_\*y^+1^_5_y^+1^_6_y^+1^_7_y^+1^_8_

0778155523333111

Fragment Matches Table

Show background peaks

| Position | Ion type | Intensity | mz Theoretical | mz Error (Th) | mz Error (ppm) | Charge | Series Number |
| --- | --- | --- | --- | --- | --- | --- | --- |
| - | - | 3.042E+04 | 120.1 | - | - | 0 | - |
| - | - | 2433 | 121.1 | - | - | 0 | - |
| - | - | 374.6 | 121.4 | - | - | 0 | - |
| - | - | 480.3 | 123 | - | - | 0 | - |
| - | - | 3175 | 129.1 | - | - | 0 | - |
| - | - | 502.3 | 130.1 | - | - | 0 | - |
| - | - | 758.5 | 132.1 | - | - | 0 | - |
| - | - | 496.8 | 132.1 | - | - | 0 | - |
| - | - | 2556 | 133.1 | - | - | 0 | - |
| - | - | 1.133E+04 | 136.1 | - | - | 0 | - |
| - | - | 802 | 137.1 | - | - | 0 | - |
| - | - | 458.3 | 138.1 | - | - | 0 | - |
| - | - | 828.5 | 146.1 | - | - | 0 | - |
| - | - | 529.5 | 147.1 | - | - | 0 | - |
| - | - | 484.5 | 149 | - | - | 0 | - |
| - | - | 774.1 | 156.1 | - | - | 0 | - |
| - | - | 1518 | 158.1 | - | - | 0 | - |
| - | - | 1086 | 159.1 | - | - | 0 | - |
| - | - | 433.1 | 159.6 | - | - | 0 | - |
| - | - | 599.8 | 160.1 | - | - | 0 | - |
| - | - | 2534 | 165.1 | - | - | 0 | - |
| - | - | 1973 | 166.1 | - | - | 0 | - |
| - | - | 2025 | 167.1 | - | - | 0 | - |
| - | - | 961.8 | 167.1 | - | - | 0 | - |
| - | - | 428.2 | 169.1 | - | - | 0 | - |
| - | - | 2066 | 173.1 | - | - | 0 | - |
| - | - | 3014 | 176.1 | - | - | 0 | - |
| - | - | 2830 | 177.1 | - | - | 0 | - |
| - | - | 599.4 | 177.1 | - | - | 0 | - |
| - | - | 7666 | 182.1 | - | - | 0 | - |
| - | - | 567.1 | 183.9 | - | - | 0 | - |
| - | - | 548.6 | 185.4 | - | - | 0 | - |
| - | - | 1261 | 188.1 | - | - | 0 | - |
| - | - | 572.3 | 191.1 | - | - | 0 | - |
| - | - | 620 | 193.1 | - | - | 0 | - |
| - | - | 559.4 | 197.1 | - | - | 0 | - |
| - | - | 610.2 | 199.1 | - | - | 0 | - |
| - | - | 916.3 | 201.1 | - | - | 0 | - |
| 2 | a | 1212 | 203.1 | 0.0001067 | 0.5252 | +1 | 2 |
| 9 | y | 926.8 | 205.1 | 0.0007616 | 3.713 | +1 | 1 |
| - | - | 602.5 | 209.1 | - | - | 0 | - |
| - | - | 612.8 | 213.6 | - | - | 0 | - |
| - | - | 567.3 | 214.1 | - | - | 0 | - |
| - | - | 688.2 | 217.1 | - | - | 0 | - |
| - | - | 951.3 | 217.1 | - | - | 0 | - |
| - | - | 2462 | 221.1 | - | - | 0 | - |
| 2 | a | 4.71E+04 | 221.1 | 4.004E-05 | 0.1811 | +1 | 2 |
| - | - | 6109 | 222.1 | - | - | 0 | - |
| - | - | 1455 | 223.1 | - | - | 0 | - |
| - | - | 510.6 | 223.1 | - | - | 0 | - |
| - | - | 801.6 | 227.1 | - | - | 0 | - |
| - | - | 655.1 | 229.2 | - | - | 0 | - |
| - | - | 831.5 | 231.1 | - | - | 0 | - |
| - | - | 811.8 | 231.1 | - | - | 0 | - |
| - | - | 1131 | 235.1 | - | - | 0 | - |
| - | - | 536.1 | 237.7 | - | - | 0 | - |
| - | - | 920.4 | 239.1 | - | - | 0 | - |
| - | - | 553.8 | 241.1 | - | - | 0 | - |
| - | - | 530 | 246.1 | - | - | 0 | - |
| 2 | b | 9096 | 249.1 | 9.002E-05 | 0.3613 | +1 | 2 |
| - | - | 1142 | 250.1 | - | - | 0 | - |
| - | - | 1058 | 251.1 | - | - | 0 | - |
| - | - | 646.6 | 251.2 | - | - | 0 | - |
| - | - | 1308 | 263.1 | - | - | 0 | - |
| - | - | 1326 | 264.2 | - | - | 0 | - |
| - | - | 571.3 | 274.1 | - | - | 0 | - |
| - | - | 1321 | 285 | - | - | 0 | - |
| - | - | 999.7 | 285.1 | - | - | 0 | - |
| - | - | 892.1 | 296.2 | - | - | 0 | - |
| - | - | 602 | 319.1 | - | - | 0 | - |
| - | - | 1194 | 324.1 | - | - | 0 | - |
| - | - | 637.8 | 324.7 | - | - | 0 | - |
| - | - | 665.4 | 325.1 | - | - | 0 | - |
| - | - | 4995 | 329.1 | - | - | 0 | - |
| - | - | 556 | 340.2 | - | - | 0 | - |
| - | - | 607.4 | 342.1 | - | - | 0 | - |
| 8 | y | 9788 | 342.2 | 0.0002146 | 0.6271 | +1 | 2 |
| - | - | 1438 | 343.2 | - | - | 0 | - |
| - | - | 545.8 | 354 | - | - | 0 | - |
| - | - | 1229 | 355.1 | - | - | 0 | - |
| - | - | 709.2 | 356.1 | - | - | 0 | - |
| - | - | 1534 | 357.2 | - | - | 0 | - |
| 3 | b | 1496 | 364.2 | 0.0004755 | 1.306 | +1 | 3 |
| - | - | 773.7 | 378.1 | - | - | 0 | - |
| - | - | 610.7 | 380.2 | - | - | 0 | - |
| - | - | 1803 | 394.1 | - | - | 0 | - |
| - | - | 612.8 | 398.2 | - | - | 0 | - |
| - | - | 2920 | 413.3 | - | - | 0 | - |
| - | - | 809.1 | 414.3 | - | - | 0 | - |
| - | - | 819.1 | 425.2 | - | - | 0 | - |
| - | - | 840.7 | 427.7 | - | - | 0 | - |
| - | - | 628.2 | 433.3 | - | - | 0 | - |
| - | - | 609.6 | 438.4 | - | - | 0 | - |
| - | - | 595.4 | 452.8 | - | - | 0 | - |
| - | - | 1069 | 465.2 | - | - | 0 | - |
| 4 | b | 758.6 | 479.2 | 8.814E-06 | 0.01839 | +1 | 4 |
| 7 | y | 1885 | 489.2 | 0.004642 | 9.489 | +1 | 3 |
| - | - | 884.8 | 490.2 | - | - | 0 | - |
| - | - | 739.7 | 496.2 | - | - | 0 | - |
| - | - | 797.1 | 527.8 | - | - | 0 | - |
| 2 | y | 1057 | 550.7 | 0.004057 | 7.366 | +2 | 8 |
| - | - | 574.4 | 551.2 | - | - | 0 | - |
| 6 | y | 3362 | 560.2 | 0.004485 | 8.006 | +1 | 4 |
| - | - | 695.1 | 561.2 | - | - | 0 | - |
| - | - | 591.7 | 565.6 | - | - | 0 | - |
| - | - | 720.5 | 584.3 | - | - | 0 | - |
| - | - | 4889 | 585.3 | - | - | 0 | - |
| - | - | 793.4 | 598.3 | - | - | 0 | - |
| 0 | Precursor | 1118 | 601.2 | 0.0001149 | 0.1911 | +2 | -1 |
| - | - | 819.6 | 601.7 | - | - | 0 | - |
| - | - | 617 | 602.3 | - | - | 0 | - |
| - | - | 683.5 | 602.3 | - | - | 0 | - |
| - | - | 611.9 | 613.2 | - | - | 0 | - |
| - | - | 667.8 | 622.4 | - | - | 0 | - |
| - | - | 816.5 | 646.9 | - | - | 0 | - |
| - | - | 694.5 | 657.3 | - | - | 0 | - |
| - | - | 1307 | 659.3 | - | - | 0 | - |
| - | - | 741.8 | 665.3 | - | - | 0 | - |
| - | - | 1102 | 685.3 | - | - | 0 | - |
| - | - | 1291 | 697.4 | - | - | 0 | - |
| - | - | 3235 | 700.4 | - | - | 0 | - |
| - | - | 1963 | 701.4 | - | - | 0 | - |
| 5 | y | 2643 | 723.3 | 0.003351 | 4.633 | +1 | 5 |
| - | - | 1348 | 724.3 | - | - | 0 | - |
| - | - | 1276 | 749.3 | - | - | 0 | - |
| - | - | 837.2 | 767.3 | - | - | 0 | - |
| - | - | 1886 | 801.4 | - | - | 0 | - |
| - | - | 806.4 | 802.4 | - | - | 0 | - |
| - | - | 1217 | 828.4 | - | - | 0 | - |
| 4 | y | 4197 | 838.3 | 0.003142 | 3.748 | +1 | 6 |
| - | - | 1299 | 839.3 | - | - | 0 | - |
| - | - | 2927 | 844.5 | - | - | 0 | - |
| - | - | 813.8 | 845.5 | - | - | 0 | - |
| - | - | 1.148E+04 | 872.4 | - | - | 0 | - |
| - | - | 4975 | 873.4 | - | - | 0 | - |
| - | - | 1496 | 874.5 | - | - | 0 | - |
| - | - | 4586 | 889.3 | - | - | 0 | - |
| - | - | 2747 | 890.4 | - | - | 0 | - |
| 3 | y | 1.826E+04 | 953.3 | 0.004092 | 4.292 | +1 | 7 |
| - | - | 9714 | 954.3 | - | - | 0 | - |
| - | - | 2206 | 955.3 | - | - | 0 | - |
| - | - | 1272 | 991.5 | - | - | 0 | - |
| - | - | 1140 | 992.5 | - | - | 0 | - |
| - | - | 6399 | 1020 | - | - | 0 | - |
| - | - | 3126 | 1021 | - | - | 0 | - |
| - | - | 768 | 1022 | - | - | 0 | - |
| 2 | y | 912.6 | 1100 | 0.002939 | 2.67 | +1 | 8 |
| - | - | 702.6 | 1101 | - | - | 0 | - |
| - | - | 652.1 | 1492 | - | - | 0 | - |
| - | - | 651.5 | 1891 | - | - | 0 | - |
| - | - | 705.8 | 3080 | - | - | 0 | - |

m/z Charge Intensity FragmentType MassShift Position
120.0809555053711 0 30417.77
121.0842514038086 0 2433.407
121.39942169189453 0 374.64426
123.04473114013672 0 480.2858
129.10235595703125 0 3174.9465
130.10597229003906 0 502.3035
132.0809326171875 0 758.506
132.10203552246094 0 496.8077
133.08609008789062 0 2556.4487
136.07586669921875 0 11334.682
137.0789337158203 0 801.978
138.06576538085938 0 458.3096
146.0601806640625 0 828.5388
147.1127166748047 0 529.5136
149.04505920410156 0 484.54834
156.076904296875 0 774.1249
158.0963897705078 0 1517.8961
159.09194946289062 0 1086.1573
159.58184814453125 0 433.1272
160.1122283935547 0 599.76404
165.05491638183594 0 2533.8362
166.08644104003906 0 1973.3484
167.05551147460938 0 2025.3088
167.08168029785156 0 961.7832
169.13401794433594 0 428.22906
173.12884521484375 0 2066.1602
176.10704040527344 0 3014.3538
177.10235595703125 0 2830.0403
177.1105499267578 0 599.3652
182.08123779296875 0 7666.1
183.92567443847656 0 567.11975
185.36537170410156 0 548.5778
188.07095336914062 0 1260.6215
191.11807250976562 0 572.34467
193.10816955566406 0 619.96924
197.1278839111328 0 559.3762
199.1077117919922 0 610.18994
201.1236114501953 0 916.33453
203.1179962158203 0 1212.0862 a Water loss 1
205.09791564941406 0 926.765 y 8
209.09132385253906 0 602.4595
213.64161682128906 0 612.8422
214.11831665039062 0 567.28357
217.08261108398438 0 688.23346
217.09669494628906 0 951.2594
221.10308837890625 0 2461.8628
221.1284942626953 0 47101.348 a 1
222.13194274902344 0 6109.3906
223.06362915039062 0 1454.5625
223.0952606201172 0 510.61752
227.10398864746094 0 801.63165
229.17100524902344 0 655.12683
231.061279296875 0 831.54333
231.12342834472656 0 811.78766
235.1080780029297 0 1131.4432
237.67791748046875 0 536.06323
239.0948486328125 0 920.35144
241.0749053955078 0 553.7565
246.12274169921875 0 530.0302
249.1234588623047 0 9095.84 b 1
250.12652587890625 0 1141.5321
251.1026153564453 0 1058.0999
251.15078735351562 0 646.64703
263.102783203125 0 1308.0845
264.1953125 0 1326.0496
274.13201904296875 0 571.26385
285.010009765625 0 1321.1356
285.1024169921875 0 999.72284
296.1508483886719 0 892.059
319.1385498046875 0 601.9959
324.1455993652344 0 1193.5916
324.7236022949219 0 637.8308
325.1464538574219 0 665.4005
329.14947509765625 0 4994.7686
340.19744873046875 0 556.0132
342.1339111328125 0 607.4383
342.1562805175781 0 9787.799 y 7
343.1588439941406 0 1438.2386
353.9761047363281 0 545.84705
355.0697937011719 0 1229.2595
356.0722351074219 0 709.2445
357.2252197265625 0 1533.5735
364.1507873535156 0 1496.2336 b 2
378.129150390625 0 773.6595
380.1933898925781 0 610.72485
394.1233825683594 0 1803.3862
398.20318603515625 0 612.7791
413.266357421875 0 2920.1672
414.2678527832031 0 809.06757
425.1933898925781 0 819.0792
427.72186279296875 0 840.67194
433.2843933105469 0 628.15137
438.3756103515625 0 609.6447
452.8394470214844 0 595.38214
465.16131591796875 0 1068.809
479.17724609375 0 758.5542 b 3
489.1911926269531 0 1885.4781 y 6
490.1946716308594 0 884.7941
496.22882080078125 0 739.6869
527.7676391601562 0 797.09894
550.7123413085938 0 1057.2015 y 1
551.2103271484375 0 574.3892
560.2281494140625 0 3361.5503 y 5
561.2329711914062 0 695.13654
565.6196899414062 0 591.69684
584.315673828125 0 720.4608
585.342529296875 0 4889.371
598.2987060546875 0 793.41406
601.2322387695312 0 1117.9465 Precursor
601.7333374023438 0 819.5993
602.2564697265625 0 616.98804
602.3267211914062 0 683.4873
613.1993408203125 0 611.92145
622.3624267578125 0 667.78577
646.9398193359375 0 816.5199
657.2598876953125 0 694.5263
659.2926635742188 0 1307.201
665.257080078125 0 741.8177
685.25830078125 0 1102.3364
697.3670043945312 0 1291.2434
700.3611450195312 0 3234.9458
701.363037109375 0 1963.1139
723.2903442382812 0 2643.2817 y 4
724.291748046875 0 1347.7787
749.2555541992188 0 1276.2585
767.2591552734375 0 837.23346
801.4110107421875 0 1886.4547
802.4093627929688 0 806.4157
828.4195556640625 0 1216.8334
838.3170776367188 0 4197.287 y 3
839.3189697265625 0 1298.7024
844.4508666992188 0 2927.0461
845.454833984375 0 813.7881
872.4462890625 0 11478.527
873.4489135742188 0 4975.4106
874.4509887695312 0 1496.2822
889.3463745117188 0 4586.171
890.351806640625 0 2747.0322
953.344970703125 0 18256.174 y 2
954.3475341796875 0 9713.962
955.3474731445312 0 2205.8308
991.5187377929688 0 1271.949
992.5194702148438 0 1140.2615
1019.5137939453125 0 6399.4165
1020.517578125 0 3126.1604
1021.517333984375 0 767.98834
1100.4122314453125 0 912.5657 y 1
1101.4239501953125 0 702.5961
1491.8173828125 0 652.10986
1890.7672119140625 0 651.5268
3080.1357421875 0 705.8161

Spectrum Details

|  |  |
| --- | --- |
| Matched peaks? Matched peaks The total absolute number of peaks matched. Additionally in brackets the total fraction of peaks matched and the total number of peaks is shown. | 15 (9.93% of 151) |
| FDR? FDR The false discovery rate estimated for this peptide. It is calculated by matching all theoretical fragments with a non-integer shift with the raw peaks for this spectrum. This is done with 40 different shifts. The resulting percentage is the average number of annotated peaks over the number of annotated peaks with the correct spectrum. | 1.27% |
| Satellite FDR? Satellite FDR See the FDR for details on its calculation. This satellite ion specific FDR only contains the satellite ions (d/w) for I/L/J positions. | - |
| PSM Score? PSM Score The PSM Score as given by Hecklib to this annotated spectrum. It is shown with three significant figures. | 160 |

#### Spectrum 11624? Spectrum 11624 The raw spectrum of this peptide as annotated by Hecklib. The fragments are coloured according to ion type (see legend). Any peaks with a star '\*' as text can be hovered over to see the full details, first the ion type second the mass shift type. By hovering over the amino acids in the peptide or ions in the legend the corresponding peaks are highlighted. By toggling the 'Unassigned' label you can turn the background (unassigned) peaks on or off in the plot. By updating the slider in the Ion legend you can update the spectrum to only show the top X% of the peaks with labels. The top X% means any peak that is within X% of the highest intensity. By dragging in the spectrum you can zoom in to a specific part of the spectrum and use 'Zoom Out' to get back to the original zoom level. The annotation of the spectrum is based on the given sequence in the peptides file and is done with different software so inconsistencies are likely. The peaks are annotated based on the given sequence, with 20 ppm tolerance.

Copy Data

##### Spectrum 11624 (TSV)

###### Preview

```
Loading example...
```

*Click on the button to copy the data to your clipboard.*

Mz MinMz MaxIntensity Max

WidthHeightPeptide font sizePeptide stroke widthSpectrum font sizeSpectrum stroke widthCompact peptide

Ion legend

wxyz

abcd

OtherUnassignedIon^Charge^_Position_Show for top:%

TFDDYAMHW

01.22e+42.43e+43.65e+44.86e+4

Zoom Out

a^+1^_2_y^+1^_1_a^+1^_2_b^+1^_2_y^+1^_2_b^+1^_3_b^+1^_4_y^+1^_3_y^+2^_8_y^+1^_4_\*y^+1^_5_y^+1^_6_y^+1^_7_y^+1^_8_

0778155523333110

Fragment Matches Table

Show background peaks

| Position | Ion type | Intensity | mz Theoretical | mz Error (Th) | mz Error (ppm) | Charge | Series Number |
| --- | --- | --- | --- | --- | --- | --- | --- |
| - | - | 3.856E+04 | 120.1 | - | - | 0 | - |
| - | - | 3019 | 121.1 | - | - | 0 | - |
| - | - | 374.7 | 121.4 | - | - | 0 | - |
| - | - | 389.5 | 122.1 | - | - | 0 | - |
| - | - | 350.6 | 122.9 | - | - | 0 | - |
| - | - | 590.1 | 123 | - | - | 0 | - |
| - | - | 715.6 | 127.1 | - | - | 0 | - |
| - | - | 3694 | 129.1 | - | - | 0 | - |
| - | - | 766.4 | 130.1 | - | - | 0 | - |
| - | - | 451.2 | 130.1 | - | - | 0 | - |
| - | - | 401.8 | 131.4 | - | - | 0 | - |
| - | - | 1066 | 132.1 | - | - | 0 | - |
| - | - | 1.223E+04 | 136.1 | - | - | 0 | - |
| - | - | 388.9 | 137.1 | - | - | 0 | - |
| - | - | 1103 | 137.1 | - | - | 0 | - |
| - | - | 2435 | 139.1 | - | - | 0 | - |
| - | - | 832.1 | 147 | - | - | 0 | - |
| - | - | 679.6 | 149 | - | - | 0 | - |
| - | - | 469.4 | 149.1 | - | - | 0 | - |
| - | - | 651.4 | 155.1 | - | - | 0 | - |
| - | - | 1535 | 156.1 | - | - | 0 | - |
| - | - | 398 | 156.1 | - | - | 0 | - |
| - | - | 1045 | 157.1 | - | - | 0 | - |
| - | - | 765.9 | 157.1 | - | - | 0 | - |
| - | - | 1789 | 158.1 | - | - | 0 | - |
| - | - | 2360 | 159.1 | - | - | 0 | - |
| - | - | 727.6 | 159.1 | - | - | 0 | - |
| - | - | 1173 | 160.1 | - | - | 0 | - |
| - | - | 601.8 | 162.1 | - | - | 0 | - |
| - | - | 4030 | 165.1 | - | - | 0 | - |
| - | - | 484.8 | 166.1 | - | - | 0 | - |
| - | - | 5807 | 166.1 | - | - | 0 | - |
| - | - | 1317 | 167.1 | - | - | 0 | - |
| - | - | 6531 | 167.1 | - | - | 0 | - |
| - | - | 788.7 | 167.1 | - | - | 0 | - |
| - | - | 1358 | 169.1 | - | - | 0 | - |
| - | - | 2578 | 173.1 | - | - | 0 | - |
| - | - | 4763 | 173.4 | - | - | 0 | - |
| - | - | 484.6 | 175.1 | - | - | 0 | - |
| - | - | 4063 | 176.1 | - | - | 0 | - |
| - | - | 492.5 | 176.7 | - | - | 0 | - |
| - | - | 2618 | 177.1 | - | - | 0 | - |
| - | - | 694.6 | 177.1 | - | - | 0 | - |
| - | - | 7967 | 182.1 | - | - | 0 | - |
| - | - | 567.1 | 183.1 | - | - | 0 | - |
| - | - | 782.1 | 185.1 | - | - | 0 | - |
| - | - | 1058 | 185.1 | - | - | 0 | - |
| - | - | 475.6 | 187.4 | - | - | 0 | - |
| - | - | 2413 | 188.1 | - | - | 0 | - |
| - | - | 626.2 | 191.1 | - | - | 0 | - |
| - | - | 921.2 | 193.1 | - | - | 0 | - |
| - | - | 485.5 | 194 | - | - | 0 | - |
| - | - | 1685 | 199.1 | - | - | 0 | - |
| - | - | 1507 | 201.1 | - | - | 0 | - |
| 2 | a | 1099 | 203.1 | 0.0003813 | 1.877 | +1 | 2 |
| - | - | 555.4 | 204.1 | - | - | 0 | - |
| 9 | y | 1548 | 205.1 | 7.491E-05 | 0.3652 | +1 | 1 |
| - | - | 615.1 | 207.1 | - | - | 0 | - |
| - | - | 576.9 | 209.1 | - | - | 0 | - |
| - | - | 613 | 211.1 | - | - | 0 | - |
| - | - | 854.9 | 212.1 | - | - | 0 | - |
| - | - | 506.2 | 219 | - | - | 0 | - |
| - | - | 3028 | 221.1 | - | - | 0 | - |
| 2 | a | 4.815E+04 | 221.1 | 0.0001316 | 0.5951 | +1 | 2 |
| - | - | 5976 | 222.1 | - | - | 0 | - |
| - | - | 1232 | 223.1 | - | - | 0 | - |
| - | - | 1354 | 224.1 | - | - | 0 | - |
| - | - | 5319 | 227.1 | - | - | 0 | - |
| - | - | 618.9 | 228.1 | - | - | 0 | - |
| - | - | 535.7 | 229.2 | - | - | 0 | - |
| - | - | 887.4 | 231.1 | - | - | 0 | - |
| - | - | 1826 | 235.1 | - | - | 0 | - |
| - | - | 1193 | 239.1 | - | - | 0 | - |
| - | - | 729.1 | 241.1 | - | - | 0 | - |
| 2 | b | 9332 | 249.1 | 0.0001511 | 0.6063 | +1 | 2 |
| - | - | 721.7 | 250.1 | - | - | 0 | - |
| - | - | 1289 | 251.1 | - | - | 0 | - |
| - | - | 626.6 | 251.1 | - | - | 0 | - |
| - | - | 1876 | 263.1 | - | - | 0 | - |
| - | - | 1983 | 264.1 | - | - | 0 | - |
| - | - | 1001 | 265.2 | - | - | 0 | - |
| - | - | 1040 | 279.1 | - | - | 0 | - |
| - | - | 3018 | 281.1 | - | - | 0 | - |
| - | - | 990.4 | 285 | - | - | 0 | - |
| - | - | 915.4 | 285.1 | - | - | 0 | - |
| - | - | 554.4 | 288.1 | - | - | 0 | - |
| - | - | 716.9 | 296.2 | - | - | 0 | - |
| - | - | 1058 | 302.2 | - | - | 0 | - |
| - | - | 661.7 | 319.1 | - | - | 0 | - |
| - | - | 635.2 | 320.4 | - | - | 0 | - |
| - | - | 1712 | 324.1 | - | - | 0 | - |
| - | - | 5269 | 329.1 | - | - | 0 | - |
| - | - | 1435 | 330.2 | - | - | 0 | - |
| - | - | 5226 | 340.2 | - | - | 0 | - |
| - | - | 818.8 | 341.2 | - | - | 0 | - |
| 8 | y | 8986 | 342.2 | 0.0002146 | 0.6271 | +1 | 2 |
| - | - | 2009 | 343.2 | - | - | 0 | - |
| - | - | 1657 | 355.1 | - | - | 0 | - |
| - | - | 727.1 | 357.2 | - | - | 0 | - |
| 3 | b | 2208 | 364.2 | 0.0003839 | 1.054 | +1 | 3 |
| - | - | 1230 | 365.2 | - | - | 0 | - |
| - | - | 1064 | 367.2 | - | - | 0 | - |
| - | - | 843.4 | 378.1 | - | - | 0 | - |
| - | - | 2334 | 380.2 | - | - | 0 | - |
| - | - | 903.6 | 392.2 | - | - | 0 | - |
| - | - | 6516 | 393.2 | - | - | 0 | - |
| - | - | 1427 | 394.1 | - | - | 0 | - |
| - | - | 1814 | 394.2 | - | - | 0 | - |
| - | - | 648.2 | 394.2 | - | - | 0 | - |
| - | - | 635.1 | 395.2 | - | - | 0 | - |
| - | - | 2323 | 398.2 | - | - | 0 | - |
| - | - | 981.7 | 413.3 | - | - | 0 | - |
| - | - | 1220 | 425.2 | - | - | 0 | - |
| - | - | 581.8 | 426.2 | - | - | 0 | - |
| - | - | 1027 | 449.2 | - | - | 0 | - |
| - | - | 826.2 | 450.2 | - | - | 0 | - |
| - | - | 773.5 | 453.3 | - | - | 0 | - |
| - | - | 584.4 | 465 | - | - | 0 | - |
| - | - | 1306 | 465.2 | - | - | 0 | - |
| 4 | b | 1475 | 479.2 | 0.00051 | 1.064 | +1 | 4 |
| - | - | 768.6 | 483.2 | - | - | 0 | - |
| 7 | y | 1811 | 489.2 | 0.004581 | 9.365 | +1 | 3 |
| - | - | 977.1 | 490.2 | - | - | 0 | - |
| - | - | 1540 | 496.2 | - | - | 0 | - |
| - | - | 5543 | 506.3 | - | - | 0 | - |
| - | - | 1105 | 507.3 | - | - | 0 | - |
| - | - | 1438 | 508.3 | - | - | 0 | - |
| 2 | y | 984.3 | 550.7 | 0.001981 | 3.598 | +2 | 8 |
| - | - | 1087 | 551.2 | - | - | 0 | - |
| 6 | y | 3610 | 560.2 | 0.004912 | 8.768 | +1 | 4 |
| - | - | 784.6 | 561.2 | - | - | 0 | - |
| - | - | 1389 | 563.3 | - | - | 0 | - |
| - | - | 824 | 564.3 | - | - | 0 | - |
| - | - | 741.2 | 570.2 | - | - | 0 | - |
| - | - | 666.4 | 573 | - | - | 0 | - |
| - | - | 2522 | 585.3 | - | - | 0 | - |
| 0 | Precursor | 2558 | 601.2 | 0.002251 | 3.744 | +2 | -1 |
| - | - | 692.5 | 602.2 | - | - | 0 | - |
| - | - | 742.5 | 602.3 | - | - | 0 | - |
| - | - | 2526 | 619.3 | - | - | 0 | - |
| - | - | 1023 | 620.3 | - | - | 0 | - |
| - | - | 1667 | 621.4 | - | - | 0 | - |
| - | - | 954.1 | 659.3 | - | - | 0 | - |
| - | - | 2205 | 676.4 | - | - | 0 | - |
| - | - | 598 | 677.4 | - | - | 0 | - |
| - | - | 1030 | 678.4 | - | - | 0 | - |
| - | - | 927.9 | 685.3 | - | - | 0 | - |
| - | - | 5250 | 700.4 | - | - | 0 | - |
| - | - | 1356 | 701.4 | - | - | 0 | - |
| - | - | 1125 | 721.3 | - | - | 0 | - |
| 5 | y | 3208 | 723.3 | 0.003229 | 4.465 | +1 | 5 |
| - | - | 1518 | 724.3 | - | - | 0 | - |
| - | - | 2420 | 733.4 | - | - | 0 | - |
| - | - | 1074 | 734.4 | - | - | 0 | - |
| - | - | 744.3 | 735.4 | - | - | 0 | - |
| - | - | 1485 | 749.3 | - | - | 0 | - |
| - | - | 800.3 | 750.3 | - | - | 0 | - |
| - | - | 844 | 757.4 | - | - | 0 | - |
| - | - | 1297 | 774.3 | - | - | 0 | - |
| - | - | 851.5 | 775.3 | - | - | 0 | - |
| - | - | 1739 | 783.4 | - | - | 0 | - |
| - | - | 672.3 | 784.4 | - | - | 0 | - |
| - | - | 1186 | 800.4 | - | - | 0 | - |
| - | - | 2221 | 801.4 | - | - | 0 | - |
| - | - | 1309 | 802.4 | - | - | 0 | - |
| - | - | 1407 | 828.4 | - | - | 0 | - |
| - | - | 1372 | 832.5 | - | - | 0 | - |
| 4 | y | 4337 | 838.3 | 0.002409 | 2.874 | +1 | 6 |
| - | - | 1703 | 839.3 | - | - | 0 | - |
| - | - | 4090 | 844.5 | - | - | 0 | - |
| - | - | 1563 | 845.5 | - | - | 0 | - |
| - | - | 400.3 | 872.4 | - | - | 0 | - |
| - | - | 1.191E+04 | 872.4 | - | - | 0 | - |
| - | - | 6719 | 873.4 | - | - | 0 | - |
| - | - | 1489 | 874.5 | - | - | 0 | - |
| - | - | 6465 | 889.3 | - | - | 0 | - |
| - | - | 3000 | 890.4 | - | - | 0 | - |
| - | - | 1235 | 901.5 | - | - | 0 | - |
| 3 | y | 1.976E+04 | 953.3 | 0.004092 | 4.292 | +1 | 7 |
| - | - | 1.012E+04 | 954.3 | - | - | 0 | - |
| - | - | 3698 | 955.3 | - | - | 0 | - |
| - | - | 2599 | 991.5 | - | - | 0 | - |
| - | - | 5257 | 1020 | - | - | 0 | - |
| - | - | 2987 | 1021 | - | - | 0 | - |
| - | - | 826.7 | 1022 | - | - | 0 | - |
| - | - | 618.4 | 1064 | - | - | 0 | - |
| 2 | y | 1599 | 1100 | 0.006845 | 6.22 | +1 | 8 |
| - | - | 611.5 | 1391 | - | - | 0 | - |
| - | - | 684 | 3079 | - | - | 0 | - |

m/z Charge Intensity FragmentType MassShift Position
120.08099365234375 0 38562.695
121.08429718017578 0 3019.261
121.37055969238281 0 374.6895
122.06739044189453 0 389.49072
122.9253921508789 0 350.6116
123.04432678222656 0 590.09705
127.08682250976562 0 715.5921
129.10247802734375 0 3693.9473
130.0655517578125 0 766.3692
130.08644104003906 0 451.24655
131.42230224609375 0 401.8315
132.0810089111328 0 1066.2811
136.07586669921875 0 12228.6455
137.0535430908203 0 388.8831
137.0792999267578 0 1103.0984
139.08676147460938 0 2435.1821
147.0444793701172 0 832.0792
149.0448760986328 0 679.6137
149.05987548828125 0 469.35178
155.08172607421875 0 651.4141
156.0770721435547 0 1534.6462
156.1215362548828 0 398.0092
157.09690856933594 0 1044.7535
157.1332244873047 0 765.8832
158.09654235839844 0 1788.6465
159.09178161621094 0 2359.691
159.11245727539062 0 727.6254
160.11256408691406 0 1173.4125
162.09210205078125 0 601.7581
165.05484008789062 0 4029.842
166.06015014648438 0 484.7621
166.08644104003906 0 5807.4946
167.05564880371094 0 1317.0739
167.0817108154297 0 6531.427
167.08953857421875 0 788.72485
169.09730529785156 0 1358.1859
173.12869262695312 0 2577.725
173.4404754638672 0 4762.7817
175.08702087402344 0 484.60828
176.10708618164062 0 4063.448
176.7496795654297 0 492.49033
177.10244750976562 0 2618.243
177.1109619140625 0 694.625
182.08128356933594 0 7966.914
183.08441162109375 0 567.1496
185.09275817871094 0 782.0727
185.12864685058594 0 1057.6953
187.39376831054688 0 475.604
188.0709991455078 0 2412.9712
191.11851501464844 0 626.19025
193.1081085205078 0 921.1523
194.00949096679688 0 485.53714
199.10789489746094 0 1684.988
201.123779296875 0 1507.0658
203.11827087402344 0 1099.1996 a Water loss 1
204.07765197753906 0 555.37933
205.09722900390625 0 1548.4558 y 8
207.1129150390625 0 615.1409
209.10279846191406 0 576.924
211.0603790283203 0 612.9784
212.10313415527344 0 854.94977
219.02828979492188 0 506.17014
221.1031951904297 0 3027.827
221.1285858154297 0 48153.707 a 1
222.1319122314453 0 5976.2515
223.06341552734375 0 1231.7429
224.10333251953125 0 1354.3663
227.1027374267578 0 5319.437
228.1053924560547 0 618.92285
229.1713409423828 0 535.69336
231.06153869628906 0 887.4328
235.1080322265625 0 1825.8292
239.09486389160156 0 1192.5746
241.07513427734375 0 729.0693
249.12351989746094 0 9331.534 b 1
250.12637329101562 0 721.6859
251.1029052734375 0 1289.2985
251.1268310546875 0 626.645
263.1025085449219 0 1876.3949
264.13409423828125 0 1982.8915
265.1544494628906 0 1000.5908
279.09771728515625 0 1040.3969
281.1244812011719 0 3017.7817
285.00994873046875 0 990.3775
285.1014099121094 0 915.39905
288.1198425292969 0 554.42303
296.15093994140625 0 716.91455
302.15032958984375 0 1058.1552
319.1401672363281 0 661.72656
320.3844299316406 0 635.2452
324.1454162597656 0 1711.7285
329.149658203125 0 5269.3545
330.153076171875 0 1435.1228
340.18743896484375 0 5226.227
341.19134521484375 0 818.75323
342.1562805175781 0 8986.292 y 7
343.1599426269531 0 2009.2582
355.07025146484375 0 1657.3322
357.2254638671875 0 727.0651
364.15069580078125 0 2207.7441 b 2
365.1808776855469 0 1229.6023
367.1965637207031 0 1064.1757
378.13006591796875 0 843.37067
380.1929016113281 0 2333.7612
392.20440673828125 0 903.6073
393.1766052246094 0 6515.9624
394.1247253417969 0 1426.7201
394.1800842285156 0 1814.4558
394.208984375 0 648.19446
395.1928405761719 0 635.0844
398.2036437988281 0 2323.4263
413.2667236328125 0 981.73035
425.1923522949219 0 1220.1277
426.194580078125 0 581.8033
449.24041748046875 0 1026.9165
450.2373962402344 0 826.18054
453.2715759277344 0 773.4992
464.9791564941406 0 584.4424
465.1619873046875 0 1306.3301
479.1777648925781 0 1475.4637 b 3
483.2193603515625 0 768.58984
489.1911315917969 0 1811.0363 y 6
490.1940002441406 0 977.1202
496.22967529296875 0 1540.3788
506.26068115234375 0 5542.7266
507.2645568847656 0 1104.7566
508.2744140625 0 1438.0529
550.7102661132812 0 984.34485 y 1
551.2127075195312 0 1087.3806
560.2285766601562 0 3609.5408 y 5
561.2291870117188 0 784.63007
563.2806396484375 0 1388.8497
564.2823486328125 0 824.04724
570.2330322265625 0 741.2018
572.9761962890625 0 666.3634
585.334716796875 0 2522.1208
601.234375 0 2557.5625 Precursor
602.2478637695312 0 692.4581
602.3181762695312 0 742.5059
619.3455810546875 0 2526.1458
620.3489990234375 0 1023.1703
621.360107421875 0 1666.5543
659.290283203125 0 954.0595
676.3675537109375 0 2205.3794
677.3652954101562 0 597.9738
678.3744506835938 0 1029.9062
685.2538452148438 0 927.8736
700.3624877929688 0 5249.949
701.3645629882812 0 1355.651
721.260498046875 0 1124.7573
723.2902221679688 0 3208.167 y 4
724.2916870117188 0 1518.4187
733.3878784179688 0 2420.2869
734.3896484375 0 1073.9692
735.4024658203125 0 744.3364
749.2564697265625 0 1485.395
750.2571411132812 0 800.3333
757.381591796875 0 844.05
774.3157348632812 0 1297.4186
775.32177734375 0 851.4585
783.396484375 0 1738.7692
784.3958129882812 0 672.2652
800.423095703125 0 1186.3978
801.409423828125 0 2220.878
802.4156494140625 0 1309.062
828.4212036132812 0 1407.0247
832.4585571289062 0 1371.5314
838.3163452148438 0 4337.485 y 3
839.3234252929688 0 1702.676
844.451416015625 0 4090.1782
845.4551391601562 0 1563.4131
872.3536987304688 0 400.3381
872.4467163085938 0 11909.5205
873.4485473632812 0 6718.741
874.4517211914062 0 1488.8584
889.3472290039062 0 6464.887
890.3501586914062 0 2999.8843
901.474609375 0 1235.2114
953.344970703125 0 19763.627 y 2
954.3475952148438 0 10123.913
955.3477783203125 0 3698.117
991.5217895507812 0 2599.22
1019.5136108398438 0 5257.262
1020.5172119140625 0 2987.3823
1021.5187377929688 0 826.73517
1063.755859375 0 618.4305
1100.4161376953125 0 1599.3372 y 1
1390.760498046875 0 611.55
3079.45751953125 0 683.99744

Spectrum Details

|  |  |
| --- | --- |
| Matched peaks? Matched peaks The total absolute number of peaks matched. Additionally in brackets the total fraction of peaks matched and the total number of peaks is shown. | 15 (7.94% of 189) |
| FDR? FDR The false discovery rate estimated for this peptide. It is calculated by matching all theoretical fragments with a non-integer shift with the raw peaks for this spectrum. This is done with 40 different shifts. The resulting percentage is the average number of annotated peaks over the number of annotated peaks with the correct spectrum. | 1.90% |
| Satellite FDR? Satellite FDR See the FDR for details on its calculation. This satellite ion specific FDR only contains the satellite ions (d/w) for I/L/J positions. | - |
| PSM Score? PSM Score The PSM Score as given by Hecklib to this annotated spectrum. It is shown with three significant figures. | 160 |

#### Spectrum 12295? Spectrum 12295 The raw spectrum of this peptide as annotated by Hecklib. The fragments are coloured according to ion type (see legend). Any peaks with a star '\*' as text can be hovered over to see the full details, first the ion type second the mass shift type. By hovering over the amino acids in the peptide or ions in the legend the corresponding peaks are highlighted. By toggling the 'Unassigned' label you can turn the background (unassigned) peaks on or off in the plot. By updating the slider in the Ion legend you can update the spectrum to only show the top X% of the peaks with labels. The top X% means any peak that is within X% of the highest intensity. By dragging in the spectrum you can zoom in to a specific part of the spectrum and use 'Zoom Out' to get back to the original zoom level. The annotation of the spectrum is based on the given sequence in the peptides file and is done with different software so inconsistencies are likely. The peaks are annotated based on the given sequence, with 20 ppm tolerance.

Copy Data

##### Spectrum 12295 (TSV)

###### Preview

```
Loading example...
```

*Click on the button to copy the data to your clipboard.*

Mz MinMz MaxIntensity Max

WidthHeightPeptide font sizePeptide stroke widthSpectrum font sizeSpectrum stroke widthCompact peptide

Ion legend

wxyz

abcd

OtherUnassignedIon^Charge^_Position_Show for top:%

TFDDYAMHW

01.00e+42.01e+43.01e+44.02e+4

Zoom Out

a^+1^_2_y^+1^_1_a^+1^_2_b^+1^_2_b^+1^_2_y^+1^_2_b^+1^_3_y^+1^_3_y^+2^_8_y^+1^_4_\*\*b^+1^_5_y^+1^_5_y^+1^_6_y^+1^_7_y^+1^_7_y^+1^_8_

0777155523323110

Fragment Matches Table

Show background peaks

| Position | Ion type | Intensity | mz Theoretical | mz Error (Th) | mz Error (ppm) | Charge | Series Number |
| --- | --- | --- | --- | --- | --- | --- | --- |
| - | - | 2.796E+04 | 120.1 | - | - | 0 | - |
| - | - | 353.7 | 121.1 | - | - | 0 | - |
| - | - | 2264 | 121.1 | - | - | 0 | - |
| - | - | 427.1 | 123 | - | - | 0 | - |
| - | - | 384.5 | 126.6 | - | - | 0 | - |
| - | - | 3319 | 129.1 | - | - | 0 | - |
| - | - | 433.6 | 130.1 | - | - | 0 | - |
| - | - | 659.4 | 132.1 | - | - | 0 | - |
| - | - | 2040 | 133.1 | - | - | 0 | - |
| - | - | 371.8 | 133.7 | - | - | 0 | - |
| - | - | 9417 | 136.1 | - | - | 0 | - |
| - | - | 584.2 | 137.1 | - | - | 0 | - |
| - | - | 393.8 | 137.3 | - | - | 0 | - |
| - | - | 486.6 | 148.9 | - | - | 0 | - |
| - | - | 1484 | 149 | - | - | 0 | - |
| - | - | 483.3 | 151 | - | - | 0 | - |
| - | - | 450.9 | 151.6 | - | - | 0 | - |
| - | - | 471.9 | 155.1 | - | - | 0 | - |
| - | - | 590.2 | 156.1 | - | - | 0 | - |
| - | - | 445.1 | 157.1 | - | - | 0 | - |
| - | - | 1422 | 158.1 | - | - | 0 | - |
| - | - | 1452 | 159.1 | - | - | 0 | - |
| - | - | 480.1 | 159.3 | - | - | 0 | - |
| - | - | 646.3 | 160.1 | - | - | 0 | - |
| - | - | 491.7 | 161.1 | - | - | 0 | - |
| - | - | 2794 | 165.1 | - | - | 0 | - |
| - | - | 772.9 | 166.1 | - | - | 0 | - |
| - | - | 3093 | 167.1 | - | - | 0 | - |
| - | - | 545.3 | 167.1 | - | - | 0 | - |
| - | - | 2302 | 173.1 | - | - | 0 | - |
| - | - | 1669 | 173.5 | - | - | 0 | - |
| - | - | 2662 | 176.1 | - | - | 0 | - |
| - | - | 1741 | 177.1 | - | - | 0 | - |
| - | - | 1465 | 177.1 | - | - | 0 | - |
| - | - | 6845 | 182.1 | - | - | 0 | - |
| - | - | 967.3 | 183.1 | - | - | 0 | - |
| - | - | 530.2 | 183.1 | - | - | 0 | - |
| - | - | 648.6 | 185.1 | - | - | 0 | - |
| - | - | 570.8 | 187.1 | - | - | 0 | - |
| - | - | 2098 | 188.1 | - | - | 0 | - |
| - | - | 425.7 | 190 | - | - | 0 | - |
| - | - | 972.5 | 193.1 | - | - | 0 | - |
| - | - | 622.5 | 197.1 | - | - | 0 | - |
| - | - | 504.7 | 199.8 | - | - | 0 | - |
| - | - | 589.7 | 200.1 | - | - | 0 | - |
| - | - | 1220 | 201.1 | - | - | 0 | - |
| 2 | a | 1174 | 203.1 | 0.0002593 | 1.276 | +1 | 2 |
| 9 | y | 1137 | 205.1 | 0.000154 | 0.7507 | +1 | 1 |
| - | - | 558.7 | 207.1 | - | - | 0 | - |
| - | - | 456.5 | 211.1 | - | - | 0 | - |
| - | - | 648.6 | 217.1 | - | - | 0 | - |
| - | - | 3014 | 221.1 | - | - | 0 | - |
| 2 | a | 3.978E+04 | 221.1 | 4.004E-05 | 0.1811 | +1 | 2 |
| - | - | 5475 | 222.1 | - | - | 0 | - |
| - | - | 1905 | 223.1 | - | - | 0 | - |
| - | - | 457.7 | 229.2 | - | - | 0 | - |
| - | - | 828.8 | 231.1 | - | - | 0 | - |
| - | - | 670.1 | 231.1 | - | - | 0 | - |
| 2 | b | 841.2 | 231.1 | 0.0003469 | 1.501 | +1 | 2 |
| - | - | 1273 | 235.1 | - | - | 0 | - |
| - | - | 1761 | 239.1 | - | - | 0 | - |
| - | - | 580.5 | 240.1 | - | - | 0 | - |
| - | - | 1652 | 241.1 | - | - | 0 | - |
| - | - | 568.2 | 248.1 | - | - | 0 | - |
| 2 | b | 6475 | 249.1 | 0.0001663 | 0.6676 | +1 | 2 |
| - | - | 945.1 | 250.1 | - | - | 0 | - |
| - | - | 1243 | 251.1 | - | - | 0 | - |
| - | - | 752.3 | 257.1 | - | - | 0 | - |
| - | - | 503.6 | 261.1 | - | - | 0 | - |
| - | - | 1644 | 263.1 | - | - | 0 | - |
| - | - | 809.2 | 279.1 | - | - | 0 | - |
| - | - | 721.8 | 283 | - | - | 0 | - |
| - | - | 1333 | 285 | - | - | 0 | - |
| - | - | 1104 | 285.1 | - | - | 0 | - |
| - | - | 576.7 | 286 | - | - | 0 | - |
| - | - | 777.6 | 296.2 | - | - | 0 | - |
| - | - | 727.7 | 299.1 | - | - | 0 | - |
| - | - | 696.9 | 309.2 | - | - | 0 | - |
| - | - | 585.9 | 310.7 | - | - | 0 | - |
| - | - | 1778 | 324.1 | - | - | 0 | - |
| - | - | 4330 | 329.1 | - | - | 0 | - |
| 8 | y | 8456 | 342.2 | 0.0003366 | 0.9839 | +1 | 2 |
| - | - | 1443 | 343.2 | - | - | 0 | - |
| - | - | 3006 | 355.1 | - | - | 0 | - |
| - | - | 1155 | 356.1 | - | - | 0 | - |
| - | - | 973.6 | 357.2 | - | - | 0 | - |
| 3 | b | 2013 | 364.2 | 1.771E-05 | 0.04865 | +1 | 3 |
| - | - | 907.8 | 378.1 | - | - | 0 | - |
| - | - | 620.9 | 380.2 | - | - | 0 | - |
| - | - | 1501 | 394.1 | - | - | 0 | - |
| - | - | 525.6 | 401.2 | - | - | 0 | - |
| - | - | 1540 | 413.3 | - | - | 0 | - |
| - | - | 706.4 | 417.1 | - | - | 0 | - |
| - | - | 1002 | 425.2 | - | - | 0 | - |
| - | - | 786.1 | 436.7 | - | - | 0 | - |
| - | - | 660.1 | 441.2 | - | - | 0 | - |
| - | - | 1175 | 465.2 | - | - | 0 | - |
| - | - | 525.3 | 479.9 | - | - | 0 | - |
| 7 | y | 1438 | 489.2 | 0.004825 | 9.864 | +1 | 3 |
| - | - | 671.7 | 496.2 | - | - | 0 | - |
| - | - | 713.2 | 545.7 | - | - | 0 | - |
| 2 | y | 1098 | 550.7 | 0.005888 | 10.69 | +2 | 8 |
| - | - | 702.6 | 551.2 | - | - | 0 | - |
| 6 | y | 3126 | 560.2 | 0.005156 | 9.204 | +1 | 4 |
| - | - | 1057 | 561.2 | - | - | 0 | - |
| - | - | 4230 | 585.3 | - | - | 0 | - |
| 0 | Precursor | 711.8 | 592.2 | 0.0008806 | 1.487 | +2 | -1 |
| 0 | Precursor | 696.1 | 601.2 | 0.008538 | 14.2 | +2 | -1 |
| - | - | 1028 | 601.7 | - | - | 0 | - |
| - | - | 648.6 | 602.2 | - | - | 0 | - |
| - | - | 616.8 | 612.2 | - | - | 0 | - |
| - | - | 1073 | 621.4 | - | - | 0 | - |
| 5 | b | 638.8 | 624.2 | 0.002113 | 3.386 | +1 | 5 |
| - | - | 736.9 | 635.8 | - | - | 0 | - |
| - | - | 798.3 | 659.3 | - | - | 0 | - |
| - | - | 700.2 | 685.3 | - | - | 0 | - |
| - | - | 3648 | 700.4 | - | - | 0 | - |
| - | - | 615.3 | 701.4 | - | - | 0 | - |
| - | - | 752.5 | 721.3 | - | - | 0 | - |
| 5 | y | 2799 | 723.3 | 0.004023 | 5.562 | +1 | 5 |
| - | - | 1006 | 724.3 | - | - | 0 | - |
| - | - | 1024 | 749.3 | - | - | 0 | - |
| - | - | 817.6 | 774.3 | - | - | 0 | - |
| - | - | 1066 | 783.4 | - | - | 0 | - |
| - | - | 648.2 | 797.4 | - | - | 0 | - |
| - | - | 2294 | 801.4 | - | - | 0 | - |
| - | - | 1111 | 802.4 | - | - | 0 | - |
| - | - | 1151 | 828.4 | - | - | 0 | - |
| 4 | y | 3687 | 838.3 | 0.006377 | 7.606 | +1 | 6 |
| - | - | 1552 | 839.3 | - | - | 0 | - |
| - | - | 3170 | 844.5 | - | - | 0 | - |
| - | - | 957.6 | 845.5 | - | - | 0 | - |
| - | - | 595.9 | 846.5 | - | - | 0 | - |
| - | - | 1E+04 | 872.4 | - | - | 0 | - |
| - | - | 5800 | 873.5 | - | - | 0 | - |
| - | - | 1503 | 874.5 | - | - | 0 | - |
| - | - | 5316 | 889.3 | - | - | 0 | - |
| - | - | 2398 | 890.4 | - | - | 0 | - |
| 3 | y | 645.1 | 935.3 | 0.006905 | 7.382 | +1 | 7 |
| 3 | y | 1.444E+04 | 953.3 | 0.006045 | 6.341 | +1 | 7 |
| - | - | 7645 | 954.3 | - | - | 0 | - |
| - | - | 3041 | 955.4 | - | - | 0 | - |
| - | - | 1972 | 991.5 | - | - | 0 | - |
| - | - | 980.8 | 992.5 | - | - | 0 | - |
| - | - | 620.7 | 994.3 | - | - | 0 | - |
| - | - | 5104 | 1020 | - | - | 0 | - |
| - | - | 2982 | 1021 | - | - | 0 | - |
| - | - | 889.6 | 1022 | - | - | 0 | - |
| 2 | y | 1146 | 1100 | 0.001352 | 1.228 | +1 | 8 |
| - | - | 620.1 | 1249 | - | - | 0 | - |
| - | - | 774.5 | 3079 | - | - | 0 | - |

m/z Charge Intensity FragmentType MassShift Position
120.08087921142578 0 27964.002
121.06486511230469 0 353.7413
121.08419036865234 0 2264.0586
123.04447937011719 0 427.12173
126.6005630493164 0 384.45087
129.10231018066406 0 3319.0813
130.06504821777344 0 433.55576
132.10191345214844 0 659.39355
133.08604431152344 0 2040.3643
133.66282653808594 0 371.82205
136.07579040527344 0 9417.165
137.0791015625 0 584.24146
137.32012939453125 0 393.7953
148.94644165039062 0 486.56354
149.04489135742188 0 1484.4745
151.02432250976562 0 483.28122
151.64002990722656 0 450.87308
155.08184814453125 0 471.89523
156.07737731933594 0 590.20166
157.1337432861328 0 445.13013
158.09671020507812 0 1422.1813
159.09161376953125 0 1451.9935
159.30174255371094 0 480.0949
160.11183166503906 0 646.28156
161.0814971923828 0 491.7024
165.05474853515625 0 2793.7693
166.08631896972656 0 772.931
167.0553741455078 0 3093.2307
167.08148193359375 0 545.25354
173.1286163330078 0 2302.2554
173.45228576660156 0 1669.4983
176.10693359375 0 2662.382
177.1022491455078 0 1740.6998
177.11172485351562 0 1465.3582
182.0812225341797 0 6845.4
183.0845947265625 0 967.3201
183.14907836914062 0 530.1642
185.1280059814453 0 648.61505
187.13279724121094 0 570.79254
188.07064819335938 0 2097.8347
189.9889678955078 0 425.71045
193.1080780029297 0 972.514
197.1275177001953 0 622.53467
199.77047729492188 0 504.72708
200.1401824951172 0 589.65656
201.1237030029297 0 1220.3824
203.11814880371094 0 1174.4219 a Water loss 1
205.0970001220703 0 1136.9993 y 8
207.1127166748047 0 558.65155
211.05960083007812 0 456.5436
217.0971221923828 0 648.5601
221.1031951904297 0 3014.4832
221.1284942626953 0 39782.766 a 1
222.1318817138672 0 5474.617
223.06407165527344 0 1904.5851
229.16964721679688 0 457.6715
231.0611572265625 0 828.8344
231.0975799560547 0 670.0832
231.11245727539062 0 841.1905 b Water loss 1
235.1077117919922 0 1272.7705
239.09547424316406 0 1760.7863
240.13404846191406 0 580.5249
241.07415771484375 0 1651.7631
248.1066131591797 0 568.24554
249.12353515625 0 6474.6675 b 1
250.12754821777344 0 945.0871
251.10208129882812 0 1243.0952
257.1057434082031 0 752.2614
261.10931396484375 0 503.64597
263.1026306152344 0 1644.2725
279.09716796875 0 809.2088
283.0306091308594 0 721.84094
285.0100402832031 0 1333.3435
285.1022644042969 0 1104.2827
286.0111389160156 0 576.68787
296.1518249511719 0 777.6401
299.06060791015625 0 727.7096
309.2027282714844 0 696.941
310.67767333984375 0 585.899
324.14569091796875 0 1777.5647
329.1497802734375 0 4330.075
342.1564025878906 0 8456.34 y 7
343.1597595214844 0 1443.055
355.0698547363281 0 3005.6567
356.0702819824219 0 1155.4467
357.22479248046875 0 973.5852
364.15032958984375 0 2013.2102 b 2
378.1290588378906 0 907.8285
380.1921081542969 0 620.89264
394.1255187988281 0 1500.8121
401.20281982421875 0 525.6344
413.26641845703125 0 1539.6444
417.0881042480469 0 706.4347
425.1934509277344 0 1001.8534
436.727783203125 0 786.13007
441.21038818359375 0 660.08057
465.1622619628906 0 1174.5867
479.879150390625 0 525.3186
489.1913757324219 0 1437.6552 y 6
496.23095703125 0 671.71
545.7279663085938 0 713.247
550.7141723632812 0 1097.5613 y 1
551.216796875 0 702.56464
560.2288208007812 0 3125.926 y 5
561.2300415039062 0 1057.3746
585.3458862304688 0 4230.048
592.2277221679688 0 711.7681 Precursor Water loss
601.2406616210938 0 696.1019 Precursor
601.7351684570312 0 1027.7537
602.2456665039062 0 648.5781
612.197509765625 0 616.80664
621.3589477539062 0 1073.277
624.2279052734375 0 638.75037 b Water loss 4
635.7685546875 0 736.88165
659.294677734375 0 798.25226
685.2548828125 0 700.175
700.36328125 0 3648.3916
701.3623046875 0 615.2598
721.2567138671875 0 752.496
723.291015625 0 2798.6392 y 4
724.2925415039062 0 1006.1654
749.2612915039062 0 1024.2329
774.3193969726562 0 817.6421
783.3992309570312 0 1066.329
797.4252319335938 0 648.22644
801.4124145507812 0 2293.6409
802.4127197265625 0 1111.3221
828.422607421875 0 1151.3336
838.3203125 0 3687 y 3
839.3225708007812 0 1552.1656
844.4525756835938 0 3169.8567
845.4573974609375 0 957.5673
846.455810546875 0 595.8627
872.448486328125 0 9999.758
873.4508056640625 0 5800.397
874.4571533203125 0 1503.061
889.348388671875 0 5315.9897
890.3515625 0 2398.0562
935.3372192382812 0 645.13153 y Water loss 2
953.346923828125 0 14441.024 y 2
954.3494873046875 0 7644.8965
955.351806640625 0 3041.1523
991.5188598632812 0 1971.7799
992.5260620117188 0 980.7891
994.28515625 0 620.7179
1019.516845703125 0 5104.4023
1020.5242309570312 0 2982.4878
1021.5232543945312 0 889.6254
1100.41064453125 0 1145.9518 y 1
1249.3287353515625 0 620.0526
3078.743896484375 0 774.52484

Spectrum Details

|  |  |
| --- | --- |
| Matched peaks? Matched peaks The total absolute number of peaks matched. Additionally in brackets the total fraction of peaks matched and the total number of peaks is shown. | 18 (11.92% of 151) |
| FDR? FDR The false discovery rate estimated for this peptide. It is calculated by matching all theoretical fragments with a non-integer shift with the raw peaks for this spectrum. This is done with 40 different shifts. The resulting percentage is the average number of annotated peaks over the number of annotated peaks with the correct spectrum. | 0.66% |
| Satellite FDR? Satellite FDR See the FDR for details on its calculation. This satellite ion specific FDR only contains the satellite ions (d/w) for I/L/J positions. | - |
| PSM Score? PSM Score The PSM Score as given by Hecklib to this annotated spectrum. It is shown with three significant figures. | 195 |

#### Spectrum 11678? Spectrum 11678 The raw spectrum of this peptide as annotated by Hecklib. The fragments are coloured according to ion type (see legend). Any peaks with a star '\*' as text can be hovered over to see the full details, first the ion type second the mass shift type. By hovering over the amino acids in the peptide or ions in the legend the corresponding peaks are highlighted. By toggling the 'Unassigned' label you can turn the background (unassigned) peaks on or off in the plot. By updating the slider in the Ion legend you can update the spectrum to only show the top X% of the peaks with labels. The top X% means any peak that is within X% of the highest intensity. By dragging in the spectrum you can zoom in to a specific part of the spectrum and use 'Zoom Out' to get back to the original zoom level. The annotation of the spectrum is based on the given sequence in the peptides file and is done with different software so inconsistencies are likely. The peaks are annotated based on the given sequence, with 20 ppm tolerance.

Copy Data

##### Spectrum 11678 (TSV)

###### Preview

```
Loading example...
```

*Click on the button to copy the data to your clipboard.*

Mz MinMz MaxIntensity Max

WidthHeightPeptide font sizePeptide stroke widthSpectrum font sizeSpectrum stroke widthCompact peptide

Ion legend

wxyz

abcd

OtherUnassignedIon^Charge^_Position_Show for top:%

TFDDYAMHW

01.28e+42.57e+43.85e+45.14e+4

Zoom Out

a^+1^_2_y^+1^_1_a^+1^_2_b^+1^_2_y^+1^_2_b^+1^_3_b^+1^_4_y^+1^_3_y^+2^_8_y^+1^_4_\*y^+1^_5_y^+1^_6_y^+1^_7_y^+1^_8_

041983812581677

Fragment Matches Table

Show background peaks

| Position | Ion type | Intensity | mz Theoretical | mz Error (Th) | mz Error (ppm) | Charge | Series Number |
| --- | --- | --- | --- | --- | --- | --- | --- |
| - | - | 3.694E+04 | 120.1 | - | - | 0 | - |
| - | - | 2835 | 121.1 | - | - | 0 | - |
| - | - | 465.4 | 127.1 | - | - | 0 | - |
| - | - | 3404 | 129.1 | - | - | 0 | - |
| - | - | 1178 | 130.1 | - | - | 0 | - |
| - | - | 424.5 | 132.1 | - | - | 0 | - |
| - | - | 871.9 | 133.1 | - | - | 0 | - |
| - | - | 1.041E+04 | 136.1 | - | - | 0 | - |
| - | - | 731.3 | 137.1 | - | - | 0 | - |
| - | - | 741 | 138.1 | - | - | 0 | - |
| - | - | 644.8 | 139.1 | - | - | 0 | - |
| - | - | 469.6 | 146.1 | - | - | 0 | - |
| - | - | 856.4 | 149 | - | - | 0 | - |
| - | - | 718.7 | 155.1 | - | - | 0 | - |
| - | - | 1182 | 156.1 | - | - | 0 | - |
| - | - | 918.1 | 157.1 | - | - | 0 | - |
| - | - | 2200 | 158.1 | - | - | 0 | - |
| - | - | 1250 | 159.1 | - | - | 0 | - |
| - | - | 684.5 | 160.1 | - | - | 0 | - |
| - | - | 1148 | 160.1 | - | - | 0 | - |
| - | - | 935.5 | 162.1 | - | - | 0 | - |
| - | - | 3347 | 165.1 | - | - | 0 | - |
| - | - | 417.8 | 166.1 | - | - | 0 | - |
| - | - | 3949 | 166.1 | - | - | 0 | - |
| - | - | 1575 | 167.1 | - | - | 0 | - |
| - | - | 2412 | 167.1 | - | - | 0 | - |
| - | - | 859.4 | 169.1 | - | - | 0 | - |
| - | - | 735.5 | 171.1 | - | - | 0 | - |
| - | - | 2530 | 173.1 | - | - | 0 | - |
| - | - | 3192 | 173.5 | - | - | 0 | - |
| - | - | 3372 | 176.1 | - | - | 0 | - |
| - | - | 2837 | 177.1 | - | - | 0 | - |
| - | - | 953 | 177.1 | - | - | 0 | - |
| - | - | 7385 | 182.1 | - | - | 0 | - |
| - | - | 1191 | 183.1 | - | - | 0 | - |
| - | - | 980.4 | 183.1 | - | - | 0 | - |
| - | - | 452 | 185.1 | - | - | 0 | - |
| - | - | 1763 | 188.1 | - | - | 0 | - |
| - | - | 1209 | 193.1 | - | - | 0 | - |
| - | - | 1073 | 199.1 | - | - | 0 | - |
| - | - | 576.6 | 200.1 | - | - | 0 | - |
| - | - | 1648 | 201.1 | - | - | 0 | - |
| - | - | 617.5 | 203.1 | - | - | 0 | - |
| 2 | a | 1183 | 203.1 | 0.0001067 | 0.5252 | +1 | 2 |
| 9 | y | 1073 | 205.1 | 0.0002428 | 1.184 | +1 | 1 |
| - | - | 704 | 207.1 | - | - | 0 | - |
| - | - | 2726 | 212.2 | - | - | 0 | - |
| - | - | 579.8 | 217.1 | - | - | 0 | - |
| - | - | 523.6 | 218.7 | - | - | 0 | - |
| - | - | 3132 | 221.1 | - | - | 0 | - |
| 2 | a | 5.085E+04 | 221.1 | 0.0001621 | 0.7331 | +1 | 2 |
| - | - | 5748 | 222.1 | - | - | 0 | - |
| - | - | 1196 | 223.1 | - | - | 0 | - |
| - | - | 461.6 | 223.1 | - | - | 0 | - |
| - | - | 769.5 | 224.1 | - | - | 0 | - |
| - | - | 3682 | 227.1 | - | - | 0 | - |
| - | - | 574.3 | 229.2 | - | - | 0 | - |
| - | - | 1477 | 231.1 | - | - | 0 | - |
| - | - | 1143 | 235.1 | - | - | 0 | - |
| - | - | 641 | 239.1 | - | - | 0 | - |
| - | - | 1103 | 241.1 | - | - | 0 | - |
| 2 | b | 8138 | 249.1 | 0.0002273 | 0.9126 | +1 | 2 |
| - | - | 726.2 | 250.1 | - | - | 0 | - |
| - | - | 955.3 | 251.1 | - | - | 0 | - |
| - | - | 1869 | 252.2 | - | - | 0 | - |
| - | - | 598.3 | 253.2 | - | - | 0 | - |
| - | - | 984.8 | 257.1 | - | - | 0 | - |
| - | - | 1725 | 263.1 | - | - | 0 | - |
| - | - | 915.3 | 264.1 | - | - | 0 | - |
| - | - | 653.2 | 265.2 | - | - | 0 | - |
| - | - | 574.8 | 269 | - | - | 0 | - |
| - | - | 1438 | 281.1 | - | - | 0 | - |
| - | - | 835.6 | 285 | - | - | 0 | - |
| - | - | 922.9 | 285.1 | - | - | 0 | - |
| - | - | 463.2 | 288.5 | - | - | 0 | - |
| - | - | 767.7 | 292.1 | - | - | 0 | - |
| - | - | 640.7 | 296.2 | - | - | 0 | - |
| - | - | 504.6 | 298.1 | - | - | 0 | - |
| - | - | 735.2 | 299.1 | - | - | 0 | - |
| - | - | 799.8 | 319.1 | - | - | 0 | - |
| - | - | 1497 | 324.1 | - | - | 0 | - |
| - | - | 496.8 | 325.1 | - | - | 0 | - |
| - | - | 4702 | 329.1 | - | - | 0 | - |
| - | - | 1011 | 330.2 | - | - | 0 | - |
| - | - | 2165 | 340.2 | - | - | 0 | - |
| - | - | 701.5 | 342.1 | - | - | 0 | - |
| 8 | y | 8707 | 342.2 | 0.0003672 | 1.073 | +1 | 2 |
| - | - | 1916 | 343.2 | - | - | 0 | - |
| - | - | 1187 | 355.1 | - | - | 0 | - |
| - | - | 1053 | 357.2 | - | - | 0 | - |
| 3 | b | 2603 | 364.2 | 0.0002569 | 0.7056 | +1 | 3 |
| - | - | 576.4 | 365.2 | - | - | 0 | - |
| - | - | 586.4 | 368.2 | - | - | 0 | - |
| - | - | 1123 | 378.1 | - | - | 0 | - |
| - | - | 1101 | 380.2 | - | - | 0 | - |
| - | - | 576.8 | 382.3 | - | - | 0 | - |
| - | - | 681.8 | 390.2 | - | - | 0 | - |
| - | - | 2425 | 393.2 | - | - | 0 | - |
| - | - | 1955 | 394.1 | - | - | 0 | - |
| - | - | 915.5 | 398.2 | - | - | 0 | - |
| - | - | 1049 | 413.3 | - | - | 0 | - |
| - | - | 576 | 413.7 | - | - | 0 | - |
| - | - | 537.1 | 415.5 | - | - | 0 | - |
| - | - | 1442 | 425.2 | - | - | 0 | - |
| - | - | 1182 | 427.7 | - | - | 0 | - |
| - | - | 548.4 | 430.9 | - | - | 0 | - |
| - | - | 587.6 | 435.6 | - | - | 0 | - |
| - | - | 1279 | 465.2 | - | - | 0 | - |
| 4 | b | 917.9 | 479.2 | 0.001639 | 3.421 | +1 | 4 |
| 7 | y | 2793 | 489.2 | 0.00513 | 10.49 | +1 | 3 |
| - | - | 568.4 | 491.2 | - | - | 0 | - |
| - | - | 729.3 | 496.2 | - | - | 0 | - |
| - | - | 2016 | 506.3 | - | - | 0 | - |
| - | - | 875.2 | 508.3 | - | - | 0 | - |
| 2 | y | 1407 | 550.7 | 0.00247 | 4.485 | +2 | 8 |
| - | - | 608.7 | 555.3 | - | - | 0 | - |
| 6 | y | 3391 | 560.2 | 0.004485 | 8.006 | +1 | 4 |
| - | - | 1002 | 561.2 | - | - | 0 | - |
| - | - | 652.3 | 569.2 | - | - | 0 | - |
| - | - | 666.7 | 583.2 | - | - | 0 | - |
| - | - | 2549 | 585.3 | - | - | 0 | - |
| - | - | 953.9 | 586.3 | - | - | 0 | - |
| 0 | Precursor | 1872 | 601.2 | 0.002617 | 4.353 | +2 | -1 |
| - | - | 590.8 | 601.8 | - | - | 0 | - |
| - | - | 1391 | 602.3 | - | - | 0 | - |
| - | - | 673.3 | 612.2 | - | - | 0 | - |
| - | - | 893.6 | 619.3 | - | - | 0 | - |
| - | - | 1099 | 621.4 | - | - | 0 | - |
| - | - | 1044 | 628.4 | - | - | 0 | - |
| - | - | 657 | 658.3 | - | - | 0 | - |
| - | - | 1698 | 659.3 | - | - | 0 | - |
| - | - | 793.6 | 676.4 | - | - | 0 | - |
| - | - | 1152 | 685.3 | - | - | 0 | - |
| - | - | 5703 | 700.4 | - | - | 0 | - |
| - | - | 1357 | 701.4 | - | - | 0 | - |
| - | - | 681.4 | 702.4 | - | - | 0 | - |
| - | - | 832.2 | 721.3 | - | - | 0 | - |
| 5 | y | 3502 | 723.3 | 0.004999 | 6.912 | +1 | 5 |
| - | - | 1352 | 724.3 | - | - | 0 | - |
| - | - | 1038 | 733.4 | - | - | 0 | - |
| - | - | 987.9 | 735.4 | - | - | 0 | - |
| - | - | 1598 | 749.3 | - | - | 0 | - |
| - | - | 764.5 | 757.4 | - | - | 0 | - |
| - | - | 1658 | 774.3 | - | - | 0 | - |
| - | - | 1008 | 775.3 | - | - | 0 | - |
| - | - | 2834 | 801.4 | - | - | 0 | - |
| - | - | 1206 | 802.4 | - | - | 0 | - |
| - | - | 752.2 | 828.4 | - | - | 0 | - |
| 4 | y | 3563 | 838.3 | 0.003569 | 4.257 | +1 | 6 |
| - | - | 1852 | 839.3 | - | - | 0 | - |
| - | - | 788.8 | 840.3 | - | - | 0 | - |
| - | - | 3952 | 844.5 | - | - | 0 | - |
| - | - | 2208 | 845.5 | - | - | 0 | - |
| - | - | 506.7 | 872.4 | - | - | 0 | - |
| - | - | 1.366E+04 | 872.4 | - | - | 0 | - |
| - | - | 6649 | 873.4 | - | - | 0 | - |
| - | - | 2259 | 874.4 | - | - | 0 | - |
| - | - | 5343 | 889.3 | - | - | 0 | - |
| - | - | 3378 | 890.3 | - | - | 0 | - |
| - | - | 564.7 | 896.5 | - | - | 0 | - |
| - | - | 684.6 | 901.5 | - | - | 0 | - |
| - | - | 922.5 | 918.5 | - | - | 0 | - |
| - | - | 1000 | 936.3 | - | - | 0 | - |
| 3 | y | 1.928E+04 | 953.3 | 0.004031 | 4.228 | +1 | 7 |
| - | - | 9556 | 954.3 | - | - | 0 | - |
| - | - | 3098 | 955.3 | - | - | 0 | - |
| - | - | 2479 | 991.5 | - | - | 0 | - |
| - | - | 644.5 | 992.5 | - | - | 0 | - |
| - | - | 786.9 | 1003 | - | - | 0 | - |
| - | - | 5433 | 1020 | - | - | 0 | - |
| - | - | 3673 | 1021 | - | - | 0 | - |
| - | - | 833.3 | 1022 | - | - | 0 | - |
| - | - | 626.9 | 1036 | - | - | 0 | - |
| - | - | 662.8 | 1036 | - | - | 0 | - |
| 2 | y | 941.2 | 1100 | 0.000131 | 0.119 | +1 | 8 |
| - | - | 703.7 | 1101 | - | - | 0 | - |
| - | - | 586.3 | 1635 | - | - | 0 | - |
| - | - | 646.2 | 1660 | - | - | 0 | - |

m/z Charge Intensity FragmentType MassShift Position
120.0810317993164 0 36938.723
121.08438873291016 0 2834.9314
127.08702850341797 0 465.36023
129.10252380371094 0 3403.8994
130.06517028808594 0 1177.7631
132.08084106445312 0 424.46143
133.08609008789062 0 871.9345
136.075927734375 0 10407.028
137.0793914794922 0 731.3005
138.0666046142578 0 741.0414
139.0867156982422 0 644.8231
146.06033325195312 0 469.55063
149.04502868652344 0 856.3714
155.0814208984375 0 718.7391
156.07705688476562 0 1182.0532
157.13365173339844 0 918.0508
158.09661865234375 0 2199.711
159.09176635742188 0 1249.8983
160.0760498046875 0 684.52484
160.11239624023438 0 1147.9583
162.0912322998047 0 935.4509
165.0547637939453 0 3347.1719
166.0613555908203 0 417.81183
166.0865478515625 0 3949.3513
167.05567932128906 0 1574.9507
167.08172607421875 0 2412.0493
169.09756469726562 0 859.38556
171.11239624023438 0 735.508
173.12864685058594 0 2529.7751
173.45086669921875 0 3191.9624
176.10707092285156 0 3372.5
177.10235595703125 0 2836.7534
177.11083984375 0 952.9568
182.08143615722656 0 7384.657
183.0849609375 0 1190.8169
183.11314392089844 0 980.4189
185.12864685058594 0 452.0022
188.07095336914062 0 1762.7306
193.10821533203125 0 1209.4647
199.10787963867188 0 1072.5293
200.1390838623047 0 576.55835
201.12359619140625 0 1647.6216
203.066650390625 0 617.475
203.1179962158203 0 1183.0757 a Water loss 1
205.09739685058594 0 1073.2556 y 8
207.11282348632812 0 703.98956
212.16488647460938 0 2725.731
217.0974578857422 0 579.7669
218.7394256591797 0 523.58234
221.10348510742188 0 3131.715
221.1286163330078 0 50847.383 a 1
222.1319580078125 0 5747.56
223.0638427734375 0 1195.8053
223.13311767578125 0 461.55615
224.10342407226562 0 769.4638
227.10293579101562 0 3682.3445
229.16909790039062 0 574.3088
231.0611114501953 0 1476.67
235.1078338623047 0 1143.1614
239.0948486328125 0 640.9737
241.07373046875 0 1103.1266
249.12359619140625 0 8137.573 b 1
250.12693786621094 0 726.2491
251.10244750976562 0 955.2804
252.15744018554688 0 1869.0753
253.1656494140625 0 598.305
257.10626220703125 0 984.75104
263.1028137207031 0 1725.2927
264.1334228515625 0 915.2749
265.15496826171875 0 653.1675
269.0217590332031 0 574.84186
281.1244201660156 0 1438.2124
285.0092468261719 0 835.60736
285.1014709472656 0 922.9388
288.5335998535156 0 463.168
292.14056396484375 0 767.70215
296.1512756347656 0 640.68365
298.1418762207031 0 504.5987
299.062255859375 0 735.1621
319.1396179199219 0 799.78467
324.14605712890625 0 1497.2307
325.146484375 0 496.84573
329.1498718261719 0 4701.7173
330.15509033203125 0 1011.0364
340.1877746582031 0 2164.6023
342.13665771484375 0 701.5112
342.15643310546875 0 8706.555 y 7
343.160400390625 0 1915.6926
355.0693664550781 0 1186.6737
357.22491455078125 0 1053.1107
364.1500549316406 0 2603.061 b 2
365.1547546386719 0 576.4441
368.19171142578125 0 586.43085
378.1295166015625 0 1122.6497
380.1917419433594 0 1100.5907
382.2534484863281 0 576.7659
390.1694030761719 0 681.7525
393.1764221191406 0 2425.0605
394.1250915527344 0 1954.701
398.20428466796875 0 915.49634
413.2672119140625 0 1048.6426
413.72198486328125 0 575.952
415.5132141113281 0 537.10364
425.19378662109375 0 1441.6324
427.72406005859375 0 1182.3965
430.9208984375 0 548.4424
435.6271057128906 0 587.6187
465.1616516113281 0 1278.7695
479.17889404296875 0 917.8829 b 3
489.1916809082031 0 2792.748 y 6
491.2010192871094 0 568.4347
496.2301940917969 0 729.3451
506.2609558105469 0 2015.6986
508.273681640625 0 875.2012
550.7107543945312 0 1407.4113 y 1
555.2780151367188 0 608.7222
560.2281494140625 0 3390.5815 y 5
561.2294311523438 0 1002.34515
569.2312622070312 0 652.2641
583.2132568359375 0 666.6662
585.336181640625 0 2548.5728
586.3372802734375 0 953.94946
601.2347412109375 0 1871.9883 Precursor
601.8273315429688 0 590.7634
602.3211669921875 0 1390.5852
612.1912841796875 0 673.31573
619.3453979492188 0 893.5914
621.3593139648438 0 1098.9703
628.3765869140625 0 1043.7319
658.2637939453125 0 657.04315
659.2930297851562 0 1697.5706
676.365234375 0 793.60034
685.2517700195312 0 1152.344
700.361328125 0 5703.1963
701.3660278320312 0 1356.5747
702.3694458007812 0 681.38055
721.259521484375 0 832.2032
723.2919921875 0 3502.1572 y 4
724.295166015625 0 1352.0829
733.3863525390625 0 1038.1097
735.4015502929688 0 987.8851
749.2562255859375 0 1597.9033
757.3839721679688 0 764.50635
774.3214721679688 0 1658.3961
775.3241577148438 0 1008.1393
801.4114379882812 0 2834.4277
802.418212890625 0 1205.8267
828.43017578125 0 752.21765
838.3175048828125 0 3562.787 y 3
839.3221435546875 0 1852.1892
840.3211669921875 0 788.804
844.4517822265625 0 3952.275
845.4548950195312 0 2208.0542
872.3583374023438 0 506.74573
872.4464721679688 0 13658.945
873.4495849609375 0 6649.406
874.449951171875 0 2259.458
889.346923828125 0 5343.266
890.3497314453125 0 3377.7744
896.5247802734375 0 564.6865
901.4791259765625 0 684.6409
918.4703979492188 0 922.47205
936.3304443359375 0 1000.4138
953.3449096679688 0 19284.902 y 2
954.3473510742188 0 9556.494
955.3463134765625 0 3097.6843
991.51904296875 0 2479.3718
992.5182495117188 0 644.5173
1003.371826171875 0 786.8503
1019.5130004882812 0 5433.3804
1020.5173950195312 0 3673.059
1021.5181884765625 0 833.32227
1036.2188720703125 0 626.901
1036.4168701171875 0 662.7559
1100.409423828125 0 941.23193 y 1
1101.4149169921875 0 703.71436
1634.6484375 0 586.299
1660.3424072265625 0 646.20746

Spectrum Details

|  |  |
| --- | --- |
| Matched peaks? Matched peaks The total absolute number of peaks matched. Additionally in brackets the total fraction of peaks matched and the total number of peaks is shown. | 15 (8.43% of 178) |
| FDR? FDR The false discovery rate estimated for this peptide. It is calculated by matching all theoretical fragments with a non-integer shift with the raw peaks for this spectrum. This is done with 40 different shifts. The resulting percentage is the average number of annotated peaks over the number of annotated peaks with the correct spectrum. | 2.22% |
| Satellite FDR? Satellite FDR See the FDR for details on its calculation. This satellite ion specific FDR only contains the satellite ions (d/w) for I/L/J positions. | - |
| PSM Score? PSM Score The PSM Score as given by Hecklib to this annotated spectrum. It is shown with three significant figures. | 160 |

#### Spectrum 12356? Spectrum 12356 The raw spectrum of this peptide as annotated by Hecklib. The fragments are coloured according to ion type (see legend). Any peaks with a star '\*' as text can be hovered over to see the full details, first the ion type second the mass shift type. By hovering over the amino acids in the peptide or ions in the legend the corresponding peaks are highlighted. By toggling the 'Unassigned' label you can turn the background (unassigned) peaks on or off in the plot. By updating the slider in the Ion legend you can update the spectrum to only show the top X% of the peaks with labels. The top X% means any peak that is within X% of the highest intensity. By dragging in the spectrum you can zoom in to a specific part of the spectrum and use 'Zoom Out' to get back to the original zoom level. The annotation of the spectrum is based on the given sequence in the peptides file and is done with different software so inconsistencies are likely. The peaks are annotated based on the given sequence, with 20 ppm tolerance.

Copy Data

##### Spectrum 12356 (TSV)

###### Preview

```
Loading example...
```

*Click on the button to copy the data to your clipboard.*

Mz MinMz MaxIntensity Max

WidthHeightPeptide font sizePeptide stroke widthSpectrum font sizeSpectrum stroke widthCompact peptide

Ion legend

wxyz

abcd

OtherUnassignedIon^Charge^_Position_Show for top:%

TFDDYAMHW

01.10e+42.21e+43.31e+44.41e+4

Zoom Out

a^+1^_2_y^+1^_1_a^+1^_2_b^+1^_2_y^+1^_2_b^+1^_3_b^+1^_4_y^+1^_3_y^+1^_4_\*y^+1^_5_y^+1^_6_y^+1^_7_y^+1^_7_y^+1^_8_

0777155423323109

Fragment Matches Table

Show background peaks

| Position | Ion type | Intensity | mz Theoretical | mz Error (Th) | mz Error (ppm) | Charge | Series Number |
| --- | --- | --- | --- | --- | --- | --- | --- |
| - | - | 2.998E+04 | 120.1 | - | - | 0 | - |
| - | - | 2736 | 121.1 | - | - | 0 | - |
| - | - | 377.3 | 122 | - | - | 0 | - |
| - | - | 491.4 | 125.9 | - | - | 0 | - |
| - | - | 3605 | 129.1 | - | - | 0 | - |
| - | - | 653.2 | 130.1 | - | - | 0 | - |
| - | - | 553.8 | 130.1 | - | - | 0 | - |
| - | - | 2062 | 133.1 | - | - | 0 | - |
| - | - | 368 | 134.1 | - | - | 0 | - |
| - | - | 1.049E+04 | 136.1 | - | - | 0 | - |
| - | - | 474.6 | 138.1 | - | - | 0 | - |
| - | - | 578.3 | 139.1 | - | - | 0 | - |
| - | - | 379.7 | 141 | - | - | 0 | - |
| - | - | 411.9 | 142.4 | - | - | 0 | - |
| - | - | 758 | 146.1 | - | - | 0 | - |
| - | - | 554.9 | 147 | - | - | 0 | - |
| - | - | 1243 | 149 | - | - | 0 | - |
| - | - | 423.9 | 155.2 | - | - | 0 | - |
| - | - | 848.7 | 156.1 | - | - | 0 | - |
| - | - | 1598 | 158.1 | - | - | 0 | - |
| - | - | 1615 | 159.1 | - | - | 0 | - |
| - | - | 718.4 | 160.1 | - | - | 0 | - |
| - | - | 654 | 162.1 | - | - | 0 | - |
| - | - | 3110 | 165.1 | - | - | 0 | - |
| - | - | 610.5 | 166.1 | - | - | 0 | - |
| - | - | 666.7 | 166.1 | - | - | 0 | - |
| - | - | 2397 | 167.1 | - | - | 0 | - |
| - | - | 416.7 | 171.6 | - | - | 0 | - |
| - | - | 1709 | 173.1 | - | - | 0 | - |
| - | - | 924.2 | 173.4 | - | - | 0 | - |
| - | - | 2665 | 176.1 | - | - | 0 | - |
| - | - | 2737 | 177.1 | - | - | 0 | - |
| - | - | 690.6 | 177.1 | - | - | 0 | - |
| - | - | 458.1 | 182 | - | - | 0 | - |
| - | - | 5542 | 182.1 | - | - | 0 | - |
| - | - | 587.9 | 183.1 | - | - | 0 | - |
| - | - | 520.6 | 185.1 | - | - | 0 | - |
| - | - | 562 | 185.1 | - | - | 0 | - |
| - | - | 522.1 | 186.1 | - | - | 0 | - |
| - | - | 505.7 | 186.1 | - | - | 0 | - |
| - | - | 1194 | 188.1 | - | - | 0 | - |
| - | - | 492.1 | 193.1 | - | - | 0 | - |
| - | - | 871.3 | 193.1 | - | - | 0 | - |
| - | - | 606 | 199.1 | - | - | 0 | - |
| - | - | 910.4 | 201.1 | - | - | 0 | - |
| - | - | 621 | 203.1 | - | - | 0 | - |
| 2 | a | 992.6 | 203.1 | 0.0001982 | 0.976 | +1 | 2 |
| 9 | y | 1058 | 205.1 | 3.19E-05 | 0.1555 | +1 | 1 |
| - | - | 1560 | 221.1 | - | - | 0 | - |
| 2 | a | 4.367E+04 | 221.1 | 7.056E-05 | 0.3191 | +1 | 2 |
| - | - | 5527 | 222.1 | - | - | 0 | - |
| - | - | 1324 | 223.1 | - | - | 0 | - |
| - | - | 750.4 | 227.1 | - | - | 0 | - |
| - | - | 905.9 | 231.1 | - | - | 0 | - |
| - | - | 1104 | 235.1 | - | - | 0 | - |
| - | - | 1875 | 239.1 | - | - | 0 | - |
| - | - | 624.6 | 240.1 | - | - | 0 | - |
| - | - | 773.8 | 241.1 | - | - | 0 | - |
| 2 | b | 7957 | 249.1 | 0.0002426 | 0.9738 | +1 | 2 |
| - | - | 797.4 | 250.1 | - | - | 0 | - |
| - | - | 736.6 | 251.1 | - | - | 0 | - |
| - | - | 557.8 | 257.2 | - | - | 0 | - |
| - | - | 1637 | 263.1 | - | - | 0 | - |
| - | - | 579.3 | 283 | - | - | 0 | - |
| - | - | 2122 | 285 | - | - | 0 | - |
| - | - | 868.9 | 285.1 | - | - | 0 | - |
| - | - | 478.2 | 285.2 | - | - | 0 | - |
| - | - | 571.2 | 286 | - | - | 0 | - |
| - | - | 547.9 | 291.2 | - | - | 0 | - |
| - | - | 766.2 | 292.1 | - | - | 0 | - |
| - | - | 933 | 296.2 | - | - | 0 | - |
| - | - | 959.7 | 303.2 | - | - | 0 | - |
| - | - | 937.8 | 309.2 | - | - | 0 | - |
| - | - | 524.2 | 315.1 | - | - | 0 | - |
| - | - | 1967 | 324.1 | - | - | 0 | - |
| - | - | 4184 | 329.1 | - | - | 0 | - |
| - | - | 962 | 330.2 | - | - | 0 | - |
| 8 | y | 6836 | 342.2 | 0.0003061 | 0.8947 | +1 | 2 |
| - | - | 1849 | 343.2 | - | - | 0 | - |
| - | - | 648 | 350.1 | - | - | 0 | - |
| - | - | 2053 | 355.1 | - | - | 0 | - |
| - | - | 721.2 | 356.1 | - | - | 0 | - |
| - | - | 683.2 | 357.2 | - | - | 0 | - |
| 3 | b | 2330 | 364.2 | 0.000318 | 0.8732 | +1 | 3 |
| - | - | 883.1 | 378.1 | - | - | 0 | - |
| - | - | 1037 | 394.1 | - | - | 0 | - |
| - | - | 720.7 | 413.3 | - | - | 0 | - |
| - | - | 1362 | 425.2 | - | - | 0 | - |
| - | - | 866.8 | 435.3 | - | - | 0 | - |
| - | - | 604.6 | 455.2 | - | - | 0 | - |
| - | - | 1041 | 465.2 | - | - | 0 | - |
| - | - | 552.5 | 465.2 | - | - | 0 | - |
| - | - | 584.2 | 471.2 | - | - | 0 | - |
| 4 | b | 1064 | 479.2 | 0.0004055 | 0.8463 | +1 | 4 |
| 7 | y | 1317 | 489.2 | 0.005985 | 12.23 | +1 | 3 |
| - | - | 618.3 | 496.2 | - | - | 0 | - |
| - | - | 761.1 | 510.3 | - | - | 0 | - |
| 6 | y | 2781 | 560.2 | 0.005401 | 9.64 | +1 | 4 |
| - | - | 865.3 | 561.2 | - | - | 0 | - |
| - | - | 2183 | 585.3 | - | - | 0 | - |
| - | - | 719.7 | 592.7 | - | - | 0 | - |
| 0 | Precursor | 1510 | 601.2 | 0.003167 | 5.267 | +2 | -1 |
| - | - | 1135 | 601.3 | - | - | 0 | - |
| - | - | 955 | 602.2 | - | - | 0 | - |
| - | - | 686.5 | 612.2 | - | - | 0 | - |
| - | - | 837.1 | 628.4 | - | - | 0 | - |
| - | - | 779.9 | 640.3 | - | - | 0 | - |
| - | - | 689.4 | 654.3 | - | - | 0 | - |
| - | - | 687.5 | 659.3 | - | - | 0 | - |
| - | - | 724.3 | 667.2 | - | - | 0 | - |
| - | - | 572.1 | 675.7 | - | - | 0 | - |
| - | - | 958.1 | 685.3 | - | - | 0 | - |
| - | - | 3690 | 700.4 | - | - | 0 | - |
| - | - | 1733 | 701.4 | - | - | 0 | - |
| - | - | 589.7 | 709.6 | - | - | 0 | - |
| - | - | 1283 | 721.3 | - | - | 0 | - |
| 5 | y | 3201 | 723.3 | 0.005366 | 7.418 | +1 | 5 |
| - | - | 1243 | 724.3 | - | - | 0 | - |
| - | - | 888.1 | 774.3 | - | - | 0 | - |
| - | - | 725.5 | 775.3 | - | - | 0 | - |
| - | - | 845.7 | 783.4 | - | - | 0 | - |
| - | - | 1963 | 801.4 | - | - | 0 | - |
| - | - | 1462 | 802.4 | - | - | 0 | - |
| - | - | 867.1 | 828.4 | - | - | 0 | - |
| 4 | y | 3526 | 838.3 | 0.006621 | 7.898 | +1 | 6 |
| - | - | 1188 | 839.3 | - | - | 0 | - |
| - | - | 1159 | 840.3 | - | - | 0 | - |
| - | - | 3042 | 844.5 | - | - | 0 | - |
| - | - | 1303 | 845.5 | - | - | 0 | - |
| - | - | 1.057E+04 | 872.4 | - | - | 0 | - |
| - | - | 5730 | 873.5 | - | - | 0 | - |
| - | - | 1193 | 874.5 | - | - | 0 | - |
| - | - | 5111 | 889.4 | - | - | 0 | - |
| - | - | 2253 | 890.4 | - | - | 0 | - |
| 3 | y | 1126 | 935.3 | 0.004341 | 4.642 | +1 | 7 |
| 3 | y | 1.456E+04 | 953.3 | 0.006167 | 6.469 | +1 | 7 |
| - | - | 8875 | 954.3 | - | - | 0 | - |
| - | - | 2754 | 955.4 | - | - | 0 | - |
| - | - | 1362 | 991.5 | - | - | 0 | - |
| - | - | 764.4 | 992.5 | - | - | 0 | - |
| - | - | 4545 | 1020 | - | - | 0 | - |
| - | - | 3178 | 1021 | - | - | 0 | - |
| - | - | 740.1 | 1022 | - | - | 0 | - |
| 2 | y | 935.7 | 1100 | 0.003915 | 3.558 | +1 | 8 |
| - | - | 653.1 | 1527 | - | - | 0 | - |
| - | - | 677.2 | 2853 | - | - | 0 | - |
| - | - | 689.6 | 2908 | - | - | 0 | - |
| - | - | 628.4 | 3078 | - | - | 0 | - |

m/z Charge Intensity FragmentType MassShift Position
120.08089447021484 0 29978.695
121.08418273925781 0 2735.7388
121.96521759033203 0 377.3168
125.86990356445312 0 491.40637
129.102294921875 0 3604.6777
130.05010986328125 0 653.15533
130.06536865234375 0 553.76526
133.08616638183594 0 2061.865
134.10353088378906 0 367.96484
136.07577514648438 0 10493.486
138.05474853515625 0 474.57382
139.0868682861328 0 578.31976
140.9649200439453 0 379.71344
142.44813537597656 0 411.87094
146.06015014648438 0 758.04956
147.0442657470703 0 554.91986
149.04483032226562 0 1242.7129
155.24989318847656 0 423.87228
156.0766143798828 0 848.70764
158.09642028808594 0 1598.4417
159.091796875 0 1615.1241
160.11264038085938 0 718.41425
162.09144592285156 0 653.9968
165.05465698242188 0 3109.768
166.05963134765625 0 610.46643
166.08631896972656 0 666.73505
167.05545043945312 0 2396.735
171.6282501220703 0 416.68625
173.1286163330078 0 1709.0421
173.43841552734375 0 924.16284
176.10702514648438 0 2664.8948
177.10231018066406 0 2736.5283
177.11082458496094 0 690.58923
182.02279663085938 0 458.0867
182.08120727539062 0 5542.1226
183.1494140625 0 587.8962
185.05567932128906 0 520.59076
185.1283721923828 0 561.9585
186.09158325195312 0 522.1333
186.12335205078125 0 505.74414
188.07081604003906 0 1193.927
193.0837860107422 0 492.10162
193.10865783691406 0 871.32837
199.07000732421875 0 605.9926
201.12339782714844 0 910.3673
203.06544494628906 0 620.99744
203.1180877685547 0 992.5942 a Water loss 1
205.0971221923828 0 1058.3468 y 8
221.1033172607422 0 1560.0928
221.12852478027344 0 43669.824 a 1
222.13194274902344 0 5527.443
223.0641326904297 0 1324.2706
227.10264587402344 0 750.4115
231.06173706054688 0 905.90875
235.10768127441406 0 1103.6587
239.0948486328125 0 1875.335
240.0977020263672 0 624.57056
241.07369995117188 0 773.79553
249.1236114501953 0 7957.22 b 1
250.12745666503906 0 797.37396
251.10231018066406 0 736.61206
257.1651306152344 0 557.7839
263.1029357910156 0 1637.3943
283.0304260253906 0 579.3004
285.0094909667969 0 2122.4727
285.1018371582031 0 868.8592
285.15850830078125 0 478.24936
286.0128173828125 0 571.1519
291.16748046875 0 547.9078
292.1417541503906 0 766.16925
296.15179443359375 0 933.0409
303.1808776855469 0 959.67706
309.2042541503906 0 937.7857
315.10546875 0 524.2318
324.1452331542969 0 1967.0986
329.1497802734375 0 4183.9517
330.15325927734375 0 961.9919
342.1563720703125 0 6835.86 y 7
343.15972900390625 0 1848.5516
350.134765625 0 647.9734
355.0695495605469 0 2052.6028
356.06976318359375 0 721.1896
357.22412109375 0 683.23883
364.1499938964844 0 2330.4802 b 2
378.12847900390625 0 883.11084
394.1244201660156 0 1036.5706
413.26446533203125 0 720.6736
425.19305419921875 0 1362.4576
435.2587890625 0 866.7729
455.2223815917969 0 604.5626
465.16241455078125 0 1040.8624
465.22857666015625 0 552.4569
471.1792907714844 0 584.1631
479.1768493652344 0 1064.4003 b 3
489.1925354003906 0 1317.22 y 6
496.2336120605469 0 618.2869
510.2655029296875 0 761.055
560.2290649414062 0 2781.4275 y 5
561.2314453125 0 865.2756
585.3414916992188 0 2182.8132
592.7318115234375 0 719.73114
601.2352905273438 0 1509.8497 Precursor
601.31884765625 0 1135.2108
602.2373657226562 0 955.02795
612.1964111328125 0 686.48315
628.3821411132812 0 837.14746
640.343505859375 0 779.9447
654.3265991210938 0 689.3594
659.2891235351562 0 687.52606
667.159912109375 0 724.3365
675.6826171875 0 572.05255
685.2584838867188 0 958.1389
700.3630981445312 0 3689.8435
701.3638916015625 0 1732.5588
709.6005859375 0 589.73376
721.2628173828125 0 1282.663
723.2923583984375 0 3200.6228 y 4
724.2944946289062 0 1243.0051
774.322998046875 0 888.0634
775.3216552734375 0 725.459
783.3991088867188 0 845.7462
801.4141845703125 0 1962.8611
802.416748046875 0 1462.2188
828.4223022460938 0 867.0673
838.320556640625 0 3525.9407 y 3
839.3181762695312 0 1187.5436
840.323486328125 0 1159.2944
844.4542846679688 0 3042.2659
845.4572143554688 0 1303.2083
872.4486083984375 0 10574.009
873.4518432617188 0 5730.0728
874.4545288085938 0 1192.5223
889.3502197265625 0 5111.4595
890.3512573242188 0 2253.13
935.3346557617188 0 1126.3379 y Water loss 2
953.3470458984375 0 14556.184 y 2
954.34912109375 0 8874.64
955.3505859375 0 2754.0771
991.52392578125 0 1361.8574
992.5169067382812 0 764.3754
1019.5169067382812 0 4545.0674
1020.5194702148438 0 3177.6577
1021.5238037109375 0 740.1128
1100.4132080078125 0 935.71484 y 1
1527.237060546875 0 653.074
2853.142333984375 0 677.2018
2907.7470703125 0 689.59265
3078.07763671875 0 628.38306

Spectrum Details

|  |  |
| --- | --- |
| Matched peaks? Matched peaks The total absolute number of peaks matched. Additionally in brackets the total fraction of peaks matched and the total number of peaks is shown. | 15 (10.14% of 148) |
| FDR? FDR The false discovery rate estimated for this peptide. It is calculated by matching all theoretical fragments with a non-integer shift with the raw peaks for this spectrum. This is done with 40 different shifts. The resulting percentage is the average number of annotated peaks over the number of annotated peaks with the correct spectrum. | 1.11% |
| Satellite FDR? Satellite FDR See the FDR for details on its calculation. This satellite ion specific FDR only contains the satellite ions (d/w) for I/L/J positions. | - |
| PSM Score? PSM Score The PSM Score as given by Hecklib to this annotated spectrum. It is shown with three significant figures. | 177 |

#### Spectrum 12909? Spectrum 12909 The raw spectrum of this peptide as annotated by Hecklib. The fragments are coloured according to ion type (see legend). Any peaks with a star '\*' as text can be hovered over to see the full details, first the ion type second the mass shift type. By hovering over the amino acids in the peptide or ions in the legend the corresponding peaks are highlighted. By toggling the 'Unassigned' label you can turn the background (unassigned) peaks on or off in the plot. By updating the slider in the Ion legend you can update the spectrum to only show the top X% of the peaks with labels. The top X% means any peak that is within X% of the highest intensity. By dragging in the spectrum you can zoom in to a specific part of the spectrum and use 'Zoom Out' to get back to the original zoom level. The annotation of the spectrum is based on the given sequence in the peptides file and is done with different software so inconsistencies are likely. The peaks are annotated based on the given sequence, with 20 ppm tolerance.

Copy Data

##### Spectrum 12909 (TSV)

###### Preview

```
Loading example...
```

*Click on the button to copy the data to your clipboard.*

Mz MinMz MaxIntensity Max

WidthHeightPeptide font sizePeptide stroke widthSpectrum font sizeSpectrum stroke widthCompact peptide

Ion legend

wxyz

abcd

OtherUnassignedIon^Charge^_Position_Show for top:%

TFDDYAMHW

08.22e+31.64e+42.47e+43.29e+4

Zoom Out

y^+1^_1_a^+1^_2_b^+1^_2_y^+1^_2_b^+1^_3_b^+1^_4_y^+1^_3_y^+2^_8_y^+1^_4_\*y^+1^_5_y^+1^_6_y^+1^_7_y^+1^_7_y^+1^_8_

040480912131617

Fragment Matches Table

Show background peaks

| Position | Ion type | Intensity | mz Theoretical | mz Error (Th) | mz Error (ppm) | Charge | Series Number |
| --- | --- | --- | --- | --- | --- | --- | --- |
| - | - | 2.282E+04 | 120.1 | - | - | 0 | - |
| - | - | 1715 | 121.1 | - | - | 0 | - |
| - | - | 506.6 | 129.1 | - | - | 0 | - |
| - | - | 3124 | 129.1 | - | - | 0 | - |
| - | - | 650.8 | 132.1 | - | - | 0 | - |
| - | - | 413.8 | 133.1 | - | - | 0 | - |
| - | - | 1136 | 133.1 | - | - | 0 | - |
| - | - | 641 | 136.1 | - | - | 0 | - |
| - | - | 8067 | 136.1 | - | - | 0 | - |
| - | - | 617.9 | 137.1 | - | - | 0 | - |
| - | - | 600.8 | 138.1 | - | - | 0 | - |
| - | - | 684.6 | 141.1 | - | - | 0 | - |
| - | - | 829.8 | 146.1 | - | - | 0 | - |
| - | - | 572.3 | 148.9 | - | - | 0 | - |
| - | - | 744.5 | 148.9 | - | - | 0 | - |
| - | - | 581.7 | 148.9 | - | - | 0 | - |
| - | - | 853.9 | 148.9 | - | - | 0 | - |
| - | - | 834.4 | 148.9 | - | - | 0 | - |
| - | - | 1145 | 148.9 | - | - | 0 | - |
| - | - | 1389 | 148.9 | - | - | 0 | - |
| - | - | 2707 | 148.9 | - | - | 0 | - |
| - | - | 4169 | 148.9 | - | - | 0 | - |
| - | - | 4220 | 149 | - | - | 0 | - |
| - | - | 2290 | 149 | - | - | 0 | - |
| - | - | 1427 | 149 | - | - | 0 | - |
| - | - | 1075 | 149 | - | - | 0 | - |
| - | - | 604.1 | 149 | - | - | 0 | - |
| - | - | 568.7 | 149 | - | - | 0 | - |
| - | - | 468.7 | 149 | - | - | 0 | - |
| - | - | 501.7 | 149 | - | - | 0 | - |
| - | - | 679.4 | 149 | - | - | 0 | - |
| - | - | 418.3 | 149.3 | - | - | 0 | - |
| - | - | 480.3 | 151 | - | - | 0 | - |
| - | - | 929.7 | 156.1 | - | - | 0 | - |
| - | - | 559.1 | 157.1 | - | - | 0 | - |
| - | - | 848.6 | 158.1 | - | - | 0 | - |
| - | - | 1349 | 159.1 | - | - | 0 | - |
| - | - | 916.1 | 160.1 | - | - | 0 | - |
| - | - | 420 | 161.3 | - | - | 0 | - |
| - | - | 558.8 | 162.1 | - | - | 0 | - |
| - | - | 2771 | 165.1 | - | - | 0 | - |
| - | - | 975.8 | 166.1 | - | - | 0 | - |
| - | - | 1811 | 167.1 | - | - | 0 | - |
| - | - | 1358 | 173.1 | - | - | 0 | - |
| - | - | 642.7 | 173.5 | - | - | 0 | - |
| - | - | 2766 | 176.1 | - | - | 0 | - |
| - | - | 577.1 | 176.2 | - | - | 0 | - |
| - | - | 1848 | 177.1 | - | - | 0 | - |
| - | - | 807.2 | 177.1 | - | - | 0 | - |
| - | - | 4894 | 182.1 | - | - | 0 | - |
| - | - | 577.3 | 185.1 | - | - | 0 | - |
| - | - | 1929 | 188.1 | - | - | 0 | - |
| - | - | 505.7 | 192.8 | - | - | 0 | - |
| - | - | 515.3 | 200.1 | - | - | 0 | - |
| - | - | 1418 | 201.1 | - | - | 0 | - |
| 9 | y | 754.5 | 205.1 | 0.0003038 | 1.481 | +1 | 1 |
| - | - | 942.6 | 215.1 | - | - | 0 | - |
| - | - | 600.8 | 217.1 | - | - | 0 | - |
| - | - | 1936 | 221.1 | - | - | 0 | - |
| 2 | a | 3.256E+04 | 221.1 | 0.0001316 | 0.5951 | +1 | 2 |
| - | - | 4863 | 222.1 | - | - | 0 | - |
| - | - | 1393 | 223.1 | - | - | 0 | - |
| - | - | 595 | 226.2 | - | - | 0 | - |
| - | - | 730.5 | 227.1 | - | - | 0 | - |
| - | - | 777.7 | 229.2 | - | - | 0 | - |
| - | - | 885 | 235.1 | - | - | 0 | - |
| - | - | 624.9 | 237.9 | - | - | 0 | - |
| - | - | 531.6 | 238 | - | - | 0 | - |
| - | - | 1117 | 239.1 | - | - | 0 | - |
| - | - | 976.8 | 241.1 | - | - | 0 | - |
| 2 | b | 6447 | 249.1 | 0.0002731 | 1.096 | +1 | 2 |
| - | - | 1185 | 263.1 | - | - | 0 | - |
| - | - | 1012 | 285 | - | - | 0 | - |
| - | - | 645.1 | 296.2 | - | - | 0 | - |
| - | - | 614.7 | 299.1 | - | - | 0 | - |
| - | - | 730.2 | 309.2 | - | - | 0 | - |
| - | - | 1160 | 319.1 | - | - | 0 | - |
| - | - | 1400 | 324.1 | - | - | 0 | - |
| - | - | 2972 | 329.2 | - | - | 0 | - |
| 8 | y | 5563 | 342.2 | 0.0005197 | 1.519 | +1 | 2 |
| - | - | 791.6 | 342.2 | - | - | 0 | - |
| - | - | 961 | 343.2 | - | - | 0 | - |
| - | - | 1172 | 355.1 | - | - | 0 | - |
| - | - | 574.2 | 356.1 | - | - | 0 | - |
| - | - | 584.7 | 358.6 | - | - | 0 | - |
| 3 | b | 1974 | 364.2 | 0.0004755 | 1.306 | +1 | 3 |
| - | - | 613.5 | 365.2 | - | - | 0 | - |
| - | - | 988.2 | 378.1 | - | - | 0 | - |
| - | - | 1167 | 394.1 | - | - | 0 | - |
| - | - | 602.7 | 398.3 | - | - | 0 | - |
| - | - | 815 | 413.3 | - | - | 0 | - |
| - | - | 529.4 | 440 | - | - | 0 | - |
| - | - | 615.3 | 455.2 | - | - | 0 | - |
| - | - | 809 | 465.2 | - | - | 0 | - |
| 4 | b | 705.8 | 479.2 | 0.004416 | 9.216 | +1 | 4 |
| 7 | y | 1563 | 489.2 | 0.006046 | 12.36 | +1 | 3 |
| - | - | 1231 | 496.2 | - | - | 0 | - |
| - | - | 615.5 | 496.6 | - | - | 0 | - |
| - | - | 528.1 | 542.2 | - | - | 0 | - |
| 2 | y | 1023 | 550.7 | 0.001066 | 1.936 | +2 | 8 |
| - | - | 1178 | 551.2 | - | - | 0 | - |
| 6 | y | 1997 | 560.2 | 0.005156 | 9.204 | +1 | 4 |
| - | - | 612.3 | 561.2 | - | - | 0 | - |
| - | - | 1674 | 585.3 | - | - | 0 | - |
| - | - | 636 | 586.3 | - | - | 0 | - |
| 0 | Precursor | 1516 | 601.2 | 0.005486 | 9.125 | +2 | -1 |
| - | - | 1339 | 601.3 | - | - | 0 | - |
| - | - | 544.5 | 601.8 | - | - | 0 | - |
| - | - | 774 | 602.2 | - | - | 0 | - |
| - | - | 1358 | 659.3 | - | - | 0 | - |
| - | - | 1000 | 678.2 | - | - | 0 | - |
| - | - | 723.8 | 685.3 | - | - | 0 | - |
| - | - | 3691 | 700.4 | - | - | 0 | - |
| - | - | 864.7 | 701.4 | - | - | 0 | - |
| - | - | 588.5 | 701.7 | - | - | 0 | - |
| - | - | 961.2 | 721.3 | - | - | 0 | - |
| 5 | y | 1674 | 723.3 | 0.006159 | 8.515 | +1 | 5 |
| - | - | 875.5 | 724.3 | - | - | 0 | - |
| - | - | 992.6 | 749.3 | - | - | 0 | - |
| - | - | 778.3 | 754.4 | - | - | 0 | - |
| - | - | 1202 | 774.3 | - | - | 0 | - |
| - | - | 1671 | 801.4 | - | - | 0 | - |
| - | - | 769.5 | 802.4 | - | - | 0 | - |
| - | - | 943.7 | 828.4 | - | - | 0 | - |
| 4 | y | 3062 | 838.3 | 0.005278 | 6.296 | +1 | 6 |
| - | - | 613.2 | 838.4 | - | - | 0 | - |
| - | - | 1606 | 839.3 | - | - | 0 | - |
| - | - | 2397 | 844.5 | - | - | 0 | - |
| - | - | 626.8 | 864.4 | - | - | 0 | - |
| - | - | 6830 | 872.4 | - | - | 0 | - |
| - | - | 627 | 872.9 | - | - | 0 | - |
| - | - | 4610 | 873.5 | - | - | 0 | - |
| - | - | 844.2 | 874.5 | - | - | 0 | - |
| - | - | 4323 | 889.4 | - | - | 0 | - |
| - | - | 2193 | 890.4 | - | - | 0 | - |
| 3 | y | 1002 | 935.3 | 0.01075 | 11.49 | +1 | 7 |
| - | - | 804.2 | 936.3 | - | - | 0 | - |
| 3 | y | 1.301E+04 | 953.3 | 0.006411 | 6.725 | +1 | 7 |
| - | - | 6177 | 954.3 | - | - | 0 | - |
| - | - | 2571 | 955.3 | - | - | 0 | - |
| - | - | 1190 | 991.5 | - | - | 0 | - |
| - | - | 652.4 | 992.5 | - | - | 0 | - |
| - | - | 2801 | 1020 | - | - | 0 | - |
| - | - | 2176 | 1021 | - | - | 0 | - |
| 2 | y | 768.8 | 1100 | 0.01344 | 12.21 | +1 | 8 |
| - | - | 845.4 | 1460 | - | - | 0 | - |
| - | - | 684.9 | 1601 | - | - | 0 | - |

m/z Charge Intensity FragmentType MassShift Position
120.08092498779297 0 22817.213
121.08419799804688 0 1714.6277
129.0655975341797 0 506.6044
129.10240173339844 0 3124.1338
132.08111572265625 0 650.7507
133.06092834472656 0 413.78287
133.08572387695312 0 1136.0754
136.07090759277344 0 641.01843
136.07582092285156 0 8067.218
137.0793914794922 0 617.949
138.06625366210938 0 600.76404
141.10263061523438 0 684.591
146.06044006347656 0 829.75055
148.8860626220703 0 572.25726
148.89266967773438 0 744.5499
148.90023803710938 0 581.66864
148.9072265625 0 853.8832
148.9142303466797 0 834.386
148.922119140625 0 1145.4005
148.92926025390625 0 1389.3447
148.9363250732422 0 2707.3442
148.94407653808594 0 4168.941
148.96067810058594 0 4219.5254
148.96859741210938 0 2290.0571
148.97576904296875 0 1426.8658
148.98292541503906 0 1074.568
148.9906463623047 0 604.1373
148.9976348876953 0 568.6575
149.0050811767578 0 468.67944
149.0269317626953 0 501.69177
149.04498291015625 0 679.4357
149.29759216308594 0 418.34592
150.98916625976562 0 480.2945
156.076904296875 0 929.7246
157.1335906982422 0 559.1481
158.0967559814453 0 848.56085
159.09178161621094 0 1349.1229
160.1120147705078 0 916.1325
161.30117797851562 0 420.00186
162.09164428710938 0 558.8116
165.0547637939453 0 2771.2676
166.08656311035156 0 975.78143
167.05540466308594 0 1810.8483
173.12847900390625 0 1358.116
173.450439453125 0 642.74176
176.10714721679688 0 2766.3877
176.2101287841797 0 577.1484
177.1024932861328 0 1847.5563
177.11109924316406 0 807.2005
182.08119201660156 0 4894.4443
185.12838745117188 0 577.3001
188.07049560546875 0 1928.8826
192.8450927734375 0 505.73688
200.13966369628906 0 515.3076
201.12322998046875 0 1417.8718
205.0974578857422 0 754.53107 y 8
215.12835693359375 0 942.6146
217.09742736816406 0 600.75507
221.10348510742188 0 1936.4363
221.1285858154297 0 32561.62 a 1
222.13198852539062 0 4863.3564
223.06378173828125 0 1393.3695
226.1545867919922 0 595.0327
227.1023406982422 0 730.4842
229.16978454589844 0 777.7018
235.10777282714844 0 885.02875
237.93112182617188 0 624.9347
237.9678192138672 0 531.61847
239.09532165527344 0 1116.9745
241.07400512695312 0 976.79926
249.12364196777344 0 6447.007 b 1
263.1031799316406 0 1185.3011
285.00885009765625 0 1012.0667
296.15057373046875 0 645.13336
299.06103515625 0 614.7477
309.20343017578125 0 730.2056
319.13958740234375 0 1159.8884
324.14630126953125 0 1399.5232
329.150146484375 0 2972.4985
342.1565856933594 0 5563.4556 y 7
342.1769104003906 0 791.56177
343.16021728515625 0 961.0366
355.0704650878906 0 1172.0198
356.0691833496094 0 574.2061
358.6365051269531 0 584.6981
364.1507873535156 0 1973.8585 b 2
365.15313720703125 0 613.4508
378.1292724609375 0 988.1871
394.12451171875 0 1166.8188
398.2809753417969 0 602.679
413.26513671875 0 815.03394
439.96856689453125 0 529.4043
455.228759765625 0 615.29724
465.1610107421875 0 809.0209
479.1816711425781 0 705.847 b 3
489.1925964355469 0 1563.452 y 6
496.2303771972656 0 1230.7628
496.60870361328125 0 615.4503
542.2203979492188 0 528.07117
550.7093505859375 0 1023.229 y 1
551.214111328125 0 1178.4258
560.2288208007812 0 1997.2009 y 5
561.2308959960938 0 612.25995
585.3399658203125 0 1674.1129
586.3391723632812 0 636.0431
601.2376098632812 0 1515.8453 Precursor
601.3206176757812 0 1339.0032
601.7510375976562 0 544.51117
602.236328125 0 774.0154
659.2932739257812 0 1358.3156
678.2261962890625 0 1000.233
685.2601318359375 0 723.83484
700.3635864257812 0 3691.0242
701.3658447265625 0 864.6883
701.6921997070312 0 588.52057
721.2620239257812 0 961.1511
723.2931518554688 0 1673.7924 y 4
724.2942504882812 0 875.4949
749.2604370117188 0 992.61096
754.374267578125 0 778.306
774.3258666992188 0 1202.3586
801.41259765625 0 1671.14
802.4097290039062 0 769.4957
828.4201049804688 0 943.6829
838.3192138671875 0 3062.4788 y 3
838.3988037109375 0 613.1801
839.3216552734375 0 1606.4508
844.4531860351562 0 2396.938
864.3976440429688 0 626.82983
872.4481811523438 0 6830.312
872.9080200195312 0 626.98944
873.4508056640625 0 4609.6333
874.4552612304688 0 844.18787
889.3501586914062 0 4323.4937
890.3521118164062 0 2193.48
935.341064453125 0 1002.0851 y Water loss 2
936.3342895507812 0 804.23944
953.3472900390625 0 13008.89 y 2
954.3497924804688 0 6177.426
955.3493041992188 0 2570.7825
991.524658203125 0 1189.6671
992.52001953125 0 652.3755
1019.5186767578125 0 2800.9668
1020.5216674804688 0 2176.395
1100.4227294921875 0 768.8318 y 1
1459.6927490234375 0 845.3935
1601.4476318359375 0 684.88776

Spectrum Details

|  |  |
| --- | --- |
| Matched peaks? Matched peaks The total absolute number of peaks matched. Additionally in brackets the total fraction of peaks matched and the total number of peaks is shown. | 15 (10.20% of 147) |
| FDR? FDR The false discovery rate estimated for this peptide. It is calculated by matching all theoretical fragments with a non-integer shift with the raw peaks for this spectrum. This is done with 40 different shifts. The resulting percentage is the average number of annotated peaks over the number of annotated peaks with the correct spectrum. | 1.43% |
| Satellite FDR? Satellite FDR See the FDR for details on its calculation. This satellite ion specific FDR only contains the satellite ions (d/w) for I/L/J positions. | - |
| PSM Score? PSM Score The PSM Score as given by Hecklib to this annotated spectrum. It is shown with three significant figures. | 160 |

#### Spectrum 6950? Spectrum 6950 The raw spectrum of this peptide as annotated by Hecklib. The fragments are coloured according to ion type (see legend). Any peaks with a star '\*' as text can be hovered over to see the full details, first the ion type second the mass shift type. By hovering over the amino acids in the peptide or ions in the legend the corresponding peaks are highlighted. By toggling the 'Unassigned' label you can turn the background (unassigned) peaks on or off in the plot. By updating the slider in the Ion legend you can update the spectrum to only show the top X% of the peaks with labels. The top X% means any peak that is within X% of the highest intensity. By dragging in the spectrum you can zoom in to a specific part of the spectrum and use 'Zoom Out' to get back to the original zoom level. The annotation of the spectrum is based on the given sequence in the peptides file and is done with different software so inconsistencies are likely. The peaks are annotated based on the given sequence, with 20 ppm tolerance.

Copy Data

##### Spectrum 6950 (TSV)

###### Preview

```
Loading example...
```

*Click on the button to copy the data to your clipboard.*

Mz MinMz MaxIntensity Max

WidthHeightPeptide font sizePeptide stroke widthSpectrum font sizeSpectrum stroke widthCompact peptide

Ion legend

wxyz

abcd

OtherUnassignedIon^Charge^_Position_Show for top:%

TFDDYAMHW

02.66e+45.32e+47.98e+41.06e+5

Zoom Out

c^+1^_2_y^+1^_2_z^+1^_3_y^+2^_7_c^+1^_4_y^+1^_3_z^+1^_4_y^+2^_8_y^+1^_4_z^+1^_5_y^+1^_5_c^+1^_6_z^+1^_6_y^+1^_6_c^+1^_7_c^+1^_7_c^+1^_7_y^+1^_7_z^+1^_7_y^+1^_7_c^+1^_8_z^+1^_8_y^+1^_8_

0702140321052807

Fragment Matches Table

Show background peaks

| Position | Ion type | Intensity | mz Theoretical | mz Error (Th) | mz Error (ppm) | Charge | Series Number |
| --- | --- | --- | --- | --- | --- | --- | --- |
| - | - | 912.5 | 120.1 | - | - | 0 | - |
| - | - | 333.5 | 120.3 | - | - | 0 | - |
| - | - | 527.4 | 127 | - | - | 0 | - |
| - | - | 373.3 | 134.9 | - | - | 0 | - |
| - | - | 407.5 | 136.5 | - | - | 0 | - |
| - | - | 414.1 | 139.6 | - | - | 0 | - |
| - | - | 376.4 | 140.5 | - | - | 0 | - |
| - | - | 373.4 | 143.8 | - | - | 0 | - |
| - | - | 402.5 | 145.9 | - | - | 0 | - |
| - | - | 588.2 | 149 | - | - | 0 | - |
| - | - | 468.2 | 149.3 | - | - | 0 | - |
| - | - | 373.2 | 154.1 | - | - | 0 | - |
| - | - | 466.1 | 162.9 | - | - | 0 | - |
| - | - | 433.2 | 168.4 | - | - | 0 | - |
| - | - | 477.2 | 169.6 | - | - | 0 | - |
| - | - | 5943 | 173.4 | - | - | 0 | - |
| - | - | 486.8 | 176.2 | - | - | 0 | - |
| - | - | 446 | 184.8 | - | - | 0 | - |
| - | - | 460.9 | 196.5 | - | - | 0 | - |
| - | - | 574.1 | 203.1 | - | - | 0 | - |
| - | - | 546.3 | 216.7 | - | - | 0 | - |
| - | - | 570.4 | 217.1 | - | - | 0 | - |
| - | - | 1648 | 219.1 | - | - | 0 | - |
| - | - | 2.954E+04 | 221.1 | - | - | 0 | - |
| - | - | 3331 | 222.1 | - | - | 0 | - |
| - | - | 644.4 | 223.1 | - | - | 0 | - |
| - | - | 449.8 | 225.4 | - | - | 0 | - |
| - | - | 469.3 | 229.2 | - | - | 0 | - |
| - | - | 504.1 | 235.9 | - | - | 0 | - |
| 2 | c | 9669 | 249.1 | 0.0002273 | 0.9126 | +1 | 2 |
| - | - | 1756 | 250.1 | - | - | 0 | - |
| - | - | 593.1 | 258.5 | - | - | 0 | - |
| - | - | 625.2 | 282.1 | - | - | 0 | - |
| - | - | 619.4 | 283.2 | - | - | 0 | - |
| - | - | 1026 | 327.1 | - | - | 0 | - |
| 8 | y | 3117 | 342.2 | 0.0001535 | 0.4487 | +1 | 2 |
| - | - | 1128 | 355.1 | - | - | 0 | - |
| - | - | 570.5 | 468.3 | - | - | 0 | - |
| 7 | z | 2097 | 473.2 | 0.004995 | 10.56 | +1 | 3 |
| - | - | 1647 | 474.2 | - | - | 0 | - |
| 3 | y | 1177 | 477.2 | 0.0005439 | 1.14 | +2 | 7 |
| 4 | c | 1041 | 479.2 | 0.001748 | 3.649 | +1 | 4 |
| 7 | y | 2169 | 489.2 | 0.00571 | 11.67 | +1 | 3 |
| - | - | 812.4 | 500.2 | - | - | 0 | - |
| - | - | 980 | 521.2 | - | - | 0 | - |
| - | - | 858.2 | 543.3 | - | - | 0 | - |
| 6 | z | 1.481E+04 | 544.2 | 0.005082 | 9.338 | +1 | 4 |
| - | - | 6951 | 545.2 | - | - | 0 | - |
| - | - | 1503 | 546.2 | - | - | 0 | - |
| 2 | y | 2549 | 550.7 | 0.002836 | 5.15 | +2 | 8 |
| - | - | 2019 | 551.2 | - | - | 0 | - |
| - | - | 670 | 551.7 | - | - | 0 | - |
| 6 | y | 4257 | 560.2 | 0.00479 | 8.551 | +1 | 4 |
| - | - | 525.3 | 560.7 | - | - | 0 | - |
| - | - | 1065 | 561.2 | - | - | 0 | - |
| - | - | 610.4 | 575.3 | - | - | 0 | - |
| - | - | 740.7 | 592.3 | - | - | 0 | - |
| - | - | 896 | 592.8 | - | - | 0 | - |
| - | - | 692.4 | 600.4 | - | - | 0 | - |
| - | - | 1770 | 601.2 | - | - | 0 | - |
| - | - | 1831 | 601.7 | - | - | 0 | - |
| - | - | 1033 | 602.2 | - | - | 0 | - |
| - | - | 923.7 | 663.3 | - | - | 0 | - |
| - | - | 1413 | 675.3 | - | - | 0 | - |
| 5 | z | 1.786E+04 | 707.3 | 0.004558 | 6.445 | +1 | 5 |
| - | - | 8993 | 708.3 | - | - | 0 | - |
| - | - | 2552 | 709.3 | - | - | 0 | - |
| - | - | 785.7 | 710.3 | - | - | 0 | - |
| - | - | 1062 | 715.3 | - | - | 0 | - |
| - | - | 669.3 | 720.4 | - | - | 0 | - |
| - | - | 564.7 | 721.3 | - | - | 0 | - |
| 5 | y | 8000 | 723.3 | 0.004267 | 5.899 | +1 | 5 |
| - | - | 2471 | 724.3 | - | - | 0 | - |
| 6 | c | 998.5 | 730.3 | 0.002611 | 3.575 | +1 | 6 |
| - | - | 701.9 | 738.3 | - | - | 0 | - |
| - | - | 548.3 | 740.3 | - | - | 0 | - |
| - | - | 1953 | 741.4 | - | - | 0 | - |
| - | - | 1450 | 742.4 | - | - | 0 | - |
| - | - | 1738 | 749.3 | - | - | 0 | - |
| - | - | 4894 | 778.3 | - | - | 0 | - |
| - | - | 1855 | 779.3 | - | - | 0 | - |
| - | - | 616 | 787.5 | - | - | 0 | - |
| - | - | 1078 | 801.9 | - | - | 0 | - |
| - | - | 2364 | 802.4 | - | - | 0 | - |
| - | - | 844.2 | 802.9 | - | - | 0 | - |
| - | - | 691.8 | 803.4 | - | - | 0 | - |
| - | - | 1008 | 814.4 | - | - | 0 | - |
| 4 | z | 8106 | 822.3 | 0.005264 | 6.402 | +1 | 6 |
| - | - | 9409 | 823.3 | - | - | 0 | - |
| - | - | 2628 | 824.3 | - | - | 0 | - |
| - | - | 827.1 | 825.3 | - | - | 0 | - |
| - | - | 6289 | 833.3 | - | - | 0 | - |
| - | - | 3095 | 834.3 | - | - | 0 | - |
| - | - | 886.8 | 835.3 | - | - | 0 | - |
| 4 | y | 1674 | 838.3 | 0.003752 | 4.476 | +1 | 6 |
| - | - | 1280 | 839.3 | - | - | 0 | - |
| - | - | 3261 | 858.3 | - | - | 0 | - |
| 7 | c | 1657 | 859.3 | 0.003854 | 4.485 | +1 | 7 |
| 7 | c | 733.6 | 860.3 | 0.01109 | 12.89 | +1 | 7 |
| - | - | 2951 | 876.3 | - | - | 0 | - |
| 7 | c | 2.439E+04 | 877.3 | 0.003892 | 4.436 | +1 | 7 |
| - | - | 1.13E+04 | 878.3 | - | - | 0 | - |
| - | - | 4188 | 879.3 | - | - | 0 | - |
| - | - | 3982 | 880.3 | - | - | 0 | - |
| - | - | 2043 | 881.3 | - | - | 0 | - |
| - | - | 3395 | 889.3 | - | - | 0 | - |
| - | - | 2454 | 890.4 | - | - | 0 | - |
| - | - | 3537 | 893.3 | - | - | 0 | - |
| - | - | 2233 | 894.3 | - | - | 0 | - |
| - | - | 1026 | 895.3 | - | - | 0 | - |
| - | - | 802.8 | 895.5 | - | - | 0 | - |
| - | - | 793.2 | 896.3 | - | - | 0 | - |
| - | - | 855 | 907.3 | - | - | 0 | - |
| 3 | y | 779.1 | 935.3 | 0.001839 | 1.966 | +1 | 7 |
| - | - | 853.2 | 936.3 | - | - | 0 | - |
| 3 | z | 2594 | 937.3 | 0.004688 | 5.002 | +1 | 7 |
| - | - | 1325 | 938.3 | - | - | 0 | - |
| - | - | 1275 | 952.4 | - | - | 0 | - |
| 3 | y | 2.623E+04 | 953.3 | 0.004275 | 4.484 | +1 | 7 |
| - | - | 1.47E+04 | 954.3 | - | - | 0 | - |
| - | - | 4788 | 955.3 | - | - | 0 | - |
| - | - | 1347 | 970.4 | - | - | 0 | - |
| - | - | 1510 | 984.5 | - | - | 0 | - |
| - | - | 1039 | 985.5 | - | - | 0 | - |
| - | - | 2254 | 1001 | - | - | 0 | - |
| - | - | 1276 | 1002 | - | - | 0 | - |
| - | - | 723.8 | 1012 | - | - | 0 | - |
| 8 | c | 2.191E+04 | 1014 | 0.003269 | 3.222 | +1 | 8 |
| - | - | 1.11E+04 | 1015 | - | - | 0 | - |
| - | - | 4357 | 1016 | - | - | 0 | - |
| - | - | 792.5 | 1017 | - | - | 0 | - |
| - | - | 702.8 | 1038 | - | - | 0 | - |
| - | - | 1565 | 1072 | - | - | 0 | - |
| - | - | 671.3 | 1073 | - | - | 0 | - |
| 2 | z | 1219 | 1084 | 0.005671 | 5.23 | +1 | 8 |
| - | - | 646.3 | 1094 | - | - | 0 | - |
| 2 | y | 926 | 1100 | 0.002799 | 2.543 | +1 | 8 |
| - | - | 1047 | 1101 | - | - | 0 | - |
| - | - | 2322 | 1103 | - | - | 0 | - |
| - | - | 1147 | 1110 | - | - | 0 | - |
| - | - | 1069 | 1111 | - | - | 0 | - |
| - | - | 684.2 | 1113 | - | - | 0 | - |
| - | - | 1904 | 1123 | - | - | 0 | - |
| - | - | 790.9 | 1124 | - | - | 0 | - |
| - | - | 2016 | 1132 | - | - | 0 | - |
| - | - | 1206 | 1133 | - | - | 0 | - |
| - | - | 2287 | 1136 | - | - | 0 | - |
| - | - | 1642 | 1137 | - | - | 0 | - |
| - | - | 1378 | 1138 | - | - | 0 | - |
| - | - | 5784 | 1140 | - | - | 0 | - |
| - | - | 1.046E+04 | 1141 | - | - | 0 | - |
| - | - | 1.018E+04 | 1142 | - | - | 0 | - |
| - | - | 1141 | 1143 | - | - | 0 | - |
| - | - | 4701 | 1143 | - | - | 0 | - |
| - | - | 1068 | 1144 | - | - | 0 | - |
| - | - | 1207 | 1144 | - | - | 0 | - |
| - | - | 2051 | 1149 | - | - | 0 | - |
| - | - | 2453 | 1150 | - | - | 0 | - |
| - | - | 2941 | 1151 | - | - | 0 | - |
| - | - | 966.4 | 1156 | - | - | 0 | - |
| - | - | 1064 | 1158 | - | - | 0 | - |
| - | - | 918.3 | 1159 | - | - | 0 | - |
| - | - | 1028 | 1159 | - | - | 0 | - |
| - | - | 1204 | 1163 | - | - | 0 | - |
| - | - | 1930 | 1167 | - | - | 0 | - |
| - | - | 1448 | 1168 | - | - | 0 | - |
| - | - | 6329 | 1184 | - | - | 0 | - |
| - | - | 3.454E+04 | 1185 | - | - | 0 | - |
| - | - | 2.006E+04 | 1186 | - | - | 0 | - |
| - | - | 3807 | 1187 | - | - | 0 | - |
| - | - | 9239 | 1187 | - | - | 0 | - |
| - | - | 2011 | 1188 | - | - | 0 | - |
| - | - | 1707 | 1188 | - | - | 0 | - |
| - | - | 1050 | 1200 | - | - | 0 | - |
| - | - | 714.1 | 1201 | - | - | 0 | - |
| - | - | 2.409E+04 | 1201 | - | - | 0 | - |
| - | - | 1.053E+05 | 1202 | - | - | 0 | - |
| - | - | 6.535E+04 | 1203 | - | - | 0 | - |
| - | - | 2.701E+04 | 1204 | - | - | 0 | - |
| - | - | 2931 | 1205 | - | - | 0 | - |
| - | - | 773.9 | 1275 | - | - | 0 | - |
| - | - | 1602 | 1588 | - | - | 0 | - |
| - | - | 1109 | 1589 | - | - | 0 | - |
| - | - | 1532 | 1604 | - | - | 0 | - |
| - | - | 2798 | 1605 | - | - | 0 | - |
| - | - | 1829 | 1606 | - | - | 0 | - |
| - | - | 892.4 | 1791 | - | - | 0 | - |
| - | - | 2404 | 1808 | - | - | 0 | - |
| - | - | 717.6 | 2779 | - | - | 0 | - |

m/z Charge Intensity FragmentType MassShift Position
120.08097076416016 0 912.4929
120.25736236572266 0 333.5259
126.98407745361328 0 527.3557
134.9166259765625 0 373.29156
136.47311401367188 0 407.51846
139.639892578125 0 414.1032
140.52664184570312 0 376.3862
143.83724975585938 0 373.3865
145.92982482910156 0 402.51877
149.04519653320312 0 588.16766
149.34291076660156 0 468.15036
154.0717010498047 0 373.19815
162.91552734375 0 466.1497
168.35520935058594 0 433.1628
169.5570831298828 0 477.19916
173.44039916992188 0 5943.3945
176.1687469482422 0 486.77826
184.80459594726562 0 445.9893
196.46658325195312 0 460.90762
203.06581115722656 0 574.14996
216.6542510986328 0 546.30365
217.07200622558594 0 570.4069
219.13417053222656 0 1647.8376
221.12855529785156 0 29542.379
222.13189697265625 0 3331.3447
223.0635223388672 0 644.3834
225.40602111816406 0 449.81088
229.15509033203125 0 469.29282
235.94264221191406 0 504.14044
249.12359619140625 0 9668.826 c Ammonia loss 1
250.1265869140625 0 1755.6681
258.4731140136719 0 593.1449
282.1473083496094 0 625.1903
283.1553955078125 0 619.4132
327.1460266113281 0 1025.5409
342.1562194824219 0 3116.5703 y 7
355.07012939453125 0 1128.2953
468.29205322265625 0 570.47485
473.1728210449219 0 2097.0754 z 6
474.1805419921875 0 1646.8815
477.17462158203125 0 1177.009 y 2
479.1755065917969 0 1040.5925 c Ammonia loss 3
489.1922607421875 0 2168.9883 y 6
500.2201843261719 0 812.381
521.2185668945312 0 979.99054
543.2531127929688 0 858.2381
544.2100219726562 0 14805.565 z 5
545.2146606445312 0 6950.792
546.2162475585938 0 1502.8989
550.7111206054688 0 2548.964 y 1
551.2129516601562 0 2018.7615
551.7164306640625 0 669.99524
560.2284545898438 0 4257.316 y 5
560.7186279296875 0 525.33435
561.2315063476562 0 1065.3486
575.2877807617188 0 610.3779
592.2522583007812 0 740.65424
592.7900390625 0 896.00476
600.4215698242188 0 692.3984
601.2345581054688 0 1769.9233
601.7372436523438 0 1831.0281
602.2329711914062 0 1032.8091
663.2833251953125 0 923.6913
675.3063354492188 0 1412.8962
707.2728271484375 0 17859.174 z 4
708.2767944335938 0 8992.7295
709.2763671875 0 2551.5212
710.276611328125 0 785.735
715.3424682617188 0 1061.5779
720.3675537109375 0 669.2913
721.2501831054688 0 564.7071
723.291259765625 0 8000.379 y 4
724.2953491210938 0 2471.4229
730.3016357421875 0 998.4794 c 5
738.3270263671875 0 701.8786
740.2620849609375 0 548.30145
741.4014282226562 0 1952.7861
742.4070434570312 0 1450.1542
749.2571411132812 0 1738.1421
778.3090209960938 0 4893.6865
779.312744140625 0 1855.2524
787.4609375 0 616.0441
801.9017333984375 0 1078.1046
802.399169921875 0 2363.7258
802.901123046875 0 844.22266
803.4046630859375 0 691.82477
814.4448852539062 0 1008.04034
822.3004760742188 0 8106.2456 z 3
823.3057250976562 0 9409.175
824.3096923828125 0 2627.5933
825.3143310546875 0 827.1267
833.324951171875 0 6289.049
834.330810546875 0 3094.932
835.3279418945312 0 886.84784
838.3176879882812 0 1673.7311 y 3
839.31982421875 0 1280.3125
858.3200073242188 0 3260.9106
859.3203125 0 1657.2019 c Water loss 6
860.3192749023438 0 733.5778 c Ammonia loss 6
876.3319091796875 0 2950.757
877.338623046875 0 24387.143 c 6
878.3416137695312 0 11299.087
879.3423461914062 0 4187.901
880.3296508789062 0 3981.501
881.33349609375 0 2042.9243
889.3466796875 0 3395.087
890.352294921875 0 2453.6445
893.335693359375 0 3536.6252
894.34130859375 0 2233.461
895.3455200195312 0 1026.3214
895.4551391601562 0 802.8302
896.341064453125 0 793.2378
907.3441772460938 0 855.0022
935.3321533203125 0 779.1499 y Water loss 2
936.3281860351562 0 853.1998
937.3268432617188 0 2593.7744 z 2
938.3279418945312 0 1325.2756
952.3718872070312 0 1275.4431
953.3451538085938 0 26227.57 y 2
954.3478393554688 0 14701.197
955.3480224609375 0 4788.428
970.385498046875 0 1347.2969
984.458251953125 0 1510.005
985.4635620117188 0 1038.5846
1001.4802856445312 0 2254.2937
1002.4769897460938 0 1275.6864
1011.5425415039062 0 723.75433
1014.3969116210938 0 21906.3 c 7
1015.4006958007812 0 11104.867
1016.3999633789062 0 4356.894
1017.396240234375 0 792.45245
1037.5672607421875 0 702.7884
1071.5311279296875 0 1565.4009
1072.5247802734375 0 671.331
1084.396240234375 0 1218.9269 z 1
1094.4747314453125 0 646.33563
1100.406494140625 0 925.9852 y 1
1101.41259765625 0 1046.534
1102.522705078125 0 2321.5854
1110.4578857421875 0 1146.6785
1111.4869384765625 0 1068.8013
1113.497314453125 0 684.16425
1123.46240234375 0 1903.7032
1124.4481201171875 0 790.8867
1132.4351806640625 0 2015.9773
1133.441162109375 0 1206.3331
1136.458740234375 0 2287.1858
1137.4818115234375 0 1642.3622
1138.4649658203125 0 1377.5946
1140.4652099609375 0 5784.409
1141.4561767578125 0 10460.59
1142.45166015625 0 10176.704
1142.579345703125 0 1140.7788
1143.4505615234375 0 4700.7417
1143.576904296875 0 1068.4492
1144.4417724609375 0 1207.4059
1149.4381103515625 0 2051.2095
1150.442626953125 0 2452.571
1151.449951171875 0 2941.39
1156.4603271484375 0 966.4353
1158.4388427734375 0 1063.8191
1158.584228515625 0 918.26483
1159.4388427734375 0 1028.276
1163.432861328125 0 1204.2422
1167.4471435546875 0 1929.8507
1168.447021484375 0 1448.3397
1184.4564208984375 0 6329.3774
1185.446044921875 0 34538.918
1186.447509765625 0 20061.787
1186.5704345703125 0 3807.4048
1187.4495849609375 0 9238.8955
1187.5626220703125 0 2011.1536
1188.4488525390625 0 1707.3475
1199.6734619140625 0 1050.4666
1200.6402587890625 0 714.0642
1201.4598388671875 0 24086.178
1202.466552734375 0 105326.59
1203.4696044921875 0 65352.76
1204.4713134765625 0 27012.062
1205.474853515625 0 2931.1416
1274.5606689453125 0 773.88055
1587.7769775390625 0 1602.092
1588.7728271484375 0 1109.4105
1603.7911376953125 0 1532.4204
1604.7896728515625 0 2797.8928
1605.78662109375 0 1828.5508
1790.893310546875 0 892.35455
1807.900146484375 0 2404.1414
2778.994384765625 0 717.60767

Spectrum Details

|  |  |
| --- | --- |
| Matched peaks? Matched peaks The total absolute number of peaks matched. Additionally in brackets the total fraction of peaks matched and the total number of peaks is shown. | 23 (12.17% of 189) |
| FDR? FDR The false discovery rate estimated for this peptide. It is calculated by matching all theoretical fragments with a non-integer shift with the raw peaks for this spectrum. This is done with 40 different shifts. The resulting percentage is the average number of annotated peaks over the number of annotated peaks with the correct spectrum. | 1.35% |
| Satellite FDR? Satellite FDR See the FDR for details on its calculation. This satellite ion specific FDR only contains the satellite ions (d/w) for I/L/J positions. | - |
| PSM Score? PSM Score The PSM Score as given by Hecklib to this annotated spectrum. It is shown with three significant figures. | 301 |

#### Spectrum 12523? Spectrum 12523 The raw spectrum of this peptide as annotated by Hecklib. The fragments are coloured according to ion type (see legend). Any peaks with a star '\*' as text can be hovered over to see the full details, first the ion type second the mass shift type. By hovering over the amino acids in the peptide or ions in the legend the corresponding peaks are highlighted. By toggling the 'Unassigned' label you can turn the background (unassigned) peaks on or off in the plot. By updating the slider in the Ion legend you can update the spectrum to only show the top X% of the peaks with labels. The top X% means any peak that is within X% of the highest intensity. By dragging in the spectrum you can zoom in to a specific part of the spectrum and use 'Zoom Out' to get back to the original zoom level. The annotation of the spectrum is based on the given sequence in the peptides file and is done with different software so inconsistencies are likely. The peaks are annotated based on the given sequence, with 20 ppm tolerance.

Copy Data

##### Spectrum 12523 (TSV)

###### Preview

```
Loading example...
```

*Click on the button to copy the data to your clipboard.*

Mz MinMz MaxIntensity Max

WidthHeightPeptide font sizePeptide stroke widthSpectrum font sizeSpectrum stroke widthCompact peptide

Ion legend

wxyz

abcd

OtherUnassignedIon^Charge^_Position_Show for top:%

TFDDYAMHW

09.80e+31.96e+42.94e+43.92e+4

Zoom Out

a^+1^_2_y^+1^_1_a^+1^_2_b^+1^_2_b^+2^_4_b^+1^_2_y^+1^_2_b^+1^_3_b^+1^_4_y^+1^_3_y^+1^_4_\*y^+1^_5_y^+1^_6_y^+1^_7_y^+1^_7_y^+1^_8_

0683136520482730

Fragment Matches Table

Show background peaks

| Position | Ion type | Intensity | mz Theoretical | mz Error (Th) | mz Error (ppm) | Charge | Series Number |
| --- | --- | --- | --- | --- | --- | --- | --- |
| - | - | 2.736E+04 | 120.1 | - | - | 0 | - |
| - | - | 2328 | 121.1 | - | - | 0 | - |
| - | - | 445.5 | 125.1 | - | - | 0 | - |
| - | - | 3589 | 129.1 | - | - | 0 | - |
| - | - | 972.4 | 130.1 | - | - | 0 | - |
| - | - | 414.6 | 130.8 | - | - | 0 | - |
| - | - | 466.5 | 132.1 | - | - | 0 | - |
| - | - | 5379 | 133.1 | - | - | 0 | - |
| - | - | 498 | 134.1 | - | - | 0 | - |
| - | - | 8700 | 136.1 | - | - | 0 | - |
| - | - | 1129 | 137.1 | - | - | 0 | - |
| - | - | 760.4 | 146.1 | - | - | 0 | - |
| - | - | 765.5 | 148.9 | - | - | 0 | - |
| - | - | 752.4 | 149 | - | - | 0 | - |
| - | - | 663.6 | 155.1 | - | - | 0 | - |
| - | - | 1176 | 156.1 | - | - | 0 | - |
| - | - | 1115 | 157.1 | - | - | 0 | - |
| - | - | 1513 | 158.1 | - | - | 0 | - |
| - | - | 1460 | 159.1 | - | - | 0 | - |
| - | - | 953.9 | 160.1 | - | - | 0 | - |
| - | - | 2225 | 165.1 | - | - | 0 | - |
| - | - | 817.1 | 166.1 | - | - | 0 | - |
| - | - | 1359 | 167.1 | - | - | 0 | - |
| - | - | 492.9 | 169.1 | - | - | 0 | - |
| - | - | 450.6 | 171.1 | - | - | 0 | - |
| - | - | 2096 | 173.1 | - | - | 0 | - |
| - | - | 2293 | 173.4 | - | - | 0 | - |
| - | - | 2372 | 176.1 | - | - | 0 | - |
| - | - | 1913 | 177.1 | - | - | 0 | - |
| - | - | 2697 | 177.1 | - | - | 0 | - |
| - | - | 4656 | 182.1 | - | - | 0 | - |
| - | - | 560.1 | 185.1 | - | - | 0 | - |
| - | - | 666.3 | 185.1 | - | - | 0 | - |
| - | - | 1876 | 188.1 | - | - | 0 | - |
| - | - | 648.9 | 193.1 | - | - | 0 | - |
| - | - | 991.1 | 193.1 | - | - | 0 | - |
| - | - | 606.7 | 197.1 | - | - | 0 | - |
| - | - | 845.6 | 200.1 | - | - | 0 | - |
| - | - | 1592 | 201.1 | - | - | 0 | - |
| 2 | a | 1031 | 203.1 | 4.565E-05 | 0.2247 | +1 | 2 |
| - | - | 487.9 | 204.1 | - | - | 0 | - |
| 9 | y | 1342 | 205.1 | 0.000154 | 0.7507 | +1 | 1 |
| - | - | 2510 | 215.1 | - | - | 0 | - |
| - | - | 497 | 217.1 | - | - | 0 | - |
| - | - | 523.4 | 217.1 | - | - | 0 | - |
| - | - | 1977 | 221.1 | - | - | 0 | - |
| 2 | a | 3.881E+04 | 221.1 | 9.524E-06 | 0.04307 | +1 | 2 |
| - | - | 5109 | 222.1 | - | - | 0 | - |
| - | - | 561.3 | 223.1 | - | - | 0 | - |
| - | - | 835.4 | 227.1 | - | - | 0 | - |
| - | - | 714.6 | 229.2 | - | - | 0 | - |
| 2 | b | 613.9 | 231.1 | 0.0006521 | 2.821 | +1 | 2 |
| - | - | 1345 | 235.1 | - | - | 0 | - |
| - | - | 1034 | 239.1 | - | - | 0 | - |
| 4 | b | 751.8 | 240.1 | 0.003254 | 13.55 | +2 | 4 |
| - | - | 937.3 | 241.1 | - | - | 0 | - |
| - | - | 557.7 | 246.2 | - | - | 0 | - |
| 2 | b | 6722 | 249.1 | 1.534E-06 | 0.006156 | +1 | 2 |
| - | - | 859.8 | 251.1 | - | - | 0 | - |
| - | - | 557.6 | 252.6 | - | - | 0 | - |
| - | - | 912 | 259.2 | - | - | 0 | - |
| - | - | 1305 | 263.1 | - | - | 0 | - |
| - | - | 611.4 | 263.7 | - | - | 0 | - |
| - | - | 972.2 | 279.1 | - | - | 0 | - |
| - | - | 880.4 | 285 | - | - | 0 | - |
| - | - | 680.6 | 285.1 | - | - | 0 | - |
| - | - | 540.2 | 286 | - | - | 0 | - |
| - | - | 754.6 | 296.2 | - | - | 0 | - |
| - | - | 680.2 | 300.2 | - | - | 0 | - |
| - | - | 873.1 | 309.2 | - | - | 0 | - |
| - | - | 583.5 | 319.1 | - | - | 0 | - |
| - | - | 1793 | 324.1 | - | - | 0 | - |
| - | - | 3799 | 329.1 | - | - | 0 | - |
| - | - | 774.9 | 330.2 | - | - | 0 | - |
| - | - | 563.4 | 341.7 | - | - | 0 | - |
| 8 | y | 6943 | 342.2 | 0.0003366 | 0.9839 | +1 | 2 |
| - | - | 1117 | 343.2 | - | - | 0 | - |
| - | - | 627 | 347.2 | - | - | 0 | - |
| - | - | 541.3 | 348.1 | - | - | 0 | - |
| - | - | 737.7 | 355.1 | - | - | 0 | - |
| - | - | 858.9 | 357.2 | - | - | 0 | - |
| 3 | b | 2067 | 364.2 | 7.384E-05 | 0.2028 | +1 | 3 |
| - | - | 574.7 | 378.1 | - | - | 0 | - |
| - | - | 1030 | 394.1 | - | - | 0 | - |
| - | - | 981.8 | 413.3 | - | - | 0 | - |
| - | - | 1050 | 425.2 | - | - | 0 | - |
| - | - | 805.9 | 465.2 | - | - | 0 | - |
| 4 | b | 788.3 | 479.2 | 0.0006931 | 1.446 | +1 | 4 |
| 7 | y | 1296 | 489.2 | 0.00571 | 11.67 | +1 | 3 |
| 6 | y | 3429 | 560.2 | 0.005523 | 9.858 | +1 | 4 |
| - | - | 1130 | 561.2 | - | - | 0 | - |
| - | - | 700.3 | 569.3 | - | - | 0 | - |
| - | - | 2649 | 585.3 | - | - | 0 | - |
| - | - | 1129 | 586.3 | - | - | 0 | - |
| - | - | 570.1 | 588.4 | - | - | 0 | - |
| 0 | Precursor | 1162 | 601.2 | 0.003289 | 5.47 | +2 | -1 |
| - | - | 802.4 | 602.4 | - | - | 0 | - |
| - | - | 957.9 | 659.3 | - | - | 0 | - |
| - | - | 2714 | 700.4 | - | - | 0 | - |
| - | - | 991.2 | 701.4 | - | - | 0 | - |
| 5 | y | 2097 | 723.3 | 0.003962 | 5.477 | +1 | 5 |
| - | - | 769.6 | 735.1 | - | - | 0 | - |
| - | - | 987.4 | 749.3 | - | - | 0 | - |
| - | - | 859.5 | 774.3 | - | - | 0 | - |
| - | - | 1291 | 783.4 | - | - | 0 | - |
| - | - | 1535 | 801.4 | - | - | 0 | - |
| - | - | 828.9 | 828.4 | - | - | 0 | - |
| 4 | y | 3178 | 838.3 | 0.005827 | 6.951 | +1 | 6 |
| - | - | 1123 | 839.3 | - | - | 0 | - |
| - | - | 2471 | 844.5 | - | - | 0 | - |
| - | - | 1311 | 845.5 | - | - | 0 | - |
| - | - | 619.6 | 854.4 | - | - | 0 | - |
| - | - | 8550 | 872.4 | - | - | 0 | - |
| - | - | 4207 | 873.5 | - | - | 0 | - |
| - | - | 981.9 | 874.5 | - | - | 0 | - |
| - | - | 4139 | 889.3 | - | - | 0 | - |
| - | - | 1700 | 890.4 | - | - | 0 | - |
| 3 | y | 761.4 | 935.3 | 0.004769 | 5.098 | +1 | 7 |
| 3 | y | 1.356E+04 | 953.3 | 0.005129 | 5.38 | +1 | 7 |
| - | - | 6462 | 954.3 | - | - | 0 | - |
| - | - | 2676 | 955.3 | - | - | 0 | - |
| - | - | 1102 | 991.5 | - | - | 0 | - |
| - | - | 1013 | 992.5 | - | - | 0 | - |
| - | - | 3459 | 1020 | - | - | 0 | - |
| - | - | 2396 | 1021 | - | - | 0 | - |
| 2 | y | 851.8 | 1100 | 0.006967 | 6.331 | +1 | 8 |
| - | - | 714.6 | 1101 | - | - | 0 | - |
| - | - | 581.6 | 1524 | - | - | 0 | - |
| - | - | 759.1 | 1641 | - | - | 0 | - |
| - | - | 607.1 | 1906 | - | - | 0 | - |
| - | - | 610.5 | 2081 | - | - | 0 | - |
| - | - | 636.1 | 2439 | - | - | 0 | - |
| - | - | 746.2 | 2703 | - | - | 0 | - |

m/z Charge Intensity FragmentType MassShift Position
120.08086395263672 0 27356.965
121.08428955078125 0 2328.29
125.09585571289062 0 445.54434
129.102294921875 0 3589.3982
130.06520080566406 0 972.365
130.84706115722656 0 414.59293
132.08143615722656 0 466.5375
133.08595275878906 0 5379.4487
134.08956909179688 0 497.97763
136.0757598876953 0 8700.486
137.07887268066406 0 1129.3917
146.05978393554688 0 760.4116
148.94705200195312 0 765.49084
149.0447998046875 0 752.43146
155.08139038085938 0 663.5872
156.07691955566406 0 1175.9824
157.13372802734375 0 1114.7236
158.09640502929688 0 1512.9834
159.09178161621094 0 1460.4802
160.11196899414062 0 953.9127
165.0546417236328 0 2225.128
166.08639526367188 0 817.14154
167.05560302734375 0 1359.0479
169.096435546875 0 492.92664
171.11273193359375 0 450.56717
173.12843322753906 0 2096.161
173.43907165527344 0 2293.4905
176.10687255859375 0 2372.3127
177.102294921875 0 1913.2423
177.11207580566406 0 2696.651
182.08114624023438 0 4656.34
185.09173583984375 0 560.0914
185.12857055664062 0 666.3019
188.0704803466797 0 1875.9272
193.09930419921875 0 648.8966
193.108642578125 0 991.0913
197.12852478027344 0 606.6508
200.1385498046875 0 845.59753
201.1232452392578 0 1591.5996
203.11793518066406 0 1031.0122 a Water loss 1
204.07537841796875 0 487.94455
205.0970001220703 0 1342.0239 y 8
215.12767028808594 0 2510.0237
217.08352661132812 0 496.9826
217.09765625 0 523.43524
221.10305786132812 0 1977.424
221.1284637451172 0 38812.023 a 1
222.13177490234375 0 5108.7266
223.06466674804688 0 561.29816
227.10267639160156 0 835.44684
229.17037963867188 0 714.62
231.11215209960938 0 613.90125 b Water loss 1
235.10772705078125 0 1344.9465
239.09494018554688 0 1034.0781
240.09552001953125 0 751.84485 b 3
241.07470703125 0 937.2619
246.1672821044922 0 557.7039
249.1233673095703 0 6721.761 b 1
251.1034393310547 0 859.7813
252.5567169189453 0 557.5585
259.15283203125 0 912.0236
263.1022644042969 0 1304.6385
263.6759948730469 0 611.3551
279.0975341796875 0 972.24286
285.0091247558594 0 880.39856
285.10247802734375 0 680.5678
286.0104675292969 0 540.1519
296.15155029296875 0 754.60065
300.1697692871094 0 680.2249
309.2045593261719 0 873.13745
319.1407470703125 0 583.5472
324.1457214355469 0 1793.0942
329.1498107910156 0 3799.0522
330.15228271484375 0 774.8781
341.7355041503906 0 563.4268
342.1564025878906 0 6943.37 y 7
343.1598205566406 0 1117.1927
347.2066650390625 0 627.0073
348.1172790527344 0 541.3355
355.0684509277344 0 737.6757
357.22344970703125 0 858.8854
364.1502380371094 0 2066.9417 b 2
378.1284484863281 0 574.69434
394.123779296875 0 1030.1483
413.2666015625 0 981.83826
425.1931457519531 0 1050.0446
465.162353515625 0 805.904
479.1779479980469 0 788.31805 b 3
489.1922607421875 0 1295.9319 y 6
560.2291870117188 0 3428.941 y 5
561.232421875 0 1129.5496
569.2553100585938 0 700.3177
585.3429565429688 0 2649.278
586.3373413085938 0 1128.9308
588.4264526367188 0 570.08624
601.2354125976562 0 1161.7721 Precursor
602.3710327148438 0 802.3906
659.29345703125 0 957.8654
700.3641357421875 0 2713.7415
701.3682861328125 0 991.18616
723.2909545898438 0 2097.3057 y 4
735.1377563476562 0 769.60266
749.2572021484375 0 987.3566
774.3214111328125 0 859.5161
783.4010620117188 0 1291.456
801.4120483398438 0 1534.6443
828.4203491210938 0 828.865
838.3197631835938 0 3178.1426 y 3
839.3201904296875 0 1123.4851
844.452392578125 0 2470.669
845.455810546875 0 1311.0271
854.4417114257812 0 619.5729
872.4481201171875 0 8549.569
873.4507446289062 0 4207.456
874.4541015625 0 981.8671
889.347900390625 0 4139.166
890.3510131835938 0 1699.5532
935.3350830078125 0 761.37866 y Water loss 2
953.3460083007812 0 13561.982 y 2
954.3496704101562 0 6461.976
955.3487548828125 0 2676.1375
991.5189819335938 0 1102.0466
992.5211181640625 0 1012.63385
1019.5155639648438 0 3459.1714
1020.5187377929688 0 2395.8691
1100.416259765625 0 851.7532 y 1
1101.423095703125 0 714.63257
1524.33447265625 0 581.62836
1640.6895751953125 0 759.1054
1905.8360595703125 0 607.1228
2080.5009765625 0 610.496
2438.941162109375 0 636.14624
2703.400146484375 0 746.2

Spectrum Details

|  |  |
| --- | --- |
| Matched peaks? Matched peaks The total absolute number of peaks matched. Additionally in brackets the total fraction of peaks matched and the total number of peaks is shown. | 17 (12.78% of 133) |
| FDR? FDR The false discovery rate estimated for this peptide. It is calculated by matching all theoretical fragments with a non-integer shift with the raw peaks for this spectrum. This is done with 40 different shifts. The resulting percentage is the average number of annotated peaks over the number of annotated peaks with the correct spectrum. | 0.70% |
| Satellite FDR? Satellite FDR See the FDR for details on its calculation. This satellite ion specific FDR only contains the satellite ions (d/w) for I/L/J positions. | - |
| PSM Score? PSM Score The PSM Score as given by Hecklib to this annotated spectrum. It is shown with three significant figures. | 195 |

#### Spectrum 12414? Spectrum 12414 The raw spectrum of this peptide as annotated by Hecklib. The fragments are coloured according to ion type (see legend). Any peaks with a star '\*' as text can be hovered over to see the full details, first the ion type second the mass shift type. By hovering over the amino acids in the peptide or ions in the legend the corresponding peaks are highlighted. By toggling the 'Unassigned' label you can turn the background (unassigned) peaks on or off in the plot. By updating the slider in the Ion legend you can update the spectrum to only show the top X% of the peaks with labels. The top X% means any peak that is within X% of the highest intensity. By dragging in the spectrum you can zoom in to a specific part of the spectrum and use 'Zoom Out' to get back to the original zoom level. The annotation of the spectrum is based on the given sequence in the peptides file and is done with different software so inconsistencies are likely. The peaks are annotated based on the given sequence, with 20 ppm tolerance.

Copy Data

##### Spectrum 12414 (TSV)

###### Preview

```
Loading example...
```

*Click on the button to copy the data to your clipboard.*

Mz MinMz MaxIntensity Max

WidthHeightPeptide font sizePeptide stroke widthSpectrum font sizeSpectrum stroke widthCompact peptide

Ion legend

wxyz

abcd

OtherUnassignedIon^Charge^_Position_Show for top:%

TFDDYAMHW

01.00e+42.01e+43.01e+44.02e+4

Zoom Out

a^+1^_2_y^+1^_1_a^+1^_2_b^+1^_2_y^+1^_2_b^+1^_3_b^+1^_4_y^+1^_3_y^+2^_8_y^+1^_4_\*y^+1^_5_y^+1^_6_y^+1^_7_y^+1^_8_

035370610591412

Fragment Matches Table

Show background peaks

| Position | Ion type | Intensity | mz Theoretical | mz Error (Th) | mz Error (ppm) | Charge | Series Number |
| --- | --- | --- | --- | --- | --- | --- | --- |
| - | - | 2.778E+04 | 120.1 | - | - | 0 | - |
| - | - | 2026 | 121.1 | - | - | 0 | - |
| - | - | 350 | 128.9 | - | - | 0 | - |
| - | - | 3474 | 129.1 | - | - | 0 | - |
| - | - | 749 | 130.1 | - | - | 0 | - |
| - | - | 394.7 | 130.9 | - | - | 0 | - |
| - | - | 486.1 | 131.9 | - | - | 0 | - |
| - | - | 661.3 | 132.1 | - | - | 0 | - |
| - | - | 551.3 | 133.1 | - | - | 0 | - |
| - | - | 5633 | 133.1 | - | - | 0 | - |
| - | - | 9284 | 136.1 | - | - | 0 | - |
| - | - | 983.2 | 137.1 | - | - | 0 | - |
| - | - | 485.9 | 138.1 | - | - | 0 | - |
| - | - | 765.9 | 143.1 | - | - | 0 | - |
| - | - | 414.7 | 147.1 | - | - | 0 | - |
| - | - | 412.1 | 148.9 | - | - | 0 | - |
| - | - | 932.9 | 149 | - | - | 0 | - |
| - | - | 429.5 | 154 | - | - | 0 | - |
| - | - | 469 | 155.1 | - | - | 0 | - |
| - | - | 495.4 | 155.1 | - | - | 0 | - |
| - | - | 1027 | 156.1 | - | - | 0 | - |
| - | - | 868 | 159.1 | - | - | 0 | - |
| - | - | 781.1 | 160.1 | - | - | 0 | - |
| - | - | 400.8 | 160.2 | - | - | 0 | - |
| - | - | 548.5 | 162.1 | - | - | 0 | - |
| - | - | 402 | 163.6 | - | - | 0 | - |
| - | - | 2569 | 165.1 | - | - | 0 | - |
| - | - | 706 | 166.1 | - | - | 0 | - |
| - | - | 2333 | 167.1 | - | - | 0 | - |
| - | - | 461.9 | 172.9 | - | - | 0 | - |
| - | - | 2007 | 173.1 | - | - | 0 | - |
| - | - | 485.2 | 175.1 | - | - | 0 | - |
| - | - | 547 | 175.1 | - | - | 0 | - |
| - | - | 2830 | 176.1 | - | - | 0 | - |
| - | - | 2339 | 177.1 | - | - | 0 | - |
| - | - | 2447 | 177.1 | - | - | 0 | - |
| - | - | 5648 | 182.1 | - | - | 0 | - |
| - | - | 512 | 183.1 | - | - | 0 | - |
| - | - | 1168 | 187.1 | - | - | 0 | - |
| - | - | 1587 | 188.1 | - | - | 0 | - |
| - | - | 722.4 | 191.1 | - | - | 0 | - |
| - | - | 978.2 | 193.1 | - | - | 0 | - |
| - | - | 1163 | 201.1 | - | - | 0 | - |
| 2 | a | 986.9 | 203.1 | 1.539E-05 | 0.07577 | +1 | 2 |
| 9 | y | 1510 | 205.1 | 0.0003066 | 1.495 | +1 | 1 |
| - | - | 511.2 | 212.2 | - | - | 0 | - |
| - | - | 588.6 | 215.1 | - | - | 0 | - |
| - | - | 610 | 217.1 | - | - | 0 | - |
| - | - | 1685 | 221.1 | - | - | 0 | - |
| 2 | a | 3.976E+04 | 221.1 | 0.0001011 | 0.4571 | +1 | 2 |
| - | - | 5116 | 222.1 | - | - | 0 | - |
| - | - | 1052 | 223.1 | - | - | 0 | - |
| - | - | 578.3 | 223.1 | - | - | 0 | - |
| - | - | 518.7 | 223.6 | - | - | 0 | - |
| - | - | 548.8 | 231.1 | - | - | 0 | - |
| - | - | 959.9 | 231.2 | - | - | 0 | - |
| - | - | 1304 | 235.1 | - | - | 0 | - |
| - | - | 927.3 | 239.1 | - | - | 0 | - |
| - | - | 1012 | 241.1 | - | - | 0 | - |
| 2 | b | 7303 | 249.1 | 0.0001968 | 0.7901 | +1 | 2 |
| - | - | 919.4 | 251.1 | - | - | 0 | - |
| - | - | 972 | 257.1 | - | - | 0 | - |
| - | - | 573.9 | 257.2 | - | - | 0 | - |
| - | - | 554.2 | 257.4 | - | - | 0 | - |
| - | - | 1588 | 263.1 | - | - | 0 | - |
| - | - | 1136 | 285 | - | - | 0 | - |
| - | - | 953.4 | 285.1 | - | - | 0 | - |
| - | - | 677.1 | 285.2 | - | - | 0 | - |
| - | - | 593.6 | 302.1 | - | - | 0 | - |
| - | - | 559.4 | 312.9 | - | - | 0 | - |
| - | - | 792.4 | 319.1 | - | - | 0 | - |
| - | - | 2000 | 324.1 | - | - | 0 | - |
| - | - | 3783 | 329.1 | - | - | 0 | - |
| - | - | 747.5 | 330.2 | - | - | 0 | - |
| 8 | y | 6290 | 342.2 | 0.0003061 | 0.8947 | +1 | 2 |
| - | - | 1763 | 343.2 | - | - | 0 | - |
| - | - | 2796 | 347.2 | - | - | 0 | - |
| - | - | 754.1 | 350.1 | - | - | 0 | - |
| - | - | 1407 | 355.1 | - | - | 0 | - |
| - | - | 986.5 | 357.2 | - | - | 0 | - |
| 3 | b | 1808 | 364.2 | 7.875E-05 | 0.2163 | +1 | 3 |
| - | - | 1117 | 378.1 | - | - | 0 | - |
| - | - | 2869 | 391.2 | - | - | 0 | - |
| - | - | 1580 | 394.1 | - | - | 0 | - |
| - | - | 626.3 | 397.2 | - | - | 0 | - |
| - | - | 1106 | 413.3 | - | - | 0 | - |
| - | - | 817.6 | 425.2 | - | - | 0 | - |
| - | - | 623.1 | 427.7 | - | - | 0 | - |
| - | - | 700.4 | 436.7 | - | - | 0 | - |
| - | - | 835 | 455.2 | - | - | 0 | - |
| - | - | 1290 | 465.2 | - | - | 0 | - |
| - | - | 687.9 | 476.2 | - | - | 0 | - |
| 4 | b | 752.2 | 479.2 | 0.0005886 | 1.228 | +1 | 4 |
| 7 | y | 1577 | 489.2 | 0.006168 | 12.61 | +1 | 3 |
| - | - | 717.6 | 490.2 | - | - | 0 | - |
| 2 | y | 836.1 | 550.7 | 0.001493 | 2.711 | +2 | 8 |
| - | - | 1610 | 551.2 | - | - | 0 | - |
| 6 | y | 3108 | 560.2 | 0.004912 | 8.768 | +1 | 4 |
| - | - | 881.7 | 561.2 | - | - | 0 | - |
| - | - | 6184 | 585.3 | - | - | 0 | - |
| - | - | 661.1 | 597.8 | - | - | 0 | - |
| 0 | Precursor | 774.9 | 601.2 | 0.007195 | 11.97 | +2 | -1 |
| - | - | 800.9 | 601.3 | - | - | 0 | - |
| - | - | 748.7 | 602.4 | - | - | 0 | - |
| - | - | 938.3 | 659.3 | - | - | 0 | - |
| - | - | 5198 | 700.4 | - | - | 0 | - |
| - | - | 1654 | 701.4 | - | - | 0 | - |
| - | - | 788.4 | 721.3 | - | - | 0 | - |
| 5 | y | 2756 | 723.3 | 0.004389 | 6.068 | +1 | 5 |
| - | - | 1034 | 724.3 | - | - | 0 | - |
| - | - | 979.5 | 749.3 | - | - | 0 | - |
| - | - | 916.5 | 774.3 | - | - | 0 | - |
| - | - | 809.1 | 783.4 | - | - | 0 | - |
| - | - | 705.6 | 800.4 | - | - | 0 | - |
| - | - | 2154 | 801.4 | - | - | 0 | - |
| - | - | 1328 | 802.4 | - | - | 0 | - |
| - | - | 785.5 | 828.4 | - | - | 0 | - |
| 4 | y | 2241 | 838.3 | 0.005644 | 6.733 | +1 | 6 |
| - | - | 1282 | 839.3 | - | - | 0 | - |
| - | - | 2210 | 844.5 | - | - | 0 | - |
| - | - | 1071 | 845.5 | - | - | 0 | - |
| - | - | 1.167E+04 | 872.4 | - | - | 0 | - |
| - | - | 4599 | 873.5 | - | - | 0 | - |
| - | - | 1168 | 874.5 | - | - | 0 | - |
| - | - | 527 | 881.3 | - | - | 0 | - |
| - | - | 5564 | 889.3 | - | - | 0 | - |
| - | - | 2280 | 890.4 | - | - | 0 | - |
| - | - | 590.8 | 907.3 | - | - | 0 | - |
| 3 | y | 1.376E+04 | 953.3 | 0.005923 | 6.213 | +1 | 7 |
| - | - | 8449 | 954.3 | - | - | 0 | - |
| - | - | 3354 | 955.4 | - | - | 0 | - |
| - | - | 1468 | 991.5 | - | - | 0 | - |
| - | - | 706.9 | 1003 | - | - | 0 | - |
| - | - | 3741 | 1020 | - | - | 0 | - |
| - | - | 2971 | 1021 | - | - | 0 | - |
| - | - | 1286 | 1022 | - | - | 0 | - |
| 2 | y | 688.5 | 1100 | 0.007333 | 6.664 | +1 | 8 |
| - | - | 740.1 | 1398 | - | - | 0 | - |

m/z Charge Intensity FragmentType MassShift Position
120.08091735839844 0 27783.47
121.0842514038086 0 2025.713
128.92051696777344 0 350.03964
129.10238647460938 0 3474.1719
130.0652618408203 0 749.0423
130.8983612060547 0 394.74954
131.86807250976562 0 486.11526
132.0810089111328 0 661.31976
133.08143615722656 0 551.317
133.0860137939453 0 5632.66
136.07582092285156 0 9283.709
137.07913208007812 0 983.21716
138.06634521484375 0 485.87344
143.1068115234375 0 765.85986
147.0636444091797 0 414.68542
148.9466552734375 0 412.0746
149.0450897216797 0 932.9434
154.0204315185547 0 429.53534
155.08131408691406 0 469.01422
155.11842346191406 0 495.37854
156.07666015625 0 1026.5756
159.0917205810547 0 867.9746
160.1121063232422 0 781.08417
160.19590759277344 0 400.79355
162.09141540527344 0 548.4509
163.6310577392578 0 402.01562
165.05465698242188 0 2568.9546
166.08624267578125 0 705.9616
167.0555877685547 0 2333.149
172.88662719726562 0 461.87375
173.12855529785156 0 2007.4967
175.09597778320312 0 485.15366
175.11859130859375 0 546.9804
176.10719299316406 0 2830.3071
177.10247802734375 0 2338.65
177.1122283935547 0 2447.1062
182.08126831054688 0 5648.052
183.08502197265625 0 512.0193
187.13340759277344 0 1167.519
188.0706329345703 0 1587.275
191.11785888671875 0 722.44946
193.10890197753906 0 978.24146
201.12347412109375 0 1162.8257
203.1178741455078 0 986.92865 a Water loss 1
205.0968475341797 0 1510.0099 y 8
212.1549072265625 0 511.23807
215.12831115722656 0 588.5536
217.0826873779297 0 610.0029
221.10337829589844 0 1685.3757
221.12855529785156 0 39760.86 a 1
222.13180541992188 0 5116.2866
223.0636444091797 0 1052.2279
223.13369750976562 0 578.3149
223.57147216796875 0 518.65326
231.12332153320312 0 548.8475
231.1590576171875 0 959.8951
235.1081085205078 0 1303.5306
239.1491241455078 0 927.3309
241.07455444335938 0 1012.4698
249.12356567382812 0 7302.953 b 1
251.1029815673828 0 919.3559
257.1075744628906 0 972.0383
257.1649475097656 0 573.92786
257.4211730957031 0 554.1831
263.102783203125 0 1587.8262
285.009521484375 0 1135.9038
285.1013488769531 0 953.4068
285.1597900390625 0 677.0567
302.1138610839844 0 593.5513
312.93280029296875 0 559.39154
319.13885498046875 0 792.39233
324.145263671875 0 1999.9307
329.1498107910156 0 3783.1992
330.1580505371094 0 747.45825
342.1563720703125 0 6290.2847 y 7
343.1591491699219 0 1763.1387
347.2069091796875 0 2795.7773
350.1348876953125 0 754.1267
355.0700988769531 0 1406.6628
357.2251281738281 0 986.5043
364.150390625 0 1808.2009 b 2
378.12969970703125 0 1116.9484
391.2330017089844 0 2868.8723
394.1250305175781 0 1580.1266
397.2085876464844 0 626.3372
413.2664794921875 0 1106.0062
425.19293212890625 0 817.59467
427.7200012207031 0 623.05194
436.7276916503906 0 700.37036
455.2278137207031 0 834.9715
465.1620788574219 0 1289.8922
476.17254638671875 0 687.88
479.1766662597656 0 752.213 b 3
489.1927185058594 0 1576.7323 y 6
490.1957092285156 0 717.5967
550.7097778320312 0 836.05164 y 1
551.2122192382812 0 1610.1389
560.2285766601562 0 3107.8457 y 5
561.2300415039062 0 881.6837
585.347412109375 0 6184.28
597.81298828125 0 661.12476
601.2393188476562 0 774.934 Precursor
601.2940063476562 0 800.8626
602.3719482421875 0 748.6825
659.2938232421875 0 938.3125
700.36279296875 0 5197.6685
701.3687133789062 0 1653.899
721.262939453125 0 788.3815
723.2913818359375 0 2756.3594 y 4
724.2962646484375 0 1033.5245
749.2560424804688 0 979.5399
774.3212890625 0 916.5256
783.4005737304688 0 809.1397
800.4267578125 0 705.5578
801.4127807617188 0 2154.3442
802.4149780273438 0 1328.4038
828.4285278320312 0 785.519
838.319580078125 0 2240.8354 y 3
839.323974609375 0 1281.9736
844.4520263671875 0 2210.3123
845.4525146484375 0 1070.8678
872.4483642578125 0 11666.553
873.4520874023438 0 4598.7026
874.4519653320312 0 1167.9385
881.3446655273438 0 527.0469
889.34765625 0 5564.218
890.3512573242188 0 2280.1892
907.3323364257812 0 590.82367
953.3468017578125 0 13759.571 y 2
954.3494873046875 0 8448.53
955.3521728515625 0 3354.4644
991.52392578125 0 1468.0874
1003.3822631835938 0 706.87537
1019.5166625976562 0 3741.2434
1020.5175170898438 0 2970.8281
1021.5221557617188 0 1285.9681
1100.4166259765625 0 688.48285 y 1
1398.30712890625 0 740.1343

Spectrum Details

|  |  |
| --- | --- |
| Matched peaks? Matched peaks The total absolute number of peaks matched. Additionally in brackets the total fraction of peaks matched and the total number of peaks is shown. | 15 (10.87% of 138) |
| FDR? FDR The false discovery rate estimated for this peptide. It is calculated by matching all theoretical fragments with a non-integer shift with the raw peaks for this spectrum. This is done with 40 different shifts. The resulting percentage is the average number of annotated peaks over the number of annotated peaks with the correct spectrum. | 0.63% |
| Satellite FDR? Satellite FDR See the FDR for details on its calculation. This satellite ion specific FDR only contains the satellite ions (d/w) for I/L/J positions. | - |
| PSM Score? PSM Score The PSM Score as given by Hecklib to this annotated spectrum. It is shown with three significant figures. | 160 |

#### Spectrum 12185? Spectrum 12185 The raw spectrum of this peptide as annotated by Hecklib. The fragments are coloured according to ion type (see legend). Any peaks with a star '\*' as text can be hovered over to see the full details, first the ion type second the mass shift type. By hovering over the amino acids in the peptide or ions in the legend the corresponding peaks are highlighted. By toggling the 'Unassigned' label you can turn the background (unassigned) peaks on or off in the plot. By updating the slider in the Ion legend you can update the spectrum to only show the top X% of the peaks with labels. The top X% means any peak that is within X% of the highest intensity. By dragging in the spectrum you can zoom in to a specific part of the spectrum and use 'Zoom Out' to get back to the original zoom level. The annotation of the spectrum is based on the given sequence in the peptides file and is done with different software so inconsistencies are likely. The peaks are annotated based on the given sequence, with 20 ppm tolerance.

Copy Data

##### Spectrum 12185 (TSV)

###### Preview

```
Loading example...
```

*Click on the button to copy the data to your clipboard.*

Mz MinMz MaxIntensity Max

WidthHeightPeptide font sizePeptide stroke widthSpectrum font sizeSpectrum stroke widthCompact peptide

Ion legend

wxyz

abcd

OtherUnassignedIon^Charge^_Position_Show for top:%

TFDDYAMHW

01.17e+42.34e+43.50e+44.67e+4

Zoom Out

a^+1^_2_y^+1^_1_a^+1^_2_b^+1^_2_y^+1^_2_b^+1^_3_b^+1^_3_b^+1^_4_y^+1^_3_y^+2^_8_y^+1^_4_\*y^+1^_5_y^+1^_6_y^+1^_7_y^+1^_7_

0783156523483130

Fragment Matches Table

Show background peaks

| Position | Ion type | Intensity | mz Theoretical | mz Error (Th) | mz Error (ppm) | Charge | Series Number |
| --- | --- | --- | --- | --- | --- | --- | --- |
| - | - | 2.947E+04 | 120.1 | - | - | 0 | - |
| - | - | 2549 | 121.1 | - | - | 0 | - |
| - | - | 378.3 | 121.6 | - | - | 0 | - |
| - | - | 392.3 | 127.6 | - | - | 0 | - |
| - | - | 3031 | 129.1 | - | - | 0 | - |
| - | - | 805.7 | 130.1 | - | - | 0 | - |
| - | - | 595 | 132.1 | - | - | 0 | - |
| - | - | 375.2 | 132.4 | - | - | 0 | - |
| - | - | 956.2 | 133.1 | - | - | 0 | - |
| - | - | 372.3 | 135.4 | - | - | 0 | - |
| - | - | 9432 | 136.1 | - | - | 0 | - |
| - | - | 669.3 | 137.1 | - | - | 0 | - |
| - | - | 465.6 | 138.1 | - | - | 0 | - |
| - | - | 698.3 | 138.1 | - | - | 0 | - |
| - | - | 383.6 | 140.3 | - | - | 0 | - |
| - | - | 449.9 | 143.1 | - | - | 0 | - |
| - | - | 568.4 | 146.1 | - | - | 0 | - |
| - | - | 724.3 | 148.9 | - | - | 0 | - |
| - | - | 450.6 | 148.9 | - | - | 0 | - |
| - | - | 661.2 | 148.9 | - | - | 0 | - |
| - | - | 766.7 | 148.9 | - | - | 0 | - |
| - | - | 855.4 | 148.9 | - | - | 0 | - |
| - | - | 1172 | 148.9 | - | - | 0 | - |
| - | - | 1522 | 148.9 | - | - | 0 | - |
| - | - | 3244 | 148.9 | - | - | 0 | - |
| - | - | 6430 | 149 | - | - | 0 | - |
| - | - | 3775 | 149 | - | - | 0 | - |
| - | - | 1697 | 149 | - | - | 0 | - |
| - | - | 1240 | 149 | - | - | 0 | - |
| - | - | 1195 | 149 | - | - | 0 | - |
| - | - | 685.7 | 149 | - | - | 0 | - |
| - | - | 837.8 | 149 | - | - | 0 | - |
| - | - | 597.4 | 149 | - | - | 0 | - |
| - | - | 611.6 | 149 | - | - | 0 | - |
| - | - | 1654 | 149 | - | - | 0 | - |
| - | - | 434.8 | 149.1 | - | - | 0 | - |
| - | - | 463 | 152 | - | - | 0 | - |
| - | - | 452.5 | 153.9 | - | - | 0 | - |
| - | - | 1252 | 156.1 | - | - | 0 | - |
| - | - | 1982 | 158.1 | - | - | 0 | - |
| - | - | 2809 | 159.1 | - | - | 0 | - |
| - | - | 567.2 | 159.1 | - | - | 0 | - |
| - | - | 724.6 | 160.1 | - | - | 0 | - |
| - | - | 3542 | 165.1 | - | - | 0 | - |
| - | - | 653.7 | 166.1 | - | - | 0 | - |
| - | - | 1376 | 166.1 | - | - | 0 | - |
| - | - | 2249 | 167.1 | - | - | 0 | - |
| - | - | 2251 | 173.1 | - | - | 0 | - |
| - | - | 2888 | 176.1 | - | - | 0 | - |
| - | - | 1442 | 177.1 | - | - | 0 | - |
| - | - | 525.2 | 178.1 | - | - | 0 | - |
| - | - | 6827 | 182.1 | - | - | 0 | - |
| - | - | 919.7 | 183.1 | - | - | 0 | - |
| - | - | 580.8 | 185.1 | - | - | 0 | - |
| - | - | 1649 | 188.1 | - | - | 0 | - |
| - | - | 544.9 | 191.1 | - | - | 0 | - |
| - | - | 1062 | 193.1 | - | - | 0 | - |
| - | - | 556.8 | 200.1 | - | - | 0 | - |
| - | - | 1106 | 201.1 | - | - | 0 | - |
| - | - | 507.8 | 201.4 | - | - | 0 | - |
| 2 | a | 1097 | 203.1 | 0.001297 | 6.385 | +1 | 2 |
| 9 | y | 1176 | 205.1 | 7.491E-05 | 0.3652 | +1 | 1 |
| - | - | 523.4 | 213.1 | - | - | 0 | - |
| - | - | 1666 | 221.1 | - | - | 0 | - |
| 2 | a | 4.626E+04 | 221.1 | 8.582E-05 | 0.3881 | +1 | 2 |
| - | - | 5717 | 222.1 | - | - | 0 | - |
| - | - | 2174 | 223.1 | - | - | 0 | - |
| - | - | 992.3 | 227.1 | - | - | 0 | - |
| - | - | 743.7 | 229.2 | - | - | 0 | - |
| - | - | 802.4 | 231.1 | - | - | 0 | - |
| - | - | 604.9 | 231.1 | - | - | 0 | - |
| - | - | 694.5 | 235.1 | - | - | 0 | - |
| - | - | 1314 | 239.1 | - | - | 0 | - |
| - | - | 855.6 | 241.1 | - | - | 0 | - |
| - | - | 619.4 | 243.1 | - | - | 0 | - |
| 2 | b | 7333 | 249.1 | 0.0001511 | 0.6063 | +1 | 2 |
| - | - | 1205 | 251.1 | - | - | 0 | - |
| - | - | 671 | 253.2 | - | - | 0 | - |
| - | - | 587.1 | 260.2 | - | - | 0 | - |
| - | - | 1876 | 263.1 | - | - | 0 | - |
| - | - | 694.3 | 269 | - | - | 0 | - |
| - | - | 880.9 | 279.1 | - | - | 0 | - |
| - | - | 1605 | 285 | - | - | 0 | - |
| - | - | 887.9 | 285.1 | - | - | 0 | - |
| - | - | 803 | 296.2 | - | - | 0 | - |
| - | - | 647.3 | 299.1 | - | - | 0 | - |
| - | - | 791.8 | 319.1 | - | - | 0 | - |
| - | - | 1844 | 324.1 | - | - | 0 | - |
| - | - | 821.7 | 325.1 | - | - | 0 | - |
| - | - | 528.9 | 327.2 | - | - | 0 | - |
| - | - | 3972 | 329.1 | - | - | 0 | - |
| - | - | 639.9 | 330.2 | - | - | 0 | - |
| - | - | 576.5 | 339.2 | - | - | 0 | - |
| - | - | 627.3 | 342.1 | - | - | 0 | - |
| 8 | y | 6267 | 342.2 | 0.0003366 | 0.9839 | +1 | 2 |
| - | - | 1510 | 343.2 | - | - | 0 | - |
| 3 | b | 1037 | 346.1 | 0.0001598 | 0.4616 | +1 | 3 |
| - | - | 601.8 | 348.1 | - | - | 0 | - |
| - | - | 568.2 | 351 | - | - | 0 | - |
| - | - | 2507 | 355.1 | - | - | 0 | - |
| - | - | 759.4 | 356.1 | - | - | 0 | - |
| - | - | 1247 | 357.2 | - | - | 0 | - |
| 3 | b | 1736 | 364.2 | 0.0005365 | 1.473 | +1 | 3 |
| - | - | 907.7 | 378.1 | - | - | 0 | - |
| - | - | 637.6 | 380.5 | - | - | 0 | - |
| - | - | 1357 | 394.1 | - | - | 0 | - |
| - | - | 1857 | 413.3 | - | - | 0 | - |
| - | - | 669 | 414.3 | - | - | 0 | - |
| - | - | 966.1 | 425.2 | - | - | 0 | - |
| - | - | 645.5 | 427.7 | - | - | 0 | - |
| - | - | 662.3 | 436.7 | - | - | 0 | - |
| - | - | 951 | 465.2 | - | - | 0 | - |
| 4 | b | 1372 | 479.2 | 0.002738 | 5.713 | +1 | 4 |
| - | - | 736.2 | 481.3 | - | - | 0 | - |
| 7 | y | 1877 | 489.2 | 0.004886 | 9.988 | +1 | 3 |
| - | - | 644.6 | 496.2 | - | - | 0 | - |
| 2 | y | 1259 | 550.7 | 0.00192 | 3.487 | +2 | 8 |
| - | - | 1463 | 551.2 | - | - | 0 | - |
| 6 | y | 3156 | 560.2 | 0.004607 | 8.224 | +1 | 4 |
| - | - | 758.2 | 561.2 | - | - | 0 | - |
| - | - | 605.2 | 583.5 | - | - | 0 | - |
| - | - | 2765 | 585.3 | - | - | 0 | - |
| - | - | 577.2 | 592.7 | - | - | 0 | - |
| - | - | 660.6 | 600.3 | - | - | 0 | - |
| - | - | 647.8 | 600.8 | - | - | 0 | - |
| 0 | Precursor | 2114 | 601.2 | 0.002556 | 4.252 | +2 | -1 |
| - | - | 774.7 | 601.7 | - | - | 0 | - |
| - | - | 545.3 | 649 | - | - | 0 | - |
| - | - | 658.7 | 649.4 | - | - | 0 | - |
| - | - | 738.2 | 657.3 | - | - | 0 | - |
| - | - | 3464 | 700.4 | - | - | 0 | - |
| - | - | 1793 | 701.4 | - | - | 0 | - |
| - | - | 866.9 | 721.3 | - | - | 0 | - |
| 5 | y | 3438 | 723.3 | 0.003412 | 4.718 | +1 | 5 |
| - | - | 1803 | 724.3 | - | - | 0 | - |
| - | - | 1121 | 749.3 | - | - | 0 | - |
| - | - | 798.4 | 774.3 | - | - | 0 | - |
| - | - | 1409 | 783.4 | - | - | 0 | - |
| - | - | 1832 | 801.4 | - | - | 0 | - |
| - | - | 1445 | 802.4 | - | - | 0 | - |
| - | - | 1144 | 828.4 | - | - | 0 | - |
| 4 | y | 3554 | 838.3 | 0.003813 | 4.549 | +1 | 6 |
| - | - | 1613 | 839.3 | - | - | 0 | - |
| - | - | 2969 | 844.5 | - | - | 0 | - |
| - | - | 1727 | 845.5 | - | - | 0 | - |
| - | - | 1205 | 871.3 | - | - | 0 | - |
| - | - | 1.052E+04 | 872.4 | - | - | 0 | - |
| - | - | 5345 | 873.4 | - | - | 0 | - |
| - | - | 1610 | 874.5 | - | - | 0 | - |
| - | - | 5601 | 889.3 | - | - | 0 | - |
| - | - | 2842 | 890.3 | - | - | 0 | - |
| - | - | 613.8 | 891.4 | - | - | 0 | - |
| - | - | 672.6 | 929.6 | - | - | 0 | - |
| 3 | y | 1087 | 935.3 | 0.002266 | 2.423 | +1 | 7 |
| 3 | y | 1.703E+04 | 953.3 | 0.004031 | 4.228 | +1 | 7 |
| - | - | 7726 | 954.3 | - | - | 0 | - |
| - | - | 2669 | 955.3 | - | - | 0 | - |
| - | - | 1357 | 991.5 | - | - | 0 | - |
| - | - | 960.5 | 992.5 | - | - | 0 | - |
| - | - | 752.4 | 993.5 | - | - | 0 | - |
| - | - | 4814 | 1020 | - | - | 0 | - |
| - | - | 2434 | 1021 | - | - | 0 | - |
| - | - | 1270 | 1022 | - | - | 0 | - |
| - | - | 626.2 | 1240 | - | - | 0 | - |
| - | - | 699.7 | 3079 | - | - | 0 | - |
| - | - | 676.1 | 3099 | - | - | 0 | - |

m/z Charge Intensity FragmentType MassShift Position
120.08096313476562 0 29467.988
121.08429718017578 0 2548.818
121.55816650390625 0 378.2711
127.58011627197266 0 392.31656
129.1024627685547 0 3031.466
130.0654754638672 0 805.65295
132.08090209960938 0 594.9928
132.4221649169922 0 375.19696
133.08612060546875 0 956.18713
135.37722778320312 0 372.26874
136.07586669921875 0 9432.057
137.0795440673828 0 669.2674
138.0550994873047 0 465.57913
138.06654357910156 0 698.3075
140.31234741210938 0 383.58942
143.10659790039062 0 449.85358
146.06053161621094 0 568.35785
148.8809814453125 0 724.26776
148.895751953125 0 450.62292
148.90318298339844 0 661.2456
148.9101104736328 0 766.7213
148.9171905517578 0 855.3585
148.92442321777344 0 1172.3363
148.93133544921875 0 1521.6918
148.9390869140625 0 3243.947
148.9556884765625 0 6429.956
148.9636688232422 0 3774.7305
148.97122192382812 0 1697.1726
148.9783477783203 0 1240.2798
148.9855194091797 0 1195.127
148.99264526367188 0 685.7481
149.00009155273438 0 837.8061
149.00701904296875 0 597.40314
149.01466369628906 0 611.60693
149.04443359375 0 1653.8046
149.13796997070312 0 434.77176
151.95838928222656 0 462.9542
153.8879852294922 0 452.47644
156.0768280029297 0 1252.1812
158.0966339111328 0 1982.2341
159.09181213378906 0 2808.7776
159.0987548828125 0 567.19855
160.11239624023438 0 724.5873
165.05487060546875 0 3541.5784
166.0608367919922 0 653.69855
166.08657836914062 0 1375.5138
167.0554962158203 0 2249.268
173.12850952148438 0 2250.5984
176.10711669921875 0 2887.8354
177.10226440429688 0 1442.1727
178.1057891845703 0 525.179
182.081298828125 0 6827.2715
183.08460998535156 0 919.7394
185.12876892089844 0 580.7924
188.070556640625 0 1649.037
191.11785888671875 0 544.8548
193.108642578125 0 1062.1615
200.13987731933594 0 556.7744
201.12327575683594 0 1106.0057
201.38587951660156 0 507.82584
203.1191864013672 0 1096.9224 a Water loss 1
205.09722900390625 0 1176.3772 y 8
213.08657836914062 0 523.38245
221.10345458984375 0 1666.3019
221.1285400390625 0 46259.766 a 1
222.1319580078125 0 5717.165
223.06378173828125 0 2174.0737
227.10328674316406 0 992.33997
229.1710662841797 0 743.68994
231.06105041503906 0 802.3732
231.0987091064453 0 604.85754
235.10792541503906 0 694.50793
239.0948028564453 0 1314.4454
241.07398986816406 0 855.5965
243.13241577148438 0 619.36975
249.12351989746094 0 7332.928 b 1
251.10311889648438 0 1204.5424
253.16693115234375 0 671.00653
260.19708251953125 0 587.12024
263.10260009765625 0 1875.5763
268.97845458984375 0 694.3366
279.0982360839844 0 880.85596
285.0102233886719 0 1605.0072
285.10235595703125 0 887.94385
296.1502685546875 0 803.03284
299.0618896484375 0 647.3228
319.1408996582031 0 791.8183
324.14605712890625 0 1843.9009
325.14666748046875 0 821.6513
327.1956481933594 0 528.8884
329.1498107910156 0 3971.5955
330.15277099609375 0 639.893
339.206298828125 0 576.4795
342.13665771484375 0 627.3499
342.1564025878906 0 6267.047 y 7
343.1598205566406 0 1510.2666
346.13958740234375 0 1036.51 b Water loss 2
348.12060546875 0 601.7624
351.0234375 0 568.18085
355.0696716308594 0 2506.8376
356.0711669921875 0 759.35706
357.224853515625 0 1247.3153
364.1508483886719 0 1735.9463 b 2
378.1288146972656 0 907.6687
380.5254211425781 0 637.6053
394.12518310546875 0 1357.1501
413.2658386230469 0 1857.198
414.2690734863281 0 668.9697
425.1947021484375 0 966.14874
427.72052001953125 0 645.5279
436.7291564941406 0 662.2807
465.16259765625 0 950.98083
479.17999267578125 0 1371.8542 b 3
481.2864685058594 0 736.1557
489.1914367675781 0 1877.2063 y 6
496.2276306152344 0 644.5803
550.710205078125 0 1258.5342 y 1
551.2129516601562 0 1462.7445
560.228271484375 0 3156.472 y 5
561.2295532226562 0 758.1599
583.5242919921875 0 605.17584
585.3411254882812 0 2764.6125
592.7343139648438 0 577.2237
600.28271484375 0 660.6446
600.8182983398438 0 647.77094
601.2346801757812 0 2113.7686 Precursor
601.736328125 0 774.74536
649.0045776367188 0 545.2587
649.396240234375 0 658.6679
657.2633056640625 0 738.1627
700.362548828125 0 3463.8096
701.3646240234375 0 1792.9915
721.2568969726562 0 866.94965
723.2904052734375 0 3437.5916 y 4
724.2960205078125 0 1802.8872
749.2562866210938 0 1121.4183
774.3199462890625 0 798.4293
783.3984985351562 0 1408.5138
801.4132690429688 0 1831.8224
802.4129028320312 0 1444.9802
828.4281616210938 0 1144.2806
838.3177490234375 0 3553.531 y 3
839.3250732421875 0 1612.6356
844.4528198242188 0 2968.9185
845.455810546875 0 1726.8864
871.3350219726562 0 1205.4984
872.4462280273438 0 10518.253
873.4495849609375 0 5344.7144
874.4530029296875 0 1610.4812
889.3468017578125 0 5600.8506
890.3477783203125 0 2842.088
891.3565063476562 0 613.7753
929.5887451171875 0 672.55707
935.3325805664062 0 1086.7372 y Water loss 2
953.3449096679688 0 17031.328 y 2
954.3471069335938 0 7726.2593
955.34814453125 0 2668.8098
991.5199584960938 0 1356.7826
992.512451171875 0 960.4659
993.52490234375 0 752.4082
1019.513916015625 0 4814.494
1020.517822265625 0 2434.3552
1021.5153198242188 0 1269.637
1240.40966796875 0 626.1537
3079.4697265625 0 699.7289
3099.37744140625 0 676.1341

Spectrum Details

|  |  |
| --- | --- |
| Matched peaks? Matched peaks The total absolute number of peaks matched. Additionally in brackets the total fraction of peaks matched and the total number of peaks is shown. | 16 (9.64% of 166) |
| FDR? FDR The false discovery rate estimated for this peptide. It is calculated by matching all theoretical fragments with a non-integer shift with the raw peaks for this spectrum. This is done with 40 different shifts. The resulting percentage is the average number of annotated peaks over the number of annotated peaks with the correct spectrum. | 1.34% |
| Satellite FDR? Satellite FDR See the FDR for details on its calculation. This satellite ion specific FDR only contains the satellite ions (d/w) for I/L/J positions. | - |
| PSM Score? PSM Score The PSM Score as given by Hecklib to this annotated spectrum. It is shown with three significant figures. | 177 |

#### Spectrum 11842? Spectrum 11842 The raw spectrum of this peptide as annotated by Hecklib. The fragments are coloured according to ion type (see legend). Any peaks with a star '\*' as text can be hovered over to see the full details, first the ion type second the mass shift type. By hovering over the amino acids in the peptide or ions in the legend the corresponding peaks are highlighted. By toggling the 'Unassigned' label you can turn the background (unassigned) peaks on or off in the plot. By updating the slider in the Ion legend you can update the spectrum to only show the top X% of the peaks with labels. The top X% means any peak that is within X% of the highest intensity. By dragging in the spectrum you can zoom in to a specific part of the spectrum and use 'Zoom Out' to get back to the original zoom level. The annotation of the spectrum is based on the given sequence in the peptides file and is done with different software so inconsistencies are likely. The peaks are annotated based on the given sequence, with 20 ppm tolerance.

Copy Data

##### Spectrum 11842 (TSV)

###### Preview

```
Loading example...
```

*Click on the button to copy the data to your clipboard.*

Mz MinMz MaxIntensity Max

WidthHeightPeptide font sizePeptide stroke widthSpectrum font sizeSpectrum stroke widthCompact peptide

Ion legend

wxyz

abcd

OtherUnassignedIon^Charge^_Position_Show for top:%

TFDDYAMHW

01.24e+42.48e+43.72e+44.95e+4

Zoom Out

a^+1^_2_y^+1^_1_a^+1^_2_b^+1^_2_a^+1^_3_y^+1^_2_b^+1^_3_y^+2^_7_b^+1^_4_y^+1^_3_y^+2^_8_y^+1^_4_\*\*y^+1^_5_y^+1^_6_y^+1^_7_

0788157523633151

Fragment Matches Table

Show background peaks

| Position | Ion type | Intensity | mz Theoretical | mz Error (Th) | mz Error (ppm) | Charge | Series Number |
| --- | --- | --- | --- | --- | --- | --- | --- |
| - | - | 3.451E+04 | 120.1 | - | - | 0 | - |
| - | - | 367.2 | 121 | - | - | 0 | - |
| - | - | 3173 | 121.1 | - | - | 0 | - |
| - | - | 416.7 | 123 | - | - | 0 | - |
| - | - | 3235 | 129.1 | - | - | 0 | - |
| - | - | 1045 | 130.1 | - | - | 0 | - |
| - | - | 939.4 | 132.1 | - | - | 0 | - |
| - | - | 463.5 | 132.1 | - | - | 0 | - |
| - | - | 1242 | 133.1 | - | - | 0 | - |
| - | - | 1.104E+04 | 136.1 | - | - | 0 | - |
| - | - | 966.5 | 137.1 | - | - | 0 | - |
| - | - | 412.4 | 138.1 | - | - | 0 | - |
| - | - | 454.6 | 141.1 | - | - | 0 | - |
| - | - | 512.8 | 141.1 | - | - | 0 | - |
| - | - | 466.4 | 146.1 | - | - | 0 | - |
| - | - | 418.3 | 148.8 | - | - | 0 | - |
| - | - | 579.4 | 148.9 | - | - | 0 | - |
| - | - | 497.6 | 148.9 | - | - | 0 | - |
| - | - | 608.6 | 148.9 | - | - | 0 | - |
| - | - | 573.7 | 148.9 | - | - | 0 | - |
| - | - | 800.3 | 148.9 | - | - | 0 | - |
| - | - | 1181 | 148.9 | - | - | 0 | - |
| - | - | 1401 | 148.9 | - | - | 0 | - |
| - | - | 1443 | 148.9 | - | - | 0 | - |
| - | - | 3344 | 148.9 | - | - | 0 | - |
| - | - | 6118 | 149 | - | - | 0 | - |
| - | - | 3384 | 149 | - | - | 0 | - |
| - | - | 1457 | 149 | - | - | 0 | - |
| - | - | 1160 | 149 | - | - | 0 | - |
| - | - | 1023 | 149 | - | - | 0 | - |
| - | - | 829.4 | 149 | - | - | 0 | - |
| - | - | 791.7 | 149 | - | - | 0 | - |
| - | - | 429.3 | 149 | - | - | 0 | - |
| - | - | 429 | 149 | - | - | 0 | - |
| - | - | 564.7 | 149 | - | - | 0 | - |
| - | - | 1418 | 149 | - | - | 0 | - |
| - | - | 472.5 | 149.1 | - | - | 0 | - |
| - | - | 1521 | 156.1 | - | - | 0 | - |
| - | - | 433.2 | 157.1 | - | - | 0 | - |
| - | - | 1221 | 158.1 | - | - | 0 | - |
| - | - | 1998 | 159.1 | - | - | 0 | - |
| - | - | 745.1 | 160.1 | - | - | 0 | - |
| - | - | 514.7 | 162.1 | - | - | 0 | - |
| - | - | 3799 | 165.1 | - | - | 0 | - |
| - | - | 1607 | 166.1 | - | - | 0 | - |
| - | - | 435.6 | 166.5 | - | - | 0 | - |
| - | - | 2117 | 167.1 | - | - | 0 | - |
| - | - | 922.7 | 167.1 | - | - | 0 | - |
| - | - | 517.9 | 172.8 | - | - | 0 | - |
| - | - | 1902 | 173.1 | - | - | 0 | - |
| - | - | 600.2 | 175.1 | - | - | 0 | - |
| - | - | 1042 | 175.1 | - | - | 0 | - |
| - | - | 3925 | 176.1 | - | - | 0 | - |
| - | - | 497.5 | 176.3 | - | - | 0 | - |
| - | - | 2821 | 177.1 | - | - | 0 | - |
| - | - | 920.6 | 177.1 | - | - | 0 | - |
| - | - | 6860 | 182.1 | - | - | 0 | - |
| - | - | 728.7 | 183.1 | - | - | 0 | - |
| - | - | 1238 | 184.1 | - | - | 0 | - |
| - | - | 640.7 | 185.1 | - | - | 0 | - |
| - | - | 1444 | 188.1 | - | - | 0 | - |
| - | - | 631.3 | 191.1 | - | - | 0 | - |
| - | - | 910.6 | 193.1 | - | - | 0 | - |
| - | - | 1846 | 201.1 | - | - | 0 | - |
| 2 | a | 1137 | 203.1 | 0.000244 | 1.201 | +1 | 2 |
| 9 | y | 1475 | 205.1 | 0.0001997 | 0.9739 | +1 | 1 |
| - | - | 680.5 | 209.1 | - | - | 0 | - |
| - | - | 3298 | 221.1 | - | - | 0 | - |
| 2 | a | 4.906E+04 | 221.1 | 0.0001469 | 0.6641 | +1 | 2 |
| - | - | 5549 | 222.1 | - | - | 0 | - |
| - | - | 1387 | 223.1 | - | - | 0 | - |
| - | - | 1083 | 227.1 | - | - | 0 | - |
| - | - | 1028 | 229.1 | - | - | 0 | - |
| - | - | 615.1 | 229.2 | - | - | 0 | - |
| - | - | 1434 | 231.1 | - | - | 0 | - |
| - | - | 1734 | 235.1 | - | - | 0 | - |
| - | - | 1282 | 239.1 | - | - | 0 | - |
| - | - | 629.8 | 240.1 | - | - | 0 | - |
| - | - | 629.7 | 243.1 | - | - | 0 | - |
| - | - | 547.9 | 246.9 | - | - | 0 | - |
| 2 | b | 8085 | 249.1 | 0.0002579 | 1.035 | +1 | 2 |
| - | - | 1004 | 250.1 | - | - | 0 | - |
| - | - | 1046 | 251.1 | - | - | 0 | - |
| - | - | 554.5 | 251.1 | - | - | 0 | - |
| - | - | 536.2 | 257.1 | - | - | 0 | - |
| - | - | 494.2 | 262.8 | - | - | 0 | - |
| - | - | 2297 | 263.1 | - | - | 0 | - |
| - | - | 592.7 | 274.1 | - | - | 0 | - |
| - | - | 519.6 | 276.1 | - | - | 0 | - |
| - | - | 636.8 | 279.1 | - | - | 0 | - |
| - | - | 1200 | 285 | - | - | 0 | - |
| - | - | 1106 | 285.1 | - | - | 0 | - |
| - | - | 758.9 | 296.2 | - | - | 0 | - |
| - | - | 556 | 301 | - | - | 0 | - |
| - | - | 758 | 309.2 | - | - | 0 | - |
| 3 | a | 520.4 | 318.1 | 0.0003701 | 1.163 | +1 | 3 |
| - | - | 1106 | 324.1 | - | - | 0 | - |
| - | - | 5006 | 329.2 | - | - | 0 | - |
| - | - | 950 | 330.2 | - | - | 0 | - |
| - | - | 501.1 | 340.2 | - | - | 0 | - |
| - | - | 687.2 | 342.1 | - | - | 0 | - |
| 8 | y | 8865 | 342.2 | 0.0004587 | 1.341 | +1 | 2 |
| - | - | 759 | 343.2 | - | - | 0 | - |
| - | - | 1901 | 355.1 | - | - | 0 | - |
| - | - | 676.7 | 359.2 | - | - | 0 | - |
| 3 | b | 1501 | 364.2 | 0.0006232 | 1.711 | +1 | 3 |
| - | - | 612.4 | 365.2 | - | - | 0 | - |
| - | - | 995.7 | 378.1 | - | - | 0 | - |
| - | - | 794 | 380.2 | - | - | 0 | - |
| - | - | 1874 | 394.1 | - | - | 0 | - |
| - | - | 576.6 | 408.7 | - | - | 0 | - |
| - | - | 2059 | 413.3 | - | - | 0 | - |
| - | - | 766.8 | 425.2 | - | - | 0 | - |
| - | - | 567.1 | 451.4 | - | - | 0 | - |
| - | - | 894.7 | 455.2 | - | - | 0 | - |
| - | - | 1102 | 465.2 | - | - | 0 | - |
| 3 | y | 565.3 | 477.2 | 0.007757 | 16.26 | +2 | 7 |
| 4 | b | 900.2 | 479.2 | 0.0009372 | 1.956 | +1 | 4 |
| - | - | 660.5 | 483.2 | - | - | 0 | - |
| 7 | y | 1950 | 489.2 | 0.005649 | 11.55 | +1 | 3 |
| - | - | 607 | 491.2 | - | - | 0 | - |
| - | - | 824.9 | 495.2 | - | - | 0 | - |
| - | - | 905.8 | 496.2 | - | - | 0 | - |
| - | - | 661.2 | 508.3 | - | - | 0 | - |
| 2 | y | 1309 | 550.7 | 0.004789 | 8.696 | +2 | 8 |
| - | - | 809.7 | 551.2 | - | - | 0 | - |
| 6 | y | 2478 | 560.2 | 0.003753 | 6.698 | +1 | 4 |
| - | - | 1322 | 561.2 | - | - | 0 | - |
| - | - | 647.4 | 584.3 | - | - | 0 | - |
| - | - | 2575 | 585.3 | - | - | 0 | - |
| 0 | Precursor | 586.6 | 592.2 | 0.005031 | 8.495 | +2 | -1 |
| 0 | Precursor | 1517 | 601.2 | 0.001519 | 2.526 | +2 | -1 |
| - | - | 2165 | 601.4 | - | - | 0 | - |
| - | - | 1262 | 601.7 | - | - | 0 | - |
| - | - | 799.3 | 612.2 | - | - | 0 | - |
| - | - | 654.6 | 614.2 | - | - | 0 | - |
| - | - | 979.9 | 621.4 | - | - | 0 | - |
| - | - | 624 | 657.3 | - | - | 0 | - |
| - | - | 1074 | 659.3 | - | - | 0 | - |
| - | - | 919.4 | 685.3 | - | - | 0 | - |
| - | - | 4955 | 700.4 | - | - | 0 | - |
| - | - | 1461 | 701.4 | - | - | 0 | - |
| 5 | y | 3314 | 723.3 | 0.005549 | 7.671 | +1 | 5 |
| - | - | 1559 | 724.3 | - | - | 0 | - |
| - | - | 1299 | 749.3 | - | - | 0 | - |
| - | - | 623.5 | 750.3 | - | - | 0 | - |
| - | - | 1629 | 774.3 | - | - | 0 | - |
| - | - | 633.1 | 783.4 | - | - | 0 | - |
| - | - | 2157 | 801.4 | - | - | 0 | - |
| - | - | 989.3 | 802.4 | - | - | 0 | - |
| 4 | y | 3271 | 838.3 | 0.005461 | 6.514 | +1 | 6 |
| - | - | 1334 | 839.3 | - | - | 0 | - |
| - | - | 4025 | 844.5 | - | - | 0 | - |
| - | - | 1488 | 845.5 | - | - | 0 | - |
| - | - | 862.4 | 871.3 | - | - | 0 | - |
| - | - | 1.327E+04 | 872.4 | - | - | 0 | - |
| - | - | 6501 | 873.4 | - | - | 0 | - |
| - | - | 1458 | 874.4 | - | - | 0 | - |
| - | - | 706.8 | 875.5 | - | - | 0 | - |
| - | - | 6409 | 889.3 | - | - | 0 | - |
| - | - | 2173 | 890.3 | - | - | 0 | - |
| - | - | 992.1 | 891.4 | - | - | 0 | - |
| 3 | y | 1.793E+04 | 953.3 | 0.004153 | 4.356 | +1 | 7 |
| - | - | 1.044E+04 | 954.3 | - | - | 0 | - |
| - | - | 4081 | 955.3 | - | - | 0 | - |
| - | - | 1919 | 991.5 | - | - | 0 | - |
| - | - | 1224 | 992.5 | - | - | 0 | - |
| - | - | 831.8 | 1003 | - | - | 0 | - |
| - | - | 5243 | 1020 | - | - | 0 | - |
| - | - | 3134 | 1021 | - | - | 0 | - |
| - | - | 740.5 | 1022 | - | - | 0 | - |
| - | - | 650.4 | 1102 | - | - | 0 | - |
| - | - | 595.8 | 1561 | - | - | 0 | - |
| - | - | 623.1 | 2341 | - | - | 0 | - |
| - | - | 764.5 | 2512 | - | - | 0 | - |
| - | - | 779.3 | 2702 | - | - | 0 | - |
| - | - | 758.1 | 2731 | - | - | 0 | - |
| - | - | 695.6 | 3120 | - | - | 0 | - |

m/z Charge Intensity FragmentType MassShift Position
120.08102416992188 0 34512.156
121.02893829345703 0 367.18307
121.08429718017578 0 3173.3367
123.0439224243164 0 416.71304
129.1024627685547 0 3234.6487
130.065185546875 0 1045.404
132.08119201660156 0 939.3678
132.10153198242188 0 463.52783
133.08639526367188 0 1242.2365
136.07589721679688 0 11042.541
137.07948303222656 0 966.50494
138.06686401367188 0 412.4447
141.0913543701172 0 454.55017
141.1024627685547 0 512.7614
146.06045532226562 0 466.3747
148.84542846679688 0 418.29395
148.86695861816406 0 579.40686
148.8887481689453 0 497.5633
148.89544677734375 0 608.5947
148.90341186523438 0 573.6922
148.91067504882812 0 800.281
148.917724609375 0 1181.2024
148.9250030517578 0 1400.9072
148.93234252929688 0 1443.348
148.9399871826172 0 3343.7896
148.95657348632812 0 6118.141
148.96444702148438 0 3384.061
148.9720458984375 0 1457.3346
148.97926330566406 0 1160.2986
148.98631286621094 0 1022.8826
148.99351501464844 0 829.3552
149.00086975097656 0 791.7432
149.00827026367188 0 429.2532
149.0225372314453 0 428.96655
149.03756713867188 0 564.70123
149.04486083984375 0 1418.0236
149.06634521484375 0 472.49887
156.07701110839844 0 1521.4122
157.09710693359375 0 433.16672
158.09671020507812 0 1221.1041
159.0918426513672 0 1997.7704
160.11207580566406 0 745.1154
162.09201049804688 0 514.688
165.0548858642578 0 3799.1892
166.08639526367188 0 1606.857
166.52413940429688 0 435.57465
167.0555419921875 0 2116.7915
167.08155822753906 0 922.73175
172.80128479003906 0 517.92236
173.1285858154297 0 1901.7101
175.0863037109375 0 600.1586
175.11932373046875 0 1042.3228
176.10726928710938 0 3924.6108
176.29345703125 0 497.4824
177.10238647460938 0 2820.965
177.1111297607422 0 920.5757
182.08135986328125 0 6860.0864
183.0848846435547 0 728.68134
184.07298278808594 0 1237.9342
185.12889099121094 0 640.6539
188.0709686279297 0 1443.5963
191.11903381347656 0 631.29504
193.1084747314453 0 910.63446
201.12359619140625 0 1845.8312
203.11813354492188 0 1136.5966 a Water loss 1
205.09695434570312 0 1474.958 y 8
209.09170532226562 0 680.5411
221.10353088378906 0 3298.0222
221.12860107421875 0 49056.652 a 1
222.1319580078125 0 5549.2905
223.06402587890625 0 1386.5686
227.102783203125 0 1083.0354
229.11880493164062 0 1027.5845
229.17074584960938 0 615.1389
231.06101989746094 0 1434.202
235.1081085205078 0 1734.4617
239.09449768066406 0 1282.2454
240.13436889648438 0 629.7601
243.13418579101562 0 629.7205
246.8527374267578 0 547.9187
249.12362670898438 0 8085.4346 b 1
250.12696838378906 0 1004.2845
251.10296630859375 0 1045.9532
251.1495819091797 0 554.4855
257.10546875 0 536.24097
262.843505859375 0 494.16315
263.1026611328125 0 2297.346
274.12945556640625 0 592.69257
276.1123046875 0 519.6262
279.0984802246094 0 636.78107
285.0093078613281 0 1200.2823
285.1012878417969 0 1105.7416
296.1512756347656 0 758.8688
301.0417175292969 0 556.02277
309.2037353515625 0 757.9513
318.14520263671875 0 520.36066 a Water loss 2
324.1455383300781 0 1105.6298
329.1500244140625 0 5005.846
330.15252685546875 0 950.0109
340.188232421875 0 501.07376
342.13690185546875 0 687.1751
342.1565246582031 0 8865.009 y 7
343.1593017578125 0 758.9792
355.0706481933594 0 1901.3701
359.1913146972656 0 676.73096
364.1496887207031 0 1501.3942 b 2
365.1523132324219 0 612.4015
378.13018798828125 0 995.7332
380.1923522949219 0 793.98254
394.12469482421875 0 1873.5778
408.7013854980469 0 576.6034
413.2667236328125 0 2059.3823
425.1920166015625 0 766.7859
451.4329528808594 0 567.06104
455.2251281738281 0 894.7461
465.16375732421875 0 1101.5397
477.16632080078125 0 565.3292 y 2
479.1781921386719 0 900.1707 b 3
483.2230529785156 0 660.4907
489.19219970703125 0 1949.8185 y 6
491.19757080078125 0 607.04584
495.18499755859375 0 824.8894
496.2309875488281 0 905.7562
508.2742004394531 0 661.1856
550.7130737304688 0 1308.9452 y 1
551.212158203125 0 809.7417
560.2274169921875 0 2478.4375 y 5
561.2320556640625 0 1321.7499
584.3024291992188 0 647.3975
585.3406982421875 0 2575.3835
592.2318725585938 0 586.5783 Precursor Water loss
601.233642578125 0 1517.2585 Precursor
601.3914184570312 0 2164.873
601.7361450195312 0 1262.0032
612.1992797851562 0 799.30817
614.241455078125 0 654.58704
621.35986328125 0 979.93195
657.25634765625 0 623.99023
659.2938842773438 0 1073.585
685.260986328125 0 919.3971
700.3623657226562 0 4954.5684
701.3623657226562 0 1461.0046
723.2925415039062 0 3313.951 y 4
724.2976684570312 0 1559.2567
749.2540283203125 0 1298.7859
750.252685546875 0 623.4686
774.3189697265625 0 1629.4181
783.4046020507812 0 633.14294
801.411865234375 0 2157.311
802.4122314453125 0 989.33496
838.3193969726562 0 3270.824 y 3
839.3191528320312 0 1334.4692
844.4513549804688 0 4025.3816
845.4526977539062 0 1487.9847
871.3330688476562 0 862.43616
872.4467163085938 0 13271.19
873.4495239257812 0 6501.027
874.4483032226562 0 1458.1084
875.4614868164062 0 706.8266
889.3457641601562 0 6408.529
890.3490600585938 0 2172.522
891.3526611328125 0 992.098
953.3450317382812 0 17926.63 y 2
954.3477172851562 0 10438.775
955.347900390625 0 4080.9604
991.5195922851562 0 1918.6329
992.5221557617188 0 1223.8896
1003.3822631835938 0 831.78986
1019.5147705078125 0 5243.2344
1020.5183715820312 0 3133.9773
1021.5228881835938 0 740.4925
1102.41943359375 0 650.415
1560.5941162109375 0 595.7763
2341.117431640625 0 623.062
2512.332275390625 0 764.4881
2701.58447265625 0 779.3423
2730.551513671875 0 758.0821
3119.68212890625 0 695.56744

Spectrum Details

|  |  |
| --- | --- |
| Matched peaks? Matched peaks The total absolute number of peaks matched. Additionally in brackets the total fraction of peaks matched and the total number of peaks is shown. | 17 (9.55% of 178) |
| FDR? FDR The false discovery rate estimated for this peptide. It is calculated by matching all theoretical fragments with a non-integer shift with the raw peaks for this spectrum. This is done with 40 different shifts. The resulting percentage is the average number of annotated peaks over the number of annotated peaks with the correct spectrum. | 0.70% |
| Satellite FDR? Satellite FDR See the FDR for details on its calculation. This satellite ion specific FDR only contains the satellite ions (d/w) for I/L/J positions. | - |
| PSM Score? PSM Score The PSM Score as given by Hecklib to this annotated spectrum. It is shown with three significant figures. | 160 |

#### Spectrum 12129? Spectrum 12129 The raw spectrum of this peptide as annotated by Hecklib. The fragments are coloured according to ion type (see legend). Any peaks with a star '\*' as text can be hovered over to see the full details, first the ion type second the mass shift type. By hovering over the amino acids in the peptide or ions in the legend the corresponding peaks are highlighted. By toggling the 'Unassigned' label you can turn the background (unassigned) peaks on or off in the plot. By updating the slider in the Ion legend you can update the spectrum to only show the top X% of the peaks with labels. The top X% means any peak that is within X% of the highest intensity. By dragging in the spectrum you can zoom in to a specific part of the spectrum and use 'Zoom Out' to get back to the original zoom level. The annotation of the spectrum is based on the given sequence in the peptides file and is done with different software so inconsistencies are likely. The peaks are annotated based on the given sequence, with 20 ppm tolerance.

Copy Data

##### Spectrum 12129 (TSV)

###### Preview

```
Loading example...
```

*Click on the button to copy the data to your clipboard.*

Mz MinMz MaxIntensity Max

WidthHeightPeptide font sizePeptide stroke widthSpectrum font sizeSpectrum stroke widthCompact peptide

Ion legend

wxyz

abcd

OtherUnassignedIon^Charge^_Position_Show for top:%

TFDDYAMHW

01.10e+42.21e+43.31e+44.42e+4

Zoom Out

a^+1^_2_y^+1^_1_a^+1^_2_b^+2^_4_b^+1^_2_y^+1^_2_b^+1^_3_b^+1^_4_y^+1^_3_y^+2^_8_y^+1^_4_\*y^+1^_5_y^+1^_6_y^+1^_7_y^+1^_7_

0505101015152019

Fragment Matches Table

Show background peaks

| Position | Ion type | Intensity | mz Theoretical | mz Error (Th) | mz Error (ppm) | Charge | Series Number |
| --- | --- | --- | --- | --- | --- | --- | --- |
| - | - | 3.057E+04 | 120.1 | - | - | 0 | - |
| - | - | 2917 | 121.1 | - | - | 0 | - |
| - | - | 643.6 | 123 | - | - | 0 | - |
| - | - | 360 | 123.2 | - | - | 0 | - |
| - | - | 509.5 | 127.1 | - | - | 0 | - |
| - | - | 545.4 | 129.1 | - | - | 0 | - |
| - | - | 3798 | 129.1 | - | - | 0 | - |
| - | - | 364.4 | 129.2 | - | - | 0 | - |
| - | - | 861.3 | 130.1 | - | - | 0 | - |
| - | - | 416.7 | 131.1 | - | - | 0 | - |
| - | - | 925.2 | 132.1 | - | - | 0 | - |
| - | - | 1007 | 132.1 | - | - | 0 | - |
| - | - | 461.8 | 133.1 | - | - | 0 | - |
| - | - | 1598 | 133.1 | - | - | 0 | - |
| - | - | 1.109E+04 | 136.1 | - | - | 0 | - |
| - | - | 441.1 | 137.1 | - | - | 0 | - |
| - | - | 477.8 | 138.1 | - | - | 0 | - |
| - | - | 613 | 138.1 | - | - | 0 | - |
| - | - | 613.8 | 141.1 | - | - | 0 | - |
| - | - | 486 | 143.1 | - | - | 0 | - |
| - | - | 897.1 | 149 | - | - | 0 | - |
| - | - | 926.2 | 156.1 | - | - | 0 | - |
| - | - | 780.8 | 157.1 | - | - | 0 | - |
| - | - | 1224 | 158.1 | - | - | 0 | - |
| - | - | 1950 | 159.1 | - | - | 0 | - |
| - | - | 424 | 159.7 | - | - | 0 | - |
| - | - | 529.7 | 160.1 | - | - | 0 | - |
| - | - | 603.8 | 160.1 | - | - | 0 | - |
| - | - | 661.5 | 162.1 | - | - | 0 | - |
| - | - | 3146 | 165.1 | - | - | 0 | - |
| - | - | 1851 | 166.1 | - | - | 0 | - |
| - | - | 2123 | 167.1 | - | - | 0 | - |
| - | - | 535.2 | 167.1 | - | - | 0 | - |
| - | - | 422.8 | 167.1 | - | - | 0 | - |
| - | - | 705.2 | 169 | - | - | 0 | - |
| - | - | 1049 | 169.1 | - | - | 0 | - |
| - | - | 540 | 173.1 | - | - | 0 | - |
| - | - | 1964 | 173.1 | - | - | 0 | - |
| - | - | 4054 | 176.1 | - | - | 0 | - |
| - | - | 2433 | 177.1 | - | - | 0 | - |
| - | - | 1746 | 177.1 | - | - | 0 | - |
| - | - | 6016 | 182.1 | - | - | 0 | - |
| - | - | 711.4 | 183.1 | - | - | 0 | - |
| - | - | 1973 | 183.1 | - | - | 0 | - |
| - | - | 511.2 | 184.2 | - | - | 0 | - |
| - | - | 569.4 | 185.1 | - | - | 0 | - |
| - | - | 1483 | 188.1 | - | - | 0 | - |
| - | - | 554.7 | 191.1 | - | - | 0 | - |
| - | - | 977 | 193.1 | - | - | 0 | - |
| - | - | 484 | 196.3 | - | - | 0 | - |
| - | - | 799.9 | 201.1 | - | - | 0 | - |
| - | - | 1189 | 201.1 | - | - | 0 | - |
| 2 | a | 994.2 | 203.1 | 0.0002443 | 1.203 | +1 | 2 |
| - | - | 674.8 | 204.1 | - | - | 0 | - |
| 9 | y | 1436 | 205.1 | 0.0001845 | 0.8995 | +1 | 1 |
| - | - | 494.5 | 206.6 | - | - | 0 | - |
| - | - | 1656 | 211.1 | - | - | 0 | - |
| - | - | 639.5 | 215.1 | - | - | 0 | - |
| - | - | 1749 | 221.1 | - | - | 0 | - |
| 2 | a | 4.374E+04 | 221.1 | 7.056E-05 | 0.3191 | +1 | 2 |
| - | - | 5304 | 222.1 | - | - | 0 | - |
| - | - | 2565 | 223.1 | - | - | 0 | - |
| - | - | 628.3 | 223.1 | - | - | 0 | - |
| - | - | 548.7 | 226.2 | - | - | 0 | - |
| - | - | 685.3 | 227.1 | - | - | 0 | - |
| - | - | 695.1 | 231.1 | - | - | 0 | - |
| - | - | 1035 | 235.1 | - | - | 0 | - |
| - | - | 721.7 | 238.2 | - | - | 0 | - |
| - | - | 931.8 | 239.1 | - | - | 0 | - |
| 4 | b | 720.4 | 240.1 | 0.004002 | 16.67 | +2 | 4 |
| - | - | 922.3 | 241.1 | - | - | 0 | - |
| 2 | b | 8289 | 249.1 | 0.0001663 | 0.6676 | +1 | 2 |
| - | - | 850.4 | 250.1 | - | - | 0 | - |
| - | - | 1095 | 251.1 | - | - | 0 | - |
| - | - | 650.6 | 251.1 | - | - | 0 | - |
| - | - | 577.2 | 253.2 | - | - | 0 | - |
| - | - | 1722 | 263.1 | - | - | 0 | - |
| - | - | 539.9 | 264.2 | - | - | 0 | - |
| - | - | 1957 | 285 | - | - | 0 | - |
| - | - | 639.6 | 287.3 | - | - | 0 | - |
| - | - | 826 | 292.1 | - | - | 0 | - |
| - | - | 623.1 | 293.2 | - | - | 0 | - |
| - | - | 639.9 | 296.2 | - | - | 0 | - |
| - | - | 1070 | 299.1 | - | - | 0 | - |
| - | - | 636.6 | 302.1 | - | - | 0 | - |
| - | - | 851.5 | 319.1 | - | - | 0 | - |
| - | - | 2080 | 324.1 | - | - | 0 | - |
| - | - | 1006 | 325.1 | - | - | 0 | - |
| - | - | 4206 | 329.1 | - | - | 0 | - |
| - | - | 1005 | 330.2 | - | - | 0 | - |
| - | - | 648.2 | 340.2 | - | - | 0 | - |
| 8 | y | 8426 | 342.2 | 0.0005197 | 1.519 | +1 | 2 |
| - | - | 1905 | 343.2 | - | - | 0 | - |
| - | - | 1499 | 355.1 | - | - | 0 | - |
| - | - | 782.5 | 356.1 | - | - | 0 | - |
| - | - | 1042 | 357.2 | - | - | 0 | - |
| 3 | b | 2218 | 364.2 | 0.0001044 | 0.2866 | +1 | 3 |
| - | - | 768.4 | 376.1 | - | - | 0 | - |
| - | - | 1583 | 378.1 | - | - | 0 | - |
| - | - | 723.4 | 380.2 | - | - | 0 | - |
| - | - | 1006 | 394.1 | - | - | 0 | - |
| - | - | 673.6 | 396.2 | - | - | 0 | - |
| - | - | 946.1 | 401.2 | - | - | 0 | - |
| - | - | 2195 | 413.3 | - | - | 0 | - |
| - | - | 1030 | 425.2 | - | - | 0 | - |
| - | - | 747.1 | 425.3 | - | - | 0 | - |
| - | - | 524.1 | 449.6 | - | - | 0 | - |
| - | - | 1139 | 465.2 | - | - | 0 | - |
| 4 | b | 980.8 | 479.2 | 0.00051 | 1.064 | +1 | 4 |
| - | - | 641.6 | 480.3 | - | - | 0 | - |
| - | - | 704.6 | 483.2 | - | - | 0 | - |
| 7 | y | 2556 | 489.2 | 0.005985 | 12.23 | +1 | 3 |
| - | - | 782.2 | 490.2 | - | - | 0 | - |
| - | - | 1269 | 496.2 | - | - | 0 | - |
| - | - | 623.8 | 529.7 | - | - | 0 | - |
| 2 | y | 1769 | 550.7 | 0.002042 | 3.709 | +2 | 8 |
| 6 | y | 2511 | 560.2 | 0.005706 | 10.18 | +1 | 4 |
| - | - | 1051 | 561.2 | - | - | 0 | - |
| - | - | 618.5 | 564.2 | - | - | 0 | - |
| - | - | 555 | 569.6 | - | - | 0 | - |
| - | - | 4675 | 585.3 | - | - | 0 | - |
| - | - | 726 | 591.5 | - | - | 0 | - |
| - | - | 568.5 | 592.9 | - | - | 0 | - |
| - | - | 625.2 | 593.4 | - | - | 0 | - |
| - | - | 593.4 | 601 | - | - | 0 | - |
| 0 | Precursor | 981.4 | 601.2 | 0.008416 | 14 | +2 | -1 |
| - | - | 1174 | 601.7 | - | - | 0 | - |
| - | - | 667.8 | 604.4 | - | - | 0 | - |
| - | - | 1125 | 621.4 | - | - | 0 | - |
| - | - | 817.3 | 657.3 | - | - | 0 | - |
| - | - | 1311 | 659.3 | - | - | 0 | - |
| - | - | 921.3 | 660.3 | - | - | 0 | - |
| - | - | 948.9 | 685.3 | - | - | 0 | - |
| - | - | 3522 | 700.4 | - | - | 0 | - |
| - | - | 1345 | 701.4 | - | - | 0 | - |
| - | - | 1617 | 721.3 | - | - | 0 | - |
| 5 | y | 3326 | 723.3 | 0.005488 | 7.587 | +1 | 5 |
| - | - | 932.9 | 724.3 | - | - | 0 | - |
| - | - | 1581 | 749.3 | - | - | 0 | - |
| - | - | 679.9 | 750.3 | - | - | 0 | - |
| - | - | 1280 | 774.3 | - | - | 0 | - |
| - | - | 1125 | 783.4 | - | - | 0 | - |
| - | - | 1550 | 801.4 | - | - | 0 | - |
| - | - | 784.3 | 828.4 | - | - | 0 | - |
| 4 | y | 3366 | 838.3 | 0.005034 | 6.005 | +1 | 6 |
| - | - | 1038 | 839.3 | - | - | 0 | - |
| - | - | 2753 | 844.5 | - | - | 0 | - |
| - | - | 795.4 | 845.5 | - | - | 0 | - |
| - | - | 1.171E+04 | 872.4 | - | - | 0 | - |
| - | - | 6207 | 873.5 | - | - | 0 | - |
| - | - | 990.1 | 874.5 | - | - | 0 | - |
| - | - | 4064 | 889.3 | - | - | 0 | - |
| - | - | 3170 | 890.4 | - | - | 0 | - |
| - | - | 672.3 | 909.7 | - | - | 0 | - |
| 3 | y | 615.7 | 935.3 | 0.01209 | 12.93 | +1 | 7 |
| - | - | 650.1 | 936.3 | - | - | 0 | - |
| 3 | y | 1.529E+04 | 953.3 | 0.006533 | 6.853 | +1 | 7 |
| - | - | 8104 | 954.3 | - | - | 0 | - |
| - | - | 2196 | 955.3 | - | - | 0 | - |
| - | - | 2418 | 991.5 | - | - | 0 | - |
| - | - | 1391 | 992.5 | - | - | 0 | - |
| - | - | 4860 | 1020 | - | - | 0 | - |
| - | - | 2883 | 1021 | - | - | 0 | - |
| - | - | 1340 | 1022 | - | - | 0 | - |
| - | - | 616.1 | 1308 | - | - | 0 | - |
| - | - | 661 | 1999 | - | - | 0 | - |

m/z Charge Intensity FragmentType MassShift Position
120.08090209960938 0 30570.934
121.08423614501953 0 2916.6226
123.04374694824219 0 643.6425
123.22562408447266 0 359.96506
127.08643341064453 0 509.50076
129.06593322753906 0 545.356
129.1023712158203 0 3798.4563
129.17787170410156 0 364.4153
130.06533813476562 0 861.33594
131.0706024169922 0 416.66718
132.0808868408203 0 925.21857
132.10195922851562 0 1006.9181
133.06137084960938 0 461.77002
133.0858612060547 0 1598.4304
136.07577514648438 0 11089.086
137.0791473388672 0 441.09787
138.05482482910156 0 477.809
138.0662078857422 0 613.00287
141.10203552246094 0 613.8126
143.07025146484375 0 486.02652
149.0448760986328 0 897.10345
156.07679748535156 0 926.1828
157.13339233398438 0 780.76685
158.09677124023438 0 1224.3671
159.09185791015625 0 1949.5468
159.68215942382812 0 424.03018
160.0756072998047 0 529.70374
160.11209106445312 0 603.7581
162.09207153320312 0 661.5394
165.0546112060547 0 3146.189
166.0863800048828 0 1851.0482
167.05543518066406 0 2122.8025
167.08921813964844 0 535.23505
167.11891174316406 0 422.80554
169.0343780517578 0 705.16
169.09768676757812 0 1049.0095
173.09109497070312 0 539.9624
173.12864685058594 0 1963.7107
176.10691833496094 0 4053.6262
177.10232543945312 0 2433.1184
177.11196899414062 0 1746.1012
182.0812225341797 0 6015.89
183.11312866210938 0 711.4473
183.14935302734375 0 1972.8547
184.15182495117188 0 511.20834
185.1284637451172 0 569.4155
188.07040405273438 0 1483.3376
191.11753845214844 0 554.6502
193.10845947265625 0 976.96594
196.26292419433594 0 484.033
201.1121826171875 0 799.8564
201.12332153320312 0 1189.3552
203.11764526367188 0 994.2326 a Water loss 1
204.0767059326172 0 674.82324
205.0969696044922 0 1436.3335 y 8
206.60467529296875 0 494.54337
211.1441192626953 0 1655.5844
215.1390838623047 0 639.5087
221.1033935546875 0 1748.9054
221.12852478027344 0 43736.113 a 1
222.13185119628906 0 5304.418
223.06390380859375 0 2565.1108
223.10882568359375 0 628.2694
226.15525817871094 0 548.6525
227.10324096679688 0 685.27435
231.06137084960938 0 695.1393
235.10816955566406 0 1035.1729
238.1776885986328 0 721.7115
239.09451293945312 0 931.75146
240.0962677001953 0 720.4067 b 3
241.07400512695312 0 922.33203
249.12353515625 0 8288.621 b 1
250.12696838378906 0 850.42377
251.1027069091797 0 1095.3618
251.14901733398438 0 650.64954
253.1669158935547 0 577.1555
263.1025695800781 0 1721.9343
264.2467346191406 0 539.87854
285.01007080078125 0 1957.4673
287.2785949707031 0 639.596
292.1393127441406 0 826.02466
293.1591796875 0 623.0912
296.15106201171875 0 639.8664
299.0616455078125 0 1069.617
302.1130676269531 0 636.59674
319.140869140625 0 851.48413
324.1459045410156 0 2079.6997
325.1456298828125 0 1005.78455
329.14996337890625 0 4206.3657
330.15216064453125 0 1004.5062
340.197265625 0 648.212
342.1565856933594 0 8425.569 y 7
343.1593933105469 0 1905.2319
355.0707702636719 0 1498.6416
356.07098388671875 0 782.52966
357.22369384765625 0 1042.3073
364.15020751953125 0 2217.7222 b 2
376.11358642578125 0 768.38605
378.1300048828125 0 1583.3148
380.19482421875 0 723.3954
394.1252136230469 0 1006.3897
396.19036865234375 0 673.63696
401.2080078125 0 946.1309
413.2671203613281 0 2195.047
425.1918640136719 0 1029.6414
425.2582092285156 0 747.1185
449.5809631347656 0 524.1191
465.162841796875 0 1139.3629
479.1777648925781 0 980.78033 b 3
480.2840576171875 0 641.63873
483.2200012207031 0 704.55493
489.1925354003906 0 2556.41 y 6
490.1929016113281 0 782.1914
496.22857666015625 0 1268.6125
529.7430419921875 0 623.80316
550.7103271484375 0 1769.2393 y 1
560.2293701171875 0 2510.6184 y 5
561.22998046875 0 1051.3297
564.204833984375 0 618.4675
569.6256713867188 0 555.0416
585.3450927734375 0 4675.477
591.4869384765625 0 725.9817
592.927978515625 0 568.5294
593.36376953125 0 625.1974
601.0025024414062 0 593.4377
601.2405395507812 0 981.38007 Precursor
601.7368774414062 0 1173.5146
604.3676147460938 0 667.8435
621.360107421875 0 1125.3536
657.2658081054688 0 817.29535
659.298828125 0 1310.6624
660.2962646484375 0 921.267
685.2584838867188 0 948.8523
700.3642578125 0 3522.4226
701.3651123046875 0 1345.3848
721.2609252929688 0 1616.6237
723.29248046875 0 3326.2153 y 4
724.298095703125 0 932.9468
749.2540893554688 0 1580.7046
750.2587280273438 0 679.90106
774.321044921875 0 1280.4668
783.4002685546875 0 1124.9961
801.4136962890625 0 1549.7351
828.42578125 0 784.25165
838.3189697265625 0 3365.8386 y 3
839.3231201171875 0 1038.3308
844.4537353515625 0 2753.1099
845.4584350585938 0 795.4098
872.4485473632812 0 11713.479
873.4517211914062 0 6207.2876
874.4534301757812 0 990.08484
889.34716796875 0 4064.0688
890.3529052734375 0 3169.9988
909.6851196289062 0 672.2612
935.3424072265625 0 615.6618 y Water loss 2
936.3345947265625 0 650.0558
953.347412109375 0 15289.762 y 2
954.3499145507812 0 8103.9907
955.3485107421875 0 2196.2344
991.5194702148438 0 2417.9707
992.525390625 0 1390.9335
1019.517822265625 0 4859.7124
1020.5233764648438 0 2883.1978
1021.5208740234375 0 1339.9154
1308.171630859375 0 616.13116
1999.4052734375 0 660.9736

Spectrum Details

|  |  |
| --- | --- |
| Matched peaks? Matched peaks The total absolute number of peaks matched. Additionally in brackets the total fraction of peaks matched and the total number of peaks is shown. | 16 (9.64% of 166) |
| FDR? FDR The false discovery rate estimated for this peptide. It is calculated by matching all theoretical fragments with a non-integer shift with the raw peaks for this spectrum. This is done with 40 different shifts. The resulting percentage is the average number of annotated peaks over the number of annotated peaks with the correct spectrum. | 0.60% |
| Satellite FDR? Satellite FDR See the FDR for details on its calculation. This satellite ion specific FDR only contains the satellite ions (d/w) for I/L/J positions. | - |
| PSM Score? PSM Score The PSM Score as given by Hecklib to this annotated spectrum. It is shown with three significant figures. | 160 |

#### Spectrum 5874? Spectrum 5874 The raw spectrum of this peptide as annotated by Hecklib. The fragments are coloured according to ion type (see legend). Any peaks with a star '\*' as text can be hovered over to see the full details, first the ion type second the mass shift type. By hovering over the amino acids in the peptide or ions in the legend the corresponding peaks are highlighted. By toggling the 'Unassigned' label you can turn the background (unassigned) peaks on or off in the plot. By updating the slider in the Ion legend you can update the spectrum to only show the top X% of the peaks with labels. The top X% means any peak that is within X% of the highest intensity. By dragging in the spectrum you can zoom in to a specific part of the spectrum and use 'Zoom Out' to get back to the original zoom level. The annotation of the spectrum is based on the given sequence in the peptides file and is done with different software so inconsistencies are likely. The peaks are annotated based on the given sequence, with 20 ppm tolerance.

Copy Data

##### Spectrum 5874 (TSV)

###### Preview

```
Loading example...
```

*Click on the button to copy the data to your clipboard.*

Mz MinMz MaxIntensity Max

WidthHeightPeptide font sizePeptide stroke widthSpectrum font sizeSpectrum stroke widthCompact peptide

Ion legend

wxyz

abcd

OtherUnassignedIon^Charge^_Position_Show for top:%

TFDDYAMHW

04.25e+48.50e+41.27e+51.70e+5

Zoom Out

y^+1^_1_c^+2^_4_c^+1^_2_z^+1^_2_y^+1^_2_c^+1^_3_c^+1^_4_y^+1^_3_y^+2^_8_y^+2^_8_z^+1^_4_y^+1^_4_c^+1^_5_z^+1^_5_y^+1^_5_z^+1^_6_y^+1^_6_c^+1^_7_c^+1^_7_y^+1^_7_z^+1^_7_y^+1^_7_c^+1^_8_c^+1^_8_y^+1^_8_

0770154123113082

Fragment Matches Table

Show background peaks

| Position | Ion type | Intensity | mz Theoretical | mz Error (Th) | mz Error (ppm) | Charge | Series Number |
| --- | --- | --- | --- | --- | --- | --- | --- |
| - | - | 1898 | 120.1 | - | - | 0 | - |
| - | - | 357.6 | 123.3 | - | - | 0 | - |
| - | - | 609.7 | 132.1 | - | - | 0 | - |
| - | - | 636.1 | 136.1 | - | - | 0 | - |
| - | - | 423.4 | 139.7 | - | - | 0 | - |
| - | - | 1689 | 149 | - | - | 0 | - |
| - | - | 417.4 | 153.5 | - | - | 0 | - |
| - | - | 408.9 | 164.8 | - | - | 0 | - |
| - | - | 419.1 | 165.3 | - | - | 0 | - |
| - | - | 771 | 167.1 | - | - | 0 | - |
| - | - | 443.7 | 170.5 | - | - | 0 | - |
| - | - | 636 | 175.1 | - | - | 0 | - |
| - | - | 485.9 | 201.9 | - | - | 0 | - |
| - | - | 2052 | 203.1 | - | - | 0 | - |
| 9 | y | 5007 | 221.1 | 0.002834 | 12.82 | +1 | 1 |
| - | - | 5.058E+04 | 221.1 | - | - | 0 | - |
| - | - | 1367 | 222.1 | - | - | 0 | - |
| - | - | 6563 | 222.1 | - | - | 0 | - |
| - | - | 640.9 | 223.1 | - | - | 0 | - |
| - | - | 2527 | 225 | - | - | 0 | - |
| - | - | 666.4 | 227 | - | - | 0 | - |
| - | - | 514.4 | 235.1 | - | - | 0 | - |
| - | - | 8281 | 239.1 | - | - | 0 | - |
| 4 | c | 1716 | 240.1 | 0.00388 | 16.16 | +2 | 4 |
| - | - | 555.2 | 241.1 | - | - | 0 | - |
| - | - | 504.5 | 241.3 | - | - | 0 | - |
| - | - | 798.9 | 246.1 | - | - | 0 | - |
| 2 | c | 1.94E+04 | 249.1 | 0.0001358 | 0.5451 | +1 | 2 |
| - | - | 1780 | 250.1 | - | - | 0 | - |
| - | - | 1216 | 263.1 | - | - | 0 | - |
| - | - | 1536 | 274.1 | - | - | 0 | - |
| - | - | 1507 | 281.1 | - | - | 0 | - |
| - | - | 503.3 | 285.1 | - | - | 0 | - |
| - | - | 2495 | 295.1 | - | - | 0 | - |
| - | - | 1862 | 296.1 | - | - | 0 | - |
| - | - | 813.5 | 297.1 | - | - | 0 | - |
| - | - | 5973 | 299.1 | - | - | 0 | - |
| - | - | 1044 | 300.1 | - | - | 0 | - |
| - | - | 691.4 | 301.1 | - | - | 0 | - |
| - | - | 634.4 | 313.1 | - | - | 0 | - |
| 8 | z | 1048 | 342.1 | 0.004555 | 13.31 | +1 | 2 |
| - | - | 1573 | 343.1 | - | - | 0 | - |
| - | - | 3.752E+04 | 355.1 | - | - | 0 | - |
| 8 | y | 6315 | 358.1 | 0.004752 | 13.27 | +1 | 2 |
| - | - | 1027 | 359.2 | - | - | 0 | - |
| 3 | c | 5389 | 364.2 | 0.0005011 | 1.376 | +1 | 3 |
| - | - | 989.8 | 365.2 | - | - | 0 | - |
| - | - | 1056 | 369.1 | - | - | 0 | - |
| - | - | 1077 | 370.1 | - | - | 0 | - |
| - | - | 1437 | 371.1 | - | - | 0 | - |
| - | - | 861.5 | 371.1 | - | - | 0 | - |
| - | - | 698.7 | 373.1 | - | - | 0 | - |
| - | - | 1931 | 378.1 | - | - | 0 | - |
| - | - | 2220 | 415 | - | - | 0 | - |
| - | - | 3014 | 415.2 | - | - | 0 | - |
| - | - | 663.6 | 455.1 | - | - | 0 | - |
| - | - | 940.2 | 458.2 | - | - | 0 | - |
| - | - | 775.6 | 461.2 | - | - | 0 | - |
| - | - | 560.5 | 470.9 | - | - | 0 | - |
| 4 | c | 3813 | 479.2 | 0.0009677 | 2.02 | +1 | 4 |
| - | - | 2044 | 489.1 | - | - | 0 | - |
| - | - | 1924 | 489.2 | - | - | 0 | - |
| - | - | 7402 | 490.2 | - | - | 0 | - |
| - | - | 1923 | 491.2 | - | - | 0 | - |
| - | - | 3301 | 504.2 | - | - | 0 | - |
| 7 | y | 5331 | 505.2 | 0.009027 | 17.87 | +1 | 3 |
| - | - | 1268 | 506.2 | - | - | 0 | - |
| - | - | 1198 | 512.2 | - | - | 0 | - |
| - | - | 718.9 | 530.3 | - | - | 0 | - |
| - | - | 742.8 | 547.2 | - | - | 0 | - |
| 2 | y | 976 | 549.7 | 0.004634 | 8.431 | +2 | 8 |
| 2 | y | 4047 | 558.7 | 0.004845 | 8.672 | +2 | 8 |
| - | - | 1800 | 559.2 | - | - | 0 | - |
| - | - | 883.9 | 559.7 | - | - | 0 | - |
| 6 | z | 1.806E+04 | 560.2 | 0.009955 | 17.77 | +1 | 4 |
| - | - | 9241 | 561.2 | - | - | 0 | - |
| - | - | 2288 | 562.2 | - | - | 0 | - |
| - | - | 543.9 | 569.2 | - | - | 0 | - |
| - | - | 1047 | 575.2 | - | - | 0 | - |
| 6 | y | 9127 | 576.2 | 0.009968 | 17.3 | +1 | 4 |
| - | - | 1205 | 577.1 | - | - | 0 | - |
| - | - | 3266 | 577.2 | - | - | 0 | - |
| - | - | 4601 | 578.2 | - | - | 0 | - |
| - | - | 1485 | 579.2 | - | - | 0 | - |
| - | - | 1152 | 588.2 | - | - | 0 | - |
| - | - | 689.7 | 591.2 | - | - | 0 | - |
| - | - | 2297 | 600.2 | - | - | 0 | - |
| - | - | 1775 | 600.7 | - | - | 0 | - |
| - | - | 887.5 | 603.2 | - | - | 0 | - |
| - | - | 1294 | 608.2 | - | - | 0 | - |
| - | - | 6293 | 609.2 | - | - | 0 | - |
| - | - | 954.4 | 609.3 | - | - | 0 | - |
| - | - | 6911 | 609.7 | - | - | 0 | - |
| - | - | 1700 | 609.8 | - | - | 0 | - |
| - | - | 7143 | 610.2 | - | - | 0 | - |
| - | - | 1782 | 610.2 | - | - | 0 | - |
| - | - | 2177 | 610.3 | - | - | 0 | - |
| - | - | 1148 | 610.3 | - | - | 0 | - |
| - | - | 1287 | 619.2 | - | - | 0 | - |
| - | - | 604.7 | 634.2 | - | - | 0 | - |
| 5 | c | 835.1 | 642.2 | 0.00374 | 5.824 | +1 | 5 |
| - | - | 7495 | 649.3 | - | - | 0 | - |
| - | - | 2692 | 650.3 | - | - | 0 | - |
| - | - | 1124 | 662.3 | - | - | 0 | - |
| - | - | 3648 | 668.3 | - | - | 0 | - |
| - | - | 1998 | 669.4 | - | - | 0 | - |
| - | - | 743.8 | 669.4 | - | - | 0 | - |
| - | - | 882.5 | 670.4 | - | - | 0 | - |
| - | - | 650.8 | 675.3 | - | - | 0 | - |
| - | - | 582.8 | 676.3 | - | - | 0 | - |
| - | - | 653.8 | 677.4 | - | - | 0 | - |
| - | - | 848.8 | 690.3 | - | - | 0 | - |
| - | - | 8379 | 693.2 | - | - | 0 | - |
| - | - | 3086 | 694.2 | - | - | 0 | - |
| - | - | 724.5 | 695.2 | - | - | 0 | - |
| - | - | 776.8 | 701.4 | - | - | 0 | - |
| 5 | z | 2.168E+04 | 723.3 | 0.009676 | 13.38 | +1 | 5 |
| - | - | 1.025E+04 | 724.3 | - | - | 0 | - |
| - | - | 3700 | 725.3 | - | - | 0 | - |
| - | - | 1477 | 731.3 | - | - | 0 | - |
| - | - | 806.4 | 731.4 | - | - | 0 | - |
| - | - | 786 | 732.3 | - | - | 0 | - |
| - | - | 767.8 | 732.4 | - | - | 0 | - |
| - | - | 920.2 | 734.2 | - | - | 0 | - |
| - | - | 2696 | 738.3 | - | - | 0 | - |
| 5 | y | 1.061E+04 | 739.3 | 0.008469 | 11.46 | +1 | 5 |
| - | - | 3118 | 740.3 | - | - | 0 | - |
| - | - | 968 | 741.3 | - | - | 0 | - |
| - | - | 1750 | 749.3 | - | - | 0 | - |
| - | - | 5001 | 764.3 | - | - | 0 | - |
| - | - | 1439 | 765.3 | - | - | 0 | - |
| - | - | 1475 | 766.3 | - | - | 0 | - |
| - | - | 1664 | 775.3 | - | - | 0 | - |
| - | - | 982.7 | 776.3 | - | - | 0 | - |
| - | - | 879.5 | 777.3 | - | - | 0 | - |
| - | - | 801.3 | 778.3 | - | - | 0 | - |
| - | - | 1014 | 783.4 | - | - | 0 | - |
| - | - | 604.2 | 790.3 | - | - | 0 | - |
| - | - | 1.169E+04 | 794.3 | - | - | 0 | - |
| - | - | 4675 | 795.3 | - | - | 0 | - |
| - | - | 1956 | 796.3 | - | - | 0 | - |
| - | - | 1358 | 798.4 | - | - | 0 | - |
| - | - | 1916 | 808.3 | - | - | 0 | - |
| - | - | 1413 | 809.3 | - | - | 0 | - |
| - | - | 788 | 813.4 | - | - | 0 | - |
| - | - | 630.1 | 823.3 | - | - | 0 | - |
| - | - | 635.6 | 830.5 | - | - | 0 | - |
| - | - | 2253 | 833.3 | - | - | 0 | - |
| - | - | 5022 | 834.3 | - | - | 0 | - |
| - | - | 1691 | 835.3 | - | - | 0 | - |
| - | - | 844.5 | 835.4 | - | - | 0 | - |
| 4 | z | 2.417E+04 | 838.3 | 0.00971 | 11.58 | +1 | 6 |
| - | - | 3.063E+04 | 839.3 | - | - | 0 | - |
| - | - | 1.154E+04 | 840.3 | - | - | 0 | - |
| - | - | 3471 | 841.3 | - | - | 0 | - |
| - | - | 7036 | 845.5 | - | - | 0 | - |
| - | - | 1975 | 846.5 | - | - | 0 | - |
| - | - | 1377 | 853.3 | - | - | 0 | - |
| 4 | y | 1.006E+04 | 854.3 | 0.009846 | 11.52 | +1 | 6 |
| - | - | 6280 | 855.3 | - | - | 0 | - |
| - | - | 1785 | 856.3 | - | - | 0 | - |
| - | - | 1072 | 858.4 | - | - | 0 | - |
| 7 | c | 685.9 | 860.3 | 0.002243 | 2.607 | +1 | 7 |
| - | - | 1865 | 865.3 | - | - | 0 | - |
| - | - | 715 | 866.3 | - | - | 0 | - |
| - | - | 810.2 | 871.5 | - | - | 0 | - |
| - | - | 1537 | 876.3 | - | - | 0 | - |
| 7 | c | 1.605E+04 | 877.3 | 0.004258 | 4.854 | +1 | 7 |
| - | - | 8141 | 878.3 | - | - | 0 | - |
| - | - | 2192 | 879.3 | - | - | 0 | - |
| - | - | 934.9 | 879.4 | - | - | 0 | - |
| - | - | 918.6 | 881.3 | - | - | 0 | - |
| - | - | 941.5 | 890.3 | - | - | 0 | - |
| - | - | 6809 | 905.3 | - | - | 0 | - |
| - | - | 3046 | 906.3 | - | - | 0 | - |
| - | - | 967.9 | 907.3 | - | - | 0 | - |
| - | - | 667.3 | 907.5 | - | - | 0 | - |
| - | - | 8383 | 909.3 | - | - | 0 | - |
| - | - | 659.5 | 909.4 | - | - | 0 | - |
| - | - | 4150 | 910.3 | - | - | 0 | - |
| - | - | 836.8 | 911.3 | - | - | 0 | - |
| - | - | 1004 | 915.5 | - | - | 0 | - |
| - | - | 1178 | 916 | - | - | 0 | - |
| - | - | 1099 | 927.4 | - | - | 0 | - |
| - | - | 792.5 | 928.4 | - | - | 0 | - |
| - | - | 1573 | 943.4 | - | - | 0 | - |
| - | - | 623.5 | 944.5 | - | - | 0 | - |
| 3 | y | 2978 | 951.3 | 0.01037 | 10.91 | +1 | 7 |
| - | - | 1736 | 952.3 | - | - | 0 | - |
| 3 | z | 3117 | 953.3 | 0.01127 | 11.82 | +1 | 7 |
| - | - | 3003 | 954.3 | - | - | 0 | - |
| - | - | 897.7 | 955.3 | - | - | 0 | - |
| 3 | y | 4.274E+04 | 969.3 | 0.009331 | 9.626 | +1 | 7 |
| - | - | 2.216E+04 | 970.3 | - | - | 0 | - |
| - | - | 6193 | 971.3 | - | - | 0 | - |
| - | - | 1047 | 972.3 | - | - | 0 | - |
| - | - | 1054 | 981.4 | - | - | 0 | - |
| - | - | 2665 | 982.4 | - | - | 0 | - |
| - | - | 858.7 | 983.4 | - | - | 0 | - |
| 8 | c | 3294 | 997.4 | 0.003085 | 3.093 | +1 | 8 |
| - | - | 1563 | 998.4 | - | - | 0 | - |
| - | - | 4793 | 999.4 | - | - | 0 | - |
| - | - | 2513 | 1000 | - | - | 0 | - |
| - | - | 797.4 | 1001 | - | - | 0 | - |
| - | - | 1606 | 1011 | - | - | 0 | - |
| 8 | c | 5.186E+04 | 1014 | 0.00394 | 3.884 | +1 | 8 |
| - | - | 2.767E+04 | 1015 | - | - | 0 | - |
| - | - | 9304 | 1016 | - | - | 0 | - |
| - | - | 1020 | 1017 | - | - | 0 | - |
| - | - | 1373 | 1034 | - | - | 0 | - |
| - | - | 841.6 | 1039 | - | - | 0 | - |
| - | - | 9671 | 1040 | - | - | 0 | - |
| - | - | 6061 | 1041 | - | - | 0 | - |
| - | - | 2523 | 1042 | - | - | 0 | - |
| - | - | 3456 | 1055 | - | - | 0 | - |
| - | - | 9533 | 1056 | - | - | 0 | - |
| - | - | 3561 | 1057 | - | - | 0 | - |
| - | - | 1455 | 1058 | - | - | 0 | - |
| - | - | 742.8 | 1059 | - | - | 0 | - |
| - | - | 3457 | 1069 | - | - | 0 | - |
| - | - | 2080 | 1070 | - | - | 0 | - |
| - | - | 5283 | 1071 | - | - | 0 | - |
| - | - | 1788 | 1072 | - | - | 0 | - |
| - | - | 1646 | 1073 | - | - | 0 | - |
| - | - | 1122 | 1079 | - | - | 0 | - |
| - | - | 800.1 | 1084 | - | - | 0 | - |
| - | - | 1023 | 1101 | - | - | 0 | - |
| 2 | y | 2046 | 1116 | 0.01001 | 8.965 | +1 | 8 |
| - | - | 1109 | 1117 | - | - | 0 | - |
| - | - | 1510 | 1138 | - | - | 0 | - |
| - | - | 2507 | 1139 | - | - | 0 | - |
| - | - | 1805 | 1140 | - | - | 0 | - |
| - | - | 1727 | 1140 | - | - | 0 | - |
| - | - | 1743 | 1154 | - | - | 0 | - |
| - | - | 1302 | 1155 | - | - | 0 | - |
| - | - | 2.316E+04 | 1156 | - | - | 0 | - |
| - | - | 1.63E+04 | 1157 | - | - | 0 | - |
| - | - | 2.307E+04 | 1158 | - | - | 0 | - |
| - | - | 1.213E+04 | 1159 | - | - | 0 | - |
| - | - | 938.2 | 1160 | - | - | 0 | - |
| - | - | 4616 | 1160 | - | - | 0 | - |
| - | - | 1655 | 1174 | - | - | 0 | - |
| - | - | 1510 | 1175 | - | - | 0 | - |
| - | - | 906.4 | 1181 | - | - | 0 | - |
| - | - | 857.8 | 1182 | - | - | 0 | - |
| - | - | 1533 | 1183 | - | - | 0 | - |
| - | - | 934.6 | 1184 | - | - | 0 | - |
| - | - | 1824 | 1190 | - | - | 0 | - |
| - | - | 1341 | 1199 | - | - | 0 | - |
| - | - | 1775 | 1200 | - | - | 0 | - |
| - | - | 1.106E+04 | 1200 | - | - | 0 | - |
| - | - | 7.838E+04 | 1201 | - | - | 0 | - |
| - | - | 5.47E+04 | 1202 | - | - | 0 | - |
| - | - | 2.506E+04 | 1203 | - | - | 0 | - |
| - | - | 1115 | 1204 | - | - | 0 | - |
| - | - | 3971 | 1204 | - | - | 0 | - |
| - | - | 3864 | 1217 | - | - | 0 | - |
| - | - | 4.783E+04 | 1217 | - | - | 0 | - |
| - | - | 1.683E+05 | 1218 | - | - | 0 | - |
| - | - | 1.089E+05 | 1219 | - | - | 0 | - |
| - | - | 6867 | 1220 | - | - | 0 | - |
| - | - | 4.475E+04 | 1220 | - | - | 0 | - |
| - | - | 4204 | 1221 | - | - | 0 | - |
| - | - | 2555 | 1221 | - | - | 0 | - |
| - | - | 5537 | 1221 | - | - | 0 | - |
| - | - | 982 | 1319 | - | - | 0 | - |
| - | - | 1085 | 1402 | - | - | 0 | - |
| - | - | 922.5 | 1521 | - | - | 0 | - |
| - | - | 1245 | 1530 | - | - | 0 | - |
| - | - | 1015 | 1714 | - | - | 0 | - |
| - | - | 824 | 1716 | - | - | 0 | - |
| - | - | 1035 | 1717 | - | - | 0 | - |
| - | - | 1058 | 1787 | - | - | 0 | - |
| - | - | 1089 | 1814 | - | - | 0 | - |
| - | - | 2956 | 1815 | - | - | 0 | - |
| - | - | 2241 | 1827 | - | - | 0 | - |
| - | - | 2078 | 1828 | - | - | 0 | - |
| - | - | 2723 | 1829 | - | - | 0 | - |
| - | - | 1184 | 1830 | - | - | 0 | - |
| - | - | 1997 | 1831 | - | - | 0 | - |
| - | - | 6036 | 1832 | - | - | 0 | - |
| - | - | 862.4 | 2438 | - | - | 0 | - |
| - | - | 728.1 | 3051 | - | - | 0 | - |

m/z Charge Intensity FragmentType MassShift Position
120.08094024658203 0 1898.1055
123.3228988647461 0 357.61163
132.1020965576172 0 609.74896
136.07615661621094 0 636.05963
139.68692016601562 0 423.36893
149.04507446289062 0 1689.3739
153.5248260498047 0 417.4344
164.8037872314453 0 408.92163
165.25682067871094 0 419.05475
167.05596923828125 0 771.03876
170.5362091064453 0 443.67316
175.08665466308594 0 636.03644
201.93124389648438 0 485.90787
203.08168029785156 0 2051.871
221.08432006835938 0 5007.481 y 8
221.1284942626953 0 50581.332
222.085205078125 0 1366.9734
222.13186645507812 0 6562.883
223.0799102783203 0 640.8508
225.04306030273438 0 2526.8987
227.04010009765625 0 666.4157
235.10885620117188 0 514.4239
239.0950927734375 0 8280.633
240.0961456298828 0 1715.559 c Ammonia loss 3
241.0919952392578 0 555.1968
241.25941467285156 0 504.53098
246.12408447265625 0 798.91223
249.12350463867188 0 19395.508 c Ammonia loss 1
250.1271514892578 0 1779.7137
263.1025695800781 0 1216.1108
274.11895751953125 0 1535.7542
281.05059814453125 0 1506.8301
285.1011962890625 0 503.26007
295.10345458984375 0 2495.4866
296.1042785644531 0 1861.9366
297.1023864746094 0 813.47723
299.06182861328125 0 5972.6836
300.0628967285156 0 1044.0394
301.05718994140625 0 691.44794
313.11614990234375 0 634.3697
342.13189697265625 0 1047.6134 z 7
343.1393127441406 0 1572.6536
355.0697937011719 0 37519.85
358.15081787109375 0 6314.824 y 7
359.15374755859375 0 1027.1931
364.1498107910156 0 5388.953 c Ammonia loss 2
365.1542053222656 0 989.75366
369.1221618652344 0 1056.0522
370.1217041015625 0 1076.8767
371.1002197265625 0 1436.5682
371.1222229003906 0 861.492
373.07989501953125 0 698.66846
378.1287841796875 0 1931.0945
415.0373840332031 0 2219.7366
415.15313720703125 0 3014.4468
455.1393127441406 0 663.5885
458.1565246582031 0 940.2233
461.1654052734375 0 775.55994
470.9030456542969 0 560.52576
479.17822265625 0 3813.4998 c Ammonia loss 3
489.0560302734375 0 2044.1897
489.1680603027344 0 1924.0742
490.1758728027344 0 7401.654
491.1778564453125 0 1923.0735
504.17919921875 0 3301.0334
505.1855773925781 0 5330.673 y 6
506.18768310546875 0 1267.9102
512.224609375 0 1198.0562
530.3172607421875 0 718.86676
547.2338256835938 0 742.7769
549.70263671875 0 975.97186 y Water loss 1
558.7081298828125 0 4047.2493 y 1
559.209716796875 0 1800.4675
559.7130126953125 0 883.87665
560.2048950195312 0 18058.53 z 5
561.2096557617188 0 9240.553
562.2111206054688 0 2288.2727
569.217529296875 0 543.9396
575.2169799804688 0 1047.1411
576.2236328125 0 9127.4375 y 5
577.1280517578125 0 1205.188
577.2274169921875 0 3266.0654
578.2169189453125 0 4601.3
579.218994140625 0 1485.3961
588.243896484375 0 1152.2808
591.223388671875 0 689.6708
600.2276611328125 0 2296.6633
600.7285766601562 0 1774.8391
603.2109375 0 887.49976
608.2484741210938 0 1294.1658
609.2321166992188 0 6293.336
609.3314208984375 0 954.3718
609.73291015625 0 6910.9824
609.8473510742188 0 1700.4224
610.1835327148438 0 7143.1836
610.2335815429688 0 1781.5424
610.2879028320312 0 2176.8813
610.3451538085938 0 1148.422
619.21923828125 0 1286.6033
634.2247314453125 0 604.7172
642.2443237304688 0 835.1062 c Ammonia loss 4
649.2525024414062 0 7495.3945
650.2551879882812 0 2691.5706
662.2584838867188 0 1124.2966
668.3482666015625 0 3648.4688
669.352783203125 0 1997.8792
669.408447265625 0 743.8421
670.3587646484375 0 882.49005
675.2892456054688 0 650.7991
676.2872924804688 0 582.7608
677.3519287109375 0 653.811
690.25634765625 0 848.77136
693.2421264648438 0 8379.4795
694.2449340820312 0 3085.6086
695.2456665039062 0 724.5493
701.3623046875 0 776.7748
723.2679443359375 0 21679.773 z 4
724.2715454101562 0 10245.0205
725.2725830078125 0 3699.7249
731.3136596679688 0 1476.614
731.4392700195312 0 806.4353
732.3178100585938 0 786.00604
732.4485473632812 0 767.84143
734.245361328125 0 920.1842
738.2782592773438 0 2696.3225
739.2854614257812 0 10608.171 y 4
740.2890014648438 0 3117.7473
741.2884521484375 0 967.97406
749.2598266601562 0 1749.5477
764.2796630859375 0 5000.6807
765.28271484375 0 1438.8612
766.2755126953125 0 1474.6183
775.3067626953125 0 1663.997
776.30517578125 0 982.6999
777.2927856445312 0 879.53253
778.2883911132812 0 801.27997
783.39794921875 0 1014.20435
790.3174438476562 0 604.17456
794.3047485351562 0 11689.119
795.3087768554688 0 4674.9995
796.3102416992188 0 1956.3195
798.4473876953125 0 1358.2701
808.26806640625 0 1915.8704
809.2720947265625 0 1412.5729
813.4107666015625 0 788.017
823.2755737304688 0 630.0829
830.4909057617188 0 635.5805
833.3282470703125 0 2253.307
834.3320922851562 0 5022.4624
835.3367309570312 0 1691.1296
835.4132690429688 0 844.49445
838.294921875 0 24165.305 z 3
839.3007202148438 0 30628.266
840.3037109375 0 11537.477
841.3072509765625 0 3471.2927
845.5086669921875 0 7035.7983
846.5130615234375 0 1975.139
853.311279296875 0 1376.7303
854.3137817382812 0 10059.371 y 3
855.3162841796875 0 6279.657
856.3162231445312 0 1785.2892
858.4379272460938 0 1072.2817
860.3104248046875 0 685.9362 c Ammonia loss 6
865.339111328125 0 1864.5359
866.3357543945312 0 715.0258
871.4794311523438 0 810.21094
876.3322143554688 0 1537.16
877.3389892578125 0 16046.02 c 6
878.3413696289062 0 8140.8354
879.3402709960938 0 2191.9004
879.4305419921875 0 934.86694
881.3134155273438 0 918.6248
890.3302612304688 0 941.542
905.3422241210938 0 6809.1025
906.345947265625 0 3045.7534
907.3451538085938 0 967.8989
907.4599609375 0 667.2587
909.3308715820312 0 8382.813
909.4227905273438 0 659.4854
910.3326416015625 0 4149.744
911.337646484375 0 836.8101
915.4716186523438 0 1004.1463
915.977783203125 0 1177.5734
927.4124755859375 0 1098.7334
928.4293823242188 0 792.527
943.4447021484375 0 1573.29
944.4505004882812 0 623.46655
951.3306884765625 0 2978.0068 y Water loss 2
952.328125 0 1735.6289
953.3234252929688 0 3117.2395 z 2
954.3231201171875 0 3002.9692
955.3389282226562 0 897.70605
969.3402099609375 0 42739.14 y 2
970.3434448242188 0 22162.65
971.3452758789062 0 6193.392
972.343505859375 0 1046.6865
981.36376953125 0 1053.7747
982.3663940429688 0 2665.239
983.3652954101562 0 858.66833
997.3701782226562 0 3294.2917 c Ammonia loss 7
998.3731689453125 0 1563.4933
999.3854370117188 0 4793.1543
1000.3904418945312 0 2513.1108
1001.3980102539062 0 797.3561
1011.4152221679688 0 1605.6401
1014.3975830078125 0 51863.723 c 7
1015.4011840820312 0 27668.459
1016.4013671875 0 9303.74
1017.400634765625 0 1019.9132
1033.509765625 0 1372.5187
1039.4029541015625 0 841.5988
1040.413818359375 0 9671.239
1041.415771484375 0 6060.5854
1042.4183349609375 0 2523.1536
1055.3994140625 0 3456.123
1056.3895263671875 0 9533.418
1057.3883056640625 0 3560.6956
1058.395263671875 0 1454.6836
1058.5179443359375 0 742.7887
1069.3941650390625 0 3456.7837
1070.3935546875 0 2080.369
1071.3948974609375 0 5283.232
1072.397705078125 0 1787.5375
1073.4019775390625 0 1646.2056
1078.5238037109375 0 1122.0826
1084.3994140625 0 800.1053
1101.392578125 0 1023.00793
1116.4093017578125 0 2046.3278 y 1
1117.4180908203125 0 1108.7664
1138.44970703125 0 1509.8965
1139.4512939453125 0 2507.3596
1139.6529541015625 0 1805.4238
1140.4483642578125 0 1727.1451
1154.465576171875 0 1743.1888
1155.468994140625 0 1301.9407
1156.4622802734375 0 23160.191
1157.4632568359375 0 16304.938
1158.4482421875 0 23067.291
1159.4464111328125 0 12128.235
1159.5765380859375 0 938.15137
1160.4461669921875 0 4615.5854
1174.4755859375 0 1655.2717
1175.4783935546875 0 1510.2373
1180.681396484375 0 906.4327
1182.445556640625 0 857.7676
1183.441650390625 0 1533.221
1184.420166015625 0 934.5923
1190.4676513671875 0 1824.439
1198.68994140625 0 1340.7305
1199.5579833984375 0 1774.5898
1200.4530029296875 0 11056.537
1201.442626953125 0 78377.836
1202.4439697265625 0 54700.42
1203.44482421875 0 25056.828
1203.6041259765625 0 1114.5264
1204.4453125 0 3970.661
1216.572265625 0 3864.032
1217.454833984375 0 47833.79
1218.46142578125 0 168275.56
1219.46435546875 0 108910.305
1219.7020263671875 0 6867.133
1220.4666748046875 0 44752.543
1220.7000732421875 0 4203.501
1221.199462890625 0 2554.9758
1221.467529296875 0 5537.24
1318.67041015625 0 982.016
1401.7093505859375 0 1084.9713
1521.336181640625 0 922.48047
1529.77294921875 0 1245.1552
1713.8785400390625 0 1014.99475
1715.84814453125 0 823.9993
1716.8919677734375 0 1035.0424
1786.930908203125 0 1058.1315
1813.914794921875 0 1088.9175
1814.9293212890625 0 2956.273
1826.799560546875 0 2240.973
1827.7950439453125 0 2077.6929
1828.792724609375 0 2722.5183
1829.787841796875 0 1184.1483
1830.9364013671875 0 1997.2202
1831.9500732421875 0 6035.5117
2438.307373046875 0 862.4217
3051.4248046875 0 728.0898

Spectrum Details

|  |  |
| --- | --- |
| Matched peaks? Matched peaks The total absolute number of peaks matched. Additionally in brackets the total fraction of peaks matched and the total number of peaks is shown. | 25 (8.83% of 283) |
| FDR? FDR The false discovery rate estimated for this peptide. It is calculated by matching all theoretical fragments with a non-integer shift with the raw peaks for this spectrum. This is done with 40 different shifts. The resulting percentage is the average number of annotated peaks over the number of annotated peaks with the correct spectrum. | 0.67% |
| Satellite FDR? Satellite FDR See the FDR for details on its calculation. This satellite ion specific FDR only contains the satellite ions (d/w) for I/L/J positions. | - |
| PSM Score? PSM Score The PSM Score as given by Hecklib to this annotated spectrum. It is shown with three significant figures. | 321 |

#### Spectrum 12469? Spectrum 12469 The raw spectrum of this peptide as annotated by Hecklib. The fragments are coloured according to ion type (see legend). Any peaks with a star '\*' as text can be hovered over to see the full details, first the ion type second the mass shift type. By hovering over the amino acids in the peptide or ions in the legend the corresponding peaks are highlighted. By toggling the 'Unassigned' label you can turn the background (unassigned) peaks on or off in the plot. By updating the slider in the Ion legend you can update the spectrum to only show the top X% of the peaks with labels. The top X% means any peak that is within X% of the highest intensity. By dragging in the spectrum you can zoom in to a specific part of the spectrum and use 'Zoom Out' to get back to the original zoom level. The annotation of the spectrum is based on the given sequence in the peptides file and is done with different software so inconsistencies are likely. The peaks are annotated based on the given sequence, with 20 ppm tolerance.

Copy Data

##### Spectrum 12469 (TSV)

###### Preview

```
Loading example...
```

*Click on the button to copy the data to your clipboard.*

Mz MinMz MaxIntensity Max

WidthHeightPeptide font sizePeptide stroke widthSpectrum font sizeSpectrum stroke widthCompact peptide

Ion legend

wxyz

abcd

OtherUnassignedIon^Charge^_Position_Show for top:%

TFDDYAMHW

01.00e+42.01e+43.01e+44.02e+4

Zoom Out

a^+1^_2_y^+1^_1_a^+1^_2_b^+1^_2_b^+1^_2_y^+1^_2_b^+1^_3_b^+1^_4_y^+1^_3_y^+2^_8_y^+1^_4_\*\*y^+1^_5_y^+1^_6_y^+1^_7_

0728145721852913

Fragment Matches Table

Show background peaks

| Position | Ion type | Intensity | mz Theoretical | mz Error (Th) | mz Error (ppm) | Charge | Series Number |
| --- | --- | --- | --- | --- | --- | --- | --- |
| - | - | 2.719E+04 | 120.1 | - | - | 0 | - |
| - | - | 315.1 | 120.8 | - | - | 0 | - |
| - | - | 308.7 | 121.1 | - | - | 0 | - |
| - | - | 2212 | 121.1 | - | - | 0 | - |
| - | - | 336.8 | 122.4 | - | - | 0 | - |
| - | - | 472 | 123 | - | - | 0 | - |
| - | - | 3246 | 129.1 | - | - | 0 | - |
| - | - | 717.2 | 130.1 | - | - | 0 | - |
| - | - | 417.4 | 130.1 | - | - | 0 | - |
| - | - | 731.9 | 132.1 | - | - | 0 | - |
| - | - | 1067 | 133.1 | - | - | 0 | - |
| - | - | 622.3 | 136.1 | - | - | 0 | - |
| - | - | 1.014E+04 | 136.1 | - | - | 0 | - |
| - | - | 392.6 | 136.4 | - | - | 0 | - |
| - | - | 994.4 | 137.1 | - | - | 0 | - |
| - | - | 527.8 | 146.1 | - | - | 0 | - |
| - | - | 1580 | 149 | - | - | 0 | - |
| - | - | 436.8 | 151.5 | - | - | 0 | - |
| - | - | 714.3 | 156.1 | - | - | 0 | - |
| - | - | 914 | 158.1 | - | - | 0 | - |
| - | - | 628.1 | 159.1 | - | - | 0 | - |
| - | - | 2825 | 165.1 | - | - | 0 | - |
| - | - | 637.7 | 166.1 | - | - | 0 | - |
| - | - | 2050 | 167.1 | - | - | 0 | - |
| - | - | 434.6 | 173.1 | - | - | 0 | - |
| - | - | 2341 | 173.1 | - | - | 0 | - |
| - | - | 1133 | 173.4 | - | - | 0 | - |
| - | - | 2568 | 176.1 | - | - | 0 | - |
| - | - | 1755 | 177.1 | - | - | 0 | - |
| - | - | 1046 | 177.1 | - | - | 0 | - |
| - | - | 6085 | 182.1 | - | - | 0 | - |
| - | - | 568.6 | 183.1 | - | - | 0 | - |
| - | - | 549.3 | 185.1 | - | - | 0 | - |
| - | - | 1392 | 188.1 | - | - | 0 | - |
| - | - | 1217 | 193.1 | - | - | 0 | - |
| - | - | 1047 | 201.1 | - | - | 0 | - |
| 2 | a | 1058 | 203.1 | 9.142E-05 | 0.4501 | +1 | 2 |
| 9 | y | 810.1 | 205.1 | 0.0001387 | 0.6763 | +1 | 1 |
| - | - | 605.1 | 209.1 | - | - | 0 | - |
| - | - | 532.6 | 217.1 | - | - | 0 | - |
| - | - | 2034 | 221.1 | - | - | 0 | - |
| 2 | a | 3.979E+04 | 221.1 | 6.677E-05 | 0.302 | +1 | 2 |
| - | - | 4691 | 222.1 | - | - | 0 | - |
| - | - | 984.2 | 223.1 | - | - | 0 | - |
| - | - | 517.2 | 224.1 | - | - | 0 | - |
| - | - | 868.9 | 227.1 | - | - | 0 | - |
| - | - | 661.3 | 231.1 | - | - | 0 | - |
| 2 | b | 610.4 | 231.1 | 0.001667 | 7.214 | +1 | 2 |
| - | - | 1105 | 235.1 | - | - | 0 | - |
| - | - | 1348 | 239.1 | - | - | 0 | - |
| - | - | 504.9 | 239.1 | - | - | 0 | - |
| - | - | 542.8 | 245.1 | - | - | 0 | - |
| - | - | 475.4 | 247.4 | - | - | 0 | - |
| 2 | b | 6704 | 249.1 | 1.679E-05 | 0.06741 | +1 | 2 |
| - | - | 488.4 | 249.6 | - | - | 0 | - |
| - | - | 776.1 | 251.1 | - | - | 0 | - |
| - | - | 668.8 | 253.2 | - | - | 0 | - |
| - | - | 1935 | 263.1 | - | - | 0 | - |
| - | - | 488.3 | 264.7 | - | - | 0 | - |
| - | - | 511.2 | 272.9 | - | - | 0 | - |
| - | - | 842 | 279.1 | - | - | 0 | - |
| - | - | 1379 | 285 | - | - | 0 | - |
| - | - | 911.6 | 285.1 | - | - | 0 | - |
| - | - | 757.3 | 296.2 | - | - | 0 | - |
| - | - | 725.5 | 299.1 | - | - | 0 | - |
| - | - | 800.5 | 324.1 | - | - | 0 | - |
| - | - | 3291 | 329.1 | - | - | 0 | - |
| 8 | y | 6458 | 342.2 | 3.146E-05 | 0.09194 | +1 | 2 |
| - | - | 823.7 | 343.2 | - | - | 0 | - |
| - | - | 1148 | 355.1 | - | - | 0 | - |
| - | - | 684.3 | 356.1 | - | - | 0 | - |
| - | - | 962.3 | 357.2 | - | - | 0 | - |
| 3 | b | 1552 | 364.2 | 0.0001044 | 0.2866 | +1 | 3 |
| - | - | 1435 | 394.1 | - | - | 0 | - |
| - | - | 1567 | 413.3 | - | - | 0 | - |
| - | - | 1150 | 425.2 | - | - | 0 | - |
| - | - | 793.6 | 436.7 | - | - | 0 | - |
| - | - | 647.5 | 441.2 | - | - | 0 | - |
| - | - | 580.7 | 455.2 | - | - | 0 | - |
| - | - | 1205 | 465.2 | - | - | 0 | - |
| 4 | b | 1150 | 479.2 | 0.0005581 | 1.165 | +1 | 4 |
| - | - | 557.4 | 481.6 | - | - | 0 | - |
| 7 | y | 1595 | 489.2 | 0.004978 | 10.18 | +1 | 3 |
| - | - | 634.3 | 513.2 | - | - | 0 | - |
| 2 | y | 1093 | 550.7 | 0.0004556 | 0.8272 | +2 | 8 |
| 6 | y | 2468 | 560.2 | 0.004363 | 7.788 | +1 | 4 |
| - | - | 630.2 | 561.2 | - | - | 0 | - |
| - | - | 780 | 569.2 | - | - | 0 | - |
| - | - | 1397 | 585.3 | - | - | 0 | - |
| 0 | Precursor | 692.8 | 592.2 | 0.003017 | 5.094 | +2 | -1 |
| - | - | 688 | 593.8 | - | - | 0 | - |
| 0 | Precursor | 2467 | 601.2 | 0.0028 | 4.658 | +2 | -1 |
| - | - | 628.4 | 601.3 | - | - | 0 | - |
| - | - | 831.1 | 601.7 | - | - | 0 | - |
| - | - | 815.7 | 602.2 | - | - | 0 | - |
| - | - | 737 | 612.2 | - | - | 0 | - |
| - | - | 666.9 | 659.3 | - | - | 0 | - |
| - | - | 918.7 | 685.3 | - | - | 0 | - |
| - | - | 3711 | 700.4 | - | - | 0 | - |
| - | - | 1331 | 701.4 | - | - | 0 | - |
| - | - | 699.6 | 702.6 | - | - | 0 | - |
| 5 | y | 3014 | 723.3 | 0.004816 | 6.659 | +1 | 5 |
| - | - | 1896 | 749.3 | - | - | 0 | - |
| - | - | 647.2 | 774.3 | - | - | 0 | - |
| - | - | 2006 | 801.4 | - | - | 0 | - |
| - | - | 769.1 | 802.4 | - | - | 0 | - |
| 4 | y | 2631 | 838.3 | 0.006621 | 7.898 | +1 | 6 |
| - | - | 1848 | 839.3 | - | - | 0 | - |
| - | - | 2648 | 844.5 | - | - | 0 | - |
| - | - | 1050 | 845.5 | - | - | 0 | - |
| - | - | 1.091E+04 | 872.4 | - | - | 0 | - |
| - | - | 4386 | 873.5 | - | - | 0 | - |
| - | - | 1497 | 874.5 | - | - | 0 | - |
| - | - | 4162 | 889.3 | - | - | 0 | - |
| - | - | 1927 | 890.4 | - | - | 0 | - |
| 3 | y | 1.159E+04 | 953.3 | 0.005129 | 5.38 | +1 | 7 |
| - | - | 6295 | 954.3 | - | - | 0 | - |
| - | - | 2223 | 955.3 | - | - | 0 | - |
| - | - | 1376 | 991.5 | - | - | 0 | - |
| - | - | 1058 | 992.5 | - | - | 0 | - |
| - | - | 3817 | 1020 | - | - | 0 | - |
| - | - | 3264 | 1021 | - | - | 0 | - |
| - | - | 985.5 | 1022 | - | - | 0 | - |
| - | - | 895.6 | 1101 | - | - | 0 | - |
| - | - | 637.9 | 2689 | - | - | 0 | - |
| - | - | 689.4 | 2884 | - | - | 0 | - |

m/z Charge Intensity FragmentType MassShift Position
120.08080291748047 0 27186.912
120.77432250976562 0 315.12497
121.07766723632812 0 308.6871
121.0842056274414 0 2212.3965
122.40233612060547 0 336.83844
123.04349517822266 0 472.0145
129.10227966308594 0 3246.09
130.0653839111328 0 717.1824
130.0859832763672 0 417.4286
132.08094787597656 0 731.89667
133.0859832763672 0 1066.7511
136.07015991210938 0 622.29114
136.0757293701172 0 10138.028
136.44821166992188 0 392.562
137.07888793945312 0 994.44824
146.0596923828125 0 527.8127
149.04510498046875 0 1579.7363
151.5146026611328 0 436.7834
156.07659912109375 0 714.2977
158.09657287597656 0 914.0158
159.09144592285156 0 628.10187
165.05455017089844 0 2824.598
166.08639526367188 0 637.7155
167.0553436279297 0 2050.4807
173.0919952392578 0 434.5851
173.1282196044922 0 2341.448
173.4387664794922 0 1133.0856
176.10693359375 0 2568.0964
177.1022186279297 0 1754.7992
177.11119079589844 0 1045.9376
182.0811309814453 0 6085.404
183.08465576171875 0 568.5786
185.1287841796875 0 549.3446
188.07054138183594 0 1392.0847
193.10861206054688 0 1217.118
201.12332153320312 0 1046.738
203.11798095703125 0 1057.5217 a Water loss 1
205.09701538085938 0 810.11646 y 8
209.09136962890625 0 605.14
217.09715270996094 0 532.5976
221.1031494140625 0 2033.7811
221.12838745117188 0 39792.844 a 1
222.1318359375 0 4690.864
223.06341552734375 0 984.1969
224.06431579589844 0 517.1923
227.10263061523438 0 868.8701
231.06118774414062 0 661.3308
231.11447143554688 0 610.4034 b Water loss 1
235.10714721679688 0 1105.2745
239.09445190429688 0 1348.1259
239.14901733398438 0 504.88254
245.078857421875 0 542.8383
247.43666076660156 0 475.4286
249.12335205078125 0 6703.786 b 1
249.58839416503906 0 488.38223
251.1029815673828 0 776.1456
253.16517639160156 0 668.7935
263.1022644042969 0 1934.5449
264.7229309082031 0 488.34647
272.9347229003906 0 511.2324
279.09765625 0 842.027
285.0087890625 0 1379.2717
285.1019287109375 0 911.5556
296.1507873535156 0 757.2738
299.0601806640625 0 725.54865
324.14422607421875 0 800.49646
329.1495361328125 0 3291.4226
342.1560974121094 0 6458.4263 y 7
343.15911865234375 0 823.6654
355.0695495605469 0 1148.0037
356.1387939453125 0 684.2932
357.22393798828125 0 962.3295
364.15020751953125 0 1551.885 b 2
394.1242980957031 0 1434.806
413.2664489746094 0 1566.666
425.1905212402344 0 1150.4261
436.7265319824219 0 793.5742
441.20855712890625 0 647.5182
455.2239685058594 0 580.71643
465.1631774902344 0 1204.5651
479.17669677734375 0 1149.868 b 3
481.5709228515625 0 557.4013
489.1915283203125 0 1595.4159 y 6
513.1661376953125 0 634.28076
550.708740234375 0 1093.2866 y 1
560.22802734375 0 2468.2026 y 5
561.2322998046875 0 630.19775
569.232177734375 0 779.98706
585.3406982421875 0 1396.5719
592.2298583984375 0 692.75287 Precursor Water loss
593.7916870117188 0 687.99
601.2349243164062 0 2466.676 Precursor
601.2859497070312 0 628.4401
601.7318725585938 0 831.1477
602.2374877929688 0 815.6576
612.1940307617188 0 736.9555
659.29833984375 0 666.933
685.2566528320312 0 918.7223
700.3634643554688 0 3711.4824
701.3672485351562 0 1331.3337
702.574462890625 0 699.60016
723.2918090820312 0 3013.9165 y 4
749.254150390625 0 1895.6445
774.3193969726562 0 647.24817
801.411376953125 0 2006.343
802.4210205078125 0 769.07654
838.320556640625 0 2631.3516 y 3
839.3206787109375 0 1847.5885
844.4532470703125 0 2647.5735
845.453125 0 1050.2948
872.4476928710938 0 10905.713
873.4503784179688 0 4386.025
874.453125 0 1497.4537
889.3474731445312 0 4161.6665
890.353271484375 0 1926.5664
953.3460083007812 0 11591.274 y 2
954.3497924804688 0 6294.999
955.3480834960938 0 2223.4658
991.5216674804688 0 1376.1865
992.5281372070312 0 1057.8911
1019.516845703125 0 3817.0366
1020.5176391601562 0 3263.725
1021.5169677734375 0 985.49036
1101.4263916015625 0 895.55457
2688.849609375 0 637.8819
2884.16259765625 0 689.41724

Spectrum Details

|  |  |
| --- | --- |
| Matched peaks? Matched peaks The total absolute number of peaks matched. Additionally in brackets the total fraction of peaks matched and the total number of peaks is shown. | 16 (12.70% of 126) |
| FDR? FDR The false discovery rate estimated for this peptide. It is calculated by matching all theoretical fragments with a non-integer shift with the raw peaks for this spectrum. This is done with 40 different shifts. The resulting percentage is the average number of annotated peaks over the number of annotated peaks with the correct spectrum. | 0.74% |
| Satellite FDR? Satellite FDR See the FDR for details on its calculation. This satellite ion specific FDR only contains the satellite ions (d/w) for I/L/J positions. | - |
| PSM Score? PSM Score The PSM Score as given by Hecklib to this annotated spectrum. It is shown with three significant figures. | 160 |

#### Spectrum 11965? Spectrum 11965 The raw spectrum of this peptide as annotated by Hecklib. The fragments are coloured according to ion type (see legend). Any peaks with a star '\*' as text can be hovered over to see the full details, first the ion type second the mass shift type. By hovering over the amino acids in the peptide or ions in the legend the corresponding peaks are highlighted. By toggling the 'Unassigned' label you can turn the background (unassigned) peaks on or off in the plot. By updating the slider in the Ion legend you can update the spectrum to only show the top X% of the peaks with labels. The top X% means any peak that is within X% of the highest intensity. By dragging in the spectrum you can zoom in to a specific part of the spectrum and use 'Zoom Out' to get back to the original zoom level. The annotation of the spectrum is based on the given sequence in the peptides file and is done with different software so inconsistencies are likely. The peaks are annotated based on the given sequence, with 20 ppm tolerance.

Copy Data

##### Spectrum 11965 (TSV)

###### Preview

```
Loading example...
```

*Click on the button to copy the data to your clipboard.*

Mz MinMz MaxIntensity Max

WidthHeightPeptide font sizePeptide stroke widthSpectrum font sizeSpectrum stroke widthCompact peptide

Ion legend

wxyz

abcd

OtherUnassignedIon^Charge^_Position_Show for top:%

TFDDYAMHW

01.20e+42.39e+43.59e+44.79e+4

Zoom Out

a^+1^_2_y^+1^_1_a^+1^_2_b^+1^_2_y^+1^_2_b^+1^_3_b^+1^_4_y^+1^_3_y^+2^_8_y^+1^_4_\*\*y^+1^_5_y^+1^_6_y^+1^_7_y^+1^_7_

0839167925183357

Fragment Matches Table

Show background peaks

| Position | Ion type | Intensity | mz Theoretical | mz Error (Th) | mz Error (ppm) | Charge | Series Number |
| --- | --- | --- | --- | --- | --- | --- | --- |
| - | - | 3.212E+04 | 120.1 | - | - | 0 | - |
| - | - | 2596 | 121.1 | - | - | 0 | - |
| - | - | 323.4 | 122.1 | - | - | 0 | - |
| - | - | 341.5 | 122.7 | - | - | 0 | - |
| - | - | 364.1 | 125.1 | - | - | 0 | - |
| - | - | 461.6 | 126.1 | - | - | 0 | - |
| - | - | 3503 | 129.1 | - | - | 0 | - |
| - | - | 581.8 | 130.1 | - | - | 0 | - |
| - | - | 363.9 | 131.6 | - | - | 0 | - |
| - | - | 362.3 | 132 | - | - | 0 | - |
| - | - | 779.7 | 132.1 | - | - | 0 | - |
| - | - | 1079 | 133.1 | - | - | 0 | - |
| - | - | 9700 | 136.1 | - | - | 0 | - |
| - | - | 732.5 | 136.1 | - | - | 0 | - |
| - | - | 705.9 | 137.1 | - | - | 0 | - |
| - | - | 464.4 | 138.1 | - | - | 0 | - |
| - | - | 545.3 | 141.1 | - | - | 0 | - |
| - | - | 496.3 | 146.1 | - | - | 0 | - |
| - | - | 1242 | 149 | - | - | 0 | - |
| - | - | 453.4 | 150.4 | - | - | 0 | - |
| - | - | 497.7 | 155.1 | - | - | 0 | - |
| - | - | 1075 | 156.1 | - | - | 0 | - |
| - | - | 2152 | 158.1 | - | - | 0 | - |
| - | - | 1563 | 159.1 | - | - | 0 | - |
| - | - | 683.9 | 160.1 | - | - | 0 | - |
| - | - | 2747 | 165.1 | - | - | 0 | - |
| - | - | 802.2 | 166.1 | - | - | 0 | - |
| - | - | 2365 | 166.1 | - | - | 0 | - |
| - | - | 1702 | 167.1 | - | - | 0 | - |
| - | - | 564.1 | 167.1 | - | - | 0 | - |
| - | - | 479.2 | 173.1 | - | - | 0 | - |
| - | - | 1824 | 173.1 | - | - | 0 | - |
| - | - | 3165 | 176.1 | - | - | 0 | - |
| - | - | 3018 | 177.1 | - | - | 0 | - |
| - | - | 624.1 | 177.1 | - | - | 0 | - |
| - | - | 7130 | 182.1 | - | - | 0 | - |
| - | - | 639.5 | 183.1 | - | - | 0 | - |
| - | - | 582.6 | 183.1 | - | - | 0 | - |
| - | - | 516 | 183.1 | - | - | 0 | - |
| - | - | 1651 | 188.1 | - | - | 0 | - |
| - | - | 553.5 | 190.2 | - | - | 0 | - |
| - | - | 1217 | 193.1 | - | - | 0 | - |
| - | - | 669.5 | 195.1 | - | - | 0 | - |
| - | - | 491.3 | 196.8 | - | - | 0 | - |
| - | - | 631.5 | 197.1 | - | - | 0 | - |
| - | - | 992.2 | 201.1 | - | - | 0 | - |
| - | - | 1319 | 201.1 | - | - | 0 | - |
| 2 | a | 891.9 | 203.1 | 1.539E-05 | 0.07577 | +1 | 2 |
| 9 | y | 1430 | 205.1 | 2.913E-05 | 0.1421 | +1 | 1 |
| - | - | 709.9 | 207.1 | - | - | 0 | - |
| - | - | 607.7 | 219.1 | - | - | 0 | - |
| - | - | 2858 | 221.1 | - | - | 0 | - |
| 2 | a | 4.738E+04 | 221.1 | 9.524E-06 | 0.04307 | +1 | 2 |
| - | - | 5688 | 222.1 | - | - | 0 | - |
| - | - | 1379 | 223.1 | - | - | 0 | - |
| - | - | 1130 | 227.1 | - | - | 0 | - |
| - | - | 586.8 | 230.1 | - | - | 0 | - |
| - | - | 636.7 | 231.1 | - | - | 0 | - |
| - | - | 575.3 | 231.1 | - | - | 0 | - |
| - | - | 1293 | 235.1 | - | - | 0 | - |
| - | - | 1743 | 239.1 | - | - | 0 | - |
| - | - | 802 | 241.1 | - | - | 0 | - |
| 2 | b | 7909 | 249.1 | 2.898E-05 | 0.1163 | +1 | 2 |
| - | - | 1004 | 250.1 | - | - | 0 | - |
| - | - | 1097 | 251.1 | - | - | 0 | - |
| - | - | 655 | 253.1 | - | - | 0 | - |
| - | - | 1704 | 263.1 | - | - | 0 | - |
| - | - | 624.8 | 279.1 | - | - | 0 | - |
| - | - | 640.7 | 282.1 | - | - | 0 | - |
| - | - | 1228 | 285 | - | - | 0 | - |
| - | - | 532.2 | 285.1 | - | - | 0 | - |
| - | - | 1171 | 296.2 | - | - | 0 | - |
| - | - | 617.7 | 309.2 | - | - | 0 | - |
| - | - | 1436 | 324.1 | - | - | 0 | - |
| - | - | 684.6 | 325.1 | - | - | 0 | - |
| - | - | 4070 | 329.1 | - | - | 0 | - |
| - | - | 771.5 | 330.2 | - | - | 0 | - |
| - | - | 616.6 | 338.1 | - | - | 0 | - |
| - | - | 793.4 | 342.1 | - | - | 0 | - |
| 8 | y | 9969 | 342.2 | 6.01E-05 | 0.1756 | +1 | 2 |
| - | - | 1833 | 343.2 | - | - | 0 | - |
| - | - | 1963 | 355.1 | - | - | 0 | - |
| - | - | 1194 | 357.2 | - | - | 0 | - |
| 3 | b | 2483 | 364.2 | 0.0008063 | 2.214 | +1 | 3 |
| - | - | 710.5 | 365.2 | - | - | 0 | - |
| - | - | 1021 | 378.1 | - | - | 0 | - |
| - | - | 1091 | 394.1 | - | - | 0 | - |
| - | - | 683.6 | 398.2 | - | - | 0 | - |
| - | - | 2788 | 413.3 | - | - | 0 | - |
| - | - | 608.5 | 415.2 | - | - | 0 | - |
| - | - | 1380 | 425.2 | - | - | 0 | - |
| - | - | 701.8 | 427.7 | - | - | 0 | - |
| - | - | 934.8 | 465.2 | - | - | 0 | - |
| 4 | b | 905.1 | 479.2 | 0.0009983 | 2.083 | +1 | 4 |
| 7 | y | 1997 | 489.2 | 0.004917 | 10.05 | +1 | 3 |
| - | - | 732.9 | 496.2 | - | - | 0 | - |
| - | - | 759.3 | 508.3 | - | - | 0 | - |
| 2 | y | 892.8 | 550.7 | 0.00107 | 1.944 | +2 | 8 |
| - | - | 635.3 | 551.2 | - | - | 0 | - |
| 6 | y | 2730 | 560.2 | 0.003631 | 6.481 | +1 | 4 |
| - | - | 1372 | 561.2 | - | - | 0 | - |
| - | - | 2007 | 585.3 | - | - | 0 | - |
| - | - | 647.3 | 591.8 | - | - | 0 | - |
| 0 | Precursor | 876.3 | 592.2 | 0.002345 | 3.96 | +2 | -1 |
| 0 | Precursor | 2368 | 601.2 | 0.0004811 | 0.8002 | +2 | -1 |
| - | - | 559 | 601.7 | - | - | 0 | - |
| - | - | 624.3 | 618.2 | - | - | 0 | - |
| - | - | 1115 | 657.3 | - | - | 0 | - |
| - | - | 814.1 | 659.3 | - | - | 0 | - |
| - | - | 999.1 | 660.3 | - | - | 0 | - |
| - | - | 4689 | 700.4 | - | - | 0 | - |
| - | - | 1326 | 701.4 | - | - | 0 | - |
| 5 | y | 2518 | 723.3 | 0.00445 | 6.152 | +1 | 5 |
| - | - | 1749 | 724.3 | - | - | 0 | - |
| - | - | 1006 | 749.3 | - | - | 0 | - |
| - | - | 874.3 | 750.3 | - | - | 0 | - |
| - | - | 754.5 | 761.4 | - | - | 0 | - |
| - | - | 1054 | 774.3 | - | - | 0 | - |
| - | - | 930.1 | 783.4 | - | - | 0 | - |
| - | - | 2086 | 801.4 | - | - | 0 | - |
| - | - | 1058 | 802.4 | - | - | 0 | - |
| - | - | 1464 | 828.4 | - | - | 0 | - |
| 4 | y | 3382 | 838.3 | 0.003874 | 4.621 | +1 | 6 |
| - | - | 1737 | 839.3 | - | - | 0 | - |
| - | - | 680.7 | 840.3 | - | - | 0 | - |
| - | - | 3082 | 844.5 | - | - | 0 | - |
| - | - | 1194 | 845.5 | - | - | 0 | - |
| - | - | 648.9 | 871.3 | - | - | 0 | - |
| - | - | 1.237E+04 | 872.4 | - | - | 0 | - |
| - | - | 5247 | 873.4 | - | - | 0 | - |
| - | - | 1318 | 874.5 | - | - | 0 | - |
| - | - | 3994 | 889.3 | - | - | 0 | - |
| - | - | 3141 | 890.3 | - | - | 0 | - |
| - | - | 592.9 | 891.4 | - | - | 0 | - |
| - | - | 783.5 | 902.5 | - | - | 0 | - |
| 3 | y | 875.6 | 935.3 | 0.01136 | 12.15 | +1 | 7 |
| 3 | y | 1.639E+04 | 953.3 | 0.003909 | 4.1 | +1 | 7 |
| - | - | 7766 | 954.3 | - | - | 0 | - |
| - | - | 2888 | 955.3 | - | - | 0 | - |
| - | - | 1501 | 991.5 | - | - | 0 | - |
| - | - | 696.6 | 992.5 | - | - | 0 | - |
| - | - | 605.4 | 1003 | - | - | 0 | - |
| - | - | 5182 | 1020 | - | - | 0 | - |
| - | - | 3377 | 1021 | - | - | 0 | - |
| - | - | 794 | 1022 | - | - | 0 | - |
| - | - | 619.3 | 1102 | - | - | 0 | - |
| - | - | 672.2 | 2788 | - | - | 0 | - |
| - | - | 680.7 | 3324 | - | - | 0 | - |

m/z Charge Intensity FragmentType MassShift Position
120.08092498779297 0 32115.207
121.08435821533203 0 2596.3867
122.12374114990234 0 323.4389
122.69454193115234 0 341.4588
125.0966567993164 0 364.09207
126.05529022216797 0 461.63843
129.10232543945312 0 3503.3313
130.06517028808594 0 581.78253
131.61666870117188 0 363.94788
132.02774047851562 0 362.31693
132.0809783935547 0 779.6712
133.0859832763672 0 1079.1493
136.07579040527344 0 9700.238
136.0817108154297 0 732.5178
137.07919311523438 0 705.91785
138.0660400390625 0 464.43896
141.10304260253906 0 545.253
146.06002807617188 0 496.26996
149.0450439453125 0 1242.2051
150.3700408935547 0 453.44556
155.08139038085938 0 497.72775
156.07745361328125 0 1075.1536
158.09646606445312 0 2151.9934
159.09169006347656 0 1563.4418
160.11175537109375 0 683.92053
165.05474853515625 0 2747.3586
166.0534210205078 0 802.2434
166.08628845214844 0 2365.178
167.0555419921875 0 1702.4259
167.08154296875 0 564.11774
173.09117126464844 0 479.16022
173.12864685058594 0 1824.3699
176.1071319580078 0 3164.5256
177.1022491455078 0 3018.4365
177.1104278564453 0 624.12494
182.08119201660156 0 7129.956
183.084228515625 0 639.54767
183.1011505126953 0 582.56903
183.11146545410156 0 516.0073
188.0706329345703 0 1651.2314
190.24996948242188 0 553.488
193.10830688476562 0 1217.3337
195.0888214111328 0 669.4684
196.7786407470703 0 491.30582
197.12908935546875 0 631.45685
201.11215209960938 0 992.18475
201.12315368652344 0 1319.222
203.1178741455078 0 891.9061 a Water loss 1
205.09718322753906 0 1429.9042 y 8
207.11318969726562 0 709.934
219.1126708984375 0 607.66364
221.10333251953125 0 2857.6008
221.1284637451172 0 47379.473 a 1
222.1318359375 0 5687.7524
223.06336975097656 0 1378.766
227.1020965576172 0 1129.7871
230.11294555664062 0 586.83264
231.061279296875 0 636.71783
231.0984649658203 0 575.34393
235.10755920410156 0 1292.6718
239.09474182128906 0 1742.6022
241.0740966796875 0 801.98206
249.12339782714844 0 7908.8545 b 1
250.126708984375 0 1003.5923
251.10255432128906 0 1096.952
253.11874389648438 0 654.9689
263.1029052734375 0 1704.4167
279.0971374511719 0 624.7784
282.14398193359375 0 640.6765
285.00946044921875 0 1228.3962
285.1018371582031 0 532.16205
296.1503601074219 0 1171.1805
309.2044982910156 0 617.7455
324.1455383300781 0 1435.667
325.14593505859375 0 684.57086
329.14959716796875 0 4070.0588
330.15277099609375 0 771.5127
338.13311767578125 0 616.64166
342.136474609375 0 793.37366
342.156005859375 0 9969.361 y 7
343.15924072265625 0 1832.8093
355.0700378417969 0 1962.6841
357.223876953125 0 1193.8635
364.1495056152344 0 2483.0479 b 2
365.1543273925781 0 710.5132
378.12896728515625 0 1020.8948
394.1248474121094 0 1091.1648
398.2054748535156 0 683.6422
413.2657165527344 0 2787.5806
415.1645202636719 0 608.5054
425.1930847167969 0 1379.7914
427.7205505371094 0 701.78815
465.1630859375 0 934.79126
479.1782531738281 0 905.1197 b 3
489.19146728515625 0 1996.7247 y 6
496.22760009765625 0 732.8545
508.27392578125 0 759.3047
550.7072143554688 0 892.8291 y 1
551.2099609375 0 635.3147
560.227294921875 0 2730.487 y 5
561.2303466796875 0 1372.2333
585.3397827148438 0 2006.948
591.7916870117188 0 647.343
592.2291870117188 0 876.2764 Precursor Water loss
601.2326049804688 0 2367.5166 Precursor
601.743408203125 0 558.96295
618.1923828125 0 624.34314
657.2598876953125 0 1114.6449
659.2952880859375 0 814.14484
660.29541015625 0 999.10693
700.3612670898438 0 4688.982
701.3667602539062 0 1326.408
723.2914428710938 0 2518.283 y 4
724.2933349609375 0 1749.3706
749.2584838867188 0 1005.9764
750.2568359375 0 874.2566
761.3777465820312 0 754.5049
774.3224487304688 0 1054.4034
783.3973999023438 0 930.075
801.4119873046875 0 2086.2446
802.4144897460938 0 1057.8269
828.42041015625 0 1463.9331
838.3178100585938 0 3382.32 y 3
839.3193359375 0 1736.694
840.3187866210938 0 680.6651
844.4509887695312 0 3081.6296
845.456787109375 0 1194.1504
871.3275146484375 0 648.9149
872.4466552734375 0 12366.735
873.4494018554688 0 5246.9272
874.453369140625 0 1318.4948
889.3461303710938 0 3994.3506
890.3480834960938 0 3141.2883
891.359375 0 592.8882
902.4788818359375 0 783.5188
935.3416748046875 0 875.5627 y Water loss 2
953.3447875976562 0 16388.07 y 2
954.3475341796875 0 7766.099
955.34716796875 0 2888.394
991.5183715820312 0 1501
992.5178833007812 0 696.64014
1003.3710327148438 0 605.38617
1019.5137939453125 0 5181.848
1020.5151977539062 0 3376.61
1021.525634765625 0 793.9607
1102.4219970703125 0 619.3135
2788.134033203125 0 672.219
3324.061767578125 0 680.72516

Spectrum Details

|  |  |
| --- | --- |
| Matched peaks? Matched peaks The total absolute number of peaks matched. Additionally in brackets the total fraction of peaks matched and the total number of peaks is shown. | 16 (10.81% of 148) |
| FDR? FDR The false discovery rate estimated for this peptide. It is calculated by matching all theoretical fragments with a non-integer shift with the raw peaks for this spectrum. This is done with 40 different shifts. The resulting percentage is the average number of annotated peaks over the number of annotated peaks with the correct spectrum. | 0.74% |
| Satellite FDR? Satellite FDR See the FDR for details on its calculation. This satellite ion specific FDR only contains the satellite ions (d/w) for I/L/J positions. | - |
| PSM Score? PSM Score The PSM Score as given by Hecklib to this annotated spectrum. It is shown with three significant figures. | 160 |

#### Spectrum 12241? Spectrum 12241 The raw spectrum of this peptide as annotated by Hecklib. The fragments are coloured according to ion type (see legend). Any peaks with a star '\*' as text can be hovered over to see the full details, first the ion type second the mass shift type. By hovering over the amino acids in the peptide or ions in the legend the corresponding peaks are highlighted. By toggling the 'Unassigned' label you can turn the background (unassigned) peaks on or off in the plot. By updating the slider in the Ion legend you can update the spectrum to only show the top X% of the peaks with labels. The top X% means any peak that is within X% of the highest intensity. By dragging in the spectrum you can zoom in to a specific part of the spectrum and use 'Zoom Out' to get back to the original zoom level. The annotation of the spectrum is based on the given sequence in the peptides file and is done with different software so inconsistencies are likely. The peaks are annotated based on the given sequence, with 20 ppm tolerance.

Copy Data

##### Spectrum 12241 (TSV)

###### Preview

```
Loading example...
```

*Click on the button to copy the data to your clipboard.*

Mz MinMz MaxIntensity Max

WidthHeightPeptide font sizePeptide stroke widthSpectrum font sizeSpectrum stroke widthCompact peptide

Ion legend

wxyz

abcd

OtherUnassignedIon^Charge^_Position_Show for top:%

TFDDYAMHW

01.11e+42.22e+43.33e+44.44e+4

Zoom Out

a^+1^_2_y^+1^_1_a^+1^_2_b^+2^_4_b^+1^_2_a^+1^_3_y^+1^_2_b^+1^_3_b^+1^_4_y^+1^_3_b^+2^_8_y^+2^_8_y^+1^_4_\*y^+1^_5_y^+1^_6_y^+1^_7_y^+1^_7_

0804160924133218

Fragment Matches Table

Show background peaks

| Position | Ion type | Intensity | mz Theoretical | mz Error (Th) | mz Error (ppm) | Charge | Series Number |
| --- | --- | --- | --- | --- | --- | --- | --- |
| - | - | 3.018E+04 | 120.1 | - | - | 0 | - |
| - | - | 2458 | 121.1 | - | - | 0 | - |
| - | - | 468.9 | 123 | - | - | 0 | - |
| - | - | 370.9 | 125 | - | - | 0 | - |
| - | - | 3272 | 129.1 | - | - | 0 | - |
| - | - | 912.7 | 130.1 | - | - | 0 | - |
| - | - | 1075 | 133.1 | - | - | 0 | - |
| - | - | 444.7 | 134 | - | - | 0 | - |
| - | - | 383.5 | 134.1 | - | - | 0 | - |
| - | - | 9405 | 136.1 | - | - | 0 | - |
| - | - | 722.1 | 137.1 | - | - | 0 | - |
| - | - | 570.4 | 138.1 | - | - | 0 | - |
| - | - | 562.4 | 147 | - | - | 0 | - |
| - | - | 1525 | 149 | - | - | 0 | - |
| - | - | 452.4 | 150.9 | - | - | 0 | - |
| - | - | 442.3 | 153.1 | - | - | 0 | - |
| - | - | 613.7 | 153.8 | - | - | 0 | - |
| - | - | 701 | 155.1 | - | - | 0 | - |
| - | - | 841.5 | 156.1 | - | - | 0 | - |
| - | - | 2114 | 158.1 | - | - | 0 | - |
| - | - | 1518 | 159.1 | - | - | 0 | - |
| - | - | 562.7 | 160.1 | - | - | 0 | - |
| - | - | 1013 | 160.1 | - | - | 0 | - |
| - | - | 486 | 160.2 | - | - | 0 | - |
| - | - | 671.6 | 162.1 | - | - | 0 | - |
| - | - | 3254 | 165.1 | - | - | 0 | - |
| - | - | 1171 | 166.1 | - | - | 0 | - |
| - | - | 2477 | 167.1 | - | - | 0 | - |
| - | - | 598.5 | 167.1 | - | - | 0 | - |
| - | - | 740.7 | 169.1 | - | - | 0 | - |
| - | - | 1584 | 173.1 | - | - | 0 | - |
| - | - | 5325 | 173.4 | - | - | 0 | - |
| - | - | 557.2 | 175.1 | - | - | 0 | - |
| - | - | 2704 | 176.1 | - | - | 0 | - |
| - | - | 1665 | 177.1 | - | - | 0 | - |
| - | - | 477.8 | 178.1 | - | - | 0 | - |
| - | - | 6273 | 182.1 | - | - | 0 | - |
| - | - | 462.3 | 185.1 | - | - | 0 | - |
| - | - | 682.4 | 185.1 | - | - | 0 | - |
| - | - | 914.7 | 187.1 | - | - | 0 | - |
| - | - | 1188 | 188.1 | - | - | 0 | - |
| - | - | 1025 | 193.1 | - | - | 0 | - |
| - | - | 504 | 199.1 | - | - | 0 | - |
| - | - | 1039 | 200.1 | - | - | 0 | - |
| - | - | 1064 | 201.1 | - | - | 0 | - |
| 2 | a | 1325 | 203.1 | 6.09E-05 | 0.2998 | +1 | 2 |
| 9 | y | 893.4 | 205.1 | 4.716E-05 | 0.2299 | +1 | 1 |
| - | - | 810.6 | 207.1 | - | - | 0 | - |
| - | - | 537.9 | 209.1 | - | - | 0 | - |
| - | - | 718.3 | 211.1 | - | - | 0 | - |
| - | - | 545.8 | 217.1 | - | - | 0 | - |
| - | - | 2526 | 221.1 | - | - | 0 | - |
| 2 | a | 4.399E+04 | 221.1 | 5.735E-06 | 0.02593 | +1 | 2 |
| - | - | 5122 | 222.1 | - | - | 0 | - |
| - | - | 1408 | 223.1 | - | - | 0 | - |
| - | - | 826.7 | 227.1 | - | - | 0 | - |
| - | - | 732.3 | 235.1 | - | - | 0 | - |
| - | - | 2161 | 239.1 | - | - | 0 | - |
| 4 | b | 528.9 | 240.1 | 0.004017 | 16.73 | +2 | 4 |
| - | - | 574.1 | 240.1 | - | - | 0 | - |
| - | - | 1134 | 241.1 | - | - | 0 | - |
| 2 | b | 6489 | 249.1 | 4.424E-05 | 0.1776 | +1 | 2 |
| - | - | 547.1 | 250.1 | - | - | 0 | - |
| - | - | 630.9 | 250.1 | - | - | 0 | - |
| - | - | 669.9 | 253.2 | - | - | 0 | - |
| - | - | 526 | 257.1 | - | - | 0 | - |
| - | - | 534.1 | 257.2 | - | - | 0 | - |
| - | - | 1883 | 263.1 | - | - | 0 | - |
| - | - | 1121 | 283 | - | - | 0 | - |
| - | - | 2095 | 285 | - | - | 0 | - |
| - | - | 949.7 | 285.1 | - | - | 0 | - |
| - | - | 664.7 | 296.2 | - | - | 0 | - |
| - | - | 657.3 | 299.1 | - | - | 0 | - |
| - | - | 733.6 | 301 | - | - | 0 | - |
| - | - | 598.8 | 302.1 | - | - | 0 | - |
| 3 | a | 553.2 | 318.1 | 0.0002098 | 0.6593 | +1 | 3 |
| - | - | 874.1 | 319.1 | - | - | 0 | - |
| - | - | 1996 | 324.1 | - | - | 0 | - |
| - | - | 3765 | 329.1 | - | - | 0 | - |
| - | - | 880.3 | 340.2 | - | - | 0 | - |
| - | - | 560.8 | 342 | - | - | 0 | - |
| - | - | 817.5 | 342.1 | - | - | 0 | - |
| 8 | y | 6917 | 342.2 | 3.146E-05 | 0.09194 | +1 | 2 |
| - | - | 1675 | 343.2 | - | - | 0 | - |
| - | - | 3047 | 355.1 | - | - | 0 | - |
| - | - | 903.9 | 356.1 | - | - | 0 | - |
| - | - | 695.1 | 357.1 | - | - | 0 | - |
| - | - | 682.5 | 357.2 | - | - | 0 | - |
| 3 | b | 1935 | 364.2 | 0.0001654 | 0.4542 | +1 | 3 |
| - | - | 651.4 | 366.1 | - | - | 0 | - |
| - | - | 1069 | 378.1 | - | - | 0 | - |
| - | - | 617.9 | 385.1 | - | - | 0 | - |
| - | - | 1106 | 394.1 | - | - | 0 | - |
| - | - | 886.9 | 413.3 | - | - | 0 | - |
| - | - | 881.8 | 415.1 | - | - | 0 | - |
| - | - | 677 | 415.2 | - | - | 0 | - |
| - | - | 1653 | 425.2 | - | - | 0 | - |
| - | - | 577.2 | 441.2 | - | - | 0 | - |
| - | - | 1017 | 465.2 | - | - | 0 | - |
| 4 | b | 1065 | 479.2 | 0.002829 | 5.905 | +1 | 4 |
| - | - | 753.6 | 483.2 | - | - | 0 | - |
| 7 | y | 1807 | 489.2 | 0.004459 | 9.115 | +1 | 3 |
| 8 | b | 599.6 | 490.2 | 0.007856 | 16.03 | +2 | 8 |
| - | - | 588.3 | 491.2 | - | - | 0 | - |
| - | - | 1196 | 496.2 | - | - | 0 | - |
| - | - | 630.6 | 527.9 | - | - | 0 | - |
| 2 | y | 1867 | 550.7 | 0.0009439 | 1.714 | +2 | 8 |
| 6 | y | 3742 | 560.2 | 0.004912 | 8.768 | +1 | 4 |
| - | - | 2392 | 585.3 | - | - | 0 | - |
| - | - | 777.1 | 592.7 | - | - | 0 | - |
| 0 | Precursor | 787.2 | 601.2 | 0.007012 | 11.66 | +2 | -1 |
| - | - | 809.9 | 602.2 | - | - | 0 | - |
| - | - | 885.4 | 606.2 | - | - | 0 | - |
| - | - | 675.7 | 614.2 | - | - | 0 | - |
| - | - | 609.2 | 629.4 | - | - | 0 | - |
| - | - | 777.1 | 648.1 | - | - | 0 | - |
| - | - | 752.8 | 657.3 | - | - | 0 | - |
| - | - | 1081 | 659.3 | - | - | 0 | - |
| - | - | 702.7 | 685.3 | - | - | 0 | - |
| - | - | 3692 | 700.4 | - | - | 0 | - |
| - | - | 1202 | 701.4 | - | - | 0 | - |
| - | - | 1130 | 721.3 | - | - | 0 | - |
| 5 | y | 2920 | 723.3 | 0.003168 | 4.38 | +1 | 5 |
| - | - | 1273 | 724.3 | - | - | 0 | - |
| - | - | 734.7 | 749.3 | - | - | 0 | - |
| - | - | 597.2 | 773.4 | - | - | 0 | - |
| - | - | 1483 | 774.3 | - | - | 0 | - |
| - | - | 2080 | 801.4 | - | - | 0 | - |
| - | - | 1018 | 828.4 | - | - | 0 | - |
| - | - | 699.7 | 829.4 | - | - | 0 | - |
| 4 | y | 3253 | 838.3 | 0.004362 | 5.204 | +1 | 6 |
| - | - | 1364 | 839.3 | - | - | 0 | - |
| - | - | 3497 | 844.5 | - | - | 0 | - |
| - | - | 1074 | 845.5 | - | - | 0 | - |
| - | - | 585.1 | 846.5 | - | - | 0 | - |
| - | - | 793.2 | 872.3 | - | - | 0 | - |
| - | - | 1.066E+04 | 872.4 | - | - | 0 | - |
| - | - | 5192 | 873.4 | - | - | 0 | - |
| - | - | 1073 | 874.5 | - | - | 0 | - |
| - | - | 4174 | 889.3 | - | - | 0 | - |
| - | - | 1895 | 890.3 | - | - | 0 | - |
| 3 | y | 864.3 | 935.3 | 0.002388 | 2.554 | +1 | 7 |
| 3 | y | 1.613E+04 | 953.3 | 0.003665 | 3.844 | +1 | 7 |
| - | - | 7947 | 954.3 | - | - | 0 | - |
| - | - | 3628 | 955.3 | - | - | 0 | - |
| - | - | 1908 | 991.5 | - | - | 0 | - |
| - | - | 5225 | 1020 | - | - | 0 | - |
| - | - | 1942 | 1021 | - | - | 0 | - |
| - | - | 637.2 | 2897 | - | - | 0 | - |
| - | - | 628 | 3186 | - | - | 0 | - |

m/z Charge Intensity FragmentType MassShift Position
120.0809326171875 0 30181.04
121.0843505859375 0 2458.2046
123.04438018798828 0 468.88782
124.98545837402344 0 370.8968
129.1023406982422 0 3272.4553
130.0652313232422 0 912.7123
133.08604431152344 0 1075.0975
134.04458618164062 0 444.72916
134.12014770507812 0 383.45065
136.0758056640625 0 9405.004
137.0791778564453 0 722.13763
138.06646728515625 0 570.36224
147.04437255859375 0 562.3942
149.04507446289062 0 1524.73
150.93258666992188 0 452.4329
153.09056091308594 0 442.31003
153.82362365722656 0 613.67664
155.08213806152344 0 700.9819
156.07736206054688 0 841.5301
158.0963592529297 0 2113.8757
159.091796875 0 1517.66
160.0756072998047 0 562.7319
160.1125030517578 0 1012.9254
160.1652374267578 0 485.98154
162.09161376953125 0 671.6401
165.05482482910156 0 3254.4636
166.0864715576172 0 1170.5748
167.0554962158203 0 2477.1643
167.08187866210938 0 598.472
169.09742736816406 0 740.66864
173.12884521484375 0 1584.2332
173.44029235839844 0 5324.6235
175.0967559814453 0 557.24396
176.1069793701172 0 2704.4224
177.10250854492188 0 1665.3219
178.10574340820312 0 477.77698
182.08119201660156 0 6272.7456
185.093505859375 0 462.34726
185.12847900390625 0 682.43677
187.13307189941406 0 914.6702
188.070556640625 0 1188.1329
193.10787963867188 0 1025.3046
199.07183837890625 0 504.01117
200.1397247314453 0 1039.0789
201.12306213378906 0 1064.2188
203.11795043945312 0 1324.9946 a Water loss 1
205.09710693359375 0 893.4428 y 8
207.11280822753906 0 810.6028
209.0928497314453 0 537.88666
211.0599822998047 0 718.3322
217.08145141601562 0 545.82825
221.1033477783203 0 2526.256
221.12844848632812 0 43993.73 a 1
222.13189697265625 0 5121.7803
223.06358337402344 0 1407.6155
227.1029052734375 0 826.71783
235.10804748535156 0 732.26855
239.0948028564453 0 2160.668
240.09628295898438 0 528.9151 b 3
240.13433837890625 0 574.0534
241.0746307373047 0 1133.5459
249.1234130859375 0 6489.278 b 1
250.08865356445312 0 547.0632
250.10414123535156 0 630.91656
253.16549682617188 0 669.86346
257.10589599609375 0 526.00366
257.1652526855469 0 534.0557
263.10296630859375 0 1883.0563
283.0307312011719 0 1120.7385
285.00958251953125 0 2094.915
285.10162353515625 0 949.69696
296.1503601074219 0 664.7192
299.06146240234375 0 657.2526
301.04010009765625 0 733.6465
302.11279296875 0 598.7676
318.1446228027344 0 553.2345 a Water loss 2
319.1410217285156 0 874.1259
324.1451110839844 0 1995.9348
329.1493225097656 0 3765.2542
340.19842529296875 0 880.25635
342.0039978027344 0 560.8469
342.1368713378906 0 817.53406
342.1560974121094 0 6916.572 y 7
343.1584167480469 0 1675.0972
355.0699157714844 0 3046.7979
356.07147216796875 0 903.9482
357.0686950683594 0 695.14667
357.2239685058594 0 682.466
364.150146484375 0 1935.4858 b 2
366.1294250488281 0 651.389
378.12908935546875 0 1068.6873
385.13104248046875 0 617.9493
394.12457275390625 0 1106.0785
413.265625 0 886.9113
415.1251525878906 0 881.8017
415.1578063964844 0 677.0286
425.19317626953125 0 1652.9646
441.2089538574219 0 577.1839
465.1629638671875 0 1016.6021
479.1800842285156 0 1065.2579 b 3
483.2221984863281 0 753.60986
489.1910095214844 0 1806.97 y 6
490.18975830078125 0 599.642 b Water loss 7
491.2121887207031 0 588.2791
496.2285461425781 0 1195.546
527.9207763671875 0 630.56116
550.709228515625 0 1867.2987 y 1
560.2285766601562 0 3741.8567 y 5
585.339111328125 0 2392.029
592.7296752929688 0 777.1139
601.2391357421875 0 787.2169 Precursor
602.2379150390625 0 809.85565
606.2251586914062 0 885.4131
614.245849609375 0 675.69617
629.3662109375 0 609.18884
648.0947875976562 0 777.1102
657.2656860351562 0 752.79346
659.290283203125 0 1080.6858
685.2612915039062 0 702.73193
700.363525390625 0 3691.9915
701.3641357421875 0 1202.4476
721.2604370117188 0 1130.2446
723.2901611328125 0 2919.5242 y 4
724.2962646484375 0 1273.0283
749.2560424804688 0 734.7388
773.416259765625 0 597.1848
774.3194580078125 0 1483.1152
801.408447265625 0 2080.031
828.4219360351562 0 1018.25256
829.4150390625 0 699.6548
838.3182983398438 0 3253.145 y 3
839.3199462890625 0 1363.8032
844.4501953125 0 3497.387
845.454345703125 0 1073.769
846.454833984375 0 585.05206
872.3428955078125 0 793.19476
872.4464111328125 0 10664.184
873.4483642578125 0 5192.232
874.4503784179688 0 1072.8617
889.3463745117188 0 4174.154
890.3458251953125 0 1894.6193
935.3327026367188 0 864.2913 y Water loss 2
953.3445434570312 0 16133.579 y 2
954.3474731445312 0 7946.5317
955.3490600585938 0 3627.9639
991.5191650390625 0 1907.6102
1019.5155029296875 0 5224.604
1020.5172119140625 0 1942.1935
2896.732177734375 0 637.15607
3185.81396484375 0 627.98206

Spectrum Details

|  |  |
| --- | --- |
| Matched peaks? Matched peaks The total absolute number of peaks matched. Additionally in brackets the total fraction of peaks matched and the total number of peaks is shown. | 18 (12.00% of 150) |
| FDR? FDR The false discovery rate estimated for this peptide. It is calculated by matching all theoretical fragments with a non-integer shift with the raw peaks for this spectrum. This is done with 40 different shifts. The resulting percentage is the average number of annotated peaks over the number of annotated peaks with the correct spectrum. | 1.06% |
| Satellite FDR? Satellite FDR See the FDR for details on its calculation. This satellite ion specific FDR only contains the satellite ions (d/w) for I/L/J positions. | - |
| PSM Score? PSM Score The PSM Score as given by Hecklib to this annotated spectrum. It is shown with three significant figures. | 177 |

#### Spectrum 8771? Spectrum 8771 The raw spectrum of this peptide as annotated by Hecklib. The fragments are coloured according to ion type (see legend). Any peaks with a star '\*' as text can be hovered over to see the full details, first the ion type second the mass shift type. By hovering over the amino acids in the peptide or ions in the legend the corresponding peaks are highlighted. By toggling the 'Unassigned' label you can turn the background (unassigned) peaks on or off in the plot. By updating the slider in the Ion legend you can update the spectrum to only show the top X% of the peaks with labels. The top X% means any peak that is within X% of the highest intensity. By dragging in the spectrum you can zoom in to a specific part of the spectrum and use 'Zoom Out' to get back to the original zoom level. The annotation of the spectrum is based on the given sequence in the peptides file and is done with different software so inconsistencies are likely. The peaks are annotated based on the given sequence, with 20 ppm tolerance.

Copy Data

##### Spectrum 8771 (TSV)

###### Preview

```
Loading example...
```

*Click on the button to copy the data to your clipboard.*

Mz MinMz MaxIntensity Max

WidthHeightPeptide font sizePeptide stroke widthSpectrum font sizeSpectrum stroke widthCompact peptide

Ion legend

wxyz

abcd

OtherUnassignedIon^Charge^_Position_Show for top:%

TFDDYAMHW

01.26e+52.52e+53.78e+55.03e+5

Zoom Out

a^+1^_2_y^+1^_1_a^+1^_2_b^+1^_2_b^+1^_2_a^+1^_3_y^+1^_2_b^+1^_3_b^+1^_3_b^+1^_4_y^+2^_7_b^+1^_4_y^+1^_3_y^+2^_8_y^+1^_4_\*\*b^+1^_5_b^+1^_6_y^+1^_5_y^+1^_6_y^+1^_7_y^+1^_7_b^+1^_8_y^+1^_8_

0732146521972930

Fragment Matches Table

Show background peaks

| Position | Ion type | Intensity | mz Theoretical | mz Error (Th) | mz Error (ppm) | Charge | Series Number |
| --- | --- | --- | --- | --- | --- | --- | --- |
| - | - | 4.984E+05 | 120.1 | - | - | 0 | - |
| - | - | 573.4 | 121.1 | - | - | 0 | - |
| - | - | 4.058E+04 | 121.1 | - | - | 0 | - |
| - | - | 965.5 | 122.1 | - | - | 0 | - |
| - | - | 1.175E+04 | 123 | - | - | 0 | - |
| - | - | 856.3 | 124 | - | - | 0 | - |
| - | - | 364.6 | 125.2 | - | - | 0 | - |
| - | - | 1402 | 127.1 | - | - | 0 | - |
| - | - | 1990 | 128.1 | - | - | 0 | - |
| - | - | 440.9 | 129.1 | - | - | 0 | - |
| - | - | 3557 | 129.1 | - | - | 0 | - |
| - | - | 3210 | 130.1 | - | - | 0 | - |
| - | - | 626.2 | 131 | - | - | 0 | - |
| - | - | 607.2 | 131.1 | - | - | 0 | - |
| - | - | 1547 | 131.1 | - | - | 0 | - |
| - | - | 2388 | 132.1 | - | - | 0 | - |
| - | - | 1163 | 132.1 | - | - | 0 | - |
| - | - | 1125 | 134 | - | - | 0 | - |
| - | - | 1.314E+05 | 136.1 | - | - | 0 | - |
| - | - | 9818 | 137.1 | - | - | 0 | - |
| - | - | 821.7 | 138.1 | - | - | 0 | - |
| - | - | 1415 | 138.1 | - | - | 0 | - |
| - | - | 433.1 | 139 | - | - | 0 | - |
| - | - | 966.1 | 140.1 | - | - | 0 | - |
| - | - | 831.9 | 141.1 | - | - | 0 | - |
| - | - | 550.3 | 143.1 | - | - | 0 | - |
| - | - | 497.2 | 144.1 | - | - | 0 | - |
| - | - | 3141 | 145.1 | - | - | 0 | - |
| - | - | 1207 | 146.1 | - | - | 0 | - |
| - | - | 9229 | 147 | - | - | 0 | - |
| - | - | 820.8 | 148 | - | - | 0 | - |
| - | - | 860 | 149 | - | - | 0 | - |
| - | - | 602.7 | 153.1 | - | - | 0 | - |
| - | - | 2708 | 155.1 | - | - | 0 | - |
| - | - | 897 | 155.1 | - | - | 0 | - |
| - | - | 4975 | 156.1 | - | - | 0 | - |
| - | - | 566 | 157.1 | - | - | 0 | - |
| - | - | 680.9 | 157.1 | - | - | 0 | - |
| - | - | 2787 | 157.1 | - | - | 0 | - |
| - | - | 1739 | 157.1 | - | - | 0 | - |
| - | - | 5611 | 158.1 | - | - | 0 | - |
| - | - | 5329 | 159.1 | - | - | 0 | - |
| - | - | 802.7 | 159.1 | - | - | 0 | - |
| - | - | 1117 | 160.1 | - | - | 0 | - |
| - | - | 2232 | 160.1 | - | - | 0 | - |
| - | - | 999.3 | 162.1 | - | - | 0 | - |
| - | - | 462 | 162.1 | - | - | 0 | - |
| - | - | 429.6 | 164.1 | - | - | 0 | - |
| - | - | 9.154E+04 | 165.1 | - | - | 0 | - |
| - | - | 8432 | 166.1 | - | - | 0 | - |
| - | - | 6435 | 166.1 | - | - | 0 | - |
| - | - | 572.8 | 166.1 | - | - | 0 | - |
| - | - | 462.6 | 166.1 | - | - | 0 | - |
| - | - | 1494 | 167.1 | - | - | 0 | - |
| - | - | 734.2 | 167.1 | - | - | 0 | - |
| - | - | 508.1 | 167.1 | - | - | 0 | - |
| - | - | 436.8 | 168.9 | - | - | 0 | - |
| - | - | 527.6 | 169.1 | - | - | 0 | - |
| - | - | 1373 | 169.1 | - | - | 0 | - |
| - | - | 1696 | 171.1 | - | - | 0 | - |
| - | - | 670.9 | 171.1 | - | - | 0 | - |
| - | - | 3050 | 173.1 | - | - | 0 | - |
| - | - | 4.042E+04 | 173.1 | - | - | 0 | - |
| - | - | 2879 | 174.1 | - | - | 0 | - |
| - | - | 539.8 | 175.1 | - | - | 0 | - |
| - | - | 1354 | 176.1 | - | - | 0 | - |
| - | - | 1.152E+04 | 176.1 | - | - | 0 | - |
| - | - | 7502 | 177.1 | - | - | 0 | - |
| - | - | 1329 | 177.1 | - | - | 0 | - |
| - | - | 528.7 | 180.1 | - | - | 0 | - |
| - | - | 980.2 | 181.1 | - | - | 0 | - |
| - | - | 1.801E+05 | 182.1 | - | - | 0 | - |
| - | - | 1.599E+04 | 183.1 | - | - | 0 | - |
| - | - | 984.6 | 183.1 | - | - | 0 | - |
| - | - | 826.7 | 183.1 | - | - | 0 | - |
| - | - | 818.1 | 184.1 | - | - | 0 | - |
| - | - | 1144 | 185.1 | - | - | 0 | - |
| - | - | 2363 | 185.1 | - | - | 0 | - |
| - | - | 938.6 | 186.1 | - | - | 0 | - |
| - | - | 615.8 | 187.1 | - | - | 0 | - |
| - | - | 5687 | 188.1 | - | - | 0 | - |
| - | - | 2665 | 189.1 | - | - | 0 | - |
| - | - | 1.46E+04 | 191.1 | - | - | 0 | - |
| - | - | 936.8 | 192.1 | - | - | 0 | - |
| - | - | 1128 | 192.1 | - | - | 0 | - |
| - | - | 567.3 | 192.1 | - | - | 0 | - |
| - | - | 3571 | 193.1 | - | - | 0 | - |
| - | - | 1123 | 195.1 | - | - | 0 | - |
| - | - | 1600 | 197.1 | - | - | 0 | - |
| - | - | 3398 | 199.1 | - | - | 0 | - |
| - | - | 2054 | 199.1 | - | - | 0 | - |
| - | - | 2.477E+04 | 201.1 | - | - | 0 | - |
| - | - | 2268 | 202.1 | - | - | 0 | - |
| - | - | 1280 | 203.1 | - | - | 0 | - |
| 2 | a | 3616 | 203.1 | 0.0003813 | 1.877 | +1 | 2 |
| - | - | 1249 | 204.1 | - | - | 0 | - |
| - | - | 561.2 | 205.1 | - | - | 0 | - |
| 9 | y | 4133 | 205.1 | 0.0003648 | 1.779 | +1 | 1 |
| - | - | 2589 | 207.1 | - | - | 0 | - |
| - | - | 698.3 | 208.1 | - | - | 0 | - |
| - | - | 1462 | 209.1 | - | - | 0 | - |
| - | - | 1278 | 210.2 | - | - | 0 | - |
| - | - | 1586 | 214.1 | - | - | 0 | - |
| - | - | 7049 | 215.1 | - | - | 0 | - |
| - | - | 742.4 | 216.1 | - | - | 0 | - |
| - | - | 2594 | 217.1 | - | - | 0 | - |
| - | - | 2596 | 217.1 | - | - | 0 | - |
| - | - | 666.1 | 218.2 | - | - | 0 | - |
| - | - | 8336 | 219.1 | - | - | 0 | - |
| - | - | 700.9 | 219.1 | - | - | 0 | - |
| - | - | 1041 | 220.1 | - | - | 0 | - |
| - | - | 9339 | 221.1 | - | - | 0 | - |
| 2 | a | 1.606E+05 | 221.1 | 0.0004673 | 2.113 | +1 | 2 |
| - | - | 819.2 | 222.1 | - | - | 0 | - |
| - | - | 1.565E+04 | 222.1 | - | - | 0 | - |
| - | - | 1131 | 223.1 | - | - | 0 | - |
| - | - | 706.3 | 223.1 | - | - | 0 | - |
| - | - | 7188 | 225.2 | - | - | 0 | - |
| - | - | 648 | 226.1 | - | - | 0 | - |
| - | - | 2116 | 226.2 | - | - | 0 | - |
| - | - | 566 | 227.1 | - | - | 0 | - |
| - | - | 892 | 227.1 | - | - | 0 | - |
| - | - | 1273 | 227.1 | - | - | 0 | - |
| - | - | 936 | 228.2 | - | - | 0 | - |
| - | - | 5102 | 229.1 | - | - | 0 | - |
| - | - | 1396 | 229.2 | - | - | 0 | - |
| - | - | 892.3 | 229.2 | - | - | 0 | - |
| - | - | 888.6 | 231 | - | - | 0 | - |
| - | - | 4285 | 231.1 | - | - | 0 | - |
| 2 | b | 1538 | 231.1 | 0.0007212 | 3.121 | +1 | 2 |
| - | - | 669.2 | 232.1 | - | - | 0 | - |
| - | - | 1245 | 233.1 | - | - | 0 | - |
| - | - | 812.3 | 233.1 | - | - | 0 | - |
| - | - | 5942 | 235.1 | - | - | 0 | - |
| - | - | 670.2 | 238.2 | - | - | 0 | - |
| - | - | 743.6 | 239.1 | - | - | 0 | - |
| - | - | 1121 | 242.1 | - | - | 0 | - |
| - | - | 2222 | 248.1 | - | - | 0 | - |
| - | - | 745.2 | 249.1 | - | - | 0 | - |
| 2 | b | 3.039E+04 | 249.1 | 0.0004257 | 1.709 | +1 | 2 |
| - | - | 2933 | 250.1 | - | - | 0 | - |
| - | - | 5026 | 251.1 | - | - | 0 | - |
| - | - | 5000 | 253.1 | - | - | 0 | - |
| - | - | 1.121E+04 | 253.2 | - | - | 0 | - |
| - | - | 805.3 | 254.1 | - | - | 0 | - |
| - | - | 702.4 | 254.1 | - | - | 0 | - |
| - | - | 679.3 | 254.2 | - | - | 0 | - |
| - | - | 733.7 | 256.1 | - | - | 0 | - |
| - | - | 1188 | 257.1 | - | - | 0 | - |
| - | - | 746.2 | 257.2 | - | - | 0 | - |
| - | - | 5334 | 263.1 | - | - | 0 | - |
| - | - | 904.4 | 263.1 | - | - | 0 | - |
| - | - | 615.3 | 264.1 | - | - | 0 | - |
| - | - | 580.8 | 269.1 | - | - | 0 | - |
| - | - | 2700 | 270.1 | - | - | 0 | - |
| - | - | 1726 | 270.2 | - | - | 0 | - |
| - | - | 882.5 | 271.1 | - | - | 0 | - |
| - | - | 1592 | 274.1 | - | - | 0 | - |
| - | - | 629.8 | 275.1 | - | - | 0 | - |
| - | - | 1307 | 275.2 | - | - | 0 | - |
| - | - | 5343 | 279.1 | - | - | 0 | - |
| - | - | 690.8 | 280.2 | - | - | 0 | - |
| - | - | 5270 | 283.1 | - | - | 0 | - |
| - | - | 758.9 | 284.1 | - | - | 0 | - |
| - | - | 1268 | 284.1 | - | - | 0 | - |
| - | - | 919.8 | 284.2 | - | - | 0 | - |
| - | - | 973.5 | 285 | - | - | 0 | - |
| - | - | 706.7 | 285.1 | - | - | 0 | - |
| - | - | 3388 | 285.1 | - | - | 0 | - |
| - | - | 1116 | 286.1 | - | - | 0 | - |
| - | - | 625.4 | 287.2 | - | - | 0 | - |
| - | - | 819.9 | 288.1 | - | - | 0 | - |
| - | - | 2057 | 292.1 | - | - | 0 | - |
| - | - | 593.5 | 295.2 | - | - | 0 | - |
| - | - | 715.5 | 296.1 | - | - | 0 | - |
| - | - | 2257 | 296.2 | - | - | 0 | - |
| - | - | 2561 | 296.2 | - | - | 0 | - |
| - | - | 1262 | 297.2 | - | - | 0 | - |
| - | - | 2193 | 302.1 | - | - | 0 | - |
| - | - | 901.1 | 303.2 | - | - | 0 | - |
| - | - | 1717 | 312.2 | - | - | 0 | - |
| - | - | 3147 | 314.1 | - | - | 0 | - |
| - | - | 743.3 | 314.2 | - | - | 0 | - |
| - | - | 1238 | 315.2 | - | - | 0 | - |
| - | - | 725.5 | 318.1 | - | - | 0 | - |
| 3 | a | 695.5 | 318.1 | 0.0003624 | 1.139 | +1 | 3 |
| - | - | 3382 | 319.1 | - | - | 0 | - |
| - | - | 673.4 | 320.1 | - | - | 0 | - |
| - | - | 1136 | 320.2 | - | - | 0 | - |
| - | - | 1285 | 322.2 | - | - | 0 | - |
| - | - | 1005 | 323.1 | - | - | 0 | - |
| - | - | 1464 | 323.2 | - | - | 0 | - |
| - | - | 6052 | 324.1 | - | - | 0 | - |
| - | - | 549.5 | 325.1 | - | - | 0 | - |
| - | - | 1040 | 326.2 | - | - | 0 | - |
| - | - | 626.1 | 328.1 | - | - | 0 | - |
| - | - | 1.306E+05 | 329.2 | - | - | 0 | - |
| - | - | 3649 | 329.2 | - | - | 0 | - |
| - | - | 2.371E+04 | 330.2 | - | - | 0 | - |
| - | - | 1001 | 330.2 | - | - | 0 | - |
| - | - | 2672 | 331.2 | - | - | 0 | - |
| - | - | 1167 | 332.1 | - | - | 0 | - |
| - | - | 1512 | 338.1 | - | - | 0 | - |
| - | - | 608.1 | 339.2 | - | - | 0 | - |
| - | - | 1286 | 339.2 | - | - | 0 | - |
| - | - | 2.479E+04 | 340.2 | - | - | 0 | - |
| - | - | 3603 | 341.2 | - | - | 0 | - |
| - | - | 614.3 | 341.7 | - | - | 0 | - |
| 8 | y | 3.279E+04 | 342.2 | 0.0006113 | 1.787 | +1 | 2 |
| - | - | 5054 | 343.2 | - | - | 0 | - |
| - | - | 2923 | 344.1 | - | - | 0 | - |
| 3 | b | 1897 | 346.1 | 7.199E-06 | 0.0208 | +1 | 3 |
| - | - | 645.7 | 346.2 | - | - | 0 | - |
| - | - | 1007 | 348.1 | - | - | 0 | - |
| - | - | 646.1 | 348.7 | - | - | 0 | - |
| - | - | 2196 | 350.1 | - | - | 0 | - |
| - | - | 802.9 | 350.7 | - | - | 0 | - |
| - | - | 1096 | 352.1 | - | - | 0 | - |
| - | - | 698.1 | 354.2 | - | - | 0 | - |
| - | - | 1174 | 355.1 | - | - | 0 | - |
| - | - | 1902 | 356.1 | - | - | 0 | - |
| - | - | 2.691E+04 | 357.2 | - | - | 0 | - |
| - | - | 993.1 | 357.7 | - | - | 0 | - |
| - | - | 4667 | 358.2 | - | - | 0 | - |
| 3 | b | 7047 | 364.2 | 0.0003534 | 0.9705 | +1 | 3 |
| - | - | 1431 | 365.2 | - | - | 0 | - |
| - | - | 1517 | 366.2 | - | - | 0 | - |
| - | - | 906.7 | 366.3 | - | - | 0 | - |
| - | - | 9064 | 368.2 | - | - | 0 | - |
| - | - | 1135 | 369.2 | - | - | 0 | - |
| - | - | 1052 | 369.7 | - | - | 0 | - |
| - | - | 950.3 | 375.2 | - | - | 0 | - |
| - | - | 929.2 | 376.1 | - | - | 0 | - |
| - | - | 4998 | 378.1 | - | - | 0 | - |
| - | - | 1021 | 378.2 | - | - | 0 | - |
| - | - | 831.1 | 378.7 | - | - | 0 | - |
| - | - | 1054 | 379.1 | - | - | 0 | - |
| - | - | 842.5 | 380.1 | - | - | 0 | - |
| - | - | 1619 | 383.2 | - | - | 0 | - |
| - | - | 868.7 | 383.7 | - | - | 0 | - |
| - | - | 1086 | 385.1 | - | - | 0 | - |
| - | - | 3229 | 385.2 | - | - | 0 | - |
| - | - | 1996 | 387.2 | - | - | 0 | - |
| - | - | 7667 | 392.2 | - | - | 0 | - |
| - | - | 4338 | 392.7 | - | - | 0 | - |
| - | - | 6265 | 394.1 | - | - | 0 | - |
| - | - | 863 | 394.2 | - | - | 0 | - |
| - | - | 1461 | 395.1 | - | - | 0 | - |
| - | - | 1576 | 398.2 | - | - | 0 | - |
| - | - | 1861 | 400.2 | - | - | 0 | - |
| - | - | 735.3 | 401.2 | - | - | 0 | - |
| - | - | 5770 | 401.2 | - | - | 0 | - |
| - | - | 2876 | 401.7 | - | - | 0 | - |
| - | - | 833.1 | 402.2 | - | - | 0 | - |
| - | - | 956.2 | 403.1 | - | - | 0 | - |
| - | - | 1975 | 405.2 | - | - | 0 | - |
| - | - | 782.1 | 405.7 | - | - | 0 | - |
| - | - | 711.1 | 406.2 | - | - | 0 | - |
| - | - | 1610 | 408.2 | - | - | 0 | - |
| - | - | 1820 | 410.3 | - | - | 0 | - |
| - | - | 2441 | 411.2 | - | - | 0 | - |
| - | - | 2307 | 413.7 | - | - | 0 | - |
| - | - | 1901 | 414.2 | - | - | 0 | - |
| - | - | 1224 | 414.7 | - | - | 0 | - |
| - | - | 1449 | 418.2 | - | - | 0 | - |
| - | - | 2633 | 418.7 | - | - | 0 | - |
| - | - | 803.7 | 419.2 | - | - | 0 | - |
| - | - | 1204 | 419.2 | - | - | 0 | - |
| - | - | 2897 | 422.7 | - | - | 0 | - |
| - | - | 2063 | 423.2 | - | - | 0 | - |
| - | - | 705.9 | 424.2 | - | - | 0 | - |
| - | - | 3462 | 425.2 | - | - | 0 | - |
| - | - | 2069 | 425.3 | - | - | 0 | - |
| - | - | 858.9 | 426.2 | - | - | 0 | - |
| - | - | 1073 | 426.2 | - | - | 0 | - |
| - | - | 889.8 | 426.3 | - | - | 0 | - |
| - | - | 1485 | 427.2 | - | - | 0 | - |
| - | - | 846.1 | 427.2 | - | - | 0 | - |
| - | - | 1.673E+04 | 427.7 | - | - | 0 | - |
| - | - | 6635 | 428.2 | - | - | 0 | - |
| - | - | 1097 | 428.3 | - | - | 0 | - |
| - | - | 1238 | 428.7 | - | - | 0 | - |
| - | - | 971.7 | 433.2 | - | - | 0 | - |
| - | - | 1.702E+04 | 436.7 | - | - | 0 | - |
| - | - | 8159 | 437.2 | - | - | 0 | - |
| - | - | 2436 | 437.7 | - | - | 0 | - |
| - | - | 3806 | 442.3 | - | - | 0 | - |
| - | - | 1748 | 443.2 | - | - | 0 | - |
| - | - | 766.9 | 443.3 | - | - | 0 | - |
| - | - | 984.7 | 443.7 | - | - | 0 | - |
| - | - | 1008 | 444.7 | - | - | 0 | - |
| - | - | 1118 | 445.2 | - | - | 0 | - |
| - | - | 818.1 | 447.2 | - | - | 0 | - |
| - | - | 1585 | 448.3 | - | - | 0 | - |
| - | - | 1993 | 448.8 | - | - | 0 | - |
| - | - | 1924 | 452.2 | - | - | 0 | - |
| - | - | 1277 | 452.7 | - | - | 0 | - |
| - | - | 3184 | 453.3 | - | - | 0 | - |
| - | - | 952.3 | 453.7 | - | - | 0 | - |
| - | - | 881.9 | 455.2 | - | - | 0 | - |
| - | - | 675.5 | 460.2 | - | - | 0 | - |
| - | - | 761.9 | 460.3 | - | - | 0 | - |
| 4 | b | 1760 | 461.2 | 0.0004851 | 1.052 | +1 | 4 |
| - | - | 5133 | 465.2 | - | - | 0 | - |
| - | - | 1206 | 465.2 | - | - | 0 | - |
| - | - | 964 | 465.7 | - | - | 0 | - |
| - | - | 827.2 | 466.2 | - | - | 0 | - |
| - | - | 699.5 | 467.2 | - | - | 0 | - |
| - | - | 1036 | 469.3 | - | - | 0 | - |
| - | - | 5030 | 470.3 | - | - | 0 | - |
| - | - | 789.7 | 471.2 | - | - | 0 | - |
| - | - | 1443 | 471.3 | - | - | 0 | - |
| - | - | 1548 | 474.7 | - | - | 0 | - |
| 3 | y | 859.2 | 477.2 | 0.005254 | 11.01 | +2 | 7 |
| - | - | 1663 | 478.7 | - | - | 0 | - |
| 4 | b | 4002 | 479.2 | 0.0006931 | 1.446 | +1 | 4 |
| - | - | 2136 | 479.2 | - | - | 0 | - |
| - | - | 941 | 481.3 | - | - | 0 | - |
| - | - | 1.235E+04 | 483.2 | - | - | 0 | - |
| - | - | 2998 | 484.2 | - | - | 0 | - |
| - | - | 744.7 | 486.2 | - | - | 0 | - |
| - | - | 2560 | 487.3 | - | - | 0 | - |
| - | - | 7221 | 487.3 | - | - | 0 | - |
| - | - | 1639 | 487.8 | - | - | 0 | - |
| - | - | 1172 | 488.3 | - | - | 0 | - |
| - | - | 2068 | 488.3 | - | - | 0 | - |
| 7 | y | 8272 | 489.2 | 0.005466 | 11.17 | +1 | 3 |
| - | - | 1870 | 490.2 | - | - | 0 | - |
| - | - | 1446 | 491.2 | - | - | 0 | - |
| - | - | 738.6 | 492.3 | - | - | 0 | - |
| - | - | 782.4 | 495.2 | - | - | 0 | - |
| - | - | 2298 | 496.2 | - | - | 0 | - |
| - | - | 5378 | 496.3 | - | - | 0 | - |
| - | - | 3696 | 496.8 | - | - | 0 | - |
| - | - | 747.1 | 497.2 | - | - | 0 | - |
| - | - | 709.4 | 498.2 | - | - | 0 | - |
| - | - | 1971 | 498.3 | - | - | 0 | - |
| - | - | 4650 | 500.2 | - | - | 0 | - |
| - | - | 6282 | 501.3 | - | - | 0 | - |
| - | - | 4768 | 501.8 | - | - | 0 | - |
| - | - | 757.9 | 502.3 | - | - | 0 | - |
| - | - | 1021 | 504.3 | - | - | 0 | - |
| - | - | 7603 | 510.3 | - | - | 0 | - |
| - | - | 4431 | 510.8 | - | - | 0 | - |
| - | - | 1646 | 511.3 | - | - | 0 | - |
| - | - | 786.5 | 513.2 | - | - | 0 | - |
| - | - | 8367 | 513.4 | - | - | 0 | - |
| - | - | 740 | 514.2 | - | - | 0 | - |
| - | - | 2156 | 514.4 | - | - | 0 | - |
| - | - | 1082 | 516.2 | - | - | 0 | - |
| - | - | 1226 | 519.2 | - | - | 0 | - |
| - | - | 849.4 | 520.2 | - | - | 0 | - |
| - | - | 612.4 | 527.6 | - | - | 0 | - |
| - | - | 976.7 | 529.3 | - | - | 0 | - |
| - | - | 677.2 | 531.3 | - | - | 0 | - |
| - | - | 1124 | 540.3 | - | - | 0 | - |
| - | - | 1499 | 548.2 | - | - | 0 | - |
| - | - | 976.1 | 550.2 | - | - | 0 | - |
| - | - | 1183 | 550.3 | - | - | 0 | - |
| 2 | y | 5873 | 550.7 | 0.003446 | 6.258 | +2 | 8 |
| - | - | 2872 | 551.2 | - | - | 0 | - |
| - | - | 776.7 | 552.2 | - | - | 0 | - |
| - | - | 1902 | 556.3 | - | - | 0 | - |
| - | - | 737.9 | 556.3 | - | - | 0 | - |
| - | - | 722.5 | 557.2 | - | - | 0 | - |
| 6 | y | 1.14E+04 | 560.2 | 0.005279 | 9.422 | +1 | 4 |
| - | - | 3720 | 561.2 | - | - | 0 | - |
| - | - | 632.8 | 564.7 | - | - | 0 | - |
| - | - | 2189 | 566.3 | - | - | 0 | - |
| - | - | 1442 | 567.3 | - | - | 0 | - |
| - | - | 2936 | 568.3 | - | - | 0 | - |
| - | - | 969 | 569.2 | - | - | 0 | - |
| - | - | 1154 | 569.3 | - | - | 0 | - |
| - | - | 929.9 | 569.7 | - | - | 0 | - |
| - | - | 1005 | 569.8 | - | - | 0 | - |
| - | - | 728.8 | 573.3 | - | - | 0 | - |
| - | - | 1479 | 575.3 | - | - | 0 | - |
| - | - | 3805 | 583.3 | - | - | 0 | - |
| - | - | 835.4 | 584.3 | - | - | 0 | - |
| - | - | 4.95E+04 | 585.3 | - | - | 0 | - |
| - | - | 1.761E+04 | 586.3 | - | - | 0 | - |
| - | - | 2903 | 587.3 | - | - | 0 | - |
| - | - | 1310 | 590.8 | - | - | 0 | - |
| - | - | 870.1 | 591.8 | - | - | 0 | - |
| 0 | Precursor | 1417 | 592.2 | 0.001613 | 2.724 | +2 | -1 |
| - | - | 1932 | 592.7 | - | - | 0 | - |
| - | - | 823.2 | 593.2 | - | - | 0 | - |
| - | - | 999.3 | 596.2 | - | - | 0 | - |
| - | - | 853 | 596.3 | - | - | 0 | - |
| - | - | 1529 | 599.9 | - | - | 0 | - |
| - | - | 1002 | 600.3 | - | - | 0 | - |
| 0 | Precursor | 6666 | 601.2 | 0.003045 | 5.064 | +2 | -1 |
| - | - | 1653 | 601.3 | - | - | 0 | - |
| - | - | 4146 | 601.7 | - | - | 0 | - |
| - | - | 1165 | 602.2 | - | - | 0 | - |
| - | - | 6475 | 602.4 | - | - | 0 | - |
| - | - | 3611 | 603.4 | - | - | 0 | - |
| - | - | 679.8 | 604.4 | - | - | 0 | - |
| - | - | 906.8 | 606.2 | - | - | 0 | - |
| - | - | 1947 | 612.2 | - | - | 0 | - |
| - | - | 2086 | 613.3 | - | - | 0 | - |
| - | - | 1127 | 614.2 | - | - | 0 | - |
| - | - | 1270 | 614.3 | - | - | 0 | - |
| - | - | 742.6 | 615.8 | - | - | 0 | - |
| 5 | b | 911.4 | 624.2 | 3.829E-05 | 0.06134 | +1 | 5 |
| - | - | 1153 | 628.3 | - | - | 0 | - |
| - | - | 1.459E+04 | 628.4 | - | - | 0 | - |
| - | - | 4292 | 629.4 | - | - | 0 | - |
| - | - | 856.8 | 630.4 | - | - | 0 | - |
| - | - | 1591 | 634.2 | - | - | 0 | - |
| - | - | 710.3 | 635.2 | - | - | 0 | - |
| - | - | 659.6 | 640.2 | - | - | 0 | - |
| - | - | 1513 | 644.3 | - | - | 0 | - |
| - | - | 806.2 | 655.3 | - | - | 0 | - |
| - | - | 1560 | 656.4 | - | - | 0 | - |
| - | - | 2906 | 657.3 | - | - | 0 | - |
| - | - | 4259 | 659.3 | - | - | 0 | - |
| - | - | 1093 | 660.3 | - | - | 0 | - |
| - | - | 816.3 | 663.3 | - | - | 0 | - |
| - | - | 1499 | 665.3 | - | - | 0 | - |
| - | - | 1786 | 670.4 | - | - | 0 | - |
| - | - | 740.5 | 673.3 | - | - | 0 | - |
| - | - | 830.9 | 676.3 | - | - | 0 | - |
| - | - | 731.3 | 678.4 | - | - | 0 | - |
| - | - | 6917 | 682.4 | - | - | 0 | - |
| - | - | 3356 | 683.3 | - | - | 0 | - |
| - | - | 4114 | 685.3 | - | - | 0 | - |
| - | - | 660.6 | 686.3 | - | - | 0 | - |
| - | - | 830.7 | 690.3 | - | - | 0 | - |
| 6 | b | 1083 | 695.3 | 4.27E-05 | 0.06142 | +1 | 6 |
| - | - | 2460 | 696.4 | - | - | 0 | - |
| - | - | 851 | 697.4 | - | - | 0 | - |
| - | - | 1.257E+05 | 700.4 | - | - | 0 | - |
| - | - | 4.344E+04 | 701.4 | - | - | 0 | - |
| - | - | 8064 | 702.4 | - | - | 0 | - |
| - | - | 775 | 703.3 | - | - | 0 | - |
| - | - | 943.8 | 712.3 | - | - | 0 | - |
| - | - | 848.5 | 714.4 | - | - | 0 | - |
| - | - | 816.6 | 718.4 | - | - | 0 | - |
| - | - | 4145 | 721.3 | - | - | 0 | - |
| - | - | 1146 | 722.3 | - | - | 0 | - |
| 5 | y | 1.151E+04 | 723.3 | 0.005121 | 7.081 | +1 | 5 |
| - | - | 4920 | 724.3 | - | - | 0 | - |
| - | - | 1030 | 725.3 | - | - | 0 | - |
| - | - | 2272 | 726.4 | - | - | 0 | - |
| - | - | 1094 | 727.4 | - | - | 0 | - |
| - | - | 1773 | 729.4 | - | - | 0 | - |
| - | - | 797.9 | 732.4 | - | - | 0 | - |
| - | - | 1002 | 739.4 | - | - | 0 | - |
| - | - | 1024 | 741.4 | - | - | 0 | - |
| - | - | 1033 | 742.4 | - | - | 0 | - |
| - | - | 5573 | 749.3 | - | - | 0 | - |
| - | - | 1433 | 750.3 | - | - | 0 | - |
| - | - | 1133 | 754.4 | - | - | 0 | - |
| - | - | 1460 | 755.4 | - | - | 0 | - |
| - | - | 2325 | 756.4 | - | - | 0 | - |
| - | - | 1.259E+04 | 757.4 | - | - | 0 | - |
| - | - | 5184 | 758.4 | - | - | 0 | - |
| - | - | 1899 | 759.4 | - | - | 0 | - |
| - | - | 1752 | 766.4 | - | - | 0 | - |
| - | - | 829.3 | 767.3 | - | - | 0 | - |
| - | - | 1036 | 767.4 | - | - | 0 | - |
| - | - | 1959 | 771.4 | - | - | 0 | - |
| - | - | 4351 | 772.4 | - | - | 0 | - |
| - | - | 1.046E+04 | 773.4 | - | - | 0 | - |
| - | - | 4352 | 774.3 | - | - | 0 | - |
| - | - | 3281 | 774.4 | - | - | 0 | - |
| - | - | 2011 | 775.3 | - | - | 0 | - |
| - | - | 1348 | 775.4 | - | - | 0 | - |
| - | - | 930 | 782.4 | - | - | 0 | - |
| - | - | 3.536E+04 | 783.4 | - | - | 0 | - |
| - | - | 1.526E+04 | 784.4 | - | - | 0 | - |
| - | - | 5489 | 785.4 | - | - | 0 | - |
| - | - | 1291 | 791.4 | - | - | 0 | - |
| - | - | 1014 | 792.4 | - | - | 0 | - |
| - | - | 1.952E+04 | 800.4 | - | - | 0 | - |
| - | - | 6.48E+04 | 801.4 | - | - | 0 | - |
| - | - | 2.557E+04 | 802.4 | - | - | 0 | - |
| - | - | 6686 | 803.4 | - | - | 0 | - |
| - | - | 1410 | 804.4 | - | - | 0 | - |
| - | - | 929.8 | 809.4 | - | - | 0 | - |
| - | - | 899.6 | 810.4 | - | - | 0 | - |
| - | - | 1489 | 814.4 | - | - | 0 | - |
| - | - | 950.7 | 819.4 | - | - | 0 | - |
| - | - | 1034 | 821.3 | - | - | 0 | - |
| - | - | 3718 | 826.4 | - | - | 0 | - |
| - | - | 1642 | 827.4 | - | - | 0 | - |
| - | - | 2.62E+04 | 828.4 | - | - | 0 | - |
| - | - | 1.111E+04 | 829.4 | - | - | 0 | - |
| - | - | 2146 | 830.4 | - | - | 0 | - |
| - | - | 1936 | 837.4 | - | - | 0 | - |
| 4 | y | 1.563E+04 | 838.3 | 0.005522 | 6.587 | +1 | 6 |
| - | - | 6470 | 839.3 | - | - | 0 | - |
| - | - | 1132 | 840.3 | - | - | 0 | - |
| - | - | 8.928E+04 | 844.5 | - | - | 0 | - |
| - | - | 4.04E+04 | 845.5 | - | - | 0 | - |
| - | - | 9667 | 846.5 | - | - | 0 | - |
| - | - | 3109 | 847.4 | - | - | 0 | - |
| - | - | 1159 | 848.4 | - | - | 0 | - |
| - | - | 1.516E+04 | 854.4 | - | - | 0 | - |
| - | - | 6664 | 855.4 | - | - | 0 | - |
| - | - | 2170 | 856.4 | - | - | 0 | - |
| - | - | 1041 | 857.5 | - | - | 0 | - |
| - | - | 1963 | 871.3 | - | - | 0 | - |
| - | - | 843 | 871.5 | - | - | 0 | - |
| - | - | 3.677E+05 | 872.4 | - | - | 0 | - |
| - | - | 1.622E+05 | 873.5 | - | - | 0 | - |
| - | - | 3.892E+04 | 874.5 | - | - | 0 | - |
| - | - | 1210 | 874.6 | - | - | 0 | - |
| - | - | 3468 | 875.5 | - | - | 0 | - |
| - | - | 707.5 | 885.4 | - | - | 0 | - |
| - | - | 1693 | 888.4 | - | - | 0 | - |
| - | - | 1.957E+04 | 889.3 | - | - | 0 | - |
| - | - | 1.011E+04 | 890.4 | - | - | 0 | - |
| - | - | 1796 | 891.4 | - | - | 0 | - |
| - | - | 2845 | 895.5 | - | - | 0 | - |
| - | - | 2122 | 896.5 | - | - | 0 | - |
| - | - | 1347 | 903.5 | - | - | 0 | - |
| - | - | 1186 | 904.5 | - | - | 0 | - |
| - | - | 1544 | 906.4 | - | - | 0 | - |
| - | - | 1323 | 907.3 | - | - | 0 | - |
| - | - | 1129 | 918.5 | - | - | 0 | - |
| - | - | 3152 | 919.5 | - | - | 0 | - |
| - | - | 2037 | 920.5 | - | - | 0 | - |
| - | - | 1100 | 930.5 | - | - | 0 | - |
| - | - | 924.9 | 932.5 | - | - | 0 | - |
| 3 | y | 4158 | 935.3 | 0.002633 | 2.815 | +1 | 7 |
| - | - | 2482 | 936.3 | - | - | 0 | - |
| - | - | 944 | 947.5 | - | - | 0 | - |
| - | - | 1997 | 948.5 | - | - | 0 | - |
| - | - | 1227 | 949.5 | - | - | 0 | - |
| 3 | y | 7.066E+04 | 953.3 | 0.005251 | 5.508 | +1 | 7 |
| - | - | 3.504E+04 | 954.3 | - | - | 0 | - |
| - | - | 8768 | 955.3 | - | - | 0 | - |
| - | - | 742.8 | 955.4 | - | - | 0 | - |
| - | - | 684.7 | 965.4 | - | - | 0 | - |
| - | - | 1963 | 973.5 | - | - | 0 | - |
| - | - | 840.4 | 974.5 | - | - | 0 | - |
| - | - | 1061 | 975.5 | - | - | 0 | - |
| - | - | 2549 | 983.4 | - | - | 0 | - |
| - | - | 2256 | 984.4 | - | - | 0 | - |
| - | - | 6.15E+04 | 991.5 | - | - | 0 | - |
| - | - | 3.334E+04 | 992.5 | - | - | 0 | - |
| - | - | 9114 | 993.5 | - | - | 0 | - |
| - | - | 1024 | 994.5 | - | - | 0 | - |
| 8 | b | 1155 | 997.4 | 0.0008873 | 0.8896 | +1 | 8 |
| - | - | 2446 | 1000 | - | - | 0 | - |
| - | - | 2558 | 1001 | - | - | 0 | - |
| - | - | 1102 | 1003 | - | - | 0 | - |
| - | - | 1.611E+05 | 1020 | - | - | 0 | - |
| - | - | 8.812E+04 | 1021 | - | - | 0 | - |
| - | - | 2.909E+04 | 1022 | - | - | 0 | - |
| - | - | 1037 | 1022 | - | - | 0 | - |
| - | - | 2749 | 1023 | - | - | 0 | - |
| - | - | 1187 | 1036 | - | - | 0 | - |
| - | - | 856 | 1037 | - | - | 0 | - |
| - | - | 762.2 | 1038 | - | - | 0 | - |
| 2 | y | 3500 | 1100 | 0.004892 | 4.445 | +1 | 8 |
| - | - | 1728 | 1101 | - | - | 0 | - |
| - | - | 640.3 | 1525 | - | - | 0 | - |
| - | - | 750.9 | 1915 | - | - | 0 | - |
| - | - | 679.5 | 2901 | - | - | 0 | - |

m/z Charge Intensity FragmentType MassShift Position
120.08121490478516 0 498354.66
121.06515502929688 0 573.436
121.08446502685547 0 40579.89
122.08771514892578 0 965.4787
123.04439544677734 0 11745.228
124.04769134521484 0 856.2674
125.15376281738281 0 364.58615
127.08690643310547 0 1402.472
128.10733032226562 0 1990.3306
129.0662841796875 0 440.86044
129.1025390625 0 3556.5122
130.06544494628906 0 3209.6033
131.04965209960938 0 626.2415
131.08169555664062 0 607.17084
131.1182861328125 0 1546.7372
132.0811767578125 0 2388.3438
132.1023406982422 0 1163.1564
134.02732849121094 0 1125.0161
136.07608032226562 0 131411.05
137.07940673828125 0 9818.034
138.05551147460938 0 821.71906
138.0666046142578 0 1414.926
139.00694274902344 0 433.1157
140.0823211669922 0 966.139
141.1027069091797 0 831.90424
143.1185302734375 0 550.2594
144.06541442871094 0 497.2014
145.0974578857422 0 3140.6667
146.0604705810547 0 1206.688
147.04440307617188 0 9229.137
148.04808044433594 0 820.8061
149.0452423095703 0 860.0201
153.066162109375 0 602.6939
155.08189392089844 0 2708.0269
155.1183319091797 0 896.9594
156.07713317871094 0 4974.737
157.06101989746094 0 565.9807
157.09730529785156 0 680.9425
157.10873413085938 0 2787.4304
157.1339569091797 0 1739.0642
158.09671020507812 0 5611.186
159.09201049804688 0 5328.5386
159.0990753173828 0 802.662
160.0760040283203 0 1116.8398
160.11248779296875 0 2231.5735
162.09207153320312 0 999.26086
162.10226440429688 0 461.97705
164.0825653076172 0 429.57013
165.0549774169922 0 91542.24
166.05841064453125 0 8432.096
166.08660888671875 0 6434.536
166.09396362304688 0 572.7938
166.09776306152344 0 462.64078
167.0560760498047 0 1493.559
167.09300231933594 0 734.1603
167.11798095703125 0 508.0989
168.92257690429688 0 436.82138
169.0975341796875 0 527.6401
169.13389587402344 0 1372.8685
171.07704162597656 0 1696.3685
171.11317443847656 0 670.87695
173.09243774414062 0 3050.4673
173.1288299560547 0 40415.01
174.13226318359375 0 2878.76
175.1190948486328 0 539.8318
176.08218383789062 0 1353.6343
176.10739135742188 0 11516.168
177.10263061523438 0 7502.3345
177.11122131347656 0 1329.2405
180.07740783691406 0 528.73706
181.09823608398438 0 980.1898
182.0816192626953 0 180133.83
183.08494567871094 0 15989.723
183.1128692626953 0 984.621
183.1494140625 0 826.6829
184.08787536621094 0 818.1384
185.05662536621094 0 1144.2794
185.1285858154297 0 2362.5544
186.091552734375 0 938.62805
187.1448516845703 0 615.829
188.07093811035156 0 5686.6157
189.08729553222656 0 2664.6152
191.11830139160156 0 14601.816
192.10284423828125 0 936.80023
192.12147521972656 0 1127.5677
192.14962768554688 0 567.3201
193.1087646484375 0 3571.0996
195.0877685546875 0 1123.2561
197.1284637451172 0 1599.773
199.07164001464844 0 3398.1465
199.14437866210938 0 2053.9468
201.12364196777344 0 24773.475
202.126953125 0 2268.2751
203.0663604736328 0 1280.1564
203.11827087402344 0 3615.6658 a Water loss 1
204.07691955566406 0 1248.899
205.08108520507812 0 561.15894
205.09751892089844 0 4132.6436 y 8
207.11305236816406 0 2588.5515
208.14456176757812 0 698.3155
209.0923614501953 0 1461.9004
210.1601104736328 0 1278.2864
214.1190643310547 0 1585.6865
215.13937377929688 0 7048.8506
216.14321899414062 0 742.3847
217.08201599121094 0 2593.9128
217.0976104736328 0 2595.5989
218.15003967285156 0 666.0787
219.11326599121094 0 8335.778
219.1345672607422 0 700.86774
220.11614990234375 0 1040.8953
221.10365295410156 0 9339.056
221.12892150878906 0 160599.58 a 1
222.1072998046875 0 819.22144
222.13221740722656 0 15654.93
223.10736083984375 0 1131.2861
223.1351776123047 0 706.3464
225.1714630126953 0 7187.782
226.11891174316406 0 647.97784
226.1553955078125 0 2116.253
227.09051513671875 0 565.9674
227.10281372070312 0 891.9921
227.138671875 0 1272.9099
228.17115783691406 0 935.9595
229.1186981201172 0 5101.693
229.15493774414062 0 1396.4843
229.16917419433594 0 892.34515
231.04331970214844 0 888.58527
231.0615692138672 0 4285.295
231.113525390625 0 1537.7262 b Water loss 1
232.11581420898438 0 669.16425
233.09219360351562 0 1244.6493
233.13156127929688 0 812.3155
235.10821533203125 0 5942.2397
238.15611267089844 0 670.1565
239.11456298828125 0 743.5756
242.11412048339844 0 1120.825
248.1075439453125 0 2221.9688
249.05393981933594 0 745.2163
249.12379455566406 0 30387.795 b 1
250.1273193359375 0 2932.6218
251.10313415527344 0 5025.765
253.11863708496094 0 5000.2104
253.1663818359375 0 11209.474
254.12265014648438 0 805.30145
254.149658203125 0 702.44727
254.1704559326172 0 679.2616
256.10906982421875 0 733.745
257.106689453125 0 1187.9128
257.1656188964844 0 746.2345
263.10296630859375 0 5333.8916
263.1282958984375 0 904.3682
264.10687255859375 0 615.29297
269.07635498046875 0 580.79675
270.1088562011719 0 2700.119
270.1929626464844 0 1726.2648
271.1100769042969 0 882.5331
274.12982177734375 0 1591.9114
275.1392822265625 0 629.8416
275.17559814453125 0 1306.7988
279.097900390625 0 5342.755
280.16668701171875 0 690.75726
283.1445617675781 0 5269.9775
284.1019287109375 0 758.85614
284.14764404296875 0 1268.3268
284.1974182128906 0 919.82684
285.0111999511719 0 973.48444
285.087158203125 0 706.6999
285.10211181640625 0 3387.8506
286.1045227050781 0 1116.1831
287.21942138671875 0 625.37994
288.1194152832031 0 819.9196
292.1406555175781 0 2056.9226
295.178466796875 0 593.4737
296.0880432128906 0 715.5475
296.1508483886719 0 2257.2666
296.20880126953125 0 2560.9841
297.1932067871094 0 1262.0189
302.11419677734375 0 2193.2937
303.1707458496094 0 901.149
312.2036437988281 0 1716.5485
314.098876953125 0 3147.296
314.2190856933594 0 743.30237
315.20306396484375 0 1237.5481
318.075439453125 0 725.5052
318.14447021484375 0 695.506 a Water loss 2
319.1409606933594 0 3382.4353
320.12652587890625 0 673.4265
320.1609191894531 0 1136.4685
322.1875 0 1285.1941
323.09881591796875 0 1004.69745
323.171630859375 0 1464.2548
324.146240234375 0 6051.541
325.1499938964844 0 549.5344
326.1712951660156 0 1039.8685
328.12884521484375 0 626.051
329.15020751953125 0 130641.03
329.22991943359375 0 3648.6465
330.15350341796875 0 23708.293
330.1756896972656 0 1001.1656
331.1560974121094 0 2672.4878
332.1079406738281 0 1167.056
338.1343078613281 0 1512.4329
339.1713562011719 0 608.14185
339.2146301269531 0 1285.5154
340.1985168457031 0 24790.793
341.2012023925781 0 3603.267
341.6800537109375 0 614.3001
342.15667724609375 0 32792.734 y 7
343.15966796875 0 5053.639
344.1456298828125 0 2922.9639
346.1397399902344 0 1897.0612 b Water loss 2
346.2253723144531 0 645.665
348.11981201171875 0 1007.3793
348.6875 0 646.09937
350.13507080078125 0 2196.0322
350.6858825683594 0 802.85065
352.1407470703125 0 1096.1506
354.1659851074219 0 698.0981
355.0699157714844 0 1174.0868
356.1396789550781 0 1902.0442
357.22491455078125 0 26912.72
357.6805419921875 0 993.1401
358.22747802734375 0 4667.179
364.1506652832031 0 7047.4077 b 2
365.1531677246094 0 1431.3306
366.21392822265625 0 1516.6439
366.2508850097656 0 906.74023
368.19342041015625 0 9063.646
369.1979675292969 0 1135.4789
369.69378662109375 0 1051.5328
375.2359924316406 0 950.2889
376.11492919921875 0 929.2238
378.1299133300781 0 4997.988
378.2062072753906 0 1021.1037
378.7042541503906 0 831.06635
379.13458251953125 0 1054.1707
380.1426696777344 0 842.49677
383.1996154785156 0 1618.5616
383.6998596191406 0 868.66425
385.1355895996094 0 1085.5856
385.2200012207031 0 3228.9019
387.21099853515625 0 1996.1151
392.203857421875 0 7666.613
392.7052307128906 0 4337.9307
394.12506103515625 0 6265.1035
394.2458801269531 0 863.0167
395.12744140625 0 1461.3657
398.2402038574219 0 1576.0621
400.1883850097656 0 1861.1345
401.18646240234375 0 735.2636
401.20965576171875 0 5769.5977
401.710693359375 0 2875.771
402.2136535644531 0 833.0981
403.1479187011719 0 956.1925
405.2120056152344 0 1974.6217
405.71240234375 0 782.1136
406.210693359375 0 711.1342
408.16644287109375 0 1609.6678
410.27685546875 0 1819.8043
411.2359313964844 0 2441.0845
413.725341796875 0 2307.0347
414.2230529785156 0 1900.8862
414.714111328125 0 1224.1353
418.208740234375 0 1449.2211
418.7176513671875 0 2632.7017
419.1710205078125 0 803.73596
419.2168273925781 0 1204.1575
422.73028564453125 0 2897.083
423.2316589355469 0 2063.3416
424.2201232910156 0 705.9261
425.19390869140625 0 3461.949
425.2883605957031 0 2069.2686
426.1675109863281 0 858.91705
426.1964111328125 0 1072.848
426.2891845703125 0 889.76685
427.18145751953125 0 1485.1641
427.2103271484375 0 846.0715
427.7226867675781 0 16728.555
428.2236633300781 0 6635.2085
428.2581481933594 0 1097.0262
428.7244873046875 0 1237.9463
433.17193603515625 0 971.6698
436.7279357910156 0 17024.656
437.2289733886719 0 8159.2837
437.7314453125 0 2435.6702
442.3143310546875 0 3806.0508
443.2279968261719 0 1747.5226
443.31585693359375 0 766.92175
443.7318420410156 0 984.6919
444.7159118652344 0 1007.8205
445.1927795410156 0 1118.1453
447.16595458984375 0 818.1466
448.2660217285156 0 1585.2106
448.7685546875 0 1992.9135
452.2313537597656 0 1923.9512
452.7340393066406 0 1277.0465
453.2829284667969 0 3184.204
453.71844482421875 0 952.2814
455.2276916503906 0 881.89014
460.23773193359375 0 675.4518
460.2679443359375 0 761.8629
461.16717529296875 0 1760.4451 b Water loss 3
465.1627197265625 0 5133.3027
465.2096252441406 0 1205.5964
465.73907470703125 0 964.00714
466.1641845703125 0 827.1914
467.17767333984375 0 699.4761
469.32415771484375 0 1036.4353
470.3094482421875 0 5029.7534
471.1817932128906 0 789.72095
471.31243896484375 0 1442.932
474.7422790527344 0 1547.7671
477.1688232421875 0 859.176 y 2
478.74664306640625 0 1663.1406
479.1779479980469 0 4002.3936 b 3
479.2476501464844 0 2135.6804
481.2776184082031 0 940.97437
483.22039794921875 0 12351.805
484.2230224609375 0 2998.053
486.1626281738281 0 744.6998
487.26043701171875 0 2560.4102
487.33587646484375 0 7221.422
487.75274658203125 0 1639.0999
488.2548828125 0 1172.4008
488.3364562988281 0 2067.6858
489.1920166015625 0 8272.227 y 6
490.19354248046875 0 1870.3027
491.2108459472656 0 1446.4436
492.2536315917969 0 738.64545
495.1845397949219 0 782.37195
496.2302551269531 0 2298.107
496.26519775390625 0 5377.9844
496.7662658691406 0 3696.132
497.2333068847656 0 747.136
498.21917724609375 0 709.4053
498.304931640625 0 1971.2562
500.2465515136719 0 4650.4165
501.25537109375 0 6281.8657
501.7584228515625 0 4767.65
502.2599792480469 0 757.8738
504.29364013671875 0 1021.3695
510.26190185546875 0 7602.7275
510.7636413574219 0 4431.0303
511.26361083984375 0 1646.4932
513.1967163085938 0 786.4917
513.3513793945312 0 8366.912
514.2094116210938 0 740.0182
514.3545532226562 0 2156.4797
516.230224609375 0 1082.1948
519.2020263671875 0 1225.9553
520.2062377929688 0 849.375
527.5525512695312 0 612.3776
529.3092041015625 0 976.72534
531.2890625 0 677.19763
540.278076171875 0 1124.049
548.1997680664062 0 1498.5612
550.216064453125 0 976.10034
550.2974243164062 0 1183.2876
550.7117309570312 0 5873.4663 y 1
551.2130126953125 0 2871.8384
552.2410278320312 0 776.6949
556.2742919921875 0 1902.1515
556.319580078125 0 737.8947
557.19775390625 0 722.51605
560.2289428710938 0 11398.551 y 5
561.2319946289062 0 3720.1282
564.7247924804688 0 632.81384
566.257080078125 0 2188.9998
567.325927734375 0 1442.2753
568.31005859375 0 2936.274
569.23828125 0 969.0456
569.3138427734375 0 1153.5681
569.7346801757812 0 929.93976
569.7813720703125 0 1005.39233
573.3247680664062 0 728.8153
575.3310546875 0 1478.9556
583.2842407226562 0 3804.516
584.2725830078125 0 835.403
585.3361206054688 0 49498.844
586.338134765625 0 17605.791
587.3394165039062 0 2903.107
590.8440551757812 0 1310.1921
591.7938842773438 0 870.14667
592.2284545898438 0 1416.891 Precursor Water loss
592.7308349609375 0 1931.9121
593.2332763671875 0 823.2426
596.2393188476562 0 999.25415
596.302978515625 0 853.04694
599.8512573242188 0 1529.3645
600.304443359375 0 1002.1524
601.2351684570312 0 6665.5986 Precursor
601.2918701171875 0 1652.8873
601.7369384765625 0 4145.64
602.2371215820312 0 1165.4108
602.3629760742188 0 6475.2056
603.3518676757812 0 3611.3118
604.3545532226562 0 679.7982
606.2318115234375 0 906.76776
612.1972045898438 0 1946.736
613.3326416015625 0 2085.6768
614.2473754882812 0 1127.2513
614.33203125 0 1269.8008
615.7504272460938 0 742.6309
624.22998046875 0 911.35175 b Water loss 4
628.3189086914062 0 1152.7273
628.3780517578125 0 14593.091
629.3812866210938 0 4292.0737
630.3848876953125 0 856.844
634.23046875 0 1591.184
635.231689453125 0 710.2762
640.2459716796875 0 659.57526
644.3379516601562 0 1513.0018
655.3048706054688 0 806.2
656.3714599609375 0 1559.5132
657.2636108398438 0 2905.7012
659.2938842773438 0 4259.1978
660.2965698242188 0 1093.1403
663.302490234375 0 816.2778
665.3256225585938 0 1499.1079
670.3518676757812 0 1786.0499
673.3494873046875 0 740.52936
676.2726440429688 0 830.8521
678.3583374023438 0 731.2854
682.3524169921875 0 6916.693
683.3456420898438 0 3356.2356
685.2579956054688 0 4113.6724
686.261474609375 0 660.58716
690.2828979492188 0 830.69604
695.26708984375 0 1083.4572 b Water loss 5
696.3673706054688 0 2459.6243
697.3721313476562 0 851.0498
700.363037109375 0 125650.016
701.3658447265625 0 43440.402
702.3682250976562 0 8064.419
703.2720947265625 0 775.01587
712.2955932617188 0 943.82886
714.3679809570312 0 848.4573
718.372802734375 0 816.57117
721.2606811523438 0 4145.128
722.2665405273438 0 1146.0293
723.2921142578125 0 11506.685 y 4
724.2955932617188 0 4920.4478
725.2921752929688 0 1030.3602
726.3788452148438 0 2272.2944
727.3793334960938 0 1094.2506
729.3894653320312 0 1773.2332
732.4066162109375 0 797.93146
739.3823852539062 0 1001.62396
741.3667602539062 0 1024.1025
742.3544311523438 0 1033.0707
749.2566528320312 0 5572.7344
750.2553100585938 0 1433.1565
754.3743896484375 0 1132.6644
755.3997802734375 0 1460.2389
756.3939208984375 0 2325.0186
757.38623046875 0 12594.04
758.388427734375 0 5183.8604
759.3764038085938 0 1899.3099
766.3733520507812 0 1752.0636
767.2694702148438 0 829.25885
767.3971557617188 0 1035.8093
771.4010620117188 0 1958.96
772.409423828125 0 4350.5776
773.4146728515625 0 10462.517
774.3202514648438 0 4351.7964
774.4160766601562 0 3280.731
775.322998046875 0 2010.8342
775.415771484375 0 1347.6825
782.41796875 0 929.9592
783.3997192382812 0 35356.24
784.4011840820312 0 15257.921
785.4074096679688 0 5488.571
791.4074096679688 0 1291.318
792.4019165039062 0 1014.393
800.427001953125 0 19515.441
801.41259765625 0 64799.26
802.4142456054688 0 25574.967
803.4179077148438 0 6686.166
804.4263305664062 0 1409.9362
809.41162109375 0 929.7805
810.406494140625 0 899.62305
814.4379272460938 0 1488.8549
819.401123046875 0 950.6511
821.31005859375 0 1033.7836
826.4412841796875 0 3718.136
827.4437255859375 0 1642.2454
828.42236328125 0 26203.434
829.4246215820312 0 11109.639
830.42919921875 0 2145.7188
837.4136352539062 0 1936.066
838.3194580078125 0 15626.035 y 3
839.3221435546875 0 6469.781
840.322021484375 0 1132.4355
844.4526977539062 0 89278.766
845.455322265625 0 40397.83
846.4578247070312 0 9667.393
847.43408203125 0 3109.2354
848.436767578125 0 1158.6685
854.4369506835938 0 15162.18
855.4392700195312 0 6664.216
856.4453125 0 2170.473
857.4506225585938 0 1040.9171
871.3374633789062 0 1963.1482
871.4638671875 0 843.0239
872.4478149414062 0 367706.56
873.4505004882812 0 162161.14
874.4529418945312 0 38918.88
874.5546264648438 0 1209.7963
875.455810546875 0 3468.1157
885.4443969726562 0 707.49475
888.4239501953125 0 1693.3429
889.3478393554688 0 19574.43
890.3501586914062 0 10106.176
891.3567504882812 0 1796.2689
895.5272216796875 0 2844.6582
896.5254516601562 0 2121.5735
903.4573364257812 0 1347.1313
904.4666137695312 0 1186.1512
906.4306030273438 0 1543.6852
907.3379516601562 0 1323.1018
918.4714965820312 0 1129.452
919.4781494140625 0 3151.8643
920.4800415039062 0 2036.8984
930.4674682617188 0 1099.7084
932.48681640625 0 924.8503
935.3329467773438 0 4158.496 y Water loss 2
936.3367309570312 0 2481.8347
947.4918823242188 0 944.04645
948.4827880859375 0 1997.4224
949.4844970703125 0 1226.5366
953.3461303710938 0 70660.63 y 2
954.348876953125 0 35042.527
955.348876953125 0 8768.496
955.4356079101562 0 742.75354
965.427490234375 0 684.67316
973.511474609375 0 1963.4048
974.5086669921875 0 840.41266
975.5166625976562 0 1061.3802
983.43505859375 0 2549.2046
984.44287109375 0 2256.2239
991.5206298828125 0 61500.56
992.5238037109375 0 33336.305
993.52685546875 0 9113.672
994.5294799804688 0 1023.9285
997.3679809570312 0 1154.9409 b 7
1000.4757690429688 0 2445.5974
1001.4933471679688 0 2557.9497
1002.5027465820312 0 1101.7849
1019.5156860351562 0 161074.22
1020.5183715820312 0 88119.984
1021.5216674804688 0 29094.457
1021.6466064453125 0 1036.7059
1022.526611328125 0 2748.6628
1035.51220703125 0 1186.7993
1037.4124755859375 0 856.04846
1037.5277099609375 0 762.1837
1100.4141845703125 0 3500.007 y 1
1101.4227294921875 0 1728.0411
1524.712158203125 0 640.34393
1915.2979736328125 0 750.91516
2900.748779296875 0 679.54034

Spectrum Details

|  |  |
| --- | --- |
| Matched peaks? Matched peaks The total absolute number of peaks matched. Additionally in brackets the total fraction of peaks matched and the total number of peaks is shown. | 25 (4.45% of 562) |
| FDR? FDR The false discovery rate estimated for this peptide. It is calculated by matching all theoretical fragments with a non-integer shift with the raw peaks for this spectrum. This is done with 40 different shifts. The resulting percentage is the average number of annotated peaks over the number of annotated peaks with the correct spectrum. | 4.10% |
| Satellite FDR? Satellite FDR See the FDR for details on its calculation. This satellite ion specific FDR only contains the satellite ions (d/w) for I/L/J positions. | - |
| PSM Score? PSM Score The PSM Score as given by Hecklib to this annotated spectrum. It is shown with three significant figures. | 310 |

#### Spectrum 12715? Spectrum 12715 The raw spectrum of this peptide as annotated by Hecklib. The fragments are coloured according to ion type (see legend). Any peaks with a star '\*' as text can be hovered over to see the full details, first the ion type second the mass shift type. By hovering over the amino acids in the peptide or ions in the legend the corresponding peaks are highlighted. By toggling the 'Unassigned' label you can turn the background (unassigned) peaks on or off in the plot. By updating the slider in the Ion legend you can update the spectrum to only show the top X% of the peaks with labels. The top X% means any peak that is within X% of the highest intensity. By dragging in the spectrum you can zoom in to a specific part of the spectrum and use 'Zoom Out' to get back to the original zoom level. The annotation of the spectrum is based on the given sequence in the peptides file and is done with different software so inconsistencies are likely. The peaks are annotated based on the given sequence, with 20 ppm tolerance.

Copy Data

##### Spectrum 12715 (TSV)

###### Preview

```
Loading example...
```

*Click on the button to copy the data to your clipboard.*

Mz MinMz MaxIntensity Max

WidthHeightPeptide font sizePeptide stroke widthSpectrum font sizeSpectrum stroke widthCompact peptide

Ion legend

wxyz

abcd

OtherUnassignedIon^Charge^_Position_Show for top:%

TFDDYAMHW

09.69e+31.94e+42.91e+43.88e+4

Zoom Out

a^+1^_2_y^+1^_1_a^+1^_2_b^+1^_2_y^+1^_2_b^+1^_3_b^+1^_4_y^+1^_3_y^+2^_8_y^+1^_4_\*y^+1^_5_y^+1^_6_y^+1^_7_y^+1^_8_

0855171125663421

Fragment Matches Table

Show background peaks

| Position | Ion type | Intensity | mz Theoretical | mz Error (Th) | mz Error (ppm) | Charge | Series Number |
| --- | --- | --- | --- | --- | --- | --- | --- |
| - | - | 2.378E+04 | 120.1 | - | - | 0 | - |
| - | - | 1254 | 120.1 | - | - | 0 | - |
| - | - | 328.7 | 120.2 | - | - | 0 | - |
| - | - | 2286 | 121.1 | - | - | 0 | - |
| - | - | 421.6 | 127.1 | - | - | 0 | - |
| - | - | 3115 | 129.1 | - | - | 0 | - |
| - | - | 524.5 | 130.1 | - | - | 0 | - |
| - | - | 485.3 | 132.1 | - | - | 0 | - |
| - | - | 2375 | 133.1 | - | - | 0 | - |
| - | - | 8621 | 136.1 | - | - | 0 | - |
| - | - | 576.3 | 138.1 | - | - | 0 | - |
| - | - | 654.7 | 138.1 | - | - | 0 | - |
| - | - | 466.1 | 138.6 | - | - | 0 | - |
| - | - | 438 | 139.9 | - | - | 0 | - |
| - | - | 867.4 | 148.9 | - | - | 0 | - |
| - | - | 1125 | 149 | - | - | 0 | - |
| - | - | 513.6 | 155.1 | - | - | 0 | - |
| - | - | 798.2 | 156.1 | - | - | 0 | - |
| - | - | 443.3 | 156.1 | - | - | 0 | - |
| - | - | 1387 | 158.1 | - | - | 0 | - |
| - | - | 1075 | 159.1 | - | - | 0 | - |
| - | - | 673.4 | 160.1 | - | - | 0 | - |
| - | - | 2526 | 165.1 | - | - | 0 | - |
| - | - | 1017 | 166.1 | - | - | 0 | - |
| - | - | 1397 | 167.1 | - | - | 0 | - |
| - | - | 407.6 | 168.2 | - | - | 0 | - |
| - | - | 1956 | 173.1 | - | - | 0 | - |
| - | - | 629.2 | 173.4 | - | - | 0 | - |
| - | - | 4830 | 173.4 | - | - | 0 | - |
| - | - | 3347 | 176.1 | - | - | 0 | - |
| - | - | 2156 | 177.1 | - | - | 0 | - |
| - | - | 1041 | 177.1 | - | - | 0 | - |
| - | - | 5212 | 182.1 | - | - | 0 | - |
| - | - | 665.9 | 183.1 | - | - | 0 | - |
| - | - | 532 | 183.1 | - | - | 0 | - |
| - | - | 801.7 | 185.1 | - | - | 0 | - |
| - | - | 1326 | 188.1 | - | - | 0 | - |
| - | - | 466.5 | 193.1 | - | - | 0 | - |
| - | - | 649.1 | 195.1 | - | - | 0 | - |
| - | - | 462.6 | 195.1 | - | - | 0 | - |
| - | - | 1471 | 201.1 | - | - | 0 | - |
| - | - | 440.5 | 201.4 | - | - | 0 | - |
| 2 | a | 620.7 | 203.1 | 1.305E-07 | 0.0006424 | +1 | 2 |
| - | - | 563.8 | 204.1 | - | - | 0 | - |
| 9 | y | 1370 | 205.1 | 0.0001845 | 0.8995 | +1 | 1 |
| - | - | 471.9 | 210.1 | - | - | 0 | - |
| - | - | 715.9 | 211.1 | - | - | 0 | - |
| - | - | 1394 | 215.1 | - | - | 0 | - |
| - | - | 2107 | 221.1 | - | - | 0 | - |
| 2 | a | 3.839E+04 | 221.1 | 7.056E-05 | 0.3191 | +1 | 2 |
| - | - | 3832 | 222.1 | - | - | 0 | - |
| - | - | 1181 | 223.1 | - | - | 0 | - |
| - | - | 736.7 | 224.1 | - | - | 0 | - |
| - | - | 497.7 | 226.2 | - | - | 0 | - |
| - | - | 867.2 | 227.1 | - | - | 0 | - |
| - | - | 620.9 | 229.2 | - | - | 0 | - |
| - | - | 554.7 | 231.1 | - | - | 0 | - |
| - | - | 563.4 | 231.1 | - | - | 0 | - |
| - | - | 928.4 | 235.1 | - | - | 0 | - |
| - | - | 1005 | 239.1 | - | - | 0 | - |
| 2 | b | 6132 | 249.1 | 0.0001816 | 0.7288 | +1 | 2 |
| - | - | 720.9 | 250.1 | - | - | 0 | - |
| - | - | 711.9 | 251.1 | - | - | 0 | - |
| - | - | 1017 | 263.1 | - | - | 0 | - |
| - | - | 674.2 | 266.1 | - | - | 0 | - |
| - | - | 643.6 | 278.1 | - | - | 0 | - |
| - | - | 780 | 283 | - | - | 0 | - |
| - | - | 1081 | 285 | - | - | 0 | - |
| - | - | 591.1 | 285.1 | - | - | 0 | - |
| - | - | 8340 | 309.2 | - | - | 0 | - |
| - | - | 991.9 | 310.2 | - | - | 0 | - |
| - | - | 959.8 | 315.2 | - | - | 0 | - |
| - | - | 611.2 | 319.1 | - | - | 0 | - |
| - | - | 1329 | 324.1 | - | - | 0 | - |
| - | - | 3607 | 329.1 | - | - | 0 | - |
| 8 | y | 6190 | 342.2 | 0.0003672 | 1.073 | +1 | 2 |
| - | - | 949.1 | 343.2 | - | - | 0 | - |
| - | - | 1180 | 355.1 | - | - | 0 | - |
| - | - | 569.8 | 357.2 | - | - | 0 | - |
| 3 | b | 2079 | 364.2 | 0.0001349 | 0.3704 | +1 | 3 |
| - | - | 536.8 | 368.2 | - | - | 0 | - |
| - | - | 1291 | 378.1 | - | - | 0 | - |
| - | - | 1186 | 394.1 | - | - | 0 | - |
| - | - | 787.3 | 413.3 | - | - | 0 | - |
| - | - | 1145 | 425.2 | - | - | 0 | - |
| - | - | 800.7 | 436.7 | - | - | 0 | - |
| - | - | 593 | 454.9 | - | - | 0 | - |
| - | - | 830 | 465.2 | - | - | 0 | - |
| 4 | b | 1194 | 479.2 | 0.0001438 | 0.3 | +1 | 4 |
| - | - | 563.8 | 489.2 | - | - | 0 | - |
| 7 | y | 1492 | 489.2 | 0.006382 | 13.05 | +1 | 3 |
| - | - | 574.5 | 491.1 | - | - | 0 | - |
| - | - | 601.1 | 491.3 | - | - | 0 | - |
| - | - | 873.5 | 496.2 | - | - | 0 | - |
| 2 | y | 1361 | 550.7 | 0.002836 | 5.15 | +2 | 8 |
| 6 | y | 2994 | 560.2 | 0.006011 | 10.73 | +1 | 4 |
| - | - | 660.8 | 561.2 | - | - | 0 | - |
| - | - | 1689 | 585.3 | - | - | 0 | - |
| - | - | 607.6 | 594.4 | - | - | 0 | - |
| - | - | 699.8 | 598.2 | - | - | 0 | - |
| 0 | Precursor | 1328 | 601.2 | 0.001458 | 2.424 | +2 | -1 |
| - | - | 604.9 | 636.2 | - | - | 0 | - |
| - | - | 631.5 | 651.7 | - | - | 0 | - |
| - | - | 1046 | 657.3 | - | - | 0 | - |
| - | - | 1333 | 659.3 | - | - | 0 | - |
| - | - | 698.1 | 685.3 | - | - | 0 | - |
| - | - | 2926 | 700.4 | - | - | 0 | - |
| - | - | 846.1 | 701.4 | - | - | 0 | - |
| - | - | 618.7 | 721.3 | - | - | 0 | - |
| 5 | y | 2077 | 723.3 | 0.005854 | 8.093 | +1 | 5 |
| - | - | 771.5 | 724.3 | - | - | 0 | - |
| - | - | 1053 | 749.3 | - | - | 0 | - |
| - | - | 567.1 | 758.4 | - | - | 0 | - |
| - | - | 823.1 | 774.3 | - | - | 0 | - |
| - | - | 740.5 | 783.4 | - | - | 0 | - |
| - | - | 1636 | 801.4 | - | - | 0 | - |
| - | - | 749.6 | 828.4 | - | - | 0 | - |
| 4 | y | 3791 | 838.3 | 0.007658 | 9.135 | +1 | 6 |
| - | - | 1629 | 839.3 | - | - | 0 | - |
| - | - | 2590 | 844.5 | - | - | 0 | - |
| - | - | 1563 | 845.5 | - | - | 0 | - |
| - | - | 860.9 | 854.4 | - | - | 0 | - |
| - | - | 491.9 | 872.4 | - | - | 0 | - |
| - | - | 9778 | 872.4 | - | - | 0 | - |
| - | - | 4158 | 873.5 | - | - | 0 | - |
| - | - | 784.5 | 874.5 | - | - | 0 | - |
| - | - | 4237 | 889.3 | - | - | 0 | - |
| - | - | 655.7 | 890.2 | - | - | 0 | - |
| - | - | 1886 | 890.3 | - | - | 0 | - |
| - | - | 840.8 | 936.3 | - | - | 0 | - |
| 3 | y | 1.376E+04 | 953.3 | 0.005862 | 6.149 | +1 | 7 |
| - | - | 7234 | 954.3 | - | - | 0 | - |
| - | - | 2653 | 955.4 | - | - | 0 | - |
| - | - | 1668 | 991.5 | - | - | 0 | - |
| - | - | 952.7 | 992.5 | - | - | 0 | - |
| - | - | 625 | 1008 | - | - | 0 | - |
| - | - | 3546 | 1020 | - | - | 0 | - |
| - | - | 2463 | 1021 | - | - | 0 | - |
| 2 | y | 1149 | 1100 | 0.005136 | 4.667 | +1 | 8 |
| - | - | 639.7 | 1419 | - | - | 0 | - |
| - | - | 736.1 | 1432 | - | - | 0 | - |
| - | - | 828.1 | 3078 | - | - | 0 | - |
| - | - | 859.4 | 3388 | - | - | 0 | - |

m/z Charge Intensity FragmentType MassShift Position
120.08089447021484 0 23775.24
120.0859375 0 1254.3997
120.248046875 0 328.65305
121.08429718017578 0 2285.5554
127.08662414550781 0 421.59283
129.10231018066406 0 3115.0715
130.06544494628906 0 524.52246
132.080322265625 0 485.2791
133.0860595703125 0 2375.3572
136.07579040527344 0 8621.041
138.0548095703125 0 576.2923
138.0663604736328 0 654.7321
138.6060791015625 0 466.14737
139.93899536132812 0 438.00266
148.9461212158203 0 867.4204
149.0449981689453 0 1125.1552
155.0819854736328 0 513.63873
156.0768585205078 0 798.22577
156.10267639160156 0 443.3348
158.0966033935547 0 1386.5121
159.09188842773438 0 1075.4078
160.11227416992188 0 673.3815
165.05467224121094 0 2525.9985
166.0864715576172 0 1016.80597
167.05552673339844 0 1397.354
168.22366333007812 0 407.60458
173.12864685058594 0 1955.5706
173.43374633789062 0 629.1791
173.4395751953125 0 4829.6357
176.1071014404297 0 3347.4888
177.1022491455078 0 2156.3987
177.11129760742188 0 1041.2311
182.0812225341797 0 5211.8857
183.0845184326172 0 665.93207
183.14859008789062 0 531.9535
185.12884521484375 0 801.66125
188.07029724121094 0 1326.3374
193.108154296875 0 466.5137
195.08795166015625 0 649.1477
195.1235809326172 0 462.59186
201.12353515625 0 1470.8275
201.35491943359375 0 440.46957
203.11788940429688 0 620.69666 a Water loss 1
204.07662963867188 0 563.7936
205.0969696044922 0 1369.6958 y 8
210.14918518066406 0 471.9118
211.0606689453125 0 715.93604
215.127685546875 0 1394.1992
221.1033172607422 0 2107.4128
221.12852478027344 0 38385.656 a 1
222.1320343017578 0 3832.49
223.0637969970703 0 1181.0289
224.103271484375 0 736.6806
226.15489196777344 0 497.7023
227.10267639160156 0 867.2367
229.1692657470703 0 620.9187
231.06234741210938 0 554.72986
231.09844970703125 0 563.40356
235.1077880859375 0 928.39343
239.09486389160156 0 1005.2411
249.12355041503906 0 6132.1787 b 1
250.12799072265625 0 720.85614
251.10292053222656 0 711.9264
263.1023864746094 0 1016.53973
266.1253356933594 0 674.17505
278.1489562988281 0 643.62335
283.0316467285156 0 779.95374
285.0098571777344 0 1080.748
285.10107421875 0 591.11035
309.2038879394531 0 8339.541
310.20770263671875 0 991.9307
315.17852783203125 0 959.75354
319.1394348144531 0 611.1913
324.1447448730469 0 1328.742
329.14990234375 0 3606.5835
342.15643310546875 0 6189.953 y 7
343.1596984863281 0 949.12067
355.0688781738281 0 1179.7637
357.22344970703125 0 569.82214
364.1501770019531 0 2078.6565 b 2
368.19439697265625 0 536.8361
378.12982177734375 0 1290.5232
394.1244201660156 0 1185.8932
413.2665100097656 0 787.2744
425.1914978027344 0 1144.5059
436.7252197265625 0 800.6517
454.92364501953125 0 592.99866
465.15966796875 0 830.0252
479.1773986816406 0 1193.6602 b 3
489.15478515625 0 563.819
489.19293212890625 0 1492.2358 y 6
491.0805969238281 0 574.5137
491.27337646484375 0 601.0616
496.2305603027344 0 873.52344
550.7111206054688 0 1360.634 y 1
560.2296752929688 0 2993.737 y 5
561.231689453125 0 660.8255
585.3423461914062 0 1688.6547
594.3579711914062 0 607.62585
598.2166137695312 0 699.8251
601.2335815429688 0 1328.1724 Precursor
636.1939697265625 0 604.88556
651.6864013671875 0 631.5235
657.2633666992188 0 1045.9434
659.2922973632812 0 1332.512
685.2587280273438 0 698.07544
700.3630981445312 0 2926.2075
701.3687744140625 0 846.05426
721.26123046875 0 618.7337
723.2928466796875 0 2077.1436 y 4
724.2979736328125 0 771.4975
749.2512817382812 0 1052.6112
758.3746948242188 0 567.0943
774.3170776367188 0 823.117
783.399169921875 0 740.52924
801.4132080078125 0 1636.2167
828.4193115234375 0 749.56604
838.3215942382812 0 3791.339 y 3
839.3247680664062 0 1629.3269
844.45361328125 0 2590.1758
845.4539184570312 0 1563.0609
854.4402465820312 0 860.89355
872.3517456054688 0 491.9062
872.4486694335938 0 9777.895
873.4514770507812 0 4157.5605
874.4530639648438 0 784.4748
889.3485717773438 0 4236.9927
890.2006225585938 0 655.7195
890.3487548828125 0 1885.7112
936.341796875 0 840.8001
953.3467407226562 0 13755.387 y 2
954.3493041992188 0 7233.987
955.3501586914062 0 2653.2686
991.5222778320312 0 1667.5331
992.5234985351562 0 952.67
1008.0990600585938 0 624.9598
1019.5177001953125 0 3545.5212
1020.5195922851562 0 2463.2556
1100.4144287109375 0 1148.7981 y 1
1418.6822509765625 0 639.6698
1432.1917724609375 0 736.0832
3078.169921875 0 828.06195
3387.515380859375 0 859.4243

Spectrum Details

|  |  |
| --- | --- |
| Matched peaks? Matched peaks The total absolute number of peaks matched. Additionally in brackets the total fraction of peaks matched and the total number of peaks is shown. | 15 (10.49% of 143) |
| FDR? FDR The false discovery rate estimated for this peptide. It is calculated by matching all theoretical fragments with a non-integer shift with the raw peaks for this spectrum. This is done with 40 different shifts. The resulting percentage is the average number of annotated peaks over the number of annotated peaks with the correct spectrum. | 1.43% |
| Satellite FDR? Satellite FDR See the FDR for details on its calculation. This satellite ion specific FDR only contains the satellite ions (d/w) for I/L/J positions. | - |
| PSM Score? PSM Score The PSM Score as given by Hecklib to this annotated spectrum. It is shown with three significant figures. | 160 |

#### Spectrum 11363? Spectrum 11363 The raw spectrum of this peptide as annotated by Hecklib. The fragments are coloured according to ion type (see legend). Any peaks with a star '\*' as text can be hovered over to see the full details, first the ion type second the mass shift type. By hovering over the amino acids in the peptide or ions in the legend the corresponding peaks are highlighted. By toggling the 'Unassigned' label you can turn the background (unassigned) peaks on or off in the plot. By updating the slider in the Ion legend you can update the spectrum to only show the top X% of the peaks with labels. The top X% means any peak that is within X% of the highest intensity. By dragging in the spectrum you can zoom in to a specific part of the spectrum and use 'Zoom Out' to get back to the original zoom level. The annotation of the spectrum is based on the given sequence in the peptides file and is done with different software so inconsistencies are likely. The peaks are annotated based on the given sequence, with 20 ppm tolerance.

Copy Data

##### Spectrum 11363 (TSV)

###### Preview

```
Loading example...
```

*Click on the button to copy the data to your clipboard.*

Mz MinMz MaxIntensity Max

WidthHeightPeptide font sizePeptide stroke widthSpectrum font sizeSpectrum stroke widthCompact peptide

Ion legend

wxyz

abcd

OtherUnassignedIon^Charge^_Position_Show for top:%

TFDDYAMHW

01.30e+42.59e+43.89e+45.18e+4

Zoom Out

a^+1^_2_y^+1^_1_a^+1^_2_b^+1^_2_b^+1^_2_y^+1^_2_b^+1^_3_b^+1^_4_b^+1^_4_y^+1^_3_y^+2^_8_y^+1^_4_\*y^+1^_5_y^+1^_6_y^+1^_7_y^+1^_7_

0874174826223496

Fragment Matches Table

Show background peaks

| Position | Ion type | Intensity | mz Theoretical | mz Error (Th) | mz Error (ppm) | Charge | Series Number |
| --- | --- | --- | --- | --- | --- | --- | --- |
| - | - | 4.22E+04 | 120.1 | - | - | 0 | - |
| - | - | 381.4 | 121 | - | - | 0 | - |
| - | - | 3227 | 121.1 | - | - | 0 | - |
| - | - | 352.8 | 127.1 | - | - | 0 | - |
| - | - | 3862 | 129.1 | - | - | 0 | - |
| - | - | 1409 | 130.1 | - | - | 0 | - |
| - | - | 1090 | 132.1 | - | - | 0 | - |
| - | - | 770.1 | 132.1 | - | - | 0 | - |
| - | - | 377.8 | 132.8 | - | - | 0 | - |
| - | - | 2842 | 133.1 | - | - | 0 | - |
| - | - | 1.296E+04 | 136.1 | - | - | 0 | - |
| - | - | 922.3 | 137.1 | - | - | 0 | - |
| - | - | 581.5 | 138.1 | - | - | 0 | - |
| - | - | 507.6 | 139.1 | - | - | 0 | - |
| - | - | 1344 | 141.1 | - | - | 0 | - |
| - | - | 467.1 | 144 | - | - | 0 | - |
| - | - | 1492 | 146.1 | - | - | 0 | - |
| - | - | 988.3 | 147 | - | - | 0 | - |
| - | - | 699.4 | 149 | - | - | 0 | - |
| - | - | 688.4 | 155.1 | - | - | 0 | - |
| - | - | 910.2 | 156.1 | - | - | 0 | - |
| - | - | 550.3 | 156.4 | - | - | 0 | - |
| - | - | 541.3 | 157.1 | - | - | 0 | - |
| - | - | 2130 | 158.1 | - | - | 0 | - |
| - | - | 1632 | 159.1 | - | - | 0 | - |
| - | - | 660 | 159.1 | - | - | 0 | - |
| - | - | 461.7 | 159.1 | - | - | 0 | - |
| - | - | 796.1 | 160.1 | - | - | 0 | - |
| - | - | 584.2 | 162.1 | - | - | 0 | - |
| - | - | 1315 | 163.1 | - | - | 0 | - |
| - | - | 4711 | 165.1 | - | - | 0 | - |
| - | - | 505.3 | 166.1 | - | - | 0 | - |
| - | - | 5923 | 166.1 | - | - | 0 | - |
| - | - | 1386 | 167.1 | - | - | 0 | - |
| - | - | 2071 | 167.1 | - | - | 0 | - |
| - | - | 1725 | 169.1 | - | - | 0 | - |
| - | - | 465.2 | 171.1 | - | - | 0 | - |
| - | - | 518.3 | 173.1 | - | - | 0 | - |
| - | - | 508.5 | 173.1 | - | - | 0 | - |
| - | - | 2925 | 173.1 | - | - | 0 | - |
| - | - | 2855 | 173.4 | - | - | 0 | - |
| - | - | 3727 | 176.1 | - | - | 0 | - |
| - | - | 2716 | 177.1 | - | - | 0 | - |
| - | - | 905.5 | 177.1 | - | - | 0 | - |
| - | - | 618.1 | 178.1 | - | - | 0 | - |
| - | - | 7674 | 182.1 | - | - | 0 | - |
| - | - | 507.6 | 182.1 | - | - | 0 | - |
| - | - | 1063 | 183.1 | - | - | 0 | - |
| - | - | 557.3 | 183.1 | - | - | 0 | - |
| - | - | 3404 | 183.1 | - | - | 0 | - |
| - | - | 559.2 | 185.1 | - | - | 0 | - |
| - | - | 729.9 | 185.1 | - | - | 0 | - |
| - | - | 571.1 | 187.1 | - | - | 0 | - |
| - | - | 2411 | 188.1 | - | - | 0 | - |
| - | - | 511 | 191.1 | - | - | 0 | - |
| - | - | 773.4 | 191.1 | - | - | 0 | - |
| - | - | 945 | 193.1 | - | - | 0 | - |
| - | - | 581.1 | 195.1 | - | - | 0 | - |
| - | - | 622 | 197.1 | - | - | 0 | - |
| - | - | 466 | 197.5 | - | - | 0 | - |
| - | - | 853.1 | 199.1 | - | - | 0 | - |
| - | - | 555.1 | 200.1 | - | - | 0 | - |
| - | - | 583.3 | 201.1 | - | - | 0 | - |
| - | - | 1374 | 201.1 | - | - | 0 | - |
| 2 | a | 1103 | 203.1 | 0.0004426 | 2.179 | +1 | 2 |
| - | - | 591.7 | 204.1 | - | - | 0 | - |
| 9 | y | 1422 | 205.1 | 0.0001207 | 0.5884 | +1 | 1 |
| - | - | 1067 | 207.1 | - | - | 0 | - |
| - | - | 894.5 | 207.1 | - | - | 0 | - |
| - | - | 559.3 | 211.1 | - | - | 0 | - |
| - | - | 3125 | 211.1 | - | - | 0 | - |
| - | - | 701.9 | 213.2 | - | - | 0 | - |
| - | - | 620.4 | 214.1 | - | - | 0 | - |
| - | - | 704.3 | 215.1 | - | - | 0 | - |
| - | - | 788.8 | 217.1 | - | - | 0 | - |
| - | - | 3011 | 221.1 | - | - | 0 | - |
| 2 | a | 5.13E+04 | 221.1 | 0.0001469 | 0.6641 | +1 | 2 |
| - | - | 6085 | 222.1 | - | - | 0 | - |
| - | - | 1376 | 223.1 | - | - | 0 | - |
| - | - | 1166 | 224.1 | - | - | 0 | - |
| - | - | 3239 | 227.1 | - | - | 0 | - |
| - | - | 563.9 | 229.2 | - | - | 0 | - |
| - | - | 709.6 | 229.2 | - | - | 0 | - |
| - | - | 837 | 231.1 | - | - | 0 | - |
| 2 | b | 776.4 | 231.1 | 0.0005147 | 2.227 | +1 | 2 |
| - | - | 519.4 | 233.1 | - | - | 0 | - |
| - | - | 1451 | 233.2 | - | - | 0 | - |
| - | - | 470 | 234.9 | - | - | 0 | - |
| - | - | 1427 | 235.1 | - | - | 0 | - |
| - | - | 1363 | 239.1 | - | - | 0 | - |
| - | - | 753.7 | 239.1 | - | - | 0 | - |
| - | - | 702.1 | 241.1 | - | - | 0 | - |
| - | - | 607.7 | 242.1 | - | - | 0 | - |
| 2 | b | 9422 | 249.1 | 0.0001663 | 0.6676 | +1 | 2 |
| - | - | 1210 | 250.1 | - | - | 0 | - |
| - | - | 1163 | 251.1 | - | - | 0 | - |
| - | - | 802.2 | 253.1 | - | - | 0 | - |
| - | - | 2190 | 263.1 | - | - | 0 | - |
| - | - | 656.6 | 263.1 | - | - | 0 | - |
| - | - | 744.8 | 265.2 | - | - | 0 | - |
| - | - | 900.6 | 266.2 | - | - | 0 | - |
| - | - | 957.2 | 279.1 | - | - | 0 | - |
| - | - | 2004 | 281.1 | - | - | 0 | - |
| - | - | 851.3 | 285 | - | - | 0 | - |
| - | - | 1084 | 285.1 | - | - | 0 | - |
| - | - | 763.9 | 292.1 | - | - | 0 | - |
| - | - | 1056 | 296.2 | - | - | 0 | - |
| - | - | 751.7 | 302.1 | - | - | 0 | - |
| - | - | 924.2 | 302.2 | - | - | 0 | - |
| - | - | 697.4 | 305.1 | - | - | 0 | - |
| - | - | 655.8 | 309.2 | - | - | 0 | - |
| - | - | 840.7 | 310.2 | - | - | 0 | - |
| - | - | 808.6 | 312.2 | - | - | 0 | - |
| - | - | 986.9 | 319.1 | - | - | 0 | - |
| - | - | 1695 | 324.1 | - | - | 0 | - |
| - | - | 5015 | 329.1 | - | - | 0 | - |
| - | - | 909.4 | 330.2 | - | - | 0 | - |
| - | - | 4201 | 340.2 | - | - | 0 | - |
| - | - | 760.3 | 341.2 | - | - | 0 | - |
| - | - | 679.2 | 342.1 | - | - | 0 | - |
| 8 | y | 9580 | 342.2 | 0.0003061 | 0.8947 | +1 | 2 |
| - | - | 2275 | 343.2 | - | - | 0 | - |
| - | - | 798.8 | 352.2 | - | - | 0 | - |
| - | - | 607.6 | 354.9 | - | - | 0 | - |
| - | - | 1662 | 355.1 | - | - | 0 | - |
| - | - | 577.4 | 356.1 | - | - | 0 | - |
| - | - | 1023 | 357.2 | - | - | 0 | - |
| 3 | b | 2433 | 364.2 | 0.0001398 | 0.3839 | +1 | 3 |
| - | - | 947.5 | 378.1 | - | - | 0 | - |
| - | - | 1807 | 380.2 | - | - | 0 | - |
| - | - | 2090 | 394.1 | - | - | 0 | - |
| - | - | 1499 | 395.2 | - | - | 0 | - |
| - | - | 628.8 | 396.2 | - | - | 0 | - |
| - | - | 1422 | 398.2 | - | - | 0 | - |
| - | - | 747.3 | 413.3 | - | - | 0 | - |
| - | - | 679.3 | 414.9 | - | - | 0 | - |
| - | - | 568.4 | 415 | - | - | 0 | - |
| - | - | 1702 | 425.2 | - | - | 0 | - |
| - | - | 576 | 437.2 | - | - | 0 | - |
| - | - | 853.9 | 449.2 | - | - | 0 | - |
| - | - | 1147 | 453.3 | - | - | 0 | - |
| 4 | b | 729 | 461.2 | 0.002865 | 6.213 | +1 | 4 |
| - | - | 1062 | 465.2 | - | - | 0 | - |
| 4 | b | 1077 | 479.2 | 0.0004055 | 0.8463 | +1 | 4 |
| - | - | 741 | 480.3 | - | - | 0 | - |
| 7 | y | 2022 | 489.2 | 0.003177 | 6.495 | +1 | 3 |
| - | - | 1096 | 496.2 | - | - | 0 | - |
| - | - | 876.8 | 500.8 | - | - | 0 | - |
| - | - | 3136 | 508.3 | - | - | 0 | - |
| - | - | 634.4 | 509.3 | - | - | 0 | - |
| - | - | 613.6 | 512.3 | - | - | 0 | - |
| - | - | 635.3 | 512.6 | - | - | 0 | - |
| - | - | 667.2 | 519.2 | - | - | 0 | - |
| - | - | 652.5 | 548.2 | - | - | 0 | - |
| 2 | y | 954.9 | 550.7 | 0.001676 | 3.044 | +2 | 8 |
| - | - | 1121 | 551.2 | - | - | 0 | - |
| 6 | y | 3820 | 560.2 | 0.004668 | 8.333 | +1 | 4 |
| - | - | 1172 | 561.2 | - | - | 0 | - |
| - | - | 1386 | 563.3 | - | - | 0 | - |
| - | - | 2198 | 585.3 | - | - | 0 | - |
| - | - | 801 | 586.3 | - | - | 0 | - |
| 0 | Precursor | 1910 | 601.2 | 0.0007863 | 1.308 | +2 | -1 |
| - | - | 683.3 | 602.2 | - | - | 0 | - |
| - | - | 1074 | 602.3 | - | - | 0 | - |
| - | - | 798.4 | 614.2 | - | - | 0 | - |
| - | - | 865.1 | 614.3 | - | - | 0 | - |
| - | - | 3745 | 621.4 | - | - | 0 | - |
| - | - | 1499 | 622.4 | - | - | 0 | - |
| - | - | 612.4 | 657.3 | - | - | 0 | - |
| - | - | 1594 | 659.3 | - | - | 0 | - |
| - | - | 1144 | 660.3 | - | - | 0 | - |
| - | - | 602.2 | 676.3 | - | - | 0 | - |
| - | - | 875.4 | 676.4 | - | - | 0 | - |
| - | - | 1266 | 678.4 | - | - | 0 | - |
| - | - | 840.3 | 685.3 | - | - | 0 | - |
| - | - | 5404 | 700.4 | - | - | 0 | - |
| - | - | 2346 | 701.4 | - | - | 0 | - |
| - | - | 678 | 702.4 | - | - | 0 | - |
| - | - | 588.7 | 719.6 | - | - | 0 | - |
| - | - | 1269 | 721.3 | - | - | 0 | - |
| 5 | y | 3202 | 723.3 | 0.003046 | 4.212 | +1 | 5 |
| - | - | 1277 | 724.3 | - | - | 0 | - |
| - | - | 604 | 729.3 | - | - | 0 | - |
| - | - | 1990 | 735.4 | - | - | 0 | - |
| - | - | 733.7 | 736.4 | - | - | 0 | - |
| - | - | 1160 | 749.3 | - | - | 0 | - |
| - | - | 849.1 | 750.3 | - | - | 0 | - |
| - | - | 1081 | 774.3 | - | - | 0 | - |
| - | - | 815.3 | 775.3 | - | - | 0 | - |
| - | - | 1402 | 783.4 | - | - | 0 | - |
| - | - | 670.1 | 784.4 | - | - | 0 | - |
| - | - | 616.3 | 800.4 | - | - | 0 | - |
| - | - | 2722 | 801.4 | - | - | 0 | - |
| - | - | 1183 | 802.4 | - | - | 0 | - |
| - | - | 986.6 | 828.4 | - | - | 0 | - |
| 4 | y | 4537 | 838.3 | 0.004057 | 4.84 | +1 | 6 |
| - | - | 2818 | 839.3 | - | - | 0 | - |
| - | - | 4561 | 844.5 | - | - | 0 | - |
| - | - | 1973 | 845.5 | - | - | 0 | - |
| - | - | 667.4 | 855.4 | - | - | 0 | - |
| - | - | 956.2 | 871.3 | - | - | 0 | - |
| - | - | 1.56E+04 | 872.4 | - | - | 0 | - |
| - | - | 7221 | 873.4 | - | - | 0 | - |
| - | - | 1483 | 874.4 | - | - | 0 | - |
| - | - | 5972 | 889.3 | - | - | 0 | - |
| - | - | 2814 | 890.3 | - | - | 0 | - |
| - | - | 1010 | 891.3 | - | - | 0 | - |
| - | - | 2555 | 901.5 | - | - | 0 | - |
| - | - | 1930 | 902.5 | - | - | 0 | - |
| - | - | 793.6 | 919.5 | - | - | 0 | - |
| 3 | y | 1420 | 935.3 | 0.008858 | 9.471 | +1 | 7 |
| - | - | 789.4 | 936.3 | - | - | 0 | - |
| 3 | y | 2.069E+04 | 953.3 | 0.002871 | 3.012 | +1 | 7 |
| - | - | 1.133E+04 | 954.3 | - | - | 0 | - |
| - | - | 4360 | 955.3 | - | - | 0 | - |
| - | - | 2905 | 991.5 | - | - | 0 | - |
| - | - | 1523 | 992.5 | - | - | 0 | - |
| - | - | 813.5 | 1003 | - | - | 0 | - |
| - | - | 840.1 | 1019 | - | - | 0 | - |
| - | - | 6344 | 1020 | - | - | 0 | - |
| - | - | 3061 | 1021 | - | - | 0 | - |
| - | - | 1667 | 1022 | - | - | 0 | - |
| - | - | 1075 | 1101 | - | - | 0 | - |
| - | - | 653.7 | 1210 | - | - | 0 | - |
| - | - | 733.2 | 2387 | - | - | 0 | - |
| - | - | 641.4 | 2894 | - | - | 0 | - |
| - | - | 991.2 | 3028 | - | - | 0 | - |
| - | - | 630.4 | 3167 | - | - | 0 | - |
| - | - | 814.6 | 3461 | - | - | 0 | - |

m/z Charge Intensity FragmentType MassShift Position
120.0810317993164 0 42198.957
121.02406311035156 0 381.4344
121.08441925048828 0 3227.2278
127.08661651611328 0 352.8362
129.10250854492188 0 3862.4812
130.06512451171875 0 1409.463
132.08108520507812 0 1090.1956
132.1023406982422 0 770.0972
132.798583984375 0 377.83572
133.08616638183594 0 2842.1606
136.075927734375 0 12955.481
137.07955932617188 0 922.25195
138.0659942626953 0 581.5129
139.08692932128906 0 507.5765
141.10267639160156 0 1344.0869
144.04806518554688 0 467.13004
146.0603485107422 0 1491.6802
147.047607421875 0 988.2856
149.0452423095703 0 699.39197
155.0818328857422 0 688.4023
156.07701110839844 0 910.2477
156.37469482421875 0 550.27856
157.0974884033203 0 541.2938
158.0966033935547 0 2130.345
159.09194946289062 0 1632.3507
159.10081481933594 0 660.0045
159.11294555664062 0 461.72015
160.11209106445312 0 796.0863
162.09153747558594 0 584.186
163.07882690429688 0 1315.1621
165.05487060546875 0 4710.975
166.0596466064453 0 505.2714
166.08645629882812 0 5922.904
167.05535888671875 0 1385.7355
167.08169555664062 0 2071.1816
169.09730529785156 0 1724.7518
171.076416015625 0 465.2337
173.07809448242188 0 518.2993
173.09242248535156 0 508.5086
173.12855529785156 0 2925.3152
173.43930053710938 0 2855.4766
176.1072540283203 0 3726.7754
177.10247802734375 0 2716.435
177.1109619140625 0 905.4606
178.10589599609375 0 618.13617
182.08132934570312 0 7673.699
182.12879943847656 0 507.59653
183.08494567871094 0 1063.1562
183.11294555664062 0 557.2869
183.14935302734375 0 3403.881
185.05587768554688 0 559.1778
185.09210205078125 0 729.9005
187.14430236816406 0 571.10223
188.07070922851562 0 2410.926
191.08163452148438 0 510.99127
191.11807250976562 0 773.38477
193.10873413085938 0 945.0208
195.0884246826172 0 581.09064
197.1285400390625 0 621.9761
197.49932861328125 0 465.97684
199.1078643798828 0 853.14777
200.13961791992188 0 555.0672
201.1131134033203 0 583.2839
201.12359619140625 0 1373.8862
203.11744689941406 0 1103.2135 a Water loss 1
204.07675170898438 0 591.66864
205.09727478027344 0 1422.3762 y 8
207.1040802001953 0 1066.7949
207.1138153076172 0 894.5225
211.05856323242188 0 559.2983
211.1441192626953 0 3125.3008
213.160888671875 0 701.9482
214.1178741455078 0 620.43994
215.13864135742188 0 704.28723
217.09765625 0 788.8377
221.1034698486328 0 3011.288
221.12860107421875 0 51297.97 a 1
222.13192749023438 0 6084.733
223.0634765625 0 1375.9484
224.10324096679688 0 1165.9587
227.10296630859375 0 3238.8977
229.15530395507812 0 563.8512
229.1702880859375 0 709.64294
231.06124877929688 0 837.03015
231.11228942871094 0 776.35156 b Water loss 1
233.1180877685547 0 519.3771
233.150146484375 0 1450.7987
234.9269561767578 0 470.0426
235.10781860351562 0 1427.2936
239.0945281982422 0 1363.2435
239.1146697998047 0 753.7288
241.07366943359375 0 702.1187
242.11341857910156 0 607.7488
249.12353515625 0 9422.175 b 1
250.12669372558594 0 1210.407
251.1026153564453 0 1163.2798
253.11727905273438 0 802.20435
263.1027526855469 0 2190.2043
263.12811279296875 0 656.622
265.154541015625 0 744.75073
266.150146484375 0 900.64355
279.09783935546875 0 957.1935
281.1244201660156 0 2003.5432
285.0091247558594 0 851.26996
285.10113525390625 0 1083.9415
292.14141845703125 0 763.8726
296.151123046875 0 1055.7202
302.11456298828125 0 751.7032
302.1507263183594 0 924.1543
305.1256408691406 0 697.43164
309.2056579589844 0 655.7898
310.2131042480469 0 840.72015
312.1911315917969 0 808.5859
319.1398620605469 0 986.89905
324.1455383300781 0 1695.1028
329.14971923828125 0 5015.46
330.1532897949219 0 909.35004
340.18719482421875 0 4201.136
341.1892395019531 0 760.28253
342.1373291015625 0 679.2202
342.1563720703125 0 9579.74 y 7
343.1589050292969 0 2274.9602
352.19781494140625 0 798.8135
354.86865234375 0 607.6008
355.0707092285156 0 1661.8966
356.14166259765625 0 577.4079
357.2252197265625 0 1022.5301
364.15045166015625 0 2433.4026 b 2
378.1291809082031 0 947.50714
380.1922912597656 0 1807.3958
394.12451171875 0 2089.8586
395.192138671875 0 1498.6809
396.2223205566406 0 628.8127
398.20440673828125 0 1421.6278
413.26861572265625 0 747.3446
414.868896484375 0 679.3158
415.009521484375 0 568.42596
425.1925964355469 0 1702.1631
437.2309265136719 0 576.02985
449.2409362792969 0 853.8567
453.2716979980469 0 1146.6735
461.1695556640625 0 728.9587 b Water loss 3
465.16229248046875 0 1062.2379
479.1768493652344 0 1077.0282 b 3
480.2832336425781 0 740.95337
489.1897277832031 0 2022.193 y 6
496.2306213378906 0 1096.316
500.776123046875 0 876.8083
508.2760009765625 0 3136.201
509.2798767089844 0 634.44305
512.2913208007812 0 613.63947
512.6204223632812 0 635.26117
519.2017822265625 0 667.1818
548.2008666992188 0 652.4809
550.7099609375 0 954.85834 y 1
551.2119750976562 0 1121.3704
560.2283325195312 0 3820.3728 y 5
561.2304077148438 0 1172.3778
563.2818603515625 0 1386.1704
585.3360595703125 0 2197.7095
586.3372192382812 0 800.9755
601.23291015625 0 1910.0317 Precursor
602.2391357421875 0 683.29156
602.3267211914062 0 1073.5555
614.2425537109375 0 798.44543
614.330322265625 0 865.0576
621.3600463867188 0 3745.0413
622.362548828125 0 1499.14
657.2647705078125 0 612.3558
659.2920532226562 0 1594.4225
660.2982788085938 0 1143.5964
676.2852783203125 0 602.2104
676.36474609375 0 875.44464
678.3820190429688 0 1265.8337
685.2557983398438 0 840.3472
700.361572265625 0 5403.6226
701.3643188476562 0 2345.5298
702.361328125 0 678.00214
719.6399536132812 0 588.7426
721.2614135742188 0 1269.2472
723.2900390625 0 3202.1125 y 4
724.2955322265625 0 1277.4935
729.3343505859375 0 603.9708
735.4022216796875 0 1990.3583
736.404296875 0 733.6684
749.2533569335938 0 1160.4803
750.2583618164062 0 849.0776
774.3193969726562 0 1081.2897
775.3231201171875 0 815.2875
783.3983154296875 0 1401.7773
784.3992919921875 0 670.12256
800.428955078125 0 616.327
801.4105224609375 0 2722.299
802.4131469726562 0 1182.7344
828.4181518554688 0 986.56055
838.3179931640625 0 4537.1465 y 3
839.3189086914062 0 2817.701
844.4500122070312 0 4561.46
845.4505004882812 0 1973.4585
855.4385375976562 0 667.4448
871.3338623046875 0 956.18634
872.445556640625 0 15600.729
873.4474487304688 0 7220.5776
874.4481811523438 0 1482.7275
889.345703125 0 5972.4473
890.3482055664062 0 2813.767
891.3473510742188 0 1009.5864
901.4771728515625 0 2555.3735
902.4791259765625 0 1929.5732
919.4894409179688 0 793.61755
935.3391723632812 0 1419.7175 y Water loss 2
936.3265991210938 0 789.4414
953.34375 0 20687.1 y 2
954.34619140625 0 11329.713
955.3469848632812 0 4360.371
991.5182495117188 0 2905.2073
992.5194091796875 0 1522.6196
1003.3748779296875 0 813.5069
1018.550537109375 0 840.14465
1019.5143432617188 0 6344.466
1020.5150146484375 0 3061.1428
1021.5155029296875 0 1666.7823
1101.416259765625 0 1074.7385
1209.6656494140625 0 653.6849
2387.129638671875 0 733.2083
2893.816162109375 0 641.4473
3027.551513671875 0 991.2393
3167.176513671875 0 630.3818
3461.294189453125 0 814.588

Spectrum Details

|  |  |
| --- | --- |
| Matched peaks? Matched peaks The total absolute number of peaks matched. Additionally in brackets the total fraction of peaks matched and the total number of peaks is shown. | 17 (7.42% of 229) |
| FDR? FDR The false discovery rate estimated for this peptide. It is calculated by matching all theoretical fragments with a non-integer shift with the raw peaks for this spectrum. This is done with 40 different shifts. The resulting percentage is the average number of annotated peaks over the number of annotated peaks with the correct spectrum. | 2.66% |
| Satellite FDR? Satellite FDR See the FDR for details on its calculation. This satellite ion specific FDR only contains the satellite ions (d/w) for I/L/J positions. | - |
| PSM Score? PSM Score The PSM Score as given by Hecklib to this annotated spectrum. It is shown with three significant figures. | 195 |

#### Spectrum 12975? Spectrum 12975 The raw spectrum of this peptide as annotated by Hecklib. The fragments are coloured according to ion type (see legend). Any peaks with a star '\*' as text can be hovered over to see the full details, first the ion type second the mass shift type. By hovering over the amino acids in the peptide or ions in the legend the corresponding peaks are highlighted. By toggling the 'Unassigned' label you can turn the background (unassigned) peaks on or off in the plot. By updating the slider in the Ion legend you can update the spectrum to only show the top X% of the peaks with labels. The top X% means any peak that is within X% of the highest intensity. By dragging in the spectrum you can zoom in to a specific part of the spectrum and use 'Zoom Out' to get back to the original zoom level. The annotation of the spectrum is based on the given sequence in the peptides file and is done with different software so inconsistencies are likely. The peaks are annotated based on the given sequence, with 20 ppm tolerance.

Copy Data

##### Spectrum 12975 (TSV)

###### Preview

```
Loading example...
```

*Click on the button to copy the data to your clipboard.*

Mz MinMz MaxIntensity Max

WidthHeightPeptide font sizePeptide stroke widthSpectrum font sizeSpectrum stroke widthCompact peptide

Ion legend

wxyz

abcd

OtherUnassignedIon^Charge^_Position_Show for top:%

TFDDYAMHW

08.43e+31.69e+42.53e+43.37e+4

Zoom Out

a^+1^_2_y^+1^_1_a^+1^_2_b^+2^_4_b^+1^_2_y^+1^_2_b^+1^_3_b^+1^_3_b^+1^_4_y^+1^_3_y^+2^_8_y^+1^_4_\*y^+1^_5_y^+1^_6_y^+1^_7_y^+1^_8_

046993914081877

Fragment Matches Table

Show background peaks

| Position | Ion type | Intensity | mz Theoretical | mz Error (Th) | mz Error (ppm) | Charge | Series Number |
| --- | --- | --- | --- | --- | --- | --- | --- |
| - | - | 2.305E+04 | 120.1 | - | - | 0 | - |
| - | - | 1683 | 121.1 | - | - | 0 | - |
| - | - | 343.5 | 122 | - | - | 0 | - |
| - | - | 339.4 | 122.1 | - | - | 0 | - |
| - | - | 400.1 | 126.3 | - | - | 0 | - |
| - | - | 4048 | 129.1 | - | - | 0 | - |
| - | - | 464.5 | 130.1 | - | - | 0 | - |
| - | - | 879.5 | 133.1 | - | - | 0 | - |
| - | - | 7542 | 136.1 | - | - | 0 | - |
| - | - | 447.5 | 137.1 | - | - | 0 | - |
| - | - | 394.6 | 138.9 | - | - | 0 | - |
| - | - | 559.9 | 146.1 | - | - | 0 | - |
| - | - | 1107 | 148.9 | - | - | 0 | - |
| - | - | 1206 | 149 | - | - | 0 | - |
| - | - | 559.9 | 155.1 | - | - | 0 | - |
| - | - | 955.2 | 156.1 | - | - | 0 | - |
| - | - | 1068 | 158.1 | - | - | 0 | - |
| - | - | 643.8 | 159.1 | - | - | 0 | - |
| - | - | 2624 | 165.1 | - | - | 0 | - |
| - | - | 1551 | 167.1 | - | - | 0 | - |
| - | - | 1394 | 173.1 | - | - | 0 | - |
| - | - | 1919 | 173.5 | - | - | 0 | - |
| - | - | 2118 | 176.1 | - | - | 0 | - |
| - | - | 490.4 | 176.2 | - | - | 0 | - |
| - | - | 1189 | 177.1 | - | - | 0 | - |
| - | - | 987.9 | 177.1 | - | - | 0 | - |
| - | - | 3896 | 182.1 | - | - | 0 | - |
| - | - | 650.1 | 183.1 | - | - | 0 | - |
| - | - | 518.3 | 183.6 | - | - | 0 | - |
| - | - | 538.3 | 185.1 | - | - | 0 | - |
| - | - | 1328 | 188.1 | - | - | 0 | - |
| - | - | 504.7 | 191.1 | - | - | 0 | - |
| - | - | 772.3 | 193.1 | - | - | 0 | - |
| - | - | 512.9 | 199.1 | - | - | 0 | - |
| - | - | 536 | 200.1 | - | - | 0 | - |
| - | - | 1540 | 201.1 | - | - | 0 | - |
| 2 | a | 653.4 | 203.1 | 4.591E-05 | 0.226 | +1 | 2 |
| 9 | y | 913.9 | 205.1 | 0.0001692 | 0.8251 | +1 | 1 |
| - | - | 1100 | 215.1 | - | - | 0 | - |
| - | - | 2174 | 221.1 | - | - | 0 | - |
| 2 | a | 3.34E+04 | 221.1 | 0.0001011 | 0.4571 | +1 | 2 |
| - | - | 4852 | 222.1 | - | - | 0 | - |
| - | - | 909.2 | 223.1 | - | - | 0 | - |
| - | - | 628 | 239.1 | - | - | 0 | - |
| 4 | b | 1045 | 240.1 | 0.003605 | 15.02 | +2 | 4 |
| - | - | 1152 | 241.1 | - | - | 0 | - |
| 2 | b | 5275 | 249.1 | 0.0002121 | 0.8513 | +1 | 2 |
| - | - | 850.7 | 250.1 | - | - | 0 | - |
| - | - | 576.8 | 251.1 | - | - | 0 | - |
| - | - | 1186 | 263.1 | - | - | 0 | - |
| - | - | 655.6 | 283 | - | - | 0 | - |
| - | - | 1186 | 285 | - | - | 0 | - |
| - | - | 656 | 299.1 | - | - | 0 | - |
| - | - | 1623 | 309.2 | - | - | 0 | - |
| - | - | 604 | 322.2 | - | - | 0 | - |
| - | - | 1595 | 324.1 | - | - | 0 | - |
| - | - | 2453 | 329.2 | - | - | 0 | - |
| 8 | y | 6032 | 342.2 | 0.0006113 | 1.787 | +1 | 2 |
| - | - | 1620 | 343.2 | - | - | 0 | - |
| 3 | b | 524.6 | 346.1 | 0.002678 | 7.738 | +1 | 3 |
| - | - | 1369 | 355.1 | - | - | 0 | - |
| 3 | b | 1568 | 364.2 | 0.001635 | 4.49 | +1 | 3 |
| - | - | 685.3 | 368.2 | - | - | 0 | - |
| - | - | 690.9 | 378.1 | - | - | 0 | - |
| - | - | 1065 | 394.1 | - | - | 0 | - |
| - | - | 583.3 | 413.3 | - | - | 0 | - |
| - | - | 526.1 | 415.9 | - | - | 0 | - |
| - | - | 625.1 | 421.8 | - | - | 0 | - |
| - | - | 556.6 | 424.6 | - | - | 0 | - |
| - | - | 906.3 | 425.2 | - | - | 0 | - |
| - | - | 612.9 | 438.9 | - | - | 0 | - |
| - | - | 808.5 | 465.2 | - | - | 0 | - |
| - | - | 534.3 | 468 | - | - | 0 | - |
| 4 | b | 1212 | 479.2 | 6.985E-05 | 0.1458 | +1 | 4 |
| 7 | y | 1940 | 489.2 | 0.005527 | 11.3 | +1 | 3 |
| 2 | y | 820.8 | 550.7 | 0.003751 | 6.812 | +2 | 8 |
| 6 | y | 2340 | 560.2 | 0.005584 | 9.967 | +1 | 4 |
| - | - | 639.1 | 561.2 | - | - | 0 | - |
| - | - | 1197 | 585.3 | - | - | 0 | - |
| 0 | Precursor | 811.3 | 601.2 | 0.0009083 | 1.511 | +2 | -1 |
| - | - | 646.3 | 602.3 | - | - | 0 | - |
| - | - | 646 | 613.3 | - | - | 0 | - |
| - | - | 677.6 | 614.2 | - | - | 0 | - |
| - | - | 575.7 | 625.7 | - | - | 0 | - |
| - | - | 653.8 | 685.3 | - | - | 0 | - |
| - | - | 605.1 | 686.3 | - | - | 0 | - |
| - | - | 2633 | 700.4 | - | - | 0 | - |
| - | - | 841.4 | 701.4 | - | - | 0 | - |
| - | - | 648.1 | 721.3 | - | - | 0 | - |
| 5 | y | 2186 | 723.3 | 0.005366 | 7.418 | +1 | 5 |
| - | - | 602.2 | 724.3 | - | - | 0 | - |
| - | - | 915.2 | 774.3 | - | - | 0 | - |
| - | - | 1482 | 801.4 | - | - | 0 | - |
| 4 | y | 2876 | 838.3 | 0.007353 | 8.771 | +1 | 6 |
| - | - | 1636 | 839.3 | - | - | 0 | - |
| - | - | 1633 | 844.5 | - | - | 0 | - |
| - | - | 1384 | 845.5 | - | - | 0 | - |
| - | - | 7397 | 872.4 | - | - | 0 | - |
| - | - | 2923 | 873.5 | - | - | 0 | - |
| - | - | 1104 | 874.5 | - | - | 0 | - |
| - | - | 3196 | 889.3 | - | - | 0 | - |
| - | - | 1605 | 890.4 | - | - | 0 | - |
| - | - | 979.5 | 891.4 | - | - | 0 | - |
| - | - | 652.6 | 922.8 | - | - | 0 | - |
| 3 | y | 1.196E+04 | 953.3 | 0.005862 | 6.149 | +1 | 7 |
| - | - | 6472 | 954.3 | - | - | 0 | - |
| - | - | 2554 | 955.3 | - | - | 0 | - |
| - | - | 972.8 | 991.5 | - | - | 0 | - |
| - | - | 2957 | 1020 | - | - | 0 | - |
| - | - | 1794 | 1021 | - | - | 0 | - |
| 2 | y | 698.1 | 1100 | 0.008798 | 7.995 | +1 | 8 |
| - | - | 607.8 | 1101 | - | - | 0 | - |
| - | - | 636.8 | 1788 | - | - | 0 | - |
| - | - | 643.5 | 1859 | - | - | 0 | - |

m/z Charge Intensity FragmentType MassShift Position
120.08091735839844 0 23050.451
121.0843505859375 0 1682.6952
121.97453308105469 0 343.5329
122.06492614746094 0 339.3878
126.33463287353516 0 400.08463
129.1024169921875 0 4047.6904
130.06517028808594 0 464.51706
133.08607482910156 0 879.50574
136.07582092285156 0 7542.396
137.0790557861328 0 447.539
138.91835021972656 0 394.6437
146.060546875 0 559.8517
148.9472198486328 0 1107.1702
149.04495239257812 0 1205.9272
155.08200073242188 0 559.9267
156.0768280029297 0 955.1914
158.0966796875 0 1068.2714
159.0918426513672 0 643.7532
165.05482482910156 0 2623.503
167.05564880371094 0 1550.6652
173.12850952148438 0 1393.935
173.4520263671875 0 1919.1523
176.10719299316406 0 2118.2017
176.24058532714844 0 490.44196
177.1024169921875 0 1188.849
177.111083984375 0 987.93604
182.08123779296875 0 3896.4019
183.08465576171875 0 650.06287
183.64031982421875 0 518.27856
185.1288299560547 0 538.28
188.0707244873047 0 1328.4014
191.1176300048828 0 504.7279
193.10792541503906 0 772.34186
199.0548553466797 0 512.9308
200.1388702392578 0 535.98285
201.12342834472656 0 1540.4952
203.1178436279297 0 653.3963 a Water loss 1
205.09698486328125 0 913.8893 y 8
215.12828063964844 0 1099.977
221.1033935546875 0 2174.458
221.12855529785156 0 33404.004 a 1
222.1320037841797 0 4851.6426
223.0631561279297 0 909.20624
239.0950469970703 0 627.9778
240.0958709716797 0 1044.708 b 3
241.07398986816406 0 1152.4971
249.1235809326172 0 5275.235 b 1
250.12692260742188 0 850.7255
251.1029510498047 0 576.7963
263.1023254394531 0 1185.8761
283.0310363769531 0 655.64514
285.009521484375 0 1185.5597
299.0608825683594 0 655.96014
309.2041015625 0 1623.1908
322.20697021484375 0 603.95087
324.1454772949219 0 1594.6768
329.1505126953125 0 2453.2085
342.15667724609375 0 6032.1177 y 7
343.1587829589844 0 1619.5554
346.1424255371094 0 524.6082 b Water loss 2
355.0696716308594 0 1368.6156
364.1519470214844 0 1567.7953 b 2
368.1901550292969 0 685.27905
378.12835693359375 0 690.9159
394.12628173828125 0 1064.8755
413.2660827636719 0 583.2538
415.9132995605469 0 526.0584
421.7803649902344 0 625.14124
424.627685546875 0 556.6484
425.1910705566406 0 906.3271
438.9053955078125 0 612.856
465.16033935546875 0 808.54663
468.04644775390625 0 534.34644
479.17718505859375 0 1211.833 b 3
489.19207763671875 0 1940.2706 y 6
550.7120361328125 0 820.77814 y 1
560.229248046875 0 2340.1274 y 5
561.2298583984375 0 639.1212
585.339599609375 0 1196.7185
601.2330322265625 0 811.3406 Precursor
602.3145141601562 0 646.2567
613.3444213867188 0 645.9654
614.2407836914062 0 677.6188
625.7219848632812 0 575.6957
685.2603149414062 0 653.8171
686.2556762695312 0 605.11694
700.3645629882812 0 2632.616
701.3670654296875 0 841.40454
721.2572021484375 0 648.07806
723.2923583984375 0 2186.3494 y 4
724.2955322265625 0 602.22363
774.3260498046875 0 915.15515
801.4122924804688 0 1482.4294
838.3212890625 0 2875.795 y 3
839.325439453125 0 1636.1659
844.4539794921875 0 1632.8617
845.45458984375 0 1383.9042
872.4481811523438 0 7396.597
873.4500732421875 0 2922.7988
874.4556884765625 0 1103.8738
889.3478393554688 0 3195.837
890.3544921875 0 1605.2029
891.3509521484375 0 979.4821
922.7581787109375 0 652.5889
953.3467407226562 0 11955.578 y 2
954.349853515625 0 6472.266
955.3483276367188 0 2553.7522
991.523681640625 0 972.8074
1019.5185546875 0 2957.2563
1020.517333984375 0 1793.712
1100.4180908203125 0 698.12244 y 1
1101.4215087890625 0 607.775
1787.7490234375 0 636.81683
1858.6639404296875 0 643.47107

Spectrum Details

|  |  |
| --- | --- |
| Matched peaks? Matched peaks The total absolute number of peaks matched. Additionally in brackets the total fraction of peaks matched and the total number of peaks is shown. | 17 (14.91% of 114) |
| FDR? FDR The false discovery rate estimated for this peptide. It is calculated by matching all theoretical fragments with a non-integer shift with the raw peaks for this spectrum. This is done with 40 different shifts. The resulting percentage is the average number of annotated peaks over the number of annotated peaks with the correct spectrum. | 0.56% |
| Satellite FDR? Satellite FDR See the FDR for details on its calculation. This satellite ion specific FDR only contains the satellite ions (d/w) for I/L/J positions. | - |
| PSM Score? PSM Score The PSM Score as given by Hecklib to this annotated spectrum. It is shown with three significant figures. | 177 |

#### Spectrum 11913? Spectrum 11913 The raw spectrum of this peptide as annotated by Hecklib. The fragments are coloured according to ion type (see legend). Any peaks with a star '\*' as text can be hovered over to see the full details, first the ion type second the mass shift type. By hovering over the amino acids in the peptide or ions in the legend the corresponding peaks are highlighted. By toggling the 'Unassigned' label you can turn the background (unassigned) peaks on or off in the plot. By updating the slider in the Ion legend you can update the spectrum to only show the top X% of the peaks with labels. The top X% means any peak that is within X% of the highest intensity. By dragging in the spectrum you can zoom in to a specific part of the spectrum and use 'Zoom Out' to get back to the original zoom level. The annotation of the spectrum is based on the given sequence in the peptides file and is done with different software so inconsistencies are likely. The peaks are annotated based on the given sequence, with 20 ppm tolerance.

Copy Data

##### Spectrum 11913 (TSV)

###### Preview

```
Loading example...
```

*Click on the button to copy the data to your clipboard.*

Mz MinMz MaxIntensity Max

WidthHeightPeptide font sizePeptide stroke widthSpectrum font sizeSpectrum stroke widthCompact peptide

Ion legend

wxyz

abcd

OtherUnassignedIon^Charge^_Position_Show for top:%

TFDDYAMHW

01.22e+42.45e+43.67e+44.89e+4

Zoom Out

a^+1^_2_y^+1^_1_a^+1^_2_b^+1^_2_y^+1^_2_b^+1^_3_b^+2^_6_b^+1^_3_y^+1^_3_y^+2^_8_y^+1^_4_\*y^+1^_5_y^+1^_6_y^+1^_7_y^+1^_7_y^+1^_8_

0848169725453394

Fragment Matches Table

Show background peaks

| Position | Ion type | Intensity | mz Theoretical | mz Error (Th) | mz Error (ppm) | Charge | Series Number |
| --- | --- | --- | --- | --- | --- | --- | --- |
| - | - | 4.093E+04 | 120.1 | - | - | 0 | - |
| - | - | 3805 | 121.1 | - | - | 0 | - |
| - | - | 316.8 | 121.5 | - | - | 0 | - |
| - | - | 388.8 | 121.9 | - | - | 0 | - |
| - | - | 614.9 | 123 | - | - | 0 | - |
| - | - | 555.2 | 127.1 | - | - | 0 | - |
| - | - | 2812 | 129.1 | - | - | 0 | - |
| - | - | 629.4 | 130.1 | - | - | 0 | - |
| - | - | 784 | 132.1 | - | - | 0 | - |
| - | - | 1472 | 133.1 | - | - | 0 | - |
| - | - | 1206 | 134 | - | - | 0 | - |
| - | - | 422.2 | 135 | - | - | 0 | - |
| - | - | 1.321E+04 | 136.1 | - | - | 0 | - |
| - | - | 882.5 | 137.1 | - | - | 0 | - |
| - | - | 492.4 | 138.1 | - | - | 0 | - |
| - | - | 596.3 | 138.1 | - | - | 0 | - |
| - | - | 990.2 | 141.1 | - | - | 0 | - |
| - | - | 469.9 | 142.4 | - | - | 0 | - |
| - | - | 661.4 | 149 | - | - | 0 | - |
| - | - | 467.1 | 154.2 | - | - | 0 | - |
| - | - | 1020 | 155.1 | - | - | 0 | - |
| - | - | 841.9 | 156.1 | - | - | 0 | - |
| - | - | 385.1 | 156.4 | - | - | 0 | - |
| - | - | 1276 | 157.1 | - | - | 0 | - |
| - | - | 491.2 | 157.1 | - | - | 0 | - |
| - | - | 1561 | 158.1 | - | - | 0 | - |
| - | - | 488.4 | 158.4 | - | - | 0 | - |
| - | - | 1373 | 159.1 | - | - | 0 | - |
| - | - | 969.2 | 160.1 | - | - | 0 | - |
| - | - | 457.2 | 161.1 | - | - | 0 | - |
| - | - | 451.1 | 161.6 | - | - | 0 | - |
| - | - | 427.2 | 161.9 | - | - | 0 | - |
| - | - | 881.6 | 162.1 | - | - | 0 | - |
| - | - | 3522 | 165.1 | - | - | 0 | - |
| - | - | 8320 | 166.1 | - | - | 0 | - |
| - | - | 1716 | 167.1 | - | - | 0 | - |
| - | - | 3189 | 167.1 | - | - | 0 | - |
| - | - | 1227 | 167.1 | - | - | 0 | - |
| - | - | 1172 | 169.1 | - | - | 0 | - |
| - | - | 1881 | 173.1 | - | - | 0 | - |
| - | - | 1246 | 173.4 | - | - | 0 | - |
| - | - | 3787 | 176.1 | - | - | 0 | - |
| - | - | 1917 | 177.1 | - | - | 0 | - |
| - | - | 833.1 | 177.1 | - | - | 0 | - |
| - | - | 6762 | 182.1 | - | - | 0 | - |
| - | - | 490.8 | 183.1 | - | - | 0 | - |
| - | - | 1104 | 185.1 | - | - | 0 | - |
| - | - | 2903 | 188.1 | - | - | 0 | - |
| - | - | 590.5 | 192.1 | - | - | 0 | - |
| - | - | 1150 | 193.1 | - | - | 0 | - |
| - | - | 605.2 | 195.1 | - | - | 0 | - |
| - | - | 894.8 | 195.1 | - | - | 0 | - |
| - | - | 583.1 | 197.1 | - | - | 0 | - |
| - | - | 1294 | 199.1 | - | - | 0 | - |
| - | - | 1295 | 201.1 | - | - | 0 | - |
| - | - | 1719 | 201.1 | - | - | 0 | - |
| 2 | a | 1376 | 203.1 | 7.616E-05 | 0.375 | +1 | 2 |
| - | - | 542.5 | 203.2 | - | - | 0 | - |
| - | - | 695.9 | 204.1 | - | - | 0 | - |
| 9 | y | 1857 | 205.1 | 9.017E-05 | 0.4396 | +1 | 1 |
| - | - | 544.7 | 207.1 | - | - | 0 | - |
| - | - | 664.4 | 209.1 | - | - | 0 | - |
| - | - | 670.1 | 211.1 | - | - | 0 | - |
| - | - | 778.3 | 217.1 | - | - | 0 | - |
| - | - | 3198 | 221.1 | - | - | 0 | - |
| 2 | a | 4.843E+04 | 221.1 | 0.0001163 | 0.5261 | +1 | 2 |
| - | - | 6116 | 222.1 | - | - | 0 | - |
| - | - | 991.6 | 223.1 | - | - | 0 | - |
| - | - | 2342 | 224.1 | - | - | 0 | - |
| - | - | 2965 | 227.1 | - | - | 0 | - |
| - | - | 539.4 | 229.2 | - | - | 0 | - |
| - | - | 729.6 | 231.1 | - | - | 0 | - |
| - | - | 534 | 233.1 | - | - | 0 | - |
| - | - | 3212 | 235.1 | - | - | 0 | - |
| - | - | 1460 | 235.1 | - | - | 0 | - |
| - | - | 1267 | 239.1 | - | - | 0 | - |
| - | - | 956 | 241.1 | - | - | 0 | - |
| - | - | 674.6 | 242.1 | - | - | 0 | - |
| - | - | 518.3 | 245 | - | - | 0 | - |
| 2 | b | 8078 | 249.1 | 0.0002121 | 0.8513 | +1 | 2 |
| - | - | 923.9 | 250.1 | - | - | 0 | - |
| - | - | 1338 | 251.1 | - | - | 0 | - |
| - | - | 649.4 | 251.1 | - | - | 0 | - |
| - | - | 607.4 | 257.2 | - | - | 0 | - |
| - | - | 2260 | 263.1 | - | - | 0 | - |
| - | - | 1385 | 263.1 | - | - | 0 | - |
| - | - | 785 | 265.2 | - | - | 0 | - |
| - | - | 555.3 | 268.1 | - | - | 0 | - |
| - | - | 1000 | 279.1 | - | - | 0 | - |
| - | - | 595.1 | 279.1 | - | - | 0 | - |
| - | - | 1774 | 281.1 | - | - | 0 | - |
| - | - | 1314 | 285 | - | - | 0 | - |
| - | - | 532.6 | 285.1 | - | - | 0 | - |
| - | - | 643.3 | 292.1 | - | - | 0 | - |
| - | - | 500.6 | 296.2 | - | - | 0 | - |
| - | - | 583.9 | 299.1 | - | - | 0 | - |
| - | - | 623.1 | 299.1 | - | - | 0 | - |
| - | - | 798.5 | 300.1 | - | - | 0 | - |
| - | - | 1211 | 302.1 | - | - | 0 | - |
| - | - | 774 | 309.2 | - | - | 0 | - |
| - | - | 593.4 | 315.2 | - | - | 0 | - |
| - | - | 572.6 | 319.4 | - | - | 0 | - |
| - | - | 571.8 | 320.1 | - | - | 0 | - |
| - | - | 1209 | 324.1 | - | - | 0 | - |
| - | - | 602.8 | 325.2 | - | - | 0 | - |
| - | - | 767.8 | 328.1 | - | - | 0 | - |
| - | - | 4226 | 329.1 | - | - | 0 | - |
| - | - | 1116 | 330.2 | - | - | 0 | - |
| - | - | 540.5 | 335 | - | - | 0 | - |
| - | - | 3078 | 340.2 | - | - | 0 | - |
| 8 | y | 9203 | 342.2 | 0.0002451 | 0.7163 | +1 | 2 |
| - | - | 2206 | 343.2 | - | - | 0 | - |
| 3 | b | 590.3 | 346.1 | 0.003228 | 9.325 | +1 | 3 |
| - | - | 638 | 347.2 | - | - | 0 | - |
| - | - | 744.4 | 352.2 | - | - | 0 | - |
| - | - | 1885 | 355.1 | - | - | 0 | - |
| - | - | 888.3 | 356.1 | - | - | 0 | - |
| - | - | 3510 | 356.1 | - | - | 0 | - |
| 6 | b | 569.9 | 357.1 | 0.005554 | 15.55 | +2 | 6 |
| - | - | 895 | 357.2 | - | - | 0 | - |
| - | - | 835.3 | 358.1 | - | - | 0 | - |
| - | - | 645.4 | 359.2 | - | - | 0 | - |
| - | - | 548.5 | 360.3 | - | - | 0 | - |
| 3 | b | 2457 | 364.2 | 0.0002313 | 0.6353 | +1 | 3 |
| - | - | 751.8 | 367.2 | - | - | 0 | - |
| - | - | 1296 | 378.1 | - | - | 0 | - |
| - | - | 1660 | 380.2 | - | - | 0 | - |
| - | - | 675.9 | 391.2 | - | - | 0 | - |
| - | - | 1569 | 394.1 | - | - | 0 | - |
| - | - | 1314 | 398.2 | - | - | 0 | - |
| - | - | 1436 | 413.3 | - | - | 0 | - |
| - | - | 594.4 | 423.2 | - | - | 0 | - |
| - | - | 1081 | 425.2 | - | - | 0 | - |
| - | - | 523.4 | 425.7 | - | - | 0 | - |
| - | - | 622.5 | 437.2 | - | - | 0 | - |
| - | - | 1145 | 465.2 | - | - | 0 | - |
| - | - | 674.4 | 483.2 | - | - | 0 | - |
| 7 | y | 1979 | 489.2 | 0.003849 | 7.867 | +1 | 3 |
| - | - | 733.3 | 493.3 | - | - | 0 | - |
| - | - | 2242 | 496.2 | - | - | 0 | - |
| - | - | 921 | 508.3 | - | - | 0 | - |
| - | - | 2540 | 524.2 | - | - | 0 | - |
| - | - | 762.9 | 525.2 | - | - | 0 | - |
| - | - | 717.6 | 548.2 | - | - | 0 | - |
| 2 | y | 1189 | 550.7 | 0.00131 | 2.379 | +2 | 8 |
| - | - | 737.6 | 551.2 | - | - | 0 | - |
| 6 | y | 3059 | 560.2 | 0.005462 | 9.749 | +1 | 4 |
| - | - | 1491 | 561.2 | - | - | 0 | - |
| - | - | 610 | 568.7 | - | - | 0 | - |
| - | - | 4521 | 585.3 | - | - | 0 | - |
| - | - | 621.4 | 586.3 | - | - | 0 | - |
| - | - | 1018 | 592.7 | - | - | 0 | - |
| 0 | Precursor | 1572 | 601.2 | 0.004509 | 7.5 | +2 | -1 |
| - | - | 829.5 | 601.7 | - | - | 0 | - |
| - | - | 596.1 | 602.3 | - | - | 0 | - |
| - | - | 936.7 | 602.3 | - | - | 0 | - |
| - | - | 649.8 | 615.3 | - | - | 0 | - |
| - | - | 1682 | 618.2 | - | - | 0 | - |
| - | - | 1409 | 621.4 | - | - | 0 | - |
| - | - | 628.7 | 629.5 | - | - | 0 | - |
| - | - | 1743 | 659.3 | - | - | 0 | - |
| - | - | 812 | 685.3 | - | - | 0 | - |
| - | - | 4357 | 700.4 | - | - | 0 | - |
| - | - | 1356 | 701.4 | - | - | 0 | - |
| - | - | 1463 | 721.3 | - | - | 0 | - |
| 5 | y | 2512 | 723.3 | 0.002497 | 3.452 | +1 | 5 |
| - | - | 1694 | 724.3 | - | - | 0 | - |
| - | - | 1357 | 733.4 | - | - | 0 | - |
| - | - | 688.1 | 734.4 | - | - | 0 | - |
| - | - | 1510 | 749.3 | - | - | 0 | - |
| - | - | 1254 | 750.3 | - | - | 0 | - |
| - | - | 1265 | 774.3 | - | - | 0 | - |
| - | - | 1533 | 783.4 | - | - | 0 | - |
| - | - | 732.6 | 784.4 | - | - | 0 | - |
| - | - | 1960 | 786.3 | - | - | 0 | - |
| - | - | 761.7 | 787.3 | - | - | 0 | - |
| - | - | 776.3 | 788.3 | - | - | 0 | - |
| - | - | 935.8 | 800.4 | - | - | 0 | - |
| - | - | 2266 | 801.4 | - | - | 0 | - |
| - | - | 744 | 802.4 | - | - | 0 | - |
| - | - | 1374 | 828.4 | - | - | 0 | - |
| - | - | 778.8 | 832.5 | - | - | 0 | - |
| - | - | 723.2 | 835.4 | - | - | 0 | - |
| 4 | y | 3728 | 838.3 | 0.002959 | 3.529 | +1 | 6 |
| - | - | 1385 | 839.3 | - | - | 0 | - |
| - | - | 3658 | 844.5 | - | - | 0 | - |
| - | - | 1074 | 845.5 | - | - | 0 | - |
| - | - | 1892 | 863.4 | - | - | 0 | - |
| - | - | 890.6 | 864.4 | - | - | 0 | - |
| - | - | 860.2 | 865.4 | - | - | 0 | - |
| - | - | 1017 | 872.3 | - | - | 0 | - |
| - | - | 1.166E+04 | 872.4 | - | - | 0 | - |
| - | - | 6207 | 873.4 | - | - | 0 | - |
| - | - | 1181 | 874.5 | - | - | 0 | - |
| - | - | 5486 | 889.3 | - | - | 0 | - |
| - | - | 2345 | 890.4 | - | - | 0 | - |
| - | - | 711.2 | 902.5 | - | - | 0 | - |
| 3 | y | 1766 | 935.3 | 0.001961 | 2.097 | +1 | 7 |
| - | - | 1011 | 936.3 | - | - | 0 | - |
| 3 | y | 1.706E+04 | 953.3 | 0.003603 | 3.78 | +1 | 7 |
| - | - | 9159 | 954.3 | - | - | 0 | - |
| - | - | 3548 | 955.3 | - | - | 0 | - |
| - | - | 872.1 | 976.5 | - | - | 0 | - |
| - | - | 690.4 | 977.5 | - | - | 0 | - |
| - | - | 1476 | 991.5 | - | - | 0 | - |
| - | - | 1058 | 992.5 | - | - | 0 | - |
| - | - | 5165 | 1020 | - | - | 0 | - |
| - | - | 2183 | 1021 | - | - | 0 | - |
| - | - | 875.3 | 1022 | - | - | 0 | - |
| - | - | 1021 | 1091 | - | - | 0 | - |
| - | - | 693.5 | 1092 | - | - | 0 | - |
| 2 | y | 1382 | 1100 | 0.003305 | 3.003 | +1 | 8 |
| - | - | 694.3 | 2016 | - | - | 0 | - |
| - | - | 703.7 | 2412 | - | - | 0 | - |
| - | - | 637 | 2807 | - | - | 0 | - |
| - | - | 670.8 | 2943 | - | - | 0 | - |
| - | - | 736.1 | 2953 | - | - | 0 | - |
| - | - | 661.3 | 3096 | - | - | 0 | - |
| - | - | 715.6 | 3135 | - | - | 0 | - |
| - | - | 714.7 | 3257 | - | - | 0 | - |
| - | - | 717.8 | 3360 | - | - | 0 | - |

m/z Charge Intensity FragmentType MassShift Position
120.08100128173828 0 40928.44
121.08432006835938 0 3805.117
121.48499298095703 0 316.82214
121.88029479980469 0 388.76553
123.04469299316406 0 614.9135
127.08695983886719 0 555.1989
129.10240173339844 0 2812.0322
130.06532287597656 0 629.4043
132.0810089111328 0 783.97626
133.08599853515625 0 1471.5593
134.02716064453125 0 1205.5123
134.9943084716797 0 422.17752
136.0758819580078 0 13209.471
137.07945251464844 0 882.4943
138.05532836914062 0 492.35635
138.06640625 0 596.25696
141.1025390625 0 990.19775
142.43565368652344 0 469.8874
149.0452423095703 0 661.4245
154.23385620117188 0 467.12857
155.08192443847656 0 1020.4192
156.07701110839844 0 841.9449
156.354248046875 0 385.07788
157.09725952148438 0 1276.2101
157.1341552734375 0 491.17157
158.09642028808594 0 1561.3951
158.44166564941406 0 488.41037
159.0918426513672 0 1372.5121
160.11245727539062 0 969.2395
161.08123779296875 0 457.19373
161.57778930664062 0 451.0537
161.86097717285156 0 427.22977
162.09194946289062 0 881.6158
165.0547332763672 0 3521.6187
166.08641052246094 0 8320.012
167.05569458007812 0 1716.3264
167.08163452148438 0 3189.4788
167.08929443359375 0 1227.1405
169.0976104736328 0 1172.4468
173.12855529785156 0 1881.1722
173.43869018554688 0 1245.5698
176.10714721679688 0 3787.085
177.10256958007812 0 1916.7705
177.1112823486328 0 833.0693
182.08132934570312 0 6761.7837
183.08482360839844 0 490.76556
185.09213256835938 0 1104.1731
188.0706787109375 0 2903.4575
192.06582641601562 0 590.51154
193.10874938964844 0 1150.2129
195.08761596679688 0 605.20844
195.11241149902344 0 894.7924
197.12867736816406 0 583.13776
199.10775756835938 0 1294.2678
201.1128387451172 0 1295.4368
201.1238250732422 0 1719.2493
203.1179656982422 0 1376.259 a Water loss 1
203.1755828857422 0 542.45374
204.07705688476562 0 695.8974
205.0972442626953 0 1856.9075 y 8
207.11196899414062 0 544.69586
209.09225463867188 0 664.41315
211.06007385253906 0 670.0552
217.0970001220703 0 778.28076
221.10353088378906 0 3197.82
221.12857055664062 0 48425.547 a 1
222.13198852539062 0 6116.1826
223.06317138671875 0 991.5845
224.10313415527344 0 2342.1453
227.10304260253906 0 2965.4316
229.1547393798828 0 539.4224
231.09799194335938 0 729.5525
233.0919647216797 0 534.0288
235.07492065429688 0 3211.5183
235.1082000732422 0 1460.2375
239.09510803222656 0 1266.8896
241.07493591308594 0 956.0336
242.11277770996094 0 674.58563
244.95416259765625 0 518.349
249.1235809326172 0 8078.1763 b 1
250.12644958496094 0 923.8932
251.1027374267578 0 1337.864
251.12982177734375 0 649.39197
257.1642761230469 0 607.39984
263.06976318359375 0 2259.8545
263.102783203125 0 1384.6411
265.1542663574219 0 785.00726
268.127197265625 0 555.31067
279.09869384765625 0 1000.4822
279.11810302734375 0 595.1444
281.1246032714844 0 1773.8613
285.01019287109375 0 1313.8687
285.1014404296875 0 532.6352
292.1419372558594 0 643.2723
296.1504821777344 0 500.5965
299.06121826171875 0 583.8811
299.1357421875 0 623.115
300.1188049316406 0 798.52026
302.114013671875 0 1211.2988
309.2060852050781 0 774.02185
315.162353515625 0 593.3856
319.3555603027344 0 572.62213
320.1229553222656 0 571.8297
324.1456298828125 0 1209.3816
325.18634033203125 0 602.7525
328.1332092285156 0 767.8149
329.1497497558594 0 4226.037
330.1534118652344 0 1115.5682
335.0032043457031 0 540.5138
340.18743896484375 0 3078.1394
342.15631103515625 0 9203.359 y 7
343.1595764160156 0 2206.301
346.1429748535156 0 590.27576 b Water loss 2
347.2057800292969 0 637.9964
352.1972351074219 0 744.37427
355.07061767578125 0 1884.775
356.07061767578125 0 888.32605
356.1273193359375 0 3509.6765
357.1369323730469 0 569.91394 b 5
357.22467041015625 0 895.02686
358.1237487792969 0 835.3107
359.19158935546875 0 645.37103
360.3264465332031 0 548.4815
364.1505432128906 0 2457.0225 b 2
367.1959228515625 0 751.798
378.1296081542969 0 1296.1222
380.194580078125 0 1659.8335
391.2330627441406 0 675.9322
394.12493896484375 0 1569.1768
398.20306396484375 0 1313.8965
413.26702880859375 0 1436.329
423.2309875488281 0 594.4116
425.191650390625 0 1081.4174
425.6763916015625 0 523.36633
437.22845458984375 0 622.52155
465.1627502441406 0 1144.797
483.2207946777344 0 674.35065
489.1903991699219 0 1979.0731 y 6
493.27923583984375 0 733.27637
496.2276916503906 0 2242.061
508.2734680175781 0 920.99304
524.2171630859375 0 2540.4468
525.221435546875 0 762.9089
548.2033081054688 0 717.6131
550.7095947265625 0 1189.4681 y 1
551.2096557617188 0 737.6281
560.2291259765625 0 3058.9136 y 5
561.2318115234375 0 1491.2083
568.6983032226562 0 610.02014
585.3435668945312 0 4521.344
586.339111328125 0 621.42084
592.7289428710938 0 1017.6919
601.2366333007812 0 1572.0062 Precursor
601.7359619140625 0 829.5081
602.266845703125 0 596.13336
602.3194580078125 0 936.7257
615.3043212890625 0 649.76715
618.1871948242188 0 1682.2402
621.3602294921875 0 1409.0792
629.5187377929688 0 628.70935
659.2925415039062 0 1742.9052
685.2552490234375 0 811.99365
700.362060546875 0 4356.892
701.3634033203125 0 1355.5168
721.2611694335938 0 1463.2178
723.2894897460938 0 2512.338 y 4
724.2938232421875 0 1693.5474
733.3881225585938 0 1357.2601
734.3955078125 0 688.09985
749.2555541992188 0 1510.3817
750.3104248046875 0 1253.5807
774.32080078125 0 1264.7301
783.3995971679688 0 1533.1357
784.4000244140625 0 732.60583
786.280029296875 0 1960.3085
787.2805786132812 0 761.6861
788.28369140625 0 776.2822
800.4266357421875 0 935.7588
801.4122924804688 0 2266.107
802.412109375 0 743.98
828.4234619140625 0 1374.3651
832.4569702148438 0 778.8448
835.40185546875 0 723.1752
838.31689453125 0 3728.3247 y 3
839.3207397460938 0 1384.9873
844.451904296875 0 3657.7798
845.4561767578125 0 1074.0134
863.3932495117188 0 1892.485
864.4000244140625 0 890.61957
865.39990234375 0 860.21515
872.3456420898438 0 1016.9001
872.4459838867188 0 11662.519
873.447998046875 0 6207.098
874.4506225585938 0 1181.3506
889.3458251953125 0 5486.303
890.3504638671875 0 2345.1355
902.4830322265625 0 711.18134
935.332275390625 0 1766.1473 y Water loss 2
936.3333129882812 0 1010.90497
953.344482421875 0 17061.146 y 2
954.3480834960938 0 9159.155
955.3472290039062 0 3547.9297
976.4818115234375 0 872.0965
977.47607421875 0 690.44324
991.5194091796875 0 1476.1221
992.5214233398438 0 1058.0293
1019.5145874023438 0 5164.855
1020.5193481445312 0 2182.8547
1021.5187377929688 0 875.31915
1090.5167236328125 0 1020.5725
1091.5360107421875 0 693.47284
1100.41259765625 0 1382.3242 y 1
2015.5303955078125 0 694.33453
2412.1884765625 0 703.6901
2807.45458984375 0 636.9927
2943.489013671875 0 670.81946
2952.840087890625 0 736.09753
3096.334716796875 0 661.3402
3134.6591796875 0 715.56177
3257.262451171875 0 714.7392
3360.232177734375 0 717.7942

Spectrum Details

|  |  |
| --- | --- |
| Matched peaks? Matched peaks The total absolute number of peaks matched. Additionally in brackets the total fraction of peaks matched and the total number of peaks is shown. | 17 (7.69% of 221) |
| FDR? FDR The false discovery rate estimated for this peptide. It is calculated by matching all theoretical fragments with a non-integer shift with the raw peaks for this spectrum. This is done with 40 different shifts. The resulting percentage is the average number of annotated peaks over the number of annotated peaks with the correct spectrum. | 3.08% |
| Satellite FDR? Satellite FDR See the FDR for details on its calculation. This satellite ion specific FDR only contains the satellite ions (d/w) for I/L/J positions. | - |
| PSM Score? PSM Score The PSM Score as given by Hecklib to this annotated spectrum. It is shown with three significant figures. | 177 |

#### Spectrum 13536? Spectrum 13536 The raw spectrum of this peptide as annotated by Hecklib. The fragments are coloured according to ion type (see legend). Any peaks with a star '\*' as text can be hovered over to see the full details, first the ion type second the mass shift type. By hovering over the amino acids in the peptide or ions in the legend the corresponding peaks are highlighted. By toggling the 'Unassigned' label you can turn the background (unassigned) peaks on or off in the plot. By updating the slider in the Ion legend you can update the spectrum to only show the top X% of the peaks with labels. The top X% means any peak that is within X% of the highest intensity. By dragging in the spectrum you can zoom in to a specific part of the spectrum and use 'Zoom Out' to get back to the original zoom level. The annotation of the spectrum is based on the given sequence in the peptides file and is done with different software so inconsistencies are likely. The peaks are annotated based on the given sequence, with 20 ppm tolerance.

Copy Data

##### Spectrum 13536 (TSV)

###### Preview

```
Loading example...
```

*Click on the button to copy the data to your clipboard.*

Mz MinMz MaxIntensity Max

WidthHeightPeptide font sizePeptide stroke widthSpectrum font sizeSpectrum stroke widthCompact peptide

Ion legend

wxyz

abcd

OtherUnassignedIon^Charge^_Position_Show for top:%

TFDDYAMHW

07.28e+31.46e+42.19e+42.91e+4

Zoom Out

a^+1^_2_y^+1^_1_a^+1^_2_b^+1^_2_y^+1^_2_b^+1^_3_y^+1^_3_b^+2^_8_y^+2^_8_y^+1^_4_\*y^+1^_5_y^+1^_6_y^+1^_7_y^+1^_7_y^+1^_8_

0568113717052273

Fragment Matches Table

Show background peaks

| Position | Ion type | Intensity | mz Theoretical | mz Error (Th) | mz Error (ppm) | Charge | Series Number |
| --- | --- | --- | --- | --- | --- | --- | --- |
| - | - | 1.944E+04 | 120.1 | - | - | 0 | - |
| - | - | 1917 | 121.1 | - | - | 0 | - |
| - | - | 349.9 | 121.9 | - | - | 0 | - |
| - | - | 478.8 | 129.1 | - | - | 0 | - |
| - | - | 3378 | 129.1 | - | - | 0 | - |
| - | - | 733.2 | 130.1 | - | - | 0 | - |
| - | - | 397.6 | 130.6 | - | - | 0 | - |
| - | - | 469.2 | 132.1 | - | - | 0 | - |
| - | - | 713.6 | 133.1 | - | - | 0 | - |
| - | - | 7535 | 136.1 | - | - | 0 | - |
| - | - | 977.2 | 137.1 | - | - | 0 | - |
| - | - | 432.1 | 143.5 | - | - | 0 | - |
| - | - | 394.4 | 145.5 | - | - | 0 | - |
| - | - | 433.5 | 147.1 | - | - | 0 | - |
| - | - | 1433 | 149 | - | - | 0 | - |
| - | - | 1109 | 156.1 | - | - | 0 | - |
| - | - | 492.1 | 157.1 | - | - | 0 | - |
| - | - | 1065 | 158.1 | - | - | 0 | - |
| - | - | 1031 | 159.1 | - | - | 0 | - |
| - | - | 2595 | 165.1 | - | - | 0 | - |
| - | - | 1942 | 167.1 | - | - | 0 | - |
| - | - | 1433 | 173.1 | - | - | 0 | - |
| - | - | 493 | 173.4 | - | - | 0 | - |
| - | - | 576.4 | 174.1 | - | - | 0 | - |
| - | - | 613 | 175.1 | - | - | 0 | - |
| - | - | 567.7 | 176.1 | - | - | 0 | - |
| - | - | 2126 | 176.1 | - | - | 0 | - |
| - | - | 1382 | 177.1 | - | - | 0 | - |
| - | - | 861 | 177.1 | - | - | 0 | - |
| - | - | 5159 | 182.1 | - | - | 0 | - |
| - | - | 609.5 | 185.1 | - | - | 0 | - |
| - | - | 1260 | 188.1 | - | - | 0 | - |
| - | - | 873.7 | 200.1 | - | - | 0 | - |
| - | - | 1128 | 201.1 | - | - | 0 | - |
| 2 | a | 1016 | 203.1 | 0.0001372 | 0.6755 | +1 | 2 |
| 9 | y | 969.9 | 205.1 | 0.0003648 | 1.779 | +1 | 1 |
| - | - | 2119 | 221.1 | - | - | 0 | - |
| 2 | a | 2.885E+04 | 221.1 | 7.056E-05 | 0.3191 | +1 | 2 |
| - | - | 3797 | 222.1 | - | - | 0 | - |
| - | - | 1495 | 223.1 | - | - | 0 | - |
| - | - | 509.7 | 226.2 | - | - | 0 | - |
| - | - | 1077 | 239.1 | - | - | 0 | - |
| - | - | 1038 | 241.1 | - | - | 0 | - |
| 2 | b | 5184 | 249.1 | 0.0001205 | 0.4838 | +1 | 2 |
| - | - | 716.3 | 250.1 | - | - | 0 | - |
| - | - | 587.4 | 253.2 | - | - | 0 | - |
| - | - | 752.9 | 257.2 | - | - | 0 | - |
| - | - | 1036 | 263.1 | - | - | 0 | - |
| - | - | 634.3 | 269 | - | - | 0 | - |
| - | - | 1086 | 285 | - | - | 0 | - |
| - | - | 601.6 | 287.3 | - | - | 0 | - |
| - | - | 607.7 | 296.2 | - | - | 0 | - |
| - | - | 684.2 | 302.1 | - | - | 0 | - |
| - | - | 517.5 | 319.1 | - | - | 0 | - |
| - | - | 977.1 | 324.1 | - | - | 0 | - |
| - | - | 3066 | 329.1 | - | - | 0 | - |
| - | - | 732.8 | 330.2 | - | - | 0 | - |
| - | - | 566.1 | 334.2 | - | - | 0 | - |
| 8 | y | 4660 | 342.2 | 0.0003366 | 0.9839 | +1 | 2 |
| - | - | 695.2 | 343.2 | - | - | 0 | - |
| - | - | 1593 | 355.1 | - | - | 0 | - |
| - | - | 856 | 356.1 | - | - | 0 | - |
| 3 | b | 1820 | 364.2 | 0.0005316 | 1.46 | +1 | 3 |
| - | - | 593.6 | 378.1 | - | - | 0 | - |
| - | - | 1020 | 394.1 | - | - | 0 | - |
| - | - | 641.8 | 413.3 | - | - | 0 | - |
| - | - | 541 | 417.1 | - | - | 0 | - |
| - | - | 561.1 | 419.8 | - | - | 0 | - |
| - | - | 825.7 | 425.2 | - | - | 0 | - |
| - | - | 612 | 436.7 | - | - | 0 | - |
| - | - | 816.5 | 465.2 | - | - | 0 | - |
| 7 | y | 1985 | 489.2 | 0.004978 | 10.18 | +1 | 3 |
| 8 | b | 696.4 | 490.2 | 0.007703 | 15.71 | +2 | 8 |
| - | - | 644.3 | 496.2 | - | - | 0 | - |
| - | - | 600.4 | 511.5 | - | - | 0 | - |
| - | - | 586.8 | 525.8 | - | - | 0 | - |
| 2 | y | 744.6 | 550.7 | 0.001192 | 2.165 | +2 | 8 |
| 6 | y | 2046 | 560.2 | 0.007659 | 13.67 | +1 | 4 |
| - | - | 669.4 | 571.3 | - | - | 0 | - |
| - | - | 564.4 | 575.6 | - | - | 0 | - |
| - | - | 1025 | 585.3 | - | - | 0 | - |
| - | - | 572.2 | 600.5 | - | - | 0 | - |
| 0 | Precursor | 842.4 | 601.2 | 0.001946 | 3.237 | +2 | -1 |
| - | - | 601.4 | 601.8 | - | - | 0 | - |
| - | - | 587.9 | 619.6 | - | - | 0 | - |
| - | - | 586.4 | 640.3 | - | - | 0 | - |
| - | - | 3459 | 700.4 | - | - | 0 | - |
| - | - | 712.9 | 701.4 | - | - | 0 | - |
| - | - | 900.9 | 715.9 | - | - | 0 | - |
| 5 | y | 1843 | 723.3 | 0.005427 | 7.503 | +1 | 5 |
| - | - | 886.4 | 724.3 | - | - | 0 | - |
| - | - | 980.4 | 738.3 | - | - | 0 | - |
| - | - | 677.1 | 749.2 | - | - | 0 | - |
| - | - | 644.1 | 783.4 | - | - | 0 | - |
| - | - | 1633 | 801.4 | - | - | 0 | - |
| 4 | y | 1681 | 838.3 | 0.002898 | 3.456 | +1 | 6 |
| - | - | 661.9 | 839.3 | - | - | 0 | - |
| - | - | 2103 | 844.5 | - | - | 0 | - |
| - | - | 638.8 | 845.4 | - | - | 0 | - |
| - | - | 779.7 | 871.3 | - | - | 0 | - |
| - | - | 7033 | 872.4 | - | - | 0 | - |
| - | - | 3150 | 873.5 | - | - | 0 | - |
| - | - | 797.1 | 874.5 | - | - | 0 | - |
| - | - | 3415 | 889.3 | - | - | 0 | - |
| - | - | 2260 | 890.4 | - | - | 0 | - |
| 3 | y | 620.5 | 935.3 | 0.00721 | 7.709 | +1 | 7 |
| 3 | y | 1.036E+04 | 953.3 | 0.005618 | 5.893 | +1 | 7 |
| - | - | 6038 | 954.3 | - | - | 0 | - |
| - | - | 2379 | 955.3 | - | - | 0 | - |
| - | - | 850.4 | 991.5 | - | - | 0 | - |
| - | - | 654.1 | 1003 | - | - | 0 | - |
| - | - | 3249 | 1020 | - | - | 0 | - |
| - | - | 2182 | 1021 | - | - | 0 | - |
| - | - | 627.8 | 1035 | - | - | 0 | - |
| 2 | y | 871.8 | 1100 | 0.01075 | 9.77 | +1 | 8 |
| - | - | 638.8 | 1465 | - | - | 0 | - |
| - | - | 781.7 | 1608 | - | - | 0 | - |
| - | - | 731.3 | 2251 | - | - | 0 | - |

m/z Charge Intensity FragmentType MassShift Position
120.0809097290039 0 19439.127
121.0842514038086 0 1917.1809
121.85750579833984 0 349.9443
129.0658721923828 0 478.81497
129.10235595703125 0 3377.5598
130.06517028808594 0 733.21674
130.566650390625 0 397.56363
132.08102416992188 0 469.2119
133.08595275878906 0 713.61
136.0758056640625 0 7535.468
137.07937622070312 0 977.17456
143.5342254638672 0 432.0762
145.50311279296875 0 394.39145
147.11019897460938 0 433.51636
149.04507446289062 0 1432.7992
156.0767364501953 0 1109.0123
157.1337432861328 0 492.14923
158.09681701660156 0 1065.4241
159.09188842773438 0 1030.8132
165.05474853515625 0 2594.828
167.05563354492188 0 1941.5791
173.12840270996094 0 1433.2332
173.404052734375 0 492.98883
174.0554656982422 0 576.391
175.1191864013672 0 612.9983
176.08216857910156 0 567.69684
176.10716247558594 0 2125.654
177.10243225097656 0 1382.0244
177.11184692382812 0 860.997
182.0812530517578 0 5159.1855
185.129150390625 0 609.50946
188.07070922851562 0 1260.3499
200.1395721435547 0 873.69366
201.12344360351562 0 1128.2343
203.11802673339844 0 1015.7012 a Water loss 1
205.09751892089844 0 969.86206 y 8
221.10337829589844 0 2118.789
221.12852478027344 0 28849.342 a 1
222.1318817138672 0 3797.1982
223.0633087158203 0 1494.6271
226.15505981445312 0 509.6991
239.0945281982422 0 1076.9375
241.07444763183594 0 1037.6122
249.1234893798828 0 5184.168 b 1
250.1263427734375 0 716.29913
253.16612243652344 0 587.4427
257.16412353515625 0 752.9283
263.10302734375 0 1035.5892
268.97900390625 0 634.3303
285.0099182128906 0 1086.2838
287.2911682128906 0 601.61597
296.15155029296875 0 607.71344
302.1139221191406 0 684.1987
319.1408996582031 0 517.4997
324.1452331542969 0 977.0596
329.1495361328125 0 3066.1902
330.1519470214844 0 732.78925
334.2149353027344 0 566.1405
342.1564025878906 0 4659.918 y 7
343.1583557128906 0 695.1991
355.069580078125 0 1593.1556
356.0704650878906 0 856.0214
364.1497802734375 0 1820.142 b 2
378.1306457519531 0 593.6171
394.1249694824219 0 1020.1358
413.2660217285156 0 641.80493
417.0854187011719 0 540.99677
419.838623046875 0 561.1183
425.1929016113281 0 825.68243
436.7276916503906 0 612.02594
465.16192626953125 0 816.5424
489.1915283203125 0 1984.82 y 6
490.1896057128906 0 696.4179 b Water loss 7
496.22930908203125 0 644.2729
511.5450439453125 0 600.3865
525.8155517578125 0 586.78125
550.7070922851562 0 744.5998 y 1
560.2313232421875 0 2046.0023 y 5
571.315185546875 0 669.3752
575.6355590820312 0 564.3718
585.3381958007812 0 1024.503
600.5475463867188 0 572.1546
601.2340698242188 0 842.42206 Precursor
601.7503051757812 0 601.3801
619.6246337890625 0 587.8877
640.3397827148438 0 586.3556
700.3632202148438 0 3459.359
701.3663330078125 0 712.895
715.8626708984375 0 900.8767
723.2924194335938 0 1842.7648 y 4
724.2919311523438 0 886.39655
738.3451538085938 0 980.3913
749.24853515625 0 677.0585
783.4046020507812 0 644.11475
801.411865234375 0 1632.6785
838.3168334960938 0 1680.6133 y 3
839.3193359375 0 661.9332
844.45361328125 0 2103.375
845.449462890625 0 638.8118
871.3294677734375 0 779.70184
872.4480590820312 0 7032.8096
873.4511108398438 0 3149.6252
874.45703125 0 797.1077
889.346435546875 0 3414.6697
890.3512573242188 0 2260.4114
935.3375244140625 0 620.53656 y Water loss 2
953.3464965820312 0 10357.566 y 2
954.3494873046875 0 6038.3623
955.3488159179688 0 2378.7961
991.525146484375 0 850.3755
1003.3829956054688 0 654.1322
1019.5160522460938 0 3248.9233
1020.5192260742188 0 2181.6716
1034.519775390625 0 627.8392
1100.4200439453125 0 871.7981 y 1
1464.702392578125 0 638.76746
1608.144287109375 0 781.74805
2250.92529296875 0 731.2677

Spectrum Details

|  |  |
| --- | --- |
| Matched peaks? Matched peaks The total absolute number of peaks matched. Additionally in brackets the total fraction of peaks matched and the total number of peaks is shown. | 16 (13.56% of 118) |
| FDR? FDR The false discovery rate estimated for this peptide. It is calculated by matching all theoretical fragments with a non-integer shift with the raw peaks for this spectrum. This is done with 40 different shifts. The resulting percentage is the average number of annotated peaks over the number of annotated peaks with the correct spectrum. | 0.89% |
| Satellite FDR? Satellite FDR See the FDR for details on its calculation. This satellite ion specific FDR only contains the satellite ions (d/w) for I/L/J positions. | - |
| PSM Score? PSM Score The PSM Score as given by Hecklib to this annotated spectrum. It is shown with three significant figures. | 160 |

#### Spectrum 10975? Spectrum 10975 The raw spectrum of this peptide as annotated by Hecklib. The fragments are coloured according to ion type (see legend). Any peaks with a star '\*' as text can be hovered over to see the full details, first the ion type second the mass shift type. By hovering over the amino acids in the peptide or ions in the legend the corresponding peaks are highlighted. By toggling the 'Unassigned' label you can turn the background (unassigned) peaks on or off in the plot. By updating the slider in the Ion legend you can update the spectrum to only show the top X% of the peaks with labels. The top X% means any peak that is within X% of the highest intensity. By dragging in the spectrum you can zoom in to a specific part of the spectrum and use 'Zoom Out' to get back to the original zoom level. The annotation of the spectrum is based on the given sequence in the peptides file and is done with different software so inconsistencies are likely. The peaks are annotated based on the given sequence, with 20 ppm tolerance.

Copy Data

##### Spectrum 10975 (TSV)

###### Preview

```
Loading example...
```

*Click on the button to copy the data to your clipboard.*

Mz MinMz MaxIntensity Max

WidthHeightPeptide font sizePeptide stroke widthSpectrum font sizeSpectrum stroke widthCompact peptide

Ion legend

wxyz

abcd

OtherUnassignedIon^Charge^_Position_Show for top:%

TFDDYAMHW

01.62e+43.25e+44.87e+46.50e+4

Zoom Out

a^+1^_2_y^+1^_1_a^+1^_2_b^+1^_2_b^+1^_2_y^+1^_2_b^+1^_3_b^+1^_3_b^+1^_4_y^+1^_3_y^+2^_8_y^+1^_4_\*\*y^+1^_5_y^+1^_6_y^+1^_7_y^+1^_7_

0629125818872516

Fragment Matches Table

Show background peaks

| Position | Ion type | Intensity | mz Theoretical | mz Error (Th) | mz Error (ppm) | Charge | Series Number |
| --- | --- | --- | --- | --- | --- | --- | --- |
| - | - | 4.674E+04 | 120.1 | - | - | 0 | - |
| - | - | 393.2 | 120.6 | - | - | 0 | - |
| - | - | 2756 | 121.1 | - | - | 0 | - |
| - | - | 395.9 | 122.7 | - | - | 0 | - |
| - | - | 738.4 | 123 | - | - | 0 | - |
| - | - | 6301 | 129.1 | - | - | 0 | - |
| - | - | 609.1 | 130 | - | - | 0 | - |
| - | - | 1245 | 130.1 | - | - | 0 | - |
| - | - | 447.5 | 130.1 | - | - | 0 | - |
| - | - | 528.5 | 132.1 | - | - | 0 | - |
| - | - | 715.4 | 133.1 | - | - | 0 | - |
| - | - | 1.559E+04 | 136.1 | - | - | 0 | - |
| - | - | 1431 | 137.1 | - | - | 0 | - |
| - | - | 494.9 | 138.1 | - | - | 0 | - |
| - | - | 813.2 | 146.1 | - | - | 0 | - |
| - | - | 810.3 | 147 | - | - | 0 | - |
| - | - | 1168 | 147.1 | - | - | 0 | - |
| - | - | 661.4 | 149 | - | - | 0 | - |
| - | - | 1752 | 149 | - | - | 0 | - |
| - | - | 503.2 | 155.1 | - | - | 0 | - |
| - | - | 575.3 | 155.1 | - | - | 0 | - |
| - | - | 1007 | 156.1 | - | - | 0 | - |
| - | - | 790.1 | 157.1 | - | - | 0 | - |
| - | - | 2694 | 158.1 | - | - | 0 | - |
| - | - | 1754 | 159.1 | - | - | 0 | - |
| - | - | 1078 | 160.1 | - | - | 0 | - |
| - | - | 443.1 | 161.5 | - | - | 0 | - |
| - | - | 654.8 | 162.1 | - | - | 0 | - |
| - | - | 5710 | 165.1 | - | - | 0 | - |
| - | - | 1507 | 166.1 | - | - | 0 | - |
| - | - | 2553 | 167.1 | - | - | 0 | - |
| - | - | 583.4 | 169 | - | - | 0 | - |
| - | - | 3211 | 173.1 | - | - | 0 | - |
| - | - | 464.3 | 174.1 | - | - | 0 | - |
| - | - | 725.6 | 176.1 | - | - | 0 | - |
| - | - | 4144 | 176.1 | - | - | 0 | - |
| - | - | 3517 | 177.1 | - | - | 0 | - |
| - | - | 1116 | 177.1 | - | - | 0 | - |
| - | - | 1.19E+04 | 182.1 | - | - | 0 | - |
| - | - | 972.2 | 183.1 | - | - | 0 | - |
| - | - | 2611 | 183.1 | - | - | 0 | - |
| - | - | 469.5 | 183.6 | - | - | 0 | - |
| - | - | 904.8 | 185.1 | - | - | 0 | - |
| - | - | 2301 | 188.1 | - | - | 0 | - |
| - | - | 716.7 | 191.1 | - | - | 0 | - |
| - | - | 1105 | 193.1 | - | - | 0 | - |
| - | - | 566.3 | 195.1 | - | - | 0 | - |
| - | - | 1026 | 197.1 | - | - | 0 | - |
| - | - | 874.8 | 201.1 | - | - | 0 | - |
| - | - | 3824 | 201.1 | - | - | 0 | - |
| - | - | 974.8 | 203.1 | - | - | 0 | - |
| 2 | a | 1835 | 203.1 | 0.0001525 | 0.7506 | +1 | 2 |
| 9 | y | 1404 | 205.1 | 0.0003218 | 1.569 | +1 | 1 |
| - | - | 671.4 | 207.4 | - | - | 0 | - |
| - | - | 651.1 | 211.1 | - | - | 0 | - |
| - | - | 712.2 | 214.1 | - | - | 0 | - |
| - | - | 462.2 | 214.3 | - | - | 0 | - |
| - | - | 954.2 | 217.1 | - | - | 0 | - |
| - | - | 3724 | 221.1 | - | - | 0 | - |
| 2 | a | 6.433E+04 | 221.1 | 0.0001316 | 0.5951 | +1 | 2 |
| - | - | 7922 | 222.1 | - | - | 0 | - |
| - | - | 1447 | 223.1 | - | - | 0 | - |
| - | - | 616.3 | 226.1 | - | - | 0 | - |
| - | - | 1119 | 229.1 | - | - | 0 | - |
| - | - | 943.7 | 231.1 | - | - | 0 | - |
| 2 | b | 715.3 | 231.1 | 0.001057 | 4.573 | +1 | 2 |
| - | - | 1264 | 235.1 | - | - | 0 | - |
| - | - | 510.4 | 235.7 | - | - | 0 | - |
| - | - | 1955 | 239.1 | - | - | 0 | - |
| - | - | 549.3 | 239.1 | - | - | 0 | - |
| - | - | 512.9 | 239.2 | - | - | 0 | - |
| - | - | 594.7 | 240.1 | - | - | 0 | - |
| - | - | 843.5 | 241.1 | - | - | 0 | - |
| 2 | b | 1.086E+04 | 249.1 | 0.0002273 | 0.9126 | +1 | 2 |
| - | - | 1347 | 250.1 | - | - | 0 | - |
| - | - | 1667 | 251.1 | - | - | 0 | - |
| - | - | 880.5 | 251.2 | - | - | 0 | - |
| - | - | 670.6 | 253.1 | - | - | 0 | - |
| - | - | 1051 | 257.1 | - | - | 0 | - |
| - | - | 684.8 | 257.2 | - | - | 0 | - |
| - | - | 2079 | 263.1 | - | - | 0 | - |
| - | - | 1177 | 279.1 | - | - | 0 | - |
| - | - | 1479 | 285 | - | - | 0 | - |
| - | - | 1426 | 285.1 | - | - | 0 | - |
| - | - | 766.9 | 292.1 | - | - | 0 | - |
| - | - | 982.1 | 296.2 | - | - | 0 | - |
| - | - | 1056 | 299.1 | - | - | 0 | - |
| - | - | 960.1 | 302.2 | - | - | 0 | - |
| - | - | 534.1 | 314.4 | - | - | 0 | - |
| - | - | 574 | 319.1 | - | - | 0 | - |
| - | - | 2325 | 324.1 | - | - | 0 | - |
| - | - | 636.5 | 326.2 | - | - | 0 | - |
| - | - | 6455 | 329.1 | - | - | 0 | - |
| - | - | 1823 | 330.2 | - | - | 0 | - |
| - | - | 488.3 | 333.5 | - | - | 0 | - |
| - | - | 1333 | 340.2 | - | - | 0 | - |
| 8 | y | 1.232E+04 | 342.2 | 0.0002146 | 0.6271 | +1 | 2 |
| - | - | 2479 | 343.2 | - | - | 0 | - |
| 3 | b | 648 | 346.1 | 0.0009999 | 2.889 | +1 | 3 |
| - | - | 1873 | 355.1 | - | - | 0 | - |
| - | - | 694.5 | 356.1 | - | - | 0 | - |
| - | - | 1200 | 357.2 | - | - | 0 | - |
| 3 | b | 3386 | 364.2 | 0.0001959 | 0.538 | +1 | 3 |
| - | - | 1014 | 368.2 | - | - | 0 | - |
| - | - | 545.5 | 376.1 | - | - | 0 | - |
| - | - | 2148 | 378.1 | - | - | 0 | - |
| - | - | 819.6 | 392.2 | - | - | 0 | - |
| - | - | 2198 | 394.1 | - | - | 0 | - |
| - | - | 635.9 | 395.1 | - | - | 0 | - |
| - | - | 610.1 | 401.7 | - | - | 0 | - |
| - | - | 918 | 413.3 | - | - | 0 | - |
| - | - | 1149 | 425.2 | - | - | 0 | - |
| - | - | 751 | 426.2 | - | - | 0 | - |
| - | - | 869.4 | 427.7 | - | - | 0 | - |
| - | - | 950.1 | 436.7 | - | - | 0 | - |
| - | - | 1209 | 465.2 | - | - | 0 | - |
| - | - | 537.3 | 471.2 | - | - | 0 | - |
| 4 | b | 1408 | 479.2 | 0.0003879 | 0.8095 | +1 | 4 |
| - | - | 839.1 | 483.2 | - | - | 0 | - |
| 7 | y | 2285 | 489.2 | 0.004093 | 8.366 | +1 | 3 |
| - | - | 1073 | 490.2 | - | - | 0 | - |
| - | - | 752.6 | 496.2 | - | - | 0 | - |
| - | - | 605.2 | 501.3 | - | - | 0 | - |
| - | - | 630.5 | 501.8 | - | - | 0 | - |
| - | - | 582.6 | 521.2 | - | - | 0 | - |
| - | - | 524.4 | 535.6 | - | - | 0 | - |
| 2 | y | 697.3 | 550.7 | 0.005094 | 9.25 | +2 | 8 |
| - | - | 1304 | 551.2 | - | - | 0 | - |
| - | - | 506.9 | 551.7 | - | - | 0 | - |
| 6 | y | 4637 | 560.2 | 0.004241 | 7.57 | +1 | 4 |
| - | - | 1294 | 561.2 | - | - | 0 | - |
| - | - | 2796 | 585.3 | - | - | 0 | - |
| - | - | 1373 | 586.3 | - | - | 0 | - |
| 0 | Precursor | 1004 | 592.2 | 0.0001482 | 0.2502 | +2 | -1 |
| - | - | 764.1 | 592.7 | - | - | 0 | - |
| - | - | 587.6 | 600.2 | - | - | 0 | - |
| 0 | Precursor | 1835 | 601.2 | 0.001336 | 2.221 | +2 | -1 |
| - | - | 627.6 | 601.3 | - | - | 0 | - |
| - | - | 1108 | 601.7 | - | - | 0 | - |
| - | - | 826.2 | 602.2 | - | - | 0 | - |
| - | - | 823.2 | 606.2 | - | - | 0 | - |
| - | - | 944.1 | 612.2 | - | - | 0 | - |
| - | - | 596.2 | 634.2 | - | - | 0 | - |
| - | - | 627.8 | 640.3 | - | - | 0 | - |
| - | - | 1177 | 657.3 | - | - | 0 | - |
| - | - | 1973 | 659.3 | - | - | 0 | - |
| - | - | 735.5 | 663.4 | - | - | 0 | - |
| - | - | 716.8 | 665.8 | - | - | 0 | - |
| - | - | 2000 | 685.3 | - | - | 0 | - |
| - | - | 6034 | 700.4 | - | - | 0 | - |
| - | - | 2757 | 701.4 | - | - | 0 | - |
| - | - | 959.1 | 705.1 | - | - | 0 | - |
| - | - | 1122 | 721.3 | - | - | 0 | - |
| 5 | y | 4558 | 723.3 | 0.004328 | 5.984 | +1 | 5 |
| - | - | 2129 | 724.3 | - | - | 0 | - |
| - | - | 2138 | 749.3 | - | - | 0 | - |
| - | - | 1078 | 750.3 | - | - | 0 | - |
| - | - | 909.1 | 757.4 | - | - | 0 | - |
| - | - | 1887 | 774.3 | - | - | 0 | - |
| - | - | 864.6 | 775.3 | - | - | 0 | - |
| - | - | 1495 | 783.4 | - | - | 0 | - |
| - | - | 976.6 | 784.4 | - | - | 0 | - |
| - | - | 2781 | 801.4 | - | - | 0 | - |
| - | - | 1453 | 802.4 | - | - | 0 | - |
| - | - | 1501 | 828.4 | - | - | 0 | - |
| 4 | y | 4975 | 838.3 | 0.005888 | 7.024 | +1 | 6 |
| - | - | 2838 | 839.3 | - | - | 0 | - |
| - | - | 787.6 | 840.3 | - | - | 0 | - |
| - | - | 4752 | 844.5 | - | - | 0 | - |
| - | - | 1767 | 845.5 | - | - | 0 | - |
| - | - | 742.5 | 854.4 | - | - | 0 | - |
| - | - | 718.4 | 871.3 | - | - | 0 | - |
| - | - | 1.867E+04 | 872.4 | - | - | 0 | - |
| - | - | 9645 | 873.5 | - | - | 0 | - |
| - | - | 3013 | 874.5 | - | - | 0 | - |
| - | - | 731.9 | 875.4 | - | - | 0 | - |
| - | - | 7981 | 889.3 | - | - | 0 | - |
| - | - | 3855 | 890.3 | - | - | 0 | - |
| 3 | y | 828.3 | 935.3 | 0.001595 | 1.705 | +1 | 7 |
| - | - | 758.9 | 936.3 | - | - | 0 | - |
| 3 | y | 2.423E+04 | 953.3 | 0.003726 | 3.908 | +1 | 7 |
| - | - | 1.359E+04 | 954.3 | - | - | 0 | - |
| - | - | 4921 | 955.3 | - | - | 0 | - |
| - | - | 737.4 | 956.3 | - | - | 0 | - |
| - | - | 741.5 | 969.4 | - | - | 0 | - |
| - | - | 3408 | 991.5 | - | - | 0 | - |
| - | - | 1685 | 992.5 | - | - | 0 | - |
| - | - | 8527 | 1020 | - | - | 0 | - |
| - | - | 5245 | 1021 | - | - | 0 | - |
| - | - | 1446 | 1022 | - | - | 0 | - |
| - | - | 606.2 | 1995 | - | - | 0 | - |
| - | - | 698.1 | 2491 | - | - | 0 | - |

m/z Charge Intensity FragmentType MassShift Position
120.08100128173828 0 46740.51
120.56090545654297 0 393.1669
121.08431243896484 0 2755.7686
122.74697875976562 0 395.8555
123.04412078857422 0 738.385
129.10243225097656 0 6301.154
130.04995727539062 0 609.0855
130.0651397705078 0 1245.0934
130.10585021972656 0 447.4767
132.08099365234375 0 528.4666
133.08592224121094 0 715.3761
136.0758819580078 0 15593.492
137.07925415039062 0 1431.3043
138.09117126464844 0 494.92267
146.05990600585938 0 813.20605
147.04408264160156 0 810.2961
147.11293029785156 0 1167.6353
148.9544677734375 0 661.4277
149.04518127441406 0 1751.7511
155.08180236816406 0 503.17816
155.0925750732422 0 575.30145
156.07679748535156 0 1006.5841
157.13414001464844 0 790.0677
158.09649658203125 0 2694.0532
159.0917205810547 0 1754.0331
160.11241149902344 0 1077.6149
161.4528045654297 0 443.08765
162.0916748046875 0 654.79816
165.05479431152344 0 5709.547
166.0863800048828 0 1506.6515
167.05560302734375 0 2552.6787
169.03480529785156 0 583.4233
173.1287078857422 0 3211.188
174.0555419921875 0 464.30237
176.08140563964844 0 725.6238
176.107177734375 0 4143.715
177.10238647460938 0 3516.6401
177.1107940673828 0 1115.6007
182.081298828125 0 11900.494
183.0849151611328 0 972.22485
183.1129608154297 0 2610.681
183.5887451171875 0 469.47165
185.12879943847656 0 904.7648
188.07073974609375 0 2300.9243
191.11846923828125 0 716.6707
193.1086883544922 0 1104.5521
195.11366271972656 0 566.2662
197.12820434570312 0 1025.851
201.06619262695312 0 874.75073
201.1234130859375 0 3824.05
203.06640625 0 974.8374
203.1180419921875 0 1834.6345 a Water loss 1
205.09683227539062 0 1404.1304 y 8
207.42982482910156 0 671.4481
211.1074981689453 0 651.0696
214.1190948486328 0 712.16644
214.31985473632812 0 462.20847
217.09738159179688 0 954.2214
221.1033935546875 0 3724.158
221.1285858154297 0 64326.234 a 1
222.1320343017578 0 7922.498
223.063720703125 0 1447.0482
226.11875915527344 0 616.3162
229.11856079101562 0 1119.41
231.06166076660156 0 943.7185
231.11386108398438 0 715.25867 b Water loss 1
235.10806274414062 0 1264.3804
235.6918487548828 0 510.43488
239.09506225585938 0 1955.398
239.11514282226562 0 549.3318
239.15087890625 0 512.94794
240.09803771972656 0 594.6757
241.07479858398438 0 843.46063
249.12359619140625 0 10858.471 b 1
250.1266326904297 0 1347.4777
251.10302734375 0 1666.8827
251.1503448486328 0 880.5416
253.1186065673828 0 670.6445
257.1070861816406 0 1050.7532
257.1660461425781 0 684.77985
263.1029052734375 0 2078.8098
279.0982666015625 0 1177.4305
285.01043701171875 0 1479.261
285.1011962890625 0 1426.2645
292.14093017578125 0 766.8946
296.1502380371094 0 982.05634
299.0608215332031 0 1056.1381
302.1510314941406 0 960.06726
314.3644714355469 0 534.08777
319.1399841308594 0 574.0252
324.1460266113281 0 2324.7764
326.17266845703125 0 636.4515
329.14959716796875 0 6455.3965
330.1524353027344 0 1822.8987
333.5063171386719 0 488.3156
340.19873046875 0 1333.343
342.1562805175781 0 12320.013 y 7
343.159912109375 0 2479.0103
346.1407470703125 0 648.0394 b Water loss 2
355.0693359375 0 1872.7556
356.1391296386719 0 694.5188
357.22344970703125 0 1200.3235
364.1501159667969 0 3385.565 b 2
368.19329833984375 0 1014.0484
376.1138000488281 0 545.4798
378.1297607421875 0 2148.2625
392.20098876953125 0 819.5587
394.12432861328125 0 2198.3872
395.1277770996094 0 635.91473
401.7090148925781 0 610.061
413.26806640625 0 917.98926
425.19219970703125 0 1149.4167
426.1963806152344 0 751.03906
427.72265625 0 869.42633
436.72705078125 0 950.1381
465.16253662109375 0 1208.5813
471.17877197265625 0 537.2725
479.1776428222656 0 1408.0588 b 3
483.2194519042969 0 839.0514
489.1906433105469 0 2284.8916 y 6
490.1966247558594 0 1073.326
496.22900390625 0 752.5939
501.2586364746094 0 605.2061
501.7550964355469 0 630.53156
521.1957397460938 0 582.5681
535.5989379882812 0 524.3951
550.71337890625 0 697.33356 y 1
551.2122192382812 0 1304.3818
551.712890625 0 506.875
560.2279052734375 0 4636.975 y 5
561.2335815429688 0 1294.362
585.3357543945312 0 2795.8323
586.3384399414062 0 1372.5186
592.2269897460938 0 1004.05084 Precursor Water loss
592.7288208007812 0 764.085
600.2341918945312 0 587.6354
601.2334594726562 0 1835.32 Precursor
601.3290405273438 0 627.6167
601.7385864257812 0 1108.4683
602.2393188476562 0 826.21844
606.2278442382812 0 823.2384
612.199462890625 0 944.13995
634.2247314453125 0 596.1634
640.3389282226562 0 627.7767
657.2630615234375 0 1176.9474
659.29150390625 0 1973.2502
663.3670654296875 0 735.4609
665.775146484375 0 716.8094
685.2573852539062 0 2000.158
700.3619384765625 0 6033.7163
701.3651733398438 0 2757.2695
705.0889892578125 0 959.0772
721.2636108398438 0 1122.2278
723.2913208007812 0 4557.7373 y 4
724.2938842773438 0 2128.7815
749.2557983398438 0 2137.8767
750.2570190429688 0 1078.4265
757.3778686523438 0 909.06494
774.320556640625 0 1886.8313
775.3211669921875 0 864.55505
783.3990478515625 0 1494.9152
784.4005126953125 0 976.55725
801.4112548828125 0 2780.8213
802.4124755859375 0 1453.1184
828.4187622070312 0 1500.5481
838.31982421875 0 4975.474 y 3
839.3209228515625 0 2838.2498
840.3252563476562 0 787.6156
844.4518432617188 0 4751.9907
845.4539794921875 0 1766.712
854.435791015625 0 742.467
871.34228515625 0 718.4388
872.44677734375 0 18674.553
873.4500122070312 0 9645.181
874.4528198242188 0 3013.2292
875.4478759765625 0 731.879
889.3461303710938 0 7980.592
890.348876953125 0 3855.1892
935.3319091796875 0 828.3349 y Water loss 2
936.3255004882812 0 758.9032
953.3446044921875 0 24227.498 y 2
954.3477172851562 0 13588.331
955.3477172851562 0 4920.8325
956.3436279296875 0 737.4179
969.371826171875 0 741.539
991.5183715820312 0 3407.66
992.5233764648438 0 1685.4426
1019.513916015625 0 8526.58
1020.5161743164062 0 5245.145
1021.525390625 0 1445.5961
1995.14892578125 0 606.2345
2491.3642578125 0 698.0795

Spectrum Details

|  |  |
| --- | --- |
| Matched peaks? Matched peaks The total absolute number of peaks matched. Additionally in brackets the total fraction of peaks matched and the total number of peaks is shown. | 18 (9.38% of 192) |
| FDR? FDR The false discovery rate estimated for this peptide. It is calculated by matching all theoretical fragments with a non-integer shift with the raw peaks for this spectrum. This is done with 40 different shifts. The resulting percentage is the average number of annotated peaks over the number of annotated peaks with the correct spectrum. | 1.32% |
| Satellite FDR? Satellite FDR See the FDR for details on its calculation. This satellite ion specific FDR only contains the satellite ions (d/w) for I/L/J positions. | - |
| PSM Score? PSM Score The PSM Score as given by Hecklib to this annotated spectrum. It is shown with three significant figures. | 195 |

#### Spectrum 13093? Spectrum 13093 The raw spectrum of this peptide as annotated by Hecklib. The fragments are coloured according to ion type (see legend). Any peaks with a star '\*' as text can be hovered over to see the full details, first the ion type second the mass shift type. By hovering over the amino acids in the peptide or ions in the legend the corresponding peaks are highlighted. By toggling the 'Unassigned' label you can turn the background (unassigned) peaks on or off in the plot. By updating the slider in the Ion legend you can update the spectrum to only show the top X% of the peaks with labels. The top X% means any peak that is within X% of the highest intensity. By dragging in the spectrum you can zoom in to a specific part of the spectrum and use 'Zoom Out' to get back to the original zoom level. The annotation of the spectrum is based on the given sequence in the peptides file and is done with different software so inconsistencies are likely. The peaks are annotated based on the given sequence, with 20 ppm tolerance.

Copy Data

##### Spectrum 13093 (TSV)

###### Preview

```
Loading example...
```

*Click on the button to copy the data to your clipboard.*

Mz MinMz MaxIntensity Max

WidthHeightPeptide font sizePeptide stroke widthSpectrum font sizeSpectrum stroke widthCompact peptide

Ion legend

wxyz

abcd

OtherUnassignedIon^Charge^_Position_Show for top:%

TFDDYAMHW

08.42e+31.68e+42.53e+43.37e+4

Zoom Out

a^+1^_2_y^+1^_1_a^+1^_2_b^+1^_2_y^+1^_2_b^+1^_3_b^+1^_3_y^+1^_3_y^+2^_8_y^+1^_4_\*y^+1^_5_y^+1^_6_y^+1^_7_

0830166024903320

Fragment Matches Table

Show background peaks

| Position | Ion type | Intensity | mz Theoretical | mz Error (Th) | mz Error (ppm) | Charge | Series Number |
| --- | --- | --- | --- | --- | --- | --- | --- |
| - | - | 2.208E+04 | 120.1 | - | - | 0 | - |
| - | - | 1577 | 121.1 | - | - | 0 | - |
| - | - | 3609 | 129.1 | - | - | 0 | - |
| - | - | 415.1 | 130 | - | - | 0 | - |
| - | - | 462.7 | 130 | - | - | 0 | - |
| - | - | 472.6 | 133.1 | - | - | 0 | - |
| - | - | 652.8 | 133.1 | - | - | 0 | - |
| - | - | 356.5 | 134.8 | - | - | 0 | - |
| - | - | 7140 | 136.1 | - | - | 0 | - |
| - | - | 451.4 | 138.3 | - | - | 0 | - |
| - | - | 1022 | 149 | - | - | 0 | - |
| - | - | 1154 | 156.1 | - | - | 0 | - |
| - | - | 836.8 | 158.1 | - | - | 0 | - |
| - | - | 995.6 | 159.1 | - | - | 0 | - |
| - | - | 2556 | 165.1 | - | - | 0 | - |
| - | - | 495.7 | 166.1 | - | - | 0 | - |
| - | - | 2774 | 167.1 | - | - | 0 | - |
| - | - | 564.4 | 169.4 | - | - | 0 | - |
| - | - | 1136 | 173.1 | - | - | 0 | - |
| - | - | 1333 | 173.4 | - | - | 0 | - |
| - | - | 567.3 | 175.1 | - | - | 0 | - |
| - | - | 2603 | 176.1 | - | - | 0 | - |
| - | - | 2313 | 177.1 | - | - | 0 | - |
| - | - | 4358 | 182.1 | - | - | 0 | - |
| - | - | 1160 | 183.1 | - | - | 0 | - |
| - | - | 606.9 | 187.1 | - | - | 0 | - |
| - | - | 951.2 | 188.1 | - | - | 0 | - |
| - | - | 1003 | 191.1 | - | - | 0 | - |
| - | - | 531.6 | 193.1 | - | - | 0 | - |
| - | - | 1473 | 201.1 | - | - | 0 | - |
| 2 | a | 1136 | 203.1 | 1.539E-05 | 0.07577 | +1 | 2 |
| 9 | y | 968.9 | 205.1 | 0.0001082 | 0.5275 | +1 | 1 |
| - | - | 496.9 | 208.2 | - | - | 0 | - |
| - | - | 653.3 | 211.1 | - | - | 0 | - |
| - | - | 879.9 | 215.1 | - | - | 0 | - |
| - | - | 871 | 221.1 | - | - | 0 | - |
| - | - | 2442 | 221.1 | - | - | 0 | - |
| 2 | a | 3.336E+04 | 221.1 | 0.0001469 | 0.6641 | +1 | 2 |
| - | - | 4706 | 222.1 | - | - | 0 | - |
| - | - | 1412 | 223.1 | - | - | 0 | - |
| - | - | 801.9 | 227.1 | - | - | 0 | - |
| - | - | 589.7 | 229.2 | - | - | 0 | - |
| - | - | 1213 | 235.1 | - | - | 0 | - |
| - | - | 1734 | 239.1 | - | - | 0 | - |
| - | - | 1377 | 241.1 | - | - | 0 | - |
| - | - | 635.5 | 247.1 | - | - | 0 | - |
| 2 | b | 5128 | 249.1 | 0.0003036 | 1.219 | +1 | 2 |
| - | - | 911.5 | 250.1 | - | - | 0 | - |
| - | - | 658.2 | 251.1 | - | - | 0 | - |
| - | - | 751.4 | 263.1 | - | - | 0 | - |
| - | - | 647 | 270.1 | - | - | 0 | - |
| - | - | 589.1 | 274.1 | - | - | 0 | - |
| - | - | 735.1 | 283 | - | - | 0 | - |
| - | - | 1894 | 285 | - | - | 0 | - |
| - | - | 681.9 | 296.1 | - | - | 0 | - |
| - | - | 544.1 | 299.1 | - | - | 0 | - |
| - | - | 600.1 | 324.1 | - | - | 0 | - |
| - | - | 3045 | 329.2 | - | - | 0 | - |
| 8 | y | 5404 | 342.2 | 0.0004892 | 1.43 | +1 | 2 |
| 3 | b | 705 | 346.1 | 0.001519 | 4.387 | +1 | 3 |
| - | - | 530.3 | 353.6 | - | - | 0 | - |
| - | - | 2330 | 355.1 | - | - | 0 | - |
| - | - | 923 | 357.2 | - | - | 0 | - |
| 3 | b | 1891 | 364.2 | 4.332E-05 | 0.119 | +1 | 3 |
| - | - | 618.5 | 378.1 | - | - | 0 | - |
| - | - | 1412 | 394.1 | - | - | 0 | - |
| - | - | 855.4 | 413.3 | - | - | 0 | - |
| - | - | 514.5 | 414 | - | - | 0 | - |
| - | - | 636.3 | 417.1 | - | - | 0 | - |
| - | - | 570.7 | 448.8 | - | - | 0 | - |
| - | - | 653.3 | 455.2 | - | - | 0 | - |
| - | - | 645.8 | 456 | - | - | 0 | - |
| 7 | y | 1766 | 489.2 | 0.004886 | 9.988 | +1 | 3 |
| 2 | y | 685.2 | 550.7 | 0.001615 | 2.933 | +2 | 8 |
| - | - | 695.1 | 551.2 | - | - | 0 | - |
| 6 | y | 2411 | 560.2 | 0.006499 | 11.6 | +1 | 4 |
| - | - | 818.7 | 592.7 | - | - | 0 | - |
| 0 | Precursor | 1211 | 601.2 | 0.006707 | 11.15 | +2 | -1 |
| - | - | 635.7 | 640.4 | - | - | 0 | - |
| - | - | 623.6 | 689.3 | - | - | 0 | - |
| - | - | 2692 | 700.4 | - | - | 0 | - |
| - | - | 920.5 | 701.4 | - | - | 0 | - |
| - | - | 890.8 | 721.3 | - | - | 0 | - |
| - | - | 723.8 | 722.3 | - | - | 0 | - |
| 5 | y | 2163 | 723.3 | 0.006159 | 8.515 | +1 | 5 |
| - | - | 842.1 | 724.3 | - | - | 0 | - |
| - | - | 1139 | 749.3 | - | - | 0 | - |
| - | - | 1005 | 774.3 | - | - | 0 | - |
| - | - | 1138 | 801.4 | - | - | 0 | - |
| 4 | y | 3077 | 838.3 | 0.005888 | 7.024 | +1 | 6 |
| - | - | 889.4 | 839.3 | - | - | 0 | - |
| - | - | 1868 | 844.5 | - | - | 0 | - |
| - | - | 729.8 | 845.4 | - | - | 0 | - |
| - | - | 7934 | 872.4 | - | - | 0 | - |
| - | - | 3775 | 873.5 | - | - | 0 | - |
| - | - | 996.1 | 874.5 | - | - | 0 | - |
| - | - | 3589 | 889.3 | - | - | 0 | - |
| - | - | 2334 | 890.4 | - | - | 0 | - |
| - | - | 763.5 | 901.5 | - | - | 0 | - |
| 3 | y | 1.276E+04 | 953.3 | 0.005801 | 6.085 | +1 | 7 |
| - | - | 5902 | 954.4 | - | - | 0 | - |
| - | - | 1380 | 955.3 | - | - | 0 | - |
| - | - | 641 | 956.3 | - | - | 0 | - |
| - | - | 1390 | 991.5 | - | - | 0 | - |
| - | - | 632.6 | 992.5 | - | - | 0 | - |
| - | - | 3628 | 1020 | - | - | 0 | - |
| - | - | 3000 | 1021 | - | - | 0 | - |
| - | - | 760.2 | 1022 | - | - | 0 | - |
| - | - | 698.8 | 1101 | - | - | 0 | - |
| - | - | 649.2 | 3287 | - | - | 0 | - |

m/z Charge Intensity FragmentType MassShift Position
120.0809326171875 0 22080.77
121.08432006835938 0 1576.9175
129.10235595703125 0 3608.9568
129.98031616210938 0 415.0951
130.0493621826172 0 462.6644
133.06057739257812 0 472.62576
133.08621215820312 0 652.7604
134.8091278076172 0 356.50604
136.0758514404297 0 7140.2417
138.29080200195312 0 451.42816
149.04498291015625 0 1022.09973
156.0770721435547 0 1153.8038
158.09657287597656 0 836.839
159.09205627441406 0 995.5878
165.05474853515625 0 2555.5134
166.08641052246094 0 495.74368
167.05552673339844 0 2773.684
169.36862182617188 0 564.3967
173.1289520263672 0 1136.257
173.43870544433594 0 1333.1235
175.119140625 0 567.29193
176.107177734375 0 2602.6653
177.1023712158203 0 2313.093
182.08126831054688 0 4358.077
183.08493041992188 0 1159.9467
187.13279724121094 0 606.888
188.07101440429688 0 951.18353
191.1177978515625 0 1003.26447
193.1085205078125 0 531.6233
201.12330627441406 0 1472.7181
203.1178741455078 0 1136.1957 a Water loss 1
205.0970458984375 0 968.9142 y 8
208.15145874023438 0 496.88928
211.06069946289062 0 653.2689
215.12820434570312 0 879.921
221.08511352539062 0 870.985
221.10345458984375 0 2442.4194
221.12860107421875 0 33356.85 a 1
222.13209533691406 0 4705.945
223.06396484375 0 1412.1677
227.1031951904297 0 801.91797
229.17063903808594 0 589.7266
235.10861206054688 0 1213.1595
239.0945281982422 0 1733.564
241.07443237304688 0 1376.5278
247.09402465820312 0 635.51447
249.12367248535156 0 5127.537 b 1
250.1273651123047 0 911.45593
251.10269165039062 0 658.2147
263.103515625 0 751.38605
270.1089782714844 0 646.9733
274.12969970703125 0 589.14087
283.0307922363281 0 735.0573
285.0096435546875 0 1894.2228
296.1497802734375 0 681.8841
299.0610046386719 0 544.1051
324.1452331542969 0 600.08905
329.15045166015625 0 3045.065
342.15655517578125 0 5403.5 y 7
346.1412658691406 0 704.9697 b Water loss 2
353.6466369628906 0 530.34827
355.0702819824219 0 2330.4224
357.2239074707031 0 923.00555
364.1502685546875 0 1890.9886 b 2
378.1302795410156 0 618.5408
394.1248474121094 0 1412.2301
413.2653503417969 0 855.3711
414.00811767578125 0 514.461
417.08380126953125 0 636.34827
448.7728271484375 0 570.73645
455.2276306152344 0 653.3155
456.01141357421875 0 645.76166
489.1914367675781 0 1765.7192 y 6
550.7098999023438 0 685.20703 y 1
551.2152099609375 0 695.092
560.2301635742188 0 2410.9182 y 5
592.7279052734375 0 818.701
601.2388305664062 0 1211.1605 Precursor
640.36083984375 0 635.7143
689.3440551757812 0 623.5568
700.3624877929688 0 2691.7996
701.3681030273438 0 920.48596
721.2626953125 0 890.7849
722.2667236328125 0 723.785
723.2931518554688 0 2162.7034 y 4
724.2942504882812 0 842.0513
749.2559814453125 0 1139.124
774.32177734375 0 1004.65936
801.41357421875 0 1138.2968
838.31982421875 0 3076.5125 y 3
839.3267822265625 0 889.41614
844.45361328125 0 1867.9467
845.44970703125 0 729.77344
872.4488525390625 0 7933.616
873.451416015625 0 3774.5327
874.4501342773438 0 996.1317
889.3480224609375 0 3588.6333
890.35205078125 0 2333.5244
901.4705810546875 0 763.4791
953.3466796875 0 12762.509 y 2
954.350341796875 0 5901.767
955.3487548828125 0 1379.6625
956.3443603515625 0 641.0159
991.5243530273438 0 1390.0103
992.5196533203125 0 632.58856
1019.5175170898438 0 3628.3887
1020.5203857421875 0 2999.8035
1021.5264282226562 0 760.2413
1101.406005859375 0 698.7982
3287.195068359375 0 649.2061

Spectrum Details

|  |  |
| --- | --- |
| Matched peaks? Matched peaks The total absolute number of peaks matched. Additionally in brackets the total fraction of peaks matched and the total number of peaks is shown. | 14 (12.73% of 110) |
| FDR? FDR The false discovery rate estimated for this peptide. It is calculated by matching all theoretical fragments with a non-integer shift with the raw peaks for this spectrum. This is done with 40 different shifts. The resulting percentage is the average number of annotated peaks over the number of annotated peaks with the correct spectrum. | 0.68% |
| Satellite FDR? Satellite FDR See the FDR for details on its calculation. This satellite ion specific FDR only contains the satellite ions (d/w) for I/L/J positions. | - |
| PSM Score? PSM Score The PSM Score as given by Hecklib to this annotated spectrum. It is shown with three significant figures. | 144 |

#### Spectrum 13277? Spectrum 13277 The raw spectrum of this peptide as annotated by Hecklib. The fragments are coloured according to ion type (see legend). Any peaks with a star '\*' as text can be hovered over to see the full details, first the ion type second the mass shift type. By hovering over the amino acids in the peptide or ions in the legend the corresponding peaks are highlighted. By toggling the 'Unassigned' label you can turn the background (unassigned) peaks on or off in the plot. By updating the slider in the Ion legend you can update the spectrum to only show the top X% of the peaks with labels. The top X% means any peak that is within X% of the highest intensity. By dragging in the spectrum you can zoom in to a specific part of the spectrum and use 'Zoom Out' to get back to the original zoom level. The annotation of the spectrum is based on the given sequence in the peptides file and is done with different software so inconsistencies are likely. The peaks are annotated based on the given sequence, with 20 ppm tolerance.

Copy Data

##### Spectrum 13277 (TSV)

###### Preview

```
Loading example...
```

*Click on the button to copy the data to your clipboard.*

Mz MinMz MaxIntensity Max

WidthHeightPeptide font sizePeptide stroke widthSpectrum font sizeSpectrum stroke widthCompact peptide

Ion legend

wxyz

abcd

OtherUnassignedIon^Charge^_Position_Show for top:%

TFDDYAMHW

08.01e+31.60e+42.40e+43.20e+4

Zoom Out

a^+1^_2_y^+1^_1_a^+1^_2_b^+1^_2_y^+1^_2_b^+1^_3_b^+1^_4_y^+1^_3_y^+1^_4_\*y^+1^_5_y^+1^_6_y^+1^_7_y^+1^_8_

0804160824123216

Fragment Matches Table

Show background peaks

| Position | Ion type | Intensity | mz Theoretical | mz Error (Th) | mz Error (ppm) | Charge | Series Number |
| --- | --- | --- | --- | --- | --- | --- | --- |
| - | - | 1.802E+04 | 120.1 | - | - | 0 | - |
| - | - | 1656 | 121.1 | - | - | 0 | - |
| - | - | 410.2 | 125.5 | - | - | 0 | - |
| - | - | 509.1 | 129.1 | - | - | 0 | - |
| - | - | 2616 | 129.1 | - | - | 0 | - |
| - | - | 695 | 130.1 | - | - | 0 | - |
| - | - | 918.5 | 133.1 | - | - | 0 | - |
| - | - | 687.2 | 136.1 | - | - | 0 | - |
| - | - | 7660 | 136.1 | - | - | 0 | - |
| - | - | 428.1 | 137.1 | - | - | 0 | - |
| - | - | 738.2 | 137.1 | - | - | 0 | - |
| - | - | 426.9 | 141.5 | - | - | 0 | - |
| - | - | 389 | 141.7 | - | - | 0 | - |
| - | - | 403.4 | 144.1 | - | - | 0 | - |
| - | - | 618.6 | 147.1 | - | - | 0 | - |
| - | - | 971.5 | 149 | - | - | 0 | - |
| - | - | 543.3 | 150 | - | - | 0 | - |
| - | - | 482.5 | 154.6 | - | - | 0 | - |
| - | - | 994.6 | 156.1 | - | - | 0 | - |
| - | - | 1126 | 158.1 | - | - | 0 | - |
| - | - | 1405 | 159.1 | - | - | 0 | - |
| - | - | 622.4 | 160.1 | - | - | 0 | - |
| - | - | 616.5 | 162.1 | - | - | 0 | - |
| - | - | 2567 | 165.1 | - | - | 0 | - |
| - | - | 785 | 166.1 | - | - | 0 | - |
| - | - | 1736 | 167.1 | - | - | 0 | - |
| - | - | 896.1 | 173.1 | - | - | 0 | - |
| - | - | 6131 | 173.4 | - | - | 0 | - |
| - | - | 2308 | 176.1 | - | - | 0 | - |
| - | - | 1753 | 177.1 | - | - | 0 | - |
| - | - | 4120 | 182.1 | - | - | 0 | - |
| - | - | 498.1 | 183.1 | - | - | 0 | - |
| - | - | 460.5 | 183.1 | - | - | 0 | - |
| - | - | 469.6 | 185.1 | - | - | 0 | - |
| - | - | 1266 | 188.1 | - | - | 0 | - |
| - | - | 1152 | 193.1 | - | - | 0 | - |
| - | - | 528.9 | 197.1 | - | - | 0 | - |
| - | - | 466.4 | 197.3 | - | - | 0 | - |
| - | - | 554.4 | 199.1 | - | - | 0 | - |
| - | - | 1068 | 201.1 | - | - | 0 | - |
| 2 | a | 793.9 | 203.1 | 0.0005339 | 2.629 | +1 | 2 |
| 9 | y | 1110 | 205.1 | 0.0001054 | 0.514 | +1 | 1 |
| - | - | 528 | 216.1 | - | - | 0 | - |
| - | - | 1457 | 221.1 | - | - | 0 | - |
| 2 | a | 3.172E+04 | 221.1 | 5.735E-06 | 0.02593 | +1 | 2 |
| - | - | 3151 | 222.1 | - | - | 0 | - |
| - | - | 1534 | 223.1 | - | - | 0 | - |
| - | - | 803.8 | 235.1 | - | - | 0 | - |
| - | - | 1367 | 239.1 | - | - | 0 | - |
| - | - | 875.8 | 241.1 | - | - | 0 | - |
| - | - | 713 | 241.1 | - | - | 0 | - |
| - | - | 577.2 | 241.7 | - | - | 0 | - |
| 2 | b | 5239 | 249.1 | 7.476E-05 | 0.3001 | +1 | 2 |
| - | - | 955.9 | 250.1 | - | - | 0 | - |
| - | - | 676 | 251.1 | - | - | 0 | - |
| - | - | 650.4 | 251.2 | - | - | 0 | - |
| - | - | 1553 | 263.1 | - | - | 0 | - |
| - | - | 1122 | 285 | - | - | 0 | - |
| - | - | 809.6 | 285.1 | - | - | 0 | - |
| - | - | 604.9 | 294.8 | - | - | 0 | - |
| - | - | 897.3 | 309.2 | - | - | 0 | - |
| - | - | 875.7 | 315.2 | - | - | 0 | - |
| - | - | 606.5 | 319.1 | - | - | 0 | - |
| - | - | 668.6 | 323.1 | - | - | 0 | - |
| - | - | 1030 | 324.1 | - | - | 0 | - |
| - | - | 3117 | 329.1 | - | - | 0 | - |
| 8 | y | 5704 | 342.2 | 0.0002146 | 0.6271 | +1 | 2 |
| - | - | 1218 | 343.2 | - | - | 0 | - |
| - | - | 1398 | 355.1 | - | - | 0 | - |
| 3 | b | 1456 | 364.2 | 0.000379 | 1.041 | +1 | 3 |
| - | - | 574.3 | 375.4 | - | - | 0 | - |
| - | - | 853.2 | 378.1 | - | - | 0 | - |
| - | - | 491.8 | 392.2 | - | - | 0 | - |
| - | - | 911.7 | 394.1 | - | - | 0 | - |
| - | - | 607.1 | 408.2 | - | - | 0 | - |
| - | - | 757 | 425.2 | - | - | 0 | - |
| - | - | 541.7 | 443.2 | - | - | 0 | - |
| 4 | b | 954.8 | 479.2 | 0.00123 | 2.566 | +1 | 4 |
| 7 | y | 1373 | 489.2 | 0.005985 | 12.23 | +1 | 3 |
| - | - | 613.1 | 496.2 | - | - | 0 | - |
| 6 | y | 2238 | 560.2 | 0.003447 | 6.154 | +1 | 4 |
| - | - | 1266 | 585.3 | - | - | 0 | - |
| 0 | Precursor | 1288 | 601.2 | 0.003777 | 6.282 | +2 | -1 |
| - | - | 831.2 | 602.2 | - | - | 0 | - |
| - | - | 859.8 | 642 | - | - | 0 | - |
| - | - | 646.7 | 654.7 | - | - | 0 | - |
| - | - | 880.6 | 659.3 | - | - | 0 | - |
| - | - | 582.9 | 696.5 | - | - | 0 | - |
| - | - | 3407 | 700.4 | - | - | 0 | - |
| - | - | 602.5 | 701.4 | - | - | 0 | - |
| - | - | 684.2 | 702.4 | - | - | 0 | - |
| - | - | 1164 | 721.3 | - | - | 0 | - |
| 5 | y | 2195 | 723.3 | 0.005427 | 7.503 | +1 | 5 |
| - | - | 984.2 | 724.3 | - | - | 0 | - |
| - | - | 736.9 | 749.3 | - | - | 0 | - |
| - | - | 800.5 | 774.3 | - | - | 0 | - |
| - | - | 1495 | 801.4 | - | - | 0 | - |
| 4 | y | 2497 | 838.3 | 0.004423 | 5.277 | +1 | 6 |
| - | - | 1292 | 839.3 | - | - | 0 | - |
| - | - | 1796 | 844.5 | - | - | 0 | - |
| - | - | 815.4 | 845.5 | - | - | 0 | - |
| - | - | 645 | 857 | - | - | 0 | - |
| - | - | 8832 | 872.4 | - | - | 0 | - |
| - | - | 370.5 | 872.5 | - | - | 0 | - |
| - | - | 3109 | 873.5 | - | - | 0 | - |
| - | - | 714 | 874.5 | - | - | 0 | - |
| - | - | 800.5 | 882.1 | - | - | 0 | - |
| - | - | 2955 | 889.4 | - | - | 0 | - |
| - | - | 1598 | 890.4 | - | - | 0 | - |
| 3 | y | 1.189E+04 | 953.3 | 0.00574 | 6.021 | +1 | 7 |
| - | - | 5956 | 954.3 | - | - | 0 | - |
| - | - | 2057 | 955.3 | - | - | 0 | - |
| - | - | 2640 | 1020 | - | - | 0 | - |
| - | - | 1577 | 1021 | - | - | 0 | - |
| - | - | 662.1 | 1036 | - | - | 0 | - |
| 2 | y | 852 | 1100 | 0.00831 | 7.551 | +1 | 8 |
| - | - | 703.7 | 2072 | - | - | 0 | - |
| - | - | 651.7 | 3184 | - | - | 0 | - |

m/z Charge Intensity FragmentType MassShift Position
120.08086395263672 0 18016.148
121.08418273925781 0 1656.2689
125.51823425292969 0 410.21967
129.06642150878906 0 509.11218
129.1023406982422 0 2615.521
130.0653076171875 0 695.0329
133.0861358642578 0 918.52167
136.07012939453125 0 687.2431
136.07574462890625 0 7659.5703
137.0596466064453 0 428.1105
137.07896423339844 0 738.17883
141.5169677734375 0 426.9046
141.73275756835938 0 388.98706
144.1453094482422 0 403.4323
147.1130828857422 0 618.5808
149.04507446289062 0 971.4944
149.96182250976562 0 543.3333
154.57781982421875 0 482.5229
156.07679748535156 0 994.5802
158.09629821777344 0 1125.9391
159.09173583984375 0 1404.5745
160.11285400390625 0 622.43097
162.09164428710938 0 616.49896
165.05467224121094 0 2567.224
166.0864715576172 0 785.01965
167.05540466308594 0 1735.8917
173.12893676757812 0 896.1074
173.44046020507812 0 6130.8965
176.1068878173828 0 2307.9954
177.1022186279297 0 1753.1863
182.08116149902344 0 4119.618
183.0846710205078 0 498.07712
183.11233520507812 0 460.46783
185.1270294189453 0 469.56232
188.07086181640625 0 1266.1243
193.1084442138672 0 1152.1913
197.12925720214844 0 528.9457
197.2881622314453 0 466.37485
199.13323974609375 0 554.3849
201.12294006347656 0 1068.2324
203.11842346191406 0 793.8783 a Water loss 1
205.09725952148438 0 1110.0808 y 8
216.09771728515625 0 527.97363
221.10302734375 0 1457.3759
221.12844848632812 0 31722.26 a 1
222.1317138671875 0 3150.9941
223.0630645751953 0 1533.656
235.1071014404297 0 803.7747
239.0943145751953 0 1366.8348
241.0749053955078 0 875.7959
241.09068298339844 0 713.0219
241.7460174560547 0 577.1501
249.12344360351562 0 5239.1943 b 1
250.12757873535156 0 955.8835
251.10232543945312 0 676.0171
251.1503143310547 0 650.38226
263.1022033691406 0 1552.9204
285.0096740722656 0 1122.4236
285.10211181640625 0 809.60767
294.7982482910156 0 604.92865
309.2038879394531 0 897.2849
315.1778564453125 0 875.6817
319.1407775878906 0 606.486
323.0975341796875 0 668.56836
324.14569091796875 0 1029.9966
329.14984130859375 0 3116.9758
342.1562805175781 0 5704.356 y 7
343.15972900390625 0 1218.2479
355.0694885253906 0 1398.451
364.1499328613281 0 1455.7572 b 2
375.35205078125 0 574.3244
378.1309814453125 0 853.1631
392.20587158203125 0 491.8096
394.12493896484375 0 911.69293
408.1693420410156 0 607.1363
425.19207763671875 0 756.9961
443.1911315917969 0 541.7169
479.176025390625 0 954.8163 b 3
489.1925354003906 0 1373.2151 y 6
496.229248046875 0 613.1113
560.2271118164062 0 2238.2852 y 5
585.3367309570312 0 1266.1764
601.2359008789062 0 1287.8115 Precursor
602.23486328125 0 831.1945
642.0023803710938 0 859.8182
654.67529296875 0 646.6747
659.2960205078125 0 880.5559
696.4611206054688 0 582.8665
700.3629150390625 0 3406.5505
701.3695678710938 0 602.50726
702.3663330078125 0 684.17303
721.261474609375 0 1164.3619
723.2924194335938 0 2194.597 y 4
724.2941284179688 0 984.15216
749.257568359375 0 736.907
774.3204956054688 0 800.4912
801.4097900390625 0 1494.6411
838.318359375 0 2497.229 y 3
839.3211669921875 0 1291.8557
844.45263671875 0 1795.8038
845.4549560546875 0 815.40796
857.0350952148438 0 645.0463
872.4472045898438 0 8831.689
872.5299682617188 0 370.45813
873.4508056640625 0 3108.7524
874.45263671875 0 713.983
882.0787963867188 0 800.54083
889.35009765625 0 2954.8923
890.3515625 0 1597.8972
953.3466186523438 0 11887.638 y 2
954.3487548828125 0 5956.43
955.3475952148438 0 2057.201
1019.5155029296875 0 2640.3884
1020.5211791992188 0 1576.7002
1036.4114990234375 0 662.09
1100.4176025390625 0 851.97797 y 1
2072.300048828125 0 703.67505
3183.8896484375 0 651.67395

Spectrum Details

|  |  |
| --- | --- |
| Matched peaks? Matched peaks The total absolute number of peaks matched. Additionally in brackets the total fraction of peaks matched and the total number of peaks is shown. | 14 (11.86% of 118) |
| FDR? FDR The false discovery rate estimated for this peptide. It is calculated by matching all theoretical fragments with a non-integer shift with the raw peaks for this spectrum. This is done with 40 different shifts. The resulting percentage is the average number of annotated peaks over the number of annotated peaks with the correct spectrum. | 1.19% |
| Satellite FDR? Satellite FDR See the FDR for details on its calculation. This satellite ion specific FDR only contains the satellite ions (d/w) for I/L/J positions. | - |
| PSM Score? PSM Score The PSM Score as given by Hecklib to this annotated spectrum. It is shown with three significant figures. | 160 |

#### Spectrum 12777? Spectrum 12777 The raw spectrum of this peptide as annotated by Hecklib. The fragments are coloured according to ion type (see legend). Any peaks with a star '\*' as text can be hovered over to see the full details, first the ion type second the mass shift type. By hovering over the amino acids in the peptide or ions in the legend the corresponding peaks are highlighted. By toggling the 'Unassigned' label you can turn the background (unassigned) peaks on or off in the plot. By updating the slider in the Ion legend you can update the spectrum to only show the top X% of the peaks with labels. The top X% means any peak that is within X% of the highest intensity. By dragging in the spectrum you can zoom in to a specific part of the spectrum and use 'Zoom Out' to get back to the original zoom level. The annotation of the spectrum is based on the given sequence in the peptides file and is done with different software so inconsistencies are likely. The peaks are annotated based on the given sequence, with 20 ppm tolerance.

Copy Data

##### Spectrum 12777 (TSV)

###### Preview

```
Loading example...
```

*Click on the button to copy the data to your clipboard.*

Mz MinMz MaxIntensity Max

WidthHeightPeptide font sizePeptide stroke widthSpectrum font sizeSpectrum stroke widthCompact peptide

Ion legend

wxyz

abcd

OtherUnassignedIon^Charge^_Position_Show for top:%

TFDDYAMHW

08.89e+31.78e+42.67e+43.56e+4

Zoom Out

a^+1^_2_y^+1^_1_a^+1^_2_b^+1^_2_y^+1^_2_b^+1^_3_b^+1^_4_y^+1^_3_y^+2^_8_y^+1^_4_\*y^+1^_5_y^+1^_6_y^+1^_7_y^+1^_7_y^+1^_8_

0742148322252967

Fragment Matches Table

Show background peaks

| Position | Ion type | Intensity | mz Theoretical | mz Error (Th) | mz Error (ppm) | Charge | Series Number |
| --- | --- | --- | --- | --- | --- | --- | --- |
| - | - | 2.544E+04 | 120.1 | - | - | 0 | - |
| - | - | 353.1 | 121.1 | - | - | 0 | - |
| - | - | 2137 | 121.1 | - | - | 0 | - |
| - | - | 486.4 | 123 | - | - | 0 | - |
| - | - | 3333 | 129.1 | - | - | 0 | - |
| - | - | 481.3 | 132.1 | - | - | 0 | - |
| - | - | 1314 | 133.1 | - | - | 0 | - |
| - | - | 8118 | 136.1 | - | - | 0 | - |
| - | - | 485.7 | 137.1 | - | - | 0 | - |
| - | - | 446.9 | 137.9 | - | - | 0 | - |
| - | - | 638.3 | 138.1 | - | - | 0 | - |
| - | - | 537.4 | 148.9 | - | - | 0 | - |
| - | - | 991 | 149 | - | - | 0 | - |
| - | - | 422.1 | 151 | - | - | 0 | - |
| - | - | 467.9 | 153.1 | - | - | 0 | - |
| - | - | 1065 | 156.1 | - | - | 0 | - |
| - | - | 837.3 | 158.1 | - | - | 0 | - |
| - | - | 873.1 | 159.1 | - | - | 0 | - |
| - | - | 2628 | 165.1 | - | - | 0 | - |
| - | - | 1244 | 166.1 | - | - | 0 | - |
| - | - | 429.9 | 166.7 | - | - | 0 | - |
| - | - | 1993 | 167.1 | - | - | 0 | - |
| - | - | 613.3 | 167.1 | - | - | 0 | - |
| - | - | 1427 | 173.1 | - | - | 0 | - |
| - | - | 1685 | 173.5 | - | - | 0 | - |
| - | - | 2854 | 176.1 | - | - | 0 | - |
| - | - | 2015 | 177.1 | - | - | 0 | - |
| - | - | 833.3 | 177.1 | - | - | 0 | - |
| - | - | 446.1 | 177.5 | - | - | 0 | - |
| - | - | 4757 | 182.1 | - | - | 0 | - |
| - | - | 666.2 | 183.1 | - | - | 0 | - |
| - | - | 504.6 | 183.2 | - | - | 0 | - |
| - | - | 741.3 | 185.1 | - | - | 0 | - |
| - | - | 1572 | 188.1 | - | - | 0 | - |
| - | - | 668.4 | 191.1 | - | - | 0 | - |
| - | - | 714.1 | 193.1 | - | - | 0 | - |
| - | - | 1031 | 201.1 | - | - | 0 | - |
| - | - | 671.4 | 203.1 | - | - | 0 | - |
| 2 | a | 729 | 203.1 | 0.0004274 | 2.104 | +1 | 2 |
| 9 | y | 756.1 | 205.1 | 1.388E-05 | 0.06766 | +1 | 1 |
| - | - | 738 | 211.1 | - | - | 0 | - |
| - | - | 640.9 | 215.1 | - | - | 0 | - |
| - | - | 2584 | 221.1 | - | - | 0 | - |
| 2 | a | 3.52E+04 | 221.1 | 4.004E-05 | 0.1811 | +1 | 2 |
| - | - | 3744 | 222.1 | - | - | 0 | - |
| - | - | 849.9 | 223.1 | - | - | 0 | - |
| - | - | 580.1 | 230.2 | - | - | 0 | - |
| - | - | 719.4 | 231.1 | - | - | 0 | - |
| - | - | 953.2 | 239.1 | - | - | 0 | - |
| - | - | 916.3 | 241.1 | - | - | 0 | - |
| 2 | b | 7692 | 249.1 | 5.95E-05 | 0.2388 | +1 | 2 |
| - | - | 974.5 | 250.1 | - | - | 0 | - |
| - | - | 713.9 | 251.1 | - | - | 0 | - |
| - | - | 696.6 | 251.2 | - | - | 0 | - |
| - | - | 519.4 | 257.2 | - | - | 0 | - |
| - | - | 608.4 | 260 | - | - | 0 | - |
| - | - | 583.8 | 262.1 | - | - | 0 | - |
| - | - | 1441 | 263.1 | - | - | 0 | - |
| - | - | 468 | 279.1 | - | - | 0 | - |
| - | - | 559.2 | 279.1 | - | - | 0 | - |
| - | - | 643.7 | 283 | - | - | 0 | - |
| - | - | 891.5 | 285 | - | - | 0 | - |
| - | - | 639.1 | 296.2 | - | - | 0 | - |
| - | - | 584.4 | 308.4 | - | - | 0 | - |
| - | - | 1811 | 309.2 | - | - | 0 | - |
| - | - | 1332 | 311.1 | - | - | 0 | - |
| - | - | 811.3 | 324.1 | - | - | 0 | - |
| - | - | 3510 | 329.1 | - | - | 0 | - |
| - | - | 633.8 | 342.1 | - | - | 0 | - |
| 8 | y | 7558 | 342.2 | 0.000184 | 0.5379 | +1 | 2 |
| - | - | 1141 | 343.2 | - | - | 0 | - |
| - | - | 589.4 | 349.7 | - | - | 0 | - |
| - | - | 1659 | 355.1 | - | - | 0 | - |
| - | - | 1008 | 357.2 | - | - | 0 | - |
| 3 | b | 1759 | 364.2 | 0.0001959 | 0.538 | +1 | 3 |
| - | - | 604.2 | 376.2 | - | - | 0 | - |
| - | - | 627.5 | 378.1 | - | - | 0 | - |
| - | - | 789.2 | 394.1 | - | - | 0 | - |
| - | - | 629.3 | 413.2 | - | - | 0 | - |
| - | - | 1106 | 413.3 | - | - | 0 | - |
| - | - | 664.2 | 416.2 | - | - | 0 | - |
| - | - | 884.5 | 465.2 | - | - | 0 | - |
| - | - | 562.4 | 475.3 | - | - | 0 | - |
| 4 | b | 1505 | 479.2 | 0.0008633 | 1.802 | +1 | 4 |
| 7 | y | 1406 | 489.2 | 0.003666 | 7.493 | +1 | 3 |
| - | - | 681.9 | 493.2 | - | - | 0 | - |
| - | - | 1120 | 496.2 | - | - | 0 | - |
| - | - | 554.6 | 529.2 | - | - | 0 | - |
| 2 | y | 977.1 | 550.7 | 0.002836 | 5.15 | +2 | 8 |
| 6 | y | 2239 | 560.2 | 0.00534 | 9.531 | +1 | 4 |
| - | - | 995.5 | 561.2 | - | - | 0 | - |
| - | - | 725.9 | 576.3 | - | - | 0 | - |
| - | - | 1406 | 585.3 | - | - | 0 | - |
| - | - | 961.2 | 586.3 | - | - | 0 | - |
| 0 | Precursor | 1179 | 601.2 | 0.003777 | 6.282 | +2 | -1 |
| - | - | 722.8 | 657.3 | - | - | 0 | - |
| - | - | 1195 | 659.3 | - | - | 0 | - |
| - | - | 1086 | 661.7 | - | - | 0 | - |
| - | - | 578.4 | 695.9 | - | - | 0 | - |
| - | - | 2903 | 700.4 | - | - | 0 | - |
| - | - | 1369 | 701.4 | - | - | 0 | - |
| - | - | 752.5 | 721.3 | - | - | 0 | - |
| 5 | y | 2545 | 723.3 | 0.004938 | 6.828 | +1 | 5 |
| - | - | 833.4 | 749.3 | - | - | 0 | - |
| - | - | 704 | 750.3 | - | - | 0 | - |
| - | - | 1131 | 774.3 | - | - | 0 | - |
| - | - | 1140 | 783.4 | - | - | 0 | - |
| - | - | 617.9 | 791.3 | - | - | 0 | - |
| - | - | 1830 | 801.4 | - | - | 0 | - |
| - | - | 720.7 | 802.4 | - | - | 0 | - |
| - | - | 852.3 | 828.4 | - | - | 0 | - |
| 4 | y | 2461 | 838.3 | 0.004301 | 5.131 | +1 | 6 |
| - | - | 1744 | 839.3 | - | - | 0 | - |
| - | - | 1895 | 844.5 | - | - | 0 | - |
| - | - | 766.1 | 845.5 | - | - | 0 | - |
| - | - | 1179 | 871.3 | - | - | 0 | - |
| - | - | 9571 | 872.4 | - | - | 0 | - |
| - | - | 4280 | 873.5 | - | - | 0 | - |
| - | - | 1161 | 874.5 | - | - | 0 | - |
| - | - | 3650 | 889.3 | - | - | 0 | - |
| - | - | 2512 | 890.4 | - | - | 0 | - |
| - | - | 936.8 | 891.4 | - | - | 0 | - |
| 3 | y | 874.9 | 935.3 | 0.00782 | 8.361 | +1 | 7 |
| 3 | y | 1.428E+04 | 953.3 | 0.005496 | 5.765 | +1 | 7 |
| - | - | 6597 | 954.3 | - | - | 0 | - |
| - | - | 2220 | 955.4 | - | - | 0 | - |
| - | - | 1215 | 991.5 | - | - | 0 | - |
| - | - | 3821 | 1020 | - | - | 0 | - |
| - | - | 2443 | 1021 | - | - | 0 | - |
| - | - | 588.5 | 1086 | - | - | 0 | - |
| 2 | y | 1403 | 1100 | 0.007577 | 6.886 | +1 | 8 |
| - | - | 1005 | 1101 | - | - | 0 | - |
| - | - | 626.8 | 1786 | - | - | 0 | - |
| - | - | 700.4 | 2333 | - | - | 0 | - |
| - | - | 738.8 | 2937 | - | - | 0 | - |

m/z Charge Intensity FragmentType MassShift Position
120.08088684082031 0 25438.506
121.06486511230469 0 353.09247
121.08419036865234 0 2136.621
123.04427337646484 0 486.3629
129.1023712158203 0 3333.1353
132.08090209960938 0 481.3178
133.08596801757812 0 1313.7412
136.07579040527344 0 8118.0977
137.07931518554688 0 485.69644
137.9230499267578 0 446.85233
138.0657958984375 0 638.2685
148.947509765625 0 537.4
149.0449676513672 0 991.0294
151.02511596679688 0 422.12634
153.073486328125 0 467.9438
156.07688903808594 0 1064.6211
158.09634399414062 0 837.25903
159.09176635742188 0 873.1423
165.0546112060547 0 2627.5044
166.0860137939453 0 1243.75
166.68809509277344 0 429.9139
167.05569458007812 0 1993.0717
167.08152770996094 0 613.25574
173.12828063964844 0 1426.951
173.45172119140625 0 1685.261
176.10708618164062 0 2854.0327
177.1020965576172 0 2014.5177
177.11041259765625 0 833.2688
177.47689819335938 0 446.12747
182.08120727539062 0 4757.321
183.08396911621094 0 666.2348
183.15013122558594 0 504.59158
185.12838745117188 0 741.33075
188.07066345214844 0 1572.001
191.11793518066406 0 668.38776
193.10887145996094 0 714.09
201.12342834472656 0 1030.8281
203.0661163330078 0 671.3797
203.11746215820312 0 729.0377 a Water loss 1
205.09716796875 0 756.0616 y 8
211.1443328857422 0 738.04474
215.12863159179688 0 640.88995
221.1033935546875 0 2583.7534
221.1284942626953 0 35202 a 1
222.13198852539062 0 3743.6526
223.0637664794922 0 849.8944
230.17291259765625 0 580.0958
231.0974884033203 0 719.3672
239.09458923339844 0 953.20953
241.07424926757812 0 916.2796
249.12342834472656 0 7691.515 b 1
250.126220703125 0 974.52924
251.10330200195312 0 713.85254
251.15135192871094 0 696.6232
257.1654968261719 0 519.39984
260.0452575683594 0 608.4061
262.11822509765625 0 583.7557
263.1019592285156 0 1441.4625
279.071533203125 0 468.0337
279.09783935546875 0 559.20935
283.0305480957031 0 643.6843
285.0101623535156 0 891.51996
296.150634765625 0 639.0827
308.37969970703125 0 584.39655
309.2038879394531 0 1811.323
311.14666748046875 0 1332.4395
324.1455383300781 0 811.34485
329.1493835449219 0 3510.4858
342.1368408203125 0 633.82117
342.15625 0 7558.269 y 7
343.1597900390625 0 1140.753
349.6751403808594 0 589.445
355.069580078125 0 1658.537
357.22381591796875 0 1007.5857
364.1501159667969 0 1759.4476 b 2
376.1643981933594 0 604.2137
378.128662109375 0 627.50836
394.1239318847656 0 789.18286
413.2382507324219 0 629.2947
413.2659912109375 0 1106.1699
416.2477111816406 0 664.1598
465.1602478027344 0 884.4755
475.30194091796875 0 562.446
479.1763916015625 0 1505.1265 b 3
489.1902160644531 0 1406.027 y 6
493.2221374511719 0 681.93066
496.2298278808594 0 1119.5216
529.248046875 0 554.58716
550.7111206054688 0 977.0682 y 1
560.22900390625 0 2239.2952 y 5
561.232666015625 0 995.53973
576.3291625976562 0 725.92883
585.3387451171875 0 1405.5477
586.3358154296875 0 961.1562
601.2359008789062 0 1179.2792 Precursor
657.2667846679688 0 722.83234
659.2933349609375 0 1194.9398
661.716796875 0 1085.6952
695.8772583007812 0 578.4397
700.3621826171875 0 2902.9463
701.3648681640625 0 1369.2532
721.2596435546875 0 752.54803
723.2919311523438 0 2545.2075 y 4
749.2551879882812 0 833.4124
750.25732421875 0 704.00574
774.3214721679688 0 1131.3047
783.3992919921875 0 1139.5741
791.2949829101562 0 617.91644
801.4110107421875 0 1830.0579
802.4151000976562 0 720.7168
828.421142578125 0 852.28326
838.3182373046875 0 2461.1262 y 3
839.3246459960938 0 1743.6882
844.4545288085938 0 1895.1727
845.460205078125 0 766.1321
871.3422241210938 0 1179.3053
872.4481811523438 0 9570.609
873.4506225585938 0 4279.8003
874.4550170898438 0 1160.5227
889.3484497070312 0 3650.2563
890.3513793945312 0 2512.0657
891.3554077148438 0 936.8184
935.338134765625 0 874.9303 y Water loss 2
953.3463745117188 0 14280.294 y 2
954.3493041992188 0 6597.0156
955.350341796875 0 2219.5688
991.5171508789062 0 1214.8608
1019.5162353515625 0 3820.6067
1020.5200805664062 0 2442.8586
1085.8895263671875 0 588.5472
1100.4168701171875 0 1403.2059 y 1
1101.4136962890625 0 1004.9681
1786.2098388671875 0 626.831
2333.375244140625 0 700.43677
2937.308837890625 0 738.838

Spectrum Details

|  |  |
| --- | --- |
| Matched peaks? Matched peaks The total absolute number of peaks matched. Additionally in brackets the total fraction of peaks matched and the total number of peaks is shown. | 16 (11.85% of 135) |
| FDR? FDR The false discovery rate estimated for this peptide. It is calculated by matching all theoretical fragments with a non-integer shift with the raw peaks for this spectrum. This is done with 40 different shifts. The resulting percentage is the average number of annotated peaks over the number of annotated peaks with the correct spectrum. | 1.34% |
| Satellite FDR? Satellite FDR See the FDR for details on its calculation. This satellite ion specific FDR only contains the satellite ions (d/w) for I/L/J positions. | - |
| PSM Score? PSM Score The PSM Score as given by Hecklib to this annotated spectrum. It is shown with three significant figures. | 177 |

#### Spectrum 10066? Spectrum 10066 The raw spectrum of this peptide as annotated by Hecklib. The fragments are coloured according to ion type (see legend). Any peaks with a star '\*' as text can be hovered over to see the full details, first the ion type second the mass shift type. By hovering over the amino acids in the peptide or ions in the legend the corresponding peaks are highlighted. By toggling the 'Unassigned' label you can turn the background (unassigned) peaks on or off in the plot. By updating the slider in the Ion legend you can update the spectrum to only show the top X% of the peaks with labels. The top X% means any peak that is within X% of the highest intensity. By dragging in the spectrum you can zoom in to a specific part of the spectrum and use 'Zoom Out' to get back to the original zoom level. The annotation of the spectrum is based on the given sequence in the peptides file and is done with different software so inconsistencies are likely. The peaks are annotated based on the given sequence, with 20 ppm tolerance.

Copy Data

##### Spectrum 10066 (TSV)

###### Preview

```
Loading example...
```

*Click on the button to copy the data to your clipboard.*

Mz MinMz MaxIntensity Max

WidthHeightPeptide font sizePeptide stroke widthSpectrum font sizeSpectrum stroke widthCompact peptide

Ion legend

wxyz

abcd

OtherUnassignedIon^Charge^_Position_Show for top:%

TFDDYAMHW

09.53e+31.91e+42.86e+43.81e+4

Zoom Out

c^+1^_2_y^+1^_2_c^+1^_3_z^+1^_3_c^+1^_4_y^+1^_3_z^+1^_4_y^+2^_8_y^+1^_4_z^+1^_5_y^+1^_5_z^+1^_6_y^+1^_6_c^+1^_7_y^+1^_7_z^+1^_7_y^+1^_7_c^+1^_8_z^+1^_8_

0778155623333111

Fragment Matches Table

Show background peaks

| Position | Ion type | Intensity | mz Theoretical | mz Error (Th) | mz Error (ppm) | Charge | Series Number |
| --- | --- | --- | --- | --- | --- | --- | --- |
| - | - | 392.4 | 120.1 | - | - | 0 | - |
| - | - | 3609 | 120.1 | - | - | 0 | - |
| - | - | 397.7 | 128.3 | - | - | 0 | - |
| - | - | 434.9 | 134.2 | - | - | 0 | - |
| - | - | 524.7 | 136.1 | - | - | 0 | - |
| - | - | 448.4 | 142.6 | - | - | 0 | - |
| - | - | 735.4 | 148.9 | - | - | 0 | - |
| - | - | 459.1 | 166.1 | - | - | 0 | - |
| - | - | 513.8 | 167.5 | - | - | 0 | - |
| - | - | 2534 | 173.5 | - | - | 0 | - |
| - | - | 3115 | 182.1 | - | - | 0 | - |
| - | - | 1163 | 185.1 | - | - | 0 | - |
| - | - | 555.3 | 210.5 | - | - | 0 | - |
| - | - | 488.1 | 213.2 | - | - | 0 | - |
| - | - | 574.6 | 214.1 | - | - | 0 | - |
| - | - | 462.7 | 216.4 | - | - | 0 | - |
| - | - | 1.088E+04 | 221.1 | - | - | 0 | - |
| - | - | 1119 | 222.1 | - | - | 0 | - |
| - | - | 600.3 | 223.1 | - | - | 0 | - |
| - | - | 515.9 | 240.3 | - | - | 0 | - |
| 2 | c | 4135 | 249.1 | 0.0002426 | 0.9738 | +1 | 2 |
| - | - | 871.2 | 250.1 | - | - | 0 | - |
| - | - | 567.4 | 264.1 | - | - | 0 | - |
| - | - | 530.6 | 282.1 | - | - | 0 | - |
| - | - | 2271 | 319.1 | - | - | 0 | - |
| - | - | 2362 | 329.2 | - | - | 0 | - |
| 8 | y | 1390 | 342.2 | 3.146E-05 | 0.09194 | +1 | 2 |
| - | - | 1051 | 355.1 | - | - | 0 | - |
| - | - | 591.2 | 357.1 | - | - | 0 | - |
| 3 | c | 1016 | 364.2 | 0.001172 | 3.22 | +1 | 3 |
| - | - | 1609 | 373.2 | - | - | 0 | - |
| - | - | 6015 | 374.3 | - | - | 0 | - |
| - | - | 626 | 375.3 | - | - | 0 | - |
| - | - | 684.5 | 398.2 | - | - | 0 | - |
| - | - | 754.4 | 413.3 | - | - | 0 | - |
| - | - | 787.6 | 418.2 | - | - | 0 | - |
| - | - | 674 | 427.7 | - | - | 0 | - |
| 7 | z | 833.4 | 473.2 | 0.0001423 | 0.3006 | +1 | 3 |
| - | - | 3187 | 474.2 | - | - | 0 | - |
| - | - | 835.2 | 475.2 | - | - | 0 | - |
| 4 | c | 790.9 | 479.2 | 0.001718 | 3.585 | +1 | 4 |
| - | - | 5086 | 486.3 | - | - | 0 | - |
| - | - | 2748 | 487.3 | - | - | 0 | - |
| - | - | 600.9 | 488.3 | - | - | 0 | - |
| 7 | y | 973.3 | 489.2 | 0.003696 | 7.555 | +1 | 3 |
| - | - | 604.3 | 504 | - | - | 0 | - |
| - | - | 514.7 | 531.3 | - | - | 0 | - |
| 6 | z | 8812 | 544.2 | 0.005814 | 10.68 | +1 | 4 |
| - | - | 740.5 | 544.3 | - | - | 0 | - |
| - | - | 5024 | 545.2 | - | - | 0 | - |
| - | - | 1134 | 546.2 | - | - | 0 | - |
| 2 | y | 651.2 | 550.7 | 0.006559 | 11.91 | +2 | 8 |
| - | - | 661.1 | 551.2 | - | - | 0 | - |
| 6 | y | 1339 | 560.2 | 0.00534 | 9.531 | +1 | 4 |
| - | - | 533.3 | 584.1 | - | - | 0 | - |
| - | - | 938.9 | 601.2 | - | - | 0 | - |
| - | - | 2023 | 601.4 | - | - | 0 | - |
| - | - | 807.5 | 601.7 | - | - | 0 | - |
| - | - | 767.7 | 602.2 | - | - | 0 | - |
| - | - | 753.3 | 602.4 | - | - | 0 | - |
| 5 | z | 4521 | 707.3 | 0.005901 | 8.344 | +1 | 5 |
| - | - | 1962 | 708.3 | - | - | 0 | - |
| - | - | 1025 | 709.3 | - | - | 0 | - |
| - | - | 2116 | 716.4 | - | - | 0 | - |
| - | - | 1391 | 717.4 | - | - | 0 | - |
| - | - | 556.5 | 718.4 | - | - | 0 | - |
| 5 | y | 2483 | 723.3 | 0.005671 | 7.84 | +1 | 5 |
| - | - | 781.6 | 724.3 | - | - | 0 | - |
| - | - | 821.6 | 732.4 | - | - | 0 | - |
| - | - | 1880 | 778.3 | - | - | 0 | - |
| - | - | 1506 | 779.3 | - | - | 0 | - |
| - | - | 634.7 | 799.4 | - | - | 0 | - |
| - | - | 1184 | 817.4 | - | - | 0 | - |
| - | - | 7525 | 818.4 | - | - | 0 | - |
| - | - | 560.2 | 819.2 | - | - | 0 | - |
| - | - | 3040 | 819.4 | - | - | 0 | - |
| - | - | 1177 | 820.4 | - | - | 0 | - |
| 4 | z | 2576 | 822.3 | 0.006241 | 7.589 | +1 | 6 |
| - | - | 3005 | 823.3 | - | - | 0 | - |
| - | - | 1421 | 824.3 | - | - | 0 | - |
| - | - | 1318 | 833.3 | - | - | 0 | - |
| - | - | 865.7 | 834.3 | - | - | 0 | - |
| 4 | y | 1740 | 838.3 | 0.005339 | 6.369 | +1 | 6 |
| - | - | 978.2 | 839.3 | - | - | 0 | - |
| - | - | 888.3 | 845.5 | - | - | 0 | - |
| - | - | 1663 | 849.4 | - | - | 0 | - |
| - | - | 836 | 850.3 | - | - | 0 | - |
| - | - | 920.3 | 860.5 | - | - | 0 | - |
| - | - | 703.5 | 870.9 | - | - | 0 | - |
| - | - | 6101 | 872.4 | - | - | 0 | - |
| - | - | 2782 | 873.5 | - | - | 0 | - |
| - | - | 898 | 874.4 | - | - | 0 | - |
| - | - | 1362 | 876.3 | - | - | 0 | - |
| 7 | c | 8424 | 877.3 | 0.005662 | 6.454 | +1 | 7 |
| - | - | 2880 | 878.3 | - | - | 0 | - |
| - | - | 1444 | 879.3 | - | - | 0 | - |
| - | - | 1228 | 889.3 | - | - | 0 | - |
| - | - | 1540 | 889.5 | - | - | 0 | - |
| - | - | 1377 | 890.4 | - | - | 0 | - |
| - | - | 978.5 | 890.5 | - | - | 0 | - |
| - | - | 3859 | 893.3 | - | - | 0 | - |
| - | - | 1533 | 894.3 | - | - | 0 | - |
| - | - | 674.5 | 895.3 | - | - | 0 | - |
| 3 | y | 824 | 935.3 | 0.002388 | 2.554 | +1 | 7 |
| 3 | z | 756.2 | 937.3 | 0.007801 | 8.323 | +1 | 7 |
| - | - | 579.4 | 952.4 | - | - | 0 | - |
| 3 | y | 8675 | 953.3 | 0.006594 | 6.917 | +1 | 7 |
| - | - | 4609 | 954.3 | - | - | 0 | - |
| - | - | 2032 | 955.4 | - | - | 0 | - |
| - | - | 1261 | 970.4 | - | - | 0 | - |
| - | - | 764.5 | 971.4 | - | - | 0 | - |
| - | - | 5080 | 984.5 | - | - | 0 | - |
| - | - | 3508 | 985.5 | - | - | 0 | - |
| - | - | 850.3 | 986.5 | - | - | 0 | - |
| - | - | 1464 | 988.5 | - | - | 0 | - |
| - | - | 1462 | 992.5 | - | - | 0 | - |
| - | - | 1062 | 993.5 | - | - | 0 | - |
| 8 | c | 1.238E+04 | 1014 | 0.006504 | 6.411 | +1 | 8 |
| - | - | 6808 | 1015 | - | - | 0 | - |
| - | - | 3740 | 1016 | - | - | 0 | - |
| - | - | 3405 | 1020 | - | - | 0 | - |
| - | - | 1141 | 1021 | - | - | 0 | - |
| - | - | 7429 | 1037 | - | - | 0 | - |
| - | - | 4203 | 1038 | - | - | 0 | - |
| - | - | 1181 | 1039 | - | - | 0 | - |
| - | - | 1223 | 1040 | - | - | 0 | - |
| - | - | 9048 | 1042 | - | - | 0 | - |
| - | - | 5756 | 1043 | - | - | 0 | - |
| - | - | 1971 | 1054 | - | - | 0 | - |
| - | - | 824.9 | 1055 | - | - | 0 | - |
| - | - | 731.4 | 1071 | - | - | 0 | - |
| 2 | z | 1120 | 1084 | 0.006404 | 5.905 | +1 | 8 |
| - | - | 768.6 | 1085 | - | - | 0 | - |
| - | - | 802.2 | 1090 | - | - | 0 | - |
| - | - | 4165 | 1098 | - | - | 0 | - |
| - | - | 3171 | 1099 | - | - | 0 | - |
| - | - | 947.3 | 1100 | - | - | 0 | - |
| - | - | 1085 | 1129 | - | - | 0 | - |
| - | - | 774.1 | 1130 | - | - | 0 | - |
| - | - | 2662 | 1140 | - | - | 0 | - |
| - | - | 3291 | 1141 | - | - | 0 | - |
| - | - | 4698 | 1142 | - | - | 0 | - |
| - | - | 3479 | 1142 | - | - | 0 | - |
| - | - | 4616 | 1143 | - | - | 0 | - |
| - | - | 2146 | 1143 | - | - | 0 | - |
| - | - | 1514 | 1144 | - | - | 0 | - |
| - | - | 897.3 | 1144 | - | - | 0 | - |
| - | - | 726.2 | 1145 | - | - | 0 | - |
| - | - | 985.6 | 1184 | - | - | 0 | - |
| - | - | 1.413E+04 | 1185 | - | - | 0 | - |
| - | - | 1.404E+04 | 1185 | - | - | 0 | - |
| - | - | 8871 | 1186 | - | - | 0 | - |
| - | - | 1.256E+04 | 1186 | - | - | 0 | - |
| - | - | 3634 | 1187 | - | - | 0 | - |
| - | - | 5857 | 1187 | - | - | 0 | - |
| - | - | 1575 | 1188 | - | - | 0 | - |
| - | - | 2297 | 1188 | - | - | 0 | - |
| - | - | 1160 | 1189 | - | - | 0 | - |
| - | - | 5462 | 1201 | - | - | 0 | - |
| - | - | 9906 | 1201 | - | - | 0 | - |
| - | - | 9003 | 1202 | - | - | 0 | - |
| - | - | 3.775E+04 | 1202 | - | - | 0 | - |
| - | - | 4732 | 1203 | - | - | 0 | - |
| - | - | 2.494E+04 | 1203 | - | - | 0 | - |
| - | - | 2291 | 1204 | - | - | 0 | - |
| - | - | 1.083E+04 | 1204 | - | - | 0 | - |
| - | - | 6104 | 1205 | - | - | 0 | - |
| - | - | 2808 | 1205 | - | - | 0 | - |
| - | - | 3341 | 1206 | - | - | 0 | - |
| - | - | 2193 | 1206 | - | - | 0 | - |
| - | - | 675.5 | 1366 | - | - | 0 | - |
| - | - | 749.6 | 1784 | - | - | 0 | - |
| - | - | 822.5 | 1785 | - | - | 0 | - |
| - | - | 764.3 | 1786 | - | - | 0 | - |
| - | - | 1881 | 1801 | - | - | 0 | - |
| - | - | 718.3 | 1804 | - | - | 0 | - |
| - | - | 836.1 | 1809 | - | - | 0 | - |
| - | - | 709.8 | 2547 | - | - | 0 | - |
| - | - | 769.5 | 3080 | - | - | 0 | - |

m/z Charge Intensity FragmentType MassShift Position
120.07720947265625 0 392.35696
120.0809555053711 0 3609.1755
128.25790405273438 0 397.73746
134.21234130859375 0 434.87387
136.07586669921875 0 524.6956
142.5833282470703 0 448.43848
148.947021484375 0 735.37756
166.0857696533203 0 459.13855
167.4925079345703 0 513.8434
173.45135498046875 0 2533.5442
182.08132934570312 0 3114.6948
185.1287384033203 0 1163.32
210.5415802001953 0 555.2503
213.22772216796875 0 488.09195
214.11965942382812 0 574.5937
216.35382080078125 0 462.70026
221.12860107421875 0 10882.924
222.13165283203125 0 1118.7128
223.06350708007812 0 600.2839
240.2763214111328 0 515.86304
249.1236114501953 0 4134.7944 c Ammonia loss 1
250.1273193359375 0 871.1849
264.13629150390625 0 567.42444
282.1479797363281 0 530.5947
319.14044189453125 0 2270.948
329.1503601074219 0 2361.6174
342.1560974121094 0 1389.7971 y 7
355.0703125 0 1050.9562
357.06842041015625 0 591.20636
364.1491394042969 0 1016.0804 c Ammonia loss 2
373.2430725097656 0 1608.9988
374.2509765625 0 6015.0864
375.2524108886719 0 625.97656
398.240966796875 0 684.5027
413.2669982910156 0 754.4061
418.2096252441406 0 787.58417
427.72235107421875 0 674.0451
473.16796875 0 833.4068 z 6
474.1810607910156 0 3186.774
475.18524169921875 0 835.17706
479.175537109375 0 790.93585 c Ammonia loss 3
486.328125 0 5085.51
487.33355712890625 0 2748.1147
488.3323669433594 0 600.9006
489.19024658203125 0 973.3041 y 6
504.0219421386719 0 604.3297
531.2598876953125 0 514.6887
544.2107543945312 0 8812.171 z 5
544.2520141601562 0 740.4521
545.2158203125 0 5023.5786
546.2130126953125 0 1133.8002
550.71484375 0 651.20856 y 1
551.2149047851562 0 661.1239
560.22900390625 0 1339.133 y 5
584.0921630859375 0 533.2862
601.2384643554688 0 938.9315
601.3538208007812 0 2023.1301
601.7352905273438 0 807.54004
602.2335205078125 0 767.6733
602.3599243164062 0 753.25085
707.274169921875 0 4520.981 z 4
708.2797241210938 0 1961.9747
709.275390625 0 1025.1722
716.3815307617188 0 2116.2559
717.3837280273438 0 1391.4183
718.387939453125 0 556.4602
723.2926635742188 0 2482.7559 y 4
724.2969360351562 0 781.60675
732.4013061523438 0 821.58374
778.309814453125 0 1879.7396
779.3140258789062 0 1505.7412
799.3650512695312 0 634.73303
817.4330444335938 0 1184.2294
818.4384765625 0 7524.753
819.1574096679688 0 560.16
819.4413452148438 0 3040.2139
820.4464111328125 0 1176.9502
822.3014526367188 0 2576.2112 z 3
823.3070678710938 0 3005.1182
824.3060302734375 0 1420.7067
833.32958984375 0 1317.8936
834.3323364257812 0 865.67444
838.3192749023438 0 1739.5522 y 3
839.3193359375 0 978.17267
845.4639282226562 0 888.3354
849.3505249023438 0 1662.9263
850.3455200195312 0 836.00104
860.4620361328125 0 920.27026
870.8924560546875 0 703.4702
872.4483032226562 0 6101.381
873.450927734375 0 2782.0671
874.4439697265625 0 897.95435
876.3325805664062 0 1362.4229
877.3403930664062 0 8424.283 c 6
878.3425903320312 0 2880.1843
879.3434448242188 0 1443.7644
889.3468627929688 0 1227.7004
889.4711303710938 0 1540.4147
890.350341796875 0 1376.7908
890.47607421875 0 978.49335
893.3385009765625 0 3858.611
894.34033203125 0 1532.5275
895.34619140625 0 674.48175
935.3327026367188 0 823.9843 y Water loss 2
937.3299560546875 0 756.19495 z 2
952.3563232421875 0 579.35297
953.3474731445312 0 8675.297 y 2
954.3495483398438 0 4608.5645
955.3538208007812 0 2031.8015
970.3858032226562 0 1261.1433
971.3904418945312 0 764.49274
984.4559936523438 0 5079.863
985.4593505859375 0 3507.574
986.46044921875 0 850.25226
988.5222778320312 0 1464.0941
992.52734375 0 1462.1648
993.53369140625 0 1061.7377
1014.400146484375 0 12378.892 c 7
1015.4027099609375 0 6808.118
1016.4041748046875 0 3740.238
1019.517822265625 0 3404.875
1020.5211181640625 0 1141.1782
1036.543701171875 0 7428.5063
1037.546630859375 0 4203.232
1038.54736328125 0 1181.4822
1040.4051513671875 0 1222.8514
1041.58349609375 0 9047.662
1042.58349609375 0 5756.3525
1054.4844970703125 0 1971.0555
1055.49072265625 0 824.9298
1071.396240234375 0 731.3513
1084.39697265625 0 1120.3229 z 1
1085.3946533203125 0 768.6358
1089.559814453125 0 802.23645
1097.5406494140625 0 4164.703
1098.5413818359375 0 3171.1917
1099.5474853515625 0 947.30286
1128.516357421875 0 1085.3485
1129.5057373046875 0 774.0529
1140.4661865234375 0 2662.2185
1141.45703125 0 3290.5073
1141.5811767578125 0 4698.106
1142.4530029296875 0 3478.9773
1142.579345703125 0 4616.184
1143.458740234375 0 2145.934
1143.580078125 0 1513.5743
1144.4676513671875 0 897.2881
1144.5908203125 0 726.247
1183.58349609375 0 985.6184
1184.57470703125 0 14125.345
1185.447509765625 0 14039.586
1185.5797119140625 0 8871.146
1186.44970703125 0 12556.988
1186.5882568359375 0 3633.9343
1187.4530029296875 0 5857.0083
1187.60107421875 0 1574.7407
1188.4552001953125 0 2297.3718
1188.5965576171875 0 1160.389
1200.5931396484375 0 5461.9575
1201.4615478515625 0 9906.155
1201.5986328125 0 9003.276
1202.47021484375 0 37745.465
1202.6043701171875 0 4732.243
1203.4739990234375 0 24935.645
1203.6175537109375 0 2291.318
1204.475830078125 0 10833.24
1204.639892578125 0 6104.27
1205.4786376953125 0 2807.7134
1205.6409912109375 0 3341.3022
1206.395263671875 0 2192.7349
1365.7352294921875 0 675.4538
1783.7559814453125 0 749.63257
1784.766845703125 0 822.46436
1785.77001953125 0 764.29407
1800.78369140625 0 1881.2198
1803.798583984375 0 718.2816
1809.4644775390625 0 836.12714
2546.85595703125 0 709.77997
3080.281494140625 0 769.5022

Spectrum Details

|  |  |
| --- | --- |
| Matched peaks? Matched peaks The total absolute number of peaks matched. Additionally in brackets the total fraction of peaks matched and the total number of peaks is shown. | 19 (10.61% of 179) |
| FDR? FDR The false discovery rate estimated for this peptide. It is calculated by matching all theoretical fragments with a non-integer shift with the raw peaks for this spectrum. This is done with 40 different shifts. The resulting percentage is the average number of annotated peaks over the number of annotated peaks with the correct spectrum. | 2.38% |
| Satellite FDR? Satellite FDR See the FDR for details on its calculation. This satellite ion specific FDR only contains the satellite ions (d/w) for I/L/J positions. | - |
| PSM Score? PSM Score The PSM Score as given by Hecklib to this annotated spectrum. It is shown with three significant figures. | 243 |

#### Spectrum 14047? Spectrum 14047 The raw spectrum of this peptide as annotated by Hecklib. The fragments are coloured according to ion type (see legend). Any peaks with a star '\*' as text can be hovered over to see the full details, first the ion type second the mass shift type. By hovering over the amino acids in the peptide or ions in the legend the corresponding peaks are highlighted. By toggling the 'Unassigned' label you can turn the background (unassigned) peaks on or off in the plot. By updating the slider in the Ion legend you can update the spectrum to only show the top X% of the peaks with labels. The top X% means any peak that is within X% of the highest intensity. By dragging in the spectrum you can zoom in to a specific part of the spectrum and use 'Zoom Out' to get back to the original zoom level. The annotation of the spectrum is based on the given sequence in the peptides file and is done with different software so inconsistencies are likely. The peaks are annotated based on the given sequence, with 20 ppm tolerance.

Copy Data

##### Spectrum 14047 (TSV)

###### Preview

```
Loading example...
```

*Click on the button to copy the data to your clipboard.*

Mz MinMz MaxIntensity Max

WidthHeightPeptide font sizePeptide stroke widthSpectrum font sizeSpectrum stroke widthCompact peptide

Ion legend

wxyz

abcd

OtherUnassignedIon^Charge^_Position_Show for top:%

TFDDYAMHW

06.52e+31.30e+41.96e+42.61e+4

Zoom Out

a^+1^_2_y^+1^_1_a^+1^_2_b^+1^_2_y^+1^_2_b^+1^_3_b^+1^_4_y^+1^_3_y^+2^_8_y^+1^_4_\*y^+1^_5_y^+1^_6_y^+1^_7_y^+1^_7_

0731146221932924

Fragment Matches Table

Show background peaks

| Position | Ion type | Intensity | mz Theoretical | mz Error (Th) | mz Error (ppm) | Charge | Series Number |
| --- | --- | --- | --- | --- | --- | --- | --- |
| - | - | 841.6 | 120.1 | - | - | 0 | - |
| - | - | 1.651E+04 | 120.1 | - | - | 0 | - |
| - | - | 1419 | 121.1 | - | - | 0 | - |
| - | - | 360.7 | 122.8 | - | - | 0 | - |
| - | - | 408.4 | 125.4 | - | - | 0 | - |
| - | - | 453 | 127.1 | - | - | 0 | - |
| - | - | 3836 | 129.1 | - | - | 0 | - |
| - | - | 465.2 | 130.1 | - | - | 0 | - |
| - | - | 397.9 | 130.3 | - | - | 0 | - |
| - | - | 491.9 | 131.3 | - | - | 0 | - |
| - | - | 470.8 | 133.1 | - | - | 0 | - |
| - | - | 5963 | 136.1 | - | - | 0 | - |
| - | - | 397 | 136.8 | - | - | 0 | - |
| - | - | 483.5 | 137.1 | - | - | 0 | - |
| - | - | 369.6 | 140 | - | - | 0 | - |
| - | - | 379.6 | 142.6 | - | - | 0 | - |
| - | - | 1088 | 149 | - | - | 0 | - |
| - | - | 413.5 | 149 | - | - | 0 | - |
| - | - | 591.2 | 149 | - | - | 0 | - |
| - | - | 554.6 | 151.1 | - | - | 0 | - |
| - | - | 663.8 | 157.1 | - | - | 0 | - |
| - | - | 632.6 | 159.1 | - | - | 0 | - |
| - | - | 1599 | 165.1 | - | - | 0 | - |
| - | - | 624.6 | 165.1 | - | - | 0 | - |
| - | - | 633.9 | 166.1 | - | - | 0 | - |
| - | - | 1079 | 167.1 | - | - | 0 | - |
| - | - | 1143 | 173.1 | - | - | 0 | - |
| - | - | 436.7 | 173.4 | - | - | 0 | - |
| - | - | 981.5 | 173.5 | - | - | 0 | - |
| - | - | 1985 | 176.1 | - | - | 0 | - |
| - | - | 1492 | 177.1 | - | - | 0 | - |
| - | - | 2963 | 182.1 | - | - | 0 | - |
| - | - | 1110 | 188.1 | - | - | 0 | - |
| - | - | 503.6 | 189.5 | - | - | 0 | - |
| - | - | 838.1 | 201.1 | - | - | 0 | - |
| 2 | a | 532.3 | 203.1 | 0.0002135 | 1.051 | +1 | 2 |
| 9 | y | 692.5 | 205.1 | 0.000215 | 1.048 | +1 | 1 |
| - | - | 640.1 | 207.1 | - | - | 0 | - |
| - | - | 1868 | 221.1 | - | - | 0 | - |
| 2 | a | 2.583E+04 | 221.1 | 9.524E-06 | 0.04307 | +1 | 2 |
| - | - | 3866 | 222.1 | - | - | 0 | - |
| - | - | 680.8 | 223.1 | - | - | 0 | - |
| - | - | 519.4 | 223.1 | - | - | 0 | - |
| - | - | 588.5 | 225.1 | - | - | 0 | - |
| - | - | 628 | 226.1 | - | - | 0 | - |
| - | - | 9311 | 226.2 | - | - | 0 | - |
| - | - | 1303 | 227.2 | - | - | 0 | - |
| - | - | 520.5 | 229.2 | - | - | 0 | - |
| - | - | 572.3 | 231.1 | - | - | 0 | - |
| - | - | 886.1 | 235.1 | - | - | 0 | - |
| - | - | 512.8 | 235.8 | - | - | 0 | - |
| - | - | 674.4 | 239.1 | - | - | 0 | - |
| - | - | 634.5 | 241.1 | - | - | 0 | - |
| - | - | 1303 | 244.2 | - | - | 0 | - |
| 2 | b | 3789 | 249.1 | 0.0001511 | 0.6063 | +1 | 2 |
| - | - | 677.9 | 251.2 | - | - | 0 | - |
| - | - | 674.8 | 257.2 | - | - | 0 | - |
| - | - | 875.4 | 263.1 | - | - | 0 | - |
| - | - | 8135 | 266.2 | - | - | 0 | - |
| - | - | 1412 | 267.2 | - | - | 0 | - |
| - | - | 555.9 | 281.9 | - | - | 0 | - |
| - | - | 586.1 | 296.2 | - | - | 0 | - |
| - | - | 688.7 | 299.1 | - | - | 0 | - |
| - | - | 712.1 | 309.2 | - | - | 0 | - |
| - | - | 559.1 | 313.6 | - | - | 0 | - |
| - | - | 693.2 | 314.2 | - | - | 0 | - |
| - | - | 2343 | 329.2 | - | - | 0 | - |
| 8 | y | 4992 | 342.2 | 0.0003061 | 0.8947 | +1 | 2 |
| - | - | 604.1 | 343.2 | - | - | 0 | - |
| - | - | 945.6 | 350.1 | - | - | 0 | - |
| - | - | 1063 | 355.1 | - | - | 0 | - |
| - | - | 624.3 | 357.1 | - | - | 0 | - |
| - | - | 1274 | 357.2 | - | - | 0 | - |
| - | - | 533.5 | 357.3 | - | - | 0 | - |
| 3 | b | 1220 | 364.2 | 0.0001654 | 0.4542 | +1 | 3 |
| - | - | 601.8 | 374.9 | - | - | 0 | - |
| - | - | 651.2 | 401 | - | - | 0 | - |
| - | - | 742.2 | 465.2 | - | - | 0 | - |
| 4 | b | 967.6 | 479.2 | 0.0005276 | 1.101 | +1 | 4 |
| - | - | 1687 | 486.1 | - | - | 0 | - |
| 7 | y | 1486 | 489.2 | 0.004551 | 9.302 | +1 | 3 |
| - | - | 876.4 | 511.3 | - | - | 0 | - |
| 2 | y | 892 | 550.7 | 0.001005 | 1.825 | +2 | 8 |
| 6 | y | 1586 | 560.2 | 0.004546 | 8.115 | +1 | 4 |
| - | - | 905.5 | 561.2 | - | - | 0 | - |
| - | - | 615.9 | 566.5 | - | - | 0 | - |
| - | - | 1788 | 584 | - | - | 0 | - |
| - | - | 684.7 | 584.3 | - | - | 0 | - |
| - | - | 875.3 | 585.3 | - | - | 0 | - |
| - | - | 670.6 | 600.2 | - | - | 0 | - |
| - | - | 643.8 | 601.2 | - | - | 0 | - |
| 0 | Precursor | 2135 | 601.2 | 0.01183 | 19.68 | +2 | -1 |
| - | - | 723.2 | 657.3 | - | - | 0 | - |
| - | - | 603.3 | 662.3 | - | - | 0 | - |
| - | - | 846.8 | 679.4 | - | - | 0 | - |
| - | - | 650.1 | 680.8 | - | - | 0 | - |
| - | - | 2086 | 700.4 | - | - | 0 | - |
| - | - | 627.4 | 718.7 | - | - | 0 | - |
| 5 | y | 1229 | 723.3 | 0.005854 | 8.093 | +1 | 5 |
| - | - | 751.6 | 749.3 | - | - | 0 | - |
| - | - | 656.7 | 767.3 | - | - | 0 | - |
| - | - | 721.4 | 774.3 | - | - | 0 | - |
| - | - | 703.6 | 801.4 | - | - | 0 | - |
| - | - | 693.9 | 802.4 | - | - | 0 | - |
| 4 | y | 1518 | 838.3 | 0.007231 | 8.626 | +1 | 6 |
| - | - | 1036 | 839.3 | - | - | 0 | - |
| - | - | 1245 | 844.4 | - | - | 0 | - |
| - | - | 886.9 | 845.5 | - | - | 0 | - |
| - | - | 5810 | 872.4 | - | - | 0 | - |
| - | - | 2839 | 873.5 | - | - | 0 | - |
| - | - | 661.6 | 874.5 | - | - | 0 | - |
| - | - | 3185 | 889.3 | - | - | 0 | - |
| - | - | 740.4 | 890.4 | - | - | 0 | - |
| 3 | y | 709.5 | 935.3 | 0.004525 | 4.837 | +1 | 7 |
| 3 | y | 9682 | 953.3 | 0.005679 | 5.957 | +1 | 7 |
| - | - | 4715 | 954.3 | - | - | 0 | - |
| - | - | 1446 | 955.4 | - | - | 0 | - |
| - | - | 771.5 | 971.3 | - | - | 0 | - |
| - | - | 2326 | 1020 | - | - | 0 | - |
| - | - | 926.1 | 1021 | - | - | 0 | - |
| - | - | 790.4 | 1022 | - | - | 0 | - |
| - | - | 683.3 | 1036 | - | - | 0 | - |
| - | - | 660.8 | 2895 | - | - | 0 | - |

m/z Charge Intensity FragmentType MassShift Position
120.07633209228516 0 841.6318
120.08084869384766 0 16508.441
121.08428192138672 0 1419.3641
122.84864044189453 0 360.66705
125.3641586303711 0 408.41376
127.07516479492188 0 452.98322
129.102294921875 0 3835.6118
130.0651092529297 0 465.1554
130.33425903320312 0 397.92383
131.34854125976562 0 491.86276
133.08604431152344 0 470.76624
136.0757598876953 0 5962.8706
136.8127899169922 0 396.97302
137.07899475097656 0 483.52948
140.0132598876953 0 369.64206
142.5841827392578 0 379.56635
148.9544677734375 0 1088.2542
149.0065155029297 0 413.5424
149.04486083984375 0 591.2234
151.07460021972656 0 554.5776
157.1333770751953 0 663.75494
159.0917205810547 0 632.6386
165.0546417236328 0 1599.324
165.07681274414062 0 624.63257
166.0866241455078 0 633.935
167.0553741455078 0 1079.0381
173.12875366210938 0 1143.0729
173.36720275878906 0 436.72415
173.45286560058594 0 981.4665
176.10693359375 0 1984.864
177.10227966308594 0 1492.2686
182.08111572265625 0 2963.3274
188.07107543945312 0 1110.4368
189.54867553710938 0 503.6037
201.1227569580078 0 838.0893
203.11810302734375 0 532.343 a Water loss 1
205.09693908691406 0 692.47815 y 8
207.1128692626953 0 640.12726
221.1033477783203 0 1867.9974
221.1284637451172 0 25829.852 a 1
222.13206481933594 0 3865.5317
223.0631103515625 0 680.82043
223.13311767578125 0 519.44946
225.12258911132812 0 588.5373
226.1172637939453 0 628.0244
226.1802215576172 0 9310.572
227.18359375 0 1302.7139
229.1702880859375 0 520.50385
231.06312561035156 0 572.2706
235.1078643798828 0 886.1249
235.849365234375 0 512.80566
239.0946502685547 0 674.4359
241.0735626220703 0 634.5361
244.19065856933594 0 1302.9109
249.12351989746094 0 3788.6487 b 1
251.1505889892578 0 677.85156
257.1661376953125 0 674.81793
263.1023254394531 0 875.37695
266.1725769042969 0 8134.6895
267.1761169433594 0 1411.5243
281.92938232421875 0 555.8931
296.15252685546875 0 586.1324
299.0602722167969 0 688.6824
309.20306396484375 0 712.0556
313.6346435546875 0 559.10785
314.15460205078125 0 693.21826
329.1500549316406 0 2343.1855
342.1563720703125 0 4992.4414 y 7
343.160400390625 0 604.1172
350.1365051269531 0 945.58417
355.0697326660156 0 1062.5426
357.068115234375 0 624.3108
357.1697692871094 0 1273.7405
357.28466796875 0 533.4891
364.150146484375 0 1219.5393 b 2
374.90301513671875 0 601.8269
400.95782470703125 0 651.1797
465.1619873046875 0 742.2343
479.1767272949219 0 967.6452 b 3
486.0781555175781 0 1687.1987
489.19110107421875 0 1485.8885 y 6
511.3334655761719 0 876.3556
550.7092895507812 0 891.97906 y 1
560.2282104492188 0 1585.7893 y 5
561.23193359375 0 905.5229
566.5447387695312 0 615.9173
584.0415649414062 0 1788.1785
584.298095703125 0 684.7223
585.3345947265625 0 875.29913
600.2298583984375 0 670.56055
601.1917114257812 0 643.7906
601.2439575195312 0 2134.7156 Precursor
657.26025390625 0 723.17554
662.3340454101562 0 603.27924
679.3641967773438 0 846.7782
680.7731323242188 0 650.1441
700.3618774414062 0 2085.7998
718.6948852539062 0 627.42194
723.2928466796875 0 1229.0833 y 4
749.255126953125 0 751.5679
767.274658203125 0 656.7169
774.3173217773438 0 721.38257
801.4096069335938 0 703.56177
802.4397583007812 0 693.937
838.3211669921875 0 1518.302 y 3
839.3213500976562 0 1036.3098
844.4498291015625 0 1244.747
845.455322265625 0 886.9131
872.4483032226562 0 5809.9727
873.4505615234375 0 2839.0747
874.4607543945312 0 661.5845
889.3496704101562 0 3184.526
890.3501586914062 0 740.4328
935.3348388671875 0 709.51306 y Water loss 2
953.3465576171875 0 9681.976 y 2
954.34912109375 0 4715.005
955.3522338867188 0 1445.964
971.3077392578125 0 771.50226
1019.5153198242188 0 2326.2297
1020.5180053710938 0 926.0928
1021.5162353515625 0 790.415
1036.4193115234375 0 683.304
2895.1708984375 0 660.8155

Spectrum Details

|  |  |
| --- | --- |
| Matched peaks? Matched peaks The total absolute number of peaks matched. Additionally in brackets the total fraction of peaks matched and the total number of peaks is shown. | 15 (12.20% of 123) |
| FDR? FDR The false discovery rate estimated for this peptide. It is calculated by matching all theoretical fragments with a non-integer shift with the raw peaks for this spectrum. This is done with 40 different shifts. The resulting percentage is the average number of annotated peaks over the number of annotated peaks with the correct spectrum. | 0.79% |
| Satellite FDR? Satellite FDR See the FDR for details on its calculation. This satellite ion specific FDR only contains the satellite ions (d/w) for I/L/J positions. | - |
| PSM Score? PSM Score The PSM Score as given by Hecklib to this annotated spectrum. It is shown with three significant figures. | 160 |

#### Spectrum 7257? Spectrum 7257 The raw spectrum of this peptide as annotated by Hecklib. The fragments are coloured according to ion type (see legend). Any peaks with a star '\*' as text can be hovered over to see the full details, first the ion type second the mass shift type. By hovering over the amino acids in the peptide or ions in the legend the corresponding peaks are highlighted. By toggling the 'Unassigned' label you can turn the background (unassigned) peaks on or off in the plot. By updating the slider in the Ion legend you can update the spectrum to only show the top X% of the peaks with labels. The top X% means any peak that is within X% of the highest intensity. By dragging in the spectrum you can zoom in to a specific part of the spectrum and use 'Zoom Out' to get back to the original zoom level. The annotation of the spectrum is based on the given sequence in the peptides file and is done with different software so inconsistencies are likely. The peaks are annotated based on the given sequence, with 20 ppm tolerance.

Copy Data

##### Spectrum 7257 (TSV)

###### Preview

```
Loading example...
```

*Click on the button to copy the data to your clipboard.*

Mz MinMz MaxIntensity Max

WidthHeightPeptide font sizePeptide stroke widthSpectrum font sizeSpectrum stroke widthCompact peptide

Ion legend

wxyz

abcd

OtherUnassignedIon^Charge^_Position_Show for top:%

TFDDYAMHW

07.88e+51.58e+62.36e+63.15e+6

Zoom Out

z^+1^_1_y^+1^_1_c^+1^_2_c^+1^_2_y^+1^_2_c^+2^_6_y^+2^_5_c^+1^_3_w^+2^_6_y^+2^_6_z^+2^_7_y^+2^_7_y^+2^_7_c^+1^_4_y^+1^_3_c^+2^_8_y^+2^_8_z^+1^_4_y^+2^_8_y^+1^_4_c^+1^_5_z^+1^_5_c^+1^_6_y^+1^_5_y^+1^_6_z^+1^_6_y^+1^_6_c^+1^_7_c^+1^_7_z^+1^_7_y^+1^_7_z^+1^_7_y^+1^_7_c^+1^_8_c^+1^_8_y^+1^_8_z^+1^_8_y^+1^_8_

03046089121217

Fragment Matches Table

Show background peaks

| Position | Ion type | Intensity | mz Theoretical | mz Error (Th) | mz Error (ppm) | Charge | Series Number |
| --- | --- | --- | --- | --- | --- | --- | --- |
| - | - | 9.469E+04 | 120.1 | - | - | 0 | - |
| - | - | 821.6 | 121.1 | - | - | 0 | - |
| - | - | 7722 | 121.1 | - | - | 0 | - |
| - | - | 6266 | 130.1 | - | - | 0 | - |
| - | - | 2182 | 132.1 | - | - | 0 | - |
| - | - | 410.7 | 132.6 | - | - | 0 | - |
| - | - | 837.4 | 133.1 | - | - | 0 | - |
| - | - | 1.993E+04 | 136.1 | - | - | 0 | - |
| - | - | 1697 | 137.1 | - | - | 0 | - |
| - | - | 2106 | 138.1 | - | - | 0 | - |
| - | - | 474.6 | 141.1 | - | - | 0 | - |
| - | - | 586.7 | 143.1 | - | - | 0 | - |
| - | - | 582.1 | 149 | - | - | 0 | - |
| - | - | 1390 | 156.1 | - | - | 0 | - |
| - | - | 4921 | 158.1 | - | - | 0 | - |
| - | - | 2.878E+04 | 159.1 | - | - | 0 | - |
| - | - | 1196 | 160.1 | - | - | 0 | - |
| - | - | 3599 | 160.1 | - | - | 0 | - |
| - | - | 4865 | 160.1 | - | - | 0 | - |
| - | - | 489.9 | 161.1 | - | - | 0 | - |
| - | - | 582.4 | 165.1 | - | - | 0 | - |
| - | - | 481.1 | 166.1 | - | - | 0 | - |
| - | - | 545.5 | 168.7 | - | - | 0 | - |
| - | - | 8205 | 171.1 | - | - | 0 | - |
| - | - | 740.4 | 171.2 | - | - | 0 | - |
| - | - | 433 | 172.1 | - | - | 0 | - |
| - | - | 479 | 173.1 | - | - | 0 | - |
| - | - | 4500 | 173.5 | - | - | 0 | - |
| - | - | 1.85E+04 | 176.1 | - | - | 0 | - |
| - | - | 3.743E+04 | 177.1 | - | - | 0 | - |
| - | - | 1680 | 177.1 | - | - | 0 | - |
| - | - | 4444 | 178.1 | - | - | 0 | - |
| - | - | 1626 | 185.1 | - | - | 0 | - |
| - | - | 2010 | 186.1 | - | - | 0 | - |
| - | - | 3.232E+04 | 188.1 | - | - | 0 | - |
| 9 | z | 3554 | 189.1 | 0.003738 | 19.77 | +1 | 1 |
| - | - | 928.4 | 189.1 | - | - | 0 | - |
| - | - | 2453 | 191.1 | - | - | 0 | - |
| - | - | 618.2 | 200.6 | - | - | 0 | - |
| - | - | 4171 | 203.1 | - | - | 0 | - |
| - | - | 3.268E+04 | 203.1 | - | - | 0 | - |
| - | - | 733.3 | 204.1 | - | - | 0 | - |
| - | - | 4009 | 204.1 | - | - | 0 | - |
| 9 | y | 6.729E+04 | 205.1 | 0.0007616 | 3.713 | +1 | 1 |
| - | - | 7622 | 206.1 | - | - | 0 | - |
| - | - | 671.1 | 206.6 | - | - | 0 | - |
| - | - | 629.6 | 214 | - | - | 0 | - |
| - | - | 1481 | 217.1 | - | - | 0 | - |
| - | - | 4193 | 217.1 | - | - | 0 | - |
| - | - | 793.9 | 218.1 | - | - | 0 | - |
| - | - | 1401 | 219.1 | - | - | 0 | - |
| - | - | 3.121E+06 | 221.1 | - | - | 0 | - |
| - | - | 3.875E+05 | 222.1 | - | - | 0 | - |
| - | - | 2.292E+04 | 223.1 | - | - | 0 | - |
| - | - | 1.545E+04 | 231.1 | - | - | 0 | - |
| - | - | 3.516E+04 | 231.1 | - | - | 0 | - |
| - | - | 875.3 | 232.1 | - | - | 0 | - |
| - | - | 4982 | 232.1 | - | - | 0 | - |
| - | - | 1066 | 233.1 | - | - | 0 | - |
| - | - | 2.401E+04 | 235.1 | - | - | 0 | - |
| - | - | 3303 | 236.1 | - | - | 0 | - |
| - | - | 764.6 | 237.1 | - | - | 0 | - |
| - | - | 1205 | 238.5 | - | - | 0 | - |
| - | - | 1084 | 245.1 | - | - | 0 | - |
| 2 | c | 1.238E+06 | 249.1 | 0.001036 | 4.159 | +1 | 2 |
| - | - | 1.724E+05 | 250.1 | - | - | 0 | - |
| - | - | 3752 | 251.1 | - | - | 0 | - |
| - | - | 1.281E+04 | 251.1 | - | - | 0 | - |
| - | - | 2447 | 257.1 | - | - | 0 | - |
| - | - | 1242 | 262.1 | - | - | 0 | - |
| - | - | 8.915E+04 | 263.1 | - | - | 0 | - |
| - | - | 1.261E+04 | 264.1 | - | - | 0 | - |
| - | - | 2457 | 265.1 | - | - | 0 | - |
| 2 | c | 864.5 | 266.1 | 0.0007168 | 2.693 | +1 | 2 |
| - | - | 1142 | 268.1 | - | - | 0 | - |
| - | - | 5118 | 279.1 | - | - | 0 | - |
| - | - | 902.2 | 280.1 | - | - | 0 | - |
| - | - | 2.086E+04 | 285.1 | - | - | 0 | - |
| - | - | 3234 | 286.1 | - | - | 0 | - |
| - | - | 4148 | 292.1 | - | - | 0 | - |
| - | - | 1.699E+04 | 292.2 | - | - | 0 | - |
| - | - | 3104 | 293.2 | - | - | 0 | - |
| - | - | 2228 | 296.2 | - | - | 0 | - |
| - | - | 1182 | 300.1 | - | - | 0 | - |
| - | - | 1177 | 301.1 | - | - | 0 | - |
| - | - | 1946 | 302.1 | - | - | 0 | - |
| - | - | 1007 | 303.1 | - | - | 0 | - |
| - | - | 1239 | 303.1 | - | - | 0 | - |
| - | - | 1125 | 314.1 | - | - | 0 | - |
| - | - | 2122 | 318.1 | - | - | 0 | - |
| - | - | 1311 | 319.1 | - | - | 0 | - |
| - | - | 2311 | 320.1 | - | - | 0 | - |
| - | - | 1.435E+04 | 324.1 | - | - | 0 | - |
| - | - | 2389 | 325.1 | - | - | 0 | - |
| - | - | 9399 | 328.1 | - | - | 0 | - |
| - | - | 1755 | 329.1 | - | - | 0 | - |
| - | - | 1860 | 329.1 | - | - | 0 | - |
| - | - | 2150 | 332.1 | - | - | 0 | - |
| - | - | 1249 | 334.1 | - | - | 0 | - |
| - | - | 2362 | 336.2 | - | - | 0 | - |
| - | - | 1832 | 338.1 | - | - | 0 | - |
| - | - | 4107 | 339.2 | - | - | 0 | - |
| 8 | y | 3.112E+05 | 342.2 | 0.001374 | 4.016 | +1 | 2 |
| - | - | 5.726E+04 | 343.2 | - | - | 0 | - |
| - | - | 5945 | 344.2 | - | - | 0 | - |
| - | - | 3.651E+04 | 346.1 | - | - | 0 | - |
| - | - | 5663 | 347.1 | - | - | 0 | - |
| - | - | 1777 | 348.1 | - | - | 0 | - |
| - | - | 1446 | 348.1 | - | - | 0 | - |
| - | - | 4984 | 350.1 | - | - | 0 | - |
| - | - | 2.357E+04 | 356.1 | - | - | 0 | - |
| 6 | c | 4693 | 357.1 | 0.0007016 | 1.965 | +2 | 6 |
| - | - | 958.5 | 358.1 | - | - | 0 | - |
| - | - | 1457 | 360.1 | - | - | 0 | - |
| 5 | y | 7922 | 362.1 | 0.003988 | 11.01 | +2 | 5 |
| - | - | 4367 | 362.7 | - | - | 0 | - |
| - | - | 1988 | 363.1 | - | - | 0 | - |
| 3 | c | 3.395E+05 | 364.2 | 0.00136 | 3.736 | +1 | 3 |
| - | - | 6.448E+04 | 365.2 | - | - | 0 | - |
| - | - | 3409 | 366.1 | - | - | 0 | - |
| - | - | 7376 | 366.2 | - | - | 0 | - |
| - | - | 1.064E+04 | 367.2 | - | - | 0 | - |
| - | - | 2754 | 368.2 | - | - | 0 | - |
| - | - | 2998 | 374.2 | - | - | 0 | - |
| - | - | 1586 | 376.1 | - | - | 0 | - |
| - | - | 5.774E+04 | 378.1 | - | - | 0 | - |
| - | - | 1.092E+04 | 379.1 | - | - | 0 | - |
| - | - | 7336 | 380.1 | - | - | 0 | - |
| - | - | 1086 | 381.1 | - | - | 0 | - |
| - | - | 3859 | 382.1 | - | - | 0 | - |
| - | - | 3402 | 388.2 | - | - | 0 | - |
| 4 | w | 2013 | 389.2 | 0.0008667 | 2.227 | +2 | 6 |
| - | - | 1.524E+04 | 394.1 | - | - | 0 | - |
| - | - | 1305 | 394.2 | - | - | 0 | - |
| - | - | 2937 | 395.1 | - | - | 0 | - |
| - | - | 1253 | 395.2 | - | - | 0 | - |
| - | - | 953.2 | 399.2 | - | - | 0 | - |
| - | - | 2062 | 406.1 | - | - | 0 | - |
| - | - | 1521 | 406.2 | - | - | 0 | - |
| - | - | 985.1 | 407.2 | - | - | 0 | - |
| - | - | 841.1 | 408.2 | - | - | 0 | - |
| - | - | 4412 | 415.2 | - | - | 0 | - |
| - | - | 1115 | 416.2 | - | - | 0 | - |
| - | - | 1292 | 417.1 | - | - | 0 | - |
| - | - | 996.1 | 417.2 | - | - | 0 | - |
| - | - | 1281 | 419.2 | - | - | 0 | - |
| 4 | y | 5300 | 419.7 | 0.004189 | 9.981 | +2 | 6 |
| - | - | 1763 | 420.2 | - | - | 0 | - |
| - | - | 1.931E+04 | 425.2 | - | - | 0 | - |
| - | - | 4262 | 426.2 | - | - | 0 | - |
| - | - | 1.638E+04 | 433.2 | - | - | 0 | - |
| - | - | 3479 | 434.2 | - | - | 0 | - |
| - | - | 991.3 | 434.7 | - | - | 0 | - |
| - | - | 947 | 435.2 | - | - | 0 | - |
| - | - | 1757 | 435.2 | - | - | 0 | - |
| - | - | 831.5 | 436.7 | - | - | 0 | - |
| - | - | 3206 | 443.2 | - | - | 0 | - |
| - | - | 739.1 | 443.2 | - | - | 0 | - |
| - | - | 1321 | 445.2 | - | - | 0 | - |
| - | - | 942.6 | 445.7 | - | - | 0 | - |
| - | - | 3062 | 447.2 | - | - | 0 | - |
| - | - | 2507 | 448.7 | - | - | 0 | - |
| - | - | 2454 | 449.2 | - | - | 0 | - |
| - | - | 3900 | 451.2 | - | - | 0 | - |
| - | - | 1284 | 452.2 | - | - | 0 | - |
| - | - | 1616 | 452.7 | - | - | 0 | - |
| - | - | 795.3 | 453.2 | - | - | 0 | - |
| - | - | 1731 | 454.2 | - | - | 0 | - |
| - | - | 942.1 | 454.7 | - | - | 0 | - |
| - | - | 2394 | 455.2 | - | - | 0 | - |
| - | - | 2694 | 459.2 | - | - | 0 | - |
| - | - | 1082 | 459.7 | - | - | 0 | - |
| 3 | z | 1108 | 460.2 | 0.0001431 | 0.311 | +2 | 7 |
| - | - | 4.428E+04 | 461.2 | - | - | 0 | - |
| - | - | 1.162E+04 | 462.2 | - | - | 0 | - |
| - | - | 1209 | 463.2 | - | - | 0 | - |
| - | - | 952.9 | 464.2 | - | - | 0 | - |
| - | - | 1.284E+04 | 465.2 | - | - | 0 | - |
| - | - | 2653 | 466.2 | - | - | 0 | - |
| - | - | 1374 | 467.7 | - | - | 0 | - |
| 3 | y | 7178 | 468.2 | 0.004544 | 9.707 | +2 | 7 |
| - | - | 3177 | 468.7 | - | - | 0 | - |
| - | - | 990.2 | 469.2 | - | - | 0 | - |
| - | - | 965.5 | 469.2 | - | - | 0 | - |
| - | - | 8180 | 471.2 | - | - | 0 | - |
| - | - | 1983 | 472.2 | - | - | 0 | - |
| - | - | 941.2 | 473.2 | - | - | 0 | - |
| - | - | 1156 | 474.2 | - | - | 0 | - |
| - | - | 2829 | 476.2 | - | - | 0 | - |
| - | - | 1292 | 476.7 | - | - | 0 | - |
| 3 | y | 2.796E+04 | 477.2 | 0.0005134 | 1.076 | +2 | 7 |
| - | - | 1.239E+04 | 477.7 | - | - | 0 | - |
| - | - | 6247 | 478.2 | - | - | 0 | - |
| - | - | 1312 | 478.2 | - | - | 0 | - |
| - | - | 1130 | 478.7 | - | - | 0 | - |
| 4 | c | 1.857E+05 | 479.2 | 0.002097 | 4.376 | +1 | 4 |
| - | - | 4.431E+04 | 480.2 | - | - | 0 | - |
| - | - | 8048 | 481.2 | - | - | 0 | - |
| - | - | 1057 | 481.7 | - | - | 0 | - |
| - | - | 4520 | 482.2 | - | - | 0 | - |
| - | - | 1925 | 483.2 | - | - | 0 | - |
| - | - | 3821 | 485.2 | - | - | 0 | - |
| - | - | 2368 | 485.7 | - | - | 0 | - |
| - | - | 713 | 486.2 | - | - | 0 | - |
| 7 | y | 1.883E+05 | 489.2 | 0.007114 | 14.54 | +1 | 3 |
| - | - | 5.268E+04 | 490.2 | - | - | 0 | - |
| - | - | 3655 | 490.7 | - | - | 0 | - |
| - | - | 1.219E+04 | 491.2 | - | - | 0 | - |
| - | - | 1292 | 492.2 | - | - | 0 | - |
| - | - | 9718 | 495.2 | - | - | 0 | - |
| - | - | 2709 | 495.7 | - | - | 0 | - |
| - | - | 2817 | 496.2 | - | - | 0 | - |
| - | - | 3223 | 496.2 | - | - | 0 | - |
| - | - | 4.12E+04 | 496.2 | - | - | 0 | - |
| - | - | 776.9 | 496.7 | - | - | 0 | - |
| - | - | 4724 | 497.2 | - | - | 0 | - |
| - | - | 1.127E+04 | 497.2 | - | - | 0 | - |
| - | - | 1106 | 498.2 | - | - | 0 | - |
| - | - | 1285 | 498.2 | - | - | 0 | - |
| 8 | c | 1.591E+04 | 499.2 | 0.004465 | 8.945 | +2 | 8 |
| - | - | 1.024E+04 | 499.7 | - | - | 0 | - |
| - | - | 4000 | 500.2 | - | - | 0 | - |
| - | - | 936.3 | 500.7 | - | - | 0 | - |
| - | - | 1170 | 505.2 | - | - | 0 | - |
| - | - | 4365 | 508.2 | - | - | 0 | - |
| - | - | 3999 | 508.7 | - | - | 0 | - |
| - | - | 3062 | 509.2 | - | - | 0 | - |
| - | - | 2264 | 509.7 | - | - | 0 | - |
| - | - | 1750 | 510.2 | - | - | 0 | - |
| - | - | 1154 | 511.2 | - | - | 0 | - |
| - | - | 1.154E+04 | 513.2 | - | - | 0 | - |
| - | - | 3183 | 514.2 | - | - | 0 | - |
| - | - | 1644 | 514.2 | - | - | 0 | - |
| - | - | 1.13E+04 | 518.7 | - | - | 0 | - |
| - | - | 2.084E+04 | 519.2 | - | - | 0 | - |
| - | - | 2704 | 519.7 | - | - | 0 | - |
| - | - | 5310 | 520.2 | - | - | 0 | - |
| - | - | 1240 | 521.2 | - | - | 0 | - |
| - | - | 3069 | 523.2 | - | - | 0 | - |
| - | - | 1240 | 524.2 | - | - | 0 | - |
| - | - | 3623 | 525.2 | - | - | 0 | - |
| - | - | 1277 | 526.2 | - | - | 0 | - |
| - | - | 1838 | 527.2 | - | - | 0 | - |
| - | - | 3217 | 527.7 | - | - | 0 | - |
| - | - | 2200 | 528.2 | - | - | 0 | - |
| - | - | 1000 | 528.7 | - | - | 0 | - |
| - | - | 1021 | 530.2 | - | - | 0 | - |
| - | - | 1795 | 532.2 | - | - | 0 | - |
| - | - | 2920 | 532.7 | - | - | 0 | - |
| - | - | 2882 | 533.2 | - | - | 0 | - |
| - | - | 1319 | 533.7 | - | - | 0 | - |
| - | - | 3302 | 537.2 | - | - | 0 | - |
| - | - | 3349 | 538.2 | - | - | 0 | - |
| - | - | 984.3 | 538.7 | - | - | 0 | - |
| - | - | 1.05E+04 | 541.2 | - | - | 0 | - |
| 2 | y | 2.527E+04 | 541.7 | 0.004578 | 8.452 | +2 | 8 |
| - | - | 2.49E+04 | 542.2 | - | - | 0 | - |
| - | - | 7877 | 542.7 | - | - | 0 | - |
| - | - | 3261 | 543.2 | - | - | 0 | - |
| 6 | z | 5045 | 544.2 | 0.007279 | 13.38 | +1 | 4 |
| - | - | 2175 | 545.2 | - | - | 0 | - |
| - | - | 1727 | 546.2 | - | - | 0 | - |
| - | - | 1275 | 546.7 | - | - | 0 | - |
| - | - | 6402 | 548.2 | - | - | 0 | - |
| - | - | 2554 | 549.2 | - | - | 0 | - |
| - | - | 2912 | 550.2 | - | - | 0 | - |
| 2 | y | 2.106E+05 | 550.7 | 0.004667 | 8.475 | +2 | 8 |
| - | - | 1.515E+05 | 551.2 | - | - | 0 | - |
| - | - | 6.147E+04 | 551.7 | - | - | 0 | - |
| - | - | 8961 | 552.2 | - | - | 0 | - |
| - | - | 4406 | 555.7 | - | - | 0 | - |
| - | - | 4389 | 556.2 | - | - | 0 | - |
| - | - | 909 | 556.7 | - | - | 0 | - |
| 6 | y | 4.455E+05 | 560.2 | 0.007293 | 13.02 | +1 | 4 |
| - | - | 1.573E+04 | 560.7 | - | - | 0 | - |
| - | - | 1.429E+05 | 561.2 | - | - | 0 | - |
| - | - | 2875 | 561.7 | - | - | 0 | - |
| - | - | 3.191E+04 | 562.2 | - | - | 0 | - |
| - | - | 2472 | 563.2 | - | - | 0 | - |
| - | - | 1034 | 563.7 | - | - | 0 | - |
| - | - | 972.6 | 565.7 | - | - | 0 | - |
| - | - | 2699 | 566.2 | - | - | 0 | - |
| - | - | 2724 | 567.2 | - | - | 0 | - |
| - | - | 5.362E+04 | 569.2 | - | - | 0 | - |
| - | - | 3.678E+04 | 569.7 | - | - | 0 | - |
| - | - | 1.528E+04 | 570.2 | - | - | 0 | - |
| - | - | 5664 | 570.7 | - | - | 0 | - |
| - | - | 3486 | 571.2 | - | - | 0 | - |
| - | - | 2102 | 574.2 | - | - | 0 | - |
| - | - | 5665 | 574.7 | - | - | 0 | - |
| - | - | 3145 | 575.2 | - | - | 0 | - |
| - | - | 828.9 | 575.7 | - | - | 0 | - |
| - | - | 2.787E+04 | 578.2 | - | - | 0 | - |
| - | - | 1.558E+04 | 578.7 | - | - | 0 | - |
| - | - | 8385 | 579.2 | - | - | 0 | - |
| - | - | 1105 | 579.7 | - | - | 0 | - |
| - | - | 1588 | 580.2 | - | - | 0 | - |
| - | - | 939.1 | 582.2 | - | - | 0 | - |
| - | - | 1.889E+04 | 583.2 | - | - | 0 | - |
| - | - | 1.225E+04 | 583.7 | - | - | 0 | - |
| - | - | 8038 | 584.2 | - | - | 0 | - |
| - | - | 1091 | 584.7 | - | - | 0 | - |
| - | - | 1.328E+05 | 592.2 | - | - | 0 | - |
| - | - | 9.906E+04 | 592.7 | - | - | 0 | - |
| - | - | 4.532E+04 | 593.2 | - | - | 0 | - |
| - | - | 7223 | 593.7 | - | - | 0 | - |
| - | - | 2369 | 594.2 | - | - | 0 | - |
| - | - | 1240 | 594.2 | - | - | 0 | - |
| - | - | 3.035E+04 | 596.2 | - | - | 0 | - |
| - | - | 1.358E+04 | 597.2 | - | - | 0 | - |
| - | - | 5570 | 598.2 | - | - | 0 | - |
| - | - | 1052 | 599.2 | - | - | 0 | - |
| - | - | 4.637E+05 | 601.2 | - | - | 0 | - |
| - | - | 3.398E+05 | 601.7 | - | - | 0 | - |
| - | - | 1.457E+05 | 602.2 | - | - | 0 | - |
| - | - | 2.167E+04 | 602.7 | - | - | 0 | - |
| - | - | 1.075E+04 | 606.2 | - | - | 0 | - |
| - | - | 2468 | 607.2 | - | - | 0 | - |
| - | - | 1252 | 608.2 | - | - | 0 | - |
| - | - | 1.878E+04 | 612.2 | - | - | 0 | - |
| - | - | 6680 | 613.2 | - | - | 0 | - |
| - | - | 6.353E+04 | 614.2 | - | - | 0 | - |
| - | - | 1011 | 615.2 | - | - | 0 | - |
| - | - | 2.208E+04 | 615.3 | - | - | 0 | - |
| - | - | 3151 | 616.3 | - | - | 0 | - |
| - | - | 4.756E+04 | 624.2 | - | - | 0 | - |
| - | - | 1.585E+04 | 625.2 | - | - | 0 | - |
| - | - | 3397 | 626.2 | - | - | 0 | - |
| - | - | 1097 | 632.3 | - | - | 0 | - |
| - | - | 2.329E+04 | 634.2 | - | - | 0 | - |
| - | - | 7732 | 635.2 | - | - | 0 | - |
| - | - | 1900 | 636.2 | - | - | 0 | - |
| - | - | 1862 | 641.3 | - | - | 0 | - |
| 5 | c | 5.169E+04 | 642.2 | 0.002581 | 4.018 | +1 | 5 |
| - | - | 1.946E+04 | 643.2 | - | - | 0 | - |
| - | - | 4322 | 644.2 | - | - | 0 | - |
| - | - | 983 | 649.3 | - | - | 0 | - |
| - | - | 1288 | 650.3 | - | - | 0 | - |
| - | - | 3488 | 652.2 | - | - | 0 | - |
| - | - | 1849 | 653.2 | - | - | 0 | - |
| - | - | 5601 | 657.3 | - | - | 0 | - |
| - | - | 1404 | 658.3 | - | - | 0 | - |
| - | - | 4.255E+04 | 659.3 | - | - | 0 | - |
| - | - | 1.612E+04 | 660.3 | - | - | 0 | - |
| - | - | 3650 | 661.3 | - | - | 0 | - |
| - | - | 1146 | 662.2 | - | - | 0 | - |
| - | - | 3866 | 667.3 | - | - | 0 | - |
| - | - | 3475 | 668.3 | - | - | 0 | - |
| - | - | 1850 | 669.3 | - | - | 0 | - |
| - | - | 946.8 | 676.3 | - | - | 0 | - |
| - | - | 3871 | 677.3 | - | - | 0 | - |
| - | - | 1126 | 678.3 | - | - | 0 | - |
| - | - | 866.7 | 679.2 | - | - | 0 | - |
| - | - | 937.8 | 680.2 | - | - | 0 | - |
| - | - | 2.263E+04 | 685.3 | - | - | 0 | - |
| - | - | 9334 | 686.3 | - | - | 0 | - |
| - | - | 1450 | 687.3 | - | - | 0 | - |
| - | - | 2.311E+04 | 695.3 | - | - | 0 | - |
| - | - | 1.009E+04 | 696.3 | - | - | 0 | - |
| - | - | 1407 | 697.3 | - | - | 0 | - |
| - | - | 2711 | 703.3 | - | - | 0 | - |
| - | - | 1333 | 704.2 | - | - | 0 | - |
| - | - | 1.045E+04 | 705.3 | - | - | 0 | - |
| - | - | 4043 | 706.3 | - | - | 0 | - |
| 5 | z | 2091 | 707.3 | 0.008648 | 12.23 | +1 | 5 |
| - | - | 2200 | 708.3 | - | - | 0 | - |
| 6 | c | 2.198E+04 | 713.3 | 0.002881 | 4.04 | +1 | 6 |
| - | - | 8481 | 714.3 | - | - | 0 | - |
| - | - | 1320 | 715.3 | - | - | 0 | - |
| - | - | 3.953E+04 | 721.3 | - | - | 0 | - |
| - | - | 1.557E+04 | 722.3 | - | - | 0 | - |
| 5 | y | 3.82E+05 | 723.3 | 0.007746 | 10.71 | +1 | 5 |
| - | - | 1.625E+05 | 724.3 | - | - | 0 | - |
| - | - | 4.852E+04 | 725.3 | - | - | 0 | - |
| - | - | 4962 | 726.3 | - | - | 0 | - |
| - | - | 6152 | 731.2 | - | - | 0 | - |
| - | - | 2914 | 732.2 | - | - | 0 | - |
| - | - | 2228 | 733.3 | - | - | 0 | - |
| - | - | 1084 | 747.3 | - | - | 0 | - |
| - | - | 1.328E+05 | 749.3 | - | - | 0 | - |
| - | - | 5.467E+04 | 750.3 | - | - | 0 | - |
| - | - | 1.704E+04 | 751.3 | - | - | 0 | - |
| - | - | 934.4 | 752.3 | - | - | 0 | - |
| - | - | 3176 | 756.3 | - | - | 0 | - |
| - | - | 3419 | 759.3 | - | - | 0 | - |
| - | - | 822.1 | 760.3 | - | - | 0 | - |
| - | - | 2.153E+04 | 767.3 | - | - | 0 | - |
| - | - | 7607 | 768.3 | - | - | 0 | - |
| - | - | 2959 | 769.3 | - | - | 0 | - |
| - | - | 6.306E+04 | 774.3 | - | - | 0 | - |
| - | - | 2.823E+04 | 775.3 | - | - | 0 | - |
| - | - | 7420 | 776.3 | - | - | 0 | - |
| - | - | 5044 | 778.3 | - | - | 0 | - |
| - | - | 2288 | 779.3 | - | - | 0 | - |
| - | - | 3510 | 792.3 | - | - | 0 | - |
| - | - | 2065 | 793.3 | - | - | 0 | - |
| - | - | 4464 | 796.3 | - | - | 0 | - |
| - | - | 1941 | 797.3 | - | - | 0 | - |
| - | - | 2291 | 802.4 | - | - | 0 | - |
| - | - | 1657 | 802.9 | - | - | 0 | - |
| - | - | 1039 | 803.3 | - | - | 0 | - |
| - | - | 1294 | 814.3 | - | - | 0 | - |
| - | - | 1241 | 815.3 | - | - | 0 | - |
| 4 | y | 1.683E+04 | 820.3 | 0.00803 | 9.789 | +1 | 6 |
| - | - | 793.4 | 820.4 | - | - | 0 | - |
| - | - | 1.051E+04 | 821.3 | - | - | 0 | - |
| - | - | 1268 | 822 | - | - | 0 | - |
| - | - | 1347 | 822.1 | - | - | 0 | - |
| 4 | z | 5172 | 822.3 | 0.01155 | 14.05 | +1 | 6 |
| - | - | 1470 | 822.8 | - | - | 0 | - |
| - | - | 2273 | 822.9 | - | - | 0 | - |
| - | - | 2843 | 823.3 | - | - | 0 | - |
| - | - | 3877 | 824.3 | - | - | 0 | - |
| - | - | 1466 | 825.3 | - | - | 0 | - |
| - | - | 3539 | 832.3 | - | - | 0 | - |
| - | - | 2619 | 833.3 | - | - | 0 | - |
| 4 | y | 5.126E+05 | 838.3 | 0.008208 | 9.791 | +1 | 6 |
| - | - | 2.515E+05 | 839.3 | - | - | 0 | - |
| - | - | 7.902E+04 | 840.3 | - | - | 0 | - |
| - | - | 8746 | 841.3 | - | - | 0 | - |
| - | - | 1.276E+04 | 842.3 | - | - | 0 | - |
| - | - | 5331 | 843.3 | - | - | 0 | - |
| - | - | 1140 | 844.3 | - | - | 0 | - |
| - | - | 1122 | 846.4 | - | - | 0 | - |
| - | - | 3233 | 848.3 | - | - | 0 | - |
| - | - | 1130 | 849.3 | - | - | 0 | - |
| - | - | 1101 | 854.3 | - | - | 0 | - |
| - | - | 2201 | 855.9 | - | - | 0 | - |
| - | - | 2148 | 855.9 | - | - | 0 | - |
| - | - | 5918 | 856 | - | - | 0 | - |
| 7 | c | 1.279E+04 | 860.3 | 0.00798 | 9.276 | +1 | 7 |
| - | - | 6695 | 861.3 | - | - | 0 | - |
| - | - | 9029 | 862.3 | - | - | 0 | - |
| - | - | 3432 | 863.3 | - | - | 0 | - |
| - | - | 1171 | 864.3 | - | - | 0 | - |
| - | - | 1363 | 868.3 | - | - | 0 | - |
| - | - | 2.117E+04 | 871.3 | - | - | 0 | - |
| - | - | 1.138E+04 | 872.3 | - | - | 0 | - |
| - | - | 3077 | 873.3 | - | - | 0 | - |
| - | - | 1557 | 874.3 | - | - | 0 | - |
| - | - | 1343 | 875.3 | - | - | 0 | - |
| 7 | c | 2483 | 877.3 | 0.008409 | 9.584 | +1 | 7 |
| - | - | 1926 | 878.3 | - | - | 0 | - |
| - | - | 1215 | 879.3 | - | - | 0 | - |
| - | - | 1206 | 887.3 | - | - | 0 | - |
| - | - | 3.753E+05 | 889.4 | - | - | 0 | - |
| - | - | 1.939E+05 | 890.4 | - | - | 0 | - |
| - | - | 5.056E+04 | 891.4 | - | - | 0 | - |
| - | - | 4144 | 892.4 | - | - | 0 | - |
| - | - | 3030 | 893.3 | - | - | 0 | - |
| - | - | 1469 | 894.3 | - | - | 0 | - |
| - | - | 1.014E+04 | 896.3 | - | - | 0 | - |
| - | - | 5523 | 897.3 | - | - | 0 | - |
| - | - | 1331 | 898.3 | - | - | 0 | - |
| - | - | 1278 | 899.3 | - | - | 0 | - |
| - | - | 1609 | 904.3 | - | - | 0 | - |
| - | - | 896.4 | 905.4 | - | - | 0 | - |
| - | - | 2.016E+04 | 907.3 | - | - | 0 | - |
| - | - | 1.277E+04 | 908.3 | - | - | 0 | - |
| - | - | 3981 | 909.3 | - | - | 0 | - |
| - | - | 2181 | 914.3 | - | - | 0 | - |
| - | - | 1651 | 914.5 | - | - | 0 | - |
| - | - | 1677 | 915.3 | - | - | 0 | - |
| - | - | 761.6 | 916.4 | - | - | 0 | - |
| - | - | 2859 | 917.3 | - | - | 0 | - |
| - | - | 4899 | 918.3 | - | - | 0 | - |
| 3 | z | 2605 | 919.3 | 0.008417 | 9.156 | +1 | 7 |
| - | - | 1078 | 920.3 | - | - | 0 | - |
| - | - | 1690 | 924.3 | - | - | 0 | - |
| - | - | 5722 | 933.4 | - | - | 0 | - |
| - | - | 2654 | 934.4 | - | - | 0 | - |
| 3 | y | 1.21E+05 | 935.3 | 0.008614 | 9.21 | +1 | 7 |
| - | - | 7.621E+04 | 936.3 | - | - | 0 | - |
| 3 | z | 3E+04 | 937.3 | 0.01665 | 17.76 | +1 | 7 |
| - | - | 1.17E+04 | 938.3 | - | - | 0 | - |
| - | - | 4023 | 939.3 | - | - | 0 | - |
| - | - | 1828 | 940.3 | - | - | 0 | - |
| - | - | 2260 | 951.4 | - | - | 0 | - |
| 3 | y | 2.546E+06 | 953.3 | 0.009463 | 9.926 | +1 | 7 |
| - | - | 1.399E+06 | 954.4 | - | - | 0 | - |
| - | - | 4.855E+05 | 955.4 | - | - | 0 | - |
| - | - | 5.659E+04 | 956.4 | - | - | 0 | - |
| - | - | 1.253E+04 | 963.3 | - | - | 0 | - |
| - | - | 6195 | 964.3 | - | - | 0 | - |
| - | - | 3081 | 965.3 | - | - | 0 | - |
| - | - | 9022 | 969.4 | - | - | 0 | - |
| - | - | 5532 | 970.4 | - | - | 0 | - |
| - | - | 1590 | 971.4 | - | - | 0 | - |
| - | - | 6120 | 979.4 | - | - | 0 | - |
| - | - | 3786 | 980.4 | - | - | 0 | - |
| - | - | 1771 | 981.4 | - | - | 0 | - |
| 8 | c | 3.39E+04 | 997.4 | 0.008395 | 8.417 | +1 | 8 |
| - | - | 1.974E+04 | 998.4 | - | - | 0 | - |
| - | - | 7388 | 999.4 | - | - | 0 | - |
| - | - | 896.1 | 1000 | - | - | 0 | - |
| 8 | c | 5364 | 1014 | 0.008701 | 8.577 | +1 | 8 |
| - | - | 1.167E+04 | 1015 | - | - | 0 | - |
| - | - | 7163 | 1016 | - | - | 0 | - |
| - | - | 2308 | 1017 | - | - | 0 | - |
| - | - | 5226 | 1018 | - | - | 0 | - |
| - | - | 2715 | 1019 | - | - | 0 | - |
| - | - | 2.024E+04 | 1036 | - | - | 0 | - |
| - | - | 1.097E+04 | 1037 | - | - | 0 | - |
| - | - | 4107 | 1038 | - | - | 0 | - |
| - | - | 1177 | 1039 | - | - | 0 | - |
| - | - | 1014 | 1046 | - | - | 0 | - |
| - | - | 2447 | 1054 | - | - | 0 | - |
| - | - | 994.4 | 1064 | - | - | 0 | - |
| - | - | 1241 | 1072 | - | - | 0 | - |
| 2 | y | 1.25E+04 | 1082 | 0.008132 | 7.513 | +1 | 8 |
| - | - | 7629 | 1083 | - | - | 0 | - |
| 2 | z | 3743 | 1084 | 0.01947 | 17.95 | +1 | 8 |
| 2 | y | 1.276E+05 | 1100 | 0.008798 | 7.995 | +1 | 8 |
| - | - | 8.564E+04 | 1101 | - | - | 0 | - |
| - | - | 3.835E+04 | 1102 | - | - | 0 | - |
| - | - | 4956 | 1103 | - | - | 0 | - |
| - | - | 2.211E+04 | 1110 | - | - | 0 | - |
| - | - | 1.415E+04 | 1111 | - | - | 0 | - |
| - | - | 6169 | 1112 | - | - | 0 | - |
| - | - | 1151 | 1140 | - | - | 0 | - |
| - | - | 1649 | 1141 | - | - | 0 | - |
| - | - | 2116 | 1142 | - | - | 0 | - |
| - | - | 7184 | 1185 | - | - | 0 | - |
| - | - | 5222 | 1186 | - | - | 0 | - |
| - | - | 1697 | 1187 | - | - | 0 | - |
| - | - | 1.435E+04 | 1201 | - | - | 0 | - |
| - | - | 2.041E+04 | 1202 | - | - | 0 | - |
| - | - | 1.035E+04 | 1203 | - | - | 0 | - |
| - | - | 3783 | 1204 | - | - | 0 | - |

m/z Charge Intensity FragmentType MassShift Position
120.08137512207031 0 94691.47
121.07966613769531 0 821.62476
121.08470916748047 0 7721.7827
130.0657501220703 0 6265.7754
132.08139038085938 0 2181.699
132.58984375 0 410.65894
133.06155395507812 0 837.41974
136.0762939453125 0 19928.459
137.07962036132812 0 1697.2734
138.06683349609375 0 2106.0093
141.06640625 0 474.57828
143.073974609375 0 586.7119
148.95411682128906 0 582.06934
156.07778930664062 0 1390.3127
158.09710693359375 0 4920.6704
159.09231567382812 0 28782.17
160.07644653320312 0 1196.3291
160.09564208984375 0 3599.1472
160.11276245117188 0 4865.123
161.11602783203125 0 489.93936
165.10275268554688 0 582.4324
166.061767578125 0 481.14673
168.7340545654297 0 545.5151
171.07708740234375 0 8205.44
171.15023803710938 0 740.42975
172.08059692382812 0 433.02985
173.09228515625 0 479.00293
173.45030212402344 0 4500.064
176.1077423095703 0 18502.28
177.10293579101562 0 37425.95
177.1109161376953 0 1679.8578
178.10646057128906 0 4444.173
185.0565948486328 0 1625.6708
186.0919647216797 0 2009.9275
188.07139587402344 0 32323.662
189.07469177246094 0 3554.3228 z 8
189.08705139160156 0 928.38416
191.1184844970703 0 2452.6257
200.62803649902344 0 618.1891
203.06686401367188 0 4171.2676
203.11863708496094 0 32681.18
204.1024932861328 0 733.3138
204.12191772460938 0 4009.2383
205.09791564941406 0 67290.36 y 8
206.10130310058594 0 7621.865
206.61672973632812 0 671.1429
214.00367736816406 0 629.615
217.08294677734375 0 1480.9269
217.0980682373047 0 4192.603
218.1023712158203 0 793.8504
219.08090209960938 0 1400.5734
221.12942504882812 0 3120549
222.1324005126953 0 387484.25
223.13497924804688 0 22921.61
231.06210327148438 0 15449.818
231.11373901367188 0 35159.56
232.06442260742188 0 875.2754
232.1168975830078 0 4981.5537
233.1289825439453 0 1066.3134
235.10861206054688 0 24012.264
236.11212158203125 0 3302.9092
237.12449645996094 0 764.61554
238.4553985595703 0 1204.6979
245.13043212890625 0 1084.0051
249.12440490722656 0 1238243.6 c Ammonia loss 1
250.1275634765625 0 172374.47
251.1036834716797 0 3751.9543
251.1300048828125 0 12810.53
257.10784912109375 0 2446.887
262.1204528808594 0 1241.6909
263.10357666015625 0 89145.164
264.10699462890625 0 12613.067
265.1189880371094 0 2456.7478
266.150634765625 0 864.47736 c 1
268.075439453125 0 1142.015
279.0985412597656 0 5117.805
280.13116455078125 0 902.1937
285.1027526855469 0 20864.426
286.1056213378906 0 3233.7776
292.14190673828125 0 4148.1445
292.16680908203125 0 16991.863
293.1698913574219 0 3103.8066
296.1519775390625 0 2227.9917
300.1353454589844 0 1181.8285
301.1198425292969 0 1177.4065
302.1146545410156 0 1945.5762
303.09637451171875 0 1007.0555
303.11553955078125 0 1239.1853
314.10009765625 0 1125.1594
318.1465759277344 0 2122.2595
319.1294860839844 0 1311.4891
320.12579345703125 0 2310.6465
324.1468200683594 0 14346.202
325.149658203125 0 2388.8896
328.1304626464844 0 9399.032
329.1138610839844 0 1755.0164
329.1349792480469 0 1859.7754
332.1100158691406 0 2150.3538
334.1420593261719 0 1249.2434
336.1572570800781 0 2361.8926
338.1297607421875 0 1831.8816
339.1715087890625 0 4106.9043
342.1574401855469 0 311244.62 y 7
343.1605224609375 0 57260.605
344.16326904296875 0 5944.573
346.1412353515625 0 36505.52
347.144287109375 0 5662.695
348.1199035644531 0 1776.639
348.1452941894531 0 1445.8854
350.1359558105469 0 4984.431
356.14007568359375 0 23572.031
357.1431884765625 0 4692.856 c Ammonia loss 5
358.134765625 0 958.5234
360.12042236328125 0 1456.5956
362.151123046875 0 7921.937 y 4
362.6522216796875 0 4366.5186
363.11944580078125 0 1988.2952
364.15167236328125 0 339496.66 c Ammonia loss 2
365.1549072265625 0 64478.24
366.13018798828125 0 3408.7612
366.15716552734375 0 7376.3247
367.16650390625 0 10642.456
368.16937255859375 0 2754.2295
374.1504821777344 0 2997.744
376.1168212890625 0 1586.0792
378.13104248046875 0 57741.33
379.1343078613281 0 10921.276
380.145751953125 0 7335.6846
381.1488952636719 0 1085.5518
382.144775390625 0 3859.4849
388.1514587402344 0 3401.6204
389.15155029296875 0 2012.7899 w 3
394.1262512207031 0 15235.497
394.17681884765625 0 1305.1417
395.1286315917969 0 2937.1965
395.1572570800781 0 1253.2163
399.17041015625 0 953.2481
406.12567138671875 0 2061.9036
406.1559143066406 0 1521.2676
407.1921081542969 0 985.1283
408.169921875 0 841.1364
415.16339111328125 0 4412.2554
416.16888427734375 0 1114.982
417.1419372558594 0 1292.16
417.17401123046875 0 996.0992
419.1576232910156 0 1280.6644
419.664794921875 0 5300.125 y 3
420.16729736328125 0 1763.0614
425.195068359375 0 19306.574
426.1977844238281 0 4262.1665
433.173583984375 0 16378.157
434.1771240234375 0 3478.7942
434.6693115234375 0 991.32666
435.15380859375 0 946.9739
435.1865234375 0 1756.5729
436.67669677734375 0 831.4689
443.15753173828125 0 3205.7646
443.1895751953125 0 739.1239
445.1726989746094 0 1320.875
445.67742919921875 0 942.63055
447.1527404785156 0 3061.7993
448.6683044433594 0 2506.833
449.1689758300781 0 2453.5874
451.1839294433594 0 3900.4316
452.18621826171875 0 1284.2467
452.6683349609375 0 1616.1108
453.1708984375 0 795.28876
454.15802001953125 0 1730.7322
454.6773681640625 0 942.0571
455.2049255371094 0 2394.4517
459.15374755859375 0 2693.568
459.6603698730469 0 1081.8429
460.1595764160156 0 1107.7683 z Water loss 2
461.1686096191406 0 44279.03
462.1720275878906 0 11621.221
463.17138671875 0 1208.6873
464.1824645996094 0 952.87054
465.1636657714844 0 12836.17
466.1673278808594 0 2653.2683
467.6932373046875 0 1373.5718
468.17333984375 0 7177.886 y Water loss 2
468.673828125 0 3176.7034
469.1751403808594 0 990.23206
469.2123718261719 0 965.5139
471.18304443359375 0 8180.4062
472.1838684082031 0 1982.5481
473.1786193847656 0 941.23773
474.1847229003906 0 1156.1438
476.19012451171875 0 2828.9736
476.69024658203125 0 1291.7894
477.1745910644531 0 27961.05 y 2
477.68023681640625 0 12389.849
478.1744079589844 0 6247.08
478.225341796875 0 1311.6725
478.6810607910156 0 1130.3684
479.1793518066406 0 185708.14 c Ammonia loss 3
480.1826477050781 0 44313.164
481.18426513671875 0 8047.5283
481.68353271484375 0 1056.7922
482.1937255859375 0 4519.781
483.1959533691406 0 1924.5045
485.1949462890625 0 3821.4092
485.6957092285156 0 2368.0288
486.1933898925781 0 712.9843
489.19366455078125 0 188286.9 y 6
490.19598388671875 0 52675.527
490.68756103515625 0 3655.117
491.1958312988281 0 12192.15
492.2091369628906 0 1291.5634
495.1794128417969 0 9717.735
495.69488525390625 0 2708.847
496.1742248535156 0 2816.532
496.18792724609375 0 3223.0305
496.232666015625 0 41201.28
496.690673828125 0 776.92224
497.1729736328125 0 4724.269
497.2353210449219 0 11270.037
498.1741027832031 0 1106.2188
498.239501953125 0 1285.4404
499.191650390625 0 15911.783 c Ammonia loss 7
499.69354248046875 0 10236.961
500.1925048828125 0 3999.8628
500.6898193359375 0 936.3165
505.1916809082031 0 1170.2549
508.1976623535156 0 4365.456
508.69781494140625 0 3998.709
509.2044677734375 0 3061.648
509.7039489746094 0 2264.0032
510.20428466796875 0 1749.88
511.1924133300781 0 1154.0231
513.2001953125 0 11539.133
514.2002563476562 0 3183.1265
514.2290649414062 0 1644.3058
518.712646484375 0 11298.141
519.20703125 0 20835.393
519.7146606445312 0 2703.7568
520.2064819335938 0 5309.605
521.2020263671875 0 1240.1946
523.18505859375 0 3068.8923
524.1913452148438 0 1240.2695
525.1932983398438 0 3623.215
526.1975708007812 0 1277.4354
527.2136840820312 0 1837.5981
527.7108764648438 0 3216.7737
528.2137451171875 0 2200.1338
528.7114868164062 0 1000.4661
530.1908569335938 0 1020.9563
532.2037963867188 0 1795.1089
532.7020263671875 0 2920.4531
533.203125 0 2881.8281
533.70654296875 0 1319.055
537.2145385742188 0 3302.1675
538.2181396484375 0 3349.3276
538.7177124023438 0 984.3286
541.1953125 0 10503.255
541.7075805664062 0 25274.314 y Water loss 1
542.2124633789062 0 24897.287
542.71044921875 0 7876.748
543.2206420898438 0 3260.5781
544.2122192382812 0 5044.7026 z 5
545.21484375 0 2175.0557
546.2324829101562 0 1727.3065
546.7268676757812 0 1275.263
548.20166015625 0 6401.916
549.203857421875 0 2553.676
550.2160034179688 0 2911.714
550.7129516601562 0 210636.34 y 1
551.2147827148438 0 151536.14
551.7152709960938 0 61465.1
552.2167358398438 0 8960.893
555.7064208984375 0 4406.3867
556.2085571289062 0 4388.847
556.715576171875 0 908.95966
560.23095703125 0 445474.9 y 5
560.7319946289062 0 15733.137
561.2335205078125 0 142946.61
561.71142578125 0 2874.65
562.23193359375 0 31906.143
563.2354125976562 0 2471.5647
563.7222290039062 0 1034.4939
565.7086181640625 0 972.61865
566.2103271484375 0 2698.594
567.2137451171875 0 2723.6475
569.2362670898438 0 53621.715
569.7372436523438 0 36782.125
570.2301025390625 0 15282.202
570.7236328125 0 5664.3574
571.2223510742188 0 3486.2473
574.2181396484375 0 2101.5273
574.7152099609375 0 5664.5923
575.2168579101562 0 3145.2761
575.7202758789062 0 828.9319
578.2330932617188 0 27870.97
578.735595703125 0 15575.587
579.2335815429688 0 8385.296
579.7307739257812 0 1105.2546
580.2064819335938 0 1587.7458
582.2177124023438 0 939.11163
583.2265625 0 18891.592
583.7276611328125 0 12250.181
584.2294921875 0 8038.4272
584.7254638671875 0 1091.0763
592.23193359375 0 132763.64
592.7332763671875 0 99057.89
593.2340087890625 0 45318.203
593.7332153320312 0 7223.3496
594.1875610351562 0 2369.1094
594.2298583984375 0 1240.3356
596.2377319335938 0 30346.633
597.23095703125 0 13576.194
598.2274169921875 0 5570.0464
599.2314453125 0 1051.6641
601.2373046875 0 463728
601.7386474609375 0 339778.5
602.2391967773438 0 145656.3
602.7398071289062 0 21673.033
606.2296142578125 0 10754.474
607.2335815429688 0 2467.6157
608.239013671875 0 1252.0172
612.1993408203125 0 18780.383
613.2035522460938 0 6679.8677
614.24951171875 0 63533.617
615.1953125 0 1011.05994
615.2514038085938 0 22084.71
616.2533569335938 0 3151.4927
624.2327880859375 0 47564.89
625.235595703125 0 15846.036
626.2353515625 0 3396.6016
632.2654418945312 0 1096.505
634.2317504882812 0 23291.275
635.2353515625 0 7732.4443
636.2360229492188 0 1900.4581
641.2841796875 0 1862.2971
642.2431640625 0 51689.555 c Ammonia loss 4
643.24658203125 0 19464.863
644.249755859375 0 4322.37
649.2633056640625 0 982.9912
650.2523803710938 0 1287.5106
652.2432861328125 0 3487.8623
653.2482299804688 0 1848.7711
657.2649536132812 0 5600.712
658.2714233398438 0 1404.2555
659.2963256835938 0 42553.16
660.2994384765625 0 16121.853
661.3021850585938 0 3650.3232
662.2323608398438 0 1146.446
667.2747192382812 0 3866.3584
668.26025390625 0 3475.2725
669.2575073242188 0 1849.7218
676.2645263671875 0 946.80066
677.2640991210938 0 3870.9675
678.267578125 0 1126.1765
679.2452392578125 0 866.7493
680.2482299804688 0 937.8065
685.2623901367188 0 22631.89
686.2644653320312 0 9333.921
687.2679443359375 0 1450.3191
695.2697143554688 0 23107.064
696.2728881835938 0 10085.081
697.279541015625 0 1407.1854
703.2603759765625 0 2710.8184
704.24658203125 0 1332.9634
705.2845458984375 0 10448.134
706.287109375 0 4043.0706
707.2769165039062 0 2090.5818 z 4
708.2752685546875 0 2199.9495
713.2805786132812 0 21975.438 c Ammonia loss 5
714.2830810546875 0 8480.729
715.293701171875 0 1320.2876
721.2639770507812 0 39532.004
722.2665405273438 0 15572.384
723.2947387695312 0 382034.94 y 4
724.2976684570312 0 162524.1
725.297607421875 0 48519.49
726.2982788085938 0 4962.3887
731.2498168945312 0 6152.219
732.2420043945312 0 2913.6514
733.2745971679688 0 2227.964
747.303955078125 0 1084.4229
749.2592163085938 0 132780.53
750.261962890625 0 54666.2
751.2621459960938 0 17036.305
752.2568359375 0 934.357
756.3114624023438 0 3176.0015
759.2659301757812 0 3419.4932
760.2837524414062 0 822.11273
767.269775390625 0 21528.545
768.2722778320312 0 7607.1284
769.2760009765625 0 2958.782
774.3237915039062 0 63063.004
775.3262939453125 0 28227.496
776.3297119140625 0 7420.215
778.3046875 0 5043.62
779.3121337890625 0 2288.0183
792.3178100585938 0 3510.2393
793.321044921875 0 2064.883
796.31884765625 0 4463.687
797.3217163085938 0 1940.5223
802.3923950195312 0 2290.7732
802.8945922851562 0 1656.5197
803.2754516601562 0 1039.3422
814.3026123046875 0 1294.0808
815.2884521484375 0 1240.6353
820.3114013671875 0 16825.705 y Water loss 3
820.4019165039062 0 793.42914
821.3114624023438 0 10505.95
822.0269165039062 0 1268.0631
822.12255859375 0 1346.5356
822.3067626953125 0 5171.717 z 3
822.8336791992188 0 1469.9645
822.932373046875 0 2273.0723
823.3037719726562 0 2843.2568
824.2977294921875 0 3876.9177
825.3011474609375 0 1465.7181
832.3257446289062 0 3538.557
833.326171875 0 2618.9246
838.3221435546875 0 512588.25 y 3
839.3250732421875 0 251541.98
840.325439453125 0 79022.21
841.3261108398438 0 8745.913
842.3057861328125 0 12762.364
843.314208984375 0 5331.091
844.2943115234375 0 1140.1842
846.3681030273438 0 1122.0632
848.3057250976562 0 3232.9084
849.3082885742188 0 1130.4823
854.326416015625 0 1100.5116
855.8878173828125 0 2201.042
855.9424438476562 0 2148.3806
856.048828125 0 5917.794
860.316162109375 0 12793.571 c Ammonia loss 6
861.3197021484375 0 6694.5435
862.3257446289062 0 9029.205
863.3285522460938 0 3432.1194
864.3364868164062 0 1171.1531
868.33251953125 0 1362.6438
871.3402709960938 0 21166.564
872.3416137695312 0 11382.648
873.3424682617188 0 3077.415
874.3174438476562 0 1557.301
875.3281860351562 0 1342.8043
877.3431396484375 0 2483.0073 c 6
878.3229370117188 0 1926.2036
879.316650390625 0 1214.6637
887.329345703125 0 1206.0127
889.3509521484375 0 375307.66
890.3536376953125 0 193933.08
891.3565063476562 0 50563.785
892.3551635742188 0 4143.627
893.337158203125 0 3030.3718
894.342041015625 0 1468.5903
896.327392578125 0 10141.698
897.3318481445312 0 5522.9697
898.3265991210938 0 1330.9275
899.338623046875 0 1278.0685
904.3431396484375 0 1609.1938
905.3675537109375 0 896.37415
907.3438720703125 0 20162.34
908.3472290039062 0 12772.653
909.3479614257812 0 3980.6392
914.3384399414062 0 2180.7449
914.5029907226562 0 1650.8752
915.3452758789062 0 1676.5276
916.3602905273438 0 761.6048
917.33251953125 0 2858.689
918.31884765625 0 4898.8857
919.3200073242188 0 2605.2222 z Water loss 2
920.3318481445312 0 1078.0728
924.3262329101562 0 1689.779
933.3765869140625 0 5722.0767
934.3784790039062 0 2654.282
935.3389282226562 0 121019 y Water loss 2
936.3386840820312 0 76206.41
937.3388061523438 0 30002.234 z 2
938.3311157226562 0 11702.162
939.3331298828125 0 4022.5786
940.3334350585938 0 1828.2234
951.3818969726562 0 2260.4626
953.350341796875 0 2545618.8 y 2
954.3525390625 0 1399154.6
955.3530883789062 0 485521.8
956.3538818359375 0 56589.21
963.3338623046875 0 12530.955
964.3369750976562 0 6195.0737
965.3380737304688 0 3080.7878
969.3807373046875 0 9021.554
970.384033203125 0 5531.9004
971.3814697265625 0 1590.0303
979.3643798828125 0 6119.8027
980.36767578125 0 3785.7588
981.373291015625 0 1771.4744
997.37548828125 0 33901.57 c Ammonia loss 7
998.3786010742188 0 19738.125
999.3800659179688 0 7388.0586
1000.387451171875 0 896.1375
1014.40234375 0 5363.8823 c 7
1015.3905029296875 0 11667.986
1016.3912963867188 0 7163.093
1017.3933715820312 0 2307.8325
1018.3868408203125 0 5226.088
1019.3818969726562 0 2714.8813
1036.417236328125 0 20236.209
1037.42041015625 0 10971.042
1038.423583984375 0 4107.025
1039.4337158203125 0 1177.1259
1046.3917236328125 0 1013.5508
1054.4080810546875 0 2446.9187
1064.4117431640625 0 994.42444
1072.4083251953125 0 1241.0488
1082.4068603515625 0 12499.828 y Water loss 1
1083.4111328125 0 7629.498
1084.4100341796875 0 3742.5227 z 1
1100.4180908203125 0 127573.63 y 1
1101.4208984375 0 85635.52
1102.422119140625 0 38354.254
1103.4224853515625 0 4955.738
1110.4024658203125 0 22108.14
1111.405029296875 0 14152.305
1112.4061279296875 0 6169.1016
1140.4630126953125 0 1151.2484
1141.4534912109375 0 1648.9711
1142.453857421875 0 2115.7683
1185.4541015625 0 7184.343
1186.455322265625 0 5221.5977
1187.44775390625 0 1696.8442
1201.4654541015625 0 14347.04
1202.4722900390625 0 20411.936
1203.4742431640625 0 10346.218
1204.478759765625 0 3782.8123

Spectrum Details

|  |  |
| --- | --- |
| Matched peaks? Matched peaks The total absolute number of peaks matched. Additionally in brackets the total fraction of peaks matched and the total number of peaks is shown. | 38 (7.18% of 529) |
| FDR? FDR The false discovery rate estimated for this peptide. It is calculated by matching all theoretical fragments with a non-integer shift with the raw peaks for this spectrum. This is done with 40 different shifts. The resulting percentage is the average number of annotated peaks over the number of annotated peaks with the correct spectrum. | 0.13% |
| Satellite FDR? Satellite FDR See the FDR for details on its calculation. This satellite ion specific FDR only contains the satellite ions (d/w) for I/L/J positions. | - |
| PSM Score? PSM Score The PSM Score as given by Hecklib to this annotated spectrum. It is shown with three significant figures. | 449 |

#### Spectrum 13395? Spectrum 13395 The raw spectrum of this peptide as annotated by Hecklib. The fragments are coloured according to ion type (see legend). Any peaks with a star '\*' as text can be hovered over to see the full details, first the ion type second the mass shift type. By hovering over the amino acids in the peptide or ions in the legend the corresponding peaks are highlighted. By toggling the 'Unassigned' label you can turn the background (unassigned) peaks on or off in the plot. By updating the slider in the Ion legend you can update the spectrum to only show the top X% of the peaks with labels. The top X% means any peak that is within X% of the highest intensity. By dragging in the spectrum you can zoom in to a specific part of the spectrum and use 'Zoom Out' to get back to the original zoom level. The annotation of the spectrum is based on the given sequence in the peptides file and is done with different software so inconsistencies are likely. The peaks are annotated based on the given sequence, with 20 ppm tolerance.

Copy Data

##### Spectrum 13395 (TSV)

###### Preview

```
Loading example...
```

*Click on the button to copy the data to your clipboard.*

Mz MinMz MaxIntensity Max

WidthHeightPeptide font sizePeptide stroke widthSpectrum font sizeSpectrum stroke widthCompact peptide

Ion legend

wxyz

abcd

OtherUnassignedIon^Charge^_Position_Show for top:%

TFDDYAMHW

08.40e+31.68e+42.52e+43.36e+4

Zoom Out

a^+1^_2_y^+1^_1_a^+1^_2_b^+1^_2_b^+1^_2_y^+1^_2_b^+1^_3_b^+1^_4_y^+1^_3_y^+2^_8_y^+1^_4_\*y^+1^_5_y^+1^_6_y^+1^_7_y^+1^_7_y^+1^_8_

0777155423313107

Fragment Matches Table

Show background peaks

| Position | Ion type | Intensity | mz Theoretical | mz Error (Th) | mz Error (ppm) | Charge | Series Number |
| --- | --- | --- | --- | --- | --- | --- | --- |
| - | - | 2.012E+04 | 120.1 | - | - | 0 | - |
| - | - | 1853 | 121.1 | - | - | 0 | - |
| - | - | 325.4 | 121.6 | - | - | 0 | - |
| - | - | 337.9 | 122.6 | - | - | 0 | - |
| - | - | 3738 | 129.1 | - | - | 0 | - |
| - | - | 440.7 | 130.1 | - | - | 0 | - |
| - | - | 502.5 | 132.1 | - | - | 0 | - |
| - | - | 1569 | 133.1 | - | - | 0 | - |
| - | - | 406.9 | 133.2 | - | - | 0 | - |
| - | - | 7769 | 136.1 | - | - | 0 | - |
| - | - | 519.3 | 137.4 | - | - | 0 | - |
| - | - | 573.2 | 138.1 | - | - | 0 | - |
| - | - | 491.3 | 139.1 | - | - | 0 | - |
| - | - | 412.3 | 142.6 | - | - | 0 | - |
| - | - | 783.2 | 148.9 | - | - | 0 | - |
| - | - | 1096 | 149 | - | - | 0 | - |
| - | - | 1399 | 158.1 | - | - | 0 | - |
| - | - | 923.5 | 159.1 | - | - | 0 | - |
| - | - | 565.4 | 162.1 | - | - | 0 | - |
| - | - | 2131 | 165.1 | - | - | 0 | - |
| - | - | 942.1 | 166.1 | - | - | 0 | - |
| - | - | 2145 | 167.1 | - | - | 0 | - |
| - | - | 483.1 | 167.1 | - | - | 0 | - |
| - | - | 1820 | 173.1 | - | - | 0 | - |
| - | - | 5261 | 173.5 | - | - | 0 | - |
| - | - | 2178 | 176.1 | - | - | 0 | - |
| - | - | 1714 | 177.1 | - | - | 0 | - |
| - | - | 1222 | 177.1 | - | - | 0 | - |
| - | - | 449.3 | 181.1 | - | - | 0 | - |
| - | - | 4476 | 182.1 | - | - | 0 | - |
| - | - | 511 | 185.1 | - | - | 0 | - |
| - | - | 1334 | 188.1 | - | - | 0 | - |
| - | - | 514.7 | 193.1 | - | - | 0 | - |
| - | - | 827.9 | 200.1 | - | - | 0 | - |
| - | - | 972.9 | 201.1 | - | - | 0 | - |
| 2 | a | 572.8 | 203.1 | 7.616E-05 | 0.375 | +1 | 2 |
| 9 | y | 1073 | 205.1 | 0.0005785 | 2.82 | +1 | 1 |
| - | - | 943.6 | 215.1 | - | - | 0 | - |
| - | - | 565.2 | 219.1 | - | - | 0 | - |
| - | - | 1789 | 221.1 | - | - | 0 | - |
| 2 | a | 3.327E+04 | 221.1 | 5.735E-06 | 0.02593 | +1 | 2 |
| - | - | 3706 | 222.1 | - | - | 0 | - |
| - | - | 1293 | 223.1 | - | - | 0 | - |
| - | - | 703.4 | 231.1 | - | - | 0 | - |
| 2 | b | 808 | 231.1 | 3.459E-05 | 0.1497 | +1 | 2 |
| - | - | 843.2 | 235.1 | - | - | 0 | - |
| - | - | 602.2 | 239.1 | - | - | 0 | - |
| - | - | 866.9 | 241.1 | - | - | 0 | - |
| - | - | 466.4 | 241.1 | - | - | 0 | - |
| - | - | 500.5 | 243.3 | - | - | 0 | - |
| 2 | b | 5160 | 249.1 | 7.476E-05 | 0.3001 | +1 | 2 |
| - | - | 678.7 | 251.1 | - | - | 0 | - |
| - | - | 1383 | 263.1 | - | - | 0 | - |
| - | - | 709.9 | 279.1 | - | - | 0 | - |
| - | - | 1196 | 285 | - | - | 0 | - |
| - | - | 582.1 | 285.1 | - | - | 0 | - |
| - | - | 500.5 | 289.2 | - | - | 0 | - |
| - | - | 620.5 | 296.2 | - | - | 0 | - |
| - | - | 669.4 | 299.2 | - | - | 0 | - |
| - | - | 591.2 | 306.6 | - | - | 0 | - |
| - | - | 892.4 | 309.2 | - | - | 0 | - |
| - | - | 924.7 | 324.1 | - | - | 0 | - |
| - | - | 2898 | 329.2 | - | - | 0 | - |
| - | - | 589.1 | 330.2 | - | - | 0 | - |
| 8 | y | 4980 | 342.2 | 0.0002756 | 0.8055 | +1 | 2 |
| - | - | 1043 | 343.2 | - | - | 0 | - |
| - | - | 1741 | 355.1 | - | - | 0 | - |
| - | - | 863.7 | 356.1 | - | - | 0 | - |
| - | - | 1063 | 357.2 | - | - | 0 | - |
| - | - | 539.7 | 359 | - | - | 0 | - |
| 3 | b | 1450 | 364.2 | 0.00102 | 2.801 | +1 | 3 |
| - | - | 533.9 | 381.1 | - | - | 0 | - |
| - | - | 1349 | 394.1 | - | - | 0 | - |
| - | - | 540.5 | 403.4 | - | - | 0 | - |
| - | - | 614.7 | 407.2 | - | - | 0 | - |
| - | - | 556.6 | 408.2 | - | - | 0 | - |
| - | - | 606 | 412.2 | - | - | 0 | - |
| - | - | 728.6 | 413.3 | - | - | 0 | - |
| - | - | 558.3 | 415.3 | - | - | 0 | - |
| - | - | 1073 | 425.2 | - | - | 0 | - |
| 4 | b | 933.5 | 479.2 | 0.0004361 | 0.91 | +1 | 4 |
| - | - | 595.5 | 486.9 | - | - | 0 | - |
| 7 | y | 1342 | 489.2 | 0.005069 | 10.36 | +1 | 3 |
| - | - | 578.9 | 495.2 | - | - | 0 | - |
| - | - | 589.7 | 508.3 | - | - | 0 | - |
| 2 | y | 810.9 | 541.7 | 0.001588 | 2.931 | +2 | 8 |
| 6 | y | 1561 | 560.2 | 0.003509 | 6.263 | +1 | 4 |
| - | - | 796 | 561.2 | - | - | 0 | - |
| - | - | 1168 | 585.3 | - | - | 0 | - |
| - | - | 723.2 | 586.3 | - | - | 0 | - |
| 0 | Precursor | 1144 | 601.2 | 0.003411 | 5.673 | +2 | -1 |
| - | - | 764.1 | 602.2 | - | - | 0 | - |
| - | - | 632.2 | 612.2 | - | - | 0 | - |
| - | - | 562.6 | 630 | - | - | 0 | - |
| - | - | 691.8 | 685.3 | - | - | 0 | - |
| - | - | 2313 | 700.4 | - | - | 0 | - |
| - | - | 1393 | 701.4 | - | - | 0 | - |
| - | - | 1005 | 715.6 | - | - | 0 | - |
| - | - | 907.7 | 721.3 | - | - | 0 | - |
| 5 | y | 1504 | 723.3 | 0.0002996 | 0.4142 | +1 | 5 |
| - | - | 722.1 | 724.3 | - | - | 0 | - |
| - | - | 634.3 | 750.3 | - | - | 0 | - |
| - | - | 617.2 | 753.6 | - | - | 0 | - |
| - | - | 1137 | 774.3 | - | - | 0 | - |
| - | - | 1617 | 801.4 | - | - | 0 | - |
| 4 | y | 2835 | 838.3 | 0.003569 | 4.257 | +1 | 6 |
| - | - | 927.4 | 839.3 | - | - | 0 | - |
| - | - | 2136 | 844.5 | - | - | 0 | - |
| - | - | 1153 | 845.5 | - | - | 0 | - |
| - | - | 633.8 | 864.5 | - | - | 0 | - |
| - | - | 7629 | 872.4 | - | - | 0 | - |
| - | - | 3468 | 873.5 | - | - | 0 | - |
| - | - | 720.5 | 874 | - | - | 0 | - |
| - | - | 856.4 | 874.5 | - | - | 0 | - |
| - | - | 4245 | 889.3 | - | - | 0 | - |
| - | - | 1155 | 890.4 | - | - | 0 | - |
| 3 | y | 975 | 935.3 | 0.006295 | 6.73 | +1 | 7 |
| 3 | y | 1.145E+04 | 953.3 | 0.005373 | 5.636 | +1 | 7 |
| - | - | 5372 | 954.4 | - | - | 0 | - |
| - | - | 2560 | 955.3 | - | - | 0 | - |
| - | - | 1514 | 991.5 | - | - | 0 | - |
| - | - | 702.2 | 992.5 | - | - | 0 | - |
| - | - | 3051 | 1020 | - | - | 0 | - |
| - | - | 1643 | 1021 | - | - | 0 | - |
| 2 | y | 844.5 | 1100 | 0.0149 | 13.54 | +1 | 8 |
| - | - | 627.8 | 2727 | - | - | 0 | - |
| - | - | 657 | 3077 | - | - | 0 | - |

m/z Charge Intensity FragmentType MassShift Position
120.08084869384766 0 20121.07
121.08415985107422 0 1853.3635
121.58132934570312 0 325.43753
122.56163787841797 0 337.86182
129.10231018066406 0 3737.652
130.0651092529297 0 440.6535
132.0806121826172 0 502.46503
133.08596801757812 0 1568.6665
133.19509887695312 0 406.93045
136.07577514648438 0 7769.115
137.3627166748047 0 519.2632
138.0555419921875 0 573.2474
139.0753631591797 0 491.27042
142.56698608398438 0 412.30264
148.94888305664062 0 783.1585
149.04489135742188 0 1095.9722
158.09646606445312 0 1399.4291
159.09176635742188 0 923.512
162.0917510986328 0 565.39825
165.0545654296875 0 2130.6106
166.0859832763672 0 942.09216
167.0554656982422 0 2144.5894
167.08189392089844 0 483.10785
173.12844848632812 0 1820.005
173.4508819580078 0 5261.3623
176.10708618164062 0 2177.9827
177.10264587402344 0 1713.5693
177.11183166503906 0 1221.987
181.09925842285156 0 449.3238
182.08116149902344 0 4475.8467
185.12881469726562 0 510.95468
188.07061767578125 0 1333.785
193.10983276367188 0 514.684
200.13961791992188 0 827.8503
201.123046875 0 972.8656
203.1179656982422 0 572.80164 a Water loss 1
205.0977325439453 0 1073.0825 y 8
215.1279296875 0 943.6486
219.1123809814453 0 565.21716
221.1034393310547 0 1789.072
221.12844848632812 0 33269.906 a 1
222.13192749023438 0 3706.467
223.0634765625 0 1292.6671
231.0618896484375 0 703.3748
231.1128387451172 0 808.0447 b Water loss 1
235.10775756835938 0 843.217
239.09523010253906 0 602.21704
241.07449340820312 0 866.87494
241.0917510986328 0 466.42902
243.28358459472656 0 500.4894
249.12344360351562 0 5160.081 b 1
251.10150146484375 0 678.70386
263.1020812988281 0 1382.5712
279.0975341796875 0 709.8619
285.0088806152344 0 1196.0009
285.1018981933594 0 582.05994
289.2032470703125 0 500.48688
296.1502685546875 0 620.4856
299.1649169921875 0 669.42444
306.63153076171875 0 591.1802
309.2040100097656 0 892.38184
324.1453857421875 0 924.66754
329.1500244140625 0 2897.5305
330.15167236328125 0 589.0576
342.1563415527344 0 4980.443 y 7
343.158203125 0 1042.9274
355.0701904296875 0 1740.7838
356.0696105957031 0 863.72375
357.2244567871094 0 1063.2944
359.0303955078125 0 539.71
364.1492919921875 0 1450.425 b 2
381.1125183105469 0 533.8712
394.1243896484375 0 1348.6525
403.43695068359375 0 540.48517
407.1815185546875 0 614.69446
408.16650390625 0 556.5555
412.2181396484375 0 605.9521
413.26690673828125 0 728.6272
415.2657775878906 0 558.3335
425.1921691894531 0 1072.5454
479.17681884765625 0 933.5083 b 3
486.8794860839844 0 595.5121
489.1916198730469 0 1341.8181 y 6
495.1860046386719 0 578.9053
508.2772521972656 0 589.67816
541.70458984375 0 810.9066 y Water loss 1
560.2271728515625 0 1561.334 y 5
561.2318115234375 0 796.0448
585.3359985351562 0 1167.9957
586.3374633789062 0 723.2476
601.2355346679688 0 1144.4445 Precursor
602.2430419921875 0 764.07166
612.1986083984375 0 632.16986
630.0360717773438 0 562.6038
685.2550659179688 0 691.8122
700.3640747070312 0 2313.4185
701.3638916015625 0 1392.7953
715.5675048828125 0 1004.7674
721.2601928710938 0 907.72943
723.2872924804688 0 1503.7023 y 4
724.2931518554688 0 722.07684
750.2642211914062 0 634.34753
753.6182250976562 0 617.21405
774.3199462890625 0 1136.5198
801.4151000976562 0 1617.3091
838.3175048828125 0 2835.4338 y 3
839.32177734375 0 927.3583
844.4532470703125 0 2135.8572
845.4600219726562 0 1153.2867
864.53564453125 0 633.7791
872.4484252929688 0 7629.275
873.4520263671875 0 3467.8357
873.9771728515625 0 720.5442
874.4542236328125 0 856.4018
889.3493041992188 0 4244.721
890.3524780273438 0 1154.5332
935.3366088867188 0 974.97 y Water loss 2
953.3462524414062 0 11453.153 y 2
954.3502807617188 0 5372.142
955.3485107421875 0 2560.0752
991.5208129882812 0 1514.4628
992.5243530273438 0 702.2077
1019.5140380859375 0 3051.0303
1020.5193481445312 0 1642.7782
1100.4241943359375 0 844.5226 y 1
2727.04443359375 0 627.7952
3076.61865234375 0 656.9955

Spectrum Details

|  |  |
| --- | --- |
| Matched peaks? Matched peaks The total absolute number of peaks matched. Additionally in brackets the total fraction of peaks matched and the total number of peaks is shown. | 17 (13.39% of 127) |
| FDR? FDR The false discovery rate estimated for this peptide. It is calculated by matching all theoretical fragments with a non-integer shift with the raw peaks for this spectrum. This is done with 40 different shifts. The resulting percentage is the average number of annotated peaks over the number of annotated peaks with the correct spectrum. | 0.84% |
| Satellite FDR? Satellite FDR See the FDR for details on its calculation. This satellite ion specific FDR only contains the satellite ions (d/w) for I/L/J positions. | - |
| PSM Score? PSM Score The PSM Score as given by Hecklib to this annotated spectrum. It is shown with three significant figures. | 195 |

#### Spectrum 11133? Spectrum 11133 The raw spectrum of this peptide as annotated by Hecklib. The fragments are coloured according to ion type (see legend). Any peaks with a star '\*' as text can be hovered over to see the full details, first the ion type second the mass shift type. By hovering over the amino acids in the peptide or ions in the legend the corresponding peaks are highlighted. By toggling the 'Unassigned' label you can turn the background (unassigned) peaks on or off in the plot. By updating the slider in the Ion legend you can update the spectrum to only show the top X% of the peaks with labels. The top X% means any peak that is within X% of the highest intensity. By dragging in the spectrum you can zoom in to a specific part of the spectrum and use 'Zoom Out' to get back to the original zoom level. The annotation of the spectrum is based on the given sequence in the peptides file and is done with different software so inconsistencies are likely. The peaks are annotated based on the given sequence, with 20 ppm tolerance.

Copy Data

##### Spectrum 11133 (TSV)

###### Preview

```
Loading example...
```

*Click on the button to copy the data to your clipboard.*

Mz MinMz MaxIntensity Max

WidthHeightPeptide font sizePeptide stroke widthSpectrum font sizeSpectrum stroke widthCompact peptide

Ion legend

wxyz

abcd

OtherUnassignedIon^Charge^_Position_Show for top:%

TFDDYAMHW

01.44e+42.87e+44.31e+45.74e+4

Zoom Out

a^+1^_2_y^+1^_1_a^+1^_2_b^+1^_2_b^+1^_2_y^+1^_2_b^+1^_3_b^+1^_4_y^+1^_3_y^+2^_8_y^+1^_4_\*y^+1^_5_y^+1^_6_y^+1^_7_y^+1^_7_y^+1^_8_

0778155723353114

Fragment Matches Table

Show background peaks

| Position | Ion type | Intensity | mz Theoretical | mz Error (Th) | mz Error (ppm) | Charge | Series Number |
| --- | --- | --- | --- | --- | --- | --- | --- |
| - | - | 5.684E+04 | 120.1 | - | - | 0 | - |
| - | - | 4181 | 121.1 | - | - | 0 | - |
| - | - | 607.6 | 123 | - | - | 0 | - |
| - | - | 1943 | 127.1 | - | - | 0 | - |
| - | - | 584.6 | 129.1 | - | - | 0 | - |
| - | - | 3885 | 129.1 | - | - | 0 | - |
| - | - | 768.3 | 130.1 | - | - | 0 | - |
| - | - | 671.8 | 131 | - | - | 0 | - |
| - | - | 1196 | 133.1 | - | - | 0 | - |
| - | - | 501.5 | 135 | - | - | 0 | - |
| - | - | 1.481E+04 | 136.1 | - | - | 0 | - |
| - | - | 1011 | 137.1 | - | - | 0 | - |
| - | - | 509.2 | 138.1 | - | - | 0 | - |
| - | - | 595.8 | 139.1 | - | - | 0 | - |
| - | - | 4730 | 141.1 | - | - | 0 | - |
| - | - | 698.4 | 147 | - | - | 0 | - |
| - | - | 999 | 149 | - | - | 0 | - |
| - | - | 663.3 | 149 | - | - | 0 | - |
| - | - | 2068 | 155.1 | - | - | 0 | - |
| - | - | 420.2 | 155.1 | - | - | 0 | - |
| - | - | 1367 | 156.1 | - | - | 0 | - |
| - | - | 2911 | 157.1 | - | - | 0 | - |
| - | - | 822.2 | 157.1 | - | - | 0 | - |
| - | - | 2283 | 158.1 | - | - | 0 | - |
| - | - | 1593 | 159.1 | - | - | 0 | - |
| - | - | 573.4 | 159.1 | - | - | 0 | - |
| - | - | 1180 | 160.1 | - | - | 0 | - |
| - | - | 4501 | 165.1 | - | - | 0 | - |
| - | - | 465.7 | 165.1 | - | - | 0 | - |
| - | - | 2.527E+04 | 166.1 | - | - | 0 | - |
| - | - | 1552 | 167.1 | - | - | 0 | - |
| - | - | 9861 | 167.1 | - | - | 0 | - |
| - | - | 2923 | 167.1 | - | - | 0 | - |
| - | - | 789.4 | 168.1 | - | - | 0 | - |
| - | - | 6771 | 169.1 | - | - | 0 | - |
| - | - | 667.1 | 171.1 | - | - | 0 | - |
| - | - | 3186 | 173.1 | - | - | 0 | - |
| - | - | 555.5 | 175.1 | - | - | 0 | - |
| - | - | 891.8 | 175.1 | - | - | 0 | - |
| - | - | 515.9 | 176.1 | - | - | 0 | - |
| - | - | 4128 | 176.1 | - | - | 0 | - |
| - | - | 3179 | 177.1 | - | - | 0 | - |
| - | - | 711.9 | 177.1 | - | - | 0 | - |
| - | - | 493.1 | 181.1 | - | - | 0 | - |
| - | - | 9590 | 182.1 | - | - | 0 | - |
| - | - | 883.5 | 183.1 | - | - | 0 | - |
| - | - | 1207 | 183.1 | - | - | 0 | - |
| - | - | 3020 | 185.1 | - | - | 0 | - |
| - | - | 565.9 | 185.1 | - | - | 0 | - |
| - | - | 578 | 186.1 | - | - | 0 | - |
| - | - | 1334 | 187.1 | - | - | 0 | - |
| - | - | 2697 | 188.1 | - | - | 0 | - |
| - | - | 500.9 | 189.1 | - | - | 0 | - |
| - | - | 683.8 | 191.1 | - | - | 0 | - |
| - | - | 1117 | 193.1 | - | - | 0 | - |
| - | - | 521.3 | 194.1 | - | - | 0 | - |
| - | - | 760.8 | 197.1 | - | - | 0 | - |
| - | - | 480.4 | 199.1 | - | - | 0 | - |
| - | - | 4960 | 199.1 | - | - | 0 | - |
| - | - | 643.2 | 199.2 | - | - | 0 | - |
| - | - | 543 | 200.1 | - | - | 0 | - |
| - | - | 1820 | 201.1 | - | - | 0 | - |
| 2 | a | 1896 | 203.1 | 6.117E-05 | 0.3011 | +1 | 2 |
| - | - | 792.3 | 204.1 | - | - | 0 | - |
| 9 | y | 1804 | 205.1 | 0.0001235 | 0.6019 | +1 | 1 |
| - | - | 750.7 | 207.1 | - | - | 0 | - |
| - | - | 511.1 | 209.1 | - | - | 0 | - |
| - | - | 1005 | 211.1 | - | - | 0 | - |
| - | - | 1368 | 212.1 | - | - | 0 | - |
| - | - | 644.2 | 214.1 | - | - | 0 | - |
| - | - | 611.9 | 215.1 | - | - | 0 | - |
| - | - | 3811 | 221.1 | - | - | 0 | - |
| 2 | a | 5.599E+04 | 221.1 | 0.0001011 | 0.4571 | +1 | 2 |
| - | - | 7195 | 222.1 | - | - | 0 | - |
| - | - | 1601 | 223.1 | - | - | 0 | - |
| - | - | 6955 | 224.1 | - | - | 0 | - |
| - | - | 1652 | 226.1 | - | - | 0 | - |
| - | - | 2.019E+04 | 227.1 | - | - | 0 | - |
| - | - | 587.7 | 227.2 | - | - | 0 | - |
| - | - | 2210 | 228.1 | - | - | 0 | - |
| - | - | 658.5 | 228.1 | - | - | 0 | - |
| - | - | 485.9 | 229.8 | - | - | 0 | - |
| - | - | 1010 | 231.1 | - | - | 0 | - |
| 2 | b | 840.2 | 231.1 | 0.000118 | 0.5106 | +1 | 2 |
| - | - | 1338 | 235.1 | - | - | 0 | - |
| - | - | 734.4 | 238.2 | - | - | 0 | - |
| - | - | 793.7 | 239.1 | - | - | 0 | - |
| - | - | 1060 | 241.1 | - | - | 0 | - |
| - | - | 2117 | 242.1 | - | - | 0 | - |
| - | - | 949.7 | 243.1 | - | - | 0 | - |
| - | - | 568.1 | 244.1 | - | - | 0 | - |
| 2 | b | 1.068E+04 | 249.1 | 0.0001053 | 0.4226 | +1 | 2 |
| - | - | 1501 | 250.1 | - | - | 0 | - |
| - | - | 1196 | 251.1 | - | - | 0 | - |
| - | - | 861.6 | 251.2 | - | - | 0 | - |
| - | - | 794.8 | 253.1 | - | - | 0 | - |
| - | - | 778.2 | 254.1 | - | - | 0 | - |
| - | - | 649.4 | 256.2 | - | - | 0 | - |
| - | - | 2358 | 263.1 | - | - | 0 | - |
| - | - | 3991 | 265.2 | - | - | 0 | - |
| - | - | 3534 | 266.2 | - | - | 0 | - |
| - | - | 606.3 | 268.2 | - | - | 0 | - |
| - | - | 604.2 | 270.1 | - | - | 0 | - |
| - | - | 739.9 | 271.1 | - | - | 0 | - |
| - | - | 870.5 | 274.1 | - | - | 0 | - |
| - | - | 692.4 | 279.1 | - | - | 0 | - |
| - | - | 664.1 | 280.1 | - | - | 0 | - |
| - | - | 1.402E+04 | 281.1 | - | - | 0 | - |
| - | - | 1320 | 282.1 | - | - | 0 | - |
| - | - | 713 | 282.2 | - | - | 0 | - |
| - | - | 1001 | 283.1 | - | - | 0 | - |
| - | - | 1223 | 284.2 | - | - | 0 | - |
| - | - | 1387 | 285 | - | - | 0 | - |
| - | - | 1205 | 285.1 | - | - | 0 | - |
| - | - | 1103 | 294.2 | - | - | 0 | - |
| - | - | 882 | 295.2 | - | - | 0 | - |
| - | - | 617.2 | 296.2 | - | - | 0 | - |
| - | - | 619.1 | 297.1 | - | - | 0 | - |
| - | - | 2004 | 298.1 | - | - | 0 | - |
| - | - | 2926 | 299.1 | - | - | 0 | - |
| - | - | 597.5 | 302.1 | - | - | 0 | - |
| - | - | 515.9 | 309.2 | - | - | 0 | - |
| - | - | 485.8 | 312.1 | - | - | 0 | - |
| - | - | 1667 | 312.2 | - | - | 0 | - |
| - | - | 727 | 313.2 | - | - | 0 | - |
| - | - | 983.6 | 322.2 | - | - | 0 | - |
| - | - | 1230 | 323.2 | - | - | 0 | - |
| - | - | 1938 | 324.1 | - | - | 0 | - |
| - | - | 532.4 | 325.2 | - | - | 0 | - |
| - | - | 518.8 | 328.1 | - | - | 0 | - |
| - | - | 5457 | 329.1 | - | - | 0 | - |
| - | - | 1345 | 330.2 | - | - | 0 | - |
| - | - | 955.7 | 335.2 | - | - | 0 | - |
| - | - | 599 | 338.2 | - | - | 0 | - |
| - | - | 585 | 339.2 | - | - | 0 | - |
| - | - | 1441 | 339.7 | - | - | 0 | - |
| - | - | 2.074E+04 | 340.2 | - | - | 0 | - |
| - | - | 4177 | 341.2 | - | - | 0 | - |
| - | - | 429.6 | 341.2 | - | - | 0 | - |
| - | - | 906.8 | 342.1 | - | - | 0 | - |
| 8 | y | 1.166E+04 | 342.2 | 0.0003061 | 0.8947 | +1 | 2 |
| - | - | 505.4 | 342.2 | - | - | 0 | - |
| - | - | 1708 | 343.2 | - | - | 0 | - |
| - | - | 508.2 | 345.1 | - | - | 0 | - |
| - | - | 953.6 | 349.2 | - | - | 0 | - |
| - | - | 3981 | 352.2 | - | - | 0 | - |
| - | - | 1903 | 355.1 | - | - | 0 | - |
| - | - | 844.2 | 356.1 | - | - | 0 | - |
| - | - | 879.4 | 357.2 | - | - | 0 | - |
| 3 | b | 3134 | 364.2 | 0.0002619 | 0.7191 | +1 | 3 |
| - | - | 486.4 | 364.5 | - | - | 0 | - |
| - | - | 536.8 | 365.2 | - | - | 0 | - |
| - | - | 915 | 366.2 | - | - | 0 | - |
| - | - | 1595 | 367.2 | - | - | 0 | - |
| - | - | 1452 | 368.2 | - | - | 0 | - |
| - | - | 2950 | 370.2 | - | - | 0 | - |
| - | - | 834.4 | 377.2 | - | - | 0 | - |
| - | - | 1679 | 378.1 | - | - | 0 | - |
| - | - | 1.367E+04 | 380.2 | - | - | 0 | - |
| - | - | 2667 | 381.2 | - | - | 0 | - |
| - | - | 1192 | 383.2 | - | - | 0 | - |
| - | - | 2641 | 394.1 | - | - | 0 | - |
| - | - | 2455 | 394.2 | - | - | 0 | - |
| - | - | 8073 | 395.2 | - | - | 0 | - |
| - | - | 2561 | 396.2 | - | - | 0 | - |
| - | - | 953.4 | 396.2 | - | - | 0 | - |
| - | - | 613.7 | 397.2 | - | - | 0 | - |
| - | - | 1.114E+04 | 398.2 | - | - | 0 | - |
| - | - | 2926 | 399.2 | - | - | 0 | - |
| - | - | 607.6 | 403.2 | - | - | 0 | - |
| - | - | 611.3 | 411.2 | - | - | 0 | - |
| - | - | 715.7 | 412.2 | - | - | 0 | - |
| - | - | 1195 | 413.3 | - | - | 0 | - |
| - | - | 1307 | 416.7 | - | - | 0 | - |
| - | - | 804.6 | 417.2 | - | - | 0 | - |
| - | - | 1903 | 425.2 | - | - | 0 | - |
| - | - | 1896 | 425.3 | - | - | 0 | - |
| - | - | 698.2 | 426.3 | - | - | 0 | - |
| - | - | 1063 | 427.7 | - | - | 0 | - |
| - | - | 924.5 | 435.3 | - | - | 0 | - |
| - | - | 628.8 | 437.2 | - | - | 0 | - |
| - | - | 836.5 | 442.2 | - | - | 0 | - |
| - | - | 1062 | 446.3 | - | - | 0 | - |
| - | - | 665.6 | 448.3 | - | - | 0 | - |
| - | - | 4639 | 449.2 | - | - | 0 | - |
| - | - | 1688 | 450.2 | - | - | 0 | - |
| - | - | 3904 | 451.2 | - | - | 0 | - |
| - | - | 1929 | 451.7 | - | - | 0 | - |
| - | - | 7351 | 453.3 | - | - | 0 | - |
| - | - | 961.6 | 454.2 | - | - | 0 | - |
| - | - | 2112 | 454.3 | - | - | 0 | - |
| - | - | 1418 | 460.2 | - | - | 0 | - |
| - | - | 839.4 | 462.3 | - | - | 0 | - |
| - | - | 1476 | 465.2 | - | - | 0 | - |
| 4 | b | 1182 | 479.2 | 0.001609 | 3.357 | +1 | 4 |
| - | - | 778.7 | 479.3 | - | - | 0 | - |
| - | - | 5506 | 480.3 | - | - | 0 | - |
| - | - | 826 | 483.3 | - | - | 0 | - |
| 7 | y | 2387 | 489.2 | 0.004886 | 9.988 | +1 | 3 |
| - | - | 1384 | 490.3 | - | - | 0 | - |
| - | - | 1880 | 493.3 | - | - | 0 | - |
| - | - | 667 | 494.3 | - | - | 0 | - |
| - | - | 730.4 | 496.2 | - | - | 0 | - |
| - | - | 6674 | 500.8 | - | - | 0 | - |
| - | - | 2830 | 501.3 | - | - | 0 | - |
| - | - | 827.7 | 501.8 | - | - | 0 | - |
| - | - | 3523 | 506.3 | - | - | 0 | - |
| - | - | 1.609E+04 | 508.3 | - | - | 0 | - |
| - | - | 4206 | 509.3 | - | - | 0 | - |
| - | - | 1185 | 510.3 | - | - | 0 | - |
| - | - | 1578 | 511.3 | - | - | 0 | - |
| - | - | 696.1 | 539.3 | - | - | 0 | - |
| - | - | 812.4 | 545.3 | - | - | 0 | - |
| 2 | y | 1722 | 550.7 | 0.001249 | 2.268 | +2 | 8 |
| 6 | y | 3623 | 560.2 | 0.004912 | 8.768 | +1 | 4 |
| - | - | 1408 | 561.2 | - | - | 0 | - |
| - | - | 7792 | 563.3 | - | - | 0 | - |
| - | - | 2802 | 564.3 | - | - | 0 | - |
| - | - | 975.1 | 565.3 | - | - | 0 | - |
| - | - | 1672 | 567.3 | - | - | 0 | - |
| - | - | 658 | 584.3 | - | - | 0 | - |
| - | - | 2444 | 585.3 | - | - | 0 | - |
| - | - | 770.2 | 586.3 | - | - | 0 | - |
| - | - | 700 | 592.7 | - | - | 0 | - |
| - | - | 5313 | 593.4 | - | - | 0 | - |
| - | - | 2450 | 594.4 | - | - | 0 | - |
| - | - | 747.4 | 595.4 | - | - | 0 | - |
| 0 | Precursor | 2096 | 601.2 | 0.003777 | 6.282 | +2 | -1 |
| - | - | 995.1 | 601.6 | - | - | 0 | - |
| - | - | 823 | 601.7 | - | - | 0 | - |
| - | - | 621.4 | 612.2 | - | - | 0 | - |
| - | - | 3.024E+04 | 621.4 | - | - | 0 | - |
| - | - | 850 | 622.3 | - | - | 0 | - |
| - | - | 1.117E+04 | 622.4 | - | - | 0 | - |
| - | - | 2128 | 623.4 | - | - | 0 | - |
| - | - | 1343 | 650.4 | - | - | 0 | - |
| - | - | 779.3 | 652.8 | - | - | 0 | - |
| - | - | 1004 | 657.3 | - | - | 0 | - |
| - | - | 2009 | 659.3 | - | - | 0 | - |
| - | - | 916.8 | 660.4 | - | - | 0 | - |
| - | - | 2872 | 676.4 | - | - | 0 | - |
| - | - | 1898 | 677.4 | - | - | 0 | - |
| - | - | 6689 | 678.4 | - | - | 0 | - |
| - | - | 2384 | 679.4 | - | - | 0 | - |
| - | - | 591.6 | 680.4 | - | - | 0 | - |
| - | - | 1112 | 685.3 | - | - | 0 | - |
| - | - | 608 | 693.4 | - | - | 0 | - |
| - | - | 5826 | 700.4 | - | - | 0 | - |
| - | - | 1981 | 701.4 | - | - | 0 | - |
| - | - | 1527 | 707.4 | - | - | 0 | - |
| - | - | 1448 | 717.4 | - | - | 0 | - |
| - | - | 1206 | 721.3 | - | - | 0 | - |
| 5 | y | 3687 | 723.3 | 0.004206 | 5.815 | +1 | 5 |
| - | - | 1718 | 724.3 | - | - | 0 | - |
| - | - | 3116 | 733.4 | - | - | 0 | - |
| - | - | 9109 | 735.4 | - | - | 0 | - |
| - | - | 3962 | 736.4 | - | - | 0 | - |
| - | - | 1625 | 749.3 | - | - | 0 | - |
| - | - | 803.2 | 751.4 | - | - | 0 | - |
| - | - | 892.5 | 773.4 | - | - | 0 | - |
| - | - | 2214 | 774.3 | - | - | 0 | - |
| - | - | 697.5 | 783.3 | - | - | 0 | - |
| - | - | 1230 | 783.4 | - | - | 0 | - |
| - | - | 653.6 | 799.6 | - | - | 0 | - |
| - | - | 726.6 | 800.4 | - | - | 0 | - |
| - | - | 2518 | 801.4 | - | - | 0 | - |
| - | - | 1749 | 802.4 | - | - | 0 | - |
| - | - | 694.1 | 804.4 | - | - | 0 | - |
| - | - | 1749 | 828.4 | - | - | 0 | - |
| - | - | 1374 | 832.5 | - | - | 0 | - |
| 4 | y | 4308 | 838.3 | 0.00302 | 3.602 | +1 | 6 |
| - | - | 2177 | 839.3 | - | - | 0 | - |
| - | - | 817.2 | 840.3 | - | - | 0 | - |
| - | - | 4292 | 844.5 | - | - | 0 | - |
| - | - | 2847 | 845.5 | - | - | 0 | - |
| - | - | 728.8 | 846.5 | - | - | 0 | - |
| - | - | 819 | 854.4 | - | - | 0 | - |
| - | - | 656.7 | 871.3 | - | - | 0 | - |
| - | - | 1.603E+04 | 872.4 | - | - | 0 | - |
| - | - | 6634 | 873.4 | - | - | 0 | - |
| - | - | 1735 | 874.5 | - | - | 0 | - |
| - | - | 836.3 | 883.5 | - | - | 0 | - |
| - | - | 6446 | 889.3 | - | - | 0 | - |
| - | - | 3652 | 890.3 | - | - | 0 | - |
| - | - | 681.5 | 892.5 | - | - | 0 | - |
| - | - | 1.841E+04 | 901.5 | - | - | 0 | - |
| - | - | 1.049E+04 | 902.5 | - | - | 0 | - |
| - | - | 3422 | 903.5 | - | - | 0 | - |
| - | - | 747.2 | 904.5 | - | - | 0 | - |
| - | - | 7722 | 919.5 | - | - | 0 | - |
| - | - | 4503 | 920.5 | - | - | 0 | - |
| - | - | 834 | 921.5 | - | - | 0 | - |
| 3 | y | 923.3 | 935.3 | 0.002327 | 2.488 | +1 | 7 |
| - | - | 838.6 | 936.3 | - | - | 0 | - |
| 3 | y | 2.078E+04 | 953.3 | 0.00397 | 4.164 | +1 | 7 |
| - | - | 1.29E+04 | 954.3 | - | - | 0 | - |
| - | - | 3485 | 955.3 | - | - | 0 | - |
| - | - | 1279 | 990.6 | - | - | 0 | - |
| - | - | 3576 | 991.5 | - | - | 0 | - |
| - | - | 987 | 992.5 | - | - | 0 | - |
| - | - | 5050 | 1001 | - | - | 0 | - |
| - | - | 2994 | 1002 | - | - | 0 | - |
| - | - | 870.6 | 1003 | - | - | 0 | - |
| - | - | 5721 | 1019 | - | - | 0 | - |
| - | - | 7304 | 1020 | - | - | 0 | - |
| - | - | 3117 | 1021 | - | - | 0 | - |
| - | - | 978.8 | 1022 | - | - | 0 | - |
| 2 | y | 1456 | 1100 | 0.002694 | 2.449 | +1 | 8 |
| - | - | 622.1 | 1298 | - | - | 0 | - |
| - | - | 668.1 | 1354 | - | - | 0 | - |
| - | - | 747.4 | 3083 | - | - | 0 | - |

m/z Charge Intensity FragmentType MassShift Position
120.08099365234375 0 56835.168
121.08433532714844 0 4181.261
123.04440307617188 0 607.5503
127.08670806884766 0 1943.3029
129.06619262695312 0 584.64667
129.10243225097656 0 3885.075
130.065185546875 0 768.26764
131.0492401123047 0 671.7907
133.08602905273438 0 1195.9183
134.99351501464844 0 501.46753
136.0758819580078 0 14809.093
137.0791015625 0 1011.3936
138.0662384033203 0 509.23505
139.0867919921875 0 595.7926
141.1024169921875 0 4730.303
147.04408264160156 0 698.40393
148.95455932617188 0 998.9878
149.04495239257812 0 663.32196
155.08169555664062 0 2067.813
155.0930633544922 0 420.18518
156.0765838623047 0 1367.3777
157.09715270996094 0 2910.7393
157.13388061523438 0 822.23975
158.0966339111328 0 2283.3638
159.09178161621094 0 1592.9485
159.1126251220703 0 573.4358
160.1121368408203 0 1179.5082
165.05474853515625 0 4501.268
165.10292053222656 0 465.72794
166.08642578125 0 25274.16
167.05535888671875 0 1552.2843
167.08160400390625 0 9861.475
167.0899658203125 0 2922.679
168.08473205566406 0 789.4434
169.09732055664062 0 6770.676
171.11270141601562 0 667.1008
173.12867736816406 0 3185.817
175.0865936279297 0 555.51526
175.11904907226562 0 891.8414
176.0821533203125 0 515.9172
176.10716247558594 0 4128.0654
177.10232543945312 0 3179.4333
177.11024475097656 0 711.8807
181.1326141357422 0 493.13885
182.08128356933594 0 9590.044
183.08465576171875 0 883.4918
183.11297607421875 0 1206.5887
185.0922088623047 0 3019.7378
185.12890625 0 565.92615
186.1234588623047 0 578.0074
187.10781860351562 0 1334.3809
188.0706329345703 0 2696.75
189.06558227539062 0 500.91385
191.117919921875 0 683.7938
193.10830688476562 0 1117.1804
194.07870483398438 0 521.2547
197.1288299560547 0 760.7945
199.0707244873047 0 480.36093
199.10781860351562 0 4959.8555
199.18040466308594 0 643.22577
200.1396026611328 0 543.0057
201.12351989746094 0 1820.3767
203.11782836914062 0 1896.0576 a Water loss 1
204.0768280029297 0 792.2868
205.09703063964844 0 1804.471 y 8
207.11358642578125 0 750.68195
209.09310913085938 0 511.06708
211.1440887451172 0 1005.20685
212.10301208496094 0 1368.0361
214.117431640625 0 644.2057
215.1387481689453 0 611.9009
221.103271484375 0 3810.615
221.12855529785156 0 55985.67 a 1
222.1320037841797 0 7195.1763
223.0635223388672 0 1601.0927
224.10305786132812 0 6954.7827
226.1190643310547 0 1651.5719
227.102783203125 0 20193.676
227.17532348632812 0 587.6767
228.106201171875 0 2209.7034
228.13388061523438 0 658.53784
229.84555053710938 0 485.8999
231.0614776611328 0 1010.4789
231.11268615722656 0 840.2095 b Water loss 1
235.1078643798828 0 1338.4536
238.1544189453125 0 734.4089
239.09475708007812 0 793.6869
241.0745391845703 0 1059.634
242.11390686035156 0 2116.5356
243.1333770751953 0 949.66205
244.12948608398438 0 568.08185
249.12347412109375 0 10675.484 b 1
250.126220703125 0 1501.1161
251.10296630859375 0 1196.0166
251.15101623535156 0 861.6181
253.1177978515625 0 794.8471
254.1133270263672 0 778.1568
256.16546630859375 0 649.41144
263.1023254394531 0 2357.6184
265.1546936035156 0 3991.3777
266.1500244140625 0 3534.306
268.1636047363281 0 606.3136
270.107421875 0 604.24713
271.1401062011719 0 739.94495
274.1308898925781 0 870.5036
279.097900390625 0 692.42303
280.1297607421875 0 664.092
281.12445068359375 0 14022.412
282.1271057128906 0 1320.2919
282.18145751953125 0 712.98785
283.14154052734375 0 1001.4883
284.1602783203125 0 1222.9276
285.01007080078125 0 1386.7098
285.1021728515625 0 1204.7983
294.1817932128906 0 1103.4403
295.1760559082031 0 882.0021
296.15045166015625 0 617.23584
297.0932312011719 0 619.1254
298.1395263671875 0 2003.9247
299.1356201171875 0 2925.5945
302.114013671875 0 597.46704
309.2048645019531 0 515.9394
312.1209716796875 0 485.79556
312.1919860839844 0 1667.0988
313.1869201660156 0 727.01154
322.1759338378906 0 983.57135
323.17108154296875 0 1229.7572
324.1466064453125 0 1938.3173
325.18707275390625 0 532.4191
328.1286315917969 0 518.7602
329.1497497558594 0 5457.2983
330.1530456542969 0 1344.6764
335.17193603515625 0 955.6893
338.2082824707031 0 598.973
339.2025451660156 0 584.957
339.6953125 0 1441.1927
340.1870422363281 0 20735.08
341.1882019042969 0 4177.3887
341.218505859375 0 429.55447
342.1364440917969 0 906.79834
342.1563720703125 0 11661.445 y 7
342.2150573730469 0 505.35397
343.1596984863281 0 1707.664
345.08746337890625 0 508.18658
349.1865234375 0 953.64844
352.19781494140625 0 3980.762
355.0690612792969 0 1902.8071
356.13885498046875 0 844.1878
357.2215881347656 0 879.3507
364.15057373046875 0 3133.797 b 2
364.5205383300781 0 486.42465
365.15289306640625 0 536.82947
366.2021484375 0 914.9657
367.19866943359375 0 1594.6056
368.20404052734375 0 1451.7941
370.2087097167969 0 2950.4167
377.1811218261719 0 834.3669
378.13031005859375 0 1679.2981
380.1927795410156 0 13667.08
381.1955871582031 0 2666.8125
383.22845458984375 0 1192.3573
394.1253356933594 0 2641.379
394.2089538574219 0 2455.1414
395.19268798828125 0 8072.9097
396.1953125 0 2561.443
396.22357177734375 0 953.4023
397.2073059082031 0 613.6737
398.203369140625 0 11136.808
399.20709228515625 0 2926.2405
403.23992919921875 0 607.5796
411.22601318359375 0 611.29694
412.22210693359375 0 715.6757
413.2660217285156 0 1195.2239
416.7315673828125 0 1306.914
417.232177734375 0 804.6233
425.19378662109375 0 1902.9395
425.2770080566406 0 1895.7444
426.2782287597656 0 698.1884
427.7227478027344 0 1063.2255
435.2607116699219 0 924.51666
437.2342529296875 0 628.8253
442.237060546875 0 836.49445
446.2523193359375 0 1061.5154
448.2552795410156 0 665.55664
449.23980712890625 0 4638.5957
450.2435607910156 0 1688.3094
451.2438049316406 0 3903.5547
451.7423400878906 0 1928.969
453.2712707519531 0 7351.3774
454.2323303222656 0 961.6297
454.2743835449219 0 2112.3164
460.24957275390625 0 1417.9012
462.2668762207031 0 839.4476
465.1619567871094 0 1475.5757
479.1788635253906 0 1182.4445 b 3
479.2851867675781 0 778.68054
480.28204345703125 0 5505.657
483.291748046875 0 825.98474
489.1914367675781 0 2387.1863 y 6
490.2650146484375 0 1384.3969
493.2775573730469 0 1879.7922
494.2808837890625 0 666.96735
496.2308654785156 0 730.373
500.7763671875 0 6673.887
501.2776794433594 0 2830.064
501.7783508300781 0 827.69666
506.260986328125 0 3523.0234
508.2765197753906 0 16092.773
509.2793273925781 0 4205.744
510.29052734375 0 1184.9988
511.2893371582031 0 1577.5287
539.3173217773438 0 696.0789
545.2728881835938 0 812.3704
550.7095336914062 0 1721.6802 y 1
560.2285766601562 0 3623.361 y 5
561.2305908203125 0 1408.3535
563.2823486328125 0 7791.6743
564.2857666015625 0 2801.756
565.2921752929688 0 975.1243
567.3110961914062 0 1671.7202
584.2675170898438 0 658.0234
585.337158203125 0 2443.6533
586.3374633789062 0 770.1543
592.7303466796875 0 699.9697
593.365234375 0 5312.506
594.3687133789062 0 2450.1907
595.3699951171875 0 747.38696
601.2359008789062 0 2095.9343 Precursor
601.6473999023438 0 995.13586
601.7386474609375 0 823.0269
612.197265625 0 621.4453
621.3600463867188 0 30240.45
622.3086547851562 0 850.0063
622.3632202148438 0 11170.264
623.36572265625 0 2127.739
650.3871459960938 0 1342.6254
652.8107299804688 0 779.25684
657.2626342773438 0 1004.0894
659.2944946289062 0 2009.3333
660.3690795898438 0 916.83466
676.36572265625 0 2872.2896
677.3687744140625 0 1897.7158
678.380615234375 0 6688.8535
679.3831176757812 0 2383.6995
680.3923950195312 0 591.61224
685.2545776367188 0 1112.3401
693.3930053710938 0 607.96466
700.3622436523438 0 5826.404
701.3641967773438 0 1980.8994
707.4089965820312 0 1527.4087
717.3926391601562 0 1448.3052
721.2603149414062 0 1205.7474
723.2911987304688 0 3687.0732 y 4
724.2938842773438 0 1718.3569
733.3863525390625 0 3115.774
735.4024047851562 0 9108.88
736.4063110351562 0 3962.141
749.256103515625 0 1625.492
751.399169921875 0 803.1572
773.4194946289062 0 892.548
774.3201904296875 0 2214.2832
783.3156127929688 0 697.488
783.3944091796875 0 1229.5615
799.6331787109375 0 653.55365
800.4217529296875 0 726.59955
801.4105224609375 0 2517.7583
802.4165649414062 0 1748.7246
804.4328002929688 0 694.0829
828.4193725585938 0 1748.5334
832.4546508789062 0 1373.6824
838.3169555664062 0 4308.437 y 3
839.320068359375 0 2177.0974
840.3181762695312 0 817.24164
844.4514770507812 0 4291.5615
845.4535522460938 0 2847.006
846.4518432617188 0 728.7631
854.440185546875 0 819.0353
871.3427124023438 0 656.6519
872.4462890625 0 16027.901
873.4485473632812 0 6634.2925
874.4522705078125 0 1735.428
883.4646606445312 0 836.2531
889.34619140625 0 6446.1963
890.3490600585938 0 3651.9434
892.4869384765625 0 681.5096
901.4765625 0 18409.443
902.4794921875 0 10493.596
903.48193359375 0 3422.3633
904.4915161132812 0 747.2281
919.4862670898438 0 7722.0957
920.4888916015625 0 4503.48
921.4940185546875 0 834.0414
935.3326416015625 0 923.34064 y Water loss 2
936.3351440429688 0 838.64886
953.3448486328125 0 20783.98 y 2
954.347412109375 0 12896.045
955.3469848632812 0 3484.7961
990.5628051757812 0 1278.5828
991.521728515625 0 3575.9297
992.5231323242188 0 987.031
1000.5439453125 0 5050.164
1001.5480346679688 0 2993.9534
1003.3732299804688 0 870.5924
1018.5549926757812 0 5721.285
1019.5233154296875 0 7304.1626
1020.5230102539062 0 3117.37
1021.5260620117188 0 978.764
1100.4119873046875 0 1455.7749 y 1
1298.481689453125 0 622.075
1353.5682373046875 0 668.1099
3082.71630859375 0 747.35425

Spectrum Details

|  |  |
| --- | --- |
| Matched peaks? Matched peaks The total absolute number of peaks matched. Additionally in brackets the total fraction of peaks matched and the total number of peaks is shown. | 17 (5.47% of 311) |
| FDR? FDR The false discovery rate estimated for this peptide. It is calculated by matching all theoretical fragments with a non-integer shift with the raw peaks for this spectrum. This is done with 40 different shifts. The resulting percentage is the average number of annotated peaks over the number of annotated peaks with the correct spectrum. | 4.48% |
| Satellite FDR? Satellite FDR See the FDR for details on its calculation. This satellite ion specific FDR only contains the satellite ions (d/w) for I/L/J positions. | - |
| PSM Score? PSM Score The PSM Score as given by Hecklib to this annotated spectrum. It is shown with three significant figures. | 195 |

#### Reverse Lookup? Reverse Lookup All places where this read could be placed.

| Group | Segment | Template | Template Part | Read Part | Score | Unique |
| --- | --- | --- | --- | --- | --- | --- |
| Homo sapiens Heavy Chain | IGHV | IGHV3-9 | [27..36] | [0..9] | 72 | True |

| Recombined | Template Part | Read Part | Score | Unique |
| --- | --- | --- | --- | --- |
| REC-0-1 | [27..36] | [0..9] | 72 | True |

#### Meta Information from Multiple reads

##### Number of combined reads

63

##### Intensity

0.6392

##### TotalArea

9.133E+09

#### Positional Score

Copy Data

##### Positional Score (TSV)

###### Preview

```
Loading example...
```

*Click on the button to copy the data to your clipboard.*

10012345678

Label Value
"0" 0.934
"1" 0.946
"2" 0.967
"3" 0.964
"4" 0.946
"5" 0.929
"6" 0.93
"7" 0.907
"8" 0.961

#### Meta Information from PEAKS

##### Scan Identifier

F3:7317

##### Original sequence

T

F

D

D

Y

A

M

+15.99

H

W

##### Posttranslational Modifications

Oxidation (M)

##### Source File

D:\separate\_stitch\_analyses\xle-disambiguation\raw\20210323\_F1\_UM1\_Peng0013\_SA\_F59\_ingel\_3ug\_chymo.raw

##### Fraction

3

##### Scan Feature

F3:8256

##### De Novo Score

99

##### ConfidenceScore

99

### m/z

601.2359

##### Mass

1200.4546

##### Charge

2

##### Retention Time

40.46

##### Predicted Retention Time

-

##### Area

1.459E+09

##### Parts Per Million

2.2

##### Fragmentation mode

HCD

##### Originating file

01 D:\separate\_stitch\_analyses\xle-disambiguation\20210325\_F59\_3ug\_DENOVO\_12.csv

#### Meta Information from PEAKS

##### Scan Identifier

F3:7379

##### Original sequence

T

F

D

D

Y

A

M

+15.99

H

W

##### Posttranslational Modifications

Oxidation (M)

##### Source File

D:\separate\_stitch\_analyses\xle-disambiguation\raw\20210323\_F1\_UM1\_Peng0013\_SA\_F59\_ingel\_3ug\_chymo.raw

##### Fraction

3

##### Scan Feature

F3:8256

##### De Novo Score

99

##### ConfidenceScore

99

### m/z

601.2359

##### Mass

1200.4546

##### Charge

2

##### Retention Time

40.46

##### Predicted Retention Time

-

##### Area

1.459E+09

##### Parts Per Million

2.2

##### Fragmentation mode

HCD

##### Originating file

01 D:\separate\_stitch\_analyses\xle-disambiguation\20210325\_F59\_3ug\_DENOVO\_12.csv

#### Meta Information from PEAKS

##### Scan Identifier

F3:7537

##### Original sequence

T

F

D

D

Y

A

M

+15.99

H

W

##### Posttranslational Modifications

Oxidation (M)

##### Source File

D:\separate\_stitch\_analyses\xle-disambiguation\raw\20210323\_F1\_UM1\_Peng0013\_SA\_F59\_ingel\_3ug\_chymo.raw

##### Fraction

3

##### Scan Feature

F3:8256

##### De Novo Score

99

##### ConfidenceScore

99

### m/z

601.2359

##### Mass

1200.4546

##### Charge

2

##### Retention Time

40.46

##### Predicted Retention Time

-

##### Area

1.459E+09

##### Parts Per Million

2.2

##### Fragmentation mode

HCD

##### Originating file

01 D:\separate\_stitch\_analyses\xle-disambiguation\20210325\_F59\_3ug\_DENOVO\_12.csv

#### Meta Information from PEAKS

##### Scan Identifier

F3:7636

##### Original sequence

T

F

D

D

Y

A

M

+15.99

H

W

##### Posttranslational Modifications

Oxidation (M)

##### Source File

D:\separate\_stitch\_analyses\xle-disambiguation\raw\20210323\_F1\_UM1\_Peng0013\_SA\_F59\_ingel\_3ug\_chymo.raw

##### Fraction

3

##### Scan Feature

F3:8256

##### De Novo Score

99

##### ConfidenceScore

99

### m/z

601.2359

##### Mass

1200.4546

##### Charge

2

##### Retention Time

40.46

##### Predicted Retention Time

-

##### Area

1.459E+09

##### Parts Per Million

2.2

##### Fragmentation mode

HCD

##### Originating file

01 D:\separate\_stitch\_analyses\xle-disambiguation\20210325\_F59\_3ug\_DENOVO\_12.csv

#### Meta Information from PEAKS

##### Scan Identifier

F3:9198

##### Original sequence

T

F

D

D

Y

A

M

H

W

##### Posttranslational Modifications

##### Source File

D:\separate\_stitch\_analyses\xle-disambiguation\raw\20210323\_F1\_UM1\_Peng0013\_SA\_F59\_ingel\_3ug\_chymo.raw

##### Fraction

3

##### Scan Feature

F3:7951

##### De Novo Score

99

##### ConfidenceScore

99

### m/z

593.2379

##### Mass

1184.4597

##### Charge

2

##### Retention Time

51.24

##### Predicted Retention Time

-

##### Area

7.327E+06

##### Parts Per Million

1.2

##### Fragmentation mode

HCD

##### Originating file

01 D:\separate\_stitch\_analyses\xle-disambiguation\20210325\_F59\_3ug\_DENOVO\_12.csv

#### Meta Information from PEAKS

##### Scan Identifier

F3:8296

##### Original sequence

T

F

D

D

Y

A

M

+15.99

H

W

##### Posttranslational Modifications

Oxidation (M)

##### Source File

D:\separate\_stitch\_analyses\xle-disambiguation\raw\20210323\_F1\_UM1\_Peng0013\_SA\_F59\_ingel\_3ug\_chymo.raw

##### Fraction

3

##### Scan Feature

-

##### De Novo Score

99

##### ConfidenceScore

99

### m/z

601.2349

##### Mass

1200.4546

##### Charge

2

##### Retention Time

46.32

##### Predicted Retention Time

-

##### Area

0

##### Parts Per Million

0.5

##### Fragmentation mode

HCD

##### Originating file

01 D:\separate\_stitch\_analyses\xle-disambiguation\20210325\_F59\_3ug\_DENOVO\_12.csv

#### Meta Information from PEAKS

##### Scan Identifier

F3:10292

##### Original sequence

T

F

D

D

Y

A

M

+15.99

H

W

##### Posttranslational Modifications

Oxidation (M)

##### Source File

D:\separate\_stitch\_analyses\xle-disambiguation\raw\20210323\_F1\_UM1\_Peng0013\_SA\_F59\_ingel\_3ug\_chymo.raw

##### Fraction

3

##### Scan Feature

F3:8254

##### De Novo Score

98

##### ConfidenceScore

98

### m/z

601.2358

##### Mass

1200.4546

##### Charge

2

##### Retention Time

55.46

##### Predicted Retention Time

-

##### Area

9.853E+06

##### Parts Per Million

2.1

##### Fragmentation mode

HCD

##### Originating file

01 D:\separate\_stitch\_analyses\xle-disambiguation\20210325\_F59\_3ug\_DENOVO\_12.csv

#### Meta Information from PEAKS

##### Scan Identifier

F3:10402

##### Original sequence

T

F

D

D

Y

A

M

+15.99

H

W

##### Posttranslational Modifications

Oxidation (M)

##### Source File

D:\separate\_stitch\_analyses\xle-disambiguation\raw\20210323\_F1\_UM1\_Peng0013\_SA\_F59\_ingel\_3ug\_chymo.raw

##### Fraction

3

##### Scan Feature

F3:8254

##### De Novo Score

98

##### ConfidenceScore

98

### m/z

601.2358

##### Mass

1200.4546

##### Charge

2

##### Retention Time

55.46

##### Predicted Retention Time

-

##### Area

9.853E+06

##### Parts Per Million

2.1

##### Fragmentation mode

HCD

##### Originating file

01 D:\separate\_stitch\_analyses\xle-disambiguation\20210325\_F59\_3ug\_DENOVO\_12.csv

#### Meta Information from PEAKS

##### Scan Identifier

F3:10187

##### Original sequence

T

F

D

D

Y

A

M

+15.99

H

W

##### Posttranslational Modifications

Oxidation (M)

##### Source File

D:\separate\_stitch\_analyses\xle-disambiguation\raw\20210323\_F1\_UM1\_Peng0013\_SA\_F59\_ingel\_3ug\_chymo.raw

##### Fraction

3

##### Scan Feature

F3:8254

##### De Novo Score

98

##### ConfidenceScore

98

### m/z

601.2358

##### Mass

1200.4546

##### Charge

2

##### Retention Time

55.46

##### Predicted Retention Time

-

##### Area

9.853E+06

##### Parts Per Million

2.1

##### Fragmentation mode

HCD

##### Originating file

01 D:\separate\_stitch\_analyses\xle-disambiguation\20210325\_F59\_3ug\_DENOVO\_12.csv

#### Meta Information from PEAKS

##### Scan Identifier

F3:10126

##### Original sequence

T

F

D

D

Y

A

M

+15.99

H

W

##### Posttranslational Modifications

Oxidation (M)

##### Source File

D:\separate\_stitch\_analyses\xle-disambiguation\raw\20210323\_F1\_UM1\_Peng0013\_SA\_F59\_ingel\_3ug\_chymo.raw

##### Fraction

3

##### Scan Feature

F3:8254

##### De Novo Score

98

##### ConfidenceScore

98

### m/z

601.2358

##### Mass

1200.4546

##### Charge

2

##### Retention Time

55.46

##### Predicted Retention Time

-

##### Area

9.853E+06

##### Parts Per Million

2.1

##### Fragmentation mode

HCD

##### Originating file

01 D:\separate\_stitch\_analyses\xle-disambiguation\20210325\_F59\_3ug\_DENOVO\_12.csv

#### Meta Information from PEAKS

##### Scan Identifier

F3:7474

##### Original sequence

T

F

D

D

Y

A

M

+15.99

H

W

##### Posttranslational Modifications

Oxidation (M)

##### Source File

D:\separate\_stitch\_analyses\xle-disambiguation\raw\20210323\_F1\_UM1\_Peng0013\_SA\_F59\_ingel\_3ug\_chymo.raw

##### Fraction

3

##### Scan Feature

F3:8256

##### De Novo Score

98

##### ConfidenceScore

98

### m/z

601.2359

##### Mass

1200.4546

##### Charge

2

##### Retention Time

40.46

##### Predicted Retention Time

-

##### Area

1.459E+09

##### Parts Per Million

2.2

##### Fragmentation mode

ETHCD

##### Originating file

01 D:\separate\_stitch\_analyses\xle-disambiguation\20210325\_F59\_3ug\_DENOVO\_12.csv

#### Meta Information from PEAKS

##### Scan Identifier

F3:10919

##### Original sequence

T

F

D

D

Y

A

M

+15.99

H

W

##### Posttranslational Modifications

Oxidation (M)

##### Source File

D:\separate\_stitch\_analyses\xle-disambiguation\raw\20210323\_F1\_UM1\_Peng0013\_SA\_F59\_ingel\_3ug\_chymo.raw

##### Fraction

3

##### Scan Feature

F3:8250

##### De Novo Score

98

##### ConfidenceScore

98

### m/z

601.2349

##### Mass

1200.4546

##### Charge

2

##### Retention Time

62.13

##### Predicted Retention Time

-

##### Area

1.739E+06

##### Parts Per Million

0.5

##### Fragmentation mode

HCD

##### Originating file

01 D:\separate\_stitch\_analyses\xle-disambiguation\20210325\_F59\_3ug\_DENOVO\_12.csv

#### Meta Information from PEAKS

##### Scan Identifier

F3:9794

##### Original sequence

T

F

D

D

Y

A

M

+15.99

H

W

##### Posttranslational Modifications

Oxidation (M)

##### Source File

D:\separate\_stitch\_analyses\xle-disambiguation\raw\20210323\_F1\_UM1\_Peng0013\_SA\_F59\_ingel\_3ug\_chymo.raw

##### Fraction

3

##### Scan Feature

F3:8254

##### De Novo Score

98

##### ConfidenceScore

98

### m/z

601.2358

##### Mass

1200.4546

##### Charge

2

##### Retention Time

55.46

##### Predicted Retention Time

-

##### Area

9.853E+06

##### Parts Per Million

2.1

##### Fragmentation mode

HCD

##### Originating file

01 D:\separate\_stitch\_analyses\xle-disambiguation\20210325\_F59\_3ug\_DENOVO\_12.csv

#### Meta Information from PEAKS

##### Scan Identifier

F3:10004

##### Original sequence

T

F

D

D

Y

A

M

+15.99

H

W

##### Posttranslational Modifications

Oxidation (M)

##### Source File

D:\separate\_stitch\_analyses\xle-disambiguation\raw\20210323\_F1\_UM1\_Peng0013\_SA\_F59\_ingel\_3ug\_chymo.raw

##### Fraction

3

##### Scan Feature

F3:8254

##### De Novo Score

98

##### ConfidenceScore

98

### m/z

601.2358

##### Mass

1200.4546

##### Charge

2

##### Retention Time

55.46

##### Predicted Retention Time

-

##### Area

9.853E+06

##### Parts Per Million

2.1

##### Fragmentation mode

HCD

##### Originating file

01 D:\separate\_stitch\_analyses\xle-disambiguation\20210325\_F59\_3ug\_DENOVO\_12.csv

#### Meta Information from PEAKS

##### Scan Identifier

F3:9585

##### Original sequence

T

F

D

D

Y

A

M

+15.99

H

W

##### Posttranslational Modifications

Oxidation (M)

##### Source File

D:\separate\_stitch\_analyses\xle-disambiguation\raw\20210323\_F1\_UM1\_Peng0013\_SA\_F59\_ingel\_3ug\_chymo.raw

##### Fraction

3

##### Scan Feature

F3:8254

##### De Novo Score

98

##### ConfidenceScore

98

### m/z

601.2358

##### Mass

1200.4546

##### Charge

2

##### Retention Time

55.46

##### Predicted Retention Time

-

##### Area

9.853E+06

##### Parts Per Million

2.1

##### Fragmentation mode

HCD

##### Originating file

01 D:\separate\_stitch\_analyses\xle-disambiguation\20210325\_F59\_3ug\_DENOVO\_12.csv

#### Meta Information from PEAKS

##### Scan Identifier

F3:9669

##### Original sequence

T

F

D

D

Y

A

M

+15.99

H

W

##### Posttranslational Modifications

Oxidation (M)

##### Source File

D:\separate\_stitch\_analyses\xle-disambiguation\raw\20210323\_F1\_UM1\_Peng0013\_SA\_F59\_ingel\_3ug\_chymo.raw

##### Fraction

3

##### Scan Feature

F3:8254

##### De Novo Score

98

##### ConfidenceScore

98

### m/z

601.2358

##### Mass

1200.4546

##### Charge

2

##### Retention Time

55.46

##### Predicted Retention Time

-

##### Area

9.853E+06

##### Parts Per Million

2.1

##### Fragmentation mode

HCD

##### Originating file

01 D:\separate\_stitch\_analyses\xle-disambiguation\20210325\_F59\_3ug\_DENOVO\_12.csv

#### Meta Information from PEAKS

##### Scan Identifier

F3:6163

##### Original sequence

T

F

D

D

Y

A

M

+15.99

H

W

+15.99

##### Posttranslational Modifications

Oxidation (M); Oxidation (HW)

##### Source File

D:\separate\_stitch\_analyses\xle-disambiguation\raw\20210323\_F1\_UM1\_Peng0013\_SA\_F59\_ingel\_3ug\_chymo.raw

##### Fraction

3

##### Scan Feature

F3:8552

##### De Novo Score

98

##### ConfidenceScore

98

### m/z

609.2325

##### Mass

1216.4495

##### Charge

2

##### Retention Time

34.29

##### Predicted Retention Time

-

##### Area

5.968E+07

##### Parts Per Million

0.9

##### Fragmentation mode

ETHCD

##### Originating file

01 D:\separate\_stitch\_analyses\xle-disambiguation\20210325\_F59\_3ug\_DENOVO\_12.csv

#### Meta Information from PEAKS

##### Scan Identifier

F3:11784

##### Original sequence

T

F

D

D

Y

A

M

+15.99

H

W

##### Posttranslational Modifications

Oxidation (M)

##### Source File

D:\separate\_stitch\_analyses\xle-disambiguation\raw\20210323\_F1\_UM1\_Peng0013\_SA\_F59\_ingel\_3ug\_chymo.raw

##### Fraction

3

##### Scan Feature

F3:8253

##### De Novo Score

98

##### ConfidenceScore

98

### m/z

601.2356

##### Mass

1200.4546

##### Charge

2

##### Retention Time

69.12

##### Predicted Retention Time

-

##### Area

4.502E+06

##### Parts Per Million

1.7

##### Fragmentation mode

HCD

##### Originating file

01 D:\separate\_stitch\_analyses\xle-disambiguation\20210325\_F59\_3ug\_DENOVO\_12.csv

#### Meta Information from PEAKS

##### Scan Identifier

F3:9943

##### Original sequence

T

F

D

D

Y

A

M

+15.99

H

W

##### Posttranslational Modifications

Oxidation (M)

##### Source File

D:\separate\_stitch\_analyses\xle-disambiguation\raw\20210323\_F1\_UM1\_Peng0013\_SA\_F59\_ingel\_3ug\_chymo.raw

##### Fraction

3

##### Scan Feature

F3:8254

##### De Novo Score

98

##### ConfidenceScore

98

### m/z

601.2358

##### Mass

1200.4546

##### Charge

2

##### Retention Time

55.46

##### Predicted Retention Time

-

##### Area

9.853E+06

##### Parts Per Million

2.1

##### Fragmentation mode

HCD

##### Originating file

01 D:\separate\_stitch\_analyses\xle-disambiguation\20210325\_F59\_3ug\_DENOVO\_12.csv

#### Meta Information from PEAKS

##### Scan Identifier

F3:10780

##### Original sequence

T

F

D

D

Y

A

M

+15.99

H

W

##### Posttranslational Modifications

Oxidation (M)

##### Source File

D:\separate\_stitch\_analyses\xle-disambiguation\raw\20210323\_F1\_UM1\_Peng0013\_SA\_F59\_ingel\_3ug\_chymo.raw

##### Fraction

3

##### Scan Feature

F3:8260

##### De Novo Score

98

##### ConfidenceScore

98

### m/z

601.2388

##### Mass

1200.4546

##### Charge

2

##### Retention Time

60.68

##### Predicted Retention Time

-

##### Area

1.126E+06

##### Parts Per Million

7

##### Fragmentation mode

HCD

##### Originating file

01 D:\separate\_stitch\_analyses\xle-disambiguation\20210325\_F59\_3ug\_DENOVO\_12.csv

#### Meta Information from PEAKS

##### Scan Identifier

F3:9135

##### Original sequence

T

F

D

D

Y

A

M

H

W

##### Posttranslational Modifications

##### Source File

D:\separate\_stitch\_analyses\xle-disambiguation\raw\20210323\_F1\_UM1\_Peng0013\_SA\_F59\_ingel\_3ug\_chymo.raw

##### Fraction

3

##### Scan Feature

F3:7951

##### De Novo Score

98

##### ConfidenceScore

98

### m/z

593.2379

##### Mass

1184.4597

##### Charge

2

##### Retention Time

51.24

##### Predicted Retention Time

-

##### Area

7.327E+06

##### Parts Per Million

1.2

##### Fragmentation mode

ETHCD

##### Originating file

01 D:\separate\_stitch\_analyses\xle-disambiguation\20210325\_F59\_3ug\_DENOVO\_12.csv

#### Meta Information from PEAKS

##### Scan Identifier

F3:7880

##### Original sequence

T

F

D

D

Y

A

M

+15.99

H

W

##### Posttranslational Modifications

Oxidation (M)

##### Source File

D:\separate\_stitch\_analyses\xle-disambiguation\raw\20210323\_F1\_UM1\_Peng0013\_SA\_F59\_ingel\_3ug\_chymo.raw

##### Fraction

3

##### Scan Feature

-

##### De Novo Score

98

##### ConfidenceScore

98

### m/z

601.235

##### Mass

1200.4546

##### Charge

2

##### Retention Time

43.91

##### Predicted Retention Time

-

##### Area

0

##### Parts Per Million

0.8

##### Fragmentation mode

ETHCD

##### Originating file

01 D:\separate\_stitch\_analyses\xle-disambiguation\20210325\_F59\_3ug\_DENOVO\_12.csv

#### Meta Information from PEAKS

##### Scan Identifier

F3:9729

##### Original sequence

T

F

D

D

Y

A

M

+15.99

H

W

##### Posttranslational Modifications

Oxidation (M)

##### Source File

D:\separate\_stitch\_analyses\xle-disambiguation\raw\20210323\_F1\_UM1\_Peng0013\_SA\_F59\_ingel\_3ug\_chymo.raw

##### Fraction

3

##### Scan Feature

F3:8254

##### De Novo Score

98

##### ConfidenceScore

98

### m/z

601.2358

##### Mass

1200.4546

##### Charge

2

##### Retention Time

55.46

##### Predicted Retention Time

-

##### Area

9.853E+06

##### Parts Per Million

2.1

##### Fragmentation mode

HCD

##### Originating file

01 D:\separate\_stitch\_analyses\xle-disambiguation\20210325\_F59\_3ug\_DENOVO\_12.csv

#### Meta Information from PEAKS

##### Scan Identifier

F3:11731

##### Original sequence

T

F

D

D

Y

A

M

+15.99

H

W

##### Posttranslational Modifications

Oxidation (M)

##### Source File

D:\separate\_stitch\_analyses\xle-disambiguation\raw\20210323\_F1\_UM1\_Peng0013\_SA\_F59\_ingel\_3ug\_chymo.raw

##### Fraction

3

##### Scan Feature

F3:8253

##### De Novo Score

98

##### ConfidenceScore

98

### m/z

601.2356

##### Mass

1200.4546

##### Charge

2

##### Retention Time

69.12

##### Predicted Retention Time

-

##### Area

4.502E+06

##### Parts Per Million

1.7

##### Fragmentation mode

HCD

##### Originating file

01 D:\separate\_stitch\_analyses\xle-disambiguation\20210325\_F59\_3ug\_DENOVO\_12.csv

#### Meta Information from PEAKS

##### Scan Identifier

F3:10660

##### Original sequence

T

F

D

D

Y

A

M

+15.99

H

W

##### Posttranslational Modifications

Oxidation (M)

##### Source File

D:\separate\_stitch\_analyses\xle-disambiguation\raw\20210323\_F1\_UM1\_Peng0013\_SA\_F59\_ingel\_3ug\_chymo.raw

##### Fraction

3

##### Scan Feature

F3:8254

##### De Novo Score

97

##### ConfidenceScore

97

### m/z

601.2358

##### Mass

1200.4546

##### Charge

2

##### Retention Time

55.46

##### Predicted Retention Time

-

##### Area

9.853E+06

##### Parts Per Million

2.1

##### Fragmentation mode

HCD

##### Originating file

01 D:\separate\_stitch\_analyses\xle-disambiguation\20210325\_F59\_3ug\_DENOVO\_12.csv

#### Meta Information from PEAKS

##### Scan Identifier

F3:11253

##### Original sequence

T

F

D

D

Y

A

M

+15.99

H

W

##### Posttranslational Modifications

Oxidation (M)

##### Source File

D:\separate\_stitch\_analyses\xle-disambiguation\raw\20210323\_F1\_UM1\_Peng0013\_SA\_F59\_ingel\_3ug\_chymo.raw

##### Fraction

3

##### Scan Feature

F3:8251

##### De Novo Score

97

##### ConfidenceScore

97

### m/z

601.2349

##### Mass

1200.4546

##### Charge

2

##### Retention Time

63.77

##### Predicted Retention Time

-

##### Area

9.887E+05

##### Parts Per Million

0.5

##### Fragmentation mode

HCD

##### Originating file

01 D:\separate\_stitch\_analyses\xle-disambiguation\20210325\_F59\_3ug\_DENOVO\_12.csv

#### Meta Information from PEAKS

##### Scan Identifier

F3:11490

##### Original sequence

T

F

D

D

Y

A

M

+15.99

H

W

##### Posttranslational Modifications

Oxidation (M)

##### Source File

D:\separate\_stitch\_analyses\xle-disambiguation\raw\20210323\_F1\_UM1\_Peng0013\_SA\_F59\_ingel\_3ug\_chymo.raw

##### Fraction

3

##### Scan Feature

F3:8253

##### De Novo Score

97

##### ConfidenceScore

97

### m/z

601.2356

##### Mass

1200.4546

##### Charge

2

##### Retention Time

69.12

##### Predicted Retention Time

-

##### Area

4.502E+06

##### Parts Per Million

1.7

##### Fragmentation mode

HCD

##### Originating file

01 D:\separate\_stitch\_analyses\xle-disambiguation\20210325\_F59\_3ug\_DENOVO\_12.csv

#### Meta Information from PEAKS

##### Scan Identifier

F3:9421

##### Original sequence

T

F

D

D

Y

A

M

H

W

##### Posttranslational Modifications

##### Source File

D:\separate\_stitch\_analyses\xle-disambiguation\raw\20210323\_F1\_UM1\_Peng0013\_SA\_F59\_ingel\_3ug\_chymo.raw

##### Fraction

3

##### Scan Feature

F3:7950

##### De Novo Score

97

##### ConfidenceScore

97

### m/z

593.236

##### Mass

1184.4597

##### Charge

2

##### Retention Time

52.11

##### Predicted Retention Time

-

##### Area

5.776E+05

##### Fragmentation mode

HCD

##### Originating file

01 D:\separate\_stitch\_analyses\xle-disambiguation\20210325\_F59\_3ug\_DENOVO\_12.csv

#### Meta Information from PEAKS

##### Scan Identifier

F3:9404

##### Original sequence

T

F

D

D

Y

A

M

+15.99

H

W

##### Posttranslational Modifications

Oxidation (M)

##### Source File

D:\separate\_stitch\_analyses\xle-disambiguation\raw\20210323\_F1\_UM1\_Peng0013\_SA\_F59\_ingel\_3ug\_chymo.raw

##### Fraction

3

##### Scan Feature

-

##### De Novo Score

97

##### ConfidenceScore

97

### m/z

601.2349

##### Mass

1200.4546

##### Charge

2

##### Retention Time

52.74

##### Predicted Retention Time

-

##### Area

0

##### Parts Per Million

0.6

##### Fragmentation mode

HCD

##### Originating file

01 D:\separate\_stitch\_analyses\xle-disambiguation\20210325\_F59\_3ug\_DENOVO\_12.csv

#### Meta Information from PEAKS

##### Scan Identifier

F3:11192

##### Original sequence

T

F

D

D

Y

A

M

+15.99

H

W

##### Posttranslational Modifications

Oxidation (M)

##### Source File

D:\separate\_stitch\_analyses\xle-disambiguation\raw\20210323\_F1\_UM1\_Peng0013\_SA\_F59\_ingel\_3ug\_chymo.raw

##### Fraction

3

##### Scan Feature

F3:8251

##### De Novo Score

97

##### ConfidenceScore

97

### m/z

601.2349

##### Mass

1200.4546

##### Charge

2

##### Retention Time

63.77

##### Predicted Retention Time

-

##### Area

9.887E+05

##### Parts Per Million

0.5

##### Fragmentation mode

HCD

##### Originating file

01 D:\separate\_stitch\_analyses\xle-disambiguation\20210325\_F59\_3ug\_DENOVO\_12.csv

#### Meta Information from PEAKS

##### Scan Identifier

F3:11306

##### Original sequence

T

F

D

D

Y

A

M

+15.99

H

W

##### Posttranslational Modifications

Oxidation (M)

##### Source File

D:\separate\_stitch\_analyses\xle-disambiguation\raw\20210323\_F1\_UM1\_Peng0013\_SA\_F59\_ingel\_3ug\_chymo.raw

##### Fraction

3

##### Scan Feature

F3:8251

##### De Novo Score

97

##### ConfidenceScore

97

### m/z

601.2349

##### Mass

1200.4546

##### Charge

2

##### Retention Time

63.77

##### Predicted Retention Time

-

##### Area

9.887E+05

##### Parts Per Million

0.5

##### Fragmentation mode

HCD

##### Originating file

01 D:\separate\_stitch\_analyses\xle-disambiguation\20210325\_F59\_3ug\_DENOVO\_12.csv

#### Meta Information from PEAKS

##### Scan Identifier

F3:9335

##### Original sequence

T

F

D

D

Y

A

M

H

W

##### Posttranslational Modifications

##### Source File

D:\separate\_stitch\_analyses\xle-disambiguation\raw\20210323\_F1\_UM1\_Peng0013\_SA\_F59\_ingel\_3ug\_chymo.raw

##### Fraction

3

##### Scan Feature

F3:7950

##### De Novo Score

97

##### ConfidenceScore

97

### m/z

593.236

##### Mass

1184.4597

##### Charge

2

##### Retention Time

52.11

##### Predicted Retention Time

-

##### Area

5.776E+05

##### Fragmentation mode

HCD

##### Originating file

01 D:\separate\_stitch\_analyses\xle-disambiguation\20210325\_F59\_3ug\_DENOVO\_12.csv

#### Meta Information from PEAKS

##### Scan Identifier

F3:12075

##### Original sequence

T

F

D

D

Y

A

M

+15.99

H

W

##### Posttranslational Modifications

Oxidation (M)

##### Source File

D:\separate\_stitch\_analyses\xle-disambiguation\raw\20210323\_F1\_UM1\_Peng0013\_SA\_F59\_ingel\_3ug\_chymo.raw

##### Fraction

3

##### Scan Feature

F3:8253

##### De Novo Score

97

##### ConfidenceScore

97

### m/z

601.2356

##### Mass

1200.4546

##### Charge

2

##### Retention Time

69.12

##### Predicted Retention Time

-

##### Area

4.502E+06

##### Parts Per Million

1.7

##### Fragmentation mode

HCD

##### Originating file

01 D:\separate\_stitch\_analyses\xle-disambiguation\20210325\_F59\_3ug\_DENOVO\_12.csv

#### Meta Information from PEAKS

##### Scan Identifier

F3:11624

##### Original sequence

T

F

D

D

Y

A

M

+15.99

H

W

##### Posttranslational Modifications

Oxidation (M)

##### Source File

D:\separate\_stitch\_analyses\xle-disambiguation\raw\20210323\_F1\_UM1\_Peng0013\_SA\_F59\_ingel\_3ug\_chymo.raw

##### Fraction

3

##### Scan Feature

F3:8253

##### De Novo Score

97

##### ConfidenceScore

97

### m/z

601.2356

##### Mass

1200.4546

##### Charge

2

##### Retention Time

69.12

##### Predicted Retention Time

-

##### Area

4.502E+06

##### Parts Per Million

1.7

##### Fragmentation mode

HCD

##### Originating file

01 D:\separate\_stitch\_analyses\xle-disambiguation\20210325\_F59\_3ug\_DENOVO\_12.csv

#### Meta Information from PEAKS

##### Scan Identifier

F3:12295

##### Original sequence

T

F

D

D

Y

A

M

+15.99

H

W

##### Posttranslational Modifications

Oxidation (M)

##### Source File

D:\separate\_stitch\_analyses\xle-disambiguation\raw\20210323\_F1\_UM1\_Peng0013\_SA\_F59\_ingel\_3ug\_chymo.raw

##### Fraction

3

##### Scan Feature

F3:8253

##### De Novo Score

97

##### ConfidenceScore

97

### m/z

601.2356

##### Mass

1200.4546

##### Charge

2

##### Retention Time

69.12

##### Predicted Retention Time

-

##### Area

4.502E+06

##### Parts Per Million

1.7

##### Fragmentation mode

HCD

##### Originating file

01 D:\separate\_stitch\_analyses\xle-disambiguation\20210325\_F59\_3ug\_DENOVO\_12.csv

#### Meta Information from PEAKS

##### Scan Identifier

F3:11678

##### Original sequence

T

F

D

D

Y

A

M

+15.99

H

W

##### Posttranslational Modifications

Oxidation (M)

##### Source File

D:\separate\_stitch\_analyses\xle-disambiguation\raw\20210323\_F1\_UM1\_Peng0013\_SA\_F59\_ingel\_3ug\_chymo.raw

##### Fraction

3

##### Scan Feature

F3:8253

##### De Novo Score

97

##### ConfidenceScore

97

### m/z

601.2356

##### Mass

1200.4546

##### Charge

2

##### Retention Time

69.12

##### Predicted Retention Time

-

##### Area

4.502E+06

##### Parts Per Million

1.7

##### Fragmentation mode

HCD

##### Originating file

01 D:\separate\_stitch\_analyses\xle-disambiguation\20210325\_F59\_3ug\_DENOVO\_12.csv

#### Meta Information from PEAKS

##### Scan Identifier

F3:12356

##### Original sequence

T

F

D

D

Y

A

M

+15.99

H

W

##### Posttranslational Modifications

Oxidation (M)

##### Source File

D:\separate\_stitch\_analyses\xle-disambiguation\raw\20210323\_F1\_UM1\_Peng0013\_SA\_F59\_ingel\_3ug\_chymo.raw

##### Fraction

3

##### Scan Feature

F3:8253

##### De Novo Score

97

##### ConfidenceScore

97

### m/z

601.2356

##### Mass

1200.4546

##### Charge

2

##### Retention Time

69.12

##### Predicted Retention Time

-

##### Area

4.502E+06

##### Parts Per Million

1.7

##### Fragmentation mode

HCD

##### Originating file

01 D:\separate\_stitch\_analyses\xle-disambiguation\20210325\_F59\_3ug\_DENOVO\_12.csv

#### Meta Information from PEAKS

##### Scan Identifier

F3:12909

##### Original sequence

T

F

D

D

Y

A

M

+15.99

H

W

##### Posttranslational Modifications

Oxidation (M)

##### Source File

D:\separate\_stitch\_analyses\xle-disambiguation\raw\20210323\_F1\_UM1\_Peng0013\_SA\_F59\_ingel\_3ug\_chymo.raw

##### Fraction

3

##### Scan Feature

F3:8253

##### De Novo Score

97

##### ConfidenceScore

97

### m/z

601.2356

##### Mass

1200.4546

##### Charge

2

##### Retention Time

69.12

##### Predicted Retention Time

-

##### Area

4.502E+06

##### Parts Per Million

1.7

##### Fragmentation mode

HCD

##### Originating file

01 D:\separate\_stitch\_analyses\xle-disambiguation\20210325\_F59\_3ug\_DENOVO\_12.csv

#### Meta Information from PEAKS

##### Scan Identifier

F3:6950

##### Original sequence

T

F

D

D

Y

A

M

+15.99

H

W

##### Posttranslational Modifications

Oxidation (M)

##### Source File

D:\separate\_stitch\_analyses\xle-disambiguation\raw\20210323\_F1\_UM1\_Peng0013\_SA\_F59\_ingel\_3ug\_chymo.raw

##### Fraction

3

##### Scan Feature

F3:8249

##### De Novo Score

97

##### ConfidenceScore

97

### m/z

601.2349

##### Mass

1200.4546

##### Charge

2

##### Retention Time

38.54

##### Predicted Retention Time

-

##### Area

2.343E+06

##### Parts Per Million

0.5

##### Fragmentation mode

ETHCD

##### Originating file

01 D:\separate\_stitch\_analyses\xle-disambiguation\20210325\_F59\_3ug\_DENOVO\_12.csv

#### Meta Information from PEAKS

##### Scan Identifier

F3:12523

##### Original sequence

T

F

D

D

Y

A

M

+15.99

H

W

##### Posttranslational Modifications

Oxidation (M)

##### Source File

D:\separate\_stitch\_analyses\xle-disambiguation\raw\20210323\_F1\_UM1\_Peng0013\_SA\_F59\_ingel\_3ug\_chymo.raw

##### Fraction

3

##### Scan Feature

F3:8253

##### De Novo Score

97

##### ConfidenceScore

97

### m/z

601.2356

##### Mass

1200.4546

##### Charge

2

##### Retention Time

69.12

##### Predicted Retention Time

-

##### Area

4.502E+06

##### Parts Per Million

1.7

##### Fragmentation mode

HCD

##### Originating file

01 D:\separate\_stitch\_analyses\xle-disambiguation\20210325\_F59\_3ug\_DENOVO\_12.csv

#### Meta Information from PEAKS

##### Scan Identifier

F3:12414

##### Original sequence

T

F

D

D

Y

A

M

+15.99

H

W

##### Posttranslational Modifications

Oxidation (M)

##### Source File

D:\separate\_stitch\_analyses\xle-disambiguation\raw\20210323\_F1\_UM1\_Peng0013\_SA\_F59\_ingel\_3ug\_chymo.raw

##### Fraction

3

##### Scan Feature

F3:8253

##### De Novo Score

97

##### ConfidenceScore

97

### m/z

601.2356

##### Mass

1200.4546

##### Charge

2

##### Retention Time

69.12

##### Predicted Retention Time

-

##### Area

4.502E+06

##### Parts Per Million

1.7

##### Fragmentation mode

HCD

##### Originating file

01 D:\separate\_stitch\_analyses\xle-disambiguation\20210325\_F59\_3ug\_DENOVO\_12.csv

#### Meta Information from PEAKS

##### Scan Identifier

F3:12185

##### Original sequence

T

F

D

D

Y

A

M

+15.99

H

W

##### Posttranslational Modifications

Oxidation (M)

##### Source File

D:\separate\_stitch\_analyses\xle-disambiguation\raw\20210323\_F1\_UM1\_Peng0013\_SA\_F59\_ingel\_3ug\_chymo.raw

##### Fraction

3

##### Scan Feature

F3:8253

##### De Novo Score

96

##### ConfidenceScore

96

### m/z

601.2356

##### Mass

1200.4546

##### Charge

2

##### Retention Time

69.12

##### Predicted Retention Time

-

##### Area

4.502E+06

##### Parts Per Million

1.7

##### Fragmentation mode

HCD

##### Originating file

01 D:\separate\_stitch\_analyses\xle-disambiguation\20210325\_F59\_3ug\_DENOVO\_12.csv

#### Meta Information from PEAKS

##### Scan Identifier

F3:11842

##### Original sequence

T

F

D

D

Y

A

M

+15.99

H

W

##### Posttranslational Modifications

Oxidation (M)

##### Source File

D:\separate\_stitch\_analyses\xle-disambiguation\raw\20210323\_F1\_UM1\_Peng0013\_SA\_F59\_ingel\_3ug\_chymo.raw

##### Fraction

3

##### Scan Feature

F3:8253

##### De Novo Score

96

##### ConfidenceScore

96

### m/z

601.2356

##### Mass

1200.4546

##### Charge

2

##### Retention Time

69.12

##### Predicted Retention Time

-

##### Area

4.502E+06

##### Parts Per Million

1.7

##### Fragmentation mode

HCD

##### Originating file

01 D:\separate\_stitch\_analyses\xle-disambiguation\20210325\_F59\_3ug\_DENOVO\_12.csv

#### Meta Information from PEAKS

##### Scan Identifier

F3:12129

##### Original sequence

T

F

D

D

Y

A

M

+15.99

H

W

##### Posttranslational Modifications

Oxidation (M)

##### Source File

D:\separate\_stitch\_analyses\xle-disambiguation\raw\20210323\_F1\_UM1\_Peng0013\_SA\_F59\_ingel\_3ug\_chymo.raw

##### Fraction

3

##### Scan Feature

F3:8253

##### De Novo Score

96

##### ConfidenceScore

96

### m/z

601.2356

##### Mass

1200.4546

##### Charge

2

##### Retention Time

69.12

##### Predicted Retention Time

-

##### Area

4.502E+06

##### Parts Per Million

1.7

##### Fragmentation mode

HCD

##### Originating file

01 D:\separate\_stitch\_analyses\xle-disambiguation\20210325\_F59\_3ug\_DENOVO\_12.csv

#### Meta Information from PEAKS

##### Scan Identifier

F3:5874

##### Original sequence

T

F

D

D

Y

A

M

+15.99

H

W

+15.99

##### Posttranslational Modifications

Oxidation (M); Oxidation (HW)

##### Source File

D:\separate\_stitch\_analyses\xle-disambiguation\raw\20210323\_F1\_UM1\_Peng0013\_SA\_F59\_ingel\_3ug\_chymo.raw

##### Fraction

3

##### Scan Feature

F3:8552

##### De Novo Score

96

##### ConfidenceScore

96

### m/z

609.2325

##### Mass

1216.4495

##### Charge

2

##### Retention Time

34.29

##### Predicted Retention Time

-

##### Area

5.968E+07

##### Parts Per Million

0.9

##### Fragmentation mode

ETHCD

##### Originating file

01 D:\separate\_stitch\_analyses\xle-disambiguation\20210325\_F59\_3ug\_DENOVO\_12.csv

#### Meta Information from PEAKS

##### Scan Identifier

F3:12469

##### Original sequence

T

F

D

D

Y

A

M

+15.99

H

W

##### Posttranslational Modifications

Oxidation (M)

##### Source File

D:\separate\_stitch\_analyses\xle-disambiguation\raw\20210323\_F1\_UM1\_Peng0013\_SA\_F59\_ingel\_3ug\_chymo.raw

##### Fraction

3

##### Scan Feature

F3:8253

##### De Novo Score

96

##### ConfidenceScore

96

### m/z

601.2356

##### Mass

1200.4546

##### Charge

2

##### Retention Time

69.12

##### Predicted Retention Time

-

##### Area

4.502E+06

##### Parts Per Million

1.7

##### Fragmentation mode

HCD

##### Originating file

01 D:\separate\_stitch\_analyses\xle-disambiguation\20210325\_F59\_3ug\_DENOVO\_12.csv

#### Meta Information from PEAKS

##### Scan Identifier

F3:11965

##### Original sequence

T

F

D

D

Y

A

M

+15.99

H

W

##### Posttranslational Modifications

Oxidation (M)

##### Source File

D:\separate\_stitch\_analyses\xle-disambiguation\raw\20210323\_F1\_UM1\_Peng0013\_SA\_F59\_ingel\_3ug\_chymo.raw

##### Fraction

3

##### Scan Feature

F3:8253

##### De Novo Score

96

##### ConfidenceScore

96

### m/z

601.2356

##### Mass

1200.4546

##### Charge

2

##### Retention Time

69.12

##### Predicted Retention Time

-

##### Area

4.502E+06

##### Parts Per Million

1.7

##### Fragmentation mode

HCD

##### Originating file

01 D:\separate\_stitch\_analyses\xle-disambiguation\20210325\_F59\_3ug\_DENOVO\_12.csv

#### Meta Information from PEAKS

##### Scan Identifier

F3:12241

##### Original sequence

T

F

D

D

Y

A

M

+15.99

H

W

##### Posttranslational Modifications

Oxidation (M)

##### Source File

D:\separate\_stitch\_analyses\xle-disambiguation\raw\20210323\_F1\_UM1\_Peng0013\_SA\_F59\_ingel\_3ug\_chymo.raw

##### Fraction

3

##### Scan Feature

F3:8253

##### De Novo Score

96

##### ConfidenceScore

96

### m/z

601.2356

##### Mass

1200.4546

##### Charge

2

##### Retention Time

69.12

##### Predicted Retention Time

-

##### Area

4.502E+06

##### Parts Per Million

1.7

##### Fragmentation mode

HCD

##### Originating file

01 D:\separate\_stitch\_analyses\xle-disambiguation\20210325\_F59\_3ug\_DENOVO\_12.csv

#### Meta Information from PEAKS

##### Scan Identifier

F3:8771

##### Original sequence

T

F

D

D

Y

A

M

+15.99

H

W

##### Posttranslational Modifications

Oxidation (M)

##### Source File

D:\separate\_stitch\_analyses\xle-disambiguation\raw\20210323\_F1\_UM1\_Peng0013\_SA\_F59\_ingel\_3ug\_chymo.raw

##### Fraction

3

##### Scan Feature

F3:8257

##### De Novo Score

96

##### ConfidenceScore

96

### m/z

601.236

##### Mass

1200.4546

##### Charge

2

##### Retention Time

47.04

##### Predicted Retention Time

-

##### Area

1.483E+07

##### Parts Per Million

2.4

##### Fragmentation mode

HCD

##### Originating file

01 D:\separate\_stitch\_analyses\xle-disambiguation\20210325\_F59\_3ug\_DENOVO\_12.csv

#### Meta Information from PEAKS

##### Scan Identifier

F3:12715

##### Original sequence

T

F

D

D

Y

A

M

+15.99

H

W

##### Posttranslational Modifications

Oxidation (M)

##### Source File

D:\separate\_stitch\_analyses\xle-disambiguation\raw\20210323\_F1\_UM1\_Peng0013\_SA\_F59\_ingel\_3ug\_chymo.raw

##### Fraction

3

##### Scan Feature

F3:8253

##### De Novo Score

96

##### ConfidenceScore

96

### m/z

601.2356

##### Mass

1200.4546

##### Charge

2

##### Retention Time

69.12

##### Predicted Retention Time

-

##### Area

4.502E+06

##### Parts Per Million

1.7

##### Fragmentation mode

HCD

##### Originating file

01 D:\separate\_stitch\_analyses\xle-disambiguation\20210325\_F59\_3ug\_DENOVO\_12.csv

#### Meta Information from PEAKS

##### Scan Identifier

F3:11363

##### Original sequence

T

F

D

D

Y

A

M

+15.99

H

W

##### Posttranslational Modifications

Oxidation (M)

##### Source File

D:\separate\_stitch\_analyses\xle-disambiguation\raw\20210323\_F1\_UM1\_Peng0013\_SA\_F59\_ingel\_3ug\_chymo.raw

##### Fraction

3

##### Scan Feature

F3:8251

##### De Novo Score

96

##### ConfidenceScore

96

### m/z

601.2349

##### Mass

1200.4546

##### Charge

2

##### Retention Time

63.77

##### Predicted Retention Time

-

##### Area

9.887E+05

##### Parts Per Million

0.5

##### Fragmentation mode

HCD

##### Originating file

01 D:\separate\_stitch\_analyses\xle-disambiguation\20210325\_F59\_3ug\_DENOVO\_12.csv

#### Meta Information from PEAKS

##### Scan Identifier

F3:12975

##### Original sequence

T

F

D

D

Y

A

M

+15.99

H

W

##### Posttranslational Modifications

Oxidation (M)

##### Source File

D:\separate\_stitch\_analyses\xle-disambiguation\raw\20210323\_F1\_UM1\_Peng0013\_SA\_F59\_ingel\_3ug\_chymo.raw

##### Fraction

3

##### Scan Feature

F3:8253

##### De Novo Score

96

##### ConfidenceScore

96

### m/z

601.2356

##### Mass

1200.4546

##### Charge

2

##### Retention Time

69.12

##### Predicted Retention Time

-

##### Area

4.502E+06

##### Parts Per Million

1.7

##### Fragmentation mode

HCD

##### Originating file

01 D:\separate\_stitch\_analyses\xle-disambiguation\20210325\_F59\_3ug\_DENOVO\_12.csv

#### Meta Information from PEAKS

##### Scan Identifier

F3:11913

##### Original sequence

T

F

D

D

Y

A

M

+15.99

H

W

##### Posttranslational Modifications

Oxidation (M)

##### Source File

D:\separate\_stitch\_analyses\xle-disambiguation\raw\20210323\_F1\_UM1\_Peng0013\_SA\_F59\_ingel\_3ug\_chymo.raw

##### Fraction

3

##### Scan Feature

F3:8253

##### De Novo Score

96

##### ConfidenceScore

96

### m/z

601.2356

##### Mass

1200.4546

##### Charge

2

##### Retention Time

69.12

##### Predicted Retention Time

-

##### Area

4.502E+06

##### Parts Per Million

1.7

##### Fragmentation mode

HCD

##### Originating file

01 D:\separate\_stitch\_analyses\xle-disambiguation\20210325\_F59\_3ug\_DENOVO\_12.csv

#### Meta Information from PEAKS

##### Scan Identifier

F3:13536

##### Original sequence

T

F

D

D

Y

A

M

+15.99

H

W

##### Posttranslational Modifications

Oxidation (M)

##### Source File

D:\separate\_stitch\_analyses\xle-disambiguation\raw\20210323\_F1\_UM1\_Peng0013\_SA\_F59\_ingel\_3ug\_chymo.raw

##### Fraction

3

##### Scan Feature

-

##### De Novo Score

96

##### ConfidenceScore

96

### m/z

601.2353

##### Mass

1200.4546

##### Charge

2

##### Retention Time

76.16

##### Predicted Retention Time

-

##### Area

0

##### Parts Per Million

1.2

##### Fragmentation mode

HCD

##### Originating file

01 D:\separate\_stitch\_analyses\xle-disambiguation\20210325\_F59\_3ug\_DENOVO\_12.csv

#### Meta Information from PEAKS

##### Scan Identifier

F3:10975

##### Original sequence

T

F

D

D

Y

A

M

+15.99

H

W

##### Posttranslational Modifications

Oxidation (M)

##### Source File

D:\separate\_stitch\_analyses\xle-disambiguation\raw\20210323\_F1\_UM1\_Peng0013\_SA\_F59\_ingel\_3ug\_chymo.raw

##### Fraction

3

##### Scan Feature

F3:8250

##### De Novo Score

96

##### ConfidenceScore

96

### m/z

601.2349

##### Mass

1200.4546

##### Charge

2

##### Retention Time

62.13

##### Predicted Retention Time

-

##### Area

1.739E+06

##### Parts Per Million

0.5

##### Fragmentation mode

HCD

##### Originating file

01 D:\separate\_stitch\_analyses\xle-disambiguation\20210325\_F59\_3ug\_DENOVO\_12.csv

#### Meta Information from PEAKS

##### Scan Identifier

F3:13093

##### Original sequence

T

F

D

D

Y

A

M

+15.99

H

W

##### Posttranslational Modifications

Oxidation (M)

##### Source File

D:\separate\_stitch\_analyses\xle-disambiguation\raw\20210323\_F1\_UM1\_Peng0013\_SA\_F59\_ingel\_3ug\_chymo.raw

##### Fraction

3

##### Scan Feature

F3:8253

##### De Novo Score

96

##### ConfidenceScore

96

### m/z

601.2356

##### Mass

1200.4546

##### Charge

2

##### Retention Time

69.12

##### Predicted Retention Time

-

##### Area

4.502E+06

##### Parts Per Million

1.7

##### Fragmentation mode

HCD

##### Originating file

01 D:\separate\_stitch\_analyses\xle-disambiguation\20210325\_F59\_3ug\_DENOVO\_12.csv

#### Meta Information from PEAKS

##### Scan Identifier

F3:13277

##### Original sequence

T

F

D

D

Y

A

M

+15.99

H

W

##### Posttranslational Modifications

Oxidation (M)

##### Source File

D:\separate\_stitch\_analyses\xle-disambiguation\raw\20210323\_F1\_UM1\_Peng0013\_SA\_F59\_ingel\_3ug\_chymo.raw

##### Fraction

3

##### Scan Feature

-

##### De Novo Score

95

##### ConfidenceScore

95

### m/z

601.235

##### Mass

1200.4546

##### Charge

2

##### Retention Time

74.8

##### Predicted Retention Time

-

##### Area

0

##### Parts Per Million

0.8

##### Fragmentation mode

HCD

##### Originating file

01 D:\separate\_stitch\_analyses\xle-disambiguation\20210325\_F59\_3ug\_DENOVO\_12.csv

#### Meta Information from PEAKS

##### Scan Identifier

F3:12777

##### Original sequence

T

F

D

D

Y

A

M

+15.99

H

W

##### Posttranslational Modifications

Oxidation (M)

##### Source File

D:\separate\_stitch\_analyses\xle-disambiguation\raw\20210323\_F1\_UM1\_Peng0013\_SA\_F59\_ingel\_3ug\_chymo.raw

##### Fraction

3

##### Scan Feature

F3:8253

##### De Novo Score

95

##### ConfidenceScore

95

### m/z

601.2356

##### Mass

1200.4546

##### Charge

2

##### Retention Time

69.12

##### Predicted Retention Time

-

##### Area

4.502E+06

##### Parts Per Million

1.7

##### Fragmentation mode

HCD

##### Originating file

01 D:\separate\_stitch\_analyses\xle-disambiguation\20210325\_F59\_3ug\_DENOVO\_12.csv

#### Meta Information from PEAKS

##### Scan Identifier

F3:10066

##### Original sequence

T

F

D

D

Y

A

M

+15.99

H

W

##### Posttranslational Modifications

Oxidation (M)

##### Source File

D:\separate\_stitch\_analyses\xle-disambiguation\raw\20210323\_F1\_UM1\_Peng0013\_SA\_F59\_ingel\_3ug\_chymo.raw

##### Fraction

3

##### Scan Feature

F3:8254

##### De Novo Score

95

##### ConfidenceScore

95

### m/z

601.2358

##### Mass

1200.4546

##### Charge

2

##### Retention Time

55.46

##### Predicted Retention Time

-

##### Area

9.853E+06

##### Parts Per Million

2.1

##### Fragmentation mode

ETHCD

##### Originating file

01 D:\separate\_stitch\_analyses\xle-disambiguation\20210325\_F59\_3ug\_DENOVO\_12.csv

#### Meta Information from PEAKS

##### Scan Identifier

F3:14047

##### Original sequence

T

F

D

D

Y

A

M

+15.99

H

W

##### Posttranslational Modifications

Oxidation (M)

##### Source File

D:\separate\_stitch\_analyses\xle-disambiguation\raw\20210323\_F1\_UM1\_Peng0013\_SA\_F59\_ingel\_3ug\_chymo.raw

##### Fraction

3

##### Scan Feature

-

##### De Novo Score

95

##### ConfidenceScore

95

### m/z

601.2344

##### Mass

1200.4546

##### Charge

2

##### Retention Time

78.85

##### Predicted Retention Time

-

##### Area

0

##### Fragmentation mode

HCD

##### Originating file

01 D:\separate\_stitch\_analyses\xle-disambiguation\20210325\_F59\_3ug\_DENOVO\_12.csv

#### Meta Information from PEAKS

##### Scan Identifier

F3:7257

##### Original sequence

T

F

D

D

Y

A

M

+15.99

H

W

##### Posttranslational Modifications

Oxidation (M)

##### Source File

D:\separate\_stitch\_analyses\xle-disambiguation\raw\20210323\_F1\_UM1\_Peng0013\_SA\_F59\_ingel\_3ug\_chymo.raw

##### Fraction

3

##### Scan Feature

F3:8256

##### De Novo Score

95

##### ConfidenceScore

95

### m/z

601.2359

##### Mass

1200.4546

##### Charge

2

##### Retention Time

40.46

##### Predicted Retention Time

-

##### Area

1.459E+09

##### Parts Per Million

2.2

##### Fragmentation mode

ETHCD

##### Originating file

01 D:\separate\_stitch\_analyses\xle-disambiguation\20210325\_F59\_3ug\_DENOVO\_12.csv

#### Meta Information from PEAKS

##### Scan Identifier

F3:13395

##### Original sequence

T

F

D

D

Y

A

M

+15.99

H

W

##### Posttranslational Modifications

Oxidation (M)

##### Source File

D:\separate\_stitch\_analyses\xle-disambiguation\raw\20210323\_F1\_UM1\_Peng0013\_SA\_F59\_ingel\_3ug\_chymo.raw

##### Fraction

3

##### Scan Feature

-

##### De Novo Score

95

##### ConfidenceScore

95

### m/z

601.235

##### Mass

1200.4546

##### Charge

2

##### Retention Time

75.41

##### Predicted Retention Time

-

##### Area

0

##### Parts Per Million

0.7

##### Fragmentation mode

HCD

##### Originating file

01 D:\separate\_stitch\_analyses\xle-disambiguation\20210325\_F59\_3ug\_DENOVO\_12.csv

#### Meta Information from PEAKS

##### Scan Identifier

F3:11133

##### Original sequence

T

F

D

D

Y

A

M

+15.99

H

W

##### Posttranslational Modifications

Oxidation (M)

##### Source File

D:\separate\_stitch\_analyses\xle-disambiguation\raw\20210323\_F1\_UM1\_Peng0013\_SA\_F59\_ingel\_3ug\_chymo.raw

##### Fraction

3

##### Scan Feature

F3:8250

##### De Novo Score

95

##### ConfidenceScore

95

### m/z

601.2349

##### Mass

1200.4546

##### Charge

2

##### Retention Time

62.13

##### Predicted Retention Time

-

##### Area

1.739E+06

##### Parts Per Million

0.5

##### Fragmentation mode

HCD

##### Originating file

01 D:\separate\_stitch\_analyses\xle-disambiguation\20210325\_F59\_3ug\_DENOVO\_12.csv
