## Supplementary material for "A handle on mass coincidence errors in *de novo* sequencing of antibodies by bottom-up proteomics": Combined_006.html

Details Combined\_006 | Stitch OverviewUndefined

### Read Combined\_006

#### Sequence (length=14)

VVVDVSHEDPEVKF

#### Spectrum 7070? Spectrum 7070 The raw spectrum of this peptide as annotated by Hecklib. The fragments are coloured according to ion type (see legend). Any peaks with a star '\*' as text can be hovered over to see the full details, first the ion type second the mass shift type. By hovering over the amino acids in the peptide or ions in the legend the corresponding peaks are highlighted. By toggling the 'Unassigned' label you can turn the background (unassigned) peaks on or off in the plot. By updating the slider in the Ion legend you can update the spectrum to only show the top X% of the peaks with labels. The top X% means any peak that is within X% of the highest intensity. By dragging in the spectrum you can zoom in to a specific part of the spectrum and use 'Zoom Out' to get back to the original zoom level. The annotation of the spectrum is based on the given sequence in the peptides file and is done with different software so inconsistencies are likely. The peaks are annotated based on the given sequence, with 20 ppm tolerance.

Copy Data

##### Spectrum 7070 (TSV)

###### Preview

```
Loading example...
```

*Click on the button to copy the data to your clipboard.*

Mz MinMz MaxIntensity Max

WidthHeightPeptide font sizePeptide stroke widthSpectrum font sizeSpectrum stroke widthCompact peptide

Ion legend

wxyz

abcd

OtherUnassignedIon^Charge^_Position_Show for top:%

VVVDVSHEDPEVKF

02.62e+55.23e+57.85e+51.05e+6

Zoom Out

y^+1^_1_a^+1^_2_b^+1^_2_y^+1^_2_y^+1^_2_b^+1^_3_y^+1^_3_b^+1^_4_b^+1^_4_b^+2^_9_y^+2^_8_y^+2^_8_y^+1^_4_b^+1^_5_y^+1^_4_y^+2^_9_b^+2^_10_y^+2^_9_b^+1^_6_y^+2^_10_y^+2^_10_b^+2^_11_b^+1^_6_y^+1^_5_b^+2^_11_y^+1^_5_y^+2^_11_b^+2^_12_y^+2^_12_y^+2^_12_b^+2^_13_b^+2^_13_b^+1^_7_y^+1^_6_b^+1^_7_y^+2^_13_\*\*\*y^+1^_7_b^+1^_8_y^+1^_7_b^+1^_8_b^+1^_9_b^+1^_9_y^+1^_8_y^+1^_8_y^+1^_9_y^+1^_9_b^+1^_10_y^+1^_9_y^+1^_10_b^+1^_11_b^+1^_11_y^+1^_11_b^+1^_12_y^+1^_11_b^+1^_12_y^+1^_12_y^+1^_12_b^+1^_13_

036272410861448

Fragment Matches Table

Show background peaks

| Position | Ion type | Intensity | mz Theoretical | mz Error (Th) | mz Error (ppm) | Charge | Series Number |
| --- | --- | --- | --- | --- | --- | --- | --- |
| - | - | 1.444E+04 | 120.1 | - | - | 0 | - |
| - | - | 1076 | 124 | - | - | 0 | - |
| - | - | 992.4 | 125.7 | - | - | 0 | - |
| - | - | 2065 | 126.1 | - | - | 0 | - |
| - | - | 1317 | 127.1 | - | - | 0 | - |
| - | - | 5.745E+05 | 129.1 | - | - | 0 | - |
| - | - | 6446 | 130.1 | - | - | 0 | - |
| - | - | 3333 | 130.1 | - | - | 0 | - |
| - | - | 3.347E+04 | 130.1 | - | - | 0 | - |
| - | - | 1134 | 136.9 | - | - | 0 | - |
| - | - | 3552 | 138.1 | - | - | 0 | - |
| - | - | 1128 | 139.1 | - | - | 0 | - |
| - | - | 1006 | 140.4 | - | - | 0 | - |
| - | - | 1696 | 141.1 | - | - | 0 | - |
| - | - | 1080 | 143 | - | - | 0 | - |
| - | - | 1472 | 143.2 | - | - | 0 | - |
| - | - | 1889 | 146.1 | - | - | 0 | - |
| - | - | 1.842E+04 | 147.1 | - | - | 0 | - |
| - | - | 1446 | 148.9 | - | - | 0 | - |
| - | - | 1602 | 152.1 | - | - | 0 | - |
| - | - | 1603 | 159.1 | - | - | 0 | - |
| - | - | 3194 | 163.1 | - | - | 0 | - |
| - | - | 1.685E+04 | 164.1 | - | - | 0 | - |
| - | - | 4246 | 166.1 | - | - | 0 | - |
| 14 | y | 9029 | 166.1 | 0.000247 | 1.487 | +1 | 1 |
| - | - | 2230 | 167.1 | - | - | 0 | - |
| - | - | 1509 | 168.9 | - | - | 0 | - |
| - | - | 1348 | 168.9 | - | - | 0 | - |
| - | - | 6250 | 169.1 | - | - | 0 | - |
| 2 | a | 3.406E+05 | 171.1 | 0.000316 | 1.846 | +1 | 2 |
| - | - | 2.802E+04 | 172.2 | - | - | 0 | - |
| - | - | 1739 | 172.2 | - | - | 0 | - |
| - | - | 1.334E+04 | 173.5 | - | - | 0 | - |
| - | - | 1.303E+04 | 179.1 | - | - | 0 | - |
| - | - | 2267 | 180.1 | - | - | 0 | - |
| - | - | 2650 | 183.1 | - | - | 0 | - |
| - | - | 8385 | 183.1 | - | - | 0 | - |
| - | - | 2740 | 185.1 | - | - | 0 | - |
| - | - | 3.058E+04 | 187.1 | - | - | 0 | - |
| - | - | 1650 | 188.1 | - | - | 0 | - |
| - | - | 1874 | 190.1 | - | - | 0 | - |
| - | - | 1593 | 192.1 | - | - | 0 | - |
| - | - | 2336 | 194.1 | - | - | 0 | - |
| - | - | 2874 | 195.1 | - | - | 0 | - |
| - | - | 4311 | 195.1 | - | - | 0 | - |
| - | - | 2.316E+04 | 197.1 | - | - | 0 | - |
| - | - | 1575 | 198.1 | - | - | 0 | - |
| - | - | 3.12E+04 | 199.1 | - | - | 0 | - |
| 2 | b | 2.579E+05 | 199.1 | 0.0002591 | 1.301 | +1 | 2 |
| - | - | 2.472E+04 | 200.1 | - | - | 0 | - |
| - | - | 4468 | 201.1 | - | - | 0 | - |
| - | - | 1490 | 201.1 | - | - | 0 | - |
| - | - | 1.495E+04 | 207.1 | - | - | 0 | - |
| - | - | 1.491E+04 | 208.1 | - | - | 0 | - |
| - | - | 1649 | 209.1 | - | - | 0 | - |
| - | - | 2765 | 211.1 | - | - | 0 | - |
| - | - | 7156 | 213.1 | - | - | 0 | - |
| - | - | 1864 | 213.2 | - | - | 0 | - |
| - | - | 3.551E+04 | 215.1 | - | - | 0 | - |
| - | - | 3183 | 216.1 | - | - | 0 | - |
| - | - | 3478 | 221.1 | - | - | 0 | - |
| - | - | 8.945E+04 | 225.1 | - | - | 0 | - |
| - | - | 5899 | 225.2 | - | - | 0 | - |
| - | - | 8158 | 226.1 | - | - | 0 | - |
| - | - | 2936 | 227.1 | - | - | 0 | - |
| - | - | 1.154E+05 | 227.1 | - | - | 0 | - |
| - | - | 1.073E+04 | 228.1 | - | - | 0 | - |
| - | - | 9.275E+04 | 228.2 | - | - | 0 | - |
| - | - | 2278 | 229.1 | - | - | 0 | - |
| - | - | 9575 | 229.2 | - | - | 0 | - |
| - | - | 5416 | 231.1 | - | - | 0 | - |
| - | - | 1557 | 238.2 | - | - | 0 | - |
| - | - | 1.47E+04 | 239.1 | - | - | 0 | - |
| - | - | 2273 | 239.2 | - | - | 0 | - |
| - | - | 2598 | 240.1 | - | - | 0 | - |
| - | - | 3116 | 241.2 | - | - | 0 | - |
| - | - | 7617 | 246.2 | - | - | 0 | - |
| - | - | 1423 | 247 | - | - | 0 | - |
| - | - | 1.126E+04 | 249.1 | - | - | 0 | - |
| - | - | 1.284E+04 | 253.2 | - | - | 0 | - |
| - | - | 1807 | 258.2 | - | - | 0 | - |
| - | - | 3609 | 259.1 | - | - | 0 | - |
| - | - | 2978 | 260.1 | - | - | 0 | - |
| - | - | 1866 | 262.2 | - | - | 0 | - |
| - | - | 4175 | 263.2 | - | - | 0 | - |
| - | - | 2568 | 266.1 | - | - | 0 | - |
| - | - | 3.627E+04 | 267.1 | - | - | 0 | - |
| - | - | 4094 | 268.1 | - | - | 0 | - |
| - | - | 1.129E+04 | 268.2 | - | - | 0 | - |
| - | - | 3565 | 269.1 | - | - | 0 | - |
| - | - | 1871 | 269.2 | - | - | 0 | - |
| - | - | 1662 | 275.1 | - | - | 0 | - |
| - | - | 7910 | 276.2 | - | - | 0 | - |
| 13 | y | 4.223E+04 | 277.2 | 0.0002688 | 0.9698 | +1 | 2 |
| - | - | 1816 | 277.2 | - | - | 0 | - |
| - | - | 5333 | 278.2 | - | - | 0 | - |
| - | - | 2916 | 280.2 | - | - | 0 | - |
| - | - | 1711 | 280.2 | - | - | 0 | - |
| - | - | 6330 | 284.1 | - | - | 0 | - |
| - | - | 8394 | 286.2 | - | - | 0 | - |
| - | - | 1883 | 287.2 | - | - | 0 | - |
| - | - | 3204 | 288.1 | - | - | 0 | - |
| - | - | 3830 | 291.2 | - | - | 0 | - |
| 13 | y | 2.687E+05 | 294.2 | 0.0004836 | 1.644 | +1 | 2 |
| - | - | 2434 | 295.1 | - | - | 0 | - |
| - | - | 3.773E+04 | 295.2 | - | - | 0 | - |
| - | - | 2804 | 296.1 | - | - | 0 | - |
| - | - | 3006 | 296.2 | - | - | 0 | - |
| - | - | 6260 | 298.2 | - | - | 0 | - |
| 3 | b | 2.654E+04 | 298.2 | 0.0004945 | 1.658 | +1 | 3 |
| - | - | 3621 | 299.2 | - | - | 0 | - |
| - | - | 9518 | 302.1 | - | - | 0 | - |
| - | - | 2863 | 304.2 | - | - | 0 | - |
| - | - | 9805 | 306.2 | - | - | 0 | - |
| - | - | 2054 | 307.1 | - | - | 0 | - |
| - | - | 6617 | 307.1 | - | - | 0 | - |
| - | - | 1693 | 307.2 | - | - | 0 | - |
| - | - | 2175 | 308.2 | - | - | 0 | - |
| - | - | 3273 | 310.2 | - | - | 0 | - |
| - | - | 7.729E+04 | 314.2 | - | - | 0 | - |
| - | - | 9414 | 315.2 | - | - | 0 | - |
| - | - | 4453 | 318.1 | - | - | 0 | - |
| - | - | 2121 | 319.1 | - | - | 0 | - |
| - | - | 4592 | 320.1 | - | - | 0 | - |
| - | - | 1.116E+04 | 324.1 | - | - | 0 | - |
| - | - | 1.165E+04 | 324.2 | - | - | 0 | - |
| - | - | 2637 | 325.1 | - | - | 0 | - |
| - | - | 1981 | 325.2 | - | - | 0 | - |
| - | - | 5.206E+04 | 326.2 | - | - | 0 | - |
| - | - | 7398 | 327.2 | - | - | 0 | - |
| - | - | 2872 | 334.2 | - | - | 0 | - |
| - | - | 9635 | 336.1 | - | - | 0 | - |
| - | - | 1580 | 337.3 | - | - | 0 | - |
| - | - | 2.621E+04 | 342.1 | - | - | 0 | - |
| - | - | 1826 | 342.2 | - | - | 0 | - |
| - | - | 3271 | 343.1 | - | - | 0 | - |
| - | - | 6232 | 346.1 | - | - | 0 | - |
| - | - | 2524 | 352.3 | - | - | 0 | - |
| - | - | 3.666E+04 | 354.1 | - | - | 0 | - |
| - | - | 5701 | 355.1 | - | - | 0 | - |
| - | - | 1556 | 356.2 | - | - | 0 | - |
| - | - | 3250 | 357.2 | - | - | 0 | - |
| - | - | 1573 | 359.2 | - | - | 0 | - |
| - | - | 4.402E+04 | 364.1 | - | - | 0 | - |
| - | - | 6527 | 365.1 | - | - | 0 | - |
| - | - | 4818 | 367.2 | - | - | 0 | - |
| - | - | 2438 | 368.2 | - | - | 0 | - |
| - | - | 3635 | 375.2 | - | - | 0 | - |
| - | - | 8.794E+04 | 382.1 | - | - | 0 | - |
| - | - | 1.367E+04 | 383.1 | - | - | 0 | - |
| - | - | 1.956E+04 | 383.2 | - | - | 0 | - |
| - | - | 1836 | 384.1 | - | - | 0 | - |
| - | - | 2444 | 384.2 | - | - | 0 | - |
| - | - | 1.275E+04 | 385.2 | - | - | 0 | - |
| - | - | 3760 | 386.2 | - | - | 0 | - |
| - | - | 3854 | 392.1 | - | - | 0 | - |
| - | - | 2852 | 393.2 | - | - | 0 | - |
| 12 | y | 9.056E+04 | 393.2 | 0.0004596 | 1.169 | +1 | 3 |
| - | - | 1.898E+04 | 394.3 | - | - | 0 | - |
| 4 | b | 3270 | 395.2 | 0.001999 | 5.059 | +1 | 4 |
| - | - | 2213 | 395.3 | - | - | 0 | - |
| - | - | 2361 | 397.2 | - | - | 0 | - |
| - | - | 1.531E+04 | 401.2 | - | - | 0 | - |
| - | - | 1933 | 402.2 | - | - | 0 | - |
| - | - | 1760 | 404.1 | - | - | 0 | - |
| - | - | 2412 | 404.2 | - | - | 0 | - |
| - | - | 7956 | 406.2 | - | - | 0 | - |
| - | - | 7741 | 410.1 | - | - | 0 | - |
| - | - | 8347 | 411.2 | - | - | 0 | - |
| 4 | b | 1.208E+05 | 413.2 | 0.0006816 | 1.649 | +1 | 4 |
| - | - | 2.426E+04 | 414.2 | - | - | 0 | - |
| - | - | 1566 | 415.2 | - | - | 0 | - |
| - | - | 9866 | 421.2 | - | - | 0 | - |
| - | - | 1917 | 423.2 | - | - | 0 | - |
| - | - | 3091 | 425.2 | - | - | 0 | - |
| - | - | 2561 | 433.1 | - | - | 0 | - |
| - | - | 4522 | 434.1 | - | - | 0 | - |
| - | - | 2516 | 435.2 | - | - | 0 | - |
| - | - | 1905 | 436.2 | - | - | 0 | - |
| - | - | 3821 | 436.3 | - | - | 0 | - |
| - | - | 2807 | 439.2 | - | - | 0 | - |
| - | - | 1.57E+04 | 439.2 | - | - | 0 | - |
| - | - | 2728 | 440.2 | - | - | 0 | - |
| - | - | 8965 | 441.2 | - | - | 0 | - |
| - | - | 2908 | 441.2 | - | - | 0 | - |
| - | - | 2262 | 442.2 | - | - | 0 | - |
| - | - | 2703 | 449.2 | - | - | 0 | - |
| - | - | 1.972E+04 | 451.2 | - | - | 0 | - |
| - | - | 2104 | 452.2 | - | - | 0 | - |
| - | - | 4304 | 453.2 | - | - | 0 | - |
| - | - | 1.075E+04 | 453.2 | - | - | 0 | - |
| - | - | 1659 | 454.2 | - | - | 0 | - |
| - | - | 1967 | 454.2 | - | - | 0 | - |
| - | - | 3.769E+04 | 454.3 | - | - | 0 | - |
| - | - | 8127 | 455.3 | - | - | 0 | - |
| - | - | 2235 | 463.2 | - | - | 0 | - |
| - | - | 2184 | 467.2 | - | - | 0 | - |
| - | - | 2733 | 467.3 | - | - | 0 | - |
| - | - | 1.083E+05 | 469.2 | - | - | 0 | - |
| - | - | 2.114E+04 | 470.2 | - | - | 0 | - |
| - | - | 8975 | 471.2 | - | - | 0 | - |
| - | - | 9586 | 472.3 | - | - | 0 | - |
| - | - | 1852 | 473.3 | - | - | 0 | - |
| - | - | 2048 | 475.2 | - | - | 0 | - |
| - | - | 3573 | 479.2 | - | - | 0 | - |
| - | - | 1503 | 479.2 | - | - | 0 | - |
| - | - | 2265 | 481.2 | - | - | 0 | - |
| - | - | 1.524E+04 | 482.3 | - | - | 0 | - |
| - | - | 2614 | 483.3 | - | - | 0 | - |
| - | - | 1.676E+04 | 484.3 | - | - | 0 | - |
| - | - | 6085 | 485.3 | - | - | 0 | - |
| 9 | b | 4305 | 490.7 | 0.002676 | 5.453 | +2 | 9 |
| 7 | y | 1725 | 491.7 | 0.0025 | 5.084 | +2 | 8 |
| - | - | 6143 | 492.3 | - | - | 0 | - |
| - | - | 2033 | 497.2 | - | - | 0 | - |
| - | - | 1.071E+04 | 500.3 | - | - | 0 | - |
| 7 | y | 3940 | 500.7 | 0.0004575 | 0.9137 | +2 | 8 |
| - | - | 3631 | 501.3 | - | - | 0 | - |
| 11 | y | 4087 | 504.3 | 0.001268 | 2.515 | +1 | 4 |
| - | - | 3859 | 505.3 | - | - | 0 | - |
| - | - | 1621 | 506.3 | - | - | 0 | - |
| - | - | 2994 | 507.2 | - | - | 0 | - |
| - | - | 1.678E+04 | 510.3 | - | - | 0 | - |
| - | - | 2908 | 511.3 | - | - | 0 | - |
| 5 | b | 2.196E+04 | 512.3 | 0.0007796 | 1.522 | +1 | 5 |
| - | - | 6760 | 513.3 | - | - | 0 | - |
| - | - | 9526 | 520.3 | - | - | 0 | - |
| - | - | 3829 | 521.3 | - | - | 0 | - |
| 11 | y | 6070 | 522.3 | 0.002209 | 4.228 | +1 | 4 |
| - | - | 1737 | 533.2 | - | - | 0 | - |
| 6 | y | 1445 | 535.7 | 0.008674 | 16.19 | +2 | 9 |
| - | - | 1641 | 536.2 | - | - | 0 | - |
| - | - | 1.45E+04 | 538.3 | - | - | 0 | - |
| 10 | b | 6240 | 539.3 | 0.0006356 | 1.179 | +2 | 10 |
| - | - | 5540 | 540.2 | - | - | 0 | - |
| - | - | 1962 | 542.2 | - | - | 0 | - |
| 6 | y | 4745 | 544.3 | 0.0006481 | 1.191 | +2 | 9 |
| - | - | 2948 | 544.8 | - | - | 0 | - |
| - | - | 2237 | 548.2 | - | - | 0 | - |
| - | - | 5169 | 550.2 | - | - | 0 | - |
| - | - | 2022 | 551.2 | - | - | 0 | - |
| - | - | 1360 | 551.3 | - | - | 0 | - |
| - | - | 2433 | 552.2 | - | - | 0 | - |
| - | - | 2159 | 554.2 | - | - | 0 | - |
| - | - | 2465 | 554.8 | - | - | 0 | - |
| - | - | 4570 | 566.2 | - | - | 0 | - |
| - | - | 5.51E+04 | 568.2 | - | - | 0 | - |
| - | - | 1.335E+04 | 569.2 | - | - | 0 | - |
| - | - | 2309 | 569.3 | - | - | 0 | - |
| - | - | 7894 | 570.2 | - | - | 0 | - |
| - | - | 2328 | 571.2 | - | - | 0 | - |
| - | - | 4606 | 578.2 | - | - | 0 | - |
| - | - | 6982 | 580.2 | - | - | 0 | - |
| - | - | 1519 | 580.8 | - | - | 0 | - |
| 6 | b | 1.101E+04 | 581.3 | 0.0001152 | 0.1982 | +1 | 6 |
| - | - | 2872 | 582.3 | - | - | 0 | - |
| - | - | 6527 | 583.3 | - | - | 0 | - |
| - | - | 2254 | 584.3 | - | - | 0 | - |
| 5 | y | 1644 | 584.8 | 0.00252 | 4.309 | +2 | 10 |
| - | - | 4055 | 589.8 | - | - | 0 | - |
| - | - | 1940 | 590.3 | - | - | 0 | - |
| 5 | y | 1.259E+04 | 593.8 | 0.000804 | 1.354 | +2 | 10 |
| - | - | 7909 | 594.3 | - | - | 0 | - |
| 11 | b | 2703 | 594.8 | 0.007632 | 12.83 | +2 | 11 |
| - | - | 2831 | 596.2 | - | - | 0 | - |
| 6 | b | 1.491E+04 | 599.3 | 0.0001843 | 0.3076 | +1 | 6 |
| - | - | 2230 | 600.3 | - | - | 0 | - |
| 10 | y | 6053 | 601.3 | 0.002673 | 4.445 | +1 | 5 |
| - | - | 2470 | 602.3 | - | - | 0 | - |
| 11 | b | 1.577E+04 | 603.8 | 0.0009455 | 1.566 | +2 | 11 |
| - | - | 8189 | 604.3 | - | - | 0 | - |
| - | - | 3795 | 604.3 | - | - | 0 | - |
| - | - | 2289 | 604.8 | - | - | 0 | - |
| - | - | 2.08E+04 | 608.2 | - | - | 0 | - |
| - | - | 5162 | 609.2 | - | - | 0 | - |
| - | - | 1823 | 609.3 | - | - | 0 | - |
| 10 | y | 2.864E+05 | 619.3 | 0.000409 | 0.6604 | +1 | 5 |
| - | - | 9.203E+04 | 620.3 | - | - | 0 | - |
| - | - | 2584 | 620.4 | - | - | 0 | - |
| - | - | 1.381E+04 | 621.4 | - | - | 0 | - |
| - | - | 1661 | 622.3 | - | - | 0 | - |
| - | - | 6548 | 637.3 | - | - | 0 | - |
| - | - | 5435 | 639.3 | - | - | 0 | - |
| - | - | 1844 | 649.3 | - | - | 0 | - |
| 4 | y | 1.678E+04 | 651.3 | 0.0002719 | 0.4175 | +2 | 11 |
| - | - | 1.458E+04 | 651.8 | - | - | 0 | - |
| - | - | 4117 | 652.3 | - | - | 0 | - |
| 12 | b | 6152 | 653.3 | 0.0009793 | 1.499 | +2 | 12 |
| - | - | 3202 | 653.8 | - | - | 0 | - |
| - | - | 5670 | 665.3 | - | - | 0 | - |
| - | - | 3160 | 666.2 | - | - | 0 | - |
| - | - | 1.952E+04 | 667.3 | - | - | 0 | - |
| - | - | 6319 | 668.3 | - | - | 0 | - |
| - | - | 3017 | 673.4 | - | - | 0 | - |
| - | - | 3961 | 677.3 | - | - | 0 | - |
| - | - | 6637 | 679.3 | - | - | 0 | - |
| - | - | 1960 | 680.3 | - | - | 0 | - |
| - | - | 5.254E+04 | 683.3 | - | - | 0 | - |
| - | - | 1.486E+04 | 684.3 | - | - | 0 | - |
| - | - | 2344 | 684.3 | - | - | 0 | - |
| - | - | 6601 | 690.4 | - | - | 0 | - |
| - | - | 2678 | 691.4 | - | - | 0 | - |
| 3 | y | 2133 | 691.8 | 0.003384 | 4.892 | +2 | 12 |
| - | - | 2.417E+04 | 695.3 | - | - | 0 | - |
| - | - | 1.007E+04 | 696.3 | - | - | 0 | - |
| 3 | y | 6.365E+04 | 700.8 | 0.0006109 | 0.8716 | +2 | 12 |
| - | - | 4.151E+04 | 701.3 | - | - | 0 | - |
| - | - | 1.705E+04 | 701.8 | - | - | 0 | - |
| - | - | 2570 | 702.3 | - | - | 0 | - |
| - | - | 1.899E+04 | 707.3 | - | - | 0 | - |
| - | - | 4690 | 708.3 | - | - | 0 | - |
| - | - | 1.212E+04 | 708.4 | - | - | 0 | - |
| 13 | b | 2824 | 708.9 | 0.009314 | 13.14 | +2 | 13 |
| - | - | 1837 | 709.3 | - | - | 0 | - |
| - | - | 5127 | 709.4 | - | - | 0 | - |
| 13 | b | 5931 | 717.4 | 0.0003728 | 0.5196 | +2 | 13 |
| - | - | 2579 | 717.9 | - | - | 0 | - |
| 7 | b | 2.587E+04 | 718.4 | 0.0003601 | 0.5013 | +1 | 7 |
| - | - | 7743 | 719.4 | - | - | 0 | - |
| - | - | 2526 | 720.4 | - | - | 0 | - |
| - | - | 1.152E+04 | 726.4 | - | - | 0 | - |
| - | - | 1.305E+04 | 726.9 | - | - | 0 | - |
| - | - | 2609 | 727.4 | - | - | 0 | - |
| 9 | y | 2.348E+04 | 734.4 | 0.0005656 | 0.7702 | +1 | 6 |
| - | - | 9003 | 735.4 | - | - | 0 | - |
| - | - | 1106 | 736.3 | - | - | 0 | - |
| 7 | b | 2.964E+04 | 736.4 | 1.171E-05 | 0.01591 | +1 | 7 |
| - | - | 9538 | 737.4 | - | - | 0 | - |
| 2 | y | 1.185E+04 | 750.4 | 0.001072 | 1.428 | +2 | 13 |
| - | - | 5980 | 750.9 | - | - | 0 | - |
| - | - | 2828 | 751.4 | - | - | 0 | - |
| - | - | 2269 | 755.9 | - | - | 0 | - |
| - | - | 8762 | 764.3 | - | - | 0 | - |
| - | - | 2858 | 765.3 | - | - | 0 | - |
| - | - | 5915 | 766.3 | - | - | 0 | - |
| - | - | 4935 | 766.4 | - | - | 0 | - |
| - | - | 2833 | 767.3 | - | - | 0 | - |
| - | - | 1934 | 770.4 | - | - | 0 | - |
| - | - | 6492 | 776.3 | - | - | 0 | - |
| - | - | 2162 | 776.9 | - | - | 0 | - |
| - | - | 2234 | 777.1 | - | - | 0 | - |
| - | - | 2356 | 777.3 | - | - | 0 | - |
| - | - | 2692 | 777.4 | - | - | 0 | - |
| - | - | 3835 | 778.4 | - | - | 0 | - |
| - | - | 2162 | 778.9 | - | - | 0 | - |
| - | - | 2772 | 780.3 | - | - | 0 | - |
| - | - | 2317 | 781.3 | - | - | 0 | - |
| - | - | 3012 | 781.9 | - | - | 0 | - |
| - | - | 7.312E+04 | 782.3 | - | - | 0 | - |
| - | - | 2.93E+04 | 783.3 | - | - | 0 | - |
| - | - | 5635 | 784.3 | - | - | 0 | - |
| 0 | Precursor | 2.138E+04 | 790.9 | 0.001139 | 1.44 | +2 | -1 |
| 0 | Precursor | 2.411E+04 | 791.4 | 0.009253 | 11.69 | +2 | -1 |
| - | - | 1.065E+04 | 791.9 | - | - | 0 | - |
| - | - | 4167 | 792.4 | - | - | 0 | - |
| - | - | 3.386E+04 | 794.3 | - | - | 0 | - |
| - | - | 1.146E+04 | 795.3 | - | - | 0 | - |
| - | - | 2913 | 796.3 | - | - | 0 | - |
| - | - | 1.343E+04 | 798.9 | - | - | 0 | - |
| - | - | 1.464E+04 | 799.4 | - | - | 0 | - |
| 0 | Precursor | 1.036E+06 | 799.9 | 0.0007394 | 0.9244 | +2 | -1 |
| - | - | 9.538E+05 | 800.4 | - | - | 0 | - |
| - | - | 4.13E+05 | 800.9 | - | - | 0 | - |
| - | - | 5.554E+04 | 801.4 | - | - | 0 | - |
| - | - | 1876 | 834.3 | - | - | 0 | - |
| - | - | 7173 | 835.4 | - | - | 0 | - |
| - | - | 4712 | 836.4 | - | - | 0 | - |
| - | - | 7310 | 837.4 | - | - | 0 | - |
| - | - | 3468 | 838.4 | - | - | 0 | - |
| 8 | y | 5753 | 845.4 | 0.0004586 | 0.5424 | +1 | 7 |
| 8 | b | 2177 | 847.4 | 0.005246 | 6.19 | +1 | 8 |
| - | - | 2014 | 853.4 | - | - | 0 | - |
| 8 | y | 2.124E+04 | 863.4 | 8.675E-05 | 0.1005 | +1 | 7 |
| - | - | 1.078E+04 | 864.4 | - | - | 0 | - |
| 8 | b | 6.982E+04 | 865.4 | 0.001223 | 1.413 | +1 | 8 |
| - | - | 2.425E+04 | 866.4 | - | - | 0 | - |
| - | - | 7544 | 867.4 | - | - | 0 | - |
| - | - | 2253 | 875.4 | - | - | 0 | - |
| - | - | 4973 | 879.4 | - | - | 0 | - |
| - | - | 2635 | 880.4 | - | - | 0 | - |
| - | - | 1.211E+04 | 881.4 | - | - | 0 | - |
| - | - | 7337 | 882.4 | - | - | 0 | - |
| - | - | 1983 | 883.4 | - | - | 0 | - |
| - | - | 2760 | 891.4 | - | - | 0 | - |
| - | - | 1.067E+04 | 893.4 | - | - | 0 | - |
| - | - | 3815 | 894.4 | - | - | 0 | - |
| - | - | 2838 | 908.5 | - | - | 0 | - |
| - | - | 2.077E+04 | 909.4 | - | - | 0 | - |
| - | - | 9702 | 910.4 | - | - | 0 | - |
| - | - | 1.04E+04 | 922.4 | - | - | 0 | - |
| - | - | 5912 | 923.4 | - | - | 0 | - |
| - | - | 1873 | 937.6 | - | - | 0 | - |
| - | - | 1957 | 940.4 | - | - | 0 | - |
| 9 | b | 1.17E+04 | 962.5 | 0.0006946 | 0.7217 | +1 | 9 |
| - | - | 8025 | 963.5 | - | - | 0 | - |
| - | - | 2722 | 964.5 | - | - | 0 | - |
| - | - | 2883 | 978.5 | - | - | 0 | - |
| 9 | b | 3.51E+05 | 980.5 | 0.000456 | 0.4651 | +1 | 9 |
| - | - | 1.791E+05 | 981.5 | - | - | 0 | - |
| 7 | y | 4.141E+04 | 982.5 | 0.01125 | 11.45 | +1 | 8 |
| - | - | 4301 | 983.5 | - | - | 0 | - |
| - | - | 4877 | 990.4 | - | - | 0 | - |
| - | - | 1944 | 991.4 | - | - | 0 | - |
| 7 | y | 4.151E+04 | 1000 | 0.0005011 | 0.5008 | +1 | 8 |
| - | - | 2.3E+04 | 1001 | - | - | 0 | - |
| - | - | 5142 | 1002 | - | - | 0 | - |
| - | - | 5.493E+04 | 1008 | - | - | 0 | - |
| - | - | 3.136E+04 | 1009 | - | - | 0 | - |
| - | - | 8144 | 1010 | - | - | 0 | - |
| - | - | 3145 | 1021 | - | - | 0 | - |
| 6 | y | 4620 | 1069 | 0.002957 | 2.765 | +1 | 9 |
| 6 | y | 2578 | 1070 | 0.01498 | 13.99 | +1 | 9 |
| 10 | b | 8395 | 1078 | 0.0002469 | 0.2291 | +1 | 10 |
| - | - | 4222 | 1079 | - | - | 0 | - |
| - | - | 1.232E+04 | 1079 | - | - | 0 | - |
| - | - | 5015 | 1081 | - | - | 0 | - |
| 6 | y | 8.376E+04 | 1088 | 0.0005161 | 0.4746 | +1 | 9 |
| - | - | 5.195E+04 | 1089 | - | - | 0 | - |
| - | - | 1.697E+04 | 1090 | - | - | 0 | - |
| - | - | 4.46E+04 | 1107 | - | - | 0 | - |
| - | - | 2.408E+04 | 1108 | - | - | 0 | - |
| - | - | 6424 | 1109 | - | - | 0 | - |
| - | - | 1.084E+04 | 1137 | - | - | 0 | - |
| - | - | 4328 | 1138 | - | - | 0 | - |
| - | - | 2423 | 1155 | - | - | 0 | - |
| - | - | 1.804E+04 | 1179 | - | - | 0 | - |
| - | - | 1.199E+04 | 1180 | - | - | 0 | - |
| - | - | 3268 | 1181 | - | - | 0 | - |
| 5 | y | 2.6E+04 | 1187 | 2.672E-05 | 0.02252 | +1 | 10 |
| - | - | 1.608E+04 | 1188 | - | - | 0 | - |
| 11 | b | 1.129E+04 | 1189 | 0.008135 | 6.845 | +1 | 11 |
| - | - | 5117 | 1190 | - | - | 0 | - |
| - | - | 2451 | 1191 | - | - | 0 | - |
| 11 | b | 1.849E+05 | 1207 | 0.0001343 | 0.1113 | +1 | 11 |
| - | - | 1.115E+05 | 1208 | - | - | 0 | - |
| - | - | 3.329E+04 | 1209 | - | - | 0 | - |
| - | - | 3436 | 1210 | - | - | 0 | - |
| - | - | 1.436E+04 | 1236 | - | - | 0 | - |
| - | - | 7808 | 1237 | - | - | 0 | - |
| - | - | 4019 | 1238 | - | - | 0 | - |
| - | - | 2971 | 1254 | - | - | 0 | - |
| - | - | 2857 | 1260 | - | - | 0 | - |
| - | - | 2.311E+04 | 1278 | - | - | 0 | - |
| - | - | 1.737E+04 | 1279 | - | - | 0 | - |
| - | - | 5677 | 1280 | - | - | 0 | - |
| 4 | y | 2287 | 1284 | 0.001295 | 1.009 | +1 | 11 |
| - | - | 2396 | 1286 | - | - | 0 | - |
| 12 | b | 7553 | 1288 | 2.43E-05 | 0.01887 | +1 | 12 |
| - | - | 3228 | 1289 | - | - | 0 | - |
| 4 | y | 4.829E+04 | 1302 | 0.0007246 | 0.5567 | +1 | 11 |
| - | - | 3.495E+04 | 1303 | - | - | 0 | - |
| - | - | 1.153E+04 | 1304 | - | - | 0 | - |
| 12 | b | 2.045E+05 | 1306 | 0.0004086 | 0.3129 | +1 | 12 |
| - | - | 1.449E+05 | 1307 | - | - | 0 | - |
| - | - | 4.722E+04 | 1308 | - | - | 0 | - |
| - | - | 6224 | 1309 | - | - | 0 | - |
| 3 | y | 2541 | 1383 | 0.002788 | 2.016 | +1 | 12 |
| 3 | y | 5.277E+04 | 1401 | 0.00261 | 1.864 | +1 | 12 |
| - | - | 4.484E+04 | 1402 | - | - | 0 | - |
| - | - | 1.35E+04 | 1403 | - | - | 0 | - |
| 13 | b | 3055 | 1434 | 0.000645 | 0.4499 | +1 | 13 |

m/z Charge Intensity FragmentType MassShift Position
120.08103942871094 0 14436.393
124.02106475830078 0 1075.6091
125.73689270019531 0 992.39233
126.1281967163086 0 2065.019
127.086669921875 0 1316.8269
129.10255432128906 0 574464.75
130.08656311035156 0 6445.569
130.09976196289062 0 3332.751
130.1058349609375 0 33469.92
136.87176513671875 0 1134.0619
138.06643676757812 0 3551.8413
139.08694458007812 0 1128.4615
140.37918090820312 0 1005.7521
141.1022491455078 0 1695.5175
143.0118865966797 0 1079.5679
143.15428161621094 0 1472.1425
146.0712127685547 0 1888.7344
147.1130828857422 0 18415.348
148.94769287109375 0 1446.3867
152.08204650878906 0 1601.5155
159.11375427246094 0 1603.131
163.09799194335938 0 3194.052
164.11849975585938 0 16853.227
166.0614013671875 0 4246.3535
166.0865020751953 0 9029.388 y 13
167.0928955078125 0 2230.424
168.90420532226562 0 1509.1638
168.91212463378906 0 1347.8534
169.09754943847656 0 6250.271
171.14950561523438 0 340647.72 a 1
172.15286254882812 0 28018.176
172.1613006591797 0 1738.9719
173.45030212402344 0 13340.942
179.0930633544922 0 13025.173
180.07705688476562 0 2267.4487
183.11361694335938 0 2650.3467
183.14952087402344 0 8385.0625
185.09268188476562 0 2740.1548
187.10797119140625 0 30580.201
188.1114044189453 0 1649.8716
190.06146240234375 0 1873.9198
192.11306762695312 0 1593.4819
194.09275817871094 0 2335.7637
195.0774383544922 0 2874.367
195.0880889892578 0 4311.257
197.1035614013672 0 23162.355
198.10659790039062 0 1574.7793
199.10797119140625 0 31199.996
199.1443634033203 0 257890.55 b 1
200.147705078125 0 24718.717
201.1232147216797 0 4468.451
201.14987182617188 0 1489.8435
207.0879669189453 0 14948.694
208.07191467285156 0 14914.896
209.07545471191406 0 1649.2238
211.14466857910156 0 2764.6602
213.0870361328125 0 7156.3716
213.1591339111328 0 1864.2573
215.1029052734375 0 35506.15
216.10646057128906 0 3183.061
221.10299682617188 0 3478.4797
225.09844970703125 0 89454.88
225.19650268554688 0 5898.95
226.10203552246094 0 8157.501
227.06712341308594 0 2936.2896
227.1029510498047 0 115389.266
228.1060791015625 0 10732.66
228.17092895507812 0 92754.3
229.118896484375 0 2277.8267
229.17405700683594 0 9574.76
231.1497344970703 0 5416.082
238.1547088623047 0 1557.0619
239.1142578125 0 14700.3125
239.15017700195312 0 2273.1697
240.11801147460938 0 2597.5828
241.1549530029297 0 3116.0764
246.18167114257812 0 7617.457
247.01097106933594 0 1423.3425
249.09866333007812 0 11259.495
253.19149780273438 0 12836.959
258.1595764160156 0 1806.6111
259.14453125 0 3608.6858
260.12750244140625 0 2977.5942
262.16668701171875 0 1866.2745
263.1868896484375 0 4175.268
266.12567138671875 0 2567.798
267.10894775390625 0 36266.734
268.112060546875 0 4094.0237
268.1658020019531 0 11287.765
269.1498718261719 0 3564.7659
269.167236328125 0 1870.73
275.13836669921875 0 1662.3711
276.1713562011719 0 7909.853
277.1549377441406 0 42227.137 y Ammonia loss 12
277.17120361328125 0 1815.7327
278.1587829589844 0 5332.6094
280.16546630859375 0 2916.1973
280.2018737792969 0 1710.7545
284.1243896484375 0 6330.376
286.1766052246094 0 8394.443
287.1810302734375 0 1882.9281
288.1454162597656 0 3204.071
291.1820373535156 0 3830.018
294.18170166015625 0 268693.1 y 12
295.1038818359375 0 2434.4958
295.1848449707031 0 37726.387
296.1236572265625 0 2804.2664
296.1875915527344 0 3006.0938
298.17669677734375 0 6260.079
298.2130126953125 0 26540.607 b 2
299.2164001464844 0 3620.8245
302.1348876953125 0 9518.186
304.16571044921875 0 2863.3245
306.1564025878906 0 9804.865
307.09197998046875 0 2054.3438
307.1404113769531 0 6616.825
307.15869140625 0 1693.0801
308.1618957519531 0 2175.4558
310.1514587402344 0 3273.282
314.1715087890625 0 77287.13
315.17486572265625 0 9414.055
318.1199645996094 0 4452.509
319.1040954589844 0 2120.7937
320.1357116699219 0 4591.588
324.11962890625 0 11161.571
324.167236328125 0 11650.511
325.10345458984375 0 2636.6724
325.1690979003906 0 1981.1432
326.1715087890625 0 52055.844
327.1748962402344 0 7397.5903
334.1507873535156 0 2872.2305
336.1312561035156 0 9635.475
337.2958984375 0 1579.5896
342.13006591796875 0 26211.525
342.16558837890625 0 1826.4177
343.1332702636719 0 3270.6357
346.1150817871094 0 6231.5645
352.2590026855469 0 2524.0276
354.14111328125 0 36661.12
355.14434814453125 0 5700.7334
356.18145751953125 0 1556.4484
357.2117614746094 0 3249.568
359.15606689453125 0 1573.4954
364.1253967285156 0 44022.215
365.1290588378906 0 6526.6724
367.23406982421875 0 4817.7925
368.2176513671875 0 2438.3433
375.2398681640625 0 3635.3486
382.1361389160156 0 87942.56
383.1391296386719 0 13665.853
383.1927185058594 0 19562.057
384.1414489746094 0 1836.0486
384.1973571777344 0 2443.6008
385.2449951171875 0 12752.97
386.2470397949219 0 3760.1167
392.1217346191406 0 3853.835
393.18914794921875 0 2851.9087
393.2500915527344 0 90562.55 y 11
394.2535400390625 0 18976.469
395.23089599609375 0 3270.1357 b Water loss 3
395.255126953125 0 2213.453
397.1808776855469 0 2361.4656
401.2034606933594 0 15311.24
402.2080993652344 0 1933.1481
404.12255859375 0 1759.9138
404.1555480957031 0 2412.3186
406.2088928222656 0 7956.2153
410.13092041015625 0 7740.9067
411.19927978515625 0 8347.444
413.2401428222656 0 120830.61 b 3
414.24310302734375 0 24261.059
415.2444763183594 0 1566.1893
421.1836853027344 0 9866.453
423.1874084472656 0 1916.6606
425.2160949707031 0 3091.4912
433.146728515625 0 2560.7798
434.1322326660156 0 4522.1357
435.1986083984375 0 2515.591
436.18255615234375 0 1904.8016
436.25604248046875 0 3821.4658
439.1618957519531 0 2806.8853
439.1943359375 0 15696.959
440.19281005859375 0 2728.4138
441.19854736328125 0 8964.832
441.2313537597656 0 2907.7102
442.19964599609375 0 2261.9026
449.1792907714844 0 2703.3247
451.1578369140625 0 19719.98
452.160888671875 0 2104.1257
453.1634826660156 0 4304.0137
453.2101135253906 0 10752.776
454.16424560546875 0 1659.1919
454.21307373046875 0 1967.243
454.26678466796875 0 37688.086
455.2688293457031 0 8127.293
463.19586181640625 0 2235.324
467.1896057128906 0 2183.8494
467.2850646972656 0 2732.9316
469.16851806640625 0 108290.14
470.17138671875 0 21142.682
471.1733703613281 0 8974.669
472.2772521972656 0 9585.663
473.28076171875 0 1851.7041
475.2304992675781 0 2047.5765
479.1535949707031 0 3573.1123
479.1909484863281 0 1502.5045
481.2048034667969 0 2264.564
482.2615661621094 0 15240.564
483.26666259765625 0 2614.3
484.3138427734375 0 16764.629
485.3169860839844 0 6085.0757
490.7351379394531 0 4304.706 b 8
491.7325744628906 0 1725.4822 y Water loss 6
492.2573547363281 0 6142.6196
497.1631164550781 0 2032.6937
500.27215576171875 0 10712.523
500.7408142089844 0 3940.0256 y 6
501.2784118652344 0 3630.5366
504.2829284667969 0 4086.8157 y Water loss 10
505.2789611816406 0 3858.562
506.28143310546875 0 1621.2386
507.1824645996094 0 2993.7336
510.26812744140625 0 16779.707
511.2702331542969 0 2908.389
512.3086547851562 0 21959.502 b 4
513.310302734375 0 6759.5586
520.2528076171875 0 9525.834
521.2557373046875 0 3829.0632
522.29443359375 0 6070.2886 y 10
533.2057495117188 0 1736.9014
535.7517700195312 0 1445.1614 y Ammonia loss 5
536.1797485351562 0 1641.3411
538.2627563476562 0 14497.475
539.2648315429688 0 6239.5337 b 9
540.2474365234375 0 5540.05
542.2445068359375 0 1961.7511
544.2570190429688 0 4745.1895 y 5
544.7589721679688 0 2947.6396
548.2092895507812 0 2236.8945
550.2254028320312 0 5168.5957
551.22705078125 0 2022.3712
551.2805786132812 0 1359.9385
552.232666015625 0 2433.027
554.2487182617188 0 2159.3064
554.7528076171875 0 2464.6267
566.2207641601562 0 4569.7603
568.236572265625 0 55096.88
569.239013671875 0 13353.533
569.2887573242188 0 2308.6152
570.2412719726562 0 7893.57
571.2440185546875 0 2328.3362
578.2210693359375 0 4605.61
580.23583984375 0 6981.5913
580.7866821289062 0 1519.1874
581.3292236328125 0 11009.982 b Water loss 5
582.3336791992188 0 2871.7246
583.3262939453125 0 6527.4194
584.3251342773438 0 2254.1711
584.7827758789062 0 1644.2435 y Water loss 4
589.7889404296875 0 4055.0151
590.29150390625 0 1940.1953
593.7913818359375 0 12585.489 y 4
594.292724609375 0 7909.411
594.787841796875 0 2702.7788 b Water loss 10
596.2301025390625 0 2831.472
599.340087890625 0 14905.838 b 5
600.3453979492188 0 2230.444
601.3370971679688 0 6052.6963 y Water loss 9
602.3369750976562 0 2469.7876
603.7864379882812 0 15772.135 b 10
604.2874145507812 0 8189.346
604.34716796875 0 3794.604
604.7897338867188 0 2288.7463
608.2318725585938 0 20797.432
609.234130859375 0 5161.9697
609.33935546875 0 1823.042
619.3453979492188 0 286400.78 y 9
620.3485107421875 0 92026.81
620.4077758789062 0 2584.214
621.3507690429688 0 13812.211
622.2805786132812 0 1661.3654
637.3290405273438 0 6547.9556
639.3181762695312 0 5435.171
649.3051147460938 0 1843.5853
651.3043212890625 0 16782.104 y 3
651.8060913085938 0 14576.81
652.3064575195312 0 4116.549
653.3206787109375 0 6151.695 b 11
653.8211059570312 0 3202.005
665.2527465820312 0 5670.4087
666.24267578125 0 3159.848
667.3045043945312 0 19519.676
668.3084106445312 0 6319.3643
673.367431640625 0 3017.3157
677.252197265625 0 3960.787
679.3047485351562 0 6637.4277
680.3073120117188 0 1959.6421
683.263671875 0 52541.824
684.2660522460938 0 14858.086
684.3273315429688 0 2344.0745
690.3953857421875 0 6601.035
691.3901977539062 0 2678.0107
691.82958984375 0 2132.6091 y Water loss 2
695.263916015625 0 24173.219
696.2666015625 0 10072.52
700.8388671875 0 63649.24 y 2
701.3399047851562 0 41513.457
701.841796875 0 17052.29
702.3418579101562 0 2569.848
707.3008422851562 0 18990.762
708.300537109375 0 4690.3623
708.4038696289062 0 12120.576
708.8632202148438 0 2823.8728 b Ammonia loss 12
709.307373046875 0 1837.4897
709.4085083007812 0 5126.7915
717.3675537109375 0 5931.2773 b 12
717.8692626953125 0 2579.3499
718.3886108398438 0 25874.215 b Water loss 6
719.391845703125 0 7743.4043
720.3917846679688 0 2525.6638
726.373291015625 0 11523.363
726.8736572265625 0 13049.231
727.3760986328125 0 2608.623
734.3724975585938 0 23476.693 y 8
735.3751831054688 0 9003.407
736.32861328125 0 1105.6975
736.3988037109375 0 29641.373 b 6
737.4025268554688 0 9538.266
750.37353515625 0 11845.963 y 1
750.8746948242188 0 5980.1157
751.3739013671875 0 2827.6213
755.8656005859375 0 2269.0693
764.322265625 0 8761.684
765.3204345703125 0 2858.4526
766.3377075195312 0 5914.594
766.37109375 0 4935.361
767.339599609375 0 2832.8896
770.4022827148438 0 1934.0872
776.3218383789062 0 6491.5986
776.9009399414062 0 2161.8208
777.0802001953125 0 2234.2039
777.3246459960938 0 2356.095
777.406005859375 0 2692.1252
778.3795776367188 0 3834.6172
778.8806762695312 0 2162.0112
780.3179931640625 0 2771.6057
781.3206787109375 0 2317.3147
781.8939819335938 0 3011.5813
782.3322143554688 0 73123.58
783.3349609375 0 29296.512
784.336669921875 0 5634.9927
790.9025268554688 0 21383.137 Precursor Water loss
791.4026489257812 0 24109.117 Precursor Ammonia loss
791.9033203125 0 10647.42
792.3970947265625 0 4166.537
794.3324584960938 0 33857.848
795.335693359375 0 11462.04
796.340087890625 0 2913.1118
798.9016723632812 0 13425.877
799.4033813476562 0 14636.385
799.9074096679688 0 1036222.75 Precursor
800.4089965820312 0 953833.8
800.9100341796875 0 413013.75
801.4118041992188 0 55543.082
834.2508544921875 0 1875.6514
835.3974609375 0 7172.7305
836.3956298828125 0 4711.6943
837.4463500976562 0 7310.191
838.4483642578125 0 3467.5413
845.4044189453125 0 5752.6475 y Water loss 7
847.4255981445312 0 2177.3171 b Water loss 7
853.4075927734375 0 2014.3418
863.4146118164062 0 21237.54 y 7
864.4168701171875 0 10783.438
865.440185546875 0 69818.41 b 7
866.444580078125 0 24249.023
867.4466552734375 0 7543.6514
875.3907470703125 0 2253.4114
879.3836669921875 0 4972.857
880.3861694335938 0 2634.9287
881.3944091796875 0 12108.329
882.3970336914062 0 7336.926
883.44970703125 0 1983.4199
891.3717651367188 0 2760.271
893.39892578125 0 10665.729
894.4066162109375 0 3815.114
908.4844970703125 0 2838.473
909.3594360351562 0 20766.482
910.3629760742188 0 9702.29
922.4270629882812 0 10400.079
923.4314575195312 0 5911.9844
937.64501953125 0 1872.8922
940.4407958984375 0 1957.2228
962.4570922851562 0 11695.281 b Water loss 8
963.4598388671875 0 8025.368
964.4627075195312 0 2722.052
978.4563598632812 0 2883.165
980.4678955078125 0 350963.47 b 8
981.4713134765625 0 179125.38
982.47412109375 0 41414.652 y Water loss 6
983.4734497070312 0 4300.741
990.416015625 0 4877.3467
991.4238891601562 0 1944.0946
1000.4739379882812 0 41511.457 y 6
1001.4765625 0 23000.23
1002.4813232421875 0 5142.191
1008.4271850585938 0 54925.508
1009.430908203125 0 31364.246
1010.4331665039062 0 8143.7773
1021.4906616210938 0 3145.0564
1069.491943359375 0 4619.569 y Water loss 5
1070.493896484375 0 2577.7915 y Ammonia loss 5
1077.5213623046875 0 8395.085 b 9
1078.5224609375 0 4221.9434
1079.4996337890625 0 12318.93
1080.5057373046875 0 5014.9434
1087.5059814453125 0 83762.14 y 5
1088.5089111328125 0 51945.004
1089.505859375 0 16974.99
1107.495849609375 0 44604.668
1108.4986572265625 0 24084.166
1109.4974365234375 0 6424.2817
1136.5224609375 0 10843.624
1137.523193359375 0 4328.0483
1154.5379638671875 0 2422.801
1178.5689697265625 0 18044.635
1179.571533203125 0 11988.657
1180.5723876953125 0 3267.5837
1186.5738525390625 0 26000.295 y 4
1187.5753173828125 0 16080.11
1188.561279296875 0 11291.46 b Water loss 10
1189.562255859375 0 5117.3916
1190.5582275390625 0 2451.4329
1206.5638427734375 0 184857.52 b 10
1207.566650390625 0 111476.37
1208.5689697265625 0 33292.66
1209.57421875 0 3435.5574
1235.5894775390625 0 14356.142
1236.59228515625 0 7808.2183
1237.591552734375 0 4018.9832
1253.611572265625 0 2970.851
1259.625732421875 0 2856.6504
1277.63623046875 0 23106.756
1278.6414794921875 0 17371.242
1279.642822265625 0 5676.8003
1283.591552734375 0 2286.6475 y Water loss 3
1285.5855712890625 0 2396.1785
1287.62158203125 0 7553.1616 b Water loss 11
1288.6199951171875 0 3228.2083
1301.60009765625 0 48288.18 y 3
1302.6033935546875 0 34951.145
1303.6080322265625 0 11526.173
1305.6317138671875 0 204508.17 b 11
1306.6337890625 0 144890.73
1307.6370849609375 0 47218.684
1308.6365966796875 0 6223.515
1382.6558837890625 0 2541.3894 y Water loss 2
1400.6666259765625 0 52772.06 y 2
1401.671875 0 44837.97
1402.6756591796875 0 13503.042
1433.7264404296875 0 3054.9678 b 12

Spectrum Details

|  |  |
| --- | --- |
| Matched peaks? Matched peaks The total absolute number of peaks matched. Additionally in brackets the total fraction of peaks matched and the total number of peaks is shown. | 61 (13.23% of 461) |
| FDR? FDR The false discovery rate estimated for this peptide. It is calculated by matching all theoretical fragments with a non-integer shift with the raw peaks for this spectrum. This is done with 40 different shifts. The resulting percentage is the average number of annotated peaks over the number of annotated peaks with the correct spectrum. | 1.01% |
| Satellite FDR? Satellite FDR See the FDR for details on its calculation. This satellite ion specific FDR only contains the satellite ions (d/w) for I/L/J positions. | - |
| PSM Score? PSM Score The PSM Score as given by Hecklib to this annotated spectrum. It is shown with three significant figures. | 654 |

#### Spectrum 7048? Spectrum 7048 The raw spectrum of this peptide as annotated by Hecklib. The fragments are coloured according to ion type (see legend). Any peaks with a star '\*' as text can be hovered over to see the full details, first the ion type second the mass shift type. By hovering over the amino acids in the peptide or ions in the legend the corresponding peaks are highlighted. By toggling the 'Unassigned' label you can turn the background (unassigned) peaks on or off in the plot. By updating the slider in the Ion legend you can update the spectrum to only show the top X% of the peaks with labels. The top X% means any peak that is within X% of the highest intensity. By dragging in the spectrum you can zoom in to a specific part of the spectrum and use 'Zoom Out' to get back to the original zoom level. The annotation of the spectrum is based on the given sequence in the peptides file and is done with different software so inconsistencies are likely. The peaks are annotated based on the given sequence, with 20 ppm tolerance.

Copy Data

##### Spectrum 7048 (TSV)

###### Preview

```
Loading example...
```

*Click on the button to copy the data to your clipboard.*

Mz MinMz MaxIntensity Max

WidthHeightPeptide font sizePeptide stroke widthSpectrum font sizeSpectrum stroke widthCompact peptide

Ion legend

wxyz

abcd

OtherUnassignedIon^Charge^_Position_Show for top:%

VVVDVSHEDPEVKF

02.02e+54.03e+56.05e+58.07e+5

Zoom Out

y^+1^_1_y^+2^_3_c^+1^_2_y^+1^_2_z^+1^_2_y^+1^_2_y^+2^_5_c^+1^_3_w^+1^_3_y^+1^_3_z^+1^_3_y^+1^_3_c^+1^_4_w^+1^_4_y^+3^_12_y^+3^_13_y^+2^_8_y^+1^_4_z^+1^_4_y^+1^_4_c^+1^_5_y^+2^_9_y^+2^_9_z^+2^_9_y^+2^_9_w^+1^_5_w^+2^_10_y^+2^_10_z^+2^_10_y^+2^_10_y^+1^_5_c^+2^_11_c^+1^_6_y^+1^_5_y^+2^_11_y^+2^_11_z^+2^_11_y^+2^_11_w^+2^_12_y^+2^_12_y^+2^_12_z^+2^_12_z^+1^_6_y^+2^_12_c^+2^_13_z^+1^_6_c^+2^_13_z^+2^_13_y^+1^_6_w^+2^_13_z^+2^_13_y^+2^_13_c^+1^_7_w^+1^_7_y^+1^_7_y^+1^_7_z^+1^_7_y^+1^_7_c^+1^_8_z^+1^_8_y^+1^_8_y^+1^_8_z^+1^_8_y^+1^_8_y^+1^_9_z^+1^_9_y^+1^_9_c^+1^_10_w^+1^_10_y^+1^_10_z^+1^_10_y^+1^_10_c^+1^_11_w^+1^_11_y^+1^_11_z^+1^_11_y^+1^_11_c^+1^_12_z^+1^_12_c^+1^_13_z^+1^_13_

0776155123273103

Fragment Matches Table

Show background peaks

| Position | Ion type | Intensity | mz Theoretical | mz Error (Th) | mz Error (ppm) | Charge | Series Number |
| --- | --- | --- | --- | --- | --- | --- | --- |
| - | - | 5499 | 120.1 | - | - | 0 | - |
| - | - | 404.4 | 122.1 | - | - | 0 | - |
| - | - | 368.4 | 124 | - | - | 0 | - |
| - | - | 2805 | 127.1 | - | - | 0 | - |
| - | - | 6276 | 128.1 | - | - | 0 | - |
| - | - | 5.381E+04 | 129.1 | - | - | 0 | - |
| - | - | 3636 | 130.1 | - | - | 0 | - |
| - | - | 459.4 | 131.1 | - | - | 0 | - |
| - | - | 560.5 | 133.1 | - | - | 0 | - |
| - | - | 881.7 | 147 | - | - | 0 | - |
| - | - | 1411 | 147.1 | - | - | 0 | - |
| - | - | 1335 | 148.1 | - | - | 0 | - |
| - | - | 985.4 | 149 | - | - | 0 | - |
| - | - | 1716 | 149 | - | - | 0 | - |
| - | - | 543.2 | 156.1 | - | - | 0 | - |
| 14 | y | 6922 | 166.1 | 0.0005217 | 3.141 | +1 | 1 |
| - | - | 839.9 | 167.1 | - | - | 0 | - |
| - | - | 2658 | 170.1 | - | - | 0 | - |
| - | - | 2.006E+05 | 171.1 | - | - | 0 | - |
| - | - | 596.3 | 172.1 | - | - | 0 | - |
| - | - | 1.983E+04 | 172.2 | - | - | 0 | - |
| - | - | 451.8 | 172.2 | - | - | 0 | - |
| - | - | 630.1 | 173.2 | - | - | 0 | - |
| - | - | 1565 | 173.5 | - | - | 0 | - |
| - | - | 533.7 | 174.1 | - | - | 0 | - |
| - | - | 1762 | 179.1 | - | - | 0 | - |
| - | - | 909.3 | 183.1 | - | - | 0 | - |
| - | - | 691.6 | 185.2 | - | - | 0 | - |
| - | - | 2887 | 187.1 | - | - | 0 | - |
| - | - | 937 | 189.1 | - | - | 0 | - |
| 12 | y | 1.297E+04 | 197.1 | 0.0006046 | 3.067 | +2 | 3 |
| - | - | 1.478E+04 | 198.1 | - | - | 0 | - |
| - | - | 1.664E+05 | 199.1 | - | - | 0 | - |
| - | - | 1.846E+04 | 200.1 | - | - | 0 | - |
| - | - | 1152 | 201.2 | - | - | 0 | - |
| - | - | 959 | 211.1 | - | - | 0 | - |
| - | - | 2923 | 212.2 | - | - | 0 | - |
| - | - | 1187 | 213.2 | - | - | 0 | - |
| - | - | 7304 | 215.1 | - | - | 0 | - |
| - | - | 535.5 | 215.1 | - | - | 0 | - |
| - | - | 3637 | 216.1 | - | - | 0 | - |
| - | - | 989.4 | 216.2 | - | - | 0 | - |
| 2 | c | 1337 | 216.2 | 0.001222 | 5.651 | +1 | 2 |
| - | - | 761.4 | 217.1 | - | - | 0 | - |
| - | - | 7784 | 221.1 | - | - | 0 | - |
| - | - | 774.4 | 222.1 | - | - | 0 | - |
| - | - | 1943 | 225 | - | - | 0 | - |
| - | - | 931 | 225.1 | - | - | 0 | - |
| - | - | 1765 | 227.1 | - | - | 0 | - |
| - | - | 1.386E+04 | 228.2 | - | - | 0 | - |
| - | - | 1539 | 229.2 | - | - | 0 | - |
| - | - | 1.06E+04 | 232.1 | - | - | 0 | - |
| - | - | 1444 | 233.1 | - | - | 0 | - |
| - | - | 7862 | 233.2 | - | - | 0 | - |
| - | - | 1006 | 234.2 | - | - | 0 | - |
| - | - | 532.8 | 235.9 | - | - | 0 | - |
| - | - | 8546 | 239.1 | - | - | 0 | - |
| - | - | 1523 | 240.1 | - | - | 0 | - |
| - | - | 616 | 241.2 | - | - | 0 | - |
| - | - | 3740 | 253.2 | - | - | 0 | - |
| - | - | 1288 | 260.2 | - | - | 0 | - |
| - | - | 1.308E+04 | 270.2 | - | - | 0 | - |
| - | - | 2799 | 271.2 | - | - | 0 | - |
| - | - | 1758 | 272.2 | - | - | 0 | - |
| - | - | 3882 | 275.1 | - | - | 0 | - |
| - | - | 778.6 | 276.2 | - | - | 0 | - |
| 13 | y | 1994 | 277.2 | 0.000635 | 2.291 | +1 | 2 |
| 13 | z | 1.742E+04 | 278.2 | 0.0007445 | 2.677 | +1 | 2 |
| - | - | 3793 | 279.2 | - | - | 0 | - |
| - | - | 709.3 | 281.1 | - | - | 0 | - |
| - | - | 1104 | 286.2 | - | - | 0 | - |
| 13 | y | 4.227E+04 | 294.2 | 0.0009719 | 3.304 | +1 | 2 |
| - | - | 5627 | 295.1 | - | - | 0 | - |
| - | - | 6780 | 295.2 | - | - | 0 | - |
| - | - | 1442 | 296.1 | - | - | 0 | - |
| - | - | 1564 | 297.2 | - | - | 0 | - |
| - | - | 2.219E+04 | 298.2 | - | - | 0 | - |
| - | - | 2609 | 299.1 | - | - | 0 | - |
| - | - | 4020 | 299.2 | - | - | 0 | - |
| - | - | 736.2 | 302.1 | - | - | 0 | - |
| 10 | y | 2207 | 310.2 | 0.001266 | 4.082 | +2 | 5 |
| - | - | 7182 | 314.2 | - | - | 0 | - |
| - | - | 1.306E+04 | 314.2 | - | - | 0 | - |
| - | - | 1105 | 315.2 | - | - | 0 | - |
| 3 | c | 3.141E+04 | 315.2 | 0.0008619 | 2.734 | +1 | 3 |
| - | - | 5781 | 316.2 | - | - | 0 | - |
| - | - | 712.2 | 317.2 | - | - | 0 | - |
| - | - | 2117 | 326.2 | - | - | 0 | - |
| - | - | 4310 | 331.2 | - | - | 0 | - |
| - | - | 1095 | 352.2 | - | - | 0 | - |
| - | - | 741.4 | 354.2 | - | - | 0 | - |
| - | - | 1167 | 355.1 | - | - | 0 | - |
| 12 | w | 9051 | 362.2 | 0.001124 | 3.104 | +1 | 3 |
| - | - | 1562 | 363.2 | - | - | 0 | - |
| - | - | 3583 | 368.2 | - | - | 0 | - |
| - | - | 5461 | 369.1 | - | - | 0 | - |
| - | - | 698.7 | 369.2 | - | - | 0 | - |
| - | - | 2290 | 370.1 | - | - | 0 | - |
| - | - | 924.8 | 370.2 | - | - | 0 | - |
| - | - | 1221 | 371.2 | - | - | 0 | - |
| 12 | y | 1781 | 376.2 | 0.00171 | 4.544 | +1 | 3 |
| - | - | 1098 | 377.2 | - | - | 0 | - |
| 12 | z | 3.259E+04 | 377.2 | 0.001026 | 2.719 | +1 | 3 |
| - | - | 9263 | 378.2 | - | - | 0 | - |
| - | - | 935.5 | 379.2 | - | - | 0 | - |
| - | - | 1137 | 381.2 | - | - | 0 | - |
| - | - | 1150 | 382.1 | - | - | 0 | - |
| - | - | 1052 | 382.2 | - | - | 0 | - |
| - | - | 1010 | 383.2 | - | - | 0 | - |
| - | - | 4384 | 385.2 | - | - | 0 | - |
| - | - | 1.275E+04 | 386.3 | - | - | 0 | - |
| - | - | 4564 | 387.3 | - | - | 0 | - |
| - | - | 758.6 | 388.3 | - | - | 0 | - |
| - | - | 813.3 | 391.7 | - | - | 0 | - |
| - | - | 558.1 | 392.2 | - | - | 0 | - |
| 12 | y | 5.756E+04 | 393.2 | 0.001314 | 3.342 | +1 | 3 |
| - | - | 1.435E+04 | 394.3 | - | - | 0 | - |
| - | - | 1121 | 395.2 | - | - | 0 | - |
| - | - | 1844 | 395.3 | - | - | 0 | - |
| - | - | 592.7 | 399.2 | - | - | 0 | - |
| - | - | 1954 | 401.2 | - | - | 0 | - |
| - | - | 537.2 | 411.5 | - | - | 0 | - |
| - | - | 795.5 | 412.2 | - | - | 0 | - |
| - | - | 3.318E+04 | 413.2 | - | - | 0 | - |
| - | - | 7363 | 414.2 | - | - | 0 | - |
| - | - | 1071 | 415.3 | - | - | 0 | - |
| - | - | 1271 | 418.2 | - | - | 0 | - |
| - | - | 880.2 | 419.2 | - | - | 0 | - |
| 4 | c | 1.73E+04 | 430.3 | 0.001507 | 3.502 | +1 | 4 |
| - | - | 4055 | 431.3 | - | - | 0 | - |
| - | - | 1057 | 432.3 | - | - | 0 | - |
| - | - | 808.3 | 434.2 | - | - | 0 | - |
| - | - | 642.1 | 441.2 | - | - | 0 | - |
| - | - | 669.9 | 441.2 | - | - | 0 | - |
| - | - | 1390 | 441.3 | - | - | 0 | - |
| 11 | w | 6.003E+04 | 447.3 | 0.001736 | 3.881 | +1 | 4 |
| - | - | 1.635E+04 | 448.3 | - | - | 0 | - |
| - | - | 2502 | 449.3 | - | - | 0 | - |
| - | - | 2828 | 454.3 | - | - | 0 | - |
| - | - | 1754 | 455.3 | - | - | 0 | - |
| - | - | 664.9 | 466.3 | - | - | 0 | - |
| 3 | y | 3727 | 467.6 | 0.001939 | 4.147 | +3 | 12 |
| - | - | 3725 | 467.9 | - | - | 0 | - |
| - | - | 2154 | 468.2 | - | - | 0 | - |
| - | - | 1553 | 469.2 | - | - | 0 | - |
| - | - | 1993 | 469.3 | - | - | 0 | - |
| - | - | 2433 | 470.3 | - | - | 0 | - |
| - | - | 1840 | 471.3 | - | - | 0 | - |
| - | - | 587.4 | 472.3 | - | - | 0 | - |
| - | - | 1776 | 482.3 | - | - | 0 | - |
| - | - | 831.4 | 482.3 | - | - | 0 | - |
| - | - | 1743 | 484.3 | - | - | 0 | - |
| - | - | 9102 | 484.3 | - | - | 0 | - |
| - | - | 1.436E+04 | 485.3 | - | - | 0 | - |
| - | - | 5673 | 486.3 | - | - | 0 | - |
| - | - | 990.2 | 487.3 | - | - | 0 | - |
| - | - | 2211 | 490.7 | - | - | 0 | - |
| - | - | 1002 | 494.3 | - | - | 0 | - |
| - | - | 1467 | 500.3 | - | - | 0 | - |
| 2 | y | 2646 | 500.6 | 0.001137 | 2.272 | +3 | 13 |
| 7 | y | 4829 | 500.7 | 0.001129 | 2.254 | +2 | 8 |
| - | - | 1715 | 500.9 | - | - | 0 | - |
| - | - | 3553 | 501.2 | - | - | 0 | - |
| 11 | y | 698.7 | 504.3 | 0.0003494 | 0.6928 | +1 | 4 |
| - | - | 1578 | 504.7 | - | - | 0 | - |
| - | - | 1604 | 505.2 | - | - | 0 | - |
| 11 | z | 2828 | 506.3 | 0.001951 | 3.853 | +1 | 4 |
| - | - | 736.4 | 507.3 | - | - | 0 | - |
| - | - | 1274 | 509.3 | - | - | 0 | - |
| - | - | 2191 | 511.3 | - | - | 0 | - |
| - | - | 4.42E+04 | 512.3 | - | - | 0 | - |
| - | - | 1.453E+04 | 513.3 | - | - | 0 | - |
| - | - | 657.3 | 513.4 | - | - | 0 | - |
| - | - | 2450 | 514.3 | - | - | 0 | - |
| - | - | 1014 | 518.3 | - | - | 0 | - |
| - | - | 1381 | 520.3 | - | - | 0 | - |
| 11 | y | 3400 | 522.3 | 0.001293 | 2.476 | +1 | 4 |
| - | - | 1142 | 523.3 | - | - | 0 | - |
| - | - | 1355 | 528.3 | - | - | 0 | - |
| 5 | c | 8844 | 529.3 | 0.001574 | 2.974 | +1 | 5 |
| - | - | 3595 | 530.3 | - | - | 0 | - |
| - | - | 1394 | 532.3 | - | - | 0 | - |
| - | - | 1554 | 533.3 | - | - | 0 | - |
| - | - | 1.635E+04 | 533.6 | - | - | 0 | - |
| - | - | 1.505E+04 | 533.9 | - | - | 0 | - |
| - | - | 1139 | 534.2 | - | - | 0 | - |
| - | - | 9360 | 534.3 | - | - | 0 | - |
| - | - | 831.5 | 534.3 | - | - | 0 | - |
| - | - | 2900 | 534.6 | - | - | 0 | - |
| 6 | y | 1351 | 535.3 | 0.009511 | 17.77 | +2 | 9 |
| 6 | y | 2812 | 535.7 | 0.002143 | 4 | +2 | 9 |
| 6 | z | 4208 | 536.2 | 0.001404 | 2.619 | +2 | 9 |
| - | - | 3364 | 536.7 | - | - | 0 | - |
| - | - | 1160 | 537.3 | - | - | 0 | - |
| - | - | 3512 | 538.3 | - | - | 0 | - |
| - | - | 1269 | 539.3 | - | - | 0 | - |
| 6 | y | 9309 | 544.3 | 0.001747 | 3.209 | +2 | 9 |
| - | - | 4606 | 544.8 | - | - | 0 | - |
| - | - | 1756 | 545.3 | - | - | 0 | - |
| - | - | 783.2 | 545.8 | - | - | 0 | - |
| - | - | 1202 | 548.2 | - | - | 0 | - |
| - | - | 1127 | 548.7 | - | - | 0 | - |
| - | - | 922.9 | 549.3 | - | - | 0 | - |
| - | - | 1177 | 550.3 | - | - | 0 | - |
| - | - | 1191 | 554.3 | - | - | 0 | - |
| - | - | 694.5 | 554.8 | - | - | 0 | - |
| - | - | 586.9 | 555.2 | - | - | 0 | - |
| - | - | 621.3 | 560.3 | - | - | 0 | - |
| - | - | 1257 | 563.3 | - | - | 0 | - |
| - | - | 2905 | 568.2 | - | - | 0 | - |
| - | - | 861.4 | 571.3 | - | - | 0 | - |
| - | - | 740.1 | 572.3 | - | - | 0 | - |
| 10 | w | 1.002E+04 | 576.3 | 0.001898 | 3.293 | +1 | 5 |
| - | - | 5317 | 577.3 | - | - | 0 | - |
| 5 | w | 1070 | 578.3 | 0.001717 | 2.969 | +2 | 10 |
| - | - | 1693 | 578.3 | - | - | 0 | - |
| - | - | 3674 | 581.3 | - | - | 0 | - |
| - | - | 1031 | 582.3 | - | - | 0 | - |
| 5 | y | 883.8 | 585.3 | 0.01115 | 19.05 | +2 | 10 |
| 5 | z | 1.024E+04 | 585.8 | 0.002292 | 3.914 | +2 | 10 |
| - | - | 7888 | 586.3 | - | - | 0 | - |
| - | - | 2450 | 586.8 | - | - | 0 | - |
| - | - | 1881 | 589.3 | - | - | 0 | - |
| - | - | 1740 | 591.3 | - | - | 0 | - |
| 5 | y | 1.671E+04 | 593.8 | 0.001842 | 3.101 | +2 | 10 |
| - | - | 1.03E+04 | 594.3 | - | - | 0 | - |
| - | - | 3437 | 594.8 | - | - | 0 | - |
| - | - | 1290 | 595.3 | - | - | 0 | - |
| - | - | 1.481E+04 | 599.3 | - | - | 0 | - |
| - | - | 4885 | 600.3 | - | - | 0 | - |
| 10 | y | 1655 | 601.3 | 0.01073 | 17.84 | +1 | 5 |
| - | - | 2209 | 602.3 | - | - | 0 | - |
| 11 | c | 3081 | 603.3 | 0.003934 | 6.521 | +2 | 11 |
| - | - | 800.6 | 603.3 | - | - | 0 | - |
| - | - | 2773 | 603.8 | - | - | 0 | - |
| - | - | 1321 | 604.3 | - | - | 0 | - |
| - | - | 1862 | 607.3 | - | - | 0 | - |
| - | - | 1312 | 607.8 | - | - | 0 | - |
| - | - | 750.7 | 613.8 | - | - | 0 | - |
| - | - | 1604 | 614.3 | - | - | 0 | - |
| - | - | 927.1 | 614.8 | - | - | 0 | - |
| - | - | 7477 | 615.4 | - | - | 0 | - |
| 6 | c | 3.92E+04 | 616.4 | 0.001101 | 1.786 | +1 | 6 |
| - | - | 7422 | 617.3 | - | - | 0 | - |
| - | - | 1.225E+04 | 617.4 | - | - | 0 | - |
| - | - | 3255 | 618.3 | - | - | 0 | - |
| - | - | 1832 | 618.4 | - | - | 0 | - |
| - | - | 2161 | 618.8 | - | - | 0 | - |
| 10 | y | 5.572E+04 | 619.3 | 0.001935 | 3.124 | +1 | 5 |
| - | - | 2.297E+04 | 620.4 | - | - | 0 | - |
| - | - | 9858 | 621.3 | - | - | 0 | - |
| - | - | 4289 | 621.4 | - | - | 0 | - |
| - | - | 6727 | 621.8 | - | - | 0 | - |
| - | - | 2761 | 622.3 | - | - | 0 | - |
| - | - | 1171 | 622.4 | - | - | 0 | - |
| - | - | 739.5 | 622.8 | - | - | 0 | - |
| - | - | 2387 | 627.3 | - | - | 0 | - |
| - | - | 1872 | 627.8 | - | - | 0 | - |
| - | - | 1211 | 628.3 | - | - | 0 | - |
| - | - | 1186 | 628.8 | - | - | 0 | - |
| - | - | 2274 | 630.4 | - | - | 0 | - |
| - | - | 2409 | 631.4 | - | - | 0 | - |
| - | - | 3768 | 632.3 | - | - | 0 | - |
| - | - | 1267 | 633.3 | - | - | 0 | - |
| - | - | 1650 | 639.3 | - | - | 0 | - |
| - | - | 1474 | 639.8 | - | - | 0 | - |
| 4 | y | 1560 | 642.3 | 0.001587 | 2.471 | +2 | 11 |
| 4 | y | 869.4 | 642.8 | 0.006955 | 10.82 | +2 | 11 |
| 4 | z | 7253 | 643.3 | 0.002005 | 3.116 | +2 | 11 |
| - | - | 9854 | 643.8 | - | - | 0 | - |
| - | - | 4431 | 644.3 | - | - | 0 | - |
| - | - | 2441 | 644.8 | - | - | 0 | - |
| - | - | 1.312E+04 | 646.3 | - | - | 0 | - |
| - | - | 7466 | 647.3 | - | - | 0 | - |
| - | - | 1.6E+04 | 648.3 | - | - | 0 | - |
| - | - | 6445 | 649.4 | - | - | 0 | - |
| - | - | 1680 | 650.4 | - | - | 0 | - |
| - | - | 1026 | 650.8 | - | - | 0 | - |
| 4 | y | 4.547E+04 | 651.3 | 0.002103 | 3.229 | +2 | 11 |
| - | - | 3.506E+04 | 651.8 | - | - | 0 | - |
| - | - | 1.617E+04 | 652.3 | - | - | 0 | - |
| - | - | 4701 | 652.8 | - | - | 0 | - |
| - | - | 5715 | 653.3 | - | - | 0 | - |
| - | - | 3123 | 653.8 | - | - | 0 | - |
| - | - | 1135 | 654.3 | - | - | 0 | - |
| - | - | 787.6 | 656.4 | - | - | 0 | - |
| - | - | 820.4 | 663.8 | - | - | 0 | - |
| - | - | 609.3 | 664.3 | - | - | 0 | - |
| - | - | 836.2 | 665.3 | - | - | 0 | - |
| - | - | 1233 | 665.4 | - | - | 0 | - |
| - | - | 5125 | 667.3 | - | - | 0 | - |
| - | - | 2755 | 667.9 | - | - | 0 | - |
| - | - | 1292 | 668.3 | - | - | 0 | - |
| - | - | 1756 | 668.4 | - | - | 0 | - |
| - | - | 7809 | 670.8 | - | - | 0 | - |
| - | - | 5740 | 671.3 | - | - | 0 | - |
| - | - | 4040 | 671.8 | - | - | 0 | - |
| - | - | 1.219E+05 | 674.4 | - | - | 0 | - |
| - | - | 1038 | 674.9 | - | - | 0 | - |
| - | - | 5.115E+04 | 675.4 | - | - | 0 | - |
| - | - | 1.342E+04 | 676.4 | - | - | 0 | - |
| - | - | 1882 | 677.4 | - | - | 0 | - |
| - | - | 1236 | 677.8 | - | - | 0 | - |
| - | - | 1230 | 678.8 | - | - | 0 | - |
| - | - | 949.1 | 679.3 | - | - | 0 | - |
| - | - | 652.7 | 682.8 | - | - | 0 | - |
| - | - | 5843 | 683.3 | - | - | 0 | - |
| - | - | 902.6 | 683.3 | - | - | 0 | - |
| - | - | 2037 | 684.3 | - | - | 0 | - |
| 3 | w | 8119 | 685.3 | 0.002362 | 3.447 | +2 | 12 |
| - | - | 6997 | 685.8 | - | - | 0 | - |
| - | - | 3020 | 686.3 | - | - | 0 | - |
| - | - | 737.8 | 686.8 | - | - | 0 | - |
| - | - | 843.3 | 687.9 | - | - | 0 | - |
| - | - | 898.3 | 689.4 | - | - | 0 | - |
| 3 | y | 5483 | 691.8 | 0.00217 | 3.137 | +2 | 12 |
| 3 | y | 5221 | 692.3 | 0.008209 | 11.86 | +2 | 12 |
| 3 | z | 3.522E+04 | 692.8 | 0.002649 | 3.823 | +2 | 12 |
| - | - | 2.85E+04 | 693.3 | - | - | 0 | - |
| - | - | 1145 | 693.4 | - | - | 0 | - |
| - | - | 1.138E+04 | 693.8 | - | - | 0 | - |
| - | - | 2631 | 694.3 | - | - | 0 | - |
| - | - | 693.1 | 695.9 | - | - | 0 | - |
| - | - | 1162 | 696.4 | - | - | 0 | - |
| - | - | 910 | 696.9 | - | - | 0 | - |
| - | - | 2060 | 699.8 | - | - | 0 | - |
| 9 | z | 3308 | 700.3 | 0.006401 | 9.139 | +1 | 6 |
| 3 | y | 2.784E+05 | 700.8 | 0.00232 | 3.31 | +2 | 12 |
| - | - | 2.208E+05 | 701.3 | - | - | 0 | - |
| - | - | 1.04E+05 | 701.8 | - | - | 0 | - |
| - | - | 1263 | 701.9 | - | - | 0 | - |
| - | - | 3.1E+04 | 702.3 | - | - | 0 | - |
| - | - | 7436 | 702.8 | - | - | 0 | - |
| - | - | 964.8 | 703.4 | - | - | 0 | - |
| - | - | 1.089E+04 | 703.9 | - | - | 0 | - |
| - | - | 8707 | 704.4 | - | - | 0 | - |
| - | - | 4700 | 704.9 | - | - | 0 | - |
| - | - | 2114 | 705.4 | - | - | 0 | - |
| - | - | 1675 | 706.3 | - | - | 0 | - |
| - | - | 833.7 | 707.4 | - | - | 0 | - |
| - | - | 2113 | 708.4 | - | - | 0 | - |
| - | - | 1.758E+04 | 709.4 | - | - | 0 | - |
| - | - | 1530 | 709.9 | - | - | 0 | - |
| - | - | 8033 | 710.4 | - | - | 0 | - |
| - | - | 1944 | 711.4 | - | - | 0 | - |
| - | - | 733.6 | 714.3 | - | - | 0 | - |
| - | - | 1813 | 714.8 | - | - | 0 | - |
| 13 | c | 1498 | 717.4 | 0.001044 | 1.456 | +2 | 13 |
| - | - | 1840 | 717.9 | - | - | 0 | - |
| 9 | z | 4739 | 718.4 | 0.009707 | 13.51 | +1 | 6 |
| - | - | 1704 | 718.9 | - | - | 0 | - |
| - | - | 1705 | 719.4 | - | - | 0 | - |
| - | - | 1239 | 719.9 | - | - | 0 | - |
| - | - | 5384 | 720.4 | - | - | 0 | - |
| - | - | 4622 | 720.9 | - | - | 0 | - |
| - | - | 1584 | 721.4 | - | - | 0 | - |
| - | - | 1758 | 721.8 | - | - | 0 | - |
| - | - | 5860 | 723.9 | - | - | 0 | - |
| - | - | 6023 | 724.4 | - | - | 0 | - |
| - | - | 4327 | 724.9 | - | - | 0 | - |
| - | - | 2085 | 725.4 | - | - | 0 | - |
| 13 | c | 1.765E+05 | 725.9 | 0.002296 | 3.163 | +2 | 13 |
| - | - | 1.574E+05 | 726.4 | - | - | 0 | - |
| - | - | 7.12E+04 | 726.9 | - | - | 0 | - |
| - | - | 2.306E+04 | 727.4 | - | - | 0 | - |
| - | - | 6174 | 727.9 | - | - | 0 | - |
| - | - | 721.5 | 729.3 | - | - | 0 | - |
| 2 | z | 691.9 | 733.8 | 0.006436 | 8.77 | +2 | 13 |
| 9 | y | 1.177E+04 | 734.4 | 0.006913 | 9.414 | +1 | 6 |
| 2 | w | 5749 | 734.9 | 0.002335 | 3.178 | +2 | 13 |
| - | - | 2280 | 734.9 | - | - | 0 | - |
| - | - | 8680 | 735.4 | - | - | 0 | - |
| - | - | 2946 | 735.9 | - | - | 0 | - |
| - | - | 1050 | 736.4 | - | - | 0 | - |
| - | - | 3798 | 740.9 | - | - | 0 | - |
| - | - | 7286 | 741.4 | - | - | 0 | - |
| - | - | 7579 | 741.9 | - | - | 0 | - |
| 2 | z | 1.661E+04 | 742.4 | 0.002499 | 3.367 | +2 | 13 |
| - | - | 1.727E+04 | 742.9 | - | - | 0 | - |
| - | - | 6846 | 743.4 | - | - | 0 | - |
| - | - | 2702 | 743.9 | - | - | 0 | - |
| - | - | 1020 | 744.4 | - | - | 0 | - |
| - | - | 1982 | 747.9 | - | - | 0 | - |
| - | - | 1849 | 748.4 | - | - | 0 | - |
| - | - | 1569 | 749.4 | - | - | 0 | - |
| - | - | 1983 | 749.9 | - | - | 0 | - |
| 2 | y | 4.159E+04 | 750.4 | 0.002293 | 3.055 | +2 | 13 |
| - | - | 3.807E+04 | 750.9 | - | - | 0 | - |
| - | - | 1.747E+04 | 751.4 | - | - | 0 | - |
| - | - | 5112 | 751.9 | - | - | 0 | - |
| - | - | 1553 | 752.4 | - | - | 0 | - |
| 7 | c | 1.684E+05 | 753.4 | 0.002614 | 3.469 | +1 | 7 |
| - | - | 7.147E+04 | 754.4 | - | - | 0 | - |
| - | - | 1754 | 754.9 | - | - | 0 | - |
| - | - | 1.885E+04 | 755.4 | - | - | 0 | - |
| - | - | 924.1 | 755.9 | - | - | 0 | - |
| - | - | 2576 | 756.4 | - | - | 0 | - |
| - | - | 891.3 | 756.9 | - | - | 0 | - |
| - | - | 1202 | 759.9 | - | - | 0 | - |
| - | - | 1525 | 760.4 | - | - | 0 | - |
| - | - | 1060 | 761.9 | - | - | 0 | - |
| - | - | 2299 | 762.4 | - | - | 0 | - |
| - | - | 1235 | 762.9 | - | - | 0 | - |
| - | - | 2652 | 763.4 | - | - | 0 | - |
| - | - | 2536 | 763.9 | - | - | 0 | - |
| - | - | 943.1 | 764.3 | - | - | 0 | - |
| - | - | 7902 | 764.4 | - | - | 0 | - |
| - | - | 5455 | 764.9 | - | - | 0 | - |
| - | - | 3072 | 765.4 | - | - | 0 | - |
| - | - | 802.3 | 765.9 | - | - | 0 | - |
| - | - | 1249 | 766.4 | - | - | 0 | - |
| - | - | 4336 | 769.4 | - | - | 0 | - |
| - | - | 6767 | 769.9 | - | - | 0 | - |
| - | - | 2.327E+05 | 770.4 | - | - | 0 | - |
| - | - | 2.429E+05 | 770.9 | - | - | 0 | - |
| - | - | 1.302E+05 | 771.4 | - | - | 0 | - |
| - | - | 5.08E+04 | 771.9 | - | - | 0 | - |
| - | - | 1.287E+04 | 772.4 | - | - | 0 | - |
| - | - | 936 | 772.9 | - | - | 0 | - |
| - | - | 1511 | 775.4 | - | - | 0 | - |
| - | - | 888.7 | 776.3 | - | - | 0 | - |
| - | - | 1072 | 776.9 | - | - | 0 | - |
| - | - | 2.175E+04 | 777.4 | - | - | 0 | - |
| - | - | 2.054E+04 | 777.9 | - | - | 0 | - |
| - | - | 2.564E+04 | 778.4 | - | - | 0 | - |
| - | - | 1.956E+04 | 778.9 | - | - | 0 | - |
| - | - | 8630 | 779.4 | - | - | 0 | - |
| - | - | 2850 | 779.9 | - | - | 0 | - |
| - | - | 1563 | 780.4 | - | - | 0 | - |
| - | - | 3.662E+04 | 782.3 | - | - | 0 | - |
| - | - | 1673 | 782.9 | - | - | 0 | - |
| - | - | 1.543E+04 | 783.3 | - | - | 0 | - |
| - | - | 936.6 | 783.9 | - | - | 0 | - |
| - | - | 5.913E+04 | 784.4 | - | - | 0 | - |
| - | - | 5.323E+04 | 784.9 | - | - | 0 | - |
| - | - | 2.226E+04 | 785.4 | - | - | 0 | - |
| - | - | 1.07E+04 | 785.9 | - | - | 0 | - |
| - | - | 3273 | 786.4 | - | - | 0 | - |
| 8 | w | 4.923E+04 | 788.4 | 0.002818 | 3.575 | +1 | 7 |
| - | - | 2.125E+04 | 789.4 | - | - | 0 | - |
| - | - | 6062 | 790.4 | - | - | 0 | - |
| - | - | 2293 | 790.9 | - | - | 0 | - |
| - | - | 1.451E+04 | 791.4 | - | - | 0 | - |
| - | - | 2.081E+05 | 791.9 | - | - | 0 | - |
| - | - | 1.925E+05 | 792.4 | - | - | 0 | - |
| - | - | 1.035E+05 | 792.9 | - | - | 0 | - |
| - | - | 3.948E+04 | 793.4 | - | - | 0 | - |
| - | - | 1.044E+04 | 793.9 | - | - | 0 | - |
| - | - | 1772 | 794.3 | - | - | 0 | - |
| - | - | 1085 | 795.3 | - | - | 0 | - |
| - | - | 4371 | 798.9 | - | - | 0 | - |
| - | - | 9012 | 799.4 | - | - | 0 | - |
| - | - | 3.727E+05 | 799.9 | - | - | 0 | - |
| - | - | 7.99E+05 | 800.4 | - | - | 0 | - |
| - | - | 6.319E+05 | 800.9 | - | - | 0 | - |
| - | - | 2.886E+05 | 801.4 | - | - | 0 | - |
| - | - | 9.644E+04 | 801.9 | - | - | 0 | - |
| - | - | 2.389E+04 | 802.4 | - | - | 0 | - |
| - | - | 7288 | 803.4 | - | - | 0 | - |
| - | - | 5161 | 804.4 | - | - | 0 | - |
| - | - | 1154 | 805.4 | - | - | 0 | - |
| - | - | 1063 | 807.9 | - | - | 0 | - |
| - | - | 1766 | 837.4 | - | - | 0 | - |
| - | - | 3231 | 838.5 | - | - | 0 | - |
| - | - | 1081 | 839.5 | - | - | 0 | - |
| 8 | y | 804.4 | 845.4 | 0.000579 | 0.6849 | +1 | 7 |
| 8 | y | 847.1 | 846.4 | 0.01071 | 12.65 | +1 | 7 |
| 8 | z | 8.458E+04 | 847.4 | 0.002575 | 3.039 | +1 | 7 |
| - | - | 3.908E+04 | 848.4 | - | - | 0 | - |
| - | - | 1.209E+04 | 849.4 | - | - | 0 | - |
| - | - | 2992 | 850.4 | - | - | 0 | - |
| 8 | y | 2.848E+04 | 863.4 | 0.002528 | 2.928 | +1 | 7 |
| - | - | 1.346E+04 | 864.4 | - | - | 0 | - |
| - | - | 4356 | 865.4 | - | - | 0 | - |
| - | - | 1130 | 866.4 | - | - | 0 | - |
| - | - | 4551 | 867.4 | - | - | 0 | - |
| - | - | 2019 | 868.4 | - | - | 0 | - |
| - | - | 1884 | 880.5 | - | - | 0 | - |
| - | - | 5470 | 881.4 | - | - | 0 | - |
| 8 | c | 1.995E+05 | 882.5 | 0.002868 | 3.249 | +1 | 8 |
| - | - | 9.735E+04 | 883.5 | - | - | 0 | - |
| - | - | 1082 | 883.6 | - | - | 0 | - |
| - | - | 3.196E+04 | 884.5 | - | - | 0 | - |
| - | - | 6531 | 885.5 | - | - | 0 | - |
| - | - | 1540 | 893.4 | - | - | 0 | - |
| - | - | 1990 | 909.4 | - | - | 0 | - |
| - | - | 1326 | 909.5 | - | - | 0 | - |
| - | - | 1862 | 910.4 | - | - | 0 | - |
| - | - | 2740 | 912.4 | - | - | 0 | - |
| - | - | 864.2 | 913.4 | - | - | 0 | - |
| - | - | 1759 | 925.4 | - | - | 0 | - |
| - | - | 1511 | 940.5 | - | - | 0 | - |
| - | - | 1252 | 941.5 | - | - | 0 | - |
| - | - | 2.74E+04 | 952.5 | - | - | 0 | - |
| - | - | 2.067E+04 | 953.5 | - | - | 0 | - |
| - | - | 6874 | 954.5 | - | - | 0 | - |
| - | - | 2823 | 955.5 | - | - | 0 | - |
| - | - | 1229 | 964.5 | - | - | 0 | - |
| 7 | z | 737.1 | 966.4 | 0.01178 | 12.19 | +1 | 8 |
| - | - | 6402 | 980.5 | - | - | 0 | - |
| - | - | 5097 | 981.5 | - | - | 0 | - |
| 7 | y | 2301 | 982.5 | 0.01363 | 13.87 | +1 | 8 |
| 7 | y | 1514 | 983.4 | 0.01478 | 15.03 | +1 | 8 |
| 7 | z | 7.295E+04 | 984.5 | 0.003295 | 3.347 | +1 | 8 |
| - | - | 4.56E+04 | 985.5 | - | - | 0 | - |
| - | - | 1.463E+04 | 986.5 | - | - | 0 | - |
| - | - | 2629 | 987.5 | - | - | 0 | - |
| - | - | 1075 | 999.5 | - | - | 0 | - |
| 7 | y | 1.509E+04 | 1000 | 0.002576 | 2.575 | +1 | 8 |
| - | - | 8999 | 1001 | - | - | 0 | - |
| - | - | 3246 | 1002 | - | - | 0 | - |
| - | - | 759.1 | 1003 | - | - | 0 | - |
| - | - | 1391 | 1007 | - | - | 0 | - |
| - | - | 801.4 | 1008 | - | - | 0 | - |
| - | - | 2.361E+04 | 1008 | - | - | 0 | - |
| - | - | 1.228E+04 | 1009 | - | - | 0 | - |
| - | - | 4588 | 1010 | - | - | 0 | - |
| - | - | 913.1 | 1011 | - | - | 0 | - |
| - | - | 932.5 | 1012 | - | - | 0 | - |
| - | - | 1092 | 1023 | - | - | 0 | - |
| - | - | 3159 | 1024 | - | - | 0 | - |
| - | - | 3426 | 1025 | - | - | 0 | - |
| - | - | 1141 | 1026 | - | - | 0 | - |
| - | - | 1005 | 1027 | - | - | 0 | - |
| - | - | 1504 | 1028 | - | - | 0 | - |
| - | - | 1024 | 1049 | - | - | 0 | - |
| - | - | 2930 | 1051 | - | - | 0 | - |
| - | - | 1023 | 1052 | - | - | 0 | - |
| - | - | 949.5 | 1066 | - | - | 0 | - |
| - | - | 3098 | 1067 | - | - | 0 | - |
| - | - | 2169 | 1068 | - | - | 0 | - |
| - | - | 2151 | 1069 | - | - | 0 | - |
| 6 | y | 7460 | 1070 | 0.002407 | 2.249 | +1 | 9 |
| 6 | z | 6.804E+04 | 1071 | 0.003005 | 2.804 | +1 | 9 |
| - | - | 4.712E+04 | 1072 | - | - | 0 | - |
| - | - | 2.042E+04 | 1073 | - | - | 0 | - |
| - | - | 5127 | 1074 | - | - | 0 | - |
| - | - | 1246 | 1076 | - | - | 0 | - |
| - | - | 2088 | 1080 | - | - | 0 | - |
| - | - | 1731 | 1081 | - | - | 0 | - |
| - | - | 2827 | 1086 | - | - | 0 | - |
| 6 | y | 4.169E+04 | 1088 | 0.002835 | 2.607 | +1 | 9 |
| - | - | 2.708E+04 | 1089 | - | - | 0 | - |
| - | - | 1.174E+04 | 1090 | - | - | 0 | - |
| - | - | 2891 | 1091 | - | - | 0 | - |
| - | - | 1911 | 1094 | - | - | 0 | - |
| 10 | c | 7.146E+04 | 1095 | 0.002995 | 2.736 | +1 | 10 |
| - | - | 4.306E+04 | 1096 | - | - | 0 | - |
| - | - | 1.573E+04 | 1097 | - | - | 0 | - |
| - | - | 2883 | 1098 | - | - | 0 | - |
| - | - | 1297 | 1099 | - | - | 0 | - |
| - | - | 1041 | 1100 | - | - | 0 | - |
| - | - | 3.904E+04 | 1107 | - | - | 0 | - |
| - | - | 2.484E+04 | 1109 | - | - | 0 | - |
| - | - | 8381 | 1110 | - | - | 0 | - |
| - | - | 2239 | 1111 | - | - | 0 | - |
| - | - | 2.007E+04 | 1121 | - | - | 0 | - |
| - | - | 1.268E+04 | 1122 | - | - | 0 | - |
| - | - | 4592 | 1123 | - | - | 0 | - |
| - | - | 885 | 1124 | - | - | 0 | - |
| - | - | 1175 | 1127 | - | - | 0 | - |
| - | - | 2420 | 1138 | - | - | 0 | - |
| - | - | 3315 | 1139 | - | - | 0 | - |
| - | - | 1921 | 1140 | - | - | 0 | - |
| - | - | 1402 | 1141 | - | - | 0 | - |
| - | - | 751.1 | 1148 | - | - | 0 | - |
| - | - | 2525 | 1154 | - | - | 0 | - |
| - | - | 1615 | 1155 | - | - | 0 | - |
| 5 | w | 1276 | 1156 | 0.002866 | 2.48 | +1 | 10 |
| - | - | 846.2 | 1164 | - | - | 0 | - |
| 5 | y | 1457 | 1170 | 0.01956 | 16.73 | +1 | 10 |
| 5 | z | 4.466E+04 | 1171 | 0.003439 | 2.938 | +1 | 10 |
| - | - | 3.936E+04 | 1172 | - | - | 0 | - |
| - | - | 1.655E+04 | 1173 | - | - | 0 | - |
| - | - | 4103 | 1174 | - | - | 0 | - |
| - | - | 1458 | 1179 | - | - | 0 | - |
| - | - | 3.041E+04 | 1180 | - | - | 0 | - |
| - | - | 2.481E+04 | 1181 | - | - | 0 | - |
| - | - | 1.321E+04 | 1182 | - | - | 0 | - |
| - | - | 3751 | 1183 | - | - | 0 | - |
| - | - | 1011 | 1184 | - | - | 0 | - |
| - | - | 2649 | 1186 | - | - | 0 | - |
| 5 | y | 3.698E+04 | 1187 | 0.003025 | 2.549 | +1 | 10 |
| - | - | 2.529E+04 | 1188 | - | - | 0 | - |
| - | - | 1.108E+04 | 1189 | - | - | 0 | - |
| - | - | 2893 | 1190 | - | - | 0 | - |
| - | - | 9090 | 1207 | - | - | 0 | - |
| - | - | 6597 | 1208 | - | - | 0 | - |
| - | - | 3637 | 1209 | - | - | 0 | - |
| - | - | 1616 | 1210 | - | - | 0 | - |
| - | - | 961.2 | 1215 | - | - | 0 | - |
| - | - | 1031 | 1216 | - | - | 0 | - |
| - | - | 1453 | 1220 | - | - | 0 | - |
| - | - | 1188 | 1221 | - | - | 0 | - |
| - | - | 852.8 | 1222 | - | - | 0 | - |
| - | - | 2127 | 1223 | - | - | 0 | - |
| 11 | c | 1.027E+05 | 1224 | 0.002394 | 1.956 | +1 | 11 |
| - | - | 7.153E+04 | 1225 | - | - | 0 | - |
| - | - | 2.898E+04 | 1226 | - | - | 0 | - |
| - | - | 8674 | 1227 | - | - | 0 | - |
| - | - | 3784 | 1228 | - | - | 0 | - |
| - | - | 1366 | 1229 | - | - | 0 | - |
| - | - | 725.5 | 1235 | - | - | 0 | - |
| - | - | 875.5 | 1236 | - | - | 0 | - |
| - | - | 1310 | 1237 | - | - | 0 | - |
| - | - | 1467 | 1238 | - | - | 0 | - |
| 4 | w | 1544 | 1241 | 0.002958 | 2.385 | +1 | 11 |
| - | - | 3.964E+04 | 1242 | - | - | 0 | - |
| - | - | 2.815E+04 | 1243 | - | - | 0 | - |
| - | - | 1.254E+04 | 1244 | - | - | 0 | - |
| - | - | 3474 | 1245 | - | - | 0 | - |
| - | - | 1036 | 1253 | - | - | 0 | - |
| - | - | 1066 | 1271 | - | - | 0 | - |
| - | - | 1254 | 1278 | - | - | 0 | - |
| - | - | 2.195E+04 | 1279 | - | - | 0 | - |
| - | - | 1.995E+04 | 1280 | - | - | 0 | - |
| - | - | 1.052E+04 | 1281 | - | - | 0 | - |
| - | - | 2462 | 1282 | - | - | 0 | - |
| 4 | y | 1536 | 1285 | 0.003852 | 2.999 | +1 | 11 |
| 4 | z | 4.792E+04 | 1286 | 0.002863 | 2.227 | +1 | 11 |
| - | - | 9.47E+04 | 1287 | - | - | 0 | - |
| - | - | 5.752E+04 | 1288 | - | - | 0 | - |
| - | - | 2.325E+04 | 1289 | - | - | 0 | - |
| - | - | 6796 | 1290 | - | - | 0 | - |
| - | - | 1637 | 1291 | - | - | 0 | - |
| - | - | 2734 | 1301 | - | - | 0 | - |
| 4 | y | 1.241E+04 | 1302 | 0.001228 | 0.9438 | +1 | 11 |
| - | - | 7854 | 1303 | - | - | 0 | - |
| - | - | 3366 | 1304 | - | - | 0 | - |
| - | - | 1699 | 1305 | - | - | 0 | - |
| - | - | 2142 | 1306 | - | - | 0 | - |
| - | - | 1693 | 1307 | - | - | 0 | - |
| - | - | 1485 | 1308 | - | - | 0 | - |
| - | - | 1028 | 1321 | - | - | 0 | - |
| - | - | 1464 | 1322 | - | - | 0 | - |
| 12 | c | 6.771E+04 | 1323 | 0.002705 | 2.045 | +1 | 12 |
| - | - | 5.131E+04 | 1324 | - | - | 0 | - |
| - | - | 2.308E+04 | 1325 | - | - | 0 | - |
| - | - | 7088 | 1326 | - | - | 0 | - |
| - | - | 1694 | 1327 | - | - | 0 | - |
| 3 | z | 7302 | 1385 | 0.002564 | 1.852 | +1 | 12 |
| - | - | 1.561E+04 | 1386 | - | - | 0 | - |
| - | - | 1.205E+04 | 1387 | - | - | 0 | - |
| - | - | 5681 | 1388 | - | - | 0 | - |
| - | - | 1035 | 1391 | - | - | 0 | - |
| - | - | 1779 | 1392 | - | - | 0 | - |
| - | - | 2272 | 1393 | - | - | 0 | - |
| - | - | 2052 | 1402 | - | - | 0 | - |
| - | - | 2623 | 1407 | - | - | 0 | - |
| - | - | 1.014E+04 | 1408 | - | - | 0 | - |
| - | - | 7447 | 1409 | - | - | 0 | - |
| - | - | 3530 | 1410 | - | - | 0 | - |
| - | - | 1338 | 1418 | - | - | 0 | - |
| - | - | 826.3 | 1425 | - | - | 0 | - |
| - | - | 1962 | 1435 | - | - | 0 | - |
| - | - | 1466 | 1436 | - | - | 0 | - |
| - | - | 821.1 | 1447 | - | - | 0 | - |
| 13 | c | 1.101E+04 | 1451 | 0.001859 | 1.281 | +1 | 13 |
| - | - | 2.937E+04 | 1452 | - | - | 0 | - |
| - | - | 1.882E+04 | 1453 | - | - | 0 | - |
| - | - | 8154 | 1454 | - | - | 0 | - |
| - | - | 1735 | 1455 | - | - | 0 | - |
| - | - | 821.8 | 1482 | - | - | 0 | - |
| 2 | z | 1996 | 1484 | 0.002632 | 1.774 | +1 | 13 |
| - | - | 1.414E+04 | 1485 | - | - | 0 | - |
| - | - | 1.097E+04 | 1486 | - | - | 0 | - |
| - | - | 5627 | 1487 | - | - | 0 | - |
| - | - | 2150 | 1488 | - | - | 0 | - |
| - | - | 892.2 | 1513 | - | - | 0 | - |
| - | - | 913.7 | 1514 | - | - | 0 | - |
| - | - | 2126 | 1524 | - | - | 0 | - |
| - | - | 2375 | 1525 | - | - | 0 | - |
| - | - | 862.7 | 1526 | - | - | 0 | - |
| - | - | 1298 | 1538 | - | - | 0 | - |
| - | - | 3869 | 1539 | - | - | 0 | - |
| - | - | 1.17E+04 | 1540 | - | - | 0 | - |
| - | - | 6.272E+04 | 1541 | - | - | 0 | - |
| - | - | 5.7E+04 | 1542 | - | - | 0 | - |
| - | - | 2.698E+04 | 1543 | - | - | 0 | - |
| - | - | 9828 | 1544 | - | - | 0 | - |
| - | - | 2187 | 1545 | - | - | 0 | - |
| - | - | 2014 | 1554 | - | - | 0 | - |
| - | - | 8833 | 1555 | - | - | 0 | - |
| - | - | 1.016E+04 | 1556 | - | - | 0 | - |
| - | - | 9013 | 1557 | - | - | 0 | - |
| - | - | 3012 | 1558 | - | - | 0 | - |
| - | - | 1063 | 1566 | - | - | 0 | - |
| - | - | 1361 | 1567 | - | - | 0 | - |
| - | - | 3871 | 1572 | - | - | 0 | - |
| - | - | 1.409E+04 | 1573 | - | - | 0 | - |
| - | - | 1.141E+04 | 1574 | - | - | 0 | - |
| - | - | 5704 | 1575 | - | - | 0 | - |
| - | - | 1615 | 1576 | - | - | 0 | - |
| - | - | 1749 | 1582 | - | - | 0 | - |
| - | - | 1.761E+04 | 1583 | - | - | 0 | - |
| - | - | 8.168E+04 | 1584 | - | - | 0 | - |
| - | - | 6.944E+04 | 1585 | - | - | 0 | - |
| - | - | 3.817E+04 | 1586 | - | - | 0 | - |
| - | - | 1.231E+04 | 1587 | - | - | 0 | - |
| - | - | 2386 | 1588 | - | - | 0 | - |
| - | - | 1.546E+04 | 1599 | - | - | 0 | - |
| - | - | 6.131E+04 | 1600 | - | - | 0 | - |
| - | - | 1.643E+05 | 1601 | - | - | 0 | - |
| - | - | 1.306E+05 | 1602 | - | - | 0 | - |
| - | - | 6.5E+04 | 1603 | - | - | 0 | - |
| - | - | 2.061E+04 | 1604 | - | - | 0 | - |
| - | - | 5827 | 1605 | - | - | 0 | - |
| - | - | 1234 | 3072 | - | - | 0 | - |

m/z Charge Intensity FragmentType MassShift Position
120.08124542236328 0 5498.961
122.09708404541016 0 404.42
124.0398178100586 0 368.4411
127.0870361328125 0 2805.0215
128.09490966796875 0 6276.4473
129.10272216796875 0 53805.164
130.10604858398438 0 3635.6294
131.11827087402344 0 459.42914
133.0612030029297 0 560.51117
147.04441833496094 0 881.6501
147.1130828857422 0 1410.9663
148.05239868164062 0 1334.7734
148.95480346679688 0 985.4453
149.0456085205078 0 1715.5847
156.13916015625 0 543.2142
166.08677673339844 0 6922.28 y 13
167.0905303955078 0 839.9017
170.1420440673828 0 2658.3071
171.1497802734375 0 200568.9
172.0970916748047 0 596.32947
172.15318298339844 0 19829.506
172.20436096191406 0 451.79144
173.15545654296875 0 630.05054
173.45166015625 0 1565.299
174.0911102294922 0 533.679
179.11846923828125 0 1762.3306
183.14981079101562 0 909.3484
185.16539001464844 0 691.57324
187.1082763671875 0 2887.0095
189.12391662597656 0 937.00055
197.12905883789062 0 12966.468 y 11
198.1367645263672 0 14780.869
199.14468383789062 0 166393.72
200.14808654785156 0 18456.246
201.15121459960938 0 1151.8711
211.14418029785156 0 959.0021
212.1523895263672 0 2923.4675
213.16055297851562 0 1187.2356
215.103271484375 0 7304.0786
215.1393585205078 0 535.46484
216.13905334472656 0 3637.0322
216.16180419921875 0 989.4184
216.171875 0 1337.3306 c 1
217.14263916015625 0 761.43396
221.0850830078125 0 7783.768
222.0865936279297 0 774.3563
225.04351806640625 0 1943.4857
225.0984649658203 0 931.0355
227.10354614257812 0 1765.1176
228.17137145996094 0 13856.601
229.1748809814453 0 1539.167
232.1299591064453 0 10601.028
233.1338653564453 0 1443.94
233.16558837890625 0 7861.552
234.16929626464844 0 1005.78253
235.85926818847656 0 532.80725
239.09564208984375 0 8545.511
240.09564208984375 0 1523.4382
241.19337463378906 0 616.01056
253.19178771972656 0 3740.2234
260.1763610839844 0 1287.979
270.21826171875 0 13076.463
271.22442626953125 0 2798.887
272.2337951660156 0 1758.4348
275.1399230957031 0 3882.128
276.1719970703125 0 778.5679
277.1553039550781 0 1994.4658 y Ammonia loss 12
278.1632385253906 0 17419.521 z 12
279.168212890625 0 3793.2551
281.0508728027344 0 709.3075
286.1759338378906 0 1103.7288
294.18218994140625 0 42273.492 y 12
295.1040344238281 0 5626.7324
295.1855773925781 0 6780.4424
296.1046142578125 0 1441.8077
297.20550537109375 0 1564.2432
298.2135009765625 0 22190.648
299.0623474121094 0 2608.7537
299.2165222167969 0 4019.6208
302.1370544433594 0 736.1853
310.1773986816406 0 2206.6821 y 9
314.1722412109375 0 7181.6904
314.2323913574219 0 13057.333
315.1757507324219 0 1104.7808
315.23992919921875 0 31410.244 c 2
316.2430725097656 0 5780.8496
317.2483215332031 0 712.2353
326.1719055175781 0 2117.1104
331.19891357421875 0 4310.318
352.2244567871094 0 1094.9528
354.2015686035156 0 741.3987
355.0691223144531 0 1166.6796
362.20855712890625 0 9050.757 w 11
363.21160888671875 0 1561.7336
368.2430725097656 0 3582.959
369.1227111816406 0 5460.891
369.249267578125 0 698.7221
370.1238708496094 0 2290.4346
370.234130859375 0 924.8074
371.2427062988281 0 1220.6259
376.22479248046875 0 1781.4287 y Ammonia loss 11
377.1706237792969 0 1097.5327
377.23193359375 0 32588.291 z 11
378.23529052734375 0 9263.224
379.2402038574219 0 935.5475
381.19036865234375 0 1137.4819
382.1360778808594 0 1150.2224
382.1968078613281 0 1052.1603
383.1961975097656 0 1010.4112
385.2457275390625 0 4384.028
386.2532653808594 0 12752.022
387.25872802734375 0 4563.969
388.2613220214844 0 758.6116
391.6713562011719 0 813.26465
392.17767333984375 0 558.1068
393.2509460449219 0 57559.938 y 11
394.2540588378906 0 14345.741
395.2303771972656 0 1120.9752
395.25701904296875 0 1843.8412
399.2259216308594 0 592.66644
401.2043762207031 0 1954.4592
411.4648742675781 0 537.2355
412.23272705078125 0 795.5051
413.24078369140625 0 33177.906
414.2444763183594 0 7363.2134
415.2517395019531 0 1071.4037
418.2307434082031 0 1270.9425
419.2327575683594 0 880.2446
430.26751708984375 0 17297.732 c 3
431.2706604003906 0 4054.8254
432.2716979980469 0 1057.3389
434.1960144042969 0 808.30493
441.2010498046875 0 642.11633
441.23516845703125 0 669.9264
441.33294677734375 0 1389.5316
447.2619323730469 0 60033.895 w 10
448.26483154296875 0 16353.97
449.2669982910156 0 2501.8198
454.2677001953125 0 2827.6616
455.2726135253906 0 1753.9929
466.3040466308594 0 664.894
467.5632019042969 0 3727.3572 y 2
467.89678955078125 0 3725.1301
468.2313232421875 0 2154.0522
469.16961669921875 0 1552.7759
469.3040771484375 0 1993.0447
470.3091125488281 0 2433.14
471.2940979003906 0 1839.7716
472.2784423828125 0 587.44104
482.2624816894531 0 1776.2112
482.2989807128906 0 831.3683
484.2774353027344 0 1742.7842
484.31488037109375 0 9102.307
485.3216857910156 0 14362.199
486.3270263671875 0 5673.281
487.33056640625 0 990.18274
490.73797607421875 0 2211.028
494.3005065917969 0 1002.27765
500.27313232421875 0 1466.6139
500.585205078125 0 2646.179 y 1
500.7414855957031 0 4828.5273 y 6
500.919677734375 0 1715.1577
501.24676513671875 0 3552.8145
504.28131103515625 0 698.7328 y Water loss 10
504.718505859375 0 1578.2942
505.2201843261719 0 1604.3545
506.27545166015625 0 2828.4976 z 10
507.2787780761719 0 736.39484
509.28607177734375 0 1273.7957
511.3022155761719 0 2190.539
512.3096923828125 0 44201.96
513.3125610351562 0 14529.184
513.3565063476562 0 657.28876
514.3158569335938 0 2449.6812
518.3005981445312 0 1013.5357
520.2515869140625 0 1380.593
522.2935180664062 0 3400.0674 y 10
523.2978515625 0 1141.836
528.3283081054688 0 1354.8077
529.3359985351562 0 8844.465 c 4
530.33935546875 0 3595.1338
532.2952270507812 0 1394.2539
533.26611328125 0 1554.3285
533.6087646484375 0 16347.639
533.9426879882812 0 15051.062
534.2305297851562 0 1139.1013
534.2770385742188 0 9359.555
534.3142700195312 0 831.50024
534.611572265625 0 2899.5044
535.2415771484375 0 1351.068 y Water loss 5
535.7452392578125 0 2812.289 y Ammonia loss 5
536.2484130859375 0 4208.0405 z 5
536.749755859375 0 3364.3428
537.2523803710938 0 1159.6685
538.2635498046875 0 3511.601
539.26708984375 0 1269.2267
544.2581176757812 0 9309.252 y 5
544.7589111328125 0 4605.5586
545.259765625 0 1755.7999
545.7575073242188 0 783.16833
548.23828125 0 1201.9846
548.7432861328125 0 1126.6277
549.280029296875 0 922.90265
550.2750854492188 0 1176.9954
554.2550659179688 0 1190.9045
554.7537231445312 0 694.45593
555.248046875 0 586.86206
560.2926025390625 0 621.3472
563.2969360351562 0 1256.5991
568.2387084960938 0 2905.0874
571.3450927734375 0 861.42566
572.2601928710938 0 740.09344
576.3046875 0 10021.929 w 9
577.3092041015625 0 5316.899
578.2677612304688 0 1069.7323 w 4
578.3139038085938 0 1692.9976
581.3316040039062 0 3673.6567
582.3327026367188 0 1031.4833
585.2884521484375 0 883.77606 y Ammonia loss 4
585.7835083007812 0 10243.674 z 4
586.2850341796875 0 7887.806
586.7847290039062 0 2450.1968
589.3360595703125 0 1881.4294
591.3273315429688 0 1739.7101
593.7924194335938 0 16705.287 y 4
594.294189453125 0 10296.276
594.7952880859375 0 3437.4968
595.2962646484375 0 1289.5187
599.3419189453125 0 14805.962
600.3441772460938 0 4885.347
601.3451538085938 0 1654.9878 y Water loss 9
602.3432006835938 0 2208.9182
603.28955078125 0 3081.4963 c Water loss 10
603.340576171875 0 800.5548
603.7877197265625 0 2773.1504
604.2896728515625 0 1320.7721
607.2855224609375 0 1861.8479
607.785888671875 0 1311.563
613.7877197265625 0 750.6716
614.2945556640625 0 1604.3787
614.795166015625 0 927.13947
615.3515014648438 0 7477.212
616.3675537109375 0 39197.242 c 5
617.3292846679688 0 7421.6914
617.3720703125 0 12254.526
618.3355102539062 0 3255.3018
618.3779296875 0 1832.3512
618.8037109375 0 2160.9202
619.346923828125 0 55721.83 y 9
620.350341796875 0 22973.893
621.300537109375 0 9857.658
621.3543701171875 0 4288.7393
621.80322265625 0 6726.821
622.3033447265625 0 2761.3005
622.3555297851562 0 1170.5378
622.80419921875 0 739.46405
627.3063354492188 0 2386.869
627.8060302734375 0 1872.2642
628.3060913085938 0 1211.4114
628.8098754882812 0 1185.5565
630.3745727539062 0 2273.926
631.36328125 0 2409.0894
632.3184204101562 0 3767.614
633.3211669921875 0 1267.2898
639.3226318359375 0 1650.0913
639.8286743164062 0 1473.8208
642.3003540039062 0 1560.2242 y Water loss 3
642.7977294921875 0 869.43604 y Ammonia loss 3
643.2966918945312 0 7253.234 z 3
643.79931640625 0 9854.247
644.3016357421875 0 4430.9067
644.8032836914062 0 2440.704
646.3342895507812 0 13122.492
647.3382568359375 0 7466.3447
648.3490600585938 0 16002
649.352294921875 0 6444.934
650.3554077148438 0 1679.9534
650.8030395507812 0 1025.9115
651.30615234375 0 45465.35 y 3
651.8075561523438 0 35063.49
652.3088989257812 0 16165.045
652.8110961914062 0 4701.137
653.3196411132812 0 5714.5884
653.82275390625 0 3123.1404
654.3222045898438 0 1135.4535
656.355224609375 0 787.5752
663.8325805664062 0 820.4111
664.3253173828125 0 609.34155
665.2993774414062 0 836.1543
665.4219970703125 0 1233.0896
667.306884765625 0 5125.1694
667.8636474609375 0 2755.3367
668.3084106445312 0 1292.3165
668.3659057617188 0 1756.0151
670.8355712890625 0 7808.968
671.3372802734375 0 5740.0034
671.8374633789062 0 4040.0615
674.36572265625 0 121886.5
674.8621215820312 0 1038.3931
675.3688354492188 0 51148.934
676.3712768554688 0 13421.551
677.3746948242188 0 1881.5947
677.8351440429688 0 1235.677
678.8109741210938 0 1229.8588
679.3110961914062 0 949.13495
682.82958984375 0 652.73914
683.2651977539062 0 5843.042
683.3291625976562 0 902.5662
684.26708984375 0 2037.0198
685.3195190429688 0 8118.8765 w 2
685.8211059570312 0 6997.255
686.322509765625 0 3019.8606
686.8233642578125 0 737.846
687.8687133789062 0 843.3494
689.3640747070312 0 898.31537
691.8351440429688 0 5483.3413 y Water loss 2
692.3331909179688 0 5220.6484 y Ammonia loss 2
692.83154296875 0 35215.625 z 2
693.3323364257812 0 28498.475
693.398681640625 0 1145.0933
693.8341674804688 0 11379.514
694.3367309570312 0 2630.8503
695.8770751953125 0 693.0588
696.3671875 0 1161.7987
696.86572265625 0 910
699.83154296875 0 2059.8865
700.3362426757812 0 3308.4675 z Water loss 8
700.840576171875 0 278388.47 y 2
701.3419799804688 0 220773.28
701.8432006835938 0 104043.484
701.916015625 0 1263.4978
702.3447265625 0 30999.258
702.846435546875 0 7435.509
703.37841796875 0 964.80835
703.8760986328125 0 10893.257
704.3772583007812 0 8706.72
704.877197265625 0 4699.8164
705.3800659179688 0 2114.426
706.3497924804688 0 1675.4861
707.3682250976562 0 833.6648
708.4049072265625 0 2113.1296
709.4139404296875 0 17582.115
709.8710327148438 0 1529.8623
710.4161376953125 0 8032.7446
711.4169311523438 0 1943.9158
714.3334350585938 0 733.6428
714.8357543945312 0 1812.7208
717.3682250976562 0 1497.9797 c Ammonia loss 12
717.8724365234375 0 1840.4579
718.3629150390625 0 4739.066 z 8
718.8779296875 0 1703.6633
719.3715209960938 0 1705.3881
719.8870239257812 0 1239.4391
720.370849609375 0 5383.6255
720.8709716796875 0 4621.5854
721.3760375976562 0 1583.9166
721.8477172851562 0 1757.6898
723.8848876953125 0 5859.983
724.3861083984375 0 6023.2114
724.8832397460938 0 4326.9136
725.3842163085938 0 2084.9868
725.8827514648438 0 176527.44 c 12
726.384033203125 0 157430.6
726.8851928710938 0 71195.34
727.38671875 0 23062.105
727.8870849609375 0 6174.259
729.3390502929688 0 721.54663
733.8562622070312 0 691.9412 z Ammonia loss 1
734.3788452148438 0 11767.263 y 8
734.8536987304688 0 5749.3896 w 1
734.900390625 0 2279.7017
735.3643798828125 0 8679.575
735.8562622070312 0 2945.7297
736.3656005859375 0 1049.9542
740.8935546875 0 3798.3467
741.3984375 0 7285.9756
741.8985595703125 0 7578.5063
742.3656005859375 0 16607.281 z 1
742.8671875 0 17267.732
743.3696899414062 0 6846.2217
743.869384765625 0 2701.8982
744.3710327148438 0 1019.84595
747.9043579101562 0 1982.1932
748.4068603515625 0 1849.3091
749.38623046875 0 1568.6531
749.8737182617188 0 1983.1818
750.374755859375 0 41590.316 y 1
750.876220703125 0 38071.727
751.3781127929688 0 17473.496
751.8782958984375 0 5112.3457
752.3801879882812 0 1553.0623
753.427978515625 0 168387.02 c 6
754.4306640625 0 71468.414
754.8866577148438 0 1754.4858
755.4324951171875 0 18847.486
755.8814697265625 0 924.14795
756.4368896484375 0 2576.3088
756.87841796875 0 891.2805
759.8895874023438 0 1202.086
760.3925170898438 0 1524.8431
761.8953247070312 0 1059.6967
762.3992919921875 0 2298.713
762.8932495117188 0 1235.2803
763.3936157226562 0 2651.7278
763.892333984375 0 2536.4233
764.3301391601562 0 943.14044
764.4008178710938 0 7902.064
764.9027709960938 0 5455.0264
765.3991088867188 0 3072.1895
765.9053344726562 0 802.25305
766.3820190429688 0 1248.8418
769.4082641601562 0 4336.1997
769.9051513671875 0 6767.1016
770.4024658203125 0 232727.45
770.9039916992188 0 242944.38
771.4056396484375 0 130198.76
771.9066162109375 0 50799.082
772.4074096679688 0 12872.769
772.9103393554688 0 936.0036
775.3767700195312 0 1510.852
776.3190307617188 0 888.6917
776.8925170898438 0 1071.6129
777.4100341796875 0 21746.121
777.9118041992188 0 20536.6
778.415771484375 0 25638.264
778.9173583984375 0 19556.812
779.4228515625 0 8630.204
779.9156494140625 0 2849.5737
780.4423217773438 0 1562.9565
782.3344116210938 0 36623.926
782.8955688476562 0 1673.1709
783.3374633789062 0 15433.39
783.89453125 0 936.6295
784.3880004882812 0 59129.645
784.8898315429688 0 53234.863
785.3912353515625 0 22263.25
785.8923950195312 0 10698.642
786.3956909179688 0 3272.8118
788.3853149414062 0 49228.92 w 7
789.38818359375 0 21248.402
790.3909912109375 0 6061.8525
790.8983154296875 0 2293.1252
791.4033813476562 0 14505.025
791.9004516601562 0 208093.23
792.4019165039062 0 192475.53
792.9026489257812 0 103502.01
793.4042358398438 0 39477.566
793.904296875 0 10436.386
794.3336791992188 0 1771.9474
795.3406982421875 0 1084.8652
798.9004516601562 0 4371.308
799.404541015625 0 9011.815
799.9095458984375 0 372654.53
800.412353515625 0 798993.4
800.9141235351562 0 631928.44
801.4152221679688 0 288648.9
801.9166259765625 0 96439.45
802.4180908203125 0 23885.959
803.4080200195312 0 7287.551
804.4064331054688 0 5160.6836
805.4052124023438 0 1153.8213
807.9008178710938 0 1063.4116
837.4459228515625 0 1765.9233
838.4566040039062 0 3231.2651
839.4627685546875 0 1081.1627
845.4033813476562 0 804.3745 y Water loss 7
846.398681640625 0 847.13403 y Ammonia loss 7
847.3983764648438 0 84583.36 z 7
848.4014282226562 0 39076.02
849.4036865234375 0 12086.848
850.403564453125 0 2992.4558
863.4170532226562 0 28475.088 y 7
864.4197998046875 0 13463.332
865.4249267578125 0 4356.153
866.4344482421875 0 1130.3108
867.3855590820312 0 4551.069
868.3923950195312 0 2018.5271
880.4540405273438 0 1884.3336
881.4013671875 0 5470.3833
882.4708251953125 0 199479.89 c 7
883.473388671875 0 97354.984
883.5773315429688 0 1082.4182
884.475830078125 0 31960.578
885.477783203125 0 6531.072
893.4022827148438 0 1540.2642
909.3626708984375 0 1990.0632
909.4983520507812 0 1325.8083
910.3654174804688 0 1862.0428
912.4354858398438 0 2739.5068
913.4462280273438 0 864.16187
925.441650390625 0 1759.0571
940.4710083007812 0 1510.6663
941.480712890625 0 1252.247
952.4761962890625 0 27400.682
953.4803466796875 0 20671.828
954.4849853515625 0 6874.029
955.4898681640625 0 2822.5525
964.4586791992188 0 1228.9027
966.4559326171875 0 737.05646 z Water loss 6
980.468994140625 0 6402.1567
981.4752197265625 0 5097.324
982.4765014648438 0 2300.6057 y Water loss 6
983.461669921875 0 1513.8926 y Ammonia loss 6
984.4580078125 0 72945.414 z 6
985.4611206054688 0 45602.004
986.463623046875 0 14634.511
987.4639282226562 0 2629.4788
999.4793090820312 0 1074.7886
1000.4760131835938 0 15086.907 y 6
1001.4779052734375 0 8998.589
1002.4800415039062 0 3246.4082
1003.4771118164062 0 759.11774
1006.5054321289062 0 1390.6152
1007.51025390625 0 801.4109
1008.4295654296875 0 23608.906
1009.4324340820312 0 12276.076
1010.4338989257812 0 4587.5386
1011.4498901367188 0 913.1145
1012.4716796875 0 932.5299
1022.50537109375 0 1092.1548
1023.5111694335938 0 3159.382
1024.519287109375 0 3426.1965
1025.5255126953125 0 1141.2518
1027.49658203125 0 1005.4782
1028.4803466796875 0 1504.0212
1049.49560546875 0 1023.6123
1050.5357666015625 0 2930.2004
1051.5263671875 0 1023.3817
1065.507080078125 0 949.4873
1066.528076171875 0 3097.6687
1067.5302734375 0 2169.0103
1069.4732666015625 0 2151.073
1070.4813232421875 0 7460.089 y Ammonia loss 5
1071.48974609375 0 68044.89 z 5
1072.4931640625 0 47118.78
1073.496337890625 0 20415.68
1074.4998779296875 0 5126.8438
1075.50244140625 0 1245.9604
1079.5050048828125 0 2087.6257
1080.5054931640625 0 1731.3574
1086.4974365234375 0 2827.311
1087.50830078125 0 41693.457 y 5
1088.51123046875 0 27084.684
1089.51220703125 0 11741.602
1090.510009765625 0 2890.5837
1093.5390625 0 1911.3114
1094.5506591796875 0 71456.6 c 9
1095.553955078125 0 43061.68
1096.556396484375 0 15727.192
1097.5570068359375 0 2882.5576
1098.51416015625 0 1297.2037
1099.51220703125 0 1041.2307
1107.4984130859375 0 39037.035
1108.501708984375 0 24837.158
1109.505859375 0 8381.138
1110.508056640625 0 2239.499
1120.566650390625 0 20069.326
1121.569580078125 0 12677.138
1122.573486328125 0 4592.3506
1123.579833984375 0 884.9808
1126.5618896484375 0 1175.2759
1137.5341796875 0 2420.1587
1138.5390625 0 3315.4426
1139.5391845703125 0 1920.9774
1140.5601806640625 0 1402.2795
1147.5648193359375 0 751.087
1153.5552978515625 0 2524.9912
1154.5506591796875 0 1615.0754
1155.5345458984375 0 1276.025 w 4
1163.573486328125 0 846.1964
1169.56689453125 0 1456.6879 y Ammonia loss 4
1170.55859375 0 44655.215 z 4
1171.5623779296875 0 39363.42
1172.5654296875 0 16552.885
1173.5675048828125 0 4103.087
1178.5709228515625 0 1458.373
1179.5802001953125 0 30408.617
1180.5841064453125 0 24806.852
1181.5877685546875 0 13207.743
1182.5867919921875 0 3750.6572
1183.5904541015625 0 1010.9986
1185.5677490234375 0 2648.5544
1186.576904296875 0 36980.586 y 4
1187.58056640625 0 25294.467
1188.583251953125 0 11078.466
1189.58642578125 0 2893.0068
1206.5731201171875 0 9090.128
1207.573974609375 0 6597.1494
1208.57861328125 0 3637.319
1209.57470703125 0 1615.7252
1214.5697021484375 0 961.16895
1215.5694580078125 0 1030.9923
1219.6376953125 0 1453.4303
1220.640625 0 1188.2239
1221.5927734375 0 852.8122
1222.58447265625 0 2126.9448
1223.5926513671875 0 102714.54 c 10
1224.5960693359375 0 71533.484
1225.5982666015625 0 28980.676
1226.595458984375 0 8673.625
1227.587158203125 0 3784.1023
1228.5836181640625 0 1365.6428
1234.6650390625 0 725.51135
1235.602783203125 0 875.54956
1236.5965576171875 0 1309.9668
1237.60400390625 0 1467.1266
1240.58740234375 0 1544.1216 w 3
1241.5941162109375 0 39641.016
1242.5970458984375 0 28151.846
1243.6015625 0 12536.467
1244.602783203125 0 3473.9333
1252.61376953125 0 1036.0547
1270.58251953125 0 1065.8513
1277.63134765625 0 1253.7477
1278.64697265625 0 21949.44
1279.6510009765625 0 19949.74
1280.655029296875 0 10516.19
1281.6544189453125 0 2462.4363
1284.578125 0 1536.4886 y Ammonia loss 3
1285.5849609375 0 47918.113 z 3
1286.5908203125 0 94703.164
1287.59423828125 0 57516.844
1288.59765625 0 23248.787
1289.601318359375 0 6795.948
1290.601806640625 0 1637.2784
1300.5947265625 0 2733.9827
1301.60205078125 0 12405.769 y 3
1302.60595703125 0 7854.4414
1303.6082763671875 0 3365.8408
1304.613525390625 0 1698.7394
1305.6368408203125 0 2141.6504
1306.6431884765625 0 1692.9626
1307.642333984375 0 1484.8921
1320.6494140625 0 1028.0403
1321.6539306640625 0 1463.895
1322.661376953125 0 67713.27 c 11
1323.663818359375 0 51310.543
1324.66650390625 0 23083.604
1325.66796875 0 7088.4087
1326.661865234375 0 1693.8027
1384.653076171875 0 7301.8096 z 2
1385.6590576171875 0 15611.007
1386.6624755859375 0 12045.76
1387.6654052734375 0 5681.3574
1390.72900390625 0 1034.7474
1391.7357177734375 0 1778.6176
1392.7432861328125 0 2272.304
1401.6785888671875 0 2052.221
1406.7440185546875 0 2623.3462
1407.752197265625 0 10143.349
1408.755859375 0 7447.3506
1409.75634765625 0 3529.8904
1417.7308349609375 0 1338.2112
1424.75244140625 0 826.29443
1434.735595703125 0 1961.9651
1435.7275390625 0 1465.8967
1446.5147705078125 0 821.13416
1450.7554931640625 0 11013.318 c 12
1451.76171875 0 29374.455
1452.7647705078125 0 18819.93
1453.76806640625 0 8154.083
1454.7706298828125 0 1735.4269
1481.7880859375 0 821.841
1483.7215576171875 0 1996.3999 z 1
1484.727783203125 0 14136.967
1485.73095703125 0 10966.047
1486.7353515625 0 5627.1904
1487.7396240234375 0 2150.332
1512.823486328125 0 892.18835
1513.8077392578125 0 913.7264
1523.7728271484375 0 2126.0593
1524.77734375 0 2374.507
1525.79296875 0 862.67566
1537.818603515625 0 1297.8733
1538.8118896484375 0 3868.751
1539.80029296875 0 11696.646
1540.8017578125 0 62717.426
1541.804931640625 0 56997.26
1542.8079833984375 0 26980.18
1543.81005859375 0 9828.013
1544.8121337890625 0 2187.0947
1553.8118896484375 0 2013.6504
1554.8184814453125 0 8832.937
1555.81884765625 0 10164.592
1556.8162841796875 0 9012.596
1557.8143310546875 0 3012.0598
1565.7821044921875 0 1062.5106
1566.794677734375 0 1360.8188
1571.8128662109375 0 3871.267
1572.8265380859375 0 14093.053
1573.829833984375 0 11410.666
1574.8348388671875 0 5703.7427
1575.838623046875 0 1614.8809
1581.7823486328125 0 1748.7545
1582.795654296875 0 17613.498
1583.7977294921875 0 81678.86
1584.80029296875 0 69439.58
1585.80322265625 0 38166.99
1586.8045654296875 0 12313.296
1587.811279296875 0 2385.9219
1598.80615234375 0 15463.22
1599.813720703125 0 61310.81
1600.8221435546875 0 164295.17
1601.825439453125 0 130627.99
1602.8282470703125 0 65003.44
1603.8311767578125 0 20611.443
1604.833251953125 0 5826.572
3072.216552734375 0 1234.196

Spectrum Details

|  |  |
| --- | --- |
| Matched peaks? Matched peaks The total absolute number of peaks matched. Additionally in brackets the total fraction of peaks matched and the total number of peaks is shown. | 81 (11.44% of 708) |
| FDR? FDR The false discovery rate estimated for this peptide. It is calculated by matching all theoretical fragments with a non-integer shift with the raw peaks for this spectrum. This is done with 40 different shifts. The resulting percentage is the average number of annotated peaks over the number of annotated peaks with the correct spectrum. | 1.91% |
| Satellite FDR? Satellite FDR See the FDR for details on its calculation. This satellite ion specific FDR only contains the satellite ions (d/w) for I/L/J positions. | - |
| PSM Score? PSM Score The PSM Score as given by Hecklib to this annotated spectrum. It is shown with three significant figures. | 653 |

#### Spectrum 7099? Spectrum 7099 The raw spectrum of this peptide as annotated by Hecklib. The fragments are coloured according to ion type (see legend). Any peaks with a star '\*' as text can be hovered over to see the full details, first the ion type second the mass shift type. By hovering over the amino acids in the peptide or ions in the legend the corresponding peaks are highlighted. By toggling the 'Unassigned' label you can turn the background (unassigned) peaks on or off in the plot. By updating the slider in the Ion legend you can update the spectrum to only show the top X% of the peaks with labels. The top X% means any peak that is within X% of the highest intensity. By dragging in the spectrum you can zoom in to a specific part of the spectrum and use 'Zoom Out' to get back to the original zoom level. The annotation of the spectrum is based on the given sequence in the peptides file and is done with different software so inconsistencies are likely. The peaks are annotated based on the given sequence, with 20 ppm tolerance.

Copy Data

##### Spectrum 7099 (TSV)

###### Preview

```
Loading example...
```

*Click on the button to copy the data to your clipboard.*

Mz MinMz MaxIntensity Max

WidthHeightPeptide font sizePeptide stroke widthSpectrum font sizeSpectrum stroke widthCompact peptide

Ion legend

wxyz

abcd

OtherUnassignedIon^Charge^_Position_Show for top:%

VVVDVSHEDPEVKF

04.90e+49.80e+41.47e+51.96e+5

Zoom Out

y^+1^_1_a^+1^_2_y^+2^_3_b^+1^_2_y^+1^_2_y^+1^_2_b^+1^_3_y^+2^_5_y^+1^_3_b^+1^_4_y^+3^_12_y^+3^_13_y^+2^_8_y^+1^_4_\*y^+2^_9_b^+2^_10_y^+2^_9_y^+2^_10_y^+2^_10_y^+2^_10_y^+1^_5_b^+2^_11_y^+1^_5_y^+2^_11_y^+2^_11_y^+2^_12_y^+2^_12_y^+2^_12_b^+1^_7_y^+1^_6_y^+2^_13_y^+2^_13_y^+2^_13_y^+1^_7_y^+1^_7_y^+1^_8_y^+1^_8_y^+1^_9_y^+1^_10_b^+1^_11_y^+1^_11_b^+1^_12_

0558111616742231

Fragment Matches Table

Show background peaks

| Position | Ion type | Intensity | mz Theoretical | mz Error (Th) | mz Error (ppm) | Charge | Series Number |
| --- | --- | --- | --- | --- | --- | --- | --- |
| - | - | 9820 | 120.1 | - | - | 0 | - |
| - | - | 543.3 | 121.1 | - | - | 0 | - |
| - | - | 421 | 122.1 | - | - | 0 | - |
| - | - | 588.4 | 127.1 | - | - | 0 | - |
| - | - | 1009 | 127.1 | - | - | 0 | - |
| - | - | 1364 | 128.1 | - | - | 0 | - |
| - | - | 465.3 | 129 | - | - | 0 | - |
| - | - | 1.399E+05 | 129.1 | - | - | 0 | - |
| - | - | 1647 | 130.1 | - | - | 0 | - |
| - | - | 783.9 | 130.1 | - | - | 0 | - |
| - | - | 8963 | 130.1 | - | - | 0 | - |
| - | - | 482.1 | 131.1 | - | - | 0 | - |
| - | - | 1340 | 132.1 | - | - | 0 | - |
| - | - | 545.2 | 133.1 | - | - | 0 | - |
| - | - | 3444 | 133.1 | - | - | 0 | - |
| - | - | 2533 | 136.1 | - | - | 0 | - |
| - | - | 444.6 | 137.9 | - | - | 0 | - |
| - | - | 907.6 | 138.1 | - | - | 0 | - |
| - | - | 723 | 138.1 | - | - | 0 | - |
| - | - | 607.7 | 140.1 | - | - | 0 | - |
| - | - | 2526 | 141.1 | - | - | 0 | - |
| - | - | 690.2 | 143.1 | - | - | 0 | - |
| - | - | 420.8 | 145.1 | - | - | 0 | - |
| - | - | 883.8 | 147.1 | - | - | 0 | - |
| - | - | 5219 | 147.1 | - | - | 0 | - |
| - | - | 464.9 | 147.5 | - | - | 0 | - |
| - | - | 5016 | 149 | - | - | 0 | - |
| - | - | 566.4 | 152.1 | - | - | 0 | - |
| - | - | 8067 | 155.1 | - | - | 0 | - |
| - | - | 569.1 | 156.1 | - | - | 0 | - |
| - | - | 1348 | 163.1 | - | - | 0 | - |
| - | - | 3942 | 164.1 | - | - | 0 | - |
| - | - | 605.6 | 165.1 | - | - | 0 | - |
| 14 | y | 5316 | 166.1 | 0.000247 | 1.487 | +1 | 1 |
| - | - | 6725 | 167.1 | - | - | 0 | - |
| - | - | 1235 | 167.1 | - | - | 0 | - |
| - | - | 631.5 | 168.1 | - | - | 0 | - |
| - | - | 4797 | 169.1 | - | - | 0 | - |
| - | - | 542.6 | 169.1 | - | - | 0 | - |
| 2 | a | 1.94E+05 | 171.1 | 0.0003617 | 2.114 | +1 | 2 |
| - | - | 1.813E+04 | 172.2 | - | - | 0 | - |
| - | - | 1745 | 173.1 | - | - | 0 | - |
| - | - | 923.1 | 173.2 | - | - | 0 | - |
| - | - | 777.7 | 173.4 | - | - | 0 | - |
| - | - | 1404 | 177.1 | - | - | 0 | - |
| - | - | 2488 | 179.1 | - | - | 0 | - |
| - | - | 510.1 | 180.1 | - | - | 0 | - |
| - | - | 459.3 | 181.1 | - | - | 0 | - |
| - | - | 963.9 | 182.1 | - | - | 0 | - |
| - | - | 2009 | 183.1 | - | - | 0 | - |
| - | - | 5960 | 183.1 | - | - | 0 | - |
| - | - | 795.6 | 184.2 | - | - | 0 | - |
| - | - | 875 | 185.1 | - | - | 0 | - |
| - | - | 585.6 | 185.1 | - | - | 0 | - |
| - | - | 4.034E+04 | 185.2 | - | - | 0 | - |
| - | - | 1726 | 186.1 | - | - | 0 | - |
| - | - | 3822 | 186.2 | - | - | 0 | - |
| - | - | 962.4 | 187.1 | - | - | 0 | - |
| - | - | 1.625E+04 | 187.1 | - | - | 0 | - |
| - | - | 836.6 | 187.1 | - | - | 0 | - |
| - | - | 1894 | 188.1 | - | - | 0 | - |
| - | - | 557.8 | 191 | - | - | 0 | - |
| - | - | 503.2 | 191.1 | - | - | 0 | - |
| - | - | 563.6 | 194.1 | - | - | 0 | - |
| - | - | 450.4 | 195.1 | - | - | 0 | - |
| - | - | 807.5 | 195.1 | - | - | 0 | - |
| - | - | 565.8 | 196.1 | - | - | 0 | - |
| - | - | 4844 | 197.1 | - | - | 0 | - |
| 12 | y | 897.8 | 197.1 | 9.729E-05 | 0.4935 | +2 | 3 |
| - | - | 5288 | 199.1 | - | - | 0 | - |
| 2 | b | 8.681E+04 | 199.1 | 0.0002744 | 1.378 | +1 | 2 |
| - | - | 700.9 | 200.1 | - | - | 0 | - |
| - | - | 9138 | 200.1 | - | - | 0 | - |
| - | - | 1.165E+04 | 201.1 | - | - | 0 | - |
| - | - | 1216 | 202.1 | - | - | 0 | - |
| - | - | 1718 | 203.1 | - | - | 0 | - |
| - | - | 3039 | 207.1 | - | - | 0 | - |
| - | - | 2888 | 208.1 | - | - | 0 | - |
| - | - | 618.4 | 208.1 | - | - | 0 | - |
| - | - | 661.7 | 211.1 | - | - | 0 | - |
| - | - | 2716 | 211.1 | - | - | 0 | - |
| - | - | 1311 | 213.1 | - | - | 0 | - |
| - | - | 3.919E+04 | 213.2 | - | - | 0 | - |
| - | - | 4487 | 214.2 | - | - | 0 | - |
| - | - | 1.246E+04 | 215.1 | - | - | 0 | - |
| - | - | 995.6 | 215.1 | - | - | 0 | - |
| - | - | 1751 | 216.1 | - | - | 0 | - |
| - | - | 1736 | 216.1 | - | - | 0 | - |
| - | - | 568.4 | 217.1 | - | - | 0 | - |
| - | - | 809.4 | 218.1 | - | - | 0 | - |
| - | - | 1.048E+04 | 221.1 | - | - | 0 | - |
| - | - | 1010 | 221.1 | - | - | 0 | - |
| - | - | 1443 | 222.1 | - | - | 0 | - |
| - | - | 2761 | 223.1 | - | - | 0 | - |
| - | - | 558.2 | 224.1 | - | - | 0 | - |
| - | - | 1.431E+04 | 225 | - | - | 0 | - |
| - | - | 1.825E+04 | 225.1 | - | - | 0 | - |
| - | - | 2425 | 226 | - | - | 0 | - |
| - | - | 1248 | 226.1 | - | - | 0 | - |
| - | - | 1605 | 226.1 | - | - | 0 | - |
| - | - | 938 | 227 | - | - | 0 | - |
| - | - | 1.839E+04 | 227.1 | - | - | 0 | - |
| - | - | 2408 | 228.1 | - | - | 0 | - |
| - | - | 1.651E+04 | 228.2 | - | - | 0 | - |
| - | - | 626.9 | 229.1 | - | - | 0 | - |
| - | - | 622 | 229.2 | - | - | 0 | - |
| - | - | 2289 | 229.2 | - | - | 0 | - |
| - | - | 695.2 | 231.1 | - | - | 0 | - |
| - | - | 6.753E+04 | 233.1 | - | - | 0 | - |
| - | - | 1565 | 234.1 | - | - | 0 | - |
| - | - | 7224 | 234.1 | - | - | 0 | - |
| - | - | 6590 | 235.1 | - | - | 0 | - |
| - | - | 2.505E+04 | 239.1 | - | - | 0 | - |
| - | - | 1647 | 239.1 | - | - | 0 | - |
| - | - | 2000 | 239.1 | - | - | 0 | - |
| - | - | 3505 | 240.1 | - | - | 0 | - |
| - | - | 699.7 | 240.1 | - | - | 0 | - |
| - | - | 692.8 | 243.1 | - | - | 0 | - |
| - | - | 1035 | 245.1 | - | - | 0 | - |
| - | - | 1260 | 246.2 | - | - | 0 | - |
| - | - | 1732 | 249.1 | - | - | 0 | - |
| - | - | 1.397E+04 | 251.1 | - | - | 0 | - |
| - | - | 7498 | 251.1 | - | - | 0 | - |
| - | - | 1506 | 252.1 | - | - | 0 | - |
| - | - | 674.6 | 253.1 | - | - | 0 | - |
| - | - | 1409 | 253.2 | - | - | 0 | - |
| - | - | 766.6 | 257.1 | - | - | 0 | - |
| - | - | 743.1 | 259.2 | - | - | 0 | - |
| - | - | 714.7 | 260.1 | - | - | 0 | - |
| - | - | 679.2 | 266.1 | - | - | 0 | - |
| - | - | 7735 | 267.1 | - | - | 0 | - |
| - | - | 952.9 | 268.1 | - | - | 0 | - |
| - | - | 707.7 | 268.2 | - | - | 0 | - |
| - | - | 664.1 | 274.1 | - | - | 0 | - |
| - | - | 803.4 | 275.1 | - | - | 0 | - |
| - | - | 1201 | 276.2 | - | - | 0 | - |
| 13 | y | 7313 | 277.2 | 0.0002383 | 0.8596 | +1 | 2 |
| - | - | 1459 | 278.2 | - | - | 0 | - |
| - | - | 4150 | 279.1 | - | - | 0 | - |
| - | - | 676 | 279.1 | - | - | 0 | - |
| - | - | 941.8 | 281.1 | - | - | 0 | - |
| - | - | 843.7 | 281.1 | - | - | 0 | - |
| - | - | 1.545E+04 | 282.2 | - | - | 0 | - |
| - | - | 2086 | 283.2 | - | - | 0 | - |
| - | - | 1442 | 284.1 | - | - | 0 | - |
| - | - | 771.3 | 285 | - | - | 0 | - |
| - | - | 954.9 | 286.2 | - | - | 0 | - |
| - | - | 765.8 | 286.2 | - | - | 0 | - |
| - | - | 941.8 | 287.1 | - | - | 0 | - |
| - | - | 807.5 | 288.1 | - | - | 0 | - |
| - | - | 545 | 289.9 | - | - | 0 | - |
| 13 | y | 4.626E+04 | 294.2 | 0.000392 | 1.333 | +1 | 2 |
| - | - | 5094 | 295.1 | - | - | 0 | - |
| - | - | 7198 | 295.2 | - | - | 0 | - |
| - | - | 1076 | 296.1 | - | - | 0 | - |
| - | - | 1190 | 298.2 | - | - | 0 | - |
| 3 | b | 3890 | 298.2 | 0.0001283 | 0.4302 | +1 | 3 |
| - | - | 7622 | 299.1 | - | - | 0 | - |
| - | - | 2477 | 300.1 | - | - | 0 | - |
| - | - | 1849 | 300.2 | - | - | 0 | - |
| - | - | 3657 | 302.1 | - | - | 0 | - |
| - | - | 5706 | 303.1 | - | - | 0 | - |
| - | - | 899.2 | 304.1 | - | - | 0 | - |
| - | - | 977 | 304.2 | - | - | 0 | - |
| - | - | 2081 | 306.2 | - | - | 0 | - |
| - | - | 3528 | 307.1 | - | - | 0 | - |
| - | - | 918.1 | 307.2 | - | - | 0 | - |
| - | - | 562.2 | 310.2 | - | - | 0 | - |
| 10 | y | 2794 | 310.2 | 0.0002894 | 0.9331 | +2 | 5 |
| - | - | 1947 | 313.1 | - | - | 0 | - |
| - | - | 1054 | 314.2 | - | - | 0 | - |
| - | - | 1.057E+04 | 314.2 | - | - | 0 | - |
| - | - | 1217 | 315.2 | - | - | 0 | - |
| - | - | 4768 | 321.1 | - | - | 0 | - |
| - | - | 653.4 | 322.1 | - | - | 0 | - |
| - | - | 594.8 | 322.2 | - | - | 0 | - |
| - | - | 1756 | 324.1 | - | - | 0 | - |
| - | - | 2757 | 324.2 | - | - | 0 | - |
| - | - | 623.1 | 325.2 | - | - | 0 | - |
| - | - | 9854 | 326.2 | - | - | 0 | - |
| - | - | 1841 | 327.2 | - | - | 0 | - |
| - | - | 3218 | 332.2 | - | - | 0 | - |
| - | - | 657.8 | 334.2 | - | - | 0 | - |
| - | - | 2227 | 336.1 | - | - | 0 | - |
| - | - | 3615 | 342.1 | - | - | 0 | - |
| - | - | 876.8 | 343.1 | - | - | 0 | - |
| - | - | 868.2 | 346.1 | - | - | 0 | - |
| - | - | 7956 | 354.1 | - | - | 0 | - |
| - | - | 981.1 | 355.1 | - | - | 0 | - |
| - | - | 1297 | 355.1 | - | - | 0 | - |
| - | - | 1145 | 359 | - | - | 0 | - |
| - | - | 8378 | 364.1 | - | - | 0 | - |
| - | - | 1.011E+04 | 364.2 | - | - | 0 | - |
| - | - | 792 | 365.1 | - | - | 0 | - |
| - | - | 2210 | 365.2 | - | - | 0 | - |
| - | - | 620.6 | 367.3 | - | - | 0 | - |
| - | - | 591.1 | 368.3 | - | - | 0 | - |
| - | - | 738.4 | 369.1 | - | - | 0 | - |
| - | - | 5274 | 369.1 | - | - | 0 | - |
| - | - | 1685 | 370.1 | - | - | 0 | - |
| - | - | 1061 | 377.2 | - | - | 0 | - |
| - | - | 556.5 | 378.8 | - | - | 0 | - |
| - | - | 1.927E+04 | 382.1 | - | - | 0 | - |
| - | - | 3205 | 383.1 | - | - | 0 | - |
| - | - | 2320 | 383.2 | - | - | 0 | - |
| - | - | 699.5 | 385.2 | - | - | 0 | - |
| - | - | 652.5 | 391.7 | - | - | 0 | - |
| - | - | 1112 | 392.2 | - | - | 0 | - |
| 12 | y | 1.794E+04 | 393.2 | 0.000429 | 1.091 | +1 | 3 |
| - | - | 4789 | 394.3 | - | - | 0 | - |
| - | - | 1886 | 395.3 | - | - | 0 | - |
| - | - | 1908 | 399.2 | - | - | 0 | - |
| - | - | 2811 | 401.2 | - | - | 0 | - |
| - | - | 1339 | 402.2 | - | - | 0 | - |
| - | - | 2271 | 406.2 | - | - | 0 | - |
| - | - | 622 | 407.2 | - | - | 0 | - |
| - | - | 1258 | 410.1 | - | - | 0 | - |
| - | - | 1.114E+04 | 410.2 | - | - | 0 | - |
| - | - | 1547 | 411.2 | - | - | 0 | - |
| - | - | 953.2 | 411.2 | - | - | 0 | - |
| - | - | 979.4 | 413.2 | - | - | 0 | - |
| 4 | b | 2871 | 413.2 | 0.0008037 | 1.945 | +1 | 4 |
| - | - | 9491 | 413.3 | - | - | 0 | - |
| - | - | 682.2 | 414.2 | - | - | 0 | - |
| - | - | 2920 | 414.3 | - | - | 0 | - |
| - | - | 3528 | 421.2 | - | - | 0 | - |
| - | - | 678.7 | 422.2 | - | - | 0 | - |
| - | - | 813.2 | 423.2 | - | - | 0 | - |
| - | - | 663.5 | 425.2 | - | - | 0 | - |
| - | - | 4.362E+04 | 427.2 | - | - | 0 | - |
| - | - | 1.111E+04 | 428.2 | - | - | 0 | - |
| - | - | 646.8 | 433.1 | - | - | 0 | - |
| - | - | 5112 | 439.2 | - | - | 0 | - |
| - | - | 1165 | 440.2 | - | - | 0 | - |
| - | - | 1640 | 441.2 | - | - | 0 | - |
| - | - | 570 | 442.2 | - | - | 0 | - |
| - | - | 1.068E+04 | 445.2 | - | - | 0 | - |
| - | - | 2216 | 446.3 | - | - | 0 | - |
| - | - | 657.3 | 446.3 | - | - | 0 | - |
| - | - | 1062 | 449.2 | - | - | 0 | - |
| - | - | 4106 | 451.2 | - | - | 0 | - |
| - | - | 835.6 | 453.2 | - | - | 0 | - |
| - | - | 2983 | 453.2 | - | - | 0 | - |
| - | - | 5197 | 454.3 | - | - | 0 | - |
| - | - | 1845 | 455.3 | - | - | 0 | - |
| - | - | 890.8 | 463.2 | - | - | 0 | - |
| 3 | y | 1692 | 467.6 | 0.001054 | 2.254 | +3 | 12 |
| - | - | 710.2 | 467.9 | - | - | 0 | - |
| - | - | 1871 | 468.2 | - | - | 0 | - |
| - | - | 1.954E+04 | 469.2 | - | - | 0 | - |
| - | - | 4677 | 470.2 | - | - | 0 | - |
| - | - | 560.7 | 470.7 | - | - | 0 | - |
| - | - | 2879 | 471.2 | - | - | 0 | - |
| - | - | 1994 | 472.3 | - | - | 0 | - |
| - | - | 831.8 | 479.2 | - | - | 0 | - |
| - | - | 619.2 | 479.2 | - | - | 0 | - |
| - | - | 663.9 | 489.8 | - | - | 0 | - |
| - | - | 593.1 | 490.7 | - | - | 0 | - |
| - | - | 2742 | 492.3 | - | - | 0 | - |
| 2 | y | 813.1 | 500.6 | 0.0001914 | 0.3823 | +3 | 13 |
| 7 | y | 5398 | 500.7 | 0.000427 | 0.8527 | +2 | 8 |
| - | - | 1111 | 500.9 | - | - | 0 | - |
| - | - | 2704 | 501.2 | - | - | 0 | - |
| - | - | 584.1 | 501.7 | - | - | 0 | - |
| - | - | 2196 | 504.7 | - | - | 0 | - |
| - | - | 779.5 | 505.2 | - | - | 0 | - |
| - | - | 7350 | 510.3 | - | - | 0 | - |
| - | - | 2892 | 511.3 | - | - | 0 | - |
| - | - | 696.8 | 512.3 | - | - | 0 | - |
| - | - | 3623 | 517.3 | - | - | 0 | - |
| - | - | 5173 | 520.3 | - | - | 0 | - |
| - | - | 1535 | 521.3 | - | - | 0 | - |
| 11 | y | 1077 | 522.3 | 0.0008047 | 1.541 | +1 | 4 |
| - | - | 641.1 | 532.2 | - | - | 0 | - |
| - | - | 660 | 532.9 | - | - | 0 | - |
| - | - | 720.5 | 533.2 | - | - | 0 | - |
| - | - | 710 | 533.3 | - | - | 0 | - |
| - | - | 7074 | 533.3 | - | - | 0 | - |
| 0 | Precursor | 4040 | 533.6 | 0.0007327 | 1.373 | +3 | -1 |
| - | - | 686.6 | 533.8 | - | - | 0 | - |
| - | - | 3229 | 533.9 | - | - | 0 | - |
| - | - | 739.5 | 534.2 | - | - | 0 | - |
| - | - | 2151 | 534.3 | - | - | 0 | - |
| - | - | 3489 | 534.3 | - | - | 0 | - |
| 6 | y | 970.8 | 535.3 | 0.002329 | 4.352 | +2 | 9 |
| - | - | 7451 | 538.3 | - | - | 0 | - |
| 10 | b | 2894 | 539.3 | 0.0008798 | 1.631 | +2 | 10 |
| - | - | 1942 | 540.2 | - | - | 0 | - |
| - | - | 698.3 | 540.8 | - | - | 0 | - |
| 6 | y | 6866 | 544.3 | 0.0002819 | 0.518 | +2 | 9 |
| - | - | 3766 | 544.8 | - | - | 0 | - |
| - | - | 1295 | 545.3 | - | - | 0 | - |
| - | - | 1880 | 550.2 | - | - | 0 | - |
| - | - | 670.6 | 551.2 | - | - | 0 | - |
| - | - | 741.1 | 552.2 | - | - | 0 | - |
| - | - | 970.6 | 554.3 | - | - | 0 | - |
| - | - | 1.457E+04 | 568.2 | - | - | 0 | - |
| - | - | 845.4 | 568.8 | - | - | 0 | - |
| - | - | 5470 | 569.2 | - | - | 0 | - |
| - | - | 2700 | 570.2 | - | - | 0 | - |
| - | - | 767.7 | 578.2 | - | - | 0 | - |
| - | - | 2017 | 580.2 | - | - | 0 | - |
| - | - | 1033 | 583.3 | - | - | 0 | - |
| 5 | y | 996.9 | 584.8 | 0.0007152 | 1.223 | +2 | 10 |
| 5 | y | 642 | 585.3 | 0.003825 | 6.535 | +2 | 10 |
| - | - | 698.9 | 590.3 | - | - | 0 | - |
| 5 | y | 1.181E+04 | 593.8 | 0.0001936 | 0.3261 | +2 | 10 |
| - | - | 4559 | 594.3 | - | - | 0 | - |
| - | - | 3002 | 594.8 | - | - | 0 | - |
| - | - | 777.5 | 595.3 | - | - | 0 | - |
| 10 | y | 979.6 | 601.3 | 0.001269 | 2.11 | +1 | 5 |
| 11 | b | 727 | 603.8 | 0.0007024 | 1.163 | +2 | 11 |
| - | - | 5107 | 608.2 | - | - | 0 | - |
| - | - | 1101 | 609.2 | - | - | 0 | - |
| - | - | 1183 | 618.3 | - | - | 0 | - |
| - | - | 1188 | 618.8 | - | - | 0 | - |
| 10 | y | 4.062E+04 | 619.3 | 0.0001403 | 0.2265 | +1 | 5 |
| - | - | 1.493E+04 | 620.3 | - | - | 0 | - |
| - | - | 2904 | 621.4 | - | - | 0 | - |
| - | - | 703.3 | 622.4 | - | - | 0 | - |
| - | - | 2576 | 627.3 | - | - | 0 | - |
| - | - | 1651 | 627.8 | - | - | 0 | - |
| - | - | 1213 | 637.3 | - | - | 0 | - |
| - | - | 2194 | 639.3 | - | - | 0 | - |
| - | - | 1190 | 640.3 | - | - | 0 | - |
| 4 | y | 1167 | 642.8 | 0.00372 | 5.787 | +2 | 11 |
| - | - | 686.5 | 643.3 | - | - | 0 | - |
| - | - | 973.1 | 649.3 | - | - | 0 | - |
| 4 | y | 2.446E+04 | 651.3 | 2.778E-05 | 0.04265 | +2 | 11 |
| - | - | 1.794E+04 | 651.8 | - | - | 0 | - |
| - | - | 6227 | 652.3 | - | - | 0 | - |
| - | - | 2586 | 652.8 | - | - | 0 | - |
| - | - | 1480 | 665.3 | - | - | 0 | - |
| - | - | 791.8 | 666.2 | - | - | 0 | - |
| - | - | 716.8 | 667.3 | - | - | 0 | - |
| - | - | 1.219E+04 | 667.3 | - | - | 0 | - |
| - | - | 3827 | 668.3 | - | - | 0 | - |
| - | - | 745.4 | 669.8 | - | - | 0 | - |
| - | - | 742.8 | 677.8 | - | - | 0 | - |
| - | - | 1101 | 679.3 | - | - | 0 | - |
| - | - | 834.8 | 680.3 | - | - | 0 | - |
| - | - | 693.9 | 681.8 | - | - | 0 | - |
| - | - | 659.9 | 682.8 | - | - | 0 | - |
| - | - | 1.834E+04 | 683.3 | - | - | 0 | - |
| - | - | 797.6 | 683.3 | - | - | 0 | - |
| - | - | 5880 | 684.3 | - | - | 0 | - |
| - | - | 1180 | 685.3 | - | - | 0 | - |
| - | - | 1128 | 688.8 | - | - | 0 | - |
| 3 | y | 2378 | 691.8 | 0.002048 | 2.96 | +2 | 12 |
| 3 | y | 4003 | 692.3 | 0.006561 | 9.477 | +2 | 12 |
| - | - | 2124 | 692.8 | - | - | 0 | - |
| - | - | 1006 | 693.2 | - | - | 0 | - |
| - | - | 670.7 | 693.3 | - | - | 0 | - |
| - | - | 5081 | 695.3 | - | - | 0 | - |
| - | - | 1183 | 696.3 | - | - | 0 | - |
| - | - | 726 | 700.3 | - | - | 0 | - |
| 3 | y | 1.328E+05 | 700.8 | 6.154E-05 | 0.08781 | +2 | 12 |
| - | - | 1.138E+05 | 701.3 | - | - | 0 | - |
| - | - | 5.103E+04 | 701.8 | - | - | 0 | - |
| - | - | 1.555E+04 | 702.3 | - | - | 0 | - |
| - | - | 2409 | 702.8 | - | - | 0 | - |
| - | - | 3361 | 707.3 | - | - | 0 | - |
| - | - | 1604 | 708.3 | - | - | 0 | - |
| 7 | b | 613.4 | 718.4 | 0.01386 | 19.29 | +1 | 7 |
| 9 | y | 3619 | 734.4 | 0.000533 | 0.7259 | +1 | 6 |
| - | - | 1678 | 735.4 | - | - | 0 | - |
| 2 | y | 840.5 | 741.4 | 0.00097 | 1.308 | +2 | 13 |
| 2 | y | 732.4 | 741.9 | 0.009647 | 13 | +2 | 13 |
| 2 | y | 2.295E+04 | 750.4 | 3.427E-05 | 0.04567 | +2 | 13 |
| - | - | 2.201E+04 | 750.9 | - | - | 0 | - |
| - | - | 1.034E+04 | 751.4 | - | - | 0 | - |
| - | - | 3246 | 751.9 | - | - | 0 | - |
| - | - | 729.3 | 752.4 | - | - | 0 | - |
| - | - | 3357 | 764.3 | - | - | 0 | - |
| - | - | 1622 | 765.3 | - | - | 0 | - |
| - | - | 1707 | 766.3 | - | - | 0 | - |
| - | - | 1868 | 766.4 | - | - | 0 | - |
| - | - | 828.4 | 767.3 | - | - | 0 | - |
| - | - | 688.7 | 776.3 | - | - | 0 | - |
| - | - | 948.2 | 777.3 | - | - | 0 | - |
| - | - | 1652 | 780.3 | - | - | 0 | - |
| - | - | 5.232E+04 | 782.3 | - | - | 0 | - |
| - | - | 2.267E+04 | 783.3 | - | - | 0 | - |
| - | - | 5681 | 784.3 | - | - | 0 | - |
| - | - | 8079 | 794.3 | - | - | 0 | - |
| - | - | 3996 | 795.3 | - | - | 0 | - |
| - | - | 790.6 | 796.3 | - | - | 0 | - |
| - | - | 950 | 835.4 | - | - | 0 | - |
| - | - | 1022 | 836.4 | - | - | 0 | - |
| 8 | y | 782.5 | 845.4 | 0.001434 | 1.696 | +1 | 7 |
| 8 | y | 6962 | 863.4 | 0.003514 | 4.07 | +1 | 7 |
| - | - | 3095 | 864.4 | - | - | 0 | - |
| - | - | 1345 | 865.4 | - | - | 0 | - |
| - | - | 2717 | 879.4 | - | - | 0 | - |
| - | - | 998.8 | 880.4 | - | - | 0 | - |
| - | - | 9037 | 881.4 | - | - | 0 | - |
| - | - | 3980 | 882.4 | - | - | 0 | - |
| - | - | 762.9 | 883.4 | - | - | 0 | - |
| - | - | 848.3 | 891.3 | - | - | 0 | - |
| - | - | 3069 | 893.4 | - | - | 0 | - |
| - | - | 1976 | 894.4 | - | - | 0 | - |
| - | - | 8635 | 909.4 | - | - | 0 | - |
| - | - | 3680 | 910.4 | - | - | 0 | - |
| - | - | 1472 | 911.4 | - | - | 0 | - |
| - | - | 882.6 | 922.4 | - | - | 0 | - |
| - | - | 911.1 | 962.4 | - | - | 0 | - |
| - | - | 747.2 | 979.5 | - | - | 0 | - |
| - | - | 5624 | 980.4 | - | - | 0 | - |
| - | - | 2856 | 981.4 | - | - | 0 | - |
| 7 | y | 918.6 | 982.5 | 0.008588 | 8.741 | +1 | 8 |
| - | - | 3074 | 990.4 | - | - | 0 | - |
| - | - | 1230 | 991.4 | - | - | 0 | - |
| - | - | 888.8 | 992.4 | - | - | 0 | - |
| 7 | y | 5863 | 1000 | 0.001269 | 1.268 | +1 | 8 |
| - | - | 3494 | 1001 | - | - | 0 | - |
| - | - | 1502 | 1002 | - | - | 0 | - |
| - | - | 3.498E+04 | 1008 | - | - | 0 | - |
| - | - | 1.756E+04 | 1009 | - | - | 0 | - |
| - | - | 6226 | 1010 | - | - | 0 | - |
| - | - | 982.9 | 1011 | - | - | 0 | - |
| - | - | 708.1 | 1021 | - | - | 0 | - |
| - | - | 5953 | 1079 | - | - | 0 | - |
| - | - | 3710 | 1081 | - | - | 0 | - |
| - | - | 979.9 | 1081 | - | - | 0 | - |
| 6 | y | 7577 | 1088 | 0.002047 | 1.883 | +1 | 9 |
| - | - | 699 | 1088 | - | - | 0 | - |
| - | - | 4524 | 1089 | - | - | 0 | - |
| - | - | 3994 | 1089 | - | - | 0 | - |
| - | - | 1855 | 1090 | - | - | 0 | - |
| - | - | 4.08E+04 | 1107 | - | - | 0 | - |
| - | - | 2.262E+04 | 1108 | - | - | 0 | - |
| - | - | 8084 | 1109 | - | - | 0 | - |
| - | - | 1942 | 1111 | - | - | 0 | - |
| - | - | 808.4 | 1137 | - | - | 0 | - |
| - | - | 1009 | 1138 | - | - | 0 | - |
| - | - | 1177 | 1179 | - | - | 0 | - |
| - | - | 654.7 | 1180 | - | - | 0 | - |
| 5 | y | 2221 | 1187 | 0.0003395 | 0.2861 | +1 | 10 |
| - | - | 994 | 1188 | - | - | 0 | - |
| 11 | b | 7236 | 1207 | 0.001941 | 1.609 | +1 | 11 |
| - | - | 5108 | 1208 | - | - | 0 | - |
| - | - | 2420 | 1209 | - | - | 0 | - |
| - | - | 1175 | 1237 | - | - | 0 | - |
| 4 | y | 1850 | 1302 | 0.003044 | 2.339 | +1 | 11 |
| - | - | 1164 | 1303 | - | - | 0 | - |
| 12 | b | 923.5 | 1306 | 0.004437 | 3.398 | +1 | 12 |
| - | - | 808.3 | 1808 | - | - | 0 | - |
| - | - | 715.7 | 2209 | - | - | 0 | - |

m/z Charge Intensity FragmentType MassShift Position
120.08104705810547 0 9820.002
121.08454895019531 0 543.33673
122.07193756103516 0 420.97745
127.07526397705078 0 588.40173
127.08683013916016 0 1008.8964
128.10726928710938 0 1363.9568
128.98464965820312 0 465.26837
129.10255432128906 0 139858.84
130.08648681640625 0 1646.7078
130.1000213623047 0 783.90704
130.10586547851562 0 8962.821
131.0702362060547 0 482.13766
132.10218811035156 0 1340.3931
133.06109619140625 0 545.20593
133.08619689941406 0 3443.948
136.0759735107422 0 2533.0996
137.8849639892578 0 444.55057
138.06666564941406 0 907.63226
138.091796875 0 722.96625
140.10707092285156 0 607.69305
141.10243225097656 0 2525.709
143.11843872070312 0 690.16046
145.06141662597656 0 420.8248
147.1019744873047 0 883.76984
147.11312866210938 0 5218.521
147.45620727539062 0 464.90198
149.04519653320312 0 5016.253
152.08189392089844 0 566.3804
155.11819458007812 0 8067.467
156.07708740234375 0 569.1373
163.07156372070312 0 1348.4146
164.11851501464844 0 3942.0408
165.0545654296875 0 605.64655
166.0865020751953 0 5316.3457 y 13
167.05577087402344 0 6724.5933
167.09243774414062 0 1234.9325
168.10218811035156 0 631.5269
169.097412109375 0 4796.5474
169.1337890625 0 542.642
171.14955139160156 0 194031.88 a 1
172.15286254882812 0 18126.34
173.12879943847656 0 1744.8534
173.15576171875 0 923.1167
173.4381866455078 0 777.70544
177.1121368408203 0 1404.0889
179.0930633544922 0 2488.2954
180.0767059326172 0 510.10544
181.06130981445312 0 459.34268
182.0811004638672 0 963.87823
183.11312866210938 0 2009.4006
183.1494903564453 0 5960.229
184.15252685546875 0 795.57794
185.09230041503906 0 875.0426
185.12832641601562 0 585.64905
185.16513061523438 0 40337.484
186.11277770996094 0 1726.1829
186.16864013671875 0 3822.0964
187.0908966064453 0 962.3637
187.1079864501953 0 16245.06
187.1444091796875 0 836.6162
188.11138916015625 0 1894.4747
190.96888732910156 0 557.8411
191.12742614746094 0 503.17358
194.09303283691406 0 563.58594
195.0775909423828 0 450.44196
195.08787536621094 0 807.4601
196.0609130859375 0 565.7861
197.10353088378906 0 4844.144
197.12835693359375 0 897.8006 y 11
199.10809326171875 0 5288.0625
199.14437866210938 0 86811.266 b 1
200.1122589111328 0 700.8602
200.14773559570312 0 9137.823
201.12351989746094 0 11652.217
202.12677001953125 0 1216.2318
203.10276794433594 0 1718.0387
207.0880584716797 0 3039.4846
208.0722198486328 0 2887.8386
208.09068298339844 0 618.41925
211.10791015625 0 661.7018
211.14437866210938 0 2716.3923
213.08738708496094 0 1310.7887
213.16000366210938 0 39187.04
214.1633758544922 0 4486.64
215.10289001464844 0 12463.404
215.13954162597656 0 995.5625
216.06898498535156 0 1750.6398
216.10670471191406 0 1736.1494
217.07298278808594 0 568.35394
218.08517456054688 0 809.4199
221.0846710205078 0 10479.812
221.10336303710938 0 1010.27997
222.0854949951172 0 1442.7828
223.12667846679688 0 2761.1292
224.129150390625 0 558.20215
225.043212890625 0 14306.378
225.09848022460938 0 18250.865
226.043212890625 0 2424.9011
226.064697265625 0 1247.8191
226.1017608642578 0 1604.6184
227.02249145507812 0 938.04254
227.1029052734375 0 18385.266
228.10618591308594 0 2408.4268
228.1708984375 0 16506.625
229.1191864013672 0 626.8991
229.1552276611328 0 622.0033
229.17422485351562 0 2288.8245
231.14955139160156 0 695.2372
233.09573364257812 0 67528.055
234.079345703125 0 1564.6029
234.09902954101562 0 7223.8447
235.11143493652344 0 6589.653
239.09524536132812 0 25051.69
239.11376953125 0 1647.4225
239.1393280029297 0 1999.5856
240.09559631347656 0 3504.5955
240.14402770996094 0 699.7296
243.1346435546875 0 692.82477
245.0771484375 0 1034.9208
246.1811981201172 0 1259.6714
249.09837341308594 0 1732.4095
251.1062469482422 0 13968.793
251.1214599609375 0 7497.925
252.11012268066406 0 1506.4329
253.09365844726562 0 674.6323
253.19117736816406 0 1408.8528
257.1494445800781 0 766.55695
259.18011474609375 0 743.14557
260.1274719238281 0 714.717
266.1247863769531 0 679.163
267.1090393066406 0 7734.6313
268.1117248535156 0 952.8513
268.1648254394531 0 707.71246
274.1290283203125 0 664.06055
275.14068603515625 0 803.3561
276.171142578125 0 1200.7213
277.1549072265625 0 7313.091 y Ammonia loss 12
278.15826416015625 0 1459.0944
279.1164855957031 0 4149.5054
279.1432800292969 0 675.99097
281.0519714355469 0 941.76404
281.080078125 0 843.67114
282.18145751953125 0 15447.981
283.18487548828125 0 2086.4963
284.1243591308594 0 1442.327
285.0099182128906 0 771.26984
286.1587829589844 0 954.8825
286.17572021484375 0 765.83026
287.14337158203125 0 941.8126
288.1453857421875 0 807.48193
289.92431640625 0 545.04803
294.1816101074219 0 46259.008 y 12
295.103515625 0 5094.1865
295.18475341796875 0 7198.1924
296.1043395996094 0 1075.9794
298.1756286621094 0 1189.6544
298.212646484375 0 3890.4006 b 2
299.0621032714844 0 7622.1
300.0624694824219 0 2477.038
300.19195556640625 0 1849.2874
302.1350402832031 0 3656.958
303.1374206542969 0 5706.409
304.14056396484375 0 899.1584
304.1650695800781 0 976.97815
306.1568603515625 0 2080.5605
307.14056396484375 0 3527.7197
307.15924072265625 0 918.10095
310.15576171875 0 562.2046
310.1764221191406 0 2793.8918 y 9
313.11419677734375 0 1946.6779
314.153076171875 0 1053.6576
314.17138671875 0 10567.198
315.1756896972656 0 1216.7683
321.1483459472656 0 4768.386
322.1119384765625 0 653.3973
322.1509094238281 0 594.7853
324.11932373046875 0 1756.3079
324.16668701171875 0 2757.289
325.1690979003906 0 623.08716
326.1713562011719 0 9853.754
327.1748962402344 0 1840.5016
332.1642761230469 0 3217.6343
334.15179443359375 0 657.81006
336.1305847167969 0 2227.312
342.1297607421875 0 3614.8066
343.1322021484375 0 876.7569
346.1152038574219 0 868.24554
354.1409606933594 0 7955.9272
355.07110595703125 0 981.1391
355.1441650390625 0 1296.8827
359.0281982421875 0 1144.9043
364.12548828125 0 8377.818
364.20556640625 0 10113.01
365.1268615722656 0 792.04
365.2085876464844 0 2210.0369
367.2704772949219 0 620.5561
368.271728515625 0 591.0963
369.0971374511719 0 738.39514
369.122314453125 0 5273.6255
370.1227722167969 0 1684.7386
377.1659851074219 0 1061.0502
378.81048583984375 0 556.49164
382.135986328125 0 19274.244
383.1386413574219 0 3205.143
383.1929626464844 0 2319.6792
385.24468994140625 0 699.496
391.6690979003906 0 652.4561
392.1999816894531 0 1112.0618
393.25006103515625 0 17941.91 y 11
394.2527160644531 0 4788.8228
395.2640075683594 0 1886.0319
399.242919921875 0 1907.8048
401.2041015625 0 2810.822
402.2068786621094 0 1339.132
406.208740234375 0 2271.1436
407.2104187011719 0 621.98846
410.1317443847656 0 1257.7374
410.2113342285156 0 11135.405
411.1977233886719 0 1547.003
411.2188415527344 0 953.2444
413.2068176269531 0 979.35657
413.2402648925781 0 2870.8535 b 3
413.2762756347656 0 9491.21
414.2410583496094 0 682.17487
414.2795104980469 0 2919.773
421.1839294433594 0 3527.9187
422.1846008300781 0 678.69086
423.19189453125 0 813.15564
425.2119140625 0 663.5042
427.23779296875 0 43616.336
428.2407531738281 0 11114.268
433.14892578125 0 646.75433
439.1937561035156 0 5111.878
440.1967468261719 0 1164.6364
441.1990051269531 0 1640.1984
442.2007141113281 0 570.0096
445.2483825683594 0 10676.528
446.2506408691406 0 2216.182
446.2825012207031 0 657.3319
449.1784973144531 0 1061.5553
451.15765380859375 0 4105.8823
453.1603698730469 0 835.5783
453.2093505859375 0 2983.3555
454.2667236328125 0 5197.0107
455.2681884765625 0 1845.3871
463.1956481933594 0 890.7568
467.56231689453125 0 1691.5483 y 2
467.8961181640625 0 710.2336
468.2294006347656 0 1870.9353
469.1683044433594 0 19540.498
470.17132568359375 0 4676.758
470.7226867675781 0 560.6956
471.174072265625 0 2878.5234
472.2770080566406 0 1993.8091
479.1508483886719 0 831.78546
479.18585205078125 0 619.17346
489.77130126953125 0 663.9365
490.72467041015625 0 593.0583
492.2576599121094 0 2742.1814
500.5842590332031 0 813.1312 y 1
500.74078369140625 0 5398.1123 y 6
500.9180908203125 0 1110.5681
501.2434997558594 0 2703.8833
501.7455139160156 0 584.10645
504.7177429199219 0 2195.6804
505.21759033203125 0 779.48346
510.2675476074219 0 7350.101
511.26953125 0 2891.8599
512.2731323242188 0 696.7856
517.3219604492188 0 3622.92
520.2515258789062 0 5172.6396
521.2540283203125 0 1535.3792
522.2930297851562 0 1077.2126 y 10
532.2134399414062 0 641.0836
532.9246826171875 0 660.0127
533.209228515625 0 720.50214
533.26513671875 0 710.04694
533.3003540039062 0 7073.627
533.6076049804688 0 4039.5222 Precursor
533.8011474609375 0 686.58734
533.9418334960938 0 3229.3193
534.23583984375 0 739.50964
534.3050537109375 0 2150.861
534.3497314453125 0 3488.5156
535.25341796875 0 970.8249 y Water loss 5
538.2622680664062 0 7450.805
539.2650756835938 0 2894.0056 b 9
540.2428588867188 0 1941.9076
540.7549438476562 0 698.2697
544.2566528320312 0 6866.171 y 5
544.7576904296875 0 3765.5618
545.258056640625 0 1294.5764
550.2245483398438 0 1879.5529
551.226806640625 0 670.6004
552.2291870117188 0 741.1262
554.2504272460938 0 970.57794
568.2363891601562 0 14572.961
568.808349609375 0 845.4447
569.2393798828125 0 5470.322
570.2404174804688 0 2699.7188
578.2189331054688 0 767.68896
580.2350463867188 0 2016.9159
583.3242797851562 0 1033.4183
584.7860107421875 0 996.9403 y Water loss 4
585.2811279296875 0 641.95404 y Ammonia loss 4
590.2837524414062 0 698.89764
593.790771484375 0 11807.747 y 4
594.2920532226562 0 4558.516
594.7922973632812 0 3002.4983
595.2924194335938 0 777.5237
601.335693359375 0 979.5509 y Water loss 9
603.7847900390625 0 727.00977 b 10
608.23095703125 0 5107.491
609.2333374023438 0 1101.4277
618.2998657226562 0 1182.8136
618.8012084960938 0 1187.986
619.3448486328125 0 40616.152 y 9
620.3477783203125 0 14934.031
621.3509521484375 0 2904.2053
622.3515625 0 703.26025
627.3046875 0 2575.9817
627.80517578125 0 1651.4492
637.3319091796875 0 1213.0715
639.309814453125 0 2194.314
640.3120727539062 0 1189.9473
642.7944946289062 0 1167.3186 y Ammonia loss 3
643.2964477539062 0 686.54865
649.2965087890625 0 973.12415
651.3040771484375 0 24463.902 y 3
651.8057861328125 0 17941.312
652.3074951171875 0 6227.1978
652.8095703125 0 2586.4531
665.2520751953125 0 1480.278
666.25 0 791.78876
667.2527465820312 0 716.7775
667.3045654296875 0 12190.549
668.3076782226562 0 3826.6318
669.8455200195312 0 745.40204
677.8360595703125 0 742.83716
679.3104858398438 0 1100.8469
680.3110961914062 0 834.7502
681.8314208984375 0 693.8958
682.83056640625 0 659.9487
683.2627563476562 0 18340.625
683.3215942382812 0 797.59125
684.2666625976562 0 5880.1104
685.2676391601562 0 1179.6691
688.8248901367188 0 1128.3726
691.8350219726562 0 2377.5554 y Water loss 2
692.33154296875 0 4002.5693 y Ammonia loss 2
692.8323364257812 0 2123.77
693.2484741210938 0 1006.19806
693.3194580078125 0 670.7469
695.2642822265625 0 5081.0737
696.2650756835938 0 1183.2141
700.3359375 0 725.9929
700.8383178710938 0 132787.42 y 2
701.339599609375 0 113754.4
701.8409423828125 0 51026.758
702.3421630859375 0 15552.213
702.8439331054688 0 2409.225
707.3004150390625 0 3360.9873
708.3006591796875 0 1604.4086
718.3743896484375 0 613.36786 b Water loss 6
734.3713989257812 0 3618.849 y 8
735.374267578125 0 1678.4507
741.3662109375 0 840.5137 y Water loss 1
741.8688354492188 0 732.4222 y Ammonia loss 1
750.3724975585938 0 22952.816 y 1
750.873779296875 0 22005.28
751.3748168945312 0 10341.686
751.876708984375 0 3246.2688
752.3784790039062 0 729.3225
764.3199462890625 0 3357.2844
765.32373046875 0 1622.2567
766.34423828125 0 1706.5476
766.3695068359375 0 1867.83
767.3384399414062 0 828.38324
776.3203735351562 0 688.7234
777.3225708007812 0 948.2246
780.3139038085938 0 1652.4529
782.331298828125 0 52322.016
783.3341064453125 0 22671.2
784.3378295898438 0 5680.8447
794.3311767578125 0 8078.9604
795.3348388671875 0 3996.4695
796.3391723632812 0 790.554
835.3955078125 0 949.9799
836.3893432617188 0 1021.65564
845.4025268554688 0 782.4779 y Water loss 7
863.4110107421875 0 6961.534 y 7
864.4141845703125 0 3094.759
865.41015625 0 1345.4437
879.3863525390625 0 2717.0588
880.3834228515625 0 998.768
881.3965454101562 0 9037.059
882.3994750976562 0 3979.5225
883.403564453125 0 762.93695
891.3499145507812 0 848.3496
893.40087890625 0 3068.7197
894.4015502929688 0 1975.6615
909.3585205078125 0 8635.449
910.3612670898438 0 3679.5076
911.3642578125 0 1472.484
922.4212036132812 0 882.6476
962.41943359375 0 911.1067
979.4746704101562 0 747.2318
980.439208984375 0 5624.115
981.44140625 0 2855.779
982.4542846679688 0 918.5619 y Water loss 6
990.4154663085938 0 3074.4695
991.4171142578125 0 1230.388
992.4171142578125 0 888.7923
1000.47216796875 0 5862.5757 y 6
1001.4752197265625 0 3493.758
1002.4775390625 0 1502.4039
1008.4258422851562 0 34982.58
1009.4291381835938 0 17556.717
1010.4315185546875 0 6225.754
1011.4410400390625 0 982.8861
1021.4912109375 0 708.0819
1079.4984130859375 0 5952.9077
1080.5035400390625 0 3710.3274
1081.4969482421875 0 979.8959
1087.50341796875 0 7576.893 y 5
1087.970458984375 0 699.03156
1088.505859375 0 4524.1733
1089.494873046875 0 3994.1353
1090.4959716796875 0 1854.6147
1107.493896484375 0 40804.145
1108.4971923828125 0 22618.28
1109.4991455078125 0 8083.9336
1110.504150390625 0 1942.2322
1136.521728515625 0 808.36304
1137.522216796875 0 1008.91724
1178.5684814453125 0 1177.4037
1179.587158203125 0 654.7202
1186.57421875 0 2220.6187 y 4
1187.5806884765625 0 994.0151
1206.561767578125 0 7235.5312 b 10
1207.5645751953125 0 5108.0513
1208.5660400390625 0 2419.8088
1236.583984375 0 1175.4982
1301.5977783203125 0 1850.004 y 3
1302.6136474609375 0 1163.757
1305.627685546875 0 923.51263 b 11
1807.9559326171875 0 808.3285
2209.265380859375 0 715.65436

Spectrum Details

|  |  |
| --- | --- |
| Matched peaks? Matched peaks The total absolute number of peaks matched. Additionally in brackets the total fraction of peaks matched and the total number of peaks is shown. | 43 (9.60% of 448) |
| FDR? FDR The false discovery rate estimated for this peptide. It is calculated by matching all theoretical fragments with a non-integer shift with the raw peaks for this spectrum. This is done with 40 different shifts. The resulting percentage is the average number of annotated peaks over the number of annotated peaks with the correct spectrum. | 2.27% |
| Satellite FDR? Satellite FDR See the FDR for details on its calculation. This satellite ion specific FDR only contains the satellite ions (d/w) for I/L/J positions. | - |
| PSM Score? PSM Score The PSM Score as given by Hecklib to this annotated spectrum. It is shown with three significant figures. | 274 |

#### Spectrum 7148? Spectrum 7148 The raw spectrum of this peptide as annotated by Hecklib. The fragments are coloured according to ion type (see legend). Any peaks with a star '\*' as text can be hovered over to see the full details, first the ion type second the mass shift type. By hovering over the amino acids in the peptide or ions in the legend the corresponding peaks are highlighted. By toggling the 'Unassigned' label you can turn the background (unassigned) peaks on or off in the plot. By updating the slider in the Ion legend you can update the spectrum to only show the top X% of the peaks with labels. The top X% means any peak that is within X% of the highest intensity. By dragging in the spectrum you can zoom in to a specific part of the spectrum and use 'Zoom Out' to get back to the original zoom level. The annotation of the spectrum is based on the given sequence in the peptides file and is done with different software so inconsistencies are likely. The peaks are annotated based on the given sequence, with 20 ppm tolerance.

Copy Data

##### Spectrum 7148 (TSV)

###### Preview

```
Loading example...
```

*Click on the button to copy the data to your clipboard.*

Mz MinMz MaxIntensity Max

WidthHeightPeptide font sizePeptide stroke widthSpectrum font sizeSpectrum stroke widthCompact peptide

Ion legend

wxyz

abcd

OtherUnassignedIon^Charge^_Position_Show for top:%

VVVDVSHEDPEVKF

01.18e+42.36e+43.53e+44.71e+4

Zoom Out

z^+2^_2_y^+2^_3_z^+1^_2_y^+1^_2_c^+1^_3_w^+1^_3_z^+1^_3_y^+1^_3_c^+1^_4_w^+1^_4_c^+1^_5_z^+2^_10_y^+2^_10_c^+1^_6_y^+1^_5_z^+2^_11_y^+2^_11_z^+2^_12_y^+2^_12_z^+1^_6_c^+2^_13_y^+1^_6_y^+2^_13_c^+1^_7_w^+1^_7_z^+1^_7_y^+1^_7_c^+1^_8_z^+1^_8_y^+1^_8_z^+1^_9_y^+1^_9_c^+1^_10_z^+1^_10_y^+1^_10_c^+1^_11_z^+1^_11_y^+1^_11_c^+1^_12_c^+1^_13_

0802160424063208

Fragment Matches Table

Show background peaks

| Position | Ion type | Intensity | mz Theoretical | mz Error (Th) | mz Error (ppm) | Charge | Series Number |
| --- | --- | --- | --- | --- | --- | --- | --- |
| - | - | 708 | 120.1 | - | - | 0 | - |
| - | - | 363.6 | 121.9 | - | - | 0 | - |
| - | - | 374.7 | 122.9 | - | - | 0 | - |
| - | - | 377 | 127.1 | - | - | 0 | - |
| - | - | 458 | 128.1 | - | - | 0 | - |
| - | - | 2366 | 129.1 | - | - | 0 | - |
| - | - | 402.9 | 130.8 | - | - | 0 | - |
| 13 | z | 1566 | 131.1 | 0.001237 | 9.439 | +2 | 2 |
| - | - | 424.1 | 131.7 | - | - | 0 | - |
| - | - | 1.504E+04 | 133.1 | - | - | 0 | - |
| - | - | 2206 | 134.1 | - | - | 0 | - |
| - | - | 418.4 | 135.2 | - | - | 0 | - |
| - | - | 7026 | 147.1 | - | - | 0 | - |
| - | - | 450.2 | 147.4 | - | - | 0 | - |
| - | - | 3524 | 149 | - | - | 0 | - |
| - | - | 424.2 | 150.7 | - | - | 0 | - |
| - | - | 1162 | 161.1 | - | - | 0 | - |
| - | - | 764.1 | 167.1 | - | - | 0 | - |
| - | - | 7192 | 171.1 | - | - | 0 | - |
| - | - | 553 | 172.2 | - | - | 0 | - |
| - | - | 1667 | 173.5 | - | - | 0 | - |
| - | - | 1643 | 175.1 | - | - | 0 | - |
| - | - | 9614 | 177.1 | - | - | 0 | - |
| - | - | 1508 | 178.1 | - | - | 0 | - |
| - | - | 454.8 | 181.3 | - | - | 0 | - |
| - | - | 794.9 | 185.2 | - | - | 0 | - |
| - | - | 1426 | 187.1 | - | - | 0 | - |
| - | - | 1151 | 191.1 | - | - | 0 | - |
| 12 | y | 860.9 | 197.1 | 0.0003147 | 1.596 | +2 | 3 |
| - | - | 979.6 | 198.1 | - | - | 0 | - |
| - | - | 7560 | 199.1 | - | - | 0 | - |
| - | - | 1035 | 201.1 | - | - | 0 | - |
| - | - | 5041 | 205.1 | - | - | 0 | - |
| - | - | 1758 | 206.1 | - | - | 0 | - |
| - | - | 5340 | 213.2 | - | - | 0 | - |
| - | - | 729 | 215.1 | - | - | 0 | - |
| - | - | 552.7 | 216.7 | - | - | 0 | - |
| - | - | 1221 | 219.1 | - | - | 0 | - |
| - | - | 545.2 | 219.6 | - | - | 0 | - |
| - | - | 1.221E+04 | 221.1 | - | - | 0 | - |
| - | - | 2774 | 221.1 | - | - | 0 | - |
| - | - | 2257 | 222.1 | - | - | 0 | - |
| - | - | 4182 | 225 | - | - | 0 | - |
| - | - | 637.9 | 227 | - | - | 0 | - |
| - | - | 725.9 | 228.2 | - | - | 0 | - |
| - | - | 790 | 232.1 | - | - | 0 | - |
| - | - | 4649 | 233.1 | - | - | 0 | - |
| - | - | 725.9 | 233.2 | - | - | 0 | - |
| - | - | 739 | 235.2 | - | - | 0 | - |
| - | - | 1.958E+04 | 239.1 | - | - | 0 | - |
| - | - | 1030 | 239.1 | - | - | 0 | - |
| - | - | 2545 | 240.1 | - | - | 0 | - |
| - | - | 1186 | 249.1 | - | - | 0 | - |
| - | - | 1059 | 251.1 | - | - | 0 | - |
| - | - | 1622 | 257.1 | - | - | 0 | - |
| - | - | 2352 | 265.2 | - | - | 0 | - |
| - | - | 887.4 | 276.2 | - | - | 0 | - |
| 13 | z | 823.1 | 278.2 | 0.000714 | 2.567 | +1 | 2 |
| - | - | 1332 | 281.1 | - | - | 0 | - |
| - | - | 1831 | 282.2 | - | - | 0 | - |
| - | - | 1169 | 283.2 | - | - | 0 | - |
| - | - | 554.7 | 292.1 | - | - | 0 | - |
| 13 | y | 1775 | 294.2 | 0.0008897 | 3.024 | +1 | 2 |
| - | - | 9175 | 295.1 | - | - | 0 | - |
| - | - | 2271 | 296.1 | - | - | 0 | - |
| - | - | 1242 | 298.2 | - | - | 0 | - |
| - | - | 3640 | 299.1 | - | - | 0 | - |
| - | - | 1309 | 300.1 | - | - | 0 | - |
| - | - | 627.6 | 303.1 | - | - | 0 | - |
| - | - | 2035 | 309.2 | - | - | 0 | - |
| - | - | 1703 | 313.1 | - | - | 0 | - |
| - | - | 535.8 | 313.6 | - | - | 0 | - |
| - | - | 578.2 | 314.1 | - | - | 0 | - |
| - | - | 539.2 | 314.2 | - | - | 0 | - |
| - | - | 1443 | 314.2 | - | - | 0 | - |
| 3 | c | 2060 | 315.2 | 0.0005419 | 1.719 | +1 | 3 |
| - | - | 970.3 | 321.1 | - | - | 0 | - |
| - | - | 2883 | 352.2 | - | - | 0 | - |
| - | - | 940.7 | 353.2 | - | - | 0 | - |
| - | - | 1637 | 355.1 | - | - | 0 | - |
| - | - | 521 | 356.1 | - | - | 0 | - |
| - | - | 848.2 | 359 | - | - | 0 | - |
| 12 | w | 1048 | 362.2 | 0.0007678 | 2.12 | +1 | 3 |
| - | - | 503.1 | 364.8 | - | - | 0 | - |
| - | - | 543.1 | 368.9 | - | - | 0 | - |
| - | - | 9237 | 369.1 | - | - | 0 | - |
| - | - | 2912 | 370.1 | - | - | 0 | - |
| - | - | 975.2 | 370.2 | - | - | 0 | - |
| - | - | 832.2 | 371.2 | - | - | 0 | - |
| - | - | 602.4 | 372.1 | - | - | 0 | - |
| - | - | 773.9 | 372.2 | - | - | 0 | - |
| - | - | 803.5 | 377.2 | - | - | 0 | - |
| 12 | z | 1311 | 377.2 | 0.001019 | 2.701 | +1 | 3 |
| - | - | 685.1 | 378.2 | - | - | 0 | - |
| - | - | 720.5 | 386.3 | - | - | 0 | - |
| 12 | y | 3024 | 393.2 | 0.0005475 | 1.392 | +1 | 3 |
| - | - | 577.2 | 394.3 | - | - | 0 | - |
| - | - | 2259 | 413.2 | - | - | 0 | - |
| - | - | 2331 | 413.3 | - | - | 0 | - |
| - | - | 922.7 | 414.2 | - | - | 0 | - |
| - | - | 587.6 | 414.3 | - | - | 0 | - |
| - | - | 8571 | 427.2 | - | - | 0 | - |
| - | - | 2431 | 428.2 | - | - | 0 | - |
| - | - | 1477 | 429.3 | - | - | 0 | - |
| 4 | c | 1155 | 430.3 | 1.139E-05 | 0.02648 | +1 | 4 |
| - | - | 793.4 | 443.2 | - | - | 0 | - |
| - | - | 3074 | 445.2 | - | - | 0 | - |
| - | - | 825.4 | 446.3 | - | - | 0 | - |
| 11 | w | 5322 | 447.3 | 0.0001183 | 0.2644 | +1 | 4 |
| - | - | 1298 | 448.3 | - | - | 0 | - |
| - | - | 1183 | 457.3 | - | - | 0 | - |
| - | - | 4150 | 459.3 | - | - | 0 | - |
| - | - | 1685 | 460.3 | - | - | 0 | - |
| - | - | 1691 | 473.3 | - | - | 0 | - |
| - | - | 834.2 | 476.2 | - | - | 0 | - |
| - | - | 616.1 | 481.3 | - | - | 0 | - |
| - | - | 537.9 | 483.1 | - | - | 0 | - |
| - | - | 4682 | 487.3 | - | - | 0 | - |
| - | - | 1373 | 488.3 | - | - | 0 | - |
| - | - | 757.4 | 489.3 | - | - | 0 | - |
| - | - | 1964 | 504.3 | - | - | 0 | - |
| - | - | 3405 | 512.3 | - | - | 0 | - |
| - | - | 1143 | 513.3 | - | - | 0 | - |
| - | - | 737 | 516.3 | - | - | 0 | - |
| - | - | 3.424E+04 | 517.3 | - | - | 0 | - |
| - | - | 595.4 | 518.5 | - | - | 0 | - |
| 5 | c | 1179 | 529.3 | 0.000684 | 1.292 | +1 | 5 |
| - | - | 1.355E+04 | 532.3 | - | - | 0 | - |
| - | - | 2759 | 532.3 | - | - | 0 | - |
| - | - | 8670 | 533.3 | - | - | 0 | - |
| - | - | 776.6 | 533.3 | - | - | 0 | - |
| - | - | 2922 | 534.3 | - | - | 0 | - |
| - | - | 3.859E+04 | 534.3 | - | - | 0 | - |
| - | - | 608.3 | 534.8 | - | - | 0 | - |
| 5 | z | 952.4 | 585.8 | 0.001187 | 2.026 | +2 | 10 |
| 5 | y | 882.6 | 593.8 | 0.0001936 | 0.3261 | +2 | 10 |
| - | - | 1170 | 599.3 | - | - | 0 | - |
| - | - | 815.5 | 615.3 | - | - | 0 | - |
| 6 | c | 2924 | 616.4 | 0.000791 | 1.283 | +1 | 6 |
| 10 | y | 2756 | 619.3 | 0.001727 | 2.789 | +1 | 5 |
| 4 | z | 713.5 | 643.3 | 0.000742 | 1.153 | +2 | 11 |
| - | - | 1380 | 646.3 | - | - | 0 | - |
| - | - | 573.2 | 648.3 | - | - | 0 | - |
| 4 | y | 1588 | 651.3 | 0.0007046 | 1.082 | +2 | 11 |
| - | - | 1103 | 651.8 | - | - | 0 | - |
| - | - | 1.002E+04 | 674.4 | - | - | 0 | - |
| - | - | 4638 | 675.4 | - | - | 0 | - |
| - | - | 786.4 | 676.4 | - | - | 0 | - |
| 3 | z | 1878 | 692.8 | 0.0004514 | 0.6515 | +2 | 12 |
| - | - | 989.6 | 693.3 | - | - | 0 | - |
| - | - | 974.4 | 693.8 | - | - | 0 | - |
| 3 | y | 8269 | 700.8 | 0.0004878 | 0.696 | +2 | 12 |
| 9 | z | 5433 | 701.3 | 0.01196 | 17.06 | +1 | 6 |
| - | - | 2071 | 701.8 | - | - | 0 | - |
| - | - | 1239 | 702.3 | - | - | 0 | - |
| - | - | 890.2 | 703.9 | - | - | 0 | - |
| - | - | 980.8 | 709.4 | - | - | 0 | - |
| - | - | 820.8 | 710.4 | - | - | 0 | - |
| 13 | c | 1.225E+04 | 725.9 | 0.0008168 | 1.125 | +2 | 13 |
| - | - | 8719 | 726.4 | - | - | 0 | - |
| - | - | 3442 | 726.9 | - | - | 0 | - |
| - | - | 724.8 | 727.4 | - | - | 0 | - |
| 9 | y | 685.3 | 734.4 | 0.002608 | 3.552 | +1 | 6 |
| - | - | 851.6 | 742.9 | - | - | 0 | - |
| 2 | y | 815.5 | 750.4 | 0.001499 | 1.998 | +2 | 13 |
| - | - | 1517 | 750.9 | - | - | 0 | - |
| 7 | c | 1.449E+04 | 753.4 | 0.001353 | 1.796 | +1 | 7 |
| - | - | 6616 | 754.4 | - | - | 0 | - |
| - | - | 969.4 | 755.4 | - | - | 0 | - |
| - | - | 636.5 | 764.4 | - | - | 0 | - |
| - | - | 792.5 | 764.9 | - | - | 0 | - |
| - | - | 1.352E+04 | 770.4 | - | - | 0 | - |
| - | - | 1.212E+04 | 770.9 | - | - | 0 | - |
| - | - | 8550 | 771.4 | - | - | 0 | - |
| - | - | 3051 | 771.9 | - | - | 0 | - |
| - | - | 1007 | 777.4 | - | - | 0 | - |
| - | - | 2190 | 778.4 | - | - | 0 | - |
| - | - | 1109 | 778.9 | - | - | 0 | - |
| - | - | 1045 | 782.3 | - | - | 0 | - |
| - | - | 2562 | 784.4 | - | - | 0 | - |
| - | - | 2698 | 784.9 | - | - | 0 | - |
| - | - | 2095 | 785.4 | - | - | 0 | - |
| 8 | w | 4370 | 788.4 | 0.0006607 | 0.8381 | +1 | 7 |
| - | - | 1942 | 789.4 | - | - | 0 | - |
| - | - | 737.5 | 790.4 | - | - | 0 | - |
| - | - | 956.3 | 791.4 | - | - | 0 | - |
| - | - | 1.162E+04 | 791.9 | - | - | 0 | - |
| - | - | 9719 | 792.4 | - | - | 0 | - |
| - | - | 4642 | 792.9 | - | - | 0 | - |
| - | - | 1941 | 793.4 | - | - | 0 | - |
| - | - | 1550 | 793.9 | - | - | 0 | - |
| - | - | 1040 | 799.4 | - | - | 0 | - |
| - | - | 2.186E+04 | 799.9 | - | - | 0 | - |
| - | - | 4.667E+04 | 800.4 | - | - | 0 | - |
| - | - | 3.606E+04 | 800.9 | - | - | 0 | - |
| - | - | 1.519E+04 | 801.4 | - | - | 0 | - |
| - | - | 4591 | 801.9 | - | - | 0 | - |
| - | - | 2087 | 802.4 | - | - | 0 | - |
| - | - | 1009 | 803.4 | - | - | 0 | - |
| 8 | z | 7371 | 847.4 | 0.0003542 | 0.418 | +1 | 7 |
| - | - | 3423 | 848.4 | - | - | 0 | - |
| - | - | 860.2 | 849.4 | - | - | 0 | - |
| 8 | y | 2474 | 863.4 | 0.0003405 | 0.3944 | +1 | 7 |
| - | - | 692.5 | 864.4 | - | - | 0 | - |
| 8 | c | 1.671E+04 | 882.5 | 0.001283 | 1.454 | +1 | 8 |
| - | - | 8112 | 883.5 | - | - | 0 | - |
| - | - | 2115 | 884.5 | - | - | 0 | - |
| - | - | 1924 | 952.5 | - | - | 0 | - |
| - | - | 1017 | 953.5 | - | - | 0 | - |
| 7 | z | 5532 | 984.5 | 0.002198 | 2.233 | +1 | 8 |
| - | - | 4651 | 985.5 | - | - | 0 | - |
| - | - | 1113 | 986.5 | - | - | 0 | - |
| 7 | y | 1069 | 1000 | 0.002612 | 2.61 | +1 | 8 |
| - | - | 863.2 | 1047 | - | - | 0 | - |
| - | - | 749.9 | 1065 | - | - | 0 | - |
| - | - | 787.1 | 1066 | - | - | 0 | - |
| - | - | 1120 | 1067 | - | - | 0 | - |
| - | - | 732.9 | 1067 | - | - | 0 | - |
| 6 | z | 4088 | 1071 | 0.001023 | 0.9552 | +1 | 9 |
| - | - | 3312 | 1072 | - | - | 0 | - |
| - | - | 1341 | 1073 | - | - | 0 | - |
| 6 | y | 2249 | 1088 | 0.0008823 | 0.8113 | +1 | 9 |
| - | - | 2082 | 1089 | - | - | 0 | - |
| - | - | 758.8 | 1090 | - | - | 0 | - |
| 10 | c | 4859 | 1095 | 0.003475 | 3.175 | +1 | 10 |
| - | - | 3462 | 1096 | - | - | 0 | - |
| - | - | 746.1 | 1097 | - | - | 0 | - |
| - | - | 1213 | 1107 | - | - | 0 | - |
| - | - | 1092 | 1121 | - | - | 0 | - |
| - | - | 1036 | 1122 | - | - | 0 | - |
| 5 | z | 2203 | 1171 | 0.003153 | 2.694 | +1 | 10 |
| - | - | 1904 | 1172 | - | - | 0 | - |
| - | - | 1503 | 1173 | - | - | 0 | - |
| - | - | 749.4 | 1176 | - | - | 0 | - |
| - | - | 2868 | 1180 | - | - | 0 | - |
| - | - | 1656 | 1181 | - | - | 0 | - |
| - | - | 970 | 1182 | - | - | 0 | - |
| 5 | y | 2893 | 1187 | 0.0005836 | 0.4919 | +1 | 10 |
| - | - | 1505 | 1188 | - | - | 0 | - |
| 11 | c | 7732 | 1224 | 0.003099 | 2.533 | +1 | 11 |
| - | - | 4563 | 1225 | - | - | 0 | - |
| - | - | 2307 | 1226 | - | - | 0 | - |
| - | - | 2574 | 1242 | - | - | 0 | - |
| - | - | 2080 | 1243 | - | - | 0 | - |
| - | - | 703 | 1264 | - | - | 0 | - |
| - | - | 2179 | 1279 | - | - | 0 | - |
| - | - | 2193 | 1280 | - | - | 0 | - |
| - | - | 857.4 | 1281 | - | - | 0 | - |
| 4 | z | 4076 | 1286 | 0.005072 | 3.945 | +1 | 11 |
| - | - | 1.039E+04 | 1287 | - | - | 0 | - |
| - | - | 4348 | 1288 | - | - | 0 | - |
| - | - | 2582 | 1289 | - | - | 0 | - |
| 4 | y | 1118 | 1302 | 0.002449 | 1.882 | +1 | 11 |
| - | - | 988.3 | 1303 | - | - | 0 | - |
| 12 | c | 5976 | 1323 | 0.004863 | 3.677 | +1 | 12 |
| - | - | 3396 | 1324 | - | - | 0 | - |
| - | - | 1556 | 1325 | - | - | 0 | - |
| - | - | 1838 | 1386 | - | - | 0 | - |
| - | - | 1445 | 1387 | - | - | 0 | - |
| - | - | 1494 | 1408 | - | - | 0 | - |
| 13 | c | 1196 | 1451 | 0.0005159 | 0.3556 | +1 | 13 |
| - | - | 4819 | 1452 | - | - | 0 | - |
| - | - | 3089 | 1453 | - | - | 0 | - |
| - | - | 1341 | 1454 | - | - | 0 | - |
| - | - | 2066 | 1485 | - | - | 0 | - |
| - | - | 964.7 | 1486 | - | - | 0 | - |
| - | - | 1087 | 1487 | - | - | 0 | - |
| - | - | 1494 | 1540 | - | - | 0 | - |
| - | - | 8456 | 1541 | - | - | 0 | - |
| - | - | 8801 | 1542 | - | - | 0 | - |
| - | - | 3700 | 1543 | - | - | 0 | - |
| - | - | 1076 | 1544 | - | - | 0 | - |
| - | - | 1988 | 1555 | - | - | 0 | - |
| - | - | 1442 | 1556 | - | - | 0 | - |
| - | - | 780.4 | 1558 | - | - | 0 | - |
| - | - | 1997 | 1573 | - | - | 0 | - |
| - | - | 1503 | 1574 | - | - | 0 | - |
| - | - | 836 | 1575 | - | - | 0 | - |
| - | - | 2549 | 1583 | - | - | 0 | - |
| - | - | 1.358E+04 | 1584 | - | - | 0 | - |
| - | - | 1.117E+04 | 1585 | - | - | 0 | - |
| - | - | 5961 | 1586 | - | - | 0 | - |
| - | - | 2512 | 1587 | - | - | 0 | - |
| - | - | 2960 | 1599 | - | - | 0 | - |
| - | - | 9962 | 1600 | - | - | 0 | - |
| - | - | 2.777E+04 | 1601 | - | - | 0 | - |
| - | - | 1.985E+04 | 1602 | - | - | 0 | - |
| - | - | 9502 | 1603 | - | - | 0 | - |
| - | - | 3880 | 1604 | - | - | 0 | - |
| - | - | 930.3 | 1605 | - | - | 0 | - |
| - | - | 624.9 | 3176 | - | - | 0 | - |

m/z Charge Intensity FragmentType MassShift Position
120.08084106445312 0 707.98987
121.85839080810547 0 363.57242
122.8731918334961 0 374.71463
127.08665466308594 0 377.00916
128.09469604492188 0 458.02014
129.10243225097656 0 2365.6448
130.779296875 0 402.93195
131.07037353515625 0 1565.9849 z Ammonia loss 12
131.6971893310547 0 424.09875
133.08596801757812 0 15037.098
134.0894317626953 0 2206.1526
135.192626953125 0 418.4181
147.10162353515625 0 7026.3833
147.44261169433594 0 450.21823
149.04489135742188 0 3524.271
150.72976684570312 0 424.1674
161.08065795898438 0 1161.72
167.05543518066406 0 764.1258
171.14918518066406 0 7192.3047
172.15309143066406 0 552.97314
173.45269775390625 0 1667.4668
175.09664916992188 0 1642.7019
177.1121826171875 0 9613.794
178.1154327392578 0 1507.8202
181.34312438964844 0 454.7873
185.1644744873047 0 794.8541
187.10787963867188 0 1425.859
191.12721252441406 0 1151.2642
197.12876892089844 0 860.9008 y 11
198.13661193847656 0 979.5831
199.1439971923828 0 7559.736
201.12371826171875 0 1034.8483
205.10704040527344 0 5041.243
206.11058044433594 0 1758.2893
213.15966796875 0 5339.6514
215.1026611328125 0 729.0266
216.71621704101562 0 552.6953
219.122314453125 0 1221.0397
219.61622619628906 0 545.203
221.08432006835938 0 12205.452
221.13829040527344 0 2774.425
222.0854034423828 0 2257.06
225.04295349121094 0 4182.2812
227.0224151611328 0 637.8526
228.17047119140625 0 725.94965
232.1293182373047 0 790.0095
233.09548950195312 0 4648.9106
233.16482543945312 0 725.8604
235.15382385253906 0 739.0309
239.09490966796875 0 19576.982
239.10873413085938 0 1030.2205
240.09547424316406 0 2544.683
249.1332244873047 0 1185.9169
251.10592651367188 0 1058.815
257.1496276855469 0 1622.4843
265.1643371582031 0 2352.3394
276.1554260253906 0 887.4163
278.1632080078125 0 823.09766 z 12
281.05120849609375 0 1332.2874
282.1810607910156 0 1831.4099
283.1759948730469 0 1169.0991
292.0936584472656 0 554.72015
294.1803283691406 0 1775.0685 y 12
295.1031188964844 0 9174.732
296.1043701171875 0 2270.7554
298.21246337890625 0 1242.3872
299.0616760253906 0 3640.126
300.0625 0 1308.9178
303.1373596191406 0 627.584
309.1911315917969 0 2035.3981
313.1139221191406 0 1702.9446
313.58203125 0 535.8338
314.1139831542969 0 578.1503
314.17010498046875 0 539.2224
314.2312316894531 0 1443.0796
315.238525390625 0 2060.1348 c 2
321.14752197265625 0 970.3017
352.22265625 0 2883.4905
353.21844482421875 0 940.6605
355.0697937011719 0 1637.18
356.0697326660156 0 521.0221
359.0282287597656 0 848.2043
362.2066650390625 0 1047.6332 w 11
364.77056884765625 0 503.11932
368.8622741699219 0 543.08875
369.12164306640625 0 9236.782
370.1216125488281 0 2912.1406
370.2334899902344 0 975.2407
371.2274169921875 0 832.2368
372.08203125 0 602.3594
372.2346496582031 0 773.92804
377.1659851074219 0 803.4927
377.2298889160156 0 1310.686 z 11
378.2330322265625 0 685.13293
386.2500305175781 0 720.5469
393.24908447265625 0 3024.0342 y 11
394.2527770996094 0 577.18823
413.2391357421875 0 2259.2583
413.2758483886719 0 2331.0425
414.2425231933594 0 922.6576
414.280517578125 0 587.60156
427.2373962402344 0 8571.085
428.2408142089844 0 2430.8596
429.2688903808594 0 1477.1481
430.2660217285156 0 1155.0653 c 3
443.2491455078125 0 793.42773
445.2476501464844 0 3073.6218
446.2519226074219 0 825.4
447.26031494140625 0 5321.5415 w 10
448.2626647949219 0 1298.2917
457.26531982421875 0 1183.0795
459.27984619140625 0 4150.3086
460.2809143066406 0 1684.9725
473.296630859375 0 1690.8118
476.2344970703125 0 834.1982
481.2641296386719 0 616.1156
483.05517578125 0 537.9248
487.2748107910156 0 4682.025
488.279052734375 0 1373.0304
489.2784729003906 0 757.4353
504.30950927734375 0 1963.7007
512.3072509765625 0 3404.7812
513.311279296875 0 1143.3926
516.309326171875 0 737.00476
517.3217163085938 0 34241.48
518.4591064453125 0 595.3644
529.333740234375 0 1178.7098 c 4
532.2962036132812 0 13545.873
532.3329467773438 0 2759.4226
533.2996215820312 0 8669.957
533.3406982421875 0 776.6159
534.3073120117188 0 2922.2732
534.3482055664062 0 38590.66
534.801513671875 0 608.3498
585.780029296875 0 952.4463 z 4
593.790771484375 0 882.57245 y 4
599.33740234375 0 1169.9135
615.3470458984375 0 815.4715
616.3656616210938 0 2923.7207 c 5
619.34326171875 0 2756.3027 y 9
643.2939453125 0 713.54193 z 3
646.331787109375 0 1380.2147
648.3488159179688 0 573.23016
651.3033447265625 0 1587.8044 y 3
651.8057250976562 0 1103.1697
674.3631591796875 0 10019.999
675.365234375 0 4637.89
676.3680419921875 0 786.3772
692.829345703125 0 1877.7695 z 2
693.3311767578125 0 989.5629
693.83251953125 0 974.38074
700.8377685546875 0 8268.611 y 2
701.338623046875 0 5433.286 z Ammonia loss 8
701.8408203125 0 2070.864
702.338623046875 0 1238.5026
703.873291015625 0 890.159
709.4140625 0 980.7753
710.4144287109375 0 820.8497
725.879638671875 0 12248.113 c 12
726.3806762695312 0 8718.818
726.8822021484375 0 3442.0059
727.3850708007812 0 724.8391
734.3693237304688 0 685.34265 y 8
742.8644409179688 0 851.6025
750.3739624023438 0 815.46344 y 1
750.873046875 0 1516.8025
753.4240112304688 0 14488.247 c 6
754.42822265625 0 6616.381
755.4301147460938 0 969.4405
764.3948364257812 0 636.50256
764.8904418945312 0 792.45105
770.3988647460938 0 13520.018
770.89990234375 0 12124.206
771.40185546875 0 8549.828
771.903076171875 0 3051.0747
777.4061279296875 0 1006.81946
778.4124145507812 0 2190.3733
778.9192504882812 0 1108.8237
782.3295288085938 0 1045.3394
784.385009765625 0 2561.5122
784.8880004882812 0 2697.5876
785.390625 0 2095.0164
788.3818359375 0 4369.9966 w 7
789.384033203125 0 1941.7587
790.3909912109375 0 737.4626
791.3906860351562 0 956.2592
791.8964233398438 0 11622.416
792.3980712890625 0 9719.24
792.899658203125 0 4642.2134
793.4016723632812 0 1940.6113
793.9031982421875 0 1550.1356
799.4033813476562 0 1040.1252
799.9055786132812 0 21863.822
800.4085693359375 0 46665.387
800.910400390625 0 36060.5
801.4122924804688 0 15191.324
801.9132080078125 0 4590.9575
802.412353515625 0 2087.13
803.4072265625 0 1008.6355
847.3954467773438 0 7370.5366 z 7
848.3983154296875 0 3422.5603
849.4010620117188 0 860.184
863.4141845703125 0 2474.134 y 7
864.4136352539062 0 692.45636
882.4666748046875 0 16709.27 c 7
883.46923828125 0 8112.4014
884.4705810546875 0 2114.5107
952.4713745117188 0 1924.476
953.47900390625 0 1017.2334
984.4525146484375 0 5531.927 z 6
985.4583129882812 0 4650.9556
986.459716796875 0 1112.8927
1000.4708251953125 0 1069.0298 y 6
1046.5240478515625 0 863.22327
1064.5296630859375 0 749.9369
1065.5081787109375 0 787.10223
1066.5152587890625 0 1120.0071
1067.4901123046875 0 732.9405
1071.4857177734375 0 4087.9211 z 5
1072.4881591796875 0 3312.4944
1073.48486328125 0 1340.5212
1087.50634765625 0 2248.8977 y 5
1088.5076904296875 0 2081.7573
1089.5196533203125 0 758.82837
1094.544189453125 0 4858.7285 c 9
1095.5474853515625 0 3461.838
1096.552001953125 0 746.1205
1107.494140625 0 1212.8636
1120.568115234375 0 1091.9283
1121.5662841796875 0 1035.595
1170.552001953125 0 2202.906 z 4
1171.561279296875 0 1903.7825
1172.564453125 0 1502.7219
1176.3001708984375 0 749.4492
1179.5732421875 0 2867.6335
1180.5821533203125 0 1655.5024
1181.5855712890625 0 969.9588
1186.574462890625 0 2893.065 y 4
1187.5780029296875 0 1504.6097
1223.587158203125 0 7732.3096 c 10
1224.5911865234375 0 4562.538
1225.5965576171875 0 2307.1833
1241.5888671875 0 2574.2202
1242.591552734375 0 2079.8157
1263.64404296875 0 703.03485
1278.645751953125 0 2179.0134
1279.64892578125 0 2193.4807
1280.636962890625 0 857.4334
1285.5770263671875 0 4075.5347 z 3
1286.58447265625 0 10385.483
1287.5892333984375 0 4347.708
1288.5908203125 0 2581.518
1301.603271484375 0 1117.64 y 3
1302.5997314453125 0 988.3493
1322.65380859375 0 5975.8745 c 11
1323.6588134765625 0 3395.8105
1324.6602783203125 0 1555.576
1385.6552734375 0 1838.094
1386.6513671875 0 1445.2644
1407.74365234375 0 1493.7429
1450.754150390625 0 1195.7855 c 12
1451.753662109375 0 4818.666
1452.758056640625 0 3089.2632
1453.759521484375 0 1341.1747
1484.72216796875 0 2066.2866
1485.736083984375 0 964.7387
1486.7176513671875 0 1086.5177
1539.7940673828125 0 1493.9211
1540.79296875 0 8456.131
1541.797607421875 0 8800.54
1542.8006591796875 0 3700.3118
1543.8082275390625 0 1075.9016
1554.8016357421875 0 1987.7678
1555.8121337890625 0 1442.2397
1557.8072509765625 0 780.43317
1572.8177490234375 0 1996.6223
1573.820068359375 0 1503.1135
1574.8310546875 0 835.95306
1582.787353515625 0 2549.1558
1583.7901611328125 0 13575.248
1584.7926025390625 0 11167.894
1585.795654296875 0 5961.481
1586.796142578125 0 2512.3267
1598.8009033203125 0 2960.3804
1599.8048095703125 0 9962.07
1600.81396484375 0 27769.166
1601.81787109375 0 19852.164
1602.8233642578125 0 9502.368
1603.828125 0 3880.2498
1604.8228759765625 0 930.27026
3176.46240234375 0 624.92303

Spectrum Details

|  |  |
| --- | --- |
| Matched peaks? Matched peaks The total absolute number of peaks matched. Additionally in brackets the total fraction of peaks matched and the total number of peaks is shown. | 40 (13.75% of 291) |
| FDR? FDR The false discovery rate estimated for this peptide. It is calculated by matching all theoretical fragments with a non-integer shift with the raw peaks for this spectrum. This is done with 40 different shifts. The resulting percentage is the average number of annotated peaks over the number of annotated peaks with the correct spectrum. | 1.49% |
| Satellite FDR? Satellite FDR See the FDR for details on its calculation. This satellite ion specific FDR only contains the satellite ions (d/w) for I/L/J positions. | - |
| PSM Score? PSM Score The PSM Score as given by Hecklib to this annotated spectrum. It is shown with three significant figures. | 310 |

#### Reverse Lookup? Reverse Lookup All places where this read could be placed.

| Group | Segment | Template | Template Part | Read Part | Score | Unique |
| --- | --- | --- | --- | --- | --- | --- |
| Homo sapiens Heavy Chain | IGHC | IGHG1 | [144..158] | [0..14] | 112 | False |
| Homo sapiens Heavy Chain | IGHC | IGHG3 | [191..205] | [0..14] | 103 | False |
| Homo sapiens Heavy Chain | IGHC | IGHG2 | [140..154] | [0..14] | 103 | False |

| Recombined | Template Part | Read Part | Score | Unique |
| --- | --- | --- | --- | --- |
| REC-0-1 | [267..281] | [0..14] | 112 | True |

#### Meta Information from Multiple reads

##### Number of combined reads

4

##### Intensity

0.9039

##### TotalArea

1.189E+07

#### Positional Score

Copy Data

##### Positional Score (TSV)

###### Preview

```
Loading example...
```

*Click on the button to copy the data to your clipboard.*

10012345678910111213

Label Value
"0" 0.5
"1" 0.5
"2" 0.5
"3" 0.5
"4" 0.497
"5" 0.497
"6" 0.497
"7" 0.497
"8" 0.478
"9" 0.465
"10" 0.487
"11" 0.492
"12" 0.492
"13" 0.492

#### Meta Information from PEAKS

##### Scan Identifier

F2:7070

##### Original sequence

V

V

V

D

V

S

H

E

D

P

E

V

K

F

##### Posttranslational Modifications

##### Source File

D:\separate\_stitch\_analyses\xle-disambiguation\raw\20210323\_F1\_UM1\_Peng0013\_SA\_F59\_ingel\_3ug\_TL.raw

##### Fraction

2

##### Scan Feature

-

##### De Novo Score

99

##### ConfidenceScore

99

### m/z

799.9069

##### Mass

1597.7988

##### Charge

2

##### Retention Time

39.34

##### Predicted Retention Time

-

##### Area

0

##### Parts Per Million

0.2

##### Fragmentation mode

HCD

##### Originating file

01 D:\separate\_stitch\_analyses\xle-disambiguation\20210325\_F59\_3ug\_DENOVO\_12.csv

#### Meta Information from PEAKS

##### Scan Identifier

F2:7048

##### Original sequence

V

V

V

D

V

S

H

E

D

P

E

V

K

F

##### Posttranslational Modifications

##### Source File

D:\separate\_stitch\_analyses\xle-disambiguation\raw\20210323\_F1\_UM1\_Peng0013\_SA\_F59\_ingel\_3ug\_TL.raw

##### Fraction

2

##### Scan Feature

F2:5530

##### De Novo Score

99

##### ConfidenceScore

99

### m/z

533.6078

##### Mass

1597.7988

##### Charge

3

##### Retention Time

39.25

##### Predicted Retention Time

-

##### Area

1.189E+07

##### Parts Per Million

1.7

##### Fragmentation mode

ETHCD

##### Originating file

01 D:\separate\_stitch\_analyses\xle-disambiguation\20210325\_F59\_3ug\_DENOVO\_12.csv

#### Meta Information from PEAKS

##### Scan Identifier

F2:7099

##### Original sequence

V

V

V

D

V

S

H

E

D

P

E

V

K

F

##### Posttranslational Modifications

##### Source File

D:\separate\_stitch\_analyses\xle-disambiguation\raw\20210323\_F1\_UM1\_Peng0013\_SA\_F59\_ingel\_3ug\_TL.raw

##### Fraction

2

##### Scan Feature

-

##### De Novo Score

99

##### ConfidenceScore

99

### m/z

533.6078

##### Mass

1597.7988

##### Charge

3

##### Retention Time

39.53

##### Predicted Retention Time

-

##### Area

0

##### Parts Per Million

1.8

##### Fragmentation mode

HCD

##### Originating file

01 D:\separate\_stitch\_analyses\xle-disambiguation\20210325\_F59\_3ug\_DENOVO\_12.csv

#### Meta Information from PEAKS

##### Scan Identifier

F2:7148

##### Original sequence

V

V

V

D

V

S

H

E

D

P

E

V

K

F

##### Posttranslational Modifications

##### Source File

D:\separate\_stitch\_analyses\xle-disambiguation\raw\20210323\_F1\_UM1\_Peng0013\_SA\_F59\_ingel\_3ug\_TL.raw

##### Fraction

2

##### Scan Feature

-

##### De Novo Score

97

##### ConfidenceScore

97

### m/z

533.6076

##### Mass

1597.7988

##### Charge

3

##### Retention Time

39.83

##### Predicted Retention Time

-

##### Area

0

##### Parts Per Million

1.4

##### Fragmentation mode

ETHCD

##### Originating file

01 D:\separate\_stitch\_analyses\xle-disambiguation\20210325\_F59\_3ug\_DENOVO\_12.csv
