## Supplementary material for "A handle on mass coincidence errors in *de novo* sequencing of antibodies by bottom-up proteomics": Combined_007.html

Details Combined\_007 | Stitch OverviewUndefined

### Read Combined\_007

#### Sequence (length=12)

SFVVFGGGTKJT

#### Spectrum 8995? Spectrum 8995 The raw spectrum of this peptide as annotated by Hecklib. The fragments are coloured according to ion type (see legend). Any peaks with a star '\*' as text can be hovered over to see the full details, first the ion type second the mass shift type. By hovering over the amino acids in the peptide or ions in the legend the corresponding peaks are highlighted. By toggling the 'Unassigned' label you can turn the background (unassigned) peaks on or off in the plot. By updating the slider in the Ion legend you can update the spectrum to only show the top X% of the peaks with labels. The top X% means any peak that is within X% of the highest intensity. By dragging in the spectrum you can zoom in to a specific part of the spectrum and use 'Zoom Out' to get back to the original zoom level. The annotation of the spectrum is based on the given sequence in the peptides file and is done with different software so inconsistencies are likely. The peaks are annotated based on the given sequence, with 20 ppm tolerance.

Copy Data

##### Spectrum 8995 (TSV)

###### Preview

```
Loading example...
```

*Click on the button to copy the data to your clipboard.*

Mz MinMz MaxIntensity Max

WidthHeightPeptide font sizePeptide stroke widthSpectrum font sizeSpectrum stroke widthCompact peptide

Ion legend

wxyz

abcd

OtherUnassignedIon^Charge^_Position_Show for top:%

SFVVFGGGTKJT

03.29e+56.58e+59.87e+51.32e+6

Zoom Out

y^+1^_1_y^+1^_2_y^+1^_2_z^+1^_3_y^+1^_3_y^+2^_8_y^+2^_9_y^+1^_4_z^+1^_4_y^+1^_4_y^+2^_10_y^+2^_10_c^+2^_10_z^+1^_5_y^+1^_5_y^+1^_6_z^+1^_6_y^+2^_11_y^+1^_6_y^+1^_7_z^+1^_7_y^+1^_7_c^+1^_6_c^+1^_7_c^+1^_8_y^+1^_8_y^+1^_8_z^+1^_8_c^+1^_8_y^+1^_8_z^+1^_9_w^+1^_9_c^+1^_9_y^+1^_9_z^+1^_9_c^+1^_9_y^+1^_9_z^+1^_10_w^+1^_10_z^+1^_10_y^+1^_10_c^+1^_10_c^+1^_10_c^+1^_11_y^+1^_11_z^+1^_11_c^+1^_11_

0767153423013068

Fragment Matches Table

Show background peaks

| Position | Ion type | Intensity | mz Theoretical | mz Error (Th) | mz Error (ppm) | Charge | Series Number |
| --- | --- | --- | --- | --- | --- | --- | --- |
| - | - | 4069 | 120.1 | - | - | 0 | - |
| 12 | y | 5.613E+04 | 120.1 | 0.0002305 | 1.92 | +1 | 1 |
| - | - | 6189 | 120.1 | - | - | 0 | - |
| - | - | 2339 | 121.1 | - | - | 0 | - |
| - | - | 2025 | 124.6 | - | - | 0 | - |
| - | - | 2525 | 129.1 | - | - | 0 | - |
| - | - | 2.958E+04 | 129.1 | - | - | 0 | - |
| - | - | 2087 | 143 | - | - | 0 | - |
| - | - | 5662 | 149 | - | - | 0 | - |
| - | - | 2282 | 152.9 | - | - | 0 | - |
| - | - | 2393 | 162.5 | - | - | 0 | - |
| - | - | 2680 | 163.5 | - | - | 0 | - |
| - | - | 2720 | 164.3 | - | - | 0 | - |
| - | - | 2.2E+04 | 171.1 | - | - | 0 | - |
| - | - | 6478 | 173.5 | - | - | 0 | - |
| - | - | 2.856E+04 | 199.1 | - | - | 0 | - |
| - | - | 5271 | 200.1 | - | - | 0 | - |
| - | - | 1.427E+05 | 207.1 | - | - | 0 | - |
| - | - | 1.609E+04 | 208.1 | - | - | 0 | - |
| 11 | y | 8819 | 215.1 | 0.0006448 | 2.997 | +1 | 2 |
| - | - | 9516 | 219.1 | - | - | 0 | - |
| - | - | 2869 | 226.6 | - | - | 0 | - |
| - | - | 2827 | 227.2 | - | - | 0 | - |
| - | - | 2427 | 229.7 | - | - | 0 | - |
| - | - | 5019 | 230.2 | - | - | 0 | - |
| 11 | y | 6.886E+04 | 233.1 | 0.0002272 | 0.9745 | +1 | 2 |
| - | - | 2986 | 234.2 | - | - | 0 | - |
| - | - | 2.752E+05 | 235.1 | - | - | 0 | - |
| - | - | 3.206E+04 | 236.1 | - | - | 0 | - |
| - | - | 5510 | 239.1 | - | - | 0 | - |
| - | - | 1.183E+04 | 247.1 | - | - | 0 | - |
| - | - | 6721 | 273.1 | - | - | 0 | - |
| - | - | 2.084E+04 | 289.2 | - | - | 0 | - |
| - | - | 3627 | 295.1 | - | - | 0 | - |
| - | - | 3770 | 301.2 | - | - | 0 | - |
| - | - | 1.231E+04 | 316.2 | - | - | 0 | - |
| - | - | 4870 | 326.2 | - | - | 0 | - |
| - | - | 3.5E+05 | 334.2 | - | - | 0 | - |
| - | - | 6.272E+04 | 335.2 | - | - | 0 | - |
| - | - | 5153 | 336.2 | - | - | 0 | - |
| - | - | 3215 | 344.2 | - | - | 0 | - |
| 10 | z | 3993 | 345.2 | 0.0001451 | 0.4202 | +1 | 3 |
| - | - | 6533 | 346.2 | - | - | 0 | - |
| - | - | 6590 | 346.2 | - | - | 0 | - |
| 10 | y | 2.128E+04 | 361.2 | 0.0001008 | 0.2791 | +1 | 3 |
| - | - | 4444 | 362.2 | - | - | 0 | - |
| - | - | 3706 | 382.2 | - | - | 0 | - |
| - | - | 3491 | 383.2 | - | - | 0 | - |
| - | - | 4764 | 388.2 | - | - | 0 | - |
| 5 | y | 9330 | 390.7 | 0.0004299 | 1.1 | +2 | 8 |
| - | - | 5421 | 391.2 | - | - | 0 | - |
| - | - | 4615 | 400.3 | - | - | 0 | - |
| - | - | 3.641E+04 | 401.2 | - | - | 0 | - |
| - | - | 3432 | 402.2 | - | - | 0 | - |
| - | - | 3849 | 402.2 | - | - | 0 | - |
| - | - | 3.157E+04 | 405.2 | - | - | 0 | - |
| - | - | 6679 | 406.3 | - | - | 0 | - |
| - | - | 1.228E+04 | 415.2 | - | - | 0 | - |
| - | - | 4306 | 416.2 | - | - | 0 | - |
| - | - | 9077 | 420.2 | - | - | 0 | - |
| - | - | 1.418E+05 | 433.2 | - | - | 0 | - |
| - | - | 3.646E+04 | 434.2 | - | - | 0 | - |
| - | - | 9911 | 436.2 | - | - | 0 | - |
| - | - | 3329 | 437.2 | - | - | 0 | - |
| 4 | y | 5357 | 440.3 | 0.0002382 | 0.5411 | +2 | 9 |
| - | - | 6542 | 440.8 | - | - | 0 | - |
| - | - | 1.361E+04 | 444.3 | - | - | 0 | - |
| 9 | y | 3.723E+04 | 445.3 | 0.0003152 | 0.708 | +1 | 4 |
| 9 | z | 9172 | 446.3 | 0.004336 | 9.716 | +1 | 4 |
| - | - | 1.578E+04 | 447.3 | - | - | 0 | - |
| - | - | 6644 | 453.2 | - | - | 0 | - |
| - | - | 1.3E+04 | 457.3 | - | - | 0 | - |
| - | - | 7457 | 459.3 | - | - | 0 | - |
| 9 | y | 1.162E+04 | 462.3 | 0.001781 | 3.853 | +1 | 4 |
| - | - | 1.022E+04 | 463.2 | - | - | 0 | - |
| - | - | 4902 | 470.3 | - | - | 0 | - |
| 3 | y | 1.152E+04 | 480.8 | 0.0002561 | 0.5327 | +2 | 10 |
| - | - | 2.631E+04 | 481.2 | - | - | 0 | - |
| - | - | 4668 | 481.8 | - | - | 0 | - |
| - | - | 9323 | 482.2 | - | - | 0 | - |
| 3 | y | 9493 | 489.8 | 0.0001618 | 0.3303 | +2 | 10 |
| - | - | 8524 | 490.3 | - | - | 0 | - |
| 10 | c | 8019 | 490.8 | 0.0005733 | 1.168 | +2 | 10 |
| - | - | 1.127E+04 | 491.3 | - | - | 0 | - |
| - | - | 4442 | 496.3 | - | - | 0 | - |
| - | - | 3583 | 498.3 | - | - | 0 | - |
| - | - | 3883 | 501.2 | - | - | 0 | - |
| 8 | z | 5685 | 503.3 | 7.338E-05 | 0.1458 | +1 | 5 |
| - | - | 9.506E+04 | 504.3 | - | - | 0 | - |
| - | - | 8.284E+04 | 504.3 | - | - | 0 | - |
| - | - | 3.181E+04 | 505.3 | - | - | 0 | - |
| - | - | 1.436E+04 | 505.3 | - | - | 0 | - |
| - | - | 5854 | 506.3 | - | - | 0 | - |
| - | - | 2.697E+04 | 514.3 | - | - | 0 | - |
| - | - | 8346 | 515.3 | - | - | 0 | - |
| - | - | 1.552E+04 | 516.3 | - | - | 0 | - |
| - | - | 4337 | 517.3 | - | - | 0 | - |
| - | - | 2.797E+04 | 518.3 | - | - | 0 | - |
| - | - | 3406 | 519.3 | - | - | 0 | - |
| 8 | y | 3.618E+04 | 519.3 | 0.0006395 | 1.231 | +1 | 5 |
| - | - | 8012 | 520.3 | - | - | 0 | - |
| - | - | 6175 | 530.3 | - | - | 0 | - |
| - | - | 9124 | 533.3 | - | - | 0 | - |
| - | - | 4496 | 534.3 | - | - | 0 | - |
| - | - | 8919 | 535.3 | - | - | 0 | - |
| - | - | 3274 | 536.3 | - | - | 0 | - |
| - | - | 1.077E+04 | 538.3 | - | - | 0 | - |
| - | - | 2934 | 545.1 | - | - | 0 | - |
| - | - | 4236 | 547.8 | - | - | 0 | - |
| - | - | 4.961E+04 | 548.3 | - | - | 0 | - |
| - | - | 1.414E+04 | 549.3 | - | - | 0 | - |
| - | - | 1.906E+04 | 552.3 | - | - | 0 | - |
| - | - | 6727 | 553.3 | - | - | 0 | - |
| 7 | y | 2975 | 558.3 | 0.0006132 | 1.098 | +1 | 6 |
| 7 | z | 4.757E+04 | 560.3 | 3.882E-05 | 0.06928 | +1 | 6 |
| - | - | 1.094E+04 | 561.3 | - | - | 0 | - |
| - | - | 1.858E+05 | 561.3 | - | - | 0 | - |
| - | - | 4.586E+04 | 562.3 | - | - | 0 | - |
| 2 | y | 8688 | 563.3 | 0.005384 | 9.557 | +2 | 11 |
| - | - | 3500 | 563.8 | - | - | 0 | - |
| - | - | 2.361E+04 | 575.3 | - | - | 0 | - |
| 7 | y | 1.933E+05 | 576.3 | 8.491E-06 | 0.01473 | +1 | 6 |
| - | - | 5.405E+04 | 577.3 | - | - | 0 | - |
| - | - | 1.108E+04 | 578.3 | - | - | 0 | - |
| - | - | 4.235E+04 | 580.3 | - | - | 0 | - |
| - | - | 1.626E+04 | 581.3 | - | - | 0 | - |
| - | - | 3998 | 582.3 | - | - | 0 | - |
| - | - | 2.286E+04 | 597.8 | - | - | 0 | - |
| - | - | 2.849E+04 | 598.3 | - | - | 0 | - |
| - | - | 3583 | 598.8 | - | - | 0 | - |
| - | - | 3849 | 603.3 | - | - | 0 | - |
| - | - | 6518 | 609.3 | - | - | 0 | - |
| 6 | y | 1.515E+04 | 615.3 | 1.776E-05 | 0.02887 | +1 | 7 |
| 6 | z | 1E+05 | 617.3 | 0.0001205 | 0.1952 | +1 | 7 |
| - | - | 2.528E+05 | 618.3 | - | - | 0 | - |
| - | - | 7.827E+04 | 619.3 | - | - | 0 | - |
| - | - | 5460 | 620.4 | - | - | 0 | - |
| - | - | 5283 | 631.3 | - | - | 0 | - |
| - | - | 1.007E+05 | 632.3 | - | - | 0 | - |
| 6 | y | 4.456E+05 | 633.4 | 0.0001342 | 0.2119 | +1 | 7 |
| - | - | 1.364E+05 | 634.4 | - | - | 0 | - |
| - | - | 2.403E+04 | 635.4 | - | - | 0 | - |
| - | - | 1.397E+04 | 637.3 | - | - | 0 | - |
| - | - | 7220 | 638.3 | - | - | 0 | - |
| - | - | 6909 | 643.4 | - | - | 0 | - |
| - | - | 3809 | 646.3 | - | - | 0 | - |
| - | - | 2.397E+04 | 647.4 | - | - | 0 | - |
| - | - | 7972 | 648.4 | - | - | 0 | - |
| 6 | c | 2.628E+04 | 654.4 | 1.144E-05 | 0.01748 | +1 | 6 |
| - | - | 1.179E+04 | 655.4 | - | - | 0 | - |
| - | - | 6929 | 660.3 | - | - | 0 | - |
| - | - | 3.53E+04 | 661.4 | - | - | 0 | - |
| - | - | 9100 | 662.4 | - | - | 0 | - |
| - | - | 1.877E+04 | 663.4 | - | - | 0 | - |
| - | - | 6239 | 664.4 | - | - | 0 | - |
| - | - | 1.017E+04 | 668.4 | - | - | 0 | - |
| - | - | 3842 | 669.4 | - | - | 0 | - |
| - | - | 8170 | 678.4 | - | - | 0 | - |
| - | - | 4352 | 692.4 | - | - | 0 | - |
| - | - | 8732 | 694.4 | - | - | 0 | - |
| - | - | 8.269E+04 | 710.4 | - | - | 0 | - |
| 7 | c | 7.41E+04 | 711.4 | 0.002493 | 3.505 | +1 | 7 |
| - | - | 2.397E+04 | 712.4 | - | - | 0 | - |
| - | - | 4319 | 718.4 | - | - | 0 | - |
| - | - | 5589 | 725.4 | - | - | 0 | - |
| - | - | 4922 | 728.4 | - | - | 0 | - |
| - | - | 4681 | 734.4 | - | - | 0 | - |
| - | - | 1.415E+04 | 746.4 | - | - | 0 | - |
| - | - | 6445 | 747.4 | - | - | 0 | - |
| - | - | 4901 | 749.4 | - | - | 0 | - |
| 8 | c | 4357 | 750.4 | 0.007106 | 9.469 | +1 | 8 |
| - | - | 1.362E+04 | 751.4 | - | - | 0 | - |
| - | - | 3477 | 753.4 | - | - | 0 | - |
| - | - | 2.564E+04 | 760.4 | - | - | 0 | - |
| - | - | 1.505E+04 | 761.4 | - | - | 0 | - |
| 5 | y | 1.458E+04 | 762.4 | 0.002527 | 3.314 | +1 | 8 |
| 5 | y | 9168 | 763.4 | 0.0121 | 15.85 | +1 | 8 |
| 5 | z | 2.151E+05 | 764.4 | 0.0002491 | 0.3258 | +1 | 8 |
| - | - | 3.175E+05 | 765.4 | - | - | 0 | - |
| - | - | 1.214E+05 | 766.4 | - | - | 0 | - |
| - | - | 5.951E+04 | 767.4 | - | - | 0 | - |
| 8 | c | 3.642E+04 | 768.4 | 0.002595 | 3.377 | +1 | 8 |
| - | - | 8031 | 769.4 | - | - | 0 | - |
| - | - | 4620 | 770.4 | - | - | 0 | - |
| - | - | 3695 | 778.4 | - | - | 0 | - |
| - | - | 1.47E+05 | 779.4 | - | - | 0 | - |
| 5 | y | 6.02E+05 | 780.4 | 0.0002865 | 0.3671 | +1 | 8 |
| - | - | 2.413E+05 | 781.4 | - | - | 0 | - |
| - | - | 5.439E+04 | 782.4 | - | - | 0 | - |
| - | - | 7209 | 783.4 | - | - | 0 | - |
| - | - | 1.406E+04 | 807.4 | - | - | 0 | - |
| - | - | 6379 | 808.4 | - | - | 0 | - |
| - | - | 4469 | 817.5 | - | - | 0 | - |
| - | - | 3875 | 818.5 | - | - | 0 | - |
| - | - | 4402 | 826.4 | - | - | 0 | - |
| - | - | 1.141E+04 | 834.4 | - | - | 0 | - |
| - | - | 4666 | 835.4 | - | - | 0 | - |
| 4 | z | 5680 | 845.5 | 0.004228 | 5.001 | +1 | 9 |
| - | - | 3591 | 846.5 | - | - | 0 | - |
| 4 | w | 1.111E+04 | 848.5 | 0.0009035 | 1.065 | +1 | 9 |
| - | - | 7129 | 849.5 | - | - | 0 | - |
| - | - | 1.92E+04 | 850.4 | - | - | 0 | - |
| 9 | c | 7016 | 851.4 | 0.009374 | 11.01 | +1 | 9 |
| - | - | 4.117E+04 | 852.4 | - | - | 0 | - |
| - | - | 1.956E+04 | 853.4 | - | - | 0 | - |
| - | - | 2.483E+04 | 859.5 | - | - | 0 | - |
| - | - | 1.311E+04 | 860.5 | - | - | 0 | - |
| 4 | y | 1.185E+04 | 861.5 | 0.007599 | 8.821 | +1 | 9 |
| - | - | 4345 | 862.5 | - | - | 0 | - |
| 4 | z | 2.499E+05 | 863.5 | 0.0001335 | 0.1546 | +1 | 9 |
| - | - | 2.189E+05 | 864.5 | - | - | 0 | - |
| - | - | 7.641E+04 | 865.5 | - | - | 0 | - |
| - | - | 2.086E+04 | 866.5 | - | - | 0 | - |
| - | - | 9.393E+04 | 868.4 | - | - | 0 | - |
| 9 | c | 6.261E+05 | 869.5 | 0.0001022 | 0.1176 | +1 | 9 |
| - | - | 3.002E+05 | 870.5 | - | - | 0 | - |
| - | - | 6.793E+04 | 871.5 | - | - | 0 | - |
| - | - | 9316 | 872.5 | - | - | 0 | - |
| - | - | 2.033E+04 | 878.5 | - | - | 0 | - |
| 4 | y | 2.889E+05 | 879.5 | 0.000219 | 0.249 | +1 | 9 |
| - | - | 1.403E+05 | 880.5 | - | - | 0 | - |
| - | - | 3.385E+04 | 881.5 | - | - | 0 | - |
| - | - | 3425 | 882.5 | - | - | 0 | - |
| - | - | 3881 | 906.5 | - | - | 0 | - |
| 3 | z | 4289 | 945.5 | 0.01589 | 16.8 | +1 | 10 |
| 3 | w | 1.291E+04 | 947.5 | 0.001215 | 1.282 | +1 | 10 |
| - | - | 5206 | 948.5 | - | - | 0 | - |
| - | - | 5219 | 953.5 | - | - | 0 | - |
| - | - | 4289 | 954.5 | - | - | 0 | - |
| 3 | z | 2.768E+05 | 962.5 | 0.001447 | 1.503 | +1 | 10 |
| - | - | 1.461E+05 | 963.5 | - | - | 0 | - |
| - | - | 3.5E+04 | 964.5 | - | - | 0 | - |
| - | - | 4808 | 965.6 | - | - | 0 | - |
| 3 | y | 1.738E+05 | 978.6 | 2.939E-05 | 0.03004 | +1 | 10 |
| - | - | 9.658E+04 | 979.6 | - | - | 0 | - |
| 10 | c | 6.932E+04 | 980.5 | 0.004055 | 4.135 | +1 | 10 |
| - | - | 4.944E+04 | 981.5 | - | - | 0 | - |
| - | - | 1.205E+04 | 982.5 | - | - | 0 | - |
| - | - | 4919 | 994.5 | - | - | 0 | - |
| - | - | 5546 | 995.5 | - | - | 0 | - |
| 10 | c | 3.866E+05 | 997.5 | 0.0002166 | 0.2171 | +1 | 10 |
| - | - | 2.326E+05 | 998.5 | - | - | 0 | - |
| - | - | 6.909E+04 | 999.6 | - | - | 0 | - |
| - | - | 7966 | 1001 | - | - | 0 | - |
| - | - | 6424 | 1009 | - | - | 0 | - |
| - | - | 5600 | 1010 | - | - | 0 | - |
| - | - | 5090 | 1052 | - | - | 0 | - |
| - | - | 4670 | 1053 | - | - | 0 | - |
| - | - | 4350 | 1066 | - | - | 0 | - |
| - | - | 2.264E+04 | 1067 | - | - | 0 | - |
| - | - | 1.247E+04 | 1068 | - | - | 0 | - |
| - | - | 4314 | 1069 | - | - | 0 | - |
| - | - | 6588 | 1070 | - | - | 0 | - |
| - | - | 4028 | 1071 | - | - | 0 | - |
| - | - | 5042 | 1076 | - | - | 0 | - |
| 11 | c | 6.978E+04 | 1094 | 0.0004195 | 0.3836 | +1 | 11 |
| - | - | 5.169E+04 | 1095 | - | - | 0 | - |
| - | - | 4.114E+04 | 1096 | - | - | 0 | - |
| - | - | 2.007E+04 | 1097 | - | - | 0 | - |
| - | - | 6023 | 1098 | - | - | 0 | - |
| 2 | y | 4669 | 1109 | 0.003747 | 3.38 | +1 | 11 |
| 2 | z | 2.206E+05 | 1110 | 9.766E-05 | 0.08801 | +1 | 11 |
| 11 | c | 7.404E+05 | 1111 | 0.003531 | 3.179 | +1 | 11 |
| - | - | 4.295E+05 | 1112 | - | - | 0 | - |
| - | - | 1.44E+05 | 1113 | - | - | 0 | - |
| - | - | 2.804E+04 | 1114 | - | - | 0 | - |
| - | - | 6951 | 1135 | - | - | 0 | - |
| - | - | 1.042E+04 | 1136 | - | - | 0 | - |
| - | - | 6811 | 1137 | - | - | 0 | - |
| - | - | 7645 | 1141 | - | - | 0 | - |
| - | - | 5912 | 1142 | - | - | 0 | - |
| - | - | 4195 | 1143 | - | - | 0 | - |
| - | - | 3973 | 1152 | - | - | 0 | - |
| - | - | 1.691E+04 | 1154 | - | - | 0 | - |
| - | - | 9608 | 1155 | - | - | 0 | - |
| - | - | 4179 | 1156 | - | - | 0 | - |
| - | - | 2.262E+05 | 1158 | - | - | 0 | - |
| - | - | 1.5E+05 | 1159 | - | - | 0 | - |
| - | - | 5.629E+04 | 1160 | - | - | 0 | - |
| - | - | 6049 | 1161 | - | - | 0 | - |
| - | - | 1.559E+04 | 1168 | - | - | 0 | - |
| - | - | 7686 | 1169 | - | - | 0 | - |
| - | - | 6671 | 1170 | - | - | 0 | - |
| - | - | 7470 | 1179 | - | - | 0 | - |
| - | - | 7253 | 1180 | - | - | 0 | - |
| - | - | 7.852E+04 | 1186 | - | - | 0 | - |
| - | - | 4.962E+04 | 1187 | - | - | 0 | - |
| - | - | 2.221E+04 | 1188 | - | - | 0 | - |
| - | - | 5945 | 1195 | - | - | 0 | - |
| - | - | 4.368E+04 | 1196 | - | - | 0 | - |
| - | - | 1.147E+06 | 1197 | - | - | 0 | - |
| - | - | 7.795E+05 | 1198 | - | - | 0 | - |
| - | - | 2.923E+05 | 1199 | - | - | 0 | - |
| - | - | 4.493E+04 | 1200 | - | - | 0 | - |
| - | - | 5216 | 1211 | - | - | 0 | - |
| - | - | 2.216E+04 | 1212 | - | - | 0 | - |
| - | - | 6.004E+05 | 1213 | - | - | 0 | - |
| - | - | 1.304E+06 | 1214 | - | - | 0 | - |
| - | - | 7.574E+05 | 1215 | - | - | 0 | - |
| - | - | 2.583E+05 | 1216 | - | - | 0 | - |
| - | - | 3.121E+04 | 1217 | - | - | 0 | - |
| - | - | 5154 | 3037 | - | - | 0 | - |

m/z Charge Intensity FragmentType MassShift Position
120.06135559082031 0 4069.114
120.06575012207031 0 56132.215 y 11
120.08106994628906 0 6189.463
121.06903839111328 0 2338.6677
124.5621566772461 0 2024.7637
129.09835815429688 0 2524.977
129.1024627685547 0 29580.146
142.95350646972656 0 2087.1125
148.9536590576172 0 5662.366
152.94227600097656 0 2281.8196
162.53575134277344 0 2393.0066
163.51829528808594 0 2679.731
164.2591552734375 0 2720.4792
171.14942932128906 0 21997.145
173.45147705078125 0 6477.935
199.14427185058594 0 28558.152
200.1473846435547 0 5270.5244
207.1129913330078 0 142734.14
208.11642456054688 0 16094.391
215.13966369628906 0 8818.741 y Water loss 10
219.14955139160156 0 9515.776
226.62161254882812 0 2868.7341
227.17672729492188 0 2826.7173
229.65829467773438 0 2426.828
230.15040588378906 0 5018.823
233.14981079101562 0 68863.89 y 10
234.153076171875 0 2986.066
235.10797119140625 0 275239.12
236.11138916015625 0 32059.86
239.09475708007812 0 5509.5283
247.144287109375 0 11830.123
273.11956787109375 0 6721.0234
289.1549377441406 0 20838.69
295.1038818359375 0 3626.5452
301.19256591796875 0 3770.2146
316.1664123535156 0 12309.368
326.1824951171875 0 4869.976
334.17657470703125 0 350037.25
335.1797180175781 0 62719.695
336.18231201171875 0 5153.234
344.19488525390625 0 3215.3347
345.2256774902344 0 3992.55 z 9
346.2124938964844 0 6533.0356
346.23297119140625 0 6590.0815
361.24444580078125 0 21280.615 y 9
362.24810791015625 0 4443.9634
382.1759033203125 0 3706.1558
383.2031555175781 0 3490.7942
388.2237243652344 0 4764.0967
390.7165832519531 0 9330.41 y 4
391.217529296875 0 5421.0605
400.25543212890625 0 4614.7656
401.2147521972656 0 36410.11
402.1778564453125 0 3432.0378
402.21759033203125 0 3849.3633
405.2497863769531 0 31570.947
406.2539978027344 0 6679.4927
415.2344665527344 0 12277.593
416.2377624511719 0 4305.845
420.18829345703125 0 9076.959
433.24493408203125 0 141815.98
434.2479553222656 0 36459.004
436.2225341796875 0 9910.574
437.2245178222656 0 3328.9026
440.2501220703125 0 5357.4243 y 3
440.751708984375 0 6541.5093
444.2575988769531 0 13611.49
445.2659912109375 0 37230.367 y Ammonia loss 8
446.2691650390625 0 9172.159 z 8
447.2811584472656 0 15782.826
453.2491760253906 0 6643.703
457.2765197753906 0 12998.2295
459.29425048828125 0 7456.589
462.29400634765625 0 11621.581 y 8
463.2351379394531 0 10222.643
470.26165771484375 0 4901.9717
480.779541015625 0 11519.368 y Water loss 2
481.2449645996094 0 26310.21
481.7566223144531 0 4667.7847
482.2486572265625 0 9323.059
489.78472900390625 0 9492.819 y 2
490.2865295410156 0 8523.755
490.7630615234375 0 8019.387 c Ammonia loss 9
491.26629638671875 0 11270.556
496.2872314453125 0 4442.287
498.2767639160156 0 3583.4106
501.2460021972656 0 3883.2761
503.2948913574219 0 5684.8745 z 7
504.2543029785156 0 95062.42
504.30291748046875 0 82835.6
505.25848388671875 0 31810.125
505.3061218261719 0 14358.36
506.2617492675781 0 5853.96
514.298828125 0 26965.309
515.3020629882812 0 8345.518
516.3130493164062 0 15515.728
517.3182373046875 0 4336.6353
518.3060913085938 0 27974.52
519.2639770507812 0 3405.7983
519.3130493164062 0 36179.707 y 7
520.3169555664062 0 8012.456
530.2731323242188 0 6175.063
533.3092651367188 0 9124.497
534.3048706054688 0 4495.938
535.2913208007812 0 8919.493
536.2902221679688 0 3274.3623
538.2664184570312 0 10771.767
545.0595703125 0 2933.7969
547.81201171875 0 4235.679
548.2825927734375 0 49607.1
549.2852172851562 0 14140.872
552.318115234375 0 19059.06
553.3219604492188 0 6727.335
558.323974609375 0 2974.9443 y Water loss 6
560.3164672851562 0 47569.29 z 6
561.2774658203125 0 10940.816
561.3243408203125 0 185783.64
562.326416015625 0 45855.855
563.3241577148438 0 8688.025 y 1
563.8212890625 0 3500.3699
575.3268432617188 0 23614.576
576.3351440429688 0 193328.55 y 6
577.3384399414062 0 54054.707
578.3411254882812 0 11075.433
580.31298828125 0 42354.758
581.3175048828125 0 16255.019
582.3209228515625 0 3998.378
597.8302612304688 0 22858.807
598.3313598632812 0 28493.373
598.8330078125 0 3582.8987
603.3243408203125 0 3848.9397
609.3364868164062 0 6518.0454
615.3460693359375 0 15146.052 y Water loss 5
617.3380126953125 0 100009.36 z 5
618.3453979492188 0 252824.33
619.3480224609375 0 78272.484
620.3513793945312 0 5459.743
631.3414306640625 0 5282.709
632.3489379882812 0 100740.98
633.3567504882812 0 445575.53 y 5
634.3594360351562 0 136426.1
635.3616333007812 0 24027.287
637.333251953125 0 13968.848
638.3375244140625 0 7219.9395
643.3550415039062 0 6909.361
646.3442993164062 0 3808.737
647.350341796875 0 23972.14
648.353759765625 0 7971.7534
654.3609619140625 0 26281.504 c 5
655.364501953125 0 11785.467
660.3450317382812 0 6928.852
661.3660888671875 0 35299.88
662.3679809570312 0 9100.31
663.3812866210938 0 18767.402
664.3827514648438 0 6238.8955
668.376708984375 0 10171.404
669.37548828125 0 3841.7966
678.397216796875 0 8170.179
692.3612060546875 0 4351.6523
694.3563842773438 0 8732.157
710.3745727539062 0 82687.23
711.3799438476562 0 74100.125 c 6
712.384765625 0 23969.248
718.4298706054688 0 4318.909
725.3935546875 0 5589.4814
728.4022827148438 0 4922.284
734.4205932617188 0 4681.103
746.416748046875 0 14150.529
747.4175415039062 0 6444.999
749.3848876953125 0 4901.318
750.38623046875 0 4356.64 c Water loss 7
751.3775024414062 0 13615.009
753.3650512695312 0 3477.065
760.4362182617188 0 25637.441
761.4387817382812 0 15046.241
762.4169921875 0 14580.655 y Water loss 4
763.4105834960938 0 9167.738 y Ammonia loss 4
764.4065551757812 0 215140.75 z 4
765.4132080078125 0 317495.47
766.41650390625 0 121367.59
767.4021606445312 0 59509.566
768.4013061523438 0 36420.62 c 7
769.4049072265625 0 8031.005
770.40576171875 0 4619.975
778.408203125 0 3695.048
779.4172973632812 0 147022.17
780.4247436523438 0 602007 y 4
781.4277954101562 0 241297.03
782.4304809570312 0 54392.62
783.4359130859375 0 7208.5557
807.4143676757812 0 14063.23
808.4178466796875 0 6379.337
817.4968872070312 0 4468.8647
818.5064086914062 0 3875.0906
826.446044921875 0 4402.029
834.4140625 0 11409.639
835.4186401367188 0 4666.3086
845.4683837890625 0 5679.959 z Water loss 3
846.4698486328125 0 3591.303
848.4521484375 0 11106.481 w 3
849.4569091796875 0 7129.471
850.4327392578125 0 19200.191
851.431640625 0 7016.039 c Water loss 8
852.4262084960938 0 41167.59
853.4302368164062 0 19557.275
859.504150390625 0 24828.453
860.50732421875 0 13111.809
861.490478515625 0 11851.069 y Water loss 3
862.489501953125 0 4345.0728
863.474853515625 0 249860.5 z 3
864.4797973632812 0 218937.62
865.483642578125 0 76407.13
866.4884033203125 0 20858.055
868.4434814453125 0 93932.87
869.4514770507812 0 626069.3 c 8
870.4541625976562 0 300203.38
871.45703125 0 67934.58
872.4603881835938 0 9316.218
878.4854125976562 0 20331.68
879.4932250976562 0 288874.7 y 3
880.4962158203125 0 140329.23
881.4989624023438 0 33847.598
882.4959106445312 0 3425.4055
906.479248046875 0 3880.5396
945.532470703125 0 4288.9385 z Ammonia loss 2
947.5208740234375 0 12908.139 w 2
948.5223999023438 0 5206.0283
953.5332641601562 0 5218.738
954.522705078125 0 4288.65
962.5416870117188 0 276764.62 z 2
963.5443115234375 0 146087.77
964.5479125976562 0 35001.023
965.552490234375 0 4807.6797
978.5618286132812 0 173770.67 y 2
979.5650634765625 0 96578.38
980.5240478515625 0 69317.98 c Ammonia loss 9
981.5242919921875 0 49436.883
982.5263061523438 0 12048.259
994.5286865234375 0 4919.0996
995.53466796875 0 5546.157
997.5463256835938 0 386647.06 c 9
998.5491943359375 0 232613.03
999.5520629882812 0 69094.47
1000.5571899414062 0 7965.503
1008.5836181640625 0 6424.451
1009.5961303710938 0 5599.9976
1051.60791015625 0 5089.7417
1052.60498046875 0 4670.0728
1065.603271484375 0 4349.524
1066.6136474609375 0 22643.91
1067.6163330078125 0 12470.116
1068.62060546875 0 4314.3086
1069.6112060546875 0 6588.0396
1070.6192626953125 0 4027.7224
1075.5933837890625 0 5042.0693
1093.6036376953125 0 69783.64 c Ammonia loss 10
1094.6063232421875 0 51691.945
1095.6160888671875 0 41142.406
1096.622314453125 0 20073.812
1097.6171875 0 6023.0933
1108.5999755859375 0 4668.834 y Ammonia loss 1
1109.6114501953125 0 220635.05 z 1
1110.6270751953125 0 740403.5 c 10
1111.6318359375 0 429495.2
1112.6358642578125 0 144043.08
1113.64111328125 0 28042.242
1134.6444091796875 0 6950.772
1135.621826171875 0 10420.964
1136.6304931640625 0 6810.6367
1140.5804443359375 0 7644.7554
1141.584716796875 0 5912.417
1142.5919189453125 0 4195.1543
1151.638427734375 0 3973.244
1153.632080078125 0 16906.303
1154.6392822265625 0 9608.018
1155.6231689453125 0 4178.747
1157.6068115234375 0 226152.27
1158.609619140625 0 150020.16
1159.611328125 0 56292.96
1160.6103515625 0 6049.197
1167.665283203125 0 15589.461
1168.6650390625 0 7685.88
1169.666259765625 0 6671.261
1178.6378173828125 0 7469.937
1179.636474609375 0 7252.6123
1185.6734619140625 0 78523.35
1186.6771240234375 0 49615.934
1187.6800537109375 0 22214.236
1194.636962890625 0 5945.454
1195.654052734375 0 43681.715
1196.6435546875 0 1146565.9
1197.6461181640625 0 779481.1
1198.6483154296875 0 292325.34
1199.651123046875 0 44931.05
1210.656494140625 0 5216.453
1211.64990234375 0 22158.611
1212.6612548828125 0 600405.4
1213.6678466796875 0 1303541.6
1214.67138671875 0 757405.6
1215.6741943359375 0 258319.53
1216.676513671875 0 31213.686
3037.41845703125 0 5154.3203

Spectrum Details

|  |  |
| --- | --- |
| Matched peaks? Matched peaks The total absolute number of peaks matched. Additionally in brackets the total fraction of peaks matched and the total number of peaks is shown. | 47 (15.56% of 302) |
| FDR? FDR The false discovery rate estimated for this peptide. It is calculated by matching all theoretical fragments with a non-integer shift with the raw peaks for this spectrum. This is done with 40 different shifts. The resulting percentage is the average number of annotated peaks over the number of annotated peaks with the correct spectrum. | 0.00% |
| Satellite FDR? Satellite FDR See the FDR for details on its calculation. This satellite ion specific FDR only contains the satellite ions (d/w) for I/L/J positions. | - |
| PSM Score? PSM Score The PSM Score as given by Hecklib to this annotated spectrum. It is shown with three significant figures. | 541 |

#### Spectrum 9056? Spectrum 9056 The raw spectrum of this peptide as annotated by Hecklib. The fragments are coloured according to ion type (see legend). Any peaks with a star '\*' as text can be hovered over to see the full details, first the ion type second the mass shift type. By hovering over the amino acids in the peptide or ions in the legend the corresponding peaks are highlighted. By toggling the 'Unassigned' label you can turn the background (unassigned) peaks on or off in the plot. By updating the slider in the Ion legend you can update the spectrum to only show the top X% of the peaks with labels. The top X% means any peak that is within X% of the highest intensity. By dragging in the spectrum you can zoom in to a specific part of the spectrum and use 'Zoom Out' to get back to the original zoom level. The annotation of the spectrum is based on the given sequence in the peptides file and is done with different software so inconsistencies are likely. The peaks are annotated based on the given sequence, with 20 ppm tolerance.

Copy Data

##### Spectrum 9056 (TSV)

###### Preview

```
Loading example...
```

*Click on the button to copy the data to your clipboard.*

Mz MinMz MaxIntensity Max

WidthHeightPeptide font sizePeptide stroke widthSpectrum font sizeSpectrum stroke widthCompact peptide

Ion legend

wxyz

abcd

OtherUnassignedIon^Charge^_Position_Show for top:%

SFVVFGGGTKJT

01.07e+52.14e+53.20e+54.27e+5

Zoom Out

y^+1^_1_y^+1^_2_y^+1^_2_z^+1^_3_y^+1^_3_y^+2^_8_y^+2^_9_y^+1^_4_z^+1^_4_y^+1^_4_y^+2^_10_y^+2^_10_c^+2^_10_z^+1^_5_y^+1^_5_c^+2^_11_y^+1^_6_z^+1^_6_y^+1^_6_z^+1^_7_y^+1^_7_z^+1^_7_y^+1^_7_c^+1^_6_c^+1^_7_c^+1^_8_y^+1^_8_z^+1^_8_c^+1^_8_y^+1^_8_z^+1^_9_w^+1^_9_c^+1^_9_y^+1^_9_y^+1^_9_z^+1^_9_c^+1^_9_y^+1^_9_z^+1^_10_w^+1^_10_y^+1^_10_y^+1^_10_z^+1^_10_y^+1^_10_c^+1^_10_c^+1^_10_z^+1^_11_c^+1^_11_y^+1^_11_z^+1^_11_c^+1^_11_

0765153122963062

Fragment Matches Table

Show background peaks

| Position | Ion type | Intensity | mz Theoretical | mz Error (Th) | mz Error (ppm) | Charge | Series Number |
| --- | --- | --- | --- | --- | --- | --- | --- |
| 12 | y | 1.941E+04 | 120.1 | 0.0002763 | 2.301 | +1 | 1 |
| - | - | 2140 | 120.1 | - | - | 0 | - |
| - | - | 563.6 | 120.4 | - | - | 0 | - |
| - | - | 933.7 | 121.1 | - | - | 0 | - |
| - | - | 7773 | 129.1 | - | - | 0 | - |
| - | - | 531.5 | 140.8 | - | - | 0 | - |
| - | - | 562.6 | 147.7 | - | - | 0 | - |
| - | - | 877.3 | 149 | - | - | 0 | - |
| - | - | 696.8 | 150.4 | - | - | 0 | - |
| - | - | 654.7 | 160 | - | - | 0 | - |
| - | - | 5709 | 171.1 | - | - | 0 | - |
| - | - | 811.6 | 172.2 | - | - | 0 | - |
| - | - | 2604 | 173.4 | - | - | 0 | - |
| - | - | 1559 | 177.1 | - | - | 0 | - |
| - | - | 550.5 | 186.1 | - | - | 0 | - |
| - | - | 696.8 | 193.1 | - | - | 0 | - |
| - | - | 9118 | 199.1 | - | - | 0 | - |
| - | - | 923.3 | 200.1 | - | - | 0 | - |
| - | - | 682.8 | 201.6 | - | - | 0 | - |
| - | - | 1756 | 203.1 | - | - | 0 | - |
| - | - | 4.481E+04 | 207.1 | - | - | 0 | - |
| - | - | 4633 | 208.1 | - | - | 0 | - |
| 11 | y | 2595 | 215.1 | 0.0003701 | 1.72 | +1 | 2 |
| - | - | 1680 | 219.1 | - | - | 0 | - |
| - | - | 3417 | 221.1 | - | - | 0 | - |
| - | - | 2832 | 225 | - | - | 0 | - |
| - | - | 1432 | 230.2 | - | - | 0 | - |
| 11 | y | 2.261E+04 | 233.1 | 0.0003493 | 1.498 | +1 | 2 |
| - | - | 2120 | 234.2 | - | - | 0 | - |
| - | - | 8.546E+04 | 235.1 | - | - | 0 | - |
| - | - | 1.115E+04 | 236.1 | - | - | 0 | - |
| - | - | 5225 | 239.1 | - | - | 0 | - |
| - | - | 1076 | 240.1 | - | - | 0 | - |
| - | - | 3413 | 247.1 | - | - | 0 | - |
| - | - | 784.7 | 248.1 | - | - | 0 | - |
| - | - | 653.3 | 256.8 | - | - | 0 | - |
| - | - | 1733 | 273.1 | - | - | 0 | - |
| - | - | 780.6 | 279.7 | - | - | 0 | - |
| - | - | 702.9 | 287.2 | - | - | 0 | - |
| - | - | 8157 | 289.2 | - | - | 0 | - |
| - | - | 1298 | 290.2 | - | - | 0 | - |
| - | - | 3850 | 295.1 | - | - | 0 | - |
| - | - | 2168 | 299.1 | - | - | 0 | - |
| - | - | 1650 | 301.2 | - | - | 0 | - |
| - | - | 1030 | 306.2 | - | - | 0 | - |
| - | - | 840.8 | 313.1 | - | - | 0 | - |
| - | - | 2430 | 316.2 | - | - | 0 | - |
| - | - | 735.9 | 317.2 | - | - | 0 | - |
| - | - | 1663 | 326.2 | - | - | 0 | - |
| - | - | 1139 | 332.2 | - | - | 0 | - |
| - | - | 1.199E+05 | 334.2 | - | - | 0 | - |
| - | - | 2.205E+04 | 335.2 | - | - | 0 | - |
| - | - | 2985 | 336.2 | - | - | 0 | - |
| - | - | 1504 | 344.2 | - | - | 0 | - |
| 10 | z | 1932 | 345.2 | 0.0007705 | 2.232 | +1 | 3 |
| - | - | 3455 | 346.2 | - | - | 0 | - |
| - | - | 1582 | 346.2 | - | - | 0 | - |
| 10 | y | 5687 | 361.2 | 0.0005095 | 1.411 | +1 | 3 |
| - | - | 1430 | 362.2 | - | - | 0 | - |
| - | - | 822.5 | 364.2 | - | - | 0 | - |
| - | - | 2364 | 369.1 | - | - | 0 | - |
| - | - | 1033 | 382.2 | - | - | 0 | - |
| - | - | 2794 | 383.2 | - | - | 0 | - |
| - | - | 820.4 | 387.2 | - | - | 0 | - |
| - | - | 1593 | 388.2 | - | - | 0 | - |
| 5 | y | 4272 | 390.7 | 0.000333 | 0.8523 | +2 | 8 |
| - | - | 1.167E+04 | 401.2 | - | - | 0 | - |
| - | - | 1428 | 402.2 | - | - | 0 | - |
| - | - | 869.3 | 403.2 | - | - | 0 | - |
| - | - | 9231 | 405.3 | - | - | 0 | - |
| - | - | 2579 | 406.3 | - | - | 0 | - |
| - | - | 5656 | 415.2 | - | - | 0 | - |
| - | - | 1353 | 416.2 | - | - | 0 | - |
| - | - | 1562 | 420.2 | - | - | 0 | - |
| - | - | 4.243E+04 | 433.2 | - | - | 0 | - |
| - | - | 1.421E+04 | 434.2 | - | - | 0 | - |
| - | - | 1010 | 435.3 | - | - | 0 | - |
| - | - | 3135 | 436.2 | - | - | 0 | - |
| - | - | 1216 | 439.3 | - | - | 0 | - |
| 4 | y | 2304 | 440.3 | 6.697E-05 | 0.1521 | +2 | 9 |
| - | - | 1500 | 440.8 | - | - | 0 | - |
| - | - | 5351 | 444.3 | - | - | 0 | - |
| 9 | y | 1.002E+04 | 445.3 | 0.0004678 | 1.051 | +1 | 4 |
| 9 | z | 2155 | 446.3 | 0.004214 | 9.442 | +1 | 4 |
| - | - | 3868 | 447.3 | - | - | 0 | - |
| - | - | 2011 | 453.2 | - | - | 0 | - |
| - | - | 3087 | 457.3 | - | - | 0 | - |
| - | - | 2436 | 459.3 | - | - | 0 | - |
| 9 | y | 2648 | 462.3 | 0.001903 | 4.117 | +1 | 4 |
| - | - | 2266 | 463.2 | - | - | 0 | - |
| - | - | 851.8 | 470.3 | - | - | 0 | - |
| 3 | y | 1597 | 480.8 | 0.0006834 | 1.421 | +2 | 10 |
| - | - | 9295 | 481.2 | - | - | 0 | - |
| - | - | 1249 | 481.8 | - | - | 0 | - |
| - | - | 2817 | 482.3 | - | - | 0 | - |
| 3 | y | 3302 | 489.8 | 0.000589 | 1.203 | +2 | 10 |
| - | - | 1291 | 490.3 | - | - | 0 | - |
| 10 | c | 2201 | 490.8 | 0.0002681 | 0.5463 | +2 | 10 |
| - | - | 2731 | 491.3 | - | - | 0 | - |
| - | - | 1115 | 491.8 | - | - | 0 | - |
| - | - | 2162 | 496.3 | - | - | 0 | - |
| 8 | z | 1990 | 503.3 | 0.0001402 | 0.2786 | +1 | 5 |
| - | - | 2.817E+04 | 504.3 | - | - | 0 | - |
| - | - | 2.65E+04 | 504.3 | - | - | 0 | - |
| - | - | 9518 | 505.3 | - | - | 0 | - |
| - | - | 7511 | 505.3 | - | - | 0 | - |
| - | - | 1167 | 506.3 | - | - | 0 | - |
| - | - | 1185 | 506.3 | - | - | 0 | - |
| - | - | 868.8 | 512.6 | - | - | 0 | - |
| - | - | 1.015E+04 | 514.3 | - | - | 0 | - |
| - | - | 2253 | 515.3 | - | - | 0 | - |
| - | - | 7130 | 516.3 | - | - | 0 | - |
| - | - | 1629 | 517.3 | - | - | 0 | - |
| - | - | 8364 | 518.3 | - | - | 0 | - |
| 8 | y | 9975 | 519.3 | 0.0003343 | 0.6438 | +1 | 5 |
| - | - | 1663 | 520.3 | - | - | 0 | - |
| - | - | 2210 | 530.3 | - | - | 0 | - |
| - | - | 2918 | 533.3 | - | - | 0 | - |
| - | - | 1688 | 533.8 | - | - | 0 | - |
| - | - | 1516 | 534.3 | - | - | 0 | - |
| - | - | 2747 | 535.3 | - | - | 0 | - |
| - | - | 1235 | 536.3 | - | - | 0 | - |
| - | - | 1646 | 538.3 | - | - | 0 | - |
| 11 | c | 1028 | 547.3 | 0.002627 | 4.8 | +2 | 11 |
| - | - | 1.783E+04 | 548.3 | - | - | 0 | - |
| - | - | 4377 | 549.3 | - | - | 0 | - |
| - | - | 7137 | 552.3 | - | - | 0 | - |
| - | - | 2178 | 553.3 | - | - | 0 | - |
| 7 | y | 1082 | 558.3 | 0.003665 | 6.564 | +1 | 6 |
| 7 | z | 1.411E+04 | 560.3 | 0.0007102 | 1.268 | +1 | 6 |
| - | - | 2969 | 561.3 | - | - | 0 | - |
| - | - | 6.121E+04 | 561.3 | - | - | 0 | - |
| - | - | 1.404E+04 | 562.3 | - | - | 0 | - |
| - | - | 2058 | 563.3 | - | - | 0 | - |
| - | - | 813.3 | 566.3 | - | - | 0 | - |
| - | - | 2351 | 571.3 | - | - | 0 | - |
| - | - | 939.1 | 572.3 | - | - | 0 | - |
| - | - | 4787 | 575.3 | - | - | 0 | - |
| - | - | 2365 | 575.4 | - | - | 0 | - |
| 7 | y | 5.861E+04 | 576.3 | 0.0001746 | 0.303 | +1 | 6 |
| - | - | 1.795E+04 | 577.3 | - | - | 0 | - |
| - | - | 2772 | 578.3 | - | - | 0 | - |
| - | - | 1.422E+04 | 580.3 | - | - | 0 | - |
| - | - | 5849 | 581.3 | - | - | 0 | - |
| - | - | 1021 | 587.4 | - | - | 0 | - |
| - | - | 1994 | 589.3 | - | - | 0 | - |
| - | - | 1.059E+04 | 597.8 | - | - | 0 | - |
| - | - | 5804 | 598.3 | - | - | 0 | - |
| - | - | 3707 | 598.8 | - | - | 0 | - |
| 6 | z | 1129 | 599.3 | 0.003056 | 5.099 | +1 | 7 |
| - | - | 1589 | 604.3 | - | - | 0 | - |
| - | - | 1316 | 606.8 | - | - | 0 | - |
| - | - | 980.8 | 608.3 | - | - | 0 | - |
| - | - | 1261 | 609.3 | - | - | 0 | - |
| 6 | y | 4434 | 615.3 | 0.0008112 | 1.318 | +1 | 7 |
| - | - | 1207 | 616.3 | - | - | 0 | - |
| 6 | z | 3.233E+04 | 617.3 | 0.0005478 | 0.8873 | +1 | 7 |
| - | - | 7.43E+04 | 618.3 | - | - | 0 | - |
| - | - | 2.245E+04 | 619.3 | - | - | 0 | - |
| - | - | 3821 | 620.4 | - | - | 0 | - |
| - | - | 1892 | 629.3 | - | - | 0 | - |
| - | - | 1121 | 631.3 | - | - | 0 | - |
| - | - | 3.214E+04 | 632.3 | - | - | 0 | - |
| 6 | y | 1.497E+05 | 633.4 | 0.0005004 | 0.7901 | +1 | 7 |
| - | - | 4.63E+04 | 634.4 | - | - | 0 | - |
| - | - | 7732 | 635.4 | - | - | 0 | - |
| - | - | 5388 | 637.3 | - | - | 0 | - |
| - | - | 1132 | 638.3 | - | - | 0 | - |
| - | - | 3240 | 643.4 | - | - | 0 | - |
| - | - | 1556 | 644.4 | - | - | 0 | - |
| - | - | 7603 | 647.4 | - | - | 0 | - |
| - | - | 3175 | 648.4 | - | - | 0 | - |
| 6 | c | 7176 | 654.4 | 1.144E-05 | 0.01748 | +1 | 6 |
| - | - | 3016 | 655.4 | - | - | 0 | - |
| - | - | 1027 | 659.4 | - | - | 0 | - |
| - | - | 4662 | 660.3 | - | - | 0 | - |
| - | - | 1.365E+04 | 661.4 | - | - | 0 | - |
| - | - | 5267 | 662.4 | - | - | 0 | - |
| - | - | 6066 | 663.4 | - | - | 0 | - |
| - | - | 1956 | 664.4 | - | - | 0 | - |
| - | - | 1020 | 667.4 | - | - | 0 | - |
| - | - | 3033 | 668.4 | - | - | 0 | - |
| - | - | 1666 | 678.4 | - | - | 0 | - |
| - | - | 1189 | 679.4 | - | - | 0 | - |
| - | - | 1349 | 692.4 | - | - | 0 | - |
| - | - | 2977 | 694.4 | - | - | 0 | - |
| - | - | 1903 | 695.4 | - | - | 0 | - |
| - | - | 901.3 | 708.3 | - | - | 0 | - |
| - | - | 2.532E+04 | 710.4 | - | - | 0 | - |
| 7 | c | 2.478E+04 | 711.4 | 0.001578 | 2.218 | +1 | 7 |
| - | - | 8807 | 712.4 | - | - | 0 | - |
| - | - | 1981 | 713.4 | - | - | 0 | - |
| - | - | 1309 | 718.4 | - | - | 0 | - |
| - | - | 1531 | 724.4 | - | - | 0 | - |
| - | - | 3990 | 725.4 | - | - | 0 | - |
| - | - | 1676 | 726.4 | - | - | 0 | - |
| - | - | 955.9 | 728.4 | - | - | 0 | - |
| - | - | 4916 | 746.4 | - | - | 0 | - |
| - | - | 1218 | 747.4 | - | - | 0 | - |
| - | - | 2544 | 749.4 | - | - | 0 | - |
| 8 | c | 1172 | 750.4 | 0.008632 | 11.5 | +1 | 8 |
| - | - | 2451 | 751.4 | - | - | 0 | - |
| - | - | 1284 | 752.4 | - | - | 0 | - |
| - | - | 8580 | 760.4 | - | - | 0 | - |
| - | - | 2852 | 761.4 | - | - | 0 | - |
| 5 | y | 5141 | 762.4 | 0.003259 | 4.275 | +1 | 8 |
| - | - | 3860 | 763.4 | - | - | 0 | - |
| 5 | z | 6.583E+04 | 764.4 | 0.0004932 | 0.6452 | +1 | 8 |
| - | - | 1.057E+05 | 765.4 | - | - | 0 | - |
| - | - | 3.972E+04 | 766.4 | - | - | 0 | - |
| - | - | 1.814E+04 | 767.4 | - | - | 0 | - |
| 8 | c | 9626 | 768.4 | 0.00174 | 2.265 | +1 | 8 |
| - | - | 2093 | 769.4 | - | - | 0 | - |
| - | - | 1169 | 778.4 | - | - | 0 | - |
| - | - | 4.318E+04 | 779.4 | - | - | 0 | - |
| 5 | y | 1.962E+05 | 780.4 | 0.0001407 | 0.1803 | +1 | 8 |
| - | - | 7.998E+04 | 781.4 | - | - | 0 | - |
| - | - | 1.843E+04 | 782.4 | - | - | 0 | - |
| - | - | 2771 | 783.4 | - | - | 0 | - |
| - | - | 886.7 | 792.4 | - | - | 0 | - |
| - | - | 1206 | 806.4 | - | - | 0 | - |
| - | - | 5125 | 807.4 | - | - | 0 | - |
| - | - | 1938 | 808.4 | - | - | 0 | - |
| - | - | 1654 | 817.5 | - | - | 0 | - |
| - | - | 1080 | 824.4 | - | - | 0 | - |
| - | - | 1813 | 826.4 | - | - | 0 | - |
| - | - | 3387 | 834.4 | - | - | 0 | - |
| - | - | 2529 | 835.4 | - | - | 0 | - |
| 4 | z | 1573 | 845.5 | 0.003618 | 4.279 | +1 | 9 |
| 4 | w | 2793 | 848.5 | 0.002124 | 2.504 | +1 | 9 |
| - | - | 2602 | 849.5 | - | - | 0 | - |
| - | - | 6369 | 850.4 | - | - | 0 | - |
| 9 | c | 2627 | 851.4 | 0.009313 | 10.94 | +1 | 9 |
| - | - | 1.095E+04 | 852.4 | - | - | 0 | - |
| - | - | 6743 | 853.4 | - | - | 0 | - |
| - | - | 1621 | 854.4 | - | - | 0 | - |
| - | - | 5490 | 859.5 | - | - | 0 | - |
| - | - | 3335 | 860.5 | - | - | 0 | - |
| 4 | y | 2753 | 861.5 | 0.01706 | 19.8 | +1 | 9 |
| 4 | y | 1514 | 862.5 | 0.005883 | 6.822 | +1 | 9 |
| 4 | z | 8.05E+04 | 863.5 | 0.0004997 | 0.5787 | +1 | 9 |
| - | - | 6.863E+04 | 864.5 | - | - | 0 | - |
| - | - | 2.536E+04 | 865.5 | - | - | 0 | - |
| - | - | 4475 | 866.5 | - | - | 0 | - |
| - | - | 3.349E+04 | 868.4 | - | - | 0 | - |
| 9 | c | 1.966E+05 | 869.5 | 0.000264 | 0.3036 | +1 | 9 |
| - | - | 9.182E+04 | 870.5 | - | - | 0 | - |
| - | - | 2.557E+04 | 871.5 | - | - | 0 | - |
| - | - | 3186 | 872.5 | - | - | 0 | - |
| - | - | 7214 | 878.5 | - | - | 0 | - |
| 4 | y | 9.41E+04 | 879.5 | 0.0004524 | 0.5144 | +1 | 9 |
| - | - | 4.168E+04 | 880.5 | - | - | 0 | - |
| - | - | 1.196E+04 | 881.5 | - | - | 0 | - |
| - | - | 951.5 | 900.9 | - | - | 0 | - |
| - | - | 1202 | 906.5 | - | - | 0 | - |
| 3 | z | 1515 | 944.5 | 0.003014 | 3.191 | +1 | 10 |
| 3 | w | 3204 | 947.5 | 0.003107 | 3.279 | +1 | 10 |
| - | - | 2244 | 948.5 | - | - | 0 | - |
| - | - | 2406 | 953.5 | - | - | 0 | - |
| - | - | 1357 | 954.5 | - | - | 0 | - |
| 3 | y | 1534 | 960.6 | 0.002282 | 2.376 | +1 | 10 |
| 3 | y | 2251 | 961.5 | 0.00296 | 3.079 | +1 | 10 |
| 3 | z | 8.378E+04 | 962.5 | 0.0005314 | 0.5521 | +1 | 10 |
| - | - | 4.508E+04 | 963.5 | - | - | 0 | - |
| - | - | 1.283E+04 | 964.5 | - | - | 0 | - |
| - | - | 1226 | 965.6 | - | - | 0 | - |
| 3 | y | 5.33E+04 | 978.6 | 0.0003979 | 0.4066 | +1 | 10 |
| - | - | 2.663E+04 | 979.6 | - | - | 0 | - |
| 10 | c | 2.347E+04 | 980.5 | 0.00613 | 6.252 | +1 | 10 |
| - | - | 1.573E+04 | 981.5 | - | - | 0 | - |
| - | - | 4961 | 982.5 | - | - | 0 | - |
| - | - | 1253 | 994.5 | - | - | 0 | - |
| - | - | 2378 | 995.5 | - | - | 0 | - |
| - | - | 1554 | 996.5 | - | - | 0 | - |
| 10 | c | 1.221E+05 | 997.5 | 0.0002717 | 0.2723 | +1 | 10 |
| - | - | 6.638E+04 | 998.6 | - | - | 0 | - |
| - | - | 1.921E+04 | 999.6 | - | - | 0 | - |
| - | - | 2907 | 1001 | - | - | 0 | - |
| - | - | 2208 | 1009 | - | - | 0 | - |
| - | - | 1155 | 1010 | - | - | 0 | - |
| - | - | 1231 | 1011 | - | - | 0 | - |
| - | - | 1115 | 1039 | - | - | 0 | - |
| - | - | 2287 | 1052 | - | - | 0 | - |
| - | - | 1869 | 1053 | - | - | 0 | - |
| - | - | 9440 | 1067 | - | - | 0 | - |
| - | - | 3800 | 1068 | - | - | 0 | - |
| - | - | 2532 | 1069 | - | - | 0 | - |
| - | - | 2207 | 1070 | - | - | 0 | - |
| - | - | 3024 | 1076 | - | - | 0 | - |
| - | - | 1122 | 1077 | - | - | 0 | - |
| 2 | z | 1156 | 1092 | 0.005706 | 5.227 | +1 | 11 |
| 11 | c | 2.591E+04 | 1094 | 0.00129 | 1.179 | +1 | 11 |
| - | - | 1.63E+04 | 1095 | - | - | 0 | - |
| - | - | 1.341E+04 | 1096 | - | - | 0 | - |
| - | - | 6975 | 1097 | - | - | 0 | - |
| - | - | 2897 | 1098 | - | - | 0 | - |
| 2 | y | 2118 | 1109 | 0.01053 | 9.503 | +1 | 11 |
| 2 | z | 7.309E+04 | 1110 | 0.0003906 | 0.352 | +1 | 11 |
| 11 | c | 2.355E+05 | 1111 | 0.003043 | 2.74 | +1 | 11 |
| - | - | 1.377E+05 | 1112 | - | - | 0 | - |
| - | - | 4.889E+04 | 1113 | - | - | 0 | - |
| - | - | 8104 | 1114 | - | - | 0 | - |
| - | - | 1266 | 1135 | - | - | 0 | - |
| - | - | 5317 | 1136 | - | - | 0 | - |
| - | - | 2585 | 1137 | - | - | 0 | - |
| - | - | 2877 | 1141 | - | - | 0 | - |
| - | - | 2113 | 1142 | - | - | 0 | - |
| - | - | 1679 | 1143 | - | - | 0 | - |
| - | - | 1317 | 1151 | - | - | 0 | - |
| - | - | 1298 | 1152 | - | - | 0 | - |
| - | - | 3760 | 1154 | - | - | 0 | - |
| - | - | 3642 | 1155 | - | - | 0 | - |
| - | - | 1500 | 1156 | - | - | 0 | - |
| - | - | 7.379E+04 | 1158 | - | - | 0 | - |
| - | - | 4.659E+04 | 1159 | - | - | 0 | - |
| - | - | 1.518E+04 | 1160 | - | - | 0 | - |
| - | - | 3001 | 1161 | - | - | 0 | - |
| - | - | 4348 | 1168 | - | - | 0 | - |
| - | - | 2591 | 1169 | - | - | 0 | - |
| - | - | 2204 | 1170 | - | - | 0 | - |
| - | - | 1291 | 1171 | - | - | 0 | - |
| - | - | 2225 | 1179 | - | - | 0 | - |
| - | - | 1629 | 1180 | - | - | 0 | - |
| - | - | 2.529E+04 | 1186 | - | - | 0 | - |
| - | - | 1.93E+04 | 1187 | - | - | 0 | - |
| - | - | 7299 | 1188 | - | - | 0 | - |
| - | - | 1715 | 1189 | - | - | 0 | - |
| - | - | 4731 | 1195 | - | - | 0 | - |
| - | - | 1.756E+04 | 1196 | - | - | 0 | - |
| - | - | 3.654E+05 | 1197 | - | - | 0 | - |
| - | - | 2.505E+05 | 1198 | - | - | 0 | - |
| - | - | 9.408E+04 | 1199 | - | - | 0 | - |
| - | - | 1.25E+04 | 1200 | - | - | 0 | - |
| - | - | 3373 | 1211 | - | - | 0 | - |
| - | - | 8038 | 1212 | - | - | 0 | - |
| - | - | 1.932E+05 | 1213 | - | - | 0 | - |
| - | - | 4.23E+05 | 1214 | - | - | 0 | - |
| - | - | 2.486E+05 | 1215 | - | - | 0 | - |
| - | - | 8.172E+04 | 1216 | - | - | 0 | - |
| - | - | 1.189E+04 | 1217 | - | - | 0 | - |
| - | - | 1085 | 2355 | - | - | 0 | - |
| - | - | 1022 | 2615 | - | - | 0 | - |
| - | - | 926.4 | 2812 | - | - | 0 | - |
| - | - | 897.9 | 3031 | - | - | 0 | - |

m/z Charge Intensity FragmentType MassShift Position
120.0657958984375 0 19411.246 y 11
120.08109283447266 0 2139.5977
120.36032104492188 0 563.633
121.0693359375 0 933.69934
129.1024932861328 0 7772.9756
140.82196044921875 0 531.50836
147.7397918701172 0 562.5603
149.04539489746094 0 877.3276
150.44775390625 0 696.7925
159.975341796875 0 654.6883
171.14942932128906 0 5709.4595
172.15281677246094 0 811.5558
173.4390869140625 0 2604.451
177.11280822753906 0 1559.209
186.14944458007812 0 550.5399
193.1420440673828 0 696.7633
199.14431762695312 0 9117.83
200.1475372314453 0 923.32935
201.5834197998047 0 682.83563
203.10292053222656 0 1756.0745
207.11309814453125 0 44809.758
208.11642456054688 0 4632.9897
215.13938903808594 0 2594.9036 y Water loss 10
219.14913940429688 0 1680.4548
221.08457946777344 0 3416.824
225.04312133789062 0 2832.3008
230.15028381347656 0 1432.1877
233.14993286132812 0 22614.158 y 10
234.153076171875 0 2120.4653
235.1080780029297 0 85463.21
236.11134338378906 0 11151.244
239.09535217285156 0 5224.666
240.09649658203125 0 1075.6357
247.144287109375 0 3412.8462
248.14842224121094 0 784.66693
256.835693359375 0 653.337
273.1193542480469 0 1733.3555
279.7364196777344 0 780.577
287.17144775390625 0 702.92236
289.1549987792969 0 8156.5874
290.1589660644531 0 1297.9463
295.1036682128906 0 3849.522
299.0625915527344 0 2167.5073
301.1918640136719 0 1650.0027
306.1817626953125 0 1029.6112
313.11407470703125 0 840.75946
316.16632080078125 0 2430.287
317.168212890625 0 735.9419
326.1827087402344 0 1662.6122
332.16363525390625 0 1138.8116
334.1767272949219 0 119883.18
335.17987060546875 0 22050.64
336.1819152832031 0 2985.304
344.19390869140625 0 1503.9194
345.2265930175781 0 1931.8464 z 9
346.2127685546875 0 3454.668
346.23455810546875 0 1581.9156
361.24505615234375 0 5687.396 y 9
362.2486877441406 0 1429.8612
364.16558837890625 0 822.47595
369.12298583984375 0 2363.5864
382.1783142089844 0 1032.7809
383.20428466796875 0 2794.4138
387.2404479980469 0 820.417
388.2239685058594 0 1592.523
390.7158203125 0 4272.319 y 4
401.21484375 0 11673.455
402.21832275390625 0 1427.8024
403.23248291015625 0 869.2986
405.2500305175781 0 9231.289
406.2538757324219 0 2578.8132
415.23419189453125 0 5655.9424
416.2354736328125 0 1353.4995
420.1884460449219 0 1562.2645
433.2451477050781 0 42430.793
434.2483215332031 0 14210.476
435.25042724609375 0 1010.3161
436.2239074707031 0 3134.7344
439.2676086425781 0 1216.0487
440.25042724609375 0 2303.8103 y 3
440.7519226074219 0 1499.9712
444.2586975097656 0 5351.0933
445.2661437988281 0 10022.8545 y Ammonia loss 8
446.269287109375 0 2154.7056 z 8
447.28253173828125 0 3867.9182
453.24981689453125 0 2010.9197
457.27764892578125 0 3086.8408
459.2926025390625 0 2435.518
462.29412841796875 0 2648.373 y 8
463.2347106933594 0 2265.8572
470.26177978515625 0 851.7763
480.77996826171875 0 1596.8003 y Water loss 2
481.2446594238281 0 9295.2295
481.7561340332031 0 1248.7699
482.25262451171875 0 2816.805
489.78515625 0 3302.27 y 2
490.287353515625 0 1290.9316
490.76336669921875 0 2201.173 c Ammonia loss 9
491.2669982910156 0 2731.1633
491.76885986328125 0 1114.9475
496.28875732421875 0 2161.5122
503.29510498046875 0 1989.8547 z 7
504.2542724609375 0 28165.236
504.30316162109375 0 26498.852
505.2587890625 0 9517.887
505.3066101074219 0 7511.3945
506.2662353515625 0 1167.2578
506.3080749511719 0 1185.1793
512.5986328125 0 868.7823
514.298583984375 0 10152.438
515.3018188476562 0 2252.923
516.3146362304688 0 7129.5547
517.3182373046875 0 1628.9009
518.306396484375 0 8363.81
519.3133544921875 0 9974.615 y 7
520.3162231445312 0 1662.9229
530.2711181640625 0 2209.9197
533.308837890625 0 2917.527
533.81201171875 0 1688.1737
534.3091430664062 0 1516.1025
535.2930297851562 0 2746.8545
536.2944946289062 0 1235.1936
538.2638549804688 0 1645.8524
547.3030395507812 0 1027.6896 c Ammonia loss 10
548.2836303710938 0 17832.402
549.2861938476562 0 4376.828
552.3184204101562 0 7137.043
553.321044921875 0 2177.8079
558.3209228515625 0 1081.5359 y Water loss 6
560.317138671875 0 14113.234 z 6
561.277587890625 0 2968.9333
561.3245239257812 0 61210.54
562.3263549804688 0 14043.246
563.330078125 0 2057.7922
566.2890014648438 0 813.26184
571.33203125 0 2351.0935
572.3369140625 0 939.0914
575.3270874023438 0 4787.01
575.3658447265625 0 2365.209
576.3353271484375 0 58606.543 y 6
577.33837890625 0 17953.639
578.3412475585938 0 2771.8435
580.3132934570312 0 14221.874
581.3162841796875 0 5848.557
587.3535766601562 0 1020.8079
589.3407592773438 0 1993.6136
597.8302001953125 0 10590.67
598.3325805664062 0 5804.2764
598.8339233398438 0 3707.3074
599.3303833007812 0 1128.6416 z Water loss 5
604.33056640625 0 1589.4984
606.8344116210938 0 1315.7865
608.3040161132812 0 980.7904
609.3377685546875 0 1261.1584
615.3468627929688 0 4433.7373 y Water loss 5
616.349365234375 0 1206.558
617.3384399414062 0 32334.3 z 5
618.3455810546875 0 74300.31
619.3486328125 0 22450.537
620.3524169921875 0 3821.0872
629.3419189453125 0 1891.7461
631.3397216796875 0 1120.5117
632.3494873046875 0 32142.291
633.3571166992188 0 149712.1 y 5
634.3601684570312 0 46303.758
635.36328125 0 7732.2876
637.3345947265625 0 5388.4805
638.3367919921875 0 1131.806
643.3573608398438 0 3240.2346
644.3573608398438 0 1556.1437
647.3521118164062 0 7603.4663
648.35400390625 0 3175.0034
654.3609619140625 0 7175.51 c 5
655.3633422851562 0 3015.9216
659.35546875 0 1026.649
660.3446655273438 0 4661.8833
661.3661499023438 0 13653.444
662.3697509765625 0 5266.524
663.3831176757812 0 6065.8574
664.3869018554688 0 1955.9697
667.3681030273438 0 1020.1113
668.3755493164062 0 3033.0034
678.3966674804688 0 1666.4806
679.396240234375 0 1189.2443
692.3681640625 0 1348.7896
694.3593139648438 0 2977.293
695.35986328125 0 1902.9736
708.3419189453125 0 901.2729
710.375244140625 0 25323.941
711.380859375 0 24781.793 c 6
712.3848266601562 0 8806.911
713.38623046875 0 1981.3395
718.4326171875 0 1309.1708
724.3878784179688 0 1530.8164
725.396728515625 0 3990.0542
726.3971557617188 0 1675.6031
728.4129638671875 0 955.94104
746.416748046875 0 4915.601
747.416748046875 0 1217.8066
749.3871459960938 0 2544.2136
750.3847045898438 0 1171.522 c Water loss 7
751.3796997070312 0 2450.9404
752.3810424804688 0 1283.8481
760.4356689453125 0 8580.4795
761.4358520507812 0 2852.244
762.417724609375 0 5141.4316 y Water loss 4
763.4141235351562 0 3860.3545
764.4067993164062 0 65833.81 z 4
765.4135131835938 0 105652.64
766.4168090820312 0 39723.03
767.4036865234375 0 18143.508
768.4021606445312 0 9625.997 c 7
769.4057006835938 0 2093.3467
778.4078979492188 0 1168.5868
779.4176635742188 0 43182.383
780.4251708984375 0 196171.44 y 4
781.4285278320312 0 79977.4
782.4310913085938 0 18426.084
783.431884765625 0 2771.1328
792.39697265625 0 886.71936
806.411865234375 0 1206.3141
807.416748046875 0 5125.179
808.4124755859375 0 1938.0537
817.5006103515625 0 1654.1823
824.4281005859375 0 1080.2101
826.4470825195312 0 1812.9192
834.4173583984375 0 3386.6084
835.4173583984375 0 2528.878
845.4677734375 0 1573.4747 z Water loss 3
848.453369140625 0 2793.2715 w 3
849.4546508789062 0 2602.106
850.4353637695312 0 6368.6763
851.4317016601562 0 2626.9175 c Water loss 8
852.4270629882812 0 10951.613
853.428955078125 0 6742.6626
854.43359375 0 1620.7788
859.50439453125 0 5490.269
860.5068969726562 0 3335.4006
861.4999389648438 0 2753.0618 y Water loss 3
862.4727783203125 0 1513.7039 y Ammonia loss 3
863.4752197265625 0 80503.39 z 3
864.4804077148438 0 68629.875
865.4839477539062 0 25364.709
866.4888305664062 0 4474.6455
868.444091796875 0 33491.953
869.4518432617188 0 196630.06 c 8
870.4550170898438 0 91821.305
871.4573974609375 0 25573.025
872.4595336914062 0 3186.098
878.4855346679688 0 7213.7935
879.493896484375 0 94095.42 y 3
880.4970092773438 0 41681.71
881.4993286132812 0 11963.602
900.8659057617188 0 951.51013
906.4852294921875 0 1201.8322
944.5355834960938 0 1515.2599 z Water loss 2
947.5227661132812 0 3204.246 w 2
948.5198974609375 0 2244.128
953.5333862304688 0 2405.819
954.5305786132812 0 1356.6462
960.5490112304688 0 1533.5029 y Water loss 2
961.5382690429688 0 2251.2996 y Ammonia loss 2
962.5426025390625 0 83778.69 z 2
963.54541015625 0 45079.09
964.548095703125 0 12826.914
965.5513916015625 0 1225.6467
978.562255859375 0 53301.156 y 2
979.5653686523438 0 26633
980.526123046875 0 23474.781 c Ammonia loss 9
981.5233764648438 0 15734.709
982.526123046875 0 4961.1465
994.5419921875 0 1253.4424
995.5288696289062 0 2377.806
996.5383911132812 0 1554.08
997.5468139648438 0 122059.24 c 9
998.5501708984375 0 66379.1
999.552490234375 0 19211.012
1000.5543823242188 0 2906.5288
1008.5866088867188 0 2207.6455
1009.5892333984375 0 1155.1707
1010.6008911132812 0 1231.0665
1038.5418701171875 0 1114.6333
1051.6065673828125 0 2286.948
1052.6107177734375 0 1868.6045
1066.61328125 0 9439.527
1067.61376953125 0 3800.0378
1068.61865234375 0 2532.0103
1069.619140625 0 2207.3645
1075.594970703125 0 3023.9421
1076.58935546875 0 1122.1229
1091.606689453125 0 1155.5558 z Water loss 1
1093.6053466796875 0 25912.547 c Ammonia loss 10
1094.60693359375 0 16303.468
1095.61572265625 0 13407.238
1096.62158203125 0 6974.502
1097.620361328125 0 2896.529
1108.6142578125 0 2117.735 y Ammonia loss 1
1109.6119384765625 0 73088.59 z 1
1110.6275634765625 0 235471.89 c 10
1111.632080078125 0 137712.28
1112.63623046875 0 48888.617
1113.6414794921875 0 8103.8496
1134.6446533203125 0 1266.4637
1135.630859375 0 5317.2676
1136.62841796875 0 2585.265
1140.5748291015625 0 2876.8862
1141.5859375 0 2113.3613
1142.593994140625 0 1679.2998
1150.642822265625 0 1316.5018
1151.6375732421875 0 1298.1763
1153.630615234375 0 3760.442
1154.6329345703125 0 3642.271
1155.6253662109375 0 1499.9043
1157.607666015625 0 73792.42
1158.6099853515625 0 46591.414
1159.61328125 0 15180.112
1160.6173095703125 0 3000.5256
1167.6632080078125 0 4348.2476
1168.6617431640625 0 2590.7844
1169.667236328125 0 2203.9966
1170.6475830078125 0 1291.2891
1178.6317138671875 0 2224.6296
1179.6461181640625 0 1629.0028
1185.6759033203125 0 25287.605
1186.6783447265625 0 19297.852
1187.680908203125 0 7298.833
1188.6737060546875 0 1714.9784
1194.63134765625 0 4731.3247
1195.6524658203125 0 17556.326
1196.6441650390625 0 365361.22
1197.646728515625 0 250515.31
1198.6495361328125 0 94075.84
1199.6510009765625 0 12500.173
1210.6561279296875 0 3373.256
1211.6558837890625 0 8038.425
1212.6622314453125 0 193162.25
1213.668701171875 0 422950.97
1214.6724853515625 0 248630.1
1215.675048828125 0 81720.97
1216.6778564453125 0 11893.817
2355.045166015625 0 1085.4609
2615.30126953125 0 1022.1301
2812.3203125 0 926.37994
3031.22509765625 0 897.8656

Spectrum Details

|  |  |
| --- | --- |
| Matched peaks? Matched peaks The total absolute number of peaks matched. Additionally in brackets the total fraction of peaks matched and the total number of peaks is shown. | 51 (14.83% of 344) |
| FDR? FDR The false discovery rate estimated for this peptide. It is calculated by matching all theoretical fragments with a non-integer shift with the raw peaks for this spectrum. This is done with 40 different shifts. The resulting percentage is the average number of annotated peaks over the number of annotated peaks with the correct spectrum. | 0.19% |
| Satellite FDR? Satellite FDR See the FDR for details on its calculation. This satellite ion specific FDR only contains the satellite ions (d/w) for I/L/J positions. | - |
| PSM Score? PSM Score The PSM Score as given by Hecklib to this annotated spectrum. It is shown with three significant figures. | 620 |

#### Spectrum 9114? Spectrum 9114 The raw spectrum of this peptide as annotated by Hecklib. The fragments are coloured according to ion type (see legend). Any peaks with a star '\*' as text can be hovered over to see the full details, first the ion type second the mass shift type. By hovering over the amino acids in the peptide or ions in the legend the corresponding peaks are highlighted. By toggling the 'Unassigned' label you can turn the background (unassigned) peaks on or off in the plot. By updating the slider in the Ion legend you can update the spectrum to only show the top X% of the peaks with labels. The top X% means any peak that is within X% of the highest intensity. By dragging in the spectrum you can zoom in to a specific part of the spectrum and use 'Zoom Out' to get back to the original zoom level. The annotation of the spectrum is based on the given sequence in the peptides file and is done with different software so inconsistencies are likely. The peaks are annotated based on the given sequence, with 20 ppm tolerance.

Copy Data

##### Spectrum 9114 (TSV)

###### Preview

```
Loading example...
```

*Click on the button to copy the data to your clipboard.*

Mz MinMz MaxIntensity Max

WidthHeightPeptide font sizePeptide stroke widthSpectrum font sizeSpectrum stroke widthCompact peptide

Ion legend

wxyz

abcd

OtherUnassignedIon^Charge^_Position_Show for top:%

SFVVFGGGTKJT

01.08e+52.15e+53.23e+54.30e+5

Zoom Out

y^+1^_1_a^+1^_2_a^+1^_2_b^+2^_4_y^+1^_2_b^+1^_2_y^+1^_2_b^+1^_2_y^+2^_5_a^+1^_3_a^+1^_3_y^+2^_7_b^+2^_6_b^+1^_3_b^+1^_3_y^+1^_3_y^+1^_3_y^+2^_8_y^+2^_8_y^+2^_8_b^+1^_4_b^+1^_4_y^+2^_9_y^+1^_4_y^+1^_4_y^+2^_10_y^+2^_10_b^+2^_10_b^+2^_10_y^+2^_10_b^+2^_10_y^+1^_5_y^+1^_5_b^+2^_11_y^+1^_6_b^+1^_5_y^+1^_6_b^+1^_5_\*\*\*y^+1^_7_b^+1^_6_y^+1^_7_b^+1^_6_b^+1^_7_b^+1^_7_b^+1^_8_b^+1^_8_y^+1^_8_y^+1^_8_b^+1^_9_b^+1^_9_y^+1^_9_y^+1^_9_y^+1^_10_b^+1^_10_b^+1^_10_y^+1^_10_b^+1^_10_b^+1^_11_b^+1^_11_b^+1^_11_y^+1^_11_

046793414011869

Fragment Matches Table

Show background peaks

| Position | Ion type | Intensity | mz Theoretical | mz Error (Th) | mz Error (ppm) | Charge | Series Number |
| --- | --- | --- | --- | --- | --- | --- | --- |
| 12 | y | 3.515E+04 | 120.1 | 0.0003602 | 3 | +1 | 1 |
| - | - | 9.503E+04 | 120.1 | - | - | 0 | - |
| - | - | 868.5 | 121.1 | - | - | 0 | - |
| - | - | 8525 | 121.1 | - | - | 0 | - |
| - | - | 1538 | 127.1 | - | - | 0 | - |
| - | - | 917.1 | 128.1 | - | - | 0 | - |
| - | - | 1.729E+05 | 129.1 | - | - | 0 | - |
| - | - | 521.8 | 130 | - | - | 0 | - |
| - | - | 815 | 130.1 | - | - | 0 | - |
| - | - | 9972 | 130.1 | - | - | 0 | - |
| - | - | 426.6 | 131.1 | - | - | 0 | - |
| - | - | 5114 | 131.1 | - | - | 0 | - |
| - | - | 1352 | 132.1 | - | - | 0 | - |
| - | - | 1137 | 132.1 | - | - | 0 | - |
| - | - | 509.9 | 133.1 | - | - | 0 | - |
| - | - | 2819 | 133.1 | - | - | 0 | - |
| - | - | 2091 | 136.1 | - | - | 0 | - |
| - | - | 415.1 | 138.1 | - | - | 0 | - |
| - | - | 1421 | 139.1 | - | - | 0 | - |
| - | - | 394.9 | 139.9 | - | - | 0 | - |
| - | - | 689.8 | 140.1 | - | - | 0 | - |
| - | - | 3150 | 141.1 | - | - | 0 | - |
| - | - | 1199 | 141.1 | - | - | 0 | - |
| - | - | 521.9 | 142.1 | - | - | 0 | - |
| - | - | 629.5 | 143.1 | - | - | 0 | - |
| - | - | 1089 | 143.1 | - | - | 0 | - |
| - | - | 463.5 | 144.1 | - | - | 0 | - |
| - | - | 6654 | 144.1 | - | - | 0 | - |
| - | - | 454.1 | 148.6 | - | - | 0 | - |
| - | - | 488.2 | 148.9 | - | - | 0 | - |
| - | - | 1773 | 149 | - | - | 0 | - |
| - | - | 464.8 | 150.9 | - | - | 0 | - |
| - | - | 913.2 | 151.1 | - | - | 0 | - |
| - | - | 1690 | 152.1 | - | - | 0 | - |
| - | - | 986 | 153.1 | - | - | 0 | - |
| - | - | 586.4 | 153.1 | - | - | 0 | - |
| - | - | 816 | 154.1 | - | - | 0 | - |
| - | - | 1298 | 155.1 | - | - | 0 | - |
| - | - | 3940 | 155.1 | - | - | 0 | - |
| - | - | 752.8 | 156.1 | - | - | 0 | - |
| - | - | 1845 | 158.1 | - | - | 0 | - |
| - | - | 4451 | 159.1 | - | - | 0 | - |
| - | - | 1200 | 159.1 | - | - | 0 | - |
| - | - | 542.1 | 159.1 | - | - | 0 | - |
| - | - | 2.06E+04 | 162.1 | - | - | 0 | - |
| - | - | 2367 | 163.1 | - | - | 0 | - |
| - | - | 3292 | 167.1 | - | - | 0 | - |
| - | - | 1129 | 167.1 | - | - | 0 | - |
| - | - | 1248 | 169.1 | - | - | 0 | - |
| - | - | 962.5 | 169.1 | - | - | 0 | - |
| - | - | 2978 | 170.1 | - | - | 0 | - |
| - | - | 1816 | 171.1 | - | - | 0 | - |
| - | - | 691.2 | 171.1 | - | - | 0 | - |
| - | - | 4.506E+04 | 171.1 | - | - | 0 | - |
| - | - | 6209 | 172.1 | - | - | 0 | - |
| - | - | 3420 | 172.2 | - | - | 0 | - |
| - | - | 896.3 | 173.1 | - | - | 0 | - |
| - | - | 1377 | 174.1 | - | - | 0 | - |
| - | - | 5534 | 176.1 | - | - | 0 | - |
| - | - | 480.1 | 176.7 | - | - | 0 | - |
| - | - | 5112 | 177.1 | - | - | 0 | - |
| - | - | 1458 | 177.1 | - | - | 0 | - |
| - | - | 577.2 | 181.2 | - | - | 0 | - |
| - | - | 501.9 | 182.1 | - | - | 0 | - |
| - | - | 1130 | 183.1 | - | - | 0 | - |
| - | - | 513.9 | 183.1 | - | - | 0 | - |
| - | - | 1031 | 183.1 | - | - | 0 | - |
| - | - | 872.4 | 185.1 | - | - | 0 | - |
| - | - | 2570 | 185.1 | - | - | 0 | - |
| - | - | 496.7 | 185.9 | - | - | 0 | - |
| - | - | 1716 | 186.1 | - | - | 0 | - |
| - | - | 1146 | 187.1 | - | - | 0 | - |
| - | - | 739.3 | 187.1 | - | - | 0 | - |
| - | - | 1852 | 188.1 | - | - | 0 | - |
| 2 | a | 2068 | 189.1 | 0.0001775 | 0.9388 | +1 | 2 |
| - | - | 629.1 | 189.1 | - | - | 0 | - |
| - | - | 798.2 | 194.1 | - | - | 0 | - |
| - | - | 703.2 | 195.1 | - | - | 0 | - |
| - | - | 915.5 | 196.1 | - | - | 0 | - |
| - | - | 536.7 | 197.1 | - | - | 0 | - |
| - | - | 6273 | 197.2 | - | - | 0 | - |
| - | - | 5636 | 198.1 | - | - | 0 | - |
| - | - | 577.2 | 199.1 | - | - | 0 | - |
| - | - | 2.671E+04 | 199.1 | - | - | 0 | - |
| - | - | 2059 | 200.1 | - | - | 0 | - |
| - | - | 632.3 | 201.1 | - | - | 0 | - |
| - | - | 940.2 | 201.1 | - | - | 0 | - |
| - | - | 1821 | 203.1 | - | - | 0 | - |
| - | - | 1.232E+04 | 205.1 | - | - | 0 | - |
| - | - | 1419 | 206.1 | - | - | 0 | - |
| 2 | a | 4.262E+05 | 207.1 | 0.0004618 | 2.23 | +1 | 2 |
| 4 | b | 4.93E+04 | 208.1 | 0.004113 | 19.76 | +2 | 4 |
| - | - | 1269 | 209.1 | - | - | 0 | - |
| - | - | 3044 | 209.1 | - | - | 0 | - |
| - | - | 1256 | 210.1 | - | - | 0 | - |
| - | - | 587.7 | 211.1 | - | - | 0 | - |
| - | - | 2.02E+04 | 212.1 | - | - | 0 | - |
| - | - | 1829 | 213.1 | - | - | 0 | - |
| - | - | 602.2 | 214.1 | - | - | 0 | - |
| - | - | 1592 | 214.2 | - | - | 0 | - |
| - | - | 610.6 | 215.1 | - | - | 0 | - |
| - | - | 2502 | 215.1 | - | - | 0 | - |
| 11 | y | 1.162E+04 | 215.1 | 0.0003091 | 1.437 | +1 | 2 |
| - | - | 4972 | 216.1 | - | - | 0 | - |
| - | - | 1731 | 216.1 | - | - | 0 | - |
| 2 | b | 4509 | 217.1 | 0.0005327 | 2.454 | +1 | 2 |
| - | - | 622.9 | 218.1 | - | - | 0 | - |
| - | - | 2.89E+04 | 219.1 | - | - | 0 | - |
| - | - | 3925 | 220.2 | - | - | 0 | - |
| - | - | 3752 | 221.1 | - | - | 0 | - |
| - | - | 1319 | 221.1 | - | - | 0 | - |
| - | - | 509.1 | 222.1 | - | - | 0 | - |
| - | - | 898.8 | 222.1 | - | - | 0 | - |
| - | - | 916.4 | 223.1 | - | - | 0 | - |
| - | - | 531.2 | 224.1 | - | - | 0 | - |
| - | - | 5535 | 224.2 | - | - | 0 | - |
| - | - | 8180 | 225 | - | - | 0 | - |
| - | - | 2104 | 225.1 | - | - | 0 | - |
| - | - | 1487 | 225.2 | - | - | 0 | - |
| - | - | 858 | 226 | - | - | 0 | - |
| - | - | 1224 | 226.1 | - | - | 0 | - |
| - | - | 500.8 | 226.2 | - | - | 0 | - |
| - | - | 1138 | 227 | - | - | 0 | - |
| - | - | 709.8 | 227.1 | - | - | 0 | - |
| - | - | 8460 | 227.1 | - | - | 0 | - |
| - | - | 3884 | 228.1 | - | - | 0 | - |
| - | - | 941.1 | 228.1 | - | - | 0 | - |
| - | - | 548.7 | 228.1 | - | - | 0 | - |
| - | - | 1291 | 229.1 | - | - | 0 | - |
| - | - | 1876 | 229.1 | - | - | 0 | - |
| - | - | 3.536E+04 | 230.2 | - | - | 0 | - |
| - | - | 1335 | 231.1 | - | - | 0 | - |
| - | - | 4106 | 231.2 | - | - | 0 | - |
| - | - | 1593 | 233.1 | - | - | 0 | - |
| 11 | y | 3.069E+04 | 233.1 | 0.0004408 | 1.891 | +1 | 2 |
| - | - | 981.1 | 234.1 | - | - | 0 | - |
| - | - | 3551 | 234.2 | - | - | 0 | - |
| 2 | b | 3.91E+05 | 235.1 | 0.0005118 | 2.177 | +1 | 2 |
| - | - | 4.812E+04 | 236.1 | - | - | 0 | - |
| - | - | 1700 | 237.1 | - | - | 0 | - |
| - | - | 4128 | 237.1 | - | - | 0 | - |
| - | - | 537.8 | 237.2 | - | - | 0 | - |
| - | - | 1.048E+04 | 239.1 | - | - | 0 | - |
| - | - | 1245 | 240.1 | - | - | 0 | - |
| - | - | 1562 | 240.1 | - | - | 0 | - |
| - | - | 665.2 | 240.2 | - | - | 0 | - |
| - | - | 547.9 | 241.1 | - | - | 0 | - |
| - | - | 1341 | 241.2 | - | - | 0 | - |
| - | - | 7274 | 242.2 | - | - | 0 | - |
| - | - | 2583 | 243.1 | - | - | 0 | - |
| - | - | 2136 | 243.1 | - | - | 0 | - |
| - | - | 715.3 | 243.2 | - | - | 0 | - |
| - | - | 6846 | 245.1 | - | - | 0 | - |
| - | - | 554.9 | 245.5 | - | - | 0 | - |
| - | - | 735.3 | 246.1 | - | - | 0 | - |
| - | - | 1482 | 247.1 | - | - | 0 | - |
| - | - | 1.972E+04 | 247.1 | - | - | 0 | - |
| - | - | 3054 | 248.1 | - | - | 0 | - |
| 8 | y | 1588 | 251.2 | 0.004367 | 17.39 | +2 | 5 |
| - | - | 847.4 | 254.1 | - | - | 0 | - |
| - | - | 2.09E+04 | 255.1 | - | - | 0 | - |
| - | - | 637.2 | 255.1 | - | - | 0 | - |
| - | - | 2489 | 256.1 | - | - | 0 | - |
| - | - | 619.3 | 257.1 | - | - | 0 | - |
| - | - | 2230 | 259.1 | - | - | 0 | - |
| - | - | 1082 | 260.2 | - | - | 0 | - |
| - | - | 2849 | 261.1 | - | - | 0 | - |
| - | - | 3345 | 261.2 | - | - | 0 | - |
| - | - | 7195 | 262.1 | - | - | 0 | - |
| - | - | 663.6 | 262.2 | - | - | 0 | - |
| - | - | 574.2 | 263.1 | - | - | 0 | - |
| - | - | 639.2 | 265.2 | - | - | 0 | - |
| - | - | 1426 | 268.2 | - | - | 0 | - |
| - | - | 2.185E+04 | 269.2 | - | - | 0 | - |
| - | - | 3218 | 270.2 | - | - | 0 | - |
| - | - | 1903 | 270.2 | - | - | 0 | - |
| - | - | 1549 | 271.1 | - | - | 0 | - |
| - | - | 2724 | 272.1 | - | - | 0 | - |
| - | - | 2.539E+04 | 273.1 | - | - | 0 | - |
| - | - | 1079 | 273.2 | - | - | 0 | - |
| - | - | 1983 | 273.2 | - | - | 0 | - |
| - | - | 8121 | 274.1 | - | - | 0 | - |
| - | - | 1430 | 275.1 | - | - | 0 | - |
| - | - | 1660 | 275.2 | - | - | 0 | - |
| - | - | 624.7 | 279.1 | - | - | 0 | - |
| - | - | 1013 | 281.2 | - | - | 0 | - |
| - | - | 2300 | 282.2 | - | - | 0 | - |
| - | - | 819.3 | 285.2 | - | - | 0 | - |
| - | - | 1059 | 286.2 | - | - | 0 | - |
| - | - | 7469 | 287.2 | - | - | 0 | - |
| - | - | 1428 | 288.1 | - | - | 0 | - |
| 3 | a | 2938 | 288.2 | 0.001466 | 5.086 | +1 | 3 |
| - | - | 2.879E+04 | 289.2 | - | - | 0 | - |
| - | - | 6180 | 290.2 | - | - | 0 | - |
| - | - | 4113 | 291.1 | - | - | 0 | - |
| - | - | 766.3 | 291.2 | - | - | 0 | - |
| - | - | 1366 | 292.1 | - | - | 0 | - |
| - | - | 1013 | 292.1 | - | - | 0 | - |
| - | - | 2952 | 295.1 | - | - | 0 | - |
| - | - | 891.8 | 297.2 | - | - | 0 | - |
| - | - | 4655 | 299.1 | - | - | 0 | - |
| - | - | 1377 | 300.1 | - | - | 0 | - |
| - | - | 2281 | 300.1 | - | - | 0 | - |
| - | - | 950.7 | 300.2 | - | - | 0 | - |
| - | - | 4900 | 301.2 | - | - | 0 | - |
| - | - | 869.7 | 302.2 | - | - | 0 | - |
| - | - | 1449 | 303.2 | - | - | 0 | - |
| - | - | 2946 | 304.2 | - | - | 0 | - |
| 3 | a | 1271 | 306.2 | 0.0009719 | 3.174 | +1 | 3 |
| 6 | y | 654.6 | 308.2 | 0.002287 | 7.42 | +2 | 7 |
| - | - | 1288 | 308.2 | - | - | 0 | - |
| - | - | 4495 | 309.2 | - | - | 0 | - |
| 6 | b | 842.9 | 310.2 | 0.001597 | 5.149 | +2 | 6 |
| - | - | 722 | 313.2 | - | - | 0 | - |
| - | - | 600.3 | 315.2 | - | - | 0 | - |
| - | - | 1031 | 315.2 | - | - | 0 | - |
| 3 | b | 1.678E+04 | 316.2 | 0.0006307 | 1.995 | +1 | 3 |
| - | - | 2298 | 317.2 | - | - | 0 | - |
| - | - | 2628 | 318.1 | - | - | 0 | - |
| - | - | 8772 | 319.1 | - | - | 0 | - |
| - | - | 1233 | 320.1 | - | - | 0 | - |
| - | - | 1008 | 321.2 | - | - | 0 | - |
| - | - | 3826 | 325.2 | - | - | 0 | - |
| - | - | 2.04E+04 | 326.2 | - | - | 0 | - |
| - | - | 3210 | 327.2 | - | - | 0 | - |
| - | - | 696.8 | 328.1 | - | - | 0 | - |
| - | - | 704.8 | 328.2 | - | - | 0 | - |
| - | - | 673.5 | 328.2 | - | - | 0 | - |
| - | - | 669.3 | 330.2 | - | - | 0 | - |
| - | - | 1164 | 331.1 | - | - | 0 | - |
| - | - | 1016 | 332.2 | - | - | 0 | - |
| - | - | 695.5 | 333.2 | - | - | 0 | - |
| - | - | 1174 | 333.2 | - | - | 0 | - |
| 3 | b | 2.553E+05 | 334.2 | 0.0007167 | 2.145 | +1 | 3 |
| - | - | 5.079E+04 | 335.2 | - | - | 0 | - |
| - | - | 3856 | 336.2 | - | - | 0 | - |
| - | - | 7005 | 337.2 | - | - | 0 | - |
| - | - | 1184 | 338.2 | - | - | 0 | - |
| - | - | 2189 | 339.2 | - | - | 0 | - |
| - | - | 786.4 | 340.2 | - | - | 0 | - |
| - | - | 925.1 | 341.2 | - | - | 0 | - |
| 10 | y | 8556 | 343.2 | 0.0006677 | 1.945 | +1 | 3 |
| - | - | 6296 | 344.2 | - | - | 0 | - |
| - | - | 1172 | 344.2 | - | - | 0 | - |
| - | - | 683 | 345.2 | - | - | 0 | - |
| - | - | 1487 | 345.2 | - | - | 0 | - |
| - | - | 1989 | 346.2 | - | - | 0 | - |
| - | - | 6750 | 346.2 | - | - | 0 | - |
| - | - | 930 | 347.2 | - | - | 0 | - |
| - | - | 4167 | 348.2 | - | - | 0 | - |
| - | - | 3087 | 354.2 | - | - | 0 | - |
| - | - | 5668 | 357.2 | - | - | 0 | - |
| - | - | 795.4 | 358.2 | - | - | 0 | - |
| - | - | 682.5 | 358.2 | - | - | 0 | - |
| - | - | 927.8 | 359 | - | - | 0 | - |
| - | - | 1523 | 361.2 | - | - | 0 | - |
| 10 | y | 2.714E+04 | 361.2 | 0.0003875 | 1.073 | +1 | 3 |
| - | - | 757.8 | 362.2 | - | - | 0 | - |
| - | - | 5759 | 362.2 | - | - | 0 | - |
| - | - | 6881 | 364.2 | - | - | 0 | - |
| - | - | 1320 | 365.2 | - | - | 0 | - |
| - | - | 3811 | 365.2 | - | - | 0 | - |
| - | - | 1208 | 366.2 | - | - | 0 | - |
| - | - | 1113 | 368.2 | - | - | 0 | - |
| - | - | 1850 | 369.1 | - | - | 0 | - |
| - | - | 1726 | 372.2 | - | - | 0 | - |
| - | - | 2814 | 373.2 | - | - | 0 | - |
| - | - | 6728 | 374.2 | - | - | 0 | - |
| - | - | 2150 | 374.2 | - | - | 0 | - |
| - | - | 3161 | 375.2 | - | - | 0 | - |
| - | - | 1824 | 376.2 | - | - | 0 | - |
| 5 | y | 1131 | 381.7 | 0.0005853 | 1.533 | +2 | 8 |
| - | - | 5059 | 382.2 | - | - | 0 | - |
| 5 | y | 745.5 | 382.2 | 0.0004293 | 1.123 | +2 | 8 |
| - | - | 1057 | 382.2 | - | - | 0 | - |
| - | - | 5083 | 382.2 | - | - | 0 | - |
| - | - | 1092 | 383.2 | - | - | 0 | - |
| - | - | 2.37E+04 | 383.2 | - | - | 0 | - |
| - | - | 1321 | 384.2 | - | - | 0 | - |
| - | - | 4580 | 384.2 | - | - | 0 | - |
| - | - | 1382 | 385.2 | - | - | 0 | - |
| - | - | 721.7 | 386.2 | - | - | 0 | - |
| - | - | 1175 | 387.2 | - | - | 0 | - |
| - | - | 3785 | 388.2 | - | - | 0 | - |
| - | - | 1421 | 389.2 | - | - | 0 | - |
| - | - | 4085 | 390.2 | - | - | 0 | - |
| 5 | y | 8361 | 390.7 | 0.0007046 | 1.803 | +2 | 8 |
| - | - | 1129 | 391.2 | - | - | 0 | - |
| - | - | 3031 | 391.2 | - | - | 0 | - |
| - | - | 732.4 | 391.7 | - | - | 0 | - |
| - | - | 7956 | 392.2 | - | - | 0 | - |
| - | - | 2250 | 393.2 | - | - | 0 | - |
| - | - | 7112 | 400.3 | - | - | 0 | - |
| - | - | 2.79E+04 | 401.2 | - | - | 0 | - |
| - | - | 1792 | 401.3 | - | - | 0 | - |
| - | - | 2.498E+04 | 402.2 | - | - | 0 | - |
| - | - | 5653 | 402.2 | - | - | 0 | - |
| - | - | 5425 | 403.2 | - | - | 0 | - |
| - | - | 945.7 | 403.2 | - | - | 0 | - |
| - | - | 971.8 | 404.2 | - | - | 0 | - |
| - | - | 1.737E+04 | 405.3 | - | - | 0 | - |
| - | - | 3330 | 406.3 | - | - | 0 | - |
| - | - | 791.5 | 411.2 | - | - | 0 | - |
| 4 | b | 9773 | 415.2 | 0.0007593 | 1.829 | +1 | 4 |
| - | - | 1798 | 416.2 | - | - | 0 | - |
| - | - | 2928 | 418.2 | - | - | 0 | - |
| - | - | 1501 | 419.2 | - | - | 0 | - |
| - | - | 1.678E+04 | 420.2 | - | - | 0 | - |
| - | - | 5469 | 421.2 | - | - | 0 | - |
| - | - | 1232 | 421.3 | - | - | 0 | - |
| - | - | 2597 | 429.3 | - | - | 0 | - |
| - | - | 630.1 | 430.2 | - | - | 0 | - |
| 4 | b | 6.989E+04 | 433.2 | 0.0006926 | 1.599 | +1 | 4 |
| - | - | 1.56E+04 | 434.2 | - | - | 0 | - |
| - | - | 2488 | 435.2 | - | - | 0 | - |
| - | - | 6834 | 436.2 | - | - | 0 | - |
| - | - | 595.4 | 436.2 | - | - | 0 | - |
| - | - | 2784 | 437.2 | - | - | 0 | - |
| - | - | 551.7 | 438.2 | - | - | 0 | - |
| - | - | 644.7 | 439.2 | - | - | 0 | - |
| - | - | 7524 | 439.3 | - | - | 0 | - |
| 4 | y | 2258 | 440.3 | 0.0008909 | 2.024 | +2 | 9 |
| - | - | 1254 | 440.8 | - | - | 0 | - |
| - | - | 1017 | 441.3 | - | - | 0 | - |
| - | - | 732.7 | 444.2 | - | - | 0 | - |
| 9 | y | 894.5 | 444.3 | 0.0001663 | 0.3742 | +1 | 4 |
| - | - | 1219 | 447.2 | - | - | 0 | - |
| - | - | 611.2 | 450.2 | - | - | 0 | - |
| - | - | 1812 | 452.3 | - | - | 0 | - |
| - | - | 5854 | 453.3 | - | - | 0 | - |
| - | - | 2217 | 454.3 | - | - | 0 | - |
| - | - | 1580 | 456.2 | - | - | 0 | - |
| - | - | 672.9 | 457.2 | - | - | 0 | - |
| - | - | 1.371E+04 | 457.3 | - | - | 0 | - |
| - | - | 3274 | 458.3 | - | - | 0 | - |
| - | - | 780.8 | 460.3 | - | - | 0 | - |
| - | - | 655.9 | 461.3 | - | - | 0 | - |
| 9 | y | 6820 | 462.3 | 0.001171 | 2.533 | +1 | 4 |
| - | - | 6191 | 463.2 | - | - | 0 | - |
| - | - | 1489 | 463.3 | - | - | 0 | - |
| - | - | 1768 | 464.2 | - | - | 0 | - |
| - | - | 851.6 | 468.3 | - | - | 0 | - |
| - | - | 892.3 | 469.3 | - | - | 0 | - |
| - | - | 1990 | 470.3 | - | - | 0 | - |
| - | - | 656.8 | 471.3 | - | - | 0 | - |
| - | - | 1653 | 472.3 | - | - | 0 | - |
| - | - | 3521 | 473.3 | - | - | 0 | - |
| - | - | 1671 | 474.2 | - | - | 0 | - |
| - | - | 1364 | 475.2 | - | - | 0 | - |
| - | - | 1800 | 478.3 | - | - | 0 | - |
| 3 | y | 1366 | 480.8 | 0.000958 | 1.993 | +2 | 10 |
| - | - | 1.325E+04 | 481.2 | - | - | 0 | - |
| 3 | y | 1195 | 481.3 | 0.008401 | 17.46 | +2 | 10 |
| 10 | b | 2013 | 481.8 | 0.0004366 | 0.9062 | +2 | 10 |
| 10 | b | 5099 | 482.3 | 0.0002501 | 0.5186 | +2 | 10 |
| - | - | 775.7 | 483.3 | - | - | 0 | - |
| - | - | 1168 | 486.2 | - | - | 0 | - |
| - | - | 8621 | 486.3 | - | - | 0 | - |
| - | - | 2309 | 487.3 | - | - | 0 | - |
| - | - | 1047 | 489.3 | - | - | 0 | - |
| 3 | y | 3434 | 489.8 | 0.00123 | 2.511 | +2 | 10 |
| - | - | 919.7 | 490.3 | - | - | 0 | - |
| - | - | 1046 | 490.3 | - | - | 0 | - |
| 10 | b | 3080 | 490.8 | 0.0008915 | 1.817 | +2 | 10 |
| - | - | 5089 | 491.3 | - | - | 0 | - |
| - | - | 1672 | 492.3 | - | - | 0 | - |
| - | - | 1231 | 493.2 | - | - | 0 | - |
| - | - | 1195 | 495.2 | - | - | 0 | - |
| - | - | 1.943E+04 | 496.3 | - | - | 0 | - |
| - | - | 5162 | 497.3 | - | - | 0 | - |
| - | - | 9391 | 501.2 | - | - | 0 | - |
| 8 | y | 1465 | 501.3 | 0.001075 | 2.145 | +1 | 5 |
| - | - | 2520 | 502.3 | - | - | 0 | - |
| - | - | 686.1 | 503.3 | - | - | 0 | - |
| - | - | 839.5 | 504.3 | - | - | 0 | - |
| - | - | 838.7 | 506.7 | - | - | 0 | - |
| - | - | 1610 | 510.3 | - | - | 0 | - |
| - | - | 4128 | 512.3 | - | - | 0 | - |
| - | - | 713.1 | 513.3 | - | - | 0 | - |
| - | - | 4.587E+04 | 514.3 | - | - | 0 | - |
| - | - | 444.8 | 515.3 | - | - | 0 | - |
| - | - | 1.236E+04 | 515.3 | - | - | 0 | - |
| - | - | 1802 | 516.3 | - | - | 0 | - |
| - | - | 880.6 | 517.3 | - | - | 0 | - |
| - | - | 4806 | 519.3 | - | - | 0 | - |
| 8 | y | 1.165E+04 | 519.3 | 0.0005202 | 1.002 | +1 | 5 |
| - | - | 2954 | 520.3 | - | - | 0 | - |
| - | - | 1979 | 520.3 | - | - | 0 | - |
| - | - | 818.7 | 521.3 | - | - | 0 | - |
| - | - | 1.756E+04 | 530.3 | - | - | 0 | - |
| - | - | 5272 | 531.3 | - | - | 0 | - |
| - | - | 2550 | 533.3 | - | - | 0 | - |
| - | - | 2048 | 533.8 | - | - | 0 | - |
| - | - | 3098 | 534.3 | - | - | 0 | - |
| - | - | 6195 | 535.3 | - | - | 0 | - |
| - | - | 2688 | 536.3 | - | - | 0 | - |
| - | - | 3784 | 538.3 | - | - | 0 | - |
| - | - | 644.9 | 538.9 | - | - | 0 | - |
| - | - | 1118 | 539.3 | - | - | 0 | - |
| - | - | 866.5 | 540.3 | - | - | 0 | - |
| 11 | b | 1166 | 547.3 | 0.003238 | 5.916 | +2 | 11 |
| - | - | 1475 | 547.8 | - | - | 0 | - |
| - | - | 4.039E+04 | 548.3 | - | - | 0 | - |
| - | - | 1.375E+04 | 549.3 | - | - | 0 | - |
| - | - | 2267 | 550.3 | - | - | 0 | - |
| - | - | 9204 | 552.3 | - | - | 0 | - |
| - | - | 3029 | 553.3 | - | - | 0 | - |
| 7 | y | 2613 | 558.3 | 0.0007906 | 1.416 | +1 | 6 |
| 5 | b | 7066 | 562.3 | 3.336E-05 | 0.05932 | +1 | 5 |
| - | - | 3236 | 563.3 | - | - | 0 | - |
| - | - | 1023 | 564.3 | - | - | 0 | - |
| - | - | 695.4 | 567.3 | - | - | 0 | - |
| - | - | 755.4 | 571.4 | - | - | 0 | - |
| - | - | 1316 | 574.3 | - | - | 0 | - |
| - | - | 822.7 | 575.8 | - | - | 0 | - |
| 7 | y | 2.851E+04 | 576.3 | 0.0002967 | 0.5148 | +1 | 6 |
| - | - | 742.4 | 577.3 | - | - | 0 | - |
| - | - | 8173 | 577.3 | - | - | 0 | - |
| - | - | 1516 | 578.3 | - | - | 0 | - |
| 5 | b | 1.924E+04 | 580.3 | 0.000516 | 0.8892 | +1 | 5 |
| - | - | 6776 | 581.3 | - | - | 0 | - |
| - | - | 1152 | 582.3 | - | - | 0 | - |
| - | - | 3008 | 589.3 | - | - | 0 | - |
| - | - | 994.6 | 590.3 | - | - | 0 | - |
| - | - | 838.6 | 591.3 | - | - | 0 | - |
| - | - | 1441 | 592.3 | - | - | 0 | - |
| - | - | 1125 | 595.3 | - | - | 0 | - |
| - | - | 759.4 | 596.3 | - | - | 0 | - |
| - | - | 1799 | 597.3 | - | - | 0 | - |
| 0 | Precursor | 1.225E+04 | 597.8 | 0.0006331 | 1.059 | +2 | -1 |
| 0 | Precursor | 9559 | 598.3 | 0.01021 | 17.07 | +2 | -1 |
| - | - | 2523 | 598.8 | - | - | 0 | - |
| - | - | 1692 | 599.3 | - | - | 0 | - |
| - | - | 3239 | 600.3 | - | - | 0 | - |
| - | - | 1020 | 601.3 | - | - | 0 | - |
| - | - | 1167 | 606.3 | - | - | 0 | - |
| 0 | Precursor | 995.6 | 606.8 | 0.0004378 | 0.7215 | +2 | -1 |
| - | - | 685.8 | 607.3 | - | - | 0 | - |
| - | - | 982.4 | 607.3 | - | - | 0 | - |
| - | - | 2474 | 609.3 | - | - | 0 | - |
| - | - | 1442 | 611.3 | - | - | 0 | - |
| 6 | y | 9496 | 615.3 | 0.001544 | 2.509 | +1 | 7 |
| - | - | 4349 | 616.3 | - | - | 0 | - |
| - | - | 1424 | 617.3 | - | - | 0 | - |
| - | - | 3054 | 618.3 | - | - | 0 | - |
| 6 | b | 4414 | 619.3 | 0.001458 | 2.354 | +1 | 6 |
| - | - | 1699 | 620.3 | - | - | 0 | - |
| - | - | 2812 | 625.3 | - | - | 0 | - |
| - | - | 756.2 | 626.3 | - | - | 0 | - |
| - | - | 8153 | 629.3 | - | - | 0 | - |
| - | - | 2842 | 630.3 | - | - | 0 | - |
| - | - | 1086 | 631.3 | - | - | 0 | - |
| 6 | y | 1.363E+05 | 633.4 | 0.001416 | 2.236 | +1 | 7 |
| - | - | 4.518E+04 | 634.4 | - | - | 0 | - |
| - | - | 8468 | 635.4 | - | - | 0 | - |
| - | - | 742.5 | 636.4 | - | - | 0 | - |
| 6 | b | 5834 | 637.3 | 0.0005977 | 0.9378 | +1 | 6 |
| - | - | 2789 | 638.3 | - | - | 0 | - |
| - | - | 2.365E+04 | 643.4 | - | - | 0 | - |
| - | - | 1.091E+04 | 644.4 | - | - | 0 | - |
| - | - | 1467 | 645.4 | - | - | 0 | - |
| - | - | 1.753E+04 | 647.4 | - | - | 0 | - |
| - | - | 7539 | 648.4 | - | - | 0 | - |
| - | - | 1544 | 649.3 | - | - | 0 | - |
| - | - | 1232 | 652.3 | - | - | 0 | - |
| - | - | 4.821E+04 | 661.4 | - | - | 0 | - |
| - | - | 1.74E+04 | 662.4 | - | - | 0 | - |
| - | - | 3248 | 663.4 | - | - | 0 | - |
| - | - | 1094 | 666.4 | - | - | 0 | - |
| 7 | b | 1700 | 676.3 | 0.001662 | 2.457 | +1 | 7 |
| - | - | 1386 | 679.4 | - | - | 0 | - |
| 7 | b | 3077 | 694.4 | 0.0001751 | 0.2522 | +1 | 7 |
| - | - | 1126 | 695.4 | - | - | 0 | - |
| - | - | 670.7 | 705.4 | - | - | 0 | - |
| - | - | 689.7 | 714.4 | - | - | 0 | - |
| - | - | 2951 | 718.4 | - | - | 0 | - |
| - | - | 1189 | 719.4 | - | - | 0 | - |
| - | - | 862.8 | 723.4 | - | - | 0 | - |
| - | - | 3430 | 728.4 | - | - | 0 | - |
| - | - | 1623 | 729.4 | - | - | 0 | - |
| - | - | 4631 | 732.4 | - | - | 0 | - |
| 8 | b | 971.9 | 733.4 | 0.007175 | 9.784 | +1 | 8 |
| - | - | 2160 | 733.4 | - | - | 0 | - |
| - | - | 688.2 | 734.4 | - | - | 0 | - |
| - | - | 905.2 | 734.4 | - | - | 0 | - |
| - | - | 743.7 | 735.4 | - | - | 0 | - |
| - | - | 748.7 | 736.4 | - | - | 0 | - |
| - | - | 596.7 | 737.4 | - | - | 0 | - |
| - | - | 9392 | 742.4 | - | - | 0 | - |
| - | - | 4045 | 743.4 | - | - | 0 | - |
| - | - | 2069 | 744.4 | - | - | 0 | - |
| - | - | 1057 | 745.4 | - | - | 0 | - |
| - | - | 8657 | 746.4 | - | - | 0 | - |
| - | - | 3267 | 747.4 | - | - | 0 | - |
| - | - | 925.6 | 748.4 | - | - | 0 | - |
| 8 | b | 4711 | 751.4 | 0.0008832 | 1.175 | +1 | 8 |
| - | - | 1121 | 752.4 | - | - | 0 | - |
| - | - | 1312 | 753.4 | - | - | 0 | - |
| - | - | 1027 | 754.4 | - | - | 0 | - |
| - | - | 2.749E+04 | 760.4 | - | - | 0 | - |
| - | - | 1.104E+04 | 761.4 | - | - | 0 | - |
| 5 | y | 1.423E+04 | 762.4 | 0.002466 | 3.234 | +1 | 8 |
| - | - | 6643 | 763.4 | - | - | 0 | - |
| - | - | 812.1 | 764.4 | - | - | 0 | - |
| - | - | 1248 | 778.4 | - | - | 0 | - |
| 5 | y | 1.866E+05 | 780.4 | 0.0005069 | 0.6496 | +1 | 8 |
| - | - | 7.948E+04 | 781.4 | - | - | 0 | - |
| - | - | 1.783E+04 | 782.4 | - | - | 0 | - |
| - | - | 1569 | 783.4 | - | - | 0 | - |
| - | - | 1774 | 807.4 | - | - | 0 | - |
| - | - | 762.2 | 808.4 | - | - | 0 | - |
| - | - | 1112 | 816.4 | - | - | 0 | - |
| - | - | 985.5 | 817.5 | - | - | 0 | - |
| - | - | 1255 | 824.4 | - | - | 0 | - |
| - | - | 1185 | 825.4 | - | - | 0 | - |
| - | - | 1427 | 831.5 | - | - | 0 | - |
| - | - | 1132 | 832.5 | - | - | 0 | - |
| - | - | 804.8 | 833.5 | - | - | 0 | - |
| 9 | b | 7575 | 834.4 | 2.426E-05 | 0.02907 | +1 | 9 |
| - | - | 2179 | 835.4 | - | - | 0 | - |
| - | - | 959.1 | 836.4 | - | - | 0 | - |
| - | - | 3520 | 841.5 | - | - | 0 | - |
| - | - | 1676 | 842.5 | - | - | 0 | - |
| - | - | 790 | 844.5 | - | - | 0 | - |
| - | - | 737.4 | 847.5 | - | - | 0 | - |
| 9 | b | 1.57E+04 | 852.4 | 4.238E-05 | 0.04972 | +1 | 9 |
| - | - | 8860 | 853.4 | - | - | 0 | - |
| - | - | 2091 | 854.4 | - | - | 0 | - |
| - | - | 1.514E+04 | 859.5 | - | - | 0 | - |
| - | - | 7334 | 860.5 | - | - | 0 | - |
| 4 | y | 8827 | 861.5 | 0.005402 | 6.27 | +1 | 9 |
| - | - | 2761 | 862.5 | - | - | 0 | - |
| - | - | 988 | 863.5 | - | - | 0 | - |
| 4 | y | 1.061E+05 | 879.5 | 0.0002083 | 0.2368 | +1 | 9 |
| - | - | 5.137E+04 | 880.5 | - | - | 0 | - |
| - | - | 1.426E+04 | 881.5 | - | - | 0 | - |
| - | - | 1782 | 882.5 | - | - | 0 | - |
| 3 | y | 2576 | 960.6 | 0.0008917 | 0.9284 | +1 | 10 |
| - | - | 933.5 | 961.6 | - | - | 0 | - |
| 10 | b | 1.03E+04 | 962.5 | 0.0007644 | 0.7941 | +1 | 10 |
| 10 | b | 5438 | 963.5 | 0.0176 | 18.27 | +1 | 10 |
| - | - | 1725 | 964.5 | - | - | 0 | - |
| - | - | 726.2 | 976.5 | - | - | 0 | - |
| 3 | y | 5.195E+04 | 978.6 | 0.0003368 | 0.3442 | +1 | 10 |
| - | - | 2.925E+04 | 979.6 | - | - | 0 | - |
| 10 | b | 2.985E+04 | 980.5 | 0.0032 | 3.264 | +1 | 10 |
| - | - | 1.877E+04 | 981.5 | - | - | 0 | - |
| - | - | 5449 | 982.5 | - | - | 0 | - |
| - | - | 1357 | 994.5 | - | - | 0 | - |
| - | - | 733.4 | 995.5 | - | - | 0 | - |
| - | - | 1593 | 1066 | - | - | 0 | - |
| 11 | b | 3680 | 1076 | 0.002306 | 2.144 | +1 | 11 |
| 11 | b | 3503 | 1077 | 0.01942 | 18.03 | +1 | 11 |
| - | - | 686.7 | 1077 | - | - | 0 | - |
| - | - | 1078 | 1078 | - | - | 0 | - |
| 11 | b | 3.225E+04 | 1094 | 0.0002974 | 0.272 | +1 | 11 |
| - | - | 1126 | 1094 | - | - | 0 | - |
| - | - | 2.13E+04 | 1095 | - | - | 0 | - |
| - | - | 7857 | 1096 | - | - | 0 | - |
| - | - | 999 | 1097 | - | - | 0 | - |
| 2 | y | 859.7 | 1126 | 0.002464 | 2.189 | +1 | 11 |
| - | - | 650 | 1720 | - | - | 0 | - |
| - | - | 626.1 | 1850 | - | - | 0 | - |

m/z Charge Intensity FragmentType MassShift Position
120.06587982177734 0 35147.773 y 11
120.0811538696289 0 95030.55
121.06922912597656 0 868.54254
121.08447265625 0 8524.6
127.05076599121094 0 1537.7279
128.107177734375 0 917.0667
129.10263061523438 0 172918.17
130.0499267578125 0 521.8136
130.10008239746094 0 815.0418
130.1059112548828 0 9971.654
131.0701141357422 0 426.58737
131.08180236816406 0 5113.7954
132.0811767578125 0 1351.9532
132.10223388671875 0 1137.3099
133.06109619140625 0 509.86127
133.08615112304688 0 2819.0232
136.07606506347656 0 2090.7832
138.0556182861328 0 415.05322
139.08726501464844 0 1420.6389
139.90699768066406 0 394.88913
140.0822296142578 0 689.8213
141.06625366210938 0 3150.162
141.1024627685547 0 1198.9044
142.1230926513672 0 521.92755
143.08189392089844 0 629.53503
143.11849975585938 0 1088.9336
144.0759735107422 0 463.47043
144.08116149902344 0 6653.735
148.59017944335938 0 454.05298
148.94699096679688 0 488.20038
149.04515075683594 0 1773.2445
150.91412353515625 0 464.7542
151.08697509765625 0 913.2304
152.14370727539062 0 1690.143
153.06634521484375 0 986.0259
153.1024627685547 0 586.39374
154.06153869628906 0 815.99194
155.08192443847656 0 1297.6864
155.11827087402344 0 3940.3076
156.07699584960938 0 752.83734
158.09298706054688 0 1845.1344
159.07667541503906 0 4450.864
159.09210205078125 0 1199.769
159.11338806152344 0 542.0788
162.0916748046875 0 20598.596
163.0951385498047 0 2366.655
167.05580139160156 0 3292.175
167.11813354492188 0 1129.0109
169.09738159179688 0 1248.1567
169.13365173339844 0 962.4791
170.09274291992188 0 2978.4106
171.07701110839844 0 1815.684
171.08753967285156 0 691.19525
171.14955139160156 0 45060.07
172.07205200195312 0 6209.421
172.1528778076172 0 3420.3608
173.12881469726562 0 896.2623
174.1280517578125 0 1377.3336
176.10733032226562 0 5534.24
176.65689086914062 0 480.14975
177.10256958007812 0 5112.3086
177.1115264892578 0 1458.3112
181.17063903808594 0 577.2036
182.08078002929688 0 501.8507
183.11354064941406 0 1130.4434
183.12393188476562 0 513.9147
183.1494598388672 0 1031.4154
185.09291076660156 0 872.4327
185.12879943847656 0 2570.3557
185.88490295410156 0 496.69217
186.12411499023438 0 1715.8282
187.1079864501953 0 1145.8092
187.1444854736328 0 739.28906
188.10340881347656 0 1851.8956
189.1024169921875 0 2068.3489 a Water loss 1
189.12332153320312 0 629.1114
194.12908935546875 0 798.1717
195.14935302734375 0 703.2136
196.10806274414062 0 915.5221
197.10496520996094 0 536.748
197.16519165039062 0 6273.05
198.08758544921875 0 5636.266
199.10800170898438 0 577.1738
199.14439392089844 0 26706.385
200.14793395996094 0 2059.0378
201.09805297851562 0 632.30743
201.1239776611328 0 940.174
203.1181640625 0 1820.7975
205.0975799560547 0 12324.633
206.10101318359375 0 1418.9958
207.11326599121094 0 426154.7 a 1
208.11651611328125 0 49297.95 b Water loss 3
209.1035614013672 0 1269.228
209.1190185546875 0 3044.2378
210.08753967285156 0 1256.2802
211.14398193359375 0 587.66113
212.13970947265625 0 20195.076
213.14312744140625 0 1828.8638
214.1188507080078 0 602.22736
214.19180297851562 0 1592.284
215.10345458984375 0 610.6253
215.11451721191406 0 2501.7463
215.1393280029297 0 11619.076 y Water loss 10
216.09814453125 0 4972.4985
216.14244079589844 0 1730.8611
217.09768676757812 0 4509.4277 b Water loss 1
218.10140991210938 0 622.88306
219.14956665039062 0 28897.25
220.1528778076172 0 3924.7314
221.08462524414062 0 3751.6506
221.12869262695312 0 1319.3269
222.08523559570312 0 509.12625
222.12403869628906 0 898.81384
223.10850524902344 0 916.36725
224.13978576660156 0 531.1708
224.1761016845703 0 5534.9824
225.0432586669922 0 8179.65
225.1350555419922 0 2103.7192
225.1604461669922 0 1487.0295
226.04327392578125 0 857.9613
226.11859130859375 0 1223.9321
226.22250366210938 0 500.77176
227.02296447753906 0 1138.3141
227.1023712158203 0 709.81824
227.1141815185547 0 8460.264
228.09815979003906 0 3884.4797
228.1085662841797 0 941.0624
228.11936950683594 0 548.6766
229.093505859375 0 1291.2676
229.13345336914062 0 1875.6191
230.15032958984375 0 35356.715
231.1129913330078 0 1335.1465
231.15281677246094 0 4106.1733
233.12855529785156 0 1592.746
233.1500244140625 0 30687.822 y 10
234.1235809326172 0 981.0856
234.1532440185547 0 3550.5933
235.1082305908203 0 390962.88 b 1
236.11148071289062 0 48118.43
237.09820556640625 0 1699.5797
237.11363220214844 0 4127.5005
237.15957641601562 0 537.809
239.09536743164062 0 10476.346
240.09634399414062 0 1244.6971
240.134765625 0 1561.9673
240.17066955566406 0 665.1986
241.13836669921875 0 547.9356
241.1550750732422 0 1341.3519
242.18675231933594 0 7274.495
243.11317443847656 0 2582.5823
243.1461181640625 0 2135.5593
243.18978881835938 0 715.2824
245.1248779296875 0 6845.819
245.49044799804688 0 554.8782
246.12509155273438 0 735.3264
247.10813903808594 0 1482.1526
247.14454650878906 0 19720.58
248.14788818359375 0 3053.5261
251.1508331298828 0 1588.0314 y Water loss 7
254.125732421875 0 847.43915
255.10919189453125 0 20902.068
255.14793395996094 0 637.2152
256.1121826171875 0 2488.5928
257.12860107421875 0 619.3341
259.1446228027344 0 2230.1343
260.1962890625 0 1082.0118
261.1235656738281 0 2849.3518
261.159912109375 0 3345.269
262.119140625 0 7195.1597
262.1625671386719 0 663.5796
263.12255859375 0 574.2008
265.15509033203125 0 639.2043
268.166259765625 0 1425.6779
269.16119384765625 0 21850.414
270.16400146484375 0 3217.8342
270.1814880371094 0 1903.0732
271.1412048339844 0 1548.5392
272.13555908203125 0 2724.3445
273.1196594238281 0 25394.822
273.15936279296875 0 1078.6626
273.1961975097656 0 1983.2197
274.11993408203125 0 8120.7305
275.12347412109375 0 1429.9384
275.176025390625 0 1659.6637
279.1467590332031 0 624.72394
281.19769287109375 0 1012.6806
282.1568298339844 0 2299.935
285.1585693359375 0 819.34454
286.17626953125 0 1058.7092
287.17193603515625 0 7468.957
288.134033203125 0 1428.3136
288.172119140625 0 2938.1306 a Water loss 2
289.1551208496094 0 28787.654
290.1585388183594 0 6179.561
291.1457824707031 0 4113.0703
291.1621398925781 0 766.30615
292.1293029785156 0 1366.0337
292.1485900878906 0 1013.2787
295.1034851074219 0 2951.7034
297.156494140625 0 891.84937
299.06201171875 0 4654.648
300.062744140625 0 1376.9956
300.13427734375 0 2281.0356
300.16729736328125 0 950.6537
301.1917724609375 0 4900.043
302.19464111328125 0 869.6698
303.1709899902344 0 1448.6342
304.16644287109375 0 2945.5469
306.18218994140625 0 1271.3698 a 2
308.17437744140625 0 654.5858 y Water loss 5
308.19744873046875 0 1287.51
309.1598815917969 0 4495.367
310.1639709472656 0 842.8684 b Water loss 5
313.1874084472656 0 721.9532
315.1644287109375 0 600.29156
315.2389831542969 0 1030.9272
316.16619873046875 0 16782.516 b Water loss 2
317.16827392578125 0 2298.383
318.1453552246094 0 2627.8623
319.14093017578125 0 8772.389
320.14398193359375 0 1233.1367
321.1956481933594 0 1007.80927
325.2240905761719 0 3826.4758
326.18292236328125 0 20403.432
327.1861877441406 0 3210.0793
328.13116455078125 0 696.845
328.164794921875 0 704.7911
328.2038269042969 0 673.5349
330.1825866699219 0 669.311
331.1404724121094 0 1164.2394
332.162109375 0 1016.3899
333.1614685058594 0 695.542
333.1920471191406 0 1174.0922
334.1768493652344 0 255313.73 b 2
335.1798400878906 0 50787.55
336.1822204589844 0 3855.5571
337.1551208496094 0 7004.5723
338.1594543457031 0 1183.5693
339.1781005859375 0 2189.4338
340.1995544433594 0 786.35046
341.1837158203125 0 925.12067
343.2346496582031 0 8556.442 y Water loss 9
344.1932678222656 0 6295.7207
344.2384948730469 0 1171.6619
345.15740966796875 0 682.9739
345.19439697265625 0 1487.1406
346.177001953125 0 1989.3052
346.2129821777344 0 6750.209
347.2168273925781 0 930.0384
348.1674499511719 0 4166.7427
354.1780700683594 0 3086.5098
357.1559753417969 0 5667.9365
358.15679931640625 0 795.4278
358.213134765625 0 682.46735
359.02764892578125 0 927.7657
361.1888122558594 0 1523.2235
361.24493408203125 0 27138.912 y 9
362.1917419433594 0 757.75476
362.24749755859375 0 5759.3936
364.16607666015625 0 6881.02
365.1692199707031 0 1320.0776
365.1937255859375 0 3810.7925
366.1766662597656 0 1208.0574
368.1936950683594 0 1113.3251
369.12225341796875 0 1849.8401
372.18829345703125 0 1725.7825
373.18731689453125 0 2813.5195
374.18316650390625 0 6728.4004
374.2090759277344 0 2150.267
375.16754150390625 0 3160.9385
376.16522216796875 0 1823.6909
381.7114562988281 0 1131.3214 y Water loss 4
382.1764221191406 0 5058.9136
382.20330810546875 0 745.4721 y Ammonia loss 4
382.2167053222656 0 1056.6041
382.24542236328125 0 5083.474
383.1797790527344 0 1091.5436
383.2041931152344 0 23698.531
384.1663513183594 0 1320.5293
384.2073059082031 0 4580.3037
385.1524658203125 0 1382.0482
386.20184326171875 0 721.725
387.2039794921875 0 1175.3312
388.2236022949219 0 3785.4812
389.22650146484375 0 1420.7529
390.2135314941406 0 4084.8464
390.71685791015625 0 8360.881 y 4
391.19464111328125 0 1128.7518
391.2184753417969 0 3031.3865
391.71795654296875 0 732.4467
392.1937255859375 0 7955.666
393.1941223144531 0 2249.9375
400.2559509277344 0 7111.5024
401.2149353027344 0 27897.979
401.2613525390625 0 1791.864
402.17779541015625 0 24980.541
402.2173156738281 0 5653.3325
403.18096923828125 0 5424.924
403.234130859375 0 945.6593
404.1831359863281 0 971.8093
405.2502746582031 0 17369.184
406.2542724609375 0 3330.008
411.20098876953125 0 791.5425
415.2347412109375 0 9773.211 b Water loss 3
416.2364196777344 0 1798.0751
418.2093200683594 0 2927.5542
419.20660400390625 0 1500.872
420.1884765625 0 16775.79
421.1905212402344 0 5468.6904
421.2550964355469 0 1231.6697
429.2828369140625 0 2597.325
430.20904541015625 0 630.10535
433.2452392578125 0 69885.42 b 3
434.24853515625 0 15603.99
435.2462463378906 0 2487.8699
436.2236022949219 0 6834.3657
436.24920654296875 0 595.42615
437.22467041015625 0 2783.883
438.21990966796875 0 551.6591
439.1964111328125 0 644.7051
439.2670593261719 0 7524.2495
440.2512512207031 0 2258.3538 y 3
440.7522888183594 0 1253.9711
441.25103759765625 0 1017.21295
444.22515869140625 0 732.6673
444.281494140625 0 894.5103 y Water loss 8
447.24041748046875 0 1219.1343
450.2214050292969 0 611.19415
452.26312255859375 0 1811.8306
453.2500305175781 0 5853.5815
454.2528076171875 0 2216.5422
456.2249755859375 0 1579.906
457.2254638671875 0 672.9042
457.27783203125 0 13708.943
458.2803039550781 0 3274.2563
460.253662109375 0 780.823
461.2543640136719 0 655.88916
462.29339599609375 0 6820.357 y 8
463.23431396484375 0 6190.7886
463.2969970703125 0 1488.7545
464.23876953125 0 1767.8779
468.2958679199219 0 851.6349
469.2754211425781 0 892.2755
470.27276611328125 0 1990.3584
471.27215576171875 0 656.76587
472.25604248046875 0 1652.594
473.2520751953125 0 3521.0234
474.2369384765625 0 1671.1964
475.2335510253906 0 1363.6815
478.27825927734375 0 1800.3928
480.7802429199219 0 1365.7119 y Water loss 2
481.2453918457031 0 13253.189
481.2796936035156 0 1195.0583 y Ammonia loss 2
481.7587890625 0 2013.0171 b Water loss 9
482.2506103515625 0 5098.769 b Ammonia loss 9
483.25042724609375 0 775.6782
486.2450866699219 0 1168.0985
486.3042907714844 0 8621.3955
487.3069152832031 0 2308.8518
489.28277587890625 0 1046.7686
489.7857971191406 0 3433.533 y 2
490.2604064941406 0 919.7444
490.2955017089844 0 1046.4946
490.7645263671875 0 3080.1155 b 9
491.26409912109375 0 5088.699
492.26422119140625 0 1671.6083
493.24481201171875 0 1231.3058
495.2347106933594 0 1194.9633
496.2883605957031 0 19432.73
497.2909851074219 0 5162.483
501.2462158203125 0 9390.665
501.30419921875 0 1464.7595 y Water loss 7
502.2504577636719 0 2519.827
503.2565002441406 0 686.11914
504.25445556640625 0 839.45386
506.7110595703125 0 838.7364
510.2721862792969 0 1609.5823
512.2615966796875 0 4128.4736
513.2651977539062 0 713.1353
514.2988891601562 0 45870.688
515.26416015625 0 444.76746
515.3018798828125 0 12359.931
516.3048095703125 0 1802.325
517.2759399414062 0 880.58734
519.2571411132812 0 4805.821
519.314208984375 0 11652.889 y 7
520.257568359375 0 2954.2695
520.3157958984375 0 1978.505
521.262451171875 0 818.72723
530.2728271484375 0 17560.74
531.2748413085938 0 5271.99
533.3078002929688 0 2549.83
533.8091430664062 0 2048.204
534.3084106445312 0 3097.7476
535.2932739257812 0 6194.9624
536.2930908203125 0 2687.7454
538.2664794921875 0 3784.4255
538.9472045898438 0 644.91266
539.268798828125 0 1118.4513
540.315185546875 0 866.47906
547.3024291992188 0 1165.7695 b 10
547.80712890625 0 1474.6313
548.2832641601562 0 40389.32
549.2859497070312 0 13747.865
550.2887573242188 0 2266.9531
552.3184814453125 0 9204.407
553.3206787109375 0 3029.03
558.3253784179688 0 2613.1677 y Water loss 6
562.3024291992188 0 7066.156 b Water loss 4
563.3071899414062 0 3236.059
564.3125610351562 0 1022.9725
567.2953491210938 0 695.38556
571.3595581054688 0 755.3704
574.30322265625 0 1315.7397
575.81982421875 0 822.6738
576.33544921875 0 28507.346 y 6
577.2828369140625 0 742.44885
577.338623046875 0 8172.509
578.3423461914062 0 1516.2311
580.3134765625 0 19235.281 b 4
581.3167724609375 0 6776.15
582.3182373046875 0 1152.4508
589.3411254882812 0 3007.8098
590.341796875 0 994.58264
591.3339233398438 0 838.59045
592.3132934570312 0 1440.6787
595.2901000976562 0 1124.9214
596.2879638671875 0 759.3821
597.3352661132812 0 1799.1299
597.8301391601562 0 12249.058 Precursor Water loss
598.3317260742188 0 9558.508 Precursor Ammonia loss
598.8329467773438 0 2522.5479
599.3311157226562 0 1692.0242
600.31591796875 0 3239.4863
601.3173828125 0 1019.79816
606.279052734375 0 1167.1398
606.8343505859375 0 995.6088 Precursor
607.2838134765625 0 685.8037
607.333251953125 0 982.4094
609.3390502929688 0 2474.2346
611.3319091796875 0 1442.2329
615.3475952148438 0 9496.212 y Water loss 5
616.3492431640625 0 4349.2637
617.3458862304688 0 1424.4092
618.3262329101562 0 3054.089
619.3253173828125 0 4414.119 b Water loss 5
620.3298950195312 0 1698.791
625.3449096679688 0 2812.046
626.3456420898438 0 756.228
629.3411865234375 0 8153.1055
630.343017578125 0 2841.7485
631.3351440429688 0 1086.0194
633.3580322265625 0 136254.56 y 5
634.3607788085938 0 45180.49
635.3635864257812 0 8467.873
636.3630981445312 0 742.4861
637.3350219726562 0 5833.8193 b 5
638.3391723632812 0 2788.8162
643.3563842773438 0 23651.875
644.3593139648438 0 10912.077
645.3623657226562 0 1467.2617
647.3517456054688 0 17533.73
648.3544311523438 0 7538.9297
649.3482666015625 0 1544.4507
652.3058471679688 0 1231.5583
661.3671875 0 48207.613
662.3699951171875 0 17402.334
663.3724975585938 0 3247.5486
666.3613891601562 0 1094.2565
676.3469848632812 0 1699.5356 b Water loss 6
679.375732421875 0 1386.1974
694.355712890625 0 3076.9526 b 6
695.35791015625 0 1126.359
705.3545532226562 0 670.6752
714.431884765625 0 689.7006
718.4243774414062 0 2950.6284
719.4259033203125 0 1189.2723
723.3787841796875 0 862.77014
728.4092407226562 0 3430.018
729.4140625 0 1622.5166
732.4396362304688 0 4631.3755
733.3739624023438 0 971.86066 b Water loss 7
733.4447021484375 0 2159.71
734.3613891601562 0 688.1655
734.4251708984375 0 905.18207
735.41796875 0 743.689
736.4014892578125 0 748.7322
737.4134521484375 0 596.6973
742.425048828125 0 9391.533
743.428466796875 0 4045.1711
744.4103393554688 0 2068.5364
745.3995971679688 0 1057.1501
746.418701171875 0 8657.038
747.4237060546875 0 3267.2656
748.4135131835938 0 925.5887
751.3782348632812 0 4710.8813 b 7
752.3818969726562 0 1120.7662
753.3633422851562 0 1312.442
754.3619995117188 0 1026.5897
760.4356079101562 0 27491.586
761.4382934570312 0 11042.97
762.4169311523438 0 14229.879 y Water loss 4
763.4181518554688 0 6642.9136
764.4243774414062 0 812.07513
778.405029296875 0 1248.2714
780.425537109375 0 186552.83 y 4
781.4284057617188 0 79483.945
782.43115234375 0 17834.693
783.4338989257812 0 1569.3584
807.40771484375 0 1773.6451
808.4095458984375 0 762.2485
816.4047241210938 0 1111.8054
817.4910278320312 0 985.5401
824.4315185546875 0 1255.0914
825.4358520507812 0 1185.0967
831.5073852539062 0 1427.4708
832.5120849609375 0 1132.2401
833.4912719726562 0 804.7871
834.4144897460938 0 7575.1357 b Water loss 8
835.4185180664062 0 2178.7864
836.421142578125 0 959.0795
841.4929809570312 0 3520.042
842.4976196289062 0 1675.9015
844.466796875 0 789.98517
847.4654541015625 0 737.37506
852.4249877929688 0 15696.787 b 8
853.4286499023438 0 8859.731
854.4302978515625 0 2091.1067
859.503662109375 0 15136.564
860.5057373046875 0 7333.692
861.48828125 0 8826.646 y Water loss 3
862.4881591796875 0 2761.2593
863.4887084960938 0 987.96515
879.49365234375 0 106078.92 y 3
880.4967651367188 0 51368.406
881.4991455078125 0 14255.539
882.4998779296875 0 1781.6666
960.5521850585938 0 2576.3118 y Water loss 2
961.5595703125 0 933.4678
962.5101928710938 0 10301.917 b Water loss 9
963.5110473632812 0 5438.297 b Ammonia loss 9
964.515625 0 1724.7949
976.5421142578125 0 726.16547
978.5621948242188 0 51949.77 y 2
979.5643920898438 0 29247.615
980.523193359375 0 29845.324 b 9
981.5225830078125 0 18770.504
982.5267333984375 0 5449.1284
994.5299072265625 0 1356.6582
995.5324096679688 0 733.4193
1065.6160888671875 0 1593.41
1075.5911865234375 0 3680.3823 b Water loss 10
1076.596923828125 0 3503.158 b Ammonia loss 10
1076.7177734375 0 686.73303
1077.6019287109375 0 1078.1924
1093.603759765625 0 32253.781 b 10
1094.471923828125 0 1126.1581
1094.6065673828125 0 21297.967
1095.609619140625 0 7857.4727
1096.6165771484375 0 999.0271
1125.6278076171875 0 859.7444 y 1
1719.936767578125 0 649.96967
1850.0477294921875 0 626.1025

Spectrum Details

|  |  |
| --- | --- |
| Matched peaks? Matched peaks The total absolute number of peaks matched. Additionally in brackets the total fraction of peaks matched and the total number of peaks is shown. | 64 (11.37% of 563) |
| FDR? FDR The false discovery rate estimated for this peptide. It is calculated by matching all theoretical fragments with a non-integer shift with the raw peaks for this spectrum. This is done with 40 different shifts. The resulting percentage is the average number of annotated peaks over the number of annotated peaks with the correct spectrum. | 0.45% |
| Satellite FDR? Satellite FDR See the FDR for details on its calculation. This satellite ion specific FDR only contains the satellite ions (d/w) for I/L/J positions. | - |
| PSM Score? PSM Score The PSM Score as given by Hecklib to this annotated spectrum. It is shown with three significant figures. | 788 |

#### Spectrum 9175? Spectrum 9175 The raw spectrum of this peptide as annotated by Hecklib. The fragments are coloured according to ion type (see legend). Any peaks with a star '\*' as text can be hovered over to see the full details, first the ion type second the mass shift type. By hovering over the amino acids in the peptide or ions in the legend the corresponding peaks are highlighted. By toggling the 'Unassigned' label you can turn the background (unassigned) peaks on or off in the plot. By updating the slider in the Ion legend you can update the spectrum to only show the top X% of the peaks with labels. The top X% means any peak that is within X% of the highest intensity. By dragging in the spectrum you can zoom in to a specific part of the spectrum and use 'Zoom Out' to get back to the original zoom level. The annotation of the spectrum is based on the given sequence in the peptides file and is done with different software so inconsistencies are likely. The peaks are annotated based on the given sequence, with 20 ppm tolerance.

Copy Data

##### Spectrum 9175 (TSV)

###### Preview

```
Loading example...
```

*Click on the button to copy the data to your clipboard.*

Mz MinMz MaxIntensity Max

WidthHeightPeptide font sizePeptide stroke widthSpectrum font sizeSpectrum stroke widthCompact peptide

Ion legend

wxyz

abcd

OtherUnassignedIon^Charge^_Position_Show for top:%

SFVVFGGGTKJT

03.86e+47.73e+41.16e+51.55e+5

Zoom Out

y^+1^_1_y^+1^_2_y^+1^_2_y^+1^_3_y^+2^_8_y^+2^_9_y^+1^_4_z^+1^_4_y^+1^_4_y^+2^_10_y^+2^_10_y^+2^_10_c^+2^_10_y^+1^_5_z^+1^_6_z^+1^_6_y^+1^_6_y^+1^_7_z^+1^_7_y^+1^_7_c^+1^_6_c^+1^_7_z^+1^_8_y^+1^_8_z^+1^_8_c^+1^_8_y^+1^_8_w^+1^_9_c^+1^_9_y^+1^_9_z^+1^_9_c^+1^_9_y^+1^_9_w^+1^_10_z^+1^_10_y^+1^_10_c^+1^_10_c^+1^_10_c^+1^_11_z^+1^_11_c^+1^_11_

0821164324643286

Fragment Matches Table

Show background peaks

| Position | Ion type | Intensity | mz Theoretical | mz Error (Th) | mz Error (ppm) | Charge | Series Number |
| --- | --- | --- | --- | --- | --- | --- | --- |
| - | - | 579.1 | 120.1 | - | - | 0 | - |
| 12 | y | 8210 | 120.1 | 0.0001847 | 1.539 | +1 | 1 |
| - | - | 1219 | 120.1 | - | - | 0 | - |
| - | - | 364.5 | 122.6 | - | - | 0 | - |
| - | - | 2769 | 129.1 | - | - | 0 | - |
| - | - | 446.6 | 129.4 | - | - | 0 | - |
| - | - | 459.1 | 134.2 | - | - | 0 | - |
| - | - | 1531 | 149 | - | - | 0 | - |
| - | - | 728.2 | 167.1 | - | - | 0 | - |
| - | - | 2273 | 171.1 | - | - | 0 | - |
| - | - | 3591 | 173.5 | - | - | 0 | - |
| - | - | 1960 | 177.1 | - | - | 0 | - |
| - | - | 451.2 | 184.4 | - | - | 0 | - |
| - | - | 478 | 189.6 | - | - | 0 | - |
| - | - | 3770 | 199.1 | - | - | 0 | - |
| - | - | 540.2 | 200.1 | - | - | 0 | - |
| - | - | 548 | 206.3 | - | - | 0 | - |
| - | - | 1.523E+04 | 207.1 | - | - | 0 | - |
| - | - | 1629 | 208.1 | - | - | 0 | - |
| - | - | 476.8 | 209.3 | - | - | 0 | - |
| 11 | y | 1522 | 215.1 | 0.0001487 | 0.691 | +1 | 2 |
| - | - | 894.8 | 219.1 | - | - | 0 | - |
| - | - | 3708 | 221.1 | - | - | 0 | - |
| - | - | 2465 | 225 | - | - | 0 | - |
| - | - | 940.9 | 230.2 | - | - | 0 | - |
| - | - | 647.2 | 231.2 | - | - | 0 | - |
| 11 | y | 7071 | 233.1 | 5.935E-05 | 0.2546 | +1 | 2 |
| - | - | 958.6 | 234.2 | - | - | 0 | - |
| - | - | 3.277E+04 | 235.1 | - | - | 0 | - |
| - | - | 3806 | 236.1 | - | - | 0 | - |
| - | - | 4876 | 239.1 | - | - | 0 | - |
| - | - | 1433 | 247.1 | - | - | 0 | - |
| - | - | 525.8 | 263.5 | - | - | 0 | - |
| - | - | 572.7 | 273.1 | - | - | 0 | - |
| - | - | 3847 | 289.2 | - | - | 0 | - |
| - | - | 4736 | 295.1 | - | - | 0 | - |
| - | - | 740.1 | 296.1 | - | - | 0 | - |
| - | - | 1749 | 299.1 | - | - | 0 | - |
| - | - | 868.7 | 313.1 | - | - | 0 | - |
| - | - | 4.17E+04 | 334.2 | - | - | 0 | - |
| - | - | 6734 | 335.2 | - | - | 0 | - |
| - | - | 635.5 | 336.2 | - | - | 0 | - |
| - | - | 1286 | 346.2 | - | - | 0 | - |
| 10 | y | 3004 | 361.2 | 7.029E-05 | 0.1946 | +1 | 3 |
| - | - | 2447 | 369.1 | - | - | 0 | - |
| 5 | y | 1065 | 390.7 | 0.000394 | 1.008 | +2 | 8 |
| - | - | 883.1 | 391.2 | - | - | 0 | - |
| - | - | 3443 | 401.2 | - | - | 0 | - |
| - | - | 1012 | 402.2 | - | - | 0 | - |
| - | - | 627.3 | 402.2 | - | - | 0 | - |
| - | - | 4153 | 405.2 | - | - | 0 | - |
| - | - | 1052 | 406.3 | - | - | 0 | - |
| - | - | 1453 | 415.2 | - | - | 0 | - |
| - | - | 1.661E+04 | 433.2 | - | - | 0 | - |
| - | - | 4825 | 434.2 | - | - | 0 | - |
| - | - | 650.4 | 436.2 | - | - | 0 | - |
| 4 | y | 1689 | 440.3 | 9.749E-05 | 0.2214 | +2 | 9 |
| - | - | 1879 | 444.3 | - | - | 0 | - |
| 9 | y | 4920 | 445.3 | 0.0004678 | 1.051 | +1 | 4 |
| 9 | z | 1003 | 446.3 | 0.005343 | 11.97 | +1 | 4 |
| - | - | 1197 | 447.3 | - | - | 0 | - |
| - | - | 957.5 | 453.2 | - | - | 0 | - |
| - | - | 1065 | 457.3 | - | - | 0 | - |
| - | - | 722.6 | 459.3 | - | - | 0 | - |
| 9 | y | 931.2 | 462.3 | 0.0003469 | 0.7505 | +1 | 4 |
| - | - | 1195 | 463.2 | - | - | 0 | - |
| - | - | 1001 | 470.3 | - | - | 0 | - |
| 3 | y | 593.5 | 480.8 | 0.001331 | 2.768 | +2 | 10 |
| - | - | 2699 | 481.2 | - | - | 0 | - |
| 3 | y | 652.7 | 481.3 | 0.009286 | 19.29 | +2 | 10 |
| - | - | 967.9 | 481.8 | - | - | 0 | - |
| - | - | 836.1 | 482.3 | - | - | 0 | - |
| 3 | y | 1431 | 489.8 | 0.000589 | 1.203 | +2 | 10 |
| 10 | c | 1510 | 490.8 | 0.0006779 | 1.381 | +2 | 10 |
| - | - | 643.1 | 491.3 | - | - | 0 | - |
| - | - | 1.176E+04 | 504.3 | - | - | 0 | - |
| - | - | 9310 | 504.3 | - | - | 0 | - |
| - | - | 4349 | 505.3 | - | - | 0 | - |
| - | - | 2597 | 505.3 | - | - | 0 | - |
| - | - | 592.5 | 506.3 | - | - | 0 | - |
| - | - | 3445 | 514.3 | - | - | 0 | - |
| - | - | 814.8 | 515.3 | - | - | 0 | - |
| - | - | 2682 | 516.3 | - | - | 0 | - |
| - | - | 2485 | 518.3 | - | - | 0 | - |
| 8 | y | 4467 | 519.3 | 0.0008226 | 1.584 | +1 | 5 |
| - | - | 874.1 | 520.3 | - | - | 0 | - |
| - | - | 932.7 | 530.3 | - | - | 0 | - |
| - | - | 1222 | 533.3 | - | - | 0 | - |
| - | - | 965.9 | 535.3 | - | - | 0 | - |
| - | - | 828 | 538.3 | - | - | 0 | - |
| 7 | z | 767.9 | 542.3 | 0.00258 | 4.758 | +1 | 6 |
| - | - | 4884 | 548.3 | - | - | 0 | - |
| - | - | 1383 | 549.3 | - | - | 0 | - |
| - | - | 2018 | 552.3 | - | - | 0 | - |
| - | - | 824.2 | 553.3 | - | - | 0 | - |
| 7 | z | 5943 | 560.3 | 0.000405 | 0.7229 | +1 | 6 |
| - | - | 936.7 | 561.3 | - | - | 0 | - |
| - | - | 2.376E+04 | 561.3 | - | - | 0 | - |
| - | - | 5593 | 562.3 | - | - | 0 | - |
| - | - | 1068 | 563.3 | - | - | 0 | - |
| - | - | 2970 | 575.3 | - | - | 0 | - |
| 7 | y | 2.148E+04 | 576.3 | 0.0004968 | 0.862 | +1 | 6 |
| - | - | 5912 | 577.3 | - | - | 0 | - |
| - | - | 1048 | 578.3 | - | - | 0 | - |
| - | - | 5195 | 580.3 | - | - | 0 | - |
| - | - | 1457 | 581.3 | - | - | 0 | - |
| - | - | 1696 | 589.3 | - | - | 0 | - |
| - | - | 1051 | 591.4 | - | - | 0 | - |
| - | - | 2651 | 597.8 | - | - | 0 | - |
| - | - | 2351 | 598.3 | - | - | 0 | - |
| - | - | 888.4 | 598.8 | - | - | 0 | - |
| - | - | 712.8 | 607.3 | - | - | 0 | - |
| - | - | 826.8 | 609.3 | - | - | 0 | - |
| 6 | y | 1297 | 615.3 | 0.0004095 | 0.6654 | +1 | 7 |
| 6 | z | 1.357E+04 | 617.3 | 5.947E-05 | 0.09634 | +1 | 7 |
| - | - | 3.044E+04 | 618.3 | - | - | 0 | - |
| - | - | 8619 | 619.3 | - | - | 0 | - |
| - | - | 1588 | 620.3 | - | - | 0 | - |
| - | - | 1.2E+04 | 632.3 | - | - | 0 | - |
| 6 | y | 5.426E+04 | 633.4 | 0.0001099 | 0.1735 | +1 | 7 |
| - | - | 1.627E+04 | 634.4 | - | - | 0 | - |
| - | - | 3604 | 635.4 | - | - | 0 | - |
| - | - | 1239 | 637.3 | - | - | 0 | - |
| - | - | 591.8 | 638.3 | - | - | 0 | - |
| - | - | 2472 | 647.4 | - | - | 0 | - |
| - | - | 953.8 | 648.4 | - | - | 0 | - |
| 6 | c | 2751 | 654.4 | 0.001659 | 2.536 | +1 | 6 |
| - | - | 956.4 | 655.4 | - | - | 0 | - |
| - | - | 1161 | 660.3 | - | - | 0 | - |
| - | - | 4093 | 661.4 | - | - | 0 | - |
| - | - | 2177 | 662.4 | - | - | 0 | - |
| - | - | 2166 | 663.4 | - | - | 0 | - |
| - | - | 1395 | 664.4 | - | - | 0 | - |
| - | - | 1079 | 668.4 | - | - | 0 | - |
| - | - | 786 | 679.4 | - | - | 0 | - |
| - | - | 810.7 | 692.4 | - | - | 0 | - |
| - | - | 1091 | 694.4 | - | - | 0 | - |
| - | - | 9955 | 710.4 | - | - | 0 | - |
| 7 | c | 8801 | 711.4 | 0.002188 | 3.076 | +1 | 7 |
| - | - | 3673 | 712.4 | - | - | 0 | - |
| - | - | 710 | 718.4 | - | - | 0 | - |
| - | - | 750.9 | 724.4 | - | - | 0 | - |
| - | - | 1661 | 725.4 | - | - | 0 | - |
| - | - | 689 | 742.4 | - | - | 0 | - |
| 5 | z | 1909 | 746.4 | 0.01258 | 16.86 | +1 | 8 |
| - | - | 617.9 | 749.4 | - | - | 0 | - |
| - | - | 1521 | 751.4 | - | - | 0 | - |
| - | - | 710.2 | 752.4 | - | - | 0 | - |
| - | - | 2257 | 760.4 | - | - | 0 | - |
| - | - | 1322 | 761.4 | - | - | 0 | - |
| 5 | y | 1487 | 762.4 | 0.004614 | 6.052 | +1 | 8 |
| 5 | z | 2.669E+04 | 764.4 | 0.0001782 | 0.2331 | +1 | 8 |
| - | - | 4.137E+04 | 765.4 | - | - | 0 | - |
| - | - | 1.469E+04 | 766.4 | - | - | 0 | - |
| - | - | 6480 | 767.4 | - | - | 0 | - |
| 8 | c | 3852 | 768.4 | 0.001679 | 2.185 | +1 | 8 |
| - | - | 1062 | 769.4 | - | - | 0 | - |
| - | - | 1.7E+04 | 779.4 | - | - | 0 | - |
| 5 | y | 6.782E+04 | 780.4 | 0.0007748 | 0.9928 | +1 | 8 |
| - | - | 2.783E+04 | 781.4 | - | - | 0 | - |
| - | - | 7190 | 782.4 | - | - | 0 | - |
| - | - | 1981 | 807.4 | - | - | 0 | - |
| - | - | 1489 | 808.4 | - | - | 0 | - |
| - | - | 1236 | 834.4 | - | - | 0 | - |
| 4 | w | 2648 | 848.5 | 0.0002321 | 0.2736 | +1 | 9 |
| - | - | 950.9 | 849.5 | - | - | 0 | - |
| - | - | 2062 | 850.4 | - | - | 0 | - |
| 9 | c | 1056 | 851.4 | 0.00504 | 5.92 | +1 | 9 |
| - | - | 4599 | 852.4 | - | - | 0 | - |
| - | - | 1723 | 853.4 | - | - | 0 | - |
| - | - | 697.6 | 854.4 | - | - | 0 | - |
| - | - | 2283 | 859.5 | - | - | 0 | - |
| - | - | 1337 | 860.5 | - | - | 0 | - |
| - | - | 1096 | 861.5 | - | - | 0 | - |
| 4 | y | 905.1 | 862.5 | 0.005212 | 6.043 | +1 | 9 |
| 4 | z | 3.225E+04 | 863.5 | 0.0001717 | 0.1988 | +1 | 9 |
| - | - | 2.655E+04 | 864.5 | - | - | 0 | - |
| - | - | 1.022E+04 | 865.5 | - | - | 0 | - |
| - | - | 2271 | 866.5 | - | - | 0 | - |
| - | - | 1.18E+04 | 868.4 | - | - | 0 | - |
| 9 | c | 7.234E+04 | 869.5 | 0.0006515 | 0.7494 | +1 | 9 |
| - | - | 3.572E+04 | 870.5 | - | - | 0 | - |
| - | - | 8875 | 871.5 | - | - | 0 | - |
| - | - | 1145 | 872.5 | - | - | 0 | - |
| - | - | 2472 | 878.5 | - | - | 0 | - |
| 4 | y | 3.262E+04 | 879.5 | 0.0007073 | 0.8042 | +1 | 9 |
| - | - | 1.545E+04 | 880.5 | - | - | 0 | - |
| - | - | 3375 | 881.5 | - | - | 0 | - |
| - | - | 816.9 | 882.5 | - | - | 0 | - |
| - | - | 971.8 | 906.5 | - | - | 0 | - |
| 3 | w | 1453 | 947.5 | 0.003423 | 3.613 | +1 | 10 |
| 3 | z | 3.26E+04 | 962.5 | 0.00163 | 1.693 | +1 | 10 |
| - | - | 1.851E+04 | 963.5 | - | - | 0 | - |
| - | - | 5372 | 964.5 | - | - | 0 | - |
| 3 | y | 1.888E+04 | 978.6 | 0.0008839 | 0.9033 | +1 | 10 |
| - | - | 905.1 | 978.7 | - | - | 0 | - |
| - | - | 9734 | 979.6 | - | - | 0 | - |
| 10 | c | 8123 | 980.5 | 0.004238 | 4.322 | +1 | 10 |
| - | - | 5207 | 981.5 | - | - | 0 | - |
| - | - | 2561 | 982.5 | - | - | 0 | - |
| 10 | c | 4.656E+04 | 997.5 | 0.000949 | 0.9514 | +1 | 10 |
| - | - | 2.867E+04 | 998.5 | - | - | 0 | - |
| - | - | 8352 | 999.6 | - | - | 0 | - |
| - | - | 1574 | 1001 | - | - | 0 | - |
| - | - | 926.3 | 1009 | - | - | 0 | - |
| - | - | 857.3 | 1052 | - | - | 0 | - |
| - | - | 3308 | 1067 | - | - | 0 | - |
| - | - | 1661 | 1068 | - | - | 0 | - |
| - | - | 890.2 | 1069 | - | - | 0 | - |
| 11 | c | 8630 | 1094 | 0.00103 | 0.9417 | +1 | 11 |
| - | - | 6114 | 1095 | - | - | 0 | - |
| - | - | 4327 | 1096 | - | - | 0 | - |
| - | - | 1920 | 1097 | - | - | 0 | - |
| - | - | 1200 | 1098 | - | - | 0 | - |
| 2 | z | 2.69E+04 | 1110 | 0.000708 | 0.6381 | +1 | 11 |
| 11 | c | 9.09E+04 | 1111 | 0.004019 | 3.619 | +1 | 11 |
| - | - | 5.327E+04 | 1112 | - | - | 0 | - |
| - | - | 1.917E+04 | 1113 | - | - | 0 | - |
| - | - | 2990 | 1114 | - | - | 0 | - |
| - | - | 1571 | 1136 | - | - | 0 | - |
| - | - | 1063 | 1137 | - | - | 0 | - |
| - | - | 853 | 1141 | - | - | 0 | - |
| - | - | 750.5 | 1143 | - | - | 0 | - |
| - | - | 836 | 1151 | - | - | 0 | - |
| - | - | 843.9 | 1152 | - | - | 0 | - |
| - | - | 732.5 | 1155 | - | - | 0 | - |
| - | - | 2.788E+04 | 1158 | - | - | 0 | - |
| - | - | 1.828E+04 | 1159 | - | - | 0 | - |
| - | - | 6577 | 1160 | - | - | 0 | - |
| - | - | 1091 | 1161 | - | - | 0 | - |
| - | - | 2663 | 1168 | - | - | 0 | - |
| - | - | 958.8 | 1169 | - | - | 0 | - |
| - | - | 9664 | 1186 | - | - | 0 | - |
| - | - | 6646 | 1187 | - | - | 0 | - |
| - | - | 3252 | 1188 | - | - | 0 | - |
| - | - | 2012 | 1195 | - | - | 0 | - |
| - | - | 7111 | 1196 | - | - | 0 | - |
| - | - | 1.427E+05 | 1197 | - | - | 0 | - |
| - | - | 9.481E+04 | 1198 | - | - | 0 | - |
| - | - | 3.756E+04 | 1199 | - | - | 0 | - |
| - | - | 5995 | 1200 | - | - | 0 | - |
| - | - | 2869 | 1211 | - | - | 0 | - |
| - | - | 3219 | 1212 | - | - | 0 | - |
| - | - | 7.497E+04 | 1213 | - | - | 0 | - |
| - | - | 1.53E+05 | 1214 | - | - | 0 | - |
| - | - | 9.268E+04 | 1215 | - | - | 0 | - |
| - | - | 3.243E+04 | 1216 | - | - | 0 | - |
| - | - | 3395 | 1217 | - | - | 0 | - |
| - | - | 647.6 | 3253 | - | - | 0 | - |

m/z Charge Intensity FragmentType MassShift Position
120.06149291992188 0 579.1328
120.06570434570312 0 8209.722 y 11
120.08097839355469 0 1218.7012
122.55304718017578 0 364.4779
129.102294921875 0 2769.1306
129.42604064941406 0 446.5931
134.2092742919922 0 459.12518
149.04493713378906 0 1531.3947
167.05564880371094 0 728.1618
171.14939880371094 0 2273.108
173.45025634765625 0 3590.7878
177.1123809814453 0 1959.9204
184.3584442138672 0 451.2443
189.645263671875 0 477.95343
199.1443328857422 0 3769.944
200.14768981933594 0 540.2346
206.28302001953125 0 547.9591
207.11285400390625 0 15230.876
208.11618041992188 0 1628.6627
209.29415893554688 0 476.76263
215.1388702392578 0 1521.9149 y Water loss 10
219.14944458007812 0 894.81464
221.0845489501953 0 3707.903
225.04281616210938 0 2464.5662
230.15023803710938 0 940.9317
231.1553497314453 0 647.2043
233.14964294433594 0 7070.6025 y 10
234.1525115966797 0 958.59827
235.1078643798828 0 32769.14
236.1110076904297 0 3805.929
239.09521484375 0 4876.4673
247.1442108154297 0 1433.15
263.4625549316406 0 525.81464
273.1189880371094 0 572.74554
289.1550598144531 0 3847.2368
295.10333251953125 0 4736
296.1048889160156 0 740.148
299.0616455078125 0 1749.1486
313.11541748046875 0 868.7017
334.17645263671875 0 41701.344
335.1796569824219 0 6734.415
336.1827392578125 0 635.45667
346.2123107910156 0 1285.8196
361.2444763183594 0 3004.2405 y 9
369.1210632324219 0 2447.0117
390.71575927734375 0 1065.1483 y 4
391.21832275390625 0 883.0954
401.2143249511719 0 3442.961
402.1781921386719 0 1011.8326
402.21868896484375 0 627.2988
405.24981689453125 0 4152.7095
406.2518615722656 0 1051.7703
415.2341613769531 0 1453.0662
433.244873046875 0 16614.664
434.2480773925781 0 4824.6807
436.222412109375 0 650.3591
440.2504577636719 0 1689.285 y 3
444.2586975097656 0 1879.4729
445.2661437988281 0 4920.338 y Ammonia loss 8
446.2681579589844 0 1002.7047 z 8
447.2798767089844 0 1196.9524
453.2495422363281 0 957.4552
457.2771301269531 0 1065.465
459.2924499511719 0 722.58777
462.2925720214844 0 931.17914 y 8
463.2331848144531 0 1195.0768
470.260986328125 0 1000.5261
480.7779541015625 0 593.4512 y Water loss 2
481.2442626953125 0 2698.5427
481.28057861328125 0 652.7261 y Ammonia loss 2
481.7565612792969 0 967.8741
482.2507629394531 0 836.0563
489.78515625 0 1431.2478 y 2
490.7643127441406 0 1509.9468 c Ammonia loss 9
491.2649230957031 0 643.1467
504.2539367675781 0 11762.781
504.3028259277344 0 9310.024
505.2580871582031 0 4349.14
505.30633544921875 0 2597.2566
506.2630615234375 0 592.458
514.2981567382812 0 3444.6785
515.3026733398438 0 814.76935
516.3131103515625 0 2682.2144
518.3057861328125 0 2485.3706
519.3128662109375 0 4466.962 y 7
520.3170776367188 0 874.0685
530.2719116210938 0 932.7212
533.3059692382812 0 1221.6515
535.293701171875 0 965.94434
538.2649536132812 0 828.0098
542.3032836914062 0 767.9385 z Water loss 6
548.2837524414062 0 4884.1636
549.2838745117188 0 1383.3356
552.317626953125 0 2018.0038
553.3220825195312 0 824.2044
560.3168334960938 0 5942.5435 z 6
561.2778930664062 0 936.71387
561.32373046875 0 23762.7
562.325927734375 0 5593.153
563.3303833007812 0 1067.6442
575.3260498046875 0 2969.9243
576.3346557617188 0 21479.373 y 6
577.337890625 0 5911.711
578.3399658203125 0 1048.3154
580.312744140625 0 5194.7188
581.3164672851562 0 1456.9362
589.3416748046875 0 1695.7415
591.3587646484375 0 1050.6768
597.8295288085938 0 2651.2935
598.3310546875 0 2351.062
598.8327026367188 0 888.3716
607.3333740234375 0 712.81946
609.3402709960938 0 826.7517
615.3456420898438 0 1296.6509 y Water loss 5
617.3379516601562 0 13566.272 z 5
618.3448486328125 0 30444.52
619.347900390625 0 8618.563
620.3497924804688 0 1587.692
632.3489379882812 0 11996.58
633.3565063476562 0 54258.76 y 5
634.3594360351562 0 16272.474
635.3603515625 0 3603.9631
637.3328247070312 0 1239.1785
638.3350830078125 0 591.7598
647.3511352539062 0 2471.7964
648.3536376953125 0 953.7974
654.3593139648438 0 2751.22 c 5
655.3662109375 0 956.4121
660.3439331054688 0 1161.3618
661.3660888671875 0 4093.2122
662.3694458007812 0 2177.1672
663.3792114257812 0 2165.8804
664.3828735351562 0 1394.8413
668.3741455078125 0 1079.0526
679.400634765625 0 786.02966
692.3624267578125 0 810.7155
694.3510131835938 0 1091.4171
710.374267578125 0 9955.036
711.3802490234375 0 8800.961 c 6
712.3853149414062 0 3672.7644
718.429443359375 0 709.991
724.3912963867188 0 750.8706
725.3973999023438 0 1660.727
742.426513671875 0 689.04333
746.4083251953125 0 1909.3573 z Water loss 4
749.3841552734375 0 617.8789
751.3785400390625 0 1521.2498
752.3892211914062 0 710.2064
760.43408203125 0 2257.015
761.4404296875 0 1321.848
762.4098510742188 0 1486.8645 y Water loss 4
764.4061279296875 0 26685.875 z 4
765.4126586914062 0 41368.457
766.4161987304688 0 14693.424
767.4030151367188 0 6480.0425
768.4022216796875 0 3852.3806 c 7
769.4053955078125 0 1062.172
779.416748046875 0 16995.16
780.4242553710938 0 67815.055 y 4
781.4271850585938 0 27826.031
782.4310302734375 0 7189.899
807.4154663085938 0 1980.9502
808.4187622070312 0 1488.994
834.4116821289062 0 1235.5828
848.4514770507812 0 2648.3967 w 3
849.4539794921875 0 950.87427
850.4315795898438 0 2062.1643
851.4359741210938 0 1055.6915 c Water loss 8
852.4251098632812 0 4599.299
853.4246215820312 0 1722.5093
854.4326171875 0 697.5569
859.5047607421875 0 2282.5723
860.5052490234375 0 1336.8049
861.5037231445312 0 1096.2183
862.4721069335938 0 905.14453 y Ammonia loss 3
863.4745483398438 0 32251.422 z 3
864.4793701171875 0 26547.047
865.4823608398438 0 10215.1455
866.484130859375 0 2271.1196
868.4426879882812 0 11796.649
869.450927734375 0 72342.54 c 8
870.45361328125 0 35721.21
871.4571533203125 0 8875.069
872.4579467773438 0 1144.6809
878.4862060546875 0 2471.8196
879.4927368164062 0 32620.2 y 3
880.4960327148438 0 15453.751
881.4994506835938 0 3374.9397
882.4960327148438 0 816.89795
906.4808349609375 0 971.7986
947.5162353515625 0 1452.978 w 2
962.54150390625 0 32597.533 z 2
963.5446166992188 0 18507.934
964.5477294921875 0 5371.526
978.5609741210938 0 18883.082 y 2
978.6793823242188 0 905.1125
979.564453125 0 9733.656
980.5242309570312 0 8122.513 c Ammonia loss 9
981.5216064453125 0 5206.743
982.5283813476562 0 2560.6118
997.5455932617188 0 46563.75 c 9
998.5486450195312 0 28672.408
999.55224609375 0 8351.998
1000.5560913085938 0 1573.9469
1008.5806274414062 0 926.3158
1051.6090087890625 0 857.3277
1066.6126708984375 0 3308.3174
1067.6119384765625 0 1660.6737
1068.616943359375 0 890.1815
1093.60302734375 0 8630.3955 c Ammonia loss 10
1094.6068115234375 0 6114.139
1095.613525390625 0 4327.0938
1096.61572265625 0 1919.5195
1097.632568359375 0 1200.4803
1109.61083984375 0 26899.045 z 1
1110.6265869140625 0 90904.22 c 10
1111.6309814453125 0 53266.508
1112.634033203125 0 19166.213
1113.6370849609375 0 2990.4253
1135.6302490234375 0 1571.0704
1136.627685546875 0 1062.7666
1140.5888671875 0 852.9776
1142.58984375 0 750.4977
1150.6480712890625 0 835.9912
1151.6505126953125 0 843.85864
1154.6312255859375 0 732.504
1157.6065673828125 0 27883.762
1158.6083984375 0 18278.838
1159.611083984375 0 6577.3145
1160.6185302734375 0 1091.4163
1167.662353515625 0 2662.6333
1168.6700439453125 0 958.7509
1185.6734619140625 0 9664.325
1186.6759033203125 0 6645.9634
1187.6802978515625 0 3252.3728
1194.6336669921875 0 2011.892
1195.6541748046875 0 7110.9375
1196.6427001953125 0 142747.31
1197.645263671875 0 94810.805
1198.6474609375 0 37556.46
1199.6497802734375 0 5994.557
1210.6549072265625 0 2869.2507
1211.6531982421875 0 3218.6182
1212.6605224609375 0 74972.67
1213.6671142578125 0 152982.64
1214.6702880859375 0 92681.234
1215.673095703125 0 32427.49
1216.6690673828125 0 3395.339
3253.27001953125 0 647.59106

Spectrum Details

|  |  |
| --- | --- |
| Matched peaks? Matched peaks The total absolute number of peaks matched. Additionally in brackets the total fraction of peaks matched and the total number of peaks is shown. | 41 (16.47% of 249) |
| FDR? FDR The false discovery rate estimated for this peptide. It is calculated by matching all theoretical fragments with a non-integer shift with the raw peaks for this spectrum. This is done with 40 different shifts. The resulting percentage is the average number of annotated peaks over the number of annotated peaks with the correct spectrum. | 0.17% |
| Satellite FDR? Satellite FDR See the FDR for details on its calculation. This satellite ion specific FDR only contains the satellite ions (d/w) for I/L/J positions. | - |
| PSM Score? PSM Score The PSM Score as given by Hecklib to this annotated spectrum. It is shown with three significant figures. | 428 |

#### Spectrum 9363? Spectrum 9363 The raw spectrum of this peptide as annotated by Hecklib. The fragments are coloured according to ion type (see legend). Any peaks with a star '\*' as text can be hovered over to see the full details, first the ion type second the mass shift type. By hovering over the amino acids in the peptide or ions in the legend the corresponding peaks are highlighted. By toggling the 'Unassigned' label you can turn the background (unassigned) peaks on or off in the plot. By updating the slider in the Ion legend you can update the spectrum to only show the top X% of the peaks with labels. The top X% means any peak that is within X% of the highest intensity. By dragging in the spectrum you can zoom in to a specific part of the spectrum and use 'Zoom Out' to get back to the original zoom level. The annotation of the spectrum is based on the given sequence in the peptides file and is done with different software so inconsistencies are likely. The peaks are annotated based on the given sequence, with 20 ppm tolerance.

Copy Data

##### Spectrum 9363 (TSV)

###### Preview

```
Loading example...
```

*Click on the button to copy the data to your clipboard.*

Mz MinMz MaxIntensity Max

WidthHeightPeptide font sizePeptide stroke widthSpectrum font sizeSpectrum stroke widthCompact peptide

Ion legend

wxyz

abcd

OtherUnassignedIon^Charge^_Position_Show for top:%

SFVVFGGGTKJT

01.83e+43.66e+45.49e+47.31e+4

Zoom Out

y^+1^_1_y^+1^_2_y^+1^_2_z^+1^_3_y^+1^_3_y^+2^_8_y^+1^_4_c^+2^_10_y^+1^_5_z^+1^_6_y^+1^_6_y^+1^_7_z^+1^_7_y^+1^_7_c^+1^_6_c^+1^_7_z^+1^_8_c^+1^_8_y^+1^_8_c^+1^_9_z^+1^_9_c^+1^_9_y^+1^_9_z^+1^_10_y^+1^_10_c^+1^_10_c^+1^_10_c^+1^_11_z^+1^_11_c^+1^_11_

046092113811841

Fragment Matches Table

Show background peaks

| Position | Ion type | Intensity | mz Theoretical | mz Error (Th) | mz Error (ppm) | Charge | Series Number |
| --- | --- | --- | --- | --- | --- | --- | --- |
| 12 | y | 3157 | 120.1 | 0.000139 | 1.157 | +1 | 1 |
| - | - | 358.6 | 121.8 | - | - | 0 | - |
| - | - | 356 | 128 | - | - | 0 | - |
| - | - | 1568 | 129.1 | - | - | 0 | - |
| - | - | 389.9 | 140.8 | - | - | 0 | - |
| - | - | 587.3 | 143.1 | - | - | 0 | - |
| - | - | 1576 | 149 | - | - | 0 | - |
| - | - | 437.1 | 159.3 | - | - | 0 | - |
| - | - | 500.4 | 161.2 | - | - | 0 | - |
| - | - | 535.6 | 162 | - | - | 0 | - |
| - | - | 594.4 | 167.1 | - | - | 0 | - |
| - | - | 1140 | 171.1 | - | - | 0 | - |
| - | - | 2051 | 173.4 | - | - | 0 | - |
| - | - | 417.4 | 174.4 | - | - | 0 | - |
| - | - | 445.8 | 176.2 | - | - | 0 | - |
| - | - | 1495 | 177.1 | - | - | 0 | - |
| - | - | 592.2 | 188.1 | - | - | 0 | - |
| - | - | 845.6 | 191.1 | - | - | 0 | - |
| - | - | 820.4 | 199.1 | - | - | 0 | - |
| - | - | 7120 | 207.1 | - | - | 0 | - |
| - | - | 1185 | 208.1 | - | - | 0 | - |
| 11 | y | 926 | 215.1 | 3.92E-06 | 0.01822 | +1 | 2 |
| - | - | 739.3 | 219.1 | - | - | 0 | - |
| - | - | 4117 | 221.1 | - | - | 0 | - |
| - | - | 769.1 | 221.1 | - | - | 0 | - |
| - | - | 776.3 | 222.1 | - | - | 0 | - |
| - | - | 3139 | 225 | - | - | 0 | - |
| - | - | 610.9 | 227 | - | - | 0 | - |
| 11 | y | 4031 | 233.1 | 9.324E-05 | 0.3999 | +1 | 2 |
| - | - | 518.2 | 234.2 | - | - | 0 | - |
| - | - | 906.7 | 235.1 | - | - | 0 | - |
| - | - | 1.487E+04 | 235.1 | - | - | 0 | - |
| - | - | 1465 | 236.1 | - | - | 0 | - |
| - | - | 6782 | 239.1 | - | - | 0 | - |
| - | - | 491.7 | 240.1 | - | - | 0 | - |
| - | - | 439.6 | 245.6 | - | - | 0 | - |
| - | - | 901.8 | 247.1 | - | - | 0 | - |
| - | - | 926.1 | 289.2 | - | - | 0 | - |
| - | - | 4411 | 295.1 | - | - | 0 | - |
| - | - | 808.9 | 296.1 | - | - | 0 | - |
| - | - | 1732 | 299.1 | - | - | 0 | - |
| - | - | 1271 | 313.1 | - | - | 0 | - |
| - | - | 1.98E+04 | 334.2 | - | - | 0 | - |
| - | - | 3126 | 335.2 | - | - | 0 | - |
| 10 | z | 684 | 345.2 | 0.002281 | 6.608 | +1 | 3 |
| - | - | 655.8 | 346.2 | - | - | 0 | - |
| - | - | 651.6 | 355.1 | - | - | 0 | - |
| 10 | y | 572.3 | 361.2 | 0.0004975 | 1.377 | +1 | 3 |
| - | - | 3567 | 369.1 | - | - | 0 | - |
| 5 | y | 789.7 | 390.7 | 0.0005161 | 1.321 | +2 | 8 |
| - | - | 1497 | 401.2 | - | - | 0 | - |
| - | - | 1428 | 405.3 | - | - | 0 | - |
| - | - | 786.5 | 406.3 | - | - | 0 | - |
| - | - | 523.4 | 413.1 | - | - | 0 | - |
| - | - | 7099 | 433.2 | - | - | 0 | - |
| - | - | 1963 | 434.2 | - | - | 0 | - |
| - | - | 632.5 | 436.2 | - | - | 0 | - |
| - | - | 1110 | 444.3 | - | - | 0 | - |
| 9 | y | 2188 | 445.3 | 7.109E-05 | 0.1596 | +1 | 4 |
| - | - | 1022 | 447.3 | - | - | 0 | - |
| - | - | 541.5 | 457.3 | - | - | 0 | - |
| - | - | 702.2 | 463.2 | - | - | 0 | - |
| - | - | 1302 | 481.2 | - | - | 0 | - |
| - | - | 605.2 | 481.8 | - | - | 0 | - |
| 10 | c | 708 | 490.8 | 0.003778 | 7.697 | +2 | 10 |
| - | - | 675.1 | 501.3 | - | - | 0 | - |
| - | - | 6062 | 504.3 | - | - | 0 | - |
| - | - | 5438 | 504.3 | - | - | 0 | - |
| - | - | 1546 | 505.3 | - | - | 0 | - |
| - | - | 1062 | 505.3 | - | - | 0 | - |
| - | - | 2155 | 514.3 | - | - | 0 | - |
| - | - | 1116 | 516.3 | - | - | 0 | - |
| - | - | 1364 | 518.3 | - | - | 0 | - |
| 8 | y | 2427 | 519.3 | 0.001006 | 1.937 | +1 | 5 |
| - | - | 2769 | 548.3 | - | - | 0 | - |
| - | - | 907.1 | 549.3 | - | - | 0 | - |
| - | - | 1358 | 552.3 | - | - | 0 | - |
| 7 | z | 2935 | 560.3 | 8.325E-05 | 0.1486 | +1 | 6 |
| - | - | 1165 | 561.3 | - | - | 0 | - |
| - | - | 1.216E+04 | 561.3 | - | - | 0 | - |
| - | - | 2669 | 562.3 | - | - | 0 | - |
| - | - | 742.9 | 571.3 | - | - | 0 | - |
| - | - | 938.1 | 575.3 | - | - | 0 | - |
| 7 | y | 9585 | 576.3 | 0.0006188 | 1.074 | +1 | 6 |
| - | - | 2807 | 577.3 | - | - | 0 | - |
| - | - | 2382 | 580.3 | - | - | 0 | - |
| - | - | 2047 | 589.3 | - | - | 0 | - |
| - | - | 699 | 589.4 | - | - | 0 | - |
| - | - | 723.5 | 590.3 | - | - | 0 | - |
| - | - | 1201 | 597.8 | - | - | 0 | - |
| - | - | 1164 | 598.3 | - | - | 0 | - |
| - | - | 808.9 | 606.3 | - | - | 0 | - |
| - | - | 1189 | 606.4 | - | - | 0 | - |
| 6 | y | 1013 | 615.3 | 7.88E-05 | 0.1281 | +1 | 7 |
| 6 | z | 5731 | 617.3 | 0.0008561 | 1.387 | +1 | 7 |
| - | - | 1.255E+04 | 618.3 | - | - | 0 | - |
| - | - | 3917 | 619.3 | - | - | 0 | - |
| - | - | 6096 | 632.3 | - | - | 0 | - |
| 6 | y | 2.643E+04 | 633.4 | 0.001025 | 1.619 | +1 | 7 |
| - | - | 7387 | 634.4 | - | - | 0 | - |
| - | - | 1817 | 635.4 | - | - | 0 | - |
| - | - | 1553 | 647.3 | - | - | 0 | - |
| 6 | c | 1059 | 654.4 | 0.00227 | 3.469 | +1 | 6 |
| - | - | 2462 | 661.4 | - | - | 0 | - |
| - | - | 854.1 | 662.4 | - | - | 0 | - |
| - | - | 739 | 663.4 | - | - | 0 | - |
| - | - | 706.6 | 682.5 | - | - | 0 | - |
| - | - | 4379 | 710.4 | - | - | 0 | - |
| 7 | c | 4107 | 711.4 | 0.003653 | 5.135 | +1 | 7 |
| - | - | 1090 | 712.4 | - | - | 0 | - |
| - | - | 588.2 | 746.4 | - | - | 0 | - |
| - | - | 1197 | 760.4 | - | - | 0 | - |
| - | - | 900.8 | 761.4 | - | - | 0 | - |
| 5 | z | 1.12E+04 | 764.4 | 0.0008496 | 1.111 | +1 | 8 |
| - | - | 1.868E+04 | 765.4 | - | - | 0 | - |
| - | - | 6838 | 766.4 | - | - | 0 | - |
| - | - | 3734 | 767.4 | - | - | 0 | - |
| 8 | c | 1708 | 768.4 | 0.004365 | 5.68 | +1 | 8 |
| - | - | 6346 | 779.4 | - | - | 0 | - |
| 5 | y | 3.246E+04 | 780.4 | 0.001568 | 2.009 | +1 | 8 |
| - | - | 1.222E+04 | 781.4 | - | - | 0 | - |
| - | - | 3075 | 782.4 | - | - | 0 | - |
| - | - | 1051 | 807.4 | - | - | 0 | - |
| - | - | 1322 | 850.4 | - | - | 0 | - |
| 9 | c | 982.7 | 851.4 | 0.009069 | 10.65 | +1 | 9 |
| - | - | 2201 | 852.4 | - | - | 0 | - |
| - | - | 1262 | 853.4 | - | - | 0 | - |
| - | - | 1017 | 859.5 | - | - | 0 | - |
| 4 | z | 1.301E+04 | 863.5 | 0.001209 | 1.4 | +1 | 9 |
| - | - | 1.072E+04 | 864.5 | - | - | 0 | - |
| - | - | 3716 | 865.5 | - | - | 0 | - |
| - | - | 787.1 | 866.5 | - | - | 0 | - |
| - | - | 5973 | 868.4 | - | - | 0 | - |
| 9 | c | 3.502E+04 | 869.5 | 0.001384 | 1.592 | +1 | 9 |
| - | - | 1.73E+04 | 870.5 | - | - | 0 | - |
| - | - | 5424 | 871.5 | - | - | 0 | - |
| - | - | 1495 | 878.5 | - | - | 0 | - |
| 4 | y | 1.714E+04 | 879.5 | 0.001989 | 2.262 | +1 | 9 |
| - | - | 8116 | 880.5 | - | - | 0 | - |
| - | - | 2100 | 881.5 | - | - | 0 | - |
| - | - | 689.3 | 910.5 | - | - | 0 | - |
| 3 | z | 1.379E+04 | 962.5 | 0.002912 | 3.025 | +1 | 10 |
| - | - | 9468 | 963.5 | - | - | 0 | - |
| - | - | 2170 | 964.5 | - | - | 0 | - |
| 3 | y | 9594 | 978.6 | 0.00241 | 2.463 | +1 | 10 |
| - | - | 5495 | 979.6 | - | - | 0 | - |
| 10 | c | 3430 | 980.5 | 0.0006977 | 0.7116 | +1 | 10 |
| - | - | 2124 | 981.5 | - | - | 0 | - |
| 10 | c | 2.213E+04 | 997.5 | 0.001437 | 1.441 | +1 | 10 |
| - | - | 1.418E+04 | 998.5 | - | - | 0 | - |
| - | - | 5538 | 999.6 | - | - | 0 | - |
| - | - | 822.3 | 1067 | - | - | 0 | - |
| - | - | 896.3 | 1068 | - | - | 0 | - |
| - | - | 826.7 | 1070 | - | - | 0 | - |
| 11 | c | 3378 | 1094 | 0.002617 | 2.393 | +1 | 11 |
| - | - | 2647 | 1095 | - | - | 0 | - |
| - | - | 1905 | 1096 | - | - | 0 | - |
| - | - | 1235 | 1097 | - | - | 0 | - |
| 2 | z | 1.238E+04 | 1110 | 0.002051 | 1.848 | +1 | 11 |
| 11 | c | 3.844E+04 | 1111 | 0.00524 | 4.718 | +1 | 11 |
| - | - | 2.273E+04 | 1112 | - | - | 0 | - |
| - | - | 8124 | 1113 | - | - | 0 | - |
| - | - | 1700 | 1114 | - | - | 0 | - |
| - | - | 799.1 | 1154 | - | - | 0 | - |
| - | - | 1.359E+04 | 1158 | - | - | 0 | - |
| - | - | 7229 | 1159 | - | - | 0 | - |
| - | - | 3216 | 1160 | - | - | 0 | - |
| - | - | 1214 | 1168 | - | - | 0 | - |
| - | - | 4272 | 1186 | - | - | 0 | - |
| - | - | 3661 | 1187 | - | - | 0 | - |
| - | - | 1335 | 1188 | - | - | 0 | - |
| - | - | 866.5 | 1194 | - | - | 0 | - |
| - | - | 1754 | 1195 | - | - | 0 | - |
| - | - | 3391 | 1196 | - | - | 0 | - |
| - | - | 6.845E+04 | 1197 | - | - | 0 | - |
| - | - | 4.53E+04 | 1198 | - | - | 0 | - |
| - | - | 1.568E+04 | 1199 | - | - | 0 | - |
| - | - | 2111 | 1200 | - | - | 0 | - |
| - | - | 3089 | 1211 | - | - | 0 | - |
| - | - | 1192 | 1212 | - | - | 0 | - |
| - | - | 3.438E+04 | 1213 | - | - | 0 | - |
| - | - | 7.241E+04 | 1214 | - | - | 0 | - |
| - | - | 4.642E+04 | 1215 | - | - | 0 | - |
| - | - | 1.464E+04 | 1216 | - | - | 0 | - |
| - | - | 1772 | 1217 | - | - | 0 | - |
| - | - | 691 | 1819 | - | - | 0 | - |
| - | - | 682.8 | 1821 | - | - | 0 | - |
| - | - | 982.2 | 1823 | - | - | 0 | - |

m/z Charge Intensity FragmentType MassShift Position
120.06565856933594 0 3156.7654 y 11
121.84554290771484 0 358.59378
127.95999145507812 0 355.96494
129.102294921875 0 1568.4163
140.83384704589844 0 389.9333
143.10702514648438 0 587.267
149.04505920410156 0 1575.9608
159.32664489746094 0 437.0753
161.1990966796875 0 500.36188
161.99839782714844 0 535.60925
167.055419921875 0 594.4476
171.14926147460938 0 1139.5607
173.43885803222656 0 2051.0469
174.4185791015625 0 417.43378
176.1614990234375 0 445.82803
177.1123046875 0 1494.5557
188.0667724609375 0 592.1805
191.1280975341797 0 845.6237
199.14453125 0 820.35736
207.11273193359375 0 7119.725
208.116455078125 0 1185.1763
215.13902282714844 0 925.9866 y Water loss 10
219.06492614746094 0 739.32825
221.08432006835938 0 4116.509
221.1386260986328 0 769.1154
222.08401489257812 0 776.2723
225.0428009033203 0 3138.502
227.02212524414062 0 610.8589
233.1494903564453 0 4030.9673 y 10
234.15037536621094 0 518.234
235.09544372558594 0 906.74164
235.10769653320312 0 14872.152
236.11134338378906 0 1465.2693
239.0951385498047 0 6781.7886
240.09671020507812 0 491.71655
245.58969116210938 0 439.6413
247.14532470703125 0 901.7652
289.1541442871094 0 926.06
295.1030578613281 0 4411.4614
296.1046142578125 0 808.8697
299.06207275390625 0 1731.9589
313.11370849609375 0 1271.0482
334.17620849609375 0 19797.104
335.17919921875 0 3125.511
345.2235412597656 0 683.9552 z 9
346.2110595703125 0 655.7712
355.0709533691406 0 651.5753
361.2440490722656 0 572.2621 y 9
369.1214904785156 0 3567.0654
390.71563720703125 0 789.7422 y 4
401.2137145996094 0 1497.4806
405.25006103515625 0 1427.8843
406.2525939941406 0 786.5129
413.14483642578125 0 523.4323
433.24456787109375 0 7099.462
434.2469177246094 0 1962.7152
436.22247314453125 0 632.5315
444.2572937011719 0 1109.9426
445.2657470703125 0 2188.1282 y Ammonia loss 8
447.28216552734375 0 1022.0198
457.2762756347656 0 541.49
463.23052978515625 0 702.23975
481.2447814941406 0 1302.3047
481.7586364746094 0 605.22656
490.7598571777344 0 707.99774 c Ammonia loss 9
501.28802490234375 0 675.10455
504.2533874511719 0 6061.6685
504.3021545410156 0 5438.0728
505.2589111328125 0 1545.8058
505.3038330078125 0 1061.5134
514.2977294921875 0 2155.471
516.3128051757812 0 1116.462
518.3047485351562 0 1364.005
519.3126831054688 0 2427.4636 y 7
548.2822265625 0 2769.2483
549.2879638671875 0 907.10675
552.3191528320312 0 1358.4613
560.3163452148438 0 2935.2227 z 6
561.2750244140625 0 1164.7367
561.3235473632812 0 12160.813
562.3251342773438 0 2668.609
571.3321533203125 0 742.91833
575.3255004882812 0 938.05756
576.3345336914062 0 9584.77 y 6
577.3377685546875 0 2806.9458
580.3121948242188 0 2381.9941
589.3411865234375 0 2046.971
589.3854370117188 0 699.031
590.3413696289062 0 723.48486
597.8284301757812 0 1200.5024
598.3334350585938 0 1164.1226
606.2769165039062 0 808.9323
606.4091186523438 0 1189.1547
615.3461303710938 0 1013.09247 y Water loss 5
617.3370361328125 0 5730.888 z 5
618.3440551757812 0 12549.402
619.3469848632812 0 3917.0034
632.3482666015625 0 6095.899
633.3555908203125 0 26434.076 y 5
634.359130859375 0 7386.9805
635.3610229492188 0 1816.8126
647.3490600585938 0 1553.278
654.3587036132812 0 1058.8186 c 5
661.3640747070312 0 2462.2344
662.3673095703125 0 854.0613
663.3805541992188 0 739.04895
682.4808959960938 0 706.6482
710.3734741210938 0 4378.964
711.3787841796875 0 4106.6826 c 6
712.384521484375 0 1089.8264
746.4177856445312 0 588.2468
760.435546875 0 1196.8229
761.43115234375 0 900.7712
764.4054565429688 0 11196.528 z 4
765.4116821289062 0 18682.217
766.4156494140625 0 6838.1465
767.4011840820312 0 3733.6418
768.3995361328125 0 1707.6724 c 7
779.4165649414062 0 6345.854
780.4234619140625 0 32456.26 y 4
781.427001953125 0 12217.983
782.4292602539062 0 3075.3206
807.4070434570312 0 1050.6702
850.42529296875 0 1321.8271
851.4319458007812 0 982.65375 c Water loss 8
852.4229736328125 0 2201.058
853.4290771484375 0 1261.8643
859.49853515625 0 1016.90247
863.4735107421875 0 13006.3 z 3
864.478759765625 0 10718.516
865.4827880859375 0 3715.972
866.4837036132812 0 787.1224
868.4427490234375 0 5972.587
869.4501953125 0 35019.266 c 8
870.45263671875 0 17301.818
871.4564208984375 0 5423.6265
878.4804077148438 0 1495.456
879.491455078125 0 17138.361 y 3
880.4944458007812 0 8115.6353
881.4959716796875 0 2100.182
910.4806518554688 0 689.2818
962.5402221679688 0 13785.547 z 2
963.5429077148438 0 9468.2
964.5494995117188 0 2170.3462
978.5594482421875 0 9594.324 y 2
979.5645141601562 0 5495.355
980.5206909179688 0 3429.8967 c Ammonia loss 9
981.5205688476562 0 2123.8086
997.5451049804688 0 22125.494 c 9
998.5482788085938 0 14182.62
999.5513916015625 0 5538.252
1066.614013671875 0 822.3475
1067.6170654296875 0 896.29333
1069.612548828125 0 826.7087
1093.6014404296875 0 3378.4666 c Ammonia loss 10
1094.6082763671875 0 2646.6558
1095.614501953125 0 1904.6931
1096.6278076171875 0 1235.3687
1109.6094970703125 0 12379.849 z 1
1110.6253662109375 0 38440.523 c 10
1111.6304931640625 0 22733.557
1112.6353759765625 0 8124.284
1113.642333984375 0 1699.8198
1153.6458740234375 0 799.0653
1157.6053466796875 0 13587.68
1158.6064453125 0 7228.862
1159.609130859375 0 3215.5632
1167.6605224609375 0 1213.6107
1185.6732177734375 0 4272.129
1186.67529296875 0 3661.138
1187.6707763671875 0 1334.9069
1193.6434326171875 0 866.4958
1194.6929931640625 0 1753.5267
1195.6595458984375 0 3391.1865
1196.641845703125 0 68451.445
1197.6441650390625 0 45304.31
1198.6458740234375 0 15677.902
1199.6497802734375 0 2110.6663
1210.6651611328125 0 3088.6152
1211.6636962890625 0 1192.3448
1212.6593017578125 0 34384.598
1213.6661376953125 0 72411.01
1214.6688232421875 0 46415.09
1215.672119140625 0 14636.942
1216.6697998046875 0 1771.5388
1818.986083984375 0 690.9613
1820.9571533203125 0 682.7604
1822.9368896484375 0 982.1627

Spectrum Details

|  |  |
| --- | --- |
| Matched peaks? Matched peaks The total absolute number of peaks matched. Additionally in brackets the total fraction of peaks matched and the total number of peaks is shown. | 30 (15.96% of 188) |
| FDR? FDR The false discovery rate estimated for this peptide. It is calculated by matching all theoretical fragments with a non-integer shift with the raw peaks for this spectrum. This is done with 40 different shifts. The resulting percentage is the average number of annotated peaks over the number of annotated peaks with the correct spectrum. | 0.79% |
| Satellite FDR? Satellite FDR See the FDR for details on its calculation. This satellite ion specific FDR only contains the satellite ions (d/w) for I/L/J positions. | - |
| PSM Score? PSM Score The PSM Score as given by Hecklib to this annotated spectrum. It is shown with three significant figures. | 341 |

#### Spectrum 9545? Spectrum 9545 The raw spectrum of this peptide as annotated by Hecklib. The fragments are coloured according to ion type (see legend). Any peaks with a star '\*' as text can be hovered over to see the full details, first the ion type second the mass shift type. By hovering over the amino acids in the peptide or ions in the legend the corresponding peaks are highlighted. By toggling the 'Unassigned' label you can turn the background (unassigned) peaks on or off in the plot. By updating the slider in the Ion legend you can update the spectrum to only show the top X% of the peaks with labels. The top X% means any peak that is within X% of the highest intensity. By dragging in the spectrum you can zoom in to a specific part of the spectrum and use 'Zoom Out' to get back to the original zoom level. The annotation of the spectrum is based on the given sequence in the peptides file and is done with different software so inconsistencies are likely. The peaks are annotated based on the given sequence, with 20 ppm tolerance.

Copy Data

##### Spectrum 9545 (TSV)

###### Preview

```
Loading example...
```

*Click on the button to copy the data to your clipboard.*

Mz MinMz MaxIntensity Max

WidthHeightPeptide font sizePeptide stroke widthSpectrum font sizeSpectrum stroke widthCompact peptide

Ion legend

wxyz

abcd

OtherUnassignedIon^Charge^_Position_Show for top:%

SFVVFGGGTKJT

09.66e+31.93e+42.90e+43.86e+4

Zoom Out

y^+1^_1_y^+1^_2_y^+1^_3_y^+1^_4_y^+1^_5_z^+1^_6_y^+1^_6_z^+1^_7_y^+1^_7_c^+1^_7_z^+1^_8_c^+1^_8_y^+1^_8_z^+1^_9_c^+1^_9_y^+1^_9_z^+1^_10_y^+1^_10_c^+1^_10_c^+1^_10_c^+1^_11_z^+1^_11_c^+1^_11_

0765153022963061

Fragment Matches Table

Show background peaks

| Position | Ion type | Intensity | mz Theoretical | mz Error (Th) | mz Error (ppm) | Charge | Series Number |
| --- | --- | --- | --- | --- | --- | --- | --- |
| 12 | y | 1439 | 120.1 | 0.0004441 | 3.699 | +1 | 1 |
| - | - | 506.8 | 128 | - | - | 0 | - |
| - | - | 1359 | 129.1 | - | - | 0 | - |
| - | - | 743.6 | 133.1 | - | - | 0 | - |
| - | - | 2860 | 133.1 | - | - | 0 | - |
| - | - | 429.1 | 135.7 | - | - | 0 | - |
| - | - | 377.3 | 137 | - | - | 0 | - |
| - | - | 414.5 | 142.6 | - | - | 0 | - |
| - | - | 424.8 | 142.7 | - | - | 0 | - |
| - | - | 1075 | 149 | - | - | 0 | - |
| - | - | 830.1 | 167.1 | - | - | 0 | - |
| - | - | 423.7 | 167.2 | - | - | 0 | - |
| - | - | 910.2 | 171.1 | - | - | 0 | - |
| - | - | 683.1 | 173.4 | - | - | 0 | - |
| - | - | 1429 | 177.1 | - | - | 0 | - |
| - | - | 468.6 | 178.2 | - | - | 0 | - |
| - | - | 406.3 | 178.8 | - | - | 0 | - |
| - | - | 558.6 | 179.5 | - | - | 0 | - |
| - | - | 470.9 | 186.1 | - | - | 0 | - |
| - | - | 481.7 | 194.6 | - | - | 0 | - |
| - | - | 985.6 | 199.1 | - | - | 0 | - |
| - | - | 4058 | 207.1 | - | - | 0 | - |
| - | - | 654.1 | 208.1 | - | - | 0 | - |
| - | - | 535 | 219.1 | - | - | 0 | - |
| - | - | 3974 | 221.1 | - | - | 0 | - |
| - | - | 471.9 | 221.8 | - | - | 0 | - |
| - | - | 2938 | 225 | - | - | 0 | - |
| 11 | y | 2236 | 233.1 | 0.0002577 | 1.105 | +1 | 2 |
| - | - | 8184 | 235.1 | - | - | 0 | - |
| - | - | 545.1 | 235.8 | - | - | 0 | - |
| - | - | 1332 | 236.1 | - | - | 0 | - |
| - | - | 593.6 | 238.6 | - | - | 0 | - |
| - | - | 5222 | 239.1 | - | - | 0 | - |
| - | - | 822 | 240.1 | - | - | 0 | - |
| - | - | 994.3 | 247.1 | - | - | 0 | - |
| - | - | 553.9 | 281.1 | - | - | 0 | - |
| - | - | 5144 | 295.1 | - | - | 0 | - |
| - | - | 1164 | 296.1 | - | - | 0 | - |
| - | - | 1594 | 299.1 | - | - | 0 | - |
| - | - | 979.5 | 313.1 | - | - | 0 | - |
| - | - | 1.164E+04 | 334.2 | - | - | 0 | - |
| - | - | 2560 | 335.2 | - | - | 0 | - |
| - | - | 525 | 336.2 | - | - | 0 | - |
| 10 | y | 565.9 | 361.2 | 0.002084 | 5.77 | +1 | 3 |
| - | - | 3100 | 369.1 | - | - | 0 | - |
| - | - | 1262 | 401.2 | - | - | 0 | - |
| - | - | 1082 | 405.3 | - | - | 0 | - |
| - | - | 726.4 | 415.2 | - | - | 0 | - |
| - | - | 3737 | 433.2 | - | - | 0 | - |
| - | - | 1138 | 434.2 | - | - | 0 | - |
| 9 | y | 1179 | 445.3 | 0.001058 | 2.376 | +1 | 4 |
| - | - | 826 | 447.3 | - | - | 0 | - |
| - | - | 683.1 | 463.2 | - | - | 0 | - |
| - | - | 1537 | 475.3 | - | - | 0 | - |
| - | - | 679.6 | 481.2 | - | - | 0 | - |
| - | - | 3647 | 504.3 | - | - | 0 | - |
| - | - | 1734 | 504.3 | - | - | 0 | - |
| - | - | 1062 | 505.3 | - | - | 0 | - |
| - | - | 950.9 | 505.3 | - | - | 0 | - |
| - | - | 1013 | 514.3 | - | - | 0 | - |
| - | - | 678.9 | 516.3 | - | - | 0 | - |
| - | - | 828.5 | 518.3 | - | - | 0 | - |
| 8 | y | 804.1 | 519.3 | 0.001924 | 3.705 | +1 | 5 |
| - | - | 1893 | 532.3 | - | - | 0 | - |
| - | - | 774 | 533.3 | - | - | 0 | - |
| - | - | 701.8 | 546.8 | - | - | 0 | - |
| - | - | 1459 | 548.3 | - | - | 0 | - |
| 7 | z | 1422 | 560.3 | 0.002297 | 4.1 | +1 | 6 |
| - | - | 5725 | 561.3 | - | - | 0 | - |
| - | - | 1616 | 562.3 | - | - | 0 | - |
| - | - | 767.5 | 575.3 | - | - | 0 | - |
| 7 | y | 5174 | 576.3 | 0.0001136 | 0.1971 | +1 | 6 |
| - | - | 1501 | 577.3 | - | - | 0 | - |
| - | - | 1190 | 580.3 | - | - | 0 | - |
| - | - | 3276 | 589.3 | - | - | 0 | - |
| - | - | 1205 | 590.3 | - | - | 0 | - |
| - | - | 1051 | 597.8 | - | - | 0 | - |
| - | - | 578.8 | 602.8 | - | - | 0 | - |
| - | - | 665.1 | 604.7 | - | - | 0 | - |
| 6 | z | 3554 | 617.3 | 0.0005478 | 0.8873 | +1 | 7 |
| - | - | 7507 | 618.3 | - | - | 0 | - |
| - | - | 1507 | 619.3 | - | - | 0 | - |
| - | - | 2930 | 632.3 | - | - | 0 | - |
| 6 | y | 1.446E+04 | 633.4 | 0.0003541 | 0.559 | +1 | 7 |
| - | - | 5218 | 634.4 | - | - | 0 | - |
| - | - | 757 | 635.4 | - | - | 0 | - |
| - | - | 736.5 | 647.3 | - | - | 0 | - |
| - | - | 621.3 | 663.4 | - | - | 0 | - |
| - | - | 3524 | 710.4 | - | - | 0 | - |
| 7 | c | 2153 | 711.4 | 0.0007232 | 1.017 | +1 | 7 |
| - | - | 695.7 | 712.4 | - | - | 0 | - |
| - | - | 772.1 | 746.4 | - | - | 0 | - |
| 5 | z | 7194 | 764.4 | 0.0005444 | 0.7122 | +1 | 8 |
| - | - | 1.029E+04 | 765.4 | - | - | 0 | - |
| - | - | 3098 | 766.4 | - | - | 0 | - |
| - | - | 1606 | 767.4 | - | - | 0 | - |
| 8 | c | 1501 | 768.4 | 0.002471 | 3.216 | +1 | 8 |
| - | - | 4849 | 779.4 | - | - | 0 | - |
| 5 | y | 1.68E+04 | 780.4 | 0.0008358 | 1.071 | +1 | 8 |
| - | - | 7456 | 781.4 | - | - | 0 | - |
| - | - | 1209 | 782.4 | - | - | 0 | - |
| - | - | 694.7 | 832.7 | - | - | 0 | - |
| - | - | 746.4 | 850.4 | - | - | 0 | - |
| - | - | 1204 | 852.4 | - | - | 0 | - |
| 4 | z | 8066 | 863.5 | 0.0002327 | 0.2695 | +1 | 9 |
| - | - | 8403 | 864.5 | - | - | 0 | - |
| - | - | 2843 | 865.5 | - | - | 0 | - |
| - | - | 582.2 | 866.5 | - | - | 0 | - |
| - | - | 3050 | 868.4 | - | - | 0 | - |
| 9 | c | 1.877E+04 | 869.5 | 0.0004684 | 0.5388 | +1 | 9 |
| - | - | 8929 | 870.5 | - | - | 0 | - |
| - | - | 1375 | 871.5 | - | - | 0 | - |
| - | - | 688.9 | 875.5 | - | - | 0 | - |
| - | - | 792.8 | 877.5 | - | - | 0 | - |
| - | - | 1139 | 878.5 | - | - | 0 | - |
| 4 | y | 9227 | 879.5 | 0.0004631 | 0.5266 | +1 | 9 |
| - | - | 4136 | 880.5 | - | - | 0 | - |
| - | - | 1058 | 881.5 | - | - | 0 | - |
| - | - | 820 | 883.5 | - | - | 0 | - |
| 3 | z | 8793 | 962.5 | 0.001264 | 1.313 | +1 | 10 |
| - | - | 4443 | 963.5 | - | - | 0 | - |
| - | - | 1611 | 964.5 | - | - | 0 | - |
| 3 | y | 5058 | 978.6 | 0.0001515 | 0.1548 | +1 | 10 |
| - | - | 2489 | 979.6 | - | - | 0 | - |
| 10 | c | 2002 | 980.5 | 0.005764 | 5.878 | +1 | 10 |
| - | - | 1189 | 981.5 | - | - | 0 | - |
| - | - | 696 | 982.5 | - | - | 0 | - |
| - | - | 873.9 | 995.5 | - | - | 0 | - |
| 10 | c | 1.219E+04 | 997.5 | 0.000949 | 0.9514 | +1 | 10 |
| - | - | 8256 | 998.5 | - | - | 0 | - |
| - | - | 2403 | 999.6 | - | - | 0 | - |
| 11 | c | 1901 | 1094 | 0.0003129 | 0.2862 | +1 | 11 |
| - | - | 1198 | 1095 | - | - | 0 | - |
| - | - | 1530 | 1096 | - | - | 0 | - |
| 2 | z | 8195 | 1110 | 0.00144 | 1.298 | +1 | 11 |
| 11 | c | 2.121E+04 | 1111 | 0.004264 | 3.839 | +1 | 11 |
| - | - | 1.398E+04 | 1112 | - | - | 0 | - |
| - | - | 4226 | 1113 | - | - | 0 | - |
| - | - | 1281 | 1114 | - | - | 0 | - |
| - | - | 1117 | 1137 | - | - | 0 | - |
| - | - | 918.5 | 1141 | - | - | 0 | - |
| - | - | 7570 | 1158 | - | - | 0 | - |
| - | - | 5812 | 1159 | - | - | 0 | - |
| - | - | 2741 | 1160 | - | - | 0 | - |
| - | - | 1603 | 1161 | - | - | 0 | - |
| - | - | 1224 | 1162 | - | - | 0 | - |
| - | - | 1761 | 1163 | - | - | 0 | - |
| - | - | 734.7 | 1168 | - | - | 0 | - |
| - | - | 998.1 | 1177 | - | - | 0 | - |
| - | - | 3932 | 1178 | - | - | 0 | - |
| - | - | 3126 | 1179 | - | - | 0 | - |
| - | - | 2194 | 1180 | - | - | 0 | - |
| - | - | 2414 | 1186 | - | - | 0 | - |
| - | - | 1583 | 1187 | - | - | 0 | - |
| - | - | 1825 | 1194 | - | - | 0 | - |
| - | - | 1836 | 1195 | - | - | 0 | - |
| - | - | 7320 | 1195 | - | - | 0 | - |
| - | - | 3827 | 1196 | - | - | 0 | - |
| - | - | 3494 | 1196 | - | - | 0 | - |
| - | - | 3.64E+04 | 1197 | - | - | 0 | - |
| - | - | 2.512E+04 | 1198 | - | - | 0 | - |
| - | - | 9601 | 1199 | - | - | 0 | - |
| - | - | 1500 | 1200 | - | - | 0 | - |
| - | - | 2504 | 1211 | - | - | 0 | - |
| - | - | 3384 | 1212 | - | - | 0 | - |
| - | - | 2.011E+04 | 1213 | - | - | 0 | - |
| - | - | 3.825E+04 | 1214 | - | - | 0 | - |
| - | - | 2.3E+04 | 1215 | - | - | 0 | - |
| - | - | 9066 | 1216 | - | - | 0 | - |
| - | - | 1106 | 1217 | - | - | 0 | - |
| - | - | 883.5 | 1766 | - | - | 0 | - |
| - | - | 868 | 1800 | - | - | 0 | - |
| - | - | 1069 | 1806 | - | - | 0 | - |
| - | - | 1246 | 1822 | - | - | 0 | - |
| - | - | 1651 | 1823 | - | - | 0 | - |
| - | - | 853.2 | 1824 | - | - | 0 | - |
| - | - | 655.6 | 2485 | - | - | 0 | - |
| - | - | 887.8 | 3030 | - | - | 0 | - |

m/z Charge Intensity FragmentType MassShift Position
120.06596374511719 0 1438.5326 y 11
128.04660034179688 0 506.75232
129.10232543945312 0 1359.2825
133.06947326660156 0 743.5662
133.08621215820312 0 2860.2678
135.66061401367188 0 429.05402
136.96563720703125 0 377.30206
142.63063049316406 0 414.50467
142.7060089111328 0 424.8359
149.04522705078125 0 1074.9545
167.05563354492188 0 830.09955
167.24050903320312 0 423.6792
171.1494903564453 0 910.2302
173.4373016357422 0 683.07794
177.11264038085938 0 1429.2739
178.23043823242188 0 468.6021
178.75155639648438 0 406.2764
179.52703857421875 0 558.5606
186.08721923828125 0 470.87027
194.5599365234375 0 481.6511
199.14434814453125 0 985.5767
207.1132049560547 0 4057.8806
208.11683654785156 0 654.0664
219.14999389648438 0 535.0118
221.0849609375 0 3973.9858
221.7697296142578 0 471.90872
225.043212890625 0 2938.1072
233.14984130859375 0 2235.7583 y 10
235.10818481445312 0 8184.038
235.82742309570312 0 545.13855
236.11146545410156 0 1331.8162
238.61770629882812 0 593.5638
239.09542846679688 0 5222.1226
240.09542846679688 0 821.97546
247.14414978027344 0 994.3192
281.05169677734375 0 553.9224
295.1036376953125 0 5143.9536
296.1038818359375 0 1164.0432
299.0617370605469 0 1593.8373
313.11468505859375 0 979.48505
334.1768493652344 0 11641.169
335.1795349121094 0 2559.8933
336.1842956542969 0 524.99994
361.2424621582031 0 565.9268 y 9
369.12213134765625 0 3099.7603
401.2155456542969 0 1262.1122
405.2501525878906 0 1082.1646
415.23260498046875 0 726.35834
433.2449035644531 0 3736.506
434.2491455078125 0 1138.2576
445.2646179199219 0 1179.3268 y Ammonia loss 8
447.2795715332031 0 825.98816
463.2364196777344 0 683.1319
475.2865295410156 0 1537.356
481.2457580566406 0 679.61646
504.2544250488281 0 3647.2786
504.3036804199219 0 1734.0646
505.2603454589844 0 1062.3331
505.3063659667969 0 950.928
514.2981567382812 0 1013.1907
516.3126220703125 0 678.87244
518.30615234375 0 828.50433
519.3156127929688 0 804.0724 y 7
532.3103637695312 0 1893.0671
533.3114624023438 0 774.04297
546.7783203125 0 701.81714
548.285400390625 0 1459.0741
560.3187255859375 0 1421.5039 z 6
561.3242797851562 0 5724.95
562.3278198242188 0 1616.4658
575.3256225585938 0 767.45624
576.3352661132812 0 5173.981 y 6
577.3389892578125 0 1501.1353
580.3128662109375 0 1190.0126
589.3417358398438 0 3276.1973
590.3453369140625 0 1205.1565
597.8284912109375 0 1050.6594
602.8065185546875 0 578.7914
604.7078857421875 0 665.0684
617.3384399414062 0 3553.9185 z 5
618.34521484375 0 7507.2134
619.3446655273438 0 1507.2113
632.3483276367188 0 2930.2112
633.3562622070312 0 14462.402 y 5
634.3598022460938 0 5217.609
635.3615112304688 0 756.99615
647.3467407226562 0 736.4642
663.3780517578125 0 621.31696
710.3741455078125 0 3523.9922
711.3817138671875 0 2153.4165 c 6
712.3792114257812 0 695.71436
746.4235229492188 0 772.0727
764.40576171875 0 7193.5947 z 4
765.4130859375 0 10285.997
766.415283203125 0 3098.4373
767.40380859375 0 1605.5035
768.4063720703125 0 1500.5161 c 7
779.4181518554688 0 4848.976
780.4241943359375 0 16798.676 y 4
781.4283447265625 0 7456.2485
782.4302368164062 0 1208.9246
832.7168579101562 0 694.73065
850.4365844726562 0 746.405
852.4254760742188 0 1204.3973
863.4744873046875 0 8065.6133 z 3
864.4797973632812 0 8403.34
865.4819946289062 0 2842.7551
866.47705078125 0 582.2079
868.443603515625 0 3049.7834
869.4511108398438 0 18768.797 c 8
870.4534301757812 0 8929.09
871.4591674804688 0 1375.2405
875.4841918945312 0 688.9042
877.5137329101562 0 792.7991
878.4909057617188 0 1138.8381
879.4929809570312 0 9227.202 y 3
880.495849609375 0 4136.111
881.4969482421875 0 1057.661
883.48095703125 0 819.96783
962.5418701171875 0 8792.756 z 2
963.5410766601562 0 4442.9697
964.54736328125 0 1610.7574
978.5617065429688 0 5057.575 y 2
979.5655517578125 0 2489.4827
980.5257568359375 0 2002.3579 c Ammonia loss 9
981.5266723632812 0 1189.0819
982.529541015625 0 696.01245
995.5198364257812 0 873.94763
997.5455932617188 0 12193.622 c 9
998.5482788085938 0 8256.433
999.5511474609375 0 2402.5566
1093.6043701171875 0 1901.0924 c Ammonia loss 10
1094.607177734375 0 1197.8235
1095.606201171875 0 1529.9944
1109.610107421875 0 8194.521 z 1
1110.6263427734375 0 21211.29 c 10
1111.6307373046875 0 13983.299
1112.630615234375 0 4226.377
1113.6300048828125 0 1280.5273
1136.5908203125 0 1117.0981
1140.5855712890625 0 918.49927
1157.605712890625 0 7570.0684
1158.607177734375 0 5811.8447
1159.6053466796875 0 2740.995
1160.6005859375 0 1602.5092
1161.61328125 0 1223.6654
1162.6075439453125 0 1761.1818
1167.655517578125 0 734.7063
1176.6033935546875 0 998.05096
1177.6094970703125 0 3932.488
1178.603759765625 0 3126.0867
1179.604736328125 0 2194.05
1185.6768798828125 0 2413.906
1186.672607421875 0 1583.4153
1193.612060546875 0 1825.454
1194.5802001953125 0 1835.8765
1194.7039794921875 0 7320.2104
1195.6171875 0 3826.7617
1195.7119140625 0 3493.9077
1196.6402587890625 0 36396.39
1197.6434326171875 0 25121.072
1198.64404296875 0 9601.151
1199.645263671875 0 1499.6609
1210.658935546875 0 2504.2273
1211.628662109375 0 3384.2378
1212.65576171875 0 20112.844
1213.6656494140625 0 38254.273
1214.6680908203125 0 23003.154
1215.6683349609375 0 9066.472
1216.6824951171875 0 1106.1803
1765.9573974609375 0 883.5275
1799.96923828125 0 868.003
1805.982177734375 0 1068.8763
1821.9334716796875 0 1246.3575
1822.946533203125 0 1651.1552
1823.9715576171875 0 853.1978
2484.59375 0 655.58636
3030.402099609375 0 887.84924

Spectrum Details

|  |  |
| --- | --- |
| Matched peaks? Matched peaks The total absolute number of peaks matched. Additionally in brackets the total fraction of peaks matched and the total number of peaks is shown. | 23 (12.92% of 178) |
| FDR? FDR The false discovery rate estimated for this peptide. It is calculated by matching all theoretical fragments with a non-integer shift with the raw peaks for this spectrum. This is done with 40 different shifts. The resulting percentage is the average number of annotated peaks over the number of annotated peaks with the correct spectrum. | 0.41% |
| Satellite FDR? Satellite FDR See the FDR for details on its calculation. This satellite ion specific FDR only contains the satellite ions (d/w) for I/L/J positions. | - |
| PSM Score? PSM Score The PSM Score as given by Hecklib to this annotated spectrum. It is shown with three significant figures. | 259 |

#### Spectrum 8930? Spectrum 8930 The raw spectrum of this peptide as annotated by Hecklib. The fragments are coloured according to ion type (see legend). Any peaks with a star '\*' as text can be hovered over to see the full details, first the ion type second the mass shift type. By hovering over the amino acids in the peptide or ions in the legend the corresponding peaks are highlighted. By toggling the 'Unassigned' label you can turn the background (unassigned) peaks on or off in the plot. By updating the slider in the Ion legend you can update the spectrum to only show the top X% of the peaks with labels. The top X% means any peak that is within X% of the highest intensity. By dragging in the spectrum you can zoom in to a specific part of the spectrum and use 'Zoom Out' to get back to the original zoom level. The annotation of the spectrum is based on the given sequence in the peptides file and is done with different software so inconsistencies are likely. The peaks are annotated based on the given sequence, with 20 ppm tolerance.

Copy Data

##### Spectrum 8930 (TSV)

###### Preview

```
Loading example...
```

*Click on the button to copy the data to your clipboard.*

Mz MinMz MaxIntensity Max

WidthHeightPeptide font sizePeptide stroke widthSpectrum font sizeSpectrum stroke widthCompact peptide

Ion legend

wxyz

abcd

OtherUnassignedIon^Charge^_Position_Show for top:%

SFVVFGGGTKJT

01.23e+62.46e+63.69e+64.92e+6

Zoom Out

y^+1^_1_y^+1^_2_y^+1^_2_y^+1^_3_c^+2^_7_y^+1^_3_y^+2^_8_y^+2^_8_y^+2^_9_z^+2^_9_y^+2^_9_y^+1^_4_y^+1^_4_z^+2^_10_y^+2^_10_y^+2^_10_c^+2^_10_c^+2^_10_y^+1^_5_y^+1^_5_z^+2^_11_c^+2^_11_y^+2^_11_y^+2^_11_y^+1^_6_y^+2^_11_y^+1^_6_c^+1^_5_z^+1^_7_z^+1^_7_y^+1^_7_y^+1^_7_y^+1^_8_z^+1^_8_y^+1^_8_z^+1^_9_y^+1^_9_z^+1^_9_c^+1^_9_y^+1^_9_z^+1^_10_w^+1^_10_y^+1^_10_y^+1^_10_c^+1^_10_c^+1^_10_z^+1^_11_z^+1^_11_c^+1^_11_z^+1^_11_c^+1^_11_y^+1^_11_

03076149211228

Fragment Matches Table

Show background peaks

| Position | Ion type | Intensity | mz Theoretical | mz Error (Th) | mz Error (ppm) | Charge | Series Number |
| --- | --- | --- | --- | --- | --- | --- | --- |
| 12 | y | 8.231E+05 | 120.1 | 0.000612 | 5.097 | +1 | 1 |
| - | - | 1.009E+05 | 120.1 | - | - | 0 | - |
| - | - | 3163 | 121.1 | - | - | 0 | - |
| - | - | 3.607E+04 | 121.1 | - | - | 0 | - |
| - | - | 7330 | 121.1 | - | - | 0 | - |
| - | - | 3560 | 122.1 | - | - | 0 | - |
| - | - | 1342 | 122.1 | - | - | 0 | - |
| - | - | 1458 | 127.9 | - | - | 0 | - |
| - | - | 3.137E+05 | 129.1 | - | - | 0 | - |
| - | - | 3212 | 130.1 | - | - | 0 | - |
| - | - | 1.955E+04 | 130.1 | - | - | 0 | - |
| - | - | 2190 | 131.1 | - | - | 0 | - |
| - | - | 1402 | 143.3 | - | - | 0 | - |
| - | - | 2132 | 159.1 | - | - | 0 | - |
| - | - | 1.149E+04 | 162.1 | - | - | 0 | - |
| - | - | 2.524E+05 | 171.1 | - | - | 0 | - |
| - | - | 2.324E+04 | 172.2 | - | - | 0 | - |
| - | - | 1952 | 173.4 | - | - | 0 | - |
| - | - | 6709 | 173.4 | - | - | 0 | - |
| - | - | 8323 | 187.1 | - | - | 0 | - |
| - | - | 7285 | 187.1 | - | - | 0 | - |
| - | - | 1994 | 190.3 | - | - | 0 | - |
| - | - | 9175 | 197.2 | - | - | 0 | - |
| - | - | 3.406E+05 | 199.1 | - | - | 0 | - |
| - | - | 4.3E+04 | 200.1 | - | - | 0 | - |
| - | - | 1.34E+04 | 205.1 | - | - | 0 | - |
| - | - | 1.814E+06 | 207.1 | - | - | 0 | - |
| - | - | 2.168E+05 | 208.1 | - | - | 0 | - |
| - | - | 1.097E+04 | 209.1 | - | - | 0 | - |
| - | - | 1.13E+04 | 212.1 | - | - | 0 | - |
| - | - | 2460 | 214.2 | - | - | 0 | - |
| 11 | y | 1.525E+05 | 215.1 | 0.0008279 | 3.848 | +1 | 2 |
| - | - | 1.739E+04 | 216.1 | - | - | 0 | - |
| - | - | 2597 | 217.2 | - | - | 0 | - |
| - | - | 1.01E+05 | 219.2 | - | - | 0 | - |
| - | - | 1.608E+04 | 220.2 | - | - | 0 | - |
| - | - | 2094 | 222.1 | - | - | 0 | - |
| - | - | 3054 | 228.1 | - | - | 0 | - |
| - | - | 7.311E+04 | 230.2 | - | - | 0 | - |
| - | - | 7640 | 231.2 | - | - | 0 | - |
| 11 | y | 9.522E+05 | 233.1 | 0.001005 | 4.312 | +1 | 2 |
| - | - | 9.481E+04 | 234.2 | - | - | 0 | - |
| - | - | 3.586E+06 | 235.1 | - | - | 0 | - |
| - | - | 4.655E+05 | 236.1 | - | - | 0 | - |
| - | - | 3.366E+04 | 237.1 | - | - | 0 | - |
| - | - | 1.411E+04 | 242.2 | - | - | 0 | - |
| - | - | 1.669E+05 | 247.1 | - | - | 0 | - |
| - | - | 2.633E+04 | 248.1 | - | - | 0 | - |
| - | - | 7679 | 255.1 | - | - | 0 | - |
| - | - | 2532 | 257.1 | - | - | 0 | - |
| - | - | 4181 | 261.2 | - | - | 0 | - |
| - | - | 2979 | 262.1 | - | - | 0 | - |
| - | - | 3023 | 264.2 | - | - | 0 | - |
| - | - | 3340 | 265.2 | - | - | 0 | - |
| - | - | 8386 | 268.2 | - | - | 0 | - |
| - | - | 1.531E+04 | 269.2 | - | - | 0 | - |
| - | - | 7.294E+04 | 273.1 | - | - | 0 | - |
| - | - | 2421 | 273.2 | - | - | 0 | - |
| - | - | 9747 | 274.1 | - | - | 0 | - |
| - | - | 1.12E+04 | 286.2 | - | - | 0 | - |
| - | - | 4.267E+04 | 287.2 | - | - | 0 | - |
| - | - | 9410 | 288.2 | - | - | 0 | - |
| - | - | 3.275E+05 | 289.2 | - | - | 0 | - |
| - | - | 5.623E+04 | 290.2 | - | - | 0 | - |
| - | - | 5199 | 291.2 | - | - | 0 | - |
| - | - | 6891 | 292.1 | - | - | 0 | - |
| - | - | 2250 | 294.9 | - | - | 0 | - |
| - | - | 4.55E+04 | 301.2 | - | - | 0 | - |
| - | - | 9197 | 302.2 | - | - | 0 | - |
| - | - | 5746 | 303.2 | - | - | 0 | - |
| - | - | 2.015E+04 | 304.2 | - | - | 0 | - |
| - | - | 4216 | 305.2 | - | - | 0 | - |
| - | - | 1.948E+04 | 306.2 | - | - | 0 | - |
| - | - | 4369 | 307.2 | - | - | 0 | - |
| - | - | 3752 | 309.2 | - | - | 0 | - |
| - | - | 9.392E+04 | 316.2 | - | - | 0 | - |
| - | - | 1.773E+04 | 317.2 | - | - | 0 | - |
| - | - | 7618 | 318.2 | - | - | 0 | - |
| - | - | 5980 | 319.1 | - | - | 0 | - |
| - | - | 6.368E+04 | 326.2 | - | - | 0 | - |
| - | - | 4287 | 326.2 | - | - | 0 | - |
| - | - | 8807 | 327.2 | - | - | 0 | - |
| - | - | 3240 | 329.2 | - | - | 0 | - |
| - | - | 1.149E+04 | 332.2 | - | - | 0 | - |
| - | - | 4998 | 333.2 | - | - | 0 | - |
| - | - | 4.873E+06 | 334.2 | - | - | 0 | - |
| - | - | 9.506E+05 | 335.2 | - | - | 0 | - |
| - | - | 9.359E+04 | 336.2 | - | - | 0 | - |
| - | - | 2.214E+04 | 337.2 | - | - | 0 | - |
| - | - | 2533 | 337.2 | - | - | 0 | - |
| - | - | 2882 | 337.2 | - | - | 0 | - |
| - | - | 4813 | 338.2 | - | - | 0 | - |
| - | - | 3184 | 342.2 | - | - | 0 | - |
| 10 | y | 1.967E+04 | 343.2 | 0.001644 | 4.791 | +1 | 3 |
| - | - | 6.229E+04 | 344.2 | - | - | 0 | - |
| - | - | 3025 | 344.2 | - | - | 0 | - |
| - | - | 2338 | 345.1 | - | - | 0 | - |
| - | - | 8909 | 345.2 | - | - | 0 | - |
| - | - | 8529 | 346.2 | - | - | 0 | - |
| - | - | 1.341E+05 | 346.2 | - | - | 0 | - |
| - | - | 2.85E+04 | 347.2 | - | - | 0 | - |
| - | - | 4028 | 349.2 | - | - | 0 | - |
| - | - | 5310 | 354.2 | - | - | 0 | - |
| 7 | c | 3786 | 356.2 | 0.003263 | 9.162 | +2 | 7 |
| - | - | 2666 | 357.2 | - | - | 0 | - |
| - | - | 3173 | 358.2 | - | - | 0 | - |
| - | - | 9070 | 361.2 | - | - | 0 | - |
| 10 | y | 2.525E+05 | 361.2 | 0.001334 | 3.691 | +1 | 3 |
| - | - | 4.499E+04 | 362.2 | - | - | 0 | - |
| - | - | 4469 | 363.2 | - | - | 0 | - |
| - | - | 3.844E+04 | 364.2 | - | - | 0 | - |
| - | - | 8062 | 364.7 | - | - | 0 | - |
| - | - | 7409 | 365.2 | - | - | 0 | - |
| - | - | 3399 | 365.2 | - | - | 0 | - |
| - | - | 2832 | 365.2 | - | - | 0 | - |
| - | - | 3817 | 366.2 | - | - | 0 | - |
| - | - | 3605 | 366.7 | - | - | 0 | - |
| - | - | 4322 | 368.2 | - | - | 0 | - |
| - | - | 3023 | 370.2 | - | - | 0 | - |
| - | - | 2007 | 372.2 | - | - | 0 | - |
| - | - | 2795 | 372.3 | - | - | 0 | - |
| - | - | 9504 | 373.2 | - | - | 0 | - |
| - | - | 2.721E+04 | 373.7 | - | - | 0 | - |
| - | - | 3139 | 374.2 | - | - | 0 | - |
| - | - | 2.779E+04 | 374.2 | - | - | 0 | - |
| - | - | 3939 | 374.7 | - | - | 0 | - |
| - | - | 4197 | 375.2 | - | - | 0 | - |
| - | - | 4425 | 375.2 | - | - | 0 | - |
| - | - | 6033 | 380.7 | - | - | 0 | - |
| - | - | 3117 | 381.2 | - | - | 0 | - |
| 5 | y | 1.788E+04 | 381.7 | 0.001013 | 2.653 | +2 | 8 |
| - | - | 4.612E+04 | 382.2 | - | - | 0 | - |
| - | - | 8272 | 382.2 | - | - | 0 | - |
| - | - | 1.164E+04 | 382.2 | - | - | 0 | - |
| - | - | 9341 | 383.2 | - | - | 0 | - |
| - | - | 8.959E+04 | 383.2 | - | - | 0 | - |
| - | - | 1.399E+04 | 384.2 | - | - | 0 | - |
| - | - | 4687 | 386.2 | - | - | 0 | - |
| - | - | 1.378E+04 | 387.2 | - | - | 0 | - |
| - | - | 6.583E+04 | 388.2 | - | - | 0 | - |
| - | - | 1.669E+04 | 389.2 | - | - | 0 | - |
| 5 | y | 1.999E+05 | 390.7 | 0.001468 | 3.756 | +2 | 8 |
| - | - | 1.238E+04 | 391.2 | - | - | 0 | - |
| - | - | 8.067E+04 | 391.2 | - | - | 0 | - |
| - | - | 1.79E+04 | 391.7 | - | - | 0 | - |
| - | - | 1.872E+04 | 392.2 | - | - | 0 | - |
| - | - | 7445 | 393.2 | - | - | 0 | - |
| - | - | 4.572E+04 | 400.3 | - | - | 0 | - |
| - | - | 5.061E+05 | 401.2 | - | - | 0 | - |
| - | - | 5.402E+04 | 402.2 | - | - | 0 | - |
| - | - | 8.849E+04 | 402.2 | - | - | 0 | - |
| - | - | 1.038E+04 | 403.2 | - | - | 0 | - |
| - | - | 2.209E+04 | 403.2 | - | - | 0 | - |
| - | - | 5612 | 404.2 | - | - | 0 | - |
| - | - | 4.347E+05 | 405.3 | - | - | 0 | - |
| - | - | 3000 | 406.2 | - | - | 0 | - |
| - | - | 1.069E+05 | 406.3 | - | - | 0 | - |
| - | - | 1.401E+04 | 407.3 | - | - | 0 | - |
| - | - | 2777 | 410.2 | - | - | 0 | - |
| - | - | 3783 | 411.2 | - | - | 0 | - |
| - | - | 2.187E+05 | 415.2 | - | - | 0 | - |
| - | - | 5.227E+04 | 416.2 | - | - | 0 | - |
| - | - | 5546 | 416.8 | - | - | 0 | - |
| - | - | 8311 | 417.2 | - | - | 0 | - |
| - | - | 1.144E+04 | 418.2 | - | - | 0 | - |
| - | - | 2689 | 419.2 | - | - | 0 | - |
| - | - | 8.032E+04 | 420.2 | - | - | 0 | - |
| - | - | 2.743E+04 | 421.2 | - | - | 0 | - |
| - | - | 3855 | 421.3 | - | - | 0 | - |
| - | - | 3665 | 421.8 | - | - | 0 | - |
| - | - | 3371 | 422.2 | - | - | 0 | - |
| - | - | 2305 | 426.7 | - | - | 0 | - |
| - | - | 9283 | 429.3 | - | - | 0 | - |
| - | - | 2549 | 430.2 | - | - | 0 | - |
| - | - | 2.585E+04 | 430.3 | - | - | 0 | - |
| - | - | 1.44E+04 | 430.8 | - | - | 0 | - |
| 4 | y | 1.73E+04 | 431.2 | 0.001992 | 4.62 | +2 | 9 |
| - | - | 1.061E+04 | 431.7 | - | - | 0 | - |
| 4 | z | 4181 | 432.2 | 0.004182 | 9.675 | +2 | 9 |
| - | - | 2.075E+06 | 433.2 | - | - | 0 | - |
| - | - | 5.378E+05 | 434.2 | - | - | 0 | - |
| - | - | 7.48E+04 | 435.3 | - | - | 0 | - |
| - | - | 1.154E+05 | 436.2 | - | - | 0 | - |
| - | - | 3.384E+04 | 437.2 | - | - | 0 | - |
| - | - | 4294 | 438.2 | - | - | 0 | - |
| - | - | 8900 | 439.2 | - | - | 0 | - |
| - | - | 3878 | 439.2 | - | - | 0 | - |
| - | - | 2.97E+04 | 439.3 | - | - | 0 | - |
| 4 | y | 1.237E+05 | 440.3 | 0.001684 | 3.826 | +2 | 9 |
| - | - | 6.23E+04 | 440.8 | - | - | 0 | - |
| - | - | 1.556E+04 | 441.3 | - | - | 0 | - |
| - | - | 7094 | 443.2 | - | - | 0 | - |
| - | - | 3048 | 444.2 | - | - | 0 | - |
| 9 | y | 7781 | 444.3 | 0.002794 | 6.289 | +1 | 4 |
| - | - | 1.516E+04 | 445.2 | - | - | 0 | - |
| - | - | 2671 | 446.2 | - | - | 0 | - |
| - | - | 5783 | 447.2 | - | - | 0 | - |
| - | - | 2443 | 447.8 | - | - | 0 | - |
| - | - | 3891 | 448.2 | - | - | 0 | - |
| - | - | 6587 | 448.3 | - | - | 0 | - |
| - | - | 2361 | 449.2 | - | - | 0 | - |
| - | - | 2539 | 451.2 | - | - | 0 | - |
| - | - | 7.787E+04 | 453.3 | - | - | 0 | - |
| - | - | 2.379E+04 | 454.3 | - | - | 0 | - |
| - | - | 2954 | 455.3 | - | - | 0 | - |
| - | - | 1.14E+05 | 457.3 | - | - | 0 | - |
| - | - | 2.561E+04 | 458.3 | - | - | 0 | - |
| - | - | 3021 | 459.3 | - | - | 0 | - |
| - | - | 9281 | 460.3 | - | - | 0 | - |
| - | - | 5580 | 461.2 | - | - | 0 | - |
| 9 | y | 1.072E+05 | 462.3 | 0.001964 | 4.249 | +1 | 4 |
| - | - | 1.171E+05 | 463.2 | - | - | 0 | - |
| - | - | 2.155E+04 | 463.3 | - | - | 0 | - |
| - | - | 3.768E+04 | 464.2 | - | - | 0 | - |
| - | - | 3208 | 464.3 | - | - | 0 | - |
| - | - | 3336 | 464.8 | - | - | 0 | - |
| - | - | 5712 | 465.2 | - | - | 0 | - |
| - | - | 4786 | 465.3 | - | - | 0 | - |
| - | - | 2.196E+04 | 472.3 | - | - | 0 | - |
| 3 | z | 8929 | 473.3 | 0.004973 | 10.51 | +2 | 10 |
| - | - | 6103 | 473.8 | - | - | 0 | - |
| - | - | 3373 | 474.8 | - | - | 0 | - |
| - | - | 4357 | 475.2 | - | - | 0 | - |
| - | - | 3142 | 476.8 | - | - | 0 | - |
| - | - | 3123 | 478.2 | - | - | 0 | - |
| - | - | 5511 | 478.3 | - | - | 0 | - |
| 3 | y | 8.044E+04 | 480.8 | 0.001904 | 3.96 | +2 | 10 |
| - | - | 3.657E+05 | 481.2 | - | - | 0 | - |
| - | - | 2.985E+04 | 481.3 | - | - | 0 | - |
| - | - | 9.696E+04 | 481.8 | - | - | 0 | - |
| - | - | 1.27E+05 | 482.3 | - | - | 0 | - |
| - | - | 1.837E+04 | 482.8 | - | - | 0 | - |
| - | - | 1.325E+04 | 483.3 | - | - | 0 | - |
| - | - | 1.076E+04 | 483.8 | - | - | 0 | - |
| - | - | 3679 | 484.3 | - | - | 0 | - |
| - | - | 3.38E+04 | 486.3 | - | - | 0 | - |
| - | - | 8128 | 487.3 | - | - | 0 | - |
| - | - | 2437 | 489.2 | - | - | 0 | - |
| - | - | 7576 | 489.3 | - | - | 0 | - |
| 3 | y | 1.811E+05 | 489.8 | 0.002054 | 4.193 | +2 | 10 |
| - | - | 8.934E+04 | 490.3 | - | - | 0 | - |
| 10 | c | 1.616E+05 | 490.8 | 0.001929 | 3.931 | +2 | 10 |
| - | - | 1.222E+05 | 491.3 | - | - | 0 | - |
| - | - | 2.999E+04 | 491.8 | - | - | 0 | - |
| - | - | 8906 | 492.3 | - | - | 0 | - |
| - | - | 1.518E+04 | 493.2 | - | - | 0 | - |
| - | - | 1.421E+04 | 493.3 | - | - | 0 | - |
| - | - | 4101 | 494.2 | - | - | 0 | - |
| - | - | 6033 | 494.3 | - | - | 0 | - |
| - | - | 8.085E+04 | 496.3 | - | - | 0 | - |
| - | - | 1.973E+04 | 497.3 | - | - | 0 | - |
| - | - | 1.63E+04 | 497.8 | - | - | 0 | - |
| - | - | 1.421E+04 | 498.3 | - | - | 0 | - |
| - | - | 4004 | 498.8 | - | - | 0 | - |
| 10 | c | 3555 | 499.3 | 0.007744 | 15.51 | +2 | 10 |
| - | - | 2.997E+04 | 501.2 | - | - | 0 | - |
| 8 | y | 1.819E+04 | 501.3 | 0.002631 | 5.249 | +1 | 5 |
| - | - | 1.037E+04 | 502.3 | - | - | 0 | - |
| - | - | 4919 | 502.3 | - | - | 0 | - |
| - | - | 6748 | 503.8 | - | - | 0 | - |
| - | - | 4651 | 504.3 | - | - | 0 | - |
| - | - | 2253 | 504.3 | - | - | 0 | - |
| - | - | 2940 | 507.2 | - | - | 0 | - |
| - | - | 4839 | 507.3 | - | - | 0 | - |
| - | - | 3172 | 508.3 | - | - | 0 | - |
| - | - | 1.439E+04 | 509.2 | - | - | 0 | - |
| - | - | 1.874E+04 | 510.3 | - | - | 0 | - |
| - | - | 6708 | 511.3 | - | - | 0 | - |
| - | - | 7279 | 512.3 | - | - | 0 | - |
| - | - | 4.277E+05 | 514.3 | - | - | 0 | - |
| - | - | 1.159E+05 | 515.3 | - | - | 0 | - |
| - | - | 1.679E+04 | 516.3 | - | - | 0 | - |
| - | - | 1.71E+04 | 517.3 | - | - | 0 | - |
| - | - | 8033 | 518.3 | - | - | 0 | - |
| - | - | 3.663E+04 | 519.3 | - | - | 0 | - |
| 8 | y | 2.167E+05 | 519.3 | 0.001863 | 3.587 | +1 | 5 |
| - | - | 4.676E+04 | 520.3 | - | - | 0 | - |
| - | - | 5.663E+04 | 520.3 | - | - | 0 | - |
| - | - | 1.604E+04 | 521.3 | - | - | 0 | - |
| - | - | 8139 | 521.3 | - | - | 0 | - |
| - | - | 4801 | 522.3 | - | - | 0 | - |
| - | - | 8363 | 524.3 | - | - | 0 | - |
| - | - | 4667 | 524.8 | - | - | 0 | - |
| - | - | 6762 | 525.3 | - | - | 0 | - |
| - | - | 2378 | 525.8 | - | - | 0 | - |
| - | - | 5628 | 529.3 | - | - | 0 | - |
| - | - | 3544 | 529.8 | - | - | 0 | - |
| - | - | 7.905E+04 | 530.3 | - | - | 0 | - |
| - | - | 2.426E+04 | 531.3 | - | - | 0 | - |
| - | - | 6553 | 532.3 | - | - | 0 | - |
| - | - | 1.363E+05 | 533.3 | - | - | 0 | - |
| - | - | 8.977E+04 | 533.8 | - | - | 0 | - |
| - | - | 6.537E+04 | 534.3 | - | - | 0 | - |
| - | - | 3560 | 534.8 | - | - | 0 | - |
| - | - | 1.403E+05 | 535.3 | - | - | 0 | - |
| - | - | 4.855E+04 | 536.3 | - | - | 0 | - |
| - | - | 9060 | 537.3 | - | - | 0 | - |
| - | - | 8.401E+04 | 538.3 | - | - | 0 | - |
| - | - | 1.866E+04 | 538.3 | - | - | 0 | - |
| - | - | 1.962E+04 | 538.8 | - | - | 0 | - |
| - | - | 2.614E+04 | 539.3 | - | - | 0 | - |
| - | - | 2359 | 539.3 | - | - | 0 | - |
| - | - | 4530 | 540.3 | - | - | 0 | - |
| 2 | z | 2307 | 546.3 | 0.008231 | 15.07 | +2 | 11 |
| 11 | c | 5.833E+04 | 547.3 | 0.0001859 | 0.3396 | +2 | 11 |
| - | - | 4.848E+04 | 547.8 | - | - | 0 | - |
| - | - | 7.02E+05 | 548.3 | - | - | 0 | - |
| - | - | 3053 | 548.8 | - | - | 0 | - |
| - | - | 2.144E+05 | 549.3 | - | - | 0 | - |
| - | - | 2.951E+04 | 550.3 | - | - | 0 | - |
| - | - | 2.725E+05 | 552.3 | - | - | 0 | - |
| - | - | 1.089E+05 | 553.3 | - | - | 0 | - |
| 2 | y | 1.997E+04 | 554.3 | 0.00914 | 16.49 | +2 | 11 |
| 2 | y | 3292 | 554.8 | 0.009686 | 17.46 | +2 | 11 |
| 7 | y | 4.888E+04 | 558.3 | 0.001706 | 3.056 | +1 | 6 |
| - | - | 1.665E+04 | 559.3 | - | - | 0 | - |
| - | - | 2890 | 560.3 | - | - | 0 | - |
| - | - | 5927 | 561.3 | - | - | 0 | - |
| - | - | 1.92E+05 | 562.3 | - | - | 0 | - |
| 2 | y | 7.852E+04 | 563.3 | 0.009936 | 17.64 | +2 | 11 |
| - | - | 1.224E+04 | 563.8 | - | - | 0 | - |
| - | - | 1.939E+04 | 564.3 | - | - | 0 | - |
| - | - | 4158 | 565.3 | - | - | 0 | - |
| - | - | 2220 | 566.8 | - | - | 0 | - |
| - | - | 4762 | 567.3 | - | - | 0 | - |
| - | - | 3933 | 568.3 | - | - | 0 | - |
| - | - | 4566 | 571.3 | - | - | 0 | - |
| - | - | 2.111E+04 | 571.4 | - | - | 0 | - |
| - | - | 7494 | 572.4 | - | - | 0 | - |
| - | - | 2377 | 573.3 | - | - | 0 | - |
| - | - | 2393 | 573.4 | - | - | 0 | - |
| - | - | 8906 | 574.3 | - | - | 0 | - |
| - | - | 3670 | 574.8 | - | - | 0 | - |
| - | - | 1.022E+04 | 575.3 | - | - | 0 | - |
| - | - | 1.312E+04 | 575.8 | - | - | 0 | - |
| 7 | y | 7.233E+05 | 576.3 | 0.002067 | 3.586 | +1 | 6 |
| - | - | 4298 | 576.8 | - | - | 0 | - |
| - | - | 7376 | 577.3 | - | - | 0 | - |
| - | - | 2.121E+05 | 577.3 | - | - | 0 | - |
| - | - | 9767 | 578.3 | - | - | 0 | - |
| - | - | 3.447E+04 | 578.3 | - | - | 0 | - |
| - | - | 3786 | 579.3 | - | - | 0 | - |
| - | - | 6.481E+05 | 580.3 | - | - | 0 | - |
| - | - | 2.505E+05 | 581.3 | - | - | 0 | - |
| - | - | 4.652E+04 | 582.3 | - | - | 0 | - |
| - | - | 4978 | 583.3 | - | - | 0 | - |
| - | - | 2265 | 586.3 | - | - | 0 | - |
| - | - | 1.607E+04 | 587.4 | - | - | 0 | - |
| - | - | 4303 | 588.4 | - | - | 0 | - |
| - | - | 1.536E+04 | 588.8 | - | - | 0 | - |
| - | - | 1.725E+04 | 589.3 | - | - | 0 | - |
| - | - | 5234 | 589.8 | - | - | 0 | - |
| - | - | 4432 | 590.3 | - | - | 0 | - |
| - | - | 1.664E+04 | 591.3 | - | - | 0 | - |
| - | - | 3.772E+04 | 592.3 | - | - | 0 | - |
| - | - | 1.475E+04 | 593.3 | - | - | 0 | - |
| - | - | 3114 | 594.3 | - | - | 0 | - |
| - | - | 2.833E+04 | 595.3 | - | - | 0 | - |
| - | - | 8958 | 596.3 | - | - | 0 | - |
| - | - | 2245 | 596.8 | - | - | 0 | - |
| 5 | c | 1.695E+04 | 597.3 | 0.004854 | 8.126 | +1 | 5 |
| - | - | 4.572E+05 | 597.8 | - | - | 0 | - |
| - | - | 3.473E+05 | 598.3 | - | - | 0 | - |
| - | - | 1.205E+05 | 598.8 | - | - | 0 | - |
| - | - | 4207 | 598.9 | - | - | 0 | - |
| 6 | z | 1.827E+04 | 599.3 | 0.008305 | 13.86 | +1 | 7 |
| 6 | z | 1.274E+04 | 600.3 | 0.006589 | 10.98 | +1 | 7 |
| - | - | 7743 | 601.3 | - | - | 0 | - |
| - | - | 8539 | 604.3 | - | - | 0 | - |
| - | - | 4518 | 606.3 | - | - | 0 | - |
| - | - | 3537 | 606.8 | - | - | 0 | - |
| - | - | 6.782E+04 | 606.8 | - | - | 0 | - |
| - | - | 3544 | 607.3 | - | - | 0 | - |
| - | - | 4.609E+04 | 607.3 | - | - | 0 | - |
| - | - | 1.548E+04 | 607.8 | - | - | 0 | - |
| - | - | 6.931E+04 | 609.3 | - | - | 0 | - |
| - | - | 3.032E+04 | 610.3 | - | - | 0 | - |
| - | - | 6808 | 611.3 | - | - | 0 | - |
| 6 | y | 1.712E+05 | 615.3 | 0.002337 | 3.798 | +1 | 7 |
| - | - | 5.551E+04 | 616.4 | - | - | 0 | - |
| - | - | 1.102E+04 | 617.4 | - | - | 0 | - |
| - | - | 3.588E+04 | 618.3 | - | - | 0 | - |
| - | - | 1.216E+05 | 619.3 | - | - | 0 | - |
| - | - | 4.929E+04 | 620.3 | - | - | 0 | - |
| - | - | 7798 | 621.3 | - | - | 0 | - |
| - | - | 6009 | 625.3 | - | - | 0 | - |
| - | - | 4.748E+04 | 629.3 | - | - | 0 | - |
| - | - | 2.116E+04 | 630.3 | - | - | 0 | - |
| - | - | 1.576E+04 | 631.3 | - | - | 0 | - |
| - | - | 9622 | 632.3 | - | - | 0 | - |
| 6 | y | 3.186E+06 | 633.4 | 0.002576 | 4.067 | +1 | 7 |
| - | - | 1.042E+06 | 634.4 | - | - | 0 | - |
| - | - | 1.916E+05 | 635.4 | - | - | 0 | - |
| - | - | 1.019E+04 | 636.4 | - | - | 0 | - |
| - | - | 1.888E+05 | 637.3 | - | - | 0 | - |
| - | - | 7.647E+04 | 638.3 | - | - | 0 | - |
| - | - | 1.414E+04 | 639.3 | - | - | 0 | - |
| - | - | 9.497E+04 | 643.4 | - | - | 0 | - |
| - | - | 3.758E+04 | 644.4 | - | - | 0 | - |
| - | - | 7408 | 645.4 | - | - | 0 | - |
| - | - | 2.841E+05 | 647.4 | - | - | 0 | - |
| - | - | 1.034E+05 | 648.4 | - | - | 0 | - |
| - | - | 2.322E+04 | 649.4 | - | - | 0 | - |
| - | - | 3653 | 650.3 | - | - | 0 | - |
| - | - | 1.482E+04 | 652.3 | - | - | 0 | - |
| - | - | 5282 | 653.3 | - | - | 0 | - |
| - | - | 2551 | 654.3 | - | - | 0 | - |
| - | - | 5377 | 658.3 | - | - | 0 | - |
| - | - | 4146 | 659.3 | - | - | 0 | - |
| - | - | 5.039E+05 | 661.4 | - | - | 0 | - |
| - | - | 1.936E+05 | 662.4 | - | - | 0 | - |
| - | - | 4.429E+04 | 663.4 | - | - | 0 | - |
| - | - | 3477 | 664.4 | - | - | 0 | - |
| - | - | 2.854E+04 | 666.4 | - | - | 0 | - |
| - | - | 1.069E+04 | 667.4 | - | - | 0 | - |
| - | - | 3009 | 668.4 | - | - | 0 | - |
| - | - | 3.979E+04 | 676.3 | - | - | 0 | - |
| - | - | 1.914E+04 | 677.4 | - | - | 0 | - |
| - | - | 5150 | 678.4 | - | - | 0 | - |
| - | - | 6928 | 679.4 | - | - | 0 | - |
| - | - | 5385 | 687.3 | - | - | 0 | - |
| - | - | 3344 | 688.4 | - | - | 0 | - |
| - | - | 1.103E+05 | 694.4 | - | - | 0 | - |
| - | - | 4.794E+04 | 695.4 | - | - | 0 | - |
| - | - | 8878 | 696.4 | - | - | 0 | - |
| - | - | 1.493E+04 | 705.4 | - | - | 0 | - |
| - | - | 1.196E+04 | 706.4 | - | - | 0 | - |
| - | - | 2976 | 707.4 | - | - | 0 | - |
| - | - | 4144 | 716.4 | - | - | 0 | - |
| - | - | 4.123E+04 | 718.4 | - | - | 0 | - |
| - | - | 1.513E+04 | 719.4 | - | - | 0 | - |
| - | - | 1.834E+04 | 723.4 | - | - | 0 | - |
| - | - | 2940 | 724.4 | - | - | 0 | - |
| - | - | 4.073E+04 | 728.4 | - | - | 0 | - |
| - | - | 1.79E+04 | 729.4 | - | - | 0 | - |
| - | - | 3718 | 730.4 | - | - | 0 | - |
| - | - | 3.061E+04 | 732.4 | - | - | 0 | - |
| - | - | 3.466E+04 | 733.4 | - | - | 0 | - |
| - | - | 1.552E+04 | 733.4 | - | - | 0 | - |
| - | - | 9625 | 734.4 | - | - | 0 | - |
| - | - | 1.864E+04 | 734.4 | - | - | 0 | - |
| - | - | 9356 | 735.4 | - | - | 0 | - |
| - | - | 9288 | 735.4 | - | - | 0 | - |
| - | - | 3326 | 736.3 | - | - | 0 | - |
| - | - | 8343 | 736.4 | - | - | 0 | - |
| - | - | 4.777E+04 | 742.4 | - | - | 0 | - |
| - | - | 1.91E+04 | 743.4 | - | - | 0 | - |
| - | - | 1.855E+04 | 744.4 | - | - | 0 | - |
| - | - | 1.096E+04 | 745.4 | - | - | 0 | - |
| - | - | 1.695E+05 | 746.4 | - | - | 0 | - |
| - | - | 7.434E+04 | 747.4 | - | - | 0 | - |
| - | - | 1.524E+04 | 748.4 | - | - | 0 | - |
| - | - | 1.255E+05 | 751.4 | - | - | 0 | - |
| - | - | 5.866E+04 | 752.4 | - | - | 0 | - |
| - | - | 2.544E+04 | 753.4 | - | - | 0 | - |
| - | - | 9283 | 754.4 | - | - | 0 | - |
| - | - | 3.484E+05 | 760.4 | - | - | 0 | - |
| - | - | 1.504E+05 | 761.4 | - | - | 0 | - |
| 5 | y | 2.268E+05 | 762.4 | 0.004053 | 5.315 | +1 | 8 |
| - | - | 9.662E+04 | 763.4 | - | - | 0 | - |
| 5 | z | 2.555E+04 | 764.4 | 0.01459 | 19.09 | +1 | 8 |
| - | - | 1.058E+04 | 765.4 | - | - | 0 | - |
| - | - | 2.075E+04 | 778.4 | - | - | 0 | - |
| - | - | 1.138E+04 | 779.4 | - | - | 0 | - |
| 5 | y | 4.477E+06 | 780.4 | 0.002826 | 3.621 | +1 | 8 |
| - | - | 1.957E+06 | 781.4 | - | - | 0 | - |
| - | - | 4.578E+05 | 782.4 | - | - | 0 | - |
| - | - | 3.809E+04 | 783.4 | - | - | 0 | - |
| - | - | 5705 | 789.4 | - | - | 0 | - |
| - | - | 1.173E+04 | 790.4 | - | - | 0 | - |
| - | - | 3398 | 791.4 | - | - | 0 | - |
| - | - | 1.469E+04 | 806.4 | - | - | 0 | - |
| - | - | 2.795E+04 | 807.4 | - | - | 0 | - |
| - | - | 1.997E+04 | 808.4 | - | - | 0 | - |
| - | - | 6892 | 809.4 | - | - | 0 | - |
| - | - | 2.6E+04 | 816.4 | - | - | 0 | - |
| - | - | 1.341E+04 | 817.4 | - | - | 0 | - |
| - | - | 2.791E+04 | 817.5 | - | - | 0 | - |
| - | - | 3163 | 818.4 | - | - | 0 | - |
| - | - | 8607 | 818.5 | - | - | 0 | - |
| - | - | 3174 | 819.5 | - | - | 0 | - |
| - | - | 3.855E+04 | 824.4 | - | - | 0 | - |
| - | - | 1.987E+04 | 825.4 | - | - | 0 | - |
| - | - | 6853 | 826.4 | - | - | 0 | - |
| - | - | 2.648E+04 | 831.5 | - | - | 0 | - |
| - | - | 1.234E+04 | 832.5 | - | - | 0 | - |
| - | - | 1.455E+04 | 833.5 | - | - | 0 | - |
| - | - | 1.528E+05 | 834.4 | - | - | 0 | - |
| - | - | 8.052E+04 | 835.4 | - | - | 0 | - |
| - | - | 2.055E+04 | 836.4 | - | - | 0 | - |
| - | - | 2.4E+04 | 841.5 | - | - | 0 | - |
| - | - | 9401 | 842.5 | - | - | 0 | - |
| - | - | 1.114E+04 | 843.5 | - | - | 0 | - |
| - | - | 7012 | 844.5 | - | - | 0 | - |
| 4 | z | 3728 | 845.5 | 0.005693 | 6.734 | +1 | 9 |
| - | - | 1.162E+04 | 847.5 | - | - | 0 | - |
| - | - | 6658 | 848.5 | - | - | 0 | - |
| - | - | 2802 | 849.5 | - | - | 0 | - |
| - | - | 4.579E+05 | 852.4 | - | - | 0 | - |
| - | - | 2.301E+05 | 853.4 | - | - | 0 | - |
| - | - | 4199 | 853.5 | - | - | 0 | - |
| - | - | 6.094E+04 | 854.4 | - | - | 0 | - |
| - | - | 5051 | 855.4 | - | - | 0 | - |
| - | - | 2.615E+05 | 859.5 | - | - | 0 | - |
| - | - | 1.338E+05 | 860.5 | - | - | 0 | - |
| 4 | y | 1.293E+05 | 861.5 | 0.006684 | 7.758 | +1 | 9 |
| - | - | 6.38E+04 | 862.5 | - | - | 0 | - |
| 4 | z | 1.968E+04 | 863.5 | 0.01228 | 14.22 | +1 | 9 |
| - | - | 5772 | 864.5 | - | - | 0 | - |
| - | - | 3815 | 865.5 | - | - | 0 | - |
| 9 | c | 2.114E+04 | 869.5 | 0.002217 | 2.55 | +1 | 9 |
| - | - | 1.037E+04 | 870.5 | - | - | 0 | - |
| - | - | 1.405E+04 | 877.5 | - | - | 0 | - |
| - | - | 8824 | 878.5 | - | - | 0 | - |
| 4 | y | 2.678E+06 | 879.5 | 0.002528 | 2.874 | +1 | 9 |
| - | - | 1.342E+06 | 880.5 | - | - | 0 | - |
| - | - | 3.606E+05 | 881.5 | - | - | 0 | - |
| - | - | 4574 | 881.6 | - | - | 0 | - |
| - | - | 3.838E+04 | 882.5 | - | - | 0 | - |
| - | - | 3384 | 889.5 | - | - | 0 | - |
| - | - | 1.093E+04 | 893.5 | - | - | 0 | - |
| - | - | 3820 | 894.5 | - | - | 0 | - |
| - | - | 1.169E+04 | 895.5 | - | - | 0 | - |
| - | - | 4080 | 896.5 | - | - | 0 | - |
| - | - | 8206 | 916.6 | - | - | 0 | - |
| - | - | 4212 | 917.6 | - | - | 0 | - |
| - | - | 7426 | 918.5 | - | - | 0 | - |
| - | - | 5214 | 928.5 | - | - | 0 | - |
| - | - | 4323 | 932.6 | - | - | 0 | - |
| - | - | 3928 | 933.6 | - | - | 0 | - |
| - | - | 2555 | 936.5 | - | - | 0 | - |
| - | - | 5569 | 942.5 | - | - | 0 | - |
| - | - | 2603 | 943.5 | - | - | 0 | - |
| - | - | 1.523E+04 | 944.5 | - | - | 0 | - |
| 3 | z | 5778 | 945.5 | 0.008528 | 9.02 | +1 | 10 |
| - | - | 1.673E+04 | 946.5 | - | - | 0 | - |
| 3 | w | 1.066E+04 | 947.5 | 0.01745 | 18.42 | +1 | 10 |
| - | - | 4917 | 948.5 | - | - | 0 | - |
| 3 | y | 4.716E+04 | 960.6 | 0.002296 | 2.39 | +1 | 10 |
| - | - | 2.646E+04 | 961.6 | - | - | 0 | - |
| - | - | 2.52E+05 | 962.5 | - | - | 0 | - |
| - | - | 1.511E+05 | 963.5 | - | - | 0 | - |
| - | - | 4.435E+04 | 964.5 | - | - | 0 | - |
| - | - | 6741 | 965.5 | - | - | 0 | - |
| - | - | 1.227E+04 | 976.5 | - | - | 0 | - |
| - | - | 6485 | 977.5 | - | - | 0 | - |
| 3 | y | 1.2E+06 | 978.6 | 0.003083 | 3.151 | +1 | 10 |
| - | - | 6.992E+05 | 979.6 | - | - | 0 | - |
| 10 | c | 1.051E+06 | 980.5 | 0.005336 | 5.442 | +1 | 10 |
| - | - | 6.151E+05 | 981.5 | - | - | 0 | - |
| - | - | 1.941E+05 | 982.5 | - | - | 0 | - |
| - | - | 1902 | 982.6 | - | - | 0 | - |
| - | - | 1.926E+04 | 983.5 | - | - | 0 | - |
| - | - | 4.034E+04 | 994.5 | - | - | 0 | - |
| - | - | 2.36E+04 | 995.5 | - | - | 0 | - |
| - | - | 8578 | 996.5 | - | - | 0 | - |
| 10 | c | 9648 | 997.5 | 0.0003387 | 0.3395 | +1 | 10 |
| - | - | 2999 | 998.5 | - | - | 0 | - |
| - | - | 9224 | 1007 | - | - | 0 | - |
| - | - | 3656 | 1008 | - | - | 0 | - |
| - | - | 3383 | 1050 | - | - | 0 | - |
| - | - | 2.832E+04 | 1066 | - | - | 0 | - |
| - | - | 1.845E+04 | 1067 | - | - | 0 | - |
| - | - | 6779 | 1068 | - | - | 0 | - |
| - | - | 7.834E+04 | 1076 | - | - | 0 | - |
| - | - | 5.668E+04 | 1077 | - | - | 0 | - |
| - | - | 1.72E+04 | 1078 | - | - | 0 | - |
| 2 | z | 6235 | 1092 | 0.01016 | 9.31 | +1 | 11 |
| 2 | z | 5439 | 1093 | 0.008873 | 8.121 | +1 | 11 |
| 11 | c | 9.348E+05 | 1094 | 0.002998 | 2.742 | +1 | 11 |
| - | - | 6.114E+05 | 1095 | - | - | 0 | - |
| - | - | 2.115E+05 | 1096 | - | - | 0 | - |
| - | - | 2.951E+04 | 1097 | - | - | 0 | - |
| 2 | z | 5347 | 1110 | 0.002051 | 1.848 | +1 | 11 |
| 11 | c | 1.499E+04 | 1111 | 0.003653 | 3.289 | +1 | 11 |
| - | - | 1.576E+04 | 1112 | - | - | 0 | - |
| - | - | 1.174E+04 | 1113 | - | - | 0 | - |
| 2 | y | 1.031E+04 | 1126 | 0.003151 | 2.799 | +1 | 11 |
| - | - | 5804 | 1127 | - | - | 0 | - |
| - | - | 3594 | 1136 | - | - | 0 | - |
| - | - | 5318 | 1158 | - | - | 0 | - |
| - | - | 4233 | 1159 | - | - | 0 | - |
| - | - | 2.153E+04 | 1197 | - | - | 0 | - |
| - | - | 2.025E+04 | 1198 | - | - | 0 | - |
| - | - | 4875 | 1199 | - | - | 0 | - |
| - | - | 2.179E+04 | 1213 | - | - | 0 | - |
| - | - | 3.052E+04 | 1214 | - | - | 0 | - |
| - | - | 1.899E+04 | 1215 | - | - | 0 | - |
| - | - | 5160 | 1216 | - | - | 0 | - |

m/z Charge Intensity FragmentType MassShift Position
120.06613159179688 0 823101.06 y 11
120.08148193359375 0 100888.54
121.06463623046875 0 3162.826
121.0694808959961 0 36070.64
121.08476257324219 0 7329.5854
122.0704574584961 0 3560.003
122.09748077392578 0 1341.8265
127.89823913574219 0 1457.7036
129.1028594970703 0 313714.06
130.10025024414062 0 3211.798
130.10617065429688 0 19546.004
131.0823516845703 0 2189.8472
143.3053741455078 0 1401.521
159.11322021484375 0 2131.554
162.09205627441406 0 11488.769
171.14991760253906 0 252427.42
172.15335083007812 0 23236.58
173.39242553710938 0 1951.7379
173.4391632080078 0 6708.887
187.10841369628906 0 8322.859
187.14483642578125 0 7285.329
190.2730712890625 0 1993.8265
197.16575622558594 0 9174.656
199.1448516845703 0 340560.66
200.14825439453125 0 42996.996
205.097900390625 0 13397.484
207.11358642578125 0 1814070.5
208.11697387695312 0 216775.95
209.1193084716797 0 10968.747
212.14016723632812 0 11304.133
214.19224548339844 0 2460.2498
215.1398468017578 0 152524.4 y Water loss 10
216.1432342529297 0 17388.332
217.15524291992188 0 2596.5208
219.15005493164062 0 100981.164
220.153564453125 0 16079.937
222.12452697753906 0 2093.7217
228.09963989257812 0 3054.453
230.15086364746094 0 73107.46
231.1543426513672 0 7640.44
233.1505889892578 0 952246.3 y 10
234.15386962890625 0 94806.43
235.10867309570312 0 3585977
236.1118927001953 0 465537.47
237.11412048339844 0 33661.508
242.18734741210938 0 14114.098
247.14505004882812 0 166926.22
248.14842224121094 0 26326.25
255.1095733642578 0 7678.8755
257.1298522949219 0 2532.3977
261.16143798828125 0 4180.7456
262.1190185546875 0 2978.8506
264.17169189453125 0 3023.1724
265.156494140625 0 3339.9067
268.166748046875 0 8385.605
269.16192626953125 0 15309.381
273.1203918457031 0 72935.805
273.19818115234375 0 2421.4106
274.12164306640625 0 9747.345
286.1769714355469 0 11195.049
287.1725769042969 0 42665.887
288.1729736328125 0 9410.19
289.15576171875 0 327452.2
290.1590576171875 0 56225.293
291.1615295410156 0 5198.996
292.1303405761719 0 6890.5264
294.9287109375 0 2249.814
301.1923522949219 0 45497.7
302.19561767578125 0 9197.076
303.17108154296875 0 5746.045
304.1667785644531 0 20145.709
305.17083740234375 0 4216.4487
306.1824951171875 0 19480.021
307.1861267089844 0 4369.281
309.160400390625 0 3751.5781
316.1669006347656 0 93920.05
317.1700744628906 0 17730.996
318.21832275390625 0 7618.333
319.14215087890625 0 5980.2695
326.1836242675781 0 63677.46
326.2053527832031 0 4286.6226
327.186767578125 0 8806.599
329.1848449707031 0 3240.2178
332.1619567871094 0 11492.818
333.16851806640625 0 4998.1426
334.1775207519531 0 4873465
335.1805419921875 0 950565.6
336.1831970214844 0 93585.016
337.15582275390625 0 22138.82
337.1804504394531 0 2532.7917
337.1873779296875 0 2882.2383
338.15985107421875 0 4813.29
342.204345703125 0 3184.3928
343.2356262207031 0 19671.725 y Water loss 9
344.194091796875 0 62286.38
344.2386169433594 0 3025.2874
345.11700439453125 0 2338.11
345.1976623535156 0 8909.161
346.1768798828125 0 8528.973
346.21392822265625 0 134143.48
347.21710205078125 0 28498.756
349.15167236328125 0 4027.5696
354.180419921875 0 5310.2153
356.1981201171875 0 3786.4744 c 6
357.1876525878906 0 2665.513
358.21429443359375 0 3173.37
361.19012451171875 0 9070.122
361.2458801269531 0 252527.98 y 9
362.24884033203125 0 44994.414
363.2044982910156 0 4468.975
364.16680908203125 0 38439.516
364.7093505859375 0 8062.4883
365.17010498046875 0 7409.0684
365.1915283203125 0 3399.1216
365.21514892578125 0 2832.0103
366.1774597167969 0 3817.021
366.7242126464844 0 3605.3599
368.19268798828125 0 4321.709
370.2146911621094 0 3023.2007
372.1895446777344 0 2006.9366
372.2608947753906 0 2794.6768
373.18817138671875 0 9504.371
373.71466064453125 0 27210.385
374.183837890625 0 3139.1072
374.2107849121094 0 27790.115
374.7184143066406 0 3939.4592
375.21282958984375 0 4196.6533
375.24072265625 0 4425.134
380.7221984863281 0 6032.7617
381.2239685058594 0 3117.2158
381.7118835449219 0 17875.217 y Water loss 4
382.17767333984375 0 46120.32
382.2134704589844 0 8272.269
382.2465515136719 0 11636.36
383.18023681640625 0 9340.657
383.2052917480469 0 89592.51
384.2083435058594 0 13992.103
386.2083435058594 0 4686.879
387.2405700683594 0 13782.507
388.2245788574219 0 65829.83
389.22772216796875 0 16694.463
390.7176208496094 0 199882.48 y 4
391.19976806640625 0 12378.15
391.2193298339844 0 80666.664
391.72003173828125 0 17895.387
392.1956481933594 0 18717.977
393.1966247558594 0 7445.3813
400.2572326660156 0 45720.33
401.2158508300781 0 506109.62
402.1791076660156 0 54015.67
402.21875 0 88488.734
403.1818542480469 0 10376.249
403.2349853515625 0 22089.844
404.2388000488281 0 5611.5166
405.2513122558594 0 434670.7
406.17413330078125 0 2999.6006
406.25439453125 0 106915.13
407.2569580078125 0 14011.182
410.1725769042969 0 2776.7932
411.2046813964844 0 3782.533
415.2356872558594 0 218731.39
416.2386779785156 0 52270.42
416.76092529296875 0 5546.407
417.24151611328125 0 8310.971
418.21124267578125 0 11441.922
419.2127685546875 0 2689.3005
420.1893310546875 0 80315.76
421.19146728515625 0 27430.559
421.2560119628906 0 3855.2898
421.75518798828125 0 3664.7173
422.19268798828125 0 3371.1995
426.7156677246094 0 2305.0063
429.2836608886719 0 9283.397
430.2090148925781 0 2549.4172
430.25701904296875 0 25850.871
430.75823974609375 0 14403.873
431.2470703125 0 17296.498 y Water loss 3
431.7488098144531 0 10607.055
432.23681640625 0 4181.015 z 3
433.2464294433594 0 2074711.8
434.24932861328125 0 537768.56
435.2504577636719 0 74796.305
436.224853515625 0 115389.76
437.2275695800781 0 33838.37
438.22882080078125 0 4293.8496
439.1985168457031 0 8900.183
439.23565673828125 0 3877.5117
439.26800537109375 0 29696.627
440.2520446777344 0 123666.71 y 3
440.75360107421875 0 62297.38
441.25506591796875 0 15560.55
443.2315673828125 0 7094.376
444.232421875 0 3047.6624
444.2844543457031 0 7780.607 y Water loss 8
445.2460021972656 0 15163.02
446.24951171875 0 2670.6907
447.24920654296875 0 5783.2915
447.7533874511719 0 2443.3877
448.2213439941406 0 3890.9019
448.2617492675781 0 6587.211
449.223388671875 0 2361.3623
451.23406982421875 0 2538.6624
453.25140380859375 0 77868.72
454.2546081542969 0 23785.395
455.2594909667969 0 2954.157
457.27874755859375 0 114034.26
458.2812805175781 0 25611.957
459.2864074707031 0 3021.3064
460.2564697265625 0 9280.817
461.24224853515625 0 5579.6943
462.294189453125 0 107162.94 y 8
463.23583984375 0 117119.17
463.296875 0 21552.348
464.2391052246094 0 37675.305
464.2967529296875 0 3208.1243
464.767578125 0 3335.6296
465.2437438964844 0 5712.112
465.2886047363281 0 4786.356
472.2574157714844 0 21962.719
473.2569580078125 0 8928.696 z Ammonia loss 2
473.7738342285156 0 6103.0483
474.77301025390625 0 3372.6821
475.23370361328125 0 4356.5957
476.76666259765625 0 3141.9583
478.2082824707031 0 3123.0808
478.2792663574219 0 5510.736
480.78118896484375 0 80438.24 y Water loss 2
481.2464599609375 0 365665
481.2823486328125 0 29845.26
481.7601623535156 0 96960.69
482.2528381347656 0 126969.555
482.7628479003906 0 18367.031
483.25201416015625 0 13246.491
483.7762451171875 0 10755.263
484.2796630859375 0 3679.0515
486.3052978515625 0 33803.68
487.3083190917969 0 8127.5977
489.21112060546875 0 2437.4888
489.2748107910156 0 7575.9053
489.78662109375 0 181148.97 y 2
490.2866516113281 0 89342.516
490.76556396484375 0 161569.53 c Ammonia loss 9
491.2674255371094 0 122213.43
491.7689514160156 0 29985.41
492.2672424316406 0 8905.668
493.24615478515625 0 15183.045
493.2834777832031 0 14208.785
494.24859619140625 0 4101.1343
494.2861328125 0 6032.8003
496.289794921875 0 80852.086
497.2923889160156 0 19730.39
497.7735900878906 0 16301.887
498.2760925292969 0 14214.111
498.77679443359375 0 4003.5066
499.2691650390625 0 3554.7244 c 9
501.2477722167969 0 29970.78
501.3057556152344 0 18192.484 y Water loss 7
502.25103759765625 0 10374.962
502.30902099609375 0 4918.625
503.79119873046875 0 6747.9883
504.2586364746094 0 4651.426
504.3006896972656 0 2253.4082
507.2214660644531 0 2939.8755
507.29974365234375 0 4838.5503
508.3014831542969 0 3171.5776
509.2418212890625 0 14390.673
510.27325439453125 0 18743.127
511.2757263183594 0 6707.703
512.2655639648438 0 7278.856
514.3004760742188 0 427659.8
515.30322265625 0 115944.81
516.3048095703125 0 16789.502
517.2821044921875 0 17101.076
518.284912109375 0 8033.419
519.2589721679688 0 36629.953
519.3155517578125 0 216652.36 y 7
520.2579956054688 0 46755.484
520.318603515625 0 56626.844
521.2706909179688 0 16036.301
521.3218383789062 0 8138.549
522.2770385742188 0 4801.182
524.3028564453125 0 8363.2705
524.8036499023438 0 4666.996
525.3001708984375 0 6762.3584
525.7943725585938 0 2377.7595
529.28125 0 5627.647
529.7974243164062 0 3544.4106
530.2744140625 0 79046.414
531.27734375 0 24262.564
532.3075561523438 0 6552.8555
533.3099975585938 0 136268.23
533.8116455078125 0 89768.34
534.3106689453125 0 65372.797
534.8119506835938 0 3560.3325
535.2939453125 0 140345.95
536.296630859375 0 48551.93
537.29833984375 0 9060.207
538.2678833007812 0 84005.08
538.30322265625 0 18663.285
538.8027954101562 0 19624.66
539.2708740234375 0 26138.03
539.30859375 0 2358.8213
540.2733764648438 0 4529.588
546.2958984375 0 2307.4475 z Water loss 1
547.3054809570312 0 58328.902 c Ammonia loss 10
547.80908203125 0 48483
548.2848510742188 0 701985.8
548.8072509765625 0 3052.7776
549.2877197265625 0 214434.69
550.2899169921875 0 29509.133
552.31982421875 0 272535.56
553.3231201171875 0 108912.336
554.3226318359375 0 19974.418 y Water loss 1
554.815185546875 0 3291.7842 y Ammonia loss 1
558.3262939453125 0 48883.188 y Water loss 6
559.3290405273438 0 16648.154
560.3314819335938 0 2890.081
561.3258666992188 0 5927.0215
562.3043212890625 0 191976.23
563.308837890625 0 78520.47 y 1
563.8220825195312 0 12238.38
564.31494140625 0 19393.268
565.3124389648438 0 4157.98
566.8117065429688 0 2220.0671
567.2965698242188 0 4762.4463
568.30517578125 0 3932.5432
571.3197021484375 0 4565.9663
571.3584594726562 0 21109.69
572.3616943359375 0 7493.811
573.3077392578125 0 2376.9329
573.3644409179688 0 2392.998
574.306884765625 0 8906.298
574.830322265625 0 3670.2434
575.3235473632812 0 10221.598
575.8184204101562 0 13118.461
576.3372192382812 0 723275.75 y 6
576.8208618164062 0 4298.391
577.2850341796875 0 7376.368
577.3401489257812 0 212078
578.2947998046875 0 9766.756
578.3428955078125 0 34467.32
579.3004150390625 0 3786.1428
580.31494140625 0 648051.5
581.3180541992188 0 250521.33
582.3207397460938 0 46523.61
583.3240356445312 0 4978.064
586.3018798828125 0 2264.723
587.353759765625 0 16069.374
588.3568725585938 0 4302.576
588.82568359375 0 15361.541
589.3290405273438 0 17250.229
589.83056640625 0 5233.994
590.3336791992188 0 4432.394
591.330810546875 0 16636.613
592.316650390625 0 37717.26
593.3187866210938 0 14745.791
594.3192749023438 0 3114.4285
595.2899780273438 0 28328.182
596.2930297851562 0 8958.329
596.8225708007812 0 2244.872
597.3346557617188 0 16949.549 c 4
597.8316650390625 0 457186.78
598.3331909179688 0 347293.44
598.8344116210938 0 120518.56
598.8922729492188 0 4207.344
599.3356323242188 0 18267.275 z Water loss 5
600.3179321289062 0 12737.952 z Ammonia loss 5
601.31787109375 0 7743.3794
604.311767578125 0 8539.469
606.2875366210938 0 4518.253
606.7799072265625 0 3536.7812
606.837158203125 0 67820.48
607.285400390625 0 3543.951
607.337890625 0 46093.96
607.839111328125 0 15477.806
609.3419189453125 0 69312.125
610.3443603515625 0 30317.848
611.3424072265625 0 6808.3574
615.348388671875 0 171229.33 y Water loss 5
616.3505859375 0 55512.527
617.3519897460938 0 11021.659
618.328369140625 0 35878.703
619.326416015625 0 121553.95
620.3295288085938 0 49290.363
621.3341064453125 0 7797.9497
625.345458984375 0 6009.4575
629.3427124023438 0 47484.984
630.3453369140625 0 21160.379
631.3428344726562 0 15759.054
632.3478393554688 0 9621.925
633.3591918945312 0 3185817.2 y 5
634.36181640625 0 1041925.4
635.3641967773438 0 191604.23
636.364501953125 0 10185.249
637.3366088867188 0 188820.9
638.3397216796875 0 76465.44
639.3423461914062 0 14135.442
643.35888671875 0 94966.76
644.3616333007812 0 37575.5
645.3616943359375 0 7407.931
647.3535766601562 0 284148.88
648.3560180664062 0 103433.91
649.3533935546875 0 23215.75
650.347900390625 0 3653.1646
652.3109741210938 0 14822.972
653.3126831054688 0 5281.9907
654.2880249023438 0 2551.0044
658.3375244140625 0 5377.1606
659.3494873046875 0 4146.308
661.3690795898438 0 503939.28
662.3721313476562 0 193629.84
663.374267578125 0 44287.43
664.381591796875 0 3477.264
666.3634643554688 0 28541.65
667.36669921875 0 10685.607
668.370361328125 0 3009.394
676.3470458984375 0 39790.414
677.3516845703125 0 19141.18
678.3553466796875 0 5149.946
679.3800048828125 0 6927.7373
687.3493041992188 0 5384.976
688.3529663085938 0 3344.497
694.3585815429688 0 110334.34
695.361572265625 0 47939.95
696.3637084960938 0 8878.13
705.3616943359375 0 14927.513
706.3602905273438 0 11956.443
707.3621826171875 0 2976.1636
716.41845703125 0 4143.858
718.4263305664062 0 41226.52
719.429931640625 0 15130.348
723.383544921875 0 18341.08
724.3939208984375 0 2939.9744
728.4111938476562 0 40725.977
729.4143676757812 0 17903.865
730.4129638671875 0 3718.1624
732.4431762695312 0 30610.066
733.369140625 0 34659.273
733.445068359375 0 15515.982
734.3699951171875 0 9625.195
734.4248657226562 0 18636.668
735.3524780273438 0 9355.749
735.4247436523438 0 9287.642
736.3421020507812 0 3325.5454
736.4058227539062 0 8343.005
742.4271240234375 0 47774.336
743.4307250976562 0 19098.71
744.4098510742188 0 18551.812
745.4044799804688 0 10960.991
746.4217529296875 0 169526.1
747.4248657226562 0 74337.59
748.4244384765625 0 15235.773
751.3797607421875 0 125548.79
752.3828735351562 0 58655.49
753.367431640625 0 25439.8
754.3616333007812 0 9283.174
760.4378662109375 0 348376.62
761.4407958984375 0 150424.45
762.4185180664062 0 226797.34 y Water loss 4
763.4205932617188 0 96619.89
764.4208984375 0 25551.256 z 4
765.4140014648438 0 10575.427
778.41357421875 0 20753.086
779.4187622070312 0 11382.154
780.4278564453125 0 4477257.5 y 4
781.4306640625 0 1956828.6
782.4329833984375 0 457841.16
783.435791015625 0 38088.9
789.3974609375 0 5704.7476
790.3941650390625 0 11728.33
791.3942260742188 0 3397.6743
806.4223022460938 0 14687.376
807.408935546875 0 27945.107
808.40576171875 0 19969.557
809.4082641601562 0 6891.6987
816.4065551757812 0 25998.027
817.4097290039062 0 13405.899
817.4959106445312 0 27911.256
818.410400390625 0 3163.253
818.4993896484375 0 8607.369
819.4991455078125 0 3174.0146
824.4322509765625 0 38552.594
825.4361572265625 0 19871.334
826.4400634765625 0 6852.57
831.5105590820312 0 26476.21
832.5145263671875 0 12337.674
833.4949340820312 0 14553.008
834.417236328125 0 152794.94
835.4200439453125 0 80519.266
836.4229736328125 0 20548.102
841.49560546875 0 24001.088
842.4972534179688 0 9400.506
843.4788208007812 0 11139.008
844.4667358398438 0 7012.054
845.4698486328125 0 3728.2314 z Water loss 3
847.4702758789062 0 11615.914
848.4722290039062 0 6658.4756
849.4607543945312 0 2802.1155
852.4274291992188 0 457880.97
853.4303588867188 0 230066.8
853.5313720703125 0 4199.3623
854.4335327148438 0 60942.254
855.4360961914062 0 5050.653
859.5062255859375 0 261491.58
860.509033203125 0 133776.75
861.4895629882812 0 129335.95 y Water loss 3
862.4887084960938 0 63799.234
863.4869995117188 0 19675.31 z 3
864.482177734375 0 5771.9746
865.4862670898438 0 3814.539
869.4537963867188 0 21139.965 c 8
870.4576416015625 0 10366.582
877.4820556640625 0 14052.668
878.486572265625 0 8824.251
879.4959716796875 0 2678192.5 y 3
880.4989013671875 0 1341995
881.5013427734375 0 360636.66
881.6061401367188 0 4574.44
882.5028076171875 0 38380.06
889.4844360351562 0 3384.0657
893.4921875 0 10934.918
894.4915771484375 0 3820.4758
895.4710693359375 0 11694.24
896.4808349609375 0 4080.467
916.56005859375 0 8205.984
917.55859375 0 4212.358
918.488525390625 0 7425.8013
928.5214233398438 0 5213.7236
932.5590209960938 0 4322.6562
933.5518188476562 0 3927.6848
936.5045166015625 0 2555.277
942.5402221679688 0 5569.3604
943.5352783203125 0 2602.5417
944.5054321289062 0 15231.615
945.508056640625 0 5778.437 z Ammonia loss 2
946.5369262695312 0 16727.082
947.537109375 0 10662.665 w 2
948.5283203125 0 4917.428
960.5535888671875 0 47156.77 y Water loss 2
961.5569458007812 0 26461.674
962.5130615234375 0 252038.4
963.515625 0 151122.28
964.5192260742188 0 44350.727
965.5176391601562 0 6741.044
976.5441284179688 0 12270.085
977.5463256835938 0 6484.8677
978.56494140625 0 1200260.1 y 2
979.5677490234375 0 699193.56
980.5253295898438 0 1050702.9 c Ammonia loss 9
981.5261840820312 0 615081.6
982.5284423828125 0 194093.83
982.61865234375 0 1902.1122
983.5302124023438 0 19264.404
994.538330078125 0 40340.492
995.5407104492188 0 23602.947
996.5374145507812 0 8577.929
997.5462036132812 0 9648.186 c 9
998.546875 0 2998.5535
1006.576416015625 0 9224.497
1007.5860595703125 0 3655.7378
1049.5897216796875 0 3382.8782
1065.6124267578125 0 28322.479
1066.6153564453125 0 18453.75
1067.6199951171875 0 6779.2646
1075.595458984375 0 78339.05
1076.5985107421875 0 56684.246
1077.601318359375 0 17195.604
1091.5908203125 0 6235.15 z Water loss 1
1092.5938720703125 0 5438.737 z Ammonia loss 1
1093.6070556640625 0 934784.06 c Ammonia loss 10
1094.609375 0 611420.06
1095.6121826171875 0 211519.94
1096.6129150390625 0 29507.361
1109.6094970703125 0 5346.901 z 1
1110.626953125 0 14993.517 c 10
1111.6248779296875 0 15764.921
1112.6243896484375 0 11736.272
1125.6334228515625 0 10311.917 y 1
1126.6336669921875 0 5804.3916
1135.6214599609375 0 3594.4827
1157.6146240234375 0 5317.9917
1158.6199951171875 0 4232.8604
1196.6446533203125 0 21532.703
1197.648681640625 0 20254.402
1198.6510009765625 0 4874.858
1212.6646728515625 0 21788.672
1213.6690673828125 0 30518.291
1214.6739501953125 0 18986.56
1215.67626953125 0 5159.848

Spectrum Details

|  |  |
| --- | --- |
| Matched peaks? Matched peaks The total absolute number of peaks matched. Additionally in brackets the total fraction of peaks matched and the total number of peaks is shown. | 52 (8.83% of 589) |
| FDR? FDR The false discovery rate estimated for this peptide. It is calculated by matching all theoretical fragments with a non-integer shift with the raw peaks for this spectrum. This is done with 40 different shifts. The resulting percentage is the average number of annotated peaks over the number of annotated peaks with the correct spectrum. | 0.14% |
| Satellite FDR? Satellite FDR See the FDR for details on its calculation. This satellite ion specific FDR only contains the satellite ions (d/w) for I/L/J positions. | - |
| PSM Score? PSM Score The PSM Score as given by Hecklib to this annotated spectrum. It is shown with three significant figures. | 465 |

#### Spectrum 9485? Spectrum 9485 The raw spectrum of this peptide as annotated by Hecklib. The fragments are coloured according to ion type (see legend). Any peaks with a star '\*' as text can be hovered over to see the full details, first the ion type second the mass shift type. By hovering over the amino acids in the peptide or ions in the legend the corresponding peaks are highlighted. By toggling the 'Unassigned' label you can turn the background (unassigned) peaks on or off in the plot. By updating the slider in the Ion legend you can update the spectrum to only show the top X% of the peaks with labels. The top X% means any peak that is within X% of the highest intensity. By dragging in the spectrum you can zoom in to a specific part of the spectrum and use 'Zoom Out' to get back to the original zoom level. The annotation of the spectrum is based on the given sequence in the peptides file and is done with different software so inconsistencies are likely. The peaks are annotated based on the given sequence, with 20 ppm tolerance.

Copy Data

##### Spectrum 9485 (TSV)

###### Preview

```
Loading example...
```

*Click on the button to copy the data to your clipboard.*

Mz MinMz MaxIntensity Max

WidthHeightPeptide font sizePeptide stroke widthSpectrum font sizeSpectrum stroke widthCompact peptide

Ion legend

wxyz

abcd

OtherUnassignedIon^Charge^_Position_Show for top:%

SFVVFGGGTKJT

01.06e+42.12e+43.18e+44.24e+4

Zoom Out

y^+1^_1_y^+1^_2_y^+1^_2_y^+1^_4_y^+1^_5_z^+1^_6_y^+1^_6_z^+1^_7_y^+1^_7_c^+1^_7_z^+1^_8_c^+1^_8_y^+1^_8_c^+1^_9_z^+1^_9_c^+1^_9_y^+1^_9_z^+1^_10_y^+1^_10_c^+1^_10_c^+1^_10_c^+1^_11_z^+1^_11_c^+1^_11_

0763152622903053

Fragment Matches Table

Show background peaks

| Position | Ion type | Intensity | mz Theoretical | mz Error (Th) | mz Error (ppm) | Charge | Series Number |
| --- | --- | --- | --- | --- | --- | --- | --- |
| 12 | y | 2394 | 120.1 | 0.0001618 | 1.348 | +1 | 1 |
| - | - | 962.3 | 129.1 | - | - | 0 | - |
| - | - | 450.6 | 133.1 | - | - | 0 | - |
| - | - | 861.2 | 133.1 | - | - | 0 | - |
| - | - | 425.2 | 133.1 | - | - | 0 | - |
| - | - | 4283 | 133.1 | - | - | 0 | - |
| - | - | 454.8 | 134 | - | - | 0 | - |
| - | - | 449.7 | 134 | - | - | 0 | - |
| - | - | 429.3 | 143.1 | - | - | 0 | - |
| - | - | 572.3 | 149 | - | - | 0 | - |
| - | - | 1282 | 149 | - | - | 0 | - |
| - | - | 393.2 | 159 | - | - | 0 | - |
| - | - | 457.1 | 166.9 | - | - | 0 | - |
| - | - | 667.5 | 167.1 | - | - | 0 | - |
| - | - | 686.7 | 171.1 | - | - | 0 | - |
| - | - | 1187 | 173.4 | - | - | 0 | - |
| - | - | 601 | 175.1 | - | - | 0 | - |
| - | - | 482 | 177.1 | - | - | 0 | - |
| - | - | 2952 | 177.1 | - | - | 0 | - |
| - | - | 484.3 | 184.2 | - | - | 0 | - |
| - | - | 515.8 | 185.5 | - | - | 0 | - |
| - | - | 696.4 | 187.1 | - | - | 0 | - |
| - | - | 802 | 191.1 | - | - | 0 | - |
| - | - | 427.4 | 192.1 | - | - | 0 | - |
| - | - | 1092 | 199.1 | - | - | 0 | - |
| - | - | 694.7 | 205.1 | - | - | 0 | - |
| - | - | 4031 | 207.1 | - | - | 0 | - |
| - | - | 644.3 | 208.1 | - | - | 0 | - |
| 11 | y | 1070 | 215.1 | 0.0008279 | 3.848 | +1 | 2 |
| - | - | 3554 | 221.1 | - | - | 0 | - |
| - | - | 621 | 221.1 | - | - | 0 | - |
| - | - | 3116 | 225 | - | - | 0 | - |
| - | - | 615.2 | 227 | - | - | 0 | - |
| 11 | y | 2436 | 233.1 | 0.0002425 | 1.04 | +1 | 2 |
| - | - | 9698 | 235.1 | - | - | 0 | - |
| - | - | 764.2 | 235.1 | - | - | 0 | - |
| - | - | 741.1 | 236.1 | - | - | 0 | - |
| - | - | 7501 | 239.1 | - | - | 0 | - |
| - | - | 613.1 | 239.1 | - | - | 0 | - |
| - | - | 659 | 240.1 | - | - | 0 | - |
| - | - | 983.4 | 247.1 | - | - | 0 | - |
| - | - | 525.7 | 270.4 | - | - | 0 | - |
| - | - | 544.6 | 283.2 | - | - | 0 | - |
| - | - | 5997 | 295.1 | - | - | 0 | - |
| - | - | 999.4 | 296.1 | - | - | 0 | - |
| - | - | 2951 | 299.1 | - | - | 0 | - |
| - | - | 815 | 313.1 | - | - | 0 | - |
| - | - | 1030 | 334.2 | - | - | 0 | - |
| - | - | 1.361E+04 | 334.2 | - | - | 0 | - |
| - | - | 1353 | 335.2 | - | - | 0 | - |
| - | - | 956.4 | 337.2 | - | - | 0 | - |
| - | - | 950.4 | 355.1 | - | - | 0 | - |
| - | - | 3218 | 369.1 | - | - | 0 | - |
| - | - | 552.7 | 399.2 | - | - | 0 | - |
| - | - | 1334 | 401.2 | - | - | 0 | - |
| - | - | 732.6 | 405.3 | - | - | 0 | - |
| - | - | 647.5 | 406.3 | - | - | 0 | - |
| - | - | 5063 | 433.2 | - | - | 0 | - |
| - | - | 1222 | 434.2 | - | - | 0 | - |
| - | - | 667.3 | 444.2 | - | - | 0 | - |
| 9 | y | 1277 | 445.3 | 0.0003562 | 0.7999 | +1 | 4 |
| - | - | 581.3 | 448.4 | - | - | 0 | - |
| - | - | 588.3 | 481.2 | - | - | 0 | - |
| - | - | 3486 | 504.3 | - | - | 0 | - |
| - | - | 2903 | 504.3 | - | - | 0 | - |
| - | - | 1111 | 505.3 | - | - | 0 | - |
| - | - | 989 | 505.3 | - | - | 0 | - |
| - | - | 719.2 | 508.5 | - | - | 0 | - |
| - | - | 1244 | 514.3 | - | - | 0 | - |
| 8 | y | 1704 | 519.3 | 0.0004564 | 0.8788 | +1 | 5 |
| - | - | 828.6 | 535.3 | - | - | 0 | - |
| - | - | 600.7 | 540.4 | - | - | 0 | - |
| - | - | 590.9 | 545.3 | - | - | 0 | - |
| - | - | 1213 | 548.3 | - | - | 0 | - |
| - | - | 665.8 | 552.3 | - | - | 0 | - |
| 7 | z | 1830 | 560.3 | 9.986E-05 | 0.1782 | +1 | 6 |
| - | - | 7327 | 561.3 | - | - | 0 | - |
| - | - | 1162 | 562.3 | - | - | 0 | - |
| - | - | 611.7 | 566 | - | - | 0 | - |
| - | - | 971.8 | 575.3 | - | - | 0 | - |
| 7 | y | 7235 | 576.3 | 0.0002356 | 0.4089 | +1 | 6 |
| - | - | 1327 | 577.3 | - | - | 0 | - |
| - | - | 1546 | 580.3 | - | - | 0 | - |
| - | - | 582.8 | 585.9 | - | - | 0 | - |
| - | - | 7279 | 589.3 | - | - | 0 | - |
| - | - | 707.4 | 589.4 | - | - | 0 | - |
| - | - | 2281 | 590.3 | - | - | 0 | - |
| - | - | 1204 | 591.4 | - | - | 0 | - |
| - | - | 612.7 | 606.3 | - | - | 0 | - |
| - | - | 782.2 | 606.4 | - | - | 0 | - |
| - | - | 981 | 606.8 | - | - | 0 | - |
| - | - | 827 | 607.3 | - | - | 0 | - |
| 6 | z | 3876 | 617.3 | 5.947E-05 | 0.09634 | +1 | 7 |
| - | - | 9297 | 618.3 | - | - | 0 | - |
| - | - | 1933 | 619.3 | - | - | 0 | - |
| - | - | 3348 | 632.3 | - | - | 0 | - |
| 6 | y | 1.553E+04 | 633.4 | 0.0005615 | 0.8865 | +1 | 7 |
| - | - | 4163 | 634.4 | - | - | 0 | - |
| - | - | 622.4 | 635.4 | - | - | 0 | - |
| - | - | 782.5 | 647.4 | - | - | 0 | - |
| - | - | 938.1 | 661.4 | - | - | 0 | - |
| - | - | 660.8 | 663.4 | - | - | 0 | - |
| - | - | 917.6 | 668.4 | - | - | 0 | - |
| - | - | 3792 | 710.4 | - | - | 0 | - |
| 7 | c | 2670 | 711.4 | 0.002615 | 3.676 | +1 | 7 |
| - | - | 1257 | 712.4 | - | - | 0 | - |
| 5 | z | 7414 | 764.4 | 0.0004932 | 0.6452 | +1 | 8 |
| - | - | 9854 | 765.4 | - | - | 0 | - |
| - | - | 4270 | 766.4 | - | - | 0 | - |
| - | - | 2073 | 767.4 | - | - | 0 | - |
| 8 | c | 1718 | 768.4 | 0.001313 | 1.709 | +1 | 8 |
| - | - | 4078 | 779.4 | - | - | 0 | - |
| 5 | y | 1.944E+04 | 780.4 | 0.0003476 | 0.4453 | +1 | 8 |
| - | - | 6574 | 781.4 | - | - | 0 | - |
| - | - | 1547 | 782.4 | - | - | 0 | - |
| - | - | 595.5 | 785.3 | - | - | 0 | - |
| - | - | 804.5 | 850.4 | - | - | 0 | - |
| 9 | c | 690.3 | 851.4 | 0.006627 | 7.784 | +1 | 9 |
| - | - | 1493 | 859.5 | - | - | 0 | - |
| 4 | z | 8866 | 863.5 | 0.0001335 | 0.1546 | +1 | 9 |
| - | - | 7690 | 864.5 | - | - | 0 | - |
| - | - | 2980 | 865.5 | - | - | 0 | - |
| - | - | 848.8 | 866.5 | - | - | 0 | - |
| - | - | 3517 | 868.4 | - | - | 0 | - |
| 9 | c | 2.181E+04 | 869.5 | 0.0001022 | 0.1176 | +1 | 9 |
| - | - | 1.109E+04 | 870.5 | - | - | 0 | - |
| - | - | 2844 | 871.5 | - | - | 0 | - |
| - | - | 791.7 | 878.5 | - | - | 0 | - |
| 4 | y | 9478 | 879.5 | 0.000219 | 0.249 | +1 | 9 |
| - | - | 4368 | 880.5 | - | - | 0 | - |
| - | - | 1806 | 881.5 | - | - | 0 | - |
| - | - | 645.3 | 903.5 | - | - | 0 | - |
| - | - | 903.2 | 910.5 | - | - | 0 | - |
| 3 | z | 9866 | 962.5 | 0.0001652 | 0.1716 | +1 | 10 |
| - | - | 5061 | 963.5 | - | - | 0 | - |
| - | - | 1124 | 964.5 | - | - | 0 | - |
| 3 | y | 4819 | 978.6 | 0.0002735 | 0.2795 | +1 | 10 |
| - | - | 2371 | 979.6 | - | - | 0 | - |
| 10 | c | 1927 | 980.5 | 0.01016 | 10.36 | +1 | 10 |
| - | - | 1615 | 981.5 | - | - | 0 | - |
| 10 | c | 1.248E+04 | 997.5 | 0.0002776 | 0.2783 | +1 | 10 |
| - | - | 7000 | 998.5 | - | - | 0 | - |
| - | - | 2214 | 999.6 | - | - | 0 | - |
| - | - | 1135 | 1067 | - | - | 0 | - |
| 11 | c | 2688 | 1094 | 0.002632 | 2.407 | +1 | 11 |
| - | - | 1955 | 1095 | - | - | 0 | - |
| - | - | 1373 | 1096 | - | - | 0 | - |
| 2 | z | 8145 | 1110 | 0.0005859 | 0.5281 | +1 | 11 |
| 11 | c | 2.311E+04 | 1111 | 0.003165 | 2.85 | +1 | 11 |
| - | - | 1.434E+04 | 1112 | - | - | 0 | - |
| - | - | 4891 | 1113 | - | - | 0 | - |
| - | - | 6754 | 1158 | - | - | 0 | - |
| - | - | 5368 | 1159 | - | - | 0 | - |
| - | - | 2157 | 1160 | - | - | 0 | - |
| - | - | 836.4 | 1183 | - | - | 0 | - |
| - | - | 1013 | 1184 | - | - | 0 | - |
| - | - | 2622 | 1186 | - | - | 0 | - |
| - | - | 2103 | 1187 | - | - | 0 | - |
| - | - | 829.4 | 1188 | - | - | 0 | - |
| - | - | 5341 | 1195 | - | - | 0 | - |
| - | - | 4316 | 1196 | - | - | 0 | - |
| - | - | 4.063E+04 | 1197 | - | - | 0 | - |
| - | - | 2.536E+04 | 1198 | - | - | 0 | - |
| - | - | 1.06E+04 | 1199 | - | - | 0 | - |
| - | - | 1754 | 1200 | - | - | 0 | - |
| - | - | 1884 | 1211 | - | - | 0 | - |
| - | - | 1085 | 1212 | - | - | 0 | - |
| - | - | 2.021E+04 | 1213 | - | - | 0 | - |
| - | - | 4.193E+04 | 1214 | - | - | 0 | - |
| - | - | 2.496E+04 | 1215 | - | - | 0 | - |
| - | - | 8511 | 1216 | - | - | 0 | - |
| - | - | 1141 | 1217 | - | - | 0 | - |
| - | - | 974.7 | 1766 | - | - | 0 | - |
| - | - | 628.7 | 1774 | - | - | 0 | - |
| - | - | 790.2 | 1782 | - | - | 0 | - |
| - | - | 860.4 | 1800 | - | - | 0 | - |
| - | - | 755.6 | 1802 | - | - | 0 | - |
| - | - | 741.7 | 1819 | - | - | 0 | - |
| - | - | 753.4 | 1820 | - | - | 0 | - |
| - | - | 709.3 | 1821 | - | - | 0 | - |
| - | - | 764.8 | 1822 | - | - | 0 | - |
| - | - | 847.3 | 1823 | - | - | 0 | - |
| - | - | 735.5 | 3023 | - | - | 0 | - |

m/z Charge Intensity FragmentType MassShift Position
120.06568145751953 0 2394.144 y 11
129.10240173339844 0 962.30365
133.06155395507812 0 450.61517
133.06922912597656 0 861.1501
133.07412719726562 0 425.19363
133.08612060546875 0 4283.1133
133.98985290527344 0 454.81253
134.0139617919922 0 449.6913
143.10655212402344 0 429.25806
148.95443725585938 0 572.26807
149.0451202392578 0 1281.7926
158.98585510253906 0 393.22427
166.85940551757812 0 457.14478
167.055908203125 0 667.5023
171.14950561523438 0 686.71643
173.43858337402344 0 1187.1154
175.09681701660156 0 601.01
177.10455322265625 0 482.0244
177.1121368408203 0 2952.4849
184.1797332763672 0 484.26093
185.52523803710938 0 515.7565
187.13311767578125 0 696.3854
191.09194946289062 0 801.9577
192.13230895996094 0 427.3746
199.14442443847656 0 1091.8514
205.10708618164062 0 694.65784
207.11280822753906 0 4031.0212
208.1164093017578 0 644.32385
215.1398468017578 0 1069.9508 y Water loss 10
221.08468627929688 0 3554.0425
221.13816833496094 0 621.0232
225.0427703857422 0 3116.218
227.0228271484375 0 615.24274
233.1498260498047 0 2435.582 y 10
235.10794067382812 0 9697.658
235.11978149414062 0 764.194
236.11102294921875 0 741.11365
239.09521484375 0 7501.439
239.14915466308594 0 613.1055
240.0980682373047 0 659.02374
247.14395141601562 0 983.3567
270.4299621582031 0 525.6587
283.17864990234375 0 544.5757
295.1034240722656 0 5996.527
296.1041564941406 0 999.38165
299.0621337890625 0 2950.774
313.1142578125 0 815.0293
334.1550598144531 0 1030.2709
334.1765441894531 0 13610.455
335.1792297363281 0 1353.0544
337.2217102050781 0 956.4019
355.0707092285156 0 950.4267
369.12176513671875 0 3217.9275
399.1939392089844 0 552.7179
401.2138366699219 0 1334.3347
405.2508850097656 0 732.6283
406.2528076171875 0 647.4557
433.24462890625 0 5062.5244
434.2498474121094 0 1222.3679
444.2495422363281 0 667.3381
445.26531982421875 0 1276.5902 y Ammonia loss 8
448.36822509765625 0 581.2846
481.24273681640625 0 588.3412
504.2537536621094 0 3486.0903
504.3031005859375 0 2903.0303
505.25909423828125 0 1110.7355
505.3073425292969 0 989.0354
508.48388671875 0 719.2002
514.298583984375 0 1244.1526
519.313232421875 0 1704.2104 y 7
535.2905883789062 0 828.55145
540.42724609375 0 600.7258
545.316650390625 0 590.8652
548.2823486328125 0 1213.406
552.318603515625 0 665.81396
560.3165283203125 0 1829.9202 z 6
561.3236694335938 0 7326.816
562.3311157226562 0 1162.2023
565.9561157226562 0 611.7457
575.3269653320312 0 971.82135
576.3353881835938 0 7235.2144 y 6
577.3372192382812 0 1326.9056
580.31298828125 0 1545.5638
585.923095703125 0 582.8171
589.3429565429688 0 7278.83
589.3831787109375 0 707.4159
590.3458862304688 0 2281.3035
591.3562622070312 0 1204.3944
606.2774047851562 0 612.7397
606.3681640625 0 782.2028
606.7791748046875 0 980.956
607.3128662109375 0 826.9758
617.3379516601562 0 3875.6577 z 5
618.34521484375 0 9296.848
619.3469848632812 0 1933.4584
632.3486328125 0 3347.588
633.357177734375 0 15533.174 y 5
634.3591918945312 0 4162.567
635.3582763671875 0 622.37164
647.3519287109375 0 782.5376
661.3648071289062 0 938.09784
663.3798217773438 0 660.7666
668.3731079101562 0 917.55457
710.3745727539062 0 3792.0366
711.3798217773438 0 2669.9272 c 6
712.3829956054688 0 1256.6821
764.4067993164062 0 7413.884 z 4
765.4132690429688 0 9853.929
766.4161987304688 0 4269.966
767.400390625 0 2073.4539
768.402587890625 0 1717.8553 c 7
779.4182739257812 0 4077.7747
780.4246826171875 0 19436.49 y 4
781.4287109375 0 6573.848
782.4283447265625 0 1546.8345
785.3273315429688 0 595.4629
850.4381713867188 0 804.5376
851.4343872070312 0 690.3344 c Water loss 8
859.4999389648438 0 1492.7289
863.474853515625 0 8865.647 z 3
864.4800415039062 0 7689.8916
865.4822387695312 0 2979.8242
866.487060546875 0 848.75116
868.4435424804688 0 3517.1094
869.4514770507812 0 21805.562 c 8
870.454345703125 0 11090.435
871.4584350585938 0 2844.0977
878.4885864257812 0 791.7174
879.4932250976562 0 9478.073 y 3
880.4957885742188 0 4368.4165
881.5004272460938 0 1805.6149
903.4680786132812 0 645.25806
910.4699096679688 0 903.17163
962.54296875 0 9866.398 z 2
963.5445556640625 0 5061.397
964.5481567382812 0 1124.0465
978.5615844726562 0 4818.771 y 2
979.565673828125 0 2370.9067
980.5301513671875 0 1927.1799 c Ammonia loss 9
981.5182495117188 0 1615.0557
997.5462646484375 0 12476.844 c 9
998.5493774414062 0 7000.243
999.5536499023438 0 2214.076
1066.6165771484375 0 1135.2866
1093.606689453125 0 2688.0703 c Ammonia loss 10
1094.6075439453125 0 1954.6603
1095.618408203125 0 1372.9197
1109.6109619140625 0 8144.604 z 1
1110.62744140625 0 23114.898 c 10
1111.631103515625 0 14336.38
1112.634521484375 0 4890.7476
1157.6063232421875 0 6754.184
1158.60791015625 0 5368.0054
1159.6116943359375 0 2156.8481
1182.600830078125 0 836.4035
1183.613525390625 0 1012.61566
1185.6768798828125 0 2622.0344
1186.67919921875 0 2103.2688
1187.67822265625 0 829.3597
1194.703857421875 0 5341.278
1195.6932373046875 0 4315.564
1196.6439208984375 0 40632.742
1197.6458740234375 0 25358.996
1198.6455078125 0 10601.993
1199.6376953125 0 1753.6415
1210.6580810546875 0 1883.912
1211.660400390625 0 1085.0702
1212.6619873046875 0 20214.373
1213.667236328125 0 41933.492
1214.6705322265625 0 24958.62
1215.672607421875 0 8510.898
1216.561767578125 0 1141.4896
1765.9708251953125 0 974.66504
1774.365234375 0 628.65497
1781.9327392578125 0 790.24713
1799.7451171875 0 860.43933
1801.9586181640625 0 755.62775
1819.0697021484375 0 741.6924
1819.9493408203125 0 753.3967
1820.9534912109375 0 709.29865
1821.9576416015625 0 764.81384
1822.9530029296875 0 847.3443
3022.677978515625 0 735.49554

Spectrum Details

|  |  |
| --- | --- |
| Matched peaks? Matched peaks The total absolute number of peaks matched. Additionally in brackets the total fraction of peaks matched and the total number of peaks is shown. | 24 (13.11% of 183) |
| FDR? FDR The false discovery rate estimated for this peptide. It is calculated by matching all theoretical fragments with a non-integer shift with the raw peaks for this spectrum. This is done with 40 different shifts. The resulting percentage is the average number of annotated peaks over the number of annotated peaks with the correct spectrum. | 0.89% |
| Satellite FDR? Satellite FDR See the FDR for details on its calculation. This satellite ion specific FDR only contains the satellite ions (d/w) for I/L/J positions. | ∞ |
| PSM Score? PSM Score The PSM Score as given by Hecklib to this annotated spectrum. It is shown with three significant figures. | 274 |

#### Spectrum 9661? Spectrum 9661 The raw spectrum of this peptide as annotated by Hecklib. The fragments are coloured according to ion type (see legend). Any peaks with a star '\*' as text can be hovered over to see the full details, first the ion type second the mass shift type. By hovering over the amino acids in the peptide or ions in the legend the corresponding peaks are highlighted. By toggling the 'Unassigned' label you can turn the background (unassigned) peaks on or off in the plot. By updating the slider in the Ion legend you can update the spectrum to only show the top X% of the peaks with labels. The top X% means any peak that is within X% of the highest intensity. By dragging in the spectrum you can zoom in to a specific part of the spectrum and use 'Zoom Out' to get back to the original zoom level. The annotation of the spectrum is based on the given sequence in the peptides file and is done with different software so inconsistencies are likely. The peaks are annotated based on the given sequence, with 20 ppm tolerance.

Copy Data

##### Spectrum 9661 (TSV)

###### Preview

```
Loading example...
```

*Click on the button to copy the data to your clipboard.*

Mz MinMz MaxIntensity Max

WidthHeightPeptide font sizePeptide stroke widthSpectrum font sizeSpectrum stroke widthCompact peptide

Ion legend

wxyz

abcd

OtherUnassignedIon^Charge^_Position_Show for top:%

SFVVFGGGTKJT

01.74e+43.48e+45.23e+46.97e+4

Zoom Out

y^+1^_1_a^+1^_2_b^+2^_4_y^+1^_2_b^+1^_2_y^+1^_2_b^+1^_2_b^+1^_3_b^+1^_3_y^+1^_3_y^+1^_3_y^+2^_8_b^+1^_4_b^+1^_4_y^+1^_4_y^+2^_10_b^+2^_10_y^+1^_5_b^+1^_5_y^+1^_6_b^+1^_5_\*y^+1^_7_b^+1^_6_y^+1^_7_b^+1^_8_y^+1^_8_y^+1^_8_b^+1^_9_b^+1^_9_y^+1^_9_b^+1^_10_b^+1^_10_y^+1^_10_b^+1^_10_b^+1^_11_b^+1^_11_

0765153122963062

Fragment Matches Table

Show background peaks

| Position | Ion type | Intensity | mz Theoretical | mz Error (Th) | mz Error (ppm) | Charge | Series Number |
| --- | --- | --- | --- | --- | --- | --- | --- |
| - | - | 673 | 120.1 | - | - | 0 | - |
| 12 | y | 6035 | 120.1 | 0.0003907 | 3.254 | +1 | 1 |
| - | - | 2.177E+04 | 120.1 | - | - | 0 | - |
| - | - | 1666 | 121.1 | - | - | 0 | - |
| - | - | 542.9 | 126.1 | - | - | 0 | - |
| - | - | 778.8 | 127.1 | - | - | 0 | - |
| - | - | 352.1 | 127.1 | - | - | 0 | - |
| - | - | 719.3 | 127.1 | - | - | 0 | - |
| - | - | 469.2 | 128 | - | - | 0 | - |
| - | - | 786.4 | 129.1 | - | - | 0 | - |
| - | - | 3.399E+04 | 129.1 | - | - | 0 | - |
| - | - | 570.8 | 130.1 | - | - | 0 | - |
| - | - | 387.1 | 130.1 | - | - | 0 | - |
| - | - | 547.9 | 130.1 | - | - | 0 | - |
| - | - | 2185 | 130.1 | - | - | 0 | - |
| - | - | 1286 | 131 | - | - | 0 | - |
| - | - | 410.1 | 131.1 | - | - | 0 | - |
| - | - | 980 | 131.1 | - | - | 0 | - |
| - | - | 402.4 | 132.1 | - | - | 0 | - |
| - | - | 680.2 | 132.1 | - | - | 0 | - |
| - | - | 834.2 | 133.1 | - | - | 0 | - |
| - | - | 3760 | 133.1 | - | - | 0 | - |
| - | - | 4027 | 136.1 | - | - | 0 | - |
| - | - | 412.1 | 137.1 | - | - | 0 | - |
| - | - | 412.3 | 139.7 | - | - | 0 | - |
| - | - | 547.2 | 140.1 | - | - | 0 | - |
| - | - | 802.1 | 141.1 | - | - | 0 | - |
| - | - | 788.3 | 141.1 | - | - | 0 | - |
| - | - | 432.6 | 142.1 | - | - | 0 | - |
| - | - | 1164 | 144.1 | - | - | 0 | - |
| - | - | 429.4 | 144.1 | - | - | 0 | - |
| - | - | 598.1 | 148.9 | - | - | 0 | - |
| - | - | 2181 | 149 | - | - | 0 | - |
| - | - | 425.5 | 151.1 | - | - | 0 | - |
| - | - | 815.6 | 152.1 | - | - | 0 | - |
| - | - | 1555 | 155.1 | - | - | 0 | - |
| - | - | 445.7 | 155.6 | - | - | 0 | - |
| - | - | 933.8 | 156.1 | - | - | 0 | - |
| - | - | 620.7 | 157.1 | - | - | 0 | - |
| - | - | 1078 | 157.1 | - | - | 0 | - |
| - | - | 581.6 | 158.1 | - | - | 0 | - |
| - | - | 1203 | 159.1 | - | - | 0 | - |
| - | - | 734.1 | 159.1 | - | - | 0 | - |
| - | - | 1426 | 159.1 | - | - | 0 | - |
| - | - | 3104 | 162.1 | - | - | 0 | - |
| - | - | 590.6 | 163.1 | - | - | 0 | - |
| - | - | 1213 | 165.1 | - | - | 0 | - |
| - | - | 4196 | 167.1 | - | - | 0 | - |
| - | - | 1139 | 167.1 | - | - | 0 | - |
| - | - | 995.5 | 169.1 | - | - | 0 | - |
| - | - | 814.1 | 170.1 | - | - | 0 | - |
| - | - | 6415 | 171.1 | - | - | 0 | - |
| - | - | 417.9 | 171.2 | - | - | 0 | - |
| - | - | 1686 | 172.1 | - | - | 0 | - |
| - | - | 783.5 | 172.2 | - | - | 0 | - |
| - | - | 1066 | 173.1 | - | - | 0 | - |
| - | - | 903.2 | 173.1 | - | - | 0 | - |
| - | - | 1117 | 173.4 | - | - | 0 | - |
| - | - | 580.9 | 175.1 | - | - | 0 | - |
| - | - | 1604 | 175.1 | - | - | 0 | - |
| - | - | 841.3 | 176.1 | - | - | 0 | - |
| - | - | 987.3 | 177.1 | - | - | 0 | - |
| - | - | 2080 | 177.1 | - | - | 0 | - |
| - | - | 558.5 | 183.1 | - | - | 0 | - |
| - | - | 910.2 | 183.1 | - | - | 0 | - |
| - | - | 1395 | 185.1 | - | - | 0 | - |
| - | - | 606.9 | 185.1 | - | - | 0 | - |
| - | - | 562.1 | 187.1 | - | - | 0 | - |
| - | - | 780.4 | 187.1 | - | - | 0 | - |
| - | - | 560.4 | 187.1 | - | - | 0 | - |
| - | - | 522 | 190.1 | - | - | 0 | - |
| - | - | 569.9 | 193.1 | - | - | 0 | - |
| - | - | 1518 | 197.2 | - | - | 0 | - |
| - | - | 1157 | 198.1 | - | - | 0 | - |
| - | - | 472.2 | 199.1 | - | - | 0 | - |
| - | - | 3866 | 199.1 | - | - | 0 | - |
| - | - | 548.4 | 199.2 | - | - | 0 | - |
| - | - | 511.6 | 199.2 | - | - | 0 | - |
| - | - | 748.2 | 200.1 | - | - | 0 | - |
| - | - | 897.8 | 201.1 | - | - | 0 | - |
| - | - | 1246 | 203.1 | - | - | 0 | - |
| - | - | 1677 | 205.1 | - | - | 0 | - |
| 2 | a | 6.899E+04 | 207.1 | 0.0003245 | 1.567 | +1 | 2 |
| 4 | b | 7753 | 208.1 | 0.004128 | 19.84 | +2 | 4 |
| - | - | 922.9 | 212.1 | - | - | 0 | - |
| - | - | 2829 | 212.1 | - | - | 0 | - |
| - | - | 624.2 | 214.1 | - | - | 0 | - |
| 11 | y | 3060 | 215.1 | 0.0004922 | 2.288 | +1 | 2 |
| - | - | 927.6 | 216.1 | - | - | 0 | - |
| - | - | 1358 | 216.1 | - | - | 0 | - |
| 2 | b | 778.3 | 217.1 | 0.0001817 | 0.8371 | +1 | 2 |
| - | - | 7077 | 219.1 | - | - | 0 | - |
| - | - | 545.8 | 220.2 | - | - | 0 | - |
| - | - | 4165 | 221.1 | - | - | 0 | - |
| - | - | 730.4 | 223.1 | - | - | 0 | - |
| - | - | 612.7 | 224.1 | - | - | 0 | - |
| - | - | 953 | 224.1 | - | - | 0 | - |
| - | - | 649.3 | 224.2 | - | - | 0 | - |
| - | - | 1.027E+04 | 225 | - | - | 0 | - |
| - | - | 1561 | 226 | - | - | 0 | - |
| - | - | 807.2 | 226.1 | - | - | 0 | - |
| - | - | 3255 | 226.2 | - | - | 0 | - |
| - | - | 861.9 | 227 | - | - | 0 | - |
| - | - | 901.2 | 228.1 | - | - | 0 | - |
| - | - | 6355 | 230.2 | - | - | 0 | - |
| - | - | 816.2 | 233.1 | - | - | 0 | - |
| 11 | y | 6096 | 233.1 | 0.000395 | 1.694 | +1 | 2 |
| - | - | 1501 | 233.2 | - | - | 0 | - |
| - | - | 631.1 | 234.2 | - | - | 0 | - |
| 2 | b | 5.653E+04 | 235.1 | 0.000344 | 1.463 | +1 | 2 |
| - | - | 8553 | 236.1 | - | - | 0 | - |
| - | - | 883.1 | 237.1 | - | - | 0 | - |
| - | - | 1.375E+04 | 239.1 | - | - | 0 | - |
| - | - | 731.8 | 239.1 | - | - | 0 | - |
| - | - | 1472 | 240.1 | - | - | 0 | - |
| - | - | 816.5 | 240.1 | - | - | 0 | - |
| - | - | 529.2 | 241.2 | - | - | 0 | - |
| - | - | 635.8 | 242.2 | - | - | 0 | - |
| - | - | 831.1 | 242.2 | - | - | 0 | - |
| - | - | 1610 | 245.1 | - | - | 0 | - |
| - | - | 546.3 | 246.1 | - | - | 0 | - |
| - | - | 5716 | 247.1 | - | - | 0 | - |
| - | - | 992.9 | 248.1 | - | - | 0 | - |
| - | - | 508.6 | 249.2 | - | - | 0 | - |
| - | - | 2138 | 251.1 | - | - | 0 | - |
| - | - | 925.2 | 254.1 | - | - | 0 | - |
| - | - | 3845 | 255.1 | - | - | 0 | - |
| - | - | 1104 | 261.1 | - | - | 0 | - |
| - | - | 1365 | 261.2 | - | - | 0 | - |
| - | - | 929.1 | 262.1 | - | - | 0 | - |
| - | - | 595.3 | 263.1 | - | - | 0 | - |
| - | - | 809.7 | 268.2 | - | - | 0 | - |
| - | - | 3790 | 269.2 | - | - | 0 | - |
| - | - | 652.1 | 272.1 | - | - | 0 | - |
| - | - | 4005 | 273.1 | - | - | 0 | - |
| - | - | 1185 | 274.1 | - | - | 0 | - |
| - | - | 601.6 | 277.2 | - | - | 0 | - |
| - | - | 646.5 | 279.1 | - | - | 0 | - |
| - | - | 1143 | 281.1 | - | - | 0 | - |
| - | - | 1205 | 287.2 | - | - | 0 | - |
| - | - | 4109 | 289.2 | - | - | 0 | - |
| - | - | 872.9 | 290.2 | - | - | 0 | - |
| - | - | 760.3 | 291.1 | - | - | 0 | - |
| - | - | 3578 | 295.1 | - | - | 0 | - |
| - | - | 633.2 | 296.1 | - | - | 0 | - |
| - | - | 6717 | 299.1 | - | - | 0 | - |
| - | - | 545.9 | 299.1 | - | - | 0 | - |
| - | - | 1093 | 300.1 | - | - | 0 | - |
| - | - | 780.5 | 301.2 | - | - | 0 | - |
| - | - | 730.4 | 309.2 | - | - | 0 | - |
| - | - | 840.1 | 311.2 | - | - | 0 | - |
| - | - | 1182 | 313.1 | - | - | 0 | - |
| - | - | 603.1 | 315.7 | - | - | 0 | - |
| 3 | b | 2327 | 316.2 | 0.0005392 | 1.705 | +1 | 3 |
| - | - | 995.2 | 319.1 | - | - | 0 | - |
| - | - | 767.7 | 323.2 | - | - | 0 | - |
| - | - | 2614 | 323.2 | - | - | 0 | - |
| - | - | 681.1 | 324.2 | - | - | 0 | - |
| - | - | 747.8 | 325.2 | - | - | 0 | - |
| - | - | 2772 | 326.2 | - | - | 0 | - |
| 3 | b | 3.709E+04 | 334.2 | 0.0004115 | 1.231 | +1 | 3 |
| - | - | 6970 | 335.2 | - | - | 0 | - |
| - | - | 1093 | 337.2 | - | - | 0 | - |
| - | - | 737.7 | 341.2 | - | - | 0 | - |
| 10 | y | 690.4 | 343.2 | 0.0004004 | 1.167 | +1 | 3 |
| - | - | 685.9 | 346.2 | - | - | 0 | - |
| - | - | 954.8 | 352.2 | - | - | 0 | - |
| - | - | 708.2 | 354.2 | - | - | 0 | - |
| - | - | 691.3 | 357.2 | - | - | 0 | - |
| - | - | 650.3 | 359 | - | - | 0 | - |
| - | - | 1010 | 360.2 | - | - | 0 | - |
| 10 | y | 3885 | 361.2 | 0.0001618 | 0.448 | +1 | 3 |
| - | - | 681.7 | 364.2 | - | - | 0 | - |
| - | - | 2243 | 369.1 | - | - | 0 | - |
| - | - | 787.7 | 374.2 | - | - | 0 | - |
| - | - | 911 | 380.2 | - | - | 0 | - |
| - | - | 1028 | 382.2 | - | - | 0 | - |
| - | - | 3788 | 383.2 | - | - | 0 | - |
| 5 | y | 918.8 | 390.7 | 0.000394 | 1.008 | +2 | 8 |
| - | - | 1094 | 392.2 | - | - | 0 | - |
| - | - | 1087 | 400.3 | - | - | 0 | - |
| - | - | 4936 | 401.2 | - | - | 0 | - |
| - | - | 4436 | 402.2 | - | - | 0 | - |
| - | - | 704.3 | 402.2 | - | - | 0 | - |
| - | - | 965.7 | 403.2 | - | - | 0 | - |
| - | - | 2408 | 405.2 | - | - | 0 | - |
| - | - | 2223 | 408.3 | - | - | 0 | - |
| 4 | b | 1397 | 415.2 | 0.0005457 | 1.314 | +1 | 4 |
| - | - | 592.7 | 416.2 | - | - | 0 | - |
| - | - | 1823 | 420.2 | - | - | 0 | - |
| - | - | 1799 | 428.8 | - | - | 0 | - |
| 4 | b | 9163 | 433.2 | 0.0001433 | 0.3308 | +1 | 4 |
| - | - | 2638 | 434.2 | - | - | 0 | - |
| - | - | 1813 | 436.2 | - | - | 0 | - |
| - | - | 1148 | 439.3 | - | - | 0 | - |
| - | - | 605.2 | 448.1 | - | - | 0 | - |
| - | - | 927 | 453.3 | - | - | 0 | - |
| - | - | 594.4 | 453.5 | - | - | 0 | - |
| - | - | 702.2 | 456.2 | - | - | 0 | - |
| - | - | 2100 | 457.3 | - | - | 0 | - |
| 9 | y | 768.2 | 462.3 | 0.0002329 | 0.5038 | +1 | 4 |
| - | - | 1026 | 463.2 | - | - | 0 | - |
| 3 | y | 797.7 | 480.8 | 0.0003782 | 0.7866 | +2 | 10 |
| - | - | 1752 | 481.2 | - | - | 0 | - |
| 10 | b | 805.1 | 482.3 | 0.005304 | 11 | +2 | 10 |
| - | - | 1220 | 486.3 | - | - | 0 | - |
| - | - | 533 | 490.1 | - | - | 0 | - |
| - | - | 2385 | 496.3 | - | - | 0 | - |
| - | - | 909.3 | 497.3 | - | - | 0 | - |
| - | - | 761.4 | 502.2 | - | - | 0 | - |
| - | - | 701.9 | 504.2 | - | - | 0 | - |
| - | - | 563.9 | 511.3 | - | - | 0 | - |
| - | - | 781.6 | 512.8 | - | - | 0 | - |
| - | - | 5886 | 514.3 | - | - | 0 | - |
| - | - | 1282 | 515.3 | - | - | 0 | - |
| - | - | 1076 | 519.3 | - | - | 0 | - |
| 8 | y | 1479 | 519.3 | 0.0008864 | 1.707 | +1 | 5 |
| - | - | 985.3 | 520.3 | - | - | 0 | - |
| - | - | 2604 | 530.3 | - | - | 0 | - |
| - | - | 673.2 | 531.3 | - | - | 0 | - |
| - | - | 856.9 | 532.3 | - | - | 0 | - |
| - | - | 875.2 | 533.3 | - | - | 0 | - |
| - | - | 753 | 535.3 | - | - | 0 | - |
| - | - | 618.2 | 537.6 | - | - | 0 | - |
| - | - | 4985 | 548.3 | - | - | 0 | - |
| - | - | 1724 | 549.3 | - | - | 0 | - |
| - | - | 699.6 | 552.3 | - | - | 0 | - |
| 5 | b | 1145 | 562.3 | 0.0006991 | 1.243 | +1 | 5 |
| - | - | 697 | 563.3 | - | - | 0 | - |
| - | - | 1635 | 573.3 | - | - | 0 | - |
| 7 | y | 3847 | 576.3 | 0.0005578 | 0.9679 | +1 | 6 |
| - | - | 1230 | 577.3 | - | - | 0 | - |
| 5 | b | 2304 | 580.3 | 0.0006436 | 1.109 | +1 | 5 |
| - | - | 757.6 | 580.7 | - | - | 0 | - |
| - | - | 1016 | 581.3 | - | - | 0 | - |
| - | - | 966.4 | 585.8 | - | - | 0 | - |
| - | - | 667 | 586.3 | - | - | 0 | - |
| - | - | 662.4 | 587.3 | - | - | 0 | - |
| 0 | Precursor | 1491 | 597.8 | 0.0009383 | 1.569 | +2 | -1 |
| - | - | 1163 | 598.3 | - | - | 0 | - |
| - | - | 2740 | 606.3 | - | - | 0 | - |
| - | - | 1694 | 606.5 | - | - | 0 | - |
| - | - | 1528 | 606.7 | - | - | 0 | - |
| - | - | 1134 | 606.8 | - | - | 0 | - |
| - | - | 945.7 | 606.9 | - | - | 0 | - |
| - | - | 1490 | 607.1 | - | - | 0 | - |
| - | - | 1382 | 607.3 | - | - | 0 | - |
| - | - | 635.7 | 609.3 | - | - | 0 | - |
| 6 | y | 1090 | 615.3 | 0.002398 | 3.897 | +1 | 7 |
| 6 | b | 1058 | 619.3 | 0.0002371 | 0.3829 | +1 | 6 |
| - | - | 681.1 | 622.3 | - | - | 0 | - |
| - | - | 1543 | 629.3 | - | - | 0 | - |
| 6 | y | 1.807E+04 | 633.4 | 0.0003784 | 0.5974 | +1 | 7 |
| - | - | 7379 | 634.4 | - | - | 0 | - |
| - | - | 863.5 | 635.4 | - | - | 0 | - |
| - | - | 1078 | 642.4 | - | - | 0 | - |
| - | - | 2876 | 643.4 | - | - | 0 | - |
| - | - | 1415 | 644.4 | - | - | 0 | - |
| - | - | 1848 | 647.4 | - | - | 0 | - |
| - | - | 724 | 648.4 | - | - | 0 | - |
| - | - | 6142 | 661.4 | - | - | 0 | - |
| - | - | 1803 | 662.4 | - | - | 0 | - |
| - | - | 1470 | 679.4 | - | - | 0 | - |
| - | - | 1335 | 697.9 | - | - | 0 | - |
| - | - | 1550 | 698.1 | - | - | 0 | - |
| - | - | 1572 | 742.4 | - | - | 0 | - |
| - | - | 1223 | 743.4 | - | - | 0 | - |
| - | - | 1001 | 746.4 | - | - | 0 | - |
| - | - | 914.1 | 747.4 | - | - | 0 | - |
| 8 | b | 880.2 | 751.4 | 0.002718 | 3.617 | +1 | 8 |
| - | - | 753.2 | 756.4 | - | - | 0 | - |
| - | - | 4470 | 760.4 | - | - | 0 | - |
| - | - | 1961 | 761.4 | - | - | 0 | - |
| 5 | y | 2120 | 762.4 | 0.001624 | 2.13 | +1 | 8 |
| 5 | y | 2.528E+04 | 780.4 | 0.0007748 | 0.9928 | +1 | 8 |
| - | - | 1.095E+04 | 781.4 | - | - | 0 | - |
| - | - | 2324 | 782.4 | - | - | 0 | - |
| - | - | 1101 | 815.5 | - | - | 0 | - |
| - | - | 763.7 | 815.9 | - | - | 0 | - |
| 9 | b | 1457 | 834.4 | 0.001685 | 2.019 | +1 | 9 |
| - | - | 699.2 | 841.7 | - | - | 0 | - |
| 9 | b | 2472 | 852.4 | 0.001019 | 1.195 | +1 | 9 |
| - | - | 962.5 | 853.4 | - | - | 0 | - |
| - | - | 1869 | 859.5 | - | - | 0 | - |
| - | - | 1559 | 860.5 | - | - | 0 | - |
| 4 | y | 1.532E+04 | 879.5 | 0.001562 | 1.776 | +1 | 9 |
| - | - | 7752 | 880.5 | - | - | 0 | - |
| - | - | 1772 | 881.5 | - | - | 0 | - |
| - | - | 922 | 895.5 | - | - | 0 | - |
| 10 | b | 1024 | 962.5 | 0.003084 | 3.204 | +1 | 10 |
| 10 | b | 826.6 | 963.5 | 0.01882 | 19.54 | +1 | 10 |
| - | - | 680.1 | 977.5 | - | - | 0 | - |
| 3 | y | 7302 | 978.6 | 0.00241 | 2.463 | +1 | 10 |
| - | - | 4223 | 979.6 | - | - | 0 | - |
| 10 | b | 3815 | 980.5 | 0.002468 | 2.517 | +1 | 10 |
| - | - | 2070 | 981.5 | - | - | 0 | - |
| 11 | b | 984.9 | 1076 | 0.002916 | 2.711 | +1 | 11 |
| 11 | b | 3348 | 1094 | 0.004082 | 3.732 | +1 | 11 |
| - | - | 2383 | 1095 | - | - | 0 | - |
| - | - | 663.5 | 1096 | - | - | 0 | - |
| - | - | 610.8 | 1889 | - | - | 0 | - |
| - | - | 672.7 | 2450 | - | - | 0 | - |
| - | - | 617.1 | 2464 | - | - | 0 | - |
| - | - | 810.7 | 2725 | - | - | 0 | - |
| - | - | 695.2 | 3031 | - | - | 0 | - |

m/z Charge Intensity FragmentType MassShift Position
120.06188201904297 0 672.9735
120.06591033935547 0 6035.075 y 11
120.08116149902344 0 21771.406
121.08452606201172 0 1666.2638
126.05540466308594 0 542.85614
127.05068969726562 0 778.76294
127.0762939453125 0 352.07285
127.08677673339844 0 719.32715
128.0457763671875 0 469.16968
129.06622314453125 0 786.3627
129.10260009765625 0 33988.953
130.05027770996094 0 570.8297
130.06117248535156 0 387.122
130.08663940429688 0 547.92206
130.10595703125 0 2184.7744
131.04554748535156 0 1285.9916
131.0704345703125 0 410.1383
131.08169555664062 0 980.0355
132.08143615722656 0 402.35223
132.1017303466797 0 680.18176
133.0611572265625 0 834.1754
133.08624267578125 0 3759.8267
136.07611083984375 0 4027.455
137.08106994628906 0 412.0851
139.6734161376953 0 412.2514
140.081787109375 0 547.1924
141.06600952148438 0 802.09607
141.10255432128906 0 788.2614
142.06121826171875 0 432.59103
144.08128356933594 0 1163.7428
144.10279846191406 0 429.40384
148.94686889648438 0 598.09406
149.04518127441406 0 2180.7263
151.0866241455078 0 425.53918
152.0708465576172 0 815.61255
155.08175659179688 0 1555.2725
155.6284942626953 0 445.6542
156.0770263671875 0 933.7993
157.0614776611328 0 620.6933
157.09732055664062 0 1077.9178
158.0929412841797 0 581.5963
159.0767822265625 0 1203.4171
159.0923309326172 0 734.05255
159.11309814453125 0 1425.7826
162.09173583984375 0 3104.3662
163.09556579589844 0 590.6109
165.1028289794922 0 1212.7083
167.0558624267578 0 4195.5903
167.08175659179688 0 1139.3947
169.09751892089844 0 995.51746
170.09291076660156 0 814.0924
171.14950561523438 0 6415.4194
171.22776794433594 0 417.8696
172.072021484375 0 1686.0779
172.15293884277344 0 783.4813
173.09255981445312 0 1065.8445
173.12872314453125 0 903.2209
173.4386444091797 0 1117.4111
175.0966796875 0 580.87024
175.11940002441406 0 1603.9492
176.10733032226562 0 841.27704
177.10284423828125 0 987.3392
177.1122589111328 0 2080.1052
183.07704162597656 0 558.518
183.11309814453125 0 910.18445
185.09254455566406 0 1394.5054
185.12937927246094 0 606.855
187.07101440429688 0 562.0784
187.10787963867188 0 780.38965
187.14512634277344 0 560.3556
190.0684356689453 0 521.96375
193.09738159179688 0 569.8724
197.16513061523438 0 1518.3518
198.0875244140625 0 1156.7172
199.10784912109375 0 472.17465
199.14447021484375 0 3866.0415
199.1698455810547 0 548.4484
199.1803741455078 0 511.6387
200.13941955566406 0 748.16437
201.12330627441406 0 897.8
203.10299682617188 0 1245.9985
205.09718322753906 0 1677.2477
207.11312866210938 0 68994.055 a 1
208.1165008544922 0 7753.128 b Water loss 3
212.10308837890625 0 922.9119
212.1396026611328 0 2829.3416
214.11898803710938 0 624.1966
215.13951110839844 0 3060.0984 y Water loss 10
216.09828186035156 0 927.57825
216.14312744140625 0 1358.3281
217.0973358154297 0 778.2721 b Water loss 1
219.14959716796875 0 7077.4624
220.15309143066406 0 545.82605
221.0847930908203 0 4165.1035
223.1077117919922 0 730.3835
224.10357666015625 0 612.73706
224.1390838623047 0 953.03094
224.17628479003906 0 649.27454
225.04322814941406 0 10265.936
226.04396057128906 0 1560.8494
226.11935424804688 0 807.234
226.1553192138672 0 3254.9087
227.02256774902344 0 861.9235
228.09776306152344 0 901.2025
230.1502227783203 0 6354.5684
233.12977600097656 0 816.22644
233.1499786376953 0 6096.3374 y 10
233.16506958007812 0 1500.5275
234.15330505371094 0 631.0876
235.10806274414062 0 56527.34 b 1
236.11135864257812 0 8552.604
237.1128692626953 0 883.06683
239.09527587890625 0 13754.456
239.14987182617188 0 731.77625
240.09703063964844 0 1472.0004
240.13482666015625 0 816.5208
241.15415954589844 0 529.2082
242.1509246826172 0 635.84735
242.18663024902344 0 831.0638
245.12493896484375 0 1609.9531
246.14845275878906 0 546.25793
247.1444091796875 0 5716.438
248.1481475830078 0 992.88257
249.15916442871094 0 508.55142
251.10325622558594 0 2138.0312
254.1138153076172 0 925.16437
255.10906982421875 0 3845.0032
261.12408447265625 0 1103.6123
261.1599426269531 0 1365.0795
262.11883544921875 0 929.1356
263.1387634277344 0 595.30853
268.16497802734375 0 809.70996
269.16119384765625 0 3789.831
272.13726806640625 0 652.11316
273.1197204589844 0 4004.7942
274.11883544921875 0 1184.9539
277.1546936035156 0 601.6284
279.0984802246094 0 646.51074
281.12530517578125 0 1143.3534
287.1723327636719 0 1205.3494
289.155029296875 0 4108.954
290.1588134765625 0 872.86224
291.14599609375 0 760.2769
295.1036071777344 0 3578.4343
296.10430908203125 0 633.1965
299.0620422363281 0 6716.5327
299.1371765136719 0 545.91644
300.0615539550781 0 1092.5674
301.19140625 0 780.4909
309.159423828125 0 730.417
311.2090148925781 0 840.0716
313.11358642578125 0 1182.2885
315.669921875 0 603.1151
316.1661071777344 0 2326.8684 b Water loss 2
319.139892578125 0 995.19965
323.1720275878906 0 767.6649
323.2082824707031 0 2614.1733
324.212890625 0 681.1152
325.2228088378906 0 747.75507
326.1828918457031 0 2772.0166
334.1765441894531 0 37093.83 b 2
335.17938232421875 0 6969.6997
337.1545715332031 0 1093.0356
341.180419921875 0 737.7315
343.23358154296875 0 690.3851 y Water loss 9
346.2126159667969 0 685.8631
352.1976013183594 0 954.78796
354.1786804199219 0 708.24994
357.15484619140625 0 691.34656
359.02850341796875 0 650.26697
360.2283935546875 0 1010.2682
361.244384765625 0 3885.1304 y 9
364.1648864746094 0 681.6615
369.12152099609375 0 2243.148
374.1813049316406 0 787.7337
380.1922302246094 0 910.97394
382.2446594238281 0 1027.69
383.2034912109375 0 3787.7407
390.71575927734375 0 918.8246 y 4
392.1934509277344 0 1094.0548
400.2564392089844 0 1086.7555
401.2151184082031 0 4935.6987
402.17742919921875 0 4436.057
402.217041015625 0 704.30304
403.17938232421875 0 965.7184
405.24993896484375 0 2407.9158
408.2608337402344 0 2223.3545
415.2345275878906 0 1396.7214 b Water loss 3
416.2369689941406 0 592.6776
420.1880798339844 0 1822.9142
428.753173828125 0 1798.9878
433.24468994140625 0 9162.93 b 3
434.24822998046875 0 2637.7078
436.2234802246094 0 1813.4205
439.2669677734375 0 1148.3041
448.072998046875 0 605.15356
453.2507629394531 0 927.0006
453.5410461425781 0 594.4278
456.2240905761719 0 702.23474
457.2776184082031 0 2100.1077
462.2919921875 0 768.1592 y 8
463.23260498046875 0 1025.5548
480.7796630859375 0 797.6655 y Water loss 2
481.2456970214844 0 1751.5365
482.24505615234375 0 805.0696 b Ammonia loss 9
486.3014221191406 0 1220.5
490.1416320800781 0 532.99365
496.2872009277344 0 2385.2102
497.2924499511719 0 909.29346
502.24932861328125 0 761.40857
504.2428283691406 0 701.8542
511.2843933105469 0 563.863
512.7767944335938 0 781.58405
514.2982788085938 0 5885.5664
515.3025512695312 0 1281.7986
519.2540283203125 0 1075.574
519.3145751953125 0 1479.2726 y 7
520.3172607421875 0 985.28503
530.271728515625 0 2603.8445
531.2747802734375 0 673.1624
532.3109741210938 0 856.933
533.311767578125 0 875.2404
535.2953491210938 0 752.99414
537.6278076171875 0 618.2233
548.282470703125 0 4984.8047
549.285888671875 0 1723.945
552.3187255859375 0 699.58527
562.3016967773438 0 1145.4037 b Water loss 4
563.3043212890625 0 697.04126
573.32568359375 0 1635.3585
576.3345947265625 0 3846.9897 y 6
577.3370971679688 0 1230.1759
580.3123168945312 0 2303.8806 b 4
580.7053833007812 0 757.55524
581.3150024414062 0 1016.285
585.8318481445312 0 966.3647
586.3340454101562 0 666.95416
587.3021850585938 0 662.4332
597.8304443359375 0 1491.3923 Precursor Water loss
598.3340454101562 0 1162.5034
606.2760620117188 0 2740.1165
606.526611328125 0 1693.5089
606.7263793945312 0 1527.8169
606.7796020507812 0 1134.3204
606.9269409179688 0 945.7423
607.1280517578125 0 1490.0917
607.3312377929688 0 1382.2174
609.3421630859375 0 635.746
615.3484497070312 0 1090.2893 y Water loss 5
619.3240966796875 0 1057.6968 b Water loss 5
622.3416748046875 0 681.0979
629.34033203125 0 1542.9467
633.3569946289062 0 18069.352 y 5
634.360595703125 0 7378.655
635.3644409179688 0 863.5345
642.3743286132812 0 1078.293
643.355224609375 0 2876.3965
644.3560180664062 0 1415.4843
647.3531494140625 0 1848.027
648.35302734375 0 723.96497
661.3662719726562 0 6141.7393
662.3697509765625 0 1803.4647
679.3744506835938 0 1470.277
697.8658447265625 0 1334.7618
698.1138916015625 0 1549.6403
742.4241943359375 0 1572.2106
743.42724609375 0 1223.3823
746.4147338867188 0 1001.28687
747.4257202148438 0 914.14636
751.3746337890625 0 880.1501 b 7
756.4376220703125 0 753.1794
760.4336547851562 0 4470.042
761.4379272460938 0 1961.4297
762.412841796875 0 2120.408 y Water loss 4
780.4242553710938 0 25284.516 y 4
781.4268798828125 0 10945.498
782.4323120117188 0 2324.222
815.4830322265625 0 1100.7092
815.9192504882812 0 763.6936
834.4127807617188 0 1456.9244 b Water loss 8
841.7075805664062 0 699.2454
852.4240112304688 0 2471.5217 b 8
853.4296264648438 0 962.50616
859.4991455078125 0 1868.8892
860.5089721679688 0 1559.2214
879.4918823242188 0 15316.652 y 3
880.4953002929688 0 7752.1606
881.4990844726562 0 1772.2559
895.5260009765625 0 921.96674
962.5125122070312 0 1024.0785 b Water loss 9
963.5122680664062 0 826.60175 b Ammonia loss 9
977.5040893554688 0 680.12164
978.5594482421875 0 7302.1895 y 2
979.5623779296875 0 4222.532
980.5224609375 0 3815.2676 b 9
981.5186157226562 0 2069.9973
1075.590576171875 0 984.85724 b Water loss 10
1093.5999755859375 0 3347.695 b 10
1094.6015625 0 2383.0815
1095.6131591796875 0 663.5456
1889.0115966796875 0 610.8067
2450.051513671875 0 672.6921
2464.10546875 0 617.0569
2724.686767578125 0 810.74054
3031.49853515625 0 695.21246

Spectrum Details

|  |  |
| --- | --- |
| Matched peaks? Matched peaks The total absolute number of peaks matched. Additionally in brackets the total fraction of peaks matched and the total number of peaks is shown. | 37 (12.13% of 305) |
| FDR? FDR The false discovery rate estimated for this peptide. It is calculated by matching all theoretical fragments with a non-integer shift with the raw peaks for this spectrum. This is done with 40 different shifts. The resulting percentage is the average number of annotated peaks over the number of annotated peaks with the correct spectrum. | 0.58% |
| Satellite FDR? Satellite FDR See the FDR for details on its calculation. This satellite ion specific FDR only contains the satellite ions (d/w) for I/L/J positions. | - |
| PSM Score? PSM Score The PSM Score as given by Hecklib to this annotated spectrum. It is shown with three significant figures. | 471 |

#### Reverse Lookup? Reverse Lookup All places where this read could be placed.

| Group | Segment | Template | Template Part | Read Part | Score | Unique |
| --- | --- | --- | --- | --- | --- | --- |
| Homo sapiens Light Chain | IGLJ | IGLJ2 | [0..10] | [2..12] | 80 | True |

| Recombined | Template Part | Read Part | Score | Unique |
| --- | --- | --- | --- | --- |
| REC-0-1\_002 | [98..109] | [0..12] | 87 | True |

#### Meta Information from Multiple reads

##### Number of combined reads

9

##### Intensity

0.9425

##### TotalArea

2.139E+09

##### Changes to the peptide sequence

SFVVFGGGTKJT

L→JNo support for either Leucine or Isoleucine based on side chain ions (Position: 11)

#### Positional Score

Copy Data

##### Positional Score (TSV)

###### Preview

```
Loading example...
```

*Click on the button to copy the data to your clipboard.*

1001234567891011

Label Value
"0" 0.738
"1" 0.752
"2" 0.777
"3" 0.774
"4" 0.773
"5" 0.771
"6" 0.744
"7" 0.716
"8" 0.738
"9" 0.774
"10" 0.778
"11" 0.778

#### Meta Information from PEAKS

##### Scan Identifier

F1:8995

##### Original sequence

S

F

V

V

F

G

G

G

T

K

L

T

##### Posttranslational Modifications

##### Source File

D:\separate\_stitch\_analyses\xle-disambiguation\raw\20210323\_F1\_UM1\_Peng0013\_SA\_F59\_ingel\_3ug\_ELA.raw

##### Fraction

1

##### Scan Feature

F1:8634

##### De Novo Score

99

##### ConfidenceScore

99

### m/z

606.8351

##### Mass

1211.655

##### Charge

2

##### Retention Time

49.18

##### Predicted Retention Time

-

##### Area

5.347E+08

##### Parts Per Million

0.5

##### Fragmentation mode

ETHCD

##### Originating file

01 D:\separate\_stitch\_analyses\xle-disambiguation\20210325\_F59\_3ug\_DENOVO\_12.csv

#### Meta Information from PEAKS

##### Scan Identifier

F1:9056

##### Original sequence

S

F

V

V

F

G

G

G

T

K

L

T

##### Posttranslational Modifications

##### Source File

D:\separate\_stitch\_analyses\xle-disambiguation\raw\20210323\_F1\_UM1\_Peng0013\_SA\_F59\_ingel\_3ug\_ELA.raw

##### Fraction

1

##### Scan Feature

F1:8634

##### De Novo Score

99

##### ConfidenceScore

99

### m/z

606.8351

##### Mass

1211.655

##### Charge

2

##### Retention Time

49.18

##### Predicted Retention Time

-

##### Area

5.347E+08

##### Parts Per Million

0.5

##### Fragmentation mode

ETHCD

##### Originating file

01 D:\separate\_stitch\_analyses\xle-disambiguation\20210325\_F59\_3ug\_DENOVO\_12.csv

#### Meta Information from PEAKS

##### Scan Identifier

F1:9114

##### Original sequence

S

F

V

V

F

G

G

G

T

K

L

T

##### Posttranslational Modifications

##### Source File

D:\separate\_stitch\_analyses\xle-disambiguation\raw\20210323\_F1\_UM1\_Peng0013\_SA\_F59\_ingel\_3ug\_ELA.raw

##### Fraction

1

##### Scan Feature

F1:8634

##### De Novo Score

99

##### ConfidenceScore

99

### m/z

606.8351

##### Mass

1211.655

##### Charge

2

##### Retention Time

49.18

##### Predicted Retention Time

-

##### Area

5.347E+08

##### Parts Per Million

0.5

##### Fragmentation mode

HCD

##### Originating file

01 D:\separate\_stitch\_analyses\xle-disambiguation\20210325\_F59\_3ug\_DENOVO\_12.csv

#### Meta Information from PEAKS

##### Scan Identifier

F1:9175

##### Original sequence

S

F

V

V

F

G

G

G

T

K

L

T

##### Posttranslational Modifications

##### Source File

D:\separate\_stitch\_analyses\xle-disambiguation\raw\20210323\_F1\_UM1\_Peng0013\_SA\_F59\_ingel\_3ug\_ELA.raw

##### Fraction

1

##### Scan Feature

-

##### De Novo Score

99

##### ConfidenceScore

99

### m/z

606.8361

##### Mass

1211.655

##### Charge

2

##### Retention Time

50.62

##### Predicted Retention Time

-

##### Area

0

##### Parts Per Million

2.2

##### Fragmentation mode

ETHCD

##### Originating file

01 D:\separate\_stitch\_analyses\xle-disambiguation\20210325\_F59\_3ug\_DENOVO\_12.csv

#### Meta Information from PEAKS

##### Scan Identifier

F1:9363

##### Original sequence

S

F

V

V

F

G

G

G

T

K

L

T

##### Posttranslational Modifications

##### Source File

D:\separate\_stitch\_analyses\xle-disambiguation\raw\20210323\_F1\_UM1\_Peng0013\_SA\_F59\_ingel\_3ug\_ELA.raw

##### Fraction

1

##### Scan Feature

-

##### De Novo Score

97

##### ConfidenceScore

97

### m/z

606.836

##### Mass

1211.655

##### Charge

2

##### Retention Time

51.74

##### Predicted Retention Time

-

##### Area

0

##### Parts Per Million

2

##### Fragmentation mode

ETHCD

##### Originating file

01 D:\separate\_stitch\_analyses\xle-disambiguation\20210325\_F59\_3ug\_DENOVO\_12.csv

#### Meta Information from PEAKS

##### Scan Identifier

F1:9545

##### Original sequence

S

F

V

V

F

G

G

G

T

K

L

T

##### Posttranslational Modifications

##### Source File

D:\separate\_stitch\_analyses\xle-disambiguation\raw\20210323\_F1\_UM1\_Peng0013\_SA\_F59\_ingel\_3ug\_ELA.raw

##### Fraction

1

##### Scan Feature

-

##### De Novo Score

97

##### ConfidenceScore

97

### m/z

606.8358

##### Mass

1211.655

##### Charge

2

##### Retention Time

52.84

##### Predicted Retention Time

-

##### Area

0

##### Parts Per Million

1.7

##### Fragmentation mode

ETHCD

##### Originating file

01 D:\separate\_stitch\_analyses\xle-disambiguation\20210325\_F59\_3ug\_DENOVO\_12.csv

#### Meta Information from PEAKS

##### Scan Identifier

F1:8930

##### Original sequence

S

F

V

V

F

G

G

G

T

K

L

T

##### Posttranslational Modifications

##### Source File

D:\separate\_stitch\_analyses\xle-disambiguation\raw\20210323\_F1\_UM1\_Peng0013\_SA\_F59\_ingel\_3ug\_ELA.raw

##### Fraction

1

##### Scan Feature

F1:8634

##### De Novo Score

97

##### ConfidenceScore

97

### m/z

606.8351

##### Mass

1211.655

##### Charge

2

##### Retention Time

49.18

##### Predicted Retention Time

-

##### Area

5.347E+08

##### Parts Per Million

0.5

##### Fragmentation mode

ETHCD

##### Originating file

01 D:\separate\_stitch\_analyses\xle-disambiguation\20210325\_F59\_3ug\_DENOVO\_12.csv

#### Meta Information from PEAKS

##### Scan Identifier

F1:9485

##### Original sequence

S

F

V

V

F

G

G

G

T

K

L

T

##### Posttranslational Modifications

##### Source File

D:\separate\_stitch\_analyses\xle-disambiguation\raw\20210323\_F1\_UM1\_Peng0013\_SA\_F59\_ingel\_3ug\_ELA.raw

##### Fraction

1

##### Scan Feature

-

##### De Novo Score

97

##### ConfidenceScore

97

### m/z

606.8356

##### Mass

1211.655

##### Charge

2

##### Retention Time

52.48

##### Predicted Retention Time

-

##### Area

0

##### Parts Per Million

1.4

##### Fragmentation mode

ETHCD

##### Originating file

01 D:\separate\_stitch\_analyses\xle-disambiguation\20210325\_F59\_3ug\_DENOVO\_12.csv

#### Meta Information from PEAKS

##### Scan Identifier

F1:9661

##### Original sequence

S

F

V

V

F

G

G

G

T

K

L

T

##### Posttranslational Modifications

##### Source File

D:\separate\_stitch\_analyses\xle-disambiguation\raw\20210323\_F1\_UM1\_Peng0013\_SA\_F59\_ingel\_3ug\_ELA.raw

##### Fraction

1

##### Scan Feature

-

##### De Novo Score

95

##### ConfidenceScore

95

### m/z

606.8349

##### Mass

1211.655

##### Charge

2

##### Retention Time

53.55

##### Predicted Retention Time

-

##### Area

0

##### Parts Per Million

0.2

##### Fragmentation mode

HCD

##### Originating file

01 D:\separate\_stitch\_analyses\xle-disambiguation\20210325\_F59\_3ug\_DENOVO\_12.csv
