## Supplementary material for "A handle on mass coincidence errors in *de novo* sequencing of antibodies by bottom-up proteomics": Combined_008.html

Details Combined\_008 | Stitch OverviewUndefined

### Read Combined\_008

#### Sequence (length=15)

JHQDWJDGKEYKCKV

#### Spectrum 5478? Spectrum 5478 The raw spectrum of this peptide as annotated by Hecklib. The fragments are coloured according to ion type (see legend). Any peaks with a star '\*' as text can be hovered over to see the full details, first the ion type second the mass shift type. By hovering over the amino acids in the peptide or ions in the legend the corresponding peaks are highlighted. By toggling the 'Unassigned' label you can turn the background (unassigned) peaks on or off in the plot. By updating the slider in the Ion legend you can update the spectrum to only show the top X% of the peaks with labels. The top X% means any peak that is within X% of the highest intensity. By dragging in the spectrum you can zoom in to a specific part of the spectrum and use 'Zoom Out' to get back to the original zoom level. The annotation of the spectrum is based on the given sequence in the peptides file and is done with different software so inconsistencies are likely. The peaks are annotated based on the given sequence, with 20 ppm tolerance.

Copy Data

##### Spectrum 5478 (TSV)

###### Preview

```
Loading example...
```

*Click on the button to copy the data to your clipboard.*

Mz MinMz MaxIntensity Max

WidthHeightPeptide font sizePeptide stroke widthSpectrum font sizeSpectrum stroke widthCompact peptide

Ion legend

wxyz

abcd

OtherUnassignedIon^Charge^_Position_Show for top:%

JHQDWJDGKEYKCKV

01.18e+52.35e+53.53e+54.71e+5

Zoom Out

z^+1^_2_y^+1^_2_c^+1^_2_c^+1^_3_c^+1^_3_y^+1^_3_c^+3^_10_w^+1^_4_c^+1^_4_y^+2^_8_c^+1^_4_z^+1^_4_y^+1^_4_y^+2^_9_c^+3^_13_y^+3^_14_c^+3^_14_y^+3^_14_z^+2^_10_y^+2^_10_c^+1^_5_y^+1^_5_z^+1^_5_c^+1^_5_y^+1^_5_y^+2^_11_w^+1^_6_z^+2^_12_c^+2^_12_y^+2^_12_c^+1^_6_c^+1^_6_z^+1^_6_y^+2^_13_y^+2^_13_y^+1^_6_y^+2^_13_c^+2^_13_c^+2^_13_c^+2^_13_y^+2^_14_z^+2^_14_c^+2^_14_c^+2^_14_y^+2^_14_c^+1^_7_c^+2^_14_z^+1^_7_c^+1^_7_y^+1^_7_y^+1^_7_z^+1^_7_y^+1^_7_c^+1^_8_c^+1^_8_z^+1^_8_y^+1^_8_w^+1^_9_c^+1^_9_y^+1^_9_c^+1^_10_y^+1^_10_y^+1^_10_z^+1^_10_c^+1^_10_y^+1^_10_c^+1^_11_y^+1^_11_z^+1^_11_y^+1^_11_y^+1^_12_z^+1^_12_c^+1^_12_y^+1^_12_z^+1^_13_y^+1^_13_c^+1^_13_z^+1^_14_y^+1^_14_c^+1^_14_

0536107316092145

Fragment Matches Table

Show background peaks

| Position | Ion type | Intensity | mz Theoretical | mz Error (Th) | mz Error (ppm) | Charge | Series Number |
| --- | --- | --- | --- | --- | --- | --- | --- |
| - | - | 1536 | 128.1 | - | - | 0 | - |
| - | - | 5548 | 129.1 | - | - | 0 | - |
| - | - | 2517 | 146.1 | - | - | 0 | - |
| - | - | 1231 | 147.5 | - | - | 0 | - |
| - | - | 1360 | 151.6 | - | - | 0 | - |
| - | - | 5039 | 155.1 | - | - | 0 | - |
| - | - | 1.28E+04 | 166.1 | - | - | 0 | - |
| - | - | 4162 | 178.1 | - | - | 0 | - |
| - | - | 2189 | 183.1 | - | - | 0 | - |
| - | - | 8643 | 185.2 | - | - | 0 | - |
| - | - | 2107 | 207.2 | - | - | 0 | - |
| - | - | 1679 | 222.1 | - | - | 0 | - |
| - | - | 1.373E+04 | 223.2 | - | - | 0 | - |
| 14 | z | 4.514E+04 | 230.2 | 8.839E-05 | 0.384 | +1 | 2 |
| - | - | 7934 | 231.2 | - | - | 0 | - |
| - | - | 2.117E+04 | 234.1 | - | - | 0 | - |
| 14 | y | 3528 | 246.2 | 0.0001784 | 0.7247 | +1 | 2 |
| - | - | 4084 | 250.1 | - | - | 0 | - |
| - | - | 8.668E+04 | 251.2 | - | - | 0 | - |
| - | - | 1.252E+04 | 252.2 | - | - | 0 | - |
| - | - | 3460 | 266.1 | - | - | 0 | - |
| 2 | c | 4.159E+04 | 268.2 | 1.307E-05 | 0.04873 | +1 | 2 |
| - | - | 4803 | 269.2 | - | - | 0 | - |
| - | - | 6233 | 283.2 | - | - | 0 | - |
| - | - | 1469 | 286.2 | - | - | 0 | - |
| - | - | 6.919E+04 | 300.2 | - | - | 0 | - |
| - | - | 1.077E+04 | 301.2 | - | - | 0 | - |
| - | - | 1767 | 331.9 | - | - | 0 | - |
| - | - | 2154 | 351.2 | - | - | 0 | - |
| - | - | 6309 | 352.2 | - | - | 0 | - |
| - | - | 1554 | 357.1 | - | - | 0 | - |
| 3 | c | 1.566E+04 | 379.2 | 0.0003377 | 0.8905 | +1 | 3 |
| - | - | 2823 | 380.2 | - | - | 0 | - |
| - | - | 5613 | 395.2 | - | - | 0 | - |
| 3 | c | 8.75E+04 | 396.2 | 0.0003083 | 0.7782 | +1 | 3 |
| - | - | 1.893E+04 | 397.2 | - | - | 0 | - |
| - | - | 2206 | 398.2 | - | - | 0 | - |
| 13 | y | 8545 | 407.2 | 0.004175 | 10.25 | +1 | 3 |
| 10 | c | 2188 | 408.2 | 0.0005477 | 1.342 | +3 | 10 |
| - | - | 3317 | 409.2 | - | - | 0 | - |
| - | - | 7701 | 412.1 | - | - | 0 | - |
| - | - | 1651 | 415.6 | - | - | 0 | - |
| - | - | 1838 | 416.1 | - | - | 0 | - |
| - | - | 2675 | 428.3 | - | - | 0 | - |
| - | - | 1833 | 448.9 | - | - | 0 | - |
| - | - | 1622 | 451.2 | - | - | 0 | - |
| - | - | 2897 | 460.3 | - | - | 0 | - |
| 12 | w | 3370 | 461.2 | 0.006591 | 14.29 | +1 | 4 |
| - | - | 5.159E+04 | 467.2 | - | - | 0 | - |
| - | - | 1.887E+04 | 468.3 | - | - | 0 | - |
| - | - | 3545 | 469.3 | - | - | 0 | - |
| - | - | 3693 | 475.3 | - | - | 0 | - |
| - | - | 2437 | 479.2 | - | - | 0 | - |
| 4 | c | 1.043E+04 | 494.2 | 0.0002806 | 0.5678 | +1 | 4 |
| - | - | 2039 | 506.2 | - | - | 0 | - |
| 8 | y | 4.333E+04 | 506.8 | 0.001722 | 3.399 | +2 | 8 |
| - | - | 2.021E+04 | 507.3 | - | - | 0 | - |
| - | - | 7507 | 507.8 | - | - | 0 | - |
| - | - | 3247 | 508.3 | - | - | 0 | - |
| 4 | c | 7.852E+04 | 511.3 | 0.0001903 | 0.3721 | +1 | 4 |
| - | - | 1.894E+04 | 512.3 | - | - | 0 | - |
| - | - | 2458 | 513.3 | - | - | 0 | - |
| 12 | z | 3.924E+04 | 519.3 | 0.004242 | 8.168 | +1 | 4 |
| - | - | 1.366E+04 | 520.3 | - | - | 0 | - |
| - | - | 4246 | 521.3 | - | - | 0 | - |
| - | - | 2390 | 522.2 | - | - | 0 | - |
| - | - | 4497 | 525.2 | - | - | 0 | - |
| 12 | y | 1.273E+04 | 535.3 | 0.004106 | 7.67 | +1 | 4 |
| - | - | 4797 | 536.3 | - | - | 0 | - |
| - | - | 3158 | 563.9 | - | - | 0 | - |
| 7 | y | 1.849E+04 | 564.3 | 0.002834 | 5.023 | +2 | 9 |
| 13 | c | 2479 | 564.6 | 0.003269 | 5.79 | +3 | 13 |
| - | - | 5591 | 564.8 | - | - | 0 | - |
| - | - | 2262 | 565.3 | - | - | 0 | - |
| - | - | 4214 | 578.2 | - | - | 0 | - |
| - | - | 1.331E+04 | 584.4 | - | - | 0 | - |
| - | - | 4112 | 585.4 | - | - | 0 | - |
| - | - | 3606 | 591.3 | - | - | 0 | - |
| - | - | 2407 | 592.4 | - | - | 0 | - |
| - | - | 3980 | 596.3 | - | - | 0 | - |
| 2 | y | 4748 | 597.3 | 0.005305 | 8.881 | +3 | 14 |
| 14 | c | 2.57E+04 | 601.6 | 0.0004997 | 0.8306 | +3 | 14 |
| - | - | 1.906E+04 | 602 | - | - | 0 | - |
| - | - | 1.946E+04 | 602.3 | - | - | 0 | - |
| - | - | 9787 | 602.6 | - | - | 0 | - |
| 2 | y | 3.401E+04 | 603 | 0.000595 | 0.9867 | +3 | 14 |
| - | - | 3.241E+04 | 603.3 | - | - | 0 | - |
| - | - | 1.671E+04 | 603.6 | - | - | 0 | - |
| - | - | 7226 | 604 | - | - | 0 | - |
| 6 | z | 4019 | 604.3 | 0.006055 | 10.02 | +2 | 10 |
| - | - | 7420 | 607.6 | - | - | 0 | - |
| - | - | 7057 | 608 | - | - | 0 | - |
| - | - | 4572 | 608.3 | - | - | 0 | - |
| - | - | 2487 | 610.3 | - | - | 0 | - |
| 6 | y | 3028 | 620.8 | 6.658E-05 | 0.1072 | +2 | 10 |
| - | - | 5156 | 622.2 | - | - | 0 | - |
| - | - | 3182 | 623.3 | - | - | 0 | - |
| - | - | 3726 | 624.2 | - | - | 0 | - |
| - | - | 6112 | 624.3 | - | - | 0 | - |
| - | - | 2289 | 625.6 | - | - | 0 | - |
| - | - | 3175 | 626 | - | - | 0 | - |
| - | - | 1.213E+04 | 629 | - | - | 0 | - |
| - | - | 1.224E+04 | 629.3 | - | - | 0 | - |
| - | - | 4369 | 629.6 | - | - | 0 | - |
| - | - | 4428 | 630 | - | - | 0 | - |
| - | - | 2064 | 632 | - | - | 0 | - |
| - | - | 2423 | 634.3 | - | - | 0 | - |
| - | - | 2.781E+04 | 634.6 | - | - | 0 | - |
| - | - | 4.294E+04 | 635 | - | - | 0 | - |
| - | - | 2.864E+04 | 635.3 | - | - | 0 | - |
| - | - | 1.529E+04 | 635.6 | - | - | 0 | - |
| - | - | 9122 | 636 | - | - | 0 | - |
| - | - | 2040 | 636.3 | - | - | 0 | - |
| - | - | 2961 | 637.3 | - | - | 0 | - |
| - | - | 2404 | 638.3 | - | - | 0 | - |
| - | - | 2040 | 639.3 | - | - | 0 | - |
| - | - | 2348 | 640 | - | - | 0 | - |
| - | - | 2.108E+04 | 640.3 | - | - | 0 | - |
| - | - | 4.303E+05 | 640.6 | - | - | 0 | - |
| - | - | 4.662E+05 | 641 | - | - | 0 | - |
| - | - | 2.897E+05 | 641.3 | - | - | 0 | - |
| - | - | 1.466E+05 | 641.7 | - | - | 0 | - |
| - | - | 3130 | 641.8 | - | - | 0 | - |
| - | - | 4.82E+04 | 642 | - | - | 0 | - |
| - | - | 5004 | 642.2 | - | - | 0 | - |
| - | - | 1.666E+04 | 642.3 | - | - | 0 | - |
| - | - | 3686 | 651.4 | - | - | 0 | - |
| - | - | 5641 | 653.3 | - | - | 0 | - |
| - | - | 2537 | 658.3 | - | - | 0 | - |
| 5 | c | 9670 | 680.3 | 0.001198 | 1.762 | +1 | 5 |
| 11 | y | 5178 | 681.3 | 0.01128 | 16.56 | +1 | 5 |
| 11 | z | 9.468E+04 | 682.3 | 0.004094 | 5.999 | +1 | 5 |
| - | - | 4.246E+04 | 683.3 | - | - | 0 | - |
| - | - | 1.394E+04 | 684.3 | - | - | 0 | - |
| - | - | 3259 | 685.3 | - | - | 0 | - |
| 5 | c | 6.295E+04 | 697.3 | 0.0004061 | 0.5824 | +1 | 5 |
| 11 | y | 4.789E+04 | 698.4 | 0.008719 | 12.48 | +1 | 5 |
| - | - | 1.52E+04 | 699.4 | - | - | 0 | - |
| - | - | 3135 | 700.4 | - | - | 0 | - |
| 5 | y | 1.016E+04 | 713.9 | 0.0004991 | 0.6992 | +2 | 11 |
| - | - | 7658 | 714.4 | - | - | 0 | - |
| - | - | 3235 | 714.9 | - | - | 0 | - |
| - | - | 6842 | 720.4 | - | - | 0 | - |
| - | - | 4225 | 721.4 | - | - | 0 | - |
| 10 | w | 3.012E+04 | 752.4 | 0.003231 | 4.294 | +1 | 6 |
| - | - | 1.31E+04 | 753.4 | - | - | 0 | - |
| 4 | z | 4496 | 754.4 | 0.01327 | 17.59 | +2 | 12 |
| - | - | 5213 | 765.4 | - | - | 0 | - |
| 12 | c | 2.398E+04 | 765.9 | 0.00201 | 2.625 | +2 | 12 |
| - | - | 2.224E+04 | 766.4 | - | - | 0 | - |
| - | - | 7990 | 766.9 | - | - | 0 | - |
| - | - | 8428 | 767.4 | - | - | 0 | - |
| 4 | y | 1.024E+04 | 771.4 | 0.0003381 | 0.4383 | +2 | 12 |
| - | - | 6893 | 771.9 | - | - | 0 | - |
| - | - | 3441 | 772.4 | - | - | 0 | - |
| - | - | 2753 | 775.4 | - | - | 0 | - |
| - | - | 4157 | 777.4 | - | - | 0 | - |
| - | - | 4095 | 779.4 | - | - | 0 | - |
| - | - | 2599 | 787.4 | - | - | 0 | - |
| - | - | 2615 | 787.7 | - | - | 0 | - |
| - | - | 2387 | 789.4 | - | - | 0 | - |
| - | - | 2201 | 789.9 | - | - | 0 | - |
| 6 | c | 7570 | 793.4 | 0.002665 | 3.359 | +1 | 6 |
| - | - | 4718 | 794.4 | - | - | 0 | - |
| - | - | 2165 | 799.4 | - | - | 0 | - |
| - | - | 3439 | 809.4 | - | - | 0 | - |
| 6 | c | 9.381E+04 | 810.4 | 0.0009979 | 1.231 | +1 | 6 |
| 10 | z | 1.338E+05 | 811.4 | 0.004511 | 5.56 | +1 | 6 |
| - | - | 5.921E+04 | 812.4 | - | - | 0 | - |
| - | - | 2.213E+04 | 813.4 | - | - | 0 | - |
| - | - | 5793 | 814.4 | - | - | 0 | - |
| - | - | 2254 | 815.4 | - | - | 0 | - |
| - | - | 2654 | 817.4 | - | - | 0 | - |
| 3 | y | 9951 | 826.4 | 0.002677 | 3.239 | +2 | 13 |
| 3 | y | 2.34E+04 | 826.9 | 0.002874 | 3.475 | +2 | 13 |
| 10 | y | 3.599E+04 | 827.4 | 0.009075 | 10.97 | +1 | 6 |
| - | - | 1.377E+04 | 827.9 | - | - | 0 | - |
| - | - | 1.107E+04 | 828.4 | - | - | 0 | - |
| - | - | 3208 | 829.4 | - | - | 0 | - |
| - | - | 2822 | 834.9 | - | - | 0 | - |
| 3 | y | 5.335E+04 | 835.4 | 0.001429 | 1.71 | +2 | 13 |
| - | - | 4.794E+04 | 835.9 | - | - | 0 | - |
| - | - | 2.277E+04 | 836.4 | - | - | 0 | - |
| - | - | 1.221E+04 | 836.9 | - | - | 0 | - |
| 13 | c | 4065 | 837.4 | 0.01381 | 16.5 | +2 | 13 |
| 13 | c | 2351 | 837.9 | 0.00789 | 9.416 | +2 | 13 |
| - | - | 3512 | 838.4 | - | - | 0 | - |
| - | - | 2548 | 838.9 | - | - | 0 | - |
| - | - | 2927 | 841.9 | - | - | 0 | - |
| - | - | 7429 | 842.9 | - | - | 0 | - |
| - | - | 6634 | 843.4 | - | - | 0 | - |
| - | - | 3300 | 844 | - | - | 0 | - |
| - | - | 6364 | 845.9 | - | - | 0 | - |
| 13 | c | 3.436E+04 | 846.4 | 0.001051 | 1.242 | +2 | 13 |
| - | - | 2.448E+04 | 846.9 | - | - | 0 | - |
| - | - | 1.505E+04 | 847.4 | - | - | 0 | - |
| - | - | 1.152E+04 | 847.9 | - | - | 0 | - |
| - | - | 6765 | 848.5 | - | - | 0 | - |
| - | - | 2696 | 849.5 | - | - | 0 | - |
| - | - | 4983 | 850.4 | - | - | 0 | - |
| - | - | 2194 | 858.9 | - | - | 0 | - |
| - | - | 3048 | 859.4 | - | - | 0 | - |
| - | - | 2600 | 860.4 | - | - | 0 | - |
| - | - | 7886 | 866.4 | - | - | 0 | - |
| - | - | 4448 | 866.9 | - | - | 0 | - |
| - | - | 6.323E+04 | 867.4 | - | - | 0 | - |
| - | - | 3.083E+04 | 868.4 | - | - | 0 | - |
| - | - | 8270 | 869.4 | - | - | 0 | - |
| - | - | 5059 | 873.9 | - | - | 0 | - |
| - | - | 3519 | 874.4 | - | - | 0 | - |
| - | - | 4838 | 874.9 | - | - | 0 | - |
| - | - | 2444 | 875.4 | - | - | 0 | - |
| - | - | 5260 | 880.4 | - | - | 0 | - |
| - | - | 1.472E+04 | 881.4 | - | - | 0 | - |
| - | - | 6085 | 882.5 | - | - | 0 | - |
| - | - | 2564 | 884 | - | - | 0 | - |
| - | - | 3517 | 885.5 | - | - | 0 | - |
| - | - | 4111 | 887.9 | - | - | 0 | - |
| - | - | 1.012E+04 | 888.4 | - | - | 0 | - |
| - | - | 9643 | 888.9 | - | - | 0 | - |
| - | - | 4462 | 889.4 | - | - | 0 | - |
| - | - | 4446 | 893.5 | - | - | 0 | - |
| - | - | 2866 | 894.4 | - | - | 0 | - |
| 2 | y | 3487 | 894.9 | 0.004488 | 5.015 | +2 | 14 |
| - | - | 1.038E+04 | 895.4 | - | - | 0 | - |
| 2 | z | 3.627E+04 | 895.9 | 0.002159 | 2.41 | +2 | 14 |
| - | - | 3.418E+04 | 896.4 | - | - | 0 | - |
| - | - | 1.79E+04 | 896.9 | - | - | 0 | - |
| - | - | 9695 | 897.4 | - | - | 0 | - |
| - | - | 2647 | 897.9 | - | - | 0 | - |
| 14 | c | 5067 | 901.4 | 0.00692 | 7.677 | +2 | 14 |
| 14 | c | 5416 | 901.9 | 0.007039 | 7.804 | +2 | 14 |
| - | - | 9442 | 902.4 | - | - | 0 | - |
| - | - | 5240 | 902.9 | - | - | 0 | - |
| - | - | 3491 | 903.4 | - | - | 0 | - |
| 2 | y | 2496 | 903.9 | 0.0008537 | 0.9444 | +2 | 14 |
| - | - | 2228 | 904.4 | - | - | 0 | - |
| - | - | 2720 | 906.5 | - | - | 0 | - |
| - | - | 7699 | 907 | - | - | 0 | - |
| - | - | 1.326E+04 | 907.5 | - | - | 0 | - |
| - | - | 8494 | 908 | - | - | 0 | - |
| 7 | c | 2.087E+04 | 908.4 | 0.003839 | 4.226 | +1 | 7 |
| - | - | 8595 | 909 | - | - | 0 | - |
| - | - | 1.481E+04 | 909.4 | - | - | 0 | - |
| - | - | 8663 | 910 | - | - | 0 | - |
| 14 | c | 1.597E+05 | 910.4 | 0.001719 | 1.888 | +2 | 14 |
| - | - | 1.539E+05 | 910.9 | - | - | 0 | - |
| - | - | 9.284E+04 | 911.4 | - | - | 0 | - |
| - | - | 4.071E+04 | 911.9 | - | - | 0 | - |
| - | - | 1.63E+04 | 912.4 | - | - | 0 | - |
| - | - | 6133 | 913 | - | - | 0 | - |
| - | - | 2953 | 915 | - | - | 0 | - |
| - | - | 2460 | 915.5 | - | - | 0 | - |
| - | - | 7583 | 916 | - | - | 0 | - |
| - | - | 1.149E+04 | 916.5 | - | - | 0 | - |
| - | - | 1.036E+04 | 917 | - | - | 0 | - |
| - | - | 5332 | 917.5 | - | - | 0 | - |
| - | - | 2302 | 918 | - | - | 0 | - |
| 9 | z | 1.981E+04 | 922.5 | 0.01246 | 13.51 | +1 | 7 |
| - | - | 1.71E+04 | 923 | - | - | 0 | - |
| - | - | 1.174E+04 | 923.5 | - | - | 0 | - |
| - | - | 8069 | 924 | - | - | 0 | - |
| - | - | 2.625E+04 | 924.4 | - | - | 0 | - |
| - | - | 1.039E+04 | 924.9 | - | - | 0 | - |
| 7 | c | 4.851E+04 | 925.5 | 0.001287 | 1.391 | +1 | 7 |
| - | - | 4308 | 926 | - | - | 0 | - |
| - | - | 2.421E+04 | 926.5 | - | - | 0 | - |
| - | - | 5807 | 927.5 | - | - | 0 | - |
| - | - | 3190 | 930 | - | - | 0 | - |
| - | - | 2.899E+04 | 930.5 | - | - | 0 | - |
| - | - | 8.225E+04 | 931 | - | - | 0 | - |
| - | - | 2.049E+05 | 931.5 | - | - | 0 | - |
| - | - | 1.978E+05 | 932 | - | - | 0 | - |
| - | - | 1.064E+05 | 932.5 | - | - | 0 | - |
| - | - | 4.931E+04 | 933 | - | - | 0 | - |
| - | - | 1.857E+04 | 933.5 | - | - | 0 | - |
| - | - | 7925 | 934 | - | - | 0 | - |
| 9 | y | 4357 | 937.5 | 0.01413 | 15.07 | +1 | 7 |
| - | - | 1.965E+04 | 938 | - | - | 0 | - |
| 9 | y | 3.316E+04 | 938.5 | 0.0002187 | 0.233 | +1 | 7 |
| - | - | 2.916E+04 | 939 | - | - | 0 | - |
| 9 | z | 8.941E+04 | 939.5 | 0.005663 | 6.028 | +1 | 7 |
| - | - | 8515 | 940 | - | - | 0 | - |
| - | - | 5.473E+04 | 940.5 | - | - | 0 | - |
| - | - | 2.062E+04 | 941.5 | - | - | 0 | - |
| - | - | 7903 | 942.5 | - | - | 0 | - |
| - | - | 4425 | 943 | - | - | 0 | - |
| - | - | 5598 | 943.5 | - | - | 0 | - |
| - | - | 2439 | 944 | - | - | 0 | - |
| - | - | 2160 | 944.4 | - | - | 0 | - |
| - | - | 2522 | 946.5 | - | - | 0 | - |
| - | - | 6077 | 951.5 | - | - | 0 | - |
| - | - | 1.624E+04 | 952 | - | - | 0 | - |
| - | - | 6.757E+04 | 952.5 | - | - | 0 | - |
| - | - | 6.244E+04 | 953 | - | - | 0 | - |
| - | - | 4.198E+04 | 953.5 | - | - | 0 | - |
| - | - | 2.269E+04 | 954 | - | - | 0 | - |
| - | - | 9481 | 954.5 | - | - | 0 | - |
| - | - | 4452 | 955 | - | - | 0 | - |
| 9 | y | 5105 | 955.5 | 0.006809 | 7.126 | +1 | 7 |
| - | - | 2795 | 956.5 | - | - | 0 | - |
| - | - | 1.195E+04 | 960 | - | - | 0 | - |
| - | - | 2.166E+05 | 960.5 | - | - | 0 | - |
| - | - | 3.192E+05 | 961 | - | - | 0 | - |
| - | - | 2.514E+05 | 961.5 | - | - | 0 | - |
| - | - | 1.467E+05 | 962 | - | - | 0 | - |
| - | - | 5.441E+04 | 962.5 | - | - | 0 | - |
| - | - | 2.257E+04 | 963 | - | - | 0 | - |
| - | - | 6040 | 963.5 | - | - | 0 | - |
| 8 | c | 3059 | 965.4 | 0.01131 | 11.71 | +1 | 8 |
| - | - | 1900 | 966.5 | - | - | 0 | - |
| - | - | 2124 | 976.5 | - | - | 0 | - |
| - | - | 1.714E+04 | 981.5 | - | - | 0 | - |
| 8 | c | 9.71E+04 | 982.5 | 0.00087 | 0.8855 | +1 | 8 |
| - | - | 5.381E+04 | 983.5 | - | - | 0 | - |
| - | - | 1.631E+04 | 984.5 | - | - | 0 | - |
| - | - | 4222 | 985.5 | - | - | 0 | - |
| - | - | 2540 | 991.5 | - | - | 0 | - |
| 8 | z | 2.773E+04 | 996.5 | 0.0043 | 4.315 | +1 | 8 |
| - | - | 3.126E+04 | 997.5 | - | - | 0 | - |
| - | - | 1.468E+04 | 998.5 | - | - | 0 | - |
| - | - | 4534 | 999.5 | - | - | 0 | - |
| - | - | 2971 | 1012 | - | - | 0 | - |
| 8 | y | 2.142E+04 | 1013 | 0.00508 | 5.017 | +1 | 8 |
| - | - | 1.441E+04 | 1014 | - | - | 0 | - |
| - | - | 3124 | 1015 | - | - | 0 | - |
| - | - | 3104 | 1016 | - | - | 0 | - |
| - | - | 5012 | 1021 | - | - | 0 | - |
| - | - | 3442 | 1025 | - | - | 0 | - |
| - | - | 3500 | 1028 | - | - | 0 | - |
| - | - | 4684 | 1031 | - | - | 0 | - |
| - | - | 3192 | 1032 | - | - | 0 | - |
| - | - | 3538 | 1033 | - | - | 0 | - |
| - | - | 3682 | 1034 | - | - | 0 | - |
| - | - | 4746 | 1040 | - | - | 0 | - |
| - | - | 3151 | 1041 | - | - | 0 | - |
| - | - | 2713 | 1043 | - | - | 0 | - |
| - | - | 2344 | 1044 | - | - | 0 | - |
| - | - | 3456 | 1044 | - | - | 0 | - |
| - | - | 2459 | 1045 | - | - | 0 | - |
| - | - | 5637 | 1045 | - | - | 0 | - |
| - | - | 2565 | 1046 | - | - | 0 | - |
| - | - | 4596 | 1048 | - | - | 0 | - |
| - | - | 3743 | 1048 | - | - | 0 | - |
| - | - | 4691 | 1049 | - | - | 0 | - |
| - | - | 5554 | 1049 | - | - | 0 | - |
| - | - | 6847 | 1050 | - | - | 0 | - |
| - | - | 5077 | 1050 | - | - | 0 | - |
| - | - | 3473 | 1050 | - | - | 0 | - |
| - | - | 4361 | 1051 | - | - | 0 | - |
| - | - | 3488 | 1053 | - | - | 0 | - |
| - | - | 4668 | 1054 | - | - | 0 | - |
| - | - | 4433 | 1054 | - | - | 0 | - |
| - | - | 4252 | 1054 | - | - | 0 | - |
| - | - | 7.052E+04 | 1055 | - | - | 0 | - |
| - | - | 3661 | 1055 | - | - | 0 | - |
| - | - | 3.834E+04 | 1056 | - | - | 0 | - |
| - | - | 1.812E+04 | 1057 | - | - | 0 | - |
| - | - | 5685 | 1058 | - | - | 0 | - |
| - | - | 4476 | 1059 | - | - | 0 | - |
| - | - | 3662 | 1060 | - | - | 0 | - |
| - | - | 2346 | 1060 | - | - | 0 | - |
| - | - | 2911 | 1060 | - | - | 0 | - |
| - | - | 7260 | 1061 | - | - | 0 | - |
| - | - | 7606 | 1061 | - | - | 0 | - |
| - | - | 2448 | 1061 | - | - | 0 | - |
| - | - | 3674 | 1062 | - | - | 0 | - |
| - | - | 3106 | 1062 | - | - | 0 | - |
| - | - | 6128 | 1063 | - | - | 0 | - |
| - | - | 5317 | 1063 | - | - | 0 | - |
| - | - | 7402 | 1063 | - | - | 0 | - |
| - | - | 6362 | 1064 | - | - | 0 | - |
| - | - | 3108 | 1064 | - | - | 0 | - |
| - | - | 3350 | 1065 | - | - | 0 | - |
| - | - | 3337 | 1066 | - | - | 0 | - |
| - | - | 7585 | 1066 | - | - | 0 | - |
| - | - | 6211 | 1066 | - | - | 0 | - |
| 7 | w | 1.306E+04 | 1067 | 0.01945 | 18.24 | +1 | 9 |
| - | - | 8012 | 1068 | - | - | 0 | - |
| - | - | 4375 | 1068 | - | - | 0 | - |
| - | - | 3605 | 1068 | - | - | 0 | - |
| - | - | 1.088E+04 | 1069 | - | - | 0 | - |
| - | - | 5534 | 1069 | - | - | 0 | - |
| - | - | 2882 | 1069 | - | - | 0 | - |
| - | - | 6350 | 1070 | - | - | 0 | - |
| - | - | 2662 | 1078 | - | - | 0 | - |
| - | - | 5545 | 1110 | - | - | 0 | - |
| 9 | c | 2.027E+05 | 1111 | 0.0002673 | 0.2407 | +1 | 9 |
| - | - | 1.284E+05 | 1112 | - | - | 0 | - |
| - | - | 4.886E+04 | 1113 | - | - | 0 | - |
| - | - | 1.862E+04 | 1114 | - | - | 0 | - |
| - | - | 5237 | 1115 | - | - | 0 | - |
| - | - | 2628 | 1116 | - | - | 0 | - |
| 7 | y | 1.666E+04 | 1128 | 0.004374 | 3.879 | +1 | 9 |
| - | - | 1.201E+04 | 1129 | - | - | 0 | - |
| - | - | 4041 | 1130 | - | - | 0 | - |
| - | - | 7907 | 1137 | - | - | 0 | - |
| - | - | 5539 | 1138 | - | - | 0 | - |
| - | - | 3231 | 1139 | - | - | 0 | - |
| - | - | 2634 | 1153 | - | - | 0 | - |
| - | - | 2182 | 1172 | - | - | 0 | - |
| - | - | 2907 | 1173 | - | - | 0 | - |
| - | - | 7601 | 1181 | - | - | 0 | - |
| - | - | 4757 | 1182 | - | - | 0 | - |
| - | - | 3161 | 1183 | - | - | 0 | - |
| - | - | 8308 | 1196 | - | - | 0 | - |
| - | - | 8061 | 1197 | - | - | 0 | - |
| - | - | 3207 | 1198 | - | - | 0 | - |
| 10 | c | 2740 | 1222 | 0.0003434 | 0.2811 | +1 | 10 |
| 6 | y | 2342 | 1223 | 0.0008623 | 0.7053 | +1 | 10 |
| 6 | y | 3161 | 1224 | 0.002931 | 2.395 | +1 | 10 |
| 6 | z | 3.723E+04 | 1225 | 0.00343 | 2.8 | +1 | 10 |
| - | - | 2.779E+04 | 1226 | - | - | 0 | - |
| - | - | 1.306E+04 | 1227 | - | - | 0 | - |
| - | - | 5236 | 1228 | - | - | 0 | - |
| - | - | 9291 | 1239 | - | - | 0 | - |
| 10 | c | 1.182E+05 | 1240 | 8.948E-05 | 0.07219 | +1 | 10 |
| 6 | y | 9.468E+04 | 1241 | 0.01336 | 10.77 | +1 | 10 |
| - | - | 3.767E+04 | 1242 | - | - | 0 | - |
| - | - | 9693 | 1243 | - | - | 0 | - |
| - | - | 2960 | 1263 | - | - | 0 | - |
| - | - | 3422 | 1264 | - | - | 0 | - |
| - | - | 2331 | 1265 | - | - | 0 | - |
| - | - | 2973 | 1275 | - | - | 0 | - |
| - | - | 3128 | 1280 | - | - | 0 | - |
| - | - | 4024 | 1281 | - | - | 0 | - |
| - | - | 2704 | 1302 | - | - | 0 | - |
| - | - | 3090 | 1318 | - | - | 0 | - |
| - | - | 2535 | 1321 | - | - | 0 | - |
| - | - | 2089 | 1340 | - | - | 0 | - |
| - | - | 2971 | 1355 | - | - | 0 | - |
| - | - | 2918 | 1356 | - | - | 0 | - |
| - | - | 7769 | 1359 | - | - | 0 | - |
| - | - | 4712 | 1360 | - | - | 0 | - |
| - | - | 3136 | 1367 | - | - | 0 | - |
| - | - | 2306 | 1368 | - | - | 0 | - |
| - | - | 6402 | 1402 | - | - | 0 | - |
| 11 | c | 5.659E+04 | 1403 | 0.0001806 | 0.1288 | +1 | 11 |
| - | - | 3.743E+04 | 1404 | - | - | 0 | - |
| - | - | 1.917E+04 | 1405 | - | - | 0 | - |
| - | - | 5583 | 1406 | - | - | 0 | - |
| 5 | y | 2834 | 1410 | 0.008457 | 5.999 | +1 | 11 |
| 5 | z | 4.721E+04 | 1411 | 0.003641 | 2.581 | +1 | 11 |
| - | - | 4.368E+04 | 1412 | - | - | 0 | - |
| - | - | 2.675E+04 | 1413 | - | - | 0 | - |
| - | - | 1.105E+04 | 1414 | - | - | 0 | - |
| - | - | 3168 | 1415 | - | - | 0 | - |
| - | - | 3980 | 1426 | - | - | 0 | - |
| 5 | y | 5.148E+04 | 1427 | 0.008083 | 5.665 | +1 | 11 |
| - | - | 4.31E+04 | 1428 | - | - | 0 | - |
| - | - | 1.829E+04 | 1429 | - | - | 0 | - |
| - | - | 8112 | 1430 | - | - | 0 | - |
| - | - | 3190 | 1431 | - | - | 0 | - |
| - | - | 5134 | 1437 | - | - | 0 | - |
| - | - | 2377 | 1438 | - | - | 0 | - |
| - | - | 4895 | 1464 | - | - | 0 | - |
| - | - | 2775 | 1465 | - | - | 0 | - |
| - | - | 2542 | 1466 | - | - | 0 | - |
| - | - | 4173 | 1467 | - | - | 0 | - |
| - | - | 2533 | 1468 | - | - | 0 | - |
| - | - | 2363 | 1471 | - | - | 0 | - |
| - | - | 2455 | 1480 | - | - | 0 | - |
| - | - | 3.558E+04 | 1482 | - | - | 0 | - |
| - | - | 2.859E+04 | 1483 | - | - | 0 | - |
| - | - | 1.735E+04 | 1484 | - | - | 0 | - |
| - | - | 8232 | 1485 | - | - | 0 | - |
| - | - | 5851 | 1486 | - | - | 0 | - |
| - | - | 2629 | 1486 | - | - | 0 | - |
| - | - | 7163 | 1487 | - | - | 0 | - |
| - | - | 1.07E+04 | 1488 | - | - | 0 | - |
| - | - | 4190 | 1489 | - | - | 0 | - |
| - | - | 2855 | 1499 | - | - | 0 | - |
| - | - | 3873 | 1501 | - | - | 0 | - |
| - | - | 2743 | 1502 | - | - | 0 | - |
| - | - | 2611 | 1515 | - | - | 0 | - |
| - | - | 6062 | 1516 | - | - | 0 | - |
| - | - | 2903 | 1517 | - | - | 0 | - |
| - | - | 2831 | 1518 | - | - | 0 | - |
| 4 | y | 3437 | 1525 | 0.007515 | 4.929 | +1 | 12 |
| 4 | z | 6.813E+04 | 1526 | 0.004949 | 3.244 | +1 | 12 |
| - | - | 8.212E+04 | 1527 | - | - | 0 | - |
| - | - | 6.48E+04 | 1528 | - | - | 0 | - |
| - | - | 2755 | 1528 | - | - | 0 | - |
| - | - | 2.647E+04 | 1529 | - | - | 0 | - |
| - | - | 1.384E+04 | 1530 | - | - | 0 | - |
| 12 | c | 8.742E+04 | 1531 | 0.001765 | 1.153 | +1 | 12 |
| - | - | 8.792E+04 | 1532 | - | - | 0 | - |
| - | - | 4.201E+04 | 1533 | - | - | 0 | - |
| - | - | 1.526E+04 | 1534 | - | - | 0 | - |
| - | - | 6237 | 1535 | - | - | 0 | - |
| - | - | 2733 | 1537 | - | - | 0 | - |
| - | - | 4152 | 1540 | - | - | 0 | - |
| 4 | y | 1.485E+04 | 1542 | 0.003532 | 2.291 | +1 | 12 |
| - | - | 1.25E+04 | 1543 | - | - | 0 | - |
| - | - | 7759 | 1544 | - | - | 0 | - |
| - | - | 2676 | 1549 | - | - | 0 | - |
| - | - | 2440 | 1550 | - | - | 0 | - |
| - | - | 4786 | 1550 | - | - | 0 | - |
| - | - | 4482 | 1551 | - | - | 0 | - |
| - | - | 3967 | 1551 | - | - | 0 | - |
| - | - | 4124 | 1552 | - | - | 0 | - |
| - | - | 2777 | 1552 | - | - | 0 | - |
| - | - | 2901 | 1553 | - | - | 0 | - |
| - | - | 2695 | 1554 | - | - | 0 | - |
| - | - | 3055 | 1556 | - | - | 0 | - |
| - | - | 2452 | 1556 | - | - | 0 | - |
| - | - | 4643 | 1557 | - | - | 0 | - |
| - | - | 5579 | 1558 | - | - | 0 | - |
| - | - | 6660 | 1559 | - | - | 0 | - |
| - | - | 5550 | 1559 | - | - | 0 | - |
| - | - | 2918 | 1560 | - | - | 0 | - |
| - | - | 3701 | 1560 | - | - | 0 | - |
| - | - | 2981 | 1564 | - | - | 0 | - |
| - | - | 5171 | 1565 | - | - | 0 | - |
| - | - | 6212 | 1565 | - | - | 0 | - |
| - | - | 6533 | 1566 | - | - | 0 | - |
| - | - | 6821 | 1566 | - | - | 0 | - |
| - | - | 8878 | 1567 | - | - | 0 | - |
| - | - | 6689 | 1567 | - | - | 0 | - |
| - | - | 5366 | 1568 | - | - | 0 | - |
| - | - | 5407 | 1568 | - | - | 0 | - |
| - | - | 2554 | 1569 | - | - | 0 | - |
| - | - | 3704 | 1570 | - | - | 0 | - |
| - | - | 2990 | 1571 | - | - | 0 | - |
| - | - | 4740 | 1572 | - | - | 0 | - |
| - | - | 6510 | 1572 | - | - | 0 | - |
| - | - | 4791 | 1573 | - | - | 0 | - |
| - | - | 6015 | 1573 | - | - | 0 | - |
| - | - | 8170 | 1574 | - | - | 0 | - |
| - | - | 1.127E+04 | 1574 | - | - | 0 | - |
| - | - | 1.035E+04 | 1575 | - | - | 0 | - |
| - | - | 5648 | 1575 | - | - | 0 | - |
| - | - | 5110 | 1576 | - | - | 0 | - |
| - | - | 3194 | 1576 | - | - | 0 | - |
| - | - | 3059 | 1577 | - | - | 0 | - |
| - | - | 2885 | 1578 | - | - | 0 | - |
| - | - | 3356 | 1579 | - | - | 0 | - |
| - | - | 3535 | 1579 | - | - | 0 | - |
| - | - | 3126 | 1580 | - | - | 0 | - |
| - | - | 4316 | 1580 | - | - | 0 | - |
| - | - | 2515 | 1581 | - | - | 0 | - |
| - | - | 6565 | 1581 | - | - | 0 | - |
| - | - | 5503 | 1582 | - | - | 0 | - |
| - | - | 4180 | 1582 | - | - | 0 | - |
| - | - | 5178 | 1583 | - | - | 0 | - |
| - | - | 7684 | 1583 | - | - | 0 | - |
| - | - | 2928 | 1585 | - | - | 0 | - |
| - | - | 2876 | 1586 | - | - | 0 | - |
| - | - | 3436 | 1586 | - | - | 0 | - |
| - | - | 3743 | 1587 | - | - | 0 | - |
| - | - | 4106 | 1588 | - | - | 0 | - |
| - | - | 3023 | 1589 | - | - | 0 | - |
| - | - | 6467 | 1590 | - | - | 0 | - |
| - | - | 6895 | 1590 | - | - | 0 | - |
| - | - | 7418 | 1591 | - | - | 0 | - |
| - | - | 6265 | 1591 | - | - | 0 | - |
| - | - | 4707 | 1592 | - | - | 0 | - |
| - | - | 2468 | 1592 | - | - | 0 | - |
| - | - | 4098 | 1593 | - | - | 0 | - |
| - | - | 3757 | 1593 | - | - | 0 | - |
| - | - | 7016 | 1594 | - | - | 0 | - |
| - | - | 5763 | 1594 | - | - | 0 | - |
| - | - | 8912 | 1595 | - | - | 0 | - |
| - | - | 6585 | 1595 | - | - | 0 | - |
| - | - | 6690 | 1596 | - | - | 0 | - |
| - | - | 5062 | 1597 | - | - | 0 | - |
| - | - | 3775 | 1598 | - | - | 0 | - |
| - | - | 7291 | 1599 | - | - | 0 | - |
| - | - | 3444 | 1599 | - | - | 0 | - |
| - | - | 2904 | 1600 | - | - | 0 | - |
| - | - | 3571 | 1600 | - | - | 0 | - |
| - | - | 3506 | 1601 | - | - | 0 | - |
| - | - | 5129 | 1602 | - | - | 0 | - |
| - | - | 8346 | 1602 | - | - | 0 | - |
| - | - | 8767 | 1603 | - | - | 0 | - |
| - | - | 2541 | 1603 | - | - | 0 | - |
| - | - | 3085 | 1604 | - | - | 0 | - |
| - | - | 4259 | 1610 | - | - | 0 | - |
| - | - | 2404 | 1635 | - | - | 0 | - |
| 3 | z | 6.413E+04 | 1654 | 0.004567 | 2.761 | +1 | 13 |
| - | - | 7.212E+04 | 1655 | - | - | 0 | - |
| - | - | 4.467E+04 | 1656 | - | - | 0 | - |
| - | - | 2.46E+04 | 1657 | - | - | 0 | - |
| - | - | 6332 | 1658 | - | - | 0 | - |
| 3 | y | 4327 | 1670 | 0.006689 | 4.006 | +1 | 13 |
| - | - | 4694 | 1671 | - | - | 0 | - |
| - | - | 2405 | 1677 | - | - | 0 | - |
| - | - | 6303 | 1691 | - | - | 0 | - |
| 13 | c | 9.682E+04 | 1692 | 0.005202 | 3.075 | +1 | 13 |
| - | - | 7.601E+04 | 1693 | - | - | 0 | - |
| - | - | 4.598E+04 | 1694 | - | - | 0 | - |
| - | - | 1.945E+04 | 1695 | - | - | 0 | - |
| - | - | 6048 | 1696 | - | - | 0 | - |
| - | - | 2480 | 1734 | - | - | 0 | - |
| - | - | 4497 | 1761 | - | - | 0 | - |
| - | - | 4485 | 1762 | - | - | 0 | - |
| - | - | 4018 | 1763 | - | - | 0 | - |
| - | - | 3264 | 1764 | - | - | 0 | - |
| - | - | 4388 | 1776 | - | - | 0 | - |
| - | - | 9918 | 1777 | - | - | 0 | - |
| - | - | 9405 | 1778 | - | - | 0 | - |
| - | - | 5853 | 1779 | - | - | 0 | - |
| 2 | z | 6347 | 1791 | 0.001829 | 1.021 | +1 | 14 |
| - | - | 2.661E+04 | 1792 | - | - | 0 | - |
| - | - | 2.242E+04 | 1793 | - | - | 0 | - |
| - | - | 1.852E+04 | 1794 | - | - | 0 | - |
| - | - | 6052 | 1795 | - | - | 0 | - |
| - | - | 2517 | 1796 | - | - | 0 | - |
| - | - | 4870 | 1804 | - | - | 0 | - |
| - | - | 5699 | 1805 | - | - | 0 | - |
| - | - | 3748 | 1806 | - | - | 0 | - |
| 2 | y | 3349 | 1807 | 0.02815 | 15.58 | +1 | 14 |
| - | - | 2785 | 1814 | - | - | 0 | - |
| - | - | 4494 | 1818 | - | - | 0 | - |
| - | - | 4833 | 1819 | - | - | 0 | - |
| 14 | c | 1.71E+04 | 1820 | 0.002984 | 1.64 | +1 | 14 |
| - | - | 3.87E+04 | 1821 | - | - | 0 | - |
| - | - | 3.408E+04 | 1822 | - | - | 0 | - |
| - | - | 1.474E+04 | 1823 | - | - | 0 | - |
| - | - | 7505 | 1824 | - | - | 0 | - |
| - | - | 3134 | 1825 | - | - | 0 | - |
| - | - | 1.074E+04 | 1831 | - | - | 0 | - |
| - | - | 1.098E+04 | 1832 | - | - | 0 | - |
| - | - | 7514 | 1833 | - | - | 0 | - |
| - | - | 4205 | 1834 | - | - | 0 | - |
| - | - | 5990 | 1835 | - | - | 0 | - |
| - | - | 5055 | 1836 | - | - | 0 | - |
| - | - | 2677 | 1837 | - | - | 0 | - |
| - | - | 1.015E+04 | 1845 | - | - | 0 | - |
| - | - | 1.448E+04 | 1846 | - | - | 0 | - |
| - | - | 1.281E+04 | 1847 | - | - | 0 | - |
| - | - | 7187 | 1848 | - | - | 0 | - |
| - | - | 4188 | 1849 | - | - | 0 | - |
| - | - | 3271 | 1859 | - | - | 0 | - |
| - | - | 6419 | 1860 | - | - | 0 | - |
| - | - | 9604 | 1861 | - | - | 0 | - |
| - | - | 4.32E+04 | 1862 | - | - | 0 | - |
| - | - | 8.291E+04 | 1863 | - | - | 0 | - |
| - | - | 6.323E+04 | 1864 | - | - | 0 | - |
| - | - | 2795 | 1864 | - | - | 0 | - |
| - | - | 3.847E+04 | 1865 | - | - | 0 | - |
| - | - | 1.504E+04 | 1866 | - | - | 0 | - |
| - | - | 4412 | 1867 | - | - | 0 | - |
| - | - | 3339 | 1875 | - | - | 0 | - |
| - | - | 1.712E+04 | 1876 | - | - | 0 | - |
| - | - | 3.58E+04 | 1877 | - | - | 0 | - |
| - | - | 3.29E+04 | 1878 | - | - | 0 | - |
| - | - | 1.775E+04 | 1879 | - | - | 0 | - |
| - | - | 7382 | 1880 | - | - | 0 | - |
| - | - | 2910 | 1881 | - | - | 0 | - |
| - | - | 4543 | 1887 | - | - | 0 | - |
| - | - | 3515 | 1888 | - | - | 0 | - |
| - | - | 2437 | 1891 | - | - | 0 | - |
| - | - | 1.492E+04 | 1893 | - | - | 0 | - |
| - | - | 4.202E+04 | 1894 | - | - | 0 | - |
| - | - | 4.518E+04 | 1895 | - | - | 0 | - |
| - | - | 2.371E+04 | 1896 | - | - | 0 | - |
| - | - | 8249 | 1897 | - | - | 0 | - |
| - | - | 3343 | 1898 | - | - | 0 | - |
| - | - | 4443 | 1903 | - | - | 0 | - |
| - | - | 3.644E+04 | 1904 | - | - | 0 | - |
| - | - | 9.766E+04 | 1905 | - | - | 0 | - |
| - | - | 9.852E+04 | 1906 | - | - | 0 | - |
| - | - | 5.776E+04 | 1907 | - | - | 0 | - |
| - | - | 2.737E+04 | 1908 | - | - | 0 | - |
| - | - | 9314 | 1909 | - | - | 0 | - |
| - | - | 3230 | 1910 | - | - | 0 | - |
| - | - | 4423 | 1919 | - | - | 0 | - |
| - | - | 2.36E+04 | 1920 | - | - | 0 | - |
| - | - | 1.208E+05 | 1921 | - | - | 0 | - |
| - | - | 4.025E+05 | 1922 | - | - | 0 | - |
| - | - | 3.719E+05 | 1923 | - | - | 0 | - |
| - | - | 2.06E+05 | 1924 | - | - | 0 | - |
| - | - | 1.093E+05 | 1925 | - | - | 0 | - |
| - | - | 3.353E+04 | 1926 | - | - | 0 | - |
| - | - | 9033 | 1927 | - | - | 0 | - |
| - | - | 2743 | 2124 | - | - | 0 | - |

m/z Charge Intensity FragmentType MassShift Position
128.0819549560547 0 1536.171
129.1024169921875 0 5547.814
146.0928192138672 0 2517.3796
147.5020294189453 0 1231.0679
151.6353759765625 0 1359.6807
155.0928192138672 0 5039.008
166.0612335205078 0 12795.595
178.1342010498047 0 4162.4404
183.11293029785156 0 2188.7754
185.1649932861328 0 8643.277
207.16073608398438 0 2107.1018
222.1477508544922 0 1678.9928
223.155517578125 0 13729.022
230.16258239746094 0 45143.945 z 13
231.16864013671875 0 7934.157
234.12403869628906 0 21172.72
246.181396484375 0 3528.3066 y 13
250.1425323486328 0 4083.5647
251.15042114257812 0 86684.33
252.15380859375 0 12520.279
266.1251220703125 0 3459.9626
268.1767883300781 0 41589.09 c 1
269.1803283691406 0 4802.7544
283.15142822265625 0 6233.492
286.17633056640625 0 1468.8859
300.1920471191406 0 69190.34
301.1952819824219 0 10766.403
331.8556213378906 0 1767.3711
351.2154235839844 0 2153.6062
352.2218933105469 0 6308.935
357.1310119628906 0 1553.6213
379.20916748046875 0 15663.073 c Ammonia loss 2
380.2131652832031 0 2822.7937
395.2278747558594 0 5613.2705
396.2356872558594 0 87503.01 c 2
397.2388610839844 0 18928.523
398.2417907714844 0 2206.3735
407.19622802734375 0 8545.492 y 12
408.1993408203125 0 2187.882 c Ammonia loss 9
409.2200927734375 0 3317.2297
412.13519287109375 0 7700.8223
415.5943603515625 0 1650.6333
416.1307067871094 0 1838.1388
428.2882385253906 0 2675.2217
448.8985900878906 0 1833.3866
451.22119140625 0 1622.0586
460.25775146484375 0 2897.1174
461.2043762207031 0 3369.6821 w 11
467.2491455078125 0 51592.508
468.2536926269531 0 18870.518
469.2558898925781 0 3545.3333
475.2820129394531 0 3692.6628
479.1514892578125 0 2436.785
494.2360534667969 0 10425.1 c Ammonia loss 3
506.2276306152344 0 2038.6947
506.7607727050781 0 43328.117 y 7
507.2620544433594 0 20212.996
507.7627868652344 0 7506.9956
508.2624206542969 0 3246.8564
511.26251220703125 0 78518.79 c 3
512.2652587890625 0 18935.225
513.2685546875 0 2457.5232
519.2723999023438 0 39244.234 z 11
520.2772827148438 0 13660.305
521.2717895507812 0 4245.7656
522.1572875976562 0 2390.0264
525.2216796875 0 4496.665
535.291259765625 0 12730.082 y 11
536.29296875 0 4796.553
563.9287109375 0 3157.6648
564.2731323242188 0 18492.844 y 6
564.5979614257812 0 2479.2346 c 12
564.776123046875 0 5590.861
565.2754516601562 0 2262.3577
578.2220458984375 0 4214.0195
584.3631591796875 0 13309.11
585.364501953125 0 4111.998
591.3486938476562 0 3605.6238
592.3560180664062 0 2406.6274
596.284912109375 0 3980.2395
597.2854614257812 0 4747.5547 y Ammonia loss 1
601.62353515625 0 25703.93 c Ammonia loss 13
601.9573364257812 0 19057.521
602.291748046875 0 19458.8
602.6242065429688 0 9786.779
602.955078125 0 34014.45 y 1
603.289306640625 0 32411.19
603.6229248046875 0 16707.756
603.9551391601562 0 7225.9297
604.289306640625 0 4018.9495 z Ammonia loss 5
607.6271362304688 0 7419.903
607.960205078125 0 7056.7974
608.293701171875 0 4572.2427
610.3170776367188 0 2486.5986
620.8179321289062 0 3027.8157 y 5
622.218017578125 0 5156.2417
623.3072509765625 0 3182.0437
624.2202758789062 0 3725.9944
624.2711181640625 0 6111.7754
625.6455078125 0 2289.2202
625.9760131835938 0 3175.2734
628.970703125 0 12128.2705
629.3048706054688 0 12238.82
629.6353759765625 0 4369.2227
629.972900390625 0 4427.8613
631.9825439453125 0 2064.1812
634.2984008789062 0 2422.5354
634.645263671875 0 27807.6
634.9785766601562 0 42944.004
635.3116455078125 0 28639.453
635.6453857421875 0 15285.594
635.979248046875 0 9121.941
636.315673828125 0 2039.9835
637.3071899414062 0 2960.9553
638.3495483398438 0 2404.182
639.2998657226562 0 2039.8397
639.9788818359375 0 2347.705
640.3189086914062 0 21077.484
640.6494140625 0 430346.72
640.9835815429688 0 466212.28
641.317626953125 0 289729.06
641.6511840820312 0 146624.56
641.82421875 0 3130.0935
641.9853515625 0 48195.934
642.2291259765625 0 5003.9067
642.3177490234375 0 16663.555
651.3509521484375 0 3685.666
653.3263549804688 0 5640.8496
658.283447265625 0 2537.3896
680.3162841796875 0 9669.6455 c Ammonia loss 4
681.3208618164062 0 5177.7188 y Ammonia loss 10
682.3358764648438 0 94680.414 z 10
683.33984375 0 42459.453
684.3400268554688 0 13942.501
685.34619140625 0 3259.0032
697.342041015625 0 62946.117 c 4
698.3499755859375 0 47893.44 y 10
699.354248046875 0 15199.663
700.3616333007812 0 3135.0747
713.858154296875 0 10156.204 y 4
714.3566284179688 0 7658.3643
714.8652954101562 0 3235.218
720.395263671875 0 6841.6904
721.3975219726562 0 4224.747
752.3660278320312 0 30117.297 w 9
753.3681030273438 0 13101.511
754.3697509765625 0 4496.3057 z Water loss 3
765.4005737304688 0 5212.963
765.890625 0 23983.004 c 11
766.3935546875 0 22235.977
766.891357421875 0 7990.1113
767.404296875 0 8427.56
771.3707885742188 0 10244.708 y 3
771.8729248046875 0 6892.788
772.3727416992188 0 3440.5452
775.4007568359375 0 2753.4866
777.3571166992188 0 4156.5063
779.4006958007812 0 4094.6013
787.4019775390625 0 2599.0254
787.65771484375 0 2614.6475
789.4473266601562 0 2387.432
789.9047241210938 0 2200.635
793.396484375 0 7569.5664 c Ammonia loss 5
794.4036865234375 0 4718.2026
799.3936157226562 0 2165.3503
809.4179077148438 0 3439.4312
810.4266967773438 0 93808.99 c 5
811.3780517578125 0 133840.17 z 9
812.3812866210938 0 59210.164
813.3819580078125 0 22131.41
814.3850708007812 0 5793.0063
815.38525390625 0 2254.3184
817.3807373046875 0 2653.5654
826.3924560546875 0 9950.663 y Water loss 2
826.8900146484375 0 23397.463 y Ammonia loss 2
827.3922119140625 0 35989.785 y 9
827.891357421875 0 13773.892
828.3963012695312 0 11069.354
829.3952026367188 0 3207.7588
834.90625 0 2822.2034
835.3989868164062 0 53349.32 y 2
835.9005737304688 0 47936.68
836.40234375 0 22770.586
836.9019165039062 0 12210.237
837.40673828125 0 4065.471 c Water loss 12
837.892822265625 0 2350.5444 c Ammonia loss 12
838.3983764648438 0 3511.5337
838.8900146484375 0 2547.8884
841.9364013671875 0 2926.819
842.9444580078125 0 7429.342
843.4450073242188 0 6633.685
843.9527587890625 0 3300.4167
845.9020385742188 0 6363.68
846.3971557617188 0 34358.04 c 12
846.89794921875 0 24482.56
847.3986206054688 0 15054.435
847.9012451171875 0 11519.823
848.4845581054688 0 6765.0547
849.4950561523438 0 2695.7402
850.4241943359375 0 4982.642
858.9320068359375 0 2194.1597
859.4176025390625 0 3047.8977
860.4070434570312 0 2600.3384
866.4296264648438 0 7886.4893
866.933837890625 0 4448.256
867.4241943359375 0 63234.105
868.426025390625 0 30825.402
869.4298095703125 0 8269.975
873.92333984375 0 5058.97
874.418212890625 0 3519.2131
874.9207763671875 0 4838.377
875.414306640625 0 2444.198
880.4442138671875 0 5260
881.4398193359375 0 14721.28
882.4530639648438 0 6084.591
883.9683837890625 0 2564.051
885.4742431640625 0 3516.8425
887.9284057617188 0 4111.0776
888.4347534179688 0 10116.439
888.938720703125 0 9642.878
889.4330444335938 0 4461.718
893.4840698242188 0 4445.8374
894.43994140625 0 2865.7708
894.9290771484375 0 3487.4612 y Water loss 1
895.4405517578125 0 10378.671
895.9226684570312 0 36274.047 z 1
896.4241333007812 0 34180.59
896.9227294921875 0 17896.264
897.4268798828125 0 9695.205
897.9190063476562 0 2646.503
901.4473266601562 0 5067.3506 c Water loss 13
901.939453125 0 5416.0073 c Ammonia loss 13
902.4412841796875 0 9441.726
902.9425048828125 0 5239.7026
903.4385986328125 0 3491.3125
903.9307250976562 0 2496.1265 y 1
904.4329223632812 0 2228.2407
906.5078125 0 2720.462
906.9640502929688 0 7698.709
907.4651489257812 0 13258.484
907.967529296875 0 8493.643
908.429931640625 0 20867.885 c Ammonia loss 6
908.9650268554688 0 8594.596
909.4398193359375 0 14805.879
909.9521484375 0 8662.708
910.4439697265625 0 159650.66 c 13
910.9452514648438 0 153933.86
911.4463500976562 0 92837.61
911.9476928710938 0 40710.035
912.448974609375 0 16300.46
912.9534912109375 0 6132.621
914.9766235351562 0 2952.5815
915.4642333984375 0 2460.046
915.9700927734375 0 7582.785
916.4701538085938 0 11492.242
916.9708862304688 0 10362.341
917.46630859375 0 5331.649
917.9620361328125 0 2301.7012
922.4634399414062 0 19811.135 z Ammonia loss 8
922.965576171875 0 17097.234
923.4581298828125 0 11742.695
923.9588012695312 0 8068.5215
924.44775390625 0 26245.791
924.947509765625 0 10394.451
925.4513549804688 0 48505.035 c 6
925.9517822265625 0 4308.2886
926.4555053710938 0 24214.79
927.4566040039062 0 5807.1704
929.9630737304688 0 3190.1301
930.466552734375 0 28985.342
930.9525146484375 0 82246.98
931.462890625 0 204945.08
931.9662475585938 0 197763.3
932.466796875 0 106380.07
932.9673461914062 0 49308.883
933.4682006835938 0 18566.959
933.972412109375 0 7924.789
937.4715576171875 0 4357.1924 y Water loss 8
937.9688110351562 0 19651.074
938.469482421875 0 33157.22 y Ammonia loss 8
938.96826171875 0 29159.473
939.4718627929688 0 89407.12 z 8
939.9683837890625 0 8515.467
940.4768676757812 0 54726.305
941.4799194335938 0 20623.574
942.4810791015625 0 7902.967
942.9688720703125 0 4425.12
943.4717407226562 0 5597.5254
943.9642333984375 0 2439.3396
944.446533203125 0 2159.696
946.4619750976562 0 2522.3374
951.4664916992188 0 6076.9683
951.966796875 0 16235.027
952.4630737304688 0 67565.63
952.9635620117188 0 62442.137
953.46435546875 0 41982.977
953.9645385742188 0 22690.504
954.4656982421875 0 9480.607
954.9580078125 0 4452.481
955.4894409179688 0 5105.2334 y 8
956.4973754882812 0 2794.8086
959.9707641601562 0 11948.132
960.4706420898438 0 216594.53
960.9727172851562 0 319214.16
961.4736328125 0 251425.98
961.9749145507812 0 146711.58
962.47509765625 0 54410.355
962.9730834960938 0 22571.81
963.46728515625 0 6040.123
965.4588623046875 0 3058.7712 c Ammonia loss 7
966.4705810546875 0 1899.7146
976.485107421875 0 2124.313
981.47119140625 0 17141.85
982.4749755859375 0 97100.05 c 7
983.4778442382812 0 53811.344
984.4783935546875 0 16306.716
985.4837646484375 0 4222.341
991.5325317382812 0 2540.2268
996.4946899414062 0 27734.654 z 7
997.4990844726562 0 31260.115
998.5006103515625 0 14684.662
999.5005493164062 0 4534.007
1011.5148315429688 0 2970.8867
1012.5126342773438 0 21415.396 y 7
1013.5143432617188 0 14408.133
1014.5201416015625 0 3124.053
1015.519775390625 0 3103.7693
1020.53564453125 0 5012.2935
1024.5152587890625 0 3441.909
1028.196044921875 0 3499.739
1030.526611328125 0 4684.199
1031.52392578125 0 3191.8606
1032.885498046875 0 3538.3516
1033.8719482421875 0 3682.4548
1039.5118408203125 0 4745.5576
1040.5028076171875 0 3150.9397
1042.533447265625 0 2713.1575
1043.52734375 0 2343.9602
1044.20263671875 0 3455.973
1044.5330810546875 0 2458.8396
1045.2025146484375 0 5637.372
1045.53955078125 0 2565.4795
1047.5386962890625 0 4595.569
1047.865966796875 0 3742.6638
1048.5264892578125 0 4691.289
1049.1986083984375 0 5554.11
1049.5333251953125 0 6847.085
1049.864990234375 0 5076.5977
1050.20947265625 0 3473.2505
1050.527587890625 0 4360.924
1053.2093505859375 0 3488.326
1053.5396728515625 0 4667.5576
1053.85595703125 0 4432.8213
1054.2027587890625 0 4251.8423
1054.524658203125 0 70524.99
1054.86474609375 0 3660.7065
1055.5272216796875 0 38337.21
1056.528564453125 0 18120.527
1057.5322265625 0 5685.172
1058.533447265625 0 4475.7305
1059.5255126953125 0 3662.0364
1059.8441162109375 0 2345.6365
1060.185791015625 0 2911.1853
1060.5299072265625 0 7259.688
1060.8677978515625 0 7605.644
1061.1905517578125 0 2447.7754
1061.5308837890625 0 3674.0066
1062.18994140625 0 3106.2236
1062.5321044921875 0 6128.2393
1062.865966796875 0 5317.4307
1063.2049560546875 0 7401.9854
1063.5406494140625 0 6362.376
1063.8726806640625 0 3108.4753
1064.5238037109375 0 3350.2117
1065.534912109375 0 3337.3481
1065.8787841796875 0 7585.4253
1066.203369140625 0 6211.259
1066.5477294921875 0 13061.25 w 6
1067.53662109375 0 8012.2783
1067.8702392578125 0 4375.3213
1068.210693359375 0 3604.8242
1068.544189453125 0 10875.338
1068.8817138671875 0 5533.89
1069.20751953125 0 2882.2202
1069.5457763671875 0 6350.421
1077.548095703125 0 2661.5618
1109.57080078125 0 5544.957
1110.5693359375 0 202701.72 c 8
1111.5611572265625 0 128438.516
1112.54736328125 0 48859.793
1113.5401611328125 0 18621.707
1114.53515625 0 5236.957
1115.5421142578125 0 2627.8496
1127.540283203125 0 16658.38 y 6
1128.542724609375 0 12007.891
1129.5482177734375 0 4041.022
1136.586181640625 0 7906.995
1137.587646484375 0 5538.7964
1138.5810546875 0 3230.8613
1152.579833984375 0 2634.1309
1171.6046142578125 0 2181.9863
1172.6072998046875 0 2907.1782
1180.6124267578125 0 7600.5093
1181.604248046875 0 4756.9165
1182.606689453125 0 3161.4548
1195.5994873046875 0 8307.767
1196.6005859375 0 8060.5596
1197.6085205078125 0 3206.9106
1221.6014404296875 0 2740.362 c Water loss 9
1222.6190185546875 0 2342.382 y Water loss 5
1223.6051025390625 0 3160.5425 y Ammonia loss 5
1224.6065673828125 0 37227.465 z 5
1225.6080322265625 0 27791.354
1226.6097412109375 0 13055.042
1227.6114501953125 0 5235.7485
1238.61279296875 0 9291.188
1239.611572265625 0 118172.26 c 9
1240.6153564453125 0 94684.836 y 5
1241.6175537109375 0 37666.363
1242.6199951171875 0 9692.978
1262.6231689453125 0 2959.5027
1263.64501953125 0 3422.4895
1264.6370849609375 0 2331.1106
1274.630859375 0 2973.1802
1279.613525390625 0 3128.1367
1280.6290283203125 0 4024.14
1301.7027587890625 0 2703.737
1317.6885986328125 0 3089.8022
1320.7039794921875 0 2535.1968
1339.6456298828125 0 2088.645
1354.642578125 0 2971.485
1355.6217041015625 0 2917.5671
1358.663818359375 0 7768.7466
1359.6583251953125 0 4711.971
1366.6876220703125 0 3136.0547
1367.685302734375 0 2305.7305
1401.6854248046875 0 6401.5444
1402.6751708984375 0 56588.074 c 10
1403.6788330078125 0 37434.414
1404.6795654296875 0 19169.557
1405.68115234375 0 5582.9453
1409.68994140625 0 2833.5757 y Ammonia loss 4
1410.6856689453125 0 47208.285 z 4
1411.687744140625 0 43683.93
1412.690673828125 0 26754.432
1413.693603515625 0 11053.795
1414.6988525390625 0 3167.962
1425.6876220703125 0 3980.3872
1426.699951171875 0 51478.05 y 4
1427.7044677734375 0 43098.15
1428.7042236328125 0 18287.248
1429.71044921875 0 8111.8457
1430.7222900390625 0 3189.7036
1436.71826171875 0 5133.9907
1437.717529296875 0 2377.1704
1463.71630859375 0 4895.4053
1464.71240234375 0 2774.7512
1465.747314453125 0 2541.607
1466.7032470703125 0 4172.924
1467.744384765625 0 2532.761
1470.7325439453125 0 2362.9844
1480.19970703125 0 2454.9421
1481.720947265625 0 35580.984
1482.72265625 0 28588.896
1483.7264404296875 0 17348.074
1484.72607421875 0 8231.912
1485.747314453125 0 5850.6377
1486.2435302734375 0 2628.9866
1486.7513427734375 0 7162.698
1487.754638671875 0 10697.098
1488.7598876953125 0 4190.3164
1498.7313232421875 0 2854.908
1501.234619140625 0 3873.231
1501.7410888671875 0 2743.4265
1514.75732421875 0 2610.9895
1515.7532958984375 0 6062.143
1516.7735595703125 0 2903.3718
1517.758056640625 0 2831.167
1524.7159423828125 0 3437.3677 y Ammonia loss 3
1525.7113037109375 0 68130.79 z 3
1526.71630859375 0 82115.805
1527.718505859375 0 64801.145
1528.299560546875 0 2755.115
1528.720458984375 0 26473.207
1529.73779296875 0 13839.771
1530.7681884765625 0 87420.734 c 11
1531.7720947265625 0 87918.05
1532.7738037109375 0 42008.85
1533.7750244140625 0 15262.434
1534.7747802734375 0 6236.579
1537.275146484375 0 2732.943
1539.770263671875 0 4152.492
1541.7314453125 0 14849.182 y 3
1542.7342529296875 0 12498.133
1543.741943359375 0 7759.0317
1549.3184814453125 0 2676.4507
1549.812744140625 0 2439.8064
1550.28515625 0 4786.137
1550.7900390625 0 4481.505
1551.2889404296875 0 3966.8188
1551.7783203125 0 4124.076
1552.28955078125 0 2777.288
1552.7857666015625 0 2901.1726
1553.7216796875 0 2695.1875
1555.7547607421875 0 3055.3845
1556.295166015625 0 2451.6736
1556.7685546875 0 4643.357
1557.793212890625 0 5579.2695
1558.7943115234375 0 6659.7173
1559.286865234375 0 5550.4614
1559.8179931640625 0 2918.0872
1560.2918701171875 0 3700.7358
1563.7857666015625 0 2981.4563
1564.7957763671875 0 5171.4536
1565.2706298828125 0 6211.9595
1565.7935791015625 0 6532.863
1566.294677734375 0 6821.227
1566.8001708984375 0 8878.082
1567.2978515625 0 6688.7124
1567.7977294921875 0 5366.406
1568.2816162109375 0 5407.011
1568.777587890625 0 2554.1604
1569.78515625 0 3704.487
1571.30517578125 0 2989.6309
1571.793701171875 0 4740.305
1572.3031005859375 0 6509.905
1572.7984619140625 0 4791.1006
1573.284912109375 0 6014.7837
1573.79345703125 0 8170.4
1574.293701171875 0 11273.4375
1574.8060302734375 0 10349.094
1575.293701171875 0 5647.8213
1575.8055419921875 0 5110.46
1576.2938232421875 0 3194.0957
1577.31201171875 0 3058.7231
1577.77099609375 0 2885.2935
1578.770751953125 0 3355.8108
1579.2955322265625 0 3534.8994
1579.793212890625 0 3126.032
1580.296630859375 0 4316.3965
1580.7806396484375 0 2515.099
1581.291748046875 0 6564.91
1581.7965087890625 0 5503.4365
1582.2752685546875 0 4179.7246
1582.783203125 0 5178.283
1583.292724609375 0 7683.812
1585.305419921875 0 2927.742
1585.7845458984375 0 2876.06
1586.3035888671875 0 3436.075
1586.80322265625 0 3742.694
1588.279541015625 0 4106.2344
1589.2923583984375 0 3022.633
1589.793212890625 0 6466.8677
1590.2940673828125 0 6895.446
1590.7994384765625 0 7418.1743
1591.305419921875 0 6265.097
1591.7908935546875 0 4706.736
1592.3026123046875 0 2467.6204
1592.8004150390625 0 4098.4136
1593.3101806640625 0 3756.7393
1593.7994384765625 0 7015.6953
1594.3033447265625 0 5762.867
1594.7972412109375 0 8912.328
1595.2999267578125 0 6584.7363
1595.7801513671875 0 6689.5273
1596.7781982421875 0 5061.7144
1597.7471923828125 0 3775.3586
1598.7984619140625 0 7290.8647
1599.29443359375 0 3443.5596
1599.7823486328125 0 2903.5906
1600.3096923828125 0 3571.058
1600.79638671875 0 3506.2476
1601.8048095703125 0 5129.2974
1602.30615234375 0 8346.185
1602.815673828125 0 8767.116
1603.3116455078125 0 2540.5967
1603.82421875 0 3085.2942
1609.7791748046875 0 4258.622
1634.7808837890625 0 2404.242
1653.770263671875 0 64125.78 z 2
1654.7745361328125 0 72123.23
1655.7791748046875 0 44666.9
1656.7806396484375 0 24602.066
1657.773193359375 0 6332.1025
1669.786865234375 0 4327.3228 y 2
1670.7901611328125 0 4693.812
1676.78125 0 2404.5981
1690.7926025390625 0 6303.305
1691.783935546875 0 96819.39 c 12
1692.787841796875 0 76012.055
1693.7894287109375 0 45983.223
1694.791015625 0 19451.736
1695.80126953125 0 6048.4004
1733.8460693359375 0 2479.9663
1760.8367919921875 0 4497.404
1761.8687744140625 0 4485.372
1762.8602294921875 0 4017.5244
1763.868896484375 0 3264.455
1775.8726806640625 0 4388.4272
1776.875244140625 0 9918.185
1777.86962890625 0 9404.61
1778.8795166015625 0 5853.4575
1790.8355712890625 0 6346.912 z 1
1791.8375244140625 0 26611.701
1792.847412109375 0 22418.807
1793.8560791015625 0 18522.49
1794.8619384765625 0 6052.378
1795.88720703125 0 2516.861
1803.8839111328125 0 4869.6284
1804.8824462890625 0 5698.6587
1805.88720703125 0 3748.2656
1806.880615234375 0 3349.2532 y 1
1813.9371337890625 0 2785.4167
1817.9063720703125 0 4494.054
1818.9100341796875 0 4833.071
1819.8870849609375 0 17102.54 c 13
1820.8843994140625 0 38701.54
1821.8875732421875 0 34084.22
1822.88916015625 0 14736.756
1823.8839111328125 0 7505.316
1824.9073486328125 0 3133.5452
1830.955078125 0 10740.938
1831.951171875 0 10977.864
1832.9532470703125 0 7513.6655
1833.9376220703125 0 4204.684
1834.9278564453125 0 5990.3555
1835.9464111328125 0 5055.009
1836.9375 0 2677.3281
1844.9185791015625 0 10153.28
1845.9139404296875 0 14477.621
1846.9154052734375 0 12807.05
1847.92431640625 0 7186.953
1848.926513671875 0 4188.279
1858.9302978515625 0 3271.4597
1859.9232177734375 0 6419.1377
1860.9287109375 0 9603.528
1861.9268798828125 0 43204.098
1862.93212890625 0 82910.52
1863.9327392578125 0 63231.312
1864.2457275390625 0 2794.7988
1864.9371337890625 0 38474.023
1865.940673828125 0 15037.313
1866.93310546875 0 4411.988
1874.9393310546875 0 3339.013
1875.932861328125 0 17119.605
1876.9302978515625 0 35799.637
1877.9359130859375 0 32901.383
1878.9326171875 0 17754.977
1879.9337158203125 0 7381.7456
1880.9346923828125 0 2909.704
1886.931640625 0 4543.416
1887.903564453125 0 3514.9062
1890.901123046875 0 2437.3882
1892.9466552734375 0 14924.877
1893.950439453125 0 42016.285
1894.95263671875 0 45179.938
1895.9580078125 0 23708.531
1896.956298828125 0 8248.751
1897.95654296875 0 3342.5269
1902.9183349609375 0 4443.1353
1903.9268798828125 0 36436.81
1904.9254150390625 0 97658.336
1905.9248046875 0 98522.914
1906.925537109375 0 57759.223
1907.92529296875 0 27372.414
1908.930419921875 0 9313.51
1909.936767578125 0 3230.448
1918.9185791015625 0 4422.901
1919.93359375 0 23595.41
1920.939697265625 0 120820.63
1921.94677734375 0 402528.72
1922.94921875 0 371866.3
1923.950927734375 0 206027.23
1924.953125 0 109348.83
1925.95068359375 0 33533.1
1926.95068359375 0 9033.112
2124.000244140625 0 2743.3208

Spectrum Details

|  |  |
| --- | --- |
| Matched peaks? Matched peaks The total absolute number of peaks matched. Additionally in brackets the total fraction of peaks matched and the total number of peaks is shown. | 80 (11.82% of 677) |
| FDR? FDR The false discovery rate estimated for this peptide. It is calculated by matching all theoretical fragments with a non-integer shift with the raw peaks for this spectrum. This is done with 40 different shifts. The resulting percentage is the average number of annotated peaks over the number of annotated peaks with the correct spectrum. | 1.10% |
| Satellite FDR? Satellite FDR See the FDR for details on its calculation. This satellite ion specific FDR only contains the satellite ions (d/w) for I/L/J positions. | - |
| PSM Score? PSM Score The PSM Score as given by Hecklib to this annotated spectrum. It is shown with three significant figures. | 729 |

#### Spectrum 5773? Spectrum 5773 The raw spectrum of this peptide as annotated by Hecklib. The fragments are coloured according to ion type (see legend). Any peaks with a star '\*' as text can be hovered over to see the full details, first the ion type second the mass shift type. By hovering over the amino acids in the peptide or ions in the legend the corresponding peaks are highlighted. By toggling the 'Unassigned' label you can turn the background (unassigned) peaks on or off in the plot. By updating the slider in the Ion legend you can update the spectrum to only show the top X% of the peaks with labels. The top X% means any peak that is within X% of the highest intensity. By dragging in the spectrum you can zoom in to a specific part of the spectrum and use 'Zoom Out' to get back to the original zoom level. The annotation of the spectrum is based on the given sequence in the peptides file and is done with different software so inconsistencies are likely. The peaks are annotated based on the given sequence, with 20 ppm tolerance.

Copy Data

##### Spectrum 5773 (TSV)

###### Preview

```
Loading example...
```

*Click on the button to copy the data to your clipboard.*

Mz MinMz MaxIntensity Max

WidthHeightPeptide font sizePeptide stroke widthSpectrum font sizeSpectrum stroke widthCompact peptide

Ion legend

wxyz

abcd

OtherUnassignedIon^Charge^_Position_Show for top:%

JHQDWJDGKEYKCKV

01.36e+42.73e+44.09e+45.45e+4

Zoom Out

z^+1^_2_y^+1^_2_c^+1^_2_c^+1^_3_c^+1^_3_y^+1^_3_c^+3^_10_w^+1^_4_c^+1^_4_y^+2^_8_c^+1^_4_z^+1^_4_y^+1^_4_c^+3^_14_y^+3^_14_c^+1^_5_z^+1^_5_c^+1^_5_y^+1^_5_y^+2^_11_w^+1^_6_c^+2^_12_y^+2^_12_c^+1^_6_z^+1^_6_y^+2^_13_y^+2^_13_y^+1^_6_y^+2^_13_c^+2^_13_z^+2^_14_c^+1^_7_c^+2^_14_c^+1^_7_y^+1^_7_y^+1^_7_z^+1^_7_y^+1^_7_c^+1^_8_z^+1^_8_y^+1^_8_c^+1^_9_w^+1^_10_z^+1^_10_c^+1^_10_y^+1^_10_c^+1^_11_y^+1^_11_z^+1^_11_y^+1^_11_y^+1^_12_y^+1^_12_z^+1^_12_c^+1^_12_y^+1^_12_z^+1^_13_c^+1^_13_z^+1^_14_c^+1^_14_

0515103015452060

Fragment Matches Table

Show background peaks

| Position | Ion type | Intensity | mz Theoretical | mz Error (Th) | mz Error (ppm) | Charge | Series Number |
| --- | --- | --- | --- | --- | --- | --- | --- |
| - | - | 362.2 | 121 | - | - | 0 | - |
| - | - | 1592 | 129.1 | - | - | 0 | - |
| - | - | 439.5 | 133 | - | - | 0 | - |
| - | - | 458.7 | 136.4 | - | - | 0 | - |
| - | - | 418.2 | 141.1 | - | - | 0 | - |
| - | - | 660 | 149 | - | - | 0 | - |
| - | - | 549.6 | 153.1 | - | - | 0 | - |
| - | - | 427.4 | 153.7 | - | - | 0 | - |
| - | - | 729.3 | 155.1 | - | - | 0 | - |
| - | - | 474.7 | 156.9 | - | - | 0 | - |
| - | - | 457.4 | 157.1 | - | - | 0 | - |
| - | - | 441.7 | 165.1 | - | - | 0 | - |
| - | - | 852.5 | 166.1 | - | - | 0 | - |
| - | - | 1890 | 173.4 | - | - | 0 | - |
| - | - | 458.8 | 173.7 | - | - | 0 | - |
| - | - | 528.3 | 177 | - | - | 0 | - |
| - | - | 675.9 | 183.1 | - | - | 0 | - |
| - | - | 743.2 | 185.2 | - | - | 0 | - |
| - | - | 530 | 186.1 | - | - | 0 | - |
| - | - | 1182 | 223.2 | - | - | 0 | - |
| - | - | 520.8 | 225.4 | - | - | 0 | - |
| 14 | z | 3651 | 230.2 | 0.0002015 | 0.8756 | +1 | 2 |
| - | - | 4397 | 231.2 | - | - | 0 | - |
| - | - | 1502 | 234.1 | - | - | 0 | - |
| - | - | 540.9 | 243.3 | - | - | 0 | - |
| 14 | y | 1126 | 246.2 | 0.0003768 | 1.53 | +1 | 2 |
| - | - | 1052 | 250.1 | - | - | 0 | - |
| - | - | 8163 | 251.2 | - | - | 0 | - |
| - | - | 1436 | 252.2 | - | - | 0 | - |
| 2 | c | 4847 | 268.2 | 0.0002572 | 0.9591 | +1 | 2 |
| - | - | 889.3 | 269.2 | - | - | 0 | - |
| - | - | 703.4 | 292.9 | - | - | 0 | - |
| - | - | 7840 | 300.2 | - | - | 0 | - |
| - | - | 1067 | 301.2 | - | - | 0 | - |
| - | - | 681.6 | 302.2 | - | - | 0 | - |
| - | - | 577.7 | 318.1 | - | - | 0 | - |
| - | - | 607.9 | 333.3 | - | - | 0 | - |
| - | - | 1823 | 352.2 | - | - | 0 | - |
| 3 | c | 1052 | 379.2 | 0.0005168 | 1.363 | +1 | 3 |
| - | - | 4111 | 395.2 | - | - | 0 | - |
| - | - | 807.1 | 396.2 | - | - | 0 | - |
| 3 | c | 1.42E+04 | 396.2 | 0.0002715 | 0.6852 | +1 | 3 |
| - | - | 3472 | 397.2 | - | - | 0 | - |
| 13 | y | 4014 | 407.2 | 0.004052 | 9.952 | +1 | 3 |
| 10 | c | 1109 | 408.2 | 0.001677 | 4.108 | +3 | 10 |
| - | - | 577.5 | 409.4 | - | - | 0 | - |
| - | - | 970.6 | 427.3 | - | - | 0 | - |
| - | - | 733.6 | 428.3 | - | - | 0 | - |
| 12 | w | 738.4 | 461.2 | 0.002746 | 5.953 | +1 | 4 |
| - | - | 5486 | 467.2 | - | - | 0 | - |
| - | - | 2430 | 468.3 | - | - | 0 | - |
| - | - | 670.8 | 475.3 | - | - | 0 | - |
| - | - | 851 | 488.3 | - | - | 0 | - |
| 4 | c | 745.4 | 494.2 | 0.000891 | 1.803 | +1 | 4 |
| 8 | y | 2852 | 506.8 | 0.002424 | 4.784 | +2 | 8 |
| - | - | 1128 | 507.3 | - | - | 0 | - |
| - | - | 689.4 | 507.8 | - | - | 0 | - |
| 4 | c | 2.174E+04 | 511.3 | 6.82E-05 | 0.1334 | +1 | 4 |
| - | - | 5520 | 512.3 | - | - | 0 | - |
| - | - | 614.7 | 513.3 | - | - | 0 | - |
| 12 | z | 5606 | 519.3 | 0.004425 | 8.521 | +1 | 4 |
| - | - | 2207 | 520.3 | - | - | 0 | - |
| - | - | 558.5 | 521.3 | - | - | 0 | - |
| 12 | y | 1846 | 535.3 | 0.003679 | 6.872 | +1 | 4 |
| - | - | 832.4 | 536.3 | - | - | 0 | - |
| - | - | 618.9 | 564.3 | - | - | 0 | - |
| - | - | 4251 | 584.4 | - | - | 0 | - |
| - | - | 1237 | 585.4 | - | - | 0 | - |
| - | - | 1765 | 590.9 | - | - | 0 | - |
| - | - | 554.8 | 598.3 | - | - | 0 | - |
| 14 | c | 2778 | 601.6 | 0.001232 | 2.048 | +3 | 14 |
| - | - | 2063 | 602 | - | - | 0 | - |
| - | - | 1848 | 602.3 | - | - | 0 | - |
| 2 | y | 5113 | 603 | 0.000717 | 1.189 | +3 | 14 |
| - | - | 3829 | 603.3 | - | - | 0 | - |
| - | - | 2682 | 603.6 | - | - | 0 | - |
| - | - | 644.8 | 606.3 | - | - | 0 | - |
| - | - | 1393 | 607.6 | - | - | 0 | - |
| - | - | 1326 | 608 | - | - | 0 | - |
| - | - | 838.5 | 618.9 | - | - | 0 | - |
| - | - | 667.8 | 619.4 | - | - | 0 | - |
| - | - | 711.5 | 625.3 | - | - | 0 | - |
| - | - | 1070 | 629 | - | - | 0 | - |
| - | - | 1102 | 629.3 | - | - | 0 | - |
| - | - | 1225 | 629.6 | - | - | 0 | - |
| - | - | 550.4 | 633.3 | - | - | 0 | - |
| - | - | 2168 | 634.6 | - | - | 0 | - |
| - | - | 2652 | 635 | - | - | 0 | - |
| - | - | 2773 | 635.3 | - | - | 0 | - |
| - | - | 1374 | 635.6 | - | - | 0 | - |
| - | - | 1234 | 639.3 | - | - | 0 | - |
| - | - | 1682 | 639.9 | - | - | 0 | - |
| - | - | 4130 | 640.3 | - | - | 0 | - |
| - | - | 1523 | 640.4 | - | - | 0 | - |
| - | - | 4.78E+04 | 640.6 | - | - | 0 | - |
| - | - | 1951 | 640.9 | - | - | 0 | - |
| - | - | 4.783E+04 | 641 | - | - | 0 | - |
| - | - | 3.742E+04 | 641.3 | - | - | 0 | - |
| - | - | 1.235E+04 | 641.7 | - | - | 0 | - |
| - | - | 2142 | 641.9 | - | - | 0 | - |
| - | - | 3971 | 642 | - | - | 0 | - |
| - | - | 1762 | 651.4 | - | - | 0 | - |
| - | - | 1400 | 651.9 | - | - | 0 | - |
| 5 | c | 972.8 | 680.3 | 0.00344 | 5.057 | +1 | 5 |
| 11 | z | 1.306E+04 | 682.3 | 0.004155 | 6.089 | +1 | 5 |
| - | - | 5270 | 683.3 | - | - | 0 | - |
| - | - | 1691 | 684.3 | - | - | 0 | - |
| - | - | 809.4 | 685.3 | - | - | 0 | - |
| 5 | c | 6203 | 697.3 | 0.0004672 | 0.6699 | +1 | 5 |
| 11 | y | 4094 | 698.4 | 0.01067 | 15.28 | +1 | 5 |
| - | - | 1310 | 699.3 | - | - | 0 | - |
| 5 | y | 864.8 | 713.9 | 0.001149 | 1.609 | +2 | 11 |
| 10 | w | 2287 | 752.4 | 0.003658 | 4.862 | +1 | 6 |
| - | - | 607 | 753.9 | - | - | 0 | - |
| - | - | 859.4 | 760.9 | - | - | 0 | - |
| - | - | 700.5 | 761.4 | - | - | 0 | - |
| - | - | 1422 | 765.4 | - | - | 0 | - |
| 12 | c | 4659 | 765.9 | 0.00201 | 2.625 | +2 | 12 |
| - | - | 4856 | 766.4 | - | - | 0 | - |
| - | - | 2253 | 766.9 | - | - | 0 | - |
| - | - | 778 | 767.4 | - | - | 0 | - |
| - | - | 655.1 | 768.3 | - | - | 0 | - |
| - | - | 857.9 | 769.1 | - | - | 0 | - |
| 4 | y | 1240 | 771.4 | 0.003207 | 4.157 | +2 | 12 |
| - | - | 1470 | 771.9 | - | - | 0 | - |
| - | - | 642.1 | 772.4 | - | - | 0 | - |
| - | - | 854.4 | 776.1 | - | - | 0 | - |
| - | - | 918.8 | 779.4 | - | - | 0 | - |
| - | - | 716.2 | 780.9 | - | - | 0 | - |
| - | - | 819.8 | 787.1 | - | - | 0 | - |
| - | - | 1187 | 787.4 | - | - | 0 | - |
| - | - | 663 | 794.4 | - | - | 0 | - |
| - | - | 614 | 795.2 | - | - | 0 | - |
| - | - | 835.2 | 799.2 | - | - | 0 | - |
| - | - | 669.4 | 801.9 | - | - | 0 | - |
| - | - | 1185 | 809.4 | - | - | 0 | - |
| 6 | c | 6203 | 810.4 | 0.001425 | 1.759 | +1 | 6 |
| 10 | z | 6285 | 811.4 | 0.004389 | 5.41 | +1 | 6 |
| - | - | 2912 | 812.4 | - | - | 0 | - |
| - | - | 1417 | 813.4 | - | - | 0 | - |
| - | - | 1046 | 817.4 | - | - | 0 | - |
| - | - | 1042 | 825.4 | - | - | 0 | - |
| 3 | y | 1711 | 826.4 | 0.006278 | 7.597 | +2 | 13 |
| 3 | y | 1996 | 826.9 | 0.005498 | 6.649 | +2 | 13 |
| 10 | y | 4585 | 827.4 | 0.008831 | 10.67 | +1 | 6 |
| - | - | 1875 | 827.9 | - | - | 0 | - |
| - | - | 1224 | 828.4 | - | - | 0 | - |
| 3 | y | 5848 | 835.4 | 0.002344 | 2.806 | +2 | 13 |
| - | - | 5583 | 835.9 | - | - | 0 | - |
| - | - | 3053 | 836.4 | - | - | 0 | - |
| - | - | 1042 | 836.9 | - | - | 0 | - |
| - | - | 2080 | 842.4 | - | - | 0 | - |
| - | - | 1044 | 843.4 | - | - | 0 | - |
| - | - | 3077 | 845.9 | - | - | 0 | - |
| 13 | c | 7235 | 846.4 | 0.0008072 | 0.9537 | +2 | 13 |
| - | - | 7842 | 846.9 | - | - | 0 | - |
| - | - | 4526 | 847.4 | - | - | 0 | - |
| - | - | 1499 | 847.9 | - | - | 0 | - |
| - | - | 864.2 | 848.5 | - | - | 0 | - |
| - | - | 990.3 | 867.4 | - | - | 0 | - |
| - | - | 775 | 875.5 | - | - | 0 | - |
| - | - | 1615 | 880.5 | - | - | 0 | - |
| - | - | 4476 | 881.4 | - | - | 0 | - |
| - | - | 2814 | 882.4 | - | - | 0 | - |
| - | - | 1449 | 883.5 | - | - | 0 | - |
| - | - | 799.3 | 888.4 | - | - | 0 | - |
| - | - | 751.6 | 888.9 | - | - | 0 | - |
| - | - | 1622 | 893.9 | - | - | 0 | - |
| - | - | 624.7 | 894.5 | - | - | 0 | - |
| - | - | 1930 | 895.4 | - | - | 0 | - |
| 2 | z | 2446 | 895.9 | 0.005455 | 6.089 | +2 | 14 |
| - | - | 2451 | 896.4 | - | - | 0 | - |
| - | - | 1811 | 896.9 | - | - | 0 | - |
| - | - | 1769 | 902.4 | - | - | 0 | - |
| - | - | 967.6 | 902.9 | - | - | 0 | - |
| - | - | 1055 | 903.4 | - | - | 0 | - |
| 7 | c | 2128 | 908.4 | 0.00744 | 8.19 | +1 | 7 |
| - | - | 1237 | 909 | - | - | 0 | - |
| - | - | 2095 | 909.4 | - | - | 0 | - |
| - | - | 1490 | 909.9 | - | - | 0 | - |
| 14 | c | 1.561E+04 | 910.4 | 0.001231 | 1.352 | +2 | 14 |
| - | - | 1.837E+04 | 910.9 | - | - | 0 | - |
| - | - | 1.243E+04 | 911.4 | - | - | 0 | - |
| - | - | 4227 | 911.9 | - | - | 0 | - |
| - | - | 1644 | 912.4 | - | - | 0 | - |
| - | - | 990.3 | 915.5 | - | - | 0 | - |
| - | - | 1566 | 916.5 | - | - | 0 | - |
| - | - | 1518 | 917 | - | - | 0 | - |
| - | - | 820.7 | 917.5 | - | - | 0 | - |
| - | - | 1098 | 918 | - | - | 0 | - |
| - | - | 838.7 | 921.4 | - | - | 0 | - |
| - | - | 1399 | 923 | - | - | 0 | - |
| - | - | 1184 | 923.5 | - | - | 0 | - |
| - | - | 1469 | 924 | - | - | 0 | - |
| - | - | 4540 | 924.4 | - | - | 0 | - |
| - | - | 3105 | 924.9 | - | - | 0 | - |
| 7 | c | 7781 | 925.5 | 0.0004935 | 0.5332 | +1 | 7 |
| - | - | 1465 | 925.9 | - | - | 0 | - |
| - | - | 3077 | 926.5 | - | - | 0 | - |
| - | - | 854.7 | 927.5 | - | - | 0 | - |
| - | - | 1191 | 929.5 | - | - | 0 | - |
| - | - | 2805 | 930.5 | - | - | 0 | - |
| - | - | 1.066E+04 | 931 | - | - | 0 | - |
| - | - | 2.62E+04 | 931.5 | - | - | 0 | - |
| - | - | 2.172E+04 | 932 | - | - | 0 | - |
| - | - | 1.29E+04 | 932.5 | - | - | 0 | - |
| - | - | 5327 | 933 | - | - | 0 | - |
| - | - | 1851 | 933.5 | - | - | 0 | - |
| 9 | y | 781.4 | 937.5 | 0.01114 | 11.88 | +1 | 7 |
| - | - | 3755 | 938 | - | - | 0 | - |
| 9 | y | 6499 | 938.5 | 0.001246 | 1.328 | +1 | 7 |
| - | - | 4496 | 939 | - | - | 0 | - |
| 9 | z | 1.558E+04 | 939.5 | 0.005236 | 5.573 | +1 | 7 |
| - | - | 728.4 | 940 | - | - | 0 | - |
| - | - | 9615 | 940.5 | - | - | 0 | - |
| - | - | 3417 | 941.5 | - | - | 0 | - |
| - | - | 1418 | 942.5 | - | - | 0 | - |
| - | - | 970.9 | 943 | - | - | 0 | - |
| - | - | 835.4 | 949.2 | - | - | 0 | - |
| - | - | 827.2 | 949.9 | - | - | 0 | - |
| - | - | 761.4 | 951 | - | - | 0 | - |
| - | - | 3342 | 952 | - | - | 0 | - |
| - | - | 6383 | 952.5 | - | - | 0 | - |
| - | - | 6511 | 953 | - | - | 0 | - |
| - | - | 3576 | 953.5 | - | - | 0 | - |
| - | - | 2344 | 954 | - | - | 0 | - |
| - | - | 1569 | 954.5 | - | - | 0 | - |
| 9 | y | 2080 | 955.5 | 0.00272 | 2.847 | +1 | 7 |
| - | - | 851.2 | 956.5 | - | - | 0 | - |
| - | - | 737.8 | 958.5 | - | - | 0 | - |
| - | - | 823.7 | 959.5 | - | - | 0 | - |
| - | - | 3486 | 960 | - | - | 0 | - |
| - | - | 2.409E+04 | 960.5 | - | - | 0 | - |
| - | - | 3.881E+04 | 961 | - | - | 0 | - |
| - | - | 2.901E+04 | 961.5 | - | - | 0 | - |
| - | - | 1.643E+04 | 962 | - | - | 0 | - |
| - | - | 6138 | 962.5 | - | - | 0 | - |
| - | - | 2719 | 963 | - | - | 0 | - |
| - | - | 742 | 978.2 | - | - | 0 | - |
| - | - | 3872 | 981.5 | - | - | 0 | - |
| 8 | c | 1.826E+04 | 982.5 | 0.001053 | 1.072 | +1 | 8 |
| - | - | 8078 | 983.5 | - | - | 0 | - |
| - | - | 2585 | 984.5 | - | - | 0 | - |
| - | - | 1047 | 986.5 | - | - | 0 | - |
| 8 | z | 4140 | 996.5 | 0.001309 | 1.314 | +1 | 8 |
| - | - | 3121 | 997.5 | - | - | 0 | - |
| - | - | 1532 | 998.5 | - | - | 0 | - |
| - | - | 799.1 | 999.5 | - | - | 0 | - |
| 8 | y | 6649 | 1013 | 0.004103 | 4.052 | +1 | 8 |
| - | - | 3502 | 1014 | - | - | 0 | - |
| - | - | 1907 | 1014 | - | - | 0 | - |
| - | - | 3240 | 1015 | - | - | 0 | - |
| - | - | 1111 | 1015 | - | - | 0 | - |
| - | - | 1381 | 1020 | - | - | 0 | - |
| - | - | 1134 | 1020 | - | - | 0 | - |
| - | - | 1456 | 1026 | - | - | 0 | - |
| - | - | 1405 | 1028 | - | - | 0 | - |
| - | - | 1060 | 1032 | - | - | 0 | - |
| - | - | 995.6 | 1032 | - | - | 0 | - |
| - | - | 1192 | 1033 | - | - | 0 | - |
| - | - | 908.7 | 1034 | - | - | 0 | - |
| - | - | 778.2 | 1035 | - | - | 0 | - |
| - | - | 1220 | 1035 | - | - | 0 | - |
| - | - | 864.2 | 1041 | - | - | 0 | - |
| - | - | 844.3 | 1044 | - | - | 0 | - |
| - | - | 1781 | 1044 | - | - | 0 | - |
| - | - | 1019 | 1045 | - | - | 0 | - |
| - | - | 796.4 | 1045 | - | - | 0 | - |
| - | - | 1472 | 1045 | - | - | 0 | - |
| - | - | 1046 | 1046 | - | - | 0 | - |
| - | - | 714.3 | 1046 | - | - | 0 | - |
| - | - | 782.9 | 1047 | - | - | 0 | - |
| - | - | 1799 | 1048 | - | - | 0 | - |
| - | - | 1060 | 1048 | - | - | 0 | - |
| - | - | 1458 | 1048 | - | - | 0 | - |
| - | - | 1002 | 1049 | - | - | 0 | - |
| - | - | 1537 | 1049 | - | - | 0 | - |
| - | - | 1414 | 1049 | - | - | 0 | - |
| - | - | 1176 | 1050 | - | - | 0 | - |
| - | - | 1481 | 1050 | - | - | 0 | - |
| - | - | 1095 | 1050 | - | - | 0 | - |
| - | - | 1014 | 1053 | - | - | 0 | - |
| - | - | 2555 | 1054 | - | - | 0 | - |
| - | - | 3577 | 1054 | - | - | 0 | - |
| - | - | 1723 | 1054 | - | - | 0 | - |
| - | - | 2506 | 1055 | - | - | 0 | - |
| - | - | 1898 | 1055 | - | - | 0 | - |
| - | - | 906.2 | 1055 | - | - | 0 | - |
| - | - | 2088 | 1056 | - | - | 0 | - |
| - | - | 834.4 | 1058 | - | - | 0 | - |
| - | - | 2302 | 1060 | - | - | 0 | - |
| - | - | 1479 | 1060 | - | - | 0 | - |
| - | - | 2974 | 1060 | - | - | 0 | - |
| - | - | 2083 | 1061 | - | - | 0 | - |
| - | - | 862.5 | 1061 | - | - | 0 | - |
| - | - | 1267 | 1061 | - | - | 0 | - |
| - | - | 1498 | 1062 | - | - | 0 | - |
| - | - | 1670 | 1062 | - | - | 0 | - |
| - | - | 1710 | 1062 | - | - | 0 | - |
| - | - | 1188 | 1063 | - | - | 0 | - |
| - | - | 3020 | 1063 | - | - | 0 | - |
| - | - | 3263 | 1063 | - | - | 0 | - |
| - | - | 3989 | 1064 | - | - | 0 | - |
| - | - | 1649 | 1064 | - | - | 0 | - |
| - | - | 1207 | 1064 | - | - | 0 | - |
| - | - | 894.6 | 1065 | - | - | 0 | - |
| - | - | 1691 | 1065 | - | - | 0 | - |
| - | - | 2060 | 1066 | - | - | 0 | - |
| - | - | 1537 | 1066 | - | - | 0 | - |
| - | - | 1392 | 1066 | - | - | 0 | - |
| - | - | 1154 | 1067 | - | - | 0 | - |
| - | - | 990.3 | 1067 | - | - | 0 | - |
| - | - | 1628 | 1068 | - | - | 0 | - |
| - | - | 4473 | 1069 | - | - | 0 | - |
| - | - | 5931 | 1069 | - | - | 0 | - |
| - | - | 4689 | 1069 | - | - | 0 | - |
| - | - | 4242 | 1070 | - | - | 0 | - |
| - | - | 2457 | 1070 | - | - | 0 | - |
| - | - | 863.7 | 1070 | - | - | 0 | - |
| - | - | 781 | 1095 | - | - | 0 | - |
| - | - | 1360 | 1110 | - | - | 0 | - |
| 9 | c | 1.149E+04 | 1111 | 0.0005114 | 0.4605 | +1 | 9 |
| - | - | 7015 | 1112 | - | - | 0 | - |
| - | - | 3645 | 1113 | - | - | 0 | - |
| - | - | 996.3 | 1114 | - | - | 0 | - |
| - | - | 922 | 1138 | - | - | 0 | - |
| - | - | 1703 | 1143 | - | - | 0 | - |
| - | - | 1015 | 1144 | - | - | 0 | - |
| - | - | 1438 | 1167 | - | - | 0 | - |
| - | - | 1070 | 1182 | - | - | 0 | - |
| - | - | 1025 | 1182 | - | - | 0 | - |
| - | - | 821.4 | 1190 | - | - | 0 | - |
| 6 | w | 904.8 | 1196 | 0.0158 | 13.21 | +1 | 10 |
| - | - | 853.7 | 1221 | - | - | 0 | - |
| 6 | z | 3772 | 1225 | 1.154E-05 | 0.009423 | +1 | 10 |
| - | - | 2429 | 1226 | - | - | 0 | - |
| - | - | 1997 | 1227 | - | - | 0 | - |
| - | - | 2420 | 1239 | - | - | 0 | - |
| 10 | c | 1.533E+04 | 1240 | 0.00223 | 1.799 | +1 | 10 |
| 6 | y | 9575 | 1241 | 0.01251 | 10.08 | +1 | 10 |
| - | - | 3990 | 1242 | - | - | 0 | - |
| - | - | 1151 | 1243 | - | - | 0 | - |
| - | - | 825.6 | 1244 | - | - | 0 | - |
| - | - | 905.2 | 1267 | - | - | 0 | - |
| - | - | 1118 | 1272 | - | - | 0 | - |
| - | - | 934.3 | 1272 | - | - | 0 | - |
| - | - | 1234 | 1273 | - | - | 0 | - |
| - | - | 877.2 | 1280 | - | - | 0 | - |
| - | - | 834.7 | 1280 | - | - | 0 | - |
| - | - | 1670 | 1281 | - | - | 0 | - |
| - | - | 1647 | 1281 | - | - | 0 | - |
| - | - | 2302 | 1282 | - | - | 0 | - |
| - | - | 2424 | 1283 | - | - | 0 | - |
| - | - | 1638 | 1284 | - | - | 0 | - |
| - | - | 908.1 | 1302 | - | - | 0 | - |
| - | - | 892.3 | 1356 | - | - | 0 | - |
| - | - | 1013 | 1367 | - | - | 0 | - |
| - | - | 2347 | 1402 | - | - | 0 | - |
| 11 | c | 9452 | 1403 | 0.003232 | 2.304 | +1 | 11 |
| - | - | 6888 | 1404 | - | - | 0 | - |
| - | - | 2711 | 1405 | - | - | 0 | - |
| - | - | 1121 | 1406 | - | - | 0 | - |
| 5 | y | 953.2 | 1410 | 0.0109 | 7.731 | +1 | 11 |
| 5 | z | 1.171E+04 | 1411 | 0.002664 | 1.889 | +1 | 11 |
| - | - | 9084 | 1412 | - | - | 0 | - |
| - | - | 5350 | 1413 | - | - | 0 | - |
| - | - | 1188 | 1414 | - | - | 0 | - |
| - | - | 1661 | 1422 | - | - | 0 | - |
| - | - | 1109 | 1423 | - | - | 0 | - |
| - | - | 1528 | 1426 | - | - | 0 | - |
| 5 | y | 6015 | 1427 | 0.003444 | 2.414 | +1 | 11 |
| - | - | 4145 | 1428 | - | - | 0 | - |
| - | - | 2462 | 1429 | - | - | 0 | - |
| - | - | 1019 | 1430 | - | - | 0 | - |
| - | - | 1193 | 1436 | - | - | 0 | - |
| - | - | 1424 | 1437 | - | - | 0 | - |
| - | - | 790.3 | 1439 | - | - | 0 | - |
| - | - | 923.3 | 1466 | - | - | 0 | - |
| - | - | 878.5 | 1475 | - | - | 0 | - |
| - | - | 3525 | 1482 | - | - | 0 | - |
| - | - | 2533 | 1483 | - | - | 0 | - |
| - | - | 1656 | 1484 | - | - | 0 | - |
| - | - | 1040 | 1485 | - | - | 0 | - |
| - | - | 939.7 | 1486 | - | - | 0 | - |
| - | - | 3288 | 1487 | - | - | 0 | - |
| - | - | 1603 | 1487 | - | - | 0 | - |
| - | - | 4171 | 1488 | - | - | 0 | - |
| - | - | 1659 | 1489 | - | - | 0 | - |
| - | - | 939.6 | 1496 | - | - | 0 | - |
| - | - | 1026 | 1501 | - | - | 0 | - |
| - | - | 1183 | 1502 | - | - | 0 | - |
| - | - | 839.1 | 1510 | - | - | 0 | - |
| - | - | 931.6 | 1515 | - | - | 0 | - |
| - | - | 1054 | 1516 | - | - | 0 | - |
| - | - | 806.5 | 1521 | - | - | 0 | - |
| - | - | 1694 | 1521 | - | - | 0 | - |
| - | - | 1071 | 1522 | - | - | 0 | - |
| 4 | y | 1027 | 1524 | 0.01265 | 8.301 | +1 | 12 |
| 4 | y | 1148 | 1525 | 0.02558 | 16.78 | +1 | 12 |
| 4 | z | 9164 | 1526 | 0.002874 | 1.884 | +1 | 12 |
| - | - | 1.551E+04 | 1527 | - | - | 0 | - |
| - | - | 1056 | 1527 | - | - | 0 | - |
| - | - | 9888 | 1528 | - | - | 0 | - |
| - | - | 1376 | 1528 | - | - | 0 | - |
| - | - | 4842 | 1529 | - | - | 0 | - |
| - | - | 4030 | 1530 | - | - | 0 | - |
| 12 | c | 1.088E+04 | 1531 | 0.0005545 | 0.3622 | +1 | 12 |
| - | - | 8956 | 1532 | - | - | 0 | - |
| - | - | 4019 | 1533 | - | - | 0 | - |
| - | - | 2138 | 1534 | - | - | 0 | - |
| - | - | 1035 | 1535 | - | - | 0 | - |
| - | - | 818.6 | 1540 | - | - | 0 | - |
| 4 | y | 3230 | 1542 | 0.002433 | 1.578 | +1 | 12 |
| - | - | 2805 | 1543 | - | - | 0 | - |
| - | - | 1178 | 1543 | - | - | 0 | - |
| - | - | 1151 | 1544 | - | - | 0 | - |
| - | - | 995.4 | 1545 | - | - | 0 | - |
| - | - | 1039 | 1545 | - | - | 0 | - |
| - | - | 784.2 | 1547 | - | - | 0 | - |
| - | - | 1060 | 1548 | - | - | 0 | - |
| - | - | 1773 | 1552 | - | - | 0 | - |
| - | - | 2227 | 1552 | - | - | 0 | - |
| - | - | 995.3 | 1553 | - | - | 0 | - |
| - | - | 823.8 | 1555 | - | - | 0 | - |
| - | - | 1036 | 1556 | - | - | 0 | - |
| - | - | 1427 | 1557 | - | - | 0 | - |
| - | - | 1887 | 1558 | - | - | 0 | - |
| - | - | 1421 | 1558 | - | - | 0 | - |
| - | - | 1767 | 1559 | - | - | 0 | - |
| - | - | 1218 | 1559 | - | - | 0 | - |
| - | - | 1900 | 1560 | - | - | 0 | - |
| - | - | 1025 | 1560 | - | - | 0 | - |
| - | - | 1097 | 1561 | - | - | 0 | - |
| - | - | 1613 | 1563 | - | - | 0 | - |
| - | - | 921.9 | 1563 | - | - | 0 | - |
| - | - | 1215 | 1565 | - | - | 0 | - |
| - | - | 2261 | 1565 | - | - | 0 | - |
| - | - | 2055 | 1566 | - | - | 0 | - |
| - | - | 1961 | 1566 | - | - | 0 | - |
| - | - | 2252 | 1567 | - | - | 0 | - |
| - | - | 2121 | 1567 | - | - | 0 | - |
| - | - | 1487 | 1568 | - | - | 0 | - |
| - | - | 783.8 | 1568 | - | - | 0 | - |
| - | - | 919.5 | 1570 | - | - | 0 | - |
| - | - | 936.8 | 1571 | - | - | 0 | - |
| - | - | 875.2 | 1571 | - | - | 0 | - |
| - | - | 1728 | 1572 | - | - | 0 | - |
| - | - | 1576 | 1572 | - | - | 0 | - |
| - | - | 2723 | 1573 | - | - | 0 | - |
| - | - | 3713 | 1573 | - | - | 0 | - |
| - | - | 2999 | 1574 | - | - | 0 | - |
| - | - | 3350 | 1574 | - | - | 0 | - |
| - | - | 3347 | 1575 | - | - | 0 | - |
| - | - | 2906 | 1575 | - | - | 0 | - |
| - | - | 1074 | 1576 | - | - | 0 | - |
| - | - | 1001 | 1576 | - | - | 0 | - |
| - | - | 1347 | 1577 | - | - | 0 | - |
| - | - | 1988 | 1579 | - | - | 0 | - |
| - | - | 1356 | 1580 | - | - | 0 | - |
| - | - | 3894 | 1580 | - | - | 0 | - |
| - | - | 4839 | 1581 | - | - | 0 | - |
| - | - | 3052 | 1581 | - | - | 0 | - |
| - | - | 2308 | 1582 | - | - | 0 | - |
| - | - | 2516 | 1582 | - | - | 0 | - |
| - | - | 2518 | 1583 | - | - | 0 | - |
| - | - | 1984 | 1583 | - | - | 0 | - |
| - | - | 1207 | 1584 | - | - | 0 | - |
| - | - | 1104 | 1585 | - | - | 0 | - |
| - | - | 1159 | 1586 | - | - | 0 | - |
| - | - | 1146 | 1587 | - | - | 0 | - |
| - | - | 1417 | 1588 | - | - | 0 | - |
| - | - | 1105 | 1588 | - | - | 0 | - |
| - | - | 2969 | 1589 | - | - | 0 | - |
| - | - | 3111 | 1589 | - | - | 0 | - |
| - | - | 1589 | 1590 | - | - | 0 | - |
| - | - | 2642 | 1590 | - | - | 0 | - |
| - | - | 2670 | 1591 | - | - | 0 | - |
| - | - | 2307 | 1591 | - | - | 0 | - |
| - | - | 2107 | 1592 | - | - | 0 | - |
| - | - | 1380 | 1592 | - | - | 0 | - |
| - | - | 1675 | 1593 | - | - | 0 | - |
| - | - | 1609 | 1594 | - | - | 0 | - |
| - | - | 4121 | 1594 | - | - | 0 | - |
| - | - | 3712 | 1595 | - | - | 0 | - |
| - | - | 3834 | 1595 | - | - | 0 | - |
| - | - | 2520 | 1596 | - | - | 0 | - |
| - | - | 1761 | 1596 | - | - | 0 | - |
| - | - | 998.6 | 1597 | - | - | 0 | - |
| - | - | 1711 | 1597 | - | - | 0 | - |
| - | - | 2042 | 1598 | - | - | 0 | - |
| - | - | 1124 | 1598 | - | - | 0 | - |
| - | - | 964.7 | 1599 | - | - | 0 | - |
| - | - | 1513 | 1599 | - | - | 0 | - |
| - | - | 2467 | 1600 | - | - | 0 | - |
| - | - | 1856 | 1600 | - | - | 0 | - |
| - | - | 1523 | 1601 | - | - | 0 | - |
| - | - | 1538 | 1601 | - | - | 0 | - |
| - | - | 1398 | 1602 | - | - | 0 | - |
| - | - | 4005 | 1602 | - | - | 0 | - |
| - | - | 6080 | 1603 | - | - | 0 | - |
| - | - | 9312 | 1603 | - | - | 0 | - |
| - | - | 4531 | 1604 | - | - | 0 | - |
| - | - | 4044 | 1604 | - | - | 0 | - |
| - | - | 2098 | 1605 | - | - | 0 | - |
| - | - | 1146 | 1605 | - | - | 0 | - |
| - | - | 767.1 | 1610 | - | - | 0 | - |
| - | - | 860.8 | 1633 | - | - | 0 | - |
| - | - | 861.3 | 1634 | - | - | 0 | - |
| 3 | z | 5062 | 1654 | 0.002736 | 1.654 | +1 | 13 |
| - | - | 7818 | 1655 | - | - | 0 | - |
| - | - | 6045 | 1656 | - | - | 0 | - |
| - | - | 3171 | 1657 | - | - | 0 | - |
| - | - | 3384 | 1691 | - | - | 0 | - |
| 13 | c | 1.598E+04 | 1692 | 0.002395 | 1.415 | +1 | 13 |
| - | - | 1.569E+04 | 1693 | - | - | 0 | - |
| - | - | 7864 | 1694 | - | - | 0 | - |
| - | - | 3432 | 1695 | - | - | 0 | - |
| - | - | 947.3 | 1696 | - | - | 0 | - |
| - | - | 1101 | 1777 | - | - | 0 | - |
| 2 | z | 989.2 | 1791 | 0.01428 | 7.974 | +1 | 14 |
| - | - | 2402 | 1792 | - | - | 0 | - |
| - | - | 1902 | 1793 | - | - | 0 | - |
| - | - | 1587 | 1794 | - | - | 0 | - |
| - | - | 1038 | 1795 | - | - | 0 | - |
| - | - | 1038 | 1819 | - | - | 0 | - |
| 14 | c | 2850 | 1820 | 0.007379 | 4.054 | +1 | 14 |
| - | - | 4541 | 1821 | - | - | 0 | - |
| - | - | 4214 | 1822 | - | - | 0 | - |
| - | - | 2318 | 1823 | - | - | 0 | - |
| - | - | 1296 | 1824 | - | - | 0 | - |
| - | - | 1983 | 1831 | - | - | 0 | - |
| - | - | 2347 | 1832 | - | - | 0 | - |
| - | - | 987.7 | 1833 | - | - | 0 | - |
| - | - | 1138 | 1835 | - | - | 0 | - |
| - | - | 1031 | 1845 | - | - | 0 | - |
| - | - | 1177 | 1846 | - | - | 0 | - |
| - | - | 1703 | 1847 | - | - | 0 | - |
| - | - | 958.9 | 1848 | - | - | 0 | - |
| - | - | 1040 | 1849 | - | - | 0 | - |
| - | - | 2211 | 1861 | - | - | 0 | - |
| - | - | 7485 | 1862 | - | - | 0 | - |
| - | - | 9603 | 1863 | - | - | 0 | - |
| - | - | 9144 | 1864 | - | - | 0 | - |
| - | - | 4978 | 1865 | - | - | 0 | - |
| - | - | 2001 | 1866 | - | - | 0 | - |
| - | - | 2814 | 1876 | - | - | 0 | - |
| - | - | 6287 | 1877 | - | - | 0 | - |
| - | - | 5339 | 1878 | - | - | 0 | - |
| - | - | 3596 | 1879 | - | - | 0 | - |
| - | - | 1796 | 1880 | - | - | 0 | - |
| - | - | 3219 | 1893 | - | - | 0 | - |
| - | - | 7920 | 1894 | - | - | 0 | - |
| - | - | 6946 | 1895 | - | - | 0 | - |
| - | - | 4258 | 1896 | - | - | 0 | - |
| - | - | 1538 | 1897 | - | - | 0 | - |
| - | - | 890.3 | 1903 | - | - | 0 | - |
| - | - | 5196 | 1904 | - | - | 0 | - |
| - | - | 1.149E+04 | 1905 | - | - | 0 | - |
| - | - | 1.115E+04 | 1906 | - | - | 0 | - |
| - | - | 6002 | 1907 | - | - | 0 | - |
| - | - | 3943 | 1908 | - | - | 0 | - |
| - | - | 5218 | 1920 | - | - | 0 | - |
| - | - | 1.829E+04 | 1921 | - | - | 0 | - |
| - | - | 5.397E+04 | 1922 | - | - | 0 | - |
| - | - | 4.966E+04 | 1923 | - | - | 0 | - |
| - | - | 2.929E+04 | 1924 | - | - | 0 | - |
| - | - | 1.287E+04 | 1925 | - | - | 0 | - |
| - | - | 4773 | 1926 | - | - | 0 | - |
| - | - | 955.5 | 2040 | - | - | 0 | - |

m/z Charge Intensity FragmentType MassShift Position
121.01114654541016 0 362.24066
129.1021728515625 0 1591.722
132.95233154296875 0 439.53796
136.41448974609375 0 458.74744
141.14561462402344 0 418.16064
148.95443725585938 0 659.9974
153.13702392578125 0 549.58887
153.65750122070312 0 427.44025
155.0928955078125 0 729.288
156.90858459472656 0 474.73975
157.13308715820312 0 457.37683
165.10147094726562 0 441.6555
166.0610809326172 0 852.5235
173.4387969970703 0 1889.9125
173.70831298828125 0 458.8428
177.040283203125 0 528.2872
183.1123809814453 0 675.8766
185.1644287109375 0 743.16815
186.12396240234375 0 529.9749
223.15516662597656 0 1182.2533
225.36428833007812 0 520.7979
230.16229248046875 0 3650.7007 z 13
231.16989135742188 0 4396.9844
234.1234130859375 0 1501.95
243.25466918945312 0 540.9293
246.1815948486328 0 1125.6843 y 13
250.14263916015625 0 1051.7563
251.15011596679688 0 8163.084
252.15316772460938 0 1436.3341
268.1765441894531 0 4847.032 c 1
269.18023681640625 0 889.2566
292.9272155761719 0 703.4383
300.19183349609375 0 7840.476
301.1952209472656 0 1067.0345
302.1705017089844 0 681.5515
318.1410217285156 0 577.7359
333.3406982421875 0 607.88855
352.2212829589844 0 1822.9457
379.20831298828125 0 1051.9011 c Ammonia loss 2
395.2269592285156 0 4110.6094
396.2066650390625 0 807.09503
396.235107421875 0 14201.79 c 2
397.2382507324219 0 3471.9587
407.19635009765625 0 4014.355 y 12
408.1982116699219 0 1108.684 c Ammonia loss 9
409.3603210449219 0 577.471
427.26654052734375 0 970.5706
428.2640075683594 0 733.56757
461.2082214355469 0 738.3848 w 11
467.24932861328125 0 5486.465
468.2534484863281 0 2429.513
475.27874755859375 0 670.8108
488.3190002441406 0 850.9567
494.2366638183594 0 745.36743 c Ammonia loss 3
506.76007080078125 0 2852.0955 y 7
507.2604064941406 0 1128.2573
507.7631530761719 0 689.44336
511.26239013671875 0 21741.973 c 3
512.2655639648438 0 5520.1333
513.2685546875 0 614.67145
519.272216796875 0 5606.095 z 11
520.2775268554688 0 2207.003
521.2731323242188 0 558.5486
535.2916870117188 0 1845.653 y 11
536.2930297851562 0 832.3729
564.2645874023438 0 618.9392
584.362548828125 0 4250.6187
585.3668212890625 0 1236.5853
590.8561401367188 0 1764.5717
598.3446044921875 0 554.8201
601.622802734375 0 2778.4463 c Ammonia loss 13
601.9573974609375 0 2062.5752
602.29443359375 0 1848.1936
602.9549560546875 0 5112.6753 y 1
603.2899169921875 0 3828.5298
603.6240234375 0 2682.1707
606.3243408203125 0 644.78326
607.6270751953125 0 1393.3785
607.9584350585938 0 1326.2295
618.8707275390625 0 838.45874
619.3637084960938 0 667.7756
625.2781372070312 0 711.4991
628.9700927734375 0 1069.7234
629.3027954101562 0 1101.9606
629.6366577148438 0 1225.2253
633.30908203125 0 550.39716
634.6474609375 0 2167.6123
634.978271484375 0 2651.9246
635.3106689453125 0 2772.609
635.6446533203125 0 1374.3181
639.3353881835938 0 1234.3986
639.87646484375 0 1681.6272
640.32080078125 0 4130.027
640.3772583007812 0 1523.2025
640.6495971679688 0 47797.44
640.8822631835938 0 1950.6461
640.983642578125 0 47831.234
641.317626953125 0 37424.15
641.6517944335938 0 12348.154
641.8623046875 0 2142.0864
641.98583984375 0 3971.1611
651.353271484375 0 1762.1061
651.8541259765625 0 1400.4492
680.3116455078125 0 972.8174 c Ammonia loss 4
682.3358154296875 0 13062.256 z 10
683.339599609375 0 5270.413
684.340087890625 0 1691.3596
685.3382568359375 0 809.43146
697.3421020507812 0 6202.623 c 4
698.3480224609375 0 4094.4785 y 10
699.3499145507812 0 1309.8374
713.8565063476562 0 864.7748 y 4
752.3656005859375 0 2286.5942 w 9
753.8645629882812 0 607.0039
760.9014282226562 0 859.444
761.39892578125 0 700.4823
765.4010009765625 0 1422.4158
765.890625 0 4658.858 c 11
766.3909912109375 0 4856.1675
766.8909301757812 0 2252.9397
767.3902587890625 0 777.97186
768.3475952148438 0 655.1472
769.0875854492188 0 857.9069
771.367919921875 0 1240.3282 y 3
771.8734130859375 0 1469.5554
772.3682250976562 0 642.0901
776.1499633789062 0 854.36304
779.3955078125 0 918.79114
780.9130249023438 0 716.2036
787.1487426757812 0 819.7676
787.4033813476562 0 1186.7039
794.3986206054688 0 663.0441
795.1544189453125 0 614.03156
799.1505126953125 0 835.1879
801.9102783203125 0 669.3754
809.4354858398438 0 1184.6019
810.4271240234375 0 6202.8022 c 5
811.378173828125 0 6284.8066 z 9
812.3834228515625 0 2911.987
813.3788452148438 0 1417.099
817.3958129882812 0 1045.8934
825.3795776367188 0 1041.744
826.3888549804688 0 1710.5596 y Water loss 2
826.8926391601562 0 1996.3982 y Ammonia loss 2
827.3924560546875 0 4584.538 y 9
827.888427734375 0 1874.6144
828.3897094726562 0 1223.5415
835.3980712890625 0 5847.6753 y 2
835.9000244140625 0 5582.628
836.4005737304688 0 3053.3464
836.9072265625 0 1041.7151
842.4202270507812 0 2080.0544
843.4210815429688 0 1044.0834
845.8939819335938 0 3076.5356
846.3973999023438 0 7235.0493 c 12
846.8989868164062 0 7841.623
847.3984985351562 0 4525.993
847.8983154296875 0 1499.1235
848.4902954101562 0 864.18286
867.4183349609375 0 990.3162
875.475830078125 0 774.9977
880.4520874023438 0 1615.1804
881.441162109375 0 4475.6943
882.4459228515625 0 2813.9048
883.450439453125 0 1448.607
888.4371337890625 0 799.2724
888.9393310546875 0 751.6198
893.9420166015625 0 1622.4309
894.456787109375 0 624.7249
895.4374389648438 0 1929.9658
895.9259643554688 0 2445.5051 z 1
896.425048828125 0 2450.7122
896.9202880859375 0 1811.3754
902.4432983398438 0 1768.9893
902.9279174804688 0 967.59607
903.4434204101562 0 1054.9391
908.4335327148438 0 2127.5044 c Ammonia loss 6
908.9603881835938 0 1236.6157
909.4459838867188 0 2095.0647
909.9495849609375 0 1490.1694
910.4444580078125 0 15609.762 c 13
910.9456787109375 0 18373.766
911.4454345703125 0 12432.967
911.9443359375 0 4226.759
912.4412231445312 0 1644.4353
915.4635009765625 0 990.26544
916.4664916992188 0 1565.9526
916.9658203125 0 1518.2815
917.4847412109375 0 820.6724
917.9711303710938 0 1098.3569
921.4428100585938 0 838.6561
922.9641723632812 0 1398.9615
923.455810546875 0 1184.2189
923.9503784179688 0 1469.3726
924.4490356445312 0 4540.376
924.9474487304688 0 3104.8425
925.4521484375 0 7781.4443 c 6
925.9459228515625 0 1465.0427
926.4548950195312 0 3076.7273
927.4595947265625 0 854.68555
929.4575805664062 0 1190.8777
930.46142578125 0 2804.7095
930.9588012695312 0 10656.335
931.4656982421875 0 26203.523
931.9659423828125 0 21715.908
932.4683837890625 0 12902.279
932.9676513671875 0 5327.066
933.4711303710938 0 1850.5984
937.4745483398438 0 781.4391 y Water loss 8
937.9732666015625 0 3754.7139
938.470947265625 0 6499.054 y Ammonia loss 8
938.9683227539062 0 4496.366
939.4722900390625 0 15584.309 z 8
939.9669799804688 0 728.37994
940.478271484375 0 9615.378
941.4819946289062 0 3417.2715
942.4857177734375 0 1418.1407
942.9667358398438 0 970.9467
949.1539306640625 0 835.3791
949.9437255859375 0 827.1539
950.9572143554688 0 761.40436
951.9688110351562 0 3341.8801
952.465576171875 0 6382.731
952.964599609375 0 6510.9326
953.464599609375 0 3576.1738
953.9614868164062 0 2344.3962
954.470703125 0 1569.4724
955.4935302734375 0 2079.8342 y 8
956.50244140625 0 851.16504
958.4854125976562 0 737.77484
959.4995727539062 0 823.6815
959.9729614257812 0 3486.1226
960.470947265625 0 24085.531
960.972900390625 0 38807.01
961.473876953125 0 29014.287
961.975341796875 0 16433.021
962.47509765625 0 6138.245
962.9695434570312 0 2718.6333
978.1808471679688 0 741.9859
981.4732055664062 0 3872.19
982.4751586914062 0 18264.176 c 7
983.4782104492188 0 8078.196
984.4813842773438 0 2584.5376
986.49609375 0 1047.1307
996.4976806640625 0 4139.6006 z 7
997.4982299804688 0 3120.5735
998.4982299804688 0 1531.9128
999.5105590820312 0 799.0743
1012.5136108398438 0 6649.3447 y 7
1013.5182495117188 0 3501.5967
1014.198486328125 0 1906.5957
1014.5235595703125 0 3239.8289
1014.8566284179688 0 1111.4418
1019.9845581054688 0 1380.9005
1020.4962158203125 0 1133.8411
1025.5206298828125 0 1456.0199
1027.5367431640625 0 1405.2396
1031.543701171875 0 1060.1157
1032.0419921875 0 995.58887
1032.540771484375 0 1191.8335
1033.8778076171875 0 908.74817
1034.53759765625 0 778.1585
1034.8641357421875 0 1219.6779
1040.505126953125 0 864.21216
1043.8773193359375 0 844.3208
1044.2083740234375 0 1781.0259
1044.5164794921875 0 1018.7305
1044.8680419921875 0 796.42566
1045.1983642578125 0 1472.2672
1045.5252685546875 0 1045.8077
1046.186767578125 0 714.26666
1046.8626708984375 0 782.8877
1047.533447265625 0 1798.571
1047.867431640625 0 1060.1937
1048.2005615234375 0 1458.2073
1048.540771484375 0 1002.2864
1048.87158203125 0 1536.684
1049.2061767578125 0 1413.8657
1049.537841796875 0 1175.9402
1049.8685302734375 0 1480.5273
1050.202880859375 0 1095.3594
1053.209716796875 0 1014.12946
1053.5369873046875 0 2555.4265
1053.876220703125 0 3576.7927
1054.202392578125 0 1722.7513
1054.5352783203125 0 2505.9358
1054.8729248046875 0 1897.6564
1055.1953125 0 906.1721
1055.5286865234375 0 2087.867
1057.5308837890625 0 834.42126
1059.533203125 0 2302.3276
1059.8681640625 0 1478.5985
1060.2017822265625 0 2974.4756
1060.53271484375 0 2083.3901
1060.86083984375 0 862.50494
1061.2093505859375 0 1266.8285
1061.5115966796875 0 1498.2773
1061.8558349609375 0 1669.6913
1062.2022705078125 0 1709.9294
1062.5390625 0 1188.1268
1062.8717041015625 0 3019.73
1063.2034912109375 0 3262.7268
1063.541015625 0 3989.0898
1063.874755859375 0 1648.691
1064.1949462890625 0 1207.1176
1064.5364990234375 0 894.63226
1065.211669921875 0 1691.3887
1065.5401611328125 0 2059.969
1065.8763427734375 0 1536.5704
1066.212890625 0 1392.3513
1066.8680419921875 0 1154.2943
1067.185791015625 0 990.329
1067.535400390625 0 1627.5753
1068.548095703125 0 4473.371
1068.882568359375 0 5930.8887
1069.2142333984375 0 4688.759
1069.5506591796875 0 4242.0054
1069.8843994140625 0 2457.0986
1070.2191162109375 0 863.7108
1094.530517578125 0 781.03784
1109.591796875 0 1359.9974
1110.569580078125 0 11493.059 c 8
1111.5570068359375 0 7015.2236
1112.54345703125 0 3644.77
1113.5394287109375 0 996.3376
1137.566650390625 0 922.04205
1142.5625 0 1702.7489
1143.5631103515625 0 1014.62427
1166.672607421875 0 1438.4001
1181.618408203125 0 1069.9009
1182.1234130859375 0 1025.4705
1189.61328125 0 821.427
1195.586669921875 0 904.7873 w 5
1220.5736083984375 0 853.68
1224.6099853515625 0 3771.9724 z 5
1225.611083984375 0 2429.1326
1226.60986328125 0 1996.6891
1238.6192626953125 0 2420.154
1239.6138916015625 0 15331.498 c 9
1240.6162109375 0 9574.918 y 5
1241.61767578125 0 3990.3997
1242.632568359375 0 1150.6256
1243.6385498046875 0 825.6265
1266.66552734375 0 905.16327
1271.6402587890625 0 1117.5049
1272.15966796875 0 934.2592
1272.6328125 0 1233.9796
1279.7293701171875 0 877.24194
1280.1611328125 0 834.707
1280.5924072265625 0 1670.0225
1280.7393798828125 0 1646.9207
1281.7498779296875 0 2301.6406
1282.736572265625 0 2423.7485
1283.7197265625 0 1638.2388
1301.6759033203125 0 908.09296
1355.6273193359375 0 892.34326
1366.6932373046875 0 1013.1501
1401.6834716796875 0 2346.7864
1402.67822265625 0 9452.215 c 10
1403.6802978515625 0 6888.0195
1404.681884765625 0 2711.1077
1405.6961669921875 0 1120.5819
1409.6923828125 0 953.2428 y Ammonia loss 4
1410.6866455078125 0 11711.009 z 4
1411.6881103515625 0 9083.944
1412.6920166015625 0 5349.5615
1413.6859130859375 0 1188.2941
1422.232421875 0 1661.3838
1423.23681640625 0 1109.3195
1425.70703125 0 1527.666
1426.70458984375 0 6015.206 y 4
1427.708251953125 0 4144.595
1428.7047119140625 0 2461.8826
1429.7071533203125 0 1018.72217
1435.700439453125 0 1193.0026
1436.7071533203125 0 1423.6609
1438.7235107421875 0 790.2755
1465.732666015625 0 923.33234
1475.2086181640625 0 878.505
1481.723876953125 0 3524.6404
1482.7264404296875 0 2532.8035
1483.7254638671875 0 1656.321
1484.7369384765625 0 1039.5933
1485.7569580078125 0 939.71954
1486.7532958984375 0 3287.6013
1487.2486572265625 0 1602.6854
1487.756591796875 0 4171.333
1488.7557373046875 0 1659.0048
1496.238037109375 0 939.64343
1500.73046875 0 1025.537
1501.7156982421875 0 1182.6185
1509.709716796875 0 839.1436
1515.2430419921875 0 931.5627
1516.2354736328125 0 1053.5938
1520.7791748046875 0 806.52094
1521.28466796875 0 1694.2931
1522.272705078125 0 1071.3467
1523.737060546875 0 1027.354 y Water loss 3
1524.7340087890625 0 1148.138 y Ammonia loss 3
1525.71337890625 0 9164.07 z 3
1526.7176513671875 0 15506.993
1527.2484130859375 0 1055.9164
1527.724609375 0 9888.238
1528.2637939453125 0 1376.4568
1528.730224609375 0 4841.9717
1529.7506103515625 0 4029.9219
1530.7705078125 0 10881.687 c 11
1531.77392578125 0 8956.298
1532.781005859375 0 4018.7778
1533.783447265625 0 2138.328
1534.7940673828125 0 1034.648
1539.767822265625 0 818.6016
1541.7325439453125 0 3229.7244 y 3
1542.73681640625 0 2804.5903
1543.2718505859375 0 1177.8702
1543.738525390625 0 1150.7888
1544.76025390625 0 995.35864
1545.2781982421875 0 1039.0109
1546.775634765625 0 784.15955
1547.7960205078125 0 1060.1
1551.7906494140625 0 1772.7067
1552.2794189453125 0 2226.9294
1552.7730712890625 0 995.34607
1554.7376708984375 0 823.77734
1555.726318359375 0 1035.8846
1557.2969970703125 0 1426.9857
1557.8021240234375 0 1887.4321
1558.2880859375 0 1421.0612
1558.79248046875 0 1766.797
1559.2933349609375 0 1218.3467
1559.7867431640625 0 1900.4484
1560.286376953125 0 1025.3743
1561.2723388671875 0 1097.4626
1562.771484375 0 1613.2091
1563.2557373046875 0 921.8985
1564.802978515625 0 1215.1095
1565.3045654296875 0 2261.1558
1565.7889404296875 0 2055.1377
1566.2943115234375 0 1961.1268
1566.802001953125 0 2252.499
1567.2913818359375 0 2121.147
1567.80615234375 0 1487.1776
1568.2911376953125 0 783.7626
1569.765625 0 919.47675
1570.7542724609375 0 936.8138
1571.2969970703125 0 875.2425
1571.786865234375 0 1728.1929
1572.3035888671875 0 1576.3314
1572.8057861328125 0 2722.6035
1573.29833984375 0 3712.8882
1573.791259765625 0 2999.0176
1574.3092041015625 0 3349.5957
1574.8056640625 0 3346.6213
1575.302490234375 0 2905.7163
1575.8104248046875 0 1073.8281
1576.26611328125 0 1000.5056
1576.79345703125 0 1347.1835
1579.2957763671875 0 1987.6058
1579.81494140625 0 1356.0702
1580.3023681640625 0 3893.6262
1580.7999267578125 0 4839.363
1581.307373046875 0 3052.402
1581.7928466796875 0 2307.6511
1582.290771484375 0 2515.9758
1582.802001953125 0 2517.9585
1583.294921875 0 1984.3297
1584.3026123046875 0 1206.5255
1585.299560546875 0 1103.9818
1586.2843017578125 0 1159.1774
1587.307861328125 0 1146.2247
1587.818115234375 0 1416.5103
1588.2882080078125 0 1105.4009
1588.8082275390625 0 2969.1284
1589.300048828125 0 3110.7727
1589.8076171875 0 1588.7207
1590.2879638671875 0 2642.2708
1590.7874755859375 0 2669.719
1591.3011474609375 0 2307.4026
1591.79443359375 0 2107.2583
1592.29345703125 0 1379.7067
1593.2889404296875 0 1675.322
1593.814697265625 0 1609.3031
1594.302490234375 0 4120.68
1594.7984619140625 0 3711.6333
1595.315673828125 0 3834.184
1595.796875 0 2519.6294
1596.30224609375 0 1760.5167
1596.7884521484375 0 998.5671
1597.311767578125 0 1711.2632
1597.8013916015625 0 2042.0485
1598.3106689453125 0 1123.636
1598.79541015625 0 964.66156
1599.2962646484375 0 1512.742
1599.79541015625 0 2467.1748
1600.3017578125 0 1855.9199
1600.782958984375 0 1522.7557
1601.28857421875 0 1537.8325
1601.8026123046875 0 1398.3864
1602.3177490234375 0 4004.9736
1602.8201904296875 0 6079.868
1603.320068359375 0 9312.056
1603.8226318359375 0 4531.104
1604.3209228515625 0 4044.3936
1604.8203125 0 2097.9546
1605.3212890625 0 1146.428
1609.8021240234375 0 767.0986
1632.7928466796875 0 860.7744
1633.785400390625 0 861.2769
1653.7720947265625 0 5061.6636 z 2
1654.777587890625 0 7817.628
1655.78125 0 6045.1978
1656.7845458984375 0 3171.4978
1690.785400390625 0 3383.6714
1691.7867431640625 0 15980.4795 c 12
1692.7874755859375 0 15686.411
1693.791748046875 0 7863.689
1694.7974853515625 0 3431.8186
1695.7760009765625 0 947.2656
1776.877685546875 0 1100.5055
1790.8480224609375 0 989.18524 z 1
1791.8438720703125 0 2402.2197
1792.8564453125 0 1901.6088
1793.8477783203125 0 1586.8796
1794.856689453125 0 1038.0919
1818.92919921875 0 1037.9188
1819.8914794921875 0 2850.2673 c 13
1820.8868408203125 0 4541.3765
1821.8909912109375 0 4214.339
1822.89453125 0 2318.4336
1823.89306640625 0 1296.0188
1830.950927734375 0 1982.7234
1831.9530029296875 0 2346.5415
1832.958984375 0 987.7029
1834.9376220703125 0 1138.0325
1844.917236328125 0 1030.9459
1845.916015625 0 1177.1183
1846.914794921875 0 1703.2488
1847.946533203125 0 958.8891
1848.8975830078125 0 1039.5656
1860.9207763671875 0 2210.535
1861.927734375 0 7484.763
1862.934326171875 0 9603.4375
1863.9339599609375 0 9144.3955
1864.9322509765625 0 4978.4683
1865.923583984375 0 2001.2747
1875.935302734375 0 2813.5938
1876.9354248046875 0 6287.1978
1877.9368896484375 0 5339.459
1878.93994140625 0 3596.2815
1879.9307861328125 0 1795.9495
1892.9501953125 0 3218.502
1893.9541015625 0 7919.6963
1894.9525146484375 0 6945.6333
1895.9599609375 0 4257.745
1896.960693359375 0 1538.1348
1902.9356689453125 0 890.33997
1903.92724609375 0 5196.2246
1904.925537109375 0 11489.714
1905.928955078125 0 11153.585
1906.9234619140625 0 6001.885
1907.92529296875 0 3942.5398
1919.9405517578125 0 5217.88
1920.9449462890625 0 18285.373
1921.9503173828125 0 53972.785
1922.9527587890625 0 49657.746
1923.9534912109375 0 29291.186
1924.9510498046875 0 12871.952
1925.949951171875 0 4773.4473
2039.955078125 0 955.5345

Spectrum Details

|  |  |
| --- | --- |
| Matched peaks? Matched peaks The total absolute number of peaks matched. Additionally in brackets the total fraction of peaks matched and the total number of peaks is shown. | 59 (10.37% of 569) |
| FDR? FDR The false discovery rate estimated for this peptide. It is calculated by matching all theoretical fragments with a non-integer shift with the raw peaks for this spectrum. This is done with 40 different shifts. The resulting percentage is the average number of annotated peaks over the number of annotated peaks with the correct spectrum. | 1.17% |
| Satellite FDR? Satellite FDR See the FDR for details on its calculation. This satellite ion specific FDR only contains the satellite ions (d/w) for I/L/J positions. | 0.00% |
| PSM Score? PSM Score The PSM Score as given by Hecklib to this annotated spectrum. It is shown with three significant figures. | 559 |

#### Spectrum 5773? Spectrum 5773 The raw spectrum of this peptide as annotated by Hecklib. The fragments are coloured according to ion type (see legend). Any peaks with a star '\*' as text can be hovered over to see the full details, first the ion type second the mass shift type. By hovering over the amino acids in the peptide or ions in the legend the corresponding peaks are highlighted. By toggling the 'Unassigned' label you can turn the background (unassigned) peaks on or off in the plot. By updating the slider in the Ion legend you can update the spectrum to only show the top X% of the peaks with labels. The top X% means any peak that is within X% of the highest intensity. By dragging in the spectrum you can zoom in to a specific part of the spectrum and use 'Zoom Out' to get back to the original zoom level. The annotation of the spectrum is based on the given sequence in the peptides file and is done with different software so inconsistencies are likely. The peaks are annotated based on the given sequence, with 20 ppm tolerance.

Copy Data

##### Spectrum 5773 (TSV)

###### Preview

```
Loading example...
```

*Click on the button to copy the data to your clipboard.*

Mz MinMz MaxIntensity Max

WidthHeightPeptide font sizePeptide stroke widthSpectrum font sizeSpectrum stroke widthCompact peptide

Ion legend

wxyz

abcd

OtherUnassignedIon^Charge^_Position_Show for top:%

JHQDWJDGKEYKCKV

01.36e+42.73e+44.09e+45.45e+4

Zoom Out

z^+1^_2_y^+1^_2_c^+1^_2_c^+1^_3_c^+1^_3_y^+1^_3_c^+3^_10_w^+1^_4_c^+1^_4_y^+2^_8_c^+1^_4_z^+1^_4_y^+1^_4_c^+3^_14_y^+3^_14_c^+1^_5_z^+1^_5_c^+1^_5_y^+1^_5_y^+2^_11_w^+1^_6_c^+2^_12_y^+2^_12_c^+1^_6_z^+1^_6_y^+2^_13_y^+2^_13_y^+1^_6_y^+2^_13_c^+2^_13_z^+2^_14_c^+1^_7_c^+2^_14_c^+1^_7_y^+1^_7_y^+1^_7_z^+1^_7_y^+1^_7_c^+1^_8_z^+1^_8_y^+1^_8_c^+1^_9_w^+1^_10_z^+1^_10_c^+1^_10_y^+1^_10_c^+1^_11_y^+1^_11_z^+1^_11_y^+1^_11_y^+1^_12_y^+1^_12_z^+1^_12_c^+1^_12_y^+1^_12_z^+1^_13_c^+1^_13_z^+1^_14_c^+1^_14_

0515103015452060

Fragment Matches Table

Show background peaks

| Position | Ion type | Intensity | mz Theoretical | mz Error (Th) | mz Error (ppm) | Charge | Series Number |
| --- | --- | --- | --- | --- | --- | --- | --- |
| - | - | 362.2 | 121 | - | - | 0 | - |
| - | - | 1592 | 129.1 | - | - | 0 | - |
| - | - | 439.5 | 133 | - | - | 0 | - |
| - | - | 458.7 | 136.4 | - | - | 0 | - |
| - | - | 418.2 | 141.1 | - | - | 0 | - |
| - | - | 660 | 149 | - | - | 0 | - |
| - | - | 549.6 | 153.1 | - | - | 0 | - |
| - | - | 427.4 | 153.7 | - | - | 0 | - |
| - | - | 729.3 | 155.1 | - | - | 0 | - |
| - | - | 474.7 | 156.9 | - | - | 0 | - |
| - | - | 457.4 | 157.1 | - | - | 0 | - |
| - | - | 441.7 | 165.1 | - | - | 0 | - |
| - | - | 852.5 | 166.1 | - | - | 0 | - |
| - | - | 1890 | 173.4 | - | - | 0 | - |
| - | - | 458.8 | 173.7 | - | - | 0 | - |
| - | - | 528.3 | 177 | - | - | 0 | - |
| - | - | 675.9 | 183.1 | - | - | 0 | - |
| - | - | 743.2 | 185.2 | - | - | 0 | - |
| - | - | 530 | 186.1 | - | - | 0 | - |
| - | - | 1182 | 223.2 | - | - | 0 | - |
| - | - | 520.8 | 225.4 | - | - | 0 | - |
| 14 | z | 3651 | 230.2 | 0.0002015 | 0.8756 | +1 | 2 |
| - | - | 4397 | 231.2 | - | - | 0 | - |
| - | - | 1502 | 234.1 | - | - | 0 | - |
| - | - | 540.9 | 243.3 | - | - | 0 | - |
| 14 | y | 1126 | 246.2 | 0.0003768 | 1.53 | +1 | 2 |
| - | - | 1052 | 250.1 | - | - | 0 | - |
| - | - | 8163 | 251.2 | - | - | 0 | - |
| - | - | 1436 | 252.2 | - | - | 0 | - |
| 2 | c | 4847 | 268.2 | 0.0002572 | 0.9591 | +1 | 2 |
| - | - | 889.3 | 269.2 | - | - | 0 | - |
| - | - | 703.4 | 292.9 | - | - | 0 | - |
| - | - | 7840 | 300.2 | - | - | 0 | - |
| - | - | 1067 | 301.2 | - | - | 0 | - |
| - | - | 681.6 | 302.2 | - | - | 0 | - |
| - | - | 577.7 | 318.1 | - | - | 0 | - |
| - | - | 607.9 | 333.3 | - | - | 0 | - |
| - | - | 1823 | 352.2 | - | - | 0 | - |
| 3 | c | 1052 | 379.2 | 0.0005168 | 1.363 | +1 | 3 |
| - | - | 4111 | 395.2 | - | - | 0 | - |
| - | - | 807.1 | 396.2 | - | - | 0 | - |
| 3 | c | 1.42E+04 | 396.2 | 0.0002715 | 0.6852 | +1 | 3 |
| - | - | 3472 | 397.2 | - | - | 0 | - |
| 13 | y | 4014 | 407.2 | 0.004052 | 9.952 | +1 | 3 |
| 10 | c | 1109 | 408.2 | 0.001677 | 4.108 | +3 | 10 |
| - | - | 577.5 | 409.4 | - | - | 0 | - |
| - | - | 970.6 | 427.3 | - | - | 0 | - |
| - | - | 733.6 | 428.3 | - | - | 0 | - |
| 12 | w | 738.4 | 461.2 | 0.002746 | 5.953 | +1 | 4 |
| - | - | 5486 | 467.2 | - | - | 0 | - |
| - | - | 2430 | 468.3 | - | - | 0 | - |
| - | - | 670.8 | 475.3 | - | - | 0 | - |
| - | - | 851 | 488.3 | - | - | 0 | - |
| 4 | c | 745.4 | 494.2 | 0.000891 | 1.803 | +1 | 4 |
| 8 | y | 2852 | 506.8 | 0.002424 | 4.784 | +2 | 8 |
| - | - | 1128 | 507.3 | - | - | 0 | - |
| - | - | 689.4 | 507.8 | - | - | 0 | - |
| 4 | c | 2.174E+04 | 511.3 | 6.82E-05 | 0.1334 | +1 | 4 |
| - | - | 5520 | 512.3 | - | - | 0 | - |
| - | - | 614.7 | 513.3 | - | - | 0 | - |
| 12 | z | 5606 | 519.3 | 0.004425 | 8.521 | +1 | 4 |
| - | - | 2207 | 520.3 | - | - | 0 | - |
| - | - | 558.5 | 521.3 | - | - | 0 | - |
| 12 | y | 1846 | 535.3 | 0.003679 | 6.872 | +1 | 4 |
| - | - | 832.4 | 536.3 | - | - | 0 | - |
| - | - | 618.9 | 564.3 | - | - | 0 | - |
| - | - | 4251 | 584.4 | - | - | 0 | - |
| - | - | 1237 | 585.4 | - | - | 0 | - |
| - | - | 1765 | 590.9 | - | - | 0 | - |
| - | - | 554.8 | 598.3 | - | - | 0 | - |
| 14 | c | 2778 | 601.6 | 0.001232 | 2.048 | +3 | 14 |
| - | - | 2063 | 602 | - | - | 0 | - |
| - | - | 1848 | 602.3 | - | - | 0 | - |
| 2 | y | 5113 | 603 | 0.000717 | 1.189 | +3 | 14 |
| - | - | 3829 | 603.3 | - | - | 0 | - |
| - | - | 2682 | 603.6 | - | - | 0 | - |
| - | - | 644.8 | 606.3 | - | - | 0 | - |
| - | - | 1393 | 607.6 | - | - | 0 | - |
| - | - | 1326 | 608 | - | - | 0 | - |
| - | - | 838.5 | 618.9 | - | - | 0 | - |
| - | - | 667.8 | 619.4 | - | - | 0 | - |
| - | - | 711.5 | 625.3 | - | - | 0 | - |
| - | - | 1070 | 629 | - | - | 0 | - |
| - | - | 1102 | 629.3 | - | - | 0 | - |
| - | - | 1225 | 629.6 | - | - | 0 | - |
| - | - | 550.4 | 633.3 | - | - | 0 | - |
| - | - | 2168 | 634.6 | - | - | 0 | - |
| - | - | 2652 | 635 | - | - | 0 | - |
| - | - | 2773 | 635.3 | - | - | 0 | - |
| - | - | 1374 | 635.6 | - | - | 0 | - |
| - | - | 1234 | 639.3 | - | - | 0 | - |
| - | - | 1682 | 639.9 | - | - | 0 | - |
| - | - | 4130 | 640.3 | - | - | 0 | - |
| - | - | 1523 | 640.4 | - | - | 0 | - |
| - | - | 4.78E+04 | 640.6 | - | - | 0 | - |
| - | - | 1951 | 640.9 | - | - | 0 | - |
| - | - | 4.783E+04 | 641 | - | - | 0 | - |
| - | - | 3.742E+04 | 641.3 | - | - | 0 | - |
| - | - | 1.235E+04 | 641.7 | - | - | 0 | - |
| - | - | 2142 | 641.9 | - | - | 0 | - |
| - | - | 3971 | 642 | - | - | 0 | - |
| - | - | 1762 | 651.4 | - | - | 0 | - |
| - | - | 1400 | 651.9 | - | - | 0 | - |
| 5 | c | 972.8 | 680.3 | 0.00344 | 5.057 | +1 | 5 |
| 11 | z | 1.306E+04 | 682.3 | 0.004155 | 6.089 | +1 | 5 |
| - | - | 5270 | 683.3 | - | - | 0 | - |
| - | - | 1691 | 684.3 | - | - | 0 | - |
| - | - | 809.4 | 685.3 | - | - | 0 | - |
| 5 | c | 6203 | 697.3 | 0.0004672 | 0.6699 | +1 | 5 |
| 11 | y | 4094 | 698.4 | 0.01067 | 15.28 | +1 | 5 |
| - | - | 1310 | 699.3 | - | - | 0 | - |
| 5 | y | 864.8 | 713.9 | 0.001149 | 1.609 | +2 | 11 |
| 10 | w | 2287 | 752.4 | 0.003658 | 4.862 | +1 | 6 |
| - | - | 607 | 753.9 | - | - | 0 | - |
| - | - | 859.4 | 760.9 | - | - | 0 | - |
| - | - | 700.5 | 761.4 | - | - | 0 | - |
| - | - | 1422 | 765.4 | - | - | 0 | - |
| 12 | c | 4659 | 765.9 | 0.00201 | 2.625 | +2 | 12 |
| - | - | 4856 | 766.4 | - | - | 0 | - |
| - | - | 2253 | 766.9 | - | - | 0 | - |
| - | - | 778 | 767.4 | - | - | 0 | - |
| - | - | 655.1 | 768.3 | - | - | 0 | - |
| - | - | 857.9 | 769.1 | - | - | 0 | - |
| 4 | y | 1240 | 771.4 | 0.003207 | 4.157 | +2 | 12 |
| - | - | 1470 | 771.9 | - | - | 0 | - |
| - | - | 642.1 | 772.4 | - | - | 0 | - |
| - | - | 854.4 | 776.1 | - | - | 0 | - |
| - | - | 918.8 | 779.4 | - | - | 0 | - |
| - | - | 716.2 | 780.9 | - | - | 0 | - |
| - | - | 819.8 | 787.1 | - | - | 0 | - |
| - | - | 1187 | 787.4 | - | - | 0 | - |
| - | - | 663 | 794.4 | - | - | 0 | - |
| - | - | 614 | 795.2 | - | - | 0 | - |
| - | - | 835.2 | 799.2 | - | - | 0 | - |
| - | - | 669.4 | 801.9 | - | - | 0 | - |
| - | - | 1185 | 809.4 | - | - | 0 | - |
| 6 | c | 6203 | 810.4 | 0.001425 | 1.759 | +1 | 6 |
| 10 | z | 6285 | 811.4 | 0.004389 | 5.41 | +1 | 6 |
| - | - | 2912 | 812.4 | - | - | 0 | - |
| - | - | 1417 | 813.4 | - | - | 0 | - |
| - | - | 1046 | 817.4 | - | - | 0 | - |
| - | - | 1042 | 825.4 | - | - | 0 | - |
| 3 | y | 1711 | 826.4 | 0.006278 | 7.597 | +2 | 13 |
| 3 | y | 1996 | 826.9 | 0.005498 | 6.649 | +2 | 13 |
| 10 | y | 4585 | 827.4 | 0.008831 | 10.67 | +1 | 6 |
| - | - | 1875 | 827.9 | - | - | 0 | - |
| - | - | 1224 | 828.4 | - | - | 0 | - |
| 3 | y | 5848 | 835.4 | 0.002344 | 2.806 | +2 | 13 |
| - | - | 5583 | 835.9 | - | - | 0 | - |
| - | - | 3053 | 836.4 | - | - | 0 | - |
| - | - | 1042 | 836.9 | - | - | 0 | - |
| - | - | 2080 | 842.4 | - | - | 0 | - |
| - | - | 1044 | 843.4 | - | - | 0 | - |
| - | - | 3077 | 845.9 | - | - | 0 | - |
| 13 | c | 7235 | 846.4 | 0.0008072 | 0.9537 | +2 | 13 |
| - | - | 7842 | 846.9 | - | - | 0 | - |
| - | - | 4526 | 847.4 | - | - | 0 | - |
| - | - | 1499 | 847.9 | - | - | 0 | - |
| - | - | 864.2 | 848.5 | - | - | 0 | - |
| - | - | 990.3 | 867.4 | - | - | 0 | - |
| - | - | 775 | 875.5 | - | - | 0 | - |
| - | - | 1615 | 880.5 | - | - | 0 | - |
| - | - | 4476 | 881.4 | - | - | 0 | - |
| - | - | 2814 | 882.4 | - | - | 0 | - |
| - | - | 1449 | 883.5 | - | - | 0 | - |
| - | - | 799.3 | 888.4 | - | - | 0 | - |
| - | - | 751.6 | 888.9 | - | - | 0 | - |
| - | - | 1622 | 893.9 | - | - | 0 | - |
| - | - | 624.7 | 894.5 | - | - | 0 | - |
| - | - | 1930 | 895.4 | - | - | 0 | - |
| 2 | z | 2446 | 895.9 | 0.005455 | 6.089 | +2 | 14 |
| - | - | 2451 | 896.4 | - | - | 0 | - |
| - | - | 1811 | 896.9 | - | - | 0 | - |
| - | - | 1769 | 902.4 | - | - | 0 | - |
| - | - | 967.6 | 902.9 | - | - | 0 | - |
| - | - | 1055 | 903.4 | - | - | 0 | - |
| 7 | c | 2128 | 908.4 | 0.00744 | 8.19 | +1 | 7 |
| - | - | 1237 | 909 | - | - | 0 | - |
| - | - | 2095 | 909.4 | - | - | 0 | - |
| - | - | 1490 | 909.9 | - | - | 0 | - |
| 14 | c | 1.561E+04 | 910.4 | 0.001231 | 1.352 | +2 | 14 |
| - | - | 1.837E+04 | 910.9 | - | - | 0 | - |
| - | - | 1.243E+04 | 911.4 | - | - | 0 | - |
| - | - | 4227 | 911.9 | - | - | 0 | - |
| - | - | 1644 | 912.4 | - | - | 0 | - |
| - | - | 990.3 | 915.5 | - | - | 0 | - |
| - | - | 1566 | 916.5 | - | - | 0 | - |
| - | - | 1518 | 917 | - | - | 0 | - |
| - | - | 820.7 | 917.5 | - | - | 0 | - |
| - | - | 1098 | 918 | - | - | 0 | - |
| - | - | 838.7 | 921.4 | - | - | 0 | - |
| - | - | 1399 | 923 | - | - | 0 | - |
| - | - | 1184 | 923.5 | - | - | 0 | - |
| - | - | 1469 | 924 | - | - | 0 | - |
| - | - | 4540 | 924.4 | - | - | 0 | - |
| - | - | 3105 | 924.9 | - | - | 0 | - |
| 7 | c | 7781 | 925.5 | 0.0004935 | 0.5332 | +1 | 7 |
| - | - | 1465 | 925.9 | - | - | 0 | - |
| - | - | 3077 | 926.5 | - | - | 0 | - |
| - | - | 854.7 | 927.5 | - | - | 0 | - |
| - | - | 1191 | 929.5 | - | - | 0 | - |
| - | - | 2805 | 930.5 | - | - | 0 | - |
| - | - | 1.066E+04 | 931 | - | - | 0 | - |
| - | - | 2.62E+04 | 931.5 | - | - | 0 | - |
| - | - | 2.172E+04 | 932 | - | - | 0 | - |
| - | - | 1.29E+04 | 932.5 | - | - | 0 | - |
| - | - | 5327 | 933 | - | - | 0 | - |
| - | - | 1851 | 933.5 | - | - | 0 | - |
| 9 | y | 781.4 | 937.5 | 0.01114 | 11.88 | +1 | 7 |
| - | - | 3755 | 938 | - | - | 0 | - |
| 9 | y | 6499 | 938.5 | 0.001246 | 1.328 | +1 | 7 |
| - | - | 4496 | 939 | - | - | 0 | - |
| 9 | z | 1.558E+04 | 939.5 | 0.005236 | 5.573 | +1 | 7 |
| - | - | 728.4 | 940 | - | - | 0 | - |
| - | - | 9615 | 940.5 | - | - | 0 | - |
| - | - | 3417 | 941.5 | - | - | 0 | - |
| - | - | 1418 | 942.5 | - | - | 0 | - |
| - | - | 970.9 | 943 | - | - | 0 | - |
| - | - | 835.4 | 949.2 | - | - | 0 | - |
| - | - | 827.2 | 949.9 | - | - | 0 | - |
| - | - | 761.4 | 951 | - | - | 0 | - |
| - | - | 3342 | 952 | - | - | 0 | - |
| - | - | 6383 | 952.5 | - | - | 0 | - |
| - | - | 6511 | 953 | - | - | 0 | - |
| - | - | 3576 | 953.5 | - | - | 0 | - |
| - | - | 2344 | 954 | - | - | 0 | - |
| - | - | 1569 | 954.5 | - | - | 0 | - |
| 9 | y | 2080 | 955.5 | 0.00272 | 2.847 | +1 | 7 |
| - | - | 851.2 | 956.5 | - | - | 0 | - |
| - | - | 737.8 | 958.5 | - | - | 0 | - |
| - | - | 823.7 | 959.5 | - | - | 0 | - |
| - | - | 3486 | 960 | - | - | 0 | - |
| - | - | 2.409E+04 | 960.5 | - | - | 0 | - |
| - | - | 3.881E+04 | 961 | - | - | 0 | - |
| - | - | 2.901E+04 | 961.5 | - | - | 0 | - |
| - | - | 1.643E+04 | 962 | - | - | 0 | - |
| - | - | 6138 | 962.5 | - | - | 0 | - |
| - | - | 2719 | 963 | - | - | 0 | - |
| - | - | 742 | 978.2 | - | - | 0 | - |
| - | - | 3872 | 981.5 | - | - | 0 | - |
| 8 | c | 1.826E+04 | 982.5 | 0.001053 | 1.072 | +1 | 8 |
| - | - | 8078 | 983.5 | - | - | 0 | - |
| - | - | 2585 | 984.5 | - | - | 0 | - |
| - | - | 1047 | 986.5 | - | - | 0 | - |
| 8 | z | 4140 | 996.5 | 0.001309 | 1.314 | +1 | 8 |
| - | - | 3121 | 997.5 | - | - | 0 | - |
| - | - | 1532 | 998.5 | - | - | 0 | - |
| - | - | 799.1 | 999.5 | - | - | 0 | - |
| 8 | y | 6649 | 1013 | 0.004103 | 4.052 | +1 | 8 |
| - | - | 3502 | 1014 | - | - | 0 | - |
| - | - | 1907 | 1014 | - | - | 0 | - |
| - | - | 3240 | 1015 | - | - | 0 | - |
| - | - | 1111 | 1015 | - | - | 0 | - |
| - | - | 1381 | 1020 | - | - | 0 | - |
| - | - | 1134 | 1020 | - | - | 0 | - |
| - | - | 1456 | 1026 | - | - | 0 | - |
| - | - | 1405 | 1028 | - | - | 0 | - |
| - | - | 1060 | 1032 | - | - | 0 | - |
| - | - | 995.6 | 1032 | - | - | 0 | - |
| - | - | 1192 | 1033 | - | - | 0 | - |
| - | - | 908.7 | 1034 | - | - | 0 | - |
| - | - | 778.2 | 1035 | - | - | 0 | - |
| - | - | 1220 | 1035 | - | - | 0 | - |
| - | - | 864.2 | 1041 | - | - | 0 | - |
| - | - | 844.3 | 1044 | - | - | 0 | - |
| - | - | 1781 | 1044 | - | - | 0 | - |
| - | - | 1019 | 1045 | - | - | 0 | - |
| - | - | 796.4 | 1045 | - | - | 0 | - |
| - | - | 1472 | 1045 | - | - | 0 | - |
| - | - | 1046 | 1046 | - | - | 0 | - |
| - | - | 714.3 | 1046 | - | - | 0 | - |
| - | - | 782.9 | 1047 | - | - | 0 | - |
| - | - | 1799 | 1048 | - | - | 0 | - |
| - | - | 1060 | 1048 | - | - | 0 | - |
| - | - | 1458 | 1048 | - | - | 0 | - |
| - | - | 1002 | 1049 | - | - | 0 | - |
| - | - | 1537 | 1049 | - | - | 0 | - |
| - | - | 1414 | 1049 | - | - | 0 | - |
| - | - | 1176 | 1050 | - | - | 0 | - |
| - | - | 1481 | 1050 | - | - | 0 | - |
| - | - | 1095 | 1050 | - | - | 0 | - |
| - | - | 1014 | 1053 | - | - | 0 | - |
| - | - | 2555 | 1054 | - | - | 0 | - |
| - | - | 3577 | 1054 | - | - | 0 | - |
| - | - | 1723 | 1054 | - | - | 0 | - |
| - | - | 2506 | 1055 | - | - | 0 | - |
| - | - | 1898 | 1055 | - | - | 0 | - |
| - | - | 906.2 | 1055 | - | - | 0 | - |
| - | - | 2088 | 1056 | - | - | 0 | - |
| - | - | 834.4 | 1058 | - | - | 0 | - |
| - | - | 2302 | 1060 | - | - | 0 | - |
| - | - | 1479 | 1060 | - | - | 0 | - |
| - | - | 2974 | 1060 | - | - | 0 | - |
| - | - | 2083 | 1061 | - | - | 0 | - |
| - | - | 862.5 | 1061 | - | - | 0 | - |
| - | - | 1267 | 1061 | - | - | 0 | - |
| - | - | 1498 | 1062 | - | - | 0 | - |
| - | - | 1670 | 1062 | - | - | 0 | - |
| - | - | 1710 | 1062 | - | - | 0 | - |
| - | - | 1188 | 1063 | - | - | 0 | - |
| - | - | 3020 | 1063 | - | - | 0 | - |
| - | - | 3263 | 1063 | - | - | 0 | - |
| - | - | 3989 | 1064 | - | - | 0 | - |
| - | - | 1649 | 1064 | - | - | 0 | - |
| - | - | 1207 | 1064 | - | - | 0 | - |
| - | - | 894.6 | 1065 | - | - | 0 | - |
| - | - | 1691 | 1065 | - | - | 0 | - |
| - | - | 2060 | 1066 | - | - | 0 | - |
| - | - | 1537 | 1066 | - | - | 0 | - |
| - | - | 1392 | 1066 | - | - | 0 | - |
| - | - | 1154 | 1067 | - | - | 0 | - |
| - | - | 990.3 | 1067 | - | - | 0 | - |
| - | - | 1628 | 1068 | - | - | 0 | - |
| - | - | 4473 | 1069 | - | - | 0 | - |
| - | - | 5931 | 1069 | - | - | 0 | - |
| - | - | 4689 | 1069 | - | - | 0 | - |
| - | - | 4242 | 1070 | - | - | 0 | - |
| - | - | 2457 | 1070 | - | - | 0 | - |
| - | - | 863.7 | 1070 | - | - | 0 | - |
| - | - | 781 | 1095 | - | - | 0 | - |
| - | - | 1360 | 1110 | - | - | 0 | - |
| 9 | c | 1.149E+04 | 1111 | 0.0005114 | 0.4605 | +1 | 9 |
| - | - | 7015 | 1112 | - | - | 0 | - |
| - | - | 3645 | 1113 | - | - | 0 | - |
| - | - | 996.3 | 1114 | - | - | 0 | - |
| - | - | 922 | 1138 | - | - | 0 | - |
| - | - | 1703 | 1143 | - | - | 0 | - |
| - | - | 1015 | 1144 | - | - | 0 | - |
| - | - | 1438 | 1167 | - | - | 0 | - |
| - | - | 1070 | 1182 | - | - | 0 | - |
| - | - | 1025 | 1182 | - | - | 0 | - |
| - | - | 821.4 | 1190 | - | - | 0 | - |
| 6 | w | 904.8 | 1196 | 0.0158 | 13.21 | +1 | 10 |
| - | - | 853.7 | 1221 | - | - | 0 | - |
| 6 | z | 3772 | 1225 | 1.154E-05 | 0.009423 | +1 | 10 |
| - | - | 2429 | 1226 | - | - | 0 | - |
| - | - | 1997 | 1227 | - | - | 0 | - |
| - | - | 2420 | 1239 | - | - | 0 | - |
| 10 | c | 1.533E+04 | 1240 | 0.00223 | 1.799 | +1 | 10 |
| 6 | y | 9575 | 1241 | 0.01251 | 10.08 | +1 | 10 |
| - | - | 3990 | 1242 | - | - | 0 | - |
| - | - | 1151 | 1243 | - | - | 0 | - |
| - | - | 825.6 | 1244 | - | - | 0 | - |
| - | - | 905.2 | 1267 | - | - | 0 | - |
| - | - | 1118 | 1272 | - | - | 0 | - |
| - | - | 934.3 | 1272 | - | - | 0 | - |
| - | - | 1234 | 1273 | - | - | 0 | - |
| - | - | 877.2 | 1280 | - | - | 0 | - |
| - | - | 834.7 | 1280 | - | - | 0 | - |
| - | - | 1670 | 1281 | - | - | 0 | - |
| - | - | 1647 | 1281 | - | - | 0 | - |
| - | - | 2302 | 1282 | - | - | 0 | - |
| - | - | 2424 | 1283 | - | - | 0 | - |
| - | - | 1638 | 1284 | - | - | 0 | - |
| - | - | 908.1 | 1302 | - | - | 0 | - |
| - | - | 892.3 | 1356 | - | - | 0 | - |
| - | - | 1013 | 1367 | - | - | 0 | - |
| - | - | 2347 | 1402 | - | - | 0 | - |
| 11 | c | 9452 | 1403 | 0.003232 | 2.304 | +1 | 11 |
| - | - | 6888 | 1404 | - | - | 0 | - |
| - | - | 2711 | 1405 | - | - | 0 | - |
| - | - | 1121 | 1406 | - | - | 0 | - |
| 5 | y | 953.2 | 1410 | 0.0109 | 7.731 | +1 | 11 |
| 5 | z | 1.171E+04 | 1411 | 0.002664 | 1.889 | +1 | 11 |
| - | - | 9084 | 1412 | - | - | 0 | - |
| - | - | 5350 | 1413 | - | - | 0 | - |
| - | - | 1188 | 1414 | - | - | 0 | - |
| - | - | 1661 | 1422 | - | - | 0 | - |
| - | - | 1109 | 1423 | - | - | 0 | - |
| - | - | 1528 | 1426 | - | - | 0 | - |
| 5 | y | 6015 | 1427 | 0.003444 | 2.414 | +1 | 11 |
| - | - | 4145 | 1428 | - | - | 0 | - |
| - | - | 2462 | 1429 | - | - | 0 | - |
| - | - | 1019 | 1430 | - | - | 0 | - |
| - | - | 1193 | 1436 | - | - | 0 | - |
| - | - | 1424 | 1437 | - | - | 0 | - |
| - | - | 790.3 | 1439 | - | - | 0 | - |
| - | - | 923.3 | 1466 | - | - | 0 | - |
| - | - | 878.5 | 1475 | - | - | 0 | - |
| - | - | 3525 | 1482 | - | - | 0 | - |
| - | - | 2533 | 1483 | - | - | 0 | - |
| - | - | 1656 | 1484 | - | - | 0 | - |
| - | - | 1040 | 1485 | - | - | 0 | - |
| - | - | 939.7 | 1486 | - | - | 0 | - |
| - | - | 3288 | 1487 | - | - | 0 | - |
| - | - | 1603 | 1487 | - | - | 0 | - |
| - | - | 4171 | 1488 | - | - | 0 | - |
| - | - | 1659 | 1489 | - | - | 0 | - |
| - | - | 939.6 | 1496 | - | - | 0 | - |
| - | - | 1026 | 1501 | - | - | 0 | - |
| - | - | 1183 | 1502 | - | - | 0 | - |
| - | - | 839.1 | 1510 | - | - | 0 | - |
| - | - | 931.6 | 1515 | - | - | 0 | - |
| - | - | 1054 | 1516 | - | - | 0 | - |
| - | - | 806.5 | 1521 | - | - | 0 | - |
| - | - | 1694 | 1521 | - | - | 0 | - |
| - | - | 1071 | 1522 | - | - | 0 | - |
| 4 | y | 1027 | 1524 | 0.01265 | 8.301 | +1 | 12 |
| 4 | y | 1148 | 1525 | 0.02558 | 16.78 | +1 | 12 |
| 4 | z | 9164 | 1526 | 0.002874 | 1.884 | +1 | 12 |
| - | - | 1.551E+04 | 1527 | - | - | 0 | - |
| - | - | 1056 | 1527 | - | - | 0 | - |
| - | - | 9888 | 1528 | - | - | 0 | - |
| - | - | 1376 | 1528 | - | - | 0 | - |
| - | - | 4842 | 1529 | - | - | 0 | - |
| - | - | 4030 | 1530 | - | - | 0 | - |
| 12 | c | 1.088E+04 | 1531 | 0.0005545 | 0.3622 | +1 | 12 |
| - | - | 8956 | 1532 | - | - | 0 | - |
| - | - | 4019 | 1533 | - | - | 0 | - |
| - | - | 2138 | 1534 | - | - | 0 | - |
| - | - | 1035 | 1535 | - | - | 0 | - |
| - | - | 818.6 | 1540 | - | - | 0 | - |
| 4 | y | 3230 | 1542 | 0.002433 | 1.578 | +1 | 12 |
| - | - | 2805 | 1543 | - | - | 0 | - |
| - | - | 1178 | 1543 | - | - | 0 | - |
| - | - | 1151 | 1544 | - | - | 0 | - |
| - | - | 995.4 | 1545 | - | - | 0 | - |
| - | - | 1039 | 1545 | - | - | 0 | - |
| - | - | 784.2 | 1547 | - | - | 0 | - |
| - | - | 1060 | 1548 | - | - | 0 | - |
| - | - | 1773 | 1552 | - | - | 0 | - |
| - | - | 2227 | 1552 | - | - | 0 | - |
| - | - | 995.3 | 1553 | - | - | 0 | - |
| - | - | 823.8 | 1555 | - | - | 0 | - |
| - | - | 1036 | 1556 | - | - | 0 | - |
| - | - | 1427 | 1557 | - | - | 0 | - |
| - | - | 1887 | 1558 | - | - | 0 | - |
| - | - | 1421 | 1558 | - | - | 0 | - |
| - | - | 1767 | 1559 | - | - | 0 | - |
| - | - | 1218 | 1559 | - | - | 0 | - |
| - | - | 1900 | 1560 | - | - | 0 | - |
| - | - | 1025 | 1560 | - | - | 0 | - |
| - | - | 1097 | 1561 | - | - | 0 | - |
| - | - | 1613 | 1563 | - | - | 0 | - |
| - | - | 921.9 | 1563 | - | - | 0 | - |
| - | - | 1215 | 1565 | - | - | 0 | - |
| - | - | 2261 | 1565 | - | - | 0 | - |
| - | - | 2055 | 1566 | - | - | 0 | - |
| - | - | 1961 | 1566 | - | - | 0 | - |
| - | - | 2252 | 1567 | - | - | 0 | - |
| - | - | 2121 | 1567 | - | - | 0 | - |
| - | - | 1487 | 1568 | - | - | 0 | - |
| - | - | 783.8 | 1568 | - | - | 0 | - |
| - | - | 919.5 | 1570 | - | - | 0 | - |
| - | - | 936.8 | 1571 | - | - | 0 | - |
| - | - | 875.2 | 1571 | - | - | 0 | - |
| - | - | 1728 | 1572 | - | - | 0 | - |
| - | - | 1576 | 1572 | - | - | 0 | - |
| - | - | 2723 | 1573 | - | - | 0 | - |
| - | - | 3713 | 1573 | - | - | 0 | - |
| - | - | 2999 | 1574 | - | - | 0 | - |
| - | - | 3350 | 1574 | - | - | 0 | - |
| - | - | 3347 | 1575 | - | - | 0 | - |
| - | - | 2906 | 1575 | - | - | 0 | - |
| - | - | 1074 | 1576 | - | - | 0 | - |
| - | - | 1001 | 1576 | - | - | 0 | - |
| - | - | 1347 | 1577 | - | - | 0 | - |
| - | - | 1988 | 1579 | - | - | 0 | - |
| - | - | 1356 | 1580 | - | - | 0 | - |
| - | - | 3894 | 1580 | - | - | 0 | - |
| - | - | 4839 | 1581 | - | - | 0 | - |
| - | - | 3052 | 1581 | - | - | 0 | - |
| - | - | 2308 | 1582 | - | - | 0 | - |
| - | - | 2516 | 1582 | - | - | 0 | - |
| - | - | 2518 | 1583 | - | - | 0 | - |
| - | - | 1984 | 1583 | - | - | 0 | - |
| - | - | 1207 | 1584 | - | - | 0 | - |
| - | - | 1104 | 1585 | - | - | 0 | - |
| - | - | 1159 | 1586 | - | - | 0 | - |
| - | - | 1146 | 1587 | - | - | 0 | - |
| - | - | 1417 | 1588 | - | - | 0 | - |
| - | - | 1105 | 1588 | - | - | 0 | - |
| - | - | 2969 | 1589 | - | - | 0 | - |
| - | - | 3111 | 1589 | - | - | 0 | - |
| - | - | 1589 | 1590 | - | - | 0 | - |
| - | - | 2642 | 1590 | - | - | 0 | - |
| - | - | 2670 | 1591 | - | - | 0 | - |
| - | - | 2307 | 1591 | - | - | 0 | - |
| - | - | 2107 | 1592 | - | - | 0 | - |
| - | - | 1380 | 1592 | - | - | 0 | - |
| - | - | 1675 | 1593 | - | - | 0 | - |
| - | - | 1609 | 1594 | - | - | 0 | - |
| - | - | 4121 | 1594 | - | - | 0 | - |
| - | - | 3712 | 1595 | - | - | 0 | - |
| - | - | 3834 | 1595 | - | - | 0 | - |
| - | - | 2520 | 1596 | - | - | 0 | - |
| - | - | 1761 | 1596 | - | - | 0 | - |
| - | - | 998.6 | 1597 | - | - | 0 | - |
| - | - | 1711 | 1597 | - | - | 0 | - |
| - | - | 2042 | 1598 | - | - | 0 | - |
| - | - | 1124 | 1598 | - | - | 0 | - |
| - | - | 964.7 | 1599 | - | - | 0 | - |
| - | - | 1513 | 1599 | - | - | 0 | - |
| - | - | 2467 | 1600 | - | - | 0 | - |
| - | - | 1856 | 1600 | - | - | 0 | - |
| - | - | 1523 | 1601 | - | - | 0 | - |
| - | - | 1538 | 1601 | - | - | 0 | - |
| - | - | 1398 | 1602 | - | - | 0 | - |
| - | - | 4005 | 1602 | - | - | 0 | - |
| - | - | 6080 | 1603 | - | - | 0 | - |
| - | - | 9312 | 1603 | - | - | 0 | - |
| - | - | 4531 | 1604 | - | - | 0 | - |
| - | - | 4044 | 1604 | - | - | 0 | - |
| - | - | 2098 | 1605 | - | - | 0 | - |
| - | - | 1146 | 1605 | - | - | 0 | - |
| - | - | 767.1 | 1610 | - | - | 0 | - |
| - | - | 860.8 | 1633 | - | - | 0 | - |
| - | - | 861.3 | 1634 | - | - | 0 | - |
| 3 | z | 5062 | 1654 | 0.002736 | 1.654 | +1 | 13 |
| - | - | 7818 | 1655 | - | - | 0 | - |
| - | - | 6045 | 1656 | - | - | 0 | - |
| - | - | 3171 | 1657 | - | - | 0 | - |
| - | - | 3384 | 1691 | - | - | 0 | - |
| 13 | c | 1.598E+04 | 1692 | 0.002395 | 1.415 | +1 | 13 |
| - | - | 1.569E+04 | 1693 | - | - | 0 | - |
| - | - | 7864 | 1694 | - | - | 0 | - |
| - | - | 3432 | 1695 | - | - | 0 | - |
| - | - | 947.3 | 1696 | - | - | 0 | - |
| - | - | 1101 | 1777 | - | - | 0 | - |
| 2 | z | 989.2 | 1791 | 0.01428 | 7.974 | +1 | 14 |
| - | - | 2402 | 1792 | - | - | 0 | - |
| - | - | 1902 | 1793 | - | - | 0 | - |
| - | - | 1587 | 1794 | - | - | 0 | - |
| - | - | 1038 | 1795 | - | - | 0 | - |
| - | - | 1038 | 1819 | - | - | 0 | - |
| 14 | c | 2850 | 1820 | 0.007379 | 4.054 | +1 | 14 |
| - | - | 4541 | 1821 | - | - | 0 | - |
| - | - | 4214 | 1822 | - | - | 0 | - |
| - | - | 2318 | 1823 | - | - | 0 | - |
| - | - | 1296 | 1824 | - | - | 0 | - |
| - | - | 1983 | 1831 | - | - | 0 | - |
| - | - | 2347 | 1832 | - | - | 0 | - |
| - | - | 987.7 | 1833 | - | - | 0 | - |
| - | - | 1138 | 1835 | - | - | 0 | - |
| - | - | 1031 | 1845 | - | - | 0 | - |
| - | - | 1177 | 1846 | - | - | 0 | - |
| - | - | 1703 | 1847 | - | - | 0 | - |
| - | - | 958.9 | 1848 | - | - | 0 | - |
| - | - | 1040 | 1849 | - | - | 0 | - |
| - | - | 2211 | 1861 | - | - | 0 | - |
| - | - | 7485 | 1862 | - | - | 0 | - |
| - | - | 9603 | 1863 | - | - | 0 | - |
| - | - | 9144 | 1864 | - | - | 0 | - |
| - | - | 4978 | 1865 | - | - | 0 | - |
| - | - | 2001 | 1866 | - | - | 0 | - |
| - | - | 2814 | 1876 | - | - | 0 | - |
| - | - | 6287 | 1877 | - | - | 0 | - |
| - | - | 5339 | 1878 | - | - | 0 | - |
| - | - | 3596 | 1879 | - | - | 0 | - |
| - | - | 1796 | 1880 | - | - | 0 | - |
| - | - | 3219 | 1893 | - | - | 0 | - |
| - | - | 7920 | 1894 | - | - | 0 | - |
| - | - | 6946 | 1895 | - | - | 0 | - |
| - | - | 4258 | 1896 | - | - | 0 | - |
| - | - | 1538 | 1897 | - | - | 0 | - |
| - | - | 890.3 | 1903 | - | - | 0 | - |
| - | - | 5196 | 1904 | - | - | 0 | - |
| - | - | 1.149E+04 | 1905 | - | - | 0 | - |
| - | - | 1.115E+04 | 1906 | - | - | 0 | - |
| - | - | 6002 | 1907 | - | - | 0 | - |
| - | - | 3943 | 1908 | - | - | 0 | - |
| - | - | 5218 | 1920 | - | - | 0 | - |
| - | - | 1.829E+04 | 1921 | - | - | 0 | - |
| - | - | 5.397E+04 | 1922 | - | - | 0 | - |
| - | - | 4.966E+04 | 1923 | - | - | 0 | - |
| - | - | 2.929E+04 | 1924 | - | - | 0 | - |
| - | - | 1.287E+04 | 1925 | - | - | 0 | - |
| - | - | 4773 | 1926 | - | - | 0 | - |
| - | - | 955.5 | 2040 | - | - | 0 | - |

m/z Charge Intensity FragmentType MassShift Position
121.01114654541016 0 362.24066
129.1021728515625 0 1591.722
132.95233154296875 0 439.53796
136.41448974609375 0 458.74744
141.14561462402344 0 418.16064
148.95443725585938 0 659.9974
153.13702392578125 0 549.58887
153.65750122070312 0 427.44025
155.0928955078125 0 729.288
156.90858459472656 0 474.73975
157.13308715820312 0 457.37683
165.10147094726562 0 441.6555
166.0610809326172 0 852.5235
173.4387969970703 0 1889.9125
173.70831298828125 0 458.8428
177.040283203125 0 528.2872
183.1123809814453 0 675.8766
185.1644287109375 0 743.16815
186.12396240234375 0 529.9749
223.15516662597656 0 1182.2533
225.36428833007812 0 520.7979
230.16229248046875 0 3650.7007 z 13
231.16989135742188 0 4396.9844
234.1234130859375 0 1501.95
243.25466918945312 0 540.9293
246.1815948486328 0 1125.6843 y 13
250.14263916015625 0 1051.7563
251.15011596679688 0 8163.084
252.15316772460938 0 1436.3341
268.1765441894531 0 4847.032 c 1
269.18023681640625 0 889.2566
292.9272155761719 0 703.4383
300.19183349609375 0 7840.476
301.1952209472656 0 1067.0345
302.1705017089844 0 681.5515
318.1410217285156 0 577.7359
333.3406982421875 0 607.88855
352.2212829589844 0 1822.9457
379.20831298828125 0 1051.9011 c Ammonia loss 2
395.2269592285156 0 4110.6094
396.2066650390625 0 807.09503
396.235107421875 0 14201.79 c 2
397.2382507324219 0 3471.9587
407.19635009765625 0 4014.355 y 12
408.1982116699219 0 1108.684 c Ammonia loss 9
409.3603210449219 0 577.471
427.26654052734375 0 970.5706
428.2640075683594 0 733.56757
461.2082214355469 0 738.3848 w 11
467.24932861328125 0 5486.465
468.2534484863281 0 2429.513
475.27874755859375 0 670.8108
488.3190002441406 0 850.9567
494.2366638183594 0 745.36743 c Ammonia loss 3
506.76007080078125 0 2852.0955 y 7
507.2604064941406 0 1128.2573
507.7631530761719 0 689.44336
511.26239013671875 0 21741.973 c 3
512.2655639648438 0 5520.1333
513.2685546875 0 614.67145
519.272216796875 0 5606.095 z 11
520.2775268554688 0 2207.003
521.2731323242188 0 558.5486
535.2916870117188 0 1845.653 y 11
536.2930297851562 0 832.3729
564.2645874023438 0 618.9392
584.362548828125 0 4250.6187
585.3668212890625 0 1236.5853
590.8561401367188 0 1764.5717
598.3446044921875 0 554.8201
601.622802734375 0 2778.4463 c Ammonia loss 13
601.9573974609375 0 2062.5752
602.29443359375 0 1848.1936
602.9549560546875 0 5112.6753 y 1
603.2899169921875 0 3828.5298
603.6240234375 0 2682.1707
606.3243408203125 0 644.78326
607.6270751953125 0 1393.3785
607.9584350585938 0 1326.2295
618.8707275390625 0 838.45874
619.3637084960938 0 667.7756
625.2781372070312 0 711.4991
628.9700927734375 0 1069.7234
629.3027954101562 0 1101.9606
629.6366577148438 0 1225.2253
633.30908203125 0 550.39716
634.6474609375 0 2167.6123
634.978271484375 0 2651.9246
635.3106689453125 0 2772.609
635.6446533203125 0 1374.3181
639.3353881835938 0 1234.3986
639.87646484375 0 1681.6272
640.32080078125 0 4130.027
640.3772583007812 0 1523.2025
640.6495971679688 0 47797.44
640.8822631835938 0 1950.6461
640.983642578125 0 47831.234
641.317626953125 0 37424.15
641.6517944335938 0 12348.154
641.8623046875 0 2142.0864
641.98583984375 0 3971.1611
651.353271484375 0 1762.1061
651.8541259765625 0 1400.4492
680.3116455078125 0 972.8174 c Ammonia loss 4
682.3358154296875 0 13062.256 z 10
683.339599609375 0 5270.413
684.340087890625 0 1691.3596
685.3382568359375 0 809.43146
697.3421020507812 0 6202.623 c 4
698.3480224609375 0 4094.4785 y 10
699.3499145507812 0 1309.8374
713.8565063476562 0 864.7748 y 4
752.3656005859375 0 2286.5942 w 9
753.8645629882812 0 607.0039
760.9014282226562 0 859.444
761.39892578125 0 700.4823
765.4010009765625 0 1422.4158
765.890625 0 4658.858 c 11
766.3909912109375 0 4856.1675
766.8909301757812 0 2252.9397
767.3902587890625 0 777.97186
768.3475952148438 0 655.1472
769.0875854492188 0 857.9069
771.367919921875 0 1240.3282 y 3
771.8734130859375 0 1469.5554
772.3682250976562 0 642.0901
776.1499633789062 0 854.36304
779.3955078125 0 918.79114
780.9130249023438 0 716.2036
787.1487426757812 0 819.7676
787.4033813476562 0 1186.7039
794.3986206054688 0 663.0441
795.1544189453125 0 614.03156
799.1505126953125 0 835.1879
801.9102783203125 0 669.3754
809.4354858398438 0 1184.6019
810.4271240234375 0 6202.8022 c 5
811.378173828125 0 6284.8066 z 9
812.3834228515625 0 2911.987
813.3788452148438 0 1417.099
817.3958129882812 0 1045.8934
825.3795776367188 0 1041.744
826.3888549804688 0 1710.5596 y Water loss 2
826.8926391601562 0 1996.3982 y Ammonia loss 2
827.3924560546875 0 4584.538 y 9
827.888427734375 0 1874.6144
828.3897094726562 0 1223.5415
835.3980712890625 0 5847.6753 y 2
835.9000244140625 0 5582.628
836.4005737304688 0 3053.3464
836.9072265625 0 1041.7151
842.4202270507812 0 2080.0544
843.4210815429688 0 1044.0834
845.8939819335938 0 3076.5356
846.3973999023438 0 7235.0493 c 12
846.8989868164062 0 7841.623
847.3984985351562 0 4525.993
847.8983154296875 0 1499.1235
848.4902954101562 0 864.18286
867.4183349609375 0 990.3162
875.475830078125 0 774.9977
880.4520874023438 0 1615.1804
881.441162109375 0 4475.6943
882.4459228515625 0 2813.9048
883.450439453125 0 1448.607
888.4371337890625 0 799.2724
888.9393310546875 0 751.6198
893.9420166015625 0 1622.4309
894.456787109375 0 624.7249
895.4374389648438 0 1929.9658
895.9259643554688 0 2445.5051 z 1
896.425048828125 0 2450.7122
896.9202880859375 0 1811.3754
902.4432983398438 0 1768.9893
902.9279174804688 0 967.59607
903.4434204101562 0 1054.9391
908.4335327148438 0 2127.5044 c Ammonia loss 6
908.9603881835938 0 1236.6157
909.4459838867188 0 2095.0647
909.9495849609375 0 1490.1694
910.4444580078125 0 15609.762 c 13
910.9456787109375 0 18373.766
911.4454345703125 0 12432.967
911.9443359375 0 4226.759
912.4412231445312 0 1644.4353
915.4635009765625 0 990.26544
916.4664916992188 0 1565.9526
916.9658203125 0 1518.2815
917.4847412109375 0 820.6724
917.9711303710938 0 1098.3569
921.4428100585938 0 838.6561
922.9641723632812 0 1398.9615
923.455810546875 0 1184.2189
923.9503784179688 0 1469.3726
924.4490356445312 0 4540.376
924.9474487304688 0 3104.8425
925.4521484375 0 7781.4443 c 6
925.9459228515625 0 1465.0427
926.4548950195312 0 3076.7273
927.4595947265625 0 854.68555
929.4575805664062 0 1190.8777
930.46142578125 0 2804.7095
930.9588012695312 0 10656.335
931.4656982421875 0 26203.523
931.9659423828125 0 21715.908
932.4683837890625 0 12902.279
932.9676513671875 0 5327.066
933.4711303710938 0 1850.5984
937.4745483398438 0 781.4391 y Water loss 8
937.9732666015625 0 3754.7139
938.470947265625 0 6499.054 y Ammonia loss 8
938.9683227539062 0 4496.366
939.4722900390625 0 15584.309 z 8
939.9669799804688 0 728.37994
940.478271484375 0 9615.378
941.4819946289062 0 3417.2715
942.4857177734375 0 1418.1407
942.9667358398438 0 970.9467
949.1539306640625 0 835.3791
949.9437255859375 0 827.1539
950.9572143554688 0 761.40436
951.9688110351562 0 3341.8801
952.465576171875 0 6382.731
952.964599609375 0 6510.9326
953.464599609375 0 3576.1738
953.9614868164062 0 2344.3962
954.470703125 0 1569.4724
955.4935302734375 0 2079.8342 y 8
956.50244140625 0 851.16504
958.4854125976562 0 737.77484
959.4995727539062 0 823.6815
959.9729614257812 0 3486.1226
960.470947265625 0 24085.531
960.972900390625 0 38807.01
961.473876953125 0 29014.287
961.975341796875 0 16433.021
962.47509765625 0 6138.245
962.9695434570312 0 2718.6333
978.1808471679688 0 741.9859
981.4732055664062 0 3872.19
982.4751586914062 0 18264.176 c 7
983.4782104492188 0 8078.196
984.4813842773438 0 2584.5376
986.49609375 0 1047.1307
996.4976806640625 0 4139.6006 z 7
997.4982299804688 0 3120.5735
998.4982299804688 0 1531.9128
999.5105590820312 0 799.0743
1012.5136108398438 0 6649.3447 y 7
1013.5182495117188 0 3501.5967
1014.198486328125 0 1906.5957
1014.5235595703125 0 3239.8289
1014.8566284179688 0 1111.4418
1019.9845581054688 0 1380.9005
1020.4962158203125 0 1133.8411
1025.5206298828125 0 1456.0199
1027.5367431640625 0 1405.2396
1031.543701171875 0 1060.1157
1032.0419921875 0 995.58887
1032.540771484375 0 1191.8335
1033.8778076171875 0 908.74817
1034.53759765625 0 778.1585
1034.8641357421875 0 1219.6779
1040.505126953125 0 864.21216
1043.8773193359375 0 844.3208
1044.2083740234375 0 1781.0259
1044.5164794921875 0 1018.7305
1044.8680419921875 0 796.42566
1045.1983642578125 0 1472.2672
1045.5252685546875 0 1045.8077
1046.186767578125 0 714.26666
1046.8626708984375 0 782.8877
1047.533447265625 0 1798.571
1047.867431640625 0 1060.1937
1048.2005615234375 0 1458.2073
1048.540771484375 0 1002.2864
1048.87158203125 0 1536.684
1049.2061767578125 0 1413.8657
1049.537841796875 0 1175.9402
1049.8685302734375 0 1480.5273
1050.202880859375 0 1095.3594
1053.209716796875 0 1014.12946
1053.5369873046875 0 2555.4265
1053.876220703125 0 3576.7927
1054.202392578125 0 1722.7513
1054.5352783203125 0 2505.9358
1054.8729248046875 0 1897.6564
1055.1953125 0 906.1721
1055.5286865234375 0 2087.867
1057.5308837890625 0 834.42126
1059.533203125 0 2302.3276
1059.8681640625 0 1478.5985
1060.2017822265625 0 2974.4756
1060.53271484375 0 2083.3901
1060.86083984375 0 862.50494
1061.2093505859375 0 1266.8285
1061.5115966796875 0 1498.2773
1061.8558349609375 0 1669.6913
1062.2022705078125 0 1709.9294
1062.5390625 0 1188.1268
1062.8717041015625 0 3019.73
1063.2034912109375 0 3262.7268
1063.541015625 0 3989.0898
1063.874755859375 0 1648.691
1064.1949462890625 0 1207.1176
1064.5364990234375 0 894.63226
1065.211669921875 0 1691.3887
1065.5401611328125 0 2059.969
1065.8763427734375 0 1536.5704
1066.212890625 0 1392.3513
1066.8680419921875 0 1154.2943
1067.185791015625 0 990.329
1067.535400390625 0 1627.5753
1068.548095703125 0 4473.371
1068.882568359375 0 5930.8887
1069.2142333984375 0 4688.759
1069.5506591796875 0 4242.0054
1069.8843994140625 0 2457.0986
1070.2191162109375 0 863.7108
1094.530517578125 0 781.03784
1109.591796875 0 1359.9974
1110.569580078125 0 11493.059 c 8
1111.5570068359375 0 7015.2236
1112.54345703125 0 3644.77
1113.5394287109375 0 996.3376
1137.566650390625 0 922.04205
1142.5625 0 1702.7489
1143.5631103515625 0 1014.62427
1166.672607421875 0 1438.4001
1181.618408203125 0 1069.9009
1182.1234130859375 0 1025.4705
1189.61328125 0 821.427
1195.586669921875 0 904.7873 w 5
1220.5736083984375 0 853.68
1224.6099853515625 0 3771.9724 z 5
1225.611083984375 0 2429.1326
1226.60986328125 0 1996.6891
1238.6192626953125 0 2420.154
1239.6138916015625 0 15331.498 c 9
1240.6162109375 0 9574.918 y 5
1241.61767578125 0 3990.3997
1242.632568359375 0 1150.6256
1243.6385498046875 0 825.6265
1266.66552734375 0 905.16327
1271.6402587890625 0 1117.5049
1272.15966796875 0 934.2592
1272.6328125 0 1233.9796
1279.7293701171875 0 877.24194
1280.1611328125 0 834.707
1280.5924072265625 0 1670.0225
1280.7393798828125 0 1646.9207
1281.7498779296875 0 2301.6406
1282.736572265625 0 2423.7485
1283.7197265625 0 1638.2388
1301.6759033203125 0 908.09296
1355.6273193359375 0 892.34326
1366.6932373046875 0 1013.1501
1401.6834716796875 0 2346.7864
1402.67822265625 0 9452.215 c 10
1403.6802978515625 0 6888.0195
1404.681884765625 0 2711.1077
1405.6961669921875 0 1120.5819
1409.6923828125 0 953.2428 y Ammonia loss 4
1410.6866455078125 0 11711.009 z 4
1411.6881103515625 0 9083.944
1412.6920166015625 0 5349.5615
1413.6859130859375 0 1188.2941
1422.232421875 0 1661.3838
1423.23681640625 0 1109.3195
1425.70703125 0 1527.666
1426.70458984375 0 6015.206 y 4
1427.708251953125 0 4144.595
1428.7047119140625 0 2461.8826
1429.7071533203125 0 1018.72217
1435.700439453125 0 1193.0026
1436.7071533203125 0 1423.6609
1438.7235107421875 0 790.2755
1465.732666015625 0 923.33234
1475.2086181640625 0 878.505
1481.723876953125 0 3524.6404
1482.7264404296875 0 2532.8035
1483.7254638671875 0 1656.321
1484.7369384765625 0 1039.5933
1485.7569580078125 0 939.71954
1486.7532958984375 0 3287.6013
1487.2486572265625 0 1602.6854
1487.756591796875 0 4171.333
1488.7557373046875 0 1659.0048
1496.238037109375 0 939.64343
1500.73046875 0 1025.537
1501.7156982421875 0 1182.6185
1509.709716796875 0 839.1436
1515.2430419921875 0 931.5627
1516.2354736328125 0 1053.5938
1520.7791748046875 0 806.52094
1521.28466796875 0 1694.2931
1522.272705078125 0 1071.3467
1523.737060546875 0 1027.354 y Water loss 3
1524.7340087890625 0 1148.138 y Ammonia loss 3
1525.71337890625 0 9164.07 z 3
1526.7176513671875 0 15506.993
1527.2484130859375 0 1055.9164
1527.724609375 0 9888.238
1528.2637939453125 0 1376.4568
1528.730224609375 0 4841.9717
1529.7506103515625 0 4029.9219
1530.7705078125 0 10881.687 c 11
1531.77392578125 0 8956.298
1532.781005859375 0 4018.7778
1533.783447265625 0 2138.328
1534.7940673828125 0 1034.648
1539.767822265625 0 818.6016
1541.7325439453125 0 3229.7244 y 3
1542.73681640625 0 2804.5903
1543.2718505859375 0 1177.8702
1543.738525390625 0 1150.7888
1544.76025390625 0 995.35864
1545.2781982421875 0 1039.0109
1546.775634765625 0 784.15955
1547.7960205078125 0 1060.1
1551.7906494140625 0 1772.7067
1552.2794189453125 0 2226.9294
1552.7730712890625 0 995.34607
1554.7376708984375 0 823.77734
1555.726318359375 0 1035.8846
1557.2969970703125 0 1426.9857
1557.8021240234375 0 1887.4321
1558.2880859375 0 1421.0612
1558.79248046875 0 1766.797
1559.2933349609375 0 1218.3467
1559.7867431640625 0 1900.4484
1560.286376953125 0 1025.3743
1561.2723388671875 0 1097.4626
1562.771484375 0 1613.2091
1563.2557373046875 0 921.8985
1564.802978515625 0 1215.1095
1565.3045654296875 0 2261.1558
1565.7889404296875 0 2055.1377
1566.2943115234375 0 1961.1268
1566.802001953125 0 2252.499
1567.2913818359375 0 2121.147
1567.80615234375 0 1487.1776
1568.2911376953125 0 783.7626
1569.765625 0 919.47675
1570.7542724609375 0 936.8138
1571.2969970703125 0 875.2425
1571.786865234375 0 1728.1929
1572.3035888671875 0 1576.3314
1572.8057861328125 0 2722.6035
1573.29833984375 0 3712.8882
1573.791259765625 0 2999.0176
1574.3092041015625 0 3349.5957
1574.8056640625 0 3346.6213
1575.302490234375 0 2905.7163
1575.8104248046875 0 1073.8281
1576.26611328125 0 1000.5056
1576.79345703125 0 1347.1835
1579.2957763671875 0 1987.6058
1579.81494140625 0 1356.0702
1580.3023681640625 0 3893.6262
1580.7999267578125 0 4839.363
1581.307373046875 0 3052.402
1581.7928466796875 0 2307.6511
1582.290771484375 0 2515.9758
1582.802001953125 0 2517.9585
1583.294921875 0 1984.3297
1584.3026123046875 0 1206.5255
1585.299560546875 0 1103.9818
1586.2843017578125 0 1159.1774
1587.307861328125 0 1146.2247
1587.818115234375 0 1416.5103
1588.2882080078125 0 1105.4009
1588.8082275390625 0 2969.1284
1589.300048828125 0 3110.7727
1589.8076171875 0 1588.7207
1590.2879638671875 0 2642.2708
1590.7874755859375 0 2669.719
1591.3011474609375 0 2307.4026
1591.79443359375 0 2107.2583
1592.29345703125 0 1379.7067
1593.2889404296875 0 1675.322
1593.814697265625 0 1609.3031
1594.302490234375 0 4120.68
1594.7984619140625 0 3711.6333
1595.315673828125 0 3834.184
1595.796875 0 2519.6294
1596.30224609375 0 1760.5167
1596.7884521484375 0 998.5671
1597.311767578125 0 1711.2632
1597.8013916015625 0 2042.0485
1598.3106689453125 0 1123.636
1598.79541015625 0 964.66156
1599.2962646484375 0 1512.742
1599.79541015625 0 2467.1748
1600.3017578125 0 1855.9199
1600.782958984375 0 1522.7557
1601.28857421875 0 1537.8325
1601.8026123046875 0 1398.3864
1602.3177490234375 0 4004.9736
1602.8201904296875 0 6079.868
1603.320068359375 0 9312.056
1603.8226318359375 0 4531.104
1604.3209228515625 0 4044.3936
1604.8203125 0 2097.9546
1605.3212890625 0 1146.428
1609.8021240234375 0 767.0986
1632.7928466796875 0 860.7744
1633.785400390625 0 861.2769
1653.7720947265625 0 5061.6636 z 2
1654.777587890625 0 7817.628
1655.78125 0 6045.1978
1656.7845458984375 0 3171.4978
1690.785400390625 0 3383.6714
1691.7867431640625 0 15980.4795 c 12
1692.7874755859375 0 15686.411
1693.791748046875 0 7863.689
1694.7974853515625 0 3431.8186
1695.7760009765625 0 947.2656
1776.877685546875 0 1100.5055
1790.8480224609375 0 989.18524 z 1
1791.8438720703125 0 2402.2197
1792.8564453125 0 1901.6088
1793.8477783203125 0 1586.8796
1794.856689453125 0 1038.0919
1818.92919921875 0 1037.9188
1819.8914794921875 0 2850.2673 c 13
1820.8868408203125 0 4541.3765
1821.8909912109375 0 4214.339
1822.89453125 0 2318.4336
1823.89306640625 0 1296.0188
1830.950927734375 0 1982.7234
1831.9530029296875 0 2346.5415
1832.958984375 0 987.7029
1834.9376220703125 0 1138.0325
1844.917236328125 0 1030.9459
1845.916015625 0 1177.1183
1846.914794921875 0 1703.2488
1847.946533203125 0 958.8891
1848.8975830078125 0 1039.5656
1860.9207763671875 0 2210.535
1861.927734375 0 7484.763
1862.934326171875 0 9603.4375
1863.9339599609375 0 9144.3955
1864.9322509765625 0 4978.4683
1865.923583984375 0 2001.2747
1875.935302734375 0 2813.5938
1876.9354248046875 0 6287.1978
1877.9368896484375 0 5339.459
1878.93994140625 0 3596.2815
1879.9307861328125 0 1795.9495
1892.9501953125 0 3218.502
1893.9541015625 0 7919.6963
1894.9525146484375 0 6945.6333
1895.9599609375 0 4257.745
1896.960693359375 0 1538.1348
1902.9356689453125 0 890.33997
1903.92724609375 0 5196.2246
1904.925537109375 0 11489.714
1905.928955078125 0 11153.585
1906.9234619140625 0 6001.885
1907.92529296875 0 3942.5398
1919.9405517578125 0 5217.88
1920.9449462890625 0 18285.373
1921.9503173828125 0 53972.785
1922.9527587890625 0 49657.746
1923.9534912109375 0 29291.186
1924.9510498046875 0 12871.952
1925.949951171875 0 4773.4473
2039.955078125 0 955.5345

Spectrum Details

|  |  |
| --- | --- |
| Matched peaks? Matched peaks The total absolute number of peaks matched. Additionally in brackets the total fraction of peaks matched and the total number of peaks is shown. | 59 (10.37% of 569) |
| FDR? FDR The false discovery rate estimated for this peptide. It is calculated by matching all theoretical fragments with a non-integer shift with the raw peaks for this spectrum. This is done with 40 different shifts. The resulting percentage is the average number of annotated peaks over the number of annotated peaks with the correct spectrum. | 1.17% |
| Satellite FDR? Satellite FDR See the FDR for details on its calculation. This satellite ion specific FDR only contains the satellite ions (d/w) for I/L/J positions. | 0.00% |
| PSM Score? PSM Score The PSM Score as given by Hecklib to this annotated spectrum. It is shown with three significant figures. | 559 |

#### Spectrum 5764? Spectrum 5764 The raw spectrum of this peptide as annotated by Hecklib. The fragments are coloured according to ion type (see legend). Any peaks with a star '\*' as text can be hovered over to see the full details, first the ion type second the mass shift type. By hovering over the amino acids in the peptide or ions in the legend the corresponding peaks are highlighted. By toggling the 'Unassigned' label you can turn the background (unassigned) peaks on or off in the plot. By updating the slider in the Ion legend you can update the spectrum to only show the top X% of the peaks with labels. The top X% means any peak that is within X% of the highest intensity. By dragging in the spectrum you can zoom in to a specific part of the spectrum and use 'Zoom Out' to get back to the original zoom level. The annotation of the spectrum is based on the given sequence in the peptides file and is done with different software so inconsistencies are likely. The peaks are annotated based on the given sequence, with 20 ppm tolerance.

Copy Data

##### Spectrum 5764 (TSV)

###### Preview

```
Loading example...
```

*Click on the button to copy the data to your clipboard.*

Mz MinMz MaxIntensity Max

WidthHeightPeptide font sizePeptide stroke widthSpectrum font sizeSpectrum stroke widthCompact peptide

Ion legend

wxyz

abcd

OtherUnassignedIon^Charge^_Position_Show for top:%

JHQDWJDGKEYKCKV

04.03e+48.07e+41.21e+51.61e+5

Zoom Out

w^+1^_2_y^+1^_2_z^+1^_2_y^+1^_2_c^+1^_2_z^+3^_8_y^+3^_8_c^+2^_5_y^+2^_5_y^+2^_5_y^+3^_9_w^+2^_6_z^+4^_12_c^+1^_3_y^+4^_12_z^+4^_12_c^+1^_3_c^+2^_6_z^+3^_10_y^+1^_3_c^+3^_10_y^+2^_6_w^+2^_7_y^+4^_14_w^+1^_4_y^+2^_7_y^+3^_11_z^+3^_11_y^+3^_11_y^+2^_7_z^+2^_8_c^+1^_4_c^+1^_4_z^+2^_8_z^+3^_12_y^+2^_8_c^+1^_4_z^+1^_4_w^+2^_9_y^+1^_4_c^+2^_9_y^+3^_13_y^+3^_13_z^+3^_13_c^+2^_9_y^+3^_13_y^+2^_9_w^+2^_10_z^+3^_14_c^+3^_14_c^+3^_14_y^+3^_14_c^+3^_14_c^+2^_10_c^+2^_10_y^+2^_10_z^+2^_10_c^+2^_10_y^+2^_10_c^+1^_5_c^+1^_5_y^+1^_5_z^+1^_5_c^+2^_11_c^+1^_5_y^+1^_5_c^+2^_11_y^+2^_11_z^+2^_11_y^+2^_11_w^+1^_6_y^+2^_12_z^+2^_12_c^+2^_12_y^+2^_12_c^+1^_6_w^+2^_13_c^+1^_6_z^+1^_6_y^+2^_13_y^+1^_6_y^+2^_13_c^+2^_13_y^+2^_14_z^+2^_14_y^+2^_14_c^+1^_7_c^+2^_14_z^+1^_7_c^+1^_7_y^+1^_7_z^+1^_7_c^+1^_8_c^+1^_8_z^+1^_8_y^+1^_8_c^+1^_9_y^+1^_10_z^+1^_10_c^+1^_10_y^+1^_10_c^+1^_11_y^+1^_11_z^+1^_11_y^+1^_11_z^+1^_12_c^+1^_12_y^+1^_12_z^+1^_13_c^+1^_13_

048697214581944

Fragment Matches Table

Show background peaks

| Position | Ion type | Intensity | mz Theoretical | mz Error (Th) | mz Error (ppm) | Charge | Series Number |
| --- | --- | --- | --- | --- | --- | --- | --- |
| - | - | 4550 | 120.1 | - | - | 0 | - |
| - | - | 570.8 | 126.1 | - | - | 0 | - |
| - | - | 424.3 | 126.6 | - | - | 0 | - |
| - | - | 683 | 129.1 | - | - | 0 | - |
| - | - | 3581 | 129.1 | - | - | 0 | - |
| - | - | 619.7 | 130.1 | - | - | 0 | - |
| - | - | 4930 | 130.1 | - | - | 0 | - |
| - | - | 2348 | 132.1 | - | - | 0 | - |
| - | - | 1768 | 136.1 | - | - | 0 | - |
| - | - | 566.3 | 138.1 | - | - | 0 | - |
| - | - | 1053 | 146.1 | - | - | 0 | - |
| - | - | 515.2 | 148.9 | - | - | 0 | - |
| - | - | 497.2 | 149 | - | - | 0 | - |
| - | - | 9240 | 155.1 | - | - | 0 | - |
| - | - | 708.8 | 157.1 | - | - | 0 | - |
| - | - | 2450 | 159.1 | - | - | 0 | - |
| - | - | 488.3 | 163.1 | - | - | 0 | - |
| - | - | 1901 | 164.1 | - | - | 0 | - |
| - | - | 1867 | 165.1 | - | - | 0 | - |
| - | - | 6750 | 166.1 | - | - | 0 | - |
| - | - | 572.5 | 167.1 | - | - | 0 | - |
| - | - | 3742 | 168.1 | - | - | 0 | - |
| - | - | 467.4 | 170.5 | - | - | 0 | - |
| 14 | w | 1682 | 172.1 | 0.0002414 | 1.403 | +1 | 2 |
| - | - | 467.9 | 177.3 | - | - | 0 | - |
| - | - | 971.6 | 178.1 | - | - | 0 | - |
| - | - | 2860 | 183.1 | - | - | 0 | - |
| - | - | 6167 | 185.2 | - | - | 0 | - |
| - | - | 617 | 185.3 | - | - | 0 | - |
| - | - | 650.5 | 186.1 | - | - | 0 | - |
| - | - | 586.9 | 187.1 | - | - | 0 | - |
| - | - | 1763 | 190.1 | - | - | 0 | - |
| - | - | 456.2 | 194.7 | - | - | 0 | - |
| - | - | 1029 | 201.1 | - | - | 0 | - |
| - | - | 1013 | 203.2 | - | - | 0 | - |
| - | - | 2951 | 215.1 | - | - | 0 | - |
| - | - | 729.4 | 219.1 | - | - | 0 | - |
| - | - | 7051 | 223.2 | - | - | 0 | - |
| - | - | 782.4 | 224.2 | - | - | 0 | - |
| - | - | 884.4 | 227.1 | - | - | 0 | - |
| - | - | 587.5 | 229 | - | - | 0 | - |
| - | - | 722.1 | 229.1 | - | - | 0 | - |
| 14 | y | 2458 | 229.2 | 0.0004977 | 2.172 | +1 | 2 |
| 14 | z | 2599 | 230.2 | 4.262E-05 | 0.1852 | +1 | 2 |
| - | - | 1097 | 231.2 | - | - | 0 | - |
| - | - | 1648 | 232.1 | - | - | 0 | - |
| - | - | 6103 | 233.2 | - | - | 0 | - |
| - | - | 6432 | 234.1 | - | - | 0 | - |
| - | - | 899.1 | 243.1 | - | - | 0 | - |
| 14 | y | 2022 | 246.2 | 0.0004988 | 2.026 | +1 | 2 |
| - | - | 1582 | 250.1 | - | - | 0 | - |
| - | - | 3.36E+04 | 251.2 | - | - | 0 | - |
| - | - | 3835 | 252.2 | - | - | 0 | - |
| - | - | 845.8 | 253.1 | - | - | 0 | - |
| - | - | 3589 | 261.1 | - | - | 0 | - |
| - | - | 771 | 263.1 | - | - | 0 | - |
| 2 | c | 3876 | 268.2 | 0.0003226 | 1.203 | +1 | 2 |
| - | - | 718.1 | 269.2 | - | - | 0 | - |
| - | - | 716.8 | 270.1 | - | - | 0 | - |
| - | - | 775.4 | 274.1 | - | - | 0 | - |
| - | - | 2083 | 283.2 | - | - | 0 | - |
| - | - | 506.5 | 287.7 | - | - | 0 | - |
| - | - | 752.3 | 288.1 | - | - | 0 | - |
| - | - | 1131 | 290.1 | - | - | 0 | - |
| - | - | 1303 | 292.1 | - | - | 0 | - |
| - | - | 830.2 | 293.1 | - | - | 0 | - |
| - | - | 506.7 | 294.1 | - | - | 0 | - |
| - | - | 1335 | 294.2 | - | - | 0 | - |
| - | - | 657.3 | 295.2 | - | - | 0 | - |
| - | - | 547.2 | 297.1 | - | - | 0 | - |
| - | - | 629 | 298.5 | - | - | 0 | - |
| - | - | 859.5 | 299.1 | - | - | 0 | - |
| - | - | 4981 | 300.2 | - | - | 0 | - |
| - | - | 716.6 | 301.2 | - | - | 0 | - |
| - | - | 736.9 | 318.1 | - | - | 0 | - |
| - | - | 8382 | 326.7 | - | - | 0 | - |
| 8 | z | 2974 | 327.2 | 0.003532 | 10.8 | +3 | 8 |
| - | - | 635.4 | 327.7 | - | - | 0 | - |
| - | - | 850.6 | 329.2 | - | - | 0 | - |
| - | - | 1343 | 331.7 | - | - | 0 | - |
| - | - | 869.4 | 332.2 | - | - | 0 | - |
| 8 | y | 885.8 | 338.2 | 0.003075 | 9.094 | +3 | 8 |
| - | - | 713.5 | 339.2 | - | - | 0 | - |
| 5 | c | 5220 | 340.7 | 0.0007757 | 2.277 | +2 | 5 |
| 11 | y | 1592 | 341.2 | 0.006014 | 17.63 | +2 | 5 |
| - | - | 1042 | 341.7 | - | - | 0 | - |
| - | - | 1212 | 349.2 | - | - | 0 | - |
| 11 | y | 5304 | 349.7 | 0.001619 | 4.631 | +2 | 5 |
| - | - | 1137 | 350.2 | - | - | 0 | - |
| - | - | 1472 | 351.2 | - | - | 0 | - |
| - | - | 955.1 | 352.2 | - | - | 0 | - |
| - | - | 662.1 | 355.6 | - | - | 0 | - |
| - | - | 900 | 361.2 | - | - | 0 | - |
| - | - | 725.6 | 367.7 | - | - | 0 | - |
| - | - | 739.1 | 370.7 | - | - | 0 | - |
| - | - | 1053 | 372.2 | - | - | 0 | - |
| - | - | 1076 | 374.7 | - | - | 0 | - |
| - | - | 2015 | 375.2 | - | - | 0 | - |
| - | - | 1200 | 375.2 | - | - | 0 | - |
| 7 | y | 1661 | 376.5 | 0.00164 | 4.355 | +3 | 9 |
| 10 | w | 1.756E+04 | 376.7 | 0.001927 | 5.116 | +2 | 6 |
| - | - | 1173 | 376.9 | - | - | 0 | - |
| - | - | 8231 | 377.2 | - | - | 0 | - |
| 4 | z | 1900 | 377.7 | 0.0062 | 16.42 | +4 | 12 |
| - | - | 2130 | 378.2 | - | - | 0 | - |
| 3 | c | 9259 | 379.2 | 0.0003377 | 0.8905 | +1 | 3 |
| - | - | 2049 | 380.2 | - | - | 0 | - |
| - | - | 672.9 | 381.2 | - | - | 0 | - |
| - | - | 1072 | 381.2 | - | - | 0 | - |
| 4 | y | 793.6 | 381.7 | 0.001349 | 3.535 | +4 | 12 |
| 4 | z | 704.3 | 382.2 | 0.001659 | 4.341 | +4 | 12 |
| - | - | 2062 | 383.2 | - | - | 0 | - |
| - | - | 1509 | 383.7 | - | - | 0 | - |
| - | - | 1300 | 384.3 | - | - | 0 | - |
| - | - | 834.8 | 387.2 | - | - | 0 | - |
| - | - | 1094 | 387.7 | - | - | 0 | - |
| - | - | 1072 | 388.2 | - | - | 0 | - |
| - | - | 756.7 | 390.2 | - | - | 0 | - |
| - | - | 2411 | 395.2 | - | - | 0 | - |
| - | - | 987.1 | 396.2 | - | - | 0 | - |
| 3 | c | 1.493E+04 | 396.2 | 0.0004915 | 1.24 | +1 | 3 |
| 6 | c | 668.2 | 397.2 | 0.006748 | 16.99 | +2 | 6 |
| - | - | 2648 | 397.2 | - | - | 0 | - |
| - | - | 1215 | 397.7 | - | - | 0 | - |
| - | - | 778.6 | 399.2 | - | - | 0 | - |
| 6 | z | 1060 | 403.2 | 0.00193 | 4.786 | +3 | 10 |
| - | - | 981 | 404.2 | - | - | 0 | - |
| 13 | y | 3757 | 407.2 | 0.005182 | 12.72 | +1 | 3 |
| - | - | 1324 | 408.2 | - | - | 0 | - |
| 10 | c | 701 | 408.2 | 0.0002763 | 0.6768 | +3 | 10 |
| - | - | 976.6 | 408.7 | - | - | 0 | - |
| - | - | 2016 | 409.2 | - | - | 0 | - |
| 10 | y | 2604 | 414.2 | 0.008212 | 19.83 | +2 | 6 |
| - | - | 1045 | 416.3 | - | - | 0 | - |
| - | - | 6660 | 417.2 | - | - | 0 | - |
| - | - | 2522 | 417.7 | - | - | 0 | - |
| - | - | 978.1 | 418.2 | - | - | 0 | - |
| - | - | 983.1 | 419.2 | - | - | 0 | - |
| - | - | 1039 | 424.8 | - | - | 0 | - |
| - | - | 1358 | 425.3 | - | - | 0 | - |
| - | - | 5136 | 428.3 | - | - | 0 | - |
| - | - | 2678 | 429.3 | - | - | 0 | - |
| - | - | 3958 | 440.7 | - | - | 0 | - |
| 9 | w | 3805 | 441.2 | 0.001678 | 3.804 | +2 | 7 |
| - | - | 1277 | 441.2 | - | - | 0 | - |
| - | - | 1467 | 441.7 | - | - | 0 | - |
| - | - | 1169 | 445.2 | - | - | 0 | - |
| - | - | 2159 | 448.2 | - | - | 0 | - |
| - | - | 1330 | 449.2 | - | - | 0 | - |
| - | - | 687.5 | 450.2 | - | - | 0 | - |
| 2 | y | 1512 | 452.5 | 0.0004813 | 1.064 | +4 | 14 |
| - | - | 2609 | 452.7 | - | - | 0 | - |
| - | - | 1113 | 453 | - | - | 0 | - |
| - | - | 850.7 | 453.2 | - | - | 0 | - |
| - | - | 3813 | 453.3 | - | - | 0 | - |
| - | - | 2397 | 453.8 | - | - | 0 | - |
| - | - | 1342 | 454.3 | - | - | 0 | - |
| - | - | 1126 | 456.2 | - | - | 0 | - |
| - | - | 899.3 | 457.2 | - | - | 0 | - |
| - | - | 4959 | 460.3 | - | - | 0 | - |
| 12 | w | 4792 | 461.2 | 0.003844 | 8.336 | +1 | 4 |
| - | - | 1827 | 461.3 | - | - | 0 | - |
| - | - | 978.3 | 462.2 | - | - | 0 | - |
| - | - | 1666 | 466.2 | - | - | 0 | - |
| - | - | 793.4 | 467.2 | - | - | 0 | - |
| - | - | 3213 | 467.2 | - | - | 0 | - |
| - | - | 820.7 | 468.3 | - | - | 0 | - |
| 9 | y | 692.2 | 469.2 | 0.002268 | 4.833 | +2 | 7 |
| 5 | y | 1928 | 470.2 | 0.003809 | 8.101 | +3 | 11 |
| - | - | 1108 | 470.7 | - | - | 0 | - |
| 5 | z | 780.8 | 470.9 | 0.002911 | 6.182 | +3 | 11 |
| - | - | 652.9 | 472.2 | - | - | 0 | - |
| - | - | 2035 | 474.3 | - | - | 0 | - |
| - | - | 3360 | 475.3 | - | - | 0 | - |
| 5 | y | 3648 | 476.2 | 0.005999 | 12.6 | +3 | 11 |
| - | - | 1823 | 476.5 | - | - | 0 | - |
| - | - | 960 | 476.7 | - | - | 0 | - |
| 9 | y | 1.143E+04 | 478.3 | 0.001519 | 3.177 | +2 | 7 |
| - | - | 6671 | 478.8 | - | - | 0 | - |
| - | - | 2379 | 479.3 | - | - | 0 | - |
| - | - | 1843 | 480.5 | - | - | 0 | - |
| - | - | 3318 | 480.7 | - | - | 0 | - |
| - | - | 2.395E+04 | 480.7 | - | - | 0 | - |
| - | - | 2.077E+04 | 481 | - | - | 0 | - |
| - | - | 1475 | 481.2 | - | - | 0 | - |
| - | - | 1.461E+04 | 481.2 | - | - | 0 | - |
| - | - | 2484 | 481.3 | - | - | 0 | - |
| - | - | 4360 | 481.5 | - | - | 0 | - |
| - | - | 1057 | 481.7 | - | - | 0 | - |
| - | - | 2521 | 481.7 | - | - | 0 | - |
| - | - | 860.6 | 482 | - | - | 0 | - |
| - | - | 1300 | 482.2 | - | - | 0 | - |
| - | - | 2370 | 488.8 | - | - | 0 | - |
| - | - | 841.2 | 489.3 | - | - | 0 | - |
| 8 | z | 1620 | 490.2 | 0.006113 | 12.47 | +2 | 8 |
| - | - | 791.8 | 491.3 | - | - | 0 | - |
| 4 | c | 1058 | 493.3 | 0.0002924 | 0.5928 | +1 | 4 |
| 4 | c | 7786 | 494.2 | 0.0003722 | 0.753 | +1 | 4 |
| - | - | 2802 | 495.2 | - | - | 0 | - |
| - | - | 1590 | 496.3 | - | - | 0 | - |
| - | - | 800 | 498.3 | - | - | 0 | - |
| 8 | z | 1736 | 498.8 | 0.002187 | 4.385 | +2 | 8 |
| - | - | 2027 | 499.3 | - | - | 0 | - |
| - | - | 831.2 | 499.8 | - | - | 0 | - |
| 4 | z | 2740 | 503.2 | 0.003694 | 7.341 | +3 | 12 |
| - | - | 763 | 504.2 | - | - | 0 | - |
| - | - | 1787 | 504.8 | - | - | 0 | - |
| - | - | 3791 | 505.2 | - | - | 0 | - |
| - | - | 4173 | 505.7 | - | - | 0 | - |
| - | - | 6712 | 506.3 | - | - | 0 | - |
| 8 | y | 1.786E+04 | 506.8 | 0.00157 | 3.098 | +2 | 8 |
| - | - | 7803 | 507.3 | - | - | 0 | - |
| - | - | 3010 | 507.8 | - | - | 0 | - |
| - | - | 1304 | 508.3 | - | - | 0 | - |
| - | - | 841.1 | 509.3 | - | - | 0 | - |
| - | - | 5206 | 510.8 | - | - | 0 | - |
| 4 | c | 6.211E+04 | 511.3 | 0.0009227 | 1.805 | +1 | 4 |
| - | - | 1.571E+04 | 512.3 | - | - | 0 | - |
| - | - | 3174 | 513.3 | - | - | 0 | - |
| - | - | 1251 | 514.7 | - | - | 0 | - |
| - | - | 778.8 | 515.7 | - | - | 0 | - |
| 12 | z | 1.938E+04 | 519.3 | 0.003814 | 7.346 | +1 | 4 |
| - | - | 770.3 | 519.8 | - | - | 0 | - |
| - | - | 6185 | 520.3 | - | - | 0 | - |
| - | - | 1180 | 521.3 | - | - | 0 | - |
| - | - | 1056 | 525.2 | - | - | 0 | - |
| - | - | 2709 | 526.8 | - | - | 0 | - |
| - | - | 5959 | 527.2 | - | - | 0 | - |
| - | - | 1334 | 527.7 | - | - | 0 | - |
| - | - | 4026 | 527.8 | - | - | 0 | - |
| - | - | 2149 | 528.3 | - | - | 0 | - |
| - | - | 938.9 | 528.8 | - | - | 0 | - |
| - | - | 1086 | 530.2 | - | - | 0 | - |
| - | - | 771.2 | 530.3 | - | - | 0 | - |
| 7 | w | 5313 | 533.8 | 0.001603 | 3.004 | +2 | 9 |
| - | - | 1.247E+04 | 534.3 | - | - | 0 | - |
| - | - | 5326 | 534.8 | - | - | 0 | - |
| 12 | y | 1.12E+04 | 535.3 | 0.004777 | 8.924 | +1 | 4 |
| - | - | 2837 | 536.3 | - | - | 0 | - |
| - | - | 1046 | 537.3 | - | - | 0 | - |
| - | - | 901.2 | 538.3 | - | - | 0 | - |
| - | - | 1198 | 544.3 | - | - | 0 | - |
| - | - | 1597 | 546.2 | - | - | 0 | - |
| 9 | c | 2195 | 547.3 | 0.004277 | 7.815 | +2 | 9 |
| - | - | 1807 | 547.8 | - | - | 0 | - |
| - | - | 741 | 549.3 | - | - | 0 | - |
| - | - | 1122 | 549.6 | - | - | 0 | - |
| - | - | 844 | 549.9 | - | - | 0 | - |
| - | - | 1.34E+04 | 550.3 | - | - | 0 | - |
| 3 | y | 5638 | 551.3 | 0.007486 | 13.58 | +3 | 13 |
| 3 | y | 2130 | 551.6 | 0.003254 | 5.9 | +3 | 13 |
| 3 | z | 1278 | 551.9 | 0.00029 | 0.5255 | +3 | 13 |
| - | - | 3216 | 552.3 | - | - | 0 | - |
| - | - | 3205 | 555.3 | - | - | 0 | - |
| 9 | c | 5.448E+04 | 555.8 | 0.0006458 | 1.162 | +2 | 9 |
| - | - | 2.565E+04 | 556.3 | - | - | 0 | - |
| - | - | 6093 | 556.8 | - | - | 0 | - |
| - | - | 7853 | 556.8 | - | - | 0 | - |
| 3 | y | 4299 | 557.3 | 0.001444 | 2.591 | +3 | 13 |
| - | - | 1762 | 557.8 | - | - | 0 | - |
| - | - | 820.1 | 559.6 | - | - | 0 | - |
| - | - | 1835 | 562.3 | - | - | 0 | - |
| - | - | 1270 | 562.6 | - | - | 0 | - |
| - | - | 1214 | 563 | - | - | 0 | - |
| - | - | 765.6 | 563.6 | - | - | 0 | - |
| - | - | 1998 | 563.8 | - | - | 0 | - |
| 7 | y | 3.448E+04 | 564.3 | 0.001431 | 2.535 | +2 | 9 |
| - | - | 2.128E+04 | 564.8 | - | - | 0 | - |
| - | - | 9420 | 565.3 | - | - | 0 | - |
| - | - | 2081 | 565.8 | - | - | 0 | - |
| - | - | 820.8 | 567.2 | - | - | 0 | - |
| - | - | 1154 | 567.3 | - | - | 0 | - |
| - | - | 4782 | 568.8 | - | - | 0 | - |
| - | - | 4049 | 569.3 | - | - | 0 | - |
| - | - | 1204 | 569.8 | - | - | 0 | - |
| - | - | 909.9 | 569.9 | - | - | 0 | - |
| - | - | 1483 | 573 | - | - | 0 | - |
| - | - | 1264 | 573.3 | - | - | 0 | - |
| - | - | 1864 | 573.9 | - | - | 0 | - |
| - | - | 1684 | 574.3 | - | - | 0 | - |
| - | - | 1143 | 574.9 | - | - | 0 | - |
| - | - | 767.9 | 575.3 | - | - | 0 | - |
| - | - | 770.1 | 577.3 | - | - | 0 | - |
| - | - | 3220 | 578 | - | - | 0 | - |
| - | - | 3082 | 578.3 | - | - | 0 | - |
| - | - | 1311 | 578.6 | - | - | 0 | - |
| - | - | 1955 | 578.9 | - | - | 0 | - |
| - | - | 759.6 | 579.3 | - | - | 0 | - |
| - | - | 843.2 | 582.3 | - | - | 0 | - |
| - | - | 1277 | 582.6 | - | - | 0 | - |
| - | - | 2661 | 583 | - | - | 0 | - |
| - | - | 2380 | 583.3 | - | - | 0 | - |
| - | - | 1593 | 583.6 | - | - | 0 | - |
| - | - | 840.7 | 585.3 | - | - | 0 | - |
| - | - | 842.4 | 586.3 | - | - | 0 | - |
| - | - | 1810 | 590.8 | - | - | 0 | - |
| 6 | w | 3.446E+04 | 591.3 | 0.00122 | 2.063 | +2 | 10 |
| - | - | 1299 | 591.4 | - | - | 0 | - |
| - | - | 2.052E+04 | 591.8 | - | - | 0 | - |
| - | - | 9577 | 592.3 | - | - | 0 | - |
| - | - | 1419 | 592.4 | - | - | 0 | - |
| - | - | 1701 | 592.6 | - | - | 0 | - |
| - | - | 3588 | 592.8 | - | - | 0 | - |
| - | - | 2491 | 593 | - | - | 0 | - |
| - | - | 1954 | 593.3 | - | - | 0 | - |
| - | - | 877 | 595.3 | - | - | 0 | - |
| - | - | 7724 | 597.3 | - | - | 0 | - |
| 2 | z | 7635 | 597.6 | 0.01104 | 18.47 | +3 | 14 |
| - | - | 4499 | 598 | - | - | 0 | - |
| - | - | 3079 | 598.3 | - | - | 0 | - |
| - | - | 880 | 599.2 | - | - | 0 | - |
| - | - | 757.7 | 600.3 | - | - | 0 | - |
| 14 | c | 2228 | 601.3 | 0.005362 | 8.917 | +3 | 14 |
| 14 | c | 2632 | 601.6 | 0.005238 | 8.706 | +3 | 14 |
| - | - | 6749 | 602 | - | - | 0 | - |
| - | - | 3039 | 602.3 | - | - | 0 | - |
| - | - | 1249 | 602.6 | - | - | 0 | - |
| 2 | y | 3672 | 603 | 0.002091 | 3.467 | +3 | 14 |
| - | - | 3302 | 603.3 | - | - | 0 | - |
| - | - | 1174 | 603.6 | - | - | 0 | - |
| - | - | 1010 | 605.6 | - | - | 0 | - |
| - | - | 891.7 | 606 | - | - | 0 | - |
| - | - | 1214 | 606.3 | - | - | 0 | - |
| - | - | 1693 | 606.6 | - | - | 0 | - |
| - | - | 4727 | 607 | - | - | 0 | - |
| 14 | c | 5.5E+04 | 607.3 | 0.000479 | 0.7887 | +3 | 14 |
| - | - | 6.002E+04 | 607.6 | - | - | 0 | - |
| - | - | 3.127E+04 | 608 | - | - | 0 | - |
| - | - | 1.765E+04 | 608.3 | - | - | 0 | - |
| - | - | 4825 | 608.6 | - | - | 0 | - |
| - | - | 1565 | 609 | - | - | 0 | - |
| - | - | 1252 | 610 | - | - | 0 | - |
| - | - | 906.9 | 610.3 | - | - | 0 | - |
| - | - | 886.2 | 611 | - | - | 0 | - |
| 10 | c | 2239 | 611.3 | 0.008496 | 13.9 | +2 | 10 |
| - | - | 1026 | 611.6 | - | - | 0 | - |
| 10 | c | 5002 | 611.8 | 0.004159 | 6.799 | +2 | 10 |
| 6 | y | 3737 | 612.3 | 0.004248 | 6.938 | +2 | 10 |
| 6 | z | 7790 | 612.8 | 0.002973 | 4.851 | +2 | 10 |
| - | - | 4871 | 613.3 | - | - | 0 | - |
| - | - | 1813 | 613.8 | - | - | 0 | - |
| - | - | 903.8 | 614.3 | - | - | 0 | - |
| - | - | 782.1 | 614.8 | - | - | 0 | - |
| - | - | 1656 | 615 | - | - | 0 | - |
| - | - | 2925 | 615.3 | - | - | 0 | - |
| - | - | 2048 | 615.6 | - | - | 0 | - |
| - | - | 771.6 | 616 | - | - | 0 | - |
| - | - | 1147 | 616.3 | - | - | 0 | - |
| - | - | 6527 | 616.6 | - | - | 0 | - |
| - | - | 6483 | 617 | - | - | 0 | - |
| - | - | 3104 | 617.3 | - | - | 0 | - |
| - | - | 2306 | 617.6 | - | - | 0 | - |
| - | - | 4518 | 619.8 | - | - | 0 | - |
| 10 | c | 6.287E+04 | 620.3 | 0.0009557 | 1.541 | +2 | 10 |
| - | - | 4332 | 620.6 | - | - | 0 | - |
| 6 | y | 5.892E+04 | 620.8 | 0.004766 | 7.677 | +2 | 10 |
| - | - | 5.177E+04 | 621 | - | - | 0 | - |
| - | - | 7.828E+04 | 621.3 | - | - | 0 | - |
| - | - | 3.681E+04 | 621.6 | - | - | 0 | - |
| - | - | 1.052E+04 | 621.8 | - | - | 0 | - |
| - | - | 2.125E+04 | 622 | - | - | 0 | - |
| - | - | 9332 | 622.3 | - | - | 0 | - |
| - | - | 2825 | 622.6 | - | - | 0 | - |
| - | - | 2159 | 624.3 | - | - | 0 | - |
| - | - | 3063 | 625.7 | - | - | 0 | - |
| - | - | 4595 | 626 | - | - | 0 | - |
| - | - | 2141 | 626.3 | - | - | 0 | - |
| - | - | 2104 | 626.6 | - | - | 0 | - |
| - | - | 1883 | 627 | - | - | 0 | - |
| - | - | 1580 | 629.3 | - | - | 0 | - |
| - | - | 1432 | 629.6 | - | - | 0 | - |
| - | - | 816 | 630 | - | - | 0 | - |
| - | - | 900.9 | 631.3 | - | - | 0 | - |
| - | - | 1161 | 634.3 | - | - | 0 | - |
| - | - | 806.7 | 634.7 | - | - | 0 | - |
| - | - | 796.2 | 634.8 | - | - | 0 | - |
| - | - | 3485 | 635 | - | - | 0 | - |
| - | - | 5495 | 635.3 | - | - | 0 | - |
| - | - | 6910 | 635.6 | - | - | 0 | - |
| - | - | 3156 | 636 | - | - | 0 | - |
| - | - | 1783 | 636.3 | - | - | 0 | - |
| - | - | 1686 | 638.3 | - | - | 0 | - |
| - | - | 1697 | 640 | - | - | 0 | - |
| - | - | 917.3 | 640.3 | - | - | 0 | - |
| - | - | 8238 | 640.3 | - | - | 0 | - |
| - | - | 1.115E+05 | 640.7 | - | - | 0 | - |
| - | - | 1.598E+05 | 641 | - | - | 0 | - |
| - | - | 1.088E+05 | 641.3 | - | - | 0 | - |
| - | - | 5.457E+04 | 641.7 | - | - | 0 | - |
| - | - | 2.351E+04 | 642 | - | - | 0 | - |
| - | - | 8270 | 642.3 | - | - | 0 | - |
| - | - | 2474 | 642.6 | - | - | 0 | - |
| - | - | 1510 | 643 | - | - | 0 | - |
| - | - | 847.9 | 647.8 | - | - | 0 | - |
| - | - | 2061 | 648.3 | - | - | 0 | - |
| - | - | 1407 | 648.8 | - | - | 0 | - |
| - | - | 2891 | 652.3 | - | - | 0 | - |
| - | - | 3145 | 653.3 | - | - | 0 | - |
| - | - | 1019 | 654.3 | - | - | 0 | - |
| - | - | 1569 | 655.8 | - | - | 0 | - |
| - | - | 1040 | 656.3 | - | - | 0 | - |
| - | - | 1080 | 657.3 | - | - | 0 | - |
| - | - | 2133 | 660.4 | - | - | 0 | - |
| - | - | 2153 | 660.9 | - | - | 0 | - |
| - | - | 1209 | 661.4 | - | - | 0 | - |
| - | - | 2182 | 664.3 | - | - | 0 | - |
| - | - | 1330 | 669.8 | - | - | 0 | - |
| - | - | 2258 | 670.3 | - | - | 0 | - |
| - | - | 1167 | 671.8 | - | - | 0 | - |
| - | - | 1855 | 676.3 | - | - | 0 | - |
| - | - | 4487 | 677.8 | - | - | 0 | - |
| - | - | 3128 | 678.3 | - | - | 0 | - |
| - | - | 888.8 | 678.8 | - | - | 0 | - |
| 5 | c | 1130 | 679.3 | 0.002731 | 4.02 | +1 | 5 |
| - | - | 3663 | 679.8 | - | - | 0 | - |
| 5 | c | 1.455E+04 | 680.3 | 0.002724 | 4.004 | +1 | 5 |
| - | - | 1316 | 680.8 | - | - | 0 | - |
| 11 | y | 7954 | 681.3 | 0.01025 | 15.04 | +1 | 5 |
| 11 | z | 5.025E+04 | 682.3 | 0.004155 | 6.089 | +1 | 5 |
| - | - | 988.1 | 682.8 | - | - | 0 | - |
| - | - | 1.812E+04 | 683.3 | - | - | 0 | - |
| - | - | 1152 | 683.4 | - | - | 0 | - |
| - | - | 4155 | 683.9 | - | - | 0 | - |
| - | - | 7514 | 684.3 | - | - | 0 | - |
| - | - | 639.4 | 684.8 | - | - | 0 | - |
| - | - | 840.9 | 684.9 | - | - | 0 | - |
| - | - | 1691 | 685.3 | - | - | 0 | - |
| 11 | c | 1117 | 693.3 | 0.0008155 | 1.176 | +2 | 11 |
| - | - | 814.9 | 694.3 | - | - | 0 | - |
| 5 | c | 4.9E+04 | 697.3 | 0.0005282 | 0.7574 | +1 | 5 |
| 11 | y | 2.671E+04 | 698.4 | 0.01037 | 14.84 | +1 | 5 |
| - | - | 7378 | 699.4 | - | - | 0 | - |
| - | - | 1328 | 700.4 | - | - | 0 | - |
| - | - | 2750 | 701.3 | - | - | 0 | - |
| 11 | c | 4.561E+04 | 701.8 | 0.0009687 | 1.38 | +2 | 11 |
| - | - | 3.587E+04 | 702.3 | - | - | 0 | - |
| - | - | 1.572E+04 | 702.8 | - | - | 0 | - |
| - | - | 3777 | 703.3 | - | - | 0 | - |
| 5 | y | 2727 | 705.3 | 0.003642 | 5.163 | +2 | 11 |
| 5 | z | 3.217E+04 | 705.8 | 0.00143 | 2.026 | +2 | 11 |
| - | - | 2.382E+04 | 706.3 | - | - | 0 | - |
| - | - | 9619 | 706.8 | - | - | 0 | - |
| - | - | 3271 | 707.4 | - | - | 0 | - |
| - | - | 2623 | 711.3 | - | - | 0 | - |
| - | - | 3054 | 712.3 | - | - | 0 | - |
| - | - | 788.5 | 713.4 | - | - | 0 | - |
| 5 | y | 1.139E+04 | 713.9 | 0.001576 | 2.208 | +2 | 11 |
| - | - | 9283 | 714.4 | - | - | 0 | - |
| - | - | 4977 | 714.9 | - | - | 0 | - |
| - | - | 959.9 | 715.4 | - | - | 0 | - |
| - | - | 1175 | 717.4 | - | - | 0 | - |
| - | - | 864.9 | 718.9 | - | - | 0 | - |
| - | - | 2047 | 719.4 | - | - | 0 | - |
| - | - | 1255 | 720.4 | - | - | 0 | - |
| - | - | 1202 | 721.4 | - | - | 0 | - |
| - | - | 903.4 | 722.3 | - | - | 0 | - |
| - | - | 820.7 | 723.4 | - | - | 0 | - |
| - | - | 1548 | 727.3 | - | - | 0 | - |
| - | - | 955.4 | 727.8 | - | - | 0 | - |
| - | - | 862.7 | 728.3 | - | - | 0 | - |
| - | - | 2008 | 730.3 | - | - | 0 | - |
| - | - | 1485 | 731.4 | - | - | 0 | - |
| - | - | 860 | 733.4 | - | - | 0 | - |
| - | - | 852.2 | 733.9 | - | - | 0 | - |
| - | - | 1625 | 740.3 | - | - | 0 | - |
| - | - | 7156 | 741.4 | - | - | 0 | - |
| - | - | 3615 | 741.9 | - | - | 0 | - |
| - | - | 1990 | 742.4 | - | - | 0 | - |
| - | - | 852.8 | 743.4 | - | - | 0 | - |
| - | - | 968.4 | 743.9 | - | - | 0 | - |
| - | - | 973.3 | 744.4 | - | - | 0 | - |
| - | - | 2191 | 747.4 | - | - | 0 | - |
| 10 | w | 9795 | 752.4 | 0.00378 | 5.024 | +1 | 6 |
| - | - | 4131 | 753.4 | - | - | 0 | - |
| - | - | 1568 | 754.4 | - | - | 0 | - |
| 4 | y | 1186 | 762.9 | 0.01349 | 17.68 | +2 | 12 |
| 4 | z | 2.442E+04 | 763.4 | 0.001108 | 1.451 | +2 | 12 |
| - | - | 1.9E+04 | 763.9 | - | - | 0 | - |
| - | - | 8879 | 764.4 | - | - | 0 | - |
| - | - | 4498 | 764.9 | - | - | 0 | - |
| - | - | 1070 | 765.4 | - | - | 0 | - |
| 12 | c | 1.075E+04 | 765.9 | 0.001217 | 1.589 | +2 | 12 |
| - | - | 1.201E+04 | 766.4 | - | - | 0 | - |
| - | - | 5074 | 766.9 | - | - | 0 | - |
| - | - | 1320 | 767.4 | - | - | 0 | - |
| 4 | y | 3723 | 771.4 | 0.001071 | 1.388 | +2 | 12 |
| - | - | 3802 | 771.9 | - | - | 0 | - |
| - | - | 1286 | 772.4 | - | - | 0 | - |
| - | - | 1615 | 772.9 | - | - | 0 | - |
| - | - | 1659 | 777.4 | - | - | 0 | - |
| - | - | 2152 | 778.9 | - | - | 0 | - |
| - | - | 2167 | 779.4 | - | - | 0 | - |
| - | - | 1176 | 785.4 | - | - | 0 | - |
| - | - | 996.6 | 786.4 | - | - | 0 | - |
| - | - | 1107 | 792.4 | - | - | 0 | - |
| 6 | c | 2202 | 793.4 | 0.001973 | 2.487 | +1 | 6 |
| - | - | 1192 | 794.4 | - | - | 0 | - |
| - | - | 964.5 | 797.9 | - | - | 0 | - |
| 3 | w | 5317 | 798.4 | 0.001093 | 1.37 | +2 | 13 |
| - | - | 6904 | 798.9 | - | - | 0 | - |
| - | - | 1934 | 799.4 | - | - | 0 | - |
| - | - | 1127 | 799.9 | - | - | 0 | - |
| - | - | 3595 | 805.4 | - | - | 0 | - |
| - | - | 2513 | 805.9 | - | - | 0 | - |
| - | - | 2345 | 806.4 | - | - | 0 | - |
| - | - | 1658 | 806.9 | - | - | 0 | - |
| - | - | 2096 | 809.4 | - | - | 0 | - |
| - | - | 936.3 | 809.9 | - | - | 0 | - |
| 6 | c | 9.789E+04 | 810.4 | 0.0008758 | 1.081 | +1 | 6 |
| 10 | z | 3.454E+04 | 811.4 | 0.007807 | 9.622 | +1 | 6 |
| - | - | 2.889E+04 | 811.4 | - | - | 0 | - |
| - | - | 2.514E+04 | 812.4 | - | - | 0 | - |
| - | - | 1.064E+04 | 813.4 | - | - | 0 | - |
| - | - | 3762 | 814.4 | - | - | 0 | - |
| - | - | 1372 | 825.4 | - | - | 0 | - |
| 3 | y | 3150 | 826.9 | 0.008672 | 10.49 | +2 | 13 |
| 10 | y | 3.641E+04 | 827.4 | 0.01023 | 12.37 | +1 | 6 |
| - | - | 2.648E+04 | 827.9 | - | - | 0 | - |
| - | - | 1.634E+04 | 828.4 | - | - | 0 | - |
| - | - | 3178 | 828.9 | - | - | 0 | - |
| - | - | 2005 | 829.4 | - | - | 0 | - |
| - | - | 2890 | 831.4 | - | - | 0 | - |
| 3 | y | 4566 | 835.4 | 0.0007573 | 0.9065 | +2 | 13 |
| - | - | 4222 | 835.9 | - | - | 0 | - |
| - | - | 2633 | 836.4 | - | - | 0 | - |
| - | - | 812.9 | 836.9 | - | - | 0 | - |
| - | - | 2918 | 839.4 | - | - | 0 | - |
| - | - | 1766 | 840.4 | - | - | 0 | - |
| - | - | 1397 | 844.4 | - | - | 0 | - |
| - | - | 1637 | 845.4 | - | - | 0 | - |
| - | - | 4098 | 845.9 | - | - | 0 | - |
| 13 | c | 4.871E+04 | 846.4 | 0.0009293 | 1.098 | +2 | 13 |
| - | - | 3.761E+04 | 846.9 | - | - | 0 | - |
| - | - | 2.377E+04 | 847.4 | - | - | 0 | - |
| - | - | 9736 | 847.9 | - | - | 0 | - |
| - | - | 4084 | 848.4 | - | - | 0 | - |
| - | - | 965.2 | 848.9 | - | - | 0 | - |
| - | - | 934.2 | 850.4 | - | - | 0 | - |
| - | - | 2279 | 860.4 | - | - | 0 | - |
| - | - | 978.1 | 861.4 | - | - | 0 | - |
| - | - | 2241 | 866.4 | - | - | 0 | - |
| - | - | 1986 | 866.9 | - | - | 0 | - |
| - | - | 5786 | 867.4 | - | - | 0 | - |
| - | - | 1689 | 868.4 | - | - | 0 | - |
| - | - | 951.5 | 873.9 | - | - | 0 | - |
| - | - | 946.7 | 879.9 | - | - | 0 | - |
| - | - | 3900 | 880.4 | - | - | 0 | - |
| - | - | 3797 | 880.9 | - | - | 0 | - |
| - | - | 1.003E+04 | 881.4 | - | - | 0 | - |
| - | - | 3968 | 881.9 | - | - | 0 | - |
| - | - | 5578 | 882.4 | - | - | 0 | - |
| - | - | 1850 | 883.4 | - | - | 0 | - |
| - | - | 1187 | 887.4 | - | - | 0 | - |
| - | - | 3301 | 888.4 | - | - | 0 | - |
| - | - | 6592 | 888.9 | - | - | 0 | - |
| - | - | 2088 | 889.3 | - | - | 0 | - |
| - | - | 6791 | 889.4 | - | - | 0 | - |
| - | - | 3827 | 889.9 | - | - | 0 | - |
| 2 | y | 1340 | 895.4 | 0.005766 | 6.44 | +2 | 14 |
| 2 | z | 1.507E+04 | 895.9 | 0.001015 | 1.133 | +2 | 14 |
| - | - | 1.707E+04 | 896.4 | - | - | 0 | - |
| - | - | 1.105E+04 | 896.9 | - | - | 0 | - |
| - | - | 6180 | 897.4 | - | - | 0 | - |
| - | - | 2273 | 897.9 | - | - | 0 | - |
| - | - | 1107 | 902 | - | - | 0 | - |
| - | - | 3344 | 902.4 | - | - | 0 | - |
| - | - | 3417 | 902.9 | - | - | 0 | - |
| - | - | 1882 | 903.4 | - | - | 0 | - |
| 2 | y | 1661 | 903.9 | 0.009399 | 10.4 | +2 | 14 |
| 7 | c | 3585 | 908.4 | 0.0008481 | 0.9336 | +1 | 7 |
| - | - | 1009 | 909 | - | - | 0 | - |
| - | - | 3106 | 909.5 | - | - | 0 | - |
| - | - | 2120 | 910 | - | - | 0 | - |
| 14 | c | 9407 | 910.4 | 0.00178 | 1.955 | +2 | 14 |
| - | - | 1.075E+04 | 910.9 | - | - | 0 | - |
| - | - | 7295 | 911.4 | - | - | 0 | - |
| - | - | 3755 | 911.9 | - | - | 0 | - |
| - | - | 2938 | 912.5 | - | - | 0 | - |
| - | - | 1063 | 913.5 | - | - | 0 | - |
| - | - | 1344 | 914.5 | - | - | 0 | - |
| - | - | 955.6 | 915 | - | - | 0 | - |
| - | - | 1273 | 917.3 | - | - | 0 | - |
| 9 | z | 2094 | 922.5 | 0.007397 | 8.019 | +1 | 7 |
| - | - | 6248 | 923 | - | - | 0 | - |
| - | - | 8569 | 923.5 | - | - | 0 | - |
| - | - | 5579 | 924 | - | - | 0 | - |
| - | - | 5118 | 924.5 | - | - | 0 | - |
| - | - | 2849 | 925 | - | - | 0 | - |
| 7 | c | 1.434E+04 | 925.5 | 0.001277 | 1.379 | +1 | 7 |
| - | - | 6776 | 926.5 | - | - | 0 | - |
| - | - | 1355 | 927.5 | - | - | 0 | - |
| - | - | 1041 | 930 | - | - | 0 | - |
| - | - | 3713 | 930.5 | - | - | 0 | - |
| - | - | 7724 | 931 | - | - | 0 | - |
| - | - | 4.248E+04 | 931.5 | - | - | 0 | - |
| - | - | 4.259E+04 | 932 | - | - | 0 | - |
| - | - | 2.861E+04 | 932.5 | - | - | 0 | - |
| - | - | 1.178E+04 | 933 | - | - | 0 | - |
| - | - | 5453 | 933.5 | - | - | 0 | - |
| - | - | 1522 | 934 | - | - | 0 | - |
| - | - | 1131 | 938 | - | - | 0 | - |
| 9 | y | 1.7E+04 | 938.5 | 0.002172 | 2.314 | +1 | 7 |
| - | - | 1.346E+04 | 939 | - | - | 0 | - |
| 9 | z | 2.233E+04 | 939.5 | 0.006579 | 7.003 | +1 | 7 |
| - | - | 4973 | 940 | - | - | 0 | - |
| - | - | 1.784E+04 | 940.5 | - | - | 0 | - |
| - | - | 7123 | 941.5 | - | - | 0 | - |
| - | - | 2326 | 942.5 | - | - | 0 | - |
| - | - | 2422 | 943.5 | - | - | 0 | - |
| - | - | 1760 | 944 | - | - | 0 | - |
| - | - | 1632 | 944.5 | - | - | 0 | - |
| - | - | 977.1 | 946 | - | - | 0 | - |
| - | - | 1652 | 952 | - | - | 0 | - |
| - | - | 1.363E+04 | 952.5 | - | - | 0 | - |
| - | - | 4.306E+04 | 953 | - | - | 0 | - |
| - | - | 4.274E+04 | 953.5 | - | - | 0 | - |
| - | - | 2.594E+04 | 954 | - | - | 0 | - |
| - | - | 1.159E+04 | 954.5 | - | - | 0 | - |
| - | - | 4390 | 955 | - | - | 0 | - |
| - | - | 2921 | 955.5 | - | - | 0 | - |
| - | - | 1572 | 956.5 | - | - | 0 | - |
| - | - | 818.4 | 957.5 | - | - | 0 | - |
| - | - | 1104 | 960 | - | - | 0 | - |
| - | - | 8374 | 960.5 | - | - | 0 | - |
| - | - | 1.624E+04 | 961 | - | - | 0 | - |
| - | - | 4.461E+04 | 961.5 | - | - | 0 | - |
| - | - | 3.86E+04 | 962 | - | - | 0 | - |
| - | - | 2.536E+04 | 962.5 | - | - | 0 | - |
| - | - | 9914 | 963 | - | - | 0 | - |
| - | - | 8552 | 963.5 | - | - | 0 | - |
| - | - | 1679 | 964 | - | - | 0 | - |
| 8 | c | 3254 | 964.5 | 0.004416 | 4.578 | +1 | 8 |
| - | - | 1285 | 967.5 | - | - | 0 | - |
| - | - | 1173 | 973.4 | - | - | 0 | - |
| - | - | 3827 | 981.5 | - | - | 0 | - |
| 8 | c | 4.602E+04 | 982.5 | 0.000931 | 0.9476 | +1 | 8 |
| - | - | 2.654E+04 | 983.5 | - | - | 0 | - |
| - | - | 7650 | 984.5 | - | - | 0 | - |
| - | - | 1751 | 985.5 | - | - | 0 | - |
| 8 | z | 2364 | 996.5 | 0.001248 | 1.253 | +1 | 8 |
| - | - | 3972 | 997.5 | - | - | 0 | - |
| - | - | 2222 | 998.5 | - | - | 0 | - |
| - | - | 1971 | 1011 | - | - | 0 | - |
| 8 | y | 3863 | 1013 | 0.01637 | 16.17 | +1 | 8 |
| - | - | 1915 | 1014 | - | - | 0 | - |
| - | - | 1520 | 1052 | - | - | 0 | - |
| - | - | 2759 | 1054 | - | - | 0 | - |
| - | - | 1867 | 1055 | - | - | 0 | - |
| - | - | 893.4 | 1056 | - | - | 0 | - |
| - | - | 787.3 | 1066 | - | - | 0 | - |
| - | - | 1554 | 1067 | - | - | 0 | - |
| - | - | 1961 | 1068 | - | - | 0 | - |
| - | - | 935.1 | 1069 | - | - | 0 | - |
| - | - | 2385 | 1095 | - | - | 0 | - |
| - | - | 1282 | 1096 | - | - | 0 | - |
| - | - | 1813 | 1110 | - | - | 0 | - |
| 9 | c | 3.482E+04 | 1111 | 0.0003894 | 0.3506 | +1 | 9 |
| - | - | 2.59E+04 | 1112 | - | - | 0 | - |
| - | - | 2.734E+04 | 1113 | - | - | 0 | - |
| - | - | 1.329E+04 | 1114 | - | - | 0 | - |
| - | - | 6178 | 1115 | - | - | 0 | - |
| - | - | 1300 | 1116 | - | - | 0 | - |
| - | - | 1346 | 1129 | - | - | 0 | - |
| - | - | 1081 | 1168 | - | - | 0 | - |
| - | - | 949.3 | 1181 | - | - | 0 | - |
| - | - | 1052 | 1183 | - | - | 0 | - |
| - | - | 1271 | 1196 | - | - | 0 | - |
| - | - | 3560 | 1197 | - | - | 0 | - |
| - | - | 1215 | 1198 | - | - | 0 | - |
| - | - | 925.2 | 1199 | - | - | 0 | - |
| - | - | 1426 | 1213 | - | - | 0 | - |
| 6 | y | 2802 | 1224 | 0.004027 | 3.291 | +1 | 10 |
| 6 | z | 4677 | 1225 | 0.00343 | 2.8 | +1 | 10 |
| - | - | 1.294E+04 | 1226 | - | - | 0 | - |
| - | - | 7828 | 1227 | - | - | 0 | - |
| - | - | 2973 | 1228 | - | - | 0 | - |
| - | - | 1062 | 1229 | - | - | 0 | - |
| - | - | 1125 | 1239 | - | - | 0 | - |
| 10 | c | 1.191E+04 | 1240 | 0.000765 | 0.6171 | +1 | 10 |
| 6 | y | 1.042E+04 | 1241 | 0.01092 | 8.804 | +1 | 10 |
| - | - | 6258 | 1242 | - | - | 0 | - |
| - | - | 2875 | 1243 | - | - | 0 | - |
| - | - | 1153 | 1244 | - | - | 0 | - |
| - | - | 2139 | 1311 | - | - | 0 | - |
| - | - | 1797 | 1312 | - | - | 0 | - |
| - | - | 1242 | 1344 | - | - | 0 | - |
| - | - | 1080 | 1353 | - | - | 0 | - |
| - | - | 948.3 | 1354 | - | - | 0 | - |
| - | - | 908 | 1359 | - | - | 0 | - |
| - | - | 2008 | 1360 | - | - | 0 | - |
| - | - | 1495 | 1361 | - | - | 0 | - |
| - | - | 739.7 | 1362 | - | - | 0 | - |
| - | - | 1231 | 1387 | - | - | 0 | - |
| - | - | 1359 | 1402 | - | - | 0 | - |
| 11 | c | 6715 | 1403 | 0.001523 | 1.086 | +1 | 11 |
| - | - | 7725 | 1404 | - | - | 0 | - |
| - | - | 4684 | 1405 | - | - | 0 | - |
| - | - | 1573 | 1406 | - | - | 0 | - |
| - | - | 750.1 | 1407 | - | - | 0 | - |
| 5 | y | 846.5 | 1410 | 0.005649 | 4.007 | +1 | 11 |
| 5 | z | 5631 | 1411 | 0.002054 | 1.456 | +1 | 11 |
| - | - | 1.659E+04 | 1412 | - | - | 0 | - |
| - | - | 1.191E+04 | 1413 | - | - | 0 | - |
| - | - | 6479 | 1414 | - | - | 0 | - |
| - | - | 2117 | 1415 | - | - | 0 | - |
| - | - | 913.1 | 1426 | - | - | 0 | - |
| 5 | y | 3186 | 1427 | 0.01712 | 12 | +1 | 11 |
| - | - | 3652 | 1428 | - | - | 0 | - |
| - | - | 1038 | 1469 | - | - | 0 | - |
| - | - | 1047 | 1472 | - | - | 0 | - |
| - | - | 1043 | 1482 | - | - | 0 | - |
| - | - | 1072 | 1487 | - | - | 0 | - |
| - | - | 1244 | 1488 | - | - | 0 | - |
| - | - | 1033 | 1497 | - | - | 0 | - |
| - | - | 969.7 | 1498 | - | - | 0 | - |
| - | - | 810.2 | 1515 | - | - | 0 | - |
| 4 | z | 3416 | 1526 | 0.000544 | 0.3566 | +1 | 12 |
| - | - | 1.231E+04 | 1527 | - | - | 0 | - |
| - | - | 7510 | 1528 | - | - | 0 | - |
| - | - | 3350 | 1529 | - | - | 0 | - |
| - | - | 2279 | 1530 | - | - | 0 | - |
| 12 | c | 3146 | 1531 | 0.003596 | 2.349 | +1 | 12 |
| - | - | 3433 | 1532 | - | - | 0 | - |
| - | - | 2576 | 1533 | - | - | 0 | - |
| - | - | 960.1 | 1534 | - | - | 0 | - |
| 4 | y | 774 | 1542 | 0.001456 | 0.9447 | +1 | 12 |
| - | - | 1283 | 1554 | - | - | 0 | - |
| - | - | 860 | 1555 | - | - | 0 | - |
| - | - | 790.7 | 1597 | - | - | 0 | - |
| - | - | 1361 | 1634 | - | - | 0 | - |
| 3 | z | 2561 | 1654 | 5.012E-05 | 0.0303 | +1 | 13 |
| - | - | 9419 | 1655 | - | - | 0 | - |
| - | - | 6994 | 1656 | - | - | 0 | - |
| - | - | 4089 | 1657 | - | - | 0 | - |
| - | - | 1340 | 1658 | - | - | 0 | - |
| - | - | 1002 | 1676 | - | - | 0 | - |
| - | - | 1834 | 1677 | - | - | 0 | - |
| - | - | 1237 | 1691 | - | - | 0 | - |
| 13 | c | 4032 | 1692 | 0.004714 | 2.786 | +1 | 13 |
| - | - | 6496 | 1693 | - | - | 0 | - |
| - | - | 5762 | 1694 | - | - | 0 | - |
| - | - | 3722 | 1695 | - | - | 0 | - |
| - | - | 1686 | 1696 | - | - | 0 | - |
| - | - | 1125 | 1777 | - | - | 0 | - |
| - | - | 3408 | 1792 | - | - | 0 | - |
| - | - | 4031 | 1793 | - | - | 0 | - |
| - | - | 2070 | 1794 | - | - | 0 | - |
| - | - | 1152 | 1795 | - | - | 0 | - |
| - | - | 1038 | 1805 | - | - | 0 | - |
| - | - | 1051 | 1806 | - | - | 0 | - |
| - | - | 1833 | 1821 | - | - | 0 | - |
| - | - | 2652 | 1822 | - | - | 0 | - |
| - | - | 2422 | 1823 | - | - | 0 | - |
| - | - | 1119 | 1846 | - | - | 0 | - |
| - | - | 1384 | 1847 | - | - | 0 | - |
| - | - | 1181 | 1848 | - | - | 0 | - |
| - | - | 2211 | 1862 | - | - | 0 | - |
| - | - | 4613 | 1863 | - | - | 0 | - |
| - | - | 3821 | 1864 | - | - | 0 | - |
| - | - | 2472 | 1865 | - | - | 0 | - |
| - | - | 1557 | 1866 | - | - | 0 | - |
| - | - | 1111 | 1877 | - | - | 0 | - |
| - | - | 1656 | 1878 | - | - | 0 | - |
| - | - | 1909 | 1879 | - | - | 0 | - |
| - | - | 1309 | 1880 | - | - | 0 | - |
| - | - | 1148 | 1895 | - | - | 0 | - |
| - | - | 913.4 | 1904 | - | - | 0 | - |
| - | - | 2397 | 1905 | - | - | 0 | - |
| - | - | 4214 | 1906 | - | - | 0 | - |
| - | - | 3993 | 1907 | - | - | 0 | - |
| - | - | 1351 | 1908 | - | - | 0 | - |
| - | - | 1504 | 1921 | - | - | 0 | - |
| - | - | 4489 | 1922 | - | - | 0 | - |
| - | - | 4871 | 1923 | - | - | 0 | - |
| - | - | 4872 | 1924 | - | - | 0 | - |
| - | - | 2022 | 1925 | - | - | 0 | - |

m/z Charge Intensity FragmentType MassShift Position
120.06582641601562 0 4549.803
126.09190368652344 0 570.8417
126.57588195800781 0 424.34073
129.066162109375 0 682.9878
129.1025390625 0 3581.3762
130.0611114501953 0 619.6942
130.06544494628906 0 4929.7134
132.10214233398438 0 2348.185
136.07601928710938 0 1768.3832
138.0664825439453 0 566.2513
146.09259033203125 0 1052.5292
148.94717407226562 0 515.2497
149.02369689941406 0 497.20462
155.09300231933594 0 9240.334
157.13369750976562 0 708.8434
159.09193420410156 0 2450.1177
163.06178283691406 0 488.27124
164.0822296142578 0 1901.3286
165.1024932861328 0 1867.1267
166.0613250732422 0 6749.8047
167.0648651123047 0 572.47955
168.13848876953125 0 3742.1042
170.47900390625 0 467.38223
172.09706115722656 0 1682.3281 w 13
177.3397979736328 0 467.90988
178.13401794433594 0 971.6311
183.11305236816406 0 2860.0098
185.1650390625 0 6166.6064
185.33541870117188 0 616.9569
186.12379455566406 0 650.4835
187.0872344970703 0 586.89886
190.08627319335938 0 1763.2234
194.71609497070312 0 456.2221
201.1232147216797 0 1029.0917
203.15078735351562 0 1012.5226
215.13946533203125 0 2950.998
219.1331787109375 0 729.44617
223.1556854248047 0 7051.0557
224.159912109375 0 782.42914
227.13885498046875 0 884.39294
228.97096252441406 0 587.5379
229.1186065673828 0 722.0617
229.15516662597656 0 2458.0369 y Ammonia loss 13
230.16253662109375 0 2598.5027 z 13
231.1708526611328 0 1096.8438
232.0936737060547 0 1648.3867
233.15013122558594 0 6102.823
234.1240234375 0 6432.113
243.13412475585938 0 899.05096
246.1817169189453 0 2021.5842 y 13
250.1428680419922 0 1581.853
251.15065002441406 0 33598.28
252.15370178222656 0 3834.89
253.11883544921875 0 845.75995
261.11968994140625 0 3588.6697
263.1393127441406 0 771.0413
268.1771240234375 0 3875.5645 c 1
269.1796875 0 718.1126
270.12091064453125 0 716.7552
274.1187744140625 0 775.4057
283.15167236328125 0 2083.2017
287.69781494140625 0 506.46625
288.10101318359375 0 752.3076
290.1164855957031 0 1131.3773
292.13287353515625 0 1302.6982
293.13592529296875 0 830.1786
294.1124267578125 0 506.67838
294.1934509277344 0 1335.0793
295.1651611328125 0 657.2556
297.08270263671875 0 547.1876
298.45343017578125 0 628.9602
299.1349182128906 0 859.4524
300.1922302246094 0 4981.324
301.1968994140625 0 716.62537
318.14898681640625 0 736.8742
326.66424560546875 0 8382.163
327.1658630371094 0 2973.6165 z Ammonia loss 7
327.6668701171875 0 635.4288
329.18255615234375 0 850.58105
331.65704345703125 0 1342.807
332.1600646972656 0 869.37213
338.1743469238281 0 885.8173 y 7
339.1786804199219 0 713.5273
340.6619567871094 0 5220.0005 c Ammonia loss 4
341.1636962890625 0 1591.9978 y Ammonia loss 10
341.6631164550781 0 1042.328
349.19036865234375 0 1212.1716
349.6813659667969 0 5304.31 y 10
350.1810607910156 0 1137.373
351.21502685546875 0 1471.7135
352.2203674316406 0 955.14197
355.61468505859375 0 662.1175
361.19903564453125 0 899.958
367.6794738769531 0 725.5573
370.6517333984375 0 739.09375
372.2291259765625 0 1053.3228
374.6923828125 0 1076.0302
375.1695251464844 0 2014.5336
375.2073669433594 0 1200.0254
376.5180969238281 0 1660.7173 y 6
376.68634033203125 0 17555.93 w 9
376.853515625 0 1172.9966
377.18780517578125 0 8230.838
377.6880798339844 0 1899.6107 z Water loss 3
378.2028503417969 0 2129.7527
379.20916748046875 0 9259.147 c Ammonia loss 2
380.212158203125 0 2049.017
381.1519775390625 0 672.9395
381.21160888671875 0 1072.0437
381.6852111816406 0 793.5976 y Water loss 3
382.182861328125 0 704.26196 z 3
383.2054443359375 0 2061.9497
383.7078552246094 0 1509.1301
384.2972106933594 0 1300.137
387.1834716796875 0 834.81024
387.71331787109375 0 1093.8474
388.19061279296875 0 1072.1274
390.1952209472656 0 756.72455
395.22784423828125 0 2411.2969
396.20770263671875 0 987.115
396.2358703613281 0 14929.906 c 2
397.2099609375 0 668.17993 c Ammonia loss 5
397.2397766113281 0 2648.1555
397.70550537109375 0 1215.0999
399.16168212890625 0 778.62946
403.2012634277344 0 1060.2673 z Ammonia loss 5
404.208740234375 0 981.0324
407.1952209472656 0 3757.0342 y 12
408.1679992675781 0 1323.6807
408.2001647949219 0 700.98444 c Ammonia loss 9
408.6693420410156 0 976.62054
409.2186279296875 0 2015.8098
414.2124938964844 0 2604.408 y 9
416.2720031738281 0 1045.4077
417.2196044921875 0 6660.4043
417.7211608886719 0 2522.1738
418.2213439941406 0 978.08795
419.2183532714844 0 983.1219
424.7506103515625 0 1038.8082
425.25042724609375 0 1358.431
428.28759765625 0 5136.112
429.29296875 0 2677.655
440.7338562011719 0 3957.7317
441.2078857421875 0 3804.953 w 8
441.23675537109375 0 1276.5931
441.70904541015625 0 1467.3447
445.16082763671875 0 1168.7117
448.1996154785156 0 2159.361
449.19989013671875 0 1329.5056
450.1701354980469 0 687.4756
452.46905517578125 0 1511.8435 y 1
452.71978759765625 0 2609.4314
452.9687194824219 0 1113.2501
453.2200927734375 0 850.7386
453.2591857910156 0 3812.9502
453.76153564453125 0 2397.3794
454.2632751464844 0 1341.8965
456.2200622558594 0 1125.5283
457.2277526855469 0 899.26263
460.2593688964844 0 4958.9956
461.2071228027344 0 4791.8354 w 11
461.2630615234375 0 1826.5365
462.20843505859375 0 978.33746
466.2420349121094 0 1665.9237
467.2167053222656 0 793.393
467.2489929199219 0 3213.4558
468.2552490234375 0 820.7013
469.2487487792969 0 692.1951 y Water loss 8
470.24114990234375 0 1927.8687 y Water loss 4
470.74163818359375 0 1107.8922
470.89837646484375 0 780.8011 z 4
472.22784423828125 0 652.9407
474.2752990722656 0 2034.8402
475.2821960449219 0 3360.065
476.23486328125 0 3648.2122 y 4
476.4866638183594 0 1823.2579
476.7351989746094 0 960.03143
478.250244140625 0 11426.972 y 8
478.7521667480469 0 6671.2485
479.2521057128906 0 2379.042
480.4938659667969 0 1843.0734
480.69305419921875 0 3317.8567
480.7394104003906 0 23946.19
480.9901123046875 0 20772.02
481.1954650878906 0 1475.2732
481.2408752441406 0 14614.893
481.3145446777344 0 2484.2966
481.49066162109375 0 4359.991
481.6941223144531 0 1056.7919
481.7408142089844 0 2520.866
481.9916687011719 0 860.59064
482.23492431640625 0 1299.6304
488.7770080566406 0 2369.6348
489.28033447265625 0 841.22363
490.2459716796875 0 1619.8723 z Ammonia loss 7
491.2508544921875 0 791.7697
493.25146484375 0 1057.8364 c Water loss 3
494.23614501953125 0 7786.041 c Ammonia loss 3
495.240478515625 0 2802.415
496.2505187988281 0 1589.864
498.2609558105469 0 799.95825
498.7509460449219 0 1735.8363 z 7
499.2519226074219 0 2027.3617
499.7523498535156 0 831.2345
503.2437744140625 0 2739.6946 z Water loss 3
504.24676513671875 0 763.03577
504.76348876953125 0 1786.724
505.23675537109375 0 3791.3828
505.7361145019531 0 4173.266
506.2606506347656 0 6712.2876
506.76092529296875 0 17856.078 y 7
507.26324462890625 0 7802.7397
507.76214599609375 0 3010.2776
508.2652893066406 0 1303.8933
509.25946044921875 0 841.0896
510.7719421386719 0 5206.102
511.26324462890625 0 62110.188 c 3
512.2661743164062 0 15708.417
513.269287109375 0 3173.5752
514.7393798828125 0 1251.4087
515.7406616210938 0 778.75287
519.2728271484375 0 19377.098 z 11
519.7581787109375 0 770.29034
520.2759399414062 0 6184.85
521.2764282226562 0 1180.0479
525.2313232421875 0 1056.36
526.7587890625 0 2709.0383
527.2324829101562 0 5958.9976
527.728271484375 0 1333.6754
527.7658081054688 0 4025.659
528.2672729492188 0 2148.9314
528.769287109375 0 938.8828
530.2203369140625 0 1086.2397
530.2960815429688 0 771.202
533.7661743164062 0 5312.662 w 6
534.2697143554688 0 12471.787
534.7694702148438 0 5326.3716
535.2905883789062 0 11204.904 y 11
536.293701171875 0 2836.9634
537.2791748046875 0 1046.2327
538.2731323242188 0 901.17035
544.2731323242188 0 1198.0043
546.2415161132812 0 1596.5211
547.2791748046875 0 2194.8118 c Ammonia loss 8
547.7759399414062 0 1806.8721
549.2639770507812 0 741.01495
549.6060180664062 0 1122.3152
549.9439086914062 0 843.95654
550.2516479492188 0 13398.437
551.2583618164062 0 5637.5615 y Water loss 2
551.5971069335938 0 2130.14 y Ammonia loss 2
551.9295043945312 0 1278.3273 z 2
552.2612915039062 0 3216.1018
555.2965698242188 0 3204.5718
555.788818359375 0 54477.676 c 8
556.2892456054688 0 25653.81
556.7642822265625 0 6093.085
556.7933349609375 0 7853.2944
557.2708129882812 0 4298.8726 y 2
557.7684936523438 0 1761.8097
559.5968017578125 0 820.109
562.278564453125 0 1834.7611
562.6336059570312 0 1270.217
562.965576171875 0 1214.1343
563.6135864257812 0 765.57745
563.7830200195312 0 1997.6262
564.2745361328125 0 34481.152 y 6
564.7759399414062 0 21280.355
565.27587890625 0 9420.3545
565.7764892578125 0 2080.6033
567.2295532226562 0 820.8216
567.3142700195312 0 1154.0046
568.796630859375 0 4781.5107
569.296142578125 0 4048.8
569.7960815429688 0 1203.845
569.934814453125 0 909.8992
572.95458984375 0 1482.6487
573.2825927734375 0 1263.6084
573.935302734375 0 1863.5021
574.2722778320312 0 1684.248
574.9404907226562 0 1143.2383
575.269775390625 0 767.9009
577.2719116210938 0 770.1108
577.958740234375 0 3220.2441
578.2902221679688 0 3081.888
578.60693359375 0 1310.6567
578.9443969726562 0 1955.187
579.2764282226562 0 759.5909
582.2811279296875 0 843.21704
582.623779296875 0 1276.6244
582.9541015625 0 2661.4854
583.291259765625 0 2380.086
583.6185302734375 0 1593.1622
585.2616577148438 0 840.7314
586.2890014648438 0 842.38715
590.8088989257812 0 1809.6426
591.280029296875 0 34463.285 w 5
591.3510131835938 0 1298.5012
591.7810668945312 0 20520.018
592.2824096679688 0 9576.554
592.355712890625 0 1418.8397
592.6256103515625 0 1700.6188
592.7822265625 0 3588.3992
592.9598388671875 0 2490.5723
593.2817993164062 0 1953.503
595.3208618164062 0 877.0239
597.2940063476562 0 7724.469
597.6271362304688 0 7634.6265 z 1
597.9633178710938 0 4499.35
598.2959594726562 0 3079.486
599.2435913085938 0 879.9924
600.2959594726562 0 757.70044
601.3013916015625 0 2228.11 c Water loss 13
601.6292724609375 0 2632.4897 c Ammonia loss 13
601.9625854492188 0 6748.861
602.2954711914062 0 3039.3403
602.6253051757812 0 1249.0295
602.957763671875 0 3672.3408 y 1
603.2915649414062 0 3302.443
603.62353515625 0 1174.1184
605.646240234375 0 1010.28564
605.961181640625 0 891.69336
606.318115234375 0 1214.2319
606.6372680664062 0 1692.5529
606.9698486328125 0 4726.6396
607.299072265625 0 55002.38 c 13
607.6329345703125 0 60016.11
607.9667358398438 0 31271.318
608.301025390625 0 17653.205
608.6346435546875 0 4824.9844
608.9666748046875 0 1565.0359
609.9839477539062 0 1251.6593
610.3214111328125 0 906.85834
610.9876708984375 0 886.1876
611.3126831054688 0 2238.8037 c Water loss 9
611.6260375976562 0 1026.2001
611.8003540039062 0 5001.576 c Ammonia loss 9
612.3004760742188 0 3736.6414 y Ammonia loss 5
612.8056640625 0 7789.93 z 5
613.3089599609375 0 4870.5737
613.8048706054688 0 1813.2255
614.3106079101562 0 903.7724
614.80517578125 0 782.07074
614.95849609375 0 1656.0219
615.2935180664062 0 2925.1191
615.631591796875 0 2048.2288
615.9636840820312 0 771.59033
616.304443359375 0 1146.7131
616.6356811523438 0 6527.358
616.9680786132812 0 6483.301
617.301513671875 0 3103.522
617.636962890625 0 2305.9626
619.8173828125 0 4517.9272
620.3104248046875 0 62866.12 c 9
620.641357421875 0 4331.5645
620.813232421875 0 58917.86 y 5
620.9623413085938 0 51770.773
621.3089599609375 0 78277.25
621.6416015625 0 36807.844
621.816650390625 0 10520.665
621.9769897460938 0 21248.637
622.3121948242188 0 9332.089
622.6448364257812 0 2825.1533
624.2693481445312 0 2158.9426
625.6514282226562 0 3063.1511
625.9828491210938 0 4594.525
626.3182983398438 0 2140.654
626.6478881835938 0 2104.268
626.9805908203125 0 1882.8839
629.3043823242188 0 1580.2496
629.642578125 0 1432.1523
629.974853515625 0 816.0147
631.3046875 0 900.9398
634.3090209960938 0 1161.2173
634.6514892578125 0 806.6877
634.81884765625 0 796.24915
634.9788208007812 0 3484.9514
635.3139038085938 0 5494.748
635.6453247070312 0 6910.3984
635.9779052734375 0 3155.927
636.3115234375 0 1783.2925
638.3489990234375 0 1686.1462
639.97900390625 0 1697.0148
640.258544921875 0 917.31525
640.3203735351562 0 8238.062
640.6500854492188 0 111473.58
640.9845581054688 0 159780.67
641.3185424804688 0 108781.05
641.6526489257812 0 54572.13
641.9866333007812 0 23514.578
642.3199462890625 0 8270.056
642.6490478515625 0 2473.918
642.9813232421875 0 1509.9237
647.8213500976562 0 847.9124
648.3233032226562 0 2061.0073
648.8269653320312 0 1406.7028
652.3198852539062 0 2891.113
653.3253784179688 0 3144.5662
654.3279418945312 0 1018.9855
655.8209228515625 0 1568.8684
656.3216552734375 0 1039.5897
657.3323364257812 0 1079.9679
660.3543090820312 0 2132.6162
660.856201171875 0 2153.4724
661.3555297851562 0 1208.6982
664.34130859375 0 2182.4045
669.8366088867188 0 1330.2711
670.3344116210938 0 2257.5862
671.8212890625 0 1166.5908
676.3418579101562 0 1854.6986
677.815185546875 0 4486.974
678.3167114257812 0 3128.3193
678.8189086914062 0 888.7932
679.3338012695312 0 1129.5164 c Water loss 4
679.8349609375 0 3663.292
680.3178100585938 0 14547.186 c Ammonia loss 4
680.8370971679688 0 1316.1786
681.3218994140625 0 7954.447 y Ammonia loss 10
682.3358154296875 0 50246.043 z 10
682.8447875976562 0 988.0746
683.3397216796875 0 18124.26
683.4027709960938 0 1151.5724
683.85107421875 0 4154.6133
684.3417358398438 0 7514.174
684.8194580078125 0 639.35486
684.8569946289062 0 840.8549
685.3383178710938 0 1690.7406
693.3286743164062 0 1117.1694 c Ammonia loss 10
694.3203125 0 814.94946
697.3421630859375 0 49004.66 c 4
698.3483276367188 0 26705.828 y 10
699.3505249023438 0 7378.3184
700.3541259765625 0 1327.917
701.3494262695312 0 2750.057
701.8421020507812 0 45605.426 c 10
702.3433837890625 0 35871.656
702.8445434570312 0 15716.554
703.346923828125 0 3777.438
705.3480224609375 0 2726.5647 y Ammonia loss 4
705.8468627929688 0 32166.697 z 4
706.34814453125 0 23822.184
706.8491821289062 0 9618.788
707.3505859375 0 3270.8633
711.3257446289062 0 2622.876
712.326171875 0 3053.5906
713.3665771484375 0 788.5394
713.8560791015625 0 11390.178 y 4
714.359130859375 0 9283.186
714.858154296875 0 4977.312
715.3629760742188 0 959.8518
717.3568115234375 0 1175.095
718.8607788085938 0 864.94257
719.3656616210938 0 2047.372
720.3942260742188 0 1255.001
721.35205078125 0 1201.9489
722.3402099609375 0 903.3997
723.3750610351562 0 820.6808
727.3487548828125 0 1547.9381
727.8432006835938 0 955.41077
728.334716796875 0 862.6707
730.3395385742188 0 2008.3511
731.3938598632812 0 1485.2733
733.36962890625 0 860.02277
733.8555908203125 0 852.2072
740.2991333007812 0 1625.4606
741.3659057617188 0 7155.797
741.8651123046875 0 3614.6487
742.3660888671875 0 1989.8209
743.3720703125 0 852.8343
743.8798217773438 0 968.43854
744.384033203125 0 973.291
747.3627319335938 0 2191.3645
752.365478515625 0 9794.62 w 9
753.3683471679688 0 4130.962
754.3729248046875 0 1568.4119
762.871337890625 0 1186.2839 y Ammonia loss 3
763.3606567382812 0 24419.164 z 3
763.8617553710938 0 18995.193
764.3629760742188 0 8878.855
764.8617553710938 0 4497.726
765.3778076171875 0 1069.7764
765.8898315429688 0 10746.912 c 11
766.3936767578125 0 12009.862
766.8926391601562 0 5073.875
767.405029296875 0 1320.0316
771.3700561523438 0 3722.8975 y 3
771.8719482421875 0 3801.5168
772.3670043945312 0 1286.1428
772.873046875 0 1614.8234
777.359375 0 1659.3812
778.897705078125 0 2151.7307
779.4008178710938 0 2166.7253
785.3707275390625 0 1176.0248
786.3720092773438 0 996.5852
792.3567504882812 0 1107.2311
793.401123046875 0 2201.9746 c Ammonia loss 5
794.4035034179688 0 1191.7095
797.881591796875 0 964.49365
798.3775024414062 0 5317.171 w 2
798.8771362304688 0 6904.0044
799.3765869140625 0 1934.3776
799.8779907226562 0 1127.3055
805.3932495117188 0 3595.4756
805.8956909179688 0 2513.2502
806.39501953125 0 2344.9648
806.8958129882812 0 1657.6421
809.4155883789062 0 2095.6523
809.8936157226562 0 936.3
810.4265747070312 0 97891.22 c 5
811.374755859375 0 34535.336 z 9
811.435302734375 0 28890.002
812.3828735351562 0 25139.78
813.3824462890625 0 10644.498
814.3848876953125 0 3762.0913
825.38671875 0 1372.1953
826.8958129882812 0 3149.7034 y Ammonia loss 2
827.3910522460938 0 36410.523 y 9
827.89111328125 0 26482.922
828.392578125 0 16341.965
828.8935546875 0 3178.3542
829.3910522460938 0 2005.4462
831.3856811523438 0 2889.7056
835.399658203125 0 4566.3574 y 2
835.900634765625 0 4221.7324
836.402587890625 0 2633.4717
836.9000244140625 0 812.90295
839.4205932617188 0 2917.8247
840.4242553710938 0 1765.5056
844.40966796875 0 1396.6196
845.4066162109375 0 1636.5884
845.9000854492188 0 4097.8135
846.3972778320312 0 48713.586 c 12
846.8980712890625 0 37606.805
847.3989868164062 0 23768.338
847.900146484375 0 9735.597
848.4031982421875 0 4084.0503
848.8981323242188 0 965.1595
850.402099609375 0 934.19366
860.42626953125 0 2278.623
861.4268188476562 0 978.0655
866.4310913085938 0 2241.3674
866.9166259765625 0 1986.1284
867.4239501953125 0 5785.6206
868.4185791015625 0 1688.9955
873.924560546875 0 951.4791
879.9430541992188 0 946.6762
880.4390258789062 0 3899.8875
880.935302734375 0 3796.5256
881.4411010742188 0 10026.783
881.9395751953125 0 3968.313
882.44580078125 0 5577.551
883.4454345703125 0 1849.5992
887.4281616210938 0 1186.6006
888.4371948242188 0 3301.4497
888.9390258789062 0 6592.1025
889.3187255859375 0 2088.2031
889.4425659179688 0 6790.6553
889.9439086914062 0 3826.7104
895.42236328125 0 1339.7406 y Ammonia loss 1
895.9194946289062 0 15071.304 z 1
896.4223022460938 0 17073.828
896.9234008789062 0 11046.872
897.4255981445312 0 6180.422
897.926025390625 0 2272.8838
901.9611206054688 0 1107.1704
902.4405517578125 0 3343.5625
902.9409790039062 0 3417.0598
903.4359741210938 0 1882.3798
903.9392700195312 0 1661.3126 y 1
908.4269409179688 0 3585.2693 c Ammonia loss 6
908.9611206054688 0 1008.82776
909.4505615234375 0 3106.1858
909.957275390625 0 2119.5203
910.4439086914062 0 9407.151 c 13
910.9474487304688 0 10751.783
911.4483032226562 0 7294.852
911.9473876953125 0 3755.2803
912.458251953125 0 2937.8733
913.4625854492188 0 1062.9064
914.4619140625 0 1343.8202
914.9596557617188 0 955.58563
917.3107299804688 0 1273.187
922.4583740234375 0 2094.0046 z Ammonia loss 8
922.962158203125 0 6247.713
923.4610595703125 0 8568.721
923.9612426757812 0 5578.8716
924.45849609375 0 5118.234
924.9576416015625 0 2849.039
925.4539184570312 0 14343.058 c 6
926.4547119140625 0 6776.306
927.4552612304688 0 1355.0574
929.9610595703125 0 1041.1799
930.4673461914062 0 3713.1863
930.9632568359375 0 7724.26
931.466552734375 0 42480.664
931.968505859375 0 42594.934
932.4701538085938 0 28613.043
932.9710693359375 0 11783.04
933.4713745117188 0 5453.023
933.9710083007812 0 1521.622
937.9727172851562 0 1130.6765
938.467529296875 0 16995.379 y Ammonia loss 8
938.9713745117188 0 13455.368
939.470947265625 0 22331.545 z 8
939.9705810546875 0 4972.612
940.4773559570312 0 17835.773
941.4826049804688 0 7123.091
942.4812622070312 0 2326.4797
943.4685668945312 0 2422.3462
943.9667358398438 0 1760.3436
944.4694213867188 0 1632.183
945.9583740234375 0 977.10004
951.969970703125 0 1651.8278
952.4692993164062 0 13627.2
952.9662475585938 0 43059.95
953.4667358398438 0 42743.914
953.9677734375 0 25939.213
954.468017578125 0 11590.87
954.9697265625 0 4390.2197
955.4730224609375 0 2920.6047
956.49462890625 0 1571.5989
957.5013427734375 0 818.4263
959.9691772460938 0 1103.9572
960.470703125 0 8373.971
960.974365234375 0 16237.634
961.4779052734375 0 44611.09
961.97900390625 0 38598.56
962.4799194335938 0 25356.121
962.981201171875 0 9914.273
963.4830932617188 0 8551.809
963.974609375 0 1678.9675
964.4679565429688 0 3254.1794 c Water loss 7
967.46337890625 0 1285.0662
973.447265625 0 1173.2399
981.4856567382812 0 3826.9448
982.4750366210938 0 46018.652 c 7
983.4779052734375 0 26536.193
984.480224609375 0 7650.1772
985.4823608398438 0 1750.6263
996.4977416992188 0 2364.0547 z 7
997.5023803710938 0 3971.626
998.5052490234375 0 2222.16
1011.4718017578125 0 1971.4554
1012.5013427734375 0 3863.3516 y 7
1013.5057373046875 0 1915.4991
1051.5576171875 0 1519.9003
1053.5220947265625 0 2759.2632
1054.52587890625 0 1866.9509
1055.5303955078125 0 893.3577
1065.56640625 0 787.306
1066.560302734375 0 1554.0613
1067.552734375 0 1960.8895
1068.5462646484375 0 935.1199
1094.5487060546875 0 2384.5464
1095.5556640625 0 1281.5013
1109.575927734375 0 1812.8527
1110.5694580078125 0 34817.02 c 8
1111.5706787109375 0 25895.072
1112.5380859375 0 27342.666
1113.536376953125 0 13293.144
1114.534912109375 0 6178.3184
1115.548095703125 0 1300.1528
1128.5421142578125 0 1345.7881
1167.6075439453125 0 1080.8196
1180.607421875 0 949.31854
1182.6082763671875 0 1051.7861
1195.5989990234375 0 1271.0247
1196.6053466796875 0 3559.659
1197.6068115234375 0 1215.0269
1198.612060546875 0 925.1531
1212.62841796875 0 1425.8218
1223.59814453125 0 2801.7483 y Ammonia loss 5
1224.6065673828125 0 4677.154 z 5
1225.6123046875 0 12938.424
1226.6153564453125 0 7827.852
1227.6156005859375 0 2973.0981
1228.6163330078125 0 1061.9143
1238.60009765625 0 1124.893
1239.6124267578125 0 11905.106 c 9
1240.6177978515625 0 10418.299 y 5
1241.622314453125 0 6258.097
1242.6224365234375 0 2874.6348
1243.6319580078125 0 1153.4423
1310.6357421875 0 2139.0808
1311.638671875 0 1797.4431
1343.66015625 0 1242.0586
1352.679443359375 0 1080.222
1353.693359375 0 948.2799
1358.6729736328125 0 907.9848
1359.6658935546875 0 2008.487
1360.675537109375 0 1494.598
1361.677001953125 0 739.73193
1386.6632080078125 0 1230.6816
1401.6727294921875 0 1358.5322
1402.676513671875 0 6715.26 c 10
1403.6796875 0 7724.993
1404.6820068359375 0 4684.2837
1405.687255859375 0 1573.3254
1406.6810302734375 0 750.10175
1409.6871337890625 0 846.4635 y Ammonia loss 4
1410.687255859375 0 5631.095 z 4
1411.6898193359375 0 16588.465
1412.6943359375 0 11908.762
1413.697265625 0 6479.413
1414.6988525390625 0 2116.7312
1425.66015625 0 913.0689
1426.69091796875 0 3186.1123 y 4
1427.6990966796875 0 3652.231
1468.6998291015625 0 1038.0824
1471.731689453125 0 1047.2483
1481.7225341796875 0 1042.6975
1486.7659912109375 0 1071.5483
1487.759765625 0 1244.1854
1496.74365234375 0 1032.5594
1497.7373046875 0 969.7282
1514.7506103515625 0 810.2389
1525.716796875 0 3416.4778 z 3
1526.718017578125 0 12309.742
1527.717529296875 0 7510.3276
1528.7183837890625 0 3350.2703
1529.7322998046875 0 2279.2893
1530.766357421875 0 3145.6382 c 11
1531.7713623046875 0 3433.1877
1532.7803955078125 0 2576.4087
1533.7843017578125 0 960.0643
1541.7335205078125 0 773.98944 y 3
1553.71826171875 0 1282.7638
1554.718017578125 0 860.0016
1596.751708984375 0 790.69574
1633.7720947265625 0 1361.1697
1653.7747802734375 0 2560.6992 z 2
1654.77734375 0 9418.94
1655.7781982421875 0 6993.62
1656.781005859375 0 4088.524
1657.7762451171875 0 1340.0116
1675.7669677734375 0 1002.3952
1676.7672119140625 0 1834.2892
1690.7708740234375 0 1236.6825
1691.784423828125 0 4032.382 c 12
1692.7889404296875 0 6495.7793
1693.7943115234375 0 5762.4946
1694.8004150390625 0 3722.2444
1695.786865234375 0 1685.637
1776.8896484375 0 1125.0155
1791.8353271484375 0 3408.2732
1792.842041015625 0 4030.7974
1793.8419189453125 0 2069.9097
1794.8446044921875 0 1151.5435
1804.8873291015625 0 1038.0878
1805.86767578125 0 1051.151
1820.892822265625 0 1833.0986
1821.891845703125 0 2651.7026
1822.8892822265625 0 2422.069
1845.925537109375 0 1119.1343
1846.9306640625 0 1384.4714
1847.919677734375 0 1181.1064
1861.9246826171875 0 2210.6306
1862.930908203125 0 4612.557
1863.9346923828125 0 3820.842
1864.9364013671875 0 2471.5112
1865.9346923828125 0 1556.5503
1876.931884765625 0 1111.279
1877.924072265625 0 1655.9729
1878.928955078125 0 1908.587
1879.94091796875 0 1308.6188
1894.9521484375 0 1147.8499
1903.9422607421875 0 913.4287
1904.9246826171875 0 2397.3884
1905.9266357421875 0 4214.0435
1906.9300537109375 0 3993.2896
1907.93017578125 0 1351.3945
1920.93798828125 0 1503.5486
1921.94677734375 0 4488.6743
1922.949951171875 0 4870.786
1923.9508056640625 0 4872.3086
1924.9527587890625 0 2021.9691

Spectrum Details

|  |  |
| --- | --- |
| Matched peaks? Matched peaks The total absolute number of peaks matched. Additionally in brackets the total fraction of peaks matched and the total number of peaks is shown. | 110 (14.18% of 776) |
| FDR? FDR The false discovery rate estimated for this peptide. It is calculated by matching all theoretical fragments with a non-integer shift with the raw peaks for this spectrum. This is done with 40 different shifts. The resulting percentage is the average number of annotated peaks over the number of annotated peaks with the correct spectrum. | 0.32% |
| Satellite FDR? Satellite FDR See the FDR for details on its calculation. This satellite ion specific FDR only contains the satellite ions (d/w) for I/L/J positions. | 0.00% |
| PSM Score? PSM Score The PSM Score as given by Hecklib to this annotated spectrum. It is shown with three significant figures. | 595 |

#### Spectrum 5829? Spectrum 5829 The raw spectrum of this peptide as annotated by Hecklib. The fragments are coloured according to ion type (see legend). Any peaks with a star '\*' as text can be hovered over to see the full details, first the ion type second the mass shift type. By hovering over the amino acids in the peptide or ions in the legend the corresponding peaks are highlighted. By toggling the 'Unassigned' label you can turn the background (unassigned) peaks on or off in the plot. By updating the slider in the Ion legend you can update the spectrum to only show the top X% of the peaks with labels. The top X% means any peak that is within X% of the highest intensity. By dragging in the spectrum you can zoom in to a specific part of the spectrum and use 'Zoom Out' to get back to the original zoom level. The annotation of the spectrum is based on the given sequence in the peptides file and is done with different software so inconsistencies are likely. The peaks are annotated based on the given sequence, with 20 ppm tolerance.

Copy Data

##### Spectrum 5829 (TSV)

###### Preview

```
Loading example...
```

*Click on the button to copy the data to your clipboard.*

Mz MinMz MaxIntensity Max

WidthHeightPeptide font sizePeptide stroke widthSpectrum font sizeSpectrum stroke widthCompact peptide

Ion legend

wxyz

abcd

OtherUnassignedIon^Charge^_Position_Show for top:%

JHQDWJDGKEYKCKV

01.52e+43.05e+44.57e+46.10e+4

Zoom Out

w^+1^_2_y^+1^_2_z^+1^_2_y^+1^_2_c^+1^_2_z^+3^_8_c^+2^_5_y^+2^_5_w^+2^_6_z^+4^_12_c^+1^_3_c^+1^_3_y^+1^_3_y^+2^_6_w^+2^_7_y^+4^_14_w^+1^_4_y^+2^_7_y^+3^_11_y^+2^_7_c^+1^_4_z^+2^_8_z^+3^_12_y^+2^_8_c^+1^_4_z^+1^_4_w^+2^_9_y^+1^_4_c^+2^_9_y^+3^_13_z^+3^_13_c^+2^_9_z^+2^_9_y^+3^_13_y^+2^_9_w^+2^_10_z^+3^_14_c^+3^_14_c^+3^_14_y^+3^_14_c^+3^_14_c^+2^_10_c^+2^_10_y^+2^_10_z^+2^_10_c^+2^_10_y^+2^_10_c^+1^_5_y^+1^_5_z^+1^_5_c^+1^_5_y^+1^_5_c^+2^_11_y^+2^_11_z^+2^_11_y^+2^_11_w^+2^_12_w^+1^_6_y^+2^_12_z^+2^_12_c^+2^_12_y^+2^_12_c^+1^_6_w^+2^_13_c^+1^_6_z^+1^_6_y^+2^_13_y^+1^_6_y^+2^_13_c^+2^_13_y^+2^_14_z^+2^_14_c^+2^_14_c^+2^_14_z^+1^_7_z^+1^_7_c^+1^_7_y^+1^_7_y^+1^_7_z^+1^_7_y^+1^_7_c^+1^_8_c^+1^_8_z^+1^_8_y^+1^_8_c^+1^_9_y^+1^_10_z^+1^_10_c^+1^_10_y^+1^_10_c^+1^_11_z^+1^_11_y^+1^_11_z^+1^_12_c^+1^_12_z^+1^_13_c^+1^_13_

0769153823083077

Fragment Matches Table

Show background peaks

| Position | Ion type | Intensity | mz Theoretical | mz Error (Th) | mz Error (ppm) | Charge | Series Number |
| --- | --- | --- | --- | --- | --- | --- | --- |
| - | - | 486.5 | 120.1 | - | - | 0 | - |
| - | - | 351.4 | 126.1 | - | - | 0 | - |
| - | - | 2545 | 129.1 | - | - | 0 | - |
| - | - | 2806 | 130.1 | - | - | 0 | - |
| - | - | 623.5 | 130.1 | - | - | 0 | - |
| - | - | 863.2 | 132.1 | - | - | 0 | - |
| - | - | 377.9 | 135.3 | - | - | 0 | - |
| - | - | 808.7 | 136.1 | - | - | 0 | - |
| - | - | 440.5 | 137.1 | - | - | 0 | - |
| - | - | 622.9 | 138.1 | - | - | 0 | - |
| - | - | 440 | 139.2 | - | - | 0 | - |
| - | - | 499.5 | 140.1 | - | - | 0 | - |
| - | - | 552.7 | 146.1 | - | - | 0 | - |
| - | - | 604.4 | 147.1 | - | - | 0 | - |
| - | - | 719.4 | 148.9 | - | - | 0 | - |
| - | - | 457 | 149 | - | - | 0 | - |
| - | - | 4519 | 155.1 | - | - | 0 | - |
| - | - | 493.6 | 155.8 | - | - | 0 | - |
| - | - | 668.2 | 157.1 | - | - | 0 | - |
| - | - | 689.9 | 159.1 | - | - | 0 | - |
| - | - | 2999 | 159.1 | - | - | 0 | - |
| - | - | 1029 | 164.1 | - | - | 0 | - |
| - | - | 556.3 | 165.1 | - | - | 0 | - |
| - | - | 3021 | 166.1 | - | - | 0 | - |
| - | - | 457.3 | 166.7 | - | - | 0 | - |
| - | - | 484.2 | 168.1 | - | - | 0 | - |
| - | - | 2143 | 168.1 | - | - | 0 | - |
| 14 | w | 1011 | 172.1 | 0.0001706 | 0.9911 | +1 | 2 |
| - | - | 504 | 178.1 | - | - | 0 | - |
| - | - | 965 | 183.1 | - | - | 0 | - |
| - | - | 2778 | 185.2 | - | - | 0 | - |
| - | - | 536.5 | 187.1 | - | - | 0 | - |
| - | - | 703.4 | 187.1 | - | - | 0 | - |
| - | - | 787 | 195.1 | - | - | 0 | - |
| - | - | 559.2 | 201.1 | - | - | 0 | - |
| - | - | 723.5 | 203.1 | - | - | 0 | - |
| - | - | 774.4 | 207.1 | - | - | 0 | - |
| - | - | 925.8 | 215.1 | - | - | 0 | - |
| - | - | 2136 | 219.1 | - | - | 0 | - |
| - | - | 486.9 | 219.1 | - | - | 0 | - |
| - | - | 957.7 | 223.1 | - | - | 0 | - |
| - | - | 2299 | 223.2 | - | - | 0 | - |
| - | - | 604.4 | 224.2 | - | - | 0 | - |
| 14 | y | 5432 | 229.2 | 0.0002653 | 1.158 | +1 | 2 |
| 14 | z | 1070 | 230.2 | 0.001285 | 5.583 | +1 | 2 |
| - | - | 943.8 | 233.1 | - | - | 0 | - |
| - | - | 1297 | 233.1 | - | - | 0 | - |
| - | - | 2786 | 234.1 | - | - | 0 | - |
| 14 | y | 962.8 | 246.2 | 0.000493 | 2.002 | +1 | 2 |
| - | - | 916.7 | 250.1 | - | - | 0 | - |
| - | - | 1.455E+04 | 251.1 | - | - | 0 | - |
| - | - | 1164 | 252.2 | - | - | 0 | - |
| - | - | 960.4 | 261.1 | - | - | 0 | - |
| 2 | c | 1124 | 268.2 | 0.0003226 | 1.203 | +1 | 2 |
| - | - | 509.7 | 270.2 | - | - | 0 | - |
| - | - | 490.2 | 272.5 | - | - | 0 | - |
| - | - | 614.9 | 278.2 | - | - | 0 | - |
| - | - | 1001 | 283.2 | - | - | 0 | - |
| - | - | 735.6 | 285.2 | - | - | 0 | - |
| - | - | 645.9 | 294.2 | - | - | 0 | - |
| - | - | 1859 | 295.2 | - | - | 0 | - |
| - | - | 596.4 | 298.9 | - | - | 0 | - |
| - | - | 2197 | 300.2 | - | - | 0 | - |
| - | - | 1200 | 310.1 | - | - | 0 | - |
| - | - | 1259 | 312.2 | - | - | 0 | - |
| - | - | 619.2 | 315.2 | - | - | 0 | - |
| - | - | 4322 | 326.7 | - | - | 0 | - |
| 8 | z | 1430 | 327.2 | 0.001914 | 5.851 | +3 | 8 |
| - | - | 819 | 328.2 | - | - | 0 | - |
| - | - | 549.3 | 332.2 | - | - | 0 | - |
| 5 | c | 1797 | 340.7 | 0.000323 | 0.9481 | +2 | 5 |
| - | - | 857.5 | 341.2 | - | - | 0 | - |
| - | - | 1268 | 345.2 | - | - | 0 | - |
| 11 | y | 1489 | 349.7 | 0.003115 | 8.907 | +2 | 5 |
| - | - | 703.5 | 350.2 | - | - | 0 | - |
| - | - | 653.6 | 350.7 | - | - | 0 | - |
| - | - | 814.5 | 356.2 | - | - | 0 | - |
| - | - | 572 | 364.2 | - | - | 0 | - |
| - | - | 830 | 365.1 | - | - | 0 | - |
| - | - | 644.4 | 367.2 | - | - | 0 | - |
| - | - | 785.9 | 371.7 | - | - | 0 | - |
| - | - | 641.4 | 375.2 | - | - | 0 | - |
| 10 | w | 6740 | 376.7 | 0.00269 | 7.142 | +2 | 6 |
| - | - | 2269 | 377.2 | - | - | 0 | - |
| 4 | z | 986.1 | 377.7 | 0.005987 | 15.85 | +4 | 12 |
| - | - | 628.8 | 378.2 | - | - | 0 | - |
| 3 | c | 4284 | 379.2 | 0.000761 | 2.007 | +1 | 3 |
| - | - | 755.2 | 380.2 | - | - | 0 | - |
| - | - | 1356 | 383.2 | - | - | 0 | - |
| - | - | 660.7 | 387.2 | - | - | 0 | - |
| - | - | 610.6 | 387.2 | - | - | 0 | - |
| - | - | 724.6 | 394.2 | - | - | 0 | - |
| - | - | 845.2 | 396.2 | - | - | 0 | - |
| 3 | c | 6893 | 396.2 | 0.0004546 | 1.147 | +1 | 3 |
| - | - | 787 | 397.2 | - | - | 0 | - |
| - | - | 695.3 | 402.2 | - | - | 0 | - |
| - | - | 2194 | 402.7 | - | - | 0 | - |
| - | - | 799.3 | 403.2 | - | - | 0 | - |
| - | - | 571.6 | 406.2 | - | - | 0 | - |
| 13 | y | 2299 | 407.2 | 0.005487 | 13.47 | +1 | 3 |
| 10 | y | 608.3 | 414.2 | 0.007998 | 19.31 | +2 | 6 |
| - | - | 695.9 | 414.5 | - | - | 0 | - |
| - | - | 875.7 | 416.2 | - | - | 0 | - |
| - | - | 2239 | 417.2 | - | - | 0 | - |
| - | - | 943.5 | 417.7 | - | - | 0 | - |
| - | - | 571.9 | 425.2 | - | - | 0 | - |
| - | - | 2350 | 428.3 | - | - | 0 | - |
| - | - | 777.9 | 429.3 | - | - | 0 | - |
| - | - | 1182 | 431.1 | - | - | 0 | - |
| - | - | 931.5 | 440.7 | - | - | 0 | - |
| 9 | w | 1071 | 441.2 | 0.003418 | 7.747 | +2 | 7 |
| - | - | 809 | 441.2 | - | - | 0 | - |
| - | - | 824 | 448.2 | - | - | 0 | - |
| - | - | 698.7 | 448.3 | - | - | 0 | - |
| 2 | y | 1029 | 452.5 | 2.349E-05 | 0.05192 | +4 | 14 |
| - | - | 2217 | 453.3 | - | - | 0 | - |
| - | - | 950.9 | 453.8 | - | - | 0 | - |
| - | - | 2451 | 460.3 | - | - | 0 | - |
| 12 | w | 2392 | 461.2 | 0.003905 | 8.468 | +1 | 4 |
| - | - | 849.9 | 461.3 | - | - | 0 | - |
| - | - | 890.1 | 464.2 | - | - | 0 | - |
| - | - | 1240 | 466.2 | - | - | 0 | - |
| - | - | 1610 | 467.2 | - | - | 0 | - |
| - | - | 559.4 | 467.7 | - | - | 0 | - |
| - | - | 835.6 | 468.3 | - | - | 0 | - |
| 9 | y | 1862 | 469.2 | 0.005503 | 11.73 | +2 | 7 |
| 5 | y | 1205 | 470.2 | 0.00442 | 9.399 | +3 | 11 |
| - | - | 1147 | 472.8 | - | - | 0 | - |
| - | - | 3918 | 474.3 | - | - | 0 | - |
| - | - | 1888 | 475.3 | - | - | 0 | - |
| - | - | 689.1 | 476.2 | - | - | 0 | - |
| - | - | 1248 | 476.2 | - | - | 0 | - |
| - | - | 1407 | 476.5 | - | - | 0 | - |
| 9 | y | 4169 | 478.3 | 0.003106 | 6.495 | +2 | 7 |
| - | - | 2239 | 478.8 | - | - | 0 | - |
| - | - | 927.9 | 479.2 | - | - | 0 | - |
| - | - | 881.8 | 480.3 | - | - | 0 | - |
| - | - | 2605 | 480.5 | - | - | 0 | - |
| - | - | 3261 | 480.7 | - | - | 0 | - |
| - | - | 7715 | 480.7 | - | - | 0 | - |
| - | - | 9181 | 481 | - | - | 0 | - |
| - | - | 1475 | 481.2 | - | - | 0 | - |
| - | - | 4684 | 481.2 | - | - | 0 | - |
| - | - | 1.647E+04 | 481.3 | - | - | 0 | - |
| - | - | 856.3 | 481.3 | - | - | 0 | - |
| - | - | 2183 | 481.5 | - | - | 0 | - |
| - | - | 1070 | 481.7 | - | - | 0 | - |
| - | - | 7791 | 481.8 | - | - | 0 | - |
| - | - | 680.4 | 482.2 | - | - | 0 | - |
| - | - | 1365 | 482.3 | - | - | 0 | - |
| - | - | 1176 | 484.3 | - | - | 0 | - |
| 4 | c | 3499 | 494.2 | 0.0002687 | 0.5436 | +1 | 4 |
| 8 | z | 1824 | 498.8 | 0.002035 | 4.079 | +2 | 8 |
| - | - | 930.3 | 499.3 | - | - | 0 | - |
| 4 | z | 849.1 | 503.2 | 0.007539 | 14.98 | +3 | 12 |
| - | - | 1996 | 505.7 | - | - | 0 | - |
| - | - | 2139 | 506.3 | - | - | 0 | - |
| 8 | y | 4922 | 506.8 | 0.003096 | 6.109 | +2 | 8 |
| - | - | 3286 | 507.3 | - | - | 0 | - |
| - | - | 1526 | 507.8 | - | - | 0 | - |
| - | - | 2204 | 510.8 | - | - | 0 | - |
| 4 | c | 2.175E+04 | 511.3 | 0.0001759 | 0.3441 | +1 | 4 |
| - | - | 5267 | 512.3 | - | - | 0 | - |
| - | - | 793.1 | 514.7 | - | - | 0 | - |
| 12 | z | 8061 | 519.3 | 0.005096 | 9.814 | +1 | 4 |
| - | - | 3286 | 520.3 | - | - | 0 | - |
| - | - | 1668 | 526.8 | - | - | 0 | - |
| - | - | 2482 | 527.2 | - | - | 0 | - |
| - | - | 4807 | 527.8 | - | - | 0 | - |
| - | - | 705.9 | 528.2 | - | - | 0 | - |
| - | - | 2927 | 528.3 | - | - | 0 | - |
| 7 | w | 3149 | 533.8 | 0.002336 | 4.376 | +2 | 9 |
| - | - | 3873 | 534.3 | - | - | 0 | - |
| - | - | 2441 | 534.8 | - | - | 0 | - |
| 12 | y | 4709 | 535.3 | 0.005937 | 11.09 | +1 | 4 |
| - | - | 786.8 | 536.3 | - | - | 0 | - |
| - | - | 785.9 | 537.3 | - | - | 0 | - |
| - | - | 590.6 | 538.3 | - | - | 0 | - |
| - | - | 807.6 | 542.3 | - | - | 0 | - |
| 9 | c | 686.2 | 547.3 | 0.001957 | 3.577 | +2 | 9 |
| - | - | 564.6 | 549.9 | - | - | 0 | - |
| - | - | 4002 | 550.3 | - | - | 0 | - |
| 3 | y | 1835 | 551.3 | 0.009378 | 17.01 | +3 | 13 |
| 3 | z | 1110 | 551.9 | 0.0005342 | 0.9678 | +3 | 13 |
| - | - | 1231 | 552.3 | - | - | 0 | - |
| - | - | 3955 | 555.3 | - | - | 0 | - |
| 9 | c | 2.182E+04 | 555.8 | 0.0001575 | 0.2834 | +2 | 9 |
| 7 | z | 2243 | 556.3 | 0.009586 | 17.23 | +2 | 9 |
| - | - | 9590 | 556.3 | - | - | 0 | - |
| - | - | 2274 | 556.8 | - | - | 0 | - |
| - | - | 1842 | 556.8 | - | - | 0 | - |
| 3 | y | 1903 | 557.3 | 0.001242 | 2.228 | +3 | 13 |
| - | - | 1147 | 557.8 | - | - | 0 | - |
| - | - | 1294 | 561.3 | - | - | 0 | - |
| - | - | 1655 | 563.8 | - | - | 0 | - |
| 7 | y | 1.504E+04 | 564.3 | 0.002224 | 3.941 | +2 | 9 |
| - | - | 7537 | 564.8 | - | - | 0 | - |
| - | - | 2920 | 565.3 | - | - | 0 | - |
| - | - | 1465 | 565.8 | - | - | 0 | - |
| - | - | 795.7 | 567.8 | - | - | 0 | - |
| - | - | 2683 | 568.8 | - | - | 0 | - |
| - | - | 1855 | 569.3 | - | - | 0 | - |
| - | - | 695.9 | 571.3 | - | - | 0 | - |
| - | - | 600.8 | 574.3 | - | - | 0 | - |
| - | - | 664.3 | 576.8 | - | - | 0 | - |
| - | - | 1103 | 578 | - | - | 0 | - |
| - | - | 778.7 | 578.6 | - | - | 0 | - |
| - | - | 739.5 | 578.9 | - | - | 0 | - |
| - | - | 1027 | 582.3 | - | - | 0 | - |
| - | - | 746 | 588.3 | - | - | 0 | - |
| - | - | 2211 | 590.8 | - | - | 0 | - |
| 6 | w | 1.193E+04 | 591.3 | 0.001891 | 3.198 | +2 | 10 |
| - | - | 8939 | 591.8 | - | - | 0 | - |
| - | - | 2850 | 592.3 | - | - | 0 | - |
| - | - | 746.3 | 592.4 | - | - | 0 | - |
| - | - | 1135 | 592.8 | - | - | 0 | - |
| - | - | 646.9 | 593 | - | - | 0 | - |
| - | - | 883.5 | 595.3 | - | - | 0 | - |
| - | - | 805.8 | 595.7 | - | - | 0 | - |
| - | - | 2216 | 597.3 | - | - | 0 | - |
| 2 | z | 3212 | 597.6 | 0.01147 | 19.18 | +3 | 14 |
| - | - | 1374 | 598 | - | - | 0 | - |
| - | - | 897.2 | 598.3 | - | - | 0 | - |
| 14 | c | 1582 | 601.3 | 0.005789 | 9.628 | +3 | 14 |
| 14 | c | 1393 | 601.6 | 0.002186 | 3.633 | +3 | 14 |
| - | - | 1475 | 602 | - | - | 0 | - |
| - | - | 1218 | 602.3 | - | - | 0 | - |
| 2 | y | 1120 | 603 | 0.0005339 | 0.8855 | +3 | 14 |
| - | - | 1062 | 603.3 | - | - | 0 | - |
| - | - | 859.4 | 603.6 | - | - | 0 | - |
| - | - | 601.1 | 606.3 | - | - | 0 | - |
| - | - | 766.5 | 606.6 | - | - | 0 | - |
| - | - | 3714 | 607 | - | - | 0 | - |
| 14 | c | 1.929E+04 | 607.3 | 0.001394 | 2.296 | +3 | 14 |
| - | - | 2.134E+04 | 607.6 | - | - | 0 | - |
| - | - | 1.328E+04 | 608 | - | - | 0 | - |
| - | - | 6283 | 608.3 | - | - | 0 | - |
| - | - | 2998 | 608.6 | - | - | 0 | - |
| 10 | c | 1246 | 611.3 | 0.003858 | 6.311 | +2 | 10 |
| 10 | c | 1292 | 611.8 | 0.002756 | 4.504 | +2 | 10 |
| 6 | y | 805 | 612.3 | 0.004663 | 7.616 | +2 | 10 |
| 6 | z | 2362 | 612.8 | 0.002912 | 4.751 | +2 | 10 |
| - | - | 1491 | 613.3 | - | - | 0 | - |
| - | - | 905.5 | 615.3 | - | - | 0 | - |
| - | - | 1354 | 615.6 | - | - | 0 | - |
| - | - | 3269 | 616.6 | - | - | 0 | - |
| - | - | 2113 | 617 | - | - | 0 | - |
| - | - | 1187 | 617.3 | - | - | 0 | - |
| - | - | 876.6 | 618.3 | - | - | 0 | - |
| - | - | 6495 | 618.4 | - | - | 0 | - |
| - | - | 1198 | 619.4 | - | - | 0 | - |
| - | - | 3416 | 619.8 | - | - | 0 | - |
| 10 | c | 2.144E+04 | 620.3 | 0.001139 | 1.836 | +2 | 10 |
| - | - | 1837 | 620.6 | - | - | 0 | - |
| 6 | y | 2.349E+04 | 620.8 | 0.005804 | 9.349 | +2 | 10 |
| - | - | 1.148E+04 | 621 | - | - | 0 | - |
| - | - | 2.363E+04 | 621.3 | - | - | 0 | - |
| - | - | 1.418E+04 | 621.6 | - | - | 0 | - |
| - | - | 3850 | 621.8 | - | - | 0 | - |
| - | - | 6487 | 622 | - | - | 0 | - |
| - | - | 4185 | 622.3 | - | - | 0 | - |
| - | - | 1012 | 622.6 | - | - | 0 | - |
| - | - | 2003 | 625.6 | - | - | 0 | - |
| - | - | 2069 | 626 | - | - | 0 | - |
| - | - | 1732 | 626.3 | - | - | 0 | - |
| - | - | 765.8 | 626.6 | - | - | 0 | - |
| - | - | 656.5 | 629.6 | - | - | 0 | - |
| - | - | 919.3 | 635 | - | - | 0 | - |
| - | - | 2444 | 635.3 | - | - | 0 | - |
| - | - | 1595 | 635.6 | - | - | 0 | - |
| - | - | 1453 | 636 | - | - | 0 | - |
| - | - | 825.7 | 636.6 | - | - | 0 | - |
| - | - | 6451 | 640.3 | - | - | 0 | - |
| - | - | 4.078E+04 | 640.6 | - | - | 0 | - |
| - | - | 6.037E+04 | 641 | - | - | 0 | - |
| - | - | 4.223E+04 | 641.3 | - | - | 0 | - |
| - | - | 2.071E+04 | 641.7 | - | - | 0 | - |
| - | - | 6612 | 642 | - | - | 0 | - |
| - | - | 3008 | 642.3 | - | - | 0 | - |
| - | - | 1176 | 642.7 | - | - | 0 | - |
| - | - | 704.8 | 648.8 | - | - | 0 | - |
| - | - | 1390 | 652.3 | - | - | 0 | - |
| - | - | 1503 | 653.3 | - | - | 0 | - |
| - | - | 690.1 | 660.9 | - | - | 0 | - |
| - | - | 559.6 | 670.3 | - | - | 0 | - |
| - | - | 927.1 | 676.3 | - | - | 0 | - |
| - | - | 725.3 | 677.3 | - | - | 0 | - |
| - | - | 1768 | 677.8 | - | - | 0 | - |
| - | - | 1603 | 678.3 | - | - | 0 | - |
| - | - | 1563 | 679.8 | - | - | 0 | - |
| 5 | c | 6483 | 680.3 | 0.001565 | 2.3 | +1 | 5 |
| 11 | y | 3353 | 681.3 | 0.01171 | 17.19 | +1 | 5 |
| 11 | z | 1.796E+04 | 682.3 | 0.005375 | 7.878 | +1 | 5 |
| - | - | 8865 | 683.3 | - | - | 0 | - |
| - | - | 914.5 | 683.9 | - | - | 0 | - |
| - | - | 3018 | 684.3 | - | - | 0 | - |
| 5 | c | 2.013E+04 | 697.3 | 0.0008146 | 1.168 | +1 | 5 |
| 11 | y | 1.333E+04 | 698.4 | 0.01159 | 16.59 | +1 | 5 |
| - | - | 2935 | 699.4 | - | - | 0 | - |
| - | - | 892.2 | 700.4 | - | - | 0 | - |
| - | - | 3500 | 701.3 | - | - | 0 | - |
| 11 | c | 1.901E+04 | 701.8 | 0.0002363 | 0.3366 | +2 | 11 |
| - | - | 1.471E+04 | 702.3 | - | - | 0 | - |
| - | - | 6085 | 702.8 | - | - | 0 | - |
| - | - | 1896 | 703.3 | - | - | 0 | - |
| 5 | y | 1113 | 705.3 | 0.006144 | 8.711 | +2 | 11 |
| 5 | z | 9959 | 705.8 | 0.002468 | 3.496 | +2 | 11 |
| - | - | 6795 | 706.3 | - | - | 0 | - |
| - | - | 4232 | 706.8 | - | - | 0 | - |
| - | - | 1726 | 707.3 | - | - | 0 | - |
| - | - | 974.1 | 712.3 | - | - | 0 | - |
| 5 | y | 4749 | 713.9 | 0.002431 | 3.405 | +2 | 11 |
| - | - | 2752 | 714.4 | - | - | 0 | - |
| - | - | 1534 | 714.9 | - | - | 0 | - |
| - | - | 886.9 | 717.9 | - | - | 0 | - |
| - | - | 850.8 | 719.4 | - | - | 0 | - |
| - | - | 842.2 | 727.4 | - | - | 0 | - |
| - | - | 805.5 | 731.4 | - | - | 0 | - |
| - | - | 2887 | 733.4 | - | - | 0 | - |
| - | - | 977.5 | 734.4 | - | - | 0 | - |
| 4 | w | 700.2 | 740.9 | 0.003762 | 5.077 | +2 | 12 |
| - | - | 4528 | 741.4 | - | - | 0 | - |
| - | - | 1253 | 741.9 | - | - | 0 | - |
| - | - | 1614 | 742.4 | - | - | 0 | - |
| 10 | w | 3759 | 752.4 | 0.005001 | 6.647 | +1 | 6 |
| - | - | 1594 | 753.4 | - | - | 0 | - |
| 4 | y | 2030 | 762.9 | 0.007687 | 10.08 | +2 | 12 |
| 4 | z | 9624 | 763.4 | 0.0006196 | 0.8117 | +2 | 12 |
| - | - | 5982 | 763.9 | - | - | 0 | - |
| - | - | 3913 | 764.4 | - | - | 0 | - |
| - | - | 2020 | 764.9 | - | - | 0 | - |
| 12 | c | 5780 | 765.9 | 0.0014 | 1.828 | +2 | 12 |
| - | - | 3990 | 766.4 | - | - | 0 | - |
| - | - | 1583 | 766.9 | - | - | 0 | - |
| - | - | 677.1 | 767.4 | - | - | 0 | - |
| 4 | y | 1132 | 771.4 | 0.0008264 | 1.071 | +2 | 12 |
| - | - | 1234 | 771.9 | - | - | 0 | - |
| - | - | 901.8 | 778.9 | - | - | 0 | - |
| - | - | 759.7 | 786.4 | - | - | 0 | - |
| - | - | 2623 | 789.4 | - | - | 0 | - |
| 6 | c | 920 | 793.4 | 0.004985 | 6.283 | +1 | 6 |
| - | - | 889.3 | 794.4 | - | - | 0 | - |
| - | - | 1514 | 797.9 | - | - | 0 | - |
| 3 | w | 3828 | 798.4 | 0.002386 | 2.988 | +2 | 13 |
| - | - | 1180 | 798.9 | - | - | 0 | - |
| - | - | 937.8 | 799.4 | - | - | 0 | - |
| - | - | 1420 | 804.4 | - | - | 0 | - |
| - | - | 903.9 | 806.4 | - | - | 0 | - |
| 6 | c | 3.538E+04 | 810.4 | 0.000589 | 0.7268 | +1 | 6 |
| 10 | z | 1.634E+04 | 811.4 | 0.007441 | 9.171 | +1 | 6 |
| - | - | 1.048E+04 | 811.4 | - | - | 0 | - |
| - | - | 1.219E+04 | 812.4 | - | - | 0 | - |
| - | - | 2842 | 813.4 | - | - | 0 | - |
| - | - | 908.5 | 814.4 | - | - | 0 | - |
| 3 | y | 3412 | 826.9 | 0.00855 | 10.34 | +2 | 13 |
| 10 | y | 1.37E+04 | 827.4 | 0.01054 | 12.74 | +1 | 6 |
| - | - | 9733 | 827.9 | - | - | 0 | - |
| - | - | 6078 | 828.4 | - | - | 0 | - |
| - | - | 1907 | 828.9 | - | - | 0 | - |
| - | - | 2079 | 829.4 | - | - | 0 | - |
| 3 | y | 1078 | 835.4 | 0.007532 | 9.016 | +2 | 13 |
| - | - | 1563 | 835.9 | - | - | 0 | - |
| - | - | 1045 | 839.4 | - | - | 0 | - |
| - | - | 1602 | 844.4 | - | - | 0 | - |
| - | - | 806.8 | 845.4 | - | - | 0 | - |
| - | - | 3081 | 845.9 | - | - | 0 | - |
| 13 | c | 1.758E+04 | 846.4 | 0.001845 | 2.18 | +2 | 13 |
| - | - | 1.45E+04 | 846.9 | - | - | 0 | - |
| - | - | 8775 | 847.4 | - | - | 0 | - |
| - | - | 3231 | 847.9 | - | - | 0 | - |
| - | - | 799.4 | 848.4 | - | - | 0 | - |
| - | - | 1133 | 860.4 | - | - | 0 | - |
| - | - | 1102 | 866.4 | - | - | 0 | - |
| - | - | 4189 | 867.4 | - | - | 0 | - |
| - | - | 2247 | 868.4 | - | - | 0 | - |
| - | - | 641.9 | 871.4 | - | - | 0 | - |
| - | - | 756.1 | 874.4 | - | - | 0 | - |
| - | - | 827.9 | 876.5 | - | - | 0 | - |
| - | - | 1052 | 880.4 | - | - | 0 | - |
| - | - | 1171 | 880.9 | - | - | 0 | - |
| - | - | 4075 | 881.4 | - | - | 0 | - |
| - | - | 1667 | 881.9 | - | - | 0 | - |
| - | - | 1680 | 882.5 | - | - | 0 | - |
| - | - | 2269 | 888.4 | - | - | 0 | - |
| - | - | 2752 | 888.9 | - | - | 0 | - |
| - | - | 2985 | 889.4 | - | - | 0 | - |
| - | - | 1747 | 889.9 | - | - | 0 | - |
| - | - | 1602 | 894.5 | - | - | 0 | - |
| 2 | y | 1323 | 895.4 | 0.007109 | 7.94 | +2 | 14 |
| - | - | 1120 | 895.5 | - | - | 0 | - |
| 2 | z | 5963 | 895.9 | 0.0005722 | 0.6387 | +2 | 14 |
| - | - | 8108 | 896.4 | - | - | 0 | - |
| - | - | 4789 | 896.9 | - | - | 0 | - |
| - | - | 754.5 | 901.5 | - | - | 0 | - |
| 14 | c | 1237 | 901.9 | 0.01754 | 19.44 | +2 | 14 |
| - | - | 2171 | 902.9 | - | - | 0 | - |
| - | - | 785.7 | 904.4 | - | - | 0 | - |
| - | - | 895.7 | 909.5 | - | - | 0 | - |
| - | - | 1340 | 910 | - | - | 0 | - |
| 14 | c | 2893 | 910.4 | 0.002065 | 2.268 | +2 | 14 |
| - | - | 4428 | 910.9 | - | - | 0 | - |
| - | - | 3601 | 911.4 | - | - | 0 | - |
| - | - | 1211 | 912 | - | - | 0 | - |
| - | - | 1102 | 912.5 | - | - | 0 | - |
| - | - | 1543 | 919.5 | - | - | 0 | - |
| - | - | 901.2 | 920.5 | - | - | 0 | - |
| 9 | z | 782.1 | 921.5 | 4.252E-05 | 0.04615 | +1 | 7 |
| 9 | z | 2091 | 922.5 | 0.01149 | 12.45 | +1 | 7 |
| - | - | 2614 | 923 | - | - | 0 | - |
| - | - | 4467 | 923.5 | - | - | 0 | - |
| - | - | 3383 | 924 | - | - | 0 | - |
| - | - | 3314 | 924.5 | - | - | 0 | - |
| 7 | c | 5897 | 925.5 | 0.0003104 | 0.3354 | +1 | 7 |
| - | - | 2525 | 926.5 | - | - | 0 | - |
| - | - | 787.5 | 927.5 | - | - | 0 | - |
| - | - | 1283 | 930.5 | - | - | 0 | - |
| - | - | 5801 | 931 | - | - | 0 | - |
| - | - | 1.862E+04 | 931.5 | - | - | 0 | - |
| - | - | 1.606E+04 | 932 | - | - | 0 | - |
| - | - | 9177 | 932.5 | - | - | 0 | - |
| - | - | 4098 | 933 | - | - | 0 | - |
| - | - | 1484 | 933.5 | - | - | 0 | - |
| 9 | y | 889 | 937.5 | 0.01443 | 15.4 | +1 | 7 |
| - | - | 1452 | 938 | - | - | 0 | - |
| 9 | y | 9056 | 938.5 | 0.001684 | 1.794 | +1 | 7 |
| - | - | 7549 | 939 | - | - | 0 | - |
| 9 | z | 1.041E+04 | 939.5 | 0.008471 | 9.017 | +1 | 7 |
| - | - | 2514 | 940 | - | - | 0 | - |
| - | - | 7399 | 940.5 | - | - | 0 | - |
| - | - | 2816 | 941.5 | - | - | 0 | - |
| - | - | 844.1 | 942.5 | - | - | 0 | - |
| - | - | 2613 | 943.5 | - | - | 0 | - |
| - | - | 2506 | 944.5 | - | - | 0 | - |
| - | - | 1509 | 945.5 | - | - | 0 | - |
| - | - | 2522 | 946.5 | - | - | 0 | - |
| - | - | 1563 | 947.5 | - | - | 0 | - |
| - | - | 1070 | 952 | - | - | 0 | - |
| - | - | 7533 | 952.5 | - | - | 0 | - |
| - | - | 2.029E+04 | 953 | - | - | 0 | - |
| - | - | 1.554E+04 | 953.5 | - | - | 0 | - |
| - | - | 9415 | 954 | - | - | 0 | - |
| - | - | 4822 | 954.5 | - | - | 0 | - |
| - | - | 1576 | 955 | - | - | 0 | - |
| 9 | y | 1478 | 955.5 | 0.0148 | 15.49 | +1 | 7 |
| - | - | 720.3 | 956.5 | - | - | 0 | - |
| - | - | 2097 | 958.5 | - | - | 0 | - |
| - | - | 2061 | 959.5 | - | - | 0 | - |
| - | - | 1286 | 960 | - | - | 0 | - |
| - | - | 5340 | 960.5 | - | - | 0 | - |
| - | - | 7888 | 961 | - | - | 0 | - |
| - | - | 1264 | 961.4 | - | - | 0 | - |
| - | - | 2.437E+04 | 961.5 | - | - | 0 | - |
| - | - | 1.738E+04 | 962 | - | - | 0 | - |
| - | - | 1.055E+04 | 962.5 | - | - | 0 | - |
| - | - | 1137 | 962.6 | - | - | 0 | - |
| - | - | 4668 | 963 | - | - | 0 | - |
| - | - | 2943 | 963.5 | - | - | 0 | - |
| - | - | 903.8 | 964 | - | - | 0 | - |
| 8 | c | 1126 | 964.5 | 0.0009555 | 0.9907 | +1 | 8 |
| - | - | 2434 | 981.5 | - | - | 0 | - |
| 8 | c | 1.665E+04 | 982.5 | 0.0003817 | 0.3885 | +1 | 8 |
| - | - | 8877 | 983.5 | - | - | 0 | - |
| - | - | 2507 | 984.5 | - | - | 0 | - |
| 8 | z | 705.8 | 996.5 | 3.356E-05 | 0.03368 | +1 | 8 |
| - | - | 1875 | 997.5 | - | - | 0 | - |
| - | - | 1112 | 998.5 | - | - | 0 | - |
| 8 | y | 1776 | 1013 | 0.008742 | 8.634 | +1 | 8 |
| - | - | 691.3 | 1014 | - | - | 0 | - |
| - | - | 860.9 | 1052 | - | - | 0 | - |
| - | - | 924.1 | 1054 | - | - | 0 | - |
| - | - | 1006 | 1055 | - | - | 0 | - |
| - | - | 1392 | 1067 | - | - | 0 | - |
| - | - | 1523 | 1095 | - | - | 0 | - |
| - | - | 1522 | 1110 | - | - | 0 | - |
| 9 | c | 1.254E+04 | 1111 | 0.0008776 | 0.7903 | +1 | 9 |
| - | - | 1.044E+04 | 1112 | - | - | 0 | - |
| - | - | 1.102E+04 | 1113 | - | - | 0 | - |
| - | - | 5355 | 1114 | - | - | 0 | - |
| - | - | 1771 | 1115 | - | - | 0 | - |
| - | - | 835.3 | 1129 | - | - | 0 | - |
| - | - | 608.4 | 1181 | - | - | 0 | - |
| - | - | 797.4 | 1197 | - | - | 0 | - |
| 6 | y | 1040 | 1224 | 0.006959 | 5.687 | +1 | 10 |
| 6 | z | 2277 | 1225 | 0.000965 | 0.788 | +1 | 10 |
| - | - | 6342 | 1226 | - | - | 0 | - |
| - | - | 3837 | 1227 | - | - | 0 | - |
| - | - | 699.7 | 1228 | - | - | 0 | - |
| - | - | 672 | 1239 | - | - | 0 | - |
| 10 | c | 3682 | 1240 | 0.001864 | 1.503 | +1 | 10 |
| 6 | y | 4917 | 1241 | 0.01178 | 9.493 | +1 | 10 |
| - | - | 2633 | 1242 | - | - | 0 | - |
| - | - | 682.3 | 1243 | - | - | 0 | - |
| - | - | 780.1 | 1250 | - | - | 0 | - |
| - | - | 855.3 | 1311 | - | - | 0 | - |
| - | - | 1094 | 1312 | - | - | 0 | - |
| - | - | 976 | 1359 | - | - | 0 | - |
| - | - | 863.6 | 1360 | - | - | 0 | - |
| - | - | 731.1 | 1368 | - | - | 0 | - |
| 11 | c | 2647 | 1403 | 0.007383 | 5.263 | +1 | 11 |
| - | - | 3519 | 1404 | - | - | 0 | - |
| - | - | 2405 | 1405 | - | - | 0 | - |
| 5 | z | 2617 | 1411 | 0.002585 | 1.832 | +1 | 11 |
| - | - | 5457 | 1412 | - | - | 0 | - |
| - | - | 3965 | 1413 | - | - | 0 | - |
| - | - | 2142 | 1414 | - | - | 0 | - |
| - | - | 1168 | 1415 | - | - | 0 | - |
| 5 | y | 1390 | 1427 | 0.02224 | 15.59 | +1 | 11 |
| - | - | 988.2 | 1428 | - | - | 0 | - |
| - | - | 734.3 | 1429 | - | - | 0 | - |
| 4 | z | 2142 | 1526 | 0.001032 | 0.6766 | +1 | 12 |
| - | - | 4791 | 1527 | - | - | 0 | - |
| - | - | 2902 | 1528 | - | - | 0 | - |
| - | - | 1003 | 1529 | - | - | 0 | - |
| 12 | c | 1647 | 1531 | 0.001653 | 1.08 | +1 | 12 |
| - | - | 2509 | 1532 | - | - | 0 | - |
| 3 | z | 831.2 | 1654 | 0.008495 | 5.137 | +1 | 13 |
| - | - | 3907 | 1655 | - | - | 0 | - |
| - | - | 2664 | 1656 | - | - | 0 | - |
| - | - | 1757 | 1657 | - | - | 0 | - |
| 13 | c | 1877 | 1692 | 0.002028 | 1.199 | +1 | 13 |
| - | - | 2921 | 1693 | - | - | 0 | - |
| - | - | 2174 | 1694 | - | - | 0 | - |
| - | - | 720 | 1696 | - | - | 0 | - |
| - | - | 1717 | 1792 | - | - | 0 | - |
| - | - | 1861 | 1793 | - | - | 0 | - |
| - | - | 1519 | 1794 | - | - | 0 | - |
| - | - | 889.4 | 1821 | - | - | 0 | - |
| - | - | 1155 | 1822 | - | - | 0 | - |
| - | - | 1148 | 1823 | - | - | 0 | - |
| - | - | 964.4 | 1846 | - | - | 0 | - |
| - | - | 1167 | 1848 | - | - | 0 | - |
| - | - | 1984 | 1863 | - | - | 0 | - |
| - | - | 2378 | 1864 | - | - | 0 | - |
| - | - | 1408 | 1865 | - | - | 0 | - |
| - | - | 1486 | 1877 | - | - | 0 | - |
| - | - | 1204 | 1878 | - | - | 0 | - |
| - | - | 751.6 | 1880 | - | - | 0 | - |
| - | - | 1450 | 1905 | - | - | 0 | - |
| - | - | 1650 | 1906 | - | - | 0 | - |
| - | - | 1791 | 1907 | - | - | 0 | - |
| - | - | 2871 | 1922 | - | - | 0 | - |
| - | - | 3194 | 1923 | - | - | 0 | - |
| - | - | 1954 | 1924 | - | - | 0 | - |
| - | - | 1019 | 1925 | - | - | 0 | - |
| - | - | 700.4 | 2578 | - | - | 0 | - |
| - | - | 693.8 | 3046 | - | - | 0 | - |
| - | - | 962 | 3047 | - | - | 0 | - |

m/z Charge Intensity FragmentType MassShift Position
120.06541442871094 0 486.45807
126.09095001220703 0 351.41098
129.10198974609375 0 2545.2615
130.06494140625 0 2806.4785
130.06956481933594 0 623.49316
132.10220336914062 0 863.1779
135.2641143798828 0 377.91916
136.0760040283203 0 808.68915
137.0819091796875 0 440.54782
138.0663604736328 0 622.86346
139.2493438720703 0 439.9635
140.08238220214844 0 499.49106
146.0916290283203 0 552.707
147.076171875 0 604.397
148.9477081298828 0 719.41675
149.0442657470703 0 457.01187
155.09268188476562 0 4518.5947
155.81890869140625 0 493.62476
157.13369750976562 0 668.23987
159.0764617919922 0 689.86566
159.0913848876953 0 2999.163
164.0816650390625 0 1028.94
165.1022186279297 0 556.2939
166.06089782714844 0 3020.8608
166.688232421875 0 457.29315
168.06788635253906 0 484.21017
168.13803100585938 0 2142.73
172.09664916992188 0 1011.39984 w 13
178.13356018066406 0 503.9745
183.11282348632812 0 965.0356
185.1646270751953 0 2777.7703
187.0868377685547 0 536.5492
187.107177734375 0 703.4236
195.0878143310547 0 787.0457
201.1227264404297 0 559.1802
203.13917541503906 0 723.4951
207.0973358154297 0 774.4048
215.13890075683594 0 925.7909
219.133544921875 0 2136.475
219.14886474609375 0 486.88745
223.06324768066406 0 957.70483
223.15501403808594 0 2299.146
224.1592559814453 0 604.4084
229.15440368652344 0 5432.3037 y Ammonia loss 13
230.1612091064453 0 1069.7257 z 13
233.1387939453125 0 943.7661
233.14993286132812 0 1296.8005
234.1233673095703 0 2786.3098
246.18072509765625 0 962.8315 y 13
250.1422576904297 0 916.7135
251.14987182617188 0 14553.763
252.15399169921875 0 1163.5747
261.1192626953125 0 960.39014
268.1771240234375 0 1123.8525 c 1
270.1608581542969 0 509.66208
272.5189208984375 0 490.2213
278.15777587890625 0 614.92474
283.1518859863281 0 1001.23737
285.15460205078125 0 735.5797
294.1932678222656 0 645.92993
295.1646728515625 0 1858.6263
298.94439697265625 0 596.4377
300.19122314453125 0 2197.3115
310.1391906738281 0 1200.3215
312.1915588378906 0 1258.6042
315.16839599609375 0 619.19226
326.6635437011719 0 4322.128
327.16424560546875 0 1430.2915 z Ammonia loss 7
328.1975402832031 0 818.9568
332.1601257324219 0 549.3082
340.6608581542969 0 1797.2947 c Ammonia loss 4
341.1614074707031 0 857.54114
345.15533447265625 0 1267.9758
349.67987060546875 0 1488.6194 y 10
350.181396484375 0 703.45435
350.71533203125 0 653.6071
356.1920166015625 0 814.48706
364.1842346191406 0 572.0148
365.0837097167969 0 830.03766
367.202880859375 0 644.3914
371.7259216308594 0 785.92236
375.1668701171875 0 641.4463
376.6855773925781 0 6739.6855 w 9
377.18548583984375 0 2269.2087
377.6878662109375 0 986.1165 z Water loss 3
378.2013244628906 0 628.7832
379.20806884765625 0 4283.983 c Ammonia loss 2
380.21270751953125 0 755.215
383.2046813964844 0 1355.5107
387.1828918457031 0 660.6714
387.2237548828125 0 610.5515
394.2060241699219 0 724.60236
396.1876220703125 0 845.2347
396.23492431640625 0 6893.1265 c 2
397.23846435546875 0 787.01843
402.177001953125 0 695.3139
402.7212219238281 0 2194.0242
403.2243957519531 0 799.3132
406.20465087890625 0 571.60315
407.1949157714844 0 2298.6802 y 12
414.2122802734375 0 608.29987 y 9
414.5493469238281 0 695.85956
416.2303466796875 0 875.68176
417.2177734375 0 2239.273
417.72119140625 0 943.5028
425.2467346191406 0 571.85455
428.2865295410156 0 2349.9666
429.29254150390625 0 777.90607
431.0858154296875 0 1182.4813
440.73095703125 0 931.48
441.2061462402344 0 1071.2878 w 8
441.2383728027344 0 808.95123
448.1960144042969 0 823.95874
448.25250244140625 0 698.7217
452.4685974121094 0 1029.249 y 1
453.25799560546875 0 2216.7544
453.76129150390625 0 950.86707
460.2586669921875 0 2450.847
461.2070617675781 0 2392.321 w 11
461.2607421875 0 849.8957
464.23907470703125 0 890.12305
466.23992919921875 0 1239.7322
467.24737548828125 0 1610.0055
467.7052001953125 0 559.3764
468.2519836425781 0 835.62317
469.2519836425781 0 1862.3541 y Water loss 8
470.24176025390625 0 1205.0654 y Water loss 4
472.7514343261719 0 1146.7847
474.2563171386719 0 3917.5923
475.2592468261719 0 1888.2197
476.1907653808594 0 689.1455
476.22955322265625 0 1248.0431
476.48590087890625 0 1406.9258
478.2486572265625 0 4169.134 y 8
478.7503356933594 0 2238.6216
479.249755859375 0 927.9202
480.2528991699219 0 881.8182
480.49200439453125 0 2605.1667
480.6902770996094 0 3261.4717
480.7389221191406 0 7715.2446
480.9896240234375 0 9181.234
481.1925048828125 0 1474.6598
481.23919677734375 0 4684.302
481.2763977050781 0 16467.21
481.3153991699219 0 856.2621
481.48919677734375 0 2182.5242
481.6950378417969 0 1069.9657
481.7781677246094 0 7791.2095
482.19775390625 0 680.3547
482.2813415527344 0 1365.1985
484.2518615722656 0 1175.5668
494.2355041503906 0 3499.1992 c Ammonia loss 3
498.7510986328125 0 1823.8497 z 7
499.254150390625 0 930.2863
503.24761962890625 0 849.12726 z Water loss 3
505.7376403808594 0 1996.0455
506.2596130371094 0 2139.1218
506.7593994140625 0 4921.9897 y 7
507.26043701171875 0 3285.686
507.76123046875 0 1525.6316
510.7730407714844 0 2204.1155
511.26214599609375 0 21750.2 c 3
512.2649536132812 0 5266.9707
514.7398071289062 0 793.0862
519.2715454101562 0 8061.121 z 11
520.2757568359375 0 3285.7932
526.758056640625 0 1667.6965
527.2311401367188 0 2482.1206
527.7649536132812 0 4807.4673
528.228515625 0 705.899
528.267578125 0 2927.32
533.7654418945312 0 3148.938 w 6
534.2681884765625 0 3873.498
534.76806640625 0 2440.7473
535.2894287109375 0 4709.336 y 11
536.29443359375 0 786.817
537.2857666015625 0 785.93274
538.2772827148438 0 590.5684
542.2704467773438 0 807.60547
547.27685546875 0 686.24713 c Ammonia loss 8
549.9410400390625 0 564.621
550.2510375976562 0 4001.978
551.2564697265625 0 1835.4232 y Water loss 2
551.9292602539062 0 1109.517 z 2
552.2601928710938 0 1230.7162
555.2958374023438 0 3955.3875
555.788330078125 0 21824.29 c 8
556.2570190429688 0 2242.7588 z 6
556.2887573242188 0 9590.499
556.765380859375 0 2273.8572
556.791748046875 0 1841.8938
557.2681274414062 0 1902.6814 y 2
557.7638549804688 0 1146.9324
561.3344116210938 0 1294.1719
563.7830200195312 0 1654.9825
564.2737426757812 0 15039.841 y 6
564.7744140625 0 7537.132
565.2742919921875 0 2920.1282
565.7742309570312 0 1465.0701
567.81640625 0 795.6652
568.7958374023438 0 2683.3318
569.29248046875 0 1855.0283
571.28076171875 0 695.8886
574.2667846679688 0 600.818
576.7929077148438 0 664.3479
577.9630737304688 0 1102.9417
578.6148071289062 0 778.6572
578.9442749023438 0 739.5162
582.2805786132812 0 1026.6422
588.2882080078125 0 746.0135
590.7879638671875 0 2211.0999
591.2793579101562 0 11926.653 w 5
591.7808227539062 0 8939.472
592.2843017578125 0 2850.1
592.3550415039062 0 746.2619
592.7838745117188 0 1135.4513
592.96044921875 0 646.93726
595.3198852539062 0 883.4848
595.6520385742188 0 805.7727
597.2926635742188 0 2216.2563
597.6275634765625 0 3211.6672 z 1
597.9574584960938 0 1374.1484
598.29833984375 0 897.1841
601.3018188476562 0 1581.7037 c Water loss 13
601.626220703125 0 1393.1937 c Ammonia loss 13
601.9634399414062 0 1475.4757
602.2947387695312 0 1218.3595
602.9551391601562 0 1120.0732 y 1
603.2892456054688 0 1061.5411
603.6233520507812 0 859.4179
606.294189453125 0 601.0505
606.6345825195312 0 766.50964
606.969482421875 0 3714.2085
607.2981567382812 0 19287.201 c 13
607.6321411132812 0 21338.281
607.9663696289062 0 13278.526
608.2986450195312 0 6282.6987
608.6322631835938 0 2998.328
611.3080444335938 0 1246.0282 c Water loss 9
611.7989501953125 0 1291.7238 c Ammonia loss 9
612.3093872070312 0 804.9626 y Ammonia loss 5
612.8057250976562 0 2362.471 z 5
613.3076782226562 0 1491.4772
615.3074951171875 0 905.47003
615.6395874023438 0 1353.7616
616.6343994140625 0 3268.8335
616.96630859375 0 2112.8965
617.3032836914062 0 1186.7318
618.3037719726562 0 876.6083
618.3563232421875 0 6495.261
619.357666015625 0 1197.6371
619.81689453125 0 3415.574
620.3106079101562 0 21439.871 c 9
620.63671875 0 1836.881
620.8121948242188 0 23491.387 y 5
620.9664916992188 0 11480.258
621.3060913085938 0 23630.25
621.6420288085938 0 14180.688
621.8169555664062 0 3850.0105
621.976318359375 0 6486.7476
622.3158569335938 0 4185.123
622.642333984375 0 1011.5266
625.6499633789062 0 2003.2794
625.9822998046875 0 2068.625
626.3171997070312 0 1731.6367
626.6484375 0 765.7655
629.637939453125 0 656.5051
634.9771728515625 0 919.2699
635.311279296875 0 2443.6555
635.6459350585938 0 1594.9453
635.9789428710938 0 1453.4104
636.6453247070312 0 825.74646
640.3198852539062 0 6451.231
640.6498413085938 0 40778.2
640.9840698242188 0 60368.176
641.3179321289062 0 42228.88
641.6524047851562 0 20712.36
641.98583984375 0 6612.274
642.32080078125 0 3008.0151
642.653076171875 0 1175.961
648.824951171875 0 704.83905
652.3171997070312 0 1390.0095
653.3242797851562 0 1503.1548
660.8606567382812 0 690.0705
670.3208618164062 0 559.6256
676.3370361328125 0 927.12164
677.3162841796875 0 725.3059
677.815185546875 0 1767.7627
678.3157958984375 0 1603.0283
679.8338623046875 0 1563.4647
680.316650390625 0 6482.6343 c Ammonia loss 4
681.3204345703125 0 3353.0728 y Ammonia loss 10
682.3345947265625 0 17956.754 z 10
683.3380126953125 0 8864.78
683.8515014648438 0 914.5494
684.3363647460938 0 3017.606
697.3408203125 0 20129.92 c 4
698.3471069335938 0 13325.35 y 10
699.3504638671875 0 2935.4578
700.3521118164062 0 892.21747
701.3480834960938 0 3499.8704
701.8413696289062 0 19010.78 c 10
702.34228515625 0 14710.719
702.8441772460938 0 6084.7495
703.3463134765625 0 1896.3121
705.3505249023438 0 1113.3525 y Ammonia loss 4
705.8458251953125 0 9959.193 z 4
706.3489990234375 0 6794.534
706.8486938476562 0 4232.4316
707.34765625 0 1726.0244
712.3316650390625 0 974.1489
713.855224609375 0 4749.361 y 4
714.3592529296875 0 2751.665
714.86083984375 0 1534.1349
717.8726806640625 0 886.9036
719.3705444335938 0 850.78687
727.3510131835938 0 842.1906
731.3941650390625 0 805.463
733.3987426757812 0 2886.589
734.4048461914062 0 977.54663
740.86669921875 0 700.1695 w 3
741.3651733398438 0 4527.746
741.868408203125 0 1252.5038
742.3607788085938 0 1614.2739
752.3642578125 0 3759.4548 w 9
753.3677368164062 0 1594.0474
762.8655395507812 0 2029.627 y Ammonia loss 3
763.3611450195312 0 9623.957 z 3
763.8604125976562 0 5982.1016
764.3626098632812 0 3913.3481
764.8628540039062 0 2019.9622
765.8900146484375 0 5780.258 c 11
766.395263671875 0 3990.012
766.8887329101562 0 1583.0299
767.40771484375 0 677.0607
771.3703002929688 0 1132.3085 y 3
771.8743896484375 0 1233.9574
778.8999633789062 0 901.8058
786.3793334960938 0 759.73535
789.4330444335938 0 2623.3625
793.3941650390625 0 919.99725 c Ammonia loss 5
794.4005126953125 0 889.34296
797.8860473632812 0 1514.4674
798.3740234375 0 3827.7278 w 2
798.874755859375 0 1179.8746
799.374267578125 0 937.77423
804.4368286132812 0 1419.8402
806.39501953125 0 903.90967
810.4251098632812 0 35381.977 c 5
811.3751220703125 0 16339.842 z 9
811.4363403320312 0 10480.455
812.3819580078125 0 12190.388
813.3817138671875 0 2842.4197
814.3919067382812 0 908.47516
826.8956909179688 0 3411.7766 y Ammonia loss 2
827.3907470703125 0 13704.777 y 9
827.890625 0 9733.177
828.3931274414062 0 6077.6855
828.8922119140625 0 1907.2598
829.394287109375 0 2079.1426
835.3928833007812 0 1078.4 y 2
835.90234375 0 1563.2412
839.413818359375 0 1044.9298
844.4169921875 0 1602.3143
845.4193725585938 0 806.78986
845.900634765625 0 3081.3596
846.3963623046875 0 17579.723 c 12
846.8971557617188 0 14498.781
847.3980102539062 0 8775.025
847.8987426757812 0 3231.2634
848.3987426757812 0 799.4177
860.4275512695312 0 1133.3011
866.42724609375 0 1102.4742
867.4251098632812 0 4189.115
868.422119140625 0 2246.7974
871.4249877929688 0 641.9372
874.4336547851562 0 756.1444
876.453857421875 0 827.93066
880.4412231445312 0 1052.3602
880.9342041015625 0 1170.5614
881.4426879882812 0 4075.193
881.9420776367188 0 1666.7743
882.4508666992188 0 1679.9347
888.4340209960938 0 2268.7576
888.9384765625 0 2752.0823
889.44140625 0 2985.016
889.9434814453125 0 1747.0396
894.51708984375 0 1602.2914
895.4237060546875 0 1322.9747 y Ammonia loss 1
895.5197143554688 0 1119.7805
895.9210815429688 0 5963.068 z 1
896.4215087890625 0 8108.4985
896.922119140625 0 4788.91
901.470947265625 0 754.5387
901.949951171875 0 1236.7399 c Ammonia loss 13
902.9456787109375 0 2170.9124
904.4365844726562 0 785.74274
909.4562377929688 0 895.7204
909.9617919921875 0 1339.6266
910.44775390625 0 2892.8115 c 13
910.9462280273438 0 4427.809
911.4483032226562 0 3600.6042
911.9515991210938 0 1210.7821
912.4517211914062 0 1102.39
919.46240234375 0 1543.0298
920.4771728515625 0 901.16565
921.4669189453125 0 782.0692 z Water loss 8
922.4624633789062 0 2091.4465 z Ammonia loss 8
922.9618530273438 0 2613.7134
923.4595336914062 0 4466.8345
923.96240234375 0 3383.3743
924.4589233398438 0 3314.0437
925.4523315429688 0 5897.233 c 6
926.4561767578125 0 2525.142
927.459228515625 0 787.5463
930.4697265625 0 1283.143
930.9674072265625 0 5801.497
931.4672241210938 0 18621.225
931.9686889648438 0 16060.214
932.4686279296875 0 9176.853
932.9716186523438 0 4098.229
933.475830078125 0 1484.0101
937.4712524414062 0 888.95953 y Water loss 8
937.9703979492188 0 1452.1698
938.468017578125 0 9056.018 y Ammonia loss 8
938.9725341796875 0 7549.465
939.4690551757812 0 10406.957 z 8
939.9702758789062 0 2514.2083
940.4774169921875 0 7398.5576
941.478759765625 0 2816.0256
942.475341796875 0 844.06793
943.472900390625 0 2612.6003
944.4717407226562 0 2506.4507
945.4747314453125 0 1509.0428
946.4825439453125 0 2521.9453
947.48583984375 0 1562.7874
951.9691162109375 0 1070.4126
952.4685668945312 0 7532.7188
952.9649658203125 0 20289.195
953.466064453125 0 15543.437
953.9671020507812 0 9415.126
954.4680786132812 0 4822.1597
954.9655151367188 0 1575.7363
955.4814453125 0 1477.7124 y 8
956.4896850585938 0 720.2699
958.5028686523438 0 2097.0664
959.500732421875 0 2061.0208
959.985595703125 0 1285.9882
960.4764404296875 0 5339.991
960.97607421875 0 7887.5513
961.3690185546875 0 1263.6134
961.478271484375 0 24366.395
961.9784545898438 0 17377.658
962.4808349609375 0 10552.898
962.57470703125 0 1136.7832
962.9791259765625 0 4668.206
963.4826049804688 0 2942.8618
963.9793090820312 0 903.79913
964.4625854492188 0 1125.9948 c Water loss 7
981.4833374023438 0 2434.2473
982.4744873046875 0 16650.08 c 7
983.47705078125 0 8877.318
984.4747314453125 0 2507.1067
996.4990234375 0 705.7635 z 7
997.50537109375 0 1875.2981
998.4994506835938 0 1111.7332
1012.5089721679688 0 1775.6685 y 7
1013.5120849609375 0 691.3005
1051.552734375 0 860.8727
1053.51806640625 0 924.13293
1054.52294921875 0 1006.2344
1066.55859375 0 1392.1301
1094.5400390625 0 1523.1112
1109.574951171875 0 1521.8583
1110.5699462890625 0 12535.323 c 8
1111.5684814453125 0 10439.11
1112.5379638671875 0 11018.874
1113.5357666015625 0 5355.092
1114.5357666015625 0 1771.0648
1128.5433349609375 0 835.2657
1180.5869140625 0 608.39136
1196.6190185546875 0 797.3945
1223.609130859375 0 1039.9137 y Ammonia loss 5
1224.6109619140625 0 2276.7942 z 5
1225.6119384765625 0 6342.3994
1226.615966796875 0 3837.2668
1227.6287841796875 0 699.69226
1238.625732421875 0 672.01166
1239.613525390625 0 3681.9368 c 9
1240.616943359375 0 4916.95 y 5
1241.6195068359375 0 2633.295
1242.6241455078125 0 682.2516
1249.8873291015625 0 780.1383
1310.6466064453125 0 855.3042
1311.6353759765625 0 1093.5173
1358.670166015625 0 975.969
1359.67578125 0 863.60175
1367.67041015625 0 731.1023
1402.682373046875 0 2646.7446 c 10
1403.6763916015625 0 3519.4717
1404.685546875 0 2405.2573
1410.69189453125 0 2616.5679 z 4
1411.690673828125 0 5456.8057
1412.6929931640625 0 3964.6772
1413.6893310546875 0 2141.5913
1414.6968994140625 0 1167.5148
1426.685791015625 0 1390.1494 y 4
1427.7005615234375 0 988.23865
1428.6864013671875 0 734.2906
1525.71728515625 0 2141.5935 z 3
1526.71728515625 0 4791.2983
1527.719970703125 0 2901.6516
1528.7203369140625 0 1002.7071
1530.7716064453125 0 1646.7084 c 11
1531.7747802734375 0 2508.6914
1653.7833251953125 0 831.1929 z 2
1654.77587890625 0 3906.9067
1655.779296875 0 2664.2056
1656.7752685546875 0 1756.5353
1691.787109375 0 1876.639 c 12
1692.790283203125 0 2920.8145
1693.79541015625 0 2173.7173
1695.803955078125 0 720.0077
1791.840087890625 0 1716.9216
1792.848876953125 0 1860.7145
1793.8326416015625 0 1519.3575
1820.915771484375 0 889.44885
1821.8828125 0 1154.8689
1822.9075927734375 0 1147.5444
1845.9202880859375 0 964.444
1847.9349365234375 0 1166.7389
1862.9324951171875 0 1983.5051
1863.9288330078125 0 2377.7961
1864.9393310546875 0 1408.3539
1876.9395751953125 0 1486.0181
1877.938720703125 0 1203.8401
1879.946533203125 0 751.60443
1904.9293212890625 0 1449.7538
1905.931884765625 0 1649.5817
1906.93212890625 0 1791.3878
1921.9517822265625 0 2870.9695
1922.952880859375 0 3193.56
1923.957763671875 0 1954.4353
1924.950439453125 0 1018.7744
2578.44677734375 0 700.3657
3045.826904296875 0 693.7646
3046.514892578125 0 961.9582

Spectrum Details

|  |  |
| --- | --- |
| Matched peaks? Matched peaks The total absolute number of peaks matched. Additionally in brackets the total fraction of peaks matched and the total number of peaks is shown. | 97 (17.73% of 547) |
| FDR? FDR The false discovery rate estimated for this peptide. It is calculated by matching all theoretical fragments with a non-integer shift with the raw peaks for this spectrum. This is done with 40 different shifts. The resulting percentage is the average number of annotated peaks over the number of annotated peaks with the correct spectrum. | 0.44% |
| Satellite FDR? Satellite FDR See the FDR for details on its calculation. This satellite ion specific FDR only contains the satellite ions (d/w) for I/L/J positions. | 0.00% |
| PSM Score? PSM Score The PSM Score as given by Hecklib to this annotated spectrum. It is shown with three significant figures. | 559 |

#### Spectrum 5464? Spectrum 5464 The raw spectrum of this peptide as annotated by Hecklib. The fragments are coloured according to ion type (see legend). Any peaks with a star '\*' as text can be hovered over to see the full details, first the ion type second the mass shift type. By hovering over the amino acids in the peptide or ions in the legend the corresponding peaks are highlighted. By toggling the 'Unassigned' label you can turn the background (unassigned) peaks on or off in the plot. By updating the slider in the Ion legend you can update the spectrum to only show the top X% of the peaks with labels. The top X% means any peak that is within X% of the highest intensity. By dragging in the spectrum you can zoom in to a specific part of the spectrum and use 'Zoom Out' to get back to the original zoom level. The annotation of the spectrum is based on the given sequence in the peptides file and is done with different software so inconsistencies are likely. The peaks are annotated based on the given sequence, with 20 ppm tolerance.

Copy Data

##### Spectrum 5464 (TSV)

###### Preview

```
Loading example...
```

*Click on the button to copy the data to your clipboard.*

Mz MinMz MaxIntensity Max

WidthHeightPeptide font sizePeptide stroke widthSpectrum font sizeSpectrum stroke widthCompact peptide

Ion legend

wxyz

abcd

OtherUnassignedIon^Charge^_Position_Show for top:%

JHQDWJDGKEYKCKV

06.84e+41.37e+52.05e+52.74e+5

Zoom Out

z^+3^_3_w^+1^_2_z^+5^_9_z^+3^_5_y^+1^_2_z^+1^_2_y^+4^_7_y^+1^_2_c^+2^_4_y^+4^_8_z^+2^_4_c^+2^_4_y^+2^_4_z^+2^_4_z^+3^_6_y^+2^_4_c^+1^_2_y^+3^_7_y^+3^_7_z^+3^_7_y^+3^_7_z^+3^_8_c^+3^_8_y^+3^_8_z^+3^_8_y^+3^_8_c^+2^_5_y^+2^_5_z^+2^_5_y^+2^_5_y^+4^_11_z^+4^_11_w^+3^_9_c^+5^_14_y^+3^_9_y^+3^_9_z^+3^_9_y^+3^_9_w^+2^_6_z^+4^_12_c^+1^_3_y^+4^_12_z^+4^_12_y^+4^_12_w^+3^_10_c^+1^_3_c^+2^_6_z^+3^_10_y^+2^_6_z^+2^_6_y^+1^_3_c^+3^_10_y^+3^_10_z^+3^_10_z^+4^_13_y^+4^_13_y^+4^_13_y^+3^_10_w^+2^_7_z^+4^_14_y^+4^_14_y^+4^_14_z^+4^_14_c^+4^_14_c^+4^_14_y^+4^_14_c^+2^_7_c^+2^_7_c^+4^_14_w^+1^_4_c^+3^_11_c^+2^_7_z^+3^_11_y^+2^_7_y^+3^_11_y^+3^_11_z^+3^_11_y^+3^_11_y^+2^_7_c^+2^_8_c^+2^_8_z^+2^_8_c^+2^_8_c^+1^_4_c^+1^_4_y^+2^_8_z^+2^_8_z^+3^_12_c^+3^_12_y^+2^_8_z^+3^_12_c^+3^_12_c^+1^_4_y^+3^_12_z^+1^_4_w^+3^_13_w^+2^_9_y^+1^_4_z^+3^_13_z^+3^_13_c^+2^_9_y^+3^_13_y^+3^_13_z^+3^_13_y^+2^_9_c^+2^_9_y^+3^_13_c^+3^_13_y^+2^_9_c^+3^_13_w^+2^_10_y^+3^_14_z^+3^_14_c^+3^_14_y^+3^_14_z^+2^_10_c^+3^_14_c^+2^_10_y^+2^_10_z^+2^_10_c^+2^_10_y^+2^_10_c^+1^_5_c^+1^_5_y^+1^_5_z^+1^_5_c^+2^_11_c^+1^_5_y^+1^_5_c^+2^_11_y^+2^_11_y^+2^_11_z^+2^_11_y^+2^_11_w^+2^_12_w^+1^_6_c^+2^_12_y^+2^_12_y^+2^_12_z^+2^_12_c^+2^_12_y^+2^_12_c^+1^_6_w^+2^_13_y^+1^_6_c^+1^_6_z^+1^_6_z^+2^_13_y^+2^_13_y^+1^_6_y^+2^_13_c^+2^_13_z^+2^_14_y^+2^_14_c^+1^_7_c^+2^_14_z^+1^_7_c^+1^_7_y^+1^_7_z^+1^_7_y^+1^_7_c^+1^_8_c^+1^_8_z^+1^_8_y^+1^_8_c^+1^_9_y^+1^_9_z^+1^_10_c^+1^_10_y^+1^_10_c^+1^_11_

0770154023103080

Fragment Matches Table

Show background peaks

| Position | Ion type | Intensity | mz Theoretical | mz Error (Th) | mz Error (ppm) | Charge | Series Number |
| --- | --- | --- | --- | --- | --- | --- | --- |
| - | - | 666 | 120.1 | - | - | 0 | - |
| - | - | 391.2 | 121.3 | - | - | 0 | - |
| - | - | 629.1 | 122 | - | - | 0 | - |
| - | - | 641.1 | 126.1 | - | - | 0 | - |
| - | - | 406 | 127.1 | - | - | 0 | - |
| - | - | 621.8 | 128.1 | - | - | 0 | - |
| - | - | 765.2 | 128.1 | - | - | 0 | - |
| - | - | 716 | 129.1 | - | - | 0 | - |
| - | - | 3.51E+04 | 129.1 | - | - | 0 | - |
| - | - | 9.843E+04 | 130.1 | - | - | 0 | - |
| - | - | 1940 | 130.1 | - | - | 0 | - |
| 13 | z | 1101 | 131.1 | 0.001537 | 11.73 | +3 | 3 |
| - | - | 9560 | 131.1 | - | - | 0 | - |
| - | - | 3084 | 131.1 | - | - | 0 | - |
| - | - | 392.7 | 131.8 | - | - | 0 | - |
| - | - | 3027 | 132.1 | - | - | 0 | - |
| - | - | 1164 | 132.1 | - | - | 0 | - |
| - | - | 1.289E+04 | 133.1 | - | - | 0 | - |
| - | - | 554.6 | 134 | - | - | 0 | - |
| - | - | 7059 | 136.1 | - | - | 0 | - |
| - | - | 1205 | 138.1 | - | - | 0 | - |
| - | - | 448.9 | 139.1 | - | - | 0 | - |
| - | - | 545.6 | 140.1 | - | - | 0 | - |
| - | - | 1357 | 141.1 | - | - | 0 | - |
| - | - | 421.6 | 142.3 | - | - | 0 | - |
| - | - | 2520 | 144.1 | - | - | 0 | - |
| - | - | 1606 | 145.1 | - | - | 0 | - |
| - | - | 499.8 | 145.1 | - | - | 0 | - |
| - | - | 546.6 | 146.1 | - | - | 0 | - |
| - | - | 1929 | 146.1 | - | - | 0 | - |
| - | - | 444.1 | 146.1 | - | - | 0 | - |
| - | - | 1095 | 147 | - | - | 0 | - |
| - | - | 574.8 | 147.1 | - | - | 0 | - |
| - | - | 651.2 | 147.1 | - | - | 0 | - |
| - | - | 416.6 | 148.1 | - | - | 0 | - |
| - | - | 472.4 | 148.9 | - | - | 0 | - |
| - | - | 506.4 | 154.1 | - | - | 0 | - |
| - | - | 1.642E+04 | 155.1 | - | - | 0 | - |
| - | - | 844.7 | 156.1 | - | - | 0 | - |
| - | - | 626.7 | 156.1 | - | - | 0 | - |
| - | - | 790.4 | 157.1 | - | - | 0 | - |
| - | - | 1347 | 157.1 | - | - | 0 | - |
| - | - | 497.7 | 158 | - | - | 0 | - |
| - | - | 1881 | 158.1 | - | - | 0 | - |
| - | - | 601.3 | 158.1 | - | - | 0 | - |
| - | - | 4.229E+04 | 159.1 | - | - | 0 | - |
| - | - | 3681 | 160.1 | - | - | 0 | - |
| - | - | 2391 | 164.1 | - | - | 0 | - |
| - | - | 4560 | 165.1 | - | - | 0 | - |
| - | - | 1.633E+04 | 166.1 | - | - | 0 | - |
| - | - | 1436 | 167.1 | - | - | 0 | - |
| - | - | 4139 | 168.1 | - | - | 0 | - |
| - | - | 590.2 | 169.1 | - | - | 0 | - |
| - | - | 2.05E+04 | 170.1 | - | - | 0 | - |
| - | - | 850.8 | 170.1 | - | - | 0 | - |
| - | - | 1071 | 171.1 | - | - | 0 | - |
| - | - | 1402 | 171.1 | - | - | 0 | - |
| - | - | 3738 | 172.1 | - | - | 0 | - |
| 14 | w | 3749 | 172.1 | 0.0008212 | 4.772 | +1 | 2 |
| - | - | 2340 | 173.1 | - | - | 0 | - |
| - | - | 793.9 | 173.4 | - | - | 0 | - |
| - | - | 553.8 | 175.1 | - | - | 0 | - |
| - | - | 488.6 | 175.7 | - | - | 0 | - |
| - | - | 1237 | 176.1 | - | - | 0 | - |
| - | - | 2469 | 178.1 | - | - | 0 | - |
| - | - | 752.7 | 181.1 | - | - | 0 | - |
| - | - | 1.597E+04 | 183.1 | - | - | 0 | - |
| - | - | 725.7 | 184.1 | - | - | 0 | - |
| - | - | 941.9 | 184.1 | - | - | 0 | - |
| - | - | 661.4 | 185.1 | - | - | 0 | - |
| - | - | 1.773E+04 | 185.2 | - | - | 0 | - |
| - | - | 805.4 | 186.1 | - | - | 0 | - |
| - | - | 3460 | 186.1 | - | - | 0 | - |
| - | - | 1886 | 186.2 | - | - | 0 | - |
| - | - | 7379 | 187.1 | - | - | 0 | - |
| - | - | 1888 | 187.1 | - | - | 0 | - |
| - | - | 1038 | 188.1 | - | - | 0 | - |
| - | - | 1275 | 188.1 | - | - | 0 | - |
| - | - | 3.061E+04 | 190.1 | - | - | 0 | - |
| - | - | 1330 | 191.1 | - | - | 0 | - |
| - | - | 1704 | 192.1 | - | - | 0 | - |
| - | - | 1421 | 195.1 | - | - | 0 | - |
| - | - | 565.6 | 198.1 | - | - | 0 | - |
| - | - | 1.031E+04 | 198.1 | - | - | 0 | - |
| - | - | 708.9 | 199.1 | - | - | 0 | - |
| - | - | 1050 | 199.1 | - | - | 0 | - |
| - | - | 858.7 | 199.2 | - | - | 0 | - |
| - | - | 715.7 | 200.1 | - | - | 0 | - |
| - | - | 2270 | 201.1 | - | - | 0 | - |
| - | - | 587.4 | 201.1 | - | - | 0 | - |
| - | - | 3504 | 203.1 | - | - | 0 | - |
| - | - | 1406 | 203.2 | - | - | 0 | - |
| - | - | 756.4 | 203.6 | - | - | 0 | - |
| - | - | 648.3 | 204.1 | - | - | 0 | - |
| - | - | 2205 | 207.2 | - | - | 0 | - |
| - | - | 3035 | 210.1 | - | - | 0 | - |
| - | - | 4047 | 211.6 | - | - | 0 | - |
| - | - | 1169 | 212.1 | - | - | 0 | - |
| - | - | 539.4 | 214.2 | - | - | 0 | - |
| - | - | 1350 | 214.6 | - | - | 0 | - |
| - | - | 3734 | 215.1 | - | - | 0 | - |
| - | - | 1613 | 218.1 | - | - | 0 | - |
| - | - | 1434 | 221.6 | - | - | 0 | - |
| - | - | 1323 | 222.1 | - | - | 0 | - |
| 7 | z | 959.8 | 223.1 | 0.002213 | 9.917 | +5 | 9 |
| - | - | 5744 | 223.1 | - | - | 0 | - |
| - | - | 1.671E+04 | 223.2 | - | - | 0 | - |
| - | - | 1896 | 224.2 | - | - | 0 | - |
| - | - | 628.2 | 225 | - | - | 0 | - |
| - | - | 1299 | 227.1 | - | - | 0 | - |
| - | - | 1277 | 227.1 | - | - | 0 | - |
| - | - | 567.9 | 227.2 | - | - | 0 | - |
| 11 | z | 909.1 | 228.1 | 0.002309 | 10.12 | +3 | 5 |
| - | - | 815.9 | 228.1 | - | - | 0 | - |
| - | - | 935 | 229.1 | - | - | 0 | - |
| - | - | 3623 | 229.1 | - | - | 0 | - |
| 14 | y | 2653 | 229.2 | 0.001093 | 4.769 | +1 | 2 |
| 14 | z | 4007 | 230.2 | 0.0009429 | 4.097 | +1 | 2 |
| - | - | 9573 | 230.6 | - | - | 0 | - |
| - | - | 3203 | 231.1 | - | - | 0 | - |
| - | - | 705.3 | 231.2 | - | - | 0 | - |
| - | - | 5307 | 233.1 | - | - | 0 | - |
| - | - | 1517 | 233.2 | - | - | 0 | - |
| - | - | 1.917E+04 | 234.1 | - | - | 0 | - |
| - | - | 818.6 | 234.1 | - | - | 0 | - |
| 9 | y | 2133 | 235.1 | 0.001265 | 5.378 | +4 | 7 |
| - | - | 1759 | 236.6 | - | - | 0 | - |
| - | - | 549.9 | 237.1 | - | - | 0 | - |
| - | - | 703.1 | 239.1 | - | - | 0 | - |
| - | - | 1577 | 239.1 | - | - | 0 | - |
| - | - | 1294 | 241.1 | - | - | 0 | - |
| - | - | 643.7 | 243.1 | - | - | 0 | - |
| - | - | 1360 | 243.1 | - | - | 0 | - |
| - | - | 1144 | 243.1 | - | - | 0 | - |
| - | - | 2044 | 244.1 | - | - | 0 | - |
| - | - | 1040 | 245.1 | - | - | 0 | - |
| - | - | 1010 | 246.1 | - | - | 0 | - |
| - | - | 1230 | 246.1 | - | - | 0 | - |
| 14 | y | 6492 | 246.2 | 0.001048 | 4.258 | +1 | 2 |
| - | - | 625.5 | 247.2 | - | - | 0 | - |
| 4 | c | 6038 | 247.6 | 0.0008966 | 3.621 | +2 | 4 |
| - | - | 743.8 | 248.1 | - | - | 0 | - |
| - | - | 488.3 | 248.6 | - | - | 0 | - |
| - | - | 679 | 249.1 | - | - | 0 | - |
| - | - | 833.8 | 249.1 | - | - | 0 | - |
| 8 | y | 594.1 | 249.6 | 0.002412 | 9.664 | +4 | 8 |
| - | - | 736.2 | 249.8 | - | - | 0 | - |
| - | - | 619 | 250.1 | - | - | 0 | - |
| - | - | 3808 | 250.1 | - | - | 0 | - |
| - | - | 1138 | 250.1 | - | - | 0 | - |
| - | - | 1158 | 250.1 | - | - | 0 | - |
| - | - | 2265 | 250.2 | - | - | 0 | - |
| - | - | 9.064E+04 | 251.2 | - | - | 0 | - |
| 12 | z | 623 | 251.6 | 0.001701 | 6.759 | +2 | 4 |
| - | - | 1.057E+04 | 252.2 | - | - | 0 | - |
| - | - | 2668 | 252.6 | - | - | 0 | - |
| - | - | 785.3 | 253.1 | - | - | 0 | - |
| - | - | 995.6 | 253.2 | - | - | 0 | - |
| - | - | 526.9 | 253.4 | - | - | 0 | - |
| - | - | 684 | 254.2 | - | - | 0 | - |
| - | - | 2301 | 255.1 | - | - | 0 | - |
| - | - | 3498 | 255.8 | - | - | 0 | - |
| 4 | c | 913.9 | 256.1 | 0.004453 | 17.38 | +2 | 4 |
| - | - | 625.5 | 256.2 | - | - | 0 | - |
| - | - | 1024 | 257.2 | - | - | 0 | - |
| - | - | 1904 | 258.6 | - | - | 0 | - |
| 12 | y | 4763 | 259.6 | 0.00148 | 5.701 | +2 | 4 |
| 12 | z | 1218 | 260.1 | 0.003257 | 12.52 | +2 | 4 |
| - | - | 678.3 | 260.6 | - | - | 0 | - |
| - | - | 3428 | 261.1 | - | - | 0 | - |
| - | - | 723.3 | 261.2 | - | - | 0 | - |
| - | - | 1714 | 261.6 | - | - | 0 | - |
| - | - | 890.3 | 262.1 | - | - | 0 | - |
| 10 | z | 663.3 | 265.1 | 0.0006051 | 2.282 | +3 | 6 |
| - | - | 4116 | 266.1 | - | - | 0 | - |
| - | - | 1087 | 266.6 | - | - | 0 | - |
| 12 | y | 4557 | 268.2 | 0.001113 | 4.152 | +2 | 4 |
| 2 | c | 1.371E+04 | 268.2 | 0.0009635 | 3.593 | +1 | 2 |
| - | - | 893.7 | 268.7 | - | - | 0 | - |
| - | - | 736 | 269.1 | - | - | 0 | - |
| - | - | 1021 | 269.2 | - | - | 0 | - |
| - | - | 888 | 270.1 | - | - | 0 | - |
| - | - | 1.525E+04 | 270.1 | - | - | 0 | - |
| - | - | 1076 | 270.2 | - | - | 0 | - |
| - | - | 4372 | 270.6 | - | - | 0 | - |
| - | - | 1141 | 271.1 | - | - | 0 | - |
| - | - | 7771 | 272.1 | - | - | 0 | - |
| - | - | 981.3 | 272.2 | - | - | 0 | - |
| - | - | 4302 | 272.2 | - | - | 0 | - |
| - | - | 3113 | 273.1 | - | - | 0 | - |
| - | - | 1099 | 273.1 | - | - | 0 | - |
| - | - | 947.4 | 273.2 | - | - | 0 | - |
| - | - | 684.2 | 273.5 | - | - | 0 | - |
| - | - | 2977 | 274.1 | - | - | 0 | - |
| - | - | 1610 | 275.1 | - | - | 0 | - |
| - | - | 825.4 | 277.2 | - | - | 0 | - |
| - | - | 885.6 | 282.6 | - | - | 0 | - |
| - | - | 5721 | 283.1 | - | - | 0 | - |
| - | - | 1558 | 284.1 | - | - | 0 | - |
| - | - | 3651 | 284.1 | - | - | 0 | - |
| - | - | 600.6 | 284.1 | - | - | 0 | - |
| - | - | 1722 | 284.6 | - | - | 0 | - |
| - | - | 1278 | 285.2 | - | - | 0 | - |
| - | - | 1423 | 286.1 | - | - | 0 | - |
| - | - | 639.5 | 287.8 | - | - | 0 | - |
| - | - | 630.7 | 289.2 | - | - | 0 | - |
| - | - | 6589 | 290.1 | - | - | 0 | - |
| - | - | 696 | 291.1 | - | - | 0 | - |
| - | - | 770.5 | 292.1 | - | - | 0 | - |
| - | - | 1696 | 292.2 | - | - | 0 | - |
| - | - | 750 | 293.1 | - | - | 0 | - |
| - | - | 796.9 | 294.1 | - | - | 0 | - |
| - | - | 678.7 | 294.2 | - | - | 0 | - |
| - | - | 1717 | 295.1 | - | - | 0 | - |
| - | - | 1918 | 296.2 | - | - | 0 | - |
| - | - | 961.8 | 296.7 | - | - | 0 | - |
| - | - | 2973 | 297.1 | - | - | 0 | - |
| - | - | 1348 | 297.2 | - | - | 0 | - |
| - | - | 1847 | 297.5 | - | - | 0 | - |
| - | - | 1815 | 297.6 | - | - | 0 | - |
| - | - | 595.1 | 297.8 | - | - | 0 | - |
| - | - | 665.3 | 298.2 | - | - | 0 | - |
| - | - | 1611 | 299.7 | - | - | 0 | - |
| - | - | 1007 | 300.2 | - | - | 0 | - |
| - | - | 1492 | 300.2 | - | - | 0 | - |
| - | - | 1.133E+04 | 300.2 | - | - | 0 | - |
| - | - | 581.3 | 300.6 | - | - | 0 | - |
| - | - | 1011 | 301.1 | - | - | 0 | - |
| - | - | 3413 | 301.2 | - | - | 0 | - |
| - | - | 1756 | 301.2 | - | - | 0 | - |
| - | - | 3190 | 302.1 | - | - | 0 | - |
| - | - | 714.7 | 302.2 | - | - | 0 | - |
| - | - | 1222 | 302.5 | - | - | 0 | - |
| - | - | 5308 | 303.2 | - | - | 0 | - |
| - | - | 1566 | 304.8 | - | - | 0 | - |
| - | - | 734.8 | 305.2 | - | - | 0 | - |
| - | - | 1561 | 305.6 | - | - | 0 | - |
| - | - | 599.3 | 307.1 | - | - | 0 | - |
| - | - | 560.5 | 307.5 | - | - | 0 | - |
| - | - | 2792 | 308.1 | - | - | 0 | - |
| - | - | 4678 | 309.1 | - | - | 0 | - |
| - | - | 2068 | 309.6 | - | - | 0 | - |
| - | - | 3446 | 311.1 | - | - | 0 | - |
| - | - | 774.7 | 312.2 | - | - | 0 | - |
| - | - | 995.3 | 312.6 | - | - | 0 | - |
| - | - | 924.4 | 312.8 | - | - | 0 | - |
| - | - | 793.4 | 313.1 | - | - | 0 | - |
| 9 | y | 2359 | 313.2 | 0.0007916 | 2.528 | +3 | 7 |
| 9 | y | 1356 | 313.5 | 3.015E-05 | 0.09618 | +3 | 7 |
| 9 | z | 1624 | 313.8 | 0.0006759 | 2.154 | +3 | 7 |
| - | - | 908.9 | 314.2 | - | - | 0 | - |
| - | - | 998.9 | 314.5 | - | - | 0 | - |
| - | - | 1861 | 315.2 | - | - | 0 | - |
| - | - | 1811 | 315.2 | - | - | 0 | - |
| - | - | 1193 | 316.2 | - | - | 0 | - |
| - | - | 5558 | 317.7 | - | - | 0 | - |
| - | - | 1.729E+04 | 318.2 | - | - | 0 | - |
| - | - | 2885 | 318.2 | - | - | 0 | - |
| - | - | 7142 | 318.7 | - | - | 0 | - |
| - | - | 1791 | 318.8 | - | - | 0 | - |
| - | - | 1946 | 319.2 | - | - | 0 | - |
| 9 | y | 896.6 | 319.2 | 0.003347 | 10.49 | +3 | 7 |
| - | - | 1807 | 320.7 | - | - | 0 | - |
| - | - | 597.7 | 322.8 | - | - | 0 | - |
| - | - | 4339 | 323.1 | - | - | 0 | - |
| - | - | 1557 | 323.6 | - | - | 0 | - |
| - | - | 1575 | 324.7 | - | - | 0 | - |
| - | - | 8.423E+04 | 326.7 | - | - | 0 | - |
| 8 | z | 3.154E+04 | 327.2 | 0.004264 | 13.03 | +3 | 8 |
| - | - | 5723 | 327.7 | - | - | 0 | - |
| - | - | 1772 | 327.8 | - | - | 0 | - |
| 8 | c | 1670 | 328.2 | 0.004503 | 13.72 | +3 | 8 |
| - | - | 680.7 | 330.7 | - | - | 0 | - |
| - | - | 1.331E+04 | 331.7 | - | - | 0 | - |
| - | - | 4683 | 332.2 | - | - | 0 | - |
| - | - | 1291 | 332.2 | - | - | 0 | - |
| 8 | y | 2711 | 332.5 | 0.0008712 | 2.62 | +3 | 8 |
| - | - | 984.8 | 332.7 | - | - | 0 | - |
| 8 | z | 3523 | 332.8 | 0.0003232 | 0.971 | +3 | 8 |
| - | - | 1068 | 333.2 | - | - | 0 | - |
| - | - | 1153 | 337.2 | - | - | 0 | - |
| - | - | 1190 | 337.8 | - | - | 0 | - |
| - | - | 1712 | 338.1 | - | - | 0 | - |
| 8 | y | 3.733E+04 | 338.2 | 0.0002067 | 0.6112 | +3 | 8 |
| - | - | 1.86E+04 | 338.5 | - | - | 0 | - |
| - | - | 1721 | 338.7 | - | - | 0 | - |
| - | - | 5439 | 338.8 | - | - | 0 | - |
| - | - | 2095 | 339.2 | - | - | 0 | - |
| - | - | 1401 | 339.5 | - | - | 0 | - |
| - | - | 848.7 | 340.3 | - | - | 0 | - |
| 5 | c | 4.154E+04 | 340.7 | 0.001355 | 3.979 | +2 | 5 |
| 11 | y | 2.227E+04 | 341.2 | 0.004611 | 13.51 | +2 | 5 |
| 11 | z | 7274 | 341.7 | 0.005502 | 16.1 | +2 | 5 |
| - | - | 2128 | 342.2 | - | - | 0 | - |
| - | - | 957.7 | 344.2 | - | - | 0 | - |
| - | - | 860.1 | 345.2 | - | - | 0 | - |
| - | - | 1134 | 346.1 | - | - | 0 | - |
| - | - | 914.4 | 346.2 | - | - | 0 | - |
| - | - | 1186 | 347.2 | - | - | 0 | - |
| - | - | 765.8 | 349.2 | - | - | 0 | - |
| 11 | y | 5.127E+04 | 349.7 | 0.0009479 | 2.711 | +2 | 5 |
| - | - | 1.87E+04 | 350.2 | - | - | 0 | - |
| - | - | 6986 | 350.7 | - | - | 0 | - |
| - | - | 1131 | 351.2 | - | - | 0 | - |
| - | - | 1215 | 351.2 | - | - | 0 | - |
| - | - | 9944 | 352.2 | - | - | 0 | - |
| - | - | 4395 | 352.5 | - | - | 0 | - |
| - | - | 2257 | 352.8 | - | - | 0 | - |
| 5 | y | 7364 | 353.2 | 0.000502 | 1.421 | +4 | 11 |
| 5 | z | 4855 | 353.4 | 0.00121 | 3.424 | +4 | 11 |
| - | - | 4907 | 353.7 | - | - | 0 | - |
| - | - | 1000 | 353.9 | - | - | 0 | - |
| - | - | 1675 | 354.2 | - | - | 0 | - |
| - | - | 1243 | 355.2 | - | - | 0 | - |
| 7 | w | 2120 | 356.2 | 0.0003103 | 0.8712 | +3 | 9 |
| - | - | 2527 | 356.5 | - | - | 0 | - |
| - | - | 862.2 | 356.9 | - | - | 0 | - |
| - | - | 1230 | 357.2 | - | - | 0 | - |
| - | - | 5184 | 360.7 | - | - | 0 | - |
| 14 | c | 1293 | 361.2 | 0.0002577 | 0.7136 | +5 | 14 |
| - | - | 1811 | 361.2 | - | - | 0 | - |
| - | - | 1591 | 361.5 | - | - | 0 | - |
| - | - | 837.1 | 361.9 | - | - | 0 | - |
| - | - | 3168 | 362.2 | - | - | 0 | - |
| - | - | 845.3 | 362.7 | - | - | 0 | - |
| - | - | 1840 | 363.1 | - | - | 0 | - |
| - | - | 888 | 363.2 | - | - | 0 | - |
| - | - | 4619 | 363.7 | - | - | 0 | - |
| - | - | 1173 | 364.1 | - | - | 0 | - |
| - | - | 761.6 | 364.3 | - | - | 0 | - |
| - | - | 657.1 | 365.6 | - | - | 0 | - |
| - | - | 1810 | 365.7 | - | - | 0 | - |
| - | - | 970.1 | 366.2 | - | - | 0 | - |
| - | - | 609.2 | 367.1 | - | - | 0 | - |
| - | - | 1891 | 367.3 | - | - | 0 | - |
| - | - | 816 | 367.7 | - | - | 0 | - |
| - | - | 1631 | 368.2 | - | - | 0 | - |
| - | - | 1192 | 368.7 | - | - | 0 | - |
| - | - | 1355 | 370.1 | - | - | 0 | - |
| 7 | y | 1570 | 370.5 | 0.001384 | 3.734 | +3 | 9 |
| 7 | y | 2676 | 370.8 | 0.0031 | 8.36 | +3 | 9 |
| 7 | z | 2048 | 371.2 | 0.001204 | 3.244 | +3 | 9 |
| - | - | 1255 | 371.5 | - | - | 0 | - |
| - | - | 1562 | 371.7 | - | - | 0 | - |
| - | - | 1255 | 372.2 | - | - | 0 | - |
| - | - | 1761 | 372.2 | - | - | 0 | - |
| - | - | 768.8 | 372.6 | - | - | 0 | - |
| - | - | 3452 | 374.2 | - | - | 0 | - |
| - | - | 1709 | 374.4 | - | - | 0 | - |
| - | - | 1.198E+04 | 374.7 | - | - | 0 | - |
| - | - | 6135 | 375.2 | - | - | 0 | - |
| - | - | 5880 | 375.2 | - | - | 0 | - |
| - | - | 1674 | 375.7 | - | - | 0 | - |
| 7 | y | 2.896E+04 | 376.5 | 0.0001749 | 0.4644 | +3 | 9 |
| 10 | w | 2.147E+05 | 376.7 | 0.001012 | 2.686 | +2 | 6 |
| - | - | 1.405E+04 | 376.9 | - | - | 0 | - |
| - | - | 8.758E+04 | 377.2 | - | - | 0 | - |
| - | - | 1588 | 377.4 | - | - | 0 | - |
| - | - | 1976 | 377.5 | - | - | 0 | - |
| - | - | 2565 | 377.6 | - | - | 0 | - |
| 4 | z | 2.765E+04 | 377.7 | 0.00678 | 17.95 | +4 | 12 |
| - | - | 5961 | 377.8 | - | - | 0 | - |
| - | - | 5837 | 378 | - | - | 0 | - |
| - | - | 8220 | 378.2 | - | - | 0 | - |
| - | - | 3408 | 378.2 | - | - | 0 | - |
| - | - | 1039 | 378.4 | - | - | 0 | - |
| - | - | 1896 | 378.7 | - | - | 0 | - |
| 3 | c | 2.797E+04 | 379.2 | 0.001253 | 3.305 | +1 | 3 |
| - | - | 1869 | 379.9 | - | - | 0 | - |
| - | - | 4268 | 380.2 | - | - | 0 | - |
| - | - | 1152 | 380.5 | - | - | 0 | - |
| - | - | 1895 | 380.9 | - | - | 0 | - |
| - | - | 7379 | 381.2 | - | - | 0 | - |
| - | - | 9008 | 381.2 | - | - | 0 | - |
| - | - | 1.41E+04 | 381.4 | - | - | 0 | - |
| - | - | 1.516E+04 | 381.6 | - | - | 0 | - |
| 4 | y | 1021 | 381.7 | 0.002387 | 6.253 | +4 | 12 |
| - | - | 8572 | 381.8 | - | - | 0 | - |
| - | - | 2157 | 382 | - | - | 0 | - |
| - | - | 956.4 | 382.2 | - | - | 0 | - |
| 4 | z | 2076 | 382.2 | 0.00713 | 18.66 | +4 | 12 |
| - | - | 3.92E+04 | 383.2 | - | - | 0 | - |
| - | - | 1.854E+04 | 383.7 | - | - | 0 | - |
| - | - | 1174 | 384.2 | - | - | 0 | - |
| - | - | 5601 | 384.2 | - | - | 0 | - |
| - | - | 1738 | 384.3 | - | - | 0 | - |
| - | - | 937 | 384.6 | - | - | 0 | - |
| - | - | 1487 | 384.8 | - | - | 0 | - |
| - | - | 1.03E+04 | 384.8 | - | - | 0 | - |
| - | - | 1.311E+04 | 385 | - | - | 0 | - |
| - | - | 2761 | 385.2 | - | - | 0 | - |
| - | - | 8413 | 385.2 | - | - | 0 | - |
| - | - | 908.6 | 385.2 | - | - | 0 | - |
| - | - | 1624 | 385.3 | - | - | 0 | - |
| - | - | 3289 | 385.4 | - | - | 0 | - |
| - | - | 1024 | 385.6 | - | - | 0 | - |
| - | - | 860.4 | 386.2 | - | - | 0 | - |
| 4 | y | 993.3 | 386.2 | 0.003151 | 8.158 | +4 | 12 |
| - | - | 2768 | 387.2 | - | - | 0 | - |
| - | - | 2356 | 387.7 | - | - | 0 | - |
| - | - | 853.8 | 388.2 | - | - | 0 | - |
| - | - | 3342 | 388.7 | - | - | 0 | - |
| - | - | 2256 | 389.2 | - | - | 0 | - |
| - | - | 1330 | 390.2 | - | - | 0 | - |
| - | - | 955.4 | 392.7 | - | - | 0 | - |
| 6 | w | 1.664E+04 | 394.5 | 3.431E-05 | 0.08697 | +3 | 10 |
| - | - | 9539 | 394.9 | - | - | 0 | - |
| - | - | 1259 | 395.1 | - | - | 0 | - |
| - | - | 2382 | 395.2 | - | - | 0 | - |
| - | - | 806.6 | 395.2 | - | - | 0 | - |
| - | - | 866.9 | 395.5 | - | - | 0 | - |
| 3 | c | 4.03E+04 | 396.2 | 0.001346 | 3.397 | +1 | 3 |
| 6 | c | 2.539E+04 | 397.2 | 0.001529 | 3.85 | +2 | 6 |
| - | - | 7646 | 397.2 | - | - | 0 | - |
| - | - | 1.162E+04 | 397.7 | - | - | 0 | - |
| - | - | 1890 | 398.2 | - | - | 0 | - |
| - | - | 4094 | 398.2 | - | - | 0 | - |
| - | - | 1179 | 398.2 | - | - | 0 | - |
| - | - | 1760 | 402.2 | - | - | 0 | - |
| 6 | z | 2601 | 403.2 | 0.003181 | 7.89 | +3 | 10 |
| - | - | 766.4 | 403.7 | - | - | 0 | - |
| 10 | y | 6808 | 405.2 | 0.0004826 | 1.191 | +2 | 6 |
| - | - | 2049 | 405.7 | - | - | 0 | - |
| 10 | z | 1302 | 406.2 | 0.0009501 | 2.339 | +2 | 6 |
| 13 | y | 2.194E+04 | 407.2 | 0.002984 | 7.329 | +1 | 3 |
| 10 | c | 4548 | 408.2 | 0.005129 | 12.56 | +3 | 10 |
| 6 | y | 3800 | 408.5 | 0.004518 | 11.06 | +3 | 10 |
| 6 | z | 1427 | 408.9 | 0.0009435 | 2.308 | +3 | 10 |
| - | - | 1803 | 409.1 | - | - | 0 | - |
| - | - | 1321 | 409.2 | - | - | 0 | - |
| 3 | z | 959.7 | 409.7 | 0.003439 | 8.394 | +4 | 13 |
| - | - | 817.9 | 410.2 | - | - | 0 | - |
| - | - | 2790 | 412.2 | - | - | 0 | - |
| - | - | 994.2 | 412.2 | - | - | 0 | - |
| - | - | 1116 | 412.7 | - | - | 0 | - |
| 3 | y | 1987 | 413.7 | 0.0007911 | 1.912 | +4 | 13 |
| - | - | 1342 | 413.9 | - | - | 0 | - |
| 3 | y | 1893 | 413.9 | 0.00494 | 11.93 | +4 | 13 |
| 6 | y | 3.42E+04 | 414.2 | 0.0004659 | 1.125 | +3 | 10 |
| - | - | 1027 | 414.5 | - | - | 0 | - |
| - | - | 2.428E+04 | 414.5 | - | - | 0 | - |
| - | - | 2141 | 414.7 | - | - | 0 | - |
| - | - | 9430 | 414.9 | - | - | 0 | - |
| - | - | 1722 | 415.2 | - | - | 0 | - |
| - | - | 762.8 | 416.2 | - | - | 0 | - |
| - | - | 3.892E+04 | 417.2 | - | - | 0 | - |
| - | - | 1.53E+04 | 417.7 | - | - | 0 | - |
| - | - | 7617 | 418.2 | - | - | 0 | - |
| - | - | 716.4 | 418.7 | - | - | 0 | - |
| - | - | 2587 | 419.2 | - | - | 0 | - |
| - | - | 2192 | 419.7 | - | - | 0 | - |
| - | - | 1186 | 420.2 | - | - | 0 | - |
| - | - | 1014 | 421.2 | - | - | 0 | - |
| - | - | 957 | 422.3 | - | - | 0 | - |
| - | - | 1.513E+04 | 424.7 | - | - | 0 | - |
| - | - | 1.035E+04 | 425.3 | - | - | 0 | - |
| - | - | 3189 | 425.8 | - | - | 0 | - |
| - | - | 1331 | 426.2 | - | - | 0 | - |
| - | - | 1081 | 426.7 | - | - | 0 | - |
| - | - | 1027 | 426.7 | - | - | 0 | - |
| - | - | 812.3 | 427.5 | - | - | 0 | - |
| - | - | 8202 | 428.3 | - | - | 0 | - |
| - | - | 3603 | 429.3 | - | - | 0 | - |
| - | - | 6993 | 430.2 | - | - | 0 | - |
| - | - | 638.9 | 430.3 | - | - | 0 | - |
| - | - | 1137 | 431 | - | - | 0 | - |
| - | - | 2023 | 431.2 | - | - | 0 | - |
| - | - | 1185 | 432.7 | - | - | 0 | - |
| - | - | 1843 | 433.7 | - | - | 0 | - |
| - | - | 1608 | 434.2 | - | - | 0 | - |
| - | - | 3801 | 434.7 | - | - | 0 | - |
| - | - | 1663 | 435.2 | - | - | 0 | - |
| - | - | 2807 | 435.2 | - | - | 0 | - |
| - | - | 856.6 | 436.2 | - | - | 0 | - |
| - | - | 3210 | 436.2 | - | - | 0 | - |
| - | - | 1492 | 436.9 | - | - | 0 | - |
| - | - | 785.3 | 437.5 | - | - | 0 | - |
| - | - | 2596 | 437.5 | - | - | 0 | - |
| - | - | 674.6 | 437.7 | - | - | 0 | - |
| - | - | 1943 | 437.9 | - | - | 0 | - |
| - | - | 1157 | 438 | - | - | 0 | - |
| - | - | 882 | 438.2 | - | - | 0 | - |
| - | - | 877.3 | 439.2 | - | - | 0 | - |
| - | - | 1038 | 439.5 | - | - | 0 | - |
| - | - | 1078 | 439.7 | - | - | 0 | - |
| - | - | 1.951E+04 | 440.7 | - | - | 0 | - |
| 9 | w | 1.204E+04 | 441.2 | 0.000885 | 2.006 | +2 | 7 |
| - | - | 1.09E+04 | 441.2 | - | - | 0 | - |
| - | - | 5714 | 441.7 | - | - | 0 | - |
| - | - | 1957 | 441.7 | - | - | 0 | - |
| - | - | 3579 | 442.2 | - | - | 0 | - |
| - | - | 875.5 | 443.2 | - | - | 0 | - |
| - | - | 904.1 | 443.5 | - | - | 0 | - |
| - | - | 6746 | 443.7 | - | - | 0 | - |
| 2 | z | 5462 | 444 | 0.0001741 | 0.392 | +4 | 14 |
| - | - | 4212 | 444.2 | - | - | 0 | - |
| - | - | 1351 | 444.5 | - | - | 0 | - |
| - | - | 819.4 | 444.7 | - | - | 0 | - |
| - | - | 943.8 | 445 | - | - | 0 | - |
| - | - | 1685 | 445.7 | - | - | 0 | - |
| - | - | 1250 | 445.7 | - | - | 0 | - |
| - | - | 908 | 446.2 | - | - | 0 | - |
| - | - | 8672 | 447.2 | - | - | 0 | - |
| - | - | 1124 | 447.3 | - | - | 0 | - |
| - | - | 882.3 | 447.5 | - | - | 0 | - |
| - | - | 2204 | 447.7 | - | - | 0 | - |
| 2 | y | 4844 | 448 | 0.0008946 | 1.997 | +4 | 14 |
| 2 | y | 9761 | 448.2 | 0.007576 | 16.9 | +4 | 14 |
| - | - | 2834 | 448.3 | - | - | 0 | - |
| 2 | z | 5891 | 448.5 | 0.004766 | 10.63 | +4 | 14 |
| - | - | 6686 | 448.7 | - | - | 0 | - |
| - | - | 1872 | 449 | - | - | 0 | - |
| - | - | 1762 | 449.2 | - | - | 0 | - |
| - | - | 869.4 | 450.9 | - | - | 0 | - |
| 14 | c | 3183 | 451.2 | 0.002111 | 4.678 | +4 | 14 |
| 14 | c | 3935 | 451.5 | 0.00391 | 8.66 | +4 | 14 |
| - | - | 710.7 | 451.5 | - | - | 0 | - |
| - | - | 4111 | 451.7 | - | - | 0 | - |
| - | - | 757.3 | 451.9 | - | - | 0 | - |
| - | - | 1873 | 452 | - | - | 0 | - |
| - | - | 3063 | 452.2 | - | - | 0 | - |
| 2 | y | 7392 | 452.5 | 0.000878 | 1.94 | +4 | 14 |
| - | - | 7195 | 452.7 | - | - | 0 | - |
| - | - | 3318 | 453 | - | - | 0 | - |
| - | - | 2358 | 453.2 | - | - | 0 | - |
| - | - | 1.555E+04 | 453.3 | - | - | 0 | - |
| - | - | 1.155E+04 | 453.8 | - | - | 0 | - |
| - | - | 873.9 | 454 | - | - | 0 | - |
| 7 | c | 2735 | 454.2 | 0.00155 | 3.412 | +2 | 7 |
| - | - | 4215 | 454.3 | - | - | 0 | - |
| 7 | c | 6243 | 454.7 | 0.003286 | 7.227 | +2 | 7 |
| - | - | 1222 | 454.8 | - | - | 0 | - |
| - | - | 2023 | 455.2 | - | - | 0 | - |
| - | - | 1987 | 455.6 | - | - | 0 | - |
| 14 | c | 2.71E+04 | 455.7 | 0.0008734 | 1.917 | +4 | 14 |
| - | - | 3205 | 455.9 | - | - | 0 | - |
| - | - | 2.589E+04 | 456 | - | - | 0 | - |
| - | - | 1.462E+04 | 456.2 | - | - | 0 | - |
| - | - | 4178 | 456.5 | - | - | 0 | - |
| - | - | 1232 | 456.6 | - | - | 0 | - |
| - | - | 3659 | 456.7 | - | - | 0 | - |
| - | - | 727.1 | 456.9 | - | - | 0 | - |
| - | - | 7049 | 457.2 | - | - | 0 | - |
| - | - | 2467 | 457.7 | - | - | 0 | - |
| - | - | 888.6 | 458.2 | - | - | 0 | - |
| - | - | 1106 | 459.2 | - | - | 0 | - |
| - | - | 1007 | 460.2 | - | - | 0 | - |
| - | - | 6946 | 460.3 | - | - | 0 | - |
| 12 | w | 5281 | 461.2 | 0.002898 | 6.284 | +1 | 4 |
| - | - | 1541 | 461.3 | - | - | 0 | - |
| - | - | 1988 | 461.5 | - | - | 0 | - |
| - | - | 1124 | 461.6 | - | - | 0 | - |
| - | - | 3647 | 461.7 | - | - | 0 | - |
| - | - | 935.6 | 462 | - | - | 0 | - |
| 11 | c | 3641 | 462.2 | 0.003929 | 8.5 | +3 | 11 |
| - | - | 934.1 | 462.5 | - | - | 0 | - |
| - | - | 6776 | 462.7 | - | - | 0 | - |
| - | - | 4282 | 463 | - | - | 0 | - |
| 7 | c | 3667 | 463.2 | 0.0005401 | 1.166 | +2 | 7 |
| - | - | 1202 | 463.5 | - | - | 0 | - |
| - | - | 1354 | 463.7 | - | - | 0 | - |
| 5 | z | 1591 | 465.2 | 0.000643 | 1.382 | +3 | 11 |
| - | - | 5248 | 465.7 | - | - | 0 | - |
| - | - | 1.099E+05 | 466 | - | - | 0 | - |
| - | - | 9.866E+04 | 466.2 | - | - | 0 | - |
| - | - | 5.043E+04 | 466.5 | - | - | 0 | - |
| - | - | 2.299E+04 | 466.7 | - | - | 0 | - |
| - | - | 7494 | 467 | - | - | 0 | - |
| - | - | 2848 | 467.2 | - | - | 0 | - |
| - | - | 2749 | 467.2 | - | - | 0 | - |
| - | - | 1328 | 468.6 | - | - | 0 | - |
| 9 | y | 2924 | 469.2 | 0.0004788 | 1.02 | +2 | 7 |
| - | - | 1.049E+04 | 469.7 | - | - | 0 | - |
| - | - | 2239 | 470 | - | - | 0 | - |
| 5 | y | 2.357E+04 | 470.2 | 0.001337 | 2.844 | +3 | 11 |
| 5 | y | 1.572E+04 | 470.6 | 5.326E-05 | 0.1132 | +3 | 11 |
| - | - | 1.207E+04 | 470.7 | - | - | 0 | - |
| 5 | z | 1.408E+04 | 470.9 | 0.00108 | 2.294 | +3 | 11 |
| - | - | 8833 | 471.2 | - | - | 0 | - |
| - | - | 3405 | 471.6 | - | - | 0 | - |
| - | - | 1026 | 471.7 | - | - | 0 | - |
| - | - | 1618 | 471.9 | - | - | 0 | - |
| - | - | 1764 | 472 | - | - | 0 | - |
| - | - | 1176 | 472.2 | - | - | 0 | - |
| - | - | 955.8 | 473.7 | - | - | 0 | - |
| - | - | 1730 | 474.2 | - | - | 0 | - |
| - | - | 1555 | 474.3 | - | - | 0 | - |
| - | - | 862.4 | 474.7 | - | - | 0 | - |
| - | - | 6487 | 475.3 | - | - | 0 | - |
| - | - | 1166 | 475.9 | - | - | 0 | - |
| 5 | y | 1.768E+04 | 476.2 | 0.001086 | 2.28 | +3 | 11 |
| - | - | 1947 | 476.3 | - | - | 0 | - |
| - | - | 1.142E+04 | 476.5 | - | - | 0 | - |
| - | - | 9866 | 476.6 | - | - | 0 | - |
| - | - | 1.096E+04 | 476.7 | - | - | 0 | - |
| - | - | 3680 | 476.9 | - | - | 0 | - |
| - | - | 3808 | 477 | - | - | 0 | - |
| - | - | 2240 | 477.2 | - | - | 0 | - |
| - | - | 5337 | 477.2 | - | - | 0 | - |
| - | - | 1313 | 477.3 | - | - | 0 | - |
| - | - | 1958 | 477.5 | - | - | 0 | - |
| - | - | 2996 | 477.7 | - | - | 0 | - |
| 9 | y | 2.916E+04 | 478.3 | 0.0009394 | 1.964 | +2 | 7 |
| - | - | 1.346E+04 | 478.8 | - | - | 0 | - |
| - | - | 5657 | 479.3 | - | - | 0 | - |
| - | - | 3580 | 479.6 | - | - | 0 | - |
| - | - | 3058 | 479.9 | - | - | 0 | - |
| - | - | 3102 | 480.2 | - | - | 0 | - |
| - | - | 7834 | 480.5 | - | - | 0 | - |
| - | - | 1.591E+05 | 480.7 | - | - | 0 | - |
| - | - | 2.049E+05 | 481 | - | - | 0 | - |
| - | - | 1.499E+05 | 481.2 | - | - | 0 | - |
| - | - | 7.671E+04 | 481.5 | - | - | 0 | - |
| - | - | 2.941E+04 | 481.7 | - | - | 0 | - |
| - | - | 9575 | 482 | - | - | 0 | - |
| - | - | 4444 | 482.2 | - | - | 0 | - |
| 8 | c | 6335 | 482.7 | 0.001835 | 3.801 | +2 | 8 |
| 8 | c | 1.275E+04 | 483.2 | 0.003693 | 7.643 | +2 | 8 |
| - | - | 5455 | 483.7 | - | - | 0 | - |
| - | - | 3453 | 484.2 | - | - | 0 | - |
| - | - | 984.2 | 484.7 | - | - | 0 | - |
| - | - | 5310 | 485.2 | - | - | 0 | - |
| - | - | 5333 | 485.6 | - | - | 0 | - |
| - | - | 2807 | 485.9 | - | - | 0 | - |
| - | - | 891.4 | 489.2 | - | - | 0 | - |
| 8 | z | 2820 | 490.2 | 0.007151 | 14.59 | +2 | 8 |
| - | - | 1934 | 491.2 | - | - | 0 | - |
| 8 | c | 2.69E+04 | 491.7 | 0.00168 | 3.416 | +2 | 8 |
| - | - | 1.759E+04 | 492.2 | - | - | 0 | - |
| - | - | 4588 | 492.7 | - | - | 0 | - |
| 4 | c | 1852 | 493.3 | 0.002246 | 4.553 | +1 | 4 |
| 4 | c | 4.404E+04 | 494.2 | 0.002051 | 4.149 | +1 | 4 |
| - | - | 1577 | 494.6 | - | - | 0 | - |
| - | - | 1.268E+04 | 495.2 | - | - | 0 | - |
| - | - | 3772 | 496.3 | - | - | 0 | - |
| - | - | 1381 | 496.7 | - | - | 0 | - |
| - | - | 1764 | 497.2 | - | - | 0 | - |
| - | - | 965.7 | 497.7 | - | - | 0 | - |
| 8 | y | 1933 | 497.8 | 0.00233 | 4.681 | +2 | 8 |
| - | - | 1179 | 498.2 | - | - | 0 | - |
| 8 | z | 2.058E+04 | 498.8 | 0.0003764 | 0.7546 | +2 | 8 |
| - | - | 1.157E+04 | 499.3 | - | - | 0 | - |
| - | - | 4278 | 499.8 | - | - | 0 | - |
| - | - | 1766 | 500.3 | - | - | 0 | - |
| 4 | z | 2881 | 503.2 | 0.008119 | 16.13 | +3 | 12 |
| - | - | 1083 | 504.2 | - | - | 0 | - |
| 12 | c | 2883 | 505.3 | 0.003263 | 6.458 | +3 | 12 |
| - | - | 2.259E+04 | 505.7 | - | - | 0 | - |
| - | - | 1.604E+04 | 506.2 | - | - | 0 | - |
| 8 | y | 6.362E+04 | 506.8 | 0.0005018 | 0.9902 | +2 | 8 |
| - | - | 3.6E+04 | 507.3 | - | - | 0 | - |
| - | - | 1.207E+04 | 507.8 | - | - | 0 | - |
| - | - | 3317 | 508.3 | - | - | 0 | - |
| 4 | z | 2043 | 509.2 | 0.0007828 | 1.537 | +3 | 12 |
| - | - | 1041 | 509.8 | - | - | 0 | - |
| - | - | 910.9 | 510.6 | - | - | 0 | - |
| - | - | 4861 | 510.8 | - | - | 0 | - |
| 12 | c | 3.931E+04 | 510.9 | 0.001977 | 3.869 | +3 | 12 |
| 4 | c | 1.226E+05 | 511.3 | 0.002265 | 4.431 | +1 | 4 |
| - | - | 1.62E+04 | 511.6 | - | - | 0 | - |
| - | - | 1615 | 511.8 | - | - | 0 | - |
| - | - | 5585 | 511.9 | - | - | 0 | - |
| - | - | 2.057E+04 | 512.3 | - | - | 0 | - |
| - | - | 883.7 | 512.6 | - | - | 0 | - |
| - | - | 2800 | 513.3 | - | - | 0 | - |
| - | - | 1327 | 513.8 | - | - | 0 | - |
| - | - | 2.694E+04 | 514.2 | - | - | 0 | - |
| 4 | y | 3489 | 514.6 | 0.0001688 | 0.328 | +3 | 12 |
| - | - | 2.91E+04 | 514.7 | - | - | 0 | - |
| - | - | 2210 | 514.9 | - | - | 0 | - |
| - | - | 1.418E+04 | 515.2 | - | - | 0 | - |
| - | - | 4620 | 515.7 | - | - | 0 | - |
| - | - | 1913 | 516.3 | - | - | 0 | - |
| - | - | 2265 | 518.2 | - | - | 0 | - |
| - | - | 1010 | 518.3 | - | - | 0 | - |
| - | - | 1697 | 518.6 | - | - | 0 | - |
| 12 | z | 3.936E+04 | 519.3 | 0.002655 | 5.112 | +1 | 4 |
| - | - | 1383 | 519.6 | - | - | 0 | - |
| - | - | 975.9 | 519.7 | - | - | 0 | - |
| - | - | 2834 | 519.9 | - | - | 0 | - |
| - | - | 1.14E+04 | 520.3 | - | - | 0 | - |
| - | - | 1119 | 520.7 | - | - | 0 | - |
| - | - | 1066 | 521.2 | - | - | 0 | - |
| - | - | 3089 | 521.3 | - | - | 0 | - |
| - | - | 1369 | 524.6 | - | - | 0 | - |
| - | - | 1517 | 525.3 | - | - | 0 | - |
| - | - | 5194 | 526.3 | - | - | 0 | - |
| - | - | 913.3 | 526.6 | - | - | 0 | - |
| - | - | 7254 | 526.8 | - | - | 0 | - |
| - | - | 3716 | 527.2 | - | - | 0 | - |
| - | - | 4288 | 527.3 | - | - | 0 | - |
| - | - | 5.852E+04 | 527.8 | - | - | 0 | - |
| - | - | 3.203E+04 | 528.3 | - | - | 0 | - |
| - | - | 1.068E+04 | 528.8 | - | - | 0 | - |
| - | - | 2863 | 529.3 | - | - | 0 | - |
| - | - | 864.9 | 530.6 | - | - | 0 | - |
| - | - | 1089 | 531.3 | - | - | 0 | - |
| - | - | 1008 | 531.6 | - | - | 0 | - |
| - | - | 3320 | 532.2 | - | - | 0 | - |
| - | - | 3474 | 532.3 | - | - | 0 | - |
| 3 | w | 6.577E+04 | 532.6 | 0.0006431 | 1.208 | +3 | 13 |
| - | - | 5116 | 532.8 | - | - | 0 | - |
| - | - | 5.714E+04 | 532.9 | - | - | 0 | - |
| - | - | 3.356E+04 | 533.3 | - | - | 0 | - |
| - | - | 1.018E+04 | 533.6 | - | - | 0 | - |
| 7 | w | 2766 | 533.8 | 0.005721 | 10.72 | +2 | 9 |
| - | - | 4948 | 533.9 | - | - | 0 | - |
| - | - | 6731 | 534.3 | - | - | 0 | - |
| - | - | 4115 | 534.8 | - | - | 0 | - |
| - | - | 1181 | 534.9 | - | - | 0 | - |
| 12 | y | 2.435E+04 | 535.3 | 0.00496 | 9.266 | +1 | 4 |
| - | - | 1.195E+04 | 535.8 | - | - | 0 | - |
| - | - | 4726 | 536.3 | - | - | 0 | - |
| - | - | 1016 | 536.9 | - | - | 0 | - |
| - | - | 4507 | 537.3 | - | - | 0 | - |
| - | - | 2358 | 537.6 | - | - | 0 | - |
| - | - | 975.1 | 537.9 | - | - | 0 | - |
| - | - | 2348 | 538.3 | - | - | 0 | - |
| - | - | 2881 | 539.2 | - | - | 0 | - |
| - | - | 1126 | 539.3 | - | - | 0 | - |
| - | - | 1189 | 539.6 | - | - | 0 | - |
| - | - | 1032 | 540.3 | - | - | 0 | - |
| - | - | 1729 | 541.3 | - | - | 0 | - |
| - | - | 1120 | 541.8 | - | - | 0 | - |
| - | - | 3308 | 542.3 | - | - | 0 | - |
| - | - | 867.8 | 542.8 | - | - | 0 | - |
| - | - | 2042 | 543.3 | - | - | 0 | - |
| - | - | 994.1 | 544.3 | - | - | 0 | - |
| - | - | 1037 | 544.6 | - | - | 0 | - |
| - | - | 962.5 | 544.9 | - | - | 0 | - |
| - | - | 2083 | 545.6 | - | - | 0 | - |
| 3 | z | 1333 | 545.9 | 0.003232 | 5.919 | +3 | 13 |
| 3 | z | 1978 | 546.3 | 0.007807 | 14.29 | +3 | 13 |
| - | - | 1319 | 546.8 | - | - | 0 | - |
| 9 | c | 4232 | 547.3 | 0.002385 | 4.357 | +2 | 9 |
| - | - | 3253 | 547.8 | - | - | 0 | - |
| - | - | 1070 | 548.3 | - | - | 0 | - |
| - | - | 1111 | 548.9 | - | - | 0 | - |
| - | - | 1842 | 549.2 | - | - | 0 | - |
| - | - | 2309 | 549.3 | - | - | 0 | - |
| - | - | 8131 | 549.6 | - | - | 0 | - |
| - | - | 7690 | 549.9 | - | - | 0 | - |
| - | - | 2848 | 550.2 | - | - | 0 | - |
| - | - | 1.057E+04 | 550.3 | - | - | 0 | - |
| - | - | 2698 | 550.6 | - | - | 0 | - |
| 3 | y | 1.327E+04 | 551.3 | 0.005411 | 9.815 | +3 | 13 |
| 3 | y | 6891 | 551.6 | 0.003865 | 7.006 | +3 | 13 |
| 3 | z | 1.48E+04 | 551.9 | 0.001541 | 2.792 | +3 | 13 |
| - | - | 1.51E+04 | 552.3 | - | - | 0 | - |
| - | - | 7024 | 552.6 | - | - | 0 | - |
| - | - | 2446 | 552.9 | - | - | 0 | - |
| - | - | 2145 | 553.3 | - | - | 0 | - |
| - | - | 1774 | 554.3 | - | - | 0 | - |
| - | - | 1246 | 554.6 | - | - | 0 | - |
| - | - | 1657 | 554.8 | - | - | 0 | - |
| - | - | 2246 | 554.9 | - | - | 0 | - |
| 7 | y | 1.241E+04 | 555.3 | 0.000909 | 1.637 | +2 | 9 |
| 9 | c | 2.709E+05 | 555.8 | 0.001867 | 3.358 | +2 | 9 |
| - | - | 1.43E+05 | 556.3 | - | - | 0 | - |
| - | - | 4.361E+04 | 556.8 | - | - | 0 | - |
| - | - | 3.437E+04 | 556.8 | - | - | 0 | - |
| 3 | y | 2.928E+04 | 557.3 | 0.001871 | 3.358 | +3 | 13 |
| - | - | 7438 | 557.6 | - | - | 0 | - |
| - | - | 9781 | 557.8 | - | - | 0 | - |
| - | - | 6132 | 557.9 | - | - | 0 | - |
| - | - | 5441 | 558.3 | - | - | 0 | - |
| 13 | c | 2690 | 558.6 | 0.003915 | 7.008 | +3 | 13 |
| - | - | 2411 | 558.9 | - | - | 0 | - |
| - | - | 5774 | 559.3 | - | - | 0 | - |
| - | - | 4035 | 559.6 | - | - | 0 | - |
| - | - | 1471 | 559.9 | - | - | 0 | - |
| - | - | 1866 | 562 | - | - | 0 | - |
| - | - | 5260 | 562.3 | - | - | 0 | - |
| - | - | 5492 | 562.6 | - | - | 0 | - |
| - | - | 4872 | 562.8 | - | - | 0 | - |
| - | - | 4507 | 563 | - | - | 0 | - |
| - | - | 1540 | 563.3 | - | - | 0 | - |
| - | - | 2988 | 563.3 | - | - | 0 | - |
| - | - | 2632 | 563.6 | - | - | 0 | - |
| - | - | 4704 | 563.8 | - | - | 0 | - |
| - | - | 5452 | 563.9 | - | - | 0 | - |
| 7 | y | 2.619E+05 | 564.3 | 0.000332 | 0.5883 | +2 | 9 |
| 13 | c | 6.87E+04 | 564.6 | 0.0008816 | 1.561 | +3 | 13 |
| - | - | 1.516E+05 | 564.8 | - | - | 0 | - |
| - | - | 6.747E+04 | 564.9 | - | - | 0 | - |
| - | - | 8.799E+04 | 565.3 | - | - | 0 | - |
| - | - | 1.017E+04 | 565.6 | - | - | 0 | - |
| - | - | 1.701E+04 | 565.8 | - | - | 0 | - |
| - | - | 5105 | 565.9 | - | - | 0 | - |
| - | - | 7886 | 566.3 | - | - | 0 | - |
| - | - | 961.3 | 566.8 | - | - | 0 | - |
| - | - | 1.323E+04 | 567.2 | - | - | 0 | - |
| - | - | 2796 | 567.3 | - | - | 0 | - |
| - | - | 902.4 | 568 | - | - | 0 | - |
| - | - | 3586 | 568.2 | - | - | 0 | - |
| - | - | 4137 | 568.3 | - | - | 0 | - |
| - | - | 1901 | 568.6 | - | - | 0 | - |
| - | - | 1.631E+04 | 568.8 | - | - | 0 | - |
| - | - | 1.419E+04 | 569.3 | - | - | 0 | - |
| - | - | 1014 | 569.6 | - | - | 0 | - |
| - | - | 7073 | 569.8 | - | - | 0 | - |
| - | - | 1888 | 569.9 | - | - | 0 | - |
| - | - | 3748 | 570.3 | - | - | 0 | - |
| - | - | 8412 | 570.8 | - | - | 0 | - |
| - | - | 8988 | 571.3 | - | - | 0 | - |
| - | - | 5919 | 571.8 | - | - | 0 | - |
| - | - | 1884 | 572.3 | - | - | 0 | - |
| - | - | 1196 | 572.6 | - | - | 0 | - |
| - | - | 3054 | 573 | - | - | 0 | - |
| - | - | 3544 | 573.3 | - | - | 0 | - |
| - | - | 2458 | 573.6 | - | - | 0 | - |
| - | - | 2.251E+04 | 573.9 | - | - | 0 | - |
| - | - | 2.042E+04 | 574.3 | - | - | 0 | - |
| - | - | 1.005E+04 | 574.6 | - | - | 0 | - |
| - | - | 1195 | 574.8 | - | - | 0 | - |
| - | - | 3100 | 574.9 | - | - | 0 | - |
| - | - | 1263 | 576.4 | - | - | 0 | - |
| - | - | 1723 | 576.8 | - | - | 0 | - |
| - | - | 1326 | 577.6 | - | - | 0 | - |
| - | - | 1.223E+04 | 578 | - | - | 0 | - |
| - | - | 1.237E+04 | 578.3 | - | - | 0 | - |
| - | - | 9838 | 578.6 | - | - | 0 | - |
| - | - | 1011 | 578.8 | - | - | 0 | - |
| - | - | 6110 | 578.9 | - | - | 0 | - |
| - | - | 2184 | 579.3 | - | - | 0 | - |
| - | - | 1361 | 579.6 | - | - | 0 | - |
| - | - | 2939 | 581.3 | - | - | 0 | - |
| - | - | 2983 | 582.3 | - | - | 0 | - |
| - | - | 3044 | 582.6 | - | - | 0 | - |
| - | - | 1677 | 582.8 | - | - | 0 | - |
| - | - | 1054 | 582.8 | - | - | 0 | - |
| - | - | 1.718E+04 | 583 | - | - | 0 | - |
| - | - | 1.581E+04 | 583.3 | - | - | 0 | - |
| - | - | 1.145E+04 | 583.6 | - | - | 0 | - |
| - | - | 2236 | 583.8 | - | - | 0 | - |
| - | - | 3845 | 584 | - | - | 0 | - |
| - | - | 2880 | 584.3 | - | - | 0 | - |
| - | - | 2074 | 587.3 | - | - | 0 | - |
| - | - | 8882 | 587.6 | - | - | 0 | - |
| - | - | 8792 | 587.9 | - | - | 0 | - |
| - | - | 6278 | 588.3 | - | - | 0 | - |
| - | - | 2487 | 588.6 | - | - | 0 | - |
| - | - | 2062 | 588.9 | - | - | 0 | - |
| - | - | 1193 | 589.3 | - | - | 0 | - |
| - | - | 993.4 | 589.8 | - | - | 0 | - |
| - | - | 3979 | 590.3 | - | - | 0 | - |
| - | - | 737.2 | 590.8 | - | - | 0 | - |
| - | - | 1.064E+04 | 590.8 | - | - | 0 | - |
| 6 | w | 2.42E+04 | 591.3 | 6.009E-05 | 0.1016 | +2 | 10 |
| - | - | 2861 | 591.3 | - | - | 0 | - |
| - | - | 1.341E+04 | 591.8 | - | - | 0 | - |
| - | - | 4638 | 591.8 | - | - | 0 | - |
| - | - | 1329 | 592 | - | - | 0 | - |
| - | - | 1.032E+04 | 592.3 | - | - | 0 | - |
| - | - | 2.424E+04 | 592.6 | - | - | 0 | - |
| - | - | 2755 | 592.8 | - | - | 0 | - |
| - | - | 2.126E+04 | 593 | - | - | 0 | - |
| - | - | 4700 | 593.3 | - | - | 0 | - |
| - | - | 1.091E+04 | 593.3 | - | - | 0 | - |
| - | - | 5790 | 593.6 | - | - | 0 | - |
| - | - | 1051 | 594 | - | - | 0 | - |
| - | - | 3802 | 594.3 | - | - | 0 | - |
| - | - | 1308 | 595.3 | - | - | 0 | - |
| - | - | 1220 | 596.6 | - | - | 0 | - |
| 2 | y | 1075 | 597 | 0.00488 | 8.174 | +3 | 14 |
| - | - | 6851 | 597.3 | - | - | 0 | - |
| 2 | z | 2.568E+04 | 597.6 | 0.003775 | 6.316 | +3 | 14 |
| - | - | 1104 | 597.8 | - | - | 0 | - |
| - | - | 3.014E+04 | 598 | - | - | 0 | - |
| - | - | 1.507E+04 | 598.3 | - | - | 0 | - |
| - | - | 5549 | 598.6 | - | - | 0 | - |
| - | - | 7410 | 598.8 | - | - | 0 | - |
| - | - | 2719 | 599 | - | - | 0 | - |
| - | - | 4067 | 599.3 | - | - | 0 | - |
| - | - | 2021 | 601.3 | - | - | 0 | - |
| 14 | c | 7363 | 601.6 | 0.003651 | 6.068 | +3 | 14 |
| - | - | 1.015E+04 | 602 | - | - | 0 | - |
| - | - | 5494 | 602.3 | - | - | 0 | - |
| - | - | 1870 | 602.6 | - | - | 0 | - |
| - | - | 1656 | 602.8 | - | - | 0 | - |
| 2 | y | 4214 | 603 | 0.001969 | 3.265 | +3 | 14 |
| - | - | 6880 | 603.3 | - | - | 0 | - |
| - | - | 2705 | 603.6 | - | - | 0 | - |
| 6 | z | 1211 | 603.8 | 0.00176 | 2.916 | +2 | 10 |
| - | - | 1739 | 604 | - | - | 0 | - |
| - | - | 1072 | 604.8 | - | - | 0 | - |
| - | - | 4461 | 606.3 | - | - | 0 | - |
| - | - | 3042 | 606.6 | - | - | 0 | - |
| - | - | 3630 | 607 | - | - | 0 | - |
| 14 | c | 2.454E+04 | 607.3 | 0.001474 | 2.427 | +3 | 14 |
| - | - | 3.077E+04 | 607.6 | - | - | 0 | - |
| - | - | 1.986E+04 | 608 | - | - | 0 | - |
| - | - | 1.076E+04 | 608.3 | - | - | 0 | - |
| - | - | 4757 | 608.6 | - | - | 0 | - |
| - | - | 2037 | 609 | - | - | 0 | - |
| - | - | 2.184E+04 | 610.3 | - | - | 0 | - |
| - | - | 6147 | 611.3 | - | - | 0 | - |
| 10 | c | 1.539E+04 | 611.8 | 0.006784 | 11.09 | +2 | 10 |
| 6 | y | 1.197E+04 | 612.3 | 0.002356 | 3.848 | +2 | 10 |
| 6 | z | 3.097E+04 | 612.8 | 0.0007143 | 1.166 | +2 | 10 |
| - | - | 2.406E+04 | 613.3 | - | - | 0 | - |
| - | - | 1.078E+04 | 613.8 | - | - | 0 | - |
| - | - | 3839 | 614.3 | - | - | 0 | - |
| - | - | 1123 | 614.6 | - | - | 0 | - |
| - | - | 6716 | 615.3 | - | - | 0 | - |
| - | - | 8417 | 615.6 | - | - | 0 | - |
| - | - | 1608 | 615.8 | - | - | 0 | - |
| - | - | 7994 | 616 | - | - | 0 | - |
| - | - | 8521 | 616.3 | - | - | 0 | - |
| - | - | 6422 | 616.6 | - | - | 0 | - |
| - | - | 5763 | 617 | - | - | 0 | - |
| - | - | 3824 | 617.3 | - | - | 0 | - |
| - | - | 1092 | 617.6 | - | - | 0 | - |
| - | - | 3584 | 619.8 | - | - | 0 | - |
| 10 | c | 9.519E+04 | 620.3 | 0.002604 | 4.197 | +2 | 10 |
| - | - | 5659 | 620.6 | - | - | 0 | - |
| 6 | y | 1.131E+05 | 620.8 | 0.002447 | 3.941 | +2 | 10 |
| - | - | 2.09E+04 | 621 | - | - | 0 | - |
| - | - | 6693 | 621.2 | - | - | 0 | - |
| - | - | 1.132E+05 | 621.3 | - | - | 0 | - |
| - | - | 5.792E+04 | 621.6 | - | - | 0 | - |
| - | - | 2.302E+04 | 621.8 | - | - | 0 | - |
| - | - | 3.56E+04 | 622 | - | - | 0 | - |
| - | - | 4047 | 622.2 | - | - | 0 | - |
| - | - | 1.927E+04 | 622.3 | - | - | 0 | - |
| - | - | 6273 | 622.7 | - | - | 0 | - |
| - | - | 1621 | 622.8 | - | - | 0 | - |
| - | - | 1183 | 623 | - | - | 0 | - |
| - | - | 1072 | 623.3 | - | - | 0 | - |
| - | - | 3765 | 624.3 | - | - | 0 | - |
| - | - | 1550 | 624.3 | - | - | 0 | - |
| - | - | 1305 | 625.3 | - | - | 0 | - |
| - | - | 4925 | 625.7 | - | - | 0 | - |
| - | - | 8127 | 626 | - | - | 0 | - |
| - | - | 1.396E+04 | 626.3 | - | - | 0 | - |
| - | - | 7662 | 626.7 | - | - | 0 | - |
| - | - | 1854 | 626.8 | - | - | 0 | - |
| - | - | 4926 | 627 | - | - | 0 | - |
| - | - | 2232 | 627.3 | - | - | 0 | - |
| - | - | 2327 | 629.3 | - | - | 0 | - |
| - | - | 5072 | 629.6 | - | - | 0 | - |
| - | - | 1.028E+04 | 630 | - | - | 0 | - |
| - | - | 9299 | 630.3 | - | - | 0 | - |
| - | - | 2616 | 630.6 | - | - | 0 | - |
| - | - | 2202 | 631 | - | - | 0 | - |
| - | - | 1373 | 632 | - | - | 0 | - |
| - | - | 1593 | 632.3 | - | - | 0 | - |
| - | - | 1673 | 633.3 | - | - | 0 | - |
| - | - | 1141 | 633.8 | - | - | 0 | - |
| - | - | 1010 | 634.3 | - | - | 0 | - |
| - | - | 2189 | 635 | - | - | 0 | - |
| - | - | 1.59E+04 | 635.3 | - | - | 0 | - |
| - | - | 1.01E+05 | 635.6 | - | - | 0 | - |
| - | - | 1.004E+05 | 636 | - | - | 0 | - |
| - | - | 5.878E+04 | 636.3 | - | - | 0 | - |
| - | - | 2.375E+04 | 636.6 | - | - | 0 | - |
| - | - | 8664 | 637 | - | - | 0 | - |
| - | - | 4145 | 637.3 | - | - | 0 | - |
| - | - | 1426 | 637.6 | - | - | 0 | - |
| - | - | 1643 | 639.8 | - | - | 0 | - |
| - | - | 2208 | 640.3 | - | - | 0 | - |
| - | - | 2.827E+04 | 640.7 | - | - | 0 | - |
| - | - | 1966 | 640.8 | - | - | 0 | - |
| - | - | 5.549E+04 | 641 | - | - | 0 | - |
| - | - | 6.674E+04 | 641.3 | - | - | 0 | - |
| - | - | 4.425E+04 | 641.7 | - | - | 0 | - |
| - | - | 2.03E+04 | 642 | - | - | 0 | - |
| - | - | 1.027E+04 | 642.3 | - | - | 0 | - |
| - | - | 4543 | 642.7 | - | - | 0 | - |
| - | - | 4602 | 643.8 | - | - | 0 | - |
| - | - | 3411 | 644.3 | - | - | 0 | - |
| - | - | 1637 | 644.8 | - | - | 0 | - |
| - | - | 1114 | 646.3 | - | - | 0 | - |
| - | - | 3409 | 647.3 | - | - | 0 | - |
| - | - | 3.811E+04 | 647.8 | - | - | 0 | - |
| - | - | 2.855E+04 | 648.3 | - | - | 0 | - |
| - | - | 1.329E+04 | 648.8 | - | - | 0 | - |
| - | - | 5098 | 649.3 | - | - | 0 | - |
| - | - | 2037 | 649.8 | - | - | 0 | - |
| - | - | 1151 | 650.3 | - | - | 0 | - |
| - | - | 1335 | 650.8 | - | - | 0 | - |
| - | - | 1021 | 651.3 | - | - | 0 | - |
| - | - | 1.546E+04 | 652.3 | - | - | 0 | - |
| - | - | 5561 | 653.3 | - | - | 0 | - |
| - | - | 1105 | 654.3 | - | - | 0 | - |
| - | - | 1327 | 655.3 | - | - | 0 | - |
| - | - | 2992 | 655.8 | - | - | 0 | - |
| - | - | 1666 | 656.3 | - | - | 0 | - |
| - | - | 2581 | 657.8 | - | - | 0 | - |
| - | - | 2818 | 658.3 | - | - | 0 | - |
| - | - | 1015 | 658.8 | - | - | 0 | - |
| - | - | 2850 | 662.3 | - | - | 0 | - |
| - | - | 7450 | 663.3 | - | - | 0 | - |
| - | - | 2365 | 663.3 | - | - | 0 | - |
| - | - | 1.792E+04 | 664.3 | - | - | 0 | - |
| - | - | 8115 | 665.3 | - | - | 0 | - |
| - | - | 2295 | 666.4 | - | - | 0 | - |
| - | - | 1736 | 670.8 | - | - | 0 | - |
| - | - | 1208 | 671.3 | - | - | 0 | - |
| - | - | 1768 | 671.8 | - | - | 0 | - |
| - | - | 2262 | 672.3 | - | - | 0 | - |
| - | - | 1131 | 672.8 | - | - | 0 | - |
| - | - | 3071 | 676.3 | - | - | 0 | - |
| - | - | 4014 | 676.8 | - | - | 0 | - |
| - | - | 3657 | 677.3 | - | - | 0 | - |
| - | - | 2202 | 677.8 | - | - | 0 | - |
| - | - | 850.4 | 678.3 | - | - | 0 | - |
| 5 | c | 2905 | 679.3 | 0.004928 | 7.255 | +1 | 5 |
| - | - | 2.632E+04 | 679.8 | - | - | 0 | - |
| 5 | c | 1.255E+05 | 680.3 | 0.004555 | 6.696 | +1 | 5 |
| - | - | 1.091E+04 | 680.8 | - | - | 0 | - |
| 11 | y | 5.565E+04 | 681.3 | 0.009696 | 14.23 | +1 | 5 |
| - | - | 1072 | 681.8 | - | - | 0 | - |
| 11 | z | 4.222E+04 | 682.3 | 0.006047 | 8.862 | +1 | 5 |
| - | - | 2412 | 682.8 | - | - | 0 | - |
| - | - | 2.16E+04 | 683.3 | - | - | 0 | - |
| - | - | 2391 | 683.8 | - | - | 0 | - |
| - | - | 8049 | 684.3 | - | - | 0 | - |
| - | - | 1925 | 685.3 | - | - | 0 | - |
| - | - | 1279 | 691.3 | - | - | 0 | - |
| - | - | 1032 | 691.8 | - | - | 0 | - |
| 11 | c | 1570 | 693.3 | 0.001853 | 2.673 | +2 | 11 |
| - | - | 3334 | 693.8 | - | - | 0 | - |
| - | - | 2025 | 694.3 | - | - | 0 | - |
| - | - | 983.7 | 694.8 | - | - | 0 | - |
| - | - | 1804 | 695.3 | - | - | 0 | - |
| - | - | 1901 | 696.3 | - | - | 0 | - |
| 5 | c | 3.047E+04 | 697.3 | 0.002481 | 3.558 | +1 | 5 |
| 11 | y | 3.008E+04 | 698.4 | 0.006399 | 9.163 | +1 | 5 |
| - | - | 4698 | 698.8 | - | - | 0 | - |
| - | - | 9818 | 699.4 | - | - | 0 | - |
| - | - | 2209 | 699.8 | - | - | 0 | - |
| - | - | 2332 | 700.4 | - | - | 0 | - |
| - | - | 2416 | 701.3 | - | - | 0 | - |
| 11 | c | 4.089E+04 | 701.8 | 0.002372 | 3.38 | +2 | 11 |
| - | - | 4.067E+04 | 702.3 | - | - | 0 | - |
| - | - | 2.162E+04 | 702.8 | - | - | 0 | - |
| - | - | 6816 | 703.3 | - | - | 0 | - |
| - | - | 1789 | 703.9 | - | - | 0 | - |
| - | - | 1091 | 704.3 | - | - | 0 | - |
| 5 | y | 1224 | 704.9 | 0.007524 | 10.67 | +2 | 11 |
| 5 | y | 3083 | 705.3 | 0.00179 | 2.538 | +2 | 11 |
| 5 | z | 9500 | 705.8 | 0.001003 | 1.421 | +2 | 11 |
| - | - | 1.998E+04 | 706.4 | - | - | 0 | - |
| - | - | 1.322E+04 | 706.9 | - | - | 0 | - |
| - | - | 8062 | 707.4 | - | - | 0 | - |
| - | - | 2903 | 710.3 | - | - | 0 | - |
| - | - | 6547 | 711.3 | - | - | 0 | - |
| - | - | 4053 | 712.3 | - | - | 0 | - |
| - | - | 1799 | 712.8 | - | - | 0 | - |
| - | - | 3106 | 713.3 | - | - | 0 | - |
| 5 | y | 7201 | 713.9 | 0.001332 | 1.866 | +2 | 11 |
| - | - | 5276 | 714.4 | - | - | 0 | - |
| - | - | 3154 | 714.9 | - | - | 0 | - |
| - | - | 987.6 | 719.4 | - | - | 0 | - |
| - | - | 4110 | 721.4 | - | - | 0 | - |
| - | - | 2137 | 722.4 | - | - | 0 | - |
| - | - | 3598 | 723.4 | - | - | 0 | - |
| - | - | 1100 | 724.4 | - | - | 0 | - |
| - | - | 3093 | 727.4 | - | - | 0 | - |
| - | - | 1841 | 727.9 | - | - | 0 | - |
| - | - | 2293 | 728.3 | - | - | 0 | - |
| - | - | 1128 | 729.3 | - | - | 0 | - |
| - | - | 1852 | 733.9 | - | - | 0 | - |
| - | - | 2642 | 734.4 | - | - | 0 | - |
| - | - | 1952 | 734.9 | - | - | 0 | - |
| - | - | 1797 | 735.3 | - | - | 0 | - |
| - | - | 1920 | 738.4 | - | - | 0 | - |
| 4 | w | 1694 | 740.9 | 0.002053 | 2.771 | +2 | 12 |
| - | - | 5246 | 741.4 | - | - | 0 | - |
| - | - | 6340 | 741.9 | - | - | 0 | - |
| - | - | 1941 | 742.4 | - | - | 0 | - |
| - | - | 1062 | 743.4 | - | - | 0 | - |
| - | - | 1064 | 743.9 | - | - | 0 | - |
| - | - | 1676 | 744.4 | - | - | 0 | - |
| - | - | 1189 | 744.9 | - | - | 0 | - |
| - | - | 958.4 | 745.4 | - | - | 0 | - |
| - | - | 1285 | 749.4 | - | - | 0 | - |
| 10 | w | 7198 | 752.4 | 0.0014 | 1.861 | +1 | 6 |
| - | - | 5031 | 753.4 | - | - | 0 | - |
| - | - | 2694 | 754.4 | - | - | 0 | - |
| - | - | 1097 | 755.4 | - | - | 0 | - |
| - | - | 1136 | 756.4 | - | - | 0 | - |
| 12 | c | 1038 | 757.4 | 0.008449 | 11.16 | +2 | 12 |
| - | - | 2371 | 757.9 | - | - | 0 | - |
| - | - | 1352 | 758.4 | - | - | 0 | - |
| 4 | y | 955.8 | 762.4 | 0.003967 | 5.203 | +2 | 12 |
| 4 | y | 4250 | 762.9 | 0.003598 | 4.717 | +2 | 12 |
| 4 | z | 1.476E+04 | 763.4 | 0.0008452 | 1.107 | +2 | 12 |
| - | - | 3.06E+04 | 763.9 | - | - | 0 | - |
| - | - | 1.557E+04 | 764.4 | - | - | 0 | - |
| - | - | 8957 | 764.9 | - | - | 0 | - |
| - | - | 1975 | 765.4 | - | - | 0 | - |
| 12 | c | 6294 | 765.9 | 0.001827 | 2.385 | +2 | 12 |
| - | - | 7940 | 766.4 | - | - | 0 | - |
| - | - | 3593 | 766.9 | - | - | 0 | - |
| - | - | 2635 | 767.4 | - | - | 0 | - |
| - | - | 2951 | 769.4 | - | - | 0 | - |
| - | - | 1996 | 769.9 | - | - | 0 | - |
| - | - | 2025 | 770.4 | - | - | 0 | - |
| - | - | 1086 | 770.9 | - | - | 0 | - |
| 4 | y | 3224 | 771.4 | 0.002286 | 2.964 | +2 | 12 |
| - | - | 3321 | 771.9 | - | - | 0 | - |
| - | - | 1968 | 772.4 | - | - | 0 | - |
| - | - | 1211 | 776.4 | - | - | 0 | - |
| - | - | 2182 | 777.4 | - | - | 0 | - |
| - | - | 1766 | 777.9 | - | - | 0 | - |
| - | - | 1322 | 778.4 | - | - | 0 | - |
| - | - | 1077 | 778.9 | - | - | 0 | - |
| - | - | 1820 | 779.4 | - | - | 0 | - |
| - | - | 973.8 | 786.4 | - | - | 0 | - |
| 6 | c | 6.248E+04 | 793.4 | 0.002339 | 2.949 | +1 | 6 |
| - | - | 3.049E+04 | 794.4 | - | - | 0 | - |
| - | - | 8425 | 795.4 | - | - | 0 | - |
| - | - | 1380 | 796.4 | - | - | 0 | - |
| 3 | w | 4754 | 798.4 | 0.009455 | 11.84 | +2 | 13 |
| - | - | 3405 | 798.9 | - | - | 0 | - |
| - | - | 1101 | 799.4 | - | - | 0 | - |
| 10 | y | 2801 | 809.4 | 0.009241 | 11.42 | +1 | 6 |
| 6 | c | 9.301E+04 | 810.4 | 0.002524 | 3.114 | +1 | 6 |
| 10 | z | 1.846E+04 | 811.4 | 0.01129 | 13.91 | +1 | 6 |
| - | - | 3.576E+04 | 811.4 | - | - | 0 | - |
| - | - | 2.395E+04 | 812.4 | - | - | 0 | - |
| - | - | 3021 | 812.4 | - | - | 0 | - |
| - | - | 1.112E+04 | 813.4 | - | - | 0 | - |
| - | - | 3002 | 814.4 | - | - | 0 | - |
| - | - | 993.9 | 815.4 | - | - | 0 | - |
| - | - | 3307 | 816.9 | - | - | 0 | - |
| - | - | 4356 | 817.4 | - | - | 0 | - |
| - | - | 3649 | 817.9 | - | - | 0 | - |
| 3 | z | 1851 | 818.4 | 0.009554 | 11.67 | +2 | 13 |
| - | - | 2190 | 822.4 | - | - | 0 | - |
| - | - | 1147 | 823.4 | - | - | 0 | - |
| - | - | 3938 | 823.9 | - | - | 0 | - |
| - | - | 4792 | 824.4 | - | - | 0 | - |
| - | - | 2676 | 824.9 | - | - | 0 | - |
| - | - | 1118 | 825.4 | - | - | 0 | - |
| - | - | 945.9 | 825.9 | - | - | 0 | - |
| 3 | y | 1751 | 826.4 | 0.01647 | 19.93 | +2 | 13 |
| 10 | y | 2.051E+04 | 827.4 | 0.005352 | 6.469 | +1 | 6 |
| - | - | 3.326E+04 | 827.9 | - | - | 0 | - |
| - | - | 2.878E+04 | 828.4 | - | - | 0 | - |
| - | - | 1.235E+04 | 828.9 | - | - | 0 | - |
| - | - | 6429 | 829.4 | - | - | 0 | - |
| - | - | 1653 | 829.9 | - | - | 0 | - |
| - | - | 1015 | 830.4 | - | - | 0 | - |
| - | - | 1345 | 831.4 | - | - | 0 | - |
| 3 | y | 3426 | 835.4 | 0.001551 | 1.856 | +2 | 13 |
| - | - | 3456 | 835.9 | - | - | 0 | - |
| - | - | 1081 | 836.4 | - | - | 0 | - |
| - | - | 1242 | 836.9 | - | - | 0 | - |
| - | - | 4223 | 838.4 | - | - | 0 | - |
| - | - | 4701 | 838.9 | - | - | 0 | - |
| - | - | 2078 | 839.4 | - | - | 0 | - |
| - | - | 1141 | 839.9 | - | - | 0 | - |
| - | - | 2695 | 844.4 | - | - | 0 | - |
| - | - | 1063 | 845.4 | - | - | 0 | - |
| - | - | 2192 | 845.9 | - | - | 0 | - |
| 13 | c | 1.716E+04 | 846.4 | 0.0002304 | 0.2722 | +2 | 13 |
| - | - | 1.807E+04 | 846.9 | - | - | 0 | - |
| - | - | 1.269E+04 | 847.4 | - | - | 0 | - |
| - | - | 4776 | 847.9 | - | - | 0 | - |
| - | - | 3226 | 848.4 | - | - | 0 | - |
| - | - | 1875 | 851.4 | - | - | 0 | - |
| - | - | 1340 | 852.4 | - | - | 0 | - |
| - | - | 1422 | 853.4 | - | - | 0 | - |
| - | - | 1397 | 854.4 | - | - | 0 | - |
| - | - | 1814 | 860.4 | - | - | 0 | - |
| - | - | 3098 | 866.9 | - | - | 0 | - |
| - | - | 3.503E+04 | 867.4 | - | - | 0 | - |
| - | - | 1185 | 867.9 | - | - | 0 | - |
| - | - | 2.121E+04 | 868.4 | - | - | 0 | - |
| - | - | 6727 | 869.4 | - | - | 0 | - |
| - | - | 925.8 | 870.4 | - | - | 0 | - |
| - | - | 1768 | 880.4 | - | - | 0 | - |
| - | - | 1768 | 880.9 | - | - | 0 | - |
| - | - | 4306 | 881.4 | - | - | 0 | - |
| - | - | 1079 | 881.9 | - | - | 0 | - |
| - | - | 1362 | 882.4 | - | - | 0 | - |
| - | - | 2477 | 882.5 | - | - | 0 | - |
| - | - | 1027 | 883.4 | - | - | 0 | - |
| - | - | 1790 | 888.4 | - | - | 0 | - |
| - | - | 1.343E+04 | 888.9 | - | - | 0 | - |
| - | - | 8344 | 889.4 | - | - | 0 | - |
| - | - | 4934 | 889.9 | - | - | 0 | - |
| - | - | 2166 | 890.4 | - | - | 0 | - |
| - | - | 1756 | 893.4 | - | - | 0 | - |
| 2 | z | 4852 | 895.9 | 0.001671 | 1.865 | +2 | 14 |
| - | - | 2.392E+04 | 896.4 | - | - | 0 | - |
| - | - | 1.911E+04 | 896.9 | - | - | 0 | - |
| - | - | 9447 | 897.4 | - | - | 0 | - |
| - | - | 3995 | 897.9 | - | - | 0 | - |
| - | - | 2425 | 898.4 | - | - | 0 | - |
| - | - | 1776 | 902.9 | - | - | 0 | - |
| - | - | 1229 | 903.4 | - | - | 0 | - |
| 2 | y | 1306 | 903.9 | 0.01678 | 18.57 | +2 | 14 |
| 7 | c | 7277 | 908.4 | 0.001703 | 1.874 | +1 | 7 |
| - | - | 5094 | 909.4 | - | - | 0 | - |
| 14 | c | 2891 | 910.4 | 0.007517 | 8.257 | +2 | 14 |
| - | - | 2001 | 911.4 | - | - | 0 | - |
| - | - | 1899 | 912.5 | - | - | 0 | - |
| - | - | 1023 | 920.4 | - | - | 0 | - |
| 9 | z | 1079 | 922.5 | 0.005566 | 6.034 | +1 | 7 |
| - | - | 2878 | 923.5 | - | - | 0 | - |
| - | - | 2693 | 924 | - | - | 0 | - |
| - | - | 2613 | 924.5 | - | - | 0 | - |
| - | - | 1124 | 925 | - | - | 0 | - |
| 7 | c | 6390 | 925.5 | 0.001704 | 1.841 | +1 | 7 |
| - | - | 5030 | 926.5 | - | - | 0 | - |
| - | - | 1719 | 927.5 | - | - | 0 | - |
| - | - | 3126 | 931.5 | - | - | 0 | - |
| - | - | 3150 | 932 | - | - | 0 | - |
| - | - | 2332 | 932.5 | - | - | 0 | - |
| - | - | 1407 | 933 | - | - | 0 | - |
| 9 | y | 4520 | 938.5 | 0.005163 | 5.501 | +1 | 7 |
| 9 | z | 6866 | 939.5 | 0.005846 | 6.223 | +1 | 7 |
| - | - | 1.365E+04 | 940.5 | - | - | 0 | - |
| - | - | 5731 | 941.5 | - | - | 0 | - |
| - | - | 2123 | 942.5 | - | - | 0 | - |
| - | - | 1304 | 952.5 | - | - | 0 | - |
| - | - | 5222 | 953 | - | - | 0 | - |
| - | - | 4900 | 953.5 | - | - | 0 | - |
| - | - | 2732 | 954 | - | - | 0 | - |
| - | - | 2063 | 954.5 | - | - | 0 | - |
| 9 | y | 1627 | 955.5 | 0.007664 | 8.021 | +1 | 7 |
| - | - | 1247 | 961 | - | - | 0 | - |
| - | - | 3182 | 961.5 | - | - | 0 | - |
| - | - | 3814 | 962 | - | - | 0 | - |
| - | - | 2193 | 962.5 | - | - | 0 | - |
| 8 | c | 1052 | 965.4 | 0.009658 | 10 | +1 | 8 |
| - | - | 2712 | 966.5 | - | - | 0 | - |
| - | - | 2784 | 967.5 | - | - | 0 | - |
| - | - | 1168 | 968.5 | - | - | 0 | - |
| - | - | 1244 | 981.5 | - | - | 0 | - |
| 8 | c | 1.548E+04 | 982.5 | 0.003067 | 3.122 | +1 | 8 |
| - | - | 1.236E+04 | 983.5 | - | - | 0 | - |
| - | - | 5318 | 984.5 | - | - | 0 | - |
| - | - | 857.3 | 985.5 | - | - | 0 | - |
| 8 | z | 1753 | 996.5 | 0.005582 | 5.601 | +1 | 8 |
| - | - | 4489 | 997.5 | - | - | 0 | - |
| - | - | 2221 | 998.5 | - | - | 0 | - |
| - | - | 1038 | 999.5 | - | - | 0 | - |
| - | - | 954.7 | 1011 | - | - | 0 | - |
| 8 | y | 2642 | 1013 | 0.009962 | 9.839 | +1 | 8 |
| - | - | 1364 | 1014 | - | - | 0 | - |
| - | - | 894.4 | 1015 | - | - | 0 | - |
| - | - | 1415 | 1028 | - | - | 0 | - |
| - | - | 1059 | 1055 | - | - | 0 | - |
| - | - | 1582 | 1067 | - | - | 0 | - |
| - | - | 1329 | 1068 | - | - | 0 | - |
| - | - | 2404 | 1095 | - | - | 0 | - |
| - | - | 2546 | 1096 | - | - | 0 | - |
| 9 | c | 1.009E+04 | 1111 | 0.002953 | 2.659 | +1 | 9 |
| - | - | 1.182E+04 | 1112 | - | - | 0 | - |
| - | - | 7138 | 1113 | - | - | 0 | - |
| - | - | 4345 | 1114 | - | - | 0 | - |
| - | - | 1710 | 1115 | - | - | 0 | - |
| 7 | y | 1643 | 1128 | 0.00535 | 4.745 | +1 | 9 |
| - | - | 1026 | 1137 | - | - | 0 | - |
| - | - | 982.7 | 1138 | - | - | 0 | - |
| - | - | 714.1 | 1196 | - | - | 0 | - |
| - | - | 1301 | 1197 | - | - | 0 | - |
| - | - | 871.4 | 1198 | - | - | 0 | - |
| 6 | z | 1132 | 1225 | 0.003674 | 3 | +1 | 10 |
| - | - | 4409 | 1226 | - | - | 0 | - |
| - | - | 2156 | 1227 | - | - | 0 | - |
| 10 | c | 2155 | 1240 | 0.0002767 | 0.2232 | +1 | 10 |
| 6 | y | 3293 | 1241 | 0.007505 | 6.049 | +1 | 10 |
| - | - | 2234 | 1242 | - | - | 0 | - |
| - | - | 898.8 | 1266 | - | - | 0 | - |
| - | - | 2131 | 1360 | - | - | 0 | - |
| - | - | 1945 | 1361 | - | - | 0 | - |
| - | - | 771.2 | 1362 | - | - | 0 | - |
| 11 | c | 1150 | 1403 | 0.003599 | 2.566 | +1 | 11 |
| - | - | 2010 | 1404 | - | - | 0 | - |
| - | - | 1533 | 1405 | - | - | 0 | - |
| - | - | 920.3 | 1406 | - | - | 0 | - |
| - | - | 663.3 | 3008 | - | - | 0 | - |
| - | - | 1076 | 3050 | - | - | 0 | - |

m/z Charge Intensity FragmentType MassShift Position
120.05738830566406 0 665.9623
121.33567810058594 0 391.15915
122.04839324951172 0 629.051
126.07923889160156 0 641.11566
127.0510025024414 0 406.0198
128.0824737548828 0 621.7898
128.09532165527344 0 765.2001
129.06649780273438 0 715.99817
129.10287475585938 0 35103.7
130.06578063964844 0 98425.45
130.10609436035156 0 1940.1355
131.06387329101562 0 1100.7821 z 12
131.069091796875 0 9560.28
131.11846923828125 0 3084.005
131.75698852539062 0 392.66452
132.08143615722656 0 3027.441
132.10267639160156 0 1164.0581
133.0614013671875 0 12886.342
134.02780151367188 0 554.6042
136.07640075683594 0 7058.995
138.06671142578125 0 1204.9191
139.05093383789062 0 448.85623
140.08334350585938 0 545.639
141.10299682617188 0 1356.8507
142.2604522705078 0 421.6101
144.0814971923828 0 2520.2124
145.06149291992188 0 1605.5054
145.08499145507812 0 499.7789
146.06944274902344 0 546.5928
146.09304809570312 0 1928.9349
146.12940979003906 0 444.0773
147.04489135742188 0 1094.6536
147.0775604248047 0 574.80884
147.1136474609375 0 651.2416
148.1068115234375 0 416.63345
148.94540405273438 0 472.43185
154.08724975585938 0 506.43982
155.09339904785156 0 16422.223
156.09666442871094 0 844.70557
156.13856506347656 0 626.7382
157.09780883789062 0 790.38635
157.1343536376953 0 1346.5978
158.04525756835938 0 497.66223
158.0608367919922 0 1880.9257
158.08485412597656 0 601.2539
159.09234619140625 0 42290.14
160.09584045410156 0 3681.4377
164.08250427246094 0 2390.9873
165.1029510498047 0 4560.439
166.06179809570312 0 16328.537
167.0652313232422 0 1436.2715
168.13905334472656 0 4138.7764
169.09860229492188 0 590.1667
170.060791015625 0 20501.723
170.10519409179688 0 850.83905
171.06304931640625 0 1070.7408
171.06983947753906 0 1401.6082
172.07237243652344 0 3737.5754
172.09764099121094 0 3748.6736 w 13
173.05648803710938 0 2339.957
173.43850708007812 0 793.9048
175.0858917236328 0 553.7514
175.7394561767578 0 488.58228
176.11126708984375 0 1237.3131
178.13449096679688 0 2468.8445
181.09835815429688 0 752.70496
183.1135711669922 0 15967.446
184.08534240722656 0 725.7059
184.11720275878906 0 941.8966
185.07119750976562 0 661.3854
185.16561889648438 0 17728.61
186.0795440673828 0 805.3602
186.12452697753906 0 3460.4639
186.16952514648438 0 1886.4777
187.0873260498047 0 7378.882
187.10838317871094 0 1888.0664
188.09080505371094 0 1038.2267
188.11643981933594 0 1275.307
190.08311462402344 0 30609.443
191.08663940429688 0 1329.7081
192.0771484375 0 1703.826
195.11349487304688 0 1421.4633
198.1134490966797 0 565.60156
198.12843322753906 0 10305.413
199.07196044921875 0 708.85016
199.13223266601562 0 1050.4034
199.16993713378906 0 858.7215
200.1434326171875 0 715.68365
201.12420654296875 0 2269.7664
201.136474609375 0 587.4391
203.1166534423828 0 3503.949
203.15078735351562 0 1406.2002
203.61807250976562 0 756.37805
204.11427307128906 0 648.2827
207.16134643554688 0 2204.7703
210.1284942626953 0 3034.856
211.6299285888672 0 4047.3892
212.1309814453125 0 1168.8123
214.15591430664062 0 539.4399
214.64781188964844 0 1350.3354
215.13990783691406 0 3734.1194
218.0820770263672 0 1612.7904
221.62860107421875 0 1433.7958
222.12078857421875 0 1323.2477
223.10879516601562 0 959.75665 z 6
223.1197509765625 0 5744.314
223.15618896484375 0 16711.3
224.1598663330078 0 1895.5325
225.04493713378906 0 628.16583
227.06707763671875 0 1298.5458
227.13992309570312 0 1276.7238
227.15452575683594 0 567.8742
228.1204833984375 0 909.14685 z 10
228.13548278808594 0 815.9143
229.0826873779297 0 934.9911
229.119140625 0 3622.855
229.15576171875 0 2652.6382 y Ammonia loss 13
230.16343688964844 0 4006.8262 z 13
230.63394165039062 0 9573.463
231.1354522705078 0 3202.8318
231.16990661621094 0 705.31726
233.14060974121094 0 5306.5337
233.1660919189453 0 1517.2787
234.12466430664062 0 19173.307
234.144287109375 0 818.55505
235.12814331054688 0 2133.047 y Water loss 8
236.63417053222656 0 1759.3748
237.1357879638672 0 549.9472
239.09637451171875 0 703.058
239.1092071533203 0 1577.1713
241.11924743652344 0 1294.2568
243.1107635498047 0 643.69116
243.13507080078125 0 1360.3936
243.14866638183594 0 1143.8973
244.09368896484375 0 2044.2179
245.09609985351562 0 1040.2153
246.1094207763672 0 1009.865
246.1460723876953 0 1229.5598
246.18226623535156 0 6491.651 y 13
247.18557739257812 0 625.5103
247.62242126464844 0 6037.6675 c Ammonia loss 3
248.12466430664062 0 743.79596
248.62762451171875 0 488.3043
249.09913635253906 0 679.04694
249.1306915283203 0 833.7734
249.6306610107422 0 594.1407 y Ammonia loss 7
249.80349731445312 0 736.2422
250.0955810546875 0 618.9678
250.10816955566406 0 3807.9978
250.13111877441406 0 1137.8899
250.1444854736328 0 1157.924
250.16744995117188 0 2264.5808
251.15122985839844 0 90639.4
251.62698364257812 0 622.9888 z Ammonia loss 11
252.15452575683594 0 10568.84
252.6042938232422 0 2667.7153
253.10531616210938 0 785.2968
253.1573486328125 0 995.59924
253.44692993164062 0 526.9305
254.15078735351562 0 683.98236
255.14990234375 0 2300.6301
255.80718994140625 0 3497.6492
256.1392517089844 0 913.9406 c 3
256.15631103515625 0 625.5363
257.1630554199219 0 1024.2933
258.6288146972656 0 1904.3625
259.6365661621094 0 4763.3667 y Ammonia loss 11
260.1387023925781 0 1218.2771 z 11
260.6368713378906 0 678.3452
261.1191711425781 0 3428.4343
261.1604309082031 0 723.2551
261.60919189453125 0 1714.3688
262.11029052734375 0 890.28375
265.12945556640625 0 663.2514 z Water loss 9
266.12548828125 0 4116.2744
266.6013488769531 0 1086.8644
268.15020751953125 0 4557.181 y 11
268.1777648925781 0 13705.508 c 1
268.6502380371094 0 893.69226
269.11346435546875 0 735.9683
269.1805419921875 0 1020.8157
270.10595703125 0 887.9615
270.12255859375 0 15251.979
270.1819152832031 0 1075.8275
270.6243896484375 0 4372.4673
271.1245422363281 0 1141.452
272.1072692871094 0 7770.917
272.161865234375 0 981.26514
272.17681884765625 0 4302.453
273.09124755859375 0 3112.893
273.1079406738281 0 1099.2379
273.18035888671875 0 947.43225
273.4820251464844 0 684.2358
274.1197204589844 0 2976.937
275.11572265625 0 1610.1034
277.1555480957031 0 825.4133
282.63482666015625 0 885.60315
283.14129638671875 0 5721.3374
284.1041259765625 0 1557.6786
284.12030029296875 0 3650.6338
284.1457824707031 0 600.5666
284.62164306640625 0 1721.5828
285.15771484375 0 1277.8344
286.1410827636719 0 1423.2501
287.8470764160156 0 639.5288
289.17022705078125 0 630.6587
290.1179504394531 0 6589.2515
291.14306640625 0 696.0188
292.13787841796875 0 770.4547
292.16680908203125 0 1695.9698
293.1375732421875 0 750.0071
294.1207580566406 0 796.9072
294.1935119628906 0 678.684
295.1041259765625 0 1717.1239
296.1797790527344 0 1917.611
296.6831970214844 0 961.8227
297.1285705566406 0 2973.3728
297.15570068359375 0 1348.4108
297.4905090332031 0 1847.0677
297.62969970703125 0 1814.8837
297.8236083984375 0 595.0592
298.1781005859375 0 665.29816
299.6561584472656 0 1611.3885
300.1567077636719 0 1006.5856
300.1728820800781 0 1492.0259
300.19305419921875 0 11331.118
300.6337585449219 0 581.2744
301.10797119140625 0 1011.1039
301.1528625488281 0 3412.585
301.1970520019531 0 1755.762
302.115478515625 0 3189.857
302.1553039550781 0 714.7232
302.509033203125 0 1222.1501
303.1673583984375 0 5307.798
304.8262023925781 0 1565.6041
305.1560974121094 0 734.8168
305.64630126953125 0 1561.4114
307.1448669433594 0 599.33496
307.4872741699219 0 560.5465
308.1289367675781 0 2791.55
309.14630126953125 0 4678.017
309.64788818359375 0 2068.0718
311.1258850097656 0 3445.733
312.16546630859375 0 774.6605
312.6305236816406 0 995.2533
312.82220458984375 0 924.4052
313.1159973144531 0 793.42505
313.16595458984375 0 2358.8928 y Water loss 8
313.4947814941406 0 1356.0037 y Ammonia loss 8
313.83001708984375 0 1623.808 z 8
314.163818359375 0 908.9204
314.50091552734375 0 998.9205
315.1673889160156 0 1861.2285
315.2039489746094 0 1810.7988
316.1748046875 0 1193.1145
317.6597900390625 0 5557.919
318.1516418457031 0 17289.986
318.1781005859375 0 2884.966
318.6534729003906 0 7142.361
318.84210205078125 0 1791.2565
319.1537780761719 0 1946.164
319.1736145019531 0 896.5557 y 8
320.6785583496094 0 1806.6703
322.8415832519531 0 597.72217
323.14385986328125 0 4339.465
323.6446533203125 0 1557.3633
324.67889404296875 0 1575.2305
326.6650390625 0 84225.96
327.1665954589844 0 31543.52 z Ammonia loss 7
327.6685485839844 0 5723.477
327.83026123046875 0 1771.532
328.1673889160156 0 1669.6565 c 7
330.69097900390625 0 680.69446
331.6574401855469 0 13305.648
332.1585998535156 0 4683.0547
332.1939697265625 0 1291.2073
332.5027770996094 0 2711.1465 y Ammonia loss 7
332.6538391113281 0 984.7837
332.8375244140625 0 3522.6775 z 7
333.171630859375 0 1068.192
337.1816711425781 0 1153.2842
337.84014892578125 0 1189.5139
338.1487121582031 0 1711.5874
338.1772155761719 0 37334.76 y 7
338.51171875 0 18595.498
338.67620849609375 0 1720.6274
338.845458984375 0 5438.923
339.177734375 0 2095.385
339.5107421875 0 1401.1356
340.2603454589844 0 848.6555
340.66253662109375 0 41541.934 c Ammonia loss 4
341.16510009765625 0 22269.988 y Ammonia loss 10
341.6681213378906 0 7273.7183 z 10
342.169189453125 0 2128.2637
344.19476318359375 0 957.66797
345.1562805175781 0 860.07495
346.1151123046875 0 1134.1194
346.1794738769531 0 914.36053
347.1763610839844 0 1185.869
349.19268798828125 0 765.7915
349.6820373535156 0 51270.94 y 10
350.1834716796875 0 18702.777
350.68310546875 0 6985.6045
351.18365478515625 0 1130.8851
351.2154541015625 0 1215.153
352.18060302734375 0 9944.314
352.5152282714844 0 4395.272
352.84844970703125 0 2257.0935
353.17633056640625 0 7363.5474 y Ammonia loss 4
353.42657470703125 0 4854.802 z 4
353.6718444824219 0 4907.385
353.9268798828125 0 1000.3275
354.1711120605469 0 1674.6423
355.17303466796875 0 1242.9036
356.1806335449219 0 2119.9526 w 6
356.5157470703125 0 2527.236
356.85009765625 0 862.18164
357.178466796875 0 1230.2502
360.6978454589844 0 5183.983
361.1807861328125 0 1293.3528 c Water loss 13
361.2001647949219 0 1811.3755
361.5180969238281 0 1591.315
361.8548889160156 0 837.113
362.1865539550781 0 3168.3403
362.6897888183594 0 845.32684
363.1421813964844 0 1839.5579
363.2049560546875 0 887.99536
363.74859619140625 0 4619.0337
364.1259765625 0 1173.238
364.2507629394531 0 761.55786
365.578857421875 0 657.10156
365.68902587890625 0 1809.9255
366.1878662109375 0 970.1194
367.1417236328125 0 609.1919
367.2701110839844 0 1891.4788
367.66650390625 0 815.96423
368.1708068847656 0 1630.8218
368.67584228515625 0 1191.8027
370.145751953125 0 1355.303
370.51483154296875 0 1570.0581 y Water loss 6
370.8473205566406 0 2675.615 y Ammonia loss 6
371.1813659667969 0 2048.1948 z 6
371.51275634765625 0 1254.8844
371.72760009765625 0 1561.9995
372.1893310546875 0 1255.4517
372.23028564453125 0 1760.5425
372.58929443359375 0 768.8037
374.2017517089844 0 3451.878
374.3845520019531 0 1708.5591
374.694091796875 0 11977.632
375.1710510253906 0 6135.2686
375.19659423828125 0 5880.06
375.69818115234375 0 1673.978
376.5195617675781 0 28962.465 y 6
376.687255859375 0 214730.12 w 9
376.8536682128906 0 14048.169
377.18865966796875 0 87582.35
377.4306640625 0 1587.5127
377.5225830078125 0 1975.5212
377.5897216796875 0 2564.931
377.68865966796875 0 27647.357 z Water loss 3
377.78729248046875 0 5961.1353
377.98797607421875 0 5837.028
378.1876220703125 0 8220.451
378.20538330078125 0 3408.1067
378.3858337402344 0 1038.7526
378.69036865234375 0 1895.5554
379.2100830078125 0 27971.71 c Ammonia loss 2
379.86004638671875 0 1869.3833
380.2131042480469 0 4268.101
380.5281066894531 0 1152.1918
380.863525390625 0 1895.4639
381.1530456542969 0 7379.27
381.19158935546875 0 9008.002
381.3896484375 0 14101.136
381.5897521972656 0 15160.677
381.6841735839844 0 1021.21155 y Water loss 3
381.7895202636719 0 8572.083
381.98974609375 0 2156.5042
382.1563415527344 0 956.35876
382.191650390625 0 2076.09 z 3
383.2070007324219 0 39198.94
383.7088623046875 0 18538.9
384.16522216796875 0 1173.8073
384.2105712890625 0 5601.2793
384.2994384765625 0 1738.2339
384.583251953125 0 936.96094
384.77301025390625 0 1486.8181
384.794189453125 0 10300.689
384.994140625 0 13111.651
385.151123046875 0 2761.354
385.1947937011719 0 8412.693
385.2486572265625 0 908.58936
385.2748718261719 0 1623.5403
385.3948974609375 0 3289.4104
385.5938415527344 0 1024.046
386.1558837890625 0 860.4045
386.1923522949219 0 993.34674 y 3
387.1845397949219 0 2768.4849
387.7144470214844 0 2356.416
388.1866760253906 0 853.7546
388.6916809082031 0 3341.576
389.18841552734375 0 2255.8562
390.1937255859375 0 1330.3494
392.6877746582031 0 955.43
394.5232238769531 0 16636.713 w 5
394.8572998046875 0 9539.245
395.1362609863281 0 1258.8802
395.19171142578125 0 2382.1104
395.2188720703125 0 806.6079
395.5263977050781 0 866.9128
396.2367248535156 0 40295.78 c 2
397.2047424316406 0 25388.025 c Ammonia loss 5
397.239501953125 0 7646.172
397.7067565917969 0 11622.275
398.1780700683594 0 1890.0546
398.208251953125 0 4094.4434
398.2392272949219 0 1179.3313
402.1775817871094 0 1760.1805
403.2025146484375 0 2600.5312 z Ammonia loss 5
403.7330017089844 0 766.4371
405.1985168457031 0 6808.0884 y Water loss 9
405.699462890625 0 2049.077
406.1939697265625 0 1301.9114 z 9
407.1974182128906 0 21943.19 y 12
408.20501708984375 0 4547.8823 c Ammonia loss 9
408.5434265136719 0 3800.0972 y Ammonia loss 5
408.87579345703125 0 1426.5165 z 5
409.1469421386719 0 1802.9939
409.19451904296875 0 1320.63
409.6930847167969 0 959.6858 z Water loss 2
410.2356872558594 0 817.89764
412.1845703125 0 2789.5752
412.2103271484375 0 994.2437
412.70806884765625 0 1115.9323
413.7019958496094 0 1987.4869 y Water loss 2
413.8872985839844 0 1342.3586
413.9521484375 0 1892.7125 y Ammonia loss 2
414.2139587402344 0 34198.195 y 5
414.4500427246094 0 1027.3256
414.5489807128906 0 24275.594
414.705322265625 0 2140.7122
414.882568359375 0 9429.65
415.2178955078125 0 1722.3892
416.2037048339844 0 762.8345
417.2207946777344 0 38918.37
417.7223815917969 0 15300.876
418.220947265625 0 7617.0654
418.7216491699219 0 716.433
419.2174987792969 0 2587.4883
419.713134765625 0 2191.6594
420.2125244140625 0 1186.4839
421.2110595703125 0 1013.65607
422.2553405761719 0 956.9661
424.74896240234375 0 15129.017
425.25115966796875 0 10352.329
425.75213623046875 0 3189.3708
426.2193298339844 0 1331.4147
426.68707275390625 0 1080.8096
426.72314453125 0 1026.7703
427.4552307128906 0 812.2629
428.2884216308594 0 8202.343
429.29339599609375 0 3602.967
430.1950378417969 0 6993.0054
430.29681396484375 0 638.8664
430.9563293457031 0 1136.8741
431.199462890625 0 2022.7058
432.7206726074219 0 1184.8806
433.7130432128906 0 1843.3862
434.2335510253906 0 1607.8772
434.705810546875 0 3800.8933
435.16412353515625 0 1663.3376
435.2294006347656 0 2806.7002
436.17095947265625 0 856.6461
436.2268981933594 0 3209.6675
436.8804016113281 0 1492.3364
437.4629211425781 0 785.31476
437.5490417480469 0 2595.9338
437.7179260253906 0 674.57404
437.8809509277344 0 1942.5847
437.9659118652344 0 1156.6167
438.2179260253906 0 881.9778
439.2076721191406 0 877.2664
439.4600830078125 0 1037.9045
439.7169494628906 0 1077.9039
440.7353210449219 0 19513.1
441.20867919921875 0 12044.27 w 8
441.23773193359375 0 10895.583
441.71112060546875 0 5713.534
441.737548828125 0 1956.5889
442.21173095703125 0 3578.8423
443.2274169921875 0 875.49066
443.46441650390625 0 904.0928
443.7114562988281 0 6745.99
443.96142578125 0 5462.157 z Water loss 1
444.2172546386719 0 4212.1655
444.4669494628906 0 1351.006
444.7160949707031 0 819.3927
444.9730529785156 0 943.78973
445.7106018066406 0 1684.8044
445.7364501953125 0 1249.6367
446.2074279785156 0 908.00385
447.2214050292969 0 8672.157
447.2553405761719 0 1124.1498
447.47039794921875 0 882.34467
447.7156677246094 0 2203.9958
447.9668273925781 0 4844.1304 y Water loss 1
448.2195129394531 0 9761.368 y Ammonia loss 1
448.2682189941406 0 2833.6636
448.4686584472656 0 5891.324 z 1
448.7204895019531 0 6686.1978
448.9686279296875 0 1871.582
449.219970703125 0 1761.6963
450.89459228515625 0 869.3555
451.2259521484375 0 3182.863 c Water loss 13
451.4737548828125 0 3935.032 c Ammonia loss 13
451.5372619628906 0 710.7301
451.7200012207031 0 4111.4473
451.8815612792969 0 757.28186
451.97137451171875 0 1873.0547
452.2167663574219 0 3062.7224
452.4694519042969 0 7392.1255 y 1
452.72003173828125 0 7194.8813
452.9710388183594 0 3317.845
453.2231140136719 0 2357.962
453.2601318359375 0 15552.117
453.7618713378906 0 11548.378
453.9786682128906 0 873.89154
454.2262268066406 0 2735.143 c Water loss 6
454.26385498046875 0 4215
454.719970703125 0 6243.481 c Ammonia loss 6
454.7633361816406 0 1222.0542
455.2212219238281 0 2023.0093
455.566650390625 0 1987.1357
455.72735595703125 0 27100.613 c 13
455.899658203125 0 3204.987
455.9776611328125 0 25893.693
456.2286682128906 0 14622.727
456.479248046875 0 4178.187
456.5705871582031 0 1231.5088
456.7313232421875 0 3658.6729
456.9057922363281 0 727.1393
457.2294921875 0 7049.3857
457.73095703125 0 2467.1504
458.2266845703125 0 888.55896
459.19549560546875 0 1105.7308
460.22808837890625 0 1006.67566
460.2607116699219 0 6945.8306
461.20806884765625 0 5281.282 w 11
461.2656555175781 0 1540.5947
461.4698486328125 0 1987.5017
461.56915283203125 0 1124.1335
461.71966552734375 0 3647.2224
461.99188232421875 0 935.57104
462.2302551269531 0 3640.626 c Water loss 10
462.48370361328125 0 934.1101
462.7314147949219 0 6775.974
462.97906494140625 0 4282.1533
463.2304992675781 0 3667.1504 c 6
463.4794006347656 0 1202.047
463.7330017089844 0 1353.8087
465.2251281738281 0 1590.9363 z Ammonia loss 4
465.72747802734375 0 5248.077
465.9722595214844 0 109916.01
466.2239990234375 0 98660.32
466.4759521484375 0 50433.023
466.72540283203125 0 22992.13
466.9752502441406 0 7494.0396
467.22271728515625 0 2847.8457
467.2486877441406 0 2749.2637
468.5668029785156 0 1328.3016
469.2460021972656 0 2923.5588 y Water loss 8
469.7219543457031 0 10493.048
469.98699951171875 0 2239.2568
470.2386779785156 0 23574.117 y Water loss 4
470.5653991699219 0 15720.279 y Ammonia loss 4
470.74310302734375 0 12073.275
470.90020751953125 0 14078.103 z 4
471.23712158203125 0 8833.1875
471.5691223144531 0 3404.8792
471.7454833984375 0 1026.2443
471.9017333984375 0 1617.7202
471.9809875488281 0 1764.4883
472.2309265136719 0 1176.3171
473.7299499511719 0 955.7505
474.2267761230469 0 1729.6189
474.2767028808594 0 1554.8695
474.7241516113281 0 862.43115
475.283447265625 0 6486.531
475.9121398925781 0 1165.9568
476.2397766113281 0 17681.816 y 4
476.2854309082031 0 1946.5122
476.4885559082031 0 11424.969
476.5762634277344 0 9865.749
476.73773193359375 0 10956.689
476.90972900390625 0 3680.4714
476.98895263671875 0 3807.9324
477.2079162597656 0 2239.631
477.23895263671875 0 5336.8535
477.28729248046875 0 1312.9851
477.4871826171875 0 1957.9653
477.7409973144531 0 2996.4873
478.2508239746094 0 29162.547 y 8
478.7524719238281 0 13464.024
479.2524108886719 0 5657.032
479.5695495605469 0 3579.714
479.9036865234375 0 3057.8684
480.2394714355469 0 3102.0725
480.4925842285156 0 7834.218
480.74078369140625 0 159098.19
480.9914855957031 0 204879.53
481.2421875 0 149923.64
481.4925231933594 0 76710.875
481.7429504394531 0 29409.104
481.9936218261719 0 9574.547
482.24249267578125 0 4444.3535
482.73724365234375 0 6334.7485 c Water loss 7
483.2311096191406 0 12753.627 c Ammonia loss 7
483.7324523925781 0 5454.601
484.23516845703125 0 3453.3938
484.7359924316406 0 984.2024
485.2365417480469 0 5309.7217
485.57086181640625 0 5333.232
485.90655517578125 0 2806.828
489.2460021972656 0 891.4244
490.24700927734375 0 2819.5225 z Ammonia loss 7
491.2475891113281 0 1934.3381
491.74237060546875 0 26901.307 c 7
492.244384765625 0 17593.648
492.7445373535156 0 4588.2495
493.24951171875 0 1852.438 c Water loss 3
494.2378234863281 0 44035.043 c Ammonia loss 3
494.58184814453125 0 1576.7861
495.2414855957031 0 12683.05
496.2501525878906 0 3771.8064
496.7298889160156 0 1381.1694
497.22613525390625 0 1764.1646
497.71966552734375 0 965.73456
497.7548828125 0 1933.0486 y Water loss 7
498.2266845703125 0 1179.4515
498.7535095214844 0 20575.43 z 7
499.2541809082031 0 11569.297
499.7549133300781 0 4277.717
500.2559814453125 0 1766.3827
503.2481994628906 0 2881.2244 z Water loss 3
504.2002868652344 0 1082.7333
505.2493896484375 0 2883.2322 c Ammonia loss 11
505.7368469238281 0 22586.91
506.2380065917969 0 16036.265
506.7619934082031 0 63620.68 y 7
507.2635192871094 0 36004.76
507.7641296386719 0 12073.449
508.2629089355469 0 3317.4253
509.244384765625 0 2043.3276 z 3
509.7630310058594 0 1041.1873
510.6038513183594 0 910.8906
510.773193359375 0 4861.2925
510.9301452636719 0 39312 c 11
511.26458740234375 0 122617.69 c 3
511.5984191894531 0 16203.821
511.77740478515625 0 1614.5165
511.9331359863281 0 5584.947
512.2672119140625 0 20573.902
512.59912109375 0 883.72784
513.2689208984375 0 2800.1382
513.7545776367188 0 1326.5616
514.2496337890625 0 26937.227
514.5830078125 0 3488.766 y 3
514.7457885742188 0 29103.738
514.9170532226562 0 2210.2942
515.2467041015625 0 14175.158
515.7461547851562 0 4619.71
516.2520751953125 0 1912.7294
518.2457885742188 0 2264.5334
518.2805786132812 0 1010.04333
518.5769653320312 0 1696.5038
519.2739868164062 0 39363.15 z 11
519.6022338867188 0 1383.3052
519.7459106445312 0 975.8546
519.9345703125 0 2833.5605
520.2763671875 0 11397.332
520.726318359375 0 1119.0692
521.228759765625 0 1065.9141
521.2747192382812 0 3088.603
524.5861206054688 0 1369.4939
525.2706909179688 0 1516.828
526.2560424804688 0 5193.8994
526.5834350585938 0 913.311
526.7587890625 0 7254.061
527.2312622070312 0 3715.6567
527.2640380859375 0 4287.9927
527.767333984375 0 58524.637
528.2685546875 0 32026.727
528.7691040039062 0 10675.353
529.2692260742188 0 2863.2935
530.597412109375 0 864.94556
531.2586669921875 0 1088.9225
531.5861206054688 0 1008.0956
532.196044921875 0 3320.3667
532.2593994140625 0 3474.0195
532.5873413085938 0 65773.984 w 2
532.7596435546875 0 5116.4585
532.921630859375 0 57144.01
533.25537109375 0 33562.223
533.5891723632812 0 10182.404
533.7734985351562 0 2765.7983 w 6
533.9229125976562 0 4947.587
534.2705688476562 0 6730.5947
534.7724609375 0 4115.183
534.9366455078125 0 1181.054
535.2904052734375 0 24345.68 y 11
535.77392578125 0 11949.51
536.2940063476562 0 4725.876
536.929931640625 0 1015.57214
537.27099609375 0 4506.7954
537.5994262695312 0 2357.7788
537.9369506835938 0 975.0823
538.2725219726562 0 2348.2183
539.238037109375 0 2881.171
539.2809448242188 0 1125.7014
539.6013793945312 0 1189.0411
540.276611328125 0 1032.1241
541.2672119140625 0 1728.6837
541.781494140625 0 1120.3679
542.2796020507812 0 3308.3247
542.7756958007812 0 867.7572
543.276123046875 0 2041.9786
544.284423828125 0 994.13214
544.6084594726562 0 1036.7946
544.9437866210938 0 962.46094
545.60009765625 0 2082.6025
545.9295043945312 0 1333.499 z Water loss 2
546.2620849609375 0 1978.4213 z Ammonia loss 2
546.7570190429688 0 1318.9899
547.2772827148438 0 4232.334 c Ammonia loss 8
547.7782592773438 0 3252.5605
548.2659301757812 0 1070.2262
548.9435424804688 0 1110.8174
549.2222900390625 0 1842.3903
549.2716064453125 0 2308.7925
549.6100463867188 0 8130.8823
549.9427490234375 0 7690.408
550.2064208984375 0 2848.48
550.2548828125 0 10566.621
550.6060791015625 0 2698.44
551.2604370117188 0 13274.672 y Water loss 2
551.5977172851562 0 6890.663 y Ammonia loss 2
551.9313354492188 0 14798.955 z 2
552.2633056640625 0 15100.088
552.5991821289062 0 7023.957
552.93115234375 0 2446.0723
553.2681274414062 0 2145.0166
554.2700805664062 0 1774.1788
554.601806640625 0 1246.2256
554.7813720703125 0 1657.0219
554.93505859375 0 2246.3735
555.269775390625 0 12410.441 y Water loss 6
555.7900390625 0 270883.25 c 8
556.2903442382812 0 143010.98
556.7693481445312 0 43610.79
556.7938232421875 0 34371.137
557.271240234375 0 29278.566 y 2
557.6048583984375 0 7438.3
557.771728515625 0 9781.384
557.9397583007812 0 6132.281
558.26953125 0 5440.566
558.6016235351562 0 2689.523 c Water loss 12
558.9423217773438 0 2411.4424
559.2722778320312 0 5774.404
559.6085815429688 0 4035.1743
559.9415893554688 0 1471.2494
561.9645385742188 0 1865.7511
562.2944946289062 0 5260.1313
562.634033203125 0 5491.7876
562.781005859375 0 4872.3843
562.9683837890625 0 4507.4653
563.2570190429688 0 1540.1069
563.2974243164062 0 2988.313
563.6149291992188 0 2632.1016
563.7801513671875 0 4703.678
563.9425048828125 0 5451.968
564.275634765625 0 261907.48 y 6
564.6021118164062 0 68697.484 c 12
564.7769165039062 0 151613.78
564.9359130859375 0 67473.34
565.2748413085938 0 87985.06
565.6039428710938 0 10171.452
565.7782592773438 0 17014.174
565.9376220703125 0 5104.695
566.275146484375 0 7886.0264
566.78173828125 0 961.25146
567.2327270507812 0 13229.812
567.2814331054688 0 2796.2275
567.9552612304688 0 902.4134
568.2344360351562 0 3585.6865
568.2862548828125 0 4136.8516
568.6161499023438 0 1901.0192
568.7975463867188 0 16313.506
569.2911987304688 0 14188.859
569.6103515625 0 1013.72723
569.7909545898438 0 7073.491
569.936767578125 0 1888.1375
570.2824096679688 0 3748.2886
570.7911987304688 0 8412.328
571.2876586914062 0 8987.593
571.7869262695312 0 5919.087
572.2888793945312 0 1883.9354
572.6281127929688 0 1195.9287
572.9566040039062 0 3054.3203
573.2879028320312 0 3544.4717
573.6135864257812 0 2457.5015
573.9379272460938 0 22514.486
574.2720947265625 0 20417.104
574.6055297851562 0 10045.187
574.8212280273438 0 1194.7666
574.9397583007812 0 3100.4253
576.3701171875 0 1263.0425
576.8003540039062 0 1723.2013
577.62353515625 0 1326.4424
577.9609375 0 12231.668
578.28857421875 0 12369.191
578.615234375 0 9838.323
578.77587890625 0 1010.8874
578.9429321289062 0 6110.343
579.2721557617188 0 2184.0723
579.602294921875 0 1361.3807
581.2791748046875 0 2939.0435
582.2838745117188 0 2983.147
582.6266479492188 0 3043.7876
582.763427734375 0 1676.5806
582.8104858398438 0 1054.3098
582.9532470703125 0 17176.895
583.2869262695312 0 15813.82
583.620849609375 0 11447.147
583.8046264648438 0 2236.1792
583.9530029296875 0 3845.1277
584.2923583984375 0 2879.6465
587.2893676757812 0 2074.4187
587.6168823242188 0 8882.034
587.9465942382812 0 8792.272
588.2844848632812 0 6278.05
588.6140747070312 0 2487.492
588.9498291015625 0 2062.3718
589.2850341796875 0 1192.8806
589.7898559570312 0 993.394
590.3068237304688 0 3978.701
590.7671508789062 0 737.16266
590.81298828125 0 10639.861
591.2811889648438 0 24201.162 w 5
591.3243408203125 0 2860.697
591.7824096679688 0 13413.209
591.8223876953125 0 4638.355
591.9577026367188 0 1329.1844
592.28515625 0 10318.764
592.6288452148438 0 24239.361
592.7818603515625 0 2754.52
592.963134765625 0 21259.338
593.2554931640625 0 4699.996
593.298095703125 0 10907.234
593.6314697265625 0 5790.247
593.9595947265625 0 1051.4552
594.265869140625 0 3801.8528
595.270263671875 0 1307.6433
596.6217651367188 0 1219.5337
596.95703125 0 1075.348 y Water loss 1
597.2935180664062 0 6850.728
597.619873046875 0 25676.035 z 1
597.8049926757812 0 1103.9598
597.953125 0 30139.082
598.2919311523438 0 15074.563
598.621337890625 0 5548.8936
598.8094482421875 0 7410.363
598.9534301757812 0 2719.4685
599.2908325195312 0 4066.665
601.3099975585938 0 2020.8054
601.627685546875 0 7363.1343 c Ammonia loss 13
601.9614868164062 0 10146.216
602.2940063476562 0 5493.701
602.629638671875 0 1869.7185
602.7947998046875 0 1656.2112
602.9576416015625 0 4214.2065 y 1
603.2916870117188 0 6880.2744
603.625 0 2705.3525
603.8051147460938 0 1210.5945 z Water loss 5
603.9593505859375 0 1739.398
604.815673828125 0 1071.5349
606.3162231445312 0 4460.9106
606.649169921875 0 3042.2478
606.9730224609375 0 3629.8877
607.301025390625 0 24539.223 c 13
607.6342163085938 0 30767.414
607.9678344726562 0 19856.646
608.3017578125 0 10761.438
608.6340942382812 0 4757.004
608.9710693359375 0 2037.0625
610.2849731445312 0 21838.193
611.2902221679688 0 6146.534
611.802978515625 0 15392.006 c Ammonia loss 9
612.3023681640625 0 11968.985 y Ammonia loss 5
612.8079223632812 0 30968.732 z 5
613.310302734375 0 24058.668
613.8117065429688 0 10782.658
614.3114013671875 0 3839.195
614.6480102539062 0 1123.4624
615.3106689453125 0 6716.4795
615.6437377929688 0 8416.972
615.8081665039062 0 1607.822
615.97705078125 0 7994.299
616.3101196289062 0 8521.324
616.6395263671875 0 6422.3643
616.97314453125 0 5763.0415
617.3038330078125 0 3823.5188
617.6397094726562 0 1092.2537
619.8173217773438 0 3584.1548
620.3120727539062 0 95188.26 c 9
620.6497802734375 0 5659.4053
620.8155517578125 0 113101.414 y 5
620.9696044921875 0 20895.139
621.2437133789062 0 6692.731
621.3152465820312 0 113223.516
621.6484985351562 0 57919.215
621.8187866210938 0 23020.834
621.9833374023438 0 35600.94
622.246337890625 0 4047.1772
622.3176879882812 0 19268.363
622.6514282226562 0 6273.287
622.8218994140625 0 1621.3069
622.9876098632812 0 1183.136
623.253662109375 0 1071.938
624.2705078125 0 3764.673
624.330078125 0 1549.6243
625.2755126953125 0 1305.1077
625.6509399414062 0 4925.108
625.984619140625 0 8126.831
626.3176879882812 0 13956.392
626.651611328125 0 7661.9697
626.819091796875 0 1854.2087
626.9842529296875 0 4925.6265
627.3209228515625 0 2231.657
629.3089599609375 0 2326.6753
629.6453857421875 0 5071.851
629.97607421875 0 10280.958
630.3097534179688 0 9298.739
630.6443481445312 0 2616.43
630.9745483398438 0 2202.3623
631.9793701171875 0 1373.0155
632.3073120117188 0 1593.3286
633.3147583007812 0 1672.6089
633.821533203125 0 1141.3357
634.3126831054688 0 1010.3629
634.9802856445312 0 2189.2878
635.314208984375 0 15903.196
635.6482543945312 0 100975
635.9818115234375 0 100426.89
636.3162841796875 0 58783.754
636.6498413085938 0 23745.896
636.9837036132812 0 8663.833
637.3179931640625 0 4145.335
637.6487426757812 0 1425.8928
639.80810546875 0 1643.1914
640.3128662109375 0 2207.559
640.651611328125 0 28271.83
640.8175048828125 0 1966.4973
640.9863891601562 0 55492.21
641.321533203125 0 66744.7
641.6558227539062 0 44247.613
641.9899291992188 0 20296.236
642.3238525390625 0 10267.256
642.65673828125 0 4543.3667
643.826171875 0 4601.6313
644.3268432617188 0 3410.6045
644.824951171875 0 1636.762
646.3328247070312 0 1113.7264
647.3241577148438 0 3409.1077
647.823486328125 0 38113.586
648.3243408203125 0 28554.654
648.8253784179688 0 13290.996
649.3248901367188 0 5097.56
649.82958984375 0 2037.0483
650.3306274414062 0 1150.5466
650.8302001953125 0 1334.9053
651.3184814453125 0 1021.1784
652.322265625 0 15459.602
653.3216552734375 0 5560.9165
654.3287963867188 0 1104.6144
655.3163452148438 0 1327.126
655.8180541992188 0 2992.0254
656.3211059570312 0 1666.3496
657.8430786132812 0 2580.5908
658.3426513671875 0 2817.7969
658.849853515625 0 1015.197
662.3165893554688 0 2850.041
663.2908325195312 0 7450.3525
663.3389282226562 0 2365.1086
664.3428344726562 0 17924.562
665.3478393554688 0 8115.159
666.3510131835938 0 2294.9976
670.8017578125 0 1735.5687
671.3346557617188 0 1207.8622
671.82470703125 0 1767.809
672.3179321289062 0 2262.363
672.8095703125 0 1130.8135
676.3424682617188 0 3070.8044
676.8406372070312 0 4014.1772
677.3424072265625 0 3657.468
677.821533203125 0 2201.5881
678.3248291015625 0 850.4143
679.3359985351562 0 2905.086 c Water loss 4
679.8372802734375 0 26315
680.3196411132812 0 125533.89 c Ammonia loss 4
680.840087890625 0 10911.759
681.3224487304688 0 55651.246 y Ammonia loss 10
681.840576171875 0 1071.9916
682.3339233398438 0 42216.26 z 10
682.8464965820312 0 2412.2456
683.3405151367188 0 21599.24
683.8480224609375 0 2391.0583
684.3427734375 0 8049.2275
685.3357543945312 0 1924.8398
691.3357543945312 0 1278.5848
691.8344116210938 0 1031.9861
693.3297119140625 0 1569.9307 c Ammonia loss 10
693.83251953125 0 3334.159
694.335205078125 0 2025.189
694.83642578125 0 983.7222
695.3197631835938 0 1804.204
696.3197021484375 0 1900.7211
697.3441162109375 0 30471.887 c 4
698.352294921875 0 30084.762 y 10
698.8333129882812 0 4697.8574
699.3508911132812 0 9818.087
699.831787109375 0 2209.4011
700.35009765625 0 2331.9258
701.3493041992188 0 2416.3547
701.843505859375 0 40892.668 c 10
702.3456420898438 0 40667.805
702.8472900390625 0 21624.42
703.3473510742188 0 6815.902
703.8505249023438 0 1789.4323
704.345947265625 0 1090.6643
704.8448486328125 0 1224.3553 y Water loss 4
705.3425903320312 0 3083.2278 y Ammonia loss 4
705.8472900390625 0 9499.983 z 4
706.3511352539062 0 19981.191
706.8531494140625 0 13215.809
707.353271484375 0 8061.9985
710.3223876953125 0 2902.8494
711.3277587890625 0 6547.369
712.3329467773438 0 4052.7578
712.840087890625 0 1798.6514
713.3359985351562 0 3105.6533
713.8563232421875 0 7200.929 y 4
714.35546875 0 5276.1743
714.8592529296875 0 3154.3113
719.3571166992188 0 987.6337
721.352294921875 0 4110.4985
722.3544311523438 0 2137.2234
723.3679809570312 0 3597.934
724.3679809570312 0 1100.3503
727.3507080078125 0 3093.3098
727.8516235351562 0 1841.2698
728.3439331054688 0 2292.698
729.3384399414062 0 1128.4542
733.8541259765625 0 1851.6486
734.356201171875 0 2642.1074
734.8538208007812 0 1951.9568
735.3381958007812 0 1796.5813
738.3720703125 0 1920.0278
740.864990234375 0 1694.4325 w 3
741.3656005859375 0 5246.303
741.868408203125 0 6339.901
742.36767578125 0 1941.2184
743.388427734375 0 1061.8412
743.8831787109375 0 1064.3892
744.386962890625 0 1675.9849
744.88720703125 0 1189.2295
745.3887329101562 0 958.3846
749.3654174804688 0 1284.6396
752.3678588867188 0 7198.016 w 9
753.3727416992188 0 5031.3384
754.3724975585938 0 2693.9146
755.3701782226562 0 1096.8169
756.35986328125 0 1136.3949
757.3837890625 0 1037.9854 c Ammonia loss 11
757.8812866210938 0 2370.9207
758.3793334960938 0 1351.9247
762.3618774414062 0 955.7938 y Water loss 3
762.8614501953125 0 4250.455 y Ammonia loss 3
763.3626098632812 0 14760.194 z 3
763.8653564453125 0 30601.125
764.3668823242188 0 15574.345
764.8680419921875 0 8957.004
765.3743286132812 0 1975.3197
765.8904418945312 0 6293.8994 c 11
766.3998413085938 0 7939.568
766.8948364257812 0 3592.8171
767.4085083007812 0 2634.5417
769.369140625 0 2951.037
769.8682250976562 0 1995.7719
770.3665161132812 0 2025.4891
770.8677368164062 0 1085.8108
771.3734130859375 0 3224.4504 y 3
771.8726196289062 0 3321.355
772.3700561523438 0 1968.0719
776.3851928710938 0 1210.8372
777.3650512695312 0 2182.3726
777.8666381835938 0 1765.8973
778.367919921875 0 1321.5579
778.8973388671875 0 1076.6074
779.3998413085938 0 1819.9103
786.3640747070312 0 973.83417
793.4014892578125 0 62476.61 c Ammonia loss 5
794.40478515625 0 30492.428
795.406982421875 0 8424.828
796.4180908203125 0 1379.7319
798.3858642578125 0 4754.328 w 2
798.8876953125 0 3405.2844
799.379638671875 0 1100.9572
809.3999633789062 0 2801.4568 y Water loss 9
810.42822265625 0 93014.49 c 5
811.3712768554688 0 18463.992 z 9
811.4329223632812 0 35762.89
812.3867797851562 0 23952.463
812.44970703125 0 3020.8105
813.3881225585938 0 11124.528
814.3890991210938 0 3001.6265
815.3950805664062 0 993.92957
816.89111328125 0 3307.1763
817.3935546875 0 4356.02
817.8960571289062 0 3649.469
818.3953247070312 0 1850.5781 z Water loss 2
822.4113159179688 0 2189.8445
823.4189453125 0 1146.5299
823.908447265625 0 3937.6233
824.404296875 0 4792.1934
824.9039916992188 0 2675.8628
825.394287109375 0 1118.267
825.8978271484375 0 945.9291
826.378662109375 0 1750.736 y Water loss 2
827.3959350585938 0 20512.285 y 9
827.8950805664062 0 33256.42
828.3967895507812 0 28782.713
828.8980102539062 0 12350.683
829.3990478515625 0 6428.7666
829.8999633789062 0 1652.563
830.4022827148438 0 1014.79626
831.3911743164062 0 1344.7395
835.3988647460938 0 3426.3845 y 2
835.9024047851562 0 3456.397
836.3952026367188 0 1081.1802
836.9066162109375 0 1242.4994
838.390380859375 0 4223.1084
838.891845703125 0 4700.9946
839.3982543945312 0 2078.311
839.8919067382812 0 1141.4603
844.4102783203125 0 2694.9468
845.402587890625 0 1062.785
845.8988647460938 0 2191.517
846.3984375 0 17156.15 c 12
846.9006958007812 0 18067.525
847.402099609375 0 12692.802
847.90234375 0 4775.5605
848.4048461914062 0 3226.2334
851.4035034179688 0 1874.8782
852.3648071289062 0 1339.6299
853.3771362304688 0 1422.019
854.3782958984375 0 1397.0502
860.4321899414062 0 1814.2527
866.9199829101562 0 3097.7666
867.4254150390625 0 35027.734
867.9278564453125 0 1185.4083
868.4288330078125 0 21205.871
869.4327392578125 0 6726.843
870.43408203125 0 925.8242
880.440673828125 0 1768.04
880.9291381835938 0 1768.147
881.4486694335938 0 4305.993
881.9411010742188 0 1078.7507
882.3635864257812 0 1362.0117
882.4515380859375 0 2476.611
883.3735961914062 0 1027.0948
888.4415893554688 0 1790.4844
888.9424438476562 0 13433.125
889.4444580078125 0 8344.057
889.9451904296875 0 4933.6484
890.4421997070312 0 2165.828
893.42919921875 0 1756.384
895.9221801757812 0 4852.3174 z 1
896.4246826171875 0 23918.492
896.9259643554688 0 19105.93
897.4271240234375 0 9446.936
897.9279174804688 0 3994.627
898.4293212890625 0 2424.656
902.9439697265625 0 1775.6938
903.4462890625 0 1228.9718
903.9466552734375 0 1306.0842 y 1
908.4277954101562 0 7277.317 c Ammonia loss 6
909.4346923828125 0 5094.4307
910.4381713867188 0 2890.5632 c 13
911.4483032226562 0 2001.4026
912.4572143554688 0 1898.71
920.4494018554688 0 1023.25696
922.45654296875 0 1079.0067 z Ammonia loss 8
923.4622192382812 0 2878.1184
923.9642944335938 0 2692.8362
924.4599609375 0 2613.4792
924.9641723632812 0 1123.8175
925.454345703125 0 6390.066 c 6
926.4588623046875 0 5030.484
927.4669189453125 0 1718.5994
931.4691162109375 0 3125.525
931.9727172851562 0 3150.4592
932.474365234375 0 2332.371
932.9793701171875 0 1406.7579
938.4645385742188 0 4519.5503 y Ammonia loss 8
939.4716796875 0 6866.227 z 8
940.4810180664062 0 13646.357
941.4847412109375 0 5731.1963
942.4864501953125 0 2122.6023
952.4725341796875 0 1303.7922
952.9696655273438 0 5221.9155
953.4707641601562 0 4899.743
953.9705810546875 0 2732.4116
954.478271484375 0 2063.2043
955.4885864257812 0 1626.8918 y 8
960.9774780273438 0 1246.9142
961.4791259765625 0 3181.8533
961.9827270507812 0 3814.1384
962.4814453125 0 2192.659
965.4572143554688 0 1052.3357 c Ammonia loss 7
966.45654296875 0 2711.8442
967.4633178710938 0 2784.2073
968.4624633789062 0 1167.5518
981.4760131835938 0 1244.042
982.4771728515625 0 15484.254 c 7
983.4812622070312 0 12358.48
984.4852905273438 0 5317.57
985.4904174804688 0 857.2744
996.493408203125 0 1752.9653 z 7
997.5018920898438 0 4489.476
998.5031127929688 0 2220.5737
999.51220703125 0 1037.6344
1011.4644165039062 0 954.7288
1012.5077514648438 0 2642.057 y 7
1013.5206298828125 0 1364.4241
1014.5022583007812 0 894.413
1028.47265625 0 1414.9873
1054.52685546875 0 1058.9175
1066.5543212890625 0 1581.7255
1067.5621337890625 0 1328.6707
1094.55078125 0 2404.3271
1095.55712890625 0 2546.0054
1110.572021484375 0 10093.473 c 8
1111.5762939453125 0 11822.191
1112.558349609375 0 7137.7812
1113.5435791015625 0 4345.129
1114.53857421875 0 1709.5645
1127.539306640625 0 1643.2467 y 6
1136.5257568359375 0 1025.701
1137.5347900390625 0 982.6569
1195.5994873046875 0 714.07996
1196.611083984375 0 1300.5181
1197.616455078125 0 871.4398
1224.6063232421875 0 1131.6421 z 5
1225.6162109375 0 4408.761
1226.6156005859375 0 2156.4446
1239.6119384765625 0 2154.9944 c 9
1240.6212158203125 0 3293.0115 y 5
1241.6243896484375 0 2233.9285
1265.6011962890625 0 898.8099
1359.672119140625 0 2131.219
1360.6729736328125 0 1945.0535
1361.6844482421875 0 771.24927
1402.6785888671875 0 1150.012 c 10
1403.6834716796875 0 2010.2605
1404.682861328125 0 1532.9827
1405.68798828125 0 920.30475
3008.23583984375 0 663.3393
3049.968994140625 0 1076.4574

Spectrum Details

|  |  |
| --- | --- |
| Matched peaks? Matched peaks The total absolute number of peaks matched. Additionally in brackets the total fraction of peaks matched and the total number of peaks is shown. | 171 (13.37% of 1279) |
| FDR? FDR The false discovery rate estimated for this peptide. It is calculated by matching all theoretical fragments with a non-integer shift with the raw peaks for this spectrum. This is done with 40 different shifts. The resulting percentage is the average number of annotated peaks over the number of annotated peaks with the correct spectrum. | 1.50% |
| Satellite FDR? Satellite FDR See the FDR for details on its calculation. This satellite ion specific FDR only contains the satellite ions (d/w) for I/L/J positions. | 0.00% |
| PSM Score? PSM Score The PSM Score as given by Hecklib to this annotated spectrum. It is shown with three significant figures. | 487 |

#### Spectrum 5664? Spectrum 5664 The raw spectrum of this peptide as annotated by Hecklib. The fragments are coloured according to ion type (see legend). Any peaks with a star '\*' as text can be hovered over to see the full details, first the ion type second the mass shift type. By hovering over the amino acids in the peptide or ions in the legend the corresponding peaks are highlighted. By toggling the 'Unassigned' label you can turn the background (unassigned) peaks on or off in the plot. By updating the slider in the Ion legend you can update the spectrum to only show the top X% of the peaks with labels. The top X% means any peak that is within X% of the highest intensity. By dragging in the spectrum you can zoom in to a specific part of the spectrum and use 'Zoom Out' to get back to the original zoom level. The annotation of the spectrum is based on the given sequence in the peptides file and is done with different software so inconsistencies are likely. The peaks are annotated based on the given sequence, with 20 ppm tolerance.

Copy Data

##### Spectrum 5664 (TSV)

###### Preview

```
Loading example...
```

*Click on the button to copy the data to your clipboard.*

Mz MinMz MaxIntensity Max

WidthHeightPeptide font sizePeptide stroke widthSpectrum font sizeSpectrum stroke widthCompact peptide

Ion legend

wxyz

abcd

OtherUnassignedIon^Charge^_Position_Show for top:%

JHQDWJDGKEYKCKV

06.05e+31.21e+41.82e+42.42e+4

Zoom Out

y^+1^_2_z^+1^_2_c^+1^_2_c^+1^_3_c^+1^_3_y^+1^_3_c^+1^_4_y^+2^_8_c^+1^_4_z^+1^_4_y^+1^_4_c^+3^_14_y^+3^_14_z^+1^_5_c^+1^_5_y^+1^_5_y^+2^_11_w^+1^_6_c^+2^_12_c^+1^_6_c^+1^_6_z^+1^_6_y^+2^_13_y^+1^_6_y^+2^_13_c^+2^_13_y^+2^_14_z^+2^_14_c^+2^_14_c^+2^_14_z^+1^_7_c^+1^_7_y^+1^_7_y^+1^_7_z^+1^_7_c^+1^_8_z^+1^_8_y^+1^_8_w^+1^_9_c^+1^_9_y^+1^_9_y^+1^_10_z^+1^_10_c^+1^_10_y^+1^_10_c^+1^_11_y^+1^_11_z^+1^_11_y^+1^_11_w^+1^_12_y^+1^_12_z^+1^_12_c^+1^_12_y^+1^_12_z^+1^_13_c^+1^_13_c^+1^_14_

0799159823963195

Fragment Matches Table

Show background peaks

| Position | Ion type | Intensity | mz Theoretical | mz Error (Th) | mz Error (ppm) | Charge | Series Number |
| --- | --- | --- | --- | --- | --- | --- | --- |
| - | - | 530.4 | 120.1 | - | - | 0 | - |
| - | - | 349 | 122.1 | - | - | 0 | - |
| - | - | 390 | 127.8 | - | - | 0 | - |
| - | - | 2745 | 129.1 | - | - | 0 | - |
| - | - | 481.2 | 148.3 | - | - | 0 | - |
| - | - | 569.7 | 148.9 | - | - | 0 | - |
| - | - | 742.6 | 155.1 | - | - | 0 | - |
| - | - | 770.5 | 166.1 | - | - | 0 | - |
| - | - | 642.2 | 169.1 | - | - | 0 | - |
| - | - | 660.2 | 177 | - | - | 0 | - |
| - | - | 825.6 | 183.1 | - | - | 0 | - |
| - | - | 541 | 193.1 | - | - | 0 | - |
| - | - | 737.1 | 203.1 | - | - | 0 | - |
| - | - | 464.4 | 215.9 | - | - | 0 | - |
| - | - | 467.9 | 219.7 | - | - | 0 | - |
| - | - | 1023 | 223.2 | - | - | 0 | - |
| - | - | 502.4 | 225 | - | - | 0 | - |
| - | - | 678 | 227.1 | - | - | 0 | - |
| 14 | y | 1.147E+04 | 229.2 | 0.0001467 | 0.6402 | +1 | 2 |
| 14 | z | 2487 | 230.2 | 0.0007814 | 3.395 | +1 | 2 |
| - | - | 1185 | 234.1 | - | - | 0 | - |
| - | - | 789.4 | 243.1 | - | - | 0 | - |
| - | - | 485.8 | 244.9 | - | - | 0 | - |
| - | - | 4866 | 251.2 | - | - | 0 | - |
| - | - | 735 | 252.2 | - | - | 0 | - |
| 2 | c | 1973 | 268.2 | 0.0005057 | 1.886 | +1 | 2 |
| - | - | 598.7 | 274.5 | - | - | 0 | - |
| - | - | 640.4 | 293.5 | - | - | 0 | - |
| - | - | 571.9 | 297.2 | - | - | 0 | - |
| - | - | 3289 | 300.2 | - | - | 0 | - |
| - | - | 702.6 | 314.2 | - | - | 0 | - |
| - | - | 572.1 | 321.7 | - | - | 0 | - |
| - | - | 578 | 330.2 | - | - | 0 | - |
| - | - | 1104 | 342.2 | - | - | 0 | - |
| - | - | 1223 | 353.2 | - | - | 0 | - |
| - | - | 1211 | 355.1 | - | - | 0 | - |
| - | - | 657.5 | 358.5 | - | - | 0 | - |
| 3 | c | 594.4 | 379.2 | 0.0006734 | 1.776 | +1 | 3 |
| 3 | c | 4786 | 396.2 | 0.0005156 | 1.301 | +1 | 3 |
| - | - | 6845 | 397.2 | - | - | 0 | - |
| - | - | 558.1 | 398.2 | - | - | 0 | - |
| - | - | 1001 | 404.1 | - | - | 0 | - |
| - | - | 737.1 | 405.1 | - | - | 0 | - |
| 13 | y | 1100 | 407.2 | 0.005639 | 13.85 | +1 | 3 |
| - | - | 985.6 | 411.2 | - | - | 0 | - |
| - | - | 804.4 | 412.2 | - | - | 0 | - |
| - | - | 946.7 | 413.2 | - | - | 0 | - |
| - | - | 582.4 | 414.9 | - | - | 0 | - |
| - | - | 840.9 | 418.2 | - | - | 0 | - |
| - | - | 1205 | 428.3 | - | - | 0 | - |
| - | - | 1024 | 455.8 | - | - | 0 | - |
| - | - | 767 | 456.3 | - | - | 0 | - |
| - | - | 2403 | 467.3 | - | - | 0 | - |
| - | - | 1183 | 468.3 | - | - | 0 | - |
| - | - | 703.1 | 471.3 | - | - | 0 | - |
| - | - | 849.9 | 488.3 | - | - | 0 | - |
| - | - | 560.3 | 488.5 | - | - | 0 | - |
| 4 | c | 600.5 | 494.2 | 0.002405 | 4.866 | +1 | 4 |
| - | - | 574.8 | 496.8 | - | - | 0 | - |
| 8 | y | 1477 | 506.8 | 0.001539 | 3.038 | +2 | 8 |
| 4 | c | 5543 | 511.3 | 0.0001149 | 0.2248 | +1 | 4 |
| - | - | 1718 | 512.3 | - | - | 0 | - |
| - | - | 821.7 | 514.3 | - | - | 0 | - |
| 12 | z | 2300 | 519.3 | 0.003936 | 7.581 | +1 | 4 |
| - | - | 1113 | 520.3 | - | - | 0 | - |
| - | - | 635.3 | 527.3 | - | - | 0 | - |
| 12 | y | 1080 | 535.3 | 0.003495 | 6.53 | +1 | 4 |
| - | - | 595.7 | 560.3 | - | - | 0 | - |
| - | - | 639.7 | 568.3 | - | - | 0 | - |
| - | - | 659.5 | 573.9 | - | - | 0 | - |
| - | - | 1113 | 581.4 | - | - | 0 | - |
| - | - | 6078 | 584.4 | - | - | 0 | - |
| - | - | 1331 | 585.4 | - | - | 0 | - |
| - | - | 792.2 | 599.3 | - | - | 0 | - |
| 14 | c | 1328 | 601.6 | 0.001087 | 1.807 | +3 | 14 |
| - | - | 1038 | 602 | - | - | 0 | - |
| - | - | 674.8 | 602.3 | - | - | 0 | - |
| 2 | y | 2135 | 603 | 1.54E-05 | 0.02554 | +3 | 14 |
| - | - | 1017 | 603.3 | - | - | 0 | - |
| - | - | 1581 | 634.6 | - | - | 0 | - |
| - | - | 1892 | 635 | - | - | 0 | - |
| - | - | 699.2 | 635.3 | - | - | 0 | - |
| - | - | 842 | 635.6 | - | - | 0 | - |
| - | - | 1198 | 636 | - | - | 0 | - |
| - | - | 5605 | 640.3 | - | - | 0 | - |
| - | - | 2.126E+04 | 640.7 | - | - | 0 | - |
| - | - | 6460 | 640.8 | - | - | 0 | - |
| - | - | 1.874E+04 | 641 | - | - | 0 | - |
| - | - | 1.411E+04 | 641.3 | - | - | 0 | - |
| - | - | 5720 | 641.7 | - | - | 0 | - |
| - | - | 3978 | 641.8 | - | - | 0 | - |
| - | - | 1488 | 642 | - | - | 0 | - |
| - | - | 1629 | 642.3 | - | - | 0 | - |
| - | - | 1915 | 651.4 | - | - | 0 | - |
| - | - | 1537 | 651.9 | - | - | 0 | - |
| - | - | 833.1 | 659.3 | - | - | 0 | - |
| - | - | 660 | 674.4 | - | - | 0 | - |
| 11 | z | 3520 | 682.3 | 0.005619 | 8.236 | +1 | 5 |
| - | - | 1219 | 682.4 | - | - | 0 | - |
| - | - | 1541 | 683.3 | - | - | 0 | - |
| - | - | 694.6 | 684.3 | - | - | 0 | - |
| - | - | 719.1 | 685.3 | - | - | 0 | - |
| - | - | 656.3 | 688.8 | - | - | 0 | - |
| 5 | c | 4402 | 697.3 | 0.0008146 | 1.168 | +1 | 5 |
| 11 | y | 2536 | 698.4 | 0.01159 | 16.59 | +1 | 5 |
| 5 | y | 1189 | 713.9 | 0.001842 | 2.58 | +2 | 11 |
| - | - | 931.6 | 743.5 | - | - | 0 | - |
| 10 | w | 1483 | 752.4 | 0.006954 | 9.243 | +1 | 6 |
| - | - | 620.7 | 761.9 | - | - | 0 | - |
| 12 | c | 1203 | 765.9 | 0.0001791 | 0.2338 | +2 | 12 |
| - | - | 733.1 | 766.4 | - | - | 0 | - |
| - | - | 950.9 | 766.9 | - | - | 0 | - |
| - | - | 789.5 | 771.9 | - | - | 0 | - |
| - | - | 805.8 | 774.4 | - | - | 0 | - |
| - | - | 752.1 | 783.4 | - | - | 0 | - |
| - | - | 665.6 | 789.4 | - | - | 0 | - |
| 6 | c | 951.9 | 793.4 | 0.003154 | 3.975 | +1 | 6 |
| - | - | 727.2 | 794.4 | - | - | 0 | - |
| - | - | 605.7 | 794.9 | - | - | 0 | - |
| - | - | 836.9 | 797.4 | - | - | 0 | - |
| - | - | 839.7 | 802.7 | - | - | 0 | - |
| 6 | c | 4737 | 810.4 | 0.004477 | 5.524 | +1 | 6 |
| 10 | z | 5024 | 811.4 | 0.003657 | 4.507 | +1 | 6 |
| - | - | 1429 | 812.4 | - | - | 0 | - |
| - | - | 603.1 | 813.4 | - | - | 0 | - |
| - | - | 1375 | 817.4 | - | - | 0 | - |
| 3 | y | 790.6 | 826.4 | 0.002067 | 2.501 | +2 | 13 |
| 10 | y | 1643 | 827.4 | 0.01023 | 12.37 | +1 | 6 |
| - | - | 774.2 | 834.9 | - | - | 0 | - |
| 3 | y | 2845 | 835.4 | 0.001562 | 1.87 | +2 | 13 |
| - | - | 769.2 | 835.7 | - | - | 0 | - |
| - | - | 2307 | 835.9 | - | - | 0 | - |
| - | - | 1240 | 836.4 | - | - | 0 | - |
| - | - | 1470 | 845.9 | - | - | 0 | - |
| 13 | c | 2398 | 846.4 | 0.002061 | 2.436 | +2 | 13 |
| - | - | 2453 | 846.9 | - | - | 0 | - |
| - | - | 698.6 | 850 | - | - | 0 | - |
| - | - | 601.2 | 859.4 | - | - | 0 | - |
| - | - | 655.5 | 866.4 | - | - | 0 | - |
| - | - | 2547 | 867.4 | - | - | 0 | - |
| - | - | 1461 | 868.4 | - | - | 0 | - |
| - | - | 978.1 | 873.4 | - | - | 0 | - |
| - | - | 699 | 873.9 | - | - | 0 | - |
| - | - | 2053 | 874.4 | - | - | 0 | - |
| - | - | 1568 | 875.4 | - | - | 0 | - |
| - | - | 1719 | 880.5 | - | - | 0 | - |
| - | - | 818.4 | 881.4 | - | - | 0 | - |
| - | - | 853.1 | 882.5 | - | - | 0 | - |
| - | - | 1012 | 887.9 | - | - | 0 | - |
| - | - | 911.5 | 888.4 | - | - | 0 | - |
| - | - | 1458 | 893.9 | - | - | 0 | - |
| - | - | 928 | 894.4 | - | - | 0 | - |
| 2 | y | 913.8 | 895.4 | 0.01736 | 19.39 | +2 | 14 |
| 2 | z | 3077 | 895.9 | 0.004967 | 5.544 | +2 | 14 |
| - | - | 1758 | 896.4 | - | - | 0 | - |
| - | - | 902.5 | 896.9 | - | - | 0 | - |
| - | - | 911.3 | 897.4 | - | - | 0 | - |
| 14 | c | 635.1 | 901.9 | 0.004476 | 4.962 | +2 | 14 |
| - | - | 1126 | 908.4 | - | - | 0 | - |
| - | - | 2262 | 910 | - | - | 0 | - |
| 14 | c | 8042 | 910.4 | 0.002451 | 2.692 | +2 | 14 |
| - | - | 6010 | 910.5 | - | - | 0 | - |
| - | - | 6398 | 910.9 | - | - | 0 | - |
| - | - | 4643 | 911.4 | - | - | 0 | - |
| - | - | 3680 | 911.5 | - | - | 0 | - |
| - | - | 2683 | 911.9 | - | - | 0 | - |
| - | - | 1250 | 912.5 | - | - | 0 | - |
| - | - | 753.6 | 916.5 | - | - | 0 | - |
| - | - | 842.4 | 918.5 | - | - | 0 | - |
| 9 | z | 757.5 | 922.5 | 0.01081 | 11.72 | +1 | 7 |
| - | - | 841 | 923 | - | - | 0 | - |
| - | - | 1537 | 923.5 | - | - | 0 | - |
| - | - | 1049 | 924.4 | - | - | 0 | - |
| 7 | c | 1784 | 925.5 | 0.0006155 | 0.6651 | +1 | 7 |
| - | - | 1942 | 930.5 | - | - | 0 | - |
| - | - | 5512 | 931 | - | - | 0 | - |
| - | - | 1.076E+04 | 931.5 | - | - | 0 | - |
| - | - | 1.08E+04 | 932 | - | - | 0 | - |
| - | - | 7838 | 932.5 | - | - | 0 | - |
| - | - | 1495 | 933 | - | - | 0 | - |
| - | - | 1021 | 933.5 | - | - | 0 | - |
| - | - | 1619 | 934.5 | - | - | 0 | - |
| - | - | 819.3 | 934.8 | - | - | 0 | - |
| - | - | 1233 | 935.5 | - | - | 0 | - |
| 9 | y | 637.2 | 937.5 | 0.00833 | 8.885 | +1 | 7 |
| - | - | 1736 | 938 | - | - | 0 | - |
| 9 | y | 2778 | 938.5 | 0.001368 | 1.458 | +1 | 7 |
| - | - | 2540 | 939 | - | - | 0 | - |
| 9 | z | 6008 | 939.5 | 0.005908 | 6.288 | +1 | 7 |
| - | - | 3113 | 940.5 | - | - | 0 | - |
| - | - | 731.3 | 943 | - | - | 0 | - |
| - | - | 671 | 947.4 | - | - | 0 | - |
| - | - | 1034 | 948.5 | - | - | 0 | - |
| - | - | 799.1 | 951.5 | - | - | 0 | - |
| - | - | 1601 | 952 | - | - | 0 | - |
| - | - | 2668 | 952.5 | - | - | 0 | - |
| - | - | 3179 | 953 | - | - | 0 | - |
| - | - | 2087 | 953.5 | - | - | 0 | - |
| - | - | 908 | 954 | - | - | 0 | - |
| - | - | 1291 | 956.5 | - | - | 0 | - |
| - | - | 1043 | 959.4 | - | - | 0 | - |
| - | - | 2992 | 960 | - | - | 0 | - |
| - | - | 1.421E+04 | 960.5 | - | - | 0 | - |
| - | - | 1.745E+04 | 961 | - | - | 0 | - |
| - | - | 1.419E+04 | 961.5 | - | - | 0 | - |
| - | - | 8045 | 962 | - | - | 0 | - |
| - | - | 3987 | 962.5 | - | - | 0 | - |
| - | - | 2134 | 963 | - | - | 0 | - |
| - | - | 1221 | 963.5 | - | - | 0 | - |
| - | - | 1259 | 965.5 | - | - | 0 | - |
| - | - | 826.7 | 967.2 | - | - | 0 | - |
| - | - | 703.7 | 977.1 | - | - | 0 | - |
| - | - | 2948 | 981.5 | - | - | 0 | - |
| 8 | c | 4065 | 982.5 | 0.0005949 | 0.6055 | +1 | 8 |
| - | - | 2063 | 983.5 | - | - | 0 | - |
| - | - | 1726 | 984.5 | - | - | 0 | - |
| 8 | z | 2261 | 996.5 | 0.005154 | 5.173 | +1 | 8 |
| - | - | 2129 | 997.5 | - | - | 0 | - |
| - | - | 903.9 | 1003 | - | - | 0 | - |
| 8 | y | 1891 | 1013 | 0.007155 | 7.066 | +1 | 8 |
| - | - | 851.7 | 1020 | - | - | 0 | - |
| - | - | 771.8 | 1020 | - | - | 0 | - |
| - | - | 812.1 | 1025 | - | - | 0 | - |
| - | - | 885 | 1026 | - | - | 0 | - |
| - | - | 893.1 | 1027 | - | - | 0 | - |
| - | - | 1494 | 1027 | - | - | 0 | - |
| - | - | 1468 | 1029 | - | - | 0 | - |
| - | - | 1155 | 1030 | - | - | 0 | - |
| - | - | 1594 | 1033 | - | - | 0 | - |
| - | - | 1782 | 1033 | - | - | 0 | - |
| - | - | 818.5 | 1033 | - | - | 0 | - |
| - | - | 1037 | 1034 | - | - | 0 | - |
| - | - | 875.5 | 1035 | - | - | 0 | - |
| - | - | 740.1 | 1035 | - | - | 0 | - |
| - | - | 772.2 | 1036 | - | - | 0 | - |
| - | - | 910.7 | 1038 | - | - | 0 | - |
| - | - | 933.1 | 1038 | - | - | 0 | - |
| - | - | 1057 | 1039 | - | - | 0 | - |
| - | - | 1006 | 1039 | - | - | 0 | - |
| - | - | 1224 | 1040 | - | - | 0 | - |
| - | - | 937.7 | 1042 | - | - | 0 | - |
| - | - | 987.5 | 1043 | - | - | 0 | - |
| - | - | 1005 | 1044 | - | - | 0 | - |
| - | - | 1216 | 1044 | - | - | 0 | - |
| - | - | 2500 | 1044 | - | - | 0 | - |
| - | - | 2046 | 1045 | - | - | 0 | - |
| - | - | 923.8 | 1045 | - | - | 0 | - |
| - | - | 796.6 | 1046 | - | - | 0 | - |
| - | - | 1094 | 1046 | - | - | 0 | - |
| - | - | 1080 | 1048 | - | - | 0 | - |
| - | - | 1062 | 1048 | - | - | 0 | - |
| - | - | 2136 | 1048 | - | - | 0 | - |
| - | - | 1364 | 1049 | - | - | 0 | - |
| - | - | 1470 | 1049 | - | - | 0 | - |
| - | - | 1996 | 1049 | - | - | 0 | - |
| - | - | 1999 | 1050 | - | - | 0 | - |
| - | - | 2143 | 1050 | - | - | 0 | - |
| - | - | 1462 | 1050 | - | - | 0 | - |
| - | - | 1625 | 1051 | - | - | 0 | - |
| - | - | 1281 | 1052 | - | - | 0 | - |
| - | - | 1122 | 1053 | - | - | 0 | - |
| - | - | 1646 | 1054 | - | - | 0 | - |
| - | - | 2163 | 1054 | - | - | 0 | - |
| - | - | 2680 | 1054 | - | - | 0 | - |
| - | - | 5462 | 1055 | - | - | 0 | - |
| - | - | 2236 | 1055 | - | - | 0 | - |
| - | - | 3694 | 1056 | - | - | 0 | - |
| - | - | 818.2 | 1056 | - | - | 0 | - |
| - | - | 814.4 | 1056 | - | - | 0 | - |
| - | - | 964.4 | 1057 | - | - | 0 | - |
| - | - | 1935 | 1058 | - | - | 0 | - |
| - | - | 1164 | 1059 | - | - | 0 | - |
| - | - | 2340 | 1060 | - | - | 0 | - |
| - | - | 4278 | 1060 | - | - | 0 | - |
| - | - | 2929 | 1061 | - | - | 0 | - |
| - | - | 2937 | 1061 | - | - | 0 | - |
| - | - | 1218 | 1061 | - | - | 0 | - |
| - | - | 2075 | 1062 | - | - | 0 | - |
| - | - | 1123 | 1062 | - | - | 0 | - |
| - | - | 1831 | 1063 | - | - | 0 | - |
| - | - | 2208 | 1063 | - | - | 0 | - |
| - | - | 2153 | 1063 | - | - | 0 | - |
| - | - | 4373 | 1064 | - | - | 0 | - |
| - | - | 2969 | 1064 | - | - | 0 | - |
| - | - | 2546 | 1064 | - | - | 0 | - |
| - | - | 1143 | 1065 | - | - | 0 | - |
| - | - | 873.6 | 1065 | - | - | 0 | - |
| - | - | 1061 | 1065 | - | - | 0 | - |
| - | - | 1073 | 1066 | - | - | 0 | - |
| - | - | 1806 | 1066 | - | - | 0 | - |
| - | - | 1691 | 1066 | - | - | 0 | - |
| 7 | w | 1896 | 1067 | 0.01164 | 10.91 | +1 | 9 |
| - | - | 1168 | 1067 | - | - | 0 | - |
| - | - | 1118 | 1067 | - | - | 0 | - |
| - | - | 2099 | 1068 | - | - | 0 | - |
| - | - | 1556 | 1068 | - | - | 0 | - |
| - | - | 1947 | 1068 | - | - | 0 | - |
| - | - | 3494 | 1069 | - | - | 0 | - |
| - | - | 2737 | 1069 | - | - | 0 | - |
| - | - | 2469 | 1069 | - | - | 0 | - |
| - | - | 3017 | 1070 | - | - | 0 | - |
| - | - | 995 | 1070 | - | - | 0 | - |
| - | - | 1496 | 1070 | - | - | 0 | - |
| - | - | 902.4 | 1071 | - | - | 0 | - |
| - | - | 2220 | 1110 | - | - | 0 | - |
| 9 | c | 7467 | 1111 | 0.0004651 | 0.4188 | +1 | 9 |
| - | - | 4615 | 1112 | - | - | 0 | - |
| - | - | 2069 | 1113 | - | - | 0 | - |
| - | - | 911.5 | 1125 | - | - | 0 | - |
| - | - | 1665 | 1126 | - | - | 0 | - |
| - | - | 1820 | 1127 | - | - | 0 | - |
| 7 | y | 1639 | 1128 | 0.006857 | 6.081 | +1 | 9 |
| - | - | 2873 | 1143 | - | - | 0 | - |
| - | - | 2579 | 1144 | - | - | 0 | - |
| - | - | 1049 | 1145 | - | - | 0 | - |
| - | - | 962.8 | 1151 | - | - | 0 | - |
| - | - | 1173 | 1152 | - | - | 0 | - |
| - | - | 1069 | 1162 | - | - | 0 | - |
| - | - | 796.2 | 1191 | - | - | 0 | - |
| 6 | y | 1025 | 1224 | 0.01733 | 14.17 | +1 | 10 |
| 6 | z | 2072 | 1225 | 0.005138 | 4.196 | +1 | 10 |
| - | - | 1057 | 1226 | - | - | 0 | - |
| - | - | 1154 | 1227 | - | - | 0 | - |
| - | - | 1864 | 1239 | - | - | 0 | - |
| 10 | c | 5074 | 1240 | 0.0003988 | 0.3217 | +1 | 10 |
| 6 | y | 2814 | 1241 | 0.01324 | 10.67 | +1 | 10 |
| - | - | 1640 | 1242 | - | - | 0 | - |
| - | - | 796.6 | 1243 | - | - | 0 | - |
| - | - | 2137 | 1254 | - | - | 0 | - |
| - | - | 1096 | 1255 | - | - | 0 | - |
| - | - | 696.8 | 1265 | - | - | 0 | - |
| - | - | 999.4 | 1274 | - | - | 0 | - |
| - | - | 730.2 | 1275 | - | - | 0 | - |
| - | - | 807.4 | 1277 | - | - | 0 | - |
| - | - | 3227 | 1279 | - | - | 0 | - |
| - | - | 4505 | 1280 | - | - | 0 | - |
| - | - | 1488 | 1281 | - | - | 0 | - |
| - | - | 993.1 | 1282 | - | - | 0 | - |
| - | - | 1561 | 1282 | - | - | 0 | - |
| - | - | 870 | 1284 | - | - | 0 | - |
| - | - | 935 | 1285 | - | - | 0 | - |
| - | - | 819.7 | 1312 | - | - | 0 | - |
| - | - | 832.4 | 1358 | - | - | 0 | - |
| - | - | 1056 | 1359 | - | - | 0 | - |
| - | - | 956.4 | 1375 | - | - | 0 | - |
| - | - | 1022 | 1389 | - | - | 0 | - |
| - | - | 843.4 | 1393 | - | - | 0 | - |
| - | - | 1060 | 1401 | - | - | 0 | - |
| - | - | 1103 | 1402 | - | - | 0 | - |
| 11 | c | 3996 | 1403 | 0.002988 | 2.13 | +1 | 11 |
| - | - | 3875 | 1404 | - | - | 0 | - |
| - | - | 1334 | 1405 | - | - | 0 | - |
| 5 | y | 1675 | 1410 | 0.02042 | 14.49 | +1 | 11 |
| 5 | z | 2052 | 1411 | 0.00181 | 1.283 | +1 | 11 |
| - | - | 2382 | 1412 | - | - | 0 | - |
| - | - | 1028 | 1413 | - | - | 0 | - |
| - | - | 893 | 1414 | - | - | 0 | - |
| - | - | 980.8 | 1426 | - | - | 0 | - |
| 5 | y | 2587 | 1427 | 0.005885 | 4.125 | +1 | 11 |
| - | - | 2090 | 1428 | - | - | 0 | - |
| - | - | 1242 | 1437 | - | - | 0 | - |
| - | - | 1891 | 1438 | - | - | 0 | - |
| - | - | 1002 | 1439 | - | - | 0 | - |
| - | - | 844.9 | 1466 | - | - | 0 | - |
| - | - | 1119 | 1473 | - | - | 0 | - |
| - | - | 1072 | 1473 | - | - | 0 | - |
| 4 | w | 1413 | 1481 | 0.02725 | 18.4 | +1 | 12 |
| - | - | 968.3 | 1481 | - | - | 0 | - |
| - | - | 1827 | 1482 | - | - | 0 | - |
| - | - | 1354 | 1482 | - | - | 0 | - |
| - | - | 1085 | 1483 | - | - | 0 | - |
| - | - | 1018 | 1483 | - | - | 0 | - |
| - | - | 917.9 | 1485 | - | - | 0 | - |
| - | - | 1187 | 1487 | - | - | 0 | - |
| - | - | 776.8 | 1489 | - | - | 0 | - |
| - | - | 748.9 | 1491 | - | - | 0 | - |
| - | - | 1418 | 1491 | - | - | 0 | - |
| - | - | 1037 | 1492 | - | - | 0 | - |
| - | - | 884.2 | 1493 | - | - | 0 | - |
| - | - | 1031 | 1494 | - | - | 0 | - |
| - | - | 1574 | 1495 | - | - | 0 | - |
| - | - | 857.3 | 1496 | - | - | 0 | - |
| - | - | 1380 | 1497 | - | - | 0 | - |
| - | - | 950.1 | 1499 | - | - | 0 | - |
| - | - | 1132 | 1502 | - | - | 0 | - |
| - | - | 1086 | 1502 | - | - | 0 | - |
| - | - | 920.5 | 1503 | - | - | 0 | - |
| - | - | 1091 | 1507 | - | - | 0 | - |
| - | - | 1067 | 1516 | - | - | 0 | - |
| - | - | 897 | 1520 | - | - | 0 | - |
| - | - | 1106 | 1522 | - | - | 0 | - |
| - | - | 907.3 | 1523 | - | - | 0 | - |
| - | - | 953.4 | 1524 | - | - | 0 | - |
| 4 | y | 1543 | 1525 | 0.0196 | 12.85 | +1 | 12 |
| 4 | z | 4484 | 1526 | 0.003596 | 2.357 | +1 | 12 |
| - | - | 4983 | 1527 | - | - | 0 | - |
| - | - | 935.9 | 1527 | - | - | 0 | - |
| - | - | 3232 | 1528 | - | - | 0 | - |
| - | - | 1256 | 1528 | - | - | 0 | - |
| - | - | 2656 | 1529 | - | - | 0 | - |
| - | - | 2022 | 1530 | - | - | 0 | - |
| - | - | 1011 | 1530 | - | - | 0 | - |
| 12 | c | 4999 | 1531 | 0.003474 | 2.269 | +1 | 12 |
| - | - | 1564 | 1531 | - | - | 0 | - |
| - | - | 3146 | 1532 | - | - | 0 | - |
| - | - | 2056 | 1533 | - | - | 0 | - |
| - | - | 1114 | 1533 | - | - | 0 | - |
| - | - | 1157 | 1536 | - | - | 0 | - |
| - | - | 948.4 | 1536 | - | - | 0 | - |
| - | - | 1994 | 1538 | - | - | 0 | - |
| - | - | 1149 | 1538 | - | - | 0 | - |
| - | - | 1062 | 1539 | - | - | 0 | - |
| - | - | 1255 | 1540 | - | - | 0 | - |
| 4 | y | 2360 | 1542 | 0.006112 | 3.964 | +1 | 12 |
| - | - | 2112 | 1543 | - | - | 0 | - |
| - | - | 882.9 | 1543 | - | - | 0 | - |
| - | - | 1431 | 1544 | - | - | 0 | - |
| - | - | 1492 | 1545 | - | - | 0 | - |
| - | - | 1187 | 1545 | - | - | 0 | - |
| - | - | 1688 | 1546 | - | - | 0 | - |
| - | - | 1130 | 1546 | - | - | 0 | - |
| - | - | 1059 | 1547 | - | - | 0 | - |
| - | - | 1694 | 1547 | - | - | 0 | - |
| - | - | 1209 | 1549 | - | - | 0 | - |
| - | - | 2424 | 1550 | - | - | 0 | - |
| - | - | 1746 | 1550 | - | - | 0 | - |
| - | - | 1773 | 1551 | - | - | 0 | - |
| - | - | 1856 | 1551 | - | - | 0 | - |
| - | - | 2063 | 1552 | - | - | 0 | - |
| - | - | 2653 | 1552 | - | - | 0 | - |
| - | - | 2260 | 1553 | - | - | 0 | - |
| - | - | 1170 | 1553 | - | - | 0 | - |
| - | - | 944.5 | 1554 | - | - | 0 | - |
| - | - | 1060 | 1555 | - | - | 0 | - |
| - | - | 825.9 | 1556 | - | - | 0 | - |
| - | - | 940.2 | 1557 | - | - | 0 | - |
| - | - | 1244 | 1557 | - | - | 0 | - |
| - | - | 1199 | 1558 | - | - | 0 | - |
| - | - | 2065 | 1558 | - | - | 0 | - |
| - | - | 1677 | 1559 | - | - | 0 | - |
| - | - | 2500 | 1559 | - | - | 0 | - |
| - | - | 2192 | 1560 | - | - | 0 | - |
| - | - | 2693 | 1560 | - | - | 0 | - |
| - | - | 1972 | 1561 | - | - | 0 | - |
| - | - | 1290 | 1561 | - | - | 0 | - |
| - | - | 1777 | 1562 | - | - | 0 | - |
| - | - | 906.3 | 1562 | - | - | 0 | - |
| - | - | 1048 | 1563 | - | - | 0 | - |
| - | - | 988 | 1564 | - | - | 0 | - |
| - | - | 2071 | 1565 | - | - | 0 | - |
| - | - | 2361 | 1565 | - | - | 0 | - |
| - | - | 3072 | 1566 | - | - | 0 | - |
| - | - | 4324 | 1566 | - | - | 0 | - |
| - | - | 4300 | 1567 | - | - | 0 | - |
| - | - | 2415 | 1567 | - | - | 0 | - |
| - | - | 2596 | 1568 | - | - | 0 | - |
| - | - | 2085 | 1568 | - | - | 0 | - |
| - | - | 1761 | 1569 | - | - | 0 | - |
| - | - | 1708 | 1569 | - | - | 0 | - |
| - | - | 1003 | 1570 | - | - | 0 | - |
| - | - | 1601 | 1571 | - | - | 0 | - |
| - | - | 1751 | 1571 | - | - | 0 | - |
| - | - | 1838 | 1572 | - | - | 0 | - |
| - | - | 2766 | 1572 | - | - | 0 | - |
| - | - | 2953 | 1573 | - | - | 0 | - |
| - | - | 4090 | 1573 | - | - | 0 | - |
| - | - | 3870 | 1574 | - | - | 0 | - |
| - | - | 6115 | 1574 | - | - | 0 | - |
| - | - | 3795 | 1575 | - | - | 0 | - |
| - | - | 3348 | 1575 | - | - | 0 | - |
| - | - | 2156 | 1576 | - | - | 0 | - |
| - | - | 1579 | 1576 | - | - | 0 | - |
| - | - | 1902 | 1577 | - | - | 0 | - |
| - | - | 820 | 1578 | - | - | 0 | - |
| - | - | 943.2 | 1578 | - | - | 0 | - |
| - | - | 1789 | 1579 | - | - | 0 | - |
| - | - | 1575 | 1580 | - | - | 0 | - |
| - | - | 2756 | 1580 | - | - | 0 | - |
| - | - | 3568 | 1581 | - | - | 0 | - |
| - | - | 3632 | 1581 | - | - | 0 | - |
| - | - | 3434 | 1582 | - | - | 0 | - |
| - | - | 4101 | 1582 | - | - | 0 | - |
| - | - | 3238 | 1583 | - | - | 0 | - |
| - | - | 3128 | 1583 | - | - | 0 | - |
| - | - | 2100 | 1584 | - | - | 0 | - |
| - | - | 1570 | 1584 | - | - | 0 | - |
| - | - | 1209 | 1585 | - | - | 0 | - |
| - | - | 1435 | 1586 | - | - | 0 | - |
| - | - | 1742 | 1586 | - | - | 0 | - |
| - | - | 2290 | 1587 | - | - | 0 | - |
| - | - | 1255 | 1587 | - | - | 0 | - |
| - | - | 2199 | 1588 | - | - | 0 | - |
| - | - | 2487 | 1588 | - | - | 0 | - |
| - | - | 844.7 | 1589 | - | - | 0 | - |
| - | - | 2044 | 1589 | - | - | 0 | - |
| - | - | 3278 | 1590 | - | - | 0 | - |
| - | - | 5398 | 1590 | - | - | 0 | - |
| - | - | 2895 | 1591 | - | - | 0 | - |
| - | - | 3481 | 1591 | - | - | 0 | - |
| - | - | 3046 | 1592 | - | - | 0 | - |
| - | - | 2173 | 1592 | - | - | 0 | - |
| - | - | 1485 | 1593 | - | - | 0 | - |
| - | - | 3360 | 1593 | - | - | 0 | - |
| - | - | 3516 | 1594 | - | - | 0 | - |
| - | - | 2600 | 1594 | - | - | 0 | - |
| - | - | 3908 | 1595 | - | - | 0 | - |
| - | - | 2473 | 1595 | - | - | 0 | - |
| - | - | 3177 | 1596 | - | - | 0 | - |
| - | - | 3189 | 1596 | - | - | 0 | - |
| - | - | 1526 | 1597 | - | - | 0 | - |
| - | - | 2067 | 1599 | - | - | 0 | - |
| - | - | 2733 | 1599 | - | - | 0 | - |
| - | - | 2001 | 1600 | - | - | 0 | - |
| - | - | 1451 | 1600 | - | - | 0 | - |
| - | - | 1633 | 1601 | - | - | 0 | - |
| - | - | 1609 | 1601 | - | - | 0 | - |
| - | - | 1892 | 1602 | - | - | 0 | - |
| - | - | 4236 | 1602 | - | - | 0 | - |
| - | - | 4468 | 1603 | - | - | 0 | - |
| - | - | 3856 | 1603 | - | - | 0 | - |
| - | - | 3229 | 1604 | - | - | 0 | - |
| - | - | 2085 | 1604 | - | - | 0 | - |
| - | - | 1476 | 1605 | - | - | 0 | - |
| - | - | 1760 | 1605 | - | - | 0 | - |
| - | - | 1142 | 1606 | - | - | 0 | - |
| - | - | 807.8 | 1606 | - | - | 0 | - |
| 3 | z | 2749 | 1654 | 0.006032 | 3.647 | +1 | 13 |
| - | - | 3739 | 1655 | - | - | 0 | - |
| - | - | 2465 | 1656 | - | - | 0 | - |
| - | - | 1108 | 1657 | - | - | 0 | - |
| 13 | c | 4841 | 1692 | 0.006179 | 3.652 | +1 | 13 |
| - | - | 3433 | 1693 | - | - | 0 | - |
| - | - | 3101 | 1694 | - | - | 0 | - |
| - | - | 1078 | 1695 | - | - | 0 | - |
| - | - | 950.7 | 1723 | - | - | 0 | - |
| - | - | 961.2 | 1792 | - | - | 0 | - |
| - | - | 1306 | 1793 | - | - | 0 | - |
| - | - | 972.7 | 1794 | - | - | 0 | - |
| 14 | c | 1050 | 1820 | 0.004083 | 2.243 | +1 | 14 |
| - | - | 2271 | 1821 | - | - | 0 | - |
| - | - | 1829 | 1822 | - | - | 0 | - |
| - | - | 1578 | 1823 | - | - | 0 | - |
| - | - | 1629 | 1831 | - | - | 0 | - |
| - | - | 1817 | 1832 | - | - | 0 | - |
| - | - | 1505 | 1833 | - | - | 0 | - |
| - | - | 1086 | 1846 | - | - | 0 | - |
| - | - | 1176 | 1861 | - | - | 0 | - |
| - | - | 2307 | 1862 | - | - | 0 | - |
| - | - | 4503 | 1863 | - | - | 0 | - |
| - | - | 4224 | 1864 | - | - | 0 | - |
| - | - | 1899 | 1865 | - | - | 0 | - |
| - | - | 1122 | 1866 | - | - | 0 | - |
| - | - | 2130 | 1876 | - | - | 0 | - |
| - | - | 3603 | 1877 | - | - | 0 | - |
| - | - | 2145 | 1878 | - | - | 0 | - |
| - | - | 1933 | 1879 | - | - | 0 | - |
| - | - | 1440 | 1893 | - | - | 0 | - |
| - | - | 2518 | 1894 | - | - | 0 | - |
| - | - | 3190 | 1895 | - | - | 0 | - |
| - | - | 1444 | 1896 | - | - | 0 | - |
| - | - | 1679 | 1903 | - | - | 0 | - |
| - | - | 3027 | 1904 | - | - | 0 | - |
| - | - | 6597 | 1905 | - | - | 0 | - |
| - | - | 5950 | 1906 | - | - | 0 | - |
| - | - | 3059 | 1907 | - | - | 0 | - |
| - | - | 2057 | 1908 | - | - | 0 | - |
| - | - | 3763 | 1920 | - | - | 0 | - |
| - | - | 1.11E+04 | 1921 | - | - | 0 | - |
| - | - | 2.397E+04 | 1922 | - | - | 0 | - |
| - | - | 2.111E+04 | 1923 | - | - | 0 | - |
| - | - | 1.281E+04 | 1924 | - | - | 0 | - |
| - | - | 6272 | 1925 | - | - | 0 | - |
| - | - | 3262 | 1926 | - | - | 0 | - |
| - | - | 1458 | 1927 | - | - | 0 | - |
| - | - | 743.4 | 3048 | - | - | 0 | - |
| - | - | 1021 | 3149 | - | - | 0 | - |
| - | - | 901.4 | 3164 | - | - | 0 | - |

m/z Charge Intensity FragmentType MassShift Position
120.08110809326172 0 530.40955
122.06483459472656 0 348.98233
127.79460906982422 0 390.04117
129.1023712158203 0 2744.8184
148.32858276367188 0 481.22736
148.94789123535156 0 569.74365
155.0931396484375 0 742.63477
166.0612030029297 0 770.4866
169.0967254638672 0 642.1718
177.03948974609375 0 660.2414
183.113037109375 0 825.61316
193.1332244873047 0 541.0357
203.10340881347656 0 737.07996
215.92208862304688 0 464.44806
219.69964599609375 0 467.94186
223.15560913085938 0 1023.4114
225.0282440185547 0 502.40277
227.1386260986328 0 677.9804
229.15481567382812 0 11473.188 y Ammonia loss 13
230.16171264648438 0 2486.8352 z 13
234.12374877929688 0 1185.0327
243.13352966308594 0 789.4232
244.90684509277344 0 485.75888
251.15048217773438 0 4865.994
252.15338134765625 0 734.98047
268.17730712890625 0 1973.4907 c 1
274.5060729980469 0 598.71454
293.5097351074219 0 640.3947
297.1548156738281 0 571.9331
300.19219970703125 0 3288.6567
314.1824645996094 0 702.61176
321.6656494140625 0 572.12225
330.1773376464844 0 578.0354
342.1670227050781 0 1103.9153
353.218505859375 0 1222.5918
355.0687561035156 0 1210.8457
358.5284423828125 0 657.49896
379.2095031738281 0 594.3915 c Ammonia loss 2
396.23486328125 0 4785.5225 c 2
397.244384765625 0 6844.5938
398.246337890625 0 558.0792
404.1138916015625 0 1001.2051
405.1162414550781 0 737.1
407.19476318359375 0 1100.1439 y 12
411.2247314453125 0 985.6059
412.2284240722656 0 804.44684
413.2024841308594 0 946.67914
414.8685607910156 0 582.4328
418.2403564453125 0 840.9496
428.26153564453125 0 1204.7423
455.770751953125 0 1023.973
456.274169921875 0 766.9936
467.2502136230469 0 2403.0635
468.2535400390625 0 1182.7938
471.2821350097656 0 703.0872
488.3192443847656 0 849.89825
488.538818359375 0 560.28033
494.2333679199219 0 600.48706 c Ammonia loss 3
496.79083251953125 0 574.80536
506.7609558105469 0 1477.3053 y 7
511.26220703125 0 5543.273 c 3
512.2647705078125 0 1718.0797
514.30029296875 0 821.74036
519.272705078125 0 2300.4548 z 11
520.2767944335938 0 1113.3354
527.3043212890625 0 635.3109
535.2918701171875 0 1079.7687 y 11
560.2916259765625 0 595.65204
568.3070678710938 0 639.7208
573.9281616210938 0 659.4852
581.3662109375 0 1112.6365
584.3626098632812 0 6077.8794
585.366943359375 0 1331.2188
599.3477172851562 0 792.191
601.6251220703125 0 1327.941 c Ammonia loss 13
601.9603881835938 0 1037.9912
602.2971801757812 0 674.7984
602.9556884765625 0 2135.4897 y 1
603.2896118164062 0 1017.382
634.649169921875 0 1581.3157
634.97900390625 0 1892.3362
635.3082275390625 0 699.15283
635.6492919921875 0 841.9641
635.9739990234375 0 1197.7205
640.3199462890625 0 5604.7197
640.6505126953125 0 21260.098
640.8397216796875 0 6459.592
640.9839477539062 0 18743.832
641.3203125 0 14105.92
641.6510620117188 0 5719.912
641.8421630859375 0 3978.1338
641.9822387695312 0 1488.2263
642.3497314453125 0 1628.9816
651.3507690429688 0 1915.1936
651.8549194335938 0 1537.1974
659.3406982421875 0 833.1485
674.3541259765625 0 660.0294
682.3343505859375 0 3519.913 z 10
682.3909301757812 0 1219.0261
683.3399047851562 0 1541.2446
684.3472900390625 0 694.6196
685.343505859375 0 719.06964
688.8477783203125 0 656.33105
697.3408203125 0 4402.3613 c 4
698.3471069335938 0 2535.5151 y 10
713.8594970703125 0 1189.164 y 4
743.490478515625 0 931.5903
752.3623046875 0 1482.5831 w 9
761.8931274414062 0 620.7018
765.8887939453125 0 1202.968 c 11
766.3930053710938 0 733.1115
766.8948974609375 0 950.8649
771.8737182617188 0 789.5486
774.4403076171875 0 805.7714
783.3942260742188 0 752.06116
789.4420776367188 0 665.6303
793.39599609375 0 951.906 c Ammonia loss 5
794.401123046875 0 727.2398
794.8924560546875 0 605.6596
797.3983154296875 0 836.9084
802.6611328125 0 839.6888
810.43017578125 0 4736.965 c 5
811.37890625 0 5023.8486 z 9
812.3795166015625 0 1429.3713
813.3855590820312 0 603.1022
817.3978271484375 0 1375.2222
826.39306640625 0 790.63556 y Water loss 2
827.3910522460938 0 1642.5333 y 9
834.9068603515625 0 774.2131
835.4019775390625 0 2845.4097 y 2
835.7230224609375 0 769.2461
835.901123046875 0 2307.3093
836.4029541015625 0 1240.0388
845.905029296875 0 1469.9985
846.4002685546875 0 2397.6301 c 12
846.9000244140625 0 2452.602
849.955322265625 0 698.5576
859.4166870117188 0 601.1852
866.4197998046875 0 655.4653
867.4263305664062 0 2546.696
868.4239501953125 0 1461.3336
873.4315185546875 0 978.0878
873.9215698242188 0 699.04254
874.4381713867188 0 2052.9033
875.4402465820312 0 1567.5504
880.4512329101562 0 1719.1167
881.4308471679688 0 818.44037
882.4608154296875 0 853.1269
887.93701171875 0 1011.5036
888.4386596679688 0 911.4783
893.9341430664062 0 1458.2646
894.4343872070312 0 927.9781
895.4339599609375 0 913.791 y Ammonia loss 1
895.9254760742188 0 3077.4456 z 1
896.423828125 0 1757.8021
896.9124145507812 0 902.48004
897.4243774414062 0 911.27386
901.9368896484375 0 635.07654 c Ammonia loss 13
908.4447021484375 0 1125.9291
909.9503173828125 0 2262.4187
910.4432373046875 0 8041.6494 c 13
910.5367431640625 0 6010.405
910.9454956054688 0 6397.8584
911.4473266601562 0 4642.9707
911.53857421875 0 3679.8782
911.9487915039062 0 2683.0056
912.5428466796875 0 1250.0234
916.478271484375 0 753.551
918.4566040039062 0 842.3883
922.4617919921875 0 757.4964 z Ammonia loss 8
922.96630859375 0 841.0429
923.4569702148438 0 1536.5854
924.4439086914062 0 1048.6998
925.4520263671875 0 1783.8894 c 6
930.4629516601562 0 1941.8292
930.95849609375 0 5511.942
931.4636840820312 0 10761.894
931.96728515625 0 10804.396
932.4677124023438 0 7838.4863
932.9647827148438 0 1495.4282
933.4710693359375 0 1020.82196
934.4864501953125 0 1618.6519
934.80419921875 0 819.29724
935.4950561523438 0 1232.9586
937.4773559570312 0 637.19244 y Water loss 8
937.9727172851562 0 1735.7374
938.4710693359375 0 2778.4392 y Ammonia loss 8
938.9718627929688 0 2540.3762
939.4716186523438 0 6007.9673 z 8
940.4810180664062 0 3113.0513
942.9806518554688 0 731.2613
947.4387817382812 0 671.0363
948.4710693359375 0 1034.2295
951.4597778320312 0 799.11194
951.9656982421875 0 1601.2405
952.4644775390625 0 2668.4636
952.9644775390625 0 3179.2446
953.462646484375 0 2086.7341
953.9661865234375 0 907.9892
956.5086669921875 0 1290.501
959.4492797851562 0 1042.6239
959.9761352539062 0 2991.7017
960.4718627929688 0 14214.34
960.972900390625 0 17453.666
961.4737548828125 0 14193.065
961.9736938476562 0 8045.373
962.4739990234375 0 3987.2256
962.970947265625 0 2134.0825
963.4675903320312 0 1220.7849
965.4688110351562 0 1258.7177
967.1517333984375 0 826.6738
977.1458740234375 0 703.67035
981.4859619140625 0 2948.4646
982.4735107421875 0 4064.8857 c 7
983.48193359375 0 2062.9658
984.4844360351562 0 1726.1168
996.4938354492188 0 2260.649 z 7
997.49609375 0 2128.8213
1003.4907836914062 0 903.85846
1012.5105590820312 0 1891.0905 y 7
1019.83154296875 0 851.7273
1020.1976928710938 0 771.8342
1024.5301513671875 0 812.0848
1025.85107421875 0 885.0054
1026.858154296875 0 893.09045
1027.1871337890625 0 1494.1234
1028.512939453125 0 1468.4513
1030.1875 0 1154.55
1032.5423583984375 0 1594.2604
1032.8682861328125 0 1781.8898
1033.197265625 0 818.4908
1033.85400390625 0 1036.6022
1034.531005859375 0 875.54254
1035.1751708984375 0 740.1185
1035.5185546875 0 772.21564
1037.5179443359375 0 910.6836
1038.1822509765625 0 933.0659
1038.5283203125 0 1056.9998
1038.859375 0 1005.6709
1039.5223388671875 0 1224.0608
1042.2174072265625 0 937.7184
1042.521240234375 0 987.4702
1043.5303955078125 0 1004.5294
1043.87646484375 0 1216.106
1044.1922607421875 0 2499.6592
1044.863037109375 0 2045.5469
1045.206298828125 0 923.82367
1045.5362548828125 0 796.62177
1046.2005615234375 0 1094.2241
1047.5087890625 0 1080.2617
1047.857666015625 0 1062.221
1048.201171875 0 2136.2998
1048.5355224609375 0 1363.8818
1048.8665771484375 0 1469.6812
1049.1956787109375 0 1995.5396
1049.5321044921875 0 1999.2783
1049.867919921875 0 2143.3435
1050.194580078125 0 1462.1647
1050.530029296875 0 1625.367
1051.8704833984375 0 1281.0873
1052.538330078125 0 1121.8062
1053.5362548828125 0 1645.868
1053.8685302734375 0 2162.735
1054.196044921875 0 2679.7192
1054.52783203125 0 5461.9185
1054.86865234375 0 2235.5972
1055.529052734375 0 3694.2185
1055.8560791015625 0 818.15533
1056.17822265625 0 814.3728
1056.85107421875 0 964.37317
1058.197021484375 0 1935.1879
1058.5194091796875 0 1164.006
1059.8673095703125 0 2339.626
1060.1925048828125 0 4277.848
1060.524169921875 0 2929.4124
1060.861083984375 0 2937.1868
1061.1990966796875 0 1218.3978
1061.5159912109375 0 2074.8877
1062.1966552734375 0 1123.0325
1062.52001953125 0 1830.7219
1062.8699951171875 0 2208.18
1063.2030029296875 0 2152.7983
1063.5360107421875 0 4372.959
1063.8697509765625 0 2969.1838
1064.193359375 0 2546.2385
1064.5263671875 0 1143.1425
1064.857177734375 0 873.5516
1065.1956787109375 0 1060.7147
1065.52392578125 0 1072.556
1065.8685302734375 0 1805.9078
1066.1966552734375 0 1690.6353
1066.5399169921875 0 1895.5548 w 6
1066.875244140625 0 1168.2494
1067.189697265625 0 1117.6311
1067.54052734375 0 2099.3433
1067.8663330078125 0 1555.5115
1068.1982421875 0 1947.3165
1068.542724609375 0 3494.3313
1068.88134765625 0 2737.142
1069.2120361328125 0 2468.6606
1069.5416259765625 0 3017.3674
1069.865478515625 0 994.95605
1070.2071533203125 0 1496.2579
1070.5413818359375 0 902.4365
1109.576904296875 0 2220.496
1110.568603515625 0 7466.8286 c 8
1111.5589599609375 0 4614.9395
1112.5396728515625 0 2069.303
1124.560791015625 0 911.5026
1125.568359375 0 1665.0845
1126.5540771484375 0 1819.9988
1127.551513671875 0 1638.8949 y 6
1143.4801025390625 0 2872.9807
1144.4803466796875 0 2579.1707
1145.4779052734375 0 1048.8925
1150.5552978515625 0 962.769
1151.5606689453125 0 1173.3865
1162.067138671875 0 1069.2885
1190.599609375 0 796.1902
1223.6195068359375 0 1024.8783 y Ammonia loss 5
1224.6048583984375 0 2071.5784 z 5
1225.603515625 0 1056.9365
1226.60986328125 0 1154.2546
1238.6185302734375 0 1864.4475
1239.612060546875 0 5073.831 c 9
1240.615478515625 0 2814.1038 y 5
1241.6187744140625 0 1640.0713
1242.60400390625 0 796.5951
1253.6871337890625 0 2137.4648
1254.692138671875 0 1096.0034
1264.628662109375 0 696.79034
1274.1285400390625 0 999.38763
1274.6453857421875 0 730.21893
1276.6104736328125 0 807.396
1278.548828125 0 3227.0144
1279.5615234375 0 4505.4814
1280.5465087890625 0 1487.898
1281.5335693359375 0 993.10425
1281.66845703125 0 1560.5986
1283.6719970703125 0 870.0202
1284.6446533203125 0 935.01746
1311.6165771484375 0 819.6837
1357.657470703125 0 832.44745
1358.6536865234375 0 1056.2599
1375.1763916015625 0 956.38324
1388.6829833984375 0 1021.98535
1392.6490478515625 0 843.35693
1400.697021484375 0 1059.6848
1401.6873779296875 0 1102.6503
1402.677978515625 0 3995.951 c 10
1403.677490234375 0 3874.7585
1404.6859130859375 0 1333.6545
1409.701904296875 0 1674.955 y Ammonia loss 4
1410.6875 0 2051.923 z 4
1411.686767578125 0 2382.2568
1412.6827392578125 0 1028.4008
1413.6922607421875 0 893.0393
1425.711181640625 0 980.7842
1426.7021484375 0 2586.7559 y 4
1427.701171875 0 2090.1365
1436.7081298828125 0 1242.3564
1437.7061767578125 0 1891.0681
1438.7037353515625 0 1002.27344
1466.232177734375 0 844.9226
1472.714111328125 0 1118.5775
1473.218505859375 0 1071.5962
1480.745849609375 0 1413.2261 w 3
1481.2423095703125 0 968.28986
1481.735107421875 0 1826.9967
1482.223388671875 0 1353.8213
1482.72998046875 0 1085.015
1483.212890625 0 1018.06604
1484.738525390625 0 917.90717
1486.7606201171875 0 1187.4791
1488.734375 0 776.78015
1490.7344970703125 0 748.92126
1491.21435546875 0 1418.4078
1492.2288818359375 0 1037.0881
1492.7381591796875 0 884.1689
1494.245361328125 0 1030.8365
1494.728515625 0 1574.0161
1496.2142333984375 0 857.2564
1497.2215576171875 0 1380.1101
1498.7276611328125 0 950.0997
1501.7586669921875 0 1132.0795
1502.21337890625 0 1086.3883
1503.243408203125 0 920.4995
1506.72265625 0 1090.8558
1515.751953125 0 1066.7317
1520.24462890625 0 896.9626
1522.239013671875 0 1106.0272
1523.251953125 0 907.25854
1524.2403564453125 0 953.36743
1524.72802734375 0 1542.9723 y Ammonia loss 3
1525.7198486328125 0 4483.5835 z 3
1526.7196044921875 0 4982.858
1527.2523193359375 0 935.90155
1527.727294921875 0 3231.8228
1528.2430419921875 0 1256.2931
1528.732177734375 0 2656.2395
1529.767822265625 0 2021.7794
1530.2740478515625 0 1011.21063
1530.7664794921875 0 4998.652 c 11
1531.2686767578125 0 1563.6084
1531.766357421875 0 3145.602
1532.763671875 0 2055.6724
1533.231689453125 0 1114.4238
1535.752685546875 0 1157.404
1536.2767333984375 0 948.44666
1537.7593994140625 0 1994.0236
1538.247802734375 0 1148.5989
1538.7510986328125 0 1061.8239
1539.766357421875 0 1255.2551
1541.7410888671875 0 2360.0583 y 3
1542.75048828125 0 2111.5417
1543.2554931640625 0 882.8973
1544.275146484375 0 1430.5846
1544.759033203125 0 1492.4523
1545.2685546875 0 1186.9049
1545.764892578125 0 1688.2261
1546.2489013671875 0 1129.7253
1546.7513427734375 0 1059.0942
1547.2742919921875 0 1694.3215
1549.2769775390625 0 1209.3123
1549.7655029296875 0 2424.1672
1550.2840576171875 0 1746.2079
1550.774169921875 0 1773.0615
1551.286376953125 0 1856.4036
1551.7760009765625 0 2062.9167
1552.2744140625 0 2652.7402
1552.7611083984375 0 2259.51
1553.28076171875 0 1169.6271
1553.7645263671875 0 944.51086
1554.7337646484375 0 1059.9606
1555.738525390625 0 825.8623
1556.7718505859375 0 940.1802
1557.25 0 1244.0597
1557.805908203125 0 1198.9679
1558.274169921875 0 2065.0442
1558.787841796875 0 1677.0481
1559.273681640625 0 2500.3992
1559.7862548828125 0 2192.33
1560.2801513671875 0 2693.3914
1560.77880859375 0 1972.391
1561.275146484375 0 1290.4406
1561.756591796875 0 1776.986
1562.2318115234375 0 906.3147
1562.7628173828125 0 1047.6821
1563.7647705078125 0 988.01556
1564.77685546875 0 2071.1
1565.280517578125 0 2360.9338
1565.7783203125 0 3072.4038
1566.287841796875 0 4324.225
1566.7872314453125 0 4299.7446
1567.2901611328125 0 2415.423
1567.7802734375 0 2596.3218
1568.294677734375 0 2084.644
1568.792236328125 0 1761.4391
1569.2684326171875 0 1708.4781
1570.2691650390625 0 1002.6724
1570.753662109375 0 1600.7496
1571.27490234375 0 1751.2188
1571.778076171875 0 1838.2861
1572.273681640625 0 2766.0164
1572.7889404296875 0 2952.5266
1573.283935546875 0 4090.0413
1573.7906494140625 0 3869.7446
1574.2921142578125 0 6114.69
1574.788818359375 0 3795.4011
1575.28515625 0 3347.67
1575.77734375 0 2156.4497
1576.2919921875 0 1579.3279
1576.7818603515625 0 1902.4542
1577.77392578125 0 819.97766
1578.2838134765625 0 943.21234
1579.2734375 0 1788.6832
1579.8052978515625 0 1575.3999
1580.2969970703125 0 2755.5781
1580.7957763671875 0 3567.5942
1581.29638671875 0 3631.8723
1581.7830810546875 0 3434.357
1582.28857421875 0 4100.8994
1582.783203125 0 3238.103
1583.294921875 0 3127.9253
1583.7772216796875 0 2099.8293
1584.29150390625 0 1569.925
1584.7772216796875 0 1209.1853
1585.7821044921875 0 1434.5942
1586.2923583984375 0 1742.307
1586.7945556640625 0 2290.4895
1587.2791748046875 0 1255.4146
1587.7969970703125 0 2199.2322
1588.2730712890625 0 2487.4885
1588.787353515625 0 844.74115
1589.2904052734375 0 2044.389
1589.796630859375 0 3278.1033
1590.287109375 0 5398.1406
1590.786376953125 0 2895.4536
1591.2874755859375 0 3480.5745
1591.783447265625 0 3046.386
1592.2884521484375 0 2172.6963
1592.796875 0 1485.4156
1593.2957763671875 0 3359.724
1593.79248046875 0 3516.3313
1594.2979736328125 0 2599.951
1594.802978515625 0 3907.5422
1595.3016357421875 0 2473.2332
1595.7852783203125 0 3177.1946
1596.2960205078125 0 3189.2522
1596.7811279296875 0 1526.2825
1598.79248046875 0 2067.4692
1599.3028564453125 0 2733.3823
1599.787841796875 0 2001.3531
1600.2935791015625 0 1451.2125
1600.7884521484375 0 1632.8772
1601.291748046875 0 1609.3088
1601.8037109375 0 1891.7941
1602.30322265625 0 4235.668
1602.817626953125 0 4468.4717
1603.311767578125 0 3855.939
1603.8114013671875 0 3228.7214
1604.302001953125 0 2084.546
1604.80517578125 0 1476.3539
1605.2923583984375 0 1760.376
1605.8067626953125 0 1141.5033
1606.2855224609375 0 807.7657
1653.768798828125 0 2749.2554 z 2
1654.7708740234375 0 3739.0234
1655.779052734375 0 2464.611
1656.7874755859375 0 1108.1162
1691.782958984375 0 4840.8335 c 12
1692.78759765625 0 3432.679
1693.7923583984375 0 3100.5295
1694.814453125 0 1078.0715
1722.7711181640625 0 950.6535
1791.8275146484375 0 961.1993
1792.833740234375 0 1306.4249
1793.8746337890625 0 972.72614
1819.88818359375 0 1049.7827 c 13
1820.8765869140625 0 2271.1848
1821.8779296875 0 1828.7903
1822.897216796875 0 1578.2311
1830.9298095703125 0 1629.3478
1831.9404296875 0 1816.8401
1832.9212646484375 0 1504.959
1845.903564453125 0 1086.2837
1860.9395751953125 0 1175.7827
1861.922119140625 0 2307.0198
1862.9324951171875 0 4502.853
1863.9326171875 0 4223.949
1864.9249267578125 0 1899.3887
1865.922607421875 0 1121.6481
1875.9296875 0 2129.7688
1876.936767578125 0 3602.7764
1877.9329833984375 0 2144.747
1878.9266357421875 0 1932.5839
1892.94970703125 0 1440.3531
1893.9461669921875 0 2517.6248
1894.9527587890625 0 3190.1965
1895.952392578125 0 1444.1462
1902.9224853515625 0 1678.9801
1903.9180908203125 0 3026.5244
1904.921630859375 0 6597.1025
1905.9283447265625 0 5950.0454
1906.921875 0 3058.5386
1907.923828125 0 2056.809
1919.936767578125 0 3763.2422
1920.9486083984375 0 11101.898
1921.9453125 0 23969.088
1922.9488525390625 0 21105.02
1923.947509765625 0 12811.549
1924.94482421875 0 6272.122
1925.9359130859375 0 3262.0364
1926.921875 0 1458.4628
3047.595458984375 0 743.4204
3148.585693359375 0 1020.5684
3163.593994140625 0 901.4182

Spectrum Details

|  |  |
| --- | --- |
| Matched peaks? Matched peaks The total absolute number of peaks matched. Additionally in brackets the total fraction of peaks matched and the total number of peaks is shown. | 57 (9.88% of 577) |
| FDR? FDR The false discovery rate estimated for this peptide. It is calculated by matching all theoretical fragments with a non-integer shift with the raw peaks for this spectrum. This is done with 40 different shifts. The resulting percentage is the average number of annotated peaks over the number of annotated peaks with the correct spectrum. | 0.84% |
| Satellite FDR? Satellite FDR See the FDR for details on its calculation. This satellite ion specific FDR only contains the satellite ions (d/w) for I/L/J positions. | - |
| PSM Score? PSM Score The PSM Score as given by Hecklib to this annotated spectrum. It is shown with three significant figures. | 540 |

#### Spectrum 5764? Spectrum 5764 The raw spectrum of this peptide as annotated by Hecklib. The fragments are coloured according to ion type (see legend). Any peaks with a star '\*' as text can be hovered over to see the full details, first the ion type second the mass shift type. By hovering over the amino acids in the peptide or ions in the legend the corresponding peaks are highlighted. By toggling the 'Unassigned' label you can turn the background (unassigned) peaks on or off in the plot. By updating the slider in the Ion legend you can update the spectrum to only show the top X% of the peaks with labels. The top X% means any peak that is within X% of the highest intensity. By dragging in the spectrum you can zoom in to a specific part of the spectrum and use 'Zoom Out' to get back to the original zoom level. The annotation of the spectrum is based on the given sequence in the peptides file and is done with different software so inconsistencies are likely. The peaks are annotated based on the given sequence, with 20 ppm tolerance.

Copy Data

##### Spectrum 5764 (TSV)

###### Preview

```
Loading example...
```

*Click on the button to copy the data to your clipboard.*

Mz MinMz MaxIntensity Max

WidthHeightPeptide font sizePeptide stroke widthSpectrum font sizeSpectrum stroke widthCompact peptide

Ion legend

wxyz

abcd

OtherUnassignedIon^Charge^_Position_Show for top:%

JHQDWJDGKEYKCKV

04.03e+48.07e+41.21e+51.61e+5

Zoom Out

w^+1^_2_y^+1^_2_z^+1^_2_y^+1^_2_c^+1^_2_z^+3^_8_y^+3^_8_c^+2^_5_y^+2^_5_y^+2^_5_y^+3^_9_w^+2^_6_z^+4^_12_c^+1^_3_y^+4^_12_z^+4^_12_c^+1^_3_c^+2^_6_z^+3^_10_y^+1^_3_c^+3^_10_y^+2^_6_w^+2^_7_y^+4^_14_w^+1^_4_y^+2^_7_y^+3^_11_z^+3^_11_y^+3^_11_y^+2^_7_z^+2^_8_c^+1^_4_c^+1^_4_z^+2^_8_z^+3^_12_y^+2^_8_c^+1^_4_z^+1^_4_w^+2^_9_y^+1^_4_c^+2^_9_y^+3^_13_y^+3^_13_z^+3^_13_c^+2^_9_y^+3^_13_y^+2^_9_w^+2^_10_z^+3^_14_c^+3^_14_c^+3^_14_y^+3^_14_c^+3^_14_c^+2^_10_c^+2^_10_y^+2^_10_z^+2^_10_c^+2^_10_y^+2^_10_c^+1^_5_c^+1^_5_y^+1^_5_z^+1^_5_c^+2^_11_c^+1^_5_y^+1^_5_c^+2^_11_y^+2^_11_z^+2^_11_y^+2^_11_w^+1^_6_y^+2^_12_z^+2^_12_c^+2^_12_y^+2^_12_c^+1^_6_w^+2^_13_c^+1^_6_z^+1^_6_y^+2^_13_y^+1^_6_y^+2^_13_c^+2^_13_y^+2^_14_z^+2^_14_y^+2^_14_c^+1^_7_c^+2^_14_z^+1^_7_c^+1^_7_y^+1^_7_z^+1^_7_c^+1^_8_c^+1^_8_z^+1^_8_y^+1^_8_c^+1^_9_y^+1^_10_z^+1^_10_c^+1^_10_y^+1^_10_c^+1^_11_y^+1^_11_z^+1^_11_y^+1^_11_z^+1^_12_c^+1^_12_y^+1^_12_z^+1^_13_c^+1^_13_

048697214581944

Fragment Matches Table

Show background peaks

| Position | Ion type | Intensity | mz Theoretical | mz Error (Th) | mz Error (ppm) | Charge | Series Number |
| --- | --- | --- | --- | --- | --- | --- | --- |
| - | - | 4550 | 120.1 | - | - | 0 | - |
| - | - | 570.8 | 126.1 | - | - | 0 | - |
| - | - | 424.3 | 126.6 | - | - | 0 | - |
| - | - | 683 | 129.1 | - | - | 0 | - |
| - | - | 3581 | 129.1 | - | - | 0 | - |
| - | - | 619.7 | 130.1 | - | - | 0 | - |
| - | - | 4930 | 130.1 | - | - | 0 | - |
| - | - | 2348 | 132.1 | - | - | 0 | - |
| - | - | 1768 | 136.1 | - | - | 0 | - |
| - | - | 566.3 | 138.1 | - | - | 0 | - |
| - | - | 1053 | 146.1 | - | - | 0 | - |
| - | - | 515.2 | 148.9 | - | - | 0 | - |
| - | - | 497.2 | 149 | - | - | 0 | - |
| - | - | 9240 | 155.1 | - | - | 0 | - |
| - | - | 708.8 | 157.1 | - | - | 0 | - |
| - | - | 2450 | 159.1 | - | - | 0 | - |
| - | - | 488.3 | 163.1 | - | - | 0 | - |
| - | - | 1901 | 164.1 | - | - | 0 | - |
| - | - | 1867 | 165.1 | - | - | 0 | - |
| - | - | 6750 | 166.1 | - | - | 0 | - |
| - | - | 572.5 | 167.1 | - | - | 0 | - |
| - | - | 3742 | 168.1 | - | - | 0 | - |
| - | - | 467.4 | 170.5 | - | - | 0 | - |
| 14 | w | 1682 | 172.1 | 0.0002414 | 1.403 | +1 | 2 |
| - | - | 467.9 | 177.3 | - | - | 0 | - |
| - | - | 971.6 | 178.1 | - | - | 0 | - |
| - | - | 2860 | 183.1 | - | - | 0 | - |
| - | - | 6167 | 185.2 | - | - | 0 | - |
| - | - | 617 | 185.3 | - | - | 0 | - |
| - | - | 650.5 | 186.1 | - | - | 0 | - |
| - | - | 586.9 | 187.1 | - | - | 0 | - |
| - | - | 1763 | 190.1 | - | - | 0 | - |
| - | - | 456.2 | 194.7 | - | - | 0 | - |
| - | - | 1029 | 201.1 | - | - | 0 | - |
| - | - | 1013 | 203.2 | - | - | 0 | - |
| - | - | 2951 | 215.1 | - | - | 0 | - |
| - | - | 729.4 | 219.1 | - | - | 0 | - |
| - | - | 7051 | 223.2 | - | - | 0 | - |
| - | - | 782.4 | 224.2 | - | - | 0 | - |
| - | - | 884.4 | 227.1 | - | - | 0 | - |
| - | - | 587.5 | 229 | - | - | 0 | - |
| - | - | 722.1 | 229.1 | - | - | 0 | - |
| 14 | y | 2458 | 229.2 | 0.0004977 | 2.172 | +1 | 2 |
| 14 | z | 2599 | 230.2 | 4.262E-05 | 0.1852 | +1 | 2 |
| - | - | 1097 | 231.2 | - | - | 0 | - |
| - | - | 1648 | 232.1 | - | - | 0 | - |
| - | - | 6103 | 233.2 | - | - | 0 | - |
| - | - | 6432 | 234.1 | - | - | 0 | - |
| - | - | 899.1 | 243.1 | - | - | 0 | - |
| 14 | y | 2022 | 246.2 | 0.0004988 | 2.026 | +1 | 2 |
| - | - | 1582 | 250.1 | - | - | 0 | - |
| - | - | 3.36E+04 | 251.2 | - | - | 0 | - |
| - | - | 3835 | 252.2 | - | - | 0 | - |
| - | - | 845.8 | 253.1 | - | - | 0 | - |
| - | - | 3589 | 261.1 | - | - | 0 | - |
| - | - | 771 | 263.1 | - | - | 0 | - |
| 2 | c | 3876 | 268.2 | 0.0003226 | 1.203 | +1 | 2 |
| - | - | 718.1 | 269.2 | - | - | 0 | - |
| - | - | 716.8 | 270.1 | - | - | 0 | - |
| - | - | 775.4 | 274.1 | - | - | 0 | - |
| - | - | 2083 | 283.2 | - | - | 0 | - |
| - | - | 506.5 | 287.7 | - | - | 0 | - |
| - | - | 752.3 | 288.1 | - | - | 0 | - |
| - | - | 1131 | 290.1 | - | - | 0 | - |
| - | - | 1303 | 292.1 | - | - | 0 | - |
| - | - | 830.2 | 293.1 | - | - | 0 | - |
| - | - | 506.7 | 294.1 | - | - | 0 | - |
| - | - | 1335 | 294.2 | - | - | 0 | - |
| - | - | 657.3 | 295.2 | - | - | 0 | - |
| - | - | 547.2 | 297.1 | - | - | 0 | - |
| - | - | 629 | 298.5 | - | - | 0 | - |
| - | - | 859.5 | 299.1 | - | - | 0 | - |
| - | - | 4981 | 300.2 | - | - | 0 | - |
| - | - | 716.6 | 301.2 | - | - | 0 | - |
| - | - | 736.9 | 318.1 | - | - | 0 | - |
| - | - | 8382 | 326.7 | - | - | 0 | - |
| 8 | z | 2974 | 327.2 | 0.003532 | 10.8 | +3 | 8 |
| - | - | 635.4 | 327.7 | - | - | 0 | - |
| - | - | 850.6 | 329.2 | - | - | 0 | - |
| - | - | 1343 | 331.7 | - | - | 0 | - |
| - | - | 869.4 | 332.2 | - | - | 0 | - |
| 8 | y | 885.8 | 338.2 | 0.003075 | 9.094 | +3 | 8 |
| - | - | 713.5 | 339.2 | - | - | 0 | - |
| 5 | c | 5220 | 340.7 | 0.0007757 | 2.277 | +2 | 5 |
| 11 | y | 1592 | 341.2 | 0.006014 | 17.63 | +2 | 5 |
| - | - | 1042 | 341.7 | - | - | 0 | - |
| - | - | 1212 | 349.2 | - | - | 0 | - |
| 11 | y | 5304 | 349.7 | 0.001619 | 4.631 | +2 | 5 |
| - | - | 1137 | 350.2 | - | - | 0 | - |
| - | - | 1472 | 351.2 | - | - | 0 | - |
| - | - | 955.1 | 352.2 | - | - | 0 | - |
| - | - | 662.1 | 355.6 | - | - | 0 | - |
| - | - | 900 | 361.2 | - | - | 0 | - |
| - | - | 725.6 | 367.7 | - | - | 0 | - |
| - | - | 739.1 | 370.7 | - | - | 0 | - |
| - | - | 1053 | 372.2 | - | - | 0 | - |
| - | - | 1076 | 374.7 | - | - | 0 | - |
| - | - | 2015 | 375.2 | - | - | 0 | - |
| - | - | 1200 | 375.2 | - | - | 0 | - |
| 7 | y | 1661 | 376.5 | 0.00164 | 4.355 | +3 | 9 |
| 10 | w | 1.756E+04 | 376.7 | 0.001927 | 5.116 | +2 | 6 |
| - | - | 1173 | 376.9 | - | - | 0 | - |
| - | - | 8231 | 377.2 | - | - | 0 | - |
| 4 | z | 1900 | 377.7 | 0.0062 | 16.42 | +4 | 12 |
| - | - | 2130 | 378.2 | - | - | 0 | - |
| 3 | c | 9259 | 379.2 | 0.0003377 | 0.8905 | +1 | 3 |
| - | - | 2049 | 380.2 | - | - | 0 | - |
| - | - | 672.9 | 381.2 | - | - | 0 | - |
| - | - | 1072 | 381.2 | - | - | 0 | - |
| 4 | y | 793.6 | 381.7 | 0.001349 | 3.535 | +4 | 12 |
| 4 | z | 704.3 | 382.2 | 0.001659 | 4.341 | +4 | 12 |
| - | - | 2062 | 383.2 | - | - | 0 | - |
| - | - | 1509 | 383.7 | - | - | 0 | - |
| - | - | 1300 | 384.3 | - | - | 0 | - |
| - | - | 834.8 | 387.2 | - | - | 0 | - |
| - | - | 1094 | 387.7 | - | - | 0 | - |
| - | - | 1072 | 388.2 | - | - | 0 | - |
| - | - | 756.7 | 390.2 | - | - | 0 | - |
| - | - | 2411 | 395.2 | - | - | 0 | - |
| - | - | 987.1 | 396.2 | - | - | 0 | - |
| 3 | c | 1.493E+04 | 396.2 | 0.0004915 | 1.24 | +1 | 3 |
| 6 | c | 668.2 | 397.2 | 0.006748 | 16.99 | +2 | 6 |
| - | - | 2648 | 397.2 | - | - | 0 | - |
| - | - | 1215 | 397.7 | - | - | 0 | - |
| - | - | 778.6 | 399.2 | - | - | 0 | - |
| 6 | z | 1060 | 403.2 | 0.00193 | 4.786 | +3 | 10 |
| - | - | 981 | 404.2 | - | - | 0 | - |
| 13 | y | 3757 | 407.2 | 0.005182 | 12.72 | +1 | 3 |
| - | - | 1324 | 408.2 | - | - | 0 | - |
| 10 | c | 701 | 408.2 | 0.0002763 | 0.6768 | +3 | 10 |
| - | - | 976.6 | 408.7 | - | - | 0 | - |
| - | - | 2016 | 409.2 | - | - | 0 | - |
| 10 | y | 2604 | 414.2 | 0.008212 | 19.83 | +2 | 6 |
| - | - | 1045 | 416.3 | - | - | 0 | - |
| - | - | 6660 | 417.2 | - | - | 0 | - |
| - | - | 2522 | 417.7 | - | - | 0 | - |
| - | - | 978.1 | 418.2 | - | - | 0 | - |
| - | - | 983.1 | 419.2 | - | - | 0 | - |
| - | - | 1039 | 424.8 | - | - | 0 | - |
| - | - | 1358 | 425.3 | - | - | 0 | - |
| - | - | 5136 | 428.3 | - | - | 0 | - |
| - | - | 2678 | 429.3 | - | - | 0 | - |
| - | - | 3958 | 440.7 | - | - | 0 | - |
| 9 | w | 3805 | 441.2 | 0.001678 | 3.804 | +2 | 7 |
| - | - | 1277 | 441.2 | - | - | 0 | - |
| - | - | 1467 | 441.7 | - | - | 0 | - |
| - | - | 1169 | 445.2 | - | - | 0 | - |
| - | - | 2159 | 448.2 | - | - | 0 | - |
| - | - | 1330 | 449.2 | - | - | 0 | - |
| - | - | 687.5 | 450.2 | - | - | 0 | - |
| 2 | y | 1512 | 452.5 | 0.0004813 | 1.064 | +4 | 14 |
| - | - | 2609 | 452.7 | - | - | 0 | - |
| - | - | 1113 | 453 | - | - | 0 | - |
| - | - | 850.7 | 453.2 | - | - | 0 | - |
| - | - | 3813 | 453.3 | - | - | 0 | - |
| - | - | 2397 | 453.8 | - | - | 0 | - |
| - | - | 1342 | 454.3 | - | - | 0 | - |
| - | - | 1126 | 456.2 | - | - | 0 | - |
| - | - | 899.3 | 457.2 | - | - | 0 | - |
| - | - | 4959 | 460.3 | - | - | 0 | - |
| 12 | w | 4792 | 461.2 | 0.003844 | 8.336 | +1 | 4 |
| - | - | 1827 | 461.3 | - | - | 0 | - |
| - | - | 978.3 | 462.2 | - | - | 0 | - |
| - | - | 1666 | 466.2 | - | - | 0 | - |
| - | - | 793.4 | 467.2 | - | - | 0 | - |
| - | - | 3213 | 467.2 | - | - | 0 | - |
| - | - | 820.7 | 468.3 | - | - | 0 | - |
| 9 | y | 692.2 | 469.2 | 0.002268 | 4.833 | +2 | 7 |
| 5 | y | 1928 | 470.2 | 0.003809 | 8.101 | +3 | 11 |
| - | - | 1108 | 470.7 | - | - | 0 | - |
| 5 | z | 780.8 | 470.9 | 0.002911 | 6.182 | +3 | 11 |
| - | - | 652.9 | 472.2 | - | - | 0 | - |
| - | - | 2035 | 474.3 | - | - | 0 | - |
| - | - | 3360 | 475.3 | - | - | 0 | - |
| 5 | y | 3648 | 476.2 | 0.005999 | 12.6 | +3 | 11 |
| - | - | 1823 | 476.5 | - | - | 0 | - |
| - | - | 960 | 476.7 | - | - | 0 | - |
| 9 | y | 1.143E+04 | 478.3 | 0.001519 | 3.177 | +2 | 7 |
| - | - | 6671 | 478.8 | - | - | 0 | - |
| - | - | 2379 | 479.3 | - | - | 0 | - |
| - | - | 1843 | 480.5 | - | - | 0 | - |
| - | - | 3318 | 480.7 | - | - | 0 | - |
| - | - | 2.395E+04 | 480.7 | - | - | 0 | - |
| - | - | 2.077E+04 | 481 | - | - | 0 | - |
| - | - | 1475 | 481.2 | - | - | 0 | - |
| - | - | 1.461E+04 | 481.2 | - | - | 0 | - |
| - | - | 2484 | 481.3 | - | - | 0 | - |
| - | - | 4360 | 481.5 | - | - | 0 | - |
| - | - | 1057 | 481.7 | - | - | 0 | - |
| - | - | 2521 | 481.7 | - | - | 0 | - |
| - | - | 860.6 | 482 | - | - | 0 | - |
| - | - | 1300 | 482.2 | - | - | 0 | - |
| - | - | 2370 | 488.8 | - | - | 0 | - |
| - | - | 841.2 | 489.3 | - | - | 0 | - |
| 8 | z | 1620 | 490.2 | 0.006113 | 12.47 | +2 | 8 |
| - | - | 791.8 | 491.3 | - | - | 0 | - |
| 4 | c | 1058 | 493.3 | 0.0002924 | 0.5928 | +1 | 4 |
| 4 | c | 7786 | 494.2 | 0.0003722 | 0.753 | +1 | 4 |
| - | - | 2802 | 495.2 | - | - | 0 | - |
| - | - | 1590 | 496.3 | - | - | 0 | - |
| - | - | 800 | 498.3 | - | - | 0 | - |
| 8 | z | 1736 | 498.8 | 0.002187 | 4.385 | +2 | 8 |
| - | - | 2027 | 499.3 | - | - | 0 | - |
| - | - | 831.2 | 499.8 | - | - | 0 | - |
| 4 | z | 2740 | 503.2 | 0.003694 | 7.341 | +3 | 12 |
| - | - | 763 | 504.2 | - | - | 0 | - |
| - | - | 1787 | 504.8 | - | - | 0 | - |
| - | - | 3791 | 505.2 | - | - | 0 | - |
| - | - | 4173 | 505.7 | - | - | 0 | - |
| - | - | 6712 | 506.3 | - | - | 0 | - |
| 8 | y | 1.786E+04 | 506.8 | 0.00157 | 3.098 | +2 | 8 |
| - | - | 7803 | 507.3 | - | - | 0 | - |
| - | - | 3010 | 507.8 | - | - | 0 | - |
| - | - | 1304 | 508.3 | - | - | 0 | - |
| - | - | 841.1 | 509.3 | - | - | 0 | - |
| - | - | 5206 | 510.8 | - | - | 0 | - |
| 4 | c | 6.211E+04 | 511.3 | 0.0009227 | 1.805 | +1 | 4 |
| - | - | 1.571E+04 | 512.3 | - | - | 0 | - |
| - | - | 3174 | 513.3 | - | - | 0 | - |
| - | - | 1251 | 514.7 | - | - | 0 | - |
| - | - | 778.8 | 515.7 | - | - | 0 | - |
| 12 | z | 1.938E+04 | 519.3 | 0.003814 | 7.346 | +1 | 4 |
| - | - | 770.3 | 519.8 | - | - | 0 | - |
| - | - | 6185 | 520.3 | - | - | 0 | - |
| - | - | 1180 | 521.3 | - | - | 0 | - |
| - | - | 1056 | 525.2 | - | - | 0 | - |
| - | - | 2709 | 526.8 | - | - | 0 | - |
| - | - | 5959 | 527.2 | - | - | 0 | - |
| - | - | 1334 | 527.7 | - | - | 0 | - |
| - | - | 4026 | 527.8 | - | - | 0 | - |
| - | - | 2149 | 528.3 | - | - | 0 | - |
| - | - | 938.9 | 528.8 | - | - | 0 | - |
| - | - | 1086 | 530.2 | - | - | 0 | - |
| - | - | 771.2 | 530.3 | - | - | 0 | - |
| 7 | w | 5313 | 533.8 | 0.001603 | 3.004 | +2 | 9 |
| - | - | 1.247E+04 | 534.3 | - | - | 0 | - |
| - | - | 5326 | 534.8 | - | - | 0 | - |
| 12 | y | 1.12E+04 | 535.3 | 0.004777 | 8.924 | +1 | 4 |
| - | - | 2837 | 536.3 | - | - | 0 | - |
| - | - | 1046 | 537.3 | - | - | 0 | - |
| - | - | 901.2 | 538.3 | - | - | 0 | - |
| - | - | 1198 | 544.3 | - | - | 0 | - |
| - | - | 1597 | 546.2 | - | - | 0 | - |
| 9 | c | 2195 | 547.3 | 0.004277 | 7.815 | +2 | 9 |
| - | - | 1807 | 547.8 | - | - | 0 | - |
| - | - | 741 | 549.3 | - | - | 0 | - |
| - | - | 1122 | 549.6 | - | - | 0 | - |
| - | - | 844 | 549.9 | - | - | 0 | - |
| - | - | 1.34E+04 | 550.3 | - | - | 0 | - |
| 3 | y | 5638 | 551.3 | 0.007486 | 13.58 | +3 | 13 |
| 3 | y | 2130 | 551.6 | 0.003254 | 5.9 | +3 | 13 |
| 3 | z | 1278 | 551.9 | 0.00029 | 0.5255 | +3 | 13 |
| - | - | 3216 | 552.3 | - | - | 0 | - |
| - | - | 3205 | 555.3 | - | - | 0 | - |
| 9 | c | 5.448E+04 | 555.8 | 0.0006458 | 1.162 | +2 | 9 |
| - | - | 2.565E+04 | 556.3 | - | - | 0 | - |
| - | - | 6093 | 556.8 | - | - | 0 | - |
| - | - | 7853 | 556.8 | - | - | 0 | - |
| 3 | y | 4299 | 557.3 | 0.001444 | 2.591 | +3 | 13 |
| - | - | 1762 | 557.8 | - | - | 0 | - |
| - | - | 820.1 | 559.6 | - | - | 0 | - |
| - | - | 1835 | 562.3 | - | - | 0 | - |
| - | - | 1270 | 562.6 | - | - | 0 | - |
| - | - | 1214 | 563 | - | - | 0 | - |
| - | - | 765.6 | 563.6 | - | - | 0 | - |
| - | - | 1998 | 563.8 | - | - | 0 | - |
| 7 | y | 3.448E+04 | 564.3 | 0.001431 | 2.535 | +2 | 9 |
| - | - | 2.128E+04 | 564.8 | - | - | 0 | - |
| - | - | 9420 | 565.3 | - | - | 0 | - |
| - | - | 2081 | 565.8 | - | - | 0 | - |
| - | - | 820.8 | 567.2 | - | - | 0 | - |
| - | - | 1154 | 567.3 | - | - | 0 | - |
| - | - | 4782 | 568.8 | - | - | 0 | - |
| - | - | 4049 | 569.3 | - | - | 0 | - |
| - | - | 1204 | 569.8 | - | - | 0 | - |
| - | - | 909.9 | 569.9 | - | - | 0 | - |
| - | - | 1483 | 573 | - | - | 0 | - |
| - | - | 1264 | 573.3 | - | - | 0 | - |
| - | - | 1864 | 573.9 | - | - | 0 | - |
| - | - | 1684 | 574.3 | - | - | 0 | - |
| - | - | 1143 | 574.9 | - | - | 0 | - |
| - | - | 767.9 | 575.3 | - | - | 0 | - |
| - | - | 770.1 | 577.3 | - | - | 0 | - |
| - | - | 3220 | 578 | - | - | 0 | - |
| - | - | 3082 | 578.3 | - | - | 0 | - |
| - | - | 1311 | 578.6 | - | - | 0 | - |
| - | - | 1955 | 578.9 | - | - | 0 | - |
| - | - | 759.6 | 579.3 | - | - | 0 | - |
| - | - | 843.2 | 582.3 | - | - | 0 | - |
| - | - | 1277 | 582.6 | - | - | 0 | - |
| - | - | 2661 | 583 | - | - | 0 | - |
| - | - | 2380 | 583.3 | - | - | 0 | - |
| - | - | 1593 | 583.6 | - | - | 0 | - |
| - | - | 840.7 | 585.3 | - | - | 0 | - |
| - | - | 842.4 | 586.3 | - | - | 0 | - |
| - | - | 1810 | 590.8 | - | - | 0 | - |
| 6 | w | 3.446E+04 | 591.3 | 0.00122 | 2.063 | +2 | 10 |
| - | - | 1299 | 591.4 | - | - | 0 | - |
| - | - | 2.052E+04 | 591.8 | - | - | 0 | - |
| - | - | 9577 | 592.3 | - | - | 0 | - |
| - | - | 1419 | 592.4 | - | - | 0 | - |
| - | - | 1701 | 592.6 | - | - | 0 | - |
| - | - | 3588 | 592.8 | - | - | 0 | - |
| - | - | 2491 | 593 | - | - | 0 | - |
| - | - | 1954 | 593.3 | - | - | 0 | - |
| - | - | 877 | 595.3 | - | - | 0 | - |
| - | - | 7724 | 597.3 | - | - | 0 | - |
| 2 | z | 7635 | 597.6 | 0.01104 | 18.47 | +3 | 14 |
| - | - | 4499 | 598 | - | - | 0 | - |
| - | - | 3079 | 598.3 | - | - | 0 | - |
| - | - | 880 | 599.2 | - | - | 0 | - |
| - | - | 757.7 | 600.3 | - | - | 0 | - |
| 14 | c | 2228 | 601.3 | 0.005362 | 8.917 | +3 | 14 |
| 14 | c | 2632 | 601.6 | 0.005238 | 8.706 | +3 | 14 |
| - | - | 6749 | 602 | - | - | 0 | - |
| - | - | 3039 | 602.3 | - | - | 0 | - |
| - | - | 1249 | 602.6 | - | - | 0 | - |
| 2 | y | 3672 | 603 | 0.002091 | 3.467 | +3 | 14 |
| - | - | 3302 | 603.3 | - | - | 0 | - |
| - | - | 1174 | 603.6 | - | - | 0 | - |
| - | - | 1010 | 605.6 | - | - | 0 | - |
| - | - | 891.7 | 606 | - | - | 0 | - |
| - | - | 1214 | 606.3 | - | - | 0 | - |
| - | - | 1693 | 606.6 | - | - | 0 | - |
| - | - | 4727 | 607 | - | - | 0 | - |
| 14 | c | 5.5E+04 | 607.3 | 0.000479 | 0.7887 | +3 | 14 |
| - | - | 6.002E+04 | 607.6 | - | - | 0 | - |
| - | - | 3.127E+04 | 608 | - | - | 0 | - |
| - | - | 1.765E+04 | 608.3 | - | - | 0 | - |
| - | - | 4825 | 608.6 | - | - | 0 | - |
| - | - | 1565 | 609 | - | - | 0 | - |
| - | - | 1252 | 610 | - | - | 0 | - |
| - | - | 906.9 | 610.3 | - | - | 0 | - |
| - | - | 886.2 | 611 | - | - | 0 | - |
| 10 | c | 2239 | 611.3 | 0.008496 | 13.9 | +2 | 10 |
| - | - | 1026 | 611.6 | - | - | 0 | - |
| 10 | c | 5002 | 611.8 | 0.004159 | 6.799 | +2 | 10 |
| 6 | y | 3737 | 612.3 | 0.004248 | 6.938 | +2 | 10 |
| 6 | z | 7790 | 612.8 | 0.002973 | 4.851 | +2 | 10 |
| - | - | 4871 | 613.3 | - | - | 0 | - |
| - | - | 1813 | 613.8 | - | - | 0 | - |
| - | - | 903.8 | 614.3 | - | - | 0 | - |
| - | - | 782.1 | 614.8 | - | - | 0 | - |
| - | - | 1656 | 615 | - | - | 0 | - |
| - | - | 2925 | 615.3 | - | - | 0 | - |
| - | - | 2048 | 615.6 | - | - | 0 | - |
| - | - | 771.6 | 616 | - | - | 0 | - |
| - | - | 1147 | 616.3 | - | - | 0 | - |
| - | - | 6527 | 616.6 | - | - | 0 | - |
| - | - | 6483 | 617 | - | - | 0 | - |
| - | - | 3104 | 617.3 | - | - | 0 | - |
| - | - | 2306 | 617.6 | - | - | 0 | - |
| - | - | 4518 | 619.8 | - | - | 0 | - |
| 10 | c | 6.287E+04 | 620.3 | 0.0009557 | 1.541 | +2 | 10 |
| - | - | 4332 | 620.6 | - | - | 0 | - |
| 6 | y | 5.892E+04 | 620.8 | 0.004766 | 7.677 | +2 | 10 |
| - | - | 5.177E+04 | 621 | - | - | 0 | - |
| - | - | 7.828E+04 | 621.3 | - | - | 0 | - |
| - | - | 3.681E+04 | 621.6 | - | - | 0 | - |
| - | - | 1.052E+04 | 621.8 | - | - | 0 | - |
| - | - | 2.125E+04 | 622 | - | - | 0 | - |
| - | - | 9332 | 622.3 | - | - | 0 | - |
| - | - | 2825 | 622.6 | - | - | 0 | - |
| - | - | 2159 | 624.3 | - | - | 0 | - |
| - | - | 3063 | 625.7 | - | - | 0 | - |
| - | - | 4595 | 626 | - | - | 0 | - |
| - | - | 2141 | 626.3 | - | - | 0 | - |
| - | - | 2104 | 626.6 | - | - | 0 | - |
| - | - | 1883 | 627 | - | - | 0 | - |
| - | - | 1580 | 629.3 | - | - | 0 | - |
| - | - | 1432 | 629.6 | - | - | 0 | - |
| - | - | 816 | 630 | - | - | 0 | - |
| - | - | 900.9 | 631.3 | - | - | 0 | - |
| - | - | 1161 | 634.3 | - | - | 0 | - |
| - | - | 806.7 | 634.7 | - | - | 0 | - |
| - | - | 796.2 | 634.8 | - | - | 0 | - |
| - | - | 3485 | 635 | - | - | 0 | - |
| - | - | 5495 | 635.3 | - | - | 0 | - |
| - | - | 6910 | 635.6 | - | - | 0 | - |
| - | - | 3156 | 636 | - | - | 0 | - |
| - | - | 1783 | 636.3 | - | - | 0 | - |
| - | - | 1686 | 638.3 | - | - | 0 | - |
| - | - | 1697 | 640 | - | - | 0 | - |
| - | - | 917.3 | 640.3 | - | - | 0 | - |
| - | - | 8238 | 640.3 | - | - | 0 | - |
| - | - | 1.115E+05 | 640.7 | - | - | 0 | - |
| - | - | 1.598E+05 | 641 | - | - | 0 | - |
| - | - | 1.088E+05 | 641.3 | - | - | 0 | - |
| - | - | 5.457E+04 | 641.7 | - | - | 0 | - |
| - | - | 2.351E+04 | 642 | - | - | 0 | - |
| - | - | 8270 | 642.3 | - | - | 0 | - |
| - | - | 2474 | 642.6 | - | - | 0 | - |
| - | - | 1510 | 643 | - | - | 0 | - |
| - | - | 847.9 | 647.8 | - | - | 0 | - |
| - | - | 2061 | 648.3 | - | - | 0 | - |
| - | - | 1407 | 648.8 | - | - | 0 | - |
| - | - | 2891 | 652.3 | - | - | 0 | - |
| - | - | 3145 | 653.3 | - | - | 0 | - |
| - | - | 1019 | 654.3 | - | - | 0 | - |
| - | - | 1569 | 655.8 | - | - | 0 | - |
| - | - | 1040 | 656.3 | - | - | 0 | - |
| - | - | 1080 | 657.3 | - | - | 0 | - |
| - | - | 2133 | 660.4 | - | - | 0 | - |
| - | - | 2153 | 660.9 | - | - | 0 | - |
| - | - | 1209 | 661.4 | - | - | 0 | - |
| - | - | 2182 | 664.3 | - | - | 0 | - |
| - | - | 1330 | 669.8 | - | - | 0 | - |
| - | - | 2258 | 670.3 | - | - | 0 | - |
| - | - | 1167 | 671.8 | - | - | 0 | - |
| - | - | 1855 | 676.3 | - | - | 0 | - |
| - | - | 4487 | 677.8 | - | - | 0 | - |
| - | - | 3128 | 678.3 | - | - | 0 | - |
| - | - | 888.8 | 678.8 | - | - | 0 | - |
| 5 | c | 1130 | 679.3 | 0.002731 | 4.02 | +1 | 5 |
| - | - | 3663 | 679.8 | - | - | 0 | - |
| 5 | c | 1.455E+04 | 680.3 | 0.002724 | 4.004 | +1 | 5 |
| - | - | 1316 | 680.8 | - | - | 0 | - |
| 11 | y | 7954 | 681.3 | 0.01025 | 15.04 | +1 | 5 |
| 11 | z | 5.025E+04 | 682.3 | 0.004155 | 6.089 | +1 | 5 |
| - | - | 988.1 | 682.8 | - | - | 0 | - |
| - | - | 1.812E+04 | 683.3 | - | - | 0 | - |
| - | - | 1152 | 683.4 | - | - | 0 | - |
| - | - | 4155 | 683.9 | - | - | 0 | - |
| - | - | 7514 | 684.3 | - | - | 0 | - |
| - | - | 639.4 | 684.8 | - | - | 0 | - |
| - | - | 840.9 | 684.9 | - | - | 0 | - |
| - | - | 1691 | 685.3 | - | - | 0 | - |
| 11 | c | 1117 | 693.3 | 0.0008155 | 1.176 | +2 | 11 |
| - | - | 814.9 | 694.3 | - | - | 0 | - |
| 5 | c | 4.9E+04 | 697.3 | 0.0005282 | 0.7574 | +1 | 5 |
| 11 | y | 2.671E+04 | 698.4 | 0.01037 | 14.84 | +1 | 5 |
| - | - | 7378 | 699.4 | - | - | 0 | - |
| - | - | 1328 | 700.4 | - | - | 0 | - |
| - | - | 2750 | 701.3 | - | - | 0 | - |
| 11 | c | 4.561E+04 | 701.8 | 0.0009687 | 1.38 | +2 | 11 |
| - | - | 3.587E+04 | 702.3 | - | - | 0 | - |
| - | - | 1.572E+04 | 702.8 | - | - | 0 | - |
| - | - | 3777 | 703.3 | - | - | 0 | - |
| 5 | y | 2727 | 705.3 | 0.003642 | 5.163 | +2 | 11 |
| 5 | z | 3.217E+04 | 705.8 | 0.00143 | 2.026 | +2 | 11 |
| - | - | 2.382E+04 | 706.3 | - | - | 0 | - |
| - | - | 9619 | 706.8 | - | - | 0 | - |
| - | - | 3271 | 707.4 | - | - | 0 | - |
| - | - | 2623 | 711.3 | - | - | 0 | - |
| - | - | 3054 | 712.3 | - | - | 0 | - |
| - | - | 788.5 | 713.4 | - | - | 0 | - |
| 5 | y | 1.139E+04 | 713.9 | 0.001576 | 2.208 | +2 | 11 |
| - | - | 9283 | 714.4 | - | - | 0 | - |
| - | - | 4977 | 714.9 | - | - | 0 | - |
| - | - | 959.9 | 715.4 | - | - | 0 | - |
| - | - | 1175 | 717.4 | - | - | 0 | - |
| - | - | 864.9 | 718.9 | - | - | 0 | - |
| - | - | 2047 | 719.4 | - | - | 0 | - |
| - | - | 1255 | 720.4 | - | - | 0 | - |
| - | - | 1202 | 721.4 | - | - | 0 | - |
| - | - | 903.4 | 722.3 | - | - | 0 | - |
| - | - | 820.7 | 723.4 | - | - | 0 | - |
| - | - | 1548 | 727.3 | - | - | 0 | - |
| - | - | 955.4 | 727.8 | - | - | 0 | - |
| - | - | 862.7 | 728.3 | - | - | 0 | - |
| - | - | 2008 | 730.3 | - | - | 0 | - |
| - | - | 1485 | 731.4 | - | - | 0 | - |
| - | - | 860 | 733.4 | - | - | 0 | - |
| - | - | 852.2 | 733.9 | - | - | 0 | - |
| - | - | 1625 | 740.3 | - | - | 0 | - |
| - | - | 7156 | 741.4 | - | - | 0 | - |
| - | - | 3615 | 741.9 | - | - | 0 | - |
| - | - | 1990 | 742.4 | - | - | 0 | - |
| - | - | 852.8 | 743.4 | - | - | 0 | - |
| - | - | 968.4 | 743.9 | - | - | 0 | - |
| - | - | 973.3 | 744.4 | - | - | 0 | - |
| - | - | 2191 | 747.4 | - | - | 0 | - |
| 10 | w | 9795 | 752.4 | 0.00378 | 5.024 | +1 | 6 |
| - | - | 4131 | 753.4 | - | - | 0 | - |
| - | - | 1568 | 754.4 | - | - | 0 | - |
| 4 | y | 1186 | 762.9 | 0.01349 | 17.68 | +2 | 12 |
| 4 | z | 2.442E+04 | 763.4 | 0.001108 | 1.451 | +2 | 12 |
| - | - | 1.9E+04 | 763.9 | - | - | 0 | - |
| - | - | 8879 | 764.4 | - | - | 0 | - |
| - | - | 4498 | 764.9 | - | - | 0 | - |
| - | - | 1070 | 765.4 | - | - | 0 | - |
| 12 | c | 1.075E+04 | 765.9 | 0.001217 | 1.589 | +2 | 12 |
| - | - | 1.201E+04 | 766.4 | - | - | 0 | - |
| - | - | 5074 | 766.9 | - | - | 0 | - |
| - | - | 1320 | 767.4 | - | - | 0 | - |
| 4 | y | 3723 | 771.4 | 0.001071 | 1.388 | +2 | 12 |
| - | - | 3802 | 771.9 | - | - | 0 | - |
| - | - | 1286 | 772.4 | - | - | 0 | - |
| - | - | 1615 | 772.9 | - | - | 0 | - |
| - | - | 1659 | 777.4 | - | - | 0 | - |
| - | - | 2152 | 778.9 | - | - | 0 | - |
| - | - | 2167 | 779.4 | - | - | 0 | - |
| - | - | 1176 | 785.4 | - | - | 0 | - |
| - | - | 996.6 | 786.4 | - | - | 0 | - |
| - | - | 1107 | 792.4 | - | - | 0 | - |
| 6 | c | 2202 | 793.4 | 0.001973 | 2.487 | +1 | 6 |
| - | - | 1192 | 794.4 | - | - | 0 | - |
| - | - | 964.5 | 797.9 | - | - | 0 | - |
| 3 | w | 5317 | 798.4 | 0.001093 | 1.37 | +2 | 13 |
| - | - | 6904 | 798.9 | - | - | 0 | - |
| - | - | 1934 | 799.4 | - | - | 0 | - |
| - | - | 1127 | 799.9 | - | - | 0 | - |
| - | - | 3595 | 805.4 | - | - | 0 | - |
| - | - | 2513 | 805.9 | - | - | 0 | - |
| - | - | 2345 | 806.4 | - | - | 0 | - |
| - | - | 1658 | 806.9 | - | - | 0 | - |
| - | - | 2096 | 809.4 | - | - | 0 | - |
| - | - | 936.3 | 809.9 | - | - | 0 | - |
| 6 | c | 9.789E+04 | 810.4 | 0.0008758 | 1.081 | +1 | 6 |
| 10 | z | 3.454E+04 | 811.4 | 0.007807 | 9.622 | +1 | 6 |
| - | - | 2.889E+04 | 811.4 | - | - | 0 | - |
| - | - | 2.514E+04 | 812.4 | - | - | 0 | - |
| - | - | 1.064E+04 | 813.4 | - | - | 0 | - |
| - | - | 3762 | 814.4 | - | - | 0 | - |
| - | - | 1372 | 825.4 | - | - | 0 | - |
| 3 | y | 3150 | 826.9 | 0.008672 | 10.49 | +2 | 13 |
| 10 | y | 3.641E+04 | 827.4 | 0.01023 | 12.37 | +1 | 6 |
| - | - | 2.648E+04 | 827.9 | - | - | 0 | - |
| - | - | 1.634E+04 | 828.4 | - | - | 0 | - |
| - | - | 3178 | 828.9 | - | - | 0 | - |
| - | - | 2005 | 829.4 | - | - | 0 | - |
| - | - | 2890 | 831.4 | - | - | 0 | - |
| 3 | y | 4566 | 835.4 | 0.0007573 | 0.9065 | +2 | 13 |
| - | - | 4222 | 835.9 | - | - | 0 | - |
| - | - | 2633 | 836.4 | - | - | 0 | - |
| - | - | 812.9 | 836.9 | - | - | 0 | - |
| - | - | 2918 | 839.4 | - | - | 0 | - |
| - | - | 1766 | 840.4 | - | - | 0 | - |
| - | - | 1397 | 844.4 | - | - | 0 | - |
| - | - | 1637 | 845.4 | - | - | 0 | - |
| - | - | 4098 | 845.9 | - | - | 0 | - |
| 13 | c | 4.871E+04 | 846.4 | 0.0009293 | 1.098 | +2 | 13 |
| - | - | 3.761E+04 | 846.9 | - | - | 0 | - |
| - | - | 2.377E+04 | 847.4 | - | - | 0 | - |
| - | - | 9736 | 847.9 | - | - | 0 | - |
| - | - | 4084 | 848.4 | - | - | 0 | - |
| - | - | 965.2 | 848.9 | - | - | 0 | - |
| - | - | 934.2 | 850.4 | - | - | 0 | - |
| - | - | 2279 | 860.4 | - | - | 0 | - |
| - | - | 978.1 | 861.4 | - | - | 0 | - |
| - | - | 2241 | 866.4 | - | - | 0 | - |
| - | - | 1986 | 866.9 | - | - | 0 | - |
| - | - | 5786 | 867.4 | - | - | 0 | - |
| - | - | 1689 | 868.4 | - | - | 0 | - |
| - | - | 951.5 | 873.9 | - | - | 0 | - |
| - | - | 946.7 | 879.9 | - | - | 0 | - |
| - | - | 3900 | 880.4 | - | - | 0 | - |
| - | - | 3797 | 880.9 | - | - | 0 | - |
| - | - | 1.003E+04 | 881.4 | - | - | 0 | - |
| - | - | 3968 | 881.9 | - | - | 0 | - |
| - | - | 5578 | 882.4 | - | - | 0 | - |
| - | - | 1850 | 883.4 | - | - | 0 | - |
| - | - | 1187 | 887.4 | - | - | 0 | - |
| - | - | 3301 | 888.4 | - | - | 0 | - |
| - | - | 6592 | 888.9 | - | - | 0 | - |
| - | - | 2088 | 889.3 | - | - | 0 | - |
| - | - | 6791 | 889.4 | - | - | 0 | - |
| - | - | 3827 | 889.9 | - | - | 0 | - |
| 2 | y | 1340 | 895.4 | 0.005766 | 6.44 | +2 | 14 |
| 2 | z | 1.507E+04 | 895.9 | 0.001015 | 1.133 | +2 | 14 |
| - | - | 1.707E+04 | 896.4 | - | - | 0 | - |
| - | - | 1.105E+04 | 896.9 | - | - | 0 | - |
| - | - | 6180 | 897.4 | - | - | 0 | - |
| - | - | 2273 | 897.9 | - | - | 0 | - |
| - | - | 1107 | 902 | - | - | 0 | - |
| - | - | 3344 | 902.4 | - | - | 0 | - |
| - | - | 3417 | 902.9 | - | - | 0 | - |
| - | - | 1882 | 903.4 | - | - | 0 | - |
| 2 | y | 1661 | 903.9 | 0.009399 | 10.4 | +2 | 14 |
| 7 | c | 3585 | 908.4 | 0.0008481 | 0.9336 | +1 | 7 |
| - | - | 1009 | 909 | - | - | 0 | - |
| - | - | 3106 | 909.5 | - | - | 0 | - |
| - | - | 2120 | 910 | - | - | 0 | - |
| 14 | c | 9407 | 910.4 | 0.00178 | 1.955 | +2 | 14 |
| - | - | 1.075E+04 | 910.9 | - | - | 0 | - |
| - | - | 7295 | 911.4 | - | - | 0 | - |
| - | - | 3755 | 911.9 | - | - | 0 | - |
| - | - | 2938 | 912.5 | - | - | 0 | - |
| - | - | 1063 | 913.5 | - | - | 0 | - |
| - | - | 1344 | 914.5 | - | - | 0 | - |
| - | - | 955.6 | 915 | - | - | 0 | - |
| - | - | 1273 | 917.3 | - | - | 0 | - |
| 9 | z | 2094 | 922.5 | 0.007397 | 8.019 | +1 | 7 |
| - | - | 6248 | 923 | - | - | 0 | - |
| - | - | 8569 | 923.5 | - | - | 0 | - |
| - | - | 5579 | 924 | - | - | 0 | - |
| - | - | 5118 | 924.5 | - | - | 0 | - |
| - | - | 2849 | 925 | - | - | 0 | - |
| 7 | c | 1.434E+04 | 925.5 | 0.001277 | 1.379 | +1 | 7 |
| - | - | 6776 | 926.5 | - | - | 0 | - |
| - | - | 1355 | 927.5 | - | - | 0 | - |
| - | - | 1041 | 930 | - | - | 0 | - |
| - | - | 3713 | 930.5 | - | - | 0 | - |
| - | - | 7724 | 931 | - | - | 0 | - |
| - | - | 4.248E+04 | 931.5 | - | - | 0 | - |
| - | - | 4.259E+04 | 932 | - | - | 0 | - |
| - | - | 2.861E+04 | 932.5 | - | - | 0 | - |
| - | - | 1.178E+04 | 933 | - | - | 0 | - |
| - | - | 5453 | 933.5 | - | - | 0 | - |
| - | - | 1522 | 934 | - | - | 0 | - |
| - | - | 1131 | 938 | - | - | 0 | - |
| 9 | y | 1.7E+04 | 938.5 | 0.002172 | 2.314 | +1 | 7 |
| - | - | 1.346E+04 | 939 | - | - | 0 | - |
| 9 | z | 2.233E+04 | 939.5 | 0.006579 | 7.003 | +1 | 7 |
| - | - | 4973 | 940 | - | - | 0 | - |
| - | - | 1.784E+04 | 940.5 | - | - | 0 | - |
| - | - | 7123 | 941.5 | - | - | 0 | - |
| - | - | 2326 | 942.5 | - | - | 0 | - |
| - | - | 2422 | 943.5 | - | - | 0 | - |
| - | - | 1760 | 944 | - | - | 0 | - |
| - | - | 1632 | 944.5 | - | - | 0 | - |
| - | - | 977.1 | 946 | - | - | 0 | - |
| - | - | 1652 | 952 | - | - | 0 | - |
| - | - | 1.363E+04 | 952.5 | - | - | 0 | - |
| - | - | 4.306E+04 | 953 | - | - | 0 | - |
| - | - | 4.274E+04 | 953.5 | - | - | 0 | - |
| - | - | 2.594E+04 | 954 | - | - | 0 | - |
| - | - | 1.159E+04 | 954.5 | - | - | 0 | - |
| - | - | 4390 | 955 | - | - | 0 | - |
| - | - | 2921 | 955.5 | - | - | 0 | - |
| - | - | 1572 | 956.5 | - | - | 0 | - |
| - | - | 818.4 | 957.5 | - | - | 0 | - |
| - | - | 1104 | 960 | - | - | 0 | - |
| - | - | 8374 | 960.5 | - | - | 0 | - |
| - | - | 1.624E+04 | 961 | - | - | 0 | - |
| - | - | 4.461E+04 | 961.5 | - | - | 0 | - |
| - | - | 3.86E+04 | 962 | - | - | 0 | - |
| - | - | 2.536E+04 | 962.5 | - | - | 0 | - |
| - | - | 9914 | 963 | - | - | 0 | - |
| - | - | 8552 | 963.5 | - | - | 0 | - |
| - | - | 1679 | 964 | - | - | 0 | - |
| 8 | c | 3254 | 964.5 | 0.004416 | 4.578 | +1 | 8 |
| - | - | 1285 | 967.5 | - | - | 0 | - |
| - | - | 1173 | 973.4 | - | - | 0 | - |
| - | - | 3827 | 981.5 | - | - | 0 | - |
| 8 | c | 4.602E+04 | 982.5 | 0.000931 | 0.9476 | +1 | 8 |
| - | - | 2.654E+04 | 983.5 | - | - | 0 | - |
| - | - | 7650 | 984.5 | - | - | 0 | - |
| - | - | 1751 | 985.5 | - | - | 0 | - |
| 8 | z | 2364 | 996.5 | 0.001248 | 1.253 | +1 | 8 |
| - | - | 3972 | 997.5 | - | - | 0 | - |
| - | - | 2222 | 998.5 | - | - | 0 | - |
| - | - | 1971 | 1011 | - | - | 0 | - |
| 8 | y | 3863 | 1013 | 0.01637 | 16.17 | +1 | 8 |
| - | - | 1915 | 1014 | - | - | 0 | - |
| - | - | 1520 | 1052 | - | - | 0 | - |
| - | - | 2759 | 1054 | - | - | 0 | - |
| - | - | 1867 | 1055 | - | - | 0 | - |
| - | - | 893.4 | 1056 | - | - | 0 | - |
| - | - | 787.3 | 1066 | - | - | 0 | - |
| - | - | 1554 | 1067 | - | - | 0 | - |
| - | - | 1961 | 1068 | - | - | 0 | - |
| - | - | 935.1 | 1069 | - | - | 0 | - |
| - | - | 2385 | 1095 | - | - | 0 | - |
| - | - | 1282 | 1096 | - | - | 0 | - |
| - | - | 1813 | 1110 | - | - | 0 | - |
| 9 | c | 3.482E+04 | 1111 | 0.0003894 | 0.3506 | +1 | 9 |
| - | - | 2.59E+04 | 1112 | - | - | 0 | - |
| - | - | 2.734E+04 | 1113 | - | - | 0 | - |
| - | - | 1.329E+04 | 1114 | - | - | 0 | - |
| - | - | 6178 | 1115 | - | - | 0 | - |
| - | - | 1300 | 1116 | - | - | 0 | - |
| - | - | 1346 | 1129 | - | - | 0 | - |
| - | - | 1081 | 1168 | - | - | 0 | - |
| - | - | 949.3 | 1181 | - | - | 0 | - |
| - | - | 1052 | 1183 | - | - | 0 | - |
| - | - | 1271 | 1196 | - | - | 0 | - |
| - | - | 3560 | 1197 | - | - | 0 | - |
| - | - | 1215 | 1198 | - | - | 0 | - |
| - | - | 925.2 | 1199 | - | - | 0 | - |
| - | - | 1426 | 1213 | - | - | 0 | - |
| 6 | y | 2802 | 1224 | 0.004027 | 3.291 | +1 | 10 |
| 6 | z | 4677 | 1225 | 0.00343 | 2.8 | +1 | 10 |
| - | - | 1.294E+04 | 1226 | - | - | 0 | - |
| - | - | 7828 | 1227 | - | - | 0 | - |
| - | - | 2973 | 1228 | - | - | 0 | - |
| - | - | 1062 | 1229 | - | - | 0 | - |
| - | - | 1125 | 1239 | - | - | 0 | - |
| 10 | c | 1.191E+04 | 1240 | 0.000765 | 0.6171 | +1 | 10 |
| 6 | y | 1.042E+04 | 1241 | 0.01092 | 8.804 | +1 | 10 |
| - | - | 6258 | 1242 | - | - | 0 | - |
| - | - | 2875 | 1243 | - | - | 0 | - |
| - | - | 1153 | 1244 | - | - | 0 | - |
| - | - | 2139 | 1311 | - | - | 0 | - |
| - | - | 1797 | 1312 | - | - | 0 | - |
| - | - | 1242 | 1344 | - | - | 0 | - |
| - | - | 1080 | 1353 | - | - | 0 | - |
| - | - | 948.3 | 1354 | - | - | 0 | - |
| - | - | 908 | 1359 | - | - | 0 | - |
| - | - | 2008 | 1360 | - | - | 0 | - |
| - | - | 1495 | 1361 | - | - | 0 | - |
| - | - | 739.7 | 1362 | - | - | 0 | - |
| - | - | 1231 | 1387 | - | - | 0 | - |
| - | - | 1359 | 1402 | - | - | 0 | - |
| 11 | c | 6715 | 1403 | 0.001523 | 1.086 | +1 | 11 |
| - | - | 7725 | 1404 | - | - | 0 | - |
| - | - | 4684 | 1405 | - | - | 0 | - |
| - | - | 1573 | 1406 | - | - | 0 | - |
| - | - | 750.1 | 1407 | - | - | 0 | - |
| 5 | y | 846.5 | 1410 | 0.005649 | 4.007 | +1 | 11 |
| 5 | z | 5631 | 1411 | 0.002054 | 1.456 | +1 | 11 |
| - | - | 1.659E+04 | 1412 | - | - | 0 | - |
| - | - | 1.191E+04 | 1413 | - | - | 0 | - |
| - | - | 6479 | 1414 | - | - | 0 | - |
| - | - | 2117 | 1415 | - | - | 0 | - |
| - | - | 913.1 | 1426 | - | - | 0 | - |
| 5 | y | 3186 | 1427 | 0.01712 | 12 | +1 | 11 |
| - | - | 3652 | 1428 | - | - | 0 | - |
| - | - | 1038 | 1469 | - | - | 0 | - |
| - | - | 1047 | 1472 | - | - | 0 | - |
| - | - | 1043 | 1482 | - | - | 0 | - |
| - | - | 1072 | 1487 | - | - | 0 | - |
| - | - | 1244 | 1488 | - | - | 0 | - |
| - | - | 1033 | 1497 | - | - | 0 | - |
| - | - | 969.7 | 1498 | - | - | 0 | - |
| - | - | 810.2 | 1515 | - | - | 0 | - |
| 4 | z | 3416 | 1526 | 0.000544 | 0.3566 | +1 | 12 |
| - | - | 1.231E+04 | 1527 | - | - | 0 | - |
| - | - | 7510 | 1528 | - | - | 0 | - |
| - | - | 3350 | 1529 | - | - | 0 | - |
| - | - | 2279 | 1530 | - | - | 0 | - |
| 12 | c | 3146 | 1531 | 0.003596 | 2.349 | +1 | 12 |
| - | - | 3433 | 1532 | - | - | 0 | - |
| - | - | 2576 | 1533 | - | - | 0 | - |
| - | - | 960.1 | 1534 | - | - | 0 | - |
| 4 | y | 774 | 1542 | 0.001456 | 0.9447 | +1 | 12 |
| - | - | 1283 | 1554 | - | - | 0 | - |
| - | - | 860 | 1555 | - | - | 0 | - |
| - | - | 790.7 | 1597 | - | - | 0 | - |
| - | - | 1361 | 1634 | - | - | 0 | - |
| 3 | z | 2561 | 1654 | 5.012E-05 | 0.0303 | +1 | 13 |
| - | - | 9419 | 1655 | - | - | 0 | - |
| - | - | 6994 | 1656 | - | - | 0 | - |
| - | - | 4089 | 1657 | - | - | 0 | - |
| - | - | 1340 | 1658 | - | - | 0 | - |
| - | - | 1002 | 1676 | - | - | 0 | - |
| - | - | 1834 | 1677 | - | - | 0 | - |
| - | - | 1237 | 1691 | - | - | 0 | - |
| 13 | c | 4032 | 1692 | 0.004714 | 2.786 | +1 | 13 |
| - | - | 6496 | 1693 | - | - | 0 | - |
| - | - | 5762 | 1694 | - | - | 0 | - |
| - | - | 3722 | 1695 | - | - | 0 | - |
| - | - | 1686 | 1696 | - | - | 0 | - |
| - | - | 1125 | 1777 | - | - | 0 | - |
| - | - | 3408 | 1792 | - | - | 0 | - |
| - | - | 4031 | 1793 | - | - | 0 | - |
| - | - | 2070 | 1794 | - | - | 0 | - |
| - | - | 1152 | 1795 | - | - | 0 | - |
| - | - | 1038 | 1805 | - | - | 0 | - |
| - | - | 1051 | 1806 | - | - | 0 | - |
| - | - | 1833 | 1821 | - | - | 0 | - |
| - | - | 2652 | 1822 | - | - | 0 | - |
| - | - | 2422 | 1823 | - | - | 0 | - |
| - | - | 1119 | 1846 | - | - | 0 | - |
| - | - | 1384 | 1847 | - | - | 0 | - |
| - | - | 1181 | 1848 | - | - | 0 | - |
| - | - | 2211 | 1862 | - | - | 0 | - |
| - | - | 4613 | 1863 | - | - | 0 | - |
| - | - | 3821 | 1864 | - | - | 0 | - |
| - | - | 2472 | 1865 | - | - | 0 | - |
| - | - | 1557 | 1866 | - | - | 0 | - |
| - | - | 1111 | 1877 | - | - | 0 | - |
| - | - | 1656 | 1878 | - | - | 0 | - |
| - | - | 1909 | 1879 | - | - | 0 | - |
| - | - | 1309 | 1880 | - | - | 0 | - |
| - | - | 1148 | 1895 | - | - | 0 | - |
| - | - | 913.4 | 1904 | - | - | 0 | - |
| - | - | 2397 | 1905 | - | - | 0 | - |
| - | - | 4214 | 1906 | - | - | 0 | - |
| - | - | 3993 | 1907 | - | - | 0 | - |
| - | - | 1351 | 1908 | - | - | 0 | - |
| - | - | 1504 | 1921 | - | - | 0 | - |
| - | - | 4489 | 1922 | - | - | 0 | - |
| - | - | 4871 | 1923 | - | - | 0 | - |
| - | - | 4872 | 1924 | - | - | 0 | - |
| - | - | 2022 | 1925 | - | - | 0 | - |

m/z Charge Intensity FragmentType MassShift Position
120.06582641601562 0 4549.803
126.09190368652344 0 570.8417
126.57588195800781 0 424.34073
129.066162109375 0 682.9878
129.1025390625 0 3581.3762
130.0611114501953 0 619.6942
130.06544494628906 0 4929.7134
132.10214233398438 0 2348.185
136.07601928710938 0 1768.3832
138.0664825439453 0 566.2513
146.09259033203125 0 1052.5292
148.94717407226562 0 515.2497
149.02369689941406 0 497.20462
155.09300231933594 0 9240.334
157.13369750976562 0 708.8434
159.09193420410156 0 2450.1177
163.06178283691406 0 488.27124
164.0822296142578 0 1901.3286
165.1024932861328 0 1867.1267
166.0613250732422 0 6749.8047
167.0648651123047 0 572.47955
168.13848876953125 0 3742.1042
170.47900390625 0 467.38223
172.09706115722656 0 1682.3281 w 13
177.3397979736328 0 467.90988
178.13401794433594 0 971.6311
183.11305236816406 0 2860.0098
185.1650390625 0 6166.6064
185.33541870117188 0 616.9569
186.12379455566406 0 650.4835
187.0872344970703 0 586.89886
190.08627319335938 0 1763.2234
194.71609497070312 0 456.2221
201.1232147216797 0 1029.0917
203.15078735351562 0 1012.5226
215.13946533203125 0 2950.998
219.1331787109375 0 729.44617
223.1556854248047 0 7051.0557
224.159912109375 0 782.42914
227.13885498046875 0 884.39294
228.97096252441406 0 587.5379
229.1186065673828 0 722.0617
229.15516662597656 0 2458.0369 y Ammonia loss 13
230.16253662109375 0 2598.5027 z 13
231.1708526611328 0 1096.8438
232.0936737060547 0 1648.3867
233.15013122558594 0 6102.823
234.1240234375 0 6432.113
243.13412475585938 0 899.05096
246.1817169189453 0 2021.5842 y 13
250.1428680419922 0 1581.853
251.15065002441406 0 33598.28
252.15370178222656 0 3834.89
253.11883544921875 0 845.75995
261.11968994140625 0 3588.6697
263.1393127441406 0 771.0413
268.1771240234375 0 3875.5645 c 1
269.1796875 0 718.1126
270.12091064453125 0 716.7552
274.1187744140625 0 775.4057
283.15167236328125 0 2083.2017
287.69781494140625 0 506.46625
288.10101318359375 0 752.3076
290.1164855957031 0 1131.3773
292.13287353515625 0 1302.6982
293.13592529296875 0 830.1786
294.1124267578125 0 506.67838
294.1934509277344 0 1335.0793
295.1651611328125 0 657.2556
297.08270263671875 0 547.1876
298.45343017578125 0 628.9602
299.1349182128906 0 859.4524
300.1922302246094 0 4981.324
301.1968994140625 0 716.62537
318.14898681640625 0 736.8742
326.66424560546875 0 8382.163
327.1658630371094 0 2973.6165 z Ammonia loss 7
327.6668701171875 0 635.4288
329.18255615234375 0 850.58105
331.65704345703125 0 1342.807
332.1600646972656 0 869.37213
338.1743469238281 0 885.8173 y 7
339.1786804199219 0 713.5273
340.6619567871094 0 5220.0005 c Ammonia loss 4
341.1636962890625 0 1591.9978 y Ammonia loss 10
341.6631164550781 0 1042.328
349.19036865234375 0 1212.1716
349.6813659667969 0 5304.31 y 10
350.1810607910156 0 1137.373
351.21502685546875 0 1471.7135
352.2203674316406 0 955.14197
355.61468505859375 0 662.1175
361.19903564453125 0 899.958
367.6794738769531 0 725.5573
370.6517333984375 0 739.09375
372.2291259765625 0 1053.3228
374.6923828125 0 1076.0302
375.1695251464844 0 2014.5336
375.2073669433594 0 1200.0254
376.5180969238281 0 1660.7173 y 6
376.68634033203125 0 17555.93 w 9
376.853515625 0 1172.9966
377.18780517578125 0 8230.838
377.6880798339844 0 1899.6107 z Water loss 3
378.2028503417969 0 2129.7527
379.20916748046875 0 9259.147 c Ammonia loss 2
380.212158203125 0 2049.017
381.1519775390625 0 672.9395
381.21160888671875 0 1072.0437
381.6852111816406 0 793.5976 y Water loss 3
382.182861328125 0 704.26196 z 3
383.2054443359375 0 2061.9497
383.7078552246094 0 1509.1301
384.2972106933594 0 1300.137
387.1834716796875 0 834.81024
387.71331787109375 0 1093.8474
388.19061279296875 0 1072.1274
390.1952209472656 0 756.72455
395.22784423828125 0 2411.2969
396.20770263671875 0 987.115
396.2358703613281 0 14929.906 c 2
397.2099609375 0 668.17993 c Ammonia loss 5
397.2397766113281 0 2648.1555
397.70550537109375 0 1215.0999
399.16168212890625 0 778.62946
403.2012634277344 0 1060.2673 z Ammonia loss 5
404.208740234375 0 981.0324
407.1952209472656 0 3757.0342 y 12
408.1679992675781 0 1323.6807
408.2001647949219 0 700.98444 c Ammonia loss 9
408.6693420410156 0 976.62054
409.2186279296875 0 2015.8098
414.2124938964844 0 2604.408 y 9
416.2720031738281 0 1045.4077
417.2196044921875 0 6660.4043
417.7211608886719 0 2522.1738
418.2213439941406 0 978.08795
419.2183532714844 0 983.1219
424.7506103515625 0 1038.8082
425.25042724609375 0 1358.431
428.28759765625 0 5136.112
429.29296875 0 2677.655
440.7338562011719 0 3957.7317
441.2078857421875 0 3804.953 w 8
441.23675537109375 0 1276.5931
441.70904541015625 0 1467.3447
445.16082763671875 0 1168.7117
448.1996154785156 0 2159.361
449.19989013671875 0 1329.5056
450.1701354980469 0 687.4756
452.46905517578125 0 1511.8435 y 1
452.71978759765625 0 2609.4314
452.9687194824219 0 1113.2501
453.2200927734375 0 850.7386
453.2591857910156 0 3812.9502
453.76153564453125 0 2397.3794
454.2632751464844 0 1341.8965
456.2200622558594 0 1125.5283
457.2277526855469 0 899.26263
460.2593688964844 0 4958.9956
461.2071228027344 0 4791.8354 w 11
461.2630615234375 0 1826.5365
462.20843505859375 0 978.33746
466.2420349121094 0 1665.9237
467.2167053222656 0 793.393
467.2489929199219 0 3213.4558
468.2552490234375 0 820.7013
469.2487487792969 0 692.1951 y Water loss 8
470.24114990234375 0 1927.8687 y Water loss 4
470.74163818359375 0 1107.8922
470.89837646484375 0 780.8011 z 4
472.22784423828125 0 652.9407
474.2752990722656 0 2034.8402
475.2821960449219 0 3360.065
476.23486328125 0 3648.2122 y 4
476.4866638183594 0 1823.2579
476.7351989746094 0 960.03143
478.250244140625 0 11426.972 y 8
478.7521667480469 0 6671.2485
479.2521057128906 0 2379.042
480.4938659667969 0 1843.0734
480.69305419921875 0 3317.8567
480.7394104003906 0 23946.19
480.9901123046875 0 20772.02
481.1954650878906 0 1475.2732
481.2408752441406 0 14614.893
481.3145446777344 0 2484.2966
481.49066162109375 0 4359.991
481.6941223144531 0 1056.7919
481.7408142089844 0 2520.866
481.9916687011719 0 860.59064
482.23492431640625 0 1299.6304
488.7770080566406 0 2369.6348
489.28033447265625 0 841.22363
490.2459716796875 0 1619.8723 z Ammonia loss 7
491.2508544921875 0 791.7697
493.25146484375 0 1057.8364 c Water loss 3
494.23614501953125 0 7786.041 c Ammonia loss 3
495.240478515625 0 2802.415
496.2505187988281 0 1589.864
498.2609558105469 0 799.95825
498.7509460449219 0 1735.8363 z 7
499.2519226074219 0 2027.3617
499.7523498535156 0 831.2345
503.2437744140625 0 2739.6946 z Water loss 3
504.24676513671875 0 763.03577
504.76348876953125 0 1786.724
505.23675537109375 0 3791.3828
505.7361145019531 0 4173.266
506.2606506347656 0 6712.2876
506.76092529296875 0 17856.078 y 7
507.26324462890625 0 7802.7397
507.76214599609375 0 3010.2776
508.2652893066406 0 1303.8933
509.25946044921875 0 841.0896
510.7719421386719 0 5206.102
511.26324462890625 0 62110.188 c 3
512.2661743164062 0 15708.417
513.269287109375 0 3173.5752
514.7393798828125 0 1251.4087
515.7406616210938 0 778.75287
519.2728271484375 0 19377.098 z 11
519.7581787109375 0 770.29034
520.2759399414062 0 6184.85
521.2764282226562 0 1180.0479
525.2313232421875 0 1056.36
526.7587890625 0 2709.0383
527.2324829101562 0 5958.9976
527.728271484375 0 1333.6754
527.7658081054688 0 4025.659
528.2672729492188 0 2148.9314
528.769287109375 0 938.8828
530.2203369140625 0 1086.2397
530.2960815429688 0 771.202
533.7661743164062 0 5312.662 w 6
534.2697143554688 0 12471.787
534.7694702148438 0 5326.3716
535.2905883789062 0 11204.904 y 11
536.293701171875 0 2836.9634
537.2791748046875 0 1046.2327
538.2731323242188 0 901.17035
544.2731323242188 0 1198.0043
546.2415161132812 0 1596.5211
547.2791748046875 0 2194.8118 c Ammonia loss 8
547.7759399414062 0 1806.8721
549.2639770507812 0 741.01495
549.6060180664062 0 1122.3152
549.9439086914062 0 843.95654
550.2516479492188 0 13398.437
551.2583618164062 0 5637.5615 y Water loss 2
551.5971069335938 0 2130.14 y Ammonia loss 2
551.9295043945312 0 1278.3273 z 2
552.2612915039062 0 3216.1018
555.2965698242188 0 3204.5718
555.788818359375 0 54477.676 c 8
556.2892456054688 0 25653.81
556.7642822265625 0 6093.085
556.7933349609375 0 7853.2944
557.2708129882812 0 4298.8726 y 2
557.7684936523438 0 1761.8097
559.5968017578125 0 820.109
562.278564453125 0 1834.7611
562.6336059570312 0 1270.217
562.965576171875 0 1214.1343
563.6135864257812 0 765.57745
563.7830200195312 0 1997.6262
564.2745361328125 0 34481.152 y 6
564.7759399414062 0 21280.355
565.27587890625 0 9420.3545
565.7764892578125 0 2080.6033
567.2295532226562 0 820.8216
567.3142700195312 0 1154.0046
568.796630859375 0 4781.5107
569.296142578125 0 4048.8
569.7960815429688 0 1203.845
569.934814453125 0 909.8992
572.95458984375 0 1482.6487
573.2825927734375 0 1263.6084
573.935302734375 0 1863.5021
574.2722778320312 0 1684.248
574.9404907226562 0 1143.2383
575.269775390625 0 767.9009
577.2719116210938 0 770.1108
577.958740234375 0 3220.2441
578.2902221679688 0 3081.888
578.60693359375 0 1310.6567
578.9443969726562 0 1955.187
579.2764282226562 0 759.5909
582.2811279296875 0 843.21704
582.623779296875 0 1276.6244
582.9541015625 0 2661.4854
583.291259765625 0 2380.086
583.6185302734375 0 1593.1622
585.2616577148438 0 840.7314
586.2890014648438 0 842.38715
590.8088989257812 0 1809.6426
591.280029296875 0 34463.285 w 5
591.3510131835938 0 1298.5012
591.7810668945312 0 20520.018
592.2824096679688 0 9576.554
592.355712890625 0 1418.8397
592.6256103515625 0 1700.6188
592.7822265625 0 3588.3992
592.9598388671875 0 2490.5723
593.2817993164062 0 1953.503
595.3208618164062 0 877.0239
597.2940063476562 0 7724.469
597.6271362304688 0 7634.6265 z 1
597.9633178710938 0 4499.35
598.2959594726562 0 3079.486
599.2435913085938 0 879.9924
600.2959594726562 0 757.70044
601.3013916015625 0 2228.11 c Water loss 13
601.6292724609375 0 2632.4897 c Ammonia loss 13
601.9625854492188 0 6748.861
602.2954711914062 0 3039.3403
602.6253051757812 0 1249.0295
602.957763671875 0 3672.3408 y 1
603.2915649414062 0 3302.443
603.62353515625 0 1174.1184
605.646240234375 0 1010.28564
605.961181640625 0 891.69336
606.318115234375 0 1214.2319
606.6372680664062 0 1692.5529
606.9698486328125 0 4726.6396
607.299072265625 0 55002.38 c 13
607.6329345703125 0 60016.11
607.9667358398438 0 31271.318
608.301025390625 0 17653.205
608.6346435546875 0 4824.9844
608.9666748046875 0 1565.0359
609.9839477539062 0 1251.6593
610.3214111328125 0 906.85834
610.9876708984375 0 886.1876
611.3126831054688 0 2238.8037 c Water loss 9
611.6260375976562 0 1026.2001
611.8003540039062 0 5001.576 c Ammonia loss 9
612.3004760742188 0 3736.6414 y Ammonia loss 5
612.8056640625 0 7789.93 z 5
613.3089599609375 0 4870.5737
613.8048706054688 0 1813.2255
614.3106079101562 0 903.7724
614.80517578125 0 782.07074
614.95849609375 0 1656.0219
615.2935180664062 0 2925.1191
615.631591796875 0 2048.2288
615.9636840820312 0 771.59033
616.304443359375 0 1146.7131
616.6356811523438 0 6527.358
616.9680786132812 0 6483.301
617.301513671875 0 3103.522
617.636962890625 0 2305.9626
619.8173828125 0 4517.9272
620.3104248046875 0 62866.12 c 9
620.641357421875 0 4331.5645
620.813232421875 0 58917.86 y 5
620.9623413085938 0 51770.773
621.3089599609375 0 78277.25
621.6416015625 0 36807.844
621.816650390625 0 10520.665
621.9769897460938 0 21248.637
622.3121948242188 0 9332.089
622.6448364257812 0 2825.1533
624.2693481445312 0 2158.9426
625.6514282226562 0 3063.1511
625.9828491210938 0 4594.525
626.3182983398438 0 2140.654
626.6478881835938 0 2104.268
626.9805908203125 0 1882.8839
629.3043823242188 0 1580.2496
629.642578125 0 1432.1523
629.974853515625 0 816.0147
631.3046875 0 900.9398
634.3090209960938 0 1161.2173
634.6514892578125 0 806.6877
634.81884765625 0 796.24915
634.9788208007812 0 3484.9514
635.3139038085938 0 5494.748
635.6453247070312 0 6910.3984
635.9779052734375 0 3155.927
636.3115234375 0 1783.2925
638.3489990234375 0 1686.1462
639.97900390625 0 1697.0148
640.258544921875 0 917.31525
640.3203735351562 0 8238.062
640.6500854492188 0 111473.58
640.9845581054688 0 159780.67
641.3185424804688 0 108781.05
641.6526489257812 0 54572.13
641.9866333007812 0 23514.578
642.3199462890625 0 8270.056
642.6490478515625 0 2473.918
642.9813232421875 0 1509.9237
647.8213500976562 0 847.9124
648.3233032226562 0 2061.0073
648.8269653320312 0 1406.7028
652.3198852539062 0 2891.113
653.3253784179688 0 3144.5662
654.3279418945312 0 1018.9855
655.8209228515625 0 1568.8684
656.3216552734375 0 1039.5897
657.3323364257812 0 1079.9679
660.3543090820312 0 2132.6162
660.856201171875 0 2153.4724
661.3555297851562 0 1208.6982
664.34130859375 0 2182.4045
669.8366088867188 0 1330.2711
670.3344116210938 0 2257.5862
671.8212890625 0 1166.5908
676.3418579101562 0 1854.6986
677.815185546875 0 4486.974
678.3167114257812 0 3128.3193
678.8189086914062 0 888.7932
679.3338012695312 0 1129.5164 c Water loss 4
679.8349609375 0 3663.292
680.3178100585938 0 14547.186 c Ammonia loss 4
680.8370971679688 0 1316.1786
681.3218994140625 0 7954.447 y Ammonia loss 10
682.3358154296875 0 50246.043 z 10
682.8447875976562 0 988.0746
683.3397216796875 0 18124.26
683.4027709960938 0 1151.5724
683.85107421875 0 4154.6133
684.3417358398438 0 7514.174
684.8194580078125 0 639.35486
684.8569946289062 0 840.8549
685.3383178710938 0 1690.7406
693.3286743164062 0 1117.1694 c Ammonia loss 10
694.3203125 0 814.94946
697.3421630859375 0 49004.66 c 4
698.3483276367188 0 26705.828 y 10
699.3505249023438 0 7378.3184
700.3541259765625 0 1327.917
701.3494262695312 0 2750.057
701.8421020507812 0 45605.426 c 10
702.3433837890625 0 35871.656
702.8445434570312 0 15716.554
703.346923828125 0 3777.438
705.3480224609375 0 2726.5647 y Ammonia loss 4
705.8468627929688 0 32166.697 z 4
706.34814453125 0 23822.184
706.8491821289062 0 9618.788
707.3505859375 0 3270.8633
711.3257446289062 0 2622.876
712.326171875 0 3053.5906
713.3665771484375 0 788.5394
713.8560791015625 0 11390.178 y 4
714.359130859375 0 9283.186
714.858154296875 0 4977.312
715.3629760742188 0 959.8518
717.3568115234375 0 1175.095
718.8607788085938 0 864.94257
719.3656616210938 0 2047.372
720.3942260742188 0 1255.001
721.35205078125 0 1201.9489
722.3402099609375 0 903.3997
723.3750610351562 0 820.6808
727.3487548828125 0 1547.9381
727.8432006835938 0 955.41077
728.334716796875 0 862.6707
730.3395385742188 0 2008.3511
731.3938598632812 0 1485.2733
733.36962890625 0 860.02277
733.8555908203125 0 852.2072
740.2991333007812 0 1625.4606
741.3659057617188 0 7155.797
741.8651123046875 0 3614.6487
742.3660888671875 0 1989.8209
743.3720703125 0 852.8343
743.8798217773438 0 968.43854
744.384033203125 0 973.291
747.3627319335938 0 2191.3645
752.365478515625 0 9794.62 w 9
753.3683471679688 0 4130.962
754.3729248046875 0 1568.4119
762.871337890625 0 1186.2839 y Ammonia loss 3
763.3606567382812 0 24419.164 z 3
763.8617553710938 0 18995.193
764.3629760742188 0 8878.855
764.8617553710938 0 4497.726
765.3778076171875 0 1069.7764
765.8898315429688 0 10746.912 c 11
766.3936767578125 0 12009.862
766.8926391601562 0 5073.875
767.405029296875 0 1320.0316
771.3700561523438 0 3722.8975 y 3
771.8719482421875 0 3801.5168
772.3670043945312 0 1286.1428
772.873046875 0 1614.8234
777.359375 0 1659.3812
778.897705078125 0 2151.7307
779.4008178710938 0 2166.7253
785.3707275390625 0 1176.0248
786.3720092773438 0 996.5852
792.3567504882812 0 1107.2311
793.401123046875 0 2201.9746 c Ammonia loss 5
794.4035034179688 0 1191.7095
797.881591796875 0 964.49365
798.3775024414062 0 5317.171 w 2
798.8771362304688 0 6904.0044
799.3765869140625 0 1934.3776
799.8779907226562 0 1127.3055
805.3932495117188 0 3595.4756
805.8956909179688 0 2513.2502
806.39501953125 0 2344.9648
806.8958129882812 0 1657.6421
809.4155883789062 0 2095.6523
809.8936157226562 0 936.3
810.4265747070312 0 97891.22 c 5
811.374755859375 0 34535.336 z 9
811.435302734375 0 28890.002
812.3828735351562 0 25139.78
813.3824462890625 0 10644.498
814.3848876953125 0 3762.0913
825.38671875 0 1372.1953
826.8958129882812 0 3149.7034 y Ammonia loss 2
827.3910522460938 0 36410.523 y 9
827.89111328125 0 26482.922
828.392578125 0 16341.965
828.8935546875 0 3178.3542
829.3910522460938 0 2005.4462
831.3856811523438 0 2889.7056
835.399658203125 0 4566.3574 y 2
835.900634765625 0 4221.7324
836.402587890625 0 2633.4717
836.9000244140625 0 812.90295
839.4205932617188 0 2917.8247
840.4242553710938 0 1765.5056
844.40966796875 0 1396.6196
845.4066162109375 0 1636.5884
845.9000854492188 0 4097.8135
846.3972778320312 0 48713.586 c 12
846.8980712890625 0 37606.805
847.3989868164062 0 23768.338
847.900146484375 0 9735.597
848.4031982421875 0 4084.0503
848.8981323242188 0 965.1595
850.402099609375 0 934.19366
860.42626953125 0 2278.623
861.4268188476562 0 978.0655
866.4310913085938 0 2241.3674
866.9166259765625 0 1986.1284
867.4239501953125 0 5785.6206
868.4185791015625 0 1688.9955
873.924560546875 0 951.4791
879.9430541992188 0 946.6762
880.4390258789062 0 3899.8875
880.935302734375 0 3796.5256
881.4411010742188 0 10026.783
881.9395751953125 0 3968.313
882.44580078125 0 5577.551
883.4454345703125 0 1849.5992
887.4281616210938 0 1186.6006
888.4371948242188 0 3301.4497
888.9390258789062 0 6592.1025
889.3187255859375 0 2088.2031
889.4425659179688 0 6790.6553
889.9439086914062 0 3826.7104
895.42236328125 0 1339.7406 y Ammonia loss 1
895.9194946289062 0 15071.304 z 1
896.4223022460938 0 17073.828
896.9234008789062 0 11046.872
897.4255981445312 0 6180.422
897.926025390625 0 2272.8838
901.9611206054688 0 1107.1704
902.4405517578125 0 3343.5625
902.9409790039062 0 3417.0598
903.4359741210938 0 1882.3798
903.9392700195312 0 1661.3126 y 1
908.4269409179688 0 3585.2693 c Ammonia loss 6
908.9611206054688 0 1008.82776
909.4505615234375 0 3106.1858
909.957275390625 0 2119.5203
910.4439086914062 0 9407.151 c 13
910.9474487304688 0 10751.783
911.4483032226562 0 7294.852
911.9473876953125 0 3755.2803
912.458251953125 0 2937.8733
913.4625854492188 0 1062.9064
914.4619140625 0 1343.8202
914.9596557617188 0 955.58563
917.3107299804688 0 1273.187
922.4583740234375 0 2094.0046 z Ammonia loss 8
922.962158203125 0 6247.713
923.4610595703125 0 8568.721
923.9612426757812 0 5578.8716
924.45849609375 0 5118.234
924.9576416015625 0 2849.039
925.4539184570312 0 14343.058 c 6
926.4547119140625 0 6776.306
927.4552612304688 0 1355.0574
929.9610595703125 0 1041.1799
930.4673461914062 0 3713.1863
930.9632568359375 0 7724.26
931.466552734375 0 42480.664
931.968505859375 0 42594.934
932.4701538085938 0 28613.043
932.9710693359375 0 11783.04
933.4713745117188 0 5453.023
933.9710083007812 0 1521.622
937.9727172851562 0 1130.6765
938.467529296875 0 16995.379 y Ammonia loss 8
938.9713745117188 0 13455.368
939.470947265625 0 22331.545 z 8
939.9705810546875 0 4972.612
940.4773559570312 0 17835.773
941.4826049804688 0 7123.091
942.4812622070312 0 2326.4797
943.4685668945312 0 2422.3462
943.9667358398438 0 1760.3436
944.4694213867188 0 1632.183
945.9583740234375 0 977.10004
951.969970703125 0 1651.8278
952.4692993164062 0 13627.2
952.9662475585938 0 43059.95
953.4667358398438 0 42743.914
953.9677734375 0 25939.213
954.468017578125 0 11590.87
954.9697265625 0 4390.2197
955.4730224609375 0 2920.6047
956.49462890625 0 1571.5989
957.5013427734375 0 818.4263
959.9691772460938 0 1103.9572
960.470703125 0 8373.971
960.974365234375 0 16237.634
961.4779052734375 0 44611.09
961.97900390625 0 38598.56
962.4799194335938 0 25356.121
962.981201171875 0 9914.273
963.4830932617188 0 8551.809
963.974609375 0 1678.9675
964.4679565429688 0 3254.1794 c Water loss 7
967.46337890625 0 1285.0662
973.447265625 0 1173.2399
981.4856567382812 0 3826.9448
982.4750366210938 0 46018.652 c 7
983.4779052734375 0 26536.193
984.480224609375 0 7650.1772
985.4823608398438 0 1750.6263
996.4977416992188 0 2364.0547 z 7
997.5023803710938 0 3971.626
998.5052490234375 0 2222.16
1011.4718017578125 0 1971.4554
1012.5013427734375 0 3863.3516 y 7
1013.5057373046875 0 1915.4991
1051.5576171875 0 1519.9003
1053.5220947265625 0 2759.2632
1054.52587890625 0 1866.9509
1055.5303955078125 0 893.3577
1065.56640625 0 787.306
1066.560302734375 0 1554.0613
1067.552734375 0 1960.8895
1068.5462646484375 0 935.1199
1094.5487060546875 0 2384.5464
1095.5556640625 0 1281.5013
1109.575927734375 0 1812.8527
1110.5694580078125 0 34817.02 c 8
1111.5706787109375 0 25895.072
1112.5380859375 0 27342.666
1113.536376953125 0 13293.144
1114.534912109375 0 6178.3184
1115.548095703125 0 1300.1528
1128.5421142578125 0 1345.7881
1167.6075439453125 0 1080.8196
1180.607421875 0 949.31854
1182.6082763671875 0 1051.7861
1195.5989990234375 0 1271.0247
1196.6053466796875 0 3559.659
1197.6068115234375 0 1215.0269
1198.612060546875 0 925.1531
1212.62841796875 0 1425.8218
1223.59814453125 0 2801.7483 y Ammonia loss 5
1224.6065673828125 0 4677.154 z 5
1225.6123046875 0 12938.424
1226.6153564453125 0 7827.852
1227.6156005859375 0 2973.0981
1228.6163330078125 0 1061.9143
1238.60009765625 0 1124.893
1239.6124267578125 0 11905.106 c 9
1240.6177978515625 0 10418.299 y 5
1241.622314453125 0 6258.097
1242.6224365234375 0 2874.6348
1243.6319580078125 0 1153.4423
1310.6357421875 0 2139.0808
1311.638671875 0 1797.4431
1343.66015625 0 1242.0586
1352.679443359375 0 1080.222
1353.693359375 0 948.2799
1358.6729736328125 0 907.9848
1359.6658935546875 0 2008.487
1360.675537109375 0 1494.598
1361.677001953125 0 739.73193
1386.6632080078125 0 1230.6816
1401.6727294921875 0 1358.5322
1402.676513671875 0 6715.26 c 10
1403.6796875 0 7724.993
1404.6820068359375 0 4684.2837
1405.687255859375 0 1573.3254
1406.6810302734375 0 750.10175
1409.6871337890625 0 846.4635 y Ammonia loss 4
1410.687255859375 0 5631.095 z 4
1411.6898193359375 0 16588.465
1412.6943359375 0 11908.762
1413.697265625 0 6479.413
1414.6988525390625 0 2116.7312
1425.66015625 0 913.0689
1426.69091796875 0 3186.1123 y 4
1427.6990966796875 0 3652.231
1468.6998291015625 0 1038.0824
1471.731689453125 0 1047.2483
1481.7225341796875 0 1042.6975
1486.7659912109375 0 1071.5483
1487.759765625 0 1244.1854
1496.74365234375 0 1032.5594
1497.7373046875 0 969.7282
1514.7506103515625 0 810.2389
1525.716796875 0 3416.4778 z 3
1526.718017578125 0 12309.742
1527.717529296875 0 7510.3276
1528.7183837890625 0 3350.2703
1529.7322998046875 0 2279.2893
1530.766357421875 0 3145.6382 c 11
1531.7713623046875 0 3433.1877
1532.7803955078125 0 2576.4087
1533.7843017578125 0 960.0643
1541.7335205078125 0 773.98944 y 3
1553.71826171875 0 1282.7638
1554.718017578125 0 860.0016
1596.751708984375 0 790.69574
1633.7720947265625 0 1361.1697
1653.7747802734375 0 2560.6992 z 2
1654.77734375 0 9418.94
1655.7781982421875 0 6993.62
1656.781005859375 0 4088.524
1657.7762451171875 0 1340.0116
1675.7669677734375 0 1002.3952
1676.7672119140625 0 1834.2892
1690.7708740234375 0 1236.6825
1691.784423828125 0 4032.382 c 12
1692.7889404296875 0 6495.7793
1693.7943115234375 0 5762.4946
1694.8004150390625 0 3722.2444
1695.786865234375 0 1685.637
1776.8896484375 0 1125.0155
1791.8353271484375 0 3408.2732
1792.842041015625 0 4030.7974
1793.8419189453125 0 2069.9097
1794.8446044921875 0 1151.5435
1804.8873291015625 0 1038.0878
1805.86767578125 0 1051.151
1820.892822265625 0 1833.0986
1821.891845703125 0 2651.7026
1822.8892822265625 0 2422.069
1845.925537109375 0 1119.1343
1846.9306640625 0 1384.4714
1847.919677734375 0 1181.1064
1861.9246826171875 0 2210.6306
1862.930908203125 0 4612.557
1863.9346923828125 0 3820.842
1864.9364013671875 0 2471.5112
1865.9346923828125 0 1556.5503
1876.931884765625 0 1111.279
1877.924072265625 0 1655.9729
1878.928955078125 0 1908.587
1879.94091796875 0 1308.6188
1894.9521484375 0 1147.8499
1903.9422607421875 0 913.4287
1904.9246826171875 0 2397.3884
1905.9266357421875 0 4214.0435
1906.9300537109375 0 3993.2896
1907.93017578125 0 1351.3945
1920.93798828125 0 1503.5486
1921.94677734375 0 4488.6743
1922.949951171875 0 4870.786
1923.9508056640625 0 4872.3086
1924.9527587890625 0 2021.9691

Spectrum Details

|  |  |
| --- | --- |
| Matched peaks? Matched peaks The total absolute number of peaks matched. Additionally in brackets the total fraction of peaks matched and the total number of peaks is shown. | 110 (14.18% of 776) |
| FDR? FDR The false discovery rate estimated for this peptide. It is calculated by matching all theoretical fragments with a non-integer shift with the raw peaks for this spectrum. This is done with 40 different shifts. The resulting percentage is the average number of annotated peaks over the number of annotated peaks with the correct spectrum. | 0.32% |
| Satellite FDR? Satellite FDR See the FDR for details on its calculation. This satellite ion specific FDR only contains the satellite ions (d/w) for I/L/J positions. | 0.00% |
| PSM Score? PSM Score The PSM Score as given by Hecklib to this annotated spectrum. It is shown with three significant figures. | 595 |

#### Reverse Lookup? Reverse Lookup All places where this read could be placed.

| Group | Segment | Template | Template Part | Read Part | Score | Unique |
| --- | --- | --- | --- | --- | --- | --- |
| Homo sapiens Heavy Chain | IGHC | IGHG1 | [191..206] | [0..15] | 115 | False |
| Homo sapiens Heavy Chain | IGHC | IGHG3 | [238..253] | [0..15] | 115 | False |
| Homo sapiens Heavy Chain | IGHC | IGHG2 | [188..202] | [0..15] | 106 | False |
| Homo sapiens Heavy Chain | IGHC | IGHG4 | [188..203] | [0..15] | 115 | False |

| Recombined | Template Part | Read Part | Score | Unique |
| --- | --- | --- | --- | --- |
| REC-0-1 | [316..331] | [0..15] | 120 | True |

#### Meta Information from Multiple reads

##### Number of combined reads

8

##### Intensity

0.6513

##### TotalArea

2.138E+08

##### Changes to the peptide sequence

JHQDWJDGKEYKCKV

L→JNo support for either Leucine or Isoleucine based on side chain ions (Position: 6)

J→LSupport for Leucine based on side chain ions (2 for L 1 for I) (Position: 6)

I→JEqual support for both Leucine and Isoleucine based on side chain ions (1 ions for both) (Position: 6)

J→ISupport for Isoleucine based on side chain ions (1 for I 0 for L) (Position: 6)

L→JNo support for either Leucine or Isoleucine based on side chain ions (Position: 6)

L→JNo support for either Leucine or Isoleucine based on side chain ions (Position: 1)

#### Positional Score

Copy Data

##### Positional Score (TSV)

###### Preview

```
Loading example...
```

*Click on the button to copy the data to your clipboard.*

1001234567891011121314

Label Value
"0" 0.748
"1" 0.738
"2" 0.727
"3" 0.746
"4" 0.744
"5" 0.741
"6" 0.736
"7" 0.74
"8" 0.741
"9" 0.744
"10" 0.745
"11" 0.733
"12" 0.728
"13" 0.735
"14" 0.712

#### Meta Information from PEAKS

##### Scan Identifier

F1:5478

##### Original sequence

L

H

Q

D

W

L

D

G

K

E

Y

K

C

+58.01

K

V

##### Posttranslational Modifications

Carboxymethyl

##### Source File

D:\separate\_stitch\_analyses\xle-disambiguation\raw\20210323\_F1\_UM1\_Peng0013\_SA\_F59\_ingel\_3ug\_ELA.raw

##### Fraction

1

##### Scan Feature

F1:10242

##### De Novo Score

99

##### ConfidenceScore

99

### m/z

640.6482

##### Mass

1918.9248

##### Charge

3

##### Retention Time

29.26

##### Predicted Retention Time

-

##### Area

2.499E+07

##### Fragmentation mode

ETHCD

##### Originating file

01 D:\separate\_stitch\_analyses\xle-disambiguation\20210325\_F59\_3ug\_DENOVO\_12.csv

#### Meta Information from PEAKS

##### Scan Identifier

F1:5773

##### Original sequence

L

H

Q

D

W

L

D

G

K

E

Y

K

C

+58.01

K

V

##### Posttranslational Modifications

Carboxymethyl

##### Source File

D:\separate\_stitch\_analyses\xle-disambiguation\raw\20210323\_F1\_UM1\_Peng0013\_SA\_F59\_ingel\_3ug\_ELA.raw

##### Fraction

1

##### Scan Feature

F1:10242

##### De Novo Score

99

##### ConfidenceScore

99

### m/z

640.6482

##### Mass

1918.9248

##### Charge

3

##### Retention Time

29.26

##### Predicted Retention Time

-

##### Area

2.499E+07

##### Fragmentation mode

ETHCD

##### Originating file

01 D:\separate\_stitch\_analyses\xle-disambiguation\20210325\_F59\_3ug\_DENOVO\_12.csv

#### Meta Information from PEAKS

##### Scan Identifier

F1:5773

##### Original sequence

L

H

Q

D

W

L

D

G

K

E

Y

K

C

+58.01

K

V

##### Posttranslational Modifications

Carboxymethyl

##### Source File

D:\separate\_stitch\_analyses\xle-disambiguation\raw\20210323\_F1\_UM1\_Peng0013\_SA\_F59\_ingel\_3ug\_ELA.raw

##### Fraction

1

##### Scan Feature

F1:10243

##### De Novo Score

98

##### ConfidenceScore

98

### m/z

640.6497

##### Mass

1918.9248

##### Charge

3

##### Retention Time

30.72

##### Predicted Retention Time

-

##### Area

9.457E+06

##### Parts Per Million

1.2

##### Fragmentation mode

ETHCD

##### Originating file

01 D:\separate\_stitch\_analyses\xle-disambiguation\20210325\_F59\_3ug\_DENOVO\_12.csv

#### Meta Information from PEAKS

##### Scan Identifier

F1:5764

##### Original sequence

L

H

Q

D

W

L

D

G

K

E

Y

K

C

+58.01

K

V

##### Posttranslational Modifications

Carboxymethyl

##### Source File

D:\separate\_stitch\_analyses\xle-disambiguation\raw\20210323\_F1\_UM1\_Peng0013\_SA\_F59\_ingel\_3ug\_ELA.raw

##### Fraction

1

##### Scan Feature

F1:3308

##### De Novo Score

98

##### ConfidenceScore

98

### m/z

480.7385

##### Mass

1918.9248

##### Charge

4

##### Retention Time

29.26

##### Predicted Retention Time

-

##### Area

5.587E+07

##### Parts Per Million

0.1

##### Fragmentation mode

ETHCD

##### Originating file

01 D:\separate\_stitch\_analyses\xle-disambiguation\20210325\_F59\_3ug\_DENOVO\_12.csv

#### Meta Information from PEAKS

##### Scan Identifier

F1:5829

##### Original sequence

L

H

Q

D

W

L

D

G

K

E

Y

K

C

+58.01

K

V

##### Posttranslational Modifications

Carboxymethyl

##### Source File

D:\separate\_stitch\_analyses\xle-disambiguation\raw\20210323\_F1\_UM1\_Peng0013\_SA\_F59\_ingel\_3ug\_ELA.raw

##### Fraction

1

##### Scan Feature

F1:3308

##### De Novo Score

98

##### ConfidenceScore

98

### m/z

480.7385

##### Mass

1918.9248

##### Charge

4

##### Retention Time

29.26

##### Predicted Retention Time

-

##### Area

5.587E+07

##### Parts Per Million

0.1

##### Fragmentation mode

ETHCD

##### Originating file

01 D:\separate\_stitch\_analyses\xle-disambiguation\20210325\_F59\_3ug\_DENOVO\_12.csv

#### Meta Information from PEAKS

##### Scan Identifier

F1:5464

##### Original sequence

L

H

Q

D

W

L

D

G

K

E

Y

K

C

+58.01

K

V

##### Posttranslational Modifications

Carboxymethyl

##### Source File

D:\separate\_stitch\_analyses\xle-disambiguation\raw\20210323\_F1\_UM1\_Peng0013\_SA\_F59\_ingel\_3ug\_ELA.raw

##### Fraction

1

##### Scan Feature

F1:515

##### De Novo Score

98

##### ConfidenceScore

98

### m/z

384.7922

##### Mass

1918.9248

##### Charge

5

##### Retention Time

29.26

##### Predicted Retention Time

-

##### Area

4.772E+06

##### Fragmentation mode

ETHCD

##### Originating file

01 D:\separate\_stitch\_analyses\xle-disambiguation\20210325\_F59\_3ug\_DENOVO\_12.csv

#### Meta Information from PEAKS

##### Scan Identifier

F1:5664

##### Original sequence

L

H

Q

D

W

L

D

G

K

E

Y

K

C

+58.01

K

V

##### Posttranslational Modifications

Carboxymethyl

##### Source File

D:\separate\_stitch\_analyses\xle-disambiguation\raw\20210323\_F1\_UM1\_Peng0013\_SA\_F59\_ingel\_3ug\_ELA.raw

##### Fraction

1

##### Scan Feature

F1:10242

##### De Novo Score

97

##### ConfidenceScore

97

### m/z

640.6482

##### Mass

1918.9248

##### Charge

3

##### Retention Time

29.26

##### Predicted Retention Time

-

##### Area

2.499E+07

##### Fragmentation mode

ETHCD

##### Originating file

01 D:\separate\_stitch\_analyses\xle-disambiguation\20210325\_F59\_3ug\_DENOVO\_12.csv

#### Meta Information from PEAKS

##### Scan Identifier

F1:5764

##### Original sequence

L

H

Q

D

W

L

D

G

K

E

Y

K

C

+58.01

K

V

##### Posttranslational Modifications

Carboxymethyl

##### Source File

D:\separate\_stitch\_analyses\xle-disambiguation\raw\20210323\_F1\_UM1\_Peng0013\_SA\_F59\_ingel\_3ug\_ELA.raw

##### Fraction

1

##### Scan Feature

F1:3309

##### De Novo Score

97

##### ConfidenceScore

97

### m/z

480.7393

##### Mass

1918.9248

##### Charge

4

##### Retention Time

30.72

##### Predicted Retention Time

-

##### Area

1.283E+07

##### Parts Per Million

1.8

##### Fragmentation mode

ETHCD

##### Originating file

01 D:\separate\_stitch\_analyses\xle-disambiguation\20210325\_F59\_3ug\_DENOVO\_12.csv
