## Supplementary material for "A handle on mass coincidence errors in *de novo* sequencing of antibodies by bottom-up proteomics": Combined_009.html

Details Combined\_009 | Stitch OverviewUndefined

### Read Combined\_009

#### Sequence (length=15)

SVMHEAJHNHYTQKS

#### Spectrum 2869? Spectrum 2869 The raw spectrum of this peptide as annotated by Hecklib. The fragments are coloured according to ion type (see legend). Any peaks with a star '\*' as text can be hovered over to see the full details, first the ion type second the mass shift type. By hovering over the amino acids in the peptide or ions in the legend the corresponding peaks are highlighted. By toggling the 'Unassigned' label you can turn the background (unassigned) peaks on or off in the plot. By updating the slider in the Ion legend you can update the spectrum to only show the top X% of the peaks with labels. The top X% means any peak that is within X% of the highest intensity. By dragging in the spectrum you can zoom in to a specific part of the spectrum and use 'Zoom Out' to get back to the original zoom level. The annotation of the spectrum is based on the given sequence in the peptides file and is done with different software so inconsistencies are likely. The peaks are annotated based on the given sequence, with 20 ppm tolerance.

Copy Data

##### Spectrum 2869 (TSV)

###### Preview

```
Loading example...
```

*Click on the button to copy the data to your clipboard.*

Mz MinMz MaxIntensity Max

WidthHeightPeptide font sizePeptide stroke widthSpectrum font sizeSpectrum stroke widthCompact peptide

Ion legend

wxyz

abcd

OtherUnassignedIon^Charge^_Position_Show for top:%

SVMHEAJHNHYTQKS

08.24e+51.65e+62.47e+63.30e+6

Zoom Out

z^+1^_2_y^+1^_2_w^+1^_3_z^+1^_3_y^+1^_3_z^+1^_4_w^+1^_4_z^+3^_11_y^+2^_7_z^+1^_4_y^+1^_4_c^+1^_4_y^+3^_12_c^+2^_9_y^+3^_13_z^+3^_13_y^+3^_13_z^+3^_14_z^+3^_14_c^+3^_14_y^+3^_14_z^+3^_14_y^+3^_14_c^+2^_10_c^+1^_5_z^+1^_5_y^+2^_10_c^+1^_5_z^+1^_5_y^+1^_5_c^+1^_6_c^+2^_11_y^+2^_11_c^+1^_6_z^+2^_12_y^+2^_12_z^+1^_6_w^+2^_13_y^+1^_6_c^+2^_13_c^+2^_13_c^+1^_7_y^+2^_13_z^+2^_13_y^+2^_13_w^+1^_7_z^+2^_14_c^+2^_14_y^+2^_14_z^+2^_14_c^+2^_14_y^+2^_14_z^+1^_7_y^+1^_7_c^+1^_8_z^+1^_8_y^+1^_8_c^+1^_9_w^+1^_9_z^+1^_9_c^+1^_10_z^+1^_10_y^+1^_10_z^+1^_11_z^+1^_11_y^+1^_11_c^+1^_11_c^+1^_12_z^+1^_12_c^+1^_13_y^+1^_14_z^+1^_14_c^+1^_14_y^+1^_14_

0712142421362847

Fragment Matches Table

Show background peaks

| Position | Ion type | Intensity | mz Theoretical | mz Error (Th) | mz Error (ppm) | Charge | Series Number |
| --- | --- | --- | --- | --- | --- | --- | --- |
| - | - | 3605 | 128 | - | - | 0 | - |
| - | - | 4.871E+04 | 129.1 | - | - | 0 | - |
| - | - | 3769 | 145.1 | - | - | 0 | - |
| - | - | 3959 | 147.4 | - | - | 0 | - |
| - | - | 4.207E+04 | 155.1 | - | - | 0 | - |
| - | - | 4.685E+04 | 159.1 | - | - | 0 | - |
| - | - | 5118 | 165.1 | - | - | 0 | - |
| - | - | 4350 | 176 | - | - | 0 | - |
| - | - | 4014 | 177.2 | - | - | 0 | - |
| - | - | 1.101E+04 | 183.1 | - | - | 0 | - |
| - | - | 3.494E+04 | 187.1 | - | - | 0 | - |
| - | - | 4291 | 188.1 | - | - | 0 | - |
| 14 | z | 1.14E+04 | 218.1 | 0.0003608 | 1.654 | +1 | 2 |
| 14 | y | 2.647E+04 | 234.1 | 6.489E-05 | 0.2772 | +1 | 2 |
| - | - | 4386 | 236.8 | - | - | 0 | - |
| - | - | 9594 | 239.1 | - | - | 0 | - |
| - | - | 7666 | 240.1 | - | - | 0 | - |
| - | - | 8126 | 241.1 | - | - | 0 | - |
| - | - | 5111 | 251.1 | - | - | 0 | - |
| - | - | 7980 | 269.1 | - | - | 0 | - |
| - | - | 5196 | 272.3 | - | - | 0 | - |
| - | - | 6492 | 284.1 | - | - | 0 | - |
| - | - | 2.035E+04 | 286.1 | - | - | 0 | - |
| 13 | w | 5.813E+04 | 288.2 | 2.877E-05 | 0.09985 | +1 | 3 |
| - | - | 8458 | 289.2 | - | - | 0 | - |
| - | - | 6252 | 299.1 | - | - | 0 | - |
| - | - | 5670 | 315.2 | - | - | 0 | - |
| 13 | z | 1.284E+05 | 346.2 | 3.689E-05 | 0.1066 | +1 | 3 |
| - | - | 3.219E+04 | 347.2 | - | - | 0 | - |
| - | - | 5.049E+04 | 355.1 | - | - | 0 | - |
| - | - | 3.2E+04 | 356.1 | - | - | 0 | - |
| - | - | 1.589E+04 | 357.1 | - | - | 0 | - |
| 13 | y | 1.556E+04 | 362.2 | 0.000661 | 1.825 | +1 | 3 |
| - | - | 6582 | 373.2 | - | - | 0 | - |
| - | - | 1.086E+04 | 410.2 | - | - | 0 | - |
| - | - | 1.004E+04 | 412.2 | - | - | 0 | - |
| - | - | 9103 | 417.2 | - | - | 0 | - |
| - | - | 3.655E+04 | 427.2 | - | - | 0 | - |
| - | - | 8618 | 428.2 | - | - | 0 | - |
| 12 | z | 8427 | 429.2 | 0.002474 | 5.764 | +1 | 4 |
| 12 | w | 1.955E+04 | 430.2 | 5.046E-05 | 0.1173 | +1 | 4 |
| - | - | 5645 | 431.2 | - | - | 0 | - |
| 5 | z | 9716 | 432.2 | 0.007069 | 16.36 | +3 | 11 |
| 9 | y | 9781 | 439.2 | 0.0006359 | 1.448 | +2 | 7 |
| - | - | 5383 | 445.2 | - | - | 0 | - |
| 12 | z | 9.184E+04 | 447.2 | 0.0002173 | 0.4858 | +1 | 4 |
| - | - | 1.56E+05 | 448.2 | - | - | 0 | - |
| - | - | 3.492E+04 | 449.2 | - | - | 0 | - |
| - | - | 1.182E+04 | 453.2 | - | - | 0 | - |
| - | - | 4.28E+04 | 455.2 | - | - | 0 | - |
| - | - | 1.227E+04 | 456.2 | - | - | 0 | - |
| - | - | 5821 | 457.2 | - | - | 0 | - |
| 12 | y | 1.076E+04 | 463.3 | 0.0003458 | 0.7464 | +1 | 4 |
| - | - | 6.46E+04 | 471.2 | - | - | 0 | - |
| 4 | c | 1.019E+05 | 472.2 | 0.000449 | 0.9508 | +1 | 4 |
| - | - | 2.398E+04 | 473.2 | - | - | 0 | - |
| - | - | 5282 | 479.3 | - | - | 0 | - |
| 4 | y | 2.245E+04 | 488.9 | 0.001108 | 2.267 | +3 | 12 |
| - | - | 1.505E+04 | 489.2 | - | - | 0 | - |
| - | - | 6475 | 498.2 | - | - | 0 | - |
| 9 | c | 8587 | 510.2 | 0.0001328 | 0.2602 | +2 | 9 |
| - | - | 5957 | 510.7 | - | - | 0 | - |
| 3 | y | 1.184E+04 | 526.9 | 3.368E-05 | 0.06391 | +3 | 13 |
| 3 | z | 6409 | 527.2 | 0.000825 | 1.565 | +3 | 13 |
| 3 | y | 4.958E+04 | 532.6 | 0.0003596 | 0.6752 | +3 | 13 |
| - | - | 6.931E+04 | 532.9 | - | - | 0 | - |
| - | - | 3.213E+04 | 533.3 | - | - | 0 | - |
| - | - | 6522 | 533.6 | - | - | 0 | - |
| - | - | 6519 | 547.9 | - | - | 0 | - |
| - | - | 6189 | 548.6 | - | - | 0 | - |
| - | - | 3.411E+04 | 553.9 | - | - | 0 | - |
| 2 | z | 2.224E+04 | 554.3 | 0.003323 | 5.996 | +3 | 14 |
| 2 | z | 2.977E+04 | 554.6 | 0.004548 | 8.2 | +3 | 14 |
| 14 | c | 6.3E+04 | 559.6 | 0.0001857 | 0.3318 | +3 | 14 |
| 2 | y | 6.153E+04 | 559.9 | 0.006526 | 11.65 | +3 | 14 |
| 2 | z | 3.219E+04 | 560.3 | 0.004324 | 7.719 | +3 | 14 |
| - | - | 2.15E+04 | 560.6 | - | - | 0 | - |
| - | - | 1.01E+04 | 564.8 | - | - | 0 | - |
| - | - | 8995 | 565.3 | - | - | 0 | - |
| 2 | y | 2.814E+04 | 565.6 | 0.0008996 | 1.591 | +3 | 14 |
| - | - | 9433 | 565.9 | - | - | 0 | - |
| - | - | 7177 | 566.3 | - | - | 0 | - |
| - | - | 8952 | 576.9 | - | - | 0 | - |
| - | - | 7275 | 577.3 | - | - | 0 | - |
| 10 | c | 1.492E+04 | 578.8 | 0.001513 | 2.613 | +2 | 10 |
| - | - | 7749 | 579.3 | - | - | 0 | - |
| - | - | 8853 | 582.9 | - | - | 0 | - |
| 5 | c | 1.566E+04 | 583.3 | 0.009698 | 16.63 | +1 | 5 |
| - | - | 2.686E+04 | 584.2 | - | - | 0 | - |
| - | - | 2.887E+04 | 588.6 | - | - | 0 | - |
| - | - | 1.014E+05 | 588.9 | - | - | 0 | - |
| - | - | 6.53E+04 | 589.3 | - | - | 0 | - |
| - | - | 4.376E+04 | 589.6 | - | - | 0 | - |
| - | - | 1.117E+04 | 589.9 | - | - | 0 | - |
| 11 | z | 1.829E+04 | 592.3 | 0.0006383 | 1.078 | +1 | 5 |
| - | - | 7128 | 593.3 | - | - | 0 | - |
| - | - | 8199 | 594.3 | - | - | 0 | - |
| - | - | 9.082E+05 | 594.6 | - | - | 0 | - |
| - | - | 9.372E+05 | 595 | - | - | 0 | - |
| - | - | 5.142E+05 | 595.3 | - | - | 0 | - |
| - | - | 2.352E+05 | 595.6 | - | - | 0 | - |
| - | - | 8.096E+04 | 596 | - | - | 0 | - |
| 6 | y | 9934 | 599.8 | 0.000618 | 1.03 | +2 | 10 |
| - | - | 6220 | 600.8 | - | - | 0 | - |
| 5 | c | 2.426E+05 | 601.3 | 1.233E-05 | 0.02051 | +1 | 5 |
| - | - | 8.275E+04 | 602.3 | - | - | 0 | - |
| - | - | 2.695E+04 | 603.3 | - | - | 0 | - |
| - | - | 8105 | 604.3 | - | - | 0 | - |
| 11 | z | 2.198E+05 | 610.3 | 2.231E-05 | 0.03656 | +1 | 5 |
| - | - | 8.204E+04 | 611.3 | - | - | 0 | - |
| - | - | 1.248E+04 | 612.3 | - | - | 0 | - |
| 11 | y | 3.495E+04 | 626.3 | 0.0003912 | 0.6246 | +1 | 5 |
| - | - | 8853 | 627.3 | - | - | 0 | - |
| - | - | 1.574E+04 | 653.3 | - | - | 0 | - |
| 6 | c | 7412 | 654.3 | 0.003917 | 5.987 | +1 | 6 |
| - | - | 9935 | 655.3 | - | - | 0 | - |
| 11 | c | 9902 | 660.3 | 0.0008892 | 1.347 | +2 | 11 |
| - | - | 6829 | 661.3 | - | - | 0 | - |
| 5 | y | 3.911E+04 | 664.3 | 0.000598 | 0.9002 | +2 | 11 |
| - | - | 1.294E+04 | 664.8 | - | - | 0 | - |
| 6 | c | 1.283E+05 | 672.3 | 7.778E-05 | 0.1157 | +1 | 6 |
| - | - | 5.481E+04 | 673.3 | - | - | 0 | - |
| - | - | 1.822E+04 | 674.3 | - | - | 0 | - |
| - | - | 8310 | 680.8 | - | - | 0 | - |
| - | - | 6300 | 689.3 | - | - | 0 | - |
| - | - | 6110 | 698.3 | - | - | 0 | - |
| - | - | 8720 | 702.3 | - | - | 0 | - |
| - | - | 1.011E+04 | 711.3 | - | - | 0 | - |
| - | - | 5480 | 711.8 | - | - | 0 | - |
| 4 | z | 4.184E+04 | 724.8 | 0.0001821 | 0.2512 | +2 | 12 |
| - | - | 5.119E+04 | 725.3 | - | - | 0 | - |
| - | - | 2.267E+04 | 725.8 | - | - | 0 | - |
| - | - | 6014 | 726.3 | - | - | 0 | - |
| - | - | 2.114E+04 | 730.4 | - | - | 0 | - |
| - | - | 7659 | 731.4 | - | - | 0 | - |
| 4 | y | 5.813E+04 | 732.9 | 0.0001467 | 0.2002 | +2 | 12 |
| - | - | 2.657E+04 | 733.4 | - | - | 0 | - |
| - | - | 2.665E+04 | 733.9 | - | - | 0 | - |
| - | - | 6447 | 737.3 | - | - | 0 | - |
| - | - | 6514 | 737.8 | - | - | 0 | - |
| - | - | 1.985E+04 | 741.4 | - | - | 0 | - |
| - | - | 1.183E+04 | 742.4 | - | - | 0 | - |
| - | - | 6929 | 743.4 | - | - | 0 | - |
| - | - | 9466 | 745.9 | - | - | 0 | - |
| 10 | z | 1.062E+05 | 747.4 | 0.0002535 | 0.3392 | +1 | 6 |
| - | - | 4.568E+04 | 748.4 | - | - | 0 | - |
| - | - | 1.173E+04 | 749.4 | - | - | 0 | - |
| - | - | 8193 | 753.3 | - | - | 0 | - |
| 3 | w | 3.753E+04 | 759.9 | 0.0003693 | 0.486 | +2 | 13 |
| - | - | 3.232E+04 | 760.4 | - | - | 0 | - |
| - | - | 2.618E+04 | 760.9 | - | - | 0 | - |
| - | - | 8995 | 761.4 | - | - | 0 | - |
| 10 | y | 2.144E+04 | 763.4 | 0.001061 | 1.389 | +1 | 6 |
| - | - | 1.437E+04 | 764.4 | - | - | 0 | - |
| - | - | 2.363E+04 | 765.8 | - | - | 0 | - |
| - | - | 1.639E+04 | 766.4 | - | - | 0 | - |
| - | - | 1.335E+04 | 766.9 | - | - | 0 | - |
| - | - | 8169 | 767.4 | - | - | 0 | - |
| - | - | 1.049E+04 | 768.4 | - | - | 0 | - |
| - | - | 5.827E+04 | 774.4 | - | - | 0 | - |
| 13 | c | 4.538E+04 | 774.9 | 0.0004872 | 0.6288 | +2 | 13 |
| - | - | 4.084E+04 | 775.4 | - | - | 0 | - |
| - | - | 2.911E+04 | 775.9 | - | - | 0 | - |
| - | - | 6520 | 776.4 | - | - | 0 | - |
| - | - | 7859 | 782.9 | - | - | 0 | - |
| 13 | c | 1.805E+05 | 783.4 | 3.003E-05 | 0.03833 | +2 | 13 |
| - | - | 1.34E+05 | 783.9 | - | - | 0 | - |
| - | - | 8.432E+04 | 784.4 | - | - | 0 | - |
| - | - | 2.161E+04 | 784.9 | - | - | 0 | - |
| 7 | c | 9.754E+04 | 785.4 | 0.0003405 | 0.4335 | +1 | 7 |
| - | - | 3.503E+04 | 786.4 | - | - | 0 | - |
| - | - | 1.334E+04 | 787.4 | - | - | 0 | - |
| - | - | 6602 | 788.7 | - | - | 0 | - |
| 3 | y | 1.367E+04 | 789.9 | 0.005764 | 7.297 | +2 | 13 |
| 3 | z | 2.59E+04 | 790.4 | 0.001628 | 2.059 | +2 | 13 |
| - | - | 1.499E+04 | 790.9 | - | - | 0 | - |
| - | - | 7503 | 791.4 | - | - | 0 | - |
| 3 | y | 8.754E+04 | 798.4 | 0.0007967 | 0.998 | +2 | 13 |
| - | - | 6.916E+04 | 798.9 | - | - | 0 | - |
| - | - | 3.229E+04 | 799.4 | - | - | 0 | - |
| - | - | 1.153E+04 | 799.9 | - | - | 0 | - |
| - | - | 9203 | 809.9 | - | - | 0 | - |
| - | - | 6172 | 816.4 | - | - | 0 | - |
| 9 | w | 6.045E+04 | 817.4 | 0.001421 | 1.739 | +1 | 7 |
| - | - | 3.679E+04 | 818.4 | - | - | 0 | - |
| - | - | 7530 | 819.4 | - | - | 0 | - |
| - | - | 2.565E+04 | 825.4 | - | - | 0 | - |
| - | - | 1.491E+04 | 825.9 | - | - | 0 | - |
| - | - | 1.004E+04 | 826.4 | - | - | 0 | - |
| 2 | z | 1.041E+04 | 830.9 | 0.007106 | 8.553 | +2 | 14 |
| - | - | 1.322E+04 | 833.4 | - | - | 0 | - |
| - | - | 6094 | 833.9 | - | - | 0 | - |
| 14 | c | 1.472E+04 | 838.9 | 0.005252 | 6.26 | +2 | 14 |
| 2 | y | 2.27E+04 | 839.4 | 0.009338 | 11.12 | +2 | 14 |
| 2 | z | 5.05E+04 | 839.9 | 0.007745 | 9.221 | +2 | 14 |
| - | - | 4.507E+04 | 840.4 | - | - | 0 | - |
| - | - | 2.149E+04 | 840.9 | - | - | 0 | - |
| - | - | 8514 | 841.4 | - | - | 0 | - |
| - | - | 5384 | 844.4 | - | - | 0 | - |
| - | - | 1.671E+04 | 846.4 | - | - | 0 | - |
| - | - | 7102 | 846.9 | - | - | 0 | - |
| 14 | c | 4.424E+05 | 847.4 | 2.716E-05 | 0.03206 | +2 | 14 |
| 2 | y | 4.133E+05 | 847.9 | 0.008697 | 10.26 | +2 | 14 |
| - | - | 2.123E+05 | 848.4 | - | - | 0 | - |
| - | - | 8.298E+04 | 848.9 | - | - | 0 | - |
| - | - | 2.541E+04 | 849.4 | - | - | 0 | - |
| - | - | 2.357E+04 | 850.9 | - | - | 0 | - |
| - | - | 2.156E+04 | 851.4 | - | - | 0 | - |
| - | - | 1.497E+04 | 851.9 | - | - | 0 | - |
| - | - | 1.134E+05 | 852.9 | - | - | 0 | - |
| - | - | 1.053E+05 | 853.4 | - | - | 0 | - |
| - | - | 6.115E+04 | 853.9 | - | - | 0 | - |
| - | - | 2.388E+04 | 854.4 | - | - | 0 | - |
| - | - | 1.147E+04 | 854.9 | - | - | 0 | - |
| - | - | 1.58E+04 | 855.9 | - | - | 0 | - |
| - | - | 1.221E+04 | 859.9 | - | - | 0 | - |
| - | - | 2.692E+04 | 860.4 | - | - | 0 | - |
| - | - | 2.725E+04 | 860.9 | - | - | 0 | - |
| 9 | z | 3.158E+05 | 861.4 | 0.0007221 | 0.8382 | +1 | 7 |
| - | - | 9184 | 861.9 | - | - | 0 | - |
| - | - | 1.532E+05 | 862.4 | - | - | 0 | - |
| - | - | 2.16E+04 | 862.9 | - | - | 0 | - |
| - | - | 4.636E+04 | 863.4 | - | - | 0 | - |
| - | - | 1.056E+04 | 864.4 | - | - | 0 | - |
| - | - | 2.655E+05 | 869.4 | - | - | 0 | - |
| - | - | 2.553E+05 | 869.9 | - | - | 0 | - |
| - | - | 1.249E+05 | 870.4 | - | - | 0 | - |
| - | - | 5.849E+04 | 870.9 | - | - | 0 | - |
| - | - | 2.013E+04 | 871.4 | - | - | 0 | - |
| - | - | 2.906E+04 | 876.4 | - | - | 0 | - |
| 9 | y | 7.773E+04 | 877.4 | 0.0003696 | 0.4212 | +1 | 7 |
| - | - | 3.962E+04 | 878.4 | - | - | 0 | - |
| - | - | 2.878E+04 | 879.4 | - | - | 0 | - |
| - | - | 1.493E+05 | 883.4 | - | - | 0 | - |
| - | - | 1.61E+05 | 883.9 | - | - | 0 | - |
| - | - | 9.278E+04 | 884.4 | - | - | 0 | - |
| - | - | 4.294E+04 | 884.9 | - | - | 0 | - |
| - | - | 2.157E+04 | 885.4 | - | - | 0 | - |
| - | - | 1.599E+04 | 890.9 | - | - | 0 | - |
| - | - | 6.047E+05 | 891.4 | - | - | 0 | - |
| - | - | 3.265E+06 | 891.9 | - | - | 0 | - |
| - | - | 2.992E+06 | 892.4 | - | - | 0 | - |
| - | - | 1.591E+06 | 892.9 | - | - | 0 | - |
| - | - | 6.43E+05 | 893.4 | - | - | 0 | - |
| - | - | 1.945E+05 | 893.9 | - | - | 0 | - |
| - | - | 8183 | 905.4 | - | - | 0 | - |
| 8 | c | 5.481E+05 | 922.5 | 0.0002924 | 0.317 | +1 | 8 |
| - | - | 2.527E+05 | 923.5 | - | - | 0 | - |
| - | - | 1.022E+05 | 924.5 | - | - | 0 | - |
| - | - | 2.386E+04 | 925.5 | - | - | 0 | - |
| 8 | z | 7.665E+04 | 998.5 | 0.0006947 | 0.6958 | +1 | 8 |
| - | - | 4.297E+04 | 999.5 | - | - | 0 | - |
| - | - | 1.565E+04 | 1000 | - | - | 0 | - |
| 8 | y | 2.681E+04 | 1014 | 0.001963 | 1.935 | +1 | 8 |
| - | - | 1.565E+04 | 1015 | - | - | 0 | - |
| - | - | 8966 | 1016 | - | - | 0 | - |
| 9 | c | 1.786E+05 | 1036 | 0.0004813 | 0.4644 | +1 | 9 |
| - | - | 1.108E+05 | 1038 | - | - | 0 | - |
| - | - | 3.984E+04 | 1039 | - | - | 0 | - |
| - | - | 9403 | 1040 | - | - | 0 | - |
| 7 | w | 2.638E+04 | 1068 | 0.00145 | 1.357 | +1 | 9 |
| - | - | 8971 | 1069 | - | - | 0 | - |
| - | - | 8256 | 1070 | - | - | 0 | - |
| - | - | 6316 | 1096 | - | - | 0 | - |
| 7 | z | 8.402E+04 | 1112 | 0.0007133 | 0.6417 | +1 | 9 |
| - | - | 6.698E+04 | 1113 | - | - | 0 | - |
| - | - | 3.132E+04 | 1114 | - | - | 0 | - |
| - | - | 1.122E+04 | 1115 | - | - | 0 | - |
| - | - | 3.074E+04 | 1127 | - | - | 0 | - |
| - | - | 2.307E+04 | 1128 | - | - | 0 | - |
| - | - | 1.707E+04 | 1129 | - | - | 0 | - |
| - | - | 2.708E+04 | 1130 | - | - | 0 | - |
| - | - | 1.254E+04 | 1131 | - | - | 0 | - |
| - | - | 6993 | 1132 | - | - | 0 | - |
| 10 | c | 3.471E+05 | 1174 | 0.0002853 | 0.2431 | +1 | 10 |
| - | - | 2.335E+05 | 1175 | - | - | 0 | - |
| - | - | 8.547E+04 | 1176 | - | - | 0 | - |
| - | - | 2.653E+04 | 1177 | - | - | 0 | - |
| 6 | z | 1.928E+05 | 1183 | 0.0004735 | 0.4004 | +1 | 10 |
| - | - | 1.767E+05 | 1184 | - | - | 0 | - |
| - | - | 6.471E+04 | 1185 | - | - | 0 | - |
| - | - | 2.045E+04 | 1186 | - | - | 0 | - |
| 6 | y | 1.198E+04 | 1199 | 0.003141 | 2.621 | +1 | 10 |
| - | - | 1.345E+04 | 1200 | - | - | 0 | - |
| - | - | 8457 | 1240 | - | - | 0 | - |
| - | - | 8293 | 1292 | - | - | 0 | - |
| - | - | 9209 | 1293 | - | - | 0 | - |
| 5 | z | 6892 | 1294 | 0.002044 | 1.58 | +1 | 11 |
| 5 | z | 2.066E+05 | 1312 | 0.001074 | 0.8192 | +1 | 11 |
| - | - | 2.727E+05 | 1313 | - | - | 0 | - |
| - | - | 1.479E+05 | 1314 | - | - | 0 | - |
| - | - | 4.556E+04 | 1315 | - | - | 0 | - |
| - | - | 1.056E+04 | 1316 | - | - | 0 | - |
| - | - | 7089 | 1317 | - | - | 0 | - |
| - | - | 8124 | 1327 | - | - | 0 | - |
| 5 | y | 1.802E+04 | 1328 | 0.0009534 | 0.7182 | +1 | 11 |
| - | - | 1.673E+04 | 1329 | - | - | 0 | - |
| - | - | 1.087E+05 | 1336 | - | - | 0 | - |
| 11 | c | 1.883E+05 | 1337 | 0.002741 | 2.05 | +1 | 11 |
| - | - | 1.217E+05 | 1338 | - | - | 0 | - |
| - | - | 5.267E+04 | 1339 | - | - | 0 | - |
| - | - | 1.058E+04 | 1340 | - | - | 0 | - |
| - | - | 1.561E+04 | 1437 | - | - | 0 | - |
| 12 | c | 2.441E+05 | 1438 | 0.001835 | 1.276 | +1 | 12 |
| - | - | 1.814E+05 | 1439 | - | - | 0 | - |
| - | - | 8.724E+04 | 1440 | - | - | 0 | - |
| - | - | 2.227E+04 | 1441 | - | - | 0 | - |
| - | - | 6793 | 1442 | - | - | 0 | - |
| 4 | z | 9295 | 1449 | 0.00774 | 5.343 | +1 | 12 |
| - | - | 6.752E+04 | 1450 | - | - | 0 | - |
| - | - | 6.101E+04 | 1451 | - | - | 0 | - |
| - | - | 2.174E+04 | 1452 | - | - | 0 | - |
| - | - | 7254 | 1453 | - | - | 0 | - |
| - | - | 2.335E+04 | 1523 | - | - | 0 | - |
| - | - | 1.07E+04 | 1524 | - | - | 0 | - |
| 13 | c | 1.234E+05 | 1566 | 0.003162 | 2.019 | +1 | 13 |
| - | - | 1.172E+05 | 1567 | - | - | 0 | - |
| - | - | 6.051E+04 | 1568 | - | - | 0 | - |
| - | - | 2.147E+04 | 1569 | - | - | 0 | - |
| - | - | 4.76E+04 | 1581 | - | - | 0 | - |
| - | - | 4.197E+04 | 1582 | - | - | 0 | - |
| - | - | 1.746E+04 | 1583 | - | - | 0 | - |
| - | - | 8262 | 1650 | - | - | 0 | - |
| - | - | 1.665E+04 | 1651 | - | - | 0 | - |
| - | - | 1.719E+04 | 1652 | - | - | 0 | - |
| - | - | 9259 | 1662 | - | - | 0 | - |
| 2 | y | 3.424E+04 | 1678 | 0.02192 | 13.07 | +1 | 14 |
| 2 | z | 4.764E+04 | 1679 | 0.018 | 10.72 | +1 | 14 |
| - | - | 4.236E+04 | 1680 | - | - | 0 | - |
| - | - | 3.051E+04 | 1681 | - | - | 0 | - |
| - | - | 2.023E+04 | 1682 | - | - | 0 | - |
| 14 | c | 4.583E+04 | 1694 | 0.001241 | 0.7325 | +1 | 14 |
| 2 | y | 1.134E+05 | 1695 | 0.02015 | 11.89 | +1 | 14 |
| - | - | 7.899E+04 | 1696 | - | - | 0 | - |
| - | - | 5.023E+04 | 1697 | - | - | 0 | - |
| - | - | 9684 | 1698 | - | - | 0 | - |
| - | - | 8647 | 1722 | - | - | 0 | - |
| - | - | 1.42E+04 | 1723 | - | - | 0 | - |
| - | - | 1.741E+04 | 1725 | - | - | 0 | - |
| - | - | 8778 | 1726 | - | - | 0 | - |
| - | - | 3.761E+04 | 1738 | - | - | 0 | - |
| - | - | 1.766E+05 | 1739 | - | - | 0 | - |
| - | - | 1.411E+05 | 1740 | - | - | 0 | - |
| - | - | 8.75E+04 | 1741 | - | - | 0 | - |
| - | - | 3.665E+04 | 1742 | - | - | 0 | - |
| - | - | 1.159E+04 | 1743 | - | - | 0 | - |
| - | - | 3.187E+04 | 1749 | - | - | 0 | - |
| - | - | 2.541E+04 | 1750 | - | - | 0 | - |
| - | - | 1.048E+04 | 1751 | - | - | 0 | - |
| - | - | 7734 | 1752 | - | - | 0 | - |
| - | - | 2.021E+04 | 1755 | - | - | 0 | - |
| - | - | 8.231E+04 | 1756 | - | - | 0 | - |
| - | - | 6.853E+04 | 1757 | - | - | 0 | - |
| - | - | 4.022E+04 | 1758 | - | - | 0 | - |
| - | - | 1.744E+04 | 1759 | - | - | 0 | - |
| - | - | 5.973E+04 | 1766 | - | - | 0 | - |
| - | - | 2.443E+05 | 1767 | - | - | 0 | - |
| - | - | 2.398E+05 | 1768 | - | - | 0 | - |
| - | - | 1.134E+05 | 1769 | - | - | 0 | - |
| - | - | 5.194E+04 | 1770 | - | - | 0 | - |
| - | - | 1.804E+04 | 1771 | - | - | 0 | - |
| - | - | 4.655E+04 | 1782 | - | - | 0 | - |
| - | - | 4.411E+05 | 1783 | - | - | 0 | - |
| - | - | 1.776E+06 | 1784 | - | - | 0 | - |
| - | - | 1.557E+06 | 1785 | - | - | 0 | - |
| - | - | 8.475E+05 | 1786 | - | - | 0 | - |
| - | - | 3.124E+05 | 1787 | - | - | 0 | - |
| - | - | 9.252E+04 | 1788 | - | - | 0 | - |
| - | - | 6867 | 2819 | - | - | 0 | - |

m/z Charge Intensity FragmentType MassShift Position
128.00518798828125 0 3605.3206
129.102294921875 0 48713.89
145.06858825683594 0 3768.9102
147.40994262695312 0 3959.4128
155.09274291992188 0 42066.855
159.11279296875 0 46848.53
165.10226440429688 0 5118.115
176.00711059570312 0 4349.524
177.2118682861328 0 4013.8127
183.11276245117188 0 11006.354
187.10760498046875 0 34938.555
188.11053466796875 0 4290.953
218.12574768066406 0 11400.206 z 13
234.1448974609375 0 26468.822 y 13
236.8113250732422 0 4386.2754
239.09548950195312 0 9593.512
240.13465881347656 0 7665.52
241.11264038085938 0 8125.8228
251.13450622558594 0 5111.1104
269.1062316894531 0 7979.9053
272.2820129394531 0 5195.91
284.1347961425781 0 6492.287
286.1332702636719 0 20354.709
288.1554260253906 0 58132.39 w 12
289.1585998535156 0 8458.142
299.0616149902344 0 6251.8916
315.1525573730469 0 5669.563
346.1847229003906 0 128355.6 z 12
347.18902587890625 0 32188.287
355.0699157714844 0 50488.297
356.0704345703125 0 31995.904
357.06866455078125 0 15886.03
362.2040710449219 0 15561.945 y 12
373.17401123046875 0 6581.783
410.21624755859375 0 10860.512
412.2002258300781 0 10038.442
417.18621826171875 0 9103.33
427.21240234375 0 36547.32
428.21820068359375 0 8618.21
429.2242736816406 0 8427.096 z Water loss 11
430.22967529296875 0 19554.385 w 11
431.2294616699219 0 5645.1445
432.2098083496094 0 9715.722 z Ammonia loss 4
439.21240234375 0 9781.336 y 8
445.2173156738281 0 5383.273
447.2321472167969 0 91839.24 z 11
448.2397155761719 0 155973.9
449.24273681640625 0 34922.24
453.21856689453125 0 11818.016
455.2071533203125 0 42795.715
456.2087097167969 0 12274.751
457.20880126953125 0 5820.936
463.2514343261719 0 10761.737 y 11
471.2248229980469 0 64598.062
472.23321533203125 0 101937.734 c 3
473.23583984375 0 23975.93
479.2756652832031 0 5282.059
488.9052429199219 0 22453.357 y 3
489.2371520996094 0 15053.943
498.24945068359375 0 6474.917
510.2398681640625 0 8587.422 c Ammonia loss 8
510.7379150390625 0 5956.8696
526.9088134765625 0 11837.942 y Ammonia loss 2
527.243896484375 0 6409.3022 z 2
532.5846557617188 0 49582.484 y 2
532.9187622070312 0 69312.94
533.252685546875 0 32133.16
533.5833740234375 0 6521.533
547.9249877929688 0 6519.498
548.5918579101562 0 6189.045
553.9283447265625 0 34112.555
554.2606811523438 0 22236.61 z Water loss 1
554.5965576171875 0 29766.016 z Ammonia loss 1
559.6033935546875 0 63002.336 c Ammonia loss 13
559.9381103515625 0 61533.402 y Ammonia loss 1
560.2718505859375 0 32191.7 z 1
560.6061401367188 0 21504.271
564.7759399414062 0 10104.622
565.2667236328125 0 8994.704
565.606201171875 0 28136.402 y 1
565.9388427734375 0 9433.218
566.2734375 0 7177.39
576.935302734375 0 8952.011
577.2714233398438 0 7274.8564
578.7679443359375 0 14915.724 c Ammonia loss 9
579.2694702148438 0 7749.23
582.9400634765625 0 8853.363
583.275390625 0 15657.739 c Water loss 4
584.2498168945312 0 26859.1
588.6148681640625 0 28870.932
588.9445190429688 0 101407.65
589.2784423828125 0 65297.934
589.611083984375 0 43755.723
589.9454345703125 0 11166.047
592.2857666015625 0 18292.377 z Water loss 10
593.2870483398438 0 7127.5537
594.282958984375 0 8198.853
594.617919921875 0 908184.94
594.9520263671875 0 937151.8
595.2860107421875 0 514198.7
595.619873046875 0 235222.2
595.9534301757812 0 80962.375
599.8024291992188 0 9933.701 y 5
600.8021850585938 0 6220.386
601.2762451171875 0 242566.39 c 4
602.2786865234375 0 82747.3
603.2797241210938 0 26950.648
604.28271484375 0 8104.943
610.2957153320312 0 219796.97 z 10
611.2987060546875 0 82036.87
612.2999267578125 0 12484.695
626.3140258789062 0 34946.316 y 10
627.3203735351562 0 8853.025
653.3009643554688 0 15744.895
654.2988891601562 0 7411.6304 c Water loss 5
655.2871704101562 0 9934.93
660.3002319335938 0 9902.104 c Ammonia loss 10
661.3054809570312 0 6828.7544
664.322509765625 0 39110.41 y 4
664.8236083984375 0 12942.415
672.3132934570312 0 128341.69 c 5
673.3160400390625 0 54814.867
674.3171997070312 0 18217.568
680.8319091796875 0 8310.499
689.3245849609375 0 6299.508
698.3273315429688 0 6110.0537
702.3219604492188 0 8719.647
711.3243408203125 0 10109.629
711.8370971679688 0 5479.5386
724.8433837890625 0 41835.01 z 3
725.3450927734375 0 51189.52
725.8465576171875 0 22666.965
726.3469848632812 0 6013.7573
730.3734130859375 0 21139.201
731.3826904296875 0 7658.577
732.8524169921875 0 58127.906 y 3
733.3531494140625 0 26573.365
733.854248046875 0 26653.643
737.3392944335938 0 6446.707
737.836181640625 0 6514.2686
741.3804931640625 0 19850.355
742.385498046875 0 11834.048
743.3904418945312 0 6929.453
745.851318359375 0 9465.57
747.3548583984375 0 106183.4 z 9
748.3580932617188 0 45675.36
749.3587036132812 0 11729.208
753.3494873046875 0 8193.289
759.8582153320312 0 37527.78 w 2
760.3587646484375 0 32316.057
760.8597412109375 0 26178.248
761.3665161132812 0 8995.48
763.3743896484375 0 21438.1 y 9
764.376953125 0 14374.765
765.8478393554688 0 23633.238
766.3521728515625 0 16388.2
766.8519287109375 0 13348.303
767.3646240234375 0 8169.0884
768.36962890625 0 10485.316
774.3889770507812 0 58274.156
774.854736328125 0 45382.41 c Ammonia loss 12
775.3604736328125 0 40843.29
775.8563232421875 0 29110.127
776.3724975585938 0 6520.175
782.867919921875 0 7859.2197
783.3675537109375 0 180515.16 c 12
783.868408203125 0 134032.7
784.3707885742188 0 84315.4
784.8693237304688 0 21613.354
785.3970947265625 0 97542.16 c 6
786.4002685546875 0 35029.168
787.3999633789062 0 13340.439
788.6596069335938 0 6602.044
789.8652954101562 0 13674.736 y Ammonia loss 2
790.36181640625 0 25896.547 z 2
790.8609619140625 0 14985.543
791.3707885742188 0 7503.3354
798.3720092773438 0 87540.234 y 2
798.8743286132812 0 69155.164
799.3779296875 0 32288.408
799.8756103515625 0 11525.984
809.891357421875 0 9202.537
816.3959350585938 0 6172.003
817.3853149414062 0 60445.92 w 8
818.3868408203125 0 36785.395
819.3928833007812 0 7529.8755
825.4092407226562 0 25652.557
825.9078979492188 0 14911.772
826.410888671875 0 10036.626
830.8994750976562 0 10405.412 z Water loss 1
833.4032592773438 0 13224.549
833.8912963867188 0 6093.901
838.906982421875 0 14718.675 c Ammonia loss 13
839.403076171875 0 22699.303 y Ammonia loss 1
839.9053955078125 0 50500.824 z 1
840.406982421875 0 45067.77
840.9110717773438 0 21485.062
841.3834228515625 0 8514.355
844.3985595703125 0 5384.024
846.4046020507812 0 16709.197
846.90673828125 0 7101.9575
847.4149780273438 0 442403.2 c 13
847.9157104492188 0 413285.9 y 1
848.4166870117188 0 212281.78
848.9176025390625 0 82976.23
849.4190063476562 0 25414.62
850.8993530273438 0 23574.576
851.4009399414062 0 21564.854
851.8992919921875 0 14965.937
852.9078369140625 0 113433.14
853.4091796875 0 105324.625
853.9099731445312 0 61153.555
854.4131469726562 0 23875.387
854.9114379882812 0 11472.717
855.890625 0 15802.498
859.9183349609375 0 12205.179
860.4168090820312 0 26915.895
860.9185791015625 0 27253.525
861.3982543945312 0 315756.75 z 8
861.9224853515625 0 9184.432
862.401611328125 0 153180.56
862.9082641601562 0 21601.246
863.4047241210938 0 46357.184
864.4055786132812 0 10556.804
869.4163208007812 0 265469.78
869.9179077148438 0 255342.72
870.4177856445312 0 124868.945
870.9183959960938 0 58492.723
871.4183349609375 0 20128.568
876.4070434570312 0 29057.88
877.4166259765625 0 77732.93 y 8
878.4312133789062 0 39622.87
879.4412841796875 0 28784.998
883.41357421875 0 149273.55
883.916015625 0 160958.89
884.41552734375 0 92784.77
884.9149169921875 0 42943.24
885.4173583984375 0 21570.09
890.9183959960938 0 15985.082
891.4230346679688 0 604726.8
891.927001953125 0 3264687.8
892.4282836914062 0 2991920
892.9292602539062 0 1590973.6
893.4298095703125 0 642970.4
893.9310302734375 0 194545.42
905.4344482421875 0 8182.557
922.4560546875 0 548138.9 c 7
923.4591674804688 0 252705.39
924.4598999023438 0 102177.46
925.4616088867188 0 23861.719
998.4557495117188 0 76647.28 z 7
999.4597778320312 0 42974.81
1000.4659423828125 0 15650.154
1014.4732055664062 0 26807.686 y 7
1015.4739379882812 0 15651.477
1016.4957885742188 0 8965.545
1036.499755859375 0 178630.9 c 8
1037.5023193359375 0 110808.5
1038.503173828125 0 39842.98
1039.50537109375 0 9402.817
1068.4871826171875 0 26379.885 w 6
1069.48974609375 0 8970.763
1070.494384765625 0 8255.515
1095.57470703125 0 6316.0625
1111.539794921875 0 84024.43 z 6
1112.5426025390625 0 66976.805
1113.5469970703125 0 31321.758
1114.5479736328125 0 11224.787
1126.516845703125 0 30735.12
1127.5296630859375 0 23069.645
1128.53662109375 0 17067.764
1129.541748046875 0 27082.342
1130.545654296875 0 12543.315
1131.5394287109375 0 6992.635
1173.5584716796875 0 347119.16 c 9
1174.560546875 0 233487.38
1175.56201171875 0 85465.05
1176.562255859375 0 26528.17
1182.5771484375 0 192809.48 z 5
1183.5811767578125 0 176694.17
1184.5845947265625 0 64710.566
1185.5845947265625 0 20446.082
1198.5994873046875 0 11977.112 y 5
1199.6044921875 0 13451.658
1239.6209716796875 0 8456.866
1291.5982666015625 0 8292.801
1292.610107421875 0 9209.169
1293.6116943359375 0 6891.7954 z Water loss 4
1311.619140625 0 206606.66 z 4
1312.6248779296875 0 272696.28
1313.6263427734375 0 147934.06
1314.632080078125 0 45559.64
1315.635009765625 0 10557.342
1316.6268310546875 0 7089.308
1326.6278076171875 0 8124.3535
1327.639892578125 0 18019.865 y 4
1328.6361083984375 0 16733.28
1335.61279296875 0 108654.9
1336.6187744140625 0 188300.61 c 10
1337.621337890625 0 121701.97
1338.6248779296875 0 52666.594
1339.6197509765625 0 10578.218
1436.66357421875 0 15611.285
1437.6673583984375 0 244074.61 c 11
1438.6700439453125 0 181395.06
1439.6732177734375 0 87236.7
1440.6712646484375 0 22271.654
1441.6717529296875 0 6793.3257
1448.67138671875 0 9295.14 z 3
1449.6846923828125 0 67523.79
1450.68701171875 0 61013.21
1451.688720703125 0 21736.557
1452.69970703125 0 7253.895
1522.722900390625 0 23352.016
1523.7239990234375 0 10698.049
1565.724609375 0 123350.53 c 12
1566.728515625 0 117172.69
1567.7303466796875 0 60512.28
1568.7305908203125 0 21468.906
1580.7225341796875 0 47596.047
1581.728271484375 0 41974.254
1582.7353515625 0 17461.953
1649.80517578125 0 8261.766
1650.8084716796875 0 16654.39
1651.820068359375 0 17186.541
1661.797607421875 0 9258.964
1677.8021240234375 0 34244.22 y Ammonia loss 1
1678.8060302734375 0 47643.02 z 1
1679.80078125 0 42360.35
1680.7978515625 0 30514.873
1681.7984619140625 0 20227.92
1693.823974609375 0 45834.88 c 13
1694.826904296875 0 113434.38 y 1
1695.8292236328125 0 78992.65
1696.830322265625 0 50225.21
1697.830078125 0 9683.644
1721.8131103515625 0 8646.555
1722.8092041015625 0 14199.351
1724.8275146484375 0 17412.885
1725.815673828125 0 8778.101
1737.8328857421875 0 37606.703
1738.8336181640625 0 176597.44
1739.8353271484375 0 141076.12
1740.838134765625 0 87501.37
1741.8394775390625 0 36650.984
1742.837158203125 0 11591.935
1748.822265625 0 31873.588
1749.8221435546875 0 25406.861
1750.820556640625 0 10476.057
1751.8172607421875 0 7733.652
1754.8421630859375 0 20214.922
1755.857666015625 0 82313.68
1756.861572265625 0 68528.19
1757.85888671875 0 40216.957
1758.869384765625 0 17437.785
1765.8271484375 0 59726.89
1766.82763671875 0 244334.25
1767.8287353515625 0 239803.39
1768.832275390625 0 113363.74
1769.8341064453125 0 51936.88
1770.83203125 0 18040.045
1781.8369140625 0 46545.312
1782.8446044921875 0 441061.5
1783.8529052734375 0 1775605.8
1784.8551025390625 0 1556933
1785.8568115234375 0 847538.7
1786.859130859375 0 312418.22
1787.8597412109375 0 92523.96
2819.239501953125 0 6867.2837

Spectrum Details

|  |  |
| --- | --- |
| Matched peaks? Matched peaks The total absolute number of peaks matched. Additionally in brackets the total fraction of peaks matched and the total number of peaks is shown. | 74 (20.05% of 369) |
| FDR? FDR The false discovery rate estimated for this peptide. It is calculated by matching all theoretical fragments with a non-integer shift with the raw peaks for this spectrum. This is done with 40 different shifts. The resulting percentage is the average number of annotated peaks over the number of annotated peaks with the correct spectrum. | 0.19% |
| Satellite FDR? Satellite FDR See the FDR for details on its calculation. This satellite ion specific FDR only contains the satellite ions (d/w) for I/L/J positions. | 0.00% |
| PSM Score? PSM Score The PSM Score as given by Hecklib to this annotated spectrum. It is shown with three significant figures. | 476 |

#### Spectrum 2472? Spectrum 2472 The raw spectrum of this peptide as annotated by Hecklib. The fragments are coloured according to ion type (see legend). Any peaks with a star '\*' as text can be hovered over to see the full details, first the ion type second the mass shift type. By hovering over the amino acids in the peptide or ions in the legend the corresponding peaks are highlighted. By toggling the 'Unassigned' label you can turn the background (unassigned) peaks on or off in the plot. By updating the slider in the Ion legend you can update the spectrum to only show the top X% of the peaks with labels. The top X% means any peak that is within X% of the highest intensity. By dragging in the spectrum you can zoom in to a specific part of the spectrum and use 'Zoom Out' to get back to the original zoom level. The annotation of the spectrum is based on the given sequence in the peptides file and is done with different software so inconsistencies are likely. The peaks are annotated based on the given sequence, with 20 ppm tolerance.

Copy Data

##### Spectrum 2472 (TSV)

###### Preview

```
Loading example...
```

*Click on the button to copy the data to your clipboard.*

Mz MinMz MaxIntensity Max

WidthHeightPeptide font sizePeptide stroke widthSpectrum font sizeSpectrum stroke widthCompact peptide

Ion legend

wxyz

abcd

OtherUnassignedIon^Charge^_Position_Show for top:%

SVMHEAJHNHYTQKS

04.14e+48.28e+41.24e+51.66e+5

Zoom Out

z^+1^_2_y^+1^_2_w^+1^_3_z^+1^_3_c^+1^_3_w^+1^_4_y^+1^_4_z^+1^_4_y^+1^_4_c^+1^_4_y^+3^_13_z^+3^_14_z^+3^_14_c^+3^_14_y^+3^_14_z^+3^_14_y^+3^_14_c^+2^_10_z^+1^_5_c^+1^_5_y^+1^_5_c^+1^_6_z^+2^_12_y^+2^_12_z^+1^_6_y^+1^_6_c^+2^_13_c^+2^_13_z^+2^_13_c^+1^_7_y^+2^_13_w^+1^_7_c^+2^_14_z^+2^_14_c^+2^_14_y^+2^_14_y^+1^_7_y^+1^_7_z^+1^_7_y^+1^_7_c^+1^_8_z^+1^_8_y^+1^_8_c^+1^_9_w^+1^_9_z^+1^_9_z^+1^_10_c^+1^_10_y^+1^_10_z^+1^_11_y^+1^_11_c^+1^_11_z^+1^_12_c^+1^_12_c^+1^_13_z^+1^_14_c^+1^_14_y^+1^_14_

045691113671823

Fragment Matches Table

Show background peaks

| Position | Ion type | Intensity | mz Theoretical | mz Error (Th) | mz Error (ppm) | Charge | Series Number |
| --- | --- | --- | --- | --- | --- | --- | --- |
| - | - | 367.5 | 123.4 | - | - | 0 | - |
| - | - | 493.2 | 128 | - | - | 0 | - |
| - | - | 3607 | 129.1 | - | - | 0 | - |
| - | - | 366.9 | 133.8 | - | - | 0 | - |
| - | - | 500.4 | 148.9 | - | - | 0 | - |
| - | - | 498.8 | 148.9 | - | - | 0 | - |
| - | - | 761.6 | 148.9 | - | - | 0 | - |
| - | - | 1151 | 148.9 | - | - | 0 | - |
| - | - | 1319 | 148.9 | - | - | 0 | - |
| - | - | 2297 | 148.9 | - | - | 0 | - |
| - | - | 3805 | 148.9 | - | - | 0 | - |
| - | - | 3706 | 149 | - | - | 0 | - |
| - | - | 1959 | 149 | - | - | 0 | - |
| - | - | 945.2 | 149 | - | - | 0 | - |
| - | - | 942.2 | 149 | - | - | 0 | - |
| - | - | 697.4 | 149 | - | - | 0 | - |
| - | - | 701.3 | 149 | - | - | 0 | - |
| - | - | 675.6 | 149 | - | - | 0 | - |
| - | - | 552.9 | 149 | - | - | 0 | - |
| - | - | 443.4 | 149 | - | - | 0 | - |
| - | - | 460.8 | 149 | - | - | 0 | - |
| - | - | 502.7 | 149 | - | - | 0 | - |
| - | - | 410.4 | 149.1 | - | - | 0 | - |
| - | - | 1554 | 155.1 | - | - | 0 | - |
| - | - | 2115 | 159.1 | - | - | 0 | - |
| - | - | 494.1 | 169.6 | - | - | 0 | - |
| - | - | 621.3 | 173.4 | - | - | 0 | - |
| - | - | 754.1 | 183.1 | - | - | 0 | - |
| - | - | 478 | 183.9 | - | - | 0 | - |
| - | - | 973.6 | 187.1 | - | - | 0 | - |
| - | - | 468.6 | 208 | - | - | 0 | - |
| 14 | z | 805.2 | 218.1 | 3.591E-05 | 0.1646 | +1 | 2 |
| 14 | y | 1414 | 234.1 | 0.0002022 | 0.8637 | +1 | 2 |
| - | - | 498.5 | 235.4 | - | - | 0 | - |
| - | - | 518.6 | 256.8 | - | - | 0 | - |
| 13 | w | 3354 | 288.2 | 0.0007002 | 2.43 | +1 | 3 |
| - | - | 575.1 | 302.1 | - | - | 0 | - |
| - | - | 822.6 | 334.1 | - | - | 0 | - |
| 13 | z | 7355 | 346.2 | 0.0004641 | 1.341 | +1 | 3 |
| - | - | 1882 | 347.2 | - | - | 0 | - |
| 3 | c | 1000 | 351.2 | 0.005597 | 15.94 | +1 | 3 |
| - | - | 896.8 | 355.1 | - | - | 0 | - |
| - | - | 580.7 | 367.4 | - | - | 0 | - |
| - | - | 610 | 373.2 | - | - | 0 | - |
| - | - | 683.4 | 395.7 | - | - | 0 | - |
| - | - | 910.5 | 424.2 | - | - | 0 | - |
| - | - | 689.1 | 428.2 | - | - | 0 | - |
| 12 | w | 672 | 430.2 | 0.000804 | 1.869 | +1 | 4 |
| - | - | 729.7 | 439.7 | - | - | 0 | - |
| - | - | 1216 | 443.2 | - | - | 0 | - |
| - | - | 711 | 444.2 | - | - | 0 | - |
| - | - | 579 | 445.2 | - | - | 0 | - |
| 12 | y | 1360 | 446.2 | 5.215E-05 | 0.1169 | +1 | 4 |
| 12 | z | 4977 | 447.2 | 0.0001184 | 0.2648 | +1 | 4 |
| - | - | 9335 | 448.2 | - | - | 0 | - |
| - | - | 2296 | 449.2 | - | - | 0 | - |
| 12 | y | 715.6 | 463.3 | 0.001455 | 3.14 | +1 | 4 |
| - | - | 561.2 | 471.2 | - | - | 0 | - |
| 4 | c | 7705 | 488.2 | 0.005828 | 11.94 | +1 | 4 |
| - | - | 2230 | 489.2 | - | - | 0 | - |
| 3 | y | 1197 | 537.9 | 0.003978 | 7.395 | +3 | 13 |
| - | - | 1925 | 538.2 | - | - | 0 | - |
| - | - | 805 | 538.6 | - | - | 0 | - |
| - | - | 1382 | 559.3 | - | - | 0 | - |
| 2 | z | 1937 | 559.6 | 0.001149 | 2.054 | +3 | 14 |
| 2 | z | 1226 | 559.9 | 0.003222 | 5.755 | +3 | 14 |
| 14 | c | 3966 | 564.9 | 0.002029 | 3.591 | +3 | 14 |
| 2 | y | 3254 | 565.3 | 0.005566 | 9.848 | +3 | 14 |
| 2 | z | 1762 | 565.6 | 0.004342 | 7.676 | +3 | 14 |
| - | - | 1390 | 565.9 | - | - | 0 | - |
| 2 | y | 1874 | 570.9 | 0.001437 | 2.517 | +3 | 14 |
| - | - | 856.6 | 571.3 | - | - | 0 | - |
| - | - | 1016 | 571.9 | - | - | 0 | - |
| - | - | 962.7 | 572.3 | - | - | 0 | - |
| - | - | 1068 | 572.9 | - | - | 0 | - |
| - | - | 1193 | 573.3 | - | - | 0 | - |
| - | - | 2566 | 578.6 | - | - | 0 | - |
| - | - | 5187 | 579 | - | - | 0 | - |
| - | - | 2675 | 579.3 | - | - | 0 | - |
| - | - | 835.6 | 579.6 | - | - | 0 | - |
| - | - | 762.7 | 582.3 | - | - | 0 | - |
| - | - | 945.6 | 588.6 | - | - | 0 | - |
| - | - | 2413 | 593.9 | - | - | 0 | - |
| - | - | 878.6 | 594 | - | - | 0 | - |
| - | - | 5214 | 594.3 | - | - | 0 | - |
| - | - | 5038 | 594.6 | - | - | 0 | - |
| - | - | 2921 | 594.9 | - | - | 0 | - |
| 10 | c | 1822 | 595.3 | 0.0002837 | 0.4766 | +2 | 10 |
| - | - | 1025 | 599.3 | - | - | 0 | - |
| - | - | 1201 | 599.7 | - | - | 0 | - |
| - | - | 4.101E+04 | 599.9 | - | - | 0 | - |
| - | - | 6.638E+04 | 600.3 | - | - | 0 | - |
| - | - | 3.448E+04 | 600.6 | - | - | 0 | - |
| - | - | 1.401E+04 | 601 | - | - | 0 | - |
| - | - | 1.208E+04 | 601.3 | - | - | 0 | - |
| - | - | 2036 | 601.6 | - | - | 0 | - |
| 11 | z | 1.23E+04 | 610.3 | 0.0006327 | 1.037 | +1 | 5 |
| - | - | 3819 | 611.3 | - | - | 0 | - |
| - | - | 779 | 612.3 | - | - | 0 | - |
| 5 | c | 1.533E+04 | 617.3 | 0.005288 | 8.567 | +1 | 5 |
| - | - | 4515 | 618.3 | - | - | 0 | - |
| - | - | 1640 | 619.3 | - | - | 0 | - |
| - | - | 591 | 625.1 | - | - | 0 | - |
| 11 | y | 1395 | 626.3 | 0.0002081 | 0.3323 | +1 | 5 |
| - | - | 825.2 | 653.3 | - | - | 0 | - |
| 6 | c | 6345 | 688.3 | 0.005161 | 7.499 | +1 | 6 |
| - | - | 2484 | 689.3 | - | - | 0 | - |
| - | - | 764.9 | 690.3 | - | - | 0 | - |
| 4 | z | 2830 | 724.8 | 0.0001211 | 0.167 | +2 | 12 |
| - | - | 2641 | 725.3 | - | - | 0 | - |
| - | - | 1010 | 725.8 | - | - | 0 | - |
| - | - | 1098 | 730.4 | - | - | 0 | - |
| 4 | y | 1801 | 732.9 | 0.001306 | 1.783 | +2 | 12 |
| - | - | 2098 | 733.4 | - | - | 0 | - |
| - | - | 1454 | 733.9 | - | - | 0 | - |
| 10 | z | 6417 | 747.4 | 9.373E-06 | 0.01254 | +1 | 6 |
| - | - | 2777 | 748.4 | - | - | 0 | - |
| - | - | 814.2 | 749.4 | - | - | 0 | - |
| - | - | 1402 | 757.4 | - | - | 0 | - |
| 10 | y | 1139 | 763.4 | 0.002662 | 3.488 | +1 | 6 |
| - | - | 671.2 | 764.4 | - | - | 0 | - |
| - | - | 1799 | 767.4 | - | - | 0 | - |
| - | - | 1738 | 767.9 | - | - | 0 | - |
| - | - | 1128 | 773.8 | - | - | 0 | - |
| - | - | 2045 | 774.4 | - | - | 0 | - |
| - | - | 1095 | 775.4 | - | - | 0 | - |
| - | - | 905.9 | 775.5 | - | - | 0 | - |
| - | - | 859.8 | 775.8 | - | - | 0 | - |
| 13 | c | 1562 | 782.8 | 0.003717 | 4.748 | +2 | 13 |
| - | - | 2493 | 783.4 | - | - | 0 | - |
| - | - | 1392 | 783.9 | - | - | 0 | - |
| - | - | 752.1 | 784.4 | - | - | 0 | - |
| - | - | 593.3 | 785.4 | - | - | 0 | - |
| 13 | c | 9924 | 791.4 | 0.002222 | 2.808 | +2 | 13 |
| - | - | 1.093E+04 | 791.9 | - | - | 0 | - |
| - | - | 5364 | 792.4 | - | - | 0 | - |
| - | - | 2971 | 792.9 | - | - | 0 | - |
| 3 | z | 718.7 | 798.4 | 0.008438 | 10.57 | +2 | 13 |
| 7 | c | 5202 | 801.4 | 0.005143 | 6.418 | +1 | 7 |
| - | - | 1831 | 802.4 | - | - | 0 | - |
| - | - | 1434 | 803.4 | - | - | 0 | - |
| 3 | y | 2422 | 806.4 | 0.007133 | 8.846 | +2 | 13 |
| - | - | 2586 | 806.9 | - | - | 0 | - |
| - | - | 1262 | 807.4 | - | - | 0 | - |
| - | - | 902.9 | 807.9 | - | - | 0 | - |
| 9 | w | 2290 | 817.4 | 0.0002006 | 0.2454 | +1 | 7 |
| - | - | 1669 | 818.4 | - | - | 0 | - |
| - | - | 1490 | 833.4 | - | - | 0 | - |
| - | - | 1089 | 833.9 | - | - | 0 | - |
| - | - | 731.9 | 834.4 | - | - | 0 | - |
| - | - | 604.3 | 839.4 | - | - | 0 | - |
| - | - | 680.7 | 839.9 | - | - | 0 | - |
| - | - | 640.7 | 845.9 | - | - | 0 | - |
| 14 | c | 713.6 | 846.4 | 0.0007311 | 0.8638 | +2 | 14 |
| 2 | z | 3105 | 847.9 | 0.01171 | 13.81 | +2 | 14 |
| - | - | 1291 | 848.4 | - | - | 0 | - |
| - | - | 1944 | 848.9 | - | - | 0 | - |
| - | - | 1358 | 853.9 | - | - | 0 | - |
| - | - | 1446 | 854.4 | - | - | 0 | - |
| - | - | 916.2 | 854.9 | - | - | 0 | - |
| 14 | c | 1.633E+04 | 855.4 | 0.001677 | 1.96 | +2 | 14 |
| 2 | y | 1.952E+04 | 855.9 | 0.009242 | 10.8 | +2 | 14 |
| - | - | 1.25E+04 | 856.4 | - | - | 0 | - |
| - | - | 6110 | 856.9 | - | - | 0 | - |
| - | - | 2605 | 857.4 | - | - | 0 | - |
| - | - | 801.7 | 857.9 | - | - | 0 | - |
| 9 | y | 1453 | 859.4 | 0.001944 | 2.262 | +1 | 7 |
| - | - | 860.9 | 859.9 | - | - | 0 | - |
| 9 | y | 1215 | 860.4 | 0.01142 | 13.27 | +1 | 7 |
| - | - | 745.9 | 860.9 | - | - | 0 | - |
| 9 | z | 1.121E+04 | 861.4 | 0.0001935 | 0.2246 | +1 | 7 |
| - | - | 1.264E+04 | 862.4 | - | - | 0 | - |
| - | - | 5627 | 863.4 | - | - | 0 | - |
| - | - | 1169 | 863.9 | - | - | 0 | - |
| - | - | 3322 | 864.4 | - | - | 0 | - |
| - | - | 1371 | 867.9 | - | - | 0 | - |
| - | - | 1149 | 868.4 | - | - | 0 | - |
| - | - | 2625 | 868.9 | - | - | 0 | - |
| - | - | 2151 | 869.4 | - | - | 0 | - |
| - | - | 1537 | 870.4 | - | - | 0 | - |
| - | - | 1125 | 870.9 | - | - | 0 | - |
| - | - | 1593 | 876.4 | - | - | 0 | - |
| 9 | y | 1.208E+04 | 877.4 | 0.001523 | 1.735 | +1 | 7 |
| - | - | 1.157E+04 | 877.9 | - | - | 0 | - |
| - | - | 8670 | 878.4 | - | - | 0 | - |
| - | - | 3658 | 878.9 | - | - | 0 | - |
| - | - | 2655 | 879.4 | - | - | 0 | - |
| - | - | 1416 | 883.4 | - | - | 0 | - |
| - | - | 1079 | 890.5 | - | - | 0 | - |
| - | - | 1.078E+04 | 891.4 | - | - | 0 | - |
| - | - | 1.277E+04 | 891.9 | - | - | 0 | - |
| - | - | 9363 | 892.4 | - | - | 0 | - |
| - | - | 6154 | 892.9 | - | - | 0 | - |
| - | - | 2918 | 893.4 | - | - | 0 | - |
| - | - | 1192 | 893.9 | - | - | 0 | - |
| - | - | 2461 | 894.4 | - | - | 0 | - |
| - | - | 2153 | 895.4 | - | - | 0 | - |
| - | - | 858.9 | 896.4 | - | - | 0 | - |
| - | - | 1304 | 899 | - | - | 0 | - |
| - | - | 2.263E+04 | 899.4 | - | - | 0 | - |
| - | - | 1503 | 899.5 | - | - | 0 | - |
| - | - | 1.532E+05 | 899.9 | - | - | 0 | - |
| - | - | 1.64E+05 | 900.4 | - | - | 0 | - |
| - | - | 9.962E+04 | 900.9 | - | - | 0 | - |
| - | - | 4.814E+04 | 901.4 | - | - | 0 | - |
| - | - | 1.566E+04 | 901.9 | - | - | 0 | - |
| - | - | 4349 | 902.4 | - | - | 0 | - |
| 8 | c | 3.126E+04 | 938.4 | 0.00513 | 5.466 | +1 | 8 |
| - | - | 1.548E+04 | 939.5 | - | - | 0 | - |
| - | - | 4945 | 940.5 | - | - | 0 | - |
| - | - | 1925 | 941.5 | - | - | 0 | - |
| - | - | 815.5 | 942.4 | - | - | 0 | - |
| 8 | z | 3233 | 998.5 | 0.0003429 | 0.3434 | +1 | 8 |
| - | - | 2014 | 999.5 | - | - | 0 | - |
| 8 | y | 2139 | 1014 | 0.00123 | 1.213 | +1 | 8 |
| - | - | 1318 | 1015 | - | - | 0 | - |
| - | - | 830 | 1049 | - | - | 0 | - |
| 9 | c | 9314 | 1052 | 0.0045 | 4.275 | +1 | 9 |
| - | - | 6558 | 1053 | - | - | 0 | - |
| - | - | 2035 | 1054 | - | - | 0 | - |
| 7 | w | 1592 | 1068 | 0.005844 | 5.47 | +1 | 9 |
| - | - | 832 | 1069 | - | - | 0 | - |
| 7 | z | 4036 | 1112 | 0.001446 | 1.301 | +1 | 9 |
| - | - | 4287 | 1113 | - | - | 0 | - |
| - | - | 1671 | 1114 | - | - | 0 | - |
| - | - | 1057 | 1127 | - | - | 0 | - |
| - | - | 969.4 | 1147 | - | - | 0 | - |
| 6 | z | 8427 | 1183 | 0.001816 | 1.536 | +1 | 10 |
| - | - | 8532 | 1184 | - | - | 0 | - |
| - | - | 4437 | 1185 | - | - | 0 | - |
| - | - | 748.9 | 1186 | - | - | 0 | - |
| 10 | c | 1.174E+04 | 1190 | 0.003449 | 2.9 | +1 | 10 |
| - | - | 1.078E+04 | 1191 | - | - | 0 | - |
| - | - | 4988 | 1192 | - | - | 0 | - |
| - | - | 1619 | 1193 | - | - | 0 | - |
| - | - | 866.9 | 1194 | - | - | 0 | - |
| 6 | y | 1092 | 1199 | 0.001131 | 0.9438 | +1 | 10 |
| - | - | 1771 | 1204 | - | - | 0 | - |
| 5 | z | 1.136E+04 | 1312 | 0.002539 | 1.936 | +1 | 11 |
| - | - | 1.147E+04 | 1313 | - | - | 0 | - |
| - | - | 4764 | 1314 | - | - | 0 | - |
| - | - | 2082 | 1315 | - | - | 0 | - |
| 5 | y | 1504 | 1328 | 0.002098 | 1.58 | +1 | 11 |
| - | - | 1428 | 1329 | - | - | 0 | - |
| - | - | 5704 | 1352 | - | - | 0 | - |
| 11 | c | 8272 | 1353 | 0.001774 | 1.311 | +1 | 11 |
| - | - | 5907 | 1354 | - | - | 0 | - |
| - | - | 4308 | 1355 | - | - | 0 | - |
| - | - | 977.6 | 1356 | - | - | 0 | - |
| - | - | 741.5 | 1397 | - | - | 0 | - |
| 4 | z | 1247 | 1449 | 0.007984 | 5.511 | +1 | 12 |
| - | - | 3445 | 1450 | - | - | 0 | - |
| - | - | 3906 | 1451 | - | - | 0 | - |
| - | - | 1748 | 1452 | - | - | 0 | - |
| - | - | 1211 | 1453 | - | - | 0 | - |
| 12 | c | 9193 | 1454 | 0.0002304 | 0.1585 | +1 | 12 |
| - | - | 1.063E+04 | 1455 | - | - | 0 | - |
| - | - | 6230 | 1456 | - | - | 0 | - |
| - | - | 2211 | 1457 | - | - | 0 | - |
| - | - | 930 | 1539 | - | - | 0 | - |
| - | - | 607.6 | 1541 | - | - | 0 | - |
| - | - | 831 | 1569 | - | - | 0 | - |
| 13 | c | 5900 | 1582 | 0.000857 | 0.5418 | +1 | 13 |
| - | - | 6419 | 1583 | - | - | 0 | - |
| - | - | 4011 | 1584 | - | - | 0 | - |
| - | - | 1419 | 1585 | - | - | 0 | - |
| - | - | 725.7 | 1586 | - | - | 0 | - |
| - | - | 1950 | 1597 | - | - | 0 | - |
| - | - | 2395 | 1598 | - | - | 0 | - |
| - | - | 1637 | 1599 | - | - | 0 | - |
| - | - | 820.9 | 1668 | - | - | 0 | - |
| 2 | z | 1815 | 1695 | 0.01531 | 9.033 | +1 | 14 |
| - | - | 2607 | 1696 | - | - | 0 | - |
| - | - | 1963 | 1697 | - | - | 0 | - |
| - | - | 996.2 | 1698 | - | - | 0 | - |
| 14 | c | 2587 | 1710 | 0.002207 | 1.291 | +1 | 14 |
| 2 | y | 3827 | 1711 | 0.02283 | 13.34 | +1 | 14 |
| - | - | 6020 | 1712 | - | - | 0 | - |
| - | - | 3646 | 1713 | - | - | 0 | - |
| - | - | 1509 | 1714 | - | - | 0 | - |
| - | - | 1739 | 1736 | - | - | 0 | - |
| - | - | 1745 | 1737 | - | - | 0 | - |
| - | - | 1484 | 1738 | - | - | 0 | - |
| - | - | 1015 | 1739 | - | - | 0 | - |
| - | - | 1438 | 1741 | - | - | 0 | - |
| - | - | 1310 | 1754 | - | - | 0 | - |
| - | - | 6912 | 1755 | - | - | 0 | - |
| - | - | 7421 | 1756 | - | - | 0 | - |
| - | - | 4129 | 1757 | - | - | 0 | - |
| - | - | 1407 | 1758 | - | - | 0 | - |
| - | - | 1436 | 1765 | - | - | 0 | - |
| - | - | 955.9 | 1766 | - | - | 0 | - |
| - | - | 2768 | 1772 | - | - | 0 | - |
| - | - | 3597 | 1773 | - | - | 0 | - |
| - | - | 3293 | 1774 | - | - | 0 | - |
| - | - | 1142 | 1775 | - | - | 0 | - |
| - | - | 756.8 | 1776 | - | - | 0 | - |
| - | - | 3764 | 1782 | - | - | 0 | - |
| - | - | 1.277E+04 | 1783 | - | - | 0 | - |
| - | - | 1.566E+04 | 1784 | - | - | 0 | - |
| - | - | 9661 | 1785 | - | - | 0 | - |
| - | - | 5566 | 1786 | - | - | 0 | - |
| - | - | 1760 | 1787 | - | - | 0 | - |
| - | - | 827.3 | 1788 | - | - | 0 | - |
| - | - | 1915 | 1798 | - | - | 0 | - |
| - | - | 2.096E+04 | 1799 | - | - | 0 | - |
| - | - | 7.256E+04 | 1800 | - | - | 0 | - |
| - | - | 7.836E+04 | 1801 | - | - | 0 | - |
| - | - | 5.203E+04 | 1802 | - | - | 0 | - |
| - | - | 2.494E+04 | 1803 | - | - | 0 | - |
| - | - | 8354 | 1804 | - | - | 0 | - |
| - | - | 2984 | 1805 | - | - | 0 | - |

m/z Charge Intensity FragmentType MassShift Position
123.40428161621094 0 367.5267
128.02517700195312 0 493.19827
129.10252380371094 0 3606.7769
133.7906494140625 0 366.8963
148.899658203125 0 500.40134
148.90682983398438 0 498.81436
148.91444396972656 0 761.6039
148.92132568359375 0 1151.3102
148.92868041992188 0 1319.2971
148.9359130859375 0 2297.314
148.9436798095703 0 3804.6252
148.9602813720703 0 3706.4165
148.968017578125 0 1959.2568
148.97511291503906 0 945.17035
148.98294067382812 0 942.2254
148.99002075195312 0 697.4468
148.99671936035156 0 701.30133
149.00437927246094 0 675.6319
149.01153564453125 0 552.9345
149.01893615722656 0 443.4082
149.02670288085938 0 460.8179
149.03384399414062 0 502.73572
149.0772247314453 0 410.37247
155.09312438964844 0 1553.9688
159.113037109375 0 2115.013
169.60659790039062 0 494.0994
173.4496612548828 0 621.28204
183.11312866210938 0 754.1102
183.93887329101562 0 478.0158
187.1077117919922 0 973.6022
208.01268005371094 0 468.60547
218.1261444091797 0 805.1902 z 13
234.14503479003906 0 1413.8223 y 13
235.3890380859375 0 498.4797
256.75390625 0 518.64453
288.1560974121094 0 3354.3093 w 12
302.1297912597656 0 575.07745
334.1436462402344 0 822.6025
346.1851501464844 0 7354.9053 z 12
347.1893310546875 0 1881.722
351.17034912109375 0 1000.33606 c 2
355.07000732421875 0 896.81195
367.4265441894531 0 580.712
373.17669677734375 0 609.96765
395.66595458984375 0 683.3813
424.231201171875 0 910.48444
428.1971130371094 0 689.1249
430.22882080078125 0 671.9738 w 11
439.7117919921875 0 729.6669
443.20867919921875 0 1215.7665
444.21551513671875 0 711.0104
445.21746826171875 0 578.97284
446.2244873046875 0 1359.6249 y Ammonia loss 11
447.23248291015625 0 4976.783 z 11
448.2401428222656 0 9334.935
449.243408203125 0 2295.7395
463.2496337890625 0 715.64124 y 11
471.21966552734375 0 561.1539
488.2294921875 0 7705.2075 c 3
489.23101806640625 0 2230.0613
537.9182739257812 0 1196.5986 y 2
538.2496337890625 0 1924.6382
538.5831298828125 0 804.9576
559.25927734375 0 1382.1338
559.5951538085938 0 1937.4402 z Water loss 1
559.9252319335938 0 1226.0281 z Ammonia loss 1
564.9356079101562 0 3965.611 c Ammonia loss 13
565.2671508789062 0 3253.6558 y Ammonia loss 1
565.6018676757812 0 1761.5864 z 1
565.9340209960938 0 1389.9731
570.9385375976562 0 1873.5378 y 1
571.2734375 0 856.571
571.9347534179688 0 1016.1091
572.280029296875 0 962.706
572.9446411132812 0 1067.8695
573.275146484375 0 1193.3752
578.6173706054688 0 2566.3113
578.9503784179688 0 5186.6963
579.2830810546875 0 2675.0713
579.6190185546875 0 835.6228
582.2650756835938 0 762.6511
588.605712890625 0 945.63257
593.9444580078125 0 2413.1365
593.9982299804688 0 878.64844
594.2774658203125 0 5214.176
594.6090698242188 0 5037.847
594.9439697265625 0 2921.3833
595.2780151367188 0 1821.8113 c 9
599.337158203125 0 1024.5695
599.6729736328125 0 1200.716
599.9498291015625 0 41013.32
600.2803955078125 0 66380.46
600.6160278320312 0 34477.84
600.9503784179688 0 14008.23
601.280029296875 0 12082.32
601.6168823242188 0 2036.0873
610.2963256835938 0 12299.37 z 10
611.3007202148438 0 3818.7288
612.300048828125 0 779.02966
617.2715454101562 0 15334.979 c 4
618.2738647460938 0 4515.4985
619.2702026367188 0 1640.3484
625.0728759765625 0 590.956
626.314208984375 0 1394.7292 y 10
653.3011474609375 0 825.20026
688.3085327148438 0 6345.453 c 5
689.3098754882812 0 2484.2346
690.3091430664062 0 764.91327
724.8433227539062 0 2829.7031 z 3
725.3403930664062 0 2640.72
725.8453369140625 0 1009.9899
730.3768310546875 0 1098.3638
732.8512573242188 0 1801.1774 y 3
733.3511962890625 0 2097.8262
733.8514404296875 0 1453.5944
747.3546142578125 0 6416.516 z 9
748.3618774414062 0 2776.6995
749.3638305664062 0 814.24774
757.3778076171875 0 1402.4728
763.3706665039062 0 1139.2113 y 9
764.3709106445312 0 671.17255
767.3682250976562 0 1799.4178
767.8699951171875 0 1737.5024
773.8461303710938 0 1128.1989
774.383544921875 0 2045.2382
775.3868408203125 0 1095.144
775.462890625 0 905.93304
775.8435668945312 0 859.84125
782.8529663085938 0 1561.9775 c Ammonia loss 12
783.35205078125 0 2493.3428
783.8540649414062 0 1391.9742
784.3629150390625 0 752.05194
785.3637084960938 0 593.3178
791.36474609375 0 9923.632 c 12
791.86376953125 0 10931.23
792.364013671875 0 5363.8296
792.8641967773438 0 2970.617
798.3668823242188 0 718.66376 z 2
801.392578125 0 5201.8027 c 6
802.3977661132812 0 1830.679
803.3897705078125 0 1434.3407
806.3749389648438 0 2421.9512 y 2
806.8722534179688 0 2586.4934
807.3709106445312 0 1261.8586
807.8666381835938 0 902.90955
817.3840942382812 0 2289.543 w 8
818.3867797851562 0 1668.7384
833.4048461914062 0 1489.839
833.9066162109375 0 1089.0802
834.4054565429688 0 731.8548
839.399169921875 0 604.25946
839.9000854492188 0 680.66876
845.8961181640625 0 640.7352
846.4039916992188 0 713.60077 c Water loss 13
847.9043579101562 0 3104.647 z 1
848.402587890625 0 1291.1333
848.9058837890625 0 1944.0629
853.9116821289062 0 1358.4666
854.412109375 0 1446.1139
854.9108276367188 0 916.22266
855.4116821289062 0 16329.299 c 13
855.9112548828125 0 19523.752 y 1
856.41064453125 0 12504.495
856.9129638671875 0 6110.3267
857.4124145507812 0 2605.1287
857.9051513671875 0 801.7067
859.4037475585938 0 1452.693 y Water loss 8
859.905517578125 0 860.89246
860.401123046875 0 1214.5259 y Ammonia loss 8
860.9061279296875 0 745.9427
861.3973388671875 0 11206.219 z 8
862.3917846679688 0 12637.551
863.3888549804688 0 5627.0938
863.8876953125 0 1169.4111
864.3897705078125 0 3321.5984
867.9240112304688 0 1370.5077
868.4061279296875 0 1149.4294
868.9126586914062 0 2625.329
869.4141845703125 0 2150.5625
870.4042358398438 0 1536.7905
870.900146484375 0 1124.8547
876.4144287109375 0 1593.4875
877.4147338867188 0 12079.733 y 8
877.9151000976562 0 11574.058
878.4132690429688 0 8670.095
878.9140014648438 0 3657.9395
879.4130249023438 0 2654.8474
883.4039916992188 0 1416.4886
890.492431640625 0 1078.873
891.4127197265625 0 10777.128
891.912841796875 0 12771.757
892.41162109375 0 9363.049
892.9122924804688 0 6153.9062
893.4120483398438 0 2917.9998
893.9000854492188 0 1191.6888
894.4357299804688 0 2461.363
895.447509765625 0 2153.3062
896.445556640625 0 858.87537
899.0105590820312 0 1304.2506
899.4197998046875 0 22627.387
899.5062866210938 0 1502.6893
899.9234008789062 0 153240.95
900.4238891601562 0 164025.31
900.9235229492188 0 99617.02
901.42431640625 0 48143.66
901.9237060546875 0 15658.598
902.4219360351562 0 4348.6025
938.4514770507812 0 31258.043 c 7
939.4534301757812 0 15478.734
940.4539184570312 0 4945.1045
941.4542236328125 0 1924.5112
942.4462280273438 0 815.4569
998.456787109375 0 3233.039 z 7
999.4596557617188 0 2013.7856
1014.4739379882812 0 2138.9204 y 7
1015.4718017578125 0 1318.1501
1048.5196533203125 0 830.0393
1052.4937744140625 0 9314.492 c 8
1053.4915771484375 0 6557.735
1054.4893798828125 0 2035.1667
1068.4915771484375 0 1592.283 w 6
1069.486328125 0 832.01416
1111.5390625 0 4036.3494 z 6
1112.539306640625 0 4287.4854
1113.5467529296875 0 1671.4895
1126.515625 0 1057.061
1146.535400390625 0 969.37354
1182.5758056640625 0 8427.449 z 5
1183.5758056640625 0 8531.897
1184.5751953125 0 4436.5894
1185.5792236328125 0 748.88477
1189.5516357421875 0 11740.47 c 9
1190.55029296875 0 10776.281
1191.548828125 0 4987.5635
1192.5452880859375 0 1618.8369
1193.54541015625 0 866.9428
1198.59521484375 0 1091.819 y 5
1203.521728515625 0 1770.7694
1311.61767578125 0 11360.074 z 4
1312.6165771484375 0 11469.383
1313.6201171875 0 4764.343
1314.6160888671875 0 2081.7932
1327.6368408203125 0 1503.6447 y 4
1328.633056640625 0 1427.5936
1351.6033935546875 0 5703.6777
1352.6097412109375 0 8272.225 c 10
1353.6097412109375 0 5906.825
1354.610107421875 0 4308.1777
1355.6104736328125 0 977.60864
1396.6207275390625 0 741.45685
1448.671142578125 0 1247.2692 z 3
1449.67822265625 0 3444.842
1450.6846923828125 0 3905.8794
1451.6842041015625 0 1747.9902
1452.66357421875 0 1211.1866
1453.659423828125 0 9193.069 c 11
1454.6585693359375 0 10633.602
1455.6610107421875 0 6229.8823
1456.6605224609375 0 2210.6167
1538.71337890625 0 930.02094
1541.1448974609375 0 607.59644
1568.720947265625 0 831.01605
1581.7186279296875 0 5900.3096 c 12
1582.715576171875 0 6419.326
1583.7127685546875 0 4010.508
1584.71533203125 0 1419.0103
1585.6849365234375 0 725.7452
1596.718505859375 0 1949.8334
1597.7152099609375 0 2395.4233
1598.710693359375 0 1637.0648
1667.7994384765625 0 820.8648
1694.7933349609375 0 1815.121 z 1
1695.7877197265625 0 2606.7107
1696.7919921875 0 1963.0093
1697.77880859375 0 996.1663
1709.81494140625 0 2587.099 c 13
1710.819580078125 0 3826.9937 y 1
1711.8165283203125 0 6019.656
1712.81982421875 0 3646.387
1713.816650390625 0 1509.297
1735.843017578125 0 1739.1461
1736.842529296875 0 1744.9366
1737.8333740234375 0 1484.267
1738.83154296875 0 1015.35803
1740.8218994140625 0 1437.9196
1753.8209228515625 0 1310.2286
1754.8240966796875 0 6911.6133
1755.824951171875 0 7421.181
1756.82080078125 0 4128.84
1757.8240966796875 0 1407.0461
1764.8023681640625 0 1436.005
1765.81005859375 0 955.86816
1771.8468017578125 0 2768.1228
1772.8475341796875 0 3596.77
1773.84619140625 0 3293.0894
1774.8514404296875 0 1141.8699
1775.8507080078125 0 756.7547
1781.8238525390625 0 3764.4656
1782.8201904296875 0 12772.59
1783.8189697265625 0 15660.624
1784.81884765625 0 9661.218
1785.8187255859375 0 5565.6855
1786.8072509765625 0 1760.0242
1787.817626953125 0 827.31146
1797.8262939453125 0 1915.3134
1798.8377685546875 0 20958.486
1799.8416748046875 0 72558.3
1800.842529296875 0 78363
1801.8431396484375 0 52027.39
1802.8428955078125 0 24937.646
1803.8426513671875 0 8353.695
1804.839599609375 0 2983.858

Spectrum Details

|  |  |
| --- | --- |
| Matched peaks? Matched peaks The total absolute number of peaks matched. Additionally in brackets the total fraction of peaks matched and the total number of peaks is shown. | 58 (18.59% of 312) |
| FDR? FDR The false discovery rate estimated for this peptide. It is calculated by matching all theoretical fragments with a non-integer shift with the raw peaks for this spectrum. This is done with 40 different shifts. The resulting percentage is the average number of annotated peaks over the number of annotated peaks with the correct spectrum. | 0.45% |
| Satellite FDR? Satellite FDR See the FDR for details on its calculation. This satellite ion specific FDR only contains the satellite ions (d/w) for I/L/J positions. | 0.00% |
| PSM Score? PSM Score The PSM Score as given by Hecklib to this annotated spectrum. It is shown with three significant figures. | 426 |

#### Spectrum 3013? Spectrum 3013 The raw spectrum of this peptide as annotated by Hecklib. The fragments are coloured according to ion type (see legend). Any peaks with a star '\*' as text can be hovered over to see the full details, first the ion type second the mass shift type. By hovering over the amino acids in the peptide or ions in the legend the corresponding peaks are highlighted. By toggling the 'Unassigned' label you can turn the background (unassigned) peaks on or off in the plot. By updating the slider in the Ion legend you can update the spectrum to only show the top X% of the peaks with labels. The top X% means any peak that is within X% of the highest intensity. By dragging in the spectrum you can zoom in to a specific part of the spectrum and use 'Zoom Out' to get back to the original zoom level. The annotation of the spectrum is based on the given sequence in the peptides file and is done with different software so inconsistencies are likely. The peaks are annotated based on the given sequence, with 20 ppm tolerance.

Copy Data

##### Spectrum 3013 (TSV)

###### Preview

```
Loading example...
```

*Click on the button to copy the data to your clipboard.*

Mz MinMz MaxIntensity Max

WidthHeightPeptide font sizePeptide stroke widthSpectrum font sizeSpectrum stroke widthCompact peptide

Ion legend

wxyz

abcd

OtherUnassignedIon^Charge^_Position_Show for top:%

SVMHEAJHNHYTQKS

02.23e+44.46e+46.69e+48.92e+4

Zoom Out

z^+1^_2_y^+1^_2_w^+1^_3_z^+1^_3_z^+1^_4_c^+1^_4_y^+3^_12_y^+3^_13_z^+3^_14_z^+3^_14_c^+3^_14_y^+3^_14_z^+3^_14_c^+1^_5_z^+1^_5_y^+1^_5_c^+1^_6_z^+2^_12_y^+2^_12_z^+1^_6_y^+1^_6_c^+2^_13_c^+2^_13_c^+1^_7_y^+2^_13_w^+1^_7_w^+2^_14_c^+2^_14_z^+2^_14_c^+2^_14_y^+2^_14_z^+1^_7_y^+1^_7_c^+1^_8_z^+1^_8_y^+1^_8_c^+1^_9_z^+1^_9_c^+1^_10_z^+1^_10_y^+1^_10_z^+1^_11_y^+1^_11_c^+1^_11_c^+1^_12_c^+1^_13_y^+1^_14_z^+1^_14_c^+1^_14_y^+1^_14_

0767153323003066

Fragment Matches Table

Show background peaks

| Position | Ion type | Intensity | mz Theoretical | mz Error (Th) | mz Error (ppm) | Charge | Series Number |
| --- | --- | --- | --- | --- | --- | --- | --- |
| - | - | 1448 | 129.1 | - | - | 0 | - |
| - | - | 421.3 | 135.8 | - | - | 0 | - |
| - | - | 502.8 | 138.4 | - | - | 0 | - |
| - | - | 449.4 | 148.9 | - | - | 0 | - |
| - | - | 691.9 | 148.9 | - | - | 0 | - |
| - | - | 765.8 | 148.9 | - | - | 0 | - |
| - | - | 905.5 | 148.9 | - | - | 0 | - |
| - | - | 1248 | 148.9 | - | - | 0 | - |
| - | - | 2044 | 148.9 | - | - | 0 | - |
| - | - | 3431 | 148.9 | - | - | 0 | - |
| - | - | 3982 | 149 | - | - | 0 | - |
| - | - | 2402 | 149 | - | - | 0 | - |
| - | - | 1278 | 149 | - | - | 0 | - |
| - | - | 995.9 | 149 | - | - | 0 | - |
| - | - | 669.1 | 149 | - | - | 0 | - |
| - | - | 623.2 | 149 | - | - | 0 | - |
| - | - | 497.2 | 149 | - | - | 0 | - |
| - | - | 550.8 | 149 | - | - | 0 | - |
| - | - | 488.2 | 149 | - | - | 0 | - |
| - | - | 1657 | 149 | - | - | 0 | - |
| - | - | 567.6 | 155.1 | - | - | 0 | - |
| - | - | 482.5 | 158.8 | - | - | 0 | - |
| - | - | 1557 | 159.1 | - | - | 0 | - |
| - | - | 2194 | 187.1 | - | - | 0 | - |
| - | - | 424 | 188.3 | - | - | 0 | - |
| 14 | z | 784.5 | 218.1 | 0.0004784 | 2.193 | +1 | 2 |
| - | - | 3302 | 221.1 | - | - | 0 | - |
| - | - | 777.6 | 222.1 | - | - | 0 | - |
| 14 | y | 1282 | 234.1 | 1.14E-05 | 0.04869 | +1 | 2 |
| - | - | 3593 | 239.1 | - | - | 0 | - |
| - | - | 1238 | 240.1 | - | - | 0 | - |
| - | - | 1021 | 241.1 | - | - | 0 | - |
| - | - | 2460 | 285 | - | - | 0 | - |
| - | - | 1582 | 286 | - | - | 0 | - |
| - | - | 794.3 | 286.1 | - | - | 0 | - |
| - | - | 570.3 | 287 | - | - | 0 | - |
| 13 | w | 2348 | 288.2 | 0.0002764 | 0.9592 | +1 | 3 |
| - | - | 854.4 | 297.1 | - | - | 0 | - |
| - | - | 5085 | 299.1 | - | - | 0 | - |
| - | - | 2804 | 300.1 | - | - | 0 | - |
| - | - | 1369 | 301.1 | - | - | 0 | - |
| - | - | 614.4 | 301.9 | - | - | 0 | - |
| - | - | 1.01E+04 | 309.2 | - | - | 0 | - |
| - | - | 1058 | 310.2 | - | - | 0 | - |
| 13 | z | 3373 | 346.2 | 0.0001767 | 0.5105 | +1 | 3 |
| - | - | 637.2 | 347.2 | - | - | 0 | - |
| - | - | 3.129E+04 | 355.1 | - | - | 0 | - |
| - | - | 2.891E+04 | 356.1 | - | - | 0 | - |
| - | - | 1.812E+04 | 357.1 | - | - | 0 | - |
| - | - | 3503 | 358.1 | - | - | 0 | - |
| - | - | 1198 | 371.1 | - | - | 0 | - |
| - | - | 1267 | 372.1 | - | - | 0 | - |
| - | - | 908.8 | 373.1 | - | - | 0 | - |
| - | - | 839.2 | 415 | - | - | 0 | - |
| - | - | 1041 | 416 | - | - | 0 | - |
| - | - | 915.1 | 417 | - | - | 0 | - |
| - | - | 1210 | 419 | - | - | 0 | - |
| - | - | 560.7 | 420 | - | - | 0 | - |
| - | - | 1015 | 427.2 | - | - | 0 | - |
| 12 | z | 3010 | 447.2 | 0.0007361 | 1.646 | +1 | 4 |
| - | - | 5044 | 448.2 | - | - | 0 | - |
| - | - | 825.3 | 449.2 | - | - | 0 | - |
| - | - | 1285 | 455.2 | - | - | 0 | - |
| - | - | 1213 | 471.2 | - | - | 0 | - |
| 4 | c | 3218 | 472.2 | 0.0004185 | 0.8862 | +1 | 4 |
| - | - | 631.8 | 473.2 | - | - | 0 | - |
| - | - | 1292 | 475 | - | - | 0 | - |
| 4 | y | 853.4 | 488.9 | 0.001871 | 3.827 | +3 | 12 |
| - | - | 908.2 | 489.2 | - | - | 0 | - |
| - | - | 822.9 | 490.1 | - | - | 0 | - |
| - | - | 562.6 | 510.7 | - | - | 0 | - |
| 3 | y | 1191 | 532.6 | 0.001044 | 1.961 | +3 | 13 |
| - | - | 1151 | 533.3 | - | - | 0 | - |
| - | - | 692 | 539.3 | - | - | 0 | - |
| - | - | 663.5 | 553.9 | - | - | 0 | - |
| 2 | z | 1147 | 554.3 | 0.004239 | 7.648 | +3 | 14 |
| 2 | z | 1033 | 554.6 | 0.004975 | 8.971 | +3 | 14 |
| 14 | c | 1757 | 559.6 | 0.001401 | 2.504 | +3 | 14 |
| 2 | y | 1468 | 559.9 | 0.004695 | 8.385 | +3 | 14 |
| 2 | z | 1165 | 560.3 | 0.004263 | 7.61 | +3 | 14 |
| - | - | 1606 | 563.1 | - | - | 0 | - |
| - | - | 832.3 | 579.1 | - | - | 0 | - |
| - | - | 1729 | 584.3 | - | - | 0 | - |
| - | - | 1417 | 588.9 | - | - | 0 | - |
| - | - | 3091 | 589.3 | - | - | 0 | - |
| - | - | 1000 | 589.6 | - | - | 0 | - |
| - | - | 1071 | 589.9 | - | - | 0 | - |
| - | - | 1069 | 590.3 | - | - | 0 | - |
| - | - | 756.9 | 593.9 | - | - | 0 | - |
| - | - | 1542 | 594.3 | - | - | 0 | - |
| - | - | 2.637E+04 | 594.6 | - | - | 0 | - |
| - | - | 2.986E+04 | 595 | - | - | 0 | - |
| - | - | 1.558E+04 | 595.3 | - | - | 0 | - |
| - | - | 1.023E+04 | 595.6 | - | - | 0 | - |
| - | - | 1565 | 596 | - | - | 0 | - |
| - | - | 768.5 | 596.3 | - | - | 0 | - |
| - | - | 644.9 | 596.5 | - | - | 0 | - |
| 5 | c | 7351 | 601.3 | 4.871E-05 | 0.081 | +1 | 5 |
| - | - | 2185 | 602.3 | - | - | 0 | - |
| 11 | z | 5911 | 610.3 | 0.0002665 | 0.4366 | +1 | 5 |
| - | - | 2030 | 611.3 | - | - | 0 | - |
| 11 | y | 1031 | 626.3 | 0.001246 | 1.989 | +1 | 5 |
| - | - | 727.1 | 653.3 | - | - | 0 | - |
| - | - | 635.1 | 664.8 | - | - | 0 | - |
| 6 | c | 4585 | 672.3 | 0.000505 | 0.7512 | +1 | 6 |
| - | - | 1177 | 673.3 | - | - | 0 | - |
| 4 | z | 1508 | 724.8 | 0.0005483 | 0.7565 | +2 | 12 |
| - | - | 1643 | 725.3 | - | - | 0 | - |
| 4 | y | 1707 | 732.9 | 0.0004636 | 0.6326 | +2 | 12 |
| - | - | 1143 | 733.4 | - | - | 0 | - |
| 10 | z | 3202 | 747.4 | 0.00115 | 1.539 | +1 | 6 |
| - | - | 2062 | 748.4 | - | - | 0 | - |
| - | - | 617.8 | 760.9 | - | - | 0 | - |
| 10 | y | 726.2 | 763.4 | 0.0002672 | 0.3501 | +1 | 6 |
| - | - | 847.5 | 765.9 | - | - | 0 | - |
| - | - | 545.9 | 767.1 | - | - | 0 | - |
| - | - | 1017 | 774.4 | - | - | 0 | - |
| 13 | c | 1204 | 774.9 | 0.003236 | 4.176 | +2 | 13 |
| - | - | 671.4 | 775.3 | - | - | 0 | - |
| - | - | 992.7 | 775.9 | - | - | 0 | - |
| 13 | c | 4265 | 783.4 | 0.002228 | 2.844 | +2 | 13 |
| - | - | 4131 | 783.9 | - | - | 0 | - |
| - | - | 1738 | 784.4 | - | - | 0 | - |
| - | - | 1785 | 784.9 | - | - | 0 | - |
| 7 | c | 2081 | 785.4 | 0.0006457 | 0.8221 | +1 | 7 |
| - | - | 794.5 | 786.4 | - | - | 0 | - |
| - | - | 735 | 791.4 | - | - | 0 | - |
| 3 | y | 2177 | 798.4 | 0.002201 | 2.756 | +2 | 13 |
| - | - | 1370 | 798.9 | - | - | 0 | - |
| - | - | 1533 | 799.4 | - | - | 0 | - |
| - | - | 806 | 799.9 | - | - | 0 | - |
| 9 | w | 1364 | 817.4 | 0.00252 | 3.083 | +1 | 7 |
| - | - | 1520 | 818.4 | - | - | 0 | - |
| - | - | 729.8 | 819.4 | - | - | 0 | - |
| - | - | 588.7 | 826.4 | - | - | 0 | - |
| 2 | w | 1023 | 832.4 | 0.01112 | 13.36 | +2 | 14 |
| 14 | c | 1169 | 838.9 | 0.004519 | 5.387 | +2 | 14 |
| 2 | z | 819.8 | 839.9 | 0.01025 | 12.2 | +2 | 14 |
| - | - | 1696 | 840.4 | - | - | 0 | - |
| - | - | 1097 | 840.9 | - | - | 0 | - |
| 14 | c | 9567 | 847.4 | 0.001431 | 1.689 | +2 | 14 |
| 2 | y | 1.033E+04 | 847.9 | 0.007538 | 8.89 | +2 | 14 |
| - | - | 6947 | 848.4 | - | - | 0 | - |
| - | - | 2166 | 848.9 | - | - | 0 | - |
| - | - | 789.6 | 849.4 | - | - | 0 | - |
| - | - | 889.6 | 850.9 | - | - | 0 | - |
| - | - | 986.8 | 851.9 | - | - | 0 | - |
| - | - | 2736 | 852.9 | - | - | 0 | - |
| - | - | 2610 | 853.4 | - | - | 0 | - |
| - | - | 2973 | 853.9 | - | - | 0 | - |
| - | - | 1273 | 854.4 | - | - | 0 | - |
| - | - | 1511 | 855.9 | - | - | 0 | - |
| - | - | 732.7 | 860.4 | - | - | 0 | - |
| - | - | 897.8 | 860.9 | - | - | 0 | - |
| 9 | z | 8440 | 861.4 | 0.0008441 | 0.98 | +1 | 7 |
| - | - | 6819 | 862.4 | - | - | 0 | - |
| - | - | 1102 | 862.9 | - | - | 0 | - |
| - | - | 3355 | 863.4 | - | - | 0 | - |
| - | - | 5784 | 869.4 | - | - | 0 | - |
| - | - | 6961 | 869.9 | - | - | 0 | - |
| - | - | 5066 | 870.4 | - | - | 0 | - |
| - | - | 2521 | 870.9 | - | - | 0 | - |
| - | - | 848.3 | 871.4 | - | - | 0 | - |
| - | - | 721.1 | 875.9 | - | - | 0 | - |
| 9 | y | 1790 | 877.4 | 0.0006137 | 0.6995 | +1 | 7 |
| - | - | 1975 | 878.4 | - | - | 0 | - |
| - | - | 4416 | 883.4 | - | - | 0 | - |
| - | - | 5051 | 883.9 | - | - | 0 | - |
| - | - | 3366 | 884.4 | - | - | 0 | - |
| - | - | 2683 | 884.9 | - | - | 0 | - |
| - | - | 2154 | 885.4 | - | - | 0 | - |
| - | - | 2093 | 890.4 | - | - | 0 | - |
| - | - | 4164 | 890.9 | - | - | 0 | - |
| - | - | 1.789E+04 | 891.4 | - | - | 0 | - |
| - | - | 8.654E+04 | 891.9 | - | - | 0 | - |
| - | - | 8.828E+04 | 892.4 | - | - | 0 | - |
| - | - | 5.648E+04 | 892.9 | - | - | 0 | - |
| - | - | 2.873E+04 | 893.4 | - | - | 0 | - |
| - | - | 9547 | 893.9 | - | - | 0 | - |
| - | - | 2330 | 894.4 | - | - | 0 | - |
| 8 | c | 1.994E+04 | 922.5 | 0.0005976 | 0.6478 | +1 | 8 |
| - | - | 1.052E+04 | 923.5 | - | - | 0 | - |
| - | - | 3364 | 924.5 | - | - | 0 | - |
| - | - | 812.8 | 925.5 | - | - | 0 | - |
| 8 | z | 1205 | 998.5 | 0.001991 | 1.994 | +1 | 8 |
| - | - | 993.8 | 999.4 | - | - | 0 | - |
| 8 | y | 1054 | 1014 | 0.0002346 | 0.2312 | +1 | 8 |
| - | - | 1245 | 1015 | - | - | 0 | - |
| 9 | c | 5752 | 1036 | 0.000129 | 0.1245 | +1 | 9 |
| - | - | 4676 | 1037 | - | - | 0 | - |
| - | - | 1654 | 1038 | - | - | 0 | - |
| 7 | z | 2454 | 1112 | 0.0003854 | 0.3467 | +1 | 9 |
| - | - | 2256 | 1113 | - | - | 0 | - |
| - | - | 1174 | 1114 | - | - | 0 | - |
| 10 | c | 9041 | 1174 | 0.0009354 | 0.7971 | +1 | 10 |
| - | - | 7115 | 1175 | - | - | 0 | - |
| - | - | 4042 | 1176 | - | - | 0 | - |
| - | - | 1219 | 1177 | - | - | 0 | - |
| - | - | 677.6 | 1178 | - | - | 0 | - |
| 6 | z | 5515 | 1183 | 0.0003515 | 0.2972 | +1 | 10 |
| - | - | 4589 | 1184 | - | - | 0 | - |
| - | - | 2615 | 1185 | - | - | 0 | - |
| - | - | 1137 | 1186 | - | - | 0 | - |
| - | - | 845.5 | 1189 | - | - | 0 | - |
| - | - | 738 | 1192 | - | - | 0 | - |
| 6 | y | 915.2 | 1199 | 0.004728 | 3.945 | +1 | 10 |
| 5 | z | 6449 | 1312 | 0.002173 | 1.657 | +1 | 11 |
| - | - | 6847 | 1313 | - | - | 0 | - |
| - | - | 4207 | 1314 | - | - | 0 | - |
| - | - | 1448 | 1315 | - | - | 0 | - |
| 5 | y | 780.6 | 1328 | 0.01626 | 12.25 | +1 | 11 |
| - | - | 2116 | 1336 | - | - | 0 | - |
| 11 | c | 5255 | 1337 | 0.003473 | 2.598 | +1 | 11 |
| - | - | 4041 | 1338 | - | - | 0 | - |
| - | - | 2238 | 1339 | - | - | 0 | - |
| - | - | 686 | 1368 | - | - | 0 | - |
| - | - | 758.5 | 1382 | - | - | 0 | - |
| - | - | 722.2 | 1437 | - | - | 0 | - |
| 12 | c | 5407 | 1438 | 0.002323 | 1.616 | +1 | 12 |
| - | - | 7007 | 1439 | - | - | 0 | - |
| - | - | 3150 | 1440 | - | - | 0 | - |
| - | - | 852 | 1441 | - | - | 0 | - |
| - | - | 1097 | 1450 | - | - | 0 | - |
| - | - | 1678 | 1451 | - | - | 0 | - |
| - | - | 771.1 | 1452 | - | - | 0 | - |
| 13 | c | 2915 | 1566 | 0.005237 | 3.345 | +1 | 13 |
| - | - | 3669 | 1567 | - | - | 0 | - |
| - | - | 2787 | 1568 | - | - | 0 | - |
| - | - | 1167 | 1569 | - | - | 0 | - |
| - | - | 1108 | 1581 | - | - | 0 | - |
| - | - | 847.1 | 1582 | - | - | 0 | - |
| - | - | 796.3 | 1583 | - | - | 0 | - |
| - | - | 615.6 | 1584 | - | - | 0 | - |
| - | - | 717.3 | 1652 | - | - | 0 | - |
| 2 | y | 970.5 | 1678 | 0.01716 | 10.23 | +1 | 14 |
| 2 | z | 1144 | 1679 | 0.001647 | 0.9813 | +1 | 14 |
| - | - | 1618 | 1680 | - | - | 0 | - |
| - | - | 660.3 | 1682 | - | - | 0 | - |
| 14 | c | 1231 | 1694 | 0.01573 | 9.285 | +1 | 14 |
| 2 | y | 2641 | 1695 | 0.01539 | 9.083 | +1 | 14 |
| - | - | 2219 | 1696 | - | - | 0 | - |
| - | - | 1112 | 1697 | - | - | 0 | - |
| - | - | 766.9 | 1726 | - | - | 0 | - |
| - | - | 963 | 1738 | - | - | 0 | - |
| - | - | 4037 | 1739 | - | - | 0 | - |
| - | - | 4900 | 1740 | - | - | 0 | - |
| - | - | 3268 | 1741 | - | - | 0 | - |
| - | - | 1451 | 1742 | - | - | 0 | - |
| - | - | 1029 | 1750 | - | - | 0 | - |
| - | - | 1905 | 1756 | - | - | 0 | - |
| - | - | 2574 | 1757 | - | - | 0 | - |
| - | - | 1457 | 1758 | - | - | 0 | - |
| - | - | 973.1 | 1766 | - | - | 0 | - |
| - | - | 5704 | 1767 | - | - | 0 | - |
| - | - | 7690 | 1768 | - | - | 0 | - |
| - | - | 5165 | 1769 | - | - | 0 | - |
| - | - | 2532 | 1770 | - | - | 0 | - |
| - | - | 1812 | 1771 | - | - | 0 | - |
| - | - | 718.4 | 1781 | - | - | 0 | - |
| - | - | 4316 | 1782 | - | - | 0 | - |
| - | - | 1.273E+04 | 1783 | - | - | 0 | - |
| - | - | 4.67E+04 | 1784 | - | - | 0 | - |
| - | - | 4.843E+04 | 1785 | - | - | 0 | - |
| - | - | 2.985E+04 | 1786 | - | - | 0 | - |
| - | - | 1.493E+04 | 1787 | - | - | 0 | - |
| - | - | 5337 | 1788 | - | - | 0 | - |
| - | - | 1725 | 1789 | - | - | 0 | - |
| - | - | 757.2 | 1863 | - | - | 0 | - |
| - | - | 656 | 3036 | - | - | 0 | - |

m/z Charge Intensity FragmentType MassShift Position
129.10214233398438 0 1448.3218
135.7659454345703 0 421.34137
138.4300537109375 0 502.81793
148.86170959472656 0 449.44186
148.89926147460938 0 691.8728
148.9058074951172 0 765.8392
148.92030334472656 0 905.51196
148.9276580810547 0 1247.6993
148.9348602294922 0 2044.1531
148.94259643554688 0 3430.9304
148.9593048095703 0 3981.8237
148.96717834472656 0 2402.1016
148.9744110107422 0 1278.4109
148.98171997070312 0 995.93646
148.9889373779297 0 669.1125
148.9959716796875 0 623.16925
149.00360107421875 0 497.16998
149.01089477539062 0 550.83325
149.0251922607422 0 488.17902
149.04505920410156 0 1656.6871
155.09283447265625 0 567.5802
158.8022918701172 0 482.53732
159.11244201660156 0 1557.4227
187.10772705078125 0 2193.594
188.2796173095703 0 423.96564
218.1265869140625 0 784.51886 z 13
221.08413696289062 0 3301.841
222.08615112304688 0 777.6271
234.1448211669922 0 1282.0817 y 13
239.09481811523438 0 3592.9248
240.09506225585938 0 1237.6361
241.0923614501953 0 1021.3377
285.0096740722656 0 2459.5388
286.009521484375 0 1582.0876
286.1335754394531 0 794.2555
287.0076904296875 0 570.3042
288.1551208496094 0 2347.7078 w 12
297.0826721191406 0 854.4329
299.0617370605469 0 5084.62
300.06207275390625 0 2803.995
301.0595397949219 0 1369.4651
301.8958435058594 0 614.3548
309.20355224609375 0 10102.946
310.2064514160156 0 1057.6604
346.18450927734375 0 3372.7244 z 12
347.1892395019531 0 637.20825
355.0696105957031 0 31288.576
356.0701904296875 0 28911.162
357.06756591796875 0 18121.965
358.0677795410156 0 3502.9478
371.1005554199219 0 1197.8262
372.10211181640625 0 1266.7257
373.10284423828125 0 908.84564
415.0364685058594 0 839.19653
416.0371398925781 0 1041.2887
417.03466796875 0 915.1033
418.99456787109375 0 1210.0264
419.99560546875 0 560.6958
427.2143859863281 0 1014.78204
447.23162841796875 0 3009.7954 z 11
448.2396240234375 0 5044.018
449.24029541015625 0 825.2621
455.2055969238281 0 1285.0894
471.2226257324219 0 1212.5509
472.2332458496094 0 3217.8975 c 3
473.2335510253906 0 631.8475
475.0032958984375 0 1291.9578
488.906005859375 0 853.3595 y 3
489.238037109375 0 908.16016
490.05609130859375 0 822.90497
510.7396545410156 0 562.5989
532.583251953125 0 1190.5038 y 2
533.2503662109375 0 1151.1907
539.2517700195312 0 692.0098
553.9282836914062 0 663.50055
554.259765625 0 1146.5997 z Water loss 1
554.5969848632812 0 1032.9442 z Ammonia loss 1
559.60498046875 0 1757.2039 c Ammonia loss 13
559.936279296875 0 1467.7854 y Ammonia loss 1
560.2717895507812 0 1165.2825 z 1
563.0751342773438 0 1606.1825
579.1093139648438 0 832.3035
584.251708984375 0 1729.0833
588.9454956054688 0 1416.9385
589.2783203125 0 3090.7192
589.6112670898438 0 1000.18463
589.9492797851562 0 1070.754
590.26123046875 0 1068.9619
593.9442138671875 0 756.89667
594.280029296875 0 1542.2247
594.6173095703125 0 26371.855
594.9507446289062 0 29858.775
595.2842407226562 0 15580.049
595.6171264648438 0 10229.992
595.9524536132812 0 1565.0736
596.2792358398438 0 768.4875
596.4716796875 0 644.9173
601.2763061523438 0 7350.5083 c 4
602.2786254882812 0 2184.998
610.2959594726562 0 5910.627 z 10
611.2937622070312 0 2030.3485
626.3131713867188 0 1030.5402 y 10
653.301025390625 0 727.13306
664.8226318359375 0 635.1073
672.3128662109375 0 4585.0034 c 5
673.3165893554688 0 1177.4318
724.84375 0 1507.7727 z 3
725.343994140625 0 1642.6736
732.85302734375 0 1707.3524 y 3
733.35009765625 0 1142.7173
747.3534545898438 0 3201.6223 z 9
748.3554077148438 0 2061.5874
760.85693359375 0 617.8056
763.3735961914062 0 726.1669 y 9
765.8530883789062 0 847.5496
767.0591430664062 0 545.941
774.3883056640625 0 1016.67236
774.8510131835938 0 1203.8608 c Ammonia loss 12
775.346435546875 0 671.37225
775.8517456054688 0 992.6915
783.3652954101562 0 4265.1997 c 12
783.8661499023438 0 4130.998
784.3684692382812 0 1738.0496
784.8655395507812 0 1785.0547
785.3967895507812 0 2081.3408 c 6
786.3978271484375 0 794.4904
791.3610229492188 0 735.01373
798.37060546875 0 2177.242 y 2
798.8743896484375 0 1369.6681
799.372314453125 0 1533.3679
799.870849609375 0 806.04443
817.3864135742188 0 1364.4075 w 8
818.3840942382812 0 1520.2498
819.3786010742188 0 729.7786
826.405029296875 0 588.6875
832.3970336914062 0 1023.2607 w 1
838.90625 0 1168.5011 c Ammonia loss 13
839.9078979492188 0 819.824 z 1
840.4066162109375 0 1695.5205
840.9146728515625 0 1097.0809
847.41357421875 0 9567.19 c 13
847.91455078125 0 10329.995 y 1
848.4129638671875 0 6947.2153
848.912353515625 0 2165.787
849.4234008789062 0 789.5621
850.8999633789062 0 889.61176
851.897705078125 0 986.755
852.906982421875 0 2735.7307
853.4110717773438 0 2610.3203
853.9058227539062 0 2973.0552
854.403564453125 0 1272.7253
855.8880004882812 0 1511.0468
860.41748046875 0 732.69495
860.9026489257812 0 897.82477
861.3983764648438 0 8439.945 z 8
862.3936157226562 0 6819.3677
862.8912353515625 0 1101.5614
863.3973999023438 0 3354.847
869.4158325195312 0 5783.8477
869.9150390625 0 6960.5586
870.4172973632812 0 5065.9473
870.918701171875 0 2521.438
871.4100952148438 0 848.3362
875.904541015625 0 721.07434
877.4168701171875 0 1789.9935 y 8
878.4276123046875 0 1974.84
883.4107666015625 0 4416.3154
883.913330078125 0 5051.327
884.4098510742188 0 3366.377
884.9058837890625 0 2682.8455
885.3973388671875 0 2154.118
890.4078369140625 0 2093.158
890.9179077148438 0 4163.684
891.42138671875 0 17886.656
891.9256591796875 0 86540.9
892.4259643554688 0 88277.05
892.9257202148438 0 56483.445
893.4259643554688 0 28733.52
893.9240112304688 0 9546.796
894.421630859375 0 2330.2327
922.4557495117188 0 19935.412 c 7
923.4592895507812 0 10520.11
924.4606323242188 0 3364.1624
925.4517211914062 0 812.8198
998.4584350585938 0 1204.9668 z 7
999.4462280273438 0 993.7735
1014.4754028320312 0 1054.1614 y 7
1015.4690551757812 0 1245.3926
1036.4991455078125 0 5752.4673 c 8
1037.49658203125 0 4675.5493
1038.4993896484375 0 1654.1189
1111.5408935546875 0 2453.739 z 6
1112.5391845703125 0 2255.981
1113.5447998046875 0 1173.8469
1173.5572509765625 0 9041.087 c 9
1174.5555419921875 0 7115.003
1175.552978515625 0 4041.7957
1176.5498046875 0 1219.0116
1178.4957275390625 0 677.5946
1182.5772705078125 0 5514.7393 z 5
1183.576416015625 0 4588.709
1184.5797119140625 0 2614.9392
1185.5823974609375 0 1137.2052
1188.51806640625 0 845.5066
1191.5587158203125 0 738.0408
1198.60107421875 0 915.20264 y 5
1311.6180419921875 0 6448.921 z 4
1312.623291015625 0 6847.1055
1313.623291015625 0 4207.3403
1314.6234130859375 0 1448.2275
1327.6226806640625 0 780.58203 y 4
1335.6099853515625 0 2116.0903
1336.6180419921875 0 5255.0376 c 10
1337.618408203125 0 4041.0618
1338.61572265625 0 2238.4587
1367.5858154296875 0 685.9845
1381.75048828125 0 758.4754
1436.6566162109375 0 722.2194
1437.6668701171875 0 5407.3013 c 11
1438.66650390625 0 7007.2275
1439.6705322265625 0 3150.083
1440.6727294921875 0 851.99475
1449.687255859375 0 1097.3525
1450.676513671875 0 1677.8026
1451.68310546875 0 771.0927
1565.7225341796875 0 2914.798 c 12
1566.724365234375 0 3668.5393
1567.7230224609375 0 2786.8015
1568.7119140625 0 1167.0659
1580.7174072265625 0 1107.7852
1581.7197265625 0 847.113
1582.7423095703125 0 796.3378
1583.6802978515625 0 615.5788
1651.8017578125 0 717.2944
1677.79736328125 0 970.5397 y Ammonia loss 1
1678.7896728515625 0 1144.0863 z 1
1679.799560546875 0 1617.7343
1681.8009033203125 0 660.3116
1693.8070068359375 0 1231.4525 c 13
1694.8221435546875 0 2641.2136 y 1
1695.81884765625 0 2219.1057
1696.8094482421875 0 1111.972
1725.8280029296875 0 766.93445
1737.821533203125 0 963.0113
1738.832275390625 0 4036.9773
1739.830810546875 0 4899.9565
1740.8251953125 0 3267.666
1741.8282470703125 0 1450.9065
1749.82421875 0 1028.5281
1755.8563232421875 0 1905.3112
1756.8548583984375 0 2574.1426
1757.8570556640625 0 1457.0928
1765.817626953125 0 973.0803
1766.827392578125 0 5704.009
1767.8248291015625 0 7689.8677
1768.8214111328125 0 5165.2983
1769.8206787109375 0 2531.6729
1770.8221435546875 0 1811.5857
1780.8487548828125 0 718.3944
1781.840087890625 0 4315.9
1782.8419189453125 0 12732.794
1783.850341796875 0 46701.96
1784.84912109375 0 48432.65
1785.851318359375 0 29846.613
1786.8525390625 0 14928.007
1787.84423828125 0 5337.2393
1788.827392578125 0 1724.8607
1862.928466796875 0 757.18054
3035.81396484375 0 656.00055

Spectrum Details

|  |  |
| --- | --- |
| Matched peaks? Matched peaks The total absolute number of peaks matched. Additionally in brackets the total fraction of peaks matched and the total number of peaks is shown. | 50 (18.59% of 269) |
| FDR? FDR The false discovery rate estimated for this peptide. It is calculated by matching all theoretical fragments with a non-integer shift with the raw peaks for this spectrum. This is done with 40 different shifts. The resulting percentage is the average number of annotated peaks over the number of annotated peaks with the correct spectrum. | 0.90% |
| Satellite FDR? Satellite FDR See the FDR for details on its calculation. This satellite ion specific FDR only contains the satellite ions (d/w) for I/L/J positions. | - |
| PSM Score? PSM Score The PSM Score as given by Hecklib to this annotated spectrum. It is shown with three significant figures. | 345 |

#### Spectrum 2369? Spectrum 2369 The raw spectrum of this peptide as annotated by Hecklib. The fragments are coloured according to ion type (see legend). Any peaks with a star '\*' as text can be hovered over to see the full details, first the ion type second the mass shift type. By hovering over the amino acids in the peptide or ions in the legend the corresponding peaks are highlighted. By toggling the 'Unassigned' label you can turn the background (unassigned) peaks on or off in the plot. By updating the slider in the Ion legend you can update the spectrum to only show the top X% of the peaks with labels. The top X% means any peak that is within X% of the highest intensity. By dragging in the spectrum you can zoom in to a specific part of the spectrum and use 'Zoom Out' to get back to the original zoom level. The annotation of the spectrum is based on the given sequence in the peptides file and is done with different software so inconsistencies are likely. The peaks are annotated based on the given sequence, with 20 ppm tolerance.

Copy Data

##### Spectrum 2369 (TSV)

###### Preview

```
Loading example...
```

*Click on the button to copy the data to your clipboard.*

Mz MinMz MaxIntensity Max

WidthHeightPeptide font sizePeptide stroke widthSpectrum font sizeSpectrum stroke widthCompact peptide

Ion legend

wxyz

abcd

OtherUnassignedIon^Charge^_Position_Show for top:%

SVMHEAJHNHYTQKS

04.46e+58.92e+51.34e+61.78e+6

Zoom Out

a^+1^_2_y^+3^_4_a^+1^_2_b^+1^_2_y^+2^_3_b^+1^_2_y^+1^_2_y^+1^_2_y^+1^_2_b^+2^_5_a^+1^_3_y^+2^_5_b^+2^_6_b^+2^_6_y^+1^_3_y^+1^_3_y^+1^_3_y^+2^_6_y^+2^_6_y^+2^_6_b^+3^_10_y^+2^_7_y^+2^_7_y^+2^_7_y^+3^_11_y^+1^_4_b^+3^_11_y^+1^_4_b^+2^_8_y^+1^_4_b^+1^_4_b^+3^_12_b^+3^_12_y^+3^_12_y^+3^_12_y^+3^_12_y^+2^_8_y^+2^_8_y^+2^_8_b^+3^_13_b^+2^_9_b^+3^_13_y^+3^_13_y^+3^_13_y^+2^_9_y^+2^_9_b^+3^_14_b^+3^_14_y^+2^_9_b^+3^_14_y^+3^_14_y^+3^_14_b^+2^_10_b^+1^_5_b^+2^_10_y^+2^_10_y^+2^_10_\*\*y^+2^_10_\*y^+1^_5_y^+1^_5_y^+1^_5_y^+2^_11_y^+2^_11_b^+2^_11_b^+2^_11_y^+2^_11_b^+2^_11_b^+1^_6_b^+2^_12_b^+2^_12_y^+2^_12_y^+2^_12_y^+2^_12_y^+1^_6_y^+1^_6_b^+1^_7_b^+2^_13_b^+2^_13_b^+1^_7_y^+2^_13_y^+2^_13_y^+2^_13_b^+2^_14_y^+2^_14_y^+2^_14_y^+1^_7_y^+1^_7_y^+1^_7_b^+1^_8_b^+1^_8_y^+1^_8_y^+1^_8_y^+1^_8_b^+1^_9_b^+1^_9_y^+1^_9_y^+1^_9_y^+1^_9_b^+1^_10_b^+1^_10_y^+1^_10_y^+1^_10_y^+1^_11_y^+1^_11_y^+1^_11_b^+1^_12_

0766153122973063

Fragment Matches Table

Show background peaks

| Position | Ion type | Intensity | mz Theoretical | mz Error (Th) | mz Error (ppm) | Charge | Series Number |
| --- | --- | --- | --- | --- | --- | --- | --- |
| - | - | 2490 | 120.1 | - | - | 0 | - |
| - | - | 500.5 | 120.1 | - | - | 0 | - |
| - | - | 3512 | 121 | - | - | 0 | - |
| - | - | 5122 | 122.1 | - | - | 0 | - |
| - | - | 595.9 | 122.1 | - | - | 0 | - |
| - | - | 3655 | 123.1 | - | - | 0 | - |
| - | - | 463.8 | 123.1 | - | - | 0 | - |
| - | - | 811.2 | 124.1 | - | - | 0 | - |
| - | - | 485.9 | 125.7 | - | - | 0 | - |
| - | - | 1012 | 126.1 | - | - | 0 | - |
| - | - | 3492 | 127.1 | - | - | 0 | - |
| - | - | 1120 | 128.1 | - | - | 0 | - |
| - | - | 2542 | 128.1 | - | - | 0 | - |
| - | - | 5038 | 129.1 | - | - | 0 | - |
| - | - | 7.132E+05 | 129.1 | - | - | 0 | - |
| - | - | 2252 | 130.1 | - | - | 0 | - |
| - | - | 8226 | 130.1 | - | - | 0 | - |
| - | - | 5182 | 130.1 | - | - | 0 | - |
| - | - | 4.675E+04 | 130.1 | - | - | 0 | - |
| - | - | 2115 | 131 | - | - | 0 | - |
| - | - | 1032 | 131.1 | - | - | 0 | - |
| - | - | 1916 | 136.1 | - | - | 0 | - |
| - | - | 1.457E+05 | 136.1 | - | - | 0 | - |
| - | - | 675.4 | 137.1 | - | - | 0 | - |
| - | - | 1.185E+04 | 137.1 | - | - | 0 | - |
| - | - | 9497 | 138.1 | - | - | 0 | - |
| - | - | 627.9 | 138.1 | - | - | 0 | - |
| - | - | 658.3 | 138.1 | - | - | 0 | - |
| - | - | 613.7 | 139.1 | - | - | 0 | - |
| - | - | 1.061E+04 | 139.1 | - | - | 0 | - |
| - | - | 868.8 | 140.1 | - | - | 0 | - |
| - | - | 1279 | 140.1 | - | - | 0 | - |
| 2 | a | 6327 | 141.1 | 0.0008642 | 6.124 | +1 | 2 |
| - | - | 1090 | 142.1 | - | - | 0 | - |
| - | - | 1580 | 143.1 | - | - | 0 | - |
| - | - | 1024 | 143.1 | - | - | 0 | - |
| - | - | 690.9 | 146.1 | - | - | 0 | - |
| - | - | 1376 | 146.1 | - | - | 0 | - |
| - | - | 1161 | 147 | - | - | 0 | - |
| - | - | 1.165E+04 | 147.1 | - | - | 0 | - |
| - | - | 1388 | 148 | - | - | 0 | - |
| - | - | 724.5 | 148.1 | - | - | 0 | - |
| - | - | 813.5 | 148.1 | - | - | 0 | - |
| - | - | 1.249E+04 | 148.1 | - | - | 0 | - |
| - | - | 630.7 | 148.1 | - | - | 0 | - |
| - | - | 461.1 | 148.9 | - | - | 0 | - |
| - | - | 832.4 | 148.9 | - | - | 0 | - |
| - | - | 702.8 | 148.9 | - | - | 0 | - |
| - | - | 1154 | 148.9 | - | - | 0 | - |
| - | - | 1007 | 148.9 | - | - | 0 | - |
| - | - | 2615 | 148.9 | - | - | 0 | - |
| - | - | 4063 | 148.9 | - | - | 0 | - |
| - | - | 3671 | 149 | - | - | 0 | - |
| - | - | 1551 | 149 | - | - | 0 | - |
| - | - | 938 | 149 | - | - | 0 | - |
| - | - | 988 | 149 | - | - | 0 | - |
| - | - | 630.6 | 149 | - | - | 0 | - |
| - | - | 629.1 | 149 | - | - | 0 | - |
| - | - | 455.3 | 149 | - | - | 0 | - |
| - | - | 2955 | 149 | - | - | 0 | - |
| - | - | 689.8 | 149 | - | - | 0 | - |
| - | - | 872.8 | 149.1 | - | - | 0 | - |
| 12 | y | 1116 | 149.1 | 0.001438 | 9.645 | +3 | 4 |
| - | - | 888.8 | 149.1 | - | - | 0 | - |
| - | - | 571.3 | 149.1 | - | - | 0 | - |
| - | - | 671.2 | 150 | - | - | 0 | - |
| - | - | 649.9 | 150 | - | - | 0 | - |
| - | - | 495.3 | 150.1 | - | - | 0 | - |
| - | - | 609.8 | 150.1 | - | - | 0 | - |
| - | - | 700 | 151.1 | - | - | 0 | - |
| - | - | 2148 | 151.1 | - | - | 0 | - |
| - | - | 1.194E+04 | 152.1 | - | - | 0 | - |
| - | - | 571.6 | 153.1 | - | - | 0 | - |
| - | - | 2069 | 153.1 | - | - | 0 | - |
| - | - | 1390 | 154.1 | - | - | 0 | - |
| - | - | 3477 | 154.1 | - | - | 0 | - |
| - | - | 424.7 | 155.1 | - | - | 0 | - |
| - | - | 4269 | 155.1 | - | - | 0 | - |
| - | - | 1.266E+04 | 155.1 | - | - | 0 | - |
| - | - | 5034 | 155.1 | - | - | 0 | - |
| - | - | 1.019E+04 | 156.1 | - | - | 0 | - |
| - | - | 736.8 | 157.1 | - | - | 0 | - |
| - | - | 3400 | 157.1 | - | - | 0 | - |
| - | - | 4219 | 157.1 | - | - | 0 | - |
| - | - | 516.1 | 158.1 | - | - | 0 | - |
| - | - | 2304 | 159.1 | - | - | 0 | - |
| 2 | a | 7.935E+05 | 159.1 | 0.0008891 | 5.588 | +1 | 2 |
| - | - | 922 | 160.1 | - | - | 0 | - |
| - | - | 1161 | 160.1 | - | - | 0 | - |
| - | - | 4465 | 160.1 | - | - | 0 | - |
| - | - | 5.805E+04 | 160.1 | - | - | 0 | - |
| - | - | 2334 | 161.1 | - | - | 0 | - |
| - | - | 2784 | 162.1 | - | - | 0 | - |
| - | - | 1542 | 163.1 | - | - | 0 | - |
| - | - | 1051 | 164.1 | - | - | 0 | - |
| - | - | 767.3 | 164.1 | - | - | 0 | - |
| - | - | 882.7 | 165.1 | - | - | 0 | - |
| - | - | 1.283E+04 | 165.1 | - | - | 0 | - |
| - | - | 1011 | 165.1 | - | - | 0 | - |
| - | - | 645.3 | 165.1 | - | - | 0 | - |
| - | - | 1057 | 166.1 | - | - | 0 | - |
| - | - | 2.91E+04 | 166.1 | - | - | 0 | - |
| - | - | 3.025E+04 | 166.1 | - | - | 0 | - |
| - | - | 517.1 | 167 | - | - | 0 | - |
| - | - | 2448 | 167.1 | - | - | 0 | - |
| - | - | 9275 | 167.1 | - | - | 0 | - |
| - | - | 4998 | 167.1 | - | - | 0 | - |
| - | - | 1764 | 168.1 | - | - | 0 | - |
| - | - | 582.4 | 168.1 | - | - | 0 | - |
| - | - | 3385 | 169.1 | - | - | 0 | - |
| - | - | 1002 | 169.1 | - | - | 0 | - |
| 2 | b | 2114 | 169.1 | 0.0009447 | 5.587 | +1 | 2 |
| - | - | 1054 | 171.1 | - | - | 0 | - |
| - | - | 1.258E+04 | 171.1 | - | - | 0 | - |
| - | - | 963.2 | 171.2 | - | - | 0 | - |
| - | - | 1929 | 172.1 | - | - | 0 | - |
| - | - | 1327 | 172.1 | - | - | 0 | - |
| 13 | y | 7123 | 173.1 | 0.0009794 | 5.658 | +2 | 3 |
| - | - | 1028 | 173.4 | - | - | 0 | - |
| - | - | 537.7 | 174 | - | - | 0 | - |
| - | - | 4317 | 174.1 | - | - | 0 | - |
| - | - | 820.4 | 175.1 | - | - | 0 | - |
| - | - | 2475 | 175.1 | - | - | 0 | - |
| - | - | 3.099E+04 | 176.1 | - | - | 0 | - |
| - | - | 1017 | 177.1 | - | - | 0 | - |
| - | - | 2340 | 177.1 | - | - | 0 | - |
| - | - | 2339 | 177.1 | - | - | 0 | - |
| - | - | 4640 | 177.1 | - | - | 0 | - |
| - | - | 1308 | 178.1 | - | - | 0 | - |
| - | - | 1923 | 178.1 | - | - | 0 | - |
| - | - | 795.4 | 178.1 | - | - | 0 | - |
| - | - | 801.3 | 178.1 | - | - | 0 | - |
| - | - | 2147 | 178.1 | - | - | 0 | - |
| - | - | 715.9 | 179.1 | - | - | 0 | - |
| - | - | 5959 | 179.1 | - | - | 0 | - |
| - | - | 8483 | 180.1 | - | - | 0 | - |
| - | - | 1.089E+04 | 180.1 | - | - | 0 | - |
| - | - | 676.2 | 181.1 | - | - | 0 | - |
| - | - | 1860 | 181.1 | - | - | 0 | - |
| - | - | 1267 | 181.1 | - | - | 0 | - |
| - | - | 4409 | 183.1 | - | - | 0 | - |
| - | - | 9849 | 183.1 | - | - | 0 | - |
| - | - | 1038 | 184.1 | - | - | 0 | - |
| - | - | 1276 | 185.1 | - | - | 0 | - |
| - | - | 2872 | 185.1 | - | - | 0 | - |
| - | - | 1973 | 186.1 | - | - | 0 | - |
| - | - | 1191 | 186.1 | - | - | 0 | - |
| - | - | 4412 | 186.1 | - | - | 0 | - |
| 2 | b | 2.281E+05 | 187.1 | 0.001046 | 5.59 | +1 | 2 |
| - | - | 1.812E+04 | 188.1 | - | - | 0 | - |
| - | - | 9979 | 189.1 | - | - | 0 | - |
| - | - | 1164 | 189.1 | - | - | 0 | - |
| - | - | 1673 | 189.1 | - | - | 0 | - |
| - | - | 1.283E+04 | 190.1 | - | - | 0 | - |
| - | - | 793.8 | 190.1 | - | - | 0 | - |
| - | - | 1472 | 190.1 | - | - | 0 | - |
| - | - | 1119 | 191.1 | - | - | 0 | - |
| - | - | 8865 | 191.1 | - | - | 0 | - |
| - | - | 2291 | 192.1 | - | - | 0 | - |
| - | - | 8509 | 192.1 | - | - | 0 | - |
| - | - | 4546 | 193.1 | - | - | 0 | - |
| - | - | 5.909E+04 | 193.1 | - | - | 0 | - |
| - | - | 5598 | 194.1 | - | - | 0 | - |
| - | - | 1.224E+04 | 194.1 | - | - | 0 | - |
| - | - | 4795 | 194.1 | - | - | 0 | - |
| - | - | 2308 | 194.1 | - | - | 0 | - |
| - | - | 9266 | 195.1 | - | - | 0 | - |
| - | - | 4385 | 195.1 | - | - | 0 | - |
| - | - | 1258 | 195.1 | - | - | 0 | - |
| - | - | 1.78E+04 | 195.1 | - | - | 0 | - |
| - | - | 1539 | 196.1 | - | - | 0 | - |
| - | - | 1233 | 197.1 | - | - | 0 | - |
| - | - | 839.6 | 197.1 | - | - | 0 | - |
| - | - | 1994 | 198.1 | - | - | 0 | - |
| - | - | 882.4 | 199.1 | - | - | 0 | - |
| - | - | 8588 | 199.1 | - | - | 0 | - |
| - | - | 2283 | 200.1 | - | - | 0 | - |
| - | - | 904.8 | 200.1 | - | - | 0 | - |
| - | - | 1486 | 200.1 | - | - | 0 | - |
| - | - | 1145 | 201.1 | - | - | 0 | - |
| - | - | 1.029E+04 | 201.1 | - | - | 0 | - |
| - | - | 3028 | 201.1 | - | - | 0 | - |
| - | - | 4400 | 202.1 | - | - | 0 | - |
| - | - | 4749 | 202.1 | - | - | 0 | - |
| - | - | 2732 | 202.1 | - | - | 0 | - |
| - | - | 667 | 203.1 | - | - | 0 | - |
| - | - | 3898 | 203.1 | - | - | 0 | - |
| - | - | 833.1 | 203.1 | - | - | 0 | - |
| - | - | 586.1 | 203.1 | - | - | 0 | - |
| - | - | 3.642E+04 | 204.1 | - | - | 0 | - |
| - | - | 1030 | 204.1 | - | - | 0 | - |
| - | - | 665.6 | 204.1 | - | - | 0 | - |
| - | - | 3524 | 205.1 | - | - | 0 | - |
| - | - | 2215 | 205.1 | - | - | 0 | - |
| - | - | 1781 | 205.1 | - | - | 0 | - |
| - | - | 500.1 | 205.1 | - | - | 0 | - |
| - | - | 883.9 | 206.1 | - | - | 0 | - |
| - | - | 4511 | 206.1 | - | - | 0 | - |
| - | - | 1083 | 206.1 | - | - | 0 | - |
| - | - | 4.062E+04 | 207.1 | - | - | 0 | - |
| - | - | 1399 | 207.2 | - | - | 0 | - |
| - | - | 608.2 | 208.1 | - | - | 0 | - |
| - | - | 3265 | 208.1 | - | - | 0 | - |
| - | - | 1728 | 208.1 | - | - | 0 | - |
| - | - | 1249 | 209.1 | - | - | 0 | - |
| - | - | 1114 | 209.1 | - | - | 0 | - |
| - | - | 1.31E+04 | 211.1 | - | - | 0 | - |
| - | - | 7001 | 212.1 | - | - | 0 | - |
| - | - | 1772 | 212.1 | - | - | 0 | - |
| - | - | 2442 | 213.1 | - | - | 0 | - |
| - | - | 588.4 | 213.1 | - | - | 0 | - |
| - | - | 776.7 | 213.1 | - | - | 0 | - |
| - | - | 2868 | 215.1 | - | - | 0 | - |
| 14 | y | 2.913E+04 | 216.1 | 0.001169 | 5.409 | +1 | 2 |
| - | - | 1.074E+04 | 217.1 | - | - | 0 | - |
| - | - | 1647 | 217.1 | - | - | 0 | - |
| 14 | y | 3.918E+04 | 217.1 | 0.001208 | 5.564 | +1 | 2 |
| - | - | 2602 | 217.1 | - | - | 0 | - |
| - | - | 1.143E+04 | 218.1 | - | - | 0 | - |
| - | - | 902.6 | 218.1 | - | - | 0 | - |
| - | - | 3020 | 218.1 | - | - | 0 | - |
| - | - | 1147 | 219.1 | - | - | 0 | - |
| - | - | 2977 | 219.1 | - | - | 0 | - |
| - | - | 1939 | 219.1 | - | - | 0 | - |
| - | - | 1029 | 219.1 | - | - | 0 | - |
| - | - | 982.5 | 220.1 | - | - | 0 | - |
| - | - | 1.604E+05 | 221.1 | - | - | 0 | - |
| - | - | 2418 | 221.1 | - | - | 0 | - |
| - | - | 1098 | 222.1 | - | - | 0 | - |
| - | - | 4176 | 222.1 | - | - | 0 | - |
| - | - | 1.444E+04 | 222.1 | - | - | 0 | - |
| - | - | 3.105E+04 | 222.1 | - | - | 0 | - |
| - | - | 7302 | 223.1 | - | - | 0 | - |
| - | - | 2168 | 223.1 | - | - | 0 | - |
| - | - | 9011 | 223.2 | - | - | 0 | - |
| - | - | 4.725E+04 | 224.1 | - | - | 0 | - |
| - | - | 1476 | 224.1 | - | - | 0 | - |
| - | - | 1263 | 224.2 | - | - | 0 | - |
| - | - | 2245 | 225.1 | - | - | 0 | - |
| - | - | 3020 | 225.1 | - | - | 0 | - |
| - | - | 1118 | 226.1 | - | - | 0 | - |
| - | - | 2576 | 226.2 | - | - | 0 | - |
| - | - | 880.6 | 228.1 | - | - | 0 | - |
| - | - | 5991 | 228.1 | - | - | 0 | - |
| - | - | 1081 | 229.1 | - | - | 0 | - |
| - | - | 562.6 | 229.1 | - | - | 0 | - |
| - | - | 7921 | 230.1 | - | - | 0 | - |
| - | - | 5.127E+04 | 230.1 | - | - | 0 | - |
| - | - | 1862 | 231.1 | - | - | 0 | - |
| - | - | 4388 | 231.1 | - | - | 0 | - |
| - | - | 7436 | 233.1 | - | - | 0 | - |
| - | - | 1.498E+04 | 234.1 | - | - | 0 | - |
| - | - | 7486 | 234.1 | - | - | 0 | - |
| 14 | y | 1.77E+05 | 234.1 | 0.001286 | 5.491 | +1 | 2 |
| - | - | 6.815E+04 | 235.1 | - | - | 0 | - |
| - | - | 621.8 | 235.1 | - | - | 0 | - |
| - | - | 1984 | 235.1 | - | - | 0 | - |
| - | - | 744.1 | 235.1 | - | - | 0 | - |
| - | - | 1.689E+04 | 235.1 | - | - | 0 | - |
| - | - | 1285 | 236.1 | - | - | 0 | - |
| - | - | 7985 | 236.1 | - | - | 0 | - |
| - | - | 2658 | 236.1 | - | - | 0 | - |
| - | - | 1933 | 236.2 | - | - | 0 | - |
| - | - | 1270 | 237.1 | - | - | 0 | - |
| - | - | 1.37E+04 | 237.1 | - | - | 0 | - |
| - | - | 1032 | 238.1 | - | - | 0 | - |
| - | - | 2730 | 238.1 | - | - | 0 | - |
| - | - | 944.2 | 239.1 | - | - | 0 | - |
| - | - | 4.04E+04 | 239.1 | - | - | 0 | - |
| - | - | 8.99E+04 | 239.2 | - | - | 0 | - |
| - | - | 1923 | 240.1 | - | - | 0 | - |
| - | - | 1.895E+05 | 240.1 | - | - | 0 | - |
| - | - | 8741 | 240.2 | - | - | 0 | - |
| - | - | 1836 | 241.1 | - | - | 0 | - |
| - | - | 1402 | 241.1 | - | - | 0 | - |
| - | - | 2.164E+04 | 241.1 | - | - | 0 | - |
| - | - | 1186 | 241.2 | - | - | 0 | - |
| - | - | 1676 | 242.1 | - | - | 0 | - |
| - | - | 1655 | 242.1 | - | - | 0 | - |
| - | - | 1040 | 243.1 | - | - | 0 | - |
| - | - | 4850 | 244.1 | - | - | 0 | - |
| - | - | 1.74E+04 | 247.1 | - | - | 0 | - |
| - | - | 1050 | 247.1 | - | - | 0 | - |
| - | - | 2864 | 248.1 | - | - | 0 | - |
| - | - | 2.919E+04 | 249.1 | - | - | 0 | - |
| - | - | 4685 | 249.1 | - | - | 0 | - |
| - | - | 568.9 | 249.2 | - | - | 0 | - |
| - | - | 705.8 | 250.1 | - | - | 0 | - |
| - | - | 3664 | 250.1 | - | - | 0 | - |
| - | - | 2980 | 250.1 | - | - | 0 | - |
| - | - | 1813 | 250.1 | - | - | 0 | - |
| - | - | 1924 | 250.2 | - | - | 0 | - |
| - | - | 1169 | 251.1 | - | - | 0 | - |
| - | - | 1.964E+04 | 251.2 | - | - | 0 | - |
| - | - | 1.61E+05 | 252.1 | - | - | 0 | - |
| - | - | 9748 | 252.1 | - | - | 0 | - |
| - | - | 899.7 | 253.1 | - | - | 0 | - |
| - | - | 4289 | 253.1 | - | - | 0 | - |
| - | - | 1.582E+04 | 253.1 | - | - | 0 | - |
| - | - | 1219 | 253.1 | - | - | 0 | - |
| - | - | 666.2 | 254.1 | - | - | 0 | - |
| - | - | 2174 | 254.1 | - | - | 0 | - |
| - | - | 2197 | 256.1 | - | - | 0 | - |
| - | - | 2.434E+04 | 257.1 | - | - | 0 | - |
| - | - | 1797 | 257.1 | - | - | 0 | - |
| - | - | 2.47E+04 | 257.2 | - | - | 0 | - |
| - | - | 4790 | 258.1 | - | - | 0 | - |
| - | - | 7300 | 258.1 | - | - | 0 | - |
| - | - | 2123 | 258.2 | - | - | 0 | - |
| - | - | 1292 | 259.1 | - | - | 0 | - |
| - | - | 1462 | 259.2 | - | - | 0 | - |
| - | - | 1112 | 260.1 | - | - | 0 | - |
| - | - | 722.3 | 261.1 | - | - | 0 | - |
| - | - | 3741 | 261.1 | - | - | 0 | - |
| - | - | 1514 | 261.2 | - | - | 0 | - |
| - | - | 7747 | 262.1 | - | - | 0 | - |
| - | - | 1427 | 262.1 | - | - | 0 | - |
| - | - | 2724 | 263.1 | - | - | 0 | - |
| - | - | 1136 | 263.1 | - | - | 0 | - |
| - | - | 7086 | 264.1 | - | - | 0 | - |
| - | - | 1.433E+04 | 265.1 | - | - | 0 | - |
| - | - | 2259 | 266.1 | - | - | 0 | - |
| - | - | 7142 | 267.1 | - | - | 0 | - |
| - | - | 9.117E+04 | 267.1 | - | - | 0 | - |
| - | - | 3790 | 268.1 | - | - | 0 | - |
| - | - | 1210 | 268.1 | - | - | 0 | - |
| - | - | 1.048E+04 | 268.1 | - | - | 0 | - |
| - | - | 3341 | 268.1 | - | - | 0 | - |
| - | - | 2647 | 268.2 | - | - | 0 | - |
| - | - | 1115 | 269.1 | - | - | 0 | - |
| - | - | 668.5 | 269.1 | - | - | 0 | - |
| - | - | 7464 | 269.1 | - | - | 0 | - |
| - | - | 619.2 | 269.2 | - | - | 0 | - |
| - | - | 1215 | 270.1 | - | - | 0 | - |
| - | - | 1946 | 270.1 | - | - | 0 | - |
| - | - | 3.246E+04 | 270.1 | - | - | 0 | - |
| - | - | 3799 | 271.1 | - | - | 0 | - |
| - | - | 596.9 | 271.5 | - | - | 0 | - |
| - | - | 4.351E+04 | 273.1 | - | - | 0 | - |
| - | - | 4966 | 274.1 | - | - | 0 | - |
| - | - | 872.7 | 274.2 | - | - | 0 | - |
| - | - | 3.683E+04 | 275.1 | - | - | 0 | - |
| - | - | 1988 | 275.2 | - | - | 0 | - |
| - | - | 6497 | 276.1 | - | - | 0 | - |
| - | - | 1635 | 276.2 | - | - | 0 | - |
| - | - | 4672 | 277.1 | - | - | 0 | - |
| - | - | 1323 | 277.2 | - | - | 0 | - |
| - | - | 898.1 | 278.1 | - | - | 0 | - |
| - | - | 820.2 | 278.1 | - | - | 0 | - |
| - | - | 1327 | 278.2 | - | - | 0 | - |
| - | - | 1.766E+04 | 279.1 | - | - | 0 | - |
| - | - | 9922 | 280.1 | - | - | 0 | - |
| - | - | 2.775E+04 | 280.1 | - | - | 0 | - |
| - | - | 3392 | 280.1 | - | - | 0 | - |
| - | - | 2254 | 280.7 | - | - | 0 | - |
| - | - | 1440 | 281.1 | - | - | 0 | - |
| - | - | 2931 | 281.1 | - | - | 0 | - |
| - | - | 1059 | 281.2 | - | - | 0 | - |
| - | - | 2134 | 282.1 | - | - | 0 | - |
| - | - | 2062 | 283.1 | - | - | 0 | - |
| - | - | 653.2 | 283.1 | - | - | 0 | - |
| - | - | 2162 | 284.1 | - | - | 0 | - |
| - | - | 2038 | 284.1 | - | - | 0 | - |
| - | - | 6.684E+04 | 285.1 | - | - | 0 | - |
| - | - | 893.1 | 285.6 | - | - | 0 | - |
| - | - | 9812 | 286.1 | - | - | 0 | - |
| - | - | 1541 | 286.1 | - | - | 0 | - |
| - | - | 2867 | 287.1 | - | - | 0 | - |
| - | - | 1891 | 287.2 | - | - | 0 | - |
| - | - | 641.4 | 287.2 | - | - | 0 | - |
| - | - | 1126 | 291.1 | - | - | 0 | - |
| - | - | 1022 | 292.1 | - | - | 0 | - |
| - | - | 1395 | 292.2 | - | - | 0 | - |
| - | - | 2165 | 293.1 | - | - | 0 | - |
| - | - | 2480 | 294.2 | - | - | 0 | - |
| - | - | 2672 | 294.6 | - | - | 0 | - |
| - | - | 1.167E+04 | 295.1 | - | - | 0 | - |
| - | - | 1892 | 296.2 | - | - | 0 | - |
| - | - | 1.321E+04 | 297.2 | - | - | 0 | - |
| - | - | 1183 | 298.1 | - | - | 0 | - |
| - | - | 2245 | 298.2 | - | - | 0 | - |
| - | - | 823.2 | 299.6 | - | - | 0 | - |
| - | - | 1259 | 300.1 | - | - | 0 | - |
| 5 | b | 694.9 | 300.6 | 0.003796 | 12.63 | +2 | 5 |
| - | - | 7.191E+04 | 301.1 | - | - | 0 | - |
| - | - | 1.259E+04 | 302.1 | - | - | 0 | - |
| - | - | 2788 | 302.2 | - | - | 0 | - |
| - | - | 2124 | 303.1 | - | - | 0 | - |
| - | - | 694.5 | 303.1 | - | - | 0 | - |
| - | - | 1029 | 303.2 | - | - | 0 | - |
| - | - | 3409 | 304.1 | - | - | 0 | - |
| - | - | 3215 | 304.1 | - | - | 0 | - |
| - | - | 7020 | 304.2 | - | - | 0 | - |
| - | - | 2035 | 305.1 | - | - | 0 | - |
| - | - | 1889 | 305.1 | - | - | 0 | - |
| - | - | 6350 | 305.2 | - | - | 0 | - |
| - | - | 1208 | 305.2 | - | - | 0 | - |
| - | - | 1239 | 306.1 | - | - | 0 | - |
| 3 | a | 5134 | 306.1 | 0.00347 | 11.34 | +1 | 3 |
| - | - | 917.8 | 306.2 | - | - | 0 | - |
| - | - | 1432 | 307.1 | - | - | 0 | - |
| - | - | 564.7 | 307.1 | - | - | 0 | - |
| - | - | 643.2 | 309.1 | - | - | 0 | - |
| - | - | 927.4 | 309.1 | - | - | 0 | - |
| - | - | 887.9 | 309.2 | - | - | 0 | - |
| - | - | 703.8 | 309.6 | - | - | 0 | - |
| - | - | 1680 | 310.1 | - | - | 0 | - |
| - | - | 2456 | 311.1 | - | - | 0 | - |
| - | - | 1088 | 312.1 | - | - | 0 | - |
| - | - | 1418 | 312.2 | - | - | 0 | - |
| - | - | 1164 | 313.1 | - | - | 0 | - |
| - | - | 1055 | 313.1 | - | - | 0 | - |
| - | - | 669 | 313.2 | - | - | 0 | - |
| 11 | y | 1097 | 313.7 | 0.002247 | 7.164 | +2 | 5 |
| - | - | 1024 | 314.1 | - | - | 0 | - |
| - | - | 2257 | 314.2 | - | - | 0 | - |
| - | - | 1478 | 314.6 | - | - | 0 | - |
| - | - | 683.6 | 315.1 | - | - | 0 | - |
| - | - | 803 | 315.1 | - | - | 0 | - |
| - | - | 1.1E+04 | 316.1 | - | - | 0 | - |
| - | - | 801.5 | 317.1 | - | - | 0 | - |
| - | - | 1392 | 317.1 | - | - | 0 | - |
| - | - | 734.7 | 318.1 | - | - | 0 | - |
| - | - | 3848 | 318.1 | - | - | 0 | - |
| - | - | 1063 | 318.2 | - | - | 0 | - |
| - | - | 995.4 | 318.6 | - | - | 0 | - |
| - | - | 1427 | 319.1 | - | - | 0 | - |
| - | - | 3012 | 319.1 | - | - | 0 | - |
| - | - | 4.796E+04 | 320.1 | - | - | 0 | - |
| - | - | 7001 | 320.2 | - | - | 0 | - |
| - | - | 886.5 | 321.1 | - | - | 0 | - |
| - | - | 7868 | 321.1 | - | - | 0 | - |
| - | - | 1260 | 321.2 | - | - | 0 | - |
| - | - | 1106 | 321.2 | - | - | 0 | - |
| - | - | 3754 | 322.1 | - | - | 0 | - |
| - | - | 6285 | 322.2 | - | - | 0 | - |
| - | - | 2533 | 322.2 | - | - | 0 | - |
| - | - | 2.188E+04 | 322.2 | - | - | 0 | - |
| - | - | 1502 | 322.7 | - | - | 0 | - |
| - | - | 865.7 | 322.7 | - | - | 0 | - |
| - | - | 739.8 | 323.1 | - | - | 0 | - |
| - | - | 1657 | 323.2 | - | - | 0 | - |
| - | - | 1.268E+04 | 323.2 | - | - | 0 | - |
| - | - | 3533 | 323.2 | - | - | 0 | - |
| - | - | 990.6 | 324.1 | - | - | 0 | - |
| - | - | 1991 | 324.2 | - | - | 0 | - |
| - | - | 897.5 | 325.2 | - | - | 0 | - |
| - | - | 5566 | 326.1 | - | - | 0 | - |
| - | - | 1134 | 326.2 | - | - | 0 | - |
| - | - | 1499 | 326.2 | - | - | 0 | - |
| - | - | 904.7 | 326.7 | - | - | 0 | - |
| 6 | b | 2916 | 327.1 | 0.006513 | 19.91 | +2 | 6 |
| - | - | 7607 | 327.2 | - | - | 0 | - |
| - | - | 950.3 | 327.2 | - | - | 0 | - |
| - | - | 824.9 | 327.7 | - | - | 0 | - |
| - | - | 4898 | 328.2 | - | - | 0 | - |
| - | - | 804.5 | 328.2 | - | - | 0 | - |
| - | - | 2088 | 328.7 | - | - | 0 | - |
| - | - | 2911 | 329.1 | - | - | 0 | - |
| - | - | 765.4 | 330.1 | - | - | 0 | - |
| - | - | 2828 | 330.2 | - | - | 0 | - |
| - | - | 1966 | 331.1 | - | - | 0 | - |
| - | - | 660.4 | 331.1 | - | - | 0 | - |
| - | - | 1.564E+04 | 332.1 | - | - | 0 | - |
| - | - | 889 | 332.2 | - | - | 0 | - |
| - | - | 3026 | 333.1 | - | - | 0 | - |
| - | - | 1463 | 333.1 | - | - | 0 | - |
| - | - | 7.723E+04 | 334.1 | - | - | 0 | - |
| - | - | 2105 | 335.1 | - | - | 0 | - |
| - | - | 1.125E+04 | 335.1 | - | - | 0 | - |
| - | - | 603 | 335.7 | - | - | 0 | - |
| 6 | b | 5050 | 336.1 | 0.002177 | 6.476 | +2 | 6 |
| - | - | 4117 | 336.2 | - | - | 0 | - |
| - | - | 949.6 | 336.7 | - | - | 0 | - |
| - | - | 819.8 | 337.1 | - | - | 0 | - |
| - | - | 1280 | 337.1 | - | - | 0 | - |
| - | - | 695.2 | 337.2 | - | - | 0 | - |
| - | - | 1405 | 337.2 | - | - | 0 | - |
| - | - | 1259 | 337.7 | - | - | 0 | - |
| - | - | 4.624E+04 | 338.1 | - | - | 0 | - |
| - | - | 1775 | 338.7 | - | - | 0 | - |
| - | - | 8093 | 339.2 | - | - | 0 | - |
| - | - | 1010 | 339.2 | - | - | 0 | - |
| - | - | 3522 | 339.2 | - | - | 0 | - |
| - | - | 2.515E+04 | 340.1 | - | - | 0 | - |
| - | - | 3730 | 340.2 | - | - | 0 | - |
| - | - | 4900 | 341.1 | - | - | 0 | - |
| - | - | 3.022E+04 | 341.2 | - | - | 0 | - |
| - | - | 5912 | 342.2 | - | - | 0 | - |
| - | - | 1303 | 342.7 | - | - | 0 | - |
| - | - | 3282 | 343.2 | - | - | 0 | - |
| - | - | 704.3 | 343.2 | - | - | 0 | - |
| - | - | 1.029E+04 | 344.1 | - | - | 0 | - |
| 13 | y | 2.316E+04 | 344.2 | 0.001948 | 5.661 | +1 | 3 |
| - | - | 955.1 | 344.7 | - | - | 0 | - |
| - | - | 1444 | 345.2 | - | - | 0 | - |
| 13 | y | 2.957E+04 | 345.2 | 0.001942 | 5.625 | +1 | 3 |
| - | - | 2554 | 345.2 | - | - | 0 | - |
| - | - | 4805 | 346.2 | - | - | 0 | - |
| - | - | 1464 | 346.6 | - | - | 0 | - |
| - | - | 757.4 | 346.7 | - | - | 0 | - |
| - | - | 1335 | 347.1 | - | - | 0 | - |
| - | - | 2435 | 347.2 | - | - | 0 | - |
| - | - | 5794 | 348.1 | - | - | 0 | - |
| - | - | 4997 | 348.2 | - | - | 0 | - |
| - | - | 1609 | 349.1 | - | - | 0 | - |
| - | - | 1148 | 349.2 | - | - | 0 | - |
| - | - | 692.1 | 349.2 | - | - | 0 | - |
| - | - | 6.251E+04 | 350.1 | - | - | 0 | - |
| - | - | 2250 | 350.2 | - | - | 0 | - |
| - | - | 1602 | 350.7 | - | - | 0 | - |
| - | - | 1.013E+04 | 351.2 | - | - | 0 | - |
| - | - | 5565 | 351.7 | - | - | 0 | - |
| - | - | 4217 | 352.1 | - | - | 0 | - |
| - | - | 2802 | 352.2 | - | - | 0 | - |
| - | - | 918.9 | 352.7 | - | - | 0 | - |
| - | - | 3124 | 353.1 | - | - | 0 | - |
| - | - | 1670 | 353.1 | - | - | 0 | - |
| - | - | 969.4 | 353.2 | - | - | 0 | - |
| - | - | 3621 | 353.2 | - | - | 0 | - |
| - | - | 7284 | 354.1 | - | - | 0 | - |
| - | - | 3877 | 354.2 | - | - | 0 | - |
| - | - | 991.4 | 354.7 | - | - | 0 | - |
| - | - | 1146 | 355.1 | - | - | 0 | - |
| - | - | 1242 | 355.2 | - | - | 0 | - |
| - | - | 1208 | 356.1 | - | - | 0 | - |
| - | - | 2104 | 356.2 | - | - | 0 | - |
| - | - | 2873 | 358.2 | - | - | 0 | - |
| - | - | 1147 | 358.2 | - | - | 0 | - |
| - | - | 1.613E+04 | 358.2 | - | - | 0 | - |
| - | - | 1340 | 359.1 | - | - | 0 | - |
| - | - | 1411 | 359.2 | - | - | 0 | - |
| - | - | 2275 | 359.2 | - | - | 0 | - |
| - | - | 1250 | 360.2 | - | - | 0 | - |
| - | - | 899.5 | 360.9 | - | - | 0 | - |
| - | - | 1.235E+04 | 361.2 | - | - | 0 | - |
| - | - | 2145 | 361.2 | - | - | 0 | - |
| - | - | 3274 | 362.2 | - | - | 0 | - |
| 13 | y | 2.004E+04 | 362.2 | 0.002004 | 5.532 | +1 | 3 |
| - | - | 1422 | 362.5 | - | - | 0 | - |
| - | - | 821.5 | 363.1 | - | - | 0 | - |
| - | - | 830.9 | 363.2 | - | - | 0 | - |
| - | - | 3980 | 363.2 | - | - | 0 | - |
| - | - | 880.8 | 364.1 | - | - | 0 | - |
| - | - | 1033 | 364.2 | - | - | 0 | - |
| - | - | 1.414E+04 | 365.2 | - | - | 0 | - |
| - | - | 1.055E+04 | 366.1 | - | - | 0 | - |
| - | - | 1.607E+04 | 366.2 | - | - | 0 | - |
| - | - | 2629 | 366.2 | - | - | 0 | - |
| - | - | 2120 | 367.1 | - | - | 0 | - |
| - | - | 4321 | 367.2 | - | - | 0 | - |
| - | - | 977 | 368.1 | - | - | 0 | - |
| - | - | 6252 | 368.2 | - | - | 0 | - |
| - | - | 1995 | 368.7 | - | - | 0 | - |
| - | - | 3293 | 369.2 | - | - | 0 | - |
| - | - | 920.7 | 369.7 | - | - | 0 | - |
| - | - | 5013 | 370.2 | - | - | 0 | - |
| - | - | 593.4 | 370.7 | - | - | 0 | - |
| - | - | 4.333E+04 | 371.2 | - | - | 0 | - |
| - | - | 2.001E+04 | 372.1 | - | - | 0 | - |
| - | - | 5625 | 372.2 | - | - | 0 | - |
| - | - | 3903 | 373.1 | - | - | 0 | - |
| 10 | y | 1669 | 373.2 | 0.002297 | 6.154 | +2 | 6 |
| 10 | y | 5486 | 373.7 | 0.002873 | 7.688 | +2 | 6 |
| - | - | 859.4 | 374.1 | - | - | 0 | - |
| - | - | 3352 | 374.2 | - | - | 0 | - |
| - | - | 1.274E+04 | 375.2 | - | - | 0 | - |
| - | - | 1134 | 376.1 | - | - | 0 | - |
| - | - | 3683 | 376.2 | - | - | 0 | - |
| - | - | 557.6 | 376.2 | - | - | 0 | - |
| - | - | 3058 | 376.2 | - | - | 0 | - |
| - | - | 1175 | 376.7 | - | - | 0 | - |
| - | - | 585 | 377.2 | - | - | 0 | - |
| - | - | 814.2 | 377.2 | - | - | 0 | - |
| - | - | 1143 | 378.1 | - | - | 0 | - |
| - | - | 2615 | 378.2 | - | - | 0 | - |
| - | - | 2443 | 378.7 | - | - | 0 | - |
| - | - | 741.5 | 379.1 | - | - | 0 | - |
| - | - | 868.1 | 379.2 | - | - | 0 | - |
| - | - | 2.022E+04 | 379.2 | - | - | 0 | - |
| - | - | 1085 | 379.7 | - | - | 0 | - |
| - | - | 4930 | 380.1 | - | - | 0 | - |
| - | - | 558.2 | 380.2 | - | - | 0 | - |
| - | - | 1760 | 380.2 | - | - | 0 | - |
| - | - | 4377 | 380.2 | - | - | 0 | - |
| - | - | 4407 | 381.1 | - | - | 0 | - |
| - | - | 794.7 | 381.1 | - | - | 0 | - |
| - | - | 885 | 381.2 | - | - | 0 | - |
| - | - | 1447 | 382.1 | - | - | 0 | - |
| - | - | 6240 | 382.2 | - | - | 0 | - |
| 10 | y | 1.848E+04 | 382.2 | 0.001989 | 5.203 | +2 | 6 |
| - | - | 1918 | 382.2 | - | - | 0 | - |
| - | - | 2215 | 382.7 | - | - | 0 | - |
| - | - | 9859 | 382.7 | - | - | 0 | - |
| - | - | 1751 | 383.2 | - | - | 0 | - |
| - | - | 1.673E+04 | 384.2 | - | - | 0 | - |
| - | - | 1104 | 384.2 | - | - | 0 | - |
| - | - | 672.7 | 384.5 | - | - | 0 | - |
| - | - | 2015 | 384.7 | - | - | 0 | - |
| - | - | 4138 | 385.2 | - | - | 0 | - |
| - | - | 4938 | 386.1 | - | - | 0 | - |
| - | - | 868 | 386.7 | - | - | 0 | - |
| - | - | 1420 | 387.2 | - | - | 0 | - |
| - | - | 5494 | 387.2 | - | - | 0 | - |
| - | - | 1657 | 387.2 | - | - | 0 | - |
| - | - | 1184 | 388.2 | - | - | 0 | - |
| - | - | 1240 | 388.7 | - | - | 0 | - |
| - | - | 1.587E+04 | 389.2 | - | - | 0 | - |
| - | - | 2929 | 389.2 | - | - | 0 | - |
| - | - | 2890 | 390.2 | - | - | 0 | - |
| - | - | 999 | 390.5 | - | - | 0 | - |
| - | - | 1871 | 391.2 | - | - | 0 | - |
| - | - | 2078 | 391.2 | - | - | 0 | - |
| 10 | b | 1727 | 391.5 | 0.004264 | 10.89 | +3 | 10 |
| - | - | 992.6 | 391.7 | - | - | 0 | - |
| - | - | 985.9 | 392.2 | - | - | 0 | - |
| - | - | 1968 | 393.2 | - | - | 0 | - |
| - | - | 1.539E+04 | 393.2 | - | - | 0 | - |
| - | - | 2603 | 393.7 | - | - | 0 | - |
| - | - | 4056 | 394.2 | - | - | 0 | - |
| - | - | 3344 | 394.2 | - | - | 0 | - |
| - | - | 1111 | 395.2 | - | - | 0 | - |
| - | - | 1395 | 395.7 | - | - | 0 | - |
| - | - | 4331 | 396.1 | - | - | 0 | - |
| - | - | 968.9 | 396.2 | - | - | 0 | - |
| - | - | 751.6 | 396.5 | - | - | 0 | - |
| - | - | 776.4 | 397.1 | - | - | 0 | - |
| - | - | 6817 | 397.2 | - | - | 0 | - |
| - | - | 2290 | 397.2 | - | - | 0 | - |
| - | - | 9030 | 397.7 | - | - | 0 | - |
| - | - | 1.328E+04 | 398.1 | - | - | 0 | - |
| - | - | 1868 | 398.2 | - | - | 0 | - |
| - | - | 4473 | 399.2 | - | - | 0 | - |
| - | - | 1240 | 400.1 | - | - | 0 | - |
| - | - | 4679 | 401.2 | - | - | 0 | - |
| - | - | 2251 | 402.1 | - | - | 0 | - |
| - | - | 1.02E+04 | 402.2 | - | - | 0 | - |
| - | - | 2151 | 402.7 | - | - | 0 | - |
| - | - | 4699 | 403.2 | - | - | 0 | - |
| - | - | 4304 | 404.2 | - | - | 0 | - |
| - | - | 1084 | 404.7 | - | - | 0 | - |
| - | - | 1394 | 405.2 | - | - | 0 | - |
| - | - | 1.132E+04 | 405.2 | - | - | 0 | - |
| - | - | 2.204E+04 | 406.2 | - | - | 0 | - |
| - | - | 9556 | 406.7 | - | - | 0 | - |
| - | - | 2.63E+04 | 407.2 | - | - | 0 | - |
| - | - | 4531 | 408.2 | - | - | 0 | - |
| - | - | 3616 | 408.2 | - | - | 0 | - |
| - | - | 7683 | 410.2 | - | - | 0 | - |
| - | - | 604.4 | 410.7 | - | - | 0 | - |
| - | - | 5069 | 411.2 | - | - | 0 | - |
| - | - | 1712 | 411.5 | - | - | 0 | - |
| - | - | 3823 | 411.7 | - | - | 0 | - |
| - | - | 884.8 | 411.8 | - | - | 0 | - |
| - | - | 1.038E+04 | 412.2 | - | - | 0 | - |
| - | - | 738.1 | 412.7 | - | - | 0 | - |
| - | - | 3813 | 413.2 | - | - | 0 | - |
| - | - | 3866 | 413.7 | - | - | 0 | - |
| - | - | 4.137E+04 | 414.1 | - | - | 0 | - |
| - | - | 782.5 | 414.7 | - | - | 0 | - |
| - | - | 5806 | 415.1 | - | - | 0 | - |
| - | - | 3.416E+04 | 415.2 | - | - | 0 | - |
| - | - | 8513 | 415.2 | - | - | 0 | - |
| - | - | 755.4 | 415.5 | - | - | 0 | - |
| - | - | 4028 | 415.7 | - | - | 0 | - |
| - | - | 1950 | 416.1 | - | - | 0 | - |
| - | - | 8006 | 416.2 | - | - | 0 | - |
| - | - | 1130 | 416.2 | - | - | 0 | - |
| - | - | 1456 | 416.7 | - | - | 0 | - |
| - | - | 2628 | 417.2 | - | - | 0 | - |
| - | - | 1091 | 417.2 | - | - | 0 | - |
| - | - | 1223 | 417.7 | - | - | 0 | - |
| - | - | 947.8 | 417.9 | - | - | 0 | - |
| - | - | 865.7 | 418.2 | - | - | 0 | - |
| - | - | 748.6 | 418.2 | - | - | 0 | - |
| - | - | 1498 | 419.2 | - | - | 0 | - |
| - | - | 7157 | 419.2 | - | - | 0 | - |
| - | - | 741.5 | 420.1 | - | - | 0 | - |
| - | - | 1306 | 420.2 | - | - | 0 | - |
| - | - | 1.469E+04 | 420.2 | - | - | 0 | - |
| - | - | 6115 | 420.7 | - | - | 0 | - |
| - | - | 4.199E+04 | 421.2 | - | - | 0 | - |
| - | - | 2735 | 421.7 | - | - | 0 | - |
| - | - | 1.526E+04 | 422.2 | - | - | 0 | - |
| - | - | 2686 | 422.7 | - | - | 0 | - |
| - | - | 2080 | 423.2 | - | - | 0 | - |
| - | - | 1.857E+04 | 423.2 | - | - | 0 | - |
| - | - | 857.1 | 423.5 | - | - | 0 | - |
| - | - | 1490 | 424.2 | - | - | 0 | - |
| - | - | 5290 | 424.2 | - | - | 0 | - |
| - | - | 2691 | 424.7 | - | - | 0 | - |
| - | - | 734.1 | 424.9 | - | - | 0 | - |
| - | - | 1499 | 425.2 | - | - | 0 | - |
| - | - | 1.263E+04 | 425.2 | - | - | 0 | - |
| - | - | 4082 | 425.7 | - | - | 0 | - |
| - | - | 1155 | 426.2 | - | - | 0 | - |
| - | - | 6302 | 426.2 | - | - | 0 | - |
| - | - | 1231 | 427.2 | - | - | 0 | - |
| - | - | 1171 | 428.2 | - | - | 0 | - |
| - | - | 2722 | 428.2 | - | - | 0 | - |
| - | - | 928.8 | 429.2 | - | - | 0 | - |
| - | - | 9030 | 429.2 | - | - | 0 | - |
| - | - | 959.6 | 429.7 | - | - | 0 | - |
| - | - | 5607 | 429.7 | - | - | 0 | - |
| - | - | 3223 | 430.2 | - | - | 0 | - |
| 9 | y | 5984 | 430.2 | 0.0003641 | 0.8462 | +2 | 7 |
| 9 | y | 1.179E+04 | 430.7 | 0.002711 | 6.293 | +2 | 7 |
| - | - | 2234 | 431.2 | - | - | 0 | - |
| - | - | 4615 | 431.2 | - | - | 0 | - |
| - | - | 2122 | 431.7 | - | - | 0 | - |
| - | - | 1024 | 432.2 | - | - | 0 | - |
| - | - | 855.1 | 433.2 | - | - | 0 | - |
| - | - | 1.527E+04 | 433.2 | - | - | 0 | - |
| - | - | 1770 | 433.5 | - | - | 0 | - |
| - | - | 1557 | 433.7 | - | - | 0 | - |
| - | - | 1717 | 433.9 | - | - | 0 | - |
| - | - | 1486 | 434.2 | - | - | 0 | - |
| - | - | 2148 | 434.2 | - | - | 0 | - |
| - | - | 2576 | 436.2 | - | - | 0 | - |
| - | - | 1.843E+04 | 436.2 | - | - | 0 | - |
| - | - | 2272 | 436.5 | - | - | 0 | - |
| - | - | 1140 | 436.9 | - | - | 0 | - |
| - | - | 910.2 | 437.2 | - | - | 0 | - |
| - | - | 4006 | 437.2 | - | - | 0 | - |
| - | - | 1027 | 437.7 | - | - | 0 | - |
| - | - | 1130 | 437.7 | - | - | 0 | - |
| - | - | 672.2 | 437.9 | - | - | 0 | - |
| - | - | 1293 | 438.1 | - | - | 0 | - |
| - | - | 2153 | 438.2 | - | - | 0 | - |
| - | - | 5545 | 438.7 | - | - | 0 | - |
| 9 | y | 4.197E+04 | 439.2 | 0.001887 | 4.297 | +2 | 7 |
| - | - | 2.075E+04 | 439.7 | - | - | 0 | - |
| - | - | 4660 | 440.2 | - | - | 0 | - |
| - | - | 767.6 | 440.7 | - | - | 0 | - |
| - | - | 1633 | 442.1 | - | - | 0 | - |
| - | - | 3188 | 442.2 | - | - | 0 | - |
| - | - | 1085 | 442.5 | - | - | 0 | - |
| - | - | 1737 | 442.9 | - | - | 0 | - |
| - | - | 1.085E+04 | 443.2 | - | - | 0 | - |
| 5 | y | 3.457E+04 | 443.2 | 0.007748 | 17.48 | +3 | 11 |
| - | - | 768.3 | 443.6 | - | - | 0 | - |
| - | - | 2088 | 444.2 | - | - | 0 | - |
| - | - | 8572 | 444.2 | - | - | 0 | - |
| - | - | 1748 | 444.7 | - | - | 0 | - |
| - | - | 2711 | 445.2 | - | - | 0 | - |
| 12 | y | 7372 | 445.2 | 0.003098 | 6.958 | +1 | 4 |
| - | - | 681.5 | 445.7 | - | - | 0 | - |
| 11 | b | 3420 | 445.9 | 0.004374 | 9.81 | +3 | 11 |
| - | - | 3178 | 446.1 | - | - | 0 | - |
| 12 | y | 4968 | 446.2 | 0.001321 | 2.961 | +1 | 4 |
| - | - | 1048 | 446.5 | - | - | 0 | - |
| - | - | 3558 | 446.7 | - | - | 0 | - |
| - | - | 2.16E+04 | 447.2 | - | - | 0 | - |
| - | - | 2.589E+04 | 447.7 | - | - | 0 | - |
| - | - | 1.253E+04 | 448.2 | - | - | 0 | - |
| - | - | 2136 | 448.6 | - | - | 0 | - |
| - | - | 3273 | 448.7 | - | - | 0 | - |
| - | - | 1671 | 448.9 | - | - | 0 | - |
| - | - | 4857 | 449.2 | - | - | 0 | - |
| - | - | 1106 | 450.1 | - | - | 0 | - |
| - | - | 1243 | 450.2 | - | - | 0 | - |
| - | - | 1112 | 450.2 | - | - | 0 | - |
| - | - | 2791 | 450.2 | - | - | 0 | - |
| - | - | 835.8 | 450.9 | - | - | 0 | - |
| - | - | 4.942E+04 | 451.2 | - | - | 0 | - |
| - | - | 1.142E+04 | 452.2 | - | - | 0 | - |
| - | - | 1784 | 452.6 | - | - | 0 | - |
| - | - | 2626 | 452.7 | - | - | 0 | - |
| - | - | 1407 | 452.9 | - | - | 0 | - |
| - | - | 3706 | 453.2 | - | - | 0 | - |
| - | - | 1161 | 453.7 | - | - | 0 | - |
| - | - | 1030 | 453.9 | - | - | 0 | - |
| - | - | 2121 | 455.2 | - | - | 0 | - |
| - | - | 5590 | 455.2 | - | - | 0 | - |
| - | - | 1383 | 455.5 | - | - | 0 | - |
| - | - | 2960 | 455.7 | - | - | 0 | - |
| - | - | 1455 | 455.9 | - | - | 0 | - |
| - | - | 730.8 | 456.2 | - | - | 0 | - |
| - | - | 1030 | 456.2 | - | - | 0 | - |
| - | - | 2009 | 457.2 | - | - | 0 | - |
| - | - | 3370 | 457.2 | - | - | 0 | - |
| - | - | 740.3 | 457.9 | - | - | 0 | - |
| - | - | 2028 | 458.2 | - | - | 0 | - |
| - | - | 2467 | 458.6 | - | - | 0 | - |
| - | - | 1011 | 458.9 | - | - | 0 | - |
| - | - | 1238 | 459.2 | - | - | 0 | - |
| - | - | 1597 | 459.9 | - | - | 0 | - |
| - | - | 4458 | 460.2 | - | - | 0 | - |
| - | - | 838.6 | 460.5 | - | - | 0 | - |
| - | - | 1087 | 460.7 | - | - | 0 | - |
| - | - | 1196 | 460.9 | - | - | 0 | - |
| 8 | b | 4.707E+04 | 461.2 | 0.005335 | 11.57 | +2 | 8 |
| - | - | 1845 | 461.6 | - | - | 0 | - |
| - | - | 2.868E+04 | 461.7 | - | - | 0 | - |
| - | - | 1137 | 461.9 | - | - | 0 | - |
| - | - | 1.236E+04 | 462.2 | - | - | 0 | - |
| - | - | 3028 | 462.7 | - | - | 0 | - |
| - | - | 1996 | 463.2 | - | - | 0 | - |
| 12 | y | 4.692E+04 | 463.3 | 0.002696 | 5.819 | +1 | 4 |
| - | - | 5306 | 464.2 | - | - | 0 | - |
| - | - | 5722 | 464.2 | - | - | 0 | - |
| - | - | 9097 | 464.3 | - | - | 0 | - |
| - | - | 3362 | 464.6 | - | - | 0 | - |
| - | - | 920 | 464.7 | - | - | 0 | - |
| - | - | 3099 | 464.9 | - | - | 0 | - |
| - | - | 1327 | 465.2 | - | - | 0 | - |
| - | - | 2246 | 465.2 | - | - | 0 | - |
| - | - | 1393 | 465.3 | - | - | 0 | - |
| - | - | 754.3 | 465.5 | - | - | 0 | - |
| - | - | 2204 | 465.7 | - | - | 0 | - |
| - | - | 1221 | 466.2 | - | - | 0 | - |
| - | - | 1149 | 466.7 | - | - | 0 | - |
| - | - | 779.9 | 466.9 | - | - | 0 | - |
| - | - | 2039 | 467.2 | - | - | 0 | - |
| - | - | 3060 | 467.2 | - | - | 0 | - |
| - | - | 805.3 | 467.5 | - | - | 0 | - |
| - | - | 1134 | 467.9 | - | - | 0 | - |
| - | - | 984.3 | 468.2 | - | - | 0 | - |
| - | - | 1036 | 468.2 | - | - | 0 | - |
| - | - | 3509 | 468.3 | - | - | 0 | - |
| - | - | 742.9 | 469.2 | - | - | 0 | - |
| - | - | 3630 | 469.7 | - | - | 0 | - |
| - | - | 2901 | 469.9 | - | - | 0 | - |
| - | - | 6142 | 470.2 | - | - | 0 | - |
| - | - | 3265 | 470.6 | - | - | 0 | - |
| - | - | 2618 | 470.7 | - | - | 0 | - |
| - | - | 842 | 470.9 | - | - | 0 | - |
| 4 | b | 3.804E+04 | 471.2 | 0.008299 | 17.61 | +1 | 4 |
| - | - | 1233 | 471.7 | - | - | 0 | - |
| - | - | 9321 | 472.2 | - | - | 0 | - |
| - | - | 982.3 | 472.2 | - | - | 0 | - |
| - | - | 738.2 | 472.6 | - | - | 0 | - |
| - | - | 2942 | 473.2 | - | - | 0 | - |
| - | - | 1224 | 473.2 | - | - | 0 | - |
| 12 | b | 6930 | 473.5 | 0.004657 | 9.835 | +3 | 12 |
| - | - | 5529 | 473.9 | - | - | 0 | - |
| - | - | 2587 | 474.2 | - | - | 0 | - |
| - | - | 6035 | 474.3 | - | - | 0 | - |
| - | - | 3862 | 474.7 | - | - | 0 | - |
| - | - | 3087 | 475.2 | - | - | 0 | - |
| - | - | 1246 | 475.3 | - | - | 0 | - |
| - | - | 697.2 | 475.6 | - | - | 0 | - |
| - | - | 5917 | 475.9 | - | - | 0 | - |
| - | - | 4644 | 476.2 | - | - | 0 | - |
| - | - | 2971 | 476.6 | - | - | 0 | - |
| - | - | 1115 | 476.9 | - | - | 0 | - |
| - | - | 2270 | 477.2 | - | - | 0 | - |
| - | - | 1835 | 477.2 | - | - | 0 | - |
| - | - | 2334 | 477.6 | - | - | 0 | - |
| - | - | 1073 | 477.9 | - | - | 0 | - |
| - | - | 2077 | 478.2 | - | - | 0 | - |
| - | - | 1.198E+04 | 479.2 | - | - | 0 | - |
| 12 | b | 8829 | 479.5 | 0.005286 | 11.02 | +3 | 12 |
| - | - | 1.81E+04 | 479.7 | - | - | 0 | - |
| - | - | 4937 | 479.9 | - | - | 0 | - |
| - | - | 1.404E+04 | 480.2 | - | - | 0 | - |
| - | - | 2103 | 480.6 | - | - | 0 | - |
| - | - | 3163 | 480.7 | - | - | 0 | - |
| - | - | 945.7 | 480.9 | - | - | 0 | - |
| - | - | 1.079E+04 | 481.2 | - | - | 0 | - |
| - | - | 2297 | 481.2 | - | - | 0 | - |
| - | - | 3294 | 481.6 | - | - | 0 | - |
| - | - | 1024 | 481.7 | - | - | 0 | - |
| - | - | 3384 | 481.9 | - | - | 0 | - |
| - | - | 1981 | 482.2 | - | - | 0 | - |
| - | - | 1264 | 482.2 | - | - | 0 | - |
| - | - | 2292 | 482.6 | - | - | 0 | - |
| 4 | y | 3435 | 482.9 | 0.005136 | 10.64 | +3 | 12 |
| - | - | 1113 | 483.2 | - | - | 0 | - |
| - | - | 392.6 | 483.2 | - | - | 0 | - |
| 4 | y | 7996 | 483.2 | 0.004994 | 10.33 | +3 | 12 |
| - | - | 6120 | 483.6 | - | - | 0 | - |
| - | - | 2127 | 483.7 | - | - | 0 | - |
| - | - | 3148 | 483.9 | - | - | 0 | - |
| - | - | 3301 | 484.2 | - | - | 0 | - |
| - | - | 1511 | 484.7 | - | - | 0 | - |
| - | - | 2.667E+04 | 485.2 | - | - | 0 | - |
| - | - | 6038 | 485.2 | - | - | 0 | - |
| - | - | 1622 | 485.6 | - | - | 0 | - |
| - | - | 2126 | 485.9 | - | - | 0 | - |
| - | - | 5842 | 486.2 | - | - | 0 | - |
| - | - | 1.869E+04 | 486.2 | - | - | 0 | - |
| - | - | 8129 | 486.7 | - | - | 0 | - |
| - | - | 1149 | 487.2 | - | - | 0 | - |
| - | - | 3363 | 487.2 | - | - | 0 | - |
| - | - | 2741 | 487.6 | - | - | 0 | - |
| - | - | 7409 | 487.7 | - | - | 0 | - |
| - | - | 2220 | 487.9 | - | - | 0 | - |
| - | - | 9476 | 488.2 | - | - | 0 | - |
| - | - | 1096 | 488.6 | - | - | 0 | - |
| - | - | 2946 | 488.7 | - | - | 0 | - |
| 4 | y | 1.505E+04 | 488.9 | 0.002573 | 5.263 | +3 | 12 |
| - | - | 3975 | 489.2 | - | - | 0 | - |
| - | - | 1.263E+04 | 489.2 | - | - | 0 | - |
| - | - | 5258 | 489.6 | - | - | 0 | - |
| - | - | 1673 | 489.9 | - | - | 0 | - |
| - | - | 2204 | 490.2 | - | - | 0 | - |
| - | - | 3959 | 490.2 | - | - | 0 | - |
| - | - | 2016 | 490.7 | - | - | 0 | - |
| - | - | 5634 | 491.2 | - | - | 0 | - |
| - | - | 3170 | 491.6 | - | - | 0 | - |
| - | - | 822.3 | 491.9 | - | - | 0 | - |
| - | - | 650 | 491.9 | - | - | 0 | - |
| - | - | 897.3 | 492.3 | - | - | 0 | - |
| - | - | 1383 | 492.6 | - | - | 0 | - |
| - | - | 2057 | 492.7 | - | - | 0 | - |
| - | - | 1890 | 493.2 | - | - | 0 | - |
| - | - | 971.3 | 493.6 | - | - | 0 | - |
| - | - | 1.638E+04 | 493.7 | - | - | 0 | - |
| - | - | 774 | 493.9 | - | - | 0 | - |
| - | - | 9077 | 494.2 | - | - | 0 | - |
| - | - | 1080 | 494.6 | - | - | 0 | - |
| - | - | 2588 | 494.7 | - | - | 0 | - |
| - | - | 2626 | 495.2 | - | - | 0 | - |
| - | - | 1203 | 495.2 | - | - | 0 | - |
| - | - | 1953 | 495.6 | - | - | 0 | - |
| - | - | 1067 | 495.7 | - | - | 0 | - |
| - | - | 983.9 | 495.9 | - | - | 0 | - |
| - | - | 905.2 | 496.2 | - | - | 0 | - |
| - | - | 9355 | 497.2 | - | - | 0 | - |
| - | - | 7475 | 497.6 | - | - | 0 | - |
| - | - | 4142 | 497.9 | - | - | 0 | - |
| - | - | 1.463E+04 | 498.2 | - | - | 0 | - |
| - | - | 1682 | 498.6 | - | - | 0 | - |
| 8 | y | 4847 | 498.7 | 0.002707 | 5.429 | +2 | 8 |
| - | - | 1178 | 498.9 | - | - | 0 | - |
| - | - | 3666 | 499.2 | - | - | 0 | - |
| 8 | y | 7964 | 499.2 | 0.00246 | 4.927 | +2 | 8 |
| - | - | 2399 | 499.6 | - | - | 0 | - |
| - | - | 3519 | 499.7 | - | - | 0 | - |
| - | - | 1066 | 499.9 | - | - | 0 | - |
| - | - | 2490 | 500.2 | - | - | 0 | - |
| - | - | 1248 | 500.2 | - | - | 0 | - |
| - | - | 1647 | 500.9 | - | - | 0 | - |
| - | - | 1235 | 501.2 | - | - | 0 | - |
| - | - | 2993 | 501.6 | - | - | 0 | - |
| - | - | 7483 | 501.7 | - | - | 0 | - |
| - | - | 3130 | 501.9 | - | - | 0 | - |
| - | - | 1E+04 | 502.2 | - | - | 0 | - |
| - | - | 1079 | 502.6 | - | - | 0 | - |
| - | - | 1336 | 502.7 | - | - | 0 | - |
| - | - | 8456 | 502.9 | - | - | 0 | - |
| - | - | 7956 | 503.2 | - | - | 0 | - |
| - | - | 3054 | 503.6 | - | - | 0 | - |
| - | - | 1489 | 503.9 | - | - | 0 | - |
| - | - | 8587 | 504.2 | - | - | 0 | - |
| - | - | 2835 | 504.7 | - | - | 0 | - |
| - | - | 3876 | 504.9 | - | - | 0 | - |
| - | - | 7121 | 505.2 | - | - | 0 | - |
| - | - | 3382 | 505.6 | - | - | 0 | - |
| - | - | 1610 | 505.9 | - | - | 0 | - |
| - | - | 2480 | 506.2 | - | - | 0 | - |
| - | - | 7364 | 506.3 | - | - | 0 | - |
| - | - | 807.8 | 506.6 | - | - | 0 | - |
| - | - | 2464 | 506.9 | - | - | 0 | - |
| - | - | 8481 | 507.2 | - | - | 0 | - |
| - | - | 3466 | 507.2 | - | - | 0 | - |
| - | - | 930.1 | 507.3 | - | - | 0 | - |
| - | - | 4777 | 507.6 | - | - | 0 | - |
| 8 | y | 2.479E+04 | 507.7 | 0.002766 | 5.447 | +2 | 8 |
| - | - | 4175 | 507.9 | - | - | 0 | - |
| - | - | 2.667E+04 | 508.2 | - | - | 0 | - |
| - | - | 4929 | 508.7 | - | - | 0 | - |
| - | - | 4181 | 508.9 | - | - | 0 | - |
| - | - | 769.5 | 509.2 | - | - | 0 | - |
| - | - | 6157 | 509.2 | - | - | 0 | - |
| - | - | 2230 | 509.6 | - | - | 0 | - |
| - | - | 1054 | 509.9 | - | - | 0 | - |
| - | - | 1690 | 510.2 | - | - | 0 | - |
| - | - | 621.8 | 510.3 | - | - | 0 | - |
| - | - | 4875 | 510.6 | - | - | 0 | - |
| - | - | 1137 | 510.7 | - | - | 0 | - |
| - | - | 1.515E+04 | 510.9 | - | - | 0 | - |
| - | - | 1.019E+04 | 511.2 | - | - | 0 | - |
| - | - | 4564 | 511.6 | - | - | 0 | - |
| - | - | 2622 | 511.8 | - | - | 0 | - |
| - | - | 1344 | 511.9 | - | - | 0 | - |
| - | - | 7230 | 512.2 | - | - | 0 | - |
| - | - | 3491 | 512.9 | - | - | 0 | - |
| - | - | 3697 | 513.2 | - | - | 0 | - |
| - | - | 3273 | 513.2 | - | - | 0 | - |
| - | - | 2818 | 513.6 | - | - | 0 | - |
| - | - | 1067 | 513.9 | - | - | 0 | - |
| - | - | 1290 | 514.2 | - | - | 0 | - |
| - | - | 2099 | 515.2 | - | - | 0 | - |
| - | - | 1167 | 515.6 | - | - | 0 | - |
| 13 | b | 1.551E+04 | 516.2 | 0.006212 | 12.03 | +3 | 13 |
| - | - | 3.291E+04 | 516.6 | - | - | 0 | - |
| - | - | 2.876E+04 | 516.9 | - | - | 0 | - |
| - | - | 1253 | 517.2 | - | - | 0 | - |
| - | - | 1.542E+04 | 517.3 | - | - | 0 | - |
| - | - | 4185 | 517.6 | - | - | 0 | - |
| - | - | 927.8 | 517.9 | - | - | 0 | - |
| 9 | b | 5.804E+04 | 518.2 | 0.005356 | 10.33 | +2 | 9 |
| - | - | 2.664E+04 | 518.7 | - | - | 0 | - |
| - | - | 1.271E+04 | 519.2 | - | - | 0 | - |
| - | - | 4202 | 519.7 | - | - | 0 | - |
| - | - | 9808 | 520.3 | - | - | 0 | - |
| - | - | 1023 | 520.6 | - | - | 0 | - |
| - | - | 3863 | 520.7 | - | - | 0 | - |
| - | - | 1005 | 521.2 | - | - | 0 | - |
| - | - | 5115 | 521.3 | - | - | 0 | - |
| - | - | 1290 | 521.7 | - | - | 0 | - |
| 13 | b | 6123 | 522.2 | 0.005098 | 9.762 | +3 | 13 |
| - | - | 1023 | 522.3 | - | - | 0 | - |
| - | - | 6976 | 522.6 | - | - | 0 | - |
| - | - | 3019 | 522.9 | - | - | 0 | - |
| - | - | 1310 | 523.2 | - | - | 0 | - |
| - | - | 7080 | 523.3 | - | - | 0 | - |
| - | - | 4174 | 523.6 | - | - | 0 | - |
| - | - | 1002 | 523.8 | - | - | 0 | - |
| - | - | 2379 | 523.9 | - | - | 0 | - |
| - | - | 8093 | 524.2 | - | - | 0 | - |
| - | - | 2889 | 525.2 | - | - | 0 | - |
| - | - | 1843 | 525.7 | - | - | 0 | - |
| - | - | 1282 | 525.9 | - | - | 0 | - |
| - | - | 4499 | 526.2 | - | - | 0 | - |
| - | - | 5907 | 526.6 | - | - | 0 | - |
| - | - | 3585 | 526.9 | - | - | 0 | - |
| - | - | 1776 | 527.2 | - | - | 0 | - |
| - | - | 1459 | 527.3 | - | - | 0 | - |
| - | - | 5264 | 528.3 | - | - | 0 | - |
| - | - | 1049 | 528.9 | - | - | 0 | - |
| - | - | 1.234E+04 | 529.3 | - | - | 0 | - |
| - | - | 2462 | 529.3 | - | - | 0 | - |
| - | - | 8217 | 529.8 | - | - | 0 | - |
| - | - | 1.953E+04 | 530.2 | - | - | 0 | - |
| - | - | 1142 | 530.6 | - | - | 0 | - |
| - | - | 2271 | 530.8 | - | - | 0 | - |
| - | - | 4462 | 531.2 | - | - | 0 | - |
| - | - | 3788 | 531.8 | - | - | 0 | - |
| 3 | y | 1.112E+04 | 531.9 | 0.007866 | 14.79 | +3 | 13 |
| - | - | 2.025E+04 | 532.3 | - | - | 0 | - |
| - | - | 1.752E+04 | 532.6 | - | - | 0 | - |
| - | - | 3827 | 532.8 | - | - | 0 | - |
| - | - | 6831 | 532.9 | - | - | 0 | - |
| - | - | 7608 | 533.2 | - | - | 0 | - |
| - | - | 1887 | 533.6 | - | - | 0 | - |
| - | - | 1758 | 534.2 | - | - | 0 | - |
| - | - | 2.63E+04 | 534.3 | - | - | 0 | - |
| - | - | 6630 | 534.7 | - | - | 0 | - |
| - | - | 4083 | 535.2 | - | - | 0 | - |
| - | - | 1236 | 535.2 | - | - | 0 | - |
| - | - | 6677 | 535.3 | - | - | 0 | - |
| - | - | 2020 | 535.7 | - | - | 0 | - |
| - | - | 1045 | 535.9 | - | - | 0 | - |
| - | - | 4.404E+04 | 536.2 | - | - | 0 | - |
| - | - | 917.9 | 536.6 | - | - | 0 | - |
| - | - | 1.197E+04 | 537.3 | - | - | 0 | - |
| - | - | 1169 | 537.6 | - | - | 0 | - |
| 3 | y | 1.16E+05 | 537.9 | 0.00587 | 10.91 | +3 | 13 |
| - | - | 1.185E+05 | 538.3 | - | - | 0 | - |
| - | - | 5.924E+04 | 538.6 | - | - | 0 | - |
| - | - | 3238 | 538.8 | - | - | 0 | - |
| - | - | 2.362E+04 | 538.9 | - | - | 0 | - |
| - | - | 8999 | 539.3 | - | - | 0 | - |
| - | - | 1193 | 539.9 | - | - | 0 | - |
| - | - | 1093 | 540.2 | - | - | 0 | - |
| - | - | 938.1 | 540.3 | - | - | 0 | - |
| - | - | 3.366E+04 | 540.8 | - | - | 0 | - |
| - | - | 5432 | 541.2 | - | - | 0 | - |
| - | - | 1.863E+04 | 541.3 | - | - | 0 | - |
| - | - | 6680 | 541.8 | - | - | 0 | - |
| - | - | 881.5 | 541.9 | - | - | 0 | - |
| - | - | 3250 | 542.3 | - | - | 0 | - |
| - | - | 1.957E+04 | 543.3 | - | - | 0 | - |
| - | - | 1.655E+04 | 543.6 | - | - | 0 | - |
| - | - | 1.385E+04 | 543.8 | - | - | 0 | - |
| - | - | 1.445E+04 | 543.9 | - | - | 0 | - |
| - | - | 1068 | 544.2 | - | - | 0 | - |
| - | - | 8231 | 544.3 | - | - | 0 | - |
| - | - | 1.565E+04 | 544.6 | - | - | 0 | - |
| - | - | 1778 | 544.8 | - | - | 0 | - |
| - | - | 1.601E+04 | 544.9 | - | - | 0 | - |
| - | - | 8743 | 545.3 | - | - | 0 | - |
| - | - | 1.301E+04 | 545.3 | - | - | 0 | - |
| - | - | 3628 | 545.6 | - | - | 0 | - |
| - | - | 3949 | 545.8 | - | - | 0 | - |
| - | - | 5523 | 546.3 | - | - | 0 | - |
| - | - | 4014 | 546.3 | - | - | 0 | - |
| - | - | 4064 | 546.8 | - | - | 0 | - |
| - | - | 1.58E+04 | 547.3 | - | - | 0 | - |
| - | - | 1820 | 547.6 | - | - | 0 | - |
| - | - | 1887 | 547.8 | - | - | 0 | - |
| - | - | 1034 | 547.8 | - | - | 0 | - |
| - | - | 1561 | 547.9 | - | - | 0 | - |
| - | - | 9836 | 548.3 | - | - | 0 | - |
| - | - | 1138 | 548.7 | - | - | 0 | - |
| - | - | 2882 | 549.3 | - | - | 0 | - |
| - | - | 1.364E+04 | 549.6 | - | - | 0 | - |
| - | - | 1.323E+04 | 549.9 | - | - | 0 | - |
| - | - | 5636 | 550.3 | - | - | 0 | - |
| - | - | 3060 | 550.6 | - | - | 0 | - |
| - | - | 1170 | 550.9 | - | - | 0 | - |
| - | - | 940.4 | 551.3 | - | - | 0 | - |
| - | - | 2556 | 551.8 | - | - | 0 | - |
| - | - | 6749 | 552.2 | - | - | 0 | - |
| - | - | 4778 | 552.3 | - | - | 0 | - |
| - | - | 4452 | 552.8 | - | - | 0 | - |
| - | - | 951.9 | 552.9 | - | - | 0 | - |
| - | - | 2.018E+04 | 553.3 | - | - | 0 | - |
| - | - | 1.87E+04 | 553.6 | - | - | 0 | - |
| - | - | 9444 | 553.9 | - | - | 0 | - |
| - | - | 6147 | 554.3 | - | - | 0 | - |
| - | - | 2691 | 554.6 | - | - | 0 | - |
| - | - | 2.987E+04 | 554.8 | - | - | 0 | - |
| - | - | 951.4 | 554.9 | - | - | 0 | - |
| 7 | y | 1.957E+04 | 555.3 | 0.005023 | 9.046 | +2 | 9 |
| 7 | y | 8009 | 555.8 | 0.002664 | 4.794 | +2 | 9 |
| - | - | 5640 | 556.3 | - | - | 0 | - |
| - | - | 1813 | 557.3 | - | - | 0 | - |
| - | - | 1261 | 557.8 | - | - | 0 | - |
| - | - | 1084 | 557.9 | - | - | 0 | - |
| - | - | 941.9 | 558.2 | - | - | 0 | - |
| 14 | b | 1.092E+04 | 558.9 | 0.003902 | 6.982 | +3 | 14 |
| 14 | b | 9.101E+04 | 559.3 | 0.00567 | 10.14 | +3 | 14 |
| - | - | 8.957E+04 | 559.6 | - | - | 0 | - |
| - | - | 4.816E+04 | 559.9 | - | - | 0 | - |
| - | - | 1.417E+04 | 560.3 | - | - | 0 | - |
| - | - | 9202 | 560.3 | - | - | 0 | - |
| - | - | 6377 | 560.6 | - | - | 0 | - |
| - | - | 3223 | 560.8 | - | - | 0 | - |
| - | - | 2970 | 560.9 | - | - | 0 | - |
| - | - | 9147 | 561.3 | - | - | 0 | - |
| - | - | 2413 | 561.6 | - | - | 0 | - |
| - | - | 5940 | 561.8 | - | - | 0 | - |
| - | - | 4860 | 562.3 | - | - | 0 | - |
| - | - | 1246 | 562.8 | - | - | 0 | - |
| - | - | 2106 | 563.3 | - | - | 0 | - |
| - | - | 7536 | 563.8 | - | - | 0 | - |
| - | - | 2358 | 563.9 | - | - | 0 | - |
| 7 | y | 1.057E+04 | 564.3 | 0.0008447 | 1.497 | +2 | 9 |
| - | - | 1936 | 564.6 | - | - | 0 | - |
| - | - | 5257 | 564.8 | - | - | 0 | - |
| - | - | 4098 | 564.8 | - | - | 0 | - |
| 14 | b | 1.451E+05 | 564.9 | 0.004836 | 8.561 | +3 | 14 |
| 2 | y | 1.638E+05 | 565.3 | 0.01124 | 19.89 | +3 | 14 |
| - | - | 7.996E+04 | 565.6 | - | - | 0 | - |
| - | - | 1108 | 565.8 | - | - | 0 | - |
| - | - | 3.189E+04 | 565.9 | - | - | 0 | - |
| - | - | 1.404E+04 | 566.3 | - | - | 0 | - |
| - | - | 1540 | 566.6 | - | - | 0 | - |
| - | - | 2947 | 566.7 | - | - | 0 | - |
| - | - | 5390 | 566.9 | - | - | 0 | - |
| - | - | 8180 | 567.3 | - | - | 0 | - |
| - | - | 5351 | 567.6 | - | - | 0 | - |
| - | - | 1084 | 567.8 | - | - | 0 | - |
| - | - | 2328 | 567.9 | - | - | 0 | - |
| - | - | 1624 | 568.3 | - | - | 0 | - |
| - | - | 1570 | 569.3 | - | - | 0 | - |
| - | - | 3611 | 569.6 | - | - | 0 | - |
| - | - | 2422 | 569.9 | - | - | 0 | - |
| - | - | 9985 | 570.3 | - | - | 0 | - |
| 2 | y | 5.83E+04 | 570.9 | 0.005221 | 9.145 | +3 | 14 |
| - | - | 5.533E+04 | 571.3 | - | - | 0 | - |
| - | - | 3.469E+04 | 571.6 | - | - | 0 | - |
| - | - | 2662 | 571.8 | - | - | 0 | - |
| - | - | 1.356E+04 | 571.9 | - | - | 0 | - |
| - | - | 1.457E+04 | 572.3 | - | - | 0 | - |
| - | - | 1.696E+04 | 572.6 | - | - | 0 | - |
| - | - | 7.237E+04 | 572.8 | - | - | 0 | - |
| - | - | 3.819E+04 | 572.9 | - | - | 0 | - |
| - | - | 6.913E+04 | 573.3 | - | - | 0 | - |
| - | - | 1.295E+04 | 573.6 | - | - | 0 | - |
| - | - | 1.848E+04 | 573.8 | - | - | 0 | - |
| - | - | 5424 | 573.9 | - | - | 0 | - |
| - | - | 1360 | 574.2 | - | - | 0 | - |
| - | - | 6976 | 574.3 | - | - | 0 | - |
| - | - | 2417 | 574.6 | - | - | 0 | - |
| - | - | 983.5 | 574.8 | - | - | 0 | - |
| - | - | 1728 | 574.9 | - | - | 0 | - |
| - | - | 1.052E+04 | 575.3 | - | - | 0 | - |
| - | - | 899.8 | 575.3 | - | - | 0 | - |
| - | - | 7259 | 575.8 | - | - | 0 | - |
| - | - | 973.7 | 575.9 | - | - | 0 | - |
| - | - | 7644 | 576.3 | - | - | 0 | - |
| - | - | 1403 | 576.6 | - | - | 0 | - |
| - | - | 4956 | 576.8 | - | - | 0 | - |
| - | - | 1791 | 577.3 | - | - | 0 | - |
| 10 | b | 5711 | 577.8 | 0.00584 | 10.11 | +2 | 10 |
| - | - | 8420 | 578.3 | - | - | 0 | - |
| - | - | 2.249E+05 | 578.6 | - | - | 0 | - |
| - | - | 4669 | 578.8 | - | - | 0 | - |
| - | - | 2.24E+05 | 579 | - | - | 0 | - |
| - | - | 1.288E+05 | 579.3 | - | - | 0 | - |
| - | - | 3.959E+04 | 579.6 | - | - | 0 | - |
| - | - | 1864 | 579.8 | - | - | 0 | - |
| - | - | 1.002E+04 | 580 | - | - | 0 | - |
| - | - | 3137 | 580.3 | - | - | 0 | - |
| - | - | 1907 | 580.8 | - | - | 0 | - |
| - | - | 3429 | 581.3 | - | - | 0 | - |
| - | - | 878.6 | 581.6 | - | - | 0 | - |
| - | - | 1293 | 581.8 | - | - | 0 | - |
| - | - | 4307 | 581.9 | - | - | 0 | - |
| 5 | b | 2975 | 582.2 | 0.004255 | 7.308 | +1 | 5 |
| - | - | 8220 | 582.3 | - | - | 0 | - |
| - | - | 8977 | 582.6 | - | - | 0 | - |
| - | - | 1963 | 582.8 | - | - | 0 | - |
| - | - | 4890 | 582.9 | - | - | 0 | - |
| - | - | 1536 | 583.2 | - | - | 0 | - |
| - | - | 6049 | 583.3 | - | - | 0 | - |
| - | - | 1446 | 583.6 | - | - | 0 | - |
| - | - | 2087 | 584.2 | - | - | 0 | - |
| - | - | 1443 | 584.6 | - | - | 0 | - |
| - | - | 1.307E+04 | 584.8 | - | - | 0 | - |
| - | - | 2128 | 584.9 | - | - | 0 | - |
| - | - | 1.351E+04 | 585.3 | - | - | 0 | - |
| - | - | 6326 | 585.6 | - | - | 0 | - |
| - | - | 6122 | 585.8 | - | - | 0 | - |
| - | - | 3508 | 585.9 | - | - | 0 | - |
| - | - | 7344 | 586.3 | - | - | 0 | - |
| 10 | b | 7.571E+04 | 586.8 | 0.00599 | 10.21 | +2 | 10 |
| - | - | 4.875E+04 | 587.3 | - | - | 0 | - |
| - | - | 2.011E+04 | 587.8 | - | - | 0 | - |
| - | - | 6510 | 587.9 | - | - | 0 | - |
| - | - | 3.162E+04 | 588.3 | - | - | 0 | - |
| - | - | 2.121E+04 | 588.6 | - | - | 0 | - |
| - | - | 1841 | 588.8 | - | - | 0 | - |
| - | - | 1.621E+04 | 588.9 | - | - | 0 | - |
| - | - | 1.58E+04 | 589.3 | - | - | 0 | - |
| - | - | 5699 | 589.6 | - | - | 0 | - |
| - | - | 3271 | 589.9 | - | - | 0 | - |
| - | - | 5903 | 590.3 | - | - | 0 | - |
| - | - | 1356 | 590.6 | - | - | 0 | - |
| 6 | y | 3261 | 590.8 | 0.003276 | 5.545 | +2 | 10 |
| 6 | y | 1.038E+04 | 591.3 | 0.002357 | 3.986 | +2 | 10 |
| - | - | 3283 | 591.8 | - | - | 0 | - |
| - | - | 3644 | 592.2 | - | - | 0 | - |
| - | - | 1788 | 592.3 | - | - | 0 | - |
| - | - | 1219 | 592.8 | - | - | 0 | - |
| - | - | 1227 | 593.2 | - | - | 0 | - |
| - | - | 4848 | 593.3 | - | - | 0 | - |
| - | - | 9419 | 593.8 | - | - | 0 | - |
| 0 | Precursor | 1.053E+05 | 593.9 | 0.005879 | 9.898 | +3 | -1 |
| 0 | Precursor | 2.183E+05 | 594.3 | 0.008623 | 14.51 | +3 | -1 |
| - | - | 1.672E+05 | 594.6 | - | - | 0 | - |
| - | - | 5051 | 594.8 | - | - | 0 | - |
| - | - | 1.037E+05 | 594.9 | - | - | 0 | - |
| - | - | 4.989E+04 | 595.3 | - | - | 0 | - |
| - | - | 1.484E+04 | 595.6 | - | - | 0 | - |
| - | - | 1389 | 595.8 | - | - | 0 | - |
| - | - | 2068 | 595.9 | - | - | 0 | - |
| - | - | 2267 | 596.3 | - | - | 0 | - |
| - | - | 1341 | 597.3 | - | - | 0 | - |
| - | - | 2.238E+04 | 598.3 | - | - | 0 | - |
| - | - | 8821 | 598.6 | - | - | 0 | - |
| - | - | 1409 | 598.8 | - | - | 0 | - |
| - | - | 5289 | 598.8 | - | - | 0 | - |
| - | - | 2083 | 599.1 | - | - | 0 | - |
| - | - | 8019 | 599.3 | - | - | 0 | - |
| 6 | y | 2.999E+04 | 599.8 | 0.004768 | 7.95 | +2 | 10 |
| 0 | Precursor | 1.758E+06 | 599.9 | 0.005592 | 9.321 | +3 | -1 |
| - | - | 1.766E+06 | 600.3 | - | - | 0 | - |
| - | - | 1.041E+06 | 600.6 | - | - | 0 | - |
| - | - | 4.359E+05 | 601 | - | - | 0 | - |
| - | - | 1.354E+05 | 601.3 | - | - | 0 | - |
| - | - | 2981 | 601.6 | - | - | 0 | - |
| - | - | 4968 | 602.2 | - | - | 0 | - |
| - | - | 3985 | 602.3 | - | - | 0 | - |
| - | - | 6229 | 602.8 | - | - | 0 | - |
| - | - | 9133 | 603.3 | - | - | 0 | - |
| - | - | 2442 | 603.8 | - | - | 0 | - |
| - | - | 980.2 | 603.8 | - | - | 0 | - |
| - | - | 1195 | 604.3 | - | - | 0 | - |
| - | - | 2074 | 604.3 | - | - | 0 | - |
| - | - | 2761 | 604.8 | - | - | 0 | - |
| - | - | 3570 | 605.3 | - | - | 0 | - |
| - | - | 1368 | 606.3 | - | - | 0 | - |
| - | - | 4.466E+04 | 607.3 | - | - | 0 | - |
| - | - | 7407 | 607.8 | - | - | 0 | - |
| 11 | y | 1.321E+04 | 608.3 | 0.008869 | 14.58 | +1 | 5 |
| - | - | 2121 | 608.8 | - | - | 0 | - |
| - | - | 3506 | 609.2 | - | - | 0 | - |
| 11 | y | 1.021E+04 | 609.3 | 0.006749 | 11.08 | +1 | 5 |
| - | - | 2159 | 610.2 | - | - | 0 | - |
| - | - | 2627 | 610.3 | - | - | 0 | - |
| - | - | 1368 | 610.8 | - | - | 0 | - |
| - | - | 998 | 611.2 | - | - | 0 | - |
| - | - | 2211 | 611.3 | - | - | 0 | - |
| - | - | 7291 | 611.8 | - | - | 0 | - |
| - | - | 4125 | 612.2 | - | - | 0 | - |
| - | - | 5490 | 612.3 | - | - | 0 | - |
| - | - | 1881 | 612.8 | - | - | 0 | - |
| - | - | 1202 | 613.2 | - | - | 0 | - |
| - | - | 2273 | 613.3 | - | - | 0 | - |
| - | - | 6533 | 613.8 | - | - | 0 | - |
| - | - | 3615 | 614.3 | - | - | 0 | - |
| - | - | 1843 | 614.8 | - | - | 0 | - |
| - | - | 1117 | 615.3 | - | - | 0 | - |
| - | - | 1.516E+04 | 616.3 | - | - | 0 | - |
| - | - | 1.819E+04 | 616.8 | - | - | 0 | - |
| - | - | 1.267E+04 | 617.3 | - | - | 0 | - |
| - | - | 5724 | 617.8 | - | - | 0 | - |
| - | - | 6083 | 618.2 | - | - | 0 | - |
| - | - | 1646 | 619.2 | - | - | 0 | - |
| - | - | 3570 | 620.3 | - | - | 0 | - |
| - | - | 2143 | 620.8 | - | - | 0 | - |
| - | - | 3614 | 621.3 | - | - | 0 | - |
| - | - | 1.754E+04 | 622.3 | - | - | 0 | - |
| - | - | 1.182E+04 | 622.8 | - | - | 0 | - |
| - | - | 5348 | 623.3 | - | - | 0 | - |
| - | - | 1415 | 623.8 | - | - | 0 | - |
| - | - | 1276 | 624.3 | - | - | 0 | - |
| - | - | 2755 | 624.8 | - | - | 0 | - |
| - | - | 5674 | 625.3 | - | - | 0 | - |
| - | - | 7634 | 625.8 | - | - | 0 | - |
| - | - | 9093 | 626.3 | - | - | 0 | - |
| 11 | y | 4.725E+04 | 626.3 | 0.003271 | 5.222 | +1 | 5 |
| - | - | 2791 | 626.8 | - | - | 0 | - |
| - | - | 5844 | 627.3 | - | - | 0 | - |
| - | - | 1.72E+04 | 627.3 | - | - | 0 | - |
| - | - | 3407 | 627.8 | - | - | 0 | - |
| - | - | 2012 | 628.3 | - | - | 0 | - |
| - | - | 1064 | 628.8 | - | - | 0 | - |
| - | - | 932.3 | 629.3 | - | - | 0 | - |
| - | - | 1784 | 630.3 | - | - | 0 | - |
| - | - | 1331 | 631.3 | - | - | 0 | - |
| - | - | 2770 | 633.3 | - | - | 0 | - |
| - | - | 4366 | 634.3 | - | - | 0 | - |
| - | - | 2501 | 634.8 | - | - | 0 | - |
| - | - | 1.114E+04 | 635.3 | - | - | 0 | - |
| - | - | 3118 | 635.8 | - | - | 0 | - |
| - | - | 1.569E+04 | 636.3 | - | - | 0 | - |
| - | - | 1.064E+04 | 636.8 | - | - | 0 | - |
| - | - | 6997 | 637.3 | - | - | 0 | - |
| - | - | 3218 | 637.8 | - | - | 0 | - |
| - | - | 3056 | 638.3 | - | - | 0 | - |
| - | - | 4766 | 639.3 | - | - | 0 | - |
| - | - | 3773 | 640.3 | - | - | 0 | - |
| - | - | 2098 | 640.3 | - | - | 0 | - |
| - | - | 2009 | 640.8 | - | - | 0 | - |
| - | - | 3333 | 641.3 | - | - | 0 | - |
| - | - | 1686 | 642.3 | - | - | 0 | - |
| - | - | 5370 | 643.3 | - | - | 0 | - |
| - | - | 6038 | 643.3 | - | - | 0 | - |
| - | - | 2810 | 643.8 | - | - | 0 | - |
| - | - | 1.383E+04 | 644.3 | - | - | 0 | - |
| - | - | 2563 | 644.8 | - | - | 0 | - |
| - | - | 7422 | 645.3 | - | - | 0 | - |
| - | - | 8979 | 645.8 | - | - | 0 | - |
| - | - | 5755 | 646.3 | - | - | 0 | - |
| - | - | 6631 | 646.8 | - | - | 0 | - |
| - | - | 6141 | 647.3 | - | - | 0 | - |
| - | - | 3164 | 647.8 | - | - | 0 | - |
| - | - | 3763 | 648.3 | - | - | 0 | - |
| - | - | 6278 | 648.8 | - | - | 0 | - |
| - | - | 8283 | 649.3 | - | - | 0 | - |
| - | - | 8229 | 649.8 | - | - | 0 | - |
| - | - | 4291 | 650.3 | - | - | 0 | - |
| - | - | 951 | 650.8 | - | - | 0 | - |
| - | - | 1069 | 651.8 | - | - | 0 | - |
| - | - | 1803 | 652.3 | - | - | 0 | - |
| - | - | 1.059E+04 | 653.3 | - | - | 0 | - |
| - | - | 3.84E+04 | 654.3 | - | - | 0 | - |
| - | - | 2.314E+04 | 654.8 | - | - | 0 | - |
| 5 | y | 2.232E+04 | 655.3 | 0.002762 | 4.215 | +2 | 11 |
| 5 | y | 2.098E+04 | 655.8 | 0.007 | 10.67 | +2 | 11 |
| - | - | 1.734E+04 | 656.3 | - | - | 0 | - |
| - | - | 5083 | 656.8 | - | - | 0 | - |
| - | - | 1.157E+04 | 657.3 | - | - | 0 | - |
| - | - | 1.103E+04 | 657.8 | - | - | 0 | - |
| - | - | 1.256E+04 | 658.3 | - | - | 0 | - |
| - | - | 3651 | 658.8 | - | - | 0 | - |
| 11 | b | 2610 | 659.3 | 0.001642 | 2.49 | +2 | 11 |
| - | - | 2673 | 659.3 | - | - | 0 | - |
| 11 | b | 5512 | 659.8 | 0.01177 | 17.84 | +2 | 11 |
| - | - | 3677 | 660.3 | - | - | 0 | - |
| - | - | 3440 | 660.8 | - | - | 0 | - |
| - | - | 1843 | 661.3 | - | - | 0 | - |
| - | - | 3933 | 662.8 | - | - | 0 | - |
| - | - | 5552 | 663.3 | - | - | 0 | - |
| - | - | 2662 | 663.8 | - | - | 0 | - |
| 5 | y | 4.412E+04 | 664.3 | 0.003308 | 4.98 | +2 | 11 |
| - | - | 3.509E+04 | 664.8 | - | - | 0 | - |
| - | - | 2.303E+04 | 665.3 | - | - | 0 | - |
| - | - | 4445 | 665.8 | - | - | 0 | - |
| - | - | 8072 | 666.3 | - | - | 0 | - |
| - | - | 2365 | 666.8 | - | - | 0 | - |
| - | - | 8503 | 667.3 | - | - | 0 | - |
| - | - | 3219 | 667.8 | - | - | 0 | - |
| 11 | b | 4.538E+04 | 668.3 | 0.006247 | 9.348 | +2 | 11 |
| - | - | 3.299E+04 | 668.8 | - | - | 0 | - |
| - | - | 2.194E+04 | 669.3 | - | - | 0 | - |
| - | - | 9545 | 669.8 | - | - | 0 | - |
| - | - | 6372 | 670.3 | - | - | 0 | - |
| - | - | 1130 | 670.8 | - | - | 0 | - |
| 6 | b | 6.168E+04 | 671.3 | 0.008517 | 12.69 | +1 | 6 |
| - | - | 1.101E+04 | 671.8 | - | - | 0 | - |
| - | - | 1.862E+04 | 672.3 | - | - | 0 | - |
| - | - | 5912 | 672.3 | - | - | 0 | - |
| - | - | 4536 | 672.8 | - | - | 0 | - |
| - | - | 5705 | 673.3 | - | - | 0 | - |
| - | - | 3703 | 673.3 | - | - | 0 | - |
| - | - | 2122 | 673.8 | - | - | 0 | - |
| - | - | 1.454E+04 | 674.3 | - | - | 0 | - |
| - | - | 4655 | 675.3 | - | - | 0 | - |
| - | - | 2526 | 675.8 | - | - | 0 | - |
| - | - | 1277 | 676.3 | - | - | 0 | - |
| - | - | 3036 | 676.3 | - | - | 0 | - |
| - | - | 1092 | 676.8 | - | - | 0 | - |
| - | - | 2764 | 677.3 | - | - | 0 | - |
| - | - | 2.783E+04 | 677.8 | - | - | 0 | - |
| - | - | 2.25E+04 | 678.3 | - | - | 0 | - |
| - | - | 8503 | 678.8 | - | - | 0 | - |
| - | - | 4342 | 679.3 | - | - | 0 | - |
| - | - | 1.836E+04 | 680.3 | - | - | 0 | - |
| - | - | 1.363E+04 | 680.8 | - | - | 0 | - |
| - | - | 3412 | 681.3 | - | - | 0 | - |
| - | - | 4800 | 681.3 | - | - | 0 | - |
| - | - | 3834 | 681.8 | - | - | 0 | - |
| - | - | 975.2 | 682.3 | - | - | 0 | - |
| - | - | 2444 | 682.8 | - | - | 0 | - |
| - | - | 1819 | 683.3 | - | - | 0 | - |
| - | - | 1.127E+04 | 684.3 | - | - | 0 | - |
| - | - | 8576 | 685.3 | - | - | 0 | - |
| - | - | 6519 | 686.3 | - | - | 0 | - |
| - | - | 1.459E+04 | 686.8 | - | - | 0 | - |
| - | - | 1.121E+04 | 687.3 | - | - | 0 | - |
| - | - | 4510 | 687.8 | - | - | 0 | - |
| - | - | 2325 | 688.3 | - | - | 0 | - |
| - | - | 1344 | 688.8 | - | - | 0 | - |
| - | - | 5190 | 689.3 | - | - | 0 | - |
| - | - | 1.326E+04 | 689.8 | - | - | 0 | - |
| - | - | 1.14E+04 | 690.3 | - | - | 0 | - |
| - | - | 4007 | 690.8 | - | - | 0 | - |
| - | - | 8722 | 691.3 | - | - | 0 | - |
| - | - | 4496 | 691.8 | - | - | 0 | - |
| - | - | 2850 | 692.3 | - | - | 0 | - |
| - | - | 5700 | 692.4 | - | - | 0 | - |
| - | - | 2062 | 693.3 | - | - | 0 | - |
| - | - | 1873 | 693.4 | - | - | 0 | - |
| - | - | 1821 | 694.3 | - | - | 0 | - |
| - | - | 6866 | 695.3 | - | - | 0 | - |
| - | - | 8007 | 695.8 | - | - | 0 | - |
| - | - | 4390 | 696.3 | - | - | 0 | - |
| - | - | 1.342E+04 | 696.8 | - | - | 0 | - |
| - | - | 8584 | 697.3 | - | - | 0 | - |
| - | - | 4531 | 697.8 | - | - | 0 | - |
| - | - | 4002 | 698.3 | - | - | 0 | - |
| - | - | 1094 | 698.8 | - | - | 0 | - |
| - | - | 3017 | 699.3 | - | - | 0 | - |
| - | - | 976.1 | 700.3 | - | - | 0 | - |
| - | - | 4021 | 700.8 | - | - | 0 | - |
| - | - | 8994 | 701.3 | - | - | 0 | - |
| - | - | 6408 | 701.8 | - | - | 0 | - |
| - | - | 3.647E+04 | 702.3 | - | - | 0 | - |
| - | - | 1272 | 702.8 | - | - | 0 | - |
| - | - | 1.453E+04 | 703.3 | - | - | 0 | - |
| - | - | 7920 | 704.3 | - | - | 0 | - |
| - | - | 8573 | 704.8 | - | - | 0 | - |
| - | - | 6290 | 705.3 | - | - | 0 | - |
| - | - | 4922 | 705.8 | - | - | 0 | - |
| - | - | 5177 | 706.3 | - | - | 0 | - |
| - | - | 3076 | 706.8 | - | - | 0 | - |
| - | - | 7008 | 707.3 | - | - | 0 | - |
| - | - | 1066 | 707.8 | - | - | 0 | - |
| - | - | 7238 | 708.4 | - | - | 0 | - |
| - | - | 1467 | 708.8 | - | - | 0 | - |
| - | - | 3688 | 709.4 | - | - | 0 | - |
| 12 | b | 5.08E+04 | 709.8 | 0.006794 | 9.572 | +2 | 12 |
| - | - | 4.637E+04 | 710.3 | - | - | 0 | - |
| - | - | 2.054E+04 | 710.8 | - | - | 0 | - |
| - | - | 1.095E+04 | 711.3 | - | - | 0 | - |
| - | - | 3120 | 711.8 | - | - | 0 | - |
| - | - | 5079 | 712.3 | - | - | 0 | - |
| - | - | 1579 | 712.8 | - | - | 0 | - |
| - | - | 9780 | 713.3 | - | - | 0 | - |
| - | - | 8796 | 713.8 | - | - | 0 | - |
| - | - | 4109 | 714.3 | - | - | 0 | - |
| - | - | 2747 | 714.8 | - | - | 0 | - |
| - | - | 6428 | 715.3 | - | - | 0 | - |
| - | - | 7099 | 715.8 | - | - | 0 | - |
| - | - | 3160 | 716.3 | - | - | 0 | - |
| - | - | 967.8 | 717.3 | - | - | 0 | - |
| - | - | 7017 | 718.3 | - | - | 0 | - |
| 12 | b | 3.527E+04 | 718.8 | 0.008104 | 11.27 | +2 | 12 |
| - | - | 2.996E+04 | 719.3 | - | - | 0 | - |
| - | - | 1.48E+04 | 719.8 | - | - | 0 | - |
| - | - | 3.126E+04 | 720.4 | - | - | 0 | - |
| - | - | 3350 | 720.8 | - | - | 0 | - |
| - | - | 1.142E+04 | 721.4 | - | - | 0 | - |
| - | - | 1.02E+04 | 721.9 | - | - | 0 | - |
| - | - | 1.03E+04 | 722.4 | - | - | 0 | - |
| - | - | 5269 | 722.9 | - | - | 0 | - |
| - | - | 4328 | 723.3 | - | - | 0 | - |
| 4 | y | 1.085E+04 | 723.8 | 0.001657 | 2.289 | +2 | 12 |
| 4 | y | 2.217E+04 | 724.3 | 0.00617 | 8.518 | +2 | 12 |
| - | - | 1.505E+04 | 724.8 | - | - | 0 | - |
| - | - | 8047 | 725.3 | - | - | 0 | - |
| - | - | 2096 | 725.8 | - | - | 0 | - |
| - | - | 1.312E+04 | 727.3 | - | - | 0 | - |
| - | - | 1.084E+04 | 727.8 | - | - | 0 | - |
| - | - | 9347 | 728.3 | - | - | 0 | - |
| - | - | 5982 | 728.8 | - | - | 0 | - |
| - | - | 5813 | 729.3 | - | - | 0 | - |
| - | - | 2624 | 729.8 | - | - | 0 | - |
| - | - | 3794 | 730.3 | - | - | 0 | - |
| - | - | 4296 | 730.9 | - | - | 0 | - |
| - | - | 3569 | 731.3 | - | - | 0 | - |
| - | - | 2141 | 731.9 | - | - | 0 | - |
| - | - | 1969 | 732.3 | - | - | 0 | - |
| 4 | y | 1.179E+05 | 732.9 | 0.004248 | 5.796 | +2 | 12 |
| - | - | 9.551E+04 | 733.4 | - | - | 0 | - |
| - | - | 4.267E+04 | 733.9 | - | - | 0 | - |
| - | - | 1.263E+04 | 734.4 | - | - | 0 | - |
| - | - | 3457 | 734.9 | - | - | 0 | - |
| - | - | 9993 | 735.3 | - | - | 0 | - |
| - | - | 1.215E+04 | 736.3 | - | - | 0 | - |
| - | - | 6039 | 736.8 | - | - | 0 | - |
| - | - | 9312 | 737.3 | - | - | 0 | - |
| - | - | 2428 | 737.8 | - | - | 0 | - |
| - | - | 4719 | 738.3 | - | - | 0 | - |
| - | - | 1183 | 738.8 | - | - | 0 | - |
| - | - | 4126 | 739.3 | - | - | 0 | - |
| - | - | 1596 | 739.8 | - | - | 0 | - |
| - | - | 3654 | 740.3 | - | - | 0 | - |
| - | - | 1983 | 741.4 | - | - | 0 | - |
| - | - | 1.357E+04 | 741.9 | - | - | 0 | - |
| - | - | 1.448E+04 | 742.4 | - | - | 0 | - |
| - | - | 9481 | 742.9 | - | - | 0 | - |
| - | - | 5061 | 743.3 | - | - | 0 | - |
| - | - | 2418 | 743.8 | - | - | 0 | - |
| - | - | 2265 | 744.3 | - | - | 0 | - |
| - | - | 2036 | 744.8 | - | - | 0 | - |
| - | - | 1.227E+04 | 745.3 | - | - | 0 | - |
| - | - | 1.055E+04 | 745.8 | - | - | 0 | - |
| 10 | y | 7554 | 746.3 | 0.003257 | 4.364 | +1 | 6 |
| - | - | 3141 | 746.8 | - | - | 0 | - |
| - | - | 3918 | 747.4 | - | - | 0 | - |
| - | - | 3532 | 747.9 | - | - | 0 | - |
| - | - | 6062 | 748.4 | - | - | 0 | - |
| - | - | 1224 | 748.9 | - | - | 0 | - |
| - | - | 2141 | 749.4 | - | - | 0 | - |
| - | - | 3030 | 750.3 | - | - | 0 | - |
| - | - | 2.922E+04 | 750.9 | - | - | 0 | - |
| - | - | 2.43E+04 | 751.4 | - | - | 0 | - |
| - | - | 1.3E+04 | 751.9 | - | - | 0 | - |
| - | - | 4601 | 752.4 | - | - | 0 | - |
| - | - | 2023 | 752.9 | - | - | 0 | - |
| - | - | 3660 | 753.3 | - | - | 0 | - |
| - | - | 1.838E+04 | 753.9 | - | - | 0 | - |
| - | - | 1.768E+04 | 754.4 | - | - | 0 | - |
| - | - | 8854 | 754.9 | - | - | 0 | - |
| - | - | 8641 | 755.4 | - | - | 0 | - |
| - | - | 1698 | 755.8 | - | - | 0 | - |
| - | - | 1.101E+04 | 756.4 | - | - | 0 | - |
| - | - | 3459 | 756.9 | - | - | 0 | - |
| - | - | 7612 | 757.4 | - | - | 0 | - |
| - | - | 2308 | 757.9 | - | - | 0 | - |
| - | - | 3298 | 758.4 | - | - | 0 | - |
| - | - | 3228 | 759.4 | - | - | 0 | - |
| - | - | 5869 | 759.9 | - | - | 0 | - |
| - | - | 6607 | 760.4 | - | - | 0 | - |
| - | - | 1.517E+04 | 760.8 | - | - | 0 | - |
| - | - | 1.102E+04 | 761.4 | - | - | 0 | - |
| - | - | 6881 | 761.8 | - | - | 0 | - |
| - | - | 2744 | 762.3 | - | - | 0 | - |
| - | - | 3778 | 762.9 | - | - | 0 | - |
| 10 | y | 4.181E+04 | 763.4 | 0.003441 | 4.508 | +1 | 6 |
| - | - | 2005 | 763.9 | - | - | 0 | - |
| - | - | 3178 | 764.3 | - | - | 0 | - |
| - | - | 1.526E+04 | 764.4 | - | - | 0 | - |
| - | - | 2439 | 764.8 | - | - | 0 | - |
| - | - | 4739 | 765.3 | - | - | 0 | - |
| - | - | 6012 | 765.4 | - | - | 0 | - |
| - | - | 1.592E+04 | 765.9 | - | - | 0 | - |
| 7 | b | 1.921E+04 | 766.4 | 0.0118 | 15.4 | +1 | 7 |
| - | - | 4864 | 766.9 | - | - | 0 | - |
| - | - | 7156 | 767.4 | - | - | 0 | - |
| - | - | 5508 | 768.4 | - | - | 0 | - |
| - | - | 1.294E+04 | 768.9 | - | - | 0 | - |
| - | - | 1.253E+04 | 769.4 | - | - | 0 | - |
| - | - | 8717 | 769.9 | - | - | 0 | - |
| - | - | 2622 | 770.4 | - | - | 0 | - |
| - | - | 8455 | 772.4 | - | - | 0 | - |
| - | - | 3957 | 773.4 | - | - | 0 | - |
| 13 | b | 4.594E+04 | 773.8 | 0.007291 | 9.421 | +2 | 13 |
| - | - | 6.579E+04 | 774.4 | - | - | 0 | - |
| - | - | 4.435E+04 | 774.9 | - | - | 0 | - |
| - | - | 1.935E+04 | 775.4 | - | - | 0 | - |
| - | - | 5993 | 775.9 | - | - | 0 | - |
| - | - | 2320 | 776.4 | - | - | 0 | - |
| - | - | 1379 | 779.8 | - | - | 0 | - |
| - | - | 1372 | 780.4 | - | - | 0 | - |
| - | - | 1194 | 780.9 | - | - | 0 | - |
| - | - | 1.011E+04 | 781.3 | - | - | 0 | - |
| - | - | 6039 | 782.3 | - | - | 0 | - |
| 13 | b | 1.109E+05 | 782.8 | 0.006952 | 8.88 | +2 | 13 |
| - | - | 9.404E+04 | 783.4 | - | - | 0 | - |
| - | - | 5.354E+04 | 783.9 | - | - | 0 | - |
| 7 | b | 7.426E+04 | 784.4 | 0.00624 | 7.956 | +1 | 7 |
| - | - | 6079 | 784.9 | - | - | 0 | - |
| - | - | 3.739E+04 | 785.4 | - | - | 0 | - |
| - | - | 1.569E+04 | 786.4 | - | - | 0 | - |
| - | - | 4437 | 787.4 | - | - | 0 | - |
| - | - | 2113 | 788.4 | - | - | 0 | - |
| - | - | 2139 | 788.9 | - | - | 0 | - |
| - | - | 2842 | 789.4 | - | - | 0 | - |
| - | - | 1115 | 789.9 | - | - | 0 | - |
| - | - | 1846 | 790.4 | - | - | 0 | - |
| - | - | 7496 | 791.4 | - | - | 0 | - |
| - | - | 6643 | 791.9 | - | - | 0 | - |
| - | - | 5550 | 792.4 | - | - | 0 | - |
| - | - | 2081 | 792.9 | - | - | 0 | - |
| - | - | 2806 | 793.4 | - | - | 0 | - |
| - | - | 3772 | 794.4 | - | - | 0 | - |
| - | - | 1093 | 794.9 | - | - | 0 | - |
| - | - | 2755 | 795.4 | - | - | 0 | - |
| 3 | y | 6931 | 797.4 | 0.006556 | 8.222 | +2 | 13 |
| 3 | y | 1.463E+04 | 797.9 | 0.00997 | 12.5 | +2 | 13 |
| - | - | 1.353E+04 | 798.4 | - | - | 0 | - |
| - | - | 6076 | 798.9 | - | - | 0 | - |
| - | - | 3240 | 799.4 | - | - | 0 | - |
| - | - | 1407 | 801.4 | - | - | 0 | - |
| - | - | 5840 | 802.4 | - | - | 0 | - |
| - | - | 4347 | 803.4 | - | - | 0 | - |
| - | - | 2510 | 804.4 | - | - | 0 | - |
| - | - | 1111 | 805.4 | - | - | 0 | - |
| 3 | y | 9.257E+04 | 806.4 | 0.00695 | 8.619 | +2 | 13 |
| - | - | 8.139E+04 | 806.9 | - | - | 0 | - |
| - | - | 4.663E+04 | 807.4 | - | - | 0 | - |
| - | - | 1.735E+04 | 807.9 | - | - | 0 | - |
| - | - | 4281 | 808.4 | - | - | 0 | - |
| - | - | 1.035E+04 | 811.4 | - | - | 0 | - |
| - | - | 5662 | 812.4 | - | - | 0 | - |
| - | - | 986.9 | 813.4 | - | - | 0 | - |
| - | - | 2349 | 814.9 | - | - | 0 | - |
| - | - | 2544 | 815.4 | - | - | 0 | - |
| - | - | 1598 | 815.9 | - | - | 0 | - |
| - | - | 1062 | 816.4 | - | - | 0 | - |
| - | - | 1.196E+04 | 819.4 | - | - | 0 | - |
| - | - | 9600 | 820.4 | - | - | 0 | - |
| - | - | 7602 | 821.4 | - | - | 0 | - |
| - | - | 5232 | 822.4 | - | - | 0 | - |
| - | - | 2967 | 823.4 | - | - | 0 | - |
| - | - | 1396 | 824.4 | - | - | 0 | - |
| - | - | 2312 | 825.4 | - | - | 0 | - |
| - | - | 1180 | 826.4 | - | - | 0 | - |
| - | - | 5760 | 829.4 | - | - | 0 | - |
| - | - | 1523 | 830.4 | - | - | 0 | - |
| - | - | 2919 | 830.4 | - | - | 0 | - |
| - | - | 1929 | 831.4 | - | - | 0 | - |
| - | - | 1.01E+04 | 837.4 | - | - | 0 | - |
| - | - | 7929 | 838.4 | - | - | 0 | - |
| - | - | 2158 | 838.9 | - | - | 0 | - |
| - | - | 1.679E+04 | 839.4 | - | - | 0 | - |
| - | - | 7005 | 840.4 | - | - | 0 | - |
| - | - | 2408 | 841.4 | - | - | 0 | - |
| - | - | 2273 | 842.4 | - | - | 0 | - |
| - | - | 3348 | 843.4 | - | - | 0 | - |
| - | - | 1039 | 844.4 | - | - | 0 | - |
| 14 | b | 9787 | 846.9 | 0.007261 | 8.574 | +2 | 14 |
| 2 | y | 1.237E+04 | 847.4 | 0.01068 | 12.6 | +2 | 14 |
| - | - | 5816 | 847.9 | - | - | 0 | - |
| - | - | 6092 | 848.4 | - | - | 0 | - |
| - | - | 1.235E+04 | 849.4 | - | - | 0 | - |
| - | - | 6834 | 850.4 | - | - | 0 | - |
| - | - | 2235 | 851.4 | - | - | 0 | - |
| - | - | 2986 | 854.4 | - | - | 0 | - |
| 2 | y | 2114 | 855.9 | 0.009547 | 11.15 | +2 | 14 |
| - | - | 1968 | 856.4 | - | - | 0 | - |
| - | - | 1527 | 856.9 | - | - | 0 | - |
| - | - | 1.39E+04 | 857.4 | - | - | 0 | - |
| - | - | 6822 | 858.4 | - | - | 0 | - |
| 9 | y | 3605 | 859.4 | 0.01588 | 18.48 | +1 | 7 |
| 9 | y | 9375 | 860.4 | 0.006594 | 7.664 | +1 | 7 |
| - | - | 4713 | 861.4 | - | - | 0 | - |
| - | - | 2737 | 862.4 | - | - | 0 | - |
| - | - | 8430 | 865.4 | - | - | 0 | - |
| - | - | 6511 | 866.4 | - | - | 0 | - |
| - | - | 1727 | 867.4 | - | - | 0 | - |
| - | - | 1988 | 874.5 | - | - | 0 | - |
| - | - | 1005 | 875.4 | - | - | 0 | - |
| - | - | 3615 | 876.4 | - | - | 0 | - |
| 9 | y | 3.446E+04 | 877.4 | 0.003421 | 3.899 | +1 | 7 |
| - | - | 1.626E+04 | 878.4 | - | - | 0 | - |
| - | - | 5839 | 879.4 | - | - | 0 | - |
| - | - | 1270 | 887.4 | - | - | 0 | - |
| - | - | 1672 | 888.4 | - | - | 0 | - |
| - | - | 1352 | 891.4 | - | - | 0 | - |
| - | - | 3741 | 892.4 | - | - | 0 | - |
| - | - | 9159 | 893.4 | - | - | 0 | - |
| - | - | 1.354E+04 | 894.4 | - | - | 0 | - |
| - | - | 6796 | 895.4 | - | - | 0 | - |
| - | - | 1857 | 896.4 | - | - | 0 | - |
| 8 | b | 2401 | 903.4 | 0.006416 | 7.102 | +1 | 8 |
| - | - | 3177 | 904.4 | - | - | 0 | - |
| - | - | 4385 | 905.4 | - | - | 0 | - |
| - | - | 2546 | 906.4 | - | - | 0 | - |
| - | - | 7116 | 909.4 | - | - | 0 | - |
| - | - | 3267 | 910.4 | - | - | 0 | - |
| - | - | 1763 | 911.4 | - | - | 0 | - |
| - | - | 1794 | 913.4 | - | - | 0 | - |
| - | - | 2135 | 920.4 | - | - | 0 | - |
| 8 | b | 2.342E+04 | 921.4 | 0.009828 | 10.67 | +1 | 8 |
| - | - | 2.38E+04 | 922.4 | - | - | 0 | - |
| - | - | 9213 | 923.4 | - | - | 0 | - |
| - | - | 2950 | 924.4 | - | - | 0 | - |
| - | - | 2089 | 926.5 | - | - | 0 | - |
| - | - | 1742 | 927.5 | - | - | 0 | - |
| - | - | 1188 | 928.5 | - | - | 0 | - |
| - | - | 4548 | 930.4 | - | - | 0 | - |
| - | - | 4973 | 931.4 | - | - | 0 | - |
| - | - | 2552 | 932.4 | - | - | 0 | - |
| - | - | 1723 | 937.5 | - | - | 0 | - |
| - | - | 4888 | 938.5 | - | - | 0 | - |
| - | - | 2594 | 939.5 | - | - | 0 | - |
| - | - | 1124 | 940.4 | - | - | 0 | - |
| - | - | 4148 | 943.5 | - | - | 0 | - |
| - | - | 1182 | 944.5 | - | - | 0 | - |
| - | - | 3814 | 947.5 | - | - | 0 | - |
| - | - | 1.256E+04 | 948.4 | - | - | 0 | - |
| - | - | 8352 | 949.4 | - | - | 0 | - |
| - | - | 2024 | 950.4 | - | - | 0 | - |
| - | - | 2102 | 953.5 | - | - | 0 | - |
| - | - | 1416 | 954.5 | - | - | 0 | - |
| - | - | 2763 | 957.4 | - | - | 0 | - |
| - | - | 4433 | 958.4 | - | - | 0 | - |
| - | - | 1579 | 959.4 | - | - | 0 | - |
| - | - | 1277 | 961.4 | - | - | 0 | - |
| - | - | 8923 | 965.5 | - | - | 0 | - |
| - | - | 9089 | 966.5 | - | - | 0 | - |
| - | - | 3526 | 967.5 | - | - | 0 | - |
| - | - | 1216 | 968.4 | - | - | 0 | - |
| - | - | 1971 | 969.4 | - | - | 0 | - |
| - | - | 2.002E+04 | 971.5 | - | - | 0 | - |
| - | - | 9739 | 972.5 | - | - | 0 | - |
| - | - | 3199 | 973.5 | - | - | 0 | - |
| - | - | 6524 | 974.5 | - | - | 0 | - |
| - | - | 3558 | 975.5 | - | - | 0 | - |
| - | - | 1293 | 979.4 | - | - | 0 | - |
| - | - | 2426 | 983.5 | - | - | 0 | - |
| - | - | 2070 | 984.5 | - | - | 0 | - |
| - | - | 2250 | 985.4 | - | - | 0 | - |
| - | - | 7396 | 986.4 | - | - | 0 | - |
| - | - | 3621 | 987.4 | - | - | 0 | - |
| - | - | 1852 | 988.5 | - | - | 0 | - |
| - | - | 1429 | 990.4 | - | - | 0 | - |
| 8 | y | 1307 | 996.5 | 0.002865 | 2.875 | +1 | 8 |
| 8 | y | 4963 | 997.4 | 0.009267 | 9.29 | +1 | 8 |
| - | - | 2947 | 998.5 | - | - | 0 | - |
| - | - | 8957 | 1002 | - | - | 0 | - |
| - | - | 5589 | 1003 | - | - | 0 | - |
| - | - | 1834 | 1004 | - | - | 0 | - |
| - | - | 1186 | 1005 | - | - | 0 | - |
| - | - | 1192 | 1006 | - | - | 0 | - |
| - | - | 3873 | 1007 | - | - | 0 | - |
| - | - | 1215 | 1008 | - | - | 0 | - |
| 8 | y | 1.544E+04 | 1014 | 0.004568 | 4.503 | +1 | 8 |
| - | - | 8540 | 1015 | - | - | 0 | - |
| - | - | 2628 | 1016 | - | - | 0 | - |
| 9 | b | 2313 | 1017 | 0.01134 | 11.15 | +1 | 9 |
| - | - | 2082 | 1018 | - | - | 0 | - |
| - | - | 1674 | 1019 | - | - | 0 | - |
| - | - | 1079 | 1020 | - | - | 0 | - |
| - | - | 2146 | 1023 | - | - | 0 | - |
| - | - | 1063 | 1024 | - | - | 0 | - |
| - | - | 1197 | 1031 | - | - | 0 | - |
| - | - | 1697 | 1032 | - | - | 0 | - |
| 9 | b | 2.585E+04 | 1035 | 0.01005 | 9.709 | +1 | 9 |
| - | - | 1.549E+04 | 1036 | - | - | 0 | - |
| - | - | 5631 | 1037 | - | - | 0 | - |
| - | - | 2363 | 1038 | - | - | 0 | - |
| - | - | 1581 | 1040 | - | - | 0 | - |
| - | - | 1778 | 1041 | - | - | 0 | - |
| - | - | 1057 | 1049 | - | - | 0 | - |
| - | - | 2539 | 1050 | - | - | 0 | - |
| - | - | 3631 | 1057 | - | - | 0 | - |
| - | - | 4701 | 1059 | - | - | 0 | - |
| - | - | 4431 | 1059 | - | - | 0 | - |
| - | - | 2288 | 1060 | - | - | 0 | - |
| - | - | 1854 | 1067 | - | - | 0 | - |
| - | - | 1704 | 1067 | - | - | 0 | - |
| - | - | 3436 | 1068 | - | - | 0 | - |
| - | - | 2119 | 1069 | - | - | 0 | - |
| - | - | 1113 | 1075 | - | - | 0 | - |
| - | - | 1377 | 1076 | - | - | 0 | - |
| - | - | 7351 | 1077 | - | - | 0 | - |
| - | - | 4864 | 1078 | - | - | 0 | - |
| - | - | 1814 | 1079 | - | - | 0 | - |
| - | - | 1.399E+04 | 1085 | - | - | 0 | - |
| - | - | 8711 | 1086 | - | - | 0 | - |
| - | - | 2951 | 1087 | - | - | 0 | - |
| - | - | 2144 | 1092 | - | - | 0 | - |
| - | - | 1134 | 1093 | - | - | 0 | - |
| - | - | 4783 | 1094 | - | - | 0 | - |
| - | - | 8095 | 1095 | - | - | 0 | - |
| - | - | 5173 | 1096 | - | - | 0 | - |
| - | - | 1385 | 1097 | - | - | 0 | - |
| - | - | 1036 | 1103 | - | - | 0 | - |
| - | - | 4318 | 1104 | - | - | 0 | - |
| - | - | 3643 | 1105 | - | - | 0 | - |
| - | - | 1142 | 1105 | - | - | 0 | - |
| - | - | 4639 | 1109 | - | - | 0 | - |
| 7 | y | 1818 | 1110 | 0.00814 | 7.337 | +1 | 9 |
| 7 | y | 2661 | 1111 | 0.003572 | 3.216 | +1 | 9 |
| - | - | 2155 | 1112 | - | - | 0 | - |
| - | - | 1738 | 1121 | - | - | 0 | - |
| - | - | 1939 | 1122 | - | - | 0 | - |
| - | - | 1667 | 1123 | - | - | 0 | - |
| 7 | y | 6256 | 1128 | 0.002535 | 2.249 | +1 | 9 |
| - | - | 4090 | 1129 | - | - | 0 | - |
| - | - | 1203 | 1145 | - | - | 0 | - |
| - | - | 1176 | 1146 | - | - | 0 | - |
| - | - | 1023 | 1147 | - | - | 0 | - |
| - | - | 5502 | 1149 | - | - | 0 | - |
| - | - | 3490 | 1150 | - | - | 0 | - |
| - | - | 2186 | 1152 | - | - | 0 | - |
| - | - | 1033 | 1153 | - | - | 0 | - |
| 10 | b | 941.2 | 1155 | 0.01565 | 13.55 | +1 | 10 |
| - | - | 1245 | 1164 | - | - | 0 | - |
| - | - | 1154 | 1165 | - | - | 0 | - |
| - | - | 2984 | 1169 | - | - | 0 | - |
| - | - | 2402 | 1170 | - | - | 0 | - |
| - | - | 1118 | 1171 | - | - | 0 | - |
| 10 | b | 4237 | 1173 | 0.00827 | 7.053 | +1 | 10 |
| - | - | 3185 | 1174 | - | - | 0 | - |
| - | - | 1457 | 1175 | - | - | 0 | - |
| 6 | y | 3548 | 1182 | 0.007718 | 6.532 | +1 | 10 |
| - | - | 2103 | 1183 | - | - | 0 | - |
| - | - | 2668 | 1187 | - | - | 0 | - |
| - | - | 4138 | 1188 | - | - | 0 | - |
| - | - | 1296 | 1189 | - | - | 0 | - |
| 6 | y | 1.007E+04 | 1199 | 0.004972 | 4.148 | +1 | 10 |
| - | - | 7857 | 1200 | - | - | 0 | - |
| - | - | 2484 | 1201 | - | - | 0 | - |
| - | - | 1501 | 1204 | - | - | 0 | - |
| - | - | 1825 | 1205 | - | - | 0 | - |
| - | - | 2850 | 1206 | - | - | 0 | - |
| - | - | 1204 | 1207 | - | - | 0 | - |
| - | - | 2243 | 1214 | - | - | 0 | - |
| - | - | 1951 | 1215 | - | - | 0 | - |
| - | - | 1795 | 1216 | - | - | 0 | - |
| - | - | 2897 | 1223 | - | - | 0 | - |
| - | - | 1821 | 1224 | - | - | 0 | - |
| - | - | 6565 | 1232 | - | - | 0 | - |
| - | - | 5346 | 1233 | - | - | 0 | - |
| - | - | 3047 | 1234 | - | - | 0 | - |
| - | - | 1049 | 1249 | - | - | 0 | - |
| - | - | 1329 | 1251 | - | - | 0 | - |
| - | - | 1813 | 1272 | - | - | 0 | - |
| 5 | y | 1477 | 1310 | 0.0007759 | 0.5925 | +1 | 11 |
| 5 | y | 3528 | 1311 | 0.01383 | 10.55 | +1 | 11 |
| - | - | 1469 | 1312 | - | - | 0 | - |
| - | - | 2264 | 1315 | - | - | 0 | - |
| - | - | 1786 | 1316 | - | - | 0 | - |
| - | - | 1114 | 1317 | - | - | 0 | - |
| 5 | y | 6183 | 1328 | 0.004249 | 3.201 | +1 | 11 |
| - | - | 5125 | 1329 | - | - | 0 | - |
| - | - | 2145 | 1330 | - | - | 0 | - |
| - | - | 900.5 | 1337 | - | - | 0 | - |
| - | - | 1211 | 1356 | - | - | 0 | - |
| - | - | 1433 | 1379 | - | - | 0 | - |
| 12 | b | 1024 | 1420 | 0.02464 | 17.36 | +1 | 12 |
| - | - | 682.7 | 2274 | - | - | 0 | - |
| - | - | 875.1 | 3032 | - | - | 0 | - |

m/z Charge Intensity FragmentType MassShift Position
120.05638885498047 0 2490.3835
120.08515167236328 0 500.48578
121.04039764404297 0 3512.1106
122.07205963134766 0 5122.414
122.0970458984375 0 595.8909
123.05603790283203 0 3655.065
123.07530975341797 0 463.79037
124.07603454589844 0 811.172
125.67017364501953 0 485.93042
126.09198760986328 0 1012.4643
127.0874252319336 0 3491.6292
128.0716094970703 0 1120.1649
128.08250427246094 0 2542.313
129.06671142578125 0 5037.592
129.10302734375 0 713195
130.0504913330078 0 2251.9585
130.0869903564453 0 8226.165
130.10037231445312 0 5181.6406
130.1063232421875 0 46754.543
131.01695251464844 0 2114.571
131.10763549804688 0 1031.8945
136.0511474609375 0 1915.6826
136.07650756835938 0 145695.98
137.07427978515625 0 675.42664
137.0798797607422 0 11852.58
138.06698608398438 0 9496.933
138.08311462402344 0 627.918
138.09242248535156 0 658.26794
139.07037353515625 0 613.7167
139.08737182617188 0 10608.051
140.07164001464844 0 868.8226
140.0907440185547 0 1279.1874
141.1031036376953 0 6326.9277 a Water loss 1
142.0873565673828 0 1089.7615
143.09475708007812 0 1579.8347
143.11891174316406 0 1024.4478
146.07249450683594 0 690.93854
146.0931854248047 0 1376.3125
147.0447998046875 0 1160.9103
147.11363220214844 0 11645.145
148.04315185546875 0 1387.8793
148.0519561767578 0 724.46686
148.07696533203125 0 813.4611
148.08779907226562 0 12493.048
148.11767578125 0 630.72644
148.87242126464844 0 461.13144
148.9081573486328 0 832.3526
148.9155731201172 0 702.75305
148.92266845703125 0 1153.8186
148.929931640625 0 1006.9325
148.93701171875 0 2614.6626
148.94471740722656 0 4063.2593
148.96142578125 0 3671.2256
148.96925354003906 0 1551.0121
148.97610473632812 0 938.01074
148.98304748535156 0 988.02405
148.99061584472656 0 630.57117
149.00534057617188 0 629.1022
149.0211181640625 0 455.27283
149.02732849121094 0 2955.4033
149.03321838378906 0 689.77136
149.0717315673828 0 872.7557
149.08358764648438 0 1116.2413 y Water loss 11
149.0912628173828 0 888.78436
149.10731506347656 0 571.29553
150.02708435058594 0 671.20886
150.03302001953125 0 649.8763
150.10301208496094 0 495.27057
150.11036682128906 0 609.7568
151.07492065429688 0 699.97675
151.08750915527344 0 2147.8162
152.0826873779297 0 11935.977
153.08645629882812 0 571.55835
153.10308837890625 0 2068.784
154.0511474609375 0 1389.5271
154.06195068359375 0 3477.2117
155.07740783691406 0 424.65118
155.08241271972656 0 4269.391
155.0936279296875 0 12661.165
155.1187286376953 0 5033.868
156.07762145996094 0 10186.877
157.0812530517578 0 736.8035
157.09808349609375 0 3400.2444
157.13453674316406 0 4218.6646
158.061279296875 0 516.0797
159.0563201904297 0 2303.8855
159.1136932373047 0 793534.94 a 1
160.07643127441406 0 922.0034
160.0878448486328 0 1161.3772
160.11077880859375 0 4465.3486
160.1170196533203 0 58053.016
161.11859130859375 0 2334.4382
162.0670928955078 0 2784.1562
163.098876953125 0 1542.257
164.08265686035156 0 1050.507
164.11964416503906 0 767.252
165.07012939453125 0 882.7208
165.07798767089844 0 12825.553
165.10336303710938 0 1010.66235
165.11459350585938 0 645.2669
166.0559539794922 0 1056.9955
166.06201171875 0 29104.592
166.08689880371094 0 30251.744
167.0443878173828 0 517.0712
167.0653076171875 0 2448.3428
167.09339904785156 0 9275.111
167.1188507080078 0 4997.527
168.11410522460938 0 1763.8911
168.1215362548828 0 582.4295
169.06158447265625 0 3384.5588
169.0742950439453 0 1001.8508
169.0980987548828 0 2113.7236 b Water loss 1
171.07705688476562 0 1054.461
171.1138153076172 0 12581.237
171.15028381347656 0 963.20935
172.05136108398438 0 1929.4858
172.11758422851562 0 1326.8502
173.09304809570312 0 7123.1655 y Ammonia loss 12
173.4490203857422 0 1027.8711
174.0015869140625 0 537.6796
174.06707763671875 0 4316.728
175.08790588378906 0 820.43005
175.09866333007812 0 2475.016
176.08282470703125 0 30992.354
177.07827758789062 0 1016.5489
177.08645629882812 0 2340.4797
177.1031951904297 0 2339.3967
177.1144256591797 0 4640.431
178.0511016845703 0 1308.306
178.06219482421875 0 1922.6296
178.08792114257812 0 795.44116
178.0982208251953 0 801.2565
178.134765625 0 2147.0789
179.0699920654297 0 715.8836
179.09371948242188 0 5958.589
180.0776824951172 0 8482.852
180.1140899658203 0 10885.432
181.0734405517578 0 676.21655
181.09814453125 0 1860.4126
181.11753845214844 0 1266.7812
183.0774383544922 0 4408.971
183.1138458251953 0 9848.773
184.11734008789062 0 1037.5522
185.09327697753906 0 1275.6854
185.1295623779297 0 2872.4253
186.0884552001953 0 1972.9778
186.10113525390625 0 1191.1525
186.1248016357422 0 4411.63
187.1087646484375 0 228104.62 b 1
188.11219787597656 0 18122.926
189.07810974121094 0 9979.2705
189.1038055419922 0 1164.2347
189.1138916015625 0 1672.564
190.0621795654297 0 12828.012
190.08193969726562 0 793.8177
190.13516235351562 0 1471.6162
191.06590270996094 0 1118.9786
191.09384155273438 0 8864.666
192.07769775390625 0 2291.2249
192.10301208496094 0 8508.704
193.072998046875 0 4546.0933
193.10940551757812 0 59085.36
194.0809326171875 0 5598.1533
194.0934600830078 0 12236.343
194.11288452148438 0 4795.132
194.129638671875 0 2308.287
195.07740783691406 0 9265.73
195.08834838867188 0 4384.7866
195.0977783203125 0 1257.5481
195.11383056640625 0 17804.293
196.1172332763672 0 1538.9003
197.09295654296875 0 1232.6807
197.12904357910156 0 839.58466
198.1248016357422 0 1994.2208
199.0717315673828 0 882.4438
199.10873413085938 0 8587.806
200.09300231933594 0 2282.9534
200.11251831054688 0 904.7822
200.14083862304688 0 1485.6802
201.07766723632812 0 1145.1597
201.0880584716797 0 10292.503
201.10655212402344 0 3027.7156
202.0876007080078 0 4399.828
202.10968017578125 0 4749.1763
202.1198272705078 0 2732.4124
203.08270263671875 0 666.98267
203.09332275390625 0 3897.8245
203.10333251953125 0 833.10846
203.11325073242188 0 586.1316
204.07781982421875 0 36418.92
204.11502075195312 0 1029.5905
204.13580322265625 0 665.58356
205.08116149902344 0 3524.3513
205.09835815429688 0 2215.1558
205.10972595214844 0 1780.5643
205.14553833007812 0 500.1175
206.0944366455078 0 883.92786
206.1046600341797 0 4511.2324
206.12960815429688 0 1083.0005
207.0886993408203 0 40624.703
207.16201782226562 0 1398.9551
208.0738525390625 0 608.2499
208.09231567382812 0 3265.1287
208.12022399902344 0 1728.0452
209.09336853027344 0 1249.2983
209.10418701171875 0 1114.3248
211.10887145996094 0 13098.933
212.10403442382812 0 7001.247
212.1403045654297 0 1772.4269
213.08828735351562 0 2441.6416
213.10079956054688 0 588.36584
213.1246337890625 0 776.6798
215.103759765625 0 2867.5874
216.13543701171875 0 29131.635 y Water loss 13
217.07321166992188 0 10743.717
217.09893798828125 0 1646.8386
217.11949157714844 0 39184.008 y Ammonia loss 13
217.1388397216797 0 2602.1304
218.05718994140625 0 11432.518
218.07608032226562 0 902.5652
218.12271118164062 0 3020.1763
219.0608673095703 0 1146.9548
219.08901977539062 0 2977.4275
219.1137237548828 0 1939.3435
219.12490844726562 0 1029.3185
220.07302856445312 0 982.5034
221.10452270507812 0 160404.72
221.1404571533203 0 2417.8545
222.07095336914062 0 1097.714
222.08836364746094 0 4175.674
222.10806274414062 0 14439.259
222.12484741210938 0 31049.48
223.10902404785156 0 7302.0166
223.12855529785156 0 2167.8125
223.15660095214844 0 9011.419
224.11534118652344 0 47245.062
224.1409149169922 0 1476.1531
224.1603546142578 0 1263.4084
225.09906005859375 0 2244.7803
225.12001037597656 0 3019.971
226.11947631835938 0 1118.4344
226.15634155273438 0 2576.4175
228.1018524169922 0 880.6271
228.11431884765625 0 5991.2344
229.11846923828125 0 1081.0991
229.130859375 0 562.576
230.0833282470703 0 7921.353
230.11480712890625 0 51269.133
231.08753967285156 0 1861.8513
231.11769104003906 0 4388.205
233.1409149169922 0 7436.1733
234.09988403320312 0 14983.396
234.12460327148438 0 7485.669
234.1461181640625 0 176991.36 y 13
235.0838623046875 0 68147.19
235.09963989257812 0 621.7524
235.10623168945312 0 1984.3229
235.1298370361328 0 744.12787
235.1494598388672 0 16893.93
236.06814575195312 0 1284.6442
236.08724975585938 0 7985.375
236.1156768798828 0 2658.264
236.15086364746094 0 1932.6547
237.1006622314453 0 1269.8759
237.12460327148438 0 13704.813
238.11822509765625 0 1032.1522
238.12989807128906 0 2730.462
239.10328674316406 0 944.1886
239.1151885986328 0 40395.51
239.1515350341797 0 89899.57
240.08160400390625 0 1923.4028
240.13560485839844 0 189544.55
240.15499877929688 0 8740.648
241.08311462402344 0 1835.6356
241.11907958984375 0 1401.5968
241.13893127441406 0 21637.756
241.15476989746094 0 1186.2736
242.11465454101562 0 1676.2072
242.14117431640625 0 1655.4183
243.12222290039062 0 1040.2783
244.1305389404297 0 4850.461
247.11033630371094 0 17400.447
247.1411895751953 0 1050.1901
248.11383056640625 0 2864.248
249.09962463378906 0 29191.129
249.13600158691406 0 4685.457
249.17201232910156 0 568.85474
250.068359375 0 705.7529
250.08355712890625 0 3664.1812
250.10276794433594 0 2980.3467
250.11984252929688 0 1812.8296
250.1678924560547 0 1923.8329
251.1272430419922 0 1168.7012
251.15164184570312 0 19637.738
252.11056518554688 0 160980.39
252.13531494140625 0 9747.745
253.08047485351562 0 899.6523
253.09461975097656 0 4289.236
253.11367797851562 0 15821.62
253.13809204101562 0 1219.0264
254.0995330810547 0 666.2377
254.11550903320312 0 2173.692
256.109130859375 0 2197.1377
257.1081848144531 0 24342.879
257.1412353515625 0 1796.5013
257.1620788574219 0 24700.332
258.1107177734375 0 4790.0923
258.14599609375 0 7299.865
258.16510009765625 0 2123.229
259.1050109863281 0 1291.6289
259.1581726074219 0 1462.2689
260.11541748046875 0 1112.163
261.0989685058594 0 722.34076
261.1357421875 0 3741.376
261.17156982421875 0 1513.937
262.0948181152344 0 7746.6772
262.1195373535156 0 1427.4094
263.0790710449219 0 2723.6064
263.102294921875 0 1136.2988
264.137451171875 0 7086.488
265.1196594238281 0 14326.669
266.1232604980469 0 2258.8428
267.09320068359375 0 7142.327
267.1101379394531 0 91165.266
268.076416015625 0 3789.8833
268.094970703125 0 1210.1458
268.1133117675781 0 10476.23
268.130615234375 0 3340.9219
268.178466796875 0 2647.3315
269.1133728027344 0 1115.0471
269.1203918457031 0 668.46356
269.1371765136719 0 7463.9043
269.1806335449219 0 619.1824
270.0807800292969 0 1215.0026
270.1205749511719 0 1946.0049
270.14617919921875 0 32456.32
271.1496887207031 0 3799.469
271.47418212890625 0 596.9249
273.13604736328125 0 43509.215
274.1385192871094 0 4966.1025
274.1678161621094 0 872.696
275.107421875 0 36828.58
275.1734924316406 0 1988.251
276.1104736328125 0 6496.9854
276.18304443359375 0 1634.5039
277.0947265625 0 4672.004
277.16802978515625 0 1323.1704
278.0980224609375 0 898.1193
278.1493835449219 0 820.2213
278.1991271972656 0 1326.7991
279.1466979980469 0 17664.797
280.10540771484375 0 9921.962
280.1307373046875 0 27747.662
280.1497497558594 0 3392.0874
280.6524353027344 0 2253.879
281.1103210449219 0 1440.2631
281.1330871582031 0 2930.5789
281.1510009765625 0 1059.0828
282.1468505859375 0 2134.3928
283.12042236328125 0 2061.8416
283.1478271484375 0 653.23364
284.10443115234375 0 2161.8223
284.1366271972656 0 2038.3525
285.10321044921875 0 66835.31
285.6427307128906 0 893.1064
286.106201171875 0 9812.131
286.1241455078125 0 1541.147
287.09930419921875 0 2866.6643
287.15191650390625 0 1891.061
287.1703796386719 0 641.35614
291.1470031738281 0 1125.8179
292.1434326171875 0 1021.74066
292.1786193847656 0 1395.1539
293.1268310546875 0 2164.8574
294.19390869140625 0 2479.9133
294.64971923828125 0 2671.5698
295.10540771484375 0 11669.041
296.1625061035156 0 1891.5652
297.1575012207031 0 13207.252
298.12481689453125 0 1183.2025
298.1603698730469 0 2245.184
299.6418151855469 0 823.1744
300.1451416015625 0 1259.168
300.6272888183594 0 694.87354 b 4
301.1312561035156 0 71912.1
302.1330871582031 0 12591.39
302.1629333496094 0 2788.3958
303.1106872558594 0 2124.1296
303.1300354003906 0 694.4848
303.1650695800781 0 1028.6044
304.1058044433594 0 3408.9797
304.142333984375 0 3215.2808
304.178466796875 0 7019.6587
305.08984375 0 2035.3735
305.1268615722656 0 1888.5912
305.1627502441406 0 6349.935
305.18115234375 0 1208.2654
306.1268005371094 0 1239.1261
306.1467590332031 0 5134.354 a 2
306.1643371582031 0 917.7735
307.1209411621094 0 1432.0786
307.1451110839844 0 564.6848
309.1200866699219 0 643.2183
309.1380920410156 0 927.42224
309.15875244140625 0 887.898
309.62701416015625 0 703.819
310.14080810546875 0 1680.1517
311.1160583496094 0 2455.8162
312.120361328125 0 1088.2069
312.1515197753906 0 1417.7327
313.0981750488281 0 1164.3127
313.1446228027344 0 1054.9752
313.1892395019531 0 669.0118
313.6585998535156 0 1096.6062 y 10
314.1148681640625 0 1023.92535
314.1726379394531 0 2256.6492
314.6462097167969 0 1478.479
315.0968933105469 0 683.5566
315.1297912597656 0 803.002
316.13446044921875 0 10998.684
317.1188659667969 0 801.4793
317.1397705078125 0 1392.3771
318.12152099609375 0 734.67566
318.139404296875 0 3847.9243
318.1578369140625 0 1062.9653
318.6407470703125 0 995.35504
319.10589599609375 0 1426.7375
319.1416931152344 0 3011.8438
320.13726806640625 0 47956.55
320.1733093261719 0 7000.5425
321.12060546875 0 886.5287
321.1398010253906 0 7867.524
321.1579895019531 0 1260.2528
321.175537109375 0 1106.103
322.1164855957031 0 3753.9932
322.1506042480469 0 6285.04
322.1710510253906 0 2533.0867
322.1893005371094 0 21884.611
322.6512451171875 0 1502.3217
322.6715087890625 0 865.65857
323.1216735839844 0 739.8307
323.1513671875 0 1657.3894
323.1731262207031 0 12681.841
323.19268798828125 0 3532.7483
324.1318054199219 0 990.6411
324.1746826171875 0 1990.8455
325.15301513671875 0 897.45605
326.13763427734375 0 5566.224
326.1594543457031 0 1133.9624
326.1842956542969 0 1499.3872
326.6540832519531 0 904.73724
327.1432800292969 0 2915.8098 b Water loss 5
327.16796875 0 7607.1177
327.1891174316406 0 950.3442
327.65509033203125 0 824.8508
328.1523132324219 0 4898.336
328.1707458496094 0 804.51404
328.6683044433594 0 2088.17
329.1262512207031 0 2911.468
330.1267395019531 0 765.3845
330.158203125 0 2827.9636
331.10516357421875 0 1966.3496
331.1414794921875 0 660.41174
332.1372375488281 0 15638.854
332.17236328125 0 888.9984
333.1213684082031 0 3026.3909
333.1401062011719 0 1463.3232
334.1451416015625 0 77227.85
335.1155700683594 0 2104.9915
335.14825439453125 0 11245.112
335.6580505371094 0 602.9769
336.14422607421875 0 5050.1987 b 5
336.1688537597656 0 4117.2603
336.6680908203125 0 949.5985
337.1076354980469 0 819.8384
337.1298828125 0 1279.5627
337.1686706542969 0 695.24054
337.199951171875 0 1404.5958
337.6737060546875 0 1259.2751
338.1477355957031 0 46244.91
338.67584228515625 0 1774.7881
339.15045166015625 0 8092.859
339.1745910644531 0 1010.3148
339.21551513671875 0 3521.5422
340.14251708984375 0 25150.83
340.1998596191406 0 3730.1792
341.1452331542969 0 4900.303
341.1838073730469 0 30217.82
342.1870422363281 0 5911.998
342.66510009765625 0 1302.7477
343.1642150878906 0 3282.2947
343.1884765625 0 704.32874
344.1484069824219 0 10294.838
344.1947937011719 0 23155.35 y Water loss 12
344.68182373046875 0 955.1114
345.15313720703125 0 1444.4343
345.1788024902344 0 29568.785 y Ammonia loss 12
345.1972351074219 0 2553.8904
346.181396484375 0 4804.715
346.6459045410156 0 1463.9214
346.6920471191406 0 757.44415
347.1477355957031 0 1334.8788
347.18408203125 0 2434.6775
348.1319274902344 0 5794.4346
348.1684265136719 0 4996.5825
349.1329345703125 0 1608.755
349.15301513671875 0 1148.4856
349.19842529296875 0 692.1382
350.14788818359375 0 62514.51
350.18609619140625 0 2249.9536
350.6673278808594 0 1601.5062
351.15057373046875 0 10129.344
351.6713562011719 0 5564.749
352.1433410644531 0 4217.0103
352.17242431640625 0 2802.3132
352.6751403808594 0 918.89624
353.1263427734375 0 3124.2053
353.14764404296875 0 1669.6012
353.1716003417969 0 969.44824
353.1950988769531 0 3620.6365
354.1324768066406 0 7284.374
354.17010498046875 0 3876.5813
354.6708068847656 0 991.37646
355.13916015625 0 1145.9047
355.17431640625 0 1241.6049
356.1369323730469 0 1208.2054
356.172607421875 0 2104.1118
358.1534729003906 0 2873.3926
358.1893005371094 0 1147.2662
358.2104797363281 0 16125.615
359.1484680175781 0 1340.1971
359.1936950683594 0 1410.613
359.2144775390625 0 2275.017
360.1580810546875 0 1249.9519
360.8503723144531 0 899.4912
361.175048828125 0 12350.832
361.2008361816406 0 2145.43
362.17950439453125 0 3273.6577
362.2054138183594 0 20036.965 y 12
362.50372314453125 0 1422.2644
363.1112365722656 0 821.50397
363.1847839355469 0 830.8839
363.2091979980469 0 3980.33
364.1269226074219 0 880.84607
364.2106628417969 0 1032.598
365.1950988769531 0 14143.2705
366.1404113769531 0 10547.826
366.1587829589844 0 16070.292
366.1992492675781 0 2629.0615
367.14166259765625 0 2119.7368
367.1623840332031 0 4320.744
368.14373779296875 0 976.95734
368.16827392578125 0 6252.2866
368.6707458496094 0 1995.0774
369.17156982421875 0 3292.5754
369.6673278808594 0 920.71564
370.15313720703125 0 5012.6147
370.66619873046875 0 593.4396
371.1594543457031 0 43325.312
372.143310546875 0 20014.564
372.1635437011719 0 5625.2144
373.146484375 0 3903.1052
373.18731689453125 0 1669.3217 y Water loss 9
373.6799011230469 0 5486.3623 y Ammonia loss 9
374.14544677734375 0 859.3839
374.1822204589844 0 3351.7305
375.169189453125 0 12743.973
376.1264343261719 0 1134.0155
376.1676025390625 0 3683.4219
376.19537353515625 0 557.5718
376.2212829589844 0 3057.5845
376.6820373535156 0 1175.1201
377.1628723144531 0 585.0104
377.2255554199219 0 814.2268
378.1443176269531 0 1143.2263
378.17864990234375 0 2614.947
378.69000244140625 0 2442.8457
379.1075744628906 0 741.46564
379.17022705078125 0 868.05176
379.21099853515625 0 20221.01
379.675048828125 0 1084.8658
380.1371765136719 0 4929.6465
380.1571960449219 0 558.1974
380.1824645996094 0 1759.9639
380.2144470214844 0 4377.4624
381.1210632324219 0 4406.7617
381.1436767578125 0 794.65137
381.1886901855469 0 884.97986
382.1248779296875 0 1446.946
382.16546630859375 0 6239.806
382.1922912597656 0 18475.66 y 9
382.2221374511719 0 1917.7937
382.6652526855469 0 2214.903
382.69390869140625 0 9858.84
383.19464111328125 0 1751.2028
384.16949462890625 0 16728.09
384.1939392089844 0 1104.3756
384.5071716308594 0 672.656
384.6775817871094 0 2015.2552
385.17242431640625 0 4137.7993
386.1493225097656 0 4938.2524
386.6944580078125 0 867.9917
387.1522521972656 0 1420.3358
387.181396484375 0 5493.989
387.2156982421875 0 1656.5061
388.1933898925781 0 1183.9949
388.68487548828125 0 1239.6058
389.17010498046875 0 15867.003
389.19561767578125 0 2928.6677
390.1747131347656 0 2890.4863
390.5163269042969 0 999.028
391.17578125 0 1870.933
391.2107849121094 0 2077.6235
391.5163269042969 0 1726.971 b 9
391.6770935058594 0 992.5501
392.1968688964844 0 985.94324
393.1524353027344 0 1968.3307
393.18389892578125 0 15388.284
393.69049072265625 0 2602.5034
394.1566162109375 0 4055.5525
394.1872863769531 0 3344.157
395.16748046875 0 1111.0433
395.69891357421875 0 1395.037
396.1365661621094 0 4330.6675
396.198486328125 0 968.9078
396.52197265625 0 751.55786
397.1183166503906 0 776.41974
397.16448974609375 0 6817.0405
397.19879150390625 0 2290.0312
397.69110107421875 0 9030.479
398.1480712890625 0 13279.989
398.1941223144531 0 1867.7585
399.1523742675781 0 4472.6113
400.1371765136719 0 1240.0387
401.19580078125 0 4678.5425
402.1431579589844 0 2250.9626
402.1801452636719 0 10201.538
402.683349609375 0 2150.559
403.1791076660156 0 4699.1255
404.1588439941406 0 4303.5312
404.6685791015625 0 1084.4056
405.1631164550781 0 1394.4963
405.22711181640625 0 11318.694
406.2044372558594 0 22038.338
406.7055358886719 0 9556.162
407.2060852050781 0 26304.686
408.2082214355469 0 4530.524
408.2381591796875 0 3615.6667
410.2042236328125 0 7683.3516
410.7036437988281 0 604.40375
411.1944580078125 0 5068.7407
411.5154724121094 0 1712.4053
411.6908264160156 0 3822.514
411.8492431640625 0 884.84894
412.16680908203125 0 10380.113
412.6964416503906 0 738.1031
413.1798400878906 0 3813.0767
413.6802978515625 0 3866.0105
414.14666748046875 0 41367.527
414.6755676269531 0 782.52686
415.1485595703125 0 5805.579
415.1752624511719 0 34156.54
415.2206726074219 0 8513.467
415.5399169921875 0 755.39856
415.7232971191406 0 4027.9011
416.1485290527344 0 1949.8922
416.1785888671875 0 8006.136
416.2107238769531 0 1129.6527
416.6824645996094 0 1456.2396
417.165771484375 0 2628.4194
417.19329833984375 0 1091.4353
417.7015075683594 0 1223.2117
417.8544616699219 0 947.8053
418.160400390625 0 865.7351
418.2225646972656 0 748.55994
419.1703796386719 0 1497.9755
419.2062072753906 0 7157.3276
420.1334533691406 0 741.48444
420.16851806640625 0 1305.5854
420.20233154296875 0 14694.034
420.7031555175781 0 6114.8496
421.1854248046875 0 41986.703
421.6950988769531 0 2735.4321
422.18914794921875 0 15260.514
422.6910705566406 0 2686.3018
423.19085693359375 0 2080.1086
423.2375183105469 0 18572.088
423.5387268066406 0 857.1119
424.2001953125 0 1489.5752
424.23944091796875 0 5289.934
424.6979064941406 0 2691.4866
424.87335205078125 0 734.14325
425.1587219238281 0 1499.2054
425.19158935546875 0 12631.658
425.691650390625 0 4081.7324
426.15069580078125 0 1154.5917
426.1846008300781 0 6302.2666
427.18402099609375 0 1230.999
428.1867370605469 0 1171.4103
428.2165832519531 0 2722.1226
429.1927490234375 0 928.7825
429.219482421875 0 9029.963
429.69232177734375 0 959.5632
429.72113037109375 0 5606.974
430.1759033203125 0 3223.341
430.20684814453125 0 5984.0757 y Water loss 8
430.7012023925781 0 11794.449 y Ammonia loss 8
431.1734313964844 0 2234.0474
431.20281982421875 0 4615.1816
431.70111083984375 0 2122.068
432.2025146484375 0 1023.6451
433.1925964355469 0 855.1003
433.22174072265625 0 15270.002
433.540771484375 0 1769.5938
433.7071838378906 0 1557.1776
433.87451171875 0 1716.823
434.2049255371094 0 1486.0869
434.2236328125 0 2147.648
436.1647644042969 0 2575.7468
436.2328796386719 0 18434.186
436.5379333496094 0 2272.0203
436.8744812011719 0 1139.8279
437.15972900390625 0 910.22485
437.2358093261719 0 4005.5796
437.7040100097656 0 1026.886
437.73333740234375 0 1130.3455
437.87384033203125 0 672.19794
438.14495849609375 0 1293.2603
438.209716796875 0 2152.994
438.7115783691406 0 5545.0576
439.2136535644531 0 41972.223 y 8
439.7149963378906 0 20752.318
440.2164306640625 0 4660.2847
440.71832275390625 0 767.6412
442.14111328125 0 1633.4303
442.2131042480469 0 3187.8113
442.5479736328125 0 1084.5339
442.88330078125 0 1737.1995
443.1706237792969 0 10854.27
443.2100830078125 0 34565.023 y 4
443.55352783203125 0 768.3375
444.17230224609375 0 2088.406
444.2117919921875 0 8571.686
444.69744873046875 0 1747.6075
445.20806884765625 0 2711.488
445.2436218261719 0 7371.692 y Water loss 11
445.6953125 0 681.51624
445.8708801269531 0 3419.9854 b 10
446.14862060546875 0 3177.707
446.2258605957031 0 4967.6504 y Ammonia loss 11
446.5385437011719 0 1048.423
446.710205078125 0 3558.0398
447.2210693359375 0 21601.062
447.72247314453125 0 25891.037
448.2225341796875 0 12530.328
448.55279541015625 0 2135.9026
448.7247619628906 0 3273.1758
448.88604736328125 0 1670.6902
449.2177734375 0 4857.0396
450.1482849121094 0 1106.0463
450.18524169921875 0 1242.5527
450.2184143066406 0 1112.3779
450.24835205078125 0 2790.753
450.8763427734375 0 835.8309
451.2326965332031 0 49417.04
452.2353820800781 0 11417.889
452.55126953125 0 1783.8632
452.71337890625 0 2626.1565
452.8841247558594 0 1406.8174
453.2024230957031 0 3706.4392
453.7106018066406 0 1161.4291
453.8917236328125 0 1030.2675
455.1714172363281 0 2121.4102
455.2217712402344 0 5590.351
455.5449523925781 0 1383.0718
455.7229919433594 0 2959.972
455.8801574707031 0 1455.311
456.1809387207031 0 730.8436
456.21832275390625 0 1030.0582
457.1891174316406 0 2008.5029
457.2323303222656 0 3370.1868
457.8917541503906 0 740.27045
458.2185974121094 0 2028.1797
458.5518493652344 0 2467.2087
458.88525390625 0 1011.0005
459.2037048339844 0 1237.9783
459.8968200683594 0 1596.8384
460.2243957519531 0 4457.928
460.5478515625 0 838.61346
460.7270202636719 0 1087.3322
460.8856201171875 0 1195.6665
461.2188720703125 0 47067.207 b 7
461.55419921875 0 1845.3113
461.72021484375 0 28680.436
461.88604736328125 0 1136.839
462.2206115722656 0 12355.92
462.72149658203125 0 3028.16
463.1759948730469 0 1996.1887
463.2537841796875 0 46916.133 y 11
464.15948486328125 0 5306.1753
464.2249755859375 0 5721.8657
464.257568359375 0 9096.895
464.5541076660156 0 3362.4888
464.7292785644531 0 920.0484
464.8833312988281 0 3098.8245
465.1629943847656 0 1326.7208
465.2177734375 0 2245.6018
465.25994873046875 0 1392.7845
465.5469970703125 0 754.2985
465.7168273925781 0 2203.9814
466.2186584472656 0 1220.702
466.720703125 0 1149.4905
466.89007568359375 0 779.8609
467.1727294921875 0 2039.4672
467.2190246582031 0 3059.5303
467.54986572265625 0 805.3151
467.8768615722656 0 1134.1583
468.1571960449219 0 984.3105
468.2248840332031 0 1036.1016
468.2587585449219 0 3508.8826
469.1847839355469 0 742.8878
469.73187255859375 0 3629.6135
469.8919982910156 0 2901.1482
470.22637939453125 0 6141.724
470.5580139160156 0 3264.9822
470.7186279296875 0 2617.649
470.8888244628906 0 842.0084
471.2054138183594 0 38044.004 b 3
471.7171630859375 0 1233.0925
472.20550537109375 0 9321.232
472.2394104003906 0 982.2505
472.5634765625 0 738.2279
473.2037048339844 0 2942.2942
473.24066162109375 0 1223.6527
473.5502014160156 0 6930.176 b Water loss 11
473.88397216796875 0 5528.6577
474.2179260253906 0 2587.2808
474.2593688964844 0 6035.337
474.724365234375 0 3861.722
475.2242431640625 0 3087.1067
475.26544189453125 0 1246.3721
475.5675048828125 0 697.1606
475.897216796875 0 5917.2593
476.2300109863281 0 4644.438
476.5643615722656 0 2970.721
476.8970947265625 0 1115.0883
477.21112060546875 0 2269.8076
477.23388671875 0 1834.5219
477.55780029296875 0 2334.157
477.88916015625 0 1072.643
478.24237060546875 0 2077.378
479.2262268066406 0 11980.664
479.5543518066406 0 8828.989 b 11
479.7225036621094 0 18102.121
479.8892822265625 0 4937.1963
480.2237548828125 0 14036.326
480.55584716796875 0 2103.0344
480.7244873046875 0 3162.5315
480.8891906738281 0 945.709
481.18585205078125 0 10792.942
481.22418212890625 0 2297.3057
481.5713195800781 0 3293.7078
481.71380615234375 0 1024.1865
481.9015808105469 0 3383.721
482.1833801269531 0 1980.6901
482.2359619140625 0 1263.7083
482.55426025390625 0 2292.412
482.8954772949219 0 3434.5056 y Water loss 3
483.1700439453125 0 1112.9153
483.2052917480469 0 392.59662
483.2336120605469 0 7996.1123 y Ammonia loss 3
483.5670166015625 0 6119.844
483.7298278808594 0 2126.7659
483.8979187011719 0 3147.9023
484.236083984375 0 3300.549
484.7136535644531 0 1510.9861
485.1839599609375 0 26665.613
485.22564697265625 0 6037.8535
485.5600891113281 0 1622.307
485.8981628417969 0 2125.7837
486.187255859375 0 5841.9785
486.2403259277344 0 18690.672
486.74237060546875 0 8128.617
487.1838073730469 0 1148.6924
487.2406921386719 0 3363.3203
487.5732727050781 0 2740.9697
487.7364807128906 0 7409.141
487.90875244140625 0 2219.6094
488.2363586425781 0 9475.554
488.5726623535156 0 1095.66
488.74072265625 0 2946.1052
488.9067077636719 0 15050.952 y 3
489.2035827636719 0 3975.291
489.2430114746094 0 12631.066
489.5755310058594 0 5258.0337
489.9066467285156 0 1673.4081
490.20745849609375 0 2203.6145
490.2423095703125 0 3958.7695
490.7244567871094 0 2015.6984
491.2250061035156 0 5634.397
491.5607604980469 0 3169.8665
491.88568115234375 0 822.2731
491.905517578125 0 650.00867
492.25726318359375 0 897.263
492.56781005859375 0 1382.9615
492.7297058105469 0 2057.0007
493.23052978515625 0 1889.8041
493.56707763671875 0 971.3419
493.7200622558594 0 16383.54
493.8953552246094 0 773.9841
494.2217102050781 0 9076.533
494.577392578125 0 1080.1875
494.7196350097656 0 2587.6235
495.2071533203125 0 2625.7024
495.2409362792969 0 1202.6703
495.56622314453125 0 1953.0538
495.7322692871094 0 1066.5883
495.8984069824219 0 983.8702
496.2354736328125 0 905.22705
497.229248046875 0 9354.645
497.5644836425781 0 7475.2837
497.8979797363281 0 4141.9033
498.2131042480469 0 14625.163
498.56671142578125 0 1681.9608
498.7386474609375 0 4847.19 y Water loss 7
498.8984375 0 1177.9137
499.1963806152344 0 3666.302
499.23040771484375 0 7964.448 y Ammonia loss 7
499.5648193359375 0 2399.2756
499.73370361328125 0 3519.3542
499.8957214355469 0 1066.0222
500.197021484375 0 2490.275
500.2351379394531 0 1248.264
500.90545654296875 0 1646.7075
501.2392578125 0 1234.7191
501.5692443847656 0 2993.2322
501.7340393066406 0 7482.572
501.9034729003906 0 3129.9072
502.2497253417969 0 9999.98
502.573486328125 0 1079.1099
502.73388671875 0 1335.9686
502.90576171875 0 8455.814
503.23944091796875 0 7956.4326
503.5718078613281 0 3053.7688
503.9064636230469 0 1489.2336
504.2452087402344 0 8587.089
504.7442626953125 0 2835.4219
504.9075622558594 0 3875.766
505.24090576171875 0 7120.6704
505.5697326660156 0 3382.048
505.9058532714844 0 1609.7098
506.23712158203125 0 2479.5356
506.2752990722656 0 7364.2056
506.56378173828125 0 807.84155
506.9052429199219 0 2464.0002
507.212890625 0 8480.641
507.2445983886719 0 3465.7678
507.2799987792969 0 930.088
507.57525634765625 0 4777.1807
507.7439880371094 0 24788.385 y 7
507.91217041015625 0 4174.8896
508.2498474121094 0 26668.414
508.7461853027344 0 4928.855
508.9096984863281 0 4181.477
509.1799621582031 0 769.47687
509.2496032714844 0 6157.142
509.5767517089844 0 2229.7166
509.9118957519531 0 1054.138
510.23651123046875 0 1689.8811
510.2664489746094 0 621.7749
510.5804443359375 0 4874.7163
510.74114990234375 0 1137.3502
510.9123840332031 0 15150.639
511.2463073730469 0 10188.925
511.57861328125 0 4564.421
511.7623596191406 0 2622.0945
511.9154357910156 0 1343.9744
512.2305908203125 0 7230.129
512.9085083007812 0 3491.2573
513.2161254882812 0 3697.3464
513.2440795898438 0 3272.8354
513.5757446289062 0 2817.9678
513.9113159179688 0 1067.1311
514.2175903320312 0 1289.8556
515.2354736328125 0 2099.3115
515.578369140625 0 1166.6045
516.2255249023438 0 15514.134 b Water loss 12
516.5852661132812 0 32905.51
516.9197998046875 0 28761.559
517.1919555664062 0 1253.3356
517.2528686523438 0 15416
517.5867309570312 0 4184.8813
517.9155883789062 0 927.8485
518.2403564453125 0 58039.582 b 8
518.7415161132812 0 26644.873
519.241455078125 0 12708.1
519.7426147460938 0 4202.4727
520.2523803710938 0 9807.963
520.5729370117188 0 1022.68744
520.7417602539062 0 3863.1494
521.2359619140625 0 1005.4993
521.2733154296875 0 5115.405
521.741455078125 0 1290.0005
522.2403564453125 0 6122.8423 b 12
522.2825317382812 0 1022.5423
522.5744018554688 0 6976.2534
522.9098510742188 0 3018.8528
523.2008056640625 0 1310.3262
523.2533569335938 0 7080.3574
523.5866088867188 0 4174.481
523.7570190429688 0 1001.95056
523.92138671875 0 2379.1453
524.2406005859375 0 8093.4453
525.2448120117188 0 2888.6865
525.7401123046875 0 1842.6539
525.9183959960938 0 1281.656
526.24365234375 0 4499.477
526.5730590820312 0 5906.9766
526.9061889648438 0 3584.8262
527.2400512695312 0 1776.2927
527.2866821289062 0 1459.4452
528.269775390625 0 5264.422
528.92236328125 0 1048.729
529.2554931640625 0 12342.262
529.3024291992188 0 2462.1792
529.7559814453125 0 8217.023
530.2427368164062 0 19527.555
530.5864868164062 0 1141.772
530.7515869140625 0 2271.1724
531.2448120117188 0 4462.3794
531.768798828125 0 3787.732
531.9186401367188 0 11116.643 y Water loss 2
532.2514038085938 0 20250.457
532.5800170898438 0 17518.59
532.7638549804688 0 3827.4878
532.9138793945312 0 6830.926
533.2498779296875 0 7607.8867
533.5777587890625 0 1887.115
534.2333374023438 0 1757.8372
534.2695922851562 0 26300.666
534.7418212890625 0 6630.054
535.2069091796875 0 4082.6553
535.230712890625 0 1236.0343
535.2716674804688 0 6676.5376
535.7447509765625 0 2020.3909
535.9275512695312 0 1044.7836
536.2495727539062 0 44036.926
536.5963134765625 0 917.9068
537.2529907226562 0 11968.149
537.6033935546875 0 1169.2303
537.920166015625 0 115954.1 y 2
538.2545776367188 0 118548.86
538.587890625 0 59242.41
538.7564086914062 0 3237.5532
538.9209594726562 0 23619.04
539.2544555664062 0 8998.803
539.9153442382812 0 1192.9247
540.2210083007812 0 1092.5494
540.26611328125 0 938.0715
540.7733764648438 0 33662.094
541.2114868164062 0 5431.782
541.2743530273438 0 18633.396
541.77587890625 0 6680.192
541.9273681640625 0 881.4931
542.2775268554688 0 3249.56
543.2532958984375 0 19569.85
543.6052856445312 0 16549.62
543.7531127929688 0 13853.839
543.9385375976562 0 14447.012
544.2156982421875 0 1068.2828
544.2662353515625 0 8230.931
544.5886840820312 0 15652.9795
544.7567138671875 0 1777.6935
544.9227294921875 0 16008.909
545.2554321289062 0 8742.875
545.2977294921875 0 13005.181
545.5899047851562 0 3628.273
545.7648315429688 0 3948.5266
546.2582397460938 0 5522.507
546.30029296875 0 4013.9702
546.7620239257812 0 4064.184
547.265380859375 0 15802.913
547.587158203125 0 1819.9341
547.7554321289062 0 1887.1305
547.7930297851562 0 1033.8547
547.918212890625 0 1561.4164
548.2523193359375 0 9836.404
548.747802734375 0 1137.6632
549.2528076171875 0 2882.186
549.608642578125 0 13642.233
549.9425659179688 0 13231.753
550.2758178710938 0 5635.7456
550.6061401367188 0 3060.0708
550.9352416992188 0 1169.534
551.2601318359375 0 940.38477
551.7651977539062 0 2556.2256
552.234619140625 0 6748.8545
552.2633666992188 0 4778.135
552.7526245117188 0 4452.1997
552.9363403320312 0 951.9051
553.25830078125 0 20183.746
553.593017578125 0 18696.354
553.9254150390625 0 9443.953
554.2537231445312 0 6146.9814
554.593505859375 0 2690.9067
554.7706909179688 0 29870.582
554.9334716796875 0 951.42413
555.27294921875 0 19569.697 y Water loss 6
555.7726440429688 0 8008.5137 y Ammonia loss 6
556.270751953125 0 5640.3477
557.2593994140625 0 1813.4779
557.75537109375 0 1260.6862
557.9375610351562 0 1084.4717
558.2344360351562 0 941.9261
558.9339599609375 0 10916.41 b Water loss 13
559.2637329101562 0 91006.164 b Ammonia loss 13
559.59765625 0 89567.18
559.931396484375 0 48155.26
560.2632446289062 0 14167.198
560.2991943359375 0 9201.703
560.6010131835938 0 6377.3784
560.7699584960938 0 3222.5647
560.9381713867188 0 2969.5264
561.2595825195312 0 9146.792
561.6016845703125 0 2413.2427
561.7567138671875 0 5940.021
562.2584228515625 0 4859.9634
562.7598266601562 0 1246.4557
563.251953125 0 2105.6353
563.7671508789062 0 7536.128
563.9434204101562 0 2358.3586
564.2824096679688 0 10572.61 y 6
564.6107788085938 0 1936.1082
564.7627563476562 0 5256.5264
564.7849731445312 0 4097.586
564.9384155273438 0 145101.83 b 13
565.2728271484375 0 163830.38 y Ammonia loss 1
565.6062622070312 0 79957.41
565.7686157226562 0 1108.4867
565.9397583007812 0 31894.357
566.2754516601562 0 14038.936
566.6092529296875 0 1539.6361
566.7391357421875 0 2946.9314
566.9404296875 0 5390.1826
567.2724609375 0 8179.6436
567.6065063476562 0 5351.324
567.7523193359375 0 1083.6769
567.941162109375 0 2327.6245
568.279052734375 0 1624.1963
569.2540893554688 0 1569.794
569.61376953125 0 3610.6582
569.9483642578125 0 2422.0808
570.2767333984375 0 9985.468
570.9423217773438 0 58296.535 y 1
571.276123046875 0 55332.715
571.6102294921875 0 34687.71
571.77294921875 0 2661.9023
571.9442749023438 0 13560.99
572.2562866210938 0 14569.888
572.6163330078125 0 16957.889
572.772705078125 0 72369.45
572.946533203125 0 38193.746
573.2783203125 0 69134.23
573.6144409179688 0 12954.621
573.774658203125 0 18482.592
573.9462280273438 0 5424.009
574.2062377929688 0 1360.006
574.2830810546875 0 6975.73
574.6047973632812 0 2416.506
574.7807006835938 0 983.52924
574.9382934570312 0 1728.0404
575.2523803710938 0 10516.165
575.2964477539062 0 899.84033
575.7544555664062 0 7259.067
575.9364624023438 0 973.6639
576.2578735351562 0 7643.9067
576.5977172851562 0 1403.0197
576.759521484375 0 4955.5195
577.2666015625 0 1790.9623
577.7650146484375 0 5711.3867 b Water loss 9
578.2633666992188 0 8420.161
578.6199340820312 0 224918.5
578.760498046875 0 4668.791
578.9542236328125 0 223968.02
579.2880859375 0 128815.35
579.6223754882812 0 39587.965
579.7698974609375 0 1863.5952
579.9558715820312 0 10024.781
580.2694702148438 0 3137.0142
580.7753295898438 0 1906.6221
581.2754516601562 0 3428.558
581.59912109375 0 878.58844
581.7813110351562 0 1292.7957
581.9415283203125 0 4307.2314
582.2333984375 0 2974.6501 b Water loss 4
582.2745971679688 0 8219.842
582.6041870117188 0 8977.1875
582.7869262695312 0 1963.0663
582.9384765625 0 4890.3823
583.2340698242188 0 1535.7063
583.2767944335938 0 6049.468
583.6055908203125 0 1446.3627
584.248046875 0 2087.006
584.6134033203125 0 1442.6091
584.7711181640625 0 13072.701
584.9496459960938 0 2128.2703
585.2731323242188 0 13510.357
585.6107177734375 0 6325.7114
585.7728271484375 0 6122.3076
585.9423828125 0 3507.7344
586.2757568359375 0 7344.0747
586.7704467773438 0 75713.67 b 9
587.2720947265625 0 48750.785
587.7723388671875 0 20105.89
587.9464111328125 0 6510.3105
588.2800903320312 0 31618.318
588.608154296875 0 21213.998
588.7733764648438 0 1841.3722
588.941650390625 0 16207.88
589.2779541015625 0 15799.681
589.6096801757812 0 5698.674
589.9466552734375 0 3270.983
590.2821655273438 0 5903.1787
590.6159057617188 0 1355.536
590.7998046875 0 3260.8625 y Water loss 5
591.2908935546875 0 10379.712 y Ammonia loss 5
591.7958374023438 0 3282.625
592.2210693359375 0 3644.0215
592.293701171875 0 1787.6924
592.7880249023438 0 1219.0132
593.2232666015625 0 1227.239
593.2792358398438 0 4848.2305
593.7789916992188 0 9419.255
593.9501342773438 0 105335.62 Precursor Water loss
594.2808837890625 0 218313 Precursor Ammonia loss
594.614501953125 0 167245.88
594.7825927734375 0 5050.55
594.9473266601562 0 103689.98
595.2814331054688 0 49888.9
595.6139526367188 0 14843.965
595.784423828125 0 1388.8074
595.9492797851562 0 2068.2842
596.2815551757812 0 2266.7385
597.28125 0 1341.0944
598.2709350585938 0 22379.447
598.5645751953125 0 8820.563
598.7686767578125 0 1409.3552
598.8150024414062 0 5289.0483
599.0656127929688 0 2082.6946
599.2719116210938 0 8018.524
599.8065795898438 0 29987.059 y 5
599.953369140625 0 1757622.6 Precursor
600.287109375 0 1766051.1
600.6203002929688 0 1040751.56
600.9542846679688 0 435859.16
601.2882690429688 0 135353.23
601.6218872070312 0 2981.0425
602.2481689453125 0 4968.041
602.2899169921875 0 3984.9207
602.7927856445312 0 6228.755
603.293212890625 0 9133.248
603.763916015625 0 2442.2256
603.7987060546875 0 980.1922
604.258056640625 0 1194.7562
604.2999877929688 0 2073.8096
604.7808227539062 0 2760.8906
605.3196411132812 0 3569.7307
606.3209838867188 0 1368.3661
607.2857666015625 0 44655.953
607.779541015625 0 7407.351
608.2949829101562 0 13210.072 y Water loss 10
608.7631225585938 0 2120.9624
609.2483520507812 0 3506.4084
609.2946166992188 0 10214.119 y Ammonia loss 10
610.2312622070312 0 2158.789
610.2959594726562 0 2626.7354
610.7877197265625 0 1368.4619
611.2334594726562 0 998.0318
611.29296875 0 2211.0994
611.8036499023438 0 7290.798
612.2474975585938 0 4124.917
612.3045043945312 0 5489.52
612.8049926757812 0 1880.5636
613.2471923828125 0 1202.03
613.3009643554688 0 2273.1123
613.792724609375 0 6532.9688
614.2962646484375 0 3614.5942
614.7958374023438 0 1843.3726
615.2986450195312 0 1116.9503
616.2867431640625 0 15158.6
616.775390625 0 18191.836
617.2730712890625 0 12669.374
617.7737426757812 0 5724.463
618.2471923828125 0 6082.5127
619.248046875 0 1645.7836
620.2987670898438 0 3570.0889
620.8087158203125 0 2142.98
621.3023681640625 0 3614.132
622.3049926757812 0 17537.309
622.8060302734375 0 11815.967
623.3048095703125 0 5348.4185
623.8051147460938 0 1414.5736
624.3070678710938 0 1275.7699
624.7958374023438 0 2755.3884
625.286376953125 0 5674.037
625.7789916992188 0 7633.7095
626.2730102539062 0 9092.799
626.3176879882812 0 47245.414 y 10
626.8121948242188 0 2791.0537
627.2603149414062 0 5843.65
627.31787109375 0 17199.154
627.7944946289062 0 3407.0674
628.3262329101562 0 2011.7596
628.8018188476562 0 1064.1676
629.2918090820312 0 932.3422
630.2884521484375 0 1783.7134
631.2890625 0 1330.8391
633.3394165039062 0 2770.4954
634.292724609375 0 4366.442
634.794677734375 0 2501.3208
635.27392578125 0 11142.26
635.7939453125 0 3118.2947
636.30078125 0 15688.9
636.801513671875 0 10643.779
637.3092041015625 0 6997.3047
637.8021240234375 0 3218.1338
638.3135375976562 0 3055.713
639.3005981445312 0 4765.745
640.2901000976562 0 3772.787
640.3342895507812 0 2098.334
640.7892456054688 0 2009.305
641.3123168945312 0 3333.0896
642.3107299804688 0 1686.0226
643.2907104492188 0 5370.0283
643.333984375 0 6037.7275
643.8070068359375 0 2810.101
644.2846069335938 0 13834.364
644.8057861328125 0 2562.8103
645.2888793945312 0 7422.2983
645.7941284179688 0 8978.649
646.2998046875 0 5754.5684
646.8060302734375 0 6631.487
647.308349609375 0 6141.1636
647.806396484375 0 3163.946
648.3004150390625 0 3763.0576
648.801513671875 0 6278.2725
649.2972412109375 0 8282.862
649.8049926757812 0 8229.009
650.3037719726562 0 4291.221
650.801025390625 0 950.9847
651.8187866210938 0 1069.3567
652.298095703125 0 1803.4679
653.279541015625 0 10593.562
654.3041381835938 0 38401.027
654.80615234375 0 23142.182
655.3150634765625 0 22320.066 y Water loss 4
655.8168334960938 0 20981.908 y Ammonia loss 4
656.322021484375 0 17336.637
656.8140869140625 0 5083.1772
657.3167114257812 0 11567.838
657.8053588867188 0 11034.385
658.33154296875 0 12562.48
658.80712890625 0 3651.1357
659.29248046875 0 2610.1333 b Water loss 10
659.341064453125 0 2673.3604
659.7946166992188 0 5512.2437 b Ammonia loss 10
660.2965698242188 0 3676.7515
660.8013305664062 0 3440.2656
661.3084716796875 0 1843.3484
662.8175659179688 0 3932.6611
663.3167114257812 0 5552.2856
663.8167724609375 0 2661.9136
664.326416015625 0 44117.793 y 4
664.82763671875 0 35085.477
665.3265991210938 0 23033.191
665.828125 0 4444.7207
666.3157958984375 0 8071.6777
666.80712890625 0 2365.474
667.3012084960938 0 8503.021
667.7999877929688 0 3218.9927
668.3023681640625 0 45377.836 b 10
668.8046875 0 32985.418
669.3063354492188 0 21940.895
669.808349609375 0 9545.309
670.3125 0 6372.393
670.814453125 0 1130.0037
671.2853393554688 0 61677.082 b 5
671.8223266601562 0 11011.265
672.2886962890625 0 18620.33
672.3370971679688 0 5912.162
672.8253173828125 0 4536.055
673.2855224609375 0 5705.0244
673.3327026367188 0 3702.9058
673.8302001953125 0 2122.4028
674.3416137695312 0 14541.298
675.3418579101562 0 4654.8647
675.8117065429688 0 2526.1494
676.2966918945312 0 1277.3912
676.3482666015625 0 3035.6216
676.81005859375 0 1091.8021
677.3170776367188 0 2763.914
677.82177734375 0 27834.273
678.3229370117188 0 22502.879
678.822998046875 0 8503.068
679.323974609375 0 4341.6177
680.3349609375 0 18356.53
680.8346557617188 0 13633.9
681.2941284179688 0 3412.1025
681.3370361328125 0 4799.998
681.8007202148438 0 3833.5884
682.3241577148438 0 975.16943
682.8161010742188 0 2444.3394
683.3232421875 0 1819.144
684.3257446289062 0 11273.752
685.3179931640625 0 8576.176
686.33056640625 0 6519.387
686.8289184570312 0 14593.931
687.3292236328125 0 11211.517
687.8282470703125 0 4509.565
688.3087158203125 0 2325.4155
688.8170776367188 0 1344.3966
689.3375854492188 0 5189.895
689.80712890625 0 13261.5625
690.3072509765625 0 11398.752
690.8078002929688 0 4006.6655
691.3301391601562 0 8722.254
691.8271484375 0 4496.203
692.3202514648438 0 2849.9395
692.3780517578125 0 5699.98
693.3171997070312 0 2061.8218
693.3795166015625 0 1873.0093
694.3109741210938 0 1821.4021
695.3392333984375 0 6866.1304
695.8369140625 0 8007.0254
696.3272094726562 0 4389.9614
696.8201904296875 0 13419.6875
697.3233642578125 0 8583.666
697.8212890625 0 4530.645
698.3272094726562 0 4001.7402
698.8231811523438 0 1093.5144
699.3285522460938 0 3016.8599
700.325439453125 0 976.107
700.8197021484375 0 4021.1128
701.31494140625 0 8993.736
701.8131713867188 0 6407.8115
702.3369750976562 0 36465.336
702.8070678710938 0 1271.565
703.3402709960938 0 14528.93
704.3387451171875 0 7919.9165
704.8326416015625 0 8573.303
705.3316650390625 0 6290.1724
705.8336791992188 0 4921.5757
706.3347778320312 0 5177.3706
706.8328247070312 0 3076.1812
707.3341064453125 0 7007.7817
707.836669921875 0 1065.5496
708.3549194335938 0 7238.331
708.8413696289062 0 1467.2627
709.3527221679688 0 3687.5908
709.8214721679688 0 50804.832 b Water loss 11
710.3228149414062 0 46369.82
710.8229370117188 0 20542.752
711.3253784179688 0 10945.165
711.826904296875 0 3120.393
712.3229370117188 0 5078.672
712.8444213867188 0 1579.4618
713.3391723632812 0 9779.653
713.8427124023438 0 8795.86
714.340087890625 0 4109.212
714.845458984375 0 2747.476
715.3399658203125 0 6428.477
715.8379516601562 0 7099.333
716.3370361328125 0 3160.272
717.3291625976562 0 967.8081
718.3348999023438 0 7017.2983
718.8280639648438 0 35270.527 b 11
719.3292846679688 0 29959.572
719.8299560546875 0 14796.415
720.371337890625 0 31263.488
720.8321533203125 0 3350.0303
721.373779296875 0 11415.9
721.853759765625 0 10201.009
722.35888671875 0 10301.133
722.8552856445312 0 5269.33
723.3359985351562 0 4327.5156
723.8489379882812 0 10849.16 y Water loss 3
724.345458984375 0 22173.83 y Ammonia loss 3
724.84619140625 0 15054.562
725.344970703125 0 8046.8755
725.8477172851562 0 2096.4194
727.3405151367188 0 13124.067
727.8418579101562 0 10840.976
728.3436889648438 0 9346.775
728.8427124023438 0 5982.1855
729.342041015625 0 5813.4507
729.8428344726562 0 2623.898
730.3385620117188 0 3794.099
730.8569946289062 0 4296.137
731.3427734375 0 3568.789
731.8566284179688 0 2141.394
732.34814453125 0 1969.3553
732.8568115234375 0 117861.234 y 3
733.35791015625 0 95505.2
733.8593139648438 0 42670.125
734.3597412109375 0 12632.698
734.85595703125 0 3457.067
735.3297729492188 0 9993.345
736.3475952148438 0 12148.567
736.8479614257812 0 6039.226
737.3469848632812 0 9312.348
737.8486328125 0 2428.066
738.3439331054688 0 4718.6543
738.8494262695312 0 1182.5914
739.3431396484375 0 4126.3423
739.8457641601562 0 1595.5203
740.3480834960938 0 3653.9155
741.3529052734375 0 1982.767
741.8511962890625 0 13570.983
742.35009765625 0 14483.958
742.8505859375 0 9481.1045
743.347900390625 0 5061.451
743.848876953125 0 2417.9644
744.3493041992188 0 2265.3901
744.8453979492188 0 2036.445
745.3461303710938 0 12273.489
745.8428344726562 0 10549.418
746.3500366210938 0 7553.579 y Ammonia loss 9
746.8465576171875 0 3140.9429
747.3529663085938 0 3918.3826
747.8501586914062 0 3531.681
748.3548583984375 0 6061.724
748.8599243164062 0 1223.7875
749.3556518554688 0 2140.8591
750.3426513671875 0 3029.6055
750.85693359375 0 29222.76
751.35791015625 0 24300.084
751.8567504882812 0 12998.772
752.3527221679688 0 4600.7563
752.8555908203125 0 2022.5469
753.3494873046875 0 3660.1458
753.853515625 0 18375.715
754.3545532226562 0 17680.934
754.853759765625 0 8854.046
755.353271484375 0 8640.87
755.84765625 0 1697.9856
756.3701782226562 0 11005.424
756.8544311523438 0 3459.2612
757.367919921875 0 7611.878
757.8529052734375 0 2308.3564
758.3662719726562 0 3298.165
759.3673095703125 0 3227.9883
759.85986328125 0 5868.505
760.3575439453125 0 6606.645
760.84619140625 0 15174.388
761.3505859375 0 11019.636
761.8482666015625 0 6881.0864
762.3494873046875 0 2744.2815
762.857421875 0 3778.4373
763.3767700195312 0 41808.758 y 9
763.8602294921875 0 2004.876
764.3186645507812 0 3178.1436
764.3811645507812 0 15257.827
764.8491821289062 0 2439.3652
765.3429565429688 0 4739.026
765.37548828125 0 6011.7876
765.8616333007812 0 15918.695
766.3621215820312 0 19206.465 b Water loss 6
766.862060546875 0 4864
767.3619995117188 0 7155.7837
768.3518676757812 0 5507.6523
768.8588256835938 0 12937.46
769.3587036132812 0 12529.266
769.860595703125 0 8716.6045
770.3589477539062 0 2622.2285
772.377685546875 0 8454.63
773.3748779296875 0 3956.922
773.8512573242188 0 45942.535 b Water loss 12
774.3599243164062 0 65785.29
774.8662109375 0 44349.54
775.3673095703125 0 19352.508
775.86376953125 0 5992.6636
776.3742065429688 0 2320.3755
779.8496704101562 0 1379.2411
780.35791015625 0 1371.7623
780.850830078125 0 1193.7993
781.3422241210938 0 10105.023
782.34716796875 0 6039.034
782.856201171875 0 110850.76 b 12
783.3574829101562 0 94040.61
783.8585815429688 0 53543.906
784.3671264648438 0 74256.99 b 6
784.8590698242188 0 6079.0347
785.373291015625 0 37390.63
786.3728637695312 0 15689.127
787.3745727539062 0 4437.4116
788.3677978515625 0 2113.013
788.85986328125 0 2139.4548
789.3582153320312 0 2842.4905
789.8582153320312 0 1114.9999
790.374755859375 0 1845.7882
791.369873046875 0 7495.675
791.8684692382812 0 6642.78
792.3714599609375 0 5550.3955
792.8659057617188 0 2080.5762
793.3847045898438 0 2806.1582
794.3762817382812 0 3772.089
794.8759765625 0 1093.0549
795.3905639648438 0 2755.312
797.3690795898438 0 6930.679 y Water loss 2
797.864501953125 0 14633.323 y Ammonia loss 2
798.36572265625 0 13527.322
798.8668823242188 0 6075.788
799.3650512695312 0 3240.336
801.3809814453125 0 1407.3057
802.3685913085938 0 5839.635
803.3731689453125 0 4347.011
804.3631591796875 0 2509.8774
805.37158203125 0 1111.3442
806.374755859375 0 92573.23 y 2
806.8761596679688 0 81389.96
807.3768310546875 0 46631.836
807.877685546875 0 17351.607
808.3775024414062 0 4281.156
811.4010009765625 0 10345.083
812.404541015625 0 5662.258
813.4230346679688 0 986.8827
814.9041137695312 0 2349.061
815.4036254882812 0 2543.887
815.906005859375 0 1598.4606
816.4021606445312 0 1062.463
819.3947143554688 0 11960.619
820.3859252929688 0 9599.953
821.383544921875 0 7601.5024
822.3778076171875 0 5231.9326
823.3812866210938 0 2967.316
824.3898315429688 0 1395.9573
825.3656616210938 0 2312.095
826.365234375 0 1179.8735
829.4348754882812 0 5760.0117
830.3612670898438 0 1522.5525
830.4370727539062 0 2919.275
831.36328125 0 1928.5444
837.40576171875 0 10102.682
838.40087890625 0 7928.964
838.8896484375 0 2158.1248
839.3984375 0 16790.2
840.4014892578125 0 7004.89
841.4014282226562 0 2407.9343
842.384521484375 0 2273.4668
843.3746337890625 0 3348.104
844.3729858398438 0 1039.164
846.9039916992188 0 9786.885 b 13
847.3994140625 0 12372.425 y Ammonia loss 1
847.905029296875 0 5816.4404
848.3909912109375 0 6091.609
849.3738403320312 0 12354.181
850.377685546875 0 6834.4556
851.3795166015625 0 2234.6055
854.4298706054688 0 2986.218
855.9115600585938 0 2113.7898 y 1
856.415771484375 0 1967.7598
856.9089965820312 0 1527.3519
857.429931640625 0 13900.195
858.4326782226562 0 6821.8804
859.4215698242188 0 3604.7844 y Water loss 8
860.3963012695312 0 9374.61 y Ammonia loss 8
861.3978271484375 0 4713.0547
862.4006958007812 0 2736.9705
865.3994750976562 0 8430.16
866.4041748046875 0 6510.922
867.408935546875 0 1727.2196
874.4566650390625 0 1987.5503
875.3822631835938 0 1005.33716
876.4146118164062 0 3614.7273
877.419677734375 0 34458.625 y 8
878.4232788085938 0 16260.913
879.4258422851562 0 5839.4116
887.3977661132812 0 1270.4478
888.3887939453125 0 1672.2458
891.4248046875 0 1351.6107
892.412841796875 0 3740.8435
893.4290161132812 0 9158.9795
894.431396484375 0 13543.626
895.4302978515625 0 6796.4707
896.4270629882812 0 1857.4886
903.4156494140625 0 2400.9114 b Water loss 7
904.418701171875 0 3177.3916
905.4088134765625 0 4385.4883
906.4147338867188 0 2545.9834
909.4369506835938 0 7115.6724
910.440185546875 0 3267.3333
911.4432983398438 0 1763.1888
913.4072265625 0 1794.2324
920.4429321289062 0 2135.3682
921.4296264648438 0 23420.018 b 7
922.4329833984375 0 23795.926
923.432861328125 0 9212.722
924.4364624023438 0 2950.1128
926.4545288085938 0 2088.8792
927.4503784179688 0 1741.7408
928.4603881835938 0 1188.1821
930.42626953125 0 4548.1763
931.41796875 0 4973.278
932.4210205078125 0 2551.5854
937.465087890625 0 1723.4443
938.4561767578125 0 4887.5635
939.451904296875 0 2594.2363
940.4485473632812 0 1124.282
943.4774780273438 0 4148.439
944.482666015625 0 1181.6127
947.45361328125 0 3813.5603
948.4391479492188 0 12564.953
949.4403076171875 0 8351.863
950.43798828125 0 2023.7778
953.4639282226562 0 2102.011
954.4599609375 0 1416.2235
957.4393310546875 0 2763.361
958.4368896484375 0 4433.4385
959.4363403320312 0 1578.7155
961.4487915039062 0 1277.3992
965.462646484375 0 8922.596
966.4583129882812 0 9089.428
967.4583129882812 0 3526.3354
968.4361572265625 0 1215.7903
969.407958984375 0 1970.9797
971.4738159179688 0 20022.37
972.4758911132812 0 9739.137
973.4801025390625 0 3199.0962
974.4657592773438 0 6524.426
975.4645385742188 0 3558.1047
979.4454956054688 0 1292.8794
983.4766845703125 0 2425.834
984.4638061523438 0 2069.8894
985.44189453125 0 2249.769
986.430908203125 0 7395.9897
987.4352416992188 0 3621.2712
988.4506225585938 0 1852.2252
990.4476928710938 0 1428.6998
996.4674682617188 0 1306.9423 y Water loss 7
997.4578857421875 0 4962.521 y Ammonia loss 7
998.4576416015625 0 2946.6252
1002.458251953125 0 8957.117
1003.46142578125 0 5589.002
1004.4707641601562 0 1833.8523
1005.4918823242188 0 1186.2566
1006.487060546875 0 1192.4996
1007.4806518554688 0 3872.901
1008.480712890625 0 1215.1748
1014.479736328125 0 15443.633 y 7
1015.4818725585938 0 8539.529
1016.4849853515625 0 2628.0984
1017.4635009765625 0 2312.5764 b Water loss 8
1018.4610595703125 0 2082.1294
1019.4721069335938 0 1673.9373
1020.4728393554688 0 1078.7622
1022.5120239257812 0 2145.797
1023.5314331054688 0 1062.8617
1031.4764404296875 0 1196.6241
1032.4803466796875 0 1697.4214
1035.4727783203125 0 25849.752 b 8
1036.474365234375 0 15487.628
1037.4744873046875 0 5631.1855
1038.4796142578125 0 2362.6057
1040.473388671875 0 1580.9329
1041.4783935546875 0 1777.594
1049.485595703125 0 1056.9594
1050.4808349609375 0 2538.7893
1057.4990234375 0 3630.832
1058.50048828125 0 4701.1724
1059.49169921875 0 4430.923
1060.4901123046875 0 2288.0332
1066.5098876953125 0 1853.7458
1067.4876708984375 0 1704.4082
1068.47802734375 0 3435.7654
1069.4769287109375 0 2119.3333
1074.514404296875 0 1113.2972
1075.5169677734375 0 1377.3638
1076.5023193359375 0 7351.3174
1077.5003662109375 0 4864.3276
1078.5006103515625 0 1814.0149
1085.49658203125 0 13988.749
1086.498779296875 0 8711.239
1087.497802734375 0 2951.37
1091.5084228515625 0 2143.9868
1092.51611328125 0 1134.0112
1093.5538330078125 0 4782.6733
1094.51318359375 0 8094.885
1095.5120849609375 0 5172.8384
1096.50439453125 0 1385.435
1102.52490234375 0 1035.7769
1103.5084228515625 0 4318.1494
1104.50634765625 0 3643.0005
1105.4951171875 0 1142.1433
1108.5303955078125 0 4638.8936
1109.54052734375 0 1818.4053 y Water loss 6
1110.5362548828125 0 2661.439 y Ammonia loss 6
1111.5430908203125 0 2155.216
1120.5372314453125 0 1738.0789
1121.5048828125 0 1938.593
1122.5084228515625 0 1667.2502
1127.561767578125 0 6255.7793 y 6
1128.5628662109375 0 4090.149
1144.5423583984375 0 1202.6039
1145.5419921875 0 1176.4486
1146.5379638671875 0 1023.0876
1149.49560546875 0 5502.2617
1150.499267578125 0 3489.9695
1151.50439453125 0 2186.4863
1152.5081787109375 0 1032.7294
1155.5107421875 0 941.19977 b Ammonia loss 9
1163.5643310546875 0 1244.6661
1164.55419921875 0 1154.1833
1168.534423828125 0 2984.4932
1169.5338134765625 0 2401.9453
1170.5323486328125 0 1118.295
1172.5299072265625 0 4237.074 b 9
1173.533935546875 0 3184.605
1174.5322265625 0 1456.9458
1181.5775146484375 0 3547.847 y Ammonia loss 5
1182.5858154296875 0 2102.839
1186.545654296875 0 2667.5833
1187.55224609375 0 4138.314
1188.555908203125 0 1295.5222
1198.601318359375 0 10072.17 y 5
1199.603759765625 0 7857.3403
1200.60498046875 0 2483.9292
1203.5660400390625 0 1501.0975
1204.57568359375 0 1824.9113
1205.57666015625 0 2850.3962
1206.5726318359375 0 1204.2428
1213.5560302734375 0 2243.487
1214.5474853515625 0 1951.4854
1215.5498046875 0 1794.5387
1222.597412109375 0 2897.3438
1223.6046142578125 0 1820.5333
1231.5653076171875 0 6565.1704
1232.554443359375 0 5346.4453
1233.5499267578125 0 3046.838
1248.59033203125 0 1048.8425
1250.5482177734375 0 1328.8834
1271.5987548828125 0 1813.2103
1309.629150390625 0 1476.8337 y Water loss 4
1310.626220703125 0 3528.29 y Ammonia loss 4
1311.621337890625 0 1468.7153
1314.5994873046875 0 2264.1528
1315.60107421875 0 1786.2397
1316.6068115234375 0 1113.9093
1327.6431884765625 0 6182.679 y 4
1328.6484375 0 5124.6987
1329.650146484375 0 2145.309
1336.59423828125 0 900.46875
1355.6256103515625 0 1211.3022
1378.6068115234375 0 1433.4119
1419.6307373046875 0 1023.6285 b Ammonia loss 11
2274.173828125 0 682.69495
3032.267333984375 0 875.1444

Spectrum Details

|  |  |
| --- | --- |
| Matched peaks? Matched peaks The total absolute number of peaks matched. Additionally in brackets the total fraction of peaks matched and the total number of peaks is shown. | 109 (5.87% of 1856) |
| FDR? FDR The false discovery rate estimated for this peptide. It is calculated by matching all theoretical fragments with a non-integer shift with the raw peaks for this spectrum. This is done with 40 different shifts. The resulting percentage is the average number of annotated peaks over the number of annotated peaks with the correct spectrum. | 0.35% |
| Satellite FDR? Satellite FDR See the FDR for details on its calculation. This satellite ion specific FDR only contains the satellite ions (d/w) for I/L/J positions. | - |
| PSM Score? PSM Score The PSM Score as given by Hecklib to this annotated spectrum. It is shown with three significant figures. | 674 |

#### Spectrum 2860? Spectrum 2860 The raw spectrum of this peptide as annotated by Hecklib. The fragments are coloured according to ion type (see legend). Any peaks with a star '\*' as text can be hovered over to see the full details, first the ion type second the mass shift type. By hovering over the amino acids in the peptide or ions in the legend the corresponding peaks are highlighted. By toggling the 'Unassigned' label you can turn the background (unassigned) peaks on or off in the plot. By updating the slider in the Ion legend you can update the spectrum to only show the top X% of the peaks with labels. The top X% means any peak that is within X% of the highest intensity. By dragging in the spectrum you can zoom in to a specific part of the spectrum and use 'Zoom Out' to get back to the original zoom level. The annotation of the spectrum is based on the given sequence in the peptides file and is done with different software so inconsistencies are likely. The peaks are annotated based on the given sequence, with 20 ppm tolerance.

Copy Data

##### Spectrum 2860 (TSV)

###### Preview

```
Loading example...
```

*Click on the button to copy the data to your clipboard.*

Mz MinMz MaxIntensity Max

WidthHeightPeptide font sizePeptide stroke widthSpectrum font sizeSpectrum stroke widthCompact peptide

Ion legend

wxyz

abcd

OtherUnassignedIon^Charge^_Position_Show for top:%

SVMHEAJHNHYTQKS

02.98e+65.97e+68.95e+61.19e+7

Zoom Out

a^+1^_2_y^+3^_4_a^+1^_2_b^+4^_6_b^+1^_2_y^+2^_3_b^+1^_2_y^+1^_2_y^+1^_2_y^+2^_4_b^+2^_4_y^+2^_4_y^+1^_2_y^+4^_8_b^+2^_5_b^+1^_3_y^+2^_5_y^+2^_5_y^+2^_5_b^+2^_6_b^+2^_6_y^+3^_8_y^+3^_8_b^+3^_9_y^+3^_8_y^+1^_3_y^+1^_3_y^+1^_3_y^+4^_12_y^+4^_12_y^+3^_9_y^+2^_6_y^+2^_6_y^+3^_9_b^+3^_10_b^+3^_10_y^+2^_6_b^+3^_10_y^+3^_10_y^+4^_13_y^+4^_13_y^+4^_13_b^+4^_14_b^+4^_14_y^+4^_14_y^+4^_14_y^+2^_7_y^+2^_7_y^+3^_11_y^+3^_11_y^+2^_7_b^+3^_11_\*\*y^+3^_11_b^+2^_8_y^+1^_4_\*b^+2^_8_b^+1^_4_y^+1^_4_b^+3^_12_b^+3^_12_b^+3^_12_y^+3^_12_y^+3^_12_y^+3^_12_y^+2^_8_y^+2^_8_b^+2^_9_b^+2^_9_y^+2^_8_b^+2^_9_b^+3^_13_b^+3^_13_y^+3^_13_y^+3^_13_y^+3^_13_b^+3^_14_b^+3^_14_y^+2^_9_y^+2^_9_b^+3^_14_y^+3^_14_y^+2^_9_y^+3^_14_b^+2^_10_b^+2^_10_b^+2^_10_b^+1^_5_y^+2^_10_y^+2^_10_y^+2^_10_y^+1^_5_y^+1^_5_b^+1^_6_b^+2^_11_b^+2^_11_b^+1^_6_y^+2^_11_y^+2^_11_b^+2^_11_y^+2^_11_b^+2^_12_b^+2^_12_y^+2^_12_y^+2^_12_y^+2^_12_y^+1^_6_y^+1^_6_b^+1^_7_y^+1^_6_b^+2^_13_b^+1^_7_b^+2^_13_y^+2^_13_y^+2^_13_y^+2^_14_y^+1^_7_y^+1^_7_y^+1^_7_b^+1^_8_b^+1^_8_y^+1^_8_y^+1^_8_b^+1^_9_b^+1^_9_y^+1^_8_b^+1^_9_y^+1^_9_y^+1^_10_y^+1^_10_y^+1^_11_

041482812421656

Fragment Matches Table

Show background peaks

| Position | Ion type | Intensity | mz Theoretical | mz Error (Th) | mz Error (ppm) | Charge | Series Number |
| --- | --- | --- | --- | --- | --- | --- | --- |
| - | - | 2.456E+04 | 120.1 | - | - | 0 | - |
| - | - | 2.67E+04 | 121 | - | - | 0 | - |
| - | - | 3.705E+04 | 122.1 | - | - | 0 | - |
| - | - | 3.061E+04 | 123.1 | - | - | 0 | - |
| - | - | 1.618E+04 | 124.1 | - | - | 0 | - |
| - | - | 8089 | 126.1 | - | - | 0 | - |
| - | - | 9732 | 127.1 | - | - | 0 | - |
| - | - | 1.046E+04 | 128.1 | - | - | 0 | - |
| - | - | 1.741E+04 | 128.1 | - | - | 0 | - |
| - | - | 6.854E+04 | 129.1 | - | - | 0 | - |
| - | - | 4.395E+06 | 129.1 | - | - | 0 | - |
| - | - | 2.24E+04 | 130.1 | - | - | 0 | - |
| - | - | 8.432E+04 | 130.1 | - | - | 0 | - |
| - | - | 2.872E+04 | 130.1 | - | - | 0 | - |
| - | - | 2.721E+05 | 130.1 | - | - | 0 | - |
| - | - | 5831 | 131.1 | - | - | 0 | - |
| - | - | 9569 | 132 | - | - | 0 | - |
| - | - | 6858 | 134.1 | - | - | 0 | - |
| - | - | 2.332E+06 | 136.1 | - | - | 0 | - |
| - | - | 1.623E+04 | 137.1 | - | - | 0 | - |
| - | - | 2.107E+05 | 137.1 | - | - | 0 | - |
| - | - | 1.165E+05 | 138.1 | - | - | 0 | - |
| - | - | 5696 | 138.2 | - | - | 0 | - |
| - | - | 8.43E+04 | 139.1 | - | - | 0 | - |
| - | - | 7032 | 140.1 | - | - | 0 | - |
| - | - | 1.768E+04 | 141 | - | - | 0 | - |
| - | - | 1.126E+04 | 141.1 | - | - | 0 | - |
| 2 | a | 1.23E+05 | 141.1 | 0.00062 | 4.394 | +1 | 2 |
| - | - | 2.507E+04 | 142.1 | - | - | 0 | - |
| - | - | 1.022E+04 | 142.1 | - | - | 0 | - |
| - | - | 1.812E+04 | 146.1 | - | - | 0 | - |
| - | - | 2.476E+04 | 147 | - | - | 0 | - |
| - | - | 8.011E+04 | 147.1 | - | - | 0 | - |
| - | - | 3.174E+05 | 148.1 | - | - | 0 | - |
| - | - | 1.465E+04 | 149 | - | - | 0 | - |
| - | - | 1.075E+04 | 149 | - | - | 0 | - |
| - | - | 6.676E+04 | 149 | - | - | 0 | - |
| 12 | y | 7400 | 149.1 | 0.0009039 | 6.063 | +3 | 4 |
| - | - | 2.737E+04 | 149.1 | - | - | 0 | - |
| - | - | 1.038E+04 | 151.1 | - | - | 0 | - |
| - | - | 3.35E+04 | 152.1 | - | - | 0 | - |
| - | - | 1.733E+04 | 153.1 | - | - | 0 | - |
| - | - | 3.136E+04 | 154.1 | - | - | 0 | - |
| - | - | 7.196E+04 | 155.1 | - | - | 0 | - |
| - | - | 1.048E+05 | 155.1 | - | - | 0 | - |
| - | - | 5.144E+05 | 155.1 | - | - | 0 | - |
| - | - | 9.526E+04 | 156.1 | - | - | 0 | - |
| - | - | 8146 | 156.1 | - | - | 0 | - |
| - | - | 4.399E+04 | 156.1 | - | - | 0 | - |
| - | - | 7.321E+04 | 157.1 | - | - | 0 | - |
| - | - | 7569 | 158.1 | - | - | 0 | - |
| - | - | 3.035E+04 | 158.1 | - | - | 0 | - |
| - | - | 7613 | 158.1 | - | - | 0 | - |
| 2 | a | 1.182E+07 | 159.1 | 0.0006602 | 4.149 | +1 | 2 |
| 6 | b | 8057 | 160.1 | 0.002383 | 14.88 | +4 | 6 |
| - | - | 3.432E+04 | 160.1 | - | - | 0 | - |
| - | - | 5.713E+04 | 160.1 | - | - | 0 | - |
| - | - | 8.94E+05 | 160.1 | - | - | 0 | - |
| - | - | 6856 | 161.1 | - | - | 0 | - |
| - | - | 4.897E+04 | 161.1 | - | - | 0 | - |
| - | - | 2.058E+04 | 162.1 | - | - | 0 | - |
| - | - | 8644 | 163.1 | - | - | 0 | - |
| - | - | 1.06E+04 | 164.1 | - | - | 0 | - |
| - | - | 7.211E+04 | 165.1 | - | - | 0 | - |
| - | - | 8.442E+05 | 166.1 | - | - | 0 | - |
| - | - | 1.566E+04 | 167.1 | - | - | 0 | - |
| - | - | 6.164E+04 | 167.1 | - | - | 0 | - |
| - | - | 1.769E+04 | 167.1 | - | - | 0 | - |
| - | - | 3.109E+04 | 167.1 | - | - | 0 | - |
| - | - | 3.462E+04 | 167.1 | - | - | 0 | - |
| - | - | 8023 | 168.1 | - | - | 0 | - |
| - | - | 2.505E+04 | 169.1 | - | - | 0 | - |
| 2 | b | 1.573E+04 | 169.1 | 0.0008531 | 5.045 | +1 | 2 |
| - | - | 7320 | 170 | - | - | 0 | - |
| - | - | 4.055E+04 | 171.1 | - | - | 0 | - |
| - | - | 1.101E+04 | 172.1 | - | - | 0 | - |
| 13 | y | 1.362E+05 | 173.1 | 0.0006895 | 3.983 | +2 | 3 |
| - | - | 2.675E+04 | 173.5 | - | - | 0 | - |
| - | - | 1.093E+04 | 174.1 | - | - | 0 | - |
| - | - | 1.036E+04 | 175.1 | - | - | 0 | - |
| - | - | 2.958E+04 | 175.1 | - | - | 0 | - |
| - | - | 1.084E+05 | 176.1 | - | - | 0 | - |
| - | - | 1.392E+04 | 177.1 | - | - | 0 | - |
| - | - | 6.12E+04 | 177.1 | - | - | 0 | - |
| - | - | 1.362E+04 | 178.1 | - | - | 0 | - |
| - | - | 7188 | 178.1 | - | - | 0 | - |
| - | - | 2.93E+04 | 178.1 | - | - | 0 | - |
| - | - | 4.307E+04 | 179.1 | - | - | 0 | - |
| - | - | 2.11E+04 | 180.1 | - | - | 0 | - |
| - | - | 5.407E+04 | 180.1 | - | - | 0 | - |
| - | - | 3.362E+04 | 181.1 | - | - | 0 | - |
| - | - | 1.128E+04 | 181.1 | - | - | 0 | - |
| - | - | 6.78E+04 | 183.1 | - | - | 0 | - |
| - | - | 3.006E+04 | 183.1 | - | - | 0 | - |
| - | - | 7348 | 184.1 | - | - | 0 | - |
| - | - | 1.116E+04 | 185.1 | - | - | 0 | - |
| - | - | 4.805E+04 | 185.1 | - | - | 0 | - |
| - | - | 1.511E+04 | 186.1 | - | - | 0 | - |
| 2 | b | 1.665E+06 | 187.1 | 0.0007407 | 3.959 | +1 | 2 |
| - | - | 1.521E+05 | 188.1 | - | - | 0 | - |
| - | - | 7.046E+04 | 189.1 | - | - | 0 | - |
| - | - | 7165 | 189.1 | - | - | 0 | - |
| - | - | 1.131E+05 | 190.1 | - | - | 0 | - |
| - | - | 9766 | 190.1 | - | - | 0 | - |
| - | - | 9874 | 190.1 | - | - | 0 | - |
| - | - | 9389 | 191.1 | - | - | 0 | - |
| - | - | 6.045E+04 | 191.1 | - | - | 0 | - |
| - | - | 1.079E+04 | 192.1 | - | - | 0 | - |
| - | - | 1.038E+05 | 192.1 | - | - | 0 | - |
| - | - | 9023 | 193.1 | - | - | 0 | - |
| - | - | 1.214E+04 | 193.1 | - | - | 0 | - |
| - | - | 1.619E+05 | 193.1 | - | - | 0 | - |
| - | - | 1.478E+05 | 194.1 | - | - | 0 | - |
| - | - | 1.027E+04 | 194.1 | - | - | 0 | - |
| - | - | 1.146E+04 | 194.1 | - | - | 0 | - |
| - | - | 1.175E+05 | 195.1 | - | - | 0 | - |
| - | - | 2.016E+04 | 195.1 | - | - | 0 | - |
| - | - | 1.318E+04 | 195.1 | - | - | 0 | - |
| - | - | 1.308E+05 | 195.1 | - | - | 0 | - |
| - | - | 3.869E+05 | 196.1 | - | - | 0 | - |
| - | - | 1.361E+04 | 196.1 | - | - | 0 | - |
| - | - | 4.625E+04 | 197.1 | - | - | 0 | - |
| - | - | 8925 | 198.1 | - | - | 0 | - |
| - | - | 1.573E+04 | 198.1 | - | - | 0 | - |
| - | - | 4.259E+04 | 199.1 | - | - | 0 | - |
| - | - | 1.372E+04 | 200.1 | - | - | 0 | - |
| - | - | 1.776E+05 | 201.1 | - | - | 0 | - |
| - | - | 4.273E+04 | 202.1 | - | - | 0 | - |
| - | - | 4.678E+04 | 202.1 | - | - | 0 | - |
| - | - | 2.589E+04 | 202.1 | - | - | 0 | - |
| - | - | 3.191E+04 | 203.1 | - | - | 0 | - |
| - | - | 3.698E+05 | 203.1 | - | - | 0 | - |
| - | - | 6597 | 204.1 | - | - | 0 | - |
| - | - | 2.821E+04 | 204.1 | - | - | 0 | - |
| - | - | 3.13E+04 | 204.1 | - | - | 0 | - |
| - | - | 1.893E+04 | 205.1 | - | - | 0 | - |
| - | - | 1.151E+04 | 205.1 | - | - | 0 | - |
| - | - | 1.721E+04 | 205.1 | - | - | 0 | - |
| - | - | 3.473E+04 | 206.1 | - | - | 0 | - |
| - | - | 3.324E+05 | 207.1 | - | - | 0 | - |
| - | - | 1.01E+04 | 207.2 | - | - | 0 | - |
| - | - | 4.176E+04 | 208.1 | - | - | 0 | - |
| - | - | 1.918E+04 | 208.1 | - | - | 0 | - |
| - | - | 6.656E+04 | 212.1 | - | - | 0 | - |
| - | - | 1.394E+04 | 212.1 | - | - | 0 | - |
| - | - | 1.275E+04 | 213.1 | - | - | 0 | - |
| - | - | 1.095E+04 | 213.1 | - | - | 0 | - |
| - | - | 1.854E+04 | 213.1 | - | - | 0 | - |
| - | - | 1.349E+04 | 214.1 | - | - | 0 | - |
| - | - | 9283 | 214.1 | - | - | 0 | - |
| - | - | 1.024E+04 | 215.1 | - | - | 0 | - |
| 14 | y | 1.651E+05 | 216.1 | 0.0007724 | 3.574 | +1 | 2 |
| - | - | 1.017E+05 | 217.1 | - | - | 0 | - |
| 14 | y | 2.488E+05 | 217.1 | 0.0007503 | 3.456 | +1 | 2 |
| - | - | 1.58E+04 | 217.1 | - | - | 0 | - |
| - | - | 9.347E+04 | 218.1 | - | - | 0 | - |
| - | - | 7446 | 218.1 | - | - | 0 | - |
| - | - | 1.197E+04 | 218.1 | - | - | 0 | - |
| - | - | 8407 | 219.1 | - | - | 0 | - |
| - | - | 1.574E+04 | 219.1 | - | - | 0 | - |
| - | - | 1.678E+04 | 221.1 | - | - | 0 | - |
| - | - | 3.826E+05 | 221.1 | - | - | 0 | - |
| - | - | 1.457E+04 | 221.1 | - | - | 0 | - |
| - | - | 6.352E+04 | 222.1 | - | - | 0 | - |
| - | - | 3.515E+04 | 222.1 | - | - | 0 | - |
| - | - | 2.068E+05 | 222.1 | - | - | 0 | - |
| - | - | 1.682E+04 | 223.1 | - | - | 0 | - |
| - | - | 5.233E+04 | 223.1 | - | - | 0 | - |
| 12 | y | 2.561E+04 | 223.1 | 0.003984 | 17.85 | +2 | 4 |
| - | - | 7.617E+04 | 223.2 | - | - | 0 | - |
| - | - | 4.319E+04 | 224.1 | - | - | 0 | - |
| - | - | 3.527E+05 | 224.1 | - | - | 0 | - |
| - | - | 1.043E+04 | 224.2 | - | - | 0 | - |
| - | - | 2.325E+04 | 225 | - | - | 0 | - |
| - | - | 1.602E+04 | 225.1 | - | - | 0 | - |
| - | - | 5.266E+04 | 225.1 | - | - | 0 | - |
| - | - | 1.095E+04 | 226.1 | - | - | 0 | - |
| - | - | 9581 | 227.2 | - | - | 0 | - |
| 4 | b | 2.977E+04 | 228.1 | 0.001355 | 5.941 | +2 | 4 |
| - | - | 3.691E+04 | 228.1 | - | - | 0 | - |
| - | - | 1.411E+04 | 228.6 | - | - | 0 | - |
| - | - | 1.218E+04 | 229.1 | - | - | 0 | - |
| - | - | 5.744E+04 | 230.1 | - | - | 0 | - |
| - | - | 4.086E+05 | 230.1 | - | - | 0 | - |
| - | - | 9.483E+04 | 231.1 | - | - | 0 | - |
| 12 | y | 1.107E+04 | 232.1 | 0.001189 | 5.12 | +2 | 4 |
| - | - | 3.6E+04 | 233.1 | - | - | 0 | - |
| - | - | 1.234E+05 | 234.1 | - | - | 0 | - |
| - | - | 6.882E+04 | 234.1 | - | - | 0 | - |
| 14 | y | 1.049E+06 | 234.1 | 0.0008126 | 3.47 | +1 | 2 |
| - | - | 5.617E+05 | 235.1 | - | - | 0 | - |
| - | - | 9252 | 235.1 | - | - | 0 | - |
| - | - | 1.167E+05 | 235.1 | - | - | 0 | - |
| - | - | 6.212E+04 | 236.1 | - | - | 0 | - |
| - | - | 1.722E+04 | 236.1 | - | - | 0 | - |
| - | - | 1.084E+04 | 236.2 | - | - | 0 | - |
| - | - | 1.547E+05 | 237.1 | - | - | 0 | - |
| - | - | 7680 | 238.1 | - | - | 0 | - |
| - | - | 4.115E+05 | 239.1 | - | - | 0 | - |
| - | - | 5.426E+05 | 239.2 | - | - | 0 | - |
| - | - | 2.026E+04 | 240.1 | - | - | 0 | - |
| - | - | 1.281E+06 | 240.1 | - | - | 0 | - |
| - | - | 4.266E+04 | 240.2 | - | - | 0 | - |
| - | - | 7.551E+05 | 241.1 | - | - | 0 | - |
| - | - | 1.314E+05 | 241.1 | - | - | 0 | - |
| - | - | 6.817E+04 | 242.1 | - | - | 0 | - |
| - | - | 1.071E+04 | 242.1 | - | - | 0 | - |
| - | - | 2.821E+04 | 243.1 | - | - | 0 | - |
| - | - | 3.61E+04 | 244.1 | - | - | 0 | - |
| - | - | 9.405E+04 | 247.1 | - | - | 0 | - |
| - | - | 1.261E+04 | 247.1 | - | - | 0 | - |
| - | - | 1.926E+04 | 248.1 | - | - | 0 | - |
| - | - | 6561 | 248.1 | - | - | 0 | - |
| - | - | 3.308E+05 | 249.1 | - | - | 0 | - |
| - | - | 1.534E+04 | 249.1 | - | - | 0 | - |
| - | - | 4.016E+04 | 250.1 | - | - | 0 | - |
| - | - | 3.458E+04 | 250.1 | - | - | 0 | - |
| 8 | y | 1.126E+04 | 250.1 | 0.0004551 | 1.82 | +4 | 8 |
| - | - | 2.586E+04 | 250.2 | - | - | 0 | - |
| - | - | 4.214E+04 | 251.1 | - | - | 0 | - |
| - | - | 1.078E+04 | 251.1 | - | - | 0 | - |
| - | - | 2.051E+05 | 251.2 | - | - | 0 | - |
| - | - | 8.851E+05 | 252.1 | - | - | 0 | - |
| - | - | 1.236E+06 | 252.1 | - | - | 0 | - |
| - | - | 2.592E+04 | 252.2 | - | - | 0 | - |
| - | - | 9435 | 252.6 | - | - | 0 | - |
| - | - | 9.194E+04 | 253.1 | - | - | 0 | - |
| - | - | 1.402E+05 | 253.1 | - | - | 0 | - |
| - | - | 3.603E+04 | 254.1 | - | - | 0 | - |
| - | - | 1.253E+04 | 254.1 | - | - | 0 | - |
| - | - | 1.767E+04 | 256.1 | - | - | 0 | - |
| - | - | 1.303E+05 | 257.2 | - | - | 0 | - |
| - | - | 7.275E+04 | 258.1 | - | - | 0 | - |
| - | - | 1.52E+04 | 258.2 | - | - | 0 | - |
| - | - | 1.04E+04 | 259.2 | - | - | 0 | - |
| - | - | 2.197E+04 | 261.1 | - | - | 0 | - |
| - | - | 7.125E+04 | 262.1 | - | - | 0 | - |
| - | - | 1.382E+04 | 262.1 | - | - | 0 | - |
| - | - | 1.032E+04 | 262.6 | - | - | 0 | - |
| - | - | 1.506E+04 | 263.1 | - | - | 0 | - |
| - | - | 6942 | 263.1 | - | - | 0 | - |
| - | - | 3.655E+04 | 264.1 | - | - | 0 | - |
| - | - | 1.232E+05 | 265.1 | - | - | 0 | - |
| - | - | 9115 | 265.1 | - | - | 0 | - |
| - | - | 1.337E+04 | 266.1 | - | - | 0 | - |
| - | - | 9.525E+05 | 267.1 | - | - | 0 | - |
| - | - | 6888 | 267.6 | - | - | 0 | - |
| - | - | 1.166E+05 | 268.1 | - | - | 0 | - |
| - | - | 1.976E+04 | 268.2 | - | - | 0 | - |
| - | - | 3.067E+06 | 269.1 | - | - | 0 | - |
| - | - | 3.579E+05 | 270.1 | - | - | 0 | - |
| - | - | 1.361E+04 | 270.6 | - | - | 0 | - |
| - | - | 1.179E+05 | 271.1 | - | - | 0 | - |
| - | - | 1.906E+04 | 271.1 | - | - | 0 | - |
| - | - | 9261 | 271.5 | - | - | 0 | - |
| - | - | 8392 | 272.7 | - | - | 0 | - |
| - | - | 2.452E+05 | 273.1 | - | - | 0 | - |
| - | - | 8215 | 274.1 | - | - | 0 | - |
| - | - | 2.315E+04 | 274.1 | - | - | 0 | - |
| - | - | 2.443E+04 | 274.1 | - | - | 0 | - |
| - | - | 1.213E+04 | 275.2 | - | - | 0 | - |
| - | - | 1.447E+04 | 277.1 | - | - | 0 | - |
| - | - | 2.045E+04 | 278.6 | - | - | 0 | - |
| - | - | 1.22E+04 | 279.1 | - | - | 0 | - |
| - | - | 1.078E+05 | 279.1 | - | - | 0 | - |
| - | - | 7.464E+04 | 280.1 | - | - | 0 | - |
| - | - | 1.727E+05 | 280.1 | - | - | 0 | - |
| - | - | 1.23E+04 | 280.1 | - | - | 0 | - |
| - | - | 7986 | 280.5 | - | - | 0 | - |
| - | - | 7.685E+04 | 280.7 | - | - | 0 | - |
| - | - | 1.086E+04 | 281.1 | - | - | 0 | - |
| - | - | 1.198E+04 | 281.1 | - | - | 0 | - |
| - | - | 1.95E+04 | 281.1 | - | - | 0 | - |
| - | - | 3.408E+04 | 281.2 | - | - | 0 | - |
| - | - | 2.064E+04 | 282.1 | - | - | 0 | - |
| - | - | 1.591E+04 | 283.1 | - | - | 0 | - |
| - | - | 1.061E+04 | 284.1 | - | - | 0 | - |
| - | - | 1.561E+04 | 284.1 | - | - | 0 | - |
| - | - | 8046 | 286.1 | - | - | 0 | - |
| - | - | 1.222E+04 | 287.2 | - | - | 0 | - |
| - | - | 1.91E+04 | 291.2 | - | - | 0 | - |
| - | - | 1.307E+04 | 292.1 | - | - | 0 | - |
| 5 | b | 2.755E+04 | 292.6 | 0.000658 | 2.249 | +2 | 5 |
| - | - | 2.57E+04 | 293.1 | - | - | 0 | - |
| - | - | 1.865E+04 | 294.2 | - | - | 0 | - |
| - | - | 1.552E+05 | 294.6 | - | - | 0 | - |
| - | - | 7.131E+04 | 295.1 | - | - | 0 | - |
| - | - | 3.123E+04 | 295.2 | - | - | 0 | - |
| - | - | 1.086E+04 | 295.7 | - | - | 0 | - |
| - | - | 8334 | 296.1 | - | - | 0 | - |
| - | - | 2.558E+04 | 296.2 | - | - | 0 | - |
| - | - | 3.083E+04 | 297.1 | - | - | 0 | - |
| - | - | 8.646E+04 | 297.2 | - | - | 0 | - |
| - | - | 8494 | 298.1 | - | - | 0 | - |
| - | - | 1.138E+04 | 298.2 | - | - | 0 | - |
| - | - | 1.036E+04 | 299.1 | - | - | 0 | - |
| 3 | b | 2.473E+04 | 300.1 | 0.001552 | 5.171 | +1 | 3 |
| - | - | 4.321E+05 | 301.1 | - | - | 0 | - |
| - | - | 6.876E+04 | 302.1 | - | - | 0 | - |
| - | - | 8576 | 302.2 | - | - | 0 | - |
| - | - | 7642 | 303.1 | - | - | 0 | - |
| - | - | 2.918E+04 | 303.2 | - | - | 0 | - |
| - | - | 1.154E+04 | 304.1 | - | - | 0 | - |
| - | - | 6.885E+04 | 304.2 | - | - | 0 | - |
| 11 | y | 4.983E+04 | 304.7 | 0.0009297 | 3.052 | +2 | 5 |
| - | - | 1.201E+04 | 305.1 | - | - | 0 | - |
| 11 | y | 2.179E+04 | 305.1 | 0.001576 | 5.165 | +2 | 5 |
| - | - | 4.992E+04 | 305.2 | - | - | 0 | - |
| - | - | 1.281E+04 | 305.6 | - | - | 0 | - |
| - | - | 2.654E+04 | 306.1 | - | - | 0 | - |
| - | - | 8160 | 307.1 | - | - | 0 | - |
| - | - | 7659 | 309.1 | - | - | 0 | - |
| - | - | 1.014E+04 | 309.2 | - | - | 0 | - |
| - | - | 8195 | 309.6 | - | - | 0 | - |
| - | - | 1.612E+04 | 310.1 | - | - | 0 | - |
| - | - | 7313 | 310.6 | - | - | 0 | - |
| - | - | 1.363E+04 | 311.1 | - | - | 0 | - |
| - | - | 7640 | 311.1 | - | - | 0 | - |
| - | - | 1.715E+04 | 312.1 | - | - | 0 | - |
| - | - | 9004 | 312.1 | - | - | 0 | - |
| - | - | 1.105E+04 | 313.6 | - | - | 0 | - |
| 11 | y | 1.682E+05 | 313.7 | 0.001049 | 3.344 | +2 | 5 |
| - | - | 7513 | 314.1 | - | - | 0 | - |
| - | - | 9.253E+04 | 314.2 | - | - | 0 | - |
| - | - | 1.753E+04 | 314.2 | - | - | 0 | - |
| - | - | 3.547E+04 | 314.7 | - | - | 0 | - |
| - | - | 3.674E+04 | 314.8 | - | - | 0 | - |
| - | - | 1.426E+04 | 315.2 | - | - | 0 | - |
| - | - | 3.442E+04 | 318.1 | - | - | 0 | - |
| - | - | 7951 | 318.6 | - | - | 0 | - |
| - | - | 1.886E+04 | 319.1 | - | - | 0 | - |
| 6 | b | 2.889E+04 | 319.1 | 0.0003534 | 1.107 | +2 | 6 |
| - | - | 9001 | 319.7 | - | - | 0 | - |
| - | - | 5.734E+05 | 320.1 | - | - | 0 | - |
| - | - | 7.877E+04 | 321.1 | - | - | 0 | - |
| - | - | 1.312E+04 | 321.7 | - | - | 0 | - |
| - | - | 1.993E+05 | 322.2 | - | - | 0 | - |
| - | - | 7.123E+04 | 323.2 | - | - | 0 | - |
| - | - | 3.441E+04 | 323.2 | - | - | 0 | - |
| - | - | 1.06E+04 | 323.7 | - | - | 0 | - |
| - | - | 1.231E+04 | 324.2 | - | - | 0 | - |
| - | - | 9304 | 324.2 | - | - | 0 | - |
| - | - | 2.301E+04 | 325 | - | - | 0 | - |
| - | - | 1.167E+04 | 325.2 | - | - | 0 | - |
| - | - | 1.314E+04 | 325.8 | - | - | 0 | - |
| - | - | 1.139E+04 | 326 | - | - | 0 | - |
| - | - | 5.902E+04 | 326.1 | - | - | 0 | - |
| - | - | 2.024E+04 | 326.2 | - | - | 0 | - |
| - | - | 1.767E+04 | 326.7 | - | - | 0 | - |
| - | - | 3.007E+04 | 327.1 | - | - | 0 | - |
| - | - | 7.362E+04 | 327.2 | - | - | 0 | - |
| - | - | 9001 | 327.6 | - | - | 0 | - |
| 6 | b | 6.626E+04 | 328.1 | 0.003433 | 10.46 | +2 | 6 |
| - | - | 1.228E+04 | 328.2 | - | - | 0 | - |
| - | - | 8621 | 328.6 | - | - | 0 | - |
| - | - | 4.815E+04 | 328.7 | - | - | 0 | - |
| - | - | 2.221E+04 | 329.1 | - | - | 0 | - |
| - | - | 9216 | 329.2 | - | - | 0 | - |
| - | - | 1.325E+04 | 329.2 | - | - | 0 | - |
| - | - | 1.09E+04 | 329.7 | - | - | 0 | - |
| - | - | 1.732E+04 | 330.2 | - | - | 0 | - |
| - | - | 1.08E+04 | 332.1 | - | - | 0 | - |
| 8 | y | 2.866E+04 | 332.8 | 0.0006714 | 2.017 | +3 | 8 |
| 8 | y | 3.04E+04 | 333.2 | 0.004697 | 14.1 | +3 | 8 |
| - | - | 7211 | 333.7 | - | - | 0 | - |
| 9 | b | 1.096E+04 | 334.8 | 0.00422 | 12.6 | +3 | 9 |
| - | - | 1.438E+04 | 335.1 | - | - | 0 | - |
| - | - | 7775 | 336.2 | - | - | 0 | - |
| - | - | 1.033E+04 | 337.1 | - | - | 0 | - |
| - | - | 1.515E+04 | 337.2 | - | - | 0 | - |
| - | - | 7694 | 337.2 | - | - | 0 | - |
| - | - | 1.074E+04 | 337.7 | - | - | 0 | - |
| - | - | 1.499E+04 | 337.7 | - | - | 0 | - |
| - | - | 4.789E+05 | 338.1 | - | - | 0 | - |
| - | - | 2.372E+04 | 338.7 | - | - | 0 | - |
| 8 | y | 1.321E+05 | 338.8 | 0.001117 | 3.297 | +3 | 8 |
| - | - | 6.145E+04 | 339.1 | - | - | 0 | - |
| - | - | 3.261E+04 | 339.2 | - | - | 0 | - |
| - | - | 4.035E+04 | 339.2 | - | - | 0 | - |
| - | - | 3.221E+04 | 339.5 | - | - | 0 | - |
| - | - | 1.861E+05 | 340.1 | - | - | 0 | - |
| - | - | 1.432E+04 | 340.2 | - | - | 0 | - |
| - | - | 3.8E+04 | 340.2 | - | - | 0 | - |
| - | - | 4.794E+04 | 341 | - | - | 0 | - |
| - | - | 3.668E+04 | 341.1 | - | - | 0 | - |
| - | - | 1.715E+05 | 341.2 | - | - | 0 | - |
| - | - | 2.293E+04 | 342 | - | - | 0 | - |
| - | - | 2.423E+04 | 342.2 | - | - | 0 | - |
| - | - | 1.366E+04 | 342.7 | - | - | 0 | - |
| - | - | 1.392E+04 | 343 | - | - | 0 | - |
| - | - | 3.015E+04 | 343.2 | - | - | 0 | - |
| - | - | 7393 | 343.2 | - | - | 0 | - |
| - | - | 1.029E+05 | 344.1 | - | - | 0 | - |
| 13 | y | 2.194E+05 | 344.2 | 0.001277 | 3.71 | +1 | 3 |
| - | - | 1.5E+04 | 345.2 | - | - | 0 | - |
| 13 | y | 2.398E+05 | 345.2 | 0.001209 | 3.503 | +1 | 3 |
| - | - | 2.29E+04 | 345.2 | - | - | 0 | - |
| - | - | 8.502E+04 | 346.2 | - | - | 0 | - |
| - | - | 2.302E+04 | 346.7 | - | - | 0 | - |
| - | - | 1.683E+04 | 347.2 | - | - | 0 | - |
| - | - | 7627 | 347.8 | - | - | 0 | - |
| - | - | 3.949E+04 | 348.1 | - | - | 0 | - |
| - | - | 1.111E+04 | 348.2 | - | - | 0 | - |
| - | - | 1.304E+04 | 349.1 | - | - | 0 | - |
| - | - | 3.746E+04 | 350.2 | - | - | 0 | - |
| - | - | 1.685E+04 | 350.2 | - | - | 0 | - |
| - | - | 2.35E+04 | 351.2 | - | - | 0 | - |
| - | - | 1.657E+05 | 351.7 | - | - | 0 | - |
| - | - | 4.378E+04 | 352.1 | - | - | 0 | - |
| - | - | 6.44E+04 | 352.2 | - | - | 0 | - |
| - | - | 1.714E+04 | 352.7 | - | - | 0 | - |
| - | - | 3.106E+04 | 353.1 | - | - | 0 | - |
| - | - | 1.972E+04 | 353.1 | - | - | 0 | - |
| - | - | 2.452E+04 | 353.2 | - | - | 0 | - |
| - | - | 9077 | 353.5 | - | - | 0 | - |
| - | - | 6.018E+04 | 354.1 | - | - | 0 | - |
| - | - | 9842 | 354.7 | - | - | 0 | - |
| - | - | 1.026E+04 | 355.1 | - | - | 0 | - |
| - | - | 1.331E+04 | 355.2 | - | - | 0 | - |
| - | - | 1.084E+04 | 356.1 | - | - | 0 | - |
| - | - | 1.387E+04 | 356.2 | - | - | 0 | - |
| - | - | 1.064E+04 | 357.2 | - | - | 0 | - |
| - | - | 7553 | 358.2 | - | - | 0 | - |
| - | - | 1.302E+04 | 358.2 | - | - | 0 | - |
| - | - | 1.051E+05 | 358.2 | - | - | 0 | - |
| - | - | 2.35E+05 | 359 | - | - | 0 | - |
| - | - | 2.753E+04 | 359.1 | - | - | 0 | - |
| - | - | 2.055E+04 | 359.2 | - | - | 0 | - |
| - | - | 9.138E+04 | 360 | - | - | 0 | - |
| - | - | 1.425E+05 | 360.2 | - | - | 0 | - |
| - | - | 3.765E+04 | 360.7 | - | - | 0 | - |
| - | - | 5.081E+04 | 361 | - | - | 0 | - |
| - | - | 1.008E+04 | 361.1 | - | - | 0 | - |
| - | - | 1.186E+05 | 361.2 | - | - | 0 | - |
| - | - | 1.596E+04 | 362.1 | - | - | 0 | - |
| - | - | 2.204E+04 | 362.2 | - | - | 0 | - |
| 13 | y | 1.846E+05 | 362.2 | 0.00118 | 3.257 | +1 | 3 |
| - | - | 3.729E+04 | 362.5 | - | - | 0 | - |
| 4 | y | 1.739E+04 | 362.7 | 0.006405 | 17.66 | +4 | 12 |
| - | - | 1.496E+04 | 362.8 | - | - | 0 | - |
| - | - | 2.264E+04 | 363.1 | - | - | 0 | - |
| - | - | 1.031E+04 | 363.1 | - | - | 0 | - |
| - | - | 3.147E+04 | 363.2 | - | - | 0 | - |
| - | - | 1.414E+04 | 365.2 | - | - | 0 | - |
| - | - | 1.234E+05 | 365.2 | - | - | 0 | - |
| - | - | 1.18E+05 | 366.1 | - | - | 0 | - |
| - | - | 9.381E+04 | 366.2 | - | - | 0 | - |
| - | - | 2.111E+04 | 366.2 | - | - | 0 | - |
| 4 | y | 3.178E+04 | 366.9 | 0.001049 | 2.859 | +4 | 12 |
| - | - | 2.152E+04 | 367.1 | - | - | 0 | - |
| - | - | 1.473E+04 | 367.2 | - | - | 0 | - |
| - | - | 1.892E+04 | 367.2 | - | - | 0 | - |
| - | - | 1.937E+05 | 368.2 | - | - | 0 | - |
| - | - | 2.7E+04 | 368.5 | - | - | 0 | - |
| - | - | 1.842E+04 | 368.7 | - | - | 0 | - |
| - | - | 2.364E+04 | 368.8 | - | - | 0 | - |
| - | - | 5.649E+04 | 369.2 | - | - | 0 | - |
| - | - | 1.077E+04 | 369.7 | - | - | 0 | - |
| - | - | 5.637E+04 | 370.2 | - | - | 0 | - |
| - | - | 4.339E+04 | 370.7 | - | - | 0 | - |
| 7 | y | 3.048E+04 | 370.8 | 4.914E-05 | 0.1325 | +3 | 9 |
| - | - | 4.232E+05 | 371.2 | - | - | 0 | - |
| - | - | 2.146E+04 | 371.2 | - | - | 0 | - |
| - | - | 1.924E+05 | 372.1 | - | - | 0 | - |
| - | - | 4.113E+04 | 372.2 | - | - | 0 | - |
| - | - | 4.263E+04 | 373.1 | - | - | 0 | - |
| 10 | y | 1.911E+04 | 373.2 | 0.002052 | 5.5 | +2 | 6 |
| - | - | 8642 | 373.4 | - | - | 0 | - |
| 10 | y | 3.915E+04 | 373.7 | 0.002415 | 6.463 | +2 | 6 |
| - | - | 2.662E+04 | 374.2 | - | - | 0 | - |
| - | - | 1.18E+05 | 375.2 | - | - | 0 | - |
| - | - | 2.122E+04 | 376.2 | - | - | 0 | - |
| - | - | 4.496E+04 | 376.2 | - | - | 0 | - |
| 7 | y | 2.187E+04 | 376.5 | 0.001345 | 3.572 | +3 | 9 |
| - | - | 1.321E+05 | 376.9 | - | - | 0 | - |
| - | - | 6.071E+04 | 377.2 | - | - | 0 | - |
| - | - | 2.631E+04 | 377.5 | - | - | 0 | - |
| - | - | 1.31E+04 | 377.9 | - | - | 0 | - |
| - | - | 1.169E+04 | 378.2 | - | - | 0 | - |
| - | - | 1.766E+04 | 378.5 | - | - | 0 | - |
| - | - | 8044 | 378.7 | - | - | 0 | - |
| - | - | 2.573E+04 | 379.7 | - | - | 0 | - |
| - | - | 1.526E+05 | 380.1 | - | - | 0 | - |
| 10 | b | 1.886E+04 | 380.2 | 0.000871 | 2.291 | +3 | 10 |
| 10 | b | 1.206E+04 | 380.5 | 0.006942 | 18.24 | +3 | 10 |
| - | - | 4.039E+04 | 381.1 | - | - | 0 | - |
| - | - | 2.275E+04 | 381.1 | - | - | 0 | - |
| - | - | 1.174E+04 | 381.2 | - | - | 0 | - |
| - | - | 2.033E+04 | 382.1 | - | - | 0 | - |
| - | - | 1.178E+04 | 382.2 | - | - | 0 | - |
| 10 | y | 1.325E+05 | 382.2 | 0.001348 | 3.526 | +2 | 6 |
| - | - | 8681 | 382.2 | - | - | 0 | - |
| - | - | 9568 | 382.7 | - | - | 0 | - |
| - | - | 7.126E+04 | 382.7 | - | - | 0 | - |
| - | - | 1.758E+04 | 383.2 | - | - | 0 | - |
| - | - | 1.304E+04 | 383.4 | - | - | 0 | - |
| - | - | 1.147E+04 | 383.7 | - | - | 0 | - |
| - | - | 7.754E+04 | 384.2 | - | - | 0 | - |
| - | - | 1.047E+04 | 384.2 | - | - | 0 | - |
| - | - | 2.233E+04 | 385.2 | - | - | 0 | - |
| - | - | 1.025E+04 | 386.2 | - | - | 0 | - |
| 10 | b | 9.671E+04 | 386.2 | 0.001195 | 3.094 | +3 | 10 |
| - | - | 3.071E+04 | 386.5 | - | - | 0 | - |
| - | - | 1.592E+04 | 386.8 | - | - | 0 | - |
| - | - | 5.481E+04 | 387.2 | - | - | 0 | - |
| - | - | 2.971E+04 | 387.2 | - | - | 0 | - |
| - | - | 1.211E+04 | 387.7 | - | - | 0 | - |
| - | - | 3.527E+04 | 388.2 | - | - | 0 | - |
| - | - | 2.856E+04 | 388.7 | - | - | 0 | - |
| - | - | 1.425E+05 | 389.2 | - | - | 0 | - |
| - | - | 9397 | 389.7 | - | - | 0 | - |
| - | - | 2.101E+04 | 390.2 | - | - | 0 | - |
| - | - | 1.057E+04 | 390.9 | - | - | 0 | - |
| - | - | 2.942E+04 | 391.2 | - | - | 0 | - |
| - | - | 1.689E+04 | 391.2 | - | - | 0 | - |
| - | - | 9074 | 392.2 | - | - | 0 | - |
| - | - | 8.938E+04 | 393.2 | - | - | 0 | - |
| - | - | 1.525E+04 | 394.2 | - | - | 0 | - |
| - | - | 1.067E+04 | 394.2 | - | - | 0 | - |
| 6 | y | 1.006E+04 | 394.5 | 0.0035 | 8.87 | +3 | 10 |
| - | - | 7952 | 394.7 | - | - | 0 | - |
| 3 | y | 8.135E+04 | 395.2 | 0.001107 | 2.801 | +4 | 13 |
| 3 | y | 6.286E+04 | 395.4 | 0.003943 | 9.972 | +4 | 13 |
| - | - | 4.967E+04 | 395.7 | - | - | 0 | - |
| - | - | 1.432E+04 | 395.9 | - | - | 0 | - |
| - | - | 1.587E+04 | 396.2 | - | - | 0 | - |
| - | - | 1.124E+04 | 396.5 | - | - | 0 | - |
| - | - | 1.323E+04 | 396.9 | - | - | 0 | - |
| - | - | 5.397E+04 | 397.2 | - | - | 0 | - |
| - | - | 4.248E+04 | 397.2 | - | - | 0 | - |
| - | - | 1.624E+04 | 397.5 | - | - | 0 | - |
| - | - | 1.584E+05 | 397.7 | - | - | 0 | - |
| - | - | 1.099E+06 | 398.2 | - | - | 0 | - |
| - | - | 5.029E+04 | 398.2 | - | - | 0 | - |
| - | - | 2.289E+04 | 398.7 | - | - | 0 | - |
| - | - | 2.013E+05 | 399.2 | - | - | 0 | - |
| 3 | y | 3.149E+05 | 399.7 | 0.001243 | 3.11 | +4 | 13 |
| - | - | 2.463E+05 | 399.9 | - | - | 0 | - |
| - | - | 5.029E+04 | 400.1 | - | - | 0 | - |
| - | - | 1.326E+05 | 400.2 | - | - | 0 | - |
| - | - | 6.211E+04 | 400.4 | - | - | 0 | - |
| - | - | 2.718E+04 | 400.5 | - | - | 0 | - |
| - | - | 1.933E+04 | 400.7 | - | - | 0 | - |
| - | - | 1.822E+04 | 400.9 | - | - | 0 | - |
| - | - | 3.909E+04 | 401.2 | - | - | 0 | - |
| - | - | 1.028E+04 | 401.7 | - | - | 0 | - |
| - | - | 7.082E+04 | 402.2 | - | - | 0 | - |
| - | - | 1.049E+04 | 402.4 | - | - | 0 | - |
| - | - | 1.236E+04 | 402.7 | - | - | 0 | - |
| - | - | 8521 | 403.2 | - | - | 0 | - |
| - | - | 6996 | 403.2 | - | - | 0 | - |
| - | - | 1.724E+04 | 404.7 | - | - | 0 | - |
| - | - | 1.333E+05 | 405.2 | - | - | 0 | - |
| - | - | 9512 | 405.5 | - | - | 0 | - |
| - | - | 4.809E+05 | 406.2 | - | - | 0 | - |
| - | - | 1.839E+04 | 406.2 | - | - | 0 | - |
| - | - | 3.701E+04 | 406.5 | - | - | 0 | - |
| - | - | 1.953E+05 | 406.7 | - | - | 0 | - |
| - | - | 1.629E+04 | 406.9 | - | - | 0 | - |
| - | - | 5.889E+04 | 407.2 | - | - | 0 | - |
| - | - | 9694 | 407.7 | - | - | 0 | - |
| - | - | 1.279E+04 | 408.1 | - | - | 0 | - |
| - | - | 1.354E+04 | 408.2 | - | - | 0 | - |
| - | - | 3.407E+04 | 408.2 | - | - | 0 | - |
| - | - | 3.306E+04 | 409.2 | - | - | 0 | - |
| - | - | 2.718E+04 | 409.7 | - | - | 0 | - |
| - | - | 8.708E+04 | 410.2 | - | - | 0 | - |
| - | - | 1.908E+04 | 410.7 | - | - | 0 | - |
| - | - | 7.201E+04 | 411.2 | - | - | 0 | - |
| - | - | 1.242E+04 | 411.4 | - | - | 0 | - |
| - | - | 6.067E+04 | 411.7 | - | - | 0 | - |
| - | - | 9005 | 411.9 | - | - | 0 | - |
| - | - | 1.385E+04 | 412.2 | - | - | 0 | - |
| - | - | 6.461E+04 | 412.2 | - | - | 0 | - |
| - | - | 1.246E+04 | 412.5 | - | - | 0 | - |
| - | - | 1.613E+04 | 412.7 | - | - | 0 | - |
| - | - | 4.363E+04 | 413.2 | - | - | 0 | - |
| - | - | 3.181E+04 | 413.7 | - | - | 0 | - |
| - | - | 1.004E+04 | 414.2 | - | - | 0 | - |
| - | - | 2.772E+05 | 415.2 | - | - | 0 | - |
| - | - | 3.98E+04 | 415.2 | - | - | 0 | - |
| 14 | b | 2.806E+04 | 415.7 | 0.00239 | 5.75 | +4 | 14 |
| - | - | 1.29E+04 | 415.9 | - | - | 0 | - |
| - | - | 5.093E+04 | 416.2 | - | - | 0 | - |
| - | - | 8482 | 416.2 | - | - | 0 | - |
| - | - | 9703 | 416.4 | - | - | 0 | - |
| - | - | 1.447E+04 | 417.2 | - | - | 0 | - |
| - | - | 1.726E+05 | 417.2 | - | - | 0 | - |
| - | - | 8.409E+04 | 417.7 | - | - | 0 | - |
| - | - | 1.459E+04 | 417.9 | - | - | 0 | - |
| - | - | 8260 | 418.2 | - | - | 0 | - |
| - | - | 3.69E+04 | 418.2 | - | - | 0 | - |
| - | - | 6.803E+04 | 419.2 | - | - | 0 | - |
| - | - | 1.313E+04 | 419.7 | - | - | 0 | - |
| 14 | b | 5.392E+04 | 420 | 0.002192 | 5.22 | +4 | 14 |
| - | - | 1.847E+04 | 420.2 | - | - | 0 | - |
| 2 | y | 3.886E+05 | 420.2 | 0.0006949 | 1.654 | +4 | 14 |
| - | - | 2.979E+04 | 420.5 | - | - | 0 | - |
| - | - | 1.91E+05 | 420.7 | - | - | 0 | - |
| - | - | 1.549E+04 | 421 | - | - | 0 | - |
| - | - | 4.966E+04 | 421.2 | - | - | 0 | - |
| - | - | 3.874E+04 | 421.7 | - | - | 0 | - |
| - | - | 5.135E+04 | 422.2 | - | - | 0 | - |
| - | - | 1.238E+04 | 422.7 | - | - | 0 | - |
| - | - | 1.533E+04 | 423.2 | - | - | 0 | - |
| - | - | 2.678E+05 | 423.2 | - | - | 0 | - |
| - | - | 1.515E+04 | 424.2 | - | - | 0 | - |
| - | - | 1.03E+04 | 424.2 | - | - | 0 | - |
| - | - | 4.028E+04 | 424.2 | - | - | 0 | - |
| 2 | y | 2.181E+05 | 424.5 | 0.001473 | 3.471 | +4 | 14 |
| - | - | 1.545E+05 | 424.7 | - | - | 0 | - |
| - | - | 8.831E+04 | 425 | - | - | 0 | - |
| - | - | 1.525E+04 | 425.2 | - | - | 0 | - |
| - | - | 4.448E+04 | 425.2 | - | - | 0 | - |
| - | - | 2.556E+04 | 425.5 | - | - | 0 | - |
| - | - | 1.078E+04 | 425.9 | - | - | 0 | - |
| - | - | 4.321E+04 | 426.1 | - | - | 0 | - |
| - | - | 1.369E+05 | 427.2 | - | - | 0 | - |
| - | - | 2.166E+04 | 427.5 | - | - | 0 | - |
| - | - | 2.168E+04 | 427.9 | - | - | 0 | - |
| - | - | 5.551E+04 | 428.2 | - | - | 0 | - |
| - | - | 1.113E+04 | 428.5 | - | - | 0 | - |
| - | - | 1.129E+04 | 428.7 | - | - | 0 | - |
| - | - | 5.956E+04 | 429.1 | - | - | 0 | - |
| - | - | 1.244E+04 | 429.2 | - | - | 0 | - |
| - | - | 1.377E+04 | 430 | - | - | 0 | - |
| - | - | 3.211E+04 | 430.1 | - | - | 0 | - |
| - | - | 1.733E+04 | 430.2 | - | - | 0 | - |
| 9 | y | 3.717E+04 | 430.2 | 0.0025 | 5.812 | +2 | 7 |
| 9 | y | 1.087E+05 | 430.7 | 0.002955 | 6.86 | +2 | 7 |
| - | - | 2.597E+04 | 431.1 | - | - | 0 | - |
| - | - | 8.837E+04 | 431.2 | - | - | 0 | - |
| - | - | 5.931E+04 | 431.5 | - | - | 0 | - |
| - | - | 4.18E+04 | 431.9 | - | - | 0 | - |
| - | - | 1.551E+04 | 432.2 | - | - | 0 | - |
| - | - | 5.048E+04 | 433 | - | - | 0 | - |
| - | - | 1.561E+05 | 433.2 | - | - | 0 | - |
| - | - | 1.718E+04 | 433.5 | - | - | 0 | - |
| - | - | 1.072E+05 | 433.5 | - | - | 0 | - |
| - | - | 2.283E+04 | 433.7 | - | - | 0 | - |
| - | - | 4.182E+04 | 433.9 | - | - | 0 | - |
| - | - | 2.377E+04 | 434.2 | - | - | 0 | - |
| - | - | 3.194E+04 | 434.2 | - | - | 0 | - |
| - | - | 2.138E+04 | 434.7 | - | - | 0 | - |
| - | - | 1.849E+04 | 435 | - | - | 0 | - |
| - | - | 1.429E+04 | 436.2 | - | - | 0 | - |
| - | - | 1.6E+05 | 436.2 | - | - | 0 | - |
| - | - | 2.141E+04 | 436.5 | - | - | 0 | - |
| 5 | y | 1.106E+05 | 437.2 | 0.0007776 | 1.779 | +3 | 11 |
| - | - | 8.757E+04 | 437.5 | - | - | 0 | - |
| 5 | y | 7.55E+04 | 437.5 | 0.00688 | 15.72 | +3 | 11 |
| - | - | 8.263E+04 | 437.7 | - | - | 0 | - |
| - | - | 4.165E+04 | 437.9 | - | - | 0 | - |
| - | - | 6.423E+04 | 438 | - | - | 0 | - |
| - | - | 4.329E+04 | 438.2 | - | - | 0 | - |
| - | - | 1.035E+04 | 438.5 | - | - | 0 | - |
| - | - | 1.407E+04 | 438.7 | - | - | 0 | - |
| 9 | y | 3.005E+05 | 439.2 | 0.002803 | 6.381 | +2 | 7 |
| - | - | 1.103E+05 | 439.7 | - | - | 0 | - |
| - | - | 1.497E+04 | 439.9 | - | - | 0 | - |
| - | - | 3.346E+04 | 440.2 | - | - | 0 | - |
| 11 | b | 5.391E+04 | 440.5 | 0.0001149 | 0.2607 | +3 | 11 |
| - | - | 1.474E+04 | 440.9 | - | - | 0 | - |
| - | - | 3.44E+04 | 441.2 | - | - | 0 | - |
| 0 | Precursor | 4.509E+05 | 441.7 | 0.001509 | 3.417 | +4 | -1 |
| 0 | Precursor | 4.245E+05 | 442 | 0.004956 | 11.21 | +4 | -1 |
| - | - | 4.108E+05 | 442.2 | - | - | 0 | - |
| - | - | 1.729E+05 | 442.5 | - | - | 0 | - |
| - | - | 6.908E+04 | 442.5 | - | - | 0 | - |
| - | - | 5.964E+04 | 442.7 | - | - | 0 | - |
| - | - | 5.024E+04 | 442.9 | - | - | 0 | - |
| - | - | 3.031E+04 | 443 | - | - | 0 | - |
| - | - | 9.666E+04 | 443.2 | - | - | 0 | - |
| 5 | y | 4.339E+04 | 443.2 | 0.0002098 | 0.4734 | +3 | 11 |
| - | - | 2.731E+04 | 443.6 | - | - | 0 | - |
| - | - | 9458 | 443.9 | - | - | 0 | - |
| - | - | 1.871E+04 | 444.2 | - | - | 0 | - |
| 8 | b | 2.3E+04 | 444.2 | 0.001101 | 2.478 | +2 | 8 |
| - | - | 9994 | 445.2 | - | - | 0 | - |
| 12 | y | 8.413E+04 | 445.2 | 0.001755 | 3.942 | +1 | 4 |
| - | - | 2.533E+04 | 445.5 | - | - | 0 | - |
| - | - | 1.373E+04 | 445.9 | - | - | 0 | - |
| 0 | Precursor | 1.748E+06 | 446.2 | 0.001645 | 3.687 | +4 | -1 |
| - | - | 1.342E+06 | 446.5 | - | - | 0 | - |
| - | - | 8.506E+05 | 446.7 | - | - | 0 | - |
| - | - | 4.192E+05 | 447 | - | - | 0 | - |
| - | - | 1.078E+05 | 447.2 | - | - | 0 | - |
| - | - | 2.393E+04 | 447.5 | - | - | 0 | - |
| - | - | 2.274E+04 | 447.7 | - | - | 0 | - |
| - | - | 1.737E+05 | 448.2 | - | - | 0 | - |
| - | - | 1.245E+05 | 448.6 | - | - | 0 | - |
| - | - | 4.113E+04 | 448.9 | - | - | 0 | - |
| - | - | 1.965E+04 | 449.2 | - | - | 0 | - |
| - | - | 1.291E+04 | 449.5 | - | - | 0 | - |
| - | - | 1.109E+04 | 450.2 | - | - | 0 | - |
| - | - | 2.992E+04 | 450.2 | - | - | 0 | - |
| - | - | 3.623E+04 | 451.2 | - | - | 0 | - |
| - | - | 6.302E+05 | 451.2 | - | - | 0 | - |
| - | - | 8828 | 451.6 | - | - | 0 | - |
| - | - | 2.132E+04 | 452.2 | - | - | 0 | - |
| - | - | 1.774E+04 | 452.2 | - | - | 0 | - |
| - | - | 1.517E+05 | 452.2 | - | - | 0 | - |
| - | - | 9994 | 452.7 | - | - | 0 | - |
| 8 | b | 9.971E+04 | 453.2 | 0.002654 | 5.856 | +2 | 8 |
| - | - | 2.423E+04 | 453.7 | - | - | 0 | - |
| - | - | 9.759E+04 | 453.9 | - | - | 0 | - |
| - | - | 9.059E+04 | 454.2 | - | - | 0 | - |
| - | - | 3.55E+04 | 454.6 | - | - | 0 | - |
| - | - | 9872 | 454.7 | - | - | 0 | - |
| - | - | 2.946E+04 | 454.9 | - | - | 0 | - |
| - | - | 1.65E+04 | 455.2 | - | - | 0 | - |
| 4 | b | 2.142E+05 | 455.2 | 0.005592 | 12.29 | +1 | 4 |
| - | - | 9206 | 455.5 | - | - | 0 | - |
| - | - | 3.833E+04 | 455.7 | - | - | 0 | - |
| - | - | 4.152E+04 | 456.2 | - | - | 0 | - |
| - | - | 2.479E+04 | 457.2 | - | - | 0 | - |
| - | - | 1.51E+04 | 458.2 | - | - | 0 | - |
| - | - | 1.819E+04 | 458.9 | - | - | 0 | - |
| - | - | 1.687E+04 | 459.2 | - | - | 0 | - |
| - | - | 1.172E+04 | 459.6 | - | - | 0 | - |
| - | - | 1.157E+05 | 459.9 | - | - | 0 | - |
| - | - | 1.061E+05 | 460.2 | - | - | 0 | - |
| - | - | 4.612E+04 | 460.6 | - | - | 0 | - |
| - | - | 9432 | 460.9 | - | - | 0 | - |
| - | - | 5.484E+04 | 461.2 | - | - | 0 | - |
| - | - | 1.423E+04 | 461.7 | - | - | 0 | - |
| - | - | 8940 | 461.9 | - | - | 0 | - |
| - | - | 2.444E+04 | 462.2 | - | - | 0 | - |
| - | - | 5.381E+04 | 462.7 | - | - | 0 | - |
| - | - | 1.381E+04 | 463.2 | - | - | 0 | - |
| - | - | 1.098E+05 | 463.2 | - | - | 0 | - |
| 12 | y | 7.079E+05 | 463.3 | 0.001658 | 3.579 | +1 | 4 |
| - | - | 4.822E+04 | 463.7 | - | - | 0 | - |
| - | - | 3.687E+04 | 464.2 | - | - | 0 | - |
| - | - | 4.88E+04 | 464.2 | - | - | 0 | - |
| - | - | 1.139E+05 | 464.3 | - | - | 0 | - |
| - | - | 1.125E+04 | 464.5 | - | - | 0 | - |
| - | - | 1.483E+04 | 464.7 | - | - | 0 | - |
| - | - | 1.667E+04 | 464.9 | - | - | 0 | - |
| - | - | 1.233E+04 | 465.2 | - | - | 0 | - |
| - | - | 2.178E+04 | 465.2 | - | - | 0 | - |
| - | - | 2.01E+04 | 465.3 | - | - | 0 | - |
| - | - | 1.175E+04 | 465.6 | - | - | 0 | - |
| - | - | 5.598E+04 | 465.7 | - | - | 0 | - |
| - | - | 3.249E+04 | 466.2 | - | - | 0 | - |
| - | - | 3.315E+04 | 466.7 | - | - | 0 | - |
| - | - | 2.633E+04 | 467.2 | - | - | 0 | - |
| - | - | 1.087E+04 | 467.9 | - | - | 0 | - |
| 12 | b | 4.172E+04 | 468.2 | 0.003633 | 7.759 | +3 | 12 |
| - | - | 4.209E+04 | 468.3 | - | - | 0 | - |
| 12 | b | 5.017E+04 | 468.5 | 0.008849 | 18.89 | +3 | 12 |
| - | - | 1.089E+04 | 468.7 | - | - | 0 | - |
| - | - | 1.5E+04 | 468.9 | - | - | 0 | - |
| - | - | 4.854E+05 | 469.2 | - | - | 0 | - |
| - | - | 1.039E+05 | 470.2 | - | - | 0 | - |
| - | - | 1.987E+04 | 470.2 | - | - | 0 | - |
| - | - | 2.908E+04 | 471.2 | - | - | 0 | - |
| - | - | 3.487E+04 | 471.2 | - | - | 0 | - |
| - | - | 4.726E+04 | 471.6 | - | - | 0 | - |
| - | - | 5.065E+05 | 471.7 | - | - | 0 | - |
| - | - | 3.077E+04 | 471.9 | - | - | 0 | - |
| - | - | 2.713E+05 | 472.2 | - | - | 0 | - |
| - | - | 1.049E+04 | 472.6 | - | - | 0 | - |
| - | - | 9.736E+04 | 472.7 | - | - | 0 | - |
| - | - | 3.818E+04 | 473.2 | - | - | 0 | - |
| - | - | 9878 | 473.6 | - | - | 0 | - |
| - | - | 1.616E+04 | 473.9 | - | - | 0 | - |
| 12 | b | 6.719E+04 | 474.2 | 0.00591 | 12.46 | +3 | 12 |
| - | - | 4.686E+04 | 474.3 | - | - | 0 | - |
| - | - | 2.57E+04 | 474.6 | - | - | 0 | - |
| - | - | 5.544E+04 | 474.7 | - | - | 0 | - |
| - | - | 1.533E+04 | 474.9 | - | - | 0 | - |
| - | - | 3.39E+04 | 475.2 | - | - | 0 | - |
| - | - | 1.728E+04 | 475.3 | - | - | 0 | - |
| - | - | 2.614E+04 | 476.7 | - | - | 0 | - |
| - | - | 5.248E+04 | 476.9 | - | - | 0 | - |
| - | - | 2.513E+05 | 477.2 | - | - | 0 | - |
| - | - | 2.22E+05 | 477.6 | - | - | 0 | - |
| - | - | 2.604E+04 | 477.7 | - | - | 0 | - |
| - | - | 9.474E+04 | 477.9 | - | - | 0 | - |
| - | - | 4.83E+04 | 478.2 | - | - | 0 | - |
| - | - | 1.313E+04 | 478.6 | - | - | 0 | - |
| - | - | 1.931E+04 | 478.7 | - | - | 0 | - |
| - | - | 1.138E+04 | 479.2 | - | - | 0 | - |
| - | - | 1.3E+05 | 479.2 | - | - | 0 | - |
| - | - | 2.559E+04 | 479.7 | - | - | 0 | - |
| - | - | 4.704E+04 | 480.2 | - | - | 0 | - |
| - | - | 3.459E+04 | 480.6 | - | - | 0 | - |
| - | - | 1.187E+04 | 480.9 | - | - | 0 | - |
| - | - | 9.077E+04 | 481.2 | - | - | 0 | - |
| - | - | 3.522E+04 | 481.2 | - | - | 0 | - |
| - | - | 1.01E+04 | 481.9 | - | - | 0 | - |
| - | - | 1.446E+04 | 482.2 | - | - | 0 | - |
| 4 | y | 2.358E+05 | 482.9 | 0.001609 | 3.331 | +3 | 12 |
| 4 | y | 4.947E+05 | 483.2 | 0.003926 | 8.124 | +3 | 12 |
| - | - | 3.132E+05 | 483.6 | - | - | 0 | - |
| - | - | 4.826E+04 | 483.7 | - | - | 0 | - |
| - | - | 1.288E+05 | 483.9 | - | - | 0 | - |
| - | - | 6.215E+04 | 484.2 | - | - | 0 | - |
| - | - | 9513 | 484.6 | - | - | 0 | - |
| - | - | 3.052E+04 | 485.2 | - | - | 0 | - |
| - | - | 3.854E+05 | 485.7 | - | - | 0 | - |
| - | - | 1.495E+05 | 485.9 | - | - | 0 | - |
| - | - | 3.303E+05 | 486.2 | - | - | 0 | - |
| - | - | 9.52E+04 | 486.6 | - | - | 0 | - |
| - | - | 8.004E+04 | 486.7 | - | - | 0 | - |
| - | - | 4.653E+04 | 486.9 | - | - | 0 | - |
| - | - | 2.639E+04 | 487.2 | - | - | 0 | - |
| - | - | 1.686E+05 | 487.7 | - | - | 0 | - |
| - | - | 1.1E+05 | 488.2 | - | - | 0 | - |
| - | - | 3.08E+04 | 488.7 | - | - | 0 | - |
| 4 | y | 1.579E+06 | 488.9 | 0.001688 | 3.453 | +3 | 12 |
| - | - | 1.146E+06 | 489.2 | - | - | 0 | - |
| - | - | 4.68E+05 | 489.6 | - | - | 0 | - |
| - | - | 1.401E+05 | 489.9 | - | - | 0 | - |
| - | - | 1.149E+04 | 490.2 | - | - | 0 | - |
| - | - | 4.515E+04 | 490.2 | - | - | 0 | - |
| - | - | 4.06E+04 | 490.7 | - | - | 0 | - |
| - | - | 1.866E+04 | 491.2 | - | - | 0 | - |
| - | - | 3.533E+04 | 491.6 | - | - | 0 | - |
| - | - | 4.245E+05 | 491.9 | - | - | 0 | - |
| - | - | 2.762E+05 | 492.2 | - | - | 0 | - |
| - | - | 1.87E+05 | 492.6 | - | - | 0 | - |
| - | - | 2.445E+04 | 492.7 | - | - | 0 | - |
| - | - | 5.601E+04 | 492.9 | - | - | 0 | - |
| - | - | 4.869E+04 | 493.2 | - | - | 0 | - |
| - | - | 1.649E+04 | 493.7 | - | - | 0 | - |
| - | - | 1.071E+04 | 494.2 | - | - | 0 | - |
| - | - | 1.264E+04 | 496.2 | - | - | 0 | - |
| - | - | 1.897E+04 | 496.2 | - | - | 0 | - |
| - | - | 2.742E+04 | 497.2 | - | - | 0 | - |
| - | - | 8.238E+04 | 497.2 | - | - | 0 | - |
| - | - | 3.056E+05 | 497.6 | - | - | 0 | - |
| - | - | 2.255E+05 | 497.9 | - | - | 0 | - |
| - | - | 6.238E+04 | 498.2 | - | - | 0 | - |
| - | - | 1.228E+05 | 498.2 | - | - | 0 | - |
| - | - | 6.242E+04 | 498.6 | - | - | 0 | - |
| 8 | y | 6.527E+04 | 498.7 | 0.001273 | 2.553 | +2 | 8 |
| - | - | 2.182E+04 | 498.9 | - | - | 0 | - |
| - | - | 1.689E+04 | 499.2 | - | - | 0 | - |
| 8 | y | 1.315E+05 | 499.2 | 0.002857 | 5.722 | +2 | 8 |
| - | - | 6.251E+04 | 499.7 | - | - | 0 | - |
| - | - | 1.923E+04 | 500.2 | - | - | 0 | - |
| 9 | b | 2.629E+04 | 501.2 | 0.0007856 | 1.567 | +2 | 9 |
| - | - | 1.587E+04 | 501.6 | - | - | 0 | - |
| 9 | b | 1.958E+05 | 501.7 | 0.005787 | 11.53 | +2 | 9 |
| - | - | 1.037E+05 | 502.2 | - | - | 0 | - |
| - | - | 3.822E+04 | 502.7 | - | - | 0 | - |
| - | - | 5.746E+04 | 503.3 | - | - | 0 | - |
| - | - | 2.67E+05 | 503.6 | - | - | 0 | - |
| - | - | 1.691E+04 | 503.7 | - | - | 0 | - |
| - | - | 1.943E+05 | 503.9 | - | - | 0 | - |
| - | - | 1.161E+05 | 504.2 | - | - | 0 | - |
| - | - | 4.486E+04 | 504.6 | - | - | 0 | - |
| - | - | 1.587E+04 | 504.9 | - | - | 0 | - |
| - | - | 3.17E+04 | 505.2 | - | - | 0 | - |
| - | - | 2.05E+04 | 505.6 | - | - | 0 | - |
| - | - | 1.754E+04 | 506.2 | - | - | 0 | - |
| - | - | 1.099E+04 | 506.3 | - | - | 0 | - |
| - | - | 8.811E+04 | 507.2 | - | - | 0 | - |
| - | - | 1.491E+04 | 507.6 | - | - | 0 | - |
| 8 | y | 4.369E+05 | 507.7 | 0.001575 | 3.103 | +2 | 8 |
| - | - | 1.649E+04 | 507.9 | - | - | 0 | - |
| - | - | 2.326E+05 | 508.2 | - | - | 0 | - |
| - | - | 8.252E+04 | 508.7 | - | - | 0 | - |
| - | - | 1.876E+04 | 509.2 | - | - | 0 | - |
| - | - | 9312 | 509.6 | - | - | 0 | - |
| - | - | 1.134E+04 | 509.7 | - | - | 0 | - |
| 9 | b | 1.437E+05 | 510.2 | 0.002553 | 5.003 | +2 | 9 |
| - | - | 4.288E+04 | 510.6 | - | - | 0 | - |
| - | - | 5.654E+04 | 510.7 | - | - | 0 | - |
| 13 | b | 5.328E+04 | 510.9 | 0.007412 | 14.51 | +3 | 13 |
| - | - | 5.664E+04 | 511.2 | - | - | 0 | - |
| - | - | 2.611E+04 | 511.6 | - | - | 0 | - |
| - | - | 3.644E+04 | 511.8 | - | - | 0 | - |
| - | - | 2.272E+04 | 511.9 | - | - | 0 | - |
| - | - | 5.305E+04 | 512.2 | - | - | 0 | - |
| - | - | 9880 | 512.6 | - | - | 0 | - |
| - | - | 2.032E+04 | 512.7 | - | - | 0 | - |
| - | - | 1.284E+04 | 513.2 | - | - | 0 | - |
| - | - | 9769 | 514.2 | - | - | 0 | - |
| - | - | 1.099E+04 | 514.6 | - | - | 0 | - |
| - | - | 4.018E+04 | 514.9 | - | - | 0 | - |
| - | - | 7.942E+04 | 515.2 | - | - | 0 | - |
| - | - | 5.581E+04 | 515.6 | - | - | 0 | - |
| - | - | 2.64E+04 | 515.9 | - | - | 0 | - |
| - | - | 9.924E+04 | 516.2 | - | - | 0 | - |
| - | - | 2.483E+04 | 516.6 | - | - | 0 | - |
| 13 | b | 3.759E+04 | 516.9 | 0.00618 | 11.96 | +3 | 13 |
| - | - | 1.752E+04 | 517.2 | - | - | 0 | - |
| - | - | 3.983E+04 | 517.2 | - | - | 0 | - |
| - | - | 2.973E+04 | 517.6 | - | - | 0 | - |
| - | - | 1.964E+04 | 517.9 | - | - | 0 | - |
| - | - | 3.091E+04 | 518.2 | - | - | 0 | - |
| - | - | 2.205E+04 | 518.9 | - | - | 0 | - |
| - | - | 3.056E+04 | 519.3 | - | - | 0 | - |
| - | - | 1.331E+04 | 519.6 | - | - | 0 | - |
| - | - | 5.643E+04 | 520.3 | - | - | 0 | - |
| - | - | 5.595E+04 | 520.6 | - | - | 0 | - |
| - | - | 2.967E+05 | 520.9 | - | - | 0 | - |
| - | - | 1.894E+05 | 521.2 | - | - | 0 | - |
| - | - | 1.787E+05 | 521.6 | - | - | 0 | - |
| - | - | 3.365E+04 | 521.8 | - | - | 0 | - |
| - | - | 7.66E+04 | 521.9 | - | - | 0 | - |
| - | - | 2.586E+04 | 522.2 | - | - | 0 | - |
| - | - | 4.756E+04 | 524.2 | - | - | 0 | - |
| - | - | 1.606E+04 | 524.3 | - | - | 0 | - |
| - | - | 3.935E+04 | 524.9 | - | - | 0 | - |
| - | - | 4.338E+04 | 525.2 | - | - | 0 | - |
| - | - | 3.1E+04 | 525.6 | - | - | 0 | - |
| - | - | 2.028E+04 | 525.7 | - | - | 0 | - |
| - | - | 1.693E+04 | 526.2 | - | - | 0 | - |
| 3 | y | 5.839E+05 | 526.6 | 0.001684 | 3.198 | +3 | 13 |
| 3 | y | 1.134E+06 | 526.9 | 0.00455 | 8.636 | +3 | 13 |
| - | - | 8.678E+05 | 527.2 | - | - | 0 | - |
| - | - | 4.262E+05 | 527.6 | - | - | 0 | - |
| - | - | 1.722E+05 | 527.9 | - | - | 0 | - |
| - | - | 3.613E+04 | 528.2 | - | - | 0 | - |
| - | - | 2.776E+04 | 528.3 | - | - | 0 | - |
| - | - | 2.271E+04 | 529.3 | - | - | 0 | - |
| - | - | 1.776E+04 | 529.3 | - | - | 0 | - |
| - | - | 2.706E+04 | 529.8 | - | - | 0 | - |
| - | - | 1.839E+05 | 530.2 | - | - | 0 | - |
| - | - | 3.443E+04 | 530.6 | - | - | 0 | - |
| - | - | 3.24E+04 | 530.7 | - | - | 0 | - |
| - | - | 3.1E+04 | 530.9 | - | - | 0 | - |
| - | - | 3.497E+04 | 531.2 | - | - | 0 | - |
| - | - | 2.739E+04 | 532.2 | - | - | 0 | - |
| 3 | y | 4.904E+06 | 532.6 | 0.00158 | 2.967 | +3 | 13 |
| - | - | 3.773E+06 | 532.9 | - | - | 0 | - |
| - | - | 2.077E+06 | 533.3 | - | - | 0 | - |
| - | - | 8.936E+05 | 533.6 | - | - | 0 | - |
| - | - | 2.495E+05 | 533.9 | - | - | 0 | - |
| - | - | 7.854E+04 | 534.3 | - | - | 0 | - |
| - | - | 1.245E+04 | 534.6 | - | - | 0 | - |
| - | - | 7.821E+04 | 534.7 | - | - | 0 | - |
| - | - | 3.499E+04 | 535.2 | - | - | 0 | - |
| - | - | 7.976E+04 | 535.2 | - | - | 0 | - |
| - | - | 3.514E+04 | 535.7 | - | - | 0 | - |
| - | - | 1.887E+04 | 535.9 | - | - | 0 | - |
| - | - | 1.304E+04 | 536.2 | - | - | 0 | - |
| - | - | 2.782E+04 | 536.2 | - | - | 0 | - |
| - | - | 2.719E+04 | 536.6 | - | - | 0 | - |
| - | - | 9843 | 536.7 | - | - | 0 | - |
| - | - | 2.515E+04 | 536.9 | - | - | 0 | - |
| - | - | 4.105E+04 | 537.3 | - | - | 0 | - |
| - | - | 5.27E+04 | 538.3 | - | - | 0 | - |
| - | - | 8.243E+04 | 538.8 | - | - | 0 | - |
| - | - | 3.134E+04 | 539.2 | - | - | 0 | - |
| - | - | 2.911E+04 | 539.3 | - | - | 0 | - |
| - | - | 1.098E+04 | 539.6 | - | - | 0 | - |
| - | - | 1.711E+04 | 539.8 | - | - | 0 | - |
| - | - | 2.377E+04 | 540.3 | - | - | 0 | - |
| - | - | 1.012E+04 | 542.3 | - | - | 0 | - |
| - | - | 2.697E+05 | 543.3 | - | - | 0 | - |
| - | - | 1.659E+05 | 543.8 | - | - | 0 | - |
| - | - | 6.221E+04 | 544.3 | - | - | 0 | - |
| - | - | 6.512E+04 | 544.7 | - | - | 0 | - |
| - | - | 2.211E+04 | 545.2 | - | - | 0 | - |
| - | - | 1.47E+05 | 545.3 | - | - | 0 | - |
| - | - | 4.099E+04 | 546.3 | - | - | 0 | - |
| - | - | 2.678E+04 | 546.8 | - | - | 0 | - |
| - | - | 1.827E+05 | 547.3 | - | - | 0 | - |
| - | - | 1.772E+04 | 547.8 | - | - | 0 | - |
| - | - | 1.319E+04 | 547.8 | - | - | 0 | - |
| - | - | 1.95E+04 | 547.9 | - | - | 0 | - |
| - | - | 7.89E+04 | 548.3 | - | - | 0 | - |
| - | - | 1.653E+04 | 549.3 | - | - | 0 | - |
| - | - | 9757 | 550.3 | - | - | 0 | - |
| - | - | 3.474E+04 | 551.8 | - | - | 0 | - |
| - | - | 1.209E+05 | 552.3 | - | - | 0 | - |
| - | - | 9.551E+04 | 552.8 | - | - | 0 | - |
| - | - | 2.248E+05 | 553.3 | - | - | 0 | - |
| 14 | b | 2.006E+04 | 553.6 | 0.001322 | 2.387 | +3 | 14 |
| - | - | 1.423E+05 | 553.8 | - | - | 0 | - |
| 14 | b | 9.465E+04 | 553.9 | 0.001869 | 3.374 | +3 | 14 |
| - | - | 1.541E+05 | 554.3 | - | - | 0 | - |
| - | - | 2.68E+04 | 554.6 | - | - | 0 | - |
| - | - | 1.333E+04 | 554.8 | - | - | 0 | - |
| - | - | 1.053E+04 | 554.9 | - | - | 0 | - |
| 7 | y | 8.831E+04 | 555.3 | 0.0007145 | 1.287 | +2 | 9 |
| 7 | y | 8.615E+04 | 555.8 | 0.0002228 | 0.401 | +2 | 9 |
| - | - | 1.736E+05 | 556.3 | - | - | 0 | - |
| - | - | 5.331E+04 | 556.8 | - | - | 0 | - |
| - | - | 3.705E+04 | 557.3 | - | - | 0 | - |
| - | - | 1.672E+04 | 558.2 | - | - | 0 | - |
| - | - | 3.127E+04 | 558.7 | - | - | 0 | - |
| - | - | 1.976E+04 | 559.2 | - | - | 0 | - |
| 14 | b | 1.499E+05 | 559.6 | 0.00134 | 2.395 | +3 | 14 |
| - | - | 1.205E+04 | 559.7 | - | - | 0 | - |
| 2 | y | 1.98E+05 | 559.9 | 0.005122 | 9.148 | +3 | 14 |
| - | - | 7.647E+04 | 560.3 | - | - | 0 | - |
| - | - | 1.128E+05 | 560.3 | - | - | 0 | - |
| - | - | 8.381E+04 | 560.6 | - | - | 0 | - |
| - | - | 5.25E+04 | 560.8 | - | - | 0 | - |
| - | - | 3.117E+04 | 560.9 | - | - | 0 | - |
| - | - | 1.399E+04 | 561.2 | - | - | 0 | - |
| - | - | 2.251E+04 | 561.3 | - | - | 0 | - |
| - | - | 3.517E+04 | 561.3 | - | - | 0 | - |
| - | - | 1.241E+04 | 561.8 | - | - | 0 | - |
| - | - | 1.164E+04 | 562.3 | - | - | 0 | - |
| - | - | 1.313E+04 | 563.8 | - | - | 0 | - |
| 7 | y | 1.48E+05 | 564.3 | 0.0006202 | 1.099 | +2 | 9 |
| - | - | 4.608E+05 | 564.8 | - | - | 0 | - |
| - | - | 5.43E+05 | 565.3 | - | - | 0 | - |
| 2 | y | 7.187E+05 | 565.6 | 0.001664 | 2.942 | +3 | 14 |
| - | - | 1.054E+05 | 565.8 | - | - | 0 | - |
| - | - | 5.827E+05 | 565.9 | - | - | 0 | - |
| - | - | 4.39E+05 | 566.3 | - | - | 0 | - |
| - | - | 1.567E+05 | 566.6 | - | - | 0 | - |
| - | - | 5.07E+04 | 566.9 | - | - | 0 | - |
| - | - | 2.213E+05 | 567.3 | - | - | 0 | - |
| - | - | 1.416E+05 | 567.8 | - | - | 0 | - |
| - | - | 1.309E+05 | 568.3 | - | - | 0 | - |
| - | - | 1.557E+04 | 568.8 | - | - | 0 | - |
| - | - | 2.328E+04 | 568.9 | - | - | 0 | - |
| - | - | 4.798E+04 | 569.3 | - | - | 0 | - |
| 10 | b | 4.884E+04 | 569.8 | 0.001939 | 3.403 | +2 | 10 |
| 10 | b | 6.888E+04 | 570.3 | 0.01109 | 19.45 | +2 | 10 |
| - | - | 2.256E+04 | 570.8 | - | - | 0 | - |
| - | - | 1.748E+04 | 571.3 | - | - | 0 | - |
| - | - | 1.067E+04 | 571.8 | - | - | 0 | - |
| - | - | 1.584E+04 | 572.3 | - | - | 0 | - |
| - | - | 1.3E+05 | 573.3 | - | - | 0 | - |
| - | - | 2.851E+04 | 574.3 | - | - | 0 | - |
| - | - | 2.35E+04 | 575.3 | - | - | 0 | - |
| - | - | 2.963E+04 | 576.3 | - | - | 0 | - |
| - | - | 1.663E+04 | 576.8 | - | - | 0 | - |
| 10 | b | 3.567E+05 | 578.8 | 0.001783 | 3.081 | +2 | 10 |
| - | - | 2.186E+05 | 579.3 | - | - | 0 | - |
| - | - | 9.688E+04 | 579.8 | - | - | 0 | - |
| - | - | 1.375E+04 | 580.2 | - | - | 0 | - |
| - | - | 2.679E+04 | 580.3 | - | - | 0 | - |
| - | - | 2.196E+04 | 581.3 | - | - | 0 | - |
| - | - | 4.067E+05 | 582.3 | - | - | 0 | - |
| - | - | 1.608E+04 | 582.8 | - | - | 0 | - |
| - | - | 1.248E+05 | 583.3 | - | - | 0 | - |
| 5 | b | 2.702E+05 | 584.2 | 0.002367 | 4.051 | +1 | 5 |
| - | - | 1.304E+04 | 584.8 | - | - | 0 | - |
| - | - | 8.017E+04 | 585.3 | - | - | 0 | - |
| - | - | 2.495E+04 | 585.8 | - | - | 0 | - |
| - | - | 3.516E+04 | 586.3 | - | - | 0 | - |
| - | - | 1.317E+04 | 586.8 | - | - | 0 | - |
| - | - | 1.17E+04 | 587.3 | - | - | 0 | - |
| - | - | 1.262E+04 | 587.8 | - | - | 0 | - |
| - | - | 2.381E+05 | 588.3 | - | - | 0 | - |
| - | - | 1.296E+04 | 589.2 | - | - | 0 | - |
| - | - | 5.793E+04 | 589.3 | - | - | 0 | - |
| - | - | 1.472E+04 | 590.2 | - | - | 0 | - |
| - | - | 3.342E+04 | 590.3 | - | - | 0 | - |
| 6 | y | 3.441E+04 | 590.8 | 0.0006513 | 1.102 | +2 | 10 |
| 6 | y | 8.995E+04 | 591.3 | 0.00138 | 2.334 | +2 | 10 |
| - | - | 2.965E+04 | 591.8 | - | - | 0 | - |
| - | - | 1.867E+04 | 592.2 | - | - | 0 | - |
| - | - | 4.363E+04 | 593.3 | - | - | 0 | - |
| - | - | 2.448E+04 | 593.8 | - | - | 0 | - |
| - | - | 1.116E+05 | 594.3 | - | - | 0 | - |
| - | - | 8.857E+04 | 594.8 | - | - | 0 | - |
| - | - | 4.208E+04 | 595.3 | - | - | 0 | - |
| - | - | 7.385E+04 | 595.8 | - | - | 0 | - |
| - | - | 3.226E+04 | 596.3 | - | - | 0 | - |
| - | - | 1.372E+04 | 597.3 | - | - | 0 | - |
| - | - | 1.742E+04 | 598.3 | - | - | 0 | - |
| - | - | 2.303E+04 | 598.8 | - | - | 0 | - |
| - | - | 2.349E+04 | 599.3 | - | - | 0 | - |
| 6 | y | 3.218E+05 | 599.8 | 0.002144 | 3.574 | +2 | 10 |
| - | - | 5.544E+04 | 600.3 | - | - | 0 | - |
| - | - | 1.776E+05 | 600.3 | - | - | 0 | - |
| - | - | 3.628E+04 | 600.8 | - | - | 0 | - |
| - | - | 5.278E+04 | 600.8 | - | - | 0 | - |
| - | - | 1.256E+04 | 601.3 | - | - | 0 | - |
| - | - | 5.065E+04 | 602.3 | - | - | 0 | - |
| - | - | 9.49E+04 | 602.8 | - | - | 0 | - |
| - | - | 1.384E+05 | 603.3 | - | - | 0 | - |
| - | - | 6.907E+04 | 603.8 | - | - | 0 | - |
| - | - | 4.753E+04 | 604.3 | - | - | 0 | - |
| - | - | 2.381E+04 | 604.8 | - | - | 0 | - |
| - | - | 4.379E+04 | 605.3 | - | - | 0 | - |
| - | - | 1.31E+04 | 606.3 | - | - | 0 | - |
| - | - | 1.095E+05 | 607.3 | - | - | 0 | - |
| - | - | 1.094E+05 | 607.8 | - | - | 0 | - |
| 11 | y | 7.977E+04 | 608.3 | 0.0008739 | 1.437 | +1 | 5 |
| - | - | 2.695E+05 | 608.8 | - | - | 0 | - |
| - | - | 1.911E+05 | 609.3 | - | - | 0 | - |
| - | - | 1.067E+05 | 609.8 | - | - | 0 | - |
| - | - | 1.38E+04 | 610.2 | - | - | 0 | - |
| - | - | 3.062E+04 | 610.3 | - | - | 0 | - |
| - | - | 1.82E+05 | 611.8 | - | - | 0 | - |
| - | - | 9.703E+04 | 612.3 | - | - | 0 | - |
| - | - | 4.981E+04 | 612.8 | - | - | 0 | - |
| - | - | 2.495E+05 | 616.3 | - | - | 0 | - |
| - | - | 1.637E+05 | 616.8 | - | - | 0 | - |
| - | - | 1.25E+05 | 617.3 | - | - | 0 | - |
| - | - | 2.55E+05 | 617.8 | - | - | 0 | - |
| - | - | 2.597E+04 | 618.2 | - | - | 0 | - |
| - | - | 1.06E+05 | 618.3 | - | - | 0 | - |
| - | - | 5.85E+04 | 618.8 | - | - | 0 | - |
| - | - | 2.075E+04 | 619.2 | - | - | 0 | - |
| - | - | 2.931E+04 | 620.3 | - | - | 0 | - |
| - | - | 7.045E+04 | 620.8 | - | - | 0 | - |
| - | - | 5.869E+04 | 621.3 | - | - | 0 | - |
| - | - | 1.49E+04 | 621.8 | - | - | 0 | - |
| - | - | 3.364E+04 | 624.8 | - | - | 0 | - |
| - | - | 4.327E+04 | 625.3 | - | - | 0 | - |
| - | - | 1.52E+04 | 625.8 | - | - | 0 | - |
| 11 | y | 3.941E+05 | 626.3 | 0.0009516 | 1.519 | +1 | 5 |
| - | - | 4.679E+04 | 626.8 | - | - | 0 | - |
| - | - | 1.204E+05 | 627.3 | - | - | 0 | - |
| - | - | 2.313E+04 | 628.3 | - | - | 0 | - |
| - | - | 2.211E+04 | 628.8 | - | - | 0 | - |
| - | - | 1.578E+04 | 629.3 | - | - | 0 | - |
| - | - | 1.101E+05 | 635.3 | - | - | 0 | - |
| - | - | 2.794E+04 | 636.3 | - | - | 0 | - |
| 6 | b | 1.053E+05 | 637.3 | 0.002185 | 3.429 | +1 | 6 |
| - | - | 2.24E+04 | 637.3 | - | - | 0 | - |
| - | - | 4.097E+04 | 637.8 | - | - | 0 | - |
| - | - | 5.287E+04 | 638.3 | - | - | 0 | - |
| - | - | 2.796E+04 | 638.8 | - | - | 0 | - |
| - | - | 5.906E+04 | 639.3 | - | - | 0 | - |
| - | - | 2.63E+04 | 640.3 | - | - | 0 | - |
| - | - | 3.13E+04 | 641.3 | - | - | 0 | - |
| - | - | 9.706E+04 | 644.3 | - | - | 0 | - |
| - | - | 2.932E+04 | 645.3 | - | - | 0 | - |
| - | - | 1.374E+04 | 645.8 | - | - | 0 | - |
| - | - | 9.558E+04 | 646.3 | - | - | 0 | - |
| - | - | 1.237E+05 | 646.8 | - | - | 0 | - |
| - | - | 1.163E+05 | 647.3 | - | - | 0 | - |
| - | - | 4.334E+04 | 647.8 | - | - | 0 | - |
| - | - | 2.919E+04 | 648.3 | - | - | 0 | - |
| - | - | 2.404E+04 | 649.3 | - | - | 0 | - |
| - | - | 2.969E+04 | 649.8 | - | - | 0 | - |
| - | - | 3.361E+04 | 650.3 | - | - | 0 | - |
| - | - | 1.629E+04 | 650.8 | - | - | 0 | - |
| 11 | b | 2.319E+04 | 651.3 | 0.005067 | 7.78 | +2 | 11 |
| 11 | b | 1.427E+04 | 651.8 | 0.005916 | 9.076 | +2 | 11 |
| - | - | 2.166E+04 | 652.3 | - | - | 0 | - |
| - | - | 5.389E+04 | 653.3 | - | - | 0 | - |
| - | - | 1.625E+04 | 654.3 | - | - | 0 | - |
| 6 | b | 3.11E+05 | 655.3 | 0.002179 | 3.326 | +1 | 6 |
| 5 | y | 2.079E+05 | 655.3 | 0.003952 | 6.031 | +2 | 11 |
| 5 | y | 3.528E+05 | 655.8 | 0.007183 | 10.95 | +2 | 11 |
| - | - | 2.189E+05 | 656.3 | - | - | 0 | - |
| - | - | 6.183E+04 | 656.8 | - | - | 0 | - |
| - | - | 1.32E+05 | 657.3 | - | - | 0 | - |
| - | - | 7.767E+04 | 658.3 | - | - | 0 | - |
| - | - | 5.139E+04 | 658.8 | - | - | 0 | - |
| - | - | 4.359E+04 | 659.3 | - | - | 0 | - |
| - | - | 9.099E+04 | 659.8 | - | - | 0 | - |
| 11 | b | 8.624E+04 | 660.3 | 3.471E-05 | 0.05257 | +2 | 11 |
| - | - | 6.095E+04 | 660.8 | - | - | 0 | - |
| - | - | 2.983E+04 | 661.3 | - | - | 0 | - |
| - | - | 5.451E+04 | 662.8 | - | - | 0 | - |
| - | - | 4.714E+04 | 663.3 | - | - | 0 | - |
| - | - | 2.838E+04 | 663.8 | - | - | 0 | - |
| 5 | y | 1.043E+06 | 664.3 | 0.001965 | 2.959 | +2 | 11 |
| - | - | 6.176E+05 | 664.8 | - | - | 0 | - |
| - | - | 3.369E+05 | 665.3 | - | - | 0 | - |
| - | - | 9.201E+04 | 665.8 | - | - | 0 | - |
| - | - | 6.328E+04 | 666.3 | - | - | 0 | - |
| - | - | 8.765E+04 | 667.3 | - | - | 0 | - |
| - | - | 8.636E+04 | 667.8 | - | - | 0 | - |
| - | - | 8.574E+04 | 668.3 | - | - | 0 | - |
| - | - | 1.962E+04 | 668.8 | - | - | 0 | - |
| - | - | 4.765E+04 | 669.3 | - | - | 0 | - |
| - | - | 1.738E+04 | 669.8 | - | - | 0 | - |
| - | - | 3.652E+04 | 670.3 | - | - | 0 | - |
| - | - | 3.303E+04 | 671.3 | - | - | 0 | - |
| - | - | 1.073E+05 | 671.8 | - | - | 0 | - |
| - | - | 6.579E+04 | 672.3 | - | - | 0 | - |
| - | - | 2.411E+05 | 672.8 | - | - | 0 | - |
| - | - | 1.831E+05 | 673.3 | - | - | 0 | - |
| - | - | 1.277E+05 | 673.8 | - | - | 0 | - |
| - | - | 2.023E+05 | 674.3 | - | - | 0 | - |
| - | - | 1.655E+04 | 674.8 | - | - | 0 | - |
| - | - | 7.845E+04 | 675.3 | - | - | 0 | - |
| - | - | 4.899E+04 | 676.3 | - | - | 0 | - |
| - | - | 1.471E+05 | 680.3 | - | - | 0 | - |
| - | - | 1.242E+05 | 680.8 | - | - | 0 | - |
| - | - | 1.339E+05 | 681.3 | - | - | 0 | - |
| - | - | 4.13E+05 | 681.8 | - | - | 0 | - |
| - | - | 2.691E+05 | 682.3 | - | - | 0 | - |
| - | - | 1.597E+05 | 682.8 | - | - | 0 | - |
| - | - | 5.015E+04 | 683.3 | - | - | 0 | - |
| - | - | 1.576E+04 | 683.8 | - | - | 0 | - |
| - | - | 1.759E+05 | 684.3 | - | - | 0 | - |
| - | - | 1.028E+05 | 685.3 | - | - | 0 | - |
| - | - | 3.791E+04 | 686.3 | - | - | 0 | - |
| - | - | 1.315E+04 | 688.8 | - | - | 0 | - |
| - | - | 7.678E+04 | 689.3 | - | - | 0 | - |
| - | - | 4.562E+04 | 689.8 | - | - | 0 | - |
| - | - | 3.607E+04 | 690.3 | - | - | 0 | - |
| - | - | 2.871E+04 | 690.8 | - | - | 0 | - |
| - | - | 1.504E+05 | 691.3 | - | - | 0 | - |
| - | - | 5.021E+04 | 692.3 | - | - | 0 | - |
| - | - | 3.218E+04 | 693.3 | - | - | 0 | - |
| - | - | 3.554E+04 | 701.3 | - | - | 0 | - |
| 12 | b | 6.814E+04 | 701.8 | 0.002832 | 4.035 | +2 | 12 |
| - | - | 5.832E+05 | 702.3 | - | - | 0 | - |
| - | - | 3.576E+04 | 702.8 | - | - | 0 | - |
| - | - | 2.07E+05 | 703.3 | - | - | 0 | - |
| - | - | 4.641E+04 | 704.3 | - | - | 0 | - |
| - | - | 1.559E+04 | 705.3 | - | - | 0 | - |
| - | - | 1.154E+04 | 705.8 | - | - | 0 | - |
| - | - | 2.07E+04 | 706.3 | - | - | 0 | - |
| - | - | 3.97E+04 | 708.4 | - | - | 0 | - |
| - | - | 3.4E+04 | 709.4 | - | - | 0 | - |
| - | - | 3.717E+04 | 710.3 | - | - | 0 | - |
| 12 | b | 4.823E+04 | 710.8 | 0.004019 | 5.654 | +2 | 12 |
| - | - | 3.983E+04 | 711.3 | - | - | 0 | - |
| - | - | 1.257E+04 | 711.8 | - | - | 0 | - |
| - | - | 3.152E+04 | 712.3 | - | - | 0 | - |
| - | - | 1.411E+04 | 714.8 | - | - | 0 | - |
| - | - | 4.683E+04 | 715.3 | - | - | 0 | - |
| - | - | 3E+04 | 715.8 | - | - | 0 | - |
| - | - | 1.364E+04 | 718.3 | - | - | 0 | - |
| - | - | 2.299E+05 | 719.3 | - | - | 0 | - |
| - | - | 1.176E+04 | 719.8 | - | - | 0 | - |
| - | - | 9.57E+04 | 720.3 | - | - | 0 | - |
| - | - | 3.631E+04 | 721.3 | - | - | 0 | - |
| - | - | 3.839E+04 | 722.3 | - | - | 0 | - |
| - | - | 2.817E+04 | 722.8 | - | - | 0 | - |
| - | - | 2.17E+04 | 723.3 | - | - | 0 | - |
| 4 | y | 8.087E+04 | 723.8 | 0.000558 | 0.7708 | +2 | 12 |
| 4 | y | 1.714E+05 | 724.3 | 0.005132 | 7.085 | +2 | 12 |
| - | - | 1.204E+05 | 724.8 | - | - | 0 | - |
| - | - | 4.332E+04 | 725.3 | - | - | 0 | - |
| - | - | 3.745E+04 | 728.3 | - | - | 0 | - |
| - | - | 1.84E+04 | 728.8 | - | - | 0 | - |
| - | - | 2.261E+04 | 729.3 | - | - | 0 | - |
| - | - | 2.646E+04 | 730.3 | - | - | 0 | - |
| - | - | 7.875E+04 | 731.3 | - | - | 0 | - |
| - | - | 4.244E+04 | 731.8 | - | - | 0 | - |
| - | - | 2.555E+04 | 732.3 | - | - | 0 | - |
| 4 | y | 6.36E+05 | 732.9 | 0.001989 | 2.715 | +2 | 12 |
| - | - | 4.641E+05 | 733.4 | - | - | 0 | - |
| - | - | 2.066E+05 | 733.9 | - | - | 0 | - |
| - | - | 7.015E+04 | 734.4 | - | - | 0 | - |
| - | - | 1.684E+04 | 734.9 | - | - | 0 | - |
| - | - | 1.095E+05 | 736.4 | - | - | 0 | - |
| - | - | 1.262E+04 | 736.8 | - | - | 0 | - |
| - | - | 9.386E+04 | 737.3 | - | - | 0 | - |
| - | - | 3.466E+04 | 737.8 | - | - | 0 | - |
| - | - | 4.615E+04 | 738.3 | - | - | 0 | - |
| - | - | 3.357E+04 | 740.4 | - | - | 0 | - |
| - | - | 9874 | 741.4 | - | - | 0 | - |
| 10 | y | 3.753E+04 | 745.4 | 0.000578 | 0.7755 | +1 | 6 |
| - | - | 7.685E+04 | 745.9 | - | - | 0 | - |
| 10 | y | 1.035E+05 | 746.3 | 0.006309 | 8.453 | +1 | 6 |
| - | - | 4.187E+04 | 746.9 | - | - | 0 | - |
| - | - | 3.6E+04 | 747.4 | - | - | 0 | - |
| - | - | 4.851E+04 | 748.4 | - | - | 0 | - |
| 7 | b | 5.262E+04 | 750.4 | 0.001313 | 1.749 | +1 | 7 |
| - | - | 3.357E+04 | 751.4 | - | - | 0 | - |
| - | - | 1.156E+04 | 752.4 | - | - | 0 | - |
| - | - | 3.007E+04 | 753.3 | - | - | 0 | - |
| - | - | 1.612E+04 | 754.4 | - | - | 0 | - |
| - | - | 4.324E+04 | 754.9 | - | - | 0 | - |
| - | - | 7.463E+04 | 755.4 | - | - | 0 | - |
| - | - | 1.966E+04 | 755.9 | - | - | 0 | - |
| - | - | 3.683E+04 | 756.4 | - | - | 0 | - |
| - | - | 1.483E+04 | 758.3 | - | - | 0 | - |
| - | - | 1.599E+04 | 761.4 | - | - | 0 | - |
| 10 | y | 2.229E+05 | 763.4 | 0.001976 | 2.589 | +1 | 6 |
| - | - | 3.881E+04 | 764.3 | - | - | 0 | - |
| - | - | 7.893E+04 | 764.4 | - | - | 0 | - |
| - | - | 1.276E+04 | 765.3 | - | - | 0 | - |
| - | - | 2.307E+04 | 765.4 | - | - | 0 | - |
| 13 | b | 2.644E+04 | 765.8 | 0.002718 | 3.549 | +2 | 13 |
| - | - | 5.404E+04 | 766.4 | - | - | 0 | - |
| - | - | 1.361E+04 | 766.8 | - | - | 0 | - |
| - | - | 1.275E+04 | 767.4 | - | - | 0 | - |
| 7 | b | 1.954E+05 | 768.4 | 0.001672 | 2.177 | +1 | 7 |
| - | - | 7.607E+04 | 769.4 | - | - | 0 | - |
| - | - | 3.188E+04 | 770.4 | - | - | 0 | - |
| - | - | 4.9E+04 | 772.4 | - | - | 0 | - |
| - | - | 2.62E+04 | 773.4 | - | - | 0 | - |
| 13 | b | 3.511E+04 | 774.9 | 0.002684 | 3.464 | +2 | 13 |
| - | - | 3.655E+04 | 775.4 | - | - | 0 | - |
| - | - | 1.964E+04 | 775.9 | - | - | 0 | - |
| - | - | 1.627E+04 | 776.4 | - | - | 0 | - |
| - | - | 1.296E+05 | 781.3 | - | - | 0 | - |
| - | - | 3.832E+04 | 782.3 | - | - | 0 | - |
| - | - | 2.073E+04 | 788.4 | - | - | 0 | - |
| 3 | y | 2.719E+04 | 789.4 | 0.006989 | 8.854 | +2 | 13 |
| - | - | 2.75E+04 | 790.4 | - | - | 0 | - |
| - | - | 1.599E+04 | 791.4 | - | - | 0 | - |
| - | - | 4.266E+04 | 793.4 | - | - | 0 | - |
| - | - | 3.36E+04 | 794.4 | - | - | 0 | - |
| - | - | 2.412E+04 | 795.4 | - | - | 0 | - |
| 3 | y | 3.519E+04 | 798.4 | 0.002689 | 3.368 | +2 | 13 |
| - | - | 2.301E+04 | 798.9 | - | - | 0 | - |
| - | - | 1.541E+04 | 799.4 | - | - | 0 | - |
| - | - | 1.012E+04 | 800.3 | - | - | 0 | - |
| - | - | 5.326E+04 | 802.4 | - | - | 0 | - |
| - | - | 4.022E+04 | 803.4 | - | - | 0 | - |
| - | - | 1.98E+04 | 804.4 | - | - | 0 | - |
| - | - | 3.84E+04 | 805.4 | - | - | 0 | - |
| - | - | 1.959E+04 | 806.4 | - | - | 0 | - |
| - | - | 9.413E+04 | 811.4 | - | - | 0 | - |
| - | - | 3.31E+04 | 812.4 | - | - | 0 | - |
| - | - | 2.622E+04 | 816.4 | - | - | 0 | - |
| - | - | 1.396E+04 | 817.4 | - | - | 0 | - |
| - | - | 2.805E+04 | 818.4 | - | - | 0 | - |
| - | - | 1.439E+05 | 819.4 | - | - | 0 | - |
| - | - | 9.815E+04 | 820.4 | - | - | 0 | - |
| - | - | 6.073E+04 | 821.4 | - | - | 0 | - |
| - | - | 5.872E+04 | 822.4 | - | - | 0 | - |
| - | - | 2.69E+04 | 823.4 | - | - | 0 | - |
| - | - | 1.343E+04 | 824.4 | - | - | 0 | - |
| - | - | 1.688E+04 | 825.4 | - | - | 0 | - |
| - | - | 2.02E+04 | 830.4 | - | - | 0 | - |
| - | - | 1.434E+04 | 831.4 | - | - | 0 | - |
| - | - | 2.485E+05 | 833.4 | - | - | 0 | - |
| - | - | 1.123E+05 | 834.4 | - | - | 0 | - |
| - | - | 3.304E+04 | 835.4 | - | - | 0 | - |
| - | - | 1.086E+04 | 836.4 | - | - | 0 | - |
| - | - | 1.126E+05 | 837.4 | - | - | 0 | - |
| - | - | 6.106E+04 | 838.4 | - | - | 0 | - |
| 2 | y | 1.437E+05 | 839.4 | 0.0001227 | 0.1462 | +2 | 14 |
| - | - | 6.167E+04 | 840.4 | - | - | 0 | - |
| - | - | 2.134E+04 | 841.4 | - | - | 0 | - |
| - | - | 2.475E+04 | 842.4 | - | - | 0 | - |
| - | - | 2.272E+04 | 843.4 | - | - | 0 | - |
| - | - | 5.997E+04 | 847.4 | - | - | 0 | - |
| - | - | 4.982E+04 | 848.4 | - | - | 0 | - |
| - | - | 2.426E+04 | 849.4 | - | - | 0 | - |
| - | - | 1.243E+04 | 850.4 | - | - | 0 | - |
| - | - | 1.861E+04 | 854.4 | - | - | 0 | - |
| - | - | 1.361E+04 | 855.4 | - | - | 0 | - |
| 9 | y | 3.406E+04 | 859.4 | 0.0002352 | 0.2736 | +1 | 7 |
| 9 | y | 5.832E+04 | 860.4 | 0.004458 | 5.181 | +1 | 7 |
| - | - | 2.345E+04 | 861.4 | - | - | 0 | - |
| - | - | 1.595E+05 | 865.4 | - | - | 0 | - |
| - | - | 7.931E+04 | 866.4 | - | - | 0 | - |
| - | - | 1.252E+04 | 867.4 | - | - | 0 | - |
| - | - | 1.151E+04 | 875.4 | - | - | 0 | - |
| - | - | 1.968E+04 | 876.4 | - | - | 0 | - |
| 9 | y | 2.163E+05 | 877.4 | 0.001834 | 2.091 | +1 | 7 |
| - | - | 8.962E+04 | 878.4 | - | - | 0 | - |
| - | - | 2.806E+04 | 879.4 | - | - | 0 | - |
| 8 | b | 2.131E+04 | 887.4 | 0.0006542 | 0.7372 | +1 | 8 |
| - | - | 3.422E+04 | 892.4 | - | - | 0 | - |
| - | - | 2.391E+04 | 893.4 | - | - | 0 | - |
| - | - | 3.933E+04 | 894.4 | - | - | 0 | - |
| - | - | 2.677E+04 | 895.4 | - | - | 0 | - |
| - | - | 1.302E+04 | 896.4 | - | - | 0 | - |
| - | - | 9849 | 902.4 | - | - | 0 | - |
| - | - | 1.618E+04 | 904.4 | - | - | 0 | - |
| 8 | b | 6.464E+04 | 905.4 | 0.001514 | 1.672 | +1 | 8 |
| - | - | 1.661E+04 | 906.4 | - | - | 0 | - |
| - | - | 1.671E+04 | 907.4 | - | - | 0 | - |
| - | - | 3.984E+04 | 909.4 | - | - | 0 | - |
| - | - | 1.993E+04 | 910.4 | - | - | 0 | - |
| - | - | 1.148E+04 | 911.4 | - | - | 0 | - |
| - | - | 1.014E+04 | 912.4 | - | - | 0 | - |
| - | - | 2.49E+04 | 913.4 | - | - | 0 | - |
| - | - | 1.193E+04 | 914.4 | - | - | 0 | - |
| - | - | 1.223E+04 | 920.4 | - | - | 0 | - |
| - | - | 2.296E+04 | 921.4 | - | - | 0 | - |
| - | - | 4.653E+04 | 922.4 | - | - | 0 | - |
| - | - | 1.234E+04 | 923.4 | - | - | 0 | - |
| - | - | 1.071E+04 | 924.4 | - | - | 0 | - |
| - | - | 1.715E+04 | 927.4 | - | - | 0 | - |
| - | - | 8.201E+04 | 930.4 | - | - | 0 | - |
| - | - | 6.459E+04 | 931.4 | - | - | 0 | - |
| - | - | 6.258E+04 | 932.4 | - | - | 0 | - |
| - | - | 1.999E+04 | 933.4 | - | - | 0 | - |
| - | - | 1.142E+04 | 934.5 | - | - | 0 | - |
| - | - | 1.526E+04 | 938.5 | - | - | 0 | - |
| - | - | 1.88E+04 | 939.4 | - | - | 0 | - |
| - | - | 1.082E+04 | 940.4 | - | - | 0 | - |
| - | - | 3.85E+04 | 942.4 | - | - | 0 | - |
| - | - | 2.058E+04 | 943.4 | - | - | 0 | - |
| - | - | 4.526E+04 | 947.5 | - | - | 0 | - |
| - | - | 1.795E+05 | 948.4 | - | - | 0 | - |
| - | - | 8.565E+04 | 949.4 | - | - | 0 | - |
| - | - | 3.718E+04 | 950.4 | - | - | 0 | - |
| - | - | 1.289E+04 | 952.4 | - | - | 0 | - |
| - | - | 3.112E+04 | 953.4 | - | - | 0 | - |
| - | - | 2.714E+04 | 954.4 | - | - | 0 | - |
| - | - | 1.414E+04 | 957.4 | - | - | 0 | - |
| - | - | 9.326E+04 | 965.5 | - | - | 0 | - |
| - | - | 1.084E+05 | 966.5 | - | - | 0 | - |
| - | - | 4.914E+04 | 967.5 | - | - | 0 | - |
| - | - | 1.405E+04 | 968.5 | - | - | 0 | - |
| - | - | 1.138E+05 | 970.4 | - | - | 0 | - |
| - | - | 5.427E+04 | 971.4 | - | - | 0 | - |
| - | - | 1.287E+04 | 972.4 | - | - | 0 | - |
| - | - | 6.536E+04 | 974.5 | - | - | 0 | - |
| - | - | 3.278E+04 | 975.5 | - | - | 0 | - |
| - | - | 1.195E+04 | 976.5 | - | - | 0 | - |
| - | - | 3.613E+04 | 983.5 | - | - | 0 | - |
| - | - | 2.256E+04 | 984.5 | - | - | 0 | - |
| - | - | 1.2E+04 | 985.5 | - | - | 0 | - |
| - | - | 1.211E+04 | 991.5 | - | - | 0 | - |
| 8 | y | 1.697E+04 | 996.5 | 0.0007285 | 0.731 | +1 | 8 |
| 8 | y | 2.657E+04 | 997.4 | 0.006825 | 6.843 | +1 | 8 |
| - | - | 1.506E+04 | 998.5 | - | - | 0 | - |
| 9 | b | 2.39E+04 | 1001 | 0.001767 | 1.765 | +1 | 9 |
| 9 | b | 8.149E+04 | 1002 | 0.01006 | 10.04 | +1 | 9 |
| - | - | 5.228E+04 | 1003 | - | - | 0 | - |
| - | - | 2.212E+04 | 1004 | - | - | 0 | - |
| - | - | 1.107E+04 | 1005 | - | - | 0 | - |
| 8 | y | 6.663E+04 | 1014 | 0.002371 | 2.337 | +1 | 8 |
| - | - | 5.006E+04 | 1015 | - | - | 0 | - |
| - | - | 1.133E+04 | 1016 | - | - | 0 | - |
| 9 | b | 1.612E+05 | 1019 | 0.002983 | 2.926 | +1 | 9 |
| - | - | 7.811E+04 | 1020 | - | - | 0 | - |
| - | - | 2.537E+04 | 1021 | - | - | 0 | - |
| - | - | 1.698E+04 | 1023 | - | - | 0 | - |
| - | - | 1.359E+04 | 1049 | - | - | 0 | - |
| - | - | 2.384E+04 | 1050 | - | - | 0 | - |
| - | - | 1.143E+04 | 1051 | - | - | 0 | - |
| - | - | 3.168E+04 | 1058 | - | - | 0 | - |
| - | - | 3.824E+04 | 1059 | - | - | 0 | - |
| - | - | 2.834E+04 | 1060 | - | - | 0 | - |
| - | - | 2.049E+04 | 1067 | - | - | 0 | - |
| - | - | 1.496E+04 | 1068 | - | - | 0 | - |
| - | - | 1.21E+04 | 1068 | - | - | 0 | - |
| - | - | 2.184E+04 | 1069 | - | - | 0 | - |
| - | - | 1.129E+05 | 1076 | - | - | 0 | - |
| - | - | 8.753E+04 | 1077 | - | - | 0 | - |
| - | - | 3.035E+04 | 1078 | - | - | 0 | - |
| - | - | 5.105E+04 | 1085 | - | - | 0 | - |
| - | - | 3.806E+04 | 1086 | - | - | 0 | - |
| - | - | 2.196E+04 | 1087 | - | - | 0 | - |
| - | - | 2.113E+04 | 1094 | - | - | 0 | - |
| - | - | 1.426E+05 | 1095 | - | - | 0 | - |
| - | - | 6.384E+04 | 1096 | - | - | 0 | - |
| - | - | 2.837E+04 | 1097 | - | - | 0 | - |
| - | - | 3.321E+04 | 1104 | - | - | 0 | - |
| - | - | 2.179E+04 | 1105 | - | - | 0 | - |
| - | - | 2.296E+04 | 1105 | - | - | 0 | - |
| - | - | 1.226E+04 | 1107 | - | - | 0 | - |
| - | - | 3.129E+04 | 1112 | - | - | 0 | - |
| - | - | 1.192E+04 | 1122 | - | - | 0 | - |
| 7 | y | 2.306E+04 | 1128 | 0.002902 | 2.573 | +1 | 9 |
| - | - | 1.373E+04 | 1129 | - | - | 0 | - |
| - | - | 2.43E+04 | 1133 | - | - | 0 | - |
| - | - | 2.015E+04 | 1134 | - | - | 0 | - |
| - | - | 9310 | 1162 | - | - | 0 | - |
| 6 | y | 1.368E+04 | 1182 | 0.004666 | 3.949 | +1 | 10 |
| - | - | 1.084E+04 | 1183 | - | - | 0 | - |
| - | - | 1.313E+04 | 1189 | - | - | 0 | - |
| 6 | y | 3.91E+04 | 1199 | 0.001253 | 1.046 | +1 | 10 |
| - | - | 2.459E+04 | 1200 | - | - | 0 | - |
| - | - | 1.049E+04 | 1205 | - | - | 0 | - |
| - | - | 1.235E+04 | 1206 | - | - | 0 | - |
| - | - | 1.314E+04 | 1207 | - | - | 0 | - |
| - | - | 1.098E+04 | 1214 | - | - | 0 | - |
| - | - | 2.336E+04 | 1217 | - | - | 0 | - |
| - | - | 1.612E+04 | 1218 | - | - | 0 | - |
| - | - | 1.053E+04 | 1219 | - | - | 0 | - |
| - | - | 1.234E+04 | 1223 | - | - | 0 | - |
| - | - | 1.633E+04 | 1224 | - | - | 0 | - |
| - | - | 3.378E+04 | 1232 | - | - | 0 | - |
| - | - | 2.642E+04 | 1233 | - | - | 0 | - |
| - | - | 1.704E+04 | 1235 | - | - | 0 | - |
| - | - | 1.217E+04 | 1236 | - | - | 0 | - |
| 5 | y | 1.058E+04 | 1328 | 0.0002673 | 0.2013 | +1 | 11 |
| - | - | 8592 | 1640 | - | - | 0 | - |

m/z Charge Intensity FragmentType MassShift Position
120.05615997314453 0 24557.59
121.04019165039062 0 26698.066
122.0718994140625 0 37045.824
123.05582427978516 0 30614.494
124.07633972167969 0 16183.177
126.0918197631836 0 8088.893
127.08724212646484 0 9731.744
128.07144165039062 0 10456.169
128.0823516845703 0 17409.4
129.06642150878906 0 68535.734
129.1028289794922 0 4394940
130.05029296875 0 22402.229
130.08685302734375 0 84324.91
130.10009765625 0 28719.969
130.10614013671875 0 272089.97
131.11817932128906 0 5831.4233
132.0482177734375 0 9568.816
134.0712890625 0 6857.8623
136.07627868652344 0 2332155
137.07362365722656 0 16230.961
137.07962036132812 0 210660.25
138.06671142578125 0 116507.41
138.19842529296875 0 5696.3154
139.08714294433594 0 84303.69
140.07156372070312 0 7031.558
141.04873657226562 0 17675.08
141.0970458984375 0 11257.918
141.1028594970703 0 122981.58 a Water loss 1
142.08676147460938 0 25068.133
142.1063995361328 0 10223.202
146.09262084960938 0 18117.193
147.04470825195312 0 24762.482
147.11341857910156 0 80109.93
148.08749389648438 0 317445
148.95443725585938 0 14648.619
149.0240936279297 0 10749.149
149.04541015625 0 66758.37
149.08412170410156 0 7399.949 y Water loss 11
149.0910186767578 0 27373.285
151.08717346191406 0 10376.098
152.0823974609375 0 33502.62
153.10292053222656 0 17327.883
154.0617218017578 0 31358.613
155.0820770263672 0 71962.75
155.09332275390625 0 104758.93
155.1185302734375 0 514386.34
156.077392578125 0 95264.41
156.09669494628906 0 8145.8237
156.1219024658203 0 43993.363
157.1341552734375 0 73206.83
158.06045532226562 0 7568.878
158.10055541992188 0 30349.988
158.13833618164062 0 7613.3433
159.11346435546875 0 11820856 a 1
160.076904296875 0 8056.8184 b Water loss 5
160.08740234375 0 34317.79
160.11050415039062 0 57128.15
160.1167449951172 0 893969.06
161.08314514160156 0 6856.2246
161.11839294433594 0 48972.285
162.06678771972656 0 20582.615
163.0980987548828 0 8643.729
164.1193084716797 0 10595.322
165.0776824951172 0 72110.95
166.06170654296875 0 844197
167.05726623535156 0 15655.135
167.06512451171875 0 61642.848
167.08201599121094 0 17688.229
167.0932159423828 0 31092.129
167.11865234375 0 34620.254
168.06590270996094 0 8022.7373
169.061279296875 0 25051.928
169.09800720214844 0 15731.935 b Water loss 1
170.0459747314453 0 7320.122
171.1135711669922 0 40553.906
172.0506591796875 0 11008.493
173.09275817871094 0 136244.78 y Ammonia loss 12
173.4511260986328 0 26748.605
174.0669403076172 0 10934.428
175.08834838867188 0 10357.47
175.0986328125 0 29575.28
176.08253479003906 0 108395.555
177.10279846191406 0 13916.555
177.11416625976562 0 61200.79
178.05038452148438 0 13619.59
178.11837768554688 0 7188.064
178.13453674316406 0 29303.906
179.09332275390625 0 43067.54
180.0775909423828 0 21104.336
180.1138153076172 0 54072.734
181.09066772460938 0 33620.742
181.09835815429688 0 11275.457
183.0770263671875 0 67803.664
183.11355590820312 0 30055.568
184.10887145996094 0 7347.551
185.0924835205078 0 11159.154
185.12928771972656 0 48047.91
186.08786010742188 0 15114.725
187.10845947265625 0 1664705.2 b 1
188.1117706298828 0 152138.36
189.07772827148438 0 70457.79
189.1143798828125 0 7164.84
190.06185913085938 0 113105.625
190.0811004638672 0 9766.448
190.13455200195312 0 9874.483
191.0654296875 0 9388.983
191.0934295654297 0 60447.28
192.07675170898438 0 10788.405
192.1026153564453 0 103775.6
193.0727996826172 0 9023.357
193.0984649658203 0 12135.695
193.10894775390625 0 161910.95
194.0930633544922 0 147796.27
194.11203002929688 0 10268.862
194.1289825439453 0 11460.7705
195.07708740234375 0 117530.73
195.0882110595703 0 20156.113
195.0972137451172 0 13175.191
195.11351013183594 0 130782.07
196.0909881591797 0 386863.84
196.1166534423828 0 13608.916
197.09371948242188 0 46248.484
198.08718872070312 0 8924.746
198.1248016357422 0 15726.058
199.10824584960938 0 42591.387
200.0927734375 0 13724.653
201.08763122558594 0 177566.61
202.08712768554688 0 42725.7
202.10926818847656 0 46776.684
202.11947631835938 0 25894.008
203.09304809570312 0 31907.748
203.12197875976562 0 369845.7
204.0674591064453 0 6597.3384
204.07763671875 0 28206.559
204.1253662109375 0 31300.564
205.09799194335938 0 18926.28
205.10838317871094 0 11508.73
205.11770629882812 0 17207.863
206.10446166992188 0 34729.875
207.0883331298828 0 332439.03
207.16078186035156 0 10097.694
208.09149169921875 0 41760.117
208.12013244628906 0 19177.96
212.10369873046875 0 66560.914
212.14039611816406 0 13940.338
213.08792114257812 0 12746.602
213.10702514648438 0 10949.116
213.1242218017578 0 18539.812
214.09060668945312 0 13494.508
214.10935974121094 0 9283.33
215.1031036376953 0 10241.626
216.13504028320312 0 165126.86 y Water loss 13
217.07273864746094 0 101713.38
217.11903381347656 0 248758.28 y Ammonia loss 13
217.13821411132812 0 15799.543
218.05674743652344 0 93469.65
218.07586669921875 0 7445.8936
218.1221923828125 0 11971.309
219.0890655517578 0 8406.696
219.11300659179688 0 15744.663
221.09388732910156 0 16781.287
221.10411071777344 0 382643.6
221.1400909423828 0 14574.555
222.0880584716797 0 63519.01
222.1074676513672 0 35152.48
222.12454223632812 0 206782.44
223.06483459472656 0 16824.73
223.10848999023438 0 52330.96
223.1278839111328 0 25613.281 y Water loss 11
223.1562042236328 0 76173.516
224.08619689941406 0 43192.25
224.1149444580078 0 352720.84
224.15902709960938 0 10428.711
225.04351806640625 0 23248.973
225.0992889404297 0 16024.786
225.11807250976562 0 52660.043
226.12042236328125 0 10953.672
227.17678833007812 0 9580.647
228.10855102539062 0 29770.168 b 3
228.11351013183594 0 36909.69
228.6106414794922 0 14105.423
229.11825561523438 0 12177.464
230.08193969726562 0 57438.215
230.11439514160156 0 408612.22
231.1173095703125 0 94833.47
232.13037109375 0 11069.795 y 11
233.1405029296875 0 36003.066
234.099365234375 0 123375.9
234.124267578125 0 68818.17
234.14564514160156 0 1048766.9 y 13
235.08340454101562 0 561694.56
235.1279754638672 0 9251.502
235.14898681640625 0 116680.336
236.08663940429688 0 62121.05
236.11447143554688 0 17215.797
236.15170288085938 0 10838.269
237.12420654296875 0 154653.42
238.12811279296875 0 7680.2236
239.11473083496094 0 411545.56
239.15109252929688 0 542636.6
240.09815979003906 0 20256.96
240.13510131835938 0 1280537.9
240.1554412841797 0 42656.223
241.11265563964844 0 755123.8
241.1383056640625 0 131420.95
242.1156768798828 0 68174.16
242.1398162841797 0 10713.402
243.10873413085938 0 28211.346
244.1299591064453 0 36095.734
247.10853576660156 0 94054.75
247.14109802246094 0 12611.836
248.11294555664062 0 19261.303
248.14279174804688 0 6561.4634
249.0991668701172 0 330760.12
249.13560485839844 0 15335.704
250.08297729492188 0 40158.04
250.10275268554688 0 34580.207
250.11715698242188 0 11255.915 y Ammonia loss 7
250.16700744628906 0 25857.371
251.0972900390625 0 42137.684
251.1271514892578 0 10779.498
251.151123046875 0 205110.98
252.0810089111328 0 885130.75
252.10997009277344 0 1235845.5
252.1542510986328 0 25924.088
252.62728881835938 0 9435.393
253.08392333984375 0 91942.08
253.1133270263672 0 140186.86
254.0767822265625 0 36032.777
254.11456298828125 0 12532.188
256.1089782714844 0 17671.494
257.1616516113281 0 130331.37
258.1456298828125 0 72750.01
258.162841796875 0 15205
259.1564636230469 0 10399.679
261.1357116699219 0 21973.15
262.0942687988281 0 71246.43
262.1189270019531 0 13819.431
262.62335205078125 0 10321.88
263.07855224609375 0 15055.327
263.1016540527344 0 6942.0166
264.134765625 0 36554.824
265.11907958984375 0 123169.84
265.13385009765625 0 9115.185
266.1242370605469 0 13372.634
267.1095886230469 0 952473
267.6158447265625 0 6888.458
268.1127014160156 0 116566.7
268.1779479980469 0 19756.484
269.1075134277344 0 3067214
270.1103820800781 0 357923.62
270.63336181640625 0 13606.83
271.10333251953125 0 117938.71
271.1376037597656 0 19063.045
271.47125244140625 0 9261.182
272.6537170410156 0 8391.542
273.1352233886719 0 245245.64
274.0820617675781 0 8215.454
274.1329040527344 0 23154.941
274.13720703125 0 24431.025
275.1718444824219 0 12127.429
277.0940856933594 0 14471.005
278.631591796875 0 20448.13
279.09222412109375 0 12204.521
279.14599609375 0 107829.38
280.10498046875 0 74643.78
280.13006591796875 0 172739.06
280.1475830078125 0 12299.594
280.4683837890625 0 7985.507
280.6515197753906 0 76854.81
281.0899963378906 0 10863.297
281.115234375 0 11983.975
281.1333312988281 0 19496.805
281.1531982421875 0 34080.156
282.1460876464844 0 20643.357
283.1187438964844 0 15908.756
284.10357666015625 0 10612.511
284.135986328125 0 15606.984
286.13348388671875 0 8046.1133
287.1511535644531 0 12220.774
291.15057373046875 0 19097.514
292.1422424316406 0 13066.511
292.629150390625 0 27550.021 b 4
293.1261291503906 0 25696.28
294.19378662109375 0 18647.604
294.64910888671875 0 155191.36
295.1047668457031 0 71311.1
295.1510314941406 0 31229.357
295.6523742675781 0 10857.462
296.1100769042969 0 8333.814
296.16107177734375 0 25581.504
297.1029357910156 0 30832.377
297.1570129394531 0 86463.484
298.1405944824219 0 8493.751
298.1600036621094 0 11381.728
299.14410400390625 0 10363.916
300.1391906738281 0 24732.719 b Water loss 2
301.130615234375 0 432100.2
302.1287536621094 0 68757.49
302.1624450683594 0 8576.017
303.1327209472656 0 7642.2324
303.1625671386719 0 29184.271
304.1413879394531 0 11541.943
304.1780090332031 0 68847.57
304.656494140625 0 49830.477 y Water loss 10
305.12921142578125 0 12014.013
305.14599609375 0 21791.04 y Ammonia loss 10
305.16143798828125 0 49917.848
305.6496887207031 0 12807.966
306.1454772949219 0 26538.133
307.1199645996094 0 8160.261
309.1099853515625 0 7658.8003
309.1571960449219 0 10142.513
309.6251525878906 0 8195.471
310.1393737792969 0 16124.633
310.6311340332031 0 7312.8105
311.1136474609375 0 13628.622
311.1498107910156 0 7640.2773
312.120849609375 0 17148.314
312.1491394042969 0 9003.513
313.6426086425781 0 11052.613
313.6618957519531 0 168167.03 y 10
314.11669921875 0 7512.731
314.1506652832031 0 92525.32
314.1697082519531 0 17527.645
314.652587890625 0 35471.805
314.8179016113281 0 36740.06
315.151611328125 0 14258.176
318.1395263671875 0 34422.418
318.6407775878906 0 7951.1196
319.10443115234375 0 18859.13
319.1421203613281 0 28892.248 b Water loss 5
319.66241455078125 0 9001.049
320.13653564453125 0 573394.75
321.1393127441406 0 78771.25
321.6587829589844 0 13121.188
322.18853759765625 0 199313.69
323.1723937988281 0 71232.35
323.19207763671875 0 34407.88
323.66046142578125 0 10598.365
324.1538391113281 0 12308.754
324.18890380859375 0 9303.735
324.9879455566406 0 23008.488
325.150634765625 0 11669.535
325.8277893066406 0 13142.693
325.9888000488281 0 11394.818
326.13714599609375 0 59024.363
326.18328857421875 0 20238.088
326.660888671875 0 17673.924
327.1432189941406 0 30066.154
327.1672058105469 0 73617.84
327.6468505859375 0 9001.237
328.1504821777344 0 66257.336 b 5
328.1708984375 0 12283.218
328.64947509765625 0 8621.013
328.66802978515625 0 48154.25
329.1253662109375 0 22213.049
329.156005859375 0 9216.366
329.1732482910156 0 13254.867
329.67071533203125 0 10895.1045
330.1578369140625 0 17315.549
332.1366271972656 0 10801.612
332.8270568847656 0 28659.43 y Water loss 7
333.1590881347656 0 30397.344 y Ammonia loss 7
333.66015625 0 7211.272
334.824462890625 0 10958.951 b Ammonia loss 8
335.1149597167969 0 14375.298
336.1605529785156 0 7774.8145
337.13128662109375 0 10333.41
337.1685485839844 0 15146.179
337.1986389160156 0 7694.0166
337.65582275390625 0 10735.079
337.6757507324219 0 14994.303
338.14715576171875 0 478907.88
338.67486572265625 0 23718.082
338.8310241699219 0 132126.05 y 7
339.1492004394531 0 61445.41
339.1668395996094 0 32607.39
339.2151794433594 0 40349.562
339.49847412109375 0 32207.94
340.14190673828125 0 186051.72
340.1791687011719 0 14323.316
340.1994323730469 0 37998.38
341.0194091796875 0 47942.797
341.1448669433594 0 36679.01
341.1831359863281 0 171521.12
342.01910400390625 0 22927.166
342.1863708496094 0 24225.99
342.6685791015625 0 13656.961
343.01788330078125 0 13918.507
343.1637268066406 0 30145.877
343.1878967285156 0 7393.0957
344.14764404296875 0 102940.99
344.1941223144531 0 219442.06 y Water loss 12
345.1531982421875 0 14995.895
345.1780700683594 0 239815.52 y Ammonia loss 12
345.19732666015625 0 22901.17
346.173828125 0 85017.97
346.6734924316406 0 23016.146
347.1849060058594 0 16825.238
347.8418884277344 0 7627.4585
348.13128662109375 0 39487.97
348.16778564453125 0 11114.538
349.13140869140625 0 13037.2295
350.16558837890625 0 37455.24
350.1847229003906 0 16851.541
351.1675109863281 0 23502.234
351.67059326171875 0 165681.9
352.1427307128906 0 43775.355
352.1716003417969 0 64395.824
352.6727294921875 0 17140.441
353.12591552734375 0 31059.475
353.148193359375 0 19718.236
353.1722412109375 0 24523.781
353.50384521484375 0 9076.93
354.1316223144531 0 60176.44
354.685791015625 0 9841.851
355.1364440917969 0 10258.635
355.1729736328125 0 13312.086
356.13818359375 0 10841.173
356.1700134277344 0 13866.05
357.1718444824219 0 10640.995
358.1529541015625 0 7552.7217
358.1864318847656 0 13020.903
358.20965576171875 0 105091.984
359.02960205078125 0 234951.86
359.1475830078125 0 27526.344
359.212890625 0 20545.193
360.03021240234375 0 91381.4
360.16949462890625 0 142488.17
360.67034912109375 0 37652.758
361.02716064453125 0 50811.54
361.14874267578125 0 10079.955
361.173828125 0 118554.14
362.129150390625 0 15959.242
362.1787414550781 0 22038.312
362.20458984375 0 184625.34 y 12
362.5029296875 0 37285.668
362.6796875 0 17388.352 y Ammonia loss 3
362.8351745605469 0 14961.371
363.1116943359375 0 22636.098
363.136474609375 0 10310.128
363.2077941894531 0 31468.768
365.17120361328125 0 14136.522
365.1943359375 0 123368.74
366.14056396484375 0 117996.73
366.158935546875 0 93810.38
366.1986083984375 0 21108.795
366.93096923828125 0 31775.748 y 3
367.14166259765625 0 21523.59
367.1637268066406 0 14732.933
367.18304443359375 0 18915.15
368.17596435546875 0 193744.48
368.5068054199219 0 27003.312
368.6829833984375 0 18415.152
368.840576171875 0 23638.135
369.1765441894531 0 56488.184
369.66461181640625 0 10769.251
370.15301513671875 0 56366.66
370.69219970703125 0 43390.094
370.8490295410156 0 30480.363 y Ammonia loss 6
371.1584777832031 0 423204.16
371.18267822265625 0 21459.303
372.1425476074219 0 192441.05
372.162109375 0 41132.824
373.14593505859375 0 42629.188
373.18707275390625 0 19110.943 y Water loss 9
373.4313659667969 0 8642.101
373.679443359375 0 39145.875 y Ammonia loss 9
374.18157958984375 0 26616.814
375.1683044433594 0 118016
376.17034912109375 0 21217.969
376.2204284667969 0 44957.39
376.52593994140625 0 21874.705 y 6
376.85223388671875 0 132143.38
377.18597412109375 0 60706.113
377.5198669433594 0 26308.621
377.85345458984375 0 13104.784
378.17950439453125 0 11688.845
378.5043640136719 0 17660.38
378.6712951660156 0 8043.8003
379.6791076660156 0 25731.887
380.13916015625 0 152616.06
380.1794128417969 0 18861.68 b Water loss 9
380.51348876953125 0 12061.505 b Ammonia loss 9
381.1211242675781 0 40391.4
381.1444396972656 0 22753.281
381.20928955078125 0 11742.202
382.1275329589844 0 20327.197
382.16741943359375 0 11779.937
382.191650390625 0 132477.39 y 9
382.2181396484375 0 8681.126
382.66632080078125 0 9567.955
382.6936340332031 0 71264.27
383.1921081542969 0 17580.775
383.4366455078125 0 13042.036
383.6843566894531 0 11472.678
384.16778564453125 0 77536.03
384.18927001953125 0 10469.461
385.1719665527344 0 22332.37
386.15631103515625 0 10245.392
386.1832580566406 0 96706.15 b 9
386.51806640625 0 30709.17
386.8493957519531 0 15919.932
387.181396484375 0 54810.44
387.2151184082031 0 29709.383
387.6891174316406 0 12114.531
388.191162109375 0 35269.844
388.68536376953125 0 28557.26
389.1693420410156 0 142522.86
389.6895446777344 0 9396.739
390.1713562011719 0 21013.922
390.9342346191406 0 10573.36
391.1758117675781 0 29416.023
391.2098693847656 0 16886.39
392.19793701171875 0 9073.916
393.18035888671875 0 89378.3
394.1524353027344 0 15245.154
394.1812438964844 0 10671.276
394.5316162109375 0 10059.277 y Ammonia loss 5
394.6853942871094 0 7952.379
395.1885070800781 0 81348.65 y Water loss 2
395.4373474121094 0 62859.066 y Ammonia loss 2
395.6907043457031 0 49673.27
395.94000244140625 0 14318.794
396.170166015625 0 15874.623
396.52093505859375 0 11244.044
396.85308837890625 0 13234.344
397.1640930175781 0 53970.945
397.1969909667969 0 42476.547
397.5137939453125 0 16243.909
397.6904602050781 0 158382.9
398.1504211425781 0 1099337.9
398.1915588378906 0 50286.477
398.6932373046875 0 22894.258
399.15338134765625 0 201302.62
399.6912841796875 0 314928.22 y 2
399.94195556640625 0 246305.89
400.1475830078125 0 50290.78
400.1937561035156 0 132607.9
400.4431457519531 0 62107.16
400.5382080078125 0 27182.025
400.6917724609375 0 19329.633
400.8725891113281 0 18220.238
401.1949768066406 0 39092.13
401.68634033203125 0 10281.783
402.17974853515625 0 70818.12
402.4363098144531 0 10486.908
402.6859130859375 0 12357.631
403.18292236328125 0 8520.575
403.197021484375 0 6996.4043
404.66912841796875 0 17241.746
405.22607421875 0 133285.33
405.51800537109375 0 9511.933
406.2030944824219 0 480928.2
406.2258605957031 0 18389.611
406.5169372558594 0 37014.914
406.7040100097656 0 195346.77
406.8515930175781 0 16294.288
407.20562744140625 0 58890.047
407.70672607421875 0 9694.234
408.13446044921875 0 12785.479
408.2092590332031 0 13542.894
408.2375183105469 0 34069.844
409.20245361328125 0 33064.203
409.7030334472656 0 27176.607
410.203857421875 0 87080.66
410.70050048828125 0 19084.264
411.19464111328125 0 72012.02
411.447265625 0 12422.04
411.69000244140625 0 60674.035
411.8597412109375 0 9005.288
412.16131591796875 0 13849.86
412.1887512207031 0 64610.207
412.523193359375 0 12460.19
412.6911315917969 0 16128.823
413.18182373046875 0 43634.84
413.6795654296875 0 31806.193
414.1763000488281 0 10040.579
415.17431640625 0 277188.16
415.2091064453125 0 39796.473
415.70025634765625 0 28060.318 b Ammonia loss 13
415.9486999511719 0 12895.999
416.177978515625 0 50933.4
416.20660400390625 0 8481.648
416.4498596191406 0 9703.353
417.1651916503906 0 14466.995
417.1911926269531 0 172554.47
417.6932067871094 0 84088.17
417.8632507324219 0 14594.499
418.1544494628906 0 8259.981
418.1924743652344 0 36899.008
419.2046203613281 0 68026.76
419.7079162597656 0 13131.568
419.9566955566406 0 53921.06 b 13
420.1697998046875 0 18471.674
420.2012023925781 0 388559.6 y Ammonia loss 1
420.45458984375 0 29793.865
420.70245361328125 0 191003.3
420.95404052734375 0 15492.019
421.20355224609375 0 49657.395
421.6956787109375 0 38739.35
422.1884460449219 0 51348.47
422.68829345703125 0 12384.454
423.18646240234375 0 15329.681
423.2363586425781 0 267808.1
424.1654052734375 0 15147.685
424.1998596191406 0 10296.47
424.2391357421875 0 40283.49
424.4586181640625 0 218132.06 y 1
424.7085876464844 0 154505.48
424.9600524902344 0 88305.63
425.1593933105469 0 15248.1
425.2087707519531 0 44475.27
425.4613037109375 0 25559.32
425.8679504394531 0 10778.143
426.14532470703125 0 43213.78
427.2135314941406 0 136905.19
427.53594970703125 0 21662.79
427.869873046875 0 21680.873
428.2156066894531 0 55506.41
428.45428466796875 0 11126.696
428.697509765625 0 11293.147
429.09063720703125 0 59560.28
429.18902587890625 0 12435.251
429.9634094238281 0 13772.019
430.0904846191406 0 32107.408
430.17919921875 0 17328.59
430.208984375 0 37169.508 y Water loss 8
430.7014465332031 0 108749.15 y Ammonia loss 8
431.08795166015625 0 25974.29
431.20562744140625 0 88374.76
431.5414123535156 0 59311.48
431.8757019042969 0 41796.773
432.20684814453125 0 15506.992
432.9560546875 0 50475.86
433.21990966796875 0 156060.31
433.4564208984375 0 17177.023
433.5405578613281 0 107161.49
433.7046813964844 0 22833.896
433.8748779296875 0 41823.17
434.15093994140625 0 23772.371
434.2109375 0 31939.059
434.7158508300781 0 21377.672
434.96441650390625 0 18488.127
436.1627502441406 0 14292.83
436.23199462890625 0 160047.73
436.5384826660156 0 21409.879
437.2135314941406 0 110643.336 y Water loss 4
437.4600524902344 0 87571.914
437.5491943359375 0 75503.83 y Ammonia loss 4
437.7080993652344 0 82627.71
437.8844299316406 0 41645.473
437.959716796875 0 64229.703
438.2028503417969 0 43287.766
438.4544982910156 0 10352.68
438.7156677246094 0 14069.828
439.2145690917969 0 300463.3 y 8
439.71588134765625 0 110306.87
439.868408203125 0 14968.461
440.214599609375 0 33457.934
440.53662109375 0 53907.305 b 10
440.87286376953125 0 14742.696
441.1957092285156 0 34395.492
441.7140197753906 0 450916.75 Precursor Water loss
441.9634704589844 0 424489.53 Precursor Ammonia loss
442.2132873535156 0 410764.72
442.4637756347656 0 172854.64
442.5450744628906 0 69081.695
442.7137756347656 0 59643.504
442.8764343261719 0 50244.688
442.96533203125 0 30310.105
443.1690673828125 0 96656.48
443.2176208496094 0 43389.676 y 4
443.5526123046875 0 27309.678
443.88818359375 0 9458.192
444.1718444824219 0 18708.156
444.21435546875 0 23000.107 b Water loss 7
445.2114562988281 0 9993.582
445.2422790527344 0 84130.93 y Water loss 11
445.54266357421875 0 25325.775
445.87646484375 0 13727.681
446.216796875 0 1747653 Precursor
446.4673767089844 0 1341616.5
446.7175598144531 0 850637.9
446.96826171875 0 419221.94
447.21728515625 0 107812.91
447.4694519042969 0 23931.07
447.7152099609375 0 22736.814
448.21661376953125 0 173703.28
448.55059814453125 0 124463.414
448.8818359375 0 41130.145
449.2183837890625 0 19648.535
449.538330078125 0 12913.693
450.1879577636719 0 11092.966
450.24713134765625 0 29919.592
451.1759948730469 0 36225.195
451.2316589355469 0 630204.6
451.58953857421875 0 8828.131
452.1605224609375 0 21322.027
452.1933898925781 0 17741.262
452.2347106933594 0 151711.47
452.70989990234375 0 9993.658
453.22119140625 0 99714.98 b 7
453.72119140625 0 24231.14
453.89141845703125 0 97592.914
454.2236328125 0 90594.195
454.5572204589844 0 35497.32
454.7082214355469 0 9871.948
454.874755859375 0 29463.586
455.17059326171875 0 16498.068
455.21270751953125 0 214175.27 b 3
455.540771484375 0 9205.821
455.7209167480469 0 38330.707
456.213134765625 0 41521.57
457.2057800292969 0 24790.059
458.21282958984375 0 15100.945
458.88531494140625 0 18186.965
459.22064208984375 0 16870.65
459.55230712890625 0 11719.678
459.8945007324219 0 115669.92
460.2276916503906 0 106065.96
460.56341552734375 0 46123.508
460.8923645019531 0 9432.096
461.2168884277344 0 54840.734
461.7354736328125 0 14230.65
461.8840026855469 0 8939.619
462.2172546386719 0 24438.219
462.7184753417969 0 53811.324
463.1758117675781 0 13809.801
463.21246337890625 0 109819.06
463.25274658203125 0 707899.06 y 11
463.7139587402344 0 48221.37
464.1582946777344 0 36866.55
464.2231750488281 0 48801.69
464.2558898925781 0 113895.93
464.5481872558594 0 11249.57
464.72698974609375 0 14828.353
464.88946533203125 0 16670.479
465.15972900390625 0 12334.546
465.2237243652344 0 21779.777
465.25848388671875 0 20102.822
465.5506286621094 0 11747.598
465.7166748046875 0 55976.26
466.21478271484375 0 32492.957
466.7232666015625 0 33146.645
467.2193908691406 0 26333.229
467.8945617675781 0 10873.354
468.21917724609375 0 41718.777 b Water loss 11
468.2590026855469 0 42092
468.5523986816406 0 50166.742 b Ammonia loss 11
468.6994934082031 0 10892.273
468.8878173828125 0 14996.461
469.1879577636719 0 485443.03
470.1907653808594 0 103944.06
470.2237854003906 0 19866.19
471.1885986328125 0 29075.678
471.22283935546875 0 34873.67
471.552490234375 0 47258.062
471.7233581542969 0 506465.3
471.8839416503906 0 30766.486
472.2245788574219 0 271315.34
472.5553283691406 0 10492.05
472.7242431640625 0 97364.66
473.2283630371094 0 38179.79
473.5672912597656 0 9878.251
473.8977966308594 0 16158.21
474.2249755859375 0 67189.36 b 11
474.2593688964844 0 46861.902
474.5558776855469 0 25701.568
474.72198486328125 0 55441.39
474.8888854980469 0 15325.732
475.2216491699219 0 33896.89
475.26129150390625 0 17279.406
476.7156982421875 0 26139.76
476.8987731933594 0 52477.098
477.22222900390625 0 251276.67
477.55706787109375 0 221993.62
477.7108459472656 0 26044.611
477.8912353515625 0 94737.3
478.2235107421875 0 48303.293
478.5589294433594 0 13130.327
478.7305603027344 0 19311.584
479.1828308105469 0 11380.877
479.2246398925781 0 129983.28
479.724609375 0 25586.363
480.2249450683594 0 47040.3
480.55224609375 0 34589.35
480.88494873046875 0 11867.252
481.1845397949219 0 90773.38
481.2229919433594 0 35218.945
481.8936767578125 0 10098.905
482.1831970214844 0 14459.432
482.9022216796875 0 235818.83 y Water loss 3
483.2325439453125 0 494749.22 y Ammonia loss 3
483.5658264160156 0 313152.4
483.7284240722656 0 48258.977
483.8997497558594 0 128849.41
484.2360534667969 0 62151.06
484.5680236816406 0 9512.647
485.2286071777344 0 30521.193
485.7210693359375 0 385427.6
485.8929443359375 0 149495.86
486.2239685058594 0 330250.75
486.5583190917969 0 95200.79
486.7224426269531 0 80041.4
486.89300537109375 0 46532.504
487.2215576171875 0 26394.182
487.7345886230469 0 168633.73
488.2354736328125 0 110011.24
488.73809814453125 0 30803.87
488.90582275390625 0 1578726.2 y 3
489.2400817871094 0 1145735
489.5740966796875 0 467978.97
489.9085693359375 0 140099.92
490.1903381347656 0 11489.752
490.2311096191406 0 45146.863
490.71990966796875 0 40602.605
491.2214660644531 0 18656.035
491.5675048828125 0 35330.47
491.89666748046875 0 424453.16
492.2312316894531 0 276152.84
492.5650329589844 0 186975.06
492.7293395996094 0 24452.455
492.89892578125 0 56013.285
493.2270812988281 0 48686.36
493.7202453613281 0 16493.293
494.22027587890625 0 10707.576
496.2049560546875 0 12635.2705
496.24395751953125 0 18968.979
497.18170166015625 0 27415.703
497.22003173828125 0 82379.73
497.57159423828125 0 305559.03
497.90472412109375 0 225533.44
498.2089538574219 0 62381.992
498.2392883300781 0 122769.19
498.5734558105469 0 62422.773
498.7372131347656 0 65267.32 y Water loss 7
498.9061279296875 0 21820.39
499.1952819824219 0 16889.06
499.2308044433594 0 131508.42 y Ammonia loss 7
499.7320556640625 0 62510.914
500.1912536621094 0 19230.879
501.2355041503906 0 26286.523 b Water loss 8
501.5728759765625 0 15873.35
501.7325134277344 0 195846.19 b Ammonia loss 8
502.24072265625 0 103660.49
502.7349853515625 0 38217.934
503.2552185058594 0 57462.164
503.5753479003906 0 266953.47
503.7469177246094 0 16908.791
503.9095764160156 0 194335.44
504.24462890625 0 116082.91
504.57666015625 0 44859.13
504.90740966796875 0 15866.419
505.2391357421875 0 31699.355
505.5697937011719 0 20496.557
506.2301940917969 0 17535.678
506.27386474609375 0 10985.88
507.2113342285156 0 88109.84
507.5742492675781 0 14912.387
507.7427978515625 0 436929.94 y 7
507.9076843261719 0 16494.43
508.24420166015625 0 232635.28
508.74530029296875 0 82523.81
509.248779296875 0 18759.54
509.5576477050781 0 9312.344
509.74884033203125 0 11337.372
510.2425537109375 0 143734.44 b 8
510.57916259765625 0 42876.395
510.7416076660156 0 56544.664
510.9091491699219 0 53281.156 b Water loss 12
511.24273681640625 0 56644.145
511.5731201171875 0 26111.082
511.76226806640625 0 36444.44
511.905029296875 0 22718.95
512.2322387695312 0 53050.137
512.57421875 0 9880.36
512.7455444335938 0 20324.303
513.230224609375 0 12838.766
514.2393188476562 0 9769.099
514.5763549804688 0 10986.612
514.9068603515625 0 40176.2
515.2374877929688 0 79419.72
515.565673828125 0 55812.594
515.8994750976562 0 26397.166
516.2228393554688 0 99236.87
516.583251953125 0 24829.387
516.9114379882812 0 37594.957 b 12
517.1943359375 0 17516.79
517.2470703125 0 39834.52
517.5798950195312 0 29730.553
517.9114379882812 0 19644.25
518.24560546875 0 30910.328
518.917724609375 0 22051.846
519.2699584960938 0 30562.318
519.5819702148438 0 13307.124
520.2529296875 0 56431.375
520.5789184570312 0 55952.824
520.907958984375 0 296732.9
521.2417602539062 0 189440.53
521.574462890625 0 178725.83
521.7576904296875 0 33646.62
521.9080200195312 0 76599.13
522.2462768554688 0 25861.783
524.2381591796875 0 47559.598
524.2753295898438 0 16058.039
524.9197387695312 0 39346.96
525.2489013671875 0 43375.08
525.5875854492188 0 30995.668
525.73388671875 0 20279.156
526.244384765625 0 16933.248
526.5824584960938 0 583905.7 y Water loss 2
526.913330078125 0 1133985.9 y Ammonia loss 2
527.2467041015625 0 867780.7
527.5796508789062 0 426164.38
527.9134521484375 0 172152.8
528.2456665039062 0 36125.09
528.2699584960938 0 27761.635
529.25830078125 0 22706.912
529.3020629882812 0 17762.195
529.7523803710938 0 27057.23
530.2420654296875 0 183859.45
530.5947875976562 0 34427.64
530.749755859375 0 32404.059
530.9274291992188 0 30996.09
531.245849609375 0 34970.727
532.2478637695312 0 27386.795
532.5858764648438 0 4903720 y 2
532.9200439453125 0 3773303.2
533.2537841796875 0 2076609.4
533.5877685546875 0 893551.75
533.9215698242188 0 249530.27
534.2549438476562 0 78539.586
534.589111328125 0 12448.657
534.7395629882812 0 78209.59
535.2051391601562 0 34993.59
535.2491455078125 0 79760.836
535.7476196289062 0 35136.9
535.9144287109375 0 18871.125
536.2039794921875 0 13038.678
536.2476806640625 0 27820.322
536.593505859375 0 27194.805
536.7429809570312 0 9842.825
536.9302978515625 0 25154.605
537.2786254882812 0 41045.914
538.2528686523438 0 52700.438
538.753173828125 0 82427.94
539.249755859375 0 31343.19
539.2852783203125 0 29108.336
539.5866088867188 0 10983.133
539.7559204101562 0 17106.953
540.2556762695312 0 23768.703
542.2879638671875 0 10117.073
543.2520141601562 0 269704.62
543.7510986328125 0 165858.66
544.2515869140625 0 62209.348
544.7437744140625 0 65116.07
545.2462768554688 0 22114.201
545.2962036132812 0 146973.92
546.2972412109375 0 40986.797
546.7670288085938 0 26779.768
547.2640991210938 0 182699.9
547.7528076171875 0 17718.486
547.7877807617188 0 13186.547
547.9290161132812 0 19496.691
548.2582397460938 0 78900.57
549.2545166015625 0 16534.143
550.25341796875 0 9756.547
551.7649536132812 0 34739.2
552.2572631835938 0 120881.33
552.7576904296875 0 95507.09
553.2550048828125 0 224817.4
553.6013793945312 0 20058.764 b Water loss 13
553.7564086914062 0 142302.25
553.929931640625 0 94652.875 b Ammonia loss 13
554.270263671875 0 154072.08
554.5964965820312 0 26803.576
554.7590942382812 0 13327.158
554.9323120117188 0 10533.63
555.2786865234375 0 88306.96 y Water loss 6
555.7702026367188 0 86153.76 y Ammonia loss 6
556.2630615234375 0 173600.05
556.7682495117188 0 53307.41
557.2631225585938 0 37049.348
558.24462890625 0 16717.543
558.7422485351562 0 31274.074
559.2431640625 0 19755.719
559.6049194335938 0 149869.73 b 13
559.7398071289062 0 12045.502
559.9367065429688 0 198032.98 y Ammonia loss 1
560.2650756835938 0 76472.45
560.2966918945312 0 112783.38
560.6029052734375 0 83807.555
560.7681274414062 0 52495.258
560.9373779296875 0 31169.344
561.222900390625 0 13994.865
561.2628173828125 0 22513.713
561.298828125 0 35174.633
561.763427734375 0 12408.537
562.3026123046875 0 11635.632
563.7646484375 0 13132.021
564.2838745117188 0 148009.48 y 6
564.7755126953125 0 460780.34
565.2750854492188 0 542991.06
565.6087646484375 0 718738.94 y 1
565.7755126953125 0 105389.18
565.94287109375 0 582722
566.2769165039062 0 438994.9
566.611083984375 0 156658.48
566.9454345703125 0 50695.355
567.251953125 0 221317.28
567.7536010742188 0 141646.69
568.2554931640625 0 130893.945
568.7540893554688 0 15568.879
568.9367065429688 0 23281.805
569.2633056640625 0 47984.664
569.76611328125 0 48836.676 b Water loss 9
570.2672729492188 0 68875.98 b Ammonia loss 9
570.764404296875 0 22564.45
571.2623901367188 0 17482.754
571.763916015625 0 10669.78
572.278076171875 0 15839.084
573.2896728515625 0 130017.2
574.2910766601562 0 28508.885
575.2579956054688 0 23496.684
576.2586669921875 0 29629.967
576.761962890625 0 16630.56
578.771240234375 0 356723.2 b 9
579.2728271484375 0 218555.12
579.77294921875 0 96879.23
580.225830078125 0 13749.306
580.27294921875 0 26792.254
581.2769775390625 0 21963.078
582.2728271484375 0 406709.94
582.7828369140625 0 16084.049
583.275634765625 0 124776.45
584.2520751953125 0 270183.8 b 4
584.7687377929688 0 13039.444
585.2584838867188 0 80168.15
585.7681274414062 0 24948.607
586.261962890625 0 35162.004
586.759765625 0 13168.58
587.2606811523438 0 11699.285
587.7567138671875 0 12620.798
588.2905883789062 0 238069.22
589.2467041015625 0 12962.496
589.29345703125 0 57934.85
590.2479248046875 0 14722.385
590.2958984375 0 33419.773
590.7971801757812 0 34405.605 y Water loss 5
591.2899169921875 0 89946.09 y Ammonia loss 5
591.79345703125 0 29648.88
592.2194213867188 0 18669.148
593.2723388671875 0 43630.938
593.7840576171875 0 24484.793
594.2815551757812 0 111564.555
594.7802734375 0 88570.27
595.2749633789062 0 42080.984
595.7688598632812 0 73848.445
596.2618408203125 0 32260.846
597.2742919921875 0 13724.879
598.276611328125 0 17424.158
598.7702026367188 0 23026.709
599.2684326171875 0 23491.889
599.803955078125 0 321810.16 y 5
600.2562255859375 0 55442.14
600.3053588867188 0 177645.8
600.760498046875 0 36275.715
600.806640625 0 52782.63
601.2628784179688 0 12563.412
602.2876586914062 0 50651.77
602.7940673828125 0 94902.125
603.2919921875 0 138377.2
603.7901000976562 0 69068.914
604.2857055664062 0 47534.246
604.7844848632812 0 23807.387
605.316650390625 0 43790.566
606.320556640625 0 13103.927
607.2802734375 0 109469.31
607.7789306640625 0 109395.26
608.302978515625 0 79772.01 y Water loss 10
608.7721557617188 0 269548.8
609.275390625 0 191115.33
609.77294921875 0 106673.65
610.22314453125 0 13797.347
610.275146484375 0 30615.65
611.8037719726562 0 182024.56
612.3030395507812 0 97029.08
612.8049926757812 0 49806.598
616.2857055664062 0 249487.89
616.7865600585938 0 163691.28
617.2858276367188 0 124983.445
617.7781982421875 0 254954.75
618.2342529296875 0 25970.348
618.2799682617188 0 106030.9
618.779296875 0 58496.47
619.240966796875 0 20752.39
620.2928466796875 0 29306.137
620.8087158203125 0 70447.6
621.3064575195312 0 58693.484
621.8118896484375 0 14904.951
624.799560546875 0 33637.133
625.2953491210938 0 43268.15
625.79736328125 0 15202.741
626.3153686523438 0 394109.75 y 10
626.7911376953125 0 46793.926
627.3174438476562 0 120421.69
628.3171997070312 0 23131.736
628.795654296875 0 22108.21
629.2940063476562 0 15779.729
635.2703857421875 0 110143.89
636.265625 0 27938.559
637.2784423828125 0 105264.38 b Water loss 5
637.3211059570312 0 22396.852
637.800048828125 0 40972.78
638.2901611328125 0 52868.527
638.7972412109375 0 27957.408
639.3012084960938 0 59059.89
640.324462890625 0 26296.826
641.310791015625 0 31297.693
644.2804565429688 0 97057.914
645.2888793945312 0 29315.559
645.7962036132812 0 13737.466
646.3072509765625 0 95575.22
646.8097534179688 0 123681.484
647.3068237304688 0 116293.58
647.8078002929688 0 43337.91
648.2959594726562 0 29186.352
649.2974243164062 0 24036.688
649.8070068359375 0 29686.133
650.2990112304688 0 33605.56
650.7997436523438 0 16289.961
651.290771484375 0 23190.936 b Water loss 10
651.7937622070312 0 14271.632 b Ammonia loss 10
652.2947387695312 0 21662.986
653.2835693359375 0 53892.82
654.29638671875 0 16248.092
655.2890014648438 0 310985.8 b 5
655.32177734375 0 207864.12 y Water loss 4
655.8170166015625 0 352766.3 y Ammonia loss 4
656.3197021484375 0 218917.53
656.8178100585938 0 61829.516
657.3126831054688 0 131982.53
658.3321533203125 0 77667.61
658.8048706054688 0 51393.133
659.3073120117188 0 43587.03
659.795654296875 0 90986.69
660.3010864257812 0 86244.586 b 10
660.8031005859375 0 60948.66
661.3020629882812 0 29826.156
662.8140869140625 0 54509.33
663.318359375 0 47137.457
663.8082885742188 0 28381.307
664.3250732421875 0 1042952 y 4
664.82666015625 0 617575.06
665.326416015625 0 336874.22
665.8282470703125 0 92009.14
666.315673828125 0 63276.426
667.305908203125 0 87648.086
667.80908203125 0 86361.32
668.30908203125 0 85737.29
668.80859375 0 19621.943
669.3086547851562 0 47649.67
669.8193969726562 0 17375.312
670.307861328125 0 36522.266
671.3211059570312 0 33029.027
671.820068359375 0 107299.13
672.3206787109375 0 65789.414
672.8021850585938 0 241131.02
673.3014526367188 0 183079.16
673.800537109375 0 127703.15
674.3377075195312 0 202314.98
674.8016357421875 0 16554.834
675.3406982421875 0 78448.32
676.34033203125 0 48994.46
680.333251953125 0 147109.02
680.8326416015625 0 124188.73
681.3367919921875 0 133857.16
681.8062744140625 0 412950.47
682.30712890625 0 269083.47
682.8079223632812 0 159749.02
683.3118286132812 0 50152.586
683.811279296875 0 15760.516
684.3237915039062 0 175853.47
685.315673828125 0 102817.39
686.3258666992188 0 37912.047
688.8184204101562 0 13146.075
689.3361206054688 0 76777.086
689.8392944335938 0 45615.934
690.322265625 0 36074.402
690.820556640625 0 28706.953
691.33447265625 0 150388.3
692.3369750976562 0 50214.934
693.3260498046875 0 32183.56
701.3231811523438 0 35542.54
701.822509765625 0 68135.02 b Water loss 11
702.332275390625 0 583204.1
702.8240966796875 0 35764.46
703.33544921875 0 207001.95
704.3367309570312 0 46413.69
705.338134765625 0 15593.671
705.8399658203125 0 11544.208
706.3359985351562 0 20697.58
708.3588256835938 0 39701.418
709.3513793945312 0 33995.957
710.333984375 0 37172.58
710.8289794921875 0 48233.35 b 11
711.32763671875 0 39832.465
711.830322265625 0 12567.871
712.3235473632812 0 31523.219
714.8440551757812 0 14112.284
715.3358154296875 0 46832.8
715.8335571289062 0 29997.025
718.3450927734375 0 13637.167
719.3321533203125 0 229914.52
719.8414306640625 0 11764.269
720.3336181640625 0 95700.36
721.333251953125 0 36314.598
722.337890625 0 38385.406
722.8352661132812 0 28169.13
723.3424072265625 0 21696.674
723.8478393554688 0 80867.49 y Water loss 3
724.3444213867188 0 171374.7 y Ammonia loss 3
724.8445434570312 0 120363.48
725.3446655273438 0 43315.16
728.3346557617188 0 37451.812
728.8348388671875 0 18398.994
729.3374633789062 0 22609.56
730.3331909179688 0 26456.596
731.3369750976562 0 78754.39
731.8406372070312 0 42443.652
732.3430786132812 0 25551.176
732.8545532226562 0 635963.8 y 3
733.355712890625 0 464062.97
733.8572387695312 0 206601.69
734.3572387695312 0 70146.65
734.8590087890625 0 16843.719
736.3541870117188 0 109498.875
736.8436889648438 0 12621.372
737.3450927734375 0 93864.89
737.843505859375 0 34660.883
738.343017578125 0 46146.723
740.3756103515625 0 33567.62
741.3740844726562 0 9873.636
745.3633422851562 0 37534.074 y Water loss 9
745.8524780273438 0 76848.91
746.3530883789062 0 103531.63 y Ammonia loss 9
746.8543701171875 0 41868.86
747.3534545898438 0 35998.566
748.3538208007812 0 48508.4
750.3590087890625 0 52623 b Water loss 6
751.361083984375 0 33565.13
752.36328125 0 11555.787
753.3446655273438 0 30066.873
754.3589477539062 0 16123.574
754.8590087890625 0 43242.46
755.35302734375 0 74626.19
755.8616943359375 0 19659.697
756.3536376953125 0 36830.293
758.343994140625 0 14829.724
761.3630981445312 0 15986.706
763.3753051757812 0 222851.5 y 9
764.3155517578125 0 38810.137
764.3786010742188 0 78933.09
765.3084716796875 0 12760.451
765.3792724609375 0 23073.791
765.8516845703125 0 26437.838 b Water loss 12
766.358154296875 0 54036.207
766.8477172851562 0 13614.743
767.3680419921875 0 12746.661
768.37255859375 0 195424.61 b 6
769.3758544921875 0 76067.19
770.3727416992188 0 31882.727
772.375732421875 0 49002.11
773.3714599609375 0 26204.248
774.85693359375 0 35111.63 b 12
775.3612060546875 0 36551.523
775.8543090820312 0 19638.213
776.3571166992188 0 16273.516
781.3400268554688 0 129577.06
782.3417358398438 0 38321.066
788.3534545898438 0 20727.191
789.3605346679688 0 27194.066 y Water loss 2
790.3911743164062 0 27499.3
791.3997192382812 0 15989.288
793.382080078125 0 42658.83
794.3731689453125 0 33595.066
795.392822265625 0 24123.275
798.3701171875 0 35189.844 y 2
798.8757934570312 0 23011.947
799.37744140625 0 15412.556
800.3463134765625 0 10120.193
802.3680419921875 0 53263.586
803.36669921875 0 40218.31
804.3682861328125 0 19804.588
805.37841796875 0 38400.902
806.3806762695312 0 19585.277
811.39794921875 0 94126.164
812.4019165039062 0 33104.65
816.361572265625 0 26215.129
817.3612670898438 0 13957.583
818.3986206054688 0 28050.215
819.3927612304688 0 143897.45
820.3860473632812 0 98154.75
821.3839721679688 0 60731.156
822.3702392578125 0 58717.582
823.3749389648438 0 26903.672
824.3738403320312 0 13429.605
825.3639526367188 0 16876.975
830.3610229492188 0 20200.344
831.3607788085938 0 14335.933
833.3743896484375 0 248455.22
834.376708984375 0 112325.49
835.3753662109375 0 33036.81
836.4105224609375 0 10855.931
837.4026489257812 0 112569.586
838.4051513671875 0 61056.074
839.3936157226562 0 143660.98 y Ammonia loss 1
840.3952026367188 0 61667.926
841.393798828125 0 21339.312
842.3863525390625 0 24753.049
843.3687744140625 0 22717.031
847.38623046875 0 59969.504
848.3806762695312 0 49820.117
849.3836059570312 0 24264.732
850.39599609375 0 12434.977
854.4287719726562 0 18612.793
855.4303588867188 0 13610.594
859.4054565429688 0 34056.363 y Water loss 8
860.3941650390625 0 58324.78 y Ammonia loss 8
861.4005737304688 0 23448.7
865.39697265625 0 159538.83
866.3994750976562 0 79305.76
867.3980102539062 0 12521.664
875.387939453125 0 11512.621
876.4111938476562 0 19684.979
877.4180908203125 0 216337.34 y 8
878.4217529296875 0 89619.17
879.4240112304688 0 28058.277
887.4185791015625 0 21310.562 b Water loss 7
892.4111328125 0 34224.1
893.4035034179688 0 23906.887
894.4212036132812 0 39330.03
895.4208374023438 0 26767.936
896.4303588867188 0 13022.506
902.4186401367188 0 9848.631
904.4235229492188 0 16177.708
905.4282836914062 0 64641.773 b 7
906.432861328125 0 16614.873
907.4298095703125 0 16714.738
909.4358520507812 0 39837.246
910.4335327148438 0 19929.314
911.4310302734375 0 11476.662
912.4151000976562 0 10139.965
913.4019775390625 0 24897.502
914.4027709960938 0 11925.011
920.4386596679688 0 12226.843
921.4284057617188 0 22961.172
922.4276733398438 0 46534.477
923.43115234375 0 12335.791
924.4303588867188 0 10705.37
927.44921875 0 17150.469
930.4237670898438 0 82011.39
931.4189453125 0 64588.133
932.4312744140625 0 62579.996
933.4400024414062 0 19994.379
934.4552001953125 0 11417.443
938.4518432617188 0 15255.867
939.4471435546875 0 18800.125
940.442626953125 0 10821.809
942.4362182617188 0 38501.414
943.4403076171875 0 20582.463
947.4511108398438 0 45261.75
948.4351806640625 0 179468.53
949.4351806640625 0 85654.42
950.4326171875 0 37180.984
952.4216918945312 0 12887.691
953.4097900390625 0 31119.941
954.415283203125 0 27140.658
957.4322509765625 0 14142.524
965.4605102539062 0 93257.516
966.4514770507812 0 108449.94
967.451171875 0 49138.715
968.4536743164062 0 14053.702
970.4332885742188 0 113763.98
971.43603515625 0 54270.883
972.4435424804688 0 12868.715
974.4598999023438 0 65356.188
975.4600219726562 0 32779.977
976.4633178710938 0 11946.329
983.4697875976562 0 36127.754
984.47119140625 0 22562.973
985.4573974609375 0 12004.7705
991.47998046875 0 12114.949
996.46533203125 0 16973.104 y Water loss 7
997.4554443359375 0 26573.07 y Ammonia loss 7
998.4573974609375 0 15058.373
1001.4639282226562 0 23902.639 b Water loss 8
1002.4562377929688 0 81488.03 b Ammonia loss 8
1003.4598388671875 0 52280.55
1004.4745483398438 0 22117.834
1005.4840087890625 0 11070.808
1014.4775390625 0 66630.6 y 7
1015.477783203125 0 50061.22
1016.4826049804688 0 11329.738
1019.4757080078125 0 161235.62 b 8
1020.4765625 0 78109.12
1021.4768676757812 0 25367.2
1022.5086669921875 0 16980.857
1049.4959716796875 0 13589.908
1050.4781494140625 0 23837.514
1051.47412109375 0 11426.381
1058.487060546875 0 31675.662
1059.4765625 0 38237.633
1060.4739990234375 0 28344.264
1066.5067138671875 0 20493.684
1067.508056640625 0 14958.069
1068.473876953125 0 12101.179
1069.49560546875 0 21842.29
1076.4952392578125 0 112865.445
1077.490966796875 0 87526.234
1078.489990234375 0 30349.812
1085.49365234375 0 51052.75
1086.494140625 0 38056.53
1087.4903564453125 0 21957.537
1093.548583984375 0 21131.742
1094.5050048828125 0 142627.56
1095.506103515625 0 63838.746
1096.5078125 0 28371.348
1103.5045166015625 0 33209.97
1104.502685546875 0 21792.75
1105.49951171875 0 22955.771
1106.50537109375 0 12257.504
1111.5458984375 0 31294.428
1121.535888671875 0 11915.518
1127.5621337890625 0 23061.482 y 6
1128.560302734375 0 13728.902
1133.495361328125 0 24300.848
1134.4949951171875 0 20153.514
1161.5504150390625 0 9309.735
1181.574462890625 0 13679.925 y Ammonia loss 5
1182.576416015625 0 10840.656
1188.544677734375 0 13126.494
1198.5950927734375 0 39096.95 y 5
1199.5982666015625 0 24589.86
1204.5723876953125 0 10490.474
1205.5732421875 0 12346.789
1206.576904296875 0 13140.623
1213.551513671875 0 10983.338
1216.534912109375 0 23357.863
1217.53759765625 0 16119.441
1218.52197265625 0 10529.587
1222.599365234375 0 12336.992
1223.593017578125 0 16328.724
1231.5596923828125 0 33775.48
1232.5672607421875 0 26420.875
1234.538818359375 0 17043.11
1235.5543212890625 0 12170.214
1327.638671875 0 10575.945 y 4
1640.0467529296875 0 8591.639

Spectrum Details

|  |  |
| --- | --- |
| Matched peaks? Matched peaks The total absolute number of peaks matched. Additionally in brackets the total fraction of peaks matched and the total number of peaks is shown. | 133 (9.10% of 1461) |
| FDR? FDR The false discovery rate estimated for this peptide. It is calculated by matching all theoretical fragments with a non-integer shift with the raw peaks for this spectrum. This is done with 40 different shifts. The resulting percentage is the average number of annotated peaks over the number of annotated peaks with the correct spectrum. | 0.36% |
| Satellite FDR? Satellite FDR See the FDR for details on its calculation. This satellite ion specific FDR only contains the satellite ions (d/w) for I/L/J positions. | - |
| PSM Score? PSM Score The PSM Score as given by Hecklib to this annotated spectrum. It is shown with three significant figures. | 571 |

#### Spectrum 2615? Spectrum 2615 The raw spectrum of this peptide as annotated by Hecklib. The fragments are coloured according to ion type (see legend). Any peaks with a star '\*' as text can be hovered over to see the full details, first the ion type second the mass shift type. By hovering over the amino acids in the peptide or ions in the legend the corresponding peaks are highlighted. By toggling the 'Unassigned' label you can turn the background (unassigned) peaks on or off in the plot. By updating the slider in the Ion legend you can update the spectrum to only show the top X% of the peaks with labels. The top X% means any peak that is within X% of the highest intensity. By dragging in the spectrum you can zoom in to a specific part of the spectrum and use 'Zoom Out' to get back to the original zoom level. The annotation of the spectrum is based on the given sequence in the peptides file and is done with different software so inconsistencies are likely. The peaks are annotated based on the given sequence, with 20 ppm tolerance.

Copy Data

##### Spectrum 2615 (TSV)

###### Preview

```
Loading example...
```

*Click on the button to copy the data to your clipboard.*

Mz MinMz MaxIntensity Max

WidthHeightPeptide font sizePeptide stroke widthSpectrum font sizeSpectrum stroke widthCompact peptide

Ion legend

wxyz

abcd

OtherUnassignedIon^Charge^_Position_Show for top:%

SVMHEAJHNHYTQKS

04.52e+49.05e+41.36e+51.81e+5

Zoom Out

a^+1^_2_a^+1^_2_y^+2^_3_b^+1^_2_y^+1^_2_y^+1^_2_y^+1^_2_b^+2^_4_b^+2^_5_y^+2^_5_b^+2^_6_b^+1^_3_b^+2^_6_y^+3^_8_y^+1^_3_y^+1^_3_y^+1^_3_y^+2^_6_y^+3^_9_y^+2^_6_b^+3^_10_y^+3^_10_y^+4^_13_b^+4^_14_b^+4^_14_y^+4^_14_y^+4^_14_y^+2^_7_y^+2^_7_y^+3^_11_y^+3^_11_y^+2^_7_y^+1^_4_\*b^+3^_11_\*\*b^+2^_8_y^+1^_4_b^+1^_4_b^+3^_12_b^+3^_12_y^+3^_12_y^+3^_12_y^+3^_12_y^+2^_8_y^+2^_8_y^+2^_8_b^+3^_13_b^+2^_9_b^+3^_13_y^+3^_13_y^+3^_13_y^+3^_13_y^+2^_9_y^+2^_9_b^+3^_14_y^+2^_9_b^+3^_14_y^+3^_14_y^+3^_14_b^+2^_10_b^+2^_10_b^+1^_5_b^+2^_10_y^+2^_10_\*y^+2^_10_b^+1^_5_y^+1^_5_y^+1^_5_y^+1^_5_b^+1^_6_y^+2^_11_y^+2^_11_y^+2^_11_b^+2^_11_b^+1^_6_b^+2^_12_b^+2^_12_y^+2^_12_y^+2^_12_y^+1^_6_y^+1^_6_b^+1^_7_b^+2^_13_b^+1^_7_y^+1^_7_y^+1^_7_b^+1^_8_y^+1^_8_b^+1^_9_

0581116117422323

Fragment Matches Table

Show background peaks

| Position | Ion type | Intensity | mz Theoretical | mz Error (Th) | mz Error (ppm) | Charge | Series Number |
| --- | --- | --- | --- | --- | --- | --- | --- |
| - | - | 617.8 | 120.1 | - | - | 0 | - |
| - | - | 577.4 | 120.1 | - | - | 0 | - |
| - | - | 484.7 | 121 | - | - | 0 | - |
| - | - | 387 | 121.7 | - | - | 0 | - |
| - | - | 638.7 | 122.1 | - | - | 0 | - |
| - | - | 841.6 | 123.1 | - | - | 0 | - |
| - | - | 884.8 | 127.1 | - | - | 0 | - |
| - | - | 1552 | 129.1 | - | - | 0 | - |
| - | - | 9.173E+04 | 129.1 | - | - | 0 | - |
| - | - | 433.5 | 130.1 | - | - | 0 | - |
| - | - | 1921 | 130.1 | - | - | 0 | - |
| - | - | 5410 | 130.1 | - | - | 0 | - |
| - | - | 420.1 | 132.1 | - | - | 0 | - |
| - | - | 766.2 | 134 | - | - | 0 | - |
| - | - | 5.581E+04 | 136.1 | - | - | 0 | - |
| - | - | 4039 | 137.1 | - | - | 0 | - |
| - | - | 2150 | 138.1 | - | - | 0 | - |
| - | - | 1677 | 139.1 | - | - | 0 | - |
| - | - | 500.8 | 141.1 | - | - | 0 | - |
| 2 | a | 4021 | 141.1 | 0.0003606 | 2.556 | +1 | 2 |
| - | - | 665.5 | 147 | - | - | 0 | - |
| - | - | 1662 | 147.1 | - | - | 0 | - |
| - | - | 2544 | 148.1 | - | - | 0 | - |
| - | - | 734.7 | 149 | - | - | 0 | - |
| - | - | 510.2 | 149 | - | - | 0 | - |
| - | - | 683.4 | 149 | - | - | 0 | - |
| - | - | 478.7 | 151 | - | - | 0 | - |
| - | - | 692 | 152.1 | - | - | 0 | - |
| - | - | 512.5 | 153.1 | - | - | 0 | - |
| - | - | 518 | 154.1 | - | - | 0 | - |
| - | - | 1283 | 155.1 | - | - | 0 | - |
| - | - | 1986 | 155.1 | - | - | 0 | - |
| - | - | 2258 | 155.1 | - | - | 0 | - |
| - | - | 3550 | 156.1 | - | - | 0 | - |
| - | - | 1252 | 157.1 | - | - | 0 | - |
| 2 | a | 1.791E+05 | 159.1 | 0.0004161 | 2.615 | +1 | 2 |
| - | - | 1013 | 160.1 | - | - | 0 | - |
| - | - | 1.275E+04 | 160.1 | - | - | 0 | - |
| - | - | 538.8 | 165.1 | - | - | 0 | - |
| - | - | 2372 | 165.1 | - | - | 0 | - |
| - | - | 4953 | 166.1 | - | - | 0 | - |
| - | - | 1.041E+04 | 166.1 | - | - | 0 | - |
| - | - | 564 | 167.1 | - | - | 0 | - |
| - | - | 882.8 | 167.1 | - | - | 0 | - |
| - | - | 889 | 167.1 | - | - | 0 | - |
| - | - | 1294 | 169.1 | - | - | 0 | - |
| - | - | 752.9 | 169.1 | - | - | 0 | - |
| - | - | 2097 | 171.1 | - | - | 0 | - |
| 13 | y | 1090 | 173.1 | 0.0003843 | 2.22 | +2 | 3 |
| - | - | 5279 | 176.1 | - | - | 0 | - |
| - | - | 859.6 | 177.1 | - | - | 0 | - |
| - | - | 821.7 | 178.1 | - | - | 0 | - |
| - | - | 1055 | 179.1 | - | - | 0 | - |
| - | - | 725 | 180.1 | - | - | 0 | - |
| - | - | 1355 | 180.1 | - | - | 0 | - |
| - | - | 612.3 | 181.1 | - | - | 0 | - |
| - | - | 1471 | 183.1 | - | - | 0 | - |
| - | - | 4929 | 183.1 | - | - | 0 | - |
| - | - | 780.7 | 183.1 | - | - | 0 | - |
| - | - | 1238 | 185.1 | - | - | 0 | - |
| - | - | 553.8 | 185.2 | - | - | 0 | - |
| 2 | b | 2.644E+04 | 187.1 | 0.0003745 | 2.001 | +1 | 2 |
| - | - | 2199 | 188.1 | - | - | 0 | - |
| - | - | 1178 | 189.1 | - | - | 0 | - |
| - | - | 2206 | 190.1 | - | - | 0 | - |
| - | - | 818.1 | 191.1 | - | - | 0 | - |
| - | - | 2233 | 192.1 | - | - | 0 | - |
| - | - | 1121 | 193.1 | - | - | 0 | - |
| - | - | 8951 | 193.1 | - | - | 0 | - |
| - | - | 1053 | 194.1 | - | - | 0 | - |
| - | - | 1759 | 194.1 | - | - | 0 | - |
| - | - | 678.3 | 194.1 | - | - | 0 | - |
| - | - | 588.9 | 194.1 | - | - | 0 | - |
| - | - | 1610 | 195.1 | - | - | 0 | - |
| - | - | 594.7 | 195.1 | - | - | 0 | - |
| - | - | 3504 | 195.1 | - | - | 0 | - |
| - | - | 1794 | 199.1 | - | - | 0 | - |
| - | - | 1807 | 201.1 | - | - | 0 | - |
| - | - | 860.9 | 202.1 | - | - | 0 | - |
| - | - | 648.9 | 202.1 | - | - | 0 | - |
| - | - | 7324 | 204.1 | - | - | 0 | - |
| - | - | 1633 | 204.1 | - | - | 0 | - |
| - | - | 551.7 | 205.1 | - | - | 0 | - |
| - | - | 591.8 | 205.1 | - | - | 0 | - |
| - | - | 968.7 | 206.1 | - | - | 0 | - |
| - | - | 5879 | 207.1 | - | - | 0 | - |
| - | - | 806.8 | 208.1 | - | - | 0 | - |
| - | - | 1641 | 209 | - | - | 0 | - |
| - | - | 902.6 | 209.1 | - | - | 0 | - |
| - | - | 2679 | 211 | - | - | 0 | - |
| - | - | 755.4 | 211.1 | - | - | 0 | - |
| - | - | 564.2 | 211.1 | - | - | 0 | - |
| - | - | 993.2 | 212.1 | - | - | 0 | - |
| 14 | y | 3681 | 216.1 | 0.0004825 | 2.232 | +1 | 2 |
| - | - | 1453 | 217.1 | - | - | 0 | - |
| 14 | y | 3920 | 217.1 | 0.0003994 | 1.84 | +1 | 2 |
| - | - | 1789 | 218.1 | - | - | 0 | - |
| - | - | 512.5 | 220.1 | - | - | 0 | - |
| - | - | 2.407E+04 | 221.1 | - | - | 0 | - |
| - | - | 985.3 | 222.1 | - | - | 0 | - |
| - | - | 2514 | 222.1 | - | - | 0 | - |
| - | - | 3568 | 222.1 | - | - | 0 | - |
| - | - | 1079 | 223.1 | - | - | 0 | - |
| - | - | 1153 | 223.1 | - | - | 0 | - |
| - | - | 1894 | 223.2 | - | - | 0 | - |
| - | - | 6253 | 224.1 | - | - | 0 | - |
| - | - | 1162 | 225 | - | - | 0 | - |
| - | - | 1130 | 225.1 | - | - | 0 | - |
| - | - | 1970 | 225.1 | - | - | 0 | - |
| - | - | 719.3 | 225.1 | - | - | 0 | - |
| - | - | 601.9 | 226 | - | - | 0 | - |
| - | - | 1222 | 226.1 | - | - | 0 | - |
| - | - | 981.6 | 226.1 | - | - | 0 | - |
| - | - | 1149 | 226.2 | - | - | 0 | - |
| - | - | 1127 | 227 | - | - | 0 | - |
| - | - | 1168 | 227 | - | - | 0 | - |
| - | - | 602.3 | 228.1 | - | - | 0 | - |
| - | - | 1924 | 230.1 | - | - | 0 | - |
| - | - | 6889 | 230.1 | - | - | 0 | - |
| - | - | 649.4 | 231.1 | - | - | 0 | - |
| - | - | 957.7 | 233.1 | - | - | 0 | - |
| - | - | 1523 | 234.1 | - | - | 0 | - |
| - | - | 1375 | 234.1 | - | - | 0 | - |
| 14 | y | 1.93E+04 | 234.1 | 0.0004158 | 1.776 | +1 | 2 |
| - | - | 1.071E+04 | 235.1 | - | - | 0 | - |
| - | - | 1404 | 235.1 | - | - | 0 | - |
| - | - | 714.4 | 236.1 | - | - | 0 | - |
| - | - | 1145 | 236.1 | - | - | 0 | - |
| 4 | b | 1430 | 236.1 | 0.002937 | 12.44 | +2 | 4 |
| - | - | 2974 | 237.1 | - | - | 0 | - |
| - | - | 3180 | 239.1 | - | - | 0 | - |
| - | - | 1.002E+04 | 239.2 | - | - | 0 | - |
| - | - | 521.3 | 240.1 | - | - | 0 | - |
| - | - | 729.4 | 240.1 | - | - | 0 | - |
| - | - | 2.271E+04 | 240.1 | - | - | 0 | - |
| - | - | 2338 | 241.1 | - | - | 0 | - |
| - | - | 692.9 | 244.1 | - | - | 0 | - |
| - | - | 3855 | 247.1 | - | - | 0 | - |
| - | - | 3759 | 249.1 | - | - | 0 | - |
| - | - | 3736 | 251.2 | - | - | 0 | - |
| - | - | 2.122E+04 | 252.1 | - | - | 0 | - |
| - | - | 1240 | 252.1 | - | - | 0 | - |
| - | - | 570.8 | 252.2 | - | - | 0 | - |
| - | - | 2488 | 253.1 | - | - | 0 | - |
| - | - | 1728 | 253.1 | - | - | 0 | - |
| - | - | 3103 | 257.1 | - | - | 0 | - |
| - | - | 3206 | 257.2 | - | - | 0 | - |
| - | - | 1572 | 258.1 | - | - | 0 | - |
| - | - | 706.8 | 261.1 | - | - | 0 | - |
| - | - | 972.4 | 262.1 | - | - | 0 | - |
| - | - | 493.1 | 263.1 | - | - | 0 | - |
| - | - | 1032 | 264.1 | - | - | 0 | - |
| - | - | 2415 | 265.1 | - | - | 0 | - |
| - | - | 981.2 | 267.1 | - | - | 0 | - |
| - | - | 9688 | 267.1 | - | - | 0 | - |
| - | - | 935.7 | 268.1 | - | - | 0 | - |
| - | - | 1333 | 268.1 | - | - | 0 | - |
| - | - | 648.1 | 269.1 | - | - | 0 | - |
| - | - | 2111 | 270.1 | - | - | 0 | - |
| - | - | 5940 | 273.1 | - | - | 0 | - |
| - | - | 2333 | 275.1 | - | - | 0 | - |
| - | - | 788.8 | 276.1 | - | - | 0 | - |
| - | - | 589.7 | 276.2 | - | - | 0 | - |
| - | - | 1517 | 277.1 | - | - | 0 | - |
| - | - | 1788 | 279.1 | - | - | 0 | - |
| - | - | 1364 | 280.1 | - | - | 0 | - |
| - | - | 3819 | 280.1 | - | - | 0 | - |
| - | - | 768.6 | 280.7 | - | - | 0 | - |
| - | - | 730.8 | 281.1 | - | - | 0 | - |
| - | - | 563.4 | 282.1 | - | - | 0 | - |
| - | - | 637.3 | 284.1 | - | - | 0 | - |
| - | - | 1.341E+04 | 285.1 | - | - | 0 | - |
| - | - | 1287 | 286.1 | - | - | 0 | - |
| - | - | 1056 | 286.6 | - | - | 0 | - |
| - | - | 610.4 | 287.1 | - | - | 0 | - |
| - | - | 893.4 | 290.1 | - | - | 0 | - |
| - | - | 847.5 | 293.1 | - | - | 0 | - |
| - | - | 972 | 294.6 | - | - | 0 | - |
| - | - | 1984 | 295.1 | - | - | 0 | - |
| - | - | 1532 | 297.2 | - | - | 0 | - |
| 5 | b | 1482 | 300.6 | 0.003095 | 10.29 | +2 | 5 |
| - | - | 7677 | 301.1 | - | - | 0 | - |
| - | - | 1751 | 302.1 | - | - | 0 | - |
| - | - | 519.7 | 302.3 | - | - | 0 | - |
| - | - | 923.9 | 304.1 | - | - | 0 | - |
| - | - | 1106 | 304.2 | - | - | 0 | - |
| - | - | 698.1 | 305.1 | - | - | 0 | - |
| - | - | 986 | 305.2 | - | - | 0 | - |
| - | - | 921.7 | 309.1 | - | - | 0 | - |
| - | - | 774.3 | 310.1 | - | - | 0 | - |
| 11 | y | 4198 | 313.7 | 8.017E-05 | 0.2556 | +2 | 5 |
| - | - | 964.2 | 314.2 | - | - | 0 | - |
| - | - | 647.2 | 318.1 | - | - | 0 | - |
| - | - | 639.7 | 319.1 | - | - | 0 | - |
| - | - | 6376 | 320.1 | - | - | 0 | - |
| - | - | 2302 | 320.2 | - | - | 0 | - |
| - | - | 2933 | 320.9 | - | - | 0 | - |
| - | - | 913.5 | 321.1 | - | - | 0 | - |
| - | - | 521.8 | 321.9 | - | - | 0 | - |
| - | - | 1211 | 322.1 | - | - | 0 | - |
| - | - | 4403 | 322.1 | - | - | 0 | - |
| - | - | 2490 | 322.2 | - | - | 0 | - |
| - | - | 770.5 | 322.6 | - | - | 0 | - |
| - | - | 763.5 | 322.7 | - | - | 0 | - |
| - | - | 1635 | 322.9 | - | - | 0 | - |
| - | - | 1782 | 323.2 | - | - | 0 | - |
| - | - | 1114 | 325.1 | - | - | 0 | - |
| - | - | 1097 | 326.1 | - | - | 0 | - |
| - | - | 1464 | 326.2 | - | - | 0 | - |
| 6 | b | 952.6 | 327.1 | 0.003583 | 10.95 | +2 | 6 |
| - | - | 1108 | 327.2 | - | - | 0 | - |
| - | - | 871 | 328.2 | - | - | 0 | - |
| - | - | 895.7 | 328.7 | - | - | 0 | - |
| - | - | 1926 | 332.1 | - | - | 0 | - |
| - | - | 740.2 | 333.1 | - | - | 0 | - |
| 3 | b | 3329 | 334.1 | 0.005565 | 16.65 | +1 | 3 |
| 6 | b | 821.4 | 336.1 | 0.002909 | 8.655 | +2 | 6 |
| - | - | 1126 | 336.2 | - | - | 0 | - |
| - | - | 633.9 | 336.7 | - | - | 0 | - |
| - | - | 5300 | 338.1 | - | - | 0 | - |
| 8 | y | 1998 | 338.8 | 0.0007204 | 2.126 | +3 | 8 |
| - | - | 529.6 | 339.2 | - | - | 0 | - |
| - | - | 685.4 | 339.5 | - | - | 0 | - |
| - | - | 2538 | 340.1 | - | - | 0 | - |
| - | - | 809.4 | 340.2 | - | - | 0 | - |
| - | - | 2482 | 341.2 | - | - | 0 | - |
| - | - | 576.3 | 342.2 | - | - | 0 | - |
| - | - | 677.3 | 343.2 | - | - | 0 | - |
| - | - | 1478 | 343.2 | - | - | 0 | - |
| - | - | 2444 | 344.1 | - | - | 0 | - |
| 13 | y | 4413 | 344.2 | 0.0006361 | 1.848 | +1 | 3 |
| - | - | 869.9 | 345.2 | - | - | 0 | - |
| 13 | y | 4285 | 345.2 | 0.0003241 | 0.9389 | +1 | 3 |
| - | - | 577.9 | 345.2 | - | - | 0 | - |
| - | - | 592.8 | 346.2 | - | - | 0 | - |
| - | - | 2318 | 348.1 | - | - | 0 | - |
| - | - | 1050 | 348.2 | - | - | 0 | - |
| - | - | 7237 | 350.1 | - | - | 0 | - |
| - | - | 1451 | 351.1 | - | - | 0 | - |
| - | - | 1442 | 351.7 | - | - | 0 | - |
| - | - | 690.1 | 352.2 | - | - | 0 | - |
| - | - | 1279 | 354.1 | - | - | 0 | - |
| - | - | 1819 | 354.2 | - | - | 0 | - |
| - | - | 1106 | 354.8 | - | - | 0 | - |
| - | - | 662.2 | 356.2 | - | - | 0 | - |
| - | - | 2834 | 358.2 | - | - | 0 | - |
| - | - | 803.5 | 360 | - | - | 0 | - |
| - | - | 793.5 | 360.9 | - | - | 0 | - |
| - | - | 3889 | 361 | - | - | 0 | - |
| - | - | 2054 | 361.2 | - | - | 0 | - |
| - | - | 3579 | 362 | - | - | 0 | - |
| - | - | 989.4 | 362.2 | - | - | 0 | - |
| 13 | y | 4214 | 362.2 | 0.0002642 | 0.7295 | +1 | 3 |
| - | - | 8526 | 363 | - | - | 0 | - |
| - | - | 900.2 | 363.2 | - | - | 0 | - |
| - | - | 1866 | 364 | - | - | 0 | - |
| - | - | 1481 | 365.2 | - | - | 0 | - |
| - | - | 1639 | 366.1 | - | - | 0 | - |
| - | - | 2091 | 366.2 | - | - | 0 | - |
| - | - | 624.3 | 367.2 | - | - | 0 | - |
| - | - | 609.3 | 368.1 | - | - | 0 | - |
| - | - | 1633 | 368.2 | - | - | 0 | - |
| - | - | 1058 | 369.7 | - | - | 0 | - |
| - | - | 846.4 | 370.2 | - | - | 0 | - |
| - | - | 6030 | 371.2 | - | - | 0 | - |
| - | - | 758.7 | 371.2 | - | - | 0 | - |
| - | - | 3391 | 372.1 | - | - | 0 | - |
| 10 | y | 949.4 | 373.7 | 0.0003095 | 0.8282 | +2 | 6 |
| - | - | 1937 | 375.2 | - | - | 0 | - |
| - | - | 594.4 | 376.2 | - | - | 0 | - |
| - | - | 1079 | 376.2 | - | - | 0 | - |
| 7 | y | 936.3 | 376.5 | 0.0004294 | 1.14 | +3 | 9 |
| - | - | 951.1 | 378.7 | - | - | 0 | - |
| - | - | 1103 | 379.2 | - | - | 0 | - |
| - | - | 730.8 | 380.1 | - | - | 0 | - |
| 10 | y | 5191 | 382.2 | 0.00381 | 9.968 | +2 | 6 |
| - | - | 2082 | 382.5 | - | - | 0 | - |
| - | - | 1158 | 382.7 | - | - | 0 | - |
| - | - | 855.3 | 383.2 | - | - | 0 | - |
| - | - | 2539 | 384.2 | - | - | 0 | - |
| - | - | 1009 | 384.5 | - | - | 0 | - |
| - | - | 906.2 | 384.7 | - | - | 0 | - |
| - | - | 1405 | 385.2 | - | - | 0 | - |
| - | - | 1035 | 386.1 | - | - | 0 | - |
| - | - | 838.6 | 387.7 | - | - | 0 | - |
| - | - | 2182 | 389.2 | - | - | 0 | - |
| - | - | 660.3 | 390.2 | - | - | 0 | - |
| - | - | 774.2 | 390.5 | - | - | 0 | - |
| 10 | b | 2578 | 391.5 | 0.001303 | 3.329 | +3 | 10 |
| - | - | 785.9 | 391.8 | - | - | 0 | - |
| - | - | 761.6 | 393.2 | - | - | 0 | - |
| - | - | 1845 | 393.2 | - | - | 0 | - |
| - | - | 649.7 | 393.7 | - | - | 0 | - |
| - | - | 750.4 | 394.2 | - | - | 0 | - |
| - | - | 627.6 | 396.2 | - | - | 0 | - |
| - | - | 1226 | 397.2 | - | - | 0 | - |
| - | - | 826 | 397.7 | - | - | 0 | - |
| - | - | 1532 | 398.1 | - | - | 0 | - |
| 6 | y | 1221 | 400.2 | 0.0007127 | 1.781 | +3 | 10 |
| - | - | 762.5 | 400.9 | - | - | 0 | - |
| - | - | 1566 | 402.2 | - | - | 0 | - |
| 3 | y | 1719 | 403.7 | 0.003041 | 7.533 | +4 | 13 |
| - | - | 2566 | 403.9 | - | - | 0 | - |
| - | - | 1675 | 404.2 | - | - | 0 | - |
| - | - | 1529 | 405.2 | - | - | 0 | - |
| - | - | 3247 | 406.2 | - | - | 0 | - |
| - | - | 1798 | 406.7 | - | - | 0 | - |
| - | - | 1892 | 407.2 | - | - | 0 | - |
| - | - | 1086 | 410.2 | - | - | 0 | - |
| - | - | 904.8 | 411.2 | - | - | 0 | - |
| - | - | 981.6 | 411.5 | - | - | 0 | - |
| - | - | 934.1 | 411.7 | - | - | 0 | - |
| - | - | 884.2 | 412.2 | - | - | 0 | - |
| - | - | 849 | 412.5 | - | - | 0 | - |
| - | - | 911.3 | 412.7 | - | - | 0 | - |
| - | - | 6813 | 414.1 | - | - | 0 | - |
| - | - | 647.5 | 414.9 | - | - | 0 | - |
| - | - | 695.2 | 415 | - | - | 0 | - |
| - | - | 1229 | 415.1 | - | - | 0 | - |
| - | - | 3691 | 415.2 | - | - | 0 | - |
| - | - | 515.3 | 415.2 | - | - | 0 | - |
| - | - | 776.8 | 415.4 | - | - | 0 | - |
| - | - | 733.9 | 416.1 | - | - | 0 | - |
| - | - | 1272 | 416.2 | - | - | 0 | - |
| - | - | 610.6 | 416.3 | - | - | 0 | - |
| - | - | 646.6 | 417.5 | - | - | 0 | - |
| 14 | b | 1504 | 419.7 | 0.001411 | 3.362 | +4 | 14 |
| - | - | 1290 | 419.7 | - | - | 0 | - |
| - | - | 2142 | 419.9 | - | - | 0 | - |
| - | - | 3807 | 420.2 | - | - | 0 | - |
| - | - | 1084 | 420.7 | - | - | 0 | - |
| - | - | 876.7 | 421 | - | - | 0 | - |
| - | - | 6299 | 421.2 | - | - | 0 | - |
| - | - | 1217 | 421.7 | - | - | 0 | - |
| - | - | 2112 | 422.2 | - | - | 0 | - |
| - | - | 2228 | 423.2 | - | - | 0 | - |
| 14 | b | 2078 | 424 | 0.002129 | 5.021 | +4 | 14 |
| 2 | y | 1684 | 424.2 | 0.003409 | 8.035 | +4 | 14 |
| - | - | 1914 | 424.5 | - | - | 0 | - |
| - | - | 952.9 | 424.7 | - | - | 0 | - |
| - | - | 3295 | 425.2 | - | - | 0 | - |
| - | - | 801.4 | 425.5 | - | - | 0 | - |
| - | - | 2172 | 425.7 | - | - | 0 | - |
| - | - | 970.4 | 426.2 | - | - | 0 | - |
| - | - | 1498 | 427.7 | - | - | 0 | - |
| - | - | 797.5 | 428.2 | - | - | 0 | - |
| 2 | y | 3195 | 428.5 | 0.00144 | 3.362 | +4 | 14 |
| - | - | 4728 | 428.7 | - | - | 0 | - |
| - | - | 1872 | 429 | - | - | 0 | - |
| - | - | 799.4 | 429.2 | - | - | 0 | - |
| - | - | 3795 | 429.7 | - | - | 0 | - |
| - | - | 3623 | 430 | - | - | 0 | - |
| 9 | y | 3715 | 430.2 | 0.003446 | 8.011 | +2 | 7 |
| - | - | 1061 | 430.5 | - | - | 0 | - |
| 9 | y | 1606 | 430.7 | 0.0001887 | 0.438 | +2 | 7 |
| - | - | 1784 | 431.1 | - | - | 0 | - |
| - | - | 1489 | 431.2 | - | - | 0 | - |
| - | - | 715 | 432.1 | - | - | 0 | - |
| - | - | 3720 | 432.9 | - | - | 0 | - |
| - | - | 3859 | 433.1 | - | - | 0 | - |
| - | - | 2493 | 433.2 | - | - | 0 | - |
| - | - | 609.5 | 434.1 | - | - | 0 | - |
| - | - | 9363 | 434.2 | - | - | 0 | - |
| - | - | 6100 | 434.5 | - | - | 0 | - |
| - | - | 4890 | 434.7 | - | - | 0 | - |
| - | - | 1183 | 435 | - | - | 0 | - |
| - | - | 963.6 | 435.2 | - | - | 0 | - |
| - | - | 1171 | 436.2 | - | - | 0 | - |
| 5 | y | 1435 | 437.2 | 0.002609 | 5.967 | +3 | 11 |
| - | - | 851.7 | 437.4 | - | - | 0 | - |
| 5 | y | 1315 | 437.5 | 0.005415 | 12.38 | +3 | 11 |
| - | - | 805 | 438.2 | - | - | 0 | - |
| - | - | 814.5 | 438.7 | - | - | 0 | - |
| 9 | y | 8986 | 439.2 | 0.001378 | 3.138 | +2 | 7 |
| - | - | 3259 | 439.7 | - | - | 0 | - |
| - | - | 705 | 440 | - | - | 0 | - |
| - | - | 2709 | 441.2 | - | - | 0 | - |
| - | - | 3860 | 441.5 | - | - | 0 | - |
| - | - | 1638 | 441.7 | - | - | 0 | - |
| - | - | 1081 | 442 | - | - | 0 | - |
| - | - | 1584 | 442.2 | - | - | 0 | - |
| - | - | 765.5 | 442.5 | - | - | 0 | - |
| - | - | 1576 | 443.2 | - | - | 0 | - |
| - | - | 3781 | 443.2 | - | - | 0 | - |
| - | - | 872.8 | 445.2 | - | - | 0 | - |
| 12 | y | 2088 | 445.2 | 0.0001378 | 0.3094 | +1 | 4 |
| 0 | Precursor | 1.396E+04 | 445.7 | 0.001324 | 2.97 | +4 | -1 |
| 11 | b | 1919 | 445.9 | 0.001109 | 2.486 | +3 | 11 |
| 0 | Precursor | 1.655E+04 | 446 | 0.00474 | 10.63 | +4 | -1 |
| - | - | 1.29E+04 | 446.2 | - | - | 0 | - |
| - | - | 7252 | 446.5 | - | - | 0 | - |
| - | - | 1015 | 446.5 | - | - | 0 | - |
| - | - | 4542 | 446.7 | - | - | 0 | - |
| - | - | 3291 | 447 | - | - | 0 | - |
| - | - | 2059 | 447.2 | - | - | 0 | - |
| - | - | 3138 | 447.4 | - | - | 0 | - |
| - | - | 3412 | 447.6 | - | - | 0 | - |
| - | - | 2754 | 447.7 | - | - | 0 | - |
| - | - | 1079 | 447.8 | - | - | 0 | - |
| - | - | 2031 | 448.2 | - | - | 0 | - |
| - | - | 804.4 | 448.9 | - | - | 0 | - |
| - | - | 1034 | 449.2 | - | - | 0 | - |
| - | - | 758 | 449.7 | - | - | 0 | - |
| - | - | 1039 | 450 | - | - | 0 | - |
| 0 | Precursor | 4.739E+04 | 450.2 | 0.001765 | 3.92 | +4 | -1 |
| - | - | 3.951E+04 | 450.5 | - | - | 0 | - |
| - | - | 1138 | 450.6 | - | - | 0 | - |
| - | - | 3.071E+04 | 450.7 | - | - | 0 | - |
| - | - | 2324 | 450.8 | - | - | 0 | - |
| - | - | 1.401E+04 | 451 | - | - | 0 | - |
| - | - | 1658 | 451 | - | - | 0 | - |
| - | - | 5591 | 451.2 | - | - | 0 | - |
| - | - | 1993 | 451.5 | - | - | 0 | - |
| - | - | 1087 | 451.7 | - | - | 0 | - |
| - | - | 910.2 | 452.2 | - | - | 0 | - |
| - | - | 1615 | 453.2 | - | - | 0 | - |
| - | - | 1054 | 453.9 | - | - | 0 | - |
| - | - | 2369 | 455.2 | - | - | 0 | - |
| - | - | 721 | 455.7 | - | - | 0 | - |
| - | - | 646.5 | 455.9 | - | - | 0 | - |
| - | - | 1390 | 459.9 | - | - | 0 | - |
| - | - | 887.7 | 460.2 | - | - | 0 | - |
| 8 | b | 3637 | 461.2 | 0.002436 | 5.281 | +2 | 8 |
| - | - | 3881 | 461.7 | - | - | 0 | - |
| - | - | 1692 | 462.2 | - | - | 0 | - |
| - | - | 838.7 | 462.7 | - | - | 0 | - |
| - | - | 713.1 | 463.2 | - | - | 0 | - |
| 12 | y | 1.694E+04 | 463.3 | 0.0004373 | 0.944 | +1 | 4 |
| - | - | 885.5 | 464.2 | - | - | 0 | - |
| - | - | 2827 | 464.3 | - | - | 0 | - |
| - | - | 972.4 | 464.5 | - | - | 0 | - |
| - | - | 803.7 | 464.9 | - | - | 0 | - |
| - | - | 2431 | 465.6 | - | - | 0 | - |
| - | - | 1130 | 465.9 | - | - | 0 | - |
| - | - | 1057 | 467.2 | - | - | 0 | - |
| - | - | 982.8 | 468.3 | - | - | 0 | - |
| - | - | 1279 | 470.2 | - | - | 0 | - |
| - | - | 975 | 470.6 | - | - | 0 | - |
| - | - | 761 | 470.7 | - | - | 0 | - |
| 4 | b | 4278 | 471.2 | 0.006559 | 13.92 | +1 | 4 |
| - | - | 947.8 | 473.2 | - | - | 0 | - |
| - | - | 779.7 | 473.5 | - | - | 0 | - |
| 12 | b | 1364 | 473.5 | 0.002552 | 5.388 | +3 | 12 |
| - | - | 731.9 | 474.2 | - | - | 0 | - |
| - | - | 1227 | 474.7 | - | - | 0 | - |
| - | - | 1109 | 475.9 | - | - | 0 | - |
| - | - | 1814 | 476.2 | - | - | 0 | - |
| - | - | 1555 | 476.6 | - | - | 0 | - |
| - | - | 1097 | 477.2 | - | - | 0 | - |
| - | - | 3133 | 477.5 | - | - | 0 | - |
| - | - | 1083 | 477.6 | - | - | 0 | - |
| - | - | 741.7 | 477.7 | - | - | 0 | - |
| - | - | 1247 | 478 | - | - | 0 | - |
| - | - | 889.7 | 478.2 | - | - | 0 | - |
| - | - | 2226 | 479.2 | - | - | 0 | - |
| 12 | b | 1369 | 479.5 | 0.003333 | 6.95 | +3 | 12 |
| - | - | 6999 | 479.7 | - | - | 0 | - |
| - | - | 792.2 | 479.9 | - | - | 0 | - |
| - | - | 3401 | 480.2 | - | - | 0 | - |
| - | - | 987.5 | 480.7 | - | - | 0 | - |
| - | - | 1360 | 481.2 | - | - | 0 | - |
| - | - | 1102 | 481.2 | - | - | 0 | - |
| - | - | 876.4 | 481.6 | - | - | 0 | - |
| - | - | 1257 | 481.9 | - | - | 0 | - |
| - | - | 1268 | 482.5 | - | - | 0 | - |
| 4 | y | 1528 | 482.9 | 0.0009067 | 1.878 | +3 | 12 |
| 4 | y | 3370 | 483.2 | 0.000111 | 0.2297 | +3 | 12 |
| - | - | 2572 | 483.6 | - | - | 0 | - |
| - | - | 1793 | 483.9 | - | - | 0 | - |
| - | - | 5602 | 485.2 | - | - | 0 | - |
| - | - | 1559 | 485.2 | - | - | 0 | - |
| - | - | 1052 | 485.6 | - | - | 0 | - |
| - | - | 1200 | 486.2 | - | - | 0 | - |
| - | - | 1438 | 487.6 | - | - | 0 | - |
| - | - | 1060 | 487.9 | - | - | 0 | - |
| - | - | 1581 | 488.2 | - | - | 0 | - |
| 4 | y | 9662 | 488.9 | 0.0005285 | 1.081 | +3 | 12 |
| - | - | 7561 | 489.2 | - | - | 0 | - |
| - | - | 2692 | 489.6 | - | - | 0 | - |
| - | - | 1270 | 490.7 | - | - | 0 | - |
| - | - | 1448 | 491.2 | - | - | 0 | - |
| - | - | 2188 | 491.6 | - | - | 0 | - |
| - | - | 923.6 | 491.9 | - | - | 0 | - |
| - | - | 739.6 | 492.2 | - | - | 0 | - |
| - | - | 5008 | 493.7 | - | - | 0 | - |
| - | - | 2486 | 494.2 | - | - | 0 | - |
| - | - | 712.5 | 495.9 | - | - | 0 | - |
| - | - | 6280 | 497.2 | - | - | 0 | - |
| - | - | 3754 | 497.6 | - | - | 0 | - |
| - | - | 2014 | 497.9 | - | - | 0 | - |
| - | - | 2218 | 498.2 | - | - | 0 | - |
| 8 | y | 1565 | 498.7 | 0.000266 | 0.5334 | +2 | 8 |
| 8 | y | 3673 | 499.2 | 0.002735 | 5.478 | +2 | 8 |
| - | - | 874.4 | 499.6 | - | - | 0 | - |
| - | - | 842.2 | 499.7 | - | - | 0 | - |
| - | - | 877.9 | 500.2 | - | - | 0 | - |
| - | - | 1175 | 501.6 | - | - | 0 | - |
| - | - | 1298 | 501.7 | - | - | 0 | - |
| - | - | 2205 | 502.2 | - | - | 0 | - |
| - | - | 4038 | 502.9 | - | - | 0 | - |
| - | - | 3173 | 503.2 | - | - | 0 | - |
| - | - | 1938 | 503.6 | - | - | 0 | - |
| - | - | 801.8 | 503.9 | - | - | 0 | - |
| - | - | 689.9 | 504.2 | - | - | 0 | - |
| - | - | 2596 | 504.9 | - | - | 0 | - |
| - | - | 3270 | 505.2 | - | - | 0 | - |
| - | - | 681.6 | 505.6 | - | - | 0 | - |
| - | - | 1276 | 506.3 | - | - | 0 | - |
| - | - | 814.8 | 506.6 | - | - | 0 | - |
| - | - | 1985 | 507.2 | - | - | 0 | - |
| - | - | 855 | 507.6 | - | - | 0 | - |
| 8 | y | 1.141E+04 | 507.7 | 0.0004463 | 0.8791 | +2 | 8 |
| - | - | 2271 | 507.9 | - | - | 0 | - |
| - | - | 6951 | 508.2 | - | - | 0 | - |
| - | - | 1464 | 508.7 | - | - | 0 | - |
| - | - | 3614 | 508.9 | - | - | 0 | - |
| - | - | 3627 | 509.2 | - | - | 0 | - |
| - | - | 1115 | 509.6 | - | - | 0 | - |
| - | - | 1531 | 510.2 | - | - | 0 | - |
| - | - | 3781 | 510.6 | - | - | 0 | - |
| - | - | 7502 | 510.9 | - | - | 0 | - |
| - | - | 6932 | 511.2 | - | - | 0 | - |
| - | - | 3439 | 511.6 | - | - | 0 | - |
| - | - | 1246 | 511.9 | - | - | 0 | - |
| - | - | 4800 | 512.2 | - | - | 0 | - |
| - | - | 1433 | 512.6 | - | - | 0 | - |
| - | - | 1068 | 513.2 | - | - | 0 | - |
| - | - | 1390 | 513.3 | - | - | 0 | - |
| - | - | 694.5 | 514.2 | - | - | 0 | - |
| 13 | b | 1568 | 516.2 | 0.008836 | 17.12 | +3 | 13 |
| - | - | 1.801E+04 | 516.6 | - | - | 0 | - |
| - | - | 1.187E+04 | 516.9 | - | - | 0 | - |
| - | - | 8230 | 517.2 | - | - | 0 | - |
| - | - | 3445 | 517.6 | - | - | 0 | - |
| - | - | 1152 | 517.9 | - | - | 0 | - |
| 9 | b | 6842 | 518.2 | 0.003708 | 7.154 | +2 | 9 |
| - | - | 1157 | 518.6 | - | - | 0 | - |
| - | - | 1287 | 518.7 | - | - | 0 | - |
| - | - | 965.4 | 519.2 | - | - | 0 | - |
| - | - | 1542 | 520.2 | - | - | 0 | - |
| - | - | 784.6 | 520.6 | - | - | 0 | - |
| - | - | 1034 | 521.3 | - | - | 0 | - |
| 13 | b | 1734 | 522.2 | 0.0002764 | 0.5293 | +3 | 13 |
| - | - | 847.3 | 523.6 | - | - | 0 | - |
| - | - | 1190 | 524.2 | - | - | 0 | - |
| - | - | 4743 | 526.2 | - | - | 0 | - |
| - | - | 2304 | 526.6 | - | - | 0 | - |
| - | - | 2165 | 526.9 | - | - | 0 | - |
| - | - | 909.7 | 527.2 | - | - | 0 | - |
| - | - | 1855 | 527.9 | - | - | 0 | - |
| - | - | 1413 | 528.2 | - | - | 0 | - |
| - | - | 1182 | 528.3 | - | - | 0 | - |
| - | - | 2239 | 529.3 | - | - | 0 | - |
| - | - | 1017 | 529.8 | - | - | 0 | - |
| - | - | 3449 | 530.2 | - | - | 0 | - |
| - | - | 1289 | 531.8 | - | - | 0 | - |
| 3 | y | 8738 | 531.9 | 0.001335 | 2.509 | +3 | 13 |
| 3 | y | 1.453E+04 | 532.2 | 0.004262 | 8.008 | +3 | 13 |
| - | - | 9850 | 532.6 | - | - | 0 | - |
| - | - | 6203 | 532.9 | - | - | 0 | - |
| - | - | 2885 | 533.2 | - | - | 0 | - |
| - | - | 6020 | 533.6 | - | - | 0 | - |
| - | - | 1907 | 533.9 | - | - | 0 | - |
| - | - | 2900 | 534.3 | - | - | 0 | - |
| - | - | 835.4 | 534.7 | - | - | 0 | - |
| - | - | 2640 | 536.2 | - | - | 0 | - |
| 3 | y | 5.168E+04 | 537.9 | 0.00178 | 3.31 | +3 | 13 |
| - | - | 4.179E+04 | 538.2 | - | - | 0 | - |
| - | - | 2.377E+04 | 538.6 | - | - | 0 | - |
| - | - | 1.123E+04 | 538.9 | - | - | 0 | - |
| - | - | 2778 | 539.2 | - | - | 0 | - |
| - | - | 5169 | 539.6 | - | - | 0 | - |
| - | - | 3413 | 539.9 | - | - | 0 | - |
| - | - | 1735 | 540.8 | - | - | 0 | - |
| - | - | 1188 | 541.3 | - | - | 0 | - |
| - | - | 740.9 | 541.8 | - | - | 0 | - |
| - | - | 5419 | 543.2 | - | - | 0 | - |
| - | - | 2534 | 543.7 | - | - | 0 | - |
| - | - | 1253 | 544.3 | - | - | 0 | - |
| - | - | 791 | 545.3 | - | - | 0 | - |
| - | - | 1215 | 545.3 | - | - | 0 | - |
| - | - | 1655 | 547.3 | - | - | 0 | - |
| - | - | 907.8 | 547.8 | - | - | 0 | - |
| - | - | 1863 | 548.2 | - | - | 0 | - |
| - | - | 2443 | 549.6 | - | - | 0 | - |
| - | - | 3343 | 549.9 | - | - | 0 | - |
| - | - | 2398 | 550.3 | - | - | 0 | - |
| - | - | 818.3 | 552.3 | - | - | 0 | - |
| - | - | 1007 | 552.7 | - | - | 0 | - |
| - | - | 846.3 | 553.6 | - | - | 0 | - |
| - | - | 1865 | 554.8 | - | - | 0 | - |
| 7 | y | 2134 | 555.3 | 0.008136 | 14.65 | +2 | 9 |
| 7 | y | 1126 | 555.8 | 0.0007722 | 1.389 | +2 | 9 |
| - | - | 1581 | 556.3 | - | - | 0 | - |
| - | - | 973.3 | 557.3 | - | - | 0 | - |
| 14 | b | 1441 | 559.3 | 0.004693 | 8.392 | +3 | 14 |
| - | - | 1020 | 559.6 | - | - | 0 | - |
| - | - | 1723 | 560.3 | - | - | 0 | - |
| - | - | 3038 | 561.2 | - | - | 0 | - |
| - | - | 1235 | 561.8 | - | - | 0 | - |
| - | - | 1634 | 562.2 | - | - | 0 | - |
| - | - | 1055 | 563.8 | - | - | 0 | - |
| 7 | y | 4419 | 564.3 | 0.001577 | 2.795 | +2 | 9 |
| - | - | 2745 | 564.8 | - | - | 0 | - |
| 14 | b | 2088 | 564.9 | 0.001662 | 2.943 | +3 | 14 |
| 2 | y | 6671 | 565.3 | 0.006482 | 11.47 | +3 | 14 |
| - | - | 3160 | 565.6 | - | - | 0 | - |
| - | - | 1295 | 565.9 | - | - | 0 | - |
| - | - | 959.1 | 566.3 | - | - | 0 | - |
| - | - | 1288 | 566.6 | - | - | 0 | - |
| - | - | 1003 | 566.9 | - | - | 0 | - |
| - | - | 1064 | 567.7 | - | - | 0 | - |
| - | - | 1886 | 570.3 | - | - | 0 | - |
| 2 | y | 1.031E+04 | 570.9 | 0.001864 | 3.265 | +3 | 14 |
| - | - | 9590 | 571.3 | - | - | 0 | - |
| - | - | 5072 | 571.6 | - | - | 0 | - |
| - | - | 3559 | 571.9 | - | - | 0 | - |
| - | - | 1557 | 572.3 | - | - | 0 | - |
| - | - | 9387 | 572.8 | - | - | 0 | - |
| - | - | 3548 | 573.3 | - | - | 0 | - |
| - | - | 1596 | 573.8 | - | - | 0 | - |
| - | - | 1708 | 574.3 | - | - | 0 | - |
| - | - | 2210 | 575.3 | - | - | 0 | - |
| - | - | 2747 | 575.7 | - | - | 0 | - |
| - | - | 2154 | 576.3 | - | - | 0 | - |
| - | - | 1141 | 576.8 | - | - | 0 | - |
| 10 | b | 932.7 | 577.8 | 0.001568 | 2.713 | +2 | 10 |
| 10 | b | 1178 | 578.3 | 0.005348 | 9.249 | +2 | 10 |
| - | - | 957.1 | 578.8 | - | - | 0 | - |
| 5 | b | 836.9 | 582.2 | 0.002241 | 3.848 | +1 | 5 |
| - | - | 1379 | 584.2 | - | - | 0 | - |
| - | - | 2519 | 584.8 | - | - | 0 | - |
| - | - | 1942 | 585.3 | - | - | 0 | - |
| - | - | 1022 | 585.8 | - | - | 0 | - |
| 10 | b | 7657 | 586.8 | 0.00245 | 4.175 | +2 | 10 |
| - | - | 6066 | 587.3 | - | - | 0 | - |
| - | - | 3041 | 587.8 | - | - | 0 | - |
| - | - | 1879 | 588.3 | - | - | 0 | - |
| - | - | 1524 | 589.3 | - | - | 0 | - |
| 6 | y | 1143 | 591.3 | 0.001488 | 2.517 | +2 | 10 |
| - | - | 1016 | 593.8 | - | - | 0 | - |
| 0 | Precursor | 1572 | 594.3 | 0.002337 | 3.932 | +3 | -1 |
| - | - | 1422 | 594.8 | - | - | 0 | - |
| - | - | 5319 | 598.3 | - | - | 0 | - |
| - | - | 1453 | 599.3 | - | - | 0 | - |
| 6 | y | 3201 | 599.8 | 0.0002517 | 0.4197 | +2 | 10 |
| 5 | b | 5186 | 600.2 | 0.004188 | 6.977 | +1 | 5 |
| - | - | 3347 | 600.3 | - | - | 0 | - |
| - | - | 1858 | 600.8 | - | - | 0 | - |
| - | - | 1489 | 601.2 | - | - | 0 | - |
| - | - | 795.8 | 601.3 | - | - | 0 | - |
| - | - | 1227 | 603.3 | - | - | 0 | - |
| - | - | 851.7 | 603.8 | - | - | 0 | - |
| - | - | 4857 | 607.3 | - | - | 0 | - |
| - | - | 1061 | 607.8 | - | - | 0 | - |
| - | - | 1129 | 608.3 | - | - | 0 | - |
| 11 | y | 1295 | 608.3 | 0.002361 | 3.881 | +1 | 5 |
| 11 | y | 1286 | 609.3 | 0.005467 | 8.973 | +1 | 5 |
| - | - | 751.1 | 610.2 | - | - | 0 | - |
| - | - | 862.3 | 611.8 | - | - | 0 | - |
| - | - | 1164 | 612.3 | - | - | 0 | - |
| - | - | 1529 | 616.3 | - | - | 0 | - |
| - | - | 3949 | 616.8 | - | - | 0 | - |
| - | - | 3928 | 617.3 | - | - | 0 | - |
| - | - | 1200 | 617.8 | - | - | 0 | - |
| - | - | 918.3 | 618.2 | - | - | 0 | - |
| - | - | 888.4 | 618.8 | - | - | 0 | - |
| - | - | 960.4 | 619.2 | - | - | 0 | - |
| - | - | 970.8 | 621.8 | - | - | 0 | - |
| - | - | 918.2 | 624.8 | - | - | 0 | - |
| - | - | 2982 | 625.8 | - | - | 0 | - |
| 11 | y | 8820 | 626.3 | 0.0004522 | 0.7221 | +1 | 5 |
| - | - | 2060 | 627.3 | - | - | 0 | - |
| - | - | 1169 | 633.3 | - | - | 0 | - |
| - | - | 920.1 | 634.3 | - | - | 0 | - |
| - | - | 2003 | 635.3 | - | - | 0 | - |
| - | - | 972.9 | 636.8 | - | - | 0 | - |
| - | - | 1098 | 639.3 | - | - | 0 | - |
| - | - | 829.8 | 643.3 | - | - | 0 | - |
| - | - | 1216 | 644.3 | - | - | 0 | - |
| - | - | 1515 | 646.8 | - | - | 0 | - |
| - | - | 1241 | 648.8 | - | - | 0 | - |
| - | - | 1258 | 649.3 | - | - | 0 | - |
| - | - | 1006 | 649.8 | - | - | 0 | - |
| - | - | 986.7 | 650.3 | - | - | 0 | - |
| 6 | b | 2760 | 653.3 | 0.009744 | 14.92 | +1 | 6 |
| - | - | 1289 | 654.3 | - | - | 0 | - |
| 5 | y | 2186 | 655.3 | 0.0005037 | 0.7686 | +2 | 11 |
| 5 | y | 3703 | 655.8 | 0.005413 | 8.254 | +2 | 11 |
| - | - | 2278 | 656.3 | - | - | 0 | - |
| - | - | 1830 | 657.3 | - | - | 0 | - |
| - | - | 1388 | 657.8 | - | - | 0 | - |
| - | - | 1950 | 658.3 | - | - | 0 | - |
| - | - | 941.5 | 658.8 | - | - | 0 | - |
| 5 | y | 6331 | 664.3 | 0.001025 | 1.543 | +2 | 11 |
| - | - | 4423 | 664.8 | - | - | 0 | - |
| - | - | 3053 | 665.3 | - | - | 0 | - |
| - | - | 876.5 | 665.8 | - | - | 0 | - |
| - | - | 1204 | 666.3 | - | - | 0 | - |
| - | - | 1666 | 667.3 | - | - | 0 | - |
| - | - | 828.9 | 667.8 | - | - | 0 | - |
| 11 | b | 1709 | 668.3 | 0.005759 | 8.617 | +2 | 11 |
| - | - | 1139 | 668.8 | - | - | 0 | - |
| 6 | b | 6994 | 671.3 | 0.004367 | 6.505 | +1 | 6 |
| - | - | 1418 | 671.3 | - | - | 0 | - |
| - | - | 3632 | 672.3 | - | - | 0 | - |
| - | - | 2607 | 674.3 | - | - | 0 | - |
| - | - | 1028 | 675.3 | - | - | 0 | - |
| - | - | 1072 | 675.8 | - | - | 0 | - |
| - | - | 805.6 | 676.3 | - | - | 0 | - |
| - | - | 1935 | 680.8 | - | - | 0 | - |
| - | - | 2099 | 681.3 | - | - | 0 | - |
| - | - | 1530 | 681.8 | - | - | 0 | - |
| - | - | 961.2 | 683.3 | - | - | 0 | - |
| - | - | 1973 | 684.3 | - | - | 0 | - |
| - | - | 1251 | 685.3 | - | - | 0 | - |
| - | - | 3766 | 689.8 | - | - | 0 | - |
| - | - | 3557 | 690.3 | - | - | 0 | - |
| - | - | 2172 | 690.8 | - | - | 0 | - |
| - | - | 936.9 | 697.3 | - | - | 0 | - |
| - | - | 4310 | 702.3 | - | - | 0 | - |
| - | - | 1969 | 703.3 | - | - | 0 | - |
| - | - | 948.2 | 704.3 | - | - | 0 | - |
| - | - | 1530 | 707.3 | - | - | 0 | - |
| 12 | b | 2088 | 709.8 | 0.004048 | 5.702 | +2 | 12 |
| - | - | 2359 | 710.3 | - | - | 0 | - |
| - | - | 777 | 713.3 | - | - | 0 | - |
| - | - | 881.4 | 715.3 | - | - | 0 | - |
| - | - | 989.2 | 718.3 | - | - | 0 | - |
| 12 | b | 991.3 | 718.8 | 1.409E-05 | 0.0196 | +2 | 12 |
| - | - | 1726 | 719.3 | - | - | 0 | - |
| 4 | y | 1000 | 724.3 | 0.005926 | 8.181 | +2 | 12 |
| - | - | 1075 | 724.8 | - | - | 0 | - |
| - | - | 1481 | 726.3 | - | - | 0 | - |
| - | - | 1194 | 726.8 | - | - | 0 | - |
| 4 | y | 3982 | 732.9 | 0.0002078 | 0.2835 | +2 | 12 |
| - | - | 4328 | 733.4 | - | - | 0 | - |
| - | - | 1251 | 733.9 | - | - | 0 | - |
| - | - | 1924 | 735.3 | - | - | 0 | - |
| - | - | 1070 | 736.3 | - | - | 0 | - |
| - | - | 938.5 | 737.3 | - | - | 0 | - |
| - | - | 1500 | 739.3 | - | - | 0 | - |
| - | - | 793.2 | 742.3 | - | - | 0 | - |
| 10 | y | 930.9 | 746.3 | 0.001731 | 2.319 | +1 | 6 |
| - | - | 1151 | 754.4 | - | - | 0 | - |
| 10 | y | 3689 | 763.4 | 0.0002821 | 0.3695 | +1 | 6 |
| - | - | 1218 | 764.3 | - | - | 0 | - |
| - | - | 1695 | 764.4 | - | - | 0 | - |
| 7 | b | 1003 | 766.4 | 0.0143 | 18.66 | +1 | 7 |
| - | - | 1119 | 768.3 | - | - | 0 | - |
| - | - | 883.5 | 772.4 | - | - | 0 | - |
| - | - | 1961 | 781.3 | - | - | 0 | - |
| - | - | 744.3 | 782.3 | - | - | 0 | - |
| 13 | b | 1294 | 782.8 | 0.003473 | 4.436 | +2 | 13 |
| - | - | 1249 | 783.3 | - | - | 0 | - |
| 7 | b | 3876 | 784.4 | 0.002578 | 3.287 | +1 | 7 |
| - | - | 3511 | 785.4 | - | - | 0 | - |
| - | - | 1367 | 786.4 | - | - | 0 | - |
| - | - | 902.3 | 802.4 | - | - | 0 | - |
| - | - | 883.2 | 811.4 | - | - | 0 | - |
| - | - | 1605 | 819.4 | - | - | 0 | - |
| - | - | 1084 | 821.4 | - | - | 0 | - |
| - | - | 1493 | 837.4 | - | - | 0 | - |
| - | - | 920.7 | 838.4 | - | - | 0 | - |
| - | - | 1455 | 839.4 | - | - | 0 | - |
| - | - | 2383 | 849.4 | - | - | 0 | - |
| - | - | 1477 | 850.4 | - | - | 0 | - |
| 9 | y | 1068 | 860.4 | 0.004392 | 5.105 | +1 | 7 |
| - | - | 726.2 | 861.4 | - | - | 0 | - |
| - | - | 1744 | 865.4 | - | - | 0 | - |
| 9 | y | 3408 | 877.4 | 0.003048 | 3.474 | +1 | 7 |
| - | - | 1157 | 878.4 | - | - | 0 | - |
| - | - | 753 | 879.4 | - | - | 0 | - |
| - | - | 1046 | 894.4 | - | - | 0 | - |
| - | - | 848.5 | 895.4 | - | - | 0 | - |
| 8 | b | 1431 | 921.4 | 0.004885 | 5.301 | +1 | 8 |
| - | - | 1293 | 922.4 | - | - | 0 | - |
| - | - | 1177 | 930.4 | - | - | 0 | - |
| - | - | 2547 | 948.4 | - | - | 0 | - |
| - | - | 1282 | 949.4 | - | - | 0 | - |
| - | - | 1000 | 965.5 | - | - | 0 | - |
| - | - | 1345 | 966.5 | - | - | 0 | - |
| - | - | 715.3 | 971.5 | - | - | 0 | - |
| - | - | 690.7 | 975.5 | - | - | 0 | - |
| - | - | 853.2 | 987.4 | - | - | 0 | - |
| - | - | 804.6 | 1002 | - | - | 0 | - |
| 8 | y | 802.1 | 1014 | 0.005381 | 5.304 | +1 | 8 |
| 9 | b | 2603 | 1035 | 0.001874 | 1.81 | +1 | 9 |
| - | - | 840.2 | 1037 | - | - | 0 | - |
| - | - | 1002 | 1076 | - | - | 0 | - |
| - | - | 809.4 | 1085 | - | - | 0 | - |
| - | - | 1101 | 1086 | - | - | 0 | - |
| - | - | 945.6 | 1095 | - | - | 0 | - |
| - | - | 803.6 | 1096 | - | - | 0 | - |
| - | - | 741.3 | 2028 | - | - | 0 | - |
| - | - | 673.7 | 2300 | - | - | 0 | - |

m/z Charge Intensity FragmentType MassShift Position
120.0658950805664 0 617.79736
120.08125305175781 0 577.36456
121.04004669189453 0 484.74078
121.74261474609375 0 386.96643
122.07168579101562 0 638.67004
123.05562591552734 0 841.63824
127.08702087402344 0 884.79694
129.0662078857422 0 1551.7283
129.10264587402344 0 91733.62
130.05007934570312 0 433.48428
130.0866241455078 0 1921.2931
130.10595703125 0 5410.203
132.10223388671875 0 420.10397
134.02732849121094 0 766.2103
136.07608032226562 0 55805.535
137.07940673828125 0 4039.2585
138.06666564941406 0 2149.9128
139.0869598388672 0 1677.4648
141.06639099121094 0 500.82312
141.10260009765625 0 4021.2595 a Water loss 1
147.0447998046875 0 665.4976
147.11305236816406 0 1661.8171
148.08724975585938 0 2543.9363
148.9544219970703 0 734.6606
149.0233612060547 0 510.2209
149.0453338623047 0 683.35364
151.0423583984375 0 478.65753
152.0823974609375 0 692.01886
153.10256958007812 0 512.5121
154.0867919921875 0 518.0379
155.0817108154297 0 1283.2268
155.0931396484375 0 1986.0565
155.1183624267578 0 2257.518
156.0772247314453 0 3550.3186
157.13369750976562 0 1252.0052
159.11322021484375 0 179125.7 a 1
160.1100311279297 0 1012.8904
160.11656188964844 0 12745.278
165.07061767578125 0 538.808
165.07757568359375 0 2371.6667
166.06143188476562 0 4952.665
166.08657836914062 0 10406.764
167.055419921875 0 563.9592
167.0901641845703 0 882.75714
167.118408203125 0 888.9586
169.0529327392578 0 1293.8907
169.0607452392578 0 752.9333
171.11317443847656 0 2097.363
173.0924530029297 0 1090.1862 y Ammonia loss 12
176.0822296142578 0 5279.035
177.11375427246094 0 859.55536
178.13430786132812 0 821.69116
179.09310913085938 0 1054.7212
180.07705688476562 0 725.035
180.11386108398438 0 1355.2249
181.09027099609375 0 612.31976
183.07647705078125 0 1470.5061
183.1131591796875 0 4929.3403
183.14939880371094 0 780.70807
185.1294403076172 0 1237.6024
185.16543579101562 0 553.84705
187.10809326171875 0 26443.521 b 1
188.11151123046875 0 2199.2976
189.07757568359375 0 1178.2677
190.06158447265625 0 2205.8757
191.093505859375 0 818.1431
192.10220336914062 0 2233.1606
193.07254028320312 0 1120.7367
193.10865783691406 0 8951.405
194.08030700683594 0 1053.0632
194.0923614501953 0 1758.5117
194.11212158203125 0 678.27527
194.12838745117188 0 588.89905
195.07669067382812 0 1609.6831
195.08767700195312 0 594.66486
195.1132354736328 0 3503.8296
199.10792541503906 0 1794.2686
201.08738708496094 0 1807.07
202.10894775390625 0 860.9051
202.11880493164062 0 648.9374
204.07716369628906 0 7323.9473
204.13479614257812 0 1633.0819
205.08042907714844 0 551.73535
205.09767150878906 0 591.7857
206.10385131835938 0 968.6679
207.08802795410156 0 5878.9863
208.0714569091797 0 806.84015
208.95359802246094 0 1640.561
209.09336853027344 0 902.56476
210.95077514648438 0 2678.7676
211.1085968017578 0 755.43616
211.14486694335938 0 564.20685
212.1034698486328 0 993.2381
216.13475036621094 0 3681.1968 y Water loss 13
217.0726776123047 0 1452.7882
217.11868286132812 0 3920.0693 y Ammonia loss 13
218.05642700195312 0 1788.6616
220.0966796875 0 512.5357
221.10362243652344 0 24074.252
222.0878143310547 0 985.2955
222.10740661621094 0 2513.96
222.1240692138672 0 3567.6553
223.0634002685547 0 1079.4484
223.1078338623047 0 1152.8586
223.15541076660156 0 1894.0249
224.11456298828125 0 6253.456
225.04312133789062 0 1162.124
225.06121826171875 0 1130.4844
225.09878540039062 0 1969.7297
225.1184844970703 0 719.2835
226.04344177246094 0 601.93567
226.08287048339844 0 1222.2666
226.11817932128906 0 981.5882
226.15626525878906 0 1149.1884
227.01368713378906 0 1126.9023
227.03961181640625 0 1167.513
228.11415100097656 0 602.2805
230.08358764648438 0 1924.2491
230.11390686035156 0 6889.417
231.0866241455078 0 649.3747
233.13914489746094 0 957.6766
234.09877014160156 0 1523.2383
234.12399291992188 0 1374.5188
234.14524841308594 0 19297.617 y 13
235.08302307128906 0 10709.932
235.1485595703125 0 1404.3223
236.06710815429688 0 714.36414
236.08612060546875 0 1145.1046
236.10513305664062 0 1430.1294 b 3
237.12380981445312 0 2974.3052
239.11428833007812 0 3179.981
239.15066528320312 0 10018.395
240.08079528808594 0 521.3129
240.09835815429688 0 729.44336
240.13470458984375 0 22709.09
241.13829040527344 0 2337.667
244.1296844482422 0 692.9374
247.1101837158203 0 3854.915
249.0985107421875 0 3759.4595
251.15074157714844 0 3736.2114
252.10952758789062 0 21222.332
252.1337432861328 0 1239.9137
252.15455627441406 0 570.831
253.09347534179688 0 2488.3389
253.113037109375 0 1727.5825
257.1072692871094 0 3103.2656
257.1609802246094 0 3206.0425
258.1451110839844 0 1572.188
261.13525390625 0 706.82666
262.0941162109375 0 972.4352
263.0701904296875 0 493.0742
264.134765625 0 1032.4747
265.1182861328125 0 2415.3125
267.09222412109375 0 981.1772
267.1090393066406 0 9688.379
268.0751647949219 0 935.69354
268.1124267578125 0 1333.0409
269.1361999511719 0 648.1126
270.1451110839844 0 2110.905
273.13482666015625 0 5940.3145
275.1063537597656 0 2333.2327
276.109619140625 0 788.7676
276.1825866699219 0 589.72144
277.0935363769531 0 1516.6934
279.14556884765625 0 1787.6091
280.1036071777344 0 1363.7849
280.1297912597656 0 3818.866
280.6509094238281 0 768.6009
281.0885925292969 0 730.8313
282.1444396972656 0 563.3589
284.103271484375 0 637.29694
285.1020812988281 0 13412.757
286.1053771972656 0 1287.4187
286.62823486328125 0 1055.5245
287.1003112792969 0 610.4499
290.14874267578125 0 893.3675
293.0997009277344 0 847.53705
294.64862060546875 0 971.9705
295.1039733886719 0 1983.7961
297.15545654296875 0 1531.6646
300.6265869140625 0 1482.1326 b 4
301.12994384765625 0 7676.584
302.1311340332031 0 1750.956
302.3335876464844 0 519.67993
304.14056396484375 0 923.9211
304.1773681640625 0 1106.2526
305.12579345703125 0 698.0647
305.1607666015625 0 986.0034
309.1198425292969 0 921.69403
310.1401672363281 0 774.30115
313.6607666015625 0 4197.874 y 10
314.1625061035156 0 964.2015
318.1375732421875 0 647.1707
319.14093017578125 0 639.74115
320.1357421875 0 6376.261
320.1720886230469 0 2302.4502
320.9196472167969 0 2932.5325
321.13861083984375 0 913.5036
321.9219055175781 0 521.814
322.11492919921875 0 1210.6128
322.1478271484375 0 4402.9023
322.1878662109375 0 2489.9343
322.6494445800781 0 770.478
322.6714782714844 0 763.49567
322.91650390625 0 1635.0203
323.1714172363281 0 1782.1135
325.114990234375 0 1113.6235
326.1363525390625 0 1097.3695
326.17144775390625 0 1463.5361
327.1403503417969 0 952.5686 b Water loss 5
327.1658020019531 0 1107.6041
328.1524658203125 0 871.0066
328.6673583984375 0 895.65814
332.1357421875 0 1925.917
333.1197509765625 0 740.20355
334.1437683105469 0 3328.9707 b 2
336.14495849609375 0 821.4035 b 5
336.16668701171875 0 1125.8967
336.66741943359375 0 633.8554
338.14642333984375 0 5299.505
338.83062744140625 0 1997.6411 y 7
339.1655578613281 0 529.61835
339.49755859375 0 685.375
340.14141845703125 0 2538.004
340.1997985839844 0 809.44574
341.1825866699219 0 2481.9954
342.1871337890625 0 576.2987
343.16229248046875 0 677.302
343.1975402832031 0 1477.7493
344.14666748046875 0 2443.9045
344.1934814453125 0 4413.1333 y Water loss 12
345.1540832519531 0 869.894
345.17718505859375 0 4285.4404 y Ammonia loss 12
345.1969299316406 0 577.88605
346.1806945800781 0 592.77856
348.1302490234375 0 2317.8933
348.1663818359375 0 1050.4061
350.1461181640625 0 7236.5303
351.1498718261719 0 1450.7007
351.66864013671875 0 1442.2454
352.1702575683594 0 690.0715
354.1312561035156 0 1279.1931
354.1698913574219 0 1819.2281
354.8455810546875 0 1105.5831
356.172607421875 0 662.19684
358.2088928222656 0 2834.4917
360.0298156738281 0 803.461
360.85107421875 0 793.4672
361.0268859863281 0 3889.209
361.17376708984375 0 2053.8853
362.0268249511719 0 3579.499
362.1802978515625 0 989.3953
362.20367431640625 0 4214.21 y 12
363.0239562988281 0 8526.061
363.2066650390625 0 900.1539
364.0242919921875 0 1866.3987
365.19329833984375 0 1480.6267
366.138916015625 0 1638.8324
366.1575927734375 0 2090.7415
367.16339111328125 0 624.3323
368.1399230957031 0 609.336
368.1654052734375 0 1632.9736
369.7359313964844 0 1058.1989
370.15045166015625 0 846.42566
371.15753173828125 0 6030.027
371.181884765625 0 758.7069
372.1416015625 0 3390.9307
373.6773376464844 0 949.4439 y Ammonia loss 9
375.1676940917969 0 1937.1736
376.1695556640625 0 594.3598
376.2194519042969 0 1078.6775
376.5250244140625 0 936.27167 y 6
378.6885070800781 0 951.1287
379.2113037109375 0 1103.3093
380.1337585449219 0 730.83154
382.1864929199219 0 5190.8774 y 9
382.5156555175781 0 2082.205
382.69293212890625 0 1157.5525
383.1830139160156 0 855.2768
384.1673583984375 0 2538.864
384.5062255859375 0 1009.17377
384.6756896972656 0 906.23773
385.17144775390625 0 1405.4988
386.1462097167969 0 1035.2981
387.6893615722656 0 838.6294
389.16876220703125 0 2182.3137
390.1507873535156 0 660.2699
390.5151062011719 0 774.21606
391.51336669921875 0 2578.3152 b 9
391.8486328125 0 785.9242
393.1513671875 0 761.6297
393.1835021972656 0 1844.9446
393.68505859375 0 649.74774
394.15374755859375 0 750.3962
396.18719482421875 0 627.61194
397.1626892089844 0 1226.0864
397.6904296875 0 826.01666
398.1459655761719 0 1531.632
400.204345703125 0 1220.5046 y 5
400.86834716796875 0 762.5037
402.1775207519531 0 1566.0663
403.6905822753906 0 1718.6149 y 2
403.93817138671875 0 2565.8494
404.15667724609375 0 1674.5675
405.2246398925781 0 1529.3348
406.2024841308594 0 3246.5046
406.7015380859375 0 1797.649
407.2047424316406 0 1891.9879
410.20196533203125 0 1086.0673
411.1923828125 0 904.762
411.5149230957031 0 981.6457
411.6884765625 0 934.1208
412.1632385253906 0 884.16724
412.4573974609375 0 849.01825
412.7076721191406 0 911.2685
414.1444396972656 0 6812.8945
414.9443359375 0 647.494
415.03753662109375 0 695.19745
415.1460266113281 0 1229.2683
415.17291259765625 0 3690.9097
415.1961975097656 0 515.33325
415.44390869140625 0 776.8387
416.147216796875 0 733.9279
416.17822265625 0 1271.7699
416.2859802246094 0 610.5911
417.5163879394531 0 646.5918
419.69677734375 0 1504.369 b Ammonia loss 13
419.7427673339844 0 1290.165
419.94757080078125 0 2142.409
420.1983337402344 0 3806.6558
420.6971435546875 0 1084.0557
420.9526062011719 0 876.73096
421.1840515136719 0 6298.8228
421.6915283203125 0 1216.7129
422.1856689453125 0 2111.7825
423.2348937988281 0 2228.4714
423.9541320800781 0 2078.3997 b 13
424.201416015625 0 1683.6895 y Ammonia loss 1
424.4539489746094 0 1913.5747
424.6965637207031 0 952.9028
425.1900634765625 0 3295.1892
425.45709228515625 0 801.35565
425.6886901855469 0 2172.061
426.1794738769531 0 970.3847
427.7078857421875 0 1497.5605
428.21478271484375 0 797.49585
428.4560852050781 0 3194.7773 y 1
428.7060546875 0 4727.962
428.9555358886719 0 1872.1694
429.22186279296875 0 799.43555
429.7111511230469 0 3794.8564
429.96087646484375 0 3623.103
430.2099304199219 0 3715.3135 y Water loss 8
430.4617919921875 0 1060.9816
430.69830322265625 0 1606.4972 y Ammonia loss 8
431.0877685546875 0 1784.1699
431.20428466796875 0 1488.697
432.09051513671875 0 714.9664
432.8856201171875 0 3719.6895
433.08416748046875 0 3858.5198
433.2196960449219 0 2493.0012
434.087890625 0 609.47003
434.2143249511719 0 9362.7
434.46441650390625 0 6100.24
434.7149963378906 0 4890.234
434.9655456542969 0 1183.4888
435.21319580078125 0 963.60803
436.2312927246094 0 1171.3785
437.2117004394531 0 1435.1096 y Water loss 4
437.4497375488281 0 851.7134
437.5477294921875 0 1315.2991 y Ammonia loss 4
438.1993713378906 0 805.0483
438.7102966308594 0 814.5348
439.21038818359375 0 8985.707 y 8
439.7098388671875 0 3258.9539
439.9548034667969 0 705.0112
441.20758056640625 0 2708.7188
441.45758056640625 0 3860.1987
441.7068786621094 0 1637.8923
441.95947265625 0 1081.4316
442.2076110839844 0 1583.5502
442.5416259765625 0 765.5449
443.16815185546875 0 1575.9445
443.2084045410156 0 3781.3962
445.20623779296875 0 872.7735
445.24066162109375 0 2087.8162 y Water loss 11
445.7113342285156 0 13964.556 Precursor Water loss
445.86761474609375 0 1919.0786 b 10
445.96075439453125 0 16545.879 Precursor Ammonia loss
446.2106628417969 0 12896.8125
446.4605407714844 0 7252.0537
446.53656005859375 0 1015.0456
446.7099304199219 0 4541.7466
446.957763671875 0 3290.8608
447.232177734375 0 2058.6719
447.4339599609375 0 3137.907
447.63427734375 0 3411.6157
447.7193298339844 0 2754.4292
447.8349914550781 0 1078.6337
448.2183837890625 0 2030.6779
448.88726806640625 0 804.4059
449.2145080566406 0 1033.9375
449.7152099609375 0 758.0325
449.959716796875 0 1039.1782
450.21441650390625 0 47392.668 Precursor
450.4642639160156 0 39508.996
450.63916015625 0 1138.4214
450.7144470214844 0 30705.295
450.83935546875 0 2324.1443
450.9644470214844 0 14005.547
451.0411376953125 0 1658.3934
451.2292785644531 0 5590.634
451.462158203125 0 1992.5765
451.71026611328125 0 1087.4994
452.2376403808594 0 910.22906
453.2027282714844 0 1615.0265
453.8892822265625 0 1054.3025
455.2195739746094 0 2369.2087
455.71905517578125 0 721.02155
455.8798522949219 0 646.4675
459.8939208984375 0 1389.7263
460.2254638671875 0 887.6767
461.2159729003906 0 3636.5854 b 7
461.71795654296875 0 3880.9678
462.2135314941406 0 1691.9463
462.7172546386719 0 838.67395
463.2171325683594 0 713.1197
463.25152587890625 0 16935.176 y 11
464.1568603515625 0 885.5298
464.25518798828125 0 2826.9873
464.54888916015625 0 972.3701
464.88189697265625 0 803.67267
465.5584411621094 0 2431.1367
465.8948059082031 0 1130.3098
467.21466064453125 0 1056.8428
468.2586975097656 0 982.7624
470.2219543457031 0 1279.2722
470.5520324707031 0 975.02747
470.7131652832031 0 760.9849
471.20367431640625 0 4277.6694 b 3
473.2034912109375 0 947.8117
473.4910583496094 0 779.7141
473.548095703125 0 1364.2559 b Water loss 11
474.2169189453125 0 731.91797
474.7222900390625 0 1226.7175
475.8940734863281 0 1108.5176
476.22772216796875 0 1814.0752
476.5632019042969 0 1554.83
477.2309265136719 0 1096.7378
477.4991149902344 0 3133.0173
477.5544738769531 0 1082.7484
477.7486267089844 0 741.7119
478.0027160644531 0 1246.8641
478.2497253417969 0 889.7276
479.2239074707031 0 2225.8223
479.5523986816406 0 1369.2792 b 11
479.7197265625 0 6999.2446
479.8855895996094 0 792.1788
480.2198181152344 0 3400.5796
480.72320556640625 0 987.4937
481.18341064453125 0 1359.8254
481.2186279296875 0 1101.8397
481.56927490234375 0 876.43713
481.9000244140625 0 1256.7153
482.5498046875 0 1268.4031
482.9015197753906 0 1527.6184 y Water loss 3
483.2287292480469 0 3369.5283 y Ammonia loss 3
483.5619201660156 0 2571.7083
483.8979797363281 0 1792.5529
485.1813659667969 0 5601.974
485.219482421875 0 1558.7965
485.5511474609375 0 1052.0566
486.2333984375 0 1200.3583
487.5726623535156 0 1438.0225
487.90484619140625 0 1060.0057
488.23199462890625 0 1580.7317
488.9046630859375 0 9662.305 y 3
489.2380065917969 0 7561.4263
489.5724182128906 0 2692.1506
490.7191162109375 0 1269.5767
491.2232666015625 0 1448.4879
491.55645751953125 0 2187.7627
491.8926696777344 0 923.56915
492.2186584472656 0 739.61584
493.7166442871094 0 5007.6504
494.21728515625 0 2486.2283
495.9012145996094 0 712.5467
497.2264404296875 0 6279.9595
497.56103515625 0 3754.1917
497.89385986328125 0 2013.6744
498.21148681640625 0 2218.4622
498.7362060546875 0 1565.1011 y Water loss 7
499.2306823730469 0 3673.0837 y Ammonia loss 7
499.56011962890625 0 874.3962
499.73028564453125 0 842.2112
500.2315368652344 0 877.8884
501.56622314453125 0 1175.1981
501.7301940917969 0 1297.522
502.234375 0 2204.734
502.90203857421875 0 4037.912
503.2367858886719 0 3173.006
503.56646728515625 0 1937.6023
503.9007873535156 0 801.83325
504.2420959472656 0 689.8558
504.90380859375 0 2595.8997
505.2367248535156 0 3269.5042
505.56866455078125 0 681.62964
506.2734069824219 0 1275.7924
506.5625 0 814.8132
507.2386169433594 0 1985.41
507.5750427246094 0 855.01953
507.7416687011719 0 11411.451 y 7
507.90863037109375 0 2270.5173
508.24200439453125 0 6950.5244
508.7414245605469 0 1464.3645
508.9058837890625 0 3614.2058
509.24053955078125 0 3626.9207
509.57086181640625 0 1114.9691
510.2362976074219 0 1531.1477
510.57843017578125 0 3780.7437
510.9089660644531 0 7501.9497
511.2427673339844 0 6932.4517
511.576416015625 0 3438.9111
511.9086608886719 0 1246.2308
512.2381591796875 0 4800.17
512.5747680664062 0 1433.1515
513.2158813476562 0 1067.504
513.3154907226562 0 1389.5717
514.2269897460938 0 694.4735
516.222900390625 0 1567.5408 b Water loss 12
516.5831298828125 0 18013.258
516.916015625 0 11870.328
517.2494506835938 0 8230.266
517.5839233398438 0 3444.8
517.9174194335938 0 1151.8561
518.2387084960938 0 6842.0264 b 8
518.5760498046875 0 1157.0802
518.7372436523438 0 1287.0193
519.2345581054688 0 965.41547
520.2473754882812 0 1541.7202
520.5634765625 0 784.63635
521.2731323242188 0 1033.5172
522.2355346679688 0 1733.8389 b 12
523.572998046875 0 847.3213
524.2385864257812 0 1190.3595
526.2366943359375 0 4743.1733
526.56884765625 0 2304.2673
526.902099609375 0 2164.5022
527.23388671875 0 909.6825
527.9026489257812 0 1854.7639
528.2300415039062 0 1413.2527
528.2704467773438 0 1182.4467
529.25244140625 0 2239.2659
529.7537231445312 0 1017.4919
530.2443237304688 0 3449.189
531.7679443359375 0 1289.4915
531.912109375 0 8738.492 y Water loss 2
532.2430419921875 0 14529.101 y Ammonia loss 2
532.5758666992188 0 9850.354
532.9088745117188 0 6202.8257
533.2451171875 0 2884.633
533.5726318359375 0 6019.8193
533.9049682617188 0 1907.3286
534.2671508789062 0 2899.8938
534.7374877929688 0 835.3778
536.24755859375 0 2639.7864
537.9160766601562 0 51675.605 y 2
538.24951171875 0 41793.72
538.5828857421875 0 23766.76
538.917236328125 0 11226.134
539.2496337890625 0 2778.272
539.5758056640625 0 5169.0913
539.909423828125 0 3413.2725
540.7704467773438 0 1734.7302
541.272216796875 0 1187.6183
541.7723999023438 0 740.9227
543.2496948242188 0 5419.2
543.7467651367188 0 2534.3704
544.2579956054688 0 1253.1703
545.2564697265625 0 790.9568
545.296630859375 0 1215.3723
547.2632446289062 0 1655.385
547.7811279296875 0 907.7573
548.2476806640625 0 1862.8318
549.6065063476562 0 2442.999
549.939453125 0 3343.0354
550.2728271484375 0 2398.1702
552.2623291015625 0 818.2675
552.747802734375 0 1007.3623
553.585205078125 0 846.328
554.7665405273438 0 1864.8167
555.2698364257812 0 2133.5408 y Water loss 6
555.770751953125 0 1125.8401 y Ammonia loss 6
556.2789306640625 0 1580.8818
557.2501831054688 0 973.2722
559.2627563476562 0 1441.2234 b Ammonia loss 13
559.5933837890625 0 1019.8352
560.2952880859375 0 1723.1528
561.2494506835938 0 3038.113
561.7503051757812 0 1235.3773
562.2479858398438 0 1634.2422
563.763427734375 0 1055.177
564.2816772460938 0 4418.8457 y 6
564.7806396484375 0 2745.1123
564.9352416992188 0 2087.833 b 13
565.26806640625 0 6671.19 y Ammonia loss 1
565.6002197265625 0 3160.467
565.9317626953125 0 1294.7584
566.2737426757812 0 959.14886
566.5955810546875 0 1288.2928
566.9310302734375 0 1003.4961
567.7426147460938 0 1063.9819
570.27197265625 0 1886.0049
570.93896484375 0 10310.578 y 1
571.2725830078125 0 9590.061
571.6058349609375 0 5071.57
571.9386596679688 0 3559.42
572.2680053710938 0 1556.916
572.7694091796875 0 9387.481
573.2725830078125 0 3548.227
573.771484375 0 1595.7878
574.2733764648438 0 1708.4856
575.25244140625 0 2210.251
575.7496948242188 0 2746.9187
576.2520141601562 0 2154.0908
576.7521362304688 0 1141.141
577.7607421875 0 932.73254 b Water loss 9
578.2565307617188 0 1178.1212 b Ammonia loss 9
578.75634765625 0 957.1424
582.2313842773438 0 836.873 b Water loss 4
584.2493286132812 0 1378.8342
584.7681884765625 0 2519.291
585.26416015625 0 1941.6516
585.761474609375 0 1021.9412
586.7669067382812 0 7657.4077 b 9
587.26611328125 0 6065.9395
587.765869140625 0 3040.709
588.2874755859375 0 1879.0807
589.2740478515625 0 1523.8259
591.2870483398438 0 1143.2136 y Ammonia loss 5
593.7752685546875 0 1015.5367
594.2745971679688 0 1571.559 Precursor Ammonia loss
594.7781372070312 0 1421.9929
598.26708984375 0 5318.526
599.2686767578125 0 1452.5417
599.8020629882812 0 3200.673 y 5
600.243896484375 0 5186.4224 b 4
600.3019409179688 0 3346.9258
600.8035888671875 0 1857.8359
601.2479858398438 0 1489.1833
601.3033447265625 0 795.76294
603.2909545898438 0 1226.981
603.7616577148438 0 851.67737
607.2815551757812 0 4856.7954
607.7728271484375 0 1061.1721
608.254638671875 0 1129.1304
608.3062133789062 0 1294.7964 y Water loss 10
609.2933349609375 0 1286.2537 y Ammonia loss 10
610.238037109375 0 751.076
611.800537109375 0 862.2635
612.2957153320312 0 1163.9204
616.2857055664062 0 1528.9989
616.7691650390625 0 3949.109
617.2673950195312 0 3928.157
617.76513671875 0 1199.8503
618.2469482421875 0 918.3007
618.7658081054688 0 888.3953
619.2496948242188 0 960.35657
621.8092651367188 0 970.8042
624.7905883789062 0 918.21277
625.7737426757812 0 2981.7056
626.31396484375 0 8819.63 y 10
627.3161010742188 0 2059.7163
633.3348388671875 0 1169.361
634.2828979492188 0 920.10236
635.2725830078125 0 2002.9049
636.7916259765625 0 972.916
639.2996826171875 0 1097.7468
643.2850952148438 0 829.7936
644.2816772460938 0 1215.9166
646.805908203125 0 1515.2432
648.7978515625 0 1241.4747
649.2969360351562 0 1258.0493
649.7963256835938 0 1005.8753
650.2936401367188 0 986.7492
653.2760009765625 0 2760.3816 b Water loss 5
654.2918701171875 0 1288.6469
655.3173217773438 0 2186.255 y Water loss 4
655.8152465820312 0 3703.0142 y Ammonia loss 4
656.3126831054688 0 2278.119
657.31005859375 0 1830.3248
657.8037719726562 0 1387.6727
658.309814453125 0 1949.6663
658.801025390625 0 941.45184
664.3220825195312 0 6330.5283 y 4
664.8240356445312 0 4423.039
665.322509765625 0 3053.1995
665.8169555664062 0 876.5022
666.3135986328125 0 1204.3842
667.2987670898438 0 1666.4745
667.792236328125 0 828.8897
668.3018798828125 0 1708.7657 b 10
668.7990112304688 0 1139.4451
671.2811889648438 0 6993.847 b 5
671.3274536132812 0 1417.6536
672.28466796875 0 3631.5068
674.3372192382812 0 2606.6272
675.34130859375 0 1028.0023
675.803466796875 0 1072.48
676.29736328125 0 805.6352
680.7943725585938 0 1934.7327
681.2975463867188 0 2099.117
681.7908325195312 0 1530.37
683.2848510742188 0 961.20374
684.3206787109375 0 1973.2485
685.3060913085938 0 1251.2021
689.8017578125 0 3766.4016
690.3013305664062 0 3557.3135
690.802734375 0 2171.986
697.3300170898438 0 936.8796
702.33349609375 0 4309.671
703.3326416015625 0 1969.0667
704.3273315429688 0 948.2188
707.326904296875 0 1530.432
709.8187255859375 0 2087.615 b Water loss 11
710.3226928710938 0 2359.4077
713.3192138671875 0 777.0494
715.3289794921875 0 881.3784
718.3190307617188 0 989.17926
718.8199462890625 0 991.33185 b 11
719.3262329101562 0 1726.413
724.34521484375 0 1000.4468 y Ammonia loss 3
724.8368530273438 0 1074.8134
726.3351440429688 0 1480.5156
726.8323364257812 0 1194.0093
732.8523559570312 0 3981.6465 y 3
733.35400390625 0 4327.7285
733.8536376953125 0 1250.9229
735.3308715820312 0 1924.3938
736.341064453125 0 1069.6046
737.3326416015625 0 938.47943
739.3380737304688 0 1499.6956
742.3456420898438 0 793.17615
746.3485107421875 0 930.92285 y Ammonia loss 9
754.35107421875 0 1150.7119
763.373046875 0 3688.6438 y 9
764.3056030273438 0 1217.808
764.3753662109375 0 1695.2919
766.3646240234375 0 1002.6977 b Water loss 6
768.343994140625 0 1118.5438
772.3751831054688 0 883.5455
781.3380737304688 0 1961.3225
782.3400268554688 0 744.26495
782.8527221679688 0 1293.528 b 12
783.3490600585938 0 1248.896
784.3634643554688 0 3876.1772 b 6
785.3673706054688 0 3511.1702
786.3693237304688 0 1367.0208
802.3660888671875 0 902.2862
811.3980102539062 0 883.15015
819.3873901367188 0 1604.509
821.3792724609375 0 1084.3925
837.3995361328125 0 1493.0386
838.4026489257812 0 920.7089
839.3909912109375 0 1454.617
849.3673706054688 0 2382.7393
850.3696899414062 0 1476.7472
860.3853149414062 0 1068.2727 y Ammonia loss 8
861.39013671875 0 726.20776
865.3931884765625 0 1743.8276
877.4132080078125 0 3407.7998 y 8
878.4133911132812 0 1157.3251
879.4194946289062 0 752.95294
894.4159545898438 0 1046.0155
895.4218139648438 0 848.4582
921.4246826171875 0 1430.7747 b 7
922.4285888671875 0 1292.643
930.4221801757812 0 1177.0508
948.4307250976562 0 2547.0984
949.43408203125 0 1281.7916
965.4513549804688 0 1000.0951
966.455322265625 0 1344.7795
971.4723510742188 0 715.2768
975.45458984375 0 690.7187
987.416259765625 0 853.22375
1002.4478149414062 0 804.55115
1014.4697875976562 0 802.1458 y 7
1035.464599609375 0 2603.2827 b 8
1037.465576171875 0 840.2
1076.4908447265625 0 1002.1464
1085.49560546875 0 809.4157
1086.4791259765625 0 1100.5142
1094.5037841796875 0 945.5808
1095.5037841796875 0 803.64185
2028.1123046875 0 741.343
2299.95849609375 0 673.6653

Spectrum Details

|  |  |
| --- | --- |
| Matched peaks? Matched peaks The total absolute number of peaks matched. Additionally in brackets the total fraction of peaks matched and the total number of peaks is shown. | 92 (11.56% of 796) |
| FDR? FDR The false discovery rate estimated for this peptide. It is calculated by matching all theoretical fragments with a non-integer shift with the raw peaks for this spectrum. This is done with 40 different shifts. The resulting percentage is the average number of annotated peaks over the number of annotated peaks with the correct spectrum. | 0.83% |
| Satellite FDR? Satellite FDR See the FDR for details on its calculation. This satellite ion specific FDR only contains the satellite ions (d/w) for I/L/J positions. | - |
| PSM Score? PSM Score The PSM Score as given by Hecklib to this annotated spectrum. It is shown with three significant figures. | 365 |

#### Spectrum 2866? Spectrum 2866 The raw spectrum of this peptide as annotated by Hecklib. The fragments are coloured according to ion type (see legend). Any peaks with a star '\*' as text can be hovered over to see the full details, first the ion type second the mass shift type. By hovering over the amino acids in the peptide or ions in the legend the corresponding peaks are highlighted. By toggling the 'Unassigned' label you can turn the background (unassigned) peaks on or off in the plot. By updating the slider in the Ion legend you can update the spectrum to only show the top X% of the peaks with labels. The top X% means any peak that is within X% of the highest intensity. By dragging in the spectrum you can zoom in to a specific part of the spectrum and use 'Zoom Out' to get back to the original zoom level. The annotation of the spectrum is based on the given sequence in the peptides file and is done with different software so inconsistencies are likely. The peaks are annotated based on the given sequence, with 20 ppm tolerance.

Copy Data

##### Spectrum 2866 (TSV)

###### Preview

```
Loading example...
```

*Click on the button to copy the data to your clipboard.*

Mz MinMz MaxIntensity Max

WidthHeightPeptide font sizePeptide stroke widthSpectrum font sizeSpectrum stroke widthCompact peptide

Ion legend

wxyz

abcd

OtherUnassignedIon^Charge^_Position_Show for top:%

SVMHEAJHNHYTQKS

02.84e+65.67e+68.51e+61.13e+7

Zoom Out

a^+1^_2_a^+1^_2_b^+1^_2_y^+2^_3_b^+1^_2_y^+1^_2_y^+1^_2_y^+4^_7_y^+1^_2_b^+4^_9_b^+2^_5_b^+1^_3_y^+2^_5_y^+2^_5_b^+2^_6_b^+2^_6_y^+3^_8_y^+3^_8_y^+3^_8_y^+1^_3_y^+1^_3_y^+1^_3_y^+3^_9_y^+2^_6_y^+3^_9_y^+2^_6_b^+3^_10_y^+4^_13_y^+4^_13_y^+4^_13_b^+4^_14_b^+4^_14_y^+4^_14_y^+4^_14_y^+2^_7_y^+2^_7_y^+3^_11_y^+3^_11_y^+2^_7_b^+3^_11_\*\*y^+3^_11_y^+1^_4_\*b^+2^_8_b^+1^_4_y^+1^_4_b^+3^_12_b^+3^_12_b^+3^_12_y^+3^_12_y^+3^_12_y^+3^_12_y^+2^_8_y^+2^_8_b^+2^_9_y^+2^_8_b^+2^_9_b^+3^_13_b^+3^_13_y^+3^_13_y^+3^_13_y^+3^_13_b^+3^_14_y^+2^_9_y^+2^_9_b^+3^_14_y^+3^_14_y^+2^_9_y^+3^_14_b^+2^_10_b^+2^_10_b^+2^_10_b^+1^_5_y^+2^_10_y^+2^_10_y^+2^_10_y^+1^_5_y^+1^_5_b^+1^_6_b^+1^_6_y^+2^_11_b^+2^_11_y^+2^_11_b^+2^_12_b^+2^_12_y^+2^_12_y^+2^_12_y^+2^_12_y^+1^_6_y^+1^_6_b^+1^_7_y^+1^_6_b^+2^_13_b^+2^_13_b^+1^_7_b^+2^_13_y^+2^_14_y^+1^_7_y^+1^_7_y^+1^_7_b^+1^_8_y^+1^_8_b^+1^_9_b^+1^_9_y^+1^_8_b^+1^_9_y^+1^_10_

0785156923543139

Fragment Matches Table

Show background peaks

| Position | Ion type | Intensity | mz Theoretical | mz Error (Th) | mz Error (ppm) | Charge | Series Number |
| --- | --- | --- | --- | --- | --- | --- | --- |
| - | - | 1.383E+04 | 120.1 | - | - | 0 | - |
| - | - | 1.726E+04 | 121 | - | - | 0 | - |
| - | - | 1.646E+04 | 122.1 | - | - | 0 | - |
| - | - | 1.056E+04 | 122.8 | - | - | 0 | - |
| - | - | 2.38E+04 | 123.1 | - | - | 0 | - |
| - | - | 1.25E+04 | 125.1 | - | - | 0 | - |
| - | - | 2.661E+04 | 128.1 | - | - | 0 | - |
| - | - | 6.23E+04 | 129.1 | - | - | 0 | - |
| - | - | 4.151E+06 | 129.1 | - | - | 0 | - |
| - | - | 1.949E+04 | 130.1 | - | - | 0 | - |
| - | - | 6.033E+04 | 130.1 | - | - | 0 | - |
| - | - | 3.117E+04 | 130.1 | - | - | 0 | - |
| - | - | 2.503E+05 | 130.1 | - | - | 0 | - |
| - | - | 2.276E+06 | 136.1 | - | - | 0 | - |
| - | - | 2.856E+04 | 137.1 | - | - | 0 | - |
| - | - | 1.795E+05 | 137.1 | - | - | 0 | - |
| - | - | 1.108E+05 | 138.1 | - | - | 0 | - |
| - | - | 7.487E+04 | 139.1 | - | - | 0 | - |
| - | - | 1.186E+04 | 140.1 | - | - | 0 | - |
| - | - | 1.924E+04 | 141 | - | - | 0 | - |
| 2 | a | 1.346E+05 | 141.1 | 0.0003149 | 2.231 | +1 | 2 |
| - | - | 1.277E+04 | 142.1 | - | - | 0 | - |
| - | - | 1.896E+04 | 147 | - | - | 0 | - |
| - | - | 1.262E+04 | 147.1 | - | - | 0 | - |
| - | - | 7.438E+04 | 147.1 | - | - | 0 | - |
| - | - | 2.913E+05 | 148.1 | - | - | 0 | - |
| - | - | 8.82E+04 | 149 | - | - | 0 | - |
| - | - | 3.329E+04 | 149.1 | - | - | 0 | - |
| - | - | 2.263E+04 | 152.1 | - | - | 0 | - |
| - | - | 2.921E+04 | 153.1 | - | - | 0 | - |
| - | - | 1.961E+04 | 154.1 | - | - | 0 | - |
| - | - | 5.181E+04 | 155.1 | - | - | 0 | - |
| - | - | 1.111E+05 | 155.1 | - | - | 0 | - |
| - | - | 4.544E+05 | 155.1 | - | - | 0 | - |
| - | - | 8.367E+04 | 156.1 | - | - | 0 | - |
| - | - | 1.353E+04 | 156.1 | - | - | 0 | - |
| - | - | 3.647E+04 | 156.1 | - | - | 0 | - |
| - | - | 8.1E+04 | 157.1 | - | - | 0 | - |
| - | - | 3.001E+04 | 158.1 | - | - | 0 | - |
| 2 | a | 1.123E+07 | 159.1 | 0.0004466 | 2.807 | +1 | 2 |
| - | - | 2.269E+04 | 160.1 | - | - | 0 | - |
| - | - | 5.418E+04 | 160.1 | - | - | 0 | - |
| - | - | 8.371E+05 | 160.1 | - | - | 0 | - |
| - | - | 4.32E+04 | 161.1 | - | - | 0 | - |
| - | - | 2.306E+04 | 162.1 | - | - | 0 | - |
| - | - | 7.27E+04 | 165.1 | - | - | 0 | - |
| - | - | 7.647E+05 | 166.1 | - | - | 0 | - |
| - | - | 1.497E+04 | 167.1 | - | - | 0 | - |
| - | - | 4.013E+04 | 167.1 | - | - | 0 | - |
| - | - | 2.028E+04 | 167.1 | - | - | 0 | - |
| - | - | 3.067E+04 | 167.1 | - | - | 0 | - |
| - | - | 2.057E+04 | 169.1 | - | - | 0 | - |
| 2 | b | 1.839E+04 | 169.1 | 0.0005632 | 3.331 | +1 | 2 |
| - | - | 2.794E+04 | 171.1 | - | - | 0 | - |
| - | - | 1.742E+04 | 172.1 | - | - | 0 | - |
| 13 | y | 1.592E+05 | 173.1 | 0.0003385 | 1.956 | +2 | 3 |
| - | - | 1.784E+05 | 173.5 | - | - | 0 | - |
| - | - | 3.014E+04 | 175.1 | - | - | 0 | - |
| - | - | 1.067E+05 | 176.1 | - | - | 0 | - |
| - | - | 8.825E+04 | 177.1 | - | - | 0 | - |
| - | - | 1.496E+04 | 178.1 | - | - | 0 | - |
| - | - | 1.868E+04 | 178.1 | - | - | 0 | - |
| - | - | 2.873E+04 | 179.1 | - | - | 0 | - |
| - | - | 1.633E+04 | 180.1 | - | - | 0 | - |
| - | - | 4.471E+04 | 180.1 | - | - | 0 | - |
| - | - | 1.445E+04 | 181.1 | - | - | 0 | - |
| - | - | 7.154E+04 | 183.1 | - | - | 0 | - |
| - | - | 2.285E+04 | 183.1 | - | - | 0 | - |
| - | - | 1.472E+04 | 185.1 | - | - | 0 | - |
| - | - | 4.366E+04 | 185.1 | - | - | 0 | - |
| - | - | 1.614E+04 | 186.1 | - | - | 0 | - |
| 2 | b | 1.621E+06 | 187.1 | 0.0003897 | 2.083 | +1 | 2 |
| - | - | 1.557E+05 | 188.1 | - | - | 0 | - |
| - | - | 7.343E+04 | 189.1 | - | - | 0 | - |
| - | - | 1.226E+05 | 190.1 | - | - | 0 | - |
| - | - | 4.91E+04 | 191.1 | - | - | 0 | - |
| - | - | 1.033E+05 | 192.1 | - | - | 0 | - |
| - | - | 1.722E+04 | 193.1 | - | - | 0 | - |
| - | - | 1.201E+05 | 193.1 | - | - | 0 | - |
| - | - | 1.208E+05 | 194.1 | - | - | 0 | - |
| - | - | 9.517E+04 | 195.1 | - | - | 0 | - |
| - | - | 1.838E+05 | 195.1 | - | - | 0 | - |
| - | - | 2.852E+04 | 196.1 | - | - | 0 | - |
| - | - | 3.443E+05 | 196.1 | - | - | 0 | - |
| - | - | 6.056E+04 | 197.1 | - | - | 0 | - |
| - | - | 2.392E+04 | 198.1 | - | - | 0 | - |
| - | - | 3.886E+04 | 199.1 | - | - | 0 | - |
| - | - | 1.492E+04 | 200.1 | - | - | 0 | - |
| - | - | 1.646E+05 | 201.1 | - | - | 0 | - |
| - | - | 3.748E+04 | 202.1 | - | - | 0 | - |
| - | - | 4.361E+04 | 202.1 | - | - | 0 | - |
| - | - | 2.967E+04 | 202.1 | - | - | 0 | - |
| - | - | 3.044E+04 | 203.1 | - | - | 0 | - |
| - | - | 3.379E+05 | 203.1 | - | - | 0 | - |
| - | - | 1.438E+04 | 204.1 | - | - | 0 | - |
| - | - | 1.331E+04 | 204.1 | - | - | 0 | - |
| - | - | 1.907E+04 | 206.1 | - | - | 0 | - |
| - | - | 3.332E+05 | 207.1 | - | - | 0 | - |
| - | - | 2.819E+04 | 208.1 | - | - | 0 | - |
| - | - | 1.99E+04 | 208.1 | - | - | 0 | - |
| - | - | 6.12E+04 | 212.1 | - | - | 0 | - |
| - | - | 1.997E+04 | 213.1 | - | - | 0 | - |
| 14 | y | 1.502E+05 | 216.1 | 0.0003909 | 1.809 | +1 | 2 |
| - | - | 1.041E+05 | 217.1 | - | - | 0 | - |
| 14 | y | 1.72E+05 | 217.1 | 0.0004604 | 2.121 | +1 | 2 |
| - | - | 8.536E+04 | 218.1 | - | - | 0 | - |
| - | - | 2.186E+04 | 219.1 | - | - | 0 | - |
| 9 | y | 1.431E+04 | 220.1 | 0.0003295 | 1.497 | +4 | 7 |
| - | - | 3.181E+05 | 221.1 | - | - | 0 | - |
| - | - | 5.129E+04 | 222.1 | - | - | 0 | - |
| - | - | 2.057E+05 | 222.1 | - | - | 0 | - |
| - | - | 3.566E+04 | 223.1 | - | - | 0 | - |
| - | - | 3.323E+04 | 223.1 | - | - | 0 | - |
| - | - | 1.562E+04 | 223.1 | - | - | 0 | - |
| - | - | 7.112E+04 | 223.2 | - | - | 0 | - |
| - | - | 4.616E+04 | 224.1 | - | - | 0 | - |
| - | - | 3.117E+05 | 224.1 | - | - | 0 | - |
| - | - | 3.257E+04 | 225 | - | - | 0 | - |
| - | - | 2.071E+04 | 225.1 | - | - | 0 | - |
| - | - | 5.376E+04 | 225.1 | - | - | 0 | - |
| - | - | 1.686E+04 | 226.1 | - | - | 0 | - |
| - | - | 2.663E+04 | 228.1 | - | - | 0 | - |
| - | - | 3.458E+04 | 230.1 | - | - | 0 | - |
| - | - | 3.283E+05 | 230.1 | - | - | 0 | - |
| - | - | 2.537E+04 | 230.1 | - | - | 0 | - |
| - | - | 8.516E+04 | 231.1 | - | - | 0 | - |
| - | - | 4.252E+04 | 233.1 | - | - | 0 | - |
| - | - | 1.186E+05 | 234.1 | - | - | 0 | - |
| - | - | 5.492E+04 | 234.1 | - | - | 0 | - |
| 14 | y | 9.921E+05 | 234.1 | 0.0004769 | 2.037 | +1 | 2 |
| - | - | 5.251E+05 | 235.1 | - | - | 0 | - |
| - | - | 9.106E+04 | 235.1 | - | - | 0 | - |
| - | - | 4.375E+04 | 236.1 | - | - | 0 | - |
| - | - | 1.334E+05 | 237.1 | - | - | 0 | - |
| - | - | 3.504E+05 | 239.1 | - | - | 0 | - |
| - | - | 4.876E+05 | 239.2 | - | - | 0 | - |
| - | - | 3.241E+04 | 240.1 | - | - | 0 | - |
| - | - | 1.131E+06 | 240.1 | - | - | 0 | - |
| - | - | 4.48E+04 | 240.2 | - | - | 0 | - |
| - | - | 6.789E+05 | 241.1 | - | - | 0 | - |
| - | - | 1.183E+05 | 241.1 | - | - | 0 | - |
| - | - | 1.555E+04 | 241.4 | - | - | 0 | - |
| - | - | 5.828E+04 | 242.1 | - | - | 0 | - |
| - | - | 2.267E+04 | 243.1 | - | - | 0 | - |
| - | - | 2.532E+04 | 244.1 | - | - | 0 | - |
| - | - | 1.559E+04 | 245.2 | - | - | 0 | - |
| - | - | 8.897E+04 | 247.1 | - | - | 0 | - |
| - | - | 2.231E+05 | 249.1 | - | - | 0 | - |
| - | - | 2.302E+04 | 249.1 | - | - | 0 | - |
| - | - | 3.524E+04 | 250.1 | - | - | 0 | - |
| - | - | 2.724E+04 | 250.1 | - | - | 0 | - |
| - | - | 2.516E+04 | 251.1 | - | - | 0 | - |
| 9 | b | 2.045E+04 | 251.1 | 0.004949 | 19.71 | +4 | 9 |
| - | - | 1.884E+05 | 251.2 | - | - | 0 | - |
| - | - | 8.911E+05 | 252.1 | - | - | 0 | - |
| - | - | 1.145E+06 | 252.1 | - | - | 0 | - |
| - | - | 9.58E+04 | 253.1 | - | - | 0 | - |
| - | - | 1.223E+05 | 253.1 | - | - | 0 | - |
| - | - | 4.394E+04 | 254.1 | - | - | 0 | - |
| - | - | 1.865E+04 | 256.1 | - | - | 0 | - |
| - | - | 1.263E+05 | 257.2 | - | - | 0 | - |
| - | - | 1.821E+04 | 258.1 | - | - | 0 | - |
| - | - | 6.124E+04 | 258.1 | - | - | 0 | - |
| - | - | 1.535E+04 | 261.1 | - | - | 0 | - |
| - | - | 4.177E+04 | 262.1 | - | - | 0 | - |
| - | - | 2.825E+04 | 263.1 | - | - | 0 | - |
| - | - | 3.62E+04 | 264.1 | - | - | 0 | - |
| - | - | 1.308E+05 | 265.1 | - | - | 0 | - |
| - | - | 8.76E+05 | 267.1 | - | - | 0 | - |
| - | - | 9.626E+04 | 268.1 | - | - | 0 | - |
| - | - | 2.922E+06 | 269.1 | - | - | 0 | - |
| - | - | 3.352E+05 | 270.1 | - | - | 0 | - |
| - | - | 1.327E+05 | 271.1 | - | - | 0 | - |
| - | - | 2.1E+05 | 273.1 | - | - | 0 | - |
| - | - | 2.471E+04 | 274.1 | - | - | 0 | - |
| - | - | 2.588E+04 | 277.1 | - | - | 0 | - |
| - | - | 1.636E+04 | 279.1 | - | - | 0 | - |
| - | - | 1.059E+05 | 279.1 | - | - | 0 | - |
| - | - | 8.849E+04 | 280.1 | - | - | 0 | - |
| - | - | 1.824E+05 | 280.1 | - | - | 0 | - |
| - | - | 2.569E+04 | 280.1 | - | - | 0 | - |
| - | - | 5.591E+04 | 280.7 | - | - | 0 | - |
| - | - | 2.128E+04 | 282.1 | - | - | 0 | - |
| - | - | 1.95E+04 | 283.1 | - | - | 0 | - |
| - | - | 1.608E+04 | 286.1 | - | - | 0 | - |
| 5 | b | 2.817E+04 | 292.6 | 0.0009632 | 3.291 | +2 | 5 |
| - | - | 2.473E+04 | 293.1 | - | - | 0 | - |
| - | - | 1.936E+04 | 294.2 | - | - | 0 | - |
| - | - | 1.323E+05 | 294.6 | - | - | 0 | - |
| - | - | 7.704E+04 | 295.1 | - | - | 0 | - |
| - | - | 3.963E+04 | 295.2 | - | - | 0 | - |
| - | - | 1.796E+04 | 296.6 | - | - | 0 | - |
| - | - | 3.531E+04 | 297.1 | - | - | 0 | - |
| - | - | 9.162E+04 | 297.2 | - | - | 0 | - |
| - | - | 2.176E+04 | 298.1 | - | - | 0 | - |
| - | - | 2.303E+04 | 298.2 | - | - | 0 | - |
| 3 | b | 2.464E+04 | 300.1 | 0.001064 | 3.544 | +1 | 3 |
| - | - | 3.613E+05 | 301.1 | - | - | 0 | - |
| - | - | 5.545E+04 | 302.1 | - | - | 0 | - |
| - | - | 1.693E+04 | 303.1 | - | - | 0 | - |
| - | - | 2.944E+04 | 303.2 | - | - | 0 | - |
| - | - | 1.648E+04 | 304.1 | - | - | 0 | - |
| - | - | 6.059E+04 | 304.2 | - | - | 0 | - |
| 11 | y | 5.065E+04 | 304.7 | 0.0001057 | 0.3471 | +2 | 5 |
| - | - | 6.944E+04 | 305.2 | - | - | 0 | - |
| - | - | 1.76E+04 | 305.2 | - | - | 0 | - |
| - | - | 2.745E+04 | 306.1 | - | - | 0 | - |
| 11 | y | 1.487E+05 | 313.7 | 0.0005302 | 1.69 | +2 | 5 |
| - | - | 7.57E+04 | 314.2 | - | - | 0 | - |
| - | - | 2.326E+04 | 314.2 | - | - | 0 | - |
| - | - | 4.197E+04 | 314.7 | - | - | 0 | - |
| - | - | 2.452E+04 | 314.8 | - | - | 0 | - |
| - | - | 2.54E+04 | 318.1 | - | - | 0 | - |
| 6 | b | 1.896E+04 | 319.1 | 0.001264 | 3.961 | +2 | 6 |
| - | - | 5.154E+05 | 320.1 | - | - | 0 | - |
| - | - | 5.144E+04 | 321.1 | - | - | 0 | - |
| - | - | 2.34E+04 | 321.2 | - | - | 0 | - |
| - | - | 2.173E+05 | 322.2 | - | - | 0 | - |
| - | - | 9.35E+04 | 323.2 | - | - | 0 | - |
| - | - | 4.065E+04 | 323.2 | - | - | 0 | - |
| - | - | 5.843E+04 | 326.1 | - | - | 0 | - |
| - | - | 1.9E+04 | 327.1 | - | - | 0 | - |
| - | - | 3.644E+04 | 327.2 | - | - | 0 | - |
| 6 | b | 6.246E+04 | 328.1 | 0.002639 | 8.043 | +2 | 6 |
| - | - | 5.938E+04 | 328.7 | - | - | 0 | - |
| - | - | 2.29E+04 | 329.1 | - | - | 0 | - |
| - | - | 2.158E+04 | 330.2 | - | - | 0 | - |
| 8 | y | 4.009E+04 | 332.8 | 0.001038 | 3.118 | +3 | 8 |
| 8 | y | 4.231E+04 | 333.2 | 0.001432 | 4.299 | +3 | 8 |
| - | - | 1.821E+04 | 337.2 | - | - | 0 | - |
| - | - | 4.217E+04 | 337.7 | - | - | 0 | - |
| - | - | 4.773E+05 | 338.1 | - | - | 0 | - |
| - | - | 2.692E+04 | 338.7 | - | - | 0 | - |
| 8 | y | 6.983E+04 | 338.8 | 0.0004762 | 1.406 | +3 | 8 |
| - | - | 6.857E+04 | 339.1 | - | - | 0 | - |
| - | - | 2.612E+04 | 339.2 | - | - | 0 | - |
| - | - | 3.377E+04 | 339.2 | - | - | 0 | - |
| - | - | 1.898E+04 | 339.5 | - | - | 0 | - |
| - | - | 1.755E+05 | 340.1 | - | - | 0 | - |
| - | - | 2.985E+04 | 340.2 | - | - | 0 | - |
| - | - | 7.659E+04 | 341 | - | - | 0 | - |
| - | - | 2.608E+04 | 341.1 | - | - | 0 | - |
| - | - | 1.625E+05 | 341.2 | - | - | 0 | - |
| - | - | 4.093E+04 | 342 | - | - | 0 | - |
| - | - | 1.771E+04 | 342.2 | - | - | 0 | - |
| - | - | 1.887E+04 | 342.7 | - | - | 0 | - |
| - | - | 3.449E+04 | 343.2 | - | - | 0 | - |
| - | - | 1.16E+05 | 344.1 | - | - | 0 | - |
| 13 | y | 2.134E+05 | 344.2 | 0.0004529 | 1.316 | +1 | 3 |
| 13 | y | 2.232E+05 | 345.2 | 0.0003851 | 1.116 | +1 | 3 |
| - | - | 2.67E+04 | 345.2 | - | - | 0 | - |
| - | - | 8.349E+04 | 346.2 | - | - | 0 | - |
| - | - | 3.11E+04 | 346.7 | - | - | 0 | - |
| - | - | 3.013E+04 | 348.1 | - | - | 0 | - |
| - | - | 3.843E+04 | 350.2 | - | - | 0 | - |
| - | - | 2.424E+04 | 350.2 | - | - | 0 | - |
| - | - | 2.337E+04 | 351.2 | - | - | 0 | - |
| - | - | 2.055E+05 | 351.7 | - | - | 0 | - |
| - | - | 4.46E+04 | 352.1 | - | - | 0 | - |
| - | - | 8.266E+04 | 352.2 | - | - | 0 | - |
| - | - | 2.774E+04 | 353.1 | - | - | 0 | - |
| - | - | 2.692E+04 | 353.1 | - | - | 0 | - |
| - | - | 2.003E+04 | 353.2 | - | - | 0 | - |
| - | - | 3.367E+04 | 354.1 | - | - | 0 | - |
| - | - | 2.294E+04 | 355.1 | - | - | 0 | - |
| - | - | 9.129E+04 | 358.2 | - | - | 0 | - |
| - | - | 3.457E+05 | 359 | - | - | 0 | - |
| - | - | 2.617E+04 | 359.1 | - | - | 0 | - |
| - | - | 1.629E+05 | 360 | - | - | 0 | - |
| - | - | 1.318E+05 | 360.2 | - | - | 0 | - |
| - | - | 8.172E+04 | 360.7 | - | - | 0 | - |
| - | - | 8.139E+04 | 361 | - | - | 0 | - |
| - | - | 1.131E+05 | 361.2 | - | - | 0 | - |
| - | - | 2.4E+04 | 362.1 | - | - | 0 | - |
| - | - | 2.891E+04 | 362.2 | - | - | 0 | - |
| 13 | y | 1.762E+05 | 362.2 | 0.0005999 | 1.656 | +1 | 3 |
| - | - | 4.1E+04 | 362.5 | - | - | 0 | - |
| - | - | 2.227E+04 | 363.2 | - | - | 0 | - |
| - | - | 1.057E+05 | 365.2 | - | - | 0 | - |
| - | - | 1.207E+05 | 366.1 | - | - | 0 | - |
| - | - | 6.447E+04 | 366.2 | - | - | 0 | - |
| - | - | 2.928E+04 | 367.2 | - | - | 0 | - |
| - | - | 1.978E+05 | 368.2 | - | - | 0 | - |
| - | - | 1.75E+04 | 368.5 | - | - | 0 | - |
| - | - | 8.147E+04 | 369.2 | - | - | 0 | - |
| - | - | 1.67E+04 | 369.5 | - | - | 0 | - |
| - | - | 4.078E+04 | 370.2 | - | - | 0 | - |
| - | - | 4.563E+04 | 370.7 | - | - | 0 | - |
| 7 | y | 2.245E+04 | 370.8 | 0.001941 | 5.235 | +3 | 9 |
| - | - | 3.288E+05 | 371.2 | - | - | 0 | - |
| - | - | 1.723E+05 | 372.1 | - | - | 0 | - |
| - | - | 5.202E+04 | 372.2 | - | - | 0 | - |
| - | - | 4.285E+04 | 373.1 | - | - | 0 | - |
| 10 | y | 4.564E+04 | 373.7 | 0.000279 | 0.7466 | +2 | 6 |
| - | - | 1.87E+04 | 374.2 | - | - | 0 | - |
| - | - | 8.422E+04 | 375.2 | - | - | 0 | - |
| - | - | 1.69E+04 | 376.2 | - | - | 0 | - |
| - | - | 3.674E+04 | 376.2 | - | - | 0 | - |
| 7 | y | 1.942E+04 | 376.5 | 0.0007608 | 2.021 | +3 | 9 |
| - | - | 1.156E+05 | 376.9 | - | - | 0 | - |
| - | - | 7.563E+04 | 377.2 | - | - | 0 | - |
| - | - | 1.807E+04 | 378.7 | - | - | 0 | - |
| - | - | 2.595E+04 | 379.7 | - | - | 0 | - |
| - | - | 1.337E+05 | 380.1 | - | - | 0 | - |
| - | - | 1.995E+04 | 380.8 | - | - | 0 | - |
| - | - | 4.198E+04 | 381.1 | - | - | 0 | - |
| - | - | 2.653E+04 | 381.1 | - | - | 0 | - |
| 10 | y | 1.57E+05 | 382.2 | 0.0003711 | 0.971 | +2 | 6 |
| - | - | 4.42E+04 | 382.7 | - | - | 0 | - |
| - | - | 6.007E+04 | 384.2 | - | - | 0 | - |
| - | - | 2.129E+04 | 384.2 | - | - | 0 | - |
| 10 | b | 7.406E+04 | 386.2 | 0.0001176 | 0.3045 | +3 | 10 |
| - | - | 4.326E+04 | 386.5 | - | - | 0 | - |
| - | - | 4.601E+04 | 386.8 | - | - | 0 | - |
| - | - | 5.015E+04 | 387.2 | - | - | 0 | - |
| - | - | 3.147E+04 | 387.2 | - | - | 0 | - |
| - | - | 3.864E+04 | 388.2 | - | - | 0 | - |
| - | - | 1.807E+04 | 388.7 | - | - | 0 | - |
| - | - | 1.547E+05 | 389.2 | - | - | 0 | - |
| - | - | 1.921E+04 | 390.2 | - | - | 0 | - |
| - | - | 2.592E+04 | 390.9 | - | - | 0 | - |
| - | - | 3.266E+04 | 391.2 | - | - | 0 | - |
| - | - | 1.892E+04 | 391.2 | - | - | 0 | - |
| - | - | 1.081E+05 | 393.2 | - | - | 0 | - |
| 3 | y | 5.35E+04 | 395.2 | 0.0003273 | 0.8282 | +4 | 13 |
| 3 | y | 7.563E+04 | 395.4 | 0.00315 | 7.966 | +4 | 13 |
| - | - | 2.334E+04 | 395.7 | - | - | 0 | - |
| - | - | 3.84E+04 | 395.9 | - | - | 0 | - |
| - | - | 2.629E+04 | 396.5 | - | - | 0 | - |
| - | - | 4.451E+04 | 397.2 | - | - | 0 | - |
| - | - | 2.603E+04 | 397.2 | - | - | 0 | - |
| - | - | 1.567E+05 | 397.7 | - | - | 0 | - |
| - | - | 9.914E+05 | 398.1 | - | - | 0 | - |
| - | - | 4.739E+04 | 398.2 | - | - | 0 | - |
| - | - | 1.926E+05 | 399.2 | - | - | 0 | - |
| 3 | y | 2.718E+05 | 399.7 | 0.0006326 | 1.583 | +4 | 13 |
| - | - | 1.924E+05 | 399.9 | - | - | 0 | - |
| - | - | 5.045E+04 | 400.1 | - | - | 0 | - |
| - | - | 1.055E+05 | 400.2 | - | - | 0 | - |
| - | - | 6.317E+04 | 400.4 | - | - | 0 | - |
| - | - | 4.296E+04 | 400.5 | - | - | 0 | - |
| - | - | 2.932E+04 | 400.7 | - | - | 0 | - |
| - | - | 3.565E+04 | 401.2 | - | - | 0 | - |
| - | - | 7.015E+04 | 402.2 | - | - | 0 | - |
| - | - | 2.285E+04 | 402.7 | - | - | 0 | - |
| - | - | 1.589E+04 | 403.2 | - | - | 0 | - |
| - | - | 3.287E+04 | 405.2 | - | - | 0 | - |
| - | - | 1.169E+05 | 405.2 | - | - | 0 | - |
| - | - | 4.104E+05 | 406.2 | - | - | 0 | - |
| - | - | 1.491E+04 | 406.2 | - | - | 0 | - |
| - | - | 2.282E+05 | 406.7 | - | - | 0 | - |
| - | - | 2.806E+04 | 407.2 | - | - | 0 | - |
| - | - | 3.139E+04 | 408.2 | - | - | 0 | - |
| - | - | 3.475E+04 | 408.2 | - | - | 0 | - |
| - | - | 3.698E+04 | 409.2 | - | - | 0 | - |
| - | - | 3.01E+04 | 409.7 | - | - | 0 | - |
| - | - | 7.429E+04 | 410.2 | - | - | 0 | - |
| - | - | 3.28E+04 | 410.7 | - | - | 0 | - |
| - | - | 3.63E+04 | 411.2 | - | - | 0 | - |
| - | - | 1.893E+04 | 411.4 | - | - | 0 | - |
| - | - | 6.972E+04 | 411.7 | - | - | 0 | - |
| - | - | 5.605E+04 | 412.2 | - | - | 0 | - |
| - | - | 3.375E+04 | 412.5 | - | - | 0 | - |
| - | - | 4.55E+04 | 413.2 | - | - | 0 | - |
| - | - | 2.423E+04 | 413.7 | - | - | 0 | - |
| - | - | 2.137E+05 | 415.2 | - | - | 0 | - |
| - | - | 3.798E+04 | 415.2 | - | - | 0 | - |
| 14 | b | 4.783E+04 | 415.7 | 0.000326 | 0.7842 | +4 | 14 |
| - | - | 6.286E+04 | 416.2 | - | - | 0 | - |
| - | - | 3.195E+04 | 416.2 | - | - | 0 | - |
| - | - | 2.624E+04 | 417.2 | - | - | 0 | - |
| - | - | 2.001E+05 | 417.2 | - | - | 0 | - |
| - | - | 5.365E+04 | 417.7 | - | - | 0 | - |
| - | - | 2.837E+04 | 418.2 | - | - | 0 | - |
| - | - | 3.205E+04 | 419.2 | - | - | 0 | - |
| 14 | b | 4.924E+04 | 420 | 0.0007682 | 1.829 | +4 | 14 |
| 2 | y | 4.066E+05 | 420.2 | 0.0002206 | 0.5249 | +4 | 14 |
| - | - | 1.288E+05 | 420.7 | - | - | 0 | - |
| - | - | 3.009E+04 | 421.2 | - | - | 0 | - |
| - | - | 3.202E+04 | 421.7 | - | - | 0 | - |
| - | - | 4.598E+04 | 422.2 | - | - | 0 | - |
| - | - | 2.573E+04 | 423.2 | - | - | 0 | - |
| - | - | 2.434E+05 | 423.2 | - | - | 0 | - |
| - | - | 3.004E+04 | 424.2 | - | - | 0 | - |
| - | - | 4.252E+04 | 424.2 | - | - | 0 | - |
| 2 | y | 2.549E+05 | 424.5 | 0.0002222 | 0.5236 | +4 | 14 |
| - | - | 1.204E+05 | 424.7 | - | - | 0 | - |
| - | - | 7.014E+04 | 425 | - | - | 0 | - |
| - | - | 6.808E+04 | 425.2 | - | - | 0 | - |
| - | - | 2.983E+04 | 426.1 | - | - | 0 | - |
| - | - | 1.341E+05 | 427.2 | - | - | 0 | - |
| - | - | 2.457E+04 | 428.2 | - | - | 0 | - |
| - | - | 2.356E+04 | 428.7 | - | - | 0 | - |
| - | - | 1.025E+05 | 429.1 | - | - | 0 | - |
| - | - | 6.079E+04 | 430.1 | - | - | 0 | - |
| 9 | y | 3.587E+04 | 430.2 | 0.001432 | 3.329 | +2 | 7 |
| 9 | y | 1.097E+05 | 430.7 | 0.002741 | 6.364 | +2 | 7 |
| - | - | 3.405E+04 | 431.1 | - | - | 0 | - |
| - | - | 5.395E+04 | 431.2 | - | - | 0 | - |
| - | - | 2.227E+04 | 431.5 | - | - | 0 | - |
| - | - | 2.611E+04 | 431.9 | - | - | 0 | - |
| - | - | 1.412E+05 | 433.2 | - | - | 0 | - |
| - | - | 4.476E+04 | 433.5 | - | - | 0 | - |
| - | - | 6.205E+04 | 433.5 | - | - | 0 | - |
| - | - | 7.386E+04 | 433.9 | - | - | 0 | - |
| - | - | 2.668E+04 | 435.5 | - | - | 0 | - |
| - | - | 1.735E+05 | 436.2 | - | - | 0 | - |
| - | - | 2.701E+04 | 436.5 | - | - | 0 | - |
| 5 | y | 1.105E+05 | 437.2 | 0.004104 | 9.387 | +3 | 11 |
| - | - | 1.469E+05 | 437.5 | - | - | 0 | - |
| 5 | y | 6.461E+04 | 437.5 | 0.006117 | 13.98 | +3 | 11 |
| - | - | 1.187E+05 | 437.7 | - | - | 0 | - |
| - | - | 9.203E+04 | 438 | - | - | 0 | - |
| - | - | 6.156E+04 | 438.2 | - | - | 0 | - |
| 9 | y | 2.678E+05 | 439.2 | 0.001429 | 3.254 | +2 | 7 |
| - | - | 1.087E+05 | 439.7 | - | - | 0 | - |
| - | - | 2.135E+04 | 440.2 | - | - | 0 | - |
| 11 | b | 3.713E+04 | 440.5 | 0.0009999 | 2.27 | +3 | 11 |
| - | - | 2.149E+04 | 440.9 | - | - | 0 | - |
| - | - | 3.498E+04 | 441.2 | - | - | 0 | - |
| 0 | Precursor | 4.668E+05 | 441.7 | 0.0004715 | 1.068 | +4 | -1 |
| 0 | Precursor | 3.903E+05 | 442 | 0.004193 | 9.487 | +4 | -1 |
| - | - | 2.862E+05 | 442.2 | - | - | 0 | - |
| - | - | 1.558E+05 | 442.5 | - | - | 0 | - |
| - | - | 8.504E+04 | 442.5 | - | - | 0 | - |
| - | - | 6.162E+04 | 442.7 | - | - | 0 | - |
| - | - | 4.817E+04 | 442.9 | - | - | 0 | - |
| - | - | 5.903E+04 | 443.2 | - | - | 0 | - |
| 5 | y | 5.914E+04 | 443.2 | 0.001858 | 4.192 | +3 | 11 |
| - | - | 3.484E+04 | 443.6 | - | - | 0 | - |
| 12 | y | 6.43E+04 | 445.2 | 0.00145 | 3.257 | +1 | 4 |
| - | - | 2.653E+04 | 445.5 | - | - | 0 | - |
| - | - | 2.019E+04 | 446 | - | - | 0 | - |
| 0 | Precursor | 1.602E+06 | 446.2 | 0.0007906 | 1.772 | +4 | -1 |
| - | - | 1.171E+06 | 446.5 | - | - | 0 | - |
| - | - | 6.791E+05 | 446.7 | - | - | 0 | - |
| - | - | 3.109E+05 | 447 | - | - | 0 | - |
| - | - | 1.202E+05 | 447.2 | - | - | 0 | - |
| - | - | 2.7E+04 | 447.5 | - | - | 0 | - |
| - | - | 1.796E+04 | 447.9 | - | - | 0 | - |
| - | - | 1.534E+05 | 448.2 | - | - | 0 | - |
| - | - | 1.166E+05 | 448.5 | - | - | 0 | - |
| - | - | 5.127E+04 | 448.9 | - | - | 0 | - |
| - | - | 2.155E+04 | 449.2 | - | - | 0 | - |
| - | - | 2.44E+04 | 450.2 | - | - | 0 | - |
| - | - | 3.054E+04 | 451.2 | - | - | 0 | - |
| - | - | 6.462E+05 | 451.2 | - | - | 0 | - |
| - | - | 1.37E+05 | 452.2 | - | - | 0 | - |
| 8 | b | 7.56E+04 | 453.2 | 0.002044 | 4.51 | +2 | 8 |
| - | - | 2.191E+04 | 453.7 | - | - | 0 | - |
| - | - | 8.834E+04 | 453.9 | - | - | 0 | - |
| - | - | 1.131E+05 | 454.2 | - | - | 0 | - |
| - | - | 4.52E+04 | 454.6 | - | - | 0 | - |
| - | - | 2.007E+04 | 454.7 | - | - | 0 | - |
| 4 | b | 1.89E+05 | 455.2 | 0.003731 | 8.196 | +1 | 4 |
| - | - | 2.333E+04 | 455.5 | - | - | 0 | - |
| - | - | 3.125E+04 | 455.7 | - | - | 0 | - |
| - | - | 6.233E+04 | 456.2 | - | - | 0 | - |
| - | - | 2.626E+04 | 457.2 | - | - | 0 | - |
| - | - | 2.364E+04 | 459.5 | - | - | 0 | - |
| - | - | 1.455E+05 | 459.9 | - | - | 0 | - |
| - | - | 8.879E+04 | 460.2 | - | - | 0 | - |
| - | - | 3.423E+04 | 460.6 | - | - | 0 | - |
| - | - | 2.928E+04 | 461.2 | - | - | 0 | - |
| - | - | 2.47E+04 | 462.2 | - | - | 0 | - |
| - | - | 2.117E+04 | 462.7 | - | - | 0 | - |
| - | - | 1.21E+05 | 463.2 | - | - | 0 | - |
| 12 | y | 6.149E+05 | 463.3 | 0.0006509 | 1.405 | +1 | 4 |
| - | - | 3.49E+04 | 463.3 | - | - | 0 | - |
| - | - | 5.984E+04 | 463.7 | - | - | 0 | - |
| - | - | 3.806E+04 | 464.2 | - | - | 0 | - |
| - | - | 4.971E+04 | 464.2 | - | - | 0 | - |
| - | - | 1.335E+05 | 464.3 | - | - | 0 | - |
| - | - | 2.087E+04 | 464.7 | - | - | 0 | - |
| - | - | 3.019E+04 | 465.2 | - | - | 0 | - |
| - | - | 2.18E+04 | 465.3 | - | - | 0 | - |
| - | - | 4.46E+04 | 465.7 | - | - | 0 | - |
| - | - | 3.362E+04 | 466.2 | - | - | 0 | - |
| - | - | 3.261E+04 | 466.7 | - | - | 0 | - |
| - | - | 2.688E+04 | 467.2 | - | - | 0 | - |
| 12 | b | 2.994E+04 | 468.2 | 0.007844 | 16.75 | +3 | 12 |
| - | - | 3.905E+04 | 468.3 | - | - | 0 | - |
| 12 | b | 2.361E+04 | 468.5 | 0.00418 | 8.922 | +3 | 12 |
| - | - | 4.777E+05 | 469.2 | - | - | 0 | - |
| - | - | 9.512E+04 | 470.2 | - | - | 0 | - |
| - | - | 4.208E+04 | 470.2 | - | - | 0 | - |
| - | - | 2.906E+04 | 471.2 | - | - | 0 | - |
| - | - | 3.653E+04 | 471.2 | - | - | 0 | - |
| - | - | 2.82E+04 | 471.6 | - | - | 0 | - |
| - | - | 4.776E+05 | 471.7 | - | - | 0 | - |
| - | - | 3.035E+04 | 471.9 | - | - | 0 | - |
| - | - | 2.236E+04 | 472.2 | - | - | 0 | - |
| - | - | 2.3E+05 | 472.2 | - | - | 0 | - |
| - | - | 1.165E+05 | 472.7 | - | - | 0 | - |
| 12 | b | 4.667E+04 | 474.2 | 0.001424 | 3.002 | +3 | 12 |
| - | - | 5.926E+04 | 474.3 | - | - | 0 | - |
| - | - | 4.262E+04 | 474.6 | - | - | 0 | - |
| - | - | 5.889E+04 | 474.7 | - | - | 0 | - |
| - | - | 1.831E+04 | 474.9 | - | - | 0 | - |
| - | - | 1.748E+04 | 475.2 | - | - | 0 | - |
| - | - | 6.152E+04 | 476.7 | - | - | 0 | - |
| - | - | 5.534E+04 | 476.9 | - | - | 0 | - |
| - | - | 1.74E+05 | 477.2 | - | - | 0 | - |
| - | - | 1.925E+05 | 477.6 | - | - | 0 | - |
| - | - | 4.18E+04 | 477.7 | - | - | 0 | - |
| - | - | 1.293E+05 | 477.9 | - | - | 0 | - |
| - | - | 6.2E+04 | 478.2 | - | - | 0 | - |
| - | - | 2.884E+04 | 478.6 | - | - | 0 | - |
| - | - | 1.65E+04 | 479.2 | - | - | 0 | - |
| - | - | 1.487E+05 | 479.2 | - | - | 0 | - |
| - | - | 2.25E+04 | 479.9 | - | - | 0 | - |
| - | - | 4.481E+04 | 480.2 | - | - | 0 | - |
| - | - | 3.164E+04 | 480.6 | - | - | 0 | - |
| - | - | 9.224E+04 | 481.2 | - | - | 0 | - |
| - | - | 2.784E+04 | 481.2 | - | - | 0 | - |
| 4 | y | 2.443E+05 | 482.9 | 0.0006015 | 1.246 | +3 | 12 |
| 4 | y | 4.229E+05 | 483.2 | 0.002613 | 5.408 | +3 | 12 |
| - | - | 2.633E+05 | 483.6 | - | - | 0 | - |
| - | - | 2.647E+04 | 483.7 | - | - | 0 | - |
| - | - | 1.044E+05 | 483.9 | - | - | 0 | - |
| - | - | 6.926E+04 | 484.2 | - | - | 0 | - |
| - | - | 3.171E+05 | 485.7 | - | - | 0 | - |
| - | - | 1.802E+05 | 485.9 | - | - | 0 | - |
| - | - | 3.317E+05 | 486.2 | - | - | 0 | - |
| - | - | 5.784E+04 | 486.6 | - | - | 0 | - |
| - | - | 7.476E+04 | 486.7 | - | - | 0 | - |
| - | - | 3.921E+04 | 486.9 | - | - | 0 | - |
| - | - | 2.299E+04 | 487.2 | - | - | 0 | - |
| - | - | 1.787E+05 | 487.7 | - | - | 0 | - |
| - | - | 7.84E+04 | 488.2 | - | - | 0 | - |
| - | - | 2.384E+04 | 488.7 | - | - | 0 | - |
| 4 | y | 1.573E+06 | 488.9 | 0.0005285 | 1.081 | +3 | 12 |
| - | - | 4.536E+04 | 489.2 | - | - | 0 | - |
| - | - | 1.073E+06 | 489.2 | - | - | 0 | - |
| - | - | 5.114E+05 | 489.6 | - | - | 0 | - |
| - | - | 1.987E+05 | 489.9 | - | - | 0 | - |
| - | - | 4.369E+04 | 490.2 | - | - | 0 | - |
| - | - | 3.305E+04 | 490.7 | - | - | 0 | - |
| - | - | 3.012E+04 | 491.6 | - | - | 0 | - |
| - | - | 3.855E+05 | 491.9 | - | - | 0 | - |
| - | - | 3.523E+05 | 492.2 | - | - | 0 | - |
| - | - | 1.608E+05 | 492.6 | - | - | 0 | - |
| - | - | 2.258E+04 | 492.7 | - | - | 0 | - |
| - | - | 7.643E+04 | 492.9 | - | - | 0 | - |
| - | - | 4.602E+04 | 493.2 | - | - | 0 | - |
| - | - | 2.042E+04 | 493.7 | - | - | 0 | - |
| - | - | 7.806E+04 | 497.2 | - | - | 0 | - |
| - | - | 2.375E+05 | 497.6 | - | - | 0 | - |
| - | - | 2.657E+05 | 497.9 | - | - | 0 | - |
| - | - | 4.343E+04 | 498.2 | - | - | 0 | - |
| - | - | 1.114E+05 | 498.2 | - | - | 0 | - |
| - | - | 5.193E+04 | 498.6 | - | - | 0 | - |
| 8 | y | 7.864E+04 | 498.7 | 0.0003576 | 0.717 | +2 | 8 |
| - | - | 2.173E+04 | 498.9 | - | - | 0 | - |
| - | - | 3.226E+04 | 499.2 | - | - | 0 | - |
| 8 | y | 1.246E+05 | 499.2 | 0.002704 | 5.416 | +2 | 8 |
| - | - | 5.67E+04 | 499.7 | - | - | 0 | - |
| - | - | 2.775E+04 | 500.2 | - | - | 0 | - |
| - | - | 1.908E+04 | 501.6 | - | - | 0 | - |
| 9 | b | 1.653E+05 | 501.7 | 0.004902 | 9.77 | +2 | 9 |
| - | - | 1.189E+05 | 502.2 | - | - | 0 | - |
| - | - | 5.627E+04 | 503.3 | - | - | 0 | - |
| - | - | 2.683E+05 | 503.6 | - | - | 0 | - |
| - | - | 1.867E+05 | 503.9 | - | - | 0 | - |
| - | - | 9.755E+04 | 504.2 | - | - | 0 | - |
| - | - | 3.082E+04 | 504.6 | - | - | 0 | - |
| - | - | 3.48E+04 | 504.9 | - | - | 0 | - |
| - | - | 3.498E+04 | 505.2 | - | - | 0 | - |
| - | - | 8.333E+04 | 507.2 | - | - | 0 | - |
| - | - | 3.794E+04 | 507.6 | - | - | 0 | - |
| 8 | y | 4.307E+05 | 507.7 | 0.0005684 | 1.119 | +2 | 8 |
| - | - | 2.133E+05 | 508.2 | - | - | 0 | - |
| - | - | 6.215E+04 | 508.7 | - | - | 0 | - |
| - | - | 3.079E+04 | 509.2 | - | - | 0 | - |
| 9 | b | 1.409E+05 | 510.2 | 0.001668 | 3.269 | +2 | 9 |
| - | - | 3.143E+04 | 510.6 | - | - | 0 | - |
| - | - | 4.304E+04 | 510.7 | - | - | 0 | - |
| 13 | b | 4.383E+04 | 510.9 | 0.006466 | 12.66 | +3 | 13 |
| - | - | 6.89E+04 | 511.2 | - | - | 0 | - |
| - | - | 1.914E+04 | 511.8 | - | - | 0 | - |
| - | - | 5.122E+04 | 512.2 | - | - | 0 | - |
| - | - | 1.956E+04 | 513.2 | - | - | 0 | - |
| - | - | 4.047E+04 | 514.9 | - | - | 0 | - |
| - | - | 3.772E+04 | 515.2 | - | - | 0 | - |
| - | - | 3.152E+04 | 515.3 | - | - | 0 | - |
| - | - | 7.668E+04 | 516.2 | - | - | 0 | - |
| 13 | b | 2.109E+04 | 516.9 | 0.004105 | 7.941 | +3 | 13 |
| - | - | 2.596E+04 | 517.3 | - | - | 0 | - |
| - | - | 2.129E+04 | 518.2 | - | - | 0 | - |
| - | - | 2.555E+04 | 519.3 | - | - | 0 | - |
| - | - | 4.798E+04 | 520.3 | - | - | 0 | - |
| - | - | 4.361E+04 | 520.6 | - | - | 0 | - |
| - | - | 3.04E+05 | 520.9 | - | - | 0 | - |
| - | - | 2.472E+05 | 521.2 | - | - | 0 | - |
| - | - | 1.326E+05 | 521.6 | - | - | 0 | - |
| - | - | 7.205E+04 | 521.8 | - | - | 0 | - |
| - | - | 9.625E+04 | 521.9 | - | - | 0 | - |
| - | - | 4.62E+04 | 522.2 | - | - | 0 | - |
| - | - | 5.898E+04 | 524.2 | - | - | 0 | - |
| - | - | 4.679E+04 | 524.9 | - | - | 0 | - |
| - | - | 6.607E+04 | 525.2 | - | - | 0 | - |
| 3 | y | 6.389E+05 | 526.6 | 0.0004632 | 0.8796 | +3 | 13 |
| 3 | y | 9.742E+05 | 526.9 | 0.003024 | 5.74 | +3 | 13 |
| - | - | 7.496E+05 | 527.2 | - | - | 0 | - |
| - | - | 4.135E+05 | 527.6 | - | - | 0 | - |
| - | - | 1.887E+05 | 527.9 | - | - | 0 | - |
| - | - | 3.706E+04 | 528.2 | - | - | 0 | - |
| - | - | 3.049E+04 | 529.3 | - | - | 0 | - |
| - | - | 2.723E+04 | 529.3 | - | - | 0 | - |
| - | - | 3.225E+04 | 529.7 | - | - | 0 | - |
| - | - | 1.561E+05 | 530.2 | - | - | 0 | - |
| - | - | 3.268E+04 | 530.6 | - | - | 0 | - |
| - | - | 3.312E+04 | 530.7 | - | - | 0 | - |
| - | - | 2.811E+04 | 530.9 | - | - | 0 | - |
| - | - | 2.331E+04 | 531.2 | - | - | 0 | - |
| 3 | y | 4.545E+06 | 532.6 | 0.0003596 | 0.6752 | +3 | 13 |
| - | - | 3.608E+06 | 532.9 | - | - | 0 | - |
| - | - | 1.872E+06 | 533.3 | - | - | 0 | - |
| - | - | 7.621E+05 | 533.6 | - | - | 0 | - |
| - | - | 1.935E+05 | 533.9 | - | - | 0 | - |
| - | - | 5.666E+04 | 534.3 | - | - | 0 | - |
| - | - | 6.399E+04 | 534.7 | - | - | 0 | - |
| - | - | 7.826E+04 | 535.2 | - | - | 0 | - |
| - | - | 2.724E+04 | 535.8 | - | - | 0 | - |
| - | - | 2.599E+04 | 535.9 | - | - | 0 | - |
| - | - | 2.367E+04 | 536.3 | - | - | 0 | - |
| - | - | 2.291E+04 | 536.6 | - | - | 0 | - |
| - | - | 4.662E+04 | 536.9 | - | - | 0 | - |
| - | - | 4.701E+04 | 537.3 | - | - | 0 | - |
| - | - | 5.388E+04 | 538.3 | - | - | 0 | - |
| - | - | 7.349E+04 | 538.8 | - | - | 0 | - |
| - | - | 5.691E+04 | 539.2 | - | - | 0 | - |
| - | - | 2.798E+05 | 543.2 | - | - | 0 | - |
| - | - | 1.696E+05 | 543.8 | - | - | 0 | - |
| - | - | 5.056E+04 | 544.2 | - | - | 0 | - |
| - | - | 6.873E+04 | 544.7 | - | - | 0 | - |
| - | - | 4.496E+04 | 545.2 | - | - | 0 | - |
| - | - | 1.274E+05 | 545.3 | - | - | 0 | - |
| - | - | 1.623E+05 | 547.3 | - | - | 0 | - |
| - | - | 6.888E+04 | 547.8 | - | - | 0 | - |
| - | - | 2.696E+04 | 547.9 | - | - | 0 | - |
| - | - | 7.948E+04 | 548.3 | - | - | 0 | - |
| - | - | 5.038E+04 | 551.8 | - | - | 0 | - |
| - | - | 1.229E+05 | 552.3 | - | - | 0 | - |
| - | - | 9.521E+04 | 552.8 | - | - | 0 | - |
| - | - | 1.851E+05 | 553.3 | - | - | 0 | - |
| - | - | 1.317E+05 | 553.8 | - | - | 0 | - |
| 14 | b | 6.754E+04 | 553.9 | 8.436E-05 | 0.1523 | +3 | 14 |
| - | - | 1.331E+05 | 554.3 | - | - | 0 | - |
| 7 | y | 5.63E+04 | 555.3 | 0.002887 | 5.198 | +2 | 9 |
| 7 | y | 1.193E+05 | 555.8 | 0.001181 | 2.125 | +2 | 9 |
| - | - | 1.387E+05 | 556.3 | - | - | 0 | - |
| - | - | 4.882E+04 | 556.8 | - | - | 0 | - |
| - | - | 2.692E+04 | 558.7 | - | - | 0 | - |
| 14 | b | 8.446E+04 | 559.6 | 0.0004298 | 0.7681 | +3 | 14 |
| 2 | y | 2.058E+05 | 559.9 | 0.003474 | 6.205 | +3 | 14 |
| - | - | 5.522E+04 | 560.3 | - | - | 0 | - |
| - | - | 1.127E+05 | 560.3 | - | - | 0 | - |
| - | - | 4.846E+04 | 560.6 | - | - | 0 | - |
| - | - | 5.438E+04 | 560.8 | - | - | 0 | - |
| - | - | 3.122E+04 | 561.3 | - | - | 0 | - |
| 7 | y | 1.844E+05 | 564.3 | 0.0001733 | 0.3071 | +2 | 9 |
| - | - | 4.573E+05 | 564.8 | - | - | 0 | - |
| - | - | 4.325E+05 | 565.3 | - | - | 0 | - |
| 2 | y | 6.825E+05 | 565.6 | 0.00026 | 0.4597 | +3 | 14 |
| - | - | 9.532E+04 | 565.8 | - | - | 0 | - |
| - | - | 5.867E+05 | 565.9 | - | - | 0 | - |
| - | - | 3.666E+05 | 566.3 | - | - | 0 | - |
| - | - | 1.585E+05 | 566.6 | - | - | 0 | - |
| - | - | 3.539E+04 | 566.9 | - | - | 0 | - |
| - | - | 2.176E+05 | 567.3 | - | - | 0 | - |
| - | - | 1.566E+05 | 567.8 | - | - | 0 | - |
| - | - | 1.139E+05 | 568.3 | - | - | 0 | - |
| - | - | 3.455E+04 | 568.9 | - | - | 0 | - |
| - | - | 3.646E+04 | 569.3 | - | - | 0 | - |
| 10 | b | 4.34E+04 | 569.8 | 0.004258 | 7.473 | +2 | 10 |
| 10 | b | 5.825E+04 | 570.3 | 0.009504 | 16.67 | +2 | 10 |
| - | - | 9.467E+04 | 573.3 | - | - | 0 | - |
| - | - | 3.543E+04 | 574.3 | - | - | 0 | - |
| 10 | b | 3.741E+05 | 578.8 | 7.438E-05 | 0.1285 | +2 | 10 |
| - | - | 1.762E+05 | 579.3 | - | - | 0 | - |
| - | - | 4.583E+04 | 579.8 | - | - | 0 | - |
| - | - | 6.498E+04 | 580.3 | - | - | 0 | - |
| - | - | 4.455E+05 | 582.3 | - | - | 0 | - |
| - | - | 2.553E+04 | 582.8 | - | - | 0 | - |
| - | - | 1.092E+05 | 583.3 | - | - | 0 | - |
| 5 | b | 2.253E+05 | 584.2 | 0.001695 | 2.902 | +1 | 5 |
| - | - | 5.954E+04 | 585.3 | - | - | 0 | - |
| - | - | 3.256E+04 | 585.8 | - | - | 0 | - |
| - | - | 2.742E+04 | 586.3 | - | - | 0 | - |
| - | - | 2.054E+05 | 588.3 | - | - | 0 | - |
| - | - | 5.285E+04 | 589.3 | - | - | 0 | - |
| 6 | y | 3.786E+04 | 590.8 | 0.001201 | 2.032 | +2 | 10 |
| - | - | 2.227E+04 | 591.2 | - | - | 0 | - |
| 6 | y | 7.008E+04 | 591.3 | 0.001502 | 2.541 | +2 | 10 |
| - | - | 4.647E+04 | 591.8 | - | - | 0 | - |
| - | - | 2.881E+04 | 592.2 | - | - | 0 | - |
| - | - | 1.876E+04 | 592.3 | - | - | 0 | - |
| - | - | 4.192E+04 | 593.3 | - | - | 0 | - |
| - | - | 3.661E+04 | 593.8 | - | - | 0 | - |
| - | - | 7.601E+04 | 594.3 | - | - | 0 | - |
| - | - | 7.791E+04 | 594.8 | - | - | 0 | - |
| - | - | 5.293E+04 | 595.3 | - | - | 0 | - |
| - | - | 7.132E+04 | 595.8 | - | - | 0 | - |
| - | - | 5.721E+04 | 596.3 | - | - | 0 | - |
| - | - | 3.483E+04 | 598.3 | - | - | 0 | - |
| - | - | 2.395E+04 | 599.8 | - | - | 0 | - |
| 6 | y | 3.392E+05 | 599.8 | 0.0002517 | 0.4197 | +2 | 10 |
| - | - | 5.498E+04 | 600.3 | - | - | 0 | - |
| - | - | 1.238E+05 | 600.3 | - | - | 0 | - |
| - | - | 6.914E+04 | 600.8 | - | - | 0 | - |
| - | - | 6.039E+04 | 602.3 | - | - | 0 | - |
| - | - | 1.045E+05 | 602.8 | - | - | 0 | - |
| - | - | 1.435E+05 | 603.3 | - | - | 0 | - |
| - | - | 6.035E+04 | 603.8 | - | - | 0 | - |
| - | - | 2.483E+04 | 604.3 | - | - | 0 | - |
| - | - | 3.681E+04 | 605.3 | - | - | 0 | - |
| - | - | 1.129E+05 | 607.3 | - | - | 0 | - |
| - | - | 1.125E+05 | 607.8 | - | - | 0 | - |
| 11 | y | 7.785E+04 | 608.3 | 0.002217 | 3.644 | +1 | 5 |
| - | - | 2.893E+05 | 608.8 | - | - | 0 | - |
| - | - | 1.557E+05 | 609.3 | - | - | 0 | - |
| - | - | 8.365E+04 | 609.8 | - | - | 0 | - |
| - | - | 5.507E+04 | 610.3 | - | - | 0 | - |
| - | - | 2.192E+04 | 611.2 | - | - | 0 | - |
| - | - | 1.714E+05 | 611.8 | - | - | 0 | - |
| - | - | 7.408E+04 | 612.3 | - | - | 0 | - |
| - | - | 6.14E+04 | 612.8 | - | - | 0 | - |
| - | - | 2.073E+04 | 613.3 | - | - | 0 | - |
| - | - | 2.271E+05 | 616.3 | - | - | 0 | - |
| - | - | 1.526E+05 | 616.8 | - | - | 0 | - |
| - | - | 1.035E+05 | 617.3 | - | - | 0 | - |
| - | - | 2.302E+05 | 617.8 | - | - | 0 | - |
| - | - | 1.134E+05 | 618.3 | - | - | 0 | - |
| - | - | 6.748E+04 | 618.8 | - | - | 0 | - |
| - | - | 3.192E+04 | 620.3 | - | - | 0 | - |
| - | - | 4.859E+04 | 620.8 | - | - | 0 | - |
| - | - | 4.442E+04 | 621.3 | - | - | 0 | - |
| - | - | 2.343E+04 | 624.8 | - | - | 0 | - |
| 11 | y | 3.94E+05 | 626.3 | 0.0003912 | 0.6246 | +1 | 5 |
| - | - | 6.397E+04 | 626.8 | - | - | 0 | - |
| - | - | 9.941E+04 | 627.3 | - | - | 0 | - |
| - | - | 8.925E+04 | 635.3 | - | - | 0 | - |
| - | - | 2.343E+04 | 636.3 | - | - | 0 | - |
| 6 | b | 1.202E+05 | 637.3 | 0.0002318 | 0.3638 | +1 | 6 |
| - | - | 5.04E+04 | 637.8 | - | - | 0 | - |
| - | - | 3.518E+04 | 638.3 | - | - | 0 | - |
| - | - | 2.069E+04 | 638.8 | - | - | 0 | - |
| - | - | 5.139E+04 | 639.3 | - | - | 0 | - |
| - | - | 3.477E+04 | 640.3 | - | - | 0 | - |
| - | - | 2.2E+04 | 641.3 | - | - | 0 | - |
| - | - | 8.58E+04 | 644.3 | - | - | 0 | - |
| - | - | 2.343E+04 | 645.3 | - | - | 0 | - |
| - | - | 2.458E+04 | 645.8 | - | - | 0 | - |
| - | - | 9.717E+04 | 646.3 | - | - | 0 | - |
| - | - | 1.765E+05 | 646.8 | - | - | 0 | - |
| - | - | 1.062E+05 | 647.3 | - | - | 0 | - |
| - | - | 4.214E+04 | 647.8 | - | - | 0 | - |
| - | - | 2.312E+04 | 648.3 | - | - | 0 | - |
| - | - | 2.25E+04 | 649.8 | - | - | 0 | - |
| - | - | 2.894E+04 | 650.3 | - | - | 0 | - |
| - | - | 2.933E+04 | 652.3 | - | - | 0 | - |
| - | - | 3.423E+04 | 653.3 | - | - | 0 | - |
| - | - | 2.625E+04 | 654.3 | - | - | 0 | - |
| 6 | b | 2.583E+05 | 655.3 | 0.0008114 | 1.238 | +1 | 6 |
| 5 | y | 3.365E+05 | 655.8 | 0.00578 | 8.813 | +2 | 11 |
| - | - | 2.419E+05 | 656.3 | - | - | 0 | - |
| - | - | 7.107E+04 | 656.8 | - | - | 0 | - |
| - | - | 1.306E+05 | 657.3 | - | - | 0 | - |
| - | - | 7.651E+04 | 658.3 | - | - | 0 | - |
| - | - | 6.375E+04 | 658.8 | - | - | 0 | - |
| - | - | 7.492E+04 | 659.8 | - | - | 0 | - |
| 11 | b | 6.27E+04 | 660.3 | 0.005223 | 7.91 | +2 | 11 |
| - | - | 3.695E+04 | 660.8 | - | - | 0 | - |
| - | - | 2.643E+04 | 662.8 | - | - | 0 | - |
| - | - | 2.839E+04 | 663.3 | - | - | 0 | - |
| - | - | 3.342E+04 | 663.8 | - | - | 0 | - |
| 5 | y | 1.083E+06 | 664.3 | 7.336E-05 | 0.1104 | +2 | 11 |
| - | - | 7.188E+05 | 664.8 | - | - | 0 | - |
| - | - | 3.164E+05 | 665.3 | - | - | 0 | - |
| - | - | 1.097E+05 | 665.8 | - | - | 0 | - |
| - | - | 5.207E+04 | 666.3 | - | - | 0 | - |
| - | - | 7.681E+04 | 667.3 | - | - | 0 | - |
| - | - | 8.047E+04 | 667.8 | - | - | 0 | - |
| - | - | 7.465E+04 | 668.3 | - | - | 0 | - |
| - | - | 2.404E+04 | 668.8 | - | - | 0 | - |
| - | - | 2.716E+04 | 670.3 | - | - | 0 | - |
| - | - | 1.331E+05 | 671.8 | - | - | 0 | - |
| - | - | 9.738E+04 | 672.3 | - | - | 0 | - |
| - | - | 2.131E+05 | 672.8 | - | - | 0 | - |
| - | - | 2.272E+05 | 673.3 | - | - | 0 | - |
| - | - | 1.327E+05 | 673.8 | - | - | 0 | - |
| - | - | 2.321E+05 | 674.3 | - | - | 0 | - |
| - | - | 7.541E+04 | 675.3 | - | - | 0 | - |
| - | - | 6.249E+04 | 676.3 | - | - | 0 | - |
| - | - | 1.3E+05 | 680.3 | - | - | 0 | - |
| - | - | 1.197E+05 | 680.8 | - | - | 0 | - |
| - | - | 9.121E+04 | 681.3 | - | - | 0 | - |
| - | - | 3.738E+05 | 681.8 | - | - | 0 | - |
| - | - | 2.495E+05 | 682.3 | - | - | 0 | - |
| - | - | 1.267E+05 | 682.8 | - | - | 0 | - |
| - | - | 4.927E+04 | 683.3 | - | - | 0 | - |
| - | - | 1.641E+05 | 684.3 | - | - | 0 | - |
| - | - | 9.912E+04 | 685.3 | - | - | 0 | - |
| - | - | 3.015E+04 | 686.3 | - | - | 0 | - |
| - | - | 5.426E+04 | 689.3 | - | - | 0 | - |
| - | - | 3.171E+04 | 689.8 | - | - | 0 | - |
| - | - | 2.306E+04 | 690.3 | - | - | 0 | - |
| - | - | 1.417E+05 | 691.3 | - | - | 0 | - |
| - | - | 6.408E+04 | 692.3 | - | - | 0 | - |
| - | - | 2.751E+04 | 693.3 | - | - | 0 | - |
| 12 | b | 1.016E+05 | 701.8 | 0.0009396 | 1.339 | +2 | 12 |
| - | - | 4.964E+05 | 702.3 | - | - | 0 | - |
| - | - | 3.646E+04 | 702.8 | - | - | 0 | - |
| - | - | 1.657E+05 | 703.3 | - | - | 0 | - |
| - | - | 6.405E+04 | 704.3 | - | - | 0 | - |
| - | - | 3.651E+04 | 708.4 | - | - | 0 | - |
| 12 | b | 4.098E+04 | 710.8 | 0.002859 | 4.023 | +2 | 12 |
| - | - | 3.526E+04 | 711.3 | - | - | 0 | - |
| - | - | 3.118E+04 | 712.3 | - | - | 0 | - |
| - | - | 3.06E+04 | 715.3 | - | - | 0 | - |
| - | - | 2.702E+04 | 715.8 | - | - | 0 | - |
| - | - | 2.331E+04 | 718.3 | - | - | 0 | - |
| - | - | 1.995E+05 | 719.3 | - | - | 0 | - |
| - | - | 7.265E+04 | 720.3 | - | - | 0 | - |
| - | - | 2.864E+04 | 721.3 | - | - | 0 | - |
| - | - | 4.541E+04 | 722.3 | - | - | 0 | - |
| 4 | y | 9.311E+04 | 723.8 | 0.0004796 | 0.6626 | +2 | 12 |
| 4 | y | 1.762E+05 | 724.3 | 0.002264 | 3.125 | +2 | 12 |
| - | - | 8.389E+04 | 724.8 | - | - | 0 | - |
| - | - | 7.364E+04 | 725.3 | - | - | 0 | - |
| - | - | 7.311E+04 | 731.3 | - | - | 0 | - |
| - | - | 4.049E+04 | 731.8 | - | - | 0 | - |
| 4 | y | 6.507E+05 | 732.9 | 0.0002078 | 0.2835 | +2 | 12 |
| - | - | 4.704E+05 | 733.4 | - | - | 0 | - |
| - | - | 1.961E+05 | 733.9 | - | - | 0 | - |
| - | - | 6.627E+04 | 734.4 | - | - | 0 | - |
| - | - | 1.211E+05 | 736.4 | - | - | 0 | - |
| - | - | 1.005E+05 | 737.3 | - | - | 0 | - |
| - | - | 4.446E+04 | 737.8 | - | - | 0 | - |
| - | - | 5.654E+04 | 738.3 | - | - | 0 | - |
| - | - | 3.051E+04 | 740.4 | - | - | 0 | - |
| 10 | y | 5.879E+04 | 745.4 | 0.001009 | 1.354 | +1 | 6 |
| - | - | 6.149E+04 | 745.8 | - | - | 0 | - |
| 10 | y | 1.061E+05 | 746.3 | 0.004722 | 6.326 | +1 | 6 |
| - | - | 4.382E+04 | 746.9 | - | - | 0 | - |
| - | - | 2.781E+04 | 747.4 | - | - | 0 | - |
| - | - | 4.067E+04 | 748.3 | - | - | 0 | - |
| 7 | b | 5.001E+04 | 750.4 | 0.001617 | 2.155 | +1 | 7 |
| - | - | 5.042E+04 | 754.9 | - | - | 0 | - |
| - | - | 7.345E+04 | 755.4 | - | - | 0 | - |
| - | - | 2.944E+04 | 756.3 | - | - | 0 | - |
| 10 | y | 2.208E+05 | 763.4 | 0.0004652 | 0.6094 | +1 | 6 |
| - | - | 7.008E+04 | 764.4 | - | - | 0 | - |
| - | - | 3.407E+04 | 765.4 | - | - | 0 | - |
| 13 | b | 3.184E+04 | 765.8 | 0.0006426 | 0.8391 | +2 | 13 |
| 13 | b | 4.407E+04 | 766.3 | 0.0123 | 16.05 | +2 | 13 |
| 7 | b | 1.924E+05 | 768.4 | 0.0005858 | 0.7624 | +1 | 7 |
| - | - | 9.237E+04 | 769.4 | - | - | 0 | - |
| - | - | 6.055E+04 | 772.4 | - | - | 0 | - |
| - | - | 3.135E+04 | 773.4 | - | - | 0 | - |
| 13 | b | 4.905E+04 | 774.9 | 0.0006114 | 0.7891 | +2 | 13 |
| - | - | 1.304E+05 | 781.3 | - | - | 0 | - |
| - | - | 4.783E+04 | 782.3 | - | - | 0 | - |
| - | - | 4.061E+04 | 788.3 | - | - | 0 | - |
| - | - | 3.257E+04 | 790.4 | - | - | 0 | - |
| - | - | 5.526E+04 | 802.4 | - | - | 0 | - |
| - | - | 3.38E+04 | 803.4 | - | - | 0 | - |
| - | - | 3.13E+04 | 804.4 | - | - | 0 | - |
| - | - | 3.832E+04 | 805.4 | - | - | 0 | - |
| - | - | 2.013E+04 | 806.4 | - | - | 0 | - |
| - | - | 7.066E+04 | 811.4 | - | - | 0 | - |
| - | - | 5.504E+04 | 812.4 | - | - | 0 | - |
| - | - | 2.683E+04 | 815.4 | - | - | 0 | - |
| - | - | 3.262E+04 | 816.4 | - | - | 0 | - |
| - | - | 9.779E+04 | 819.4 | - | - | 0 | - |
| - | - | 9.87E+04 | 820.4 | - | - | 0 | - |
| - | - | 5.499E+04 | 821.4 | - | - | 0 | - |
| - | - | 6.342E+04 | 822.4 | - | - | 0 | - |
| - | - | 2.161E+04 | 824.4 | - | - | 0 | - |
| - | - | 2.135E+05 | 833.4 | - | - | 0 | - |
| - | - | 1.143E+05 | 834.4 | - | - | 0 | - |
| - | - | 5.004E+04 | 835.4 | - | - | 0 | - |
| - | - | 1.113E+05 | 837.4 | - | - | 0 | - |
| - | - | 5.305E+04 | 838.4 | - | - | 0 | - |
| 2 | y | 1.293E+05 | 839.4 | 0.002808 | 3.346 | +2 | 14 |
| - | - | 6.442E+04 | 840.4 | - | - | 0 | - |
| - | - | 2.394E+04 | 842.4 | - | - | 0 | - |
| - | - | 5.192E+04 | 847.4 | - | - | 0 | - |
| - | - | 5.075E+04 | 848.4 | - | - | 0 | - |
| - | - | 3.278E+04 | 849.4 | - | - | 0 | - |
| - | - | 2.298E+04 | 850.4 | - | - | 0 | - |
| 9 | y | 2.395E+04 | 859.4 | 0.002371 | 2.759 | +1 | 7 |
| 9 | y | 7.075E+04 | 860.4 | 0.0006125 | 0.7119 | +1 | 7 |
| - | - | 1.722E+05 | 865.4 | - | - | 0 | - |
| - | - | 6.978E+04 | 866.4 | - | - | 0 | - |
| 9 | y | 1.918E+05 | 877.4 | 0.0003018 | 0.344 | +1 | 7 |
| - | - | 7.979E+04 | 878.4 | - | - | 0 | - |
| - | - | 2.472E+04 | 893.4 | - | - | 0 | - |
| - | - | 4.848E+04 | 894.4 | - | - | 0 | - |
| 8 | b | 6.047E+04 | 905.4 | 0.007862 | 8.683 | +1 | 8 |
| - | - | 2.292E+04 | 906.4 | - | - | 0 | - |
| - | - | 4.749E+04 | 909.4 | - | - | 0 | - |
| - | - | 3.442E+04 | 910.4 | - | - | 0 | - |
| - | - | 2.68E+04 | 911.4 | - | - | 0 | - |
| - | - | 2.721E+04 | 921.4 | - | - | 0 | - |
| - | - | 3.801E+04 | 922.4 | - | - | 0 | - |
| - | - | 2.564E+04 | 927.4 | - | - | 0 | - |
| - | - | 6.853E+04 | 930.4 | - | - | 0 | - |
| - | - | 4.863E+04 | 931.4 | - | - | 0 | - |
| - | - | 5.283E+04 | 932.4 | - | - | 0 | - |
| - | - | 4.255E+04 | 942.4 | - | - | 0 | - |
| - | - | 4.103E+04 | 947.4 | - | - | 0 | - |
| - | - | 2.072E+05 | 948.4 | - | - | 0 | - |
| - | - | 8.921E+04 | 949.4 | - | - | 0 | - |
| - | - | 2.348E+04 | 950.4 | - | - | 0 | - |
| - | - | 2.412E+04 | 953.4 | - | - | 0 | - |
| - | - | 2.642E+04 | 954.4 | - | - | 0 | - |
| - | - | 9.005E+04 | 965.5 | - | - | 0 | - |
| - | - | 1.116E+05 | 966.4 | - | - | 0 | - |
| - | - | 4.875E+04 | 967.4 | - | - | 0 | - |
| - | - | 1.119E+05 | 970.4 | - | - | 0 | - |
| - | - | 4.586E+04 | 971.4 | - | - | 0 | - |
| - | - | 3.959E+04 | 974.5 | - | - | 0 | - |
| - | - | 3.979E+04 | 983.5 | - | - | 0 | - |
| - | - | 2.745E+04 | 984.5 | - | - | 0 | - |
| 8 | y | 2.58E+04 | 997.4 | 0.006581 | 6.598 | +1 | 8 |
| 9 | b | 2.836E+04 | 1001 | 0.0003077 | 0.3073 | +1 | 9 |
| 9 | b | 7.175E+04 | 1002 | 0.007925 | 7.906 | +1 | 9 |
| - | - | 5.035E+04 | 1003 | - | - | 0 | - |
| 8 | y | 7.95E+04 | 1014 | 0.0008449 | 0.8329 | +1 | 8 |
| - | - | 2.408E+04 | 1015 | - | - | 0 | - |
| 9 | b | 1.719E+05 | 1019 | 0.0009847 | 0.9659 | +1 | 9 |
| - | - | 8.034E+04 | 1020 | - | - | 0 | - |
| - | - | 1.932E+04 | 1022 | - | - | 0 | - |
| - | - | 3.14E+04 | 1050 | - | - | 0 | - |
| - | - | 2.431E+04 | 1058 | - | - | 0 | - |
| - | - | 5.252E+04 | 1059 | - | - | 0 | - |
| - | - | 2.324E+04 | 1068 | - | - | 0 | - |
| - | - | 8.405E+04 | 1076 | - | - | 0 | - |
| - | - | 7.375E+04 | 1077 | - | - | 0 | - |
| - | - | 2.745E+04 | 1078 | - | - | 0 | - |
| - | - | 5.545E+04 | 1085 | - | - | 0 | - |
| - | - | 3.024E+04 | 1086 | - | - | 0 | - |
| - | - | 3.164E+04 | 1094 | - | - | 0 | - |
| - | - | 1.559E+05 | 1095 | - | - | 0 | - |
| - | - | 7.346E+04 | 1096 | - | - | 0 | - |
| - | - | 3.489E+04 | 1103 | - | - | 0 | - |
| - | - | 1.968E+04 | 1112 | - | - | 0 | - |
| - | - | 1.824E+04 | 1113 | - | - | 0 | - |
| 6 | y | 3.582E+04 | 1199 | 0.001375 | 1.147 | +1 | 10 |
| - | - | 3.054E+04 | 1232 | - | - | 0 | - |
| - | - | 1.9E+04 | 2009 | - | - | 0 | - |
| - | - | 1.839E+04 | 3108 | - | - | 0 | - |

m/z Charge Intensity FragmentType MassShift Position
120.05621337890625 0 13826.483
121.03974914550781 0 17260.787
122.07196807861328 0 16458.332
122.75192260742188 0 10557.271
123.05587768554688 0 23797.934
125.07933044433594 0 12495.694
128.08226013183594 0 26612.715
129.06625366210938 0 62304.473
129.10267639160156 0 4150834.5
130.05015563964844 0 19488.504
130.0867156982422 0 60334.566
130.10023498535156 0 31174.146
130.1060028076172 0 250307.17
136.07608032226562 0 2275731.8
137.073974609375 0 28562.676
137.07945251464844 0 179473.98
138.06663513183594 0 110767.945
139.08692932128906 0 74868.625
140.07135009765625 0 11864.07
141.04843139648438 0 19235.715
141.10255432128906 0 134617.69 a Water loss 1
142.08663940429688 0 12770.514
147.04458618164062 0 18960.852
147.07672119140625 0 12615.781
147.11314392089844 0 74377.65
148.0873260498047 0 291254.22
149.0452423095703 0 88204.99
149.0905303955078 0 33290.227
152.08251953125 0 22630.639
153.1028594970703 0 29209.584
154.06182861328125 0 19613.441
155.08203125 0 51811.977
155.09312438964844 0 111106.72
155.11827087402344 0 454401.62
156.07716369628906 0 83670.21
156.0839080810547 0 13527.953
156.12130737304688 0 36471.066
157.1339111328125 0 80996.39
158.1004180908203 0 30008.305
159.11325073242188 0 11230081 a 1
160.08741760253906 0 22692.23
160.11019897460938 0 54181.043
160.1165771484375 0 837096.06
161.1185302734375 0 43195.086
162.06687927246094 0 23058.898
165.07748413085938 0 72696.97
166.06149291992188 0 764734.7
167.0571746826172 0 14973.884
167.06491088867188 0 40125.86
167.09339904785156 0 20277.088
167.1182403564453 0 30672.34
169.06126403808594 0 20567.338
169.09771728515625 0 18391.945 b Water loss 1
171.11355590820312 0 27938.27
172.05117797851562 0 17419.586
173.0924072265625 0 159183.52 y Ammonia loss 12
173.4502716064453 0 178358.78
175.09817504882812 0 30144.953
176.08218383789062 0 106704.43
177.1138916015625 0 88254.8
178.05087280273438 0 14961.319
178.1341552734375 0 18681.316
179.0928955078125 0 28725.777
180.07684326171875 0 16330.166
180.11343383789062 0 44713.35
181.09800720214844 0 14453.711
183.07662963867188 0 71537.52
183.1131134033203 0 22854.705
185.09182739257812 0 14719.258
185.12913513183594 0 43661.062
186.08786010742188 0 16138.393
187.1081085205078 0 1621122.5 b 1
188.11158752441406 0 155732.48
189.07748413085938 0 73430.4
190.0615692138672 0 122638.38
191.0929718017578 0 49103.66
192.10232543945312 0 103321.66
193.0714874267578 0 17216.918
193.10862731933594 0 120130.89
194.0927734375 0 120821.945
195.07691955566406 0 95166.65
195.11322021484375 0 183848.1
196.08128356933594 0 28519.506
196.0906524658203 0 344272
197.0935516357422 0 60557.195
198.12474060058594 0 23917.555
199.10765075683594 0 38863.277
200.09225463867188 0 14916.95
201.08738708496094 0 164615.38
202.0869598388672 0 37483.84
202.10882568359375 0 43614.816
202.11875915527344 0 29667.65
203.09234619140625 0 30439.766
203.1216278076172 0 337904.2
204.07672119140625 0 14384.59
204.12542724609375 0 13307.966
206.10301208496094 0 19073.053
207.0880126953125 0 333220.47
208.09144592285156 0 28192.158
208.1197967529297 0 19900.44
212.10330200195312 0 61198.6
213.08758544921875 0 19973.111
216.13465881347656 0 150206.33 y Water loss 13
217.07232666015625 0 104112.69
217.11874389648438 0 171980.45 y Ammonia loss 13
218.05638122558594 0 85361.81
219.1125946044922 0 21858.639
220.10919189453125 0 14309.995 y 8
221.1036376953125 0 318055.9
222.0878143310547 0 51293.223
222.12423706054688 0 205668.75
223.10853576660156 0 35663.246
223.128662109375 0 33232.316
223.14479064941406 0 15620.885
223.1558380126953 0 71118.9
224.0857391357422 0 46162.465
224.1145782470703 0 311659.06
225.0430450439453 0 32570.02
225.09815979003906 0 20705.172
225.11773681640625 0 53760.46
226.11968994140625 0 16861.256
228.11305236816406 0 26630.59
230.08123779296875 0 34579.598
230.11395263671875 0 328348.75
230.1260223388672 0 25368.418
231.1165313720703 0 85155.51
233.13973999023438 0 42519.516
234.09906005859375 0 118600.125
234.1238555908203 0 54918.785
234.1453094482422 0 992099.9 y 13
235.08299255371094 0 525074.06
235.14857482910156 0 91059.06
236.08627319335938 0 43746.598
237.12379455566406 0 133390.34
239.11434936523438 0 350355.34
239.15069580078125 0 487605.84
240.0980987548828 0 32413.11
240.1347198486328 0 1131212.9
240.15283203125 0 44801.07
241.1122589111328 0 678938.7
241.13807678222656 0 118314.51
241.35353088378906 0 15551.179
242.1151123046875 0 58275.004
243.1076202392578 0 22668.38
244.12994384765625 0 25319.924
245.1937255859375 0 15585.524
247.108154296875 0 88974.086
249.09872436523438 0 223118.1
249.1349639892578 0 23019.904
250.08279418945312 0 35241.406
250.10235595703125 0 27238.775
251.0968017578125 0 25162.084
251.12594604492188 0 20446.732 b Water loss 8
251.15078735351562 0 188391.67
252.08059692382812 0 891106.7
252.1095733642578 0 1144530.5
253.08349609375 0 95800.97
253.11253356933594 0 122278.06
254.07620239257812 0 43935.715
256.1078186035156 0 18648.898
257.1611328125 0 126327.8
258.1075744628906 0 18207.053
258.1447448730469 0 61235.266
261.134033203125 0 15351.335
262.0935363769531 0 41772.293
263.078125 0 28254.535
264.13470458984375 0 36203.605
265.11883544921875 0 130846.71
267.109130859375 0 875960.5
268.1121520996094 0 96258.79
269.10699462890625 0 2922109
270.1099853515625 0 335203.5
271.10296630859375 0 132702.9
273.1349182128906 0 209993.42
274.1305236816406 0 24712.281
277.0928039550781 0 25875.012
279.0916442871094 0 16355.082
279.145751953125 0 105890.22
280.1044006347656 0 88493.61
280.1297302246094 0 182421.64
280.14593505859375 0 25693.879
280.65093994140625 0 55913.777
282.1451416015625 0 21280.707
283.119140625 0 19495.137
286.1327209472656 0 16077.909
292.62945556640625 0 28165.521 b 4
293.1263732910156 0 24727.996
294.193603515625 0 19360.76
294.6482849121094 0 132344.6
295.1042785644531 0 77035.36
295.1510314941406 0 39634.61
296.6353759765625 0 17962.121
297.10198974609375 0 35312.504
297.1564025878906 0 91623.39
298.1403503417969 0 21757.121
298.1601257324219 0 23034.234
300.1387023925781 0 24641.477 b Water loss 2
301.1299743652344 0 361293.12
302.1312561035156 0 55454.84
303.1105041503906 0 16926.707
303.16156005859375 0 29441.12
304.1417236328125 0 16479.26
304.1770324707031 0 60588.484
304.6556701660156 0 50646.01 y Water loss 10
305.1606750488281 0 69439.25
305.1783447265625 0 17601.986
306.14544677734375 0 27451.938
313.661376953125 0 148735.38 y 10
314.15008544921875 0 75698.62
314.168701171875 0 23261.46
314.65087890625 0 41965.254
314.81756591796875 0 24521.932
318.1379089355469 0 25401.281
319.1405029296875 0 18958.025 b Water loss 5
320.1358947753906 0 515364.1
321.13824462890625 0 51440.58
321.15765380859375 0 23403.729
322.1879577636719 0 217265.06
323.1718444824219 0 93495.984
323.19219970703125 0 40645.652
326.1365051269531 0 58427.473
327.14483642578125 0 18998.775
327.1665954589844 0 36435.777
328.1496887207031 0 62457.043 b 5
328.6676330566406 0 59377.8
329.1248474121094 0 22898.273
330.156005859375 0 21578.605
332.8274230957031 0 40091.11 y Water loss 7
333.15582275390625 0 42306.332 y Ammonia loss 7
337.16973876953125 0 18208.23
337.6727294921875 0 42169.15
338.1463623046875 0 477293.47
338.6745910644531 0 26916.535
338.83038330078125 0 69834.12 y 7
339.1490478515625 0 68568.914
339.1681823730469 0 26115.084
339.2138977050781 0 33773.703
339.5000305175781 0 18979.326
340.1409606933594 0 175464.75
340.1983947753906 0 29851.318
341.01812744140625 0 76593.016
341.14404296875 0 26080.35
341.1825256347656 0 162536.7
342.018798828125 0 40933.562
342.1867370605469 0 17709.898
342.66497802734375 0 18874.287
343.1645202636719 0 34492.57
344.1468505859375 0 116029.67
344.19329833984375 0 213448.25 y Water loss 12
345.17724609375 0 223168.62 y Ammonia loss 12
345.19708251953125 0 26695.984
346.17333984375 0 83487.22
346.6730041503906 0 31096.625
348.1299743652344 0 30130.629
350.1644592285156 0 38427.312
350.1850891113281 0 24238.803
351.166748046875 0 23372.322
351.6698303222656 0 205488.58
352.1426086425781 0 44598.55
352.1709899902344 0 82664.01
353.12371826171875 0 27736.613
353.14788818359375 0 26923.334
353.171875 0 20029.305
354.1313781738281 0 33671.535
355.1379089355469 0 22939.188
358.209228515625 0 91293.37
359.0289611816406 0 345700.66
359.1453857421875 0 26165.982
360.0292663574219 0 162921.81
360.16839599609375 0 131821.31
360.6700134277344 0 81722.125
361.02655029296875 0 81386.61
361.1730651855469 0 113054.32
362.12847900390625 0 24001.646
362.1808776855469 0 28905.076
362.2040100097656 0 176238.44 y 12
362.50177001953125 0 41003.71
363.2073059082031 0 22266.822
365.1932373046875 0 105689.28
366.1402893066406 0 120674.41
366.1591491699219 0 64466.32
367.1614074707031 0 29284.611
368.1753234863281 0 197762.1
368.5097351074219 0 17501.16
369.17431640625 0 81472.34
369.5101318359375 0 16703.984
370.153076171875 0 40782.305
370.69183349609375 0 45625.047
370.8471374511719 0 22453.734 y Ammonia loss 6
371.1578369140625 0 328833.8
372.14129638671875 0 172306.11
372.1620178222656 0 52017.516
373.1452331542969 0 42851.69
373.67730712890625 0 45642.33 y Ammonia loss 9
374.1811218261719 0 18696.879
375.1678771972656 0 84219.23
376.16845703125 0 16903.064
376.2192687988281 0 36736.37
376.5238342285156 0 19420.133 y 6
376.8516540527344 0 115557.41
377.1841125488281 0 75634.875
378.6711730957031 0 18073.314
379.6779479980469 0 25952.777
380.1378479003906 0 133738.16
380.8485412597656 0 19946.406
381.120361328125 0 41984.047
381.1443786621094 0 26530.838
382.190673828125 0 157029.61 y 9
382.6929016113281 0 44199.867
384.1667785644531 0 60071.684
384.1910400390625 0 21288.27
386.18194580078125 0 74059.02 b 9
386.5164489746094 0 43264.504
386.8489990234375 0 46014.266
387.180908203125 0 50145.64
387.2151184082031 0 31472.232
388.18304443359375 0 38635.824
388.686279296875 0 18067.41
389.1684265136719 0 154711.92
390.1735534667969 0 19208.344
390.93292236328125 0 25924.684
391.1728515625 0 32660.637
391.2088623046875 0 18920.957
393.1784973144531 0 108122.32
395.18707275390625 0 53504.49 y Water loss 2
395.4365539550781 0 75632.695 y Ammonia loss 2
395.68670654296875 0 23338.713
395.93878173828125 0 38404.66
396.5231628417969 0 26289.008
397.1619567871094 0 44507.277
397.1933288574219 0 26028.873
397.68878173828125 0 156708.23
398.1493835449219 0 991430.8
398.1932373046875 0 47385.613
399.1527099609375 0 192555.83
399.690673828125 0 271789 y 2
399.9412536621094 0 192410.83
400.1459655761719 0 50453.926
400.19482421875 0 105456.9
400.4408264160156 0 63169.734
400.5379638671875 0 42962.16
400.6929931640625 0 29321.87
401.197021484375 0 35651.336
402.17755126953125 0 70154.4
402.6778869628906 0 22854.16
403.1886291503906 0 15889.153
405.1964416503906 0 32871.246
405.2249755859375 0 116940.88
406.2020568847656 0 410354.88
406.2290954589844 0 14907.139
406.70306396484375 0 228207.25
407.2043151855469 0 28056.705
408.2066955566406 0 31393.404
408.2353210449219 0 34750.266
409.2019348144531 0 36983.996
409.70220947265625 0 30097.143
410.2027893066406 0 74294.07
410.6999206542969 0 32795.645
411.19427490234375 0 36304.67
411.44671630859375 0 18927.03
411.6900939941406 0 69718.22
412.18927001953125 0 56045.69
412.5204772949219 0 33753.773
413.1806335449219 0 45496.305
413.67706298828125 0 24231.271
415.1734619140625 0 213748.12
415.20745849609375 0 37975.758
415.6975402832031 0 47825.668 b Ammonia loss 13
416.17572021484375 0 62857.137
416.2040710449219 0 31946.982
417.16363525390625 0 26244.762
417.1901550292969 0 200062.84
417.6925354003906 0 53646.51
418.19342041015625 0 28365.441
419.2034912109375 0 32048.09
419.9537353515625 0 49242.953 b 13
420.2002868652344 0 406646.03 y Ammonia loss 1
420.7004699707031 0 128790.85
421.2025451660156 0 30093.125
421.7005920410156 0 32023.477
422.18658447265625 0 45981.754
423.1848449707031 0 25726.996
423.23565673828125 0 243436.03
424.19390869140625 0 30038.342
424.2384948730469 0 42515.887
424.4573669433594 0 254872.17 y 1
424.70758056640625 0 120438.414
424.9587097167969 0 70136.67
425.20745849609375 0 68078.49
426.1443786621094 0 29834.791
427.2126159667969 0 134115
428.2137451171875 0 24566.79
428.70001220703125 0 23559.621
429.0894470214844 0 102499.78
430.0903625488281 0 60792.453
430.2079162597656 0 35870.97 y Water loss 8
430.70123291015625 0 109673.09 y Ammonia loss 8
431.08856201171875 0 34054.91
431.2062072753906 0 53948.055
431.54071044921875 0 22267.168
431.87322998046875 0 26113.977
433.2190246582031 0 141204.3
433.4534912109375 0 44759.3
433.5387878417969 0 62053.26
433.87371826171875 0 73864.79
435.4560852050781 0 26680.572
436.23114013671875 0 173465.16
436.53570556640625 0 27007.594
437.210205078125 0 110470.11 y Water loss 4
437.4575500488281 0 146866.9
437.5484313964844 0 64614.36 y Ammonia loss 4
437.7083435058594 0 118717.63
437.95794677734375 0 92028.19
438.20550537109375 0 61560.875
439.21319580078125 0 267818.53 y 8
439.7144775390625 0 108689.984
440.2157897949219 0 21346.217
440.5375061035156 0 37132.27 b 10
440.8728942871094 0 21491.793
441.1953125 0 34976.867
441.7129821777344 0 466804.1 Precursor Water loss
441.96270751953125 0 390288.44 Precursor Ammonia loss
442.21173095703125 0 286170.94
442.4636535644531 0 155769.5
442.543212890625 0 85044.65
442.7127990722656 0 61623.324
442.8778991699219 0 48169.13
443.1669921875 0 59026.098
443.2159729003906 0 59142.67 y 4
443.5521545410156 0 34839.707
445.2419738769531 0 64302.055 y Water loss 11
445.5398254394531 0 26525.898
445.95849609375 0 20189.21
446.2159423828125 0 1602357.4 Precursor
446.4663391113281 0 1171165.4
446.7164001464844 0 679104.06
446.9674377441406 0 310935.97
447.21661376953125 0 120242.85
447.4682312011719 0 26998.406
447.8819274902344 0 17956.941
448.2159118652344 0 153443.75
448.5491943359375 0 116641.19
448.8825988769531 0 51273.297
449.2174987792969 0 21548.78
450.2452697753906 0 24396.898
451.1755676269531 0 30542.756
451.2306213378906 0 646241.2
452.2335205078125 0 137020.47
453.2205810546875 0 75604.42 b 7
453.7167663574219 0 21912.053
453.8907775878906 0 88344.98
454.2232971191406 0 113080.46
454.557373046875 0 45198.492
454.70477294921875 0 20071.5
455.2108459472656 0 188961.17 b 3
455.54132080078125 0 23328.13
455.7224426269531 0 31248.863
456.2120666503906 0 62330.5
457.2343444824219 0 26259.61
459.5465393066406 0 23643.922
459.8938293457031 0 145454.69
460.22686767578125 0 88791.37
460.5635681152344 0 34229.63
461.2171325683594 0 29284.59
462.2163391113281 0 24703.28
462.7176818847656 0 21169.78
463.2102355957031 0 121010.625
463.2517395019531 0 614882.44 y 11
463.2885437011719 0 34904.73
463.7110290527344 0 59841.535
464.15771484375 0 38056.477
464.22283935546875 0 49711.527
464.2555236816406 0 133462.12
464.7276611328125 0 20874.203
465.2235107421875 0 30185.271
465.258544921875 0 21799.08
465.7163391113281 0 44597.004
466.21124267578125 0 33617.8
466.7249755859375 0 32605.02
467.21856689453125 0 26881.26
468.223388671875 0 29941.17 b Water loss 11
468.2581787109375 0 39048.273
468.5477294921875 0 23609.73 b Ammonia loss 11
469.18707275390625 0 477656.56
470.1893310546875 0 95123.1
470.222900390625 0 42077.72
471.1858825683594 0 29057.88
471.221923828125 0 36528.625
471.5552062988281 0 28200.72
471.7224426269531 0 477591.22
471.88433837890625 0 30350.92
472.19110107421875 0 22359.357
472.2237854003906 0 229964.1
472.72357177734375 0 116452.54
474.2204895019531 0 46670.51 b 11
474.2579345703125 0 59259.27
474.5526428222656 0 42623.3
474.72314453125 0 58891.555
474.88330078125 0 18311.805
475.21319580078125 0 17484.818
476.71527099609375 0 61517.15
476.89410400390625 0 55340.297
477.21954345703125 0 173961.55
477.5553894042969 0 192540.77
477.7097473144531 0 41798.49
477.88824462890625 0 129298.164
478.224853515625 0 62003.836
478.55657958984375 0 28839.646
479.1761779785156 0 16499.824
479.2231750488281 0 148687.39
479.8935852050781 0 22503.006
480.2253723144531 0 44808.875
480.55548095703125 0 31638.748
481.1842956542969 0 92238.31
481.22015380859375 0 27836.89
482.9012145996094 0 244328.62 y Water loss 3
483.2312316894531 0 422862.3 y Ammonia loss 3
483.56427001953125 0 263250.75
483.729248046875 0 26471.662
483.89825439453125 0 104394.09
484.23046875 0 69262.8
485.72003173828125 0 317140.1
485.8915710449219 0 180246.64
486.22259521484375 0 331683.38
486.5594177246094 0 57841.383
486.720458984375 0 74760.07
486.8890075683594 0 39210.477
487.227294921875 0 22989.893
487.73388671875 0 178744.7
488.2341613769531 0 78398.24
488.7374267578125 0 23835.469
488.9046630859375 0 1572590.9 y 3
489.203369140625 0 45355.348
489.2390441894531 0 1072724.1
489.5730285644531 0 511413.62
489.90728759765625 0 198655.38
490.2427978515625 0 43690.26
490.72406005859375 0 33046.082
491.56634521484375 0 30117.812
491.8957214355469 0 385483.97
492.22967529296875 0 352299.84
492.5646667480469 0 160801.16
492.7249755859375 0 22579.703
492.8971252441406 0 76428.52
493.22589111328125 0 46015.34
493.720947265625 0 20417.09
497.2181701660156 0 78064.586
497.5704650878906 0 237530.61
497.9042053222656 0 265688.66
498.2045593261719 0 43431.535
498.2379150390625 0 111434.67
498.5708312988281 0 51931.168
498.7362976074219 0 78638.92 y Water loss 7
498.9001770019531 0 21729.25
499.1923828125 0 32262.09
499.23065185546875 0 124647.99 y Ammonia loss 7
499.73016357421875 0 56702.31
500.19525146484375 0 27749.6
501.5714111328125 0 19084.459
501.73162841796875 0 165332.2 b Ammonia loss 8
502.23614501953125 0 118919.58
503.25543212890625 0 56273.56
503.5741271972656 0 268266.8
503.9080505371094 0 186681.89
504.2421569824219 0 97546.516
504.5767517089844 0 30815.879
504.9092102050781 0 34796.242
505.2300109863281 0 34982.32
507.2099304199219 0 83327.016
507.572509765625 0 37939.746
507.7417907714844 0 430735.8 y 7
508.2431640625 0 213338.8
508.7441711425781 0 62150.37
509.249267578125 0 30789.842
510.2416687011719 0 140878.77 b 8
510.58038330078125 0 31430.896
510.7427978515625 0 43038.934
510.908203125 0 43825.695 b Water loss 12
511.2406921386719 0 68897.06
511.7622375488281 0 19136.242
512.2249755859375 0 51223.945
513.2262573242188 0 19560.059
514.8997802734375 0 40466.117
515.2352905273438 0 37715.645
515.2739868164062 0 31519.137
516.224853515625 0 76684.81
516.9093627929688 0 21086.574 b 12
517.251953125 0 25956.924
518.2471313476562 0 21292.639
519.2711791992188 0 25552.455
520.252197265625 0 47980.16
520.57763671875 0 43606.414
520.9059448242188 0 304044.75
521.23974609375 0 247180.97
521.5730590820312 0 132586.3
521.7571411132812 0 72047.61
521.9078369140625 0 96246.4
522.2401733398438 0 46201.434
524.2369384765625 0 58975.414
524.9172973632812 0 46793.117
525.2485961914062 0 66069.6
526.5812377929688 0 638940.44 y Water loss 2
526.9118041992188 0 974151.44 y Ammonia loss 2
527.2449951171875 0 749638.06
527.5785522460938 0 413519.8
527.911376953125 0 188743.55
528.242431640625 0 37056.727
529.2521362304688 0 30493.016
529.2994995117188 0 27229.082
529.7470092773438 0 32253.65
530.239990234375 0 156149.11
530.5925903320312 0 32680.72
530.7442626953125 0 33116.016
530.9251708984375 0 28106.412
531.244873046875 0 23307.848
532.5846557617188 0 4544828 y 2
532.9189453125 0 3608450.2
533.25244140625 0 1871762.1
533.5863647460938 0 762130.2
533.9208984375 0 193451.47
534.2532348632812 0 56662.316
534.7366943359375 0 63985.484
535.2449340820312 0 78262.445
535.751953125 0 27238.494
535.911376953125 0 25994.92
536.2529296875 0 23669.904
536.5971069335938 0 22906.469
536.9290771484375 0 46621.54
537.276611328125 0 47012.43
538.2532958984375 0 53878.297
538.7521362304688 0 73493.02
539.2479248046875 0 56908.555
543.2493896484375 0 279800.5
543.7504272460938 0 169623.16
544.2498168945312 0 50560.945
544.7440185546875 0 68729.73
545.24609375 0 44960.156
545.2953491210938 0 127449.445
547.2635498046875 0 162284.81
547.755126953125 0 68882.305
547.9259033203125 0 26955.133
548.2593994140625 0 79477.91
551.7614135742188 0 50384.133
552.2537231445312 0 122905.586
552.7543334960938 0 95205.19
553.2533569335938 0 185127.53
553.7555541992188 0 131666.08
553.927978515625 0 67543.93 b Ammonia loss 13
554.2670288085938 0 133083.14
555.2750854492188 0 56298.04 y Water loss 6
555.768798828125 0 119310.56 y Ammonia loss 6
556.2619018554688 0 138728.55
556.76318359375 0 48816.92
558.7389526367188 0 26921.559
559.6031494140625 0 84464.61 b 13
559.93505859375 0 205751.05 y Ammonia loss 1
560.2606201171875 0 55220.594
560.2944946289062 0 112746.46
560.5997314453125 0 48463.418
560.766845703125 0 54376.2
561.3008422851562 0 31218.584
564.2830810546875 0 184415.28 y 6
564.7734375 0 457276.88
565.2738037109375 0 432460.9
565.6073608398438 0 682455.3 y 1
565.7740478515625 0 95315.61
565.9414672851562 0 586724
566.2753295898438 0 366607.25
566.6098022460938 0 158487.92
566.94140625 0 35393.73
567.2501831054688 0 217629.94
567.752197265625 0 156642.75
568.2542724609375 0 113851.3
568.9362182617188 0 34554.086
569.2653198242188 0 36459.305
569.7684326171875 0 43400.508 b Water loss 9
570.2656860351562 0 58249.805 b Ammonia loss 9
573.288330078125 0 94671.9
574.2945556640625 0 35426.496
578.76953125 0 374092.25 b 9
579.2708129882812 0 176221.19
579.7710571289062 0 45829.445
580.2733764648438 0 64978.465
582.27099609375 0 445530.75
582.7798461914062 0 25531.559
583.2735595703125 0 109175.055
584.2514038085938 0 225322.44 b 4
585.2545776367188 0 59544.43
585.766845703125 0 32555.232
586.2650756835938 0 27416.098
588.289306640625 0 205369.16
589.2913818359375 0 52846.49
590.7977294921875 0 37856.15 y Water loss 5
591.2422485351562 0 22273.78
591.2900390625 0 70076.61 y Ammonia loss 5
591.7920532226562 0 46474.324
592.2184448242188 0 28807.225
592.26953125 0 18763.83
593.2700805664062 0 41919.844
593.7880249023438 0 36608.402
594.2803955078125 0 76013.29
594.7796630859375 0 77910.36
595.2781982421875 0 52933.63
595.76953125 0 71324.6
596.2609252929688 0 57206.09
598.2745971679688 0 34829.617
599.7550048828125 0 23949.447
599.8020629882812 0 339161.47 y 5
600.2559814453125 0 54984.492
600.3038940429688 0 123814.92
600.8046264648438 0 69135.85
602.2855224609375 0 60387.484
602.7941284179688 0 104477.664
603.2903442382812 0 143493.83
603.7887573242188 0 60354.13
604.2911376953125 0 24833.988
605.316162109375 0 36808.363
607.2781372070312 0 112890.23
607.7763671875 0 112505.016
608.3016357421875 0 77853.94 y Water loss 10
608.7705688476562 0 289327.84
609.2734375 0 155696.72
609.7704467773438 0 83648.56
610.2747802734375 0 55070.04
611.229248046875 0 21921.662
611.801513671875 0 171417.31
612.3023681640625 0 74081.86
612.80224609375 0 61404.24
613.3034057617188 0 20730.33
616.283935546875 0 227122.31
616.7852783203125 0 152587.53
617.283935546875 0 103464.414
617.7775268554688 0 230222.44
618.2789916992188 0 113399.16
618.7764892578125 0 67478.05
620.2921752929688 0 31924.453
620.806640625 0 48588.824
621.3058471679688 0 44423.812
624.7996215820312 0 23432.754
626.3140258789062 0 393994.1 y 10
626.7896118164062 0 63967.668
627.31591796875 0 99406.72
635.2689208984375 0 89250.35
636.2619018554688 0 23434.6
637.2764892578125 0 120235.24 b Water loss 5
637.7977905273438 0 50403.117
638.2883911132812 0 35182.555
638.7938842773438 0 20690.871
639.299560546875 0 51391.594
640.3287353515625 0 34767.273
641.3015747070312 0 21995.484
644.278564453125 0 85795.055
645.2816162109375 0 23430.059
645.7952270507812 0 24579.635
646.3078002929688 0 97168.625
646.8065795898438 0 176454.6
647.3049926757812 0 106170.92
647.8009643554688 0 42135.21
648.3009033203125 0 23118.818
649.8054809570312 0 22500.316
650.2969360351562 0 28944.99
652.298583984375 0 29334.584
653.281494140625 0 34229.844
654.2980346679688 0 26246.81
655.2860107421875 0 258332.53 b 5
655.8156127929688 0 336505.72 y Ammonia loss 4
656.3143310546875 0 241924.55
656.8128662109375 0 71074.305
657.3113403320312 0 130633.32
658.3277587890625 0 76508.79
658.8030395507812 0 63753.633
659.7946166992188 0 74918.29
660.2958984375 0 62703.848 b 10
660.79833984375 0 36953.27
662.8124389648438 0 26431.885
663.3114624023438 0 28390.38
663.8065795898438 0 33419.016
664.3231811523438 0 1082796 y 4
664.8243408203125 0 718830.44
665.3245239257812 0 316360.16
665.8268432617188 0 109685
666.3155517578125 0 52074.46
667.3005981445312 0 76810.01
667.8062744140625 0 80471.44
668.306640625 0 74647.62
668.8131103515625 0 24039.67
670.3060302734375 0 27156.16
671.8180541992188 0 133149.9
672.3194580078125 0 97376.05
672.8004760742188 0 213108.4
673.29931640625 0 227179.94
673.7987060546875 0 132695.64
674.3358764648438 0 232064.39
675.337158203125 0 75407.664
676.3406372070312 0 62488.26
680.33056640625 0 130001.44
680.83056640625 0 119670.3
681.3347778320312 0 91211.586
681.8045043945312 0 373807.12
682.3053588867188 0 249494.7
682.8069458007812 0 126723.836
683.3070068359375 0 49268.51
684.3186645507812 0 164135.4
685.3143310546875 0 99116.96
686.3250732421875 0 30147.75
689.3340454101562 0 54260.01
689.8401489257812 0 31709.898
690.3239135742188 0 23056.297
691.3310546875 0 141724.31
692.3350219726562 0 64077.008
693.3167724609375 0 27512.625
701.8206176757812 0 101550.98 b Water loss 11
702.3307495117188 0 496424.38
702.8221435546875 0 36464.38
703.3331298828125 0 165725.47
704.336181640625 0 64054.03
708.3576049804688 0 36509.703
710.8278198242188 0 40980.453 b 11
711.3236694335938 0 35263.074
712.3226318359375 0 31184.908
715.3399658203125 0 30598.84
715.8339233398438 0 27020.463
718.3441772460938 0 23312.703
719.33056640625 0 199545.02
720.328857421875 0 72652.73
721.328125 0 28635.588
722.3350219726562 0 45407.434
723.8468017578125 0 93105.56 y Water loss 3
724.341552734375 0 176226.81 y Ammonia loss 3
724.84375 0 83889.66
725.3408813476562 0 73636.75
731.3348999023438 0 73108.54
731.8388671875 0 40490.516
732.8523559570312 0 650651.25 y 3
733.3536376953125 0 470435.62
733.8550415039062 0 196085.42
734.355224609375 0 66271.695
736.3526611328125 0 121133.51
737.3433227539062 0 100469.586
737.842041015625 0 44456.695
738.3432006835938 0 56535.07
740.3660278320312 0 30505.227
745.3617553710938 0 58794.504 y Water loss 9
745.848876953125 0 61494.582
746.3515014648438 0 106122.85 y Ammonia loss 9
746.8519897460938 0 43821.168
747.3533325195312 0 27810.758
748.3475341796875 0 40673.83
750.3619384765625 0 50013.344 b Water loss 6
754.8558959960938 0 50422.38
755.3525390625 0 73454.586
756.3493041992188 0 29437.54
763.3728637695312 0 220784.17 y 9
764.3760375976562 0 70082.84
765.3814086914062 0 34069.707
765.849609375 0 31840.963 b Water loss 12
766.353271484375 0 44065.695 b Ammonia loss 12
768.3703002929688 0 192372.1 b 6
769.3728637695312 0 92373.98
772.3728637695312 0 60551.12
773.3768310546875 0 31346.541
774.8536376953125 0 49050.523 b 12
781.3372192382812 0 130360.24
782.34375 0 47827.715
788.3477172851562 0 40613.707
790.394287109375 0 32574
802.36083984375 0 55260.023
803.3635864257812 0 33804.79
804.36962890625 0 31295.814
805.3799438476562 0 38317.28
806.3840942382812 0 20132.938
811.395751953125 0 70657.055
812.3973999023438 0 55037.957
815.3612060546875 0 26834.1
816.3530883789062 0 32621.75
819.390380859375 0 97794.414
820.382568359375 0 98700.69
821.3814086914062 0 54985.78
822.3676147460938 0 63415.816
824.3701171875 0 21605.393
833.3716430664062 0 213510.89
834.3743896484375 0 114289.08
835.373779296875 0 50042.55
837.4006958007812 0 111325.234
838.4003295898438 0 53051.504
839.3909301757812 0 129256.38 y Ammonia loss 1
840.391845703125 0 64417.43
842.3803100585938 0 23935.469
847.384521484375 0 51924.28
848.3834838867188 0 50753.453
849.3798828125 0 32778.105
850.3935546875 0 22975.219
859.4033203125 0 23952.742 y Water loss 8
860.3903198242188 0 70746.266 y Ammonia loss 8
865.3951416015625 0 172187.11
866.3970947265625 0 69782.95
877.4159545898438 0 191762.42 y 8
878.4177856445312 0 79787.625
893.4114990234375 0 24715.389
894.4198608398438 0 48476.92
905.4219360351562 0 60469.227 b 7
906.431640625 0 22915.416
909.4345703125 0 47491.49
910.43359375 0 34424.684
911.4402465820312 0 26799.492
921.4286499023438 0 27205.662
922.4257202148438 0 38011.57
927.4463500976562 0 25637.262
930.41943359375 0 68532.984
931.412109375 0 48630.246
932.4268798828125 0 52830.22
942.4314575195312 0 42546.746
947.44775390625 0 41034.61
948.4334106445312 0 207201.62
949.4358520507812 0 89213.2
950.4334716796875 0 23475.855
953.4096069335938 0 24121.812
954.4116821289062 0 26423.213
965.4583740234375 0 90053.47
966.4473876953125 0 111607.984
967.4483032226562 0 48746.883
970.4310913085938 0 111856.62
971.429443359375 0 45855.63
974.4586181640625 0 39590.234
983.472900390625 0 39791.062
984.4674682617188 0 27451.883
997.4552001953125 0 25802.076 y Ammonia loss 7
1001.4618530273438 0 28357.863 b Water loss 8
1002.4541015625 0 71752.38 b Ammonia loss 8
1003.458251953125 0 50354.81
1014.4760131835938 0 79501.445 y 7
1015.4706420898438 0 24083.533
1019.4717407226562 0 171919.88 b 8
1020.4718627929688 0 80335.57
1022.4900512695312 0 19319.002
1050.4813232421875 0 31401.799
1058.47314453125 0 24307.154
1059.474853515625 0 52515.035
1068.472900390625 0 23237.625
1076.4940185546875 0 84053.39
1077.49169921875 0 73745.766
1078.4833984375 0 27446.732
1085.4866943359375 0 55451.004
1086.49609375 0 30243.727
1093.5545654296875 0 31641.004
1094.5006103515625 0 155854.53
1095.5037841796875 0 73455.3
1103.499267578125 0 34887.168
1111.532958984375 0 19677.086
1112.5313720703125 0 18243.46
1198.594970703125 0 35820.117 y 5
1231.5657958984375 0 30542.46
2009.078857421875 0 19003.04
3107.63525390625 0 18389.678

Spectrum Details

|  |  |
| --- | --- |
| Matched peaks? Matched peaks The total absolute number of peaks matched. Additionally in brackets the total fraction of peaks matched and the total number of peaks is shown. | 109 (11.43% of 954) |
| FDR? FDR The false discovery rate estimated for this peptide. It is calculated by matching all theoretical fragments with a non-integer shift with the raw peaks for this spectrum. This is done with 40 different shifts. The resulting percentage is the average number of annotated peaks over the number of annotated peaks with the correct spectrum. | 0.35% |
| Satellite FDR? Satellite FDR See the FDR for details on its calculation. This satellite ion specific FDR only contains the satellite ions (d/w) for I/L/J positions. | - |
| PSM Score? PSM Score The PSM Score as given by Hecklib to this annotated spectrum. It is shown with three significant figures. | 474 |

#### Spectrum 2379? Spectrum 2379 The raw spectrum of this peptide as annotated by Hecklib. The fragments are coloured according to ion type (see legend). Any peaks with a star '\*' as text can be hovered over to see the full details, first the ion type second the mass shift type. By hovering over the amino acids in the peptide or ions in the legend the corresponding peaks are highlighted. By toggling the 'Unassigned' label you can turn the background (unassigned) peaks on or off in the plot. By updating the slider in the Ion legend you can update the spectrum to only show the top X% of the peaks with labels. The top X% means any peak that is within X% of the highest intensity. By dragging in the spectrum you can zoom in to a specific part of the spectrum and use 'Zoom Out' to get back to the original zoom level. The annotation of the spectrum is based on the given sequence in the peptides file and is done with different software so inconsistencies are likely. The peaks are annotated based on the given sequence, with 20 ppm tolerance.

Copy Data

##### Spectrum 2379 (TSV)

###### Preview

```
Loading example...
```

*Click on the button to copy the data to your clipboard.*

Mz MinMz MaxIntensity Max

WidthHeightPeptide font sizePeptide stroke widthSpectrum font sizeSpectrum stroke widthCompact peptide

Ion legend

wxyz

abcd

OtherUnassignedIon^Charge^_Position_Show for top:%

SVMHEAJHNHYTQKS

06.87e+41.37e+52.06e+52.75e+5

Zoom Out

a^+1^_2_a^+1^_2_b^+1^_2_y^+1^_2_y^+1^_2_y^+1^_2_b^+1^_3_y^+1^_3_y^+1^_3_y^+1^_3_y^+1^_4_y^+1^_4_b^+1^_4_b^+2^_9_y^+2^_9_b^+1^_5_b^+2^_10_b^+1^_5_y^+1^_5_y^+1^_5_b^+1^_6_b^+2^_11_b^+1^_6_b^+2^_12_b^+2^_12_b^+2^_12_y^+2^_12_y^+2^_12_y^+1^_6_y^+1^_6_y^+1^_6_b^+1^_7_b^+2^_13_b^+2^_13_b^+1^_7_y^+2^_13_y^+2^_13_b^+2^_14_b^+2^_14_b^+2^_14_y^+2^_14_y^+2^_14_y^+1^_7_y^+1^_7_y^+1^_7_\*\*\*b^+1^_8_y^+1^_8_y^+1^_8_b^+1^_9_y^+1^_9_y^+1^_9_y^+1^_9_b^+1^_10_y^+1^_10_y^+1^_10_y^+1^_11_y^+1^_11_y^+1^_11_b^+1^_11_b^+1^_12_b^+1^_12_b^+1^_12_y^+1^_12_y^+1^_12_b^+1^_13_

0727145321802906

Fragment Matches Table

Show background peaks

| Position | Ion type | Intensity | mz Theoretical | mz Error (Th) | mz Error (ppm) | Charge | Series Number |
| --- | --- | --- | --- | --- | --- | --- | --- |
| - | - | 422.7 | 122.1 | - | - | 0 | - |
| - | - | 495.4 | 123.1 | - | - | 0 | - |
| - | - | 540.2 | 126.1 | - | - | 0 | - |
| - | - | 368.8 | 127.1 | - | - | 0 | - |
| - | - | 2.038E+04 | 129.1 | - | - | 0 | - |
| - | - | 775.3 | 130.1 | - | - | 0 | - |
| - | - | 5331 | 136.1 | - | - | 0 | - |
| - | - | 417.6 | 136.1 | - | - | 0 | - |
| - | - | 768.4 | 138.1 | - | - | 0 | - |
| - | - | 822.7 | 138.1 | - | - | 0 | - |
| - | - | 565.8 | 139.1 | - | - | 0 | - |
| - | - | 416.5 | 140.2 | - | - | 0 | - |
| 2 | a | 410.6 | 141.1 | 0.0003149 | 2.231 | +1 | 2 |
| - | - | 446.9 | 142.8 | - | - | 0 | - |
| - | - | 397.2 | 143.5 | - | - | 0 | - |
| - | - | 837.8 | 148.1 | - | - | 0 | - |
| - | - | 565.6 | 148.9 | - | - | 0 | - |
| - | - | 576.3 | 148.9 | - | - | 0 | - |
| - | - | 631.1 | 148.9 | - | - | 0 | - |
| - | - | 701.7 | 148.9 | - | - | 0 | - |
| - | - | 942.9 | 148.9 | - | - | 0 | - |
| - | - | 1125 | 148.9 | - | - | 0 | - |
| - | - | 1383 | 148.9 | - | - | 0 | - |
| - | - | 3061 | 148.9 | - | - | 0 | - |
| - | - | 6171 | 149 | - | - | 0 | - |
| - | - | 3024 | 149 | - | - | 0 | - |
| - | - | 1310 | 149 | - | - | 0 | - |
| - | - | 1177 | 149 | - | - | 0 | - |
| - | - | 897.7 | 149 | - | - | 0 | - |
| - | - | 752.8 | 149 | - | - | 0 | - |
| - | - | 509 | 149 | - | - | 0 | - |
| - | - | 518.4 | 149 | - | - | 0 | - |
| - | - | 491.9 | 149 | - | - | 0 | - |
| - | - | 489.2 | 149 | - | - | 0 | - |
| - | - | 1195 | 152.1 | - | - | 0 | - |
| - | - | 1414 | 155.1 | - | - | 0 | - |
| - | - | 897.6 | 156.1 | - | - | 0 | - |
| 2 | a | 1.894E+04 | 159.1 | 0.000355 | 2.231 | +1 | 2 |
| - | - | 1381 | 160.1 | - | - | 0 | - |
| - | - | 1250 | 165.1 | - | - | 0 | - |
| - | - | 451.4 | 165.2 | - | - | 0 | - |
| - | - | 1813 | 166.1 | - | - | 0 | - |
| - | - | 866.5 | 166.1 | - | - | 0 | - |
| - | - | 1037 | 167.1 | - | - | 0 | - |
| - | - | 1931 | 176.1 | - | - | 0 | - |
| - | - | 1759 | 180.1 | - | - | 0 | - |
| - | - | 559.4 | 180.1 | - | - | 0 | - |
| 2 | b | 9516 | 187.1 | 0.0002982 | 1.594 | +1 | 2 |
| - | - | 583.1 | 188.1 | - | - | 0 | - |
| - | - | 783.8 | 189.1 | - | - | 0 | - |
| - | - | 1209 | 190.1 | - | - | 0 | - |
| - | - | 463.6 | 191.1 | - | - | 0 | - |
| - | - | 541.7 | 192.1 | - | - | 0 | - |
| - | - | 837.6 | 193.1 | - | - | 0 | - |
| - | - | 5032 | 193.1 | - | - | 0 | - |
| - | - | 1104 | 194.1 | - | - | 0 | - |
| - | - | 1657 | 195.1 | - | - | 0 | - |
| - | - | 560.2 | 195.1 | - | - | 0 | - |
| - | - | 672.3 | 195.1 | - | - | 0 | - |
| - | - | 504.5 | 201.1 | - | - | 0 | - |
| - | - | 567.8 | 202.1 | - | - | 0 | - |
| - | - | 2750 | 204.1 | - | - | 0 | - |
| - | - | 567.1 | 204.1 | - | - | 0 | - |
| - | - | 2724 | 207.1 | - | - | 0 | - |
| - | - | 450.4 | 209.9 | - | - | 0 | - |
| - | - | 858.6 | 211.1 | - | - | 0 | - |
| - | - | 841.3 | 212.1 | - | - | 0 | - |
| 14 | y | 1332 | 216.1 | 0.0004825 | 2.232 | +1 | 2 |
| - | - | 960.2 | 217.1 | - | - | 0 | - |
| 14 | y | 2058 | 217.1 | 0.0005367 | 2.472 | +1 | 2 |
| - | - | 1.027E+04 | 221.1 | - | - | 0 | - |
| - | - | 742.2 | 221.1 | - | - | 0 | - |
| - | - | 1068 | 222.1 | - | - | 0 | - |
| - | - | 1798 | 222.1 | - | - | 0 | - |
| - | - | 547.7 | 223.1 | - | - | 0 | - |
| - | - | 877.1 | 223.2 | - | - | 0 | - |
| - | - | 3415 | 224.1 | - | - | 0 | - |
| - | - | 561.3 | 228.1 | - | - | 0 | - |
| - | - | 7612 | 230.1 | - | - | 0 | - |
| - | - | 2026 | 233.1 | - | - | 0 | - |
| - | - | 1215 | 234.1 | - | - | 0 | - |
| - | - | 1086 | 234.1 | - | - | 0 | - |
| 14 | y | 1.384E+04 | 234.1 | 0.0004921 | 2.102 | +1 | 2 |
| - | - | 6279 | 235.1 | - | - | 0 | - |
| - | - | 2093 | 235.1 | - | - | 0 | - |
| - | - | 566.9 | 236.1 | - | - | 0 | - |
| - | - | 833.9 | 237.1 | - | - | 0 | - |
| - | - | 2615 | 239.1 | - | - | 0 | - |
| - | - | 9484 | 239.2 | - | - | 0 | - |
| - | - | 1.153E+04 | 240.1 | - | - | 0 | - |
| - | - | 1261 | 240.2 | - | - | 0 | - |
| - | - | 881.1 | 241.1 | - | - | 0 | - |
| - | - | 1212 | 247.1 | - | - | 0 | - |
| - | - | 3070 | 249.1 | - | - | 0 | - |
| - | - | 751.3 | 249.1 | - | - | 0 | - |
| - | - | 2224 | 251.2 | - | - | 0 | - |
| - | - | 1.622E+04 | 252.1 | - | - | 0 | - |
| - | - | 713.7 | 252.1 | - | - | 0 | - |
| - | - | 830.7 | 253.1 | - | - | 0 | - |
| - | - | 1613 | 253.1 | - | - | 0 | - |
| - | - | 2438 | 257.1 | - | - | 0 | - |
| - | - | 2579 | 257.2 | - | - | 0 | - |
| - | - | 590.9 | 258.1 | - | - | 0 | - |
| - | - | 497.8 | 261.1 | - | - | 0 | - |
| - | - | 985 | 262.1 | - | - | 0 | - |
| - | - | 1492 | 264.1 | - | - | 0 | - |
| - | - | 864.1 | 265.1 | - | - | 0 | - |
| - | - | 1084 | 267.1 | - | - | 0 | - |
| - | - | 7432 | 267.1 | - | - | 0 | - |
| - | - | 602.7 | 268.1 | - | - | 0 | - |
| - | - | 1216 | 269.1 | - | - | 0 | - |
| - | - | 2963 | 270.1 | - | - | 0 | - |
| - | - | 3934 | 273.1 | - | - | 0 | - |
| - | - | 2308 | 275.1 | - | - | 0 | - |
| - | - | 2206 | 279.1 | - | - | 0 | - |
| - | - | 1549 | 280.1 | - | - | 0 | - |
| - | - | 1389 | 280.1 | - | - | 0 | - |
| - | - | 3686 | 285.1 | - | - | 0 | - |
| - | - | 560.2 | 288 | - | - | 0 | - |
| - | - | 783.6 | 295.1 | - | - | 0 | - |
| - | - | 6008 | 301.1 | - | - | 0 | - |
| - | - | 913.9 | 302.1 | - | - | 0 | - |
| - | - | 635.9 | 302.2 | - | - | 0 | - |
| - | - | 608.7 | 305.2 | - | - | 0 | - |
| - | - | 5590 | 320.1 | - | - | 0 | - |
| - | - | 993.6 | 321.1 | - | - | 0 | - |
| - | - | 3368 | 322.2 | - | - | 0 | - |
| - | - | 886.6 | 323.2 | - | - | 0 | - |
| - | - | 776.7 | 326.1 | - | - | 0 | - |
| - | - | 598 | 327.2 | - | - | 0 | - |
| - | - | 789.7 | 332.1 | - | - | 0 | - |
| 3 | b | 5537 | 334.1 | 0.005534 | 16.56 | +1 | 3 |
| - | - | 4683 | 338.1 | - | - | 0 | - |
| - | - | 570.8 | 339.1 | - | - | 0 | - |
| - | - | 2227 | 340.1 | - | - | 0 | - |
| - | - | 1511 | 341.2 | - | - | 0 | - |
| - | - | 1513 | 344.1 | - | - | 0 | - |
| 13 | y | 4247 | 344.2 | 0.000575 | 1.671 | +1 | 3 |
| 13 | y | 4135 | 345.2 | 0.0005377 | 1.558 | +1 | 3 |
| - | - | 671.9 | 345.2 | - | - | 0 | - |
| - | - | 545.3 | 346.2 | - | - | 0 | - |
| - | - | 958.8 | 347.2 | - | - | 0 | - |
| - | - | 1084 | 348.2 | - | - | 0 | - |
| - | - | 6987 | 350.1 | - | - | 0 | - |
| - | - | 855 | 351.1 | - | - | 0 | - |
| - | - | 675.3 | 353.1 | - | - | 0 | - |
| - | - | 525.7 | 353.2 | - | - | 0 | - |
| - | - | 1270 | 354.1 | - | - | 0 | - |
| - | - | 2410 | 358.2 | - | - | 0 | - |
| - | - | 973.8 | 361.2 | - | - | 0 | - |
| 13 | y | 3718 | 362.2 | 0.0004779 | 1.319 | +1 | 3 |
| - | - | 1970 | 365.2 | - | - | 0 | - |
| - | - | 1813 | 366.2 | - | - | 0 | - |
| - | - | 588.4 | 370.2 | - | - | 0 | - |
| - | - | 4911 | 371.2 | - | - | 0 | - |
| - | - | 561.6 | 371.7 | - | - | 0 | - |
| - | - | 2133 | 372.1 | - | - | 0 | - |
| - | - | 774 | 373.1 | - | - | 0 | - |
| - | - | 1417 | 375.2 | - | - | 0 | - |
| - | - | 3638 | 379.2 | - | - | 0 | - |
| - | - | 681.8 | 380.2 | - | - | 0 | - |
| - | - | 1034 | 384.2 | - | - | 0 | - |
| - | - | 1984 | 389.2 | - | - | 0 | - |
| - | - | 888.8 | 389.2 | - | - | 0 | - |
| - | - | 1047 | 393.2 | - | - | 0 | - |
| - | - | 747.6 | 394.2 | - | - | 0 | - |
| - | - | 511.1 | 394.2 | - | - | 0 | - |
| - | - | 1331 | 398.1 | - | - | 0 | - |
| - | - | 1456 | 402.2 | - | - | 0 | - |
| - | - | 735.9 | 403.2 | - | - | 0 | - |
| - | - | 1271 | 405.2 | - | - | 0 | - |
| - | - | 5498 | 407.2 | - | - | 0 | - |
| - | - | 999.4 | 408.2 | - | - | 0 | - |
| - | - | 973.7 | 410.2 | - | - | 0 | - |
| - | - | 1548 | 412.2 | - | - | 0 | - |
| - | - | 2474 | 414.1 | - | - | 0 | - |
| - | - | 3031 | 415.2 | - | - | 0 | - |
| - | - | 567.9 | 416.2 | - | - | 0 | - |
| - | - | 4211 | 421.2 | - | - | 0 | - |
| - | - | 1539 | 423.2 | - | - | 0 | - |
| - | - | 880.3 | 428.2 | - | - | 0 | - |
| - | - | 867.8 | 433.2 | - | - | 0 | - |
| - | - | 1600 | 436.2 | - | - | 0 | - |
| - | - | 681.8 | 438.7 | - | - | 0 | - |
| - | - | 1314 | 443.2 | - | - | 0 | - |
| - | - | 5219 | 443.2 | - | - | 0 | - |
| - | - | 647 | 444.2 | - | - | 0 | - |
| 12 | y | 595.4 | 445.2 | 0.002091 | 4.696 | +1 | 4 |
| - | - | 3205 | 451.2 | - | - | 0 | - |
| 12 | y | 1.115E+04 | 463.3 | 0.0007425 | 1.603 | +1 | 4 |
| - | - | 1816 | 464.3 | - | - | 0 | - |
| 4 | b | 5199 | 471.2 | 0.005613 | 11.91 | +1 | 4 |
| - | - | 1254 | 472.2 | - | - | 0 | - |
| - | - | 873.5 | 474.3 | - | - | 0 | - |
| - | - | 788 | 477.2 | - | - | 0 | - |
| - | - | 745.9 | 479.2 | - | - | 0 | - |
| - | - | 1345 | 481.2 | - | - | 0 | - |
| - | - | 1595 | 485.2 | - | - | 0 | - |
| - | - | 1897 | 498.2 | - | - | 0 | - |
| - | - | 1042 | 502.2 | - | - | 0 | - |
| - | - | 720.2 | 503.3 | - | - | 0 | - |
| - | - | 606.5 | 506.3 | - | - | 0 | - |
| - | - | 847.9 | 507.2 | - | - | 0 | - |
| - | - | 3592 | 508.3 | - | - | 0 | - |
| - | - | 721.5 | 509.3 | - | - | 0 | - |
| - | - | 716.4 | 512.2 | - | - | 0 | - |
| - | - | 1131 | 516.2 | - | - | 0 | - |
| 9 | b | 2324 | 518.2 | 0.001632 | 3.15 | +2 | 9 |
| - | - | 1097 | 520.2 | - | - | 0 | - |
| - | - | 830.9 | 520.7 | - | - | 0 | - |
| - | - | 753.9 | 524.2 | - | - | 0 | - |
| - | - | 1573 | 530.2 | - | - | 0 | - |
| - | - | 3678 | 534.3 | - | - | 0 | - |
| - | - | 8892 | 536.2 | - | - | 0 | - |
| - | - | 1628 | 537.3 | - | - | 0 | - |
| - | - | 907.6 | 541.3 | - | - | 0 | - |
| - | - | 734.8 | 543.3 | - | - | 0 | - |
| - | - | 1221 | 545.3 | - | - | 0 | - |
| - | - | 694 | 546.3 | - | - | 0 | - |
| - | - | 1137 | 547.3 | - | - | 0 | - |
| - | - | 670.2 | 548.2 | - | - | 0 | - |
| - | - | 1044 | 552.2 | - | - | 0 | - |
| - | - | 1548 | 554.8 | - | - | 0 | - |
| 7 | y | 766.3 | 555.8 | 0.001321 | 2.378 | +2 | 9 |
| - | - | 1249 | 560.3 | - | - | 0 | - |
| - | - | 2246 | 565.3 | - | - | 0 | - |
| - | - | 3041 | 572.3 | - | - | 0 | - |
| - | - | 2481 | 572.8 | - | - | 0 | - |
| - | - | 1595 | 573.3 | - | - | 0 | - |
| - | - | 2047 | 579.3 | - | - | 0 | - |
| 5 | b | 876.6 | 582.2 | 0.003156 | 5.421 | +1 | 5 |
| 10 | b | 2450 | 586.8 | 0.003671 | 6.256 | +2 | 10 |
| - | - | 1660 | 587.3 | - | - | 0 | - |
| - | - | 1411 | 588.3 | - | - | 0 | - |
| - | - | 2779 | 589.3 | - | - | 0 | - |
| - | - | 1552 | 598.3 | - | - | 0 | - |
| 5 | b | 1.154E+04 | 600.2 | 0.00547 | 9.113 | +1 | 5 |
| - | - | 3780 | 601.2 | - | - | 0 | - |
| - | - | 824.2 | 603.3 | - | - | 0 | - |
| - | - | 9050 | 607.3 | - | - | 0 | - |
| - | - | 2430 | 608.3 | - | - | 0 | - |
| 11 | y | 1220 | 609.3 | 0.006444 | 10.58 | +1 | 5 |
| - | - | 957.2 | 612.2 | - | - | 0 | - |
| - | - | 634 | 612.5 | - | - | 0 | - |
| - | - | 965.3 | 613.8 | - | - | 0 | - |
| - | - | 758.4 | 616.3 | - | - | 0 | - |
| - | - | 1050 | 616.8 | - | - | 0 | - |
| - | - | 712 | 617.3 | - | - | 0 | - |
| - | - | 1061 | 618.2 | - | - | 0 | - |
| - | - | 745.5 | 620.3 | - | - | 0 | - |
| - | - | 998.3 | 622.3 | - | - | 0 | - |
| - | - | 898.6 | 626.3 | - | - | 0 | - |
| 11 | y | 7137 | 626.3 | 0.0005243 | 0.8371 | +1 | 5 |
| - | - | 690.5 | 627.3 | - | - | 0 | - |
| - | - | 2592 | 627.3 | - | - | 0 | - |
| - | - | 1253 | 635.3 | - | - | 0 | - |
| - | - | 1526 | 636.3 | - | - | 0 | - |
| - | - | 1068 | 636.8 | - | - | 0 | - |
| - | - | 652.8 | 638.6 | - | - | 0 | - |
| - | - | 715.8 | 643.3 | - | - | 0 | - |
| - | - | 929.7 | 643.3 | - | - | 0 | - |
| - | - | 876.6 | 644.3 | - | - | 0 | - |
| - | - | 1132 | 648.8 | - | - | 0 | - |
| 6 | b | 1019 | 653.3 | 0.00602 | 9.216 | +1 | 6 |
| - | - | 1129 | 654.3 | - | - | 0 | - |
| - | - | 1996 | 654.8 | - | - | 0 | - |
| - | - | 706.5 | 656.3 | - | - | 0 | - |
| - | - | 680.8 | 657.3 | - | - | 0 | - |
| - | - | 889.9 | 657.8 | - | - | 0 | - |
| - | - | 2552 | 658.3 | - | - | 0 | - |
| - | - | 737.4 | 664.1 | - | - | 0 | - |
| - | - | 935.4 | 664.8 | - | - | 0 | - |
| - | - | 1247 | 665.3 | - | - | 0 | - |
| 11 | b | 2661 | 668.3 | 0.002585 | 3.868 | +2 | 11 |
| - | - | 1167 | 668.8 | - | - | 0 | - |
| - | - | 1076 | 669.3 | - | - | 0 | - |
| - | - | 831 | 669.8 | - | - | 0 | - |
| 6 | b | 8903 | 671.3 | 0.005282 | 7.869 | +1 | 6 |
| - | - | 2492 | 672.3 | - | - | 0 | - |
| - | - | 891.3 | 673.5 | - | - | 0 | - |
| - | - | 1125 | 674.3 | - | - | 0 | - |
| - | - | 693.7 | 676.8 | - | - | 0 | - |
| - | - | 2450 | 677.8 | - | - | 0 | - |
| - | - | 2156 | 678.3 | - | - | 0 | - |
| - | - | 706.6 | 678.8 | - | - | 0 | - |
| - | - | 1439 | 680.3 | - | - | 0 | - |
| - | - | 890.8 | 680.8 | - | - | 0 | - |
| - | - | 1134 | 684.3 | - | - | 0 | - |
| - | - | 709 | 686.8 | - | - | 0 | - |
| - | - | 895.7 | 687.3 | - | - | 0 | - |
| - | - | 1224 | 691.3 | - | - | 0 | - |
| - | - | 1158 | 692.4 | - | - | 0 | - |
| - | - | 3527 | 702.3 | - | - | 0 | - |
| - | - | 1141 | 703.3 | - | - | 0 | - |
| - | - | 1754 | 703.9 | - | - | 0 | - |
| - | - | 818.1 | 708.4 | - | - | 0 | - |
| 12 | b | 2131 | 709.8 | 0.003376 | 4.756 | +2 | 12 |
| 12 | b | 1950 | 710.3 | 0.01271 | 17.9 | +2 | 12 |
| - | - | 921.3 | 713.3 | - | - | 0 | - |
| - | - | 884.8 | 718.3 | - | - | 0 | - |
| 12 | b | 829 | 718.8 | 0.002028 | 2.822 | +2 | 12 |
| - | - | 1588 | 719.3 | - | - | 0 | - |
| - | - | 1067 | 719.6 | - | - | 0 | - |
| - | - | 6029 | 720.4 | - | - | 0 | - |
| - | - | 2283 | 721.4 | - | - | 0 | - |
| - | - | 1355 | 721.8 | - | - | 0 | - |
| - | - | 666.4 | 722.4 | - | - | 0 | - |
| 4 | y | 1171 | 724.3 | 0.005804 | 8.012 | +2 | 12 |
| - | - | 1490 | 725.1 | - | - | 0 | - |
| - | - | 1189 | 726.9 | - | - | 0 | - |
| - | - | 1239 | 727.1 | - | - | 0 | - |
| - | - | 925.3 | 727.8 | - | - | 0 | - |
| - | - | 644.5 | 728.8 | - | - | 0 | - |
| - | - | 867.9 | 732.2 | - | - | 0 | - |
| - | - | 1007 | 732.3 | - | - | 0 | - |
| - | - | 769.2 | 732.7 | - | - | 0 | - |
| 4 | y | 2935 | 732.9 | 0.0004519 | 0.6166 | +2 | 12 |
| - | - | 753.8 | 733.1 | - | - | 0 | - |
| - | - | 2524 | 733.4 | - | - | 0 | - |
| - | - | 1464 | 733.9 | - | - | 0 | - |
| - | - | 1789 | 736.4 | - | - | 0 | - |
| - | - | 876.6 | 736.9 | - | - | 0 | - |
| - | - | 1867 | 737.4 | - | - | 0 | - |
| - | - | 971.4 | 739.5 | - | - | 0 | - |
| - | - | 2062 | 741.8 | - | - | 0 | - |
| - | - | 1728 | 742.3 | - | - | 0 | - |
| - | - | 860.5 | 744.6 | - | - | 0 | - |
| - | - | 1053 | 745 | - | - | 0 | - |
| 10 | y | 879.4 | 745.4 | 0.0001544 | 0.2072 | +1 | 6 |
| - | - | 850.5 | 745.7 | - | - | 0 | - |
| 10 | y | 756.9 | 746.3 | 0.007468 | 10.01 | +1 | 6 |
| - | - | 877.9 | 748.3 | - | - | 0 | - |
| - | - | 2773 | 750.9 | - | - | 0 | - |
| - | - | 677 | 751 | - | - | 0 | - |
| - | - | 2059 | 751.4 | - | - | 0 | - |
| - | - | 1759 | 751.9 | - | - | 0 | - |
| - | - | 900.3 | 753.1 | - | - | 0 | - |
| - | - | 1271 | 753.9 | - | - | 0 | - |
| - | - | 1635 | 754.4 | - | - | 0 | - |
| - | - | 786 | 754.8 | - | - | 0 | - |
| - | - | 838.6 | 755.1 | - | - | 0 | - |
| - | - | 854.8 | 755.3 | - | - | 0 | - |
| - | - | 1204 | 756.4 | - | - | 0 | - |
| - | - | 845.8 | 758.6 | - | - | 0 | - |
| - | - | 839.1 | 760.3 | - | - | 0 | - |
| - | - | 1160 | 762.9 | - | - | 0 | - |
| - | - | 689.8 | 763.3 | - | - | 0 | - |
| 10 | y | 5144 | 763.4 | 0.001061 | 1.389 | +1 | 6 |
| - | - | 2608 | 764.4 | - | - | 0 | - |
| - | - | 1011 | 765.9 | - | - | 0 | - |
| - | - | 2698 | 766.2 | - | - | 0 | - |
| 7 | b | 2152 | 766.4 | 0.006734 | 8.787 | +1 | 7 |
| - | - | 875.9 | 767.4 | - | - | 0 | - |
| - | - | 1436 | 768.4 | - | - | 0 | - |
| - | - | 986.2 | 768.5 | - | - | 0 | - |
| - | - | 1618 | 770.1 | - | - | 0 | - |
| - | - | 1566 | 772.4 | - | - | 0 | - |
| 13 | b | 1189 | 773.8 | 0.002408 | 3.111 | +2 | 13 |
| - | - | 5219 | 774.4 | - | - | 0 | - |
| - | - | 3018 | 774.9 | - | - | 0 | - |
| - | - | 1209 | 775.4 | - | - | 0 | - |
| - | - | 1326 | 775.7 | - | - | 0 | - |
| - | - | 2386 | 775.9 | - | - | 0 | - |
| - | - | 2104 | 781.3 | - | - | 0 | - |
| - | - | 1446 | 782.6 | - | - | 0 | - |
| 13 | b | 6221 | 782.8 | 0.003046 | 3.891 | +2 | 13 |
| - | - | 3934 | 783.4 | - | - | 0 | - |
| - | - | 2889 | 783.9 | - | - | 0 | - |
| 7 | b | 7709 | 784.4 | 0.006057 | 7.723 | +1 | 7 |
| - | - | 5319 | 785.4 | - | - | 0 | - |
| - | - | 1179 | 786.4 | - | - | 0 | - |
| - | - | 2450 | 787.7 | - | - | 0 | - |
| - | - | 769.3 | 788.4 | - | - | 0 | - |
| - | - | 1238 | 789.3 | - | - | 0 | - |
| - | - | 938.6 | 789.4 | - | - | 0 | - |
| - | - | 1056 | 790.6 | - | - | 0 | - |
| - | - | 1798 | 795 | - | - | 0 | - |
| - | - | 792 | 795.7 | - | - | 0 | - |
| 3 | y | 1121 | 797.4 | 0.005701 | 7.15 | +2 | 13 |
| - | - | 1557 | 799.9 | - | - | 0 | - |
| - | - | 1051 | 802.4 | - | - | 0 | - |
| - | - | 888.2 | 803.4 | - | - | 0 | - |
| - | - | 1130 | 805.7 | - | - | 0 | - |
| - | - | 775.4 | 805.9 | - | - | 0 | - |
| - | - | 976.8 | 806.3 | - | - | 0 | - |
| 3 | y | 4648 | 806.4 | 0.007072 | 8.77 | +2 | 13 |
| - | - | 1016 | 806.5 | - | - | 0 | - |
| - | - | 5180 | 806.9 | - | - | 0 | - |
| - | - | 3340 | 807.4 | - | - | 0 | - |
| - | - | 1040 | 812.4 | - | - | 0 | - |
| - | - | 6863 | 814.9 | - | - | 0 | - |
| - | - | 1137 | 815.1 | - | - | 0 | - |
| - | - | 7630 | 815.4 | - | - | 0 | - |
| - | - | 887.7 | 815.8 | - | - | 0 | - |
| - | - | 1638 | 815.9 | - | - | 0 | - |
| - | - | 1139 | 817.8 | - | - | 0 | - |
| - | - | 1285 | 819 | - | - | 0 | - |
| - | - | 1161 | 819.2 | - | - | 0 | - |
| - | - | 997 | 820.4 | - | - | 0 | - |
| - | - | 1421 | 821.1 | - | - | 0 | - |
| - | - | 1148 | 824.2 | - | - | 0 | - |
| - | - | 897.8 | 824.9 | - | - | 0 | - |
| - | - | 1094 | 828.7 | - | - | 0 | - |
| - | - | 1404 | 828.8 | - | - | 0 | - |
| - | - | 929.1 | 828.8 | - | - | 0 | - |
| - | - | 880.6 | 829.3 | - | - | 0 | - |
| - | - | 1654 | 829.4 | - | - | 0 | - |
| - | - | 782.5 | 829.6 | - | - | 0 | - |
| - | - | 1141 | 829.7 | - | - | 0 | - |
| - | - | 1031 | 829.8 | - | - | 0 | - |
| - | - | 1215 | 829.9 | - | - | 0 | - |
| - | - | 2149 | 830 | - | - | 0 | - |
| - | - | 1118 | 831.4 | - | - | 0 | - |
| - | - | 857.9 | 831.6 | - | - | 0 | - |
| - | - | 2306 | 831.7 | - | - | 0 | - |
| - | - | 1108 | 837 | - | - | 0 | - |
| - | - | 2235 | 837.4 | - | - | 0 | - |
| - | - | 976.8 | 837.7 | - | - | 0 | - |
| 14 | b | 803.2 | 837.9 | 0.006623 | 7.904 | +2 | 14 |
| 14 | b | 3106 | 838.4 | 0.006009 | 7.168 | +2 | 14 |
| - | - | 2827 | 838.9 | - | - | 0 | - |
| - | - | 2719 | 839.4 | - | - | 0 | - |
| - | - | 1148 | 840.4 | - | - | 0 | - |
| - | - | 1059 | 840.8 | - | - | 0 | - |
| - | - | 846.1 | 840.9 | - | - | 0 | - |
| - | - | 758.6 | 842.4 | - | - | 0 | - |
| - | - | 916.6 | 845.1 | - | - | 0 | - |
| 14 | b | 1.987E+04 | 846.9 | 0.003599 | 4.25 | +2 | 14 |
| - | - | 1086 | 847 | - | - | 0 | - |
| 2 | y | 2.312E+04 | 847.4 | 0.0119 | 14.04 | +2 | 14 |
| - | - | 9657 | 847.9 | - | - | 0 | - |
| - | - | 2057 | 848.4 | - | - | 0 | - |
| - | - | 1226 | 849.9 | - | - | 0 | - |
| - | - | 937.8 | 850.4 | - | - | 0 | - |
| - | - | 1278 | 851.2 | - | - | 0 | - |
| - | - | 2227 | 852 | - | - | 0 | - |
| 2 | y | 1271 | 855.9 | 8.663E-05 | 0.1012 | +2 | 14 |
| - | - | 988.3 | 856.4 | - | - | 0 | - |
| - | - | 1749 | 856.4 | - | - | 0 | - |
| - | - | 979.1 | 856.6 | - | - | 0 | - |
| - | - | 1491 | 856.7 | - | - | 0 | - |
| - | - | 976.3 | 856.9 | - | - | 0 | - |
| - | - | 3821 | 857.4 | - | - | 0 | - |
| - | - | 3490 | 858.4 | - | - | 0 | - |
| - | - | 5650 | 858.9 | - | - | 0 | - |
| 9 | y | 4505 | 859.4 | 0.007272 | 8.462 | +1 | 7 |
| - | - | 2130 | 859.9 | - | - | 0 | - |
| 9 | y | 1927 | 860.4 | 0.009341 | 10.86 | +1 | 7 |
| - | - | 1360 | 865.4 | - | - | 0 | - |
| - | - | 4.972E+04 | 867.4 | - | - | 0 | - |
| - | - | 4.694E+04 | 867.9 | - | - | 0 | - |
| - | - | 2.149E+04 | 868.4 | - | - | 0 | - |
| - | - | 4002 | 868.9 | - | - | 0 | - |
| - | - | 1457 | 873.4 | - | - | 0 | - |
| 9 | y | 7093 | 877.4 | 0.002011 | 2.292 | +1 | 7 |
| - | - | 3032 | 878.4 | - | - | 0 | - |
| - | - | 1827 | 881.9 | - | - | 0 | - |
| - | - | 1545 | 882.4 | - | - | 0 | - |
| - | - | 2547 | 882.5 | - | - | 0 | - |
| 0 | Precursor | 6566 | 890.4 | 0.003393 | 3.81 | +2 | -1 |
| 0 | Precursor | 1.566E+04 | 890.9 | 0.007967 | 8.943 | +2 | -1 |
| - | - | 1.585E+04 | 891.4 | - | - | 0 | - |
| - | - | 7395 | 891.9 | - | - | 0 | - |
| - | - | 3006 | 892.4 | - | - | 0 | - |
| - | - | 2581 | 893.4 | - | - | 0 | - |
| - | - | 2127 | 894.4 | - | - | 0 | - |
| 0 | Precursor | 2.661E+05 | 899.4 | 0.002688 | 2.989 | +2 | -1 |
| - | - | 2.72E+05 | 899.9 | - | - | 0 | - |
| - | - | 1.56E+05 | 900.4 | - | - | 0 | - |
| - | - | 3.192E+04 | 900.9 | - | - | 0 | - |
| - | - | 2226 | 906.3 | - | - | 0 | - |
| - | - | 1287 | 906.4 | - | - | 0 | - |
| - | - | 1961 | 909.4 | - | - | 0 | - |
| - | - | 1186 | 915.1 | - | - | 0 | - |
| 8 | b | 4429 | 921.4 | 0.006288 | 6.825 | +1 | 8 |
| - | - | 2871 | 922.4 | - | - | 0 | - |
| - | - | 1470 | 923.4 | - | - | 0 | - |
| - | - | 955.6 | 927.4 | - | - | 0 | - |
| - | - | 1257 | 937.7 | - | - | 0 | - |
| - | - | 1197 | 937.8 | - | - | 0 | - |
| - | - | 1258 | 937.8 | - | - | 0 | - |
| - | - | 2401 | 938 | - | - | 0 | - |
| - | - | 1431 | 948.4 | - | - | 0 | - |
| - | - | 1195 | 949.4 | - | - | 0 | - |
| - | - | 1128 | 965.5 | - | - | 0 | - |
| - | - | 3636 | 971.5 | - | - | 0 | - |
| - | - | 2084 | 972.5 | - | - | 0 | - |
| - | - | 1064 | 973.5 | - | - | 0 | - |
| - | - | 1110 | 986.4 | - | - | 0 | - |
| - | - | 925.5 | 993 | - | - | 0 | - |
| 8 | y | 1679 | 997.4 | 0.006642 | 6.659 | +1 | 8 |
| - | - | 1228 | 1002 | - | - | 0 | - |
| 8 | y | 9618 | 1014 | 0.0001316 | 0.1298 | +1 | 8 |
| - | - | 4296 | 1015 | - | - | 0 | - |
| - | - | 1217 | 1016 | - | - | 0 | - |
| - | - | 1549 | 1023 | - | - | 0 | - |
| 9 | b | 4042 | 1035 | 0.006757 | 6.526 | +1 | 9 |
| - | - | 1872 | 1036 | - | - | 0 | - |
| - | - | 907.2 | 1037 | - | - | 0 | - |
| - | - | 2014 | 1081 | - | - | 0 | - |
| - | - | 1208 | 1085 | - | - | 0 | - |
| - | - | 1990 | 1094 | - | - | 0 | - |
| - | - | 1179 | 1095 | - | - | 0 | - |
| - | - | 4700 | 1109 | - | - | 0 | - |
| 7 | y | 2969 | 1110 | 0.01327 | 11.96 | +1 | 9 |
| 7 | y | 1604 | 1111 | 0.003572 | 3.216 | +1 | 9 |
| - | - | 1031 | 1112 | - | - | 0 | - |
| 7 | y | 5066 | 1128 | 0.001127 | 0.9993 | +1 | 9 |
| - | - | 3453 | 1129 | - | - | 0 | - |
| - | - | 1053 | 1145 | - | - | 0 | - |
| - | - | 819 | 1146 | - | - | 0 | - |
| 10 | b | 4788 | 1173 | 0.006683 | 5.7 | +1 | 10 |
| - | - | 2694 | 1174 | - | - | 0 | - |
| - | - | 972 | 1175 | - | - | 0 | - |
| 6 | y | 1021 | 1182 | 0.005276 | 4.466 | +1 | 10 |
| - | - | 853.5 | 1183 | - | - | 0 | - |
| 6 | y | 8206 | 1199 | 0.001009 | 0.8419 | +1 | 10 |
| - | - | 5412 | 1200 | - | - | 0 | - |
| - | - | 2264 | 1201 | - | - | 0 | - |
| - | - | 1651 | 1223 | - | - | 0 | - |
| - | - | 922.2 | 1224 | - | - | 0 | - |
| - | - | 1124 | 1232 | - | - | 0 | - |
| - | - | 1722 | 1244 | - | - | 0 | - |
| - | - | 1711 | 1245 | - | - | 0 | - |
| - | - | 945.5 | 1251 | - | - | 0 | - |
| - | - | 3938 | 1272 | - | - | 0 | - |
| - | - | 2472 | 1273 | - | - | 0 | - |
| - | - | 1267 | 1274 | - | - | 0 | - |
| - | - | 1403 | 1308 | - | - | 0 | - |
| 5 | y | 1812 | 1310 | 0.001055 | 0.8057 | +1 | 11 |
| 5 | y | 1030 | 1311 | 0.001502 | 1.146 | +1 | 11 |
| - | - | 939 | 1312 | - | - | 0 | - |
| 5 | y | 6240 | 1328 | 0.0002673 | 0.2013 | +1 | 11 |
| - | - | 3729 | 1329 | - | - | 0 | - |
| - | - | 1632 | 1330 | - | - | 0 | - |
| 11 | b | 4747 | 1336 | 0.005 | 3.744 | +1 | 11 |
| - | - | 3583 | 1337 | - | - | 0 | - |
| - | - | 1917 | 1338 | - | - | 0 | - |
| - | - | 1581 | 1355 | - | - | 0 | - |
| - | - | 1682 | 1356 | - | - | 0 | - |
| - | - | 1370 | 1360 | - | - | 0 | - |
| - | - | 1088 | 1373 | - | - | 0 | - |
| - | - | 728.1 | 1415 | - | - | 0 | - |
| 12 | b | 1801 | 1419 | 0.006582 | 4.64 | +1 | 12 |
| 12 | b | 1566 | 1420 | 0.02818 | 19.85 | +1 | 12 |
| 12 | b | 1444 | 1437 | 0.001145 | 0.7968 | +1 | 12 |
| 4 | y | 1043 | 1448 | 0.003503 | 2.42 | +1 | 12 |
| 4 | y | 4338 | 1465 | 0.001684 | 1.15 | +1 | 12 |
| - | - | 3510 | 1466 | - | - | 0 | - |
| - | - | 1407 | 1467 | - | - | 0 | - |
| - | - | 1079 | 1501 | - | - | 0 | - |
| 13 | b | 1100 | 1565 | 0.01056 | 6.747 | +1 | 13 |
| - | - | 818.2 | 1566 | - | - | 0 | - |
| - | - | 778.2 | 2878 | - | - | 0 | - |

m/z Charge Intensity FragmentType MassShift Position
122.07170867919922 0 422.73837
123.05561065673828 0 495.4181
126.05530548095703 0 540.2263
127.08708953857422 0 368.834
129.10260009765625 0 20382.26
130.10597229003906 0 775.3441
136.07608032226562 0 5331.293
136.08689880371094 0 417.63428
138.0552978515625 0 768.3637
138.06642150878906 0 822.66583
139.08702087402344 0 565.759
140.18698120117188 0 416.50952
141.10255432128906 0 410.61093 a Water loss 1
142.8103485107422 0 446.91135
143.5092315673828 0 397.17853
148.08714294433594 0 837.7855
148.873779296875 0 565.61914
148.89483642578125 0 576.2508
148.90223693847656 0 631.1338
148.9095001220703 0 701.66345
148.91676330566406 0 942.8799
148.92416381835938 0 1125.4205
148.9311981201172 0 1383.0416
148.9387664794922 0 3061.0205
148.9553680419922 0 6170.539
148.9632568359375 0 3024.2925
148.9709930419922 0 1309.5953
148.97825622558594 0 1176.8475
148.98544311523438 0 897.67487
148.99276733398438 0 752.7748
148.99969482421875 0 508.9954
149.00735473632812 0 518.42035
149.01431274414062 0 491.90475
149.03688049316406 0 489.23184
152.0826873779297 0 1195.0985
155.09327697753906 0 1414.1306
156.07725524902344 0 897.5778
159.1131591796875 0 18944.387 a 1
160.1165771484375 0 1380.894
165.07778930664062 0 1250.4767
165.16319274902344 0 451.4199
166.06161499023438 0 1812.5217
166.0865936279297 0 866.48334
167.09291076660156 0 1037.3478
176.08221435546875 0 1930.9377
180.07705688476562 0 1759.4164
180.11297607421875 0 559.44653
187.10801696777344 0 9516.388 b 1
188.11062622070312 0 583.06854
189.076904296875 0 783.7899
190.0615997314453 0 1208.8544
191.09307861328125 0 463.5626
192.10231018066406 0 541.7441
193.09933471679688 0 837.6468
193.1087646484375 0 5031.5
194.09278869628906 0 1103.608
195.07640075683594 0 1656.7074
195.0872039794922 0 560.2292
195.11419677734375 0 672.28925
201.0883331298828 0 504.50876
202.10873413085938 0 567.80725
204.07704162597656 0 2750.253
204.08628845214844 0 567.117
207.08798217773438 0 2723.9868
209.9215545654297 0 450.35727
211.10836791992188 0 858.6268
212.1035919189453 0 841.3432
216.13475036621094 0 1331.6323 y Water loss 13
217.07252502441406 0 960.202
217.1188201904297 0 2057.75 y Ammonia loss 13
221.10366821289062 0 10267.376
221.11514282226562 0 742.1999
222.1068572998047 0 1068.0613
222.12388610839844 0 1798.2198
223.1069793701172 0 547.7318
223.1558380126953 0 877.09625
224.1145782470703 0 3415.087
228.1134033203125 0 561.28156
230.11383056640625 0 7612.153
233.14002990722656 0 2025.877
234.09927368164062 0 1215.2354
234.1243438720703 0 1085.5356
234.14532470703125 0 13836.431 y 13
235.08290100097656 0 6279.4287
235.1483917236328 0 2093.2273
236.11505126953125 0 566.9226
237.12388610839844 0 833.91266
239.11444091796875 0 2615.2446
239.15069580078125 0 9484.393
240.13475036621094 0 11529.148
240.1547393798828 0 1260.825
241.1385498046875 0 881.10315
247.1084442138672 0 1211.6633
249.09877014160156 0 3069.85
249.13528442382812 0 751.29865
251.1505584716797 0 2224.4966
252.1095733642578 0 16223.682
252.1331787109375 0 713.69745
253.09307861328125 0 830.67896
253.1132354736328 0 1613.4111
257.10711669921875 0 2438.4941
257.1614074707031 0 2579.2673
258.1439514160156 0 590.8606
261.1338195800781 0 497.78522
262.09381103515625 0 985.0203
264.1376647949219 0 1491.7441
265.11798095703125 0 864.06555
267.0927734375 0 1084.0498
267.10919189453125 0 7432.4644
268.11175537109375 0 602.7309
269.13653564453125 0 1216.0498
270.1451416015625 0 2963.1235
273.1351623535156 0 3933.8862
275.1060485839844 0 2308.1135
279.1451416015625 0 2206.1902
280.1044921875 0 1549.2566
280.1297302246094 0 1388.5227
285.1018981933594 0 3686.0508
287.9578857421875 0 560.2087
295.1050720214844 0 783.5659
301.1302795410156 0 6007.682
302.1297912597656 0 913.8934
302.1605224609375 0 635.89014
305.1606750488281 0 608.73065
320.13580322265625 0 5589.7383
321.1398010253906 0 993.6181
322.1878662109375 0 3368.4788
323.1709289550781 0 886.63135
326.1350402832031 0 776.72546
327.1668701171875 0 597.9711
332.13580322265625 0 789.66644
334.14373779296875 0 5537.1875 b 2
338.1465759277344 0 4682.9736
339.1479797363281 0 570.7537
340.141845703125 0 2226.6826
341.1827697753906 0 1511.0763
344.1484069824219 0 1513.4609
344.19342041015625 0 4247.043 y Water loss 12
345.1773986816406 0 4134.5884 y Ammonia loss 12
345.19647216796875 0 671.8672
346.1795959472656 0 545.29535
347.1808776855469 0 958.8194
348.16754150390625 0 1083.5126
350.1462707519531 0 6986.9253
351.14874267578125 0 855.02155
353.1478576660156 0 675.2662
353.1941223144531 0 525.6783
354.13134765625 0 1269.9486
358.2093200683594 0 2410.0498
361.1749267578125 0 973.7841
362.2038879394531 0 3717.7122 y 12
365.19366455078125 0 1969.5276
366.156982421875 0 1813.0027
370.1507873535156 0 588.38245
371.1578369140625 0 4911.064
371.7442321777344 0 561.58795
372.1419677734375 0 2132.8665
373.14453125 0 773.9952
375.1669921875 0 1416.5463
379.20977783203125 0 3637.884
380.2117004394531 0 681.8297
384.1661071777344 0 1033.5914
389.16888427734375 0 1984.2181
389.1952209472656 0 888.827
393.17755126953125 0 1047.4657
394.1570129394531 0 747.62604
394.1895751953125 0 511.13837
398.1456604003906 0 1330.7151
402.1773681640625 0 1456.283
403.1766357421875 0 735.86285
405.2251892089844 0 1271.4708
407.20458984375 0 5497.947
408.2085266113281 0 999.38947
410.2038269042969 0 973.6562
412.1651306152344 0 1547.8143
414.14483642578125 0 2473.9814
415.1737976074219 0 3030.5176
416.1767883300781 0 567.9237
421.1839904785156 0 4210.643
423.2354431152344 0 1539.1733
428.2141418457031 0 880.3207
433.2198791503906 0 867.8215
436.230712890625 0 1599.8733
438.7120361328125 0 681.7566
443.171630859375 0 1314.0927
443.2075500488281 0 5219.257
444.2103271484375 0 646.9948
445.24261474609375 0 595.36475 y Water loss 11
451.2309265136719 0 3205.0159
463.2518310546875 0 11150.393 y 11
464.25439453125 0 1815.7423
471.2027282714844 0 5199.1094 b 3
472.2012634277344 0 1253.5979
474.2577209472656 0 873.4641
477.2088928222656 0 787.97894
479.2218933105469 0 745.8619
481.1827087402344 0 1345.1144
485.1809387207031 0 1594.6764
498.209716796875 0 1896.5834
502.24884033203125 0 1042.4998
503.26324462890625 0 720.23883
506.2705078125 0 606.5139
507.2110595703125 0 847.8808
508.2524719238281 0 3592.3945
509.2571105957031 0 721.509
512.2259521484375 0 716.40875
516.2198486328125 0 1131.0349
518.2366333007812 0 2323.6016 b 8
520.2498168945312 0 1097.2106
520.7399291992188 0 830.8669
524.23974609375 0 753.8709
530.2399291992188 0 1573.3417
534.267822265625 0 3677.8052
536.2470092773438 0 8891.802
537.2509155273438 0 1627.9226
541.2734985351562 0 907.62354
543.2506103515625 0 734.7979
545.2938842773438 0 1220.5017
546.2977294921875 0 694.0464
547.2630004882812 0 1137.4265
548.24658203125 0 670.2206
552.2308349609375 0 1044.3737
554.7661743164062 0 1547.7157
555.7713012695312 0 766.25934 y Ammonia loss 6
560.2949829101562 0 1249.3867
565.2722778320312 0 2246.0305
572.2507934570312 0 3041.1477
572.7704467773438 0 2480.901
573.2821655273438 0 1594.677
579.2894897460938 0 2047.2712
582.2322998046875 0 876.64777 b Water loss 4
586.7681274414062 0 2449.9724 b 9
587.2669067382812 0 1660.0839
588.2906494140625 0 1411.3723
589.2753295898438 0 2779.0295
598.2674560546875 0 1551.6573
600.2451782226562 0 11542.652 b 4
601.24853515625 0 3779.777
603.2887573242188 0 824.2446
607.2836303710938 0 9049.927
608.2913818359375 0 2430.3503
609.2943115234375 0 1219.6285 y Ammonia loss 10
612.2478637695312 0 957.17444
612.4510498046875 0 633.9871
613.7898559570312 0 965.25854
616.2858276367188 0 758.3728
616.7719116210938 0 1050.4283
617.26953125 0 712.03845
618.240234375 0 1060.6185
620.2932739257812 0 745.4937
622.3040771484375 0 998.2592
626.2694091796875 0 898.56976
626.31494140625 0 7136.502 y 10
627.2565307617188 0 690.5121
627.3143310546875 0 2591.8052
635.2656860351562 0 1252.7903
636.2999267578125 0 1525.7141
636.7957153320312 0 1068.3436
638.6389770507812 0 652.7967
643.272705078125 0 715.7607
643.3299560546875 0 929.6506
644.2803955078125 0 876.6425
648.80078125 0 1132.1261
653.2722778320312 0 1018.6499 b Water loss 5
654.2999877929688 0 1129.4121
654.8003540039062 0 1995.7992
656.3251953125 0 706.5021
657.3124389648438 0 680.8227
657.8042602539062 0 889.9094
658.3297729492188 0 2552.208
664.0568237304688 0 737.4225
664.8257446289062 0 935.3937
665.3145141601562 0 1246.775
668.2987060546875 0 2660.7766 b 10
668.8001098632812 0 1166.5411
669.3048706054688 0 1075.7761
669.8076171875 0 831.00867
671.2821044921875 0 8903.001 b 5
672.2846069335938 0 2492.0366
673.545166015625 0 891.3383
674.3379516601562 0 1124.8866
676.8076171875 0 693.6819
677.8202514648438 0 2449.8953
678.3207397460938 0 2156.1653
678.8145141601562 0 706.5846
680.3314819335938 0 1438.5862
680.83349609375 0 890.7705
684.3224487304688 0 1133.7747
686.82177734375 0 709.0295
687.3152465820312 0 895.6598
691.3257446289062 0 1224.3849
692.3753051757812 0 1157.9032
702.3380126953125 0 3527.2202
703.3331909179688 0 1140.6129
703.9251098632812 0 1754.0767
708.3607177734375 0 818.0779
709.8180541992188 0 2130.6914 b Water loss 11
710.3193969726562 0 1949.6384 b Ammonia loss 11
713.3363647460938 0 921.32983
718.2774658203125 0 884.8152
718.8179321289062 0 828.9907 b 11
719.3278198242188 0 1587.5234
719.552490234375 0 1066.8997
720.3674926757812 0 6028.9194
721.3711547851562 0 2282.6077
721.8482055664062 0 1354.987
722.352783203125 0 666.3724
724.3450927734375 0 1170.9093 y Ammonia loss 3
725.1136474609375 0 1489.7178
726.923583984375 0 1188.8268
727.0571899414062 0 1238.722
727.8386840820312 0 925.3271
728.8477783203125 0 644.5149
732.20166015625 0 867.9181
732.2890014648438 0 1007.22723
732.7131958007812 0 769.19366
732.8521118164062 0 2934.6213 y 3
733.148193359375 0 753.80927
733.351318359375 0 2523.8083
733.8538208007812 0 1463.8564
736.3509521484375 0 1788.5642
736.8535766601562 0 876.6457
737.350341796875 0 1867.4094
739.4662475585938 0 971.4045
741.8486938476562 0 2062.1497
742.3447875976562 0 1728.2493
744.5693969726562 0 860.5438
745 0 1053.4224
745.3626098632812 0 879.36035 y Water loss 9
745.7410278320312 0 850.51263
746.354248046875 0 756.895 y Ammonia loss 9
748.3479614257812 0 877.91846
750.8532104492188 0 2773.244
750.9684448242188 0 676.95953
751.3538208007812 0 2058.5952
751.8546142578125 0 1759.1497
753.1254272460938 0 900.2675
753.8512573242188 0 1271.1321
754.3551025390625 0 1635.1758
754.8396606445312 0 786.0074
755.1067504882812 0 838.56586
755.3428344726562 0 854.8345
756.3652954101562 0 1204.2455
758.614990234375 0 845.769
760.2630004882812 0 839.0832
762.9358520507812 0 1160.2411
763.31396484375 0 689.7936
763.3743896484375 0 5143.976 y 9
764.3760986328125 0 2607.8904
765.8634033203125 0 1010.6041
766.20068359375 0 2698.3184
766.3570556640625 0 2152.276 b Water loss 6
767.3547973632812 0 875.87683
768.3639526367188 0 1436.2925
768.4520263671875 0 986.2309
770.1162719726562 0 1618.1489
772.3760986328125 0 1565.7021
773.8463745117188 0 1188.6355 b Water loss 12
774.3653564453125 0 5219.2236
774.8621215820312 0 3018.4954
775.3710327148438 0 1209.3016
775.715087890625 0 1326.4824
775.8707885742188 0 2386.3005
781.3381958007812 0 2104.4263
782.60546875 0 1446.1484
782.852294921875 0 6221.171 b 12
783.3540649414062 0 3934.1152
783.85498046875 0 2889.3882
784.366943359375 0 7709.39 b 6
785.3688354492188 0 5318.706
786.3692626953125 0 1178.5837
787.6753540039062 0 2449.7158
788.3543090820312 0 769.2513
789.2554321289062 0 1238.2804
789.3678588867188 0 938.5714
790.635498046875 0 1055.7194
794.9959716796875 0 1798.4038
795.7388305664062 0 791.9714
797.3682250976562 0 1121.096 y Water loss 2
799.9354248046875 0 1557.3093
802.3583374023438 0 1051.1877
803.375732421875 0 888.18085
805.67041015625 0 1129.6934
805.8941040039062 0 775.4377
806.2767333984375 0 976.7768
806.3748779296875 0 4648.474 y 2
806.4744873046875 0 1015.9876
806.877685546875 0 5180.427
807.3764038085938 0 3340.3574
812.3980102539062 0 1040.0088
814.8995971679688 0 6862.944
815.0770263671875 0 1136.8716
815.4010009765625 0 7629.9546
815.8350219726562 0 887.7075
815.9227905273438 0 1638.1555
817.811279296875 0 1138.837
819.0394897460938 0 1284.8601
819.1912841796875 0 1161.0134
820.3807373046875 0 997.02966
821.1011352539062 0 1420.7136
824.2357788085938 0 1147.5
824.899658203125 0 897.7836
828.6785278320312 0 1094.1978
828.7595825195312 0 1404.3191
828.8441162109375 0 929.1149
829.3031005859375 0 880.5577
829.4246215820312 0 1654.3867
829.5679931640625 0 782.46405
829.7115478515625 0 1140.5427
829.7987670898438 0 1030.6178
829.8963012695312 0 1214.5564
830.0188598632812 0 2148.5747
831.4053955078125 0 1117.8638
831.5761108398438 0 857.9453
831.693115234375 0 2305.644
837.0031127929688 0 1107.6503
837.39892578125 0 2235.1594
837.6736450195312 0 976.8268
837.8980712890625 0 803.2067 b Water loss 13
838.3894653320312 0 3105.9802 b Ammonia loss 13
838.8960571289062 0 2826.9658
839.3950805664062 0 2719.1165
840.3965454101562 0 1148.1265
840.7891845703125 0 1058.9038
840.9069213867188 0 846.07117
842.382080078125 0 758.6439
845.1405639648438 0 916.6159
846.9003295898438 0 19866.135 b 13
847.03564453125 0 1086.2834
847.400634765625 0 23116.762 y Ammonia loss 1
847.901123046875 0 9657.39
848.3988647460938 0 2056.5955
849.9090576171875 0 1225.7969
850.3956909179688 0 937.7996
851.169921875 0 1278.2344
851.970703125 0 2227.433
855.902099609375 0 1270.812 y 1
856.3560791015625 0 988.28033
856.4415893554688 0 1749.0532
856.550048828125 0 979.08954
856.6841430664062 0 1491.4658
856.9192504882812 0 976.3409
857.4251098632812 0 3821.47
858.4210815429688 0 3490.2375
858.91357421875 0 5650.038
859.4129638671875 0 4505.2554 y Water loss 8
859.9097900390625 0 2130.3752
860.3990478515625 0 1927.0134 y Ammonia loss 8
865.4013671875 0 1360.1879
867.4215087890625 0 49716.598
867.9232788085938 0 46938.207
868.424072265625 0 21492.018
868.9249267578125 0 4002.0593
873.4063720703125 0 1456.7662
877.4142456054688 0 7092.6123 y 8
878.4158935546875 0 3031.7183
881.9033813476562 0 1826.8779
882.402587890625 0 1544.9275
882.5343017578125 0 2546.674
890.4161376953125 0 6566.4854 Precursor Water loss
890.9127197265625 0 15662.092 Precursor Ammonia loss
891.4120483398438 0 15849.577
891.9118041992188 0 7395.4272
892.4085083007812 0 3005.7239
893.42529296875 0 2580.8254
894.427490234375 0 2126.6353
899.4207153320312 0 266135.34 Precursor
899.9224853515625 0 272037.34
900.42333984375 0 155997.05
900.9237060546875 0 31920.752
906.2582397460938 0 2225.946
906.3602905273438 0 1286.9938
909.4327392578125 0 1960.5938
915.083984375 0 1186.3341
921.4260864257812 0 4429.027 b 7
922.4274291992188 0 2870.8684
923.4306030273438 0 1469.9506
927.4398193359375 0 955.5755
937.6549072265625 0 1256.522
937.7513427734375 0 1196.6079
937.8489990234375 0 1257.5784
938.0034790039062 0 2401.1316
948.4324340820312 0 1430.6675
949.4320678710938 0 1195.1857
965.452880859375 0 1127.5361
971.4679565429688 0 3636.0437
972.47216796875 0 2084.121
973.4737548828125 0 1063.7562
986.4361572265625 0 1109.556
993.0486450195312 0 925.4551
997.4552612304688 0 1679.2859 y Ammonia loss 7
1002.4583740234375 0 1227.7546
1014.4750366210938 0 9617.955 y 7
1015.477783203125 0 4296.1855
1016.4788818359375 0 1216.5208
1022.5136108398438 0 1548.5585
1035.469482421875 0 4041.8086 b 8
1036.4703369140625 0 1871.6002
1037.471923828125 0 907.2102
1080.5318603515625 0 2014.0225
1085.4979248046875 0 1208.3552
1093.5496826171875 0 1990.3254
1094.546875 0 1178.772
1108.5294189453125 0 4699.914
1109.535400390625 0 2968.704 y Water loss 6
1110.5362548828125 0 1603.7091 y Ammonia loss 6
1111.5401611328125 0 1030.8517
1127.55810546875 0 5065.794 y 6
1128.557373046875 0 3452.548
1144.5345458984375 0 1053.2278
1145.5435791015625 0 819.0079
1172.5283203125 0 4787.8477 b 9
1173.52783203125 0 2694.0288
1174.5216064453125 0 972.0484
1181.5750732421875 0 1021.15533 y Ammonia loss 5
1182.5672607421875 0 853.51843
1198.5953369140625 0 8206.027 y 5
1199.5975341796875 0 5411.545
1200.602294921875 0 2263.5618
1222.5941162109375 0 1651.4592
1223.60009765625 0 922.1767
1231.5548095703125 0 1124.4554
1243.5947265625 0 1722.4181
1244.593505859375 0 1711.2183
1250.5382080078125 0 945.4726
1271.5924072265625 0 3938.1687
1272.5909423828125 0 2472.2227
1273.596923828125 0 1267.3599
1307.592041015625 0 1402.5939
1309.6273193359375 0 1812.2445 y Water loss 4
1310.6138916015625 0 1030.1003 y Ammonia loss 4
1311.6221923828125 0 939.027
1327.638671875 0 6240.456 y 4
1328.6363525390625 0 3728.9668
1329.6395263671875 0 1631.5544
1335.5899658203125 0 4747.2993 b 10
1336.5897216796875 0 3583.2654
1337.6036376953125 0 1916.7489
1354.622802734375 0 1580.5406
1355.6318359375 0 1682.1626
1359.6495361328125 0 1369.6273
1372.6331787109375 0 1088.4525
1414.9588623046875 0 728.14233
1418.628662109375 0 1801.2462 b Water loss 11
1419.63427734375 0 1566.3948 b Ammonia loss 11
1436.6337890625 0 1443.6387 b 11
1447.6748046875 0 1042.6536 y Ammonia loss 3
1464.6961669921875 0 4338.202 y 3
1465.6929931640625 0 3510.2126
1466.69384765625 0 1407.0066
1500.68310546875 0 1078.7657
1564.6806640625 0 1100.3779 b 12
1565.6885986328125 0 818.16034
2877.641845703125 0 778.1611

Spectrum Details

|  |  |
| --- | --- |
| Matched peaks? Matched peaks The total absolute number of peaks matched. Additionally in brackets the total fraction of peaks matched and the total number of peaks is shown. | 68 (12.27% of 554) |
| FDR? FDR The false discovery rate estimated for this peptide. It is calculated by matching all theoretical fragments with a non-integer shift with the raw peaks for this spectrum. This is done with 40 different shifts. The resulting percentage is the average number of annotated peaks over the number of annotated peaks with the correct spectrum. | 0.77% |
| Satellite FDR? Satellite FDR See the FDR for details on its calculation. This satellite ion specific FDR only contains the satellite ions (d/w) for I/L/J positions. | - |
| PSM Score? PSM Score The PSM Score as given by Hecklib to this annotated spectrum. It is shown with three significant figures. | 716 |

#### Reverse Lookup? Reverse Lookup All places where this read could be placed.

| Group | Segment | Template | Template Part | Read Part | Score | Unique |
| --- | --- | --- | --- | --- | --- | --- |
| Homo sapiens Heavy Chain | IGHC | IGHG1 | [308..323] | [0..15] | 120 | False |
| Homo sapiens Heavy Chain | IGHC | IGHG2 | [304..319] | [0..15] | 120 | False |
| Homo sapiens Heavy Chain | IGHC | IGHG4 | [305..320] | [0..15] | 120 | False |

| Recombined | Template Part | Read Part | Score | Unique |
| --- | --- | --- | --- | --- |
| REC-0-1 | [433..448] | [0..15] | 120 | True |

#### Meta Information from Multiple reads

##### Number of combined reads

8

##### Intensity

0.814

##### TotalArea

4.662E+08

##### Changes to the peptide sequence

SVMHEAJHNHYTQKS

L→JNo support for either Leucine or Isoleucine based on side chain ions (Position: 7)

#### Positional Score

Copy Data

##### Positional Score (TSV)

###### Preview

```
Loading example...
```

*Click on the button to copy the data to your clipboard.*

1001234567891011121314

Label Value
"0" 0.721
"1" 0.726
"2" 0.748
"3" 0.75
"4" 0.75
"5" 0.745
"6" 0.745
"7" 0.743
"8" 0.74
"9" 0.739
"10" 0.742
"11" 0.737
"12" 0.728
"13" 0.695
"14" 0.721

#### Meta Information from PEAKS

##### Scan Identifier

F1:2869

##### Original sequence

S

V

M

H

E

A

L

H

N

H

Y

T

Q

K

S

##### Posttranslational Modifications

##### Source File

D:\separate\_stitch\_analyses\xle-disambiguation\raw\20210323\_F1\_UM1\_Peng0013\_SA\_F59\_ingel\_3ug\_ELA.raw

##### Fraction

1

##### Scan Feature

F1:8194

##### De Novo Score

99

##### ConfidenceScore

99

### m/z

594.618

##### Mass

1780.8315

##### Charge

3

##### Retention Time

14.61

##### Predicted Retention Time

-

##### Area

8.294E+07

##### Parts Per Million

0.3

##### Fragmentation mode

ETHCD

##### Originating file

01 D:\separate\_stitch\_analyses\xle-disambiguation\20210325\_F59\_3ug\_DENOVO\_12.csv

#### Meta Information from PEAKS

##### Scan Identifier

F1:2472

##### Original sequence

S

V

M

+15.99

H

E

A

L

H

N

H

Y

T

Q

K

S

##### Posttranslational Modifications

Oxidation (M)

##### Source File

D:\separate\_stitch\_analyses\xle-disambiguation\raw\20210323\_F1\_UM1\_Peng0013\_SA\_F59\_ingel\_3ug\_ELA.raw

##### Fraction

1

##### Scan Feature

-

##### De Novo Score

99

##### ConfidenceScore

99

### m/z

599.9504

##### Mass

1796.8264

##### Charge

3

##### Retention Time

12.55

##### Predicted Retention Time

-

##### Area

0

##### Parts Per Million

1.6

##### Fragmentation mode

ETHCD

##### Originating file

01 D:\separate\_stitch\_analyses\xle-disambiguation\20210325\_F59\_3ug\_DENOVO\_12.csv

#### Meta Information from PEAKS

##### Scan Identifier

F1:3013

##### Original sequence

S

V

M

H

E

A

L

H

N

H

Y

T

Q

K

S

##### Posttranslational Modifications

##### Source File

D:\separate\_stitch\_analyses\xle-disambiguation\raw\20210323\_F1\_UM1\_Peng0013\_SA\_F59\_ingel\_3ug\_ELA.raw

##### Fraction

1

##### Scan Feature

-

##### De Novo Score

98

##### ConfidenceScore

98

### m/z

594.6177

##### Mass

1780.8315

##### Charge

3

##### Retention Time

15.42

##### Predicted Retention Time

-

##### Area

0

##### Fragmentation mode

ETHCD

##### Originating file

01 D:\separate\_stitch\_analyses\xle-disambiguation\20210325\_F59\_3ug\_DENOVO\_12.csv

#### Meta Information from PEAKS

##### Scan Identifier

F1:2369

##### Original sequence

S

V

M

+15.99

H

E

A

L

H

N

H

Y

T

Q

K

S

##### Posttranslational Modifications

Oxidation (M)

##### Source File

D:\separate\_stitch\_analyses\xle-disambiguation\raw\20210323\_F1\_UM1\_Peng0013\_SA\_F59\_ingel\_3ug\_ELA.raw

##### Fraction

1

##### Scan Feature

F1:8391

##### De Novo Score

98

##### ConfidenceScore

98

### m/z

599.9501

##### Mass

1796.8264

##### Charge

3

##### Retention Time

12.01

##### Predicted Retention Time

-

##### Area

6.205E+07

##### Parts Per Million

1.1

##### Fragmentation mode

HCD

##### Originating file

01 D:\separate\_stitch\_analyses\xle-disambiguation\20210325\_F59\_3ug\_DENOVO\_12.csv

#### Meta Information from PEAKS

##### Scan Identifier

F1:2860

##### Original sequence

S

V

M

H

E

A

L

H

N

H

Y

T

Q

K

S

##### Posttranslational Modifications

##### Source File

D:\separate\_stitch\_analyses\xle-disambiguation\raw\20210323\_F1\_UM1\_Peng0013\_SA\_F59\_ingel\_3ug\_ELA.raw

##### Fraction

1

##### Scan Feature

F1:2080

##### De Novo Score

98

##### ConfidenceScore

98

### m/z

446.2156

##### Mass

1780.8315

##### Charge

4

##### Retention Time

14.61

##### Predicted Retention Time

-

##### Area

1.6E+08

##### Parts Per Million

1.1

##### Fragmentation mode

HCD

##### Originating file

01 D:\separate\_stitch\_analyses\xle-disambiguation\20210325\_F59\_3ug\_DENOVO\_12.csv

#### Meta Information from PEAKS

##### Scan Identifier

F1:2615

##### Original sequence

S

V

M

+15.99

H

E

A

L

H

N

H

Y

T

Q

K

S

##### Posttranslational Modifications

Oxidation (M)

##### Source File

D:\separate\_stitch\_analyses\xle-disambiguation\raw\20210323\_F1\_UM1\_Peng0013\_SA\_F59\_ingel\_3ug\_ELA.raw

##### Fraction

1

##### Scan Feature

-

##### De Novo Score

98

##### ConfidenceScore

98

### m/z

450.2144

##### Mass

1796.8264

##### Charge

4

##### Retention Time

13.3

##### Predicted Retention Time

-

##### Area

0

##### Parts Per Million

1.1

##### Fragmentation mode

HCD

##### Originating file

01 D:\separate\_stitch\_analyses\xle-disambiguation\20210325\_F59\_3ug\_DENOVO\_12.csv

#### Meta Information from PEAKS

##### Scan Identifier

F1:2866

##### Original sequence

S

V

M

H

E

A

L

H

N

H

Y

T

Q

K

S

##### Posttranslational Modifications

##### Source File

D:\separate\_stitch\_analyses\xle-disambiguation\raw\20210323\_F1\_UM1\_Peng0013\_SA\_F59\_ingel\_3ug\_ELA.raw

##### Fraction

1

##### Scan Feature

F1:2080

##### De Novo Score

97

##### ConfidenceScore

97

### m/z

446.2156

##### Mass

1780.8315

##### Charge

4

##### Retention Time

14.61

##### Predicted Retention Time

-

##### Area

1.6E+08

##### Parts Per Million

1.1

##### Fragmentation mode

HCD

##### Originating file

01 D:\separate\_stitch\_analyses\xle-disambiguation\20210325\_F59\_3ug\_DENOVO\_12.csv

#### Meta Information from PEAKS

##### Scan Identifier

F1:2379

##### Original sequence

S

V

M

+15.99

H

E

A

L

H

N

H

Y

T

Q

K

S

##### Posttranslational Modifications

Oxidation (M)

##### Source File

D:\separate\_stitch\_analyses\xle-disambiguation\raw\20210323\_F1\_UM1\_Peng0013\_SA\_F59\_ingel\_3ug\_ELA.raw

##### Fraction

1

##### Scan Feature

F1:19013

##### De Novo Score

97

##### ConfidenceScore

97

### m/z

899.4222

##### Mass

1796.8264

##### Charge

2

##### Retention Time

12.03

##### Predicted Retention Time

-

##### Area

1.163E+06

##### Parts Per Million

2

##### Fragmentation mode

HCD

##### Originating file

01 D:\separate\_stitch\_analyses\xle-disambiguation\20210325\_F59\_3ug\_DENOVO\_12.csv
