## Supplementary material for "A handle on mass coincidence errors in *de novo* sequencing of antibodies by bottom-up proteomics": Combined_010.html

Details Combined\_010 | Stitch OverviewUndefined

### Read Combined\_010

#### Sequence (length=15)

JJSEVSDRPSGVSSR

#### Spectrum 4095? Spectrum 4095 The raw spectrum of this peptide as annotated by Hecklib. The fragments are coloured according to ion type (see legend). Any peaks with a star '\*' as text can be hovered over to see the full details, first the ion type second the mass shift type. By hovering over the amino acids in the peptide or ions in the legend the corresponding peaks are highlighted. By toggling the 'Unassigned' label you can turn the background (unassigned) peaks on or off in the plot. By updating the slider in the Ion legend you can update the spectrum to only show the top X% of the peaks with labels. The top X% means any peak that is within X% of the highest intensity. By dragging in the spectrum you can zoom in to a specific part of the spectrum and use 'Zoom Out' to get back to the original zoom level. The annotation of the spectrum is based on the given sequence in the peptides file and is done with different software so inconsistencies are likely. The peaks are annotated based on the given sequence, with 20 ppm tolerance.

Copy Data

##### Spectrum 4095 (TSV)

###### Preview

```
Loading example...
```

*Click on the button to copy the data to your clipboard.*

Mz MinMz MaxIntensity Max

WidthHeightPeptide font sizePeptide stroke widthSpectrum font sizeSpectrum stroke widthCompact peptide

Ion legend

wxyz

abcd

OtherUnassignedIon^Charge^_Position_Show for top:%

JJSEVSDRPSGVSSR

02.56e+45.11e+47.67e+41.02e+5

Zoom Out

z^+1^_1_y^+1^_1_y^+1^_1_w^+1^_2_y^+1^_2_z^+1^_2_y^+1^_2_w^+1^_3_c^+1^_3_y^+1^_3_z^+1^_3_y^+1^_3_z^+2^_8_w^+1^_4_y^+2^_8_y^+1^_4_y^+1^_4_c^+2^_8_y^+3^_13_c^+1^_4_c^+3^_14_z^+2^_9_y^+2^_9_z^+3^_14_z^+1^_5_y^+3^_14_c^+2^_9_y^+1^_5_y^+2^_10_z^+2^_10_y^+2^_10_z^+1^_6_w^+1^_6_c^+1^_5_z^+2^_11_y^+2^_11_y^+1^_6_y^+1^_6_z^+1^_6_y^+1^_6_w^+2^_12_c^+2^_12_z^+2^_12_y^+2^_12_c^+1^_6_z^+2^_13_y^+2^_13_y^+2^_13_z^+2^_13_y^+2^_13_y^+1^_7_w^+2^_14_y^+2^_14_z^+2^_14_y^+2^_14_w^+1^_8_c^+1^_7_z^+1^_8_y^+1^_8_z^+1^_8_y^+1^_8_c^+1^_8_z^+1^_9_z^+1^_9_y^+1^_9_c^+1^_9_z^+1^_10_y^+1^_10_c^+1^_10_z^+1^_11_y^+1^_11_c^+1^_11_c^+1^_12_y^+1^_12_w^+1^_13_c^+1^_13_y^+1^_13_c^+1^_14_z^+1^_14_

0621124218642485

Fragment Matches Table

Show background peaks

| Position | Ion type | Intensity | mz Theoretical | mz Error (Th) | mz Error (ppm) | Charge | Series Number |
| --- | --- | --- | --- | --- | --- | --- | --- |
| - | - | 373.7 | 123.9 | - | - | 0 | - |
| - | - | 452.9 | 124.1 | - | - | 0 | - |
| - | - | 376.3 | 124.4 | - | - | 0 | - |
| - | - | 1627 | 125.1 | - | - | 0 | - |
| - | - | 1909 | 126.1 | - | - | 0 | - |
| - | - | 555.4 | 130.1 | - | - | 0 | - |
| - | - | 1465 | 131.1 | - | - | 0 | - |
| - | - | 596.3 | 133.1 | - | - | 0 | - |
| - | - | 472.7 | 138.2 | - | - | 0 | - |
| 15 | z | 413.3 | 142.1 | 5.696E-06 | 0.04009 | +1 | 1 |
| - | - | 667.4 | 148.9 | - | - | 0 | - |
| 15 | y | 836.4 | 158.1 | 0.0002636 | 1.667 | +1 | 1 |
| - | - | 1800 | 163.1 | - | - | 0 | - |
| - | - | 467.3 | 164.2 | - | - | 0 | - |
| - | - | 3568 | 168.1 | - | - | 0 | - |
| - | - | 1393 | 173.5 | - | - | 0 | - |
| 15 | y | 5167 | 175.1 | 0.00028 | 1.599 | +1 | 1 |
| - | - | 538.1 | 183.1 | - | - | 0 | - |
| - | - | 1.326E+04 | 185.1 | - | - | 0 | - |
| - | - | 570.4 | 186.1 | - | - | 0 | - |
| - | - | 542.3 | 189.1 | - | - | 0 | - |
| - | - | 3.039E+04 | 199.2 | - | - | 0 | - |
| - | - | 2407 | 200.1 | - | - | 0 | - |
| - | - | 4112 | 200.2 | - | - | 0 | - |
| - | - | 514.6 | 202.1 | - | - | 0 | - |
| - | - | 677.9 | 212.1 | - | - | 0 | - |
| - | - | 1694 | 217.1 | - | - | 0 | - |
| - | - | 977.7 | 226.1 | - | - | 0 | - |
| - | - | 659 | 226.2 | - | - | 0 | - |
| - | - | 1.607E+04 | 227.2 | - | - | 0 | - |
| - | - | 1894 | 228.2 | - | - | 0 | - |
| 14 | w | 2451 | 229.1 | 0.0004728 | 2.063 | +1 | 2 |
| - | - | 793.1 | 233.2 | - | - | 0 | - |
| - | - | 592.2 | 234.1 | - | - | 0 | - |
| - | - | 641.9 | 244.1 | - | - | 0 | - |
| 14 | y | 2.489E+04 | 245.1 | 0.0003091 | 1.261 | +1 | 2 |
| 14 | z | 4112 | 246.1 | 0.0009394 | 3.817 | +1 | 2 |
| - | - | 1112 | 255.1 | - | - | 0 | - |
| - | - | 591.4 | 261.2 | - | - | 0 | - |
| 14 | y | 2447 | 262.2 | 0.0001932 | 0.7371 | +1 | 2 |
| - | - | 3627 | 270.1 | - | - | 0 | - |
| - | - | 616.2 | 272.2 | - | - | 0 | - |
| - | - | 651 | 293.3 | - | - | 0 | - |
| - | - | 929.8 | 294.2 | - | - | 0 | - |
| - | - | 585.1 | 303.2 | - | - | 0 | - |
| - | - | 1807 | 314.2 | - | - | 0 | - |
| 13 | w | 1.205E+04 | 316.2 | 0.0004726 | 1.495 | +1 | 3 |
| - | - | 2448 | 317.2 | - | - | 0 | - |
| - | - | 3226 | 331.1 | - | - | 0 | - |
| 3 | c | 873.4 | 331.2 | 0.002316 | 6.991 | +1 | 3 |
| 13 | y | 4570 | 332.2 | 0.0003089 | 0.93 | +1 | 3 |
| 13 | z | 1.021E+04 | 333.2 | 0.0001438 | 0.4316 | +1 | 3 |
| - | - | 1130 | 334.2 | - | - | 0 | - |
| 13 | y | 2762 | 349.2 | 5.611E-05 | 0.1607 | +1 | 3 |
| - | - | 6593 | 357.2 | - | - | 0 | - |
| - | - | 615.3 | 363.6 | - | - | 0 | - |
| - | - | 544.6 | 378.2 | - | - | 0 | - |
| - | - | 1347 | 391.7 | - | - | 0 | - |
| - | - | 956.8 | 392.2 | - | - | 0 | - |
| 8 | z | 2313 | 415.2 | 0.003227 | 7.772 | +2 | 8 |
| - | - | 1211 | 415.7 | - | - | 0 | - |
| 12 | w | 1.022E+04 | 417.2 | 0.0005236 | 1.255 | +1 | 4 |
| - | - | 1274 | 418.2 | - | - | 0 | - |
| 8 | y | 2110 | 423.2 | 0.0009756 | 2.305 | +2 | 8 |
| - | - | 1058 | 423.7 | - | - | 0 | - |
| - | - | 672.6 | 425.2 | - | - | 0 | - |
| - | - | 791.1 | 429.2 | - | - | 0 | - |
| 12 | y | 683.6 | 431.2 | 0.001506 | 3.491 | +1 | 4 |
| - | - | 1552 | 443.3 | - | - | 0 | - |
| - | - | 1205 | 443.3 | - | - | 0 | - |
| - | - | 721.2 | 444.3 | - | - | 0 | - |
| - | - | 1067 | 446.2 | - | - | 0 | - |
| - | - | 652.2 | 447.2 | - | - | 0 | - |
| 12 | y | 9224 | 448.3 | 0.0006218 | 1.387 | +1 | 4 |
| - | - | 1759 | 449.3 | - | - | 0 | - |
| 8 | c | 717.7 | 450.7 | 0.002218 | 4.92 | +2 | 8 |
| - | - | 803.1 | 451.2 | - | - | 0 | - |
| 3 | y | 1483 | 454.9 | 0.001283 | 2.821 | +3 | 13 |
| - | - | 626.2 | 455.3 | - | - | 0 | - |
| 4 | c | 6835 | 460.3 | 0.0006772 | 1.471 | +1 | 4 |
| - | - | 1945 | 461.3 | - | - | 0 | - |
| - | - | 606.3 | 461.9 | - | - | 0 | - |
| 14 | c | 2395 | 472.2 | 0.007897 | 16.72 | +3 | 14 |
| 7 | z | 2146 | 472.7 | 0.000162 | 0.3427 | +2 | 9 |
| - | - | 2882 | 474.2 | - | - | 0 | - |
| 7 | y | 1336 | 480.7 | 0.0004435 | 0.9226 | +2 | 9 |
| 2 | z | 1432 | 481.2 | 0.005404 | 11.23 | +3 | 14 |
| - | - | 539 | 481.6 | - | - | 0 | - |
| 11 | z | 2194 | 489.3 | 0.0004761 | 0.9731 | +1 | 5 |
| - | - | 1993 | 490.3 | - | - | 0 | - |
| 2 | y | 865.3 | 492.6 | 0.0007175 | 1.457 | +3 | 14 |
| - | - | 569.6 | 495.9 | - | - | 0 | - |
| 9 | c | 647.8 | 499.3 | 0.006098 | 12.21 | +2 | 9 |
| - | - | 2149 | 499.3 | - | - | 0 | - |
| - | - | 594.5 | 504.3 | - | - | 0 | - |
| 11 | y | 3.149E+04 | 505.3 | 0.0004593 | 0.9091 | +1 | 5 |
| - | - | 6753 | 506.3 | - | - | 0 | - |
| - | - | 1182 | 507.3 | - | - | 0 | - |
| - | - | 1145 | 515.3 | - | - | 0 | - |
| 6 | y | 1717 | 515.7 | 0.0005115 | 0.9918 | +2 | 10 |
| 6 | z | 2986 | 516.3 | 0.0004103 | 0.7947 | +2 | 10 |
| - | - | 892.3 | 516.3 | - | - | 0 | - |
| - | - | 4482 | 516.8 | - | - | 0 | - |
| - | - | 911 | 517.3 | - | - | 0 | - |
| 6 | y | 1.035E+04 | 524.3 | 0.004266 | 8.137 | +2 | 10 |
| - | - | 1894 | 524.6 | - | - | 0 | - |
| - | - | 4704 | 524.8 | - | - | 0 | - |
| - | - | 1795 | 524.9 | - | - | 0 | - |
| - | - | 2124 | 525.3 | - | - | 0 | - |
| - | - | 647.5 | 529.3 | - | - | 0 | - |
| - | - | 1381 | 529.3 | - | - | 0 | - |
| - | - | 640.1 | 530.2 | - | - | 0 | - |
| - | - | 3845 | 530.3 | - | - | 0 | - |
| - | - | 3322 | 530.6 | - | - | 0 | - |
| - | - | 1383 | 530.9 | - | - | 0 | - |
| - | - | 1478 | 541.3 | - | - | 0 | - |
| - | - | 3548 | 542.3 | - | - | 0 | - |
| - | - | 1209 | 543.3 | - | - | 0 | - |
| - | - | 748.9 | 551.3 | - | - | 0 | - |
| - | - | 857.3 | 551.8 | - | - | 0 | - |
| 10 | z | 791.2 | 558.3 | 0.0006019 | 1.078 | +1 | 6 |
| - | - | 2112 | 558.3 | - | - | 0 | - |
| 10 | w | 1523 | 559.3 | 0.0001791 | 0.3202 | +1 | 6 |
| 5 | c | 5801 | 559.3 | 0.0005065 | 0.9056 | +1 | 5 |
| - | - | 621.5 | 560.3 | - | - | 0 | - |
| - | - | 1907 | 560.3 | - | - | 0 | - |
| 5 | z | 2.294E+04 | 565.8 | 1.031E-05 | 0.01823 | +2 | 11 |
| - | - | 1.498E+04 | 566.3 | - | - | 0 | - |
| - | - | 5863 | 566.8 | - | - | 0 | - |
| - | - | 1578 | 567.3 | - | - | 0 | - |
| 5 | y | 1.321E+04 | 573.8 | 8.81E-05 | 0.1535 | +2 | 11 |
| 10 | y | 6453 | 574.3 | 0.003684 | 6.415 | +1 | 6 |
| - | - | 2527 | 574.8 | - | - | 0 | - |
| 10 | y | 956.6 | 575.3 | 0.0005647 | 0.9817 | +1 | 6 |
| 10 | z | 5914 | 576.3 | 0.0003633 | 0.6305 | +1 | 6 |
| - | - | 1307 | 577.3 | - | - | 0 | - |
| - | - | 872.1 | 586.3 | - | - | 0 | - |
| - | - | 1614 | 587.3 | - | - | 0 | - |
| 10 | y | 839.5 | 592.3 | 0.0008101 | 1.368 | +1 | 6 |
| - | - | 2579 | 600.3 | - | - | 0 | - |
| - | - | 1320 | 600.3 | - | - | 0 | - |
| 4 | w | 1.021E+04 | 600.8 | 0.0002989 | 0.4975 | +2 | 12 |
| - | - | 7019 | 601.3 | - | - | 0 | - |
| - | - | 3512 | 601.8 | - | - | 0 | - |
| - | - | 1015 | 603.4 | - | - | 0 | - |
| - | - | 789.7 | 615.3 | - | - | 0 | - |
| - | - | 950.3 | 619.3 | - | - | 0 | - |
| 12 | c | 1556 | 629.3 | 0.006528 | 10.37 | +2 | 12 |
| 4 | z | 2.289E+04 | 630.3 | 0.0001775 | 0.2816 | +2 | 12 |
| - | - | 1.744E+04 | 630.8 | - | - | 0 | - |
| - | - | 6587 | 631.3 | - | - | 0 | - |
| - | - | 2579 | 631.8 | - | - | 0 | - |
| - | - | 903 | 633.4 | - | - | 0 | - |
| - | - | 7474 | 637.8 | - | - | 0 | - |
| 4 | y | 2.711E+04 | 638.3 | 0.0003955 | 0.6195 | +2 | 12 |
| - | - | 1.36E+04 | 638.8 | - | - | 0 | - |
| - | - | 4913 | 639.3 | - | - | 0 | - |
| - | - | 1547 | 639.8 | - | - | 0 | - |
| - | - | 752.1 | 642.4 | - | - | 0 | - |
| - | - | 2082 | 644.3 | - | - | 0 | - |
| - | - | 1765 | 644.8 | - | - | 0 | - |
| - | - | 1049 | 645.4 | - | - | 0 | - |
| 6 | c | 2.196E+04 | 646.4 | 0.0004241 | 0.6561 | +1 | 6 |
| - | - | 7372 | 647.4 | - | - | 0 | - |
| - | - | 1134 | 648.4 | - | - | 0 | - |
| - | - | 1387 | 651.8 | - | - | 0 | - |
| 3 | z | 1405 | 665.3 | 0.01038 | 15.6 | +2 | 13 |
| - | - | 1020 | 665.8 | - | - | 0 | - |
| - | - | 891 | 666.3 | - | - | 0 | - |
| 3 | y | 1604 | 672.8 | 0.0008734 | 1.298 | +2 | 13 |
| 3 | y | 865 | 673.3 | 0.006508 | 9.666 | +2 | 13 |
| 3 | z | 1.587E+04 | 673.8 | 0.0002156 | 0.3199 | +2 | 13 |
| - | - | 2.034E+04 | 674.3 | - | - | 0 | - |
| - | - | 1.055E+04 | 674.8 | - | - | 0 | - |
| - | - | 3765 | 675.3 | - | - | 0 | - |
| - | - | 854.3 | 675.8 | - | - | 0 | - |
| - | - | 802.1 | 676.9 | - | - | 0 | - |
| - | - | 1050 | 680.8 | - | - | 0 | - |
| - | - | 2008 | 681.3 | - | - | 0 | - |
| 3 | y | 1.012E+05 | 681.8 | 6.983E-05 | 0.1024 | +2 | 13 |
| - | - | 6.734E+04 | 682.3 | - | - | 0 | - |
| - | - | 1115 | 682.4 | - | - | 0 | - |
| - | - | 2.779E+04 | 682.8 | - | - | 0 | - |
| - | - | 8690 | 683.3 | - | - | 0 | - |
| - | - | 1245 | 683.8 | - | - | 0 | - |
| - | - | 4339 | 684.4 | - | - | 0 | - |
| - | - | 3886 | 684.9 | - | - | 0 | - |
| - | - | 2955 | 685.4 | - | - | 0 | - |
| - | - | 1269 | 685.9 | - | - | 0 | - |
| - | - | 847 | 686.3 | - | - | 0 | - |
| - | - | 1199 | 687.3 | - | - | 0 | - |
| - | - | 722.9 | 688.3 | - | - | 0 | - |
| 9 | y | 2823 | 689.4 | 0.0007195 | 1.044 | +1 | 7 |
| - | - | 1737 | 690.4 | - | - | 0 | - |
| - | - | 1097 | 692.4 | - | - | 0 | - |
| - | - | 819.5 | 699.4 | - | - | 0 | - |
| - | - | 1737 | 700.4 | - | - | 0 | - |
| - | - | 1046 | 700.9 | - | - | 0 | - |
| - | - | 710.1 | 708.4 | - | - | 0 | - |
| 2 | w | 1752 | 715.8 | 0.001807 | 2.524 | +2 | 14 |
| - | - | 1892 | 716.4 | - | - | 0 | - |
| - | - | 962.8 | 716.9 | - | - | 0 | - |
| - | - | 1836 | 717.4 | - | - | 0 | - |
| - | - | 1846 | 718.4 | - | - | 0 | - |
| 2 | y | 765.2 | 729.4 | 0.006284 | 8.616 | +2 | 14 |
| 2 | z | 2570 | 730.4 | 0.0001758 | 0.2406 | +2 | 14 |
| - | - | 872.3 | 730.9 | - | - | 0 | - |
| - | - | 1764 | 736.9 | - | - | 0 | - |
| - | - | 714.3 | 737.4 | - | - | 0 | - |
| - | - | 1134 | 737.9 | - | - | 0 | - |
| 2 | y | 6655 | 738.4 | 0.0001521 | 0.206 | +2 | 14 |
| - | - | 4425 | 738.9 | - | - | 0 | - |
| - | - | 1950 | 739.4 | - | - | 0 | - |
| - | - | 856.4 | 742.4 | - | - | 0 | - |
| 8 | w | 1674 | 743.4 | 0.002484 | 3.341 | +1 | 8 |
| - | - | 1799 | 744.4 | - | - | 0 | - |
| - | - | 927 | 745.4 | - | - | 0 | - |
| - | - | 845.9 | 750.9 | - | - | 0 | - |
| - | - | 1153 | 751.4 | - | - | 0 | - |
| - | - | 836.9 | 751.9 | - | - | 0 | - |
| - | - | 702.3 | 756.4 | - | - | 0 | - |
| - | - | 1592 | 757.4 | - | - | 0 | - |
| - | - | 3770 | 758.4 | - | - | 0 | - |
| - | - | 4226 | 758.9 | - | - | 0 | - |
| - | - | 2829 | 759.4 | - | - | 0 | - |
| - | - | 1201 | 759.9 | - | - | 0 | - |
| - | - | 1186 | 760.4 | - | - | 0 | - |
| 7 | c | 4.115E+04 | 761.4 | 0.0001534 | 0.2015 | +1 | 7 |
| - | - | 1.672E+04 | 762.4 | - | - | 0 | - |
| - | - | 4309 | 763.4 | - | - | 0 | - |
| - | - | 2088 | 764.4 | - | - | 0 | - |
| - | - | 5182 | 764.9 | - | - | 0 | - |
| - | - | 7.38E+04 | 765.4 | - | - | 0 | - |
| - | - | 5.919E+04 | 765.9 | - | - | 0 | - |
| - | - | 2.6E+04 | 766.4 | - | - | 0 | - |
| - | - | 8434 | 766.9 | - | - | 0 | - |
| - | - | 2231 | 767.4 | - | - | 0 | - |
| - | - | 1179 | 771.9 | - | - | 0 | - |
| - | - | 3082 | 772.4 | - | - | 0 | - |
| - | - | 2611 | 772.9 | - | - | 0 | - |
| - | - | 2803 | 773.4 | - | - | 0 | - |
| - | - | 2.003E+04 | 773.9 | - | - | 0 | - |
| - | - | 1.605E+04 | 774.4 | - | - | 0 | - |
| - | - | 1.148E+04 | 774.9 | - | - | 0 | - |
| - | - | 2592 | 775.4 | - | - | 0 | - |
| - | - | 785.4 | 775.9 | - | - | 0 | - |
| - | - | 2211 | 777.9 | - | - | 0 | - |
| - | - | 2065 | 778.4 | - | - | 0 | - |
| - | - | 1036 | 778.9 | - | - | 0 | - |
| - | - | 915.1 | 780.4 | - | - | 0 | - |
| - | - | 911.5 | 780.9 | - | - | 0 | - |
| - | - | 1277 | 782.4 | - | - | 0 | - |
| - | - | 862.1 | 785.9 | - | - | 0 | - |
| - | - | 8484 | 786.4 | - | - | 0 | - |
| - | - | 7.704E+04 | 786.9 | - | - | 0 | - |
| - | - | 6.341E+04 | 787.4 | - | - | 0 | - |
| - | - | 2.913E+04 | 787.9 | - | - | 0 | - |
| - | - | 9191 | 788.4 | - | - | 0 | - |
| - | - | 2024 | 788.9 | - | - | 0 | - |
| - | - | 1253 | 793.9 | - | - | 0 | - |
| - | - | 4581 | 794.4 | - | - | 0 | - |
| - | - | 5.587E+04 | 794.9 | - | - | 0 | - |
| - | - | 6.845E+04 | 795.4 | - | - | 0 | - |
| - | - | 4.104E+04 | 795.9 | - | - | 0 | - |
| - | - | 1.497E+04 | 796.4 | - | - | 0 | - |
| - | - | 4742 | 796.9 | - | - | 0 | - |
| - | - | 699 | 797.4 | - | - | 0 | - |
| - | - | 751.7 | 798.4 | - | - | 0 | - |
| 8 | z | 2712 | 811.4 | 0.002933 | 3.615 | +1 | 8 |
| - | - | 933.4 | 812.4 | - | - | 0 | - |
| 8 | y | 1029 | 828.4 | 0.0007223 | 0.8718 | +1 | 8 |
| 8 | z | 3.213E+04 | 829.4 | 6.341E-05 | 0.07645 | +1 | 8 |
| - | - | 1.833E+04 | 830.4 | - | - | 0 | - |
| - | - | 5897 | 831.4 | - | - | 0 | - |
| - | - | 2344 | 832.5 | - | - | 0 | - |
| - | - | 1037 | 841.5 | - | - | 0 | - |
| - | - | 1543 | 844.4 | - | - | 0 | - |
| 8 | y | 5707 | 845.5 | 0.00066 | 0.7807 | +1 | 8 |
| - | - | 1703 | 846.5 | - | - | 0 | - |
| - | - | 1240 | 847.5 | - | - | 0 | - |
| - | - | 2405 | 873.5 | - | - | 0 | - |
| - | - | 1010 | 874.5 | - | - | 0 | - |
| - | - | 1180 | 876.4 | - | - | 0 | - |
| 8 | c | 4844 | 900.5 | 0.001288 | 1.431 | +1 | 8 |
| - | - | 2318 | 901.5 | - | - | 0 | - |
| - | - | 930.1 | 902.5 | - | - | 0 | - |
| 7 | z | 870.7 | 926.5 | 0.003075 | 3.319 | +1 | 9 |
| - | - | 1079 | 931.5 | - | - | 0 | - |
| - | - | 670.1 | 940.6 | - | - | 0 | - |
| 7 | z | 1.32E+04 | 944.5 | 0.0003951 | 0.4183 | +1 | 9 |
| - | - | 9444 | 945.5 | - | - | 0 | - |
| - | - | 3239 | 946.5 | - | - | 0 | - |
| - | - | 1669 | 947.5 | - | - | 0 | - |
| - | - | 1167 | 956.5 | - | - | 0 | - |
| 7 | y | 1833 | 960.5 | 0.001755 | 1.827 | +1 | 9 |
| - | - | 895.5 | 961.5 | - | - | 0 | - |
| - | - | 3563 | 970.5 | - | - | 0 | - |
| - | - | 1825 | 971.5 | - | - | 0 | - |
| - | - | 1886 | 980.5 | - | - | 0 | - |
| - | - | 1749 | 1014 | - | - | 0 | - |
| 9 | c | 3.133E+04 | 1015 | 0.0003312 | 0.3265 | +1 | 9 |
| - | - | 1.768E+04 | 1016 | - | - | 0 | - |
| - | - | 5310 | 1017 | - | - | 0 | - |
| - | - | 1683 | 1018 | - | - | 0 | - |
| - | - | 676.6 | 1022 | - | - | 0 | - |
| 6 | z | 5529 | 1031 | 0.0003801 | 0.3685 | +1 | 10 |
| - | - | 6831 | 1033 | - | - | 0 | - |
| - | - | 2808 | 1034 | - | - | 0 | - |
| - | - | 1164 | 1035 | - | - | 0 | - |
| - | - | 1602 | 1040 | - | - | 0 | - |
| - | - | 2365 | 1041 | - | - | 0 | - |
| - | - | 859.4 | 1042 | - | - | 0 | - |
| - | - | 1910 | 1043 | - | - | 0 | - |
| - | - | 631.3 | 1043 | - | - | 0 | - |
| - | - | 790.6 | 1044 | - | - | 0 | - |
| - | - | 1237 | 1045 | - | - | 0 | - |
| - | - | 919.3 | 1047 | - | - | 0 | - |
| 6 | y | 2836 | 1048 | 0.0006715 | 0.6411 | +1 | 10 |
| - | - | 2.168E+04 | 1058 | - | - | 0 | - |
| - | - | 1.331E+04 | 1059 | - | - | 0 | - |
| - | - | 5982 | 1060 | - | - | 0 | - |
| - | - | 1972 | 1061 | - | - | 0 | - |
| - | - | 869.1 | 1063 | - | - | 0 | - |
| - | - | 1020 | 1101 | - | - | 0 | - |
| 10 | c | 4.247E+04 | 1102 | 0.0003463 | 0.3144 | +1 | 10 |
| - | - | 2.401E+04 | 1103 | - | - | 0 | - |
| - | - | 9205 | 1104 | - | - | 0 | - |
| - | - | 1521 | 1105 | - | - | 0 | - |
| - | - | 6531 | 1115 | - | - | 0 | - |
| - | - | 4670 | 1116 | - | - | 0 | - |
| - | - | 2013 | 1117 | - | - | 0 | - |
| 5 | z | 6041 | 1131 | 0.001777 | 1.572 | +1 | 11 |
| - | - | 1.547E+04 | 1132 | - | - | 0 | - |
| - | - | 9187 | 1133 | - | - | 0 | - |
| - | - | 3002 | 1134 | - | - | 0 | - |
| - | - | 886.1 | 1138 | - | - | 0 | - |
| - | - | 1550 | 1139 | - | - | 0 | - |
| 5 | y | 1549 | 1147 | 0.001227 | 1.07 | +1 | 11 |
| - | - | 1074 | 1148 | - | - | 0 | - |
| - | - | 967.6 | 1149 | - | - | 0 | - |
| 11 | c | 2.317E+04 | 1159 | 0.001342 | 1.158 | +1 | 11 |
| - | - | 1.562E+04 | 1160 | - | - | 0 | - |
| - | - | 6005 | 1161 | - | - | 0 | - |
| - | - | 934.6 | 1162 | - | - | 0 | - |
| - | - | 713.1 | 1213 | - | - | 0 | - |
| - | - | 4266 | 1214 | - | - | 0 | - |
| - | - | 3322 | 1215 | - | - | 0 | - |
| - | - | 1813 | 1216 | - | - | 0 | - |
| - | - | 2491 | 1217 | - | - | 0 | - |
| - | - | 688.6 | 1257 | - | - | 0 | - |
| 12 | c | 2.441E+04 | 1258 | 0.002251 | 1.79 | +1 | 12 |
| - | - | 1.555E+04 | 1259 | - | - | 0 | - |
| - | - | 5570 | 1260 | - | - | 0 | - |
| - | - | 2.173E+04 | 1261 | - | - | 0 | - |
| - | - | 1.312E+04 | 1262 | - | - | 0 | - |
| - | - | 5351 | 1263 | - | - | 0 | - |
| - | - | 1424 | 1264 | - | - | 0 | - |
| - | - | 723.6 | 1275 | - | - | 0 | - |
| 4 | y | 7084 | 1276 | 0.003524 | 2.763 | +1 | 12 |
| - | - | 5165 | 1277 | - | - | 0 | - |
| - | - | 1948 | 1278 | - | - | 0 | - |
| - | - | 1165 | 1279 | - | - | 0 | - |
| - | - | 782.9 | 1284 | - | - | 0 | - |
| - | - | 3655 | 1301 | - | - | 0 | - |
| - | - | 3739 | 1302 | - | - | 0 | - |
| - | - | 1785 | 1303 | - | - | 0 | - |
| 3 | w | 1500 | 1330 | 0.00471 | 3.542 | +1 | 13 |
| - | - | 2143 | 1331 | - | - | 0 | - |
| - | - | 1164 | 1332 | - | - | 0 | - |
| 13 | c | 4.167E+04 | 1345 | 0.002297 | 1.708 | +1 | 13 |
| - | - | 3.123E+04 | 1346 | - | - | 0 | - |
| - | - | 1.235E+04 | 1347 | - | - | 0 | - |
| - | - | 1.83E+04 | 1348 | - | - | 0 | - |
| - | - | 1.345E+04 | 1349 | - | - | 0 | - |
| - | - | 4670 | 1350 | - | - | 0 | - |
| - | - | 1665 | 1351 | - | - | 0 | - |
| - | - | 937.6 | 1354 | - | - | 0 | - |
| 3 | y | 1065 | 1363 | 0.004425 | 3.247 | +1 | 13 |
| - | - | 1653 | 1364 | - | - | 0 | - |
| - | - | 740.1 | 1365 | - | - | 0 | - |
| - | - | 1032 | 1369 | - | - | 0 | - |
| - | - | 1518 | 1370 | - | - | 0 | - |
| - | - | 911.1 | 1372 | - | - | 0 | - |
| - | - | 3486 | 1388 | - | - | 0 | - |
| - | - | 3399 | 1389 | - | - | 0 | - |
| - | - | 1528 | 1390 | - | - | 0 | - |
| - | - | 678.9 | 1417 | - | - | 0 | - |
| - | - | 976.3 | 1430 | - | - | 0 | - |
| - | - | 1118 | 1431 | - | - | 0 | - |
| 14 | c | 5.615E+04 | 1432 | 0.002709 | 1.892 | +1 | 14 |
| - | - | 4.334E+04 | 1433 | - | - | 0 | - |
| - | - | 1.882E+04 | 1434 | - | - | 0 | - |
| - | - | 4784 | 1435 | - | - | 0 | - |
| - | - | 1341 | 1436 | - | - | 0 | - |
| 2 | z | 890.9 | 1460 | 0.02216 | 15.18 | +1 | 14 |
| - | - | 3034 | 1461 | - | - | 0 | - |
| - | - | 1918 | 1462 | - | - | 0 | - |
| - | - | 910.8 | 1463 | - | - | 0 | - |
| - | - | 825.6 | 1472 | - | - | 0 | - |
| - | - | 910.9 | 1473 | - | - | 0 | - |
| - | - | 1098 | 1474 | - | - | 0 | - |
| - | - | 803.4 | 1475 | - | - | 0 | - |
| - | - | 733.9 | 1488 | - | - | 0 | - |
| - | - | 5535 | 1489 | - | - | 0 | - |
| - | - | 4053 | 1490 | - | - | 0 | - |
| - | - | 2637 | 1491 | - | - | 0 | - |
| - | - | 1007 | 1492 | - | - | 0 | - |
| - | - | 1369 | 1513 | - | - | 0 | - |
| - | - | 2150 | 1514 | - | - | 0 | - |
| - | - | 1348 | 1515 | - | - | 0 | - |
| - | - | 757 | 1516 | - | - | 0 | - |
| - | - | 913.8 | 1517 | - | - | 0 | - |
| - | - | 886.3 | 1529 | - | - | 0 | - |
| - | - | 1690 | 1530 | - | - | 0 | - |
| - | - | 1.071E+04 | 1531 | - | - | 0 | - |
| - | - | 1.366E+04 | 1532 | - | - | 0 | - |
| - | - | 9407 | 1533 | - | - | 0 | - |
| - | - | 4066 | 1534 | - | - | 0 | - |
| - | - | 1781 | 1535 | - | - | 0 | - |
| - | - | 778.2 | 1544 | - | - | 0 | - |
| - | - | 3812 | 1545 | - | - | 0 | - |
| - | - | 4695 | 1546 | - | - | 0 | - |
| - | - | 6526 | 1547 | - | - | 0 | - |
| - | - | 7831 | 1548 | - | - | 0 | - |
| - | - | 3694 | 1549 | - | - | 0 | - |
| - | - | 1357 | 1550 | - | - | 0 | - |
| - | - | 2668 | 1556 | - | - | 0 | - |
| - | - | 6244 | 1557 | - | - | 0 | - |
| - | - | 4463 | 1558 | - | - | 0 | - |
| - | - | 2821 | 1559 | - | - | 0 | - |
| - | - | 697.1 | 1560 | - | - | 0 | - |
| - | - | 2023 | 1562 | - | - | 0 | - |
| - | - | 1.051E+04 | 1563 | - | - | 0 | - |
| - | - | 7630 | 1564 | - | - | 0 | - |
| - | - | 4113 | 1565 | - | - | 0 | - |
| - | - | 860.8 | 1566 | - | - | 0 | - |
| - | - | 2091 | 1572 | - | - | 0 | - |
| - | - | 1.444E+04 | 1573 | - | - | 0 | - |
| - | - | 8.54E+04 | 1574 | - | - | 0 | - |
| - | - | 6.072E+04 | 1575 | - | - | 0 | - |
| - | - | 3.186E+04 | 1576 | - | - | 0 | - |
| - | - | 9223 | 1577 | - | - | 0 | - |
| - | - | 2487 | 1578 | - | - | 0 | - |
| - | - | 1330 | 1588 | - | - | 0 | - |
| - | - | 7328 | 1589 | - | - | 0 | - |
| - | - | 2.483E+04 | 1590 | - | - | 0 | - |
| - | - | 8.176E+04 | 1591 | - | - | 0 | - |
| - | - | 5.976E+04 | 1592 | - | - | 0 | - |
| - | - | 2.801E+04 | 1593 | - | - | 0 | - |
| - | - | 8507 | 1594 | - | - | 0 | - |
| - | - | 1937 | 1595 | - | - | 0 | - |
| - | - | 681.4 | 1884 | - | - | 0 | - |
| - | - | 795.2 | 2460 | - | - | 0 | - |

m/z Charge Intensity FragmentType MassShift Position
123.9146499633789 0 373.69495
124.08241271972656 0 452.87756
124.38890838623047 0 376.27667
125.10768127441406 0 1627.0758
126.09161376953125 0 1908.8469
130.0615692138672 0 555.41156
131.12939453125 0 1465.1594
133.06114196777344 0 596.2612
138.1892852783203 0 472.71954
142.0736846923828 0 413.3337 z Ammonia loss 14
148.94776916503906 0 667.3628
158.09266662597656 0 836.4271 y Ammonia loss 14
163.07156372070312 0 1800.1489
164.21897888183594 0 467.25333
168.1134796142578 0 3567.813
173.4515838623047 0 1392.6841
175.11923217773438 0 5167.2266 y 14
183.11294555664062 0 538.11285
185.13990783691406 0 13257.199
186.14418029785156 0 570.4292
189.0866241455078 0 542.34534
199.18072509765625 0 30388.236
200.12725830078125 0 2406.5293
200.18408203125 0 4111.868
202.142578125 0 514.58215
212.10308837890625 0 677.94867
217.08221435546875 0 1694.0579
226.130615234375 0 977.7367
226.16812133789062 0 658.9799
227.17564392089844 0 16067.281
228.17965698242188 0 1894.3247
229.12998962402344 0 2451.1511 w 13
233.16458129882812 0 793.1449
234.10763549804688 0 592.17163
244.1167449951172 0 641.9075
245.12474060058594 0 24894.625 y Ammonia loss 13
246.13131713867188 0 4111.9873 z 13
255.10952758789062 0 1111.8522
261.1616516113281 0 591.4032
262.1507873535156 0 2447.114 y 13
270.1196594238281 0 3627.384
272.1720275878906 0 616.2013
293.3086853027344 0 650.9608
294.1743469238281 0 929.8043
303.15386962890625 0 585.07886
314.2078552246094 0 1807.2018
316.1620178222656 0 12051.422 w 12
317.1584777832031 0 2448.0625
331.1490173339844 0 3226.2144
331.2362976074219 0 873.37103 c 2
332.1567687988281 0 4569.7217 y Ammonia loss 12
333.1644287109375 0 10206.214 z 12
334.1689758300781 0 1129.965
349.1829528808594 0 2762.3992 y 12
357.1518859863281 0 6592.8564
363.5715637207031 0 615.33
378.2175598144531 0 544.5865
391.72503662109375 0 1347.0233
392.22515869140625 0 956.77185
415.2268981933594 0 2312.696 z 7
415.7243957519531 0 1210.9937
417.2097473144531 0 10224.912 w 11
418.2111511230469 0 1273.7914
423.2340087890625 0 2110.4736 y 7
423.73492431640625 0 1057.8448
425.2400817871094 0 672.629
429.23712158203125 0 791.1326
431.22637939453125 0 683.56683 y Ammonia loss 11
443.25042724609375 0 1552.1731
443.30963134765625 0 1204.6243
444.2560729980469 0 721.2014
446.2343444824219 0 1067.3217
447.2066650390625 0 652.21704
448.2520446777344 0 9223.92 y 11
449.2552185058594 0 1758.7764
450.7451171875 0 717.7113 c Ammonia loss 7
451.2428894042969 0 803.05334
454.89306640625 0 1483.3613 y 2
455.2895202636719 0 626.2378
460.2772521972656 0 6834.937 c 3
461.280517578125 0 1944.5679
461.92315673828125 0 606.285
472.2518310546875 0 2395.3223 c Ammonia loss 13
472.7373046875 0 2146.0176 z 6
474.2319641113281 0 2881.8167
480.7469482421875 0 1335.7167 y 6
481.248779296875 0 1431.5604 z Water loss 1
481.6166687011719 0 539.01416
489.254638671875 0 2193.8687 z 10
490.261474609375 0 1992.8557
492.5871887207031 0 865.30475 y 1
495.91448974609375 0 569.6013
499.26318359375 0 647.75824 c Ammonia loss 8
499.3002014160156 0 2149.4832
504.2626647949219 0 594.5403
505.2733459472656 0 31487.955 y 10
506.2760314941406 0 6753.4917
507.28057861328125 0 1182.1118
515.3294677734375 0 1145.373
515.749755859375 0 1716.7344 y Ammonia loss 5
516.2527465820312 0 2985.6575 z 5
516.336181640625 0 892.2705
516.756103515625 0 4481.73
517.2592163085938 0 910.96716
524.2667846679688 0 10350.26 y 5
524.6122436523438 0 1893.9756
524.7639770507812 0 4703.947
524.9466552734375 0 1795.1599
525.2681274414062 0 2124.1914
529.2516479492188 0 647.5391
529.2988891601562 0 1380.506
530.2221069335938 0 640.1313
530.2831420898438 0 3845.4194
530.616455078125 0 3321.8281
530.9478149414062 0 1383.2135
541.3123168945312 0 1477.6191
542.318115234375 0 3547.986
543.3218994140625 0 1209.0934
551.279052734375 0 748.8777
551.7835693359375 0 857.28235
558.2750244140625 0 791.1644 z Water loss 9
558.3368530273438 0 2112.3315
559.2836303710938 0 1523.0294 w 9
559.344482421875 0 5801.0312 c 4
560.2753295898438 0 621.53687
560.3485107421875 0 1907.487
565.787353515625 0 22942.986 z 4
566.2891845703125 0 14984.074
566.7908935546875 0 5863.3135
567.2918090820312 0 1577.5524
573.7968139648438 0 13210.699 y 4
574.2980346679688 0 6452.819 y Water loss 9
574.7979125976562 0 2526.7695
575.2789306640625 0 956.6116 y Ammonia loss 9
576.2858276367188 0 5914.4204 z 9
577.2920532226562 0 1307.3074
586.3297729492188 0 872.1142
587.3382568359375 0 1613.7103
592.3057250976562 0 839.54535 y 9
600.2742309570312 0 2579.0208
600.3306884765625 0 1320.3838
600.8023071289062 0 10209.269 w 3
601.3033447265625 0 7019.2285
601.8056640625 0 3512.4556
603.3711547851562 0 1015.32043
615.2735595703125 0 789.66815
619.342041015625 0 950.3298
629.3500366210938 0 1555.897 c 11
630.308837890625 0 22893.26 z 3
630.8106079101562 0 17444.156
631.3110961914062 0 6587.098
631.8125 0 2578.8877
633.3833618164062 0 902.97925
637.8145141601562 0 7474.4736
638.317626953125 0 27112.693 y 3
638.8193359375 0 13604.181
639.3204345703125 0 4913.406
639.8214111328125 0 1546.9144
642.356689453125 0 752.0977
644.3191528320312 0 2081.601
644.8194580078125 0 1765.4248
645.371337890625 0 1049.0807
646.37744140625 0 21955.805 c 5
647.3807373046875 0 7371.598
648.383056640625 0 1133.8412
651.82958984375 0 1386.5931
665.32177734375 0 1404.8502 z Ammonia loss 2
665.81884765625 0 1019.89655
666.318115234375 0 891.00476
672.827880859375 0 1603.7612 y Water loss 2
673.3272705078125 0 864.9809 y Ammonia loss 2
673.8248901367188 0 15871.821 z 2
674.3276977539062 0 20342.3
674.8292236328125 0 10553.859
675.3309326171875 0 3764.7607
675.83447265625 0 854.2764
676.8917846679688 0 802.11383
680.8264770507812 0 1050.422
681.3304443359375 0 2007.7329
681.8341064453125 0 101206.38 y 2
682.33544921875 0 67341.41
682.4046020507812 0 1114.5905
682.837158203125 0 27787.857
683.3380737304688 0 8690.272
683.836181640625 0 1245.1052
684.398193359375 0 4339.052
684.8995361328125 0 3885.6394
685.3994140625 0 2954.6897
685.9010009765625 0 1268.5127
686.3206176757812 0 846.9897
687.3333740234375 0 1198.8513
688.3494262695312 0 722.9357
689.3583984375 0 2823.2751 y 8
690.3611450195312 0 1736.6294
692.3788452148438 0 1096.933
699.3795776367188 0 819.5107
700.3841552734375 0 1737.1661
700.8936157226562 0 1046.3151
708.3642578125 0 710.13385
715.8453369140625 0 1751.6437 w 1
716.3518676757812 0 1891.7708
716.8515014648438 0 962.7937
717.389404296875 0 1836.1942
718.395751953125 0 1846.3954
729.364501953125 0 765.15076 y Water loss 1
730.3668823242188 0 2570.4756 z 1
730.8698120117188 0 872.332
736.9091186523438 0 1764.0702
737.411865234375 0 714.30286
737.9146118164062 0 1133.7942
738.376220703125 0 6654.5645 y 1
738.8776245117188 0 4425.191
739.3780517578125 0 1949.6361
742.3912963867188 0 856.42035
743.3707275390625 0 1674.3552 w 7
744.3736572265625 0 1799.4917
745.3739624023438 0 926.9663
750.897216796875 0 845.9343
751.4026489257812 0 1153.18
751.9058837890625 0 836.89984
756.4208374023438 0 702.3459
757.4000244140625 0 1592.012
758.3882446289062 0 3770.181
758.8837890625 0 4226.28
759.3865966796875 0 2829.369
759.8834838867188 0 1201.0005
760.3934936523438 0 1185.641
761.4041137695312 0 41154.08 c 6
762.4070434570312 0 16722.51
763.4093017578125 0 4308.705
764.4083862304688 0 2087.5222
764.9124145507812 0 5182.33
765.3822631835938 0 73799.42
765.8834838867188 0 59185.086
766.3848266601562 0 25995.455
766.8860473632812 0 8433.698
767.3867797851562 0 2231.0173
771.8961791992188 0 1179.0546
772.4133911132812 0 3081.9727
772.913330078125 0 2611.273
773.4144287109375 0 2802.5103
773.9080200195312 0 20034.3
774.4085693359375 0 16047.194
774.9093627929688 0 11479.693
775.4141845703125 0 2592.4507
775.9081420898438 0 785.4045
777.904052734375 0 2211.0806
778.4083251953125 0 2065.1611
778.8987426757812 0 1036.4651
780.4121704101562 0 915.0847
780.9205932617188 0 911.4715
782.4441528320312 0 1277.1957
785.9046020507812 0 862.1413
786.4136962890625 0 8483.63
786.9094848632812 0 77037.305
787.4108276367188 0 63407.176
787.91162109375 0 29127.43
788.4126586914062 0 9190.784
788.9156494140625 0 2023.964
793.9077758789062 0 1253.2428
794.4148559570312 0 4580.956
794.91796875 0 55868.402
795.4200439453125 0 68451.61
795.9219360351562 0 41041.27
796.4227294921875 0 14974.581
796.9215087890625 0 4742.491
797.4281616210938 0 698.9526
798.4326782226562 0 751.70087
811.4324340820312 0 2712.3354 z Water loss 7
812.431640625 0 933.4146
828.4315185546875 0 1028.5703 y Ammonia loss 7
829.4400024414062 0 32132.488 z 7
830.4434204101562 0 18334.385
831.4464111328125 0 5897.2666
832.4581298828125 0 2344.2441
841.5303344726562 0 1037.1104
844.4495849609375 0 1542.7496
845.4581298828125 0 5706.6514 y 7
846.4630737304688 0 1702.8961
847.470458984375 0 1240.0272
873.4913940429688 0 2405.3438
874.4949340820312 0 1009.66376
876.4163818359375 0 1179.5294
900.4772338867188 0 4844.407 c Ammonia loss 7
901.4800415039062 0 2318.3057
902.484375 0 930.1285
926.453369140625 0 870.7116 z Water loss 6
931.5341796875 0 1079.0609
940.55029296875 0 670.06805
944.4666137695312 0 13203.57 z 6
945.4719848632812 0 9444.137
946.4745483398438 0 3239.069
947.480712890625 0 1668.5741
956.5487060546875 0 1166.8062
960.4874877929688 0 1833.4312 y 6
961.4957885742188 0 895.47925
970.5383911132812 0 3562.8933
971.5413818359375 0 1824.6663
980.4663696289062 0 1886.4741
1013.58447265625 0 1749.3303
1014.5581665039062 0 31329.178 c 8
1015.559814453125 0 17680.252
1016.56396484375 0 5310.0654
1017.5687255859375 0 1683.4272
1022.4562377929688 0 676.56647
1031.4986572265625 0 5528.9775 z 5
1032.504638671875 0 6830.866
1033.510009765625 0 2807.9326
1034.5135498046875 0 1164.1989
1039.5672607421875 0 1602.3677
1040.5645751953125 0 2364.902
1041.5740966796875 0 859.39136
1042.5732421875 0 1909.7092
1043.412841796875 0 631.25244
1043.576416015625 0 790.60645
1044.61328125 0 1236.7784
1046.507568359375 0 919.3354
1047.51708984375 0 2835.7302 y 5
1057.576416015625 0 21682.645
1058.57958984375 0 13309.952
1059.585693359375 0 5982.1094
1060.588623046875 0 1972.4124
1062.5322265625 0 869.1199
1100.5841064453125 0 1019.88965
1101.5902099609375 0 42471.29 c 9
1102.593017578125 0 24007.54
1103.595703125 0 9205.329
1104.592041015625 0 1521.2219
1114.5946044921875 0 6531.347
1115.5997314453125 0 4670.3994
1116.603759765625 0 2012.7073
1130.565673828125 0 6041.31 z 4
1131.57373046875 0 15473.064
1132.5775146484375 0 9186.589
1133.5782470703125 0 3002.2349
1137.666259765625 0 886.12317
1138.66455078125 0 1549.8428
1146.58740234375 0 1549.3564 y 4
1147.5889892578125 0 1073.5905
1148.5947265625 0 967.6354
1158.6099853515625 0 23169.795 c 10
1159.613525390625 0 15624.081
1160.6138916015625 0 6004.608
1161.6295166015625 0 934.57556
1212.6563720703125 0 713.08435
1213.6646728515625 0 4265.5024
1214.6663818359375 0 3321.767
1215.6715087890625 0 1813.2141
1216.6668701171875 0 2491.2158
1256.6636962890625 0 688.5753
1257.677490234375 0 24407.6 c 11
1258.6788330078125 0 15547.57
1259.6751708984375 0 5569.9443
1260.6163330078125 0 21732.594
1261.6182861328125 0 13124.749
1262.6187744140625 0 5350.569
1263.6356201171875 0 1423.844
1274.619140625 0 723.6213
1275.625244140625 0 7083.743 y 3
1276.6307373046875 0 5165.0947
1277.634521484375 0 1948.2886
1278.6396484375 0 1164.5331
1283.6815185546875 0 782.88525
1300.6959228515625 0 3655.0496
1301.6986083984375 0 3739.1887
1302.699462890625 0 1785.2048
1329.64404296875 0 1499.6655 w 2
1330.6292724609375 0 2142.7375
1331.6292724609375 0 1164.0333
1344.70947265625 0 41671.46 c 12
1345.71240234375 0 31233.545
1346.70751953125 0 12354.266
1347.6519775390625 0 18303.049
1348.6494140625 0 13445.278
1349.65380859375 0 4669.9023
1350.6575927734375 0 1664.8525
1353.7760009765625 0 937.57556
1362.6563720703125 0 1064.7428 y 2
1363.6644287109375 0 1653.4307
1364.6705322265625 0 740.0645
1368.7911376953125 0 1031.6222
1369.796630859375 0 1518.1887
1371.8177490234375 0 911.078
1387.7318115234375 0 3485.756
1388.7288818359375 0 3398.798
1389.741943359375 0 1527.7659
1416.7314453125 0 678.8511
1429.7215576171875 0 976.2922
1430.734375 0 1118.3718
1431.7410888671875 0 56153.727 c 13
1432.74267578125 0 43339.73
1433.745849609375 0 18820.37
1434.748779296875 0 4784.3237
1435.74560546875 0 1340.6986
1459.7039794921875 0 890.9119 z 1
1460.7294921875 0 3033.6426
1461.73828125 0 1918.0511
1462.7335205078125 0 910.83185
1471.784423828125 0 825.5887
1472.786865234375 0 910.88666
1473.7423095703125 0 1097.6536
1474.78369140625 0 803.4156
1487.784912109375 0 733.87305
1488.7403564453125 0 5535.357
1489.744384765625 0 4053.4597
1490.743896484375 0 2637.33
1491.7545166015625 0 1007.156
1512.791748046875 0 1368.8699
1513.785888671875 0 2149.8848
1514.7843017578125 0 1347.753
1515.7916259765625 0 757.02026
1516.758056640625 0 913.7841
1528.8095703125 0 886.3369
1529.7886962890625 0 1689.905
1530.7896728515625 0 10707.874
1531.7919921875 0 13656.965
1532.79443359375 0 9407.397
1533.7960205078125 0 4066.3694
1534.8035888671875 0 1780.9235
1543.8336181640625 0 778.17975
1544.8299560546875 0 3812.2925
1545.824951171875 0 4694.7075
1546.8218994140625 0 6525.557
1547.818603515625 0 7831.0474
1548.8140869140625 0 3694.165
1549.8197021484375 0 1356.9823
1555.802978515625 0 2667.7373
1556.790283203125 0 6244.286
1557.7943115234375 0 4462.726
1558.794189453125 0 2820.9963
1559.7967529296875 0 697.099
1561.83203125 0 2023.3583
1562.8453369140625 0 10506.443
1563.8504638671875 0 7630.0166
1564.85498046875 0 4113.136
1565.86279296875 0 860.8126
1571.8055419921875 0 2090.9116
1572.8134765625 0 14442.681
1573.814697265625 0 85400.26
1574.8173828125 0 60719.715
1575.8193359375 0 31862.031
1576.8206787109375 0 9223.321
1577.8193359375 0 2486.692
1587.8231201171875 0 1330.4069
1588.8260498046875 0 7327.942
1589.83203125 0 24832.795
1590.8387451171875 0 81763.4
1591.8416748046875 0 59757.316
1592.8448486328125 0 28010.562
1593.846923828125 0 8506.956
1594.8428955078125 0 1937.3308
1884.0838623046875 0 681.36957
2460.2685546875 0 795.17426

Spectrum Details

|  |  |
| --- | --- |
| Matched peaks? Matched peaks The total absolute number of peaks matched. Additionally in brackets the total fraction of peaks matched and the total number of peaks is shown. | 79 (17.44% of 453) |
| FDR? FDR The false discovery rate estimated for this peptide. It is calculated by matching all theoretical fragments with a non-integer shift with the raw peaks for this spectrum. This is done with 40 different shifts. The resulting percentage is the average number of annotated peaks over the number of annotated peaks with the correct spectrum. | 1.33% |
| Satellite FDR? Satellite FDR See the FDR for details on its calculation. This satellite ion specific FDR only contains the satellite ions (d/w) for I/L/J positions. | 0.00% |
| PSM Score? PSM Score The PSM Score as given by Hecklib to this annotated spectrum. It is shown with three significant figures. | 600 |

#### Spectrum 3905? Spectrum 3905 The raw spectrum of this peptide as annotated by Hecklib. The fragments are coloured according to ion type (see legend). Any peaks with a star '\*' as text can be hovered over to see the full details, first the ion type second the mass shift type. By hovering over the amino acids in the peptide or ions in the legend the corresponding peaks are highlighted. By toggling the 'Unassigned' label you can turn the background (unassigned) peaks on or off in the plot. By updating the slider in the Ion legend you can update the spectrum to only show the top X% of the peaks with labels. The top X% means any peak that is within X% of the highest intensity. By dragging in the spectrum you can zoom in to a specific part of the spectrum and use 'Zoom Out' to get back to the original zoom level. The annotation of the spectrum is based on the given sequence in the peptides file and is done with different software so inconsistencies are likely. The peaks are annotated based on the given sequence, with 20 ppm tolerance.

Copy Data

##### Spectrum 3905 (TSV)

###### Preview

```
Loading example...
```

*Click on the button to copy the data to your clipboard.*

Mz MinMz MaxIntensity Max

WidthHeightPeptide font sizePeptide stroke widthSpectrum font sizeSpectrum stroke widthCompact peptide

Ion legend

wxyz

abcd

OtherUnassignedIon^Charge^_Position_Show for top:%

JJSEVSDRPSGVSSR

04.42e+48.85e+41.33e+51.77e+5

Zoom Out

y^+1^_1_y^+1^_1_w^+1^_2_y^+1^_2_z^+1^_2_y^+1^_2_z^+3^_8_w^+1^_3_y^+1^_3_z^+1^_3_y^+2^_7_y^+1^_3_c^+3^_12_z^+2^_8_w^+1^_4_y^+2^_8_y^+1^_4_y^+1^_4_y^+1^_4_y^+3^_13_c^+2^_8_y^+3^_13_c^+1^_4_c^+3^_14_z^+2^_9_y^+2^_9_y^+3^_14_y^+3^_14_z^+1^_5_y^+3^_14_y^+1^_5_y^+2^_10_z^+2^_10_y^+2^_10_z^+1^_6_w^+1^_6_c^+1^_5_z^+2^_11_y^+2^_11_y^+1^_6_y^+1^_6_z^+1^_6_y^+1^_6_w^+2^_12_z^+2^_12_c^+2^_12_y^+2^_12_z^+2^_12_y^+2^_12_c^+1^_6_z^+2^_13_y^+2^_13_y^+2^_13_z^+2^_13_y^+2^_13_y^+1^_7_w^+2^_14_y^+2^_14_z^+2^_14_y^+2^_14_w^+1^_8_c^+1^_7_z^+1^_8_z^+1^_8_y^+1^_8_z^+1^_8_y^+1^_8_w^+1^_9_c^+1^_8_z^+1^_9_y^+1^_9_z^+1^_9_y^+1^_9_c^+1^_9_y^+1^_10_z^+1^_10_y^+1^_10_c^+1^_10_z^+1^_11_y^+1^_11_c^+1^_11_c^+1^_12_y^+1^_12_w^+1^_13_c^+1^_13_y^+1^_13_c^+1^_14_y^+1^_14_

0804160824123216

Fragment Matches Table

Show background peaks

| Position | Ion type | Intensity | mz Theoretical | mz Error (Th) | mz Error (ppm) | Charge | Series Number |
| --- | --- | --- | --- | --- | --- | --- | --- |
| - | - | 1940 | 125.1 | - | - | 0 | - |
| - | - | 4750 | 126.1 | - | - | 0 | - |
| - | - | 1743 | 131.1 | - | - | 0 | - |
| - | - | 419.1 | 132 | - | - | 0 | - |
| - | - | 471.5 | 136.5 | - | - | 0 | - |
| - | - | 502 | 143.1 | - | - | 0 | - |
| - | - | 414.7 | 143.8 | - | - | 0 | - |
| - | - | 436.9 | 146.1 | - | - | 0 | - |
| - | - | 1094 | 148.9 | - | - | 0 | - |
| 15 | y | 1326 | 158.1 | 0.000172 | 1.088 | +1 | 1 |
| - | - | 2095 | 160 | - | - | 0 | - |
| - | - | 494.3 | 160.3 | - | - | 0 | - |
| - | - | 427 | 160.7 | - | - | 0 | - |
| - | - | 4905 | 168.1 | - | - | 0 | - |
| - | - | 657.7 | 169.1 | - | - | 0 | - |
| - | - | 1189 | 173.5 | - | - | 0 | - |
| 15 | y | 8719 | 175.1 | 0.0003258 | 1.86 | +1 | 1 |
| - | - | 573.1 | 175.9 | - | - | 0 | - |
| - | - | 595.9 | 176.1 | - | - | 0 | - |
| - | - | 5289 | 178.1 | - | - | 0 | - |
| - | - | 2.117E+04 | 185.1 | - | - | 0 | - |
| - | - | 2350 | 186.1 | - | - | 0 | - |
| - | - | 848.8 | 189.1 | - | - | 0 | - |
| - | - | 5.575E+04 | 199.2 | - | - | 0 | - |
| - | - | 3769 | 200.1 | - | - | 0 | - |
| - | - | 7209 | 200.2 | - | - | 0 | - |
| - | - | 1662 | 201.1 | - | - | 0 | - |
| - | - | 886.4 | 201.1 | - | - | 0 | - |
| - | - | 720.4 | 216.1 | - | - | 0 | - |
| - | - | 600 | 216.7 | - | - | 0 | - |
| - | - | 2659 | 217.1 | - | - | 0 | - |
| - | - | 898.1 | 226.1 | - | - | 0 | - |
| - | - | 777.4 | 226.2 | - | - | 0 | - |
| - | - | 2.911E+04 | 227.2 | - | - | 0 | - |
| - | - | 3527 | 228.2 | - | - | 0 | - |
| 14 | w | 4308 | 229.1 | 0.0003354 | 1.464 | +1 | 2 |
| - | - | 586.2 | 229.2 | - | - | 0 | - |
| - | - | 781 | 230.1 | - | - | 0 | - |
| - | - | 3079 | 233.2 | - | - | 0 | - |
| - | - | 1003 | 244.1 | - | - | 0 | - |
| 14 | y | 3.918E+04 | 245.1 | 0.0004312 | 1.759 | +1 | 2 |
| 14 | z | 6224 | 246.1 | 0.001534 | 6.234 | +1 | 2 |
| - | - | 647.9 | 247.1 | - | - | 0 | - |
| - | - | 1579 | 249.2 | - | - | 0 | - |
| - | - | 1182 | 255.1 | - | - | 0 | - |
| - | - | 3244 | 261.2 | - | - | 0 | - |
| 14 | y | 5641 | 262.2 | 0.0002035 | 0.7763 | +1 | 2 |
| - | - | 700.3 | 262.2 | - | - | 0 | - |
| - | - | 3831 | 270.1 | - | - | 0 | - |
| - | - | 1194 | 272.2 | - | - | 0 | - |
| 8 | z | 1927 | 277.2 | 0.003368 | 12.15 | +3 | 8 |
| - | - | 9212 | 279.1 | - | - | 0 | - |
| - | - | 891.5 | 280.1 | - | - | 0 | - |
| - | - | 567.6 | 299.2 | - | - | 0 | - |
| - | - | 1104 | 302.2 | - | - | 0 | - |
| - | - | 1047 | 303.2 | - | - | 0 | - |
| - | - | 695.8 | 304.1 | - | - | 0 | - |
| - | - | 1914 | 314.2 | - | - | 0 | - |
| 13 | w | 2.012E+04 | 316.2 | 0.0005946 | 1.881 | +1 | 3 |
| - | - | 2927 | 317.2 | - | - | 0 | - |
| - | - | 980.5 | 323.2 | - | - | 0 | - |
| - | - | 5404 | 331.1 | - | - | 0 | - |
| 13 | y | 6254 | 332.2 | 0.0001563 | 0.4706 | +1 | 3 |
| 13 | z | 1.338E+04 | 333.2 | 0.0003574 | 1.073 | +1 | 3 |
| - | - | 1783 | 334.2 | - | - | 0 | - |
| 9 | y | 634.4 | 345.2 | 0.000994 | 2.88 | +2 | 7 |
| - | - | 574.3 | 345.7 | - | - | 0 | - |
| - | - | 1759 | 347.2 | - | - | 0 | - |
| - | - | 793.6 | 349.1 | - | - | 0 | - |
| 13 | y | 4751 | 349.2 | 0.0004932 | 1.412 | +1 | 3 |
| - | - | 1050 | 350.2 | - | - | 0 | - |
| - | - | 9138 | 357.2 | - | - | 0 | - |
| - | - | 711.2 | 359.2 | - | - | 0 | - |
| - | - | 667.3 | 373.2 | - | - | 0 | - |
| - | - | 2137 | 389.7 | - | - | 0 | - |
| - | - | 799.2 | 390.2 | - | - | 0 | - |
| - | - | 1517 | 390.7 | - | - | 0 | - |
| - | - | 666.5 | 391.2 | - | - | 0 | - |
| - | - | 7797 | 391.7 | - | - | 0 | - |
| - | - | 4122 | 392.2 | - | - | 0 | - |
| - | - | 2614 | 398.7 | - | - | 0 | - |
| - | - | 1667 | 399.2 | - | - | 0 | - |
| - | - | 1876 | 399.7 | - | - | 0 | - |
| - | - | 1156 | 400.2 | - | - | 0 | - |
| 12 | c | 754.5 | 414.2 | 0.0006884 | 1.662 | +3 | 12 |
| 8 | z | 5719 | 415.2 | 0.00222 | 5.346 | +2 | 8 |
| - | - | 2710 | 415.7 | - | - | 0 | - |
| 12 | w | 1.517E+04 | 417.2 | 0.0007982 | 1.913 | +1 | 4 |
| - | - | 3072 | 418.2 | - | - | 0 | - |
| 8 | y | 3489 | 423.2 | 0.0003042 | 0.7188 | +2 | 8 |
| - | - | 1527 | 423.7 | - | - | 0 | - |
| - | - | 1544 | 425.2 | - | - | 0 | - |
| - | - | 739.4 | 426.2 | - | - | 0 | - |
| - | - | 1484 | 429.2 | - | - | 0 | - |
| 12 | y | 644.6 | 430.2 | 0.00218 | 5.067 | +1 | 4 |
| 12 | y | 1559 | 431.2 | 0.0005901 | 1.368 | +1 | 4 |
| - | - | 1087 | 433.7 | - | - | 0 | - |
| - | - | 2940 | 442.7 | - | - | 0 | - |
| - | - | 4436 | 443.2 | - | - | 0 | - |
| - | - | 1449 | 443.3 | - | - | 0 | - |
| - | - | 588.8 | 444.3 | - | - | 0 | - |
| - | - | 1232 | 446.2 | - | - | 0 | - |
| - | - | 731.2 | 447.2 | - | - | 0 | - |
| - | - | 878.6 | 447.2 | - | - | 0 | - |
| 12 | y | 1.294E+04 | 448.3 | 0.0008049 | 1.796 | +1 | 4 |
| 3 | y | 637.8 | 449.2 | 0.008159 | 18.16 | +3 | 13 |
| - | - | 3073 | 449.3 | - | - | 0 | - |
| 8 | c | 755.4 | 450.7 | 0.0005699 | 1.264 | +2 | 8 |
| 3 | y | 1041 | 454.9 | 0.0002456 | 0.5399 | +3 | 13 |
| - | - | 746.9 | 455.2 | - | - | 0 | - |
| - | - | 884 | 455.3 | - | - | 0 | - |
| - | - | 751.6 | 460.2 | - | - | 0 | - |
| 4 | c | 1.048E+04 | 460.3 | 0.0007992 | 1.736 | +1 | 4 |
| - | - | 3033 | 461.3 | - | - | 0 | - |
| - | - | 1116 | 465.3 | - | - | 0 | - |
| 14 | c | 2574 | 472.2 | 0.00808 | 17.11 | +3 | 14 |
| 7 | z | 3203 | 472.7 | 0.001017 | 2.15 | +2 | 9 |
| - | - | 930.1 | 473.2 | - | - | 0 | - |
| - | - | 4794 | 474.2 | - | - | 0 | - |
| 7 | y | 3630 | 480.7 | 0.0008403 | 1.748 | +2 | 9 |
| 2 | y | 829 | 486.6 | 0.001859 | 3.82 | +3 | 14 |
| 2 | y | 606.1 | 486.9 | 0.007319 | 15.03 | +3 | 14 |
| - | - | 608.4 | 489.1 | - | - | 0 | - |
| 11 | z | 4233 | 489.3 | 0.0005677 | 1.16 | +1 | 5 |
| - | - | 2977 | 490.3 | - | - | 0 | - |
| 2 | y | 1062 | 492.6 | 0.001053 | 2.138 | +3 | 14 |
| - | - | 641.9 | 492.9 | - | - | 0 | - |
| - | - | 811.3 | 493.3 | - | - | 0 | - |
| - | - | 1393 | 495.9 | - | - | 0 | - |
| - | - | 946.6 | 496.3 | - | - | 0 | - |
| - | - | 2433 | 499.3 | - | - | 0 | - |
| - | - | 607.6 | 504.3 | - | - | 0 | - |
| 11 | y | 4.779E+04 | 505.3 | 0.000795 | 1.573 | +1 | 5 |
| - | - | 1.12E+04 | 506.3 | - | - | 0 | - |
| - | - | 2102 | 507.3 | - | - | 0 | - |
| - | - | 784.4 | 514.3 | - | - | 0 | - |
| - | - | 1915 | 515.2 | - | - | 0 | - |
| - | - | 2093 | 515.3 | - | - | 0 | - |
| 6 | y | 3307 | 515.7 | 0.0005261 | 1.02 | +2 | 10 |
| 6 | z | 4504 | 516.3 | 0.0001051 | 0.2036 | +2 | 10 |
| - | - | 1036 | 516.3 | - | - | 0 | - |
| - | - | 5465 | 516.8 | - | - | 0 | - |
| - | - | 3630 | 517.3 | - | - | 0 | - |
| - | - | 701.6 | 517.3 | - | - | 0 | - |
| - | - | 853.2 | 520.3 | - | - | 0 | - |
| - | - | 3638 | 522.3 | - | - | 0 | - |
| - | - | 1173 | 523.8 | - | - | 0 | - |
| 6 | y | 1.884E+04 | 524.3 | 0.002374 | 4.528 | +2 | 10 |
| - | - | 6030 | 524.6 | - | - | 0 | - |
| - | - | 7524 | 524.8 | - | - | 0 | - |
| - | - | 2304 | 524.9 | - | - | 0 | - |
| - | - | 3439 | 525.3 | - | - | 0 | - |
| - | - | 3752 | 528.8 | - | - | 0 | - |
| - | - | 1.495E+04 | 529.3 | - | - | 0 | - |
| - | - | 4225 | 529.8 | - | - | 0 | - |
| - | - | 5069 | 530.3 | - | - | 0 | - |
| - | - | 4145 | 530.6 | - | - | 0 | - |
| - | - | 1412 | 530.8 | - | - | 0 | - |
| - | - | 3853 | 531 | - | - | 0 | - |
| - | - | 6745 | 531.3 | - | - | 0 | - |
| - | - | 2744 | 531.8 | - | - | 0 | - |
| - | - | 1842 | 541.3 | - | - | 0 | - |
| - | - | 6350 | 542.3 | - | - | 0 | - |
| - | - | 1261 | 543.3 | - | - | 0 | - |
| - | - | 1035 | 546.3 | - | - | 0 | - |
| 10 | z | 651.2 | 558.3 | 0.009896 | 17.73 | +1 | 6 |
| - | - | 2605 | 558.3 | - | - | 0 | - |
| 10 | w | 1837 | 559.3 | 0.001705 | 3.048 | +1 | 6 |
| 5 | c | 8314 | 559.3 | 4.28E-05 | 0.07652 | +1 | 5 |
| - | - | 2134 | 560.3 | - | - | 0 | - |
| - | - | 702.1 | 561.3 | - | - | 0 | - |
| 5 | z | 3.808E+04 | 565.8 | 0.000539 | 0.9527 | +2 | 11 |
| - | - | 2.423E+04 | 566.3 | - | - | 0 | - |
| - | - | 7802 | 566.8 | - | - | 0 | - |
| - | - | 2325 | 567.3 | - | - | 0 | - |
| - | - | 1125 | 571.3 | - | - | 0 | - |
| - | - | 732.7 | 572.3 | - | - | 0 | - |
| - | - | 4794 | 572.4 | - | - | 0 | - |
| - | - | 873.8 | 573.3 | - | - | 0 | - |
| 5 | y | 2.038E+04 | 573.8 | 0.0003322 | 0.579 | +2 | 11 |
| 10 | y | 8961 | 574.3 | 0.003928 | 6.84 | +1 | 6 |
| - | - | 4582 | 574.8 | - | - | 0 | - |
| 10 | y | 2856 | 575.3 | 0.00264 | 4.589 | +1 | 6 |
| 10 | z | 7761 | 576.3 | 0.0003691 | 0.6405 | +1 | 6 |
| - | - | 2543 | 577.3 | - | - | 0 | - |
| - | - | 1034 | 585.3 | - | - | 0 | - |
| - | - | 2421 | 586.3 | - | - | 0 | - |
| - | - | 665.1 | 587.3 | - | - | 0 | - |
| 10 | y | 1529 | 592.3 | 0.0008989 | 1.518 | +1 | 6 |
| - | - | 640.9 | 593.3 | - | - | 0 | - |
| - | - | 3382 | 600.3 | - | - | 0 | - |
| 4 | w | 1.497E+04 | 600.8 | 0.0007872 | 1.31 | +2 | 12 |
| - | - | 8685 | 601.3 | - | - | 0 | - |
| - | - | 4084 | 601.8 | - | - | 0 | - |
| - | - | 655.3 | 602.3 | - | - | 0 | - |
| - | - | 1160 | 603.4 | - | - | 0 | - |
| - | - | 738 | 608.3 | - | - | 0 | - |
| - | - | 780.2 | 611.3 | - | - | 0 | - |
| 4 | z | 668.3 | 621.3 | 0.002536 | 4.081 | +2 | 12 |
| - | - | 974 | 627.4 | - | - | 0 | - |
| 12 | c | 2197 | 629.3 | 0.006833 | 10.86 | +2 | 12 |
| 4 | y | 888.1 | 629.8 | 0.00763 | 12.11 | +2 | 12 |
| 4 | z | 3.574E+04 | 630.3 | 0.0006048 | 0.9595 | +2 | 12 |
| - | - | 2.63E+04 | 630.8 | - | - | 0 | - |
| - | - | 1.159E+04 | 631.3 | - | - | 0 | - |
| - | - | 3210 | 631.8 | - | - | 0 | - |
| - | - | 763.1 | 637.3 | - | - | 0 | - |
| - | - | 1.589E+04 | 637.8 | - | - | 0 | - |
| 4 | y | 3.628E+04 | 638.3 | 0.0002124 | 0.3327 | +2 | 12 |
| - | - | 2.213E+04 | 638.8 | - | - | 0 | - |
| - | - | 8363 | 639.3 | - | - | 0 | - |
| - | - | 1480 | 639.8 | - | - | 0 | - |
| - | - | 3681 | 644.3 | - | - | 0 | - |
| - | - | 2420 | 644.8 | - | - | 0 | - |
| - | - | 894.8 | 645.4 | - | - | 0 | - |
| 6 | c | 3.557E+04 | 646.4 | 0.0006682 | 1.034 | +1 | 6 |
| - | - | 639 | 647.3 | - | - | 0 | - |
| - | - | 1.09E+04 | 647.4 | - | - | 0 | - |
| - | - | 2238 | 648.4 | - | - | 0 | - |
| - | - | 1956 | 651.8 | - | - | 0 | - |
| - | - | 886.8 | 652.3 | - | - | 0 | - |
| 3 | z | 1951 | 665.3 | 0.00995 | 14.96 | +2 | 13 |
| - | - | 1952 | 665.8 | - | - | 0 | - |
| - | - | 1524 | 666.3 | - | - | 0 | - |
| - | - | 743.1 | 670.4 | - | - | 0 | - |
| 3 | y | 1587 | 672.8 | 0.001606 | 2.387 | +2 | 13 |
| 3 | y | 2836 | 673.3 | 0.007485 | 11.12 | +2 | 13 |
| 3 | z | 2.364E+04 | 673.8 | 0.0008259 | 1.226 | +2 | 13 |
| - | - | 3.194E+04 | 674.3 | - | - | 0 | - |
| - | - | 1.723E+04 | 674.8 | - | - | 0 | - |
| - | - | 5956 | 675.3 | - | - | 0 | - |
| - | - | 1045 | 675.8 | - | - | 0 | - |
| - | - | 2403 | 680.8 | - | - | 0 | - |
| - | - | 3202 | 681.3 | - | - | 0 | - |
| 3 | y | 1.752E+05 | 681.8 | 0.0005581 | 0.8185 | +2 | 13 |
| - | - | 1.131E+05 | 682.3 | - | - | 0 | - |
| - | - | 4.947E+04 | 682.8 | - | - | 0 | - |
| - | - | 1.379E+04 | 683.3 | - | - | 0 | - |
| - | - | 3575 | 683.8 | - | - | 0 | - |
| - | - | 1690 | 684.4 | - | - | 0 | - |
| - | - | 1819 | 685.4 | - | - | 0 | - |
| - | - | 832.9 | 686.3 | - | - | 0 | - |
| - | - | 773.1 | 686.4 | - | - | 0 | - |
| - | - | 1988 | 687.3 | - | - | 0 | - |
| - | - | 1002 | 688.3 | - | - | 0 | - |
| 9 | y | 5183 | 689.4 | 0.0009027 | 1.309 | +1 | 7 |
| - | - | 2308 | 690.4 | - | - | 0 | - |
| - | - | 659.4 | 691.4 | - | - | 0 | - |
| - | - | 716.1 | 697.4 | - | - | 0 | - |
| - | - | 1591 | 699.4 | - | - | 0 | - |
| - | - | 763.7 | 700.4 | - | - | 0 | - |
| - | - | 2092 | 701.4 | - | - | 0 | - |
| - | - | 1011 | 711.4 | - | - | 0 | - |
| - | - | 2138 | 712.4 | - | - | 0 | - |
| - | - | 2669 | 713.4 | - | - | 0 | - |
| - | - | 812.8 | 714.4 | - | - | 0 | - |
| 2 | w | 2840 | 715.8 | 0.001367 | 1.909 | +2 | 14 |
| - | - | 2476 | 716.4 | - | - | 0 | - |
| - | - | 939 | 716.8 | - | - | 0 | - |
| - | - | 3517 | 717.4 | - | - | 0 | - |
| - | - | 4852 | 718.4 | - | - | 0 | - |
| - | - | 1667 | 719.4 | - | - | 0 | - |
| 2 | y | 1412 | 729.4 | 0.0007347 | 1.007 | +2 | 14 |
| 2 | z | 3545 | 730.4 | 0.0001905 | 0.2608 | +2 | 14 |
| - | - | 1494 | 730.9 | - | - | 0 | - |
| - | - | 639.4 | 736.4 | - | - | 0 | - |
| 2 | y | 1.07E+04 | 738.4 | 0.0004573 | 0.6193 | +2 | 14 |
| - | - | 7651 | 738.9 | - | - | 0 | - |
| - | - | 4651 | 739.4 | - | - | 0 | - |
| - | - | 1031 | 739.9 | - | - | 0 | - |
| 8 | w | 3028 | 743.4 | 0.002118 | 2.849 | +1 | 8 |
| - | - | 3612 | 744.4 | - | - | 0 | - |
| - | - | 620.2 | 745.4 | - | - | 0 | - |
| - | - | 2928 | 750.9 | - | - | 0 | - |
| - | - | 2080 | 751.4 | - | - | 0 | - |
| - | - | 1250 | 751.9 | - | - | 0 | - |
| - | - | 1085 | 756.9 | - | - | 0 | - |
| - | - | 1005 | 757.4 | - | - | 0 | - |
| - | - | 3708 | 758.4 | - | - | 0 | - |
| - | - | 5751 | 758.9 | - | - | 0 | - |
| - | - | 5764 | 759.4 | - | - | 0 | - |
| - | - | 1158 | 759.9 | - | - | 0 | - |
| - | - | 1896 | 760.4 | - | - | 0 | - |
| 7 | c | 6.599E+04 | 761.4 | 0.0006417 | 0.8427 | +1 | 7 |
| - | - | 2.607E+04 | 762.4 | - | - | 0 | - |
| - | - | 7172 | 763.4 | - | - | 0 | - |
| - | - | 3135 | 764.4 | - | - | 0 | - |
| - | - | 5911 | 764.9 | - | - | 0 | - |
| - | - | 1.092E+05 | 765.4 | - | - | 0 | - |
| - | - | 9.096E+04 | 765.9 | - | - | 0 | - |
| - | - | 4.55E+04 | 766.4 | - | - | 0 | - |
| - | - | 1.311E+04 | 766.9 | - | - | 0 | - |
| - | - | 3429 | 767.4 | - | - | 0 | - |
| - | - | 1087 | 771.4 | - | - | 0 | - |
| - | - | 846.1 | 771.9 | - | - | 0 | - |
| - | - | 3261 | 772.4 | - | - | 0 | - |
| - | - | 2993 | 772.9 | - | - | 0 | - |
| - | - | 3194 | 773.4 | - | - | 0 | - |
| - | - | 3.004E+04 | 773.9 | - | - | 0 | - |
| - | - | 2.544E+04 | 774.4 | - | - | 0 | - |
| - | - | 1.263E+04 | 774.9 | - | - | 0 | - |
| - | - | 3658 | 775.4 | - | - | 0 | - |
| - | - | 1213 | 775.9 | - | - | 0 | - |
| - | - | 1500 | 777.9 | - | - | 0 | - |
| - | - | 4322 | 778.4 | - | - | 0 | - |
| - | - | 2061 | 778.9 | - | - | 0 | - |
| - | - | 1850 | 779.4 | - | - | 0 | - |
| - | - | 1212 | 779.9 | - | - | 0 | - |
| - | - | 2576 | 780.4 | - | - | 0 | - |
| - | - | 643.8 | 780.9 | - | - | 0 | - |
| - | - | 5120 | 782.4 | - | - | 0 | - |
| - | - | 1.359E+04 | 783.4 | - | - | 0 | - |
| - | - | 2531 | 784.4 | - | - | 0 | - |
| - | - | 2402 | 785.9 | - | - | 0 | - |
| - | - | 8007 | 786.4 | - | - | 0 | - |
| - | - | 1.176E+05 | 786.9 | - | - | 0 | - |
| - | - | 9.468E+04 | 787.4 | - | - | 0 | - |
| - | - | 4.777E+04 | 787.9 | - | - | 0 | - |
| - | - | 1.464E+04 | 788.4 | - | - | 0 | - |
| - | - | 3892 | 788.9 | - | - | 0 | - |
| - | - | 1048 | 793.4 | - | - | 0 | - |
| - | - | 2651 | 793.9 | - | - | 0 | - |
| - | - | 6071 | 794.4 | - | - | 0 | - |
| - | - | 7.598E+04 | 794.9 | - | - | 0 | - |
| - | - | 9.814E+04 | 795.4 | - | - | 0 | - |
| - | - | 5.846E+04 | 795.9 | - | - | 0 | - |
| - | - | 2.728E+04 | 796.4 | - | - | 0 | - |
| - | - | 7858 | 796.9 | - | - | 0 | - |
| - | - | 3407 | 797.4 | - | - | 0 | - |
| - | - | 2042 | 798.4 | - | - | 0 | - |
| - | - | 760.4 | 799.4 | - | - | 0 | - |
| - | - | 1584 | 800.5 | - | - | 0 | - |
| 8 | z | 3216 | 811.4 | 0.0001253 | 0.1544 | +1 | 8 |
| 8 | z | 1461 | 812.4 | 0.01245 | 15.32 | +1 | 8 |
| 8 | y | 1011 | 828.4 | 0.001577 | 1.903 | +1 | 8 |
| 8 | z | 5.13E+04 | 829.4 | 0.0004859 | 0.5858 | +1 | 8 |
| - | - | 2.846E+04 | 830.4 | - | - | 0 | - |
| - | - | 7261 | 831.4 | - | - | 0 | - |
| - | - | 1408 | 832.4 | - | - | 0 | - |
| - | - | 1969 | 839.5 | - | - | 0 | - |
| - | - | 1206 | 840.5 | - | - | 0 | - |
| - | - | 4309 | 841.5 | - | - | 0 | - |
| - | - | 2542 | 842.5 | - | - | 0 | - |
| - | - | 2927 | 844.5 | - | - | 0 | - |
| 8 | y | 6725 | 845.5 | 7.239E-05 | 0.08562 | +1 | 8 |
| - | - | 3153 | 846.5 | - | - | 0 | - |
| - | - | 1525 | 847.5 | - | - | 0 | - |
| - | - | 859.5 | 858.5 | - | - | 0 | - |
| - | - | 948.1 | 866.5 | - | - | 0 | - |
| - | - | 1178 | 872.5 | - | - | 0 | - |
| - | - | 3853 | 873.5 | - | - | 0 | - |
| - | - | 1620 | 874.5 | - | - | 0 | - |
| - | - | 7362 | 884.5 | - | - | 0 | - |
| - | - | 2490 | 885.5 | - | - | 0 | - |
| 7 | w | 1192 | 899.5 | 0.002264 | 2.517 | +1 | 9 |
| 8 | c | 7370 | 900.5 | 0.001288 | 1.431 | +1 | 8 |
| - | - | 5614 | 901.5 | - | - | 0 | - |
| - | - | 2418 | 902.5 | - | - | 0 | - |
| 7 | z | 2153 | 926.5 | 0.001061 | 1.145 | +1 | 9 |
| - | - | 1735 | 927.5 | - | - | 0 | - |
| - | - | 2118 | 929.5 | - | - | 0 | - |
| - | - | 3265 | 932.4 | - | - | 0 | - |
| - | - | 1091 | 933.4 | - | - | 0 | - |
| 7 | y | 802.4 | 943.5 | 0.003096 | 3.282 | +1 | 9 |
| 7 | z | 1.945E+04 | 944.5 | 0.0003373 | 0.3571 | +1 | 9 |
| - | - | 1.62E+04 | 945.5 | - | - | 0 | - |
| - | - | 6377 | 946.5 | - | - | 0 | - |
| - | - | 1211 | 947.5 | - | - | 0 | - |
| - | - | 1827 | 954.5 | - | - | 0 | - |
| - | - | 2165 | 955.5 | - | - | 0 | - |
| - | - | 3150 | 956.5 | - | - | 0 | - |
| - | - | 1467 | 957.6 | - | - | 0 | - |
| - | - | 954.1 | 958.6 | - | - | 0 | - |
| - | - | 768.1 | 959.5 | - | - | 0 | - |
| 7 | y | 2352 | 960.5 | 0.000229 | 0.2384 | +1 | 9 |
| - | - | 963.7 | 961.5 | - | - | 0 | - |
| - | - | 5153 | 970.5 | - | - | 0 | - |
| - | - | 3033 | 971.5 | - | - | 0 | - |
| - | - | 714.6 | 972.5 | - | - | 0 | - |
| - | - | 711.7 | 998.6 | - | - | 0 | - |
| - | - | 849.7 | 999.6 | - | - | 0 | - |
| - | - | 1400 | 1003 | - | - | 0 | - |
| - | - | 1.216E+04 | 1004 | - | - | 0 | - |
| - | - | 9880 | 1006 | - | - | 0 | - |
| - | - | 2058 | 1007 | - | - | 0 | - |
| - | - | 3091 | 1014 | - | - | 0 | - |
| 9 | c | 4.694E+04 | 1015 | 0.0009416 | 0.9281 | +1 | 9 |
| - | - | 2.801E+04 | 1016 | - | - | 0 | - |
| - | - | 8279 | 1017 | - | - | 0 | - |
| - | - | 1719 | 1018 | - | - | 0 | - |
| - | - | 933.8 | 1020 | - | - | 0 | - |
| 6 | y | 1319 | 1030 | 0.003663 | 3.555 | +1 | 10 |
| 6 | z | 7088 | 1031 | 0.0002303 | 0.2233 | +1 | 10 |
| - | - | 9851 | 1033 | - | - | 0 | - |
| - | - | 5548 | 1034 | - | - | 0 | - |
| - | - | 1271 | 1035 | - | - | 0 | - |
| - | - | 3548 | 1040 | - | - | 0 | - |
| - | - | 3454 | 1041 | - | - | 0 | - |
| - | - | 2998 | 1042 | - | - | 0 | - |
| - | - | 3635 | 1043 | - | - | 0 | - |
| - | - | 1535 | 1044 | - | - | 0 | - |
| - | - | 1161 | 1045 | - | - | 0 | - |
| - | - | 1.02E+04 | 1046 | - | - | 0 | - |
| - | - | 3886 | 1047 | - | - | 0 | - |
| 6 | y | 3606 | 1048 | 0.0003053 | 0.2915 | +1 | 10 |
| - | - | 1983 | 1049 | - | - | 0 | - |
| - | - | 874.6 | 1050 | - | - | 0 | - |
| - | - | 1696 | 1057 | - | - | 0 | - |
| - | - | 3.731E+04 | 1058 | - | - | 0 | - |
| - | - | 2.451E+04 | 1059 | - | - | 0 | - |
| - | - | 1.57E+04 | 1060 | - | - | 0 | - |
| - | - | 5309 | 1061 | - | - | 0 | - |
| - | - | 3958 | 1062 | - | - | 0 | - |
| - | - | 9767 | 1063 | - | - | 0 | - |
| - | - | 2996 | 1064 | - | - | 0 | - |
| - | - | 798.8 | 1071 | - | - | 0 | - |
| - | - | 791.5 | 1087 | - | - | 0 | - |
| - | - | 2467 | 1101 | - | - | 0 | - |
| 10 | c | 6.482E+04 | 1102 | 0.0002242 | 0.2035 | +1 | 10 |
| - | - | 3.734E+04 | 1103 | - | - | 0 | - |
| - | - | 1.447E+04 | 1104 | - | - | 0 | - |
| - | - | 3147 | 1105 | - | - | 0 | - |
| - | - | 9499 | 1115 | - | - | 0 | - |
| - | - | 5800 | 1116 | - | - | 0 | - |
| - | - | 3432 | 1117 | - | - | 0 | - |
| 5 | z | 9944 | 1131 | 0.0001905 | 0.1685 | +1 | 11 |
| - | - | 2.264E+04 | 1132 | - | - | 0 | - |
| - | - | 9811 | 1133 | - | - | 0 | - |
| - | - | 3694 | 1134 | - | - | 0 | - |
| - | - | 1117 | 1135 | - | - | 0 | - |
| 5 | y | 2932 | 1147 | 0.002801 | 2.443 | +1 | 11 |
| - | - | 2002 | 1148 | - | - | 0 | - |
| 11 | c | 3.665E+04 | 1159 | 0.0003655 | 0.3154 | +1 | 11 |
| - | - | 2.297E+04 | 1160 | - | - | 0 | - |
| - | - | 8294 | 1161 | - | - | 0 | - |
| - | - | 1907 | 1162 | - | - | 0 | - |
| - | - | 7468 | 1214 | - | - | 0 | - |
| - | - | 4488 | 1215 | - | - | 0 | - |
| - | - | 2990 | 1216 | - | - | 0 | - |
| - | - | 861.1 | 1217 | - | - | 0 | - |
| - | - | 837.4 | 1219 | - | - | 0 | - |
| - | - | 740.5 | 1248 | - | - | 0 | - |
| 12 | c | 3.602E+04 | 1258 | 0.001152 | 0.9163 | +1 | 12 |
| - | - | 2.306E+04 | 1259 | - | - | 0 | - |
| - | - | 7669 | 1260 | - | - | 0 | - |
| - | - | 3.114E+04 | 1261 | - | - | 0 | - |
| - | - | 1.822E+04 | 1262 | - | - | 0 | - |
| - | - | 7840 | 1263 | - | - | 0 | - |
| - | - | 1842 | 1264 | - | - | 0 | - |
| 4 | y | 1.019E+04 | 1276 | 0.001693 | 1.327 | +1 | 12 |
| - | - | 8046 | 1277 | - | - | 0 | - |
| - | - | 2559 | 1278 | - | - | 0 | - |
| - | - | 1540 | 1279 | - | - | 0 | - |
| - | - | 841.3 | 1280 | - | - | 0 | - |
| - | - | 1179 | 1284 | - | - | 0 | - |
| - | - | 756 | 1286 | - | - | 0 | - |
| - | - | 5520 | 1301 | - | - | 0 | - |
| - | - | 4731 | 1302 | - | - | 0 | - |
| - | - | 2953 | 1303 | - | - | 0 | - |
| - | - | 1072 | 1304 | - | - | 0 | - |
| - | - | 882.8 | 1305 | - | - | 0 | - |
| - | - | 969.7 | 1306 | - | - | 0 | - |
| 3 | w | 1443 | 1330 | 0.01435 | 10.8 | +1 | 13 |
| - | - | 2665 | 1331 | - | - | 0 | - |
| - | - | 1854 | 1332 | - | - | 0 | - |
| - | - | 1028 | 1333 | - | - | 0 | - |
| - | - | 1255 | 1343 | - | - | 0 | - |
| 13 | c | 6.453E+04 | 1345 | 0.001321 | 0.982 | +1 | 13 |
| - | - | 4.396E+04 | 1346 | - | - | 0 | - |
| - | - | 608.4 | 1346 | - | - | 0 | - |
| - | - | 1.895E+04 | 1347 | - | - | 0 | - |
| - | - | 2.586E+04 | 1348 | - | - | 0 | - |
| - | - | 1.722E+04 | 1349 | - | - | 0 | - |
| - | - | 7363 | 1350 | - | - | 0 | - |
| - | - | 2295 | 1351 | - | - | 0 | - |
| - | - | 762 | 1352 | - | - | 0 | - |
| 3 | y | 1658 | 1363 | 0.003266 | 2.397 | +1 | 13 |
| - | - | 1490 | 1364 | - | - | 0 | - |
| - | - | 727.5 | 1365 | - | - | 0 | - |
| - | - | 4380 | 1388 | - | - | 0 | - |
| - | - | 4696 | 1389 | - | - | 0 | - |
| - | - | 1956 | 1390 | - | - | 0 | - |
| - | - | 995.3 | 1430 | - | - | 0 | - |
| - | - | 2903 | 1431 | - | - | 0 | - |
| 14 | c | 9.159E+04 | 1432 | 0.001855 | 1.295 | +1 | 14 |
| - | - | 6.735E+04 | 1433 | - | - | 0 | - |
| - | - | 3.049E+04 | 1434 | - | - | 0 | - |
| - | - | 9825 | 1435 | - | - | 0 | - |
| - | - | 1940 | 1436 | - | - | 0 | - |
| - | - | 4219 | 1461 | - | - | 0 | - |
| - | - | 3451 | 1462 | - | - | 0 | - |
| - | - | 1551 | 1463 | - | - | 0 | - |
| - | - | 861.7 | 1474 | - | - | 0 | - |
| - | - | 853.6 | 1475 | - | - | 0 | - |
| 2 | y | 724.2 | 1476 | 0.01027 | 6.957 | +1 | 14 |
| - | - | 8030 | 1489 | - | - | 0 | - |
| - | - | 6746 | 1490 | - | - | 0 | - |
| - | - | 2560 | 1491 | - | - | 0 | - |
| - | - | 1646 | 1492 | - | - | 0 | - |
| - | - | 1259 | 1502 | - | - | 0 | - |
| - | - | 2694 | 1513 | - | - | 0 | - |
| - | - | 2135 | 1514 | - | - | 0 | - |
| - | - | 1379 | 1515 | - | - | 0 | - |
| - | - | 1240 | 1516 | - | - | 0 | - |
| - | - | 868.6 | 1519 | - | - | 0 | - |
| - | - | 735.8 | 1529 | - | - | 0 | - |
| - | - | 2117 | 1530 | - | - | 0 | - |
| - | - | 1.722E+04 | 1531 | - | - | 0 | - |
| - | - | 2.237E+04 | 1532 | - | - | 0 | - |
| - | - | 1.55E+04 | 1533 | - | - | 0 | - |
| - | - | 4465 | 1534 | - | - | 0 | - |
| - | - | 2115 | 1535 | - | - | 0 | - |
| - | - | 1361 | 1544 | - | - | 0 | - |
| - | - | 4296 | 1545 | - | - | 0 | - |
| - | - | 6853 | 1546 | - | - | 0 | - |
| - | - | 9193 | 1547 | - | - | 0 | - |
| - | - | 8693 | 1548 | - | - | 0 | - |
| - | - | 6001 | 1549 | - | - | 0 | - |
| - | - | 2486 | 1550 | - | - | 0 | - |
| - | - | 3257 | 1556 | - | - | 0 | - |
| - | - | 7839 | 1557 | - | - | 0 | - |
| - | - | 7289 | 1558 | - | - | 0 | - |
| - | - | 3241 | 1559 | - | - | 0 | - |
| - | - | 912.7 | 1560 | - | - | 0 | - |
| - | - | 3357 | 1562 | - | - | 0 | - |
| - | - | 1.355E+04 | 1563 | - | - | 0 | - |
| - | - | 1.086E+04 | 1564 | - | - | 0 | - |
| - | - | 5495 | 1565 | - | - | 0 | - |
| - | - | 1884 | 1566 | - | - | 0 | - |
| - | - | 3169 | 1572 | - | - | 0 | - |
| - | - | 1.839E+04 | 1573 | - | - | 0 | - |
| - | - | 1.221E+05 | 1574 | - | - | 0 | - |
| - | - | 9.594E+04 | 1575 | - | - | 0 | - |
| - | - | 4.545E+04 | 1576 | - | - | 0 | - |
| - | - | 1.306E+04 | 1577 | - | - | 0 | - |
| - | - | 3825 | 1578 | - | - | 0 | - |
| - | - | 889.4 | 1587 | - | - | 0 | - |
| - | - | 1963 | 1588 | - | - | 0 | - |
| - | - | 9494 | 1589 | - | - | 0 | - |
| - | - | 3.552E+04 | 1590 | - | - | 0 | - |
| - | - | 1.17E+05 | 1591 | - | - | 0 | - |
| - | - | 8.614E+04 | 1592 | - | - | 0 | - |
| - | - | 3.982E+04 | 1593 | - | - | 0 | - |
| - | - | 1.236E+04 | 1594 | - | - | 0 | - |
| - | - | 3095 | 1595 | - | - | 0 | - |
| - | - | 781.5 | 2904 | - | - | 0 | - |
| - | - | 740.8 | 3183 | - | - | 0 | - |
| - | - | 785.7 | 3184 | - | - | 0 | - |

m/z Charge Intensity FragmentType MassShift Position
125.10769653320312 0 1940.1135
126.09169006347656 0 4750.345
131.12945556640625 0 1743.439
132.0272216796875 0 419.08417
136.47999572753906 0 471.53494
143.1185302734375 0 501.95358
143.78578186035156 0 414.71024
146.0915985107422 0 436.94775
148.947265625 0 1093.8223
158.0925750732422 0 1326.3955 y Ammonia loss 14
160.04315185546875 0 2095.1604
160.27426147460938 0 494.25378
160.69541931152344 0 427.0311
168.11349487304688 0 4905.2285
169.11767578125 0 657.6754
173.45188903808594 0 1188.9608
175.11927795410156 0 8718.747 y 14
175.9103240966797 0 573.09235
176.1222686767578 0 595.8944
178.0535430908203 0 5289.161
185.14004516601562 0 21170.227
186.14337158203125 0 2349.8083
189.08738708496094 0 848.8333
199.18080139160156 0 55746.574
200.1269989013672 0 3769.2458
200.1842803955078 0 7209.0625
201.12347412109375 0 1661.5012
201.13449096679688 0 886.4029
216.14573669433594 0 720.3824
216.66163635253906 0 599.98834
217.08209228515625 0 2659.3455
226.13047790527344 0 898.0899
226.16812133789062 0 777.4442
227.1757354736328 0 29106.951
228.17906188964844 0 3526.5894
229.12985229492188 0 4307.907 w 13
229.18310546875 0 586.23846
230.1140899658203 0 781.0001
233.16513061523438 0 3078.902
244.1173553466797 0 1002.5077
245.12486267089844 0 39175.79 y Ammonia loss 13
246.13072204589844 0 6224.14 z 13
247.1354522705078 0 647.9414
249.16012573242188 0 1578.5981
255.10888671875 0 1182.2528
261.16058349609375 0 3244.3574
262.15118408203125 0 5640.817 y 13
262.16357421875 0 700.2847
270.1200866699219 0 3830.7937
272.1722412109375 0 1194.2728
277.1549072265625 0 1927.2301 z 7
279.10137939453125 0 9212.326
280.1047668457031 0 891.4635
299.1968078613281 0 567.608
302.1715393066406 0 1103.5436
303.1546325683594 0 1046.9906
304.14959716796875 0 695.7819
314.2082214355469 0 1914.1309
316.1621398925781 0 20119.203 w 12
317.1613464355469 0 2927.1768
323.2102966308594 0 980.51135
331.1489562988281 0 5404.076
332.1566162109375 0 6254.256 y Ammonia loss 12
333.1646423339844 0 13377.913 z 12
334.1675109863281 0 1782.9717
345.1834716796875 0 634.4274 y 8
345.683349609375 0 574.336
347.1672058105469 0 1759.1085
349.1149597167969 0 793.64404
349.1835021972656 0 4750.889 y 12
350.1867980957031 0 1050.2687
357.1518859863281 0 9137.837
359.1925048828125 0 711.1817
373.2430725097656 0 667.32526
389.70880126953125 0 2136.9668
390.2095642089844 0 799.1793
390.7166748046875 0 1516.5256
391.21881103515625 0 666.4529
391.7240905761719 0 7797.384
392.2254333496094 0 4121.564
398.7137756347656 0 2613.6248
399.21600341796875 0 1666.8585
399.7212219238281 0 1876.4878
400.2213134765625 0 1156.2764
414.2218933105469 0 754.53406 c Ammonia loss 11
415.22589111328125 0 5719.161 z 7
415.7258605957031 0 2710.2537
417.21002197265625 0 15169.885 w 11
418.2131042480469 0 3071.935
423.23333740234375 0 3489.2532 y 7
423.7351989746094 0 1526.5542
425.23883056640625 0 1544.2297
426.24169921875 0 739.3736
429.234619140625 0 1484.2458
430.2386779785156 0 644.5705 y Water loss 11
431.2254638671875 0 1559.0459 y Ammonia loss 11
433.74053955078125 0 1086.9081
442.7467041015625 0 2940.4731
443.2498474121094 0 4436.333
443.3101806640625 0 1448.6738
444.25457763671875 0 588.7874
446.2373962402344 0 1232.293
447.20733642578125 0 731.1601
447.24066162109375 0 878.5988
448.2522277832031 0 12938.1455 y 11
449.22442626953125 0 637.8349 y Ammonia loss 2
449.2554931640625 0 3073.4868
450.74346923828125 0 755.43506 c Ammonia loss 7
454.89202880859375 0 1041.2316 y 2
455.2291259765625 0 746.8678
455.2851257324219 0 883.9812
460.2413635253906 0 751.5774
460.2773742675781 0 10478.803 c 3
461.2803039550781 0 3033.091
465.260009765625 0 1116.1687
472.25201416015625 0 2573.7795 c Ammonia loss 13
472.7381591796875 0 3202.716 z 6
473.24267578125 0 930.069
474.2310791015625 0 4793.9604
480.7473449707031 0 3629.841 y 6
486.5848083496094 0 828.97107 y Water loss 1
486.91827392578125 0 606.05035 y Ammonia loss 1
489.12994384765625 0 608.44244
489.2547302246094 0 4233.3076 z 10
490.2609558105469 0 2976.953
492.5875244140625 0 1062.092 y 1
492.92340087890625 0 641.8604
493.2523193359375 0 811.2945
495.9151306152344 0 1392.7559
496.2511291503906 0 946.6076
499.2995300292969 0 2433.4905
504.26556396484375 0 607.58417
505.273681640625 0 47786 y 10
506.276123046875 0 11197.209
507.2796325683594 0 2102.1672
514.3232421875 0 784.351
515.2465209960938 0 1914.6603
515.3310546875 0 2092.7117
515.7487182617188 0 3306.8032 y Ammonia loss 5
516.2530517578125 0 4504.1367 z 5
516.3388061523438 0 1035.7426
516.7572021484375 0 5464.9736
517.2583618164062 0 3630.0872
517.3399047851562 0 701.62463
520.279541015625 0 853.23376
522.2637329101562 0 3637.9768
523.7589721679688 0 1172.6794
524.264892578125 0 18835.506 y 5
524.6123657226562 0 6029.556
524.7649536132812 0 7524.491
524.9464111328125 0 2303.9893
525.2670288085938 0 3438.894
528.7908325195312 0 3752.1128
529.2973022460938 0 14946.131
529.7977905273438 0 4224.5425
530.2913818359375 0 5069.3955
530.6161499023438 0 4144.9736
530.8006591796875 0 1411.734
530.9502563476562 0 3853.2705
531.2703247070312 0 6744.9224
531.7706909179688 0 2743.89
541.3128051757812 0 1842.0162
542.318115234375 0 6349.9106
543.3217163085938 0 1260.8167
546.2758178710938 0 1035.2803
558.2855224609375 0 651.15076 z Water loss 9
558.3384399414062 0 2604.7776
559.28515625 0 1837.1283 w 9
559.3450317382812 0 8314.072 c 4
560.3488159179688 0 2133.9995
561.2692260742188 0 702.1316
565.7879028320312 0 38080.617 z 4
566.2893676757812 0 24234.164
566.7911987304688 0 7802.1772
567.29248046875 0 2325.2922
571.3458251953125 0 1125.322
572.3102416992188 0 732.67145
572.351806640625 0 4793.8784
573.2877807617188 0 873.7784
573.7970581054688 0 20379.03 y 4
574.2982788085938 0 8960.967 y Water loss 9
574.7992553710938 0 4582.1543
575.281005859375 0 2856.2236 y Ammonia loss 9
576.2865600585938 0 7760.6953 z 9
577.2894287109375 0 2543.3567
585.2640380859375 0 1034.172
586.3312377929688 0 2421.2646
587.3350830078125 0 665.10693
592.3040161132812 0 1529.1438 y 9
593.3071899414062 0 640.90894
600.2733154296875 0 3381.7002
600.8027954101562 0 14969.919 w 3
601.3038330078125 0 8684.763
601.8052368164062 0 4083.925
602.3035278320312 0 655.3067
603.3743286132812 0 1160.128
608.3109741210938 0 738.0333
611.3384399414062 0 780.19525
621.3008422851562 0 668.2695 z Water loss 3
627.35595703125 0 973.9867
629.350341796875 0 2197.1506 c 11
629.8123779296875 0 888.1448 y Ammonia loss 3
630.3092651367188 0 35743.918 z 3
630.81103515625 0 26299.332
631.3118286132812 0 11588.69
631.812744140625 0 3209.7964
637.3075561523438 0 763.13354
637.8147583007812 0 15887.479
638.3178100585938 0 36280.695 y 3
638.8197021484375 0 22134.629
639.3204956054688 0 8363.434
639.8203735351562 0 1480.3673
644.3182983398438 0 3680.8716
644.8203735351562 0 2419.733
645.37548828125 0 894.8106
646.377685546875 0 35567.938 c 5
647.3213500976562 0 638.99817
647.3807983398438 0 10895.604
648.3831787109375 0 2238.2246
651.8292236328125 0 1955.9652
652.3237915039062 0 886.83905
665.3213500976562 0 1950.6807 z Ammonia loss 2
665.8247680664062 0 1951.7518
666.3259887695312 0 1524.4202
670.3534545898438 0 743.0586
672.8271484375 0 1587.4227 y Water loss 2
673.3282470703125 0 2836.4463 y Ammonia loss 2
673.8255004882812 0 23643.385 z 2
674.3279418945312 0 31938.26
674.8301391601562 0 17225.13
675.3312377929688 0 5956.3843
675.834228515625 0 1044.7172
680.8280029296875 0 2402.6804
681.3296508789062 0 3201.7441
681.8345947265625 0 175204.69 y 2
682.3358154296875 0 113144.14
682.8373413085938 0 49465.35
683.33837890625 0 13793.841
683.83984375 0 3575.2861
684.368408203125 0 1689.7577
685.3869018554688 0 1818.9343
686.3238525390625 0 832.934
686.3964233398438 0 773.12195
687.3416137695312 0 1987.9623
688.3439331054688 0 1001.92993
689.3585815429688 0 5182.824 y 8
690.3607177734375 0 2308.265
691.3699340820312 0 659.3511
697.4403686523438 0 716.0807
699.374267578125 0 1590.8431
700.3773193359375 0 763.65546
701.3939208984375 0 2092.095
711.4156494140625 0 1011.17065
712.423828125 0 2138.193
713.4339599609375 0 2668.5571
714.4410400390625 0 812.8057
715.8485107421875 0 2840.0583 w 1
716.3505859375 0 2475.775
716.84716796875 0 938.9903
717.390869140625 0 3516.8337
718.3984985351562 0 4851.6294
719.4006958007812 0 1666.6991
729.3715209960938 0 1412.3995 y Water loss 1
730.3665161132812 0 3545.444 z 1
730.8674926757812 0 1493.7749
736.406494140625 0 639.3622
738.3765258789062 0 10699.634 y 1
738.8778686523438 0 7650.524
739.3787231445312 0 4651.168
739.884521484375 0 1030.5818
743.370361328125 0 3027.9587 w 7
744.3757934570312 0 3612.3064
745.3812255859375 0 620.2006
750.897216796875 0 2927.7776
751.4017944335938 0 2079.5122
751.8945922851562 0 1250.0255
756.9012451171875 0 1084.7865
757.3950805664062 0 1004.8584
758.388427734375 0 3707.8022
758.8841552734375 0 5751.0005
759.385009765625 0 5763.5605
759.884521484375 0 1157.9652
760.3960571289062 0 1895.6384
761.4046020507812 0 65988.58 c 6
762.4071655273438 0 26073.666
763.4095458984375 0 7172.3853
764.4083862304688 0 3134.9778
764.9102783203125 0 5910.6543
765.3825073242188 0 109156.5
765.8837280273438 0 90962.086
766.3854370117188 0 45498.906
766.8876342773438 0 13108.479
767.3886108398438 0 3428.6133
771.4049682617188 0 1087.1943
771.9089965820312 0 846.1458
772.4132080078125 0 3261.2703
772.9090576171875 0 2993.3691
773.4188232421875 0 3193.973
773.9082641601562 0 30043.684
774.4091796875 0 25439.766
774.9104614257812 0 12628.8
775.4124755859375 0 3657.8733
775.9147338867188 0 1213.0315
777.9017333984375 0 1500.0464
778.4066162109375 0 4322.3784
778.9024658203125 0 2061.0303
779.4036254882812 0 1850.3705
779.904296875 0 1212.4973
780.41943359375 0 2575.855
780.91650390625 0 643.78406
782.4403076171875 0 5119.704
783.438720703125 0 13589.396
784.4413452148438 0 2531.3196
785.90283203125 0 2402.0095
786.413818359375 0 8006.7856
786.9096069335938 0 117625.016
787.4108276367188 0 94677.41
787.9120483398438 0 47770.15
788.4132690429688 0 14641.822
788.91552734375 0 3891.546
793.4025268554688 0 1048.1979
793.9072265625 0 2650.7744
794.4127807617188 0 6070.619
794.918212890625 0 75976.79
795.420654296875 0 98142.68
795.92236328125 0 58460.32
796.4235229492188 0 27279.121
796.9202880859375 0 7858.275
797.4187622070312 0 3406.8904
798.4336547851562 0 2041.5024
799.4370727539062 0 760.39374
800.458740234375 0 1583.58
811.4296264648438 0 3215.673 z Water loss 7
812.4259643554688 0 1460.6362 z Ammonia loss 7
828.4306640625 0 1011.4012 y Ammonia loss 7
829.4405517578125 0 51303.977 z 7
830.4442749023438 0 28460.21
831.4486083984375 0 7260.851
832.4485473632812 0 1407.6362
839.51123046875 0 1969.2256
840.5194091796875 0 1205.6414
841.5296020507812 0 4309.4917
842.5335693359375 0 2542.4065
844.4501342773438 0 2927.364
845.4588623046875 0 6725.1133 y 7
846.46240234375 0 3153.242
847.4631958007812 0 1525.3613
858.48486328125 0 859.5432
866.4722290039062 0 948.1236
872.492919921875 0 1178.176
873.4910888671875 0 3853.3074
874.49462890625 0 1619.8875
884.482421875 0 7361.8794
885.4844970703125 0 2490.4048
899.4716186523438 0 1192.0142 w 6
900.4772338867188 0 7370.1816 c Ammonia loss 7
901.4945068359375 0 5614.0923
902.4901123046875 0 2417.8599
926.4553833007812 0 2152.893 z Water loss 6
927.463623046875 0 1735.4949
929.4827270507812 0 2117.9014
932.4269409179688 0 3264.652
933.4264526367188 0 1090.7834
943.4622802734375 0 802.44763 y Ammonia loss 6
944.4673461914062 0 19451.35 z 6
945.4716186523438 0 16202.113
946.475341796875 0 6376.7163
947.4761352539062 0 1211.3102
954.5383911132812 0 1827.292
955.5394897460938 0 2165.4187
956.5482177734375 0 3149.602
957.5582275390625 0 1466.8088
958.5588989257812 0 954.0663
959.4825439453125 0 768.07404
960.4859619140625 0 2351.7854 y 6
961.4644165039062 0 963.68677
970.5435791015625 0 5152.794
971.5453491210938 0 3033.2717
972.5360107421875 0 714.64966
998.5574340820312 0 711.71356
999.5584106445312 0 849.7406
1003.4912109375 0 1399.7078
1004.495849609375 0 12155.253
1005.5013427734375 0 9879.515
1006.5037231445312 0 2058.3577
1013.5834350585938 0 3090.9368
1014.5587768554688 0 46935.918 c 8
1015.5619506835938 0 28006.352
1016.5638427734375 0 8279.247
1017.5696411132812 0 1718.9624
1019.5090942382812 0 933.7869
1030.487548828125 0 1319.0013 y Ammonia loss 5
1031.499267578125 0 7088.4146 z 5
1032.505615234375 0 9850.6045
1033.5079345703125 0 5548.078
1034.51904296875 0 1270.9486
1039.5660400390625 0 3547.543
1040.56396484375 0 3453.7952
1041.56884765625 0 2998.0728
1042.5692138671875 0 3635.1836
1043.572021484375 0 1534.8273
1044.5260009765625 0 1160.9906
1045.51025390625 0 10201.658
1046.51318359375 0 3885.706
1047.5174560546875 0 3606.0654 y 5
1048.522705078125 0 1983.1022
1049.532470703125 0 874.55774
1056.573486328125 0 1696.3857
1057.5772705078125 0 37308.914
1058.580810546875 0 24510.566
1059.587890625 0 15696.049
1060.59375 0 5308.848
1061.529541015625 0 3958.1584
1062.5355224609375 0 9766.88
1063.5391845703125 0 2995.8694
1070.60791015625 0 798.84906
1086.576416015625 0 791.53595
1100.57958984375 0 2466.7825
1101.590087890625 0 64815.496 c 9
1102.5931396484375 0 37340.676
1103.5948486328125 0 14471.045
1104.600341796875 0 3147.1104
1114.596435546875 0 9498.779
1115.6007080078125 0 5800.248
1116.599853515625 0 3432.1143
1130.5672607421875 0 9943.862 z 4
1131.5743408203125 0 22644.459
1132.5775146484375 0 9810.767
1133.580322265625 0 3693.8499
1134.5789794921875 0 1117.2762
1146.5833740234375 0 2931.7903 y 4
1147.5859375 0 2001.7828
1158.6109619140625 0 36651.375 c 10
1159.613525390625 0 22973.824
1160.6168212890625 0 8293.8545
1161.6192626953125 0 1907.3578
1213.6663818359375 0 7468.1313
1214.668701171875 0 4487.9253
1215.672607421875 0 2990.3845
1216.674072265625 0 861.1377
1218.59326171875 0 837.408
1247.5482177734375 0 740.50775
1257.6785888671875 0 36023.164 c 11
1258.6805419921875 0 23060.467
1259.6734619140625 0 7668.7725
1260.6175537109375 0 31141.295
1261.6199951171875 0 18224.156
1262.6197509765625 0 7839.9644
1263.621826171875 0 1842.4973
1275.6270751953125 0 10193.777 y 3
1276.6300048828125 0 8046.3
1277.634765625 0 2558.6362
1278.6383056640625 0 1540.222
1279.638916015625 0 841.30316
1283.6920166015625 0 1179.3457
1285.6708984375 0 755.95844
1300.6956787109375 0 5519.786
1301.700927734375 0 4731.085
1302.7039794921875 0 2953.4448
1303.7044677734375 0 1071.9133
1304.62158203125 0 882.84467
1305.625732421875 0 969.7096
1329.6536865234375 0 1443.4662 w 2
1330.629150390625 0 2664.7217
1331.63037109375 0 1854.2825
1332.63671875 0 1028.0265
1342.697021484375 0 1255.4824
1344.71044921875 0 64525.95 c 12
1345.713134765625 0 43955.64
1345.87060546875 0 608.41626
1346.70947265625 0 18950.582
1347.6524658203125 0 25857.084
1348.651611328125 0 17218.83
1349.654296875 0 7363.0527
1350.6572265625 0 2295.4045
1351.6551513671875 0 761.96356
1362.6640625 0 1658.2474 y 2
1363.66162109375 0 1489.5692
1364.6568603515625 0 727.49426
1387.7315673828125 0 4380.3037
1388.738525390625 0 4696.368
1389.740234375 0 1956.1814
1429.7208251953125 0 995.28674
1430.7296142578125 0 2902.8352
1431.741943359375 0 91590.03 c 13
1432.7447509765625 0 67345.266
1433.7462158203125 0 30492.707
1434.749755859375 0 9825.245
1435.75244140625 0 1940.372
1460.7298583984375 0 4219.036
1461.7347412109375 0 3450.591
1462.7352294921875 0 1550.9662
1473.729248046875 0 861.68225
1474.7177734375 0 853.59863
1475.755126953125 0 724.2234 y 1
1488.739990234375 0 8029.8237
1489.74560546875 0 6746.162
1490.75537109375 0 2560.231
1491.7559814453125 0 1645.7856
1501.7850341796875 0 1258.9233
1512.7923583984375 0 2693.6697
1513.78564453125 0 2134.6514
1514.77978515625 0 1378.8129
1515.783447265625 0 1239.9371
1518.776611328125 0 868.59766
1528.8165283203125 0 735.80536
1529.7908935546875 0 2117.3413
1530.7901611328125 0 17222.871
1531.7923583984375 0 22370.098
1532.79443359375 0 15503.7295
1533.7987060546875 0 4465.1094
1534.806396484375 0 2115.3184
1543.818359375 0 1360.5452
1544.8272705078125 0 4295.5327
1545.8271484375 0 6852.6753
1546.8216552734375 0 9193.334
1547.8187255859375 0 8693.355
1548.817626953125 0 6001.166
1549.8170166015625 0 2485.5227
1555.802001953125 0 3257.3628
1556.7926025390625 0 7839.05
1557.794921875 0 7288.922
1558.794677734375 0 3241.4485
1559.78076171875 0 912.74414
1561.835693359375 0 3357.0647
1562.8458251953125 0 13554.019
1563.846435546875 0 10859.467
1564.8502197265625 0 5494.626
1565.8492431640625 0 1883.6506
1571.803466796875 0 3169.1304
1572.813232421875 0 18388.27
1573.8154296875 0 122078.87
1574.8187255859375 0 95936.86
1575.81982421875 0 45454.547
1576.823486328125 0 13063.005
1577.8258056640625 0 3825.225
1586.8021240234375 0 889.4312
1587.8121337890625 0 1963.3413
1588.824462890625 0 9494.37
1589.8309326171875 0 35521.965
1590.83984375 0 116955.695
1591.8431396484375 0 86137.47
1592.8458251953125 0 39815.63
1593.8463134765625 0 12356.9795
1594.8392333984375 0 3095.081
2903.702392578125 0 781.45874
3183.11279296875 0 740.8437
3183.864013671875 0 785.6884

Spectrum Details

|  |  |
| --- | --- |
| Matched peaks? Matched peaks The total absolute number of peaks matched. Additionally in brackets the total fraction of peaks matched and the total number of peaks is shown. | 88 (16.06% of 548) |
| FDR? FDR The false discovery rate estimated for this peptide. It is calculated by matching all theoretical fragments with a non-integer shift with the raw peaks for this spectrum. This is done with 40 different shifts. The resulting percentage is the average number of annotated peaks over the number of annotated peaks with the correct spectrum. | 0.87% |
| Satellite FDR? Satellite FDR See the FDR for details on its calculation. This satellite ion specific FDR only contains the satellite ions (d/w) for I/L/J positions. | 0.00% |
| PSM Score? PSM Score The PSM Score as given by Hecklib to this annotated spectrum. It is shown with three significant figures. | 656 |

#### Spectrum 3840? Spectrum 3840 The raw spectrum of this peptide as annotated by Hecklib. The fragments are coloured according to ion type (see legend). Any peaks with a star '\*' as text can be hovered over to see the full details, first the ion type second the mass shift type. By hovering over the amino acids in the peptide or ions in the legend the corresponding peaks are highlighted. By toggling the 'Unassigned' label you can turn the background (unassigned) peaks on or off in the plot. By updating the slider in the Ion legend you can update the spectrum to only show the top X% of the peaks with labels. The top X% means any peak that is within X% of the highest intensity. By dragging in the spectrum you can zoom in to a specific part of the spectrum and use 'Zoom Out' to get back to the original zoom level. The annotation of the spectrum is based on the given sequence in the peptides file and is done with different software so inconsistencies are likely. The peaks are annotated based on the given sequence, with 20 ppm tolerance.

Copy Data

##### Spectrum 3840 (TSV)

###### Preview

```
Loading example...
```

*Click on the button to copy the data to your clipboard.*

Mz MinMz MaxIntensity Max

WidthHeightPeptide font sizePeptide stroke widthSpectrum font sizeSpectrum stroke widthCompact peptide

Ion legend

wxyz

abcd

OtherUnassignedIon^Charge^_Position_Show for top:%

JJSEVSDRPSGVSSR

06.20e+41.24e+51.86e+52.48e+5

Zoom Out

y^+1^_1_y^+1^_1_w^+1^_2_y^+1^_2_z^+1^_2_y^+1^_2_z^+3^_8_w^+1^_3_c^+1^_3_y^+1^_3_z^+1^_3_y^+1^_3_c^+3^_12_z^+2^_8_w^+1^_4_y^+2^_8_y^+1^_4_z^+1^_4_y^+1^_4_y^+3^_13_y^+3^_13_c^+2^_8_y^+3^_13_c^+1^_4_z^+1^_5_c^+3^_14_z^+2^_9_y^+2^_9_z^+3^_14_y^+3^_14_y^+1^_5_z^+1^_5_y^+3^_14_y^+1^_5_y^+2^_10_y^+2^_10_z^+2^_10_y^+2^_10_z^+1^_6_w^+1^_6_c^+1^_5_y^+2^_11_z^+2^_11_y^+2^_11_y^+1^_6_y^+1^_6_z^+1^_6_y^+1^_6_w^+2^_12_z^+2^_12_y^+2^_12_c^+2^_12_z^+2^_12_y^+2^_12_c^+1^_6_z^+2^_13_z^+2^_13_y^+2^_13_y^+2^_13_z^+2^_13_y^+2^_13_y^+1^_7_w^+2^_14_y^+2^_14_y^+2^_14_z^+2^_14_y^+2^_14_w^+1^_8_c^+1^_7_z^+1^_8_y^+1^_8_z^+1^_8_y^+1^_8_w^+1^_9_c^+1^_8_z^+1^_9_y^+1^_9_z^+1^_9_y^+1^_9_c^+1^_9_y^+1^_10_z^+1^_10_y^+1^_10_c^+1^_10_z^+1^_11_y^+1^_11_c^+1^_11_w^+1^_12_c^+1^_12_y^+1^_12_w^+1^_13_c^+1^_13_y^+1^_13_c^+1^_14_y^+1^_14_

0844168825323376

Fragment Matches Table

Show background peaks

| Position | Ion type | Intensity | mz Theoretical | mz Error (Th) | mz Error (ppm) | Charge | Series Number |
| --- | --- | --- | --- | --- | --- | --- | --- |
| - | - | 804.6 | 120.1 | - | - | 0 | - |
| - | - | 452.2 | 121.3 | - | - | 0 | - |
| - | - | 3033 | 125.1 | - | - | 0 | - |
| - | - | 4272 | 126.1 | - | - | 0 | - |
| - | - | 411.1 | 129.1 | - | - | 0 | - |
| - | - | 477.2 | 130.1 | - | - | 0 | - |
| - | - | 468.6 | 130.1 | - | - | 0 | - |
| - | - | 778 | 131.1 | - | - | 0 | - |
| - | - | 2241 | 131.1 | - | - | 0 | - |
| - | - | 482.2 | 142.2 | - | - | 0 | - |
| - | - | 719.8 | 143.1 | - | - | 0 | - |
| - | - | 458.8 | 147 | - | - | 0 | - |
| - | - | 401.7 | 149.3 | - | - | 0 | - |
| - | - | 507.1 | 151.3 | - | - | 0 | - |
| 15 | y | 1678 | 158.1 | 0.0004314 | 2.729 | +1 | 1 |
| - | - | 6048 | 168.1 | - | - | 0 | - |
| - | - | 458 | 172.4 | - | - | 0 | - |
| - | - | 2643 | 173.5 | - | - | 0 | - |
| 15 | y | 1.175E+04 | 175.1 | 0.0004326 | 2.47 | +1 | 1 |
| - | - | 2.82E+04 | 185.1 | - | - | 0 | - |
| - | - | 2218 | 186.1 | - | - | 0 | - |
| - | - | 873.8 | 189.1 | - | - | 0 | - |
| - | - | 480.3 | 197.8 | - | - | 0 | - |
| - | - | 7.992E+04 | 199.2 | - | - | 0 | - |
| - | - | 4922 | 200.1 | - | - | 0 | - |
| - | - | 9580 | 200.2 | - | - | 0 | - |
| - | - | 909.5 | 201.1 | - | - | 0 | - |
| - | - | 894.6 | 212.1 | - | - | 0 | - |
| - | - | 3967 | 217.1 | - | - | 0 | - |
| - | - | 1626 | 226.1 | - | - | 0 | - |
| - | - | 1166 | 226.2 | - | - | 0 | - |
| - | - | 3.979E+04 | 227.2 | - | - | 0 | - |
| - | - | 5262 | 228.2 | - | - | 0 | - |
| 14 | w | 5051 | 229.1 | 0.0006559 | 2.862 | +1 | 2 |
| - | - | 1117 | 230.1 | - | - | 0 | - |
| - | - | 4601 | 233.2 | - | - | 0 | - |
| - | - | 659.9 | 234.2 | - | - | 0 | - |
| - | - | 1644 | 244.1 | - | - | 0 | - |
| 14 | y | 4.715E+04 | 245.1 | 0.0005685 | 2.319 | +1 | 2 |
| 14 | z | 7136 | 246.1 | 0.001351 | 5.49 | +1 | 2 |
| - | - | 1196 | 248.1 | - | - | 0 | - |
| - | - | 2050 | 249.2 | - | - | 0 | - |
| - | - | 700.9 | 250.2 | - | - | 0 | - |
| - | - | 2076 | 255.1 | - | - | 0 | - |
| - | - | 727.5 | 260.1 | - | - | 0 | - |
| - | - | 947.3 | 261.1 | - | - | 0 | - |
| - | - | 4658 | 261.2 | - | - | 0 | - |
| 14 | y | 7243 | 262.2 | 0.0004782 | 1.824 | +1 | 2 |
| - | - | 521.6 | 262.2 | - | - | 0 | - |
| - | - | 1322 | 263.2 | - | - | 0 | - |
| - | - | 7496 | 270.1 | - | - | 0 | - |
| - | - | 741.9 | 271.1 | - | - | 0 | - |
| - | - | 1903 | 272.2 | - | - | 0 | - |
| 8 | z | 2303 | 277.2 | 0.003581 | 12.92 | +3 | 8 |
| - | - | 511.4 | 286 | - | - | 0 | - |
| - | - | 747.4 | 296.2 | - | - | 0 | - |
| - | - | 1040 | 299.2 | - | - | 0 | - |
| - | - | 995.7 | 303.2 | - | - | 0 | - |
| - | - | 888 | 304.2 | - | - | 0 | - |
| - | - | 828.6 | 312.2 | - | - | 0 | - |
| - | - | 860.6 | 314.1 | - | - | 0 | - |
| - | - | 4316 | 314.2 | - | - | 0 | - |
| 13 | w | 2.463E+04 | 316.2 | 0.0007777 | 2.46 | +1 | 3 |
| - | - | 4488 | 317.2 | - | - | 0 | - |
| - | - | 781.2 | 323.2 | - | - | 0 | - |
| - | - | 5750 | 331.1 | - | - | 0 | - |
| 3 | c | 997.8 | 331.2 | 5.738E-05 | 0.1732 | +1 | 3 |
| 13 | y | 1.039E+04 | 332.2 | 0.0003089 | 0.93 | +1 | 3 |
| 13 | z | 1.997E+04 | 333.2 | 0.00051 | 1.531 | +1 | 3 |
| - | - | 3377 | 334.2 | - | - | 0 | - |
| - | - | 1019 | 347.2 | - | - | 0 | - |
| - | - | 858.4 | 349.1 | - | - | 0 | - |
| 13 | y | 5793 | 349.2 | 0.0006458 | 1.849 | +1 | 3 |
| - | - | 1.289E+04 | 357.2 | - | - | 0 | - |
| - | - | 2149 | 389.7 | - | - | 0 | - |
| - | - | 2164 | 390.2 | - | - | 0 | - |
| - | - | 2778 | 390.7 | - | - | 0 | - |
| - | - | 1038 | 391.2 | - | - | 0 | - |
| - | - | 1.168E+04 | 391.7 | - | - | 0 | - |
| - | - | 5225 | 392.2 | - | - | 0 | - |
| - | - | 965 | 392.7 | - | - | 0 | - |
| - | - | 4311 | 398.7 | - | - | 0 | - |
| - | - | 1567 | 399.2 | - | - | 0 | - |
| - | - | 4756 | 399.7 | - | - | 0 | - |
| - | - | 2482 | 400.2 | - | - | 0 | - |
| 12 | c | 699.6 | 414.2 | 0.001573 | 3.798 | +3 | 12 |
| 8 | z | 6011 | 415.2 | 0.003197 | 7.698 | +2 | 8 |
| - | - | 1821 | 415.7 | - | - | 0 | - |
| - | - | 614.9 | 416.2 | - | - | 0 | - |
| 12 | w | 2.018E+04 | 417.2 | 0.001073 | 2.572 | +1 | 4 |
| - | - | 3636 | 418.2 | - | - | 0 | - |
| 8 | y | 5300 | 423.2 | 0.0007315 | 1.728 | +2 | 8 |
| - | - | 2061 | 423.7 | - | - | 0 | - |
| - | - | 785.3 | 424.2 | - | - | 0 | - |
| - | - | 2452 | 425.2 | - | - | 0 | - |
| - | - | 2088 | 429.2 | - | - | 0 | - |
| 12 | y | 1201 | 431.2 | 0.0001933 | 0.4483 | +1 | 4 |
| 12 | z | 1033 | 432.2 | 0.0008522 | 1.972 | +1 | 4 |
| - | - | 3749 | 443.3 | - | - | 0 | - |
| - | - | 1147 | 443.3 | - | - | 0 | - |
| - | - | 1031 | 444.3 | - | - | 0 | - |
| - | - | 2665 | 446.2 | - | - | 0 | - |
| - | - | 954.3 | 447.2 | - | - | 0 | - |
| 12 | y | 1.785E+04 | 448.3 | 0.001202 | 2.681 | +1 | 4 |
| 3 | y | 1162 | 448.9 | 0.0004712 | 1.05 | +3 | 13 |
| 3 | y | 1215 | 449.2 | 0.007427 | 16.53 | +3 | 13 |
| - | - | 3421 | 449.3 | - | - | 0 | - |
| 8 | c | 1513 | 450.7 | 0.001028 | 2.28 | +2 | 8 |
| 3 | y | 907.3 | 454.9 | 0.0006118 | 1.345 | +3 | 13 |
| - | - | 852.5 | 455.2 | - | - | 0 | - |
| - | - | 1962 | 455.3 | - | - | 0 | - |
| 4 | c | 1.309E+04 | 460.3 | 0.00141 | 3.062 | +1 | 4 |
| - | - | 3805 | 461.3 | - | - | 0 | - |
| - | - | 970.2 | 462.3 | - | - | 0 | - |
| - | - | 863.5 | 465.3 | - | - | 0 | - |
| - | - | 1198 | 465.8 | - | - | 0 | - |
| 11 | z | 934.7 | 471.2 | 5.447E-05 | 0.1156 | +1 | 5 |
| 14 | c | 6009 | 472.2 | 0.008416 | 17.82 | +3 | 14 |
| 7 | z | 3017 | 472.7 | 0.00184 | 3.893 | +2 | 9 |
| - | - | 2267 | 473.2 | - | - | 0 | - |
| - | - | 4729 | 474.2 | - | - | 0 | - |
| - | - | 1380 | 475.2 | - | - | 0 | - |
| 7 | y | 3714 | 480.7 | 0.001908 | 3.97 | +2 | 9 |
| 2 | z | 1112 | 481.2 | 0.006778 | 14.08 | +3 | 14 |
| 2 | y | 820.1 | 486.6 | 0.00195 | 4.008 | +3 | 14 |
| 11 | y | 891.6 | 487.3 | 0.009026 | 18.52 | +1 | 5 |
| 11 | z | 6136 | 489.3 | 0.001178 | 2.408 | +1 | 5 |
| - | - | 3805 | 490.3 | - | - | 0 | - |
| - | - | 1040 | 491.3 | - | - | 0 | - |
| 2 | y | 2202 | 492.6 | 0.001419 | 2.882 | +3 | 14 |
| - | - | 1402 | 492.9 | - | - | 0 | - |
| - | - | 2274 | 495.9 | - | - | 0 | - |
| - | - | 1627 | 496.3 | - | - | 0 | - |
| - | - | 4364 | 499.3 | - | - | 0 | - |
| - | - | 716.4 | 499.8 | - | - | 0 | - |
| - | - | 900.5 | 500.3 | - | - | 0 | - |
| - | - | 906.6 | 504.3 | - | - | 0 | - |
| 11 | y | 6.31E+04 | 505.3 | 0.001222 | 2.419 | +1 | 5 |
| - | - | 1.359E+04 | 506.3 | - | - | 0 | - |
| - | - | 785.2 | 507.2 | - | - | 0 | - |
| - | - | 2642 | 507.3 | - | - | 0 | - |
| - | - | 1265 | 514.3 | - | - | 0 | - |
| 6 | y | 1893 | 515.3 | 0.008396 | 16.3 | +2 | 10 |
| - | - | 2965 | 515.3 | - | - | 0 | - |
| 6 | y | 2072 | 515.7 | 0.0002674 | 0.5184 | +2 | 10 |
| 6 | z | 5868 | 516.3 | 0.001116 | 2.161 | +2 | 10 |
| - | - | 1884 | 516.3 | - | - | 0 | - |
| - | - | 7599 | 516.8 | - | - | 0 | - |
| - | - | 4246 | 517.3 | - | - | 0 | - |
| - | - | 1254 | 517.8 | - | - | 0 | - |
| - | - | 944 | 518.6 | - | - | 0 | - |
| - | - | 1097 | 520.8 | - | - | 0 | - |
| - | - | 1069 | 523.8 | - | - | 0 | - |
| 6 | y | 2.584E+04 | 524.3 | 0.003411 | 6.507 | +2 | 10 |
| - | - | 8581 | 524.6 | - | - | 0 | - |
| - | - | 1.278E+04 | 524.8 | - | - | 0 | - |
| - | - | 4455 | 524.9 | - | - | 0 | - |
| - | - | 4339 | 525.3 | - | - | 0 | - |
| - | - | 821 | 525.8 | - | - | 0 | - |
| - | - | 1412 | 528.3 | - | - | 0 | - |
| - | - | 5251 | 528.8 | - | - | 0 | - |
| - | - | 4116 | 529.3 | - | - | 0 | - |
| - | - | 7387 | 529.8 | - | - | 0 | - |
| - | - | 1.04E+04 | 530.3 | - | - | 0 | - |
| - | - | 1.046E+04 | 530.6 | - | - | 0 | - |
| - | - | 2534 | 530.8 | - | - | 0 | - |
| - | - | 5488 | 531 | - | - | 0 | - |
| - | - | 1086 | 531.3 | - | - | 0 | - |
| - | - | 1868 | 541.3 | - | - | 0 | - |
| - | - | 7510 | 542.3 | - | - | 0 | - |
| - | - | 1570 | 543.3 | - | - | 0 | - |
| - | - | 1406 | 550.8 | - | - | 0 | - |
| 10 | z | 1951 | 558.3 | 0.003121 | 5.591 | +1 | 6 |
| - | - | 4207 | 558.3 | - | - | 0 | - |
| - | - | 1308 | 558.8 | - | - | 0 | - |
| 10 | w | 3400 | 559.3 | 0.001156 | 2.066 | +1 | 6 |
| 5 | c | 1.1E+04 | 559.3 | 0.001019 | 1.822 | +1 | 5 |
| - | - | 681.2 | 559.8 | - | - | 0 | - |
| - | - | 1411 | 560.3 | - | - | 0 | - |
| - | - | 4303 | 560.3 | - | - | 0 | - |
| 5 | y | 738.6 | 565.3 | 0.004765 | 8.429 | +2 | 11 |
| 5 | z | 4.402E+04 | 565.8 | 0.001332 | 2.355 | +2 | 11 |
| - | - | 2.941E+04 | 566.3 | - | - | 0 | - |
| - | - | 1.033E+04 | 566.8 | - | - | 0 | - |
| - | - | 2842 | 567.3 | - | - | 0 | - |
| - | - | 2196 | 572.3 | - | - | 0 | - |
| - | - | 1801 | 573.3 | - | - | 0 | - |
| 5 | y | 2.827E+04 | 573.8 | 0.0008205 | 1.43 | +2 | 11 |
| 10 | y | 1.699E+04 | 574.3 | 0.004478 | 7.797 | +1 | 6 |
| - | - | 7856 | 574.8 | - | - | 0 | - |
| 10 | y | 3331 | 575.3 | 0.003983 | 6.923 | +1 | 6 |
| 10 | z | 9068 | 576.3 | 0.0006743 | 1.17 | +1 | 6 |
| - | - | 2754 | 577.3 | - | - | 0 | - |
| - | - | 1111 | 585.3 | - | - | 0 | - |
| - | - | 2637 | 586.3 | - | - | 0 | - |
| - | - | 885 | 587.3 | - | - | 0 | - |
| 10 | y | 2359 | 592.3 | 0.002275 | 3.841 | +1 | 6 |
| - | - | 5523 | 600.3 | - | - | 0 | - |
| 4 | w | 2.27E+04 | 600.8 | 0.00152 | 2.529 | +2 | 12 |
| - | - | 1.237E+04 | 601.3 | - | - | 0 | - |
| - | - | 513.2 | 601.4 | - | - | 0 | - |
| - | - | 5356 | 601.8 | - | - | 0 | - |
| - | - | 710.8 | 602.3 | - | - | 0 | - |
| - | - | 1720 | 603.4 | - | - | 0 | - |
| - | - | 833.6 | 614.3 | - | - | 0 | - |
| 4 | z | 715.4 | 621.3 | 0.0043 | 6.921 | +2 | 12 |
| 4 | y | 1220 | 629.3 | 0.009823 | 15.61 | +2 | 12 |
| 12 | c | 2771 | 629.3 | 0.009579 | 15.22 | +2 | 12 |
| 4 | z | 5.645E+04 | 630.3 | 0.001154 | 1.831 | +2 | 12 |
| - | - | 3.488E+04 | 630.8 | - | - | 0 | - |
| - | - | 1.635E+04 | 631.3 | - | - | 0 | - |
| - | - | 4784 | 631.8 | - | - | 0 | - |
| - | - | 999.2 | 632.3 | - | - | 0 | - |
| - | - | 1.784E+04 | 637.8 | - | - | 0 | - |
| 4 | y | 5.529E+04 | 638.3 | 0.0007642 | 1.197 | +2 | 12 |
| - | - | 2.83E+04 | 638.8 | - | - | 0 | - |
| - | - | 1.017E+04 | 639.3 | - | - | 0 | - |
| - | - | 3378 | 639.8 | - | - | 0 | - |
| - | - | 3626 | 644.3 | - | - | 0 | - |
| - | - | 2656 | 644.8 | - | - | 0 | - |
| - | - | 2070 | 645.4 | - | - | 0 | - |
| 6 | c | 4.59E+04 | 646.4 | 0.00134 | 2.072 | +1 | 6 |
| - | - | 1.795E+04 | 647.4 | - | - | 0 | - |
| - | - | 3774 | 648.4 | - | - | 0 | - |
| - | - | 1240 | 651.8 | - | - | 0 | - |
| - | - | 656 | 652.3 | - | - | 0 | - |
| - | - | 686.9 | 657.4 | - | - | 0 | - |
| - | - | 723.6 | 660.3 | - | - | 0 | - |
| 3 | z | 1884 | 664.8 | 0.005071 | 7.627 | +2 | 13 |
| 3 | z | 2703 | 665.3 | 0.009767 | 14.68 | +2 | 13 |
| - | - | 1966 | 665.8 | - | - | 0 | - |
| - | - | 1453 | 666.3 | - | - | 0 | - |
| 3 | y | 4436 | 672.8 | 0.001972 | 2.931 | +2 | 13 |
| 3 | y | 3135 | 673.3 | 0.00718 | 10.66 | +2 | 13 |
| - | - | 384.3 | 673.8 | - | - | 0 | - |
| 3 | z | 3.135E+04 | 673.8 | 0.001864 | 2.766 | +2 | 13 |
| - | - | 4.489E+04 | 674.3 | - | - | 0 | - |
| - | - | 2.258E+04 | 674.8 | - | - | 0 | - |
| - | - | 9245 | 675.3 | - | - | 0 | - |
| - | - | 3167 | 675.8 | - | - | 0 | - |
| - | - | 747.4 | 678.3 | - | - | 0 | - |
| - | - | 2660 | 680.8 | - | - | 0 | - |
| - | - | 3962 | 681.3 | - | - | 0 | - |
| 3 | y | 2.454E+05 | 681.8 | 0.001535 | 2.251 | +2 | 13 |
| - | - | 1.724E+05 | 682.3 | - | - | 0 | - |
| - | - | 7.271E+04 | 682.8 | - | - | 0 | - |
| - | - | 1.833E+04 | 683.3 | - | - | 0 | - |
| - | - | 5252 | 683.8 | - | - | 0 | - |
| - | - | 1515 | 684.4 | - | - | 0 | - |
| - | - | 2414 | 685.4 | - | - | 0 | - |
| - | - | 860.3 | 685.8 | - | - | 0 | - |
| - | - | 699.7 | 686.3 | - | - | 0 | - |
| - | - | 836.7 | 686.4 | - | - | 0 | - |
| - | - | 1149 | 686.8 | - | - | 0 | - |
| - | - | 1887 | 687.3 | - | - | 0 | - |
| 9 | y | 7128 | 689.4 | 0.001086 | 1.575 | +1 | 7 |
| - | - | 2526 | 690.4 | - | - | 0 | - |
| - | - | 674.2 | 691.4 | - | - | 0 | - |
| - | - | 784.8 | 697.4 | - | - | 0 | - |
| - | - | 1309 | 699.4 | - | - | 0 | - |
| - | - | 788.6 | 710.4 | - | - | 0 | - |
| - | - | 1262 | 711.4 | - | - | 0 | - |
| - | - | 2361 | 712.4 | - | - | 0 | - |
| - | - | 3796 | 713.4 | - | - | 0 | - |
| - | - | 1576 | 714.4 | - | - | 0 | - |
| 2 | w | 3919 | 715.8 | 0.002771 | 3.87 | +2 | 14 |
| - | - | 4244 | 716.4 | - | - | 0 | - |
| - | - | 1902 | 716.9 | - | - | 0 | - |
| - | - | 2943 | 717.4 | - | - | 0 | - |
| - | - | 6121 | 718.4 | - | - | 0 | - |
| - | - | 2228 | 719.4 | - | - | 0 | - |
| - | - | 847.2 | 723.4 | - | - | 0 | - |
| 2 | y | 1941 | 729.4 | 0.0004249 | 0.5826 | +2 | 14 |
| 2 | y | 744.4 | 729.9 | 0.009093 | 12.46 | +2 | 14 |
| 2 | z | 3127 | 730.4 | 0.001519 | 2.079 | +2 | 14 |
| - | - | 1532 | 730.9 | - | - | 0 | - |
| - | - | 1354 | 731.4 | - | - | 0 | - |
| 2 | y | 1.6E+04 | 738.4 | 0.001495 | 2.025 | +2 | 14 |
| - | - | 1.101E+04 | 738.9 | - | - | 0 | - |
| - | - | 3817 | 739.4 | - | - | 0 | - |
| - | - | 1343 | 739.9 | - | - | 0 | - |
| 8 | w | 4627 | 743.4 | 0.003827 | 5.148 | +1 | 8 |
| - | - | 3145 | 744.4 | - | - | 0 | - |
| - | - | 1731 | 745.4 | - | - | 0 | - |
| - | - | 848.7 | 749.9 | - | - | 0 | - |
| - | - | 3217 | 750.9 | - | - | 0 | - |
| - | - | 2699 | 751.4 | - | - | 0 | - |
| - | - | 934.8 | 751.9 | - | - | 0 | - |
| - | - | 1437 | 756.4 | - | - | 0 | - |
| - | - | 4745 | 758.4 | - | - | 0 | - |
| - | - | 8124 | 758.9 | - | - | 0 | - |
| - | - | 6418 | 759.4 | - | - | 0 | - |
| - | - | 1616 | 759.9 | - | - | 0 | - |
| - | - | 2045 | 760.4 | - | - | 0 | - |
| 7 | c | 8.2E+04 | 761.4 | 0.00174 | 2.286 | +1 | 7 |
| - | - | 3.542E+04 | 762.4 | - | - | 0 | - |
| - | - | 9536 | 763.4 | - | - | 0 | - |
| - | - | 4109 | 764.4 | - | - | 0 | - |
| - | - | 7874 | 764.9 | - | - | 0 | - |
| - | - | 1.48E+05 | 765.4 | - | - | 0 | - |
| - | - | 1.181E+05 | 765.9 | - | - | 0 | - |
| - | - | 5.914E+04 | 766.4 | - | - | 0 | - |
| - | - | 1.683E+04 | 766.9 | - | - | 0 | - |
| - | - | 4832 | 767.4 | - | - | 0 | - |
| - | - | 1095 | 770.9 | - | - | 0 | - |
| - | - | 1286 | 771.4 | - | - | 0 | - |
| - | - | 1700 | 771.9 | - | - | 0 | - |
| - | - | 4693 | 772.4 | - | - | 0 | - |
| - | - | 4803 | 772.9 | - | - | 0 | - |
| - | - | 4880 | 773.4 | - | - | 0 | - |
| - | - | 4.585E+04 | 773.9 | - | - | 0 | - |
| - | - | 3.535E+04 | 774.4 | - | - | 0 | - |
| - | - | 1.672E+04 | 774.9 | - | - | 0 | - |
| - | - | 5517 | 775.4 | - | - | 0 | - |
| - | - | 1682 | 775.9 | - | - | 0 | - |
| - | - | 4093 | 777.9 | - | - | 0 | - |
| - | - | 5143 | 778.4 | - | - | 0 | - |
| - | - | 2210 | 778.9 | - | - | 0 | - |
| - | - | 3064 | 779.4 | - | - | 0 | - |
| - | - | 1758 | 779.9 | - | - | 0 | - |
| - | - | 1828 | 780.4 | - | - | 0 | - |
| - | - | 1240 | 781.4 | - | - | 0 | - |
| - | - | 7989 | 782.4 | - | - | 0 | - |
| - | - | 4762 | 783.4 | - | - | 0 | - |
| - | - | 1551 | 784.4 | - | - | 0 | - |
| - | - | 3280 | 785.9 | - | - | 0 | - |
| - | - | 1.127E+04 | 786.4 | - | - | 0 | - |
| - | - | 1.504E+05 | 786.9 | - | - | 0 | - |
| - | - | 1.345E+05 | 787.4 | - | - | 0 | - |
| - | - | 6.317E+04 | 787.9 | - | - | 0 | - |
| - | - | 2.102E+04 | 788.4 | - | - | 0 | - |
| - | - | 7817 | 788.9 | - | - | 0 | - |
| - | - | 905.8 | 789.5 | - | - | 0 | - |
| - | - | 857.9 | 793.4 | - | - | 0 | - |
| - | - | 2350 | 793.9 | - | - | 0 | - |
| - | - | 7373 | 794.4 | - | - | 0 | - |
| - | - | 1.125E+05 | 794.9 | - | - | 0 | - |
| - | - | 1.288E+05 | 795.4 | - | - | 0 | - |
| - | - | 8.233E+04 | 795.9 | - | - | 0 | - |
| - | - | 3.656E+04 | 796.4 | - | - | 0 | - |
| - | - | 1.152E+04 | 796.9 | - | - | 0 | - |
| - | - | 4163 | 797.4 | - | - | 0 | - |
| - | - | 3005 | 798.4 | - | - | 0 | - |
| - | - | 1995 | 799.4 | - | - | 0 | - |
| - | - | 848.4 | 800.4 | - | - | 0 | - |
| 8 | z | 3799 | 811.4 | 0.0006746 | 0.8314 | +1 | 8 |
| - | - | 2063 | 812.4 | - | - | 0 | - |
| - | - | 795.8 | 814.4 | - | - | 0 | - |
| 8 | y | 1640 | 828.4 | 0.001475 | 1.78 | +1 | 8 |
| 8 | z | 7.194E+04 | 829.4 | 0.001401 | 1.69 | +1 | 8 |
| - | - | 3.399E+04 | 830.4 | - | - | 0 | - |
| - | - | 1.125E+04 | 831.4 | - | - | 0 | - |
| - | - | 2985 | 832.5 | - | - | 0 | - |
| - | - | 2759 | 839.5 | - | - | 0 | - |
| - | - | 3285 | 840.5 | - | - | 0 | - |
| - | - | 6824 | 841.5 | - | - | 0 | - |
| - | - | 3534 | 842.5 | - | - | 0 | - |
| - | - | 3939 | 844.5 | - | - | 0 | - |
| - | - | 1385 | 845.4 | - | - | 0 | - |
| 8 | y | 1.09E+04 | 845.5 | 0.001049 | 1.241 | +1 | 8 |
| - | - | 3384 | 846.5 | - | - | 0 | - |
| - | - | 2026 | 847.5 | - | - | 0 | - |
| - | - | 3772 | 873.5 | - | - | 0 | - |
| - | - | 2863 | 874.5 | - | - | 0 | - |
| - | - | 747.1 | 875.5 | - | - | 0 | - |
| - | - | 834 | 888.4 | - | - | 0 | - |
| 7 | w | 1736 | 899.5 | 0.001043 | 1.16 | +1 | 9 |
| 8 | c | 1.107E+04 | 900.5 | 5.435E-05 | 0.06036 | +1 | 8 |
| - | - | 4470 | 901.5 | - | - | 0 | - |
| - | - | 1779 | 902.5 | - | - | 0 | - |
| 7 | z | 3151 | 926.5 | 0.001014 | 1.095 | +1 | 9 |
| - | - | 2502 | 927.5 | - | - | 0 | - |
| - | - | 1714 | 928.5 | - | - | 0 | - |
| - | - | 2031 | 929.5 | - | - | 0 | - |
| - | - | 1252 | 930.5 | - | - | 0 | - |
| - | - | 810.8 | 943.2 | - | - | 0 | - |
| 7 | y | 1198 | 943.5 | 0.001542 | 1.635 | +1 | 9 |
| 7 | z | 2.976E+04 | 944.5 | 0.001497 | 1.585 | +1 | 9 |
| - | - | 2.337E+04 | 945.5 | - | - | 0 | - |
| - | - | 8223 | 946.5 | - | - | 0 | - |
| - | - | 2332 | 947.5 | - | - | 0 | - |
| - | - | 2607 | 954.5 | - | - | 0 | - |
| - | - | 3710 | 955.5 | - | - | 0 | - |
| - | - | 6525 | 956.6 | - | - | 0 | - |
| - | - | 3385 | 957.6 | - | - | 0 | - |
| - | - | 1093 | 958.6 | - | - | 0 | - |
| 7 | y | 3250 | 960.5 | 0.001633 | 1.7 | +1 | 9 |
| - | - | 1595 | 961.5 | - | - | 0 | - |
| - | - | 6695 | 970.5 | - | - | 0 | - |
| - | - | 2815 | 971.5 | - | - | 0 | - |
| - | - | 1221 | 987.5 | - | - | 0 | - |
| - | - | 1672 | 999.6 | - | - | 0 | - |
| - | - | 914.2 | 1001 | - | - | 0 | - |
| - | - | 3591 | 1014 | - | - | 0 | - |
| 9 | c | 6.195E+04 | 1015 | 0.002345 | 2.312 | +1 | 9 |
| - | - | 3.46E+04 | 1016 | - | - | 0 | - |
| - | - | 1.075E+04 | 1017 | - | - | 0 | - |
| - | - | 2521 | 1018 | - | - | 0 | - |
| - | - | 1094 | 1029 | - | - | 0 | - |
| 6 | y | 933.3 | 1030 | 0.01245 | 12.08 | +1 | 10 |
| 6 | z | 9795 | 1031 | 0.0003524 | 0.3416 | +1 | 10 |
| - | - | 1.427E+04 | 1033 | - | - | 0 | - |
| - | - | 6299 | 1034 | - | - | 0 | - |
| - | - | 2261 | 1035 | - | - | 0 | - |
| - | - | 2409 | 1040 | - | - | 0 | - |
| - | - | 5397 | 1041 | - | - | 0 | - |
| - | - | 3372 | 1042 | - | - | 0 | - |
| - | - | 5311 | 1043 | - | - | 0 | - |
| - | - | 3505 | 1044 | - | - | 0 | - |
| - | - | 925.8 | 1045 | - | - | 0 | - |
| - | - | 1736 | 1047 | - | - | 0 | - |
| 6 | y | 4611 | 1048 | 0.001282 | 1.223 | +1 | 10 |
| - | - | 2565 | 1049 | - | - | 0 | - |
| - | - | 924.5 | 1050 | - | - | 0 | - |
| - | - | 826.6 | 1054 | - | - | 0 | - |
| - | - | 2400 | 1057 | - | - | 0 | - |
| - | - | 4.799E+04 | 1058 | - | - | 0 | - |
| - | - | 3.596E+04 | 1059 | - | - | 0 | - |
| - | - | 2.537E+04 | 1060 | - | - | 0 | - |
| - | - | 9740 | 1061 | - | - | 0 | - |
| - | - | 973.1 | 1061 | - | - | 0 | - |
| - | - | 3161 | 1062 | - | - | 0 | - |
| - | - | 1245 | 1062 | - | - | 0 | - |
| - | - | 2492 | 1062 | - | - | 0 | - |
| - | - | 1548 | 1064 | - | - | 0 | - |
| - | - | 1127 | 1073 | - | - | 0 | - |
| - | - | 823.4 | 1090 | - | - | 0 | - |
| - | - | 1066 | 1100 | - | - | 0 | - |
| - | - | 3191 | 1101 | - | - | 0 | - |
| 10 | c | 8.455E+04 | 1102 | 0.002177 | 1.977 | +1 | 10 |
| - | - | 5.222E+04 | 1103 | - | - | 0 | - |
| - | - | 1.851E+04 | 1104 | - | - | 0 | - |
| - | - | 4204 | 1105 | - | - | 0 | - |
| - | - | 1.315E+04 | 1115 | - | - | 0 | - |
| - | - | 7991 | 1116 | - | - | 0 | - |
| - | - | 3228 | 1117 | - | - | 0 | - |
| 5 | z | 1.392E+04 | 1131 | 0.00103 | 0.9113 | +1 | 11 |
| - | - | 2.653E+04 | 1132 | - | - | 0 | - |
| - | - | 1.502E+04 | 1133 | - | - | 0 | - |
| - | - | 4596 | 1134 | - | - | 0 | - |
| - | - | 1015 | 1135 | - | - | 0 | - |
| 5 | y | 4392 | 1147 | 0.002326 | 2.028 | +1 | 11 |
| - | - | 2246 | 1148 | - | - | 0 | - |
| - | - | 1104 | 1149 | - | - | 0 | - |
| 11 | c | 5.221E+04 | 1159 | 0.001832 | 1.581 | +1 | 11 |
| - | - | 2.978E+04 | 1160 | - | - | 0 | - |
| - | - | 1.043E+04 | 1161 | - | - | 0 | - |
| - | - | 2122 | 1162 | - | - | 0 | - |
| - | - | 838.8 | 1199 | - | - | 0 | - |
| - | - | 694.3 | 1200 | - | - | 0 | - |
| 4 | w | 1184 | 1201 | 0.02399 | 19.98 | +1 | 12 |
| - | - | 1271 | 1213 | - | - | 0 | - |
| - | - | 1.026E+04 | 1214 | - | - | 0 | - |
| - | - | 5890 | 1215 | - | - | 0 | - |
| - | - | 4735 | 1216 | - | - | 0 | - |
| - | - | 1071 | 1217 | - | - | 0 | - |
| - | - | 1082 | 1218 | - | - | 0 | - |
| - | - | 859.9 | 1234 | - | - | 0 | - |
| - | - | 872.6 | 1256 | - | - | 0 | - |
| - | - | 857.4 | 1257 | - | - | 0 | - |
| 12 | c | 4.774E+04 | 1258 | 0.001533 | 1.219 | +1 | 12 |
| - | - | 3.274E+04 | 1259 | - | - | 0 | - |
| - | - | 1.11E+04 | 1260 | - | - | 0 | - |
| - | - | 4.08E+04 | 1261 | - | - | 0 | - |
| - | - | 2.405E+04 | 1262 | - | - | 0 | - |
| - | - | 8414 | 1263 | - | - | 0 | - |
| - | - | 2510 | 1264 | - | - | 0 | - |
| - | - | 847.3 | 1275 | - | - | 0 | - |
| 4 | y | 1.481E+04 | 1276 | 0.000382 | 0.2995 | +1 | 12 |
| - | - | 9422 | 1277 | - | - | 0 | - |
| - | - | 4631 | 1278 | - | - | 0 | - |
| - | - | 2116 | 1279 | - | - | 0 | - |
| - | - | 1621 | 1283 | - | - | 0 | - |
| - | - | 1187 | 1284 | - | - | 0 | - |
| - | - | 933 | 1286 | - | - | 0 | - |
| - | - | 987.4 | 1287 | - | - | 0 | - |
| - | - | 7655 | 1301 | - | - | 0 | - |
| - | - | 5649 | 1302 | - | - | 0 | - |
| - | - | 3302 | 1303 | - | - | 0 | - |
| - | - | 1261 | 1304 | - | - | 0 | - |
| - | - | 1612 | 1305 | - | - | 0 | - |
| - | - | 1377 | 1306 | - | - | 0 | - |
| 3 | w | 2668 | 1330 | 0.008006 | 6.021 | +1 | 13 |
| - | - | 3854 | 1331 | - | - | 0 | - |
| - | - | 2280 | 1332 | - | - | 0 | - |
| 13 | c | 8.206E+04 | 1345 | 0.0005106 | 0.3797 | +1 | 13 |
| - | - | 6.453E+04 | 1346 | - | - | 0 | - |
| - | - | 2.577E+04 | 1347 | - | - | 0 | - |
| - | - | 3.162E+04 | 1348 | - | - | 0 | - |
| - | - | 2.214E+04 | 1349 | - | - | 0 | - |
| - | - | 1.168E+04 | 1350 | - | - | 0 | - |
| - | - | 2868 | 1351 | - | - | 0 | - |
| 3 | y | 1157 | 1363 | 0.002533 | 1.859 | +1 | 13 |
| - | - | 1937 | 1364 | - | - | 0 | - |
| - | - | 974.9 | 1365 | - | - | 0 | - |
| - | - | 6895 | 1388 | - | - | 0 | - |
| - | - | 5528 | 1389 | - | - | 0 | - |
| - | - | 2884 | 1390 | - | - | 0 | - |
| - | - | 1566 | 1391 | - | - | 0 | - |
| - | - | 887.1 | 1417 | - | - | 0 | - |
| - | - | 1780 | 1430 | - | - | 0 | - |
| - | - | 2887 | 1431 | - | - | 0 | - |
| 14 | c | 1.182E+05 | 1432 | 0.0004646 | 0.3245 | +1 | 14 |
| - | - | 9.061E+04 | 1433 | - | - | 0 | - |
| - | - | 4.164E+04 | 1434 | - | - | 0 | - |
| - | - | 1.284E+04 | 1435 | - | - | 0 | - |
| - | - | 2189 | 1436 | - | - | 0 | - |
| - | - | 834.2 | 1459 | - | - | 0 | - |
| - | - | 4706 | 1461 | - | - | 0 | - |
| - | - | 4687 | 1462 | - | - | 0 | - |
| - | - | 1709 | 1463 | - | - | 0 | - |
| - | - | 827.6 | 1473 | - | - | 0 | - |
| - | - | 1245 | 1474 | - | - | 0 | - |
| - | - | 1196 | 1475 | - | - | 0 | - |
| 2 | y | 961.6 | 1476 | 0.02101 | 14.24 | +1 | 14 |
| - | - | 781 | 1477 | - | - | 0 | - |
| - | - | 8779 | 1489 | - | - | 0 | - |
| - | - | 8902 | 1490 | - | - | 0 | - |
| - | - | 4801 | 1491 | - | - | 0 | - |
| - | - | 1903 | 1492 | - | - | 0 | - |
| - | - | 794.1 | 1502 | - | - | 0 | - |
| - | - | 1579 | 1503 | - | - | 0 | - |
| - | - | 1308 | 1504 | - | - | 0 | - |
| - | - | 1020 | 1505 | - | - | 0 | - |
| - | - | 2493 | 1513 | - | - | 0 | - |
| - | - | 3594 | 1514 | - | - | 0 | - |
| - | - | 3081 | 1515 | - | - | 0 | - |
| - | - | 1455 | 1516 | - | - | 0 | - |
| - | - | 815.6 | 1517 | - | - | 0 | - |
| - | - | 1226 | 1520 | - | - | 0 | - |
| - | - | 1448 | 1529 | - | - | 0 | - |
| - | - | 2570 | 1530 | - | - | 0 | - |
| - | - | 2.32E+04 | 1531 | - | - | 0 | - |
| - | - | 3.06E+04 | 1532 | - | - | 0 | - |
| - | - | 1.751E+04 | 1533 | - | - | 0 | - |
| - | - | 9320 | 1534 | - | - | 0 | - |
| - | - | 2096 | 1535 | - | - | 0 | - |
| - | - | 1815 | 1544 | - | - | 0 | - |
| - | - | 4751 | 1545 | - | - | 0 | - |
| - | - | 1.025E+04 | 1546 | - | - | 0 | - |
| - | - | 1.064E+04 | 1547 | - | - | 0 | - |
| - | - | 1.174E+04 | 1548 | - | - | 0 | - |
| - | - | 6875 | 1549 | - | - | 0 | - |
| - | - | 3783 | 1550 | - | - | 0 | - |
| - | - | 4092 | 1556 | - | - | 0 | - |
| - | - | 1.022E+04 | 1557 | - | - | 0 | - |
| - | - | 6617 | 1558 | - | - | 0 | - |
| - | - | 3674 | 1559 | - | - | 0 | - |
| - | - | 1811 | 1560 | - | - | 0 | - |
| - | - | 1648 | 1561 | - | - | 0 | - |
| - | - | 3709 | 1562 | - | - | 0 | - |
| - | - | 1.588E+04 | 1563 | - | - | 0 | - |
| - | - | 1.578E+04 | 1564 | - | - | 0 | - |
| - | - | 6912 | 1565 | - | - | 0 | - |
| - | - | 1661 | 1566 | - | - | 0 | - |
| - | - | 873.4 | 1570 | - | - | 0 | - |
| - | - | 2580 | 1572 | - | - | 0 | - |
| - | - | 2.158E+04 | 1573 | - | - | 0 | - |
| - | - | 1.455E+05 | 1574 | - | - | 0 | - |
| - | - | 1.136E+05 | 1575 | - | - | 0 | - |
| - | - | 5.688E+04 | 1576 | - | - | 0 | - |
| - | - | 1.622E+04 | 1577 | - | - | 0 | - |
| - | - | 3805 | 1578 | - | - | 0 | - |
| - | - | 1229 | 1587 | - | - | 0 | - |
| - | - | 1315 | 1588 | - | - | 0 | - |
| - | - | 1.205E+04 | 1589 | - | - | 0 | - |
| - | - | 1149 | 1590 | - | - | 0 | - |
| - | - | 4.211E+04 | 1590 | - | - | 0 | - |
| - | - | 1.397E+05 | 1591 | - | - | 0 | - |
| - | - | 1.08E+05 | 1592 | - | - | 0 | - |
| - | - | 5.388E+04 | 1593 | - | - | 0 | - |
| - | - | 1.533E+04 | 1594 | - | - | 0 | - |
| - | - | 3869 | 1595 | - | - | 0 | - |
| - | - | 754.6 | 3343 | - | - | 0 | - |

m/z Charge Intensity FragmentType MassShift Position
120.08098602294922 0 804.5564
121.29129791259766 0 452.20657
125.10765075683594 0 3033.1128
126.09169006347656 0 4272.387
129.10244750976562 0 411.11755
130.0614013671875 0 477.17697
130.1219482421875 0 468.643
131.11854553222656 0 777.99054
131.12960815429688 0 2241.0662
142.21131896972656 0 482.1932
143.11875915527344 0 719.76764
146.98863220214844 0 458.75433
149.33848571777344 0 401.72006
151.3488311767578 0 507.1255
158.09283447265625 0 1677.5612 y Ammonia loss 14
168.11358642578125 0 6048.399
172.42527770996094 0 458.00302
173.45205688476562 0 2642.8987
175.119384765625 0 11745.353 y 14
185.14016723632812 0 28195.934
186.1434783935547 0 2217.6562
189.08740234375 0 873.816
197.84255981445312 0 480.32996
199.180908203125 0 79924.64
200.127197265625 0 4922.348
200.1842498779297 0 9579.798
201.12387084960938 0 909.50507
212.10342407226562 0 894.61395
217.0823211669922 0 3966.905
226.13052368164062 0 1626.1378
226.1676483154297 0 1165.7703
227.1758575439453 0 39792.145
228.17919921875 0 5261.504
229.1301727294922 0 5051.0093 w 13
230.11387634277344 0 1117.2434
233.16534423828125 0 4601.149
234.1687469482422 0 659.8661
244.11715698242188 0 1643.6061
245.125 0 47150.367 y Ammonia loss 13
246.1309051513672 0 7135.8496 z 13
248.07107543945312 0 1195.6985
249.1605224609375 0 2049.6887
250.16383361816406 0 700.8823
255.10934448242188 0 2075.9973
260.13482666015625 0 727.4608
261.14520263671875 0 947.2674
261.1603088378906 0 4657.981
262.1514587402344 0 7243.216 y 13
262.1640930175781 0 521.5974
263.1550598144531 0 1321.6498
270.12017822265625 0 7495.9067
271.12335205078125 0 741.9462
272.1718444824219 0 1903.3223
277.1551208496094 0 2303.1672 z 7
286.0365295410156 0 511.38068
296.1976013183594 0 747.36035
299.19549560546875 0 1040.448
303.154052734375 0 995.71075
304.1512145996094 0 887.95337
312.1552734375 0 828.62256
314.1462707519531 0 860.60785
314.20819091796875 0 4315.7954
316.1623229980469 0 24632.482 w 12
317.15960693359375 0 4487.892
323.2076721191406 0 781.20825
331.1494140625 0 5749.8555
331.2340393066406 0 997.78253 c 2
332.1567687988281 0 10387.215 y Ammonia loss 12
333.164794921875 0 19969.064 z 12
334.167724609375 0 3376.5295
347.16778564453125 0 1019.3971
349.1160583496094 0 858.4349
349.18365478515625 0 5792.9385 y 12
357.1523132324219 0 12886.237
389.7098693847656 0 2148.9143
390.2089538574219 0 2163.6494
390.71673583984375 0 2778.2683
391.2171630859375 0 1038.203
391.7247314453125 0 11679.612
392.2259826660156 0 5224.741
392.72845458984375 0 964.9754
398.7142639160156 0 4310.981
399.2156066894531 0 1566.5637
399.7215270996094 0 4756.1875
400.2232666015625 0 2481.9998
414.22100830078125 0 699.5824 c Ammonia loss 11
415.22686767578125 0 6010.9355 z 7
415.7264099121094 0 1821.0066
416.2316589355469 0 614.9242
417.2102966308594 0 20183.432 w 11
418.2136535644531 0 3636.4424
423.2337646484375 0 5299.922 y 7
423.7357177734375 0 2061.0686
424.23486328125 0 785.3483
425.2403564453125 0 2451.8362
429.2360534667969 0 2087.9875
431.2250671386719 0 1201.0973 y Ammonia loss 11
432.2335510253906 0 1032.6534 z 11
443.2505798339844 0 3748.8435
443.3091125488281 0 1147.2865
444.2533874511719 0 1030.5568
446.2373046875 0 2664.949
447.2081298828125 0 954.3333
448.25262451171875 0 17846.195 y 11
448.88873291015625 0 1162.2821 y Water loss 2
449.22369384765625 0 1214.5629 y Ammonia loss 2
449.2561340332031 0 3421.2112
450.7439270019531 0 1513.4515 c Ammonia loss 7
454.89239501953125 0 907.3279 y 2
455.2288818359375 0 852.54175
455.2852783203125 0 1962.4238
460.2779846191406 0 13088.703 c 3
461.28118896484375 0 3804.6672
462.283203125 0 970.1735
465.2590637207031 0 863.4919
465.757568359375 0 1197.6731
471.24365234375 0 934.6804 z Water loss 10
472.2523498535156 0 6009.205 c Ammonia loss 13
472.7389831542969 0 3017.048 z 6
473.24169921875 0 2267.0295
474.23175048828125 0 4729.4917
475.23626708984375 0 1380.0734
480.7484130859375 0 3713.7566 y 6
481.2501525878906 0 1111.9371 z Water loss 1
486.58489990234375 0 820.1314 y Water loss 1
487.2532958984375 0 891.5515 y Water loss 10
489.2553405761719 0 6136.3867 z 10
490.2618408203125 0 3805.2454
491.26409912109375 0 1040.0555
492.587890625 0 2201.6355 y 1
492.9236755371094 0 1401.5907
495.9154052734375 0 2274.1492
496.2523193359375 0 1626.6536
499.30023193359375 0 4364.376
499.7709045410156 0 716.4094
500.2703552246094 0 900.4533
504.265380859375 0 906.5661
505.27410888671875 0 63103.05 y 10
506.27734375 0 13587.479
507.1850280761719 0 785.186
507.2784423828125 0 2641.8374
514.3246459960938 0 1264.5708
515.2488403320312 0 1892.9059 y Water loss 5
515.3318481445312 0 2964.6548
515.74951171875 0 2071.503 y Ammonia loss 5
516.2542724609375 0 5867.708 z 5
516.3383178710938 0 1884.3043
516.7559814453125 0 7599.3555
517.2589721679688 0 4245.6133
517.75830078125 0 1254.0444
518.610107421875 0 943.95496
520.7921752929688 0 1096.6199
523.76025390625 0 1069.3041
524.2659301757812 0 25835.117 y 5
524.61279296875 0 8581.353
524.76513671875 0 12776.282
524.9468383789062 0 4454.7217
525.267333984375 0 4338.718
525.7691040039062 0 821.01587
528.2509155273438 0 1412.0502
528.7915649414062 0 5251.333
529.2919921875 0 4116.4956
529.7978515625 0 7387.182
530.2847290039062 0 10401.361
530.61669921875 0 10456.086
530.8019409179688 0 2533.741
530.9501953125 0 5487.703
531.291015625 0 1085.6632
541.3121337890625 0 1867.6053
542.319091796875 0 7510.1187
543.322998046875 0 1570.0277
550.7825927734375 0 1406.4647
558.2787475585938 0 1951.1029 z Water loss 9
558.3383178710938 0 4206.794
558.776611328125 0 1307.9944
559.2846069335938 0 3399.526 w 9
559.3460083007812 0 10995.46 c 4
559.7760009765625 0 681.24817
560.2908935546875 0 1410.9486
560.3485717773438 0 4303.311
565.2786865234375 0 738.5816 y Ammonia loss 4
565.7886962890625 0 44020.71 z 4
566.2899169921875 0 29411.21
566.7914428710938 0 10333.69
567.292236328125 0 2842.3865
572.3177490234375 0 2196.0354
573.2936401367188 0 1801.1256
573.7975463867188 0 28269.85 y 4
574.298828125 0 16985.793 y Water loss 9
574.801025390625 0 7856.499
575.2823486328125 0 3331.276 y Ammonia loss 9
576.286865234375 0 9067.55 z 9
577.29248046875 0 2754.4373
585.2686767578125 0 1111.1726
586.3323364257812 0 2637.2476
587.3394775390625 0 884.9655
592.3071899414062 0 2359.2024 y 9
600.274169921875 0 5522.6084
600.8035278320312 0 22702.723 w 3
601.30419921875 0 12371.696
601.3590087890625 0 513.2442
601.804931640625 0 5355.9688
602.2947387695312 0 710.78766
603.373291015625 0 1719.7303
614.3324584960938 0 833.59344
621.3076782226562 0 715.3717 z Water loss 3
629.3029174804688 0 1219.7058 y Water loss 3
629.3530883789062 0 2771.3022 c 11
630.309814453125 0 56446.008 z 3
630.8115844726562 0 34875.445
631.3130493164062 0 16349.376
631.8137817382812 0 4783.828
632.3153076171875 0 999.1537
637.8150634765625 0 17843.084
638.3187866210938 0 55289.61 y 3
638.8203735351562 0 28295.986
639.3214111328125 0 10172.317
639.8234252929688 0 3378.3333
644.3203735351562 0 3626.0833
644.8201904296875 0 2656.3762
645.3697509765625 0 2070.4028
646.3783569335938 0 45902.73 c 5
647.381591796875 0 17946.758
648.3849487304688 0 3773.5967
651.8234252929688 0 1240.055
652.3255004882812 0 656.0493
657.3699340820312 0 686.9208
660.3109741210938 0 723.6002
664.824462890625 0 1883.8934 z Water loss 2
665.3211669921875 0 2702.6284 z Ammonia loss 2
665.824951171875 0 1965.5569
666.3222045898438 0 1453.2968
672.8267822265625 0 4435.508 y Water loss 2
673.3279418945312 0 3134.5015 y Ammonia loss 2
673.7771606445312 0 384.25488
673.8265380859375 0 31351.248 z 2
674.328369140625 0 44893.6
674.8302612304688 0 22575.826
675.3321533203125 0 9245.246
675.8331909179688 0 3167.424
678.3192138671875 0 747.3768
680.8270263671875 0 2660.4941
681.3309936523438 0 3962.2192
681.8355712890625 0 245363.8 y 2
682.3367919921875 0 172404.11
682.837890625 0 72714.02
683.3394775390625 0 18334.473
683.8408203125 0 5252.042
684.3734741210938 0 1514.6704
685.3882446289062 0 2413.8486
685.8170166015625 0 860.291
686.3305053710938 0 699.6581
686.3930053710938 0 836.7497
686.8267822265625 0 1148.8728
687.342529296875 0 1887.4314
689.3587646484375 0 7127.5186 y 8
690.3637084960938 0 2525.545
691.3657836914062 0 674.1861
697.4379272460938 0 784.7597
699.3748168945312 0 1309.1924
710.4135131835938 0 788.5571
711.4189453125 0 1261.9795
712.4260864257812 0 2361.1182
713.4340209960938 0 3796.33
714.4379272460938 0 1576.1879
715.8499145507812 0 3919.405 w 1
716.3511352539062 0 4244.1777
716.85205078125 0 1901.544
717.390625 0 2942.7007
718.3980102539062 0 6121.2417
719.4016723632812 0 2228.4102
723.3610229492188 0 847.1645
729.370361328125 0 1940.737 y Water loss 1
729.8718872070312 0 744.3511 y Ammonia loss 1
730.3682250976562 0 3126.563 z 1
730.866943359375 0 1531.7347
731.367431640625 0 1353.7911
738.3775634765625 0 16004.094 y 1
738.879150390625 0 11014.0205
739.37939453125 0 3816.7903
739.8822631835938 0 1343.2225
743.3720703125 0 4626.654 w 7
744.377197265625 0 3144.843
745.3786010742188 0 1731.4733
749.8775634765625 0 848.66846
750.8971557617188 0 3216.7268
751.4006958007812 0 2698.5105
751.8927001953125 0 934.8023
756.4058227539062 0 1437.1097
758.3895874023438 0 4745.475
758.8831176757812 0 8124.1636
759.385009765625 0 6418.165
759.8816528320312 0 1615.9525
760.3927612304688 0 2044.5028
761.4057006835938 0 81996 c 6
762.4083862304688 0 35420.832
763.4103393554688 0 9536.37
764.411376953125 0 4109.084
764.913330078125 0 7874.084
765.3836059570312 0 148045.73
765.8851928710938 0 118088.266
766.386474609375 0 59135.01
766.8872680664062 0 16832.03
767.3901977539062 0 4832.3335
770.9113159179688 0 1095.2405
771.412353515625 0 1285.7786
771.9022216796875 0 1700.3763
772.4156494140625 0 4692.7095
772.916259765625 0 4803.003
773.4127807617188 0 4879.9927
773.909423828125 0 45850.6
774.4103393554688 0 35353.305
774.9110717773438 0 16720.766
775.4132690429688 0 5516.8125
775.91357421875 0 1681.7706
777.9041137695312 0 4093.4983
778.4085693359375 0 5143.149
778.9048461914062 0 2209.5024
779.4077758789062 0 3063.7034
779.8982543945312 0 1758.4346
780.4122924804688 0 1827.7463
781.4232788085938 0 1239.8323
782.4417114257812 0 7988.591
783.444580078125 0 4761.519
784.4468994140625 0 1550.7576
785.90234375 0 3280.2976
786.4147338867188 0 11265.448
786.910888671875 0 150443.67
787.4117431640625 0 134455.25
787.9132080078125 0 63169.48
788.413818359375 0 21019.256
788.9158935546875 0 7816.9165
789.4531860351562 0 905.8211
793.4160766601562 0 857.89374
793.9102172851562 0 2349.5107
794.412353515625 0 7373.4536
794.9192504882812 0 112457.19
795.4218139648438 0 128774.68
795.9236450195312 0 82334.19
796.4246826171875 0 36557.48
796.9234008789062 0 11518.162
797.4219360351562 0 4162.994
798.4349365234375 0 3004.7256
799.4344482421875 0 1995.2301
800.439453125 0 848.4365
811.43017578125 0 3799.2598 z Water loss 7
812.4315795898438 0 2062.6807
814.4243774414062 0 795.8041
828.4337158203125 0 1640.1726 y Ammonia loss 7
829.4414672851562 0 71944.78 z 7
830.4454956054688 0 33991.027
831.4481201171875 0 11252.863
832.4507446289062 0 2984.6145
839.5151977539062 0 2759.014
840.5184326171875 0 3285.299
841.5298461914062 0 6824.255
842.5333251953125 0 3533.9583
844.4522705078125 0 3939.4753
845.3850708007812 0 1385.3019
845.4598388671875 0 10899.128 y 7
846.46337890625 0 3383.8958
847.4675903320312 0 2025.5431
873.4939575195312 0 3772.2144
874.4962768554688 0 2863.421
875.5000610351562 0 747.11676
888.4468383789062 0 833.9541
899.4703979492188 0 1736.2146 w 6
900.4785766601562 0 11069.486 c Ammonia loss 7
901.4825439453125 0 4469.956
902.4852905273438 0 1779.4077
926.4574584960938 0 3150.7185 z Water loss 6
927.4651489257812 0 2501.5742
928.4749145507812 0 1713.6631
929.480712890625 0 2030.585
930.4874267578125 0 1252.3755
943.2314453125 0 810.8142
943.4576416015625 0 1197.7847 y Ammonia loss 6
944.468505859375 0 29757.514 z 6
945.473388671875 0 23368.082
946.4771118164062 0 8223.472
947.4772338867188 0 2332.176
954.54248046875 0 2606.582
955.5397338867188 0 3710.371
956.5527954101562 0 6524.985
957.5595092773438 0 3384.7573
958.5744018554688 0 1092.795
960.4873657226562 0 3250.1052 y 6
961.4906616210938 0 1594.9073
970.5420532226562 0 6694.548
971.5445556640625 0 2815.0647
987.5036010742188 0 1220.5948
999.5656127929688 0 1672.2244
1000.5726928710938 0 914.1652
1013.5866088867188 0 3590.6086
1014.5601806640625 0 61949.527 c 8
1015.56298828125 0 34598.156
1016.5657958984375 0 10750.516
1017.5643310546875 0 2521.1704
1029.477783203125 0 1093.5137
1030.503662109375 0 933.304 y Ammonia loss 5
1031.4993896484375 0 9795.338 z 5
1032.50732421875 0 14265.245
1033.5101318359375 0 6299.0386
1034.51513671875 0 2261.2869
1039.56787109375 0 2408.995
1040.563720703125 0 5397.2607
1041.565673828125 0 3372.437
1042.572509765625 0 5311.116
1043.574462890625 0 3505.3945
1045.4749755859375 0 925.79395
1046.5029296875 0 1735.6719
1047.51904296875 0 4610.934 y 5
1048.5198974609375 0 2564.6123
1049.520263671875 0 924.4747
1053.9962158203125 0 826.6082
1056.577880859375 0 2400.2383
1057.5784912109375 0 47988.95
1058.583251953125 0 35959.793
1059.5911865234375 0 25373.533
1060.595703125 0 9739.775
1061.4813232421875 0 973.05865
1061.6015625 0 3160.8936
1062.383056640625 0 1244.8351
1062.490478515625 0 2491.8506
1063.50048828125 0 1548.3778
1072.5732421875 0 1126.6261
1089.5654296875 0 823.3801
1099.5693359375 0 1065.924
1100.582275390625 0 3191.0208
1101.592041015625 0 84549.11 c 9
1102.5946044921875 0 52217.805
1103.597412109375 0 18514.836
1104.5965576171875 0 4204.124
1114.5977783203125 0 13147.484
1115.60205078125 0 7990.5146
1116.603759765625 0 3228.2563
1130.5684814453125 0 13920.154 z 4
1131.576171875 0 26531.947
1132.5791015625 0 15016.567
1133.5831298828125 0 4596.0425
1134.5828857421875 0 1014.50464
1146.5885009765625 0 4391.5015 y 4
1147.5909423828125 0 2245.7273
1148.5869140625 0 1103.6958
1158.6131591796875 0 52208.473 c 10
1159.6158447265625 0 29779.283
1160.6173095703125 0 10425.407
1161.6226806640625 0 2122.2668
1198.6485595703125 0 838.7712
1199.6556396484375 0 694.30457
1200.6207275390625 0 1184.1465 w 3
1212.6650390625 0 1270.6108
1213.6680908203125 0 10258.928
1214.6697998046875 0 5889.9277
1215.673583984375 0 4735.298
1216.6917724609375 0 1071.3539
1217.58984375 0 1081.9602
1233.61474609375 0 859.9279
1255.672119140625 0 872.6442
1256.6707763671875 0 857.36523
1257.6812744140625 0 47740.406 c 11
1258.682373046875 0 32736.621
1259.677001953125 0 11097.642
1260.6192626953125 0 40800.406
1261.6219482421875 0 24049.053
1262.6236572265625 0 8413.9795
1263.626953125 0 2509.9714
1274.614013671875 0 847.2636
1275.629150390625 0 14805.88 y 3
1276.63232421875 0 9421.521
1277.6357421875 0 4631.204
1278.63720703125 0 2115.643
1282.68896484375 0 1620.6211
1283.6890869140625 0 1186.6112
1285.693603515625 0 932.9826
1286.6783447265625 0 987.37946
1300.6978759765625 0 7655.4014
1301.701416015625 0 5648.509
1302.7069091796875 0 3301.676
1303.7030029296875 0 1261.4808
1304.6241455078125 0 1612.4689
1305.6246337890625 0 1376.9257
1329.6473388671875 0 2667.5579 w 2
1330.636962890625 0 3854.426
1331.639404296875 0 2279.6118
1344.7122802734375 0 82056.25 c 12
1345.71533203125 0 64534.734
1346.7127685546875 0 25773.928
1347.655517578125 0 31624.74
1348.655517578125 0 22144.908
1349.655517578125 0 11679.603
1350.6622314453125 0 2868.0542
1362.663330078125 0 1156.9346 y 2
1363.6632080078125 0 1936.8739
1364.6575927734375 0 974.8833
1387.7308349609375 0 6894.725
1388.7366943359375 0 5527.7314
1389.7364501953125 0 2884.1807
1390.7476806640625 0 1565.8297
1416.7266845703125 0 887.09235
1429.723876953125 0 1780.4368
1430.73095703125 0 2887.092
1431.7442626953125 0 118187.8 c 13
1432.7474365234375 0 90610.56
1433.74951171875 0 41644.137
1434.7520751953125 0 12842.337
1435.7579345703125 0 2188.8396
1458.7674560546875 0 834.19464
1460.7332763671875 0 4705.6724
1461.7342529296875 0 4686.5425
1462.730712890625 0 1709.2305
1473.2293701171875 0 827.6487
1473.74267578125 0 1245.286
1474.759765625 0 1195.7838
1475.765869140625 0 961.5968 y 1
1476.7567138671875 0 781.01624
1488.743408203125 0 8778.565
1489.7471923828125 0 8901.624
1490.753173828125 0 4800.5264
1491.7586669921875 0 1903.0743
1501.751953125 0 794.1254
1502.788818359375 0 1579.1083
1503.77880859375 0 1308.0005
1504.8031005859375 0 1020.02325
1512.7978515625 0 2492.72
1513.7933349609375 0 3594.2253
1514.78271484375 0 3081.167
1515.7813720703125 0 1455.2886
1516.781982421875 0 815.58026
1519.829833984375 0 1225.7372
1528.8070068359375 0 1448.3546
1529.8004150390625 0 2569.5852
1530.793212890625 0 23198.002
1531.795654296875 0 30602.174
1532.7977294921875 0 17507.467
1533.8011474609375 0 9320.447
1534.804931640625 0 2095.7449
1543.823486328125 0 1815.2906
1544.838134765625 0 4750.8564
1545.827880859375 0 10254.613
1546.8245849609375 0 10638.375
1547.8203125 0 11738.355
1548.8204345703125 0 6875.4297
1549.818359375 0 3783.0322
1555.8111572265625 0 4092.2024
1556.796142578125 0 10224.353
1557.797607421875 0 6616.561
1558.80078125 0 3673.6738
1559.811279296875 0 1810.6023
1560.832763671875 0 1648.0309
1561.8370361328125 0 3708.5583
1562.848388671875 0 15880.71
1563.851806640625 0 15775.028
1564.853759765625 0 6912.0815
1565.8577880859375 0 1660.515
1569.78955078125 0 873.41235
1571.809326171875 0 2579.8982
1572.817138671875 0 21579.373
1573.818115234375 0 145503.02
1574.8212890625 0 113562.83
1575.8236083984375 0 56882.555
1576.8255615234375 0 16222.507
1577.8248291015625 0 3804.664
1586.8265380859375 0 1229.3378
1587.8153076171875 0 1315.4268
1588.8255615234375 0 12047.852
1589.5904541015625 0 1149.1786
1589.8349609375 0 42111.984
1590.84326171875 0 139719.22
1591.8460693359375 0 107998.22
1592.848876953125 0 53881.664
1593.850830078125 0 15333.327
1594.8521728515625 0 3868.5867
3342.7490234375 0 754.58124

Spectrum Details

|  |  |
| --- | --- |
| Matched peaks? Matched peaks The total absolute number of peaks matched. Additionally in brackets the total fraction of peaks matched and the total number of peaks is shown. | 95 (16.55% of 574) |
| FDR? FDR The false discovery rate estimated for this peptide. It is calculated by matching all theoretical fragments with a non-integer shift with the raw peaks for this spectrum. This is done with 40 different shifts. The resulting percentage is the average number of annotated peaks over the number of annotated peaks with the correct spectrum. | 1.03% |
| Satellite FDR? Satellite FDR See the FDR for details on its calculation. This satellite ion specific FDR only contains the satellite ions (d/w) for I/L/J positions. | 0.00% |
| PSM Score? PSM Score The PSM Score as given by Hecklib to this annotated spectrum. It is shown with three significant figures. | 694 |

#### Spectrum 4284? Spectrum 4284 The raw spectrum of this peptide as annotated by Hecklib. The fragments are coloured according to ion type (see legend). Any peaks with a star '\*' as text can be hovered over to see the full details, first the ion type second the mass shift type. By hovering over the amino acids in the peptide or ions in the legend the corresponding peaks are highlighted. By toggling the 'Unassigned' label you can turn the background (unassigned) peaks on or off in the plot. By updating the slider in the Ion legend you can update the spectrum to only show the top X% of the peaks with labels. The top X% means any peak that is within X% of the highest intensity. By dragging in the spectrum you can zoom in to a specific part of the spectrum and use 'Zoom Out' to get back to the original zoom level. The annotation of the spectrum is based on the given sequence in the peptides file and is done with different software so inconsistencies are likely. The peaks are annotated based on the given sequence, with 20 ppm tolerance.

Copy Data

##### Spectrum 4284 (TSV)

###### Preview

```
Loading example...
```

*Click on the button to copy the data to your clipboard.*

Mz MinMz MaxIntensity Max

WidthHeightPeptide font sizePeptide stroke widthSpectrum font sizeSpectrum stroke widthCompact peptide

Ion legend

wxyz

abcd

OtherUnassignedIon^Charge^_Position_Show for top:%

JJSEVSDRPSGVSSR

01.93e+43.85e+45.78e+47.71e+4

Zoom Out

y^+1^_1_y^+1^_1_w^+1^_2_y^+1^_2_z^+1^_2_y^+1^_2_w^+1^_3_y^+1^_3_z^+1^_3_y^+1^_3_z^+2^_8_w^+1^_4_y^+2^_8_y^+1^_4_y^+1^_4_y^+3^_13_c^+1^_4_c^+3^_14_z^+2^_9_y^+2^_9_z^+3^_14_z^+1^_5_y^+1^_5_y^+2^_10_z^+2^_10_y^+2^_10_w^+1^_6_c^+1^_5_z^+2^_11_y^+2^_11_y^+1^_6_y^+1^_6_z^+1^_6_y^+1^_6_w^+2^_12_c^+2^_12_z^+2^_12_y^+2^_12_w^+1^_7_c^+1^_6_z^+2^_13_y^+2^_13_z^+2^_13_y^+2^_13_y^+1^_7_w^+2^_14_y^+2^_14_y^+2^_14_z^+2^_14_y^+2^_14_c^+1^_7_z^+1^_8_z^+1^_8_y^+1^_8_w^+1^_9_c^+1^_8_z^+1^_9_y^+1^_9_c^+1^_9_z^+1^_10_y^+1^_10_c^+1^_10_z^+1^_11_y^+1^_11_c^+1^_11_c^+1^_12_y^+1^_12_c^+1^_13_c^+1^_14_

0805161124163221

Fragment Matches Table

Show background peaks

| Position | Ion type | Intensity | mz Theoretical | mz Error (Th) | mz Error (ppm) | Charge | Series Number |
| --- | --- | --- | --- | --- | --- | --- | --- |
| - | - | 1209 | 125.1 | - | - | 0 | - |
| - | - | 2234 | 126.1 | - | - | 0 | - |
| - | - | 950.4 | 131.1 | - | - | 0 | - |
| - | - | 705.7 | 133.1 | - | - | 0 | - |
| - | - | 359.7 | 135.2 | - | - | 0 | - |
| - | - | 497.6 | 137.8 | - | - | 0 | - |
| - | - | 417.8 | 144.5 | - | - | 0 | - |
| - | - | 582 | 147.1 | - | - | 0 | - |
| - | - | 880.7 | 149 | - | - | 0 | - |
| - | - | 526.8 | 149.3 | - | - | 0 | - |
| - | - | 473.9 | 157.2 | - | - | 0 | - |
| 15 | y | 908.8 | 158.1 | 0.000111 | 0.7019 | +1 | 1 |
| - | - | 459.4 | 160.1 | - | - | 0 | - |
| - | - | 3434 | 168.1 | - | - | 0 | - |
| - | - | 2107 | 173.5 | - | - | 0 | - |
| 15 | y | 4085 | 175.1 | 0.0002648 | 1.512 | +1 | 1 |
| - | - | 966.3 | 185.1 | - | - | 0 | - |
| - | - | 1.051E+04 | 185.1 | - | - | 0 | - |
| - | - | 553.8 | 190.1 | - | - | 0 | - |
| - | - | 2.357E+04 | 199.2 | - | - | 0 | - |
| - | - | 1682 | 200.1 | - | - | 0 | - |
| - | - | 2908 | 200.2 | - | - | 0 | - |
| - | - | 548 | 210.9 | - | - | 0 | - |
| - | - | 988.3 | 217.1 | - | - | 0 | - |
| - | - | 1.177E+04 | 227.2 | - | - | 0 | - |
| - | - | 1594 | 228.2 | - | - | 0 | - |
| 14 | w | 1777 | 229.1 | 0.000488 | 2.13 | +1 | 2 |
| - | - | 487.8 | 230.9 | - | - | 0 | - |
| 14 | y | 1.935E+04 | 245.1 | 0.0003549 | 1.448 | +1 | 2 |
| 14 | z | 2618 | 246.1 | 0.001992 | 8.094 | +1 | 2 |
| - | - | 1198 | 255.1 | - | - | 0 | - |
| - | - | 522.2 | 257.9 | - | - | 0 | - |
| - | - | 566.9 | 261.2 | - | - | 0 | - |
| 14 | y | 1864 | 262.2 | 0.0002645 | 1.009 | +1 | 2 |
| - | - | 2648 | 270.1 | - | - | 0 | - |
| - | - | 604.7 | 277.1 | - | - | 0 | - |
| - | - | 1464 | 314.2 | - | - | 0 | - |
| 13 | w | 8810 | 316.2 | 0.0002284 | 0.7225 | +1 | 3 |
| - | - | 1841 | 317.2 | - | - | 0 | - |
| - | - | 521.9 | 318.4 | - | - | 0 | - |
| - | - | 2132 | 331.1 | - | - | 0 | - |
| 13 | y | 3299 | 332.2 | 0.0002784 | 0.8381 | +1 | 3 |
| 13 | z | 6276 | 333.2 | 0.0003574 | 1.073 | +1 | 3 |
| - | - | 890.9 | 334.2 | - | - | 0 | - |
| 13 | y | 2695 | 349.2 | 0.0005542 | 1.587 | +1 | 3 |
| - | - | 4718 | 357.2 | - | - | 0 | - |
| - | - | 1233 | 391.7 | - | - | 0 | - |
| - | - | 1169 | 392.2 | - | - | 0 | - |
| - | - | 664.8 | 395.2 | - | - | 0 | - |
| 8 | z | 2224 | 415.2 | 0.003471 | 8.36 | +2 | 8 |
| - | - | 613.8 | 415.3 | - | - | 0 | - |
| - | - | 682.2 | 415.7 | - | - | 0 | - |
| 12 | w | 7763 | 417.2 | 0.0005846 | 1.401 | +1 | 4 |
| - | - | 1239 | 418.2 | - | - | 0 | - |
| 8 | y | 1699 | 423.2 | 0.0003367 | 0.7954 | +2 | 8 |
| - | - | 1057 | 423.7 | - | - | 0 | - |
| 12 | y | 741.5 | 431.2 | 0.001353 | 3.138 | +1 | 4 |
| - | - | 1763 | 443.3 | - | - | 0 | - |
| - | - | 623.6 | 444.2 | - | - | 0 | - |
| - | - | 928.4 | 446.2 | - | - | 0 | - |
| 12 | y | 6375 | 448.3 | 0.0009269 | 2.068 | +1 | 4 |
| - | - | 848.2 | 449.3 | - | - | 0 | - |
| 3 | y | 880.2 | 454.9 | 0.0008254 | 1.815 | +3 | 13 |
| - | - | 1133 | 455.2 | - | - | 0 | - |
| - | - | 617.4 | 455.6 | - | - | 0 | - |
| 4 | c | 4348 | 460.3 | 0.0001889 | 0.4104 | +1 | 4 |
| - | - | 1416 | 461.3 | - | - | 0 | - |
| 14 | c | 1853 | 472.2 | 0.008416 | 17.82 | +3 | 14 |
| 7 | z | 874.6 | 472.7 | 0.002024 | 4.281 | +2 | 9 |
| - | - | 1050 | 473.2 | - | - | 0 | - |
| - | - | 2500 | 474.2 | - | - | 0 | - |
| - | - | 571.8 | 475.1 | - | - | 0 | - |
| 7 | y | 666.5 | 480.7 | 0.001847 | 3.843 | +2 | 9 |
| 2 | z | 739.4 | 481.2 | 0.005923 | 12.31 | +3 | 14 |
| 11 | z | 1700 | 489.3 | 0.002368 | 4.84 | +1 | 5 |
| - | - | 1498 | 490.3 | - | - | 0 | - |
| - | - | 664.3 | 496.3 | - | - | 0 | - |
| - | - | 1540 | 499.3 | - | - | 0 | - |
| 11 | y | 2.349E+04 | 505.3 | 0.0005204 | 1.03 | +1 | 5 |
| - | - | 5285 | 506.3 | - | - | 0 | - |
| - | - | 1306 | 515.2 | - | - | 0 | - |
| - | - | 918.3 | 515.3 | - | - | 0 | - |
| 6 | y | 1587 | 515.7 | 0.0002209 | 0.4283 | +2 | 10 |
| 6 | z | 1361 | 516.3 | 0.00157 | 3.041 | +2 | 10 |
| - | - | 710.4 | 516.3 | - | - | 0 | - |
| - | - | 3355 | 516.8 | - | - | 0 | - |
| - | - | 1553 | 517.3 | - | - | 0 | - |
| - | - | 680.6 | 518.6 | - | - | 0 | - |
| 6 | y | 7717 | 524.3 | 0.00335 | 6.39 | +2 | 10 |
| - | - | 2051 | 524.6 | - | - | 0 | - |
| - | - | 3769 | 524.8 | - | - | 0 | - |
| - | - | 1307 | 524.9 | - | - | 0 | - |
| - | - | 1472 | 525.3 | - | - | 0 | - |
| - | - | 865.6 | 529.3 | - | - | 0 | - |
| - | - | 1843 | 529.3 | - | - | 0 | - |
| - | - | 2035 | 530.3 | - | - | 0 | - |
| - | - | 9085 | 530.3 | - | - | 0 | - |
| - | - | 2686 | 530.6 | - | - | 0 | - |
| - | - | 1347 | 530.9 | - | - | 0 | - |
| - | - | 2465 | 531.3 | - | - | 0 | - |
| - | - | 580.4 | 537.8 | - | - | 0 | - |
| - | - | 859.1 | 540.3 | - | - | 0 | - |
| - | - | 2508 | 542.3 | - | - | 0 | - |
| - | - | 930.6 | 543.3 | - | - | 0 | - |
| - | - | 1406 | 558.3 | - | - | 0 | - |
| 10 | w | 630.1 | 559.3 | 0.002804 | 5.013 | +1 | 6 |
| 5 | c | 3708 | 559.3 | 0.0001038 | 0.1856 | +1 | 5 |
| - | - | 1524 | 560.3 | - | - | 0 | - |
| 5 | z | 1.448E+04 | 565.8 | 0.0002338 | 0.4133 | +2 | 11 |
| - | - | 1.178E+04 | 566.3 | - | - | 0 | - |
| - | - | 4656 | 566.8 | - | - | 0 | - |
| - | - | 806.4 | 573.3 | - | - | 0 | - |
| 5 | y | 1.125E+04 | 573.8 | 0.0001491 | 0.2599 | +2 | 11 |
| 10 | y | 5746 | 574.3 | 0.00344 | 5.99 | +1 | 6 |
| - | - | 3066 | 574.8 | - | - | 0 | - |
| 10 | y | 1702 | 575.3 | 0.0002898 | 0.5037 | +1 | 6 |
| 10 | z | 3640 | 576.3 | 0.001224 | 2.123 | +1 | 6 |
| - | - | 691.1 | 577.3 | - | - | 0 | - |
| - | - | 846.8 | 586.3 | - | - | 0 | - |
| 10 | y | 926.3 | 592.3 | 0.0005049 | 0.8524 | +1 | 6 |
| - | - | 2219 | 600.3 | - | - | 0 | - |
| 4 | w | 7197 | 600.8 | 0.000482 | 0.8023 | +2 | 12 |
| - | - | 5244 | 601.3 | - | - | 0 | - |
| - | - | 2601 | 601.8 | - | - | 0 | - |
| - | - | 633 | 602.3 | - | - | 0 | - |
| - | - | 886.2 | 628.8 | - | - | 0 | - |
| 12 | c | 748.2 | 629.3 | 0.01123 | 17.84 | +2 | 12 |
| 4 | z | 1.606E+04 | 630.3 | 0.0004217 | 0.669 | +2 | 12 |
| - | - | 1.265E+04 | 630.8 | - | - | 0 | - |
| - | - | 4554 | 631.3 | - | - | 0 | - |
| - | - | 1437 | 631.8 | - | - | 0 | - |
| - | - | 723 | 632.3 | - | - | 0 | - |
| - | - | 6284 | 637.8 | - | - | 0 | - |
| 4 | y | 2.104E+04 | 638.3 | 9.028E-05 | 0.1414 | +2 | 12 |
| - | - | 1.042E+04 | 638.8 | - | - | 0 | - |
| - | - | 4764 | 639.3 | - | - | 0 | - |
| - | - | 1137 | 639.8 | - | - | 0 | - |
| - | - | 1396 | 644.3 | - | - | 0 | - |
| - | - | 1151 | 644.8 | - | - | 0 | - |
| - | - | 666.6 | 645.3 | - | - | 0 | - |
| - | - | 706.4 | 645.4 | - | - | 0 | - |
| 9 | w | 614.1 | 646.3 | 0.002208 | 3.417 | +1 | 7 |
| 6 | c | 1.786E+04 | 646.4 | 0.0006682 | 1.034 | +1 | 6 |
| - | - | 5970 | 647.4 | - | - | 0 | - |
| - | - | 1563 | 648.4 | - | - | 0 | - |
| 3 | z | 1452 | 665.3 | 0.009767 | 14.68 | +2 | 13 |
| - | - | 1560 | 665.8 | - | - | 0 | - |
| 3 | y | 1006 | 673.3 | 0.005898 | 8.76 | +2 | 13 |
| 3 | z | 1.188E+04 | 673.8 | 0.0004597 | 0.6822 | +2 | 13 |
| - | - | 1.6E+04 | 674.3 | - | - | 0 | - |
| - | - | 6871 | 674.8 | - | - | 0 | - |
| - | - | 3034 | 675.3 | - | - | 0 | - |
| - | - | 1340 | 680.8 | - | - | 0 | - |
| - | - | 1564 | 681.3 | - | - | 0 | - |
| 3 | y | 7.629E+04 | 681.8 | 0.000375 | 0.55 | +2 | 13 |
| - | - | 5.27E+04 | 682.3 | - | - | 0 | - |
| - | - | 2.323E+04 | 682.8 | - | - | 0 | - |
| - | - | 6781 | 683.3 | - | - | 0 | - |
| - | - | 1616 | 683.8 | - | - | 0 | - |
| - | - | 1105 | 687.3 | - | - | 0 | - |
| 9 | y | 2655 | 689.4 | 0.0006843 | 0.9926 | +1 | 7 |
| - | - | 956 | 690.4 | - | - | 0 | - |
| - | - | 634.8 | 711.4 | - | - | 0 | - |
| 2 | w | 1406 | 715.8 | 0.0007695 | 1.075 | +2 | 14 |
| - | - | 1805 | 716.3 | - | - | 0 | - |
| - | - | 721.6 | 717.4 | - | - | 0 | - |
| - | - | 2112 | 718.4 | - | - | 0 | - |
| - | - | 973.3 | 719.4 | - | - | 0 | - |
| 2 | y | 1304 | 729.4 | 0.007566 | 10.37 | +2 | 14 |
| 2 | y | 590.7 | 729.9 | 0.01092 | 14.97 | +2 | 14 |
| 2 | z | 1045 | 730.4 | 0.001838 | 2.517 | +2 | 14 |
| 2 | y | 5609 | 738.4 | 0.0001521 | 0.206 | +2 | 14 |
| - | - | 3158 | 738.9 | - | - | 0 | - |
| - | - | 1356 | 739.4 | - | - | 0 | - |
| - | - | 802.2 | 743.9 | - | - | 0 | - |
| - | - | 1369 | 744.4 | - | - | 0 | - |
| - | - | 815.5 | 745.4 | - | - | 0 | - |
| - | - | 1455 | 750.9 | - | - | 0 | - |
| - | - | 924.9 | 757.4 | - | - | 0 | - |
| - | - | 1521 | 758.4 | - | - | 0 | - |
| - | - | 2609 | 758.9 | - | - | 0 | - |
| - | - | 2703 | 759.4 | - | - | 0 | - |
| - | - | 965.7 | 759.9 | - | - | 0 | - |
| - | - | 1147 | 760.4 | - | - | 0 | - |
| 7 | c | 3.366E+04 | 761.4 | 0.0005196 | 0.6824 | +1 | 7 |
| - | - | 1.25E+04 | 762.4 | - | - | 0 | - |
| - | - | 1080 | 762.5 | - | - | 0 | - |
| - | - | 3277 | 763.4 | - | - | 0 | - |
| - | - | 1815 | 764.4 | - | - | 0 | - |
| - | - | 4594 | 764.9 | - | - | 0 | - |
| - | - | 5.792E+04 | 765.4 | - | - | 0 | - |
| - | - | 4.389E+04 | 765.9 | - | - | 0 | - |
| - | - | 2.224E+04 | 766.4 | - | - | 0 | - |
| - | - | 5667 | 766.9 | - | - | 0 | - |
| - | - | 1784 | 767.4 | - | - | 0 | - |
| - | - | 737.3 | 771.9 | - | - | 0 | - |
| - | - | 1456 | 772.4 | - | - | 0 | - |
| - | - | 1221 | 772.9 | - | - | 0 | - |
| - | - | 2264 | 773.4 | - | - | 0 | - |
| - | - | 1.767E+04 | 773.9 | - | - | 0 | - |
| - | - | 1.284E+04 | 774.4 | - | - | 0 | - |
| - | - | 6030 | 774.9 | - | - | 0 | - |
| - | - | 1854 | 775.4 | - | - | 0 | - |
| - | - | 1490 | 778.4 | - | - | 0 | - |
| - | - | 1261 | 779.4 | - | - | 0 | - |
| - | - | 1200 | 785.9 | - | - | 0 | - |
| - | - | 3472 | 786.4 | - | - | 0 | - |
| - | - | 5.514E+04 | 786.9 | - | - | 0 | - |
| - | - | 4.513E+04 | 787.4 | - | - | 0 | - |
| - | - | 2.238E+04 | 787.9 | - | - | 0 | - |
| - | - | 8785 | 788.4 | - | - | 0 | - |
| - | - | 1518 | 788.9 | - | - | 0 | - |
| - | - | 903.8 | 793.4 | - | - | 0 | - |
| - | - | 920.4 | 793.9 | - | - | 0 | - |
| - | - | 3813 | 794.4 | - | - | 0 | - |
| - | - | 3.787E+04 | 794.9 | - | - | 0 | - |
| - | - | 4.801E+04 | 795.4 | - | - | 0 | - |
| - | - | 2.942E+04 | 795.9 | - | - | 0 | - |
| - | - | 1.215E+04 | 796.4 | - | - | 0 | - |
| - | - | 2147 | 796.9 | - | - | 0 | - |
| - | - | 1593 | 797.4 | - | - | 0 | - |
| 8 | z | 1289 | 811.4 | 0.0006746 | 0.8314 | +1 | 8 |
| 8 | z | 2.548E+04 | 829.4 | 0.0003638 | 0.4387 | +1 | 8 |
| - | - | 1.394E+04 | 830.4 | - | - | 0 | - |
| - | - | 5117 | 831.4 | - | - | 0 | - |
| - | - | 727.6 | 832.5 | - | - | 0 | - |
| - | - | 919.6 | 841.5 | - | - | 0 | - |
| - | - | 1440 | 844.5 | - | - | 0 | - |
| 8 | y | 2723 | 845.5 | 4.968E-05 | 0.05877 | +1 | 8 |
| - | - | 1615 | 846.5 | - | - | 0 | - |
| - | - | 830.9 | 847.5 | - | - | 0 | - |
| - | - | 1592 | 873.5 | - | - | 0 | - |
| 7 | w | 667 | 899.5 | 0.0006771 | 0.7528 | +1 | 9 |
| 8 | c | 3971 | 900.5 | 0.001838 | 2.041 | +1 | 8 |
| - | - | 1705 | 901.5 | - | - | 0 | - |
| 7 | z | 9109 | 944.5 | 0.000212 | 0.2245 | +1 | 9 |
| - | - | 7663 | 945.5 | - | - | 0 | - |
| - | - | 2888 | 946.5 | - | - | 0 | - |
| - | - | 870.7 | 955.5 | - | - | 0 | - |
| - | - | 1008 | 956.5 | - | - | 0 | - |
| - | - | 749.4 | 957.4 | - | - | 0 | - |
| 7 | y | 927.1 | 960.5 | 0.005966 | 6.212 | +1 | 9 |
| - | - | 2018 | 970.5 | - | - | 0 | - |
| - | - | 780.5 | 973.5 | - | - | 0 | - |
| - | - | 1556 | 1014 | - | - | 0 | - |
| 9 | c | 2.388E+04 | 1015 | 0.0006364 | 0.6273 | +1 | 9 |
| - | - | 1.137E+04 | 1016 | - | - | 0 | - |
| - | - | 3945 | 1017 | - | - | 0 | - |
| 6 | z | 2982 | 1031 | 0.0008406 | 0.815 | +1 | 10 |
| - | - | 5171 | 1033 | - | - | 0 | - |
| - | - | 2709 | 1034 | - | - | 0 | - |
| - | - | 574.8 | 1035 | - | - | 0 | - |
| - | - | 1194 | 1040 | - | - | 0 | - |
| - | - | 1105 | 1041 | - | - | 0 | - |
| - | - | 713.8 | 1042 | - | - | 0 | - |
| - | - | 729.5 | 1044 | - | - | 0 | - |
| - | - | 2036 | 1045 | - | - | 0 | - |
| - | - | 1136 | 1046 | - | - | 0 | - |
| 6 | y | 2099 | 1048 | 0.001892 | 1.806 | +1 | 10 |
| - | - | 701 | 1057 | - | - | 0 | - |
| - | - | 1.367E+04 | 1058 | - | - | 0 | - |
| - | - | 1.006E+04 | 1059 | - | - | 0 | - |
| - | - | 3814 | 1060 | - | - | 0 | - |
| - | - | 1038 | 1061 | - | - | 0 | - |
| - | - | 635.4 | 1063 | - | - | 0 | - |
| - | - | 1090 | 1101 | - | - | 0 | - |
| 10 | c | 3.024E+04 | 1102 | 0.0003463 | 0.3144 | +1 | 10 |
| - | - | 1.812E+04 | 1103 | - | - | 0 | - |
| - | - | 7137 | 1104 | - | - | 0 | - |
| - | - | 1503 | 1105 | - | - | 0 | - |
| - | - | 5296 | 1115 | - | - | 0 | - |
| - | - | 2931 | 1116 | - | - | 0 | - |
| - | - | 1120 | 1117 | - | - | 0 | - |
| 5 | z | 4831 | 1131 | 0.001045 | 0.9243 | +1 | 11 |
| - | - | 1.167E+04 | 1132 | - | - | 0 | - |
| - | - | 5599 | 1133 | - | - | 0 | - |
| - | - | 2278 | 1134 | - | - | 0 | - |
| 5 | y | 1758 | 1147 | 0.001837 | 1.603 | +1 | 11 |
| - | - | 1086 | 1148 | - | - | 0 | - |
| 11 | c | 1.748E+04 | 1159 | 0.0004876 | 0.4208 | +1 | 11 |
| - | - | 1.144E+04 | 1160 | - | - | 0 | - |
| - | - | 5617 | 1161 | - | - | 0 | - |
| - | - | 1017 | 1162 | - | - | 0 | - |
| - | - | 3274 | 1214 | - | - | 0 | - |
| - | - | 2315 | 1215 | - | - | 0 | - |
| - | - | 1538 | 1216 | - | - | 0 | - |
| 12 | c | 1.682E+04 | 1258 | 0.001397 | 1.11 | +1 | 12 |
| - | - | 1.254E+04 | 1259 | - | - | 0 | - |
| - | - | 4666 | 1260 | - | - | 0 | - |
| - | - | 1.698E+04 | 1261 | - | - | 0 | - |
| - | - | 1.071E+04 | 1262 | - | - | 0 | - |
| - | - | 4453 | 1263 | - | - | 0 | - |
| - | - | 1159 | 1264 | - | - | 0 | - |
| 4 | y | 6279 | 1276 | 0.0008387 | 0.6575 | +1 | 12 |
| - | - | 4405 | 1277 | - | - | 0 | - |
| - | - | 1462 | 1278 | - | - | 0 | - |
| - | - | 807.9 | 1283 | - | - | 0 | - |
| - | - | 710.7 | 1285 | - | - | 0 | - |
| - | - | 2472 | 1301 | - | - | 0 | - |
| - | - | 2174 | 1302 | - | - | 0 | - |
| - | - | 952 | 1303 | - | - | 0 | - |
| - | - | 1077 | 1305 | - | - | 0 | - |
| - | - | 1168 | 1331 | - | - | 0 | - |
| - | - | 1039 | 1332 | - | - | 0 | - |
| 13 | c | 3.35E+04 | 1345 | 0.001443 | 1.073 | +1 | 13 |
| - | - | 2.238E+04 | 1346 | - | - | 0 | - |
| - | - | 6894 | 1347 | - | - | 0 | - |
| - | - | 1.413E+04 | 1348 | - | - | 0 | - |
| - | - | 1.047E+04 | 1349 | - | - | 0 | - |
| - | - | 4809 | 1350 | - | - | 0 | - |
| - | - | 1242 | 1351 | - | - | 0 | - |
| - | - | 664 | 1365 | - | - | 0 | - |
| - | - | 2457 | 1388 | - | - | 0 | - |
| - | - | 2414 | 1389 | - | - | 0 | - |
| - | - | 1327 | 1390 | - | - | 0 | - |
| - | - | 1236 | 1431 | - | - | 0 | - |
| 14 | c | 4.644E+04 | 1432 | 0.001489 | 1.04 | +1 | 14 |
| - | - | 3.534E+04 | 1433 | - | - | 0 | - |
| - | - | 1067 | 1433 | - | - | 0 | - |
| - | - | 1.613E+04 | 1434 | - | - | 0 | - |
| - | - | 3780 | 1435 | - | - | 0 | - |
| - | - | 1347 | 1436 | - | - | 0 | - |
| - | - | 2718 | 1461 | - | - | 0 | - |
| - | - | 1810 | 1462 | - | - | 0 | - |
| - | - | 750.2 | 1474 | - | - | 0 | - |
| - | - | 3833 | 1489 | - | - | 0 | - |
| - | - | 2862 | 1490 | - | - | 0 | - |
| - | - | 2323 | 1491 | - | - | 0 | - |
| - | - | 1110 | 1513 | - | - | 0 | - |
| - | - | 1799 | 1514 | - | - | 0 | - |
| - | - | 884.7 | 1515 | - | - | 0 | - |
| - | - | 706.6 | 1516 | - | - | 0 | - |
| - | - | 1048 | 1517 | - | - | 0 | - |
| - | - | 1741 | 1530 | - | - | 0 | - |
| - | - | 7571 | 1531 | - | - | 0 | - |
| - | - | 1.151E+04 | 1532 | - | - | 0 | - |
| - | - | 7941 | 1533 | - | - | 0 | - |
| - | - | 3424 | 1534 | - | - | 0 | - |
| - | - | 884.6 | 1535 | - | - | 0 | - |
| - | - | 1954 | 1545 | - | - | 0 | - |
| - | - | 4166 | 1546 | - | - | 0 | - |
| - | - | 5643 | 1547 | - | - | 0 | - |
| - | - | 4880 | 1548 | - | - | 0 | - |
| - | - | 3042 | 1549 | - | - | 0 | - |
| - | - | 1564 | 1550 | - | - | 0 | - |
| - | - | 834.4 | 1551 | - | - | 0 | - |
| - | - | 2108 | 1556 | - | - | 0 | - |
| - | - | 4148 | 1557 | - | - | 0 | - |
| - | - | 3870 | 1558 | - | - | 0 | - |
| - | - | 1186 | 1559 | - | - | 0 | - |
| - | - | 1664 | 1562 | - | - | 0 | - |
| - | - | 8447 | 1563 | - | - | 0 | - |
| - | - | 5621 | 1564 | - | - | 0 | - |
| - | - | 3104 | 1565 | - | - | 0 | - |
| - | - | 1201 | 1566 | - | - | 0 | - |
| - | - | 853.1 | 1572 | - | - | 0 | - |
| - | - | 8733 | 1573 | - | - | 0 | - |
| - | - | 6.692E+04 | 1574 | - | - | 0 | - |
| - | - | 4.945E+04 | 1575 | - | - | 0 | - |
| - | - | 2.406E+04 | 1576 | - | - | 0 | - |
| - | - | 7945 | 1577 | - | - | 0 | - |
| - | - | 1563 | 1578 | - | - | 0 | - |
| - | - | 1085 | 1588 | - | - | 0 | - |
| - | - | 5714 | 1589 | - | - | 0 | - |
| - | - | 1.809E+04 | 1590 | - | - | 0 | - |
| - | - | 6.129E+04 | 1591 | - | - | 0 | - |
| - | - | 4.897E+04 | 1592 | - | - | 0 | - |
| - | - | 1070 | 1592 | - | - | 0 | - |
| - | - | 2.095E+04 | 1593 | - | - | 0 | - |
| - | - | 6153 | 1594 | - | - | 0 | - |
| - | - | 1630 | 1595 | - | - | 0 | - |
| - | - | 782.9 | 2244 | - | - | 0 | - |
| - | - | 815.9 | 2355 | - | - | 0 | - |
| - | - | 726 | 2858 | - | - | 0 | - |
| - | - | 719.8 | 3073 | - | - | 0 | - |
| - | - | 794.4 | 3190 | - | - | 0 | - |

m/z Charge Intensity FragmentType MassShift Position
125.1075439453125 0 1208.7502
126.09156799316406 0 2234.131
131.129150390625 0 950.43365
133.0611114501953 0 705.72974
135.20492553710938 0 359.71408
137.75405883789062 0 497.62973
144.4662628173828 0 417.84412
147.0771026611328 0 582.0059
148.9556884765625 0 880.73236
149.3037567138672 0 526.8447
157.15428161621094 0 473.91595
158.09251403808594 0 908.79877 y Ammonia loss 14
160.13731384277344 0 459.44702
168.11329650878906 0 3433.6973
173.45135498046875 0 2106.6895
175.1192169189453 0 4084.5688 y 14
185.1314697265625 0 966.32446
185.13995361328125 0 10509.739
190.08212280273438 0 553.76575
199.1807098388672 0 23574.447
200.12738037109375 0 1681.5221
200.18394470214844 0 2908.4412
210.87632751464844 0 548.0329
217.0816650390625 0 988.3149
227.17567443847656 0 11770.913
228.17898559570312 0 1593.6248
229.1300048828125 0 1777.4714 w 13
230.94395446777344 0 487.82516
245.12478637695312 0 19346.293 y Ammonia loss 13
246.13026428222656 0 2618.3616 z 13
255.10861206054688 0 1197.5603
257.8526611328125 0 522.23535
261.15936279296875 0 566.85175
262.1512451171875 0 1863.7562 y 13
270.11962890625 0 2648.0354
277.115234375 0 604.6759
314.2080383300781 0 1464.3318
316.1617736816406 0 8810.138 w 12
317.1617126464844 0 1841.3384
318.38824462890625 0 521.93634
331.1489562988281 0 2131.627
332.15673828125 0 3299.0671 y Ammonia loss 12
333.1646423339844 0 6275.627 z 12
334.1684265136719 0 890.85297
349.1835632324219 0 2694.609 y 12
357.1518859863281 0 4718.186
391.7236022949219 0 1233.0596
392.2251281738281 0 1168.5665
395.2273864746094 0 664.79364
415.2271423339844 0 2223.5974 z 7
415.25506591796875 0 613.8315
415.72509765625 0 682.18915
417.2098083496094 0 7762.838 w 11
418.2119140625 0 1238.9202
423.2326965332031 0 1699.1382 y 7
423.7351989746094 0 1056.7106
431.2262268066406 0 741.50195 y Ammonia loss 11
443.2504577636719 0 1763.3289
444.208251953125 0 623.5748
446.2359924316406 0 928.35394
448.2523498535156 0 6374.8765 y 11
449.2557678222656 0 848.2246
454.8926086425781 0 880.18097 y 2
455.2262878417969 0 1132.9355
455.5628967285156 0 617.421
460.2767639160156 0 4348.338 c 3
461.2804870605469 0 1416.4136
472.2523498535156 0 1853.3606 c Ammonia loss 13
472.7391662597656 0 874.5705 z 6
473.2378845214844 0 1050.2764
474.2317810058594 0 2500.244
475.1332702636719 0 571.7807
480.74835205078125 0 666.4673 y 6
481.2492980957031 0 739.37823 z Water loss 1
489.25653076171875 0 1699.849 z 10
490.26239013671875 0 1498.1565
496.25067138671875 0 664.31525
499.29937744140625 0 1540.4426
505.2734069824219 0 23493.074 y 10
506.276611328125 0 5285.1807
515.2462768554688 0 1306.1482
515.330810546875 0 918.2982
515.7490234375 0 1587.1732 y Ammonia loss 5
516.2515869140625 0 1361.2996 z 5
516.3406372070312 0 710.36115
516.7561645507812 0 3355.4448
517.258056640625 0 1552.7965
518.6083374023438 0 680.5995
524.265869140625 0 7716.977 y 5
524.6128540039062 0 2051.4246
524.7637329101562 0 3768.9565
524.9462280273438 0 1306.8414
525.2666625976562 0 1472.0934
529.2949829101562 0 865.6065
529.334716796875 0 1842.6171
530.279541015625 0 2035.3099
530.3178100585938 0 9085.429
530.6162109375 0 2686.0269
530.9486083984375 0 1347.4819
531.3193359375 0 2465.2017
537.755126953125 0 580.42676
540.3248901367188 0 859.0818
542.3172607421875 0 2507.5327
543.3209228515625 0 930.63715
558.3374633789062 0 1406.4124
559.2862548828125 0 630.138 w 9
559.3450927734375 0 3708.1736 c 4
560.3475952148438 0 1523.9193
565.78759765625 0 14480.74 z 4
566.2894287109375 0 11784.306
566.7909545898438 0 4656.069
573.2913208007812 0 806.37915
573.796875 0 11245.299 y 4
574.2977905273438 0 5746.3887 y Water loss 9
574.7991943359375 0 3065.8218
575.278076171875 0 1702.0989 y Ammonia loss 9
576.2874145507812 0 3640.1355 z 9
577.2904663085938 0 691.0897
586.3284301757812 0 846.7645
592.305419921875 0 926.34357 y 9
600.2742309570312 0 2219.364
600.802490234375 0 7197.047 w 3
601.3035888671875 0 5244.3057
601.8062133789062 0 2600.9683
602.2981567382812 0 633.03656
628.80859375 0 886.227
629.354736328125 0 748.22925 c 11
630.30908203125 0 16064.871 z 3
630.81103515625 0 12646.188
631.3120727539062 0 4553.8936
631.8134765625 0 1436.6725
632.3130493164062 0 723.00616
637.8146362304688 0 6284.101
638.3179321289062 0 21043.7 y 3
638.8194580078125 0 10420.897
639.3218383789062 0 4764.4575
639.8237915039062 0 1136.8015
644.3168334960938 0 1395.8766
644.820068359375 0 1150.6445
645.3193359375 0 666.57556
645.372314453125 0 706.3712
646.3176879882812 0 614.06445 w 8
646.377685546875 0 17863.07 c 5
647.3807373046875 0 5970.2686
648.3837890625 0 1563.4524
665.3211669921875 0 1451.8143 z Ammonia loss 2
665.824462890625 0 1559.847
673.32666015625 0 1005.8628 y Ammonia loss 2
673.8251342773438 0 11884.975 z 2
674.3278198242188 0 15999.911
674.8287353515625 0 6870.713
675.3296508789062 0 3034.1848
680.830322265625 0 1340.11
681.3291625976562 0 1564.1196
681.8344116210938 0 76294.164 y 2
682.3355712890625 0 52696.984
682.8372192382812 0 23232.498
683.3380737304688 0 6781.1533
683.8417358398438 0 1616.179
687.342529296875 0 1104.6307
689.3569946289062 0 2655.033 y 8
690.3600463867188 0 955.96954
711.4213256835938 0 634.77747
715.8463745117188 0 1406.1765 w 1
716.3484497070312 0 1804.5525
717.385009765625 0 721.56323
718.3954467773438 0 2112.2927
719.4008178710938 0 973.27313
729.3632202148438 0 1303.7963 y Water loss 1
729.8737182617188 0 590.68475 y Ammonia loss 1
730.3648681640625 0 1044.7446 z 1
738.376220703125 0 5609.2515 y 1
738.8779296875 0 3157.7275
739.3786010742188 0 1355.6895
743.8700561523438 0 802.22565
744.3720092773438 0 1369.2833
745.3755493164062 0 815.4612
750.8982543945312 0 1455.1422
757.401611328125 0 924.94104
758.3883666992188 0 1521.3817
758.8839721679688 0 2609.418
759.3816528320312 0 2702.7007
759.88671875 0 965.65314
760.3881225585938 0 1147.2856
761.4044799804688 0 33661.836 c 6
762.4075317382812 0 12499.369
762.4805297851562 0 1079.9631
763.4105834960938 0 3276.6611
764.4103393554688 0 1814.9143
764.9105224609375 0 4593.75
765.382568359375 0 57919.26
765.8837280273438 0 43889.164
766.3856811523438 0 22239.47
766.88671875 0 5666.7817
767.3875732421875 0 1783.9856
771.9075317382812 0 737.31537
772.4025268554688 0 1455.8033
772.9131469726562 0 1220.8687
773.4094848632812 0 2264.3215
773.908203125 0 17673.48
774.4088134765625 0 12838.863
774.91015625 0 6029.891
775.4083862304688 0 1853.5311
778.40771484375 0 1490.1497
779.3978271484375 0 1261.4927
785.9052734375 0 1199.9564
786.4135131835938 0 3472.2004
786.909912109375 0 55139.92
787.4109497070312 0 45128
787.9118041992188 0 22379.988
788.4127807617188 0 8785.422
788.9116821289062 0 1517.6078
793.4105834960938 0 903.84753
793.9136962890625 0 920.4067
794.4136962890625 0 3812.957
794.9180908203125 0 37868.87
795.4203491210938 0 48007.19
795.9220581054688 0 29417.453
796.4229736328125 0 12151.131
796.92431640625 0 2146.9604
797.4182739257812 0 1593.4768
811.43017578125 0 1289.3575 z Water loss 7
829.4404296875 0 25483.062 z 7
830.4437866210938 0 13938.893
831.4473266601562 0 5116.508
832.451904296875 0 727.57196
841.5297241210938 0 919.5935
844.4520874023438 0 1440.416
845.458740234375 0 2722.8303 y 7
846.4630126953125 0 1615.4132
847.4681396484375 0 830.9134
873.4888305664062 0 1591.6232
899.4700317382812 0 667.00146 w 6
900.4766845703125 0 3971.0474 c Ammonia loss 7
901.4816284179688 0 1704.6815
944.466796875 0 9109.083 z 6
945.4717407226562 0 7662.801
946.474365234375 0 2888.1548
955.53466796875 0 870.65076
956.547607421875 0 1007.73486
957.4279174804688 0 749.35974
960.49169921875 0 927.0749 y 6
970.54443359375 0 2018.2913
973.4531860351562 0 780.50275
1013.5839233398438 0 1556.3998
1014.5584716796875 0 23879.06 c 8
1015.5611572265625 0 11366.394
1016.5639038085938 0 3944.824
1031.4998779296875 0 2982.2773 z 5
1032.506591796875 0 5171.182
1033.5084228515625 0 2708.5417
1034.5162353515625 0 574.766
1039.5621337890625 0 1194.4796
1040.566650390625 0 1105.1832
1041.5721435546875 0 713.75397
1044.4808349609375 0 729.54144
1045.4970703125 0 2036.1965
1046.4931640625 0 1136.1064
1047.515869140625 0 2098.6387 y 5
1057.4608154296875 0 701.0066
1057.5772705078125 0 13672.997
1058.580078125 0 10056.594
1059.58544921875 0 3813.7656
1060.59033203125 0 1037.8503
1062.8621826171875 0 635.4369
1100.5782470703125 0 1090.4381
1101.5902099609375 0 30238.643 c 9
1102.5931396484375 0 18117.27
1103.5941162109375 0 7137.156
1104.591064453125 0 1502.7876
1114.59619140625 0 5295.9233
1115.6016845703125 0 2931.023
1116.60107421875 0 1119.8275
1130.56640625 0 4830.842 z 4
1131.57470703125 0 11666.481
1132.5782470703125 0 5598.72
1133.582275390625 0 2277.6387
1146.5880126953125 0 1757.8322 y 4
1147.590576171875 0 1085.5504
1158.61083984375 0 17483.732 c 10
1159.615234375 0 11439.918
1160.6158447265625 0 5616.9565
1161.619140625 0 1017.3306
1213.664306640625 0 3273.5942
1214.6685791015625 0 2314.664
1215.6683349609375 0 1537.779
1257.6783447265625 0 16818.848 c 11
1258.68017578125 0 12540.559
1259.6744384765625 0 4666.13
1260.61767578125 0 16983.709
1261.6197509765625 0 10714.249
1262.6187744140625 0 4452.8057
1263.6265869140625 0 1158.6244
1275.6279296875 0 6279.217 y 3
1276.630859375 0 4404.794
1277.6358642578125 0 1461.9125
1282.6707763671875 0 807.937
1284.6925048828125 0 710.7085
1300.6949462890625 0 2471.635
1301.7003173828125 0 2173.6248
1302.69482421875 0 952.0362
1304.6165771484375 0 1077.0898
1330.640869140625 0 1167.9268
1331.62841796875 0 1038.9122
1344.7103271484375 0 33498.637 c 12
1345.7135009765625 0 22384.77
1346.7066650390625 0 6894.3105
1347.6513671875 0 14131.217
1348.6522216796875 0 10465.155
1349.654541015625 0 4808.6484
1350.6551513671875 0 1242.126
1364.6588134765625 0 664.0042
1387.730224609375 0 2457.2036
1388.7337646484375 0 2413.672
1389.73779296875 0 1326.8109
1430.7247314453125 0 1235.5859
1431.7423095703125 0 46435.84 c 13
1432.7451171875 0 35341.85
1432.95654296875 0 1066.8146
1433.747802734375 0 16127.919
1434.75048828125 0 3779.5508
1435.75390625 0 1346.7804
1460.7274169921875 0 2717.5063
1461.73486328125 0 1810.017
1473.74462890625 0 750.23267
1488.73779296875 0 3833.1555
1489.7423095703125 0 2861.8813
1490.7576904296875 0 2323.2244
1512.791748046875 0 1109.9922
1513.797119140625 0 1799.0153
1514.787109375 0 884.72455
1515.79443359375 0 706.595
1516.7935791015625 0 1048.167
1529.7926025390625 0 1741.2571
1530.794921875 0 7570.6377
1531.7926025390625 0 11509.462
1532.7957763671875 0 7940.677
1533.795166015625 0 3424.4504
1534.815185546875 0 884.56335
1544.833984375 0 1954.2345
1545.8282470703125 0 4165.8047
1546.823486328125 0 5642.6436
1547.820556640625 0 4880.2954
1548.8162841796875 0 3041.8628
1549.824951171875 0 1564.098
1550.8184814453125 0 834.4211
1555.804443359375 0 2108.4392
1556.794677734375 0 4148.341
1557.7958984375 0 3869.8103
1558.7833251953125 0 1186.1329
1561.8397216796875 0 1663.5752
1562.84619140625 0 8446.742
1563.8470458984375 0 5621.026
1564.849609375 0 3104.0215
1565.8414306640625 0 1200.7899
1571.820068359375 0 853.10895
1572.8128662109375 0 8733.421
1573.81591796875 0 66920.59
1574.8187255859375 0 49445.848
1575.8203125 0 24059.422
1576.82080078125 0 7945.389
1577.8304443359375 0 1562.5739
1587.7996826171875 0 1085.4922
1588.8238525390625 0 5714.461
1589.832763671875 0 18086.713
1590.8399658203125 0 61293.316
1591.84375 0 48968.67
1592.0914306640625 0 1069.6271
1592.8465576171875 0 20954.414
1593.8458251953125 0 6152.645
1594.84814453125 0 1630.4152
2244.36474609375 0 782.9357
2355.304931640625 0 815.86743
2858.462890625 0 726.0202
3072.677490234375 0 719.7795
3189.50341796875 0 794.44495

Spectrum Details

|  |  |
| --- | --- |
| Matched peaks? Matched peaks The total absolute number of peaks matched. Additionally in brackets the total fraction of peaks matched and the total number of peaks is shown. | 69 (18.35% of 376) |
| FDR? FDR The false discovery rate estimated for this peptide. It is calculated by matching all theoretical fragments with a non-integer shift with the raw peaks for this spectrum. This is done with 40 different shifts. The resulting percentage is the average number of annotated peaks over the number of annotated peaks with the correct spectrum. | 0.79% |
| Satellite FDR? Satellite FDR See the FDR for details on its calculation. This satellite ion specific FDR only contains the satellite ions (d/w) for I/L/J positions. | 2.38% |
| PSM Score? PSM Score The PSM Score as given by Hecklib to this annotated spectrum. It is shown with three significant figures. | 476 |

#### Spectrum 3970? Spectrum 3970 The raw spectrum of this peptide as annotated by Hecklib. The fragments are coloured according to ion type (see legend). Any peaks with a star '\*' as text can be hovered over to see the full details, first the ion type second the mass shift type. By hovering over the amino acids in the peptide or ions in the legend the corresponding peaks are highlighted. By toggling the 'Unassigned' label you can turn the background (unassigned) peaks on or off in the plot. By updating the slider in the Ion legend you can update the spectrum to only show the top X% of the peaks with labels. The top X% means any peak that is within X% of the highest intensity. By dragging in the spectrum you can zoom in to a specific part of the spectrum and use 'Zoom Out' to get back to the original zoom level. The annotation of the spectrum is based on the given sequence in the peptides file and is done with different software so inconsistencies are likely. The peaks are annotated based on the given sequence, with 20 ppm tolerance.

Copy Data

##### Spectrum 3970 (TSV)

###### Preview

```
Loading example...
```

*Click on the button to copy the data to your clipboard.*

Mz MinMz MaxIntensity Max

WidthHeightPeptide font sizePeptide stroke widthSpectrum font sizeSpectrum stroke widthCompact peptide

Ion legend

wxyz

abcd

OtherUnassignedIon^Charge^_Position_Show for top:%

JJSEVSDRPSGVSSR

03.44e+46.88e+41.03e+51.38e+5

Zoom Out

y^+1^_1_y^+1^_1_w^+1^_2_y^+1^_2_z^+1^_2_y^+1^_2_w^+1^_3_c^+1^_3_y^+1^_3_z^+1^_3_y^+1^_3_z^+2^_8_w^+1^_4_y^+2^_8_y^+1^_4_y^+1^_4_y^+3^_13_c^+2^_8_y^+3^_13_c^+1^_4_z^+1^_5_c^+3^_14_z^+2^_9_y^+2^_9_z^+3^_14_z^+1^_5_y^+1^_5_y^+2^_10_y^+2^_10_z^+2^_10_y^+2^_10_z^+1^_6_w^+1^_6_c^+1^_5_z^+2^_11_y^+2^_11_y^+1^_6_y^+1^_6_z^+1^_6_y^+1^_6_w^+2^_12_c^+2^_12_y^+2^_12_z^+2^_12_y^+2^_12_c^+1^_6_z^+2^_13_y^+1^_7_y^+2^_13_y^+2^_13_z^+2^_13_y^+2^_13_y^+1^_7_w^+2^_14_w^+2^_14_y^+2^_14_z^+2^_14_y^+2^_14_w^+1^_8_c^+1^_7_z^+1^_8_y^+1^_8_z^+1^_8_y^+1^_8_c^+1^_8_z^+1^_9_z^+1^_9_y^+1^_9_c^+1^_9_z^+1^_10_y^+1^_10_c^+1^_10_y^+1^_11_z^+1^_11_y^+1^_11_c^+1^_11_c^+1^_12_y^+1^_12_w^+1^_13_c^+1^_13_y^+1^_13_c^+1^_14_z^+1^_14_y^+1^_14_

0558111616742232

Fragment Matches Table

Show background peaks

| Position | Ion type | Intensity | mz Theoretical | mz Error (Th) | mz Error (ppm) | Charge | Series Number |
| --- | --- | --- | --- | --- | --- | --- | --- |
| - | - | 367.2 | 122.9 | - | - | 0 | - |
| - | - | 378.4 | 124.9 | - | - | 0 | - |
| - | - | 2032 | 125.1 | - | - | 0 | - |
| - | - | 3691 | 126.1 | - | - | 0 | - |
| - | - | 363.9 | 126.1 | - | - | 0 | - |
| - | - | 1113 | 128.1 | - | - | 0 | - |
| - | - | 476.9 | 128.3 | - | - | 0 | - |
| - | - | 543.3 | 130.1 | - | - | 0 | - |
| - | - | 502.4 | 130.1 | - | - | 0 | - |
| - | - | 1418 | 131.1 | - | - | 0 | - |
| - | - | 415 | 137.8 | - | - | 0 | - |
| - | - | 422.6 | 143 | - | - | 0 | - |
| - | - | 489.9 | 145.1 | - | - | 0 | - |
| - | - | 891.5 | 149 | - | - | 0 | - |
| 15 | y | 804.2 | 158.1 | 0.0005077 | 3.211 | +1 | 1 |
| - | - | 441.5 | 162.9 | - | - | 0 | - |
| - | - | 4334 | 168.1 | - | - | 0 | - |
| - | - | 1039 | 173.4 | - | - | 0 | - |
| 15 | y | 6243 | 175.1 | 0.0003258 | 1.86 | +1 | 1 |
| - | - | 802.3 | 176.1 | - | - | 0 | - |
| - | - | 548.5 | 176.2 | - | - | 0 | - |
| - | - | 1.787E+04 | 185.1 | - | - | 0 | - |
| - | - | 1272 | 186.1 | - | - | 0 | - |
| - | - | 4.817E+04 | 199.2 | - | - | 0 | - |
| - | - | 2530 | 200.1 | - | - | 0 | - |
| - | - | 5406 | 200.2 | - | - | 0 | - |
| - | - | 867 | 201.1 | - | - | 0 | - |
| - | - | 4004 | 217.1 | - | - | 0 | - |
| - | - | 625.8 | 226.1 | - | - | 0 | - |
| - | - | 1107 | 226.2 | - | - | 0 | - |
| - | - | 2.238E+04 | 227.2 | - | - | 0 | - |
| - | - | 3167 | 228.2 | - | - | 0 | - |
| 14 | w | 3411 | 229.1 | 0.0004728 | 2.063 | +1 | 2 |
| - | - | 1514 | 233.2 | - | - | 0 | - |
| - | - | 551 | 234.1 | - | - | 0 | - |
| - | - | 1502 | 244.1 | - | - | 0 | - |
| 14 | y | 3.006E+04 | 245.1 | 0.0004617 | 1.884 | +1 | 2 |
| 14 | z | 3997 | 246.1 | 0.001184 | 4.808 | +1 | 2 |
| - | - | 617.3 | 249.2 | - | - | 0 | - |
| - | - | 1252 | 255.1 | - | - | 0 | - |
| - | - | 784.2 | 260.1 | - | - | 0 | - |
| - | - | 1140 | 261.2 | - | - | 0 | - |
| 14 | y | 3743 | 262.2 | 0.0004476 | 1.708 | +1 | 2 |
| - | - | 816.4 | 263.2 | - | - | 0 | - |
| - | - | 5148 | 270.1 | - | - | 0 | - |
| - | - | 1688 | 272.2 | - | - | 0 | - |
| - | - | 2716 | 304.2 | - | - | 0 | - |
| - | - | 3004 | 314.2 | - | - | 0 | - |
| 13 | w | 1.54E+04 | 316.2 | 0.0005641 | 1.784 | +1 | 3 |
| - | - | 2428 | 317.2 | - | - | 0 | - |
| - | - | 837.3 | 318.2 | - | - | 0 | - |
| - | - | 673.2 | 330.2 | - | - | 0 | - |
| - | - | 4203 | 331.1 | - | - | 0 | - |
| 3 | c | 642.6 | 331.2 | 0.0008508 | 2.569 | +1 | 3 |
| 13 | y | 4981 | 332.2 | 0.0004005 | 1.206 | +1 | 3 |
| 13 | z | 1.239E+04 | 333.2 | 0.0006321 | 1.897 | +1 | 3 |
| - | - | 2127 | 334.2 | - | - | 0 | - |
| - | - | 793 | 340.1 | - | - | 0 | - |
| - | - | 1157 | 347.2 | - | - | 0 | - |
| 13 | y | 4087 | 349.2 | 0.0003101 | 0.8881 | +1 | 3 |
| - | - | 639.3 | 350.2 | - | - | 0 | - |
| - | - | 8212 | 357.2 | - | - | 0 | - |
| - | - | 675.3 | 357.2 | - | - | 0 | - |
| - | - | 729.7 | 375.2 | - | - | 0 | - |
| - | - | 886 | 389.7 | - | - | 0 | - |
| - | - | 3538 | 391.7 | - | - | 0 | - |
| - | - | 1615 | 392.2 | - | - | 0 | - |
| - | - | 1238 | 398.7 | - | - | 0 | - |
| - | - | 641.5 | 399.7 | - | - | 0 | - |
| 8 | z | 4726 | 415.2 | 0.003288 | 7.919 | +2 | 8 |
| - | - | 1812 | 415.7 | - | - | 0 | - |
| 12 | w | 1.279E+04 | 417.2 | 0.0008898 | 2.133 | +1 | 4 |
| - | - | 2492 | 418.2 | - | - | 0 | - |
| - | - | 635.1 | 422.7 | - | - | 0 | - |
| - | - | 699.8 | 423.2 | - | - | 0 | - |
| 8 | y | 3445 | 423.2 | 0.001067 | 2.521 | +2 | 8 |
| - | - | 1554 | 423.7 | - | - | 0 | - |
| - | - | 1741 | 425.2 | - | - | 0 | - |
| - | - | 622.4 | 426.2 | - | - | 0 | - |
| 12 | y | 725.8 | 431.2 | 0.0009053 | 2.099 | +1 | 4 |
| - | - | 1627 | 441.2 | - | - | 0 | - |
| - | - | 2703 | 443.3 | - | - | 0 | - |
| - | - | 696.1 | 443.3 | - | - | 0 | - |
| - | - | 780.5 | 444.3 | - | - | 0 | - |
| - | - | 901.6 | 446.2 | - | - | 0 | - |
| - | - | 851.2 | 447.2 | - | - | 0 | - |
| - | - | 1048 | 447.2 | - | - | 0 | - |
| - | - | 414.8 | 448.2 | - | - | 0 | - |
| 12 | y | 1.308E+04 | 448.3 | 0.000988 | 2.204 | +1 | 4 |
| 3 | y | 793.1 | 449.2 | 0.008831 | 19.66 | +3 | 13 |
| - | - | 2602 | 449.3 | - | - | 0 | - |
| 8 | c | 970.9 | 450.7 | 0.00236 | 5.235 | +2 | 8 |
| - | - | 682.8 | 451.2 | - | - | 0 | - |
| 3 | y | 1235 | 454.9 | 0.001588 | 3.492 | +3 | 13 |
| - | - | 750.8 | 455.3 | - | - | 0 | - |
| - | - | 553.3 | 455.6 | - | - | 0 | - |
| 4 | c | 8320 | 460.3 | 0.001104 | 2.399 | +1 | 4 |
| - | - | 1619 | 461.3 | - | - | 0 | - |
| 11 | z | 758.8 | 471.2 | 0.003106 | 6.592 | +1 | 5 |
| 14 | c | 2395 | 472.2 | 0.007744 | 16.4 | +3 | 14 |
| 7 | z | 2182 | 472.7 | 0.003 | 6.346 | +2 | 9 |
| - | - | 1549 | 473.2 | - | - | 0 | - |
| - | - | 589.7 | 473.7 | - | - | 0 | - |
| - | - | 2678 | 474.2 | - | - | 0 | - |
| - | - | 896.2 | 475.2 | - | - | 0 | - |
| 7 | y | 2238 | 480.7 | 0.001786 | 3.716 | +2 | 9 |
| 2 | z | 1070 | 481.2 | 0.006564 | 13.64 | +3 | 14 |
| 11 | z | 2972 | 489.3 | 0.0008118 | 1.659 | +1 | 5 |
| - | - | 2530 | 490.3 | - | - | 0 | - |
| - | - | 1441 | 495.9 | - | - | 0 | - |
| - | - | 571.7 | 498.3 | - | - | 0 | - |
| - | - | 2495 | 499.3 | - | - | 0 | - |
| 11 | y | 3.927E+04 | 505.3 | 0.001009 | 1.996 | +1 | 5 |
| - | - | 1.016E+04 | 506.3 | - | - | 0 | - |
| - | - | 1057 | 507.3 | - | - | 0 | - |
| - | - | 1215 | 512.2 | - | - | 0 | - |
| - | - | 917.7 | 514.3 | - | - | 0 | - |
| 6 | y | 1745 | 515.3 | 0.008762 | 17.01 | +2 | 10 |
| - | - | 2210 | 515.3 | - | - | 0 | - |
| 6 | y | 2374 | 515.7 | 0.001122 | 2.175 | +2 | 10 |
| 6 | z | 3927 | 516.3 | 0.0006273 | 1.215 | +2 | 10 |
| - | - | 1616 | 516.3 | - | - | 0 | - |
| - | - | 5148 | 516.8 | - | - | 0 | - |
| - | - | 2041 | 517.3 | - | - | 0 | - |
| - | - | 767.8 | 517.8 | - | - | 0 | - |
| - | - | 644.2 | 523.8 | - | - | 0 | - |
| 6 | y | 1.347E+04 | 524.3 | 0.00335 | 6.39 | +2 | 10 |
| - | - | 4643 | 524.6 | - | - | 0 | - |
| - | - | 6827 | 524.8 | - | - | 0 | - |
| - | - | 2029 | 524.9 | - | - | 0 | - |
| - | - | 1775 | 525.3 | - | - | 0 | - |
| - | - | 1077 | 525.8 | - | - | 0 | - |
| - | - | 1076 | 529.3 | - | - | 0 | - |
| - | - | 1239 | 529.8 | - | - | 0 | - |
| - | - | 1634 | 530.2 | - | - | 0 | - |
| - | - | 3509 | 530.3 | - | - | 0 | - |
| - | - | 3525 | 530.3 | - | - | 0 | - |
| - | - | 3114 | 530.6 | - | - | 0 | - |
| - | - | 1813 | 530.9 | - | - | 0 | - |
| - | - | 764.3 | 531.3 | - | - | 0 | - |
| - | - | 1696 | 531.3 | - | - | 0 | - |
| - | - | 853.1 | 537.8 | - | - | 0 | - |
| - | - | 1378 | 538.3 | - | - | 0 | - |
| - | - | 711.6 | 538.8 | - | - | 0 | - |
| - | - | 1029 | 541.3 | - | - | 0 | - |
| - | - | 4076 | 542.3 | - | - | 0 | - |
| - | - | 1412 | 543.3 | - | - | 0 | - |
| 10 | z | 1636 | 558.3 | 0.0007409 | 1.327 | +1 | 6 |
| - | - | 1727 | 558.3 | - | - | 0 | - |
| 10 | w | 2004 | 559.3 | 0.0014 | 2.503 | +1 | 6 |
| 5 | c | 7128 | 559.3 | 0.0007142 | 1.277 | +1 | 5 |
| - | - | 624.4 | 560.3 | - | - | 0 | - |
| - | - | 1691 | 560.3 | - | - | 0 | - |
| 5 | z | 2.885E+04 | 565.8 | 0.0009662 | 1.708 | +2 | 11 |
| - | - | 1.905E+04 | 566.3 | - | - | 0 | - |
| - | - | 5939 | 566.8 | - | - | 0 | - |
| - | - | 2077 | 567.3 | - | - | 0 | - |
| - | - | 755 | 572.8 | - | - | 0 | - |
| 5 | y | 1.496E+04 | 573.8 | 0.0007595 | 1.324 | +2 | 11 |
| 10 | y | 7885 | 574.3 | 0.004417 | 7.691 | +1 | 6 |
| - | - | 3580 | 574.8 | - | - | 0 | - |
| 10 | y | 1698 | 575.3 | 0.004105 | 7.135 | +1 | 6 |
| 10 | z | 6510 | 576.3 | 0.0004301 | 0.7464 | +1 | 6 |
| - | - | 1459 | 577.3 | - | - | 0 | - |
| - | - | 1415 | 586.3 | - | - | 0 | - |
| - | - | 742.5 | 587.8 | - | - | 0 | - |
| 10 | y | 1816 | 592.3 | 0.0002607 | 0.4402 | +1 | 6 |
| - | - | 3079 | 600.3 | - | - | 0 | - |
| 4 | w | 1.433E+04 | 600.8 | 0.0006651 | 1.107 | +2 | 12 |
| - | - | 8022 | 601.3 | - | - | 0 | - |
| - | - | 2464 | 601.8 | - | - | 0 | - |
| - | - | 741.6 | 608.3 | - | - | 0 | - |
| - | - | 961.9 | 629.3 | - | - | 0 | - |
| 12 | c | 2184 | 629.3 | 0.006833 | 10.86 | +2 | 12 |
| 4 | y | 934.1 | 629.8 | 0.0009772 | 1.552 | +2 | 12 |
| 4 | z | 3.162E+04 | 630.3 | 0.001032 | 1.637 | +2 | 12 |
| - | - | 1.986E+04 | 630.8 | - | - | 0 | - |
| - | - | 8923 | 631.3 | - | - | 0 | - |
| - | - | 664.6 | 631.4 | - | - | 0 | - |
| - | - | 1717 | 631.8 | - | - | 0 | - |
| - | - | 9923 | 637.8 | - | - | 0 | - |
| - | - | 768 | 637.9 | - | - | 0 | - |
| 4 | y | 3.314E+04 | 638.3 | 0.0002759 | 0.4323 | +2 | 12 |
| - | - | 1.95E+04 | 638.8 | - | - | 0 | - |
| - | - | 7525 | 639.3 | - | - | 0 | - |
| - | - | 1571 | 639.8 | - | - | 0 | - |
| - | - | 3107 | 644.3 | - | - | 0 | - |
| - | - | 1728 | 644.8 | - | - | 0 | - |
| - | - | 1028 | 645.4 | - | - | 0 | - |
| 6 | c | 2.684E+04 | 646.4 | 0.0009734 | 1.506 | +1 | 6 |
| - | - | 8369 | 647.4 | - | - | 0 | - |
| - | - | 1876 | 648.4 | - | - | 0 | - |
| - | - | 1213 | 651.8 | - | - | 0 | - |
| - | - | 778.5 | 652.3 | - | - | 0 | - |
| 3 | z | 1389 | 665.3 | 0.01087 | 16.33 | +2 | 13 |
| - | - | 964.6 | 665.8 | - | - | 0 | - |
| - | - | 1162 | 666.3 | - | - | 0 | - |
| 9 | y | 972.2 | 672.3 | 0.006118 | 9.099 | +1 | 7 |
| 3 | y | 3058 | 672.8 | 0.001972 | 2.931 | +2 | 13 |
| 3 | y | 2641 | 673.3 | 0.006814 | 10.12 | +2 | 13 |
| 3 | z | 2.079E+04 | 673.8 | 0.001375 | 2.041 | +2 | 13 |
| - | - | 2.488E+04 | 674.3 | - | - | 0 | - |
| - | - | 1.607E+04 | 674.8 | - | - | 0 | - |
| - | - | 4424 | 675.3 | - | - | 0 | - |
| - | - | 1461 | 675.8 | - | - | 0 | - |
| - | - | 717.7 | 680.3 | - | - | 0 | - |
| - | - | 3925 | 680.8 | - | - | 0 | - |
| - | - | 4295 | 681.3 | - | - | 0 | - |
| 3 | y | 1.363E+05 | 681.8 | 0.001046 | 1.535 | +2 | 13 |
| - | - | 9.179E+04 | 682.3 | - | - | 0 | - |
| - | - | 3.624E+04 | 682.8 | - | - | 0 | - |
| - | - | 1.115E+04 | 683.3 | - | - | 0 | - |
| - | - | 1711 | 683.8 | - | - | 0 | - |
| - | - | 923.3 | 685.4 | - | - | 0 | - |
| - | - | 660.9 | 685.8 | - | - | 0 | - |
| - | - | 1688 | 687.3 | - | - | 0 | - |
| 9 | y | 3828 | 689.4 | 0.001025 | 1.486 | +1 | 7 |
| - | - | 1122 | 690.4 | - | - | 0 | - |
| - | - | 1161 | 711.4 | - | - | 0 | - |
| - | - | 1359 | 713.4 | - | - | 0 | - |
| - | - | 881.8 | 714.4 | - | - | 0 | - |
| 2 | w | 3572 | 715.8 | 0.001367 | 1.909 | +2 | 14 |
| - | - | 2192 | 716.3 | - | - | 0 | - |
| - | - | 793.7 | 716.9 | - | - | 0 | - |
| - | - | 1904 | 717.4 | - | - | 0 | - |
| - | - | 4221 | 718.4 | - | - | 0 | - |
| 2 | w | 797.8 | 722.9 | 0.002941 | 4.069 | +2 | 14 |
| 2 | y | 1398 | 729.4 | 0.002256 | 3.093 | +2 | 14 |
| 2 | z | 2750 | 730.4 | 0.0002515 | 0.3443 | +2 | 14 |
| - | - | 1359 | 730.9 | - | - | 0 | - |
| - | - | 697 | 737.4 | - | - | 0 | - |
| 2 | y | 6598 | 738.4 | 0.0008845 | 1.198 | +2 | 14 |
| - | - | 5649 | 738.9 | - | - | 0 | - |
| - | - | 2784 | 739.4 | - | - | 0 | - |
| - | - | 699.2 | 742.4 | - | - | 0 | - |
| 8 | w | 2944 | 743.4 | 0.005353 | 7.2 | +1 | 8 |
| - | - | 3189 | 744.4 | - | - | 0 | - |
| - | - | 2088 | 750.9 | - | - | 0 | - |
| - | - | 2005 | 751.4 | - | - | 0 | - |
| - | - | 1479 | 751.9 | - | - | 0 | - |
| - | - | 1211 | 756.4 | - | - | 0 | - |
| - | - | 1581 | 757.9 | - | - | 0 | - |
| - | - | 4718 | 758.4 | - | - | 0 | - |
| - | - | 4239 | 758.9 | - | - | 0 | - |
| - | - | 4205 | 759.4 | - | - | 0 | - |
| - | - | 1207 | 759.9 | - | - | 0 | - |
| - | - | 1430 | 760.4 | - | - | 0 | - |
| 7 | c | 5.417E+04 | 761.4 | 0.001252 | 1.644 | +1 | 7 |
| - | - | 2.068E+04 | 762.4 | - | - | 0 | - |
| - | - | 5320 | 763.4 | - | - | 0 | - |
| - | - | 2988 | 764.4 | - | - | 0 | - |
| - | - | 7001 | 764.9 | - | - | 0 | - |
| - | - | 9.289E+04 | 765.4 | - | - | 0 | - |
| - | - | 7.62E+04 | 765.9 | - | - | 0 | - |
| - | - | 3.547E+04 | 766.4 | - | - | 0 | - |
| - | - | 1.063E+04 | 766.9 | - | - | 0 | - |
| - | - | 1544 | 767.4 | - | - | 0 | - |
| - | - | 887.9 | 771.9 | - | - | 0 | - |
| - | - | 4191 | 772.4 | - | - | 0 | - |
| - | - | 3604 | 772.9 | - | - | 0 | - |
| - | - | 2112 | 773.4 | - | - | 0 | - |
| - | - | 2.572E+04 | 773.9 | - | - | 0 | - |
| - | - | 2.019E+04 | 774.4 | - | - | 0 | - |
| - | - | 9759 | 774.9 | - | - | 0 | - |
| - | - | 2695 | 775.4 | - | - | 0 | - |
| - | - | 1611 | 777.9 | - | - | 0 | - |
| - | - | 3188 | 778.4 | - | - | 0 | - |
| - | - | 1606 | 778.9 | - | - | 0 | - |
| - | - | 1581 | 779.4 | - | - | 0 | - |
| - | - | 1082 | 779.9 | - | - | 0 | - |
| - | - | 1377 | 780.4 | - | - | 0 | - |
| - | - | 717 | 781.4 | - | - | 0 | - |
| - | - | 2636 | 782.4 | - | - | 0 | - |
| - | - | 2494 | 783.4 | - | - | 0 | - |
| - | - | 3359 | 785.4 | - | - | 0 | - |
| - | - | 4521 | 785.9 | - | - | 0 | - |
| - | - | 9958 | 786.4 | - | - | 0 | - |
| - | - | 9.998E+04 | 786.9 | - | - | 0 | - |
| - | - | 8.002E+04 | 787.4 | - | - | 0 | - |
| - | - | 4.19E+04 | 787.9 | - | - | 0 | - |
| - | - | 1.185E+04 | 788.4 | - | - | 0 | - |
| - | - | 3711 | 788.9 | - | - | 0 | - |
| - | - | 949.3 | 792.9 | - | - | 0 | - |
| - | - | 4915 | 793.4 | - | - | 0 | - |
| - | - | 1.882E+04 | 793.9 | - | - | 0 | - |
| - | - | 1.841E+04 | 794.4 | - | - | 0 | - |
| - | - | 7.009E+04 | 794.9 | - | - | 0 | - |
| - | - | 7.765E+04 | 795.4 | - | - | 0 | - |
| - | - | 4.835E+04 | 795.9 | - | - | 0 | - |
| - | - | 1.932E+04 | 796.4 | - | - | 0 | - |
| - | - | 4726 | 796.9 | - | - | 0 | - |
| - | - | 2165 | 797.4 | - | - | 0 | - |
| - | - | 733.5 | 798.4 | - | - | 0 | - |
| - | - | 4708 | 803.4 | - | - | 0 | - |
| - | - | 2351 | 804.4 | - | - | 0 | - |
| 8 | z | 2728 | 811.4 | 0.0007292 | 0.8986 | +1 | 8 |
| 8 | y | 955.7 | 828.4 | 0.0004171 | 0.5035 | +1 | 8 |
| 8 | z | 4.655E+04 | 829.4 | 0.0008521 | 1.027 | +1 | 8 |
| - | - | 2.162E+04 | 830.4 | - | - | 0 | - |
| - | - | 7293 | 831.4 | - | - | 0 | - |
| - | - | 878.4 | 832.5 | - | - | 0 | - |
| - | - | 1216 | 839.5 | - | - | 0 | - |
| - | - | 925 | 840.5 | - | - | 0 | - |
| - | - | 2086 | 841.5 | - | - | 0 | - |
| - | - | 1161 | 842.5 | - | - | 0 | - |
| - | - | 2022 | 844.4 | - | - | 0 | - |
| - | - | 575.7 | 845.4 | - | - | 0 | - |
| 8 | y | 7001 | 845.5 | 0.001171 | 1.385 | +1 | 8 |
| - | - | 1936 | 846.5 | - | - | 0 | - |
| - | - | 1273 | 847.5 | - | - | 0 | - |
| - | - | 772.3 | 857.4 | - | - | 0 | - |
| - | - | 682.7 | 861.4 | - | - | 0 | - |
| - | - | 2251 | 873.5 | - | - | 0 | - |
| - | - | 895.2 | 874.5 | - | - | 0 | - |
| 8 | c | 6682 | 900.5 | 0.0001764 | 0.1959 | +1 | 8 |
| - | - | 2748 | 901.5 | - | - | 0 | - |
| - | - | 1391 | 902.5 | - | - | 0 | - |
| 7 | z | 2229 | 926.5 | 0.0006481 | 0.6995 | +1 | 9 |
| - | - | 1367 | 927.5 | - | - | 0 | - |
| - | - | 1097 | 928.5 | - | - | 0 | - |
| - | - | 1057 | 929.5 | - | - | 0 | - |
| 7 | z | 1.701E+04 | 944.5 | 0.001009 | 1.068 | +1 | 9 |
| - | - | 1.23E+04 | 945.5 | - | - | 0 | - |
| - | - | 4740 | 946.5 | - | - | 0 | - |
| - | - | 1177 | 947.5 | - | - | 0 | - |
| - | - | 1034 | 954.5 | - | - | 0 | - |
| - | - | 1712 | 955.5 | - | - | 0 | - |
| - | - | 2029 | 956.6 | - | - | 0 | - |
| - | - | 791.2 | 957.5 | - | - | 0 | - |
| - | - | 1038 | 959.5 | - | - | 0 | - |
| 7 | y | 2083 | 960.5 | 0.0007783 | 0.8103 | +1 | 9 |
| - | - | 3535 | 970.5 | - | - | 0 | - |
| - | - | 2106 | 971.5 | - | - | 0 | - |
| - | - | 958.8 | 999.6 | - | - | 0 | - |
| - | - | 725.4 | 1004 | - | - | 0 | - |
| - | - | 855.2 | 1006 | - | - | 0 | - |
| - | - | 755.4 | 1013 | - | - | 0 | - |
| - | - | 2527 | 1014 | - | - | 0 | - |
| 9 | c | 4.266E+04 | 1015 | 0.001613 | 1.59 | +1 | 9 |
| - | - | 2.141E+04 | 1016 | - | - | 0 | - |
| - | - | 6479 | 1017 | - | - | 0 | - |
| - | - | 1951 | 1018 | - | - | 0 | - |
| 6 | z | 6787 | 1031 | 0.001207 | 1.17 | +1 | 10 |
| - | - | 9130 | 1033 | - | - | 0 | - |
| - | - | 4314 | 1034 | - | - | 0 | - |
| - | - | 1182 | 1035 | - | - | 0 | - |
| - | - | 2598 | 1040 | - | - | 0 | - |
| - | - | 1587 | 1041 | - | - | 0 | - |
| - | - | 1628 | 1042 | - | - | 0 | - |
| - | - | 2791 | 1043 | - | - | 0 | - |
| - | - | 1008 | 1047 | - | - | 0 | - |
| 6 | y | 2707 | 1048 | 0.000305 | 0.2912 | +1 | 10 |
| - | - | 2150 | 1049 | - | - | 0 | - |
| - | - | 2.819E+04 | 1058 | - | - | 0 | - |
| - | - | 1.984E+04 | 1059 | - | - | 0 | - |
| - | - | 1.024E+04 | 1060 | - | - | 0 | - |
| - | - | 3899 | 1061 | - | - | 0 | - |
| - | - | 712.1 | 1062 | - | - | 0 | - |
| - | - | 2451 | 1101 | - | - | 0 | - |
| 10 | c | 5.598E+04 | 1102 | 0.001567 | 1.422 | +1 | 10 |
| - | - | 3.246E+04 | 1103 | - | - | 0 | - |
| - | - | 1.101E+04 | 1104 | - | - | 0 | - |
| - | - | 2061 | 1105 | - | - | 0 | - |
| - | - | 8608 | 1115 | - | - | 0 | - |
| - | - | 6665 | 1116 | - | - | 0 | - |
| - | - | 2321 | 1117 | - | - | 0 | - |
| - | - | 758.2 | 1118 | - | - | 0 | - |
| 5 | y | 623.6 | 1130 | 0.0007883 | 0.6979 | +1 | 11 |
| 5 | z | 8029 | 1131 | 0.0009082 | 0.8033 | +1 | 11 |
| - | - | 1.734E+04 | 1132 | - | - | 0 | - |
| - | - | 9509 | 1133 | - | - | 0 | - |
| - | - | 3107 | 1134 | - | - | 0 | - |
| 5 | y | 2748 | 1147 | 0.002814 | 2.454 | +1 | 11 |
| - | - | 1096 | 1148 | - | - | 0 | - |
| - | - | 1213 | 1149 | - | - | 0 | - |
| 11 | c | 3.243E+04 | 1159 | 0.0008552 | 0.7381 | +1 | 11 |
| - | - | 1.896E+04 | 1160 | - | - | 0 | - |
| - | - | 8091 | 1161 | - | - | 0 | - |
| - | - | 1494 | 1162 | - | - | 0 | - |
| - | - | 681.8 | 1204 | - | - | 0 | - |
| - | - | 6407 | 1214 | - | - | 0 | - |
| - | - | 4153 | 1215 | - | - | 0 | - |
| - | - | 2910 | 1216 | - | - | 0 | - |
| - | - | 1240 | 1217 | - | - | 0 | - |
| - | - | 775.8 | 1218 | - | - | 0 | - |
| - | - | 731.8 | 1248 | - | - | 0 | - |
| 12 | c | 3.113E+04 | 1258 | 0.0006786 | 0.5396 | +1 | 12 |
| - | - | 2.059E+04 | 1259 | - | - | 0 | - |
| - | - | 7752 | 1260 | - | - | 0 | - |
| - | - | 2.838E+04 | 1261 | - | - | 0 | - |
| - | - | 1.442E+04 | 1262 | - | - | 0 | - |
| - | - | 5681 | 1263 | - | - | 0 | - |
| - | - | 1282 | 1264 | - | - | 0 | - |
| - | - | 918.1 | 1275 | - | - | 0 | - |
| 4 | y | 1.037E+04 | 1276 | 0.0008387 | 0.6575 | +1 | 12 |
| - | - | 6159 | 1277 | - | - | 0 | - |
| - | - | 2458 | 1278 | - | - | 0 | - |
| - | - | 1384 | 1283 | - | - | 0 | - |
| - | - | 837 | 1285 | - | - | 0 | - |
| - | - | 5546 | 1301 | - | - | 0 | - |
| - | - | 3997 | 1302 | - | - | 0 | - |
| - | - | 2752 | 1303 | - | - | 0 | - |
| - | - | 1570 | 1305 | - | - | 0 | - |
| 3 | w | 2095 | 1330 | 0.01228 | 9.234 | +1 | 13 |
| - | - | 2601 | 1331 | - | - | 0 | - |
| - | - | 1594 | 1332 | - | - | 0 | - |
| - | - | 770.9 | 1344 | - | - | 0 | - |
| 13 | c | 5.226E+04 | 1345 | 0.0005881 | 0.4373 | +1 | 13 |
| - | - | 3.646E+04 | 1346 | - | - | 0 | - |
| - | - | 1.746E+04 | 1347 | - | - | 0 | - |
| - | - | 2.382E+04 | 1348 | - | - | 0 | - |
| - | - | 1.427E+04 | 1349 | - | - | 0 | - |
| - | - | 7120 | 1350 | - | - | 0 | - |
| - | - | 2383 | 1351 | - | - | 0 | - |
| 3 | y | 1538 | 1363 | 0.002227 | 1.635 | +1 | 13 |
| - | - | 2393 | 1364 | - | - | 0 | - |
| - | - | 5757 | 1388 | - | - | 0 | - |
| - | - | 4496 | 1389 | - | - | 0 | - |
| - | - | 1243 | 1390 | - | - | 0 | - |
| - | - | 938.4 | 1391 | - | - | 0 | - |
| - | - | 2031 | 1430 | - | - | 0 | - |
| - | - | 1601 | 1431 | - | - | 0 | - |
| 14 | c | 7.458E+04 | 1432 | 0.0006341 | 0.4429 | +1 | 14 |
| - | - | 6.148E+04 | 1433 | - | - | 0 | - |
| - | - | 2.668E+04 | 1434 | - | - | 0 | - |
| - | - | 7206 | 1435 | - | - | 0 | - |
| - | - | 1347 | 1436 | - | - | 0 | - |
| - | - | 755.5 | 1450 | - | - | 0 | - |
| 2 | z | 884.2 | 1460 | 0.005797 | 3.971 | +1 | 14 |
| - | - | 3801 | 1461 | - | - | 0 | - |
| - | - | 2944 | 1462 | - | - | 0 | - |
| - | - | 1371 | 1463 | - | - | 0 | - |
| - | - | 689.6 | 1474 | - | - | 0 | - |
| - | - | 695.2 | 1475 | - | - | 0 | - |
| 2 | y | 658.2 | 1476 | 0.0001344 | 0.09104 | +1 | 14 |
| - | - | 943.6 | 1488 | - | - | 0 | - |
| - | - | 6952 | 1489 | - | - | 0 | - |
| - | - | 6572 | 1490 | - | - | 0 | - |
| - | - | 3868 | 1491 | - | - | 0 | - |
| - | - | 1771 | 1492 | - | - | 0 | - |
| - | - | 1483 | 1513 | - | - | 0 | - |
| - | - | 1674 | 1514 | - | - | 0 | - |
| - | - | 2139 | 1515 | - | - | 0 | - |
| - | - | 987.1 | 1516 | - | - | 0 | - |
| - | - | 969.5 | 1517 | - | - | 0 | - |
| - | - | 1209 | 1529 | - | - | 0 | - |
| - | - | 2108 | 1530 | - | - | 0 | - |
| - | - | 1.614E+04 | 1531 | - | - | 0 | - |
| - | - | 2.058E+04 | 1532 | - | - | 0 | - |
| - | - | 1.327E+04 | 1533 | - | - | 0 | - |
| - | - | 4383 | 1534 | - | - | 0 | - |
| - | - | 971.8 | 1535 | - | - | 0 | - |
| - | - | 1651 | 1544 | - | - | 0 | - |
| - | - | 3727 | 1545 | - | - | 0 | - |
| - | - | 7159 | 1546 | - | - | 0 | - |
| - | - | 7140 | 1547 | - | - | 0 | - |
| - | - | 8737 | 1548 | - | - | 0 | - |
| - | - | 4784 | 1549 | - | - | 0 | - |
| - | - | 2463 | 1550 | - | - | 0 | - |
| - | - | 2076 | 1556 | - | - | 0 | - |
| - | - | 8358 | 1557 | - | - | 0 | - |
| - | - | 5227 | 1558 | - | - | 0 | - |
| - | - | 2767 | 1559 | - | - | 0 | - |
| - | - | 1793 | 1560 | - | - | 0 | - |
| - | - | 861.1 | 1561 | - | - | 0 | - |
| - | - | 3418 | 1562 | - | - | 0 | - |
| - | - | 1.182E+04 | 1563 | - | - | 0 | - |
| - | - | 8559 | 1564 | - | - | 0 | - |
| - | - | 5706 | 1565 | - | - | 0 | - |
| - | - | 1085 | 1566 | - | - | 0 | - |
| - | - | 1216 | 1570 | - | - | 0 | - |
| - | - | 2034 | 1571 | - | - | 0 | - |
| - | - | 4716 | 1572 | - | - | 0 | - |
| - | - | 1.767E+04 | 1573 | - | - | 0 | - |
| - | - | 1.023E+05 | 1574 | - | - | 0 | - |
| - | - | 7.936E+04 | 1575 | - | - | 0 | - |
| - | - | 3.934E+04 | 1576 | - | - | 0 | - |
| - | - | 1.322E+04 | 1577 | - | - | 0 | - |
| - | - | 3583 | 1578 | - | - | 0 | - |
| - | - | 4284 | 1587 | - | - | 0 | - |
| - | - | 7711 | 1588 | - | - | 0 | - |
| - | - | 1.396E+04 | 1589 | - | - | 0 | - |
| - | - | 1093 | 1590 | - | - | 0 | - |
| - | - | 3.473E+04 | 1590 | - | - | 0 | - |
| - | - | 1.022E+05 | 1591 | - | - | 0 | - |
| - | - | 7.681E+04 | 1592 | - | - | 0 | - |
| - | - | 3.865E+04 | 1593 | - | - | 0 | - |
| - | - | 1.042E+04 | 1594 | - | - | 0 | - |
| - | - | 2902 | 1595 | - | - | 0 | - |
| - | - | 635.5 | 2210 | - | - | 0 | - |

m/z Charge Intensity FragmentType MassShift Position
122.85774993896484 0 367.15152
124.91972351074219 0 378.3758
125.10771942138672 0 2031.951
126.09163665771484 0 3690.8892
126.10456848144531 0 363.92114
128.0821075439453 0 1113.4352
128.31515502929688 0 476.9033
130.06167602539062 0 543.29114
130.1215362548828 0 502.3937
131.12950134277344 0 1418.2523
137.79541015625 0 414.9647
143.02391052246094 0 422.63754
145.10855102539062 0 489.85794
148.95480346679688 0 891.5044
158.09291076660156 0 804.18274 y Ammonia loss 14
162.8594207763672 0 441.4897
168.1134796142578 0 4333.814
173.4387664794922 0 1039.0602
175.11927795410156 0 6243.0835 y 14
176.12295532226562 0 802.32227
176.15106201171875 0 548.5143
185.14007568359375 0 17871.49
186.1433868408203 0 1271.658
199.18084716796875 0 48174.547
200.12704467773438 0 2530.0261
200.18414306640625 0 5406.3604
201.1228790283203 0 866.9515
217.08221435546875 0 4003.7688
226.1300506591797 0 625.7795
226.16798400878906 0 1107.3647
227.17576599121094 0 22377.137
228.17919921875 0 3167.3428
229.12998962402344 0 3410.667 w 13
233.1651611328125 0 1514.1793
234.10911560058594 0 550.99786
244.11729431152344 0 1501.7122
245.12489318847656 0 30058.629 y Ammonia loss 13
246.13107299804688 0 3997.1902 z 13
249.16075134277344 0 617.28625
255.10948181152344 0 1251.8398
260.1359558105469 0 784.2109
261.1601867675781 0 1139.5377
262.15142822265625 0 3742.6897 y 13
263.15386962890625 0 816.36096
270.1201171875 0 5148.4893
272.1716003417969 0 1687.6649
304.1509704589844 0 2715.7546
314.2083435058594 0 3004.1484
316.162109375 0 15404.202 w 12
317.1602783203125 0 2427.8782
318.166015625 0 837.29156
330.1653137207031 0 673.17615
331.1491394042969 0 4202.7363
331.2348327636719 0 642.62256 c 2
332.1568603515625 0 4981.279 y Ammonia loss 12
333.1649169921875 0 12385.206 z 12
334.1679992675781 0 2127.4597
340.115966796875 0 793.0369
347.1678161621094 0 1157.0209
349.1833190917969 0 4087.451 y 12
350.18695068359375 0 639.264
357.1520690917969 0 8211.971
357.2486267089844 0 675.256
375.1615295410156 0 729.70135
389.7081604003906 0 885.95105
391.724365234375 0 3537.903
392.22509765625 0 1615.231
398.7146911621094 0 1237.9014
399.7210693359375 0 641.49744
415.2269592285156 0 4725.876 z 7
415.7247009277344 0 1812.499
417.2101135253906 0 12786.362 w 11
418.2123107910156 0 2491.5938
422.7301025390625 0 635.1457
423.1503601074219 0 699.8488
423.2341003417969 0 3444.8062 y 7
423.7347717285156 0 1553.7158
425.24066162109375 0 1740.8545
426.2421569824219 0 622.3697
431.2239685058594 0 725.8272 y Ammonia loss 11
441.1630859375 0 1627.3165
443.25067138671875 0 2702.9504
443.309814453125 0 696.13855
444.2547912597656 0 780.49774
446.2361755371094 0 901.59357
447.2068786621094 0 851.17566
447.2406005859375 0 1047.6027
448.2166442871094 0 414.80945
448.2524108886719 0 13077.065 y 11
449.22509765625 0 793.0998 y Ammonia loss 2
449.2557678222656 0 2601.5718
450.74053955078125 0 970.9486 c Ammonia loss 7
451.2463073730469 0 682.826
454.89337158203125 0 1234.6893 y 2
455.2864685058594 0 750.8223
455.5610046386719 0 553.30133
460.2776794433594 0 8319.849 c 3
461.2798156738281 0 1619.1385
471.2467041015625 0 758.7558 z Water loss 10
472.2516784667969 0 2395.2249 c Ammonia loss 13
472.7401428222656 0 2182.3645 z 6
473.2444152832031 0 1548.7688
473.7429504394531 0 589.68536
474.2312316894531 0 2677.747
475.2341613769531 0 896.1511
480.748291015625 0 2238.1672 y 6
481.24993896484375 0 1070.2123 z Water loss 1
489.2549743652344 0 2972.2136 z 10
490.2613525390625 0 2529.6375
495.91522216796875 0 1440.7943
498.2901611328125 0 571.66626
499.3000183105469 0 2494.8555
505.2738952636719 0 39274.094 y 10
506.2765808105469 0 10161.34
507.2798767089844 0 1057.2538
512.200439453125 0 1214.5823
514.3242797851562 0 917.74304
515.2484741210938 0 1744.5902 y Water loss 5
515.3316040039062 0 2209.7483
515.7503662109375 0 2374.467 y Ammonia loss 5
516.2537841796875 0 3927.2368 z 5
516.3366088867188 0 1616.4813
516.7567138671875 0 5148.427
517.2593383789062 0 2041.2905
517.756591796875 0 767.8077
523.7548217773438 0 644.2255
524.265869140625 0 13469.617 y 5
524.613037109375 0 4643.2744
524.7643432617188 0 6827.4805
524.9465942382812 0 2029.4315
525.2688598632812 0 1775.4342
525.7657470703125 0 1076.8999
529.2719116210938 0 1075.829
529.7965087890625 0 1238.5315
530.2081298828125 0 1633.7461
530.2842407226562 0 3509.124
530.3191528320312 0 3524.912
530.6168823242188 0 3114.4902
530.9494018554688 0 1813.1613
531.28515625 0 764.3199
531.3228759765625 0 1695.8995
537.7596435546875 0 853.1097
538.2594604492188 0 1378.1735
538.7628784179688 0 711.57324
541.3109741210938 0 1028.8967
542.3189697265625 0 4075.5781
543.32177734375 0 1412.1174
558.2763671875 0 1636.118 z Water loss 9
558.338134765625 0 1727.4645
559.2848510742188 0 2003.7513 w 9
559.345703125 0 7128.0444 c 4
560.2888793945312 0 624.36285
560.3482055664062 0 1690.6149
565.788330078125 0 28854.232 z 4
566.2899780273438 0 19051.938
566.791748046875 0 5939.2017
567.2921752929688 0 2077.4365
572.7882080078125 0 755.00507
573.7974853515625 0 14958.626 y 4
574.2987670898438 0 7885.2393 y Water loss 9
574.8010864257812 0 3580.0698
575.282470703125 0 1697.573 y Ammonia loss 9
576.28662109375 0 6510.0195 z 9
577.29052734375 0 1458.7576
586.3316650390625 0 1414.5369
587.793701171875 0 742.5308
592.30517578125 0 1815.7312 y 9
600.2748413085938 0 3079.3074
600.8026733398438 0 14327.516 w 3
601.303955078125 0 8021.9316
601.8070678710938 0 2463.729
608.3133544921875 0 741.58044
629.2994384765625 0 961.86633
629.350341796875 0 2184.1038 c 11
629.8057250976562 0 934.0562 y Ammonia loss 3
630.3096923828125 0 31615.66 z 3
630.81103515625 0 19861.963
631.3123779296875 0 8923.478
631.3684692382812 0 664.58594
631.8123168945312 0 1717.3873
637.8153076171875 0 9923.305
637.8670043945312 0 768.0165
638.3182983398438 0 33138.438 y 3
638.8199462890625 0 19504.727
639.322021484375 0 7524.7607
639.8218383789062 0 1571.3425
644.3193969726562 0 3106.8523
644.818359375 0 1727.6913
645.369873046875 0 1028.1595
646.3779907226562 0 26836.088 c 5
647.38134765625 0 8368.885
648.38525390625 0 1876.3004
651.8267822265625 0 1213.2518
652.3141479492188 0 778.50934
665.322265625 0 1388.5981 z Ammonia loss 2
665.8247680664062 0 964.6102
666.3236083984375 0 1162.2927
672.3250122070312 0 972.22186 y Ammonia loss 8
672.8267822265625 0 3057.65 y Water loss 2
673.3275756835938 0 2641.4495 y Ammonia loss 2
673.8260498046875 0 20792.557 z 2
674.3280029296875 0 24878.396
674.829833984375 0 16070.822
675.329345703125 0 4423.838
675.8308715820312 0 1461.334
680.3257446289062 0 717.67053
680.8252563476562 0 3924.552
681.3297729492188 0 4295.1943
681.8350830078125 0 136276.78 y 2
682.33642578125 0 91787.93
682.8379516601562 0 36244.363
683.339111328125 0 11154.784
683.8386840820312 0 1711.2802
685.3868408203125 0 923.31525
685.8206176757812 0 660.9096
687.3382568359375 0 1688.0173
689.3587036132812 0 3828.163 y 8
690.3639526367188 0 1122.2258
711.4237060546875 0 1160.7336
713.4305419921875 0 1359.0817
714.380859375 0 881.7509
715.8485107421875 0 3571.9329 w 1
716.3495483398438 0 2192.0984
716.8504638671875 0 793.6867
717.3887329101562 0 1904.2256
718.3975219726562 0 4221.2725
722.85791015625 0 797.75604 w 1
729.3685302734375 0 1397.8749 y Water loss 1
730.366455078125 0 2749.9712 z 1
730.8650512695312 0 1358.874
737.3715209960938 0 696.9746
738.376953125 0 6598.093 y 1
738.878662109375 0 5649.1304
739.3783569335938 0 2784.482
742.3970336914062 0 699.1597
743.3735961914062 0 2943.8086 w 7
744.3701171875 0 3188.5242
750.8948974609375 0 2087.6794
751.3970947265625 0 2004.5077
751.9047241210938 0 1478.8124
756.4065551757812 0 1210.5801
757.8973388671875 0 1580.8806
758.3885498046875 0 4718.0264
758.8844604492188 0 4239.252
759.38330078125 0 4205.444
759.8883056640625 0 1207.4473
760.392333984375 0 1429.739
761.4052124023438 0 54172.805 c 6
762.4077758789062 0 20678.463
763.4093017578125 0 5320.2437
764.4133911132812 0 2987.7888
764.910400390625 0 7000.7896
765.38330078125 0 92888.75
765.8843994140625 0 76198.336
766.3859252929688 0 35470.31
766.8867797851562 0 10627.407
767.3856201171875 0 1544.4504
771.908447265625 0 887.89154
772.4126586914062 0 4190.816
772.91064453125 0 3603.8372
773.4111938476562 0 2112.4062
773.9085693359375 0 25717.592
774.4097290039062 0 20192.312
774.9110107421875 0 9758.549
775.4129028320312 0 2694.971
777.9055786132812 0 1611.3452
778.4060668945312 0 3188.4575
778.905029296875 0 1605.7146
779.406494140625 0 1580.9348
779.9059448242188 0 1082.4738
780.4161376953125 0 1376.5923
781.4204711914062 0 716.95953
782.441650390625 0 2636.3662
783.44140625 0 2494.272
785.3975830078125 0 3358.625
785.9012451171875 0 4520.5996
786.4113159179688 0 9957.923
786.9102783203125 0 99984.75
787.4114990234375 0 80018.95
787.9124755859375 0 41895.477
788.4124145507812 0 11845.088
788.91455078125 0 3711.4387
792.8963623046875 0 949.2896
793.4061889648438 0 4915.146
793.9105224609375 0 18818.096
794.4114379882812 0 18407.34
794.918212890625 0 70091.625
795.4208374023438 0 77646.48
795.92236328125 0 48351.73
796.42333984375 0 19323.266
796.9234619140625 0 4726.0923
797.4193725585938 0 2165.0173
798.4315185546875 0 733.4808
803.4262084960938 0 4708.214
804.4280395507812 0 2351.0925
811.4287719726562 0 2728.3315 z Water loss 7
828.4318237304688 0 955.67377 y Ammonia loss 7
829.44091796875 0 46547.1 z 7
830.4447631835938 0 21617.982
831.44873046875 0 7293.2554
832.451904296875 0 878.3961
839.5147705078125 0 1215.5819
840.5286254882812 0 925.04315
841.5309448242188 0 2086.4973
842.5321655273438 0 1160.7238
844.449951171875 0 2022.0812
845.3846435546875 0 575.6875
845.4599609375 0 7001.317 y 7
846.4642333984375 0 1936.0444
847.4664916992188 0 1273.3292
857.4308471679688 0 772.33673
861.4312133789062 0 682.7247
873.490234375 0 2250.7517
874.4951171875 0 895.15643
900.4786987304688 0 6681.5635 c Ammonia loss 7
901.47998046875 0 2747.9106
902.4844970703125 0 1390.5739
926.4570922851562 0 2228.706 z Water loss 6
927.4598388671875 0 1367.017
928.4693603515625 0 1097.1993
929.4895629882812 0 1056.7506
944.468017578125 0 17006.07 z 6
945.472412109375 0 12296.853
946.4782104492188 0 4739.6045
947.4777221679688 0 1177.3715
954.5410766601562 0 1034.2607
955.5360107421875 0 1711.8217
956.553466796875 0 2028.9009
957.5497436523438 0 791.1722
959.47705078125 0 1037.6477
960.4865112304688 0 2082.6118 y 6
970.5432739257812 0 3534.8398
971.5457153320312 0 2106.3967
999.5662231445312 0 958.8327
1004.49462890625 0 725.3558
1005.5076293945312 0 855.2119
1012.5466918945312 0 755.3883
1013.5849609375 0 2527.4128
1014.5594482421875 0 42660.496 c 8
1015.5628051757812 0 21408.316
1016.5661010742188 0 6479
1017.5750122070312 0 1950.7916
1031.500244140625 0 6787.0566 z 5
1032.5064697265625 0 9130.205
1033.5106201171875 0 4314.1807
1034.51171875 0 1182.243
1039.5667724609375 0 2597.503
1040.565673828125 0 1586.5322
1041.5660400390625 0 1627.8433
1042.5733642578125 0 2790.6118
1046.5111083984375 0 1008.30646
1047.51806640625 0 2706.7078 y 5
1048.5216064453125 0 2149.7986
1057.5777587890625 0 28191.646
1058.580810546875 0 19840.363
1059.588134765625 0 10238.146
1060.592529296875 0 3899.037
1061.6146240234375 0 712.073
1100.580078125 0 2451.425
1101.5914306640625 0 55980.004 c 9
1102.593994140625 0 32463.625
1103.5968017578125 0 11012.446
1104.5968017578125 0 2060.751
1114.5980224609375 0 8607.899
1115.60107421875 0 6665.244
1116.6005859375 0 2320.702
1117.60107421875 0 758.2415
1129.558837890625 0 623.6449 y Ammonia loss 4
1130.568359375 0 8029.242 z 4
1131.5753173828125 0 17340.045
1132.578125 0 9508.834
1133.5811767578125 0 3106.8
1146.5889892578125 0 2747.8435 y 4
1147.59228515625 0 1096.177
1148.58935546875 0 1212.8643
1158.6121826171875 0 32425.36 c 10
1159.61474609375 0 18963.227
1160.6181640625 0 8091.1016
1161.62109375 0 1493.565
1203.6134033203125 0 681.75696
1213.666259765625 0 6407.138
1214.6676025390625 0 4152.664
1215.6746826171875 0 2910.0881
1216.674072265625 0 1240.4282
1217.58251953125 0 775.82434
1247.5672607421875 0 731.7912
1257.680419921875 0 31125.318 c 11
1258.681396484375 0 20594.117
1259.6783447265625 0 7752.3013
1260.6187744140625 0 28383.598
1261.6204833984375 0 14419.094
1262.6199951171875 0 5681.112
1263.634033203125 0 1282.0784
1274.61669921875 0 918.12616
1275.6279296875 0 10368.731 y 3
1276.6319580078125 0 6159.3867
1277.6324462890625 0 2458.447
1282.6854248046875 0 1383.6978
1284.6934814453125 0 836.96027
1300.6982421875 0 5545.627
1301.700439453125 0 3997.1663
1302.700927734375 0 2751.9766
1304.631103515625 0 1569.8967
1329.651611328125 0 2095.0342 w 2
1330.631591796875 0 2601.3975
1331.6368408203125 0 1594.2432
1343.699462890625 0 770.9416
1344.711181640625 0 52261.48 c 12
1345.714599609375 0 36461.89
1346.7095947265625 0 17459.38
1347.6533203125 0 23817.617
1348.6553955078125 0 14269.196
1349.654541015625 0 7120.448
1350.6563720703125 0 2382.6567
1362.6585693359375 0 1538.0071 y 2
1363.6671142578125 0 2393.3975
1387.7305908203125 0 5757.3633
1388.7381591796875 0 4495.9653
1389.735107421875 0 1243.0885
1390.74365234375 0 938.4046
1429.72021484375 0 2031.2435
1430.72900390625 0 1600.6558
1431.7431640625 0 74579.87 c 13
1432.74609375 0 61479.523
1433.7476806640625 0 26684.25
1434.750244140625 0 7205.5986
1435.76123046875 0 1346.9144
1449.732421875 0 755.4655
1459.73193359375 0 884.156 z 1
1460.7296142578125 0 3801.3157
1461.7391357421875 0 2944.081
1462.7491455078125 0 1371.2255
1473.73486328125 0 689.59033
1474.742919921875 0 695.1521
1475.7449951171875 0 658.1723 y 1
1487.74853515625 0 943.5632
1488.74169921875 0 6952.349
1489.745849609375 0 6571.6733
1490.7525634765625 0 3868.2046
1491.7567138671875 0 1770.8824
1512.7860107421875 0 1483.2006
1513.7950439453125 0 1674.4878
1514.7864990234375 0 2139.1523
1515.781982421875 0 987.1038
1516.74365234375 0 969.48987
1528.7781982421875 0 1209.083
1529.78662109375 0 2108.3765
1530.792236328125 0 16138.669
1531.794677734375 0 20578.182
1532.7974853515625 0 13268.669
1533.795166015625 0 4383.442
1534.8060302734375 0 971.8458
1543.8074951171875 0 1650.5387
1544.8258056640625 0 3726.6902
1545.8251953125 0 7158.8154
1546.82373046875 0 7139.7305
1547.8218994140625 0 8736.544
1548.81982421875 0 4783.7476
1549.818359375 0 2462.615
1555.81494140625 0 2075.77
1556.7957763671875 0 8358.168
1557.79736328125 0 5226.5557
1558.7938232421875 0 2766.6653
1559.8062744140625 0 1792.5579
1560.824462890625 0 861.12024
1561.833984375 0 3417.6077
1562.8465576171875 0 11818.269
1563.8492431640625 0 8558.93
1564.8553466796875 0 5706.2104
1565.849853515625 0 1084.5669
1569.7979736328125 0 1216.4431
1570.7950439453125 0 2033.9335
1571.799560546875 0 4715.6353
1572.81494140625 0 17667.723
1573.8167724609375 0 102266.836
1574.81982421875 0 79364.7
1575.821044921875 0 39339.89
1576.82275390625 0 13223.442
1577.8271484375 0 3583.4824
1586.805419921875 0 4284.4326
1587.8126220703125 0 7710.6724
1588.823486328125 0 13958.646
1589.59033203125 0 1092.9174
1589.83203125 0 34727.113
1590.8411865234375 0 102228.79
1591.843994140625 0 76811.69
1592.8470458984375 0 38654.516
1593.849853515625 0 10424.187
1594.8428955078125 0 2902.4844
2209.994384765625 0 635.48724

Spectrum Details

|  |  |
| --- | --- |
| Matched peaks? Matched peaks The total absolute number of peaks matched. Additionally in brackets the total fraction of peaks matched and the total number of peaks is shown. | 84 (17.14% of 490) |
| FDR? FDR The false discovery rate estimated for this peptide. It is calculated by matching all theoretical fragments with a non-integer shift with the raw peaks for this spectrum. This is done with 40 different shifts. The resulting percentage is the average number of annotated peaks over the number of annotated peaks with the correct spectrum. | 0.88% |
| Satellite FDR? Satellite FDR See the FDR for details on its calculation. This satellite ion specific FDR only contains the satellite ions (d/w) for I/L/J positions. | 0.00% |
| PSM Score? PSM Score The PSM Score as given by Hecklib to this annotated spectrum. It is shown with three significant figures. | 656 |

#### Spectrum 4542? Spectrum 4542 The raw spectrum of this peptide as annotated by Hecklib. The fragments are coloured according to ion type (see legend). Any peaks with a star '\*' as text can be hovered over to see the full details, first the ion type second the mass shift type. By hovering over the amino acids in the peptide or ions in the legend the corresponding peaks are highlighted. By toggling the 'Unassigned' label you can turn the background (unassigned) peaks on or off in the plot. By updating the slider in the Ion legend you can update the spectrum to only show the top X% of the peaks with labels. The top X% means any peak that is within X% of the highest intensity. By dragging in the spectrum you can zoom in to a specific part of the spectrum and use 'Zoom Out' to get back to the original zoom level. The annotation of the spectrum is based on the given sequence in the peptides file and is done with different software so inconsistencies are likely. The peaks are annotated based on the given sequence, with 20 ppm tolerance.

Copy Data

##### Spectrum 4542 (TSV)

###### Preview

```
Loading example...
```

*Click on the button to copy the data to your clipboard.*

Mz MinMz MaxIntensity Max

WidthHeightPeptide font sizePeptide stroke widthSpectrum font sizeSpectrum stroke widthCompact peptide

Ion legend

wxyz

abcd

OtherUnassignedIon^Charge^_Position_Show for top:%

JJSEVSDRPSGVSSR

01.36e+42.73e+44.09e+45.45e+4

Zoom Out

y^+1^_1_w^+1^_2_y^+1^_2_z^+1^_2_y^+1^_2_w^+1^_3_y^+1^_3_z^+1^_3_y^+1^_3_z^+2^_8_w^+1^_4_y^+2^_8_z^+1^_4_y^+1^_4_c^+1^_4_c^+3^_14_z^+2^_9_z^+1^_5_y^+1^_5_y^+2^_10_z^+2^_10_y^+2^_10_c^+1^_5_z^+2^_11_y^+2^_11_y^+1^_6_y^+1^_6_z^+1^_6_w^+2^_12_z^+2^_12_y^+2^_12_c^+1^_6_w^+2^_13_y^+2^_13_y^+2^_13_z^+2^_13_y^+2^_13_y^+1^_7_z^+2^_14_y^+2^_14_c^+1^_7_z^+1^_8_z^+1^_8_y^+1^_8_c^+1^_8_z^+1^_9_z^+1^_9_c^+1^_9_z^+1^_10_y^+1^_10_c^+1^_10_z^+1^_11_y^+1^_11_c^+1^_11_c^+1^_12_z^+1^_12_y^+1^_12_z^+1^_13_c^+1^_13_c^+1^_14_

0864172925933458

Fragment Matches Table

Show background peaks

| Position | Ion type | Intensity | mz Theoretical | mz Error (Th) | mz Error (ppm) | Charge | Series Number |
| --- | --- | --- | --- | --- | --- | --- | --- |
| - | - | 1064 | 120.1 | - | - | 0 | - |
| - | - | 785.6 | 125.1 | - | - | 0 | - |
| - | - | 336.2 | 125.8 | - | - | 0 | - |
| - | - | 1807 | 126.1 | - | - | 0 | - |
| - | - | 790.1 | 131.1 | - | - | 0 | - |
| - | - | 395.7 | 132.6 | - | - | 0 | - |
| - | - | 558.1 | 133.1 | - | - | 0 | - |
| - | - | 392.6 | 134.7 | - | - | 0 | - |
| - | - | 682.5 | 136.1 | - | - | 0 | - |
| - | - | 446.5 | 143.1 | - | - | 0 | - |
| - | - | 419.9 | 153.5 | - | - | 0 | - |
| - | - | 1858 | 168.1 | - | - | 0 | - |
| - | - | 469.7 | 168.5 | - | - | 0 | - |
| - | - | 438.3 | 172.5 | - | - | 0 | - |
| - | - | 528.8 | 174.1 | - | - | 0 | - |
| - | - | 484.7 | 174.6 | - | - | 0 | - |
| 15 | y | 3195 | 175.1 | 0.000219 | 1.25 | +1 | 1 |
| - | - | 485.7 | 178.2 | - | - | 0 | - |
| - | - | 7348 | 185.1 | - | - | 0 | - |
| - | - | 679.2 | 186.1 | - | - | 0 | - |
| - | - | 1.795E+04 | 199.2 | - | - | 0 | - |
| - | - | 1262 | 200.1 | - | - | 0 | - |
| - | - | 1363 | 200.2 | - | - | 0 | - |
| - | - | 552.6 | 212.1 | - | - | 0 | - |
| - | - | 626.4 | 215.1 | - | - | 0 | - |
| - | - | 916.9 | 217.1 | - | - | 0 | - |
| - | - | 471.2 | 223 | - | - | 0 | - |
| - | - | 8423 | 227.2 | - | - | 0 | - |
| - | - | 1385 | 228.2 | - | - | 0 | - |
| 14 | w | 1970 | 229.1 | 0.0006101 | 2.663 | +1 | 2 |
| - | - | 640.9 | 234.1 | - | - | 0 | - |
| 14 | y | 1.204E+04 | 245.1 | 0.0003549 | 1.448 | +1 | 2 |
| 14 | z | 1787 | 246.1 | 0.001595 | 6.482 | +1 | 2 |
| - | - | 462.7 | 255.1 | - | - | 0 | - |
| - | - | 677.6 | 261.2 | - | - | 0 | - |
| 14 | y | 2297 | 262.2 | 0.0003561 | 1.358 | +1 | 2 |
| - | - | 2433 | 270.1 | - | - | 0 | - |
| - | - | 816.8 | 272.2 | - | - | 0 | - |
| - | - | 514.4 | 300.1 | - | - | 0 | - |
| - | - | 885.7 | 304.2 | - | - | 0 | - |
| - | - | 1020 | 314.2 | - | - | 0 | - |
| 13 | w | 6985 | 316.2 | 0.00032 | 1.012 | +1 | 3 |
| - | - | 1071 | 317.2 | - | - | 0 | - |
| - | - | 976.5 | 331.1 | - | - | 0 | - |
| 13 | y | 2322 | 332.2 | 0.0004615 | 1.389 | +1 | 3 |
| 13 | z | 4723 | 333.2 | 0.0001743 | 0.5232 | +1 | 3 |
| - | - | 612.9 | 334.2 | - | - | 0 | - |
| 13 | y | 1993 | 349.2 | 0.0004627 | 1.325 | +1 | 3 |
| - | - | 3705 | 357.2 | - | - | 0 | - |
| - | - | 690.7 | 391.7 | - | - | 0 | - |
| 8 | z | 1883 | 415.2 | 0.003502 | 8.433 | +2 | 8 |
| 12 | w | 4558 | 417.2 | 0.0007677 | 1.84 | +1 | 4 |
| - | - | 1096 | 418.2 | - | - | 0 | - |
| 8 | y | 1141 | 423.2 | 0.0002146 | 0.507 | +2 | 8 |
| - | - | 903.5 | 425.2 | - | - | 0 | - |
| 12 | z | 743.8 | 432.2 | 0.002047 | 4.736 | +1 | 4 |
| - | - | 1432 | 443.2 | - | - | 0 | - |
| 12 | y | 4712 | 448.3 | 0.0007438 | 1.659 | +1 | 4 |
| - | - | 1184 | 449.3 | - | - | 0 | - |
| - | - | 553.9 | 453.1 | - | - | 0 | - |
| - | - | 602.7 | 453.3 | - | - | 0 | - |
| 4 | c | 3755 | 460.3 | 0.0009213 | 2.002 | +1 | 4 |
| - | - | 844.9 | 461.3 | - | - | 0 | - |
| 14 | c | 1346 | 472.2 | 0.008233 | 17.43 | +3 | 14 |
| 7 | z | 1034 | 472.7 | 0.000986 | 2.086 | +2 | 9 |
| - | - | 1084 | 474.2 | - | - | 0 | - |
| - | - | 722.2 | 475.2 | - | - | 0 | - |
| 11 | z | 1481 | 489.3 | 0.001117 | 2.283 | +1 | 5 |
| - | - | 1205 | 490.3 | - | - | 0 | - |
| - | - | 1102 | 495.9 | - | - | 0 | - |
| - | - | 684.7 | 499.3 | - | - | 0 | - |
| 11 | y | 1.658E+04 | 505.3 | 0.0005814 | 1.151 | +1 | 5 |
| - | - | 4431 | 506.3 | - | - | 0 | - |
| - | - | 852.9 | 507.3 | - | - | 0 | - |
| - | - | 670 | 515.3 | - | - | 0 | - |
| 6 | y | 1000 | 515.7 | 0.001564 | 3.032 | +2 | 10 |
| 6 | z | 1858 | 516.3 | 0.001665 | 3.225 | +2 | 10 |
| - | - | 736.1 | 516.3 | - | - | 0 | - |
| - | - | 1438 | 516.8 | - | - | 0 | - |
| - | - | 693.4 | 517.3 | - | - | 0 | - |
| - | - | 648.8 | 517.8 | - | - | 0 | - |
| 6 | y | 6563 | 524.3 | 0.002557 | 4.877 | +2 | 10 |
| - | - | 1202 | 524.6 | - | - | 0 | - |
| - | - | 2784 | 524.8 | - | - | 0 | - |
| - | - | 729.6 | 525.3 | - | - | 0 | - |
| - | - | 1023 | 528.7 | - | - | 0 | - |
| - | - | 842 | 529.2 | - | - | 0 | - |
| - | - | 1085 | 529.3 | - | - | 0 | - |
| - | - | 687.9 | 529.7 | - | - | 0 | - |
| - | - | 1063 | 530.3 | - | - | 0 | - |
| - | - | 8802 | 530.3 | - | - | 0 | - |
| - | - | 1776 | 530.6 | - | - | 0 | - |
| - | - | 951 | 530.7 | - | - | 0 | - |
| - | - | 3172 | 531.3 | - | - | 0 | - |
| - | - | 617.2 | 542.3 | - | - | 0 | - |
| - | - | 1334 | 542.3 | - | - | 0 | - |
| - | - | 620.2 | 543.3 | - | - | 0 | - |
| - | - | 1058 | 558.3 | - | - | 0 | - |
| 5 | c | 3184 | 559.3 | 0.0002869 | 0.513 | +1 | 5 |
| - | - | 1143 | 560.4 | - | - | 0 | - |
| 5 | z | 1.167E+04 | 565.8 | 0.0002949 | 0.5212 | +2 | 11 |
| - | - | 7370 | 566.3 | - | - | 0 | - |
| - | - | 2407 | 566.8 | - | - | 0 | - |
| - | - | 732.9 | 567.3 | - | - | 0 | - |
| 5 | y | 6383 | 573.8 | 0.0001491 | 0.2599 | +2 | 11 |
| 10 | y | 2998 | 574.3 | 0.003806 | 6.628 | +1 | 6 |
| - | - | 1678 | 574.8 | - | - | 0 | - |
| 10 | y | 1398 | 575.3 | 7.645E-05 | 0.1329 | +1 | 6 |
| 10 | z | 3538 | 576.3 | 0.0009126 | 1.584 | +1 | 6 |
| - | - | 660.2 | 586.3 | - | - | 0 | - |
| - | - | 1298 | 600.3 | - | - | 0 | - |
| 4 | w | 4981 | 600.8 | 0.0006651 | 1.107 | +2 | 12 |
| - | - | 3207 | 601.3 | - | - | 0 | - |
| - | - | 1407 | 601.8 | - | - | 0 | - |
| 4 | z | 1.14E+04 | 630.3 | 0.0002996 | 0.4753 | +2 | 12 |
| - | - | 9628 | 630.8 | - | - | 0 | - |
| - | - | 4076 | 631.3 | - | - | 0 | - |
| - | - | 1563 | 631.8 | - | - | 0 | - |
| - | - | 697.7 | 632.3 | - | - | 0 | - |
| - | - | 6018 | 637.8 | - | - | 0 | - |
| 4 | y | 1.119E+04 | 638.3 | 0.0005786 | 0.9064 | +2 | 12 |
| - | - | 8144 | 638.8 | - | - | 0 | - |
| - | - | 2597 | 639.3 | - | - | 0 | - |
| - | - | 1095 | 639.8 | - | - | 0 | - |
| - | - | 1284 | 644.3 | - | - | 0 | - |
| 6 | c | 1.167E+04 | 646.4 | 0.0006682 | 1.034 | +1 | 6 |
| - | - | 791.6 | 647.3 | - | - | 0 | - |
| - | - | 4528 | 647.4 | - | - | 0 | - |
| - | - | 661.5 | 648.4 | - | - | 0 | - |
| 3 | w | 1140 | 665.3 | 0.002623 | 3.942 | +2 | 13 |
| 3 | y | 1466 | 672.8 | 0.0006903 | 1.026 | +2 | 13 |
| 3 | y | 1007 | 673.3 | 0.008401 | 12.48 | +2 | 13 |
| 3 | z | 9011 | 673.8 | 0.0003987 | 0.5916 | +2 | 13 |
| - | - | 8275 | 674.3 | - | - | 0 | - |
| - | - | 4739 | 674.8 | - | - | 0 | - |
| - | - | 1683 | 675.3 | - | - | 0 | - |
| - | - | 941.4 | 681.3 | - | - | 0 | - |
| 3 | y | 5.396E+04 | 681.8 | 0.000314 | 0.4605 | +2 | 13 |
| - | - | 3.465E+04 | 682.3 | - | - | 0 | - |
| - | - | 1.692E+04 | 682.8 | - | - | 0 | - |
| - | - | 4081 | 683.3 | - | - | 0 | - |
| - | - | 1128 | 683.8 | - | - | 0 | - |
| 9 | y | 1817 | 689.4 | 0.0001349 | 0.1958 | +1 | 7 |
| - | - | 827 | 701.4 | - | - | 0 | - |
| - | - | 1089 | 716.3 | - | - | 0 | - |
| - | - | 1029 | 717.4 | - | - | 0 | - |
| - | - | 1866 | 718.4 | - | - | 0 | - |
| - | - | 652.3 | 726.3 | - | - | 0 | - |
| 2 | z | 883.7 | 730.4 | 0.001716 | 2.35 | +2 | 14 |
| 2 | y | 3077 | 738.4 | 0.0005193 | 0.7033 | +2 | 14 |
| - | - | 3038 | 738.9 | - | - | 0 | - |
| - | - | 1349 | 739.4 | - | - | 0 | - |
| - | - | 933.3 | 739.9 | - | - | 0 | - |
| - | - | 1044 | 744.4 | - | - | 0 | - |
| - | - | 867.7 | 745.4 | - | - | 0 | - |
| - | - | 1662 | 750.9 | - | - | 0 | - |
| - | - | 1504 | 758.4 | - | - | 0 | - |
| - | - | 1220 | 758.9 | - | - | 0 | - |
| - | - | 1045 | 760.4 | - | - | 0 | - |
| 7 | c | 2.144E+04 | 761.4 | 0.0004586 | 0.6023 | +1 | 7 |
| - | - | 7743 | 762.4 | - | - | 0 | - |
| - | - | 2403 | 763.4 | - | - | 0 | - |
| - | - | 1138 | 764.4 | - | - | 0 | - |
| - | - | 1499 | 764.9 | - | - | 0 | - |
| - | - | 3.673E+04 | 765.4 | - | - | 0 | - |
| - | - | 3.16E+04 | 765.9 | - | - | 0 | - |
| - | - | 1.692E+04 | 766.4 | - | - | 0 | - |
| - | - | 5391 | 766.9 | - | - | 0 | - |
| - | - | 1454 | 767.4 | - | - | 0 | - |
| - | - | 1778 | 772.4 | - | - | 0 | - |
| - | - | 915.9 | 772.9 | - | - | 0 | - |
| - | - | 838.9 | 773.4 | - | - | 0 | - |
| - | - | 1.116E+04 | 773.9 | - | - | 0 | - |
| - | - | 1.016E+04 | 774.4 | - | - | 0 | - |
| - | - | 4187 | 774.9 | - | - | 0 | - |
| - | - | 1383 | 775.4 | - | - | 0 | - |
| - | - | 1167 | 777.9 | - | - | 0 | - |
| - | - | 1177 | 778.4 | - | - | 0 | - |
| - | - | 769.7 | 778.9 | - | - | 0 | - |
| - | - | 686 | 780.4 | - | - | 0 | - |
| - | - | 832.2 | 782.4 | - | - | 0 | - |
| - | - | 776.9 | 785.4 | - | - | 0 | - |
| - | - | 962.3 | 785.9 | - | - | 0 | - |
| - | - | 4458 | 786.4 | - | - | 0 | - |
| - | - | 4.249E+04 | 786.9 | - | - | 0 | - |
| - | - | 3.604E+04 | 787.4 | - | - | 0 | - |
| - | - | 1.609E+04 | 787.9 | - | - | 0 | - |
| - | - | 5013 | 788.4 | - | - | 0 | - |
| - | - | 2260 | 788.9 | - | - | 0 | - |
| - | - | 624.3 | 792.9 | - | - | 0 | - |
| - | - | 1807 | 793.9 | - | - | 0 | - |
| - | - | 2872 | 794.4 | - | - | 0 | - |
| - | - | 2.536E+04 | 794.9 | - | - | 0 | - |
| - | - | 3.255E+04 | 795.4 | - | - | 0 | - |
| - | - | 2.197E+04 | 795.9 | - | - | 0 | - |
| - | - | 8829 | 796.4 | - | - | 0 | - |
| - | - | 2732 | 796.9 | - | - | 0 | - |
| - | - | 806.3 | 797.4 | - | - | 0 | - |
| 8 | z | 994.7 | 811.4 | 0.003299 | 4.066 | +1 | 8 |
| 8 | z | 1.827E+04 | 829.4 | 0.0002465 | 0.2972 | +1 | 8 |
| - | - | 9942 | 830.4 | - | - | 0 | - |
| - | - | 2659 | 831.4 | - | - | 0 | - |
| - | - | 1053 | 844.5 | - | - | 0 | - |
| 8 | y | 2701 | 845.5 | 0.0009042 | 1.069 | +1 | 8 |
| - | - | 1196 | 846.5 | - | - | 0 | - |
| - | - | 1862 | 873.5 | - | - | 0 | - |
| 8 | c | 2365 | 900.5 | 0.0006781 | 0.753 | +1 | 8 |
| - | - | 1539 | 901.5 | - | - | 0 | - |
| 7 | z | 754.7 | 926.5 | 0.006324 | 6.826 | +1 | 9 |
| 7 | z | 6618 | 944.5 | 0.0007003 | 0.7415 | +1 | 9 |
| - | - | 6335 | 945.5 | - | - | 0 | - |
| - | - | 2328 | 946.5 | - | - | 0 | - |
| - | - | 1616 | 970.5 | - | - | 0 | - |
| - | - | 640.8 | 1014 | - | - | 0 | - |
| 9 | c | 1.53E+04 | 1015 | 0.0005754 | 0.5671 | +1 | 9 |
| - | - | 8937 | 1016 | - | - | 0 | - |
| - | - | 901 | 1016 | - | - | 0 | - |
| - | - | 2622 | 1017 | - | - | 0 | - |
| 6 | z | 2238 | 1031 | 0.006728 | 6.522 | +1 | 10 |
| - | - | 3699 | 1033 | - | - | 0 | - |
| - | - | 1954 | 1034 | - | - | 0 | - |
| - | - | 878.6 | 1035 | - | - | 0 | - |
| - | - | 926.5 | 1040 | - | - | 0 | - |
| - | - | 707.1 | 1043 | - | - | 0 | - |
| 6 | y | 1200 | 1048 | 0.001526 | 1.457 | +1 | 10 |
| - | - | 803.3 | 1049 | - | - | 0 | - |
| - | - | 682 | 1057 | - | - | 0 | - |
| - | - | 1.202E+04 | 1058 | - | - | 0 | - |
| - | - | 685 | 1058 | - | - | 0 | - |
| - | - | 6605 | 1059 | - | - | 0 | - |
| - | - | 3226 | 1060 | - | - | 0 | - |
| - | - | 726.8 | 1061 | - | - | 0 | - |
| - | - | 681 | 1062 | - | - | 0 | - |
| - | - | 765.6 | 1088 | - | - | 0 | - |
| - | - | 840.6 | 1101 | - | - | 0 | - |
| 10 | c | 2.127E+04 | 1102 | 0.000142 | 0.1289 | +1 | 10 |
| - | - | 1.361E+04 | 1103 | - | - | 0 | - |
| - | - | 5487 | 1104 | - | - | 0 | - |
| - | - | 1072 | 1105 | - | - | 0 | - |
| - | - | 3892 | 1115 | - | - | 0 | - |
| - | - | 1884 | 1116 | - | - | 0 | - |
| - | - | 1105 | 1117 | - | - | 0 | - |
| 5 | z | 3877 | 1131 | 0.0001757 | 0.1554 | +1 | 11 |
| - | - | 7682 | 1132 | - | - | 0 | - |
| - | - | 3914 | 1133 | - | - | 0 | - |
| - | - | 1003 | 1134 | - | - | 0 | - |
| 5 | y | 1601 | 1147 | 0.004876 | 4.253 | +1 | 11 |
| 11 | c | 1.351E+04 | 1159 | 0.0003669 | 0.3167 | +1 | 11 |
| - | - | 8508 | 1160 | - | - | 0 | - |
| - | - | 2746 | 1161 | - | - | 0 | - |
| - | - | 850.8 | 1162 | - | - | 0 | - |
| - | - | 1895 | 1214 | - | - | 0 | - |
| - | - | 2222 | 1215 | - | - | 0 | - |
| - | - | 1145 | 1216 | - | - | 0 | - |
| 12 | c | 1.348E+04 | 1258 | 0.002617 | 2.081 | +1 | 12 |
| - | - | 7244 | 1259 | - | - | 0 | - |
| 4 | z | 2675 | 1260 | 0.005678 | 4.508 | +1 | 12 |
| - | - | 2596 | 1260 | - | - | 0 | - |
| - | - | 1.155E+04 | 1261 | - | - | 0 | - |
| - | - | 7166 | 1262 | - | - | 0 | - |
| - | - | 3072 | 1263 | - | - | 0 | - |
| - | - | 992.8 | 1264 | - | - | 0 | - |
| 4 | y | 4337 | 1276 | 0.001937 | 1.519 | +1 | 12 |
| - | - | 3055 | 1277 | - | - | 0 | - |
| - | - | 1091 | 1278 | - | - | 0 | - |
| - | - | 1866 | 1301 | - | - | 0 | - |
| - | - | 1663 | 1302 | - | - | 0 | - |
| - | - | 1564 | 1303 | - | - | 0 | - |
| - | - | 698.6 | 1305 | - | - | 0 | - |
| 3 | z | 962.5 | 1330 | 0.01436 | 10.8 | +1 | 13 |
| - | - | 1540 | 1331 | - | - | 0 | - |
| 13 | c | 2.284E+04 | 1345 | 0.001931 | 1.436 | +1 | 13 |
| - | - | 1.719E+04 | 1346 | - | - | 0 | - |
| - | - | 7177 | 1347 | - | - | 0 | - |
| - | - | 1.083E+04 | 1348 | - | - | 0 | - |
| - | - | 7765 | 1349 | - | - | 0 | - |
| - | - | 3446 | 1350 | - | - | 0 | - |
| - | - | 1483 | 1351 | - | - | 0 | - |
| - | - | 921.3 | 1364 | - | - | 0 | - |
| - | - | 631.2 | 1373 | - | - | 0 | - |
| - | - | 1959 | 1388 | - | - | 0 | - |
| - | - | 1404 | 1389 | - | - | 0 | - |
| 14 | c | 3.006E+04 | 1432 | 0.001977 | 1.381 | +1 | 14 |
| - | - | 2.523E+04 | 1433 | - | - | 0 | - |
| - | - | 1.254E+04 | 1434 | - | - | 0 | - |
| - | - | 3596 | 1435 | - | - | 0 | - |
| - | - | 705.5 | 1436 | - | - | 0 | - |
| - | - | 1085 | 1461 | - | - | 0 | - |
| - | - | 1177 | 1462 | - | - | 0 | - |
| - | - | 2936 | 1489 | - | - | 0 | - |
| - | - | 2630 | 1490 | - | - | 0 | - |
| - | - | 775.5 | 1491 | - | - | 0 | - |
| - | - | 892.5 | 1492 | - | - | 0 | - |
| - | - | 661.1 | 1513 | - | - | 0 | - |
| - | - | 1731 | 1514 | - | - | 0 | - |
| - | - | 1033 | 1515 | - | - | 0 | - |
| - | - | 1374 | 1530 | - | - | 0 | - |
| - | - | 6939 | 1531 | - | - | 0 | - |
| - | - | 8404 | 1532 | - | - | 0 | - |
| - | - | 6219 | 1533 | - | - | 0 | - |
| - | - | 2136 | 1534 | - | - | 0 | - |
| - | - | 1500 | 1545 | - | - | 0 | - |
| - | - | 3829 | 1546 | - | - | 0 | - |
| - | - | 3730 | 1547 | - | - | 0 | - |
| - | - | 4690 | 1548 | - | - | 0 | - |
| - | - | 2173 | 1549 | - | - | 0 | - |
| - | - | 1157 | 1550 | - | - | 0 | - |
| - | - | 792 | 1556 | - | - | 0 | - |
| - | - | 2753 | 1557 | - | - | 0 | - |
| - | - | 2460 | 1558 | - | - | 0 | - |
| - | - | 1742 | 1559 | - | - | 0 | - |
| - | - | 736.7 | 1560 | - | - | 0 | - |
| - | - | 973.4 | 1562 | - | - | 0 | - |
| - | - | 5060 | 1563 | - | - | 0 | - |
| - | - | 4883 | 1564 | - | - | 0 | - |
| - | - | 2073 | 1565 | - | - | 0 | - |
| - | - | 857.9 | 1571 | - | - | 0 | - |
| - | - | 2301 | 1572 | - | - | 0 | - |
| - | - | 9472 | 1573 | - | - | 0 | - |
| - | - | 4.499E+04 | 1574 | - | - | 0 | - |
| - | - | 3.597E+04 | 1575 | - | - | 0 | - |
| - | - | 1.849E+04 | 1576 | - | - | 0 | - |
| - | - | 5373 | 1577 | - | - | 0 | - |
| - | - | 1012 | 1578 | - | - | 0 | - |
| - | - | 1160 | 1587 | - | - | 0 | - |
| - | - | 1467 | 1588 | - | - | 0 | - |
| - | - | 5104 | 1589 | - | - | 0 | - |
| - | - | 1.265E+04 | 1590 | - | - | 0 | - |
| - | - | 4.687E+04 | 1591 | - | - | 0 | - |
| - | - | 3.624E+04 | 1592 | - | - | 0 | - |
| - | - | 1.709E+04 | 1593 | - | - | 0 | - |
| - | - | 6880 | 1594 | - | - | 0 | - |
| - | - | 1595 | 1595 | - | - | 0 | - |
| - | - | 912.1 | 2686 | - | - | 0 | - |
| - | - | 679.8 | 2960 | - | - | 0 | - |
| - | - | 644.3 | 3423 | - | - | 0 | - |

m/z Charge Intensity FragmentType MassShift Position
120.08113861083984 0 1063.5834
125.1076889038086 0 785.59265
125.81410217285156 0 336.20172
126.09162139892578 0 1806.8712
131.1295928955078 0 790.1039
132.63450622558594 0 395.67914
133.06106567382812 0 558.0703
134.6832733154297 0 392.56274
136.07565307617188 0 682.51044
143.1181182861328 0 446.5395
153.5372772216797 0 419.8835
168.1134490966797 0 1857.7833
168.52410888671875 0 469.6549
172.45437622070312 0 438.3173
174.05638122558594 0 528.7767
174.59815979003906 0 484.7445
175.11917114257812 0 3194.9915 y 14
178.2040557861328 0 485.73953
185.13990783691406 0 7347.7974
186.14328002929688 0 679.1589
199.18072509765625 0 17952.836
200.12681579589844 0 1261.7703
200.18405151367188 0 1362.9314
212.10348510742188 0 552.5956
215.1394500732422 0 626.44055
217.08204650878906 0 916.9089
223.01121520996094 0 471.206
227.1755828857422 0 8423.145
228.17921447753906 0 1384.8506
229.130126953125 0 1970.0186 w 13
234.11013793945312 0 640.91614
245.12478637695312 0 12036.955 y Ammonia loss 13
246.1306610107422 0 1786.536 z 13
255.11033630371094 0 462.6819
261.1585998535156 0 677.59766
262.1513366699219 0 2296.624 y 13
270.1196594238281 0 2433.3413
272.1726379394531 0 816.8088
300.1042785644531 0 514.3711
304.15106201171875 0 885.6721
314.2077941894531 0 1020.2051
316.161865234375 0 6984.5195 w 12
317.1592712402344 0 1071.4617
331.14923095703125 0 976.48016
332.15692138671875 0 2321.6445 y Ammonia loss 12
333.1644592285156 0 4722.9834 z 12
334.1680908203125 0 612.9287
349.1834716796875 0 1993.1423 y 12
357.1522216796875 0 3704.9634
391.7240295410156 0 690.69354
415.2271728515625 0 1883.4296 z 7
417.2099914550781 0 4557.7505 w 11
418.21087646484375 0 1095.8864
423.2328186035156 0 1140.9114 y 7
425.2386169433594 0 903.46466
432.23065185546875 0 743.8173 z 11
443.25 0 1431.9674
448.2521667480469 0 4711.899 y 11
449.25604248046875 0 1183.6289
453.0549621582031 0 553.92926
453.3111572265625 0 602.65405
460.2774963378906 0 3754.882 c 3
461.2821960449219 0 844.86194
472.2521667480469 0 1346.406 c Ammonia loss 13
472.7381286621094 0 1033.9417 z 6
474.2333984375 0 1083.5828
475.2345886230469 0 722.1969
489.2552795410156 0 1481.169 z 10
490.2593994140625 0 1205.4421
495.9162902832031 0 1101.6194
499.29736328125 0 684.7206
505.2734680175781 0 16579.162 y 10
506.276611328125 0 4430.575
507.2797546386719 0 852.9177
515.3319091796875 0 669.9666
515.7476806640625 0 1000.3957 y Ammonia loss 5
516.2548217773438 0 1858.029 z 5
516.3421020507812 0 736.0524
516.75537109375 0 1437.825
517.2600708007812 0 693.4367
517.758544921875 0 648.76904
524.2650756835938 0 6562.906 y 5
524.6126098632812 0 1202.4779
524.7653198242188 0 2784.0515
525.269775390625 0 729.6492
528.7394409179688 0 1023.0707
529.2383422851562 0 841.9943
529.295166015625 0 1084.7207
529.74072265625 0 687.86035
530.2783813476562 0 1062.9757
530.3176879882812 0 8801.727
530.6156005859375 0 1776.4585
530.7276000976562 0 950.9782
531.3212280273438 0 3172.0764
542.2679443359375 0 617.18933
542.3201293945312 0 1334.1582
543.3198852539062 0 620.1592
558.3333740234375 0 1057.5874
559.3452758789062 0 3184.4658 c 4
560.3504028320312 0 1143.4869
565.7876586914062 0 11666.435 z 4
566.2890625 0 7370.444
566.7897338867188 0 2407.0442
567.2885131835938 0 732.8707
573.796875 0 6383.3384 y 4
574.2981567382812 0 2998.487 y Water loss 9
574.7977294921875 0 1677.5276
575.2784423828125 0 1398.154 y Ammonia loss 9
576.2852783203125 0 3537.6729 z 9
586.32861328125 0 660.1767
600.2760009765625 0 1297.9772
600.8026733398438 0 4980.6665 w 3
601.3030395507812 0 3206.8716
601.8067016601562 0 1406.7598
630.3089599609375 0 11395.949 z 3
630.8107299804688 0 9628.229
631.3115234375 0 4075.8098
631.816162109375 0 1562.9332
632.3164672851562 0 697.7388
637.8145141601562 0 6018.336
638.3174438476562 0 11186.349 y 3
638.8192138671875 0 8143.794
639.3197021484375 0 2596.9512
639.8226318359375 0 1094.5508
644.3170776367188 0 1284.068
646.377685546875 0 11674.01 c 5
647.3226318359375 0 791.6261
647.3810424804688 0 4528.0396
648.384765625 0 661.4692
665.325927734375 0 1139.7994 w 2
672.8280639648438 0 1466.1256 y Water loss 2
673.3291625976562 0 1006.8237 y Ammonia loss 2
673.8250732421875 0 9011.227 z 2
674.3279418945312 0 8274.712
674.8285522460938 0 4739.0215
675.33154296875 0 1683.1346
681.3338623046875 0 941.35767
681.8343505859375 0 53963.434 y 2
682.335693359375 0 34652.66
682.837158203125 0 16922.514
683.3390502929688 0 4080.9492
683.8446655273438 0 1128.337
689.3575439453125 0 1817.388 y 8
701.3571166992188 0 827.0466
716.3495483398438 0 1088.5117
717.3902587890625 0 1028.7279
718.3983764648438 0 1865.7914
726.3145751953125 0 652.31305
730.364990234375 0 883.65643 z 1
738.3755493164062 0 3076.5542 y 1
738.8782348632812 0 3038.0703
739.3807983398438 0 1348.7874
739.8802490234375 0 933.2531
744.368408203125 0 1044.4829
745.3787231445312 0 867.6767
750.8987426757812 0 1661.9097
758.3870849609375 0 1503.7246
758.880615234375 0 1220.3901
760.3862915039062 0 1044.7915
761.4044189453125 0 21440.012 c 6
762.4068603515625 0 7743.231
763.4087524414062 0 2402.7273
764.4144287109375 0 1138.2214
764.9066162109375 0 1499.4388
765.3822021484375 0 36730.77
765.8834838867188 0 31599.809
766.384521484375 0 16916.047
766.8858032226562 0 5390.962
767.3851318359375 0 1454.3445
772.4082641601562 0 1778.4147
772.920654296875 0 915.8799
773.4042358398438 0 838.8854
773.9072875976562 0 11161.767
774.408203125 0 10157.737
774.9095458984375 0 4187.1
775.4092407226562 0 1383.165
777.9007568359375 0 1166.7135
778.4090576171875 0 1177.2521
778.9083862304688 0 769.67584
780.4071655273438 0 685.99274
782.4390869140625 0 832.15186
785.4033203125 0 776.8548
785.9026489257812 0 962.3309
786.4116821289062 0 4457.772
786.9096069335938 0 42486.01
787.4107055664062 0 36040.94
787.9110107421875 0 16086.465
788.4116821289062 0 5013.014
788.91064453125 0 2260.102
792.8966064453125 0 624.3255
793.908203125 0 1807.1106
794.4107055664062 0 2872.2976
794.917724609375 0 25361.164
795.4202270507812 0 32547.658
795.9217529296875 0 21972.367
796.418701171875 0 8828.599
796.905517578125 0 2732.4492
797.4158935546875 0 806.27374
811.4328002929688 0 994.71735 z Water loss 7
829.4398193359375 0 18267.615 z 7
830.4442749023438 0 9941.894
831.4451904296875 0 2658.8215
844.4520874023438 0 1053.383
845.4578857421875 0 2700.8342 y 7
846.4578857421875 0 1196.3347
873.4921264648438 0 1862.1086
900.4778442382812 0 2365.0537 c Ammonia loss 7
901.475830078125 0 1538.7393
926.4627685546875 0 754.7267 z Water loss 6
944.46630859375 0 6617.5835 z 6
945.4724731445312 0 6335.17
946.4762573242188 0 2327.89
970.5435791015625 0 1616.092
1013.5791625976562 0 640.8491
1014.5584106445312 0 15297.14 c 8
1015.5621337890625 0 8936.721
1016.4609985351562 0 900.9564
1016.5651245117188 0 2622.4316
1031.4923095703125 0 2238.095 z 5
1032.5037841796875 0 3698.9363
1033.5096435546875 0 1953.7096
1034.5054931640625 0 878.56824
1039.568359375 0 926.52405
1042.5694580078125 0 707.1015
1047.519287109375 0 1199.9957 y 5
1048.5262451171875 0 803.27454
1057.4683837890625 0 682.02716
1057.5767822265625 0 12017.541
1058.4774169921875 0 684.9558
1058.578369140625 0 6605.4053
1059.5833740234375 0 3225.5881
1060.602783203125 0 726.76807
1062.4951171875 0 680.9609
1087.579833984375 0 765.5924
1100.5760498046875 0 840.5711
1101.5897216796875 0 21267.092 c 9
1102.593505859375 0 13613.798
1103.5947265625 0 5486.956
1104.596923828125 0 1071.9567
1114.596435546875 0 3892.4067
1115.6005859375 0 1883.5349
1116.6043701171875 0 1104.5033
1130.567626953125 0 3877.1511 z 4
1131.574462890625 0 7681.8374
1132.5804443359375 0 3913.52
1133.5911865234375 0 1002.875
1146.581298828125 0 1600.7458 y 4
1158.6116943359375 0 13514.403 c 10
1159.61376953125 0 8508.491
1160.6162109375 0 2745.516
1161.628662109375 0 850.8474
1213.6634521484375 0 1894.9866
1214.66748046875 0 2222.0312
1215.6788330078125 0 1145.1431
1257.6771240234375 0 13483.689 c 11
1258.680419921875 0 7243.986
1259.61572265625 0 2674.9683 z 3
1259.671875 0 2596.1655
1260.6170654296875 0 11554.451
1261.6192626953125 0 7166.0425
1262.6202392578125 0 3071.6606
1263.6240234375 0 992.7738
1275.6268310546875 0 4336.8086 y 3
1276.630126953125 0 3055.3123
1277.631591796875 0 1091.0448
1300.7000732421875 0 1865.6403
1301.70068359375 0 1663.0846
1302.7041015625 0 1563.519
1304.6209716796875 0 698.63873
1329.6298828125 0 962.53864 z Ammonia loss 2
1330.6307373046875 0 1540.2406
1344.7098388671875 0 22836.85 c 12
1345.7135009765625 0 17190.695
1346.7052001953125 0 7176.888
1347.6517333984375 0 10828.547
1348.65234375 0 7764.556
1349.65478515625 0 3446.355
1350.662109375 0 1483.1418
1363.66357421875 0 921.3224
1372.7242431640625 0 631.23083
1387.730224609375 0 1958.8832
1388.7354736328125 0 1404.4846
1431.7418212890625 0 30062.322 c 13
1432.74462890625 0 25232.066
1433.7467041015625 0 12539.822
1434.749755859375 0 3595.6948
1435.7567138671875 0 705.4524
1460.7213134765625 0 1084.6458
1461.7413330078125 0 1177.3954
1488.738525390625 0 2936.2595
1489.7454833984375 0 2630.239
1490.74658203125 0 775.4705
1491.76123046875 0 892.5222
1512.7965087890625 0 661.1061
1513.79052734375 0 1730.5127
1514.7977294921875 0 1033.1779
1529.78955078125 0 1374.4369
1530.7899169921875 0 6938.527
1531.79296875 0 8404.23
1532.7958984375 0 6218.503
1533.7958984375 0 2136.0586
1544.82666015625 0 1499.6625
1545.828369140625 0 3828.6807
1546.826416015625 0 3730.4824
1547.8148193359375 0 4689.924
1548.811767578125 0 2173.4077
1549.8046875 0 1156.839
1555.8077392578125 0 792.0335
1556.7982177734375 0 2752.5308
1557.79443359375 0 2459.541
1558.7978515625 0 1742.3625
1559.79443359375 0 736.6543
1561.83544921875 0 973.3684
1562.8446044921875 0 5060.2197
1563.8507080078125 0 4882.6357
1564.852294921875 0 2072.9792
1570.806396484375 0 857.85284
1571.800048828125 0 2300.6567
1572.8135986328125 0 9471.573
1573.8154296875 0 44985.277
1574.8179931640625 0 35965.062
1575.8194580078125 0 18486.502
1576.822021484375 0 5373.1855
1577.820556640625 0 1011.9313
1586.829833984375 0 1160.3743
1587.8134765625 0 1466.7974
1588.82373046875 0 5104.231
1589.8302001953125 0 12653.743
1590.8397216796875 0 46866.465
1591.84228515625 0 36241.23
1592.842041015625 0 17093.73
1593.83984375 0 6880.4893
1594.828125 0 1595.2063
2685.77294921875 0 912.0725
2960.20556640625 0 679.83636
3423.3251953125 0 644.2529

Spectrum Details

|  |  |
| --- | --- |
| Matched peaks? Matched peaks The total absolute number of peaks matched. Additionally in brackets the total fraction of peaks matched and the total number of peaks is shown. | 60 (17.86% of 336) |
| FDR? FDR The false discovery rate estimated for this peptide. It is calculated by matching all theoretical fragments with a non-integer shift with the raw peaks for this spectrum. This is done with 40 different shifts. The resulting percentage is the average number of annotated peaks over the number of annotated peaks with the correct spectrum. | 0.99% |
| Satellite FDR? Satellite FDR See the FDR for details on its calculation. This satellite ion specific FDR only contains the satellite ions (d/w) for I/L/J positions. | ∞ |
| PSM Score? PSM Score The PSM Score as given by Hecklib to this annotated spectrum. It is shown with three significant figures. | 476 |

#### Spectrum 3632? Spectrum 3632 The raw spectrum of this peptide as annotated by Hecklib. The fragments are coloured according to ion type (see legend). Any peaks with a star '\*' as text can be hovered over to see the full details, first the ion type second the mass shift type. By hovering over the amino acids in the peptide or ions in the legend the corresponding peaks are highlighted. By toggling the 'Unassigned' label you can turn the background (unassigned) peaks on or off in the plot. By updating the slider in the Ion legend you can update the spectrum to only show the top X% of the peaks with labels. The top X% means any peak that is within X% of the highest intensity. By dragging in the spectrum you can zoom in to a specific part of the spectrum and use 'Zoom Out' to get back to the original zoom level. The annotation of the spectrum is based on the given sequence in the peptides file and is done with different software so inconsistencies are likely. The peaks are annotated based on the given sequence, with 20 ppm tolerance.

Copy Data

##### Spectrum 3632 (TSV)

###### Preview

```
Loading example...
```

*Click on the button to copy the data to your clipboard.*

Mz MinMz MaxIntensity Max

WidthHeightPeptide font sizePeptide stroke widthSpectrum font sizeSpectrum stroke widthCompact peptide

Ion legend

wxyz

abcd

OtherUnassignedIon^Charge^_Position_Show for top:%

JJSEVSDRPSGVSSR

04.46e+78.93e+71.34e+81.79e+8

Zoom Out

y^+1^_1_y^+3^_5_y^+1^_1_a^+1^_2_b^+1^_2_y^+1^_2_y^+1^_2_y^+1^_2_b^+1^_3_b^+1^_3_y^+1^_3_y^+1^_3_y^+2^_7_y^+2^_7_y^+1^_3_b^+3^_12_y^+2^_8_y^+2^_8_b^+1^_4_y^+1^_4_y^+1^_4_b^+1^_4_y^+1^_4_y^+3^_13_y^+3^_13_b^+2^_8_y^+3^_13_y^+2^_9_b^+3^_14_y^+2^_9_y^+3^_14_y^+3^_14_y^+1^_5_y^+1^_5_y^+3^_14_y^+1^_5_y^+2^_10_y^+2^_10_y^+2^_10_\*\*b^+1^_5_b^+2^_11_y^+2^_11_y^+2^_11_y^+2^_11_y^+1^_6_y^+1^_6_y^+2^_12_y^+2^_12_y^+2^_12_y^+1^_7_y^+1^_7_y^+2^_13_y^+2^_13_y^+2^_13_y^+1^_7_b^+2^_14_y^+2^_14_y^+2^_14_y^+2^_14_b^+1^_7_\*y^+1^_8_b^+1^_8_b^+1^_8_y^+1^_9_y^+1^_9_y^+1^_10_y^+1^_10_y^+1^_11_y^+1^_13_

034068110211361

Fragment Matches Table

Show background peaks

| Position | Ion type | Intensity | mz Theoretical | mz Error (Th) | mz Error (ppm) | Charge | Series Number |
| --- | --- | --- | --- | --- | --- | --- | --- |
| - | - | 3.613E+04 | 125.1 | - | - | 0 | - |
| - | - | 1.116E+06 | 126.1 | - | - | 0 | - |
| - | - | 7.714E+04 | 127.1 | - | - | 0 | - |
| - | - | 3.79E+04 | 127.1 | - | - | 0 | - |
| - | - | 4.904E+04 | 128.1 | - | - | 0 | - |
| - | - | 1.35E+05 | 129.1 | - | - | 0 | - |
| - | - | 1.941E+05 | 129.1 | - | - | 0 | - |
| - | - | 1.09E+05 | 129.1 | - | - | 0 | - |
| - | - | 7.074E+04 | 130.1 | - | - | 0 | - |
| - | - | 6.153E+05 | 130.1 | - | - | 0 | - |
| - | - | 3.4E+05 | 130.1 | - | - | 0 | - |
| - | - | 4.479E+04 | 131.1 | - | - | 0 | - |
| - | - | 5.32E+04 | 133.1 | - | - | 0 | - |
| - | - | 1.041E+05 | 133.1 | - | - | 0 | - |
| - | - | 4.079E+04 | 136 | - | - | 0 | - |
| - | - | 5.162E+04 | 138.1 | - | - | 0 | - |
| - | - | 7.81E+04 | 139.1 | - | - | 0 | - |
| - | - | 1.282E+05 | 140.1 | - | - | 0 | - |
| - | - | 5.742E+04 | 140.1 | - | - | 0 | - |
| - | - | 5.969E+04 | 141.1 | - | - | 0 | - |
| - | - | 1.387E+05 | 141.1 | - | - | 0 | - |
| - | - | 4.742E+04 | 143.1 | - | - | 0 | - |
| - | - | 3.766E+04 | 143.2 | - | - | 0 | - |
| - | - | 3.453E+05 | 144.1 | - | - | 0 | - |
| - | - | 3.763E+05 | 147.1 | - | - | 0 | - |
| - | - | 4.057E+04 | 150.7 | - | - | 0 | - |
| - | - | 1.033E+05 | 153.1 | - | - | 0 | - |
| - | - | 1.145E+06 | 154.1 | - | - | 0 | - |
| - | - | 8.463E+04 | 155.1 | - | - | 0 | - |
| - | - | 1.204E+05 | 155.1 | - | - | 0 | - |
| - | - | 9.637E+04 | 156.1 | - | - | 0 | - |
| - | - | 4.893E+05 | 157.1 | - | - | 0 | - |
| - | - | 5.35E+05 | 157.1 | - | - | 0 | - |
| - | - | 6.268E+05 | 157.1 | - | - | 0 | - |
| 15 | y | 3.008E+06 | 158.1 | 0.0003856 | 2.439 | +1 | 1 |
| - | - | 7.529E+04 | 158.1 | - | - | 0 | - |
| - | - | 1.92E+05 | 159.1 | - | - | 0 | - |
| - | - | 4.648E+05 | 159.1 | - | - | 0 | - |
| - | - | 4.289E+04 | 165.2 | - | - | 0 | - |
| - | - | 2.004E+05 | 167.1 | - | - | 0 | - |
| - | - | 4.23E+04 | 168.7 | - | - | 0 | - |
| 11 | y | 9.663E+04 | 169.1 | 0.001416 | 8.373 | +3 | 5 |
| - | - | 1.294E+05 | 170 | - | - | 0 | - |
| - | - | 6.831E+04 | 170.5 | - | - | 0 | - |
| - | - | 6.896E+04 | 170.5 | - | - | 0 | - |
| - | - | 1.413E+06 | 171.1 | - | - | 0 | - |
| - | - | 6.247E+04 | 171.2 | - | - | 0 | - |
| - | - | 1.623E+05 | 172.1 | - | - | 0 | - |
| - | - | 1.589E+05 | 172.1 | - | - | 0 | - |
| - | - | 4.51E+05 | 173.1 | - | - | 0 | - |
| - | - | 4.518E+04 | 174.1 | - | - | 0 | - |
| - | - | 8.624E+05 | 175.1 | - | - | 0 | - |
| 15 | y | 5.924E+06 | 175.1 | 0.0004021 | 2.296 | +1 | 1 |
| - | - | 4.323E+04 | 176.1 | - | - | 0 | - |
| - | - | 3.754E+05 | 176.1 | - | - | 0 | - |
| - | - | 1.074E+05 | 182.1 | - | - | 0 | - |
| - | - | 4.842E+04 | 182.2 | - | - | 0 | - |
| - | - | 1.294E+06 | 183.1 | - | - | 0 | - |
| - | - | 1.586E+05 | 184.1 | - | - | 0 | - |
| - | - | 1.209E+05 | 184.1 | - | - | 0 | - |
| - | - | 8.596E+04 | 185.1 | - | - | 0 | - |
| - | - | 6.782E+05 | 185.1 | - | - | 0 | - |
| - | - | 1.168E+05 | 185.1 | - | - | 0 | - |
| - | - | 4.687E+04 | 186.1 | - | - | 0 | - |
| - | - | 6.912E+04 | 187.1 | - | - | 0 | - |
| - | - | 9.052E+05 | 187.1 | - | - | 0 | - |
| - | - | 1.012E+05 | 188.1 | - | - | 0 | - |
| - | - | 7.231E+06 | 189.1 | - | - | 0 | - |
| - | - | 4.881E+05 | 190.1 | - | - | 0 | - |
| - | - | 6.981E+04 | 191.1 | - | - | 0 | - |
| - | - | 7.838E+04 | 192.1 | - | - | 0 | - |
| - | - | 6.509E+04 | 193.1 | - | - | 0 | - |
| - | - | 5.212E+04 | 193.1 | - | - | 0 | - |
| - | - | 1.619E+05 | 194.1 | - | - | 0 | - |
| - | - | 6.994E+04 | 195.1 | - | - | 0 | - |
| - | - | 1.736E+05 | 195.1 | - | - | 0 | - |
| - | - | 5.83E+04 | 197.1 | - | - | 0 | - |
| - | - | 1.431E+05 | 197.2 | - | - | 0 | - |
| - | - | 6.073E+05 | 199.1 | - | - | 0 | - |
| 2 | a | 6.912E+07 | 199.2 | 0.0004032 | 2.024 | +1 | 2 |
| - | - | 6.287E+04 | 200.1 | - | - | 0 | - |
| - | - | 5.321E+04 | 200.1 | - | - | 0 | - |
| - | - | 8.555E+06 | 200.2 | - | - | 0 | - |
| - | - | 2.554E+06 | 201.1 | - | - | 0 | - |
| - | - | 5.889E+04 | 201.1 | - | - | 0 | - |
| - | - | 4.105E+05 | 201.2 | - | - | 0 | - |
| - | - | 3.224E+05 | 202.1 | - | - | 0 | - |
| - | - | 1.955E+05 | 202.1 | - | - | 0 | - |
| - | - | 9.404E+05 | 203.1 | - | - | 0 | - |
| - | - | 1.497E+05 | 203.1 | - | - | 0 | - |
| - | - | 1.001E+05 | 204.1 | - | - | 0 | - |
| - | - | 6.705E+04 | 209.1 | - | - | 0 | - |
| - | - | 1.133E+05 | 209.1 | - | - | 0 | - |
| - | - | 1.46E+05 | 210.1 | - | - | 0 | - |
| - | - | 4.97E+05 | 211.1 | - | - | 0 | - |
| - | - | 4.69E+05 | 212.1 | - | - | 0 | - |
| - | - | 1.101E+05 | 213.2 | - | - | 0 | - |
| - | - | 5.142E+04 | 215.1 | - | - | 0 | - |
| - | - | 1.198E+05 | 216.1 | - | - | 0 | - |
| - | - | 1.236E+07 | 217.1 | - | - | 0 | - |
| - | - | 1.181E+06 | 218.1 | - | - | 0 | - |
| - | - | 1.267E+05 | 219.1 | - | - | 0 | - |
| - | - | 5.199E+04 | 220.1 | - | - | 0 | - |
| - | - | 1.066E+05 | 220.1 | - | - | 0 | - |
| - | - | 1.142E+05 | 220.1 | - | - | 0 | - |
| - | - | 5.816E+05 | 224.1 | - | - | 0 | - |
| - | - | 6.504E+04 | 225.1 | - | - | 0 | - |
| - | - | 1.285E+05 | 226.1 | - | - | 0 | - |
| - | - | 4.149E+05 | 227.1 | - | - | 0 | - |
| 2 | b | 2.08E+07 | 227.2 | 0.0004379 | 1.927 | +1 | 2 |
| - | - | 9.506E+04 | 228.1 | - | - | 0 | - |
| - | - | 2.775E+06 | 228.2 | - | - | 0 | - |
| - | - | 4.905E+05 | 229.1 | - | - | 0 | - |
| - | - | 2.201E+05 | 229.2 | - | - | 0 | - |
| - | - | 3.031E+05 | 230.1 | - | - | 0 | - |
| - | - | 1.287E+05 | 237.1 | - | - | 0 | - |
| - | - | 3.246E+05 | 237.1 | - | - | 0 | - |
| - | - | 7.845E+04 | 241.1 | - | - | 0 | - |
| - | - | 1.18E+05 | 242.1 | - | - | 0 | - |
| - | - | 1.34E+06 | 242.1 | - | - | 0 | - |
| - | - | 1.539E+05 | 243.1 | - | - | 0 | - |
| - | - | 3.186E+05 | 244.1 | - | - | 0 | - |
| 14 | y | 1.633E+05 | 244.1 | 0.0006211 | 2.544 | +1 | 2 |
| - | - | 1.179E+05 | 245.1 | - | - | 0 | - |
| 14 | y | 1.893E+06 | 245.1 | 0.0005075 | 2.07 | +1 | 2 |
| - | - | 6.471E+04 | 245.2 | - | - | 0 | - |
| - | - | 1.58E+05 | 246.1 | - | - | 0 | - |
| - | - | 1.018E+05 | 252.1 | - | - | 0 | - |
| - | - | 1.349E+05 | 253.2 | - | - | 0 | - |
| - | - | 1.649E+05 | 254.1 | - | - | 0 | - |
| - | - | 1.338E+05 | 254.2 | - | - | 0 | - |
| - | - | 8.609E+04 | 255.1 | - | - | 0 | - |
| - | - | 1.374E+06 | 255.1 | - | - | 0 | - |
| - | - | 6.875E+04 | 255.1 | - | - | 0 | - |
| - | - | 1.741E+05 | 256.1 | - | - | 0 | - |
| - | - | 9.325E+04 | 256.1 | - | - | 0 | - |
| - | - | 2.154E+05 | 259.1 | - | - | 0 | - |
| - | - | 6.763E+04 | 260.1 | - | - | 0 | - |
| 14 | y | 1.139E+06 | 262.2 | 0.0003866 | 1.475 | +1 | 2 |
| - | - | 1.311E+05 | 263.2 | - | - | 0 | - |
| - | - | 5.863E+04 | 268.1 | - | - | 0 | - |
| - | - | 2.785E+05 | 270.1 | - | - | 0 | - |
| - | - | 1.171E+05 | 271.1 | - | - | 0 | - |
| - | - | 2.145E+05 | 272.1 | - | - | 0 | - |
| - | - | 1.467E+05 | 279.1 | - | - | 0 | - |
| - | - | 1.67E+05 | 280.1 | - | - | 0 | - |
| - | - | 1.31E+05 | 281.1 | - | - | 0 | - |
| - | - | 2.964E+05 | 284.1 | - | - | 0 | - |
| - | - | 1.6E+05 | 284.2 | - | - | 0 | - |
| - | - | 6.382E+04 | 285.1 | - | - | 0 | - |
| - | - | 2.622E+05 | 288.2 | - | - | 0 | - |
| - | - | 7.31E+04 | 289.2 | - | - | 0 | - |
| - | - | 8.767E+04 | 294.1 | - | - | 0 | - |
| - | - | 9.582E+04 | 295.2 | - | - | 0 | - |
| - | - | 2.462E+05 | 296.1 | - | - | 0 | - |
| - | - | 9.401E+04 | 296.2 | - | - | 0 | - |
| 3 | b | 1.016E+06 | 296.2 | 0.000489 | 1.651 | +1 | 3 |
| - | - | 2.435E+05 | 297.1 | - | - | 0 | - |
| - | - | 1.725E+05 | 297.2 | - | - | 0 | - |
| - | - | 6.366E+05 | 298.1 | - | - | 0 | - |
| - | - | 3.201E+05 | 299.1 | - | - | 0 | - |
| - | - | 7.921E+04 | 301.2 | - | - | 0 | - |
| - | - | 2.411E+05 | 302.1 | - | - | 0 | - |
| - | - | 8.068E+04 | 302.2 | - | - | 0 | - |
| - | - | 6.868E+04 | 306.2 | - | - | 0 | - |
| - | - | 6.131E+04 | 307.2 | - | - | 0 | - |
| - | - | 7.251E+04 | 311.1 | - | - | 0 | - |
| - | - | 1.023E+06 | 312.2 | - | - | 0 | - |
| - | - | 2.46E+05 | 313.2 | - | - | 0 | - |
| - | - | 1.864E+06 | 313.2 | - | - | 0 | - |
| - | - | 6.193E+05 | 314.1 | - | - | 0 | - |
| - | - | 2.375E+05 | 314.2 | - | - | 0 | - |
| 3 | b | 2.402E+06 | 314.2 | 0.0006665 | 2.121 | +1 | 3 |
| - | - | 6.108E+04 | 315.1 | - | - | 0 | - |
| - | - | 2.988E+05 | 315.2 | - | - | 0 | - |
| - | - | 2.532E+06 | 316.2 | - | - | 0 | - |
| - | - | 3.186E+05 | 317.2 | - | - | 0 | - |
| - | - | 5.915E+04 | 323.1 | - | - | 0 | - |
| - | - | 9.339E+04 | 323.2 | - | - | 0 | - |
| - | - | 2.108E+05 | 324.1 | - | - | 0 | - |
| - | - | 8.831E+04 | 324.2 | - | - | 0 | - |
| - | - | 7.762E+04 | 325.1 | - | - | 0 | - |
| - | - | 6.195E+04 | 325.1 | - | - | 0 | - |
| - | - | 7.353E+04 | 326.1 | - | - | 0 | - |
| - | - | 9.097E+04 | 329.1 | - | - | 0 | - |
| - | - | 5.924E+05 | 330.2 | - | - | 0 | - |
| 13 | y | 5.588E+05 | 331.2 | 0.0007829 | 2.364 | +1 | 3 |
| 13 | y | 1.73E+06 | 332.2 | 0.000492 | 1.481 | +1 | 3 |
| - | - | 2.744E+05 | 333.2 | - | - | 0 | - |
| 9 | y | 1.235E+05 | 336.2 | 0.0007222 | 2.148 | +2 | 7 |
| - | - | 1.578E+05 | 340.2 | - | - | 0 | - |
| - | - | 5.219E+04 | 341.2 | - | - | 0 | - |
| - | - | 1.006E+06 | 341.2 | - | - | 0 | - |
| - | - | 1.863E+06 | 342.1 | - | - | 0 | - |
| - | - | 1.633E+05 | 342.2 | - | - | 0 | - |
| - | - | 3.288E+05 | 343.1 | - | - | 0 | - |
| - | - | 7.571E+04 | 344.1 | - | - | 0 | - |
| 9 | y | 9.066E+05 | 345.2 | 0.0006888 | 1.996 | +2 | 7 |
| - | - | 2.792E+05 | 345.7 | - | - | 0 | - |
| 13 | y | 1.551E+06 | 349.2 | 0.0004322 | 1.238 | +1 | 3 |
| - | - | 2.206E+05 | 350.2 | - | - | 0 | - |
| - | - | 7.245E+04 | 354.1 | - | - | 0 | - |
| - | - | 1.501E+05 | 358.2 | - | - | 0 | - |
| - | - | 9.476E+05 | 359.2 | - | - | 0 | - |
| - | - | 2.119E+05 | 360.2 | - | - | 0 | - |
| - | - | 8.859E+04 | 365.2 | - | - | 0 | - |
| - | - | 1.02E+05 | 367.2 | - | - | 0 | - |
| - | - | 6.26E+04 | 369.2 | - | - | 0 | - |
| - | - | 6.472E+04 | 379.2 | - | - | 0 | - |
| - | - | 1.261E+05 | 380.2 | - | - | 0 | - |
| - | - | 9.117E+04 | 381.2 | - | - | 0 | - |
| - | - | 8.521E+04 | 382.2 | - | - | 0 | - |
| - | - | 1.078E+05 | 382.2 | - | - | 0 | - |
| - | - | 8.749E+04 | 383.2 | - | - | 0 | - |
| - | - | 2.353E+05 | 385.2 | - | - | 0 | - |
| - | - | 8.03E+04 | 393.2 | - | - | 0 | - |
| - | - | 2.9E+05 | 393.7 | - | - | 0 | - |
| - | - | 9.561E+04 | 394.2 | - | - | 0 | - |
| - | - | 1.464E+05 | 396.2 | - | - | 0 | - |
| - | - | 3.151E+05 | 397.2 | - | - | 0 | - |
| - | - | 1.038E+05 | 398.2 | - | - | 0 | - |
| - | - | 2.56E+05 | 398.2 | - | - | 0 | - |
| - | - | 5.809E+04 | 398.2 | - | - | 0 | - |
| - | - | 6.813E+04 | 399.2 | - | - | 0 | - |
| - | - | 6.094E+04 | 399.7 | - | - | 0 | - |
| - | - | 2.744E+05 | 400.2 | - | - | 0 | - |
| - | - | 1.968E+05 | 401.2 | - | - | 0 | - |
| - | - | 3.292E+05 | 403.2 | - | - | 0 | - |
| - | - | 1.108E+05 | 405.2 | - | - | 0 | - |
| - | - | 1.231E+05 | 405.7 | - | - | 0 | - |
| - | - | 1.03E+05 | 407.2 | - | - | 0 | - |
| - | - | 2.228E+05 | 408.2 | - | - | 0 | - |
| - | - | 8.786E+04 | 408.7 | - | - | 0 | - |
| - | - | 2.414E+05 | 410.2 | - | - | 0 | - |
| - | - | 1.737E+05 | 411.2 | - | - | 0 | - |
| - | - | 7.153E+04 | 413.2 | - | - | 0 | - |
| - | - | 6.076E+04 | 414.2 | - | - | 0 | - |
| 12 | b | 7.493E+05 | 414.2 | 0.006209 | 14.99 | +3 | 12 |
| 8 | y | 1.053E+06 | 414.7 | 0.002196 | 5.294 | +2 | 8 |
| - | - | 6.149E+05 | 415.2 | - | - | 0 | - |
| - | - | 2.423E+05 | 415.3 | - | - | 0 | - |
| - | - | 1.284E+05 | 415.7 | - | - | 0 | - |
| - | - | 1.039E+05 | 416.2 | - | - | 0 | - |
| - | - | 8.258E+04 | 416.3 | - | - | 0 | - |
| - | - | 8.201E+04 | 418.2 | - | - | 0 | - |
| - | - | 1.3E+05 | 422.2 | - | - | 0 | - |
| 8 | y | 1.522E+07 | 423.2 | 0.000762 | 1.8 | +2 | 8 |
| - | - | 6.572E+06 | 423.7 | - | - | 0 | - |
| - | - | 1.81E+06 | 424.2 | - | - | 0 | - |
| - | - | 2.638E+05 | 424.7 | - | - | 0 | - |
| 4 | b | 1.174E+06 | 425.2 | 0.0006816 | 1.603 | +1 | 4 |
| - | - | 3.324E+05 | 426.2 | - | - | 0 | - |
| - | - | 4.157E+05 | 428.2 | - | - | 0 | - |
| - | - | 7.625E+05 | 429.2 | - | - | 0 | - |
| 12 | y | 4.201E+05 | 430.2 | 0.000868 | 2.017 | +1 | 4 |
| 12 | y | 1.428E+05 | 431.2 | 0.002492 | 5.779 | +1 | 4 |
| - | - | 7.669E+04 | 439.2 | - | - | 0 | - |
| - | - | 1.038E+05 | 440.2 | - | - | 0 | - |
| - | - | 8.424E+05 | 441.2 | - | - | 0 | - |
| - | - | 1.513E+05 | 442.2 | - | - | 0 | - |
| - | - | 1.278E+05 | 443.2 | - | - | 0 | - |
| 4 | b | 1.282E+06 | 443.3 | 0.0008591 | 1.938 | +1 | 4 |
| - | - | 2.857E+05 | 444.3 | - | - | 0 | - |
| 12 | y | 5.383E+05 | 448.3 | 0.0004692 | 1.047 | +1 | 4 |
| 3 | y | 3.256E+05 | 448.9 | 0.0007154 | 1.594 | +3 | 13 |
| 3 | y | 2.604E+05 | 449.2 | 0.005565 | 12.39 | +3 | 13 |
| - | - | 1.366E+05 | 449.3 | - | - | 0 | - |
| - | - | 1.885E+05 | 449.6 | - | - | 0 | - |
| 8 | b | 1.567E+05 | 450.7 | 4.049E-05 | 0.08984 | +2 | 8 |
| - | - | 8.727E+04 | 451.2 | - | - | 0 | - |
| - | - | 1.143E+05 | 452.2 | - | - | 0 | - |
| - | - | 3.677E+05 | 452.3 | - | - | 0 | - |
| - | - | 8.184E+04 | 453.3 | - | - | 0 | - |
| 3 | y | 6.568E+05 | 454.9 | 0.0007339 | 1.613 | +3 | 13 |
| - | - | 4.028E+05 | 455.2 | - | - | 0 | - |
| - | - | 1.839E+05 | 455.6 | - | - | 0 | - |
| - | - | 7.568E+04 | 455.9 | - | - | 0 | - |
| - | - | 7.324E+05 | 458.2 | - | - | 0 | - |
| - | - | 7.909E+04 | 459.2 | - | - | 0 | - |
| - | - | 9.607E+04 | 460.2 | - | - | 0 | - |
| - | - | 8.469E+04 | 463.2 | - | - | 0 | - |
| - | - | 3.306E+05 | 469.3 | - | - | 0 | - |
| - | - | 1.798E+05 | 470.2 | - | - | 0 | - |
| - | - | 9.962E+04 | 470.3 | - | - | 0 | - |
| 7 | y | 1.421E+05 | 471.7 | 0.0004692 | 0.9946 | +2 | 9 |
| - | - | 8.55E+04 | 471.9 | - | - | 0 | - |
| 14 | b | 2.264E+05 | 472.2 | 0.005866 | 12.42 | +3 | 14 |
| - | - | 1.576E+05 | 472.7 | - | - | 0 | - |
| - | - | 9.455E+04 | 478.2 | - | - | 0 | - |
| - | - | 1.328E+05 | 480.3 | - | - | 0 | - |
| 7 | y | 2.851E+06 | 480.7 | 0.0005961 | 1.24 | +2 | 9 |
| - | - | 1.319E+06 | 481.2 | - | - | 0 | - |
| - | - | 3.635E+05 | 481.7 | - | - | 0 | - |
| - | - | 8.389E+04 | 482.2 | - | - | 0 | - |
| - | - | 5.986E+04 | 486.2 | - | - | 0 | - |
| 2 | y | 6.159E+05 | 486.6 | 0.0005159 | 1.06 | +3 | 14 |
| 2 | y | 3.438E+05 | 486.9 | 0.007472 | 15.35 | +3 | 14 |
| 11 | y | 2.283E+05 | 487.3 | 0.007836 | 16.08 | +1 | 5 |
| 11 | y | 1.811E+05 | 488.2 | 0.001587 | 3.251 | +1 | 5 |
| 2 | y | 5.165E+05 | 492.6 | 4.611E-05 | 0.09361 | +3 | 14 |
| - | - | 4.627E+05 | 492.9 | - | - | 0 | - |
| - | - | 2.542E+05 | 493.3 | - | - | 0 | - |
| - | - | 1.2E+05 | 494.7 | - | - | 0 | - |
| - | - | 1.667E+05 | 495.2 | - | - | 0 | - |
| - | - | 8.516E+05 | 495.9 | - | - | 0 | - |
| - | - | 1.224E+05 | 496.2 | - | - | 0 | - |
| - | - | 7.404E+05 | 496.2 | - | - | 0 | - |
| - | - | 9.316E+04 | 496.3 | - | - | 0 | - |
| - | - | 3.506E+05 | 496.6 | - | - | 0 | - |
| - | - | 2.201E+05 | 497.3 | - | - | 0 | - |
| - | - | 1.263E+05 | 498.2 | - | - | 0 | - |
| - | - | 1.932E+05 | 500.2 | - | - | 0 | - |
| 11 | y | 1.752E+06 | 505.3 | 0.0006119 | 1.211 | +1 | 5 |
| - | - | 3.598E+05 | 506.3 | - | - | 0 | - |
| - | - | 8.616E+04 | 507.8 | - | - | 0 | - |
| - | - | 1.636E+05 | 509.3 | - | - | 0 | - |
| - | - | 3.726E+05 | 514.3 | - | - | 0 | - |
| 6 | y | 5.158E+05 | 515.3 | 0.0005837 | 1.133 | +2 | 10 |
| - | - | 8.825E+04 | 515.3 | - | - | 0 | - |
| 6 | y | 3.608E+05 | 515.7 | 0.004357 | 8.447 | +2 | 10 |
| - | - | 3.097E+05 | 516.3 | - | - | 0 | - |
| - | - | 2.306E+05 | 518.2 | - | - | 0 | - |
| - | - | 1.711E+05 | 518.3 | - | - | 0 | - |
| - | - | 2.034E+05 | 518.6 | - | - | 0 | - |
| - | - | 1.189E+05 | 518.9 | - | - | 0 | - |
| - | - | 8.098E+04 | 519.3 | - | - | 0 | - |
| 6 | y | 1.667E+07 | 524.3 | 0.002007 | 3.829 | +2 | 10 |
| 0 | Precursor | 2.881E+06 | 524.6 | 0.006723 | 12.82 | +3 | -1 |
| - | - | 8.372E+06 | 524.8 | - | - | 0 | - |
| - | - | 1.647E+06 | 524.9 | - | - | 0 | - |
| - | - | 2.996E+06 | 525.3 | - | - | 0 | - |
| - | - | 4.895E+05 | 525.8 | - | - | 0 | - |
| - | - | 8.278E+04 | 526.3 | - | - | 0 | - |
| - | - | 1.093E+05 | 529.3 | - | - | 0 | - |
| 0 | Precursor | 1.871E+06 | 530.3 | 0.000457 | 0.8618 | +3 | -1 |
| - | - | 1.897E+06 | 530.6 | - | - | 0 | - |
| - | - | 9.148E+05 | 530.9 | - | - | 0 | - |
| - | - | 2.447E+05 | 531.3 | - | - | 0 | - |
| - | - | 1.496E+05 | 531.6 | - | - | 0 | - |
| - | - | 1.927E+05 | 532.3 | - | - | 0 | - |
| - | - | 1.606E+05 | 533.3 | - | - | 0 | - |
| - | - | 1.107E+05 | 538.3 | - | - | 0 | - |
| - | - | 9.762E+04 | 539.3 | - | - | 0 | - |
| - | - | 6.833E+04 | 542.3 | - | - | 0 | - |
| - | - | 8.146E+04 | 542.3 | - | - | 0 | - |
| 5 | b | 5.076E+05 | 542.3 | 0.001262 | 2.328 | +1 | 5 |
| - | - | 9.922E+04 | 543.3 | - | - | 0 | - |
| - | - | 1.082E+05 | 543.3 | - | - | 0 | - |
| - | - | 1.551E+05 | 552.8 | - | - | 0 | - |
| - | - | 1.433E+05 | 556.3 | - | - | 0 | - |
| - | - | 1.247E+05 | 556.3 | - | - | 0 | - |
| - | - | 1.112E+05 | 556.8 | - | - | 0 | - |
| - | - | 1.123E+05 | 557.3 | - | - | 0 | - |
| - | - | 2.345E+05 | 558.8 | - | - | 0 | - |
| - | - | 2.18E+05 | 559.3 | - | - | 0 | - |
| 11 | b | 7.962E+04 | 562.3 | 0.0004618 | 0.8213 | +2 | 11 |
| 5 | y | 8.68E+05 | 564.8 | 0.0001227 | 0.2173 | +2 | 11 |
| - | - | 1.366E+05 | 565.2 | - | - | 0 | - |
| 5 | y | 9.622E+05 | 565.3 | 0.004757 | 8.415 | +2 | 11 |
| - | - | 4.003E+05 | 565.8 | - | - | 0 | - |
| - | - | 1.799E+05 | 566.3 | - | - | 0 | - |
| - | - | 1.72E+05 | 567.3 | - | - | 0 | - |
| - | - | 1.01E+05 | 569.3 | - | - | 0 | - |
| - | - | 1.076E+05 | 572.3 | - | - | 0 | - |
| - | - | 1.881E+05 | 572.8 | - | - | 0 | - |
| - | - | 1.196E+05 | 573.3 | - | - | 0 | - |
| 5 | y | 2.342E+07 | 573.8 | 0.0003933 | 0.6854 | +2 | 11 |
| 10 | y | 1.389E+07 | 574.3 | 0.003989 | 6.947 | +1 | 6 |
| - | - | 4.48E+06 | 574.8 | - | - | 0 | - |
| - | - | 1.104E+06 | 575.3 | - | - | 0 | - |
| - | - | 1.414E+05 | 575.8 | - | - | 0 | - |
| - | - | 1.802E+05 | 583.2 | - | - | 0 | - |
| - | - | 2.917E+05 | 584.3 | - | - | 0 | - |
| - | - | 2E+05 | 587.3 | - | - | 0 | - |
| 10 | y | 1.859E+06 | 592.3 | 0.0003828 | 0.6463 | +1 | 6 |
| - | - | 4.896E+05 | 593.3 | - | - | 0 | - |
| - | - | 1.626E+05 | 594.3 | - | - | 0 | - |
| - | - | 3.691E+05 | 600.3 | - | - | 0 | - |
| - | - | 1.938E+05 | 602.3 | - | - | 0 | - |
| - | - | 1.062E+05 | 611.3 | - | - | 0 | - |
| - | - | 1.304E+05 | 617.3 | - | - | 0 | - |
| - | - | 1.053E+05 | 621.3 | - | - | 0 | - |
| - | - | 8.632E+04 | 623.3 | - | - | 0 | - |
| - | - | 1.073E+05 | 624.3 | - | - | 0 | - |
| - | - | 1.889E+05 | 626.3 | - | - | 0 | - |
| - | - | 1.081E+05 | 627.3 | - | - | 0 | - |
| 4 | y | 1.535E+06 | 629.3 | 0.001774 | 2.819 | +2 | 12 |
| 4 | y | 1.028E+06 | 629.8 | 0.008301 | 13.18 | +2 | 12 |
| - | - | 6.142E+05 | 630.3 | - | - | 0 | - |
| - | - | 1.194E+05 | 630.8 | - | - | 0 | - |
| - | - | 2.149E+05 | 637.3 | - | - | 0 | - |
| - | - | 1.025E+05 | 637.8 | - | - | 0 | - |
| 4 | y | 1.13E+07 | 638.3 | 0.0002149 | 0.3367 | +2 | 12 |
| - | - | 8.258E+06 | 638.8 | - | - | 0 | - |
| - | - | 3.347E+06 | 639.3 | - | - | 0 | - |
| - | - | 7.39E+05 | 639.8 | - | - | 0 | - |
| - | - | 1.88E+05 | 640.3 | - | - | 0 | - |
| - | - | 3.335E+05 | 641.3 | - | - | 0 | - |
| - | - | 1.177E+05 | 642.3 | - | - | 0 | - |
| - | - | 1.091E+05 | 642.8 | - | - | 0 | - |
| - | - | 8.092E+04 | 643.3 | - | - | 0 | - |
| - | - | 2.271E+05 | 643.4 | - | - | 0 | - |
| - | - | 1.48E+05 | 643.8 | - | - | 0 | - |
| - | - | 7.376E+05 | 646.3 | - | - | 0 | - |
| - | - | 2.319E+05 | 647.3 | - | - | 0 | - |
| - | - | 3.428E+05 | 648.8 | - | - | 0 | - |
| - | - | 2.254E+05 | 649.3 | - | - | 0 | - |
| - | - | 1.001E+05 | 649.8 | - | - | 0 | - |
| - | - | 1.339E+05 | 651.8 | - | - | 0 | - |
| - | - | 2.489E+05 | 652.3 | - | - | 0 | - |
| - | - | 1.242E+05 | 652.8 | - | - | 0 | - |
| - | - | 3.245E+05 | 653.3 | - | - | 0 | - |
| - | - | 3.404E+05 | 654.3 | - | - | 0 | - |
| - | - | 3.579E+05 | 654.8 | - | - | 0 | - |
| - | - | 4.338E+05 | 655.3 | - | - | 0 | - |
| - | - | 1.293E+05 | 655.8 | - | - | 0 | - |
| - | - | 2.019E+05 | 656.3 | - | - | 0 | - |
| - | - | 3.733E+05 | 657.3 | - | - | 0 | - |
| - | - | 4.014E+05 | 657.8 | - | - | 0 | - |
| - | - | 2.274E+05 | 658.3 | - | - | 0 | - |
| - | - | 1.527E+05 | 658.8 | - | - | 0 | - |
| - | - | 4.375E+05 | 659.3 | - | - | 0 | - |
| - | - | 1.806E+05 | 659.8 | - | - | 0 | - |
| - | - | 1.243E+05 | 660.3 | - | - | 0 | - |
| - | - | 1.281E+05 | 660.4 | - | - | 0 | - |
| - | - | 5.62E+05 | 660.8 | - | - | 0 | - |
| - | - | 4.039E+05 | 661.3 | - | - | 0 | - |
| - | - | 1.691E+05 | 661.8 | - | - | 0 | - |
| - | - | 9.916E+05 | 663.8 | - | - | 0 | - |
| - | - | 7.539E+05 | 664.3 | - | - | 0 | - |
| - | - | 3.884E+05 | 664.8 | - | - | 0 | - |
| - | - | 1.672E+05 | 665.3 | - | - | 0 | - |
| - | - | 2.261E+05 | 665.8 | - | - | 0 | - |
| - | - | 4.392E+05 | 666.3 | - | - | 0 | - |
| - | - | 1.656E+06 | 666.8 | - | - | 0 | - |
| - | - | 1.156E+06 | 667.3 | - | - | 0 | - |
| - | - | 4.656E+05 | 667.8 | - | - | 0 | - |
| - | - | 7.411E+04 | 668.3 | - | - | 0 | - |
| - | - | 1.225E+05 | 671.3 | - | - | 0 | - |
| 9 | y | 1.917E+06 | 671.3 | 0.00042 | 0.6256 | +1 | 7 |
| - | - | 1.203E+05 | 671.8 | - | - | 0 | - |
| 9 | y | 1.428E+06 | 672.3 | 0.006089 | 9.057 | +1 | 7 |
| 3 | y | 5.812E+06 | 672.8 | 0.0001642 | 0.244 | +2 | 13 |
| 3 | y | 5.542E+06 | 673.3 | 0.009499 | 14.11 | +2 | 13 |
| - | - | 2.353E+06 | 673.8 | - | - | 0 | - |
| - | - | 2.967E+06 | 674.3 | - | - | 0 | - |
| - | - | 2.624E+05 | 674.8 | - | - | 0 | - |
| - | - | 8.892E+05 | 675.3 | - | - | 0 | - |
| - | - | 2.124E+05 | 676.3 | - | - | 0 | - |
| - | - | 2.104E+05 | 677.8 | - | - | 0 | - |
| - | - | 9.628E+04 | 678.3 | - | - | 0 | - |
| - | - | 1.369E+06 | 680.8 | - | - | 0 | - |
| - | - | 1.251E+06 | 681.3 | - | - | 0 | - |
| 3 | y | 1.768E+08 | 681.8 | 0.0008633 | 1.266 | +2 | 13 |
| - | - | 1.292E+08 | 682.3 | - | - | 0 | - |
| - | - | 5.258E+07 | 682.8 | - | - | 0 | - |
| - | - | 1.367E+07 | 683.3 | - | - | 0 | - |
| - | - | 2.538E+06 | 683.8 | - | - | 0 | - |
| - | - | 8.551E+04 | 684.3 | - | - | 0 | - |
| - | - | 1.495E+06 | 686.8 | - | - | 0 | - |
| - | - | 1.143E+06 | 687.3 | - | - | 0 | - |
| - | - | 3.553E+05 | 687.8 | - | - | 0 | - |
| - | - | 9.999E+04 | 688.3 | - | - | 0 | - |
| 9 | y | 1.22E+07 | 689.4 | 0.0002923 | 0.424 | +1 | 7 |
| - | - | 4.234E+06 | 690.4 | - | - | 0 | - |
| - | - | 1.089E+06 | 691.4 | - | - | 0 | - |
| - | - | 1.599E+05 | 692.4 | - | - | 0 | - |
| 14 | b | 5.976E+05 | 699.3 | 0.007802 | 11.16 | +2 | 14 |
| - | - | 2.692E+05 | 700.3 | - | - | 0 | - |
| - | - | 1.072E+05 | 716.4 | - | - | 0 | - |
| - | - | 1.861E+05 | 717.4 | - | - | 0 | - |
| - | - | 1.504E+05 | 724.4 | - | - | 0 | - |
| - | - | 1.124E+05 | 728.4 | - | - | 0 | - |
| 2 | y | 1.722E+06 | 729.4 | 0.0001244 | 0.1705 | +2 | 14 |
| 2 | y | 1.462E+06 | 729.9 | 0.009215 | 12.63 | +2 | 14 |
| - | - | 6.836E+05 | 730.4 | - | - | 0 | - |
| - | - | 1.607E+05 | 730.9 | - | - | 0 | - |
| - | - | 1.069E+05 | 734.9 | - | - | 0 | - |
| - | - | 2.493E+05 | 737.4 | - | - | 0 | - |
| - | - | 1.717E+05 | 737.9 | - | - | 0 | - |
| 2 | y | 1.445E+07 | 738.4 | 3.002E-05 | 0.04066 | +2 | 14 |
| - | - | 1.218E+07 | 738.9 | - | - | 0 | - |
| - | - | 5.652E+06 | 739.4 | - | - | 0 | - |
| - | - | 1.374E+06 | 739.9 | - | - | 0 | - |
| - | - | 3.106E+05 | 740.4 | - | - | 0 | - |
| - | - | 9.336E+05 | 743.4 | - | - | 0 | - |
| - | - | 4.862E+05 | 743.9 | - | - | 0 | - |
| 7 | b | 3.528E+05 | 744.4 | 0.005341 | 7.175 | +1 | 7 |
| - | - | 1.018E+05 | 744.9 | - | - | 0 | - |
| - | - | 1.695E+05 | 758.4 | - | - | 0 | - |
| - | - | 2.946E+05 | 768.4 | - | - | 0 | - |
| - | - | 5.734E+05 | 770.4 | - | - | 0 | - |
| - | - | 2.295E+05 | 771.4 | - | - | 0 | - |
| - | - | 1.111E+05 | 784.4 | - | - | 0 | - |
| - | - | 2.43E+05 | 785.4 | - | - | 0 | - |
| - | - | 1.065E+05 | 786.4 | - | - | 0 | - |
| 0 | Precursor | 3.069E+05 | 786.4 | 0.01162 | 14.77 | +2 | -1 |
| - | - | 3.799E+05 | 787.4 | - | - | 0 | - |
| - | - | 1.637E+05 | 788.4 | - | - | 0 | - |
| - | - | 4.409E+05 | 798.4 | - | - | 0 | - |
| - | - | 1.996E+05 | 799.4 | - | - | 0 | - |
| - | - | 2.347E+05 | 803.4 | - | - | 0 | - |
| - | - | 1.199E+05 | 804.4 | - | - | 0 | - |
| - | - | 1.363E+05 | 810.4 | - | - | 0 | - |
| - | - | 3.522E+05 | 811.4 | - | - | 0 | - |
| - | - | 1.214E+05 | 812.4 | - | - | 0 | - |
| 8 | y | 1.903E+06 | 828.4 | 0.000295 | 0.3561 | +1 | 8 |
| - | - | 7.959E+05 | 829.4 | - | - | 0 | - |
| - | - | 1.954E+05 | 830.4 | - | - | 0 | - |
| - | - | 1.611E+05 | 855.4 | - | - | 0 | - |
| - | - | 2.336E+05 | 857.4 | - | - | 0 | - |
| - | - | 2.242E+05 | 858.4 | - | - | 0 | - |
| - | - | 1.256E+05 | 859.4 | - | - | 0 | - |
| - | - | 1.648E+05 | 867.4 | - | - | 0 | - |
| - | - | 1.837E+05 | 873.4 | - | - | 0 | - |
| - | - | 1.106E+05 | 881.4 | - | - | 0 | - |
| 8 | b | 7.822E+04 | 883.5 | 0.004448 | 5.034 | +1 | 8 |
| - | - | 2.013E+05 | 885.4 | - | - | 0 | - |
| - | - | 4.509E+05 | 887.4 | - | - | 0 | - |
| - | - | 1.925E+05 | 888.4 | - | - | 0 | - |
| - | - | 1.052E+05 | 889.4 | - | - | 0 | - |
| - | - | 2.549E+05 | 891.4 | - | - | 0 | - |
| - | - | 1.099E+05 | 892.4 | - | - | 0 | - |
| - | - | 1.841E+05 | 897.4 | - | - | 0 | - |
| - | - | 1.586E+05 | 899.5 | - | - | 0 | - |
| 8 | b | 4.72E+05 | 900.5 | 0.0102 | 11.33 | +1 | 8 |
| - | - | 1.069E+05 | 901.5 | - | - | 0 | - |
| - | - | 1.139E+05 | 909.5 | - | - | 0 | - |
| - | - | 9.186E+05 | 915.4 | - | - | 0 | - |
| - | - | 3.08E+05 | 916.4 | - | - | 0 | - |
| - | - | 1.658E+05 | 917.4 | - | - | 0 | - |
| - | - | 8.736E+04 | 918.5 | - | - | 0 | - |
| - | - | 1.568E+05 | 925.5 | - | - | 0 | - |
| - | - | 2.036E+05 | 926.4 | - | - | 0 | - |
| - | - | 1.237E+05 | 927.4 | - | - | 0 | - |
| 7 | y | 1.691E+06 | 943.5 | 0.0004436 | 0.4702 | +1 | 9 |
| - | - | 7.927E+05 | 944.5 | - | - | 0 | - |
| - | - | 4.084E+05 | 945.5 | - | - | 0 | - |
| - | - | 8.488E+04 | 951.5 | - | - | 0 | - |
| - | - | 1.406E+05 | 954.5 | - | - | 0 | - |
| - | - | 1.171E+05 | 957.5 | - | - | 0 | - |
| 7 | y | 7.24E+04 | 960.5 | 0.006607 | 6.879 | +1 | 9 |
| - | - | 1.647E+05 | 968.5 | - | - | 0 | - |
| - | - | 2.127E+05 | 969.5 | - | - | 0 | - |
| - | - | 1.124E+05 | 970.5 | - | - | 0 | - |
| - | - | 2.396E+05 | 972.5 | - | - | 0 | - |
| - | - | 1.422E+05 | 973.5 | - | - | 0 | - |
| - | - | 1.397E+06 | 986.5 | - | - | 0 | - |
| - | - | 1.61E+06 | 987.5 | - | - | 0 | - |
| - | - | 7.058E+05 | 988.5 | - | - | 0 | - |
| - | - | 2.572E+05 | 989.5 | - | - | 0 | - |
| - | - | 2.784E+05 | 990.5 | - | - | 0 | - |
| - | - | 1.326E+05 | 991.5 | - | - | 0 | - |
| - | - | 2.106E+05 | 996.5 | - | - | 0 | - |
| - | - | 1.46E+05 | 997.5 | - | - | 0 | - |
| - | - | 2.025E+05 | 1000 | - | - | 0 | - |
| - | - | 1.944E+05 | 1005 | - | - | 0 | - |
| - | - | 5.926E+05 | 1012 | - | - | 0 | - |
| - | - | 3.69E+05 | 1013 | - | - | 0 | - |
| - | - | 1.382E+06 | 1014 | - | - | 0 | - |
| - | - | 7.891E+05 | 1015 | - | - | 0 | - |
| - | - | 2.525E+05 | 1016 | - | - | 0 | - |
| 6 | y | 4.757E+06 | 1030 | 0.0007337 | 0.712 | +1 | 10 |
| - | - | 2.625E+06 | 1031 | - | - | 0 | - |
| - | - | 1.015E+06 | 1032 | - | - | 0 | - |
| - | - | 1.745E+05 | 1033 | - | - | 0 | - |
| - | - | 1.178E+05 | 1040 | - | - | 0 | - |
| 6 | y | 1.366E+05 | 1048 | 0.003235 | 3.088 | +1 | 10 |
| - | - | 1.243E+05 | 1058 | - | - | 0 | - |
| - | - | 5.169E+05 | 1074 | - | - | 0 | - |
| - | - | 2.348E+05 | 1075 | - | - | 0 | - |
| - | - | 1.309E+05 | 1076 | - | - | 0 | - |
| - | - | 3.779E+05 | 1084 | - | - | 0 | - |
| - | - | 2.388E+05 | 1085 | - | - | 0 | - |
| - | - | 3.202E+05 | 1087 | - | - | 0 | - |
| - | - | 1.809E+05 | 1088 | - | - | 0 | - |
| - | - | 1.057E+05 | 1089 | - | - | 0 | - |
| - | - | 7.431E+05 | 1102 | - | - | 0 | - |
| - | - | 4.703E+05 | 1103 | - | - | 0 | - |
| - | - | 1.937E+05 | 1104 | - | - | 0 | - |
| - | - | 1.521E+05 | 1112 | - | - | 0 | - |
| - | - | 1.813E+05 | 1113 | - | - | 0 | - |
| - | - | 1.097E+05 | 1114 | - | - | 0 | - |
| - | - | 1.839E+05 | 1120 | - | - | 0 | - |
| - | - | 1.169E+05 | 1128 | - | - | 0 | - |
| 5 | y | 1.582E+06 | 1130 | 0.002131 | 1.887 | +1 | 11 |
| - | - | 9.351E+05 | 1131 | - | - | 0 | - |
| - | - | 3.809E+05 | 1132 | - | - | 0 | - |
| - | - | 1.139E+05 | 1133 | - | - | 0 | - |
| - | - | 1.711E+05 | 1143 | - | - | 0 | - |
| - | - | 1.087E+05 | 1144 | - | - | 0 | - |
| - | - | 1.457E+05 | 1145 | - | - | 0 | - |
| - | - | 1.222E+05 | 1159 | - | - | 0 | - |
| - | - | 8.428E+05 | 1161 | - | - | 0 | - |
| - | - | 4.411E+05 | 1162 | - | - | 0 | - |
| - | - | 1.948E+05 | 1163 | - | - | 0 | - |
| - | - | 2.787E+05 | 1171 | - | - | 0 | - |
| - | - | 1.685E+05 | 1172 | - | - | 0 | - |
| - | - | 1.189E+06 | 1189 | - | - | 0 | - |
| - | - | 5.742E+05 | 1190 | - | - | 0 | - |
| - | - | 1.6E+05 | 1191 | - | - | 0 | - |
| - | - | 1.14E+05 | 1206 | - | - | 0 | - |
| - | - | 9.267E+05 | 1207 | - | - | 0 | - |
| - | - | 5.712E+05 | 1208 | - | - | 0 | - |
| - | - | 2.128E+05 | 1209 | - | - | 0 | - |
| - | - | 1.109E+05 | 1216 | - | - | 0 | - |
| - | - | 8.395E+04 | 1241 | - | - | 0 | - |
| - | - | 1.158E+05 | 1274 | - | - | 0 | - |
| - | - | 1.202E+05 | 1303 | - | - | 0 | - |
| 3 | y | 1.735E+05 | 1346 | 0.001617 | 1.201 | +1 | 13 |
| - | - | 1.372E+05 | 1347 | - | - | 0 | - |
| - | - | 9.037E+04 | 1348 | - | - | 0 | - |

m/z Charge Intensity FragmentType MassShift Position
125.10782623291016 0 36132.473
126.05530548095703 0 1116264.1
127.05870056152344 0 77140.46
127.0866928100586 0 37903.957
128.08221435546875 0 49043.953
129.0661163330078 0 135009
129.1027069091797 0 194097.8
129.11386108398438 0 108999.35
130.05003356933594 0 70742.9
130.0614776611328 0 615268.75
130.0978240966797 0 340005
131.08200073242188 0 44790.23
133.0605926513672 0 53204.367
133.09756469726562 0 104138.21
136.03924560546875 0 40786.22
138.0918426513672 0 51622.164
139.08685302734375 0 78098.76
140.08216857910156 0 128198.05
140.14390563964844 0 57418.746
141.0661163330078 0 59693.734
141.102783203125 0 138668.25
143.08187866210938 0 47419.21
143.19224548339844 0 37655.02
144.0658416748047 0 345340.7
147.07672119140625 0 376316.44
150.68080139160156 0 40571.895
153.06622314453125 0 103311.82
154.05023193359375 0 1145199.9
155.05357360839844 0 84633.11
155.11825561523438 0 120434.766
156.1024169921875 0 96374.94
157.0611572265625 0 489328.88
157.09759521484375 0 535018.06
157.1087646484375 0 626794.2
158.09278869628906 0 3007740.5 y Ammonia loss 14
158.11256408691406 0 75285.95
159.09622192382812 0 191963.25
159.1131134033203 0 464811.03
165.22463989257812 0 42886.824
167.0818328857422 0 200443.7
168.658935546875 0 42298.094
169.09722900390625 0 96629.734 y 10
170.0449981689453 0 129432.47
170.51834106445312 0 68306.69
170.52516174316406 0 68960.484
171.07679748535156 0 1412562.1
171.18577575683594 0 62471.31
172.0607147216797 0 162328.08
172.0802459716797 0 158936.72
173.12887573242188 0 450959.44
174.13186645507812 0 45178.59
175.07167053222656 0 862409
175.11935424804688 0 5924248 y 14
176.07516479492188 0 43227.727
176.12277221679688 0 375380.1
182.0928955078125 0 107382.98
182.1548614501953 0 48420.332
183.11312866210938 0 1294315.4
184.108642578125 0 158610.72
184.11695861816406 0 120888.22
185.0561981201172 0 85962.33
185.0924072265625 0 678223.44
185.10256958007812 0 116813.586
186.06121826171875 0 46874.95
187.07139587402344 0 69117.32
187.1080780029297 0 905164
188.11143493652344 0 101247.625
189.08737182617188 0 7230604.5
190.09085083007812 0 488071.62
191.09201049804688 0 69809.305
192.1134490966797 0 78379.54
193.0823974609375 0 65093.254
193.09779357910156 0 52123.027
194.0924835205078 0 161893.66
195.07601928710938 0 69936.87
195.149658203125 0 173639.48
197.0927734375 0 58303.46
197.1652069091797 0 143105.44
199.07177734375 0 607349.8
199.18089294433594 0 69119864 a 1
200.0755157470703 0 62865.99
200.10385131835938 0 53211.184
200.18421936035156 0 8554672
201.12364196777344 0 2554141.8
201.13490295410156 0 58886.75
201.18707275390625 0 410545.47
202.11880493164062 0 322414.75
202.1273651123047 0 195539.88
203.06649780273438 0 940374.6
203.1027069091797 0 149681.53
204.0703582763672 0 100072.234
209.10353088378906 0 67051.57
209.1398468017578 0 113277.54
210.08792114257812 0 145991.06
211.10813903808594 0 496970.03
212.10333251953125 0 468975.4
213.16006469726562 0 110127.6
215.11363220214844 0 51423.83
216.13473510742188 0 119764.55
217.082275390625 0 12361109
218.085693359375 0 1181461.4
219.08712768554688 0 126690.97
220.0723876953125 0 51991.375
220.09291076660156 0 106633.195
220.12918090820312 0 114223.79
224.10325622558594 0 581604.6
225.10702514648438 0 65041.355
226.1192169189453 0 128521.01
227.1141357421875 0 414901.56
227.17584228515625 0 20798856 b 1
228.09783935546875 0 95062.805
228.1791534423828 0 2775234
229.11871337890625 0 490461.1
229.1812744140625 0 220095.06
230.11398315429688 0 303075.97
237.09896850585938 0 128687.89
237.13504028320312 0 324567.2
241.1309051513672 0 78449.16
242.1006317138672 0 117968.43
242.11404418945312 0 1340115.5
243.11720275878906 0 153861.34
244.1294403076172 0 318626.72
244.1410369873047 0 163326.8 y Water loss 13
245.1112823486328 0 117931.57
245.12493896484375 0 1893300.1 y Ammonia loss 13
245.1864776611328 0 64705.23
246.1285400390625 0 157967.98
252.1346893310547 0 101828.3
253.22865295410156 0 134904.25
254.11354064941406 0 164924.3
254.16171264648438 0 133826.64
255.09521484375 0 86091.88
255.1092529296875 0 1373742.2
255.14463806152344 0 68753.805
256.1124572753906 0 174095.06
256.127685546875 0 93254.97
259.1402893066406 0 215426.88
260.1443786621094 0 67627.695
262.1513671875 0 1138628.2 y 13
263.1537780761719 0 131115
268.0921936035156 0 58626.145
270.1451110839844 0 278472.4
271.1416015625 0 117060.8
272.1357421875 0 214518.47
279.10943603515625 0 146708.1
280.12945556640625 0 166989.84
281.1253662109375 0 131017.98
284.1243896484375 0 296379.94
284.1612854003906 0 160001.89
285.1293640136719 0 63822.016
288.1555480957031 0 262247.44
289.15155029296875 0 73096.21
294.145751953125 0 87672.95
295.1759948730469 0 95821.3
296.1357421875 0 246225.45
296.17706298828125 0 94007.76
296.1973571777344 0 1016211.06 b Water loss 2
297.11962890625 0 243513
297.2008972167969 0 172471.19
298.1405334472656 0 636551.06
299.1358642578125 0 320141.6
301.1504821777344 0 79211.914
302.1352233886719 0 241147.6
302.17138671875 0 80682.9
306.1583251953125 0 68676.28
307.1605224609375 0 61307.082
311.1350402832031 0 72511.17
312.1558837890625 0 1022576.3
313.1527099609375 0 246048.78
313.1875915527344 0 1864399.1
314.14666748046875 0 619336.2
314.19049072265625 0 237530.44
314.2080993652344 0 2401857 b 2
315.1497802734375 0 61080.934
315.21112060546875 0 298819.66
316.1508483886719 0 2531817.2
317.15185546875 0 318633.28
323.13775634765625 0 59154.625
323.1707763671875 0 93393.984
324.13055419921875 0 210799.38
324.167724609375 0 88311.07
325.1153564453125 0 77615.81
325.1331481933594 0 61947.277
326.13555908203125 0 73533.61
329.145751953125 0 90969.93
330.16632080078125 0 592410.5
331.1716613769531 0 558814 y Water loss 12
332.1569519042969 0 1729581.8 y Ammonia loss 12
333.1594543457031 0 274371.8
336.17791748046875 0 123517.95 y Water loss 8
340.1507263183594 0 157757.8
341.1588439941406 0 52188.047
341.18255615234375 0 1006043
342.14141845703125 0 1863294
342.1836853027344 0 163258.5
343.14447021484375 0 328769.16
344.1449279785156 0 75712.76
345.18316650390625 0 906574.6 y 8
345.6845703125 0 279241.34
349.1834411621094 0 1551484.5 y 12
350.1859130859375 0 220633.42
354.1427917480469 0 72449.66
358.16046142578125 0 150058.84
359.1675109863281 0 947641.1
360.1693420410156 0 211946.53
365.1806640625 0 88590.625
367.1622619628906 0 102008.19
369.22491455078125 0 62600.75
379.2344055175781 0 64719.914
380.1565246582031 0 126094.49
381.1892395019531 0 91166.42
382.17236328125 0 85207.99
382.2088317871094 0 107845.766
383.1982421875 0 87485.445
385.17218017578125 0 235296.12
393.22015380859375 0 80302.17
393.70941162109375 0 290043.8
394.2100524902344 0 95613.06
396.1870422363281 0 146359.44
397.24554443359375 0 315090.62
398.1702575683594 0 103786.16
398.2133483886719 0 255958.06
398.2413330078125 0 58089.82
399.2238464355469 0 68129.59
399.7130126953125 0 60944.484
400.218505859375 0 274373.12
401.24041748046875 0 196847.27
403.18267822265625 0 329152
405.22381591796875 0 110771.31
405.7159423828125 0 123084.664
407.2304992675781 0 102986.03
408.22845458984375 0 222763.36
408.73223876953125 0 87855.17
410.20361328125 0 241351.53
411.2233581542969 0 173672.9
413.2127685546875 0 71528.41
414.2012939453125 0 60758.035
414.2287902832031 0 749282.4 b 11
414.7219543457031 0 1053319.4 y Ammonia loss 7
415.2227478027344 0 614880.8
415.255615234375 0 242309.88
415.72381591796875 0 128381.55
416.2174987792969 0 103869.68
416.2559509277344 0 82579.29
418.1951599121094 0 82014.02
422.22662353515625 0 129975.62
423.2337951660156 0 15215620 y 7
423.7350769042969 0 6572338.5
424.2361755371094 0 1810305.9
424.7368469238281 0 263773.16
425.2401428222656 0 1173706.9 b Water loss 3
426.2431945800781 0 332393.16
428.2164611816406 0 415687.56
429.2348327636719 0 762522.56
430.239990234375 0 420063.5 y Water loss 11
431.2223815917969 0 142808.56 y Ammonia loss 11
439.19268798828125 0 76692.836
440.2232971191406 0 103786.58
441.210205078125 0 842370.4
442.21307373046875 0 151261.97
443.2181396484375 0 127841.4
443.2508850097656 0 1281621.6 b 3
444.2544250488281 0 285698.4
448.25189208984375 0 538347.1 y 11
448.88897705078125 0 325614.1 y Water loss 2
449.2218322753906 0 260420.33 y Ammonia loss 2
449.25543212890625 0 136615.61
449.556396484375 0 188520.25
450.74285888671875 0 156665.78 b 7
451.2411193847656 0 87273.5
452.2265625 0 114288.35
452.2625427246094 0 367674.62
453.26397705078125 0 81843.6
454.89251708984375 0 656779.94 y 2
455.2262878417969 0 402810.47
455.5613098144531 0 183895.31
455.8984069824219 0 75684.33
458.236083984375 0 732442.56
459.2370300292969 0 79089.41
460.2416687011719 0 96072.54
463.2257995605469 0 84688.016
469.2891845703125 0 330640.06
470.2386474609375 0 179777.22
470.2889099121094 0 99619.18
471.7407531738281 0 142110.67 y Water loss 6
471.908447265625 0 85502.96
472.2380676269531 0 226376.39 b 13
472.7353820800781 0 157647.28
478.20355224609375 0 94547.414
480.2523193359375 0 132848.02
480.7471008300781 0 2850922.8 y 6
481.24853515625 0 1319078.1
481.74945068359375 0 363509.22
482.2479248046875 0 83893.87
486.24371337890625 0 59861.953
486.5834655761719 0 615874.7 y Water loss 1
486.9184265136719 0 343831.56 y Ammonia loss 1
487.2544860839844 0 228349.06 y Water loss 10
488.2479248046875 0 181071.05 y Ammonia loss 10
492.5865173339844 0 516523.28 y 1
492.922119140625 0 462675.66
493.25592041015625 0 254180.75
494.7413635253906 0 120029.16
495.2427673339844 0 166669.56
495.9149475097656 0 851595.8
496.2142028808594 0 122359.99
496.2499084472656 0 740424.44
496.3101501464844 0 93164.08
496.5836181640625 0 350555.44
497.2850341796875 0 220095.39
498.23602294921875 0 126279.83
500.19915771484375 0 193167.83
505.27349853515625 0 1752090.6 y 10
506.27618408203125 0 359804.6
507.7521057128906 0 86161.26
509.2569580078125 0 163609.64
514.3237915039062 0 372559.6
515.2566528320312 0 515759.28 y Water loss 5
515.330810546875 0 88251.055
515.7536010742188 0 360822.25 y Ammonia loss 5
516.257080078125 0 309718.28
518.2094116210938 0 230600.31
518.2734375 0 171127.86
518.609619140625 0 203382.39
518.9439086914062 0 118942.06
519.2738037109375 0 80982.74
524.2645263671875 0 16665974 y 5
524.6123657226562 0 2880941 Precursor Ammonia loss
524.764404296875 0 8371530
524.9464111328125 0 1647311.5
525.2669067382812 0 2995762.2
525.7669067382812 0 489509.16
526.2640380859375 0 82780.2
529.2531127929688 0 109284.24
530.2816162109375 0 1871490.1 Precursor
530.6155395507812 0 1897216.1
530.949951171875 0 914804.25
531.2828369140625 0 244716.97
531.6165161132812 0 149638.19
532.2720947265625 0 192690.58
533.2543334960938 0 160607.58
538.3104858398438 0 110717.586
539.2998657226562 0 97624.67
542.2543334960938 0 68326.13
542.2727661132812 0 81463.05
542.3197021484375 0 507558.4 b 4
543.2512817382812 0 99219.7
543.3235473632812 0 108230.375
552.7857666015625 0 155119.75
556.2814331054688 0 143340.81
556.3207397460938 0 124724.57
556.7730712890625 0 111197.51
557.2742919921875 0 112317.74
558.7901611328125 0 234462.53
559.2886962890625 0 217950.05
562.290283203125 0 79622.54 b Water loss 10
564.7913208007812 0 867999.4 y Water loss 4
565.2384643554688 0 136648.98
565.2882080078125 0 962229.7 y Ammonia loss 4
565.7866821289062 0 400346.97
566.2880859375 0 179914.9
567.2938232421875 0 172008.42
569.2691650390625 0 101024.26
572.2736206054688 0 107587.33
572.7908935546875 0 188130.44
573.2916259765625 0 119551.46
573.797119140625 0 23417830 y 4
574.29833984375 0 13893403 y Water loss 9
574.7998046875 0 4479748
575.3002319335938 0 1103859.9
575.8069458007812 0 141368.02
583.2459106445312 0 180200.52
584.3162841796875 0 291656.4
587.2783813476562 0 200031.98
592.3052978515625 0 1859304.4 y 9
593.3082885742188 0 489566.34
594.3101196289062 0 162617.78
600.2730102539062 0 369065.2
602.2904052734375 0 193810.23
611.3163452148438 0 106154.02
617.2994384765625 0 130415.94
621.3056030273438 0 105301.44
623.3145751953125 0 86318.58
624.3096313476562 0 107310.67
626.325927734375 0 188912.16
627.3301391601562 0 108114.77
629.3145141601562 0 1534811.1 y Water loss 3
629.8130493164062 0 1028207.3 y Ammonia loss 3
630.31591796875 0 614187.25
630.8152465820312 0 119384.37
637.3060913085938 0 214915.77
637.8224487304688 0 102468.19
638.3182373046875 0 11299613 y 3
638.81982421875 0 8258296.5
639.32080078125 0 3346826.2
639.8216552734375 0 739033.94
640.3220825195312 0 188000.12
641.3360595703125 0 333502.88
642.3245849609375 0 117656.02
642.8150634765625 0 109124.21
643.3084106445312 0 80922.66
643.355712890625 0 227068.31
643.8179931640625 0 147998.14
646.3163452148438 0 737593.7
647.3201904296875 0 231878.55
648.8193359375 0 342847.25
649.318359375 0 225403.22
649.8207397460938 0 100107.484
651.8198852539062 0 133891.47
652.3148803710938 0 248880.8
652.818359375 0 124216.54
653.3339233398438 0 324501.75
654.32470703125 0 340440.16
654.8181762695312 0 357858.78
655.3211059570312 0 433834
655.8121948242188 0 129342.17
656.3064575195312 0 201888.86
657.2859497070312 0 373298.12
657.8255004882812 0 401386
658.3260498046875 0 227383.25
658.822998046875 0 152650.53
659.34375 0 437530.56
659.8331909179688 0 180569.5
660.2991943359375 0 124252.1
660.3526000976562 0 128051.29
660.8201293945312 0 562029.7
661.3231811523438 0 403944.34
661.8258666992188 0 169122.45
663.8232421875 0 991611.75
664.320556640625 0 753900.8
664.81689453125 0 388392.12
665.3234252929688 0 167180.28
665.8240966796875 0 226063.89
666.3238525390625 0 439221.16
666.829345703125 0 1656205
667.3302001953125 0 1156372.9
667.832763671875 0 465634.66
668.3333740234375 0 74109.73
671.2847900390625 0 122525.984
671.3475341796875 0 1916769.2 y Water loss 8
671.8168334960938 0 120283.93
672.3372192382812 0 1427679.1 y Ammonia loss 8
672.8289184570312 0 5811809.5 y Water loss 2
673.3302612304688 0 5542163 y Ammonia loss 2
673.82958984375 0 2352663.5
674.314208984375 0 2967246
674.8273315429688 0 262375.8
675.3137817382812 0 889190.44
676.3187255859375 0 212354.02
677.8199462890625 0 210418.27
678.3295288085938 0 96282.445
680.8268432617188 0 1368830
681.3287963867188 0 1250789.4
681.8348999023438 0 176759600 y 2
682.3359375 0 129233640
682.8372192382812 0 52579640
683.3380737304688 0 13668467
683.839111328125 0 2538401.5
684.3281860351562 0 85511.79
686.8263549804688 0 1495138.8
687.3289794921875 0 1143247.5
687.8290405273438 0 355250.7
688.340087890625 0 99988.38
689.3579711914062 0 12197581 y 8
690.3607788085938 0 4233635
691.3626708984375 0 1089073.5
692.3634033203125 0 159863.81
699.3411865234375 0 597591 b Ammonia loss 13
700.3447265625 0 269172.47
716.3635864257812 0 107245.69
717.3528442382812 0 186143.42
724.37353515625 0 150358.84
728.3660888671875 0 112442.055
729.3709106445312 0 1721839.5 y Water loss 1
729.8720092773438 0 1462102.9 y Ammonia loss 1
730.3711547851562 0 683643.06
730.8700561523438 0 160668.39
734.8665771484375 0 106874.766
737.3679809570312 0 249302
737.8715209960938 0 171701.89
738.3760986328125 0 14451505 y 1
738.877685546875 0 12175850
739.3788452148438 0 5652146.5
739.8802490234375 0 1373693.6
740.3790893554688 0 310620.62
743.3687133789062 0 933573.8
743.8696899414062 0 486227.56
744.3720703125 0 352790.16 b 6
744.864013671875 0 101794.234
758.38037109375 0 169460.12
768.3994750976562 0 294608.78
770.4159545898438 0 573437
771.420654296875 0 229498.38
784.388427734375 0 111077.555
785.4268188476562 0 242999.92
786.3612060546875 0 106533.8
786.4164428710938 0 306893.8 Precursor Ammonia loss
787.396240234375 0 379922.6
788.4010009765625 0 163680.02
798.4114990234375 0 440864.06
799.413818359375 0 199582.75
803.4351806640625 0 234690.3
804.4412231445312 0 119916.414
810.4329833984375 0 136328.27
811.4052734375 0 352155.84
812.4009399414062 0 121352.66
828.4319458007812 0 1902642.1 y Ammonia loss 7
829.4345703125 0 795870.7
830.4303588867188 0 195352.61
855.396240234375 0 161107.1
857.4420166015625 0 233554.34
858.397705078125 0 224230.92
859.3887939453125 0 125597.63
867.4349975585938 0 164769.69
873.407470703125 0 183703.52
881.4452514648438 0 110601.74
883.4564208984375 0 78222.664 b Ammonia loss 7
885.4349975585938 0 201282.3
887.4240112304688 0 450940
888.4268188476562 0 192465.61
889.4216918945312 0 105181.39
891.4152221679688 0 254926
892.4271240234375 0 109933.29
897.4036865234375 0 184143.67
899.4666748046875 0 158607.7
900.4683227539062 0 472026.84 b 7
901.4588623046875 0 106882.95
909.4527587890625 0 113896.05
915.4166870117188 0 918571.3
916.418701171875 0 308004.97
917.4175415039062 0 165847.4
918.4646606445312 0 87356.21
925.4501342773438 0 156796.97
926.4468994140625 0 203627.39
927.4476318359375 0 123742.69
943.458740234375 0 1690727.2 y Ammonia loss 6
944.466064453125 0 792650.5
945.4714965820312 0 408436.88
951.4555053710938 0 84881.23
954.46044921875 0 140642.9
957.4699096679688 0 117121.18
960.4791259765625 0 72398.516 y 6
968.4810791015625 0 164666.25
969.4717407226562 0 212707.77
970.4695434570312 0 112352.51
972.470947265625 0 239598.98
973.4830322265625 0 142247.06
986.4896240234375 0 1396708.8
987.4891967773438 0 1609574.2
988.4868774414062 0 705763.1
989.4933471679688 0 257163.75
990.4873657226562 0 278426.34
991.4887084960938 0 132613.33
996.468994140625 0 210550.75
997.4750366210938 0 146019.4
1000.480712890625 0 202501.9
1005.49755859375 0 194447.8
1012.4793701171875 0 592643.1
1013.4859619140625 0 368971.97
1014.4842529296875 0 1382367.2
1015.4869995117188 0 789116.9
1016.49072265625 0 252502.86
1030.490478515625 0 4757102.5 y Ammonia loss 5
1031.4931640625 0 2625424.2
1032.495361328125 0 1015090.5
1033.489501953125 0 174537.08
1040.4833984375 0 117838.414
1047.52099609375 0 136644.78 y 5
1057.510498046875 0 124336.45
1073.5211181640625 0 516908.38
1074.5263671875 0 234810.2
1075.5302734375 0 130917.94
1083.5091552734375 0 377853.56
1084.511474609375 0 238819.92
1086.5516357421875 0 320199.9
1087.551025390625 0 180875.38
1088.5501708984375 0 105681.68
1101.5167236328125 0 743073.3
1102.5205078125 0 470308.22
1103.519775390625 0 193725.03
1111.545654296875 0 152093.4
1112.5443115234375 0 181296.69
1113.5440673828125 0 109696.74
1119.5340576171875 0 183854.48
1127.566650390625 0 116857.24
1129.5574951171875 0 1581567.2 y Ammonia loss 4
1130.560791015625 0 935130.06
1131.5589599609375 0 380854.97
1132.5467529296875 0 113936.25
1142.5374755859375 0 171100.56
1143.5372314453125 0 108687.06
1144.5386962890625 0 145687.08
1158.531494140625 0 122231.57
1160.5531005859375 0 842797.44
1161.556884765625 0 441135.5
1162.5621337890625 0 194839.12
1170.5347900390625 0 278674.62
1171.5421142578125 0 168538.44
1188.5487060546875 0 1188509.9
1189.5494384765625 0 574174.8
1190.5596923828125 0 160014.56
1205.5614013671875 0 114029.76
1206.557861328125 0 926686.8
1207.5596923828125 0 571249.4
1208.5623779296875 0 212755.17
1215.5968017578125 0 110927.07
1240.593994140625 0 83950.23
1273.6466064453125 0 115755.34
1302.6341552734375 0 120233.055
1345.6358642578125 0 173463.73 y Ammonia loss 2
1346.6387939453125 0 137183.14
1347.62744140625 0 90374.69

Spectrum Details

|  |  |
| --- | --- |
| Matched peaks? Matched peaks The total absolute number of peaks matched. Additionally in brackets the total fraction of peaks matched and the total number of peaks is shown. | 72 (11.76% of 612) |
| FDR? FDR The false discovery rate estimated for this peptide. It is calculated by matching all theoretical fragments with a non-integer shift with the raw peaks for this spectrum. This is done with 40 different shifts. The resulting percentage is the average number of annotated peaks over the number of annotated peaks with the correct spectrum. | 0.66% |
| Satellite FDR? Satellite FDR See the FDR for details on its calculation. This satellite ion specific FDR only contains the satellite ions (d/w) for I/L/J positions. | - |
| PSM Score? PSM Score The PSM Score as given by Hecklib to this annotated spectrum. It is shown with three significant figures. | 496 |

#### Spectrum 3703? Spectrum 3703 The raw spectrum of this peptide as annotated by Hecklib. The fragments are coloured according to ion type (see legend). Any peaks with a star '\*' as text can be hovered over to see the full details, first the ion type second the mass shift type. By hovering over the amino acids in the peptide or ions in the legend the corresponding peaks are highlighted. By toggling the 'Unassigned' label you can turn the background (unassigned) peaks on or off in the plot. By updating the slider in the Ion legend you can update the spectrum to only show the top X% of the peaks with labels. The top X% means any peak that is within X% of the highest intensity. By dragging in the spectrum you can zoom in to a specific part of the spectrum and use 'Zoom Out' to get back to the original zoom level. The annotation of the spectrum is based on the given sequence in the peptides file and is done with different software so inconsistencies are likely. The peaks are annotated based on the given sequence, with 20 ppm tolerance.

Copy Data

##### Spectrum 3703 (TSV)

###### Preview

```
Loading example...
```

*Click on the button to copy the data to your clipboard.*

Mz MinMz MaxIntensity Max

WidthHeightPeptide font sizePeptide stroke widthSpectrum font sizeSpectrum stroke widthCompact peptide

Ion legend

wxyz

abcd

OtherUnassignedIon^Charge^_Position_Show for top:%

JJSEVSDRPSGVSSR

01.92e+63.83e+65.75e+67.66e+6

Zoom Out

y^+1^_1_y^+3^_5_y^+1^_1_a^+1^_2_b^+1^_2_y^+1^_2_y^+1^_2_y^+1^_2_b^+1^_3_b^+1^_3_y^+1^_3_y^+1^_3_y^+2^_7_y^+2^_7_y^+1^_3_b^+3^_12_y^+2^_8_y^+2^_8_b^+1^_4_y^+1^_4_b^+1^_4_y^+1^_4_y^+3^_13_y^+3^_13_b^+2^_8_y^+3^_13_y^+2^_9_b^+3^_14_y^+2^_9_y^+3^_14_y^+3^_14_y^+1^_5_y^+1^_5_y^+3^_14_y^+1^_5_y^+2^_10_y^+2^_10_y^+2^_10_\*\*b^+1^_5_y^+2^_11_y^+2^_11_y^+2^_11_y^+1^_6_y^+1^_6_b^+2^_12_y^+2^_12_y^+2^_12_y^+2^_12_y^+1^_7_y^+1^_7_y^+2^_13_y^+2^_13_y^+2^_13_y^+1^_7_b^+2^_14_y^+2^_14_y^+2^_14_y^+2^_14_b^+1^_7_y^+1^_8_b^+1^_8_b^+1^_8_y^+1^_9_y^+1^_10_y^+1^_11_

0733146722002933

Fragment Matches Table

Show background peaks

| Position | Ion type | Intensity | mz Theoretical | mz Error (Th) | mz Error (ppm) | Charge | Series Number |
| --- | --- | --- | --- | --- | --- | --- | --- |
| - | - | 2.278E+04 | 120.1 | - | - | 0 | - |
| - | - | 2950 | 120.1 | - | - | 0 | - |
| - | - | 3620 | 121.1 | - | - | 0 | - |
| - | - | 2518 | 122.2 | - | - | 0 | - |
| - | - | 2652 | 122.3 | - | - | 0 | - |
| - | - | 6.575E+04 | 126.1 | - | - | 0 | - |
| - | - | 2522 | 127.7 | - | - | 0 | - |
| - | - | 2567 | 128.6 | - | - | 0 | - |
| - | - | 8369 | 129.1 | - | - | 0 | - |
| - | - | 4.051E+04 | 129.1 | - | - | 0 | - |
| - | - | 4427 | 129.1 | - | - | 0 | - |
| - | - | 5763 | 130.1 | - | - | 0 | - |
| - | - | 3.32E+04 | 130.1 | - | - | 0 | - |
| - | - | 1.72E+04 | 130.1 | - | - | 0 | - |
| - | - | 3852 | 130.1 | - | - | 0 | - |
| - | - | 4713 | 131.1 | - | - | 0 | - |
| - | - | 6859 | 131.1 | - | - | 0 | - |
| - | - | 5103 | 133.1 | - | - | 0 | - |
| - | - | 5425 | 133.1 | - | - | 0 | - |
| - | - | 1.753E+04 | 136.1 | - | - | 0 | - |
| - | - | 5271 | 138.1 | - | - | 0 | - |
| - | - | 6069 | 138.1 | - | - | 0 | - |
| - | - | 7534 | 139.1 | - | - | 0 | - |
| - | - | 1.04E+04 | 140.1 | - | - | 0 | - |
| - | - | 3072 | 140.4 | - | - | 0 | - |
| - | - | 3314 | 141.1 | - | - | 0 | - |
| - | - | 8017 | 141.1 | - | - | 0 | - |
| - | - | 1.48E+04 | 144.1 | - | - | 0 | - |
| - | - | 3460 | 145.1 | - | - | 0 | - |
| - | - | 2.081E+04 | 147.1 | - | - | 0 | - |
| - | - | 4476 | 153.1 | - | - | 0 | - |
| - | - | 6.919E+04 | 154.1 | - | - | 0 | - |
| - | - | 3726 | 154.6 | - | - | 0 | - |
| - | - | 3733 | 155.1 | - | - | 0 | - |
| - | - | 7504 | 155.1 | - | - | 0 | - |
| - | - | 4955 | 156.1 | - | - | 0 | - |
| - | - | 2.788E+04 | 157.1 | - | - | 0 | - |
| - | - | 3.68E+04 | 157.1 | - | - | 0 | - |
| - | - | 3.779E+04 | 157.1 | - | - | 0 | - |
| 15 | y | 1.392E+05 | 158.1 | 0.0005077 | 3.211 | +1 | 1 |
| - | - | 8369 | 159.1 | - | - | 0 | - |
| - | - | 1.958E+04 | 159.1 | - | - | 0 | - |
| - | - | 3028 | 159.1 | - | - | 0 | - |
| - | - | 3443 | 162.5 | - | - | 0 | - |
| - | - | 4121 | 165.1 | - | - | 0 | - |
| - | - | 1.006E+04 | 167.1 | - | - | 0 | - |
| - | - | 3407 | 167.1 | - | - | 0 | - |
| 11 | y | 3103 | 169.1 | 0.00169 | 9.997 | +3 | 5 |
| - | - | 7808 | 170 | - | - | 0 | - |
| - | - | 8.203E+04 | 171.1 | - | - | 0 | - |
| - | - | 1.353E+04 | 172.1 | - | - | 0 | - |
| - | - | 4302 | 172.1 | - | - | 0 | - |
| - | - | 2.748E+04 | 173.1 | - | - | 0 | - |
| - | - | 5.775E+04 | 175.1 | - | - | 0 | - |
| 15 | y | 2.773E+05 | 175.1 | 0.0005699 | 3.255 | +1 | 1 |
| - | - | 2.117E+04 | 176.1 | - | - | 0 | - |
| - | - | 6776 | 182.1 | - | - | 0 | - |
| - | - | 6.385E+04 | 183.1 | - | - | 0 | - |
| - | - | 8707 | 184.1 | - | - | 0 | - |
| - | - | 5316 | 184.1 | - | - | 0 | - |
| - | - | 5679 | 185.1 | - | - | 0 | - |
| - | - | 5.197E+04 | 185.1 | - | - | 0 | - |
| - | - | 7770 | 185.1 | - | - | 0 | - |
| - | - | 5644 | 186.1 | - | - | 0 | - |
| - | - | 4213 | 187.1 | - | - | 0 | - |
| - | - | 5.544E+04 | 187.1 | - | - | 0 | - |
| - | - | 6832 | 188.1 | - | - | 0 | - |
| - | - | 3.797E+05 | 189.1 | - | - | 0 | - |
| - | - | 3.483E+04 | 190.1 | - | - | 0 | - |
| - | - | 6614 | 192.1 | - | - | 0 | - |
| - | - | 4683 | 193.1 | - | - | 0 | - |
| - | - | 9397 | 194.1 | - | - | 0 | - |
| - | - | 6145 | 195.1 | - | - | 0 | - |
| - | - | 5199 | 195.1 | - | - | 0 | - |
| - | - | 1.142E+04 | 195.1 | - | - | 0 | - |
| - | - | 5844 | 197.2 | - | - | 0 | - |
| - | - | 2.883E+04 | 199.1 | - | - | 0 | - |
| 2 | a | 2.957E+06 | 199.2 | 0.0006015 | 3.02 | +1 | 2 |
| - | - | 3.619E+05 | 200.2 | - | - | 0 | - |
| - | - | 1.148E+05 | 201.1 | - | - | 0 | - |
| - | - | 2.349E+04 | 201.2 | - | - | 0 | - |
| - | - | 2.262E+04 | 202.1 | - | - | 0 | - |
| - | - | 1.022E+04 | 202.1 | - | - | 0 | - |
| - | - | 5.57E+04 | 203.1 | - | - | 0 | - |
| - | - | 9457 | 203.1 | - | - | 0 | - |
| - | - | 4690 | 209.1 | - | - | 0 | - |
| - | - | 7124 | 209.1 | - | - | 0 | - |
| - | - | 6537 | 209.1 | - | - | 0 | - |
| - | - | 1.21E+04 | 210.1 | - | - | 0 | - |
| - | - | 1.674E+04 | 211.1 | - | - | 0 | - |
| - | - | 2.823E+04 | 212.1 | - | - | 0 | - |
| - | - | 5759 | 212.1 | - | - | 0 | - |
| - | - | 5805 | 213.1 | - | - | 0 | - |
| - | - | 1.111E+04 | 216.1 | - | - | 0 | - |
| - | - | 6.135E+05 | 217.1 | - | - | 0 | - |
| - | - | 5.002E+04 | 218.1 | - | - | 0 | - |
| - | - | 5746 | 219.1 | - | - | 0 | - |
| - | - | 4148 | 223.1 | - | - | 0 | - |
| - | - | 4.18E+04 | 224.1 | - | - | 0 | - |
| - | - | 6373 | 225.1 | - | - | 0 | - |
| - | - | 1.295E+04 | 226.1 | - | - | 0 | - |
| - | - | 3.225E+04 | 226.2 | - | - | 0 | - |
| - | - | 4713 | 227.1 | - | - | 0 | - |
| - | - | 2.6E+04 | 227.1 | - | - | 0 | - |
| 2 | b | 8.517E+05 | 227.2 | 0.000621 | 2.733 | +1 | 2 |
| - | - | 3984 | 228.1 | - | - | 0 | - |
| - | - | 1.136E+05 | 228.2 | - | - | 0 | - |
| - | - | 2.965E+04 | 229.1 | - | - | 0 | - |
| - | - | 1.363E+04 | 229.2 | - | - | 0 | - |
| - | - | 1.82E+04 | 230.1 | - | - | 0 | - |
| - | - | 3.925E+04 | 233.2 | - | - | 0 | - |
| - | - | 7450 | 234.2 | - | - | 0 | - |
| - | - | 4641 | 237.1 | - | - | 0 | - |
| - | - | 2.118E+04 | 237.1 | - | - | 0 | - |
| - | - | 5017 | 238.1 | - | - | 0 | - |
| - | - | 3988 | 241.2 | - | - | 0 | - |
| - | - | 8.207E+04 | 242.1 | - | - | 0 | - |
| - | - | 7600 | 243.1 | - | - | 0 | - |
| - | - | 1.427E+04 | 244.1 | - | - | 0 | - |
| 14 | y | 9382 | 244.1 | 0.001521 | 6.232 | +1 | 2 |
| 14 | y | 8.766E+04 | 245.1 | 0.0007364 | 3.004 | +1 | 2 |
| - | - | 7778 | 246.1 | - | - | 0 | - |
| - | - | 1.872E+04 | 249.2 | - | - | 0 | - |
| - | - | 4192 | 251.1 | - | - | 0 | - |
| - | - | 5114 | 253.2 | - | - | 0 | - |
| - | - | 8626 | 254.1 | - | - | 0 | - |
| - | - | 7785 | 254.2 | - | - | 0 | - |
| - | - | 7.403E+04 | 255.1 | - | - | 0 | - |
| - | - | 4035 | 255.1 | - | - | 0 | - |
| - | - | 6233 | 256.1 | - | - | 0 | - |
| - | - | 7556 | 259.1 | - | - | 0 | - |
| - | - | 1.388E+04 | 261.2 | - | - | 0 | - |
| 14 | y | 5.931E+04 | 262.2 | 0.0006308 | 2.406 | +1 | 2 |
| - | - | 6763 | 262.2 | - | - | 0 | - |
| - | - | 7543 | 263.2 | - | - | 0 | - |
| - | - | 3990 | 266.1 | - | - | 0 | - |
| - | - | 1.628E+04 | 270.1 | - | - | 0 | - |
| - | - | 8256 | 271.1 | - | - | 0 | - |
| - | - | 7215 | 272.1 | - | - | 0 | - |
| - | - | 7427 | 277.2 | - | - | 0 | - |
| - | - | 5531 | 279.1 | - | - | 0 | - |
| - | - | 4935 | 280.1 | - | - | 0 | - |
| - | - | 6160 | 281.1 | - | - | 0 | - |
| - | - | 1.824E+04 | 284.1 | - | - | 0 | - |
| - | - | 1.034E+04 | 284.2 | - | - | 0 | - |
| - | - | 1.238E+04 | 288.2 | - | - | 0 | - |
| - | - | 6752 | 289.2 | - | - | 0 | - |
| - | - | 5984 | 295.2 | - | - | 0 | - |
| - | - | 1.312E+04 | 296.1 | - | - | 0 | - |
| 3 | b | 4.318E+04 | 296.2 | 0.0008858 | 2.99 | +1 | 3 |
| - | - | 1.378E+04 | 297.1 | - | - | 0 | - |
| - | - | 8692 | 297.2 | - | - | 0 | - |
| - | - | 3.138E+04 | 298.1 | - | - | 0 | - |
| - | - | 1.284E+04 | 299.1 | - | - | 0 | - |
| - | - | 5026 | 301.2 | - | - | 0 | - |
| - | - | 1.362E+04 | 302.1 | - | - | 0 | - |
| - | - | 4147 | 302.2 | - | - | 0 | - |
| - | - | 5068 | 307.2 | - | - | 0 | - |
| - | - | 5673 | 311.1 | - | - | 0 | - |
| - | - | 4.438E+04 | 312.2 | - | - | 0 | - |
| - | - | 1.453E+04 | 313.2 | - | - | 0 | - |
| - | - | 9.794E+04 | 313.2 | - | - | 0 | - |
| - | - | 3.329E+04 | 314.1 | - | - | 0 | - |
| - | - | 1.154E+04 | 314.2 | - | - | 0 | - |
| 3 | b | 9.368E+04 | 314.2 | 0.001002 | 3.19 | +1 | 3 |
| - | - | 1.211E+04 | 315.2 | - | - | 0 | - |
| - | - | 1.202E+05 | 316.2 | - | - | 0 | - |
| - | - | 1.474E+04 | 317.2 | - | - | 0 | - |
| - | - | 3987 | 319.2 | - | - | 0 | - |
| - | - | 7052 | 323.2 | - | - | 0 | - |
| - | - | 1.647E+04 | 324.1 | - | - | 0 | - |
| - | - | 4469 | 325.1 | - | - | 0 | - |
| - | - | 5605 | 329.1 | - | - | 0 | - |
| - | - | 2.297E+04 | 330.2 | - | - | 0 | - |
| 13 | y | 2.231E+04 | 331.2 | 0.0001421 | 0.4289 | +1 | 3 |
| 13 | y | 8.234E+04 | 332.2 | 0.0009803 | 2.951 | +1 | 3 |
| - | - | 1.167E+04 | 333.2 | - | - | 0 | - |
| 9 | y | 6606 | 336.2 | 0.002004 | 5.961 | +2 | 7 |
| - | - | 5392 | 338.1 | - | - | 0 | - |
| - | - | 4717 | 339.2 | - | - | 0 | - |
| - | - | 9383 | 340.1 | - | - | 0 | - |
| - | - | 6.691E+04 | 341.2 | - | - | 0 | - |
| - | - | 9.793E+04 | 342.1 | - | - | 0 | - |
| - | - | 8652 | 342.2 | - | - | 0 | - |
| - | - | 1.493E+04 | 343.1 | - | - | 0 | - |
| - | - | 4372 | 343.2 | - | - | 0 | - |
| - | - | 5837 | 344.1 | - | - | 0 | - |
| 9 | y | 3.283E+04 | 345.2 | 0.0006278 | 1.819 | +2 | 7 |
| - | - | 1.839E+04 | 345.7 | - | - | 0 | - |
| 13 | y | 6.693E+04 | 349.2 | 0.000951 | 2.723 | +1 | 3 |
| - | - | 8275 | 350.2 | - | - | 0 | - |
| - | - | 4304 | 353.2 | - | - | 0 | - |
| - | - | 4.55E+04 | 359.2 | - | - | 0 | - |
| - | - | 6144 | 360.2 | - | - | 0 | - |
| - | - | 4100 | 362.1 | - | - | 0 | - |
| - | - | 7419 | 365.2 | - | - | 0 | - |
| - | - | 5070 | 367.2 | - | - | 0 | - |
| - | - | 4637 | 368.9 | - | - | 0 | - |
| - | - | 5451 | 379.2 | - | - | 0 | - |
| - | - | 1.094E+04 | 380.2 | - | - | 0 | - |
| - | - | 1.44E+04 | 385.2 | - | - | 0 | - |
| - | - | 7138 | 385.7 | - | - | 0 | - |
| - | - | 1.591E+04 | 389.7 | - | - | 0 | - |
| - | - | 1.007E+04 | 390.2 | - | - | 0 | - |
| - | - | 1.417E+04 | 390.7 | - | - | 0 | - |
| - | - | 8654 | 391.2 | - | - | 0 | - |
| - | - | 5.303E+04 | 391.7 | - | - | 0 | - |
| - | - | 1.395E+04 | 392.2 | - | - | 0 | - |
| - | - | 8446 | 392.7 | - | - | 0 | - |
| - | - | 1.31E+04 | 393.7 | - | - | 0 | - |
| - | - | 4888 | 395.2 | - | - | 0 | - |
| - | - | 1.203E+04 | 397.2 | - | - | 0 | - |
| - | - | 6330 | 398.2 | - | - | 0 | - |
| - | - | 1.543E+04 | 398.7 | - | - | 0 | - |
| - | - | 1.188E+04 | 399.2 | - | - | 0 | - |
| - | - | 1.717E+04 | 399.7 | - | - | 0 | - |
| - | - | 2.162E+04 | 400.2 | - | - | 0 | - |
| - | - | 5417 | 401.2 | - | - | 0 | - |
| - | - | 1.807E+04 | 403.2 | - | - | 0 | - |
| - | - | 5224 | 404.2 | - | - | 0 | - |
| - | - | 9892 | 405.2 | - | - | 0 | - |
| - | - | 1.225E+04 | 408.2 | - | - | 0 | - |
| - | - | 1.403E+04 | 410.2 | - | - | 0 | - |
| - | - | 9732 | 411.2 | - | - | 0 | - |
| - | - | 5440 | 413.2 | - | - | 0 | - |
| 12 | b | 3.471E+04 | 414.2 | 0.006422 | 15.5 | +3 | 12 |
| 8 | y | 5.241E+04 | 414.7 | 0.002501 | 6.03 | +2 | 8 |
| - | - | 2.805E+04 | 415.2 | - | - | 0 | - |
| - | - | 1.782E+04 | 415.3 | - | - | 0 | - |
| 8 | y | 6.64E+05 | 423.2 | 0.00122 | 2.882 | +2 | 8 |
| - | - | 2.849E+05 | 423.7 | - | - | 0 | - |
| - | - | 7.399E+04 | 424.2 | - | - | 0 | - |
| - | - | 1.318E+04 | 424.7 | - | - | 0 | - |
| 4 | b | 5.088E+04 | 425.2 | 0.0007731 | 1.818 | +1 | 4 |
| - | - | 1.123E+04 | 426.2 | - | - | 0 | - |
| - | - | 2.496E+04 | 428.2 | - | - | 0 | - |
| - | - | 3.431E+04 | 429.2 | - | - | 0 | - |
| 12 | y | 2.214E+04 | 430.2 | 0.0005018 | 1.166 | +1 | 4 |
| - | - | 9029 | 439.2 | - | - | 0 | - |
| - | - | 5068 | 440.2 | - | - | 0 | - |
| - | - | 4.272E+04 | 441.2 | - | - | 0 | - |
| 4 | b | 4.739E+04 | 443.3 | 0.001408 | 3.177 | +1 | 4 |
| - | - | 1.478E+04 | 444.3 | - | - | 0 | - |
| 12 | y | 2.539E+04 | 448.3 | 0.00111 | 2.476 | +1 | 4 |
| 3 | y | 1.387E+04 | 448.9 | 0.0007459 | 1.662 | +3 | 13 |
| 3 | y | 9504 | 449.2 | 0.0088 | 19.59 | +3 | 13 |
| - | - | 5381 | 449.6 | - | - | 0 | - |
| 8 | b | 1.152E+04 | 450.7 | 0.0009361 | 2.077 | +2 | 8 |
| - | - | 5026 | 451.2 | - | - | 0 | - |
| - | - | 7255 | 452.2 | - | - | 0 | - |
| - | - | 2.126E+04 | 452.3 | - | - | 0 | - |
| 3 | y | 2.469E+04 | 454.9 | 0.0008559 | 1.882 | +3 | 13 |
| - | - | 1.938E+04 | 455.2 | - | - | 0 | - |
| - | - | 1.063E+04 | 455.6 | - | - | 0 | - |
| - | - | 3.824E+04 | 458.2 | - | - | 0 | - |
| - | - | 2.132E+04 | 460.2 | - | - | 0 | - |
| - | - | 6056 | 465.3 | - | - | 0 | - |
| - | - | 1.645E+04 | 469.3 | - | - | 0 | - |
| - | - | 4556 | 470.2 | - | - | 0 | - |
| 7 | y | 7162 | 471.7 | 0.001087 | 2.305 | +2 | 9 |
| 14 | b | 1.681E+04 | 472.2 | 0.008094 | 17.14 | +3 | 14 |
| - | - | 7316 | 472.7 | - | - | 0 | - |
| - | - | 5197 | 478.2 | - | - | 0 | - |
| - | - | 4547 | 479.2 | - | - | 0 | - |
| - | - | 7971 | 480.3 | - | - | 0 | - |
| 7 | y | 1.117E+05 | 480.7 | 0.001298 | 2.7 | +2 | 9 |
| - | - | 4.941E+04 | 481.2 | - | - | 0 | - |
| - | - | 1.62E+04 | 481.8 | - | - | 0 | - |
| 2 | y | 1.059E+04 | 486.6 | 0.0003938 | 0.8094 | +3 | 14 |
| 2 | y | 2.414E+04 | 486.9 | 0.008113 | 16.66 | +3 | 14 |
| 11 | y | 7660 | 487.3 | 0.002678 | 5.497 | +1 | 5 |
| 11 | y | 8311 | 488.2 | 0.002197 | 4.5 | +1 | 5 |
| 2 | y | 2.644E+04 | 492.6 | 0.0008701 | 1.766 | +3 | 14 |
| - | - | 1.634E+04 | 492.9 | - | - | 0 | - |
| - | - | 1.127E+04 | 493.3 | - | - | 0 | - |
| - | - | 3.773E+04 | 495.9 | - | - | 0 | - |
| - | - | 8102 | 496.2 | - | - | 0 | - |
| - | - | 2.077E+04 | 496.3 | - | - | 0 | - |
| - | - | 1.299E+04 | 496.6 | - | - | 0 | - |
| - | - | 8593 | 497.2 | - | - | 0 | - |
| - | - | 1.418E+04 | 497.3 | - | - | 0 | - |
| 11 | y | 8.073E+04 | 505.3 | 0.001131 | 2.238 | +1 | 5 |
| - | - | 1.655E+04 | 506.3 | - | - | 0 | - |
| - | - | 5080 | 513.9 | - | - | 0 | - |
| - | - | 5359 | 514.3 | - | - | 0 | - |
| - | - | 1.909E+04 | 514.3 | - | - | 0 | - |
| 6 | y | 1.799E+04 | 515.3 | 0.001011 | 1.962 | +2 | 10 |
| 6 | y | 1.246E+04 | 515.7 | 0.005455 | 10.58 | +2 | 10 |
| - | - | 1.077E+04 | 516.3 | - | - | 0 | - |
| - | - | 6105 | 516.8 | - | - | 0 | - |
| - | - | 5261 | 517.3 | - | - | 0 | - |
| - | - | 8755 | 518.2 | - | - | 0 | - |
| - | - | 7143 | 518.3 | - | - | 0 | - |
| - | - | 8586 | 518.6 | - | - | 0 | - |
| - | - | 6455 | 518.9 | - | - | 0 | - |
| 6 | y | 7.423E+05 | 524.3 | 0.002618 | 4.993 | +2 | 10 |
| 0 | Precursor | 1.458E+05 | 524.6 | 0.00715 | 13.63 | +3 | -1 |
| - | - | 3.907E+05 | 524.8 | - | - | 0 | - |
| - | - | 6.247E+04 | 524.9 | - | - | 0 | - |
| - | - | 1.202E+05 | 525.3 | - | - | 0 | - |
| - | - | 7617 | 525.6 | - | - | 0 | - |
| - | - | 3.666E+04 | 525.8 | - | - | 0 | - |
| - | - | 1.209E+04 | 528.8 | - | - | 0 | - |
| - | - | 9351 | 529.3 | - | - | 0 | - |
| - | - | 1.972E+04 | 529.8 | - | - | 0 | - |
| 0 | Precursor | 8.976E+04 | 530.3 | 0.001128 | 2.128 | +3 | -1 |
| - | - | 5.192E+04 | 530.6 | - | - | 0 | - |
| - | - | 3.275E+04 | 531 | - | - | 0 | - |
| - | - | 1.355E+04 | 532.3 | - | - | 0 | - |
| - | - | 5724 | 533.3 | - | - | 0 | - |
| - | - | 8106 | 539.3 | - | - | 0 | - |
| 5 | b | 1.414E+04 | 542.3 | 0.0023 | 4.241 | +1 | 5 |
| - | - | 5650 | 543.3 | - | - | 0 | - |
| - | - | 1.047E+04 | 556.3 | - | - | 0 | - |
| - | - | 6432 | 556.8 | - | - | 0 | - |
| - | - | 5506 | 558.8 | - | - | 0 | - |
| - | - | 1.205E+04 | 559.3 | - | - | 0 | - |
| 5 | y | 3.502E+04 | 564.8 | 0.0006707 | 1.188 | +2 | 11 |
| - | - | 7286 | 565.2 | - | - | 0 | - |
| 5 | y | 3.352E+04 | 565.3 | 0.005916 | 10.47 | +2 | 11 |
| - | - | 2.135E+04 | 565.8 | - | - | 0 | - |
| - | - | 8273 | 566.3 | - | - | 0 | - |
| - | - | 7583 | 567.3 | - | - | 0 | - |
| - | - | 8924 | 572.3 | - | - | 0 | - |
| - | - | 6936 | 572.8 | - | - | 0 | - |
| - | - | 1.162E+04 | 573.3 | - | - | 0 | - |
| 5 | y | 9.816E+05 | 573.8 | 0.001187 | 2.068 | +2 | 11 |
| 10 | y | 5.855E+05 | 574.3 | 0.004783 | 8.328 | +1 | 6 |
| - | - | 1.894E+05 | 574.8 | - | - | 0 | - |
| - | - | 3.65E+04 | 575.3 | - | - | 0 | - |
| - | - | 1.352E+04 | 583.2 | - | - | 0 | - |
| - | - | 1.326E+04 | 584.3 | - | - | 0 | - |
| - | - | 5969 | 585.3 | - | - | 0 | - |
| - | - | 1.106E+04 | 587.3 | - | - | 0 | - |
| 10 | y | 1.054E+05 | 592.3 | 0.00142 | 2.398 | +1 | 6 |
| - | - | 3.093E+04 | 593.3 | - | - | 0 | - |
| - | - | 1.011E+04 | 594.3 | - | - | 0 | - |
| - | - | 5844 | 596.3 | - | - | 0 | - |
| - | - | 2.26E+04 | 600.3 | - | - | 0 | - |
| - | - | 7381 | 601.3 | - | - | 0 | - |
| - | - | 7728 | 602.3 | - | - | 0 | - |
| - | - | 9810 | 608.3 | - | - | 0 | - |
| 12 | b | 5777 | 612.3 | 0.0001873 | 0.3059 | +2 | 12 |
| - | - | 5813 | 617.3 | - | - | 0 | - |
| - | - | 5938 | 620.3 | - | - | 0 | - |
| - | - | 5256 | 624.3 | - | - | 0 | - |
| - | - | 1.584E+04 | 626.3 | - | - | 0 | - |
| 4 | y | 8.014E+04 | 629.3 | 0.002812 | 4.468 | +2 | 12 |
| 4 | y | 5E+04 | 629.8 | 0.008729 | 13.86 | +2 | 12 |
| - | - | 2.666E+04 | 630.3 | - | - | 0 | - |
| - | - | 9794 | 636.3 | - | - | 0 | - |
| 4 | y | 4.794E+05 | 638.3 | 0.001191 | 1.867 | +2 | 12 |
| - | - | 3.131E+05 | 638.8 | - | - | 0 | - |
| - | - | 1.055E+05 | 639.3 | - | - | 0 | - |
| - | - | 3.387E+04 | 639.8 | - | - | 0 | - |
| - | - | 1.041E+04 | 640.3 | - | - | 0 | - |
| - | - | 9247 | 641.3 | - | - | 0 | - |
| - | - | 7969 | 642.3 | - | - | 0 | - |
| - | - | 7197 | 642.8 | - | - | 0 | - |
| - | - | 1.353E+04 | 643.4 | - | - | 0 | - |
| - | - | 5963 | 644.4 | - | - | 0 | - |
| - | - | 5287 | 645.4 | - | - | 0 | - |
| - | - | 3.451E+04 | 646.3 | - | - | 0 | - |
| - | - | 8422 | 647.3 | - | - | 0 | - |
| - | - | 1.148E+04 | 648.8 | - | - | 0 | - |
| - | - | 1.034E+04 | 649.3 | - | - | 0 | - |
| - | - | 6021 | 649.8 | - | - | 0 | - |
| - | - | 6877 | 651.3 | - | - | 0 | - |
| - | - | 1.192E+04 | 652.3 | - | - | 0 | - |
| - | - | 1.506E+04 | 653.3 | - | - | 0 | - |
| - | - | 2.037E+04 | 654.3 | - | - | 0 | - |
| - | - | 7703 | 654.8 | - | - | 0 | - |
| - | - | 1.253E+04 | 655.3 | - | - | 0 | - |
| - | - | 1.686E+04 | 657.3 | - | - | 0 | - |
| - | - | 6668 | 657.8 | - | - | 0 | - |
| - | - | 2.59E+04 | 659.3 | - | - | 0 | - |
| - | - | 5606 | 659.8 | - | - | 0 | - |
| - | - | 8112 | 660.3 | - | - | 0 | - |
| - | - | 1.197E+04 | 660.3 | - | - | 0 | - |
| - | - | 2.325E+04 | 660.8 | - | - | 0 | - |
| - | - | 1.587E+04 | 661.3 | - | - | 0 | - |
| - | - | 4.12E+04 | 663.8 | - | - | 0 | - |
| - | - | 3.507E+04 | 664.3 | - | - | 0 | - |
| - | - | 2.125E+04 | 664.8 | - | - | 0 | - |
| - | - | 7348 | 665.3 | - | - | 0 | - |
| - | - | 6834 | 665.8 | - | - | 0 | - |
| - | - | 1.978E+04 | 666.3 | - | - | 0 | - |
| - | - | 8.617E+04 | 666.8 | - | - | 0 | - |
| - | - | 5.136E+04 | 667.3 | - | - | 0 | - |
| - | - | 2.093E+04 | 667.8 | - | - | 0 | - |
| - | - | 6859 | 668.3 | - | - | 0 | - |
| 9 | y | 1.074E+05 | 671.3 | 0.001519 | 2.262 | +1 | 7 |
| 9 | y | 7.573E+04 | 672.3 | 0.00615 | 9.148 | +1 | 7 |
| 3 | y | 2.526E+05 | 672.8 | 0.001263 | 1.877 | +2 | 13 |
| 3 | y | 2.417E+05 | 673.3 | 0.01005 | 14.92 | +2 | 13 |
| - | - | 8.987E+04 | 673.8 | - | - | 0 | - |
| - | - | 1.386E+05 | 674.3 | - | - | 0 | - |
| - | - | 1.324E+04 | 674.8 | - | - | 0 | - |
| - | - | 3.992E+04 | 675.3 | - | - | 0 | - |
| - | - | 6113 | 676.3 | - | - | 0 | - |
| - | - | 5811 | 678.3 | - | - | 0 | - |
| - | - | 7.901E+04 | 680.8 | - | - | 0 | - |
| - | - | 7.155E+04 | 681.3 | - | - | 0 | - |
| 3 | y | 7.584E+06 | 681.8 | 0.00184 | 2.698 | +2 | 13 |
| - | - | 5.303E+06 | 682.3 | - | - | 0 | - |
| - | - | 2.137E+06 | 682.8 | - | - | 0 | - |
| - | - | 5.683E+05 | 683.3 | - | - | 0 | - |
| - | - | 1.273E+05 | 683.8 | - | - | 0 | - |
| - | - | 1.121E+04 | 685.4 | - | - | 0 | - |
| - | - | 5.665E+04 | 686.8 | - | - | 0 | - |
| - | - | 4.305E+04 | 687.3 | - | - | 0 | - |
| - | - | 1.83E+04 | 687.8 | - | - | 0 | - |
| 9 | y | 6.107E+05 | 689.4 | 0.001391 | 2.018 | +1 | 7 |
| - | - | 2.148E+05 | 690.4 | - | - | 0 | - |
| - | - | 4.798E+04 | 691.4 | - | - | 0 | - |
| - | - | 8602 | 692.4 | - | - | 0 | - |
| 14 | b | 3.837E+04 | 699.3 | 0.004384 | 6.268 | +2 | 14 |
| - | - | 1.127E+04 | 700.3 | - | - | 0 | - |
| - | - | 1.056E+04 | 728.4 | - | - | 0 | - |
| 2 | y | 7.571E+04 | 729.4 | 0.001162 | 1.593 | +2 | 14 |
| 2 | y | 3.979E+04 | 729.9 | 0.009765 | 13.38 | +2 | 14 |
| - | - | 1.982E+04 | 730.4 | - | - | 0 | - |
| - | - | 8960 | 730.9 | - | - | 0 | - |
| - | - | 6592 | 737.4 | - | - | 0 | - |
| - | - | 8126 | 737.9 | - | - | 0 | - |
| 2 | y | 6.266E+05 | 738.4 | 0.001007 | 1.363 | +2 | 14 |
| - | - | 4.708E+05 | 738.9 | - | - | 0 | - |
| - | - | 2.006E+05 | 739.4 | - | - | 0 | - |
| - | - | 5.847E+04 | 739.9 | - | - | 0 | - |
| - | - | 1.884E+04 | 740.4 | - | - | 0 | - |
| - | - | 3.438E+04 | 743.4 | - | - | 0 | - |
| - | - | 2.189E+04 | 743.9 | - | - | 0 | - |
| 7 | b | 1.515E+04 | 744.4 | 0.003815 | 5.125 | +1 | 7 |
| - | - | 8331 | 758.4 | - | - | 0 | - |
| - | - | 6033 | 767.4 | - | - | 0 | - |
| - | - | 5793 | 768.4 | - | - | 0 | - |
| - | - | 1.386E+04 | 768.4 | - | - | 0 | - |
| - | - | 7650 | 769.4 | - | - | 0 | - |
| - | - | 2.931E+04 | 770.4 | - | - | 0 | - |
| - | - | 7523 | 771.4 | - | - | 0 | - |
| - | - | 1.103E+04 | 778.4 | - | - | 0 | - |
| - | - | 7168 | 779.4 | - | - | 0 | - |
| - | - | 6856 | 780.4 | - | - | 0 | - |
| - | - | 3.463E+04 | 782.4 | - | - | 0 | - |
| - | - | 1.871E+04 | 783.4 | - | - | 0 | - |
| - | - | 6731 | 784.4 | - | - | 0 | - |
| - | - | 1.482E+04 | 785.4 | - | - | 0 | - |
| - | - | 1.106E+04 | 786.4 | - | - | 0 | - |
| - | - | 7408 | 786.4 | - | - | 0 | - |
| - | - | 7059 | 787.4 | - | - | 0 | - |
| - | - | 1.921E+04 | 796.4 | - | - | 0 | - |
| - | - | 6733 | 797.4 | - | - | 0 | - |
| - | - | 2.328E+04 | 798.4 | - | - | 0 | - |
| - | - | 8375 | 799.4 | - | - | 0 | - |
| - | - | 1.496E+04 | 803.4 | - | - | 0 | - |
| - | - | 6822 | 810.4 | - | - | 0 | - |
| - | - | 2.031E+04 | 811.4 | - | - | 0 | - |
| - | - | 8488 | 812.4 | - | - | 0 | - |
| 8 | y | 8.981E+04 | 828.4 | 0.0009867 | 1.191 | +1 | 8 |
| - | - | 3.497E+04 | 829.4 | - | - | 0 | - |
| - | - | 1.36E+04 | 830.4 | - | - | 0 | - |
| - | - | 6077 | 855.4 | - | - | 0 | - |
| - | - | 1.118E+04 | 857.4 | - | - | 0 | - |
| - | - | 7851 | 858.4 | - | - | 0 | - |
| - | - | 6360 | 868.4 | - | - | 0 | - |
| - | - | 1.147E+04 | 873.4 | - | - | 0 | - |
| - | - | 6149 | 874.4 | - | - | 0 | - |
| 8 | b | 4681 | 883.5 | 0.01044 | 11.82 | +1 | 8 |
| - | - | 1.397E+04 | 885.4 | - | - | 0 | - |
| - | - | 2.149E+04 | 887.4 | - | - | 0 | - |
| - | - | 9808 | 888.4 | - | - | 0 | - |
| - | - | 6285 | 889.4 | - | - | 0 | - |
| - | - | 1.514E+04 | 891.4 | - | - | 0 | - |
| - | - | 6420 | 892.4 | - | - | 0 | - |
| - | - | 7308 | 899.5 | - | - | 0 | - |
| 8 | b | 1.891E+04 | 900.5 | 0.009528 | 10.58 | +1 | 8 |
| - | - | 1.158E+04 | 901.5 | - | - | 0 | - |
| - | - | 1.171E+04 | 909.4 | - | - | 0 | - |
| - | - | 5619 | 914.4 | - | - | 0 | - |
| - | - | 4.478E+04 | 915.4 | - | - | 0 | - |
| - | - | 2.349E+04 | 916.4 | - | - | 0 | - |
| - | - | 6523 | 917.4 | - | - | 0 | - |
| - | - | 5879 | 925.5 | - | - | 0 | - |
| - | - | 7351 | 926.4 | - | - | 0 | - |
| - | - | 1.254E+04 | 927.4 | - | - | 0 | - |
| 7 | y | 6.984E+04 | 943.5 | 0.001326 | 1.406 | +1 | 9 |
| - | - | 3.621E+04 | 944.5 | - | - | 0 | - |
| - | - | 1.288E+04 | 945.5 | - | - | 0 | - |
| - | - | 6055 | 946.5 | - | - | 0 | - |
| - | - | 6155 | 952.5 | - | - | 0 | - |
| - | - | 6720 | 953.5 | - | - | 0 | - |
| - | - | 6229 | 957.5 | - | - | 0 | - |
| - | - | 6503 | 968.5 | - | - | 0 | - |
| - | - | 1.325E+04 | 969.5 | - | - | 0 | - |
| - | - | 7073 | 970.5 | - | - | 0 | - |
| - | - | 1.118E+04 | 972.5 | - | - | 0 | - |
| - | - | 6.476E+04 | 986.5 | - | - | 0 | - |
| - | - | 6.904E+04 | 987.5 | - | - | 0 | - |
| - | - | 3.266E+04 | 988.5 | - | - | 0 | - |
| - | - | 1.163E+04 | 989.5 | - | - | 0 | - |
| - | - | 1.459E+04 | 990.5 | - | - | 0 | - |
| - | - | 6785 | 995.5 | - | - | 0 | - |
| - | - | 1.274E+04 | 996.5 | - | - | 0 | - |
| - | - | 6931 | 997.5 | - | - | 0 | - |
| - | - | 9152 | 1000 | - | - | 0 | - |
| - | - | 7910 | 1006 | - | - | 0 | - |
| - | - | 3.453E+04 | 1012 | - | - | 0 | - |
| - | - | 2.332E+04 | 1013 | - | - | 0 | - |
| - | - | 5.393E+04 | 1014 | - | - | 0 | - |
| - | - | 2.522E+04 | 1015 | - | - | 0 | - |
| 6 | y | 2.344E+05 | 1030 | 0.001097 | 1.065 | +1 | 10 |
| - | - | 1.061E+05 | 1031 | - | - | 0 | - |
| - | - | 4.055E+04 | 1032 | - | - | 0 | - |
| - | - | 8617 | 1033 | - | - | 0 | - |
| - | - | 8959 | 1056 | - | - | 0 | - |
| - | - | 7115 | 1057 | - | - | 0 | - |
| - | - | 1.034E+04 | 1058 | - | - | 0 | - |
| - | - | 7258 | 1059 | - | - | 0 | - |
| - | - | 2.3E+04 | 1074 | - | - | 0 | - |
| - | - | 8624 | 1075 | - | - | 0 | - |
| - | - | 1.147E+04 | 1084 | - | - | 0 | - |
| - | - | 1.028E+04 | 1085 | - | - | 0 | - |
| - | - | 1.557E+04 | 1087 | - | - | 0 | - |
| - | - | 6850 | 1088 | - | - | 0 | - |
| - | - | 3.454E+04 | 1102 | - | - | 0 | - |
| - | - | 2.161E+04 | 1103 | - | - | 0 | - |
| - | - | 1.105E+04 | 1112 | - | - | 0 | - |
| - | - | 6471 | 1113 | - | - | 0 | - |
| - | - | 1.427E+04 | 1120 | - | - | 0 | - |
| 5 | y | 6.364E+04 | 1130 | 0.001409 | 1.247 | +1 | 11 |
| - | - | 3.398E+04 | 1131 | - | - | 0 | - |
| - | - | 7768 | 1132 | - | - | 0 | - |
| - | - | 5647 | 1143 | - | - | 0 | - |
| - | - | 6966 | 1144 | - | - | 0 | - |
| - | - | 7645 | 1145 | - | - | 0 | - |
| - | - | 3.534E+04 | 1161 | - | - | 0 | - |
| - | - | 1.709E+04 | 1162 | - | - | 0 | - |
| - | - | 1.426E+04 | 1171 | - | - | 0 | - |
| - | - | 5810 | 1188 | - | - | 0 | - |
| - | - | 3.782E+04 | 1189 | - | - | 0 | - |
| - | - | 2.909E+04 | 1190 | - | - | 0 | - |
| - | - | 5.023E+04 | 1207 | - | - | 0 | - |
| - | - | 3.006E+04 | 1208 | - | - | 0 | - |
| - | - | 1.111E+04 | 1209 | - | - | 0 | - |
| - | - | 6484 | 1216 | - | - | 0 | - |
| - | - | 6945 | 1302 | - | - | 0 | - |
| - | - | 5654 | 2373 | - | - | 0 | - |
| - | - | 5475 | 2904 | - | - | 0 | - |

m/z Charge Intensity FragmentType MassShift Position
120.08124542236328 0 22778.979
120.08514404296875 0 2949.5374
121.0844955444336 0 3619.571
122.19901275634766 0 2517.7878
122.28245544433594 0 2652.356
126.05543518066406 0 65749.38
127.69400787353516 0 2521.8867
128.62628173828125 0 2567.382
129.06631469726562 0 8369.367
129.10267639160156 0 40513.01
129.11419677734375 0 4427.4243
130.05059814453125 0 5763.468
130.0615692138672 0 33204.645
130.0980224609375 0 17204.357
130.10609436035156 0 3851.5051
131.08192443847656 0 4713.1143
131.118408203125 0 6858.5527
133.0614013671875 0 5102.672
133.09756469726562 0 5424.7046
136.07615661621094 0 17534.412
138.06663513183594 0 5271.1953
138.0915985107422 0 6068.7515
139.08726501464844 0 7533.897
140.08245849609375 0 10400.259
140.39524841308594 0 3071.8674
141.0664520263672 0 3313.963
141.10281372070312 0 8017.4707
144.06590270996094 0 14803.255
145.0609130859375 0 3459.9802
147.0768280029297 0 20808.814
153.06605529785156 0 4475.634
154.05035400390625 0 69191.2
154.6253204345703 0 3726.0967
155.05377197265625 0 3733.0562
155.11822509765625 0 7503.593
156.10198974609375 0 4954.6514
157.06124877929688 0 27877.875
157.09759521484375 0 36801.68
157.10888671875 0 37793.67
158.09291076660156 0 139194.42 y Ammonia loss 14
159.09654235839844 0 8369.1045
159.1132354736328 0 19576.883
159.11927795410156 0 3028.4219
162.53790283203125 0 3442.5337
165.1025848388672 0 4120.6587
167.08212280273438 0 10063.839
167.11903381347656 0 3407.0847
169.09750366210938 0 3103.471 y 10
170.04542541503906 0 7808.184
171.07691955566406 0 82026.305
172.06094360351562 0 13533.684
172.0801544189453 0 4302.467
173.1290283203125 0 27483.836
175.07186889648438 0 57751.332
175.11952209472656 0 277334.56 y 14
176.12301635742188 0 21169.338
182.09307861328125 0 6775.643
183.11341857910156 0 63845.36
184.1087646484375 0 8707.118
184.11685180664062 0 5316.419
185.0558319091797 0 5679.499
185.09263610839844 0 51969.918
185.1031494140625 0 7769.515
186.0956573486328 0 5643.963
187.07147216796875 0 4212.997
187.10826110839844 0 55443.766
188.11190795898438 0 6831.594
189.08758544921875 0 379698.1
190.09091186523438 0 34827.625
192.1138153076172 0 6614.211
193.0980987548828 0 4682.588
194.093017578125 0 9397.38
195.07675170898438 0 6144.8
195.1138153076172 0 5198.7217
195.14967346191406 0 11421.175
197.16464233398438 0 5843.576
199.07183837890625 0 28830.879
199.18109130859375 0 2957183.8 a 1
200.1844482421875 0 361858.7
201.1238250732422 0 114791.125
201.18724060058594 0 23494.38
202.1192169189453 0 22623.15
202.12777709960938 0 10216.11
203.0668182373047 0 55696.766
203.10328674316406 0 9457.247
209.093505859375 0 4689.542
209.10369873046875 0 7124.039
209.13999938964844 0 6536.9766
210.08790588378906 0 12099.921
211.10848999023438 0 16742.234
212.10354614257812 0 28225.266
212.13980102539062 0 5759.4175
213.0878448486328 0 5804.656
216.13525390625 0 11111.215
217.08248901367188 0 613514.75
218.08583068847656 0 50021.11
219.0873565673828 0 5746.0947
223.10821533203125 0 4147.971
224.10360717773438 0 41795.43
225.1071319580078 0 6372.9336
226.1194610595703 0 12947.635
226.15557861328125 0 32245.03
227.0656280517578 0 4712.5103
227.11463928222656 0 26004.96
227.176025390625 0 851699.56 b 1
228.098876953125 0 3984.325
228.17935180664062 0 113587.49
229.11911010742188 0 29647.48
229.1818084716797 0 13630.56
230.1142578125 0 18199.32
233.16543579101562 0 39250.38
234.16900634765625 0 7450.011
237.09930419921875 0 4641.269
237.13523864746094 0 21178.16
238.11915588378906 0 5016.6143
241.19122314453125 0 3987.6929
242.1143035888672 0 82065.98
243.1175994873047 0 7599.986
244.12974548339844 0 14272.454
244.14193725585938 0 9382.261 y Water loss 13
245.1251678466797 0 87661.88 y Ammonia loss 13
246.12908935546875 0 7778.445
249.16030883789062 0 18721.602
251.1146240234375 0 4192.08
253.2286376953125 0 5113.7847
254.11407470703125 0 8626.061
254.16204833984375 0 7785.0107
255.1093292236328 0 74027.73
255.14576721191406 0 4034.8142
256.1121520996094 0 6232.671
259.1413269042969 0 7556.223
261.1605529785156 0 13881.969
262.151611328125 0 59307.24 y 13
262.1656188964844 0 6763.238
263.1544189453125 0 7542.6304
266.1136474609375 0 3990.292
270.1455993652344 0 16280.187
271.1291198730469 0 8256.361
272.1369323730469 0 7215.206
277.1555480957031 0 7426.902
279.1091613769531 0 5530.829
280.13006591796875 0 4935.147
281.1255187988281 0 6160.3286
284.1246643066406 0 18241.158
284.1612548828125 0 10340.446
288.15570068359375 0 12380.867
289.151611328125 0 6752.042
295.17803955078125 0 5983.8765
296.13629150390625 0 13118.611
296.19775390625 0 43179.324 b Water loss 2
297.11993408203125 0 13781.118
297.2012634277344 0 8692.093
298.1405944824219 0 31379.742
299.13690185546875 0 12841.992
301.1518859863281 0 5025.6177
302.1359558105469 0 13615.86
302.17156982421875 0 4147.262
307.1632080078125 0 5068.416
311.13543701171875 0 5673.2085
312.15625 0 44383.906
313.153076171875 0 14527.543
313.1880187988281 0 97937.766
314.1470642089844 0 33289.723
314.1903991699219 0 11543.882
314.20843505859375 0 93682.336 b 2
315.2108154296875 0 12106.86
316.1511535644531 0 120230.41
317.152099609375 0 14741.033
319.1802062988281 0 3987.2683
323.2091064453125 0 7052.2607
324.131591796875 0 16470.209
325.1152038574219 0 4468.577
329.1459655761719 0 5604.75
330.1669921875 0 22967.518
331.17230224609375 0 22307.705 y Water loss 12
332.1574401855469 0 82344.625 y Ammonia loss 12
333.15997314453125 0 11666.993
336.17919921875 0 6605.782 y Water loss 8
338.1361389160156 0 5391.927
339.1659851074219 0 4717.048
340.1496276855469 0 9382.675
341.1830139160156 0 66910.42
342.1418151855469 0 97926.53
342.18310546875 0 8652.018
343.1448059082031 0 14929.862
343.1991271972656 0 4372.287
344.145263671875 0 5837.2183
345.18310546875 0 32830.008 y 8
345.6849060058594 0 18391.85
349.1839599609375 0 66928.55 y 12
350.1858825683594 0 8275.452
353.1949462890625 0 4303.9707
359.16796875 0 45498.316
360.17230224609375 0 6143.5273
362.1482238769531 0 4100.469
365.1826171875 0 7419.3145
367.1622314453125 0 5070.3804
368.89666748046875 0 4636.974
379.2350158691406 0 5451.0557
380.15777587890625 0 10940.195
385.17291259765625 0 14404.069
385.7113952636719 0 7138.2954
389.70928955078125 0 15911.186
390.2098693847656 0 10070.386
390.71490478515625 0 14173.643
391.2178039550781 0 8653.778
391.7249755859375 0 53028.895
392.2270202636719 0 13949.553
392.7288513183594 0 8446.232
393.7102966308594 0 13099.405
395.15478515625 0 4888.214
397.2456359863281 0 12029.836
398.21533203125 0 6329.842
398.71453857421875 0 15432.571
399.2176818847656 0 11880.904
399.7208557128906 0 17170.736
400.22149658203125 0 21624.504
401.24139404296875 0 5417.007
403.1831359863281 0 18065.924
404.18896484375 0 5223.7476
405.22552490234375 0 9892.28
408.2275390625 0 12254.591
410.20526123046875 0 14027.358
411.2250061035156 0 9731.716
413.16986083984375 0 5440.374
414.22900390625 0 34706.008 b 11
414.7222595214844 0 52414.5 y Ammonia loss 7
415.2222595214844 0 28048.668
415.2558898925781 0 17820.955
423.2342529296875 0 663981.3 y 7
423.7355041503906 0 284927.53
424.2366638183594 0 73994.84
424.736328125 0 13178.529
425.240234375 0 50884.195 b Water loss 3
426.2433776855469 0 11226.575
428.21588134765625 0 24957.941
429.2353820800781 0 34310.01
430.2403564453125 0 22143.06 y Water loss 11
439.1935119628906 0 9029.377
440.2255554199219 0 5068.1777
441.2104187011719 0 42718.3
443.2514343261719 0 47385.188 b 3
444.2547912597656 0 14784.42
448.2525329589844 0 25389.156 y 11
448.8890075683594 0 13869.611 y Water loss 2
449.2250671386719 0 9504.065 y Ammonia loss 2
449.55694580078125 0 5380.814
450.74383544921875 0 11521.236 b 7
451.1967468261719 0 5025.7524
452.2269592285156 0 7255.242
452.2626647949219 0 21260.1
454.89263916015625 0 24693.3 y 2
455.2268371582031 0 19380.582
455.5616455078125 0 10631.741
458.2362060546875 0 38241.047
460.2417297363281 0 21316.932
465.2621765136719 0 6056.482
469.2898864746094 0 16448.479
470.23614501953125 0 4556.128
471.7423095703125 0 7161.8804 y Water loss 6
472.23583984375 0 16812.13 b 13
472.7350769042969 0 7316.4663
478.2032775878906 0 5197.185
479.2290954589844 0 4547.364
480.25628662109375 0 7971.1104
480.747802734375 0 111738.14 y 6
481.2495422363281 0 49413.656
481.75030517578125 0 16195.298
486.5833435058594 0 10585.407 y Water loss 1
486.9190673828125 0 24137.254 y Ammonia loss 1
487.2596435546875 0 7660.1704 y Water loss 10
488.244140625 0 8311.065 y Ammonia loss 10
492.58734130859375 0 26437.314 y 1
492.9229431152344 0 16340.905
493.25628662109375 0 11272.453
495.9148864746094 0 37726.156
496.2147216796875 0 8101.7
496.25164794921875 0 20768.057
496.5841369628906 0 12985.791
497.2359924316406 0 8593.204
497.284423828125 0 14175.045
505.2740173339844 0 80728.984 y 10
506.27691650390625 0 16551.76
513.9320068359375 0 5079.7866
514.2599487304688 0 5358.507
514.3251342773438 0 19091.717
515.2562255859375 0 17994.203 y Water loss 5
515.7546997070312 0 12463.135 y Ammonia loss 5
516.2575073242188 0 10769.559
516.757568359375 0 6104.9795
517.27001953125 0 5261.254
518.210205078125 0 8754.914
518.2760009765625 0 7143.241
518.61181640625 0 8585.731
518.9437866210938 0 6455.147
524.26513671875 0 742300.56 y 5
524.61279296875 0 145812.77 Precursor Ammonia loss
524.7650146484375 0 390743.97
524.94677734375 0 62469.406
525.2672729492188 0 120196.664
525.614501953125 0 7616.8223
525.767578125 0 36662.723
528.7910766601562 0 12091.521
529.292236328125 0 9351.173
529.7985229492188 0 19715.842
530.2822875976562 0 89755.26 Precursor
530.6165771484375 0 51924.81
530.950927734375 0 32752.064
532.2727661132812 0 13546.694
533.2543334960938 0 5723.828
539.2965698242188 0 8106.3213
542.3207397460938 0 14143.963 b 4
543.3243408203125 0 5649.568
556.320556640625 0 10465.793
556.7741088867188 0 6432.328
558.7938232421875 0 5506.2153
559.2899780273438 0 12049.292
564.7921142578125 0 35017.195 y Water loss 4
565.2421875 0 7286.4307
565.2893676757812 0 33516.58 y Ammonia loss 4
565.7901611328125 0 21345.027
566.2902221679688 0 8272.973
567.2938842773438 0 7582.9224
572.2828979492188 0 8923.7295
572.7903442382812 0 6935.7036
573.288818359375 0 11619.207
573.7979125976562 0 981592.56 y 4
574.2991333007812 0 585538.44 y Water loss 9
574.8006591796875 0 189378.11
575.3009643554688 0 36498.23
583.2463989257812 0 13523.82
584.3157958984375 0 13255.514
585.3069458007812 0 5969.221
587.2809448242188 0 11060.523
592.3063354492188 0 105402.25 y 9
593.3086547851562 0 30929.096
594.3126831054688 0 10107.252
596.3126831054688 0 5844.309
600.2747192382812 0 22601.629
601.2809448242188 0 7380.661
602.2879638671875 0 7727.8765
608.3098754882812 0 9809.591
612.3167724609375 0 5776.784 b Ammonia loss 11
617.3017578125 0 5813.1035
620.3060302734375 0 5937.514
624.3126831054688 0 5255.955
626.3255004882812 0 15836.668
629.3155517578125 0 80140.83 y Water loss 3
629.8134765625 0 49999.92 y Ammonia loss 3
630.3176879882812 0 26656.443
636.3097534179688 0 9793.5
638.3192138671875 0 479438.97 y 3
638.8206787109375 0 313128.9
639.321533203125 0 105521.445
639.8222045898438 0 33874.67
640.3242797851562 0 10413.212
641.3380126953125 0 9246.601
642.3211669921875 0 7968.9478
642.8142700195312 0 7197.2896
643.35400390625 0 13525.871
644.3514404296875 0 5962.7217
645.3626708984375 0 5287.2734
646.3165283203125 0 34507.43
647.3230590820312 0 8421.869
648.8213500976562 0 11482.218
649.31884765625 0 10341.841
649.8214721679688 0 6021.0747
651.3133544921875 0 6876.94
652.315673828125 0 11917.436
653.3363647460938 0 15060.681
654.3258666992188 0 20372.943
654.82568359375 0 7703.346
655.3178100585938 0 12530.027
657.2882690429688 0 16857.576
657.82373046875 0 6667.764
659.34521484375 0 25903.805
659.8317260742188 0 5605.7944
660.2974243164062 0 8111.7847
660.3488159179688 0 11966.877
660.8189086914062 0 23249.342
661.3262939453125 0 15866.8545
663.82373046875 0 41199.19
664.3217163085938 0 35068.42
664.8220825195312 0 21247.105
665.3212890625 0 7347.994
665.8244018554688 0 6833.821
666.325927734375 0 19778.23
666.8302612304688 0 86174.9
667.3313598632812 0 51362.92
667.8322143554688 0 20928.926
668.3248901367188 0 6859.27
671.3486328125 0 107395.336 y Water loss 8
672.3372802734375 0 75725.9 y Ammonia loss 8
672.8300170898438 0 252550.31 y Water loss 2
673.330810546875 0 241697.47 y Ammonia loss 2
673.8292236328125 0 89874.984
674.3158569335938 0 138551.61
674.82861328125 0 13239.202
675.314453125 0 39916.586
676.3168334960938 0 6113.164
678.3317260742188 0 5811.4824
680.8270263671875 0 79013.57
681.3300170898438 0 71554.89
681.8358764648438 0 7584216.5 y 2
682.3370971679688 0 5302956.5
682.8380126953125 0 2137048.2
683.3389892578125 0 568302.2
683.8400268554688 0 127256.086
685.3885498046875 0 11212.249
686.8283081054688 0 56653.285
687.3309326171875 0 43045.457
687.830322265625 0 18302.533
689.3590698242188 0 610651.3 y 8
690.3616943359375 0 214829.28
691.3643798828125 0 47984.16
692.363525390625 0 8602.467
699.3446044921875 0 38369.97 b Ammonia loss 13
700.34521484375 0 11273.154
728.3687744140625 0 10560.787
729.3719482421875 0 75709.305 y Water loss 1
729.87255859375 0 39785.316 y Ammonia loss 1
730.3736572265625 0 19824.639
730.8716430664062 0 8960.089
737.3690185546875 0 6591.5693
737.8750610351562 0 8126.4023
738.3770751953125 0 626569.44 y 1
738.8784790039062 0 470820.25
739.3798828125 0 200551.88
739.88134765625 0 58467.566
740.381103515625 0 18842.037
743.370361328125 0 34380.902
743.8705444335938 0 21887.828
744.3735961914062 0 15152.827 b 6
758.3833618164062 0 8331.376
767.4130249023438 0 6032.5933
768.3526000976562 0 5792.7847
768.40234375 0 13858.147
769.4072875976562 0 7649.9443
770.4158935546875 0 29305.045
771.4249267578125 0 7522.7197
778.4117431640625 0 11031.782
779.4102783203125 0 7167.5474
780.4329833984375 0 6856.276
782.4419555664062 0 34625.094
783.4435424804688 0 18714.678
784.3884887695312 0 6731.237
785.4259033203125 0 14817.608
786.36962890625 0 11064.375
786.4310302734375 0 7407.6313
787.3911743164062 0 7059.3047
796.4207763671875 0 19211.885
797.423095703125 0 6733.2983
798.4208984375 0 23284.152
799.4190673828125 0 8375.175
803.4395751953125 0 14964.436
810.4336547851562 0 6821.611
811.4072875976562 0 20309.768
812.4107055664062 0 8488.11
828.4332275390625 0 89807.9 y Ammonia loss 7
829.4345092773438 0 34966.31
830.405029296875 0 13595.819
855.3949584960938 0 6077.154
857.4454956054688 0 11184.967
858.3956298828125 0 7850.543
868.4337158203125 0 6359.7114
873.4068603515625 0 11468.358
874.4008178710938 0 6149.1494
883.4415283203125 0 4680.6187 b Ammonia loss 7
885.4415893554688 0 13969.439
887.4223022460938 0 21488.566
888.428466796875 0 9808.367
889.427490234375 0 6284.9355
891.4172973632812 0 15137.476
892.4227294921875 0 6419.7246
899.4700927734375 0 7308.341
900.468994140625 0 18913.396 b 7
901.4523315429688 0 11576.356
909.44189453125 0 11713.26
914.448974609375 0 5619.1865
915.4191284179688 0 44776.668
916.421875 0 23486.05
917.4136962890625 0 6523.053
925.455078125 0 5879.102
926.4258422851562 0 7350.568
927.4495239257812 0 12541.031
943.4605102539062 0 69838.17 y Ammonia loss 6
944.4680786132812 0 36210.316
945.4724731445312 0 12876.82
946.4723510742188 0 6055.473
952.46435546875 0 6154.745
953.4570922851562 0 6719.781
957.4688110351562 0 6229.0396
968.4849853515625 0 6503.3687
969.4754028320312 0 13253.384
970.4752197265625 0 7072.673
972.4776000976562 0 11180.26
986.4920043945312 0 64763.64
987.489013671875 0 69035.03
988.4887084960938 0 32661.857
989.4908447265625 0 11634.13
990.4902954101562 0 14589.525
995.4662475585938 0 6785.2144
996.4734497070312 0 12735.612
997.4696655273438 0 6930.91
1000.4842529296875 0 9152.303
1006.4990844726562 0 7910.098
1012.48046875 0 34532.26
1013.4822998046875 0 23319.502
1014.4856567382812 0 53932.9
1015.4891357421875 0 25224.908
1030.4923095703125 0 234442.14 y Ammonia loss 5
1031.4951171875 0 106127.81
1032.4981689453125 0 40546.29
1033.4959716796875 0 8616.876
1055.511962890625 0 8959.273
1056.5228271484375 0 7115.1694
1057.5184326171875 0 10336.273
1058.53466796875 0 7258.3496
1073.5206298828125 0 23004.482
1074.5301513671875 0 8624.339
1083.5133056640625 0 11474.936
1084.5101318359375 0 10284.317
1086.55224609375 0 15567.071
1087.5660400390625 0 6849.7397
1101.52001953125 0 34538.836
1102.52294921875 0 21606.082
1111.5494384765625 0 11054.265
1112.539794921875 0 6471.069
1119.5316162109375 0 14271.518
1129.56103515625 0 63643.164 y Ammonia loss 4
1130.5615234375 0 33977.24
1131.5616455078125 0 7768.152
1142.546630859375 0 5646.7783
1143.538330078125 0 6966.171
1144.560302734375 0 7644.5825
1160.55322265625 0 35335.89
1161.55615234375 0 17094.883
1170.5418701171875 0 14263.678
1187.55859375 0 5809.861
1188.5511474609375 0 37822.61
1189.552001953125 0 29092.002
1206.560546875 0 50233.39
1207.563232421875 0 30064.393
1208.56591796875 0 11108.366
1215.5977783203125 0 6483.5684
1301.63623046875 0 6945.493
2373.021728515625 0 5653.9414
2904.10595703125 0 5474.5225

Spectrum Details

|  |  |
| --- | --- |
| Matched peaks? Matched peaks The total absolute number of peaks matched. Additionally in brackets the total fraction of peaks matched and the total number of peaks is shown. | 67 (12.23% of 548) |
| FDR? FDR The false discovery rate estimated for this peptide. It is calculated by matching all theoretical fragments with a non-integer shift with the raw peaks for this spectrum. This is done with 40 different shifts. The resulting percentage is the average number of annotated peaks over the number of annotated peaks with the correct spectrum. | 0.39% |
| Satellite FDR? Satellite FDR See the FDR for details on its calculation. This satellite ion specific FDR only contains the satellite ions (d/w) for I/L/J positions. | - |
| PSM Score? PSM Score The PSM Score as given by Hecklib to this annotated spectrum. It is shown with three significant figures. | 421 |

#### Spectrum 4798? Spectrum 4798 The raw spectrum of this peptide as annotated by Hecklib. The fragments are coloured according to ion type (see legend). Any peaks with a star '\*' as text can be hovered over to see the full details, first the ion type second the mass shift type. By hovering over the amino acids in the peptide or ions in the legend the corresponding peaks are highlighted. By toggling the 'Unassigned' label you can turn the background (unassigned) peaks on or off in the plot. By updating the slider in the Ion legend you can update the spectrum to only show the top X% of the peaks with labels. The top X% means any peak that is within X% of the highest intensity. By dragging in the spectrum you can zoom in to a specific part of the spectrum and use 'Zoom Out' to get back to the original zoom level. The annotation of the spectrum is based on the given sequence in the peptides file and is done with different software so inconsistencies are likely. The peaks are annotated based on the given sequence, with 20 ppm tolerance.

Copy Data

##### Spectrum 4798 (TSV)

###### Preview

```
Loading example...
```

*Click on the button to copy the data to your clipboard.*

Mz MinMz MaxIntensity Max

WidthHeightPeptide font sizePeptide stroke widthSpectrum font sizeSpectrum stroke widthCompact peptide

Ion legend

wxyz

abcd

OtherUnassignedIon^Charge^_Position_Show for top:%

JJSEVSDRPSGVSSR

01.07e+42.14e+43.21e+44.28e+4

Zoom Out

y^+1^_1_w^+1^_2_y^+1^_2_z^+1^_2_y^+1^_2_w^+1^_3_y^+1^_3_z^+1^_3_y^+1^_3_z^+2^_8_w^+1^_4_y^+1^_4_c^+1^_4_z^+2^_9_y^+2^_9_c^+3^_14_z^+2^_9_y^+1^_5_z^+2^_10_y^+2^_10_w^+1^_6_c^+1^_5_z^+2^_11_y^+2^_11_y^+1^_6_y^+1^_6_z^+1^_6_w^+2^_12_c^+2^_12_z^+2^_12_y^+2^_12_c^+1^_6_y^+1^_7_y^+2^_13_z^+2^_13_y^+2^_13_y^+1^_7_w^+2^_14_z^+2^_14_y^+2^_14_w^+1^_8_c^+1^_7_z^+1^_8_c^+1^_8_y^+1^_9_z^+1^_9_y^+1^_9_c^+1^_9_z^+1^_10_y^+1^_10_c^+1^_10_z^+1^_11_c^+1^_11_c^+1^_12_y^+1^_12_c^+1^_13_c^+1^_14_

0843168625293373

Fragment Matches Table

Show background peaks

| Position | Ion type | Intensity | mz Theoretical | mz Error (Th) | mz Error (ppm) | Charge | Series Number |
| --- | --- | --- | --- | --- | --- | --- | --- |
| - | - | 570.3 | 120.1 | - | - | 0 | - |
| - | - | 410.6 | 123 | - | - | 0 | - |
| - | - | 899.6 | 125.1 | - | - | 0 | - |
| - | - | 1216 | 126.1 | - | - | 0 | - |
| - | - | 1191 | 133.1 | - | - | 0 | - |
| - | - | 490.6 | 136.1 | - | - | 0 | - |
| - | - | 412.9 | 139.1 | - | - | 0 | - |
| - | - | 824.4 | 149 | - | - | 0 | - |
| - | - | 495.8 | 160.4 | - | - | 0 | - |
| - | - | 511 | 163.1 | - | - | 0 | - |
| - | - | 490.4 | 166.9 | - | - | 0 | - |
| - | - | 1369 | 168.1 | - | - | 0 | - |
| - | - | 1027 | 171.1 | - | - | 0 | - |
| - | - | 1148 | 173.4 | - | - | 0 | - |
| 15 | y | 2615 | 175.1 | 9.907E-06 | 0.05657 | +1 | 1 |
| - | - | 620.3 | 177.1 | - | - | 0 | - |
| - | - | 541.9 | 179.1 | - | - | 0 | - |
| - | - | 439.2 | 179.2 | - | - | 0 | - |
| - | - | 711.3 | 185.1 | - | - | 0 | - |
| - | - | 6141 | 185.1 | - | - | 0 | - |
| - | - | 609.7 | 186.1 | - | - | 0 | - |
| - | - | 1.286E+04 | 187.1 | - | - | 0 | - |
| - | - | 952.5 | 188.1 | - | - | 0 | - |
| - | - | 542.9 | 189.1 | - | - | 0 | - |
| - | - | 739.7 | 199.2 | - | - | 0 | - |
| - | - | 1.145E+04 | 199.2 | - | - | 0 | - |
| - | - | 810.5 | 200.1 | - | - | 0 | - |
| - | - | 1731 | 200.2 | - | - | 0 | - |
| - | - | 483.9 | 201.1 | - | - | 0 | - |
| - | - | 1772 | 214.2 | - | - | 0 | - |
| - | - | 497 | 214.9 | - | - | 0 | - |
| - | - | 679.6 | 215.1 | - | - | 0 | - |
| - | - | 5779 | 227.2 | - | - | 0 | - |
| - | - | 961.8 | 228.2 | - | - | 0 | - |
| 14 | w | 953.6 | 229.1 | 0.0001218 | 0.5317 | +1 | 2 |
| 14 | y | 9895 | 245.1 | 0.0001108 | 0.4519 | +1 | 2 |
| 14 | z | 1802 | 246.1 | 0.001031 | 4.188 | +1 | 2 |
| 14 | y | 931.2 | 262.2 | 0.0003256 | 1.242 | +1 | 2 |
| - | - | 696.2 | 270.1 | - | - | 0 | - |
| - | - | 608.3 | 276.9 | - | - | 0 | - |
| 13 | w | 4339 | 316.2 | 0.00032 | 1.012 | +1 | 3 |
| - | - | 871.3 | 317.2 | - | - | 0 | - |
| - | - | 957.7 | 331.1 | - | - | 0 | - |
| 13 | y | 1445 | 332.2 | 0.0006982 | 2.102 | +1 | 3 |
| - | - | 1085 | 333.1 | - | - | 0 | - |
| 13 | z | 3579 | 333.2 | 0.0001743 | 0.5232 | +1 | 3 |
| - | - | 2117 | 334.2 | - | - | 0 | - |
| - | - | 822.1 | 349.1 | - | - | 0 | - |
| 13 | y | 1227 | 349.2 | 0.0001477 | 0.4229 | +1 | 3 |
| - | - | 2478 | 357.2 | - | - | 0 | - |
| - | - | 665.4 | 399.7 | - | - | 0 | - |
| - | - | 673.1 | 404.2 | - | - | 0 | - |
| 8 | z | 1346 | 415.2 | 0.001243 | 2.995 | +2 | 8 |
| - | - | 2316 | 415.3 | - | - | 0 | - |
| 12 | w | 3123 | 417.2 | 0.0007677 | 1.84 | +1 | 4 |
| - | - | 957.1 | 418.2 | - | - | 0 | - |
| - | - | 769.8 | 430.3 | - | - | 0 | - |
| - | - | 2262 | 439.3 | - | - | 0 | - |
| - | - | 1098 | 440.2 | - | - | 0 | - |
| - | - | 1007 | 443.3 | - | - | 0 | - |
| 12 | y | 3885 | 448.3 | 0.0004387 | 0.9786 | +1 | 4 |
| - | - | 1492 | 457.7 | - | - | 0 | - |
| - | - | 1995 | 458.3 | - | - | 0 | - |
| 4 | c | 2213 | 460.3 | 0.0001889 | 0.4104 | +1 | 4 |
| - | - | 924.3 | 461.3 | - | - | 0 | - |
| - | - | 2611 | 462.7 | - | - | 0 | - |
| - | - | 1295 | 463.2 | - | - | 0 | - |
| 7 | z | 625.4 | 463.7 | 0.002393 | 5.159 | +2 | 9 |
| 7 | y | 2523 | 471.7 | 0.003643 | 7.722 | +2 | 9 |
| 14 | c | 983.2 | 472.2 | 0.004265 | 9.032 | +3 | 14 |
| 7 | z | 980.5 | 472.7 | 0.0006198 | 1.311 | +2 | 9 |
| - | - | 1459 | 474.2 | - | - | 0 | - |
| - | - | 2546 | 475.3 | - | - | 0 | - |
| - | - | 661.4 | 486.3 | - | - | 0 | - |
| - | - | 725.9 | 490.3 | - | - | 0 | - |
| - | - | 761 | 499.3 | - | - | 0 | - |
| - | - | 2104 | 501.3 | - | - | 0 | - |
| - | - | 3440 | 502.3 | - | - | 0 | - |
| - | - | 720.6 | 503.3 | - | - | 0 | - |
| - | - | 701.5 | 504.3 | - | - | 0 | - |
| 11 | y | 1.253E+04 | 505.3 | 0.0004898 | 0.9695 | +1 | 5 |
| - | - | 2844 | 506.3 | - | - | 0 | - |
| - | - | 905.1 | 507.3 | - | - | 0 | - |
| - | - | 648.7 | 513.3 | - | - | 0 | - |
| 6 | z | 1310 | 516.3 | 0.0002272 | 0.4401 | +2 | 10 |
| - | - | 1047 | 516.8 | - | - | 0 | - |
| - | - | 770.4 | 521.3 | - | - | 0 | - |
| 6 | y | 4631 | 524.3 | 0.001214 | 2.316 | +2 | 10 |
| - | - | 1787 | 524.6 | - | - | 0 | - |
| - | - | 2255 | 524.8 | - | - | 0 | - |
| - | - | 987.6 | 524.9 | - | - | 0 | - |
| - | - | 656.8 | 529.8 | - | - | 0 | - |
| - | - | 4.242E+04 | 530.3 | - | - | 0 | - |
| - | - | 1226 | 530.6 | - | - | 0 | - |
| - | - | 1012 | 530.9 | - | - | 0 | - |
| - | - | 1.125E+04 | 531.3 | - | - | 0 | - |
| - | - | 1136 | 542.3 | - | - | 0 | - |
| - | - | 696 | 558.3 | - | - | 0 | - |
| 10 | w | 700.5 | 559.3 | 0.0006673 | 1.193 | +1 | 6 |
| 5 | c | 1845 | 559.3 | 0.001019 | 1.822 | +1 | 5 |
| - | - | 601.7 | 560.3 | - | - | 0 | - |
| 5 | z | 8826 | 565.8 | 5.072E-05 | 0.08965 | +2 | 11 |
| - | - | 3332 | 566.3 | - | - | 0 | - |
| - | - | 2894 | 566.8 | - | - | 0 | - |
| - | - | 730.8 | 567.3 | - | - | 0 | - |
| - | - | 2606 | 572.3 | - | - | 0 | - |
| - | - | 1724 | 573.3 | - | - | 0 | - |
| 5 | y | 5671 | 573.8 | 0.0002102 | 0.3663 | +2 | 11 |
| 10 | y | 3500 | 574.3 | 0.004539 | 7.903 | +1 | 6 |
| - | - | 1072 | 574.8 | - | - | 0 | - |
| 10 | y | 771.3 | 575.3 | 0.001663 | 2.891 | +1 | 6 |
| 10 | z | 1573 | 576.3 | 0.0003633 | 0.6305 | +1 | 6 |
| - | - | 869.3 | 577.3 | - | - | 0 | - |
| - | - | 3719 | 585.3 | - | - | 0 | - |
| - | - | 604.2 | 586.3 | - | - | 0 | - |
| - | - | 993.1 | 600.3 | - | - | 0 | - |
| 4 | w | 2675 | 600.8 | 6.255E-06 | 0.01041 | +2 | 12 |
| - | - | 1899 | 601.3 | - | - | 0 | - |
| - | - | 1355 | 601.8 | - | - | 0 | - |
| 12 | c | 1474 | 629.3 | 0.008664 | 13.77 | +2 | 12 |
| 4 | z | 8524 | 630.3 | 0.0002996 | 0.4753 | +2 | 12 |
| - | - | 7646 | 630.8 | - | - | 0 | - |
| - | - | 3688 | 631.3 | - | - | 0 | - |
| - | - | 834.7 | 631.8 | - | - | 0 | - |
| - | - | 3044 | 637.8 | - | - | 0 | - |
| 4 | y | 8162 | 638.3 | 0.0005786 | 0.9064 | +2 | 12 |
| - | - | 5756 | 638.8 | - | - | 0 | - |
| - | - | 2020 | 639.3 | - | - | 0 | - |
| - | - | 943.6 | 643.4 | - | - | 0 | - |
| - | - | 729.5 | 644.8 | - | - | 0 | - |
| 6 | c | 9591 | 646.4 | 0.000363 | 0.5616 | +1 | 6 |
| - | - | 2796 | 647.4 | - | - | 0 | - |
| - | - | 607.1 | 648.4 | - | - | 0 | - |
| 9 | y | 659.1 | 672.3 | 0.009385 | 13.96 | +1 | 7 |
| 3 | y | 638.5 | 673.3 | 0.01127 | 16.74 | +2 | 13 |
| 3 | z | 5626 | 673.8 | 0.0008869 | 1.316 | +2 | 13 |
| - | - | 8591 | 674.3 | - | - | 0 | - |
| - | - | 4042 | 674.8 | - | - | 0 | - |
| - | - | 1485 | 675.3 | - | - | 0 | - |
| - | - | 1167 | 675.8 | - | - | 0 | - |
| - | - | 959.6 | 681.3 | - | - | 0 | - |
| 3 | y | 3.466E+04 | 681.8 | 6.983E-05 | 0.1024 | +2 | 13 |
| - | - | 2.548E+04 | 682.3 | - | - | 0 | - |
| - | - | 9365 | 682.8 | - | - | 0 | - |
| - | - | 3169 | 683.3 | - | - | 0 | - |
| - | - | 2712 | 686.4 | - | - | 0 | - |
| - | - | 3438 | 687.4 | - | - | 0 | - |
| - | - | 1888 | 688.4 | - | - | 0 | - |
| 9 | y | 1199 | 689.4 | 0.008166 | 11.85 | +1 | 7 |
| - | - | 824 | 702.4 | - | - | 0 | - |
| - | - | 1529 | 711.4 | - | - | 0 | - |
| - | - | 1431 | 711.9 | - | - | 0 | - |
| 2 | w | 1356 | 715.8 | 0.0007084 | 0.9897 | +2 | 14 |
| - | - | 861.5 | 718.4 | - | - | 0 | - |
| - | - | 682.7 | 719.4 | - | - | 0 | - |
| - | - | 1104 | 727.4 | - | - | 0 | - |
| 2 | z | 1084 | 730.4 | 0.001899 | 2.601 | +2 | 14 |
| - | - | 625.4 | 730.9 | - | - | 0 | - |
| 2 | y | 3369 | 738.4 | 0.0006414 | 0.8686 | +2 | 14 |
| - | - | 1356 | 738.9 | - | - | 0 | - |
| 8 | w | 837.6 | 743.4 | 0.003766 | 5.066 | +1 | 8 |
| - | - | 3595 | 744.4 | - | - | 0 | - |
| - | - | 1142 | 745.4 | - | - | 0 | - |
| - | - | 956.5 | 749.9 | - | - | 0 | - |
| - | - | 914.3 | 750.9 | - | - | 0 | - |
| - | - | 1003 | 758.4 | - | - | 0 | - |
| - | - | 1641 | 758.9 | - | - | 0 | - |
| - | - | 1429 | 759.4 | - | - | 0 | - |
| 7 | c | 1.633E+04 | 761.4 | 0.0005196 | 0.6824 | +1 | 7 |
| - | - | 7465 | 762.4 | - | - | 0 | - |
| - | - | 1225 | 763.4 | - | - | 0 | - |
| - | - | 1381 | 764.9 | - | - | 0 | - |
| - | - | 2.637E+04 | 765.4 | - | - | 0 | - |
| - | - | 1.874E+04 | 765.9 | - | - | 0 | - |
| - | - | 1.048E+04 | 766.4 | - | - | 0 | - |
| - | - | 2465 | 766.9 | - | - | 0 | - |
| - | - | 940.6 | 767.4 | - | - | 0 | - |
| - | - | 756.4 | 772.4 | - | - | 0 | - |
| - | - | 724.3 | 772.9 | - | - | 0 | - |
| - | - | 918.5 | 773.4 | - | - | 0 | - |
| - | - | 8012 | 773.9 | - | - | 0 | - |
| - | - | 6226 | 774.4 | - | - | 0 | - |
| - | - | 3231 | 774.9 | - | - | 0 | - |
| - | - | 5134 | 775.4 | - | - | 0 | - |
| - | - | 2059 | 776.4 | - | - | 0 | - |
| - | - | 982.8 | 777.9 | - | - | 0 | - |
| - | - | 843.4 | 779.4 | - | - | 0 | - |
| - | - | 896.2 | 779.9 | - | - | 0 | - |
| - | - | 716.3 | 782.4 | - | - | 0 | - |
| - | - | 691.1 | 785.4 | - | - | 0 | - |
| - | - | 1101 | 785.9 | - | - | 0 | - |
| - | - | 2817 | 786.4 | - | - | 0 | - |
| - | - | 3.057E+04 | 786.9 | - | - | 0 | - |
| - | - | 2.358E+04 | 787.4 | - | - | 0 | - |
| - | - | 1.078E+04 | 787.9 | - | - | 0 | - |
| - | - | 5093 | 788.4 | - | - | 0 | - |
| - | - | 693 | 788.9 | - | - | 0 | - |
| - | - | 700.3 | 793.4 | - | - | 0 | - |
| - | - | 925.7 | 793.9 | - | - | 0 | - |
| - | - | 1122 | 794.4 | - | - | 0 | - |
| - | - | 1.908E+04 | 794.9 | - | - | 0 | - |
| - | - | 2.408E+04 | 795.4 | - | - | 0 | - |
| - | - | 1.477E+04 | 795.9 | - | - | 0 | - |
| - | - | 5781 | 796.4 | - | - | 0 | - |
| - | - | 2390 | 796.9 | - | - | 0 | - |
| - | - | 1182 | 797.4 | - | - | 0 | - |
| - | - | 3067 | 827.4 | - | - | 0 | - |
| - | - | 1663 | 828.4 | - | - | 0 | - |
| 8 | z | 1.244E+04 | 829.4 | 0.0006127 | 0.7387 | +1 | 8 |
| - | - | 6037 | 830.4 | - | - | 0 | - |
| - | - | 3335 | 831.4 | - | - | 0 | - |
| - | - | 1587 | 832.4 | - | - | 0 | - |
| - | - | 2147 | 839.4 | - | - | 0 | - |
| - | - | 875.9 | 844.5 | - | - | 0 | - |
| - | - | 3661 | 845.4 | - | - | 0 | - |
| - | - | 2498 | 846.4 | - | - | 0 | - |
| - | - | 880.8 | 847.4 | - | - | 0 | - |
| - | - | 2405 | 857.4 | - | - | 0 | - |
| - | - | 861 | 858.4 | - | - | 0 | - |
| - | - | 1014 | 871.4 | - | - | 0 | - |
| - | - | 632.5 | 872.4 | - | - | 0 | - |
| - | - | 1203 | 874.4 | - | - | 0 | - |
| - | - | 1026 | 888.5 | - | - | 0 | - |
| 8 | c | 2097 | 900.5 | 0.0005426 | 0.6026 | +1 | 8 |
| - | - | 606.4 | 910.7 | - | - | 0 | - |
| - | - | 650.3 | 914.4 | - | - | 0 | - |
| - | - | 551.7 | 928.4 | - | - | 0 | - |
| - | - | 1077 | 931.4 | - | - | 0 | - |
| - | - | 1009 | 932.4 | - | - | 0 | - |
| 7 | y | 2015 | 943.5 | 0.01695 | 17.97 | +1 | 9 |
| 7 | z | 6032 | 944.5 | 0.002962 | 3.136 | +1 | 9 |
| - | - | 5141 | 945.5 | - | - | 0 | - |
| - | - | 1410 | 946.5 | - | - | 0 | - |
| 7 | y | 1123 | 960.5 | 0.003647 | 3.797 | +1 | 9 |
| - | - | 1217 | 970.5 | - | - | 0 | - |
| - | - | 719.5 | 975.5 | - | - | 0 | - |
| - | - | 3154 | 988.5 | - | - | 0 | - |
| - | - | 2384 | 989.5 | - | - | 0 | - |
| - | - | 1567 | 999.6 | - | - | 0 | - |
| - | - | 1191 | 1001 | - | - | 0 | - |
| 9 | c | 1.454E+04 | 1015 | 0.0002791 | 0.2751 | +1 | 9 |
| - | - | 9584 | 1016 | - | - | 0 | - |
| - | - | 2746 | 1017 | - | - | 0 | - |
| - | - | 1540 | 1018 | - | - | 0 | - |
| - | - | 754.9 | 1019 | - | - | 0 | - |
| 6 | z | 1980 | 1031 | 0.0008406 | 0.815 | +1 | 10 |
| - | - | 2224 | 1033 | - | - | 0 | - |
| - | - | 1212 | 1034 | - | - | 0 | - |
| - | - | 6856 | 1043 | - | - | 0 | - |
| - | - | 4146 | 1044 | - | - | 0 | - |
| - | - | 1.276E+04 | 1045 | - | - | 0 | - |
| - | - | 6451 | 1046 | - | - | 0 | - |
| - | - | 2289 | 1047 | - | - | 0 | - |
| 6 | y | 817.9 | 1048 | 0.001526 | 1.457 | +1 | 10 |
| - | - | 1008 | 1049 | - | - | 0 | - |
| - | - | 7595 | 1058 | - | - | 0 | - |
| - | - | 7685 | 1059 | - | - | 0 | - |
| - | - | 9474 | 1060 | - | - | 0 | - |
| - | - | 5363 | 1061 | - | - | 0 | - |
| - | - | 9861 | 1062 | - | - | 0 | - |
| - | - | 5216 | 1063 | - | - | 0 | - |
| - | - | 681.7 | 1064 | - | - | 0 | - |
| 10 | c | 1.549E+04 | 1102 | 0.0003463 | 0.3144 | +1 | 10 |
| - | - | 8813 | 1103 | - | - | 0 | - |
| - | - | 3456 | 1104 | - | - | 0 | - |
| - | - | 815.3 | 1105 | - | - | 0 | - |
| - | - | 2851 | 1115 | - | - | 0 | - |
| - | - | 2032 | 1116 | - | - | 0 | - |
| - | - | 1181 | 1117 | - | - | 0 | - |
| 5 | z | 2244 | 1131 | 0.001274 | 1.127 | +1 | 11 |
| - | - | 5951 | 1132 | - | - | 0 | - |
| - | - | 3093 | 1133 | - | - | 0 | - |
| - | - | 1105 | 1134 | - | - | 0 | - |
| 11 | c | 9115 | 1159 | 0.0007332 | 0.6328 | +1 | 11 |
| - | - | 7018 | 1160 | - | - | 0 | - |
| - | - | 1368 | 1161 | - | - | 0 | - |
| - | - | 1727 | 1214 | - | - | 0 | - |
| - | - | 1017 | 1215 | - | - | 0 | - |
| 12 | c | 8918 | 1258 | 0.00103 | 0.8193 | +1 | 12 |
| - | - | 5564 | 1259 | - | - | 0 | - |
| - | - | 2051 | 1260 | - | - | 0 | - |
| - | - | 7500 | 1261 | - | - | 0 | - |
| - | - | 3848 | 1262 | - | - | 0 | - |
| - | - | 2077 | 1263 | - | - | 0 | - |
| 4 | y | 2794 | 1276 | 0.0009608 | 0.7532 | +1 | 12 |
| - | - | 1667 | 1277 | - | - | 0 | - |
| - | - | 908.6 | 1278 | - | - | 0 | - |
| - | - | 584.8 | 1281 | - | - | 0 | - |
| - | - | 1189 | 1301 | - | - | 0 | - |
| - | - | 912 | 1302 | - | - | 0 | - |
| - | - | 851.4 | 1303 | - | - | 0 | - |
| - | - | 725.4 | 1304 | - | - | 0 | - |
| - | - | 915.4 | 1331 | - | - | 0 | - |
| 13 | c | 1.447E+04 | 1345 | 0.0008322 | 0.6189 | +1 | 13 |
| - | - | 1.051E+04 | 1346 | - | - | 0 | - |
| - | - | 4684 | 1347 | - | - | 0 | - |
| - | - | 6504 | 1348 | - | - | 0 | - |
| - | - | 5223 | 1349 | - | - | 0 | - |
| - | - | 1902 | 1350 | - | - | 0 | - |
| - | - | 672.3 | 1351 | - | - | 0 | - |
| - | - | 1322 | 1388 | - | - | 0 | - |
| - | - | 1081 | 1389 | - | - | 0 | - |
| - | - | 666.2 | 1416 | - | - | 0 | - |
| 14 | c | 2.29E+04 | 1432 | 0.001 | 0.6986 | +1 | 14 |
| - | - | 1.843E+04 | 1433 | - | - | 0 | - |
| - | - | 8669 | 1434 | - | - | 0 | - |
| - | - | 1418 | 1435 | - | - | 0 | - |
| - | - | 824.8 | 1461 | - | - | 0 | - |
| - | - | 745.3 | 1462 | - | - | 0 | - |
| - | - | 2313 | 1489 | - | - | 0 | - |
| - | - | 1550 | 1490 | - | - | 0 | - |
| - | - | 1010 | 1491 | - | - | 0 | - |
| - | - | 818.8 | 1515 | - | - | 0 | - |
| - | - | 728.1 | 1516 | - | - | 0 | - |
| - | - | 4901 | 1531 | - | - | 0 | - |
| - | - | 6243 | 1532 | - | - | 0 | - |
| - | - | 4129 | 1533 | - | - | 0 | - |
| - | - | 1320 | 1534 | - | - | 0 | - |
| - | - | 710 | 1535 | - | - | 0 | - |
| - | - | 1356 | 1545 | - | - | 0 | - |
| - | - | 3052 | 1546 | - | - | 0 | - |
| - | - | 2127 | 1547 | - | - | 0 | - |
| - | - | 1853 | 1548 | - | - | 0 | - |
| - | - | 1900 | 1549 | - | - | 0 | - |
| - | - | 1006 | 1556 | - | - | 0 | - |
| - | - | 2330 | 1557 | - | - | 0 | - |
| - | - | 2029 | 1558 | - | - | 0 | - |
| - | - | 923.3 | 1559 | - | - | 0 | - |
| - | - | 776.4 | 1562 | - | - | 0 | - |
| - | - | 4330 | 1563 | - | - | 0 | - |
| - | - | 3715 | 1564 | - | - | 0 | - |
| - | - | 1304 | 1565 | - | - | 0 | - |
| - | - | 804 | 1571 | - | - | 0 | - |
| - | - | 1175 | 1572 | - | - | 0 | - |
| - | - | 5299 | 1573 | - | - | 0 | - |
| - | - | 3.279E+04 | 1574 | - | - | 0 | - |
| - | - | 2.597E+04 | 1575 | - | - | 0 | - |
| - | - | 1.308E+04 | 1576 | - | - | 0 | - |
| - | - | 3996 | 1577 | - | - | 0 | - |
| - | - | 1465 | 1578 | - | - | 0 | - |
| - | - | 645.1 | 1587 | - | - | 0 | - |
| - | - | 1878 | 1588 | - | - | 0 | - |
| - | - | 3087 | 1589 | - | - | 0 | - |
| - | - | 9374 | 1590 | - | - | 0 | - |
| - | - | 3.04E+04 | 1591 | - | - | 0 | - |
| - | - | 2.484E+04 | 1592 | - | - | 0 | - |
| - | - | 1.107E+04 | 1593 | - | - | 0 | - |
| - | - | 3091 | 1594 | - | - | 0 | - |
| - | - | 985.9 | 1595 | - | - | 0 | - |
| - | - | 597.2 | 1701 | - | - | 0 | - |
| - | - | 720.3 | 2634 | - | - | 0 | - |
| - | - | 702.5 | 2655 | - | - | 0 | - |
| - | - | 707.8 | 3075 | - | - | 0 | - |
| - | - | 746.9 | 3339 | - | - | 0 | - |

m/z Charge Intensity FragmentType MassShift Position
120.08097076416016 0 570.29694
122.95699310302734 0 410.59894
125.10733795166016 0 899.55853
126.0914077758789 0 1215.9297
133.08596801757812 0 1191.2871
136.0762481689453 0 490.6035
139.1114501953125 0 412.88312
148.95480346679688 0 824.44574
160.3935546875 0 495.81256
163.07086181640625 0 510.97116
166.8554229736328 0 490.3758
168.11349487304688 0 1368.6914
171.1135711669922 0 1027.1707
173.43824768066406 0 1148.1074
175.1189422607422 0 2615.4534 y 14
177.11212158203125 0 620.26764
179.0931854248047 0 541.9393
179.19149780273438 0 439.20718
185.1312255859375 0 711.2796
185.1397705078125 0 6141.113
186.14305114746094 0 609.6917
187.10780334472656 0 12860.603
188.11167907714844 0 952.4884
189.0869598388672 0 542.89703
199.1702423095703 0 739.6529
199.18045043945312 0 11453.598
200.12710571289062 0 810.4628
200.1836395263672 0 1731.063
201.11251831054688 0 483.9115
214.1548614501953 0 1771.572
214.91427612304688 0 497.0028
215.1385498046875 0 679.6459
227.17543029785156 0 5778.7427
228.17892456054688 0 961.79877
229.129638671875 0 953.5686 w 13
245.12454223632812 0 9894.684 y Ammonia loss 13
246.1312255859375 0 1801.9875 z 13
262.15130615234375 0 931.23737 y 13
270.1194152832031 0 696.2015
276.8674011230469 0 608.2751
316.161865234375 0 4338.9243 w 12
317.1597595214844 0 871.25275
331.1484375 0 957.6834
332.15576171875 0 1445.3718 y Ammonia loss 12
333.1192626953125 0 1084.6588
333.1644592285156 0 3579.4824 z 12
334.1756286621094 0 2117.0479
349.11444091796875 0 822.05695
349.182861328125 0 1227.1147 y 12
357.1513366699219 0 2477.7026
399.72418212890625 0 665.3786
404.21392822265625 0 673.0616
415.22491455078125 0 1345.9108 z 7
415.2546691894531 0 2316.3594
417.2099914550781 0 3123.3557 w 11
418.2128601074219 0 957.1009
430.2679138183594 0 769.753
439.2544250488281 0 2261.634
440.2488098144531 0 1098.2413
443.25054931640625 0 1006.95087
448.2518615722656 0 3884.7134 y 11
457.7405700683594 0 1491.9541
458.2613525390625 0 1995.371
460.2767639160156 0 2212.9478 c 3
461.27825927734375 0 924.29474
462.73236083984375 0 2611.4802
463.23583984375 0 1294.5879
463.7342529296875 0 625.37787 z Water loss 6
471.7375793457031 0 2523.4504 y Water loss 6
472.2481994628906 0 983.1984 c Ammonia loss 13
472.7377624511719 0 980.49445 z 6
474.23175048828125 0 1458.863
475.2881774902344 0 2546.441
486.2515869140625 0 661.4372
490.2598876953125 0 725.9089
499.2973327636719 0 761.0159
501.30572509765625 0 2104.17
502.313232421875 0 3440.132
503.318359375 0 720.61914
504.26715087890625 0 701.4971
505.27337646484375 0 12528.872 y 10
506.27587890625 0 2844.0803
507.2770080566406 0 905.13806
513.2749633789062 0 648.651
516.2529296875 0 1310.0428 z 5
516.7553100585938 0 1046.5219
521.2685546875 0 770.44916
524.2637329101562 0 4630.7764 y 5
524.6110229492188 0 1786.7555
524.7634887695312 0 2255.3533
524.945556640625 0 987.5844
529.794189453125 0 656.84875
530.3173828125 0 42424.934
530.6162719726562 0 1226.1466
530.9485473632812 0 1011.9296
531.3206176757812 0 11246.565
542.3186645507812 0 1136.3931
558.331298828125 0 696.00183
559.2841186523438 0 700.5018 w 9
559.3460083007812 0 1845.3231 c 4
560.3464965820312 0 601.6928
565.7874145507812 0 8825.936 z 4
566.289794921875 0 3332.397
566.791748046875 0 2893.6519
567.2958374023438 0 730.77356
572.34326171875 0 2605.7278
573.3477783203125 0 1723.6953
573.7969360351562 0 5671.1323 y 4
574.2988891601562 0 3500.0017 y Water loss 9
574.8010864257812 0 1071.7156
575.280029296875 0 771.32007 y Ammonia loss 9
576.2858276367188 0 1572.9569 z 9
577.2904052734375 0 869.2787
585.2560424804688 0 3718.6997
586.2589111328125 0 604.1807
600.2733154296875 0 993.1254
600.802001953125 0 2675.3047 w 3
601.3034057617188 0 1899.0735
601.80517578125 0 1355.2847
629.3521728515625 0 1473.7281 c 11
630.3089599609375 0 8524.096 z 3
630.8113403320312 0 7646.344
631.3121337890625 0 3687.9058
631.8140869140625 0 834.72327
637.815185546875 0 3043.7842
638.3174438476562 0 8161.524 y 3
638.81982421875 0 5756.029
639.3203735351562 0 2019.8044
643.3525390625 0 943.56006
644.8211669921875 0 729.46216
646.3773803710938 0 9591.306 c 5
647.38037109375 0 2796.2961
648.3827514648438 0 607.10187
672.3405151367188 0 659.1118 y Ammonia loss 8
673.33203125 0 638.50476 y Ammonia loss 2
673.8255615234375 0 5626.1787 z 2
674.3279418945312 0 8591.074
674.829833984375 0 4041.8875
675.3305053710938 0 1485.4235
675.8296508789062 0 1166.9434
681.3318481445312 0 959.63574
681.8341064453125 0 34656.668 y 2
682.3355102539062 0 25480.791
682.8369750976562 0 9364.878
683.3363647460938 0 3168.6943
686.3555297851562 0 2711.8247
687.3622436523438 0 3438.429
688.36767578125 0 1887.9208
689.3658447265625 0 1198.9814 y 8
702.35693359375 0 823.9995
711.3551635742188 0 1528.7252
711.860595703125 0 1430.7427
715.846435546875 0 1355.5934 w 1
718.3950805664062 0 861.4777
719.4039916992188 0 682.67584
727.3736572265625 0 1104.0223
730.3648071289062 0 1083.824 z 1
730.8697509765625 0 625.40137
738.3754272460938 0 3368.6208 y 1
738.8770141601562 0 1356.1555
743.3720092773438 0 837.56226 w 7
744.3936767578125 0 3595.4592
745.3956909179688 0 1141.5018
749.8759155273438 0 956.4564
750.8997802734375 0 914.2503
758.3807373046875 0 1003.43176
758.8807373046875 0 1640.9718
759.3832397460938 0 1428.6844
761.4044799804688 0 16326.344 c 6
762.4073486328125 0 7465.307
763.4078979492188 0 1225.1327
764.9139404296875 0 1380.7214
765.3826293945312 0 26369.625
765.8839111328125 0 18739.586
766.3851318359375 0 10479.434
766.888427734375 0 2465.0566
767.3824462890625 0 940.56506
772.4053344726562 0 756.38074
772.9121704101562 0 724.25024
773.4078979492188 0 918.54205
773.9063720703125 0 8011.7686
774.4089965820312 0 6226.372
774.9102172851562 0 3231.426
775.41064453125 0 5134.4233
776.4102783203125 0 2058.9836
777.8975830078125 0 982.8454
779.4142456054688 0 843.4318
779.9028930664062 0 896.1723
782.4412841796875 0 716.2875
785.3765258789062 0 691.08545
785.88818359375 0 1101.0752
786.412109375 0 2817.0427
786.9092407226562 0 30565.47
787.4105224609375 0 23584.752
787.9119873046875 0 10780.416
788.413330078125 0 5093.1685
788.9115600585938 0 693.02936
793.40478515625 0 700.2901
793.8974609375 0 925.7369
794.4044799804688 0 1122.4556
794.917236328125 0 19075.385
795.420166015625 0 24082.424
795.922119140625 0 14767.473
796.4210205078125 0 5781.448
796.9232788085938 0 2390.2812
797.4185791015625 0 1182.1404
827.405517578125 0 3067.1213
828.4111938476562 0 1663.4845
829.439453125 0 12444.168 z 7
830.442626953125 0 6037.198
831.4028930664062 0 3334.7676
832.4044799804688 0 1586.865
839.4065551757812 0 2147.4958
844.4510498046875 0 875.8668
845.4168090820312 0 3661.36
846.4202880859375 0 2498.3997
847.4164428710938 0 880.75214
857.4173583984375 0 2405.203
858.4231567382812 0 861.027
871.4337158203125 0 1013.6103
872.4412231445312 0 632.50354
874.4423828125 0 1203.3185
888.4589233398438 0 1025.568
900.4790649414062 0 2096.5305 c Ammonia loss 7
910.6825561523438 0 606.4185
914.4113159179688 0 650.27185
928.4395751953125 0 551.7491
931.4464111328125 0 1077.4926
932.4440307617188 0 1008.9127
943.4761352539062 0 2014.6724 y Ammonia loss 6
944.469970703125 0 6032.04 z 6
945.47119140625 0 5140.59
946.4752807617188 0 1410.2231
960.4893798828125 0 1122.9824 y 6
970.5458984375 0 1217.2617
975.4781494140625 0 719.4858
988.5221557617188 0 3154.4841
989.5236206054688 0 2384.2446
999.5519409179688 0 1566.7085
1000.5498046875 0 1190.6522
1014.5575561523438 0 14543.408 c 8
1015.5597534179688 0 9583.815
1016.5615844726562 0 2745.5671
1017.560302734375 0 1539.7158
1018.9257202148438 0 754.8972
1031.4998779296875 0 1980.3494 z 5
1032.5029296875 0 2223.619
1033.51171875 0 1211.528
1042.5460205078125 0 6855.6616
1043.545166015625 0 4146.2754
1044.5263671875 0 12755.511
1045.52734375 0 6450.85
1046.5281982421875 0 2289.1133
1047.5162353515625 0 817.93866 y 5
1048.5333251953125 0 1008.20044
1057.576416015625 0 7595.203
1058.572998046875 0 7684.967
1059.5728759765625 0 9474.304
1060.56396484375 0 5363.3325
1061.550537109375 0 9860.831
1062.552978515625 0 5215.549
1063.5579833984375 0 681.7349
1101.5902099609375 0 15493.942 c 9
1102.5936279296875 0 8813.3545
1103.59375 0 3456.1055
1104.5946044921875 0 815.33386
1114.597900390625 0 2851.0688
1115.599365234375 0 2032.0383
1116.6063232421875 0 1180.5872
1130.5687255859375 0 2243.7546 z 4
1131.5748291015625 0 5950.9844
1132.5762939453125 0 3092.529
1133.56884765625 0 1104.9182
1158.612060546875 0 9114.906 c 10
1159.6136474609375 0 7018.24
1160.6109619140625 0 1368.04
1213.66357421875 0 1727.3137
1214.66748046875 0 1017.1937
1257.6787109375 0 8917.731 c 11
1258.681396484375 0 5563.743
1259.674560546875 0 2051.2942
1260.616943359375 0 7499.8677
1261.6209716796875 0 3848.1038
1262.62548828125 0 2076.6423
1275.6278076171875 0 2794.0105 y 3
1276.62744140625 0 1667.4822
1277.6365966796875 0 908.5828
1281.0447998046875 0 584.75055
1300.6968994140625 0 1188.5579
1301.7042236328125 0 911.9971
1302.6903076171875 0 851.39526
1303.6724853515625 0 725.3962
1330.6336669921875 0 915.4441
1344.7109375 0 14474.144 c 12
1345.7130126953125 0 10506.652
1346.7093505859375 0 4683.826
1347.6575927734375 0 6504.1294
1348.6512451171875 0 5223.0264
1349.6556396484375 0 1901.531
1350.6571044921875 0 672.2959
1387.734130859375 0 1321.8796
1388.7452392578125 0 1080.6503
1415.727294921875 0 666.2204
1431.7427978515625 0 22899.105 c 13
1432.746337890625 0 18425.814
1433.7462158203125 0 8668.805
1434.7474365234375 0 1417.8008
1460.7403564453125 0 824.83234
1461.73095703125 0 745.28156
1488.7457275390625 0 2313.164
1489.74462890625 0 1550.0519
1490.7593994140625 0 1009.5891
1514.7911376953125 0 818.7837
1515.7762451171875 0 728.1112
1530.7904052734375 0 4900.64
1531.7926025390625 0 6243.0674
1532.7967529296875 0 4128.606
1533.7921142578125 0 1320.0979
1534.803955078125 0 710.02997
1544.83056640625 0 1356.3391
1545.8314208984375 0 3051.6177
1546.8172607421875 0 2127.0823
1547.82763671875 0 1852.8575
1548.819580078125 0 1899.6016
1555.8238525390625 0 1006.284
1556.7960205078125 0 2330.0195
1557.7894287109375 0 2029.452
1558.796630859375 0 923.3063
1561.818603515625 0 776.3557
1562.843505859375 0 4330.475
1563.84814453125 0 3714.6328
1564.8585205078125 0 1303.8942
1570.77197265625 0 804.00916
1571.770751953125 0 1174.5438
1572.8094482421875 0 5299.2544
1573.816162109375 0 32792.65
1574.81982421875 0 25969.863
1575.8221435546875 0 13080.85
1576.8238525390625 0 3996.0383
1577.8187255859375 0 1464.669
1586.7666015625 0 645.1139
1587.789306640625 0 1877.7761
1588.80517578125 0 3086.7725
1589.8287353515625 0 9373.645
1590.840576171875 0 30400.135
1591.84423828125 0 24844.762
1592.8480224609375 0 11073.557
1593.8468017578125 0 3090.8674
1594.8485107421875 0 985.91833
1700.57470703125 0 597.1866
2633.66357421875 0 720.31305
2654.6083984375 0 702.54517
3074.62158203125 0 707.8372
3339.248779296875 0 746.85596

Spectrum Details

|  |  |
| --- | --- |
| Matched peaks? Matched peaks The total absolute number of peaks matched. Additionally in brackets the total fraction of peaks matched and the total number of peaks is shown. | 57 (16.10% of 354) |
| FDR? FDR The false discovery rate estimated for this peptide. It is calculated by matching all theoretical fragments with a non-integer shift with the raw peaks for this spectrum. This is done with 40 different shifts. The resulting percentage is the average number of annotated peaks over the number of annotated peaks with the correct spectrum. | 0.67% |
| Satellite FDR? Satellite FDR See the FDR for details on its calculation. This satellite ion specific FDR only contains the satellite ions (d/w) for I/L/J positions. | 0.00% |
| PSM Score? PSM Score The PSM Score as given by Hecklib to this annotated spectrum. It is shown with three significant figures. | 393 |

#### Spectrum 4735? Spectrum 4735 The raw spectrum of this peptide as annotated by Hecklib. The fragments are coloured according to ion type (see legend). Any peaks with a star '\*' as text can be hovered over to see the full details, first the ion type second the mass shift type. By hovering over the amino acids in the peptide or ions in the legend the corresponding peaks are highlighted. By toggling the 'Unassigned' label you can turn the background (unassigned) peaks on or off in the plot. By updating the slider in the Ion legend you can update the spectrum to only show the top X% of the peaks with labels. The top X% means any peak that is within X% of the highest intensity. By dragging in the spectrum you can zoom in to a specific part of the spectrum and use 'Zoom Out' to get back to the original zoom level. The annotation of the spectrum is based on the given sequence in the peptides file and is done with different software so inconsistencies are likely. The peaks are annotated based on the given sequence, with 20 ppm tolerance.

Copy Data

##### Spectrum 4735 (TSV)

###### Preview

```
Loading example...
```

*Click on the button to copy the data to your clipboard.*

Mz MinMz MaxIntensity Max

WidthHeightPeptide font sizePeptide stroke widthSpectrum font sizeSpectrum stroke widthCompact peptide

Ion legend

wxyz

abcd

OtherUnassignedIon^Charge^_Position_Show for top:%

JJSEVSDRPSGVSSR

01.78e+43.57e+45.35e+47.13e+4

Zoom Out

y^+1^_1_y^+3^_5_y^+1^_1_w^+1^_2_y^+1^_2_z^+1^_2_y^+1^_2_w^+1^_3_y^+1^_3_z^+1^_3_y^+1^_3_z^+2^_8_w^+1^_4_y^+2^_8_y^+1^_4_c^+1^_4_y^+2^_9_z^+1^_5_y^+1^_5_y^+2^_10_w^+1^_6_c^+1^_5_z^+2^_11_y^+2^_11_y^+1^_6_z^+1^_6_w^+2^_12_z^+2^_12_y^+2^_12_c^+1^_6_z^+1^_7_z^+2^_13_y^+2^_13_y^+1^_7_w^+2^_14_y^+2^_14_z^+2^_14_y^+2^_14_w^+1^_8_c^+1^_7_z^+1^_8_y^+1^_8_c^+1^_8_z^+1^_9_y^+1^_9_z^+1^_9_y^+1^_9_c^+1^_9_z^+1^_10_y^+1^_10_c^+1^_10_z^+1^_11_y^+1^_11_c^+1^_11_c^+1^_12_y^+1^_12_c^+1^_13_c^+1^_14_

040380512081611

Fragment Matches Table

Show background peaks

| Position | Ion type | Intensity | mz Theoretical | mz Error (Th) | mz Error (ppm) | Charge | Series Number |
| --- | --- | --- | --- | --- | --- | --- | --- |
| - | - | 468.5 | 121.5 | - | - | 0 | - |
| - | - | 1186 | 126.1 | - | - | 0 | - |
| - | - | 415 | 128.4 | - | - | 0 | - |
| - | - | 563.2 | 131.1 | - | - | 0 | - |
| - | - | 360 | 131.9 | - | - | 0 | - |
| - | - | 1617 | 132.1 | - | - | 0 | - |
| - | - | 2288 | 133.1 | - | - | 0 | - |
| - | - | 1744 | 133.1 | - | - | 0 | - |
| - | - | 456.5 | 143.1 | - | - | 0 | - |
| - | - | 405.6 | 149 | - | - | 0 | - |
| - | - | 2023 | 157.1 | - | - | 0 | - |
| - | - | 516.7 | 157.1 | - | - | 0 | - |
| 15 | y | 516.4 | 158.1 | 3.468E-05 | 0.2194 | +1 | 1 |
| - | - | 447.4 | 158.1 | - | - | 0 | - |
| - | - | 1524 | 168.1 | - | - | 0 | - |
| 11 | y | 855.2 | 169.1 | 0.001355 | 8.012 | +3 | 5 |
| - | - | 439.2 | 171.1 | - | - | 0 | - |
| - | - | 612.1 | 174.1 | - | - | 0 | - |
| 15 | y | 2754 | 175.1 | 0.000132 | 0.7536 | +1 | 1 |
| - | - | 1208 | 177.1 | - | - | 0 | - |
| - | - | 2042 | 185.1 | - | - | 0 | - |
| - | - | 5848 | 185.1 | - | - | 0 | - |
| - | - | 631 | 187.1 | - | - | 0 | - |
| - | - | 777 | 199.2 | - | - | 0 | - |
| - | - | 1.287E+04 | 199.2 | - | - | 0 | - |
| - | - | 1123 | 200.1 | - | - | 0 | - |
| - | - | 1840 | 200.2 | - | - | 0 | - |
| - | - | 1187 | 201.1 | - | - | 0 | - |
| - | - | 1489 | 217.1 | - | - | 0 | - |
| - | - | 546.5 | 226.2 | - | - | 0 | - |
| - | - | 6594 | 227.2 | - | - | 0 | - |
| - | - | 666.3 | 228.2 | - | - | 0 | - |
| 14 | w | 1536 | 229.1 | 0.0004275 | 1.866 | +1 | 2 |
| 14 | y | 9762 | 245.1 | 0.0002707 | 1.104 | +1 | 2 |
| 14 | z | 1171 | 246.1 | 0.0005732 | 2.329 | +1 | 2 |
| - | - | 596.8 | 254.1 | - | - | 0 | - |
| - | - | 678.7 | 255.1 | - | - | 0 | - |
| - | - | 646.5 | 256.1 | - | - | 0 | - |
| - | - | 559.1 | 259.6 | - | - | 0 | - |
| - | - | 501.3 | 261.7 | - | - | 0 | - |
| 14 | y | 1389 | 262.2 | 5.092E-05 | 0.1942 | +1 | 2 |
| - | - | 713.7 | 265.2 | - | - | 0 | - |
| - | - | 1101 | 270.1 | - | - | 0 | - |
| - | - | 626.3 | 272.1 | - | - | 0 | - |
| - | - | 542 | 305.8 | - | - | 0 | - |
| - | - | 1185 | 314.2 | - | - | 0 | - |
| 13 | w | 5409 | 316.2 | 4.531E-05 | 0.1433 | +1 | 3 |
| - | - | 849.5 | 317.2 | - | - | 0 | - |
| - | - | 1134 | 323.1 | - | - | 0 | - |
| - | - | 530.9 | 323.4 | - | - | 0 | - |
| - | - | 1373 | 331.1 | - | - | 0 | - |
| 13 | y | 2041 | 332.2 | 0.0004845 | 1.459 | +1 | 3 |
| - | - | 3953 | 333.1 | - | - | 0 | - |
| 13 | z | 4466 | 333.2 | 0.0005581 | 1.675 | +1 | 3 |
| - | - | 1457 | 349.1 | - | - | 0 | - |
| 13 | y | 1243 | 349.2 | 0.0007275 | 2.083 | +1 | 3 |
| - | - | 692.3 | 353.2 | - | - | 0 | - |
| - | - | 3374 | 357.2 | - | - | 0 | - |
| - | - | 895.9 | 378.2 | - | - | 0 | - |
| - | - | 632.2 | 378.7 | - | - | 0 | - |
| - | - | 675.5 | 395.2 | - | - | 0 | - |
| 8 | z | 1721 | 415.2 | 0.0007246 | 1.745 | +2 | 8 |
| - | - | 7659 | 415.3 | - | - | 0 | - |
| - | - | 544.7 | 416.2 | - | - | 0 | - |
| 12 | w | 3451 | 417.2 | 0.0001783 | 0.4274 | +1 | 4 |
| - | - | 970.4 | 418.2 | - | - | 0 | - |
| 8 | y | 1107 | 423.2 | 0.0005484 | 1.296 | +2 | 8 |
| 12 | y | 3995 | 448.3 | 0.000225 | 0.5021 | +1 | 4 |
| - | - | 1127 | 449.3 | - | - | 0 | - |
| - | - | 1321 | 460.2 | - | - | 0 | - |
| 4 | c | 2925 | 460.3 | 0.0001468 | 0.3189 | +1 | 4 |
| - | - | 686.3 | 460.7 | - | - | 0 | - |
| - | - | 836.3 | 461.2 | - | - | 0 | - |
| - | - | 673.5 | 466.2 | - | - | 0 | - |
| - | - | 1084 | 474.2 | - | - | 0 | - |
| 7 | y | 874.1 | 480.7 | 0.001603 | 3.335 | +2 | 9 |
| - | - | 1112 | 482.3 | - | - | 0 | - |
| 11 | z | 1716 | 489.3 | 0.0003784 | 0.7734 | +1 | 5 |
| - | - | 722.3 | 499.3 | - | - | 0 | - |
| 11 | y | 1.373E+04 | 505.3 | 0.0001815 | 0.3593 | +1 | 5 |
| - | - | 3303 | 506.3 | - | - | 0 | - |
| - | - | 1072 | 512.2 | - | - | 0 | - |
| - | - | 919.7 | 515.3 | - | - | 0 | - |
| - | - | 1572 | 516.8 | - | - | 0 | - |
| 6 | y | 4358 | 524.3 | 0.002374 | 4.528 | +2 | 10 |
| - | - | 973.5 | 524.6 | - | - | 0 | - |
| - | - | 1894 | 524.8 | - | - | 0 | - |
| - | - | 618.8 | 529.2 | - | - | 0 | - |
| - | - | 3422 | 530.2 | - | - | 0 | - |
| - | - | 7.062E+04 | 530.3 | - | - | 0 | - |
| - | - | 1304 | 530.6 | - | - | 0 | - |
| - | - | 779.9 | 531.2 | - | - | 0 | - |
| - | - | 2.061E+04 | 531.3 | - | - | 0 | - |
| - | - | 688.5 | 531.7 | - | - | 0 | - |
| - | - | 1550 | 542.3 | - | - | 0 | - |
| - | - | 948.2 | 558.3 | - | - | 0 | - |
| 10 | w | 565.1 | 559.3 | 0.005611 | 10.03 | +1 | 6 |
| 5 | c | 2860 | 559.3 | 0.001666 | 2.979 | +1 | 5 |
| - | - | 867.4 | 560.3 | - | - | 0 | - |
| 5 | z | 8625 | 565.8 | 0.0005596 | 0.9891 | +2 | 11 |
| - | - | 7163 | 566.3 | - | - | 0 | - |
| - | - | 2160 | 566.8 | - | - | 0 | - |
| - | - | 767.1 | 567.3 | - | - | 0 | - |
| 5 | y | 4675 | 573.8 | 0.001194 | 2.08 | +2 | 11 |
| 10 | y | 2152 | 574.3 | 0.004478 | 7.797 | +1 | 6 |
| - | - | 777.5 | 574.8 | - | - | 0 | - |
| 10 | z | 2089 | 576.3 | 2.879E-06 | 0.004996 | +1 | 6 |
| - | - | 1088 | 586.3 | - | - | 0 | - |
| 4 | w | 3767 | 600.8 | 0.0007387 | 1.229 | +2 | 12 |
| - | - | 2032 | 601.3 | - | - | 0 | - |
| - | - | 924.1 | 601.8 | - | - | 0 | - |
| 4 | z | 1.016E+04 | 630.3 | 0.0004328 | 0.6867 | +2 | 12 |
| - | - | 7534 | 630.8 | - | - | 0 | - |
| - | - | 2174 | 631.3 | - | - | 0 | - |
| - | - | 1007 | 631.8 | - | - | 0 | - |
| - | - | 3053 | 637.8 | - | - | 0 | - |
| 4 | y | 9525 | 638.3 | 0.001311 | 2.054 | +2 | 12 |
| - | - | 7552 | 638.8 | - | - | 0 | - |
| - | - | 3301 | 639.3 | - | - | 0 | - |
| - | - | 653.9 | 644.3 | - | - | 0 | - |
| - | - | 737.7 | 644.8 | - | - | 0 | - |
| 6 | c | 1.042E+04 | 646.4 | 0.0001253 | 0.1938 | +1 | 6 |
| - | - | 3218 | 647.4 | - | - | 0 | - |
| 9 | z | 647.8 | 673.3 | 0.004726 | 7.019 | +1 | 7 |
| 3 | z | 5041 | 673.8 | 0.0002117 | 0.3142 | +2 | 13 |
| - | - | 8282 | 674.3 | - | - | 0 | - |
| - | - | 3652 | 674.8 | - | - | 0 | - |
| - | - | 1861 | 675.3 | - | - | 0 | - |
| 3 | y | 3.879E+04 | 681.8 | 0.0005405 | 0.7928 | +2 | 13 |
| - | - | 2.84E+04 | 682.3 | - | - | 0 | - |
| - | - | 1.081E+04 | 682.8 | - | - | 0 | - |
| - | - | 3729 | 683.3 | - | - | 0 | - |
| 9 | y | 835.5 | 689.4 | 0.00105 | 1.524 | +1 | 7 |
| - | - | 608.3 | 701.4 | - | - | 0 | - |
| - | - | 1275 | 711.9 | - | - | 0 | - |
| 2 | w | 787.8 | 715.8 | 2.397E-05 | 0.03349 | +2 | 14 |
| - | - | 622.2 | 716.9 | - | - | 0 | - |
| - | - | 930.2 | 717.4 | - | - | 0 | - |
| - | - | 861.3 | 718.4 | - | - | 0 | - |
| 2 | y | 747.7 | 729.4 | 0.005491 | 7.528 | +2 | 14 |
| 2 | z | 1205 | 730.4 | 0.002144 | 2.935 | +2 | 14 |
| - | - | 694.1 | 738.3 | - | - | 0 | - |
| 2 | y | 3268 | 738.4 | 0.001496 | 2.026 | +2 | 14 |
| - | - | 1688 | 738.9 | - | - | 0 | - |
| - | - | 1071 | 739.4 | - | - | 0 | - |
| 8 | w | 843.5 | 743.4 | 0.00362 | 4.869 | +1 | 8 |
| - | - | 735 | 744.4 | - | - | 0 | - |
| - | - | 749.2 | 755.4 | - | - | 0 | - |
| - | - | 1149 | 758.4 | - | - | 0 | - |
| - | - | 1557 | 758.9 | - | - | 0 | - |
| - | - | 997.1 | 759.4 | - | - | 0 | - |
| - | - | 795.4 | 760.4 | - | - | 0 | - |
| 7 | c | 1.886E+04 | 761.4 | 0.0008232 | 1.081 | +1 | 7 |
| - | - | 7484 | 762.4 | - | - | 0 | - |
| - | - | 1346 | 763.4 | - | - | 0 | - |
| - | - | 1101 | 764.9 | - | - | 0 | - |
| - | - | 3.182E+04 | 765.4 | - | - | 0 | - |
| - | - | 2.46E+04 | 765.9 | - | - | 0 | - |
| - | - | 1.384E+04 | 766.4 | - | - | 0 | - |
| - | - | 3909 | 766.9 | - | - | 0 | - |
| - | - | 1413 | 767.4 | - | - | 0 | - |
| - | - | 891.6 | 771.9 | - | - | 0 | - |
| - | - | 1076 | 772.4 | - | - | 0 | - |
| - | - | 724.8 | 772.9 | - | - | 0 | - |
| - | - | 1057 | 773.4 | - | - | 0 | - |
| - | - | 9209 | 773.9 | - | - | 0 | - |
| - | - | 6049 | 774.4 | - | - | 0 | - |
| - | - | 4460 | 774.9 | - | - | 0 | - |
| - | - | 999.1 | 775.4 | - | - | 0 | - |
| - | - | 880.5 | 778.9 | - | - | 0 | - |
| - | - | 843.3 | 779.4 | - | - | 0 | - |
| - | - | 1126 | 785.4 | - | - | 0 | - |
| - | - | 1098 | 785.9 | - | - | 0 | - |
| - | - | 3309 | 786.4 | - | - | 0 | - |
| - | - | 2.749E+04 | 786.9 | - | - | 0 | - |
| - | - | 3.026E+04 | 787.4 | - | - | 0 | - |
| - | - | 1.148E+04 | 787.9 | - | - | 0 | - |
| - | - | 3854 | 788.4 | - | - | 0 | - |
| - | - | 1021 | 793.9 | - | - | 0 | - |
| - | - | 1767 | 794.4 | - | - | 0 | - |
| - | - | 2.38E+04 | 794.9 | - | - | 0 | - |
| - | - | 2.829E+04 | 795.4 | - | - | 0 | - |
| - | - | 1.656E+04 | 795.9 | - | - | 0 | - |
| - | - | 6052 | 796.4 | - | - | 0 | - |
| - | - | 2482 | 796.9 | - | - | 0 | - |
| - | - | 949.5 | 801.4 | - | - | 0 | - |
| - | - | 693.1 | 813.4 | - | - | 0 | - |
| - | - | 972 | 828.4 | - | - | 0 | - |
| 8 | z | 1.373E+04 | 829.4 | 0.001528 | 1.843 | +1 | 8 |
| - | - | 7122 | 830.4 | - | - | 0 | - |
| - | - | 2303 | 831.4 | - | - | 0 | - |
| - | - | 675.8 | 839.5 | - | - | 0 | - |
| 8 | y | 1947 | 845.5 | 0.0007821 | 0.9251 | +1 | 8 |
| - | - | 795.3 | 873.5 | - | - | 0 | - |
| 8 | c | 2163 | 900.5 | 0.001777 | 1.973 | +1 | 8 |
| - | - | 1034 | 901.5 | - | - | 0 | - |
| - | - | 1834 | 924.5 | - | - | 0 | - |
| - | - | 1223 | 925.5 | - | - | 0 | - |
| 7 | z | 940.5 | 926.5 | 0.001869 | 2.017 | +1 | 9 |
| - | - | 2166 | 941.5 | - | - | 0 | - |
| 7 | y | 994.3 | 942.5 | 0.01726 | 18.32 | +1 | 9 |
| 7 | z | 5480 | 944.5 | 2.89E-05 | 0.0306 | +1 | 9 |
| - | - | 4434 | 945.5 | - | - | 0 | - |
| - | - | 1737 | 946.5 | - | - | 0 | - |
| - | - | 1057 | 954.5 | - | - | 0 | - |
| - | - | 782.1 | 955.5 | - | - | 0 | - |
| - | - | 911.7 | 956.5 | - | - | 0 | - |
| 7 | y | 669 | 960.5 | 0.005447 | 5.671 | +1 | 9 |
| - | - | 941.5 | 961.5 | - | - | 0 | - |
| - | - | 1735 | 964.5 | - | - | 0 | - |
| - | - | 1965 | 970.5 | - | - | 0 | - |
| - | - | 835.7 | 971.5 | - | - | 0 | - |
| - | - | 649.4 | 972.6 | - | - | 0 | - |
| - | - | 894.3 | 984.5 | - | - | 0 | - |
| - | - | 1208 | 986.5 | - | - | 0 | - |
| - | - | 1669 | 987.5 | - | - | 0 | - |
| - | - | 946.2 | 988.5 | - | - | 0 | - |
| - | - | 821.6 | 999.5 | - | - | 0 | - |
| - | - | 1234 | 1002 | - | - | 0 | - |
| - | - | 746.6 | 1003 | - | - | 0 | - |
| - | - | 2058 | 1004 | - | - | 0 | - |
| - | - | 1142 | 1005 | - | - | 0 | - |
| - | - | 886 | 1014 | - | - | 0 | - |
| 9 | c | 1.384E+04 | 1015 | 0.0005843 | 0.5759 | +1 | 9 |
| - | - | 6909 | 1016 | - | - | 0 | - |
| - | - | 1686 | 1017 | - | - | 0 | - |
| 6 | z | 2380 | 1031 | 0.003038 | 2.945 | +1 | 10 |
| - | - | 3494 | 1033 | - | - | 0 | - |
| - | - | 1439 | 1034 | - | - | 0 | - |
| - | - | 885.3 | 1040 | - | - | 0 | - |
| - | - | 2101 | 1041 | - | - | 0 | - |
| - | - | 2921 | 1042 | - | - | 0 | - |
| - | - | 2155 | 1043 | - | - | 0 | - |
| - | - | 1533 | 1044 | - | - | 0 | - |
| - | - | 1314 | 1045 | - | - | 0 | - |
| - | - | 893.5 | 1046 | - | - | 0 | - |
| 6 | y | 899.2 | 1048 | 0.00763 | 7.283 | +1 | 10 |
| - | - | 7990 | 1058 | - | - | 0 | - |
| - | - | 4909 | 1059 | - | - | 0 | - |
| - | - | 2713 | 1060 | - | - | 0 | - |
| - | - | 2554 | 1061 | - | - | 0 | - |
| - | - | 829.2 | 1101 | - | - | 0 | - |
| 10 | c | 1.815E+04 | 1102 | 0.001241 | 1.126 | +1 | 10 |
| - | - | 1.027E+04 | 1103 | - | - | 0 | - |
| - | - | 4433 | 1104 | - | - | 0 | - |
| - | - | 2774 | 1115 | - | - | 0 | - |
| - | - | 1916 | 1116 | - | - | 0 | - |
| 5 | z | 2734 | 1131 | 0.002022 | 1.788 | +1 | 11 |
| - | - | 7152 | 1132 | - | - | 0 | - |
| - | - | 3066 | 1133 | - | - | 0 | - |
| - | - | 1232 | 1134 | - | - | 0 | - |
| 5 | y | 1641 | 1147 | 0.001214 | 1.059 | +1 | 11 |
| - | - | 650.9 | 1148 | - | - | 0 | - |
| 11 | c | 9443 | 1159 | 0.001952 | 1.685 | +1 | 11 |
| - | - | 6663 | 1160 | - | - | 0 | - |
| - | - | 2452 | 1161 | - | - | 0 | - |
| - | - | 2768 | 1214 | - | - | 0 | - |
| - | - | 1010 | 1215 | - | - | 0 | - |
| 12 | c | 1.063E+04 | 1258 | 0.002251 | 1.79 | +1 | 12 |
| - | - | 7597 | 1259 | - | - | 0 | - |
| - | - | 1997 | 1260 | - | - | 0 | - |
| - | - | 2090 | 1260 | - | - | 0 | - |
| - | - | 8577 | 1261 | - | - | 0 | - |
| - | - | 5617 | 1262 | - | - | 0 | - |
| - | - | 2104 | 1263 | - | - | 0 | - |
| 4 | y | 3455 | 1276 | 0.001815 | 1.423 | +1 | 12 |
| - | - | 2079 | 1277 | - | - | 0 | - |
| - | - | 1383 | 1278 | - | - | 0 | - |
| - | - | 862.5 | 1283 | - | - | 0 | - |
| - | - | 966 | 1301 | - | - | 0 | - |
| - | - | 1156 | 1302 | - | - | 0 | - |
| - | - | 785.5 | 1331 | - | - | 0 | - |
| 13 | c | 1.808E+04 | 1345 | 0.001931 | 1.436 | +1 | 13 |
| - | - | 1.239E+04 | 1346 | - | - | 0 | - |
| - | - | 5973 | 1347 | - | - | 0 | - |
| - | - | 7095 | 1348 | - | - | 0 | - |
| - | - | 5399 | 1349 | - | - | 0 | - |
| - | - | 3311 | 1350 | - | - | 0 | - |
| - | - | 1468 | 1388 | - | - | 0 | - |
| - | - | 774.8 | 1390 | - | - | 0 | - |
| - | - | 805.6 | 1431 | - | - | 0 | - |
| 14 | c | 2.584E+04 | 1432 | 0.002587 | 1.807 | +1 | 14 |
| - | - | 1.71E+04 | 1433 | - | - | 0 | - |
| - | - | 8310 | 1434 | - | - | 0 | - |
| - | - | 2894 | 1435 | - | - | 0 | - |
| - | - | 1074 | 1461 | - | - | 0 | - |
| - | - | 1569 | 1462 | - | - | 0 | - |
| - | - | 2610 | 1489 | - | - | 0 | - |
| - | - | 1688 | 1490 | - | - | 0 | - |
| - | - | 944.9 | 1491 | - | - | 0 | - |
| - | - | 4429 | 1531 | - | - | 0 | - |
| - | - | 6595 | 1532 | - | - | 0 | - |
| - | - | 3682 | 1533 | - | - | 0 | - |
| - | - | 2506 | 1534 | - | - | 0 | - |
| - | - | 1048 | 1535 | - | - | 0 | - |
| - | - | 1466 | 1545 | - | - | 0 | - |
| - | - | 2736 | 1546 | - | - | 0 | - |
| - | - | 3141 | 1547 | - | - | 0 | - |
| - | - | 2985 | 1548 | - | - | 0 | - |
| - | - | 1678 | 1549 | - | - | 0 | - |
| - | - | 1315 | 1556 | - | - | 0 | - |
| - | - | 3213 | 1557 | - | - | 0 | - |
| - | - | 2168 | 1558 | - | - | 0 | - |
| - | - | 1001 | 1562 | - | - | 0 | - |
| - | - | 4421 | 1563 | - | - | 0 | - |
| - | - | 3431 | 1564 | - | - | 0 | - |
| - | - | 1610 | 1565 | - | - | 0 | - |
| - | - | 835 | 1566 | - | - | 0 | - |
| - | - | 1793 | 1572 | - | - | 0 | - |
| - | - | 6039 | 1573 | - | - | 0 | - |
| - | - | 3.654E+04 | 1574 | - | - | 0 | - |
| - | - | 3E+04 | 1575 | - | - | 0 | - |
| - | - | 1.639E+04 | 1576 | - | - | 0 | - |
| - | - | 4297 | 1577 | - | - | 0 | - |
| - | - | 1149 | 1578 | - | - | 0 | - |
| - | - | 922.4 | 1588 | - | - | 0 | - |
| - | - | 4068 | 1589 | - | - | 0 | - |
| - | - | 1.008E+04 | 1590 | - | - | 0 | - |
| - | - | 3.681E+04 | 1591 | - | - | 0 | - |
| - | - | 2.833E+04 | 1592 | - | - | 0 | - |
| - | - | 1.333E+04 | 1593 | - | - | 0 | - |
| - | - | 3974 | 1594 | - | - | 0 | - |
| - | - | 964.9 | 1595 | - | - | 0 | - |

m/z Charge Intensity FragmentType MassShift Position
121.49907684326172 0 468.50027
126.09146881103516 0 1186.2203
128.3719482421875 0 415.03265
131.12918090820312 0 563.21906
131.91085815429688 0 360.0447
132.07669067382812 0 1617.0621
133.06077575683594 0 2288.0083
133.08578491210938 0 1744.4414
143.11793518066406 0 456.53986
149.02354431152344 0 405.58905
157.1332550048828 0 2022.757
157.13963317871094 0 516.6948
158.09243774414062 0 516.41455 y Ammonia loss 14
158.13722229003906 0 447.36053
168.1128692626953 0 1523.7645
169.09716796875 0 855.20154 y 10
171.11265563964844 0 439.1553
174.05477905273438 0 612.0904
175.1188201904297 0 2754.056 y 14
177.11212158203125 0 1208.4832
185.1285400390625 0 2042.3936
185.13954162597656 0 5848.1978
187.1082305908203 0 630.9848
199.17047119140625 0 777.014
199.1802978515625 0 12867.673
200.1265869140625 0 1122.5265
200.183837890625 0 1839.665
201.12335205078125 0 1187.2986
217.1178741455078 0 1488.9839
226.16744995117188 0 546.5213
227.1752166748047 0 6594.4575
228.17945861816406 0 666.27625
229.12908935546875 0 1535.6533 w 13
245.12416076660156 0 9762.242 y Ammonia loss 13
246.13168334960938 0 1171.1702 z 13
254.14942932128906 0 596.7961
255.10853576660156 0 678.6712
256.1285400390625 0 646.4866
259.5525817871094 0 559.12524
261.656005859375 0 501.26086
262.1510314941406 0 1388.8119 y 13
265.1644287109375 0 713.7066
270.1197204589844 0 1101.4056
272.1234130859375 0 626.344
305.7637023925781 0 541.9622
314.2081604003906 0 1185.0417
316.1615905761719 0 5409.357 w 12
317.1603088378906 0 849.4811
323.0976257324219 0 1133.7146
323.38763427734375 0 530.9391
331.1480712890625 0 1372.7013
332.1559753417969 0 2041.2859 y Ammonia loss 12
333.1192321777344 0 3952.9976
333.1637268066406 0 4465.6973 z 12
349.1142578125 0 1457.4509
349.1822814941406 0 1242.6354 y 12
353.2198181152344 0 692.34894
357.1509704589844 0 3374.182
378.2201843261719 0 895.8686
378.7242431640625 0 632.20056
395.22821044921875 0 675.52747
415.2243957519531 0 1721.3162 z 7
415.2538146972656 0 7658.5356
416.22808837890625 0 544.7071
417.20904541015625 0 3451.1265 w 11
418.21148681640625 0 970.4281
423.23358154296875 0 1107.0663 y 7
448.25164794921875 0 3995.1304 y 11
449.2549743652344 0 1127.2499
460.2437744140625 0 1320.7719
460.27642822265625 0 2924.6848 c 3
460.74676513671875 0 686.3126
461.2465515136719 0 836.2583
466.17034912109375 0 673.4921
474.230712890625 0 1084.3002
480.74810791015625 0 874.1299 y 6
482.2611999511719 0 1112.43
489.2537841796875 0 1715.6001 z 10
499.29718017578125 0 722.2564
505.272705078125 0 13728.329 y 10
506.2754211425781 0 3302.5632
512.17578125 0 1072.1273
515.3269653320312 0 919.748
516.7577514648438 0 1571.9385
524.264892578125 0 4358.205 y 5
524.6107788085938 0 973.53265
524.76318359375 0 1893.6753
529.201904296875 0 618.8363
530.1884155273438 0 3422.498
530.3167724609375 0 70615.805
530.6141357421875 0 1304.4666
531.1917724609375 0 779.93256
531.3201293945312 0 20610.814
531.7416381835938 0 688.5072
542.3187866210938 0 1550.0411
558.3382568359375 0 948.15234
559.2890625 0 565.103 w 9
559.3433227539062 0 2860.3826 c 4
560.3465576171875 0 867.38965
565.7868041992188 0 8624.746 z 4
566.2886962890625 0 7162.724
566.7894287109375 0 2159.657
567.2925415039062 0 767.051
573.7955322265625 0 4674.9805 y 4
574.298828125 0 2151.5972 y Water loss 9
574.7976684570312 0 777.45715
576.2861938476562 0 2089.0874 z 9
586.3289794921875 0 1088.4904
600.80126953125 0 3767.027 w 3
601.3067626953125 0 2031.9575
601.8037109375 0 924.05054
630.3082275390625 0 10160.097 z 3
630.8095092773438 0 7533.7935
631.3103637695312 0 2174.1887
631.8136596679688 0 1006.7875
637.8143310546875 0 3052.8328
638.3167114257812 0 9525.067 y 3
638.8189086914062 0 7552.496
639.3189697265625 0 3301.3599
644.3157958984375 0 653.87177
644.81591796875 0 737.72253
646.3768920898438 0 10422.108 c 5
647.3792114257812 0 3218.153
673.334228515625 0 647.7929 z 8
673.824462890625 0 5040.815 z 2
674.3267211914062 0 8281.716
674.8291625976562 0 3652.1445
675.330078125 0 1861.4722
681.83349609375 0 38785.613 y 2
682.3348999023438 0 28399.143
682.836669921875 0 10805.567
683.3369750976562 0 3728.7065
689.3566284179688 0 835.4754 y 8
701.3577270507812 0 608.2849
711.8577880859375 0 1275.4371
715.84716796875 0 787.7558 w 1
716.8512573242188 0 622.22754
717.3927001953125 0 930.2143
718.3966674804688 0 861.27496
729.3652954101562 0 747.6666 y Water loss 1
730.3645629882812 0 1205.2172 z 1
738.2706298828125 0 694.06287
738.3745727539062 0 3268.0356 y 1
738.8748168945312 0 1687.9644
739.3784790039062 0 1070.7971
743.3646240234375 0 843.4556 w 7
744.3762817382812 0 735.033
755.4252319335938 0 749.17926
758.384033203125 0 1149.4729
758.8785400390625 0 1556.7053
759.3848876953125 0 997.10284
760.3896484375 0 795.3941
761.4031372070312 0 18855.574 c 6
762.4064331054688 0 7484.4053
763.408935546875 0 1346.0411
764.9194946289062 0 1100.5216
765.3818359375 0 31820.898
765.883056640625 0 24603.621
766.3839721679688 0 13836.088
766.8851928710938 0 3908.9836
767.3876342773438 0 1413.1821
771.9031372070312 0 891.59985
772.3912963867188 0 1075.8152
772.899658203125 0 724.7581
773.3934936523438 0 1056.5693
773.906982421875 0 9208.749
774.4078979492188 0 6048.9355
774.90966796875 0 4459.6
775.4037475585938 0 999.057
778.8998413085938 0 880.4938
779.4035034179688 0 843.32745
785.4036254882812 0 1126.3273
785.9068603515625 0 1097.6416
786.4097900390625 0 3308.9995
786.9083251953125 0 27490.252
787.4099731445312 0 30256.525
787.9104614257812 0 11478.799
788.41162109375 0 3853.7441
793.9010620117188 0 1021.1248
794.4057006835938 0 1767.2894
794.9169921875 0 23799.662
795.4193725585938 0 28285.285
795.9207153320312 0 16562.352
796.4227294921875 0 6051.8315
796.9209594726562 0 2482.1838
801.4022216796875 0 949.5226
813.3904418945312 0 693.05225
828.4071044921875 0 971.96857
829.4385375976562 0 13732.512 z 7
830.4423828125 0 7121.604
831.4437255859375 0 2302.7722
839.5322875976562 0 675.83105
845.4580078125 0 1946.8817 y 7
873.4851684570312 0 795.29755
900.4767456054688 0 2163.4045 c Ammonia loss 7
901.4754028320312 0 1033.9452
924.4569702148438 0 1833.8599
925.4612426757812 0 1222.9581
926.4583129882812 0 940.5247 z Water loss 6
941.4822998046875 0 2165.9817
942.492431640625 0 994.33813 y Water loss 6
944.4669799804688 0 5480.1206 z 6
945.4714965820312 0 4433.9224
946.4747314453125 0 1736.9512
954.4672241210938 0 1057.4485
955.5408935546875 0 782.13885
956.537841796875 0 911.6592
960.4802856445312 0 669.03534 y 6
961.4835205078125 0 941.5289
964.5159912109375 0 1734.5912
970.5332641601562 0 1964.9053
971.5491943359375 0 835.6826
972.5514526367188 0 649.3501
984.481201171875 0 894.3193
986.51416015625 0 1208.1564
987.526123046875 0 1669.374
988.5131225585938 0 946.1651
999.5147094726562 0 821.61053
1001.5196533203125 0 1234.1587
1002.5166625976562 0 746.5806
1003.5276489257812 0 2057.9937
1004.5305786132812 0 1142.1122
1013.5858764648438 0 885.9775
1014.5572509765625 0 13836.12 c 8
1015.5585327148438 0 6908.96
1016.5604858398438 0 1685.7375
1031.5020751953125 0 2379.785 z 5
1032.507080078125 0 3493.6418
1033.5076904296875 0 1438.8937
1039.564208984375 0 885.2982
1040.505615234375 0 2101.0984
1041.5098876953125 0 2920.747
1042.52685546875 0 2155.1265
1043.526611328125 0 1532.5027
1044.5411376953125 0 1313.5178
1046.4931640625 0 893.4928
1047.5101318359375 0 899.2463 y 5
1057.5689697265625 0 7990.092
1058.5714111328125 0 4909.0557
1059.569580078125 0 2713.4631
1060.53759765625 0 2553.9941
1100.581298828125 0 829.1626
1101.588623046875 0 18152.959 c 9
1102.5908203125 0 10269.73
1103.59375 0 4433.245
1114.593017578125 0 2773.545
1115.6004638671875 0 1915.9564
1130.5654296875 0 2733.9917 z 4
1131.573974609375 0 7152.021
1132.5782470703125 0 3065.6648
1133.5845947265625 0 1232.1232
1146.5849609375 0 1640.503 y 4
1147.5838623046875 0 650.9119
1158.609375 0 9442.898 c 10
1159.61376953125 0 6662.8545
1160.616455078125 0 2451.798
1213.6663818359375 0 2768.16
1214.6676025390625 0 1010.31177
1257.677490234375 0 10634.5205 c 11
1258.679931640625 0 7597.204
1259.6434326171875 0 1997.082
1259.669677734375 0 2090.416
1260.6170654296875 0 8576.994
1261.619140625 0 5617.0664
1262.6197509765625 0 2104.219
1275.626953125 0 3454.9944 y 3
1276.629150390625 0 2079.1526
1277.6339111328125 0 1382.8866
1282.693603515625 0 862.5206
1300.6956787109375 0 966.0135
1301.69384765625 0 1155.5414
1330.636474609375 0 785.486
1344.7098388671875 0 18077.896 c 12
1345.712646484375 0 12386.983
1346.70947265625 0 5973.2485
1347.65185546875 0 7094.924
1348.6531982421875 0 5399.3164
1349.6552734375 0 3310.754
1387.730224609375 0 1467.736
1389.7420654296875 0 774.756
1430.73486328125 0 805.6447
1431.7412109375 0 25841.709 c 13
1432.743896484375 0 17098.129
1433.7474365234375 0 8310.248
1434.75 0 2894.4324
1460.7239990234375 0 1073.7776
1461.733154296875 0 1569.4054
1488.7470703125 0 2609.9954
1489.7376708984375 0 1687.8467
1490.745361328125 0 944.9129
1530.78759765625 0 4429.2686
1531.7935791015625 0 6594.777
1532.796630859375 0 3682.241
1533.794189453125 0 2505.9233
1534.79248046875 0 1048.0806
1544.83251953125 0 1466.0118
1545.8250732421875 0 2735.556
1546.819580078125 0 3141.0515
1547.81689453125 0 2985.4968
1548.81005859375 0 1677.8617
1555.8116455078125 0 1314.6859
1556.7935791015625 0 3213.0505
1557.79150390625 0 2167.839
1561.825439453125 0 1000.9723
1562.8450927734375 0 4420.85
1563.8511962890625 0 3431.0237
1564.8519287109375 0 1609.6819
1565.8438720703125 0 834.95294
1571.798583984375 0 1792.8553
1572.8116455078125 0 6039.0713
1573.8148193359375 0 36543.527
1574.8175048828125 0 30004.28
1575.820068359375 0 16387.385
1576.818115234375 0 4297.0283
1577.8074951171875 0 1149.008
1587.805419921875 0 922.3559
1588.815673828125 0 4068.1877
1589.8310546875 0 10083.111
1590.83984375 0 36807.91
1591.8414306640625 0 28330.822
1592.8443603515625 0 13331.527
1593.840576171875 0 3973.5503
1594.8271484375 0 964.9091

Spectrum Details

|  |  |
| --- | --- |
| Matched peaks? Matched peaks The total absolute number of peaks matched. Additionally in brackets the total fraction of peaks matched and the total number of peaks is shown. | 58 (17.96% of 323) |
| FDR? FDR The false discovery rate estimated for this peptide. It is calculated by matching all theoretical fragments with a non-integer shift with the raw peaks for this spectrum. This is done with 40 different shifts. The resulting percentage is the average number of annotated peaks over the number of annotated peaks with the correct spectrum. | 0.90% |
| Satellite FDR? Satellite FDR See the FDR for details on its calculation. This satellite ion specific FDR only contains the satellite ions (d/w) for I/L/J positions. | 0.00% |
| PSM Score? PSM Score The PSM Score as given by Hecklib to this annotated spectrum. It is shown with three significant figures. | 442 |

#### Spectrum 4162? Spectrum 4162 The raw spectrum of this peptide as annotated by Hecklib. The fragments are coloured according to ion type (see legend). Any peaks with a star '\*' as text can be hovered over to see the full details, first the ion type second the mass shift type. By hovering over the amino acids in the peptide or ions in the legend the corresponding peaks are highlighted. By toggling the 'Unassigned' label you can turn the background (unassigned) peaks on or off in the plot. By updating the slider in the Ion legend you can update the spectrum to only show the top X% of the peaks with labels. The top X% means any peak that is within X% of the highest intensity. By dragging in the spectrum you can zoom in to a specific part of the spectrum and use 'Zoom Out' to get back to the original zoom level. The annotation of the spectrum is based on the given sequence in the peptides file and is done with different software so inconsistencies are likely. The peaks are annotated based on the given sequence, with 20 ppm tolerance.

Copy Data

##### Spectrum 4162 (TSV)

###### Preview

```
Loading example...
```

*Click on the button to copy the data to your clipboard.*

Mz MinMz MaxIntensity Max

WidthHeightPeptide font sizePeptide stroke widthSpectrum font sizeSpectrum stroke widthCompact peptide

Ion legend

wxyz

abcd

OtherUnassignedIon^Charge^_Position_Show for top:%

JJSEVSDRPSGVSSR

01.46e+52.92e+54.38e+55.84e+5

Zoom Out

y^+1^_1_y^+3^_5_y^+1^_1_d^+1^_2_a^+1^_2_b^+1^_2_y^+1^_2_y^+1^_2_y^+1^_2_b^+1^_3_b^+1^_3_y^+1^_3_y^+1^_3_y^+2^_7_y^+1^_3_b^+3^_12_y^+2^_8_y^+2^_8_b^+1^_4_y^+1^_4_b^+1^_4_y^+1^_4_y^+3^_13_y^+3^_13_y^+2^_9_y^+3^_14_y^+1^_5_y^+3^_14_y^+1^_5_y^+2^_10_y^+2^_10_y^+2^_10_\*\*b^+1^_5_b^+2^_11_y^+2^_11_y^+2^_11_y^+2^_11_y^+1^_6_y^+1^_6_y^+2^_12_y^+2^_12_y^+2^_12_y^+1^_7_y^+1^_7_y^+2^_13_y^+2^_13_y^+2^_13_y^+1^_7_b^+2^_14_y^+2^_14_y^+2^_14_y^+2^_14_y^+1^_8_y^+1^_9_y^+1^_10_y^+1^_11_

0776155323293106

Fragment Matches Table

Show background peaks

| Position | Ion type | Intensity | mz Theoretical | mz Error (Th) | mz Error (ppm) | Charge | Series Number |
| --- | --- | --- | --- | --- | --- | --- | --- |
| - | - | 3170 | 120.1 | - | - | 0 | - |
| - | - | 402.1 | 121.1 | - | - | 0 | - |
| - | - | 403.3 | 122.1 | - | - | 0 | - |
| - | - | 319 | 122.5 | - | - | 0 | - |
| - | - | 522.7 | 124.1 | - | - | 0 | - |
| - | - | 409.6 | 124.6 | - | - | 0 | - |
| - | - | 461.3 | 125.1 | - | - | 0 | - |
| - | - | 359.2 | 125.6 | - | - | 0 | - |
| - | - | 358.8 | 125.9 | - | - | 0 | - |
| - | - | 5631 | 126.1 | - | - | 0 | - |
| - | - | 525 | 127.1 | - | - | 0 | - |
| - | - | 561.5 | 127.1 | - | - | 0 | - |
| - | - | 1045 | 128.1 | - | - | 0 | - |
| - | - | 617.4 | 128.1 | - | - | 0 | - |
| - | - | 1540 | 129.1 | - | - | 0 | - |
| - | - | 4998 | 129.1 | - | - | 0 | - |
| - | - | 389.7 | 129.1 | - | - | 0 | - |
| - | - | 997.8 | 130.1 | - | - | 0 | - |
| - | - | 2860 | 130.1 | - | - | 0 | - |
| - | - | 1816 | 130.1 | - | - | 0 | - |
| - | - | 2.63E+04 | 133.1 | - | - | 0 | - |
| - | - | 455.9 | 133.1 | - | - | 0 | - |
| - | - | 900 | 134.1 | - | - | 0 | - |
| - | - | 2004 | 136.1 | - | - | 0 | - |
| - | - | 710.8 | 138.1 | - | - | 0 | - |
| - | - | 1410 | 140.1 | - | - | 0 | - |
| - | - | 732.4 | 141.1 | - | - | 0 | - |
| - | - | 895.9 | 141.1 | - | - | 0 | - |
| - | - | 577.9 | 143.1 | - | - | 0 | - |
| - | - | 1215 | 144.1 | - | - | 0 | - |
| - | - | 4599 | 145.1 | - | - | 0 | - |
| - | - | 446.3 | 147.1 | - | - | 0 | - |
| - | - | 2721 | 147.1 | - | - | 0 | - |
| - | - | 488 | 147.1 | - | - | 0 | - |
| - | - | 421 | 152.1 | - | - | 0 | - |
| - | - | 940.8 | 153.1 | - | - | 0 | - |
| - | - | 5893 | 154.1 | - | - | 0 | - |
| - | - | 643.7 | 155.1 | - | - | 0 | - |
| - | - | 1027 | 155.1 | - | - | 0 | - |
| - | - | 721.3 | 156.1 | - | - | 0 | - |
| - | - | 511 | 156.1 | - | - | 0 | - |
| - | - | 2421 | 157.1 | - | - | 0 | - |
| - | - | 3241 | 157.1 | - | - | 0 | - |
| - | - | 2708 | 157.1 | - | - | 0 | - |
| 15 | y | 1.214E+04 | 158.1 | 0.0002941 | 1.86 | +1 | 1 |
| - | - | 521.9 | 159.1 | - | - | 0 | - |
| - | - | 1897 | 159.1 | - | - | 0 | - |
| - | - | 1711 | 163.1 | - | - | 0 | - |
| - | - | 861.6 | 167.1 | - | - | 0 | - |
| 11 | y | 1043 | 169.1 | 0.001767 | 10.45 | +3 | 5 |
| - | - | 531.2 | 171 | - | - | 0 | - |
| - | - | 7000 | 171.1 | - | - | 0 | - |
| - | - | 818.4 | 172.1 | - | - | 0 | - |
| - | - | 549.4 | 173.1 | - | - | 0 | - |
| - | - | 3341 | 173.1 | - | - | 0 | - |
| - | - | 428 | 174.1 | - | - | 0 | - |
| - | - | 3852 | 175.1 | - | - | 0 | - |
| 15 | y | 2.411E+04 | 175.1 | 0.0002953 | 1.686 | +1 | 1 |
| - | - | 1902 | 176.1 | - | - | 0 | - |
| - | - | 2093 | 181.1 | - | - | 0 | - |
| - | - | 5665 | 181.1 | - | - | 0 | - |
| - | - | 557.9 | 181.1 | - | - | 0 | - |
| - | - | 447 | 181.6 | - | - | 0 | - |
| - | - | 567.9 | 182.1 | - | - | 0 | - |
| - | - | 802.4 | 182.1 | - | - | 0 | - |
| - | - | 549.6 | 182.2 | - | - | 0 | - |
| - | - | 5236 | 183.1 | - | - | 0 | - |
| - | - | 574.2 | 184.1 | - | - | 0 | - |
| - | - | 1084 | 184.1 | - | - | 0 | - |
| - | - | 801.5 | 184.1 | - | - | 0 | - |
| - | - | 4115 | 185.1 | - | - | 0 | - |
| - | - | 635.8 | 185.1 | - | - | 0 | - |
| - | - | 608.4 | 185.1 | - | - | 0 | - |
| 2 | d | 1406 | 185.2 | 0.0003214 | 1.736 | +1 | 2 |
| - | - | 896.6 | 187.1 | - | - | 0 | - |
| - | - | 4349 | 187.1 | - | - | 0 | - |
| - | - | 763.6 | 187.1 | - | - | 0 | - |
| - | - | 2.982E+04 | 189.1 | - | - | 0 | - |
| - | - | 7968 | 190.1 | - | - | 0 | - |
| - | - | 1910 | 190.1 | - | - | 0 | - |
| - | - | 856.3 | 194.1 | - | - | 0 | - |
| - | - | 1234 | 195.1 | - | - | 0 | - |
| - | - | 620.2 | 195.1 | - | - | 0 | - |
| - | - | 854.4 | 195.1 | - | - | 0 | - |
| - | - | 1949 | 197.1 | - | - | 0 | - |
| - | - | 1584 | 197.1 | - | - | 0 | - |
| - | - | 746.1 | 197.2 | - | - | 0 | - |
| - | - | 5236 | 199.1 | - | - | 0 | - |
| - | - | 2634 | 199.1 | - | - | 0 | - |
| 2 | a | 2.402E+05 | 199.2 | 0.0003269 | 1.641 | +1 | 2 |
| - | - | 575.5 | 199.4 | - | - | 0 | - |
| - | - | 497.8 | 199.8 | - | - | 0 | - |
| - | - | 2.768E+04 | 200.2 | - | - | 0 | - |
| - | - | 1.141E+04 | 201.1 | - | - | 0 | - |
| - | - | 1119 | 201.2 | - | - | 0 | - |
| - | - | 2256 | 202.1 | - | - | 0 | - |
| - | - | 1938 | 202.1 | - | - | 0 | - |
| - | - | 1049 | 202.1 | - | - | 0 | - |
| - | - | 466.2 | 203 | - | - | 0 | - |
| - | - | 4748 | 203.1 | - | - | 0 | - |
| - | - | 1132 | 203.1 | - | - | 0 | - |
| - | - | 642.5 | 204.1 | - | - | 0 | - |
| - | - | 551.4 | 207.1 | - | - | 0 | - |
| - | - | 8033 | 209.1 | - | - | 0 | - |
| - | - | 1012 | 209.1 | - | - | 0 | - |
| - | - | 525.4 | 209.1 | - | - | 0 | - |
| - | - | 1758 | 210.1 | - | - | 0 | - |
| - | - | 1935 | 211.1 | - | - | 0 | - |
| - | - | 2669 | 212.1 | - | - | 0 | - |
| - | - | 881.3 | 213.1 | - | - | 0 | - |
| - | - | 958.3 | 213.2 | - | - | 0 | - |
| - | - | 978 | 215.1 | - | - | 0 | - |
| - | - | 1674 | 215.1 | - | - | 0 | - |
| - | - | 671.8 | 216.1 | - | - | 0 | - |
| - | - | 4.767E+04 | 217.1 | - | - | 0 | - |
| - | - | 4620 | 218.1 | - | - | 0 | - |
| - | - | 1363 | 221.1 | - | - | 0 | - |
| - | - | 786 | 223.1 | - | - | 0 | - |
| - | - | 730.5 | 223.1 | - | - | 0 | - |
| - | - | 4360 | 224.1 | - | - | 0 | - |
| - | - | 1896 | 225.1 | - | - | 0 | - |
| - | - | 833.1 | 225.1 | - | - | 0 | - |
| - | - | 1271 | 226.1 | - | - | 0 | - |
| - | - | 1714 | 226.2 | - | - | 0 | - |
| - | - | 4061 | 227.1 | - | - | 0 | - |
| - | - | 1616 | 227.1 | - | - | 0 | - |
| 2 | b | 6.796E+04 | 227.2 | 0.00027 | 1.189 | +1 | 2 |
| - | - | 644.3 | 228.1 | - | - | 0 | - |
| - | - | 8593 | 228.2 | - | - | 0 | - |
| - | - | 4641 | 229.1 | - | - | 0 | - |
| - | - | 825.8 | 229.2 | - | - | 0 | - |
| - | - | 1553 | 230.1 | - | - | 0 | - |
| - | - | 1435 | 233.2 | - | - | 0 | - |
| - | - | 719.6 | 237.1 | - | - | 0 | - |
| - | - | 848 | 237.1 | - | - | 0 | - |
| - | - | 807.1 | 239.1 | - | - | 0 | - |
| - | - | 773.4 | 239.1 | - | - | 0 | - |
| - | - | 924.7 | 240.1 | - | - | 0 | - |
| - | - | 5459 | 242.1 | - | - | 0 | - |
| - | - | 771.7 | 243.1 | - | - | 0 | - |
| - | - | 1867 | 243.1 | - | - | 0 | - |
| - | - | 2086 | 244.1 | - | - | 0 | - |
| 14 | y | 776.7 | 244.1 | 0.001323 | 5.419 | +1 | 2 |
| 14 | y | 8371 | 245.1 | 0.0002786 | 1.137 | +1 | 2 |
| - | - | 650.3 | 249.2 | - | - | 0 | - |
| - | - | 573.8 | 252.1 | - | - | 0 | - |
| - | - | 1109 | 254.1 | - | - | 0 | - |
| - | - | 650 | 255.1 | - | - | 0 | - |
| - | - | 4415 | 255.1 | - | - | 0 | - |
| - | - | 800.1 | 256.1 | - | - | 0 | - |
| - | - | 950.1 | 256.1 | - | - | 0 | - |
| - | - | 684.7 | 259.1 | - | - | 0 | - |
| - | - | 828.9 | 259.1 | - | - | 0 | - |
| - | - | 630.9 | 261.2 | - | - | 0 | - |
| 14 | y | 4884 | 262.2 | 0.0002035 | 0.7763 | +1 | 2 |
| - | - | 612.5 | 267.1 | - | - | 0 | - |
| - | - | 1498 | 270.1 | - | - | 0 | - |
| - | - | 606.9 | 271.1 | - | - | 0 | - |
| - | - | 656.7 | 272.1 | - | - | 0 | - |
| - | - | 592.9 | 274.1 | - | - | 0 | - |
| - | - | 7033 | 277.1 | - | - | 0 | - |
| - | - | 578.2 | 278.1 | - | - | 0 | - |
| - | - | 1605 | 284.1 | - | - | 0 | - |
| - | - | 877.7 | 284.2 | - | - | 0 | - |
| - | - | 557.3 | 285.1 | - | - | 0 | - |
| - | - | 1007 | 288.2 | - | - | 0 | - |
| - | - | 746.3 | 289.2 | - | - | 0 | - |
| 3 | b | 3731 | 296.2 | 0.0002144 | 0.7238 | +1 | 3 |
| - | - | 1189 | 297.1 | - | - | 0 | - |
| - | - | 3061 | 298.1 | - | - | 0 | - |
| - | - | 1458 | 299.1 | - | - | 0 | - |
| - | - | 777.1 | 307.2 | - | - | 0 | - |
| - | - | 1847 | 310.1 | - | - | 0 | - |
| - | - | 590 | 312.1 | - | - | 0 | - |
| - | - | 3783 | 312.2 | - | - | 0 | - |
| - | - | 1908 | 313.1 | - | - | 0 | - |
| - | - | 702.2 | 313.2 | - | - | 0 | - |
| - | - | 8799 | 313.2 | - | - | 0 | - |
| - | - | 2277 | 314.1 | - | - | 0 | - |
| - | - | 943.7 | 314.2 | - | - | 0 | - |
| 3 | b | 7829 | 314.2 | 0.000514 | 1.636 | +1 | 3 |
| - | - | 1153 | 315.2 | - | - | 0 | - |
| - | - | 9335 | 316.2 | - | - | 0 | - |
| - | - | 1820 | 317.2 | - | - | 0 | - |
| - | - | 703.8 | 319.2 | - | - | 0 | - |
| - | - | 893.3 | 323.2 | - | - | 0 | - |
| - | - | 606.8 | 324.1 | - | - | 0 | - |
| - | - | 644.6 | 324.2 | - | - | 0 | - |
| - | - | 1390 | 326.1 | - | - | 0 | - |
| - | - | 1782 | 330.2 | - | - | 0 | - |
| 13 | y | 1640 | 331.2 | 0.0001115 | 0.3368 | +1 | 3 |
| 13 | y | 6962 | 332.2 | 0.0003699 | 1.114 | +1 | 3 |
| - | - | 1128 | 333.2 | - | - | 0 | - |
| - | - | 677.3 | 335.1 | - | - | 0 | - |
| - | - | 1.327E+04 | 338.1 | - | - | 0 | - |
| - | - | 1192 | 338.2 | - | - | 0 | - |
| - | - | 2428 | 339.1 | - | - | 0 | - |
| - | - | 2409 | 340.2 | - | - | 0 | - |
| - | - | 5515 | 341.2 | - | - | 0 | - |
| - | - | 7103 | 342.1 | - | - | 0 | - |
| - | - | 672.5 | 342.2 | - | - | 0 | - |
| - | - | 1531 | 343.1 | - | - | 0 | - |
| 9 | y | 3153 | 345.2 | 0.000109 | 0.3158 | +2 | 7 |
| - | - | 1355 | 345.7 | - | - | 0 | - |
| - | - | 722.4 | 347.7 | - | - | 0 | - |
| 13 | y | 6504 | 349.2 | 6.596E-05 | 0.1889 | +1 | 3 |
| - | - | 1335 | 350.2 | - | - | 0 | - |
| - | - | 795.2 | 352.1 | - | - | 0 | - |
| - | - | 628.1 | 353.1 | - | - | 0 | - |
| - | - | 686.5 | 353.2 | - | - | 0 | - |
| - | - | 1910 | 354.1 | - | - | 0 | - |
| - | - | 1128 | 356.1 | - | - | 0 | - |
| - | - | 3641 | 359.2 | - | - | 0 | - |
| - | - | 787.6 | 359.2 | - | - | 0 | - |
| - | - | 1289 | 361.7 | - | - | 0 | - |
| - | - | 984.9 | 370.1 | - | - | 0 | - |
| - | - | 903.7 | 381.1 | - | - | 0 | - |
| - | - | 1040 | 385.2 | - | - | 0 | - |
| - | - | 1772 | 388.1 | - | - | 0 | - |
| - | - | 635.2 | 390.7 | - | - | 0 | - |
| - | - | 2809 | 391.7 | - | - | 0 | - |
| - | - | 637.5 | 393.7 | - | - | 0 | - |
| - | - | 966.6 | 396.2 | - | - | 0 | - |
| - | - | 735.2 | 396.6 | - | - | 0 | - |
| - | - | 753.7 | 397.2 | - | - | 0 | - |
| - | - | 1138 | 398.2 | - | - | 0 | - |
| - | - | 631.8 | 398.7 | - | - | 0 | - |
| - | - | 1250 | 399.2 | - | - | 0 | - |
| - | - | 671.3 | 400.2 | - | - | 0 | - |
| - | - | 944.1 | 400.2 | - | - | 0 | - |
| - | - | 1890 | 403.2 | - | - | 0 | - |
| - | - | 708.7 | 404.2 | - | - | 0 | - |
| - | - | 564.7 | 405.2 | - | - | 0 | - |
| - | - | 3962 | 406.2 | - | - | 0 | - |
| - | - | 744.5 | 407.2 | - | - | 0 | - |
| - | - | 621 | 408.2 | - | - | 0 | - |
| - | - | 734.7 | 410.2 | - | - | 0 | - |
| - | - | 671 | 411.2 | - | - | 0 | - |
| - | - | 2159 | 411.2 | - | - | 0 | - |
| - | - | 1560 | 412.2 | - | - | 0 | - |
| - | - | 621.6 | 412.2 | - | - | 0 | - |
| - | - | 849.3 | 412.7 | - | - | 0 | - |
| 12 | b | 2231 | 414.2 | 0.006087 | 14.69 | +3 | 12 |
| 8 | y | 3843 | 414.7 | 0.001433 | 3.455 | +2 | 8 |
| - | - | 1232 | 415.2 | - | - | 0 | - |
| - | - | 799.8 | 415.3 | - | - | 0 | - |
| - | - | 924.1 | 419.2 | - | - | 0 | - |
| 8 | y | 5.319E+04 | 423.2 | 0.0001841 | 0.4349 | +2 | 8 |
| - | - | 2.01E+04 | 423.7 | - | - | 0 | - |
| - | - | 6011 | 424.2 | - | - | 0 | - |
| - | - | 1250 | 424.7 | - | - | 0 | - |
| 4 | b | 4276 | 425.2 | 8.135E-05 | 0.1913 | +1 | 4 |
| - | - | 1110 | 426.2 | - | - | 0 | - |
| - | - | 1276 | 428.2 | - | - | 0 | - |
| - | - | 2143 | 429.2 | - | - | 0 | - |
| 12 | y | 1682 | 430.2 | 0.001356 | 3.152 | +1 | 4 |
| - | - | 901.2 | 433.2 | - | - | 0 | - |
| - | - | 1161 | 437.2 | - | - | 0 | - |
| - | - | 811 | 440.2 | - | - | 0 | - |
| - | - | 4508 | 441.2 | - | - | 0 | - |
| 4 | b | 3747 | 443.3 | 0.000676 | 1.525 | +1 | 4 |
| - | - | 1193 | 444.3 | - | - | 0 | - |
| - | - | 1175 | 447.7 | - | - | 0 | - |
| 12 | y | 1763 | 448.3 | 0.0001106 | 0.2468 | +1 | 4 |
| 3 | y | 1708 | 448.9 | 0.000105 | 0.234 | +3 | 13 |
| - | - | 6782 | 451.2 | - | - | 0 | - |
| - | - | 1578 | 452.2 | - | - | 0 | - |
| - | - | 1404 | 452.3 | - | - | 0 | - |
| 3 | y | 2401 | 454.9 | 0.0004287 | 0.9424 | +3 | 13 |
| - | - | 1805 | 455.2 | - | - | 0 | - |
| - | - | 1142 | 455.6 | - | - | 0 | - |
| - | - | 3122 | 458.2 | - | - | 0 | - |
| - | - | 1405 | 467.2 | - | - | 0 | - |
| - | - | 2120 | 469.2 | - | - | 0 | - |
| - | - | 1346 | 469.3 | - | - | 0 | - |
| - | - | 1152 | 470.2 | - | - | 0 | - |
| - | - | 584.4 | 479.2 | - | - | 0 | - |
| 7 | y | 9415 | 480.7 | 0.000352 | 0.7322 | +2 | 9 |
| - | - | 4604 | 481.2 | - | - | 0 | - |
| - | - | 1316 | 481.8 | - | - | 0 | - |
| 2 | y | 962.5 | 486.6 | 0.001132 | 2.326 | +3 | 14 |
| - | - | 1017 | 487.2 | - | - | 0 | - |
| 11 | y | 1858 | 487.3 | 0.007531 | 15.46 | +1 | 5 |
| 2 | y | 1713 | 492.6 | 0.00148 | 3.005 | +3 | 14 |
| - | - | 2234 | 492.9 | - | - | 0 | - |
| - | - | 2642 | 495.9 | - | - | 0 | - |
| - | - | 1041 | 496.2 | - | - | 0 | - |
| - | - | 3334 | 496.2 | - | - | 0 | - |
| - | - | 1123 | 496.6 | - | - | 0 | - |
| - | - | 1131 | 500.2 | - | - | 0 | - |
| - | - | 807.2 | 502.2 | - | - | 0 | - |
| - | - | 661.3 | 504.3 | - | - | 0 | - |
| - | - | 1545 | 505.2 | - | - | 0 | - |
| 11 | y | 6199 | 505.3 | 0.0003678 | 0.7279 | +1 | 5 |
| - | - | 1351 | 506.3 | - | - | 0 | - |
| - | - | 861.1 | 512.2 | - | - | 0 | - |
| - | - | 971.8 | 514.3 | - | - | 0 | - |
| 6 | y | 1630 | 515.3 | 0.002049 | 3.976 | +2 | 10 |
| 6 | y | 1758 | 515.7 | 0.004113 | 7.974 | +2 | 10 |
| - | - | 1653 | 516.3 | - | - | 0 | - |
| - | - | 1168 | 518.2 | - | - | 0 | - |
| - | - | 855 | 518.3 | - | - | 0 | - |
| - | - | 3451 | 521.3 | - | - | 0 | - |
| - | - | 1448 | 521.8 | - | - | 0 | - |
| - | - | 753.8 | 522.2 | - | - | 0 | - |
| - | - | 1106 | 522.3 | - | - | 0 | - |
| 6 | y | 5.469E+04 | 524.3 | 0.001458 | 2.781 | +2 | 10 |
| 0 | Precursor | 1.01E+04 | 524.6 | 0.006113 | 11.65 | +3 | -1 |
| - | - | 2.99E+04 | 524.8 | - | - | 0 | - |
| - | - | 4166 | 524.9 | - | - | 0 | - |
| - | - | 1.154E+04 | 525.3 | - | - | 0 | - |
| - | - | 2149 | 525.8 | - | - | 0 | - |
| - | - | 929.2 | 529.3 | - | - | 0 | - |
| - | - | 791.5 | 529.8 | - | - | 0 | - |
| - | - | 892 | 530.2 | - | - | 0 | - |
| 0 | Precursor | 6866 | 530.3 | 0.0001534 | 0.2892 | +3 | -1 |
| - | - | 1875 | 530.3 | - | - | 0 | - |
| - | - | 4729 | 530.6 | - | - | 0 | - |
| - | - | 1919 | 531 | - | - | 0 | - |
| - | - | 1126 | 531.2 | - | - | 0 | - |
| - | - | 822.2 | 532.3 | - | - | 0 | - |
| - | - | 1643 | 540.3 | - | - | 0 | - |
| 5 | b | 1388 | 542.3 | 0.0004467 | 0.8236 | +1 | 5 |
| - | - | 6808 | 550.3 | - | - | 0 | - |
| - | - | 1835 | 551.3 | - | - | 0 | - |
| - | - | 1256 | 556.3 | - | - | 0 | - |
| - | - | 1078 | 558.8 | - | - | 0 | - |
| 11 | b | 697 | 562.3 | 0.00552 | 9.816 | +2 | 11 |
| 5 | y | 3137 | 564.8 | 0.001282 | 2.271 | +2 | 11 |
| 5 | y | 2635 | 565.3 | 0.003902 | 6.903 | +2 | 11 |
| - | - | 1123 | 565.8 | - | - | 0 | - |
| - | - | 1575 | 566.3 | - | - | 0 | - |
| - | - | 804.6 | 567.3 | - | - | 0 | - |
| - | - | 3943 | 568.3 | - | - | 0 | - |
| - | - | 910.3 | 569.3 | - | - | 0 | - |
| - | - | 627 | 572.3 | - | - | 0 | - |
| - | - | 860.5 | 573.3 | - | - | 0 | - |
| 5 | y | 7.6E+04 | 573.8 | 0.0002171 | 0.3783 | +2 | 11 |
| 10 | y | 4.742E+04 | 574.3 | 0.003501 | 6.097 | +1 | 6 |
| - | - | 1.354E+04 | 574.8 | - | - | 0 | - |
| - | - | 4222 | 575.3 | - | - | 0 | - |
| - | - | 1580 | 584.3 | - | - | 0 | - |
| - | - | 766.5 | 585.8 | - | - | 0 | - |
| - | - | 1215 | 587.3 | - | - | 0 | - |
| 10 | y | 8092 | 592.3 | 0.0002886 | 0.4872 | +1 | 6 |
| - | - | 2424 | 593.3 | - | - | 0 | - |
| - | - | 742.8 | 594.3 | - | - | 0 | - |
| - | - | 1799 | 600.3 | - | - | 0 | - |
| - | - | 1239 | 626.3 | - | - | 0 | - |
| - | - | 959.3 | 628.3 | - | - | 0 | - |
| 4 | y | 6407 | 629.3 | 0.00214 | 3.401 | +2 | 12 |
| 4 | y | 3540 | 629.8 | 0.008668 | 13.76 | +2 | 12 |
| - | - | 1633 | 630.3 | - | - | 0 | - |
| - | - | 1224 | 636.3 | - | - | 0 | - |
| 4 | y | 3.671E+04 | 638.3 | 0.0002734 | 0.4283 | +2 | 12 |
| - | - | 2.63E+04 | 638.8 | - | - | 0 | - |
| - | - | 9066 | 639.3 | - | - | 0 | - |
| - | - | 2255 | 639.8 | - | - | 0 | - |
| - | - | 917.8 | 641.3 | - | - | 0 | - |
| - | - | 1372 | 643.4 | - | - | 0 | - |
| - | - | 3004 | 646.3 | - | - | 0 | - |
| - | - | 1057 | 647.3 | - | - | 0 | - |
| - | - | 1561 | 649.3 | - | - | 0 | - |
| - | - | 967 | 652.3 | - | - | 0 | - |
| - | - | 2029 | 653.3 | - | - | 0 | - |
| - | - | 1460 | 654.3 | - | - | 0 | - |
| - | - | 1551 | 654.8 | - | - | 0 | - |
| - | - | 1131 | 655.3 | - | - | 0 | - |
| - | - | 1084 | 656.3 | - | - | 0 | - |
| - | - | 759.3 | 657.3 | - | - | 0 | - |
| - | - | 2322 | 659.3 | - | - | 0 | - |
| - | - | 987.2 | 660.8 | - | - | 0 | - |
| - | - | 855.4 | 661.3 | - | - | 0 | - |
| - | - | 1643 | 663.8 | - | - | 0 | - |
| - | - | 2457 | 664.3 | - | - | 0 | - |
| - | - | 1292 | 664.8 | - | - | 0 | - |
| - | - | 764.6 | 665.8 | - | - | 0 | - |
| - | - | 5630 | 666.8 | - | - | 0 | - |
| - | - | 3262 | 667.3 | - | - | 0 | - |
| - | - | 1827 | 667.8 | - | - | 0 | - |
| 9 | y | 1.047E+04 | 671.3 | 0.0006787 | 1.011 | +1 | 7 |
| 9 | y | 6247 | 672.3 | 0.004319 | 6.425 | +1 | 7 |
| 3 | y | 1.908E+04 | 672.8 | 0.0003241 | 0.4817 | +2 | 13 |
| 3 | y | 1.789E+04 | 673.3 | 0.008645 | 12.84 | +2 | 13 |
| - | - | 7669 | 673.8 | - | - | 0 | - |
| - | - | 1.072E+04 | 674.3 | - | - | 0 | - |
| - | - | 1020 | 674.8 | - | - | 0 | - |
| - | - | 3575 | 675.3 | - | - | 0 | - |
| - | - | 948.8 | 676.3 | - | - | 0 | - |
| - | - | 669.1 | 677.8 | - | - | 0 | - |
| - | - | 941.7 | 678.3 | - | - | 0 | - |
| - | - | 1261 | 679.3 | - | - | 0 | - |
| - | - | 8436 | 680.8 | - | - | 0 | - |
| - | - | 5843 | 681.3 | - | - | 0 | - |
| 3 | y | 5.78E+05 | 681.8 | 0.0001133 | 0.1661 | +2 | 13 |
| - | - | 4.033E+05 | 682.3 | - | - | 0 | - |
| - | - | 1.681E+05 | 682.8 | - | - | 0 | - |
| - | - | 4.741E+04 | 683.3 | - | - | 0 | - |
| - | - | 8894 | 683.8 | - | - | 0 | - |
| - | - | 4231 | 686.8 | - | - | 0 | - |
| - | - | 2768 | 687.3 | - | - | 0 | - |
| - | - | 1196 | 687.8 | - | - | 0 | - |
| 9 | y | 4.602E+04 | 689.4 | 0.000257 | 0.3728 | +1 | 7 |
| - | - | 1.467E+04 | 690.4 | - | - | 0 | - |
| - | - | 3686 | 691.4 | - | - | 0 | - |
| - | - | 2553 | 697.3 | - | - | 0 | - |
| 14 | b | 3424 | 699.3 | 0.007863 | 11.24 | +2 | 14 |
| - | - | 1694 | 700.3 | - | - | 0 | - |
| - | - | 3828 | 722.4 | - | - | 0 | - |
| - | - | 2190 | 723.4 | - | - | 0 | - |
| 2 | y | 6095 | 729.4 | 0.0001808 | 0.2479 | +2 | 14 |
| 2 | y | 3659 | 729.9 | 0.007506 | 10.28 | +2 | 14 |
| - | - | 1417 | 730.4 | - | - | 0 | - |
| - | - | 714.1 | 736.9 | - | - | 0 | - |
| - | - | 692.3 | 737.4 | - | - | 0 | - |
| - | - | 1062 | 737.9 | - | - | 0 | - |
| 2 | y | 4.6E+04 | 738.4 | 0.0007024 | 0.9513 | +2 | 14 |
| - | - | 3.639E+04 | 738.9 | - | - | 0 | - |
| - | - | 1.537E+04 | 739.4 | - | - | 0 | - |
| - | - | 4767 | 739.9 | - | - | 0 | - |
| - | - | 1801 | 743.4 | - | - | 0 | - |
| - | - | 1691 | 743.9 | - | - | 0 | - |
| - | - | 717.6 | 758.4 | - | - | 0 | - |
| - | - | 822.6 | 759.4 | - | - | 0 | - |
| - | - | 1066 | 766.4 | - | - | 0 | - |
| - | - | 1179 | 768.4 | - | - | 0 | - |
| - | - | 2232 | 770.4 | - | - | 0 | - |
| - | - | 751.8 | 779.4 | - | - | 0 | - |
| - | - | 1514 | 782.4 | - | - | 0 | - |
| - | - | 1149 | 783.4 | - | - | 0 | - |
| - | - | 888.9 | 784.4 | - | - | 0 | - |
| - | - | 1185 | 785.4 | - | - | 0 | - |
| - | - | 1040 | 786.4 | - | - | 0 | - |
| - | - | 1048 | 798.4 | - | - | 0 | - |
| - | - | 1028 | 799.4 | - | - | 0 | - |
| - | - | 970.9 | 803.4 | - | - | 0 | - |
| - | - | 798.5 | 805.4 | - | - | 0 | - |
| - | - | 1117 | 811.4 | - | - | 0 | - |
| - | - | 741.3 | 812.4 | - | - | 0 | - |
| - | - | 2109 | 821.5 | - | - | 0 | - |
| - | - | 1344 | 823.4 | - | - | 0 | - |
| - | - | 714.1 | 824.4 | - | - | 0 | - |
| 8 | y | 7201 | 828.4 | 0.0007223 | 0.8718 | +1 | 8 |
| - | - | 2637 | 829.4 | - | - | 0 | - |
| - | - | 851.8 | 830.4 | - | - | 0 | - |
| - | - | 2173 | 841.4 | - | - | 0 | - |
| - | - | 849.4 | 857.4 | - | - | 0 | - |
| - | - | 2137 | 887.4 | - | - | 0 | - |
| - | - | 1490 | 891.4 | - | - | 0 | - |
| - | - | 905.7 | 892.4 | - | - | 0 | - |
| - | - | 1190 | 898.4 | - | - | 0 | - |
| - | - | 924.9 | 899.5 | - | - | 0 | - |
| - | - | 1547 | 900.5 | - | - | 0 | - |
| - | - | 3368 | 915.4 | - | - | 0 | - |
| - | - | 1783 | 916.4 | - | - | 0 | - |
| - | - | 826 | 917.4 | - | - | 0 | - |
| 7 | y | 5153 | 943.5 | 0.0008098 | 0.8584 | +1 | 9 |
| - | - | 3058 | 944.5 | - | - | 0 | - |
| - | - | 983.4 | 945.5 | - | - | 0 | - |
| - | - | 1744 | 969.5 | - | - | 0 | - |
| - | - | 757.6 | 971.5 | - | - | 0 | - |
| - | - | 1257 | 972.5 | - | - | 0 | - |
| - | - | 3553 | 973.4 | - | - | 0 | - |
| - | - | 831.8 | 974.5 | - | - | 0 | - |
| - | - | 5094 | 986.5 | - | - | 0 | - |
| - | - | 5675 | 987.5 | - | - | 0 | - |
| - | - | 3377 | 988.5 | - | - | 0 | - |
| - | - | 1019 | 989.5 | - | - | 0 | - |
| - | - | 1710 | 990.5 | - | - | 0 | - |
| - | - | 988.9 | 991.5 | - | - | 0 | - |
| - | - | 778.5 | 1006 | - | - | 0 | - |
| - | - | 765.5 | 1006 | - | - | 0 | - |
| - | - | 2219 | 1012 | - | - | 0 | - |
| - | - | 1815 | 1013 | - | - | 0 | - |
| - | - | 5473 | 1014 | - | - | 0 | - |
| - | - | 1894 | 1015 | - | - | 0 | - |
| - | - | 1071 | 1016 | - | - | 0 | - |
| 6 | y | 1.667E+04 | 1030 | 0.001344 | 1.304 | +1 | 10 |
| - | - | 8787 | 1031 | - | - | 0 | - |
| - | - | 2698 | 1032 | - | - | 0 | - |
| - | - | 1902 | 1074 | - | - | 0 | - |
| - | - | 1370 | 1075 | - | - | 0 | - |
| - | - | 777.3 | 1084 | - | - | 0 | - |
| - | - | 1828 | 1087 | - | - | 0 | - |
| - | - | 2727 | 1102 | - | - | 0 | - |
| - | - | 1453 | 1103 | - | - | 0 | - |
| - | - | 1057 | 1120 | - | - | 0 | - |
| 5 | y | 5987 | 1130 | 0.002619 | 2.319 | +1 | 11 |
| - | - | 2930 | 1131 | - | - | 0 | - |
| - | - | 1216 | 1132 | - | - | 0 | - |
| - | - | 1052 | 1143 | - | - | 0 | - |
| - | - | 800.9 | 1144 | - | - | 0 | - |
| - | - | 2966 | 1161 | - | - | 0 | - |
| - | - | 1611 | 1162 | - | - | 0 | - |
| - | - | 1185 | 1171 | - | - | 0 | - |
| - | - | 3434 | 1189 | - | - | 0 | - |
| - | - | 1619 | 1190 | - | - | 0 | - |
| - | - | 796.9 | 1206 | - | - | 0 | - |
| - | - | 4485 | 1207 | - | - | 0 | - |
| - | - | 1442 | 1208 | - | - | 0 | - |
| - | - | 678 | 2623 | - | - | 0 | - |
| - | - | 1146 | 3075 | - | - | 0 | - |

m/z Charge Intensity FragmentType MassShift Position
120.08114624023438 0 3170.3452
121.08460235595703 0 402.14322
122.07172393798828 0 403.25937
122.53192901611328 0 319.03806
124.11250305175781 0 522.67316
124.64234924316406 0 409.59085
125.07086944580078 0 461.31982
125.56732177734375 0 359.17947
125.92237091064453 0 358.795
126.0552749633789 0 5630.68
127.05093383789062 0 524.96716
127.08738708496094 0 561.47675
128.0820770263672 0 1045.0455
128.10768127441406 0 617.4009
129.06619262695312 0 1539.6666
129.1025390625 0 4998.029
129.1139373779297 0 389.66718
130.05015563964844 0 997.8055
130.06138610839844 0 2860.3604
130.09779357910156 0 1815.6224
133.0610809326172 0 26304.555
133.097412109375 0 455.91608
134.0645751953125 0 899.9893
136.07601928710938 0 2004.4653
138.09176635742188 0 710.7834
140.0821075439453 0 1409.5548
141.06610107421875 0 732.3695
141.10256958007812 0 895.91156
143.08180236816406 0 577.8704
144.0659942626953 0 1215.3445
145.06117248535156 0 4599.351
147.065185546875 0 446.25403
147.0767364501953 0 2721.327
147.114013671875 0 487.99902
152.07069396972656 0 421.01785
153.06626892089844 0 940.8342
154.0502166748047 0 5893.375
155.0817413330078 0 643.6606
155.1182861328125 0 1027.254
156.07705688476562 0 721.3087
156.10208129882812 0 510.99094
157.06109619140625 0 2420.6448
157.0976104736328 0 3241.3997
157.10867309570312 0 2708.4368
158.0926971435547 0 12137.463 y Ammonia loss 14
159.07723999023438 0 521.9473
159.1130828857422 0 1897.2139
163.07167053222656 0 1710.6848
167.08193969726562 0 861.5545
169.0975799560547 0 1042.9767 y 10
171.03977966308594 0 531.20886
171.0767059326172 0 6999.5977
172.06060791015625 0 818.41125
173.09286499023438 0 549.3651
173.12875366210938 0 3340.9312
174.05455017089844 0 427.95078
175.07168579101562 0 3851.9502
175.11924743652344 0 24106.883 y 14
176.12277221679688 0 1901.8596
181.0610809326172 0 2093.4792
181.09739685058594 0 5664.8433
181.13375854492188 0 557.94666
181.5807647705078 0 446.96872
182.0561065673828 0 567.92676
182.09243774414062 0 802.3877
182.2284393310547 0 549.5857
183.113037109375 0 5236.3896
184.07171630859375 0 574.1697
184.1084442138672 0 1084.0096
184.11668395996094 0 801.5381
185.09239196777344 0 4114.6484
185.10116577148438 0 635.7854
185.129150390625 0 608.41113
185.1651611328125 0 1406.0043 d 1
187.07139587402344 0 896.5643
187.1080322265625 0 4348.6704
187.14414978027344 0 763.56366
189.0872802734375 0 29822.701
190.08253479003906 0 7967.8228
190.0911102294922 0 1910.0714
194.09262084960938 0 856.29535
195.0768585205078 0 1234.321
195.11361694335938 0 620.2235
195.14984130859375 0 854.41345
197.09239196777344 0 1949.1809
197.12844848632812 0 1583.7351
197.16468811035156 0 746.1499
199.07154846191406 0 5236.4004
199.1079559326172 0 2633.8857
199.18081665039062 0 240179.36 a 1
199.35330200195312 0 575.53705
199.80657958984375 0 497.7929
200.18408203125 0 27684.86
201.12356567382812 0 11410.201
201.18699645996094 0 1118.7347
202.08258056640625 0 2256.0803
202.11874389648438 0 1937.8376
202.12803649902344 0 1049.1967
203.04820251464844 0 466.16443
203.06643676757812 0 4748.344
203.10269165039062 0 1132.4053
204.06967163085938 0 642.50415
207.07736206054688 0 551.38153
209.0923614501953 0 8032.8555
209.10284423828125 0 1011.54346
209.14024353027344 0 525.4319
210.0875701904297 0 1757.5093
211.10794067382812 0 1935.1437
212.1032257080078 0 2668.946
213.08766174316406 0 881.26587
213.15965270996094 0 958.3061
215.10293579101562 0 978.0322
215.13929748535156 0 1674.1412
216.13441467285156 0 671.8456
217.0821533203125 0 47668.105
218.08555603027344 0 4620.347
221.0594482421875 0 1362.6857
223.1082000732422 0 786.0094
223.14447021484375 0 730.5378
224.1031951904297 0 4360.149
225.08670043945312 0 1896.0244
225.1243438720703 0 833.1058
226.1190643310547 0 1270.9933
226.15542602539062 0 1713.7269
227.1029052734375 0 4061.4746
227.11428833007812 0 1615.81
227.17567443847656 0 67958.44 b 1
228.09693908691406 0 644.2891
228.17892456054688 0 8592.8545
229.1185302734375 0 4641.15
229.1818389892578 0 825.8108
230.11367797851562 0 1553.2357
233.16513061523438 0 1434.9482
237.0989990234375 0 719.553
237.134765625 0 848.0498
239.06982421875 0 807.13684
239.13958740234375 0 773.4399
240.135009765625 0 924.7396
242.11390686035156 0 5458.5684
243.1183624267578 0 771.74066
243.13388061523438 0 1866.6234
244.12953186035156 0 2085.6245
244.14173889160156 0 776.6659 y Water loss 13
245.1247100830078 0 8370.883 y Ammonia loss 13
249.16043090820312 0 650.33136
252.1350555419922 0 573.8239
254.1136932373047 0 1108.9127
255.0972137451172 0 649.9575
255.10916137695312 0 4414.762
256.09381103515625 0 800.1105
256.111572265625 0 950.0865
259.1051330566406 0 684.74786
259.1403503417969 0 828.945
261.16015625 0 630.9207
262.15118408203125 0 4884.405 y 13
267.1095275878906 0 612.48627
270.144775390625 0 1498.2758
271.1280517578125 0 606.8581
272.136962890625 0 656.6649
274.1410217285156 0 592.92377
277.1143493652344 0 7033.199
278.1154479980469 0 578.1809
284.1248779296875 0 1604.8726
284.1605224609375 0 877.67084
285.11944580078125 0 557.33057
288.154541015625 0 1007.1364
289.1514587402344 0 746.31586
296.19708251953125 0 3730.811 b Water loss 2
297.1195983886719 0 1189.4397
298.1402893066406 0 3061.4373
299.1356506347656 0 1457.9807
307.1610412597656 0 777.1087
310.1408996582031 0 1846.7795
312.08154296875 0 589.97375
312.1557922363281 0 3782.7302
313.1141357421875 0 1907.8452
313.153564453125 0 702.15186
313.1872253417969 0 8798.912
314.14605712890625 0 2276.515
314.18890380859375 0 943.7435
314.20794677734375 0 7828.831 b 2
315.21197509765625 0 1153.025
316.1506042480469 0 9334.632
317.15301513671875 0 1820.1891
319.17315673828125 0 703.7624
323.1712951660156 0 893.262
324.1317443847656 0 606.8004
324.1922302246094 0 644.5543
326.1356201171875 0 1389.7372
330.16595458984375 0 1782.3318
331.1723327636719 0 1640.0757 y Water loss 12
332.1568298339844 0 6961.7764 y Ammonia loss 12
333.15887451171875 0 1127.6746
335.136962890625 0 677.2802
338.1349792480469 0 13265.749
338.1716613769531 0 1192.3135
339.1380920410156 0 2427.7712
340.1873779296875 0 2409.43
341.1823425292969 0 5514.591
342.1410827636719 0 7103.2744
342.2032165527344 0 672.45074
343.1438903808594 0 1530.5734
345.1825866699219 0 3153.249 y 8
345.6834411621094 0 1355.1206
347.69635009765625 0 722.3708
349.1830749511719 0 6504.1763 y 12
350.1861267089844 0 1334.9445
352.12518310546875 0 795.2039
353.1470642089844 0 628.1352
353.194580078125 0 686.4814
354.1293640136719 0 1909.6565
356.1455993652344 0 1127.8344
359.1666564941406 0 3640.7832
359.1897888183594 0 787.55524
361.6936950683594 0 1289.3612
370.13726806640625 0 984.9029
381.1393737792969 0 903.67554
385.1707458496094 0 1039.5793
388.1463317871094 0 1771.7883
390.71337890625 0 635.209
391.72344970703125 0 2808.9685
393.7067565917969 0 637.4929
396.1877136230469 0 966.6074
396.6410827636719 0 735.16504
397.2443542480469 0 753.7055
398.2157287597656 0 1138.1964
398.71484375 0 631.75665
399.2166748046875 0 1249.8755
400.1828308105469 0 671.25323
400.22113037109375 0 944.0887
403.1853332519531 0 1889.7994
404.1919250488281 0 708.6984
405.2142028808594 0 564.6915
406.15667724609375 0 3961.7864
407.2298889160156 0 744.52985
408.2292175292969 0 621.00934
410.2070007324219 0 734.6699
411.1893310546875 0 671.046
411.228271484375 0 2159.0623
412.19415283203125 0 1560.0049
412.2247619628906 0 621.56177
412.6983337402344 0 849.2705
414.2286682128906 0 2230.9558 b 11
414.72119140625 0 3842.8936 y Ammonia loss 7
415.2217712402344 0 1232.4999
415.2532653808594 0 799.83325
419.18634033203125 0 924.12646
423.23284912109375 0 53189.617 y 7
423.7347412109375 0 20096.45
424.2342224121094 0 6011.029
424.737548828125 0 1249.8557
425.2393798828125 0 4275.676 b Water loss 3
426.2439270019531 0 1109.9722
428.2162170410156 0 1275.7518
429.23431396484375 0 2143.2542
430.239501953125 0 1681.8895 y Water loss 11
433.2093200683594 0 901.1623
437.2416076660156 0 1161.2289
440.2261657714844 0 810.9781
441.20947265625 0 4508.2275
443.2507019042969 0 3746.8318 b 3
444.2556457519531 0 1192.7103
447.6972961425781 0 1174.9268
448.2513122558594 0 1762.9421 y 11
448.88836669921875 0 1707.5027 y Water loss 2
451.21942138671875 0 6782.301
452.2218017578125 0 1577.9314
452.2619323730469 0 1404.1847
454.8922119140625 0 2400.5745 y 2
455.2257995605469 0 1805.0421
455.56298828125 0 1142.3511
458.2356872558594 0 3121.6506
467.2123718261719 0 1404.5326
469.2294616699219 0 2119.6602
469.28863525390625 0 1346.1998
470.2366943359375 0 1151.8062
479.2301940917969 0 584.4105
480.7468566894531 0 9414.85 y 6
481.2484436035156 0 4603.982
481.75054931640625 0 1316.0497
486.5818176269531 0 962.5221 y Water loss 1
487.2162780761719 0 1017.39624
487.2547912597656 0 1857.6495 y Water loss 10
492.58795166015625 0 1712.7366 y 1
492.9207763671875 0 2233.851
495.9158630371094 0 2641.836
496.2118225097656 0 1040.6724
496.25 0 3333.8413
496.5838623046875 0 1122.8011
500.1986389160156 0 1130.5411
502.1819763183594 0 807.1614
504.25726318359375 0 661.2664
505.22540283203125 0 1544.9531
505.27325439453125 0 6198.687 y 10
506.27801513671875 0 1350.9326
512.25 0 861.0951
514.3239135742188 0 971.8075
515.2551879882812 0 1629.5432 y Water loss 5
515.7533569335938 0 1757.8411 y Ammonia loss 5
516.2526245117188 0 1652.8246
518.2074584960938 0 1167.6886
518.27099609375 0 855.02246
521.2554321289062 0 3450.9944
521.7562255859375 0 1448.2719
522.219482421875 0 753.8163
522.2933959960938 0 1105.831
524.2639770507812 0 54691.95 y 5
524.6117553710938 0 10100.272 Precursor Ammonia loss
524.7639770507812 0 29899.172
524.9459838867188 0 4165.544
525.2664794921875 0 11537.927
525.7670288085938 0 2148.6636
529.2914428710938 0 929.1984
529.7984008789062 0 791.5469
530.1943359375 0 892.0342
530.281005859375 0 6865.5444 Precursor
530.3187866210938 0 1874.6047
530.614990234375 0 4728.747
530.9505615234375 0 1919.0837
531.2494506835938 0 1126.1195
532.27294921875 0 822.2208
540.3037719726562 0 1643.2484
542.3179931640625 0 1388.4352 b 4
550.2866821289062 0 6807.564
551.2891845703125 0 1834.8485
556.3201293945312 0 1255.674
558.7881469726562 0 1077.7203
562.2962646484375 0 697.0356 b Water loss 10
564.7901611328125 0 3136.8499 y Water loss 4
565.287353515625 0 2634.9158 y Ammonia loss 4
565.7891235351562 0 1123.1006
566.2867431640625 0 1574.5763
567.2909545898438 0 804.573
568.2969970703125 0 3942.5488
569.302490234375 0 910.3173
572.2774658203125 0 627.0445
573.2932739257812 0 860.52625
573.7965087890625 0 75997.88 y 4
574.2978515625 0 47420.723 y Water loss 9
574.7991943359375 0 13543.275
575.3001098632812 0 4221.586
584.3148803710938 0 1579.8063
585.7874145507812 0 766.4664
587.2781372070312 0 1215.2773
592.3046264648438 0 8091.929 y 9
593.30712890625 0 2423.8164
594.303466796875 0 742.78723
600.2745361328125 0 1799.0055
626.3251342773438 0 1239.4982
628.3031616210938 0 959.28015
629.3148803710938 0 6407.432 y Water loss 3
629.8134155273438 0 3540.1375 y Ammonia loss 3
630.3176879882812 0 1633.3668
636.3120727539062 0 1224.0094
638.3177490234375 0 36705.477 y 3
638.8193359375 0 26296.943
639.3196411132812 0 9066.189
639.822265625 0 2255.3276
641.337158203125 0 917.75903
643.3554077148438 0 1371.7654
646.3161010742188 0 3004.0176
647.3189086914062 0 1056.9271
649.3194580078125 0 1560.8474
652.3076782226562 0 966.98224
653.3358154296875 0 2029.4049
654.3248291015625 0 1460.0635
654.8192138671875 0 1551.4572
655.3225708007812 0 1130.7843
656.3013916015625 0 1084.2953
657.2831420898438 0 759.2682
659.3431396484375 0 2321.8203
660.8125610351562 0 987.1757
661.3184814453125 0 855.43414
663.8211059570312 0 1643.3335
664.3175048828125 0 2456.8623
664.821044921875 0 1292.0328
665.8107299804688 0 764.6029
666.828369140625 0 5629.7954
667.3296508789062 0 3262.026
667.8306274414062 0 1827.4475
671.346435546875 0 10470.1875 y Water loss 8
672.33544921875 0 6247.229 y Ammonia loss 8
672.8284301757812 0 19077.594 y Water loss 2
673.3294067382812 0 17885.764 y Ammonia loss 2
673.829833984375 0 7669.3813
674.3134155273438 0 10722.463
674.8268432617188 0 1020.10913
675.31396484375 0 3574.7395
676.3184814453125 0 948.7936
677.8153686523438 0 669.1497
678.3233642578125 0 941.65845
679.3289184570312 0 1261.2379
680.824951171875 0 8436.075
681.326904296875 0 5842.7773
681.8339233398438 0 578034.56 y 2
682.3350219726562 0 403297.38
682.8364868164062 0 168127.56
683.3372802734375 0 47410.793
683.8383178710938 0 8893.987
686.826416015625 0 4231.126
687.3291015625 0 2768.2375
687.8306884765625 0 1196.1754
689.357421875 0 46016.367 y 8
690.3600463867188 0 14666.128
691.3628540039062 0 3686.146
697.341552734375 0 2552.5889
699.3411254882812 0 3424.2256 b Ammonia loss 13
700.3433837890625 0 1693.512
722.3812255859375 0 3828.023
723.3790893554688 0 2190.0034
729.37060546875 0 6095.141 y Water loss 1
729.8703002929688 0 3659.0784 y Ammonia loss 1
730.3699340820312 0 1416.8617
736.86572265625 0 714.10895
737.3617553710938 0 692.33044
737.8695678710938 0 1061.793
738.3753662109375 0 45996.516 y 1
738.8768920898438 0 36386.94
739.3779907226562 0 15368.792
739.8787841796875 0 4766.8965
743.3680419921875 0 1801.4653
743.8687744140625 0 1691.405
758.383056640625 0 717.6188
759.3861083984375 0 822.5848
766.3572387695312 0 1065.8303
768.4052124023438 0 1178.8641
770.4141845703125 0 2231.7122
779.4099731445312 0 751.77875
782.4403076171875 0 1513.892
783.4420166015625 0 1149.1332
784.3732299804688 0 888.9105
785.4277954101562 0 1184.9045
786.3706665039062 0 1039.8048
798.4153442382812 0 1047.6992
799.4152221679688 0 1028.0771
803.439697265625 0 970.85187
805.3803100585938 0 798.45667
811.404052734375 0 1117.0157
812.4005126953125 0 741.29004
821.452392578125 0 2109.0376
823.380126953125 0 1343.8783
824.3744506835938 0 714.0943
828.4315185546875 0 7200.999 y Ammonia loss 7
829.4348754882812 0 2636.903
830.4251098632812 0 851.76807
841.391357421875 0 2173.3364
857.438232421875 0 849.3693
887.4263916015625 0 2137.293
891.4159545898438 0 1489.5072
892.4196166992188 0 905.70764
898.4114990234375 0 1190.2018
899.4505004882812 0 924.90234
900.4542846679688 0 1547.4158
915.4150390625 0 3368.0845
916.4229736328125 0 1782.849
917.41455078125 0 826.03644
943.4583740234375 0 5153.1 y Ammonia loss 6
944.4653930664062 0 3057.6348
945.468017578125 0 983.3971
969.4764404296875 0 1744.3265
971.4730834960938 0 757.6455
972.4644165039062 0 1256.9469
973.4485473632812 0 3552.5112
974.4594116210938 0 831.8032
986.4885864257812 0 5094.097
987.4859008789062 0 5674.8784
988.4895629882812 0 3377.001
989.4872436523438 0 1019.17487
990.4848022460938 0 1710.3079
991.483642578125 0 988.92206
1005.5045166015625 0 778.4667
1006.4974975585938 0 765.5062
1012.4775390625 0 2219.3867
1013.477783203125 0 1814.5677
1014.481689453125 0 5472.816
1015.487060546875 0 1894.3938
1016.4916381835938 0 1071.4825
1030.4898681640625 0 16670.998 y Ammonia loss 5
1031.4927978515625 0 8786.68
1032.4949951171875 0 2698.15
1073.5235595703125 0 1902.4565
1074.52880859375 0 1370.4385
1083.5089111328125 0 777.2857
1086.5426025390625 0 1828.3384
1101.516357421875 0 2727.4297
1102.519287109375 0 1452.8296
1119.5291748046875 0 1056.606
1129.5570068359375 0 5987.121 y Ammonia loss 4
1130.5574951171875 0 2930.0754
1131.559814453125 0 1216.0483
1142.5350341796875 0 1051.7119
1143.5352783203125 0 800.8768
1160.546875 0 2965.8489
1161.55419921875 0 1611.1819
1170.546142578125 0 1185.3469
1188.548095703125 0 3433.6003
1189.5496826171875 0 1618.5907
1205.5589599609375 0 796.89276
1206.55810546875 0 4485.3955
1207.5517578125 0 1442.2582
2622.669677734375 0 678.0006
3075.23583984375 0 1146.2203

Spectrum Details

|  |  |
| --- | --- |
| Matched peaks? Matched peaks The total absolute number of peaks matched. Additionally in brackets the total fraction of peaks matched and the total number of peaks is shown. | 58 (11.55% of 502) |
| FDR? FDR The false discovery rate estimated for this peptide. It is calculated by matching all theoretical fragments with a non-integer shift with the raw peaks for this spectrum. This is done with 40 different shifts. The resulting percentage is the average number of annotated peaks over the number of annotated peaks with the correct spectrum. | 0.78% |
| Satellite FDR? Satellite FDR See the FDR for details on its calculation. This satellite ion specific FDR only contains the satellite ions (d/w) for I/L/J positions. | 2.38% |
| PSM Score? PSM Score The PSM Score as given by Hecklib to this annotated spectrum. It is shown with three significant figures. | 349 |

#### Spectrum 4605? Spectrum 4605 The raw spectrum of this peptide as annotated by Hecklib. The fragments are coloured according to ion type (see legend). Any peaks with a star '\*' as text can be hovered over to see the full details, first the ion type second the mass shift type. By hovering over the amino acids in the peptide or ions in the legend the corresponding peaks are highlighted. By toggling the 'Unassigned' label you can turn the background (unassigned) peaks on or off in the plot. By updating the slider in the Ion legend you can update the spectrum to only show the top X% of the peaks with labels. The top X% means any peak that is within X% of the highest intensity. By dragging in the spectrum you can zoom in to a specific part of the spectrum and use 'Zoom Out' to get back to the original zoom level. The annotation of the spectrum is based on the given sequence in the peptides file and is done with different software so inconsistencies are likely. The peaks are annotated based on the given sequence, with 20 ppm tolerance.

Copy Data

##### Spectrum 4605 (TSV)

###### Preview

```
Loading example...
```

*Click on the button to copy the data to your clipboard.*

Mz MinMz MaxIntensity Max

WidthHeightPeptide font sizePeptide stroke widthSpectrum font sizeSpectrum stroke widthCompact peptide

Ion legend

wxyz

abcd

OtherUnassignedIon^Charge^_Position_Show for top:%

JJSEVSDRPSGVSSR

09.23e+41.85e+52.77e+53.69e+5

Zoom Out

y^+1^_1_y^+3^_5_y^+1^_1_d^+1^_2_y^+3^_6_a^+1^_2_b^+1^_2_y^+1^_2_y^+1^_2_y^+1^_2_b^+1^_3_b^+1^_3_y^+1^_3_y^+2^_7_y^+1^_3_b^+3^_10_b^+3^_12_y^+2^_8_y^+2^_8_b^+1^_4_b^+1^_4_y^+1^_4_y^+3^_13_y^+3^_13_y^+2^_9_y^+3^_14_y^+3^_14_y^+1^_5_y^+2^_10_y^+2^_10_y^+2^_10_\*\*b^+1^_5_y^+2^_11_y^+2^_11_y^+2^_11_y^+1^_6_y^+1^_6_y^+2^_12_y^+2^_12_y^+2^_12_y^+1^_7_y^+1^_7_y^+2^_13_y^+2^_13_y^+2^_13_y^+1^_7_b^+2^_14_y^+2^_14_y^+2^_14_y^+2^_14_b^+1^_7_y^+1^_8_b^+1^_8_y^+1^_9_y^+1^_10_y^+1^_11_

0502100315052006

Fragment Matches Table

Show background peaks

| Position | Ion type | Intensity | mz Theoretical | mz Error (Th) | mz Error (ppm) | Charge | Series Number |
| --- | --- | --- | --- | --- | --- | --- | --- |
| - | - | 371.6 | 120.1 | - | - | 0 | - |
| - | - | 5044 | 120.1 | - | - | 0 | - |
| - | - | 598.9 | 121.1 | - | - | 0 | - |
| - | - | 570.3 | 122.1 | - | - | 0 | - |
| - | - | 393.8 | 123.7 | - | - | 0 | - |
| - | - | 568.8 | 124.1 | - | - | 0 | - |
| - | - | 3670 | 126.1 | - | - | 0 | - |
| - | - | 403.6 | 127.9 | - | - | 0 | - |
| - | - | 708 | 128.1 | - | - | 0 | - |
| - | - | 430.4 | 129.1 | - | - | 0 | - |
| - | - | 5188 | 129.1 | - | - | 0 | - |
| - | - | 630.9 | 130 | - | - | 0 | - |
| - | - | 1348 | 130.1 | - | - | 0 | - |
| - | - | 506.4 | 130.1 | - | - | 0 | - |
| - | - | 749.5 | 130.1 | - | - | 0 | - |
| - | - | 413.9 | 131.1 | - | - | 0 | - |
| - | - | 485.8 | 132.3 | - | - | 0 | - |
| - | - | 635.4 | 133.1 | - | - | 0 | - |
| - | - | 4824 | 136.1 | - | - | 0 | - |
| - | - | 584.3 | 137.1 | - | - | 0 | - |
| - | - | 496.7 | 139.1 | - | - | 0 | - |
| - | - | 674.2 | 139.1 | - | - | 0 | - |
| - | - | 1742 | 140.1 | - | - | 0 | - |
| - | - | 541.8 | 141.1 | - | - | 0 | - |
| - | - | 785.4 | 144.1 | - | - | 0 | - |
| - | - | 457.5 | 144.1 | - | - | 0 | - |
| - | - | 618 | 145.1 | - | - | 0 | - |
| - | - | 1730 | 147.1 | - | - | 0 | - |
| - | - | 664.6 | 152.1 | - | - | 0 | - |
| - | - | 625.2 | 153.1 | - | - | 0 | - |
| - | - | 3719 | 154.1 | - | - | 0 | - |
| - | - | 552.4 | 155.1 | - | - | 0 | - |
| - | - | 577.8 | 155.1 | - | - | 0 | - |
| - | - | 1882 | 157.1 | - | - | 0 | - |
| - | - | 1973 | 157.1 | - | - | 0 | - |
| - | - | 1518 | 157.1 | - | - | 0 | - |
| 15 | y | 7046 | 158.1 | 0.000233 | 1.474 | +1 | 1 |
| - | - | 1564 | 159.1 | - | - | 0 | - |
| - | - | 1144 | 163.1 | - | - | 0 | - |
| - | - | 506.8 | 165.1 | - | - | 0 | - |
| - | - | 526.7 | 167.1 | - | - | 0 | - |
| 11 | y | 591 | 169.1 | 0.001355 | 8.012 | +3 | 5 |
| - | - | 4215 | 171.1 | - | - | 0 | - |
| - | - | 2114 | 173.1 | - | - | 0 | - |
| - | - | 1986 | 173.5 | - | - | 0 | - |
| - | - | 2426 | 175.1 | - | - | 0 | - |
| - | - | 617.6 | 175.1 | - | - | 0 | - |
| 15 | y | 1.608E+04 | 175.1 | 0.00028 | 1.599 | +1 | 1 |
| - | - | 1082 | 176.1 | - | - | 0 | - |
| - | - | 605.1 | 178 | - | - | 0 | - |
| - | - | 3637 | 183.1 | - | - | 0 | - |
| - | - | 803.8 | 184.1 | - | - | 0 | - |
| - | - | 585.3 | 185.1 | - | - | 0 | - |
| - | - | 2681 | 185.1 | - | - | 0 | - |
| 2 | d | 663.7 | 185.2 | 0.0002737 | 1.478 | +1 | 2 |
| - | - | 3034 | 187.1 | - | - | 0 | - |
| - | - | 2.011E+04 | 189.1 | - | - | 0 | - |
| - | - | 1238 | 190.1 | - | - | 0 | - |
| 10 | y | 1712 | 192.1 | 0.0008254 | 4.297 | +3 | 6 |
| - | - | 652.7 | 193.1 | - | - | 0 | - |
| - | - | 584.3 | 195.1 | - | - | 0 | - |
| - | - | 872 | 195.1 | - | - | 0 | - |
| - | - | 2775 | 199.1 | - | - | 0 | - |
| - | - | 899.3 | 199.1 | - | - | 0 | - |
| 2 | a | 1.521E+05 | 199.2 | 0.0002811 | 1.411 | +1 | 2 |
| - | - | 600.2 | 200.1 | - | - | 0 | - |
| - | - | 577.6 | 200.1 | - | - | 0 | - |
| - | - | 1.92E+04 | 200.2 | - | - | 0 | - |
| - | - | 7361 | 201.1 | - | - | 0 | - |
| - | - | 1021 | 201.2 | - | - | 0 | - |
| - | - | 812.3 | 202.1 | - | - | 0 | - |
| - | - | 697.3 | 202.1 | - | - | 0 | - |
| - | - | 3383 | 203.1 | - | - | 0 | - |
| - | - | 1103 | 203.1 | - | - | 0 | - |
| - | - | 2857 | 204.1 | - | - | 0 | - |
| - | - | 1550 | 211.1 | - | - | 0 | - |
| - | - | 864.9 | 211.1 | - | - | 0 | - |
| - | - | 1121 | 212.1 | - | - | 0 | - |
| - | - | 843.3 | 213.2 | - | - | 0 | - |
| - | - | 599.8 | 216.1 | - | - | 0 | - |
| - | - | 3.295E+04 | 217.1 | - | - | 0 | - |
| - | - | 2922 | 218.1 | - | - | 0 | - |
| - | - | 2267 | 221.1 | - | - | 0 | - |
| - | - | 1899 | 224.1 | - | - | 0 | - |
| - | - | 669.8 | 226.1 | - | - | 0 | - |
| - | - | 837.1 | 226.2 | - | - | 0 | - |
| - | - | 809.3 | 227.1 | - | - | 0 | - |
| - | - | 1763 | 227.1 | - | - | 0 | - |
| 2 | b | 4.594E+04 | 227.2 | 0.0002548 | 1.121 | +1 | 2 |
| - | - | 5501 | 228.2 | - | - | 0 | - |
| - | - | 2003 | 229.1 | - | - | 0 | - |
| - | - | 618.8 | 230.1 | - | - | 0 | - |
| - | - | 658.4 | 233.2 | - | - | 0 | - |
| - | - | 850.9 | 235.1 | - | - | 0 | - |
| - | - | 544.5 | 237.1 | - | - | 0 | - |
| - | - | 1972 | 239.1 | - | - | 0 | - |
| - | - | 820.2 | 239.1 | - | - | 0 | - |
| - | - | 3646 | 242.1 | - | - | 0 | - |
| - | - | 1009 | 244.1 | - | - | 0 | - |
| 14 | y | 542.8 | 244.1 | 0.001476 | 6.044 | +1 | 2 |
| 14 | y | 4531 | 245.1 | 0.0003854 | 1.572 | +1 | 2 |
| - | - | 1180 | 249.2 | - | - | 0 | - |
| - | - | 880.6 | 254.1 | - | - | 0 | - |
| - | - | 4113 | 255.1 | - | - | 0 | - |
| - | - | 471.7 | 256.1 | - | - | 0 | - |
| - | - | 489.7 | 256.1 | - | - | 0 | - |
| - | - | 552.5 | 262.1 | - | - | 0 | - |
| 14 | y | 2612 | 262.2 | 0.0003256 | 1.242 | +1 | 2 |
| - | - | 772.7 | 270.1 | - | - | 0 | - |
| - | - | 492.1 | 271.6 | - | - | 0 | - |
| - | - | 617.5 | 277.2 | - | - | 0 | - |
| - | - | 772.8 | 284.1 | - | - | 0 | - |
| - | - | 666.6 | 284.2 | - | - | 0 | - |
| - | - | 615.8 | 288.2 | - | - | 0 | - |
| - | - | 592.5 | 296.1 | - | - | 0 | - |
| 3 | b | 1971 | 296.2 | 0.0006416 | 2.166 | +1 | 3 |
| - | - | 1020 | 297.1 | - | - | 0 | - |
| - | - | 1901 | 298.1 | - | - | 0 | - |
| - | - | 1167 | 299.1 | - | - | 0 | - |
| - | - | 2047 | 312.2 | - | - | 0 | - |
| - | - | 4894 | 313.2 | - | - | 0 | - |
| - | - | 1259 | 314.1 | - | - | 0 | - |
| 3 | b | 5193 | 314.2 | 0.0003919 | 1.247 | +1 | 3 |
| - | - | 551.1 | 315.2 | - | - | 0 | - |
| - | - | 583 | 315.2 | - | - | 0 | - |
| - | - | 5891 | 316.2 | - | - | 0 | - |
| - | - | 522.3 | 317.1 | - | - | 0 | - |
| - | - | 624.3 | 323.1 | - | - | 0 | - |
| - | - | 880.7 | 324.1 | - | - | 0 | - |
| - | - | 1477 | 330.2 | - | - | 0 | - |
| 13 | y | 4144 | 332.2 | 0.0007972 | 2.4 | +1 | 3 |
| - | - | 585.2 | 333.2 | - | - | 0 | - |
| - | - | 2504 | 341.2 | - | - | 0 | - |
| - | - | 5973 | 342.1 | - | - | 0 | - |
| - | - | 758.1 | 342.2 | - | - | 0 | - |
| - | - | 836.1 | 343.1 | - | - | 0 | - |
| 9 | y | 1975 | 345.2 | 0.0009635 | 2.791 | +2 | 7 |
| - | - | 723.1 | 345.7 | - | - | 0 | - |
| 13 | y | 3647 | 349.2 | 8.662E-05 | 0.2481 | +1 | 3 |
| - | - | 1888 | 359.2 | - | - | 0 | - |
| - | - | 2535 | 361.7 | - | - | 0 | - |
| 10 | b | 1115 | 362.2 | 0.004185 | 11.56 | +3 | 10 |
| - | - | 1314 | 385.2 | - | - | 0 | - |
| - | - | 1900 | 391.7 | - | - | 0 | - |
| - | - | 1142 | 393.7 | - | - | 0 | - |
| - | - | 907.4 | 395.2 | - | - | 0 | - |
| - | - | 777.2 | 396.3 | - | - | 0 | - |
| - | - | 1037 | 397.2 | - | - | 0 | - |
| - | - | 540.3 | 398.2 | - | - | 0 | - |
| - | - | 575.1 | 399.2 | - | - | 0 | - |
| - | - | 612.6 | 399.2 | - | - | 0 | - |
| - | - | 919.9 | 403.2 | - | - | 0 | - |
| - | - | 666.5 | 405.7 | - | - | 0 | - |
| - | - | 785.8 | 410.2 | - | - | 0 | - |
| - | - | 3234 | 411.2 | - | - | 0 | - |
| - | - | 915 | 411.7 | - | - | 0 | - |
| 12 | b | 1579 | 414.2 | 0.004774 | 11.53 | +3 | 12 |
| 8 | y | 3389 | 414.7 | 0.001219 | 2.94 | +2 | 8 |
| - | - | 1654 | 415.2 | - | - | 0 | - |
| - | - | 827.7 | 415.3 | - | - | 0 | - |
| 8 | y | 3.394E+04 | 423.2 | 0.0003958 | 0.9351 | +2 | 8 |
| - | - | 1.666E+04 | 423.7 | - | - | 0 | - |
| - | - | 3226 | 424.2 | - | - | 0 | - |
| 4 | b | 2795 | 425.2 | 0.0007121 | 1.675 | +1 | 4 |
| - | - | 714.4 | 426.2 | - | - | 0 | - |
| - | - | 1536 | 428.2 | - | - | 0 | - |
| - | - | 1487 | 429.2 | - | - | 0 | - |
| - | - | 864.4 | 439.7 | - | - | 0 | - |
| - | - | 1977 | 441.2 | - | - | 0 | - |
| 4 | b | 2870 | 443.3 | 0.0001175 | 0.265 | +1 | 4 |
| 12 | y | 1242 | 448.3 | 0.0005684 | 1.268 | +1 | 4 |
| 3 | y | 605.3 | 448.9 | 0.001482 | 3.301 | +3 | 13 |
| - | - | 542.6 | 452.2 | - | - | 0 | - |
| - | - | 1341 | 452.3 | - | - | 0 | - |
| - | - | 1534 | 452.7 | - | - | 0 | - |
| - | - | 678.8 | 453.3 | - | - | 0 | - |
| 3 | y | 1193 | 454.9 | 0.0008559 | 1.882 | +3 | 13 |
| - | - | 879.8 | 455.2 | - | - | 0 | - |
| - | - | 611.1 | 455.7 | - | - | 0 | - |
| - | - | 1450 | 458.2 | - | - | 0 | - |
| - | - | 2121 | 461.8 | - | - | 0 | - |
| - | - | 667.6 | 462.2 | - | - | 0 | - |
| - | - | 994.9 | 462.3 | - | - | 0 | - |
| - | - | 558.2 | 467.3 | - | - | 0 | - |
| - | - | 633.2 | 468.3 | - | - | 0 | - |
| - | - | 1295 | 469.3 | - | - | 0 | - |
| - | - | 684.7 | 477.2 | - | - | 0 | - |
| 7 | y | 5412 | 480.7 | 0.0001363 | 0.2835 | +2 | 9 |
| - | - | 2392 | 481.2 | - | - | 0 | - |
| 2 | y | 1877 | 486.6 | 0.001401 | 2.879 | +3 | 14 |
| - | - | 1492 | 487.3 | - | - | 0 | - |
| 2 | y | 1424 | 492.6 | 0.0002896 | 0.5879 | +3 | 14 |
| - | - | 954 | 495.9 | - | - | 0 | - |
| - | - | 1238 | 496.2 | - | - | 0 | - |
| - | - | 801.1 | 496.3 | - | - | 0 | - |
| - | - | 1155 | 496.6 | - | - | 0 | - |
| 11 | y | 4404 | 505.3 | 3.207E-05 | 0.06348 | +1 | 5 |
| - | - | 1411 | 512.3 | - | - | 0 | - |
| 6 | y | 1229 | 515.3 | 0.004429 | 8.596 | +2 | 10 |
| 6 | y | 1132 | 515.7 | 0.00515 | 9.986 | +2 | 10 |
| - | - | 855.4 | 518.3 | - | - | 0 | - |
| - | - | 1.039E+04 | 521.3 | - | - | 0 | - |
| - | - | 3880 | 521.8 | - | - | 0 | - |
| - | - | 1566 | 522.3 | - | - | 0 | - |
| 6 | y | 3.525E+04 | 524.3 | 0.001397 | 2.665 | +2 | 10 |
| 0 | Precursor | 5851 | 524.6 | 0.00599 | 11.42 | +3 | -1 |
| - | - | 1.827E+04 | 524.8 | - | - | 0 | - |
| - | - | 3374 | 524.9 | - | - | 0 | - |
| - | - | 5552 | 525.3 | - | - | 0 | - |
| - | - | 1648 | 530.2 | - | - | 0 | - |
| 0 | Precursor | 3487 | 530.3 | 0.0008858 | 1.67 | +3 | -1 |
| - | - | 1.084E+04 | 530.3 | - | - | 0 | - |
| - | - | 3620 | 530.6 | - | - | 0 | - |
| - | - | 2645 | 530.9 | - | - | 0 | - |
| - | - | 888.6 | 531.3 | - | - | 0 | - |
| - | - | 3148 | 531.3 | - | - | 0 | - |
| - | - | 1139 | 539.8 | - | - | 0 | - |
| - | - | 836.9 | 540.3 | - | - | 0 | - |
| 5 | b | 1124 | 542.3 | 0.000935 | 1.724 | +1 | 5 |
| - | - | 678.4 | 556.3 | - | - | 0 | - |
| - | - | 763 | 558.8 | - | - | 0 | - |
| 5 | y | 1446 | 564.8 | 0.001832 | 3.243 | +2 | 11 |
| 5 | y | 2212 | 565.3 | 0.004635 | 8.199 | +2 | 11 |
| 5 | y | 4.982E+04 | 573.8 | 0.0003391 | 0.5911 | +2 | 11 |
| 10 | y | 2.71E+04 | 574.3 | 0.00344 | 5.99 | +1 | 6 |
| - | - | 9919 | 574.8 | - | - | 0 | - |
| - | - | 1641 | 575.3 | - | - | 0 | - |
| - | - | 635.2 | 584.3 | - | - | 0 | - |
| - | - | 1112 | 585.8 | - | - | 0 | - |
| 10 | y | 4906 | 592.3 | 0.0002275 | 0.3842 | +1 | 6 |
| - | - | 2063 | 593.3 | - | - | 0 | - |
| - | - | 1097 | 600.3 | - | - | 0 | - |
| - | - | 674.2 | 604.7 | - | - | 0 | - |
| - | - | 684.9 | 626.3 | - | - | 0 | - |
| 4 | y | 3937 | 629.3 | 0.002323 | 3.692 | +2 | 12 |
| 4 | y | 3262 | 629.8 | 0.006898 | 10.95 | +2 | 12 |
| - | - | 751.9 | 630.3 | - | - | 0 | - |
| - | - | 756.5 | 636.3 | - | - | 0 | - |
| 4 | y | 2.433E+04 | 638.3 | 0.0007006 | 1.098 | +2 | 12 |
| - | - | 1.453E+04 | 638.8 | - | - | 0 | - |
| - | - | 5670 | 639.3 | - | - | 0 | - |
| - | - | 2034 | 639.8 | - | - | 0 | - |
| - | - | 612 | 642.3 | - | - | 0 | - |
| - | - | 831 | 643.4 | - | - | 0 | - |
| - | - | 732.8 | 643.8 | - | - | 0 | - |
| - | - | 1528 | 646.3 | - | - | 0 | - |
| - | - | 968.7 | 654.3 | - | - | 0 | - |
| - | - | 943.5 | 655.3 | - | - | 0 | - |
| - | - | 1205 | 659.3 | - | - | 0 | - |
| - | - | 980.3 | 660.8 | - | - | 0 | - |
| - | - | 1041 | 663.8 | - | - | 0 | - |
| - | - | 2286 | 664.3 | - | - | 0 | - |
| - | - | 820.5 | 664.8 | - | - | 0 | - |
| - | - | 771.8 | 665.3 | - | - | 0 | - |
| - | - | 895.1 | 666.3 | - | - | 0 | - |
| - | - | 4157 | 666.8 | - | - | 0 | - |
| - | - | 3593 | 667.3 | - | - | 0 | - |
| - | - | 1222 | 667.8 | - | - | 0 | - |
| 9 | y | 5413 | 671.3 | 0.001106 | 1.647 | +1 | 7 |
| 9 | y | 4552 | 672.3 | 0.005418 | 8.059 | +1 | 7 |
| 3 | y | 1.566E+04 | 672.8 | 0.0006293 | 0.9353 | +2 | 13 |
| 3 | y | 1.08E+04 | 673.3 | 0.007912 | 11.75 | +2 | 13 |
| - | - | 5301 | 673.8 | - | - | 0 | - |
| - | - | 5515 | 674.3 | - | - | 0 | - |
| - | - | 1896 | 675.3 | - | - | 0 | - |
| - | - | 3642 | 680.8 | - | - | 0 | - |
| - | - | 3775 | 681.3 | - | - | 0 | - |
| 3 | y | 3.655E+05 | 681.8 | 0.0003574 | 0.5242 | +2 | 13 |
| - | - | 2.632E+05 | 682.3 | - | - | 0 | - |
| - | - | 1.108E+05 | 682.8 | - | - | 0 | - |
| - | - | 2.866E+04 | 683.3 | - | - | 0 | - |
| - | - | 6436 | 683.8 | - | - | 0 | - |
| - | - | 3414 | 686.8 | - | - | 0 | - |
| - | - | 1840 | 687.3 | - | - | 0 | - |
| 9 | y | 2.978E+04 | 689.4 | 0.0005012 | 0.727 | +1 | 7 |
| - | - | 1.128E+04 | 690.4 | - | - | 0 | - |
| - | - | 2497 | 691.4 | - | - | 0 | - |
| - | - | 1833 | 694.4 | - | - | 0 | - |
| - | - | 1399 | 695.4 | - | - | 0 | - |
| 14 | b | 2090 | 699.3 | 0.005604 | 8.014 | +2 | 14 |
| - | - | 7750 | 722.4 | - | - | 0 | - |
| - | - | 2622 | 723.4 | - | - | 0 | - |
| - | - | 706.8 | 724.4 | - | - | 0 | - |
| 2 | y | 3455 | 729.4 | 0.001096 | 1.503 | +2 | 14 |
| 2 | y | 4049 | 729.9 | 0.008483 | 11.62 | +2 | 14 |
| - | - | 847.6 | 730.4 | - | - | 0 | - |
| 2 | y | 2.869E+04 | 738.4 | 0.001008 | 1.365 | +2 | 14 |
| - | - | 2.303E+04 | 738.9 | - | - | 0 | - |
| - | - | 9285 | 739.4 | - | - | 0 | - |
| - | - | 3051 | 739.9 | - | - | 0 | - |
| - | - | 842.3 | 740.4 | - | - | 0 | - |
| - | - | 1708 | 743.4 | - | - | 0 | - |
| - | - | 823.7 | 743.9 | - | - | 0 | - |
| 7 | b | 1286 | 744.4 | 0.006867 | 9.225 | +1 | 7 |
| - | - | 1494 | 770.4 | - | - | 0 | - |
| - | - | 750.6 | 771.4 | - | - | 0 | - |
| - | - | 910.1 | 782.4 | - | - | 0 | - |
| - | - | 665.1 | 783.4 | - | - | 0 | - |
| - | - | 1261 | 785.4 | - | - | 0 | - |
| - | - | 759.7 | 787.4 | - | - | 0 | - |
| - | - | 958.3 | 793.5 | - | - | 0 | - |
| - | - | 1399 | 798.4 | - | - | 0 | - |
| - | - | 1022 | 799.4 | - | - | 0 | - |
| - | - | 798.1 | 803.4 | - | - | 0 | - |
| - | - | 791.5 | 811.4 | - | - | 0 | - |
| - | - | 5569 | 821.5 | - | - | 0 | - |
| - | - | 2360 | 822.5 | - | - | 0 | - |
| - | - | 922.6 | 823.5 | - | - | 0 | - |
| 8 | y | 4548 | 828.4 | 0.001211 | 1.461 | +1 | 8 |
| - | - | 1472 | 829.4 | - | - | 0 | - |
| - | - | 864.2 | 845.4 | - | - | 0 | - |
| - | - | 997 | 887.4 | - | - | 0 | - |
| - | - | 880.1 | 897.4 | - | - | 0 | - |
| 8 | b | 1004 | 900.5 | 0.01429 | 15.87 | +1 | 8 |
| - | - | 2517 | 915.4 | - | - | 0 | - |
| - | - | 832.8 | 916.4 | - | - | 0 | - |
| - | - | 820.8 | 922.5 | - | - | 0 | - |
| 7 | y | 3811 | 943.5 | 0.00319 | 3.381 | +1 | 9 |
| - | - | 1755 | 944.5 | - | - | 0 | - |
| - | - | 811.1 | 945.5 | - | - | 0 | - |
| - | - | 743.3 | 972.5 | - | - | 0 | - |
| - | - | 3465 | 986.5 | - | - | 0 | - |
| - | - | 3751 | 987.5 | - | - | 0 | - |
| - | - | 1781 | 988.5 | - | - | 0 | - |
| - | - | 927.1 | 989.5 | - | - | 0 | - |
| - | - | 2109 | 1012 | - | - | 0 | - |
| - | - | 759.4 | 1013 | - | - | 0 | - |
| - | - | 2912 | 1014 | - | - | 0 | - |
| - | - | 1124 | 1015 | - | - | 0 | - |
| 6 | y | 9655 | 1030 | 0.002321 | 2.252 | +1 | 10 |
| - | - | 5992 | 1031 | - | - | 0 | - |
| - | - | 2235 | 1032 | - | - | 0 | - |
| - | - | 1399 | 1074 | - | - | 0 | - |
| - | - | 880 | 1075 | - | - | 0 | - |
| - | - | 709.7 | 1087 | - | - | 0 | - |
| - | - | 1545 | 1102 | - | - | 0 | - |
| - | - | 796.6 | 1104 | - | - | 0 | - |
| 5 | y | 2669 | 1130 | 0.002863 | 2.535 | +1 | 11 |
| - | - | 2173 | 1131 | - | - | 0 | - |
| - | - | 1029 | 1132 | - | - | 0 | - |
| - | - | 1428 | 1161 | - | - | 0 | - |
| - | - | 984.5 | 1162 | - | - | 0 | - |
| - | - | 1836 | 1189 | - | - | 0 | - |
| - | - | 1838 | 1190 | - | - | 0 | - |
| - | - | 715.3 | 1191 | - | - | 0 | - |
| - | - | 2100 | 1207 | - | - | 0 | - |
| - | - | 1339 | 1208 | - | - | 0 | - |
| - | - | 639.6 | 1986 | - | - | 0 | - |

m/z Charge Intensity FragmentType MassShift Position
120.06607818603516 0 371.5524
120.08108520507812 0 5044.168
121.08442687988281 0 598.9062
122.072021484375 0 570.2849
123.71929931640625 0 393.83728
124.11229705810547 0 568.7657
126.05528259277344 0 3670.2659
127.86293029785156 0 403.631
128.08213806152344 0 708.04755
129.06631469726562 0 430.4111
129.10250854492188 0 5187.8735
130.0499725341797 0 630.8891
130.0613250732422 0 1347.9576
130.0977783203125 0 506.4059
130.10585021972656 0 749.502
131.11814880371094 0 413.86337
132.2910919189453 0 485.83032
133.0609588623047 0 635.40485
136.07603454589844 0 4823.924
137.07884216308594 0 584.2705
139.0872344970703 0 496.67142
139.09799194335938 0 674.16925
140.0821075439453 0 1741.7583
141.10269165039062 0 541.82794
144.06590270996094 0 785.43207
144.1020050048828 0 457.50214
145.0605926513672 0 617.989
147.07662963867188 0 1729.9983
152.07090759277344 0 664.57635
153.0662384033203 0 625.18744
154.05006408691406 0 3718.9365
155.08187866210938 0 552.3995
155.11773681640625 0 577.77783
157.0611572265625 0 1881.5938
157.09719848632812 0 1972.8087
157.10867309570312 0 1517.97
158.09263610839844 0 7046.279 y Ammonia loss 14
159.11302185058594 0 1563.8546
163.0719451904297 0 1144.0989
165.10269165039062 0 506.75873
167.08175659179688 0 526.72424
169.09716796875 0 591.0028 y 10
171.07672119140625 0 4214.829
173.1287078857422 0 2113.7578
173.45199584960938 0 1986.4205
175.0716094970703 0 2426.3345
175.08636474609375 0 617.6103
175.11923217773438 0 16083.607 y 14
176.1228485107422 0 1082.3049
178.01705932617188 0 605.0955
183.11314392089844 0 3637.296
184.10800170898438 0 803.80084
185.05581665039062 0 585.2619
185.09228515625 0 2680.933
185.16456604003906 0 663.72003 d 1
187.10801696777344 0 3034.4724
189.08721923828125 0 20105.883
190.0906982421875 0 1238.0424
192.10214233398438 0 1711.6584 y Water loss 9
193.0971221923828 0 652.6828
195.07693481445312 0 584.3033
195.14952087402344 0 872.0089
199.07150268554688 0 2775.4114
199.10801696777344 0 899.26526
199.18077087402344 0 152076.73 a 1
200.10330200195312 0 600.20544
200.13917541503906 0 577.5611
200.18409729003906 0 19198.727
201.12359619140625 0 7361.0728
201.1872100830078 0 1021.43677
202.11802673339844 0 812.3282
202.1273651123047 0 697.3188
203.0664825439453 0 3382.9597
203.1028289794922 0 1103.1111
204.1230010986328 0 2856.533
211.10812377929688 0 1549.7252
211.14443969726562 0 864.86005
212.10304260253906 0 1121.4653
213.15963745117188 0 843.3014
216.1343536376953 0 599.7839
217.08212280273438 0 32952.023
218.08551025390625 0 2921.9058
221.0593719482422 0 2266.5583
224.10302734375 0 1898.9384
226.11911010742188 0 669.771
226.15542602539062 0 837.0507
227.1024627685547 0 809.3132
227.11398315429688 0 1763.2645
227.1756591796875 0 45940.266 b 1
228.17910766601562 0 5500.5566
229.11851501464844 0 2003.3069
230.11370849609375 0 618.81244
233.1636199951172 0 658.44104
235.10780334472656 0 850.89764
237.1348419189453 0 544.4502
239.0700225830078 0 1971.5775
239.13902282714844 0 820.1613
242.1138153076172 0 3646.2573
244.12905883789062 0 1008.5649
244.1418914794922 0 542.7889 y Water loss 13
245.12481689453125 0 4531.0615 y Ammonia loss 13
249.16014099121094 0 1180.3324
254.11373901367188 0 880.6067
255.10903930664062 0 4112.574
256.1131286621094 0 471.65973
256.1282958984375 0 489.71262
262.1380920410156 0 552.48944
262.15130615234375 0 2611.5938 y 13
270.1446533203125 0 772.6525
271.632568359375 0 492.0975
277.1555480957031 0 617.51514
284.12548828125 0 772.8316
284.15966796875 0 666.6265
288.15576171875 0 615.7753
296.1362609863281 0 592.5143
296.197509765625 0 1971.1526 b Water loss 2
297.11944580078125 0 1020.1397
298.1399841308594 0 1901.2117
299.13580322265625 0 1166.6681
312.1558837890625 0 2046.7485
313.1875 0 4894.138
314.1468200683594 0 1259.1559
314.20782470703125 0 5193.189 b 2
315.19232177734375 0 551.0703
315.2110595703125 0 583.04755
316.1504821777344 0 5890.901
317.1453857421875 0 522.334
323.1336669921875 0 624.3222
324.12835693359375 0 880.6567
330.1661071777344 0 1477.4469
332.1572570800781 0 4143.5435 y Ammonia loss 12
333.157958984375 0 585.23834
341.1825866699219 0 2503.977
342.1410827636719 0 5973.3477
342.1821594238281 0 758.0596
343.14398193359375 0 836.081
345.1834411621094 0 1974.5883 y 8
345.684814453125 0 723.1468
349.18292236328125 0 3646.745 y 12
359.166748046875 0 1888.1252
361.6948547363281 0 2534.87
362.1968078613281 0 1115.3702 b 9
385.17315673828125 0 1313.5374
391.7235412597656 0 1900.2443
393.70623779296875 0 1142.3274
395.2410888671875 0 907.42487
396.25933837890625 0 777.22156
397.244384765625 0 1036.9417
398.21173095703125 0 540.2732
399.1647033691406 0 575.10834
399.21551513671875 0 612.5692
403.18035888671875 0 919.89636
405.7154541015625 0 666.51025
410.2047424316406 0 785.79944
411.2293701171875 0 3234.495
411.731201171875 0 915.0152
414.22735595703125 0 1578.5752 b 11
414.7209777832031 0 3388.8728 y Ammonia loss 7
415.2217712402344 0 1654.229
415.2538146972656 0 827.72186
423.2334289550781 0 33937.227 y 7
423.73480224609375 0 16656.77
424.2352600097656 0 3225.55
425.24017333984375 0 2794.7356 b Water loss 3
426.24200439453125 0 714.4149
428.21502685546875 0 1535.9105
429.2339172363281 0 1486.8844
439.68609619140625 0 864.40594
441.2093505859375 0 1977.317
443.2499084472656 0 2869.9548 b 3
448.2508544921875 0 1241.9818 y 11
448.88677978515625 0 605.2672 y Water loss 2
452.2239074707031 0 542.5564
452.2622985839844 0 1340.8003
452.7471923828125 0 1533.6793
453.2505798339844 0 678.79877
454.89263916015625 0 1193.4135 y 2
455.1932067871094 0 879.8456
455.7203063964844 0 611.0736
458.23583984375 0 1449.8026
461.7530822753906 0 2121.0112
462.1966552734375 0 667.59143
462.2565002441406 0 994.8877
467.2670593261719 0 558.24304
468.2886962890625 0 633.2073
469.28802490234375 0 1294.9143
477.24627685546875 0 684.72473
480.7463684082031 0 5411.623 y 6
481.2488098144531 0 2392.2644
486.5843505859375 0 1877.163 y Water loss 1
487.25030517578125 0 1491.8187
492.586181640625 0 1423.7216 y 1
495.9159851074219 0 954.0399
496.2492980957031 0 1238.4672
496.2842712402344 0 801.07263
496.5855407714844 0 1154.607
505.2729187011719 0 4404.4795 y 10
512.2503662109375 0 1411.2274
515.2528076171875 0 1228.9252 y Water loss 5
515.75439453125 0 1131.5958 y Ammonia loss 5
518.2743530273438 0 855.3583
521.2550048828125 0 10391.399
521.75732421875 0 3880.2336
522.2545166015625 0 1565.8752
524.263916015625 0 35247.758 y 5
524.6116333007812 0 5850.836 Precursor Ammonia loss
524.7635498046875 0 18269.928
524.945556640625 0 3373.926
525.2664794921875 0 5551.708
530.1905517578125 0 1647.5481
530.2802734375 0 3487.3726 Precursor
530.31689453125 0 10840.394
530.615478515625 0 3620.179
530.9491577148438 0 2645.2737
531.2783203125 0 888.6474
531.3196411132812 0 3148.2302
539.8107299804688 0 1139.2094
540.3120727539062 0 836.89465
542.3175048828125 0 1124.2734 b 4
556.318603515625 0 678.3791
558.7909545898438 0 762.9984
564.7896118164062 0 1445.8251 y Water loss 4
565.2880859375 0 2212.256 y Ammonia loss 4
573.79638671875 0 49816.457 y 4
574.2977905273438 0 27098.637 y Water loss 9
574.79931640625 0 9918.587
575.2994384765625 0 1640.9769
584.316650390625 0 635.2316
585.7841796875 0 1112.3278
592.3046875 0 4906.2563 y 9
593.3065795898438 0 2063.2334
600.275146484375 0 1096.5009
604.6649780273438 0 674.2305
626.3240966796875 0 684.9248
629.3150634765625 0 3936.6653 y Water loss 3
629.8116455078125 0 3262.3906 y Ammonia loss 3
630.3184204101562 0 751.89056
636.30859375 0 756.51263
638.3173217773438 0 24325.43 y 3
638.81884765625 0 14534.729
639.3193359375 0 5669.937
639.8223266601562 0 2034.3301
642.3309326171875 0 612.0191
643.356689453125 0 830.9788
643.8142700195312 0 732.79944
646.3132934570312 0 1528.3427
654.3236083984375 0 968.681
655.3231201171875 0 943.5087
659.3466186523438 0 1204.5305
660.8136596679688 0 980.27136
663.8211059570312 0 1041.2863
664.31884765625 0 2285.9102
664.8231811523438 0 820.5219
665.3226928710938 0 771.8013
666.3217163085938 0 895.0667
666.8267211914062 0 4156.7485
667.3275756835938 0 3592.5208
667.8314819335938 0 1222.4213
671.3460083007812 0 5412.8237 y Water loss 8
672.3365478515625 0 4551.9414 y Ammonia loss 8
672.828125 0 15658.62 y Water loss 2
673.3286743164062 0 10802.304 y Ammonia loss 2
673.8287353515625 0 5301.258
674.3145141601562 0 5514.953
675.3146362304688 0 1896.1895
680.8257446289062 0 3641.531
681.327392578125 0 3775.1455
681.8336791992188 0 365457.44 y 2
682.3347778320312 0 263155.25
682.8362426757812 0 110811.08
683.3369140625 0 28659.86
683.8380126953125 0 6436.0356
686.8252563476562 0 3413.7117
687.3277587890625 0 1840.1266
689.357177734375 0 29775.375 y 8
690.3600463867188 0 11277.546
691.3640747070312 0 2497.1724
694.3861694335938 0 1833.0125
695.390625 0 1398.6364
699.3433837890625 0 2089.639 b Ammonia loss 13
722.3816528320312 0 7749.828
723.384033203125 0 2622.032
724.3909301757812 0 706.8427
729.3696899414062 0 3454.9492 y Water loss 1
729.8712768554688 0 4048.7175 y Ammonia loss 1
730.37109375 0 847.5742
738.3750610351562 0 28686.61 y 1
738.8768310546875 0 23026.463
739.3773193359375 0 9284.524
739.8781127929688 0 3051.0173
740.3807373046875 0 842.27454
743.3650512695312 0 1707.591
743.8705444335938 0 823.6723
744.3705444335938 0 1286.3732 b 6
770.4160766601562 0 1494.4908
771.4161376953125 0 750.615
782.4434814453125 0 910.1222
783.4423828125 0 665.095
785.4236450195312 0 1261.1906
787.4016723632812 0 759.6672
793.4605712890625 0 958.29596
798.4156494140625 0 1399.1226
799.4182739257812 0 1022.49524
803.4414672851562 0 798.1455
811.4019775390625 0 791.5117
821.45068359375 0 5568.635
822.4541015625 0 2360.2568
823.459716796875 0 922.5553
828.4310302734375 0 4547.6025 y Ammonia loss 7
829.4342651367188 0 1471.9937
845.4209594726562 0 864.1567
887.4219970703125 0 996.9534
897.3998413085938 0 880.0542
900.4642333984375 0 1004.1341 b 7
915.4187622070312 0 2517.025
916.4214477539062 0 832.78687
922.5011596679688 0 820.80426
943.4559936523438 0 3811.1467 y Ammonia loss 6
944.464599609375 0 1755.207
945.4754638671875 0 811.08453
972.4580688476562 0 743.26074
986.4896240234375 0 3465.4294
987.486083984375 0 3750.7253
988.489013671875 0 1780.6652
989.4951171875 0 927.07623
1012.4771118164062 0 2109.159
1013.4888305664062 0 759.3549
1014.4841918945312 0 2912.3667
1015.4857177734375 0 1123.8584
1030.4888916015625 0 9655.422 y Ammonia loss 5
1031.49267578125 0 5991.503
1032.49072265625 0 2234.7427
1073.5247802734375 0 1399.4615
1074.5255126953125 0 879.96216
1086.5595703125 0 709.7439
1101.5186767578125 0 1544.9325
1103.5283203125 0 796.6448
1129.5567626953125 0 2669.443 y Ammonia loss 4
1130.560546875 0 2172.5645
1131.5654296875 0 1028.5292
1160.5518798828125 0 1427.695
1161.561767578125 0 984.4627
1188.5472412109375 0 1835.7919
1189.5479736328125 0 1837.5485
1190.5576171875 0 715.2752
1206.554931640625 0 2099.628
1207.5550537109375 0 1339.3173
1986.1717529296875 0 639.5962

Spectrum Details

|  |  |
| --- | --- |
| Matched peaks? Matched peaks The total absolute number of peaks matched. Additionally in brackets the total fraction of peaks matched and the total number of peaks is shown. | 58 (16.67% of 348) |
| FDR? FDR The false discovery rate estimated for this peptide. It is calculated by matching all theoretical fragments with a non-integer shift with the raw peaks for this spectrum. This is done with 40 different shifts. The resulting percentage is the average number of annotated peaks over the number of annotated peaks with the correct spectrum. | 0.21% |
| Satellite FDR? Satellite FDR See the FDR for details on its calculation. This satellite ion specific FDR only contains the satellite ions (d/w) for I/L/J positions. | 0.00% |
| PSM Score? PSM Score The PSM Score as given by Hecklib to this annotated spectrum. It is shown with three significant figures. | 332 |

#### Spectrum 3639? Spectrum 3639 The raw spectrum of this peptide as annotated by Hecklib. The fragments are coloured according to ion type (see legend). Any peaks with a star '\*' as text can be hovered over to see the full details, first the ion type second the mass shift type. By hovering over the amino acids in the peptide or ions in the legend the corresponding peaks are highlighted. By toggling the 'Unassigned' label you can turn the background (unassigned) peaks on or off in the plot. By updating the slider in the Ion legend you can update the spectrum to only show the top X% of the peaks with labels. The top X% means any peak that is within X% of the highest intensity. By dragging in the spectrum you can zoom in to a specific part of the spectrum and use 'Zoom Out' to get back to the original zoom level. The annotation of the spectrum is based on the given sequence in the peptides file and is done with different software so inconsistencies are likely. The peaks are annotated based on the given sequence, with 20 ppm tolerance.

Copy Data

##### Spectrum 3639 (TSV)

###### Preview

```
Loading example...
```

*Click on the button to copy the data to your clipboard.*

Mz MinMz MaxIntensity Max

WidthHeightPeptide font sizePeptide stroke widthSpectrum font sizeSpectrum stroke widthCompact peptide

Ion legend

wxyz

abcd

OtherUnassignedIon^Charge^_Position_Show for top:%

JJSEVSDRPSGVSSR

02.45e+74.90e+77.35e+79.80e+7

Zoom Out

y^+1^_1_y^+1^_1_a^+1^_2_b^+1^_2_y^+1^_2_y^+1^_2_y^+1^_2_b^+1^_3_b^+1^_3_y^+1^_3_y^+1^_3_y^+2^_7_y^+2^_7_y^+1^_3_y^+2^_8_y^+2^_8_y^+2^_8_b^+1^_4_y^+1^_4_y^+1^_4_b^+1^_4_y^+1^_4_y^+2^_9_y^+2^_9_y^+2^_9_y^+1^_5_y^+1^_5_y^+1^_5_y^+2^_10_y^+2^_10_y^+2^_10_b^+1^_5_b^+2^_10_b^+1^_5_b^+2^_11_y^+2^_11_y^+2^_11_y^+2^_11_y^+1^_6_y^+1^_6_y^+1^_6_b^+2^_12_y^+2^_12_y^+2^_12_b^+2^_13_y^+1^_7_y^+1^_7_y^+2^_13_y^+2^_13_y^+2^_13_y^+1^_7_b^+2^_14_b^+1^_7_y^+2^_14_y^+2^_14_b^+1^_7_\*\*\*y^+1^_8_y^+1^_8_y^+1^_8_b^+1^_8_b^+1^_8_b^+1^_8_y^+1^_9_y^+1^_9_y^+1^_10_y^+1^_10_y^+1^_10_b^+1^_10_b^+1^_10_y^+1^_11_b^+1^_11_y^+1^_11_b^+1^_12_b^+1^_12_y^+1^_12_y^+1^_12_b^+1^_13_b^+1^_13_y^+1^_13_y^+1^_13_b^+1^_14_b^+1^_14_b^+1^_14_y^+1^_14_

0827165324803307

Fragment Matches Table

Show background peaks

| Position | Ion type | Intensity | mz Theoretical | mz Error (Th) | mz Error (ppm) | Charge | Series Number |
| --- | --- | --- | --- | --- | --- | --- | --- |
| - | - | 1.386E+04 | 125.1 | - | - | 0 | - |
| - | - | 1.621E+04 | 125.1 | - | - | 0 | - |
| - | - | 1.209E+05 | 126.1 | - | - | 0 | - |
| - | - | 2.114E+04 | 128.1 | - | - | 0 | - |
| - | - | 1.482E+04 | 128.1 | - | - | 0 | - |
| - | - | 5.656E+04 | 129.1 | - | - | 0 | - |
| - | - | 1.399E+05 | 129.1 | - | - | 0 | - |
| - | - | 8.338E+04 | 129.1 | - | - | 0 | - |
| - | - | 2.133E+05 | 130.1 | - | - | 0 | - |
| - | - | 1.891E+05 | 130.1 | - | - | 0 | - |
| - | - | 1.938E+04 | 130.5 | - | - | 0 | - |
| - | - | 2.106E+04 | 130.5 | - | - | 0 | - |
| - | - | 1.828E+04 | 130.5 | - | - | 0 | - |
| - | - | 1.273E+04 | 130.5 | - | - | 0 | - |
| - | - | 1.558E+04 | 131.1 | - | - | 0 | - |
| - | - | 1.808E+04 | 133.1 | - | - | 0 | - |
| - | - | 4.358E+04 | 133.1 | - | - | 0 | - |
| - | - | 2.011E+04 | 136.1 | - | - | 0 | - |
| - | - | 2.825E+04 | 138.1 | - | - | 0 | - |
| - | - | 1.69E+04 | 138.1 | - | - | 0 | - |
| - | - | 1.922E+04 | 139.1 | - | - | 0 | - |
| - | - | 5.815E+04 | 139.1 | - | - | 0 | - |
| - | - | 5.956E+04 | 140.1 | - | - | 0 | - |
| - | - | 3.415E+04 | 140.1 | - | - | 0 | - |
| - | - | 1.382E+04 | 140.7 | - | - | 0 | - |
| - | - | 2.79E+04 | 141.1 | - | - | 0 | - |
| - | - | 3.164E+04 | 141.1 | - | - | 0 | - |
| - | - | 3.039E+04 | 145.1 | - | - | 0 | - |
| - | - | 1.384E+05 | 147.1 | - | - | 0 | - |
| - | - | 3.903E+04 | 149 | - | - | 0 | - |
| - | - | 1.567E+04 | 153.1 | - | - | 0 | - |
| - | - | 8.528E+04 | 154.1 | - | - | 0 | - |
| - | - | 1.982E+04 | 155.1 | - | - | 0 | - |
| - | - | 3.823E+04 | 155.1 | - | - | 0 | - |
| - | - | 1.412E+05 | 157.1 | - | - | 0 | - |
| - | - | 3.155E+05 | 157.1 | - | - | 0 | - |
| - | - | 2.965E+05 | 157.1 | - | - | 0 | - |
| 15 | y | 1.126E+06 | 158.1 | 0.0005535 | 3.501 | +1 | 1 |
| - | - | 5.766E+04 | 159.1 | - | - | 0 | - |
| - | - | 9.858E+04 | 159.1 | - | - | 0 | - |
| - | - | 2.399E+04 | 165.1 | - | - | 0 | - |
| - | - | 1.72E+04 | 166.1 | - | - | 0 | - |
| - | - | 1.343E+05 | 167.1 | - | - | 0 | - |
| - | - | 3.201E+04 | 169.1 | - | - | 0 | - |
| - | - | 1.736E+05 | 171.1 | - | - | 0 | - |
| - | - | 3.088E+04 | 171.1 | - | - | 0 | - |
| - | - | 1.967E+04 | 172.1 | - | - | 0 | - |
| - | - | 1.536E+04 | 172.5 | - | - | 0 | - |
| - | - | 1.728E+05 | 173.1 | - | - | 0 | - |
| - | - | 3.967E+04 | 173.5 | - | - | 0 | - |
| - | - | 3.04E+05 | 175.1 | - | - | 0 | - |
| 15 | y | 3.179E+06 | 175.1 | 0.0006157 | 3.516 | +1 | 1 |
| - | - | 2.011E+05 | 176.1 | - | - | 0 | - |
| - | - | 3.502E+04 | 181.1 | - | - | 0 | - |
| - | - | 1.866E+04 | 181.1 | - | - | 0 | - |
| - | - | 5.188E+04 | 182.1 | - | - | 0 | - |
| - | - | 3.276E+04 | 182.2 | - | - | 0 | - |
| - | - | 4.452E+05 | 183.1 | - | - | 0 | - |
| - | - | 8.09E+04 | 184.1 | - | - | 0 | - |
| - | - | 3.522E+04 | 184.1 | - | - | 0 | - |
| - | - | 1.789E+04 | 185.1 | - | - | 0 | - |
| - | - | 3.971E+05 | 185.1 | - | - | 0 | - |
| - | - | 3.558E+04 | 185.1 | - | - | 0 | - |
| - | - | 3.141E+04 | 186.1 | - | - | 0 | - |
| - | - | 3.049E+05 | 187.1 | - | - | 0 | - |
| - | - | 2.657E+04 | 188.1 | - | - | 0 | - |
| - | - | 7.216E+05 | 189.1 | - | - | 0 | - |
| - | - | 5.861E+04 | 190.1 | - | - | 0 | - |
| - | - | 1.745E+04 | 192.1 | - | - | 0 | - |
| - | - | 7.989E+04 | 192.1 | - | - | 0 | - |
| - | - | 2.654E+04 | 193.1 | - | - | 0 | - |
| - | - | 5.586E+04 | 194.1 | - | - | 0 | - |
| - | - | 3.747E+04 | 194.1 | - | - | 0 | - |
| - | - | 3.833E+04 | 195.1 | - | - | 0 | - |
| - | - | 7.54E+04 | 195.1 | - | - | 0 | - |
| - | - | 2.27E+04 | 195.1 | - | - | 0 | - |
| - | - | 6.208E+04 | 198.8 | - | - | 0 | - |
| - | - | 4.323E+04 | 198.8 | - | - | 0 | - |
| - | - | 3.261E+04 | 198.9 | - | - | 0 | - |
| - | - | 2.028E+05 | 199.1 | - | - | 0 | - |
| 2 | a | 6.199E+06 | 199.2 | 0.0006015 | 3.02 | +1 | 2 |
| - | - | 2.123E+04 | 200.1 | - | - | 0 | - |
| - | - | 7.097E+05 | 200.2 | - | - | 0 | - |
| - | - | 6.559E+05 | 201.1 | - | - | 0 | - |
| - | - | 1.876E+05 | 202.1 | - | - | 0 | - |
| - | - | 3.707E+04 | 202.1 | - | - | 0 | - |
| - | - | 3.911E+05 | 203.1 | - | - | 0 | - |
| - | - | 7.858E+04 | 203.1 | - | - | 0 | - |
| - | - | 3.636E+04 | 204.1 | - | - | 0 | - |
| - | - | 2.431E+04 | 209.1 | - | - | 0 | - |
| - | - | 5.182E+04 | 209.1 | - | - | 0 | - |
| - | - | 7.539E+04 | 209.1 | - | - | 0 | - |
| - | - | 8.547E+04 | 210.1 | - | - | 0 | - |
| - | - | 2.42E+04 | 210.1 | - | - | 0 | - |
| - | - | 1.28E+05 | 211.1 | - | - | 0 | - |
| - | - | 2.828E+04 | 211.1 | - | - | 0 | - |
| - | - | 2.004E+04 | 211.1 | - | - | 0 | - |
| - | - | 2.689E+04 | 211.2 | - | - | 0 | - |
| - | - | 2.221E+05 | 212.1 | - | - | 0 | - |
| - | - | 3.015E+04 | 213.1 | - | - | 0 | - |
| - | - | 4.235E+04 | 213.2 | - | - | 0 | - |
| - | - | 7.504E+04 | 216.1 | - | - | 0 | - |
| - | - | 2.203E+06 | 217.1 | - | - | 0 | - |
| - | - | 1.766E+05 | 218.1 | - | - | 0 | - |
| - | - | 2.264E+04 | 220.1 | - | - | 0 | - |
| - | - | 2.954E+04 | 220.1 | - | - | 0 | - |
| - | - | 3.255E+04 | 220.1 | - | - | 0 | - |
| - | - | 4.777E+04 | 220.1 | - | - | 0 | - |
| - | - | 4.709E+05 | 224.1 | - | - | 0 | - |
| - | - | 1.084E+05 | 226.1 | - | - | 0 | - |
| - | - | 2.137E+04 | 227.1 | - | - | 0 | - |
| - | - | 1.441E+05 | 227.1 | - | - | 0 | - |
| 2 | b | 2.332E+06 | 227.2 | 0.0006362 | 2.801 | +1 | 2 |
| - | - | 3.646E+04 | 228.1 | - | - | 0 | - |
| - | - | 2.917E+05 | 228.2 | - | - | 0 | - |
| - | - | 1.789E+05 | 229.1 | - | - | 0 | - |
| - | - | 2.494E+04 | 229.1 | - | - | 0 | - |
| - | - | 1.209E+05 | 230.1 | - | - | 0 | - |
| - | - | 2.407E+04 | 232.1 | - | - | 0 | - |
| - | - | 1.994E+04 | 234.1 | - | - | 0 | - |
| - | - | 2.489E+04 | 236.1 | - | - | 0 | - |
| - | - | 3.647E+04 | 237.1 | - | - | 0 | - |
| - | - | 5.66E+04 | 237.1 | - | - | 0 | - |
| - | - | 2.202E+05 | 237.1 | - | - | 0 | - |
| - | - | 3.284E+04 | 238.1 | - | - | 0 | - |
| - | - | 3.903E+04 | 238.1 | - | - | 0 | - |
| - | - | 2.201E+04 | 240.1 | - | - | 0 | - |
| - | - | 3.501E+04 | 241.1 | - | - | 0 | - |
| - | - | 8.586E+05 | 242.1 | - | - | 0 | - |
| - | - | 7.09E+04 | 243.1 | - | - | 0 | - |
| - | - | 2.4E+04 | 243.1 | - | - | 0 | - |
| - | - | 1.983E+05 | 244.1 | - | - | 0 | - |
| 14 | y | 8.651E+04 | 244.1 | 0.001064 | 4.357 | +1 | 2 |
| 14 | y | 6.133E+05 | 245.1 | 0.0007364 | 3.004 | +1 | 2 |
| - | - | 5.411E+04 | 246.1 | - | - | 0 | - |
| - | - | 2.433E+04 | 252.1 | - | - | 0 | - |
| - | - | 2.48E+04 | 253.1 | - | - | 0 | - |
| - | - | 7.822E+04 | 253.2 | - | - | 0 | - |
| - | - | 1.351E+05 | 254.1 | - | - | 0 | - |
| - | - | 6.446E+04 | 254.2 | - | - | 0 | - |
| - | - | 1.855E+04 | 254.6 | - | - | 0 | - |
| - | - | 5.159E+05 | 255.1 | - | - | 0 | - |
| - | - | 3.455E+04 | 255.1 | - | - | 0 | - |
| - | - | 4.245E+04 | 255.1 | - | - | 0 | - |
| - | - | 5.185E+04 | 256.1 | - | - | 0 | - |
| - | - | 1.222E+05 | 259.1 | - | - | 0 | - |
| - | - | 2.473E+04 | 261.2 | - | - | 0 | - |
| - | - | 2.161E+04 | 262.1 | - | - | 0 | - |
| 14 | y | 5.955E+05 | 262.2 | 0.0006308 | 2.406 | +1 | 2 |
| - | - | 5.713E+04 | 263.2 | - | - | 0 | - |
| - | - | 1.943E+04 | 266.1 | - | - | 0 | - |
| - | - | 2.438E+04 | 266.2 | - | - | 0 | - |
| - | - | 2.841E+04 | 268.1 | - | - | 0 | - |
| - | - | 4.725E+04 | 269.2 | - | - | 0 | - |
| - | - | 9.011E+04 | 270.1 | - | - | 0 | - |
| - | - | 2.666E+04 | 271.1 | - | - | 0 | - |
| - | - | 3.788E+04 | 271.1 | - | - | 0 | - |
| - | - | 6.908E+04 | 272.1 | - | - | 0 | - |
| - | - | 2.08E+04 | 274.1 | - | - | 0 | - |
| - | - | 4.259E+04 | 279.1 | - | - | 0 | - |
| - | - | 5.092E+04 | 280.1 | - | - | 0 | - |
| - | - | 6.335E+04 | 281.1 | - | - | 0 | - |
| - | - | 6.487E+04 | 281.2 | - | - | 0 | - |
| - | - | 2.396E+04 | 282.1 | - | - | 0 | - |
| - | - | 1.932E+04 | 282.2 | - | - | 0 | - |
| - | - | 3.038E+04 | 283.2 | - | - | 0 | - |
| - | - | 4.141E+04 | 283.2 | - | - | 0 | - |
| - | - | 1.095E+05 | 284.1 | - | - | 0 | - |
| - | - | 9.4E+04 | 284.2 | - | - | 0 | - |
| - | - | 2.569E+04 | 284.2 | - | - | 0 | - |
| - | - | 1.158E+05 | 288.2 | - | - | 0 | - |
| - | - | 6.225E+04 | 289.2 | - | - | 0 | - |
| - | - | 2.512E+04 | 293.2 | - | - | 0 | - |
| - | - | 3.908E+04 | 294.1 | - | - | 0 | - |
| - | - | 5.851E+04 | 295.1 | - | - | 0 | - |
| - | - | 7.743E+04 | 295.2 | - | - | 0 | - |
| - | - | 8.289E+04 | 296.1 | - | - | 0 | - |
| - | - | 6.068E+04 | 296.2 | - | - | 0 | - |
| 3 | b | 6.918E+05 | 296.2 | 0.0008247 | 2.784 | +1 | 3 |
| - | - | 8.019E+04 | 297.1 | - | - | 0 | - |
| - | - | 1.057E+05 | 297.2 | - | - | 0 | - |
| - | - | 2.357E+05 | 298.1 | - | - | 0 | - |
| - | - | 1.597E+05 | 299.1 | - | - | 0 | - |
| - | - | 8.238E+04 | 302.1 | - | - | 0 | - |
| - | - | 5.172E+04 | 302.2 | - | - | 0 | - |
| - | - | 3.392E+04 | 306.2 | - | - | 0 | - |
| - | - | 3.136E+04 | 307.1 | - | - | 0 | - |
| - | - | 6.001E+04 | 307.2 | - | - | 0 | - |
| - | - | 5.54E+04 | 311.1 | - | - | 0 | - |
| - | - | 2.648E+04 | 312.1 | - | - | 0 | - |
| - | - | 4.996E+05 | 312.2 | - | - | 0 | - |
| - | - | 1.351E+05 | 313.2 | - | - | 0 | - |
| - | - | 1.2E+06 | 313.2 | - | - | 0 | - |
| - | - | 2.283E+05 | 314.1 | - | - | 0 | - |
| - | - | 1.216E+05 | 314.2 | - | - | 0 | - |
| 3 | b | 8.162E+05 | 314.2 | 0.0009717 | 3.093 | +1 | 3 |
| - | - | 2.929E+04 | 315.1 | - | - | 0 | - |
| - | - | 2.175E+04 | 315.2 | - | - | 0 | - |
| - | - | 1.231E+05 | 315.2 | - | - | 0 | - |
| - | - | 7.552E+05 | 316.2 | - | - | 0 | - |
| - | - | 1.32E+05 | 317.2 | - | - | 0 | - |
| - | - | 3.63E+04 | 323.1 | - | - | 0 | - |
| - | - | 5.68E+04 | 323.2 | - | - | 0 | - |
| - | - | 1.489E+05 | 324.1 | - | - | 0 | - |
| - | - | 7.46E+04 | 324.2 | - | - | 0 | - |
| - | - | 2.337E+04 | 325.1 | - | - | 0 | - |
| - | - | 2.984E+04 | 326.1 | - | - | 0 | - |
| - | - | 1.724E+04 | 326.2 | - | - | 0 | - |
| - | - | 6.986E+04 | 329.1 | - | - | 0 | - |
| - | - | 3E+05 | 330.2 | - | - | 0 | - |
| 13 | y | 1.687E+05 | 331.2 | 0.0008745 | 2.641 | +1 | 3 |
| 13 | y | 6.863E+05 | 332.2 | 0.0009803 | 2.951 | +1 | 3 |
| - | - | 9.545E+04 | 333.2 | - | - | 0 | - |
| - | - | 6.602E+04 | 335.1 | - | - | 0 | - |
| 9 | y | 3.347E+04 | 336.2 | 0.000417 | 1.24 | +2 | 7 |
| - | - | 8.259E+04 | 340.2 | - | - | 0 | - |
| - | - | 1.875E+04 | 340.2 | - | - | 0 | - |
| - | - | 4.267E+04 | 341.2 | - | - | 0 | - |
| - | - | 6.166E+05 | 341.2 | - | - | 0 | - |
| - | - | 8.005E+05 | 342.1 | - | - | 0 | - |
| - | - | 7.34E+04 | 342.2 | - | - | 0 | - |
| - | - | 1.197E+05 | 343.1 | - | - | 0 | - |
| - | - | 2.37E+04 | 344.1 | - | - | 0 | - |
| 9 | y | 6.818E+04 | 345.2 | 0.0005362 | 1.554 | +2 | 7 |
| - | - | 2.948E+04 | 345.7 | - | - | 0 | - |
| - | - | 3.788E+04 | 349.2 | - | - | 0 | - |
| 13 | y | 5.574E+05 | 349.2 | 0.0008899 | 2.549 | +1 | 3 |
| - | - | 8.709E+04 | 350.2 | - | - | 0 | - |
| - | - | 4.587E+04 | 352.1 | - | - | 0 | - |
| - | - | 6.756E+04 | 353.2 | - | - | 0 | - |
| - | - | 2.655E+04 | 354.1 | - | - | 0 | - |
| - | - | 2.158E+04 | 355.2 | - | - | 0 | - |
| - | - | 2.228E+04 | 357.2 | - | - | 0 | - |
| - | - | 6.074E+04 | 358.2 | - | - | 0 | - |
| - | - | 2.409E+05 | 359.2 | - | - | 0 | - |
| - | - | 2.851E+04 | 359.2 | - | - | 0 | - |
| - | - | 4.685E+04 | 365.2 | - | - | 0 | - |
| - | - | 3.423E+04 | 367.2 | - | - | 0 | - |
| - | - | 3.8E+04 | 368.3 | - | - | 0 | - |
| - | - | 2.844E+04 | 368.7 | - | - | 0 | - |
| - | - | 3.738E+04 | 368.7 | - | - | 0 | - |
| - | - | 3.219E+04 | 370.2 | - | - | 0 | - |
| - | - | 1.974E+04 | 377.1 | - | - | 0 | - |
| - | - | 2.079E+04 | 379.2 | - | - | 0 | - |
| - | - | 7.581E+04 | 380.2 | - | - | 0 | - |
| - | - | 9.641E+04 | 381.2 | - | - | 0 | - |
| - | - | 3.122E+04 | 382.2 | - | - | 0 | - |
| - | - | 2.525E+04 | 383.2 | - | - | 0 | - |
| - | - | 1.344E+05 | 385.2 | - | - | 0 | - |
| - | - | 2.716E+04 | 392.2 | - | - | 0 | - |
| - | - | 1.093E+05 | 393.7 | - | - | 0 | - |
| - | - | 4.653E+04 | 394.2 | - | - | 0 | - |
| - | - | 2.943E+04 | 395.2 | - | - | 0 | - |
| - | - | 6.354E+04 | 396.2 | - | - | 0 | - |
| - | - | 7.722E+04 | 397.2 | - | - | 0 | - |
| - | - | 2.2E+04 | 398.2 | - | - | 0 | - |
| - | - | 8.581E+04 | 398.2 | - | - | 0 | - |
| - | - | 2.235E+04 | 398.2 | - | - | 0 | - |
| - | - | 5.202E+04 | 399.2 | - | - | 0 | - |
| - | - | 2.975E+04 | 399.7 | - | - | 0 | - |
| - | - | 3.909E+04 | 400.2 | - | - | 0 | - |
| - | - | 1.421E+05 | 400.2 | - | - | 0 | - |
| - | - | 7.15E+04 | 401.2 | - | - | 0 | - |
| - | - | 1.385E+05 | 403.2 | - | - | 0 | - |
| - | - | 2.432E+04 | 406.2 | - | - | 0 | - |
| - | - | 5.301E+04 | 407.2 | - | - | 0 | - |
| - | - | 2.798E+04 | 408.2 | - | - | 0 | - |
| - | - | 6.352E+04 | 408.2 | - | - | 0 | - |
| - | - | 5.682E+04 | 408.7 | - | - | 0 | - |
| - | - | 9.465E+04 | 410.2 | - | - | 0 | - |
| - | - | 2.38E+04 | 411.2 | - | - | 0 | - |
| - | - | 6.204E+04 | 411.2 | - | - | 0 | - |
| - | - | 7.553E+04 | 411.2 | - | - | 0 | - |
| - | - | 3.307E+04 | 413.2 | - | - | 0 | - |
| - | - | 3.537E+04 | 413.2 | - | - | 0 | - |
| 8 | y | 2.846E+05 | 414.2 | 0.0008869 | 2.141 | +2 | 8 |
| 8 | y | 3.543E+05 | 414.7 | 0.003325 | 8.017 | +2 | 8 |
| - | - | 1.094E+05 | 415.2 | - | - | 0 | - |
| - | - | 1.721E+05 | 415.3 | - | - | 0 | - |
| - | - | 3.481E+04 | 416.2 | - | - | 0 | - |
| - | - | 4.168E+04 | 416.3 | - | - | 0 | - |
| - | - | 3.19E+04 | 422.2 | - | - | 0 | - |
| 8 | y | 6.469E+06 | 423.2 | 0.001189 | 2.81 | +2 | 8 |
| - | - | 2.622E+06 | 423.7 | - | - | 0 | - |
| - | - | 2.785E+04 | 424.2 | - | - | 0 | - |
| - | - | 4.805E+05 | 424.2 | - | - | 0 | - |
| 4 | b | 5.495E+05 | 425.2 | 0.001109 | 2.608 | +1 | 4 |
| - | - | 9.638E+04 | 426.2 | - | - | 0 | - |
| - | - | 2.379E+05 | 428.2 | - | - | 0 | - |
| - | - | 3.411E+05 | 429.2 | - | - | 0 | - |
| 12 | y | 1.556E+05 | 430.2 | 0.0005933 | 1.379 | +1 | 4 |
| - | - | 2.919E+04 | 431.2 | - | - | 0 | - |
| 12 | y | 6.172E+04 | 431.2 | 0.0006816 | 1.581 | +1 | 4 |
| - | - | 5.91E+04 | 439.2 | - | - | 0 | - |
| - | - | 3.362E+05 | 441.2 | - | - | 0 | - |
| - | - | 5.569E+04 | 442.2 | - | - | 0 | - |
| 4 | b | 6.227E+05 | 443.3 | 0.001073 | 2.42 | +1 | 4 |
| - | - | 1.38E+05 | 444.3 | - | - | 0 | - |
| 12 | y | 2.323E+05 | 448.3 | 0.001293 | 2.885 | +1 | 4 |
| - | - | 3.961E+04 | 449.3 | - | - | 0 | - |
| - | - | 3.2E+04 | 451.2 | - | - | 0 | - |
| - | - | 2.384E+05 | 452.3 | - | - | 0 | - |
| - | - | 3.855E+04 | 453.3 | - | - | 0 | - |
| - | - | 2.797E+04 | 455.2 | - | - | 0 | - |
| - | - | 2.836E+04 | 455.3 | - | - | 0 | - |
| - | - | 1.698E+05 | 458.2 | - | - | 0 | - |
| - | - | 2.856E+04 | 459.2 | - | - | 0 | - |
| - | - | 2.475E+04 | 469.2 | - | - | 0 | - |
| - | - | 2.784E+04 | 469.2 | - | - | 0 | - |
| - | - | 1.912E+05 | 469.3 | - | - | 0 | - |
| - | - | 6.618E+04 | 470.2 | - | - | 0 | - |
| - | - | 3.416E+04 | 470.3 | - | - | 0 | - |
| 7 | y | 3.968E+04 | 471.7 | 0.0001945 | 0.4124 | +2 | 9 |
| 7 | y | 3.7E+04 | 472.2 | 0.004593 | 9.727 | +2 | 9 |
| - | - | 3.787E+04 | 472.7 | - | - | 0 | - |
| - | - | 2.377E+04 | 473.2 | - | - | 0 | - |
| - | - | 5.134E+04 | 478.2 | - | - | 0 | - |
| - | - | 6.045E+04 | 479.2 | - | - | 0 | - |
| - | - | 6.087E+04 | 480.3 | - | - | 0 | - |
| 7 | y | 1.944E+05 | 480.7 | 0.001268 | 2.637 | +2 | 9 |
| - | - | 1.219E+05 | 481.2 | - | - | 0 | - |
| - | - | 4.205E+04 | 482.2 | - | - | 0 | - |
| 11 | y | 5.791E+04 | 487.3 | 0.00277 | 5.685 | +1 | 5 |
| 11 | y | 8.212E+04 | 488.2 | 0.002381 | 4.876 | +1 | 5 |
| - | - | 7.275E+04 | 496.2 | - | - | 0 | - |
| - | - | 3.701E+04 | 496.3 | - | - | 0 | - |
| - | - | 7.923E+04 | 497.2 | - | - | 0 | - |
| - | - | 8.98E+04 | 497.3 | - | - | 0 | - |
| - | - | 1.014E+05 | 500.2 | - | - | 0 | - |
| 11 | y | 5.995E+05 | 505.3 | 0.001344 | 2.661 | +1 | 5 |
| - | - | 1.149E+05 | 506.3 | - | - | 0 | - |
| - | - | 2.321E+04 | 507.8 | - | - | 0 | - |
| - | - | 3.143E+04 | 508.2 | - | - | 0 | - |
| - | - | 4.935E+04 | 514.3 | - | - | 0 | - |
| - | - | 1.612E+05 | 514.3 | - | - | 0 | - |
| 6 | y | 1.038E+05 | 515.3 | 0.003086 | 5.99 | +2 | 10 |
| - | - | 3.726E+04 | 515.3 | - | - | 0 | - |
| 6 | y | 5.338E+04 | 515.7 | 0.005394 | 10.46 | +2 | 10 |
| - | - | 5.725E+04 | 516.3 | - | - | 0 | - |
| - | - | 1.257E+05 | 518.2 | - | - | 0 | - |
| 6 | y | 1.846E+05 | 524.3 | 0.0003595 | 0.6857 | +2 | 10 |
| 5 | b | 9.827E+04 | 524.3 | 0.001756 | 3.35 | +1 | 5 |
| - | - | 7.108E+04 | 524.8 | - | - | 0 | - |
| - | - | 3.243E+04 | 529.3 | - | - | 0 | - |
| - | - | 1.09E+05 | 532.3 | - | - | 0 | - |
| - | - | 7.257E+04 | 533.3 | - | - | 0 | - |
| 10 | b | 2.448E+04 | 534.3 | 0.00841 | 15.74 | +2 | 10 |
| - | - | 2.535E+04 | 535.2 | - | - | 0 | - |
| - | - | 9.976E+04 | 539.3 | - | - | 0 | - |
| - | - | 3.102E+04 | 540.3 | - | - | 0 | - |
| - | - | 3.307E+04 | 542.3 | - | - | 0 | - |
| - | - | 2.744E+04 | 542.3 | - | - | 0 | - |
| 5 | b | 2.288E+05 | 542.3 | 0.001567 | 2.89 | +1 | 5 |
| - | - | 5.607E+04 | 543.3 | - | - | 0 | - |
| - | - | 4.508E+04 | 547.2 | - | - | 0 | - |
| - | - | 5.654E+04 | 549.3 | - | - | 0 | - |
| - | - | 3.231E+04 | 552.2 | - | - | 0 | - |
| - | - | 6.892E+04 | 556.3 | - | - | 0 | - |
| - | - | 3.513E+04 | 558.8 | - | - | 0 | - |
| - | - | 3.725E+04 | 559.3 | - | - | 0 | - |
| 11 | b | 2.678E+04 | 562.3 | 0.004482 | 7.971 | +2 | 11 |
| 5 | y | 1.083E+05 | 564.8 | 0.0008538 | 1.512 | +2 | 11 |
| - | - | 8.529E+04 | 565.2 | - | - | 0 | - |
| 5 | y | 1.018E+05 | 565.3 | 0.003414 | 6.039 | +2 | 11 |
| - | - | 6.172E+04 | 565.8 | - | - | 0 | - |
| - | - | 6.355E+04 | 566.3 | - | - | 0 | - |
| - | - | 1.035E+05 | 567.3 | - | - | 0 | - |
| - | - | 3.436E+04 | 569.3 | - | - | 0 | - |
| - | - | 3.334E+04 | 570.3 | - | - | 0 | - |
| - | - | 2.972E+04 | 572.3 | - | - | 0 | - |
| - | - | 2.411E+04 | 573.3 | - | - | 0 | - |
| 5 | y | 1.431E+06 | 573.8 | 0.0008816 | 1.536 | +2 | 11 |
| 10 | y | 9.677E+05 | 574.3 | 0.004051 | 7.053 | +1 | 6 |
| - | - | 2.474E+05 | 574.8 | - | - | 0 | - |
| 10 | y | 5.682E+04 | 575.3 | 0.000778 | 1.352 | +1 | 6 |
| - | - | 7.366E+04 | 583.2 | - | - | 0 | - |
| - | - | 2.061E+04 | 584.3 | - | - | 0 | - |
| - | - | 2.174E+04 | 584.3 | - | - | 0 | - |
| - | - | 1.313E+05 | 584.3 | - | - | 0 | - |
| - | - | 2.867E+04 | 585.3 | - | - | 0 | - |
| - | - | 3.244E+04 | 587.3 | - | - | 0 | - |
| 10 | y | 4.402E+05 | 592.3 | 0.00142 | 2.398 | +1 | 6 |
| - | - | 1.606E+05 | 593.3 | - | - | 0 | - |
| - | - | 3.644E+04 | 595.3 | - | - | 0 | - |
| - | - | 2.542E+04 | 596.3 | - | - | 0 | - |
| - | - | 2.952E+04 | 599.3 | - | - | 0 | - |
| - | - | 4.173E+04 | 600.3 | - | - | 0 | - |
| - | - | 3.801E+04 | 602.3 | - | - | 0 | - |
| - | - | 2.712E+04 | 603.3 | - | - | 0 | - |
| - | - | 2.767E+04 | 608.3 | - | - | 0 | - |
| - | - | 6.567E+04 | 611.3 | - | - | 0 | - |
| 12 | b | 2.166E+04 | 612.3 | 0.006962 | 11.37 | +2 | 12 |
| - | - | 2.553E+04 | 623.3 | - | - | 0 | - |
| - | - | 5.028E+04 | 624.3 | - | - | 0 | - |
| - | - | 1.313E+05 | 626.3 | - | - | 0 | - |
| - | - | 1.795E+05 | 629.3 | - | - | 0 | - |
| 4 | y | 6.832E+04 | 629.8 | 0.005982 | 9.498 | +2 | 12 |
| - | - | 6.411E+04 | 630.3 | - | - | 0 | - |
| - | - | 2.781E+04 | 631.3 | - | - | 0 | - |
| - | - | 3.819E+04 | 632.3 | - | - | 0 | - |
| - | - | 5.644E+04 | 636.3 | - | - | 0 | - |
| - | - | 5.601E+04 | 637.3 | - | - | 0 | - |
| 4 | y | 2.216E+05 | 638.3 | 0.0002734 | 0.4283 | +2 | 12 |
| - | - | 1.287E+05 | 638.8 | - | - | 0 | - |
| - | - | 6.74E+04 | 639.3 | - | - | 0 | - |
| - | - | 1.352E+05 | 641.3 | - | - | 0 | - |
| - | - | 9.676E+04 | 642.3 | - | - | 0 | - |
| - | - | 1.453E+05 | 643.4 | - | - | 0 | - |
| - | - | 4.699E+04 | 644.4 | - | - | 0 | - |
| - | - | 9.374E+04 | 646.3 | - | - | 0 | - |
| - | - | 2.139E+04 | 647.3 | - | - | 0 | - |
| - | - | 2.799E+04 | 652.3 | - | - | 0 | - |
| - | - | 1.173E+05 | 653.3 | - | - | 0 | - |
| - | - | 1.883E+05 | 654.3 | - | - | 0 | - |
| 13 | b | 4.299E+04 | 655.3 | 0.007531 | 11.49 | +2 | 13 |
| - | - | 2.678E+04 | 655.8 | - | - | 0 | - |
| - | - | 4.594E+04 | 656.3 | - | - | 0 | - |
| - | - | 8.784E+04 | 657.3 | - | - | 0 | - |
| - | - | 1.299E+05 | 659.4 | - | - | 0 | - |
| - | - | 4.913E+04 | 660.3 | - | - | 0 | - |
| - | - | 4.591E+04 | 660.8 | - | - | 0 | - |
| - | - | 3.742E+04 | 661.3 | - | - | 0 | - |
| - | - | 7.325E+04 | 664.3 | - | - | 0 | - |
| 9 | y | 6.145E+05 | 671.3 | 0.0009693 | 1.444 | +1 | 7 |
| 9 | y | 6.265E+05 | 672.3 | 0.004869 | 7.242 | +1 | 7 |
| 3 | y | 9.456E+04 | 672.8 | 0.001446 | 2.149 | +2 | 13 |
| 3 | y | 2.947E+05 | 673.3 | 0.01115 | 16.56 | +2 | 13 |
| - | - | 9.057E+04 | 673.8 | - | - | 0 | - |
| - | - | 1.703E+05 | 674.3 | - | - | 0 | - |
| - | - | 7.279E+04 | 675.3 | - | - | 0 | - |
| - | - | 2.447E+04 | 681.3 | - | - | 0 | - |
| 3 | y | 4.507E+05 | 681.8 | 0.001291 | 1.893 | +2 | 13 |
| - | - | 3.274E+05 | 682.3 | - | - | 0 | - |
| - | - | 7.925E+04 | 682.8 | - | - | 0 | - |
| - | - | 2.801E+04 | 683.3 | - | - | 0 | - |
| - | - | 4.017E+04 | 687.3 | - | - | 0 | - |
| 9 | y | 3.71E+06 | 689.4 | 0.001086 | 1.575 | +1 | 7 |
| - | - | 1.238E+06 | 690.4 | - | - | 0 | - |
| - | - | 2.22E+05 | 691.4 | - | - | 0 | - |
| 14 | b | 1.407E+05 | 699.3 | 0.004872 | 6.966 | +2 | 14 |
| - | - | 5.356E+04 | 700.3 | - | - | 0 | - |
| - | - | 4.042E+04 | 716.4 | - | - | 0 | - |
| - | - | 4.876E+04 | 717.4 | - | - | 0 | - |
| 7 | b | 1.065E+05 | 726.4 | 0.001429 | 1.967 | +1 | 7 |
| - | - | 3.511E+04 | 727.4 | - | - | 0 | - |
| - | - | 3.738E+04 | 728.4 | - | - | 0 | - |
| 2 | y | 3.919E+04 | 729.9 | 0.003846 | 5.27 | +2 | 14 |
| - | - | 3.298E+04 | 730.4 | - | - | 0 | - |
| - | - | 3.027E+04 | 730.9 | - | - | 0 | - |
| - | - | 3.204E+04 | 736.4 | - | - | 0 | - |
| - | - | 2.59E+04 | 737.4 | - | - | 0 | - |
| 2 | y | 1.986E+05 | 738.4 | 0.003875 | 5.248 | +2 | 14 |
| - | - | 1.379E+05 | 738.9 | - | - | 0 | - |
| - | - | 7.759E+04 | 739.4 | - | - | 0 | - |
| - | - | 5.562E+04 | 743.4 | - | - | 0 | - |
| 7 | b | 5.257E+05 | 744.4 | 0.0005794 | 0.7784 | +1 | 7 |
| - | - | 1.906E+05 | 745.4 | - | - | 0 | - |
| - | - | 3.716E+04 | 746.4 | - | - | 0 | - |
| - | - | 4.62E+04 | 750.4 | - | - | 0 | - |
| - | - | 4.062E+04 | 751.4 | - | - | 0 | - |
| - | - | 5.83E+04 | 752.4 | - | - | 0 | - |
| - | - | 8.789E+04 | 755.9 | - | - | 0 | - |
| - | - | 2.918E+04 | 756.4 | - | - | 0 | - |
| - | - | 3.918E+04 | 759.4 | - | - | 0 | - |
| - | - | 7.222E+04 | 761.9 | - | - | 0 | - |
| - | - | 1.044E+05 | 762.4 | - | - | 0 | - |
| - | - | 6.873E+04 | 764.9 | - | - | 0 | - |
| - | - | 6.87E+04 | 765.4 | - | - | 0 | - |
| - | - | 7.945E+04 | 765.9 | - | - | 0 | - |
| - | - | 2.441E+05 | 766.4 | - | - | 0 | - |
| - | - | 2.018E+05 | 766.9 | - | - | 0 | - |
| - | - | 1.055E+05 | 767.4 | - | - | 0 | - |
| - | - | 3.006E+04 | 767.9 | - | - | 0 | - |
| - | - | 3.031E+05 | 768.4 | - | - | 0 | - |
| - | - | 4.738E+04 | 768.9 | - | - | 0 | - |
| - | - | 1.496E+05 | 769.4 | - | - | 0 | - |
| - | - | 1.022E+05 | 770.4 | - | - | 0 | - |
| - | - | 1.231E+05 | 770.9 | - | - | 0 | - |
| - | - | 1.646E+05 | 771.4 | - | - | 0 | - |
| - | - | 9.896E+04 | 771.9 | - | - | 0 | - |
| - | - | 1.188E+05 | 772.4 | - | - | 0 | - |
| - | - | 1.057E+05 | 772.9 | - | - | 0 | - |
| - | - | 2.118E+05 | 773.4 | - | - | 0 | - |
| - | - | 3.82E+05 | 773.9 | - | - | 0 | - |
| - | - | 2.394E+05 | 774.4 | - | - | 0 | - |
| - | - | 1.058E+05 | 774.9 | - | - | 0 | - |
| - | - | 2.287E+05 | 776.9 | - | - | 0 | - |
| - | - | 4.78E+05 | 777.4 | - | - | 0 | - |
| - | - | 2.664E+05 | 777.9 | - | - | 0 | - |
| - | - | 1.227E+05 | 778.4 | - | - | 0 | - |
| - | - | 5.658E+04 | 778.9 | - | - | 0 | - |
| - | - | 1.287E+05 | 779.4 | - | - | 0 | - |
| - | - | 7.097E+05 | 779.9 | - | - | 0 | - |
| - | - | 6.997E+05 | 780.4 | - | - | 0 | - |
| - | - | 3.27E+05 | 780.9 | - | - | 0 | - |
| - | - | 3.035E+04 | 781.4 | - | - | 0 | - |
| - | - | 2.363E+05 | 785.4 | - | - | 0 | - |
| 0 | Precursor | 2.905E+06 | 785.9 | 0.000756 | 0.9619 | +2 | -1 |
| 0 | Precursor | 2.898E+06 | 786.4 | 0.009908 | 12.6 | +2 | -1 |
| - | - | 1.428E+06 | 786.9 | - | - | 0 | - |
| - | - | 4.421E+05 | 787.4 | - | - | 0 | - |
| - | - | 1.288E+05 | 787.9 | - | - | 0 | - |
| - | - | 4.085E+04 | 788.4 | - | - | 0 | - |
| - | - | 2.057E+05 | 793.4 | - | - | 0 | - |
| - | - | 1.347E+06 | 793.9 | - | - | 0 | - |
| - | - | 1.186E+06 | 794.4 | - | - | 0 | - |
| 0 | Precursor | 9.701E+07 | 794.9 | 0.001455 | 1.83 | +2 | -1 |
| - | - | 8.272E+07 | 795.4 | - | - | 0 | - |
| - | - | 3.649E+07 | 795.9 | - | - | 0 | - |
| - | - | 3.605E+06 | 796.4 | - | - | 0 | - |
| - | - | 3.384E+04 | 796.9 | - | - | 0 | - |
| - | - | 8.824E+04 | 798.4 | - | - | 0 | - |
| - | - | 4.065E+04 | 801.6 | - | - | 0 | - |
| - | - | 2.514E+05 | 803.4 | - | - | 0 | - |
| - | - | 6.908E+04 | 804.4 | - | - | 0 | - |
| - | - | 1.489E+05 | 810.4 | - | - | 0 | - |
| - | - | 2.317E+05 | 811.4 | - | - | 0 | - |
| - | - | 1.022E+05 | 812.4 | - | - | 0 | - |
| 8 | y | 7.846E+04 | 827.4 | 0.0006544 | 0.7909 | +1 | 8 |
| 8 | y | 1.128E+06 | 828.4 | 0.001292 | 1.559 | +1 | 8 |
| - | - | 4.278E+05 | 829.4 | - | - | 0 | - |
| - | - | 7.057E+04 | 830.4 | - | - | 0 | - |
| 8 | y | 9.764E+05 | 845.5 | 0.0004996 | 0.591 | +1 | 8 |
| - | - | 3.781E+05 | 846.5 | - | - | 0 | - |
| - | - | 5.358E+04 | 847.5 | - | - | 0 | - |
| - | - | 6.88E+04 | 857.4 | - | - | 0 | - |
| - | - | 5.924E+04 | 865.4 | - | - | 0 | - |
| - | - | 2.574E+05 | 872.5 | - | - | 0 | - |
| - | - | 4.488E+04 | 873.4 | - | - | 0 | - |
| - | - | 9.34E+04 | 873.5 | - | - | 0 | - |
| - | - | 3.182E+04 | 874.4 | - | - | 0 | - |
| - | - | 3.414E+04 | 874.5 | - | - | 0 | - |
| 8 | b | 6.319E+04 | 882.5 | 0.009156 | 10.38 | +1 | 8 |
| 8 | b | 1.128E+05 | 883.5 | 0.00428 | 4.845 | +1 | 8 |
| - | - | 3.578E+04 | 884.4 | - | - | 0 | - |
| - | - | 5.386E+04 | 885.4 | - | - | 0 | - |
| - | - | 5.878E+04 | 887.4 | - | - | 0 | - |
| - | - | 1.294E+05 | 891.4 | - | - | 0 | - |
| - | - | 5.291E+04 | 892.4 | - | - | 0 | - |
| - | - | 3.19E+04 | 899.5 | - | - | 0 | - |
| 8 | b | 9.545E+05 | 900.5 | 0.0009832 | 1.092 | +1 | 8 |
| - | - | 3.779E+05 | 901.5 | - | - | 0 | - |
| - | - | 9.214E+04 | 902.5 | - | - | 0 | - |
| - | - | 2.857E+04 | 910.4 | - | - | 0 | - |
| - | - | 1.15E+05 | 915.4 | - | - | 0 | - |
| - | - | 6.18E+04 | 916.4 | - | - | 0 | - |
| - | - | 5.802E+04 | 918.5 | - | - | 0 | - |
| - | - | 3.424E+04 | 919.5 | - | - | 0 | - |
| - | - | 6.338E+04 | 925.5 | - | - | 0 | - |
| - | - | 7.97E+04 | 926.4 | - | - | 0 | - |
| - | - | 6.687E+04 | 927.4 | - | - | 0 | - |
| 7 | y | 5.317E+05 | 943.5 | 0.000655 | 0.6943 | +1 | 9 |
| - | - | 2.33E+05 | 944.5 | - | - | 0 | - |
| - | - | 6.86E+04 | 945.5 | - | - | 0 | - |
| - | - | 2.842E+04 | 946.5 | - | - | 0 | - |
| - | - | 2.7E+04 | 953.5 | - | - | 0 | - |
| - | - | 2.913E+04 | 957.5 | - | - | 0 | - |
| 7 | y | 6.144E+04 | 960.5 | 0.004532 | 4.718 | +1 | 9 |
| - | - | 3.801E+04 | 968.5 | - | - | 0 | - |
| - | - | 7.159E+04 | 969.5 | - | - | 0 | - |
| - | - | 4.047E+04 | 970.5 | - | - | 0 | - |
| - | - | 3.905E+04 | 971.5 | - | - | 0 | - |
| - | - | 7.955E+04 | 972.5 | - | - | 0 | - |
| - | - | 1.596E+05 | 986.5 | - | - | 0 | - |
| - | - | 4.268E+05 | 987.5 | - | - | 0 | - |
| - | - | 1.718E+05 | 988.5 | - | - | 0 | - |
| - | - | 6.069E+04 | 989.5 | - | - | 0 | - |
| - | - | 8.853E+04 | 990.5 | - | - | 0 | - |
| - | - | 4.161E+04 | 991.5 | - | - | 0 | - |
| - | - | 4.257E+04 | 994.5 | - | - | 0 | - |
| - | - | 3.151E+04 | 995.5 | - | - | 0 | - |
| - | - | 4.167E+04 | 996.5 | - | - | 0 | - |
| - | - | 3.669E+04 | 997.5 | - | - | 0 | - |
| - | - | 6.827E+04 | 1000 | - | - | 0 | - |
| - | - | 4.64E+04 | 1003 | - | - | 0 | - |
| - | - | 1.313E+05 | 1005 | - | - | 0 | - |
| - | - | 7.974E+04 | 1007 | - | - | 0 | - |
| - | - | 2.69E+05 | 1012 | - | - | 0 | - |
| - | - | 1.503E+05 | 1013 | - | - | 0 | - |
| - | - | 1.591E+05 | 1014 | - | - | 0 | - |
| - | - | 6.002E+04 | 1015 | - | - | 0 | - |
| 6 | y | 4.884E+04 | 1030 | 0.008051 | 7.82 | +1 | 10 |
| 6 | y | 2.12E+06 | 1030 | 0.000609 | 0.591 | +1 | 10 |
| - | - | 1.077E+06 | 1031 | - | - | 0 | - |
| - | - | 3.235E+05 | 1032 | - | - | 0 | - |
| 6 | y | 1.095E+05 | 1048 | 0.0004274 | 0.408 | +1 | 10 |
| - | - | 4.233E+04 | 1049 | - | - | 0 | - |
| - | - | 3.161E+04 | 1057 | - | - | 0 | - |
| - | - | 2.681E+04 | 1058 | - | - | 0 | - |
| 10 | b | 2.931E+04 | 1068 | 0.01109 | 10.38 | +1 | 10 |
| - | - | 3.519E+04 | 1069 | - | - | 0 | - |
| - | - | 3.249E+04 | 1070 | - | - | 0 | - |
| - | - | 7.429E+04 | 1074 | - | - | 0 | - |
| - | - | 3.284E+04 | 1075 | - | - | 0 | - |
| - | - | 2.889E+04 | 1083 | - | - | 0 | - |
| - | - | 5.94E+04 | 1084 | - | - | 0 | - |
| 10 | b | 1.075E+05 | 1085 | 0.004599 | 4.24 | +1 | 10 |
| - | - | 4.243E+04 | 1086 | - | - | 0 | - |
| - | - | 2.131E+05 | 1087 | - | - | 0 | - |
| - | - | 1.138E+05 | 1088 | - | - | 0 | - |
| - | - | 4.222E+04 | 1089 | - | - | 0 | - |
| - | - | 5.674E+04 | 1100 | - | - | 0 | - |
| - | - | 3.024E+04 | 1101 | - | - | 0 | - |
| - | - | 1.387E+05 | 1102 | - | - | 0 | - |
| - | - | 4.396E+04 | 1103 | - | - | 0 | - |
| - | - | 3.811E+04 | 1105 | - | - | 0 | - |
| - | - | 3.122E+04 | 1106 | - | - | 0 | - |
| - | - | 7.42E+04 | 1112 | - | - | 0 | - |
| - | - | 6.889E+04 | 1113 | - | - | 0 | - |
| - | - | 5.962E+04 | 1114 | - | - | 0 | - |
| - | - | 5.153E+04 | 1120 | - | - | 0 | - |
| - | - | 3.33E+04 | 1121 | - | - | 0 | - |
| - | - | 4.13E+04 | 1128 | - | - | 0 | - |
| 5 | y | 8.573E+05 | 1130 | 5.586E-05 | 0.04946 | +1 | 11 |
| - | - | 4.402E+05 | 1131 | - | - | 0 | - |
| - | - | 1.358E+05 | 1132 | - | - | 0 | - |
| 11 | b | 1.331E+05 | 1142 | 0.0007939 | 0.6955 | +1 | 11 |
| - | - | 5.453E+04 | 1143 | - | - | 0 | - |
| - | - | 2.609E+04 | 1145 | - | - | 0 | - |
| 5 | y | 4.317E+04 | 1147 | 0.009515 | 8.299 | +1 | 11 |
| - | - | 3.51E+04 | 1148 | - | - | 0 | - |
| - | - | 2.752E+04 | 1158 | - | - | 0 | - |
| - | - | 5.48E+04 | 1161 | - | - | 0 | - |
| - | - | 8E+04 | 1189 | - | - | 0 | - |
| - | - | 6.468E+04 | 1190 | - | - | 0 | - |
| - | - | 4.211E+04 | 1198 | - | - | 0 | - |
| - | - | 1.836E+05 | 1207 | - | - | 0 | - |
| - | - | 8.869E+04 | 1208 | - | - | 0 | - |
| - | - | 2.884E+04 | 1209 | - | - | 0 | - |
| - | - | 2.519E+05 | 1213 | - | - | 0 | - |
| - | - | 1.755E+05 | 1214 | - | - | 0 | - |
| - | - | 6.074E+04 | 1215 | - | - | 0 | - |
| - | - | 9.922E+04 | 1216 | - | - | 0 | - |
| - | - | 7.466E+04 | 1217 | - | - | 0 | - |
| 12 | b | 3.441E+04 | 1224 | 0.008 | 6.538 | +1 | 12 |
| - | - | 3.666E+04 | 1234 | - | - | 0 | - |
| 12 | b | 2.549E+05 | 1241 | 0.008661 | 6.981 | +1 | 12 |
| - | - | 1.783E+05 | 1242 | - | - | 0 | - |
| - | - | 5.979E+04 | 1243 | - | - | 0 | - |
| - | - | 2.985E+04 | 1256 | - | - | 0 | - |
| 4 | y | 4.046E+05 | 1259 | 0.0009301 | 0.739 | +1 | 12 |
| - | - | 2.475E+05 | 1260 | - | - | 0 | - |
| - | - | 6.375E+04 | 1261 | - | - | 0 | - |
| - | - | 4.032E+04 | 1273 | - | - | 0 | - |
| - | - | 3.706E+04 | 1274 | - | - | 0 | - |
| - | - | 3.852E+04 | 1275 | - | - | 0 | - |
| 4 | y | 3.531E+04 | 1276 | 0.001237 | 0.9693 | +1 | 12 |
| - | - | 4.03E+04 | 1284 | - | - | 0 | - |
| - | - | 7.27E+04 | 1285 | - | - | 0 | - |
| - | - | 1.097E+05 | 1286 | - | - | 0 | - |
| - | - | 3.386E+04 | 1287 | - | - | 0 | - |
| - | - | 1.216E+05 | 1300 | - | - | 0 | - |
| - | - | 4.615E+04 | 1301 | - | - | 0 | - |
| - | - | 5.872E+05 | 1303 | - | - | 0 | - |
| - | - | 3.969E+05 | 1304 | - | - | 0 | - |
| - | - | 1.547E+05 | 1305 | - | - | 0 | - |
| 13 | b | 6.595E+04 | 1310 | 0.02414 | 18.43 | +1 | 13 |
| 13 | b | 6.251E+04 | 1311 | 0.004741 | 3.617 | +1 | 13 |
| - | - | 5.739E+04 | 1316 | - | - | 0 | - |
| - | - | 4.77E+04 | 1317 | - | - | 0 | - |
| - | - | 3.266E+04 | 1320 | - | - | 0 | - |
| - | - | 1.307E+05 | 1321 | - | - | 0 | - |
| - | - | 6.174E+04 | 1322 | - | - | 0 | - |
| - | - | 2.779E+04 | 1327 | - | - | 0 | - |
| - | - | 2.767E+05 | 1328 | - | - | 0 | - |
| - | - | 2.447E+05 | 1329 | - | - | 0 | - |
| - | - | 9.476E+04 | 1330 | - | - | 0 | - |
| 3 | y | 2.111E+06 | 1346 | 0.0007028 | 0.5223 | +1 | 13 |
| - | - | 1.407E+06 | 1347 | - | - | 0 | - |
| - | - | 5.117E+05 | 1348 | - | - | 0 | - |
| 3 | y | 1.07E+05 | 1363 | 0.001313 | 0.9633 | +1 | 13 |
| - | - | 5.836E+04 | 1364 | - | - | 0 | - |
| - | - | 5.214E+04 | 1369 | - | - | 0 | - |
| - | - | 4.676E+04 | 1370 | - | - | 0 | - |
| - | - | 3.923E+04 | 1371 | - | - | 0 | - |
| - | - | 5.454E+04 | 1385 | - | - | 0 | - |
| - | - | 5.108E+04 | 1386 | - | - | 0 | - |
| - | - | 3.943E+05 | 1387 | - | - | 0 | - |
| - | - | 2.894E+05 | 1388 | - | - | 0 | - |
| - | - | 8.616E+04 | 1389 | - | - | 0 | - |
| 14 | b | 9.205E+04 | 1397 | 0.002583 | 1.849 | +1 | 14 |
| 14 | b | 9.787E+04 | 1398 | 0.01548 | 11.07 | +1 | 14 |
| - | - | 3.285E+04 | 1399 | - | - | 0 | - |
| - | - | 2.778E+04 | 1403 | - | - | 0 | - |
| - | - | 4.581E+04 | 1413 | - | - | 0 | - |
| 14 | b | 4.327E+05 | 1415 | 0.003748 | 2.649 | +1 | 14 |
| - | - | 3.851E+05 | 1416 | - | - | 0 | - |
| - | - | 1.51E+05 | 1417 | - | - | 0 | - |
| - | - | 3.283E+04 | 1418 | - | - | 0 | - |
| - | - | 3.557E+04 | 1432 | - | - | 0 | - |
| - | - | 5.453E+05 | 1433 | - | - | 0 | - |
| - | - | 4.247E+05 | 1434 | - | - | 0 | - |
| - | - | 1.371E+05 | 1435 | - | - | 0 | - |
| 2 | y | 1.79E+05 | 1459 | 0.008351 | 5.725 | +1 | 14 |
| - | - | 1.147E+05 | 1460 | - | - | 0 | - |
| - | - | 2.617E+04 | 3075 | - | - | 0 | - |
| - | - | 2.941E+04 | 3274 | - | - | 0 | - |

m/z Charge Intensity FragmentType MassShift Position
125.07103729248047 0 13855.501
125.10733795166016 0 16208.673
126.05540466308594 0 120916.38
128.08238220214844 0 21135.188
128.10733032226562 0 14819.262
129.06629943847656 0 56558.715
129.10272216796875 0 139883.73
129.11395263671875 0 83379.85
130.0615692138672 0 213299.58
130.09805297851562 0 189112.3
130.4634246826172 0 19377.068
130.47079467773438 0 21057.43
130.49693298339844 0 18284.998
130.50320434570312 0 12725.096
131.12893676757812 0 15584.829
133.06138610839844 0 18076.996
133.0976104736328 0 43580.86
136.0880126953125 0 20110.668
138.06680297851562 0 28251.367
138.09165954589844 0 16904.697
139.05064392089844 0 19224.932
139.0869903564453 0 58154.42
140.08248901367188 0 59561.645
140.1439208984375 0 34154.71
140.697509765625 0 13818.562
141.06649780273438 0 27897.752
141.10255432128906 0 31636.16
145.06161499023438 0 30389.836
147.07699584960938 0 138377.97
148.9535675048828 0 39028.28
153.0660858154297 0 15670.1455
154.05030822753906 0 85284.31
155.08226013183594 0 19823.53
155.11827087402344 0 38232.12
157.06138610839844 0 141249.97
157.0977325439453 0 315485.72
157.10894775390625 0 296519.75
158.09295654296875 0 1126494 y Ammonia loss 14
159.09620666503906 0 57660.008
159.11331176757812 0 98577.336
165.1029052734375 0 23989.764
166.09844970703125 0 17197.229
167.08201599121094 0 134307.66
169.09776306152344 0 32010.025
171.07699584960938 0 173611.88
171.1133270263672 0 30880.045
172.06080627441406 0 19673.451
172.49903869628906 0 15363.491
173.1290740966797 0 172822.88
173.4527587890625 0 39667.89
175.0719451904297 0 304039.84
175.11956787109375 0 3178751 y 14
176.1229705810547 0 201141.05
181.09747314453125 0 35015.61
181.10887145996094 0 18664.926
182.09298706054688 0 51882.895
182.1544647216797 0 32756.445
183.11338806152344 0 445204.72
184.1087188720703 0 80898.86
184.11688232421875 0 35222.645
185.05538940429688 0 17887.717
185.09262084960938 0 397099.84
185.10269165039062 0 35584.78
186.0964813232422 0 31409.545
187.1083221435547 0 304894.47
188.11203002929688 0 26570.615
189.08761596679688 0 721580
190.0912322998047 0 58608.04
192.09832763671875 0 17454.324
192.11378479003906 0 79889.17
193.09750366210938 0 26540.482
194.09288024902344 0 55864.28
194.12940979003906 0 37467.715
195.07704162597656 0 38331.984
195.1134490966797 0 75398.19
195.14959716796875 0 22701.324
198.78671264648438 0 62084.215
198.79803466796875 0 43227.62
198.9130859375 0 32609.295
199.07188415527344 0 202839.38
199.18109130859375 0 6199274 a 1
200.10414123535156 0 21233.895
200.18446350097656 0 709658.4
201.1239013671875 0 655945.9
202.11917114257812 0 187634.25
202.127197265625 0 37074.582
203.06686401367188 0 391066.5
203.1034698486328 0 78581.47
204.06993103027344 0 36361.137
209.0926971435547 0 24307.572
209.10391235351562 0 51817.645
209.14047241210938 0 75389.46
210.08792114257812 0 85466.1
210.12411499023438 0 24200.367
211.10826110839844 0 128048.11
211.117919921875 0 28279.277
211.14512634277344 0 20040.342
211.15585327148438 0 26894.361
212.10353088378906 0 222130.61
213.0878143310547 0 30148.242
213.16043090820312 0 42346.156
216.13482666015625 0 75035.8
217.08250427246094 0 2202635.5
218.08602905273438 0 176574.66
220.0717315673828 0 22638.5
220.0929412841797 0 29541.047
220.10833740234375 0 32554.633
220.1297607421875 0 47769.273
224.1035919189453 0 470929.47
226.1191864013672 0 108371.7
227.104248046875 0 21367.219
227.1145477294922 0 144050.9
227.17604064941406 0 2331869 b 1
228.0991973876953 0 36459.68
228.1794891357422 0 291670.28
229.1189727783203 0 178859.12
229.13050842285156 0 24943.785
230.114013671875 0 120882.516
232.14244079589844 0 24069.45
234.10975646972656 0 19942.19
236.1036376953125 0 24886.125
237.0870361328125 0 36466.37
237.09910583496094 0 56602.38
237.1351776123047 0 220171.45
238.08226013183594 0 32841.555
238.13909912109375 0 39026.727
240.09852600097656 0 22010.357
241.13038635253906 0 35011.64
242.1142120361328 0 858559.2
243.11756896972656 0 70897.234
243.1324462890625 0 24004.922
244.12957763671875 0 198327.08
244.1414794921875 0 86507.61 y Water loss 13
245.1251678466797 0 613338.2 y Ammonia loss 13
246.1287384033203 0 54110.36
252.13536071777344 0 24334.113
253.1305694580078 0 24802.643
253.22801208496094 0 78219.43
254.11427307128906 0 135083.03
254.16163635253906 0 64460.34
254.61654663085938 0 18546.31
255.1094970703125 0 515858.66
255.12258911132812 0 34551.492
255.14527893066406 0 42452.75
256.1131591796875 0 51853.13
259.140380859375 0 122237.15
261.1603088378906 0 24728.541
262.11846923828125 0 21614.975
262.151611328125 0 595542.9 y 13
263.1540832519531 0 57130.207
266.1131286621094 0 19428.834
266.1509704589844 0 24380.455
268.12969970703125 0 28408.203
269.18695068359375 0 47248.715
270.1454162597656 0 90108.52
271.12762451171875 0 26664.664
271.14208984375 0 37876.57
272.13531494140625 0 69081.664
274.1412658691406 0 20801.504
279.10919189453125 0 42591.312
280.1297607421875 0 50924.633
281.1254577636719 0 63354.56
281.2232360839844 0 64866.59
282.109375 0 23961.934
282.1809387207031 0 19318.371
283.177490234375 0 30379.857
283.2020263671875 0 41414.742
284.1249694824219 0 109457.56
284.16143798828125 0 94001.76
284.1970520019531 0 25692.371
288.1562805175781 0 115780.2
289.1517333984375 0 62245.574
293.18682861328125 0 25118.363
294.145751953125 0 39080.27
295.14105224609375 0 58509.957
295.17755126953125 0 77427.84
296.1363220214844 0 82889.58
296.1744689941406 0 60683.848
296.19769287109375 0 691752.4 b Water loss 2
297.1199645996094 0 80188.01
297.2009582519531 0 105717.195
298.1407470703125 0 235707.45
299.1362609863281 0 159729.22
302.1352844238281 0 82378.58
302.1724548339844 0 51723.62
306.15606689453125 0 33924.164
307.1419677734375 0 31356.53
307.1617431640625 0 60013.234
311.13641357421875 0 55404.18
312.1343078613281 0 26477.945
312.15631103515625 0 499573.25
313.153564453125 0 135097.42
313.18798828125 0 1199842.2
314.14697265625 0 228303.55
314.190673828125 0 121634.32
314.2084045410156 0 816221 b 2
315.13079833984375 0 29293.09
315.150390625 0 21754.074
315.211669921875 0 123135.03
316.15118408203125 0 755172.1
317.1503601074219 0 131986.81
323.13543701171875 0 36295.48
323.171875 0 56799.89
324.13104248046875 0 148902.55
324.1676025390625 0 74596.02
325.1157531738281 0 23367.746
326.1348876953125 0 29836.27
326.2443542480469 0 17243.21
329.1470947265625 0 69861.41
330.1670227050781 0 300024.56
331.17156982421875 0 168704.53 y Water loss 12
332.1574401855469 0 686332.3 y Ammonia loss 12
333.1606750488281 0 95454.84
335.135986328125 0 66015.21
336.1776123046875 0 33470.887 y Water loss 8
340.15087890625 0 82588.82
340.2012939453125 0 18748.732
341.15643310546875 0 42671.535
341.1830139160156 0 616642.94
342.14178466796875 0 800495.9
342.18408203125 0 73401.125
343.14434814453125 0 119697.914
344.14642333984375 0 23703.602
345.1830139160156 0 68175.836 y 8
345.6856689453125 0 29476.71
349.1540832519531 0 37884.293
349.18389892578125 0 557401.7 y 12
350.18646240234375 0 87090.93
352.1250915527344 0 45869.105
353.1936950683594 0 67560.945
354.1417541503906 0 26547.703
355.16326904296875 0 21577.908
357.1799011230469 0 22281.117
358.1620788574219 0 60743.582
359.16766357421875 0 240871.66
359.1919860839844 0 28508.443
365.1822814941406 0 46848.508
367.1612548828125 0 34226.7
368.25640869140625 0 37996.074
368.6606750488281 0 28439.352
368.69091796875 0 37376.414
370.2202453613281 0 32188.19
377.1492919921875 0 19735.07
379.2347106933594 0 20789.629
380.157958984375 0 75814.99
381.1892395019531 0 96412.086
382.21099853515625 0 31224.533
383.2281188964844 0 25247.482
385.1723937988281 0 134413.28
392.1947326660156 0 27162.498
393.71044921875 0 109332.375
394.2109680175781 0 46530.96
395.15545654296875 0 29432.518
396.1893615722656 0 63543.047
397.24542236328125 0 77223.66
398.1690673828125 0 21995.844
398.2135925292969 0 85813.71
398.24737548828125 0 22346.744
399.16363525390625 0 52017.223
399.7178649902344 0 29749.266
400.1886291503906 0 39091.316
400.22021484375 0 142059.03
401.2402648925781 0 71503.27
403.1833190917969 0 138505.72
406.2145690917969 0 24322.314
407.23040771484375 0 53012.8
408.1530456542969 0 27975.951
408.2286376953125 0 63524.29
408.7304992675781 0 56820.29
410.2041320800781 0 94650.18
411.16656494140625 0 23803.158
411.2001037597656 0 62035.53
411.2254943847656 0 75533.86
413.1690368652344 0 33072.39
413.2126770019531 0 35371.758
414.2286376953125 0 284630.4 y Water loss 7
414.72308349609375 0 354256.66 y Ammonia loss 7
415.22308349609375 0 109410.625
415.256103515625 0 172077.14
416.21588134765625 0 34812.44
416.2579650878906 0 41680.605
422.22674560546875 0 31902.123
423.2342224121094 0 6468528.5 y 7
423.73541259765625 0 2621926.8
424.1885070800781 0 27852.242
424.2366638183594 0 480524.16
425.2405700683594 0 549491.6 b Water loss 3
426.2434997558594 0 96381.5
428.2160339355469 0 237929.38
429.235107421875 0 341067.84
430.2402648925781 0 155562.98 y Water loss 11
431.1732177734375 0 29185.771
431.2255554199219 0 61719.22 y Ammonia loss 11
439.1952819824219 0 59098.7
441.2103271484375 0 336206.56
442.21429443359375 0 55686.184
443.2510986328125 0 622697.6 b 3
444.2545166015625 0 137975.45
448.2527160644531 0 232262.03 y 11
449.253173828125 0 39605.06
451.2351379394531 0 32001.793
452.2628479003906 0 238405.6
453.2660827636719 0 38552.156
455.22833251953125 0 27971.65
455.2612609863281 0 28362.3
458.2369079589844 0 169793.73
459.2386474609375 0 28559.21
469.2417297363281 0 24745.469
469.2489318847656 0 27842.693
469.289306640625 0 191234.06
470.2366638183594 0 66179.08
470.2915344238281 0 34162.504
471.74102783203125 0 39683.387 y Water loss 6
472.2378234863281 0 37003.617 y Ammonia loss 6
472.7318420410156 0 37873.25
473.23773193359375 0 23774.361
478.20635986328125 0 51343.977
479.225341796875 0 60449.74
480.25433349609375 0 60874.85
480.7477722167969 0 194387.34 y 6
481.2485656738281 0 121940.9
482.18939208984375 0 42048.125
487.2595520019531 0 57907.758 y Water loss 10
488.24871826171875 0 82116.85 y Ammonia loss 10
496.21551513671875 0 72754.695
496.3104248046875 0 37005.383
497.2377014160156 0 79228.74
497.2867431640625 0 89804.64
500.2005920410156 0 101365.35
505.27423095703125 0 599497.44 y 10
506.2764892578125 0 114865.39
507.7541198730469 0 23211.084
508.21563720703125 0 31425.396
514.2631225585938 0 49346.93
514.3241577148438 0 161192.31
515.254150390625 0 103824.9 y Water loss 5
515.3280029296875 0 37262.203
515.754638671875 0 53382.35 y Ammonia loss 5
516.259765625 0 57248.305
518.210205078125 0 125653.234
524.2628784179688 0 184552.6 y 5
524.3096313476562 0 98270.734 b Water loss 4
524.7639770507812 0 71082.51
529.2564697265625 0 32427.67
532.2730102539062 0 109000.52
533.257080078125 0 72573.12
534.2636108398438 0 24483.57 b Ammonia loss 9
535.2391967773438 0 25348.479
539.294921875 0 99762.37
540.2965087890625 0 31016.252
542.2547607421875 0 33072.766
542.2720336914062 0 27439.68
542.3200073242188 0 228809.2 b 4
543.322998046875 0 56067.797
547.22412109375 0 45075.918
549.2804565429688 0 56539.22
552.2417602539062 0 32311.254
556.3204956054688 0 68918.93
558.7883911132812 0 35130.844
559.2867431640625 0 37246.883
562.2952270507812 0 26782.506 b Water loss 10
564.7922973632812 0 108335.46 y Water loss 4
565.2373046875 0 85294.84
565.286865234375 0 101812.984 y Ammonia loss 4
565.78515625 0 61716.24
566.303955078125 0 63547.055
567.290771484375 0 103539.305
569.27099609375 0 34358.367
570.25390625 0 33343.727
572.2799682617188 0 29717.4
573.2813720703125 0 24105.469
573.797607421875 0 1430817.2 y 4
574.2984008789062 0 967736 y Water loss 9
574.8004760742188 0 247443.03
575.277587890625 0 56817.844 y Ammonia loss 9
583.2491455078125 0 73656.086
584.251953125 0 20605.57
584.267578125 0 21738.217
584.3169555664062 0 131294.73
585.306884765625 0 28669.541
587.2777709960938 0 32440.166
592.3063354492188 0 440171.44 y 9
593.3089599609375 0 160590.7
595.2828979492188 0 36437.43
596.3148193359375 0 25420.705
599.3185424804688 0 29519.68
600.2733154296875 0 41725.652
602.2868041992188 0 38012.785
603.294677734375 0 27118.102
608.314453125 0 27670.938
611.3223876953125 0 65672.65
612.3099975585938 0 21664.58 b Ammonia loss 11
623.3267822265625 0 25534.338
624.3110961914062 0 50281.473
626.3270263671875 0 131264.44
629.3267822265625 0 179456.58
629.8107299804688 0 68322.37 y Ammonia loss 3
630.3217163085938 0 64110.72
631.292724609375 0 27810.568
632.2923583984375 0 38192.18
636.3109741210938 0 56442.707
637.3028564453125 0 56005.96
638.3177490234375 0 221568.05 y 3
638.8214111328125 0 128687.35
639.3193969726562 0 67401.664
641.3375854492188 0 135181.3
642.3250122070312 0 96755.61
643.3519897460938 0 145262.88
644.3563842773438 0 46987.016
646.3135375976562 0 93735.695
647.3326416015625 0 21389.658
652.3126220703125 0 27993.084
653.337890625 0 117278.6
654.3242797851562 0 188299.64
655.3334350585938 0 42990.34 b Water loss 12
655.80224609375 0 26780.846
656.3060302734375 0 45938.516
657.2866821289062 0 87838.234
659.3501586914062 0 129941.11
660.3471069335938 0 49125.863
660.8186645507812 0 45911.04
661.3252563476562 0 37417.883
664.30908203125 0 73250.35
671.3480834960938 0 614535.8 y Water loss 8
672.3359985351562 0 626460.7 y Ammonia loss 8
672.8302001953125 0 94560.89 y Water loss 2
673.3319091796875 0 294699.2 y Ammonia loss 2
673.8244018554688 0 90573.75
674.31591796875 0 170293.4
675.3143310546875 0 72790.086
681.3305053710938 0 24468.43
681.8353271484375 0 450665.62 y 2
682.3312377929688 0 327357.56
682.8362426757812 0 79252.9
683.32666015625 0 28010.531
687.3425903320312 0 40171.15
689.3587646484375 0 3709650 y 8
690.3613891601562 0 1237629.9
691.3644409179688 0 222014.8
699.3441162109375 0 140729.23 b Ammonia loss 13
700.3470458984375 0 53561.004
716.3720092773438 0 40422.03
717.3644409179688 0 48757.223
726.3654174804688 0 106494.65 b Water loss 6
727.3748779296875 0 35110.82
728.3681030273438 0 37376.562
729.8589477539062 0 39192.715 y Ammonia loss 1
730.36474609375 0 32978.707
730.8716430664062 0 30271.035
736.3721313476562 0 32040.848
737.4054565429688 0 25899.266
738.3799438476562 0 198619.05 y 1
738.8776245117188 0 137897.94
739.3825073242188 0 77587.08
743.3700561523438 0 55623.79
744.3779907226562 0 525716.1 b 6
745.3803100585938 0 190619.08
746.38671875 0 37162.56
750.3930053710938 0 46204.867
751.3888549804688 0 40615.234
752.37255859375 0 58302.434
755.9020385742188 0 87891.39
756.3941040039062 0 29183.896
759.3927001953125 0 39178.508
761.903076171875 0 72222.98
762.4042358398438 0 104379.95
764.9033813476562 0 68732.086
765.3995971679688 0 68701.86
765.8965454101562 0 79446.03
766.38671875 0 244062.1
766.8883056640625 0 201824.7
767.3980102539062 0 105543.945
767.902099609375 0 30058.602
768.4008178710938 0 303126.38
768.9033203125 0 47381.5
769.3971557617188 0 149569.12
770.412109375 0 102153.63
770.9076538085938 0 123115.445
771.4122924804688 0 164647.58
771.9075317382812 0 98959.28
772.4153442382812 0 118777.125
772.9176025390625 0 105707.81
773.39501953125 0 211798.73
773.9030151367188 0 381988.7
774.4068603515625 0 239353.39
774.9078369140625 0 105789.11
776.9078369140625 0 228728.11
777.4036254882812 0 477966.1
777.9024047851562 0 266386.3
778.4029541015625 0 122696.08
778.904541015625 0 56584.375
779.4120483398438 0 128724.35
779.9132080078125 0 709738.7
780.4139404296875 0 699715.75
780.916259765625 0 327018.12
781.3953857421875 0 30352.871
785.4258422851562 0 236263.67
785.91357421875 0 2905319 Precursor Water loss
786.4147338867188 0 2897888.5 Precursor Ammonia loss
786.9143676757812 0 1428225.2
787.4100952148438 0 442095
787.9088134765625 0 128763.01
788.4036865234375 0 40850.324
793.4027099609375 0 205702.44
793.9100952148438 0 1347048.8
794.4117431640625 0 1186200.9
794.9195556640625 0 97010920 Precursor
795.4207153320312 0 82717020
795.9223022460938 0 36492384
796.4227294921875 0 3605487
796.9251708984375 0 33844.53
798.4189453125 0 88239.16
801.6460571289062 0 40645.562
803.4371948242188 0 251353.84
804.4426879882812 0 69077.69
810.42138671875 0 148944.84
811.4083251953125 0 231724.81
812.4110107421875 0 102208.1
827.4475708007812 0 78464.23 y Water loss 7
828.4335327148438 0 1128169.6 y Ammonia loss 7
829.4348754882812 0 427780.4
830.4330444335938 0 70571.71
845.4592895507812 0 976397.4 y 7
846.4609375 0 378142.47
847.4639892578125 0 53583.6
857.446533203125 0 68800.086
865.4274291992188 0 59238.918
872.483642578125 0 257433.95
873.4071655273438 0 44884.12
873.4874267578125 0 93400.305
874.400390625 0 31815.63
874.4884033203125 0 34136.316
882.4588012695312 0 63193.926 b Water loss 7
883.4476928710938 0 112827.45 b Ammonia loss 7
884.4341430664062 0 35775.26
885.4441528320312 0 53862.04
887.4264526367188 0 58784
891.4197998046875 0 129379.97
892.4222412109375 0 52909.715
899.46142578125 0 31903.977
900.4775390625 0 954459.6 b 7
901.4813842773438 0 377944.88
902.47998046875 0 92144.83
910.4283447265625 0 28565.213
915.4166870117188 0 115003.484
916.4212646484375 0 61804.055
918.474609375 0 58020.086
919.4739379882812 0 34240
925.4501953125 0 63378.15
926.4424438476562 0 79698.85
927.444580078125 0 66869.2
943.4598388671875 0 531718.5 y Ammonia loss 6
944.4664916992188 0 233020
945.4658813476562 0 68602.6
946.4815063476562 0 28423.258
953.4546508789062 0 27001.43
957.4771728515625 0 29133.477
960.481201171875 0 61441.363 y 6
968.474365234375 0 38005.516
969.476318359375 0 71585.46
970.4635620117188 0 40470.09
971.4881591796875 0 39051.766
972.4707641601562 0 79547.6
986.489501953125 0 159597.16
987.4871215820312 0 426814.72
988.487060546875 0 171767.11
989.4895629882812 0 60692.316
990.4876708984375 0 88531.625
991.4927978515625 0 41608.652
994.4653930664062 0 42573.566
995.46533203125 0 31512.336
996.4714965820312 0 41669.016
997.4658813476562 0 36688.945
1000.4818725585938 0 68269.74
1002.5022583007812 0 46399.098
1005.4963989257812 0 131313.75
1006.5006713867188 0 79737.81
1012.4814453125 0 268969.25
1013.4815673828125 0 150309
1014.4854125976562 0 159098.92
1015.4901733398438 0 60020.285
1029.4991455078125 0 48838.46 y Water loss 5
1030.4918212890625 0 2119571 y Ammonia loss 5
1031.494140625 0 1076503.6
1032.496826171875 0 323464.66
1047.517333984375 0 109469.35 y 5
1048.5157470703125 0 42331.207
1056.5628662109375 0 31607.08
1057.5626220703125 0 26810.322
1067.5478515625 0 29309.102 b Ammonia loss 9
1068.5321044921875 0 35188.47
1069.546630859375 0 32490.045
1073.5208740234375 0 74291.63
1074.5244140625 0 32839.977
1082.5498046875 0 28893.398
1083.5181884765625 0 59396.727
1084.5587158203125 0 107502.55 b 9
1085.5533447265625 0 42428.85
1086.5543212890625 0 213143.3
1087.5546875 0 113849.63
1088.5465087890625 0 42223.973
1099.562255859375 0 56744.82
1100.5732421875 0 30244.352
1101.5208740234375 0 138673.03
1102.5247802734375 0 43961.69
1104.570556640625 0 38113.402
1105.5682373046875 0 31223.885
1111.548828125 0 74198.44
1112.5391845703125 0 68888.555
1113.57568359375 0 59616.27
1119.52783203125 0 51534.934
1120.526611328125 0 33304.832
1127.5821533203125 0 41301.332
1129.5595703125 0 857262.6 y Ammonia loss 4
1130.562255859375 0 440193.2
1131.56494140625 0 135837.38
1141.583984375 0 133065.36 b 10
1142.5855712890625 0 54528.92
1145.0875244140625 0 26092.238
1146.57666015625 0 43168.29 y 4
1147.5997314453125 0 35102.69
1157.8621826171875 0 27515.965
1160.5538330078125 0 54803.543
1188.5531005859375 0 79998.69
1189.54833984375 0 64682.168
1197.586669921875 0 42110.535
1206.561767578125 0 183612.98
1207.5615234375 0 88685.73
1208.5572509765625 0 28844.582
1212.6595458984375 0 251935.17
1213.6636962890625 0 175524.48
1214.65673828125 0 60737.035
1215.596923828125 0 99219.72
1216.5965576171875 0 74661.06
1223.6346435546875 0 34409.832 b Ammonia loss 11
1233.60791015625 0 36655.004
1240.64453125 0 254899.89 b 11
1241.646728515625 0 178306.42
1242.64453125 0 59789.633
1255.612060546875 0 29849.773
1258.6031494140625 0 404581.38 y Ammonia loss 3
1259.603515625 0 247549.73
1260.603271484375 0 63751.07
1272.61279296875 0 40317.41
1273.6319580078125 0 37062.707
1274.630859375 0 38522.82
1275.6300048828125 0 35310.19 y 3
1283.687744140625 0 40297.074
1284.6226806640625 0 72702.08
1285.624267578125 0 109652.516
1286.6258544921875 0 33864.418
1299.687744140625 0 121569.89
1300.683349609375 0 46148.824
1302.6263427734375 0 587210.6
1303.62841796875 0 396914.16
1304.6300048828125 0 154730.55
1309.6505126953125 0 65951.56 b Water loss 12
1310.6539306640625 0 62510.83 b Ammonia loss 12
1315.6158447265625 0 57388.066
1316.6270751953125 0 47700.613
1319.64697265625 0 32660.46
1320.635498046875 0 130688.19
1321.6417236328125 0 61735.062
1326.678466796875 0 27789.266
1327.6395263671875 0 276679.22
1328.6439208984375 0 244668.8
1329.6329345703125 0 94763.17
1345.633544921875 0 2111072 y Ammonia loss 2
1346.6356201171875 0 1407145.2
1347.63818359375 0 511668.62
1362.662109375 0 107036.6 y 2
1363.6595458984375 0 58361.1
1368.70458984375 0 52144.992
1369.7135009765625 0 46759.598
1370.724609375 0 39229.39
1384.7037353515625 0 54535.684
1385.7015380859375 0 51083.914
1386.7193603515625 0 394340.75
1387.7232666015625 0 289434.53
1388.7215576171875 0 86163.29
1396.7041015625 0 92053.2 b Water loss 13
1397.7061767578125 0 97867.72 b Ammonia loss 13
1398.7154541015625 0 32848.73
1402.698486328125 0 27780.178
1412.6986083984375 0 45806.223
1414.7135009765625 0 432654.1 b 13
1415.71533203125 0 385129.56
1416.7183837890625 0 151015.45
1417.7108154296875 0 32833.83
1431.744873046875 0 35573.492
1432.726806640625 0 545321.2
1433.7286376953125 0 424665.8
1434.7291259765625 0 137094.22
1458.7099609375 0 179001.5 y Ammonia loss 1
1459.720703125 0 114685.35
3074.689208984375 0 26170.428
3273.99365234375 0 29411.18

Spectrum Details

|  |  |
| --- | --- |
| Matched peaks? Matched peaks The total absolute number of peaks matched. Additionally in brackets the total fraction of peaks matched and the total number of peaks is shown. | 87 (12.48% of 697) |
| FDR? FDR The false discovery rate estimated for this peptide. It is calculated by matching all theoretical fragments with a non-integer shift with the raw peaks for this spectrum. This is done with 40 different shifts. The resulting percentage is the average number of annotated peaks over the number of annotated peaks with the correct spectrum. | 0.55% |
| Satellite FDR? Satellite FDR See the FDR for details on its calculation. This satellite ion specific FDR only contains the satellite ions (d/w) for I/L/J positions. | - |
| PSM Score? PSM Score The PSM Score as given by Hecklib to this annotated spectrum. It is shown with three significant figures. | 972 |

#### Spectrum 3773? Spectrum 3773 The raw spectrum of this peptide as annotated by Hecklib. The fragments are coloured according to ion type (see legend). Any peaks with a star '\*' as text can be hovered over to see the full details, first the ion type second the mass shift type. By hovering over the amino acids in the peptide or ions in the legend the corresponding peaks are highlighted. By toggling the 'Unassigned' label you can turn the background (unassigned) peaks on or off in the plot. By updating the slider in the Ion legend you can update the spectrum to only show the top X% of the peaks with labels. The top X% means any peak that is within X% of the highest intensity. By dragging in the spectrum you can zoom in to a specific part of the spectrum and use 'Zoom Out' to get back to the original zoom level. The annotation of the spectrum is based on the given sequence in the peptides file and is done with different software so inconsistencies are likely. The peaks are annotated based on the given sequence, with 20 ppm tolerance.

Copy Data

##### Spectrum 3773 (TSV)

###### Preview

```
Loading example...
```

*Click on the button to copy the data to your clipboard.*

Mz MinMz MaxIntensity Max

WidthHeightPeptide font sizePeptide stroke widthSpectrum font sizeSpectrum stroke widthCompact peptide

Ion legend

wxyz

abcd

OtherUnassignedIon^Charge^_Position_Show for top:%

JJSEVSDRPSGVSSR

06.80e+51.36e+62.04e+62.72e+6

Zoom Out

d^+1^_2_y^+1^_1_y^+2^_3_y^+3^_5_y^+1^_1_a^+1^_2_b^+1^_2_y^+1^_2_y^+1^_2_y^+1^_2_b^+1^_3_b^+1^_3_y^+1^_3_y^+1^_3_y^+2^_7_y^+1^_3_b^+3^_12_y^+2^_8_y^+2^_8_b^+1^_4_y^+1^_4_b^+1^_4_y^+1^_4_y^+3^_13_b^+2^_8_y^+3^_13_y^+2^_9_b^+3^_14_y^+2^_9_y^+3^_14_y^+3^_14_y^+1^_5_y^+1^_5_y^+3^_14_y^+1^_5_y^+2^_10_y^+2^_10_y^+2^_10_\*\*b^+1^_5_b^+2^_11_y^+2^_11_y^+2^_11_y^+2^_11_y^+1^_6_y^+1^_6_y^+2^_12_y^+2^_12_y^+2^_12_y^+1^_7_y^+1^_7_y^+2^_13_y^+2^_13_y^+2^_13_y^+1^_7_b^+2^_14_y^+2^_14_y^+2^_14_y^+2^_14_b^+1^_7_\*y^+1^_8_b^+1^_8_y^+1^_9_y^+1^_10_y^+1^_11_

034068010201360

Fragment Matches Table

Show background peaks

| Position | Ion type | Intensity | mz Theoretical | mz Error (Th) | mz Error (ppm) | Charge | Series Number |
| --- | --- | --- | --- | --- | --- | --- | --- |
| - | - | 2072 | 120.1 | - | - | 0 | - |
| - | - | 1.969E+04 | 120.1 | - | - | 0 | - |
| - | - | 1435 | 121.1 | - | - | 0 | - |
| - | - | 1001 | 125.1 | - | - | 0 | - |
| - | - | 1106 | 125.1 | - | - | 0 | - |
| - | - | 2.359E+04 | 126.1 | - | - | 0 | - |
| - | - | 1565 | 127.1 | - | - | 0 | - |
| - | - | 948.5 | 127.1 | - | - | 0 | - |
| - | - | 1967 | 128.1 | - | - | 0 | - |
| - | - | 3265 | 129.1 | - | - | 0 | - |
| - | - | 2.281E+04 | 129.1 | - | - | 0 | - |
| - | - | 1876 | 129.1 | - | - | 0 | - |
| - | - | 2336 | 130.1 | - | - | 0 | - |
| - | - | 1.2E+04 | 130.1 | - | - | 0 | - |
| - | - | 7417 | 130.1 | - | - | 0 | - |
| - | - | 2329 | 130.1 | - | - | 0 | - |
| - | - | 1455 | 131.1 | - | - | 0 | - |
| - | - | 3344 | 131.1 | - | - | 0 | - |
| - | - | 1009 | 133.1 | - | - | 0 | - |
| - | - | 3069 | 133.1 | - | - | 0 | - |
| - | - | 9137 | 136.1 | - | - | 0 | - |
| - | - | 1702 | 138.1 | - | - | 0 | - |
| - | - | 997.5 | 138.7 | - | - | 0 | - |
| - | - | 1286 | 139.1 | - | - | 0 | - |
| - | - | 4484 | 140.1 | - | - | 0 | - |
| - | - | 2028 | 140.1 | - | - | 0 | - |
| - | - | 1330 | 141.1 | - | - | 0 | - |
| - | - | 2746 | 141.1 | - | - | 0 | - |
| - | - | 1372 | 143.1 | - | - | 0 | - |
| - | - | 6173 | 144.1 | - | - | 0 | - |
| - | - | 1429 | 145.1 | - | - | 0 | - |
| - | - | 7189 | 147.1 | - | - | 0 | - |
| - | - | 2196 | 148.9 | - | - | 0 | - |
| - | - | 1237 | 152.1 | - | - | 0 | - |
| - | - | 4138 | 153.1 | - | - | 0 | - |
| - | - | 2.825E+04 | 154.1 | - | - | 0 | - |
| - | - | 1603 | 155.1 | - | - | 0 | - |
| - | - | 1948 | 155.1 | - | - | 0 | - |
| - | - | 1056 | 156.1 | - | - | 0 | - |
| - | - | 2087 | 156.1 | - | - | 0 | - |
| - | - | 7406 | 157.1 | - | - | 0 | - |
| - | - | 1.166E+04 | 157.1 | - | - | 0 | - |
| - | - | 1.115E+04 | 157.1 | - | - | 0 | - |
| 2 | d | 4980 | 157.1 | 0.0003563 | 2.267 | +1 | 2 |
| - | - | 1230 | 158.1 | - | - | 0 | - |
| 15 | y | 5.215E+04 | 158.1 | 0.0004009 | 2.536 | +1 | 1 |
| - | - | 1778 | 159.1 | - | - | 0 | - |
| - | - | 2805 | 159.1 | - | - | 0 | - |
| - | - | 7882 | 159.1 | - | - | 0 | - |
| - | - | 1373 | 160.1 | - | - | 0 | - |
| - | - | 2721 | 165.1 | - | - | 0 | - |
| - | - | 1161 | 165.1 | - | - | 0 | - |
| 13 | y | 1325 | 166.1 | 0.003313 | 19.94 | +2 | 3 |
| - | - | 4059 | 167.1 | - | - | 0 | - |
| 11 | y | 2463 | 169.1 | 0.001446 | 8.553 | +3 | 5 |
| - | - | 1657 | 170 | - | - | 0 | - |
| - | - | 3.209E+04 | 171.1 | - | - | 0 | - |
| - | - | 4217 | 172.1 | - | - | 0 | - |
| - | - | 1670 | 172.1 | - | - | 0 | - |
| - | - | 8116 | 173.1 | - | - | 0 | - |
| - | - | 1473 | 173.4 | - | - | 0 | - |
| - | - | 1.723E+04 | 175.1 | - | - | 0 | - |
| 15 | y | 1.149E+05 | 175.1 | 0.0004021 | 2.296 | +1 | 1 |
| - | - | 7543 | 176.1 | - | - | 0 | - |
| - | - | 2496 | 180 | - | - | 0 | - |
| - | - | 2012 | 181.1 | - | - | 0 | - |
| - | - | 1358 | 181.1 | - | - | 0 | - |
| - | - | 3600 | 182.1 | - | - | 0 | - |
| - | - | 2.302E+04 | 183.1 | - | - | 0 | - |
| - | - | 1993 | 183.1 | - | - | 0 | - |
| - | - | 3368 | 184.1 | - | - | 0 | - |
| - | - | 2723 | 184.1 | - | - | 0 | - |
| - | - | 1527 | 185.1 | - | - | 0 | - |
| - | - | 1.8E+04 | 185.1 | - | - | 0 | - |
| - | - | 2258 | 185.1 | - | - | 0 | - |
| - | - | 4168 | 185.1 | - | - | 0 | - |
| - | - | 2315 | 185.2 | - | - | 0 | - |
| - | - | 1389 | 186.1 | - | - | 0 | - |
| - | - | 1.667E+04 | 187.1 | - | - | 0 | - |
| - | - | 1.278E+05 | 189.1 | - | - | 0 | - |
| - | - | 8673 | 190.1 | - | - | 0 | - |
| - | - | 1294 | 191.6 | - | - | 0 | - |
| - | - | 1919 | 192.1 | - | - | 0 | - |
| - | - | 1623 | 192.1 | - | - | 0 | - |
| - | - | 2287 | 194.1 | - | - | 0 | - |
| - | - | 1329 | 194.1 | - | - | 0 | - |
| - | - | 1812 | 195.1 | - | - | 0 | - |
| - | - | 1678 | 195.1 | - | - | 0 | - |
| - | - | 2874 | 195.1 | - | - | 0 | - |
| - | - | 4294 | 195.1 | - | - | 0 | - |
| - | - | 2581 | 197.2 | - | - | 0 | - |
| - | - | 1.333E+04 | 199.1 | - | - | 0 | - |
| 2 | a | 1.078E+06 | 199.2 | 0.0004489 | 2.254 | +1 | 2 |
| - | - | 1.317E+05 | 200.2 | - | - | 0 | - |
| - | - | 4.022E+04 | 201.1 | - | - | 0 | - |
| - | - | 8431 | 201.2 | - | - | 0 | - |
| - | - | 7424 | 202.1 | - | - | 0 | - |
| - | - | 3637 | 202.1 | - | - | 0 | - |
| - | - | 2.147E+04 | 203.1 | - | - | 0 | - |
| - | - | 3048 | 203.1 | - | - | 0 | - |
| - | - | 2097 | 209.1 | - | - | 0 | - |
| - | - | 1437 | 209.1 | - | - | 0 | - |
| - | - | 1424 | 209.1 | - | - | 0 | - |
| - | - | 2975 | 210.1 | - | - | 0 | - |
| - | - | 8166 | 211.1 | - | - | 0 | - |
| - | - | 8806 | 212.1 | - | - | 0 | - |
| - | - | 1641 | 213.1 | - | - | 0 | - |
| - | - | 1642 | 213.2 | - | - | 0 | - |
| - | - | 1703 | 215.1 | - | - | 0 | - |
| - | - | 1706 | 215.1 | - | - | 0 | - |
| - | - | 3447 | 216.1 | - | - | 0 | - |
| - | - | 2.14E+05 | 217.1 | - | - | 0 | - |
| - | - | 1.835E+04 | 218.1 | - | - | 0 | - |
| - | - | 2205 | 219.1 | - | - | 0 | - |
| - | - | 1638 | 220.1 | - | - | 0 | - |
| - | - | 3477 | 220.1 | - | - | 0 | - |
| - | - | 1574 | 223.1 | - | - | 0 | - |
| - | - | 1.513E+04 | 224.1 | - | - | 0 | - |
| - | - | 2902 | 224.1 | - | - | 0 | - |
| - | - | 1954 | 225.1 | - | - | 0 | - |
| - | - | 5480 | 226.1 | - | - | 0 | - |
| - | - | 1.828E+04 | 226.2 | - | - | 0 | - |
| - | - | 1721 | 227.1 | - | - | 0 | - |
| - | - | 9332 | 227.1 | - | - | 0 | - |
| 2 | b | 3.163E+05 | 227.2 | 0.0004531 | 1.995 | +1 | 2 |
| - | - | 2595 | 228.1 | - | - | 0 | - |
| - | - | 4.262E+04 | 228.2 | - | - | 0 | - |
| - | - | 7254 | 229.1 | - | - | 0 | - |
| - | - | 3747 | 229.2 | - | - | 0 | - |
| - | - | 1281 | 230.1 | - | - | 0 | - |
| - | - | 6637 | 230.1 | - | - | 0 | - |
| - | - | 2.174E+04 | 233.2 | - | - | 0 | - |
| - | - | 1980 | 234.1 | - | - | 0 | - |
| - | - | 3907 | 234.2 | - | - | 0 | - |
| - | - | 1713 | 235.1 | - | - | 0 | - |
| - | - | 3642 | 237.1 | - | - | 0 | - |
| - | - | 3030 | 237.1 | - | - | 0 | - |
| - | - | 6337 | 237.1 | - | - | 0 | - |
| - | - | 1404 | 239.1 | - | - | 0 | - |
| - | - | 2.786E+04 | 242.1 | - | - | 0 | - |
| - | - | 1606 | 242.2 | - | - | 0 | - |
| - | - | 3029 | 243.1 | - | - | 0 | - |
| - | - | 1273 | 243.1 | - | - | 0 | - |
| - | - | 7686 | 244.1 | - | - | 0 | - |
| 14 | y | 2769 | 244.1 | 0.0009415 | 3.857 | +1 | 2 |
| 14 | y | 3.295E+04 | 245.1 | 0.0005838 | 2.382 | +1 | 2 |
| - | - | 2901 | 246.1 | - | - | 0 | - |
| - | - | 1.036E+04 | 249.2 | - | - | 0 | - |
| - | - | 2097 | 250.2 | - | - | 0 | - |
| - | - | 1748 | 252.1 | - | - | 0 | - |
| - | - | 1665 | 252.1 | - | - | 0 | - |
| - | - | 2553 | 253.2 | - | - | 0 | - |
| - | - | 4022 | 254.1 | - | - | 0 | - |
| - | - | 1773 | 254.2 | - | - | 0 | - |
| - | - | 2.503E+04 | 255.1 | - | - | 0 | - |
| - | - | 3834 | 256.1 | - | - | 0 | - |
| - | - | 2085 | 256.1 | - | - | 0 | - |
| - | - | 4987 | 259.1 | - | - | 0 | - |
| - | - | 9765 | 261.2 | - | - | 0 | - |
| 14 | y | 2.122E+04 | 262.2 | 0.0003561 | 1.358 | +1 | 2 |
| - | - | 1923 | 262.2 | - | - | 0 | - |
| - | - | 2296 | 263.2 | - | - | 0 | - |
| - | - | 1773 | 267.1 | - | - | 0 | - |
| - | - | 5356 | 270.1 | - | - | 0 | - |
| - | - | 1849 | 271.1 | - | - | 0 | - |
| - | - | 5017 | 272.1 | - | - | 0 | - |
| - | - | 4942 | 277.2 | - | - | 0 | - |
| - | - | 1396 | 278.1 | - | - | 0 | - |
| - | - | 1990 | 279.1 | - | - | 0 | - |
| - | - | 1217 | 280.1 | - | - | 0 | - |
| - | - | 3361 | 280.1 | - | - | 0 | - |
| - | - | 1420 | 281.1 | - | - | 0 | - |
| - | - | 1395 | 283.2 | - | - | 0 | - |
| - | - | 5178 | 284.1 | - | - | 0 | - |
| - | - | 3358 | 284.2 | - | - | 0 | - |
| - | - | 4053 | 288.2 | - | - | 0 | - |
| - | - | 1854 | 289.2 | - | - | 0 | - |
| - | - | 1342 | 292.2 | - | - | 0 | - |
| - | - | 2936 | 296.1 | - | - | 0 | - |
| 3 | b | 1.639E+04 | 296.2 | 0.0005501 | 1.857 | +1 | 3 |
| - | - | 3463 | 297.1 | - | - | 0 | - |
| - | - | 1824 | 297.2 | - | - | 0 | - |
| - | - | 2176 | 298.1 | - | - | 0 | - |
| - | - | 9879 | 298.1 | - | - | 0 | - |
| - | - | 7168 | 299.1 | - | - | 0 | - |
| - | - | 2533 | 302.1 | - | - | 0 | - |
| - | - | 1854 | 302.2 | - | - | 0 | - |
| - | - | 1780 | 307.2 | - | - | 0 | - |
| - | - | 1668 | 311.1 | - | - | 0 | - |
| - | - | 1.513E+04 | 312.2 | - | - | 0 | - |
| - | - | 3762 | 313.2 | - | - | 0 | - |
| - | - | 2034 | 313.2 | - | - | 0 | - |
| - | - | 3.659E+04 | 313.2 | - | - | 0 | - |
| - | - | 1.173E+04 | 314.1 | - | - | 0 | - |
| - | - | 5174 | 314.2 | - | - | 0 | - |
| 3 | b | 3.513E+04 | 314.2 | 0.0007886 | 2.51 | +1 | 3 |
| - | - | 7147 | 315.2 | - | - | 0 | - |
| - | - | 3.876E+04 | 316.2 | - | - | 0 | - |
| - | - | 2442 | 316.2 | - | - | 0 | - |
| - | - | 6157 | 317.1 | - | - | 0 | - |
| - | - | 2307 | 319.2 | - | - | 0 | - |
| - | - | 1760 | 323.2 | - | - | 0 | - |
| - | - | 7139 | 323.2 | - | - | 0 | - |
| - | - | 6931 | 324.1 | - | - | 0 | - |
| - | - | 1269 | 324.2 | - | - | 0 | - |
| - | - | 2049 | 326.1 | - | - | 0 | - |
| - | - | 2278 | 329.1 | - | - | 0 | - |
| - | - | 9071 | 330.2 | - | - | 0 | - |
| 13 | y | 5733 | 331.2 | 0.001241 | 3.746 | +1 | 3 |
| 13 | y | 3.041E+04 | 332.2 | 0.0007056 | 2.124 | +1 | 3 |
| - | - | 4222 | 333.2 | - | - | 0 | - |
| - | - | 1854 | 339.2 | - | - | 0 | - |
| - | - | 2507 | 340.2 | - | - | 0 | - |
| - | - | 2.521E+04 | 341.2 | - | - | 0 | - |
| - | - | 1727 | 341.2 | - | - | 0 | - |
| - | - | 3.746E+04 | 342.1 | - | - | 0 | - |
| - | - | 3305 | 342.2 | - | - | 0 | - |
| - | - | 4858 | 343.1 | - | - | 0 | - |
| - | - | 1781 | 343.2 | - | - | 0 | - |
| 9 | y | 1.525E+04 | 345.2 | 0.0003837 | 1.111 | +2 | 7 |
| - | - | 6036 | 345.7 | - | - | 0 | - |
| - | - | 2312 | 349.1 | - | - | 0 | - |
| 13 | y | 2.546E+04 | 349.2 | 0.0006458 | 1.849 | +1 | 3 |
| - | - | 4720 | 350.2 | - | - | 0 | - |
| - | - | 1606 | 352.2 | - | - | 0 | - |
| - | - | 1841 | 353.2 | - | - | 0 | - |
| - | - | 1667 | 358.2 | - | - | 0 | - |
| - | - | 1.585E+04 | 359.2 | - | - | 0 | - |
| - | - | 2528 | 359.2 | - | - | 0 | - |
| - | - | 2617 | 360.2 | - | - | 0 | - |
| - | - | 2507 | 363.2 | - | - | 0 | - |
| - | - | 2815 | 365.2 | - | - | 0 | - |
| - | - | 1765 | 366.2 | - | - | 0 | - |
| - | - | 1744 | 370.2 | - | - | 0 | - |
| - | - | 1899 | 380.2 | - | - | 0 | - |
| - | - | 3031 | 381.2 | - | - | 0 | - |
| - | - | 1476 | 382.2 | - | - | 0 | - |
| - | - | 1984 | 383.2 | - | - | 0 | - |
| - | - | 5412 | 385.2 | - | - | 0 | - |
| - | - | 1789 | 385.7 | - | - | 0 | - |
| - | - | 1406 | 386.2 | - | - | 0 | - |
| - | - | 1385 | 386.2 | - | - | 0 | - |
| - | - | 8583 | 389.7 | - | - | 0 | - |
| - | - | 6981 | 390.2 | - | - | 0 | - |
| - | - | 6509 | 390.7 | - | - | 0 | - |
| - | - | 6115 | 391.2 | - | - | 0 | - |
| - | - | 3.318E+04 | 391.7 | - | - | 0 | - |
| - | - | 1.761E+04 | 392.2 | - | - | 0 | - |
| - | - | 3726 | 392.7 | - | - | 0 | - |
| - | - | 4157 | 393.7 | - | - | 0 | - |
| - | - | 3577 | 394.2 | - | - | 0 | - |
| - | - | 3619 | 396.2 | - | - | 0 | - |
| - | - | 3834 | 397.2 | - | - | 0 | - |
| - | - | 1528 | 398.2 | - | - | 0 | - |
| - | - | 4740 | 398.2 | - | - | 0 | - |
| - | - | 1.08E+04 | 398.7 | - | - | 0 | - |
| - | - | 2366 | 399.2 | - | - | 0 | - |
| - | - | 6700 | 399.2 | - | - | 0 | - |
| - | - | 1.158E+04 | 399.7 | - | - | 0 | - |
| - | - | 1827 | 400.2 | - | - | 0 | - |
| - | - | 8619 | 400.2 | - | - | 0 | - |
| - | - | 3423 | 401.2 | - | - | 0 | - |
| - | - | 6551 | 403.2 | - | - | 0 | - |
| - | - | 1947 | 404.2 | - | - | 0 | - |
| - | - | 1911 | 405.7 | - | - | 0 | - |
| - | - | 2016 | 407.2 | - | - | 0 | - |
| - | - | 2075 | 408.2 | - | - | 0 | - |
| - | - | 2043 | 408.7 | - | - | 0 | - |
| - | - | 4519 | 410.2 | - | - | 0 | - |
| - | - | 1788 | 411.2 | - | - | 0 | - |
| - | - | 2769 | 411.2 | - | - | 0 | - |
| 12 | b | 8223 | 414.2 | 0.005446 | 13.15 | +3 | 12 |
| 8 | y | 2.247E+04 | 414.7 | 0.002653 | 6.398 | +2 | 8 |
| - | - | 7687 | 415.2 | - | - | 0 | - |
| - | - | 4563 | 415.3 | - | - | 0 | - |
| - | - | 2546 | 415.7 | - | - | 0 | - |
| - | - | 2112 | 416.2 | - | - | 0 | - |
| 8 | y | 2.327E+05 | 423.2 | 0.0008841 | 2.089 | +2 | 8 |
| - | - | 9.916E+04 | 423.7 | - | - | 0 | - |
| - | - | 2.259E+04 | 424.2 | - | - | 0 | - |
| - | - | 4987 | 424.7 | - | - | 0 | - |
| 4 | b | 1.705E+04 | 425.2 | 0.0009257 | 2.177 | +1 | 4 |
| - | - | 1863 | 426.2 | - | - | 0 | - |
| - | - | 5424 | 426.2 | - | - | 0 | - |
| - | - | 1554 | 428.2 | - | - | 0 | - |
| - | - | 6529 | 428.2 | - | - | 0 | - |
| - | - | 1.104E+04 | 429.2 | - | - | 0 | - |
| 12 | y | 7926 | 430.2 | 0.0004713 | 1.095 | +1 | 4 |
| - | - | 1710 | 431.2 | - | - | 0 | - |
| - | - | 2529 | 439.2 | - | - | 0 | - |
| - | - | 2593 | 440.2 | - | - | 0 | - |
| - | - | 1.383E+04 | 441.2 | - | - | 0 | - |
| - | - | 2351 | 442.2 | - | - | 0 | - |
| 4 | b | 2.021E+04 | 443.3 | 0.0009201 | 2.076 | +1 | 4 |
| - | - | 3661 | 444.3 | - | - | 0 | - |
| 12 | y | 7084 | 448.3 | 0.0001335 | 0.2978 | +1 | 4 |
| 3 | y | 3560 | 448.9 | 0.0005628 | 1.254 | +3 | 13 |
| - | - | 2269 | 449.6 | - | - | 0 | - |
| 8 | b | 4363 | 450.7 | 0.0006004 | 1.332 | +2 | 8 |
| - | - | 9024 | 452.3 | - | - | 0 | - |
| 3 | y | 9293 | 454.9 | 0.0005202 | 1.144 | +3 | 13 |
| - | - | 8662 | 455.2 | - | - | 0 | - |
| - | - | 1777 | 455.6 | - | - | 0 | - |
| - | - | 1.387E+04 | 458.2 | - | - | 0 | - |
| - | - | 1.293E+04 | 460.2 | - | - | 0 | - |
| - | - | 3411 | 461.2 | - | - | 0 | - |
| - | - | 3388 | 463.2 | - | - | 0 | - |
| - | - | 2121 | 465.3 | - | - | 0 | - |
| - | - | 2299 | 468.9 | - | - | 0 | - |
| - | - | 2480 | 469.2 | - | - | 0 | - |
| - | - | 6779 | 469.3 | - | - | 0 | - |
| - | - | 1645 | 471.2 | - | - | 0 | - |
| 7 | y | 2819 | 471.7 | 0.0009575 | 2.03 | +2 | 9 |
| 14 | b | 2962 | 472.2 | 0.006904 | 14.62 | +3 | 14 |
| - | - | 1751 | 472.7 | - | - | 0 | - |
| - | - | 1812 | 473.8 | - | - | 0 | - |
| - | - | 2041 | 478.2 | - | - | 0 | - |
| 7 | y | 3.988E+04 | 480.7 | 0.0008403 | 1.748 | +2 | 9 |
| - | - | 1425 | 481.2 | - | - | 0 | - |
| - | - | 2.01E+04 | 481.2 | - | - | 0 | - |
| - | - | 8250 | 481.7 | - | - | 0 | - |
| - | - | 1602 | 485.2 | - | - | 0 | - |
| - | - | 2461 | 485.7 | - | - | 0 | - |
| 2 | y | 8949 | 486.6 | 0.0003328 | 0.684 | +3 | 14 |
| 2 | y | 9398 | 486.9 | 0.007441 | 15.28 | +3 | 14 |
| 11 | y | 5694 | 487.3 | 0.006188 | 12.7 | +1 | 5 |
| 11 | y | 5396 | 488.2 | 0.00116 | 2.376 | +1 | 5 |
| 2 | y | 6915 | 492.6 | 0.0003818 | 0.7751 | +3 | 14 |
| - | - | 5623 | 492.9 | - | - | 0 | - |
| - | - | 3019 | 493.3 | - | - | 0 | - |
| - | - | 1995 | 494.7 | - | - | 0 | - |
| - | - | 1.385E+04 | 495.9 | - | - | 0 | - |
| - | - | 2213 | 496.2 | - | - | 0 | - |
| - | - | 1.163E+04 | 496.3 | - | - | 0 | - |
| - | - | 3735 | 496.6 | - | - | 0 | - |
| - | - | 1787 | 496.9 | - | - | 0 | - |
| - | - | 1812 | 497.2 | - | - | 0 | - |
| - | - | 3982 | 497.3 | - | - | 0 | - |
| - | - | 1909 | 499.8 | - | - | 0 | - |
| - | - | 2974 | 500.2 | - | - | 0 | - |
| 11 | y | 2.573E+04 | 505.3 | 0.0009781 | 1.936 | +1 | 5 |
| - | - | 5581 | 506.3 | - | - | 0 | - |
| - | - | 1565 | 513.2 | - | - | 0 | - |
| - | - | 1842 | 514.3 | - | - | 0 | - |
| - | - | 3877 | 514.3 | - | - | 0 | - |
| 6 | y | 7928 | 515.3 | 0.002049 | 3.976 | +2 | 10 |
| 6 | y | 3884 | 515.7 | 0.006615 | 12.83 | +2 | 10 |
| - | - | 2226 | 516.3 | - | - | 0 | - |
| - | - | 3785 | 518.2 | - | - | 0 | - |
| - | - | 3432 | 518.3 | - | - | 0 | - |
| 6 | y | 2.375E+05 | 524.3 | 0.002374 | 4.528 | +2 | 10 |
| 0 | Precursor | 4.64E+04 | 524.6 | 0.006845 | 13.05 | +3 | -1 |
| - | - | 1.316E+05 | 524.8 | - | - | 0 | - |
| - | - | 2.788E+04 | 524.9 | - | - | 0 | - |
| - | - | 3.809E+04 | 525.3 | - | - | 0 | - |
| - | - | 9015 | 525.8 | - | - | 0 | - |
| - | - | 2016 | 526.3 | - | - | 0 | - |
| - | - | 2132 | 528.7 | - | - | 0 | - |
| - | - | 7565 | 528.8 | - | - | 0 | - |
| - | - | 4175 | 529.3 | - | - | 0 | - |
| - | - | 1717 | 529.4 | - | - | 0 | - |
| - | - | 9616 | 529.8 | - | - | 0 | - |
| 0 | Precursor | 3.264E+04 | 530.3 | 0.001556 | 2.934 | +3 | -1 |
| - | - | 2.504E+04 | 530.6 | - | - | 0 | - |
| - | - | 2491 | 530.8 | - | - | 0 | - |
| - | - | 1.352E+04 | 531 | - | - | 0 | - |
| - | - | 2143 | 531.3 | - | - | 0 | - |
| - | - | 2581 | 532.3 | - | - | 0 | - |
| - | - | 1924 | 533.3 | - | - | 0 | - |
| - | - | 1989 | 539.3 | - | - | 0 | - |
| - | - | 2574 | 540.2 | - | - | 0 | - |
| 5 | b | 6666 | 542.3 | 0.0008351 | 1.54 | +1 | 5 |
| - | - | 1773 | 543.3 | - | - | 0 | - |
| - | - | 2094 | 544.3 | - | - | 0 | - |
| - | - | 1978 | 552.8 | - | - | 0 | - |
| - | - | 1996 | 556.3 | - | - | 0 | - |
| - | - | 2277 | 556.3 | - | - | 0 | - |
| - | - | 2437 | 558.8 | - | - | 0 | - |
| - | - | 2821 | 559.3 | - | - | 0 | - |
| 11 | b | 3173 | 562.3 | 0.002407 | 4.28 | +2 | 11 |
| 5 | y | 7783 | 564.8 | 0.0008551 | 1.514 | +2 | 11 |
| - | - | 3318 | 565.2 | - | - | 0 | - |
| 5 | y | 1.689E+04 | 565.3 | 0.005245 | 9.278 | +2 | 11 |
| - | - | 8166 | 565.8 | - | - | 0 | - |
| - | - | 4437 | 566.3 | - | - | 0 | - |
| - | - | 2499 | 567.3 | - | - | 0 | - |
| - | - | 2200 | 569.3 | - | - | 0 | - |
| - | - | 1334 | 570.3 | - | - | 0 | - |
| - | - | 2623 | 572.8 | - | - | 0 | - |
| - | - | 2548 | 573.3 | - | - | 0 | - |
| 5 | y | 3.409E+05 | 573.8 | 0.0005764 | 1.005 | +2 | 11 |
| 10 | y | 2.084E+05 | 574.3 | 0.004295 | 7.478 | +1 | 6 |
| - | - | 7.25E+04 | 574.8 | - | - | 0 | - |
| - | - | 1.531E+04 | 575.3 | - | - | 0 | - |
| - | - | 1741 | 575.8 | - | - | 0 | - |
| - | - | 2734 | 583.2 | - | - | 0 | - |
| - | - | 3913 | 584.3 | - | - | 0 | - |
| - | - | 4853 | 587.3 | - | - | 0 | - |
| 10 | y | 3.331E+04 | 592.3 | 0.0008101 | 1.368 | +1 | 6 |
| - | - | 1.053E+04 | 593.3 | - | - | 0 | - |
| - | - | 2269 | 594.3 | - | - | 0 | - |
| - | - | 7819 | 600.3 | - | - | 0 | - |
| - | - | 2590 | 601.3 | - | - | 0 | - |
| - | - | 1748 | 608.3 | - | - | 0 | - |
| - | - | 2008 | 611.3 | - | - | 0 | - |
| - | - | 2808 | 617.3 | - | - | 0 | - |
| - | - | 2410 | 618.3 | - | - | 0 | - |
| - | - | 2171 | 619.3 | - | - | 0 | - |
| - | - | 1747 | 621.3 | - | - | 0 | - |
| - | - | 2416 | 624.3 | - | - | 0 | - |
| - | - | 4156 | 626.3 | - | - | 0 | - |
| 4 | y | 2.439E+04 | 629.3 | 0.002445 | 3.886 | +2 | 12 |
| 4 | y | 1.612E+04 | 629.8 | 0.0094 | 14.93 | +2 | 12 |
| - | - | 5325 | 630.3 | - | - | 0 | - |
| - | - | 5178 | 630.8 | - | - | 0 | - |
| - | - | 3764 | 636.3 | - | - | 0 | - |
| - | - | 1803 | 637.3 | - | - | 0 | - |
| - | - | 2387 | 637.8 | - | - | 0 | - |
| 4 | y | 1.675E+05 | 638.3 | 0.0005811 | 0.9104 | +2 | 12 |
| - | - | 1.072E+05 | 638.8 | - | - | 0 | - |
| - | - | 4.836E+04 | 639.3 | - | - | 0 | - |
| - | - | 1.253E+04 | 639.8 | - | - | 0 | - |
| - | - | 2612 | 640.3 | - | - | 0 | - |
| - | - | 5344 | 641.3 | - | - | 0 | - |
| - | - | 1625 | 642.3 | - | - | 0 | - |
| - | - | 1753 | 642.8 | - | - | 0 | - |
| - | - | 4784 | 643.4 | - | - | 0 | - |
| - | - | 2434 | 644.4 | - | - | 0 | - |
| - | - | 1.364E+04 | 646.3 | - | - | 0 | - |
| - | - | 3920 | 647.3 | - | - | 0 | - |
| - | - | 5363 | 648.8 | - | - | 0 | - |
| - | - | 3072 | 649.3 | - | - | 0 | - |
| - | - | 2520 | 651.8 | - | - | 0 | - |
| - | - | 4104 | 652.3 | - | - | 0 | - |
| - | - | 1846 | 652.8 | - | - | 0 | - |
| - | - | 7414 | 653.3 | - | - | 0 | - |
| - | - | 6861 | 654.3 | - | - | 0 | - |
| - | - | 2532 | 654.8 | - | - | 0 | - |
| - | - | 7850 | 655.3 | - | - | 0 | - |
| - | - | 1906 | 655.8 | - | - | 0 | - |
| - | - | 3077 | 656.3 | - | - | 0 | - |
| - | - | 4858 | 657.3 | - | - | 0 | - |
| - | - | 3833 | 657.8 | - | - | 0 | - |
| - | - | 7980 | 659.3 | - | - | 0 | - |
| - | - | 2375 | 659.8 | - | - | 0 | - |
| - | - | 2953 | 660.3 | - | - | 0 | - |
| - | - | 2104 | 660.4 | - | - | 0 | - |
| - | - | 1.043E+04 | 660.8 | - | - | 0 | - |
| - | - | 5302 | 661.3 | - | - | 0 | - |
| - | - | 1.287E+04 | 663.8 | - | - | 0 | - |
| - | - | 1.145E+04 | 664.3 | - | - | 0 | - |
| - | - | 6325 | 664.8 | - | - | 0 | - |
| - | - | 5247 | 665.3 | - | - | 0 | - |
| - | - | 4189 | 666.3 | - | - | 0 | - |
| - | - | 3.232E+04 | 666.8 | - | - | 0 | - |
| - | - | 1.727E+04 | 667.3 | - | - | 0 | - |
| - | - | 7041 | 667.8 | - | - | 0 | - |
| - | - | 2502 | 668.3 | - | - | 0 | - |
| 9 | y | 3.62E+04 | 671.3 | 0.0006641 | 0.9892 | +1 | 7 |
| - | - | 2233 | 671.8 | - | - | 0 | - |
| 9 | y | 2.76E+04 | 672.3 | 0.006822 | 10.15 | +1 | 7 |
| 3 | y | 9.528E+04 | 672.8 | 0.0005914 | 0.879 | +2 | 13 |
| 3 | y | 8.407E+04 | 673.3 | 0.00956 | 14.2 | +2 | 13 |
| - | - | 4.207E+04 | 673.8 | - | - | 0 | - |
| - | - | 4.455E+04 | 674.3 | - | - | 0 | - |
| - | - | 4571 | 674.8 | - | - | 0 | - |
| - | - | 1.546E+04 | 675.3 | - | - | 0 | - |
| - | - | 3506 | 676.3 | - | - | 0 | - |
| - | - | 2744 | 678.3 | - | - | 0 | - |
| - | - | 2.788E+04 | 680.8 | - | - | 0 | - |
| - | - | 2.101E+04 | 681.3 | - | - | 0 | - |
| 3 | y | 2.694E+06 | 681.8 | 0.001168 | 1.714 | +2 | 13 |
| - | - | 1.897E+06 | 682.3 | - | - | 0 | - |
| - | - | 7.916E+05 | 682.8 | - | - | 0 | - |
| - | - | 2.156E+05 | 683.3 | - | - | 0 | - |
| - | - | 4.705E+04 | 683.8 | - | - | 0 | - |
| - | - | 7482 | 685.4 | - | - | 0 | - |
| - | - | 2.125E+04 | 686.8 | - | - | 0 | - |
| - | - | 1.913E+04 | 687.3 | - | - | 0 | - |
| - | - | 6222 | 687.8 | - | - | 0 | - |
| - | - | 1813 | 688.3 | - | - | 0 | - |
| 9 | y | 2.121E+05 | 689.4 | 0.0007806 | 1.132 | +1 | 7 |
| - | - | 7.71E+04 | 690.4 | - | - | 0 | - |
| - | - | 1.707E+04 | 691.4 | - | - | 0 | - |
| 14 | b | 1.369E+04 | 699.3 | 0.005055 | 7.228 | +2 | 14 |
| - | - | 4207 | 700.3 | - | - | 0 | - |
| - | - | 1597 | 701.4 | - | - | 0 | - |
| 2 | y | 2.188E+04 | 729.4 | 0.0001244 | 0.1705 | +2 | 14 |
| 2 | y | 1.703E+04 | 729.9 | 0.01044 | 14.3 | +2 | 14 |
| - | - | 5044 | 730.4 | - | - | 0 | - |
| - | - | 1764 | 730.9 | - | - | 0 | - |
| - | - | 4526 | 737.4 | - | - | 0 | - |
| - | - | 2400 | 737.9 | - | - | 0 | - |
| 2 | y | 2.156E+05 | 738.4 | 0.0005793 | 0.7846 | +2 | 14 |
| - | - | 1.524E+05 | 738.9 | - | - | 0 | - |
| - | - | 7.686E+04 | 739.4 | - | - | 0 | - |
| - | - | 2.067E+04 | 739.9 | - | - | 0 | - |
| - | - | 5560 | 740.4 | - | - | 0 | - |
| - | - | 1.439E+04 | 743.4 | - | - | 0 | - |
| - | - | 7224 | 743.9 | - | - | 0 | - |
| 7 | b | 3989 | 744.4 | 0.004364 | 5.863 | +1 | 7 |
| - | - | 3184 | 758.4 | - | - | 0 | - |
| - | - | 1879 | 768.4 | - | - | 0 | - |
| - | - | 5418 | 768.4 | - | - | 0 | - |
| - | - | 2137 | 769.4 | - | - | 0 | - |
| - | - | 9476 | 770.4 | - | - | 0 | - |
| - | - | 3949 | 771.4 | - | - | 0 | - |
| - | - | 7593 | 778.4 | - | - | 0 | - |
| - | - | 4190 | 779.4 | - | - | 0 | - |
| - | - | 4385 | 780.4 | - | - | 0 | - |
| - | - | 2246 | 781.4 | - | - | 0 | - |
| - | - | 2.68E+04 | 782.4 | - | - | 0 | - |
| - | - | 9714 | 783.4 | - | - | 0 | - |
| - | - | 2393 | 784.4 | - | - | 0 | - |
| - | - | 6477 | 785.4 | - | - | 0 | - |
| - | - | 3708 | 786.4 | - | - | 0 | - |
| 0 | Precursor | 5429 | 786.4 | 0.01223 | 15.55 | +2 | -1 |
| - | - | 5475 | 787.4 | - | - | 0 | - |
| - | - | 2090 | 788.4 | - | - | 0 | - |
| - | - | 9531 | 796.4 | - | - | 0 | - |
| - | - | 5884 | 797.4 | - | - | 0 | - |
| - | - | 1.072E+04 | 798.4 | - | - | 0 | - |
| - | - | 6160 | 799.4 | - | - | 0 | - |
| - | - | 3873 | 803.4 | - | - | 0 | - |
| - | - | 2862 | 804.4 | - | - | 0 | - |
| - | - | 3410 | 810.4 | - | - | 0 | - |
| - | - | 7781 | 811.4 | - | - | 0 | - |
| - | - | 3690 | 812.4 | - | - | 0 | - |
| 8 | y | 3.215E+04 | 828.4 | 0.0006815 | 0.8227 | +1 | 8 |
| - | - | 1.412E+04 | 829.4 | - | - | 0 | - |
| - | - | 5172 | 830.4 | - | - | 0 | - |
| - | - | 1961 | 845.4 | - | - | 0 | - |
| - | - | 1751 | 856.4 | - | - | 0 | - |
| - | - | 3630 | 857.4 | - | - | 0 | - |
| - | - | 3902 | 858.4 | - | - | 0 | - |
| - | - | 2201 | 859.4 | - | - | 0 | - |
| - | - | 3640 | 867.4 | - | - | 0 | - |
| - | - | 3259 | 873.4 | - | - | 0 | - |
| - | - | 2757 | 881.4 | - | - | 0 | - |
| - | - | 4565 | 885.4 | - | - | 0 | - |
| - | - | 7608 | 887.4 | - | - | 0 | - |
| - | - | 4729 | 888.4 | - | - | 0 | - |
| - | - | 7059 | 891.4 | - | - | 0 | - |
| 8 | b | 7317 | 900.5 | 0.01105 | 12.28 | +1 | 8 |
| - | - | 3423 | 901.5 | - | - | 0 | - |
| - | - | 1.431E+04 | 915.4 | - | - | 0 | - |
| - | - | 5976 | 916.4 | - | - | 0 | - |
| - | - | 1893 | 917.4 | - | - | 0 | - |
| - | - | 4121 | 925.5 | - | - | 0 | - |
| - | - | 3618 | 926.4 | - | - | 0 | - |
| - | - | 4172 | 927.5 | - | - | 0 | - |
| 7 | y | 2.493E+04 | 943.5 | 0.001021 | 1.082 | +1 | 9 |
| - | - | 1.656E+04 | 944.5 | - | - | 0 | - |
| - | - | 4584 | 945.5 | - | - | 0 | - |
| - | - | 3959 | 968.5 | - | - | 0 | - |
| - | - | 4677 | 969.5 | - | - | 0 | - |
| - | - | 4437 | 970.5 | - | - | 0 | - |
| - | - | 2172 | 971.5 | - | - | 0 | - |
| - | - | 4349 | 972.5 | - | - | 0 | - |
| - | - | 3249 | 973.5 | - | - | 0 | - |
| - | - | 2.781E+04 | 986.5 | - | - | 0 | - |
| - | - | 2.395E+04 | 987.5 | - | - | 0 | - |
| - | - | 1.338E+04 | 988.5 | - | - | 0 | - |
| - | - | 4263 | 989.5 | - | - | 0 | - |
| - | - | 4376 | 990.5 | - | - | 0 | - |
| - | - | 2770 | 991.5 | - | - | 0 | - |
| - | - | 2112 | 994.5 | - | - | 0 | - |
| - | - | 4297 | 996.5 | - | - | 0 | - |
| - | - | 2789 | 997.5 | - | - | 0 | - |
| - | - | 2649 | 1000 | - | - | 0 | - |
| - | - | 2489 | 1001 | - | - | 0 | - |
| - | - | 2630 | 1006 | - | - | 0 | - |
| - | - | 1.209E+04 | 1012 | - | - | 0 | - |
| - | - | 8561 | 1013 | - | - | 0 | - |
| - | - | 1.951E+04 | 1014 | - | - | 0 | - |
| - | - | 1.295E+04 | 1015 | - | - | 0 | - |
| - | - | 3827 | 1016 | - | - | 0 | - |
| 6 | y | 7.771E+04 | 1030 | 0.000487 | 0.4726 | +1 | 10 |
| - | - | 3.864E+04 | 1031 | - | - | 0 | - |
| - | - | 1.326E+04 | 1032 | - | - | 0 | - |
| - | - | 3053 | 1033 | - | - | 0 | - |
| - | - | 2244 | 1056 | - | - | 0 | - |
| - | - | 2763 | 1058 | - | - | 0 | - |
| - | - | 2812 | 1072 | - | - | 0 | - |
| - | - | 8992 | 1074 | - | - | 0 | - |
| - | - | 3445 | 1075 | - | - | 0 | - |
| - | - | 2475 | 1076 | - | - | 0 | - |
| - | - | 5124 | 1084 | - | - | 0 | - |
| - | - | 3117 | 1085 | - | - | 0 | - |
| - | - | 6233 | 1087 | - | - | 0 | - |
| - | - | 1921 | 1088 | - | - | 0 | - |
| - | - | 2016 | 1089 | - | - | 0 | - |
| - | - | 1.364E+04 | 1102 | - | - | 0 | - |
| - | - | 6653 | 1103 | - | - | 0 | - |
| - | - | 3160 | 1112 | - | - | 0 | - |
| - | - | 3772 | 1113 | - | - | 0 | - |
| - | - | 4293 | 1120 | - | - | 0 | - |
| 5 | y | 2.242E+04 | 1130 | 0.001277 | 1.13 | +1 | 11 |
| - | - | 1.491E+04 | 1131 | - | - | 0 | - |
| - | - | 6749 | 1132 | - | - | 0 | - |
| - | - | 2632 | 1143 | - | - | 0 | - |
| - | - | 3042 | 1144 | - | - | 0 | - |
| - | - | 1.271E+04 | 1161 | - | - | 0 | - |
| - | - | 8279 | 1162 | - | - | 0 | - |
| - | - | 2683 | 1163 | - | - | 0 | - |
| - | - | 4894 | 1171 | - | - | 0 | - |
| - | - | 1.745E+04 | 1189 | - | - | 0 | - |
| - | - | 1.027E+04 | 1190 | - | - | 0 | - |
| - | - | 4687 | 1191 | - | - | 0 | - |
| - | - | 2172 | 1206 | - | - | 0 | - |
| - | - | 1.734E+04 | 1207 | - | - | 0 | - |
| - | - | 9902 | 1208 | - | - | 0 | - |
| - | - | 2889 | 1209 | - | - | 0 | - |
| - | - | 2364 | 1347 | - | - | 0 | - |

m/z Charge Intensity FragmentType MassShift Position
120.06560516357422 0 2072.3179
120.0811538696289 0 19685.588
121.08446502685547 0 1435.0782
125.07130432128906 0 1001.3405
125.1077651977539 0 1106.101
126.05531311035156 0 23593.234
127.05843353271484 0 1565.2803
127.0870590209961 0 948.53674
128.10740661621094 0 1966.8638
129.06622314453125 0 3264.7766
129.10260009765625 0 22810.756
129.11383056640625 0 1875.7139
130.0502166748047 0 2336.1487
130.06149291992188 0 11997.342
130.09783935546875 0 7417.2817
130.1061553955078 0 2329.3713
131.08209228515625 0 1455.1528
131.11830139160156 0 3343.6677
133.06100463867188 0 1009.41223
133.09722900390625 0 3069.2837
136.0760040283203 0 9137.282
138.09188842773438 0 1701.9071
138.68203735351562 0 997.45667
139.08705139160156 0 1286.0641
140.08226013183594 0 4484.024
140.1436004638672 0 2028.2596
141.0664825439453 0 1329.9369
141.1026153564453 0 2746.0393
143.0816650390625 0 1371.5928
144.0658416748047 0 6172.66
145.06101989746094 0 1429.4374
147.07681274414062 0 7188.7515
148.947021484375 0 2196.1238
152.07113647460938 0 1237.3693
153.06631469726562 0 4138.34
154.0502471923828 0 28247.127
155.05323791503906 0 1602.8137
155.1185302734375 0 1947.5554
156.07740783691406 0 1056.0897
156.10238647460938 0 2086.5745
157.06117248535156 0 7405.8154
157.09751892089844 0 11664.506
157.1088104248047 0 11146.664
157.13389587402344 0 4980.268 d 1
158.0640411376953 0 1230.4425
158.09280395507812 0 52149.906 y Ammonia loss 14
159.07688903808594 0 1777.5758
159.0962371826172 0 2804.6208
159.113037109375 0 7881.5186
160.11605834960938 0 1372.8425
165.1023712158203 0 2721.464
165.1144561767578 0 1160.6139
166.0865478515625 0 1325.2415 y Water loss 12
167.08184814453125 0 4059.2122
169.09725952148438 0 2463.23 y 10
170.04490661621094 0 1657.0822
171.07679748535156 0 32093.951
172.06097412109375 0 4216.786
172.0799102783203 0 1669.5651
173.12872314453125 0 8115.7437
173.438232421875 0 1473.0529
175.07176208496094 0 17227.592
175.11935424804688 0 114901.76 y 14
176.12277221679688 0 7542.7686
180.03285217285156 0 2495.765
181.06100463867188 0 2011.9943
181.1083984375 0 1357.5018
182.09263610839844 0 3600.2163
183.11314392089844 0 23019.572
183.1217803955078 0 1993.312
184.10833740234375 0 3368.095
184.11660766601562 0 2722.5793
185.05624389648438 0 1527.3165
185.0924072265625 0 18003.26
185.1029052734375 0 2257.7988
185.1287078857422 0 4167.6665
185.1653594970703 0 2315.4927
186.09605407714844 0 1388.6996
187.1080780029297 0 16666.508
189.08738708496094 0 127821.27
190.0908660888672 0 8673.3
191.6056365966797 0 1294.1737
192.09768676757812 0 1919.1284
192.1134796142578 0 1623.1052
194.09298706054688 0 2287.1821
194.12847900390625 0 1328.7106
195.07696533203125 0 1811.5817
195.08779907226562 0 1678.3958
195.1132354736328 0 2873.6558
195.149658203125 0 4293.6265
197.1654510498047 0 2580.798
199.07171630859375 0 13327.9
199.18093872070312 0 1078412.2 a 1
200.18426513671875 0 131724.1
201.12368774414062 0 40215.387
201.18716430664062 0 8431.068
202.11888122558594 0 7423.9717
202.12742614746094 0 3637.2114
203.06654357910156 0 21468.615
203.1025848388672 0 3047.9932
209.09254455566406 0 2096.888
209.1038055419922 0 1437.2732
209.12832641601562 0 1424.3816
210.0874786376953 0 2975.449
211.1082305908203 0 8165.6694
212.1034393310547 0 8805.635
213.10707092285156 0 1641.1627
213.16001892089844 0 1641.8527
215.10330200195312 0 1702.7876
215.1139373779297 0 1706.4794
216.135009765625 0 3447.2122
217.08230590820312 0 214049.53
218.08567810058594 0 18352.54
219.08734130859375 0 2205.0078
220.09353637695312 0 1637.8883
220.12954711914062 0 3477.4495
223.14466857910156 0 1573.923
224.1034698486328 0 15132.329
224.1397705078125 0 2902.1797
225.1072540283203 0 1953.608
226.1189422607422 0 5480.055
226.15541076660156 0 18279.385
227.0670623779297 0 1720.7532
227.11441040039062 0 9331.881
227.1758575439453 0 316334.8 b 1
228.0979461669922 0 2594.6687
228.17918395996094 0 42618.758
229.11862182617188 0 7253.6367
229.1824188232422 0 3747.3196
230.10252380371094 0 1281.2341
230.1144256591797 0 6636.6665
233.1652069091797 0 21743.154
234.1240997314453 0 1980.2863
234.168701171875 0 3906.7102
235.09271240234375 0 1712.9951
237.05467224121094 0 3642.36
237.09884643554688 0 3029.5933
237.13526916503906 0 6337.2026
239.14002990722656 0 1404.2235
242.11402893066406 0 27858.086
242.1510467529297 0 1605.7291
243.11749267578125 0 3029.327
243.1324462890625 0 1273.2069
244.1295166015625 0 7686.4043
244.141357421875 0 2769.1487 y Water loss 13
245.12501525878906 0 32948.81 y Ammonia loss 13
246.12799072265625 0 2900.9504
249.1602783203125 0 10361.115
250.1636505126953 0 2096.783
252.09829711914062 0 1747.7622
252.13427734375 0 1665.072
253.22813415527344 0 2552.8198
254.11402893066406 0 4021.8118
254.1619873046875 0 1773.0626
255.10928344726562 0 25028.693
256.1122741699219 0 3833.5212
256.1280822753906 0 2084.5547
259.1406555175781 0 4987.3604
261.1601867675781 0 9764.99
262.1513366699219 0 21222.316 y 13
262.1639709472656 0 1922.6545
263.15399169921875 0 2295.8572
267.1106872558594 0 1773.3534
270.1451721191406 0 5356.403
271.12799072265625 0 1849.0778
272.13604736328125 0 5017.0215
277.1551208496094 0 4942.4062
278.1258239746094 0 1396.3408
279.10882568359375 0 1989.6967
280.1128845214844 0 1216.9631
280.1297912597656 0 3360.6873
281.12493896484375 0 1419.5132
283.2013244628906 0 1394.7732
284.1247253417969 0 5177.9414
284.16070556640625 0 3358.1252
288.1560363769531 0 4052.594
289.1512756347656 0 1854.4355
292.16705322265625 0 1342.1255
296.1360168457031 0 2936.2095
296.1974182128906 0 16389.854 b Water loss 2
297.1199951171875 0 3462.954
297.20074462890625 0 1823.8008
298.12261962890625 0 2175.5183
298.1405334472656 0 9879.123
299.1365661621094 0 7168.034
302.1348571777344 0 2533.2085
302.1708068847656 0 1854.2197
307.16180419921875 0 1780.1581
311.135009765625 0 1667.542
312.15594482421875 0 15127.623
313.15283203125 0 3762.2498
313.1640625 0 2034.1261
313.1876525878906 0 36594.91
314.14691162109375 0 11728.245
314.1903076171875 0 5174.3267
314.2082214355469 0 35132.812 b 2
315.2112731933594 0 7147.4194
316.1509094238281 0 38763.047
316.17041015625 0 2441.5999
317.1489562988281 0 6156.6836
319.17742919921875 0 2306.5474
323.1727600097656 0 1759.6199
323.2086181640625 0 7139.105
324.1310119628906 0 6931.1177
324.2110595703125 0 1268.5404
326.1351013183594 0 2049.223
329.1460876464844 0 2278.2078
330.16668701171875 0 9070.894
331.17120361328125 0 5733.021 y Water loss 12
332.15716552734375 0 30411.852 y Ammonia loss 12
333.1598205566406 0 4221.973
339.20440673828125 0 1853.8643
340.1507263183594 0 2507.217
341.1826171875 0 25205.852
341.2023010253906 0 1726.9601
342.14154052734375 0 37462.05
342.182861328125 0 3304.746
343.1441650390625 0 4857.69
343.1985168457031 0 1781.0289
345.182861328125 0 15252.028 y 8
345.684326171875 0 6035.677
349.14996337890625 0 2312.4722
349.18365478515625 0 25462.299 y 12
350.18621826171875 0 4719.823
352.16217041015625 0 1606.1024
353.1951599121094 0 1840.537
358.2111511230469 0 1666.8328
359.1673278808594 0 15845.174
359.191650390625 0 2527.8806
360.171142578125 0 2616.726
363.1873779296875 0 2506.6426
365.18292236328125 0 2814.7903
366.18280029296875 0 1765.3627
370.2203369140625 0 1743.5638
380.157958984375 0 1899.4254
381.18865966796875 0 3031.1409
382.21221923828125 0 1476.1765
383.19427490234375 0 1984.3701
385.1717529296875 0 5412.268
385.7093505859375 0 1788.5834
386.1795959472656 0 1405.7671
386.212158203125 0 1385.4574
389.7089538574219 0 8583.058
390.2102355957031 0 6980.893
390.71685791015625 0 6509.049
391.2174987792969 0 6115.3496
391.72442626953125 0 33177.777
392.2261962890625 0 17611.541
392.7273864746094 0 3725.6199
393.7095642089844 0 4156.648
394.20947265625 0 3576.9395
396.18804931640625 0 3619.1997
397.2456970214844 0 3834.4404
398.1703186035156 0 1528.1361
398.21380615234375 0 4739.8364
398.71453857421875 0 10803.782
399.16497802734375 0 2365.861
399.2159729003906 0 6700.491
399.7203063964844 0 11579.849
400.1923522949219 0 1826.628
400.22125244140625 0 8619.465
401.2414245605469 0 3423.046
403.1827087402344 0 6551.2764
404.1846618652344 0 1947.4198
405.717529296875 0 1910.7124
407.23016357421875 0 2016.1973
408.2284851074219 0 2074.7256
408.73150634765625 0 2042.7058
410.20477294921875 0 4518.7734
411.1622619628906 0 1787.9397
411.22589111328125 0 2769.2295
414.22802734375 0 8223.191 b 11
414.722412109375 0 22471.578 y Ammonia loss 7
415.2230224609375 0 7686.5347
415.255615234375 0 4562.9106
415.7241516113281 0 2546.4954
416.219970703125 0 2111.5315
423.2339172363281 0 232738.4 y 7
423.7352294921875 0 99164.65
424.23651123046875 0 22591.729
424.7381896972656 0 4986.8975
425.2403869628906 0 17050.172 b Water loss 3
426.2116394042969 0 1863.3446
426.24383544921875 0 5423.9697
428.1830749511719 0 1554.0702
428.2164001464844 0 6529.4893
429.235107421875 0 11042.116
430.2403869628906 0 7926.229 y Water loss 11
431.17864990234375 0 1709.5333
439.19403076171875 0 2528.7744
440.2241516113281 0 2592.6467
441.2103271484375 0 13829.939
442.2132568359375 0 2350.5398
443.2509460449219 0 20207.44 b 3
444.2552185058594 0 3661.0532
448.2515563964844 0 7083.5146 y 11
448.8888244628906 0 3559.8792 y Water loss 2
449.5552062988281 0 2269.381
450.7434997558594 0 4363.1143 b 7
452.2629699707031 0 9023.553
454.8923034667969 0 9293.225 y 2
455.2265625 0 8661.565
455.5603942871094 0 1776.8351
458.2347106933594 0 13869.248
460.2409362792969 0 12926.944
461.243896484375 0 3410.9148
463.2435302734375 0 3388.344
465.26171875 0 2120.583
468.8757019042969 0 2298.8567
469.2071838378906 0 2479.6406
469.2892761230469 0 6779.2285
471.22735595703125 0 1644.9546
471.7402648925781 0 2818.6724 y Water loss 6
472.2370300292969 0 2962.2354 b 13
472.7457580566406 0 1751.3707
473.75860595703125 0 1811.9023
478.20428466796875 0 2041.4735
480.7473449707031 0 39882.66 y 6
481.16259765625 0 1424.9182
481.24835205078125 0 20098.434
481.74957275390625 0 8249.849
485.2433776855469 0 1602.2323
485.7432556152344 0 2461.3806
486.5832824707031 0 8948.698 y Water loss 1
486.91839599609375 0 9397.955 y Ammonia loss 1
487.2561340332031 0 5693.5654 y Water loss 10
488.24749755859375 0 5396.129 y Ammonia loss 10
492.58685302734375 0 6915.065 y 1
492.92230224609375 0 5622.7827
493.2542724609375 0 3018.5847
494.7412109375 0 1995.4347
495.9151916503906 0 13853.859
496.2143249511719 0 2212.6816
496.25067138671875 0 11632.954
496.5863952636719 0 3734.7473
496.9209289550781 0 1786.9227
497.2366027832031 0 1811.9546
497.28497314453125 0 3982.1716
499.7664489746094 0 1908.7272
500.2003479003906 0 2973.7593
505.27386474609375 0 25729.252 y 10
506.2786560058594 0 5580.9287
513.2429809570312 0 1565.1018
514.2610473632812 0 1841.5442
514.3246459960938 0 3876.9026
515.2551879882812 0 7928.4507 y Water loss 5
515.755859375 0 3883.7903 y Ammonia loss 5
516.2584838867188 0 2225.5107
518.2102661132812 0 3785.0151
518.273681640625 0 3431.8064
524.264892578125 0 237495.8 y 5
524.6124877929688 0 46402.195 Precursor Ammonia loss
524.7647094726562 0 131553.86
524.9467163085938 0 27877.598
525.2681884765625 0 38091.777
525.7672729492188 0 9014.589
526.2645874023438 0 2015.7103
528.7484741210938 0 2131.5586
528.7894897460938 0 7564.786
529.2922973632812 0 4174.531
529.3521118164062 0 1717.2346
529.7970581054688 0 9615.927
530.28271484375 0 32640.531 Precursor
530.6160278320312 0 25044.6
530.7994995117188 0 2490.6335
530.9503784179688 0 13522.328
531.2880859375 0 2142.9858
532.2716064453125 0 2581.4639
533.2543334960938 0 1923.6903
539.2974243164062 0 1988.8376
540.244873046875 0 2573.7617
542.3192749023438 0 6666.3965 b 4
543.3246459960938 0 1772.9308
544.2815551757812 0 2094.2249
552.7861328125 0 1978.1578
556.280517578125 0 1996.356
556.3221435546875 0 2276.791
558.7863159179688 0 2437.3508
559.2886352539062 0 2820.5264
562.2931518554688 0 3172.785 b Water loss 10
564.7905883789062 0 7782.81 y Water loss 4
565.2379760742188 0 3318.0972
565.2886962890625 0 16885.674 y Ammonia loss 4
565.789306640625 0 8166.117
566.2875366210938 0 4437.4165
567.2901611328125 0 2498.932
569.2670288085938 0 2199.6577
570.2506713867188 0 1333.6173
572.791259765625 0 2622.5273
573.2898559570312 0 2548.164
573.7973022460938 0 340942.03 y 4
574.2986450195312 0 208414.92 y Water loss 9
574.7999877929688 0 72496.11
575.2996826171875 0 15309.9795
575.8004150390625 0 1740.6644
583.244873046875 0 2734.4756
584.3147583007812 0 3912.8928
587.2798461914062 0 4852.9004
592.3057250976562 0 33312.84 y 9
593.3079833984375 0 10527.841
594.3151245117188 0 2268.7134
600.2748413085938 0 7819.312
601.2800903320312 0 2590.2036
608.3114624023438 0 1748.1527
611.3218383789062 0 2007.6482
617.3049926757812 0 2808.422
618.2908325195312 0 2410.3696
619.2835693359375 0 2171.4097
621.3076171875 0 1747.1051
624.3080444335938 0 2416.1096
626.3265991210938 0 4155.98
629.315185546875 0 24385.676 y Water loss 3
629.8141479492188 0 16124.869 y Ammonia loss 3
630.3169555664062 0 5325.0874
630.816650390625 0 5178.049
636.3125 0 3764.3206
637.3074340820312 0 1802.9482
637.81591796875 0 2387.153
638.318603515625 0 167461.39 y 3
638.8200073242188 0 107184.04
639.3212280273438 0 48357.34
639.8222045898438 0 12530.344
640.3231811523438 0 2612.3376
641.3372802734375 0 5343.721
642.33544921875 0 1625.0043
642.8179931640625 0 1753.1201
643.3550415039062 0 4783.592
644.3523559570312 0 2433.9448
646.31591796875 0 13640.782
647.3175048828125 0 3920.3877
648.8214111328125 0 5362.8076
649.3204345703125 0 3072.0234
651.8157958984375 0 2519.557
652.3175048828125 0 4103.5767
652.814208984375 0 1845.8082
653.3361206054688 0 7413.591
654.3261108398438 0 6860.828
654.8289184570312 0 2531.8545
655.319091796875 0 7850.4253
655.816162109375 0 1906.1348
656.304443359375 0 3076.9084
657.2877197265625 0 4858.284
657.8232421875 0 3833.0474
659.3466796875 0 7980.062
659.8275756835938 0 2375.0793
660.299560546875 0 2952.8474
660.3546142578125 0 2103.5251
660.8223266601562 0 10429.046
661.323974609375 0 5301.879
663.8238525390625 0 12873.64
664.3209228515625 0 11450.082
664.819091796875 0 6324.982
665.3197021484375 0 5246.546
666.3231811523438 0 4188.583
666.8290405273438 0 32323.49
667.3301391601562 0 17266.475
667.8302001953125 0 7040.7295
668.33154296875 0 2502.382
671.3477783203125 0 36197.617 y Water loss 8
671.82080078125 0 2233.037
672.3379516601562 0 27596.43 y Ammonia loss 8
672.829345703125 0 95280.52 y Water loss 2
673.330322265625 0 84065.375 y Ammonia loss 2
673.82958984375 0 42073.508
674.3154907226562 0 44552.35
674.8240356445312 0 4570.9834
675.3147583007812 0 15464.463
676.3180541992188 0 3506.3862
678.325439453125 0 2743.6323
680.82568359375 0 27881.74
681.3285522460938 0 21014.85
681.835205078125 0 2693644.2 y 2
682.3363037109375 0 1896681.2
682.8375244140625 0 791558.25
683.3385620117188 0 215624.22
683.8394165039062 0 47047.645
685.3870239257812 0 7481.8325
686.8265380859375 0 21249.248
687.32958984375 0 19127.457
687.830810546875 0 6221.714
688.3336181640625 0 1812.9255
689.3584594726562 0 212063.47 y 8
690.361328125 0 77103.164
691.3633422851562 0 17070.729
699.3439331054688 0 13690.65 b Ammonia loss 13
700.3451538085938 0 4206.644
701.3607177734375 0 1596.5375
729.3709106445312 0 21879.553 y Water loss 1
729.8732299804688 0 17031.852 y Ammonia loss 1
730.3690795898438 0 5043.516
730.868408203125 0 1763.6279
737.3676147460938 0 4526.047
737.8690185546875 0 2399.8225
738.3766479492188 0 215601.9 y 1
738.8779907226562 0 152353.64
739.379150390625 0 76863.58
739.880615234375 0 20668.787
740.3820190429688 0 5560.091
743.369140625 0 14394.595
743.8692626953125 0 7223.64
744.373046875 0 3989.3933 b 6
758.3861694335938 0 3184.3064
768.3646240234375 0 1878.9182
768.4022216796875 0 5418.2095
769.3627319335938 0 2137.0881
770.4164428710938 0 9476.375
771.419189453125 0 3949.1316
778.4099731445312 0 7593.385
779.4094848632812 0 4190.258
780.4229125976562 0 4384.869
781.4234008789062 0 2245.6968
782.4409790039062 0 26804.584
783.4444580078125 0 9714.093
784.4469604492188 0 2393.4434
785.4280395507812 0 6476.7363
786.3833618164062 0 3708.3655
786.4170532226562 0 5428.9497 Precursor Ammonia loss
787.3944702148438 0 5475.3003
788.3965454101562 0 2090.288
796.4197387695312 0 9531.348
797.4240112304688 0 5884.4067
798.4229736328125 0 10716.11
799.4254150390625 0 6160.399
803.43798828125 0 3872.929
804.44140625 0 2862.3865
810.4331665039062 0 3409.5771
811.4085083007812 0 7781.055
812.40380859375 0 3689.7668
828.4329223632812 0 32146.924 y Ammonia loss 7
829.4346923828125 0 14118.395
830.4302978515625 0 5172.0996
845.4203491210938 0 1960.5605
856.4043579101562 0 1751.3623
857.4404907226562 0 3630.3699
858.3955078125 0 3901.8398
859.3883056640625 0 2200.5933
867.4318237304688 0 3640.4534
873.4043579101562 0 3259.043
881.4442138671875 0 2757.1423
885.4425659179688 0 4565.185
887.42333984375 0 7608.268
888.4305419921875 0 4729.4844
891.4187622070312 0 7059.259
900.4674682617188 0 7316.839 b 7
901.4601440429688 0 3423.4243
915.4185180664062 0 14305.656
916.4185180664062 0 5975.8555
917.4163818359375 0 1893.1545
925.4556884765625 0 4120.739
926.4342651367188 0 3617.7996
927.45654296875 0 4171.755
943.460205078125 0 24933.55 y Ammonia loss 6
944.4669189453125 0 16562.092
945.4692993164062 0 4583.725
968.4832763671875 0 3959.4924
969.4742431640625 0 4676.651
970.4692993164062 0 4437.3926
971.4752197265625 0 2172.474
972.469970703125 0 4349.191
973.4781494140625 0 3249.432
986.49072265625 0 27813.406
987.48828125 0 23950.342
988.489501953125 0 13383.229
989.4921875 0 4262.5938
990.4852294921875 0 4375.785
991.488525390625 0 2770.361
994.4620971679688 0 2112.2253
996.4705810546875 0 4297.304
997.4734497070312 0 2788.6897
1000.4832153320312 0 2648.5432
1001.48681640625 0 2488.6345
1005.5027465820312 0 2629.701
1012.4797973632812 0 12089.321
1013.4834594726562 0 8560.6
1014.4843139648438 0 19514.256
1015.486572265625 0 12945.072
1016.492431640625 0 3827.1543
1030.49169921875 0 77710.01 y Ammonia loss 5
1031.4940185546875 0 38641.86
1032.496826171875 0 13258.647
1033.4937744140625 0 3053.416
1055.506103515625 0 2244.2402
1057.521240234375 0 2763.4536
1071.5087890625 0 2811.827
1073.522216796875 0 8992.049
1074.5328369140625 0 3444.7834
1075.5244140625 0 2474.6323
1083.5098876953125 0 5123.7476
1084.5150146484375 0 3117.4888
1086.5574951171875 0 6232.9053
1087.570556640625 0 1921.2383
1088.549072265625 0 2016.0834
1101.517333984375 0 13635.979
1102.5208740234375 0 6653.412
1111.548583984375 0 3160.1316
1112.5452880859375 0 3771.8867
1119.53271484375 0 4292.636
1129.558349609375 0 22415.406 y Ammonia loss 4
1130.56298828125 0 14914.126
1131.561279296875 0 6748.5063
1142.53515625 0 2632.0654
1143.53857421875 0 3041.7817
1160.553955078125 0 12712.9375
1161.55615234375 0 8279.356
1162.5616455078125 0 2682.7368
1170.5438232421875 0 4893.7446
1188.5491943359375 0 17448.793
1189.5501708984375 0 10265.049
1190.558349609375 0 4686.836
1205.5606689453125 0 2171.9004
1206.5595703125 0 17336.113
1207.5596923828125 0 9902.432
1208.5743408203125 0 2889.4124
1346.63916015625 0 2363.687

Spectrum Details

|  |  |
| --- | --- |
| Matched peaks? Matched peaks The total absolute number of peaks matched. Additionally in brackets the total fraction of peaks matched and the total number of peaks is shown. | 67 (10.91% of 614) |
| FDR? FDR The false discovery rate estimated for this peptide. It is calculated by matching all theoretical fragments with a non-integer shift with the raw peaks for this spectrum. This is done with 40 different shifts. The resulting percentage is the average number of annotated peaks over the number of annotated peaks with the correct spectrum. | 0.71% |
| Satellite FDR? Satellite FDR See the FDR for details on its calculation. This satellite ion specific FDR only contains the satellite ions (d/w) for I/L/J positions. | 0.00% |
| PSM Score? PSM Score The PSM Score as given by Hecklib to this annotated spectrum. It is shown with three significant figures. | 403 |

#### Spectrum 6067? Spectrum 6067 The raw spectrum of this peptide as annotated by Hecklib. The fragments are coloured according to ion type (see legend). Any peaks with a star '\*' as text can be hovered over to see the full details, first the ion type second the mass shift type. By hovering over the amino acids in the peptide or ions in the legend the corresponding peaks are highlighted. By toggling the 'Unassigned' label you can turn the background (unassigned) peaks on or off in the plot. By updating the slider in the Ion legend you can update the spectrum to only show the top X% of the peaks with labels. The top X% means any peak that is within X% of the highest intensity. By dragging in the spectrum you can zoom in to a specific part of the spectrum and use 'Zoom Out' to get back to the original zoom level. The annotation of the spectrum is based on the given sequence in the peptides file and is done with different software so inconsistencies are likely. The peaks are annotated based on the given sequence, with 20 ppm tolerance.

Copy Data

##### Spectrum 6067 (TSV)

###### Preview

```
Loading example...
```

*Click on the button to copy the data to your clipboard.*

Mz MinMz MaxIntensity Max

WidthHeightPeptide font sizePeptide stroke widthSpectrum font sizeSpectrum stroke widthCompact peptide

Ion legend

wxyz

abcd

OtherUnassignedIon^Charge^_Position_Show for top:%

JJSEVSDRPSGVSSR

04.00e+38.01e+31.20e+41.60e+4

Zoom Out

y^+1^_1_w^+1^_2_y^+1^_2_y^+1^_2_w^+1^_3_y^+1^_3_z^+1^_3_y^+1^_3_z^+2^_8_w^+1^_4_y^+2^_8_y^+1^_4_c^+1^_4_z^+1^_5_y^+1^_5_y^+2^_10_c^+1^_5_z^+2^_11_y^+2^_11_y^+1^_6_z^+1^_6_w^+2^_12_z^+2^_12_y^+2^_12_c^+1^_6_z^+2^_13_y^+2^_13_y^+2^_14_w^+1^_8_c^+1^_7_z^+1^_8_y^+1^_8_c^+1^_8_z^+1^_9_c^+1^_9_z^+1^_10_y^+1^_10_c^+1^_10_z^+1^_11_c^+1^_11_c^+1^_12_y^+1^_12_c^+1^_13_c^+1^_14_

0854170825623416

Fragment Matches Table

Show background peaks

| Position | Ion type | Intensity | mz Theoretical | mz Error (Th) | mz Error (ppm) | Charge | Series Number |
| --- | --- | --- | --- | --- | --- | --- | --- |
| - | - | 460.4 | 120.1 | - | - | 0 | - |
| - | - | 342.4 | 123.9 | - | - | 0 | - |
| - | - | 374.7 | 126.1 | - | - | 0 | - |
| - | - | 436.4 | 129.3 | - | - | 0 | - |
| - | - | 418.8 | 130.8 | - | - | 0 | - |
| - | - | 508.8 | 133.1 | - | - | 0 | - |
| - | - | 1425 | 136.1 | - | - | 0 | - |
| - | - | 1150 | 140.1 | - | - | 0 | - |
| - | - | 1133 | 143.1 | - | - | 0 | - |
| - | - | 973.7 | 149 | - | - | 0 | - |
| - | - | 406.3 | 151.7 | - | - | 0 | - |
| - | - | 474.6 | 157.6 | - | - | 0 | - |
| - | - | 407.6 | 157.6 | - | - | 0 | - |
| - | - | 902.7 | 168.1 | - | - | 0 | - |
| - | - | 458.1 | 171.1 | - | - | 0 | - |
| - | - | 446.1 | 171.1 | - | - | 0 | - |
| - | - | 427.3 | 173.1 | - | - | 0 | - |
| - | - | 1569 | 173.4 | - | - | 0 | - |
| - | - | 528.8 | 174.1 | - | - | 0 | - |
| 15 | y | 1322 | 175.1 | 0.0001732 | 0.989 | +1 | 1 |
| - | - | 948.3 | 185.1 | - | - | 0 | - |
| - | - | 3161 | 185.1 | - | - | 0 | - |
| - | - | 5446 | 187.1 | - | - | 0 | - |
| - | - | 1441 | 187.1 | - | - | 0 | - |
| - | - | 440.7 | 197 | - | - | 0 | - |
| - | - | 654.3 | 199.1 | - | - | 0 | - |
| - | - | 5411 | 199.2 | - | - | 0 | - |
| - | - | 969.7 | 200.2 | - | - | 0 | - |
| - | - | 2190 | 215.1 | - | - | 0 | - |
| - | - | 2884 | 227.2 | - | - | 0 | - |
| - | - | 495.9 | 228.1 | - | - | 0 | - |
| - | - | 598.4 | 228.2 | - | - | 0 | - |
| 14 | w | 707.7 | 229.1 | 7.655E-05 | 0.3341 | +1 | 2 |
| 14 | y | 4734 | 245.1 | 0.0002481 | 1.012 | +1 | 2 |
| - | - | 630.2 | 254.1 | - | - | 0 | - |
| 14 | y | 778 | 262.2 | 0.0004374 | 1.668 | +1 | 2 |
| - | - | 494.2 | 262.6 | - | - | 0 | - |
| - | - | 666.3 | 270.1 | - | - | 0 | - |
| - | - | 1126 | 301.1 | - | - | 0 | - |
| - | - | 713.3 | 314.2 | - | - | 0 | - |
| 13 | w | 2163 | 316.2 | 0.0002284 | 0.7225 | +1 | 3 |
| - | - | 907.4 | 331.1 | - | - | 0 | - |
| 13 | y | 902.2 | 332.2 | 0.0007667 | 2.308 | +1 | 3 |
| - | - | 560.2 | 332.8 | - | - | 0 | - |
| 13 | z | 1829 | 333.2 | 0.0001614 | 0.4844 | +1 | 3 |
| - | - | 1017 | 349.1 | - | - | 0 | - |
| 13 | y | 785.8 | 349.2 | 0.0004528 | 1.297 | +1 | 3 |
| - | - | 1381 | 357.2 | - | - | 0 | - |
| - | - | 820 | 367.2 | - | - | 0 | - |
| - | - | 707.5 | 369.2 | - | - | 0 | - |
| - | - | 7078 | 377.2 | - | - | 0 | - |
| - | - | 751.2 | 404.7 | - | - | 0 | - |
| 8 | z | 762.2 | 415.2 | 0.0007856 | 1.892 | +2 | 8 |
| - | - | 602.3 | 415.7 | - | - | 0 | - |
| 12 | w | 2105 | 417.2 | 0.0008593 | 2.06 | +1 | 4 |
| - | - | 639.9 | 422 | - | - | 0 | - |
| 8 | y | 824.5 | 423.2 | 0.001832 | 4.329 | +2 | 8 |
| 12 | y | 740 | 448.3 | 0.0002938 | 0.6553 | +1 | 4 |
| - | - | 1463 | 453.3 | - | - | 0 | - |
| 4 | c | 718.5 | 460.3 | 0.000513 | 1.115 | +1 | 4 |
| - | - | 528.4 | 470.3 | - | - | 0 | - |
| - | - | 1113 | 476.2 | - | - | 0 | - |
| 11 | z | 568.6 | 489.3 | 0.001849 | 3.78 | +1 | 5 |
| - | - | 613.5 | 499.3 | - | - | 0 | - |
| 11 | y | 6212 | 505.3 | 0.001039 | 2.057 | +1 | 5 |
| - | - | 1532 | 506.3 | - | - | 0 | - |
| - | - | 534.9 | 509.8 | - | - | 0 | - |
| - | - | 686.5 | 515.3 | - | - | 0 | - |
| - | - | 1086 | 516.8 | - | - | 0 | - |
| - | - | 905.9 | 517.3 | - | - | 0 | - |
| - | - | 739.6 | 520.3 | - | - | 0 | - |
| 6 | y | 1866 | 524.3 | 0.002374 | 4.528 | +2 | 10 |
| - | - | 746.9 | 524.8 | - | - | 0 | - |
| - | - | 966.2 | 529.3 | - | - | 0 | - |
| - | - | 1133 | 529.8 | - | - | 0 | - |
| - | - | 862.4 | 530.3 | - | - | 0 | - |
| - | - | 7227 | 530.3 | - | - | 0 | - |
| - | - | 925.5 | 531.3 | - | - | 0 | - |
| - | - | 1837 | 531.3 | - | - | 0 | - |
| - | - | 1769 | 531.7 | - | - | 0 | - |
| - | - | 763.1 | 551.3 | - | - | 0 | - |
| 5 | c | 849.5 | 559.3 | 0.0005921 | 1.059 | +1 | 5 |
| 5 | z | 3651 | 565.8 | 0.0002338 | 0.4133 | +2 | 11 |
| - | - | 2440 | 566.3 | - | - | 0 | - |
| - | - | 1196 | 566.8 | - | - | 0 | - |
| - | - | 1183 | 569.3 | - | - | 0 | - |
| 5 | y | 2381 | 573.8 | 0.0002712 | 0.4727 | +2 | 11 |
| 10 | y | 1289 | 574.3 | 0.003745 | 6.522 | +1 | 6 |
| 10 | z | 940.4 | 576.3 | 0.002255 | 3.914 | +1 | 6 |
| - | - | 744.7 | 577.3 | - | - | 0 | - |
| - | - | 876.4 | 585.3 | - | - | 0 | - |
| - | - | 708.6 | 587.3 | - | - | 0 | - |
| 4 | w | 1450 | 600.8 | 0.001153 | 1.92 | +2 | 12 |
| - | - | 1291 | 601.3 | - | - | 0 | - |
| - | - | 1095 | 601.8 | - | - | 0 | - |
| - | - | 1206 | 604.4 | - | - | 0 | - |
| 4 | z | 4482 | 630.3 | 0.0004939 | 0.7835 | +2 | 12 |
| - | - | 2332 | 630.8 | - | - | 0 | - |
| - | - | 1478 | 631.3 | - | - | 0 | - |
| - | - | 779.4 | 637.8 | - | - | 0 | - |
| 4 | y | 4939 | 638.3 | 9.028E-05 | 0.1414 | +2 | 12 |
| - | - | 1616 | 638.8 | - | - | 0 | - |
| - | - | 824.8 | 639.3 | - | - | 0 | - |
| - | - | 1794 | 640.3 | - | - | 0 | - |
| 6 | c | 3696 | 646.4 | 0.0006682 | 1.034 | +1 | 6 |
| - | - | 1060 | 647.4 | - | - | 0 | - |
| - | - | 644.5 | 650.4 | - | - | 0 | - |
| - | - | 914.8 | 660.4 | - | - | 0 | - |
| - | - | 1010 | 668.4 | - | - | 0 | - |
| - | - | 961 | 670.4 | - | - | 0 | - |
| 3 | z | 2428 | 673.8 | 0.0008869 | 1.316 | +2 | 13 |
| - | - | 3763 | 674.3 | - | - | 0 | - |
| - | - | 1846 | 674.8 | - | - | 0 | - |
| - | - | 960.9 | 676.4 | - | - | 0 | - |
| 3 | y | 1.585E+04 | 681.8 | 0.000314 | 0.4605 | +2 | 13 |
| - | - | 1.309E+04 | 682.3 | - | - | 0 | - |
| - | - | 6199 | 682.8 | - | - | 0 | - |
| - | - | 888.8 | 683.3 | - | - | 0 | - |
| - | - | 1210 | 686.4 | - | - | 0 | - |
| - | - | 623.8 | 718.4 | - | - | 0 | - |
| - | - | 1003 | 719.4 | - | - | 0 | - |
| - | - | 637.2 | 720.4 | - | - | 0 | - |
| - | - | 830.3 | 727.4 | - | - | 0 | - |
| - | - | 766.8 | 728.4 | - | - | 0 | - |
| 2 | y | 1264 | 738.4 | 9.106E-05 | 0.1233 | +2 | 14 |
| - | - | 1068 | 738.9 | - | - | 0 | - |
| - | - | 680.3 | 739.4 | - | - | 0 | - |
| - | - | 2941 | 741.4 | - | - | 0 | - |
| - | - | 841.2 | 742.4 | - | - | 0 | - |
| 8 | w | 821.4 | 743.4 | 0.007611 | 10.24 | +1 | 8 |
| - | - | 767.4 | 759.4 | - | - | 0 | - |
| 7 | c | 7394 | 761.4 | 0.0005196 | 0.6824 | +1 | 7 |
| - | - | 3205 | 762.4 | - | - | 0 | - |
| - | - | 811 | 763.4 | - | - | 0 | - |
| - | - | 1.47E+04 | 765.4 | - | - | 0 | - |
| - | - | 9855 | 765.9 | - | - | 0 | - |
| - | - | 4146 | 766.4 | - | - | 0 | - |
| - | - | 1078 | 766.9 | - | - | 0 | - |
| - | - | 3983 | 773.9 | - | - | 0 | - |
| - | - | 3659 | 774.4 | - | - | 0 | - |
| - | - | 1478 | 774.9 | - | - | 0 | - |
| - | - | 1046 | 775.4 | - | - | 0 | - |
| - | - | 800.7 | 778.4 | - | - | 0 | - |
| - | - | 708.6 | 779.4 | - | - | 0 | - |
| - | - | 705.5 | 782.4 | - | - | 0 | - |
| - | - | 932.1 | 785.9 | - | - | 0 | - |
| - | - | 1.297E+04 | 786.9 | - | - | 0 | - |
| - | - | 1.035E+04 | 787.4 | - | - | 0 | - |
| - | - | 4959 | 787.9 | - | - | 0 | - |
| - | - | 1634 | 788.4 | - | - | 0 | - |
| - | - | 805.2 | 794.4 | - | - | 0 | - |
| - | - | 7818 | 794.9 | - | - | 0 | - |
| - | - | 1.182E+04 | 795.4 | - | - | 0 | - |
| - | - | 8024 | 795.9 | - | - | 0 | - |
| - | - | 3262 | 796.4 | - | - | 0 | - |
| - | - | 896.7 | 800.5 | - | - | 0 | - |
| - | - | 858.6 | 827.4 | - | - | 0 | - |
| 8 | z | 4902 | 829.4 | 0.0001807 | 0.2179 | +1 | 8 |
| - | - | 3186 | 830.4 | - | - | 0 | - |
| - | - | 1209 | 831.5 | - | - | 0 | - |
| - | - | 1965 | 843.4 | - | - | 0 | - |
| - | - | 1028 | 844.4 | - | - | 0 | - |
| 8 | y | 941.2 | 845.5 | 0.00243 | 2.874 | +1 | 8 |
| - | - | 1161 | 853.4 | - | - | 0 | - |
| - | - | 1953 | 871.4 | - | - | 0 | - |
| - | - | 1245 | 872.4 | - | - | 0 | - |
| - | - | 2634 | 887.5 | - | - | 0 | - |
| 8 | c | 1419 | 900.5 | 0.004706 | 5.227 | +1 | 8 |
| - | - | 655.1 | 901.5 | - | - | 0 | - |
| - | - | 889.6 | 915.5 | - | - | 0 | - |
| 7 | z | 2962 | 944.5 | 0.006438 | 6.816 | +1 | 9 |
| - | - | 2292 | 945.5 | - | - | 0 | - |
| - | - | 984.4 | 968.5 | - | - | 0 | - |
| - | - | 3739 | 974.6 | - | - | 0 | - |
| - | - | 1735 | 985.5 | - | - | 0 | - |
| - | - | 916.9 | 986.5 | - | - | 0 | - |
| - | - | 1072 | 987.5 | - | - | 0 | - |
| - | - | 1382 | 1013 | - | - | 0 | - |
| - | - | 914.7 | 1014 | - | - | 0 | - |
| 9 | c | 5461 | 1015 | 0.001012 | 0.997 | +1 | 9 |
| - | - | 2463 | 1016 | - | - | 0 | - |
| - | - | 661.3 | 1024 | - | - | 0 | - |
| - | - | 2367 | 1025 | - | - | 0 | - |
| - | - | 1236 | 1026 | - | - | 0 | - |
| - | - | 790.3 | 1028 | - | - | 0 | - |
| 6 | z | 1336 | 1031 | 0.002305 | 2.235 | +1 | 10 |
| - | - | 1288 | 1033 | - | - | 0 | - |
| - | - | 1974 | 1041 | - | - | 0 | - |
| - | - | 4308 | 1042 | - | - | 0 | - |
| - | - | 2500 | 1043 | - | - | 0 | - |
| - | - | 1184 | 1044 | - | - | 0 | - |
| - | - | 1649 | 1045 | - | - | 0 | - |
| - | - | 2065 | 1046 | - | - | 0 | - |
| - | - | 963.6 | 1047 | - | - | 0 | - |
| 6 | y | 668.6 | 1048 | 0.002869 | 2.738 | +1 | 10 |
| - | - | 5383 | 1058 | - | - | 0 | - |
| - | - | 1.141E+04 | 1059 | - | - | 0 | - |
| - | - | 5650 | 1060 | - | - | 0 | - |
| - | - | 2606 | 1061 | - | - | 0 | - |
| - | - | 3650 | 1062 | - | - | 0 | - |
| - | - | 3297 | 1063 | - | - | 0 | - |
| - | - | 4909 | 1064 | - | - | 0 | - |
| 10 | c | 8376 | 1102 | 0.0003861 | 0.3505 | +1 | 10 |
| - | - | 4882 | 1103 | - | - | 0 | - |
| - | - | 1232 | 1104 | - | - | 0 | - |
| - | - | 756.2 | 1115 | - | - | 0 | - |
| - | - | 693.1 | 1116 | - | - | 0 | - |
| 5 | z | 938.9 | 1131 | 0.002495 | 2.207 | +1 | 11 |
| - | - | 2883 | 1132 | - | - | 0 | - |
| - | - | 1500 | 1133 | - | - | 0 | - |
| - | - | 715.3 | 1134 | - | - | 0 | - |
| 11 | c | 4380 | 1159 | 0.0008538 | 0.7369 | +1 | 11 |
| - | - | 2943 | 1160 | - | - | 0 | - |
| - | - | 1160 | 1161 | - | - | 0 | - |
| - | - | 742.5 | 1214 | - | - | 0 | - |
| - | - | 695 | 1215 | - | - | 0 | - |
| 12 | c | 3834 | 1258 | 0.0001903 | 0.1513 | +1 | 12 |
| - | - | 3317 | 1259 | - | - | 0 | - |
| - | - | 4318 | 1261 | - | - | 0 | - |
| - | - | 2505 | 1262 | - | - | 0 | - |
| - | - | 857 | 1263 | - | - | 0 | - |
| 4 | y | 1463 | 1276 | 0.001237 | 0.9693 | +1 | 12 |
| - | - | 841.2 | 1277 | - | - | 0 | - |
| - | - | 889.5 | 1278 | - | - | 0 | - |
| - | - | 660 | 1288 | - | - | 0 | - |
| 13 | c | 7158 | 1345 | 0.003152 | 2.344 | +1 | 13 |
| - | - | 5670 | 1346 | - | - | 0 | - |
| - | - | 2097 | 1347 | - | - | 0 | - |
| - | - | 3626 | 1348 | - | - | 0 | - |
| - | - | 2093 | 1349 | - | - | 0 | - |
| 14 | c | 1.122E+04 | 1432 | 0.001977 | 1.381 | +1 | 14 |
| - | - | 7301 | 1433 | - | - | 0 | - |
| - | - | 4133 | 1434 | - | - | 0 | - |
| - | - | 1137 | 1435 | - | - | 0 | - |
| - | - | 913.6 | 1489 | - | - | 0 | - |
| - | - | 987.9 | 1490 | - | - | 0 | - |
| - | - | 2174 | 1531 | - | - | 0 | - |
| - | - | 3204 | 1532 | - | - | 0 | - |
| - | - | 1716 | 1533 | - | - | 0 | - |
| - | - | 913 | 1545 | - | - | 0 | - |
| - | - | 1336 | 1546 | - | - | 0 | - |
| - | - | 1929 | 1547 | - | - | 0 | - |
| - | - | 955.7 | 1548 | - | - | 0 | - |
| - | - | 742.1 | 1549 | - | - | 0 | - |
| - | - | 1003 | 1557 | - | - | 0 | - |
| - | - | 1614 | 1558 | - | - | 0 | - |
| - | - | 2801 | 1563 | - | - | 0 | - |
| - | - | 1280 | 1564 | - | - | 0 | - |
| - | - | 3116 | 1573 | - | - | 0 | - |
| - | - | 1.384E+04 | 1574 | - | - | 0 | - |
| - | - | 1.209E+04 | 1575 | - | - | 0 | - |
| - | - | 5270 | 1576 | - | - | 0 | - |
| - | - | 2280 | 1577 | - | - | 0 | - |
| - | - | 679.9 | 1578 | - | - | 0 | - |
| - | - | 869.5 | 1587 | - | - | 0 | - |
| - | - | 1411 | 1589 | - | - | 0 | - |
| - | - | 5554 | 1590 | - | - | 0 | - |
| - | - | 1.491E+04 | 1591 | - | - | 0 | - |
| - | - | 1.11E+04 | 1592 | - | - | 0 | - |
| - | - | 5758 | 1593 | - | - | 0 | - |
| - | - | 1788 | 1594 | - | - | 0 | - |
| - | - | 698.2 | 2191 | - | - | 0 | - |
| - | - | 762.3 | 2619 | - | - | 0 | - |
| - | - | 634.4 | 2647 | - | - | 0 | - |
| - | - | 764.4 | 2680 | - | - | 0 | - |
| - | - | 745.5 | 3382 | - | - | 0 | - |

m/z Charge Intensity FragmentType MassShift Position
120.08135986328125 0 460.377
123.89373779296875 0 342.35245
126.09164428710938 0 374.67432
129.31805419921875 0 436.44122
130.8098907470703 0 418.82224
133.0868682861328 0 508.77664
136.07589721679688 0 1425.146
140.0822296142578 0 1149.606
143.11810302734375 0 1132.8591
148.95443725585938 0 973.71716
151.67298889160156 0 406.26083
157.5751190185547 0 474.60672
157.5821990966797 0 407.57498
168.11331176757812 0 902.686
171.1136474609375 0 458.13535
171.14889526367188 0 446.15
173.08102416992188 0 427.3357
173.4392852783203 0 1569.4119
174.0557861328125 0 528.78467
175.11912536621094 0 1321.6638 y 14
185.0927734375 0 948.25146
185.139892578125 0 3161.28
187.1079559326172 0 5445.991
187.1443328857422 0 1440.7849
196.9701690673828 0 440.72665
199.14459228515625 0 654.2728
199.18069458007812 0 5410.7964
200.18435668945312 0 969.6554
215.1392364501953 0 2190.3772
227.17552185058594 0 2883.5024
228.1336212158203 0 495.88315
228.17955017089844 0 598.39557
229.1294403076172 0 707.6556 w 13
245.1246795654297 0 4733.717 y Ammonia loss 13
254.1139373779297 0 630.17456
262.1505432128906 0 778.0086 y 13
262.6312255859375 0 494.2031
270.1443786621094 0 666.32623
301.1031799316406 0 1126.2014
314.1719970703125 0 713.2955
316.1617736816406 0 2163.0825 w 12
331.148681640625 0 907.37964
332.1572265625 0 902.165 y Ammonia loss 12
332.759521484375 0 560.1787
333.16412353515625 0 1828.979 z 12
349.11553955078125 0 1017.3731
349.18255615234375 0 785.84686 y 12
357.1519775390625 0 1380.6399
367.2347106933594 0 819.97675
369.21185302734375 0 707.50476
377.1669006347656 0 7078.3506
404.7121276855469 0 751.167
415.2244567871094 0 762.1814 z 7
415.72296142578125 0 602.2622
417.2100830078125 0 2104.817 w 11
422.002197265625 0 639.9396
423.231201171875 0 824.4978 y 7
448.2511291503906 0 740.03485 y 11
453.34368896484375 0 1462.9159
460.27606201171875 0 718.51746 c 3
470.2623291015625 0 528.3832
476.2342224121094 0 1113.1449
489.2560119628906 0 568.55145 z 10
499.297607421875 0 613.5024
505.27392578125 0 6212.1284 y 10
506.2749328613281 0 1532.2513
509.80560302734375 0 534.8917
515.2800903320312 0 686.4672
516.7560424804688 0 1085.5449
517.2600708007812 0 905.9412
520.2643432617188 0 739.5638
524.264892578125 0 1865.9049 y 5
524.7664184570312 0 746.9053
529.2718505859375 0 966.16046
529.7680053710938 0 1132.6018
530.2769775390625 0 862.38934
530.3170166015625 0 7227.038
531.2659912109375 0 925.54486
531.319580078125 0 1836.8643
531.7434692382812 0 1768.7594
551.31982421875 0 763.05676
559.3455810546875 0 849.5051 c 4
565.78759765625 0 3650.9348 z 4
566.289306640625 0 2439.7275
566.7916870117188 0 1195.6598
569.3310546875 0 1182.709
573.7969970703125 0 2381.071 y 4
574.298095703125 0 1289.2366 y Water loss 9
576.283935546875 0 940.43054 z 9
577.280517578125 0 744.66364
585.2634887695312 0 876.3521
587.33935546875 0 708.56683
600.8031616210938 0 1450.1868 w 3
601.30517578125 0 1290.9062
601.8037109375 0 1095.2479
604.3673095703125 0 1206.1814
630.3081665039062 0 4481.9194 z 3
630.8118286132812 0 2331.521
631.3145141601562 0 1477.6611
637.81494140625 0 779.44666
638.3179321289062 0 4939.219 y 3
638.8179931640625 0 1615.5493
639.32763671875 0 824.7954
640.3394775390625 0 1794.3542
646.377685546875 0 3695.8098 c 5
647.3810424804688 0 1059.9037
650.384765625 0 644.47906
660.3656005859375 0 914.8388
668.400390625 0 1009.97534
670.3906860351562 0 960.9612
673.8255615234375 0 2428.229 z 2
674.3274536132812 0 3762.9565
674.8291015625 0 1845.6661
676.3506469726562 0 960.94556
681.8343505859375 0 15851.657 y 2
682.3355712890625 0 13093.221
682.837158203125 0 6198.5776
683.3372192382812 0 888.76227
686.409423828125 0 1209.6729
718.3973999023438 0 623.7671
719.3920288085938 0 1002.98737
720.3861694335938 0 637.2338
727.3756103515625 0 830.27966
728.3786010742188 0 766.8479
738.3761596679688 0 1264.134 y 1
738.8792114257812 0 1067.8466
739.3738403320312 0 680.31976
741.3882446289062 0 2940.8745
742.3900146484375 0 841.1801
743.3758544921875 0 821.3575 w 7
759.4004516601562 0 767.3969
761.4044799804688 0 7394.1934 c 6
762.4066772460938 0 3205.4758
763.40869140625 0 811.03705
765.382080078125 0 14704.399
765.88330078125 0 9855.419
766.3873291015625 0 4145.538
766.8878173828125 0 1077.5322
773.9083251953125 0 3983.1453
774.4083251953125 0 3658.7136
774.910400390625 0 1477.697
775.4122314453125 0 1046.4142
778.409423828125 0 800.65845
779.40771484375 0 708.5836
782.38037109375 0 705.483
785.9000854492188 0 932.1151
786.9102172851562 0 12965.258
787.4105224609375 0 10352.677
787.9103393554688 0 4958.5356
788.4130249023438 0 1634.3052
794.4140014648438 0 805.2165
794.9169311523438 0 7818.058
795.4197387695312 0 11816.19
795.9213256835938 0 8023.615
796.42138671875 0 3262.0342
800.486572265625 0 896.6573
827.3822021484375 0 858.5601
829.4402465820312 0 4902.43 z 7
830.44287109375 0 3186.1602
831.45361328125 0 1208.8275
843.4029541015625 0 1965.1791
844.407470703125 0 1028.296
845.4563598632812 0 941.2216 y 7
853.4247436523438 0 1161.3806
871.4258422851562 0 1952.5515
872.438232421875 0 1244.646
887.5194091796875 0 2634.3167
900.4738159179688 0 1418.8652 c Ammonia loss 7
901.480224609375 0 655.13525
915.45166015625 0 889.5945
944.4605712890625 0 2961.5469 z 6
945.4685668945312 0 2292.413
968.5079345703125 0 984.3663
974.5521850585938 0 3738.7378
985.5086669921875 0 1735.2393
986.509033203125 0 916.94446
987.5116577148438 0 1071.8348
1012.5315551757812 0 1382.2744
1013.5386352539062 0 914.6725
1014.5568237304688 0 5461.0317 c 8
1015.55810546875 0 2462.5615
1023.5098266601562 0 661.272
1024.5264892578125 0 2366.8474
1025.530517578125 0 1236.2634
1027.516357421875 0 790.30457
1031.5013427734375 0 1335.9473 z 5
1032.507080078125 0 1287.6614
1040.5263671875 0 1974.2208
1041.51904296875 0 4308.2563
1042.517578125 0 2499.595
1043.52978515625 0 1184.4723
1044.5216064453125 0 1649.2826
1045.5218505859375 0 2064.647
1046.5411376953125 0 963.6212
1047.5206298828125 0 668.6088 y 5
1057.5416259765625 0 5383.1323
1058.540771484375 0 11411.017
1059.5445556640625 0 5650.1577
1060.5377197265625 0 2606.4475
1061.536865234375 0 3649.643
1062.5379638671875 0 3297.082
1063.5728759765625 0 4909.4067
1101.5894775390625 0 8376.371 c 9
1102.5926513671875 0 4882.03
1103.5943603515625 0 1232.049
1114.587890625 0 756.2182
1115.607421875 0 693.0641
1130.5699462890625 0 938.947 z 4
1131.57275390625 0 2882.826
1132.579345703125 0 1499.671
1133.5791015625 0 715.32855
1158.6104736328125 0 4380.328 c 10
1159.61328125 0 2943.234
1160.6171875 0 1160.3772
1213.6693115234375 0 742.5369
1214.669921875 0 694.97076
1257.679931640625 0 3833.5142 c 11
1258.6820068359375 0 3317.2517
1260.6165771484375 0 4318.4424
1261.6212158203125 0 2504.911
1262.6282958984375 0 856.96185
1275.6300048828125 0 1462.8694 y 3
1276.6329345703125 0 841.2253
1277.63916015625 0 889.52814
1288.0716552734375 0 659.98236
1344.7086181640625 0 7158.148 c 12
1345.714111328125 0 5669.6235
1346.707763671875 0 2096.6802
1347.650390625 0 3625.714
1348.649658203125 0 2092.5483
1431.7418212890625 0 11215.724 c 13
1432.744873046875 0 7301.071
1433.74658203125 0 4132.6484
1434.741455078125 0 1137.3223
1488.7548828125 0 913.5788
1489.7440185546875 0 987.923
1530.792236328125 0 2174.328
1531.7930908203125 0 3204.121
1532.7939453125 0 1716.3408
1544.833740234375 0 912.9972
1545.8297119140625 0 1336.2157
1546.8297119140625 0 1929.204
1547.8128662109375 0 955.71204
1548.815673828125 0 742.0904
1556.7918701171875 0 1003.297
1557.800537109375 0 1613.8029
1562.846435546875 0 2800.6096
1563.8465576171875 0 1279.6105
1572.8106689453125 0 3116.239
1573.8162841796875 0 13840.798
1574.8170166015625 0 12088.425
1575.820556640625 0 5270.0503
1576.8145751953125 0 2280.4055
1577.8427734375 0 679.9369
1586.8134765625 0 869.45905
1588.8182373046875 0 1411.406
1589.8265380859375 0 5554.1562
1590.83740234375 0 14914.914
1591.8421630859375 0 11104.271
1592.844482421875 0 5757.7915
1593.8385009765625 0 1788.4319
2190.648193359375 0 698.1891
2618.82666015625 0 762.2989
2647.057373046875 0 634.43335
2680.270751953125 0 764.43176
3382.00732421875 0 745.4848

Spectrum Details

|  |  |
| --- | --- |
| Matched peaks? Matched peaks The total absolute number of peaks matched. Additionally in brackets the total fraction of peaks matched and the total number of peaks is shown. | 44 (16.54% of 266) |
| FDR? FDR The false discovery rate estimated for this peptide. It is calculated by matching all theoretical fragments with a non-integer shift with the raw peaks for this spectrum. This is done with 40 different shifts. The resulting percentage is the average number of annotated peaks over the number of annotated peaks with the correct spectrum. | 0.81% |
| Satellite FDR? Satellite FDR See the FDR for details on its calculation. This satellite ion specific FDR only contains the satellite ions (d/w) for I/L/J positions. | - |
| PSM Score? PSM Score The PSM Score as given by Hecklib to this annotated spectrum. It is shown with three significant figures. | 313 |

#### Spectrum 7196? Spectrum 7196 The raw spectrum of this peptide as annotated by Hecklib. The fragments are coloured according to ion type (see legend). Any peaks with a star '\*' as text can be hovered over to see the full details, first the ion type second the mass shift type. By hovering over the amino acids in the peptide or ions in the legend the corresponding peaks are highlighted. By toggling the 'Unassigned' label you can turn the background (unassigned) peaks on or off in the plot. By updating the slider in the Ion legend you can update the spectrum to only show the top X% of the peaks with labels. The top X% means any peak that is within X% of the highest intensity. By dragging in the spectrum you can zoom in to a specific part of the spectrum and use 'Zoom Out' to get back to the original zoom level. The annotation of the spectrum is based on the given sequence in the peptides file and is done with different software so inconsistencies are likely. The peaks are annotated based on the given sequence, with 20 ppm tolerance.

Copy Data

##### Spectrum 7196 (TSV)

###### Preview

```
Loading example...
```

*Click on the button to copy the data to your clipboard.*

Mz MinMz MaxIntensity Max

WidthHeightPeptide font sizePeptide stroke widthSpectrum font sizeSpectrum stroke widthCompact peptide

Ion legend

wxyz

abcd

OtherUnassignedIon^Charge^_Position_Show for top:%

JJSEVSDRPSGVSSR

03.05e+36.10e+39.14e+31.22e+4

Zoom Out

y^+1^_1_y^+1^_2_y^+1^_2_w^+1^_3_y^+1^_3_z^+1^_3_w^+1^_4_y^+1^_4_y^+1^_5_y^+2^_10_c^+1^_5_z^+2^_11_y^+2^_11_y^+1^_6_z^+1^_6_w^+2^_12_z^+2^_12_y^+2^_12_c^+1^_6_y^+2^_13_z^+2^_13_y^+2^_13_c^+1^_7_z^+1^_8_z^+1^_9_c^+1^_9_z^+1^_10_c^+1^_10_c^+1^_11_c^+1^_12_y^+1^_12_c^+1^_13_c^+1^_14_

0776155223273103

Fragment Matches Table

Show background peaks

| Position | Ion type | Intensity | mz Theoretical | mz Error (Th) | mz Error (ppm) | Charge | Series Number |
| --- | --- | --- | --- | --- | --- | --- | --- |
| - | - | 432.3 | 120.1 | - | - | 0 | - |
| - | - | 424.7 | 126.1 | - | - | 0 | - |
| - | - | 419 | 128.7 | - | - | 0 | - |
| - | - | 530.2 | 131.1 | - | - | 0 | - |
| - | - | 1.103E+04 | 133.1 | - | - | 0 | - |
| - | - | 455.4 | 133.6 | - | - | 0 | - |
| - | - | 1553 | 140.1 | - | - | 0 | - |
| - | - | 467.5 | 141.9 | - | - | 0 | - |
| - | - | 414.9 | 143 | - | - | 0 | - |
| 15 | y | 667.2 | 158.1 | 4.162E-05 | 0.2632 | +1 | 1 |
| - | - | 662.3 | 161.1 | - | - | 0 | - |
| - | - | 6485 | 177.1 | - | - | 0 | - |
| - | - | 2983 | 185.1 | - | - | 0 | - |
| - | - | 1.207E+04 | 187.1 | - | - | 0 | - |
| - | - | 717.3 | 187.1 | - | - | 0 | - |
| - | - | 497.7 | 187.2 | - | - | 0 | - |
| - | - | 549.7 | 188.1 | - | - | 0 | - |
| - | - | 468.9 | 198.8 | - | - | 0 | - |
| - | - | 4191 | 199.2 | - | - | 0 | - |
| - | - | 469.7 | 200.1 | - | - | 0 | - |
| - | - | 3790 | 205.1 | - | - | 0 | - |
| - | - | 1449 | 215.1 | - | - | 0 | - |
| - | - | 1314 | 221.1 | - | - | 0 | - |
| - | - | 1823 | 227.2 | - | - | 0 | - |
| - | - | 583.6 | 239.2 | - | - | 0 | - |
| 14 | y | 3820 | 245.1 | 0.000126 | 0.5141 | +1 | 2 |
| - | - | 645.8 | 249.1 | - | - | 0 | - |
| 14 | y | 583.6 | 262.2 | 0.00176 | 6.713 | +1 | 2 |
| - | - | 1202 | 265.2 | - | - | 0 | - |
| - | - | 632.2 | 270.1 | - | - | 0 | - |
| - | - | 1084 | 292.2 | - | - | 0 | - |
| - | - | 553.3 | 293.7 | - | - | 0 | - |
| 13 | w | 1287 | 316.2 | 0.0006862 | 2.17 | +1 | 3 |
| 13 | y | 1008 | 332.2 | 5.73E-05 | 0.1725 | +1 | 3 |
| 13 | z | 885.2 | 333.2 | 0.0001743 | 0.5232 | +1 | 3 |
| - | - | 961.4 | 336.2 | - | - | 0 | - |
| - | - | 533.7 | 353.2 | - | - | 0 | - |
| - | - | 866.8 | 357.2 | - | - | 0 | - |
| - | - | 683.6 | 376.1 | - | - | 0 | - |
| - | - | 658.3 | 415.3 | - | - | 0 | - |
| 12 | w | 1298 | 417.2 | 0.001073 | 2.572 | +1 | 4 |
| 12 | y | 873.5 | 448.3 | 0.0007744 | 1.728 | +1 | 4 |
| - | - | 1345 | 458.3 | - | - | 0 | - |
| - | - | 538.1 | 458.6 | - | - | 0 | - |
| - | - | 1090 | 459.3 | - | - | 0 | - |
| - | - | 1092 | 475.3 | - | - | 0 | - |
| - | - | 3384 | 503.3 | - | - | 0 | - |
| 11 | y | 3412 | 505.3 | 0.0004593 | 0.9091 | +1 | 5 |
| - | - | 636.5 | 506.3 | - | - | 0 | - |
| - | - | 708.9 | 512.3 | - | - | 0 | - |
| - | - | 1340 | 514.3 | - | - | 0 | - |
| 6 | y | 825.3 | 524.3 | 0.0004339 | 0.8277 | +2 | 10 |
| - | - | 706.1 | 524.6 | - | - | 0 | - |
| - | - | 1011 | 524.8 | - | - | 0 | - |
| - | - | 3478 | 530.2 | - | - | 0 | - |
| - | - | 6133 | 530.3 | - | - | 0 | - |
| - | - | 1585 | 531.2 | - | - | 0 | - |
| - | - | 1454 | 531.3 | - | - | 0 | - |
| 5 | c | 801.2 | 559.3 | 0.003314 | 5.925 | +1 | 5 |
| 5 | z | 1874 | 565.8 | 0.0007221 | 1.276 | +2 | 11 |
| - | - | 1658 | 566.3 | - | - | 0 | - |
| - | - | 837.6 | 566.8 | - | - | 0 | - |
| 5 | y | 827.8 | 573.8 | 0.001431 | 2.494 | +2 | 11 |
| 10 | y | 791.9 | 574.3 | 0.004844 | 8.435 | +1 | 6 |
| 10 | z | 947.8 | 576.3 | 0.001645 | 2.855 | +1 | 6 |
| 4 | w | 826.2 | 600.8 | 0.001471 | 2.449 | +2 | 12 |
| - | - | 1038 | 601.3 | - | - | 0 | - |
| 4 | z | 2469 | 630.3 | 0.0007991 | 1.268 | +2 | 12 |
| - | - | 1104 | 630.8 | - | - | 0 | - |
| - | - | 804.5 | 631.3 | - | - | 0 | - |
| 4 | y | 1976 | 638.3 | 0.000398 | 0.6235 | +2 | 12 |
| - | - | 2219 | 638.8 | - | - | 0 | - |
| - | - | 949 | 639.3 | - | - | 0 | - |
| - | - | 1916 | 643.3 | - | - | 0 | - |
| - | - | 4169 | 644.3 | - | - | 0 | - |
| 6 | c | 1733 | 646.4 | 0.0007292 | 1.128 | +1 | 6 |
| - | - | 1435 | 647.4 | - | - | 0 | - |
| - | - | 628.9 | 657.1 | - | - | 0 | - |
| - | - | 807.4 | 669.3 | - | - | 0 | - |
| - | - | 550.4 | 670.3 | - | - | 0 | - |
| 3 | y | 720.6 | 673.3 | 0.01084 | 16.1 | +2 | 13 |
| 3 | z | 2617 | 673.8 | 0.0001507 | 0.2236 | +2 | 13 |
| - | - | 2674 | 674.3 | - | - | 0 | - |
| - | - | 935.4 | 674.8 | - | - | 0 | - |
| 3 | y | 1.132E+04 | 681.8 | 6.983E-05 | 0.1024 | +2 | 13 |
| - | - | 7184 | 682.3 | - | - | 0 | - |
| - | - | 2967 | 682.8 | - | - | 0 | - |
| - | - | 1072 | 683.3 | - | - | 0 | - |
| - | - | 703.6 | 692.3 | - | - | 0 | - |
| - | - | 3215 | 727.4 | - | - | 0 | - |
| - | - | 835.5 | 728.4 | - | - | 0 | - |
| - | - | 2341 | 745.4 | - | - | 0 | - |
| - | - | 674.3 | 758.9 | - | - | 0 | - |
| 7 | c | 4143 | 761.4 | 0.0002128 | 0.2795 | +1 | 7 |
| - | - | 1918 | 762.4 | - | - | 0 | - |
| - | - | 740.7 | 763.4 | - | - | 0 | - |
| - | - | 608.7 | 764.9 | - | - | 0 | - |
| - | - | 7024 | 765.4 | - | - | 0 | - |
| - | - | 7244 | 765.9 | - | - | 0 | - |
| - | - | 2360 | 766.4 | - | - | 0 | - |
| - | - | 644 | 766.9 | - | - | 0 | - |
| - | - | 622.8 | 767.4 | - | - | 0 | - |
| - | - | 3553 | 773.9 | - | - | 0 | - |
| - | - | 1589 | 774.4 | - | - | 0 | - |
| - | - | 987.9 | 774.9 | - | - | 0 | - |
| - | - | 1164 | 786.4 | - | - | 0 | - |
| - | - | 8042 | 786.9 | - | - | 0 | - |
| - | - | 6013 | 787.4 | - | - | 0 | - |
| - | - | 3766 | 787.9 | - | - | 0 | - |
| - | - | 714.3 | 788.4 | - | - | 0 | - |
| - | - | 733.7 | 794.4 | - | - | 0 | - |
| - | - | 5533 | 794.9 | - | - | 0 | - |
| - | - | 7599 | 795.4 | - | - | 0 | - |
| - | - | 4266 | 795.9 | - | - | 0 | - |
| - | - | 2594 | 796.4 | - | - | 0 | - |
| - | - | 831.6 | 796.9 | - | - | 0 | - |
| - | - | 641.7 | 813.4 | - | - | 0 | - |
| - | - | 1688 | 821.4 | - | - | 0 | - |
| 8 | z | 3115 | 829.4 | 0.004519 | 5.448 | +1 | 8 |
| - | - | 2444 | 830.4 | - | - | 0 | - |
| - | - | 1121 | 831.4 | - | - | 0 | - |
| - | - | 875.6 | 832.4 | - | - | 0 | - |
| - | - | 1511 | 839.4 | - | - | 0 | - |
| - | - | 769.8 | 840.4 | - | - | 0 | - |
| - | - | 608.8 | 847.4 | - | - | 0 | - |
| - | - | 2113 | 857.4 | - | - | 0 | - |
| - | - | 948.4 | 858.4 | - | - | 0 | - |
| - | - | 620.7 | 860.4 | - | - | 0 | - |
| 7 | z | 1718 | 944.5 | 0.001128 | 1.194 | +1 | 9 |
| - | - | 1062 | 945.5 | - | - | 0 | - |
| - | - | 1386 | 971.5 | - | - | 0 | - |
| - | - | 665.1 | 976.4 | - | - | 0 | - |
| - | - | 645 | 983.4 | - | - | 0 | - |
| 9 | c | 3254 | 1015 | 0.005162 | 5.088 | +1 | 9 |
| - | - | 1894 | 1016 | - | - | 0 | - |
| - | - | 838.7 | 1017 | - | - | 0 | - |
| - | - | 1891 | 1027 | - | - | 0 | - |
| - | - | 647.4 | 1029 | - | - | 0 | - |
| 6 | z | 789 | 1031 | 0.000258 | 0.2501 | +1 | 10 |
| - | - | 791.6 | 1033 | - | - | 0 | - |
| - | - | 995.3 | 1044 | - | - | 0 | - |
| - | - | 6571 | 1045 | - | - | 0 | - |
| - | - | 1830 | 1046 | - | - | 0 | - |
| - | - | 871.4 | 1047 | - | - | 0 | - |
| - | - | 2463 | 1058 | - | - | 0 | - |
| - | - | 1809 | 1059 | - | - | 0 | - |
| - | - | 900.7 | 1060 | - | - | 0 | - |
| - | - | 1149 | 1061 | - | - | 0 | - |
| - | - | 2588 | 1062 | - | - | 0 | - |
| - | - | 6404 | 1063 | - | - | 0 | - |
| - | - | 1860 | 1064 | - | - | 0 | - |
| 10 | c | 4594 | 1102 | 0.0001021 | 0.09273 | +1 | 10 |
| - | - | 2907 | 1103 | - | - | 0 | - |
| - | - | 1022 | 1104 | - | - | 0 | - |
| - | - | 796.6 | 1115 | - | - | 0 | - |
| - | - | 1456 | 1132 | - | - | 0 | - |
| - | - | 1182 | 1133 | - | - | 0 | - |
| 11 | c | 2985 | 1159 | 0.0009773 | 0.8435 | +1 | 11 |
| - | - | 1675 | 1160 | - | - | 0 | - |
| - | - | 693 | 1214 | - | - | 0 | - |
| - | - | 681 | 1221 | - | - | 0 | - |
| 12 | c | 1685 | 1258 | 0.006157 | 4.896 | +1 | 12 |
| - | - | 1845 | 1259 | - | - | 0 | - |
| - | - | 2289 | 1261 | - | - | 0 | - |
| - | - | 1324 | 1262 | - | - | 0 | - |
| 4 | y | 1072 | 1276 | 0.007064 | 5.538 | +1 | 12 |
| 13 | c | 4449 | 1345 | 0.001565 | 1.164 | +1 | 13 |
| - | - | 3718 | 1346 | - | - | 0 | - |
| - | - | 1636 | 1347 | - | - | 0 | - |
| - | - | 2239 | 1348 | - | - | 0 | - |
| - | - | 1843 | 1349 | - | - | 0 | - |
| - | - | 576.9 | 1396 | - | - | 0 | - |
| 14 | c | 5877 | 1432 | 0.00332 | 2.319 | +1 | 14 |
| - | - | 3788 | 1433 | - | - | 0 | - |
| - | - | 2874 | 1434 | - | - | 0 | - |
| - | - | 1240 | 1531 | - | - | 0 | - |
| - | - | 1635 | 1532 | - | - | 0 | - |
| - | - | 1394 | 1533 | - | - | 0 | - |
| - | - | 770.3 | 1546 | - | - | 0 | - |
| - | - | 644.1 | 1549 | - | - | 0 | - |
| - | - | 693.5 | 1557 | - | - | 0 | - |
| - | - | 1162 | 1563 | - | - | 0 | - |
| - | - | 718.1 | 1564 | - | - | 0 | - |
| - | - | 1853 | 1573 | - | - | 0 | - |
| - | - | 9873 | 1574 | - | - | 0 | - |
| - | - | 7629 | 1575 | - | - | 0 | - |
| - | - | 4495 | 1576 | - | - | 0 | - |
| - | - | 1034 | 1577 | - | - | 0 | - |
| - | - | 1679 | 1589 | - | - | 0 | - |
| - | - | 3443 | 1590 | - | - | 0 | - |
| - | - | 9835 | 1591 | - | - | 0 | - |
| - | - | 7730 | 1592 | - | - | 0 | - |
| - | - | 3901 | 1593 | - | - | 0 | - |
| - | - | 1545 | 1594 | - | - | 0 | - |
| - | - | 709.9 | 2124 | - | - | 0 | - |
| - | - | 728 | 3072 | - | - | 0 | - |

m/z Charge Intensity FragmentType MassShift Position
120.08084869384766 0 432.28845
126.09163665771484 0 424.7318
128.7245635986328 0 418.9751
131.07046508789062 0 530.1984
133.08609008789062 0 11028.728
133.6487579345703 0 455.37366
140.08209228515625 0 1552.6644
141.9342803955078 0 467.524
143.033203125 0 414.9221
158.0923614501953 0 667.21545 y Ammonia loss 14
161.08090209960938 0 662.3192
177.11224365234375 0 6485.464
185.1399383544922 0 2983.0989
187.10781860351562 0 12072.17
187.14410400390625 0 717.32355
187.1609344482422 0 497.6909
188.06759643554688 0 549.67474
198.8339385986328 0 468.8962
199.18060302734375 0 4190.564
200.1253662109375 0 469.6659
205.1071014404297 0 3790.2502
215.13897705078125 0 1448.9185
221.13864135742188 0 1314.2993
227.17556762695312 0 1823.0084
239.15005493164062 0 583.5726
245.1245574951172 0 3819.9844 y Ammonia loss 13
249.13421630859375 0 645.808
262.1527404785156 0 583.5806 y 13
265.1645812988281 0 1202.4656
270.1202392578125 0 632.214
292.1758117675781 0 1083.8558
293.66839599609375 0 553.25055
316.1622314453125 0 1286.6912 w 12
332.1564025878906 0 1007.5799 y Ammonia loss 12
333.1644592285156 0 885.24945 z 12
336.2016296386719 0 961.4233
353.2157897949219 0 533.74835
357.1509704589844 0 866.84845
376.06463623046875 0 683.6438
415.2552795410156 0 658.25433
417.2102966308594 0 1297.9927 w 11
448.252197265625 0 873.4554 y 11
458.2615661621094 0 1344.9532
458.6198425292969 0 538.12665
459.27978515625 0 1090.1558
475.2864990234375 0 1091.627
503.306640625 0 3384.428
505.2733459472656 0 3411.7 y 10
506.2737731933594 0 636.5121
512.3062133789062 0 708.87
514.2864379882812 0 1340.2369
524.2620849609375 0 825.3425 y 5
524.6119995117188 0 706.0945
524.7647094726562 0 1010.5803
530.2395629882812 0 3477.6597
530.31689453125 0 6133.433
531.2431640625 0 1585.3795
531.3035278320312 0 1454.3643
559.3416748046875 0 801.2391 c 4
565.7880859375 0 1873.6387 z 4
566.289306640625 0 1658.2355
566.7866821289062 0 837.5724
573.7981567382812 0 827.78284 y 4
574.2991943359375 0 791.94116 y Water loss 9
576.2845458984375 0 947.7584 z 9
600.800537109375 0 826.2249 w 3
601.3029174804688 0 1038.2213
630.307861328125 0 2469.3933 z 3
630.8120727539062 0 1103.5823
631.3067016601562 0 804.49164
638.3184204101562 0 1975.5712 y 3
638.8175048828125 0 2219.406
639.3207397460938 0 948.97833
643.3282470703125 0 1915.5062
644.3342895507812 0 4168.5713
646.3777465820312 0 1733.4801 c 5
647.3786010742188 0 1434.6959
657.0740356445312 0 628.8586
669.3133544921875 0 807.39185
670.313232421875 0 550.3955
673.3316040039062 0 720.56903 y Ammonia loss 2
673.8245239257812 0 2616.669 z 2
674.3251953125 0 2673.7734
674.8282470703125 0 935.3681
681.8341064453125 0 11320.507 y 2
682.3350830078125 0 7183.9966
682.8365478515625 0 2966.6875
683.3361206054688 0 1071.5079
692.3380737304688 0 703.62354
727.3724975585938 0 3214.6982
728.374755859375 0 835.499
745.3836669921875 0 2341.3037
758.8749389648438 0 674.31134
761.4037475585938 0 4142.5903 c 6
762.4046630859375 0 1917.5625
763.405029296875 0 740.7339
764.906494140625 0 608.6568
765.3821411132812 0 7024.1226
765.8828125 0 7243.731
766.38623046875 0 2359.9255
766.8897094726562 0 644.0067
767.3847045898438 0 622.8341
773.906494140625 0 3552.9585
774.4080200195312 0 1588.7649
774.9075317382812 0 987.9232
786.400634765625 0 1163.578
786.90771484375 0 8041.5664
787.40966796875 0 6012.905
787.9103393554688 0 3765.694
788.4053955078125 0 714.29767
794.3937377929688 0 733.6515
794.9161987304688 0 5533.025
795.4188842773438 0 7599.008
795.9208374023438 0 4265.6567
796.4193725585938 0 2594.379
796.9146728515625 0 831.6118
813.365966796875 0 641.7449
821.395263671875 0 1688.0203
829.435546875 0 3114.504 z 7
830.4415893554688 0 2444.3633
831.448486328125 0 1121.214
832.3804321289062 0 875.61743
839.4027099609375 0 1511.3049
840.4142456054688 0 769.7506
847.3930053710938 0 608.7517
857.4164428710938 0 2112.845
858.4213256835938 0 948.3805
860.44482421875 0 620.7064
944.4658813476562 0 1718.0228 z 6
945.472412109375 0 1061.9102
971.494873046875 0 1385.8419
976.4390258789062 0 665.0762
983.4356079101562 0 645.02203
1014.5526733398438 0 3254.287 c 8
1015.5598754882812 0 1894.4955
1016.5552978515625 0 838.6617
1027.496337890625 0 1890.7292
1028.5020751953125 0 647.36896
1031.498779296875 0 789.0388 z 5
1032.51220703125 0 791.57227
1043.518310546875 0 995.25037
1044.521728515625 0 6571.392
1045.5191650390625 0 1829.5967
1046.5233154296875 0 871.4483
1057.576904296875 0 2462.9092
1058.557373046875 0 1808.6904
1059.5677490234375 0 900.6663
1060.5240478515625 0 1149.1555
1061.5301513671875 0 2588.3066
1062.5325927734375 0 6403.785
1063.5498046875 0 1860.0444
1101.5899658203125 0 4593.6787 c 9
1102.5928955078125 0 2906.9429
1103.58251953125 0 1021.6514
1114.59521484375 0 796.6152
1131.5750732421875 0 1455.5515
1132.5767822265625 0 1181.8169
1158.6123046875 0 2985.0498 c 10
1159.61962890625 0 1675.248
1213.6678466796875 0 693.0444
1220.960205078125 0 681.002
1257.673583984375 0 1684.5032 c 11
1258.6754150390625 0 1845.0259
1260.6173095703125 0 2289.4314
1261.61865234375 0 1324.3926
1275.6217041015625 0 1071.9502 y 3
1344.710205078125 0 4449.13 c 12
1345.71533203125 0 3718.4158
1346.715087890625 0 1636.1875
1347.6500244140625 0 2238.605
1348.66162109375 0 1842.7991
1396.28759765625 0 576.8587
1431.740478515625 0 5876.933 c 13
1432.74267578125 0 3788.3599
1433.7440185546875 0 2873.8667
1530.787841796875 0 1239.9974
1531.78759765625 0 1634.5115
1532.781494140625 0 1393.6079
1545.823974609375 0 770.25745
1548.802978515625 0 644.1394
1556.7864990234375 0 693.52716
1562.8394775390625 0 1162.0278
1563.8387451171875 0 718.1168
1572.823486328125 0 1852.8638
1573.814697265625 0 9872.692
1574.815673828125 0 7628.9795
1575.8182373046875 0 4494.553
1576.8209228515625 0 1034.4347
1588.802734375 0 1679.2233
1589.8294677734375 0 3443.212
1590.8370361328125 0 9834.901
1591.8406982421875 0 7730.2886
1592.84228515625 0 3900.719
1593.854736328125 0 1545.4076
2123.927001953125 0 709.9
3072.33935546875 0 728.017

Spectrum Details

|  |  |
| --- | --- |
| Matched peaks? Matched peaks The total absolute number of peaks matched. Additionally in brackets the total fraction of peaks matched and the total number of peaks is shown. | 33 (16.84% of 196) |
| FDR? FDR The false discovery rate estimated for this peptide. It is calculated by matching all theoretical fragments with a non-integer shift with the raw peaks for this spectrum. This is done with 40 different shifts. The resulting percentage is the average number of annotated peaks over the number of annotated peaks with the correct spectrum. | 1.08% |
| Satellite FDR? Satellite FDR See the FDR for details on its calculation. This satellite ion specific FDR only contains the satellite ions (d/w) for I/L/J positions. | - |
| PSM Score? PSM Score The PSM Score as given by Hecklib to this annotated spectrum. It is shown with three significant figures. | 225 |

#### Spectrum 6646? Spectrum 6646 The raw spectrum of this peptide as annotated by Hecklib. The fragments are coloured according to ion type (see legend). Any peaks with a star '\*' as text can be hovered over to see the full details, first the ion type second the mass shift type. By hovering over the amino acids in the peptide or ions in the legend the corresponding peaks are highlighted. By toggling the 'Unassigned' label you can turn the background (unassigned) peaks on or off in the plot. By updating the slider in the Ion legend you can update the spectrum to only show the top X% of the peaks with labels. The top X% means any peak that is within X% of the highest intensity. By dragging in the spectrum you can zoom in to a specific part of the spectrum and use 'Zoom Out' to get back to the original zoom level. The annotation of the spectrum is based on the given sequence in the peptides file and is done with different software so inconsistencies are likely. The peaks are annotated based on the given sequence, with 20 ppm tolerance.

Copy Data

##### Spectrum 6646 (TSV)

###### Preview

```
Loading example...
```

*Click on the button to copy the data to your clipboard.*

Mz MinMz MaxIntensity Max

WidthHeightPeptide font sizePeptide stroke widthSpectrum font sizeSpectrum stroke widthCompact peptide

Ion legend

wxyz

abcd

OtherUnassignedIon^Charge^_Position_Show for top:%

JJSEVSDRPSGVSSR

03.78e+37.57e+31.14e+41.51e+4

Zoom Out

y^+1^_2_z^+1^_2_w^+1^_3_z^+1^_3_y^+1^_3_z^+2^_8_w^+1^_4_y^+1^_4_c^+1^_4_y^+1^_5_y^+2^_10_c^+1^_5_z^+2^_11_y^+2^_11_y^+1^_6_y^+1^_6_w^+2^_12_z^+2^_12_y^+2^_12_w^+1^_7_c^+1^_6_z^+2^_13_y^+2^_13_y^+1^_7_y^+2^_14_c^+1^_7_z^+1^_8_z^+1^_9_c^+1^_9_z^+1^_10_c^+1^_10_z^+1^_11_c^+1^_11_c^+1^_12_y^+1^_12_c^+1^_13_c^+1^_14_

0757151522723029

Fragment Matches Table

Show background peaks

| Position | Ion type | Intensity | mz Theoretical | mz Error (Th) | mz Error (ppm) | Charge | Series Number |
| --- | --- | --- | --- | --- | --- | --- | --- |
| - | - | 1452 | 120.1 | - | - | 0 | - |
| - | - | 337.6 | 129 | - | - | 0 | - |
| - | - | 7857 | 133.1 | - | - | 0 | - |
| - | - | 850.7 | 133.1 | - | - | 0 | - |
| - | - | 586.7 | 136.1 | - | - | 0 | - |
| - | - | 1303 | 140.1 | - | - | 0 | - |
| - | - | 659.4 | 146.1 | - | - | 0 | - |
| - | - | 399.8 | 147.1 | - | - | 0 | - |
| - | - | 383.4 | 147.7 | - | - | 0 | - |
| - | - | 969.8 | 149 | - | - | 0 | - |
| - | - | 432.4 | 156.8 | - | - | 0 | - |
| - | - | 424.8 | 160.9 | - | - | 0 | - |
| - | - | 390 | 161.1 | - | - | 0 | - |
| - | - | 952 | 168.1 | - | - | 0 | - |
| - | - | 3250 | 173.4 | - | - | 0 | - |
| - | - | 4185 | 177.1 | - | - | 0 | - |
| - | - | 534.9 | 185.1 | - | - | 0 | - |
| - | - | 2188 | 185.1 | - | - | 0 | - |
| - | - | 3132 | 187.1 | - | - | 0 | - |
| - | - | 539.4 | 192.4 | - | - | 0 | - |
| - | - | 4180 | 199.2 | - | - | 0 | - |
| - | - | 3945 | 205.1 | - | - | 0 | - |
| - | - | 750.7 | 215.1 | - | - | 0 | - |
| - | - | 1757 | 221.1 | - | - | 0 | - |
| - | - | 2960 | 227.2 | - | - | 0 | - |
| - | - | 562.1 | 239.1 | - | - | 0 | - |
| 14 | y | 3022 | 245.1 | 0.0002023 | 0.8253 | +1 | 2 |
| 14 | z | 525.5 | 246.1 | 0.002252 | 9.148 | +1 | 2 |
| - | - | 1297 | 249.1 | - | - | 0 | - |
| - | - | 678 | 259.1 | - | - | 0 | - |
| - | - | 1211 | 265.2 | - | - | 0 | - |
| - | - | 549.5 | 266.7 | - | - | 0 | - |
| - | - | 584.2 | 292.1 | - | - | 0 | - |
| - | - | 575.4 | 303.9 | - | - | 0 | - |
| - | - | 793.5 | 309.2 | - | - | 0 | - |
| 13 | w | 2470 | 316.2 | 4.531E-05 | 0.1433 | +1 | 3 |
| 13 | z | 1954 | 333.2 | 2.172E-05 | 0.0652 | +1 | 3 |
| 13 | y | 691 | 349.2 | 0.0004834 | 1.384 | +1 | 3 |
| - | - | 978.1 | 357.2 | - | - | 0 | - |
| - | - | 1624 | 371.2 | - | - | 0 | - |
| - | - | 2105 | 377.2 | - | - | 0 | - |
| 8 | z | 685 | 415.2 | 0.003135 | 7.551 | +2 | 8 |
| 12 | w | 1497 | 417.2 | 5.626E-05 | 0.1348 | +1 | 4 |
| - | - | 631.9 | 423.2 | - | - | 0 | - |
| - | - | 649.8 | 441.2 | - | - | 0 | - |
| - | - | 789.4 | 443.3 | - | - | 0 | - |
| - | - | 546.4 | 444.3 | - | - | 0 | - |
| 12 | y | 1020 | 448.3 | 0.0004158 | 0.9277 | +1 | 4 |
| - | - | 9380 | 459.3 | - | - | 0 | - |
| 4 | c | 2275 | 460.3 | 0.004461 | 9.693 | +1 | 4 |
| - | - | 769.3 | 469.3 | - | - | 0 | - |
| - | - | 1453 | 473.3 | - | - | 0 | - |
| - | - | 687.7 | 476.2 | - | - | 0 | - |
| - | - | 1.121E+04 | 487.3 | - | - | 0 | - |
| - | - | 2664 | 488.3 | - | - | 0 | - |
| - | - | 606.5 | 499.7 | - | - | 0 | - |
| 11 | y | 4342 | 505.3 | 0.001097 | 2.171 | +1 | 5 |
| - | - | 753.9 | 506.3 | - | - | 0 | - |
| 6 | y | 1940 | 524.3 | 0.001166 | 2.225 | +2 | 10 |
| - | - | 1096 | 530.2 | - | - | 0 | - |
| - | - | 931.2 | 530.3 | - | - | 0 | - |
| - | - | 4283 | 530.3 | - | - | 0 | - |
| - | - | 861.5 | 531.3 | - | - | 0 | - |
| - | - | 733.7 | 542.3 | - | - | 0 | - |
| - | - | 630.4 | 548.3 | - | - | 0 | - |
| 5 | c | 694 | 559.3 | 0.0003234 | 0.5782 | +1 | 5 |
| 5 | z | 3578 | 565.8 | 0.0001934 | 0.3419 | +2 | 11 |
| - | - | 2264 | 566.3 | - | - | 0 | - |
| - | - | 1130 | 566.8 | - | - | 0 | - |
| 5 | y | 2456 | 573.8 | 0.001255 | 2.187 | +2 | 11 |
| 10 | y | 1763 | 574.3 | 0.003257 | 5.671 | +1 | 6 |
| 10 | y | 612.7 | 575.3 | 0.003372 | 5.862 | +1 | 6 |
| 4 | w | 1654 | 600.8 | 5.478E-05 | 0.09118 | +2 | 12 |
| - | - | 1031 | 601.3 | - | - | 0 | - |
| - | - | 558 | 605.9 | - | - | 0 | - |
| 4 | z | 2806 | 630.3 | 0.001104 | 1.752 | +2 | 12 |
| - | - | 2691 | 630.8 | - | - | 0 | - |
| - | - | 898.7 | 631.3 | - | - | 0 | - |
| - | - | 1170 | 637.8 | - | - | 0 | - |
| 4 | y | 3649 | 638.3 | 0.0007006 | 1.098 | +2 | 12 |
| - | - | 1531 | 638.8 | - | - | 0 | - |
| - | - | 1578 | 639.3 | - | - | 0 | - |
| 9 | w | 626 | 646.3 | 0.002452 | 3.794 | +1 | 7 |
| 6 | c | 2954 | 646.4 | 5.785E-05 | 0.08949 | +1 | 6 |
| - | - | 911.6 | 647.4 | - | - | 0 | - |
| 3 | z | 1537 | 673.8 | 0.001677 | 2.488 | +2 | 13 |
| - | - | 3627 | 674.3 | - | - | 0 | - |
| - | - | 1203 | 674.8 | - | - | 0 | - |
| - | - | 749.1 | 676.3 | - | - | 0 | - |
| - | - | 604.2 | 681.3 | - | - | 0 | - |
| 3 | y | 1.499E+04 | 681.8 | 0.0003574 | 0.5242 | +2 | 13 |
| - | - | 1.22E+04 | 682.3 | - | - | 0 | - |
| - | - | 5926 | 682.8 | - | - | 0 | - |
| - | - | 1403 | 683.3 | - | - | 0 | - |
| 9 | y | 687.8 | 689.4 | 0.001086 | 1.575 | +1 | 7 |
| - | - | 791 | 712.4 | - | - | 0 | - |
| 2 | y | 1054 | 738.4 | 0.004365 | 5.911 | +2 | 14 |
| - | - | 739.6 | 742.4 | - | - | 0 | - |
| - | - | 982.6 | 758.4 | - | - | 0 | - |
| - | - | 774.6 | 758.9 | - | - | 0 | - |
| 7 | c | 6073 | 761.4 | 0.001128 | 1.482 | +1 | 7 |
| - | - | 1888 | 762.4 | - | - | 0 | - |
| - | - | 722.1 | 764.4 | - | - | 0 | - |
| - | - | 921.2 | 764.9 | - | - | 0 | - |
| - | - | 1.03E+04 | 765.4 | - | - | 0 | - |
| - | - | 9576 | 765.9 | - | - | 0 | - |
| - | - | 3846 | 766.4 | - | - | 0 | - |
| - | - | 647.2 | 766.9 | - | - | 0 | - |
| - | - | 2729 | 773.9 | - | - | 0 | - |
| - | - | 2692 | 774.4 | - | - | 0 | - |
| - | - | 659.5 | 774.9 | - | - | 0 | - |
| - | - | 605.5 | 775.7 | - | - | 0 | - |
| - | - | 1194 | 779.4 | - | - | 0 | - |
| - | - | 1.189E+04 | 786.9 | - | - | 0 | - |
| - | - | 1.03E+04 | 787.4 | - | - | 0 | - |
| - | - | 5047 | 787.9 | - | - | 0 | - |
| - | - | 2195 | 788.4 | - | - | 0 | - |
| - | - | 940.4 | 788.9 | - | - | 0 | - |
| - | - | 1442 | 793.9 | - | - | 0 | - |
| - | - | 686.9 | 794.4 | - | - | 0 | - |
| - | - | 7881 | 794.9 | - | - | 0 | - |
| - | - | 8920 | 795.4 | - | - | 0 | - |
| - | - | 6247 | 795.9 | - | - | 0 | - |
| - | - | 5300 | 796.4 | - | - | 0 | - |
| - | - | 903.3 | 796.9 | - | - | 0 | - |
| 8 | z | 4808 | 829.4 | 0.001345 | 1.622 | +1 | 8 |
| - | - | 636.2 | 830.4 | - | - | 0 | - |
| - | - | 2324 | 830.4 | - | - | 0 | - |
| - | - | 778.5 | 831.4 | - | - | 0 | - |
| 7 | z | 1814 | 944.5 | 0.002043 | 2.163 | +1 | 9 |
| - | - | 1005 | 945.5 | - | - | 0 | - |
| - | - | 1478 | 974.5 | - | - | 0 | - |
| 9 | c | 4519 | 1015 | 0.0004622 | 0.4556 | +1 | 9 |
| - | - | 2586 | 1016 | - | - | 0 | - |
| - | - | 1021 | 1017 | - | - | 0 | - |
| 6 | z | 941.7 | 1031 | 0.001357 | 1.315 | +1 | 10 |
| - | - | 904.3 | 1042 | - | - | 0 | - |
| - | - | 1457 | 1043 | - | - | 0 | - |
| - | - | 965.5 | 1045 | - | - | 0 | - |
| - | - | 985.6 | 1046 | - | - | 0 | - |
| - | - | 3338 | 1058 | - | - | 0 | - |
| - | - | 2864 | 1059 | - | - | 0 | - |
| - | - | 1591 | 1060 | - | - | 0 | - |
| - | - | 946.7 | 1061 | - | - | 0 | - |
| - | - | 2227 | 1062 | - | - | 0 | - |
| - | - | 1733 | 1063 | - | - | 0 | - |
| - | - | 1722 | 1064 | - | - | 0 | - |
| 10 | c | 5870 | 1102 | 0.0002641 | 0.2397 | +1 | 10 |
| - | - | 3442 | 1103 | - | - | 0 | - |
| - | - | 1601 | 1104 | - | - | 0 | - |
| 5 | z | 667.4 | 1131 | 0.003853 | 3.408 | +1 | 11 |
| - | - | 2518 | 1132 | - | - | 0 | - |
| - | - | 1425 | 1133 | - | - | 0 | - |
| 11 | c | 4036 | 1159 | 0.003906 | 3.371 | +1 | 11 |
| - | - | 1532 | 1160 | - | - | 0 | - |
| - | - | 1311 | 1161 | - | - | 0 | - |
| 12 | c | 3617 | 1258 | 0.000298 | 0.2369 | +1 | 12 |
| - | - | 2290 | 1259 | - | - | 0 | - |
| - | - | 1288 | 1260 | - | - | 0 | - |
| - | - | 2864 | 1261 | - | - | 0 | - |
| - | - | 1809 | 1262 | - | - | 0 | - |
| - | - | 786.8 | 1263 | - | - | 0 | - |
| 4 | y | 1193 | 1276 | 0.005599 | 4.39 | +1 | 12 |
| - | - | 806.7 | 1277 | - | - | 0 | - |
| 13 | c | 6472 | 1345 | 0.003152 | 2.344 | +1 | 13 |
| - | - | 3843 | 1346 | - | - | 0 | - |
| - | - | 2657 | 1347 | - | - | 0 | - |
| - | - | 2495 | 1348 | - | - | 0 | - |
| - | - | 2148 | 1349 | - | - | 0 | - |
| 14 | c | 9864 | 1432 | 0.005151 | 3.597 | +1 | 14 |
| - | - | 5967 | 1433 | - | - | 0 | - |
| - | - | 2795 | 1434 | - | - | 0 | - |
| - | - | 858.4 | 1435 | - | - | 0 | - |
| - | - | 813.7 | 1461 | - | - | 0 | - |
| - | - | 630.3 | 1503 | - | - | 0 | - |
| - | - | 762.3 | 1527 | - | - | 0 | - |
| - | - | 2095 | 1531 | - | - | 0 | - |
| - | - | 1659 | 1532 | - | - | 0 | - |
| - | - | 1185 | 1533 | - | - | 0 | - |
| - | - | 712.8 | 1545 | - | - | 0 | - |
| - | - | 690.1 | 1546 | - | - | 0 | - |
| - | - | 1210 | 1548 | - | - | 0 | - |
| - | - | 1070 | 1549 | - | - | 0 | - |
| - | - | 1113 | 1557 | - | - | 0 | - |
| - | - | 1154 | 1558 | - | - | 0 | - |
| - | - | 670.8 | 1559 | - | - | 0 | - |
| - | - | 2112 | 1563 | - | - | 0 | - |
| - | - | 1276 | 1564 | - | - | 0 | - |
| - | - | 801.6 | 1572 | - | - | 0 | - |
| - | - | 2049 | 1573 | - | - | 0 | - |
| - | - | 1.441E+04 | 1574 | - | - | 0 | - |
| - | - | 1.119E+04 | 1575 | - | - | 0 | - |
| - | - | 6759 | 1576 | - | - | 0 | - |
| - | - | 2677 | 1577 | - | - | 0 | - |
| - | - | 915.6 | 1578 | - | - | 0 | - |
| - | - | 748.3 | 1587 | - | - | 0 | - |
| - | - | 1965 | 1589 | - | - | 0 | - |
| - | - | 4387 | 1590 | - | - | 0 | - |
| - | - | 1.28E+04 | 1591 | - | - | 0 | - |
| - | - | 1.235E+04 | 1592 | - | - | 0 | - |
| - | - | 5417 | 1593 | - | - | 0 | - |
| - | - | 2002 | 1594 | - | - | 0 | - |
| - | - | 769.6 | 1914 | - | - | 0 | - |
| - | - | 657.1 | 2329 | - | - | 0 | - |
| - | - | 673.8 | 2859 | - | - | 0 | - |
| - | - | 634.7 | 2999 | - | - | 0 | - |

m/z Charge Intensity FragmentType MassShift Position
120.08074951171875 0 1452.4435
129.03538513183594 0 337.63553
133.08602905273438 0 7857.4907
133.09141540527344 0 850.736
136.07608032226562 0 586.67163
140.08184814453125 0 1303.325
146.06027221679688 0 659.39624
147.1018829345703 0 399.81226
147.6551513671875 0 383.43945
148.9547882080078 0 969.8421
156.81198120117188 0 432.4417
160.8848419189453 0 424.7892
161.08013916015625 0 389.98325
168.11293029785156 0 952.0284
173.4404754638672 0 3249.606
177.11212158203125 0 4184.8345
185.09202575683594 0 534.8635
185.13958740234375 0 2188.0454
187.1078643798828 0 3131.509
192.3612823486328 0 539.38574
199.18043518066406 0 4180.116
205.10704040527344 0 3945.0283
215.13900756835938 0 750.7443
221.1383056640625 0 1756.5033
227.17535400390625 0 2960.144
239.1483154296875 0 562.07886
245.1246337890625 0 3021.803 y Ammonia loss 13
246.1300048828125 0 525.5415 z 13
249.1332244873047 0 1297.3207
259.1047668457031 0 678.01953
265.1646728515625 0 1211.1199
266.6546325683594 0 549.5007
292.12994384765625 0 584.2186
303.93487548828125 0 575.38153
309.189208984375 0 793.5045
316.1615905761719 0 2469.7996 w 12
333.164306640625 0 1953.992 z 12
349.1825256347656 0 690.9573 y 12
357.15240478515625 0 978.10004
371.2268981933594 0 1623.9675
377.16558837890625 0 2104.9685
415.226806640625 0 685.02 z 7
417.20916748046875 0 1496.6799 w 11
423.16656494140625 0 631.85016
441.1733703613281 0 649.753
443.25054931640625 0 789.419
444.2550964355469 0 546.35144
448.2510070800781 0 1020.2943 y 11
459.2802429199219 0 9380.196
460.2810363769531 0 2275.1914 c 3
469.2637023925781 0 769.3265
473.2942199707031 0 1452.6398
476.23895263671875 0 687.69403
487.2750244140625 0 11210.724
488.2784423828125 0 2664.1125
499.6811218261719 0 606.5265
505.27178955078125 0 4342.47 y 10
506.27685546875 0 753.87427
524.2613525390625 0 1940.2268 y 5
530.224609375 0 1095.9966
530.2770385742188 0 931.17584
530.31640625 0 4282.8564
531.3193359375 0 861.5149
542.318115234375 0 733.7391
548.3256225585938 0 630.44385
559.3446655273438 0 694.0357 c 4
565.7871704101562 0 3578.0854 z 4
566.2880859375 0 2264.1084
566.7942504882812 0 1130.1833
573.7954711914062 0 2456.2998 y 4
574.297607421875 0 1762.6584 y Water loss 9
575.28173828125 0 612.683 y Ammonia loss 9
600.8020629882812 0 1654.4651 w 3
601.3019409179688 0 1030.9901
605.8861694335938 0 557.9997
630.3075561523438 0 2806.2422 z 3
630.8101806640625 0 2690.7322
631.30712890625 0 898.71594
637.8121948242188 0 1170.4868
638.3173217773438 0 3648.8613 y 3
638.8169555664062 0 1530.5107
639.3206176757812 0 1578.2935
646.3179321289062 0 626.0293 w 8
646.3770751953125 0 2954.2148 c 5
647.3792724609375 0 911.5676
673.822998046875 0 1536.9384 z 2
674.326904296875 0 3626.9312
674.8297119140625 0 1203.4618
676.3477172851562 0 749.1124
681.3314208984375 0 604.22424
681.8336791992188 0 14988.922 y 2
682.3347778320312 0 12196.941
682.83642578125 0 5926.0254
683.3385009765625 0 1403.0304
689.3587646484375 0 687.7829 y 8
712.3815307617188 0 791.0096
738.3717041015625 0 1054.1016 y 1
742.385498046875 0 739.55493
758.3914794921875 0 982.57794
758.8819580078125 0 774.6086
761.40283203125 0 6073.4746 c 6
762.4073486328125 0 1887.9658
764.4100341796875 0 722.11566
764.9063110351562 0 921.17786
765.3818359375 0 10298.875
765.8826293945312 0 9575.936
766.3875122070312 0 3846.15
766.8853149414062 0 647.178
773.9049682617188 0 2728.848
774.4063110351562 0 2691.698
774.9011840820312 0 659.4882
775.6870727539062 0 605.52545
779.4046630859375 0 1193.6151
786.9076538085938 0 11889.995
787.4094848632812 0 10299.519
787.9088134765625 0 5046.9756
788.406494140625 0 2194.8552
788.9158935546875 0 940.4272
793.8997192382812 0 1442.493
794.4066162109375 0 686.9298
794.9151611328125 0 7881.0034
795.4166259765625 0 8919.612
795.9168701171875 0 6247.122
796.4161987304688 0 5300.1367
796.9197387695312 0 903.3223
829.438720703125 0 4807.6694 z 7
830.3614501953125 0 636.17584
830.4434814453125 0 2323.591
831.4461669921875 0 778.4936
944.4649658203125 0 1813.5886 z 6
945.4677124023438 0 1004.7497
974.5476684570312 0 1478.347
1014.557373046875 0 4519.135 c 8
1015.5567626953125 0 2586.2346
1016.5608520507812 0 1020.8966
1031.4976806640625 0 941.74347 z 5
1041.5113525390625 0 904.3354
1042.50537109375 0 1456.8767
1044.52685546875 0 965.5345
1045.5084228515625 0 985.56976
1057.569091796875 0 3338.381
1058.5531005859375 0 2863.9114
1059.5452880859375 0 1591.3029
1060.526611328125 0 946.7177
1061.5294189453125 0 2227.2944
1062.53759765625 0 1732.8693
1063.5662841796875 0 1721.7278
1101.589599609375 0 5869.9375 c 9
1102.5914306640625 0 3442.2947
1103.5950927734375 0 1601.4191
1130.5635986328125 0 667.4497 z 4
1131.5723876953125 0 2517.5815
1132.577392578125 0 1425.2341
1158.607421875 0 4035.6846 c 10
1159.6103515625 0 1531.9973
1160.6173095703125 0 1311.1266
1257.679443359375 0 3617.0647 c 11
1258.6773681640625 0 2289.601
1259.6817626953125 0 1287.5074
1260.6143798828125 0 2863.9734
1261.6146240234375 0 1809.0273
1262.635009765625 0 786.7757
1275.6231689453125 0 1192.9521 y 3
1276.6312255859375 0 806.7343
1344.7086181640625 0 6472.3916 c 12
1345.710693359375 0 3842.537
1346.706787109375 0 2656.9368
1347.649169921875 0 2495.3818
1348.6490478515625 0 2148.3193
1431.7386474609375 0 9864.052 c 13
1432.7427978515625 0 5966.664
1433.7486572265625 0 2795.383
1434.737548828125 0 858.3799
1460.72265625 0 813.71844
1502.7928466796875 0 630.3232
1526.79638671875 0 762.2855
1530.7889404296875 0 2094.6685
1531.7808837890625 0 1658.7572
1532.79248046875 0 1185.1155
1544.8416748046875 0 712.8284
1545.8026123046875 0 690.14874
1547.812744140625 0 1209.6024
1548.8223876953125 0 1070.3427
1556.80078125 0 1112.9552
1557.80322265625 0 1153.5404
1558.796875 0 670.81537
1562.8414306640625 0 2112.3408
1563.851318359375 0 1276.443
1571.816162109375 0 801.6298
1572.80126953125 0 2049.3293
1573.81201171875 0 14406.716
1574.8123779296875 0 11185.859
1575.8173828125 0 6759.2964
1576.8160400390625 0 2676.8208
1577.8311767578125 0 915.56305
1586.7930908203125 0 748.2759
1588.8291015625 0 1965.1556
1589.823486328125 0 4387.08
1590.836669921875 0 12797.241
1591.8402099609375 0 12349.514
1592.840087890625 0 5416.8286
1593.838134765625 0 2002.3395
1913.9481201171875 0 769.5895
2329.189208984375 0 657.1245
2858.517822265625 0 673.7632
2999.174072265625 0 634.66797

Spectrum Details

|  |  |
| --- | --- |
| Matched peaks? Matched peaks The total absolute number of peaks matched. Additionally in brackets the total fraction of peaks matched and the total number of peaks is shown. | 37 (17.96% of 206) |
| FDR? FDR The false discovery rate estimated for this peptide. It is calculated by matching all theoretical fragments with a non-integer shift with the raw peaks for this spectrum. This is done with 40 different shifts. The resulting percentage is the average number of annotated peaks over the number of annotated peaks with the correct spectrum. | 1.16% |
| Satellite FDR? Satellite FDR See the FDR for details on its calculation. This satellite ion specific FDR only contains the satellite ions (d/w) for I/L/J positions. | ∞ |
| PSM Score? PSM Score The PSM Score as given by Hecklib to this annotated spectrum. It is shown with three significant figures. | 239 |

#### Spectrum 4031? Spectrum 4031 The raw spectrum of this peptide as annotated by Hecklib. The fragments are coloured according to ion type (see legend). Any peaks with a star '\*' as text can be hovered over to see the full details, first the ion type second the mass shift type. By hovering over the amino acids in the peptide or ions in the legend the corresponding peaks are highlighted. By toggling the 'Unassigned' label you can turn the background (unassigned) peaks on or off in the plot. By updating the slider in the Ion legend you can update the spectrum to only show the top X% of the peaks with labels. The top X% means any peak that is within X% of the highest intensity. By dragging in the spectrum you can zoom in to a specific part of the spectrum and use 'Zoom Out' to get back to the original zoom level. The annotation of the spectrum is based on the given sequence in the peptides file and is done with different software so inconsistencies are likely. The peaks are annotated based on the given sequence, with 20 ppm tolerance.

Copy Data

##### Spectrum 4031 (TSV)

###### Preview

```
Loading example...
```

*Click on the button to copy the data to your clipboard.*

Mz MinMz MaxIntensity Max

WidthHeightPeptide font sizePeptide stroke widthSpectrum font sizeSpectrum stroke widthCompact peptide

Ion legend

wxyz

abcd

OtherUnassignedIon^Charge^_Position_Show for top:%

JJSEVSDRPSGVSSR

01.98e+53.96e+55.94e+57.92e+5

Zoom Out

y^+1^_1_y^+3^_5_y^+1^_1_d^+1^_2_a^+1^_2_b^+1^_2_y^+1^_2_y^+1^_2_y^+1^_2_b^+1^_3_b^+1^_3_y^+1^_3_y^+1^_3_y^+2^_7_y^+1^_3_b^+3^_12_y^+2^_8_y^+2^_8_b^+1^_4_y^+1^_4_y^+1^_4_b^+1^_4_y^+1^_4_y^+3^_13_y^+3^_13_b^+2^_8_y^+3^_13_b^+3^_14_y^+2^_9_y^+3^_14_y^+3^_14_y^+1^_5_y^+1^_5_y^+3^_14_y^+1^_5_y^+2^_10_y^+2^_10_y^+2^_10_\*\*b^+1^_5_y^+2^_11_y^+2^_11_y^+2^_11_y^+1^_6_y^+1^_6_y^+2^_12_y^+2^_12_y^+2^_12_y^+1^_7_y^+1^_7_y^+2^_13_y^+2^_13_y^+2^_13_y^+1^_7_b^+2^_14_y^+2^_14_y^+2^_14_y^+2^_14_b^+1^_7_y^+1^_8_b^+1^_8_y^+1^_9_y^+1^_10_y^+1^_11_y^+1^_13_

0510102115312042

Fragment Matches Table

Show background peaks

| Position | Ion type | Intensity | mz Theoretical | mz Error (Th) | mz Error (ppm) | Charge | Series Number |
| --- | --- | --- | --- | --- | --- | --- | --- |
| - | - | 2607 | 120.1 | - | - | 0 | - |
| - | - | 340.1 | 122.3 | - | - | 0 | - |
| - | - | 7867 | 126.1 | - | - | 0 | - |
| - | - | 436.8 | 127.1 | - | - | 0 | - |
| - | - | 1449 | 128.1 | - | - | 0 | - |
| - | - | 884.4 | 128.1 | - | - | 0 | - |
| - | - | 1205 | 129.1 | - | - | 0 | - |
| - | - | 4456 | 129.1 | - | - | 0 | - |
| - | - | 569.8 | 130.1 | - | - | 0 | - |
| - | - | 3814 | 130.1 | - | - | 0 | - |
| - | - | 2208 | 130.1 | - | - | 0 | - |
| - | - | 651.2 | 131.1 | - | - | 0 | - |
| - | - | 682.4 | 131.1 | - | - | 0 | - |
| - | - | 622.9 | 133.1 | - | - | 0 | - |
| - | - | 420.5 | 133.1 | - | - | 0 | - |
| - | - | 485.5 | 136.1 | - | - | 0 | - |
| - | - | 3453 | 136.1 | - | - | 0 | - |
| - | - | 1102 | 139.1 | - | - | 0 | - |
| - | - | 585.3 | 139.1 | - | - | 0 | - |
| - | - | 1074 | 140.1 | - | - | 0 | - |
| - | - | 634.7 | 140.1 | - | - | 0 | - |
| - | - | 788.3 | 141.1 | - | - | 0 | - |
| - | - | 441.8 | 143 | - | - | 0 | - |
| - | - | 589.7 | 143.1 | - | - | 0 | - |
| - | - | 1773 | 144.1 | - | - | 0 | - |
| - | - | 1100 | 145.1 | - | - | 0 | - |
| - | - | 2426 | 147.1 | - | - | 0 | - |
| - | - | 1008 | 149 | - | - | 0 | - |
| - | - | 406 | 152.9 | - | - | 0 | - |
| - | - | 1003 | 153.1 | - | - | 0 | - |
| - | - | 7770 | 154.1 | - | - | 0 | - |
| - | - | 605 | 155.1 | - | - | 0 | - |
| - | - | 1102 | 155.1 | - | - | 0 | - |
| - | - | 904.4 | 156.1 | - | - | 0 | - |
| - | - | 3185 | 157.1 | - | - | 0 | - |
| - | - | 3445 | 157.1 | - | - | 0 | - |
| - | - | 4838 | 157.1 | - | - | 0 | - |
| 15 | y | 1.752E+04 | 158.1 | 0.0004467 | 2.825 | +1 | 1 |
| - | - | 906.5 | 158.1 | - | - | 0 | - |
| - | - | 867.1 | 158.1 | - | - | 0 | - |
| - | - | 1015 | 159.1 | - | - | 0 | - |
| - | - | 1232 | 159.1 | - | - | 0 | - |
| - | - | 2598 | 159.1 | - | - | 0 | - |
| - | - | 474.3 | 161.1 | - | - | 0 | - |
| - | - | 5238 | 163.1 | - | - | 0 | - |
| - | - | 1555 | 167.1 | - | - | 0 | - |
| - | - | 496.9 | 167.6 | - | - | 0 | - |
| - | - | 895.8 | 169.1 | - | - | 0 | - |
| 11 | y | 1368 | 169.1 | 0.001813 | 10.72 | +3 | 5 |
| - | - | 647.1 | 170 | - | - | 0 | - |
| - | - | 475.7 | 170.1 | - | - | 0 | - |
| - | - | 9471 | 171.1 | - | - | 0 | - |
| - | - | 998.4 | 171.1 | - | - | 0 | - |
| - | - | 1888 | 172.1 | - | - | 0 | - |
| - | - | 542.1 | 172.1 | - | - | 0 | - |
| - | - | 538.7 | 173.1 | - | - | 0 | - |
| - | - | 3202 | 173.1 | - | - | 0 | - |
| - | - | 1386 | 173.5 | - | - | 0 | - |
| - | - | 6281 | 175.1 | - | - | 0 | - |
| 15 | y | 3.76E+04 | 175.1 | 0.0004631 | 2.645 | +1 | 1 |
| - | - | 553.5 | 176.1 | - | - | 0 | - |
| - | - | 2124 | 176.1 | - | - | 0 | - |
| - | - | 456.4 | 178.1 | - | - | 0 | - |
| - | - | 479.5 | 179.5 | - | - | 0 | - |
| - | - | 491.9 | 182.2 | - | - | 0 | - |
| - | - | 8503 | 183.1 | - | - | 0 | - |
| - | - | 1261 | 184.1 | - | - | 0 | - |
| - | - | 927.9 | 184.1 | - | - | 0 | - |
| - | - | 5393 | 185.1 | - | - | 0 | - |
| 2 | d | 846.2 | 185.2 | 0.0002909 | 1.571 | +1 | 2 |
| - | - | 703.7 | 186.1 | - | - | 0 | - |
| - | - | 1126 | 187.1 | - | - | 0 | - |
| - | - | 5623 | 187.1 | - | - | 0 | - |
| - | - | 754 | 187.1 | - | - | 0 | - |
| - | - | 487.9 | 188.1 | - | - | 0 | - |
| - | - | 4.066E+04 | 189.1 | - | - | 0 | - |
| - | - | 3675 | 190.1 | - | - | 0 | - |
| - | - | 547.1 | 191.1 | - | - | 0 | - |
| - | - | 1348 | 192.1 | - | - | 0 | - |
| - | - | 629.4 | 193.1 | - | - | 0 | - |
| - | - | 827.3 | 193.1 | - | - | 0 | - |
| - | - | 965.4 | 194.1 | - | - | 0 | - |
| - | - | 654.3 | 195.1 | - | - | 0 | - |
| - | - | 903.9 | 195.1 | - | - | 0 | - |
| - | - | 1882 | 195.1 | - | - | 0 | - |
| - | - | 471.7 | 197.1 | - | - | 0 | - |
| - | - | 641.8 | 197.1 | - | - | 0 | - |
| - | - | 894.2 | 197.2 | - | - | 0 | - |
| - | - | 4381 | 199.1 | - | - | 0 | - |
| 2 | a | 3.157E+05 | 199.2 | 0.00051 | 2.56 | +1 | 2 |
| - | - | 4.042E+04 | 200.2 | - | - | 0 | - |
| - | - | 1.18E+04 | 201.1 | - | - | 0 | - |
| - | - | 2188 | 201.2 | - | - | 0 | - |
| - | - | 2844 | 202.1 | - | - | 0 | - |
| - | - | 740.2 | 202.1 | - | - | 0 | - |
| - | - | 6901 | 203.1 | - | - | 0 | - |
| - | - | 1923 | 203.1 | - | - | 0 | - |
| - | - | 584 | 208.1 | - | - | 0 | - |
| - | - | 601.4 | 209.1 | - | - | 0 | - |
| - | - | 1088 | 210.1 | - | - | 0 | - |
| - | - | 2387 | 211.1 | - | - | 0 | - |
| - | - | 3950 | 212.1 | - | - | 0 | - |
| - | - | 956.9 | 213.1 | - | - | 0 | - |
| - | - | 1246 | 213.2 | - | - | 0 | - |
| - | - | 627.7 | 215.1 | - | - | 0 | - |
| - | - | 705.3 | 216.1 | - | - | 0 | - |
| - | - | 967.6 | 216.1 | - | - | 0 | - |
| - | - | 6.993E+04 | 217.1 | - | - | 0 | - |
| - | - | 4865 | 218.1 | - | - | 0 | - |
| - | - | 732.4 | 219.1 | - | - | 0 | - |
| - | - | 748.9 | 220.1 | - | - | 0 | - |
| - | - | 545.4 | 222.8 | - | - | 0 | - |
| - | - | 4472 | 224.1 | - | - | 0 | - |
| - | - | 727.7 | 225.2 | - | - | 0 | - |
| - | - | 1204 | 226.1 | - | - | 0 | - |
| - | - | 1760 | 226.2 | - | - | 0 | - |
| - | - | 817.8 | 227.1 | - | - | 0 | - |
| - | - | 2973 | 227.1 | - | - | 0 | - |
| 2 | b | 9.402E+04 | 227.2 | 0.0005294 | 2.33 | +1 | 2 |
| - | - | 969 | 228.1 | - | - | 0 | - |
| - | - | 1.286E+04 | 228.2 | - | - | 0 | - |
| - | - | 3077 | 229.1 | - | - | 0 | - |
| - | - | 1047 | 229.2 | - | - | 0 | - |
| - | - | 2161 | 230.1 | - | - | 0 | - |
| - | - | 545.6 | 231.1 | - | - | 0 | - |
| - | - | 2293 | 233.2 | - | - | 0 | - |
| - | - | 746.6 | 234.1 | - | - | 0 | - |
| - | - | 633.7 | 234.2 | - | - | 0 | - |
| - | - | 1112 | 237.1 | - | - | 0 | - |
| - | - | 2225 | 237.1 | - | - | 0 | - |
| - | - | 573.9 | 237.2 | - | - | 0 | - |
| - | - | 664.7 | 240.1 | - | - | 0 | - |
| - | - | 9829 | 242.1 | - | - | 0 | - |
| - | - | 954.3 | 243.1 | - | - | 0 | - |
| - | - | 2101 | 244.1 | - | - | 0 | - |
| 14 | y | 917.6 | 244.1 | 0.001354 | 5.544 | +1 | 2 |
| 14 | y | 9394 | 245.1 | 0.0006601 | 2.693 | +1 | 2 |
| - | - | 839.6 | 246.1 | - | - | 0 | - |
| - | - | 1040 | 249.2 | - | - | 0 | - |
| - | - | 2760 | 253.2 | - | - | 0 | - |
| - | - | 986.1 | 254.1 | - | - | 0 | - |
| - | - | 8484 | 255.1 | - | - | 0 | - |
| - | - | 649.6 | 256.1 | - | - | 0 | - |
| - | - | 907.5 | 259.1 | - | - | 0 | - |
| - | - | 1237 | 261.2 | - | - | 0 | - |
| 14 | y | 6713 | 262.2 | 0.0004476 | 1.708 | +1 | 2 |
| - | - | 590.7 | 267.1 | - | - | 0 | - |
| - | - | 818.1 | 269.2 | - | - | 0 | - |
| - | - | 699 | 269.2 | - | - | 0 | - |
| - | - | 1559 | 270.1 | - | - | 0 | - |
| - | - | 1648 | 270.2 | - | - | 0 | - |
| - | - | 1475 | 272.1 | - | - | 0 | - |
| - | - | 1463 | 280.1 | - | - | 0 | - |
| - | - | 879.5 | 283.2 | - | - | 0 | - |
| - | - | 1532 | 284.1 | - | - | 0 | - |
| - | - | 699 | 287.2 | - | - | 0 | - |
| - | - | 1749 | 288.2 | - | - | 0 | - |
| - | - | 661 | 295.2 | - | - | 0 | - |
| - | - | 1446 | 296.1 | - | - | 0 | - |
| 3 | b | 3697 | 296.2 | 0.0006721 | 2.269 | +1 | 3 |
| - | - | 1621 | 297.1 | - | - | 0 | - |
| - | - | 3304 | 298.1 | - | - | 0 | - |
| - | - | 1963 | 299.1 | - | - | 0 | - |
| - | - | 1122 | 302.1 | - | - | 0 | - |
| - | - | 504.8 | 309.2 | - | - | 0 | - |
| - | - | 4672 | 312.2 | - | - | 0 | - |
| - | - | 532.7 | 313.2 | - | - | 0 | - |
| - | - | 1.024E+04 | 313.2 | - | - | 0 | - |
| - | - | 3938 | 314.1 | - | - | 0 | - |
| - | - | 1422 | 314.2 | - | - | 0 | - |
| 3 | b | 9211 | 314.2 | 0.0008802 | 2.801 | +1 | 3 |
| - | - | 967.9 | 314.2 | - | - | 0 | - |
| - | - | 743.1 | 315.1 | - | - | 0 | - |
| - | - | 1119 | 315.2 | - | - | 0 | - |
| - | - | 403.2 | 316.1 | - | - | 0 | - |
| - | - | 1.19E+04 | 316.2 | - | - | 0 | - |
| - | - | 832.4 | 317.1 | - | - | 0 | - |
| - | - | 1717 | 317.1 | - | - | 0 | - |
| - | - | 1094 | 323.2 | - | - | 0 | - |
| - | - | 1523 | 324.1 | - | - | 0 | - |
| - | - | 573.2 | 324.2 | - | - | 0 | - |
| - | - | 610.6 | 325.6 | - | - | 0 | - |
| - | - | 667.4 | 329.1 | - | - | 0 | - |
| - | - | 2894 | 330.2 | - | - | 0 | - |
| 13 | y | 2273 | 331.2 | 0.0002547 | 0.769 | +1 | 3 |
| 13 | y | 9971 | 332.2 | 0.0007972 | 2.4 | +1 | 3 |
| - | - | 1762 | 333.2 | - | - | 0 | - |
| - | - | 623.5 | 335.1 | - | - | 0 | - |
| - | - | 1136 | 340.2 | - | - | 0 | - |
| - | - | 612.7 | 341.2 | - | - | 0 | - |
| - | - | 6892 | 341.2 | - | - | 0 | - |
| - | - | 786 | 341.2 | - | - | 0 | - |
| - | - | 1.189E+04 | 342.1 | - | - | 0 | - |
| - | - | 2281 | 343.1 | - | - | 0 | - |
| 9 | y | 5681 | 345.2 | 0.0008719 | 2.526 | +2 | 7 |
| - | - | 1876 | 345.7 | - | - | 0 | - |
| 13 | y | 8597 | 349.2 | 0.0007374 | 2.112 | +1 | 3 |
| - | - | 844 | 350.2 | - | - | 0 | - |
| - | - | 784.1 | 352.2 | - | - | 0 | - |
| - | - | 6087 | 359.2 | - | - | 0 | - |
| - | - | 787.7 | 360.2 | - | - | 0 | - |
| - | - | 699.5 | 365.2 | - | - | 0 | - |
| - | - | 1385 | 380.2 | - | - | 0 | - |
| - | - | 930.2 | 381.2 | - | - | 0 | - |
| - | - | 1367 | 385.2 | - | - | 0 | - |
| - | - | 1030 | 389.7 | - | - | 0 | - |
| - | - | 1023 | 390.7 | - | - | 0 | - |
| - | - | 4678 | 391.7 | - | - | 0 | - |
| - | - | 1167 | 392.2 | - | - | 0 | - |
| - | - | 1826 | 393.7 | - | - | 0 | - |
| - | - | 834.4 | 394.2 | - | - | 0 | - |
| - | - | 953.3 | 397.2 | - | - | 0 | - |
| - | - | 968.8 | 398.2 | - | - | 0 | - |
| - | - | 1515 | 398.7 | - | - | 0 | - |
| - | - | 1096 | 399.2 | - | - | 0 | - |
| - | - | 2679 | 399.7 | - | - | 0 | - |
| - | - | 651.4 | 400.2 | - | - | 0 | - |
| - | - | 1901 | 400.2 | - | - | 0 | - |
| - | - | 878.3 | 401.2 | - | - | 0 | - |
| - | - | 1519 | 403.2 | - | - | 0 | - |
| - | - | 968.2 | 405.7 | - | - | 0 | - |
| - | - | 1134 | 407.2 | - | - | 0 | - |
| - | - | 976.7 | 408.2 | - | - | 0 | - |
| - | - | 1845 | 410.2 | - | - | 0 | - |
| - | - | 815.7 | 411.2 | - | - | 0 | - |
| - | - | 765.8 | 414.2 | - | - | 0 | - |
| 12 | b | 2863 | 414.2 | 0.006056 | 14.62 | +3 | 12 |
| 8 | y | 6007 | 414.7 | 0.00247 | 5.957 | +2 | 8 |
| - | - | 2995 | 415.2 | - | - | 0 | - |
| - | - | 1526 | 415.3 | - | - | 0 | - |
| - | - | 817.8 | 415.7 | - | - | 0 | - |
| - | - | 782.7 | 415.8 | - | - | 0 | - |
| - | - | 721.8 | 416.2 | - | - | 0 | - |
| 8 | y | 7.367E+04 | 423.2 | 0.001006 | 2.377 | +2 | 8 |
| - | - | 3.039E+04 | 423.7 | - | - | 0 | - |
| - | - | 7751 | 424.2 | - | - | 0 | - |
| - | - | 1109 | 424.7 | - | - | 0 | - |
| 4 | b | 5078 | 425.2 | 0.0008342 | 1.962 | +1 | 4 |
| - | - | 1304 | 426.2 | - | - | 0 | - |
| - | - | 2439 | 428.2 | - | - | 0 | - |
| - | - | 3749 | 429.2 | - | - | 0 | - |
| 12 | y | 2964 | 430.2 | 0.0006238 | 1.45 | +1 | 4 |
| 12 | y | 777.6 | 431.2 | 0.0008748 | 2.029 | +1 | 4 |
| - | - | 723.9 | 440.2 | - | - | 0 | - |
| - | - | 3953 | 441.2 | - | - | 0 | - |
| - | - | 984.3 | 442.2 | - | - | 0 | - |
| 4 | b | 3953 | 443.3 | 0.0006149 | 1.387 | +1 | 4 |
| - | - | 997.2 | 444.3 | - | - | 0 | - |
| - | - | 627.6 | 445.2 | - | - | 0 | - |
| 12 | y | 1763 | 448.3 | 0.001476 | 3.293 | +1 | 4 |
| 3 | y | 2205 | 448.9 | 0.001051 | 2.342 | +3 | 13 |
| 3 | y | 958.1 | 449.2 | 0.007732 | 17.21 | +3 | 13 |
| 8 | b | 751.9 | 450.7 | 0.001424 | 3.16 | +2 | 8 |
| - | - | 719.8 | 452.2 | - | - | 0 | - |
| - | - | 2115 | 452.3 | - | - | 0 | - |
| 3 | y | 2521 | 454.9 | 0.0006728 | 1.479 | +3 | 13 |
| - | - | 1751 | 455.2 | - | - | 0 | - |
| - | - | 1465 | 455.3 | - | - | 0 | - |
| - | - | 4653 | 458.2 | - | - | 0 | - |
| - | - | 1608 | 460.2 | - | - | 0 | - |
| - | - | 628 | 463.2 | - | - | 0 | - |
| - | - | 1962 | 469.3 | - | - | 0 | - |
| 14 | b | 865.6 | 472.2 | 0.005836 | 12.36 | +3 | 14 |
| - | - | 1422 | 480.2 | - | - | 0 | - |
| - | - | 1093 | 480.3 | - | - | 0 | - |
| 7 | y | 1.284E+04 | 480.7 | 0.001176 | 2.446 | +2 | 9 |
| - | - | 4348 | 481.2 | - | - | 0 | - |
| - | - | 1074 | 481.8 | - | - | 0 | - |
| 2 | y | 2128 | 486.6 | 5.815E-05 | 0.1195 | +3 | 14 |
| 2 | y | 1146 | 486.9 | 0.008448 | 17.35 | +3 | 14 |
| 11 | y | 1656 | 487.3 | 0.0075 | 15.39 | +1 | 5 |
| 11 | y | 706.4 | 488.2 | 0.001342 | 2.749 | +1 | 5 |
| 2 | y | 1477 | 492.6 | 0.0004428 | 0.899 | +3 | 14 |
| - | - | 1986 | 492.9 | - | - | 0 | - |
| - | - | 1456 | 493.3 | - | - | 0 | - |
| - | - | 3868 | 495.9 | - | - | 0 | - |
| - | - | 3105 | 496.3 | - | - | 0 | - |
| - | - | 2122 | 496.6 | - | - | 0 | - |
| - | - | 831.7 | 497.2 | - | - | 0 | - |
| - | - | 761.4 | 497.3 | - | - | 0 | - |
| 11 | y | 1.001E+04 | 505.3 | 0.00107 | 2.117 | +1 | 5 |
| - | - | 2425 | 506.3 | - | - | 0 | - |
| - | - | 1929 | 514.3 | - | - | 0 | - |
| 6 | y | 1884 | 515.3 | 0.001133 | 2.199 | +2 | 10 |
| 6 | y | 2390 | 515.7 | 0.006676 | 12.94 | +2 | 10 |
| - | - | 1081 | 516.3 | - | - | 0 | - |
| - | - | 991.4 | 518.2 | - | - | 0 | - |
| - | - | 1509 | 518.3 | - | - | 0 | - |
| - | - | 822.2 | 518.6 | - | - | 0 | - |
| - | - | 758.3 | 520.8 | - | - | 0 | - |
| - | - | 936 | 521.3 | - | - | 0 | - |
| - | - | 902.9 | 523.3 | - | - | 0 | - |
| 6 | y | 7.323E+04 | 524.3 | 0.002374 | 4.528 | +2 | 10 |
| 0 | Precursor | 1.257E+04 | 524.6 | 0.007272 | 13.86 | +3 | -1 |
| - | - | 3.856E+04 | 524.8 | - | - | 0 | - |
| - | - | 7223 | 524.9 | - | - | 0 | - |
| - | - | 1.222E+04 | 525.3 | - | - | 0 | - |
| - | - | 2668 | 525.8 | - | - | 0 | - |
| - | - | 757.7 | 528.8 | - | - | 0 | - |
| - | - | 1455 | 529.3 | - | - | 0 | - |
| - | - | 1143 | 529.8 | - | - | 0 | - |
| 0 | Precursor | 1.154E+04 | 530.3 | 0.001311 | 2.473 | +3 | -1 |
| - | - | 6024 | 530.6 | - | - | 0 | - |
| - | - | 894.3 | 530.7 | - | - | 0 | - |
| - | - | 4215 | 531 | - | - | 0 | - |
| - | - | 765.5 | 531.2 | - | - | 0 | - |
| - | - | 842 | 531.3 | - | - | 0 | - |
| - | - | 664.2 | 531.8 | - | - | 0 | - |
| - | - | 1190 | 532.3 | - | - | 0 | - |
| - | - | 768.5 | 533.3 | - | - | 0 | - |
| - | - | 827.6 | 539.3 | - | - | 0 | - |
| - | - | 685.6 | 542.3 | - | - | 0 | - |
| 5 | b | 1952 | 542.3 | 0.002544 | 4.691 | +1 | 5 |
| - | - | 809.9 | 552.8 | - | - | 0 | - |
| - | - | 1072 | 556.3 | - | - | 0 | - |
| - | - | 754.1 | 557.3 | - | - | 0 | - |
| - | - | 1152 | 558.8 | - | - | 0 | - |
| - | - | 1485 | 559.3 | - | - | 0 | - |
| 5 | y | 3912 | 564.8 | 6.169E-05 | 0.1092 | +2 | 11 |
| - | - | 852.1 | 565.2 | - | - | 0 | - |
| 5 | y | 3903 | 565.3 | 0.005977 | 10.57 | +2 | 11 |
| - | - | 2931 | 565.8 | - | - | 0 | - |
| - | - | 738.9 | 566.3 | - | - | 0 | - |
| - | - | 743.9 | 573.3 | - | - | 0 | - |
| 5 | y | 9.748E+04 | 573.8 | 0.0008205 | 1.43 | +2 | 11 |
| 10 | y | 5.968E+04 | 574.3 | 0.004295 | 7.478 | +1 | 6 |
| - | - | 2.048E+04 | 574.8 | - | - | 0 | - |
| - | - | 7033 | 575.3 | - | - | 0 | - |
| - | - | 1724 | 575.8 | - | - | 0 | - |
| - | - | 1689 | 583.2 | - | - | 0 | - |
| - | - | 1755 | 584.3 | - | - | 0 | - |
| - | - | 880.4 | 587.3 | - | - | 0 | - |
| 10 | y | 1.084E+04 | 592.3 | 0.001115 | 1.883 | +1 | 6 |
| - | - | 4300 | 593.3 | - | - | 0 | - |
| - | - | 1049 | 595.3 | - | - | 0 | - |
| - | - | 3211 | 600.3 | - | - | 0 | - |
| - | - | 752.9 | 611.3 | - | - | 0 | - |
| - | - | 659.6 | 618.3 | - | - | 0 | - |
| - | - | 777 | 621.3 | - | - | 0 | - |
| - | - | 710 | 624.3 | - | - | 0 | - |
| - | - | 1078 | 626.3 | - | - | 0 | - |
| 4 | y | 8352 | 629.3 | 0.003361 | 5.341 | +2 | 12 |
| 4 | y | 4379 | 629.8 | 0.009522 | 15.12 | +2 | 12 |
| - | - | 2525 | 630.3 | - | - | 0 | - |
| - | - | 984.5 | 637.3 | - | - | 0 | - |
| - | - | 1238 | 637.8 | - | - | 0 | - |
| 4 | y | 5.518E+04 | 638.3 | 0.0008252 | 1.293 | +2 | 12 |
| - | - | 3.662E+04 | 638.8 | - | - | 0 | - |
| - | - | 1.275E+04 | 639.3 | - | - | 0 | - |
| - | - | 2984 | 639.8 | - | - | 0 | - |
| - | - | 705.9 | 640.3 | - | - | 0 | - |
| - | - | 2210 | 641.3 | - | - | 0 | - |
| - | - | 824.8 | 642.3 | - | - | 0 | - |
| - | - | 1611 | 643.4 | - | - | 0 | - |
| - | - | 3569 | 646.3 | - | - | 0 | - |
| - | - | 1076 | 648.8 | - | - | 0 | - |
| - | - | 1203 | 651.8 | - | - | 0 | - |
| - | - | 1656 | 652.3 | - | - | 0 | - |
| - | - | 1989 | 653.3 | - | - | 0 | - |
| - | - | 3122 | 654.3 | - | - | 0 | - |
| - | - | 1088 | 654.8 | - | - | 0 | - |
| - | - | 1701 | 655.3 | - | - | 0 | - |
| - | - | 1212 | 655.8 | - | - | 0 | - |
| - | - | 938.5 | 657.3 | - | - | 0 | - |
| - | - | 1569 | 657.8 | - | - | 0 | - |
| - | - | 1488 | 658.3 | - | - | 0 | - |
| - | - | 856.2 | 658.8 | - | - | 0 | - |
| - | - | 2563 | 659.3 | - | - | 0 | - |
| - | - | 988.5 | 660.3 | - | - | 0 | - |
| - | - | 3029 | 660.8 | - | - | 0 | - |
| - | - | 1451 | 661.3 | - | - | 0 | - |
| - | - | 4537 | 663.8 | - | - | 0 | - |
| - | - | 4032 | 664.3 | - | - | 0 | - |
| - | - | 1611 | 664.8 | - | - | 0 | - |
| - | - | 1722 | 665.3 | - | - | 0 | - |
| - | - | 1082 | 665.8 | - | - | 0 | - |
| - | - | 1683 | 666.3 | - | - | 0 | - |
| - | - | 1.103E+04 | 666.8 | - | - | 0 | - |
| - | - | 5639 | 667.3 | - | - | 0 | - |
| - | - | 2721 | 667.8 | - | - | 0 | - |
| 9 | y | 1.173E+04 | 671.3 | 0.001213 | 1.807 | +1 | 7 |
| - | - | 883.9 | 671.8 | - | - | 0 | - |
| 9 | y | 8562 | 672.3 | 0.005662 | 8.422 | +1 | 7 |
| 3 | y | 2.792E+04 | 672.8 | 0.0005914 | 0.879 | +2 | 13 |
| 3 | y | 2.546E+04 | 673.3 | 0.00956 | 14.2 | +2 | 13 |
| - | - | 1.1E+04 | 673.8 | - | - | 0 | - |
| - | - | 1.462E+04 | 674.3 | - | - | 0 | - |
| - | - | 1115 | 674.8 | - | - | 0 | - |
| - | - | 4299 | 675.3 | - | - | 0 | - |
| - | - | 751.9 | 676.3 | - | - | 0 | - |
| - | - | 934.6 | 677.8 | - | - | 0 | - |
| - | - | 1000 | 680.3 | - | - | 0 | - |
| - | - | 1.119E+04 | 680.8 | - | - | 0 | - |
| - | - | 8409 | 681.3 | - | - | 0 | - |
| 3 | y | 7.843E+05 | 681.8 | 0.001352 | 1.982 | +2 | 13 |
| - | - | 5.443E+05 | 682.3 | - | - | 0 | - |
| - | - | 2.301E+05 | 682.8 | - | - | 0 | - |
| - | - | 6.649E+04 | 683.3 | - | - | 0 | - |
| - | - | 1.252E+04 | 683.8 | - | - | 0 | - |
| - | - | 1131 | 685.4 | - | - | 0 | - |
| - | - | 6911 | 686.8 | - | - | 0 | - |
| - | - | 5031 | 687.3 | - | - | 0 | - |
| - | - | 2612 | 687.8 | - | - | 0 | - |
| 9 | y | 6.707E+04 | 689.4 | 0.001025 | 1.486 | +1 | 7 |
| - | - | 2.36E+04 | 690.4 | - | - | 0 | - |
| - | - | 6290 | 691.4 | - | - | 0 | - |
| - | - | 968.2 | 692.4 | - | - | 0 | - |
| 14 | b | 4320 | 699.3 | 0.006947 | 9.934 | +2 | 14 |
| - | - | 1452 | 700.3 | - | - | 0 | - |
| - | - | 1161 | 714.4 | - | - | 0 | - |
| - | - | 850.9 | 723.9 | - | - | 0 | - |
| 2 | y | 6003 | 729.4 | 0.0004249 | 0.5826 | +2 | 14 |
| 2 | y | 5517 | 729.9 | 0.01007 | 13.8 | +2 | 14 |
| - | - | 2173 | 730.4 | - | - | 0 | - |
| - | - | 900.1 | 730.9 | - | - | 0 | - |
| - | - | 886.4 | 737.4 | - | - | 0 | - |
| - | - | 1163 | 737.9 | - | - | 0 | - |
| 2 | y | 6.634E+04 | 738.4 | 0.0007014 | 0.9499 | +2 | 14 |
| - | - | 4.868E+04 | 738.9 | - | - | 0 | - |
| - | - | 2.287E+04 | 739.4 | - | - | 0 | - |
| - | - | 6871 | 739.9 | - | - | 0 | - |
| - | - | 1456 | 740.4 | - | - | 0 | - |
| - | - | 4624 | 743.4 | - | - | 0 | - |
| - | - | 1026 | 743.9 | - | - | 0 | - |
| 7 | b | 1642 | 744.4 | 0.005707 | 7.667 | +1 | 7 |
| - | - | 1759 | 758.4 | - | - | 0 | - |
| - | - | 1059 | 759.4 | - | - | 0 | - |
| - | - | 618.5 | 768.4 | - | - | 0 | - |
| - | - | 952.1 | 768.4 | - | - | 0 | - |
| - | - | 984.1 | 769.4 | - | - | 0 | - |
| - | - | 2725 | 770.4 | - | - | 0 | - |
| - | - | 938.4 | 771.4 | - | - | 0 | - |
| - | - | 903.2 | 778.4 | - | - | 0 | - |
| - | - | 2447 | 782.4 | - | - | 0 | - |
| - | - | 2007 | 783.4 | - | - | 0 | - |
| - | - | 2320 | 785.4 | - | - | 0 | - |
| - | - | 1015 | 786.4 | - | - | 0 | - |
| - | - | 1016 | 786.4 | - | - | 0 | - |
| - | - | 1030 | 787.4 | - | - | 0 | - |
| - | - | 941.6 | 788.4 | - | - | 0 | - |
| - | - | 1287 | 796.4 | - | - | 0 | - |
| - | - | 2855 | 798.4 | - | - | 0 | - |
| - | - | 2445 | 803.4 | - | - | 0 | - |
| - | - | 934.6 | 804.4 | - | - | 0 | - |
| - | - | 845.4 | 810.4 | - | - | 0 | - |
| - | - | 1622 | 811.4 | - | - | 0 | - |
| - | - | 1165 | 812.4 | - | - | 0 | - |
| 8 | y | 1.148E+04 | 828.4 | 0.0007426 | 0.8964 | +1 | 8 |
| - | - | 5350 | 829.4 | - | - | 0 | - |
| - | - | 890.3 | 857.4 | - | - | 0 | - |
| - | - | 737.1 | 858.4 | - | - | 0 | - |
| - | - | 1399 | 873.4 | - | - | 0 | - |
| - | - | 781.1 | 874.4 | - | - | 0 | - |
| - | - | 1458 | 885.4 | - | - | 0 | - |
| - | - | 785 | 886.5 | - | - | 0 | - |
| - | - | 2148 | 887.4 | - | - | 0 | - |
| - | - | 1215 | 888.4 | - | - | 0 | - |
| - | - | 1574 | 891.4 | - | - | 0 | - |
| - | - | 1348 | 897.4 | - | - | 0 | - |
| - | - | 921.1 | 899.5 | - | - | 0 | - |
| 8 | b | 2509 | 900.5 | 0.008368 | 9.293 | +1 | 8 |
| - | - | 966.5 | 901.5 | - | - | 0 | - |
| - | - | 845.8 | 909.5 | - | - | 0 | - |
| - | - | 881.8 | 910.4 | - | - | 0 | - |
| - | - | 4823 | 915.4 | - | - | 0 | - |
| - | - | 2081 | 916.4 | - | - | 0 | - |
| - | - | 1021 | 926.4 | - | - | 0 | - |
| - | - | 1395 | 927.4 | - | - | 0 | - |
| - | - | 767.4 | 928.5 | - | - | 0 | - |
| 7 | y | 8125 | 943.5 | 0.001265 | 1.341 | +1 | 9 |
| - | - | 4694 | 944.5 | - | - | 0 | - |
| - | - | 1564 | 945.5 | - | - | 0 | - |
| - | - | 1032 | 968.5 | - | - | 0 | - |
| - | - | 1878 | 969.5 | - | - | 0 | - |
| - | - | 887.6 | 970.5 | - | - | 0 | - |
| - | - | 1249 | 972.5 | - | - | 0 | - |
| - | - | 6608 | 986.5 | - | - | 0 | - |
| - | - | 7909 | 987.5 | - | - | 0 | - |
| - | - | 3618 | 988.5 | - | - | 0 | - |
| - | - | 1238 | 989.5 | - | - | 0 | - |
| - | - | 1528 | 990.5 | - | - | 0 | - |
| - | - | 1129 | 997.5 | - | - | 0 | - |
| - | - | 4103 | 1012 | - | - | 0 | - |
| - | - | 2516 | 1013 | - | - | 0 | - |
| - | - | 7408 | 1014 | - | - | 0 | - |
| - | - | 2996 | 1015 | - | - | 0 | - |
| - | - | 800.2 | 1016 | - | - | 0 | - |
| 6 | y | 2.229E+04 | 1030 | 0.000487 | 0.4726 | +1 | 10 |
| - | - | 1.398E+04 | 1031 | - | - | 0 | - |
| - | - | 4786 | 1032 | - | - | 0 | - |
| - | - | 1167 | 1033 | - | - | 0 | - |
| - | - | 927.4 | 1058 | - | - | 0 | - |
| - | - | 2123 | 1074 | - | - | 0 | - |
| - | - | 1322 | 1075 | - | - | 0 | - |
| - | - | 1793 | 1084 | - | - | 0 | - |
| - | - | 1712 | 1087 | - | - | 0 | - |
| - | - | 779.2 | 1088 | - | - | 0 | - |
| - | - | 4435 | 1102 | - | - | 0 | - |
| - | - | 2201 | 1103 | - | - | 0 | - |
| - | - | 798 | 1104 | - | - | 0 | - |
| - | - | 1028 | 1112 | - | - | 0 | - |
| - | - | 1118 | 1120 | - | - | 0 | - |
| 5 | y | 6444 | 1130 | 6.621E-05 | 0.05861 | +1 | 11 |
| - | - | 4579 | 1131 | - | - | 0 | - |
| - | - | 1746 | 1132 | - | - | 0 | - |
| - | - | 846 | 1144 | - | - | 0 | - |
| - | - | 1040 | 1159 | - | - | 0 | - |
| - | - | 4538 | 1161 | - | - | 0 | - |
| - | - | 2648 | 1162 | - | - | 0 | - |
| - | - | 729 | 1163 | - | - | 0 | - |
| - | - | 1796 | 1171 | - | - | 0 | - |
| - | - | 984 | 1172 | - | - | 0 | - |
| - | - | 4658 | 1189 | - | - | 0 | - |
| - | - | 3187 | 1190 | - | - | 0 | - |
| - | - | 990.6 | 1206 | - | - | 0 | - |
| - | - | 5466 | 1207 | - | - | 0 | - |
| - | - | 3063 | 1208 | - | - | 0 | - |
| - | - | 880.4 | 1209 | - | - | 0 | - |
| 3 | y | 813.3 | 1346 | 0.001679 | 1.248 | +1 | 13 |
| - | - | 740.7 | 2022 | - | - | 0 | - |

m/z Charge Intensity FragmentType MassShift Position
120.08121490478516 0 2607.3154
122.26435852050781 0 340.06445
126.05538940429688 0 7866.5303
127.0868148803711 0 436.7957
128.0821075439453 0 1448.6302
128.1073760986328 0 884.401
129.06637573242188 0 1204.5864
129.1027069091797 0 4455.8545
130.05001831054688 0 569.7595
130.06158447265625 0 3814.1555
130.09803771972656 0 2207.5444
131.08184814453125 0 651.1889
131.1180877685547 0 682.36743
133.0613555908203 0 622.8649
133.09780883789062 0 420.539
136.0712127685547 0 485.5046
136.07611083984375 0 3453.0884
139.0868682861328 0 1102.3745
139.09808349609375 0 585.2509
140.0823516845703 0 1074.1935
140.1439971923828 0 634.71954
141.10296630859375 0 788.26764
142.9599151611328 0 441.8123
143.08177185058594 0 589.6821
144.06591796875 0 1773.0845
145.06134033203125 0 1100.2312
147.07687377929688 0 2425.548
148.95477294921875 0 1008.353
152.864013671875 0 405.9588
153.06619262695312 0 1003.19507
154.05027770996094 0 7769.589
155.08192443847656 0 605.0288
155.1185302734375 0 1102.1924
156.1020965576172 0 904.3986
157.0612335205078 0 3184.933
157.0976104736328 0 3445.2373
157.10879516601562 0 4837.9556
158.0928497314453 0 17520.377 y Ammonia loss 14
158.1002960205078 0 906.51056
158.11785888671875 0 867.0685
159.07696533203125 0 1015.1228
159.09632873535156 0 1231.7688
159.11329650878906 0 2597.7783
161.0548553466797 0 474.3285
163.07177734375 0 5238.35
167.0818634033203 0 1555.3337
167.60394287109375 0 496.87714
169.0613250732422 0 895.7825
169.09762573242188 0 1368.1613 y 10
170.04531860351562 0 647.0883
170.117431640625 0 475.65558
171.07687377929688 0 9471.1
171.1134490966797 0 998.44995
172.0609893798828 0 1887.5459
172.08029174804688 0 542.1073
173.12181091308594 0 538.7118
173.12904357910156 0 3202.3394
173.45254516601562 0 1386.0764
175.0718231201172 0 6280.881
175.11941528320312 0 37598.91 y 14
176.0755157470703 0 553.5028
176.1228790283203 0 2124.3145
178.05447387695312 0 456.43817
179.48638916015625 0 479.48456
182.1543426513672 0 491.88428
183.11334228515625 0 8503.235
184.10853576660156 0 1261.246
184.11688232421875 0 927.9172
185.09255981445312 0 5393.0986
185.16513061523438 0 846.2367 d 1
186.08782958984375 0 703.6758
187.0715789794922 0 1125.944
187.1082305908203 0 5622.8765
187.14459228515625 0 754.0434
188.11190795898438 0 487.93332
189.0874786376953 0 40656.023
190.09107971191406 0 3675.053
191.0914764404297 0 547.08215
192.113525390625 0 1348.4928
193.08241271972656 0 629.3796
193.0972137451172 0 827.3029
194.09259033203125 0 965.3773
195.0771942138672 0 654.3498
195.1130828857422 0 903.9163
195.14974975585938 0 1882.2097
197.0919647216797 0 471.65454
197.12811279296875 0 641.7813
197.16505432128906 0 894.2123
199.07174682617188 0 4381.022
199.18099975585938 0 315715.44 a 1
200.18434143066406 0 40416.242
201.12374877929688 0 11803.318
201.18682861328125 0 2188.499
202.11880493164062 0 2843.986
202.1278839111328 0 740.21173
203.0667266845703 0 6900.604
203.10336303710938 0 1922.71
208.1085968017578 0 583.9791
209.12818908691406 0 601.38654
210.08729553222656 0 1087.7483
211.10821533203125 0 2387.132
212.1034698486328 0 3950.4036
213.0869598388672 0 956.9366
213.1599884033203 0 1245.997
215.13900756835938 0 627.68555
216.09674072265625 0 705.30695
216.1348114013672 0 967.6306
217.08238220214844 0 69932.586
218.08590698242188 0 4864.584
219.0879669189453 0 732.4365
220.09349060058594 0 748.8694
222.83856201171875 0 545.3959
224.10354614257812 0 4472.4346
225.16049194335938 0 727.6745
226.118896484375 0 1203.6095
226.1558380126953 0 1759.796
227.1031951904297 0 817.77295
227.1146697998047 0 2972.8335
227.17593383789062 0 94022.2 b 1
228.09803771972656 0 969.01825
228.17922973632812 0 12858.457
229.11883544921875 0 3076.5588
229.18170166015625 0 1046.825
230.11378479003906 0 2160.669
231.09877014160156 0 545.64575
233.16543579101562 0 2292.9966
234.10885620117188 0 746.5918
234.16836547851562 0 633.71796
237.09906005859375 0 1111.8647
237.13511657714844 0 2224.8635
237.1604461669922 0 573.9381
240.13421630859375 0 664.7295
242.11416625976562 0 9829.112
243.11788940429688 0 954.2588
244.12954711914062 0 2101.326
244.1417694091797 0 917.6074 y Water loss 13
245.12509155273438 0 9393.983 y Ammonia loss 13
246.1291046142578 0 839.6143
249.16064453125 0 1040.3907
253.16668701171875 0 2760.2676
254.11378479003906 0 986.1106
255.1094207763672 0 8483.614
256.1099548339844 0 649.5974
259.1406555175781 0 907.4593
261.1604919433594 0 1236.7954
262.15142822265625 0 6713.1323 y 13
267.1095275878906 0 590.66345
269.1612854003906 0 818.11505
269.1865234375 0 699.0019
270.1455383300781 0 1558.9
270.1925048828125 0 1647.9318
272.1355895996094 0 1475.2102
280.12994384765625 0 1462.7184
283.1767883300781 0 879.50696
284.1246643066406 0 1531.9144
287.21990966796875 0 699.0011
288.1563415527344 0 1749.3329
295.1770324707031 0 661.0155
296.136474609375 0 1446.3235
296.1975402832031 0 3696.641 b Water loss 2
297.1200256347656 0 1620.7107
298.140625 0 3303.7283
299.1361389160156 0 1963.1411
302.1342468261719 0 1121.5082
309.1575622558594 0 504.8431
312.1559753417969 0 4671.9897
313.1507873535156 0 532.69916
313.1878356933594 0 10240.862
314.14678955078125 0 3937.966
314.1900329589844 0 1422.3102
314.20831298828125 0 9210.755 b 2
314.2270202636719 0 967.91895
315.1487731933594 0 743.1076
315.21136474609375 0 1119.1644
316.13092041015625 0 403.22937
316.1510314941406 0 11899.668
317.1241760253906 0 832.4245
317.1488342285156 0 1717.2074
323.2088623046875 0 1094.0789
324.131591796875 0 1522.7864
324.16839599609375 0 573.1706
325.6253967285156 0 610.5689
329.1459045410156 0 667.4296
330.1666564941406 0 2894.3572
331.1726989746094 0 2273.3816 y Water loss 12
332.1572570800781 0 9970.6045 y Ammonia loss 12
333.15997314453125 0 1762.1615
335.1369934082031 0 623.55
340.15069580078125 0 1135.5886
341.15875244140625 0 612.66974
341.1827697753906 0 6892.236
341.2167663574219 0 786.01373
342.1414489746094 0 11888.611
343.1443786621094 0 2280.8677
345.183349609375 0 5681.084 y 8
345.6852722167969 0 1876.1378
349.1837463378906 0 8596.561 y 12
350.1871032714844 0 843.9803
352.1637268066406 0 784.1071
359.1676940917969 0 6087.2837
360.1715393066406 0 787.7276
365.1812438964844 0 699.49835
380.15667724609375 0 1384.7446
381.1888732910156 0 930.22125
385.17266845703125 0 1367.358
389.709716796875 0 1030.4998
390.71759033203125 0 1023.39966
391.7244873046875 0 4678.399
392.22625732421875 0 1166.6069
393.71051025390625 0 1826.4443
394.21173095703125 0 834.4194
397.24560546875 0 953.3169
398.21337890625 0 968.8196
398.7151184082031 0 1515.2885
399.2154235839844 0 1095.63
399.7207946777344 0 2678.9392
400.18353271484375 0 651.3555
400.2200927734375 0 1900.6504
401.2420654296875 0 878.3067
403.1839904785156 0 1518.7184
405.71649169921875 0 968.21826
407.2290954589844 0 1134.0598
408.22943115234375 0 976.7491
410.2034912109375 0 1845.129
411.2251892089844 0 815.74866
414.2010192871094 0 765.8289
414.2286376953125 0 2863.2266 b 11
414.72222900390625 0 6006.7915 y Ammonia loss 7
415.2218322753906 0 2995.3179
415.25531005859375 0 1526.0593
415.7257080078125 0 817.78467
415.7947692871094 0 782.74536
416.21575927734375 0 721.8496
423.2340393066406 0 73673.02 y 7
423.7351989746094 0 30388.203
424.2366027832031 0 7750.7354
424.7373046875 0 1109.2373
425.24029541015625 0 5077.6216 b Water loss 3
426.242919921875 0 1303.5227
428.2169494628906 0 2438.9143
429.2347412109375 0 3748.6028
430.240234375 0 2964.4922 y Water loss 11
431.2239990234375 0 777.5743 y Ammonia loss 11
440.2005920410156 0 723.94135
441.21087646484375 0 3952.6165
442.2122802734375 0 984.27155
443.2506408691406 0 3953.0923 b 3
444.2540588378906 0 997.17
445.2421569824219 0 627.5855
448.2528991699219 0 1762.5742 y 11
448.8893127441406 0 2204.574 y Water loss 2
449.2239990234375 0 958.0592 y Ammonia loss 2
450.74432373046875 0 751.91833 b 7
452.22686767578125 0 719.7688
452.2632141113281 0 2115.02
454.8924560546875 0 2521.172 y 2
455.22589111328125 0 1750.5779
455.29742431640625 0 1465.013
458.236572265625 0 4653.197
460.2417297363281 0 1607.9998
463.23089599609375 0 627.9523
469.2895202636719 0 1962.4675
472.23809814453125 0 865.6385 b 13
480.2135009765625 0 1422.2517
480.255859375 0 1092.9397
480.7476806640625 0 12840.557 y 6
481.249267578125 0 4348.2544
481.75079345703125 0 1074.3066
486.5830078125 0 2127.9182 y Water loss 1
486.9194030761719 0 1146.2952 y Ammonia loss 1
487.25482177734375 0 1656.2207 y Water loss 10
488.2449951171875 0 706.39874 y Ammonia loss 10
492.5869140625 0 1477.4592 y 1
492.9203796386719 0 1986.1589
493.2550354003906 0 1456.2417
495.9156188964844 0 3868.0083
496.2505187988281 0 3105.0164
496.58380126953125 0 2121.7446
497.2341003417969 0 831.7044
497.2817687988281 0 761.4086
505.2739562988281 0 10008.412 y 10
506.2784118652344 0 2425.3591
514.32470703125 0 1928.6824
515.256103515625 0 1883.8442 y Water loss 5
515.7559204101562 0 2390.3538 y Ammonia loss 5
516.2590942382812 0 1080.8387
518.2100219726562 0 991.39136
518.2767944335938 0 1508.5552
518.6084594726562 0 822.1602
520.7929077148438 0 758.28455
521.2899780273438 0 936.02496
523.2606201171875 0 902.858
524.264892578125 0 73234.52 y 5
524.6129150390625 0 12570.731 Precursor Ammonia loss
524.7646484375 0 38555.53
524.94677734375 0 7222.6094
525.2666015625 0 12215.168
525.76708984375 0 2667.9465
528.7888793945312 0 757.72003
529.2999267578125 0 1455.0569
529.80224609375 0 1142.5312
530.282470703125 0 11538.598 Precursor
530.6158447265625 0 6024.2197
530.7429809570312 0 894.26935
530.9505615234375 0 4215.2646
531.246337890625 0 765.46234
531.2880249023438 0 842.00586
531.7733764648438 0 664.19086
532.2732543945312 0 1189.6553
533.2565307617188 0 768.47687
539.2964477539062 0 827.5898
542.2731323242188 0 685.633
542.3209838867188 0 1951.6968 b 4
552.7825317382812 0 809.8824
556.3213500976562 0 1071.9504
557.2750854492188 0 754.13354
558.7918701171875 0 1151.7688
559.2901611328125 0 1485.4528
564.7913818359375 0 3912.1084 y Water loss 4
565.2398071289062 0 852.13074
565.2894287109375 0 3903.1902 y Ammonia loss 4
565.7883911132812 0 2931.2893
566.2958374023438 0 738.93677
573.29248046875 0 743.8826
573.7975463867188 0 97480.1 y 4
574.2986450195312 0 59677.863 y Water loss 9
574.8002319335938 0 20477.242
575.3010864257812 0 7032.9414
575.8032836914062 0 1723.722
583.2483520507812 0 1688.6575
584.3165893554688 0 1754.8214
587.2816772460938 0 880.4039
592.3060302734375 0 10839.508 y 9
593.308349609375 0 4299.8677
595.2823486328125 0 1049.0898
600.2750854492188 0 3211.2002
611.3213500976562 0 752.9139
618.2984619140625 0 659.5982
621.3062133789062 0 777.0238
624.310791015625 0 710.04785
626.326416015625 0 1078.0338
629.3161010742188 0 8351.525 y Water loss 3
629.8142700195312 0 4378.802 y Ammonia loss 3
630.3169555664062 0 2524.5413
637.3043212890625 0 984.5151
637.8123779296875 0 1237.7087
638.31884765625 0 55180.47 y 3
638.8202514648438 0 36618.72
639.321044921875 0 12748.364
639.822998046875 0 2984.2861
640.3228149414062 0 705.9284
641.3360595703125 0 2210.056
642.3363037109375 0 824.80585
643.3549194335938 0 1610.8878
646.3156127929688 0 3569.3672
648.8226928710938 0 1075.6954
651.8161010742188 0 1203.051
652.3170166015625 0 1655.7166
653.3370361328125 0 1989.1263
654.3264770507812 0 3122.2026
654.8250732421875 0 1088.4004
655.3191528320312 0 1701.3912
655.8153076171875 0 1212.4526
657.2796630859375 0 938.4817
657.8258666992188 0 1569.1973
658.3289794921875 0 1488.4691
658.8312377929688 0 856.23975
659.3479614257812 0 2563.2466
660.3012084960938 0 988.5139
660.8207397460938 0 3028.9412
661.3245849609375 0 1450.9552
663.8242797851562 0 4536.516
664.3211669921875 0 4032.0137
664.8204345703125 0 1610.8691
665.3201904296875 0 1721.8658
665.8265380859375 0 1081.6652
666.3201293945312 0 1683.2524
666.8292846679688 0 11033.733
667.3290405273438 0 5639.0293
667.8313598632812 0 2721.2898
671.3483276367188 0 11731.306 y Water loss 8
671.8197631835938 0 883.89166
672.3367919921875 0 8561.905 y Ammonia loss 8
672.829345703125 0 27919.951 y Water loss 2
673.330322265625 0 25464.33 y Ammonia loss 2
673.8295288085938 0 10997.757
674.3162841796875 0 14620.241
674.827880859375 0 1115.0471
675.315185546875 0 4299.105
676.32421875 0 751.92145
677.8201904296875 0 934.5554
680.3191528320312 0 1000.2135
680.827392578125 0 11192.823
681.3287353515625 0 8408.586
681.8353881835938 0 784277.94 y 2
682.33642578125 0 544285.75
682.8375854492188 0 230142.67
683.3386840820312 0 66486.24
683.839599609375 0 12522.452
685.3818359375 0 1130.738
686.8275146484375 0 6911.4844
687.3302001953125 0 5031.1416
687.8304443359375 0 2611.5806
689.3587036132812 0 67074.95 y 8
690.3615112304688 0 23601.555
691.364501953125 0 6290.3267
692.3675537109375 0 968.20795
699.342041015625 0 4320.172 b Ammonia loss 13
700.3394775390625 0 1451.906
714.3539428710938 0 1160.9987
723.8731689453125 0 850.85834
729.370361328125 0 6002.6406 y Water loss 1
729.8728637695312 0 5517.033 y Ammonia loss 1
730.373046875 0 2173.416
730.865966796875 0 900.0899
737.366455078125 0 886.3606
737.8727416992188 0 1163.3445
738.3767700195312 0 66342.03 y 1
738.8782348632812 0 48675.277
739.3792114257812 0 22869.352
739.8811645507812 0 6870.588
740.3789672851562 0 1456.1155
743.3694458007812 0 4623.9404
743.8699951171875 0 1026.295
744.3717041015625 0 1641.9869 b 6
758.3809814453125 0 1758.6631
759.373779296875 0 1059.495
768.3633422851562 0 618.4741
768.40478515625 0 952.0818
769.3577270507812 0 984.1494
770.4125366210938 0 2725.1167
771.4282836914062 0 938.4248
778.41796875 0 903.17145
782.4421997070312 0 2446.827
783.4422607421875 0 2007.2133
785.425537109375 0 2320.472
786.3675537109375 0 1014.9477
786.4248657226562 0 1016.41284
787.3938598632812 0 1030.0378
788.4058227539062 0 941.60077
796.4146118164062 0 1287.396
798.4190063476562 0 2855.0325
803.4376831054688 0 2445.496
804.381591796875 0 934.64557
810.3636474609375 0 845.41614
811.410400390625 0 1622.0693
812.3992919921875 0 1165.0869
828.4329833984375 0 11475.168 y Ammonia loss 7
829.4347534179688 0 5350.4116
857.44580078125 0 890.3247
858.3984985351562 0 737.0589
873.4049072265625 0 1398.5256
874.405029296875 0 781.14954
885.4413452148438 0 1457.5051
886.45166015625 0 785.0247
887.423828125 0 2148.24
888.4290161132812 0 1214.9642
891.4197387695312 0 1573.5662
897.4036254882812 0 1347.5813
899.4655151367188 0 921.1255
900.4701538085938 0 2508.604 b 7
901.4642944335938 0 966.53705
909.4532470703125 0 845.84515
910.4439697265625 0 881.7815
915.419921875 0 4822.8716
916.4212646484375 0 2081.424
926.4378051757812 0 1020.652
927.4456176757812 0 1394.8604
928.4623413085938 0 767.43414
943.46044921875 0 8125.2153 y Ammonia loss 6
944.4652099609375 0 4694.08
945.476318359375 0 1563.6033
968.4817504882812 0 1031.7775
969.474365234375 0 1878.0149
970.4859619140625 0 887.6478
972.468994140625 0 1249.3474
986.4909057617188 0 6608.0664
987.48828125 0 7908.903
988.4898071289062 0 3617.8267
989.492431640625 0 1237.9431
990.4855346679688 0 1528.0083
997.470458984375 0 1129.2621
1012.48095703125 0 4102.997
1013.4829711914062 0 2516.2732
1014.485107421875 0 7407.6743
1015.48828125 0 2995.7551
1016.4827270507812 0 800.1576
1030.49169921875 0 22286.584 y Ammonia loss 5
1031.4945068359375 0 13975.555
1032.496337890625 0 4786.05
1033.4974365234375 0 1167.0354
1057.512939453125 0 927.3687
1073.5220947265625 0 2122.8787
1074.5299072265625 0 1322.0139
1083.50927734375 0 1792.9757
1086.5513916015625 0 1711.7726
1087.552001953125 0 779.20184
1101.5166015625 0 4435.213
1102.5186767578125 0 2201.014
1103.517333984375 0 797.96497
1111.5528564453125 0 1027.9692
1119.522705078125 0 1118.4216
1129.5596923828125 0 6443.7285 y Ammonia loss 4
1130.5621337890625 0 4579.207
1131.5616455078125 0 1745.7151
1143.524169921875 0 846.04596
1158.530029296875 0 1040.3528
1160.554443359375 0 4538.1562
1161.5596923828125 0 2648.4058
1162.546630859375 0 729.00104
1170.5396728515625 0 1795.9751
1171.5438232421875 0 983.95044
1188.5506591796875 0 4657.823
1189.55029296875 0 3187.0293
1205.569580078125 0 990.60626
1206.5601806640625 0 5466.4653
1207.56201171875 0 3063.0886
1208.5736083984375 0 880.42523
1345.632568359375 0 813.3045 y Ammonia loss 2
2021.5308837890625 0 740.69696

Spectrum Details

|  |  |
| --- | --- |
| Matched peaks? Matched peaks The total absolute number of peaks matched. Additionally in brackets the total fraction of peaks matched and the total number of peaks is shown. | 66 (12.69% of 520) |
| FDR? FDR The false discovery rate estimated for this peptide. It is calculated by matching all theoretical fragments with a non-integer shift with the raw peaks for this spectrum. This is done with 40 different shifts. The resulting percentage is the average number of annotated peaks over the number of annotated peaks with the correct spectrum. | 0.40% |
| Satellite FDR? Satellite FDR See the FDR for details on its calculation. This satellite ion specific FDR only contains the satellite ions (d/w) for I/L/J positions. | 4.76% |
| PSM Score? PSM Score The PSM Score as given by Hecklib to this annotated spectrum. It is shown with three significant figures. | 439 |

#### Spectrum 6539? Spectrum 6539 The raw spectrum of this peptide as annotated by Hecklib. The fragments are coloured according to ion type (see legend). Any peaks with a star '\*' as text can be hovered over to see the full details, first the ion type second the mass shift type. By hovering over the amino acids in the peptide or ions in the legend the corresponding peaks are highlighted. By toggling the 'Unassigned' label you can turn the background (unassigned) peaks on or off in the plot. By updating the slider in the Ion legend you can update the spectrum to only show the top X% of the peaks with labels. The top X% means any peak that is within X% of the highest intensity. By dragging in the spectrum you can zoom in to a specific part of the spectrum and use 'Zoom Out' to get back to the original zoom level. The annotation of the spectrum is based on the given sequence in the peptides file and is done with different software so inconsistencies are likely. The peaks are annotated based on the given sequence, with 20 ppm tolerance.

Copy Data

##### Spectrum 6539 (TSV)

###### Preview

```
Loading example...
```

*Click on the button to copy the data to your clipboard.*

Mz MinMz MaxIntensity Max

WidthHeightPeptide font sizePeptide stroke widthSpectrum font sizeSpectrum stroke widthCompact peptide

Ion legend

wxyz

abcd

OtherUnassignedIon^Charge^_Position_Show for top:%

JJSEVSDRPSGVSSR

03.72e+37.44e+31.12e+41.49e+4

Zoom Out

y^+3^_5_w^+1^_2_y^+1^_2_y^+1^_2_w^+1^_3_z^+1^_3_w^+1^_4_y^+1^_4_c^+1^_4_y^+1^_5_y^+2^_10_z^+2^_11_c^+1^_5_z^+2^_11_y^+2^_11_y^+1^_6_w^+2^_12_z^+2^_12_y^+2^_12_c^+1^_6_z^+2^_13_y^+2^_13_y^+1^_7_w^+2^_14_z^+2^_14_y^+2^_14_c^+1^_7_z^+1^_8_c^+1^_8_z^+1^_9_c^+1^_9_z^+1^_10_c^+1^_10_z^+1^_11_c^+1^_11_c^+1^_12_y^+1^_12_c^+1^_13_c^+1^_14_

0883176626483531

Fragment Matches Table

Show background peaks

| Position | Ion type | Intensity | mz Theoretical | mz Error (Th) | mz Error (ppm) | Charge | Series Number |
| --- | --- | --- | --- | --- | --- | --- | --- |
| - | - | 1313 | 120.1 | - | - | 0 | - |
| - | - | 407.8 | 126.1 | - | - | 0 | - |
| - | - | 462 | 129.1 | - | - | 0 | - |
| - | - | 775.2 | 133.1 | - | - | 0 | - |
| - | - | 895.4 | 136.1 | - | - | 0 | - |
| - | - | 2047 | 140.1 | - | - | 0 | - |
| - | - | 446.1 | 140.3 | - | - | 0 | - |
| - | - | 581.9 | 147.1 | - | - | 0 | - |
| - | - | 594.6 | 163.1 | - | - | 0 | - |
| - | - | 547 | 168.1 | - | - | 0 | - |
| 11 | y | 603.6 | 169.1 | 0.0009123 | 5.395 | +3 | 5 |
| - | - | 444.7 | 171.3 | - | - | 0 | - |
| - | - | 1148 | 173.4 | - | - | 0 | - |
| - | - | 2940 | 185.1 | - | - | 0 | - |
| - | - | 1.235E+04 | 187.1 | - | - | 0 | - |
| - | - | 5076 | 199.2 | - | - | 0 | - |
| - | - | 668.7 | 200.2 | - | - | 0 | - |
| - | - | 953.3 | 215.1 | - | - | 0 | - |
| - | - | 643.1 | 225.9 | - | - | 0 | - |
| - | - | 2327 | 227.2 | - | - | 0 | - |
| 14 | w | 548.8 | 229.1 | 6.078E-05 | 0.2653 | +1 | 2 |
| 14 | y | 3639 | 245.1 | 0.0002481 | 1.012 | +1 | 2 |
| 14 | y | 687 | 262.2 | 0.001841 | 7.023 | +1 | 2 |
| 13 | w | 1838 | 316.2 | 0.0006251 | 1.977 | +1 | 3 |
| 13 | z | 962.7 | 333.2 | 0.0008457 | 2.538 | +1 | 3 |
| - | - | 514.2 | 333.2 | - | - | 0 | - |
| - | - | 736.5 | 357.2 | - | - | 0 | - |
| - | - | 706.8 | 362.2 | - | - | 0 | - |
| - | - | 1643 | 377.2 | - | - | 0 | - |
| - | - | 603.6 | 386.3 | - | - | 0 | - |
| - | - | 831.2 | 401.3 | - | - | 0 | - |
| 12 | w | 1425 | 417.2 | 0.0008593 | 2.06 | +1 | 4 |
| - | - | 521.6 | 418.2 | - | - | 0 | - |
| - | - | 1066 | 419.2 | - | - | 0 | - |
| - | - | 572.6 | 429.1 | - | - | 0 | - |
| 12 | y | 1161 | 448.3 | 0.002308 | 5.149 | +1 | 4 |
| - | - | 782.9 | 453.3 | - | - | 0 | - |
| - | - | 546 | 454.8 | - | - | 0 | - |
| 4 | c | 737.5 | 460.3 | 0.001154 | 2.507 | +1 | 4 |
| - | - | 769.3 | 475.3 | - | - | 0 | - |
| - | - | 575.4 | 504.3 | - | - | 0 | - |
| 11 | y | 5236 | 505.3 | 0.0003983 | 0.7883 | +1 | 5 |
| - | - | 1118 | 506.3 | - | - | 0 | - |
| 6 | y | 917 | 524.3 | 0.004205 | 8.02 | +2 | 10 |
| - | - | 949.9 | 524.8 | - | - | 0 | - |
| - | - | 796.2 | 529.2 | - | - | 0 | - |
| - | - | 613.8 | 529.3 | - | - | 0 | - |
| - | - | 657 | 530.3 | - | - | 0 | - |
| - | - | 2470 | 530.3 | - | - | 0 | - |
| - | - | 852.1 | 531.3 | - | - | 0 | - |
| - | - | 731.5 | 542.3 | - | - | 0 | - |
| 5 | z | 695.6 | 557.3 | 0.001934 | 3.47 | +2 | 11 |
| 5 | c | 735.2 | 559.3 | 0.002643 | 4.725 | +1 | 5 |
| 5 | z | 3450 | 565.8 | 0.0004986 | 0.8812 | +2 | 11 |
| - | - | 2379 | 566.3 | - | - | 0 | - |
| - | - | 877.5 | 566.8 | - | - | 0 | - |
| - | - | 534.2 | 567.3 | - | - | 0 | - |
| - | - | 636.8 | 569.3 | - | - | 0 | - |
| - | - | 552.1 | 572.5 | - | - | 0 | - |
| 5 | y | 1553 | 573.8 | 0.001187 | 2.068 | +2 | 11 |
| 10 | y | 723.4 | 574.3 | 0.00637 | 11.09 | +1 | 6 |
| - | - | 667.3 | 585.3 | - | - | 0 | - |
| - | - | 609 | 596.3 | - | - | 0 | - |
| 4 | w | 1217 | 600.8 | 0.00213 | 3.545 | +2 | 12 |
| - | - | 740.6 | 601.3 | - | - | 0 | - |
| - | - | 774.1 | 604.4 | - | - | 0 | - |
| - | - | 699.5 | 605.2 | - | - | 0 | - |
| 4 | z | 3002 | 630.3 | 0.0002996 | 0.4753 | +2 | 12 |
| - | - | 1898 | 630.8 | - | - | 0 | - |
| - | - | 622.8 | 631.3 | - | - | 0 | - |
| 4 | y | 3720 | 638.3 | 0.0006396 | 1.002 | +2 | 12 |
| - | - | 2005 | 638.8 | - | - | 0 | - |
| - | - | 926.5 | 639.3 | - | - | 0 | - |
| 6 | c | 3144 | 646.4 | 6.422E-05 | 0.09936 | +1 | 6 |
| - | - | 1099 | 647.4 | - | - | 0 | - |
| - | - | 602.3 | 651.9 | - | - | 0 | - |
| - | - | 804.5 | 661.8 | - | - | 0 | - |
| 3 | z | 2570 | 673.8 | 3.245E-05 | 0.04815 | +2 | 13 |
| - | - | 3026 | 674.3 | - | - | 0 | - |
| - | - | 1459 | 674.8 | - | - | 0 | - |
| - | - | 760 | 675.3 | - | - | 0 | - |
| 3 | y | 1.149E+04 | 681.8 | 0.0001743 | 0.2557 | +2 | 13 |
| - | - | 9909 | 682.3 | - | - | 0 | - |
| - | - | 4588 | 682.8 | - | - | 0 | - |
| 9 | y | 756.7 | 689.4 | 0.00105 | 1.524 | +1 | 7 |
| - | - | 750.2 | 703.3 | - | - | 0 | - |
| - | - | 1416 | 712.4 | - | - | 0 | - |
| - | - | 731.6 | 713.9 | - | - | 0 | - |
| 2 | w | 689.9 | 715.8 | 0.001245 | 1.739 | +2 | 14 |
| - | - | 551 | 723.8 | - | - | 0 | - |
| - | - | 989.5 | 728.4 | - | - | 0 | - |
| 2 | z | 682 | 730.4 | 5.368E-05 | 0.0735 | +2 | 14 |
| 2 | y | 1700 | 738.4 | 0.002228 | 3.018 | +2 | 14 |
| - | - | 859.3 | 739.4 | - | - | 0 | - |
| - | - | 3972 | 745.4 | - | - | 0 | - |
| - | - | 690.8 | 746.4 | - | - | 0 | - |
| 7 | c | 5103 | 761.4 | 2.972E-05 | 0.03903 | +1 | 7 |
| - | - | 2722 | 762.4 | - | - | 0 | - |
| - | - | 1313 | 763.4 | - | - | 0 | - |
| - | - | 1400 | 764.9 | - | - | 0 | - |
| - | - | 1.216E+04 | 765.4 | - | - | 0 | - |
| - | - | 9202 | 765.9 | - | - | 0 | - |
| - | - | 5118 | 766.4 | - | - | 0 | - |
| - | - | 1322 | 766.9 | - | - | 0 | - |
| - | - | 762 | 771.4 | - | - | 0 | - |
| - | - | 2097 | 773.9 | - | - | 0 | - |
| - | - | 1743 | 774.4 | - | - | 0 | - |
| - | - | 705.1 | 774.9 | - | - | 0 | - |
| - | - | 801.1 | 775.4 | - | - | 0 | - |
| - | - | 769.2 | 776.4 | - | - | 0 | - |
| - | - | 1058 | 778.4 | - | - | 0 | - |
| - | - | 1408 | 778.9 | - | - | 0 | - |
| - | - | 907.8 | 779.9 | - | - | 0 | - |
| - | - | 774.6 | 780.4 | - | - | 0 | - |
| - | - | 627.6 | 784.4 | - | - | 0 | - |
| - | - | 846.7 | 784.9 | - | - | 0 | - |
| - | - | 983.2 | 785.9 | - | - | 0 | - |
| - | - | 1864 | 786.4 | - | - | 0 | - |
| - | - | 1.049E+04 | 786.9 | - | - | 0 | - |
| - | - | 1.261E+04 | 787.4 | - | - | 0 | - |
| - | - | 8145 | 787.9 | - | - | 0 | - |
| - | - | 2785 | 788.4 | - | - | 0 | - |
| - | - | 1341 | 788.9 | - | - | 0 | - |
| - | - | 1445 | 794.4 | - | - | 0 | - |
| - | - | 8290 | 794.9 | - | - | 0 | - |
| - | - | 1.06E+04 | 795.4 | - | - | 0 | - |
| - | - | 7391 | 795.9 | - | - | 0 | - |
| - | - | 3339 | 796.4 | - | - | 0 | - |
| - | - | 1361 | 796.9 | - | - | 0 | - |
| 8 | z | 5165 | 829.4 | 0.001589 | 1.916 | +1 | 8 |
| - | - | 2532 | 830.4 | - | - | 0 | - |
| - | - | 710.2 | 831.5 | - | - | 0 | - |
| - | - | 1918 | 832.4 | - | - | 0 | - |
| - | - | 877.3 | 839.4 | - | - | 0 | - |
| - | - | 748.7 | 847.4 | - | - | 0 | - |
| - | - | 3521 | 857.4 | - | - | 0 | - |
| - | - | 1398 | 858.4 | - | - | 0 | - |
| - | - | 728.7 | 873.5 | - | - | 0 | - |
| - | - | 1040 | 875.4 | - | - | 0 | - |
| - | - | 726.5 | 887.5 | - | - | 0 | - |
| 8 | c | 646.2 | 900.5 | 0.003058 | 3.396 | +1 | 8 |
| - | - | 1399 | 932.4 | - | - | 0 | - |
| 7 | z | 1651 | 944.5 | 0.000151 | 0.1598 | +1 | 9 |
| - | - | 1652 | 945.5 | - | - | 0 | - |
| - | - | 801.6 | 974.6 | - | - | 0 | - |
| - | - | 2099 | 989.5 | - | - | 0 | - |
| - | - | 787.7 | 1013 | - | - | 0 | - |
| 9 | c | 4126 | 1015 | 0.001378 | 1.358 | +1 | 9 |
| - | - | 2399 | 1016 | - | - | 0 | - |
| - | - | 1077 | 1017 | - | - | 0 | - |
| - | - | 811.9 | 1025 | - | - | 0 | - |
| - | - | 1036 | 1027 | - | - | 0 | - |
| - | - | 639.7 | 1028 | - | - | 0 | - |
| 6 | z | 677.2 | 1031 | 0.003526 | 3.419 | +1 | 10 |
| - | - | 1084 | 1033 | - | - | 0 | - |
| - | - | 1120 | 1042 | - | - | 0 | - |
| - | - | 1966 | 1043 | - | - | 0 | - |
| - | - | 2136 | 1043 | - | - | 0 | - |
| - | - | 7403 | 1045 | - | - | 0 | - |
| - | - | 5753 | 1046 | - | - | 0 | - |
| - | - | 1149 | 1047 | - | - | 0 | - |
| - | - | 3714 | 1058 | - | - | 0 | - |
| - | - | 2721 | 1059 | - | - | 0 | - |
| - | - | 1118 | 1060 | - | - | 0 | - |
| - | - | 1674 | 1061 | - | - | 0 | - |
| - | - | 5080 | 1062 | - | - | 0 | - |
| - | - | 9727 | 1063 | - | - | 0 | - |
| - | - | 3542 | 1064 | - | - | 0 | - |
| 10 | c | 7293 | 1102 | 0.0007523 | 0.683 | +1 | 10 |
| - | - | 2542 | 1103 | - | - | 0 | - |
| - | - | 1442 | 1104 | - | - | 0 | - |
| - | - | 1269 | 1115 | - | - | 0 | - |
| 5 | z | 987.5 | 1131 | 0.004951 | 4.379 | +1 | 11 |
| - | - | 3105 | 1132 | - | - | 0 | - |
| - | - | 1277 | 1133 | - | - | 0 | - |
| - | - | 657.8 | 1148 | - | - | 0 | - |
| 11 | c | 3532 | 1159 | 0.001342 | 1.158 | +1 | 11 |
| - | - | 1652 | 1160 | - | - | 0 | - |
| 12 | c | 2901 | 1258 | 0.000298 | 0.2369 | +1 | 12 |
| - | - | 2321 | 1259 | - | - | 0 | - |
| - | - | 732.5 | 1260 | - | - | 0 | - |
| - | - | 3808 | 1261 | - | - | 0 | - |
| - | - | 1699 | 1262 | - | - | 0 | - |
| - | - | 888.4 | 1263 | - | - | 0 | - |
| 4 | y | 995.6 | 1276 | 0.0001379 | 0.1081 | +1 | 12 |
| - | - | 1059 | 1277 | - | - | 0 | - |
| 13 | c | 5650 | 1345 | 0.002419 | 1.799 | +1 | 13 |
| - | - | 4797 | 1346 | - | - | 0 | - |
| - | - | 1902 | 1347 | - | - | 0 | - |
| - | - | 3015 | 1348 | - | - | 0 | - |
| - | - | 2593 | 1349 | - | - | 0 | - |
| - | - | 1143 | 1350 | - | - | 0 | - |
| 14 | c | 9493 | 1432 | 0.00393 | 2.745 | +1 | 14 |
| - | - | 7028 | 1433 | - | - | 0 | - |
| - | - | 3831 | 1434 | - | - | 0 | - |
| - | - | 993.6 | 1435 | - | - | 0 | - |
| - | - | 1199 | 1490 | - | - | 0 | - |
| - | - | 2145 | 1531 | - | - | 0 | - |
| - | - | 1722 | 1532 | - | - | 0 | - |
| - | - | 1648 | 1533 | - | - | 0 | - |
| - | - | 1438 | 1547 | - | - | 0 | - |
| - | - | 700.6 | 1556 | - | - | 0 | - |
| - | - | 1882 | 1557 | - | - | 0 | - |
| - | - | 850.3 | 1558 | - | - | 0 | - |
| - | - | 858.7 | 1559 | - | - | 0 | - |
| - | - | 1907 | 1563 | - | - | 0 | - |
| - | - | 1070 | 1564 | - | - | 0 | - |
| - | - | 950.3 | 1570 | - | - | 0 | - |
| - | - | 1010 | 1571 | - | - | 0 | - |
| - | - | 2628 | 1573 | - | - | 0 | - |
| - | - | 1.474E+04 | 1574 | - | - | 0 | - |
| - | - | 1.117E+04 | 1575 | - | - | 0 | - |
| - | - | 6974 | 1576 | - | - | 0 | - |
| - | - | 1764 | 1577 | - | - | 0 | - |
| - | - | 887.1 | 1588 | - | - | 0 | - |
| - | - | 1468 | 1589 | - | - | 0 | - |
| - | - | 3821 | 1590 | - | - | 0 | - |
| - | - | 1.446E+04 | 1591 | - | - | 0 | - |
| - | - | 1.086E+04 | 1592 | - | - | 0 | - |
| - | - | 4869 | 1593 | - | - | 0 | - |
| - | - | 1575 | 1594 | - | - | 0 | - |
| - | - | 643.9 | 2294 | - | - | 0 | - |
| - | - | 636.8 | 2366 | - | - | 0 | - |
| - | - | 909.4 | 2647 | - | - | 0 | - |
| - | - | 751.6 | 3039 | - | - | 0 | - |
| - | - | 651 | 3496 | - | - | 0 | - |

m/z Charge Intensity FragmentType MassShift Position
120.08102416992188 0 1313.4127
126.09154510498047 0 407.7862
129.10157775878906 0 462.0266
133.0860137939453 0 775.19836
136.07571411132812 0 895.43646
140.0821533203125 0 2047.2616
140.3094482421875 0 446.12753
147.0767364501953 0 581.8778
163.07203674316406 0 594.6294
168.1138458251953 0 547.0195
169.0967254638672 0 603.57025 y 10
171.33741760253906 0 444.732
173.43893432617188 0 1147.6093
185.13983154296875 0 2940.253
187.1078643798828 0 12351.709
199.1806182861328 0 5075.8545
200.1838836669922 0 668.6634
215.1385955810547 0 953.2897
225.8979949951172 0 643.1252
227.17572021484375 0 2327.1965
229.12957763671875 0 548.7702 w 13
245.1246795654297 0 3638.5261 y Ammonia loss 13
262.1491394042969 0 687.0477 y 13
316.16217041015625 0 1837.9528 w 12
333.1651306152344 0 962.68945 z 12
333.20635986328125 0 514.19836
357.1507263183594 0 736.4546
362.2039489746094 0 706.7788
377.16729736328125 0 1643.2617
386.2833251953125 0 603.57166
401.26373291015625 0 831.1974
417.2100830078125 0 1424.8658 w 11
418.2186584472656 0 521.6355
419.2245178222656 0 1065.9432
429.08795166015625 0 572.5564
448.2491149902344 0 1160.885 y 11
453.343994140625 0 782.91583
454.7735290527344 0 546.023
460.2754211425781 0 737.46234 c 3
475.2866516113281 0 769.2974
504.25762939453125 0 575.40765
505.2732849121094 0 5236.3223 y 10
506.2751159667969 0 1118.1862
524.2667236328125 0 916.9553 y 5
524.7637939453125 0 949.9449
529.234130859375 0 796.2107
529.2769165039062 0 613.8193
530.2763061523438 0 656.98145
530.3158569335938 0 2470.1377
531.3181762695312 0 852.06415
542.3163452148438 0 731.47186
557.2721557617188 0 695.5548 z Ammonia loss 4
559.3423461914062 0 735.19116 c 4
565.786865234375 0 3449.5403 z 4
566.29150390625 0 2379.313
566.7908325195312 0 877.4568
567.2988891601562 0 534.2328
569.329833984375 0 636.827
572.4915161132812 0 552.09595
573.7979125976562 0 1552.7041 y 4
574.3007202148438 0 723.3783 y Water loss 9
585.257568359375 0 667.277
596.28125 0 608.95984
600.8041381835938 0 1217.4524 w 3
601.3037109375 0 740.60785
604.3670043945312 0 774.14606
605.1631469726562 0 699.47107
630.3089599609375 0 3001.5862 z 3
630.810791015625 0 1897.8853
631.3126220703125 0 622.7879
638.3173828125 0 3719.6821 y 3
638.8192138671875 0 2005.4116
639.3231201171875 0 926.4967
646.376953125 0 3144.0564 c 5
647.3816528320312 0 1098.5864
651.8612060546875 0 602.2977
661.8324584960938 0 804.502
673.82470703125 0 2569.6704 z 2
674.3275756835938 0 3026.3667
674.8292846679688 0 1459.3442
675.3280639648438 0 759.97723
681.8338623046875 0 11488.36 y 2
682.3361206054688 0 9909.27
682.8377685546875 0 4587.7607
689.3566284179688 0 756.7002 y 8
703.3359375 0 750.1554
712.3804931640625 0 1416.3434
713.8748168945312 0 731.5784
715.848388671875 0 689.8881 w 1
723.7708740234375 0 550.9915
728.3660888671875 0 989.49493
730.3667602539062 0 681.9658 z 1
738.3738403320312 0 1699.8888 y 1
739.3775024414062 0 859.30853
745.383544921875 0 3972.1367
746.3886108398438 0 690.84753
761.4039306640625 0 5103.083 c 6
762.406005859375 0 2722.1318
763.4071044921875 0 1313.497
764.9077758789062 0 1399.8528
765.3818969726562 0 12163.483
765.883056640625 0 9201.502
766.3858642578125 0 5118.421
766.8895874023438 0 1321.5386
771.4059448242188 0 761.9788
773.9067993164062 0 2097.397
774.4097900390625 0 1742.5482
774.9083251953125 0 705.0915
775.4132080078125 0 801.0739
776.4124145507812 0 769.22046
778.4094848632812 0 1057.5967
778.9034423828125 0 1407.8229
779.9042358398438 0 907.779
780.4083862304688 0 774.6118
784.4072875976562 0 627.5889
784.906982421875 0 846.67224
785.9003295898438 0 983.1816
786.4031982421875 0 1863.7571
786.9071655273438 0 10493.152
787.40966796875 0 12606.206
787.9093627929688 0 8145.063
788.4107666015625 0 2784.538
788.9140625 0 1340.5778
794.4110717773438 0 1444.8867
794.916748046875 0 8290.353
795.4179077148438 0 10602.013
795.9196166992188 0 7391.277
796.4136352539062 0 3339.0479
796.9112548828125 0 1361.2557
829.4384765625 0 5164.9873 z 7
830.4432373046875 0 2531.8228
831.4520263671875 0 710.1882
832.3841552734375 0 1917.8384
839.4038696289062 0 877.3416
847.4043579101562 0 748.7047
857.4141845703125 0 3521.168
858.4154052734375 0 1398.2686
873.4949951171875 0 728.6804
875.4302368164062 0 1039.5447
887.5166625976562 0 726.4563
900.4754638671875 0 646.16406 c Ammonia loss 7
932.4221801757812 0 1398.7639
944.4668579101562 0 1650.6472 z 6
945.4774169921875 0 1652.1038
974.557861328125 0 801.59564
989.5044555664062 0 2099.29
1012.5350952148438 0 787.6771
1014.5564575195312 0 4125.5986 c 8
1015.5621337890625 0 2398.7654
1016.5563354492188 0 1076.8092
1025.4788818359375 0 811.89966
1027.4993896484375 0 1036.1549
1028.4901123046875 0 639.6502
1031.5025634765625 0 677.2122 z 5
1032.5068359375 0 1083.8202
1041.5013427734375 0 1120.3589
1042.510498046875 0 1966.0869
1043.4974365234375 0 2135.6787
1044.5186767578125 0 7402.666
1045.51611328125 0 5753.0664
1046.5084228515625 0 1149.2649
1057.57080078125 0 3714.0178
1058.5556640625 0 2720.8743
1059.5538330078125 0 1118.0106
1060.5255126953125 0 1674.4296
1061.5238037109375 0 5080.0244
1062.5335693359375 0 9726.8125
1063.550048828125 0 3541.7368
1101.589111328125 0 7293.3867 c 9
1102.5927734375 0 2541.9355
1103.5909423828125 0 1441.8075
1114.5958251953125 0 1268.5162
1130.5625 0 987.4741 z 4
1131.57666015625 0 3104.8105
1132.572509765625 0 1276.6348
1147.5853271484375 0 657.8234
1158.6099853515625 0 3531.7993 c 10
1159.61572265625 0 1651.8336
1257.679443359375 0 2901.0127 c 11
1258.6798095703125 0 2321.4238
1259.705810546875 0 732.5465
1260.6187744140625 0 3807.6487
1261.6234130859375 0 1698.8236
1262.6241455078125 0 888.44666
1275.62890625 0 995.6084 y 3
1276.6126708984375 0 1058.5039
1344.7093505859375 0 5649.6206 c 12
1345.713134765625 0 4797.3613
1346.7060546875 0 1902.1301
1347.6484375 0 3014.7742
1348.65478515625 0 2592.8523
1349.65966796875 0 1143.3047
1431.7398681640625 0 9492.527 c 13
1432.7425537109375 0 7028.0684
1433.7467041015625 0 3830.6262
1434.7470703125 0 993.5832
1489.7515869140625 0 1199.3159
1530.7908935546875 0 2144.6836
1531.7838134765625 0 1721.9287
1532.7933349609375 0 1648.3204
1546.8231201171875 0 1437.5502
1555.80419921875 0 700.6072
1556.7955322265625 0 1881.8107
1557.8045654296875 0 850.2639
1558.8016357421875 0 858.71155
1562.84033203125 0 1906.8792
1563.8406982421875 0 1070.0533
1569.84375 0 950.32434
1570.823974609375 0 1009.901
1572.809326171875 0 2628.1682
1573.81298828125 0 14740.723
1574.814453125 0 11172.782
1575.8155517578125 0 6974.287
1576.8126220703125 0 1764.424
1587.8221435546875 0 887.072
1588.8258056640625 0 1467.5266
1589.8251953125 0 3820.5178
1590.8388671875 0 14462.521
1591.839599609375 0 10857.961
1592.843505859375 0 4868.82
1593.834228515625 0 1574.5078
2293.820068359375 0 643.94977
2365.5537109375 0 636.7735
2646.658447265625 0 909.39453
3039.330322265625 0 751.56213
3496.321533203125 0 651.0465

Spectrum Details

|  |  |
| --- | --- |
| Matched peaks? Matched peaks The total absolute number of peaks matched. Additionally in brackets the total fraction of peaks matched and the total number of peaks is shown. | 39 (17.26% of 226) |
| FDR? FDR The false discovery rate estimated for this peptide. It is calculated by matching all theoretical fragments with a non-integer shift with the raw peaks for this spectrum. This is done with 40 different shifts. The resulting percentage is the average number of annotated peaks over the number of annotated peaks with the correct spectrum. | 0.98% |
| Satellite FDR? Satellite FDR See the FDR for details on its calculation. This satellite ion specific FDR only contains the satellite ions (d/w) for I/L/J positions. | 0.00% |
| PSM Score? PSM Score The PSM Score as given by Hecklib to this annotated spectrum. It is shown with three significant figures. | 225 |

#### Spectrum 4473? Spectrum 4473 The raw spectrum of this peptide as annotated by Hecklib. The fragments are coloured according to ion type (see legend). Any peaks with a star '\*' as text can be hovered over to see the full details, first the ion type second the mass shift type. By hovering over the amino acids in the peptide or ions in the legend the corresponding peaks are highlighted. By toggling the 'Unassigned' label you can turn the background (unassigned) peaks on or off in the plot. By updating the slider in the Ion legend you can update the spectrum to only show the top X% of the peaks with labels. The top X% means any peak that is within X% of the highest intensity. By dragging in the spectrum you can zoom in to a specific part of the spectrum and use 'Zoom Out' to get back to the original zoom level. The annotation of the spectrum is based on the given sequence in the peptides file and is done with different software so inconsistencies are likely. The peaks are annotated based on the given sequence, with 20 ppm tolerance.

Copy Data

##### Spectrum 4473 (TSV)

###### Preview

```
Loading example...
```

*Click on the button to copy the data to your clipboard.*

Mz MinMz MaxIntensity Max

WidthHeightPeptide font sizePeptide stroke widthSpectrum font sizeSpectrum stroke widthCompact peptide

Ion legend

wxyz

abcd

OtherUnassignedIon^Charge^_Position_Show for top:%

JJSEVSDRPSGVSSR

01.08e+52.16e+53.25e+54.33e+5

Zoom Out

d^+1^_2_y^+1^_1_y^+3^_5_y^+1^_1_a^+1^_2_b^+1^_2_y^+1^_2_y^+1^_2_b^+1^_3_b^+1^_3_y^+1^_3_y^+1^_3_b^+3^_9_y^+2^_7_y^+1^_3_b^+3^_12_y^+2^_8_y^+2^_8_b^+1^_4_y^+1^_4_b^+1^_4_y^+1^_4_y^+3^_13_b^+2^_8_y^+3^_13_y^+2^_9_y^+3^_14_y^+3^_14_y^+1^_5_y^+2^_10_y^+2^_10_y^+2^_10_\*\*b^+1^_5_y^+2^_11_y^+2^_11_y^+2^_11_y^+1^_6_y^+1^_6_y^+2^_12_y^+2^_12_y^+2^_12_y^+1^_7_y^+1^_7_y^+2^_13_y^+2^_13_y^+2^_13_y^+1^_7_b^+2^_14_y^+2^_14_y^+2^_14_y^+2^_14_b^+1^_7_y^+1^_8_b^+1^_8_y^+1^_9_y^+1^_10_y^+1^_11_

03056109151220

Fragment Matches Table

Show background peaks

| Position | Ion type | Intensity | mz Theoretical | mz Error (Th) | mz Error (ppm) | Charge | Series Number |
| --- | --- | --- | --- | --- | --- | --- | --- |
| - | - | 1.692E+04 | 120.1 | - | - | 0 | - |
| - | - | 388.8 | 120.9 | - | - | 0 | - |
| - | - | 1711 | 121.1 | - | - | 0 | - |
| - | - | 586.1 | 122.1 | - | - | 0 | - |
| - | - | 369 | 124.1 | - | - | 0 | - |
| - | - | 3726 | 126.1 | - | - | 0 | - |
| - | - | 527.7 | 127.1 | - | - | 0 | - |
| - | - | 564.2 | 128.1 | - | - | 0 | - |
| - | - | 479.8 | 129.1 | - | - | 0 | - |
| - | - | 1.545E+04 | 129.1 | - | - | 0 | - |
| - | - | 877.4 | 130.1 | - | - | 0 | - |
| - | - | 2913 | 130.1 | - | - | 0 | - |
| - | - | 792.6 | 130.1 | - | - | 0 | - |
| - | - | 395.9 | 130.1 | - | - | 0 | - |
| - | - | 575 | 130.1 | - | - | 0 | - |
| - | - | 918.3 | 130.1 | - | - | 0 | - |
| - | - | 694.1 | 132.1 | - | - | 0 | - |
| - | - | 549.7 | 133.1 | - | - | 0 | - |
| - | - | 838.2 | 134 | - | - | 0 | - |
| - | - | 932.9 | 134.1 | - | - | 0 | - |
| - | - | 2.151E+04 | 136.1 | - | - | 0 | - |
| - | - | 1395 | 137.1 | - | - | 0 | - |
| - | - | 533.9 | 139 | - | - | 0 | - |
| - | - | 1005 | 139.1 | - | - | 0 | - |
| - | - | 1343 | 140.1 | - | - | 0 | - |
| - | - | 521.7 | 141.1 | - | - | 0 | - |
| - | - | 721.8 | 141.1 | - | - | 0 | - |
| - | - | 687.8 | 143.1 | - | - | 0 | - |
| - | - | 944.3 | 143.1 | - | - | 0 | - |
| - | - | 849 | 144.1 | - | - | 0 | - |
| - | - | 726.6 | 145.1 | - | - | 0 | - |
| - | - | 1610 | 146.1 | - | - | 0 | - |
| - | - | 810.3 | 147 | - | - | 0 | - |
| - | - | 435.8 | 147.1 | - | - | 0 | - |
| - | - | 1975 | 147.1 | - | - | 0 | - |
| - | - | 529.3 | 151.1 | - | - | 0 | - |
| - | - | 2359 | 152.1 | - | - | 0 | - |
| - | - | 616.3 | 152.1 | - | - | 0 | - |
| - | - | 661.9 | 153.1 | - | - | 0 | - |
| - | - | 3980 | 154.1 | - | - | 0 | - |
| - | - | 488.4 | 154.1 | - | - | 0 | - |
| - | - | 790.6 | 155.1 | - | - | 0 | - |
| - | - | 1216 | 155.1 | - | - | 0 | - |
| - | - | 676.7 | 156.1 | - | - | 0 | - |
| - | - | 1784 | 157.1 | - | - | 0 | - |
| - | - | 1840 | 157.1 | - | - | 0 | - |
| - | - | 1615 | 157.1 | - | - | 0 | - |
| 2 | d | 459.8 | 157.1 | 0.0004326 | 2.753 | +1 | 2 |
| 15 | y | 8623 | 158.1 | 0.0004314 | 2.729 | +1 | 1 |
| - | - | 513 | 158.6 | - | - | 0 | - |
| - | - | 1335 | 159.1 | - | - | 0 | - |
| - | - | 1927 | 159.1 | - | - | 0 | - |
| - | - | 872.7 | 162.1 | - | - | 0 | - |
| - | - | 409.2 | 163.1 | - | - | 0 | - |
| - | - | 2925 | 163.1 | - | - | 0 | - |
| - | - | 687.5 | 164.1 | - | - | 0 | - |
| - | - | 848.7 | 165.1 | - | - | 0 | - |
| - | - | 614.6 | 165.1 | - | - | 0 | - |
| - | - | 1441 | 167.1 | - | - | 0 | - |
| - | - | 490.3 | 169.1 | - | - | 0 | - |
| 11 | y | 1233 | 169.1 | 0.001614 | 9.546 | +3 | 5 |
| - | - | 4228 | 171.1 | - | - | 0 | - |
| - | - | 1668 | 171.1 | - | - | 0 | - |
| - | - | 629.1 | 172.1 | - | - | 0 | - |
| - | - | 1289 | 173.1 | - | - | 0 | - |
| - | - | 3512 | 173.1 | - | - | 0 | - |
| - | - | 3979 | 173.5 | - | - | 0 | - |
| - | - | 2583 | 175.1 | - | - | 0 | - |
| - | - | 1360 | 175.1 | - | - | 0 | - |
| 15 | y | 1.722E+04 | 175.1 | 0.0004479 | 2.557 | +1 | 1 |
| - | - | 778.6 | 176.1 | - | - | 0 | - |
| - | - | 1703 | 181.1 | - | - | 0 | - |
| - | - | 2916 | 182.1 | - | - | 0 | - |
| - | - | 734.3 | 182.1 | - | - | 0 | - |
| - | - | 505.8 | 182.1 | - | - | 0 | - |
| - | - | 3796 | 183.1 | - | - | 0 | - |
| - | - | 1052 | 183.1 | - | - | 0 | - |
| - | - | 520.2 | 184.1 | - | - | 0 | - |
| - | - | 806.6 | 184.1 | - | - | 0 | - |
| - | - | 1982 | 185.1 | - | - | 0 | - |
| - | - | 994.8 | 185.1 | - | - | 0 | - |
| - | - | 517.4 | 187.1 | - | - | 0 | - |
| - | - | 4684 | 187.1 | - | - | 0 | - |
| - | - | 1811 | 187.1 | - | - | 0 | - |
| - | - | 2.181E+04 | 189.1 | - | - | 0 | - |
| - | - | 1280 | 190.1 | - | - | 0 | - |
| - | - | 566.6 | 191.1 | - | - | 0 | - |
| - | - | 599.1 | 193.1 | - | - | 0 | - |
| - | - | 710.6 | 195.1 | - | - | 0 | - |
| - | - | 865.4 | 195.1 | - | - | 0 | - |
| - | - | 2510 | 199.1 | - | - | 0 | - |
| - | - | 802 | 199.1 | - | - | 0 | - |
| 2 | a | 1.696E+05 | 199.2 | 0.0004489 | 2.254 | +1 | 2 |
| - | - | 1353 | 200.1 | - | - | 0 | - |
| - | - | 2.16E+04 | 200.2 | - | - | 0 | - |
| - | - | 953 | 201.1 | - | - | 0 | - |
| - | - | 9094 | 201.1 | - | - | 0 | - |
| - | - | 603.4 | 201.2 | - | - | 0 | - |
| - | - | 835.8 | 202.1 | - | - | 0 | - |
| - | - | 889.2 | 202.1 | - | - | 0 | - |
| - | - | 3672 | 203.1 | - | - | 0 | - |
| - | - | 617.1 | 205.1 | - | - | 0 | - |
| - | - | 961.8 | 209.1 | - | - | 0 | - |
| - | - | 527.2 | 209.2 | - | - | 0 | - |
| - | - | 765.9 | 210.1 | - | - | 0 | - |
| - | - | 1479 | 211.1 | - | - | 0 | - |
| - | - | 618.9 | 211.1 | - | - | 0 | - |
| - | - | 1790 | 212.1 | - | - | 0 | - |
| - | - | 569.1 | 212.1 | - | - | 0 | - |
| - | - | 587 | 213.2 | - | - | 0 | - |
| - | - | 615.5 | 215.1 | - | - | 0 | - |
| - | - | 1115 | 215.1 | - | - | 0 | - |
| - | - | 580.9 | 216.1 | - | - | 0 | - |
| - | - | 3.472E+04 | 217.1 | - | - | 0 | - |
| - | - | 805.3 | 217.1 | - | - | 0 | - |
| - | - | 3016 | 218.1 | - | - | 0 | - |
| - | - | 1189 | 223.1 | - | - | 0 | - |
| - | - | 2749 | 224.1 | - | - | 0 | - |
| - | - | 829.9 | 226.1 | - | - | 0 | - |
| - | - | 1698 | 226.2 | - | - | 0 | - |
| - | - | 951.2 | 227.1 | - | - | 0 | - |
| - | - | 887.2 | 227.1 | - | - | 0 | - |
| 2 | b | 5.349E+04 | 227.2 | 0.0004531 | 1.995 | +1 | 2 |
| - | - | 647.8 | 228.1 | - | - | 0 | - |
| - | - | 5759 | 228.2 | - | - | 0 | - |
| - | - | 1441 | 229.1 | - | - | 0 | - |
| - | - | 742.1 | 229.2 | - | - | 0 | - |
| - | - | 628.2 | 233.2 | - | - | 0 | - |
| - | - | 509 | 235.1 | - | - | 0 | - |
| - | - | 853.5 | 237.1 | - | - | 0 | - |
| - | - | 1260 | 237.1 | - | - | 0 | - |
| - | - | 4016 | 242.1 | - | - | 0 | - |
| - | - | 569.7 | 242.2 | - | - | 0 | - |
| - | - | 933.9 | 243.1 | - | - | 0 | - |
| - | - | 537.6 | 243.1 | - | - | 0 | - |
| - | - | 657.6 | 243.1 | - | - | 0 | - |
| - | - | 1536 | 244.1 | - | - | 0 | - |
| - | - | 845.6 | 245.1 | - | - | 0 | - |
| 14 | y | 4644 | 245.1 | 0.0005685 | 2.319 | +1 | 2 |
| - | - | 500.8 | 246.1 | - | - | 0 | - |
| - | - | 826.6 | 246.1 | - | - | 0 | - |
| - | - | 1177 | 248.1 | - | - | 0 | - |
| - | - | 552.5 | 249.2 | - | - | 0 | - |
| - | - | 796.4 | 252.1 | - | - | 0 | - |
| - | - | 524.3 | 252.2 | - | - | 0 | - |
| - | - | 1277 | 253.2 | - | - | 0 | - |
| - | - | 458.8 | 254.1 | - | - | 0 | - |
| - | - | 3842 | 255.1 | - | - | 0 | - |
| - | - | 618.7 | 256.1 | - | - | 0 | - |
| - | - | 536.9 | 258.4 | - | - | 0 | - |
| - | - | 539.3 | 259.1 | - | - | 0 | - |
| - | - | 561.8 | 261.2 | - | - | 0 | - |
| - | - | 1473 | 262.1 | - | - | 0 | - |
| 14 | y | 2610 | 262.2 | 0.0006613 | 2.522 | +1 | 2 |
| - | - | 598.3 | 270.1 | - | - | 0 | - |
| - | - | 825.1 | 272.1 | - | - | 0 | - |
| - | - | 3527 | 283.1 | - | - | 0 | - |
| - | - | 664.7 | 284.1 | - | - | 0 | - |
| - | - | 1076 | 284.1 | - | - | 0 | - |
| - | - | 970.3 | 284.1 | - | - | 0 | - |
| - | - | 823.5 | 284.2 | - | - | 0 | - |
| - | - | 496.9 | 286.2 | - | - | 0 | - |
| - | - | 652.1 | 288.2 | - | - | 0 | - |
| - | - | 1137 | 296.1 | - | - | 0 | - |
| 3 | b | 2290 | 296.2 | 0.0003975 | 1.342 | +1 | 3 |
| - | - | 988.6 | 297.1 | - | - | 0 | - |
| - | - | 2617 | 298.1 | - | - | 0 | - |
| - | - | 3254 | 299.1 | - | - | 0 | - |
| - | - | 650.3 | 307.1 | - | - | 0 | - |
| - | - | 787.7 | 311.1 | - | - | 0 | - |
| - | - | 3401 | 312.2 | - | - | 0 | - |
| - | - | 735.9 | 312.2 | - | - | 0 | - |
| - | - | 622.6 | 313.2 | - | - | 0 | - |
| - | - | 922.5 | 313.2 | - | - | 0 | - |
| - | - | 5125 | 313.2 | - | - | 0 | - |
| - | - | 2340 | 314.1 | - | - | 0 | - |
| - | - | 682.8 | 314.2 | - | - | 0 | - |
| 3 | b | 5598 | 314.2 | 0.0007886 | 2.51 | +1 | 3 |
| - | - | 939 | 315.2 | - | - | 0 | - |
| - | - | 5968 | 316.2 | - | - | 0 | - |
| - | - | 731 | 317.2 | - | - | 0 | - |
| - | - | 1197 | 324.1 | - | - | 0 | - |
| - | - | 748 | 327.2 | - | - | 0 | - |
| - | - | 1450 | 330.2 | - | - | 0 | - |
| 13 | y | 624.1 | 331.2 | 0.000804 | 2.428 | +1 | 3 |
| 13 | y | 4371 | 332.2 | 0.0006141 | 1.849 | +1 | 3 |
| 9 | b | 1065 | 333.2 | 0.003052 | 9.161 | +3 | 9 |
| - | - | 711.3 | 341.1 | - | - | 0 | - |
| - | - | 2574 | 341.2 | - | - | 0 | - |
| - | - | 5539 | 342.1 | - | - | 0 | - |
| - | - | 786.2 | 342.2 | - | - | 0 | - |
| 9 | y | 1434 | 345.2 | 0.0005973 | 1.73 | +2 | 7 |
| 13 | y | 5129 | 349.2 | 0.001043 | 2.986 | +1 | 3 |
| - | - | 715.3 | 355.2 | - | - | 0 | - |
| - | - | 2551 | 359.2 | - | - | 0 | - |
| - | - | 628.8 | 359.2 | - | - | 0 | - |
| - | - | 942 | 383.2 | - | - | 0 | - |
| - | - | 687.5 | 383.7 | - | - | 0 | - |
| - | - | 1405 | 385.2 | - | - | 0 | - |
| - | - | 619.1 | 386.2 | - | - | 0 | - |
| - | - | 934.1 | 391.7 | - | - | 0 | - |
| - | - | 662.4 | 392.2 | - | - | 0 | - |
| - | - | 661.5 | 399.7 | - | - | 0 | - |
| - | - | 1027 | 400.2 | - | - | 0 | - |
| - | - | 838.9 | 401.2 | - | - | 0 | - |
| - | - | 1060 | 402.2 | - | - | 0 | - |
| - | - | 669.9 | 407.2 | - | - | 0 | - |
| - | - | 1233 | 412.2 | - | - | 0 | - |
| 12 | b | 1095 | 414.2 | 0.006422 | 15.5 | +3 | 12 |
| 8 | y | 3003 | 414.7 | 0.001402 | 3.381 | +2 | 8 |
| - | - | 591.5 | 415.2 | - | - | 0 | - |
| - | - | 667.9 | 415.3 | - | - | 0 | - |
| - | - | 689.8 | 415.7 | - | - | 0 | - |
| 8 | y | 3.631E+04 | 423.2 | 0.0009146 | 2.161 | +2 | 8 |
| - | - | 1.601E+04 | 423.7 | - | - | 0 | - |
| - | - | 967.4 | 424.2 | - | - | 0 | - |
| - | - | 3487 | 424.2 | - | - | 0 | - |
| 4 | b | 2886 | 425.2 | 0.0009257 | 2.177 | +1 | 4 |
| - | - | 897.7 | 426.2 | - | - | 0 | - |
| - | - | 675.9 | 428.2 | - | - | 0 | - |
| - | - | 1565 | 429.2 | - | - | 0 | - |
| 12 | y | 1089 | 430.2 | 0.0008985 | 2.088 | +1 | 4 |
| - | - | 2412 | 441.2 | - | - | 0 | - |
| 4 | b | 3474 | 443.3 | 0.0007675 | 1.732 | +1 | 4 |
| 12 | y | 1229 | 448.3 | 1.91E-05 | 0.0426 | +1 | 4 |
| 3 | y | 1261 | 449.2 | 0.006786 | 15.11 | +3 | 13 |
| - | - | 596.6 | 450.2 | - | - | 0 | - |
| 8 | b | 816.1 | 450.7 | 0.001322 | 2.933 | +2 | 8 |
| - | - | 929.8 | 452.3 | - | - | 0 | - |
| - | - | 913.8 | 454.2 | - | - | 0 | - |
| 3 | y | 1384 | 454.9 | 0.0004897 | 1.077 | +3 | 13 |
| - | - | 663 | 455.2 | - | - | 0 | - |
| - | - | 1844 | 458.2 | - | - | 0 | - |
| - | - | 2014 | 460.3 | - | - | 0 | - |
| - | - | 1307 | 469.3 | - | - | 0 | - |
| 7 | y | 6411 | 480.7 | 0.001115 | 2.319 | +2 | 9 |
| - | - | 3404 | 481.2 | - | - | 0 | - |
| 2 | y | 1101 | 486.9 | 0.007838 | 16.1 | +3 | 14 |
| - | - | 1071 | 487.3 | - | - | 0 | - |
| 2 | y | 1232 | 492.6 | 0.0007785 | 1.58 | +3 | 14 |
| - | - | 691.6 | 492.9 | - | - | 0 | - |
| - | - | 920 | 494.2 | - | - | 0 | - |
| - | - | 4051 | 495.9 | - | - | 0 | - |
| - | - | 971.9 | 496.2 | - | - | 0 | - |
| - | - | 745.3 | 496.6 | - | - | 0 | - |
| - | - | 860.2 | 500.2 | - | - | 0 | - |
| 11 | y | 4832 | 505.3 | 0.0009476 | 1.875 | +1 | 5 |
| - | - | 776.5 | 506.3 | - | - | 0 | - |
| - | - | 815.6 | 511.3 | - | - | 0 | - |
| - | - | 768.2 | 512.2 | - | - | 0 | - |
| - | - | 688.1 | 514.2 | - | - | 0 | - |
| - | - | 1512 | 514.3 | - | - | 0 | - |
| 6 | y | 844.6 | 515.3 | 0.0005759 | 1.118 | +2 | 10 |
| 6 | y | 868.3 | 515.7 | 0.004662 | 9.039 | +2 | 10 |
| - | - | 1021 | 516.3 | - | - | 0 | - |
| - | - | 1163 | 518.6 | - | - | 0 | - |
| - | - | 915.4 | 519.7 | - | - | 0 | - |
| 6 | y | 3.837E+04 | 524.3 | 0.002313 | 4.411 | +2 | 10 |
| 0 | Precursor | 9273 | 524.6 | 0.006784 | 12.93 | +3 | -1 |
| - | - | 2.016E+04 | 524.8 | - | - | 0 | - |
| - | - | 3399 | 524.9 | - | - | 0 | - |
| - | - | 8080 | 525.3 | - | - | 0 | - |
| - | - | 917.7 | 525.6 | - | - | 0 | - |
| - | - | 1451 | 525.8 | - | - | 0 | - |
| - | - | 801.4 | 528.6 | - | - | 0 | - |
| - | - | 8567 | 528.7 | - | - | 0 | - |
| - | - | 8135 | 529.2 | - | - | 0 | - |
| - | - | 952.8 | 529.3 | - | - | 0 | - |
| - | - | 3437 | 529.7 | - | - | 0 | - |
| - | - | 1813 | 530.2 | - | - | 0 | - |
| 0 | Precursor | 2037 | 530.3 | 0.001556 | 2.934 | +3 | -1 |
| - | - | 1179 | 530.3 | - | - | 0 | - |
| - | - | 4441 | 530.6 | - | - | 0 | - |
| - | - | 881.8 | 530.9 | - | - | 0 | - |
| - | - | 992.9 | 531.3 | - | - | 0 | - |
| - | - | 828.5 | 539.8 | - | - | 0 | - |
| 5 | b | 972.5 | 542.3 | 0.001506 | 2.778 | +1 | 5 |
| 5 | y | 2121 | 564.8 | 0.001769 | 3.133 | +2 | 11 |
| 5 | y | 1918 | 565.3 | 0.006771 | 11.98 | +2 | 11 |
| - | - | 1105 | 565.8 | - | - | 0 | - |
| - | - | 776.2 | 572.8 | - | - | 0 | - |
| 5 | y | 5.572E+04 | 573.8 | 0.0005764 | 1.005 | +2 | 11 |
| 10 | y | 3.398E+04 | 574.3 | 0.004112 | 7.159 | +1 | 6 |
| - | - | 1.227E+04 | 574.8 | - | - | 0 | - |
| - | - | 1996 | 575.3 | - | - | 0 | - |
| - | - | 1245 | 580.8 | - | - | 0 | - |
| - | - | 1120 | 584.3 | - | - | 0 | - |
| - | - | 1828 | 585.8 | - | - | 0 | - |
| - | - | 915 | 586.3 | - | - | 0 | - |
| - | - | 926.9 | 586.8 | - | - | 0 | - |
| 10 | y | 5704 | 592.3 | 0.0009321 | 1.574 | +1 | 6 |
| - | - | 2084 | 593.3 | - | - | 0 | - |
| - | - | 886.3 | 600.3 | - | - | 0 | - |
| - | - | 651.7 | 618.8 | - | - | 0 | - |
| - | - | 801.3 | 626.3 | - | - | 0 | - |
| 4 | y | 3897 | 629.3 | 0.003666 | 5.826 | +2 | 12 |
| 4 | y | 2110 | 629.8 | 0.007752 | 12.31 | +2 | 12 |
| - | - | 1515 | 630.3 | - | - | 0 | - |
| - | - | 704 | 631.3 | - | - | 0 | - |
| - | - | 1412 | 636.3 | - | - | 0 | - |
| - | - | 978 | 636.8 | - | - | 0 | - |
| - | - | 838.9 | 637.3 | - | - | 0 | - |
| 4 | y | 2.592E+04 | 638.3 | 0.0006421 | 1.006 | +2 | 12 |
| - | - | 1.782E+04 | 638.8 | - | - | 0 | - |
| - | - | 7351 | 639.3 | - | - | 0 | - |
| - | - | 1345 | 639.8 | - | - | 0 | - |
| - | - | 921.2 | 641.3 | - | - | 0 | - |
| - | - | 2440 | 646.3 | - | - | 0 | - |
| - | - | 981.7 | 649.3 | - | - | 0 | - |
| - | - | 1018 | 652.3 | - | - | 0 | - |
| - | - | 690.1 | 653.3 | - | - | 0 | - |
| - | - | 1098 | 654.3 | - | - | 0 | - |
| - | - | 1217 | 655.3 | - | - | 0 | - |
| - | - | 772.5 | 657.3 | - | - | 0 | - |
| - | - | 1549 | 659.3 | - | - | 0 | - |
| - | - | 1476 | 660.8 | - | - | 0 | - |
| - | - | 907.7 | 661.3 | - | - | 0 | - |
| - | - | 1668 | 663.8 | - | - | 0 | - |
| - | - | 2035 | 664.3 | - | - | 0 | - |
| - | - | 827.2 | 664.8 | - | - | 0 | - |
| - | - | 709.3 | 665.8 | - | - | 0 | - |
| - | - | 4622 | 666.8 | - | - | 0 | - |
| - | - | 2626 | 667.3 | - | - | 0 | - |
| - | - | 1139 | 667.8 | - | - | 0 | - |
| 9 | y | 5799 | 671.3 | 0.0002979 | 0.4437 | +1 | 7 |
| 9 | y | 3927 | 672.3 | 0.005845 | 8.694 | +1 | 7 |
| 3 | y | 1.256E+04 | 672.8 | 0.0007135 | 1.06 | +2 | 13 |
| 3 | y | 1.494E+04 | 673.3 | 0.009377 | 13.93 | +2 | 13 |
| - | - | 6901 | 673.8 | - | - | 0 | - |
| - | - | 7724 | 674.3 | - | - | 0 | - |
| - | - | 1171 | 674.8 | - | - | 0 | - |
| - | - | 2337 | 675.3 | - | - | 0 | - |
| - | - | 4022 | 680.8 | - | - | 0 | - |
| - | - | 3706 | 681.3 | - | - | 0 | - |
| 3 | y | 4.287E+05 | 681.8 | 0.001107 | 1.624 | +2 | 13 |
| - | - | 3.025E+05 | 682.3 | - | - | 0 | - |
| - | - | 1.234E+05 | 682.8 | - | - | 0 | - |
| - | - | 3.326E+04 | 683.3 | - | - | 0 | - |
| - | - | 8505 | 683.8 | - | - | 0 | - |
| - | - | 3016 | 686.8 | - | - | 0 | - |
| - | - | 3920 | 687.3 | - | - | 0 | - |
| 9 | y | 3.234E+04 | 689.4 | 0.0007806 | 1.132 | +1 | 7 |
| - | - | 1.111E+04 | 690.4 | - | - | 0 | - |
| - | - | 2309 | 691.4 | - | - | 0 | - |
| - | - | 839.6 | 692.9 | - | - | 0 | - |
| - | - | 923.2 | 693.4 | - | - | 0 | - |
| - | - | 1793 | 694.9 | - | - | 0 | - |
| - | - | 1170 | 695.4 | - | - | 0 | - |
| 14 | b | 3105 | 699.3 | 0.004689 | 6.705 | +2 | 14 |
| - | - | 1392 | 700.3 | - | - | 0 | - |
| - | - | 1508 | 712.4 | - | - | 0 | - |
| 2 | y | 5228 | 729.4 | 0.001711 | 2.346 | +2 | 14 |
| 2 | y | 3831 | 729.9 | 0.008483 | 11.62 | +2 | 14 |
| - | - | 869.3 | 730.4 | - | - | 0 | - |
| - | - | 1416 | 736.9 | - | - | 0 | - |
| - | - | 1112 | 737.4 | - | - | 0 | - |
| 2 | y | 3.675E+04 | 738.4 | 0.0004573 | 0.6193 | +2 | 14 |
| - | - | 2.841E+04 | 738.9 | - | - | 0 | - |
| - | - | 1.327E+04 | 739.4 | - | - | 0 | - |
| - | - | 3322 | 739.9 | - | - | 0 | - |
| - | - | 1012 | 740.4 | - | - | 0 | - |
| - | - | 1828 | 743.4 | - | - | 0 | - |
| - | - | 1802 | 743.9 | - | - | 0 | - |
| 7 | b | 970 | 744.4 | 0.004303 | 5.781 | +1 | 7 |
| - | - | 826.1 | 755.4 | - | - | 0 | - |
| - | - | 1079 | 768.4 | - | - | 0 | - |
| - | - | 1061 | 769.4 | - | - | 0 | - |
| - | - | 1173 | 770.4 | - | - | 0 | - |
| - | - | 857.2 | 782.4 | - | - | 0 | - |
| - | - | 1369 | 783.4 | - | - | 0 | - |
| - | - | 955.7 | 787.4 | - | - | 0 | - |
| - | - | 920.5 | 798.4 | - | - | 0 | - |
| - | - | 1771 | 811.4 | - | - | 0 | - |
| - | - | 843.3 | 816.4 | - | - | 0 | - |
| 8 | y | 4240 | 828.4 | 0.0001933 | 0.2333 | +1 | 8 |
| - | - | 1965 | 829.4 | - | - | 0 | - |
| - | - | 948.7 | 858.4 | - | - | 0 | - |
| - | - | 686.5 | 867.4 | - | - | 0 | - |
| - | - | 633 | 873.4 | - | - | 0 | - |
| - | - | 1011 | 878.4 | - | - | 0 | - |
| - | - | 774.7 | 879.4 | - | - | 0 | - |
| - | - | 663.4 | 885.4 | - | - | 0 | - |
| - | - | 1229 | 887.4 | - | - | 0 | - |
| - | - | 810.4 | 891.4 | - | - | 0 | - |
| 8 | b | 1028 | 900.5 | 0.01081 | 12 | +1 | 8 |
| - | - | 728.3 | 901.5 | - | - | 0 | - |
| - | - | 2919 | 915.4 | - | - | 0 | - |
| - | - | 1176 | 916.4 | - | - | 0 | - |
| - | - | 973.1 | 927.5 | - | - | 0 | - |
| 7 | y | 3274 | 943.5 | 0.001448 | 1.535 | +1 | 9 |
| - | - | 2687 | 944.5 | - | - | 0 | - |
| - | - | 1003 | 945.5 | - | - | 0 | - |
| - | - | 957.6 | 972.5 | - | - | 0 | - |
| - | - | 4178 | 986.5 | - | - | 0 | - |
| - | - | 3256 | 987.5 | - | - | 0 | - |
| - | - | 1765 | 988.5 | - | - | 0 | - |
| - | - | 754.7 | 989.5 | - | - | 0 | - |
| - | - | 2051 | 1012 | - | - | 0 | - |
| - | - | 1439 | 1013 | - | - | 0 | - |
| - | - | 3104 | 1014 | - | - | 0 | - |
| - | - | 1566 | 1015 | - | - | 0 | - |
| 6 | y | 1.151E+04 | 1030 | 0.0001207 | 0.1172 | +1 | 10 |
| - | - | 6935 | 1031 | - | - | 0 | - |
| - | - | 2403 | 1032 | - | - | 0 | - |
| - | - | 1636 | 1074 | - | - | 0 | - |
| - | - | 739.1 | 1075 | - | - | 0 | - |
| - | - | 839.8 | 1087 | - | - | 0 | - |
| - | - | 630.5 | 1088 | - | - | 0 | - |
| - | - | 2301 | 1102 | - | - | 0 | - |
| - | - | 1396 | 1103 | - | - | 0 | - |
| 5 | y | 3863 | 1130 | 0.0005441 | 0.4817 | +1 | 11 |
| - | - | 1870 | 1131 | - | - | 0 | - |
| - | - | 652 | 1143 | - | - | 0 | - |
| - | - | 2737 | 1161 | - | - | 0 | - |
| - | - | 937.4 | 1162 | - | - | 0 | - |
| - | - | 1030 | 1171 | - | - | 0 | - |
| - | - | 702.8 | 1187 | - | - | 0 | - |
| - | - | 2739 | 1189 | - | - | 0 | - |
| - | - | 1146 | 1190 | - | - | 0 | - |
| - | - | 2285 | 1207 | - | - | 0 | - |
| - | - | 1349 | 1208 | - | - | 0 | - |

m/z Charge Intensity FragmentType MassShift Position
120.08116912841797 0 16924.707
120.89129638671875 0 388.7524
121.08445739746094 0 1710.7897
122.07202911376953 0 586.1193
124.08732604980469 0 369.0421
126.05538940429688 0 3726.332
127.05012512207031 0 527.6954
128.08238220214844 0 564.1555
129.06597900390625 0 479.83246
129.10263061523438 0 15451.743
130.05023193359375 0 877.4378
130.06143188476562 0 2912.9568
130.06614685058594 0 792.6398
130.08680725097656 0 395.86346
130.09811401367188 0 575.0138
130.10610961914062 0 918.3117
132.10231018066406 0 694.0854
133.06100463867188 0 549.663
134.04531860351562 0 838.2028
134.06044006347656 0 932.93805
136.0760955810547 0 21510.996
137.07945251464844 0 1394.6948
139.04971313476562 0 533.85455
139.08718872070312 0 1004.99384
140.08236694335938 0 1342.9098
141.06626892089844 0 521.73035
141.10235595703125 0 721.7567
143.08187866210938 0 687.8188
143.11846923828125 0 944.3132
144.06588745117188 0 848.95355
145.06153869628906 0 726.64667
146.06060791015625 0 1610.4618
147.04432678222656 0 810.3405
147.06439208984375 0 435.7932
147.07693481445312 0 1974.5764
151.12353515625 0 529.2817
152.07101440429688 0 2359.336
152.10757446289062 0 616.3198
153.10243225097656 0 661.8521
154.0502471923828 0 3979.8577
154.09747314453125 0 488.43906
155.08148193359375 0 790.5857
155.11819458007812 0 1216.1309
156.10244750976562 0 676.6907
157.06112670898438 0 1784.054
157.09759521484375 0 1839.7736
157.10874938964844 0 1614.5713
157.13397216796875 0 459.78415 d 1
158.09283447265625 0 8623.171 y Ammonia loss 14
158.58436584472656 0 512.9871
159.09153747558594 0 1334.6178
159.11318969726562 0 1927.0165
162.05560302734375 0 872.7133
163.0647430419922 0 409.1948
163.0715789794922 0 2925.4778
164.0705108642578 0 687.47534
165.0549774169922 0 848.6874
165.10174560546875 0 614.5695
167.08193969726562 0 1440.776
169.06146240234375 0 490.31232
169.09742736816406 0 1232.9373 y 10
171.0767822265625 0 4228.022
171.11302185058594 0 1667.6907
172.06094360351562 0 629.082
173.05612182617188 0 1289.4807
173.1288299560547 0 3512.26
173.4509735107422 0 3978.7568
175.0717315673828 0 2583.1763
175.0867462158203 0 1360.2137
175.11940002441406 0 17220.61 y 14
176.12298583984375 0 778.6024
181.0975799560547 0 1702.8887
182.08164978027344 0 2915.839
182.09169006347656 0 734.31506
182.1292266845703 0 505.78046
183.1132049560547 0 3795.634
183.14942932128906 0 1051.8999
184.07095336914062 0 520.207
184.10841369628906 0 806.5714
185.09255981445312 0 1982.0878
185.12876892089844 0 994.7566
187.07186889648438 0 517.4439
187.1081085205078 0 4684.294
187.14443969726562 0 1810.5408
189.08741760253906 0 21808.55
190.09152221679688 0 1280.176
191.0818328857422 0 566.6127
193.0968780517578 0 599.10876
195.11367797851562 0 710.5971
195.1496124267578 0 865.3996
199.0716094970703 0 2509.9473
199.10838317871094 0 802.0218
199.18093872070312 0 169596.3 a 1
200.1394500732422 0 1352.9021
200.18435668945312 0 21602.072
201.06625366210938 0 952.97327
201.1237030029297 0 9093.644
201.18771362304688 0 603.39624
202.11837768554688 0 835.7637
202.12767028808594 0 889.16925
203.06668090820312 0 3671.8748
205.10842895507812 0 617.09515
209.09286499023438 0 961.82806
209.2410125732422 0 527.16125
210.0878143310547 0 765.87225
211.1080780029297 0 1478.6465
211.1446075439453 0 618.9121
212.1033172607422 0 1789.7764
212.13922119140625 0 569.122
213.1588134765625 0 586.9972
215.1025848388672 0 615.54395
215.138916015625 0 1115.2112
216.13418579101562 0 580.9155
217.08233642578125 0 34715.684
217.11843872070312 0 805.2553
218.08570861816406 0 3016.044
223.11935424804688 0 1189.4547
224.103515625 0 2748.967
226.11903381347656 0 829.85126
226.15518188476562 0 1697.652
227.10321044921875 0 951.1603
227.1148681640625 0 887.20526
227.1758575439453 0 53494.39 b 1
228.09780883789062 0 647.8079
228.17918395996094 0 5758.7705
229.11854553222656 0 1441.1731
229.18064880371094 0 742.06055
233.16461181640625 0 628.24524
235.10842895507812 0 509.02896
237.0980987548828 0 853.5292
237.13516235351562 0 1260.0325
242.11399841308594 0 4016.3582
242.15042114257812 0 569.6954
243.0988006591797 0 933.94763
243.11488342285156 0 537.5973
243.1341094970703 0 657.5645
244.12953186035156 0 1536.1843
245.11264038085938 0 845.59
245.125 0 4644.0674 y Ammonia loss 13
246.0841827392578 0 500.77164
246.1275177001953 0 826.5979
248.07086181640625 0 1176.5306
249.15960693359375 0 552.5386
252.1345977783203 0 796.3766
252.171630859375 0 524.2616
253.16648864746094 0 1277.1006
254.11526489257812 0 458.81223
255.1094970703125 0 3842.035
256.1293029785156 0 618.7491
258.4484558105469 0 536.9385
259.14056396484375 0 539.3195
261.1606750488281 0 561.827
262.1394348144531 0 1472.5786
262.1516418457031 0 2610.3032 y 13
270.1453552246094 0 598.2927
272.1358947753906 0 825.1155
283.14471435546875 0 3526.6436
284.0875549316406 0 664.7111
284.1244201660156 0 1076.2504
284.1468811035156 0 970.29504
284.16241455078125 0 823.5382
286.2497863769531 0 496.9052
288.1543884277344 0 652.1159
296.124267578125 0 1137.0707
296.197265625 0 2290.0088 b Water loss 2
297.1197814941406 0 988.609
298.1405334472656 0 2617.2456
299.1387634277344 0 3253.7183
307.1046142578125 0 650.3148
311.1388854980469 0 787.69696
312.1556396484375 0 3401.361
312.1931457519531 0 735.8994
313.1584167480469 0 622.57166
313.1672668457031 0 922.52026
313.18798828125 0 5125.1797
314.1466369628906 0 2339.5754
314.19000244140625 0 682.7832
314.2082214355469 0 5597.9824 b 2
315.21148681640625 0 938.9934
316.15093994140625 0 5967.7305
317.153564453125 0 730.9648
324.1326599121094 0 1196.6501
327.1683044433594 0 748.0456
330.16680908203125 0 1449.8586
331.1732482910156 0 624.1303 y Water loss 12
332.1570739746094 0 4370.8813 y Ammonia loss 12
333.17889404296875 0 1064.978 b 8
341.1466064453125 0 711.3125
341.182861328125 0 2573.7078
342.1415100097656 0 5539.4443
342.1832580566406 0 786.2243
345.1830749511719 0 1433.6464 y 8
349.1840515136719 0 5129.083 y 12
355.16400146484375 0 715.3248
359.1676330566406 0 2551.396
359.19110107421875 0 628.835
383.2023620605469 0 942.0232
383.73089599609375 0 687.4736
385.17291259765625 0 1404.6772
386.15484619140625 0 619.14526
391.7248229980469 0 934.0799
392.1563720703125 0 662.3843
399.71807861328125 0 661.45166
400.21820068359375 0 1026.748
401.2394104003906 0 838.92065
402.19915771484375 0 1060.2516
407.23272705078125 0 669.93
412.192626953125 0 1233.1367
414.22900390625 0 1095.0024 b 11
414.7211608886719 0 3003.2195 y Ammonia loss 7
415.2221984863281 0 591.51276
415.2533874511719 0 667.8669
415.72509765625 0 689.7768
423.23394775390625 0 36307.62 y 7
423.7353210449219 0 16009.983
424.204345703125 0 967.41394
424.2366638183594 0 3487.0398
425.2403869628906 0 2885.6243 b Water loss 3
426.2434387207031 0 897.7205
428.2167053222656 0 675.9089
429.235107421875 0 1564.951
430.2399597167969 0 1089.1031 y Water loss 11
441.20928955078125 0 2412.47
443.25079345703125 0 3473.7468 b 3
448.25140380859375 0 1228.952 y 11
449.2230529785156 0 1260.8663 y Ammonia loss 2
450.1761474609375 0 596.6031
450.7415771484375 0 816.0543 b 7
452.2671203613281 0 929.8194
454.19287109375 0 913.76355
454.89227294921875 0 1384.2642 y 2
455.226806640625 0 663.0032
458.23675537109375 0 1843.8999
460.2578125 0 2014.3134
469.28851318359375 0 1306.6489
480.74761962890625 0 6411.335 y 6
481.2482604980469 0 3404.437
486.9187927246094 0 1100.9008 y Ammonia loss 1
487.2518005371094 0 1070.6053
492.5872497558594 0 1231.6865 y 1
492.9225769042969 0 691.6115
494.19476318359375 0 920.0448
495.91998291015625 0 4051.2473
496.2489929199219 0 971.9209
496.58172607421875 0 745.3227
500.1959533691406 0 860.154
505.2738342285156 0 4832.195 y 10
506.275390625 0 776.5402
511.28875732421875 0 815.6089
512.178955078125 0 768.2467
514.2412109375 0 688.073
514.3251953125 0 1511.8564
515.2578125 0 844.6397 y Water loss 5
515.75390625 0 868.28186 y Ammonia loss 5
516.25048828125 0 1021.1408
518.6043701171875 0 1163.1088
519.7349243164062 0 915.3965
524.2648315429688 0 38371.742 y 5
524.6124267578125 0 9272.504 Precursor Ammonia loss
524.7645874023438 0 20159.34
524.946533203125 0 3398.6147
525.2669677734375 0 8079.796
525.6124877929688 0 917.6675
525.7666625976562 0 1451.2557
528.6268920898438 0 801.396
528.7430419921875 0 8567.289
529.2450561523438 0 8135.1714
529.294677734375 0 952.7531
529.7456665039062 0 3436.6711
530.208251953125 0 1812.8307
530.28271484375 0 2037.2355 Precursor
530.31982421875 0 1179.4592
530.6156005859375 0 4440.5786
530.9495239257812 0 881.7984
531.3215942382812 0 992.933
539.8104858398438 0 828.5193
542.3199462890625 0 972.4942 b 4
564.793212890625 0 2121.0544 y Water loss 4
565.2902221679688 0 1918.1007 y Ammonia loss 4
565.7839965820312 0 1105.194
572.7903442382812 0 776.2346
573.7973022460938 0 55718.902 y 4
574.2984619140625 0 33980.81 y Water loss 9
574.8002319335938 0 12267.589
575.3013916015625 0 1995.9435
580.8043212890625 0 1245.4441
584.3148193359375 0 1120.0602
585.7857055664062 0 1828.2467
586.2901611328125 0 914.9546
586.789306640625 0 926.94336
592.3058471679688 0 5703.9453 y 9
593.3086547851562 0 2083.949
600.2717895507812 0 886.2539
618.788818359375 0 651.7114
626.3246459960938 0 801.25275
629.31640625 0 3897.3694 y Water loss 3
629.8125 0 2109.9353 y Ammonia loss 3
630.3366088867188 0 1515.3921
631.3438110351562 0 704.0257
636.3098754882812 0 1412.02
636.8146362304688 0 978.04034
637.3091430664062 0 838.91296
638.3186645507812 0 25922.334 y 3
638.8200073242188 0 17824.732
639.3209228515625 0 7350.912
639.8231201171875 0 1344.561
641.3402709960938 0 921.2221
646.3140258789062 0 2439.992
649.3163452148438 0 981.6973
652.31005859375 0 1017.9541
653.33154296875 0 690.13434
654.3269653320312 0 1097.9385
655.319091796875 0 1217.0569
657.2573852539062 0 772.5277
659.3455200195312 0 1549.1769
660.8209838867188 0 1475.7554
661.3219604492188 0 907.6597
663.824951171875 0 1668.3743
664.3209228515625 0 2035.0544
664.8236694335938 0 827.1716
665.8344116210938 0 709.3135
666.83056640625 0 4622.1655
667.3274536132812 0 2625.9663
667.83056640625 0 1139.4418
671.347412109375 0 5798.999 y Water loss 8
672.3369750976562 0 3926.5378 y Ammonia loss 8
672.8294677734375 0 12557.113 y Water loss 2
673.3301391601562 0 14937.827 y Ammonia loss 2
673.8302612304688 0 6901.2285
674.3167724609375 0 7724.084
674.8259887695312 0 1171.1632
675.3139038085938 0 2336.681
680.8260498046875 0 4021.8396
681.3280029296875 0 3706.0786
681.8351440429688 0 428654.34 y 2
682.3361206054688 0 302472.4
682.83740234375 0 123354.08
683.3385009765625 0 33258.758
683.839599609375 0 8505.487
686.8265991210938 0 3015.6067
687.3289184570312 0 3919.615
689.3584594726562 0 32340.273 y 8
690.361083984375 0 11107.697
691.3630981445312 0 2309.1604
692.8589477539062 0 839.5597
693.3565673828125 0 923.2224
694.8603515625 0 1793.1637
695.3610229492188 0 1170.4791
699.3442993164062 0 3105.2263 b Ammonia loss 13
700.3477783203125 0 1391.8837
712.3678588867188 0 1507.5535
729.3724975585938 0 5227.9604 y Water loss 1
729.8712768554688 0 3831.0132 y Ammonia loss 1
730.3743896484375 0 869.28345
736.873046875 0 1415.6311
737.3645629882812 0 1112.4397
738.3765258789062 0 36753.37 y 1
738.8779907226562 0 28412.94
739.3792724609375 0 13267.917
739.8805541992188 0 3322.3271
740.3786010742188 0 1011.58496
743.3684692382812 0 1828.2886
743.869384765625 0 1802.2523
744.3731079101562 0 969.9885 b 6
755.4344482421875 0 826.10785
768.4016723632812 0 1078.904
769.3756103515625 0 1061.4885
770.4159545898438 0 1173.2408
782.4480590820312 0 857.19324
783.4387817382812 0 1369.1907
787.3804321289062 0 955.6696
798.41357421875 0 920.4971
811.4168701171875 0 1770.9663
816.4248046875 0 843.31396
828.4324340820312 0 4240.091 y Ammonia loss 7
829.4347534179688 0 1964.8486
858.3936157226562 0 948.74835
867.4349365234375 0 686.47455
873.4057006835938 0 632.9817
878.3923950195312 0 1010.98413
879.4052734375 0 774.7068
885.4362182617188 0 663.3611
887.4219970703125 0 1228.6709
891.4240112304688 0 810.36346
900.4677124023438 0 1027.6989 b 7
901.475830078125 0 728.2898
915.4180908203125 0 2919.435
916.4221801757812 0 1176.2013
927.4552612304688 0 973.06116
943.4606323242188 0 3274.0305 y Ammonia loss 6
944.4685668945312 0 2686.6646
945.4736328125 0 1003.4405
972.4666748046875 0 957.5979
986.4896850585938 0 4178.452
987.4877319335938 0 3256.3135
988.4856567382812 0 1765.1428
989.4898071289062 0 754.686
1012.4784545898438 0 2051.265
1013.4840087890625 0 1439.3523
1014.4840087890625 0 3103.5178
1015.4906005859375 0 1565.9332
1030.4913330078125 0 11509.721 y Ammonia loss 5
1031.49365234375 0 6935.23
1032.498046875 0 2403.0015
1073.5135498046875 0 1635.7639
1074.52294921875 0 739.09375
1086.5545654296875 0 839.7939
1087.5430908203125 0 630.4615
1101.5225830078125 0 2301.235
1102.518798828125 0 1396.447
1129.55908203125 0 3863.283 y Ammonia loss 4
1130.5634765625 0 1870.4414
1142.5518798828125 0 651.99945
1160.5521240234375 0 2737.0476
1161.5604248046875 0 937.442
1170.55419921875 0 1030.3657
1186.5399169921875 0 702.81726
1188.5501708984375 0 2739.105
1189.5523681640625 0 1146.234
1206.5596923828125 0 2284.6545
1207.5640869140625 0 1348.8196

Spectrum Details

|  |  |
| --- | --- |
| Matched peaks? Matched peaks The total absolute number of peaks matched. Additionally in brackets the total fraction of peaks matched and the total number of peaks is shown. | 59 (14.01% of 421) |
| FDR? FDR The false discovery rate estimated for this peptide. It is calculated by matching all theoretical fragments with a non-integer shift with the raw peaks for this spectrum. This is done with 40 different shifts. The resulting percentage is the average number of annotated peaks over the number of annotated peaks with the correct spectrum. | 0.48% |
| Satellite FDR? Satellite FDR See the FDR for details on its calculation. This satellite ion specific FDR only contains the satellite ions (d/w) for I/L/J positions. | 0.00% |
| PSM Score? PSM Score The PSM Score as given by Hecklib to this annotated spectrum. It is shown with three significant figures. | 349 |

#### Spectrum 3761? Spectrum 3761 The raw spectrum of this peptide as annotated by Hecklib. The fragments are coloured according to ion type (see legend). Any peaks with a star '\*' as text can be hovered over to see the full details, first the ion type second the mass shift type. By hovering over the amino acids in the peptide or ions in the legend the corresponding peaks are highlighted. By toggling the 'Unassigned' label you can turn the background (unassigned) peaks on or off in the plot. By updating the slider in the Ion legend you can update the spectrum to only show the top X% of the peaks with labels. The top X% means any peak that is within X% of the highest intensity. By dragging in the spectrum you can zoom in to a specific part of the spectrum and use 'Zoom Out' to get back to the original zoom level. The annotation of the spectrum is based on the given sequence in the peptides file and is done with different software so inconsistencies are likely. The peaks are annotated based on the given sequence, with 20 ppm tolerance.

Copy Data

##### Spectrum 3761 (TSV)

###### Preview

```
Loading example...
```

*Click on the button to copy the data to your clipboard.*

Mz MinMz MaxIntensity Max

WidthHeightPeptide font sizePeptide stroke widthSpectrum font sizeSpectrum stroke widthCompact peptide

Ion legend

wxyz

abcd

OtherUnassignedIon^Charge^_Position_Show for top:%

JJSEVSDRPSGVSSR

03.37e+46.74e+41.01e+51.35e+5

Zoom Out

y^+1^_1_y^+1^_1_w^+1^_2_y^+1^_2_y^+1^_2_w^+1^_3_c^+1^_3_y^+1^_3_z^+1^_3_y^+1^_3_c^+3^_12_z^+2^_8_w^+1^_4_y^+2^_8_z^+1^_4_y^+1^_4_c^+2^_8_y^+3^_13_c^+1^_4_c^+3^_14_z^+2^_9_y^+2^_9_z^+3^_14_y^+3^_14_z^+1^_5_y^+3^_14_y^+1^_5_z^+2^_10_y^+2^_10_z^+1^_6_w^+1^_6_c^+1^_5_z^+2^_11_y^+2^_11_y^+1^_6_y^+1^_6_z^+1^_6_y^+1^_6_w^+2^_12_c^+2^_12_z^+2^_12_y^+2^_12_w^+1^_7_c^+1^_6_z^+2^_13_y^+2^_13_y^+2^_13_z^+2^_13_y^+2^_13_y^+1^_7_c^+2^_14_w^+2^_14_y^+2^_14_z^+2^_14_y^+2^_14_w^+1^_8_c^+1^_7_z^+1^_8_z^+1^_8_y^+1^_8_c^+1^_8_z^+1^_9_z^+1^_9_y^+1^_9_c^+1^_9_y^+1^_10_z^+1^_10_y^+1^_10_c^+1^_10_z^+1^_11_y^+1^_11_c^+1^_11_w^+1^_12_c^+1^_12_y^+1^_12_z^+1^_13_c^+1^_13_c^+1^_14_

0767153523023069

Fragment Matches Table

Show background peaks

| Position | Ion type | Intensity | mz Theoretical | mz Error (Th) | mz Error (ppm) | Charge | Series Number |
| --- | --- | --- | --- | --- | --- | --- | --- |
| - | - | 1568 | 125.1 | - | - | 0 | - |
| - | - | 3451 | 126.1 | - | - | 0 | - |
| - | - | 938.1 | 129.1 | - | - | 0 | - |
| - | - | 730.2 | 131.1 | - | - | 0 | - |
| - | - | 900.1 | 153.1 | - | - | 0 | - |
| 15 | y | 1237 | 158.1 | 9.571E-05 | 0.6054 | +1 | 1 |
| - | - | 4520 | 168.1 | - | - | 0 | - |
| - | - | 515.3 | 168.8 | - | - | 0 | - |
| - | - | 1.196E+04 | 173.1 | - | - | 0 | - |
| - | - | 2828 | 173.5 | - | - | 0 | - |
| 15 | y | 6777 | 175.1 | 0.0001427 | 0.8148 | +1 | 1 |
| - | - | 502.5 | 176.9 | - | - | 0 | - |
| - | - | 1026 | 181.1 | - | - | 0 | - |
| - | - | 1.604E+04 | 185.1 | - | - | 0 | - |
| - | - | 605.7 | 186.1 | - | - | 0 | - |
| - | - | 944.2 | 186.1 | - | - | 0 | - |
| - | - | 1044 | 187.1 | - | - | 0 | - |
| - | - | 875.7 | 199.1 | - | - | 0 | - |
| - | - | 4.202E+04 | 199.2 | - | - | 0 | - |
| - | - | 3123 | 200.1 | - | - | 0 | - |
| - | - | 5178 | 200.2 | - | - | 0 | - |
| - | - | 543.2 | 214.4 | - | - | 0 | - |
| - | - | 1317 | 215.1 | - | - | 0 | - |
| - | - | 550.7 | 216.1 | - | - | 0 | - |
| - | - | 1708 | 217.1 | - | - | 0 | - |
| - | - | 594.7 | 226.2 | - | - | 0 | - |
| - | - | 2.087E+04 | 227.2 | - | - | 0 | - |
| - | - | 2151 | 228.2 | - | - | 0 | - |
| 14 | w | 2752 | 229.1 | 6.129E-05 | 0.2675 | +1 | 2 |
| - | - | 1142 | 244.1 | - | - | 0 | - |
| 14 | y | 2.919E+04 | 245.1 | 8.024E-05 | 0.3274 | +1 | 2 |
| - | - | 4462 | 246.1 | - | - | 0 | - |
| - | - | 545.3 | 254.9 | - | - | 0 | - |
| - | - | 1569 | 255.1 | - | - | 0 | - |
| 14 | y | 4815 | 262.2 | 0.0001627 | 0.6207 | +1 | 2 |
| - | - | 3968 | 270.1 | - | - | 0 | - |
| - | - | 930.1 | 272.2 | - | - | 0 | - |
| - | - | 8860 | 274.1 | - | - | 0 | - |
| - | - | 529.8 | 287.4 | - | - | 0 | - |
| - | - | 816.6 | 289.1 | - | - | 0 | - |
| - | - | 671.8 | 303.2 | - | - | 0 | - |
| - | - | 2235 | 314.2 | - | - | 0 | - |
| 13 | w | 1.455E+04 | 316.2 | 7.583E-05 | 0.2398 | +1 | 3 |
| - | - | 2357 | 317.2 | - | - | 0 | - |
| - | - | 1031 | 324.1 | - | - | 0 | - |
| - | - | 630.2 | 330.2 | - | - | 0 | - |
| - | - | 3520 | 331.1 | - | - | 0 | - |
| 3 | c | 702.4 | 331.2 | 5.738E-05 | 0.1732 | +1 | 3 |
| 13 | y | 4399 | 332.2 | 2.678E-05 | 0.08063 | +1 | 3 |
| 13 | z | 1.093E+04 | 333.2 | 3.931E-05 | 0.118 | +1 | 3 |
| - | - | 1952 | 334.2 | - | - | 0 | - |
| - | - | 791.5 | 347.2 | - | - | 0 | - |
| 13 | y | 3674 | 349.2 | 6.596E-05 | 0.1889 | +1 | 3 |
| - | - | 1142 | 350.2 | - | - | 0 | - |
| - | - | 7781 | 357.2 | - | - | 0 | - |
| - | - | 787.4 | 361.2 | - | - | 0 | - |
| - | - | 750.7 | 414.2 | - | - | 0 | - |
| 12 | c | 581.9 | 414.2 | 0.001573 | 3.798 | +3 | 12 |
| 8 | z | 3356 | 415.2 | 0.003105 | 7.478 | +2 | 8 |
| - | - | 644 | 415.6 | - | - | 0 | - |
| - | - | 1059 | 415.7 | - | - | 0 | - |
| - | - | 735.3 | 416.2 | - | - | 0 | - |
| 12 | w | 1.057E+04 | 417.2 | 0.000371 | 0.8892 | +1 | 4 |
| - | - | 1849 | 418.2 | - | - | 0 | - |
| 8 | y | 3970 | 423.2 | 0.0004892 | 1.156 | +2 | 8 |
| - | - | 638.3 | 423.7 | - | - | 0 | - |
| - | - | 950 | 425.2 | - | - | 0 | - |
| - | - | 1616 | 427.2 | - | - | 0 | - |
| - | - | 604.6 | 429.2 | - | - | 0 | - |
| 12 | z | 879.8 | 432.2 | 0.0007042 | 1.629 | +1 | 4 |
| - | - | 915 | 437.2 | - | - | 0 | - |
| - | - | 1655 | 442.2 | - | - | 0 | - |
| - | - | 753.3 | 443.2 | - | - | 0 | - |
| - | - | 1692 | 443.2 | - | - | 0 | - |
| - | - | 645.4 | 443.3 | - | - | 0 | - |
| - | - | 855 | 444.3 | - | - | 0 | - |
| - | - | 1479 | 446.2 | - | - | 0 | - |
| - | - | 819 | 447.2 | - | - | 0 | - |
| 12 | y | 1.082E+04 | 448.3 | 0.000225 | 0.5021 | +1 | 4 |
| - | - | 2815 | 449.3 | - | - | 0 | - |
| 8 | c | 576.5 | 450.7 | 0.002085 | 4.626 | +2 | 8 |
| - | - | 593.4 | 453.7 | - | - | 0 | - |
| 3 | y | 943.5 | 454.9 | 0.000792 | 1.741 | +3 | 13 |
| - | - | 612.8 | 460.2 | - | - | 0 | - |
| 4 | c | 8375 | 460.3 | 0.0001889 | 0.4104 | +1 | 4 |
| - | - | 1693 | 461.3 | - | - | 0 | - |
| 14 | c | 2016 | 472.2 | 0.007439 | 15.75 | +3 | 14 |
| 7 | z | 1978 | 472.7 | 0.0005282 | 1.117 | +2 | 9 |
| - | - | 1295 | 473.2 | - | - | 0 | - |
| - | - | 2742 | 474.2 | - | - | 0 | - |
| - | - | 881.9 | 474.7 | - | - | 0 | - |
| - | - | 881.6 | 475.2 | - | - | 0 | - |
| - | - | 772.2 | 475.2 | - | - | 0 | - |
| - | - | 893.2 | 480.2 | - | - | 0 | - |
| - | - | 902.9 | 480.7 | - | - | 0 | - |
| 7 | y | 1621 | 480.7 | 7.733E-05 | 0.1609 | +2 | 9 |
| 2 | z | 698.5 | 481.2 | 0.00336 | 6.981 | +3 | 14 |
| - | - | 939.7 | 482.2 | - | - | 0 | - |
| - | - | 1991 | 482.7 | - | - | 0 | - |
| - | - | 1128 | 483.7 | - | - | 0 | - |
| - | - | 2623 | 484.2 | - | - | 0 | - |
| - | - | 1399 | 484.7 | - | - | 0 | - |
| - | - | 885 | 485.3 | - | - | 0 | - |
| 2 | y | 701.3 | 486.6 | 0.0025 | 5.137 | +3 | 14 |
| 11 | z | 3237 | 489.3 | 0.0005615 | 1.148 | +1 | 5 |
| - | - | 3042 | 489.7 | - | - | 0 | - |
| - | - | 1240 | 490.2 | - | - | 0 | - |
| - | - | 2650 | 490.3 | - | - | 0 | - |
| 2 | y | 796.9 | 492.6 | 0.001144 | 2.323 | +3 | 14 |
| - | - | 1200 | 492.9 | - | - | 0 | - |
| - | - | 979.6 | 493.3 | - | - | 0 | - |
| - | - | 1200 | 493.3 | - | - | 0 | - |
| - | - | 568.2 | 494.3 | - | - | 0 | - |
| - | - | 782.3 | 495.9 | - | - | 0 | - |
| - | - | 2204 | 499.3 | - | - | 0 | - |
| - | - | 709.7 | 501.3 | - | - | 0 | - |
| - | - | 1810 | 504.2 | - | - | 0 | - |
| - | - | 1741 | 504.2 | - | - | 0 | - |
| - | - | 1615 | 504.7 | - | - | 0 | - |
| - | - | 2122 | 504.7 | - | - | 0 | - |
| 11 | y | 3.841E+04 | 505.3 | 6.259E-05 | 0.1239 | +1 | 5 |
| - | - | 9493 | 506.3 | - | - | 0 | - |
| - | - | 1832 | 507.3 | - | - | 0 | - |
| - | - | 1.457E+04 | 513.2 | - | - | 0 | - |
| - | - | 1436 | 513.7 | - | - | 0 | - |
| - | - | 5811 | 513.7 | - | - | 0 | - |
| - | - | 1002 | 514.2 | - | - | 0 | - |
| - | - | 622.2 | 514.3 | - | - | 0 | - |
| - | - | 1145 | 515.2 | - | - | 0 | - |
| - | - | 1396 | 515.3 | - | - | 0 | - |
| 6 | z | 3668 | 516.3 | 0.0005324 | 1.031 | +2 | 10 |
| - | - | 798.1 | 516.3 | - | - | 0 | - |
| - | - | 4785 | 516.8 | - | - | 0 | - |
| - | - | 2772 | 517.3 | - | - | 0 | - |
| - | - | 1043 | 523.8 | - | - | 0 | - |
| 6 | y | 1.243E+04 | 524.3 | 0.004449 | 8.486 | +2 | 10 |
| - | - | 2935 | 524.6 | - | - | 0 | - |
| - | - | 7827 | 524.8 | - | - | 0 | - |
| - | - | 2767 | 524.9 | - | - | 0 | - |
| - | - | 1723 | 525.3 | - | - | 0 | - |
| - | - | 1042 | 528.2 | - | - | 0 | - |
| - | - | 4044 | 530.3 | - | - | 0 | - |
| - | - | 4488 | 530.6 | - | - | 0 | - |
| - | - | 2863 | 531 | - | - | 0 | - |
| - | - | 1015 | 531.3 | - | - | 0 | - |
| - | - | 1163 | 541.3 | - | - | 0 | - |
| - | - | 4782 | 542.3 | - | - | 0 | - |
| - | - | 935.8 | 543.3 | - | - | 0 | - |
| 10 | z | 1835 | 558.3 | 6.952E-05 | 0.1245 | +1 | 6 |
| - | - | 1524 | 558.3 | - | - | 0 | - |
| - | - | 767.9 | 558.8 | - | - | 0 | - |
| 10 | w | 1027 | 559.3 | 0.008297 | 14.83 | +1 | 6 |
| 5 | c | 6607 | 559.3 | 0.0003234 | 0.5782 | +1 | 5 |
| - | - | 1444 | 560.3 | - | - | 0 | - |
| 5 | z | 2.901E+04 | 565.8 | 1.031E-05 | 0.01823 | +2 | 11 |
| - | - | 1.732E+04 | 566.3 | - | - | 0 | - |
| - | - | 6686 | 566.8 | - | - | 0 | - |
| - | - | 2889 | 567.3 | - | - | 0 | - |
| - | - | 1376 | 573.3 | - | - | 0 | - |
| 5 | y | 1.693E+04 | 573.8 | 0.0004002 | 0.6974 | +2 | 11 |
| 10 | y | 7927 | 574.3 | 0.00344 | 5.99 | +1 | 6 |
| - | - | 3181 | 574.8 | - | - | 0 | - |
| 10 | y | 1691 | 575.3 | 0.001144 | 1.989 | +1 | 6 |
| 10 | z | 5660 | 576.3 | 0.0005464 | 0.9482 | +1 | 6 |
| - | - | 1960 | 577.3 | - | - | 0 | - |
| - | - | 709.2 | 578.3 | - | - | 0 | - |
| - | - | 4411 | 580.2 | - | - | 0 | - |
| - | - | 1045 | 581.2 | - | - | 0 | - |
| - | - | 680 | 584.4 | - | - | 0 | - |
| - | - | 844.8 | 585.3 | - | - | 0 | - |
| - | - | 1236 | 586.3 | - | - | 0 | - |
| - | - | 932.9 | 589.2 | - | - | 0 | - |
| - | - | 897.4 | 592.2 | - | - | 0 | - |
| 10 | y | 673.9 | 592.3 | 0.004439 | 7.494 | +1 | 6 |
| - | - | 1541 | 594.3 | - | - | 0 | - |
| - | - | 2661 | 600.3 | - | - | 0 | - |
| 4 | w | 1.211E+04 | 600.8 | 0.0001158 | 0.1928 | +2 | 12 |
| - | - | 5615 | 601.3 | - | - | 0 | - |
| - | - | 2846 | 601.8 | - | - | 0 | - |
| - | - | 925.3 | 603.4 | - | - | 0 | - |
| - | - | 2154 | 610.3 | - | - | 0 | - |
| - | - | 1794 | 621.4 | - | - | 0 | - |
| 12 | c | 1072 | 629.3 | 0.009152 | 14.54 | +2 | 12 |
| 4 | z | 3.132E+04 | 630.3 | 5.544E-05 | 0.08796 | +2 | 12 |
| - | - | 1.926E+04 | 630.8 | - | - | 0 | - |
| - | - | 9092 | 631.3 | - | - | 0 | - |
| - | - | 3352 | 631.8 | - | - | 0 | - |
| - | - | 1055 | 632.3 | - | - | 0 | - |
| - | - | 870.5 | 635.7 | - | - | 0 | - |
| - | - | 7997 | 637.8 | - | - | 0 | - |
| 4 | y | 3.092E+04 | 638.3 | 0.0007006 | 1.098 | +2 | 12 |
| - | - | 1.768E+04 | 638.8 | - | - | 0 | - |
| - | - | 6601 | 639.3 | - | - | 0 | - |
| - | - | 1557 | 639.8 | - | - | 0 | - |
| - | - | 3474 | 644.3 | - | - | 0 | - |
| - | - | 1960 | 644.8 | - | - | 0 | - |
| - | - | 1244 | 645.4 | - | - | 0 | - |
| 9 | w | 1136 | 646.3 | 0.002574 | 3.983 | +1 | 7 |
| 6 | c | 2.703E+04 | 646.4 | 0.0001253 | 0.1938 | +1 | 6 |
| - | - | 1156 | 647.3 | - | - | 0 | - |
| - | - | 8877 | 647.4 | - | - | 0 | - |
| - | - | 1708 | 648.4 | - | - | 0 | - |
| - | - | 822.3 | 651.8 | - | - | 0 | - |
| 3 | z | 2143 | 665.3 | 0.01203 | 18.07 | +2 | 13 |
| - | - | 1818 | 665.8 | - | - | 0 | - |
| 3 | y | 2630 | 672.8 | 0.0007513 | 1.117 | +2 | 13 |
| 3 | y | 3149 | 673.3 | 0.01206 | 17.92 | +2 | 13 |
| 3 | z | 2.01E+04 | 673.8 | 0.0001545 | 0.2293 | +2 | 13 |
| - | - | 2.316E+04 | 674.3 | - | - | 0 | - |
| - | - | 1.413E+04 | 674.8 | - | - | 0 | - |
| - | - | 5571 | 675.3 | - | - | 0 | - |
| - | - | 1413 | 675.8 | - | - | 0 | - |
| - | - | 748.9 | 677.3 | - | - | 0 | - |
| - | - | 789.9 | 678.8 | - | - | 0 | - |
| - | - | 1422 | 681.3 | - | - | 0 | - |
| 3 | y | 1.335E+05 | 681.8 | 0.0002353 | 0.3452 | +2 | 13 |
| - | - | 8.951E+04 | 682.3 | - | - | 0 | - |
| - | - | 3.561E+04 | 682.8 | - | - | 0 | - |
| - | - | 1.004E+04 | 683.3 | - | - | 0 | - |
| - | - | 2537 | 683.8 | - | - | 0 | - |
| - | - | 686.9 | 686.3 | - | - | 0 | - |
| - | - | 822.1 | 686.8 | - | - | 0 | - |
| - | - | 1563 | 687.3 | - | - | 0 | - |
| 9 | y | 2694 | 689.4 | 0.003492 | 5.065 | +1 | 7 |
| - | - | 1104 | 690.4 | - | - | 0 | - |
| - | - | 677.5 | 691.4 | - | - | 0 | - |
| - | - | 1606 | 695.8 | - | - | 0 | - |
| - | - | 1008 | 696.3 | - | - | 0 | - |
| - | - | 804 | 699.4 | - | - | 0 | - |
| - | - | 725.3 | 701 | - | - | 0 | - |
| - | - | 734 | 701.7 | - | - | 0 | - |
| - | - | 2755 | 702.3 | - | - | 0 | - |
| - | - | 2145 | 706.7 | - | - | 0 | - |
| - | - | 1790 | 707 | - | - | 0 | - |
| 14 | c | 722.6 | 707.4 | 0.001175 | 1.662 | +2 | 14 |
| - | - | 717.9 | 707.7 | - | - | 0 | - |
| 2 | w | 3840 | 715.8 | 0.0006343 | 0.8861 | +2 | 14 |
| - | - | 2608 | 716.3 | - | - | 0 | - |
| - | - | 1359 | 716.8 | - | - | 0 | - |
| - | - | 2142 | 717.4 | - | - | 0 | - |
| - | - | 3833 | 718.4 | - | - | 0 | - |
| - | - | 9925 | 719.3 | - | - | 0 | - |
| - | - | 1298 | 719.4 | - | - | 0 | - |
| - | - | 2557 | 720.3 | - | - | 0 | - |
| - | - | 809.7 | 728.4 | - | - | 0 | - |
| 2 | y | 1900 | 729.4 | 0.006773 | 9.286 | +2 | 14 |
| 2 | z | 2187 | 730.4 | 0.0001905 | 0.2608 | +2 | 14 |
| - | - | 1312 | 730.9 | - | - | 0 | - |
| - | - | 1513 | 734.3 | - | - | 0 | - |
| - | - | 747.6 | 735.3 | - | - | 0 | - |
| - | - | 1114 | 735.5 | - | - | 0 | - |
| 2 | y | 8076 | 738.4 | 0.0008855 | 1.199 | +2 | 14 |
| - | - | 6729 | 738.9 | - | - | 0 | - |
| - | - | 2343 | 739.4 | - | - | 0 | - |
| 8 | w | 3092 | 743.4 | 0.0002016 | 0.2712 | +1 | 8 |
| - | - | 1424 | 743.9 | - | - | 0 | - |
| - | - | 1644 | 744.4 | - | - | 0 | - |
| - | - | 938 | 745.4 | - | - | 0 | - |
| - | - | 2325 | 750.9 | - | - | 0 | - |
| - | - | 2196 | 751.4 | - | - | 0 | - |
| - | - | 3866 | 752.3 | - | - | 0 | - |
| - | - | 821.4 | 756.4 | - | - | 0 | - |
| - | - | 3184 | 758.4 | - | - | 0 | - |
| - | - | 4159 | 758.9 | - | - | 0 | - |
| - | - | 2988 | 759.4 | - | - | 0 | - |
| - | - | 1972 | 759.9 | - | - | 0 | - |
| - | - | 3178 | 760.4 | - | - | 0 | - |
| - | - | 808.5 | 760.9 | - | - | 0 | - |
| 7 | c | 5.203E+04 | 761.4 | 0.0003349 | 0.4398 | +1 | 7 |
| - | - | 760.7 | 762.3 | - | - | 0 | - |
| - | - | 1.844E+04 | 762.4 | - | - | 0 | - |
| - | - | 5215 | 763.4 | - | - | 0 | - |
| - | - | 2393 | 764.4 | - | - | 0 | - |
| - | - | 3625 | 764.9 | - | - | 0 | - |
| - | - | 8.894E+04 | 765.4 | - | - | 0 | - |
| - | - | 7.06E+04 | 765.9 | - | - | 0 | - |
| - | - | 3.092E+04 | 766.4 | - | - | 0 | - |
| - | - | 9047 | 766.9 | - | - | 0 | - |
| - | - | 3614 | 767.4 | - | - | 0 | - |
| - | - | 2938 | 770.3 | - | - | 0 | - |
| - | - | 888.6 | 771.3 | - | - | 0 | - |
| - | - | 853.4 | 771.4 | - | - | 0 | - |
| - | - | 2417 | 772.4 | - | - | 0 | - |
| - | - | 1924 | 772.9 | - | - | 0 | - |
| - | - | 2851 | 773.4 | - | - | 0 | - |
| - | - | 2.747E+04 | 773.9 | - | - | 0 | - |
| - | - | 1.814E+04 | 774.4 | - | - | 0 | - |
| - | - | 1.186E+04 | 774.9 | - | - | 0 | - |
| - | - | 2865 | 775.4 | - | - | 0 | - |
| - | - | 2503 | 777.9 | - | - | 0 | - |
| - | - | 2488 | 778.4 | - | - | 0 | - |
| - | - | 1671 | 778.9 | - | - | 0 | - |
| - | - | 1395 | 779.4 | - | - | 0 | - |
| - | - | 5228 | 786.4 | - | - | 0 | - |
| - | - | 9.25E+04 | 786.9 | - | - | 0 | - |
| - | - | 7.558E+04 | 787.4 | - | - | 0 | - |
| - | - | 3.305E+04 | 787.9 | - | - | 0 | - |
| - | - | 1.032E+04 | 788.4 | - | - | 0 | - |
| - | - | 4480 | 788.9 | - | - | 0 | - |
| - | - | 5.8E+04 | 794.9 | - | - | 0 | - |
| - | - | 7.094E+04 | 795.4 | - | - | 0 | - |
| - | - | 4.192E+04 | 795.9 | - | - | 0 | - |
| - | - | 1.653E+04 | 796.4 | - | - | 0 | - |
| - | - | 3916 | 796.9 | - | - | 0 | - |
| - | - | 825.8 | 797.4 | - | - | 0 | - |
| 8 | z | 1981 | 811.4 | 0.001285 | 1.584 | +1 | 8 |
| - | - | 1552 | 813.4 | - | - | 0 | - |
| - | - | 767.7 | 817.4 | - | - | 0 | - |
| - | - | 1100 | 822.5 | - | - | 0 | - |
| - | - | 2802 | 827.4 | - | - | 0 | - |
| - | - | 1706 | 828.4 | - | - | 0 | - |
| - | - | 1221 | 828.5 | - | - | 0 | - |
| 8 | z | 4.048E+04 | 829.4 | 0.0006738 | 0.8123 | +1 | 8 |
| - | - | 2.172E+04 | 830.4 | - | - | 0 | - |
| - | - | 7026 | 831.4 | - | - | 0 | - |
| - | - | 1859 | 832.4 | - | - | 0 | - |
| - | - | 2549 | 844.4 | - | - | 0 | - |
| 8 | y | 6817 | 845.5 | 4.968E-05 | 0.05877 | +1 | 8 |
| - | - | 3129 | 846.5 | - | - | 0 | - |
| - | - | 874.4 | 847.1 | - | - | 0 | - |
| - | - | 963.9 | 847.5 | - | - | 0 | - |
| - | - | 1.113E+04 | 853.3 | - | - | 0 | - |
| - | - | 4158 | 854.3 | - | - | 0 | - |
| - | - | 1836 | 872.5 | - | - | 0 | - |
| - | - | 2918 | 873.5 | - | - | 0 | - |
| - | - | 1173 | 873.9 | - | - | 0 | - |
| - | - | 811.6 | 874.5 | - | - | 0 | - |
| - | - | 876.8 | 875 | - | - | 0 | - |
| 8 | c | 4395 | 900.5 | 0.003486 | 3.871 | +1 | 8 |
| - | - | 2970 | 901.5 | - | - | 0 | - |
| - | - | 1035 | 902.5 | - | - | 0 | - |
| - | - | 1086 | 915.4 | - | - | 0 | - |
| 7 | z | 1798 | 926.5 | 0.0006481 | 0.6995 | +1 | 9 |
| - | - | 987.6 | 927.5 | - | - | 0 | - |
| - | - | 2806 | 930.4 | - | - | 0 | - |
| - | - | 1225 | 931.4 | - | - | 0 | - |
| - | - | 895.8 | 934.5 | - | - | 0 | - |
| 7 | z | 1.618E+04 | 944.5 | 0.0009444 | 1 | +1 | 9 |
| - | - | 1.171E+04 | 945.5 | - | - | 0 | - |
| - | - | 3930 | 946.5 | - | - | 0 | - |
| - | - | 1334 | 947.5 | - | - | 0 | - |
| - | - | 642.7 | 949.4 | - | - | 0 | - |
| - | - | 1962 | 952 | - | - | 0 | - |
| - | - | 1327 | 952.5 | - | - | 0 | - |
| - | - | 1048 | 953 | - | - | 0 | - |
| 7 | y | 1689 | 960.5 | 0.0007783 | 0.8103 | +1 | 9 |
| - | - | 1575 | 961.5 | - | - | 0 | - |
| - | - | 1318 | 965.4 | - | - | 0 | - |
| - | - | 974 | 966.4 | - | - | 0 | - |
| - | - | 691.8 | 967.4 | - | - | 0 | - |
| - | - | 4012 | 968.4 | - | - | 0 | - |
| - | - | 3436 | 969.4 | - | - | 0 | - |
| - | - | 1490 | 970.4 | - | - | 0 | - |
| - | - | 4407 | 970.5 | - | - | 0 | - |
| - | - | 1612 | 971.5 | - | - | 0 | - |
| - | - | 796.6 | 972.5 | - | - | 0 | - |
| - | - | 1929 | 985.4 | - | - | 0 | - |
| - | - | 738.1 | 986.4 | - | - | 0 | - |
| - | - | 1091 | 988.5 | - | - | 0 | - |
| - | - | 2030 | 995.3 | - | - | 0 | - |
| - | - | 922.9 | 996.3 | - | - | 0 | - |
| - | - | 1170 | 1002 | - | - | 0 | - |
| - | - | 1912 | 1003 | - | - | 0 | - |
| - | - | 832.6 | 1005 | - | - | 0 | - |
| - | - | 1154 | 1006 | - | - | 0 | - |
| - | - | 1435 | 1007 | - | - | 0 | - |
| - | - | 933.7 | 1008 | - | - | 0 | - |
| - | - | 1993 | 1009 | - | - | 0 | - |
| - | - | 3688 | 1012 | - | - | 0 | - |
| - | - | 2358 | 1013 | - | - | 0 | - |
| - | - | 916.1 | 1014 | - | - | 0 | - |
| 9 | c | 3.563E+04 | 1015 | 0.0004622 | 0.4556 | +1 | 9 |
| - | - | 2.038E+04 | 1016 | - | - | 0 | - |
| - | - | 1544 | 1016 | - | - | 0 | - |
| - | - | 6343 | 1017 | - | - | 0 | - |
| - | - | 1304 | 1017 | - | - | 0 | - |
| - | - | 1654 | 1018 | - | - | 0 | - |
| - | - | 4668 | 1025 | - | - | 0 | - |
| - | - | 3878 | 1026 | - | - | 0 | - |
| - | - | 1.148E+04 | 1026 | - | - | 0 | - |
| - | - | 1000 | 1027 | - | - | 0 | - |
| - | - | 1.372E+04 | 1027 | - | - | 0 | - |
| - | - | 3821 | 1028 | - | - | 0 | - |
| 6 | y | 843.7 | 1030 | 0.01172 | 11.37 | +1 | 10 |
| 6 | z | 5120 | 1031 | 0.0006242 | 0.6051 | +1 | 10 |
| - | - | 9054 | 1033 | - | - | 0 | - |
| - | - | 3126 | 1034 | - | - | 0 | - |
| - | - | 3180 | 1038 | - | - | 0 | - |
| - | - | 1542 | 1039 | - | - | 0 | - |
| - | - | 1868 | 1040 | - | - | 0 | - |
| - | - | 2003 | 1041 | - | - | 0 | - |
| - | - | 788.8 | 1043 | - | - | 0 | - |
| - | - | 1086 | 1043 | - | - | 0 | - |
| - | - | 3278 | 1043 | - | - | 0 | - |
| - | - | 1079 | 1044 | - | - | 0 | - |
| - | - | 6348 | 1044 | - | - | 0 | - |
| - | - | 1.465E+04 | 1045 | - | - | 0 | - |
| - | - | 5332 | 1046 | - | - | 0 | - |
| 6 | y | 3114 | 1048 | 0.003723 | 3.554 | +1 | 10 |
| - | - | 1230 | 1049 | - | - | 0 | - |
| - | - | 2000 | 1052 | - | - | 0 | - |
| - | - | 1145 | 1052 | - | - | 0 | - |
| - | - | 1429 | 1053 | - | - | 0 | - |
| - | - | 2.378E+04 | 1058 | - | - | 0 | - |
| - | - | 1.42E+04 | 1059 | - | - | 0 | - |
| - | - | 5208 | 1060 | - | - | 0 | - |
| - | - | 1752 | 1060 | - | - | 0 | - |
| - | - | 2123 | 1061 | - | - | 0 | - |
| - | - | 5017 | 1061 | - | - | 0 | - |
| - | - | 2.441E+04 | 1062 | - | - | 0 | - |
| - | - | 1.144E+04 | 1063 | - | - | 0 | - |
| - | - | 1104 | 1064 | - | - | 0 | - |
| - | - | 1380 | 1101 | - | - | 0 | - |
| 10 | c | 5.175E+04 | 1102 | 0.0008744 | 0.7938 | +1 | 10 |
| - | - | 2.958E+04 | 1103 | - | - | 0 | - |
| - | - | 1.062E+04 | 1104 | - | - | 0 | - |
| - | - | 1859 | 1105 | - | - | 0 | - |
| - | - | 7837 | 1115 | - | - | 0 | - |
| - | - | 5913 | 1116 | - | - | 0 | - |
| - | - | 2158 | 1117 | - | - | 0 | - |
| - | - | 1700 | 1118 | - | - | 0 | - |
| - | - | 963.4 | 1119 | - | - | 0 | - |
| - | - | 907.1 | 1120 | - | - | 0 | - |
| 5 | z | 6740 | 1131 | 0.00103 | 0.9113 | +1 | 11 |
| - | - | 1.64E+04 | 1132 | - | - | 0 | - |
| - | - | 9849 | 1133 | - | - | 0 | - |
| - | - | 2994 | 1134 | - | - | 0 | - |
| 5 | y | 2100 | 1147 | 0.0007261 | 0.6332 | +1 | 11 |
| - | - | 2122 | 1148 | - | - | 0 | - |
| 11 | c | 2.708E+04 | 1159 | 0.001342 | 1.158 | +1 | 11 |
| - | - | 1.882E+04 | 1160 | - | - | 0 | - |
| - | - | 5858 | 1161 | - | - | 0 | - |
| - | - | 1705 | 1162 | - | - | 0 | - |
| 4 | w | 918.5 | 1201 | 0.01666 | 13.88 | +1 | 12 |
| - | - | 5001 | 1214 | - | - | 0 | - |
| - | - | 3722 | 1215 | - | - | 0 | - |
| - | - | 1916 | 1216 | - | - | 0 | - |
| - | - | 823.6 | 1217 | - | - | 0 | - |
| - | - | 1079 | 1218 | - | - | 0 | - |
| - | - | 1552 | 1248 | - | - | 0 | - |
| - | - | 805 | 1249 | - | - | 0 | - |
| 12 | c | 2.954E+04 | 1258 | 0.002007 | 1.596 | +1 | 12 |
| - | - | 1.855E+04 | 1259 | - | - | 0 | - |
| - | - | 6625 | 1260 | - | - | 0 | - |
| - | - | 2.449E+04 | 1261 | - | - | 0 | - |
| - | - | 1.602E+04 | 1262 | - | - | 0 | - |
| - | - | 6301 | 1263 | - | - | 0 | - |
| - | - | 1517 | 1264 | - | - | 0 | - |
| - | - | 920.2 | 1265 | - | - | 0 | - |
| - | - | 822.4 | 1275 | - | - | 0 | - |
| 4 | y | 8693 | 1276 | 0.001327 | 1.04 | +1 | 12 |
| - | - | 5407 | 1277 | - | - | 0 | - |
| - | - | 2197 | 1278 | - | - | 0 | - |
| - | - | 1104 | 1279 | - | - | 0 | - |
| - | - | 980.2 | 1283 | - | - | 0 | - |
| - | - | 901.8 | 1284 | - | - | 0 | - |
| - | - | 5438 | 1301 | - | - | 0 | - |
| - | - | 3681 | 1302 | - | - | 0 | - |
| - | - | 2458 | 1303 | - | - | 0 | - |
| - | - | 1020 | 1304 | - | - | 0 | - |
| - | - | 916.8 | 1305 | - | - | 0 | - |
| - | - | 915.1 | 1320 | - | - | 0 | - |
| 3 | z | 1138 | 1330 | 0.01973 | 14.84 | +1 | 13 |
| - | - | 2178 | 1331 | - | - | 0 | - |
| - | - | 1709 | 1332 | - | - | 0 | - |
| 13 | c | 4.922E+04 | 1345 | 0.002663 | 1.981 | +1 | 13 |
| - | - | 3.622E+04 | 1346 | - | - | 0 | - |
| - | - | 1.407E+04 | 1347 | - | - | 0 | - |
| - | - | 2.174E+04 | 1348 | - | - | 0 | - |
| - | - | 1.424E+04 | 1349 | - | - | 0 | - |
| - | - | 4953 | 1350 | - | - | 0 | - |
| - | - | 1451 | 1351 | - | - | 0 | - |
| - | - | 1594 | 1364 | - | - | 0 | - |
| - | - | 1002 | 1365 | - | - | 0 | - |
| - | - | 4106 | 1388 | - | - | 0 | - |
| - | - | 3396 | 1389 | - | - | 0 | - |
| - | - | 2161 | 1390 | - | - | 0 | - |
| - | - | 1044 | 1431 | - | - | 0 | - |
| 14 | c | 7.494E+04 | 1432 | 0.002953 | 2.063 | +1 | 14 |
| - | - | 5.621E+04 | 1433 | - | - | 0 | - |
| - | - | 2.493E+04 | 1434 | - | - | 0 | - |
| - | - | 6181 | 1435 | - | - | 0 | - |
| - | - | 1832 | 1436 | - | - | 0 | - |
| - | - | 3400 | 1461 | - | - | 0 | - |
| - | - | 2532 | 1462 | - | - | 0 | - |
| - | - | 1409 | 1463 | - | - | 0 | - |
| - | - | 1004 | 1474 | - | - | 0 | - |
| - | - | 5963 | 1489 | - | - | 0 | - |
| - | - | 4877 | 1490 | - | - | 0 | - |
| - | - | 2947 | 1491 | - | - | 0 | - |
| - | - | 1209 | 1492 | - | - | 0 | - |
| - | - | 1041 | 1502 | - | - | 0 | - |
| - | - | 1299 | 1513 | - | - | 0 | - |
| - | - | 1627 | 1514 | - | - | 0 | - |
| - | - | 1390 | 1515 | - | - | 0 | - |
| - | - | 890.7 | 1516 | - | - | 0 | - |
| - | - | 723.8 | 1518 | - | - | 0 | - |
| - | - | 1510 | 1519 | - | - | 0 | - |
| - | - | 981.9 | 1520 | - | - | 0 | - |
| - | - | 1236 | 1530 | - | - | 0 | - |
| - | - | 1.289E+04 | 1531 | - | - | 0 | - |
| - | - | 1.928E+04 | 1532 | - | - | 0 | - |
| - | - | 1.18E+04 | 1533 | - | - | 0 | - |
| - | - | 4261 | 1534 | - | - | 0 | - |
| - | - | 1605 | 1535 | - | - | 0 | - |
| - | - | 976.9 | 1544 | - | - | 0 | - |
| - | - | 3735 | 1545 | - | - | 0 | - |
| - | - | 6985 | 1546 | - | - | 0 | - |
| - | - | 6890 | 1547 | - | - | 0 | - |
| - | - | 5854 | 1548 | - | - | 0 | - |
| - | - | 3926 | 1549 | - | - | 0 | - |
| - | - | 1833 | 1550 | - | - | 0 | - |
| - | - | 2586 | 1556 | - | - | 0 | - |
| - | - | 5443 | 1557 | - | - | 0 | - |
| - | - | 5423 | 1558 | - | - | 0 | - |
| - | - | 1898 | 1559 | - | - | 0 | - |
| - | - | 2218 | 1562 | - | - | 0 | - |
| - | - | 1.051E+04 | 1563 | - | - | 0 | - |
| - | - | 8743 | 1564 | - | - | 0 | - |
| - | - | 3989 | 1565 | - | - | 0 | - |
| - | - | 1.397E+04 | 1573 | - | - | 0 | - |
| - | - | 9.316E+04 | 1574 | - | - | 0 | - |
| - | - | 7.543E+04 | 1575 | - | - | 0 | - |
| - | - | 3.772E+04 | 1576 | - | - | 0 | - |
| - | - | 1.071E+04 | 1577 | - | - | 0 | - |
| - | - | 1877 | 1578 | - | - | 0 | - |
| - | - | 4085 | 1589 | - | - | 0 | - |
| - | - | 810.4 | 1590 | - | - | 0 | - |
| - | - | 2.563E+04 | 1590 | - | - | 0 | - |
| - | - | 8.949E+04 | 1591 | - | - | 0 | - |
| - | - | 6.932E+04 | 1592 | - | - | 0 | - |
| - | - | 3.148E+04 | 1593 | - | - | 0 | - |
| - | - | 1.081E+04 | 1594 | - | - | 0 | - |
| - | - | 2495 | 1595 | - | - | 0 | - |
| - | - | 781 | 1627 | - | - | 0 | - |
| - | - | 776.5 | 3039 | - | - | 0 | - |

m/z Charge Intensity FragmentType MassShift Position
125.10738372802734 0 1568.4888
126.09146881103516 0 3450.8447
129.10247802734375 0 938.1266
131.12928771972656 0 730.1977
153.0661163330078 0 900.117
158.09249877929688 0 1236.8148 y Ammonia loss 14
168.11322021484375 0 4519.8145
168.7649688720703 0 515.29834
173.0921630859375 0 11964.966
173.4510955810547 0 2828.051
175.1190948486328 0 6776.712 y 14
176.92852783203125 0 502.50052
181.0609588623047 0 1025.7538
185.13970947265625 0 16043.458
186.13504028320312 0 605.7132
186.1433563232422 0 944.1761
187.10781860351562 0 1043.6803
199.07127380371094 0 875.6585
199.18052673339844 0 42023.59
200.1268310546875 0 3122.6716
200.18385314941406 0 5177.6406
214.4254150390625 0 543.16376
215.1390380859375 0 1316.8606
216.1455535888672 0 550.7378
217.08175659179688 0 1707.6464
226.1679229736328 0 594.7091
227.17550659179688 0 20867.14
228.17886352539062 0 2151.247
229.12945556640625 0 2752.472 w 13
244.1173858642578 0 1142.4751
245.12451171875 0 29185.117 y Ammonia loss 13
246.12405395507812 0 4462.3486
254.91415405273438 0 545.25854
255.1085205078125 0 1568.8984
262.15081787109375 0 4815.2725 y 13
270.1195373535156 0 3967.5872
272.1720886230469 0 930.0793
274.1183776855469 0 8860.011
287.3819580078125 0 529.82776
289.1034240722656 0 816.5912
303.152099609375 0 671.7638
314.2074890136719 0 2235.4983
316.16162109375 0 14549.874 w 12
317.1611328125 0 2357.032
324.1300964355469 0 1031.1957
330.1652526855469 0 630.2033
331.14892578125 0 3519.6047
331.2340393066406 0 702.4093 c 2
332.15643310546875 0 4398.8813 y Ammonia loss 12
333.16424560546875 0 10933.246 z 12
334.16668701171875 0 1951.8721
347.1687316894531 0 791.5365
349.1830749511719 0 3673.9573 y 12
350.1864929199219 0 1142.131
357.151611328125 0 7781.495
361.17132568359375 0 787.3688
414.1849365234375 0 750.69867
414.22100830078125 0 581.94305 c Ammonia loss 11
415.2267761230469 0 3355.9346 z 7
415.5655822753906 0 644.0269
415.7245178222656 0 1058.7008
416.23126220703125 0 735.2697
417.2095947265625 0 10567.253 w 11
418.2115478515625 0 1849.4418
423.2325439453125 0 3969.638 y 7
423.7304382324219 0 638.3172
425.2412414550781 0 950.0283
427.1716003417969 0 1616.4164
429.2349853515625 0 604.5596
432.23199462890625 0 879.8323 z 11
437.1829528808594 0 914.9685
442.2420959472656 0 1654.7517
443.18109130859375 0 753.31305
443.2495422363281 0 1692.0586
443.3111572265625 0 645.4127
444.2529296875 0 854.95447
446.23577880859375 0 1479.3229
447.24029541015625 0 818.9534
448.25164794921875 0 10815.209 y 11
449.2545471191406 0 2815.4436
450.7408142089844 0 576.5065 c Ammonia loss 7
453.6816711425781 0 593.436
454.8909912109375 0 943.4631 y 2
460.2151184082031 0 612.7955
460.2767639160156 0 8374.902 c 3
461.27960205078125 0 1692.6743
472.2513732910156 0 2015.7551 c Ammonia loss 13
472.7376708984375 0 1978.0396 z 6
473.23773193359375 0 1294.7625
474.2300109863281 0 2742.463
474.6532287597656 0 881.9302
475.185302734375 0 881.59644
475.23406982421875 0 772.1612
480.1572570800781 0 893.1834
480.6556091308594 0 902.9279
480.74658203125 0 1620.744 y 6
481.2467346191406 0 698.539 z Water loss 1
482.1999206542969 0 939.68414
482.6983337402344 0 1991.4065
483.69183349609375 0 1128.2678
484.1885986328125 0 2623.2573
484.6891174316406 0 1399.2162
485.2949523925781 0 885.02075
486.58544921875 0 701.3203 y Water loss 1
489.25360107421875 0 3236.824 z 10
489.65899658203125 0 3041.945
490.1605224609375 0 1239.6006
490.26116943359375 0 2650.2937
492.5853271484375 0 796.863 y 1
492.9202880859375 0 1200.3729
493.257080078125 0 979.623
493.31292724609375 0 1200.4205
494.25799560546875 0 568.21454
495.9167175292969 0 782.28326
499.2977294921875 0 2204.0564
501.2809143066406 0 709.6998
504.16943359375 0 1810.2518
504.20977783203125 0 1741.375
504.66680908203125 0 1614.8728
504.709228515625 0 2121.5518
505.27294921875 0 38412.367 y 10
506.27569580078125 0 9493.049
507.2791748046875 0 1832.4662
513.215087890625 0 14568.663
513.67236328125 0 1436.3118
513.7169189453125 0 5810.5317
514.171630859375 0 1001.8964
514.3226928710938 0 622.2419
515.243896484375 0 1145.4708
515.330078125 0 1395.7943
516.2526245117188 0 3667.8735 z 5
516.3375244140625 0 798.06976
516.7554321289062 0 4785.1914
517.25732421875 0 2772.4714
523.7582397460938 0 1043.0759
524.2669677734375 0 12428.214 y 5
524.6124267578125 0 2935.0603
524.7638549804688 0 7827.4927
524.9451904296875 0 2766.5608
525.2699584960938 0 1723.164
528.2466430664062 0 1042.0663
530.2809448242188 0 4043.661
530.615478515625 0 4488.4023
530.9503784179688 0 2862.5688
531.2904663085938 0 1015.27985
541.3114624023438 0 1163.0908
542.3177490234375 0 4782.4844
543.3203735351562 0 935.76605
558.2756958007812 0 1835.0183 z Water loss 9
558.3383178710938 0 1524.298
558.7772216796875 0 767.93353
559.291748046875 0 1027.15 w 9
559.3446655273438 0 6607.311 c 4
560.3469848632812 0 1443.992
565.787353515625 0 29012.703 z 4
566.2888793945312 0 17316.363
566.79052734375 0 6686.405
567.2909545898438 0 2889.4722
573.2919921875 0 1376.2649
573.7963256835938 0 16926.193 y 4
574.2977905273438 0 7926.6675 y Water loss 9
574.7999267578125 0 3180.804
575.2772216796875 0 1691.2424 y Ammonia loss 9
576.28564453125 0 5659.9014 z 9
577.28955078125 0 1959.5249
578.2982177734375 0 709.20056
580.2257080078125 0 4410.8555
581.2286987304688 0 1044.6646
584.3616333007812 0 680.0012
585.263427734375 0 844.82983
586.3295288085938 0 1235.8746
589.2470092773438 0 932.91473
592.2490844726562 0 897.38354
592.3004760742188 0 673.92566 y 9
594.2899169921875 0 1541.1074
600.273193359375 0 2661.1301
600.8021240234375 0 12111.169 w 3
601.3026733398438 0 5614.8613
601.804443359375 0 2846.1636
603.3740234375 0 925.3049
610.2581176757812 0 2153.983
621.4077758789062 0 1794.0784
629.3526611328125 0 1071.7848 c 11
630.3087158203125 0 31321.639 z 3
630.8102416992188 0 19256.95
631.3115234375 0 9091.548
631.8130493164062 0 3351.5132
632.31005859375 0 1055.4384
635.6661987304688 0 870.54517
637.8145141601562 0 7997.0312
638.3173217773438 0 30924.95 y 3
638.8190307617188 0 17681.434
639.3204345703125 0 6600.6616
639.823486328125 0 1556.8333
644.3173217773438 0 3474.3147
644.8195190429688 0 1960.292
645.3722534179688 0 1243.7543
646.3180541992188 0 1136.2106 w 8
646.3768920898438 0 27029.594 c 5
647.3233642578125 0 1156.0798
647.379638671875 0 8877.367
648.3823852539062 0 1708.1693
651.8362426757812 0 822.285
665.3234252929688 0 2143.3027 z Ammonia loss 2
665.8258056640625 0 1817.6744
672.8280029296875 0 2630.243 y Water loss 2
673.3328247070312 0 3148.94 y Ammonia loss 2
673.8248291015625 0 20104.438 z 2
674.3267822265625 0 23157.434
674.8286743164062 0 14125.668
675.3297729492188 0 5571.2905
675.8307495117188 0 1412.8756
677.3447875976562 0 748.8906
678.812255859375 0 789.8672
681.3303833007812 0 1421.6832
681.8338012695312 0 133510.77 y 2
682.3350830078125 0 89514.805
682.836669921875 0 35609.336
683.3380126953125 0 10042.49
683.8382568359375 0 2536.6262
686.3195190429688 0 686.90796
686.82763671875 0 822.05457
687.3402099609375 0 1563.2306
689.3541870117188 0 2693.7617 y 8
690.3572387695312 0 1104.3938
691.4456787109375 0 677.5449
695.84228515625 0 1605.9048
696.3458251953125 0 1007.54193
699.3788452148438 0 804.0367
701.0196533203125 0 725.25757
701.6929321289062 0 733.9875
702.2989501953125 0 2754.9583
706.6990356445312 0 2144.651
707.0341796875 0 1790.4775
707.3690795898438 0 722.6246 c Water loss 13
707.7022705078125 0 717.8996
715.8477783203125 0 3839.7556 w 1
716.349609375 0 2608.4226
716.8494262695312 0 1358.6027
717.3886108398438 0 2142.1458
718.3965454101562 0 3833.3508
719.325927734375 0 9925.253
719.3960571289062 0 1297.9308
720.328125 0 2556.8313
728.4151000976562 0 809.6755
729.364013671875 0 1899.5079 y Water loss 1
730.3665161132812 0 2186.7625 z 1
730.865478515625 0 1312.2023
734.3001098632812 0 1513.2764
735.2891235351562 0 747.6455
735.4552001953125 0 1113.6752
738.3751831054688 0 8075.724 y 1
738.8770141601562 0 6729.276
739.3777465820312 0 2343.3313
743.3680419921875 0 3092.3115 w 7
743.8668212890625 0 1424.3796
744.3726196289062 0 1644.1642
745.3778076171875 0 937.9916
750.89794921875 0 2325.1501
751.400634765625 0 2196.3123
752.3119506835938 0 3865.6736
756.4103393554688 0 821.42
758.3856811523438 0 3183.871
758.8812255859375 0 4159.247
759.3847045898438 0 2988.2168
759.8897094726562 0 1972.3755
760.39306640625 0 3178.2307
760.8919677734375 0 808.52014
761.4036254882812 0 52026.49 c 6
762.28759765625 0 760.72296
762.4061889648438 0 18440.355
763.4086303710938 0 5214.568
764.4094848632812 0 2392.979
764.9129638671875 0 3625.0466
765.3817138671875 0 88942.83
765.8829956054688 0 70595.22
766.3845825195312 0 30922.068
766.8861694335938 0 9046.835
767.3878173828125 0 3613.6052
770.3207397460938 0 2938.1694
771.3261108398438 0 888.6112
771.4016723632812 0 853.42053
772.4149169921875 0 2417.3337
772.91259765625 0 1923.9435
773.4143676757812 0 2851.3962
773.9071655273438 0 27468.035
774.408203125 0 18141.354
774.91015625 0 11858.033
775.408203125 0 2864.7393
777.897216796875 0 2503.3247
778.4039306640625 0 2488.0562
778.90576171875 0 1671.2368
779.4019165039062 0 1395.2175
786.41455078125 0 5228.3496
786.9088134765625 0 92504.914
787.4099731445312 0 75577.32
787.9112548828125 0 33049.668
788.411865234375 0 10319.176
788.9129638671875 0 4480.009
794.9174194335938 0 58001.54
795.419677734375 0 70939.516
795.9216918945312 0 41915.31
796.42333984375 0 16531.076
796.9237670898438 0 3916.2224
797.4258422851562 0 825.8409
811.4307861328125 0 1980.6699 z Water loss 7
813.3938598632812 0 1551.9149
817.4027099609375 0 767.73627
822.520263671875 0 1100.223
827.3583984375 0 2801.804
828.3551635742188 0 1706.3801
828.4995727539062 0 1221.2994
829.4393920898438 0 40477.086 z 7
830.4429931640625 0 21715.643
831.4465942382812 0 7025.5923
832.4481811523438 0 1858.7861
844.44921875 0 2548.7407
845.458740234375 0 6817.2007 y 7
846.460693359375 0 3129.3691
847.11962890625 0 874.4123
847.4628295898438 0 963.9027
853.33740234375 0 11125.829
854.3390502929688 0 4157.7627
872.507080078125 0 1836.306
873.4934692382812 0 2917.7476
873.947021484375 0 1173.0905
874.4926147460938 0 811.56396
874.950439453125 0 876.78894
900.4750366210938 0 4395.368 c Ammonia loss 7
901.4801635742188 0 2970.2896
902.4793090820312 0 1035.0527
915.389404296875 0 1086.112
926.4570922851562 0 1797.5533 z Water loss 6
927.4599609375 0 987.6091
930.3970336914062 0 2805.8613
931.3970336914062 0 1225.1864
934.4778442382812 0 895.80347
944.466064453125 0 16177.117 z 6
945.4710083007812 0 11708.11
946.4727172851562 0 3929.9414
947.4757690429688 0 1334.2307
949.3782348632812 0 642.669
951.9973754882812 0 1961.9755
952.4949340820312 0 1327.1458
952.9955444335938 0 1047.8696
960.4865112304688 0 1689.2567 y 6
961.488037109375 0 1574.827
965.3959350585938 0 1317.9736
966.3927612304688 0 973.99304
967.3736572265625 0 691.7861
968.376953125 0 4011.8003
969.381103515625 0 3436.4797
970.3817138671875 0 1490.4875
970.5437622070312 0 4406.9243
971.5477905273438 0 1611.805
972.4693603515625 0 796.6079
985.42578125 0 1928.5139
986.4249877929688 0 738.08844
988.5135498046875 0 1091.2253
995.3123779296875 0 2030.237
996.3110961914062 0 922.8828
1001.5599365234375 0 1170.2943
1003.437255859375 0 1912.1763
1005.314697265625 0 832.6456
1006.3170166015625 0 1154.1039
1007.32275390625 0 1434.8098
1008.3126220703125 0 933.7419
1009.4010009765625 0 1993.1875
1012.3189697265625 0 3688.4407
1013.3262939453125 0 2357.5059
1013.5919799804688 0 916.08105
1014.557373046875 0 35626.38 c 8
1015.559814453125 0 20384.203
1016.016845703125 0 1543.7651
1016.561279296875 0 6342.6577
1017.0171508789062 0 1303.8862
1017.5648193359375 0 1654.0485
1025.322021484375 0 4668.392
1026.3172607421875 0 3878.16
1026.4305419921875 0 11484.791
1027.3101806640625 0 1000.46704
1027.4207763671875 0 13719.041
1028.4169921875 0 3821.4695
1030.5029296875 0 843.6909 y Ammonia loss 5
1031.4984130859375 0 5119.851 z 5
1032.504150390625 0 9054.347
1033.5078125 0 3126.3826
1038.0423583984375 0 3179.5444
1038.5369873046875 0 1542.0273
1039.5626220703125 0 1867.6863
1040.570068359375 0 2003.1267
1042.570556640625 0 788.8073
1043.322998046875 0 1086.0977
1043.4373779296875 0 3278.4497
1044.32275390625 0 1079.0563
1044.44189453125 0 6348.3315
1045.4268798828125 0 14652.857
1046.4271240234375 0 5331.7144
1047.5140380859375 0 3114.0771 y 5
1048.51806640625 0 1230.4216
1051.5372314453125 0 1999.5154
1052.0369873046875 0 1145.2203
1052.541748046875 0 1429.3517
1057.5758056640625 0 23784.75
1058.57763671875 0 14200.224
1059.5791015625 0 5208.1772
1060.0533447265625 0 1752.3068
1060.5570068359375 0 2122.5698
1061.447509765625 0 5017.1206
1062.4505615234375 0 24408.498
1063.4542236328125 0 11444.64
1063.6397705078125 0 1103.823
1100.5810546875 0 1379.5464
1101.5889892578125 0 51745.258 c 9
1102.5919189453125 0 29581.256
1103.59423828125 0 10623.6045
1104.595458984375 0 1859.325
1114.5941162109375 0 7837.139
1115.6005859375 0 5913.141
1116.603271484375 0 2157.9783
1117.540283203125 0 1699.9814
1118.5509033203125 0 963.37695
1119.5316162109375 0 907.0573
1130.5684814453125 0 6740.4326 z 4
1131.5736083984375 0 16403.129
1132.5762939453125 0 9848.937
1133.5780029296875 0 2993.7864
1146.58544921875 0 2100.1245 y 4
1147.5855712890625 0 2121.8997
1158.6099853515625 0 27083.996 c 10
1159.613037109375 0 18817.879
1160.6165771484375 0 5857.8525
1161.61865234375 0 1704.9507
1200.6134033203125 0 918.5362 w 3
1213.6634521484375 0 5001.1553
1214.66650390625 0 3721.6382
1215.662841796875 0 1915.9615
1216.671630859375 0 823.5674
1217.595703125 0 1079.0527
1247.577880859375 0 1552.3046
1248.564208984375 0 805.0147
1257.677734375 0 29544.076 c 11
1258.6785888671875 0 18552.111
1259.6741943359375 0 6625.131
1260.6163330078125 0 24486.883
1261.619140625 0 16015.599
1262.6177978515625 0 6301.0376
1263.624755859375 0 1517.3538
1264.621826171875 0 920.19586
1274.62353515625 0 822.36334
1275.62744140625 0 8692.838 y 3
1276.6297607421875 0 5406.538
1277.63134765625 0 2196.7993
1278.634033203125 0 1103.9565
1282.6856689453125 0 980.151
1283.700927734375 0 901.8477
1300.69384765625 0 5438.4077
1301.6959228515625 0 3681.3828
1302.7025146484375 0 2457.7532
1303.708740234375 0 1020.1086
1304.6292724609375 0 916.78174
1319.6544189453125 0 915.1433
1329.63525390625 0 1138.2103 z Ammonia loss 2
1330.63330078125 0 2177.7798
1331.6329345703125 0 1709.1461
1344.7091064453125 0 49215.844 c 12
1345.7120361328125 0 36220.676
1346.707275390625 0 14065.6875
1347.652099609375 0 21738.2
1348.650634765625 0 14238.753
1349.6533203125 0 4952.926
1350.65185546875 0 1451.4701
1363.6646728515625 0 1594.0862
1364.667236328125 0 1001.67145
1387.72705078125 0 4105.523
1388.7313232421875 0 3395.8508
1389.7388916015625 0 2160.7866
1430.73681640625 0 1043.947
1431.7408447265625 0 74943.97 c 13
1432.742919921875 0 56207.246
1433.7452392578125 0 24930.371
1434.7457275390625 0 6181.1157
1435.759521484375 0 1831.6123
1460.7318115234375 0 3399.891
1461.733154296875 0 2532.461
1462.7340087890625 0 1409.2823
1473.7430419921875 0 1004.2911
1488.7374267578125 0 5963.0317
1489.7418212890625 0 4876.606
1490.74755859375 0 2947.3542
1491.754150390625 0 1208.8201
1501.7791748046875 0 1040.6367
1512.7867431640625 0 1299.0686
1513.7926025390625 0 1627.447
1514.774658203125 0 1389.692
1515.8096923828125 0 890.7043
1517.7750244140625 0 723.7816
1518.7757568359375 0 1510.2284
1519.7757568359375 0 981.925
1529.8045654296875 0 1235.9534
1530.7864990234375 0 12892.011
1531.791748046875 0 19281.207
1532.79443359375 0 11804.729
1533.79931640625 0 4261.415
1534.7969970703125 0 1605.2643
1543.8203125 0 976.8992
1544.8304443359375 0 3734.567
1545.823486328125 0 6985.2153
1546.8240966796875 0 6890.374
1547.817138671875 0 5853.704
1548.8140869140625 0 3925.569
1549.8251953125 0 1832.9495
1555.80419921875 0 2586.2507
1556.7921142578125 0 5443.1885
1557.792236328125 0 5422.837
1558.797119140625 0 1898.2107
1561.8326416015625 0 2217.563
1562.8455810546875 0 10512.726
1563.846923828125 0 8742.87
1564.854736328125 0 3989.388
1572.8125 0 13965.78
1573.8143310546875 0 93162.195
1574.8173828125 0 75426.625
1575.8192138671875 0 37724.85
1576.8231201171875 0 10709.628
1577.8292236328125 0 1877.1897
1588.820556640625 0 4084.537
1589.5927734375 0 810.3841
1589.83056640625 0 25625.34
1590.8388671875 0 89489.93
1591.8414306640625 0 69318.41
1592.8450927734375 0 31482.693
1593.8477783203125 0 10814.283
1594.84765625 0 2494.5132
1626.7740478515625 0 781.00226
3038.791259765625 0 776.4593

Spectrum Details

|  |  |
| --- | --- |
| Matched peaks? Matched peaks The total absolute number of peaks matched. Additionally in brackets the total fraction of peaks matched and the total number of peaks is shown. | 78 (14.55% of 536) |
| FDR? FDR The false discovery rate estimated for this peptide. It is calculated by matching all theoretical fragments with a non-integer shift with the raw peaks for this spectrum. This is done with 40 different shifts. The resulting percentage is the average number of annotated peaks over the number of annotated peaks with the correct spectrum. | 1.83% |
| Satellite FDR? Satellite FDR See the FDR for details on its calculation. This satellite ion specific FDR only contains the satellite ions (d/w) for I/L/J positions. | 0.00% |
| PSM Score? PSM Score The PSM Score as given by Hecklib to this annotated spectrum. It is shown with three significant figures. | 564 |

#### Spectrum 3560? Spectrum 3560 The raw spectrum of this peptide as annotated by Hecklib. The fragments are coloured according to ion type (see legend). Any peaks with a star '\*' as text can be hovered over to see the full details, first the ion type second the mass shift type. By hovering over the amino acids in the peptide or ions in the legend the corresponding peaks are highlighted. By toggling the 'Unassigned' label you can turn the background (unassigned) peaks on or off in the plot. By updating the slider in the Ion legend you can update the spectrum to only show the top X% of the peaks with labels. The top X% means any peak that is within X% of the highest intensity. By dragging in the spectrum you can zoom in to a specific part of the spectrum and use 'Zoom Out' to get back to the original zoom level. The annotation of the spectrum is based on the given sequence in the peptides file and is done with different software so inconsistencies are likely. The peaks are annotated based on the given sequence, with 20 ppm tolerance.

Copy Data

##### Spectrum 3560 (TSV)

###### Preview

```
Loading example...
```

*Click on the button to copy the data to your clipboard.*

Mz MinMz MaxIntensity Max

WidthHeightPeptide font sizePeptide stroke widthSpectrum font sizeSpectrum stroke widthCompact peptide

Ion legend

wxyz

abcd

OtherUnassignedIon^Charge^_Position_Show for top:%

JJSEVSDRPSGVSSR

03.93e+37.87e+31.18e+41.57e+4

Zoom Out

y^+1^_1_y^+1^_2_w^+1^_3_z^+1^_3_z^+2^_8_w^+1^_4_y^+2^_8_y^+1^_4_y^+3^_13_c^+1^_4_c^+3^_14_y^+1^_5_z^+2^_10_y^+2^_10_c^+1^_5_z^+2^_11_y^+2^_11_y^+1^_6_z^+1^_6_w^+2^_12_z^+2^_12_y^+2^_12_c^+1^_6_z^+1^_7_z^+2^_13_y^+2^_13_y^+2^_14_y^+2^_14_c^+1^_7_z^+1^_8_z^+1^_9_c^+1^_9_z^+1^_10_c^+1^_10_z^+1^_11_c^+1^_11_c^+1^_12_y^+1^_12_c^+1^_13_c^+1^_14_

0879175826373516

Fragment Matches Table

Show background peaks

| Position | Ion type | Intensity | mz Theoretical | mz Error (Th) | mz Error (ppm) | Charge | Series Number |
| --- | --- | --- | --- | --- | --- | --- | --- |
| - | - | 478.8 | 129.1 | - | - | 0 | - |
| - | - | 385.5 | 130 | - | - | 0 | - |
| - | - | 436.1 | 134.9 | - | - | 0 | - |
| - | - | 433.6 | 135.8 | - | - | 0 | - |
| - | - | 968.3 | 149 | - | - | 0 | - |
| - | - | 997 | 164.1 | - | - | 0 | - |
| - | - | 497.5 | 164.3 | - | - | 0 | - |
| - | - | 581.7 | 168.1 | - | - | 0 | - |
| - | - | 457.9 | 169.2 | - | - | 0 | - |
| - | - | 459.5 | 172.3 | - | - | 0 | - |
| 15 | y | 976.2 | 175.1 | 2.061E-05 | 0.1177 | +1 | 1 |
| - | - | 2278 | 185.1 | - | - | 0 | - |
| - | - | 6081 | 199.2 | - | - | 0 | - |
| - | - | 729.7 | 200.2 | - | - | 0 | - |
| - | - | 1642 | 203.1 | - | - | 0 | - |
| - | - | 500.9 | 213.2 | - | - | 0 | - |
| - | - | 1791 | 215.1 | - | - | 0 | - |
| - | - | 2556 | 227.2 | - | - | 0 | - |
| 14 | y | 3795 | 245.1 | 0.0002023 | 0.8253 | +1 | 2 |
| - | - | 586.1 | 289.3 | - | - | 0 | - |
| 13 | w | 1794 | 316.2 | 0.0002599 | 0.8219 | +1 | 3 |
| - | - | 549.8 | 331.1 | - | - | 0 | - |
| 13 | z | 1244 | 333.2 | 0.0002529 | 0.7592 | +1 | 3 |
| - | - | 702.4 | 343.2 | - | - | 0 | - |
| - | - | 486.7 | 345.8 | - | - | 0 | - |
| - | - | 663.6 | 357.2 | - | - | 0 | - |
| - | - | 2013 | 361.2 | - | - | 0 | - |
| - | - | 555.7 | 370.1 | - | - | 0 | - |
| - | - | 544.4 | 392.9 | - | - | 0 | - |
| - | - | 559.3 | 410.4 | - | - | 0 | - |
| 8 | z | 864.4 | 415.2 | 0.0009077 | 2.186 | +2 | 8 |
| 12 | w | 1627 | 417.2 | 0.0007982 | 1.913 | +1 | 4 |
| 8 | y | 555.2 | 423.2 | 0.004058 | 9.588 | +2 | 8 |
| 12 | y | 1181 | 448.3 | 0.0008659 | 1.932 | +1 | 4 |
| - | - | 617 | 449.3 | - | - | 0 | - |
| 3 | y | 569.3 | 454.9 | 0.001832 | 4.028 | +3 | 13 |
| 4 | c | 1266 | 460.3 | 0.002161 | 4.695 | +1 | 4 |
| 14 | c | 662.4 | 472.2 | 0.006127 | 12.97 | +3 | 14 |
| - | - | 621.3 | 483.2 | - | - | 0 | - |
| - | - | 697.3 | 503.3 | - | - | 0 | - |
| 11 | y | 4705 | 505.3 | 0.000734 | 1.453 | +1 | 5 |
| - | - | 1531 | 506.3 | - | - | 0 | - |
| - | - | 695.1 | 511.3 | - | - | 0 | - |
| - | - | 662.1 | 515.3 | - | - | 0 | - |
| 6 | z | 648.6 | 516.3 | 7.799E-05 | 0.1511 | +2 | 10 |
| 6 | y | 2638 | 524.3 | 0.001519 | 2.898 | +2 | 10 |
| - | - | 1197 | 524.6 | - | - | 0 | - |
| - | - | 858.4 | 529.8 | - | - | 0 | - |
| - | - | 1206 | 530.3 | - | - | 0 | - |
| - | - | 3186 | 530.3 | - | - | 0 | - |
| - | - | 805.9 | 530.6 | - | - | 0 | - |
| - | - | 780.8 | 530.8 | - | - | 0 | - |
| - | - | 1465 | 530.8 | - | - | 0 | - |
| - | - | 1028 | 531.2 | - | - | 0 | - |
| 5 | c | 784.6 | 559.3 | 0.002606 | 4.66 | +1 | 5 |
| 5 | z | 3679 | 565.8 | 0.0006817 | 1.205 | +2 | 11 |
| - | - | 2355 | 566.3 | - | - | 0 | - |
| - | - | 839.8 | 566.8 | - | - | 0 | - |
| - | - | 711.1 | 570.3 | - | - | 0 | - |
| 5 | y | 1848 | 573.8 | 0.0009426 | 1.643 | +2 | 11 |
| 10 | y | 1951 | 574.3 | 0.003074 | 5.353 | +1 | 6 |
| - | - | 779.4 | 574.8 | - | - | 0 | - |
| 10 | z | 616.3 | 576.3 | 0.001102 | 1.911 | +1 | 6 |
| 4 | w | 1548 | 600.8 | 0.002008 | 3.342 | +2 | 12 |
| - | - | 659.3 | 607.3 | - | - | 0 | - |
| 4 | z | 4537 | 630.3 | 0.0009099 | 1.444 | +2 | 12 |
| - | - | 2631 | 630.8 | - | - | 0 | - |
| - | - | 2082 | 631.3 | - | - | 0 | - |
| - | - | 617 | 631.8 | - | - | 0 | - |
| - | - | 1616 | 637.8 | - | - | 0 | - |
| 4 | y | 3555 | 638.3 | 0.001189 | 1.863 | +2 | 12 |
| - | - | 1731 | 638.8 | - | - | 0 | - |
| - | - | 1270 | 639.3 | - | - | 0 | - |
| - | - | 726.4 | 642.8 | - | - | 0 | - |
| 6 | c | 3116 | 646.4 | 0.0001799 | 0.2783 | +1 | 6 |
| - | - | 962.2 | 647.4 | - | - | 0 | - |
| - | - | 740.5 | 671.3 | - | - | 0 | - |
| 9 | z | 923.4 | 673.3 | 0.0005838 | 0.867 | +1 | 7 |
| 3 | z | 1382 | 673.8 | 0.0009441 | 1.401 | +2 | 13 |
| - | - | 4053 | 674.3 | - | - | 0 | - |
| - | - | 1848 | 674.8 | - | - | 0 | - |
| 3 | y | 1.558E+04 | 681.8 | 5.224E-05 | 0.07662 | +2 | 13 |
| - | - | 1.137E+04 | 682.3 | - | - | 0 | - |
| - | - | 4083 | 682.8 | - | - | 0 | - |
| - | - | 830.9 | 700.3 | - | - | 0 | - |
| 2 | y | 679.9 | 729.4 | 0.0003685 | 0.5053 | +2 | 14 |
| 2 | y | 1324 | 738.4 | 0.001191 | 1.613 | +2 | 14 |
| 7 | c | 6760 | 761.4 | 9.235E-05 | 0.1213 | +1 | 7 |
| - | - | 2270 | 762.4 | - | - | 0 | - |
| - | - | 749.5 | 763.4 | - | - | 0 | - |
| - | - | 1537 | 764.9 | - | - | 0 | - |
| - | - | 9507 | 765.4 | - | - | 0 | - |
| - | - | 9586 | 765.9 | - | - | 0 | - |
| - | - | 4352 | 766.4 | - | - | 0 | - |
| - | - | 1687 | 766.9 | - | - | 0 | - |
| - | - | 1363 | 773.4 | - | - | 0 | - |
| - | - | 3072 | 773.9 | - | - | 0 | - |
| - | - | 2131 | 774.4 | - | - | 0 | - |
| - | - | 940.5 | 774.9 | - | - | 0 | - |
| - | - | 723.4 | 785.9 | - | - | 0 | - |
| - | - | 684.7 | 786.4 | - | - | 0 | - |
| - | - | 1.154E+04 | 786.9 | - | - | 0 | - |
| - | - | 9944 | 787.4 | - | - | 0 | - |
| - | - | 5085 | 787.9 | - | - | 0 | - |
| - | - | 8586 | 794.9 | - | - | 0 | - |
| - | - | 9089 | 795.4 | - | - | 0 | - |
| - | - | 4841 | 795.9 | - | - | 0 | - |
| - | - | 1834 | 796.4 | - | - | 0 | - |
| - | - | 1042 | 816.4 | - | - | 0 | - |
| 8 | z | 6141 | 829.4 | 0.0006738 | 0.8123 | +1 | 8 |
| - | - | 2439 | 830.4 | - | - | 0 | - |
| - | - | 738.5 | 844.4 | - | - | 0 | - |
| - | - | 649.2 | 872.4 | - | - | 0 | - |
| 7 | z | 1993 | 944.5 | 0.002351 | 2.49 | +1 | 9 |
| - | - | 1579 | 945.5 | - | - | 0 | - |
| - | - | 2221 | 967.6 | - | - | 0 | - |
| - | - | 1788 | 968.6 | - | - | 0 | - |
| - | - | 1387 | 972.5 | - | - | 0 | - |
| - | - | 627.5 | 973.5 | - | - | 0 | - |
| 9 | c | 4273 | 1015 | 0.0004622 | 0.4556 | +1 | 9 |
| - | - | 2501 | 1016 | - | - | 0 | - |
| - | - | 866.2 | 1016 | - | - | 0 | - |
| 6 | z | 720.6 | 1031 | 0.01478 | 14.33 | +1 | 10 |
| - | - | 1534 | 1032 | - | - | 0 | - |
| - | - | 623.8 | 1048 | - | - | 0 | - |
| - | - | 3719 | 1058 | - | - | 0 | - |
| - | - | 2551 | 1059 | - | - | 0 | - |
| - | - | 2640 | 1060 | - | - | 0 | - |
| - | - | 778.6 | 1060 | - | - | 0 | - |
| - | - | 5007 | 1061 | - | - | 0 | - |
| - | - | 3963 | 1062 | - | - | 0 | - |
| - | - | 2103 | 1062 | - | - | 0 | - |
| - | - | 1777 | 1063 | - | - | 0 | - |
| - | - | 721.1 | 1063 | - | - | 0 | - |
| - | - | 830.6 | 1064 | - | - | 0 | - |
| 10 | c | 6632 | 1102 | 0.001119 | 1.015 | +1 | 10 |
| - | - | 3064 | 1103 | - | - | 0 | - |
| - | - | 1049 | 1104 | - | - | 0 | - |
| - | - | 992.4 | 1115 | - | - | 0 | - |
| 5 | z | 825.3 | 1131 | 0.005669 | 5.014 | +1 | 11 |
| - | - | 2046 | 1132 | - | - | 0 | - |
| - | - | 1732 | 1133 | - | - | 0 | - |
| 11 | c | 4158 | 1159 | 0.001832 | 1.581 | +1 | 11 |
| - | - | 2246 | 1160 | - | - | 0 | - |
| 12 | c | 3383 | 1258 | 0.002495 | 1.984 | +1 | 12 |
| - | - | 2368 | 1259 | - | - | 0 | - |
| - | - | 3489 | 1261 | - | - | 0 | - |
| - | - | 2351 | 1262 | - | - | 0 | - |
| - | - | 1123 | 1263 | - | - | 0 | - |
| 4 | y | 1060 | 1276 | 0.002823 | 2.213 | +1 | 12 |
| - | - | 795 | 1277 | - | - | 0 | - |
| 13 | c | 5526 | 1345 | 0.002053 | 1.527 | +1 | 13 |
| - | - | 4158 | 1346 | - | - | 0 | - |
| - | - | 1219 | 1347 | - | - | 0 | - |
| - | - | 1858 | 1348 | - | - | 0 | - |
| - | - | 1403 | 1349 | - | - | 0 | - |
| - | - | 819.7 | 1350 | - | - | 0 | - |
| - | - | 1246 | 1389 | - | - | 0 | - |
| 14 | c | 8539 | 1432 | 0.003198 | 2.233 | +1 | 14 |
| - | - | 7528 | 1433 | - | - | 0 | - |
| - | - | 2789 | 1434 | - | - | 0 | - |
| - | - | 1255 | 1435 | - | - | 0 | - |
| - | - | 641.4 | 1451 | - | - | 0 | - |
| - | - | 674.8 | 1489 | - | - | 0 | - |
| - | - | 2023 | 1531 | - | - | 0 | - |
| - | - | 2866 | 1532 | - | - | 0 | - |
| - | - | 1936 | 1533 | - | - | 0 | - |
| - | - | 962 | 1534 | - | - | 0 | - |
| - | - | 1084 | 1546 | - | - | 0 | - |
| - | - | 839 | 1547 | - | - | 0 | - |
| - | - | 1891 | 1548 | - | - | 0 | - |
| - | - | 741.6 | 1549 | - | - | 0 | - |
| - | - | 1626 | 1557 | - | - | 0 | - |
| - | - | 869.8 | 1558 | - | - | 0 | - |
| - | - | 1482 | 1563 | - | - | 0 | - |
| - | - | 1228 | 1564 | - | - | 0 | - |
| - | - | 706.4 | 1566 | - | - | 0 | - |
| - | - | 2278 | 1573 | - | - | 0 | - |
| - | - | 1.45E+04 | 1574 | - | - | 0 | - |
| - | - | 1.029E+04 | 1575 | - | - | 0 | - |
| - | - | 5926 | 1576 | - | - | 0 | - |
| - | - | 1513 | 1577 | - | - | 0 | - |
| - | - | 934.5 | 1589 | - | - | 0 | - |
| - | - | 3380 | 1590 | - | - | 0 | - |
| - | - | 1.411E+04 | 1591 | - | - | 0 | - |
| - | - | 8634 | 1592 | - | - | 0 | - |
| - | - | 4899 | 1593 | - | - | 0 | - |
| - | - | 1060 | 1594 | - | - | 0 | - |
| - | - | 647.7 | 1967 | - | - | 0 | - |
| - | - | 681.5 | 2050 | - | - | 0 | - |
| - | - | 606.5 | 3037 | - | - | 0 | - |
| - | - | 685.5 | 3481 | - | - | 0 | - |

m/z Charge Intensity FragmentType MassShift Position
129.10195922851562 0 478.83307
130.0006866455078 0 385.53046
134.9178009033203 0 436.1257
135.7812957763672 0 433.62402
148.95449829101562 0 968.34247
164.1179656982422 0 997.0047
164.30177307128906 0 497.48352
168.11346435546875 0 581.6869
169.2152862548828 0 457.88846
172.30258178710938 0 459.49426
175.1189727783203 0 976.17487 y 14
185.13990783691406 0 2277.7854
199.1805877685547 0 6081.4326
200.1837921142578 0 729.68115
203.10264587402344 0 1641.6272
213.1859130859375 0 500.9085
215.1388397216797 0 1790.8346
227.17562866210938 0 2556.4126
245.1246337890625 0 3794.5898 y Ammonia loss 13
289.2535095214844 0 586.1357
316.1612854003906 0 1793.5049 w 12
331.1484375 0 549.75275
333.1640319824219 0 1244.005 z 12
343.1607360839844 0 702.415
345.8105163574219 0 486.68384
357.15118408203125 0 663.5643
361.1726989746094 0 2012.9432
370.13623046875 0 555.6663
392.9202880859375 0 544.41486
410.4120178222656 0 559.26215
415.2245788574219 0 864.3901 z 7
417.21002197265625 0 1627.3163 w 11
423.2370910644531 0 555.1768 y 7
448.2522888183594 0 1180.9352 y 11
449.2584228515625 0 617.0284
454.89361572265625 0 569.32025 y 2
460.2744140625 0 1265.7982 c 3
472.25006103515625 0 662.4198 c Ammonia loss 13
483.2267150878906 0 621.2677
503.2893371582031 0 697.25995
505.27362060546875 0 4704.693 y 10
506.27642822265625 0 1531.081
511.25921630859375 0 695.1214
515.3301391601562 0 662.11743
516.2532348632812 0 648.6427 z 5
524.2640380859375 0 2637.6047 y 5
524.60986328125 0 1197.1589
529.7613525390625 0 858.4167
530.2562255859375 0 1206.4462
530.2996826171875 0 3185.9917
530.6177368164062 0 805.92615
530.758056640625 0 780.8171
530.8021240234375 0 1464.9668
531.2481079101562 0 1027.7842
559.3475952148438 0 784.56555 c 4
565.7866821289062 0 3679.4976 z 4
566.2894287109375 0 2354.5022
566.7921752929688 0 839.82135
570.3255615234375 0 711.0771
573.7976684570312 0 1847.8738 y 4
574.2974243164062 0 1950.918 y Water loss 9
574.7976684570312 0 779.4279
576.2872924804688 0 616.30774 z 9
600.8040161132812 0 1548.2859 w 3
607.3009033203125 0 659.2949
630.3095703125 0 4536.709 z 3
630.8104248046875 0 2630.993
631.3126831054688 0 2081.5845
631.8125610351562 0 616.97534
637.8130493164062 0 1615.7256
638.3168334960938 0 3554.635 y 3
638.8223876953125 0 1730.5288
639.3217163085938 0 1269.5438
642.813232421875 0 726.3766
646.377197265625 0 3115.8682 c 5
647.380615234375 0 962.24023
671.3135986328125 0 740.5146
673.3395385742188 0 923.41693 z 8
673.82373046875 0 1382.2161 z 2
674.3286743164062 0 4053.2002
674.8282470703125 0 1848.2136
681.833984375 0 15581.415 y 2
682.335205078125 0 11369.85
682.8367309570312 0 4082.502
700.328369140625 0 830.8617
729.3711547851562 0 679.9495 y Water loss 1
738.3748779296875 0 1324.4302 y 1
761.404052734375 0 6759.5737 c 6
762.4071655273438 0 2270.287
763.4130859375 0 749.48553
764.9066162109375 0 1537.2213
765.38232421875 0 9507.314
765.8831176757812 0 9586.045
766.3848266601562 0 4352.098
766.8880004882812 0 1687.3966
773.3958740234375 0 1363.4363
773.9089965820312 0 3071.9067
774.4064331054688 0 2130.6045
774.9105834960938 0 940.46484
785.8853759765625 0 723.3542
786.4049072265625 0 684.68274
786.9083251953125 0 11537.676
787.4100341796875 0 9943.978
787.9107055664062 0 5085.3037
794.91796875 0 8586.007
795.41943359375 0 9089.455
795.9214477539062 0 4841.4214
796.4224243164062 0 1833.7234
816.4198608398438 0 1042.1061
829.4393920898438 0 6141.1973 z 7
830.4483032226562 0 2438.5925
844.4151611328125 0 738.5298
872.435546875 0 649.1789
944.4693603515625 0 1993.1367 z 6
945.4751586914062 0 1578.5908
967.5526733398438 0 2221.2327
968.558349609375 0 1788.4443
972.4769897460938 0 1386.5706
973.4817504882812 0 627.53827
1014.557373046875 0 4273.0664 c 8
1015.5590209960938 0 2500.7083
1016.4689331054688 0 866.20514
1031.4842529296875 0 720.5732 z 5
1032.4949951171875 0 1533.786
1047.5394287109375 0 623.7589
1057.5614013671875 0 3719.1758
1058.567626953125 0 2551.3816
1059.59814453125 0 2639.54
1060.49169921875 0 778.6274
1060.6025390625 0 5007.0273
1061.5006103515625 0 3962.978
1061.6043701171875 0 2103.3647
1062.509033203125 0 1777.2285
1062.9884033203125 0 721.1247
1063.5054931640625 0 830.6238
1101.5887451171875 0 6631.5933 c 9
1102.5938720703125 0 3063.8518
1103.5958251953125 0 1048.8059
1114.6031494140625 0 992.3511
1130.5731201171875 0 825.31177 z 4
1131.5751953125 0 2046.095
1132.578369140625 0 1732.365
1158.6131591796875 0 4157.7363 c 10
1159.6141357421875 0 2246.1438
1257.67724609375 0 3382.8655 c 11
1258.6790771484375 0 2368.3337
1260.61474609375 0 3488.6152
1261.6151123046875 0 2351.1538
1262.6220703125 0 1123.2953
1275.631591796875 0 1059.6816 y 3
1276.636474609375 0 795.0452
1344.709716796875 0 5525.6274 c 12
1345.7120361328125 0 4157.811
1346.726806640625 0 1219.0609
1347.6549072265625 0 1857.727
1348.650146484375 0 1402.6211
1349.656494140625 0 819.7205
1388.7357177734375 0 1245.8484
1431.7406005859375 0 8539.158 c 13
1432.7435302734375 0 7527.6875
1433.7421875 0 2789.4128
1434.7479248046875 0 1255.0898
1450.7763671875 0 641.41174
1488.736572265625 0 674.7597
1530.7828369140625 0 2022.8787
1531.7935791015625 0 2865.5193
1532.7958984375 0 1936.3882
1533.7900390625 0 961.9659
1545.8297119140625 0 1084.3577
1546.8331298828125 0 838.96655
1547.8062744140625 0 1890.5774
1548.8223876953125 0 741.58923
1556.78564453125 0 1625.9658
1557.794921875 0 869.7748
1562.8531494140625 0 1481.9385
1563.837158203125 0 1227.6919
1565.85986328125 0 706.36536
1572.7996826171875 0 2278.0652
1573.8143310546875 0 14498.3955
1574.8177490234375 0 10290.89
1575.81884765625 0 5925.8228
1576.8184814453125 0 1513.042
1588.8133544921875 0 934.48413
1589.8275146484375 0 3379.7095
1590.84033203125 0 14111.904
1591.8399658203125 0 8633.9
1592.844482421875 0 4898.8735
1593.8355712890625 0 1060.3881
1967.3155517578125 0 647.68097
2049.935546875 0 681.4555
3037.084716796875 0 606.45135
3481.486328125 0 685.495

Spectrum Details

|  |  |
| --- | --- |
| Matched peaks? Matched peaks The total absolute number of peaks matched. Additionally in brackets the total fraction of peaks matched and the total number of peaks is shown. | 40 (20.83% of 192) |
| FDR? FDR The false discovery rate estimated for this peptide. It is calculated by matching all theoretical fragments with a non-integer shift with the raw peaks for this spectrum. This is done with 40 different shifts. The resulting percentage is the average number of annotated peaks over the number of annotated peaks with the correct spectrum. | 1.25% |
| Satellite FDR? Satellite FDR See the FDR for details on its calculation. This satellite ion specific FDR only contains the satellite ions (d/w) for I/L/J positions. | ∞ |
| PSM Score? PSM Score The PSM Score as given by Hecklib to this annotated spectrum. It is shown with three significant figures. | 225 |

#### Reverse Lookup? Reverse Lookup All places where this read could be placed.

| Group | Segment | Template | Template Part | Read Part | Score | Unique |
| --- | --- | --- | --- | --- | --- | --- |
| Homo sapiens Light Chain | IGLV | IGLV2-14 | [49..63] | [0..15] | 88 | True |

| Recombined | Template Part | Read Part | Score | Unique |
| --- | --- | --- | --- | --- |
| REC-0-1\_002 | [48..63] | [0..15] | 120 | True |

#### Meta Information from Multiple reads

##### Number of combined reads

22

##### Intensity

0.9233

##### TotalArea

5.67E+08

##### Changes to the peptide sequence

JJSEVSDRPSGVSSR

I→JNo support for either Leucine or Isoleucine based on side chain ions (Position: 2)

L→ISupport for Isoleucine based on side chain ions (1 for I 0 for L) (Position: 2)

I→LSupport for Leucine based on side chain ions (1 for L 0 for I) (Position: 2)

J→ISupport for Isoleucine based on side chain ions (1 for I 0 for L) (Position: 2)

I→JNo support for either Leucine or Isoleucine based on side chain ions (Position: 2)

J→ISupport for Isoleucine based on side chain ions (1 for I 0 for L) (Position: 2)

I→JNo support for either Leucine or Isoleucine based on side chain ions (Position: 2)

L→ISupport for Isoleucine based on side chain ions (1 for I 0 for L) (Position: 2)

L→JNo support for either Leucine or Isoleucine based on side chain ions (Position: 1)

#### Positional Score

Copy Data

##### Positional Score (TSV)

###### Preview

```
Loading example...
```

*Click on the button to copy the data to your clipboard.*

1001234567891011121314

Label Value
"0" 0.909
"1" 0.908
"2" 0.905
"3" 0.902
"4" 0.896
"5" 0.894
"6" 0.904
"7" 0.876
"8" 0.866
"9" 0.887
"10" 0.844
"11" 0.841
"12" 0.881
"13" 0.898
"14" 0.894

#### Meta Information from PEAKS

##### Scan Identifier

F2:4095

##### Original sequence

L

L

S

E

V

S

D

R

P

S

G

V

S

S

R

##### Posttranslational Modifications

##### Source File

D:\separate\_stitch\_analyses\xle-disambiguation\raw\20210323\_F1\_UM1\_Peng0013\_SA\_F59\_ingel\_3ug\_TL.raw

##### Fraction

2

##### Scan Feature

-

##### De Novo Score

99

##### ConfidenceScore

99

### m/z

530.2823

##### Mass

1587.8215

##### Charge

3

##### Retention Time

21.76

##### Predicted Retention Time

-

##### Area

0

##### Parts Per Million

2.2

##### Fragmentation mode

ETHCD

##### Originating file

01 D:\separate\_stitch\_analyses\xle-disambiguation\20210325\_F59\_3ug\_DENOVO\_12.csv

#### Meta Information from PEAKS

##### Scan Identifier

F2:3905

##### Original sequence

L

L

S

E

V

S

D

R

P

S

G

V

S

S

R

##### Posttranslational Modifications

##### Source File

D:\separate\_stitch\_analyses\xle-disambiguation\raw\20210323\_F1\_UM1\_Peng0013\_SA\_F59\_ingel\_3ug\_TL.raw

##### Fraction

2

##### Scan Feature

-

##### De Novo Score

99

##### ConfidenceScore

99

### m/z

530.2825

##### Mass

1587.8215

##### Charge

3

##### Retention Time

20.69

##### Predicted Retention Time

-

##### Area

0

##### Parts Per Million

2.6

##### Fragmentation mode

ETHCD

##### Originating file

01 D:\separate\_stitch\_analyses\xle-disambiguation\20210325\_F59\_3ug\_DENOVO\_12.csv

#### Meta Information from PEAKS

##### Scan Identifier

F2:3840

##### Original sequence

L

L

S

E

V

S

D

R

P

S

G

V

S

S

R

##### Posttranslational Modifications

##### Source File

D:\separate\_stitch\_analyses\xle-disambiguation\raw\20210323\_F1\_UM1\_Peng0013\_SA\_F59\_ingel\_3ug\_TL.raw

##### Fraction

2

##### Scan Feature

-

##### De Novo Score

99

##### ConfidenceScore

99

### m/z

530.2816

##### Mass

1587.8215

##### Charge

3

##### Retention Time

20.33

##### Predicted Retention Time

-

##### Area

0

##### Parts Per Million

0.9

##### Fragmentation mode

ETHCD

##### Originating file

01 D:\separate\_stitch\_analyses\xle-disambiguation\20210325\_F59\_3ug\_DENOVO\_12.csv

#### Meta Information from PEAKS

##### Scan Identifier

F2:4284

##### Original sequence

L

L

S

E

V

S

D

R

P

S

G

V

S

S

R

##### Posttranslational Modifications

##### Source File

D:\separate\_stitch\_analyses\xle-disambiguation\raw\20210323\_F1\_UM1\_Peng0013\_SA\_F59\_ingel\_3ug\_TL.raw

##### Fraction

2

##### Scan Feature

-

##### De Novo Score

99

##### ConfidenceScore

99

### m/z

530.2818

##### Mass

1587.8215

##### Charge

3

##### Retention Time

22.84

##### Predicted Retention Time

-

##### Area

0

##### Parts Per Million

1.3

##### Fragmentation mode

ETHCD

##### Originating file

01 D:\separate\_stitch\_analyses\xle-disambiguation\20210325\_F59\_3ug\_DENOVO\_12.csv

#### Meta Information from PEAKS

##### Scan Identifier

F2:3970

##### Original sequence

L

L

S

E

V

S

D

R

P

S

G

V

S

S

R

##### Posttranslational Modifications

##### Source File

D:\separate\_stitch\_analyses\xle-disambiguation\raw\20210323\_F1\_UM1\_Peng0013\_SA\_F59\_ingel\_3ug\_TL.raw

##### Fraction

2

##### Scan Feature

-

##### De Novo Score

99

##### ConfidenceScore

99

### m/z

530.2823

##### Mass

1587.8215

##### Charge

3

##### Retention Time

21.07

##### Predicted Retention Time

-

##### Area

0

##### Parts Per Million

2.2

##### Fragmentation mode

ETHCD

##### Originating file

01 D:\separate\_stitch\_analyses\xle-disambiguation\20210325\_F59\_3ug\_DENOVO\_12.csv

#### Meta Information from PEAKS

##### Scan Identifier

F2:4542

##### Original sequence

L

L

S

E

V

S

D

R

P

S

G

V

S

S

R

##### Posttranslational Modifications

##### Source File

D:\separate\_stitch\_analyses\xle-disambiguation\raw\20210323\_F1\_UM1\_Peng0013\_SA\_F59\_ingel\_3ug\_TL.raw

##### Fraction

2

##### Scan Feature

-

##### De Novo Score

99

##### ConfidenceScore

99

### m/z

530.2817

##### Mass

1587.8215

##### Charge

3

##### Retention Time

24.31

##### Predicted Retention Time

-

##### Area

0

##### Parts Per Million

1.2

##### Fragmentation mode

ETHCD

##### Originating file

01 D:\separate\_stitch\_analyses\xle-disambiguation\20210325\_F59\_3ug\_DENOVO\_12.csv

#### Meta Information from PEAKS

##### Scan Identifier

F1:3761

##### Original sequence

L

L

S

E

V

S

D

R

P

S

G

V

S

S

R

##### Posttranslational Modifications

##### Source File

D:\separate\_stitch\_analyses\xle-disambiguation\raw\20210323\_F1\_UM1\_Peng0013\_SA\_F59\_ingel\_3ug\_ELA.raw

##### Fraction

1

##### Scan Feature

F1:5302

##### De Novo Score

99

##### ConfidenceScore

99

### m/z

530.2798

##### Mass

1587.8215

##### Charge

3

##### Retention Time

19.59

##### Predicted Retention Time

-

##### Area

4.956E+06

##### Fragmentation mode

ETHCD

##### Originating file

01 D:\separate\_stitch\_analyses\xle-disambiguation\20210325\_F59\_3ug\_DENOVO\_12.csv

#### Meta Information from PEAKS

##### Scan Identifier

F2:3632

##### Original sequence

L

L

S

E

V

S

D

R

P

S

G

V

S

S

R

##### Posttranslational Modifications

##### Source File

D:\separate\_stitch\_analyses\xle-disambiguation\raw\20210323\_F1\_UM1\_Peng0013\_SA\_F59\_ingel\_3ug\_TL.raw

##### Fraction

2

##### Scan Feature

F2:5350

##### De Novo Score

98

##### ConfidenceScore

98

### m/z

530.2825

##### Mass

1587.8215

##### Charge

3

##### Retention Time

19.21

##### Predicted Retention Time

-

##### Area

3.118E+08

##### Parts Per Million

2.7

##### Fragmentation mode

HCD

##### Originating file

01 D:\separate\_stitch\_analyses\xle-disambiguation\20210325\_F59\_3ug\_DENOVO\_12.csv

#### Meta Information from PEAKS

##### Scan Identifier

F2:3703

##### Original sequence

L

L

S

E

V

S

D

R

P

S

G

V

S

S

R

##### Posttranslational Modifications

##### Source File

D:\separate\_stitch\_analyses\xle-disambiguation\raw\20210323\_F1\_UM1\_Peng0013\_SA\_F59\_ingel\_3ug\_TL.raw

##### Fraction

2

##### Scan Feature

-

##### De Novo Score

98

##### ConfidenceScore

98

### m/z

530.2823

##### Mass

1587.8215

##### Charge

3

##### Retention Time

19.59

##### Predicted Retention Time

-

##### Area

0

##### Parts Per Million

2.2

##### Fragmentation mode

HCD

##### Originating file

01 D:\separate\_stitch\_analyses\xle-disambiguation\20210325\_F59\_3ug\_DENOVO\_12.csv

#### Meta Information from PEAKS

##### Scan Identifier

F2:4798

##### Original sequence

L

L

S

E

V

S

D

R

P

S

G

V

S

S

R

##### Posttranslational Modifications

##### Source File

D:\separate\_stitch\_analyses\xle-disambiguation\raw\20210323\_F1\_UM1\_Peng0013\_SA\_F59\_ingel\_3ug\_TL.raw

##### Fraction

2

##### Scan Feature

-

##### De Novo Score

98

##### ConfidenceScore

98

### m/z

530.2825

##### Mass

1587.8215

##### Charge

3

##### Retention Time

25.75

##### Predicted Retention Time

-

##### Area

0

##### Parts Per Million

2.6

##### Fragmentation mode

ETHCD

##### Originating file

01 D:\separate\_stitch\_analyses\xle-disambiguation\20210325\_F59\_3ug\_DENOVO\_12.csv

#### Meta Information from PEAKS

##### Scan Identifier

F2:4735

##### Original sequence

L

L

S

E

V

S

D

R

P

S

G

V

S

S

R

##### Posttranslational Modifications

##### Source File

D:\separate\_stitch\_analyses\xle-disambiguation\raw\20210323\_F1\_UM1\_Peng0013\_SA\_F59\_ingel\_3ug\_TL.raw

##### Fraction

2

##### Scan Feature

-

##### De Novo Score

98

##### ConfidenceScore

98

### m/z

530.2822

##### Mass

1587.8215

##### Charge

3

##### Retention Time

25.39

##### Predicted Retention Time

-

##### Area

0

##### Parts Per Million

2.1

##### Fragmentation mode

ETHCD

##### Originating file

01 D:\separate\_stitch\_analyses\xle-disambiguation\20210325\_F59\_3ug\_DENOVO\_12.csv

#### Meta Information from PEAKS

##### Scan Identifier

F2:4162

##### Original sequence

L

L

S

E

V

S

D

R

P

S

G

V

S

S

R

##### Posttranslational Modifications

##### Source File

D:\separate\_stitch\_analyses\xle-disambiguation\raw\20210323\_F1\_UM1\_Peng0013\_SA\_F59\_ingel\_3ug\_TL.raw

##### Fraction

2

##### Scan Feature

-

##### De Novo Score

98

##### ConfidenceScore

98

### m/z

530.2823

##### Mass

1587.8215

##### Charge

3

##### Retention Time

22.14

##### Predicted Retention Time

-

##### Area

0

##### Parts Per Million

2.2

##### Fragmentation mode

HCD

##### Originating file

01 D:\separate\_stitch\_analyses\xle-disambiguation\20210325\_F59\_3ug\_DENOVO\_12.csv

#### Meta Information from PEAKS

##### Scan Identifier

F2:4605

##### Original sequence

L

L

S

E

V

S

D

R

P

S

G

V

S

S

R

##### Posttranslational Modifications

##### Source File

D:\separate\_stitch\_analyses\xle-disambiguation\raw\20210323\_F1\_UM1\_Peng0013\_SA\_F59\_ingel\_3ug\_TL.raw

##### Fraction

2

##### Scan Feature

-

##### De Novo Score

98

##### ConfidenceScore

98

### m/z

530.2817

##### Mass

1587.8215

##### Charge

3

##### Retention Time

24.66

##### Predicted Retention Time

-

##### Area

0

##### Parts Per Million

1.1

##### Fragmentation mode

HCD

##### Originating file

01 D:\separate\_stitch\_analyses\xle-disambiguation\20210325\_F59\_3ug\_DENOVO\_12.csv

#### Meta Information from PEAKS

##### Scan Identifier

F2:3639

##### Original sequence

L

L

S

E

V

S

D

R

P

S

G

V

S

S

R

##### Posttranslational Modifications

##### Source File

D:\separate\_stitch\_analyses\xle-disambiguation\raw\20210323\_F1\_UM1\_Peng0013\_SA\_F59\_ingel\_3ug\_TL.raw

##### Fraction

2

##### Scan Feature

F2:15032

##### De Novo Score

97

##### ConfidenceScore

97

### m/z

794.9191

##### Mass

1587.8215

##### Charge

2

##### Retention Time

19.21

##### Predicted Retention Time

-

##### Area

2.411E+08

##### Parts Per Million

1.4

##### Fragmentation mode

HCD

##### Originating file

01 D:\separate\_stitch\_analyses\xle-disambiguation\20210325\_F59\_3ug\_DENOVO\_12.csv

#### Meta Information from PEAKS

##### Scan Identifier

F2:3773

##### Original sequence

L

L

S

E

V

S

D

R

P

S

G

V

S

S

R

##### Posttranslational Modifications

##### Source File

D:\separate\_stitch\_analyses\xle-disambiguation\raw\20210323\_F1\_UM1\_Peng0013\_SA\_F59\_ingel\_3ug\_TL.raw

##### Fraction

2

##### Scan Feature

-

##### De Novo Score

96

##### ConfidenceScore

96

### m/z

530.2818

##### Mass

1587.8215

##### Charge

3

##### Retention Time

19.96

##### Predicted Retention Time

-

##### Area

0

##### Parts Per Million

1.3

##### Fragmentation mode

HCD

##### Originating file

01 D:\separate\_stitch\_analyses\xle-disambiguation\20210325\_F59\_3ug\_DENOVO\_12.csv

#### Meta Information from PEAKS

##### Scan Identifier

F2:6067

##### Original sequence

L

L

S

E

V

S

D

R

P

S

G

V

S

S

R

##### Posttranslational Modifications

##### Source File

D:\separate\_stitch\_analyses\xle-disambiguation\raw\20210323\_F1\_UM1\_Peng0013\_SA\_F59\_ingel\_3ug\_TL.raw

##### Fraction

2

##### Scan Feature

F2:5349

##### De Novo Score

95

##### ConfidenceScore

95

### m/z

530.2825

##### Mass

1587.8215

##### Charge

3

##### Retention Time

33.45

##### Predicted Retention Time

-

##### Area

4.216E+06

##### Parts Per Million

2.6

##### Fragmentation mode

ETHCD

##### Originating file

01 D:\separate\_stitch\_analyses\xle-disambiguation\20210325\_F59\_3ug\_DENOVO\_12.csv

#### Meta Information from PEAKS

##### Scan Identifier

F1:3560

##### Original sequence

L

L

S

E

V

S

D

R

P

S

G

V

S

S

R

##### Posttranslational Modifications

##### Source File

D:\separate\_stitch\_analyses\xle-disambiguation\raw\20210323\_F1\_UM1\_Peng0013\_SA\_F59\_ingel\_3ug\_ELA.raw

##### Fraction

1

##### Scan Feature

F1:5302

##### De Novo Score

95

##### ConfidenceScore

95

### m/z

530.2798

##### Mass

1587.8215

##### Charge

3

##### Retention Time

19.59

##### Predicted Retention Time

-

##### Area

4.956E+06

##### Fragmentation mode

ETHCD

##### Originating file

01 D:\separate\_stitch\_analyses\xle-disambiguation\20210325\_F59\_3ug\_DENOVO\_12.csv

#### Meta Information from PEAKS

##### Scan Identifier

F2:7196

##### Original sequence

L

L

S

E

V

S

D

R

P

S

G

V

S

S

R

##### Posttranslational Modifications

##### Source File

D:\separate\_stitch\_analyses\xle-disambiguation\raw\20210323\_F1\_UM1\_Peng0013\_SA\_F59\_ingel\_3ug\_TL.raw

##### Fraction

2

##### Scan Feature

-

##### De Novo Score

95

##### ConfidenceScore

95

### m/z

530.2817

##### Mass

1587.8215

##### Charge

3

##### Retention Time

40.13

##### Predicted Retention Time

-

##### Area

0

##### Parts Per Million

1.1

##### Fragmentation mode

ETHCD

##### Originating file

01 D:\separate\_stitch\_analyses\xle-disambiguation\20210325\_F59\_3ug\_DENOVO\_12.csv

#### Meta Information from PEAKS

##### Scan Identifier

F2:6646

##### Original sequence

L

L

S

E

V

S

D

R

P

S

G

V

S

S

R

##### Posttranslational Modifications

##### Source File

D:\separate\_stitch\_analyses\xle-disambiguation\raw\20210323\_F1\_UM1\_Peng0013\_SA\_F59\_ingel\_3ug\_TL.raw

##### Fraction

2

##### Scan Feature

-

##### De Novo Score

95

##### ConfidenceScore

95

### m/z

530.2811

##### Mass

1587.8215

##### Charge

3

##### Retention Time

36.72

##### Predicted Retention Time

-

##### Area

0

##### Parts Per Million

0

##### Fragmentation mode

ETHCD

##### Originating file

01 D:\separate\_stitch\_analyses\xle-disambiguation\20210325\_F59\_3ug\_DENOVO\_12.csv

#### Meta Information from PEAKS

##### Scan Identifier

F2:4031

##### Original sequence

L

L

S

E

V

S

D

R

P

S

G

V

S

S

R

##### Posttranslational Modifications

##### Source File

D:\separate\_stitch\_analyses\xle-disambiguation\raw\20210323\_F1\_UM1\_Peng0013\_SA\_F59\_ingel\_3ug\_TL.raw

##### Fraction

2

##### Scan Feature

-

##### De Novo Score

95

##### ConfidenceScore

95

### m/z

530.282

##### Mass

1587.8215

##### Charge

3

##### Retention Time

21.4

##### Predicted Retention Time

-

##### Area

0

##### Parts Per Million

1.7

##### Fragmentation mode

HCD

##### Originating file

01 D:\separate\_stitch\_analyses\xle-disambiguation\20210325\_F59\_3ug\_DENOVO\_12.csv

#### Meta Information from PEAKS

##### Scan Identifier

F2:6539

##### Original sequence

L

L

S

E

V

S

D

R

P

S

G

V

S

S

R

##### Posttranslational Modifications

##### Source File

D:\separate\_stitch\_analyses\xle-disambiguation\raw\20210323\_F1\_UM1\_Peng0013\_SA\_F59\_ingel\_3ug\_TL.raw

##### Fraction

2

##### Scan Feature

-

##### De Novo Score

95

##### ConfidenceScore

95

### m/z

530.2819

##### Mass

1587.8215

##### Charge

3

##### Retention Time

36.06

##### Predicted Retention Time

-

##### Area

0

##### Parts Per Million

1.5

##### Fragmentation mode

ETHCD

##### Originating file

01 D:\separate\_stitch\_analyses\xle-disambiguation\20210325\_F59\_3ug\_DENOVO\_12.csv

#### Meta Information from PEAKS

##### Scan Identifier

F2:4473

##### Original sequence

L

L

S

E

V

S

D

R

P

S

G

V

S

S

R

##### Posttranslational Modifications

##### Source File

D:\separate\_stitch\_analyses\xle-disambiguation\raw\20210323\_F1\_UM1\_Peng0013\_SA\_F59\_ingel\_3ug\_TL.raw

##### Fraction

2

##### Scan Feature

-

##### De Novo Score

95

##### ConfidenceScore

95

### m/z

530.2821

##### Mass

1587.8215

##### Charge

3

##### Retention Time

23.92

##### Predicted Retention Time

-

##### Area

0

##### Parts Per Million

1.9

##### Fragmentation mode

HCD

##### Originating file

01 D:\separate\_stitch\_analyses\xle-disambiguation\20210325\_F59\_3ug\_DENOVO\_12.csv
