## Supplementary material for "A handle on mass coincidence errors in *de novo* sequencing of antibodies by bottom-up proteomics": Combined_011.html

Details Combined\_011 | Stitch OverviewUndefined

### Read Combined\_011

#### Sequence (length=9)

YCAKDVRPY

#### Spectrum 3914? Spectrum 3914 The raw spectrum of this peptide as annotated by Hecklib. The fragments are coloured according to ion type (see legend). Any peaks with a star '\*' as text can be hovered over to see the full details, first the ion type second the mass shift type. By hovering over the amino acids in the peptide or ions in the legend the corresponding peaks are highlighted. By toggling the 'Unassigned' label you can turn the background (unassigned) peaks on or off in the plot. By updating the slider in the Ion legend you can update the spectrum to only show the top X% of the peaks with labels. The top X% means any peak that is within X% of the highest intensity. By dragging in the spectrum you can zoom in to a specific part of the spectrum and use 'Zoom Out' to get back to the original zoom level. The annotation of the spectrum is based on the given sequence in the peptides file and is done with different software so inconsistencies are likely. The peaks are annotated based on the given sequence, with 20 ppm tolerance.

Copy Data

##### Spectrum 3914 (TSV)

###### Preview

```
Loading example...
```

*Click on the button to copy the data to your clipboard.*

Mz MinMz MaxIntensity Max

WidthHeightPeptide font sizePeptide stroke widthSpectrum font sizeSpectrum stroke widthCompact peptide

Ion legend

wxyz

abcd

OtherUnassignedIon^Charge^_Position_Show for top:%

YCAKDVRPY

01.09e+52.18e+53.28e+54.37e+5

Zoom Out

y^+1^_1_y^+3^_5_y^+2^_4_y^+1^_2_a^+1^_2_b^+2^_5_b^+1^_2_b^+3^_8_y^+3^_8_b^+2^_6_y^+2^_6_\*\*y^+2^_6_\*b^+1^_3_y^+2^_7_y^+1^_3_y^+2^_7_y^+1^_3_b^+2^_7_b^+2^_8_b^+2^_8_b^+2^_8_y^+2^_8_y^+2^_8_y^+1^_4_b^+1^_5_y^+1^_5_y^+1^_5_b^+1^_5_y^+1^_5_y^+1^_6_y^+1^_6_y^+1^_6_y^+1^_7_

0778155723353114

Fragment Matches Table

Show background peaks

| Position | Ion type | Intensity | mz Theoretical | mz Error (Th) | mz Error (ppm) | Charge | Series Number |
| --- | --- | --- | --- | --- | --- | --- | --- |
| - | - | 5279 | 120.1 | - | - | 0 | - |
| - | - | 1788 | 120.1 | - | - | 0 | - |
| - | - | 1828 | 121 | - | - | 0 | - |
| - | - | 609.7 | 121.1 | - | - | 0 | - |
| - | - | 425.6 | 122.1 | - | - | 0 | - |
| - | - | 826 | 123 | - | - | 0 | - |
| - | - | 403.5 | 123.8 | - | - | 0 | - |
| - | - | 467.8 | 126.1 | - | - | 0 | - |
| - | - | 1532 | 127.1 | - | - | 0 | - |
| - | - | 605.5 | 129.1 | - | - | 0 | - |
| - | - | 1.206E+05 | 129.1 | - | - | 0 | - |
| - | - | 655.1 | 130 | - | - | 0 | - |
| - | - | 1020 | 130.1 | - | - | 0 | - |
| - | - | 7767 | 130.1 | - | - | 0 | - |
| - | - | 3.977E+04 | 134 | - | - | 0 | - |
| - | - | 1889 | 135 | - | - | 0 | - |
| - | - | 1004 | 136 | - | - | 0 | - |
| - | - | 3.274E+05 | 136.1 | - | - | 0 | - |
| - | - | 1188 | 137.1 | - | - | 0 | - |
| - | - | 1155 | 137.1 | - | - | 0 | - |
| - | - | 2.597E+04 | 137.1 | - | - | 0 | - |
| - | - | 924.3 | 137.1 | - | - | 0 | - |
| - | - | 1446 | 138.1 | - | - | 0 | - |
| - | - | 745.6 | 138.1 | - | - | 0 | - |
| - | - | 2822 | 139.1 | - | - | 0 | - |
| - | - | 1547 | 140.1 | - | - | 0 | - |
| - | - | 1188 | 141.1 | - | - | 0 | - |
| - | - | 521.8 | 141.1 | - | - | 0 | - |
| - | - | 563.9 | 142 | - | - | 0 | - |
| - | - | 621 | 142.1 | - | - | 0 | - |
| - | - | 483.7 | 143.1 | - | - | 0 | - |
| - | - | 426.1 | 144 | - | - | 0 | - |
| - | - | 3.04E+04 | 147 | - | - | 0 | - |
| - | - | 583.8 | 147.1 | - | - | 0 | - |
| - | - | 2723 | 148 | - | - | 0 | - |
| - | - | 4.327E+05 | 149 | - | - | 0 | - |
| - | - | 3.408E+04 | 150 | - | - | 0 | - |
| - | - | 505.9 | 153.1 | - | - | 0 | - |
| - | - | 3487 | 155.1 | - | - | 0 | - |
| - | - | 2.008E+04 | 156.1 | - | - | 0 | - |
| - | - | 581 | 157.1 | - | - | 0 | - |
| - | - | 1394 | 157.1 | - | - | 0 | - |
| - | - | 1123 | 157.1 | - | - | 0 | - |
| - | - | 4784 | 157.1 | - | - | 0 | - |
| - | - | 554.3 | 157.1 | - | - | 0 | - |
| - | - | 3916 | 158 | - | - | 0 | - |
| - | - | 2689 | 159.1 | - | - | 0 | - |
| - | - | 1.203E+04 | 159.1 | - | - | 0 | - |
| - | - | 2669 | 160 | - | - | 0 | - |
| - | - | 1545 | 160.1 | - | - | 0 | - |
| - | - | 936 | 160.1 | - | - | 0 | - |
| - | - | 1550 | 161 | - | - | 0 | - |
| - | - | 5821 | 162 | - | - | 0 | - |
| - | - | 1054 | 164.1 | - | - | 0 | - |
| - | - | 2959 | 165.1 | - | - | 0 | - |
| - | - | 1706 | 165.1 | - | - | 0 | - |
| - | - | 1242 | 166.1 | - | - | 0 | - |
| - | - | 832.3 | 166.1 | - | - | 0 | - |
| - | - | 4.996E+04 | 167 | - | - | 0 | - |
| - | - | 3832 | 168 | - | - | 0 | - |
| - | - | 1452 | 169.1 | - | - | 0 | - |
| - | - | 713.2 | 169.1 | - | - | 0 | - |
| - | - | 1305 | 173.1 | - | - | 0 | - |
| - | - | 3391 | 173.4 | - | - | 0 | - |
| - | - | 650.7 | 175.1 | - | - | 0 | - |
| - | - | 575.2 | 175.1 | - | - | 0 | - |
| - | - | 4926 | 176 | - | - | 0 | - |
| - | - | 609.8 | 178.1 | - | - | 0 | - |
| - | - | 620.5 | 180.1 | - | - | 0 | - |
| - | - | 739.5 | 181.1 | - | - | 0 | - |
| 9 | y | 3870 | 182.1 | 0.0001597 | 0.8769 | +1 | 1 |
| - | - | 811.2 | 182.1 | - | - | 0 | - |
| - | - | 2704 | 183.1 | - | - | 0 | - |
| - | - | 1521 | 183.1 | - | - | 0 | - |
| - | - | 1499 | 185 | - | - | 0 | - |
| - | - | 3689 | 185.1 | - | - | 0 | - |
| - | - | 1865 | 186 | - | - | 0 | - |
| - | - | 5680 | 187.1 | - | - | 0 | - |
| - | - | 1.71E+04 | 187.1 | - | - | 0 | - |
| - | - | 596.7 | 188 | - | - | 0 | - |
| - | - | 843.6 | 188.1 | - | - | 0 | - |
| - | - | 1628 | 188.1 | - | - | 0 | - |
| - | - | 1127 | 192 | - | - | 0 | - |
| - | - | 879.1 | 192.1 | - | - | 0 | - |
| - | - | 573.1 | 194.1 | - | - | 0 | - |
| - | - | 1468 | 195.1 | - | - | 0 | - |
| - | - | 735.9 | 197.1 | - | - | 0 | - |
| - | - | 1980 | 197.1 | - | - | 0 | - |
| - | - | 1177 | 198.1 | - | - | 0 | - |
| - | - | 3677 | 199.1 | - | - | 0 | - |
| - | - | 1.186E+04 | 200.1 | - | - | 0 | - |
| - | - | 2910 | 201.1 | - | - | 0 | - |
| - | - | 4.615E+04 | 205.1 | - | - | 0 | - |
| - | - | 2912 | 206.1 | - | - | 0 | - |
| - | - | 1984 | 207.1 | - | - | 0 | - |
| - | - | 2361 | 209.1 | - | - | 0 | - |
| - | - | 1412 | 209.1 | - | - | 0 | - |
| - | - | 1797 | 210.1 | - | - | 0 | - |
| 5 | y | 951.3 | 211.1 | 0.003217 | 15.24 | +3 | 5 |
| - | - | 605.4 | 211.1 | - | - | 0 | - |
| - | - | 748.1 | 211.2 | - | - | 0 | - |
| - | - | 936.4 | 214.2 | - | - | 0 | - |
| - | - | 2042 | 215 | - | - | 0 | - |
| - | - | 3991 | 215.1 | - | - | 0 | - |
| - | - | 617 | 219.1 | - | - | 0 | - |
| - | - | 2.134E+04 | 221.1 | - | - | 0 | - |
| - | - | 1929 | 221.1 | - | - | 0 | - |
| - | - | 1526 | 222.1 | - | - | 0 | - |
| - | - | 973.7 | 223.1 | - | - | 0 | - |
| - | - | 1054 | 223.1 | - | - | 0 | - |
| - | - | 796.1 | 224.1 | - | - | 0 | - |
| - | - | 987.3 | 224.1 | - | - | 0 | - |
| - | - | 919.5 | 225 | - | - | 0 | - |
| - | - | 618.7 | 225.1 | - | - | 0 | - |
| - | - | 1052 | 226 | - | - | 0 | - |
| - | - | 5857 | 226.1 | - | - | 0 | - |
| - | - | 2301 | 227.1 | - | - | 0 | - |
| - | - | 1041 | 228.2 | - | - | 0 | - |
| - | - | 2699 | 231 | - | - | 0 | - |
| - | - | 7476 | 233.1 | - | - | 0 | - |
| - | - | 2230 | 233.1 | - | - | 0 | - |
| - | - | 3038 | 233.1 | - | - | 0 | - |
| - | - | 2163 | 234.1 | - | - | 0 | - |
| - | - | 2435 | 237.1 | - | - | 0 | - |
| - | - | 820.9 | 238.1 | - | - | 0 | - |
| - | - | 718.5 | 238.1 | - | - | 0 | - |
| - | - | 1004 | 239.1 | - | - | 0 | - |
| - | - | 3.382E+04 | 239.2 | - | - | 0 | - |
| - | - | 547.2 | 240.1 | - | - | 0 | - |
| - | - | 3996 | 240.2 | - | - | 0 | - |
| - | - | 815.6 | 241.1 | - | - | 0 | - |
| - | - | 2405 | 244.1 | - | - | 0 | - |
| - | - | 792.7 | 245.1 | - | - | 0 | - |
| - | - | 3945 | 249.1 | - | - | 0 | - |
| - | - | 1577 | 249.1 | - | - | 0 | - |
| - | - | 1437 | 251 | - | - | 0 | - |
| - | - | 805.3 | 251.1 | - | - | 0 | - |
| - | - | 7709 | 252.1 | - | - | 0 | - |
| - | - | 841 | 252.1 | - | - | 0 | - |
| - | - | 815.6 | 252.1 | - | - | 0 | - |
| - | - | 919.3 | 253.1 | - | - | 0 | - |
| - | - | 623.8 | 254.1 | - | - | 0 | - |
| - | - | 4209 | 254.2 | - | - | 0 | - |
| - | - | 3053 | 255.1 | - | - | 0 | - |
| - | - | 1.383E+04 | 256.2 | - | - | 0 | - |
| - | - | 645.5 | 257.2 | - | - | 0 | - |
| - | - | 1105 | 262.1 | - | - | 0 | - |
| - | - | 3341 | 267.1 | - | - | 0 | - |
| 6 | y | 659.1 | 267.7 | 0.0001196 | 0.4468 | +2 | 4 |
| - | - | 4039 | 269.1 | - | - | 0 | - |
| - | - | 2132 | 269.2 | - | - | 0 | - |
| - | - | 889.3 | 270.1 | - | - | 0 | - |
| - | - | 3168 | 272.1 | - | - | 0 | - |
| - | - | 2759 | 272.2 | - | - | 0 | - |
| - | - | 562.3 | 273.1 | - | - | 0 | - |
| - | - | 4012 | 274.2 | - | - | 0 | - |
| - | - | 554.9 | 278 | - | - | 0 | - |
| 8 | y | 5.349E+04 | 279.1 | 3.864E-05 | 0.1384 | +1 | 2 |
| - | - | 4735 | 279.2 | - | - | 0 | - |
| - | - | 1639 | 280.1 | - | - | 0 | - |
| - | - | 9105 | 280.1 | - | - | 0 | - |
| - | - | 1181 | 280.2 | - | - | 0 | - |
| - | - | 819.9 | 281.1 | - | - | 0 | - |
| - | - | 2433 | 282.2 | - | - | 0 | - |
| - | - | 1816 | 284.2 | - | - | 0 | - |
| - | - | 1172 | 285 | - | - | 0 | - |
| - | - | 590.4 | 285.7 | - | - | 0 | - |
| - | - | 650 | 286 | - | - | 0 | - |
| - | - | 831.6 | 292.1 | - | - | 0 | - |
| 2 | a | 2.937E+04 | 297.1 | 0.004268 | 14.37 | +1 | 2 |
| - | - | 1621 | 297.2 | - | - | 0 | - |
| - | - | 3926 | 298.1 | - | - | 0 | - |
| - | - | 617.4 | 299.1 | - | - | 0 | - |
| - | - | 511.7 | 299.1 | - | - | 0 | - |
| - | - | 1485 | 299.2 | - | - | 0 | - |
| - | - | 1173 | 300.1 | - | - | 0 | - |
| - | - | 1137 | 301.1 | - | - | 0 | - |
| - | - | 1309 | 302.6 | - | - | 0 | - |
| - | - | 3962 | 303.7 | - | - | 0 | - |
| - | - | 2163 | 304.1 | - | - | 0 | - |
| - | - | 1650 | 304.2 | - | - | 0 | - |
| - | - | 701.4 | 307.6 | - | - | 0 | - |
| - | - | 1966 | 307.7 | - | - | 0 | - |
| - | - | 673.9 | 310.2 | - | - | 0 | - |
| - | - | 791.9 | 312.7 | - | - | 0 | - |
| - | - | 4636 | 315.2 | - | - | 0 | - |
| - | - | 675.9 | 315.2 | - | - | 0 | - |
| - | - | 763.1 | 316.1 | - | - | 0 | - |
| - | - | 1059 | 316.9 | - | - | 0 | - |
| 5 | b | 2847 | 320.1 | 0.006295 | 19.67 | +2 | 5 |
| - | - | 758.3 | 320.6 | - | - | 0 | - |
| - | - | 629.1 | 321.1 | - | - | 0 | - |
| 2 | b | 5836 | 325.1 | 0.00431 | 13.26 | +1 | 2 |
| - | - | 651 | 325.2 | - | - | 0 | - |
| - | - | 607.7 | 325.2 | - | - | 0 | - |
| 8 | b | 1605 | 325.5 | 0.001304 | 4.005 | +3 | 8 |
| - | - | 710.9 | 325.8 | - | - | 0 | - |
| - | - | 1029 | 326.1 | - | - | 0 | - |
| - | - | 1637 | 330.1 | - | - | 0 | - |
| - | - | 567.3 | 333.1 | - | - | 0 | - |
| 2 | y | 635.7 | 337.2 | 0.0003051 | 0.905 | +3 | 8 |
| - | - | 4145 | 338.1 | - | - | 0 | - |
| - | - | 2112 | 340.2 | - | - | 0 | - |
| - | - | 1268 | 343.1 | - | - | 0 | - |
| - | - | 1025 | 343.2 | - | - | 0 | - |
| - | - | 4666 | 343.2 | - | - | 0 | - |
| - | - | 536.7 | 344.2 | - | - | 0 | - |
| - | - | 574.2 | 345.2 | - | - | 0 | - |
| - | - | 729.9 | 348.2 | - | - | 0 | - |
| - | - | 992.6 | 353.2 | - | - | 0 | - |
| - | - | 715.2 | 353.2 | - | - | 0 | - |
| - | - | 1826 | 354.2 | - | - | 0 | - |
| - | - | 1086 | 354.5 | - | - | 0 | - |
| - | - | 1813 | 354.9 | - | - | 0 | - |
| - | - | 3586 | 355.1 | - | - | 0 | - |
| - | - | 1258 | 355.7 | - | - | 0 | - |
| - | - | 944 | 355.9 | - | - | 0 | - |
| - | - | 1204 | 356.1 | - | - | 0 | - |
| - | - | 620 | 356.9 | - | - | 0 | - |
| - | - | 2492 | 357.1 | - | - | 0 | - |
| - | - | 3336 | 357.2 | - | - | 0 | - |
| - | - | 1196 | 357.7 | - | - | 0 | - |
| - | - | 717.7 | 360.2 | - | - | 0 | - |
| 6 | b | 2704 | 361.1 | 0.004406 | 12.2 | +2 | 6 |
| - | - | 730.9 | 362.2 | - | - | 0 | - |
| - | - | 920.8 | 366.1 | - | - | 0 | - |
| - | - | 4107 | 366.2 | - | - | 0 | - |
| - | - | 1255 | 366.7 | - | - | 0 | - |
| - | - | 6096 | 369.2 | - | - | 0 | - |
| - | - | 2304 | 369.7 | - | - | 0 | - |
| - | - | 2413 | 370.5 | - | - | 0 | - |
| - | - | 600.2 | 371.2 | - | - | 0 | - |
| - | - | 1.673E+04 | 371.2 | - | - | 0 | - |
| - | - | 1.295E+04 | 371.2 | - | - | 0 | - |
| - | - | 2746 | 372.2 | - | - | 0 | - |
| - | - | 2058 | 372.2 | - | - | 0 | - |
| - | - | 566.7 | 373.9 | - | - | 0 | - |
| - | - | 3538 | 376.2 | - | - | 0 | - |
| - | - | 950.9 | 376.8 | - | - | 0 | - |
| - | - | 1730 | 379.2 | - | - | 0 | - |
| - | - | 6038 | 379.8 | - | - | 0 | - |
| - | - | 3395 | 380.2 | - | - | 0 | - |
| - | - | 1354 | 380.5 | - | - | 0 | - |
| 4 | y | 1.153E+04 | 380.7 | 0.000104 | 0.2733 | +2 | 6 |
| - | - | 5423 | 381.2 | - | - | 0 | - |
| - | - | 602.3 | 381.2 | - | - | 0 | - |
| - | - | 1232 | 381.7 | - | - | 0 | - |
| 0 | Precursor | 605.5 | 385.5 | 0.003631 | 9.419 | +3 | -1 |
| 0 | Precursor | 867.9 | 385.8 | 0.001284 | 3.327 | +3 | -1 |
| - | - | 1285 | 386.2 | - | - | 0 | - |
| 4 | y | 3.057E+04 | 389.2 | 1.19E-05 | 0.03056 | +2 | 6 |
| - | - | 1.192E+04 | 389.7 | - | - | 0 | - |
| - | - | 2701 | 390.2 | - | - | 0 | - |
| 0 | Precursor | 1E+04 | 391.5 | 0.001538 | 3.927 | +3 | -1 |
| - | - | 7362 | 391.9 | - | - | 0 | - |
| - | - | 1233 | 392.2 | - | - | 0 | - |
| - | - | 1107 | 392.5 | - | - | 0 | - |
| - | - | 1215 | 394.2 | - | - | 0 | - |
| 3 | b | 938.4 | 396.1 | 0.004955 | 12.51 | +1 | 3 |
| - | - | 4260 | 398.1 | - | - | 0 | - |
| - | - | 706.2 | 404.2 | - | - | 0 | - |
| - | - | 979.4 | 405.7 | - | - | 0 | - |
| - | - | 970.1 | 406.2 | - | - | 0 | - |
| - | - | 1188 | 407.2 | - | - | 0 | - |
| - | - | 2299 | 414.7 | - | - | 0 | - |
| - | - | 1567 | 415.2 | - | - | 0 | - |
| 3 | y | 718.4 | 415.7 | 0.000477 | 1.147 | +2 | 7 |
| - | - | 977 | 417.2 | - | - | 0 | - |
| 7 | y | 9726 | 418.2 | 6.167E-05 | 0.1475 | +1 | 3 |
| - | - | 2935 | 419.2 | - | - | 0 | - |
| - | - | 745.3 | 423.2 | - | - | 0 | - |
| - | - | 5718 | 423.7 | - | - | 0 | - |
| - | - | 2610 | 424.2 | - | - | 0 | - |
| 3 | y | 2.434E+04 | 424.7 | 0.0001995 | 0.4698 | +2 | 7 |
| - | - | 1.241E+04 | 425.2 | - | - | 0 | - |
| - | - | 3222 | 425.7 | - | - | 0 | - |
| - | - | 1360 | 426.2 | - | - | 0 | - |
| - | - | 1341 | 427.2 | - | - | 0 | - |
| - | - | 1544 | 429.7 | - | - | 0 | - |
| - | - | 917.3 | 430.2 | - | - | 0 | - |
| 7 | y | 3.08E+04 | 435.2 | 6.286E-05 | 0.1444 | +1 | 3 |
| - | - | 7544 | 436.2 | - | - | 0 | - |
| - | - | 632.4 | 437.2 | - | - | 0 | - |
| - | - | 683.7 | 439.3 | - | - | 0 | - |
| - | - | 1405 | 440.2 | - | - | 0 | - |
| - | - | 806.5 | 440.2 | - | - | 0 | - |
| - | - | 842 | 442.2 | - | - | 0 | - |
| 7 | b | 3272 | 447.7 | 0.002417 | 5.399 | +2 | 7 |
| - | - | 1873 | 448.2 | - | - | 0 | - |
| - | - | 4460 | 451.2 | - | - | 0 | - |
| - | - | 3685 | 451.7 | - | - | 0 | - |
| - | - | 2402 | 452.2 | - | - | 0 | - |
| - | - | 631.4 | 452.7 | - | - | 0 | - |
| - | - | 639.7 | 455.2 | - | - | 0 | - |
| - | - | 7923 | 458.2 | - | - | 0 | - |
| - | - | 2831 | 458.3 | - | - | 0 | - |
| - | - | 1059 | 459.2 | - | - | 0 | - |
| - | - | 863 | 459.2 | - | - | 0 | - |
| - | - | 6935 | 462.2 | - | - | 0 | - |
| - | - | 2997 | 462.7 | - | - | 0 | - |
| - | - | 704.1 | 463.2 | - | - | 0 | - |
| - | - | 873.5 | 463.7 | - | - | 0 | - |
| - | - | 2233 | 469.2 | - | - | 0 | - |
| - | - | 3194 | 469.2 | - | - | 0 | - |
| - | - | 1001 | 471.3 | - | - | 0 | - |
| - | - | 676.6 | 473.2 | - | - | 0 | - |
| - | - | 1.818E+04 | 476.2 | - | - | 0 | - |
| - | - | 3793 | 477.2 | - | - | 0 | - |
| - | - | 737.5 | 478.2 | - | - | 0 | - |
| - | - | 1975 | 482.2 | - | - | 0 | - |
| - | - | 1676 | 482.3 | - | - | 0 | - |
| - | - | 1798 | 482.7 | - | - | 0 | - |
| - | - | 1730 | 484.3 | - | - | 0 | - |
| - | - | 6828 | 486.3 | - | - | 0 | - |
| 8 | b | 734.3 | 487.2 | 0.000476 | 0.9769 | +2 | 8 |
| - | - | 1875 | 487.3 | - | - | 0 | - |
| 8 | b | 2197 | 487.7 | 0.0004183 | 0.8577 | +2 | 8 |
| - | - | 1624 | 488.2 | - | - | 0 | - |
| - | - | 1370 | 488.3 | - | - | 0 | - |
| 8 | b | 2.502E+04 | 496.2 | 0.002157 | 4.347 | +2 | 8 |
| 2 | y | 1.493E+04 | 496.7 | 0.00553 | 11.13 | +2 | 8 |
| - | - | 6121 | 497.2 | - | - | 0 | - |
| - | - | 1970 | 497.7 | - | - | 0 | - |
| - | - | 1010 | 498.2 | - | - | 0 | - |
| - | - | 3993 | 499.3 | - | - | 0 | - |
| - | - | 787.6 | 500.3 | - | - | 0 | - |
| - | - | 1177 | 502.2 | - | - | 0 | - |
| 2 | y | 1.15E+05 | 505.2 | 0.002435 | 4.819 | +2 | 8 |
| - | - | 5.54E+04 | 505.7 | - | - | 0 | - |
| - | - | 2.17E+04 | 506.2 | - | - | 0 | - |
| - | - | 6426 | 506.7 | - | - | 0 | - |
| - | - | 946.6 | 507.2 | - | - | 0 | - |
| - | - | 1509 | 513.3 | - | - | 0 | - |
| - | - | 970.2 | 514.3 | - | - | 0 | - |
| - | - | 755.4 | 517.3 | - | - | 0 | - |
| - | - | 1319 | 527.8 | - | - | 0 | - |
| - | - | 1397 | 528.3 | - | - | 0 | - |
| - | - | 1415 | 529.2 | - | - | 0 | - |
| - | - | 919.8 | 533.7 | - | - | 0 | - |
| 6 | y | 9.886E+04 | 534.3 | 0.0003579 | 0.6698 | +1 | 4 |
| - | - | 3.091E+04 | 535.3 | - | - | 0 | - |
| - | - | 5588 | 536.3 | - | - | 0 | - |
| - | - | 653 | 537.3 | - | - | 0 | - |
| - | - | 2541 | 542.7 | - | - | 0 | - |
| - | - | 1954 | 543.2 | - | - | 0 | - |
| - | - | 1715 | 544.3 | - | - | 0 | - |
| - | - | 630.2 | 545.3 | - | - | 0 | - |
| - | - | 1712 | 547.3 | - | - | 0 | - |
| - | - | 775.9 | 551.3 | - | - | 0 | - |
| - | - | 1441 | 553.3 | - | - | 0 | - |
| - | - | 1456 | 557.2 | - | - | 0 | - |
| - | - | 2587 | 570.3 | - | - | 0 | - |
| - | - | 779.3 | 571.3 | - | - | 0 | - |
| - | - | 1794 | 575.2 | - | - | 0 | - |
| - | - | 715.9 | 576.3 | - | - | 0 | - |
| - | - | 2369 | 582.3 | - | - | 0 | - |
| - | - | 944.3 | 584.2 | - | - | 0 | - |
| - | - | 662.5 | 596.4 | - | - | 0 | - |
| - | - | 1099 | 603.3 | - | - | 0 | - |
| - | - | 5788 | 606.3 | - | - | 0 | - |
| - | - | 1643 | 607.3 | - | - | 0 | - |
| - | - | 2538 | 614.4 | - | - | 0 | - |
| 5 | b | 1187 | 622.2 | 0.005341 | 8.584 | +1 | 5 |
| 5 | y | 1232 | 631.3 | 0.001452 | 2.3 | +1 | 5 |
| 5 | y | 2236 | 632.3 | 0.002666 | 4.217 | +1 | 5 |
| 5 | b | 1408 | 639.2 | 0.004486 | 7.017 | +1 | 5 |
| 5 | y | 1.134E+05 | 649.3 | 0.0005675 | 0.874 | +1 | 5 |
| - | - | 3.561E+04 | 650.3 | - | - | 0 | - |
| - | - | 8009 | 651.3 | - | - | 0 | - |
| - | - | 1184 | 652.3 | - | - | 0 | - |
| - | - | 1723 | 655.3 | - | - | 0 | - |
| - | - | 3403 | 659.3 | - | - | 0 | - |
| - | - | 743.5 | 667.4 | - | - | 0 | - |
| - | - | 1316 | 685.4 | - | - | 0 | - |
| - | - | 1725 | 713.3 | - | - | 0 | - |
| - | - | 2543 | 714.3 | - | - | 0 | - |
| - | - | 1206 | 715.3 | - | - | 0 | - |
| - | - | 660.3 | 717.4 | - | - | 0 | - |
| - | - | 4310 | 731.3 | - | - | 0 | - |
| - | - | 1523 | 732.4 | - | - | 0 | - |
| - | - | 620.8 | 733.4 | - | - | 0 | - |
| - | - | 4174 | 737.3 | - | - | 0 | - |
| - | - | 1929 | 738.3 | - | - | 0 | - |
| 4 | y | 955.6 | 759.4 | 0.000188 | 0.2476 | +1 | 6 |
| 4 | y | 1220 | 760.4 | 0.0008052 | 1.059 | +1 | 6 |
| - | - | 1329 | 768.4 | - | - | 0 | - |
| - | - | 718.3 | 769.4 | - | - | 0 | - |
| 4 | y | 2.37E+04 | 777.4 | 0.001292 | 1.662 | +1 | 6 |
| - | - | 1.01E+04 | 778.4 | - | - | 0 | - |
| - | - | 2766 | 779.4 | - | - | 0 | - |
| - | - | 566.6 | 789.4 | - | - | 0 | - |
| - | - | 1451 | 816.3 | - | - | 0 | - |
| - | - | 683.9 | 828.4 | - | - | 0 | - |
| 3 | y | 3886 | 848.5 | 0.001846 | 2.176 | +1 | 7 |
| - | - | 1210 | 849.5 | - | - | 0 | - |
| - | - | 666.1 | 929.4 | - | - | 0 | - |
| - | - | 578 | 1090 | - | - | 0 | - |
| - | - | 652 | 1136 | - | - | 0 | - |
| - | - | 597.7 | 1191 | - | - | 0 | - |
| - | - | 581.7 | 1211 | - | - | 0 | - |
| - | - | 624.5 | 2346 | - | - | 0 | - |
| - | - | 952.3 | 3083 | - | - | 0 | - |

m/z Charge Intensity FragmentType MassShift Position
120.05272674560547 0 5279.3003
120.0810317993164 0 1787.9248
121.02864074707031 0 1827.5371
121.06491088867188 0 609.6799
122.07140350341797 0 425.5675
123.0445785522461 0 825.9676
123.75751495361328 0 403.52682
126.05529022216797 0 467.78772
127.08684539794922 0 1532.1389
129.06651306152344 0 605.5221
129.10250854492188 0 120636.08
130.03245544433594 0 655.12445
130.10023498535156 0 1020.4187
130.10586547851562 0 7766.852
134.02725219726562 0 39772.582
135.03076171875 0 1888.8955
136.02279663085938 0 1004.1151
136.07601928710938 0 327434.25
137.05996704101562 0 1187.8582
137.07347106933594 0 1155.3846
137.07925415039062 0 25973.652
137.1075897216797 0 924.2623
138.06634521484375 0 1446.3081
138.08261108398438 0 745.64935
139.08689880371094 0 2822.0918
140.08216857910156 0 1546.8765
141.10240173339844 0 1187.7478
141.1138458251953 0 521.79126
142.0325469970703 0 563.91876
142.08680725097656 0 620.9765
143.08181762695312 0 483.6974
144.0109405517578 0 426.12915
147.04429626464844 0 30399.545
147.1131591796875 0 583.8478
148.04776000976562 0 2723.1294
149.0236358642578 0 432669.53
150.0269012451172 0 34081.645
153.10260009765625 0 505.8933
155.11805725097656 0 3487.2273
156.07693481445312 0 20076.713
157.0608673095703 0 580.97156
157.08033752441406 0 1393.9136
157.0973663330078 0 1122.8224
157.10855102539062 0 4783.699
157.1343231201172 0 554.288
158.0272216796875 0 3916.2505
159.05902099609375 0 2688.936
159.11302185058594 0 12026.762
160.04293823242188 0 2668.6616
160.07608032226562 0 1544.5138
160.11654663085938 0 936.0362
161.0382537841797 0 1549.6969
162.0220489501953 0 5821.217
164.07077026367188 0 1054.1323
165.0547637939453 0 2958.8174
165.10244750976562 0 1706.3473
166.06103515625 0 1242.0017
166.08660888671875 0 832.27704
167.03407287597656 0 49962.273
168.03744506835938 0 3832.408
169.09725952148438 0 1451.6472
169.13409423828125 0 713.2019
173.1287841796875 0 1305.4854
173.4403076171875 0 3390.975
175.0872802734375 0 650.6733
175.1189422607422 0 575.243
176.0377960205078 0 4926.3184
178.13409423828125 0 609.7783
180.1023406982422 0 620.4844
181.09693908691406 0 739.5239
182.08132934570312 0 3870.3777 y 8
182.12881469726562 0 811.2126
183.1128692626953 0 2703.527
183.14944458007812 0 1521.206
185.03810119628906 0 1498.8154
185.1033477783203 0 3689.0667
186.02220153808594 0 1865.2131
187.05372619628906 0 5680.216
187.10791015625 0 17098.182
188.03817749023438 0 596.7202
188.07061767578125 0 843.5831
188.1112060546875 0 1628.1425
192.04774475097656 0 1127.1467
192.11314392089844 0 879.07336
194.1278076171875 0 573.12
195.11288452148438 0 1468.4711
197.09237670898438 0 735.92126
197.1286163330078 0 1979.6992
198.12353515625 0 1176.7269
199.10775756835938 0 3676.9458
200.13951110839844 0 11855.594
201.12330627441406 0 2909.5654
205.0642547607422 0 46150.918
206.06741333007812 0 2911.6216
207.0599822998047 0 1984.3667
209.09239196777344 0 2361.0112
209.14012145996094 0 1412.3384
210.12393188476562 0 1796.612
211.10824584960938 0 951.29297 y Water loss 4
211.1433563232422 0 605.37085
211.154296875 0 748.059
214.15512084960938 0 936.4273
215.0484619140625 0 2041.7084
215.1028594970703 0 3990.9639
219.07998657226562 0 617.01184
221.05917358398438 0 21342.771
221.09213256835938 0 1929.3882
222.0633087158203 0 1525.6605
223.0537567138672 0 973.68036
223.0646514892578 0 1054.0931
224.06442260742188 0 796.06165
224.1395721435547 0 987.33417
225.04319763183594 0 919.47626
225.0596466064453 0 618.66284
226.04397583007812 0 1052.3263
226.1188201904297 0 5856.6284
227.10287475585938 0 2301.2026
228.1829071044922 0 1041.3297
231.0435791015625 0 2698.8315
233.0590362548828 0 7476.088
233.09228515625 0 2230.1074
233.1284942626953 0 3038.361
234.0590057373047 0 2163.109
237.13473510742188 0 2434.8242
238.11903381347656 0 820.90076
238.13816833496094 0 718.5328
239.11390686035156 0 1004.3034
239.150390625 0 33821.418
240.13449096679688 0 547.2299
240.1536407470703 0 3996.4026
241.11241149902344 0 815.57794
244.12930297851562 0 2405.352
245.132080078125 0 792.69696
249.0540313720703 0 3944.905
249.09805297851562 0 1577.1881
251.04737854003906 0 1436.5327
251.14988708496094 0 805.27454
252.06895446777344 0 7709.1235
252.08090209960938 0 840.9985
252.13455200195312 0 815.61743
253.0719451904297 0 919.2826
254.1131591796875 0 623.82275
254.1613006591797 0 4208.582
255.1450653076172 0 3052.9678
256.1768798828125 0 13832.897
257.18017578125 0 645.5182
262.052734375 0 1105.3809
267.1086730957031 0 3340.9336
267.6554870605469 0 659.11456 y 5
269.1067810058594 0 4038.731
269.16082763671875 0 2132.3682
270.1084289550781 0 889.2637
272.1240539550781 0 3167.8608
272.1718444824219 0 2758.7366
273.1270751953125 0 562.3121
274.1873779296875 0 4012.0352
278.0103454589844 0 554.94543
279.13397216796875 0 53489.883 y 7
279.16015625 0 4735.101
280.0632019042969 0 1639.3882
280.13726806640625 0 9104.678
280.1631774902344 0 1180.5479
281.1393737792969 0 819.90186
282.1561279296875 0 2433.0134
284.17169189453125 0 1815.7102
285.01007080078125 0 1171.6852
285.6706237792969 0 590.4046
286.01129150390625 0 649.9927
292.1299743652344 0 831.56415
297.0906066894531 0 29367.348 a 1
297.15545654296875 0 1620.921
298.09332275390625 0 3926.2427
299.0859680175781 0 617.41705
299.09796142578125 0 511.65445
299.1717529296875 0 1485.4823
300.062744140625 0 1172.6378
301.0595703125 0 1136.7686
302.6065673828125 0 1309.4114
303.6534423828125 0 3962.0195
304.1296691894531 0 2162.781
304.15533447265625 0 1650.0759
307.64990234375 0 701.38965
307.6854248046875 0 1966.4456
310.18719482421875 0 673.9182
312.67626953125 0 791.9445
315.1665344238281 0 4636.238
315.2024841308594 0 675.91235
316.133056640625 0 763.09973
316.8772277832031 0 1059.3003
320.13433837890625 0 2846.974 b 4
320.62896728515625 0 758.27167
321.1357116699219 0 629.054
325.0854797363281 0 5835.6875 b 1
325.1515808105469 0 651.01807
325.1870422363281 0 607.7001
325.4876708984375 0 1604.732 b Ammonia loss 7
325.8230895996094 0 710.88074
326.08905029296875 0 1028.6699
330.1119689941406 0 1637.4528
333.0890808105469 0 567.25854
337.16510009765625 0 635.7264 y 1
338.1455993652344 0 4145.2515
340.1973876953125 0 2112.2637
343.14263916015625 0 1267.6174
343.1628723144531 0 1024.8627
343.20721435546875 0 4666.2593
344.2105407714844 0 536.7408
345.1761169433594 0 574.20026
348.1683349609375 0 729.9334
353.1929931640625 0 992.62146
353.2305908203125 0 715.2309
354.17681884765625 0 1826.1531
354.50030517578125 0 1086.055
354.8848876953125 0 1812.7767
355.0699768066406 0 3585.5032
355.6629333496094 0 1257.5159
355.8858947753906 0 944.03564
356.06982421875 0 1203.8887
356.8899841308594 0 620.00806
357.06787109375 0 2492.446
357.17364501953125 0 3336.4502
357.6756896972656 0 1195.6157
360.1700439453125 0 717.668
361.15338134765625 0 2703.784 b Ammonia loss 5
362.1579284667969 0 730.88135
366.1407775878906 0 920.8436
366.1788330078125 0 4106.6694
366.6787109375 0 1254.5868
369.1736145019531 0 6095.9126
369.6748352050781 0 2303.5132
370.5069885253906 0 2412.5593
371.17852783203125 0 600.213
371.2034912109375 0 16732.13
371.23992919921875 0 12953.385
372.2065124511719 0 2745.803
372.24310302734375 0 2057.5095
373.9198303222656 0 566.6785
376.1841735839844 0 3538.1985
376.8469543457031 0 950.8529
379.1666564941406 0 1729.5176
379.839111328125 0 6037.862
380.1728210449219 0 3395.0132
380.506591796875 0 1353.9048
380.70294189453125 0 11525.874 y Ammonia loss 3
381.2042541503906 0 5423.4766
381.2275085449219 0 602.3275
381.7060241699219 0 1232.4438
385.5126953125 0 605.53864 Precursor Water loss
385.8430480957031 0 867.9018 Precursor Ammonia loss
386.24066162109375 0 1284.7122
389.21630859375 0 30567.402 y 3
389.7175598144531 0 11923.337
390.2186584472656 0 2700.8025
391.518310546875 0 10004.003 Precursor
391.8527526855469 0 7362.349
392.187255859375 0 1232.7065
392.51763916015625 0 1107.4059
394.2103271484375 0 1214.8802
396.1219482421875 0 938.3723 b 2
398.1494140625 0 4259.664
404.19622802734375 0 706.22363
405.6998596191406 0 979.4498
406.192626953125 0 970.1067
407.2161560058594 0 1188.048
414.7056579589844 0 2298.538
415.2071533203125 0 1566.6174
415.7300720214844 0 718.3975 y Water loss 2
417.224609375 0 976.9639
418.20855712890625 0 9726.304 y Ammonia loss 6
419.21173095703125 0 2935.4368
423.2345886230469 0 745.2839
423.7102966308594 0 5718.3545
424.2123718261719 0 2610.458
424.7350769042969 0 24341.713 y 2
425.2362976074219 0 12406.388
425.7378234863281 0 3221.8037
426.2337341308594 0 1360.0859
427.2117919921875 0 1340.6573
429.72760009765625 0 1543.8734
430.1909484863281 0 917.25824
435.235107421875 0 30797.72 y 6
436.2384948730469 0 7543.965
437.24273681640625 0 632.41266
439.2677917480469 0 683.69543
440.1600036621094 0 1405.1212
440.2496032714844 0 806.49164
442.2299499511719 0 842.0134
447.71038818359375 0 3272.2104 b 6
448.211669921875 0 1872.582
451.22967529296875 0 4460.3394
451.740478515625 0 3685.05
452.2381591796875 0 2401.546
452.74041748046875 0 631.4188
455.207275390625 0 639.68494
458.17083740234375 0 7922.613
458.2604675292969 0 2831.1233
459.1742248535156 0 1058.6517
459.2458801269531 0 863.0154
462.2237548828125 0 6934.587
462.7259521484375 0 2996.9443
463.2250671386719 0 704.1181
463.72613525390625 0 873.5458
469.1856994628906 0 2233.2417
469.24102783203125 0 3193.5144
471.3028564453125 0 1001.313
473.2327575683594 0 676.6098
476.18096923828125 0 18178.664
477.1846618652344 0 3792.7236
478.1806640625 0 737.4967
482.2395324707031 0 1974.5339
482.27276611328125 0 1676.1228
482.7392578125 0 1797.722
484.29315185546875 0 1730.0155
486.26708984375 0 6828.311
487.2334289550781 0 734.28503 b Water loss 7
487.27069091796875 0 1875.2361
487.7254943847656 0 2197.139 b Ammonia loss 7
488.2264709472656 0 1624.1104
488.2975769042969 0 1370.0042
496.2370300292969 0 25017.332 b 7
496.7367248535156 0 14931.804 y Ammonia loss 1
497.23773193359375 0 6120.7114
497.734619140625 0 1969.6892
498.23553466796875 0 1010.3339
499.2989807128906 0 3992.9963
500.3018493652344 0 787.5715
502.159423828125 0 1177.4412
505.2420349121094 0 114972.35 y 1
505.743408203125 0 55396.258
506.2438659667969 0 21704.287
506.7439880371094 0 6425.8276
507.2452392578125 0 946.6024
513.3143920898438 0 1508.9359
514.3181762695312 0 970.20233
517.3077392578125 0 755.3737
527.8079833984375 0 1318.8481
528.3113403320312 0 1396.6604
529.2451171875 0 1415.4792
533.7234497070312 0 919.7816
534.3031005859375 0 98860.33 y 5
535.3062133789062 0 30907.535
536.3089599609375 0 5588.36
537.3121337890625 0 652.9597
542.7307739257812 0 2540.6538
543.230224609375 0 1954.1321
544.2877807617188 0 1715.1445
545.2913208007812 0 630.18256
547.253173828125 0 1711.5762
551.328125 0 775.9173
553.3104858398438 0 1440.8108
557.2398681640625 0 1455.6654
570.3356323242188 0 2587.4236
571.340087890625 0 779.3238
575.2490844726562 0 1794.3478
576.256591796875 0 715.90186
582.2702026367188 0 2368.6414
584.25 0 944.3223
596.3515625 0 662.4554
603.3262329101562 0 1098.8048
606.2987670898438 0 5788.1914
607.3018798828125 0 1643.3967
614.3607788085938 0 2538.4988
622.2169189453125 0 1187.06 b Ammonia loss 4
631.3212890625 0 1231.8021 y Water loss 4
632.3065185546875 0 2236.4573 y Ammonia loss 4
639.2443237304688 0 1407.8513 b 4
649.329833984375 0 113400.64 y 4
650.3323974609375 0 35611.523
651.3348388671875 0 8009.2876
652.34033203125 0 1183.6375
655.284912109375 0 1723.0844
659.313720703125 0 3402.7966
667.385009765625 0 743.48346
685.3983154296875 0 1315.779
713.3388061523438 0 1725.4535
714.3240356445312 0 2542.5645
715.32763671875 0 1205.569
717.385009765625 0 660.34564
731.3496704101562 0 4309.54
732.3506469726562 0 1522.9233
733.3580932617188 0 620.8266
737.338623046875 0 4173.6406
738.3417358398438 0 1929.2401
759.4146118164062 0 955.6011 y Water loss 3
760.3980102539062 0 1220.3717 y Ammonia loss 3
768.3709106445312 0 1329.3057
769.3744506835938 0 718.2735
777.424072265625 0 23701.445 y 3
778.42724609375 0 10100.285
779.4296264648438 0 2765.9866
789.409423828125 0 566.6149
816.30078125 0 1451.3557
828.4039916992188 0 683.9104
848.4606323242188 0 3885.644 y 2
849.466064453125 0 1209.8708
929.38427734375 0 666.0876
1090.4185791015625 0 577.9712
1135.5010986328125 0 651.97
1191.3070068359375 0 597.73114
1210.9285888671875 0 581.6652
2346.248291015625 0 624.49927
3082.82470703125 0 952.2747

Spectrum Details

|  |  |
| --- | --- |
| Matched peaks? Matched peaks The total absolute number of peaks matched. Additionally in brackets the total fraction of peaks matched and the total number of peaks is shown. | 36 (8.96% of 402) |
| FDR? FDR The false discovery rate estimated for this peptide. It is calculated by matching all theoretical fragments with a non-integer shift with the raw peaks for this spectrum. This is done with 40 different shifts. The resulting percentage is the average number of annotated peaks over the number of annotated peaks with the correct spectrum. | 1.06% |
| Satellite FDR? Satellite FDR See the FDR for details on its calculation. This satellite ion specific FDR only contains the satellite ions (d/w) for I/L/J positions. | - |
| PSM Score? PSM Score The PSM Score as given by Hecklib to this annotated spectrum. It is shown with three significant figures. | 234 |

#### Spectrum 3990? Spectrum 3990 The raw spectrum of this peptide as annotated by Hecklib. The fragments are coloured according to ion type (see legend). Any peaks with a star '\*' as text can be hovered over to see the full details, first the ion type second the mass shift type. By hovering over the amino acids in the peptide or ions in the legend the corresponding peaks are highlighted. By toggling the 'Unassigned' label you can turn the background (unassigned) peaks on or off in the plot. By updating the slider in the Ion legend you can update the spectrum to only show the top X% of the peaks with labels. The top X% means any peak that is within X% of the highest intensity. By dragging in the spectrum you can zoom in to a specific part of the spectrum and use 'Zoom Out' to get back to the original zoom level. The annotation of the spectrum is based on the given sequence in the peptides file and is done with different software so inconsistencies are likely. The peaks are annotated based on the given sequence, with 20 ppm tolerance.

Copy Data

##### Spectrum 3990 (TSV)

###### Preview

```
Loading example...
```

*Click on the button to copy the data to your clipboard.*

Mz MinMz MaxIntensity Max

WidthHeightPeptide font sizePeptide stroke widthSpectrum font sizeSpectrum stroke widthCompact peptide

Ion legend

wxyz

abcd

OtherUnassignedIon^Charge^_Position_Show for top:%

YCAKDVRPY

01.16e+52.32e+53.48e+54.65e+5

Zoom Out

y^+1^_1_b^+2^_4_y^+1^_2_a^+1^_2_b^+2^_5_b^+1^_2_y^+3^_8_b^+2^_6_y^+2^_6_y^+2^_6_\*b^+1^_3_y^+1^_3_y^+2^_7_y^+1^_3_b^+2^_7_b^+2^_7_b^+2^_8_b^+2^_8_y^+2^_8_y^+2^_8_y^+1^_4_b^+1^_5_y^+1^_5_y^+1^_5_b^+1^_5_y^+1^_5_y^+1^_6_y^+1^_6_y^+1^_7_

0778155723353114

Fragment Matches Table

Show background peaks

| Position | Ion type | Intensity | mz Theoretical | mz Error (Th) | mz Error (ppm) | Charge | Series Number |
| --- | --- | --- | --- | --- | --- | --- | --- |
| - | - | 2852 | 120.1 | - | - | 0 | - |
| - | - | 978.9 | 120.1 | - | - | 0 | - |
| - | - | 1788 | 121 | - | - | 0 | - |
| - | - | 438.9 | 121.1 | - | - | 0 | - |
| - | - | 742.1 | 123 | - | - | 0 | - |
| - | - | 508.8 | 124.1 | - | - | 0 | - |
| - | - | 909.6 | 127.1 | - | - | 0 | - |
| - | - | 9.065E+04 | 129.1 | - | - | 0 | - |
| - | - | 871.1 | 130.1 | - | - | 0 | - |
| - | - | 4802 | 130.1 | - | - | 0 | - |
| - | - | 2.578E+04 | 134 | - | - | 0 | - |
| - | - | 969.9 | 135 | - | - | 0 | - |
| - | - | 443.9 | 135.1 | - | - | 0 | - |
| - | - | 907.3 | 136 | - | - | 0 | - |
| - | - | 2.397E+05 | 136.1 | - | - | 0 | - |
| - | - | 847.8 | 137.1 | - | - | 0 | - |
| - | - | 696.8 | 137.1 | - | - | 0 | - |
| - | - | 2.078E+04 | 137.1 | - | - | 0 | - |
| - | - | 646 | 138.1 | - | - | 0 | - |
| - | - | 1352 | 138.1 | - | - | 0 | - |
| - | - | 2839 | 139.1 | - | - | 0 | - |
| - | - | 961.5 | 140.1 | - | - | 0 | - |
| - | - | 422.5 | 141.1 | - | - | 0 | - |
| - | - | 725.3 | 141.1 | - | - | 0 | - |
| - | - | 2.203E+04 | 147 | - | - | 0 | - |
| - | - | 2285 | 148 | - | - | 0 | - |
| - | - | 943.6 | 149 | - | - | 0 | - |
| - | - | 4.6E+05 | 149 | - | - | 0 | - |
| - | - | 3.594E+04 | 150 | - | - | 0 | - |
| - | - | 518.1 | 155.1 | - | - | 0 | - |
| - | - | 1550 | 155.1 | - | - | 0 | - |
| - | - | 953.4 | 156.1 | - | - | 0 | - |
| - | - | 2863 | 157.1 | - | - | 0 | - |
| - | - | 494.7 | 158.1 | - | - | 0 | - |
| - | - | 1639 | 159.1 | - | - | 0 | - |
| - | - | 2380 | 160 | - | - | 0 | - |
| - | - | 926.3 | 160.1 | - | - | 0 | - |
| - | - | 466.2 | 160.1 | - | - | 0 | - |
| - | - | 4115 | 162 | - | - | 0 | - |
| - | - | 552.6 | 164.1 | - | - | 0 | - |
| - | - | 2664 | 165.1 | - | - | 0 | - |
| - | - | 891.6 | 165.1 | - | - | 0 | - |
| - | - | 835.7 | 166.1 | - | - | 0 | - |
| - | - | 5.134E+04 | 167 | - | - | 0 | - |
| - | - | 608.5 | 167.1 | - | - | 0 | - |
| - | - | 3431 | 168 | - | - | 0 | - |
| - | - | 941.6 | 173.1 | - | - | 0 | - |
| - | - | 1218 | 175.1 | - | - | 0 | - |
| - | - | 470 | 178.1 | - | - | 0 | - |
| - | - | 997.7 | 180.1 | - | - | 0 | - |
| - | - | 985.9 | 181.1 | - | - | 0 | - |
| 9 | y | 3512 | 182.1 | 0.0002817 | 1.547 | +1 | 1 |
| - | - | 559.2 | 182.1 | - | - | 0 | - |
| - | - | 1954 | 183.1 | - | - | 0 | - |
| - | - | 1112 | 183.1 | - | - | 0 | - |
| - | - | 679.5 | 185 | - | - | 0 | - |
| - | - | 2960 | 185.1 | - | - | 0 | - |
| - | - | 3531 | 187.1 | - | - | 0 | - |
| - | - | 1.041E+04 | 187.1 | - | - | 0 | - |
| - | - | 1143 | 188.1 | - | - | 0 | - |
| - | - | 806.1 | 192 | - | - | 0 | - |
| - | - | 626.1 | 194.1 | - | - | 0 | - |
| - | - | 1689 | 195.1 | - | - | 0 | - |
| - | - | 1954 | 197.1 | - | - | 0 | - |
| - | - | 2952 | 199.1 | - | - | 0 | - |
| - | - | 9553 | 200.1 | - | - | 0 | - |
| - | - | 981.3 | 201.1 | - | - | 0 | - |
| - | - | 3.416E+04 | 205.1 | - | - | 0 | - |
| - | - | 2363 | 206.1 | - | - | 0 | - |
| - | - | 1232 | 207.1 | - | - | 0 | - |
| - | - | 1702 | 209.1 | - | - | 0 | - |
| - | - | 1122 | 209.1 | - | - | 0 | - |
| - | - | 722 | 210.1 | - | - | 0 | - |
| - | - | 623.9 | 214.2 | - | - | 0 | - |
| - | - | 956 | 215 | - | - | 0 | - |
| - | - | 2708 | 215.1 | - | - | 0 | - |
| - | - | 807.4 | 216.1 | - | - | 0 | - |
| - | - | 1175 | 221.1 | - | - | 0 | - |
| - | - | 1061 | 223.1 | - | - | 0 | - |
| - | - | 831.9 | 224.1 | - | - | 0 | - |
| - | - | 816.3 | 224.1 | - | - | 0 | - |
| - | - | 1942 | 225 | - | - | 0 | - |
| - | - | 864.2 | 225.1 | - | - | 0 | - |
| - | - | 711.3 | 226 | - | - | 0 | - |
| - | - | 4912 | 226.1 | - | - | 0 | - |
| - | - | 804.3 | 227 | - | - | 0 | - |
| - | - | 1873 | 227.1 | - | - | 0 | - |
| - | - | 5302 | 233.1 | - | - | 0 | - |
| - | - | 2405 | 233.1 | - | - | 0 | - |
| - | - | 1570 | 233.1 | - | - | 0 | - |
| - | - | 1603 | 234.1 | - | - | 0 | - |
| - | - | 2068 | 237.1 | - | - | 0 | - |
| - | - | 2.37E+04 | 239.2 | - | - | 0 | - |
| - | - | 2903 | 240.2 | - | - | 0 | - |
| - | - | 776.1 | 243.1 | - | - | 0 | - |
| - | - | 2589 | 244.1 | - | - | 0 | - |
| - | - | 649.7 | 249.1 | - | - | 0 | - |
| - | - | 617.8 | 250.1 | - | - | 0 | - |
| - | - | 2472 | 251 | - | - | 0 | - |
| - | - | 4163 | 252.1 | - | - | 0 | - |
| - | - | 783.5 | 253.1 | - | - | 0 | - |
| - | - | 4293 | 254.2 | - | - | 0 | - |
| - | - | 3231 | 255.1 | - | - | 0 | - |
| - | - | 9726 | 256.2 | - | - | 0 | - |
| - | - | 1347 | 257.2 | - | - | 0 | - |
| 4 | b | 787.7 | 262.6 | 0.002755 | 10.49 | +2 | 4 |
| - | - | 887.6 | 265.1 | - | - | 0 | - |
| - | - | 1657 | 272.1 | - | - | 0 | - |
| - | - | 2007 | 272.2 | - | - | 0 | - |
| - | - | 3119 | 274.2 | - | - | 0 | - |
| 8 | y | 4.025E+04 | 279.1 | 0.0002217 | 0.7944 | +1 | 2 |
| - | - | 5091 | 279.2 | - | - | 0 | - |
| - | - | 1182 | 280.1 | - | - | 0 | - |
| - | - | 5260 | 280.1 | - | - | 0 | - |
| - | - | 727.6 | 280.2 | - | - | 0 | - |
| - | - | 971.3 | 281.1 | - | - | 0 | - |
| - | - | 1186 | 282.2 | - | - | 0 | - |
| - | - | 1116 | 284.2 | - | - | 0 | - |
| - | - | 1738 | 285 | - | - | 0 | - |
| - | - | 541 | 286 | - | - | 0 | - |
| - | - | 671.8 | 291 | - | - | 0 | - |
| 2 | a | 2.096E+04 | 297.1 | 0.004177 | 14.06 | +1 | 2 |
| - | - | 1081 | 297.2 | - | - | 0 | - |
| - | - | 2509 | 298.1 | - | - | 0 | - |
| - | - | 657.1 | 298.1 | - | - | 0 | - |
| - | - | 957.6 | 299.1 | - | - | 0 | - |
| - | - | 1157 | 299.2 | - | - | 0 | - |
| - | - | 1562 | 300.1 | - | - | 0 | - |
| - | - | 675.6 | 301.1 | - | - | 0 | - |
| - | - | 1144 | 302.6 | - | - | 0 | - |
| - | - | 1233 | 304.1 | - | - | 0 | - |
| - | - | 991.2 | 307.7 | - | - | 0 | - |
| - | - | 3692 | 315.2 | - | - | 0 | - |
| - | - | 947.3 | 315.2 | - | - | 0 | - |
| - | - | 627.2 | 318.4 | - | - | 0 | - |
| 5 | b | 839.1 | 320.1 | 0.002463 | 7.694 | +2 | 5 |
| 2 | b | 4752 | 325.1 | 0.004096 | 12.6 | +1 | 2 |
| - | - | 560.1 | 326.6 | - | - | 0 | - |
| 2 | y | 980.2 | 337.2 | 0.0001525 | 0.4524 | +3 | 8 |
| - | - | 1157 | 343.1 | - | - | 0 | - |
| - | - | 752 | 343.2 | - | - | 0 | - |
| - | - | 3179 | 343.2 | - | - | 0 | - |
| - | - | 722.2 | 344.2 | - | - | 0 | - |
| - | - | 761.3 | 353.2 | - | - | 0 | - |
| - | - | 1408 | 354.2 | - | - | 0 | - |
| - | - | 903.2 | 354.9 | - | - | 0 | - |
| - | - | 3381 | 355.1 | - | - | 0 | - |
| - | - | 1147 | 355.7 | - | - | 0 | - |
| - | - | 2273 | 356.1 | - | - | 0 | - |
| - | - | 771.2 | 356.2 | - | - | 0 | - |
| - | - | 2253 | 357.1 | - | - | 0 | - |
| - | - | 1521 | 357.2 | - | - | 0 | - |
| 6 | b | 1983 | 361.1 | 0.004803 | 13.3 | +2 | 6 |
| - | - | 2041 | 366.2 | - | - | 0 | - |
| - | - | 694.6 | 366.7 | - | - | 0 | - |
| - | - | 638.5 | 370.3 | - | - | 0 | - |
| - | - | 1300 | 370.5 | - | - | 0 | - |
| - | - | 1.23E+04 | 371.2 | - | - | 0 | - |
| - | - | 9760 | 371.2 | - | - | 0 | - |
| - | - | 2070 | 372.2 | - | - | 0 | - |
| - | - | 2219 | 372.2 | - | - | 0 | - |
| - | - | 759.9 | 373.9 | - | - | 0 | - |
| - | - | 939.4 | 376.2 | - | - | 0 | - |
| - | - | 862.6 | 376.5 | - | - | 0 | - |
| - | - | 1133 | 379.2 | - | - | 0 | - |
| - | - | 5137 | 379.8 | - | - | 0 | - |
| - | - | 3394 | 380.2 | - | - | 0 | - |
| 4 | y | 9180 | 380.7 | 4.854E-05 | 0.1275 | +2 | 6 |
| - | - | 2907 | 381.2 | - | - | 0 | - |
| - | - | 743.9 | 381.7 | - | - | 0 | - |
| - | - | 1111 | 383.2 | - | - | 0 | - |
| - | - | 584 | 386.2 | - | - | 0 | - |
| 4 | y | 2.217E+04 | 389.2 | 0.0001712 | 0.4399 | +2 | 6 |
| - | - | 9333 | 389.7 | - | - | 0 | - |
| - | - | 2529 | 390.2 | - | - | 0 | - |
| 0 | Precursor | 7633 | 391.5 | 0.001354 | 3.459 | +3 | -1 |
| - | - | 6086 | 391.9 | - | - | 0 | - |
| - | - | 2442 | 392.2 | - | - | 0 | - |
| - | - | 775 | 392.8 | - | - | 0 | - |
| - | - | 1093 | 394.2 | - | - | 0 | - |
| 3 | b | 752.7 | 396.1 | 0.005871 | 14.82 | +1 | 3 |
| - | - | 973.3 | 406.2 | - | - | 0 | - |
| - | - | 2054 | 414.7 | - | - | 0 | - |
| - | - | 1190 | 417.2 | - | - | 0 | - |
| 7 | y | 7381 | 418.2 | 0.0003058 | 0.7312 | +1 | 3 |
| - | - | 1742 | 419.2 | - | - | 0 | - |
| - | - | 2590 | 423.7 | - | - | 0 | - |
| - | - | 1167 | 424.2 | - | - | 0 | - |
| 3 | y | 1.926E+04 | 424.7 | 0.0004131 | 0.9727 | +2 | 7 |
| - | - | 8639 | 425.2 | - | - | 0 | - |
| - | - | 1604 | 425.7 | - | - | 0 | - |
| - | - | 1023 | 426.2 | - | - | 0 | - |
| - | - | 602.6 | 433.7 | - | - | 0 | - |
| 7 | y | 2.276E+04 | 435.2 | 0.0003986 | 0.9157 | +1 | 3 |
| - | - | 5471 | 436.2 | - | - | 0 | - |
| 7 | b | 1340 | 439.2 | 0.001594 | 3.629 | +2 | 7 |
| - | - | 619 | 439.3 | - | - | 0 | - |
| - | - | 1361 | 440.2 | - | - | 0 | - |
| - | - | 815.8 | 440.2 | - | - | 0 | - |
| 7 | b | 2268 | 447.7 | 0.001746 | 3.899 | +2 | 7 |
| - | - | 2654 | 451.7 | - | - | 0 | - |
| - | - | 740.8 | 452.2 | - | - | 0 | - |
| - | - | 768.3 | 457.3 | - | - | 0 | - |
| - | - | 6553 | 458.2 | - | - | 0 | - |
| - | - | 849.3 | 458.3 | - | - | 0 | - |
| - | - | 789.1 | 459.2 | - | - | 0 | - |
| - | - | 1204 | 468.3 | - | - | 0 | - |
| - | - | 1.279E+04 | 476.2 | - | - | 0 | - |
| - | - | 1988 | 477.2 | - | - | 0 | - |
| - | - | 699.5 | 478.2 | - | - | 0 | - |
| - | - | 1483 | 482.2 | - | - | 0 | - |
| - | - | 1033 | 482.3 | - | - | 0 | - |
| - | - | 1352 | 482.7 | - | - | 0 | - |
| - | - | 5719 | 486.3 | - | - | 0 | - |
| - | - | 728 | 487.3 | - | - | 0 | - |
| 8 | b | 1579 | 487.7 | 0.0007718 | 1.583 | +2 | 8 |
| - | - | 648.8 | 488.2 | - | - | 0 | - |
| - | - | 797.1 | 488.3 | - | - | 0 | - |
| 8 | b | 1.647E+04 | 496.2 | 0.001913 | 3.855 | +2 | 8 |
| 2 | y | 1.132E+04 | 496.7 | 0.00614 | 12.36 | +2 | 8 |
| - | - | 3481 | 497.2 | - | - | 0 | - |
| - | - | 3035 | 499.3 | - | - | 0 | - |
| - | - | 940.5 | 502.2 | - | - | 0 | - |
| 2 | y | 7.789E+04 | 505.2 | 0.001977 | 3.913 | +2 | 8 |
| - | - | 3.915E+04 | 505.7 | - | - | 0 | - |
| - | - | 1.409E+04 | 506.2 | - | - | 0 | - |
| - | - | 5343 | 506.7 | - | - | 0 | - |
| - | - | 609.8 | 528.3 | - | - | 0 | - |
| 6 | y | 7.134E+04 | 534.3 | 0.0001748 | 0.3271 | +1 | 4 |
| - | - | 2.094E+04 | 535.3 | - | - | 0 | - |
| - | - | 4392 | 536.3 | - | - | 0 | - |
| - | - | 627.9 | 537.3 | - | - | 0 | - |
| - | - | 648.5 | 542.3 | - | - | 0 | - |
| - | - | 1186 | 544.3 | - | - | 0 | - |
| - | - | 1091 | 547.3 | - | - | 0 | - |
| - | - | 652.9 | 557.2 | - | - | 0 | - |
| - | - | 1566 | 570.3 | - | - | 0 | - |
| - | - | 1143 | 575.3 | - | - | 0 | - |
| - | - | 840.9 | 603.3 | - | - | 0 | - |
| - | - | 852.1 | 614.4 | - | - | 0 | - |
| 5 | b | 754.9 | 622.2 | 0.002351 | 3.778 | +1 | 5 |
| 5 | y | 1134 | 631.3 | 0.0002926 | 0.4634 | +1 | 5 |
| 5 | y | 1095 | 632.3 | 0.001423 | 2.251 | +1 | 5 |
| 5 | b | 1478 | 639.2 | 0.004119 | 6.444 | +1 | 5 |
| 5 | y | 8.216E+04 | 649.3 | 0.0001403 | 0.216 | +1 | 5 |
| - | - | 2.805E+04 | 650.3 | - | - | 0 | - |
| - | - | 5430 | 651.3 | - | - | 0 | - |
| - | - | 790.5 | 652.3 | - | - | 0 | - |
| - | - | 1922 | 659.3 | - | - | 0 | - |
| - | - | 937.2 | 667.4 | - | - | 0 | - |
| - | - | 761.6 | 685.4 | - | - | 0 | - |
| - | - | 601 | 703.5 | - | - | 0 | - |
| - | - | 1415 | 713.3 | - | - | 0 | - |
| - | - | 664.3 | 714.3 | - | - | 0 | - |
| - | - | 2664 | 731.3 | - | - | 0 | - |
| - | - | 901.5 | 732.4 | - | - | 0 | - |
| 4 | y | 1425 | 760.4 | 0.0002935 | 0.3859 | +1 | 6 |
| 4 | y | 1.599E+04 | 777.4 | 0.0006209 | 0.7986 | +1 | 6 |
| - | - | 6545 | 778.4 | - | - | 0 | - |
| - | - | 1159 | 779.4 | - | - | 0 | - |
| - | - | 783.9 | 787.4 | - | - | 0 | - |
| - | - | 841.6 | 792.6 | - | - | 0 | - |
| 3 | y | 3273 | 848.5 | 0.00325 | 3.83 | +1 | 7 |
| - | - | 1360 | 849.5 | - | - | 0 | - |
| - | - | 574.8 | 1009 | - | - | 0 | - |
| - | - | 767.6 | 1711 | - | - | 0 | - |
| - | - | 700.7 | 2341 | - | - | 0 | - |
| - | - | 973.3 | 3083 | - | - | 0 | - |

m/z Charge Intensity FragmentType MassShift Position
120.05276489257812 0 2852.4602
120.08116149902344 0 978.8642
121.02876281738281 0 1787.7264
121.06507110595703 0 438.90787
123.04452514648438 0 742.08746
124.08675384521484 0 508.81122
127.08712768554688 0 909.6241
129.1025848388672 0 90651.2
130.10011291503906 0 871.1488
130.10592651367188 0 4801.8525
134.02737426757812 0 25775.13
135.03082275390625 0 969.87964
135.0690460205078 0 443.89258
136.02301025390625 0 907.3399
136.0760955810547 0 239651.03
137.05999755859375 0 847.8335
137.0736541748047 0 696.8136
137.0793914794922 0 20777.47
138.06649780273438 0 645.9888
138.08250427246094 0 1352.1907
139.08692932128906 0 2838.7734
140.08212280273438 0 961.5103
141.10244750976562 0 422.49567
141.1135711669922 0 725.2505
147.0443572998047 0 22034.082
148.04763793945312 0 2284.6887
148.95382690429688 0 943.6171
149.02377319335938 0 460043.16
150.02699279785156 0 35938.95
155.0825653076172 0 518.0696
155.1180877685547 0 1549.7832
156.0771484375 0 953.42535
157.1087646484375 0 2863.3933
158.09271240234375 0 494.70972
159.05899047851562 0 1638.8135
160.0428924560547 0 2379.7615
160.07626342773438 0 926.2676
160.13299560546875 0 466.16562
162.02224731445312 0 4115.025
164.07049560546875 0 552.6432
165.05482482910156 0 2663.8687
165.1026611328125 0 891.5889
166.0862274169922 0 835.67676
167.03421020507812 0 51341.727
167.11834716796875 0 608.4647
168.03746032714844 0 3431.0776
173.12879943847656 0 941.5522
175.11904907226562 0 1217.7078
178.13267517089844 0 470.03308
180.10218811035156 0 997.71954
181.09780883789062 0 985.9155
182.08145141601562 0 3512.0356 y 8
182.12921142578125 0 559.21906
183.11302185058594 0 1954.0479
183.1492919921875 0 1111.8536
185.0386199951172 0 679.509
185.10366821289062 0 2960.3188
187.0538330078125 0 3530.5664
187.1080322265625 0 10405.26
188.11160278320312 0 1142.5215
192.04798889160156 0 806.1264
194.12875366210938 0 626.1272
195.11312866210938 0 1688.6621
197.1287384033203 0 1954.2096
199.1079559326172 0 2952.0647
200.13958740234375 0 9553.308
201.12393188476562 0 981.287
205.06442260742188 0 34163.715
206.0679473876953 0 2363.029
207.05972290039062 0 1232.1887
209.0926513671875 0 1701.5065
209.13990783691406 0 1121.8828
210.123291015625 0 721.9701
214.1556396484375 0 623.8816
215.04898071289062 0 955.9638
215.103271484375 0 2707.7046
216.06622314453125 0 807.3772
221.09242248535156 0 1175.1965
223.0641632080078 0 1061.1608
224.06390380859375 0 831.8924
224.1391143798828 0 816.2539
225.04307556152344 0 1941.547
225.06101989746094 0 864.1653
226.04287719726562 0 711.3366
226.11880493164062 0 4911.697
227.03936767578125 0 804.331
227.10267639160156 0 1872.8477
233.0593719482422 0 5301.996
233.09249877929688 0 2404.805
233.12901306152344 0 1570.2828
234.0594940185547 0 1602.9503
237.1348419189453 0 2067.8945
239.15052795410156 0 23701.996
240.15374755859375 0 2902.996
243.0982208251953 0 776.0845
244.12950134277344 0 2589.1187
249.13543701171875 0 649.66534
250.08734130859375 0 617.7512
251.0470733642578 0 2472.051
252.069091796875 0 4162.8247
253.07247924804688 0 783.48596
254.16139221191406 0 4292.655
255.1453094482422 0 3231.4177
256.177001953125 0 9725.577
257.1800842285156 0 1347.1196
262.61181640625 0 787.6723 b 3
265.1274719238281 0 887.57477
272.1245422363281 0 1657.0557
272.171875 0 2007.4701
274.18756103515625 0 3119.0852
279.1341552734375 0 40249.805 y 7
279.1600646972656 0 5091.1685
280.06378173828125 0 1182.1348
280.13739013671875 0 5260.2314
280.16259765625 0 727.63196
281.1396484375 0 971.3194
282.1556091308594 0 1185.6056
284.1703796386719 0 1116.1871
285.00982666015625 0 1737.7173
286.01043701171875 0 540.95105
290.95953369140625 0 671.777
297.0906982421875 0 20960.055 a 1
297.1556701660156 0 1080.5088
298.0941162109375 0 2508.532
298.1211853027344 0 657.07404
299.08685302734375 0 957.6092
299.1711120605469 0 1157.4581
300.0626220703125 0 1562.4781
301.0596618652344 0 675.6293
302.6075134277344 0 1144.4718
304.1300048828125 0 1233.057
307.685546875 0 991.186
315.1666564941406 0 3692.1228
315.20343017578125 0 947.34143
318.43804931640625 0 627.21
320.1255798339844 0 839.08484 b 4
325.085693359375 0 4752.4253 b 1
326.6468505859375 0 560.13025
337.1652526855469 0 980.2472 y 1
343.14349365234375 0 1156.7999
343.16351318359375 0 752.0147
343.2081298828125 0 3179.0579
344.2086486816406 0 722.1803
353.1910400390625 0 761.3328
354.1771545410156 0 1407.6868
354.88525390625 0 903.2492
355.07025146484375 0 3380.6335
355.6627197265625 0 1146.9835
356.06982421875 0 2273.4639
356.16412353515625 0 771.18976
357.0671081542969 0 2252.5002
357.1739196777344 0 1521.238
361.1537780761719 0 1983.0239 b Ammonia loss 5
366.17877197265625 0 2041.23
366.6802978515625 0 694.6312
370.25469970703125 0 638.5084
370.50921630859375 0 1300.1765
371.2037048339844 0 12295.657
371.2401123046875 0 9759.774
372.2055358886719 0 2070.4534
372.2435607910156 0 2219.4792
373.9175720214844 0 759.9043
376.1863708496094 0 939.3867
376.5152893066406 0 862.6037
379.1669616699219 0 1132.9285
379.839599609375 0 5136.62
380.1733703613281 0 3393.9692
380.7030944824219 0 9180.209 y Ammonia loss 3
381.2049255371094 0 2907.1653
381.70654296875 0 743.9325
383.2040100097656 0 1110.7144
386.1794738769531 0 583.96936
389.21649169921875 0 22165.414 y 3
389.7179260253906 0 9332.692
390.2197570800781 0 2528.7432
391.51849365234375 0 7632.616 Precursor
391.85272216796875 0 6086.1143
392.1862487792969 0 2442.4749
392.838623046875 0 775.0225
394.2093200683594 0 1092.5989
396.12103271484375 0 752.6965 b 2
406.1960754394531 0 973.3262
414.7059020996094 0 2053.8118
417.22430419921875 0 1189.6332
418.20880126953125 0 7380.5425 y Ammonia loss 6
419.2119140625 0 1741.6011
423.7102355957031 0 2589.759
424.211669921875 0 1167.1885
424.73529052734375 0 19256.732 y 2
425.23675537109375 0 8639.047
425.73712158203125 0 1603.8159
426.2391052246094 0 1022.763
433.7136535644531 0 602.5971
435.2354431152344 0 22760.291 y 6
436.23797607421875 0 5471.4863
439.19793701171875 0 1340.4723 b Ammonia loss 6
439.26031494140625 0 618.9658
440.1598205566406 0 1360.8087
440.2494201660156 0 815.83514
447.7110595703125 0 2267.882 b 6
451.7402038574219 0 2653.9832
452.2409973144531 0 740.8049
457.27862548828125 0 768.3255
458.17095947265625 0 6553.3027
458.2608337402344 0 849.335
459.1767883300781 0 789.075
468.2571716308594 0 1203.7582
476.18133544921875 0 12787.248
477.1825866699219 0 1987.6619
478.1808776855469 0 699.46906
482.23773193359375 0 1483.2406
482.2705078125 0 1033.461
482.7395935058594 0 1352.0496
486.2674560546875 0 5718.779
487.2693786621094 0 727.9569
487.7266845703125 0 1578.9398 b Ammonia loss 7
488.2220458984375 0 648.8284
488.29937744140625 0 797.125
496.2372741699219 0 16469.639 b 7
496.7373352050781 0 11321.633 y Ammonia loss 1
497.239013671875 0 3481.416
499.2994079589844 0 3034.904
502.1601867675781 0 940.5236
505.24249267578125 0 77894.61 y 1
505.7436828613281 0 39154.133
506.2442321777344 0 14091.863
506.7449035644531 0 5342.59
528.3133544921875 0 609.7669
534.3032836914062 0 71343.83 y 5
535.306396484375 0 20943.709
536.30859375 0 4391.6846
537.3092041015625 0 627.93304
542.3388061523438 0 648.52435
544.2886352539062 0 1186.3561
547.2551879882812 0 1091.0383
557.2386474609375 0 652.8766
570.3367309570312 0 1565.5696
575.2510375976562 0 1142.665
603.32373046875 0 840.9284
614.3601684570312 0 852.09796
622.2199096679688 0 754.8959 b Ammonia loss 4
631.3201293945312 0 1134.1387 y Water loss 4
632.3024291992188 0 1095.1179 y Ammonia loss 4
639.2446899414062 0 1478.0154 b 4
649.3302612304688 0 82155.21 y 4
650.3327026367188 0 28054.725
651.335205078125 0 5430.128
652.3384399414062 0 790.52124
659.3150024414062 0 1922.3679
667.3853759765625 0 937.23413
685.3983764648438 0 761.64484
703.4602661132812 0 601.0177
713.3388061523438 0 1414.5504
714.3253173828125 0 664.3319
731.3486938476562 0 2663.9314
732.3536987304688 0 901.47705
760.3991088867188 0 1425.4315 y Ammonia loss 3
777.4247436523438 0 15985.805 y 3
778.4273071289062 0 6544.6265
779.4288330078125 0 1159.1973
787.4105224609375 0 783.9054
792.64013671875 0 841.6194
848.459228515625 0 3272.5686 y 2
849.4638061523438 0 1359.8032
1009.3682861328125 0 574.78864
1710.7623291015625 0 767.56946
2341.064697265625 0 700.73834
3083.15966796875 0 973.3026

Spectrum Details

|  |  |
| --- | --- |
| Matched peaks? Matched peaks The total absolute number of peaks matched. Additionally in brackets the total fraction of peaks matched and the total number of peaks is shown. | 30 (11.19% of 268) |
| FDR? FDR The false discovery rate estimated for this peptide. It is calculated by matching all theoretical fragments with a non-integer shift with the raw peaks for this spectrum. This is done with 40 different shifts. The resulting percentage is the average number of annotated peaks over the number of annotated peaks with the correct spectrum. | 0.87% |
| Satellite FDR? Satellite FDR See the FDR for details on its calculation. This satellite ion specific FDR only contains the satellite ions (d/w) for I/L/J positions. | - |
| PSM Score? PSM Score The PSM Score as given by Hecklib to this annotated spectrum. It is shown with three significant figures. | 215 |

#### Spectrum 3845? Spectrum 3845 The raw spectrum of this peptide as annotated by Hecklib. The fragments are coloured according to ion type (see legend). Any peaks with a star '\*' as text can be hovered over to see the full details, first the ion type second the mass shift type. By hovering over the amino acids in the peptide or ions in the legend the corresponding peaks are highlighted. By toggling the 'Unassigned' label you can turn the background (unassigned) peaks on or off in the plot. By updating the slider in the Ion legend you can update the spectrum to only show the top X% of the peaks with labels. The top X% means any peak that is within X% of the highest intensity. By dragging in the spectrum you can zoom in to a specific part of the spectrum and use 'Zoom Out' to get back to the original zoom level. The annotation of the spectrum is based on the given sequence in the peptides file and is done with different software so inconsistencies are likely. The peaks are annotated based on the given sequence, with 20 ppm tolerance.

Copy Data

##### Spectrum 3845 (TSV)

###### Preview

```
Loading example...
```

*Click on the button to copy the data to your clipboard.*

Mz MinMz MaxIntensity Max

WidthHeightPeptide font sizePeptide stroke widthSpectrum font sizeSpectrum stroke widthCompact peptide

Ion legend

wxyz

abcd

OtherUnassignedIon^Charge^_Position_Show for top:%

YCAKDVRPY

01.21e+52.42e+53.64e+54.85e+5

Zoom Out

y^+1^_1_b^+2^_4_y^+1^_2_a^+1^_2_y^+2^_5_b^+2^_5_b^+1^_2_b^+3^_8_y^+3^_8_b^+2^_6_y^+2^_6_\*\*y^+2^_6_\*b^+1^_3_y^+2^_7_y^+2^_7_y^+1^_3_y^+2^_7_y^+1^_3_b^+2^_7_b^+2^_8_b^+2^_8_b^+2^_8_y^+2^_8_y^+2^_8_y^+1^_4_b^+1^_5_b^+1^_5_y^+1^_5_y^+1^_5_b^+1^_5_y^+1^_5_y^+1^_6_y^+1^_6_y^+1^_6_y^+1^_7_

0778155623343112

Fragment Matches Table

Show background peaks

| Position | Ion type | Intensity | mz Theoretical | mz Error (Th) | mz Error (ppm) | Charge | Series Number |
| --- | --- | --- | --- | --- | --- | --- | --- |
| - | - | 6202 | 120.1 | - | - | 0 | - |
| - | - | 1052 | 120.1 | - | - | 0 | - |
| - | - | 1289 | 121 | - | - | 0 | - |
| - | - | 341.6 | 122.1 | - | - | 0 | - |
| - | - | 1076 | 123 | - | - | 0 | - |
| - | - | 538.2 | 124.1 | - | - | 0 | - |
| - | - | 1012 | 125.1 | - | - | 0 | - |
| - | - | 465.8 | 126.1 | - | - | 0 | - |
| - | - | 2819 | 127.1 | - | - | 0 | - |
| - | - | 2679 | 128.1 | - | - | 0 | - |
| - | - | 1.754E+05 | 129.1 | - | - | 0 | - |
| - | - | 433.7 | 130.1 | - | - | 0 | - |
| - | - | 1493 | 130.1 | - | - | 0 | - |
| - | - | 1.119E+04 | 130.1 | - | - | 0 | - |
| - | - | 599.6 | 131.1 | - | - | 0 | - |
| - | - | 1407 | 132.1 | - | - | 0 | - |
| - | - | 4.688E+04 | 134 | - | - | 0 | - |
| - | - | 2110 | 135 | - | - | 0 | - |
| - | - | 2234 | 136 | - | - | 0 | - |
| - | - | 4.8E+05 | 136.1 | - | - | 0 | - |
| - | - | 1912 | 137.1 | - | - | 0 | - |
| - | - | 3.923E+04 | 137.1 | - | - | 0 | - |
| - | - | 1461 | 138.1 | - | - | 0 | - |
| - | - | 1477 | 138.1 | - | - | 0 | - |
| - | - | 530.3 | 138.1 | - | - | 0 | - |
| - | - | 3897 | 139.1 | - | - | 0 | - |
| - | - | 676.6 | 139.1 | - | - | 0 | - |
| - | - | 2962 | 140.1 | - | - | 0 | - |
| - | - | 562.4 | 140.1 | - | - | 0 | - |
| - | - | 1599 | 141.1 | - | - | 0 | - |
| - | - | 591.5 | 141.1 | - | - | 0 | - |
| - | - | 1063 | 142 | - | - | 0 | - |
| - | - | 1336 | 142.1 | - | - | 0 | - |
| - | - | 742.8 | 143.1 | - | - | 0 | - |
| - | - | 602.6 | 144 | - | - | 0 | - |
| - | - | 507.8 | 146.1 | - | - | 0 | - |
| - | - | 4.332E+04 | 147 | - | - | 0 | - |
| - | - | 4872 | 148 | - | - | 0 | - |
| - | - | 3.42E+05 | 149 | - | - | 0 | - |
| - | - | 2.707E+04 | 150 | - | - | 0 | - |
| - | - | 637.5 | 153.1 | - | - | 0 | - |
| - | - | 513.4 | 154.1 | - | - | 0 | - |
| - | - | 2020 | 155.1 | - | - | 0 | - |
| - | - | 3920 | 155.1 | - | - | 0 | - |
| - | - | 1019 | 156.1 | - | - | 0 | - |
| - | - | 575 | 157.1 | - | - | 0 | - |
| - | - | 1225 | 157.1 | - | - | 0 | - |
| - | - | 7175 | 157.1 | - | - | 0 | - |
| - | - | 522.7 | 158.1 | - | - | 0 | - |
| - | - | 1196 | 158.1 | - | - | 0 | - |
| - | - | 642.4 | 158.1 | - | - | 0 | - |
| - | - | 4022 | 159.1 | - | - | 0 | - |
| - | - | 5177 | 160 | - | - | 0 | - |
| - | - | 2423 | 160.1 | - | - | 0 | - |
| - | - | 8052 | 162 | - | - | 0 | - |
| - | - | 960.8 | 164.1 | - | - | 0 | - |
| - | - | 569.9 | 164.1 | - | - | 0 | - |
| - | - | 5409 | 165.1 | - | - | 0 | - |
| - | - | 1883 | 165.1 | - | - | 0 | - |
| - | - | 1212 | 166.1 | - | - | 0 | - |
| - | - | 811.4 | 166.1 | - | - | 0 | - |
| - | - | 538 | 166.8 | - | - | 0 | - |
| - | - | 3.776E+04 | 167 | - | - | 0 | - |
| - | - | 747.3 | 167.1 | - | - | 0 | - |
| - | - | 3263 | 168 | - | - | 0 | - |
| - | - | 1558 | 169.1 | - | - | 0 | - |
| - | - | 1303 | 169.1 | - | - | 0 | - |
| - | - | 526.9 | 170.1 | - | - | 0 | - |
| - | - | 1527 | 172.1 | - | - | 0 | - |
| - | - | 3.799E+04 | 173.1 | - | - | 0 | - |
| - | - | 4515 | 174.1 | - | - | 0 | - |
| - | - | 674.2 | 175.1 | - | - | 0 | - |
| - | - | 1137 | 175.1 | - | - | 0 | - |
| - | - | 738.3 | 176 | - | - | 0 | - |
| - | - | 1287 | 178.1 | - | - | 0 | - |
| - | - | 492.5 | 179.1 | - | - | 0 | - |
| - | - | 1724 | 180.1 | - | - | 0 | - |
| - | - | 1679 | 181.1 | - | - | 0 | - |
| - | - | 953.1 | 181.1 | - | - | 0 | - |
| 9 | y | 5377 | 182.1 | 0.0004191 | 2.302 | +1 | 1 |
| - | - | 1390 | 182.1 | - | - | 0 | - |
| - | - | 4143 | 183.1 | - | - | 0 | - |
| - | - | 749.7 | 183.1 | - | - | 0 | - |
| - | - | 1486 | 183.1 | - | - | 0 | - |
| - | - | 756.4 | 185 | - | - | 0 | - |
| - | - | 7177 | 185.1 | - | - | 0 | - |
| - | - | 902.2 | 185.2 | - | - | 0 | - |
| - | - | 8510 | 187.1 | - | - | 0 | - |
| - | - | 2.279E+04 | 187.1 | - | - | 0 | - |
| - | - | 923.2 | 188.1 | - | - | 0 | - |
| - | - | 1755 | 188.1 | - | - | 0 | - |
| - | - | 1271 | 191.1 | - | - | 0 | - |
| - | - | 1936 | 192 | - | - | 0 | - |
| - | - | 1600 | 192.1 | - | - | 0 | - |
| - | - | 1186 | 194.1 | - | - | 0 | - |
| - | - | 2268 | 195.1 | - | - | 0 | - |
| - | - | 652.3 | 195.2 | - | - | 0 | - |
| - | - | 1025 | 197.1 | - | - | 0 | - |
| - | - | 2334 | 197.1 | - | - | 0 | - |
| - | - | 5072 | 197.1 | - | - | 0 | - |
| - | - | 1217 | 198.1 | - | - | 0 | - |
| - | - | 4615 | 199.1 | - | - | 0 | - |
| - | - | 3830 | 199.1 | - | - | 0 | - |
| - | - | 634.1 | 200.1 | - | - | 0 | - |
| - | - | 1.924E+04 | 200.1 | - | - | 0 | - |
| - | - | 1161 | 200.6 | - | - | 0 | - |
| - | - | 6521 | 201.1 | - | - | 0 | - |
| - | - | 1122 | 201.1 | - | - | 0 | - |
| - | - | 709.2 | 202.1 | - | - | 0 | - |
| - | - | 1396 | 203.1 | - | - | 0 | - |
| - | - | 862.4 | 204.1 | - | - | 0 | - |
| - | - | 7.176E+04 | 205.1 | - | - | 0 | - |
| - | - | 5367 | 206.1 | - | - | 0 | - |
| - | - | 2992 | 207.1 | - | - | 0 | - |
| - | - | 744.5 | 208.1 | - | - | 0 | - |
| - | - | 2544 | 209.1 | - | - | 0 | - |
| - | - | 624.1 | 209.1 | - | - | 0 | - |
| - | - | 1930 | 209.1 | - | - | 0 | - |
| - | - | 2069 | 210.1 | - | - | 0 | - |
| - | - | 1198 | 213.1 | - | - | 0 | - |
| - | - | 1042 | 214.1 | - | - | 0 | - |
| - | - | 1488 | 214.2 | - | - | 0 | - |
| - | - | 2856 | 215 | - | - | 0 | - |
| - | - | 5152 | 215.1 | - | - | 0 | - |
| - | - | 1387 | 216.1 | - | - | 0 | - |
| - | - | 881.7 | 216.1 | - | - | 0 | - |
| - | - | 755.8 | 220 | - | - | 0 | - |
| - | - | 711.4 | 220.1 | - | - | 0 | - |
| - | - | 1956 | 221.1 | - | - | 0 | - |
| - | - | 1163 | 222.1 | - | - | 0 | - |
| - | - | 993.9 | 223.1 | - | - | 0 | - |
| - | - | 992.8 | 223.2 | - | - | 0 | - |
| - | - | 720.6 | 224.1 | - | - | 0 | - |
| - | - | 1409 | 224.1 | - | - | 0 | - |
| - | - | 1593 | 225 | - | - | 0 | - |
| - | - | 574.7 | 225.1 | - | - | 0 | - |
| - | - | 585 | 226 | - | - | 0 | - |
| - | - | 8821 | 226.1 | - | - | 0 | - |
| - | - | 3687 | 227.1 | - | - | 0 | - |
| - | - | 911.5 | 227.1 | - | - | 0 | - |
| - | - | 841.6 | 227.1 | - | - | 0 | - |
| - | - | 533.1 | 227.9 | - | - | 0 | - |
| - | - | 632.5 | 231 | - | - | 0 | - |
| - | - | 1.069E+04 | 233.1 | - | - | 0 | - |
| - | - | 4906 | 233.1 | - | - | 0 | - |
| - | - | 3595 | 233.1 | - | - | 0 | - |
| - | - | 3138 | 234.1 | - | - | 0 | - |
| - | - | 685.8 | 234.1 | - | - | 0 | - |
| - | - | 683.1 | 235.1 | - | - | 0 | - |
| - | - | 944.5 | 237.1 | - | - | 0 | - |
| - | - | 4245 | 237.1 | - | - | 0 | - |
| - | - | 637.2 | 237.2 | - | - | 0 | - |
| - | - | 716.3 | 238.2 | - | - | 0 | - |
| - | - | 5.546E+04 | 239.2 | - | - | 0 | - |
| - | - | 4489 | 240.2 | - | - | 0 | - |
| - | - | 591.5 | 241.2 | - | - | 0 | - |
| - | - | 952.7 | 243.1 | - | - | 0 | - |
| - | - | 474.4 | 243.1 | - | - | 0 | - |
| - | - | 6209 | 244.1 | - | - | 0 | - |
| - | - | 535.7 | 250.1 | - | - | 0 | - |
| - | - | 1823 | 251 | - | - | 0 | - |
| - | - | 1217 | 251.1 | - | - | 0 | - |
| - | - | 557.7 | 251.2 | - | - | 0 | - |
| - | - | 9074 | 252.1 | - | - | 0 | - |
| - | - | 941.6 | 252.1 | - | - | 0 | - |
| - | - | 1248 | 253.1 | - | - | 0 | - |
| - | - | 1034 | 254.1 | - | - | 0 | - |
| - | - | 6960 | 254.2 | - | - | 0 | - |
| - | - | 5970 | 255.1 | - | - | 0 | - |
| - | - | 867.3 | 255.2 | - | - | 0 | - |
| - | - | 768.7 | 256.1 | - | - | 0 | - |
| - | - | 930.8 | 256.1 | - | - | 0 | - |
| - | - | 2.582E+04 | 256.2 | - | - | 0 | - |
| - | - | 7600 | 257.2 | - | - | 0 | - |
| - | - | 2679 | 257.2 | - | - | 0 | - |
| - | - | 2318 | 257.7 | - | - | 0 | - |
| - | - | 488.5 | 261.1 | - | - | 0 | - |
| - | - | 980 | 262.1 | - | - | 0 | - |
| - | - | 505.2 | 262.2 | - | - | 0 | - |
| 4 | b | 1335 | 262.6 | 0.001412 | 5.377 | +2 | 4 |
| - | - | 1004 | 263.1 | - | - | 0 | - |
| - | - | 1766 | 265.1 | - | - | 0 | - |
| - | - | 1428 | 266.2 | - | - | 0 | - |
| - | - | 1927 | 268.1 | - | - | 0 | - |
| - | - | 1112 | 269.2 | - | - | 0 | - |
| - | - | 592.3 | 270.1 | - | - | 0 | - |
| - | - | 5062 | 272.1 | - | - | 0 | - |
| - | - | 4305 | 272.2 | - | - | 0 | - |
| - | - | 1038 | 273.1 | - | - | 0 | - |
| - | - | 937.5 | 273.1 | - | - | 0 | - |
| - | - | 871.4 | 273.2 | - | - | 0 | - |
| - | - | 6183 | 274.2 | - | - | 0 | - |
| - | - | 561.8 | 276.1 | - | - | 0 | - |
| - | - | 629.8 | 277.1 | - | - | 0 | - |
| - | - | 589.7 | 278.7 | - | - | 0 | - |
| 8 | y | 8.3E+04 | 279.1 | 0.0002217 | 0.7944 | +1 | 2 |
| - | - | 3054 | 279.2 | - | - | 0 | - |
| - | - | 2575 | 280.1 | - | - | 0 | - |
| - | - | 1.287E+04 | 280.1 | - | - | 0 | - |
| - | - | 1365 | 280.2 | - | - | 0 | - |
| - | - | 1222 | 281.1 | - | - | 0 | - |
| - | - | 3454 | 282.2 | - | - | 0 | - |
| - | - | 2618 | 284.2 | - | - | 0 | - |
| - | - | 923.2 | 285 | - | - | 0 | - |
| - | - | 981.9 | 285.2 | - | - | 0 | - |
| - | - | 1398 | 285.7 | - | - | 0 | - |
| - | - | 498.3 | 289.4 | - | - | 0 | - |
| - | - | 533.1 | 291.9 | - | - | 0 | - |
| - | - | 634.5 | 292.1 | - | - | 0 | - |
| - | - | 650.8 | 292.7 | - | - | 0 | - |
| - | - | 3691 | 294.2 | - | - | 0 | - |
| - | - | 908.8 | 295.2 | - | - | 0 | - |
| 2 | a | 4.423E+04 | 297.1 | 0.004177 | 14.06 | +1 | 2 |
| - | - | 2142 | 297.2 | - | - | 0 | - |
| - | - | 7113 | 298.1 | - | - | 0 | - |
| - | - | 1060 | 298.1 | - | - | 0 | - |
| - | - | 1766 | 299.1 | - | - | 0 | - |
| - | - | 1530 | 299.2 | - | - | 0 | - |
| - | - | 825.8 | 300.1 | - | - | 0 | - |
| - | - | 725.7 | 301.1 | - | - | 0 | - |
| - | - | 778.1 | 302.2 | - | - | 0 | - |
| - | - | 1068 | 302.6 | - | - | 0 | - |
| - | - | 1018 | 303.2 | - | - | 0 | - |
| - | - | 4015 | 304.1 | - | - | 0 | - |
| - | - | 621.4 | 305.7 | - | - | 0 | - |
| - | - | 597.4 | 307.7 | - | - | 0 | - |
| - | - | 2486 | 307.7 | - | - | 0 | - |
| - | - | 1060 | 308.1 | - | - | 0 | - |
| - | - | 654.2 | 308.2 | - | - | 0 | - |
| - | - | 2677 | 311.2 | - | - | 0 | - |
| - | - | 845.3 | 312.2 | - | - | 0 | - |
| - | - | 663.1 | 312.7 | - | - | 0 | - |
| - | - | 697.9 | 313.2 | - | - | 0 | - |
| - | - | 582.5 | 313.4 | - | - | 0 | - |
| - | - | 2818 | 314.2 | - | - | 0 | - |
| - | - | 1333 | 314.7 | - | - | 0 | - |
| - | - | 6854 | 315.2 | - | - | 0 | - |
| - | - | 1693 | 315.2 | - | - | 0 | - |
| - | - | 959 | 316.1 | - | - | 0 | - |
| 5 | y | 834.9 | 316.2 | 0.005572 | 17.62 | +2 | 5 |
| - | - | 682.7 | 316.2 | - | - | 0 | - |
| - | - | 789 | 319.1 | - | - | 0 | - |
| - | - | 584.6 | 319.2 | - | - | 0 | - |
| 5 | b | 2792 | 320.1 | 0.002219 | 6.931 | +2 | 5 |
| - | - | 530.7 | 322.5 | - | - | 0 | - |
| 2 | b | 8569 | 325.1 | 0.004096 | 12.6 | +1 | 2 |
| 8 | b | 642.6 | 325.2 | 0.003669 | 11.28 | +3 | 8 |
| - | - | 1391 | 325.2 | - | - | 0 | - |
| - | - | 1439 | 326.1 | - | - | 0 | - |
| - | - | 639.7 | 327.1 | - | - | 0 | - |
| - | - | 623.5 | 333.2 | - | - | 0 | - |
| - | - | 1235 | 336.2 | - | - | 0 | - |
| 2 | y | 1465 | 337.2 | 0.0005188 | 1.539 | +3 | 8 |
| - | - | 734.4 | 337.2 | - | - | 0 | - |
| - | - | 1055 | 337.5 | - | - | 0 | - |
| - | - | 534.8 | 338.1 | - | - | 0 | - |
| - | - | 629.6 | 340.2 | - | - | 0 | - |
| - | - | 1498 | 342.2 | - | - | 0 | - |
| - | - | 2127 | 343.1 | - | - | 0 | - |
| - | - | 5700 | 343.2 | - | - | 0 | - |
| - | - | 1232 | 344.1 | - | - | 0 | - |
| - | - | 582.9 | 344.2 | - | - | 0 | - |
| - | - | 1538 | 353.2 | - | - | 0 | - |
| - | - | 881.4 | 353.2 | - | - | 0 | - |
| - | - | 3704 | 354.2 | - | - | 0 | - |
| - | - | 2246 | 355.1 | - | - | 0 | - |
| - | - | 636.1 | 355.2 | - | - | 0 | - |
| - | - | 2030 | 355.7 | - | - | 0 | - |
| - | - | 1406 | 356.1 | - | - | 0 | - |
| - | - | 779.1 | 356.2 | - | - | 0 | - |
| - | - | 2763 | 356.2 | - | - | 0 | - |
| - | - | 1673 | 357.1 | - | - | 0 | - |
| - | - | 3723 | 357.2 | - | - | 0 | - |
| - | - | 1794 | 357.7 | - | - | 0 | - |
| - | - | 917 | 358.2 | - | - | 0 | - |
| - | - | 810.5 | 359 | - | - | 0 | - |
| 6 | b | 4169 | 361.1 | 0.005077 | 14.06 | +2 | 6 |
| - | - | 937.2 | 364.2 | - | - | 0 | - |
| - | - | 755.3 | 364.5 | - | - | 0 | - |
| - | - | 1783 | 365.2 | - | - | 0 | - |
| - | - | 4561 | 366.2 | - | - | 0 | - |
| - | - | 1567 | 366.7 | - | - | 0 | - |
| - | - | 638.5 | 367.2 | - | - | 0 | - |
| - | - | 802.5 | 369.2 | - | - | 0 | - |
| - | - | 1025 | 370.3 | - | - | 0 | - |
| - | - | 1078 | 370.5 | - | - | 0 | - |
| - | - | 711.8 | 370.7 | - | - | 0 | - |
| - | - | 2.389E+04 | 371.2 | - | - | 0 | - |
| - | - | 2.147E+04 | 371.2 | - | - | 0 | - |
| - | - | 1023 | 371.7 | - | - | 0 | - |
| - | - | 1195 | 371.8 | - | - | 0 | - |
| - | - | 4143 | 372.2 | - | - | 0 | - |
| - | - | 3761 | 372.2 | - | - | 0 | - |
| - | - | 699.9 | 373.2 | - | - | 0 | - |
| - | - | 3564 | 376.2 | - | - | 0 | - |
| - | - | 677.9 | 376.5 | - | - | 0 | - |
| - | - | 988 | 376.8 | - | - | 0 | - |
| - | - | 607.5 | 377.2 | - | - | 0 | - |
| - | - | 1830 | 379.2 | - | - | 0 | - |
| - | - | 9003 | 379.8 | - | - | 0 | - |
| - | - | 5254 | 380.2 | - | - | 0 | - |
| - | - | 1665 | 380.5 | - | - | 0 | - |
| 4 | y | 2.131E+04 | 380.7 | 4.854E-05 | 0.1275 | +2 | 6 |
| - | - | 6003 | 381.2 | - | - | 0 | - |
| - | - | 596.1 | 381.2 | - | - | 0 | - |
| - | - | 1272 | 381.7 | - | - | 0 | - |
| - | - | 2449 | 383.2 | - | - | 0 | - |
| - | - | 1535 | 383.2 | - | - | 0 | - |
| - | - | 635.4 | 384.2 | - | - | 0 | - |
| - | - | 4512 | 384.2 | - | - | 0 | - |
| - | - | 1025 | 385.2 | - | - | 0 | - |
| 0 | Precursor | 753.9 | 385.5 | 0.0009762 | 2.532 | +3 | -1 |
| 0 | Precursor | 683.3 | 385.8 | 0.0002155 | 0.5585 | +3 | -1 |
| - | - | 824.4 | 386.2 | - | - | 0 | - |
| - | - | 1642 | 386.2 | - | - | 0 | - |
| - | - | 1516 | 386.7 | - | - | 0 | - |
| - | - | 685.9 | 387.1 | - | - | 0 | - |
| 4 | y | 4.336E+04 | 389.2 | 0.0001407 | 0.3615 | +2 | 6 |
| - | - | 1.763E+04 | 389.7 | - | - | 0 | - |
| - | - | 3485 | 390.2 | - | - | 0 | - |
| - | - | 762.3 | 390.7 | - | - | 0 | - |
| - | - | 1125 | 391.2 | - | - | 0 | - |
| 0 | Precursor | 1.535E+04 | 391.5 | 0.001446 | 3.693 | +3 | -1 |
| - | - | 7937 | 391.9 | - | - | 0 | - |
| - | - | 4042 | 392.2 | - | - | 0 | - |
| - | - | 1346 | 392.2 | - | - | 0 | - |
| - | - | 730.2 | 392.5 | - | - | 0 | - |
| - | - | 1342 | 394.2 | - | - | 0 | - |
| - | - | 829.5 | 394.7 | - | - | 0 | - |
| 3 | b | 1090 | 396.1 | 0.004345 | 10.97 | +1 | 3 |
| - | - | 1118 | 399.2 | - | - | 0 | - |
| - | - | 3521 | 400.2 | - | - | 0 | - |
| - | - | 988.9 | 401.2 | - | - | 0 | - |
| - | - | 740.3 | 401.2 | - | - | 0 | - |
| - | - | 4751 | 401.3 | - | - | 0 | - |
| - | - | 902.7 | 402.2 | - | - | 0 | - |
| - | - | 920.2 | 402.3 | - | - | 0 | - |
| - | - | 1237 | 405.7 | - | - | 0 | - |
| - | - | 916.4 | 407.2 | - | - | 0 | - |
| - | - | 755.8 | 414.2 | - | - | 0 | - |
| - | - | 3561 | 414.7 | - | - | 0 | - |
| - | - | 2070 | 415.2 | - | - | 0 | - |
| 3 | y | 1339 | 415.7 | 0.00243 | 5.845 | +2 | 7 |
| 3 | y | 1451 | 416.2 | 0.0008703 | 2.091 | +2 | 7 |
| - | - | 6534 | 416.3 | - | - | 0 | - |
| - | - | 2779 | 417.2 | - | - | 0 | - |
| - | - | 1280 | 417.3 | - | - | 0 | - |
| 7 | y | 1.532E+04 | 418.2 | 0.0002448 | 0.5853 | +1 | 3 |
| - | - | 2761 | 419.2 | - | - | 0 | - |
| - | - | 8102 | 423.7 | - | - | 0 | - |
| - | - | 3103 | 424.2 | - | - | 0 | - |
| 3 | y | 3.941E+04 | 424.7 | 0.0003216 | 0.7572 | +2 | 7 |
| - | - | 554.9 | 425.2 | - | - | 0 | - |
| - | - | 1.663E+04 | 425.2 | - | - | 0 | - |
| - | - | 5076 | 425.7 | - | - | 0 | - |
| - | - | 1198 | 426.2 | - | - | 0 | - |
| - | - | 703.1 | 426.7 | - | - | 0 | - |
| - | - | 3287 | 427.3 | - | - | 0 | - |
| - | - | 978 | 427.7 | - | - | 0 | - |
| - | - | 5304 | 427.8 | - | - | 0 | - |
| - | - | 2166 | 428.3 | - | - | 0 | - |
| - | - | 799.4 | 428.8 | - | - | 0 | - |
| - | - | 839.8 | 429.7 | - | - | 0 | - |
| - | - | 859.5 | 430.2 | - | - | 0 | - |
| - | - | 757.9 | 430.7 | - | - | 0 | - |
| - | - | 4024 | 434.8 | - | - | 0 | - |
| 7 | y | 4.523E+04 | 435.2 | 0.0002155 | 0.495 | +1 | 3 |
| - | - | 1.009E+04 | 436.2 | - | - | 0 | - |
| - | - | 1690 | 437.2 | - | - | 0 | - |
| - | - | 767.2 | 437.7 | - | - | 0 | - |
| - | - | 1537 | 438.2 | - | - | 0 | - |
| - | - | 1175 | 439.3 | - | - | 0 | - |
| - | - | 2789 | 440.2 | - | - | 0 | - |
| - | - | 1098 | 441.7 | - | - | 0 | - |
| - | - | 1119 | 442.2 | - | - | 0 | - |
| - | - | 998.5 | 445.2 | - | - | 0 | - |
| 7 | b | 4966 | 447.7 | 0.001685 | 3.763 | +2 | 7 |
| - | - | 1625 | 448.2 | - | - | 0 | - |
| - | - | 720.6 | 450.2 | - | - | 0 | - |
| - | - | 679.9 | 450.7 | - | - | 0 | - |
| - | - | 1195 | 451.2 | - | - | 0 | - |
| - | - | 3586 | 451.7 | - | - | 0 | - |
| - | - | 2134 | 452.2 | - | - | 0 | - |
| - | - | 795.8 | 452.7 | - | - | 0 | - |
| - | - | 4444 | 455.3 | - | - | 0 | - |
| - | - | 3100 | 456.3 | - | - | 0 | - |
| - | - | 823.4 | 457.3 | - | - | 0 | - |
| - | - | 1.298E+04 | 458.2 | - | - | 0 | - |
| - | - | 722.1 | 459.1 | - | - | 0 | - |
| - | - | 1082 | 459.2 | - | - | 0 | - |
| - | - | 2260 | 459.2 | - | - | 0 | - |
| - | - | 923.7 | 460.2 | - | - | 0 | - |
| - | - | 1011 | 468.3 | - | - | 0 | - |
| - | - | 1101 | 471.3 | - | - | 0 | - |
| - | - | 818.6 | 473.2 | - | - | 0 | - |
| - | - | 770.1 | 473.3 | - | - | 0 | - |
| - | - | 1533 | 473.7 | - | - | 0 | - |
| - | - | 689.8 | 474.2 | - | - | 0 | - |
| - | - | 1800 | 475.8 | - | - | 0 | - |
| - | - | 2.329E+04 | 476.2 | - | - | 0 | - |
| - | - | 784.1 | 476.2 | - | - | 0 | - |
| - | - | 2218 | 476.3 | - | - | 0 | - |
| - | - | 754.3 | 476.8 | - | - | 0 | - |
| - | - | 5181 | 477.2 | - | - | 0 | - |
| - | - | 1433 | 478.2 | - | - | 0 | - |
| - | - | 4076 | 482.2 | - | - | 0 | - |
| - | - | 2407 | 482.3 | - | - | 0 | - |
| - | - | 1943 | 482.7 | - | - | 0 | - |
| - | - | 650.7 | 483.2 | - | - | 0 | - |
| - | - | 3.223E+04 | 484.3 | - | - | 0 | - |
| - | - | 2.558E+04 | 484.8 | - | - | 0 | - |
| - | - | 7162 | 485.3 | - | - | 0 | - |
| - | - | 2435 | 485.8 | - | - | 0 | - |
| - | - | 1.052E+04 | 486.3 | - | - | 0 | - |
| 8 | b | 1199 | 487.2 | 0.001239 | 2.543 | +2 | 8 |
| - | - | 2041 | 487.3 | - | - | 0 | - |
| 8 | b | 2698 | 487.7 | 0.0009855 | 2.021 | +2 | 8 |
| - | - | 2854 | 488.3 | - | - | 0 | - |
| 8 | b | 3.426E+04 | 496.2 | 0.002096 | 4.224 | +2 | 8 |
| - | - | 1370 | 496.3 | - | - | 0 | - |
| 2 | y | 2.265E+04 | 496.7 | 0.005774 | 11.62 | +2 | 8 |
| - | - | 7100 | 497.2 | - | - | 0 | - |
| - | - | 3428 | 497.7 | - | - | 0 | - |
| - | - | 656 | 498.2 | - | - | 0 | - |
| - | - | 5007 | 499.3 | - | - | 0 | - |
| - | - | 2086 | 500.3 | - | - | 0 | - |
| - | - | 1737 | 502.2 | - | - | 0 | - |
| 2 | y | 1.606E+05 | 505.2 | 0.002099 | 4.154 | +2 | 8 |
| - | - | 8.467E+04 | 505.7 | - | - | 0 | - |
| - | - | 3.461E+04 | 506.2 | - | - | 0 | - |
| - | - | 8873 | 506.7 | - | - | 0 | - |
| - | - | 1516 | 507.2 | - | - | 0 | - |
| - | - | 2.573E+04 | 513.3 | - | - | 0 | - |
| - | - | 8783 | 514.3 | - | - | 0 | - |
| - | - | 1785 | 515.3 | - | - | 0 | - |
| - | - | 800.8 | 527.8 | - | - | 0 | - |
| - | - | 1520 | 528.3 | - | - | 0 | - |
| - | - | 967.1 | 529.2 | - | - | 0 | - |
| 6 | y | 1.507E+05 | 534.3 | 0.0001748 | 0.3271 | +1 | 4 |
| - | - | 3.996E+04 | 535.3 | - | - | 0 | - |
| - | - | 8106 | 536.3 | - | - | 0 | - |
| - | - | 1173 | 537.3 | - | - | 0 | - |
| - | - | 702.5 | 540.4 | - | - | 0 | - |
| - | - | 2390 | 540.8 | - | - | 0 | - |
| - | - | 1454 | 541.3 | - | - | 0 | - |
| - | - | 774 | 541.8 | - | - | 0 | - |
| - | - | 2975 | 544.3 | - | - | 0 | - |
| - | - | 2494 | 547.3 | - | - | 0 | - |
| - | - | 608.6 | 548.3 | - | - | 0 | - |
| - | - | 863.1 | 553.3 | - | - | 0 | - |
| - | - | 1647 | 557.2 | - | - | 0 | - |
| - | - | 3503 | 570.3 | - | - | 0 | - |
| - | - | 631.3 | 571.3 | - | - | 0 | - |
| - | - | 2826 | 575.2 | - | - | 0 | - |
| - | - | 6301 | 584.4 | - | - | 0 | - |
| - | - | 2383 | 585.4 | - | - | 0 | - |
| - | - | 792.8 | 586.4 | - | - | 0 | - |
| - | - | 1198 | 588.3 | - | - | 0 | - |
| - | - | 1438 | 590.3 | - | - | 0 | - |
| - | - | 628.2 | 594.2 | - | - | 0 | - |
| - | - | 1167 | 596.4 | - | - | 0 | - |
| - | - | 831.5 | 599.3 | - | - | 0 | - |
| - | - | 2396 | 603.3 | - | - | 0 | - |
| - | - | 699.4 | 609.3 | - | - | 0 | - |
| - | - | 701.6 | 614.3 | - | - | 0 | - |
| - | - | 2784 | 614.4 | - | - | 0 | - |
| - | - | 1120 | 615.4 | - | - | 0 | - |
| 5 | b | 776.1 | 621.2 | 0.006311 | 10.16 | +1 | 5 |
| 5 | b | 1889 | 622.2 | 0.003937 | 6.328 | +1 | 5 |
| - | - | 624.7 | 624.3 | - | - | 0 | - |
| - | - | 2162 | 627.4 | - | - | 0 | - |
| - | - | 1430 | 628.4 | - | - | 0 | - |
| 5 | y | 1531 | 631.3 | 0.00105 | 1.663 | +1 | 5 |
| 5 | y | 2494 | 632.3 | 0.001629 | 2.576 | +1 | 5 |
| - | - | 808.8 | 633.3 | - | - | 0 | - |
| 5 | b | 1683 | 639.2 | 0.004119 | 6.444 | +1 | 5 |
| - | - | 805.7 | 640.2 | - | - | 0 | - |
| 5 | y | 1.635E+05 | 649.3 | 0.0001403 | 0.216 | +1 | 5 |
| - | - | 5.786E+04 | 650.3 | - | - | 0 | - |
| - | - | 1.109E+04 | 651.3 | - | - | 0 | - |
| - | - | 2047 | 652.3 | - | - | 0 | - |
| - | - | 3460 | 655.4 | - | - | 0 | - |
| - | - | 1524 | 656.4 | - | - | 0 | - |
| - | - | 3104 | 659.3 | - | - | 0 | - |
| - | - | 1882 | 660.3 | - | - | 0 | - |
| - | - | 633.4 | 669.2 | - | - | 0 | - |
| - | - | 1183 | 685.4 | - | - | 0 | - |
| - | - | 804.2 | 689.3 | - | - | 0 | - |
| - | - | 1409 | 695.4 | - | - | 0 | - |
| - | - | 1000 | 696.3 | - | - | 0 | - |
| - | - | 906.4 | 697.3 | - | - | 0 | - |
| - | - | 795.2 | 703.4 | - | - | 0 | - |
| - | - | 1519 | 712.4 | - | - | 0 | - |
| - | - | 2590 | 713.3 | - | - | 0 | - |
| - | - | 980.1 | 713.4 | - | - | 0 | - |
| - | - | 2695 | 714.3 | - | - | 0 | - |
| - | - | 1118 | 715.3 | - | - | 0 | - |
| - | - | 757.2 | 720.4 | - | - | 0 | - |
| - | - | 6507 | 731.3 | - | - | 0 | - |
| - | - | 2756 | 732.4 | - | - | 0 | - |
| 4 | y | 1101 | 759.4 | 0.001714 | 2.257 | +1 | 6 |
| 4 | y | 2532 | 760.4 | 0.005481 | 7.209 | +1 | 6 |
| 4 | y | 3.728E+04 | 777.4 | 0.0009871 | 1.27 | +1 | 6 |
| - | - | 1.493E+04 | 778.4 | - | - | 0 | - |
| - | - | 3612 | 779.4 | - | - | 0 | - |
| - | - | 2128 | 787.4 | - | - | 0 | - |
| - | - | 1043 | 788.4 | - | - | 0 | - |
| - | - | 780.1 | 789.4 | - | - | 0 | - |
| - | - | 621.7 | 828.4 | - | - | 0 | - |
| - | - | 715 | 846.4 | - | - | 0 | - |
| 3 | y | 6134 | 848.5 | 0.001114 | 1.312 | +1 | 7 |
| - | - | 2973 | 849.5 | - | - | 0 | - |
| - | - | 1087 | 851.5 | - | - | 0 | - |
| - | - | 1050 | 858.4 | - | - | 0 | - |
| - | - | 721 | 1226 | - | - | 0 | - |
| - | - | 670.1 | 1477 | - | - | 0 | - |
| - | - | 640.3 | 2237 | - | - | 0 | - |
| - | - | 692.1 | 2438 | - | - | 0 | - |
| - | - | 691.3 | 3081 | - | - | 0 | - |

m/z Charge Intensity FragmentType MassShift Position
120.05278778076172 0 6201.7617
120.08111572265625 0 1051.5791
121.02864837646484 0 1289.0686
122.0975112915039 0 341.55756
123.04435729980469 0 1076.3394
124.0768051147461 0 538.15674
125.07123565673828 0 1011.5946
126.09175109863281 0 465.8199
127.08692932128906 0 2819.2488
128.10728454589844 0 2678.966
129.1025848388672 0 175397.5
130.08697509765625 0 433.65442
130.09901428222656 0 1493.0125
130.10592651367188 0 11192.542
131.1070098876953 0 599.63715
132.10215759277344 0 1406.8766
134.02735900878906 0 46883.73
135.03077697753906 0 2110.4302
136.02301025390625 0 2233.5713
136.07608032226562 0 479995.3
137.05979919433594 0 1912.0211
137.07936096191406 0 39231.17
138.06642150878906 0 1460.8339
138.08230590820312 0 1477.2921
138.12757873535156 0 530.3177
139.0868377685547 0 3897.4314
139.123291015625 0 676.57
140.08218383789062 0 2962.4812
140.09019470214844 0 562.38684
141.10247802734375 0 1599.3995
141.11355590820312 0 591.51776
142.03248596191406 0 1062.6326
142.08648681640625 0 1335.863
143.08181762695312 0 742.821
144.0115509033203 0 602.62885
146.06072998046875 0 507.81064
147.04437255859375 0 43321.31
148.0477294921875 0 4872.246
149.02371215820312 0 342033.75
150.0269775390625 0 27073.07
153.1026153564453 0 637.51697
154.0983123779297 0 513.44037
155.08193969726562 0 2019.98
155.11813354492188 0 3920.4412
156.07716369628906 0 1019.31586
157.0609588623047 0 575.0131
157.09751892089844 0 1224.9252
157.10870361328125 0 7175.0586
158.0646209716797 0 522.7415
158.09225463867188 0 1195.8513
158.11209106445312 0 642.4399
159.05897521972656 0 4022.3054
160.04293823242188 0 5177.4565
160.075927734375 0 2423.3684
162.02223205566406 0 8052.496
164.07066345214844 0 960.77765
164.11900329589844 0 569.8721
165.05487060546875 0 5408.7104
165.10264587402344 0 1882.5376
166.0868377685547 0 1211.6981
166.0978240966797 0 811.3718
166.81956481933594 0 537.9784
167.03414916992188 0 37755.73
167.11778259277344 0 747.31616
168.03761291503906 0 3263.0574
169.09756469726562 0 1558.2133
169.1339111328125 0 1302.5979
170.08059692382812 0 526.91974
172.10818481445312 0 1527.0985
173.1287384033203 0 37986.04
174.1321258544922 0 4515.2314
175.08689880371094 0 674.2166
175.11917114257812 0 1136.8167
176.0380096435547 0 738.3018
178.13375854492188 0 1287.0494
179.1180877685547 0 492.45145
180.10218811035156 0 1724.2202
181.09739685058594 0 1678.9963
181.13316345214844 0 953.08887
182.0815887451172 0 5376.816 y 8
182.1290283203125 0 1390.3705
183.11302185058594 0 4143.0693
183.12258911132812 0 749.71405
183.14915466308594 0 1485.6482
185.03761291503906 0 756.37683
185.10362243652344 0 7176.6523
185.16551208496094 0 902.15906
187.0538330078125 0 8509.801
187.1079864501953 0 22793.42
188.07139587402344 0 923.22534
188.11143493652344 0 1755.07
191.0817413330078 0 1270.7441
192.0477294921875 0 1936
192.11337280273438 0 1599.6315
194.12908935546875 0 1186.1102
195.11294555664062 0 2267.7417
195.15972900390625 0 652.34467
197.09320068359375 0 1025.0662
197.10360717773438 0 2334.0596
197.1285858154297 0 5072.145
198.12379455566406 0 1216.95
199.10784912109375 0 4615.409
199.14434814453125 0 3829.7412
200.07424926757812 0 634.0905
200.13954162597656 0 19237.04
200.61927795410156 0 1161.0055
201.12351989746094 0 6520.7
201.14248657226562 0 1122.098
202.1263427734375 0 709.1702
203.13946533203125 0 1396.1959
204.0694580078125 0 862.35315
205.06439208984375 0 71759.875
206.067626953125 0 5367.0537
207.06005859375 0 2992.1523
208.10816955566406 0 744.5224
209.09217834472656 0 2544.0742
209.1294403076172 0 624.1499
209.13990783691406 0 1929.6663
210.1238250732422 0 2069.1184
213.12367248535156 0 1197.9686
214.13003540039062 0 1041.595
214.1551971435547 0 1487.972
215.04885864257812 0 2856.1255
215.10293579101562 0 5151.577
216.06626892089844 0 1387.0492
216.1058807373047 0 881.68665
220.04246520996094 0 755.8243
220.10874938964844 0 711.43243
221.09243774414062 0 1956.2968
222.12350463867188 0 1162.6812
223.06373596191406 0 993.90625
223.15536499023438 0 992.80884
224.06263732910156 0 720.561
224.13967895507812 0 1409.2026
225.04307556152344 0 1593.4911
225.0866241455078 0 574.7021
226.04400634765625 0 585.0418
226.11886596679688 0 8820.589
227.1029815673828 0 3687.3953
227.122802734375 0 911.5356
227.1387939453125 0 841.63196
227.9048309326172 0 533.07574
231.04457092285156 0 632.4815
233.05934143066406 0 10685.648
233.0923309326172 0 4905.936
233.1288604736328 0 3595.3052
234.05905151367188 0 3138.2603
234.09487915039062 0 685.8353
235.06353759765625 0 683.0609
237.06959533691406 0 944.451
237.1348114013672 0 4244.809
237.16287231445312 0 637.2034
238.16598510742188 0 716.2579
239.15052795410156 0 55455.586
240.15382385253906 0 4488.6797
241.15509033203125 0 591.48975
243.0975799560547 0 952.7068
243.14735412597656 0 474.40985
244.12940979003906 0 6209.2954
250.08416748046875 0 535.6833
251.0474090576172 0 1823.3324
251.11439514160156 0 1216.6624
251.15208435058594 0 557.72986
252.06922912597656 0 9074.159
252.13458251953125 0 941.5873
253.07272338867188 0 1247.8042
254.11387634277344 0 1033.9833
254.16146850585938 0 6959.7026
255.1453399658203 0 5970.202
255.1644744873047 0 867.2948
256.11981201171875 0 768.71576
256.1489562988281 0 930.75195
256.1769714355469 0 25818.807
257.1609191894531 0 7600.4507
257.1800537109375 0 2678.5615
257.6623229980469 0 2317.6826
261.1227722167969 0 488.4625
262.0536193847656 0 979.95514
262.1698303222656 0 505.18182
262.6131591796875 0 1335.3754 b 3
263.1128845214844 0 1004.41925
265.1255798339844 0 1765.6589
266.161376953125 0 1428.4794
268.140380859375 0 1926.9895
269.1600036621094 0 1112.3011
270.1453552246094 0 592.2992
272.1242370605469 0 5061.9976
272.17181396484375 0 4305.2466
273.0918273925781 0 1038.4913
273.1280517578125 0 937.5427
273.203369140625 0 871.4329
274.1875915527344 0 6183.486
276.13397216796875 0 561.83527
277.12847900390625 0 629.80743
278.68359375 0 589.71606
279.1341552734375 0 82997.195 y 7
279.1607971191406 0 3054.0664
280.0643615722656 0 2575.4897
280.1373596191406 0 12872.479
280.164306640625 0 1364.9175
281.1402282714844 0 1221.526
282.1562194824219 0 3454.1033
284.1719665527344 0 2618.4587
285.0093994140625 0 923.17
285.1674499511719 0 981.884
285.6719665527344 0 1397.6057
289.36920166015625 0 498.27423
291.89874267578125 0 533.0822
292.1292724609375 0 634.4937
292.6789245605469 0 650.7926
294.15667724609375 0 3690.8108
295.1592102050781 0 908.782
297.0906982421875 0 44227.26 a 1
297.1558532714844 0 2142.1106
298.0940246582031 0 7113.378
298.14013671875 0 1060.1449
299.0861511230469 0 1765.7633
299.17254638671875 0 1529.5759
300.0631408691406 0 825.8413
301.0600280761719 0 725.7241
302.19293212890625 0 778.11804
302.6062927246094 0 1068.2847
303.1772766113281 0 1018.05585
304.1296081542969 0 4015.3718
305.68609619140625 0 621.38904
307.6517028808594 0 597.4249
307.6846618652344 0 2486.3232
308.05859375 0 1059.5144
308.1582946777344 0 654.22345
311.18218994140625 0 2676.6042
312.1857604980469 0 845.3124
312.67608642578125 0 663.14746
313.1617126464844 0 697.9079
313.3769836425781 0 582.48846
314.2008972167969 0 2818.2764
314.70294189453125 0 1332.9445
315.16656494140625 0 6853.995
315.2033386230469 0 1693.1431
316.1330871582031 0 959.01263
316.16912841796875 0 834.8972 y Water loss 4
316.20550537109375 0 682.6816
319.1391296386719 0 788.9584
319.17474365234375 0 584.5965
320.1258239746094 0 2791.5889 b 4
322.49969482421875 0 530.66113
325.085693359375 0 8568.999 b 1
325.1546936035156 0 642.579 b Water loss 7
325.18768310546875 0 1391.0166
326.08837890625 0 1438.806
327.0813903808594 0 639.7353
333.1542663574219 0 623.4716
336.1670837402344 0 1235.3523
337.1648864746094 0 1465.1554 y 1
337.19873046875 0 734.4062
337.4988098144531 0 1055.4431
338.14544677734375 0 534.8252
340.1630859375 0 629.6319
342.214599609375 0 1498.391
343.1435241699219 0 2127.4731
343.20703125 0 5700.131
344.1274719238281 0 1231.8336
344.21185302734375 0 582.86334
353.1933898925781 0 1538.3805
353.2303466796875 0 881.42236
354.177490234375 0 3703.632
355.07012939453125 0 2245.777
355.18109130859375 0 636.13275
355.66241455078125 0 2030.3705
356.0713806152344 0 1405.8413
356.20556640625 0 779.13574
356.2405700683594 0 2762.8843
357.0684814453125 0 1672.9941
357.173583984375 0 3722.9656
357.67059326171875 0 1793.6976
358.17236328125 0 917.0212
359.0299377441406 0 810.516
361.154052734375 0 4169.49 b Ammonia loss 5
364.19757080078125 0 937.1778
364.5051574707031 0 755.3223
365.1937561035156 0 1783.1659
366.1783447265625 0 4560.6904
366.6810302734375 0 1567.271
367.1748962402344 0 638.51776
369.1765441894531 0 802.45734
370.2588195800781 0 1025.1931
370.5066833496094 0 1077.9349
370.7108154296875 0 711.8098
371.2037658691406 0 23887.299
371.240234375 0 21473.986
371.6972961425781 0 1023.00006
371.840087890625 0 1195.1077
372.20623779296875 0 4142.9976
372.24310302734375 0 3760.5476
373.24688720703125 0 699.9352
376.1845703125 0 3563.8032
376.5137023925781 0 677.9318
376.84881591796875 0 987.985
377.185302734375 0 607.4871
379.16802978515625 0 1829.5844
379.8397521972656 0 9003.269
380.173583984375 0 5254.071
380.5067138671875 0 1665.2212
380.7030944824219 0 21309.53 y Ammonia loss 3
381.2039489746094 0 6003.461
381.2267761230469 0 596.0774
381.70538330078125 0 1272.492
383.2034912109375 0 2448.5266
383.2276611328125 0 1535.3914
384.1876220703125 0 635.39246
384.23541259765625 0 4512.297
385.2386779785156 0 1024.7389
385.5153503417969 0 753.93945 Precursor Water loss
385.8441162109375 0 683.3127 Precursor Ammonia loss
386.1817626953125 0 824.37134
386.2399597167969 0 1642.018
386.7027587890625 0 1516.0674
387.1304931640625 0 685.8561
389.2164611816406 0 43362.18 y 3
389.7179870605469 0 17629.756
390.218994140625 0 3485.4082
390.7215270996094 0 762.3067
391.1985778808594 0 1125.471
391.5184020996094 0 15347.382 Precursor
391.852783203125 0 7936.8193
392.1868896484375 0 4042.258
392.23260498046875 0 1345.9031
392.5206298828125 0 730.1783
394.2085266113281 0 1341.7985
394.71136474609375 0 829.5482
396.12255859375 0 1090.4899 b 2
399.2142028808594 0 1118.0522
400.2301940917969 0 3520.8086
401.1836242675781 0 988.87067
401.2344970703125 0 740.3044
401.2621154785156 0 4751.3774
402.23016357421875 0 902.6957
402.26373291015625 0 920.2463
405.698974609375 0 1237.4341
407.21630859375 0 916.3921
414.23419189453125 0 755.79785
414.70526123046875 0 3560.701
415.2060241699219 0 2069.7112
415.7320251464844 0 1339.449 y Water loss 2
416.22247314453125 0 1451.4556 y Ammonia loss 2
416.261962890625 0 6534.366
417.22515869140625 0 2778.8325
417.2630615234375 0 1280.0356
418.208740234375 0 15321.597 y Ammonia loss 6
419.21112060546875 0 2760.8599
423.7108154296875 0 8102.092
424.2120056152344 0 3103.456
424.7351989746094 0 39408.35 y 2
425.2039489746094 0 554.8785
425.23681640625 0 16626.93
425.7383117675781 0 5075.6016
426.24322509765625 0 1197.9578
426.74688720703125 0 703.14075
427.2781677246094 0 3287.361
427.6979675292969 0 977.9907
427.7515869140625 0 5303.6074
428.2525329589844 0 2165.69
428.7541809082031 0 799.4277
429.7278137207031 0 839.7611
430.19366455078125 0 859.4605
430.7320861816406 0 757.92944
434.7596435546875 0 4023.834
435.2352600097656 0 45229.242 y 6
436.2381286621094 0 10093.037
437.2409362792969 0 1690.1824
437.7447204589844 0 767.214
438.24407958984375 0 1536.6554
439.2681579589844 0 1175.0011
440.1604309082031 0 2788.8462
441.7287292480469 0 1097.7509
442.2278137207031 0 1118.9039
445.219482421875 0 998.5071
447.71112060546875 0 4966.41 b 6
448.2120666503906 0 1625.0414
450.2422790527344 0 720.6008
450.73388671875 0 679.9202
451.229736328125 0 1195.0068
451.74005126953125 0 3585.746
452.241455078125 0 2134.2012
452.7417297363281 0 795.8413
455.27294921875 0 4444.091
456.2923278808594 0 3099.8005
457.2752685546875 0 823.3977
458.1706848144531 0 12976.269
459.1493225097656 0 722.10645
459.177978515625 0 1082.1925
459.2469482421875 0 2259.6414
460.1681213378906 0 923.6776
468.2579345703125 0 1010.98303
471.30267333984375 0 1101.1205
473.2354431152344 0 818.6098
473.3191223144531 0 770.07574
473.73223876953125 0 1532.9835
474.2289733886719 0 689.75464
475.78082275390625 0 1799.5718
476.1810607910156 0 23286.795
476.2153625488281 0 784.0665
476.2819519042969 0 2218.396
476.7852783203125 0 754.3164
477.1836853027344 0 5181.3486
478.1814880371094 0 1433.2108
482.239013671875 0 4075.7454
482.27423095703125 0 2406.9204
482.73876953125 0 1942.5792
483.23992919921875 0 650.68036
484.29351806640625 0 32229.75
484.7950134277344 0 25580.902
485.29583740234375 0 7161.517
485.79705810546875 0 2435.253
486.26715087890625 0 10519.163
487.232666015625 0 1199.2827 b Water loss 7
487.2699279785156 0 2040.8547
487.7268981933594 0 2697.8606 b Ammonia loss 7
488.29791259765625 0 2853.5789
496.2370910644531 0 34259.516 b 7
496.2806091308594 0 1370.1915
496.7369689941406 0 22653.244 y Ammonia loss 1
497.23699951171875 0 7100.163
497.7372131347656 0 3428.4875
498.24249267578125 0 655.9557
499.2992248535156 0 5007.486
500.30078125 0 2085.6965
502.1604919433594 0 1736.99
505.24237060546875 0 160552.88 y 1
505.7436218261719 0 84666.516
506.24420166015625 0 34613.195
506.7444152832031 0 8873.112
507.2469787597656 0 1516.286
513.3143920898438 0 25734.285
514.31689453125 0 8782.944
515.3195190429688 0 1785.2053
527.810546875 0 800.79333
528.3135375976562 0 1520.1548
529.2420043945312 0 967.1285
534.3032836914062 0 150733.36 y 5
535.3062133789062 0 39958.64
536.3091430664062 0 8105.7285
537.3106079101562 0 1173.2767
540.3616943359375 0 702.48004
540.8344116210938 0 2389.8008
541.3359375 0 1453.6448
541.839111328125 0 774.0205
544.2880249023438 0 2974.6694
547.255126953125 0 2493.7305
548.2572021484375 0 608.6353
553.3092651367188 0 863.1114
557.2378540039062 0 1647.1425
570.3360595703125 0 3502.9001
571.3381958007812 0 631.33044
575.2491455078125 0 2825.6624
584.3515625 0 6300.9165
585.3538208007812 0 2383.2998
586.358154296875 0 792.7976
588.3104858398438 0 1197.7955
590.283203125 0 1438.4252
594.22509765625 0 628.1644
596.3531494140625 0 1166.7181
599.3294677734375 0 831.5343
603.323974609375 0 2395.637
609.3455200195312 0 699.43134
614.2971801757812 0 701.5923
614.362548828125 0 2783.63
615.3643798828125 0 1120.2004
621.23193359375 0 776.08954 b Water loss 4
622.2183227539062 0 1889.0575 b Ammonia loss 4
624.3463134765625 0 624.6596
627.3931884765625 0 2161.6455
628.3955078125 0 1430.1677
631.3187866210938 0 1531.4039 y Water loss 4
632.3054809570312 0 2493.6814 y Ammonia loss 4
633.3096313476562 0 808.81055
639.2446899414062 0 1683.0299 b 4
640.2473754882812 0 805.72284
649.3302612304688 0 163526.9 y 4
650.3328247070312 0 57859.402
651.3356323242188 0 11093.138
652.33740234375 0 2046.7703
655.3875732421875 0 3460.4705
656.3878784179688 0 1524.3723
659.31494140625 0 3104.4717
660.3156127929688 0 1881.6166
669.2252807617188 0 633.43854
685.400146484375 0 1183.4192
689.32763671875 0 804.2235
695.380859375 0 1409.4949
696.3138427734375 0 1000.30695
697.30908203125 0 906.3998
703.3551635742188 0 795.22723
712.4082641601562 0 1518.8044
713.340576171875 0 2590.0063
713.4100952148438 0 980.1001
714.3267822265625 0 2695.4011
715.3305053710938 0 1118.2103
720.4237670898438 0 757.2156
731.349609375 0 6506.773
732.3524169921875 0 2755.5056
759.4130859375 0 1101.0311 y Water loss 3
760.404296875 0 2532.4912 y Ammonia loss 3
777.4243774414062 0 37276.56 y 3
778.4275512695312 0 14931.999
779.4296264648438 0 3611.7573
787.4100341796875 0 2127.7568
788.4013671875 0 1043.1116
789.4076538085938 0 780.05884
828.3988647460938 0 621.66486
846.4116821289062 0 715.0455
848.4613647460938 0 6134.136 y 2
849.463134765625 0 2973.037
851.4826049804688 0 1087.0286
858.4484252929688 0 1049.5133
1226.3331298828125 0 721.0347
1477.4200439453125 0 670.06384
2237.46533203125 0 640.27893
2437.79541015625 0 692.08234
3081.17529296875 0 691.3368

Spectrum Details

|  |  |
| --- | --- |
| Matched peaks? Matched peaks The total absolute number of peaks matched. Additionally in brackets the total fraction of peaks matched and the total number of peaks is shown. | 38 (7.32% of 519) |
| FDR? FDR The false discovery rate estimated for this peptide. It is calculated by matching all theoretical fragments with a non-integer shift with the raw peaks for this spectrum. This is done with 40 different shifts. The resulting percentage is the average number of annotated peaks over the number of annotated peaks with the correct spectrum. | 1.88% |
| Satellite FDR? Satellite FDR See the FDR for details on its calculation. This satellite ion specific FDR only contains the satellite ions (d/w) for I/L/J positions. | - |
| PSM Score? PSM Score The PSM Score as given by Hecklib to this annotated spectrum. It is shown with three significant figures. | 253 |

#### Spectrum 3838? Spectrum 3838 The raw spectrum of this peptide as annotated by Hecklib. The fragments are coloured according to ion type (see legend). Any peaks with a star '\*' as text can be hovered over to see the full details, first the ion type second the mass shift type. By hovering over the amino acids in the peptide or ions in the legend the corresponding peaks are highlighted. By toggling the 'Unassigned' label you can turn the background (unassigned) peaks on or off in the plot. By updating the slider in the Ion legend you can update the spectrum to only show the top X% of the peaks with labels. The top X% means any peak that is within X% of the highest intensity. By dragging in the spectrum you can zoom in to a specific part of the spectrum and use 'Zoom Out' to get back to the original zoom level. The annotation of the spectrum is based on the given sequence in the peptides file and is done with different software so inconsistencies are likely. The peaks are annotated based on the given sequence, with 20 ppm tolerance.

Copy Data

##### Spectrum 3838 (TSV)

###### Preview

```
Loading example...
```

*Click on the button to copy the data to your clipboard.*

Mz MinMz MaxIntensity Max

WidthHeightPeptide font sizePeptide stroke widthSpectrum font sizeSpectrum stroke widthCompact peptide

Ion legend

wxyz

abcd

OtherUnassignedIon^Charge^_Position_Show for top:%

YCAKDVRPY

06.08e+41.22e+51.82e+52.43e+5

Zoom Out

y^+1^_1_y^+1^_2_a^+1^_2_b^+1^_2_b^+2^_6_y^+2^_6_y^+2^_6_y^+2^_6_b^+1^_3_y^+1^_3_y^+2^_7_y^+1^_3_b^+2^_7_b^+2^_8_y^+2^_8_y^+2^_8_b^+1^_4_y^+1^_4_\*\*\*b^+1^_5_b^+1^_5_b^+1^_5_y^+1^_5_b^+1^_6_y^+1^_6_y^+1^_7_y^+1^_7_b^+1^_7_b^+1^_7_b^+1^_8_y^+1^_8_y^+1^_8_

0737147422112948

Fragment Matches Table

Show background peaks

| Position | Ion type | Intensity | mz Theoretical | mz Error (Th) | mz Error (ppm) | Charge | Series Number |
| --- | --- | --- | --- | --- | --- | --- | --- |
| - | - | 1587 | 120.1 | - | - | 0 | - |
| - | - | 692.6 | 127.1 | - | - | 0 | - |
| - | - | 4.865E+04 | 129.1 | - | - | 0 | - |
| - | - | 451.5 | 130 | - | - | 0 | - |
| - | - | 2522 | 130.1 | - | - | 0 | - |
| - | - | 1115 | 131.1 | - | - | 0 | - |
| - | - | 385.5 | 132.6 | - | - | 0 | - |
| - | - | 1.121E+04 | 134 | - | - | 0 | - |
| - | - | 8.375E+04 | 136.1 | - | - | 0 | - |
| - | - | 6737 | 137.1 | - | - | 0 | - |
| - | - | 448.2 | 138.7 | - | - | 0 | - |
| - | - | 1275 | 139.1 | - | - | 0 | - |
| - | - | 1094 | 140.1 | - | - | 0 | - |
| - | - | 763 | 141.1 | - | - | 0 | - |
| - | - | 865 | 147 | - | - | 0 | - |
| - | - | 507.8 | 149 | - | - | 0 | - |
| - | - | 438.3 | 153.1 | - | - | 0 | - |
| - | - | 1826 | 155.1 | - | - | 0 | - |
| - | - | 425.4 | 156.9 | - | - | 0 | - |
| - | - | 4043 | 157.1 | - | - | 0 | - |
| - | - | 804.6 | 157.1 | - | - | 0 | - |
| - | - | 1836 | 157.1 | - | - | 0 | - |
| - | - | 434.6 | 158.1 | - | - | 0 | - |
| - | - | 707.9 | 158.1 | - | - | 0 | - |
| - | - | 2392 | 159.1 | - | - | 0 | - |
| - | - | 1081 | 160.1 | - | - | 0 | - |
| - | - | 609.1 | 162 | - | - | 0 | - |
| - | - | 1062 | 164.1 | - | - | 0 | - |
| - | - | 801.6 | 165.1 | - | - | 0 | - |
| - | - | 514.2 | 165.1 | - | - | 0 | - |
| - | - | 1233 | 166.1 | - | - | 0 | - |
| - | - | 734.1 | 169.1 | - | - | 0 | - |
| - | - | 730.6 | 169.1 | - | - | 0 | - |
| - | - | 425.8 | 169.6 | - | - | 0 | - |
| - | - | 478.7 | 170.1 | - | - | 0 | - |
| - | - | 2707 | 173.1 | - | - | 0 | - |
| - | - | 3978 | 175.1 | - | - | 0 | - |
| - | - | 544.5 | 175.1 | - | - | 0 | - |
| - | - | 519.9 | 180.1 | - | - | 0 | - |
| - | - | 768.7 | 181.1 | - | - | 0 | - |
| 9 | y | 1308 | 182.1 | 0.0001139 | 0.6255 | +1 | 1 |
| - | - | 3710 | 183.1 | - | - | 0 | - |
| - | - | 1537 | 183.1 | - | - | 0 | - |
| - | - | 496 | 185.1 | - | - | 0 | - |
| - | - | 553.9 | 185.1 | - | - | 0 | - |
| - | - | 2633 | 185.1 | - | - | 0 | - |
| - | - | 676.8 | 187.1 | - | - | 0 | - |
| - | - | 5357 | 187.1 | - | - | 0 | - |
| - | - | 696.7 | 189.1 | - | - | 0 | - |
| - | - | 847.7 | 192 | - | - | 0 | - |
| - | - | 996.3 | 197.1 | - | - | 0 | - |
| - | - | 1542 | 197.1 | - | - | 0 | - |
| - | - | 1572 | 199.1 | - | - | 0 | - |
| - | - | 1.395E+04 | 200.1 | - | - | 0 | - |
| - | - | 2910 | 201.1 | - | - | 0 | - |
| - | - | 512.3 | 201.1 | - | - | 0 | - |
| - | - | 6591 | 203.1 | - | - | 0 | - |
| - | - | 9343 | 205.1 | - | - | 0 | - |
| - | - | 502.4 | 206.1 | - | - | 0 | - |
| - | - | 1049 | 209.1 | - | - | 0 | - |
| - | - | 619.4 | 210.1 | - | - | 0 | - |
| - | - | 918.4 | 211.1 | - | - | 0 | - |
| - | - | 794 | 214.2 | - | - | 0 | - |
| - | - | 622.9 | 215 | - | - | 0 | - |
| - | - | 1970 | 215.1 | - | - | 0 | - |
| - | - | 706.4 | 215.1 | - | - | 0 | - |
| - | - | 1888 | 216.1 | - | - | 0 | - |
| - | - | 744.9 | 217.1 | - | - | 0 | - |
| - | - | 2643 | 226.1 | - | - | 0 | - |
| - | - | 1367 | 232.1 | - | - | 0 | - |
| - | - | 2166 | 233.1 | - | - | 0 | - |
| - | - | 1.384E+04 | 233.1 | - | - | 0 | - |
| - | - | 3255 | 234.1 | - | - | 0 | - |
| - | - | 909.7 | 234.1 | - | - | 0 | - |
| - | - | 1901 | 237.1 | - | - | 0 | - |
| - | - | 1383 | 237.1 | - | - | 0 | - |
| - | - | 2.185E+04 | 239.2 | - | - | 0 | - |
| - | - | 2407 | 240.2 | - | - | 0 | - |
| - | - | 2185 | 244.1 | - | - | 0 | - |
| - | - | 8467 | 252.1 | - | - | 0 | - |
| - | - | 645.4 | 253.1 | - | - | 0 | - |
| - | - | 614 | 254.1 | - | - | 0 | - |
| - | - | 1328 | 254.2 | - | - | 0 | - |
| - | - | 534.6 | 254.2 | - | - | 0 | - |
| - | - | 1527 | 255.1 | - | - | 0 | - |
| - | - | 9280 | 256.2 | - | - | 0 | - |
| - | - | 1312 | 257.2 | - | - | 0 | - |
| - | - | 951.6 | 262.1 | - | - | 0 | - |
| - | - | 625.1 | 262.1 | - | - | 0 | - |
| - | - | 2324 | 272.1 | - | - | 0 | - |
| - | - | 1459 | 272.1 | - | - | 0 | - |
| - | - | 600.1 | 272.2 | - | - | 0 | - |
| - | - | 1975 | 274.2 | - | - | 0 | - |
| - | - | 518.9 | 275.9 | - | - | 0 | - |
| 8 | y | 3.382E+04 | 279.1 | 0.0002055 | 0.7362 | +1 | 2 |
| - | - | 4295 | 280.1 | - | - | 0 | - |
| - | - | 702 | 282.2 | - | - | 0 | - |
| - | - | 649.2 | 284.2 | - | - | 0 | - |
| - | - | 2558 | 290.1 | - | - | 0 | - |
| 2 | a | 5.59E+04 | 297.1 | 0.004543 | 15.29 | +1 | 2 |
| - | - | 7715 | 298.1 | - | - | 0 | - |
| - | - | 1859 | 299.1 | - | - | 0 | - |
| - | - | 782.6 | 301.1 | - | - | 0 | - |
| - | - | 540.3 | 301.9 | - | - | 0 | - |
| - | - | 9011 | 304.1 | - | - | 0 | - |
| - | - | 2157 | 305.1 | - | - | 0 | - |
| - | - | 1202 | 305.2 | - | - | 0 | - |
| - | - | 1500 | 312.2 | - | - | 0 | - |
| - | - | 581.4 | 312.4 | - | - | 0 | - |
| - | - | 585.5 | 315 | - | - | 0 | - |
| - | - | 4353 | 315.2 | - | - | 0 | - |
| 2 | b | 2.048E+04 | 325.1 | 0.004554 | 14.01 | +1 | 2 |
| - | - | 653 | 325.2 | - | - | 0 | - |
| - | - | 3290 | 326.1 | - | - | 0 | - |
| - | - | 749.3 | 340.2 | - | - | 0 | - |
| - | - | 2721 | 343.2 | - | - | 0 | - |
| - | - | 1209 | 343.2 | - | - | 0 | - |
| - | - | 1003 | 347.7 | - | - | 0 | - |
| - | - | 493.1 | 351.2 | - | - | 0 | - |
| - | - | 1624 | 354.2 | - | - | 0 | - |
| - | - | 690.6 | 359.2 | - | - | 0 | - |
| - | - | 1766 | 359.2 | - | - | 0 | - |
| 6 | b | 2147 | 361.1 | 0.004681 | 12.96 | +2 | 6 |
| - | - | 1153 | 361.2 | - | - | 0 | - |
| - | - | 1474 | 366.2 | - | - | 0 | - |
| - | - | 1748 | 371.2 | - | - | 0 | - |
| - | - | 7066 | 371.2 | - | - | 0 | - |
| - | - | 8437 | 371.2 | - | - | 0 | - |
| - | - | 1384 | 372.2 | - | - | 0 | - |
| - | - | 1327 | 372.2 | - | - | 0 | - |
| - | - | 6279 | 379.2 | - | - | 0 | - |
| 4 | y | 1260 | 380.2 | 5.194E-05 | 0.1366 | +2 | 6 |
| 4 | y | 3295 | 380.7 | 0.0005008 | 1.315 | +2 | 6 |
| - | - | 1601 | 381.2 | - | - | 0 | - |
| - | - | 1333 | 383.2 | - | - | 0 | - |
| - | - | 773.5 | 386.2 | - | - | 0 | - |
| - | - | 573.2 | 388.2 | - | - | 0 | - |
| 4 | y | 9603 | 389.2 | 0.0004391 | 1.128 | +2 | 6 |
| - | - | 3337 | 389.7 | - | - | 0 | - |
| 3 | b | 3416 | 396.1 | 0.004467 | 11.28 | +1 | 3 |
| - | - | 1032 | 397.1 | - | - | 0 | - |
| - | - | 559.4 | 397.3 | - | - | 0 | - |
| - | - | 1175 | 414.2 | - | - | 0 | - |
| - | - | 557.9 | 414.8 | - | - | 0 | - |
| - | - | 812 | 417.2 | - | - | 0 | - |
| 7 | y | 6463 | 418.2 | 0.0001214 | 0.2904 | +1 | 3 |
| - | - | 1553 | 419.2 | - | - | 0 | - |
| - | - | 1376 | 423.7 | - | - | 0 | - |
| 3 | y | 1.409E+04 | 424.7 | 0.0001057 | 0.2488 | +2 | 7 |
| - | - | 6057 | 425.2 | - | - | 0 | - |
| - | - | 821.8 | 425.7 | - | - | 0 | - |
| - | - | 877.7 | 431.2 | - | - | 0 | - |
| 7 | y | 9501 | 435.2 | 0.0001239 | 0.2847 | +1 | 3 |
| - | - | 2036 | 436.2 | - | - | 0 | - |
| - | - | 609 | 436.9 | - | - | 0 | - |
| - | - | 2424 | 439.7 | - | - | 0 | - |
| - | - | 836.3 | 440.3 | - | - | 0 | - |
| 7 | b | 2209 | 447.7 | 0.001868 | 4.172 | +2 | 7 |
| - | - | 2428 | 448.7 | - | - | 0 | - |
| - | - | 1050 | 449.2 | - | - | 0 | - |
| - | - | 596.5 | 451.7 | - | - | 0 | - |
| - | - | 848.2 | 452.2 | - | - | 0 | - |
| - | - | 695.9 | 453.1 | - | - | 0 | - |
| - | - | 1029 | 454.2 | - | - | 0 | - |
| - | - | 628.4 | 456.7 | - | - | 0 | - |
| - | - | 1574 | 458.2 | - | - | 0 | - |
| - | - | 919.6 | 459.2 | - | - | 0 | - |
| - | - | 626 | 460.2 | - | - | 0 | - |
| - | - | 1797 | 460.7 | - | - | 0 | - |
| - | - | 1464 | 461.2 | - | - | 0 | - |
| - | - | 902.3 | 463.7 | - | - | 0 | - |
| - | - | 681.2 | 465.2 | - | - | 0 | - |
| - | - | 874.1 | 470.7 | - | - | 0 | - |
| - | - | 956.3 | 471.7 | - | - | 0 | - |
| - | - | 697.6 | 472.7 | - | - | 0 | - |
| - | - | 4612 | 476.2 | - | - | 0 | - |
| - | - | 794.1 | 477.2 | - | - | 0 | - |
| - | - | 3133 | 479.7 | - | - | 0 | - |
| - | - | 1012 | 480.2 | - | - | 0 | - |
| - | - | 1026 | 482.3 | - | - | 0 | - |
| - | - | 1984 | 486.3 | - | - | 0 | - |
| - | - | 1130 | 487.3 | - | - | 0 | - |
| - | - | 632.6 | 488.3 | - | - | 0 | - |
| - | - | 829.5 | 490.2 | - | - | 0 | - |
| - | - | 713.1 | 490.7 | - | - | 0 | - |
| 8 | b | 3145 | 496.2 | 0.002768 | 5.577 | +2 | 8 |
| 2 | y | 2187 | 496.7 | 0.005286 | 10.64 | +2 | 8 |
| - | - | 879.9 | 497.2 | - | - | 0 | - |
| - | - | 1474 | 499.3 | - | - | 0 | - |
| 2 | y | 2.724E+04 | 505.2 | 0.002801 | 5.544 | +2 | 8 |
| - | - | 1.424E+04 | 505.7 | - | - | 0 | - |
| - | - | 4726 | 506.2 | - | - | 0 | - |
| - | - | 1.449E+04 | 508.3 | - | - | 0 | - |
| - | - | 590 | 508.3 | - | - | 0 | - |
| - | - | 2660 | 509.3 | - | - | 0 | - |
| - | - | 629.4 | 511.6 | - | - | 0 | - |
| - | - | 804.1 | 515.3 | - | - | 0 | - |
| 4 | b | 2318 | 524.2 | 0.00513 | 9.787 | +1 | 4 |
| - | - | 769 | 525.2 | - | - | 0 | - |
| - | - | 779.4 | 528.3 | - | - | 0 | - |
| 6 | y | 7.61E+04 | 534.3 | 0.000541 | 1.013 | +1 | 4 |
| - | - | 2.211E+04 | 535.3 | - | - | 0 | - |
| - | - | 3179 | 536.3 | - | - | 0 | - |
| - | - | 2182 | 540.8 | - | - | 0 | - |
| - | - | 1495 | 541.3 | - | - | 0 | - |
| - | - | 1102 | 553.3 | - | - | 0 | - |
| - | - | 824.5 | 558.3 | - | - | 0 | - |
| - | - | 1149 | 569.3 | - | - | 0 | - |
| - | - | 926 | 569.8 | - | - | 0 | - |
| - | - | 4131 | 570.3 | - | - | 0 | - |
| - | - | 1593 | 571.3 | - | - | 0 | - |
| - | - | 841.7 | 575.2 | - | - | 0 | - |
| 0 | Precursor | 1937 | 577.8 | 0.002358 | 4.081 | +2 | -1 |
| 0 | Precursor | 1755 | 578.3 | 0.004292 | 7.421 | +2 | -1 |
| - | - | 817.2 | 578.8 | - | - | 0 | - |
| - | - | 580.6 | 585.3 | - | - | 0 | - |
| - | - | 5013 | 585.4 | - | - | 0 | - |
| - | - | 2189 | 585.9 | - | - | 0 | - |
| 0 | Precursor | 2.409E+05 | 586.8 | 0.002757 | 4.699 | +2 | -1 |
| - | - | 1.653E+05 | 587.3 | - | - | 0 | - |
| - | - | 7.147E+04 | 587.8 | - | - | 0 | - |
| - | - | 9216 | 588.3 | - | - | 0 | - |
| - | - | 4435 | 607.3 | - | - | 0 | - |
| - | - | 1009 | 614.4 | - | - | 0 | - |
| 5 | b | 1182 | 621.2 | 0.004785 | 7.703 | +1 | 5 |
| 5 | b | 896.1 | 622.2 | 0.01236 | 19.86 | +1 | 5 |
| 5 | b | 1.351E+04 | 639.2 | 0.00534 | 8.354 | +1 | 5 |
| - | - | 4502 | 640.2 | - | - | 0 | - |
| - | - | 1191 | 641.2 | - | - | 0 | - |
| 5 | y | 5.044E+04 | 649.3 | 0.0006896 | 1.062 | +1 | 5 |
| - | - | 1.691E+04 | 650.3 | - | - | 0 | - |
| - | - | 2699 | 651.3 | - | - | 0 | - |
| - | - | 2339 | 685.4 | - | - | 0 | - |
| - | - | 636.9 | 686.4 | - | - | 0 | - |
| - | - | 1.005E+04 | 694.4 | - | - | 0 | - |
| - | - | 2897 | 695.4 | - | - | 0 | - |
| - | - | 918.7 | 714.3 | - | - | 0 | - |
| - | - | 1546 | 717.4 | - | - | 0 | - |
| - | - | 1677 | 731.3 | - | - | 0 | - |
| - | - | 943 | 732.3 | - | - | 0 | - |
| 6 | b | 962.8 | 738.3 | 0.006737 | 9.125 | +1 | 6 |
| - | - | 2843 | 745.3 | - | - | 0 | - |
| - | - | 927.1 | 746.3 | - | - | 0 | - |
| 4 | y | 2.13E+04 | 777.4 | 0.00117 | 1.505 | +1 | 6 |
| - | - | 8590 | 778.4 | - | - | 0 | - |
| - | - | 1654 | 779.4 | - | - | 0 | - |
| - | - | 636.8 | 812.4 | - | - | 0 | - |
| - | - | 662.4 | 814.4 | - | - | 0 | - |
| - | - | 759.3 | 815.4 | - | - | 0 | - |
| - | - | 743.1 | 816.4 | - | - | 0 | - |
| 3 | y | 1482 | 831.4 | 0.002641 | 3.176 | +1 | 7 |
| - | - | 2127 | 844.4 | - | - | 0 | - |
| - | - | 781.9 | 846.4 | - | - | 0 | - |
| 3 | y | 4.21E+04 | 848.5 | 0.001907 | 2.248 | +1 | 7 |
| - | - | 1.862E+04 | 849.5 | - | - | 0 | - |
| - | - | 3491 | 850.5 | - | - | 0 | - |
| 7 | b | 1477 | 877.4 | 0.00232 | 2.644 | +1 | 7 |
| 7 | b | 4786 | 894.4 | 0.005553 | 6.209 | +1 | 7 |
| - | - | 3120 | 895.4 | - | - | 0 | - |
| - | - | 4226 | 896.4 | - | - | 0 | - |
| - | - | 1804 | 897.4 | - | - | 0 | - |
| 8 | b | 2592 | 973.5 | 0.006493 | 6.67 | +1 | 8 |
| 2 | y | 993.5 | 992.5 | 0.004979 | 5.017 | +1 | 8 |
| 2 | y | 1.24E+04 | 1009 | 0.007175 | 7.108 | +1 | 8 |
| - | - | 6195 | 1010 | - | - | 0 | - |
| - | - | 2613 | 1011 | - | - | 0 | - |
| - | - | 3288 | 1060 | - | - | 0 | - |
| - | - | 2066 | 1061 | - | - | 0 | - |
| - | - | 601.1 | 1345 | - | - | 0 | - |
| - | - | 678.6 | 2606 | - | - | 0 | - |
| - | - | 796.6 | 2919 | - | - | 0 | - |

m/z Charge Intensity FragmentType MassShift Position
120.0809555053711 0 1586.7792
127.08682250976562 0 692.597
129.1023712158203 0 48649.55
130.0499267578125 0 451.53186
130.10577392578125 0 2522.0544
131.08145141601562 0 1114.6761
132.62255859375 0 385.51917
134.0271453857422 0 11206.593
136.07582092285156 0 83754.42
137.0791778564453 0 6737.421
138.69232177734375 0 448.19818
139.08685302734375 0 1274.8003
140.08197021484375 0 1094.298
141.1022491455078 0 762.9782
147.0440673828125 0 865.00757
149.023681640625 0 507.8058
153.10203552246094 0 438.2824
155.11810302734375 0 1825.9148
156.9056854248047 0 425.44458
157.06082153320312 0 4043.4458
157.09732055664062 0 804.56055
157.1083526611328 0 1835.6844
158.06430053710938 0 434.55786
158.09249877929688 0 707.9199
159.11280822753906 0 2392.0356
160.07583618164062 0 1081.4525
162.02145385742188 0 609.1412
164.0706024169922 0 1062.0624
165.05459594726562 0 801.5923
165.10218811035156 0 514.20844
166.08624267578125 0 1232.5183
169.0973663330078 0 734.12115
169.13356018066406 0 730.5744
169.55162048339844 0 425.82712
170.09249877929688 0 478.74326
173.1285858154297 0 2707.0842
175.07131958007812 0 3978.0864
175.0863494873047 0 544.4843
180.10157775878906 0 519.9342
181.0970001220703 0 768.74457
182.08128356933594 0 1307.8423 y 8
183.11277770996094 0 3710.232
183.14932250976562 0 1536.7339
185.0556640625 0 496.01825
185.09437561035156 0 553.9477
185.1031494140625 0 2633.055
187.05368041992188 0 676.7951
187.10769653320312 0 5357.3525
189.0870819091797 0 696.6522
192.0478973388672 0 847.65137
197.10330200195312 0 996.3364
197.12840270996094 0 1541.667
199.1077117919922 0 1572.1155
200.1392822265625 0 13949.65
201.12339782714844 0 2910.395
201.14300537109375 0 512.3172
203.06622314453125 0 6591.319
205.06399536132812 0 9343.359
206.06651306152344 0 502.43753
209.09266662597656 0 1049.3534
210.1244354248047 0 619.3737
211.1439971923828 0 918.4398
214.15475463867188 0 793.9534
215.0482940673828 0 622.88385
215.10272216796875 0 1969.6667
215.13893127441406 0 706.42413
216.06539916992188 0 1888.3505
217.0817108154297 0 744.87976
226.11863708496094 0 2642.5127
232.14012145996094 0 1367.0376
233.0590057373047 0 2166.4082
233.092041015625 0 13836.532
234.058349609375 0 3255.4019
234.09600830078125 0 909.69196
237.0692596435547 0 1900.8502
237.13450622558594 0 1383.2246
239.1502227783203 0 21846.791
240.1531982421875 0 2406.6426
244.12905883789062 0 2185.193
252.06887817382812 0 8466.632
253.07212829589844 0 645.38696
254.11386108398438 0 614.0206
254.1613311767578 0 1327.8888
254.18724060058594 0 534.59796
255.14459228515625 0 1527.3955
256.1766662597656 0 9279.574
257.1800537109375 0 1311.7341
262.0530700683594 0 951.5876
262.1025085449219 0 625.1462
272.0874938964844 0 2323.648
272.1243591308594 0 1459.0936
272.171630859375 0 600.0808
274.1875915527344 0 1975.306
275.8654479980469 0 518.9005
279.13372802734375 0 33817.91 y 7
280.1369934082031 0 4295.2065
282.1553039550781 0 702.03253
284.1708679199219 0 649.1816
290.0984802246094 0 2558.3103
297.09033203125 0 55904.824 a 1
298.0933532714844 0 7715.249
299.08685302734375 0 1858.6082
301.1412658691406 0 782.61456
301.91912841796875 0 540.30334
304.129150390625 0 9010.868
305.1322326660156 0 2157.1975
305.161376953125 0 1202.199
312.1556396484375 0 1499.6937
312.4078063964844 0 581.41583
315.0218505859375 0 585.4575
315.1661376953125 0 4352.6963
325.0852355957031 0 20477.004 b 1
325.1516418457031 0 653.0272
326.08856201171875 0 3290.0522
340.1859436035156 0 749.329
343.1611633300781 0 2720.5842
343.2027282714844 0 1208.9607
347.6785888671875 0 1002.6925
351.16766357421875 0 493.14084
354.17706298828125 0 1624.3208
359.1674499511719 0 690.5682
359.192626953125 0 1765.9703
361.1536560058594 0 2147.0068 b Ammonia loss 5
361.1803894042969 0 1153.3801
366.17681884765625 0 1473.8586
371.1561584472656 0 1747.6604
371.2032165527344 0 7065.621
371.23968505859375 0 8437.14
372.2063903808594 0 1384.2742
372.24249267578125 0 1326.9205
379.20855712890625 0 6278.81
380.2110900878906 0 1259.5607 y Water loss 3
380.7025451660156 0 3295.4067 y Ammonia loss 3
381.205322265625 0 1601.3837
383.2039489746094 0 1332.9425
386.2397766113281 0 773.5209
388.21343994140625 0 573.2097
389.21588134765625 0 9602.8545 y 3
389.7172546386719 0 3337.4336
396.1224365234375 0 3416.4348 b 2
397.12286376953125 0 1031.9619
397.2579040527344 0 559.4213
414.2337341308594 0 1175.118
414.8311462402344 0 557.8997
417.2251892089844 0 812.0177
418.2083740234375 0 6463.187 y Ammonia loss 6
419.2101745605469 0 1552.7502
423.71063232421875 0 1375.5193
424.7347717285156 0 14089.82 y 2
425.23626708984375 0 6056.9634
425.7386169433594 0 821.8412
431.1895751953125 0 877.669
435.23516845703125 0 9500.682 y 6
436.2372131347656 0 2035.8845
436.8622741699219 0 608.99805
439.7039489746094 0 2423.7349
440.2518310546875 0 836.30927
447.7109375 0 2208.654 b 6
448.708984375 0 2427.711
449.2098388671875 0 1049.6244
451.685546875 0 596.4924
452.183837890625 0 848.17065
453.1063232421875 0 695.9036
454.2409362792969 0 1028.7866
456.7142639160156 0 628.40155
458.16839599609375 0 1574.2117
459.2455749511719 0 919.5996
460.19403076171875 0 626.0204
460.6878967285156 0 1796.8213
461.1905822753906 0 1463.863
463.6622619628906 0 902.27026
465.2413330078125 0 681.1961
470.655029296875 0 874.0809
471.67694091796875 0 956.3107
472.6663818359375 0 697.6435
476.18096923828125 0 4612.096
477.18182373046875 0 794.0944
479.6609191894531 0 3132.5796
480.1642761230469 0 1012.1186
482.27386474609375 0 1025.686
486.2672424316406 0 1984.068
487.26947021484375 0 1129.5547
488.29681396484375 0 632.6346
490.24053955078125 0 829.47687
490.6514587402344 0 713.1057
496.2364196777344 0 3145.149 b 7
496.7364807128906 0 2187.479 y Ammonia loss 1
497.23919677734375 0 879.8582
499.2983703613281 0 1473.6155
505.2416687011719 0 27242.26 y 1
505.74346923828125 0 14240.706
506.24346923828125 0 4726.046
508.2508850097656 0 14493.55
508.2923583984375 0 589.9869
509.2542724609375 0 2659.6558
511.6285705566406 0 629.40186
515.2943115234375 0 804.06647
524.2167358398438 0 2318.2039 b 3
525.2225952148438 0 768.9796
528.3131103515625 0 779.39734
534.3029174804688 0 76104.76 y 5
535.3058471679688 0 22110.531
536.3084106445312 0 3178.774
540.7763061523438 0 2181.981
541.2786254882812 0 1494.581
553.3106689453125 0 1101.8026
558.3356323242188 0 824.5254
569.2554321289062 0 1148.8315
569.7557983398438 0 925.98346
570.3342895507812 0 4130.963
571.3340454101562 0 1593.4323
575.2496337890625 0 841.6768
577.7684936523438 0 1936.8707 Precursor Water loss
578.2671508789062 0 1754.997 Precursor Ammonia loss
578.7680053710938 0 817.20087
585.2889404296875 0 580.5727
585.3534545898438 0 5013.164
585.8546752929688 0 2188.7283
586.7733764648438 0 240886.16 Precursor
587.2745971679688 0 165265.67
587.7753295898438 0 71471
588.2759399414062 0 9216.314
607.3193359375 0 4435.0957
614.3621826171875 0 1008.75037
621.2334594726562 0 1182.0458 b Water loss 4
622.234619140625 0 896.10675 b Ammonia loss 4
639.2434692382812 0 13507.487 b 4
640.2464599609375 0 4502.2197
641.2451171875 0 1190.8589
649.3297119140625 0 50435.434 y 4
650.3323974609375 0 16912.35
651.3353881835938 0 2699.4456
685.3970336914062 0 2338.5288
686.3983154296875 0 636.8598
694.3512573242188 0 10048.156
695.3533935546875 0 2897.2405
714.3228759765625 0 918.6983
717.3505249023438 0 1545.5571
731.3462524414062 0 1676.7949
732.3492431640625 0 943.0081
738.3104858398438 0 962.7508 b 5
745.3463134765625 0 2842.7966
746.3475952148438 0 927.0515
777.4241943359375 0 21301.402 y 3
778.4268188476562 0 8590.465
779.4315795898438 0 1654.4084
812.3936767578125 0 636.7881
814.4064331054688 0 662.4213
815.3880615234375 0 759.33496
816.4232788085938 0 743.14856
831.4332885742188 0 1481.922 y Ammonia loss 2
844.4135131835938 0 2126.8586
846.4080810546875 0 781.91833
848.4605712890625 0 42104.3 y 2
849.4638061523438 0 18618.041
850.4674682617188 0 3490.6477
877.3894653320312 0 1476.9277 b Ammonia loss 6
894.4127807617188 0 4786.3564 b 6
895.4141235351562 0 3120.011
896.4097290039062 0 4226.401
897.4112548828125 0 1803.654
973.4540405273438 0 2592.2883 b Water loss 7
992.4501342773438 0 993.46014 y Ammonia loss 1
1009.4744873046875 0 12399.608 y 1
1010.477294921875 0 6195.009
1011.4807739257812 0 2613.2285
1060.486328125 0 3287.585
1061.490966796875 0 2065.5605
1345.2667236328125 0 601.1255
2605.578125 0 678.6122
2919.294921875 0 796.645

Spectrum Details

|  |  |
| --- | --- |
| Matched peaks? Matched peaks The total absolute number of peaks matched. Additionally in brackets the total fraction of peaks matched and the total number of peaks is shown. | 34 (12.55% of 271) |
| FDR? FDR The false discovery rate estimated for this peptide. It is calculated by matching all theoretical fragments with a non-integer shift with the raw peaks for this spectrum. This is done with 40 different shifts. The resulting percentage is the average number of annotated peaks over the number of annotated peaks with the correct spectrum. | 0.63% |
| Satellite FDR? Satellite FDR See the FDR for details on its calculation. This satellite ion specific FDR only contains the satellite ions (d/w) for I/L/J positions. | - |
| PSM Score? PSM Score The PSM Score as given by Hecklib to this annotated spectrum. It is shown with three significant figures. | 334 |

#### Spectrum 3679? Spectrum 3679 The raw spectrum of this peptide as annotated by Hecklib. The fragments are coloured according to ion type (see legend). Any peaks with a star '\*' as text can be hovered over to see the full details, first the ion type second the mass shift type. By hovering over the amino acids in the peptide or ions in the legend the corresponding peaks are highlighted. By toggling the 'Unassigned' label you can turn the background (unassigned) peaks on or off in the plot. By updating the slider in the Ion legend you can update the spectrum to only show the top X% of the peaks with labels. The top X% means any peak that is within X% of the highest intensity. By dragging in the spectrum you can zoom in to a specific part of the spectrum and use 'Zoom Out' to get back to the original zoom level. The annotation of the spectrum is based on the given sequence in the peptides file and is done with different software so inconsistencies are likely. The peaks are annotated based on the given sequence, with 20 ppm tolerance.

Copy Data

##### Spectrum 3679 (TSV)

###### Preview

```
Loading example...
```

*Click on the button to copy the data to your clipboard.*

Mz MinMz MaxIntensity Max

WidthHeightPeptide font sizePeptide stroke widthSpectrum font sizeSpectrum stroke widthCompact peptide

Ion legend

wxyz

abcd

OtherUnassignedIon^Charge^_Position_Show for top:%

YCAKDVRPY

07.37e+51.47e+62.21e+62.95e+6

Zoom Out

y^+1^_1_y^+1^_2_a^+1^_2_b^+1^_2_b^+2^_6_y^+2^_6_y^+2^_6_b^+1^_3_y^+1^_3_y^+2^_7_y^+1^_3_b^+2^_7_b^+2^_8_b^+2^_8_y^+2^_8_y^+2^_8_b^+1^_4_y^+1^_4_\*\*\*b^+1^_5_y^+1^_5_b^+1^_5_y^+1^_5_b^+1^_6_y^+1^_6_y^+1^_6_y^+1^_7_y^+1^_7_b^+1^_7_b^+1^_7_b^+1^_8_y^+1^_8_y^+1^_8_

0779155823373116

Fragment Matches Table

Show background peaks

| Position | Ion type | Intensity | mz Theoretical | mz Error (Th) | mz Error (ppm) | Charge | Series Number |
| --- | --- | --- | --- | --- | --- | --- | --- |
| - | - | 4834 | 120.1 | - | - | 0 | - |
| - | - | 2687 | 125.1 | - | - | 0 | - |
| - | - | 2747 | 127.1 | - | - | 0 | - |
| - | - | 2834 | 127.6 | - | - | 0 | - |
| - | - | 5.135E+05 | 129.1 | - | - | 0 | - |
| - | - | 2731 | 130.1 | - | - | 0 | - |
| - | - | 2.832E+04 | 130.1 | - | - | 0 | - |
| - | - | 2528 | 130.9 | - | - | 0 | - |
| - | - | 1.255E+05 | 134 | - | - | 0 | - |
| - | - | 2988 | 134.1 | - | - | 0 | - |
| - | - | 4633 | 136 | - | - | 0 | - |
| - | - | 9.995E+05 | 136.1 | - | - | 0 | - |
| - | - | 6539 | 137.1 | - | - | 0 | - |
| - | - | 7.964E+04 | 137.1 | - | - | 0 | - |
| - | - | 3204 | 138.1 | - | - | 0 | - |
| - | - | 9161 | 139.1 | - | - | 0 | - |
| - | - | 7051 | 140.1 | - | - | 0 | - |
| - | - | 3359 | 142.1 | - | - | 0 | - |
| - | - | 4455 | 146.1 | - | - | 0 | - |
| - | - | 1.317E+04 | 147 | - | - | 0 | - |
| - | - | 1.371E+04 | 155.1 | - | - | 0 | - |
| - | - | 2.089E+04 | 157.1 | - | - | 0 | - |
| - | - | 4402 | 159.1 | - | - | 0 | - |
| - | - | 7030 | 160 | - | - | 0 | - |
| - | - | 1.625E+04 | 160.1 | - | - | 0 | - |
| - | - | 8280 | 162 | - | - | 0 | - |
| - | - | 1.124E+04 | 164.1 | - | - | 0 | - |
| - | - | 3017 | 168.6 | - | - | 0 | - |
| - | - | 3374 | 169.1 | - | - | 0 | - |
| - | - | 4723 | 169.1 | - | - | 0 | - |
| - | - | 1.777E+04 | 173.4 | - | - | 0 | - |
| - | - | 5581 | 180.1 | - | - | 0 | - |
| - | - | 7148 | 181.1 | - | - | 0 | - |
| 9 | y | 9391 | 182.1 | 0.0004343 | 2.385 | +1 | 1 |
| - | - | 1.405E+04 | 183.1 | - | - | 0 | - |
| - | - | 3893 | 183.2 | - | - | 0 | - |
| - | - | 2.645E+04 | 185.1 | - | - | 0 | - |
| - | - | 1.163E+04 | 187.1 | - | - | 0 | - |
| - | - | 5.815E+04 | 187.1 | - | - | 0 | - |
| - | - | 3928 | 188.1 | - | - | 0 | - |
| - | - | 3966 | 191.1 | - | - | 0 | - |
| - | - | 1.462E+04 | 192 | - | - | 0 | - |
| - | - | 4687 | 195.1 | - | - | 0 | - |
| - | - | 3572 | 195.2 | - | - | 0 | - |
| - | - | 3191 | 195.7 | - | - | 0 | - |
| - | - | 5930 | 197.1 | - | - | 0 | - |
| - | - | 1.638E+04 | 197.1 | - | - | 0 | - |
| - | - | 1.717E+04 | 199.1 | - | - | 0 | - |
| - | - | 1.417E+05 | 200.1 | - | - | 0 | - |
| - | - | 1.07E+04 | 201.1 | - | - | 0 | - |
| - | - | 1.05E+05 | 205.1 | - | - | 0 | - |
| - | - | 4870 | 205.1 | - | - | 0 | - |
| - | - | 6640 | 206.1 | - | - | 0 | - |
| - | - | 3550 | 207.1 | - | - | 0 | - |
| - | - | 8246 | 209.1 | - | - | 0 | - |
| - | - | 5443 | 209.1 | - | - | 0 | - |
| - | - | 4963 | 210.1 | - | - | 0 | - |
| - | - | 5359 | 211.2 | - | - | 0 | - |
| - | - | 4179 | 214 | - | - | 0 | - |
| - | - | 1.213E+04 | 214.2 | - | - | 0 | - |
| - | - | 4055 | 215 | - | - | 0 | - |
| - | - | 1.843E+04 | 215.1 | - | - | 0 | - |
| - | - | 1.705E+04 | 216.1 | - | - | 0 | - |
| - | - | 2.875E+04 | 226.1 | - | - | 0 | - |
| - | - | 7504 | 227.1 | - | - | 0 | - |
| - | - | 2.294E+04 | 233.1 | - | - | 0 | - |
| - | - | 1.539E+05 | 233.1 | - | - | 0 | - |
| - | - | 9441 | 233.1 | - | - | 0 | - |
| - | - | 3.685E+04 | 234.1 | - | - | 0 | - |
| - | - | 1.77E+04 | 234.1 | - | - | 0 | - |
| - | - | 1.96E+04 | 237.1 | - | - | 0 | - |
| - | - | 1.383E+04 | 237.1 | - | - | 0 | - |
| - | - | 2.375E+05 | 239.2 | - | - | 0 | - |
| - | - | 2.551E+04 | 240.2 | - | - | 0 | - |
| - | - | 7636 | 243.1 | - | - | 0 | - |
| - | - | 1.836E+04 | 244.1 | - | - | 0 | - |
| - | - | 4458 | 250.1 | - | - | 0 | - |
| - | - | 1.065E+05 | 252.1 | - | - | 0 | - |
| - | - | 9769 | 253.1 | - | - | 0 | - |
| - | - | 6006 | 254.1 | - | - | 0 | - |
| - | - | 1.715E+04 | 254.2 | - | - | 0 | - |
| - | - | 1.492E+04 | 255.1 | - | - | 0 | - |
| - | - | 1.15E+05 | 256.2 | - | - | 0 | - |
| - | - | 7831 | 257.2 | - | - | 0 | - |
| - | - | 8780 | 259.1 | - | - | 0 | - |
| - | - | 1.119E+04 | 262.1 | - | - | 0 | - |
| - | - | 2.967E+04 | 272.1 | - | - | 0 | - |
| - | - | 7820 | 272.2 | - | - | 0 | - |
| - | - | 2.967E+04 | 274.2 | - | - | 0 | - |
| 8 | y | 3.761E+05 | 279.1 | 0.0001607 | 0.5757 | +1 | 2 |
| - | - | 5.747E+04 | 280.1 | - | - | 0 | - |
| - | - | 1.085E+04 | 282.2 | - | - | 0 | - |
| 2 | a | 6.656E+05 | 297.1 | 0.004207 | 14.16 | +1 | 2 |
| - | - | 1.092E+04 | 297.2 | - | - | 0 | - |
| - | - | 1.01E+05 | 298.1 | - | - | 0 | - |
| - | - | 4830 | 298.1 | - | - | 0 | - |
| - | - | 1.961E+04 | 299.1 | - | - | 0 | - |
| - | - | 5230 | 299.2 | - | - | 0 | - |
| - | - | 1.259E+05 | 304.1 | - | - | 0 | - |
| - | - | 1.821E+04 | 305.1 | - | - | 0 | - |
| - | - | 6070 | 307.7 | - | - | 0 | - |
| - | - | 7901 | 308.1 | - | - | 0 | - |
| - | - | 4.482E+04 | 315.2 | - | - | 0 | - |
| - | - | 6324 | 315.2 | - | - | 0 | - |
| - | - | 4856 | 316.2 | - | - | 0 | - |
| - | - | 4379 | 319.1 | - | - | 0 | - |
| 2 | b | 2.453E+05 | 325.1 | 0.004188 | 12.88 | +1 | 2 |
| - | - | 4213 | 325.2 | - | - | 0 | - |
| - | - | 3.172E+04 | 326.1 | - | - | 0 | - |
| - | - | 5575 | 327.1 | - | - | 0 | - |
| - | - | 3894 | 333.1 | - | - | 0 | - |
| - | - | 7393 | 343.1 | - | - | 0 | - |
| - | - | 1.672E+04 | 343.2 | - | - | 0 | - |
| - | - | 3986 | 344.2 | - | - | 0 | - |
| - | - | 5002 | 353.2 | - | - | 0 | - |
| - | - | 1.549E+04 | 354.2 | - | - | 0 | - |
| 6 | b | 2.245E+04 | 361.1 | 0.005077 | 14.06 | +2 | 6 |
| - | - | 1.062E+04 | 366.2 | - | - | 0 | - |
| - | - | 6.37E+04 | 371.2 | - | - | 0 | - |
| - | - | 9.395E+04 | 371.2 | - | - | 0 | - |
| - | - | 1.229E+04 | 372.2 | - | - | 0 | - |
| - | - | 1.458E+04 | 372.2 | - | - | 0 | - |
| 4 | y | 2.539E+04 | 380.7 | 0.0002927 | 0.7688 | +2 | 6 |
| - | - | 1.274E+04 | 381.2 | - | - | 0 | - |
| - | - | 1.228E+04 | 383.2 | - | - | 0 | - |
| - | - | 1.106E+04 | 386.2 | - | - | 0 | - |
| - | - | 4718 | 386.7 | - | - | 0 | - |
| 4 | y | 9.089E+04 | 389.2 | 4.241E-05 | 0.109 | +2 | 6 |
| - | - | 4.137E+04 | 389.7 | - | - | 0 | - |
| - | - | 5378 | 390.2 | - | - | 0 | - |
| 3 | b | 4.761E+04 | 396.1 | 0.004528 | 11.43 | +1 | 3 |
| - | - | 1.072E+04 | 397.1 | - | - | 0 | - |
| - | - | 1.158E+04 | 414.2 | - | - | 0 | - |
| - | - | 4928 | 414.7 | - | - | 0 | - |
| - | - | 5241 | 417.2 | - | - | 0 | - |
| 7 | y | 7.356E+04 | 418.2 | 0.0003058 | 0.7312 | +1 | 3 |
| - | - | 1.351E+04 | 419.2 | - | - | 0 | - |
| - | - | 1.804E+04 | 423.7 | - | - | 0 | - |
| - | - | 5286 | 424.2 | - | - | 0 | - |
| 3 | y | 1.712E+05 | 424.7 | 0.0004437 | 1.045 | +2 | 7 |
| - | - | 8.247E+04 | 425.2 | - | - | 0 | - |
| - | - | 2.083E+04 | 425.7 | - | - | 0 | - |
| - | - | 4254 | 428.6 | - | - | 0 | - |
| - | - | 4903 | 429.7 | - | - | 0 | - |
| 7 | y | 1.086E+05 | 435.2 | 0.0002155 | 0.495 | +1 | 3 |
| - | - | 1.877E+04 | 436.2 | - | - | 0 | - |
| - | - | 6944 | 439.3 | - | - | 0 | - |
| 7 | b | 2.84E+04 | 447.7 | 0.001288 | 2.877 | +2 | 7 |
| - | - | 9774 | 448.2 | - | - | 0 | - |
| - | - | 7071 | 450.7 | - | - | 0 | - |
| - | - | 4446 | 451.2 | - | - | 0 | - |
| - | - | 6126 | 451.7 | - | - | 0 | - |
| - | - | 4257 | 452.2 | - | - | 0 | - |
| - | - | 6041 | 454.2 | - | - | 0 | - |
| - | - | 4624 | 457.3 | - | - | 0 | - |
| - | - | 2.485E+04 | 458.2 | - | - | 0 | - |
| - | - | 5155 | 468.3 | - | - | 0 | - |
| - | - | 6.084E+04 | 476.2 | - | - | 0 | - |
| - | - | 1.29E+04 | 477.2 | - | - | 0 | - |
| - | - | 1.382E+04 | 482.3 | - | - | 0 | - |
| - | - | 3.268E+04 | 486.3 | - | - | 0 | - |
| - | - | 7800 | 487.3 | - | - | 0 | - |
| 8 | b | 5399 | 487.7 | 0.003958 | 8.116 | +2 | 8 |
| 8 | b | 4.112E+04 | 496.2 | 0.002066 | 4.163 | +2 | 8 |
| 2 | y | 2.251E+04 | 496.7 | 0.005957 | 11.99 | +2 | 8 |
| - | - | 5184 | 497.2 | - | - | 0 | - |
| - | - | 2.489E+04 | 499.3 | - | - | 0 | - |
| - | - | 7660 | 500.3 | - | - | 0 | - |
| 2 | y | 3.637E+05 | 505.2 | 0.001946 | 3.852 | +2 | 8 |
| - | - | 1.901E+05 | 505.7 | - | - | 0 | - |
| - | - | 6.369E+04 | 506.2 | - | - | 0 | - |
| - | - | 1.146E+04 | 506.7 | - | - | 0 | - |
| - | - | 4319 | 518 | - | - | 0 | - |
| - | - | 6709 | 523.3 | - | - | 0 | - |
| 4 | b | 1.635E+04 | 524.2 | 0.004398 | 8.39 | +1 | 4 |
| - | - | 8091 | 532.3 | - | - | 0 | - |
| 6 | y | 9.269E+05 | 534.3 | 0.0003135 | 0.5867 | +1 | 4 |
| - | - | 2.511E+05 | 535.3 | - | - | 0 | - |
| - | - | 3.862E+04 | 536.3 | - | - | 0 | - |
| - | - | 2.301E+04 | 540.8 | - | - | 0 | - |
| - | - | 2.097E+04 | 541.3 | - | - | 0 | - |
| - | - | 5763 | 541.8 | - | - | 0 | - |
| - | - | 6627 | 542.3 | - | - | 0 | - |
| - | - | 1.176E+04 | 544.3 | - | - | 0 | - |
| - | - | 6746 | 547.3 | - | - | 0 | - |
| - | - | 5185 | 549.3 | - | - | 0 | - |
| - | - | 1.545E+04 | 553.3 | - | - | 0 | - |
| - | - | 4653 | 557.3 | - | - | 0 | - |
| - | - | 5823 | 564.3 | - | - | 0 | - |
| - | - | 4248 | 564.8 | - | - | 0 | - |
| - | - | 5416 | 565.8 | - | - | 0 | - |
| - | - | 1.952E+04 | 569.3 | - | - | 0 | - |
| - | - | 1.261E+04 | 569.8 | - | - | 0 | - |
| - | - | 4.195E+04 | 570.3 | - | - | 0 | - |
| - | - | 1.404E+04 | 571.3 | - | - | 0 | - |
| - | - | 1.041E+04 | 575.3 | - | - | 0 | - |
| 0 | Precursor | 1.392E+04 | 577.8 | 0.001992 | 3.447 | +2 | -1 |
| 0 | Precursor | 1.631E+04 | 578.3 | 0.001606 | 2.777 | +2 | -1 |
| - | - | 1.292E+04 | 578.8 | - | - | 0 | - |
| - | - | 5509 | 579.3 | - | - | 0 | - |
| 0 | Precursor | 2.918E+06 | 586.8 | 0.001842 | 3.139 | +2 | -1 |
| - | - | 1.883E+06 | 587.3 | - | - | 0 | - |
| - | - | 8.24E+05 | 587.8 | - | - | 0 | - |
| - | - | 1.022E+05 | 588.3 | - | - | 0 | - |
| - | - | 6270 | 590.3 | - | - | 0 | - |
| - | - | 5005 | 614.3 | - | - | 0 | - |
| - | - | 1.291E+04 | 614.4 | - | - | 0 | - |
| 5 | b | 2.542E+04 | 621.2 | 0.004358 | 7.015 | +1 | 5 |
| - | - | 4580 | 623.2 | - | - | 0 | - |
| 5 | y | 1.456E+04 | 632.3 | 0.002666 | 4.217 | +1 | 5 |
| - | - | 7815 | 633.3 | - | - | 0 | - |
| 5 | b | 1.458E+05 | 639.2 | 0.004486 | 7.017 | +1 | 5 |
| - | - | 4.838E+04 | 640.2 | - | - | 0 | - |
| - | - | 1.356E+04 | 641.2 | - | - | 0 | - |
| 5 | y | 6.278E+05 | 649.3 | 0.000348 | 0.5359 | +1 | 5 |
| - | - | 2.068E+05 | 650.3 | - | - | 0 | - |
| - | - | 2.986E+04 | 651.3 | - | - | 0 | - |
| - | - | 9507 | 659.3 | - | - | 0 | - |
| - | - | 2.099E+04 | 685.4 | - | - | 0 | - |
| - | - | 8407 | 686.4 | - | - | 0 | - |
| - | - | 1.272E+04 | 714.3 | - | - | 0 | - |
| - | - | 2.436E+04 | 731.4 | - | - | 0 | - |
| - | - | 8549 | 732.4 | - | - | 0 | - |
| 6 | b | 1.866E+04 | 738.3 | 0.004113 | 5.571 | +1 | 6 |
| - | - | 1.039E+04 | 739.3 | - | - | 0 | - |
| - | - | 5769 | 740.3 | - | - | 0 | - |
| 4 | y | 7951 | 760.4 | 0.001171 | 1.54 | +1 | 6 |
| 4 | y | 2.691E+05 | 777.4 | 0.0001936 | 0.2491 | +1 | 6 |
| - | - | 1.059E+05 | 778.4 | - | - | 0 | - |
| - | - | 2.917E+04 | 779.4 | - | - | 0 | - |
| - | - | 5793 | 787.4 | - | - | 0 | - |
| - | - | 5089 | 789.4 | - | - | 0 | - |
| 3 | y | 1.505E+04 | 831.4 | 0.001876 | 2.256 | +1 | 7 |
| - | - | 1.277E+04 | 846.4 | - | - | 0 | - |
| 3 | y | 4.994E+05 | 848.5 | 0.0003201 | 0.3773 | +1 | 7 |
| - | - | 2.309E+05 | 849.5 | - | - | 0 | - |
| - | - | 5.273E+04 | 850.5 | - | - | 0 | - |
| 7 | b | 1.254E+04 | 877.4 | 0.002808 | 3.201 | +1 | 7 |
| - | - | 8217 | 878.4 | - | - | 0 | - |
| 7 | b | 6.637E+04 | 894.4 | 0.004455 | 4.981 | +1 | 7 |
| - | - | 2.278E+04 | 895.4 | - | - | 0 | - |
| - | - | 1.05E+04 | 896.4 | - | - | 0 | - |
| 8 | b | 4683 | 991.5 | 0.002653 | 2.676 | +1 | 8 |
| 2 | y | 9687 | 992.5 | 0.004491 | 4.525 | +1 | 8 |
| 2 | y | 1.56E+05 | 1009 | 0.0051 | 5.052 | +1 | 8 |
| - | - | 7.731E+04 | 1010 | - | - | 0 | - |
| - | - | 2.966E+04 | 1011 | - | - | 0 | - |
| - | - | 6589 | 1012 | - | - | 0 | - |
| - | - | 4503 | 1568 | - | - | 0 | - |
| - | - | 4305 | 2134 | - | - | 0 | - |
| - | - | 5526 | 3084 | - | - | 0 | - |
| - | - | 4277 | 3085 | - | - | 0 | - |
| - | - | 5283 | 3086 | - | - | 0 | - |

m/z Charge Intensity FragmentType MassShift Position
120.05268859863281 0 4834.0957
125.0710220336914 0 2686.5576
127.05057525634766 0 2747.1733
127.61426544189453 0 2834.1702
129.1024627685547 0 513459.75
130.0859375 0 2731.3645
130.10581970214844 0 28319.271
130.94012451171875 0 2528.1897
134.02720642089844 0 125457.58
134.1243438720703 0 2988.0784
136.02291870117188 0 4633.187
136.07594299316406 0 999458.75
137.07330322265625 0 6538.5103
137.07923889160156 0 79637.54
138.0663604736328 0 3203.5032
139.086669921875 0 9160.989
140.08206176757812 0 7050.6016
142.06207275390625 0 3359.4639
146.06053161621094 0 4454.9634
147.0442352294922 0 13165.131
155.11802673339844 0 13709.278
157.10853576660156 0 20886.941
159.05870056152344 0 4401.552
160.04278564453125 0 7030.335
160.07566833496094 0 16251.043
162.02212524414062 0 8279.623
164.0708770751953 0 11236.775
168.5688934326172 0 3017.1738
169.09718322753906 0 3373.8213
169.1338653564453 0 4722.561
173.439697265625 0 17765.688
180.1014862060547 0 5581.1904
181.097412109375 0 7148.105
182.08160400390625 0 9391.327 y 8
183.113037109375 0 14053.276
183.1501007080078 0 3893.1104
185.1034698486328 0 26447.387
187.0537567138672 0 11634.852
187.10787963867188 0 58152.797
188.11097717285156 0 3928.0964
191.0814971923828 0 3966.1138
192.0480499267578 0 14616.862
195.11277770996094 0 4686.6553
195.1595916748047 0 3572.4534
195.74847412109375 0 3190.636
197.09222412109375 0 5930.0737
197.12860107421875 0 16375.431
199.10801696777344 0 17171.14
200.1394805908203 0 141716.92
201.1429901123047 0 10700.494
205.0642852783203 0 104966.6
205.0969696044922 0 4870.008
206.0677947998047 0 6640.341
207.1142120361328 0 3549.9927
209.0928497314453 0 8245.82
209.1400146484375 0 5442.66
210.12388610839844 0 4963.1885
211.1547393798828 0 5358.752
214.0167999267578 0 4178.7856
214.15528869628906 0 12128.004
215.04830932617188 0 4054.8616
215.1031951904297 0 18431.77
216.065673828125 0 17048.16
226.11875915527344 0 28751.008
227.10281372070312 0 7504.3804
233.0592041015625 0 22938.707
233.0922088623047 0 153935.3
233.1286163330078 0 9440.601
234.05844116210938 0 36853.96
234.09564208984375 0 17699.316
237.06932067871094 0 19600.473
237.1346893310547 0 13834.863
239.1504364013672 0 237464.11
240.15383911132812 0 25509.576
243.09727478027344 0 7635.996
244.12925720214844 0 18355.95
250.08575439453125 0 4457.95
252.06907653808594 0 106471.21
253.07273864746094 0 9769.256
254.1137237548828 0 6006.08
254.16152954101562 0 17150
255.14540100097656 0 14915.032
256.1769104003906 0 115026.47
257.1807556152344 0 7831.066
259.107666015625 0 8780.285
262.0533142089844 0 11185.69
272.1241149902344 0 29665.785
272.171875 0 7820.1074
274.1876525878906 0 29674.463
279.13409423828125 0 376064.06 y 7
280.1370849609375 0 57469.465
282.1560974121094 0 10847.132
297.0906677246094 0 665624.8 a 1
297.1560974121094 0 10924.517
298.09375 0 101000.91
298.1388244628906 0 4830.06
299.0863952636719 0 19610.428
299.17218017578125 0 5230.457
304.1294250488281 0 125862.97
305.1326599121094 0 18207.652
307.6849060058594 0 6069.7427
308.0603332519531 0 7900.604
315.1663818359375 0 44824.59
315.20330810546875 0 6324.498
316.1709289550781 0 4855.553
319.1409606933594 0 4378.846
325.0856018066406 0 245280.42 b 1
325.18707275390625 0 4213.4497
326.089111328125 0 31720.51
327.0823974609375 0 5575.274
333.0891418457031 0 3893.8083
343.1435546875 0 7393.0396
343.2072448730469 0 16721.846
344.20855712890625 0 3986.016
353.1916198730469 0 5001.892
354.1778259277344 0 15486.473
361.154052734375 0 22447.494 b Ammonia loss 5
366.1791076660156 0 10615.623
371.2031555175781 0 63695.418
371.2402648925781 0 93948.9
372.2057189941406 0 12293.106
372.2431945800781 0 14582.543
380.7033386230469 0 25390.404 y Ammonia loss 3
381.2048645019531 0 12740.123
383.20361328125 0 12278.668
386.23919677734375 0 11063.231
386.70281982421875 0 4717.693
389.2162780761719 0 90890.6 y 3
389.71807861328125 0 41372.21
390.2194519042969 0 5377.615
396.12237548828125 0 47612.87 b 2
397.1249084472656 0 10720.446
414.23419189453125 0 11578.649
414.7059326171875 0 4927.7573
417.2255859375 0 5240.52
418.20880126953125 0 73563.3 y Ammonia loss 6
419.21185302734375 0 13512.254
423.71136474609375 0 18041.945
424.2120056152344 0 5286.025
424.7353210449219 0 171150.42 y 2
425.23651123046875 0 82465.14
425.7383117675781 0 20833.312
428.63470458984375 0 4254.3657
429.7281799316406 0 4903.1836
435.2352600097656 0 108616.625 y 6
436.23760986328125 0 18766.008
439.2655029296875 0 6943.7695
447.7115173339844 0 28398.816 b 6
448.2127380371094 0 9773.835
450.7328186035156 0 7071.0317
451.23394775390625 0 4445.532
451.738525390625 0 6125.656
452.2431945800781 0 4257.192
454.2415466308594 0 6041.3223
457.27484130859375 0 4623.7603
458.1708068847656 0 24849.459
468.2591552734375 0 5155.0103
476.1816711425781 0 60840.355
477.18365478515625 0 12898.972
482.27410888671875 0 13820.135
486.2672119140625 0 32678.102
487.26922607421875 0 7800.0103
487.7219543457031 0 5399.4976 b Ammonia loss 7
496.23712158203125 0 41120.277 b 7
496.7371520996094 0 22513.236 y Ammonia loss 1
497.2389221191406 0 5183.56
499.2992248535156 0 24890.418
500.2991027832031 0 7660.08
505.2425231933594 0 363692.22 y 1
505.7438049316406 0 190087.25
506.2442321777344 0 63687.066
506.742919921875 0 11459.214
518.0057373046875 0 4319.0786
523.258544921875 0 6708.511
524.2174682617188 0 16345.11 b 3
532.2662963867188 0 8091.2397
534.3037719726562 0 926926.1 y 5
535.3065185546875 0 251092.44
536.3092041015625 0 38623.914
540.77783203125 0 23007.957
541.2803344726562 0 20969.65
541.7789306640625 0 5762.639
542.3427734375 0 6626.505
544.2870483398438 0 11760.456
547.2549438476562 0 6745.9375
549.25830078125 0 5184.549
553.31005859375 0 15450.353
557.2689819335938 0 4653.0967
564.2676391601562 0 5822.932
564.7694702148438 0 4248.0977
565.7586669921875 0 5415.643
569.2562866210938 0 19523.102
569.7576293945312 0 12605.076
570.335693359375 0 41945.062
571.3395385742188 0 14038.77
575.2503662109375 0 10408.608
577.7688598632812 0 13917.72 Precursor Water loss
578.2644653320312 0 16309.599 Precursor Ammonia loss
578.7646484375 0 12922.353
579.2617797851562 0 5508.835
586.7742919921875 0 2918367.2 Precursor
587.2755126953125 0 1882716.4
587.7760009765625 0 823988.5
588.2765502929688 0 102166.766
590.284423828125 0 6270.0884
614.2939453125 0 5005.2993
614.3623657226562 0 12906.43
621.23388671875 0 25422.926 b Water loss 4
623.2391357421875 0 4580.4624
632.3065185546875 0 14558.882 y Ammonia loss 4
633.3073120117188 0 7815.278
639.2443237304688 0 145815.6 b 4
640.2470703125 0 48375.586
641.2473754882812 0 13562.685
649.3307495117188 0 627817.5 y 4
650.3330078125 0 206802.03
651.3363037109375 0 29857.97
659.3157958984375 0 9507.136
685.3991088867188 0 20989.824
686.4005737304688 0 8407.127
714.3275756835938 0 12716.078
731.3502197265625 0 24357.596
732.3512573242188 0 8548.616
738.3131103515625 0 18662.428 b 5
739.3162841796875 0 10392.265
740.31298828125 0 5769.1274
760.3976440429688 0 7951.1597 y Ammonia loss 3
777.4251708984375 0 269069.22 y 3
778.4281005859375 0 105931.96
779.4298095703125 0 29171.102
787.4136352539062 0 5793.342
789.4315795898438 0 5089.244
831.4378051757812 0 15049.096 y Ammonia loss 2
846.4113159179688 0 12770.543
848.462158203125 0 499355.38 y 2
849.4652709960938 0 230928.47
850.4682006835938 0 52726.26
877.3889770507812 0 12535.388 b Ammonia loss 6
878.3876342773438 0 8216.814
894.4138793945312 0 66367.21 b 6
895.4160766601562 0 22775.809
896.4172973632812 0 10500.572
991.4684448242188 0 4683.075 b 7
992.4506225585938 0 9686.558 y Ammonia loss 1
1009.4765625 0 155977.62 y 1
1010.4791870117188 0 77307.56
1011.480224609375 0 29663.953
1012.4732055664062 0 6588.9536
1568.26416015625 0 4502.851
2134.30810546875 0 4304.8057
3083.957275390625 0 5525.939
3084.88818359375 0 4277.3086
3085.515869140625 0 5283.053

Spectrum Details

|  |  |
| --- | --- |
| Matched peaks? Matched peaks The total absolute number of peaks matched. Additionally in brackets the total fraction of peaks matched and the total number of peaks is shown. | 35 (13.83% of 253) |
| FDR? FDR The false discovery rate estimated for this peptide. It is calculated by matching all theoretical fragments with a non-integer shift with the raw peaks for this spectrum. This is done with 40 different shifts. The resulting percentage is the average number of annotated peaks over the number of annotated peaks with the correct spectrum. | 0.82% |
| Satellite FDR? Satellite FDR See the FDR for details on its calculation. This satellite ion specific FDR only contains the satellite ions (d/w) for I/L/J positions. | - |
| PSM Score? PSM Score The PSM Score as given by Hecklib to this annotated spectrum. It is shown with three significant figures. | 355 |

#### Spectrum 3777? Spectrum 3777 The raw spectrum of this peptide as annotated by Hecklib. The fragments are coloured according to ion type (see legend). Any peaks with a star '\*' as text can be hovered over to see the full details, first the ion type second the mass shift type. By hovering over the amino acids in the peptide or ions in the legend the corresponding peaks are highlighted. By toggling the 'Unassigned' label you can turn the background (unassigned) peaks on or off in the plot. By updating the slider in the Ion legend you can update the spectrum to only show the top X% of the peaks with labels. The top X% means any peak that is within X% of the highest intensity. By dragging in the spectrum you can zoom in to a specific part of the spectrum and use 'Zoom Out' to get back to the original zoom level. The annotation of the spectrum is based on the given sequence in the peptides file and is done with different software so inconsistencies are likely. The peaks are annotated based on the given sequence, with 20 ppm tolerance.

Copy Data

##### Spectrum 3777 (TSV)

###### Preview

```
Loading example...
```

*Click on the button to copy the data to your clipboard.*

Mz MinMz MaxIntensity Max

WidthHeightPeptide font sizePeptide stroke widthSpectrum font sizeSpectrum stroke widthCompact peptide

Ion legend

wxyz

abcd

OtherUnassignedIon^Charge^_Position_Show for top:%

YCAKDVRPY

04.47e+48.95e+41.34e+51.79e+5

Zoom Out

y^+1^_1_y^+1^_2_c^+2^_5_c^+2^_6_c^+2^_6_y^+2^_6_z^+2^_6_y^+2^_6_c^+1^_3_z^+2^_7_z^+1^_3_y^+2^_7_y^+1^_3_w^+2^_8_c^+2^_8_y^+2^_8_z^+2^_8_w^+1^_4_c^+2^_8_y^+2^_8_y^+1^_4_z^+1^_4_c^+1^_4_y^+1^_4_c^+1^_4_y^+1^_5_z^+1^_5_c^+1^_5_y^+1^_5_c^+1^_5_c^+1^_6_y^+1^_6_z^+1^_6_y^+1^_6_z^+1^_7_y^+1^_7_w^+1^_8_y^+1^_8_z^+1^_8_c^+1^_8_y^+1^_8_

02985958931190

Fragment Matches Table

Show background peaks

| Position | Ion type | Intensity | mz Theoretical | mz Error (Th) | mz Error (ppm) | Charge | Series Number |
| --- | --- | --- | --- | --- | --- | --- | --- |
| - | - | 1334 | 121 | - | - | 0 | - |
| - | - | 1.008E+04 | 129.1 | - | - | 0 | - |
| - | - | 1069 | 134 | - | - | 0 | - |
| - | - | 4.714E+04 | 136.1 | - | - | 0 | - |
| - | - | 424 | 137.1 | - | - | 0 | - |
| - | - | 3441 | 137.1 | - | - | 0 | - |
| - | - | 438.8 | 142.1 | - | - | 0 | - |
| - | - | 591.2 | 142.1 | - | - | 0 | - |
| - | - | 1022 | 146.1 | - | - | 0 | - |
| - | - | 3042 | 147 | - | - | 0 | - |
| - | - | 1.771E+05 | 149 | - | - | 0 | - |
| - | - | 945.1 | 149.1 | - | - | 0 | - |
| - | - | 1.401E+04 | 150 | - | - | 0 | - |
| - | - | 423.3 | 152.7 | - | - | 0 | - |
| - | - | 2447 | 156.1 | - | - | 0 | - |
| - | - | 1126 | 160.1 | - | - | 0 | - |
| - | - | 6.086E+04 | 167 | - | - | 0 | - |
| - | - | 3609 | 168 | - | - | 0 | - |
| - | - | 1206 | 173.5 | - | - | 0 | - |
| - | - | 953.8 | 179 | - | - | 0 | - |
| 9 | y | 1570 | 182.1 | 7.081E-06 | 0.03889 | +1 | 1 |
| - | - | 474.7 | 183.1 | - | - | 0 | - |
| - | - | 397.1 | 183.4 | - | - | 0 | - |
| - | - | 1220 | 184.1 | - | - | 0 | - |
| - | - | 555.3 | 187.1 | - | - | 0 | - |
| - | - | 470 | 188 | - | - | 0 | - |
| - | - | 3820 | 200.1 | - | - | 0 | - |
| - | - | 3065 | 205.1 | - | - | 0 | - |
| - | - | 5100 | 211.2 | - | - | 0 | - |
| - | - | 516 | 212.2 | - | - | 0 | - |
| - | - | 6029 | 212.2 | - | - | 0 | - |
| - | - | 1484 | 213.1 | - | - | 0 | - |
| - | - | 604.6 | 213.2 | - | - | 0 | - |
| - | - | 919.5 | 221.1 | - | - | 0 | - |
| - | - | 650.7 | 223.1 | - | - | 0 | - |
| - | - | 1147 | 233.1 | - | - | 0 | - |
| - | - | 1202 | 233.1 | - | - | 0 | - |
| - | - | 633.7 | 239.1 | - | - | 0 | - |
| - | - | 1386 | 241.1 | - | - | 0 | - |
| - | - | 1846 | 250.1 | - | - | 0 | - |
| - | - | 921.3 | 251 | - | - | 0 | - |
| - | - | 997.1 | 254.2 | - | - | 0 | - |
| - | - | 945.1 | 255.1 | - | - | 0 | - |
| - | - | 1432 | 255.2 | - | - | 0 | - |
| - | - | 2716 | 256.2 | - | - | 0 | - |
| - | - | 2189 | 266.1 | - | - | 0 | - |
| 8 | y | 6939 | 279.1 | 0.000236 | 0.8455 | +1 | 2 |
| - | - | 7452 | 279.2 | - | - | 0 | - |
| - | - | 1020 | 280.1 | - | - | 0 | - |
| - | - | 1495 | 280.2 | - | - | 0 | - |
| - | - | 1674 | 284.2 | - | - | 0 | - |
| - | - | 941 | 285.2 | - | - | 0 | - |
| - | - | 1.8E+04 | 297.1 | - | - | 0 | - |
| - | - | 2157 | 298.1 | - | - | 0 | - |
| - | - | 883.4 | 301.1 | - | - | 0 | - |
| - | - | 5012 | 304.1 | - | - | 0 | - |
| - | - | 639.3 | 309.2 | - | - | 0 | - |
| - | - | 8956 | 313.2 | - | - | 0 | - |
| - | - | 1588 | 314.2 | - | - | 0 | - |
| - | - | 809.2 | 315.2 | - | - | 0 | - |
| 5 | c | 1937 | 320.1 | 0.002433 | 7.599 | +2 | 5 |
| - | - | 628.7 | 320.2 | - | - | 0 | - |
| - | - | 1.346E+04 | 325.1 | - | - | 0 | - |
| - | - | 2344 | 326.1 | - | - | 0 | - |
| - | - | 855.9 | 354.2 | - | - | 0 | - |
| - | - | 926.9 | 354.9 | - | - | 0 | - |
| - | - | 2802 | 355.1 | - | - | 0 | - |
| - | - | 762.9 | 355.2 | - | - | 0 | - |
| - | - | 1167 | 356.1 | - | - | 0 | - |
| - | - | 2715 | 357.1 | - | - | 0 | - |
| - | - | 946.1 | 358.1 | - | - | 0 | - |
| - | - | 826.7 | 359 | - | - | 0 | - |
| - | - | 658.4 | 359.6 | - | - | 0 | - |
| - | - | 581.7 | 361.2 | - | - | 0 | - |
| - | - | 600.9 | 366.2 | - | - | 0 | - |
| 6 | c | 777 | 369.2 | 0.005417 | 14.67 | +2 | 6 |
| - | - | 679.1 | 369.2 | - | - | 0 | - |
| - | - | 614.3 | 370.2 | - | - | 0 | - |
| - | - | 2652 | 370.3 | - | - | 0 | - |
| - | - | 642.2 | 371.3 | - | - | 0 | - |
| - | - | 1.501E+04 | 374.2 | - | - | 0 | - |
| - | - | 4961 | 375.2 | - | - | 0 | - |
| - | - | 821.6 | 376.2 | - | - | 0 | - |
| - | - | 660 | 376.5 | - | - | 0 | - |
| 6 | c | 1782 | 378.2 | 0.005109 | 13.51 | +2 | 6 |
| - | - | 1961 | 379.8 | - | - | 0 | - |
| - | - | 1182 | 380.2 | - | - | 0 | - |
| - | - | 849.3 | 380.5 | - | - | 0 | - |
| 4 | y | 2323 | 380.7 | 7.353E-05 | 0.1931 | +2 | 6 |
| 4 | z | 915.8 | 381.2 | 0.00185 | 4.853 | +2 | 6 |
| - | - | 768.9 | 381.7 | - | - | 0 | - |
| 4 | y | 9184 | 389.2 | 0.0003476 | 0.893 | +2 | 6 |
| - | - | 3859 | 389.7 | - | - | 0 | - |
| - | - | 2601 | 390.7 | - | - | 0 | - |
| - | - | 3846 | 391.2 | - | - | 0 | - |
| - | - | 7626 | 391.5 | - | - | 0 | - |
| - | - | 4563 | 391.9 | - | - | 0 | - |
| - | - | 1032 | 392.2 | - | - | 0 | - |
| - | - | 1051 | 392.5 | - | - | 0 | - |
| - | - | 3184 | 396.1 | - | - | 0 | - |
| - | - | 609.8 | 397.1 | - | - | 0 | - |
| - | - | 744.3 | 397.2 | - | - | 0 | - |
| - | - | 573.8 | 399.5 | - | - | 0 | - |
| - | - | 824.6 | 404.2 | - | - | 0 | - |
| - | - | 710.8 | 404.7 | - | - | 0 | - |
| - | - | 749.4 | 405.7 | - | - | 0 | - |
| - | - | 606.2 | 410.7 | - | - | 0 | - |
| - | - | 1.378E+04 | 412.2 | - | - | 0 | - |
| 3 | c | 3394 | 413.2 | 0.004557 | 11.03 | +1 | 3 |
| - | - | 2810 | 413.2 | - | - | 0 | - |
| 3 | z | 1910 | 416.7 | 0.0001621 | 0.3891 | +2 | 7 |
| - | - | 2186 | 417.2 | - | - | 0 | - |
| 7 | z | 5.085E+04 | 419.2 | 0.0001645 | 0.3924 | +1 | 3 |
| - | - | 1.096E+04 | 420.2 | - | - | 0 | - |
| - | - | 1537 | 421.2 | - | - | 0 | - |
| - | - | 869 | 423.2 | - | - | 0 | - |
| - | - | 1074 | 423.7 | - | - | 0 | - |
| 3 | y | 8879 | 424.7 | 0.0002277 | 0.5362 | +2 | 7 |
| - | - | 4715 | 425.2 | - | - | 0 | - |
| - | - | 1739 | 425.3 | - | - | 0 | - |
| - | - | 1747 | 425.7 | - | - | 0 | - |
| - | - | 2911 | 426.2 | - | - | 0 | - |
| - | - | 1467 | 426.3 | - | - | 0 | - |
| - | - | 1419 | 426.7 | - | - | 0 | - |
| - | - | 5560 | 433.7 | - | - | 0 | - |
| - | - | 3810 | 434.2 | - | - | 0 | - |
| - | - | 855 | 434.7 | - | - | 0 | - |
| 7 | y | 1.061E+04 | 435.2 | 0.0002423 | 0.5567 | +1 | 3 |
| - | - | 2779 | 436.2 | - | - | 0 | - |
| - | - | 1410 | 450.7 | - | - | 0 | - |
| - | - | 782.9 | 451.2 | - | - | 0 | - |
| - | - | 1.66E+04 | 451.7 | - | - | 0 | - |
| 2 | w | 8040 | 452.2 | 0.006692 | 14.8 | +2 | 8 |
| - | - | 3502 | 452.7 | - | - | 0 | - |
| - | - | 1544 | 453.2 | - | - | 0 | - |
| - | - | 1074 | 458.2 | - | - | 0 | - |
| - | - | 758 | 459.2 | - | - | 0 | - |
| - | - | 1484 | 462.2 | - | - | 0 | - |
| - | - | 1920 | 462.7 | - | - | 0 | - |
| - | - | 1044 | 463.2 | - | - | 0 | - |
| - | - | 2031 | 469.2 | - | - | 0 | - |
| - | - | 1620 | 469.3 | - | - | 0 | - |
| - | - | 3268 | 469.7 | - | - | 0 | - |
| - | - | 806 | 470.2 | - | - | 0 | - |
| - | - | 2901 | 470.3 | - | - | 0 | - |
| - | - | 597.8 | 474.7 | - | - | 0 | - |
| - | - | 888.7 | 475.2 | - | - | 0 | - |
| - | - | 849.9 | 475.7 | - | - | 0 | - |
| - | - | 4295 | 476.2 | - | - | 0 | - |
| - | - | 3496 | 476.2 | - | - | 0 | - |
| - | - | 633.3 | 476.3 | - | - | 0 | - |
| - | - | 954.5 | 476.7 | - | - | 0 | - |
| - | - | 949.1 | 477.2 | - | - | 0 | - |
| - | - | 870 | 479.2 | - | - | 0 | - |
| - | - | 576.1 | 481.8 | - | - | 0 | - |
| - | - | 1004 | 482.2 | - | - | 0 | - |
| - | - | 942 | 482.7 | - | - | 0 | - |
| - | - | 1485 | 483.2 | - | - | 0 | - |
| - | - | 8318 | 483.3 | - | - | 0 | - |
| - | - | 869.2 | 483.7 | - | - | 0 | - |
| - | - | 1881 | 484.3 | - | - | 0 | - |
| - | - | 1149 | 485.3 | - | - | 0 | - |
| - | - | 1288 | 487.2 | - | - | 0 | - |
| - | - | 1164 | 487.7 | - | - | 0 | - |
| - | - | 4902 | 489.2 | - | - | 0 | - |
| - | - | 3607 | 489.7 | - | - | 0 | - |
| - | - | 1038 | 490.2 | - | - | 0 | - |
| - | - | 1044 | 490.7 | - | - | 0 | - |
| - | - | 642.7 | 493.2 | - | - | 0 | - |
| - | - | 774.4 | 495.2 | - | - | 0 | - |
| - | - | 7337 | 495.7 | - | - | 0 | - |
| 8 | c | 8964 | 496.2 | 0.004019 | 8.099 | +2 | 8 |
| 2 | y | 5983 | 496.7 | 0.005438 | 10.95 | +2 | 8 |
| 2 | z | 3681 | 497.2 | 0.001282 | 2.577 | +2 | 8 |
| - | - | 655 | 497.7 | - | - | 0 | - |
| 6 | w | 2502 | 503.3 | 0.001524 | 3.029 | +1 | 4 |
| - | - | 3188 | 503.7 | - | - | 0 | - |
| - | - | 2.434E+04 | 504.2 | - | - | 0 | - |
| 8 | c | 1.066E+05 | 504.8 | 0.002828 | 5.603 | +2 | 8 |
| 2 | y | 8.317E+04 | 505.2 | 0.003211 | 6.355 | +2 | 8 |
| - | - | 3.848E+04 | 505.7 | - | - | 0 | - |
| - | - | 1855 | 506.2 | - | - | 0 | - |
| - | - | 1.374E+04 | 506.3 | - | - | 0 | - |
| - | - | 2654 | 506.8 | - | - | 0 | - |
| - | - | 558.7 | 507.3 | - | - | 0 | - |
| - | - | 1286 | 510.3 | - | - | 0 | - |
| - | - | 841 | 511.8 | - | - | 0 | - |
| - | - | 9040 | 512.3 | - | - | 0 | - |
| - | - | 4760 | 512.8 | - | - | 0 | - |
| - | - | 1283 | 513.3 | - | - | 0 | - |
| - | - | 692.3 | 513.8 | - | - | 0 | - |
| - | - | 836.8 | 517.2 | - | - | 0 | - |
| 6 | y | 1369 | 517.3 | 0.001045 | 2.02 | +1 | 4 |
| 6 | z | 1.212E+04 | 518.3 | 0.0006158 | 1.188 | +1 | 4 |
| - | - | 4596 | 519.3 | - | - | 0 | - |
| - | - | 1886 | 519.8 | - | - | 0 | - |
| - | - | 1537 | 520.3 | - | - | 0 | - |
| 4 | c | 1506 | 524.2 | 0.004276 | 8.157 | +1 | 4 |
| - | - | 1087 | 524.3 | - | - | 0 | - |
| - | - | 1008 | 525.2 | - | - | 0 | - |
| - | - | 1.278E+04 | 525.7 | - | - | 0 | - |
| - | - | 7078 | 526.2 | - | - | 0 | - |
| - | - | 3304 | 526.7 | - | - | 0 | - |
| - | - | 729.9 | 527.3 | - | - | 0 | - |
| - | - | 1563 | 532.3 | - | - | 0 | - |
| - | - | 660.2 | 532.3 | - | - | 0 | - |
| - | - | 714.6 | 532.8 | - | - | 0 | - |
| - | - | 5.724E+04 | 533.3 | - | - | 0 | - |
| - | - | 3.785E+04 | 533.8 | - | - | 0 | - |
| - | - | 1.592E+04 | 534.3 | - | - | 0 | - |
| 6 | y | 1.829E+04 | 534.3 | 8.322E-06 | 0.01558 | +1 | 4 |
| - | - | 7211 | 534.8 | - | - | 0 | - |
| - | - | 3671 | 535.3 | - | - | 0 | - |
| - | - | 5790 | 535.3 | - | - | 0 | - |
| - | - | 1708 | 535.8 | - | - | 0 | - |
| - | - | 1344 | 536.3 | - | - | 0 | - |
| - | - | 2588 | 540.3 | - | - | 0 | - |
| - | - | 2002 | 540.8 | - | - | 0 | - |
| 4 | c | 1.692E+05 | 541.2 | 0.004885 | 9.026 | +1 | 4 |
| - | - | 2356 | 541.8 | - | - | 0 | - |
| - | - | 4.669E+04 | 542.2 | - | - | 0 | - |
| - | - | 1.309E+04 | 543.2 | - | - | 0 | - |
| - | - | 724.3 | 543.3 | - | - | 0 | - |
| - | - | 2663 | 544.2 | - | - | 0 | - |
| - | - | 1122 | 544.3 | - | - | 0 | - |
| - | - | 819.5 | 547.3 | - | - | 0 | - |
| - | - | 5884 | 548.8 | - | - | 0 | - |
| - | - | 6.28E+04 | 549.3 | - | - | 0 | - |
| - | - | 3.222E+04 | 549.8 | - | - | 0 | - |
| - | - | 1.42E+04 | 550.3 | - | - | 0 | - |
| - | - | 5360 | 550.8 | - | - | 0 | - |
| - | - | 1431 | 554.2 | - | - | 0 | - |
| - | - | 1797 | 556.3 | - | - | 0 | - |
| - | - | 6890 | 556.8 | - | - | 0 | - |
| - | - | 4.79E+04 | 557.3 | - | - | 0 | - |
| - | - | 9.02E+04 | 557.8 | - | - | 0 | - |
| - | - | 4.663E+04 | 558.3 | - | - | 0 | - |
| - | - | 1.884E+04 | 558.8 | - | - | 0 | - |
| - | - | 5168 | 559.3 | - | - | 0 | - |
| - | - | 1055 | 559.8 | - | - | 0 | - |
| - | - | 921.7 | 563.8 | - | - | 0 | - |
| - | - | 602.1 | 564.3 | - | - | 0 | - |
| - | - | 8246 | 564.8 | - | - | 0 | - |
| - | - | 2.719E+04 | 565.3 | - | - | 0 | - |
| - | - | 2.098E+04 | 565.8 | - | - | 0 | - |
| - | - | 1.368E+04 | 566.3 | - | - | 0 | - |
| - | - | 4892 | 566.8 | - | - | 0 | - |
| - | - | 2261 | 567.3 | - | - | 0 | - |
| - | - | 947.2 | 567.8 | - | - | 0 | - |
| - | - | 2.612E+04 | 570.3 | - | - | 0 | - |
| - | - | 1.569E+04 | 570.8 | - | - | 0 | - |
| - | - | 5591 | 571.3 | - | - | 0 | - |
| - | - | 2044 | 571.8 | - | - | 0 | - |
| - | - | 3651 | 574.3 | - | - | 0 | - |
| - | - | 1804 | 575.3 | - | - | 0 | - |
| - | - | 2168 | 578.3 | - | - | 0 | - |
| - | - | 3.867E+04 | 578.8 | - | - | 0 | - |
| - | - | 2.332E+04 | 579.3 | - | - | 0 | - |
| - | - | 9536 | 579.8 | - | - | 0 | - |
| - | - | 3167 | 580.3 | - | - | 0 | - |
| - | - | 1027 | 580.8 | - | - | 0 | - |
| - | - | 1310 | 582.3 | - | - | 0 | - |
| - | - | 1229 | 586.3 | - | - | 0 | - |
| - | - | 1.56E+05 | 586.8 | - | - | 0 | - |
| - | - | 1.191E+05 | 587.3 | - | - | 0 | - |
| - | - | 5.487E+04 | 587.8 | - | - | 0 | - |
| - | - | 1.926E+04 | 588.3 | - | - | 0 | - |
| - | - | 5600 | 588.8 | - | - | 0 | - |
| - | - | 1.132E+05 | 589.3 | - | - | 0 | - |
| - | - | 3.559E+04 | 590.3 | - | - | 0 | - |
| - | - | 7010 | 591.3 | - | - | 0 | - |
| - | - | 2349 | 592.3 | - | - | 0 | - |
| - | - | 1324 | 592.3 | - | - | 0 | - |
| - | - | 786.5 | 595.3 | - | - | 0 | - |
| - | - | 1594 | 597.4 | - | - | 0 | - |
| - | - | 2545 | 598.4 | - | - | 0 | - |
| - | - | 4331 | 611.2 | - | - | 0 | - |
| - | - | 5282 | 612.3 | - | - | 0 | - |
| - | - | 1283 | 613.3 | - | - | 0 | - |
| - | - | 1691 | 617.3 | - | - | 0 | - |
| 5 | y | 991.8 | 632.3 | 0.0008351 | 1.321 | +1 | 5 |
| 5 | z | 6598 | 633.3 | 0.001253 | 1.978 | +1 | 5 |
| - | - | 5874 | 634.3 | - | - | 0 | - |
| - | - | 2147 | 635.3 | - | - | 0 | - |
| 5 | c | 3501 | 639.2 | 0.004669 | 7.303 | +1 | 5 |
| - | - | 967.7 | 640.2 | - | - | 0 | - |
| - | - | 851.2 | 642.3 | - | - | 0 | - |
| 5 | y | 2.464E+04 | 649.3 | 0.0005675 | 0.874 | +1 | 5 |
| - | - | 8455 | 650.3 | - | - | 0 | - |
| - | - | 2788 | 651.3 | - | - | 0 | - |
| - | - | 6249 | 652.3 | - | - | 0 | - |
| - | - | 2568 | 653.3 | - | - | 0 | - |
| - | - | 787.1 | 654.3 | - | - | 0 | - |
| - | - | 1345 | 655.4 | - | - | 0 | - |
| 5 | c | 2.565E+04 | 656.3 | 0.005156 | 7.856 | +1 | 5 |
| - | - | 7979 | 657.3 | - | - | 0 | - |
| - | - | 2681 | 658.3 | - | - | 0 | - |
| - | - | 905.7 | 667.3 | - | - | 0 | - |
| - | - | 3763 | 668.4 | - | - | 0 | - |
| - | - | 1353 | 669.4 | - | - | 0 | - |
| - | - | 780.2 | 682.3 | - | - | 0 | - |
| - | - | 1992 | 684.4 | - | - | 0 | - |
| - | - | 979 | 685.4 | - | - | 0 | - |
| - | - | 4045 | 694.4 | - | - | 0 | - |
| - | - | 1436 | 695.4 | - | - | 0 | - |
| - | - | 2553 | 710.3 | - | - | 0 | - |
| - | - | 4090 | 711.3 | - | - | 0 | - |
| - | - | 667.6 | 712.3 | - | - | 0 | - |
| - | - | 1387 | 717.4 | - | - | 0 | - |
| - | - | 832.4 | 719.4 | - | - | 0 | - |
| - | - | 688.2 | 720.4 | - | - | 0 | - |
| - | - | 1209 | 731.4 | - | - | 0 | - |
| - | - | 622.5 | 732.4 | - | - | 0 | - |
| - | - | 6580 | 738.4 | - | - | 0 | - |
| - | - | 5817 | 738.4 | - | - | 0 | - |
| - | - | 3492 | 739.4 | - | - | 0 | - |
| - | - | 2058 | 739.4 | - | - | 0 | - |
| - | - | 1123 | 740.4 | - | - | 0 | - |
| - | - | 808 | 740.4 | - | - | 0 | - |
| - | - | 2526 | 752.4 | - | - | 0 | - |
| - | - | 1477 | 753.4 | - | - | 0 | - |
| 6 | c | 1.445E+05 | 755.3 | 0.00576 | 7.625 | +1 | 6 |
| - | - | 5.69E+04 | 756.3 | - | - | 0 | - |
| - | - | 1.914E+04 | 757.3 | - | - | 0 | - |
| - | - | 4063 | 758.3 | - | - | 0 | - |
| - | - | 1065 | 759.3 | - | - | 0 | - |
| 4 | y | 770.6 | 760.4 | 0.007435 | 9.777 | +1 | 6 |
| 4 | z | 2.662E+04 | 761.4 | 0.001123 | 1.475 | +1 | 6 |
| - | - | 1.314E+04 | 762.4 | - | - | 0 | - |
| - | - | 3565 | 763.4 | - | - | 0 | - |
| 4 | y | 6588 | 777.4 | 0.0007429 | 0.9556 | +1 | 6 |
| - | - | 2014 | 778.4 | - | - | 0 | - |
| - | - | 1172 | 780.3 | - | - | 0 | - |
| - | - | 2803 | 780.4 | - | - | 0 | - |
| - | - | 3459 | 781.3 | - | - | 0 | - |
| - | - | 5288 | 781.4 | - | - | 0 | - |
| - | - | 3652 | 782.3 | - | - | 0 | - |
| - | - | 2442 | 782.4 | - | - | 0 | - |
| - | - | 2788 | 783.3 | - | - | 0 | - |
| - | - | 787.4 | 783.4 | - | - | 0 | - |
| - | - | 958.1 | 784.3 | - | - | 0 | - |
| - | - | 733.3 | 785.3 | - | - | 0 | - |
| - | - | 2167 | 790.4 | - | - | 0 | - |
| - | - | 735.2 | 816.4 | - | - | 0 | - |
| 3 | z | 2.118E+04 | 832.4 | 0.00131 | 1.574 | +1 | 7 |
| - | - | 1.075E+04 | 833.4 | - | - | 0 | - |
| - | - | 2958 | 834.4 | - | - | 0 | - |
| - | - | 769.7 | 845.4 | - | - | 0 | - |
| - | - | 932.5 | 846.4 | - | - | 0 | - |
| - | - | 1369 | 847.5 | - | - | 0 | - |
| 3 | y | 5634 | 848.5 | 0.003616 | 4.262 | +1 | 7 |
| - | - | 3825 | 849.5 | - | - | 0 | - |
| - | - | 1916 | 852.4 | - | - | 0 | - |
| - | - | 927.4 | 853.4 | - | - | 0 | - |
| - | - | 5178 | 867.4 | - | - | 0 | - |
| - | - | 2437 | 868.4 | - | - | 0 | - |
| - | - | 946.2 | 869.4 | - | - | 0 | - |
| - | - | 672.6 | 879.4 | - | - | 0 | - |
| - | - | 1509 | 886.5 | - | - | 0 | - |
| - | - | 873 | 887.5 | - | - | 0 | - |
| - | - | 824.2 | 901.5 | - | - | 0 | - |
| - | - | 1.047E+04 | 902.5 | - | - | 0 | - |
| 2 | w | 1.122E+04 | 903.5 | 0.01511 | 16.72 | +1 | 8 |
| - | - | 5048 | 904.5 | - | - | 0 | - |
| - | - | 1359 | 905.5 | - | - | 0 | - |
| - | - | 750.2 | 918.5 | - | - | 0 | - |
| - | - | 2350 | 933.5 | - | - | 0 | - |
| - | - | 1563 | 934.4 | - | - | 0 | - |
| - | - | 1484 | 935.5 | - | - | 0 | - |
| - | - | 790 | 948.5 | - | - | 0 | - |
| - | - | 3416 | 949.5 | - | - | 0 | - |
| - | - | 6536 | 950.5 | - | - | 0 | - |
| - | - | 3368 | 951.5 | - | - | 0 | - |
| - | - | 1603 | 952.5 | - | - | 0 | - |
| - | - | 1044 | 964.5 | - | - | 0 | - |
| - | - | 1688 | 965.5 | - | - | 0 | - |
| - | - | 3058 | 966.5 | - | - | 0 | - |
| - | - | 1378 | 967.5 | - | - | 0 | - |
| - | - | 761.9 | 981.5 | - | - | 0 | - |
| 2 | y | 1.148E+04 | 992.5 | 0.01693 | 17.06 | +1 | 8 |
| 2 | z | 7631 | 993.5 | 0.01198 | 12.05 | +1 | 8 |
| - | - | 3267 | 994.5 | - | - | 0 | - |
| - | - | 1082 | 995.5 | - | - | 0 | - |
| 8 | c | 1.484E+04 | 1008 | 0.008267 | 8.198 | +1 | 8 |
| 2 | y | 2.76E+04 | 1009 | 0.01394 | 13.81 | +1 | 8 |
| - | - | 1.443E+04 | 1011 | - | - | 0 | - |
| - | - | 5301 | 1011 | - | - | 0 | - |
| - | - | 1456 | 1013 | - | - | 0 | - |
| - | - | 1596 | 1014 | - | - | 0 | - |
| - | - | 1193 | 1015 | - | - | 0 | - |
| - | - | 880.1 | 1024 | - | - | 0 | - |
| - | - | 745 | 1027 | - | - | 0 | - |
| - | - | 1150 | 1038 | - | - | 0 | - |
| - | - | 2154 | 1056 | - | - | 0 | - |
| - | - | 1545 | 1056 | - | - | 0 | - |
| - | - | 1122 | 1066 | - | - | 0 | - |
| - | - | 2974 | 1067 | - | - | 0 | - |
| - | - | 1054 | 1068 | - | - | 0 | - |
| - | - | 1426 | 1069 | - | - | 0 | - |
| - | - | 1786 | 1070 | - | - | 0 | - |
| - | - | 1884 | 1071 | - | - | 0 | - |
| - | - | 3127 | 1072 | - | - | 0 | - |
| - | - | 5358 | 1072 | - | - | 0 | - |
| - | - | 3032 | 1073 | - | - | 0 | - |
| - | - | 2093 | 1074 | - | - | 0 | - |
| - | - | 1427 | 1081 | - | - | 0 | - |
| - | - | 991.6 | 1082 | - | - | 0 | - |
| - | - | 1401 | 1083 | - | - | 0 | - |
| - | - | 1304 | 1084 | - | - | 0 | - |
| - | - | 924 | 1087 | - | - | 0 | - |
| - | - | 716.4 | 1087 | - | - | 0 | - |
| - | - | 3589 | 1097 | - | - | 0 | - |
| - | - | 1.167E+04 | 1098 | - | - | 0 | - |
| - | - | 1.6E+04 | 1099 | - | - | 0 | - |
| - | - | 1.037E+04 | 1100 | - | - | 0 | - |
| - | - | 4064 | 1101 | - | - | 0 | - |
| - | - | 688.8 | 1102 | - | - | 0 | - |
| - | - | 1094 | 1112 | - | - | 0 | - |
| - | - | 1169 | 1113 | - | - | 0 | - |
| - | - | 9542 | 1114 | - | - | 0 | - |
| - | - | 1.025E+05 | 1115 | - | - | 0 | - |
| - | - | 7.579E+04 | 1116 | - | - | 0 | - |
| - | - | 3.475E+04 | 1117 | - | - | 0 | - |
| - | - | 591.4 | 1117 | - | - | 0 | - |
| - | - | 1.304E+04 | 1118 | - | - | 0 | - |
| - | - | 2537 | 1119 | - | - | 0 | - |
| - | - | 1389 | 1128 | - | - | 0 | - |
| - | - | 2595 | 1129 | - | - | 0 | - |
| - | - | 3615 | 1130 | - | - | 0 | - |
| - | - | 4408 | 1131 | - | - | 0 | - |
| - | - | 6520 | 1132 | - | - | 0 | - |
| - | - | 2705 | 1133 | - | - | 0 | - |
| - | - | 1320 | 1134 | - | - | 0 | - |
| - | - | 3555 | 1140 | - | - | 0 | - |
| - | - | 9503 | 1141 | - | - | 0 | - |
| - | - | 6069 | 1142 | - | - | 0 | - |
| - | - | 1985 | 1143 | - | - | 0 | - |
| - | - | 643.2 | 1146 | - | - | 0 | - |
| - | - | 1102 | 1147 | - | - | 0 | - |
| - | - | 791.8 | 1148 | - | - | 0 | - |
| - | - | 1.864E+04 | 1157 | - | - | 0 | - |
| - | - | 1.303E+05 | 1158 | - | - | 0 | - |
| - | - | 7.867E+04 | 1159 | - | - | 0 | - |
| - | - | 3.325E+04 | 1160 | - | - | 0 | - |
| - | - | 1.13E+04 | 1161 | - | - | 0 | - |
| - | - | 2357 | 1162 | - | - | 0 | - |
| - | - | 6608 | 1173 | - | - | 0 | - |
| - | - | 3.174E+04 | 1174 | - | - | 0 | - |
| - | - | 7.151E+04 | 1175 | - | - | 0 | - |
| - | - | 4.077E+04 | 1176 | - | - | 0 | - |
| - | - | 1.522E+04 | 1177 | - | - | 0 | - |
| - | - | 5862 | 1178 | - | - | 0 | - |
| - | - | 1043 | 1179 | - | - | 0 | - |

m/z Charge Intensity FragmentType MassShift Position
121.02826690673828 0 1333.6461
129.10232543945312 0 10077.642
134.02700805664062 0 1068.5238
136.07577514648438 0 47138.61
137.05953979492188 0 423.96448
137.0791015625 0 3440.7422
142.08677673339844 0 438.7536
142.09776306152344 0 591.16296
146.1288299560547 0 1022.2765
147.04425048828125 0 3041.7537
149.0234375 0 177131.47
149.0708770751953 0 945.0802
150.02671813964844 0 14011.564
152.6886444091797 0 423.33554
156.07681274414062 0 2447.3496
160.1080780029297 0 1125.9375
167.03392028808594 0 60855.406
168.03726196289062 0 3608.53
173.45164489746094 0 1205.948
179.0486602783203 0 953.75635
182.0811767578125 0 1569.5215 y 8
183.0847625732422 0 474.73337
183.43612670898438 0 397.08347
184.13186645507812 0 1220.4259
187.10791015625 0 555.34625
188.03024291992188 0 469.9617
200.1392822265625 0 3819.834
205.06410217285156 0 3065.4128
211.15521240234375 0 5100.2397
212.153564453125 0 515.9539
212.16294860839844 0 6028.5776
213.12338256835938 0 1484.2946
213.17153930664062 0 604.5518
221.09202575683594 0 919.5094
223.06405639648438 0 650.73334
233.05921936035156 0 1146.5669
233.0918426513672 0 1202.4584
239.14942932128906 0 633.68646
241.11817932128906 0 1385.5247
250.0857696533203 0 1845.878
251.048095703125 0 921.3288
254.16107177734375 0 997.0651
255.14523315429688 0 945.0942
255.16851806640625 0 1432.1349
256.1770324707031 0 2716.0474
266.07171630859375 0 2189.2866
279.1336975097656 0 6939.4775 y 7
279.1590270996094 0 7451.861
280.1373291015625 0 1020.02313
280.16217041015625 0 1494.5293
284.20819091796875 0 1674.3752
285.2154541015625 0 941.0388
297.09027099609375 0 17996.098
298.0933532714844 0 2157.4666
301.05828857421875 0 883.362
304.1290283203125 0 5011.908
309.1563720703125 0 639.3325
313.1742858886719 0 8956.18
314.1776123046875 0 1588.1857
315.1636047363281 0 809.2118
320.1256103515625 0 1936.6132 c Ammonia loss 4
320.1817321777344 0 628.7084
325.08526611328125 0 13456.695
326.0878601074219 0 2343.9106
354.2351379394531 0 855.8851
354.88677978515625 0 926.9048
355.0697021484375 0 2802.2568
355.24542236328125 0 762.9185
356.0695495605469 0 1166.9141
357.0677795410156 0 2715.0876
358.0682678222656 0 946.0526
359.02545166015625 0 826.6513
359.5740051269531 0 658.3567
361.1537780761719 0 581.7044
366.1778869628906 0 600.94684
369.1756591796875 0 777.0149 c Water loss 5
369.21435546875 0 679.05963
370.21533203125 0 614.27576
370.2559509277344 0 2651.8164
371.2598876953125 0 642.2161
374.21844482421875 0 15013.738
375.2236328125 0 4961.1875
376.2290344238281 0 821.5632
376.5126953125 0 659.98364
378.1806335449219 0 1782.049 c 5
379.83843994140625 0 1961.2078
380.17572021484375 0 1181.8798
380.5063171386719 0 849.27814
380.7029724121094 0 2322.5823 y Ammonia loss 3
381.2051086425781 0 915.7685 z 3
381.6918640136719 0 768.8815
389.2159729003906 0 9184.44 y 3
389.71734619140625 0 3858.5027
390.697265625 0 2601.3582
391.1889953613281 0 3845.6704
391.5180358886719 0 7626.371
391.8526306152344 0 4563.112
392.1875915527344 0 1031.7075
392.5202941894531 0 1050.6299
396.12213134765625 0 3183.9692
397.1239929199219 0 609.8276
397.2082824707031 0 744.3262
399.5199279785156 0 573.8223
404.20404052734375 0 824.55963
404.70849609375 0 710.80554
405.6992492675781 0 749.43274
410.7127380371094 0 606.2109
412.2425842285156 0 13784.949
413.1488952636719 0 3393.748 c 2
413.24462890625 0 2810.4387
416.7256774902344 0 1910.1561 z 2
417.2269592285156 0 2186.285
419.2161560058594 0 50845.164 z 6
420.21917724609375 0 10960.038
421.22381591796875 0 1537.3372
423.2174987792969 0 868.99695
423.71575927734375 0 1074.0544
424.7346496582031 0 8879.366 y 2
425.23626708984375 0 4715.3447
425.2747497558594 0 1739.1239
425.7372741699219 0 1747.0475
426.2073059082031 0 2911.3677
426.2801818847656 0 1467.3077
426.7090148925781 0 1419.1671
433.712890625 0 5559.9
434.21795654296875 0 3809.7397
434.7145080566406 0 854.9717
435.23480224609375 0 10610.955 y 6
436.2391357421875 0 2779.3098
450.73272705078125 0 1409.9185
451.2307434082031 0 782.8926
451.7397766113281 0 16601.074
452.2411193847656 0 8039.7266 w 1
452.7431335449219 0 3501.9646
453.23883056640625 0 1543.5428
458.1720275878906 0 1073.7106
459.2491760253906 0 757.9643
462.22149658203125 0 1484.3558
462.7275085449219 0 1920.4867
463.2276306152344 0 1044.0756
469.2308654785156 0 2031.1028
469.2647705078125 0 1620.3318
469.735107421875 0 3267.6997
470.2384338378906 0 805.96765
470.27166748046875 0 2901.4204
474.73358154296875 0 597.81256
475.2343444824219 0 888.7233
475.74407958984375 0 849.89325
476.18096923828125 0 4295.0806
476.2388916015625 0 3496.1426
476.3088073730469 0 633.3201
476.7429504394531 0 954.4812
477.1873779296875 0 949.0716
479.2430114746094 0 870.0299
481.75787353515625 0 576.1292
482.24200439453125 0 1003.8469
482.7442626953125 0 942.0138
483.2454833984375 0 1485.0127
483.2803039550781 0 8317.685
483.7485656738281 0 869.1573
484.28228759765625 0 1880.9009
485.2838134765625 0 1149.2068
487.2300109863281 0 1287.5603
487.7251281738281 0 1163.6613
489.2295227050781 0 4901.726
489.7307434082031 0 3607.0093
490.2336120605469 0 1037.9539
490.73187255859375 0 1043.629
493.20574951171875 0 642.696
495.2305603027344 0 774.3785
495.7324523925781 0 7337.0894
496.23516845703125 0 8963.717 c Ammonia loss 7
496.73663330078125 0 5983.2866 y Ammonia loss 1
497.23638916015625 0 3681.0645 z 1
497.7427062988281 0 654.97845
503.2597351074219 0 2502.4739 w 5
503.7455749511719 0 3187.8657
504.2461242675781 0 24339.879
504.7496337890625 0 106589.08 c 7
505.2476806640625 0 83167.61 y 1
505.7486572265625 0 38478.56
506.2125549316406 0 1855.3923
506.2503662109375 0 13737.291
506.75079345703125 0 2654.2737
507.25732421875 0 558.659
510.2684631347656 0 1285.7126
511.75726318359375 0 841.0226
512.261474609375 0 9040.161
512.76220703125 0 4760.1143
513.2579956054688 0 1283.0198
513.7642211914062 0 692.3111
517.2352294921875 0 836.8492
517.2779541015625 0 1369.46 y Ammonia loss 5
518.2841186523438 0 12115.423 z 5
519.2871704101562 0 4595.667
519.7910766601562 0 1886.4568
520.2930908203125 0 1537.2543
524.2175903320312 0 1506.1895 c Ammonia loss 3
524.2591552734375 0 1086.7256
525.2461547851562 0 1007.67
525.7433471679688 0 12780.687
526.2457275390625 0 7078.238
526.746826171875 0 3303.8496
527.2578735351562 0 729.91016
532.2598266601562 0 1563.0684
532.2919311523438 0 660.1984
532.7723388671875 0 714.617
533.270263671875 0 57243.703
533.7722778320312 0 37846.297
534.2622680664062 0 15920.584
534.303466796875 0 18285.432 y 5
534.7615966796875 0 7211.432
535.2637329101562 0 3671.1758
535.306396484375 0 5790.05
535.7667236328125 0 1707.7665
536.310302734375 0 1344.1368
540.2769165039062 0 2587.6833
540.7770385742188 0 2001.6974
541.2435302734375 0 169193.72 c 3
541.7828369140625 0 2355.9902
542.2462158203125 0 46688.18
543.244384765625 0 13094.885
543.2919921875 0 724.316
544.2451171875 0 2662.909
544.3237915039062 0 1122.2577
547.2748413085938 0 819.5486
548.7551879882812 0 5884.398
549.2574462890625 0 62802.895
549.7586669921875 0 32221.785
550.2592163085938 0 14197.299
550.7590942382812 0 5360.317
554.1912841796875 0 1431.2548
556.2682495117188 0 1796.9546
556.7676391601562 0 6890.269
557.2669677734375 0 47897.28
557.7698364257812 0 90196.234
558.271240234375 0 46628.51
558.771484375 0 18844.805
559.2725830078125 0 5168.359
559.7762451171875 0 1055.276
563.772216796875 0 921.69183
564.2710571289062 0 602.1457
564.77783203125 0 8246.096
565.281494140625 0 27191.482
565.7659912109375 0 20978.07
566.265869140625 0 13682.546
566.7654418945312 0 4891.611
567.264892578125 0 2261.0469
567.7646484375 0 947.2363
570.2623291015625 0 26115.885
570.7640380859375 0 15688.791
571.2647705078125 0 5591.1597
571.7649536132812 0 2044.2963
574.2987060546875 0 3651.2437
575.3029174804688 0 1803.8518
578.2651977539062 0 2168.016
578.7639770507812 0 38674.56
579.2655639648438 0 23322.389
579.7664184570312 0 9535.602
580.2675170898438 0 3166.6196
580.7688598632812 0 1027.218
582.2792358398438 0 1309.8995
586.2695922851562 0 1228.6274
586.7733764648438 0 155964.78
587.2753295898438 0 119077.88
587.7764282226562 0 54869.67
588.2772216796875 0 19260.754
588.778564453125 0 5599.6064
589.3212280273438 0 113151.73
590.323974609375 0 35586.73
591.326904296875 0 7010.2173
592.2752075195312 0 2348.6511
592.329833984375 0 1323.9897
595.2948608398438 0 786.46985
597.3590087890625 0 1594.4459
598.365478515625 0 2544.7952
611.2490844726562 0 4331.345
612.2552490234375 0 5281.9517
613.2583618164062 0 1282.5106
617.2908325195312 0 1691.1874
632.3046875 0 991.7513 y Ammonia loss 4
633.3104248046875 0 6598.348 z 4
634.31689453125 0 5873.725
635.320068359375 0 2147.1697
639.244140625 0 3500.7063 c Ammonia loss 4
640.2435913085938 0 967.69006
642.3436889648438 0 851.1723
649.329833984375 0 24640.41 y 4
650.3324584960938 0 8455.026
651.3353881835938 0 2788.3716
652.3118896484375 0 6248.6343
653.3153076171875 0 2567.5776
654.3154296875 0 787.0759
655.3648681640625 0 1345.4966
656.2702026367188 0 25646.104 c 4
657.2730712890625 0 7978.566
658.2732543945312 0 2681.2869
667.3367919921875 0 905.6839
668.3953857421875 0 3762.671
669.3983154296875 0 1352.7133
682.2930297851562 0 780.1947
684.4136962890625 0 1992.0134
685.4164428710938 0 979.00464
694.3643188476562 0 4045.4722
695.3645629882812 0 1435.5549
710.317138671875 0 2552.7754
711.3230590820312 0 4089.7378
712.3268432617188 0 667.6234
717.414306640625 0 1386.8915
719.3854370117188 0 832.4278
720.3890991210938 0 688.20844
731.3519287109375 0 1208.8789
732.3554077148438 0 622.517
738.3515625 0 6580.2656
738.42529296875 0 5817.357
739.355224609375 0 3491.9277
739.4290771484375 0 2057.948
740.35595703125 0 1122.9347
740.4309692382812 0 807.96716
752.3836669921875 0 2525.8994
753.3877563476562 0 1476.5853
755.3380126953125 0 144494.78 c 5
756.3408813476562 0 56895.023
757.3405151367188 0 19140.76
758.3421630859375 0 4063.1135
759.3474731445312 0 1065.2574
760.40625 0 770.55536 y Ammonia loss 3
761.405517578125 0 26623.61 z 3
762.4091186523438 0 13142.069
763.4108276367188 0 3565.1306
777.4246215820312 0 6588.019 y 3
778.4290161132812 0 2014.0455
780.2734375 0 1172.139
780.3877563476562 0 2803.1516
781.2742919921875 0 3458.7563
781.3944702148438 0 5288.276
782.2756958007812 0 3651.758
782.3981323242188 0 2442.107
783.2747192382812 0 2788.1748
783.3927001953125 0 787.4366
784.2763671875 0 958.08044
785.2855224609375 0 733.33905
790.4192504882812 0 2166.741
816.4283447265625 0 735.18427
832.4424438476562 0 21182.916 z 2
833.4464111328125 0 10746.815
834.4484252929688 0 2957.5178
845.4269409179688 0 769.66736
846.4391479492188 0 932.4701
847.4513549804688 0 1368.5159
848.4588623046875 0 5633.8066 y 2
849.4630126953125 0 3825.17
852.3919067382812 0 1916.3625
853.3931884765625 0 927.37335
867.4251098632812 0 5177.9043
868.42578125 0 2436.5022
869.429443359375 0 946.2332
879.4078369140625 0 672.57794
886.457275390625 0 1509.089
887.4583129882812 0 873.0202
901.4868774414062 0 824.1743
902.4721069335938 0 10467.796
903.4766845703125 0 11221.551 w 1
904.4801025390625 0 5048.0874
905.4815673828125 0 1358.8075
918.5068969726562 0 750.1722
933.4581298828125 0 2349.763
934.4447021484375 0 1562.7429
935.4531860351562 0 1483.965
948.4761962890625 0 789.96173
949.4739990234375 0 3416.2495
950.480224609375 0 6536.407
951.4788818359375 0 3367.8528
952.4844360351562 0 1602.7842
964.4832763671875 0 1044.3826
965.4946899414062 0 1687.5858
966.4775390625 0 3058.2954
967.47607421875 0 1378.3407
981.5079956054688 0 761.86523
992.4720458984375 0 11478.27 y Ammonia loss 1
993.4749145507812 0 7630.9307 z 1
994.4769287109375 0 3266.9229
995.47265625 0 1081.9331
1008.4893798828125 0 14841.949 c 7
1009.49560546875 0 27599.617 y 1
1010.500244140625 0 14430.287
1011.4998168945312 0 5301.464
1012.5017700195312 0 1456.228
1014.4462280273438 0 1595.8512
1015.4509887695312 0 1192.9453
1023.5206298828125 0 880.05585
1027.4554443359375 0 745.02625
1037.5543212890625 0 1149.768
1055.5068359375 0 2153.5884
1056.4964599609375 0 1544.701
1065.5301513671875 0 1121.6183
1066.534423828125 0 2973.5498
1067.5426025390625 0 1054.3807
1068.5323486328125 0 1425.9618
1069.5406494140625 0 1785.8958
1070.5208740234375 0 1883.9186
1071.5245361328125 0 3126.7004
1072.45751953125 0 5357.923
1073.4571533203125 0 3032.1768
1074.4599609375 0 2093.4446
1080.5118408203125 0 1426.9636
1081.5230712890625 0 991.6024
1082.5587158203125 0 1400.738
1083.556884765625 0 1304.4563
1086.538818359375 0 924.02655
1087.4649658203125 0 716.43884
1096.5206298828125 0 3589.2285
1097.5093994140625 0 11671.563
1098.5108642578125 0 16000.222
1099.5146484375 0 10369.447
1100.514404296875 0 4063.738
1101.521484375 0 688.8292
1111.5328369140625 0 1093.5521
1112.5439453125 0 1168.612
1113.5269775390625 0 9541.94
1114.52197265625 0 102516.88
1115.5277099609375 0 75792.94
1116.53173828125 0 34748.25
1116.76220703125 0 591.3872
1117.533447265625 0 13037.477
1118.5382080078125 0 2537.315
1127.53857421875 0 1388.967
1128.545166015625 0 2594.5464
1129.5465087890625 0 3614.5068
1130.5491943359375 0 4408.4243
1131.5286865234375 0 6520.42
1132.53173828125 0 2705.2214
1133.532470703125 0 1320.2427
1139.5140380859375 0 3555.0442
1140.505126953125 0 9502.971
1141.506103515625 0 6069.266
1142.5108642578125 0 1985.2366
1145.5390625 0 643.15454
1146.5570068359375 0 1102.1487
1147.55859375 0 791.8162
1156.5225830078125 0 18644.744
1157.5263671875 0 130294.016
1158.5296630859375 0 78670.6
1159.5299072265625 0 33248.027
1160.5328369140625 0 11301.234
1161.53271484375 0 2356.6243
1172.5377197265625 0 6608.3857
1173.544677734375 0 31736.521
1174.5518798828125 0 71513.016
1175.5548095703125 0 40769.707
1176.5556640625 0 15220.268
1177.5584716796875 0 5862.17
1178.5596923828125 0 1042.5352

Spectrum Details

|  |  |
| --- | --- |
| Matched peaks? Matched peaks The total absolute number of peaks matched. Additionally in brackets the total fraction of peaks matched and the total number of peaks is shown. | 41 (9.05% of 453) |
| FDR? FDR The false discovery rate estimated for this peptide. It is calculated by matching all theoretical fragments with a non-integer shift with the raw peaks for this spectrum. This is done with 40 different shifts. The resulting percentage is the average number of annotated peaks over the number of annotated peaks with the correct spectrum. | 1.28% |
| Satellite FDR? Satellite FDR See the FDR for details on its calculation. This satellite ion specific FDR only contains the satellite ions (d/w) for I/L/J positions. | - |
| PSM Score? PSM Score The PSM Score as given by Hecklib to this annotated spectrum. It is shown with three significant figures. | 354 |

#### Spectrum 4095? Spectrum 4095 The raw spectrum of this peptide as annotated by Hecklib. The fragments are coloured according to ion type (see legend). Any peaks with a star '\*' as text can be hovered over to see the full details, first the ion type second the mass shift type. By hovering over the amino acids in the peptide or ions in the legend the corresponding peaks are highlighted. By toggling the 'Unassigned' label you can turn the background (unassigned) peaks on or off in the plot. By updating the slider in the Ion legend you can update the spectrum to only show the top X% of the peaks with labels. The top X% means any peak that is within X% of the highest intensity. By dragging in the spectrum you can zoom in to a specific part of the spectrum and use 'Zoom Out' to get back to the original zoom level. The annotation of the spectrum is based on the given sequence in the peptides file and is done with different software so inconsistencies are likely. The peaks are annotated based on the given sequence, with 20 ppm tolerance.

Copy Data

##### Spectrum 4095 (TSV)

###### Preview

```
Loading example...
```

*Click on the button to copy the data to your clipboard.*

Mz MinMz MaxIntensity Max

WidthHeightPeptide font sizePeptide stroke widthSpectrum font sizeSpectrum stroke widthCompact peptide

Ion legend

wxyz

abcd

OtherUnassignedIon^Charge^_Position_Show for top:%

YCAKDVRPY

06.31e+41.26e+51.89e+52.52e+5

Zoom Out

y^+1^_2_c^+2^_6_y^+2^_6_z^+2^_7_z^+1^_3_y^+2^_7_y^+1^_3_w^+2^_8_c^+2^_8_y^+2^_8_c^+2^_8_y^+2^_8_z^+1^_4_c^+1^_4_y^+1^_4_c^+1^_4_z^+1^_5_c^+1^_5_y^+1^_5_c^+1^_5_c^+1^_6_z^+1^_6_y^+1^_6_z^+1^_7_y^+1^_7_w^+1^_8_y^+1^_8_z^+1^_8_c^+1^_8_y^+1^_8_

0699139820972796

Fragment Matches Table

Show background peaks

| Position | Ion type | Intensity | mz Theoretical | mz Error (Th) | mz Error (ppm) | Charge | Series Number |
| --- | --- | --- | --- | --- | --- | --- | --- |
| - | - | 1036 | 120.1 | - | - | 0 | - |
| - | - | 1597 | 121 | - | - | 0 | - |
| - | - | 1070 | 121 | - | - | 0 | - |
| - | - | 394.8 | 127.6 | - | - | 0 | - |
| - | - | 2663 | 129.1 | - | - | 0 | - |
| - | - | 385.9 | 129.4 | - | - | 0 | - |
| - | - | 8184 | 133.1 | - | - | 0 | - |
| - | - | 4073 | 134.1 | - | - | 0 | - |
| - | - | 2286 | 135.1 | - | - | 0 | - |
| - | - | 1.106E+04 | 136.1 | - | - | 0 | - |
| - | - | 1303 | 137.1 | - | - | 0 | - |
| - | - | 403.6 | 142.8 | - | - | 0 | - |
| - | - | 662.7 | 147 | - | - | 0 | - |
| - | - | 822 | 149 | - | - | 0 | - |
| - | - | 2.499E+05 | 149 | - | - | 0 | - |
| - | - | 1.836E+04 | 150 | - | - | 0 | - |
| - | - | 377.9 | 150 | - | - | 0 | - |
| - | - | 510.3 | 156.5 | - | - | 0 | - |
| - | - | 1110 | 166.1 | - | - | 0 | - |
| - | - | 8.621E+04 | 167 | - | - | 0 | - |
| - | - | 6364 | 168 | - | - | 0 | - |
| - | - | 1565 | 173.4 | - | - | 0 | - |
| - | - | 3562 | 177.1 | - | - | 0 | - |
| - | - | 2762 | 178.1 | - | - | 0 | - |
| - | - | 433.5 | 178.6 | - | - | 0 | - |
| - | - | 2823 | 179.1 | - | - | 0 | - |
| - | - | 521.7 | 181.9 | - | - | 0 | - |
| - | - | 821.4 | 200.1 | - | - | 0 | - |
| - | - | 569 | 205.1 | - | - | 0 | - |
| - | - | 923.6 | 211.2 | - | - | 0 | - |
| - | - | 1231 | 212.2 | - | - | 0 | - |
| - | - | 559.4 | 219.1 | - | - | 0 | - |
| - | - | 1600 | 221.1 | - | - | 0 | - |
| - | - | 749.4 | 221.1 | - | - | 0 | - |
| - | - | 1057 | 223.1 | - | - | 0 | - |
| - | - | 1248 | 223.1 | - | - | 0 | - |
| - | - | 1101 | 223.1 | - | - | 0 | - |
| - | - | 503.4 | 224.1 | - | - | 0 | - |
| - | - | 713.3 | 249.1 | - | - | 0 | - |
| - | - | 1613 | 251 | - | - | 0 | - |
| - | - | 555.4 | 255.2 | - | - | 0 | - |
| - | - | 635.3 | 265.2 | - | - | 0 | - |
| 8 | y | 1353 | 279.1 | 0.0001302 | 0.4664 | +1 | 2 |
| - | - | 1.099E+04 | 279.2 | - | - | 0 | - |
| - | - | 1363 | 280.2 | - | - | 0 | - |
| - | - | 1022 | 285.2 | - | - | 0 | - |
| - | - | 4774 | 297.1 | - | - | 0 | - |
| - | - | 878.5 | 298.1 | - | - | 0 | - |
| - | - | 929.8 | 300.1 | - | - | 0 | - |
| - | - | 1502 | 301.1 | - | - | 0 | - |
| - | - | 1076 | 304.1 | - | - | 0 | - |
| - | - | 540.2 | 308.5 | - | - | 0 | - |
| - | - | 2154 | 313.2 | - | - | 0 | - |
| - | - | 3282 | 325.1 | - | - | 0 | - |
| - | - | 672.9 | 327.2 | - | - | 0 | - |
| - | - | 2045 | 329.2 | - | - | 0 | - |
| - | - | 1476 | 353.6 | - | - | 0 | - |
| - | - | 619 | 354.1 | - | - | 0 | - |
| - | - | 930 | 354.9 | - | - | 0 | - |
| - | - | 3670 | 355.1 | - | - | 0 | - |
| - | - | 739.5 | 355.2 | - | - | 0 | - |
| - | - | 572.8 | 355.9 | - | - | 0 | - |
| - | - | 2323 | 356.1 | - | - | 0 | - |
| - | - | 3463 | 357.1 | - | - | 0 | - |
| - | - | 526.6 | 358.1 | - | - | 0 | - |
| - | - | 604.1 | 370.3 | - | - | 0 | - |
| - | - | 646.9 | 371.7 | - | - | 0 | - |
| - | - | 1148 | 372.2 | - | - | 0 | - |
| - | - | 4.997E+04 | 373.2 | - | - | 0 | - |
| - | - | 1655 | 374.2 | - | - | 0 | - |
| - | - | 7345 | 374.2 | - | - | 0 | - |
| - | - | 607.3 | 374.9 | - | - | 0 | - |
| - | - | 566.6 | 375.1 | - | - | 0 | - |
| - | - | 740.7 | 375.2 | - | - | 0 | - |
| - | - | 717.3 | 376.6 | - | - | 0 | - |
| 6 | c | 701.4 | 378.2 | 0.00572 | 15.12 | +2 | 6 |
| - | - | 555.2 | 380.5 | - | - | 0 | - |
| - | - | 690.1 | 382.6 | - | - | 0 | - |
| 4 | y | 1982 | 389.2 | 0.0001407 | 0.3615 | +2 | 6 |
| - | - | 602.7 | 389.7 | - | - | 0 | - |
| - | - | 2119 | 390.3 | - | - | 0 | - |
| - | - | 1534 | 390.6 | - | - | 0 | - |
| - | - | 644 | 391.1 | - | - | 0 | - |
| - | - | 1139 | 391.2 | - | - | 0 | - |
| - | - | 977.5 | 391.5 | - | - | 0 | - |
| - | - | 642.1 | 391.9 | - | - | 0 | - |
| - | - | 624.7 | 392.2 | - | - | 0 | - |
| - | - | 661 | 393.2 | - | - | 0 | - |
| - | - | 1122 | 402.1 | - | - | 0 | - |
| - | - | 3520 | 412.2 | - | - | 0 | - |
| - | - | 973.2 | 413.2 | - | - | 0 | - |
| 3 | z | 1629 | 416.7 | 0.0004787 | 1.149 | +2 | 7 |
| - | - | 636.1 | 417.2 | - | - | 0 | - |
| 7 | z | 1.323E+04 | 419.2 | 0.0005069 | 1.209 | +1 | 3 |
| - | - | 2315 | 420.2 | - | - | 0 | - |
| 3 | y | 3426 | 424.7 | 0.0004131 | 0.9727 | +2 | 7 |
| - | - | 1287 | 425.2 | - | - | 0 | - |
| - | - | 701.9 | 426.7 | - | - | 0 | - |
| - | - | 853 | 433.7 | - | - | 0 | - |
| 7 | y | 2784 | 435.2 | 0.0005817 | 1.336 | +1 | 3 |
| - | - | 3857 | 451.7 | - | - | 0 | - |
| 2 | w | 1809 | 452.2 | 0.007241 | 16.01 | +2 | 8 |
| - | - | 639.9 | 458.2 | - | - | 0 | - |
| - | - | 689.3 | 462.7 | - | - | 0 | - |
| - | - | 666.5 | 463 | - | - | 0 | - |
| - | - | 827.9 | 469.2 | - | - | 0 | - |
| - | - | 688.5 | 469.7 | - | - | 0 | - |
| - | - | 888.6 | 470.3 | - | - | 0 | - |
| - | - | 661.9 | 476.2 | - | - | 0 | - |
| - | - | 720 | 476.2 | - | - | 0 | - |
| - | - | 1407 | 483.3 | - | - | 0 | - |
| - | - | 724.2 | 489.2 | - | - | 0 | - |
| - | - | 894.2 | 489.7 | - | - | 0 | - |
| - | - | 647.5 | 490.2 | - | - | 0 | - |
| - | - | 635 | 495.7 | - | - | 0 | - |
| 8 | c | 2043 | 496.2 | 0.002249 | 4.532 | +2 | 8 |
| 2 | y | 2064 | 496.7 | 0.005957 | 11.99 | +2 | 8 |
| - | - | 6012 | 504.2 | - | - | 0 | - |
| 8 | c | 2.567E+04 | 504.8 | 0.001943 | 3.849 | +2 | 8 |
| 2 | y | 2.011E+04 | 505.2 | 0.004584 | 9.073 | +2 | 8 |
| - | - | 8797 | 505.8 | - | - | 0 | - |
| - | - | 3271 | 506.3 | - | - | 0 | - |
| - | - | 606.6 | 506.7 | - | - | 0 | - |
| - | - | 2598 | 512.3 | - | - | 0 | - |
| - | - | 1361 | 512.8 | - | - | 0 | - |
| - | - | 739.6 | 513.3 | - | - | 0 | - |
| 6 | z | 3278 | 518.3 | 0.0004937 | 0.9525 | +1 | 4 |
| - | - | 628.8 | 519.3 | - | - | 0 | - |
| 4 | c | 568.2 | 524.2 | 0.008609 | 16.42 | +1 | 4 |
| - | - | 3249 | 525.7 | - | - | 0 | - |
| - | - | 1483 | 526.2 | - | - | 0 | - |
| - | - | 1.376E+04 | 533.3 | - | - | 0 | - |
| - | - | 9070 | 533.8 | - | - | 0 | - |
| - | - | 3316 | 534.3 | - | - | 0 | - |
| 6 | y | 4446 | 534.3 | 0.001046 | 1.958 | +1 | 4 |
| - | - | 2540 | 534.8 | - | - | 0 | - |
| - | - | 1651 | 535.3 | - | - | 0 | - |
| - | - | 1239 | 540.8 | - | - | 0 | - |
| 4 | c | 4.31E+04 | 541.2 | 0.00397 | 7.334 | +1 | 4 |
| - | - | 703.6 | 541.8 | - | - | 0 | - |
| - | - | 1.117E+04 | 542.2 | - | - | 0 | - |
| - | - | 3296 | 543.2 | - | - | 0 | - |
| - | - | 1037 | 548.8 | - | - | 0 | - |
| - | - | 1.322E+04 | 549.3 | - | - | 0 | - |
| - | - | 8467 | 549.8 | - | - | 0 | - |
| - | - | 3038 | 550.3 | - | - | 0 | - |
| - | - | 1190 | 550.8 | - | - | 0 | - |
| - | - | 891.9 | 556.3 | - | - | 0 | - |
| - | - | 2676 | 556.8 | - | - | 0 | - |
| - | - | 1.065E+04 | 557.3 | - | - | 0 | - |
| - | - | 1.903E+04 | 557.8 | - | - | 0 | - |
| - | - | 1.236E+04 | 558.3 | - | - | 0 | - |
| - | - | 4237 | 558.8 | - | - | 0 | - |
| - | - | 2481 | 564.8 | - | - | 0 | - |
| - | - | 5641 | 565.3 | - | - | 0 | - |
| - | - | 4761 | 565.8 | - | - | 0 | - |
| - | - | 2026 | 566.3 | - | - | 0 | - |
| - | - | 822.1 | 566.8 | - | - | 0 | - |
| - | - | 5056 | 570.3 | - | - | 0 | - |
| - | - | 3634 | 570.8 | - | - | 0 | - |
| - | - | 1201 | 571.3 | - | - | 0 | - |
| - | - | 1053 | 574.3 | - | - | 0 | - |
| - | - | 8845 | 578.8 | - | - | 0 | - |
| - | - | 4391 | 579.3 | - | - | 0 | - |
| - | - | 1513 | 579.8 | - | - | 0 | - |
| - | - | 997.9 | 580.3 | - | - | 0 | - |
| - | - | 902.5 | 586.2 | - | - | 0 | - |
| - | - | 3.186E+04 | 586.8 | - | - | 0 | - |
| - | - | 2.716E+04 | 587.3 | - | - | 0 | - |
| - | - | 1.237E+04 | 587.8 | - | - | 0 | - |
| - | - | 2688 | 588.2 | - | - | 0 | - |
| - | - | 4728 | 588.3 | - | - | 0 | - |
| - | - | 1369 | 588.8 | - | - | 0 | - |
| - | - | 867 | 589.2 | - | - | 0 | - |
| - | - | 2.694E+04 | 589.3 | - | - | 0 | - |
| - | - | 6995 | 590.3 | - | - | 0 | - |
| - | - | 2148 | 591.3 | - | - | 0 | - |
| - | - | 1322 | 612.3 | - | - | 0 | - |
| 5 | z | 1843 | 633.3 | 0.0006395 | 1.01 | +1 | 5 |
| - | - | 1953 | 634.2 | - | - | 0 | - |
| - | - | 1786 | 634.3 | - | - | 0 | - |
| 5 | c | 1431 | 639.2 | 0.002593 | 4.057 | +1 | 5 |
| 5 | y | 6122 | 649.3 | 0.0007753 | 1.194 | +1 | 5 |
| - | - | 2310 | 650.3 | - | - | 0 | - |
| - | - | 2103 | 652.3 | - | - | 0 | - |
| 5 | c | 5807 | 656.3 | 0.00363 | 5.531 | +1 | 5 |
| - | - | 1928 | 657.3 | - | - | 0 | - |
| - | - | 833.8 | 668.4 | - | - | 0 | - |
| - | - | 3605 | 692.2 | - | - | 0 | - |
| - | - | 2193 | 724.2 | - | - | 0 | - |
| - | - | 1387 | 738.4 | - | - | 0 | - |
| 6 | c | 3.613E+04 | 755.3 | 0.004234 | 5.605 | +1 | 6 |
| - | - | 1.377E+04 | 756.3 | - | - | 0 | - |
| - | - | 5039 | 757.3 | - | - | 0 | - |
| - | - | 1087 | 758.3 | - | - | 0 | - |
| 4 | z | 5927 | 761.4 | 0.0004515 | 0.593 | +1 | 6 |
| - | - | 2781 | 762.4 | - | - | 0 | - |
| - | - | 3255 | 765.2 | - | - | 0 | - |
| - | - | 2027 | 766.2 | - | - | 0 | - |
| 4 | y | 1247 | 777.4 | 0.002919 | 3.755 | +1 | 6 |
| - | - | 2523 | 782.2 | - | - | 0 | - |
| - | - | 2826 | 783.2 | - | - | 0 | - |
| - | - | 848.5 | 784.2 | - | - | 0 | - |
| 3 | z | 4996 | 832.4 | 0.0004559 | 0.5477 | +1 | 7 |
| - | - | 2810 | 833.4 | - | - | 0 | - |
| - | - | 976.8 | 834.5 | - | - | 0 | - |
| 3 | y | 1203 | 848.5 | 0.002976 | 3.507 | +1 | 7 |
| - | - | 1230 | 867.4 | - | - | 0 | - |
| - | - | 1701 | 902.5 | - | - | 0 | - |
| 2 | w | 3148 | 903.5 | 0.01688 | 18.68 | +1 | 8 |
| - | - | 2426 | 950.5 | - | - | 0 | - |
| - | - | 696.5 | 951.5 | - | - | 0 | - |
| 2 | y | 2715 | 992.5 | 0.01913 | 19.27 | +1 | 8 |
| 2 | z | 1655 | 993.5 | 0.01271 | 12.79 | +1 | 8 |
| - | - | 1045 | 994.5 | - | - | 0 | - |
| 8 | c | 4187 | 1008 | 0.006497 | 6.443 | +1 | 8 |
| 2 | y | 6898 | 1009 | 0.01736 | 17.2 | +1 | 8 |
| - | - | 3723 | 1011 | - | - | 0 | - |
| - | - | 1534 | 1011 | - | - | 0 | - |
| - | - | 1179 | 1072 | - | - | 0 | - |
| - | - | 1195 | 1097 | - | - | 0 | - |
| - | - | 3672 | 1098 | - | - | 0 | - |
| - | - | 4901 | 1099 | - | - | 0 | - |
| - | - | 2048 | 1100 | - | - | 0 | - |
| - | - | 833.4 | 1101 | - | - | 0 | - |
| - | - | 3031 | 1114 | - | - | 0 | - |
| - | - | 2.642E+04 | 1115 | - | - | 0 | - |
| - | - | 1.486E+04 | 1116 | - | - | 0 | - |
| - | - | 1.071E+04 | 1117 | - | - | 0 | - |
| - | - | 2283 | 1118 | - | - | 0 | - |
| - | - | 743.7 | 1131 | - | - | 0 | - |
| - | - | 1687 | 1132 | - | - | 0 | - |
| - | - | 1205 | 1133 | - | - | 0 | - |
| - | - | 1008 | 1140 | - | - | 0 | - |
| - | - | 2135 | 1141 | - | - | 0 | - |
| - | - | 1237 | 1142 | - | - | 0 | - |
| - | - | 6621 | 1157 | - | - | 0 | - |
| - | - | 3.376E+04 | 1158 | - | - | 0 | - |
| - | - | 1.991E+04 | 1159 | - | - | 0 | - |
| - | - | 8551 | 1160 | - | - | 0 | - |
| - | - | 2844 | 1161 | - | - | 0 | - |
| - | - | 1061 | 1162 | - | - | 0 | - |
| - | - | 1523 | 1173 | - | - | 0 | - |
| - | - | 8367 | 1174 | - | - | 0 | - |
| - | - | 1.841E+04 | 1175 | - | - | 0 | - |
| - | - | 1.023E+04 | 1176 | - | - | 0 | - |
| - | - | 4064 | 1177 | - | - | 0 | - |
| - | - | 1194 | 1178 | - | - | 0 | - |
| - | - | 617.6 | 1843 | - | - | 0 | - |
| - | - | 654.8 | 2768 | - | - | 0 | - |

m/z Charge Intensity FragmentType MassShift Position
120.08099365234375 0 1035.7422
121.02882385253906 0 1596.5504
121.0399398803711 0 1070.0768
127.62571716308594 0 394.8441
129.10256958007812 0 2662.8008
129.40452575683594 0 385.92288
133.0862274169922 0 8183.5815
134.0896759033203 0 4072.815
135.09066772460938 0 2286.2197
136.07601928710938 0 11063.495
137.07957458496094 0 1302.8435
142.75914001464844 0 403.57333
147.04432678222656 0 662.7333
148.95494079589844 0 821.99133
149.02374267578125 0 249934.25
150.02699279785156 0 18360.912
150.03268432617188 0 377.90402
156.5105438232422 0 510.34204
166.08680725097656 0 1109.5103
167.0342254638672 0 86214.195
168.03753662109375 0 6364.349
173.43994140625 0 1565.2052
177.1125030517578 0 3562.329
178.11595153808594 0 2761.7708
178.6236572265625 0 433.4603
179.11712646484375 0 2823.4622
181.94448852539062 0 521.6598
200.1395721435547 0 821.43475
205.0646209716797 0 568.9796
211.1552734375 0 923.5871
212.1636199951172 0 1230.5945
219.1332244873047 0 559.432
221.05966186523438 0 1600.2096
221.0929412841797 0 749.41943
223.06411743164062 0 1056.8096
223.10769653320312 0 1247.8517
223.14369201660156 0 1101.3176
224.06631469726562 0 503.3935
249.05438232421875 0 713.2957
251.04705810546875 0 1613.3245
255.16920471191406 0 555.3748
265.1658020019531 0 635.31683
279.1340637207031 0 1353.0634 y 7
279.15948486328125 0 10991.358
280.16265869140625 0 1363.3446
285.1807556152344 0 1021.96436
297.0908508300781 0 4774.175
298.0941467285156 0 878.4805
300.0619201660156 0 929.76666
301.0604248046875 0 1501.9108
304.1297302246094 0 1076.3845
308.52545166015625 0 540.1537
313.17486572265625 0 2154.4434
325.08612060546875 0 3281.9458
327.2015380859375 0 672.94434
329.20709228515625 0 2045.0872
353.60797119140625 0 1476.1515
354.1093444824219 0 619.003
354.8833312988281 0 930.02057
355.0702209472656 0 3669.9424
355.2238464355469 0 739.50024
355.909912109375 0 572.77826
356.0714111328125 0 2322.831
357.0676574707031 0 3462.6624
358.0685119628906 0 526.5647
370.25787353515625 0 604.10706
371.6980895996094 0 646.8986
372.2310485839844 0 1147.5066
373.2334899902344 0 49970.04
374.2142333984375 0 1655.4694
374.2361145019531 0 7344.723
374.90057373046875 0 607.26843
375.0697326660156 0 566.55865
375.24127197265625 0 740.71326
376.6123352050781 0 717.2878
378.1812438964844 0 701.4174 c 5
380.4542236328125 0 555.2373
382.6103820800781 0 690.0627
389.2164611816406 0 1981.7705 y 3
389.7159423828125 0 602.74176
390.2597351074219 0 2119.2725
390.6218566894531 0 1534.3354
391.1224060058594 0 643.96906
391.1880187988281 0 1138.5588
391.5185546875 0 977.54865
391.853271484375 0 642.13135
392.1865234375 0 624.69165
393.2101135253906 0 660.9665
402.0556945800781 0 1121.7755
412.24334716796875 0 3520.002
413.2476501464844 0 973.1625
416.72503662109375 0 1628.5066 z 2
417.2292785644531 0 636.1413
419.2168273925781 0 13232.409 z 6
420.2198791503906 0 2314.5557
424.73529052734375 0 3426.2961 y 2
425.2386779785156 0 1286.7192
426.71087646484375 0 701.9498
433.7127380371094 0 852.9994
435.2356262207031 0 2784.244 y 6
451.74066162109375 0 3856.9666
452.2416687011719 0 1809.043 w 1
458.1694641113281 0 639.925
462.72589111328125 0 689.29675
463.00274658203125 0 666.4557
469.2319030761719 0 827.8576
469.7364807128906 0 688.5007
470.27252197265625 0 888.6365
476.18243408203125 0 661.8661
476.2398681640625 0 719.9736
483.2803039550781 0 1407.1636
489.2264099121094 0 724.15753
489.7315673828125 0 894.2294
490.23175048828125 0 647.4744
495.7319030761719 0 635.01215
496.2369384765625 0 2043.0704 c Ammonia loss 7
496.7371520996094 0 2064.3318 y Ammonia loss 1
504.2469787597656 0 6012.028
504.7505187988281 0 25674.031 c 7
505.2490539550781 0 20113.03 y 1
505.7505798339844 0 8796.962
506.2516784667969 0 3270.8286
506.7470703125 0 606.6339
512.2625732421875 0 2597.6582
512.7622680664062 0 1361.3596
513.26220703125 0 739.60254
518.2842407226562 0 3278.4573 z 5
519.283935546875 0 628.7782
524.2132568359375 0 568.1599 c Ammonia loss 3
525.7433471679688 0 3248.6667
526.24658203125 0 1482.9281
533.2716674804688 0 13761.055
533.7728271484375 0 9069.9
534.2648315429688 0 3316.0374
534.3045043945312 0 4446.093 y 5
534.7611694335938 0 2540.2473
535.3067626953125 0 1651.0168
540.781005859375 0 1238.596
541.2444458007812 0 43103.55 c 3
541.7842407226562 0 703.6043
542.2472534179688 0 11165.642
543.244384765625 0 3296.293
548.755615234375 0 1036.9607
549.2582397460938 0 13217.021
549.7598876953125 0 8466.79
550.2598266601562 0 3038.4348
550.759765625 0 1189.6381
556.262451171875 0 891.8761
556.7708740234375 0 2675.7983
557.2681274414062 0 10648.909
557.7708129882812 0 19034.68
558.2723388671875 0 12356.144
558.7731323242188 0 4236.898
564.7798461914062 0 2481.4155
565.2823486328125 0 5641.066
565.7657470703125 0 4761.4346
566.2672119140625 0 2025.8834
566.7713623046875 0 822.09955
570.2630004882812 0 5056.379
570.7649536132812 0 3634.2043
571.2658081054688 0 1200.6199
574.298583984375 0 1053.301
578.76513671875 0 8845.131
579.2664794921875 0 4391.352
579.765869140625 0 1513.0828
580.2679443359375 0 997.9373
586.2017822265625 0 902.4853
586.774169921875 0 31860.504
587.2763061523438 0 27156.467
587.77783203125 0 12369.53
588.18896484375 0 2688.1008
588.2783203125 0 4727.507
588.7780151367188 0 1368.8712
589.1889038085938 0 866.973
589.3224487304688 0 26944.91
590.3251953125 0 6995.197
591.326171875 0 2147.6567
612.255615234375 0 1321.8597
633.3123168945312 0 1842.7051 z 4
634.1718139648438 0 1953.3507
634.3180541992188 0 1786.4407
639.2462158203125 0 1430.7349 c Ammonia loss 4
649.3311767578125 0 6122.104 y 4
650.3325805664062 0 2309.7302
652.3135375976562 0 2102.9485
656.271728515625 0 5807.165 c 4
657.273193359375 0 1928.3502
668.3958740234375 0 833.8064
692.2471313476562 0 3604.9763
724.2183837890625 0 2193.0806
738.429443359375 0 1386.6582
755.3395385742188 0 36130.336 c 5
756.34228515625 0 13774.599
757.342041015625 0 5039.0366
758.3426513671875 0 1087.0874
761.4061889648438 0 5926.6387 z 3
762.4119873046875 0 2781.4863
765.2229614257812 0 3255.463
766.2132568359375 0 2027.3583
777.4282836914062 0 1247.172 y 3
782.2256469726562 0 2523.263
783.2332153320312 0 2826.2727
784.2330322265625 0 848.5043
832.4432983398438 0 4995.5693 z 2
833.448486328125 0 2809.6252
834.4526977539062 0 976.8228
848.4654541015625 0 1202.7296 y 2
867.4271240234375 0 1230.1987
902.4746704101562 0 1700.9827
903.4784545898438 0 3148.4016 w 1
950.479736328125 0 2425.6528
951.4854736328125 0 696.4969
992.4742431640625 0 2714.8958 y Ammonia loss 1
993.4756469726562 0 1654.545 z 1
994.4825439453125 0 1044.6123
1008.4911499023438 0 4187.3936 c 7
1009.4990234375 0 6898.1323 y 1
1010.5006103515625 0 3723.3499
1011.49609375 0 1534.3026
1072.4571533203125 0 1179.3813
1096.5294189453125 0 1194.8716
1097.51220703125 0 3671.5757
1098.513916015625 0 4900.678
1099.5142822265625 0 2048.1765
1100.5172119140625 0 833.42816
1113.5281982421875 0 3030.9849
1114.52490234375 0 26418.713
1115.52880859375 0 14861.521
1116.5352783203125 0 10707.6045
1117.536865234375 0 2283.3066
1130.5489501953125 0 743.66406
1131.5255126953125 0 1686.6052
1132.526123046875 0 1204.9078
1139.5159912109375 0 1007.719
1140.5045166015625 0 2134.5264
1141.5162353515625 0 1236.5763
1156.5233154296875 0 6620.5176
1157.528076171875 0 33756.184
1158.5316162109375 0 19909.783
1159.5340576171875 0 8550.99
1160.5343017578125 0 2844.3862
1161.5313720703125 0 1060.9761
1172.5389404296875 0 1522.9551
1173.5452880859375 0 8367.353
1174.553955078125 0 18410.146
1175.556884765625 0 10231.041
1176.5614013671875 0 4064.2537
1177.5623779296875 0 1193.5916
1843.3634033203125 0 617.607
2768.100830078125 0 654.8309

Spectrum Details

|  |  |
| --- | --- |
| Matched peaks? Matched peaks The total absolute number of peaks matched. Additionally in brackets the total fraction of peaks matched and the total number of peaks is shown. | 30 (12.00% of 250) |
| FDR? FDR The false discovery rate estimated for this peptide. It is calculated by matching all theoretical fragments with a non-integer shift with the raw peaks for this spectrum. This is done with 40 different shifts. The resulting percentage is the average number of annotated peaks over the number of annotated peaks with the correct spectrum. | 2.06% |
| Satellite FDR? Satellite FDR See the FDR for details on its calculation. This satellite ion specific FDR only contains the satellite ions (d/w) for I/L/J positions. | - |
| PSM Score? PSM Score The PSM Score as given by Hecklib to this annotated spectrum. It is shown with three significant figures. | 260 |

#### Reverse Lookup? Reverse Lookup All places where this read could be placed.

| Group | Segment | Template | Template Part | Read Part | Score | Unique |
| --- | --- | --- | --- | --- | --- | --- |
| Homo sapiens Heavy Chain | IGHV | IGHV3-9 | [94..103] | [0..9] | 44 | False |
| Homo sapiens Heavy Chain | IGHV | IGHV3-43 | [94..103] | [0..9] | 44 | False |

| Recombined | Template Part | Read Part | Score | Unique |
| --- | --- | --- | --- | --- |
| REC-0-1 | [94..103] | [0..9] | 72 | True |

#### Meta Information from Multiple reads

##### Number of combined reads

7

##### Intensity

0.8772

##### TotalArea

6.465E+08

#### Positional Score

Copy Data

##### Positional Score (TSV)

###### Preview

```
Loading example...
```

*Click on the button to copy the data to your clipboard.*

10012345678

Label Value
"0" 0.681
"1" 0.687
"2" 0.709
"3" 0.707
"4" 0.714
"5" 0.709
"6" 0.693
"7" 0.686
"8" 0.703

#### Meta Information from PEAKS

##### Scan Identifier

F3:3914

##### Original sequence

Y

C

+58.01

A

K

D

V

R

P

Y

##### Posttranslational Modifications

Carboxymethyl

##### Source File

D:\separate\_stitch\_analyses\xle-disambiguation\raw\20210323\_F1\_UM1\_Peng0013\_SA\_F59\_ingel\_3ug\_chymo.raw

##### Fraction

3

##### Scan Feature

-

##### De Novo Score

99

##### ConfidenceScore

99

### m/z

391.5186

##### Mass

1171.5332

##### Charge

3

##### Retention Time

21.46

##### Predicted Retention Time

-

##### Area

0

##### Parts Per Million

0.8

##### Fragmentation mode

HCD

##### Originating file

01 D:\separate\_stitch\_analyses\xle-disambiguation\20210325\_F59\_3ug\_DENOVO\_12.csv

#### Meta Information from PEAKS

##### Scan Identifier

F3:3990

##### Original sequence

Y

C

+58.01

A

K

D

V

R

P

Y

##### Posttranslational Modifications

Carboxymethyl

##### Source File

D:\separate\_stitch\_analyses\xle-disambiguation\raw\20210323\_F1\_UM1\_Peng0013\_SA\_F59\_ingel\_3ug\_chymo.raw

##### Fraction

3

##### Scan Feature

-

##### De Novo Score

99

##### ConfidenceScore

99

### m/z

391.5191

##### Mass

1171.5332

##### Charge

3

##### Retention Time

21.87

##### Predicted Retention Time

-

##### Area

0

##### Parts Per Million

1.9

##### Fragmentation mode

HCD

##### Originating file

01 D:\separate\_stitch\_analyses\xle-disambiguation\20210325\_F59\_3ug\_DENOVO\_12.csv

#### Meta Information from PEAKS

##### Scan Identifier

F3:3845

##### Original sequence

Y

C

+58.01

A

K

D

V

R

P

Y

##### Posttranslational Modifications

Carboxymethyl

##### Source File

D:\separate\_stitch\_analyses\xle-disambiguation\raw\20210323\_F1\_UM1\_Peng0013\_SA\_F59\_ingel\_3ug\_chymo.raw

##### Fraction

3

##### Scan Feature

F3:561

##### De Novo Score

99

##### ConfidenceScore

99

### m/z

391.5191

##### Mass

1171.5332

##### Charge

3

##### Retention Time

19.96

##### Predicted Retention Time

-

##### Area

2.234E+08

##### Parts Per Million

1.9

##### Fragmentation mode

HCD

##### Originating file

01 D:\separate\_stitch\_analyses\xle-disambiguation\20210325\_F59\_3ug\_DENOVO\_12.csv

#### Meta Information from PEAKS

##### Scan Identifier

F3:3838

##### Original sequence

Y

C

+58.01

A

K

D

V

R

P

Y

##### Posttranslational Modifications

Carboxymethyl

##### Source File

D:\separate\_stitch\_analyses\xle-disambiguation\raw\20210323\_F1\_UM1\_Peng0013\_SA\_F59\_ingel\_3ug\_chymo.raw

##### Fraction

3

##### Scan Feature

F3:7686

##### De Novo Score

99

##### ConfidenceScore

99

### m/z

586.7742

##### Mass

1171.5332

##### Charge

2

##### Retention Time

19.96

##### Predicted Retention Time

-

##### Area

9.982E+07

##### Parts Per Million

0.5

##### Fragmentation mode

HCD

##### Originating file

01 D:\separate\_stitch\_analyses\xle-disambiguation\20210325\_F59\_3ug\_DENOVO\_12.csv

#### Meta Information from PEAKS

##### Scan Identifier

F3:3679

##### Original sequence

Y

C

+58.01

A

K

D

V

R

P

Y

##### Posttranslational Modifications

Carboxymethyl

##### Source File

D:\separate\_stitch\_analyses\xle-disambiguation\raw\20210323\_F1\_UM1\_Peng0013\_SA\_F59\_ingel\_3ug\_chymo.raw

##### Fraction

3

##### Scan Feature

F3:7686

##### De Novo Score

98

##### ConfidenceScore

98

### m/z

586.7742

##### Mass

1171.5332

##### Charge

2

##### Retention Time

19.96

##### Predicted Retention Time

-

##### Area

9.982E+07

##### Parts Per Million

0.5

##### Fragmentation mode

HCD

##### Originating file

01 D:\separate\_stitch\_analyses\xle-disambiguation\20210325\_F59\_3ug\_DENOVO\_12.csv

#### Meta Information from PEAKS

##### Scan Identifier

F3:3777

##### Original sequence

Y

C

+58.01

A

K

D

V

R

P

Y

##### Posttranslational Modifications

Carboxymethyl

##### Source File

D:\separate\_stitch\_analyses\xle-disambiguation\raw\20210323\_F1\_UM1\_Peng0013\_SA\_F59\_ingel\_3ug\_chymo.raw

##### Fraction

3

##### Scan Feature

F3:561

##### De Novo Score

96

##### ConfidenceScore

96

### m/z

391.5191

##### Mass

1171.5332

##### Charge

3

##### Retention Time

19.96

##### Predicted Retention Time

-

##### Area

2.234E+08

##### Parts Per Million

1.9

##### Fragmentation mode

ETHCD

##### Originating file

01 D:\separate\_stitch\_analyses\xle-disambiguation\20210325\_F59\_3ug\_DENOVO\_12.csv

#### Meta Information from PEAKS

##### Scan Identifier

F3:4095

##### Original sequence

Y

C

+58.01

A

K

D

V

R

P

Y

##### Posttranslational Modifications

Carboxymethyl

##### Source File

D:\separate\_stitch\_analyses\xle-disambiguation\raw\20210323\_F1\_UM1\_Peng0013\_SA\_F59\_ingel\_3ug\_chymo.raw

##### Fraction

3

##### Scan Feature

-

##### De Novo Score

95

##### ConfidenceScore

95

### m/z

391.5193

##### Mass

1171.5332

##### Charge

3

##### Retention Time

22.48

##### Predicted Retention Time

-

##### Area

0

##### Parts Per Million

2.3

##### Fragmentation mode

ETHCD

##### Originating file

01 D:\separate\_stitch\_analyses\xle-disambiguation\20210325\_F59\_3ug\_DENOVO\_12.csv
