## Supplementary material for "A handle on mass coincidence errors in *de novo* sequencing of antibodies by bottom-up proteomics": Combined_012.html

Details Combined\_012 | Stitch OverviewUndefined

### Read Combined\_012

#### Sequence (length=8)

VATVSJPR

#### Spectrum 5082? Spectrum 5082 The raw spectrum of this peptide as annotated by Hecklib. The fragments are coloured according to ion type (see legend). Any peaks with a star '\*' as text can be hovered over to see the full details, first the ion type second the mass shift type. By hovering over the amino acids in the peptide or ions in the legend the corresponding peaks are highlighted. By toggling the 'Unassigned' label you can turn the background (unassigned) peaks on or off in the plot. By updating the slider in the Ion legend you can update the spectrum to only show the top X% of the peaks with labels. The top X% means any peak that is within X% of the highest intensity. By dragging in the spectrum you can zoom in to a specific part of the spectrum and use 'Zoom Out' to get back to the original zoom level. The annotation of the spectrum is based on the given sequence in the peptides file and is done with different software so inconsistencies are likely. The peaks are annotated based on the given sequence, with 20 ppm tolerance.

Copy Data

##### Spectrum 5082 (TSV)

###### Preview

```
Loading example...
```

*Click on the button to copy the data to your clipboard.*

Mz MinMz MaxIntensity Max

WidthHeightPeptide font sizePeptide stroke widthSpectrum font sizeSpectrum stroke widthCompact peptide

Ion legend

wxyz

abcd

OtherUnassignedIon^Charge^_Position_Show for top:%

VATVSJPR

01.86e+53.72e+55.57e+57.43e+5

Zoom Out

a^+1^_2_y^+1^_1_b^+1^_2_y^+1^_1_y^+2^_3_a^+1^_3_y^+2^_4_d^+1^_3_a^+1^_3_b^+1^_3_y^+1^_2_b^+1^_3_y^+1^_2_y^+2^_5_y^+2^_5_y^+2^_6_y^+2^_6_b^+1^_4_y^+2^_7_y^+1^_3_b^+1^_4_y^+2^_7_y^+1^_3_\*\*b^+1^_5_y^+1^_4_y^+1^_4_b^+1^_5_y^+1^_4_b^+1^_6_y^+1^_5_y^+1^_5_y^+1^_6_y^+1^_6_y^+1^_6_y^+1^_7_y^+1^_7_

0757151522723029

Fragment Matches Table

Show background peaks

| Position | Ion type | Intensity | mz Theoretical | mz Error (Th) | mz Error (ppm) | Charge | Series Number |
| --- | --- | --- | --- | --- | --- | --- | --- |
| - | - | 392.8 | 120.1 | - | - | 0 | - |
| - | - | 380.3 | 125.1 | - | - | 0 | - |
| - | - | 1583 | 125.1 | - | - | 0 | - |
| - | - | 595.6 | 126.1 | - | - | 0 | - |
| - | - | 1372 | 126.1 | - | - | 0 | - |
| - | - | 7859 | 127.1 | - | - | 0 | - |
| - | - | 7876 | 127.1 | - | - | 0 | - |
| - | - | 1.412E+04 | 128.1 | - | - | 0 | - |
| - | - | 763.1 | 129 | - | - | 0 | - |
| - | - | 1.538E+04 | 129.1 | - | - | 0 | - |
| - | - | 1400 | 129.1 | - | - | 0 | - |
| - | - | 2219 | 130.1 | - | - | 0 | - |
| - | - | 3648 | 130.1 | - | - | 0 | - |
| - | - | 2239 | 130.1 | - | - | 0 | - |
| - | - | 1306 | 130.1 | - | - | 0 | - |
| - | - | 544.2 | 131.1 | - | - | 0 | - |
| - | - | 2318 | 133.1 | - | - | 0 | - |
| - | - | 704.9 | 136 | - | - | 0 | - |
| - | - | 5830 | 138.1 | - | - | 0 | - |
| - | - | 486.3 | 140 | - | - | 0 | - |
| - | - | 4.473E+04 | 140.1 | - | - | 0 | - |
| - | - | 855.6 | 141.1 | - | - | 0 | - |
| - | - | 2886 | 141.1 | - | - | 0 | - |
| - | - | 1.156E+04 | 141.1 | - | - | 0 | - |
| - | - | 429.1 | 142.1 | - | - | 0 | - |
| - | - | 1854 | 142.1 | - | - | 0 | - |
| 2 | a | 2.839E+05 | 143.1 | 0.0004729 | 3.304 | +1 | 2 |
| - | - | 534.4 | 144.1 | - | - | 0 | - |
| - | - | 1930 | 144.1 | - | - | 0 | - |
| - | - | 1.812E+04 | 144.1 | - | - | 0 | - |
| - | - | 1.631E+04 | 145.1 | - | - | 0 | - |
| - | - | 597.7 | 145.1 | - | - | 0 | - |
| - | - | 747.1 | 146.1 | - | - | 0 | - |
| - | - | 2799 | 147.1 | - | - | 0 | - |
| - | - | 920.4 | 148.1 | - | - | 0 | - |
| - | - | 1099 | 149 | - | - | 0 | - |
| - | - | 6523 | 149 | - | - | 0 | - |
| - | - | 3066 | 150 | - | - | 0 | - |
| - | - | 9691 | 155.1 | - | - | 0 | - |
| - | - | 3.528E+04 | 155.1 | - | - | 0 | - |
| - | - | 535.1 | 156.1 | - | - | 0 | - |
| - | - | 782.8 | 156.1 | - | - | 0 | - |
| - | - | 2976 | 156.1 | - | - | 0 | - |
| - | - | 3649 | 157.1 | - | - | 0 | - |
| - | - | 6280 | 157.1 | - | - | 0 | - |
| - | - | 1.977E+04 | 157.1 | - | - | 0 | - |
| - | - | 1065 | 157.1 | - | - | 0 | - |
| 8 | y | 3.682E+04 | 158.1 | 0.0004009 | 2.536 | +1 | 1 |
| - | - | 6050 | 158.1 | - | - | 0 | - |
| - | - | 1943 | 159.1 | - | - | 0 | - |
| - | - | 5953 | 159.1 | - | - | 0 | - |
| - | - | 442 | 159.1 | - | - | 0 | - |
| - | - | 474.9 | 160.1 | - | - | 0 | - |
| - | - | 1135 | 165.1 | - | - | 0 | - |
| - | - | 2230 | 167.1 | - | - | 0 | - |
| - | - | 1228 | 167.1 | - | - | 0 | - |
| - | - | 1.274E+04 | 169.1 | - | - | 0 | - |
| - | - | 1132 | 169.1 | - | - | 0 | - |
| - | - | 2377 | 170.1 | - | - | 0 | - |
| 2 | b | 2.498E+05 | 171.1 | 0.0004771 | 2.788 | +1 | 2 |
| - | - | 1209 | 172.1 | - | - | 0 | - |
| - | - | 2.031E+04 | 172.1 | - | - | 0 | - |
| - | - | 3.997E+04 | 173.1 | - | - | 0 | - |
| - | - | 2.934E+05 | 173.1 | - | - | 0 | - |
| - | - | 2572 | 174.1 | - | - | 0 | - |
| - | - | 2.271E+04 | 174.1 | - | - | 0 | - |
| 8 | y | 8.262E+04 | 175.1 | 0.0003868 | 2.209 | +1 | 1 |
| - | - | 573.8 | 176.1 | - | - | 0 | - |
| - | - | 5592 | 176.1 | - | - | 0 | - |
| - | - | 763.6 | 181.1 | - | - | 0 | - |
| - | - | 567.1 | 181.1 | - | - | 0 | - |
| - | - | 6.033E+04 | 183.1 | - | - | 0 | - |
| - | - | 1.258E+04 | 183.1 | - | - | 0 | - |
| - | - | 5075 | 184.1 | - | - | 0 | - |
| - | - | 3249 | 184.2 | - | - | 0 | - |
| - | - | 1025 | 185.1 | - | - | 0 | - |
| - | - | 6550 | 185.1 | - | - | 0 | - |
| - | - | 6616 | 185.1 | - | - | 0 | - |
| - | - | 1408 | 186.1 | - | - | 0 | - |
| - | - | 1.696E+04 | 187.1 | - | - | 0 | - |
| - | - | 577.5 | 187.1 | - | - | 0 | - |
| - | - | 1875 | 188.1 | - | - | 0 | - |
| - | - | 1236 | 189.1 | - | - | 0 | - |
| - | - | 1330 | 192.1 | - | - | 0 | - |
| - | - | 957.5 | 193.1 | - | - | 0 | - |
| 6 | y | 1458 | 193.1 | 0.002017 | 10.44 | +2 | 3 |
| - | - | 1652 | 194.1 | - | - | 0 | - |
| - | - | 5663 | 195.1 | - | - | 0 | - |
| - | - | 1013 | 197.1 | - | - | 0 | - |
| - | - | 627.9 | 197.1 | - | - | 0 | - |
| - | - | 1021 | 197.2 | - | - | 0 | - |
| - | - | 1231 | 199.1 | - | - | 0 | - |
| - | - | 4273 | 199.1 | - | - | 0 | - |
| - | - | 1.092E+05 | 201.1 | - | - | 0 | - |
| - | - | 1.064E+04 | 202.1 | - | - | 0 | - |
| - | - | 794.3 | 203.1 | - | - | 0 | - |
| - | - | 1438 | 208.1 | - | - | 0 | - |
| - | - | 1483 | 209.1 | - | - | 0 | - |
| - | - | 3015 | 209.1 | - | - | 0 | - |
| - | - | 6558 | 210.1 | - | - | 0 | - |
| - | - | 1.364E+04 | 211.1 | - | - | 0 | - |
| - | - | 862.3 | 211.1 | - | - | 0 | - |
| - | - | 7183 | 211.1 | - | - | 0 | - |
| - | - | 1828 | 211.2 | - | - | 0 | - |
| - | - | 1097 | 212.1 | - | - | 0 | - |
| - | - | 3.247E+04 | 212.1 | - | - | 0 | - |
| - | - | 6676 | 213.1 | - | - | 0 | - |
| - | - | 2443 | 213.1 | - | - | 0 | - |
| - | - | 776.8 | 213.2 | - | - | 0 | - |
| - | - | 657 | 215.1 | - | - | 0 | - |
| - | - | 1336 | 215.1 | - | - | 0 | - |
| - | - | 1377 | 217.1 | - | - | 0 | - |
| - | - | 1408 | 222.1 | - | - | 0 | - |
| - | - | 710 | 223.1 | - | - | 0 | - |
| - | - | 2053 | 224.1 | - | - | 0 | - |
| - | - | 1405 | 225.2 | - | - | 0 | - |
| - | - | 3434 | 226.1 | - | - | 0 | - |
| 3 | a | 4.143E+04 | 226.2 | 0.0003922 | 1.734 | +1 | 3 |
| - | - | 1723 | 227.1 | - | - | 0 | - |
| - | - | 1.965E+04 | 227.1 | - | - | 0 | - |
| - | - | 4582 | 227.2 | - | - | 0 | - |
| 5 | y | 1.148E+04 | 228.1 | 0.0004062 | 1.78 | +2 | 4 |
| - | - | 931.5 | 228.1 | - | - | 0 | - |
| - | - | 3806 | 229.1 | - | - | 0 | - |
| - | - | 994.7 | 229.1 | - | - | 0 | - |
| 3 | d | 1.591E+04 | 230.1 | 0.0004574 | 1.987 | +1 | 3 |
| - | - | 2004 | 231.2 | - | - | 0 | - |
| - | - | 903.2 | 234.1 | - | - | 0 | - |
| - | - | 973 | 236.1 | - | - | 0 | - |
| - | - | 3070 | 237.1 | - | - | 0 | - |
| - | - | 744.4 | 237.2 | - | - | 0 | - |
| - | - | 1838 | 238.1 | - | - | 0 | - |
| - | - | 972 | 239.1 | - | - | 0 | - |
| - | - | 2818 | 240.1 | - | - | 0 | - |
| - | - | 804.2 | 241.1 | - | - | 0 | - |
| - | - | 983.6 | 241.2 | - | - | 0 | - |
| - | - | 1965 | 242.1 | - | - | 0 | - |
| - | - | 1842 | 243.1 | - | - | 0 | - |
| - | - | 843.1 | 243.7 | - | - | 0 | - |
| - | - | 4254 | 244.1 | - | - | 0 | - |
| - | - | 1238 | 244.2 | - | - | 0 | - |
| 3 | a | 2939 | 244.2 | 0.000646 | 2.646 | +1 | 3 |
| - | - | 4278 | 252.1 | - | - | 0 | - |
| 3 | b | 1.188E+05 | 254.1 | 0.0004574 | 1.8 | +1 | 3 |
| 7 | y | 2.788E+05 | 255.1 | 0.0004782 | 1.874 | +1 | 2 |
| - | - | 3.023E+04 | 256.1 | - | - | 0 | - |
| - | - | 1993 | 257.2 | - | - | 0 | - |
| - | - | 7280 | 258.1 | - | - | 0 | - |
| - | - | 2290 | 259.1 | - | - | 0 | - |
| - | - | 729.3 | 260.2 | - | - | 0 | - |
| - | - | 1171 | 264.2 | - | - | 0 | - |
| - | - | 547.8 | 264.2 | - | - | 0 | - |
| - | - | 1991 | 266.2 | - | - | 0 | - |
| - | - | 616.9 | 267.2 | - | - | 0 | - |
| - | - | 611.8 | 268.1 | - | - | 0 | - |
| - | - | 1415 | 268.2 | - | - | 0 | - |
| - | - | 4.102E+04 | 270.1 | - | - | 0 | - |
| - | - | 5714 | 271.1 | - | - | 0 | - |
| - | - | 2348 | 271.2 | - | - | 0 | - |
| - | - | 2117 | 272.1 | - | - | 0 | - |
| 3 | b | 4.392E+04 | 272.2 | 0.000479 | 1.76 | +1 | 3 |
| 7 | y | 1.182E+05 | 272.2 | 0.0004947 | 1.818 | +1 | 2 |
| - | - | 4776 | 273.2 | - | - | 0 | - |
| - | - | 1.263E+04 | 273.2 | - | - | 0 | - |
| - | - | 1155 | 274.2 | - | - | 0 | - |
| - | - | 5139 | 276.2 | - | - | 0 | - |
| - | - | 1690 | 277.2 | - | - | 0 | - |
| 4 | y | 2396 | 277.2 | 0.0004434 | 1.6 | +2 | 5 |
| - | - | 1305 | 277.7 | - | - | 0 | - |
| - | - | 3805 | 280.1 | - | - | 0 | - |
| - | - | 2042 | 280.2 | - | - | 0 | - |
| - | - | 1196 | 282.1 | - | - | 0 | - |
| - | - | 1.126E+04 | 282.2 | - | - | 0 | - |
| - | - | 1384 | 283.2 | - | - | 0 | - |
| - | - | 928.6 | 284.2 | - | - | 0 | - |
| 4 | y | 9766 | 286.2 | 0.0003185 | 1.113 | +2 | 5 |
| - | - | 3305 | 286.7 | - | - | 0 | - |
| - | - | 3.546E+04 | 288.2 | - | - | 0 | - |
| - | - | 4827 | 289.2 | - | - | 0 | - |
| - | - | 735.3 | 290.2 | - | - | 0 | - |
| - | - | 667.4 | 294.2 | - | - | 0 | - |
| - | - | 706.6 | 294.2 | - | - | 0 | - |
| - | - | 2142 | 296.2 | - | - | 0 | - |
| - | - | 1026 | 297.2 | - | - | 0 | - |
| - | - | 1707 | 297.2 | - | - | 0 | - |
| - | - | 814 | 297.7 | - | - | 0 | - |
| - | - | 5615 | 298.1 | - | - | 0 | - |
| - | - | 1632 | 298.2 | - | - | 0 | - |
| - | - | 3178 | 299.2 | - | - | 0 | - |
| - | - | 2300 | 299.2 | - | - | 0 | - |
| - | - | 1066 | 299.7 | - | - | 0 | - |
| - | - | 2487 | 300.2 | - | - | 0 | - |
| - | - | 756 | 300.2 | - | - | 0 | - |
| - | - | 819.6 | 305.7 | - | - | 0 | - |
| - | - | 1004 | 306.1 | - | - | 0 | - |
| - | - | 1.775E+04 | 306.2 | - | - | 0 | - |
| - | - | 1351 | 306.2 | - | - | 0 | - |
| - | - | 4854 | 306.7 | - | - | 0 | - |
| - | - | 889.1 | 307.2 | - | - | 0 | - |
| - | - | 1592 | 308.2 | - | - | 0 | - |
| - | - | 1113 | 310.2 | - | - | 0 | - |
| - | - | 2017 | 310.7 | - | - | 0 | - |
| - | - | 836.8 | 311.2 | - | - | 0 | - |
| - | - | 1179 | 311.2 | - | - | 0 | - |
| - | - | 8752 | 312.2 | - | - | 0 | - |
| - | - | 1567 | 313.2 | - | - | 0 | - |
| - | - | 1324 | 314.2 | - | - | 0 | - |
| - | - | 620.2 | 315.2 | - | - | 0 | - |
| - | - | 1466 | 318.7 | - | - | 0 | - |
| - | - | 3016 | 319.7 | - | - | 0 | - |
| - | - | 815.4 | 320.2 | - | - | 0 | - |
| - | - | 1439 | 320.2 | - | - | 0 | - |
| - | - | 1108 | 321.2 | - | - | 0 | - |
| - | - | 1252 | 322.2 | - | - | 0 | - |
| - | - | 6957 | 323.2 | - | - | 0 | - |
| - | - | 1414 | 323.2 | - | - | 0 | - |
| - | - | 770.9 | 323.3 | - | - | 0 | - |
| - | - | 1471 | 324.2 | - | - | 0 | - |
| - | - | 5.241E+04 | 325.2 | - | - | 0 | - |
| - | - | 9067 | 326.2 | - | - | 0 | - |
| - | - | 1635 | 327.2 | - | - | 0 | - |
| - | - | 1600 | 327.2 | - | - | 0 | - |
| 3 | y | 1.07E+05 | 327.7 | 0.0004383 | 1.338 | +2 | 6 |
| - | - | 4.494E+04 | 328.2 | - | - | 0 | - |
| - | - | 1.207E+04 | 328.7 | - | - | 0 | - |
| - | - | 1618 | 329.2 | - | - | 0 | - |
| - | - | 702 | 331.2 | - | - | 0 | - |
| - | - | 4034 | 332.7 | - | - | 0 | - |
| - | - | 1226 | 333.2 | - | - | 0 | - |
| 3 | y | 1.339E+04 | 336.7 | 0.0005576 | 1.656 | +2 | 6 |
| - | - | 2538 | 337.2 | - | - | 0 | - |
| - | - | 6172 | 337.2 | - | - | 0 | - |
| - | - | 1159 | 337.7 | - | - | 0 | - |
| - | - | 1510 | 339.2 | - | - | 0 | - |
| - | - | 5239 | 340.2 | - | - | 0 | - |
| - | - | 1.972E+04 | 341.2 | - | - | 0 | - |
| - | - | 4737 | 341.7 | - | - | 0 | - |
| - | - | 2463 | 342.2 | - | - | 0 | - |
| - | - | 608 | 342.2 | - | - | 0 | - |
| - | - | 8700 | 343.2 | - | - | 0 | - |
| - | - | 597.1 | 343.4 | - | - | 0 | - |
| - | - | 1669 | 344.2 | - | - | 0 | - |
| - | - | 1769 | 349.2 | - | - | 0 | - |
| - | - | 1022 | 350.2 | - | - | 0 | - |
| - | - | 1684 | 351.2 | - | - | 0 | - |
| - | - | 951.5 | 351.2 | - | - | 0 | - |
| - | - | 681.3 | 352.2 | - | - | 0 | - |
| 4 | b | 3.326E+04 | 353.2 | 0.0004487 | 1.27 | +1 | 4 |
| - | - | 7026 | 354.2 | - | - | 0 | - |
| - | - | 1522 | 355.2 | - | - | 0 | - |
| - | - | 6594 | 355.2 | - | - | 0 | - |
| - | - | 889.5 | 356.2 | - | - | 0 | - |
| - | - | 838.3 | 358.2 | - | - | 0 | - |
| - | - | 7187 | 359.2 | - | - | 0 | - |
| - | - | 1068 | 360.2 | - | - | 0 | - |
| 2 | y | 1.338E+05 | 363.2 | 0.0003141 | 0.8647 | +2 | 7 |
| - | - | 4.803E+04 | 363.7 | - | - | 0 | - |
| - | - | 1.384E+04 | 364.2 | - | - | 0 | - |
| - | - | 950.5 | 364.7 | - | - | 0 | - |
| - | - | 2588 | 365.2 | - | - | 0 | - |
| - | - | 1143 | 366.2 | - | - | 0 | - |
| - | - | 1665 | 367.2 | - | - | 0 | - |
| - | - | 1972 | 367.2 | - | - | 0 | - |
| 6 | y | 1.776E+04 | 368.2 | 0.0002003 | 0.5438 | +1 | 3 |
| - | - | 2638 | 369.2 | - | - | 0 | - |
| 4 | b | 3801 | 371.2 | 4.632E-05 | 0.1248 | +1 | 4 |
| 2 | y | 2286 | 372.2 | 0.001379 | 3.706 | +2 | 7 |
| - | - | 795 | 372.7 | - | - | 0 | - |
| - | - | 1746 | 373.2 | - | - | 0 | - |
| - | - | 1193 | 377.2 | - | - | 0 | - |
| - | - | 1200 | 379.2 | - | - | 0 | - |
| - | - | 976.7 | 380.2 | - | - | 0 | - |
| - | - | 936 | 382.2 | - | - | 0 | - |
| - | - | 856.7 | 382.7 | - | - | 0 | - |
| - | - | 8610 | 383.2 | - | - | 0 | - |
| - | - | 2091 | 384.2 | - | - | 0 | - |
| 6 | y | 6.772E+04 | 385.3 | 0.000354 | 0.919 | +1 | 3 |
| - | - | 1.245E+04 | 386.3 | - | - | 0 | - |
| - | - | 1310 | 387.3 | - | - | 0 | - |
| - | - | 2172 | 389.2 | - | - | 0 | - |
| - | - | 1433 | 390.2 | - | - | 0 | - |
| - | - | 973.5 | 390.7 | - | - | 0 | - |
| - | - | 3.162E+04 | 391.2 | - | - | 0 | - |
| - | - | 1.332E+04 | 391.7 | - | - | 0 | - |
| - | - | 2573 | 392.2 | - | - | 0 | - |
| - | - | 1518 | 393.2 | - | - | 0 | - |
| - | - | 792 | 394.2 | - | - | 0 | - |
| - | - | 1114 | 395.2 | - | - | 0 | - |
| - | - | 633.1 | 397.2 | - | - | 0 | - |
| - | - | 2541 | 401.2 | - | - | 0 | - |
| - | - | 1847 | 403.7 | - | - | 0 | - |
| - | - | 1273 | 403.8 | - | - | 0 | - |
| - | - | 726.5 | 404.2 | - | - | 0 | - |
| - | - | 2289 | 408.3 | - | - | 0 | - |
| - | - | 569 | 409.3 | - | - | 0 | - |
| - | - | 678.2 | 410.2 | - | - | 0 | - |
| - | - | 1038 | 412.2 | - | - | 0 | - |
| - | - | 1888 | 412.3 | - | - | 0 | - |
| 0 | Precursor | 1.004E+05 | 412.8 | 0.0006225 | 1.508 | +2 | -1 |
| - | - | 4.281E+04 | 413.3 | - | - | 0 | - |
| - | - | 896.7 | 413.3 | - | - | 0 | - |
| - | - | 1.203E+04 | 413.8 | - | - | 0 | - |
| - | - | 1300 | 414.3 | - | - | 0 | - |
| - | - | 1042 | 420.9 | - | - | 0 | - |
| 0 | Precursor | 1492 | 421.8 | 0.0004061 | 0.9628 | +2 | -1 |
| - | - | 3307 | 422.2 | - | - | 0 | - |
| - | - | 908.6 | 423.2 | - | - | 0 | - |
| - | - | 4600 | 426.3 | - | - | 0 | - |
| - | - | 2916 | 430.3 | - | - | 0 | - |
| - | - | 7732 | 436.3 | - | - | 0 | - |
| - | - | 2747 | 437.3 | - | - | 0 | - |
| - | - | 746.6 | 439.8 | - | - | 0 | - |
| 5 | b | 3739 | 440.3 | 0.001044 | 2.37 | +1 | 5 |
| - | - | 4216 | 442.3 | - | - | 0 | - |
| - | - | 1056 | 443.2 | - | - | 0 | - |
| - | - | 1165 | 443.3 | - | - | 0 | - |
| - | - | 1362 | 444.3 | - | - | 0 | - |
| - | - | 1052 | 452.3 | - | - | 0 | - |
| 5 | y | 1.368E+04 | 454.3 | 0.003501 | 7.707 | +1 | 4 |
| 5 | y | 4915 | 455.3 | 0.003572 | 7.847 | +1 | 4 |
| 5 | b | 1223 | 458.3 | 3.349E-07 | 0.0007309 | +1 | 5 |
| - | - | 918.3 | 464.3 | - | - | 0 | - |
| 5 | y | 7.359E+05 | 472.3 | 0.0007963 | 1.686 | +1 | 4 |
| - | - | 1.762E+05 | 473.3 | - | - | 0 | - |
| - | - | 2.847E+04 | 474.3 | - | - | 0 | - |
| - | - | 1010 | 475.3 | - | - | 0 | - |
| - | - | 1418 | 480.3 | - | - | 0 | - |
| - | - | 7987 | 482.3 | - | - | 0 | - |
| - | - | 1510 | 483.3 | - | - | 0 | - |
| - | - | 1.091E+04 | 486.3 | - | - | 0 | - |
| - | - | 1.474E+04 | 487.3 | - | - | 0 | - |
| - | - | 3350 | 488.3 | - | - | 0 | - |
| - | - | 675.8 | 489.3 | - | - | 0 | - |
| - | - | 837.8 | 498.3 | - | - | 0 | - |
| - | - | 1045 | 508.3 | - | - | 0 | - |
| - | - | 1402 | 509.3 | - | - | 0 | - |
| - | - | 980.6 | 510.3 | - | - | 0 | - |
| - | - | 572.2 | 525.3 | - | - | 0 | - |
| - | - | 980.7 | 529.3 | - | - | 0 | - |
| - | - | 2606 | 535.3 | - | - | 0 | - |
| - | - | 954.9 | 536.3 | - | - | 0 | - |
| - | - | 1313 | 541.3 | - | - | 0 | - |
| 6 | b | 9463 | 553.3 | 0.009448 | 17.07 | +1 | 6 |
| 4 | y | 2975 | 554.3 | 0.009316 | 16.81 | +1 | 5 |
| - | - | 2364 | 557.3 | - | - | 0 | - |
| - | - | 4258 | 558.3 | - | - | 0 | - |
| - | - | 778.6 | 559.3 | - | - | 0 | - |
| 4 | y | 2.597E+05 | 571.4 | 0.0004671 | 0.8176 | +1 | 5 |
| - | - | 7.572E+04 | 572.4 | - | - | 0 | - |
| - | - | 947.5 | 572.4 | - | - | 0 | - |
| - | - | 659.5 | 573.3 | - | - | 0 | - |
| - | - | 1.298E+04 | 573.4 | - | - | 0 | - |
| - | - | 4495 | 581.3 | - | - | 0 | - |
| - | - | 886.1 | 582.3 | - | - | 0 | - |
| - | - | 1284 | 593.3 | - | - | 0 | - |
| - | - | 4217 | 597.3 | - | - | 0 | - |
| - | - | 976 | 597.4 | - | - | 0 | - |
| - | - | 989.6 | 598.3 | - | - | 0 | - |
| - | - | 679.7 | 610.4 | - | - | 0 | - |
| - | - | 9915 | 611.4 | - | - | 0 | - |
| - | - | 2571 | 612.4 | - | - | 0 | - |
| - | - | 939.7 | 613.4 | - | - | 0 | - |
| - | - | 2.395E+04 | 628.4 | - | - | 0 | - |
| - | - | 7188 | 629.4 | - | - | 0 | - |
| - | - | 2592 | 630.4 | - | - | 0 | - |
| - | - | 841.1 | 636.4 | - | - | 0 | - |
| - | - | 790.1 | 637.4 | - | - | 0 | - |
| - | - | 2905 | 638.4 | - | - | 0 | - |
| - | - | 1014 | 639.4 | - | - | 0 | - |
| - | - | 1414 | 642.4 | - | - | 0 | - |
| 3 | y | 2.19E+04 | 654.4 | 0.002101 | 3.21 | +1 | 6 |
| 3 | y | 2.478E+04 | 655.4 | 0.009001 | 13.73 | +1 | 6 |
| - | - | 9181 | 656.4 | - | - | 0 | - |
| - | - | 1369 | 657.4 | - | - | 0 | - |
| - | - | 3868 | 664.4 | - | - | 0 | - |
| 3 | y | 5.857E+05 | 672.4 | 0.0005792 | 0.8614 | +1 | 6 |
| - | - | 1.972E+05 | 673.4 | - | - | 0 | - |
| - | - | 4.61E+04 | 674.4 | - | - | 0 | - |
| - | - | 2593 | 675.4 | - | - | 0 | - |
| - | - | 4.463E+04 | 682.4 | - | - | 0 | - |
| - | - | 1.653E+04 | 683.4 | - | - | 0 | - |
| - | - | 3391 | 684.4 | - | - | 0 | - |
| - | - | 5920 | 699.4 | - | - | 0 | - |
| - | - | 1802 | 700.4 | - | - | 0 | - |
| - | - | 1032 | 707.4 | - | - | 0 | - |
| - | - | 1114 | 708.4 | - | - | 0 | - |
| 2 | y | 7.301E+04 | 725.4 | 2.998E-05 | 0.04133 | +1 | 7 |
| - | - | 2.579E+04 | 726.4 | - | - | 0 | - |
| - | - | 5633 | 727.4 | - | - | 0 | - |
| - | - | 765.5 | 728.4 | - | - | 0 | - |
| - | - | 2217 | 735.4 | - | - | 0 | - |
| - | - | 990.3 | 736.4 | - | - | 0 | - |
| 2 | y | 2.379E+05 | 743.4 | 3.559E-05 | 0.04787 | +1 | 7 |
| - | - | 8.842E+04 | 744.4 | - | - | 0 | - |
| - | - | 2.216E+04 | 745.4 | - | - | 0 | - |
| - | - | 1330 | 746.4 | - | - | 0 | - |
| - | - | 4460 | 753.4 | - | - | 0 | - |
| - | - | 2029 | 754.4 | - | - | 0 | - |
| - | - | 699.9 | 781.5 | - | - | 0 | - |
| - | - | 588 | 1004 | - | - | 0 | - |
| - | - | 674.1 | 1372 | - | - | 0 | - |
| - | - | 646.2 | 2229 | - | - | 0 | - |
| - | - | 702.4 | 2999 | - | - | 0 | - |

m/z Charge Intensity FragmentType MassShift Position
120.0809097290039 0 392.79962
125.07170867919922 0 380.34924
125.1077651977539 0 1582.9401
126.09170532226562 0 595.59216
126.12799072265625 0 1371.767
127.08695983886719 0 7858.9
127.12336730957031 0 7875.919
128.1073760986328 0 14121.86
129.0186309814453 0 763.0953
129.10263061523438 0 15376.019
129.11419677734375 0 1399.8833
130.05026245117188 0 2219.2246
130.08660888671875 0 3647.5178
130.097900390625 0 2239.4944
130.10595703125 0 1306.0591
131.0900421142578 0 544.19525
133.0975799560547 0 2318.267
136.02194213867188 0 704.9318
138.09178161621094 0 5829.6973
140.02865600585938 0 486.2809
140.08226013183594 0 44733.445
141.07957458496094 0 855.63025
141.08544921875 0 2885.655
141.10263061523438 0 11558.851
142.1003875732422 0 429.12378
142.10589599609375 0 1854.0898
143.1183624267578 0 283898.9 a 1
144.06558227539062 0 534.4227
144.1154022216797 0 1930.2141
144.12164306640625 0 18124.78
145.0975341796875 0 16311.938
145.1240692138672 0 597.67285
146.1014862060547 0 747.1341
147.11314392089844 0 2799.4426
148.11656188964844 0 920.36755
148.95468139648438 0 1098.8783
149.02369689941406 0 6523.177
150.02708435058594 0 3066.3616
155.08192443847656 0 9690.742
155.11825561523438 0 35283.773
156.08560180664062 0 535.08826
156.11488342285156 0 782.7624
156.12173461914062 0 2975.8635
157.0974884033203 0 3649.1372
157.1088104248047 0 6280.3516
157.13392639160156 0 19772.867
157.14117431640625 0 1065.3369
158.09280395507812 0 36821.605 y Ammonia loss 7
158.1372833251953 0 6049.8906
159.0961456298828 0 1942.5028
159.1131591796875 0 5952.513
159.1396026611328 0 441.95465
160.1163787841797 0 474.85434
165.1028594970703 0 1134.7163
167.08189392089844 0 2229.9573
167.11807250976562 0 1227.7953
169.09756469726562 0 12744.939
169.13429260253906 0 1132.3068
170.1009979248047 0 2376.959
171.11328125 0 249843.03 b 1
172.10984802246094 0 1209.2225
172.1166229248047 0 20313.498
173.09246826171875 0 39970.57
173.12889099121094 0 293439.62
174.0960693359375 0 2571.538
174.13223266601562 0 22714.898
175.1193389892578 0 82618.47 y 7
176.11618041992188 0 573.8212
176.12286376953125 0 5591.5615
181.09725952148438 0 763.5625
181.13369750976562 0 567.0927
183.1132049560547 0 60332.375
183.14959716796875 0 12576.758
184.11660766601562 0 5075.372
184.1529541015625 0 3249.436
185.0943603515625 0 1025.2163
185.103759765625 0 6550.12
185.12892150878906 0 6615.5166
186.1321258544922 0 1407.5437
187.10809326171875 0 16956.451
187.14468383789062 0 577.4672
188.111572265625 0 1875.4692
189.1240692138672 0 1235.5653
192.11361694335938 0 1330.0142
193.09683227539062 0 957.4745
193.133544921875 0 1458.266 y 5
194.12924194335938 0 1651.9541
195.11312866210938 0 5662.961
197.09243774414062 0 1013.3439
197.12893676757812 0 627.8539
197.16490173339844 0 1021.0014
199.1082000732422 0 1230.6627
199.1444549560547 0 4273.109
201.12371826171875 0 109151.68
202.12710571289062 0 10643.406
203.1285858154297 0 794.3317
208.14474487304688 0 1438.3068
209.0927734375 0 1483.0962
209.12876892089844 0 3015.397
210.1241455078125 0 6557.9224
211.10812377929688 0 13642.841
211.12705993652344 0 862.2592
211.14451599121094 0 7183.4756
211.155517578125 0 1827.8851
212.1127166748047 0 1097.2705
212.13978576660156 0 32468.373
213.12374877929688 0 6676.15
213.1432647705078 0 2442.892
213.16050720214844 0 776.7741
215.10398864746094 0 656.9535
215.13905334472656 0 1335.7635
217.0826416015625 0 1377.4326
222.1243896484375 0 1408.0679
223.1444549560547 0 710.03156
224.13983154296875 0 2052.6277
225.159912109375 0 1405.3007
226.11924743652344 0 3433.5186
226.1553955078125 0 41434.035 a Water loss 2
227.10255432128906 0 1723.259
227.13943481445312 0 19654.52
227.1590118408203 0 4581.515
228.13467407226562 0 11484.383 y Ammonia loss 4
228.146240234375 0 931.50684
229.11854553222656 0 3806.3804
229.13821411132812 0 994.69354
230.15037536621094 0 15905.741 d 2
231.1537628173828 0 2004.3094
234.14688110351562 0 903.1529
236.13938903808594 0 973.01166
237.1349639892578 0 3070.306
237.1603240966797 0 744.439
238.11936950683594 0 1838.2167
239.1392364501953 0 971.9917
240.13499450683594 0 2818.4626
241.13809204101562 0 804.234
241.1555938720703 0 983.64453
242.1141815185547 0 1964.8812
243.13430786132812 0 1841.7076
243.650390625 0 843.0845
244.12954711914062 0 4253.579
244.15187072753906 0 1237.6046
244.1662139892578 0 2938.7422 a 2
252.1347198486328 0 4278.366
254.15037536621094 0 118793.39 b Water loss 2
255.14564514160156 0 278817.8 y Ammonia loss 6
256.1488037109375 0 30229.234
257.1514587402344 0 1992.7437
258.145263671875 0 7279.8784
259.1484375 0 2289.8901
260.1612854003906 0 729.34076
264.17181396484375 0 1170.9888
264.20745849609375 0 547.79065
266.1500549316406 0 1990.595
267.15325927734375 0 616.9047
268.1294250488281 0 611.75586
268.1658935546875 0 1415.1886
270.1451721191406 0 41023.688
271.1484375 0 5713.988
271.1768798828125 0 2347.504
272.1253967285156 0 2116.8438
272.1600036621094 0 43924.613 b 2
272.1722106933594 0 118235.43 y 6
273.1626281738281 0 4775.6274
273.1754455566406 0 12629.438
274.1776123046875 0 1154.9972
276.1556091308594 0 5139.495
277.1587219238281 0 1689.9819
277.1769104003906 0 2395.8232 y Water loss 3
277.67718505859375 0 1305.097
280.1295471191406 0 3805.1375
280.16607666015625 0 2042.4138
282.14508056640625 0 1196.3845
282.1815490722656 0 11263.512
283.1850280761719 0 1383.7389
284.1615295410156 0 928.5657
286.18206787109375 0 9766.426 y 3
286.6838073730469 0 3305.0908
288.1558532714844 0 35455.535
289.15887451171875 0 4827.0127
290.1605529785156 0 735.29425
294.1823425292969 0 667.40515
294.2169189453125 0 706.58417
296.16156005859375 0 2141.678
297.155517578125 0 1025.7615
297.17401123046875 0 1707.4843
297.6750793457031 0 813.9551
298.14019775390625 0 5614.8203
298.17742919921875 0 1632.4183
299.1895751953125 0 3177.7302
299.2084655761719 0 2299.8213
299.6908264160156 0 1066.2479
300.1923522949219 0 2487.2324
300.21087646484375 0 755.97235
305.6875 0 819.5873
306.1482849121094 0 1004.01465
306.17974853515625 0 17747.348
306.2298278808594 0 1351.3828
306.68133544921875 0 4854.094
307.1834411621094 0 889.0625
308.1976013183594 0 1592.4235
310.17877197265625 0 1113.3009
310.6803894042969 0 2016.5531
311.1805725097656 0 836.7577
311.2088623046875 0 1179.2262
312.192138671875 0 8752.112
313.1901550292969 0 1566.5273
314.1718444824219 0 1323.6417
315.16900634765625 0 620.2131
318.69580078125 0 1465.7239
319.68536376953125 0 3016.474
320.1619567871094 0 815.4154
320.1862487792969 0 1438.6799
321.1922912597656 0 1108.2252
322.2131042480469 0 1252.1711
323.1720275878906 0 6957.303
323.208251953125 0 1413.977
323.2569580078125 0 770.8643
324.17254638671875 0 1471.1892
325.22406005859375 0 52410.508
326.2270812988281 0 9067.116
327.203857421875 0 1634.8916
327.2298889160156 0 1599.7656
327.70074462890625 0 107014.74 y Water loss 2
328.200439453125 0 44940.016
328.7005920410156 0 12067.068
329.1995849609375 0 1617.7078
331.1976013183594 0 702.0398
332.6927185058594 0 4033.6838
333.19390869140625 0 1225.659
336.7061462402344 0 13389.37 y 2
337.1871337890625 0 2537.8684
337.2077941894531 0 6172.237
337.7106628417969 0 1158.9019
339.20263671875 0 1509.7922
340.1873779296875 0 5239.0933
341.1830749511719 0 19724.15
341.6985168457031 0 4737.323
342.1846008300781 0 2462.68
342.2037048339844 0 607.9908
343.2344970703125 0 8700.396
343.3801574707031 0 597.1218
344.2373962402344 0 1668.8789
349.2347412109375 0 1769.478
350.21844482421875 0 1021.87286
351.2035827636719 0 1683.5009
351.23870849609375 0 951.5256
352.243408203125 0 681.28613
353.2187805175781 0 33262.75 b Water loss 3
354.22003173828125 0 7025.95
355.1983642578125 0 1522.3684
355.2342529296875 0 6594.4263
356.2368469238281 0 889.5419
358.1957092285156 0 838.2915
359.1933898925781 0 7186.5825
360.19482421875 0 1067.8455
363.21917724609375 0 133825.33 y Water loss 1
363.72064208984375 0 48029.39
364.221923828125 0 13836.163
364.7244873046875 0 950.4657
365.2188415527344 0 2588.143
366.2207336425781 0 1143.0242
367.197998046875 0 1664.9519
367.2454833984375 0 1972.0647
368.22943115234375 0 17762.121 y Ammonia loss 5
369.23248291015625 0 2637.7092
371.22894287109375 0 3800.6846 b 3
372.22552490234375 0 2285.642 y 1
372.72479248046875 0 795.0499
373.24493408203125 0 1745.5942
377.2171325683594 0 1193.219
379.2335205078125 0 1199.9545
380.23675537109375 0 976.6745
382.22491455078125 0 936.0498
382.72723388671875 0 856.7319
383.22930908203125 0 8610.443
384.2323303222656 0 2090.5178
385.2561340332031 0 67721.69 y 5
386.25921630859375 0 12451.354
387.2608642578125 0 1309.9578
389.23736572265625 0 2171.591
390.2435302734375 0 1433.3145
390.7408142089844 0 973.495
391.2325134277344 0 31618.21
391.73388671875 0 13323.2295
392.23443603515625 0 2572.7395
393.2142028808594 0 1518.0349
394.2447814941406 0 792.02496
395.2333068847656 0 1113.5481
397.2459716796875 0 633.05237
401.2406005859375 0 2540.618
403.7476806640625 0 1846.6567
403.813720703125 0 1273.4832
404.2483215332031 0 726.5238
408.2611389160156 0 2288.8982
409.2561340332031 0 569.0054
410.2398376464844 0 678.2392
412.2273864746094 0 1037.7524
412.25579833984375 0 1887.6011
412.7536926269531 0 100394.91 Precursor Water loss
413.255126953125 0 42814.36
413.2876892089844 0 896.656
413.7560119628906 0 12028.017
414.2606201171875 0 1299.8958
420.8584899902344 0 1042.3763
421.7587585449219 0 1492.107 Precursor
422.2404479980469 0 3307.204
423.2424011230469 0 908.58325
426.27191162109375 0 4599.749
430.2665710449219 0 2916.1277
436.2560729980469 0 7732.2563
437.2552490234375 0 2747.1357
439.8438415527344 0 746.61395
440.25140380859375 0 3739.288 b Water loss 4
442.27801513671875 0 4216.366
443.248779296875 0 1055.9922
443.2822570800781 0 1165.0022
444.2818603515625 0 1362.1456
452.25830078125 0 1051.6362
454.27374267578125 0 13675.484 y Water loss 4
455.26483154296875 0 4914.6636 y Ammonia loss 4
458.26092529296875 0 1223.1832 b 4
464.2602233886719 0 918.3165
472.2886047363281 0 735942.25 y 4
473.2913818359375 0 176197.34
474.2936706542969 0 28467.295
475.294677734375 0 1009.72754
480.2822570800781 0 1418.1986
482.2730712890625 0 7987.43
483.2757263183594 0 1510.085
486.2928466796875 0 10909.698
487.2959899902344 0 14735.292
488.2974548339844 0 3349.9126
489.30224609375 0 675.8352
498.29876708984375 0 837.8086
508.2775573730469 0 1045.2802
509.2803649902344 0 1401.8488
510.2867431640625 0 980.64404
525.3408203125 0 572.20966
529.3365478515625 0 980.69464
535.3255615234375 0 2605.5442
536.3240966796875 0 954.8728
541.3450927734375 0 1313.155
553.3438720703125 0 9463.26 b Water loss 5
554.3389892578125 0 2975.0967 y Ammonia loss 3
557.3292236328125 0 2363.9653
558.3323364257812 0 4257.711
559.33203125 0 778.5939
571.356689453125 0 259712.34 y 3
572.3594360351562 0 75719.03
572.4141845703125 0 947.47504
573.3109130859375 0 659.4673
573.3623657226562 0 12982.855
581.3408813476562 0 4495.176
582.340087890625 0 886.0546
593.3385620117188 0 1284.2699
597.3345947265625 0 4216.5635
597.3809204101562 0 975.98303
598.33349609375 0 989.58984
610.3653564453125 0 679.7302
611.3514404296875 0 9914.896
612.3539428710938 0 2570.8748
613.3574829101562 0 939.7022
628.3778686523438 0 23945.855
629.3812255859375 0 7187.763
630.3831787109375 0 2592.3552
636.3843994140625 0 841.1328
637.3690795898438 0 790.0901
638.3607788085938 0 2904.7556
639.3682250976562 0 1013.92975
642.39306640625 0 1414.2211
654.3912353515625 0 21901.959 y Water loss 2
655.3863525390625 0 24783.293 y Ammonia loss 2
656.3887329101562 0 9180.83
657.3897094726562 0 1369.1348
664.37744140625 0 3867.99
672.4044799804688 0 585725.1 y 2
673.406982421875 0 197199.17
674.409423828125 0 46100.523
675.4116821289062 0 2593.4172
682.3883056640625 0 44627.29
683.3914184570312 0 16530.453
684.3929443359375 0 3391.1353
699.4139404296875 0 5919.821
700.416259765625 0 1801.7294
707.4197387695312 0 1031.6882
708.4078369140625 0 1114.2362
725.430419921875 0 73005.805 y Water loss 1
726.4326782226562 0 25785.795
727.4345703125 0 5633.1875
728.4365844726562 0 765.4718
735.4127197265625 0 2217.4336
736.4182739257812 0 990.26483
743.4409790039062 0 237881.1 y 1
744.44384765625 0 88418.516
745.4466552734375 0 22156.023
746.4481201171875 0 1330.3708
753.4249267578125 0 4459.8906
754.4302368164062 0 2028.9369
781.4544677734375 0 699.85626
1004.3082275390625 0 587.98517
1371.819091796875 0 674.07806
2228.572509765625 0 646.19226
2999.07275390625 0 702.4492

Spectrum Details

|  |  |
| --- | --- |
| Matched peaks? Matched peaks The total absolute number of peaks matched. Additionally in brackets the total fraction of peaks matched and the total number of peaks is shown. | 38 (9.43% of 403) |
| FDR? FDR The false discovery rate estimated for this peptide. It is calculated by matching all theoretical fragments with a non-integer shift with the raw peaks for this spectrum. This is done with 40 different shifts. The resulting percentage is the average number of annotated peaks over the number of annotated peaks with the correct spectrum. | 0.06% |
| Satellite FDR? Satellite FDR See the FDR for details on its calculation. This satellite ion specific FDR only contains the satellite ions (d/w) for I/L/J positions. | - |
| PSM Score? PSM Score The PSM Score as given by Hecklib to this annotated spectrum. It is shown with three significant figures. | 467 |

#### Spectrum 5133? Spectrum 5133 The raw spectrum of this peptide as annotated by Hecklib. The fragments are coloured according to ion type (see legend). Any peaks with a star '\*' as text can be hovered over to see the full details, first the ion type second the mass shift type. By hovering over the amino acids in the peptide or ions in the legend the corresponding peaks are highlighted. By toggling the 'Unassigned' label you can turn the background (unassigned) peaks on or off in the plot. By updating the slider in the Ion legend you can update the spectrum to only show the top X% of the peaks with labels. The top X% means any peak that is within X% of the highest intensity. By dragging in the spectrum you can zoom in to a specific part of the spectrum and use 'Zoom Out' to get back to the original zoom level. The annotation of the spectrum is based on the given sequence in the peptides file and is done with different software so inconsistencies are likely. The peaks are annotated based on the given sequence, with 20 ppm tolerance.

Copy Data

##### Spectrum 5133 (TSV)

###### Preview

```
Loading example...
```

*Click on the button to copy the data to your clipboard.*

Mz MinMz MaxIntensity Max

WidthHeightPeptide font sizePeptide stroke widthSpectrum font sizeSpectrum stroke widthCompact peptide

Ion legend

wxyz

abcd

OtherUnassignedIon^Charge^_Position_Show for top:%

VATVSJPR

01.10e+52.20e+53.30e+54.40e+5

Zoom Out

a^+1^_2_y^+1^_1_b^+1^_2_y^+1^_1_a^+1^_3_y^+2^_4_d^+1^_3_a^+1^_3_b^+1^_3_y^+1^_2_b^+1^_3_y^+1^_2_y^+2^_5_y^+2^_5_y^+2^_6_y^+2^_6_b^+1^_4_y^+2^_7_y^+1^_3_b^+1^_4_y^+2^_7_y^+1^_3_\*b^+1^_5_y^+1^_4_y^+1^_4_y^+1^_4_b^+1^_6_y^+1^_5_y^+1^_5_y^+1^_6_y^+1^_6_y^+1^_6_y^+1^_7_y^+1^_7_

0821164224633284

Fragment Matches Table

Show background peaks

| Position | Ion type | Intensity | mz Theoretical | mz Error (Th) | mz Error (ppm) | Charge | Series Number |
| --- | --- | --- | --- | --- | --- | --- | --- |
| - | - | 579.8 | 120.1 | - | - | 0 | - |
| - | - | 347 | 122.1 | - | - | 0 | - |
| - | - | 371.1 | 122.1 | - | - | 0 | - |
| - | - | 522.8 | 125.1 | - | - | 0 | - |
| - | - | 414.4 | 126.1 | - | - | 0 | - |
| - | - | 350.9 | 126.5 | - | - | 0 | - |
| - | - | 4704 | 127.1 | - | - | 0 | - |
| - | - | 5154 | 127.1 | - | - | 0 | - |
| - | - | 8167 | 128.1 | - | - | 0 | - |
| - | - | 858.2 | 129 | - | - | 0 | - |
| - | - | 1.016E+04 | 129.1 | - | - | 0 | - |
| - | - | 972.6 | 129.1 | - | - | 0 | - |
| - | - | 1677 | 130 | - | - | 0 | - |
| - | - | 3238 | 130.1 | - | - | 0 | - |
| - | - | 1671 | 130.1 | - | - | 0 | - |
| - | - | 674.9 | 130.1 | - | - | 0 | - |
| - | - | 1477 | 133.1 | - | - | 0 | - |
| - | - | 586.2 | 136.1 | - | - | 0 | - |
| - | - | 3701 | 138.1 | - | - | 0 | - |
| - | - | 431.4 | 139.9 | - | - | 0 | - |
| - | - | 2.865E+04 | 140.1 | - | - | 0 | - |
| - | - | 1352 | 141.1 | - | - | 0 | - |
| - | - | 1.006E+04 | 141.1 | - | - | 0 | - |
| - | - | 2159 | 142.1 | - | - | 0 | - |
| 2 | a | 1.793E+05 | 143.1 | 0.0004424 | 3.091 | +1 | 2 |
| - | - | 752.4 | 144.1 | - | - | 0 | - |
| - | - | 1.181E+04 | 144.1 | - | - | 0 | - |
| - | - | 1.077E+04 | 145.1 | - | - | 0 | - |
| - | - | 494.2 | 145.1 | - | - | 0 | - |
| - | - | 449.3 | 146.6 | - | - | 0 | - |
| - | - | 2936 | 147.1 | - | - | 0 | - |
| - | - | 624.7 | 148.1 | - | - | 0 | - |
| - | - | 623 | 148.8 | - | - | 0 | - |
| - | - | 462.1 | 148.9 | - | - | 0 | - |
| - | - | 568.8 | 148.9 | - | - | 0 | - |
| - | - | 516.8 | 148.9 | - | - | 0 | - |
| - | - | 689.7 | 148.9 | - | - | 0 | - |
| - | - | 651.5 | 148.9 | - | - | 0 | - |
| - | - | 763.3 | 148.9 | - | - | 0 | - |
| - | - | 1140 | 148.9 | - | - | 0 | - |
| - | - | 1035 | 148.9 | - | - | 0 | - |
| - | - | 1347 | 148.9 | - | - | 0 | - |
| - | - | 3508 | 148.9 | - | - | 0 | - |
| - | - | 5943 | 149 | - | - | 0 | - |
| - | - | 3969 | 149 | - | - | 0 | - |
| - | - | 1488 | 149 | - | - | 0 | - |
| - | - | 1224 | 149 | - | - | 0 | - |
| - | - | 1019 | 149 | - | - | 0 | - |
| - | - | 986 | 149 | - | - | 0 | - |
| - | - | 666.9 | 149 | - | - | 0 | - |
| - | - | 554.1 | 149 | - | - | 0 | - |
| - | - | 6619 | 149 | - | - | 0 | - |
| - | - | 3786 | 150 | - | - | 0 | - |
| - | - | 447 | 150.1 | - | - | 0 | - |
| - | - | 402.4 | 151 | - | - | 0 | - |
| - | - | 669.9 | 153.1 | - | - | 0 | - |
| - | - | 487.3 | 155 | - | - | 0 | - |
| - | - | 5765 | 155.1 | - | - | 0 | - |
| - | - | 2.225E+04 | 155.1 | - | - | 0 | - |
| - | - | 479.6 | 156.1 | - | - | 0 | - |
| - | - | 1157 | 156.1 | - | - | 0 | - |
| - | - | 783 | 157 | - | - | 0 | - |
| - | - | 2713 | 157.1 | - | - | 0 | - |
| - | - | 3679 | 157.1 | - | - | 0 | - |
| - | - | 2.069E+04 | 157.1 | - | - | 0 | - |
| 8 | y | 2.357E+04 | 158.1 | 0.0003551 | 2.246 | +1 | 1 |
| - | - | 1131 | 158.1 | - | - | 0 | - |
| - | - | 797.7 | 158.1 | - | - | 0 | - |
| - | - | 6552 | 158.1 | - | - | 0 | - |
| - | - | 1375 | 159.1 | - | - | 0 | - |
| - | - | 3356 | 159.1 | - | - | 0 | - |
| - | - | 885.8 | 165.1 | - | - | 0 | - |
| - | - | 915.7 | 165.1 | - | - | 0 | - |
| - | - | 1068 | 167.1 | - | - | 0 | - |
| - | - | 988.4 | 167.1 | - | - | 0 | - |
| - | - | 1.28E+04 | 169.1 | - | - | 0 | - |
| - | - | 835.1 | 169.1 | - | - | 0 | - |
| - | - | 2607 | 170.1 | - | - | 0 | - |
| 2 | b | 1.533E+05 | 171.1 | 0.0004313 | 2.521 | +1 | 2 |
| - | - | 839.1 | 172.1 | - | - | 0 | - |
| - | - | 1.164E+04 | 172.1 | - | - | 0 | - |
| - | - | 2.494E+04 | 173.1 | - | - | 0 | - |
| - | - | 1.778E+05 | 173.1 | - | - | 0 | - |
| - | - | 1051 | 173.4 | - | - | 0 | - |
| - | - | 1328 | 174.1 | - | - | 0 | - |
| - | - | 1.449E+04 | 174.1 | - | - | 0 | - |
| 8 | y | 5.347E+04 | 175.1 | 0.0003563 | 2.035 | +1 | 1 |
| - | - | 3536 | 176.1 | - | - | 0 | - |
| - | - | 902.4 | 181.1 | - | - | 0 | - |
| - | - | 3.68E+04 | 183.1 | - | - | 0 | - |
| - | - | 1.045E+04 | 183.1 | - | - | 0 | - |
| - | - | 3539 | 184.1 | - | - | 0 | - |
| - | - | 4228 | 184.2 | - | - | 0 | - |
| - | - | 530.7 | 185.1 | - | - | 0 | - |
| - | - | 4564 | 185.1 | - | - | 0 | - |
| - | - | 4963 | 185.1 | - | - | 0 | - |
| - | - | 2081 | 186.1 | - | - | 0 | - |
| - | - | 1.097E+04 | 187.1 | - | - | 0 | - |
| - | - | 573.7 | 187.2 | - | - | 0 | - |
| - | - | 1074 | 188.1 | - | - | 0 | - |
| - | - | 1276 | 192.1 | - | - | 0 | - |
| - | - | 1121 | 194.1 | - | - | 0 | - |
| - | - | 3005 | 195.1 | - | - | 0 | - |
| - | - | 551.5 | 197.1 | - | - | 0 | - |
| - | - | 700.5 | 197.2 | - | - | 0 | - |
| - | - | 476.6 | 198.4 | - | - | 0 | - |
| - | - | 595.7 | 199.1 | - | - | 0 | - |
| - | - | 2320 | 199.1 | - | - | 0 | - |
| - | - | 6.606E+04 | 201.1 | - | - | 0 | - |
| - | - | 6368 | 202.1 | - | - | 0 | - |
| - | - | 1010 | 208.1 | - | - | 0 | - |
| - | - | 1582 | 209.1 | - | - | 0 | - |
| - | - | 2691 | 209.1 | - | - | 0 | - |
| - | - | 4342 | 210.1 | - | - | 0 | - |
| - | - | 8705 | 211.1 | - | - | 0 | - |
| - | - | 4974 | 211.1 | - | - | 0 | - |
| - | - | 1442 | 211.2 | - | - | 0 | - |
| - | - | 610.5 | 212.1 | - | - | 0 | - |
| - | - | 1.881E+04 | 212.1 | - | - | 0 | - |
| - | - | 4216 | 213.1 | - | - | 0 | - |
| - | - | 1783 | 213.1 | - | - | 0 | - |
| - | - | 956 | 215.1 | - | - | 0 | - |
| - | - | 859.6 | 223.1 | - | - | 0 | - |
| - | - | 685.1 | 224.1 | - | - | 0 | - |
| - | - | 664.9 | 225.1 | - | - | 0 | - |
| - | - | 868.8 | 225.2 | - | - | 0 | - |
| - | - | 2163 | 226.1 | - | - | 0 | - |
| 3 | a | 2.309E+04 | 226.2 | 0.0003311 | 1.464 | +1 | 3 |
| - | - | 773.3 | 227.1 | - | - | 0 | - |
| - | - | 1.058E+04 | 227.1 | - | - | 0 | - |
| - | - | 1981 | 227.2 | - | - | 0 | - |
| 5 | y | 5344 | 228.1 | 0.0004215 | 1.847 | +2 | 4 |
| - | - | 1282 | 229.1 | - | - | 0 | - |
| 3 | d | 1.126E+04 | 230.1 | 0.0004116 | 1.789 | +1 | 3 |
| - | - | 668.9 | 231.2 | - | - | 0 | - |
| - | - | 605.4 | 237.1 | - | - | 0 | - |
| - | - | 1244 | 237.1 | - | - | 0 | - |
| - | - | 1285 | 238.1 | - | - | 0 | - |
| - | - | 1997 | 240.1 | - | - | 0 | - |
| - | - | 684.3 | 241.1 | - | - | 0 | - |
| - | - | 733.9 | 241.2 | - | - | 0 | - |
| - | - | 871.1 | 243.1 | - | - | 0 | - |
| - | - | 1427 | 243.7 | - | - | 0 | - |
| - | - | 1645 | 244.1 | - | - | 0 | - |
| - | - | 943.8 | 244.2 | - | - | 0 | - |
| 3 | a | 1161 | 244.2 | 0.0006765 | 2.771 | +1 | 3 |
| - | - | 478.7 | 247.1 | - | - | 0 | - |
| - | - | 2012 | 252.1 | - | - | 0 | - |
| - | - | 529.4 | 253.9 | - | - | 0 | - |
| - | - | 1769 | 254.1 | - | - | 0 | - |
| 3 | b | 6.805E+04 | 254.1 | 0.0003506 | 1.38 | +1 | 3 |
| - | - | 1311 | 254.2 | - | - | 0 | - |
| 7 | y | 1.593E+05 | 255.1 | 0.0004172 | 1.635 | +1 | 2 |
| - | - | 984.6 | 256.1 | - | - | 0 | - |
| - | - | 1.954E+04 | 256.1 | - | - | 0 | - |
| - | - | 1713 | 257.2 | - | - | 0 | - |
| - | - | 6174 | 258.1 | - | - | 0 | - |
| - | - | 2823 | 259.1 | - | - | 0 | - |
| - | - | 1768 | 266.2 | - | - | 0 | - |
| - | - | 937.8 | 267.2 | - | - | 0 | - |
| - | - | 563.9 | 269.2 | - | - | 0 | - |
| - | - | 2.497E+04 | 270.1 | - | - | 0 | - |
| - | - | 2847 | 271.1 | - | - | 0 | - |
| - | - | 1831 | 271.2 | - | - | 0 | - |
| - | - | 1089 | 272.1 | - | - | 0 | - |
| 3 | b | 2.596E+04 | 272.2 | 0.0005095 | 1.872 | +1 | 3 |
| 7 | y | 7.275E+04 | 272.2 | 0.0004336 | 1.593 | +1 | 2 |
| - | - | 4176 | 273.2 | - | - | 0 | - |
| - | - | 7623 | 273.2 | - | - | 0 | - |
| - | - | 5789 | 276.2 | - | - | 0 | - |
| - | - | 2458 | 277.2 | - | - | 0 | - |
| 4 | y | 1485 | 277.2 | 0.0004739 | 1.71 | +2 | 5 |
| - | - | 2723 | 280.1 | - | - | 0 | - |
| - | - | 1666 | 280.2 | - | - | 0 | - |
| - | - | 692.3 | 282.1 | - | - | 0 | - |
| - | - | 7276 | 282.2 | - | - | 0 | - |
| - | - | 1081 | 283.2 | - | - | 0 | - |
| - | - | 574.7 | 284.2 | - | - | 0 | - |
| 4 | y | 6424 | 286.2 | 0.000288 | 1.006 | +2 | 5 |
| - | - | 1733 | 286.7 | - | - | 0 | - |
| - | - | 2.171E+04 | 288.2 | - | - | 0 | - |
| - | - | 3097 | 289.2 | - | - | 0 | - |
| - | - | 896 | 291 | - | - | 0 | - |
| - | - | 729.9 | 293.1 | - | - | 0 | - |
| - | - | 598.4 | 294.2 | - | - | 0 | - |
| - | - | 875.1 | 296.2 | - | - | 0 | - |
| - | - | 867.7 | 297.2 | - | - | 0 | - |
| - | - | 1753 | 297.2 | - | - | 0 | - |
| - | - | 4044 | 298.1 | - | - | 0 | - |
| - | - | 859.6 | 298.2 | - | - | 0 | - |
| - | - | 552.3 | 299.1 | - | - | 0 | - |
| - | - | 2687 | 299.2 | - | - | 0 | - |
| - | - | 908.6 | 299.2 | - | - | 0 | - |
| - | - | 1163 | 300.2 | - | - | 0 | - |
| - | - | 620 | 305.7 | - | - | 0 | - |
| - | - | 8542 | 306.2 | - | - | 0 | - |
| - | - | 2036 | 306.7 | - | - | 0 | - |
| - | - | 573.5 | 308 | - | - | 0 | - |
| - | - | 1553 | 308.2 | - | - | 0 | - |
| - | - | 1062 | 310.2 | - | - | 0 | - |
| - | - | 1319 | 310.7 | - | - | 0 | - |
| - | - | 5717 | 312.2 | - | - | 0 | - |
| - | - | 1175 | 313.2 | - | - | 0 | - |
| - | - | 881 | 314.2 | - | - | 0 | - |
| - | - | 1667 | 319.7 | - | - | 0 | - |
| - | - | 839.1 | 320.2 | - | - | 0 | - |
| - | - | 542.4 | 320.2 | - | - | 0 | - |
| - | - | 3970 | 323.2 | - | - | 0 | - |
| - | - | 732.9 | 323.2 | - | - | 0 | - |
| - | - | 3.182E+04 | 325.2 | - | - | 0 | - |
| - | - | 677.8 | 326.2 | - | - | 0 | - |
| - | - | 5697 | 326.2 | - | - | 0 | - |
| - | - | 591 | 327.2 | - | - | 0 | - |
| 3 | y | 6.288E+04 | 327.7 | 0.0001942 | 0.5926 | +2 | 6 |
| - | - | 3.68E+04 | 328.2 | - | - | 0 | - |
| - | - | 1.18E+04 | 328.7 | - | - | 0 | - |
| - | - | 1655 | 329.2 | - | - | 0 | - |
| - | - | 630.9 | 331.2 | - | - | 0 | - |
| - | - | 1913 | 332.7 | - | - | 0 | - |
| - | - | 650.5 | 332.9 | - | - | 0 | - |
| - | - | 604.2 | 333.2 | - | - | 0 | - |
| 3 | y | 9079 | 336.7 | 0.0003135 | 0.931 | +2 | 6 |
| - | - | 2014 | 337.2 | - | - | 0 | - |
| - | - | 2738 | 337.2 | - | - | 0 | - |
| - | - | 728.6 | 339.2 | - | - | 0 | - |
| - | - | 4418 | 340.2 | - | - | 0 | - |
| - | - | 1.301E+04 | 341.2 | - | - | 0 | - |
| - | - | 2613 | 341.7 | - | - | 0 | - |
| - | - | 1333 | 342.2 | - | - | 0 | - |
| - | - | 6226 | 343.2 | - | - | 0 | - |
| - | - | 1622 | 349.2 | - | - | 0 | - |
| - | - | 736 | 350.2 | - | - | 0 | - |
| - | - | 1220 | 350.9 | - | - | 0 | - |
| - | - | 1686 | 351.2 | - | - | 0 | - |
| - | - | 791.6 | 352.2 | - | - | 0 | - |
| 4 | b | 2.057E+04 | 353.2 | 0.000235 | 0.6654 | +1 | 4 |
| - | - | 4376 | 354.2 | - | - | 0 | - |
| - | - | 1165 | 355.2 | - | - | 0 | - |
| - | - | 2824 | 355.2 | - | - | 0 | - |
| - | - | 894.3 | 358.2 | - | - | 0 | - |
| - | - | 5260 | 359.2 | - | - | 0 | - |
| - | - | 986.2 | 360.2 | - | - | 0 | - |
| 2 | y | 7.858E+04 | 363.2 | 0.000192 | 0.5286 | +2 | 7 |
| - | - | 2.688E+04 | 363.7 | - | - | 0 | - |
| - | - | 7256 | 364.2 | - | - | 0 | - |
| - | - | 1550 | 365.2 | - | - | 0 | - |
| - | - | 754.3 | 367.2 | - | - | 0 | - |
| - | - | 846.5 | 367.2 | - | - | 0 | - |
| 6 | y | 1.003E+04 | 368.2 | 0.0001697 | 0.461 | +1 | 3 |
| - | - | 1988 | 369.2 | - | - | 0 | - |
| 4 | b | 1733 | 371.2 | 0.0002294 | 0.618 | +1 | 4 |
| 2 | y | 1478 | 372.2 | 0.0003113 | 0.8362 | +2 | 7 |
| - | - | 889.9 | 373.2 | - | - | 0 | - |
| - | - | 591.9 | 375.9 | - | - | 0 | - |
| - | - | 863.3 | 379.2 | - | - | 0 | - |
| - | - | 974.6 | 380.2 | - | - | 0 | - |
| - | - | 4661 | 383.2 | - | - | 0 | - |
| - | - | 1466 | 384.2 | - | - | 0 | - |
| 6 | y | 4.1E+04 | 385.3 | 0.0002625 | 0.6813 | +1 | 3 |
| - | - | 8109 | 386.3 | - | - | 0 | - |
| - | - | 945.2 | 387.3 | - | - | 0 | - |
| - | - | 826.3 | 389.2 | - | - | 0 | - |
| - | - | 688.9 | 389.7 | - | - | 0 | - |
| - | - | 1571 | 390.2 | - | - | 0 | - |
| - | - | 647.7 | 390.7 | - | - | 0 | - |
| - | - | 1.803E+04 | 391.2 | - | - | 0 | - |
| - | - | 8592 | 391.7 | - | - | 0 | - |
| - | - | 1439 | 392.2 | - | - | 0 | - |
| - | - | 853.3 | 393.2 | - | - | 0 | - |
| - | - | 634 | 395.2 | - | - | 0 | - |
| - | - | 826.7 | 401.2 | - | - | 0 | - |
| - | - | 1330 | 403.7 | - | - | 0 | - |
| - | - | 1377 | 403.8 | - | - | 0 | - |
| - | - | 614.7 | 404.2 | - | - | 0 | - |
| - | - | 1120 | 408.3 | - | - | 0 | - |
| - | - | 959.8 | 411.2 | - | - | 0 | - |
| 0 | Precursor | 6.138E+04 | 412.8 | 0.0005004 | 1.212 | +2 | -1 |
| - | - | 2.534E+04 | 413.3 | - | - | 0 | - |
| - | - | 8293 | 413.8 | - | - | 0 | - |
| - | - | 1100 | 420.9 | - | - | 0 | - |
| - | - | 1534 | 422.2 | - | - | 0 | - |
| - | - | 2434 | 426.3 | - | - | 0 | - |
| - | - | 968 | 430.3 | - | - | 0 | - |
| - | - | 3506 | 436.3 | - | - | 0 | - |
| - | - | 1825 | 437.3 | - | - | 0 | - |
| - | - | 1143 | 439.8 | - | - | 0 | - |
| 5 | b | 2477 | 440.3 | 0.0001585 | 0.3601 | +1 | 5 |
| - | - | 903.6 | 441.3 | - | - | 0 | - |
| - | - | 2643 | 442.3 | - | - | 0 | - |
| - | - | 817.1 | 444.3 | - | - | 0 | - |
| 5 | y | 8149 | 454.3 | 0.002494 | 5.49 | +1 | 4 |
| 5 | y | 2917 | 455.3 | 0.004122 | 9.053 | +1 | 4 |
| - | - | 675.8 | 456.3 | - | - | 0 | - |
| - | - | 814.2 | 468.3 | - | - | 0 | - |
| - | - | 1123 | 469.3 | - | - | 0 | - |
| 5 | y | 4.358E+05 | 472.3 | 0.0007048 | 1.492 | +1 | 4 |
| - | - | 1.024E+05 | 473.3 | - | - | 0 | - |
| - | - | 1.841E+04 | 474.3 | - | - | 0 | - |
| - | - | 755.8 | 475.3 | - | - | 0 | - |
| - | - | 5224 | 482.3 | - | - | 0 | - |
| - | - | 783.9 | 483.3 | - | - | 0 | - |
| - | - | 1.056E+04 | 486.3 | - | - | 0 | - |
| - | - | 1.642E+04 | 487.3 | - | - | 0 | - |
| - | - | 3202 | 488.3 | - | - | 0 | - |
| - | - | 756.7 | 489.3 | - | - | 0 | - |
| - | - | 890.2 | 508.3 | - | - | 0 | - |
| - | - | 1420 | 509.3 | - | - | 0 | - |
| - | - | 930 | 535.3 | - | - | 0 | - |
| - | - | 668.1 | 541.3 | - | - | 0 | - |
| 6 | b | 4286 | 553.3 | 0.007434 | 13.43 | +1 | 6 |
| 4 | y | 1955 | 554.3 | 0.009865 | 17.8 | +1 | 5 |
| - | - | 1949 | 557.3 | - | - | 0 | - |
| - | - | 4457 | 558.3 | - | - | 0 | - |
| - | - | 789.6 | 559.3 | - | - | 0 | - |
| 4 | y | 1.487E+05 | 571.4 | 0.000284 | 0.4971 | +1 | 5 |
| - | - | 4.209E+04 | 572.4 | - | - | 0 | - |
| - | - | 8845 | 573.4 | - | - | 0 | - |
| - | - | 2504 | 581.3 | - | - | 0 | - |
| - | - | 783.8 | 593.3 | - | - | 0 | - |
| - | - | 2251 | 597.3 | - | - | 0 | - |
| - | - | 776.8 | 598.3 | - | - | 0 | - |
| - | - | 591 | 610 | - | - | 0 | - |
| - | - | 5973 | 611.4 | - | - | 0 | - |
| - | - | 1986 | 612.4 | - | - | 0 | - |
| - | - | 1.332E+04 | 628.4 | - | - | 0 | - |
| - | - | 4561 | 629.4 | - | - | 0 | - |
| - | - | 1771 | 630.4 | - | - | 0 | - |
| - | - | 987.8 | 636.4 | - | - | 0 | - |
| - | - | 971.2 | 637.4 | - | - | 0 | - |
| - | - | 2602 | 638.4 | - | - | 0 | - |
| - | - | 800 | 643.4 | - | - | 0 | - |
| 3 | y | 1.535E+04 | 654.4 | 0.002833 | 4.329 | +1 | 6 |
| 3 | y | 2.357E+04 | 655.4 | 0.008635 | 13.18 | +1 | 6 |
| - | - | 9529 | 656.4 | - | - | 0 | - |
| - | - | 2084 | 657.4 | - | - | 0 | - |
| - | - | 2305 | 664.4 | - | - | 0 | - |
| - | - | 833.7 | 665.4 | - | - | 0 | - |
| 3 | y | 3.423E+05 | 672.4 | 0.0001519 | 0.226 | +1 | 6 |
| - | - | 1.172E+05 | 673.4 | - | - | 0 | - |
| - | - | 2.538E+04 | 674.4 | - | - | 0 | - |
| - | - | 1325 | 675.4 | - | - | 0 | - |
| - | - | 3.033E+04 | 682.4 | - | - | 0 | - |
| - | - | 9137 | 683.4 | - | - | 0 | - |
| - | - | 1772 | 684.4 | - | - | 0 | - |
| - | - | 2520 | 699.4 | - | - | 0 | - |
| - | - | 961.2 | 700.4 | - | - | 0 | - |
| - | - | 674.6 | 708.4 | - | - | 0 | - |
| - | - | 639.4 | 713.4 | - | - | 0 | - |
| 2 | y | 4.008E+04 | 725.4 | 0.0004572 | 0.6303 | +1 | 7 |
| - | - | 1.414E+04 | 726.4 | - | - | 0 | - |
| - | - | 4042 | 727.4 | - | - | 0 | - |
| - | - | 1091 | 735.4 | - | - | 0 | - |
| 2 | y | 1.413E+05 | 743.4 | 0.0001577 | 0.2121 | +1 | 7 |
| - | - | 5.182E+04 | 744.4 | - | - | 0 | - |
| - | - | 1.307E+04 | 745.4 | - | - | 0 | - |
| - | - | 721.4 | 746.4 | - | - | 0 | - |
| - | - | 2256 | 753.4 | - | - | 0 | - |
| - | - | 951.1 | 754.4 | - | - | 0 | - |
| - | - | 649.8 | 952.6 | - | - | 0 | - |
| - | - | 632.5 | 2255 | - | - | 0 | - |
| - | - | 655.2 | 2341 | - | - | 0 | - |
| - | - | 728.5 | 3135 | - | - | 0 | - |
| - | - | 640.7 | 3160 | - | - | 0 | - |
| - | - | 659.8 | 3252 | - | - | 0 | - |

m/z Charge Intensity FragmentType MassShift Position
120.08159637451172 0 579.7669
122.10186004638672 0 347.00305
122.1468505859375 0 371.11646
125.10757446289062 0 522.84406
126.09182739257812 0 414.3719
126.47966766357422 0 350.86972
127.08694458007812 0 4703.864
127.12332916259766 0 5154.3706
128.1073455810547 0 8167.0405
129.01870727539062 0 858.2076
129.10260009765625 0 10163.375
129.11415100097656 0 972.6091
130.04998779296875 0 1677.156
130.08668518066406 0 3238.4119
130.09791564941406 0 1671.1587
130.10614013671875 0 674.9021
133.0976104736328 0 1476.8278
136.0758056640625 0 586.1838
138.09173583984375 0 3700.9285
139.92047119140625 0 431.38678
140.0822296142578 0 28646.326
141.085693359375 0 1352.085
141.1026153564453 0 10063.632
142.10595703125 0 2159.451
143.1183319091797 0 179251.61 a 1
144.11569213867188 0 752.39386
144.12164306640625 0 11814.081
145.09751892089844 0 10771.623
145.12510681152344 0 494.19864
146.57032775878906 0 449.30292
147.11318969726562 0 2935.9934
148.11660766601562 0 624.73895
148.83572387695312 0 623.04895
148.8633270263672 0 462.06012
148.87051391601562 0 568.7698
148.88385009765625 0 516.78485
148.89825439453125 0 689.69763
148.905029296875 0 651.491
148.9119110107422 0 763.2808
148.91893005371094 0 1139.7991
148.9259490966797 0 1034.9258
148.9327392578125 0 1347.0303
148.940185546875 0 3508.4336
148.9561309814453 0 5943.489
148.96368408203125 0 3968.9583
148.97116088867188 0 1487.7993
148.97801208496094 0 1223.7618
148.98471069335938 0 1018.8072
148.9918975830078 0 986.0086
148.9986572265625 0 666.926
149.01296997070312 0 554.1042
149.0237274169922 0 6618.9736
150.0270538330078 0 3786.347
150.0762481689453 0 447.02783
151.02915954589844 0 402.40726
153.13890075683594 0 669.8853
155.0453338623047 0 487.31104
155.08181762695312 0 5765.098
155.11822509765625 0 22247.418
156.0857391357422 0 479.62506
156.12142944335938 0 1157.385
157.01321411132812 0 782.9824
157.09739685058594 0 2712.9268
157.10882568359375 0 3679.2358
157.13389587402344 0 20694.27
158.09275817871094 0 23569.838 y Ammonia loss 7
158.10023498535156 0 1131.1285
158.1305694580078 0 797.7049
158.13734436035156 0 6552.2764
159.0959930419922 0 1375.4628
159.11312866210938 0 3356.0386
165.06651306152344 0 885.807
165.10267639160156 0 915.693
167.08154296875 0 1067.564
167.118408203125 0 988.44403
169.0975341796875 0 12795.121
169.13414001464844 0 835.1475
170.10092163085938 0 2607.0815
171.1132354736328 0 153253.72 b 1
172.1092529296875 0 839.0963
172.11659240722656 0 11635.611
173.09243774414062 0 24944.285
173.12887573242188 0 177847.52
173.4403076171875 0 1050.9763
174.095947265625 0 1327.6559
174.1322479248047 0 14494.742
175.1193084716797 0 53465.375 y 7
176.12265014648438 0 3535.5908
181.09730529785156 0 902.3924
183.11317443847656 0 36800.445
183.1496124267578 0 10451.221
184.11659240722656 0 3539.1316
184.15306091308594 0 4228.1763
185.08053588867188 0 530.69794
185.10365295410156 0 4563.516
185.12881469726562 0 4962.579
186.13214111328125 0 2080.7998
187.10806274414062 0 10971.273
187.1785888671875 0 573.68463
188.11155700683594 0 1073.5132
192.11366271972656 0 1276.127
194.12890625 0 1120.755
195.11312866210938 0 3004.674
197.128662109375 0 551.52625
197.16551208496094 0 700.5306
198.35157775878906 0 476.61487
199.10848999023438 0 595.6959
199.1444549560547 0 2320.4404
201.12368774414062 0 66064.805
202.12709045410156 0 6367.6953
208.14462280273438 0 1009.7552
209.092529296875 0 1582.0815
209.128662109375 0 2691.1372
210.12403869628906 0 4342.437
211.10804748535156 0 8704.622
211.14453125 0 4973.6855
211.15602111816406 0 1441.8794
212.11183166503906 0 610.5429
212.13970947265625 0 18808.15
213.12351989746094 0 4215.783
213.14295959472656 0 1782.5714
215.13958740234375 0 955.97235
223.1439208984375 0 859.6378
224.13885498046875 0 685.0609
225.12368774414062 0 664.9488
225.16000366210938 0 868.83276
226.11891174316406 0 2163.0322
226.15533447265625 0 23085.209 a Water loss 2
227.10269165039062 0 773.2903
227.1394500732422 0 10577.015
227.15896606445312 0 1981.0057
228.1346893310547 0 5343.913 y Ammonia loss 4
229.11831665039062 0 1282.208
230.15032958984375 0 11256.129 d 2
231.15347290039062 0 668.8814
237.12322998046875 0 605.4368
237.1353759765625 0 1244.3342
238.11944580078125 0 1284.9854
240.13490295410156 0 1996.8596
241.13900756835938 0 684.26984
241.15464782714844 0 733.85004
243.13385009765625 0 871.0548
243.65003967285156 0 1426.9854
244.1289825439453 0 1644.6841
244.152099609375 0 943.79224
244.16624450683594 0 1161.1616 a 2
247.06326293945312 0 478.70132
252.13478088378906 0 2012.2054
253.90428161621094 0 529.38165
254.11439514160156 0 1768.8527
254.1502685546875 0 68049.695 b Water loss 2
254.18438720703125 0 1311.0797
255.1455841064453 0 159334.38 y Ammonia loss 6
256.1293029785156 0 984.60126
256.1487121582031 0 19544.238
257.150390625 0 1712.747
258.14520263671875 0 6174.216
259.1483459472656 0 2823.34
266.1502685546875 0 1768.0494
267.1539001464844 0 937.7878
269.1695556640625 0 563.8816
270.1451416015625 0 24969.512
271.14849853515625 0 2847.3726
271.1766052246094 0 1831.4751
272.1258239746094 0 1088.9236
272.15997314453125 0 25961.053 b 2
272.1721496582031 0 72750.94 y 6
273.1630554199219 0 4176.1846
273.175537109375 0 7623.3013
276.1557922363281 0 5788.8906
277.1585388183594 0 2457.5874
277.17694091796875 0 1484.5398 y Water loss 3
280.12969970703125 0 2723.3728
280.1656494140625 0 1665.6547
282.1448059082031 0 692.3375
282.1815185546875 0 7275.8955
283.1852111816406 0 1081.2715
284.16094970703125 0 574.6975
286.1820373535156 0 6423.74 y 3
286.68359375 0 1733.4672
288.15582275390625 0 21714.139
289.1593322753906 0 3096.986
290.9610290527344 0 896.00604
293.1007080078125 0 729.87646
294.1814270019531 0 598.3969
296.1610412597656 0 875.14557
297.1563415527344 0 867.70514
297.1747131347656 0 1753.2832
298.140380859375 0 4044.3806
298.17706298828125 0 859.6078
299.1446533203125 0 552.2564
299.1900329589844 0 2686.9436
299.2079162597656 0 908.6001
300.1913757324219 0 1162.942
305.6864013671875 0 620.049
306.1796569824219 0 8541.831
306.68115234375 0 2036.0933
307.9847106933594 0 573.5196
308.19775390625 0 1553.4497
310.177978515625 0 1062.1764
310.68084716796875 0 1319.4946
312.19232177734375 0 5716.9355
313.1917724609375 0 1175.4279
314.17205810546875 0 880.9546
319.68505859375 0 1666.6135
320.1612243652344 0 839.1458
320.1859436035156 0 542.4137
323.17169189453125 0 3969.8413
323.2086486816406 0 732.8774
325.2239990234375 0 31817.281
326.18231201171875 0 677.82324
326.2273254394531 0 5697.234
327.2057800292969 0 591.0027
327.70050048828125 0 62882 y Water loss 2
328.19952392578125 0 36795.605
328.69940185546875 0 11798.276
329.1998596191406 0 1655.1732
331.1972351074219 0 630.9355
332.69293212890625 0 1912.951
332.8725280761719 0 650.50476
333.192138671875 0 604.2003
336.7059020996094 0 9078.7295 y 2
337.18731689453125 0 2013.7855
337.2073974609375 0 2737.9768
339.2028503417969 0 728.5782
340.1873474121094 0 4417.7285
341.18341064453125 0 13010.11
341.6979064941406 0 2613.1045
342.183349609375 0 1333.0286
343.2343444824219 0 6225.74
349.23443603515625 0 1621.9984
350.2188720703125 0 736.02747
350.92742919921875 0 1219.6205
351.20263671875 0 1685.5052
352.2418212890625 0 791.58563
353.21856689453125 0 20572.873 b Water loss 3
354.22015380859375 0 4376.457
355.19775390625 0 1164.9617
355.23406982421875 0 2824.167
358.1972351074219 0 894.3196
359.19317626953125 0 5260.1875
360.1961364746094 0 986.1736
363.21905517578125 0 78581.984 y Water loss 1
363.720458984375 0 26882.467
364.2217712402344 0 7256.031
365.2185363769531 0 1550.2545
367.19720458984375 0 754.3465
367.2436828613281 0 846.4726
368.2294006347656 0 10032.073 y Ammonia loss 5
369.2330322265625 0 1988.2231
371.2291259765625 0 1732.8273 b 3
372.2244567871094 0 1478.2751 y 1
373.2456970214844 0 889.9409
375.8703308105469 0 591.9059
379.2345275878906 0 863.25226
380.2373046875 0 974.5702
383.229248046875 0 4661.2363
384.2327880859375 0 1466.0232
385.25604248046875 0 40998.86 y 5
386.2587890625 0 8108.9146
387.260986328125 0 945.22705
389.23907470703125 0 826.34
389.73638916015625 0 688.87256
390.2429504394531 0 1571.2616
390.73931884765625 0 647.6992
391.23260498046875 0 18029.83
391.7339172363281 0 8591.899
392.2357177734375 0 1438.5804
393.21514892578125 0 853.2601
395.2354736328125 0 633.9565
401.2388000488281 0 826.7319
403.7472839355469 0 1330.4625
403.81402587890625 0 1377.4133
404.2478942871094 0 614.67755
408.2589416503906 0 1119.7961
411.2252197265625 0 959.8324
412.7535705566406 0 61379.492 Precursor Water loss
413.25506591796875 0 25335.459
413.75616455078125 0 8292.768
420.85577392578125 0 1099.7874
422.2405090332031 0 1534.2307
426.27142333984375 0 2434.4004
430.2666320800781 0 967.9963
436.2559814453125 0 3506.215
437.2543029785156 0 1825.3895
439.8428955078125 0 1142.653
440.2505187988281 0 2477.3435 b Water loss 4
441.25152587890625 0 903.5697
442.27752685546875 0 2643.131
444.2822570800781 0 817.0681
454.2747497558594 0 8149.0933 y Water loss 4
455.265380859375 0 2916.9502 y Ammonia loss 4
456.2655944824219 0 675.7993
468.2867126464844 0 814.2123
469.2849426269531 0 1123.4916
472.28851318359375 0 435788.8 y 4
473.2912902832031 0 102419.66
474.2937316894531 0 18405.39
475.2957458496094 0 755.8108
482.2726135253906 0 5224.1904
483.2770690917969 0 783.9264
486.2926025390625 0 10557.263
487.2955627441406 0 16415.73
488.2985534667969 0 3202.4941
489.30206298828125 0 756.65393
508.2752380371094 0 890.1791
509.27984619140625 0 1420.0049
535.3268432617188 0 929.959
541.3473510742188 0 668.1048
553.3418579101562 0 4286.0425 b Water loss 5
554.3395385742188 0 1955.1156 y Ammonia loss 3
557.3306274414062 0 1949.171
558.3323364257812 0 4457.1016
559.33642578125 0 789.6399
571.3565063476562 0 148670.33 y 3
572.359375 0 42093.76
573.3616333007812 0 8844.721
581.3417358398438 0 2503.6086
593.341064453125 0 783.7504
597.33447265625 0 2250.8284
598.3380737304688 0 776.7779
609.9524536132812 0 590.99335
611.35107421875 0 5972.6104
612.3552856445312 0 1986.235
628.3777465820312 0 13321.856
629.3806762695312 0 4561.3022
630.3809204101562 0 1771.4631
636.3804931640625 0 987.7925
637.3690185546875 0 971.2164
638.36328125 0 2602.2004
643.3915405273438 0 800.0406
654.3905029296875 0 15348.673 y Water loss 2
655.385986328125 0 23574.812 y Ammonia loss 2
656.3881225585938 0 9528.982
657.3886108398438 0 2083.6584
664.3758544921875 0 2305.4604
665.3790283203125 0 833.65765
672.404052734375 0 342266.97 y 2
673.4069213867188 0 117150.32
674.4089965820312 0 25376.203
675.4103393554688 0 1325.1714
682.3883666992188 0 30325.926
683.390869140625 0 9136.977
684.3953247070312 0 1772.4825
699.4146728515625 0 2519.7998
700.4174194335938 0 961.2424
708.407470703125 0 674.6252
713.4331665039062 0 639.4142
725.4299926757812 0 40082.492 y Water loss 1
726.4325561523438 0 14143.116
727.4346313476562 0 4042.0325
735.4102172851562 0 1090.961
743.4408569335938 0 141300.44 y 1
744.4435424804688 0 51815.715
745.4459838867188 0 13073.161
746.4491577148438 0 721.36017
753.4240112304688 0 2256.336
754.4266967773438 0 951.1098
952.6106567382812 0 649.8183
2255.358642578125 0 632.521
2341.10693359375 0 655.188
3134.97265625 0 728.45105
3159.51416015625 0 640.71136
3251.77783203125 0 659.77783

Spectrum Details

|  |  |
| --- | --- |
| Matched peaks? Matched peaks The total absolute number of peaks matched. Additionally in brackets the total fraction of peaks matched and the total number of peaks is shown. | 35 (9.62% of 364) |
| FDR? FDR The false discovery rate estimated for this peptide. It is calculated by matching all theoretical fragments with a non-integer shift with the raw peaks for this spectrum. This is done with 40 different shifts. The resulting percentage is the average number of annotated peaks over the number of annotated peaks with the correct spectrum. | 0.27% |
| Satellite FDR? Satellite FDR See the FDR for details on its calculation. This satellite ion specific FDR only contains the satellite ions (d/w) for I/L/J positions. | - |
| PSM Score? PSM Score The PSM Score as given by Hecklib to this annotated spectrum. It is shown with three significant figures. | 442 |

#### Spectrum 5189? Spectrum 5189 The raw spectrum of this peptide as annotated by Hecklib. The fragments are coloured according to ion type (see legend). Any peaks with a star '\*' as text can be hovered over to see the full details, first the ion type second the mass shift type. By hovering over the amino acids in the peptide or ions in the legend the corresponding peaks are highlighted. By toggling the 'Unassigned' label you can turn the background (unassigned) peaks on or off in the plot. By updating the slider in the Ion legend you can update the spectrum to only show the top X% of the peaks with labels. The top X% means any peak that is within X% of the highest intensity. By dragging in the spectrum you can zoom in to a specific part of the spectrum and use 'Zoom Out' to get back to the original zoom level. The annotation of the spectrum is based on the given sequence in the peptides file and is done with different software so inconsistencies are likely. The peaks are annotated based on the given sequence, with 20 ppm tolerance.

Copy Data

##### Spectrum 5189 (TSV)

###### Preview

```
Loading example...
```

*Click on the button to copy the data to your clipboard.*

Mz MinMz MaxIntensity Max

WidthHeightPeptide font sizePeptide stroke widthSpectrum font sizeSpectrum stroke widthCompact peptide

Ion legend

wxyz

abcd

OtherUnassignedIon^Charge^_Position_Show for top:%

VATVSJPR

07.36e+41.47e+52.21e+52.94e+5

Zoom Out

a^+1^_2_y^+1^_1_b^+1^_2_y^+1^_1_a^+1^_3_y^+2^_4_d^+1^_3_a^+1^_3_b^+1^_3_y^+1^_2_b^+1^_3_y^+1^_2_b^+2^_6_b^+2^_6_y^+2^_6_y^+2^_6_y^+2^_6_b^+1^_4_y^+2^_7_y^+1^_3_b^+1^_4_y^+2^_7_y^+1^_3_\*b^+1^_5_y^+1^_4_y^+1^_4_y^+1^_4_b^+1^_6_y^+1^_5_b^+1^_6_y^+1^_6_y^+1^_6_y^+1^_6_y^+1^_7_y^+1^_7_

0826165224783304

Fragment Matches Table

Show background peaks

| Position | Ion type | Intensity | mz Theoretical | mz Error (Th) | mz Error (ppm) | Charge | Series Number |
| --- | --- | --- | --- | --- | --- | --- | --- |
| - | - | 590.6 | 120.1 | - | - | 0 | - |
| - | - | 346.9 | 121 | - | - | 0 | - |
| - | - | 466.5 | 123.1 | - | - | 0 | - |
| - | - | 355.3 | 123.5 | - | - | 0 | - |
| - | - | 503.3 | 126.1 | - | - | 0 | - |
| - | - | 345.1 | 127.1 | - | - | 0 | - |
| - | - | 2945 | 127.1 | - | - | 0 | - |
| - | - | 4002 | 127.1 | - | - | 0 | - |
| - | - | 4546 | 128.1 | - | - | 0 | - |
| - | - | 543.7 | 129 | - | - | 0 | - |
| - | - | 8627 | 129.1 | - | - | 0 | - |
| - | - | 623.6 | 129.1 | - | - | 0 | - |
| - | - | 1425 | 130.1 | - | - | 0 | - |
| - | - | 3325 | 130.1 | - | - | 0 | - |
| - | - | 817.4 | 130.1 | - | - | 0 | - |
| - | - | 1028 | 130.1 | - | - | 0 | - |
| - | - | 1108 | 133.1 | - | - | 0 | - |
| - | - | 457.6 | 136.1 | - | - | 0 | - |
| - | - | 463.6 | 137.1 | - | - | 0 | - |
| - | - | 2415 | 138.1 | - | - | 0 | - |
| - | - | 1.999E+04 | 140.1 | - | - | 0 | - |
| - | - | 1170 | 141.1 | - | - | 0 | - |
| - | - | 1.1E+04 | 141.1 | - | - | 0 | - |
| - | - | 3003 | 142.1 | - | - | 0 | - |
| 2 | a | 1.183E+05 | 143.1 | 0.0003966 | 2.771 | +1 | 2 |
| - | - | 699.5 | 144.1 | - | - | 0 | - |
| - | - | 8185 | 144.1 | - | - | 0 | - |
| - | - | 8183 | 145.1 | - | - | 0 | - |
| - | - | 442.1 | 145.6 | - | - | 0 | - |
| - | - | 3748 | 147.1 | - | - | 0 | - |
| - | - | 884.9 | 148.1 | - | - | 0 | - |
| - | - | 987.8 | 149 | - | - | 0 | - |
| - | - | 9202 | 149 | - | - | 0 | - |
| - | - | 3994 | 150 | - | - | 0 | - |
| - | - | 400.2 | 152.2 | - | - | 0 | - |
| - | - | 4158 | 155.1 | - | - | 0 | - |
| - | - | 1.433E+04 | 155.1 | - | - | 0 | - |
| - | - | 716.3 | 156.1 | - | - | 0 | - |
| - | - | 540.2 | 157 | - | - | 0 | - |
| - | - | 1235 | 157.1 | - | - | 0 | - |
| - | - | 2206 | 157.1 | - | - | 0 | - |
| - | - | 2.238E+04 | 157.1 | - | - | 0 | - |
| - | - | 417.9 | 158 | - | - | 0 | - |
| 8 | y | 1.521E+04 | 158.1 | 0.0003246 | 2.053 | +1 | 1 |
| - | - | 6620 | 158.1 | - | - | 0 | - |
| - | - | 1076 | 159.1 | - | - | 0 | - |
| - | - | 2238 | 159.1 | - | - | 0 | - |
| - | - | 398.3 | 163.4 | - | - | 0 | - |
| - | - | 524.2 | 165.1 | - | - | 0 | - |
| - | - | 1153 | 167.1 | - | - | 0 | - |
| - | - | 1.038E+04 | 169.1 | - | - | 0 | - |
| - | - | 739.9 | 169.1 | - | - | 0 | - |
| - | - | 2999 | 170.1 | - | - | 0 | - |
| 2 | b | 1.07E+05 | 171.1 | 0.000355 | 2.075 | +1 | 2 |
| - | - | 891.6 | 172.1 | - | - | 0 | - |
| - | - | 1.005E+04 | 172.1 | - | - | 0 | - |
| - | - | 542.8 | 172.2 | - | - | 0 | - |
| - | - | 1.798E+04 | 173.1 | - | - | 0 | - |
| - | - | 1.216E+05 | 173.1 | - | - | 0 | - |
| - | - | 1978 | 174.1 | - | - | 0 | - |
| - | - | 9071 | 174.1 | - | - | 0 | - |
| 8 | y | 3.335E+04 | 175.1 | 0.0003105 | 1.773 | +1 | 1 |
| - | - | 806.6 | 175.1 | - | - | 0 | - |
| - | - | 1832 | 176.1 | - | - | 0 | - |
| - | - | 684.3 | 181.1 | - | - | 0 | - |
| - | - | 2.475E+04 | 183.1 | - | - | 0 | - |
| - | - | 1.17E+04 | 183.1 | - | - | 0 | - |
| - | - | 561.4 | 184.1 | - | - | 0 | - |
| - | - | 2005 | 184.1 | - | - | 0 | - |
| - | - | 3506 | 184.2 | - | - | 0 | - |
| - | - | 718.2 | 185.1 | - | - | 0 | - |
| - | - | 2791 | 185.1 | - | - | 0 | - |
| - | - | 5530 | 185.1 | - | - | 0 | - |
| - | - | 1893 | 186.1 | - | - | 0 | - |
| - | - | 6805 | 187.1 | - | - | 0 | - |
| - | - | 914 | 188.1 | - | - | 0 | - |
| - | - | 750 | 189.1 | - | - | 0 | - |
| - | - | 718.8 | 192.1 | - | - | 0 | - |
| - | - | 1132 | 194.1 | - | - | 0 | - |
| - | - | 504 | 194.2 | - | - | 0 | - |
| - | - | 1765 | 195.1 | - | - | 0 | - |
| - | - | 545 | 197.1 | - | - | 0 | - |
| - | - | 453.7 | 197.2 | - | - | 0 | - |
| - | - | 1415 | 199.1 | - | - | 0 | - |
| - | - | 4.509E+04 | 201.1 | - | - | 0 | - |
| - | - | 3855 | 202.1 | - | - | 0 | - |
| - | - | 622.6 | 203.1 | - | - | 0 | - |
| - | - | 626.4 | 205.8 | - | - | 0 | - |
| - | - | 711.7 | 209.1 | - | - | 0 | - |
| - | - | 1577 | 209.1 | - | - | 0 | - |
| - | - | 620.3 | 209.1 | - | - | 0 | - |
| - | - | 2294 | 210.1 | - | - | 0 | - |
| - | - | 508.2 | 210.9 | - | - | 0 | - |
| - | - | 5740 | 211.1 | - | - | 0 | - |
| - | - | 5705 | 211.1 | - | - | 0 | - |
| - | - | 1068 | 211.2 | - | - | 0 | - |
| - | - | 1.197E+04 | 212.1 | - | - | 0 | - |
| - | - | 2452 | 213.1 | - | - | 0 | - |
| - | - | 757 | 213.1 | - | - | 0 | - |
| - | - | 556.2 | 215.1 | - | - | 0 | - |
| - | - | 682 | 217.1 | - | - | 0 | - |
| - | - | 640.9 | 222.1 | - | - | 0 | - |
| - | - | 920.4 | 225.2 | - | - | 0 | - |
| - | - | 1473 | 226.1 | - | - | 0 | - |
| - | - | 952.1 | 226.1 | - | - | 0 | - |
| 3 | a | 1.672E+04 | 226.2 | 0.0003311 | 1.464 | +1 | 3 |
| - | - | 918.8 | 227.1 | - | - | 0 | - |
| - | - | 7399 | 227.1 | - | - | 0 | - |
| - | - | 2073 | 227.2 | - | - | 0 | - |
| 5 | y | 4782 | 228.1 | 0.0003146 | 1.379 | +2 | 4 |
| - | - | 962.5 | 229.1 | - | - | 0 | - |
| 3 | d | 6581 | 230.1 | 0.0003201 | 1.391 | +1 | 3 |
| - | - | 566.6 | 231.2 | - | - | 0 | - |
| - | - | 750 | 237.1 | - | - | 0 | - |
| - | - | 691.3 | 237.2 | - | - | 0 | - |
| - | - | 1157 | 238.1 | - | - | 0 | - |
| - | - | 2477 | 240.1 | - | - | 0 | - |
| - | - | 725.1 | 241.2 | - | - | 0 | - |
| - | - | 541.6 | 242.1 | - | - | 0 | - |
| - | - | 1158 | 243.1 | - | - | 0 | - |
| - | - | 970.2 | 243.7 | - | - | 0 | - |
| - | - | 1198 | 244.1 | - | - | 0 | - |
| - | - | 1124 | 244.2 | - | - | 0 | - |
| 3 | a | 1092 | 244.2 | 0.0004476 | 1.833 | +1 | 3 |
| - | - | 463.5 | 245.1 | - | - | 0 | - |
| - | - | 1817 | 252.1 | - | - | 0 | - |
| - | - | 819.3 | 254.1 | - | - | 0 | - |
| 3 | b | 4.596E+04 | 254.1 | 0.0002591 | 1.019 | +1 | 3 |
| 7 | y | 1.147E+05 | 255.1 | 0.0003562 | 1.396 | +1 | 2 |
| - | - | 1.234E+04 | 256.1 | - | - | 0 | - |
| - | - | 529.7 | 257.1 | - | - | 0 | - |
| - | - | 7731 | 258.1 | - | - | 0 | - |
| - | - | 2690 | 259.1 | - | - | 0 | - |
| - | - | 2233 | 266.1 | - | - | 0 | - |
| - | - | 537.4 | 268.1 | - | - | 0 | - |
| - | - | 1.651E+04 | 270.1 | - | - | 0 | - |
| - | - | 1605 | 271.1 | - | - | 0 | - |
| - | - | 1180 | 271.2 | - | - | 0 | - |
| 3 | b | 2.132E+04 | 272.2 | 0.00054 | 1.984 | +1 | 3 |
| 7 | y | 4.554E+04 | 272.2 | 0.0004031 | 1.481 | +1 | 2 |
| - | - | 2274 | 273.2 | - | - | 0 | - |
| - | - | 5652 | 273.2 | - | - | 0 | - |
| - | - | 5762 | 276.2 | - | - | 0 | - |
| - | - | 2291 | 277.2 | - | - | 0 | - |
| 6 | b | 1223 | 277.2 | 0.004534 | 16.36 | +2 | 6 |
| - | - | 1743 | 280.1 | - | - | 0 | - |
| - | - | 916.5 | 280.2 | - | - | 0 | - |
| - | - | 4190 | 282.2 | - | - | 0 | - |
| - | - | 551.8 | 285.9 | - | - | 0 | - |
| 6 | b | 3940 | 286.2 | 0.005599 | 19.57 | +2 | 6 |
| - | - | 1579 | 286.7 | - | - | 0 | - |
| - | - | 1.285E+04 | 288.2 | - | - | 0 | - |
| - | - | 1583 | 289.2 | - | - | 0 | - |
| - | - | 815.3 | 291 | - | - | 0 | - |
| - | - | 737.6 | 291.2 | - | - | 0 | - |
| - | - | 1005 | 293.1 | - | - | 0 | - |
| - | - | 782.4 | 296.2 | - | - | 0 | - |
| - | - | 605.2 | 297.2 | - | - | 0 | - |
| - | - | 2304 | 298.1 | - | - | 0 | - |
| - | - | 2147 | 299.2 | - | - | 0 | - |
| - | - | 970.6 | 299.2 | - | - | 0 | - |
| - | - | 854.9 | 299.7 | - | - | 0 | - |
| - | - | 8062 | 306.2 | - | - | 0 | - |
| - | - | 2137 | 306.7 | - | - | 0 | - |
| - | - | 755.2 | 308 | - | - | 0 | - |
| - | - | 1155 | 308.2 | - | - | 0 | - |
| - | - | 748.9 | 310.7 | - | - | 0 | - |
| - | - | 4285 | 312.2 | - | - | 0 | - |
| - | - | 742.9 | 313.2 | - | - | 0 | - |
| - | - | 794.2 | 319.7 | - | - | 0 | - |
| - | - | 3232 | 323.2 | - | - | 0 | - |
| - | - | 2.104E+04 | 325.2 | - | - | 0 | - |
| - | - | 3709 | 326.2 | - | - | 0 | - |
| - | - | 840.9 | 327.2 | - | - | 0 | - |
| - | - | 735.4 | 327.2 | - | - | 0 | - |
| 3 | y | 4.273E+04 | 327.7 | 1.942E-05 | 0.05926 | +2 | 6 |
| 3 | y | 2.929E+04 | 328.2 | 0.006508 | 19.83 | +2 | 6 |
| - | - | 1.038E+04 | 328.7 | - | - | 0 | - |
| - | - | 1213 | 329.2 | - | - | 0 | - |
| - | - | 661.4 | 331.2 | - | - | 0 | - |
| - | - | 1431 | 332.7 | - | - | 0 | - |
| 3 | y | 5341 | 336.7 | 0.0005881 | 1.747 | +2 | 6 |
| - | - | 1443 | 337.2 | - | - | 0 | - |
| - | - | 1065 | 337.2 | - | - | 0 | - |
| - | - | 542.5 | 337.7 | - | - | 0 | - |
| - | - | 736.7 | 338.2 | - | - | 0 | - |
| - | - | 5262 | 340.2 | - | - | 0 | - |
| - | - | 8574 | 341.2 | - | - | 0 | - |
| - | - | 1815 | 341.7 | - | - | 0 | - |
| - | - | 949.9 | 342.2 | - | - | 0 | - |
| - | - | 829.1 | 342.2 | - | - | 0 | - |
| - | - | 3974 | 343.2 | - | - | 0 | - |
| - | - | 780.2 | 344.2 | - | - | 0 | - |
| - | - | 1484 | 350.9 | - | - | 0 | - |
| - | - | 1109 | 351.2 | - | - | 0 | - |
| - | - | 815.5 | 351.2 | - | - | 0 | - |
| - | - | 1168 | 352.2 | - | - | 0 | - |
| 4 | b | 1.347E+04 | 353.2 | 0.000235 | 0.6654 | +1 | 4 |
| - | - | 832 | 353.2 | - | - | 0 | - |
| - | - | 2672 | 354.2 | - | - | 0 | - |
| - | - | 941.1 | 355.2 | - | - | 0 | - |
| - | - | 1847 | 355.2 | - | - | 0 | - |
| - | - | 767.5 | 357.2 | - | - | 0 | - |
| - | - | 759.5 | 358.2 | - | - | 0 | - |
| - | - | 3305 | 359.2 | - | - | 0 | - |
| - | - | 732.1 | 360.2 | - | - | 0 | - |
| 2 | y | 5.269E+04 | 363.2 | 0.0001615 | 0.4446 | +2 | 7 |
| - | - | 2.103E+04 | 363.7 | - | - | 0 | - |
| - | - | 3790 | 364.2 | - | - | 0 | - |
| - | - | 586.4 | 364.7 | - | - | 0 | - |
| - | - | 1046 | 365.2 | - | - | 0 | - |
| - | - | 819.1 | 367.2 | - | - | 0 | - |
| - | - | 617 | 367.2 | - | - | 0 | - |
| 6 | y | 7071 | 368.2 | 0.0002918 | 0.7925 | +1 | 3 |
| - | - | 1156 | 369.2 | - | - | 0 | - |
| 4 | b | 1213 | 371.2 | 0.0007777 | 2.095 | +1 | 4 |
| 2 | y | 1012 | 372.2 | 0.000116 | 0.3116 | +2 | 7 |
| - | - | 1144 | 373.2 | - | - | 0 | - |
| - | - | 765.9 | 377.2 | - | - | 0 | - |
| - | - | 1043 | 379.2 | - | - | 0 | - |
| - | - | 1395 | 380.2 | - | - | 0 | - |
| - | - | 4389 | 383.2 | - | - | 0 | - |
| - | - | 602.9 | 384.1 | - | - | 0 | - |
| - | - | 845.8 | 384.2 | - | - | 0 | - |
| 6 | y | 2.981E+04 | 385.3 | 0.000232 | 0.6021 | +1 | 3 |
| - | - | 524.4 | 386.1 | - | - | 0 | - |
| - | - | 4787 | 386.3 | - | - | 0 | - |
| - | - | 1887 | 389.2 | - | - | 0 | - |
| - | - | 1.327E+04 | 391.2 | - | - | 0 | - |
| - | - | 5268 | 391.7 | - | - | 0 | - |
| - | - | 1149 | 392.2 | - | - | 0 | - |
| - | - | 771.2 | 401.2 | - | - | 0 | - |
| - | - | 1307 | 403.7 | - | - | 0 | - |
| - | - | 2036 | 403.8 | - | - | 0 | - |
| - | - | 1119 | 408.3 | - | - | 0 | - |
| - | - | 1157 | 411.2 | - | - | 0 | - |
| 0 | Precursor | 3.705E+04 | 412.8 | 0.0004089 | 0.9906 | +2 | -1 |
| - | - | 1.61E+04 | 413.3 | - | - | 0 | - |
| - | - | 3859 | 413.8 | - | - | 0 | - |
| - | - | 560.4 | 414.3 | - | - | 0 | - |
| - | - | 1432 | 420.9 | - | - | 0 | - |
| - | - | 1950 | 422.2 | - | - | 0 | - |
| - | - | 2240 | 426.3 | - | - | 0 | - |
| - | - | 903.6 | 430.3 | - | - | 0 | - |
| - | - | 2418 | 436.3 | - | - | 0 | - |
| - | - | 949.9 | 437.3 | - | - | 0 | - |
| - | - | 850.9 | 439.8 | - | - | 0 | - |
| 5 | b | 1777 | 440.3 | 0.0009215 | 2.093 | +1 | 5 |
| - | - | 2231 | 442.3 | - | - | 0 | - |
| 5 | y | 5377 | 454.3 | 0.003043 | 6.699 | +1 | 4 |
| 5 | y | 2008 | 455.3 | 0.003633 | 7.981 | +1 | 4 |
| 5 | y | 2.914E+05 | 472.3 | 0.0005827 | 1.234 | +1 | 4 |
| - | - | 7.179E+04 | 473.3 | - | - | 0 | - |
| - | - | 1.175E+04 | 474.3 | - | - | 0 | - |
| - | - | 3106 | 482.3 | - | - | 0 | - |
| - | - | 746.2 | 483.3 | - | - | 0 | - |
| - | - | 1.269E+04 | 486.3 | - | - | 0 | - |
| - | - | 1.618E+04 | 487.3 | - | - | 0 | - |
| - | - | 5055 | 488.3 | - | - | 0 | - |
| - | - | 698.3 | 508.3 | - | - | 0 | - |
| - | - | 1433 | 509.3 | - | - | 0 | - |
| - | - | 738 | 540.3 | - | - | 0 | - |
| 6 | b | 3693 | 553.3 | 0.0103 | 18.62 | +1 | 6 |
| 4 | y | 1130 | 554.3 | 0.007058 | 12.73 | +1 | 5 |
| - | - | 2063 | 557.3 | - | - | 0 | - |
| - | - | 4289 | 558.3 | - | - | 0 | - |
| - | - | 1179 | 559.3 | - | - | 0 | - |
| 6 | b | 9.821E+04 | 571.3 | 0.01133 | 19.84 | +1 | 6 |
| - | - | 2.995E+04 | 572.4 | - | - | 0 | - |
| - | - | 4508 | 573.4 | - | - | 0 | - |
| - | - | 1358 | 581.3 | - | - | 0 | - |
| - | - | 1231 | 593.3 | - | - | 0 | - |
| - | - | 1388 | 597.3 | - | - | 0 | - |
| - | - | 690.5 | 598.3 | - | - | 0 | - |
| - | - | 4379 | 611.4 | - | - | 0 | - |
| - | - | 948.3 | 612.4 | - | - | 0 | - |
| - | - | 8779 | 628.4 | - | - | 0 | - |
| - | - | 3439 | 629.4 | - | - | 0 | - |
| - | - | 995 | 637.4 | - | - | 0 | - |
| - | - | 1127 | 638.4 | - | - | 0 | - |
| - | - | 783 | 639.4 | - | - | 0 | - |
| - | - | 695.3 | 642.4 | - | - | 0 | - |
| 3 | y | 1.222E+04 | 654.4 | 0.004603 | 7.034 | +1 | 6 |
| 3 | y | 2.419E+04 | 655.4 | 0.008635 | 13.18 | +1 | 6 |
| - | - | 7971 | 656.4 | - | - | 0 | - |
| - | - | 1706 | 657.4 | - | - | 0 | - |
| - | - | 1468 | 664.4 | - | - | 0 | - |
| - | - | 711.6 | 665.4 | - | - | 0 | - |
| 3 | y | 2.331E+05 | 672.4 | 3.117E-05 | 0.04636 | +1 | 6 |
| - | - | 7.808E+04 | 673.4 | - | - | 0 | - |
| - | - | 1.798E+04 | 674.4 | - | - | 0 | - |
| - | - | 1193 | 675.4 | - | - | 0 | - |
| - | - | 1.86E+04 | 682.4 | - | - | 0 | - |
| - | - | 5021 | 683.4 | - | - | 0 | - |
| - | - | 1070 | 684.4 | - | - | 0 | - |
| - | - | 1671 | 699.4 | - | - | 0 | - |
| - | - | 649.6 | 701.4 | - | - | 0 | - |
| 2 | y | 2.831E+04 | 725.4 | 0.0004572 | 0.6303 | +1 | 7 |
| - | - | 1.041E+04 | 726.4 | - | - | 0 | - |
| - | - | 2446 | 727.4 | - | - | 0 | - |
| 2 | y | 9.024E+04 | 743.4 | 0.0003408 | 0.4584 | +1 | 7 |
| - | - | 3.482E+04 | 744.4 | - | - | 0 | - |
| - | - | 9313 | 745.4 | - | - | 0 | - |
| - | - | 1707 | 753.4 | - | - | 0 | - |
| - | - | 1143 | 754.4 | - | - | 0 | - |
| - | - | 762.1 | 928.3 | - | - | 0 | - |
| - | - | 598.2 | 1974 | - | - | 0 | - |
| - | - | 821.7 | 3082 | - | - | 0 | - |
| - | - | 784.9 | 3271 | - | - | 0 | - |

m/z Charge Intensity FragmentType MassShift Position
120.08129119873047 0 590.5516
121.02906036376953 0 346.93265
123.11690521240234 0 466.52975
123.48684692382812 0 355.32535
126.12824249267578 0 503.32455
127.08282470703125 0 345.09183
127.08698272705078 0 2944.6519
127.12335968017578 0 4002.0645
128.10731506347656 0 4546.2715
129.01866149902344 0 543.742
129.10255432128906 0 8627.257
129.1140899658203 0 623.61584
130.05006408691406 0 1425.4458
130.0866241455078 0 3325.2502
130.0975799560547 0 817.4394
130.10589599609375 0 1028.3046
133.09747314453125 0 1107.5688
136.07640075683594 0 457.63873
137.05990600585938 0 463.63605
138.09156799316406 0 2415.2097
140.08216857910156 0 19991.156
141.08558654785156 0 1169.7688
141.1025848388672 0 10999.792
142.1059112548828 0 3002.5488
143.1182861328125 0 118264.95 a 1
144.11549377441406 0 699.4561
144.12158203125 0 8184.6353
145.0974578857422 0 8182.9526
145.5792236328125 0 442.11002
147.11312866210938 0 3747.5437
148.1166229248047 0 884.8519
148.95355224609375 0 987.8322
149.02365112304688 0 9202.116
150.02699279785156 0 3993.7058
152.17538452148438 0 400.1511
155.0818634033203 0 4157.5117
155.11819458007812 0 14331.604
156.1219482421875 0 716.3437
157.01290893554688 0 540.24164
157.097412109375 0 1235.4108
157.10867309570312 0 2206.2664
157.13385009765625 0 22377.984
158.03550720214844 0 417.90527
158.0927276611328 0 15212.061 y Ammonia loss 7
158.13720703125 0 6620.2446
159.09596252441406 0 1076.397
159.1131134033203 0 2238.3975
163.38729858398438 0 398.3441
165.1027374267578 0 524.15924
167.08151245117188 0 1153.4283
169.0974884033203 0 10383.563
169.133544921875 0 739.8934
170.10086059570312 0 2998.6785
171.1131591796875 0 106988.06 b 1
172.1086883544922 0 891.55945
172.11660766601562 0 10045.557
172.23875427246094 0 542.825
173.09234619140625 0 17977.791
173.12881469726562 0 121607.836
174.0959930419922 0 1978.1277
174.13218688964844 0 9071.163
175.1192626953125 0 33347.996 y 7
175.1322479248047 0 806.5958
176.12266540527344 0 1831.5591
181.0977020263672 0 684.3297
183.11314392089844 0 24754.305
183.14956665039062 0 11695.155
184.10870361328125 0 561.37537
184.11663818359375 0 2004.7849
184.1529541015625 0 3506.0576
185.08091735839844 0 718.1863
185.1036834716797 0 2790.5637
185.1287841796875 0 5529.8105
186.13218688964844 0 1892.7734
187.10804748535156 0 6804.5566
188.11155700683594 0 913.95917
189.12290954589844 0 750.0318
192.11370849609375 0 718.8393
194.1291961669922 0 1132.1799
194.20236206054688 0 503.99054
195.11309814453125 0 1764.8906
197.0926513671875 0 545.0007
197.16143798828125 0 453.67517
199.14456176757812 0 1415.1096
201.12362670898438 0 45087.42
202.1272430419922 0 3854.724
203.1291046142578 0 622.55707
205.8291015625 0 626.35455
209.09225463867188 0 711.7216
209.12864685058594 0 1576.5482
209.139404296875 0 620.3071
210.12391662597656 0 2294.1301
210.9047393798828 0 508.20132
211.10800170898438 0 5739.883
211.14427185058594 0 5705.0405
211.1550750732422 0 1067.6079
212.13963317871094 0 11966.722
213.12367248535156 0 2451.568
213.1429901123047 0 756.995
215.1393280029297 0 556.2379
217.08255004882812 0 682.0254
222.12301635742188 0 640.94727
225.15989685058594 0 920.3993
226.11953735351562 0 1473.2675
226.14239501953125 0 952.11896
226.15533447265625 0 16720.44 a Water loss 2
227.10292053222656 0 918.7611
227.13934326171875 0 7398.876
227.15887451171875 0 2073.171
228.13458251953125 0 4781.994 y Ammonia loss 4
229.1187286376953 0 962.49585
230.15023803710938 0 6580.87 d 2
231.15357971191406 0 566.647
237.1351776123047 0 750.0442
237.16055297851562 0 691.25464
238.11866760253906 0 1157.3884
240.1347198486328 0 2477.1638
241.15533447265625 0 725.067
242.11471557617188 0 541.58685
243.13401794433594 0 1158.3557
243.65040588378906 0 970.21204
244.1291046142578 0 1197.5659
244.15232849121094 0 1123.5443
244.166015625 0 1092.0883 a 2
245.1007080078125 0 463.51672
252.1344757080078 0 1817.4629
254.11611938476562 0 819.3472
254.15017700195312 0 45958.277 b Water loss 2
255.14552307128906 0 114731.84 y Ammonia loss 6
256.14874267578125 0 12337.106
257.14984130859375 0 529.65857
258.1451110839844 0 7731.2544
259.1485290527344 0 2690.3267
266.1499328613281 0 2232.5166
268.128662109375 0 537.39355
270.14501953125 0 16512.473
271.1482849121094 0 1604.9012
271.1764221191406 0 1179.7417
272.1599426269531 0 21323.088 b 2
272.172119140625 0 45539.4 y 6
273.16265869140625 0 2273.7322
273.17535400390625 0 5652.1196
276.15570068359375 0 5762.295
277.158935546875 0 2290.796
277.1753845214844 0 1222.974 b Water loss 5
280.1293029785156 0 1742.966
280.1658020019531 0 916.51355
282.18157958984375 0 4190.065
285.8722839355469 0 551.7607
286.1817321777344 0 3939.7356 b 5
286.6837158203125 0 1578.703
288.1556396484375 0 12850.339
289.1594543457031 0 1583.2745
290.9609375 0 815.3296
291.1757507324219 0 737.6453
293.0997009277344 0 1004.5004
296.1610107421875 0 782.4151
297.1764221191406 0 605.1749
298.13983154296875 0 2303.625
299.1898498535156 0 2147.0015
299.2062683105469 0 970.5502
299.6929931640625 0 854.93445
306.17974853515625 0 8061.8555
306.6808776855469 0 2137.4084
307.98626708984375 0 755.16693
308.19677734375 0 1154.7965
310.67901611328125 0 748.9112
312.1921691894531 0 4284.608
313.1906433105469 0 742.9279
319.68389892578125 0 794.20306
323.1718444824219 0 3231.7131
325.22393798828125 0 21043.066
326.2268371582031 0 3708.78
327.20428466796875 0 840.91187
327.2291259765625 0 735.36646
327.7002868652344 0 42732.71 y Water loss 2
328.1988220214844 0 29289.025 y Ammonia loss 2
328.6993408203125 0 10375.32
329.19976806640625 0 1212.6313
331.19696044921875 0 661.37946
332.69317626953125 0 1431.489
336.7061767578125 0 5341.1777 y 2
337.18707275390625 0 1443.1333
337.20733642578125 0 1064.8666
337.708740234375 0 542.4606
338.2092590332031 0 736.71344
340.18695068359375 0 5261.8486
341.1842956542969 0 8573.966
341.69866943359375 0 1814.991
342.1821594238281 0 949.85284
342.20220947265625 0 829.0944
343.2344665527344 0 3974.3765
344.2361145019531 0 780.2151
350.9274597167969 0 1483.8748
351.20147705078125 0 1109.3328
351.2393798828125 0 815.54944
352.2411804199219 0 1167.6216
353.21856689453125 0 13472.438 b Water loss 3
353.24261474609375 0 832.035
354.21942138671875 0 2672.4314
355.1973876953125 0 941.14075
355.2342529296875 0 1846.8777
357.217041015625 0 767.54425
358.19781494140625 0 759.49243
359.1933898925781 0 3305.4927
360.1943054199219 0 732.11914
363.2190246582031 0 52689.6 y Water loss 1
363.72039794921875 0 21029.986
364.22174072265625 0 3789.5798
364.7235107421875 0 586.4477
365.2185363769531 0 1046.3524
367.19720458984375 0 819.0852
367.2452087402344 0 617.0252
368.2295227050781 0 7070.7495 y Ammonia loss 5
369.2325439453125 0 1156.0518
371.2281188964844 0 1213.2273 b 3
372.2240295410156 0 1012.17126 y 1
373.24493408203125 0 1144.267
377.2166442871094 0 765.9143
379.23388671875 0 1042.7778
380.2371826171875 0 1395.103
383.22943115234375 0 4389.4727
384.1437683105469 0 602.8601
384.23321533203125 0 845.84296
385.2560119628906 0 29806.31 y 5
386.0852966308594 0 524.44324
386.25872802734375 0 4787.258
389.23968505859375 0 1886.7289
391.2324523925781 0 13265.455
391.73370361328125 0 5268.42
392.23406982421875 0 1149.4773
401.2401428222656 0 771.2175
403.7478332519531 0 1307.4489
403.8135070800781 0 2036.251
408.2613525390625 0 1118.5126
411.22265625 0 1156.8044
412.75347900390625 0 37046.98 Precursor Water loss
413.2550048828125 0 16103.593
413.7559814453125 0 3858.9248
414.2554931640625 0 560.3635
420.85797119140625 0 1431.9993
422.24066162109375 0 1949.7225
426.2721252441406 0 2239.7532
430.2671203613281 0 903.55194
436.2558288574219 0 2418.0088
437.25457763671875 0 949.9179
439.8439025878906 0 850.9267
440.25128173828125 0 1776.7402 b Water loss 4
442.2783203125 0 2230.7166
454.2742004394531 0 5376.9507 y Water loss 4
455.264892578125 0 2007.6965 y Ammonia loss 4
472.28839111328125 0 291371.3 y 4
473.29107666015625 0 71794.734
474.2935791015625 0 11746.697
482.27337646484375 0 3106.251
483.274658203125 0 746.19275
486.29254150390625 0 12686.477
487.2955017089844 0 16183.547
488.29779052734375 0 5054.89
508.2731628417969 0 698.33606
509.2794494628906 0 1433.1753
540.3218994140625 0 738.0046
553.3447265625 0 3692.7476 b Water loss 5
554.3367309570312 0 1129.8319 y Ammonia loss 3
557.3291625976562 0 2062.5483
558.3321533203125 0 4289.1396
559.33349609375 0 1179.2806
571.3563232421875 0 98214.266 b 5
572.3591918945312 0 29948.928
573.3612670898438 0 4508.339
581.3405151367188 0 1358.1205
593.33984375 0 1230.9932
597.3366088867188 0 1387.8328
598.3358764648438 0 690.4501
611.3510131835938 0 4379.1787
612.3564453125 0 948.2789
628.3777465820312 0 8778.765
629.380615234375 0 3438.604
637.3701782226562 0 995.01245
638.3604125976562 0 1127.1129
639.3687133789062 0 782.9948
642.3844604492188 0 695.30444
654.3887329101562 0 12219.191 y Water loss 2
655.385986328125 0 24185.264 y Ammonia loss 2
656.3877563476562 0 7971.0503
657.3911743164062 0 1705.9775
664.376953125 0 1468.037
665.3818359375 0 711.63947
672.4038696289062 0 233110.89 y 2
673.4065551757812 0 78077.6
674.4090576171875 0 17983.094
675.413818359375 0 1193.3733
682.3880615234375 0 18596.914
683.39208984375 0 5020.783
684.3956298828125 0 1070.1702
699.414794921875 0 1670.9847
701.4174194335938 0 649.5988
725.4299926757812 0 28314.516 y Water loss 1
726.4328002929688 0 10411.753
727.434326171875 0 2445.7483
743.440673828125 0 90236.22 y 1
744.4432983398438 0 34819.78
745.4462280273438 0 9312.87
753.427734375 0 1706.7467
754.429931640625 0 1142.6958
928.2726440429688 0 762.056
1973.5010986328125 0 598.2189
3081.7021484375 0 821.72
3270.8349609375 0 784.9344

Spectrum Details

|  |  |
| --- | --- |
| Matched peaks? Matched peaks The total absolute number of peaks matched. Additionally in brackets the total fraction of peaks matched and the total number of peaks is shown. | 36 (11.65% of 309) |
| FDR? FDR The false discovery rate estimated for this peptide. It is calculated by matching all theoretical fragments with a non-integer shift with the raw peaks for this spectrum. This is done with 40 different shifts. The resulting percentage is the average number of annotated peaks over the number of annotated peaks with the correct spectrum. | 0.20% |
| Satellite FDR? Satellite FDR See the FDR for details on its calculation. This satellite ion specific FDR only contains the satellite ions (d/w) for I/L/J positions. | - |
| PSM Score? PSM Score The PSM Score as given by Hecklib to this annotated spectrum. It is shown with three significant figures. | 442 |

#### Spectrum 5298? Spectrum 5298 The raw spectrum of this peptide as annotated by Hecklib. The fragments are coloured according to ion type (see legend). Any peaks with a star '\*' as text can be hovered over to see the full details, first the ion type second the mass shift type. By hovering over the amino acids in the peptide or ions in the legend the corresponding peaks are highlighted. By toggling the 'Unassigned' label you can turn the background (unassigned) peaks on or off in the plot. By updating the slider in the Ion legend you can update the spectrum to only show the top X% of the peaks with labels. The top X% means any peak that is within X% of the highest intensity. By dragging in the spectrum you can zoom in to a specific part of the spectrum and use 'Zoom Out' to get back to the original zoom level. The annotation of the spectrum is based on the given sequence in the peptides file and is done with different software so inconsistencies are likely. The peaks are annotated based on the given sequence, with 20 ppm tolerance.

Copy Data

##### Spectrum 5298 (TSV)

###### Preview

```
Loading example...
```

*Click on the button to copy the data to your clipboard.*

Mz MinMz MaxIntensity Max

WidthHeightPeptide font sizePeptide stroke widthSpectrum font sizeSpectrum stroke widthCompact peptide

Ion legend

wxyz

abcd

OtherUnassignedIon^Charge^_Position_Show for top:%

VATVSJPR

04.59e+49.18e+41.38e+51.84e+5

Zoom Out

a^+1^_2_y^+1^_1_b^+1^_2_y^+1^_1_a^+1^_3_y^+2^_4_d^+1^_3_a^+1^_3_b^+1^_3_y^+1^_2_b^+1^_3_y^+1^_2_b^+2^_6_b^+2^_6_y^+2^_6_y^+2^_6_y^+2^_6_b^+1^_4_y^+2^_7_y^+1^_3_b^+1^_4_y^+2^_7_y^+1^_3_\*b^+1^_5_y^+1^_4_y^+1^_4_y^+1^_4_b^+1^_6_b^+1^_6_y^+1^_6_y^+1^_6_y^+1^_6_y^+1^_7_y^+1^_7_

0758151722753034

Fragment Matches Table

Show background peaks

| Position | Ion type | Intensity | mz Theoretical | mz Error (Th) | mz Error (ppm) | Charge | Series Number |
| --- | --- | --- | --- | --- | --- | --- | --- |
| - | - | 811.4 | 120.1 | - | - | 0 | - |
| - | - | 492.8 | 122.7 | - | - | 0 | - |
| - | - | 379.6 | 126.1 | - | - | 0 | - |
| - | - | 500.8 | 126.1 | - | - | 0 | - |
| - | - | 1881 | 127.1 | - | - | 0 | - |
| - | - | 1786 | 127.1 | - | - | 0 | - |
| - | - | 367.7 | 127.8 | - | - | 0 | - |
| - | - | 3766 | 128.1 | - | - | 0 | - |
| - | - | 341 | 128.7 | - | - | 0 | - |
| - | - | 829.4 | 129 | - | - | 0 | - |
| - | - | 7201 | 129.1 | - | - | 0 | - |
| - | - | 522 | 130.1 | - | - | 0 | - |
| - | - | 3188 | 130.1 | - | - | 0 | - |
| - | - | 456.4 | 130.1 | - | - | 0 | - |
| - | - | 842 | 130.1 | - | - | 0 | - |
| - | - | 457.9 | 131.1 | - | - | 0 | - |
| - | - | 865.4 | 133.1 | - | - | 0 | - |
| - | - | 457.7 | 134.5 | - | - | 0 | - |
| - | - | 528.7 | 136.1 | - | - | 0 | - |
| - | - | 1713 | 138.1 | - | - | 0 | - |
| - | - | 1.072E+04 | 140.1 | - | - | 0 | - |
| - | - | 908.4 | 141.1 | - | - | 0 | - |
| - | - | 9978 | 141.1 | - | - | 0 | - |
| - | - | 395.5 | 141.5 | - | - | 0 | - |
| - | - | 501.2 | 142.1 | - | - | 0 | - |
| - | - | 2661 | 142.1 | - | - | 0 | - |
| 2 | a | 7.559E+04 | 143.1 | 0.0001219 | 0.852 | +1 | 2 |
| - | - | 988.5 | 144.1 | - | - | 0 | - |
| - | - | 5443 | 144.1 | - | - | 0 | - |
| - | - | 374.2 | 145.1 | - | - | 0 | - |
| - | - | 4184 | 145.1 | - | - | 0 | - |
| - | - | 453.3 | 145.3 | - | - | 0 | - |
| - | - | 3645 | 147.1 | - | - | 0 | - |
| - | - | 559.5 | 148.1 | - | - | 0 | - |
| - | - | 1124 | 148.9 | - | - | 0 | - |
| - | - | 9596 | 149 | - | - | 0 | - |
| - | - | 5673 | 150 | - | - | 0 | - |
| - | - | 375.6 | 150.7 | - | - | 0 | - |
| - | - | 453.1 | 151 | - | - | 0 | - |
| - | - | 2009 | 155.1 | - | - | 0 | - |
| - | - | 9469 | 155.1 | - | - | 0 | - |
| - | - | 496.5 | 156.1 | - | - | 0 | - |
| - | - | 836.3 | 156.1 | - | - | 0 | - |
| - | - | 482.9 | 157 | - | - | 0 | - |
| - | - | 994.4 | 157.1 | - | - | 0 | - |
| - | - | 1867 | 157.1 | - | - | 0 | - |
| - | - | 2.052E+04 | 157.1 | - | - | 0 | - |
| - | - | 1088 | 157.1 | - | - | 0 | - |
| 8 | y | 1.068E+04 | 158.1 | 1.942E-05 | 0.1228 | +1 | 1 |
| - | - | 803.2 | 158.1 | - | - | 0 | - |
| - | - | 5603 | 158.1 | - | - | 0 | - |
| - | - | 481.4 | 159.1 | - | - | 0 | - |
| - | - | 1944 | 159.1 | - | - | 0 | - |
| - | - | 9993 | 169.1 | - | - | 0 | - |
| - | - | 2801 | 170.1 | - | - | 0 | - |
| 2 | b | 6.786E+04 | 171.1 | 6.511E-05 | 0.3805 | +1 | 2 |
| - | - | 2728 | 171.1 | - | - | 0 | - |
| - | - | 685.9 | 172.1 | - | - | 0 | - |
| - | - | 4949 | 172.1 | - | - | 0 | - |
| - | - | 9906 | 173.1 | - | - | 0 | - |
| - | - | 7.74E+04 | 173.1 | - | - | 0 | - |
| - | - | 1935 | 173.5 | - | - | 0 | - |
| - | - | 971.2 | 174.1 | - | - | 0 | - |
| - | - | 835.1 | 174.1 | - | - | 0 | - |
| - | - | 6032 | 174.1 | - | - | 0 | - |
| 8 | y | 2.156E+04 | 175.1 | 6.639E-05 | 0.3791 | +1 | 1 |
| - | - | 1114 | 176.1 | - | - | 0 | - |
| - | - | 421.1 | 176.7 | - | - | 0 | - |
| - | - | 1.655E+04 | 183.1 | - | - | 0 | - |
| - | - | 1.031E+04 | 183.1 | - | - | 0 | - |
| - | - | 1373 | 184.1 | - | - | 0 | - |
| - | - | 2875 | 184.2 | - | - | 0 | - |
| - | - | 1147 | 185.1 | - | - | 0 | - |
| - | - | 1782 | 185.1 | - | - | 0 | - |
| - | - | 5294 | 185.1 | - | - | 0 | - |
| - | - | 545.8 | 185.2 | - | - | 0 | - |
| - | - | 4317 | 187.1 | - | - | 0 | - |
| - | - | 1491 | 195.1 | - | - | 0 | - |
| - | - | 2428 | 199.1 | - | - | 0 | - |
| - | - | 2.859E+04 | 201.1 | - | - | 0 | - |
| - | - | 2863 | 202.1 | - | - | 0 | - |
| - | - | 564.8 | 209.1 | - | - | 0 | - |
| - | - | 676.2 | 209.1 | - | - | 0 | - |
| - | - | 1914 | 210.1 | - | - | 0 | - |
| - | - | 3518 | 211.1 | - | - | 0 | - |
| - | - | 5358 | 211.1 | - | - | 0 | - |
| - | - | 7698 | 212.1 | - | - | 0 | - |
| - | - | 988.3 | 212.1 | - | - | 0 | - |
| - | - | 1808 | 213.1 | - | - | 0 | - |
| - | - | 866.9 | 213.1 | - | - | 0 | - |
| - | - | 775.5 | 213.2 | - | - | 0 | - |
| - | - | 837.7 | 215.1 | - | - | 0 | - |
| - | - | 542.9 | 216.1 | - | - | 0 | - |
| - | - | 496.1 | 222.1 | - | - | 0 | - |
| - | - | 472.7 | 225.2 | - | - | 0 | - |
| - | - | 644.8 | 226.1 | - | - | 0 | - |
| 3 | a | 1.114E+04 | 226.2 | 4.544E-06 | 0.02009 | +1 | 3 |
| - | - | 724.4 | 227.1 | - | - | 0 | - |
| - | - | 4461 | 227.1 | - | - | 0 | - |
| - | - | 1353 | 227.2 | - | - | 0 | - |
| 5 | y | 2914 | 228.1 | 5.157E-05 | 0.2261 | +2 | 4 |
| - | - | 682.8 | 229.1 | - | - | 0 | - |
| 3 | d | 3781 | 230.1 | 6.069E-05 | 0.2637 | +1 | 3 |
| - | - | 580.4 | 238.1 | - | - | 0 | - |
| - | - | 1787 | 240.1 | - | - | 0 | - |
| - | - | 644.1 | 241.1 | - | - | 0 | - |
| - | - | 951 | 243.1 | - | - | 0 | - |
| - | - | 1053 | 243.6 | - | - | 0 | - |
| - | - | 1024 | 244.1 | - | - | 0 | - |
| - | - | 950.6 | 244.2 | - | - | 0 | - |
| 3 | a | 648.7 | 244.2 | 0.0002695 | 1.104 | +1 | 3 |
| - | - | 653.4 | 252.1 | - | - | 0 | - |
| 3 | b | 3.103E+04 | 254.1 | 0.0001072 | 0.4216 | +1 | 3 |
| 7 | y | 7.3E+04 | 255.1 | 4.058E-05 | 0.159 | +1 | 2 |
| - | - | 9536 | 256.1 | - | - | 0 | - |
| - | - | 5981 | 258.1 | - | - | 0 | - |
| - | - | 2801 | 259.1 | - | - | 0 | - |
| - | - | 651.9 | 262.1 | - | - | 0 | - |
| - | - | 2095 | 266.1 | - | - | 0 | - |
| - | - | 513.5 | 267.2 | - | - | 0 | - |
| - | - | 1.042E+04 | 270.1 | - | - | 0 | - |
| - | - | 1095 | 271.1 | - | - | 0 | - |
| - | - | 879 | 271.2 | - | - | 0 | - |
| 3 | b | 9473 | 272.2 | 0.0009367 | 3.442 | +1 | 3 |
| 7 | y | 3.358E+04 | 272.2 | 5.464E-05 | 0.2008 | +1 | 2 |
| - | - | 3100 | 273.2 | - | - | 0 | - |
| - | - | 5361 | 276.2 | - | - | 0 | - |
| - | - | 2037 | 277.2 | - | - | 0 | - |
| 6 | b | 563 | 277.2 | 0.002947 | 10.63 | +2 | 6 |
| - | - | 1101 | 280.1 | - | - | 0 | - |
| - | - | 688.5 | 280.2 | - | - | 0 | - |
| - | - | 2345 | 282.2 | - | - | 0 | - |
| 6 | b | 2915 | 286.2 | 0.005233 | 18.29 | +2 | 6 |
| - | - | 536.2 | 286.2 | - | - | 0 | - |
| - | - | 720.4 | 286.7 | - | - | 0 | - |
| - | - | 8860 | 288.2 | - | - | 0 | - |
| - | - | 1356 | 289.2 | - | - | 0 | - |
| - | - | 1153 | 293.1 | - | - | 0 | - |
| - | - | 1617 | 298.1 | - | - | 0 | - |
| - | - | 728.2 | 298.2 | - | - | 0 | - |
| - | - | 788.4 | 299.2 | - | - | 0 | - |
| - | - | 648.6 | 299.2 | - | - | 0 | - |
| - | - | 735 | 300.2 | - | - | 0 | - |
| - | - | 5157 | 306.2 | - | - | 0 | - |
| - | - | 1155 | 306.7 | - | - | 0 | - |
| - | - | 806.8 | 308.2 | - | - | 0 | - |
| - | - | 705.5 | 310.2 | - | - | 0 | - |
| - | - | 2795 | 312.2 | - | - | 0 | - |
| - | - | 573.8 | 312.2 | - | - | 0 | - |
| - | - | 1189 | 319.7 | - | - | 0 | - |
| - | - | 1700 | 323.2 | - | - | 0 | - |
| - | - | 670 | 323.2 | - | - | 0 | - |
| - | - | 1.484E+04 | 325.2 | - | - | 0 | - |
| - | - | 2006 | 326.2 | - | - | 0 | - |
| 3 | y | 2.703E+04 | 327.7 | 0.0007518 | 2.294 | +2 | 6 |
| 3 | y | 2.561E+04 | 328.2 | 0.005135 | 15.65 | +2 | 6 |
| - | - | 8382 | 328.7 | - | - | 0 | - |
| - | - | 1214 | 329.2 | - | - | 0 | - |
| - | - | 596.4 | 332.9 | - | - | 0 | - |
| 3 | y | 3891 | 336.7 | 6.933E-05 | 0.2059 | +2 | 6 |
| - | - | 1085 | 337.2 | - | - | 0 | - |
| - | - | 4045 | 340.2 | - | - | 0 | - |
| - | - | 5644 | 341.2 | - | - | 0 | - |
| - | - | 1484 | 341.7 | - | - | 0 | - |
| - | - | 1476 | 342.2 | - | - | 0 | - |
| - | - | 2585 | 343.2 | - | - | 0 | - |
| - | - | 1173 | 350.9 | - | - | 0 | - |
| - | - | 743.3 | 351.2 | - | - | 0 | - |
| - | - | 671.6 | 352.9 | - | - | 0 | - |
| 4 | b | 8020 | 353.2 | 0.0002532 | 0.717 | +1 | 4 |
| - | - | 1889 | 354.2 | - | - | 0 | - |
| - | - | 1544 | 355.2 | - | - | 0 | - |
| - | - | 814.6 | 358.2 | - | - | 0 | - |
| - | - | 2694 | 359.2 | - | - | 0 | - |
| - | - | 740.5 | 362 | - | - | 0 | - |
| 2 | y | 3.095E+04 | 363.2 | 0.0005709 | 1.572 | +2 | 7 |
| - | - | 1.187E+04 | 363.7 | - | - | 0 | - |
| - | - | 4336 | 364.2 | - | - | 0 | - |
| - | - | 672.8 | 365.2 | - | - | 0 | - |
| - | - | 822.7 | 366.2 | - | - | 0 | - |
| - | - | 586.6 | 367.2 | - | - | 0 | - |
| 6 | y | 3919 | 368.2 | 0.0004711 | 1.279 | +1 | 3 |
| - | - | 959.5 | 369.2 | - | - | 0 | - |
| 4 | b | 895.2 | 371.2 | 0.00148 | 3.986 | +1 | 4 |
| 2 | y | 1366 | 372.2 | 0.0006653 | 1.787 | +2 | 7 |
| - | - | 965.5 | 379.2 | - | - | 0 | - |
| - | - | 886.8 | 380.2 | - | - | 0 | - |
| - | - | 2825 | 383.2 | - | - | 0 | - |
| 6 | y | 1.684E+04 | 385.3 | 0.0003479 | 0.903 | +1 | 3 |
| - | - | 579 | 385.8 | - | - | 0 | - |
| - | - | 3149 | 386.3 | - | - | 0 | - |
| - | - | 619.6 | 387.3 | - | - | 0 | - |
| - | - | 1611 | 389.2 | - | - | 0 | - |
| - | - | 886.9 | 390.2 | - | - | 0 | - |
| - | - | 7994 | 391.2 | - | - | 0 | - |
| - | - | 3062 | 391.7 | - | - | 0 | - |
| - | - | 989.9 | 392.2 | - | - | 0 | - |
| - | - | 2945 | 403.8 | - | - | 0 | - |
| 0 | Precursor | 2.39E+04 | 412.8 | 0.0003846 | 0.9318 | +2 | -1 |
| - | - | 8732 | 413.3 | - | - | 0 | - |
| - | - | 2364 | 413.8 | - | - | 0 | - |
| - | - | 1571 | 420.9 | - | - | 0 | - |
| - | - | 1045 | 432.2 | - | - | 0 | - |
| - | - | 2110 | 436.3 | - | - | 0 | - |
| - | - | 922.3 | 439.8 | - | - | 0 | - |
| 5 | b | 761 | 440.3 | 9.749E-05 | 0.2214 | +1 | 5 |
| - | - | 1071 | 442.3 | - | - | 0 | - |
| 5 | y | 3567 | 454.3 | 0.004722 | 10.39 | +1 | 4 |
| 5 | y | 1153 | 455.3 | 0.000551 | 1.21 | +1 | 4 |
| 5 | y | 1.818E+05 | 472.3 | 0.0002108 | 0.4462 | +1 | 4 |
| - | - | 4.214E+04 | 473.3 | - | - | 0 | - |
| - | - | 7578 | 474.3 | - | - | 0 | - |
| - | - | 2591 | 482.3 | - | - | 0 | - |
| - | - | 1.096E+04 | 486.3 | - | - | 0 | - |
| - | - | 1.393E+04 | 487.3 | - | - | 0 | - |
| - | - | 3441 | 488.3 | - | - | 0 | - |
| - | - | 695.6 | 489.3 | - | - | 0 | - |
| - | - | 829.4 | 508.3 | - | - | 0 | - |
| - | - | 1393 | 509.3 | - | - | 0 | - |
| - | - | 751.2 | 531.3 | - | - | 0 | - |
| 6 | b | 1722 | 553.3 | 0.007922 | 14.32 | +1 | 6 |
| - | - | 2770 | 557.3 | - | - | 0 | - |
| - | - | 3953 | 558.3 | - | - | 0 | - |
| - | - | 732.4 | 559.3 | - | - | 0 | - |
| 6 | b | 6.04E+04 | 571.3 | 0.01011 | 17.7 | +1 | 6 |
| - | - | 1.78E+04 | 572.4 | - | - | 0 | - |
| - | - | 3764 | 573.4 | - | - | 0 | - |
| - | - | 1189 | 581.3 | - | - | 0 | - |
| - | - | 611 | 597.3 | - | - | 0 | - |
| - | - | 1828 | 611.4 | - | - | 0 | - |
| - | - | 980.3 | 612.4 | - | - | 0 | - |
| - | - | 6221 | 628.4 | - | - | 0 | - |
| - | - | 1866 | 629.4 | - | - | 0 | - |
| - | - | 883.8 | 638.4 | - | - | 0 | - |
| - | - | 2388 | 644.4 | - | - | 0 | - |
| - | - | 651.8 | 647.5 | - | - | 0 | - |
| 3 | y | 8791 | 654.4 | 0.008082 | 12.35 | +1 | 6 |
| 3 | y | 2.031E+04 | 655.4 | 0.006987 | 10.66 | +1 | 6 |
| - | - | 7257 | 656.4 | - | - | 0 | - |
| - | - | 1517 | 657.4 | - | - | 0 | - |
| 3 | y | 1.376E+05 | 672.4 | 0.001313 | 1.953 | +1 | 6 |
| - | - | 4.748E+04 | 673.4 | - | - | 0 | - |
| - | - | 762.7 | 673.5 | - | - | 0 | - |
| - | - | 1.135E+04 | 674.4 | - | - | 0 | - |
| - | - | 1.142E+04 | 682.4 | - | - | 0 | - |
| - | - | 3945 | 683.4 | - | - | 0 | - |
| - | - | 908.4 | 684.4 | - | - | 0 | - |
| - | - | 1361 | 699.4 | - | - | 0 | - |
| 2 | y | 1.907E+04 | 725.4 | 0.002044 | 2.818 | +1 | 7 |
| - | - | 6092 | 726.4 | - | - | 0 | - |
| - | - | 1299 | 727.4 | - | - | 0 | - |
| - | - | 1213 | 735.4 | - | - | 0 | - |
| 2 | y | 5.956E+04 | 743.4 | 0.001622 | 2.182 | +1 | 7 |
| - | - | 2.046E+04 | 744.4 | - | - | 0 | - |
| - | - | 5032 | 745.4 | - | - | 0 | - |
| - | - | 1185 | 753.4 | - | - | 0 | - |
| - | - | 617.3 | 985.1 | - | - | 0 | - |
| - | - | 637.8 | 1230 | - | - | 0 | - |
| - | - | 683.6 | 1468 | - | - | 0 | - |
| - | - | 624.2 | 2226 | - | - | 0 | - |
| - | - | 779.4 | 2413 | - | - | 0 | - |
| - | - | 661.9 | 2492 | - | - | 0 | - |
| - | - | 673.9 | 2827 | - | - | 0 | - |
| - | - | 692 | 3004 | - | - | 0 | - |

m/z Charge Intensity FragmentType MassShift Position
120.080810546875 0 811.44855
122.67640686035156 0 492.83173
126.05526733398438 0 379.63275
126.12830352783203 0 500.77435
127.08671569824219 0 1881.0231
127.1230239868164 0 1785.6232
127.76991271972656 0 367.6988
128.1071319580078 0 3765.9636
128.69775390625 0 341.0255
129.01858520507812 0 829.4374
129.10238647460938 0 7200.576
130.05010986328125 0 522.016
130.08641052246094 0 3188.1724
130.09776306152344 0 456.3807
130.1056365966797 0 842.01227
131.09010314941406 0 457.86395
133.09747314453125 0 865.37463
134.5106658935547 0 457.7433
136.07601928710938 0 528.7296
138.09132385253906 0 1712.5288
140.0819549560547 0 10720.032
141.08558654785156 0 908.38306
141.1023406982422 0 9978.422
141.52182006835938 0 395.49203
142.10009765625 0 501.1761
142.10572814941406 0 2661.2131
143.11801147460938 0 75590.414 a 1
144.11544799804688 0 988.45734
144.1213836669922 0 5443.275
145.06756591796875 0 374.2352
145.09730529785156 0 4184.3936
145.26612854003906 0 453.33228
147.1129913330078 0 3644.527
148.11598205566406 0 559.4661
148.94711303710938 0 1123.7865
149.02342224121094 0 9595.7295
150.02679443359375 0 5672.846
150.6641845703125 0 375.60938
151.02964782714844 0 453.10156
155.08145141601562 0 2009.2373
155.1179656982422 0 9468.691
156.1140594482422 0 496.52582
156.12118530273438 0 836.27795
157.0132293701172 0 482.86893
157.0973663330078 0 994.4228
157.10867309570312 0 1866.894
157.13365173339844 0 20515.225
157.14105224609375 0 1087.577
158.09242248535156 0 10680.168 y Ammonia loss 7
158.13034057617188 0 803.22815
158.1370086669922 0 5602.746
159.0957489013672 0 481.42242
159.1129608154297 0 1944.206
169.0972137451172 0 9993.288
170.10072326660156 0 2801.1553
171.1128692626953 0 67857.45 b 1
171.14938354492188 0 2728.4102
172.10926818847656 0 685.8918
172.1162567138672 0 4948.9844
173.09207153320312 0 9906.348
173.12852478027344 0 77396.04
173.45101928710938 0 1935.0858
174.09494018554688 0 971.1896
174.1245880126953 0 835.13983
174.13189697265625 0 6032.4097
175.1190185546875 0 21558.676 y 7
176.1222686767578 0 1113.539
176.68894958496094 0 421.1486
183.1128387451172 0 16550.006
183.14920043945312 0 10311.903
184.11643981933594 0 1372.9225
184.1526336669922 0 2874.5774
185.08045959472656 0 1146.5062
185.10337829589844 0 1782.0554
185.12844848632812 0 5293.6343
185.16481018066406 0 545.77325
187.10772705078125 0 4317.15
195.1129913330078 0 1491.4596
199.14402770996094 0 2428.1436
201.12338256835938 0 28593.535
202.12672424316406 0 2863.1472
209.09193420410156 0 564.8034
209.12840270996094 0 676.16235
210.12339782714844 0 1914.1212
211.1075897216797 0 3518.429
211.14413452148438 0 5357.6294
212.1392059326172 0 7698.0337
212.147705078125 0 988.2516
213.12338256835938 0 1807.7511
213.1427001953125 0 866.9043
213.16000366210938 0 775.46045
215.13868713378906 0 837.65735
216.1423797607422 0 542.862
222.12477111816406 0 496.05753
225.15951538085938 0 472.66748
226.1193084716797 0 644.8024
226.15499877929688 0 11138.369 a Water loss 2
227.1038360595703 0 724.42303
227.13885498046875 0 4460.5283
227.15895080566406 0 1352.8601
228.13421630859375 0 2914.2927 y Ammonia loss 4
229.11788940429688 0 682.77075
230.1499786376953 0 3780.6116 d 2
238.11907958984375 0 580.39526
240.13417053222656 0 1786.6893
241.13803100585938 0 644.0945
243.13404846191406 0 950.9568
243.6495361328125 0 1053.4928
244.1291961669922 0 1023.88214
244.15135192871094 0 950.62146
244.16529846191406 0 648.6515 a 2
252.1343536376953 0 653.3795
254.14981079101562 0 31034.172 b Water loss 2
255.14512634277344 0 72996.3 y Ammonia loss 6
256.14837646484375 0 9535.887
258.144775390625 0 5980.659
259.1484069824219 0 2801.0264
262.1396179199219 0 651.90765
266.1498718261719 0 2095.2407
267.1526184082031 0 513.5016
270.1445007324219 0 10422.556
271.14849853515625 0 1095.0634
271.1750793457031 0 878.98096
272.1595458984375 0 9472.976 b 2
272.1716613769531 0 33578.523 y 6
273.1744384765625 0 3100.3665
276.1551208496094 0 5360.6147
277.157958984375 0 2037.2748
277.1737976074219 0 563.00214 b Water loss 5
280.1291809082031 0 1101.1171
280.16412353515625 0 688.478
282.18109130859375 0 2344.7808
286.1813659667969 0 2915.2373 b 5
286.1973571777344 0 536.209
286.6841735839844 0 720.42303
288.1551818847656 0 8860.204
289.158447265625 0 1356.3206
293.0998840332031 0 1152.8341
298.1396484375 0 1616.5844
298.1766662597656 0 728.1868
299.1894226074219 0 788.4297
299.2074890136719 0 648.55853
300.19219970703125 0 734.9501
306.17919921875 0 5157.349
306.6803894042969 0 1154.5858
308.1979064941406 0 806.7802
310.17620849609375 0 705.5257
312.19146728515625 0 2794.8745
312.2278747558594 0 573.80035
319.6841735839844 0 1189.3695
323.1722412109375 0 1699.8197
323.2080993652344 0 669.9632
325.2234191894531 0 14844.218
326.2265625 0 2005.6897
327.6995544433594 0 27026.977 y Water loss 2
328.19744873046875 0 25608.982 y Ammonia loss 2
328.69805908203125 0 8381.848
329.20013427734375 0 1214.2025
332.8738708496094 0 596.448
336.7056579589844 0 3891.31 y 2
337.2069396972656 0 1085.3346
340.1866760253906 0 4044.76
341.1840515136719 0 5644.163
341.69818115234375 0 1484.3599
342.1856689453125 0 1475.532
343.2336730957031 0 2584.9578
350.9261474609375 0 1173.0797
351.2008056640625 0 743.29974
352.90374755859375 0 671.6017
353.21807861328125 0 8019.985 b Water loss 3
354.22021484375 0 1889.4733
355.233154296875 0 1543.6547
358.1971130371094 0 814.6047
359.1932067871094 0 2693.755
361.9769592285156 0 740.46454
363.2182922363281 0 30951.791 y Water loss 1
363.71966552734375 0 11867.757
364.2214050292969 0 4335.929
365.2185974121094 0 672.80115
366.18487548828125 0 822.7261
367.194580078125 0 586.6069
368.228759765625 0 3919.4797 y Ammonia loss 5
369.2314758300781 0 959.46466
371.2274169921875 0 895.1892 b 3
372.2234802246094 0 1365.5651 y 1
379.2340087890625 0 965.5169
380.2367858886719 0 886.8114
383.22833251953125 0 2824.711
385.25543212890625 0 16836.314 y 5
385.80560302734375 0 578.96356
386.2578125 0 3149.477
387.2589416503906 0 619.63806
389.23748779296875 0 1611.2944
390.241943359375 0 886.9238
391.2314758300781 0 7994.2036
391.7333679199219 0 3062.2405
392.2353515625 0 989.91437
403.8130187988281 0 2944.6323
412.752685546875 0 23897.305 Precursor Water loss
413.2543029785156 0 8732.141
413.7547607421875 0 2364.1003
420.8568115234375 0 1570.8728
432.24603271484375 0 1044.6768
436.2554626464844 0 2110.1296
439.8420715332031 0 922.30273
440.2504577636719 0 760.95306 b Water loss 4
442.2772216796875 0 1070.609
454.27252197265625 0 3567.1187 y Water loss 4
455.2618103027344 0 1152.5978 y Ammonia loss 4
472.28759765625 0 181792.03 y 4
473.290283203125 0 42141.152
474.2924499511719 0 7578.273
482.2712707519531 0 2591.1006
486.2917785644531 0 10961.611
487.2944641113281 0 13932.028
488.2971496582031 0 3440.5615
489.2984619140625 0 695.5822
508.27593994140625 0 829.4045
509.27899169921875 0 1392.9752
531.3116455078125 0 751.1597
553.3423461914062 0 1721.5765 b Water loss 5
557.3279418945312 0 2769.9885
558.3314208984375 0 3953.3008
559.3302612304688 0 732.41626
571.3551025390625 0 60398.64 b 5
572.358154296875 0 17800.193
573.3603515625 0 3763.7695
581.3397827148438 0 1189.1599
597.332275390625 0 610.9533
611.3519897460938 0 1828.1653
612.3543090820312 0 980.2851
628.3764038085938 0 6221.3794
629.3803100585938 0 1865.634
638.361328125 0 883.77563
644.3969116210938 0 2387.638
647.4617309570312 0 651.7629
654.38525390625 0 8790.682 y Water loss 2
655.3843383789062 0 20307.514 y Ammonia loss 2
656.3863525390625 0 7257.1123
657.390380859375 0 1517.2949
672.402587890625 0 137614.6 y 2
673.4052734375 0 47480.99
673.5003662109375 0 762.72394
674.4078979492188 0 11354.069
682.386962890625 0 11423.272
683.3900146484375 0 3944.8862
684.390380859375 0 908.4151
699.411865234375 0 1360.9652
725.4284057617188 0 19068.744 y Water loss 1
726.4309692382812 0 6091.819
727.4288940429688 0 1298.7628
735.412841796875 0 1212.91
743.4393920898438 0 59558.887 y 1
744.4420166015625 0 20456.771
745.444580078125 0 5032.2246
753.4224243164062 0 1185.3237
985.09716796875 0 617.267
1230.140869140625 0 637.84564
1468.07568359375 0 683.6171
2225.526611328125 0 624.19354
2413.307373046875 0 779.44867
2491.77783203125 0 661.9123
2826.57568359375 0 673.94293
3003.561767578125 0 692.03955

Spectrum Details

|  |  |
| --- | --- |
| Matched peaks? Matched peaks The total absolute number of peaks matched. Additionally in brackets the total fraction of peaks matched and the total number of peaks is shown. | 35 (13.26% of 264) |
| FDR? FDR The false discovery rate estimated for this peptide. It is calculated by matching all theoretical fragments with a non-integer shift with the raw peaks for this spectrum. This is done with 40 different shifts. The resulting percentage is the average number of annotated peaks over the number of annotated peaks with the correct spectrum. | 0.34% |
| Satellite FDR? Satellite FDR See the FDR for details on its calculation. This satellite ion specific FDR only contains the satellite ions (d/w) for I/L/J positions. | - |
| PSM Score? PSM Score The PSM Score as given by Hecklib to this annotated spectrum. It is shown with three significant figures. | 418 |

#### Spectrum 5587? Spectrum 5587 The raw spectrum of this peptide as annotated by Hecklib. The fragments are coloured according to ion type (see legend). Any peaks with a star '\*' as text can be hovered over to see the full details, first the ion type second the mass shift type. By hovering over the amino acids in the peptide or ions in the legend the corresponding peaks are highlighted. By toggling the 'Unassigned' label you can turn the background (unassigned) peaks on or off in the plot. By updating the slider in the Ion legend you can update the spectrum to only show the top X% of the peaks with labels. The top X% means any peak that is within X% of the highest intensity. By dragging in the spectrum you can zoom in to a specific part of the spectrum and use 'Zoom Out' to get back to the original zoom level. The annotation of the spectrum is based on the given sequence in the peptides file and is done with different software so inconsistencies are likely. The peaks are annotated based on the given sequence, with 20 ppm tolerance.

Copy Data

##### Spectrum 5587 (TSV)

###### Preview

```
Loading example...
```

*Click on the button to copy the data to your clipboard.*

Mz MinMz MaxIntensity Max

WidthHeightPeptide font sizePeptide stroke widthSpectrum font sizeSpectrum stroke widthCompact peptide

Ion legend

wxyz

abcd

OtherUnassignedIon^Charge^_Position_Show for top:%

VATVSJPR

02.10e+44.21e+46.31e+48.41e+4

Zoom Out

a^+1^_2_y^+1^_1_b^+1^_2_y^+1^_1_a^+1^_3_y^+2^_4_d^+1^_3_a^+1^_3_b^+1^_3_y^+1^_2_b^+1^_3_y^+1^_2_b^+2^_6_y^+2^_6_y^+2^_6_y^+2^_6_b^+1^_4_y^+2^_7_y^+1^_3_b^+1^_4_y^+1^_3_\*b^+1^_5_y^+1^_4_y^+1^_4_y^+1^_4_b^+1^_6_b^+1^_6_y^+1^_6_y^+1^_6_y^+1^_6_y^+1^_7_y^+1^_7_

0652130319552606

Fragment Matches Table

Show background peaks

| Position | Ion type | Intensity | mz Theoretical | mz Error (Th) | mz Error (ppm) | Charge | Series Number |
| --- | --- | --- | --- | --- | --- | --- | --- |
| - | - | 1700 | 120.1 | - | - | 0 | - |
| - | - | 551.5 | 127.1 | - | - | 0 | - |
| - | - | 1352 | 127.1 | - | - | 0 | - |
| - | - | 1672 | 128.1 | - | - | 0 | - |
| - | - | 425.2 | 128.9 | - | - | 0 | - |
| - | - | 1332 | 129 | - | - | 0 | - |
| - | - | 5063 | 129.1 | - | - | 0 | - |
| - | - | 1699 | 130.1 | - | - | 0 | - |
| - | - | 490.8 | 130.1 | - | - | 0 | - |
| - | - | 393.5 | 133.1 | - | - | 0 | - |
| - | - | 547.3 | 136.1 | - | - | 0 | - |
| - | - | 827.7 | 138.1 | - | - | 0 | - |
| - | - | 6841 | 140.1 | - | - | 0 | - |
| - | - | 6183 | 141.1 | - | - | 0 | - |
| - | - | 1505 | 142.1 | - | - | 0 | - |
| 2 | a | 3.622E+04 | 143.1 | 0.000183 | 1.278 | +1 | 2 |
| - | - | 387.6 | 143.3 | - | - | 0 | - |
| - | - | 441.3 | 144 | - | - | 0 | - |
| - | - | 2007 | 144.1 | - | - | 0 | - |
| - | - | 1445 | 145.1 | - | - | 0 | - |
| - | - | 491.9 | 147.1 | - | - | 0 | - |
| - | - | 2561 | 147.1 | - | - | 0 | - |
| - | - | 545.5 | 148.1 | - | - | 0 | - |
| - | - | 1.025E+04 | 149 | - | - | 0 | - |
| - | - | 5113 | 150 | - | - | 0 | - |
| - | - | 624.9 | 151 | - | - | 0 | - |
| - | - | 1496 | 155.1 | - | - | 0 | - |
| - | - | 4357 | 155.1 | - | - | 0 | - |
| - | - | 505.1 | 157 | - | - | 0 | - |
| - | - | 690.2 | 157.1 | - | - | 0 | - |
| - | - | 1096 | 157.1 | - | - | 0 | - |
| - | - | 1.638E+04 | 157.1 | - | - | 0 | - |
| 8 | y | 5158 | 158.1 | 0.0002636 | 1.667 | +1 | 1 |
| - | - | 587.6 | 158.1 | - | - | 0 | - |
| - | - | 3746 | 158.1 | - | - | 0 | - |
| - | - | 648.2 | 159.1 | - | - | 0 | - |
| - | - | 459.1 | 159.5 | - | - | 0 | - |
| - | - | 446.2 | 161.9 | - | - | 0 | - |
| - | - | 444 | 162.8 | - | - | 0 | - |
| - | - | 678.7 | 167 | - | - | 0 | - |
| - | - | 514.3 | 167.1 | - | - | 0 | - |
| - | - | 6413 | 169.1 | - | - | 0 | - |
| - | - | 2069 | 170.1 | - | - | 0 | - |
| 2 | b | 3.138E+04 | 171.1 | 0.0001567 | 0.9155 | +1 | 2 |
| - | - | 2002 | 172.1 | - | - | 0 | - |
| - | - | 5637 | 173.1 | - | - | 0 | - |
| - | - | 3.699E+04 | 173.1 | - | - | 0 | - |
| - | - | 919.5 | 173.5 | - | - | 0 | - |
| - | - | 593.9 | 174.1 | - | - | 0 | - |
| - | - | 2662 | 174.1 | - | - | 0 | - |
| 8 | y | 9309 | 175.1 | 0.0001274 | 0.7276 | +1 | 1 |
| - | - | 1166 | 176.1 | - | - | 0 | - |
| - | - | 459.3 | 180 | - | - | 0 | - |
| - | - | 7738 | 183.1 | - | - | 0 | - |
| - | - | 6720 | 183.1 | - | - | 0 | - |
| - | - | 586.3 | 184.1 | - | - | 0 | - |
| - | - | 2257 | 184.2 | - | - | 0 | - |
| - | - | 1103 | 185.1 | - | - | 0 | - |
| - | - | 921.4 | 185.1 | - | - | 0 | - |
| - | - | 3976 | 185.1 | - | - | 0 | - |
| - | - | 676 | 186.1 | - | - | 0 | - |
| - | - | 1973 | 187.1 | - | - | 0 | - |
| - | - | 474.1 | 195.7 | - | - | 0 | - |
| - | - | 1.418E+04 | 201.1 | - | - | 0 | - |
| - | - | 597.9 | 202.1 | - | - | 0 | - |
| - | - | 922.8 | 202.1 | - | - | 0 | - |
| - | - | 828.2 | 210.1 | - | - | 0 | - |
| - | - | 1783 | 211.1 | - | - | 0 | - |
| - | - | 2721 | 211.1 | - | - | 0 | - |
| - | - | 664.5 | 211.2 | - | - | 0 | - |
| - | - | 3705 | 212.1 | - | - | 0 | - |
| - | - | 810.3 | 212.1 | - | - | 0 | - |
| - | - | 1266 | 213.1 | - | - | 0 | - |
| 3 | a | 4621 | 226.2 | 0.0001023 | 0.4522 | +1 | 3 |
| - | - | 2626 | 227.1 | - | - | 0 | - |
| 5 | y | 1516 | 228.1 | 0.0001621 | 0.7103 | +2 | 4 |
| 3 | d | 1898 | 230.1 | 9.121E-05 | 0.3963 | +1 | 3 |
| - | - | 674.6 | 238.1 | - | - | 0 | - |
| - | - | 1503 | 240.1 | - | - | 0 | - |
| - | - | 875.9 | 244.2 | - | - | 0 | - |
| 3 | a | 858.7 | 244.2 | 0.0004832 | 1.979 | +1 | 3 |
| - | - | 589.4 | 251.2 | - | - | 0 | - |
| 3 | b | 1.351E+04 | 254.1 | 4.543E-05 | 0.1788 | +1 | 3 |
| 7 | y | 3.429E+04 | 255.1 | 8.15E-05 | 0.3194 | +1 | 2 |
| - | - | 3324 | 256.1 | - | - | 0 | - |
| - | - | 4125 | 258.1 | - | - | 0 | - |
| - | - | 2305 | 259.1 | - | - | 0 | - |
| - | - | 542 | 264.2 | - | - | 0 | - |
| - | - | 1416 | 266.2 | - | - | 0 | - |
| - | - | 836.7 | 267.2 | - | - | 0 | - |
| - | - | 554.5 | 269.8 | - | - | 0 | - |
| - | - | 5643 | 270.1 | - | - | 0 | - |
| 3 | b | 4902 | 272.2 | 0.0007536 | 2.769 | +1 | 3 |
| 7 | y | 1.422E+04 | 272.2 | 9.795E-05 | 0.3599 | +1 | 2 |
| - | - | 1464 | 273.2 | - | - | 0 | - |
| - | - | 3242 | 276.2 | - | - | 0 | - |
| - | - | 1424 | 277.2 | - | - | 0 | - |
| - | - | 507.1 | 277.9 | - | - | 0 | - |
| - | - | 711.9 | 278.2 | - | - | 0 | - |
| - | - | 810 | 280.1 | - | - | 0 | - |
| - | - | 1164 | 282.2 | - | - | 0 | - |
| 6 | b | 709.9 | 286.2 | 0.004898 | 17.11 | +2 | 6 |
| - | - | 622.6 | 286.7 | - | - | 0 | - |
| - | - | 2717 | 288.2 | - | - | 0 | - |
| - | - | 997.5 | 293.1 | - | - | 0 | - |
| - | - | 1060 | 298.1 | - | - | 0 | - |
| - | - | 1012 | 299.2 | - | - | 0 | - |
| - | - | 1795 | 306.2 | - | - | 0 | - |
| - | - | 623.2 | 306.7 | - | - | 0 | - |
| - | - | 801 | 308 | - | - | 0 | - |
| - | - | 537.9 | 309.2 | - | - | 0 | - |
| - | - | 1428 | 312.2 | - | - | 0 | - |
| - | - | 871.6 | 323.2 | - | - | 0 | - |
| - | - | 6633 | 325.2 | - | - | 0 | - |
| - | - | 833 | 326.2 | - | - | 0 | - |
| 3 | y | 1.308E+04 | 327.7 | 0.0007824 | 2.387 | +2 | 6 |
| 3 | y | 1.537E+04 | 328.2 | 0.004829 | 14.72 | +2 | 6 |
| - | - | 5455 | 328.7 | - | - | 0 | - |
| - | - | 1301 | 329.2 | - | - | 0 | - |
| - | - | 586.4 | 332.9 | - | - | 0 | - |
| 3 | y | 899.7 | 336.7 | 0.001107 | 3.288 | +2 | 6 |
| - | - | 1991 | 340.2 | - | - | 0 | - |
| - | - | 3207 | 341.2 | - | - | 0 | - |
| - | - | 651.1 | 341.7 | - | - | 0 | - |
| - | - | 614.2 | 343.2 | - | - | 0 | - |
| - | - | 1872 | 350.9 | - | - | 0 | - |
| - | - | 962 | 352.2 | - | - | 0 | - |
| 4 | b | 3641 | 353.2 | 0.0002045 | 0.579 | +1 | 4 |
| - | - | 827.6 | 354.2 | - | - | 0 | - |
| - | - | 641.5 | 355.2 | - | - | 0 | - |
| 2 | y | 1.599E+04 | 363.2 | 0.0002047 | 0.5637 | +2 | 7 |
| - | - | 6094 | 363.7 | - | - | 0 | - |
| - | - | 772.3 | 364.2 | - | - | 0 | - |
| 6 | y | 1210 | 368.2 | 0.0007191 | 1.953 | +1 | 3 |
| 4 | b | 672.5 | 371.2 | 0.001542 | 4.153 | +1 | 4 |
| - | - | 965.3 | 380.2 | - | - | 0 | - |
| - | - | 840.7 | 383.2 | - | - | 0 | - |
| 6 | y | 6941 | 385.3 | 0.0002258 | 0.5861 | +1 | 3 |
| - | - | 1637 | 386.3 | - | - | 0 | - |
| - | - | 971.7 | 390.2 | - | - | 0 | - |
| - | - | 3364 | 391.2 | - | - | 0 | - |
| - | - | 1255 | 391.7 | - | - | 0 | - |
| - | - | 1399 | 403.8 | - | - | 0 | - |
| - | - | 695.6 | 404 | - | - | 0 | - |
| 0 | Precursor | 1.033E+04 | 412.8 | 0.0001404 | 0.3403 | +2 | -1 |
| - | - | 5088 | 413.3 | - | - | 0 | - |
| - | - | 1467 | 413.8 | - | - | 0 | - |
| - | - | 948.8 | 420.9 | - | - | 0 | - |
| - | - | 1.737E+04 | 421.2 | - | - | 0 | - |
| - | - | 3385 | 422.2 | - | - | 0 | - |
| - | - | 653.5 | 439.8 | - | - | 0 | - |
| 5 | b | 588.3 | 440.3 | 0.0006163 | 1.4 | +1 | 5 |
| 5 | y | 1146 | 454.3 | 0.003928 | 8.647 | +1 | 4 |
| - | - | 610 | 454.4 | - | - | 0 | - |
| 5 | y | 711.2 | 455.3 | 0.001467 | 3.221 | +1 | 4 |
| 5 | y | 8.332E+04 | 472.3 | 2.869E-06 | 0.006074 | +1 | 4 |
| - | - | 1.925E+04 | 473.3 | - | - | 0 | - |
| - | - | 3244 | 474.3 | - | - | 0 | - |
| - | - | 1158 | 482.3 | - | - | 0 | - |
| - | - | 8601 | 486.3 | - | - | 0 | - |
| - | - | 1.025E+04 | 487.3 | - | - | 0 | - |
| - | - | 2891 | 488.3 | - | - | 0 | - |
| - | - | 816.7 | 508.3 | - | - | 0 | - |
| - | - | 934.8 | 509.3 | - | - | 0 | - |
| 6 | b | 776.6 | 553.3 | 0.002307 | 4.169 | +1 | 6 |
| - | - | 1587 | 557.3 | - | - | 0 | - |
| - | - | 2967 | 558.3 | - | - | 0 | - |
| - | - | 809.1 | 559.3 | - | - | 0 | - |
| 6 | b | 2.74E+04 | 571.3 | 0.01078 | 18.88 | +1 | 6 |
| - | - | 7522 | 572.4 | - | - | 0 | - |
| - | - | 1392 | 573.4 | - | - | 0 | - |
| - | - | 1243 | 611.3 | - | - | 0 | - |
| - | - | 865.7 | 621.3 | - | - | 0 | - |
| - | - | 2228 | 628.4 | - | - | 0 | - |
| 3 | y | 5283 | 654.4 | 0.00796 | 12.16 | +1 | 6 |
| 3 | y | 1.387E+04 | 655.4 | 0.007231 | 11.03 | +1 | 6 |
| - | - | 4995 | 656.4 | - | - | 0 | - |
| - | - | 942 | 657.4 | - | - | 0 | - |
| 3 | y | 6.221E+04 | 672.4 | 0.0009467 | 1.408 | +1 | 6 |
| - | - | 2.148E+04 | 673.4 | - | - | 0 | - |
| - | - | 4831 | 674.4 | - | - | 0 | - |
| - | - | 3797 | 682.4 | - | - | 0 | - |
| - | - | 1588 | 683.4 | - | - | 0 | - |
| 2 | y | 7510 | 725.4 | 0.001434 | 1.976 | +1 | 7 |
| - | - | 3508 | 726.4 | - | - | 0 | - |
| - | - | 582.4 | 727.4 | - | - | 0 | - |
| 2 | y | 2.706E+04 | 743.4 | 0.001134 | 1.526 | +1 | 7 |
| - | - | 9789 | 744.4 | - | - | 0 | - |
| - | - | 1404 | 745.4 | - | - | 0 | - |
| - | - | 581.4 | 753.4 | - | - | 0 | - |
| - | - | 683.5 | 812.6 | - | - | 0 | - |
| - | - | 518.1 | 814.4 | - | - | 0 | - |
| - | - | 642.3 | 963.7 | - | - | 0 | - |
| - | - | 692.6 | 2580 | - | - | 0 | - |

m/z Charge Intensity FragmentType MassShift Position
120.0809555053711 0 1700.3367
127.0868911743164 0 551.48517
127.12322998046875 0 1351.7178
128.10731506347656 0 1671.8844
128.94711303710938 0 425.1826
129.01840209960938 0 1331.8702
129.10243225097656 0 5063.103
130.08644104003906 0 1699.2479
130.10552978515625 0 490.8032
133.0972442626953 0 393.45615
136.07583618164062 0 547.3465
138.0914764404297 0 827.67194
140.08203125 0 6841.2935
141.10244750976562 0 6182.978
142.10568237304688 0 1504.5941
143.11807250976562 0 36223.277 a 1
143.3478240966797 0 387.59366
144.03463745117188 0 441.2989
144.12149047851562 0 2006.8129
145.09727478027344 0 1444.6718
147.0640869140625 0 491.86685
147.11289978027344 0 2560.8123
148.11648559570312 0 545.5382
149.02346801757812 0 10253.757
150.0269317626953 0 5112.9585
151.02792358398438 0 624.891
155.08160400390625 0 1496.0306
155.11814880371094 0 4357.3433
157.0136260986328 0 505.15
157.09718322753906 0 690.23285
157.10838317871094 0 1095.9445
157.1337127685547 0 16384.129
158.09266662597656 0 5158.2954 y Ammonia loss 7
158.1310272216797 0 587.5859
158.1371307373047 0 3745.5964
159.09132385253906 0 648.2393
159.50076293945312 0 459.08124
161.92274475097656 0 446.18613
162.840576171875 0 444.03662
167.0338592529297 0 678.7033
167.08111572265625 0 514.32654
169.09730529785156 0 6413.2505
170.1007080078125 0 2068.655
171.1129608154297 0 31384.006 b 1
172.11647033691406 0 2001.9594
173.09222412109375 0 5637.4424
173.12863159179688 0 36989.51
173.45230102539062 0 919.4612
174.09576416015625 0 593.8602
174.1320037841797 0 2662.0532
175.11907958984375 0 9308.647 y 7
176.12254333496094 0 1165.5403
180.00555419921875 0 459.32986
183.1129913330078 0 7737.913
183.14939880371094 0 6720.2666
184.1165008544922 0 586.3372
184.15289306640625 0 2256.5054
185.08099365234375 0 1103.2268
185.1031951904297 0 921.44775
185.12863159179688 0 3975.8354
186.13221740722656 0 676.04297
187.10792541503906 0 1973.0021
195.6663055419922 0 474.14548
201.12342834472656 0 14180.84
202.1063232421875 0 597.87335
202.1266632080078 0 922.7715
210.12344360351562 0 828.1711
211.10763549804688 0 1782.9454
211.14418029785156 0 2721.4092
211.15420532226562 0 664.5202
212.13929748535156 0 3705.3855
212.1493377685547 0 810.31903
213.1234588623047 0 1265.7369
226.1551055908203 0 4620.6924 a Water loss 2
227.1391143798828 0 2626.3674
228.13442993164062 0 1516.1145 y Ammonia loss 4
230.15000915527344 0 1897.8026 d 2
238.1191864013672 0 674.6311
240.1343536376953 0 1502.6512
244.15159606933594 0 875.92365
244.1650848388672 0 858.6682 a 2
251.20834350585938 0 589.36505
254.14996337890625 0 13511.07 b Water loss 2
255.14524841308594 0 34287.176 y Ammonia loss 6
256.1484375 0 3323.757
258.1448974609375 0 4124.926
259.1485900878906 0 2305.0789
264.1712341308594 0 541.9795
266.150146484375 0 1416.2554
267.1534729003906 0 836.7122
269.8431701660156 0 554.54266
270.14495849609375 0 5642.7134
272.15972900390625 0 4902.2515 b 2
272.17181396484375 0 14220.007 y 6
273.17529296875 0 1463.9922
276.1549987792969 0 3241.9802
277.1585998535156 0 1424.4332
277.9163513183594 0 507.09665
278.1604919433594 0 711.8921
280.1296081542969 0 810.03754
282.18121337890625 0 1164.3572
286.1810302734375 0 709.8518 b 5
286.6837158203125 0 622.58246
288.15521240234375 0 2717.099
293.1003112792969 0 997.54065
298.139404296875 0 1060.1241
299.18896484375 0 1011.8823
306.1790771484375 0 1794.6077
306.6797180175781 0 623.1757
307.98699951171875 0 801.0318
309.2073669433594 0 537.8815
312.19189453125 0 1428.1809
323.1720886230469 0 871.5997
325.2234191894531 0 6632.7773
326.2266540527344 0 832.95294
327.69952392578125 0 13082.28 y Water loss 2
328.1971435546875 0 15366.719 y Ammonia loss 2
328.69781494140625 0 5454.637
329.1995544433594 0 1301.179
332.872314453125 0 586.3874
336.7066955566406 0 899.72906 y 2
340.1868896484375 0 1991.1019
341.1846008300781 0 3207.1746
341.69842529296875 0 651.059
343.2319030761719 0 614.2069
350.926513671875 0 1871.9894
352.2405090332031 0 962.0267
353.2185363769531 0 3640.6636 b Water loss 3
354.2193908691406 0 827.5802
355.2338562011719 0 641.5226
363.2186584472656 0 15985.298 y Water loss 1
363.7203063964844 0 6094.427
364.22064208984375 0 772.2785
368.2299499511719 0 1210.2185 y Ammonia loss 5
371.2304382324219 0 672.53973 b 3
380.2370910644531 0 965.2598
383.2285461425781 0 840.65704
385.25555419921875 0 6940.8433 y 5
386.25714111328125 0 1637.0144
390.2424621582031 0 971.65283
391.23211669921875 0 3363.7336
391.73284912109375 0 1254.9587
403.8133850097656 0 1399.1404
404.0072021484375 0 695.5545
412.7529296875 0 10333.783 Precursor Water loss
413.2550354003906 0 5088.2783
413.755859375 0 1467.261
420.8586730957031 0 948.753
421.20452880859375 0 17365.68
422.2083435058594 0 3384.885
439.8445129394531 0 653.50793
440.2509765625 0 588.2711 b Water loss 4
454.2733154296875 0 1145.6599 y Water loss 4
454.38970947265625 0 609.9622
455.2627258300781 0 711.1525 y Ammonia loss 4
472.2878112792969 0 83315.805 y 4
473.2904968261719 0 19247.623
474.29351806640625 0 3243.8503
482.27069091796875 0 1157.602
486.2919921875 0 8601.395
487.2953186035156 0 10247.543
488.29736328125 0 2890.6316
508.27435302734375 0 816.669
509.27734375 0 934.8139
553.3367309570312 0 776.61646 b Water loss 5
557.3291015625 0 1586.6427
558.3314819335938 0 2966.8152
559.3380737304688 0 809.05444
571.3557739257812 0 27395.727 b 5
572.358154296875 0 7521.782
573.3632202148438 0 1391.6996
611.3495483398438 0 1242.5625
621.28466796875 0 865.6968
628.376953125 0 2227.7205
654.3853759765625 0 5282.537 y Water loss 2
655.3845825195312 0 13865.56 y Ammonia loss 2
656.3870849609375 0 4994.7227
657.3921508789062 0 942.02826
672.4029541015625 0 62206.48 y 2
673.40576171875 0 21482.254
674.4090576171875 0 4830.538
682.3876953125 0 3797.3828
683.391357421875 0 1587.7241
725.4290161132812 0 7509.6646 y Water loss 1
726.4317016601562 0 3508.12
727.4342651367188 0 582.39465
743.4398803710938 0 27063.846 y 1
744.4424438476562 0 9788.525
745.4442749023438 0 1403.8306
753.432373046875 0 581.3572
812.5517578125 0 683.53534
814.4218139648438 0 518.14777
963.7474975585938 0 642.301
2580.423583984375 0 692.6302

Spectrum Details

|  |  |
| --- | --- |
| Matched peaks? Matched peaks The total absolute number of peaks matched. Additionally in brackets the total fraction of peaks matched and the total number of peaks is shown. | 33 (17.01% of 194) |
| FDR? FDR The false discovery rate estimated for this peptide. It is calculated by matching all theoretical fragments with a non-integer shift with the raw peaks for this spectrum. This is done with 40 different shifts. The resulting percentage is the average number of annotated peaks over the number of annotated peaks with the correct spectrum. | 0.14% |
| Satellite FDR? Satellite FDR See the FDR for details on its calculation. This satellite ion specific FDR only contains the satellite ions (d/w) for I/L/J positions. | - |
| PSM Score? PSM Score The PSM Score as given by Hecklib to this annotated spectrum. It is shown with three significant figures. | 418 |

#### Spectrum 5424? Spectrum 5424 The raw spectrum of this peptide as annotated by Hecklib. The fragments are coloured according to ion type (see legend). Any peaks with a star '\*' as text can be hovered over to see the full details, first the ion type second the mass shift type. By hovering over the amino acids in the peptide or ions in the legend the corresponding peaks are highlighted. By toggling the 'Unassigned' label you can turn the background (unassigned) peaks on or off in the plot. By updating the slider in the Ion legend you can update the spectrum to only show the top X% of the peaks with labels. The top X% means any peak that is within X% of the highest intensity. By dragging in the spectrum you can zoom in to a specific part of the spectrum and use 'Zoom Out' to get back to the original zoom level. The annotation of the spectrum is based on the given sequence in the peptides file and is done with different software so inconsistencies are likely. The peaks are annotated based on the given sequence, with 20 ppm tolerance.

Copy Data

##### Spectrum 5424 (TSV)

###### Preview

```
Loading example...
```

*Click on the button to copy the data to your clipboard.*

Mz MinMz MaxIntensity Max

WidthHeightPeptide font sizePeptide stroke widthSpectrum font sizeSpectrum stroke widthCompact peptide

Ion legend

wxyz

abcd

OtherUnassignedIon^Charge^_Position_Show for top:%

VATVSJPR

02.90e+45.79e+48.69e+41.16e+5

Zoom Out

y^+2^_2_a^+1^_2_y^+1^_1_b^+1^_2_y^+1^_1_a^+1^_3_y^+2^_4_d^+1^_3_b^+1^_3_y^+1^_2_b^+1^_3_y^+1^_2_y^+2^_5_y^+2^_6_y^+2^_6_y^+2^_6_b^+1^_4_y^+2^_7_y^+1^_3_y^+1^_3_\*y^+1^_4_y^+1^_4_y^+1^_4_b^+1^_6_b^+1^_6_y^+1^_6_y^+1^_6_y^+1^_6_y^+1^_7_y^+1^_7_

037775511321510

Fragment Matches Table

Show background peaks

| Position | Ion type | Intensity | mz Theoretical | mz Error (Th) | mz Error (ppm) | Charge | Series Number |
| --- | --- | --- | --- | --- | --- | --- | --- |
| - | - | 7687 | 120.1 | - | - | 0 | - |
| - | - | 1977 | 121.1 | - | - | 0 | - |
| - | - | 384.6 | 122.7 | - | - | 0 | - |
| - | - | 1371 | 127.1 | - | - | 0 | - |
| - | - | 1745 | 127.1 | - | - | 0 | - |
| - | - | 2361 | 128.1 | - | - | 0 | - |
| - | - | 1595 | 129 | - | - | 0 | - |
| - | - | 5147 | 129.1 | - | - | 0 | - |
| - | - | 697.2 | 130.1 | - | - | 0 | - |
| - | - | 2006 | 130.1 | - | - | 0 | - |
| - | - | 681.3 | 130.1 | - | - | 0 | - |
| - | - | 821.6 | 130.1 | - | - | 0 | - |
| - | - | 1402 | 133.1 | - | - | 0 | - |
| - | - | 598.4 | 136 | - | - | 0 | - |
| - | - | 496.9 | 136.1 | - | - | 0 | - |
| 7 | y | 412.9 | 136.6 | 0.0005917 | 4.332 | +2 | 2 |
| - | - | 739.1 | 138.1 | - | - | 0 | - |
| - | - | 7382 | 140.1 | - | - | 0 | - |
| - | - | 454.8 | 141.1 | - | - | 0 | - |
| - | - | 9246 | 141.1 | - | - | 0 | - |
| - | - | 2043 | 142.1 | - | - | 0 | - |
| 2 | a | 4.711E+04 | 143.1 | 0.0003508 | 2.451 | +1 | 2 |
| - | - | 3648 | 144.1 | - | - | 0 | - |
| - | - | 2772 | 145.1 | - | - | 0 | - |
| - | - | 2456 | 147.1 | - | - | 0 | - |
| - | - | 610.5 | 149 | - | - | 0 | - |
| - | - | 9544 | 149 | - | - | 0 | - |
| - | - | 4922 | 150 | - | - | 0 | - |
| - | - | 1721 | 155.1 | - | - | 0 | - |
| - | - | 5657 | 155.1 | - | - | 0 | - |
| - | - | 916.6 | 156.1 | - | - | 0 | - |
| - | - | 851.7 | 157 | - | - | 0 | - |
| - | - | 1231 | 157.1 | - | - | 0 | - |
| - | - | 895.4 | 157.1 | - | - | 0 | - |
| - | - | 1.584E+04 | 157.1 | - | - | 0 | - |
| 8 | y | 7569 | 158.1 | 0.0003399 | 2.15 | +1 | 1 |
| - | - | 4697 | 158.1 | - | - | 0 | - |
| - | - | 1630 | 159.1 | - | - | 0 | - |
| - | - | 1469 | 159.1 | - | - | 0 | - |
| - | - | 692.3 | 160.1 | - | - | 0 | - |
| - | - | 588.5 | 167 | - | - | 0 | - |
| - | - | 531.8 | 167.1 | - | - | 0 | - |
| - | - | 8937 | 169.1 | - | - | 0 | - |
| - | - | 1700 | 170.1 | - | - | 0 | - |
| 2 | b | 4.257E+04 | 171.1 | 0.0003245 | 1.896 | +1 | 2 |
| - | - | 3256 | 172.1 | - | - | 0 | - |
| - | - | 6526 | 173.1 | - | - | 0 | - |
| - | - | 4.753E+04 | 173.1 | - | - | 0 | - |
| - | - | 2009 | 173.5 | - | - | 0 | - |
| - | - | 4165 | 174.1 | - | - | 0 | - |
| 8 | y | 1.533E+04 | 175.1 | 0.0003258 | 1.86 | +1 | 1 |
| - | - | 1313 | 176.1 | - | - | 0 | - |
| - | - | 1.116E+04 | 183.1 | - | - | 0 | - |
| - | - | 7320 | 183.1 | - | - | 0 | - |
| - | - | 1019 | 184.1 | - | - | 0 | - |
| - | - | 2545 | 184.2 | - | - | 0 | - |
| - | - | 797.5 | 185.1 | - | - | 0 | - |
| - | - | 926.3 | 185.1 | - | - | 0 | - |
| - | - | 735.9 | 185.1 | - | - | 0 | - |
| - | - | 4533 | 185.1 | - | - | 0 | - |
| - | - | 1276 | 186.1 | - | - | 0 | - |
| - | - | 3122 | 187.1 | - | - | 0 | - |
| - | - | 4369 | 191.1 | - | - | 0 | - |
| - | - | 1349 | 192.1 | - | - | 0 | - |
| - | - | 1137 | 195.1 | - | - | 0 | - |
| - | - | 1.873E+04 | 201.1 | - | - | 0 | - |
| - | - | 1555 | 202.1 | - | - | 0 | - |
| - | - | 1374 | 210.1 | - | - | 0 | - |
| - | - | 2153 | 211.1 | - | - | 0 | - |
| - | - | 3845 | 211.1 | - | - | 0 | - |
| - | - | 4041 | 212.1 | - | - | 0 | - |
| - | - | 680.3 | 212.1 | - | - | 0 | - |
| - | - | 1488 | 213.1 | - | - | 0 | - |
| - | - | 553 | 213.7 | - | - | 0 | - |
| - | - | 503.3 | 215.1 | - | - | 0 | - |
| - | - | 815.2 | 225.1 | - | - | 0 | - |
| 3 | a | 6538 | 226.2 | 0.0002549 | 1.127 | +1 | 3 |
| - | - | 676.8 | 226.2 | - | - | 0 | - |
| - | - | 3741 | 227.1 | - | - | 0 | - |
| - | - | 1001 | 227.2 | - | - | 0 | - |
| 5 | y | 1690 | 228.1 | 9.463E-06 | 0.04148 | +2 | 4 |
| - | - | 551.8 | 229.1 | - | - | 0 | - |
| 3 | d | 2734 | 230.1 | 0.0004422 | 1.921 | +1 | 3 |
| - | - | 664.9 | 232.1 | - | - | 0 | - |
| - | - | 1860 | 240.1 | - | - | 0 | - |
| - | - | 607.3 | 243.1 | - | - | 0 | - |
| - | - | 1621 | 252.1 | - | - | 0 | - |
| - | - | 700.6 | 254.1 | - | - | 0 | - |
| 3 | b | 1.742E+04 | 254.1 | 0.0002285 | 0.8992 | +1 | 3 |
| - | - | 1192 | 254.2 | - | - | 0 | - |
| 7 | y | 4.609E+04 | 255.1 | 0.0003104 | 1.216 | +1 | 2 |
| - | - | 4926 | 256.1 | - | - | 0 | - |
| - | - | 645.7 | 257.1 | - | - | 0 | - |
| - | - | 4250 | 258.1 | - | - | 0 | - |
| - | - | 1694 | 259.1 | - | - | 0 | - |
| - | - | 662.6 | 263.1 | - | - | 0 | - |
| - | - | 1422 | 266.2 | - | - | 0 | - |
| - | - | 7888 | 270.1 | - | - | 0 | - |
| 3 | b | 8642 | 272.2 | 0.00054 | 1.984 | +1 | 3 |
| 7 | y | 1.857E+04 | 272.2 | 0.0004336 | 1.593 | +1 | 2 |
| - | - | 1897 | 273.2 | - | - | 0 | - |
| - | - | 4461 | 276.2 | - | - | 0 | - |
| - | - | 1871 | 277.2 | - | - | 0 | - |
| - | - | 1815 | 282.2 | - | - | 0 | - |
| - | - | 1092 | 282.6 | - | - | 0 | - |
| 4 | y | 2472 | 286.2 | 0.000349 | 1.22 | +2 | 5 |
| - | - | 5016 | 288.2 | - | - | 0 | - |
| - | - | 809.4 | 293.1 | - | - | 0 | - |
| - | - | 644.1 | 294.8 | - | - | 0 | - |
| - | - | 1432 | 298.1 | - | - | 0 | - |
| - | - | 870.7 | 299.2 | - | - | 0 | - |
| - | - | 669.1 | 302.1 | - | - | 0 | - |
| - | - | 2675 | 306.2 | - | - | 0 | - |
| - | - | 966.4 | 306.7 | - | - | 0 | - |
| - | - | 1598 | 312.2 | - | - | 0 | - |
| - | - | 1045 | 312.7 | - | - | 0 | - |
| - | - | 883.9 | 313.2 | - | - | 0 | - |
| - | - | 744.2 | 319.7 | - | - | 0 | - |
| - | - | 728.6 | 323.2 | - | - | 0 | - |
| - | - | 9076 | 325.2 | - | - | 0 | - |
| - | - | 1268 | 326.2 | - | - | 0 | - |
| 3 | y | 1.794E+04 | 327.7 | 0.0003551 | 1.084 | +2 | 6 |
| 3 | y | 2.015E+04 | 328.2 | 0.005531 | 16.85 | +2 | 6 |
| - | - | 9074 | 328.7 | - | - | 0 | - |
| - | - | 1432 | 329.2 | - | - | 0 | - |
| - | - | 1571 | 334.6 | - | - | 0 | - |
| - | - | 900.1 | 335.6 | - | - | 0 | - |
| 3 | y | 2982 | 336.7 | 5.274E-05 | 0.1566 | +2 | 6 |
| - | - | 3029 | 340.2 | - | - | 0 | - |
| - | - | 3689 | 341.2 | - | - | 0 | - |
| - | - | 684.9 | 341.7 | - | - | 0 | - |
| - | - | 1006 | 342.7 | - | - | 0 | - |
| - | - | 1086 | 343.2 | - | - | 0 | - |
| - | - | 725 | 344.6 | - | - | 0 | - |
| - | - | 833.3 | 348.6 | - | - | 0 | - |
| - | - | 978.9 | 349.6 | - | - | 0 | - |
| - | - | 867 | 350.9 | - | - | 0 | - |
| - | - | 727.4 | 351.2 | - | - | 0 | - |
| - | - | 936.5 | 352.1 | - | - | 0 | - |
| 4 | b | 4966 | 353.2 | 0.000113 | 0.3198 | +1 | 4 |
| - | - | 891.4 | 354.1 | - | - | 0 | - |
| - | - | 1034 | 354.2 | - | - | 0 | - |
| - | - | 1355 | 354.6 | - | - | 0 | - |
| - | - | 865.6 | 355.1 | - | - | 0 | - |
| - | - | 984.6 | 359.2 | - | - | 0 | - |
| - | - | 1038 | 361.1 | - | - | 0 | - |
| 2 | y | 1.954E+04 | 363.2 | 0.000131 | 0.3606 | +2 | 7 |
| - | - | 8970 | 363.7 | - | - | 0 | - |
| - | - | 2266 | 364.2 | - | - | 0 | - |
| - | - | 1038 | 364.6 | - | - | 0 | - |
| - | - | 796.8 | 365.2 | - | - | 0 | - |
| - | - | 608.2 | 367.1 | - | - | 0 | - |
| - | - | 743.9 | 367.6 | - | - | 0 | - |
| 6 | y | 2448 | 368.2 | 0.0002575 | 0.6993 | +1 | 3 |
| - | - | 652 | 370.1 | - | - | 0 | - |
| - | - | 1544 | 373.6 | - | - | 0 | - |
| - | - | 698.9 | 374.6 | - | - | 0 | - |
| - | - | 766.3 | 380.6 | - | - | 0 | - |
| - | - | 1222 | 383.2 | - | - | 0 | - |
| - | - | 810.2 | 383.6 | - | - | 0 | - |
| 6 | y | 9831 | 385.3 | 1.834E-05 | 0.0476 | +1 | 3 |
| - | - | 1901 | 386.3 | - | - | 0 | - |
| - | - | 853.1 | 388.6 | - | - | 0 | - |
| - | - | 981.6 | 389.2 | - | - | 0 | - |
| - | - | 655.9 | 389.6 | - | - | 0 | - |
| - | - | 896.1 | 390.2 | - | - | 0 | - |
| - | - | 4620 | 391.2 | - | - | 0 | - |
| - | - | 2195 | 391.7 | - | - | 0 | - |
| - | - | 677.3 | 392.2 | - | - | 0 | - |
| - | - | 1139 | 402.2 | - | - | 0 | - |
| - | - | 1033 | 403.8 | - | - | 0 | - |
| 0 | Precursor | 1.647E+04 | 412.8 | 0.0003478 | 0.8427 | +2 | -1 |
| - | - | 6053 | 413.3 | - | - | 0 | - |
| - | - | 2021 | 413.8 | - | - | 0 | - |
| - | - | 699.6 | 418.2 | - | - | 0 | - |
| - | - | 824.6 | 420.9 | - | - | 0 | - |
| - | - | 727.6 | 422.2 | - | - | 0 | - |
| - | - | 898.1 | 422.2 | - | - | 0 | - |
| - | - | 1908 | 436.3 | - | - | 0 | - |
| - | - | 1001 | 439.8 | - | - | 0 | - |
| - | - | 846.3 | 442.3 | - | - | 0 | - |
| 5 | y | 1455 | 454.3 | 0.00463 | 10.19 | +1 | 4 |
| 5 | y | 1045 | 455.3 | 0.0005815 | 1.277 | +1 | 4 |
| 5 | y | 1.147E+05 | 472.3 | 0.0003996 | 0.8461 | +1 | 4 |
| - | - | 2.715E+04 | 473.3 | - | - | 0 | - |
| - | - | 987.5 | 473.3 | - | - | 0 | - |
| - | - | 3833 | 474.3 | - | - | 0 | - |
| - | - | 1073 | 482.3 | - | - | 0 | - |
| - | - | 7632 | 486.3 | - | - | 0 | - |
| - | - | 1.169E+04 | 487.3 | - | - | 0 | - |
| - | - | 3528 | 488.3 | - | - | 0 | - |
| - | - | 1173 | 505.3 | - | - | 0 | - |
| - | - | 2322 | 506.3 | - | - | 0 | - |
| - | - | 618.8 | 509.3 | - | - | 0 | - |
| 6 | b | 1297 | 553.3 | 0.01073 | 19.39 | +1 | 6 |
| - | - | 1565 | 557.3 | - | - | 0 | - |
| - | - | 3502 | 558.3 | - | - | 0 | - |
| - | - | 690.9 | 559.3 | - | - | 0 | - |
| 6 | b | 3.726E+04 | 571.3 | 0.01127 | 19.73 | +1 | 6 |
| - | - | 1.155E+04 | 572.4 | - | - | 0 | - |
| - | - | 2783 | 573.4 | - | - | 0 | - |
| - | - | 782.3 | 597.3 | - | - | 0 | - |
| - | - | 634 | 603.3 | - | - | 0 | - |
| - | - | 1674 | 611.3 | - | - | 0 | - |
| - | - | 638.1 | 612.3 | - | - | 0 | - |
| - | - | 2505 | 620.3 | - | - | 0 | - |
| - | - | 6757 | 621.3 | - | - | 0 | - |
| - | - | 1475 | 622.3 | - | - | 0 | - |
| - | - | 3144 | 628.4 | - | - | 0 | - |
| - | - | 1669 | 629.4 | - | - | 0 | - |
| 3 | y | 7409 | 654.4 | 0.007472 | 11.42 | +1 | 6 |
| 3 | y | 1.398E+04 | 655.4 | 0.007719 | 11.78 | +1 | 6 |
| - | - | 5444 | 656.4 | - | - | 0 | - |
| - | - | 1302 | 657.4 | - | - | 0 | - |
| 3 | y | 9.274E+04 | 672.4 | 0.0002753 | 0.4094 | +1 | 6 |
| - | - | 3.041E+04 | 673.4 | - | - | 0 | - |
| - | - | 7130 | 674.4 | - | - | 0 | - |
| - | - | 6677 | 682.4 | - | - | 0 | - |
| - | - | 1663 | 683.4 | - | - | 0 | - |
| - | - | 712 | 684.4 | - | - | 0 | - |
| - | - | 767.7 | 699.4 | - | - | 0 | - |
| 2 | y | 1.057E+04 | 725.4 | 0.0003352 | 0.462 | +1 | 7 |
| - | - | 3739 | 726.4 | - | - | 0 | - |
| - | - | 1209 | 727.4 | - | - | 0 | - |
| 2 | y | 3.573E+04 | 743.4 | 0.0004018 | 0.5405 | +1 | 7 |
| - | - | 1.309E+04 | 744.4 | - | - | 0 | - |
| - | - | 3172 | 745.4 | - | - | 0 | - |
| - | - | 784.9 | 753.4 | - | - | 0 | - |
| - | - | 898.1 | 768.3 | - | - | 0 | - |
| - | - | 676.1 | 1495 | - | - | 0 | - |

m/z Charge Intensity FragmentType MassShift Position
120.08113098144531 0 7686.9966
121.08447265625 0 1977.3906
122.69291687011719 0 384.635
127.08694458007812 0 1370.7466
127.12332153320312 0 1744.5656
128.1073760986328 0 2360.8828
129.01870727539062 0 1595.31
129.1025848388672 0 5146.615
130.05043029785156 0 697.1892
130.0865478515625 0 2005.7474
130.0980987548828 0 681.2566
130.10594177246094 0 821.5786
133.06105041503906 0 1401.553
136.0217742919922 0 598.43286
136.0760955810547 0 496.92728
136.590087890625 0 412.89764 y 6
138.0919189453125 0 739.1129
140.0821990966797 0 7381.8394
141.0851593017578 0 454.824
141.1025848388672 0 9246.181
142.10585021972656 0 2043.4353
143.1182403564453 0 47110.94 a 1
144.12152099609375 0 3647.5938
145.09750366210938 0 2771.7026
147.11322021484375 0 2456.4434
148.95608520507812 0 610.4609
149.02369689941406 0 9544.37
150.02699279785156 0 4921.6807
155.08163452148438 0 1720.674
155.1182098388672 0 5657.135
156.1216278076172 0 916.6079
157.01345825195312 0 851.6996
157.0974578857422 0 1230.6757
157.10926818847656 0 895.355
157.13385009765625 0 15835.566
158.09274291992188 0 7568.647 y Ammonia loss 7
158.13720703125 0 4697.0317
159.0921173095703 0 1630.2683
159.11331176757812 0 1468.7122
160.09542846679688 0 692.2555
167.03407287597656 0 588.5199
167.08291625976562 0 531.77985
169.0975341796875 0 8937.014
170.10096740722656 0 1699.807
171.11312866210938 0 42568.316 b 1
172.1165313720703 0 3256.2163
173.0923309326172 0 6525.7056
173.12879943847656 0 47526.84
173.45094299316406 0 2008.7281
174.13223266601562 0 4165.1777
175.11927795410156 0 15334.936 y 7
176.12274169921875 0 1312.745
183.1131591796875 0 11158.569
183.1496124267578 0 7320.4507
184.11672973632812 0 1019.0101
184.15298461914062 0 2545.1414
185.0814666748047 0 797.451
185.10353088378906 0 926.28314
185.1199188232422 0 735.8873
185.12876892089844 0 4533.0664
186.13204956054688 0 1276.236
187.10800170898438 0 3122.403
191.11822509765625 0 4368.535
192.12179565429688 0 1348.8796
195.11257934570312 0 1137.4426
201.12362670898438 0 18729.064
202.1269989013672 0 1554.802
210.1237335205078 0 1374.2688
211.10791015625 0 2152.9473
211.144287109375 0 3845.327
212.1397705078125 0 4041.384
212.1497039794922 0 680.2828
213.1235809326172 0 1487.9774
213.73287963867188 0 553.0087
215.1138916015625 0 503.25146
225.1234130859375 0 815.2308
226.15525817871094 0 6537.966 a Water loss 2
226.1671142578125 0 676.77905
227.139404296875 0 3741.4731
227.15904235839844 0 1001.0892
228.13427734375 0 1690.0593 y Ammonia loss 4
229.11837768554688 0 551.8254
230.15036010742188 0 2734.108 d 2
232.14125061035156 0 664.93274
240.1346893310547 0 1859.7886
243.1344757080078 0 607.287
252.13482666015625 0 1620.65
254.11402893066406 0 700.64594
254.150146484375 0 17415.705 b Water loss 2
254.1624298095703 0 1192.0264
255.14547729492188 0 46086.023 y Ammonia loss 6
256.1488037109375 0 4925.616
257.0709228515625 0 645.678
258.14495849609375 0 4249.871
259.1494140625 0 1693.5079
263.103271484375 0 662.61224
266.1503601074219 0 1422.259
270.1451721191406 0 7887.7153
272.1599426269531 0 8642.182 b 2
272.1721496582031 0 18566.393 y 6
273.1755676269531 0 1897.1448
276.1557922363281 0 4461.186
277.1590881347656 0 1870.6577
282.1812744140625 0 1815.2969
282.5570983886719 0 1092.375
286.1820983886719 0 2472.3923 y 3
288.1557312011719 0 5015.7847
293.0989685058594 0 809.3515
294.8403015136719 0 644.07336
298.14007568359375 0 1432.325
299.19036865234375 0 870.6632
302.117919921875 0 669.1285
306.1799011230469 0 2675.1453
306.6805114746094 0 966.4177
312.192138671875 0 1597.6407
312.6507568359375 0 1045.4335
313.19482421875 0 883.90857
319.6871337890625 0 744.24664
323.1719970703125 0 728.6102
325.2237854003906 0 9075.956
326.22747802734375 0 1268.3192
327.699951171875 0 17939.953 y Water loss 2
328.1978454589844 0 20152.25 y Ammonia loss 2
328.69879150390625 0 9074.04
329.19952392578125 0 1431.6794
334.5837707519531 0 1571.0094
335.58807373046875 0 900.1286
336.7055358886719 0 2981.8618 y 2
340.1868591308594 0 3029.459
341.1842956542969 0 3688.6594
341.69781494140625 0 684.9336
342.6606140136719 0 1006.36523
343.2337951660156 0 1085.8422
344.6001281738281 0 724.99396
348.63958740234375 0 833.26935
349.5891418457031 0 978.8648
350.9265441894531 0 867.04987
351.2413024902344 0 727.4135
352.1135559082031 0 936.4991
353.21844482421875 0 4965.833 b Water loss 3
354.11083984375 0 891.4395
354.2206726074219 0 1034.277
354.6399230957031 0 1354.5514
355.1417236328125 0 865.6389
359.1961669921875 0 984.57715
361.12042236328125 0 1038.1051
363.218994140625 0 19535.5 y Water loss 1
363.7205505371094 0 8970.259
364.2217102050781 0 2266.264
364.5940246582031 0 1037.5859
365.2182922363281 0 796.8094
367.1199035644531 0 608.24255
367.6213684082031 0 743.9104
368.2289733886719 0 2447.6887 y Ammonia loss 5
370.12554931640625 0 652.02045
373.5982666015625 0 1544.0695
374.59808349609375 0 698.89966
380.5879211425781 0 766.2954
383.2287902832031 0 1222.1307
383.5949401855469 0 810.1645
385.25579833984375 0 9830.621 y 5
386.2599182128906 0 1900.8439
388.6045227050781 0 853.1305
389.2391052246094 0 981.57556
389.6028137207031 0 655.8633
390.24432373046875 0 896.0739
391.2323913574219 0 4620.374
391.7337341308594 0 2194.8403
392.2351379394531 0 677.3286
402.20660400390625 0 1138.6965
403.81292724609375 0 1032.5833
412.75341796875 0 16466.178 Precursor Water loss
413.2547302246094 0 6052.8516
413.7567138671875 0 2020.8829
418.2190246582031 0 699.56586
420.8563537597656 0 824.64435
422.21185302734375 0 727.6329
422.24066162109375 0 898.1278
436.25567626953125 0 1907.9329
439.8473205566406 0 1001.38684
442.27880859375 0 846.30334
454.2726135253906 0 1455.0804 y Water loss 4
455.2618408203125 0 1044.8906 y Ammonia loss 4
472.2882080078125 0 114698.28 y 4
473.29119873046875 0 27151.49
473.33099365234375 0 987.53357
474.2930908203125 0 3833.192
482.2731018066406 0 1073.0273
486.2926025390625 0 7632.433
487.295654296875 0 11685.937
488.2989501953125 0 3528.3743
505.2514343261719 0 1173.4795
506.25616455078125 0 2321.8025
509.28021240234375 0 618.79266
553.3451538085938 0 1296.758 b Water loss 5
557.3285522460938 0 1565.4218
558.33203125 0 3501.5488
559.3385620117188 0 690.9251
571.3562622070312 0 37258.45 b 5
572.3590087890625 0 11547.117
573.3609008789062 0 2783.3013
597.334228515625 0 782.3332
603.2700805664062 0 634.04425
611.3499755859375 0 1673.529
612.3497924804688 0 638.1472
620.2786865234375 0 2504.5122
621.2813720703125 0 6756.7725
622.2833251953125 0 1475.1772
628.37744140625 0 3144.194
629.3787231445312 0 1668.8484
654.3858642578125 0 7409.0215 y Water loss 2
655.3850708007812 0 13980.539 y Ammonia loss 2
656.387939453125 0 5444.354
657.3900146484375 0 1301.6576
672.4036254882812 0 92737.875 y 2
673.4064331054688 0 30411
674.4095458984375 0 7130.1196
682.3875732421875 0 6676.998
683.3889770507812 0 1663.0779
684.393798828125 0 711.9868
699.4132690429688 0 767.6664
725.4301147460938 0 10570.479 y Water loss 1
726.4321899414062 0 3738.5305
727.4341430664062 0 1209.129
743.4406127929688 0 35734.613 y 1
744.443359375 0 13089.349
745.4452514648438 0 3172.1306
753.4232788085938 0 784.8793
768.3466796875 0 898.0889
1494.8489990234375 0 676.14575

Spectrum Details

|  |  |
| --- | --- |
| Matched peaks? Matched peaks The total absolute number of peaks matched. Additionally in brackets the total fraction of peaks matched and the total number of peaks is shown. | 31 (13.48% of 230) |
| FDR? FDR The false discovery rate estimated for this peptide. It is calculated by matching all theoretical fragments with a non-integer shift with the raw peaks for this spectrum. This is done with 40 different shifts. The resulting percentage is the average number of annotated peaks over the number of annotated peaks with the correct spectrum. | 1.00% |
| Satellite FDR? Satellite FDR See the FDR for details on its calculation. This satellite ion specific FDR only contains the satellite ions (d/w) for I/L/J positions. | - |
| PSM Score? PSM Score The PSM Score as given by Hecklib to this annotated spectrum. It is shown with three significant figures. | 349 |

#### Spectrum 5365? Spectrum 5365 The raw spectrum of this peptide as annotated by Hecklib. The fragments are coloured according to ion type (see legend). Any peaks with a star '\*' as text can be hovered over to see the full details, first the ion type second the mass shift type. By hovering over the amino acids in the peptide or ions in the legend the corresponding peaks are highlighted. By toggling the 'Unassigned' label you can turn the background (unassigned) peaks on or off in the plot. By updating the slider in the Ion legend you can update the spectrum to only show the top X% of the peaks with labels. The top X% means any peak that is within X% of the highest intensity. By dragging in the spectrum you can zoom in to a specific part of the spectrum and use 'Zoom Out' to get back to the original zoom level. The annotation of the spectrum is based on the given sequence in the peptides file and is done with different software so inconsistencies are likely. The peaks are annotated based on the given sequence, with 20 ppm tolerance.

Copy Data

##### Spectrum 5365 (TSV)

###### Preview

```
Loading example...
```

*Click on the button to copy the data to your clipboard.*

Mz MinMz MaxIntensity Max

WidthHeightPeptide font sizePeptide stroke widthSpectrum font sizeSpectrum stroke widthCompact peptide

Ion legend

wxyz

abcd

OtherUnassignedIon^Charge^_Position_Show for top:%

VATVSJPR

03.37e+46.74e+41.01e+51.35e+5

Zoom Out

a^+1^_2_y^+1^_1_b^+1^_2_y^+1^_1_a^+1^_3_y^+2^_4_d^+1^_3_b^+1^_3_y^+1^_2_b^+1^_3_y^+1^_2_b^+2^_6_y^+2^_6_y^+2^_6_y^+2^_6_b^+1^_4_y^+2^_7_y^+1^_3_y^+1^_3_\*y^+1^_4_y^+1^_4_y^+1^_4_b^+1^_6_b^+1^_6_y^+1^_6_y^+1^_6_y^+1^_6_y^+1^_7_y^+1^_7_

0837167425113349

Fragment Matches Table

Show background peaks

| Position | Ion type | Intensity | mz Theoretical | mz Error (Th) | mz Error (ppm) | Charge | Series Number |
| --- | --- | --- | --- | --- | --- | --- | --- |
| - | - | 9.678E+04 | 120.1 | - | - | 0 | - |
| - | - | 2.707E+04 | 121.1 | - | - | 0 | - |
| - | - | 799 | 122.1 | - | - | 0 | - |
| - | - | 351.6 | 123.1 | - | - | 0 | - |
| - | - | 1500 | 127.1 | - | - | 0 | - |
| - | - | 1629 | 127.1 | - | - | 0 | - |
| - | - | 434.5 | 127.2 | - | - | 0 | - |
| - | - | 2421 | 128.1 | - | - | 0 | - |
| - | - | 718.9 | 129 | - | - | 0 | - |
| - | - | 6186 | 129.1 | - | - | 0 | - |
| - | - | 505.8 | 129.1 | - | - | 0 | - |
| - | - | 1193 | 130.1 | - | - | 0 | - |
| - | - | 4005 | 130.1 | - | - | 0 | - |
| - | - | 2003 | 130.1 | - | - | 0 | - |
| - | - | 1098 | 130.1 | - | - | 0 | - |
| - | - | 1214 | 130.1 | - | - | 0 | - |
| - | - | 1171 | 131.1 | - | - | 0 | - |
| - | - | 474.7 | 131.1 | - | - | 0 | - |
| - | - | 4227 | 132.1 | - | - | 0 | - |
| - | - | 1129 | 133.1 | - | - | 0 | - |
| - | - | 478.2 | 133.1 | - | - | 0 | - |
| - | - | 972.4 | 138.1 | - | - | 0 | - |
| - | - | 8879 | 140.1 | - | - | 0 | - |
| - | - | 553 | 141.1 | - | - | 0 | - |
| - | - | 8621 | 141.1 | - | - | 0 | - |
| - | - | 1810 | 142.1 | - | - | 0 | - |
| 2 | a | 5.653E+04 | 143.1 | 0.0002745 | 1.918 | +1 | 2 |
| - | - | 667.5 | 144.1 | - | - | 0 | - |
| - | - | 3249 | 144.1 | - | - | 0 | - |
| - | - | 2894 | 145.1 | - | - | 0 | - |
| - | - | 2376 | 147.1 | - | - | 0 | - |
| - | - | 525.7 | 148.1 | - | - | 0 | - |
| - | - | 617.6 | 148.1 | - | - | 0 | - |
| - | - | 1.061E+04 | 149 | - | - | 0 | - |
| - | - | 4749 | 150 | - | - | 0 | - |
| - | - | 1476 | 155.1 | - | - | 0 | - |
| - | - | 6014 | 155.1 | - | - | 0 | - |
| - | - | 467.1 | 157 | - | - | 0 | - |
| - | - | 1098 | 157.1 | - | - | 0 | - |
| - | - | 1711 | 157.1 | - | - | 0 | - |
| - | - | 1.844E+04 | 157.1 | - | - | 0 | - |
| 8 | y | 1.293E+04 | 158.1 | 0.0002483 | 1.571 | +1 | 1 |
| - | - | 699.1 | 158.1 | - | - | 0 | - |
| - | - | 4639 | 158.1 | - | - | 0 | - |
| - | - | 2.455E+04 | 159.1 | - | - | 0 | - |
| - | - | 1063 | 159.1 | - | - | 0 | - |
| - | - | 996.9 | 160.1 | - | - | 0 | - |
| - | - | 9510 | 160.1 | - | - | 0 | - |
| - | - | 584.6 | 167.1 | - | - | 0 | - |
| - | - | 8717 | 169.1 | - | - | 0 | - |
| - | - | 752.8 | 170.1 | - | - | 0 | - |
| - | - | 1879 | 170.1 | - | - | 0 | - |
| - | - | 461.3 | 171.1 | - | - | 0 | - |
| 2 | b | 4.786E+04 | 171.1 | 0.0002482 | 1.451 | +1 | 2 |
| - | - | 1762 | 172.1 | - | - | 0 | - |
| - | - | 4569 | 172.1 | - | - | 0 | - |
| - | - | 9696 | 173.1 | - | - | 0 | - |
| - | - | 5.668E+04 | 173.1 | - | - | 0 | - |
| - | - | 1332 | 174.1 | - | - | 0 | - |
| - | - | 5193 | 174.1 | - | - | 0 | - |
| - | - | 1366 | 175.1 | - | - | 0 | - |
| 8 | y | 2.592E+04 | 175.1 | 0.0002495 | 1.425 | +1 | 1 |
| - | - | 2478 | 176.1 | - | - | 0 | - |
| - | - | 1.13E+04 | 183.1 | - | - | 0 | - |
| - | - | 8226 | 183.1 | - | - | 0 | - |
| - | - | 1191 | 184.1 | - | - | 0 | - |
| - | - | 2201 | 184.2 | - | - | 0 | - |
| - | - | 986.2 | 185.1 | - | - | 0 | - |
| - | - | 1132 | 185.1 | - | - | 0 | - |
| - | - | 4048 | 185.1 | - | - | 0 | - |
| - | - | 1261 | 186.1 | - | - | 0 | - |
| - | - | 1938 | 187.1 | - | - | 0 | - |
| - | - | 2999 | 187.1 | - | - | 0 | - |
| - | - | 713.2 | 188.1 | - | - | 0 | - |
| - | - | 686.1 | 190.1 | - | - | 0 | - |
| - | - | 434.5 | 190.4 | - | - | 0 | - |
| - | - | 4.344E+04 | 191.1 | - | - | 0 | - |
| - | - | 1.944E+04 | 192.1 | - | - | 0 | - |
| - | - | 528.1 | 193.1 | - | - | 0 | - |
| - | - | 1667 | 193.1 | - | - | 0 | - |
| - | - | 1128 | 195.1 | - | - | 0 | - |
| - | - | 826.7 | 198.1 | - | - | 0 | - |
| - | - | 574.1 | 199.1 | - | - | 0 | - |
| - | - | 2.019E+04 | 201.1 | - | - | 0 | - |
| - | - | 506.3 | 202.1 | - | - | 0 | - |
| - | - | 1480 | 202.1 | - | - | 0 | - |
| - | - | 1207 | 203.1 | - | - | 0 | - |
| - | - | 840.4 | 209.1 | - | - | 0 | - |
| - | - | 1556 | 210.1 | - | - | 0 | - |
| - | - | 875.6 | 210.9 | - | - | 0 | - |
| - | - | 1219 | 211.1 | - | - | 0 | - |
| - | - | 2058 | 211.1 | - | - | 0 | - |
| - | - | 3726 | 211.1 | - | - | 0 | - |
| - | - | 868.4 | 211.2 | - | - | 0 | - |
| - | - | 6194 | 212.1 | - | - | 0 | - |
| - | - | 972.7 | 212.1 | - | - | 0 | - |
| - | - | 1139 | 213.1 | - | - | 0 | - |
| - | - | 3472 | 215.1 | - | - | 0 | - |
| - | - | 638.9 | 216.1 | - | - | 0 | - |
| - | - | 1584 | 217.1 | - | - | 0 | - |
| - | - | 7625 | 219.1 | - | - | 0 | - |
| - | - | 4608 | 220.1 | - | - | 0 | - |
| - | - | 575.4 | 224.1 | - | - | 0 | - |
| - | - | 610.8 | 225.7 | - | - | 0 | - |
| 3 | a | 7512 | 226.2 | 0.0001938 | 0.857 | +1 | 3 |
| - | - | 3655 | 227.1 | - | - | 0 | - |
| - | - | 534.2 | 228.1 | - | - | 0 | - |
| 5 | y | 1916 | 228.1 | 9.463E-06 | 0.04148 | +2 | 4 |
| 3 | d | 3156 | 230.1 | 0.0001828 | 0.7941 | +1 | 3 |
| - | - | 663.1 | 231.2 | - | - | 0 | - |
| - | - | 5769 | 232.1 | - | - | 0 | - |
| - | - | 1864 | 233.1 | - | - | 0 | - |
| - | - | 1706 | 235.1 | - | - | 0 | - |
| - | - | 1155 | 236.1 | - | - | 0 | - |
| - | - | 1689 | 240.1 | - | - | 0 | - |
| - | - | 585.7 | 241.1 | - | - | 0 | - |
| - | - | 596.2 | 242.1 | - | - | 0 | - |
| - | - | 599.4 | 243.1 | - | - | 0 | - |
| - | - | 988.1 | 244.1 | - | - | 0 | - |
| - | - | 1261 | 244.1 | - | - | 0 | - |
| - | - | 1204 | 244.2 | - | - | 0 | - |
| - | - | 4145 | 246.1 | - | - | 0 | - |
| - | - | 2392 | 247.1 | - | - | 0 | - |
| - | - | 942.9 | 252.1 | - | - | 0 | - |
| - | - | 662.4 | 254.1 | - | - | 0 | - |
| 3 | b | 2.251E+04 | 254.1 | 0.000137 | 0.539 | +1 | 3 |
| 7 | y | 5.347E+04 | 255.1 | 0.0002036 | 0.7978 | +1 | 2 |
| - | - | 622.2 | 256.1 | - | - | 0 | - |
| - | - | 6384 | 256.1 | - | - | 0 | - |
| - | - | 5348 | 258.1 | - | - | 0 | - |
| - | - | 2399 | 259.1 | - | - | 0 | - |
| - | - | 4501 | 263.1 | - | - | 0 | - |
| - | - | 2453 | 264.1 | - | - | 0 | - |
| - | - | 604.9 | 265.1 | - | - | 0 | - |
| - | - | 1137 | 266.1 | - | - | 0 | - |
| - | - | 738.8 | 267.2 | - | - | 0 | - |
| - | - | 1195 | 269.1 | - | - | 0 | - |
| - | - | 606.5 | 270.1 | - | - | 0 | - |
| - | - | 9037 | 270.1 | - | - | 0 | - |
| - | - | 1123 | 271.1 | - | - | 0 | - |
| - | - | 852.4 | 271.2 | - | - | 0 | - |
| 3 | b | 7501 | 272.2 | 0.0006621 | 2.433 | +1 | 3 |
| 7 | y | 2.358E+04 | 272.2 | 0.0002811 | 1.033 | +1 | 2 |
| - | - | 2700 | 273.2 | - | - | 0 | - |
| - | - | 2233 | 274.1 | - | - | 0 | - |
| - | - | 1023 | 275.1 | - | - | 0 | - |
| - | - | 4362 | 276.2 | - | - | 0 | - |
| - | - | 2012 | 277.2 | - | - | 0 | - |
| - | - | 2795 | 282.2 | - | - | 0 | - |
| - | - | 642.2 | 282.6 | - | - | 0 | - |
| 6 | b | 2110 | 286.2 | 0.005172 | 18.07 | +2 | 6 |
| - | - | 7208 | 288.2 | - | - | 0 | - |
| - | - | 650.5 | 289.2 | - | - | 0 | - |
| - | - | 709.3 | 289.6 | - | - | 0 | - |
| - | - | 552.8 | 294.6 | - | - | 0 | - |
| - | - | 545.2 | 295.2 | - | - | 0 | - |
| - | - | 1334 | 298.1 | - | - | 0 | - |
| - | - | 973.5 | 299.2 | - | - | 0 | - |
| - | - | 656.3 | 299.2 | - | - | 0 | - |
| - | - | 684.7 | 300.2 | - | - | 0 | - |
| - | - | 3805 | 306.2 | - | - | 0 | - |
| - | - | 1255 | 306.7 | - | - | 0 | - |
| - | - | 1894 | 312.2 | - | - | 0 | - |
| - | - | 1084 | 312.7 | - | - | 0 | - |
| - | - | 1270 | 319.7 | - | - | 0 | - |
| - | - | 609.3 | 323.2 | - | - | 0 | - |
| - | - | 8987 | 325.2 | - | - | 0 | - |
| - | - | 1354 | 326.2 | - | - | 0 | - |
| - | - | 1430 | 327.7 | - | - | 0 | - |
| 3 | y | 2.105E+04 | 327.7 | 0.0006298 | 1.922 | +2 | 6 |
| 3 | y | 2.133E+04 | 328.2 | 0.005257 | 16.02 | +2 | 6 |
| - | - | 7135 | 328.7 | - | - | 0 | - |
| - | - | 1641 | 329.2 | - | - | 0 | - |
| - | - | 610.5 | 331.1 | - | - | 0 | - |
| - | - | 807.3 | 332.7 | - | - | 0 | - |
| - | - | 1898 | 334.1 | - | - | 0 | - |
| - | - | 3187 | 334.6 | - | - | 0 | - |
| - | - | 984.9 | 335.1 | - | - | 0 | - |
| - | - | 1006 | 335.1 | - | - | 0 | - |
| - | - | 898.4 | 335.6 | - | - | 0 | - |
| 3 | y | 2886 | 336.7 | 0.000283 | 0.8404 | +2 | 6 |
| - | - | 807.2 | 337.2 | - | - | 0 | - |
| - | - | 3191 | 340.2 | - | - | 0 | - |
| - | - | 438.7 | 341.2 | - | - | 0 | - |
| - | - | 609 | 341.2 | - | - | 0 | - |
| - | - | 4366 | 341.2 | - | - | 0 | - |
| - | - | 663.9 | 341.7 | - | - | 0 | - |
| - | - | 656.2 | 342.2 | - | - | 0 | - |
| - | - | 1767 | 342.7 | - | - | 0 | - |
| - | - | 1995 | 343.2 | - | - | 0 | - |
| - | - | 626.3 | 344.1 | - | - | 0 | - |
| - | - | 1635 | 349.6 | - | - | 0 | - |
| - | - | 1548 | 350.6 | - | - | 0 | - |
| - | - | 568 | 351.2 | - | - | 0 | - |
| - | - | 750.1 | 351.2 | - | - | 0 | - |
| - | - | 976.6 | 352.1 | - | - | 0 | - |
| - | - | 1134 | 352.6 | - | - | 0 | - |
| 4 | b | 6592 | 353.2 | 0.0002656 | 0.7518 | +1 | 4 |
| - | - | 652.3 | 353.6 | - | - | 0 | - |
| - | - | 814.1 | 354.1 | - | - | 0 | - |
| - | - | 827.1 | 354.2 | - | - | 0 | - |
| - | - | 1013 | 354.6 | - | - | 0 | - |
| - | - | 1007 | 355.1 | - | - | 0 | - |
| - | - | 875.9 | 355.2 | - | - | 0 | - |
| - | - | 632.7 | 356.1 | - | - | 0 | - |
| - | - | 620.3 | 358.2 | - | - | 0 | - |
| - | - | 1809 | 359.2 | - | - | 0 | - |
| - | - | 1670 | 361.1 | - | - | 0 | - |
| 2 | y | 2.423E+04 | 363.2 | 0.0001132 | 0.3116 | +2 | 7 |
| - | - | 8515 | 363.7 | - | - | 0 | - |
| - | - | 2807 | 364.2 | - | - | 0 | - |
| - | - | 3300 | 364.6 | - | - | 0 | - |
| - | - | 606.7 | 364.6 | - | - | 0 | - |
| - | - | 1480 | 365.1 | - | - | 0 | - |
| - | - | 1122 | 365.6 | - | - | 0 | - |
| - | - | 1171 | 366.6 | - | - | 0 | - |
| 6 | y | 3916 | 368.2 | 0.000288 | 0.7822 | +1 | 3 |
| - | - | 726.8 | 369.2 | - | - | 0 | - |
| - | - | 1384 | 371.1 | - | - | 0 | - |
| - | - | 989.2 | 372.1 | - | - | 0 | - |
| - | - | 2706 | 373.6 | - | - | 0 | - |
| - | - | 2593 | 374.1 | - | - | 0 | - |
| - | - | 1195 | 374.6 | - | - | 0 | - |
| - | - | 1383 | 375.7 | - | - | 0 | - |
| - | - | 755.8 | 376.1 | - | - | 0 | - |
| - | - | 831.6 | 376.2 | - | - | 0 | - |
| - | - | 967.9 | 379.6 | - | - | 0 | - |
| - | - | 886.9 | 380.1 | - | - | 0 | - |
| - | - | 1547 | 380.6 | - | - | 0 | - |
| - | - | 1914 | 383.2 | - | - | 0 | - |
| - | - | 834.6 | 384.2 | - | - | 0 | - |
| - | - | 1426 | 384.7 | - | - | 0 | - |
| - | - | 1936 | 385.2 | - | - | 0 | - |
| 6 | y | 1.16E+04 | 385.3 | 0.0001404 | 0.3645 | +1 | 3 |
| - | - | 1939 | 386.3 | - | - | 0 | - |
| - | - | 3210 | 388.6 | - | - | 0 | - |
| - | - | 1264 | 389.1 | - | - | 0 | - |
| - | - | 661.1 | 389.2 | - | - | 0 | - |
| - | - | 888.7 | 389.6 | - | - | 0 | - |
| - | - | 6804 | 391.2 | - | - | 0 | - |
| - | - | 2177 | 391.7 | - | - | 0 | - |
| - | - | 931.7 | 392.2 | - | - | 0 | - |
| - | - | 2151 | 401.2 | - | - | 0 | - |
| - | - | 1876 | 402.2 | - | - | 0 | - |
| - | - | 913.2 | 403.7 | - | - | 0 | - |
| 0 | Precursor | 1.746E+04 | 412.8 | 0.0003173 | 0.7688 | +2 | -1 |
| - | - | 7738 | 413.3 | - | - | 0 | - |
| - | - | 1169 | 413.8 | - | - | 0 | - |
| - | - | 4225 | 418.2 | - | - | 0 | - |
| - | - | 5094 | 419.2 | - | - | 0 | - |
| - | - | 1378 | 420.2 | - | - | 0 | - |
| - | - | 623.4 | 426.3 | - | - | 0 | - |
| - | - | 2282 | 436.3 | - | - | 0 | - |
| - | - | 691 | 439.8 | - | - | 0 | - |
| - | - | 991.8 | 442.3 | - | - | 0 | - |
| 5 | y | 2599 | 454.3 | 0.002647 | 5.826 | +1 | 4 |
| 5 | y | 1125 | 455.3 | 0.0005815 | 1.277 | +1 | 4 |
| 5 | y | 1.334E+05 | 472.3 | 0.0002775 | 0.5876 | +1 | 4 |
| - | - | 3.088E+04 | 473.3 | - | - | 0 | - |
| - | - | 5093 | 474.3 | - | - | 0 | - |
| - | - | 1423 | 482.3 | - | - | 0 | - |
| - | - | 9170 | 486.3 | - | - | 0 | - |
| - | - | 894.2 | 487.2 | - | - | 0 | - |
| - | - | 1.391E+04 | 487.3 | - | - | 0 | - |
| - | - | 3733 | 488.3 | - | - | 0 | - |
| - | - | 1.799E+04 | 505.3 | - | - | 0 | - |
| - | - | 2.855E+04 | 506.3 | - | - | 0 | - |
| - | - | 6646 | 507.3 | - | - | 0 | - |
| - | - | 1187 | 508.3 | - | - | 0 | - |
| - | - | 797.5 | 509.3 | - | - | 0 | - |
| - | - | 792.8 | 541.3 | - | - | 0 | - |
| 6 | b | 1428 | 553.3 | 0.007128 | 12.88 | +1 | 6 |
| - | - | 1842 | 557.3 | - | - | 0 | - |
| - | - | 3369 | 558.3 | - | - | 0 | - |
| 6 | b | 4.63E+04 | 571.3 | 0.01091 | 19.09 | +1 | 6 |
| - | - | 1.335E+04 | 572.4 | - | - | 0 | - |
| - | - | 2972 | 573.4 | - | - | 0 | - |
| - | - | 641.8 | 597.3 | - | - | 0 | - |
| - | - | 2002 | 602.3 | - | - | 0 | - |
| - | - | 5385 | 603.3 | - | - | 0 | - |
| - | - | 2154 | 604.3 | - | - | 0 | - |
| - | - | 1986 | 611.4 | - | - | 0 | - |
| - | - | 626.1 | 612.4 | - | - | 0 | - |
| - | - | 2.837E+04 | 620.3 | - | - | 0 | - |
| - | - | 6.581E+04 | 621.3 | - | - | 0 | - |
| - | - | 2.161E+04 | 622.3 | - | - | 0 | - |
| - | - | 4065 | 623.3 | - | - | 0 | - |
| - | - | 4127 | 628.4 | - | - | 0 | - |
| - | - | 1135 | 629.4 | - | - | 0 | - |
| - | - | 902.6 | 638.4 | - | - | 0 | - |
| 3 | y | 7716 | 654.4 | 0.007045 | 10.77 | +1 | 6 |
| 3 | y | 1.729E+04 | 655.4 | 0.007292 | 11.13 | +1 | 6 |
| - | - | 6280 | 656.4 | - | - | 0 | - |
| - | - | 885.8 | 657.4 | - | - | 0 | - |
| 3 | y | 1.042E+05 | 672.4 | 0.0005195 | 0.7725 | +1 | 6 |
| - | - | 3.306E+04 | 673.4 | - | - | 0 | - |
| - | - | 8599 | 674.4 | - | - | 0 | - |
| - | - | 8041 | 682.4 | - | - | 0 | - |
| - | - | 2809 | 683.4 | - | - | 0 | - |
| - | - | 716.2 | 684.4 | - | - | 0 | - |
| - | - | 978.6 | 699.4 | - | - | 0 | - |
| 2 | y | 1.222E+04 | 725.4 | 0.0009455 | 1.303 | +1 | 7 |
| - | - | 4582 | 726.4 | - | - | 0 | - |
| - | - | 936.9 | 727.4 | - | - | 0 | - |
| 2 | y | 4.252E+04 | 743.4 | 0.000707 | 0.9509 | +1 | 7 |
| - | - | 1.569E+04 | 744.4 | - | - | 0 | - |
| - | - | 3365 | 745.4 | - | - | 0 | - |
| - | - | 970.6 | 750.3 | - | - | 0 | - |
| - | - | 990.1 | 753.4 | - | - | 0 | - |
| - | - | 7449 | 768.3 | - | - | 0 | - |
| - | - | 2885 | 769.4 | - | - | 0 | - |
| - | - | 1026 | 770.4 | - | - | 0 | - |
| - | - | 662.4 | 811.2 | - | - | 0 | - |
| - | - | 631.3 | 1783 | - | - | 0 | - |
| - | - | 729.3 | 2733 | - | - | 0 | - |
| - | - | 691 | 3112 | - | - | 0 | - |
| - | - | 622.6 | 3315 | - | - | 0 | - |

m/z Charge Intensity FragmentType MassShift Position
120.08110809326172 0 96781.1
121.08441162109375 0 27070.771
122.08784484863281 0 798.968
123.05252075195312 0 351.59702
127.0867919921875 0 1499.8049
127.12320709228516 0 1629.4304
127.2105941772461 0 434.52155
128.1072540283203 0 2420.933
129.0183563232422 0 718.8925
129.10250854492188 0 6186.3755
129.11439514160156 0 505.8104
130.050048828125 0 1193.452
130.0653839111328 0 4005.2808
130.08653259277344 0 2003.1174
130.0977783203125 0 1098.3423
130.10589599609375 0 1213.742
131.06881713867188 0 1171.4749
131.09024047851562 0 474.67715
132.08099365234375 0 4226.8506
133.08444213867188 0 1128.9635
133.09768676757812 0 478.15723
138.09152221679688 0 972.3925
140.08212280273438 0 8879.316
141.08546447753906 0 553.0215
141.10250854492188 0 8620.669
142.10580444335938 0 1810.4817
143.1181640625 0 56529.887 a 1
144.1160430908203 0 667.5081
144.1216278076172 0 3249.0674
145.09738159179688 0 2894.0205
147.11317443847656 0 2376.4893
148.07574462890625 0 525.66943
148.11659240722656 0 617.6034
149.02359008789062 0 10607.868
150.0269775390625 0 4748.513
155.08168029785156 0 1476.0443
155.11813354492188 0 6013.6846
157.01315307617188 0 467.09326
157.09747314453125 0 1098.4137
157.1086883544922 0 1711.439
157.13377380371094 0 18436.885
158.0926513671875 0 12934.01 y Ammonia loss 7
158.13038635253906 0 699.07043
158.13714599609375 0 4639.307
159.0919189453125 0 24546.623
159.11289978027344 0 1063.1195
160.08831787109375 0 996.8996
160.09532165527344 0 9509.791
167.08164978027344 0 584.565
169.09739685058594 0 8717.178
170.060302734375 0 752.7758
170.1008758544922 0 1878.8657
171.06463623046875 0 461.33676
171.11305236816406 0 47855.22 b 1
172.10813903808594 0 1761.8766
172.11656188964844 0 4569.3633
173.09230041503906 0 9696.404
173.12872314453125 0 56677.973
174.09593200683594 0 1332.2917
174.132080078125 0 5193.165
175.0715789794922 0 1366.3619
175.11920166015625 0 25924.1 y 7
176.1225128173828 0 2478.2134
183.11305236816406 0 11296.832
183.14950561523438 0 8225.853
184.11654663085938 0 1190.8295
184.1529541015625 0 2201.067
185.08108520507812 0 986.1614
185.10350036621094 0 1132.0269
185.12872314453125 0 4048.2935
186.13229370117188 0 1260.9532
187.0868682861328 0 1937.5936
187.10800170898438 0 2999.4468
188.0906982421875 0 713.1669
190.11878967285156 0 686.12573
190.406494140625 0 434.49356
191.11817932128906 0 43436.4
192.12155151367188 0 19439.357
193.09678649902344 0 528.1299
193.12448120117188 0 1666.8414
195.11314392089844 0 1128.2747
198.08718872070312 0 826.6841
199.14476013183594 0 574.09357
201.12355041503906 0 20191.562
202.10699462890625 0 506.26096
202.12681579589844 0 1479.9022
203.0664825439453 0 1207.2396
209.12832641601562 0 840.42706
210.12368774414062 0 1556.4244
210.90487670898438 0 875.59314
211.0868377685547 0 1219.3545
211.10787963867188 0 2057.5037
211.14427185058594 0 3725.9617
211.15478515625 0 868.42566
212.1395263671875 0 6193.658
212.1481475830078 0 972.7158
213.12362670898438 0 1138.998
215.114013671875 0 3471.6072
216.11593627929688 0 638.8864
217.0975341796875 0 1584.2334
219.11305236816406 0 7625.4863
220.1165008544922 0 4607.6455
224.14120483398438 0 575.3784
225.69618225097656 0 610.8188
226.1551971435547 0 7511.6274 a Water loss 2
227.13914489746094 0 3654.6565
228.08558654785156 0 534.15717
228.13427734375 0 1916.0073 y Ammonia loss 4
230.1501007080078 0 3156.2683 d 2
231.15455627441406 0 663.08887
232.14065551757812 0 5769.1865
233.14366149902344 0 1864.3331
235.1077117919922 0 1706.1964
236.11126708984375 0 1154.9601
240.1343536376953 0 1689.453
241.13807678222656 0 585.72314
242.11306762695312 0 596.16425
243.1342315673828 0 599.4161
244.10818481445312 0 988.0709
244.12896728515625 0 1260.5916
244.15138244628906 0 1203.8749
246.1240234375 0 4145.2227
247.12713623046875 0 2392.1592
252.13467407226562 0 942.94727
254.11471557617188 0 662.3615
254.15005493164062 0 22506.256 b Water loss 2
255.14537048339844 0 53470.367 y Ammonia loss 6
256.10748291015625 0 622.2341
256.1485595703125 0 6384.094
258.1448974609375 0 5347.5854
259.1485900878906 0 2398.5098
263.1028137207031 0 4500.869
264.1058654785156 0 2452.7322
265.1085205078125 0 604.89246
266.14959716796875 0 1136.5223
267.1523132324219 0 738.80634
269.1034240722656 0 1194.6293
270.1069641113281 0 606.45374
270.1448974609375 0 9036.817
271.14892578125 0 1123.0244
271.176025390625 0 852.4338
272.1598205566406 0 7500.9897 b 2
272.1719970703125 0 23583.123 y 6
273.1749267578125 0 2700.2134
274.1191711425781 0 2233.3347
275.12213134765625 0 1022.8926
276.1556091308594 0 4362.32
277.15863037109375 0 2012.0052
282.1812438964844 0 2795.0642
282.55767822265625 0 642.1743
286.1813049316406 0 2109.7412 b 5
288.1556396484375 0 7207.668
289.15936279296875 0 650.4625
289.64703369140625 0 709.2617
294.5918273925781 0 552.7725
295.2009582519531 0 545.1884
298.1406555175781 0 1333.6554
299.19073486328125 0 973.47546
299.207763671875 0 656.2667
300.1922912597656 0 684.6735
306.1791687011719 0 3804.6392
306.6806945800781 0 1254.5586
312.1921081542969 0 1894.2821
312.6505126953125 0 1084.0618
319.6845703125 0 1270.4243
323.1720275878906 0 609.3049
325.22381591796875 0 8986.787
326.2270812988281 0 1354.1881
327.65496826171875 0 1430.3304
327.6996765136719 0 21054.385 y Water loss 2
328.19757080078125 0 21333.559 y Ammonia loss 2
328.6980895996094 0 7134.7764
329.1990051269531 0 1641.4747
331.1390380859375 0 610.54156
332.69232177734375 0 807.31354
334.139404296875 0 1897.684
334.58331298828125 0 3186.5144
335.0839538574219 0 984.8687
335.1419372558594 0 1006.3323
335.58685302734375 0 898.3572
336.70587158203125 0 2885.577 y 2
337.2071228027344 0 807.21826
340.18719482421875 0 3190.8838
341.154052734375 0 438.73987
341.16107177734375 0 608.98517
341.18426513671875 0 4366.116
341.6975402832031 0 663.93256
342.189697265625 0 656.2286
342.66070556640625 0 1766.9401
343.2335510253906 0 1994.8448
344.1444091796875 0 626.28394
349.5877380371094 0 1634.6898
350.5783386230469 0 1547.5182
351.2027282714844 0 568.04565
351.2386779785156 0 750.115
352.1153564453125 0 976.6432
352.6153869628906 0 1134.2172
353.2185974121094 0 6591.5474 b Water loss 3
353.6436462402344 0 652.3469
354.1110534667969 0 814.10406
354.217529296875 0 827.1378
354.6402282714844 0 1012.75195
355.14068603515625 0 1006.68384
355.2336120605469 0 875.9056
356.0868225097656 0 632.68164
358.19525146484375 0 620.3056
359.19384765625 0 1808.8477
361.1190185546875 0 1670.256
363.21875 0 24229.033 y Water loss 1
363.7203674316406 0 8515.417
364.2214050292969 0 2806.7903
364.5931396484375 0 3300.3367
364.6432800292969 0 606.6574
365.0948791503906 0 1480.4896
365.5931396484375 0 1122.0715
366.5891418457031 0 1170.6239
368.22894287109375 0 3915.5813 y Ammonia loss 5
369.2321472167969 0 726.75494
371.1346435546875 0 1384.4032
372.13818359375 0 989.1769
373.5992431640625 0 2706.2598
374.0992431640625 0 2592.5264
374.5975341796875 0 1194.972
375.67413330078125 0 1383.3436
376.12371826171875 0 755.8269
376.17474365234375 0 831.56525
379.59814453125 0 967.86163
380.0997619628906 0 886.90533
380.5894775390625 0 1547.2601
383.2283020019531 0 1913.5304
384.1666259765625 0 834.64417
384.67755126953125 0 1426.275
385.1709289550781 0 1935.5946
385.25592041015625 0 11596.627 y 5
386.2589416503906 0 1939.3594
388.6044616699219 0 3210.1873
389.1053161621094 0 1263.5194
389.23712158203125 0 661.093
389.6004333496094 0 888.718
391.2324523925781 0 6803.5713
391.73333740234375 0 2177.4126
392.2323913574219 0 931.66656
401.19354248046875 0 2150.5767
402.19744873046875 0 1876.1671
403.7492980957031 0 913.1895
412.7533874511719 0 17460.504 Precursor Water loss
413.2543029785156 0 7737.693
413.7567443847656 0 1168.8405
418.2204895019531 0 4224.7236
419.222412109375 0 5093.6357
420.2228088378906 0 1378.0133
426.2703857421875 0 623.41095
436.25567626953125 0 2282.1692
439.84417724609375 0 691.042
442.2779846191406 0 991.83765
454.27459716796875 0 2598.761 y Water loss 4
455.2618408203125 0 1124.6771 y Ammonia loss 4
472.2880859375 0 133377.78 y 4
473.2906188964844 0 30876.021
474.29302978515625 0 5093.407
482.272705078125 0 1423.1964
486.292236328125 0 9169.551
487.2420654296875 0 894.2036
487.2953796386719 0 13910.411
488.2976989746094 0 3732.5005
505.2518310546875 0 17986.725
506.2543029785156 0 28554.79
507.25726318359375 0 6645.5444
508.26220703125 0 1186.5007
509.2811279296875 0 797.545
541.3486938476562 0 792.8197
553.341552734375 0 1428.0337 b Water loss 5
557.3291625976562 0 1842.3713
558.3323974609375 0 3368.6223
571.3558959960938 0 46303.176 b 5
572.3588256835938 0 13351.811
573.3618774414062 0 2972.3718
597.333251953125 0 641.7629
602.2672729492188 0 2001.6464
603.27001953125 0 5385.45
604.26220703125 0 2154.4011
611.351318359375 0 1986.4364
612.3604736328125 0 626.0514
620.2783813476562 0 28366.703
621.2811279296875 0 65811.99
622.283447265625 0 21614.418
623.285888671875 0 4064.5032
628.3770141601562 0 4127.432
629.38037109375 0 1134.8191
638.362060546875 0 902.57794
654.3862915039062 0 7715.9253 y Water loss 2
655.3846435546875 0 17293.697 y Ammonia loss 2
656.3876342773438 0 6279.7188
657.3911743164062 0 885.7534
672.4033813476562 0 104192.61 y 2
673.4061889648438 0 33064.453
674.4089965820312 0 8599.25
682.3876342773438 0 8041.148
683.3923950195312 0 2809.28
684.3947143554688 0 716.1545
699.4130249023438 0 978.60614
725.4295043945312 0 12221.07 y Water loss 1
726.43310546875 0 4582.221
727.4370727539062 0 936.8886
743.4403076171875 0 42522.45 y 1
744.4430541992188 0 15686.51
745.4462890625 0 3364.515
750.3383178710938 0 970.6474
753.426025390625 0 990.1462
768.3490600585938 0 7449.1816
769.3512573242188 0 2885.0828
770.357177734375 0 1025.6232
811.24609375 0 662.36633
1783.453857421875 0 631.3309
2732.511474609375 0 729.3493
3111.87255859375 0 690.98395
3315.425537109375 0 622.63196

Spectrum Details

|  |  |
| --- | --- |
| Matched peaks? Matched peaks The total absolute number of peaks matched. Additionally in brackets the total fraction of peaks matched and the total number of peaks is shown. | 30 (9.46% of 317) |
| FDR? FDR The false discovery rate estimated for this peptide. It is calculated by matching all theoretical fragments with a non-integer shift with the raw peaks for this spectrum. This is done with 40 different shifts. The resulting percentage is the average number of annotated peaks over the number of annotated peaks with the correct spectrum. | 0.63% |
| Satellite FDR? Satellite FDR See the FDR for details on its calculation. This satellite ion specific FDR only contains the satellite ions (d/w) for I/L/J positions. | - |
| PSM Score? PSM Score The PSM Score as given by Hecklib to this annotated spectrum. It is shown with three significant figures. | 349 |

#### Spectrum 5646? Spectrum 5646 The raw spectrum of this peptide as annotated by Hecklib. The fragments are coloured according to ion type (see legend). Any peaks with a star '\*' as text can be hovered over to see the full details, first the ion type second the mass shift type. By hovering over the amino acids in the peptide or ions in the legend the corresponding peaks are highlighted. By toggling the 'Unassigned' label you can turn the background (unassigned) peaks on or off in the plot. By updating the slider in the Ion legend you can update the spectrum to only show the top X% of the peaks with labels. The top X% means any peak that is within X% of the highest intensity. By dragging in the spectrum you can zoom in to a specific part of the spectrum and use 'Zoom Out' to get back to the original zoom level. The annotation of the spectrum is based on the given sequence in the peptides file and is done with different software so inconsistencies are likely. The peaks are annotated based on the given sequence, with 20 ppm tolerance.

Copy Data

##### Spectrum 5646 (TSV)

###### Preview

```
Loading example...
```

*Click on the button to copy the data to your clipboard.*

Mz MinMz MaxIntensity Max

WidthHeightPeptide font sizePeptide stroke widthSpectrum font sizeSpectrum stroke widthCompact peptide

Ion legend

wxyz

abcd

OtherUnassignedIon^Charge^_Position_Show for top:%

VATVSJPR

01.83e+43.67e+45.50e+47.34e+4

Zoom Out

a^+1^_2_y^+1^_1_b^+1^_2_y^+1^_1_a^+1^_3_y^+2^_4_d^+1^_3_b^+1^_3_y^+1^_2_b^+1^_3_y^+1^_2_b^+2^_6_y^+2^_6_y^+2^_6_y^+2^_6_b^+1^_4_y^+2^_7_y^+1^_3_b^+1^_4_y^+1^_3_\*y^+1^_4_y^+1^_4_b^+1^_6_b^+1^_6_y^+1^_6_y^+1^_6_y^+1^_6_y^+1^_7_y^+1^_7_

0782156423463128

Fragment Matches Table

Show background peaks

| Position | Ion type | Intensity | mz Theoretical | mz Error (Th) | mz Error (ppm) | Charge | Series Number |
| --- | --- | --- | --- | --- | --- | --- | --- |
| - | - | 1884 | 120.1 | - | - | 0 | - |
| - | - | 1213 | 127.1 | - | - | 0 | - |
| - | - | 475.3 | 127.1 | - | - | 0 | - |
| - | - | 1073 | 127.1 | - | - | 0 | - |
| - | - | 1290 | 128.1 | - | - | 0 | - |
| - | - | 832.8 | 129 | - | - | 0 | - |
| - | - | 371.6 | 129.1 | - | - | 0 | - |
| - | - | 6930 | 129.1 | - | - | 0 | - |
| - | - | 537.8 | 130.1 | - | - | 0 | - |
| - | - | 1329 | 130.1 | - | - | 0 | - |
| - | - | 3175 | 130.1 | - | - | 0 | - |
| - | - | 463.5 | 131.1 | - | - | 0 | - |
| - | - | 1614 | 132.1 | - | - | 0 | - |
| - | - | 1.598E+04 | 136.1 | - | - | 0 | - |
| - | - | 393.6 | 137.1 | - | - | 0 | - |
| - | - | 2394 | 137.1 | - | - | 0 | - |
| - | - | 514.5 | 138.1 | - | - | 0 | - |
| - | - | 5207 | 140.1 | - | - | 0 | - |
| - | - | 8794 | 141.1 | - | - | 0 | - |
| - | - | 1547 | 142.1 | - | - | 0 | - |
| 2 | a | 3.014E+04 | 143.1 | 0.0001677 | 1.172 | +1 | 2 |
| - | - | 526.5 | 144.1 | - | - | 0 | - |
| - | - | 2146 | 144.1 | - | - | 0 | - |
| - | - | 1952 | 145.1 | - | - | 0 | - |
| - | - | 1211 | 147 | - | - | 0 | - |
| - | - | 3098 | 147.1 | - | - | 0 | - |
| - | - | 544.3 | 148.1 | - | - | 0 | - |
| - | - | 1.024E+04 | 149 | - | - | 0 | - |
| - | - | 5788 | 150 | - | - | 0 | - |
| - | - | 495.2 | 151.1 | - | - | 0 | - |
| - | - | 432.1 | 151.1 | - | - | 0 | - |
| - | - | 1340 | 155.1 | - | - | 0 | - |
| - | - | 3891 | 155.1 | - | - | 0 | - |
| - | - | 612.2 | 157.1 | - | - | 0 | - |
| - | - | 574.5 | 157.1 | - | - | 0 | - |
| - | - | 1.899E+04 | 157.1 | - | - | 0 | - |
| 8 | y | 3504 | 158.1 | 6.52E-05 | 0.4124 | +1 | 1 |
| - | - | 4268 | 158.1 | - | - | 0 | - |
| - | - | 5683 | 159.1 | - | - | 0 | - |
| - | - | 870.5 | 159.1 | - | - | 0 | - |
| - | - | 959.3 | 160.1 | - | - | 0 | - |
| - | - | 480.5 | 164.3 | - | - | 0 | - |
| - | - | 638.9 | 167 | - | - | 0 | - |
| - | - | 539.1 | 168 | - | - | 0 | - |
| - | - | 8496 | 169.1 | - | - | 0 | - |
| - | - | 1800 | 170.1 | - | - | 0 | - |
| 2 | b | 2.763E+04 | 171.1 | 0.0001261 | 0.7372 | +1 | 2 |
| - | - | 2774 | 172.1 | - | - | 0 | - |
| - | - | 6187 | 173.1 | - | - | 0 | - |
| - | - | 3.197E+04 | 173.1 | - | - | 0 | - |
| - | - | 475 | 174.1 | - | - | 0 | - |
| - | - | 2042 | 174.1 | - | - | 0 | - |
| 8 | y | 9420 | 175.1 | 0.0001579 | 0.9019 | +1 | 1 |
| - | - | 532.2 | 175.1 | - | - | 0 | - |
| - | - | 623.9 | 176.1 | - | - | 0 | - |
| - | - | 1003 | 181.1 | - | - | 0 | - |
| - | - | 7315 | 183.1 | - | - | 0 | - |
| - | - | 8035 | 183.1 | - | - | 0 | - |
| - | - | 2898 | 184.2 | - | - | 0 | - |
| - | - | 957.6 | 185.1 | - | - | 0 | - |
| - | - | 802.8 | 185.1 | - | - | 0 | - |
| - | - | 3910 | 185.1 | - | - | 0 | - |
| - | - | 894.6 | 186.1 | - | - | 0 | - |
| - | - | 788.6 | 187.1 | - | - | 0 | - |
| - | - | 2129 | 187.1 | - | - | 0 | - |
| - | - | 501.1 | 189.3 | - | - | 0 | - |
| - | - | 740.6 | 195.1 | - | - | 0 | - |
| - | - | 1.149E+04 | 201.1 | - | - | 0 | - |
| - | - | 778.7 | 201.1 | - | - | 0 | - |
| - | - | 524 | 202.1 | - | - | 0 | - |
| - | - | 765.3 | 202.1 | - | - | 0 | - |
| - | - | 1390 | 204.1 | - | - | 0 | - |
| - | - | 1597 | 211.1 | - | - | 0 | - |
| - | - | 4000 | 211.1 | - | - | 0 | - |
| - | - | 2586 | 212.1 | - | - | 0 | - |
| - | - | 1075 | 212.1 | - | - | 0 | - |
| - | - | 1056 | 213.1 | - | - | 0 | - |
| - | - | 503.3 | 215.4 | - | - | 0 | - |
| 3 | a | 4085 | 226.2 | 0.0001175 | 0.5197 | +1 | 3 |
| - | - | 2592 | 227.1 | - | - | 0 | - |
| 5 | y | 1469 | 228.1 | 7.05E-05 | 0.309 | +2 | 4 |
| 3 | d | 1168 | 230.1 | 0.0001682 | 0.7308 | +1 | 3 |
| - | - | 2070 | 233.1 | - | - | 0 | - |
| - | - | 914.8 | 234.1 | - | - | 0 | - |
| - | - | 1219 | 235.1 | - | - | 0 | - |
| - | - | 2363 | 240.1 | - | - | 0 | - |
| - | - | 821 | 243.1 | - | - | 0 | - |
| - | - | 772.7 | 244.2 | - | - | 0 | - |
| - | - | 1693 | 246.1 | - | - | 0 | - |
| - | - | 680.1 | 247.1 | - | - | 0 | - |
| - | - | 1081 | 250.1 | - | - | 0 | - |
| 3 | b | 1.02E+04 | 254.1 | 3.018E-05 | 0.1187 | +1 | 3 |
| - | - | 1034 | 254.2 | - | - | 0 | - |
| 7 | y | 2.883E+04 | 255.1 | 5.098E-05 | 0.1998 | +1 | 2 |
| - | - | 3610 | 256.1 | - | - | 0 | - |
| - | - | 775.1 | 258.1 | - | - | 0 | - |
| - | - | 4302 | 258.1 | - | - | 0 | - |
| - | - | 1762 | 259.1 | - | - | 0 | - |
| - | - | 828.4 | 261.1 | - | - | 0 | - |
| - | - | 534.9 | 265.1 | - | - | 0 | - |
| - | - | 1531 | 266.1 | - | - | 0 | - |
| - | - | 4488 | 270.1 | - | - | 0 | - |
| 3 | b | 3810 | 272.2 | 0.0007231 | 2.657 | +1 | 3 |
| 7 | y | 1.226E+04 | 272.2 | 9.795E-05 | 0.3599 | +1 | 2 |
| - | - | 1695 | 273.2 | - | - | 0 | - |
| - | - | 1088 | 275.1 | - | - | 0 | - |
| - | - | 2828 | 276.2 | - | - | 0 | - |
| - | - | 1766 | 277.2 | - | - | 0 | - |
| - | - | 2903 | 278.1 | - | - | 0 | - |
| - | - | 502.7 | 279.1 | - | - | 0 | - |
| - | - | 1443 | 282.2 | - | - | 0 | - |
| 6 | b | 728.4 | 286.2 | 0.005661 | 19.78 | +2 | 6 |
| - | - | 2865 | 288.2 | - | - | 0 | - |
| - | - | 632.8 | 289.2 | - | - | 0 | - |
| - | - | 948.7 | 290 | - | - | 0 | - |
| - | - | 1151 | 291 | - | - | 0 | - |
| - | - | 1213 | 293.1 | - | - | 0 | - |
| - | - | 620.1 | 295 | - | - | 0 | - |
| - | - | 786.8 | 298.1 | - | - | 0 | - |
| - | - | 3137 | 301.2 | - | - | 0 | - |
| - | - | 1153 | 306.2 | - | - | 0 | - |
| - | - | 1367 | 308 | - | - | 0 | - |
| - | - | 919.5 | 312.2 | - | - | 0 | - |
| - | - | 549.4 | 317.9 | - | - | 0 | - |
| - | - | 1132 | 323.2 | - | - | 0 | - |
| - | - | 5705 | 325.2 | - | - | 0 | - |
| - | - | 547.9 | 325.8 | - | - | 0 | - |
| - | - | 846 | 326.2 | - | - | 0 | - |
| 3 | y | 1.127E+04 | 327.7 | 0.001454 | 4.436 | +2 | 6 |
| 3 | y | 1.512E+04 | 328.2 | 0.004616 | 14.06 | +2 | 6 |
| - | - | 5423 | 328.7 | - | - | 0 | - |
| - | - | 893.9 | 329.2 | - | - | 0 | - |
| - | - | 954 | 332.2 | - | - | 0 | - |
| - | - | 699.9 | 332.9 | - | - | 0 | - |
| 3 | y | 1190 | 336.7 | 3.881E-05 | 0.1153 | +2 | 6 |
| - | - | 708.4 | 337.2 | - | - | 0 | - |
| - | - | 3276 | 340.2 | - | - | 0 | - |
| - | - | 2349 | 341.2 | - | - | 0 | - |
| - | - | 1672 | 350.9 | - | - | 0 | - |
| 4 | b | 3993 | 353.2 | 0.0001007 | 0.285 | +1 | 4 |
| - | - | 629.5 | 357.1 | - | - | 0 | - |
| - | - | 1901 | 359.2 | - | - | 0 | - |
| - | - | 2742 | 361.9 | - | - | 0 | - |
| 2 | y | 1.342E+04 | 363.2 | 0.0003573 | 0.9838 | +2 | 7 |
| - | - | 4406 | 363.7 | - | - | 0 | - |
| - | - | 2024 | 364.2 | - | - | 0 | - |
| 6 | y | 2266 | 368.2 | 0.0005627 | 1.528 | +1 | 3 |
| - | - | 788.9 | 368.9 | - | - | 0 | - |
| 4 | b | 586.4 | 371.2 | 0.0002599 | 0.7002 | +1 | 4 |
| - | - | 988.5 | 379.2 | - | - | 0 | - |
| - | - | 566.8 | 380.2 | - | - | 0 | - |
| - | - | 1223 | 383.2 | - | - | 0 | - |
| - | - | 690.7 | 384.2 | - | - | 0 | - |
| 6 | y | 6912 | 385.3 | 0.0004394 | 1.141 | +1 | 3 |
| - | - | 665.7 | 385.8 | - | - | 0 | - |
| - | - | 1554 | 386.3 | - | - | 0 | - |
| - | - | 792.3 | 389.2 | - | - | 0 | - |
| - | - | 581.9 | 390.4 | - | - | 0 | - |
| - | - | 4308 | 391.2 | - | - | 0 | - |
| - | - | 1787 | 391.7 | - | - | 0 | - |
| - | - | 2206 | 402.2 | - | - | 0 | - |
| - | - | 2898 | 403.8 | - | - | 0 | - |
| 0 | Precursor | 1.011E+04 | 412.8 | 0.0001404 | 0.3403 | +2 | -1 |
| - | - | 5042 | 413.3 | - | - | 0 | - |
| - | - | 895.4 | 413.8 | - | - | 0 | - |
| - | - | 601.9 | 416.9 | - | - | 0 | - |
| - | - | 1421 | 420.9 | - | - | 0 | - |
| - | - | 2648 | 421.2 | - | - | 0 | - |
| - | - | 969.1 | 422.2 | - | - | 0 | - |
| - | - | 731.6 | 422.2 | - | - | 0 | - |
| - | - | 729.5 | 429.2 | - | - | 0 | - |
| - | - | 762.8 | 433.7 | - | - | 0 | - |
| - | - | 669.5 | 434.2 | - | - | 0 | - |
| - | - | 852.3 | 436.3 | - | - | 0 | - |
| - | - | 873.6 | 439.8 | - | - | 0 | - |
| - | - | 762.2 | 441.2 | - | - | 0 | - |
| - | - | 900.7 | 442.3 | - | - | 0 | - |
| - | - | 885.2 | 448.3 | - | - | 0 | - |
| 5 | y | 1291 | 454.3 | 0.007529 | 16.57 | +1 | 4 |
| - | - | 9135 | 458.3 | - | - | 0 | - |
| - | - | 710.1 | 468.3 | - | - | 0 | - |
| 5 | y | 7.266E+04 | 472.3 | 5.817E-05 | 0.1232 | +1 | 4 |
| - | - | 1.487E+04 | 473.3 | - | - | 0 | - |
| - | - | 2518 | 474.3 | - | - | 0 | - |
| - | - | 700.7 | 482.3 | - | - | 0 | - |
| - | - | 6578 | 486.3 | - | - | 0 | - |
| - | - | 8854 | 487.3 | - | - | 0 | - |
| - | - | 2131 | 488.3 | - | - | 0 | - |
| - | - | 674.1 | 489.3 | - | - | 0 | - |
| - | - | 1070 | 493.8 | - | - | 0 | - |
| - | - | 1516 | 509.3 | - | - | 0 | - |
| - | - | 597.1 | 521.5 | - | - | 0 | - |
| - | - | 712.6 | 540.3 | - | - | 0 | - |
| - | - | 2517 | 542.3 | - | - | 0 | - |
| 6 | b | 987.6 | 553.3 | 0.004992 | 9.022 | +1 | 6 |
| - | - | 1366 | 557.3 | - | - | 0 | - |
| - | - | 1811 | 558.3 | - | - | 0 | - |
| - | - | 672.7 | 559.3 | - | - | 0 | - |
| - | - | 882.9 | 566.2 | - | - | 0 | - |
| - | - | 573.6 | 569.3 | - | - | 0 | - |
| - | - | 750.6 | 570.3 | - | - | 0 | - |
| - | - | 695 | 570.8 | - | - | 0 | - |
| 6 | b | 2.518E+04 | 571.3 | 0.0106 | 18.56 | +1 | 6 |
| - | - | 7083 | 572.4 | - | - | 0 | - |
| - | - | 1369 | 573.4 | - | - | 0 | - |
| - | - | 562.3 | 575.3 | - | - | 0 | - |
| - | - | 639.2 | 576.4 | - | - | 0 | - |
| - | - | 2100 | 628.4 | - | - | 0 | - |
| - | - | 703.7 | 629.4 | - | - | 0 | - |
| - | - | 645.4 | 644 | - | - | 0 | - |
| 3 | y | 4895 | 654.4 | 0.007716 | 11.79 | +1 | 6 |
| 3 | y | 1.112E+04 | 655.4 | 0.007353 | 11.22 | +1 | 6 |
| - | - | 5468 | 656.4 | - | - | 0 | - |
| - | - | 827.4 | 657.4 | - | - | 0 | - |
| - | - | 723.2 | 664.4 | - | - | 0 | - |
| 3 | y | 5.574E+04 | 672.4 | 0.001008 | 1.499 | +1 | 6 |
| - | - | 1.859E+04 | 673.4 | - | - | 0 | - |
| - | - | 3544 | 674.4 | - | - | 0 | - |
| - | - | 3683 | 682.4 | - | - | 0 | - |
| - | - | 1527 | 683.4 | - | - | 0 | - |
| - | - | 1505 | 705.4 | - | - | 0 | - |
| 2 | y | 6781 | 725.4 | 0.0009455 | 1.303 | +1 | 7 |
| - | - | 2821 | 726.4 | - | - | 0 | - |
| 2 | y | 2.039E+04 | 743.4 | 0.001439 | 1.936 | +1 | 7 |
| - | - | 7524 | 744.4 | - | - | 0 | - |
| - | - | 1851 | 745.4 | - | - | 0 | - |
| - | - | 895.7 | 753.4 | - | - | 0 | - |
| - | - | 662.9 | 818.5 | - | - | 0 | - |
| - | - | 2762 | 915.5 | - | - | 0 | - |
| - | - | 665.2 | 1046 | - | - | 0 | - |
| - | - | 648.7 | 1107 | - | - | 0 | - |
| - | - | 682 | 1291 | - | - | 0 | - |
| - | - | 695.9 | 1582 | - | - | 0 | - |
| - | - | 605.3 | 1743 | - | - | 0 | - |
| - | - | 702.5 | 2349 | - | - | 0 | - |
| - | - | 612.1 | 2477 | - | - | 0 | - |
| - | - | 703.3 | 2995 | - | - | 0 | - |
| - | - | 714 | 3000 | - | - | 0 | - |
| - | - | 758.5 | 3097 | - | - | 0 | - |

m/z Charge Intensity FragmentType MassShift Position
120.08084869384766 0 1883.7263
127.08675384521484 0 1212.983
127.11245727539062 0 475.32224
127.12303924560547 0 1072.9397
128.107177734375 0 1290.4022
129.01846313476562 0 832.835
129.07235717773438 0 371.6194
129.10243225097656 0 6930.246
130.05023193359375 0 537.7947
130.0653839111328 0 1329.1572
130.0864715576172 0 3175.1084
131.0897979736328 0 463.52164
132.0809783935547 0 1613.8391
136.07583618164062 0 15981.541
137.07278442382812 0 393.59143
137.07913208007812 0 2393.6013
138.09181213378906 0 514.54047
140.08201599121094 0 5207.3535
141.10240173339844 0 8794.2705
142.10560607910156 0 1546.6252
143.11805725097656 0 30140.326 a 1
144.11593627929688 0 526.5141
144.12156677246094 0 2146.3042
145.09735107421875 0 1951.902
147.04412841796875 0 1210.6194
147.11300659179688 0 3098.3523
148.1168212890625 0 544.3367
149.0234832763672 0 10237.98
150.02685546875 0 5788.0566
151.07566833496094 0 495.245
151.0870361328125 0 432.07733
155.08155822753906 0 1340.1688
155.11810302734375 0 3890.7026
157.09707641601562 0 612.15784
157.10879516601562 0 574.47046
157.13369750976562 0 18986.367
158.09246826171875 0 3503.7568 y Ammonia loss 7
158.1370849609375 0 4268.3877
159.09185791015625 0 5682.696
159.11282348632812 0 870.5045
160.09503173828125 0 959.29
164.2550811767578 0 480.4511
167.0343780517578 0 638.8524
168.03684997558594 0 539.1431
169.09727478027344 0 8495.751
170.10076904296875 0 1799.7236
171.11293029785156 0 27627.443 b 1
172.11642456054688 0 2773.786
173.0922088623047 0 6186.556
173.1286163330078 0 31971.047
174.0951690673828 0 474.97754
174.13201904296875 0 2042.491
175.11911010742188 0 9420.13 y 7
175.1348419189453 0 532.1924
176.12266540527344 0 623.9103
181.09767150878906 0 1003.3847
183.11285400390625 0 7314.6343
183.14932250976562 0 8035.2065
184.1527099609375 0 2898.3174
185.08143615722656 0 957.60815
185.1035614013672 0 802.7833
185.12860107421875 0 3910.159
186.1322784423828 0 894.57
187.0867156982422 0 788.5659
187.10772705078125 0 2129.1729
189.32711791992188 0 501.14066
195.11265563964844 0 740.61505
201.1233673095703 0 11494.694
201.13365173339844 0 778.6776
202.108154296875 0 524.03345
202.12696838378906 0 765.34406
204.13450622558594 0 1390.111
211.10752868652344 0 1597.1123
211.1443328857422 0 4000.3254
212.1392059326172 0 2586.2034
212.14910888671875 0 1074.6072
213.12371826171875 0 1055.9491
215.42416381835938 0 503.3012
226.15512084960938 0 4084.6328 a Water loss 2
227.1387939453125 0 2591.5552
228.13433837890625 0 1469.0698 y Ammonia loss 4
230.14974975585938 0 1168.3951 d 2
233.09219360351562 0 2070.444
234.1448211669922 0 914.76965
235.14463806152344 0 1219.0988
240.13426208496094 0 2363.309
243.1334686279297 0 820.99646
244.15028381347656 0 772.7005
246.12384033203125 0 1692.6265
247.1299285888672 0 680.1005
250.1188201904297 0 1080.6672
254.1499481201172 0 10204.604 b Water loss 2
254.16236877441406 0 1034.2358
255.1452178955078 0 28830.873 y Ammonia loss 6
256.1483154296875 0 3609.9617
258.0994567871094 0 775.05383
258.1448669433594 0 4302.3604
259.1478576660156 0 1762.346
261.087890625 0 828.4498
265.08087158203125 0 534.85596
266.1493225097656 0 1531.2609
270.1445007324219 0 4488.1274
272.1597595214844 0 3809.9126 b 2
272.17181396484375 0 12260.105 y 6
273.1750183105469 0 1695.328
275.1239929199219 0 1087.5754
276.1553955078125 0 2828.0464
277.1582336425781 0 1765.9536
278.1136169433594 0 2903.4246
279.08636474609375 0 502.70764
282.1812744140625 0 1442.6285
286.1817932128906 0 728.4265 b 5
288.1555480957031 0 2864.8987
289.1586608886719 0 632.81024
289.9755554199219 0 948.7296
290.96087646484375 0 1150.9998
293.0991516113281 0 1213.1104
294.99932861328125 0 620.0887
298.1394958496094 0 786.794
301.1871032714844 0 3136.6729
306.1802673339844 0 1153.4044
307.9864807128906 0 1367.3306
312.19146728515625 0 919.4932
317.90472412109375 0 549.44257
323.1719055175781 0 1131.8353
325.2234802246094 0 5704.582
325.8330383300781 0 547.85095
326.2252502441406 0 846.0461
327.6988525390625 0 11274.352 y Water loss 2
328.1969299316406 0 15118.85 y Ammonia loss 2
328.6986389160156 0 5422.692
329.1978454589844 0 893.91205
332.1602478027344 0 953.9849
332.8703308105469 0 699.94446
336.70562744140625 0 1189.9734 y 2
337.20770263671875 0 708.37885
340.1869201660156 0 3275.6973
341.18505859375 0 2348.704
350.9267272949219 0 1672.3086
353.2182312011719 0 3992.8074 b Water loss 3
357.0591735839844 0 629.526
359.1938171386719 0 1901.1067
361.88446044921875 0 2741.6646
363.218505859375 0 13418.839 y Water loss 1
363.71954345703125 0 4406.107
364.22039794921875 0 2023.577
368.2286682128906 0 2265.799 y Ammonia loss 5
368.9371337890625 0 788.8984
371.2291564941406 0 586.4493 b 3
379.23382568359375 0 988.5452
380.2348937988281 0 566.81134
383.22857666015625 0 1223.4939
384.1932067871094 0 690.724
385.2553405761719 0 6911.6094 y 5
385.80377197265625 0 665.67645
386.259521484375 0 1553.6163
389.23895263671875 0 792.2763
390.4233093261719 0 581.94476
391.2313537597656 0 4308.041
391.73394775390625 0 1786.5508
402.2066650390625 0 2205.5671
403.8129577636719 0 2898.2527
412.7529296875 0 10112.864 Precursor Water loss
413.25439453125 0 5042.0845
413.75677490234375 0 895.44055
416.88958740234375 0 601.8562
420.8581237792969 0 1420.9252
421.20452880859375 0 2648.27
422.2124328613281 0 969.14166
422.2417907714844 0 731.5993
429.18670654296875 0 729.4505
433.7186279296875 0 762.7733
434.2181396484375 0 669.47516
436.256591796875 0 852.32697
439.84332275390625 0 873.5556
441.176025390625 0 762.23175
442.27667236328125 0 900.6645
448.2769775390625 0 885.15955
454.26971435546875 0 1290.8966 y Water loss 4
458.27886962890625 0 9135.08
468.2796936035156 0 710.13416
472.2877502441406 0 72657.71 y 4
473.2904968261719 0 14871.158
474.292236328125 0 2518.0068
482.27142333984375 0 700.7142
486.29180908203125 0 6578.136
487.29443359375 0 8853.91
488.2974548339844 0 2131.4045
489.2980041503906 0 674.0612
493.79693603515625 0 1070.2793
509.2779235839844 0 1515.6644
521.462890625 0 597.07983
540.3219604492188 0 712.59595
542.3220825195312 0 2517.3179
553.3394165039062 0 987.60425 b Water loss 5
557.327880859375 0 1365.7783
558.3324584960938 0 1811.1309
559.3330078125 0 672.70874
566.2464599609375 0 882.8601
569.3269653320312 0 573.62726
570.2736206054688 0 750.6116
570.7705688476562 0 695.0035
571.3555908203125 0 25176.451 b 5
572.3585815429688 0 7083.2407
573.3603515625 0 1368.842
575.3091430664062 0 562.31085
576.368408203125 0 639.17126
628.3770751953125 0 2099.5483
629.3818969726562 0 703.6702
643.9911499023438 0 645.36786
654.3856201171875 0 4894.8516 y Water loss 2
655.3847045898438 0 11120.097 y Ammonia loss 2
656.3870239257812 0 5468.21
657.3895874023438 0 827.35095
664.3862915039062 0 723.21246
672.4028930664062 0 55736.543 y 2
673.4055786132812 0 18591.834
674.4077758789062 0 3544.3308
682.3875122070312 0 3682.6956
683.3925170898438 0 1526.755
705.412841796875 0 1505.0424
725.4295043945312 0 6780.907 y Water loss 1
726.4309692382812 0 2821.1443
743.4395751953125 0 20389.568 y 1
744.4423828125 0 7524.019
745.4453125 0 1851.1572
753.4227294921875 0 895.6661
818.501220703125 0 662.8825
915.549560546875 0 2762.1738
1046.074462890625 0 665.2232
1107.40966796875 0 648.7202
1291.2506103515625 0 681.98694
1581.5540771484375 0 695.8873
1743.0367431640625 0 605.2753
2349.47412109375 0 702.5414
2476.75341796875 0 612.08826
2995.239013671875 0 703.25073
3000.21923828125 0 714.00397
3096.7431640625 0 758.53876

Spectrum Details

|  |  |
| --- | --- |
| Matched peaks? Matched peaks The total absolute number of peaks matched. Additionally in brackets the total fraction of peaks matched and the total number of peaks is shown. | 30 (12.55% of 239) |
| FDR? FDR The false discovery rate estimated for this peptide. It is calculated by matching all theoretical fragments with a non-integer shift with the raw peaks for this spectrum. This is done with 40 different shifts. The resulting percentage is the average number of annotated peaks over the number of annotated peaks with the correct spectrum. | 0.32% |
| Satellite FDR? Satellite FDR See the FDR for details on its calculation. This satellite ion specific FDR only contains the satellite ions (d/w) for I/L/J positions. | - |
| PSM Score? PSM Score The PSM Score as given by Hecklib to this annotated spectrum. It is shown with three significant figures. | 349 |

#### Spectrum 6234? Spectrum 6234 The raw spectrum of this peptide as annotated by Hecklib. The fragments are coloured according to ion type (see legend). Any peaks with a star '\*' as text can be hovered over to see the full details, first the ion type second the mass shift type. By hovering over the amino acids in the peptide or ions in the legend the corresponding peaks are highlighted. By toggling the 'Unassigned' label you can turn the background (unassigned) peaks on or off in the plot. By updating the slider in the Ion legend you can update the spectrum to only show the top X% of the peaks with labels. The top X% means any peak that is within X% of the highest intensity. By dragging in the spectrum you can zoom in to a specific part of the spectrum and use 'Zoom Out' to get back to the original zoom level. The annotation of the spectrum is based on the given sequence in the peptides file and is done with different software so inconsistencies are likely. The peaks are annotated based on the given sequence, with 20 ppm tolerance.

Copy Data

##### Spectrum 6234 (TSV)

###### Preview

```
Loading example...
```

*Click on the button to copy the data to your clipboard.*

Mz MinMz MaxIntensity Max

WidthHeightPeptide font sizePeptide stroke widthSpectrum font sizeSpectrum stroke widthCompact peptide

Ion legend

wxyz

abcd

OtherUnassignedIon^Charge^_Position_Show for top:%

VATVSJPR

08.18e+31.64e+42.45e+43.27e+4

Zoom Out

a^+1^_2_y^+1^_1_b^+1^_2_y^+1^_1_a^+1^_3_b^+1^_3_y^+1^_2_b^+1^_3_y^+1^_2_b^+2^_6_y^+2^_6_y^+2^_6_y^+2^_6_b^+1^_4_y^+2^_7_y^+1^_3_y^+1^_3_\*y^+1^_4_b^+1^_6_y^+1^_6_y^+1^_6_y^+1^_6_y^+1^_7_y^+1^_7_

0778155623353113

Fragment Matches Table

Show background peaks

| Position | Ion type | Intensity | mz Theoretical | mz Error (Th) | mz Error (ppm) | Charge | Series Number |
| --- | --- | --- | --- | --- | --- | --- | --- |
| - | - | 762 | 120.1 | - | - | 0 | - |
| - | - | 360.2 | 127.1 | - | - | 0 | - |
| - | - | 1014 | 128.1 | - | - | 0 | - |
| - | - | 1331 | 129 | - | - | 0 | - |
| - | - | 2833 | 129.1 | - | - | 0 | - |
| - | - | 1406 | 130.1 | - | - | 0 | - |
| - | - | 912.7 | 136.1 | - | - | 0 | - |
| - | - | 1998 | 140.1 | - | - | 0 | - |
| - | - | 446.4 | 141.1 | - | - | 0 | - |
| - | - | 3366 | 141.1 | - | - | 0 | - |
| - | - | 953.7 | 142.1 | - | - | 0 | - |
| 2 | a | 1.392E+04 | 143.1 | 0.0001525 | 1.065 | +1 | 2 |
| - | - | 1193 | 144.1 | - | - | 0 | - |
| - | - | 1096 | 145.1 | - | - | 0 | - |
| - | - | 1224 | 147.1 | - | - | 0 | - |
| - | - | 1.153E+04 | 149 | - | - | 0 | - |
| - | - | 408 | 149.7 | - | - | 0 | - |
| - | - | 6570 | 150 | - | - | 0 | - |
| - | - | 799.5 | 151 | - | - | 0 | - |
| - | - | 2148 | 155.1 | - | - | 0 | - |
| - | - | 8983 | 157.1 | - | - | 0 | - |
| 8 | y | 2028 | 158.1 | 4.994E-05 | 0.3159 | +1 | 1 |
| - | - | 2314 | 158.1 | - | - | 0 | - |
| - | - | 499.3 | 167.4 | - | - | 0 | - |
| - | - | 4446 | 169.1 | - | - | 0 | - |
| - | - | 1318 | 170.1 | - | - | 0 | - |
| - | - | 607.8 | 171.1 | - | - | 0 | - |
| 2 | b | 1.254E+04 | 171.1 | 9.562E-05 | 0.5588 | +1 | 2 |
| - | - | 497.7 | 171.1 | - | - | 0 | - |
| - | - | 829.6 | 172.1 | - | - | 0 | - |
| - | - | 1556 | 173.1 | - | - | 0 | - |
| - | - | 1.545E+04 | 173.1 | - | - | 0 | - |
| - | - | 1135 | 174.1 | - | - | 0 | - |
| 8 | y | 4617 | 175.1 | 9.69E-05 | 0.5534 | +1 | 1 |
| - | - | 2813 | 183.1 | - | - | 0 | - |
| - | - | 2880 | 183.1 | - | - | 0 | - |
| - | - | 550.8 | 183.8 | - | - | 0 | - |
| - | - | 1380 | 184.2 | - | - | 0 | - |
| - | - | 724.5 | 185.1 | - | - | 0 | - |
| - | - | 2312 | 185.1 | - | - | 0 | - |
| - | - | 952.9 | 187.1 | - | - | 0 | - |
| - | - | 6007 | 201.1 | - | - | 0 | - |
| - | - | 520 | 209 | - | - | 0 | - |
| - | - | 557.9 | 210.1 | - | - | 0 | - |
| - | - | 2148 | 211.1 | - | - | 0 | - |
| - | - | 1438 | 212.1 | - | - | 0 | - |
| - | - | 809.2 | 212.1 | - | - | 0 | - |
| 3 | a | 1763 | 226.2 | 0.0002487 | 1.1 | +1 | 3 |
| - | - | 697.9 | 227.1 | - | - | 0 | - |
| - | - | 560.9 | 235.2 | - | - | 0 | - |
| - | - | 884.2 | 240.1 | - | - | 0 | - |
| - | - | 644.4 | 241.1 | - | - | 0 | - |
| - | - | 500.7 | 244.2 | - | - | 0 | - |
| - | - | 501.1 | 248 | - | - | 0 | - |
| 3 | b | 6057 | 254.1 | 3.086E-05 | 0.1214 | +1 | 3 |
| 7 | y | 1.252E+04 | 255.1 | 6.624E-05 | 0.2596 | +1 | 2 |
| - | - | 1715 | 256.1 | - | - | 0 | - |
| - | - | 1738 | 258.1 | - | - | 0 | - |
| - | - | 898.9 | 259.1 | - | - | 0 | - |
| - | - | 508.4 | 264.2 | - | - | 0 | - |
| - | - | 1763 | 270.1 | - | - | 0 | - |
| - | - | 685 | 272.1 | - | - | 0 | - |
| 3 | b | 1840 | 272.2 | 0.0006926 | 2.545 | +1 | 3 |
| 7 | y | 5394 | 272.2 | 0.0001285 | 0.472 | +1 | 2 |
| - | - | 544.4 | 273 | - | - | 0 | - |
| - | - | 861.9 | 273.2 | - | - | 0 | - |
| - | - | 1984 | 276.2 | - | - | 0 | - |
| - | - | 878 | 277.2 | - | - | 0 | - |
| - | - | 694.6 | 282.2 | - | - | 0 | - |
| - | - | 509.9 | 283.2 | - | - | 0 | - |
| 6 | b | 782.8 | 286.2 | 0.005111 | 17.86 | +2 | 6 |
| - | - | 1694 | 288.2 | - | - | 0 | - |
| - | - | 715.8 | 291 | - | - | 0 | - |
| - | - | 968.4 | 293.1 | - | - | 0 | - |
| - | - | 560.4 | 302.8 | - | - | 0 | - |
| - | - | 1143 | 306.2 | - | - | 0 | - |
| - | - | 671.2 | 309.5 | - | - | 0 | - |
| - | - | 2445 | 325.2 | - | - | 0 | - |
| 3 | y | 6119 | 327.7 | 0.002186 | 6.671 | +2 | 6 |
| 3 | y | 9395 | 328.2 | 0.004982 | 15.18 | +2 | 6 |
| - | - | 3364 | 328.7 | - | - | 0 | - |
| 3 | y | 738.5 | 336.7 | 6.933E-05 | 0.2059 | +2 | 6 |
| - | - | 1151 | 340.2 | - | - | 0 | - |
| - | - | 953.5 | 341.2 | - | - | 0 | - |
| - | - | 577.4 | 342.2 | - | - | 0 | - |
| - | - | 1208 | 350.9 | - | - | 0 | - |
| 4 | b | 682.7 | 353.2 | 0.001321 | 3.741 | +1 | 4 |
| - | - | 538.5 | 362.3 | - | - | 0 | - |
| 2 | y | 4126 | 363.2 | 0.000632 | 1.74 | +2 | 7 |
| - | - | 2378 | 363.7 | - | - | 0 | - |
| - | - | 2130 | 366.2 | - | - | 0 | - |
| 6 | y | 552.6 | 368.2 | 0.001204 | 3.268 | +1 | 3 |
| - | - | 837 | 368.9 | - | - | 0 | - |
| - | - | 687.6 | 380.2 | - | - | 0 | - |
| - | - | 823.1 | 384.2 | - | - | 0 | - |
| 6 | y | 3710 | 385.3 | 0.0001953 | 0.5069 | +1 | 3 |
| - | - | 1531 | 391.2 | - | - | 0 | - |
| - | - | 1819 | 402.2 | - | - | 0 | - |
| - | - | 1934 | 403.8 | - | - | 0 | - |
| 0 | Precursor | 4382 | 412.8 | 0.0002015 | 0.4881 | +2 | -1 |
| - | - | 1196 | 413.3 | - | - | 0 | - |
| - | - | 863.8 | 413.8 | - | - | 0 | - |
| - | - | 1405 | 420.9 | - | - | 0 | - |
| - | - | 930.5 | 421.2 | - | - | 0 | - |
| - | - | 826.3 | 421.3 | - | - | 0 | - |
| - | - | 1066 | 422.2 | - | - | 0 | - |
| 5 | y | 3.24E+04 | 472.3 | 8.868E-05 | 0.1878 | +1 | 4 |
| - | - | 7694 | 473.3 | - | - | 0 | - |
| - | - | 581.3 | 474.3 | - | - | 0 | - |
| - | - | 5394 | 486.3 | - | - | 0 | - |
| - | - | 5684 | 487.3 | - | - | 0 | - |
| - | - | 1089 | 488.3 | - | - | 0 | - |
| - | - | 2030 | 558.3 | - | - | 0 | - |
| 6 | b | 1.139E+04 | 571.3 | 0.01103 | 19.3 | +1 | 6 |
| - | - | 3612 | 572.4 | - | - | 0 | - |
| - | - | 817.2 | 628.4 | - | - | 0 | - |
| 3 | y | 3025 | 654.4 | 0.00857 | 13.1 | +1 | 6 |
| 3 | y | 7971 | 655.4 | 0.006498 | 9.916 | +1 | 6 |
| - | - | 3126 | 656.4 | - | - | 0 | - |
| - | - | 718.3 | 657.4 | - | - | 0 | - |
| 3 | y | 2.461E+04 | 672.4 | 0.0009467 | 1.408 | +1 | 6 |
| - | - | 8612 | 673.4 | - | - | 0 | - |
| - | - | 2223 | 674.4 | - | - | 0 | - |
| - | - | 1485 | 682.4 | - | - | 0 | - |
| - | - | 697.9 | 683.4 | - | - | 0 | - |
| 2 | y | 2670 | 725.4 | 3.105E-05 | 0.04281 | +1 | 7 |
| - | - | 1512 | 726.4 | - | - | 0 | - |
| 2 | y | 1.044E+04 | 743.4 | 0.001195 | 1.608 | +1 | 7 |
| - | - | 3909 | 744.4 | - | - | 0 | - |
| - | - | 736.1 | 745.4 | - | - | 0 | - |
| - | - | 635.5 | 2481 | - | - | 0 | - |
| - | - | 871.4 | 2963 | - | - | 0 | - |
| - | - | 822.8 | 3082 | - | - | 0 | - |

m/z Charge Intensity FragmentType MassShift Position
120.08088684082031 0 762.0292
127.1120376586914 0 360.22134
128.1072998046875 0 1013.57074
129.01837158203125 0 1331.0854
129.1024169921875 0 2832.8674
130.08636474609375 0 1406.0977
136.0759735107422 0 912.7199
140.0819549560547 0 1997.9552
141.08551025390625 0 446.3817
141.10232543945312 0 3366.0173
142.10565185546875 0 953.72943
143.1180419921875 0 13922.121 a 1
144.12135314941406 0 1193.3662
145.09718322753906 0 1095.9424
147.11273193359375 0 1224.0796
149.02345275878906 0 11527.978
149.68775939941406 0 408.0394
150.02682495117188 0 6569.528
151.0278778076172 0 799.5478
155.11788940429688 0 2148.3335
157.1336669921875 0 8983.087
158.0924530029297 0 2028.0714 y Ammonia loss 7
158.13697814941406 0 2313.6743
167.4368438720703 0 499.32098
169.0972137451172 0 4445.5464
170.1002197265625 0 1318.2653
171.10470581054688 0 607.7532
171.11289978027344 0 12543.168 b 1
171.1382293701172 0 497.7252
172.11663818359375 0 829.55743
173.09201049804688 0 1556.4557
173.1285400390625 0 15454.996
174.13206481933594 0 1134.7073
175.11904907226562 0 4617.3794 y 7
183.11277770996094 0 2813.4597
183.14910888671875 0 2880.0461
183.8260498046875 0 550.78156
184.15255737304688 0 1380.3175
185.08091735839844 0 724.4704
185.1288299560547 0 2312.2957
187.1081085205078 0 952.9093
201.12344360351562 0 6007.4604
209.00025939941406 0 519.9748
210.1229705810547 0 557.854
211.14430236816406 0 2147.7305
212.1392822265625 0 1437.5515
212.1488494873047 0 809.2413
226.15475463867188 0 1763.3197 a Water loss 2
227.1390838623047 0 697.8614
235.16481018066406 0 560.8763
240.1337890625 0 884.1608
241.1390838623047 0 644.43024
244.1514892578125 0 500.6817
248.04574584960938 0 501.07175
254.14988708496094 0 6057.056 b Water loss 2
255.14523315429688 0 12524.154 y Ammonia loss 6
256.1484375 0 1714.5022
258.1448059082031 0 1738.0056
259.1483154296875 0 898.8592
264.1708679199219 0 508.36868
270.1441955566406 0 1762.9998
272.12481689453125 0 684.98694
272.1597900390625 0 1840.0198 b 2
272.1718444824219 0 5394.0176 y 6
273.02227783203125 0 544.4408
273.1759033203125 0 861.8787
276.1553649902344 0 1983.6448
277.158935546875 0 877.95917
282.1813049316406 0 694.5918
283.1856384277344 0 509.8593
286.1812438964844 0 782.7839 b 5
288.155517578125 0 1694.1632
290.95904541015625 0 715.7968
293.1001892089844 0 968.41943
302.8329162597656 0 560.43256
306.1789855957031 0 1142.6588
309.482666015625 0 671.20166
325.2242431640625 0 2444.6794
327.6981201171875 0 6118.975 y Water loss 2
328.1972961425781 0 9395.164 y Ammonia loss 2
328.6982727050781 0 3363.8123
336.7056579589844 0 738.51776 y 2
340.1873779296875 0 1150.8347
341.1878662109375 0 953.47217
342.1863098144531 0 577.3629
350.9256591796875 0 1207.9729
353.2170104980469 0 682.69 b Water loss 3
362.2928161621094 0 538.4899
363.2182312011719 0 4126.3257 y Water loss 1
363.7200927734375 0 2378.0208
366.1853942871094 0 2130.3142
368.22802734375 0 552.6 y Ammonia loss 5
368.9377136230469 0 837.0079
380.233642578125 0 687.59436
384.1961669921875 0 823.0538
385.2555847167969 0 3709.6467 y 5
391.23358154296875 0 1530.6421
402.2060852050781 0 1818.6788
403.8131408691406 0 1933.7153
412.75286865234375 0 4382.165 Precursor Water loss
413.2573547363281 0 1196.091
413.7558288574219 0 863.7857
420.857666015625 0 1405.3075
421.24163818359375 0 930.5209
421.3273620605469 0 826.33545
422.2137451171875 0 1066.0807
472.2877197265625 0 32395.955 y 4
473.2907409667969 0 7693.9893
474.29473876953125 0 581.2776
486.2921447753906 0 5394.4233
487.29510498046875 0 5684.4507
488.29736328125 0 1089.4794
558.3319702148438 0 2029.9329
571.3560180664062 0 11390.0625 b 5
572.3587646484375 0 3611.8418
628.3775024414062 0 817.17944
654.384765625 0 3025.4172 y Water loss 2
655.3838500976562 0 7971.194 y Ammonia loss 2
656.3873291015625 0 3126.239
657.3927001953125 0 718.2846
672.4029541015625 0 24605.777 y 2
673.4057006835938 0 8611.877
674.4066772460938 0 2223.152
682.3881225585938 0 1484.903
683.3952026367188 0 697.94244
725.4304809570312 0 2670.0288 y Water loss 1
726.4330444335938 0 1512.4945
743.4398193359375 0 10442.679 y 1
744.4427490234375 0 3908.5151
745.444091796875 0 736.1117
2481.19580078125 0 635.53613
2963.1806640625 0 871.37067
3081.98388671875 0 822.75104

Spectrum Details

|  |  |
| --- | --- |
| Matched peaks? Matched peaks The total absolute number of peaks matched. Additionally in brackets the total fraction of peaks matched and the total number of peaks is shown. | 25 (18.80% of 133) |
| FDR? FDR The false discovery rate estimated for this peptide. It is calculated by matching all theoretical fragments with a non-integer shift with the raw peaks for this spectrum. This is done with 40 different shifts. The resulting percentage is the average number of annotated peaks over the number of annotated peaks with the correct spectrum. | 0.38% |
| Satellite FDR? Satellite FDR See the FDR for details on its calculation. This satellite ion specific FDR only contains the satellite ions (d/w) for I/L/J positions. | - |
| PSM Score? PSM Score The PSM Score as given by Hecklib to this annotated spectrum. It is shown with three significant figures. | 285 |

#### Spectrum 7239? Spectrum 7239 The raw spectrum of this peptide as annotated by Hecklib. The fragments are coloured according to ion type (see legend). Any peaks with a star '\*' as text can be hovered over to see the full details, first the ion type second the mass shift type. By hovering over the amino acids in the peptide or ions in the legend the corresponding peaks are highlighted. By toggling the 'Unassigned' label you can turn the background (unassigned) peaks on or off in the plot. By updating the slider in the Ion legend you can update the spectrum to only show the top X% of the peaks with labels. The top X% means any peak that is within X% of the highest intensity. By dragging in the spectrum you can zoom in to a specific part of the spectrum and use 'Zoom Out' to get back to the original zoom level. The annotation of the spectrum is based on the given sequence in the peptides file and is done with different software so inconsistencies are likely. The peaks are annotated based on the given sequence, with 20 ppm tolerance.

Copy Data

##### Spectrum 7239 (TSV)

###### Preview

```
Loading example...
```

*Click on the button to copy the data to your clipboard.*

Mz MinMz MaxIntensity Max

WidthHeightPeptide font sizePeptide stroke widthSpectrum font sizeSpectrum stroke widthCompact peptide

Ion legend

wxyz

abcd

OtherUnassignedIon^Charge^_Position_Show for top:%

VATVSJPR

04.12e+38.24e+31.24e+41.65e+4

Zoom Out

a^+1^_2_y^+1^_1_b^+1^_2_y^+1^_1_a^+1^_3_b^+1^_3_y^+1^_2_b^+1^_3_y^+1^_2_y^+2^_6_y^+2^_6_b^+1^_4_y^+2^_7_y^+1^_3_y^+1^_3_\*y^+1^_4_b^+1^_6_y^+1^_6_y^+1^_6_y^+1^_6_y^+1^_7_y^+1^_7_

0778155623343112

Fragment Matches Table

Show background peaks

| Position | Ion type | Intensity | mz Theoretical | mz Error (Th) | mz Error (ppm) | Charge | Series Number |
| --- | --- | --- | --- | --- | --- | --- | --- |
| - | - | 2581 | 120.1 | - | - | 0 | - |
| - | - | 432.2 | 123 | - | - | 0 | - |
| - | - | 822.6 | 129 | - | - | 0 | - |
| - | - | 1463 | 129.1 | - | - | 0 | - |
| - | - | 898.2 | 130.1 | - | - | 0 | - |
| - | - | 1994 | 136.1 | - | - | 0 | - |
| - | - | 1080 | 140.1 | - | - | 0 | - |
| - | - | 432 | 140.5 | - | - | 0 | - |
| - | - | 1970 | 141.1 | - | - | 0 | - |
| - | - | 433.5 | 142.1 | - | - | 0 | - |
| - | - | 596.9 | 143.1 | - | - | 0 | - |
| 2 | a | 6575 | 143.1 | 1.539E-05 | 0.1075 | +1 | 2 |
| - | - | 554.9 | 145.1 | - | - | 0 | - |
| - | - | 705.7 | 147.1 | - | - | 0 | - |
| - | - | 778 | 149 | - | - | 0 | - |
| - | - | 1.13E+04 | 149 | - | - | 0 | - |
| - | - | 5642 | 150 | - | - | 0 | - |
| - | - | 379 | 150.2 | - | - | 0 | - |
| - | - | 459.2 | 152 | - | - | 0 | - |
| - | - | 490.8 | 153.1 | - | - | 0 | - |
| - | - | 656.9 | 155.1 | - | - | 0 | - |
| - | - | 454.9 | 157 | - | - | 0 | - |
| - | - | 4676 | 157.1 | - | - | 0 | - |
| - | - | 555.8 | 158.1 | - | - | 0 | - |
| 8 | y | 715.3 | 158.1 | 0.0002025 | 1.281 | +1 | 1 |
| - | - | 1822 | 158.1 | - | - | 0 | - |
| - | - | 458.7 | 160 | - | - | 0 | - |
| - | - | 735.1 | 165.1 | - | - | 0 | - |
| - | - | 637.6 | 167 | - | - | 0 | - |
| - | - | 453.5 | 167.3 | - | - | 0 | - |
| - | - | 524.8 | 168 | - | - | 0 | - |
| - | - | 2434 | 169.1 | - | - | 0 | - |
| - | - | 596.9 | 170.1 | - | - | 0 | - |
| 2 | b | 5844 | 171.1 | 7.222E-05 | 0.4221 | +1 | 2 |
| - | - | 513.3 | 171.1 | - | - | 0 | - |
| - | - | 804 | 173.1 | - | - | 0 | - |
| - | - | 6679 | 173.1 | - | - | 0 | - |
| 8 | y | 2380 | 175.1 | 2.061E-05 | 0.1177 | +1 | 1 |
| - | - | 1819 | 182.1 | - | - | 0 | - |
| - | - | 1696 | 183.1 | - | - | 0 | - |
| - | - | 2170 | 183.1 | - | - | 0 | - |
| - | - | 471.3 | 184.2 | - | - | 0 | - |
| - | - | 445.8 | 184.2 | - | - | 0 | - |
| - | - | 642.9 | 185.1 | - | - | 0 | - |
| - | - | 1330 | 185.1 | - | - | 0 | - |
| - | - | 545.1 | 187.1 | - | - | 0 | - |
| - | - | 642.5 | 193.1 | - | - | 0 | - |
| - | - | 2596 | 201.1 | - | - | 0 | - |
| - | - | 597.8 | 202.1 | - | - | 0 | - |
| - | - | 1451 | 211.1 | - | - | 0 | - |
| - | - | 550.8 | 212.1 | - | - | 0 | - |
| - | - | 516.5 | 222.1 | - | - | 0 | - |
| 3 | a | 973.7 | 226.2 | 0.0003311 | 1.464 | +1 | 3 |
| 3 | b | 2165 | 254.1 | 4.543E-05 | 0.1788 | +1 | 3 |
| 7 | y | 5456 | 255.1 | 0.0002542 | 0.9963 | +1 | 2 |
| - | - | 625.1 | 256.2 | - | - | 0 | - |
| - | - | 1745 | 258.1 | - | - | 0 | - |
| - | - | 565.4 | 265 | - | - | 0 | - |
| - | - | 1072 | 270.1 | - | - | 0 | - |
| 3 | b | 923.5 | 272.2 | 0.002768 | 10.17 | +1 | 3 |
| 7 | y | 2793 | 272.2 | 9.795E-05 | 0.3599 | +1 | 2 |
| - | - | 1217 | 276.2 | - | - | 0 | - |
| - | - | 637.1 | 277.2 | - | - | 0 | - |
| - | - | 719.4 | 288.2 | - | - | 0 | - |
| - | - | 1195 | 293.1 | - | - | 0 | - |
| - | - | 828.8 | 302.9 | - | - | 0 | - |
| - | - | 740.2 | 304.1 | - | - | 0 | - |
| - | - | 633.6 | 306.2 | - | - | 0 | - |
| - | - | 535.6 | 306.2 | - | - | 0 | - |
| - | - | 700 | 308 | - | - | 0 | - |
| - | - | 1708 | 325.2 | - | - | 0 | - |
| 3 | y | 2184 | 327.7 | 0.004383 | 13.38 | +2 | 6 |
| 3 | y | 5045 | 328.2 | 0.005287 | 16.11 | +2 | 6 |
| - | - | 1599 | 328.7 | - | - | 0 | - |
| - | - | 1159 | 341.2 | - | - | 0 | - |
| 4 | b | 879.7 | 353.2 | 0.0005584 | 1.581 | +1 | 4 |
| 2 | y | 2308 | 363.2 | 0.0008151 | 2.244 | +2 | 7 |
| - | - | 712.4 | 363.7 | - | - | 0 | - |
| 6 | y | 655 | 368.2 | 0.0003834 | 1.041 | +1 | 3 |
| 6 | y | 1699 | 385.3 | 0.001019 | 2.646 | +1 | 3 |
| - | - | 547.2 | 390.2 | - | - | 0 | - |
| - | - | 647.1 | 391.2 | - | - | 0 | - |
| - | - | 543.1 | 402.2 | - | - | 0 | - |
| - | - | 1374 | 403.8 | - | - | 0 | - |
| - | - | 604.3 | 405 | - | - | 0 | - |
| 0 | Precursor | 2120 | 412.8 | 0.0002015 | 0.4881 | +2 | -1 |
| - | - | 1293 | 413.3 | - | - | 0 | - |
| - | - | 1371 | 420.9 | - | - | 0 | - |
| - | - | 634.4 | 421 | - | - | 0 | - |
| - | - | 762.4 | 421.9 | - | - | 0 | - |
| - | - | 1144 | 422.2 | - | - | 0 | - |
| - | - | 872.8 | 439.8 | - | - | 0 | - |
| 5 | y | 1.632E+04 | 472.3 | 0.0002108 | 0.4462 | +1 | 4 |
| - | - | 3516 | 473.3 | - | - | 0 | - |
| - | - | 2593 | 486.3 | - | - | 0 | - |
| - | - | 2958 | 487.3 | - | - | 0 | - |
| - | - | 699.2 | 488.3 | - | - | 0 | - |
| - | - | 716.5 | 558.3 | - | - | 0 | - |
| 6 | b | 4858 | 571.3 | 0.01103 | 19.3 | +1 | 6 |
| - | - | 1019 | 572.4 | - | - | 0 | - |
| - | - | 555.1 | 612.3 | - | - | 0 | - |
| - | - | 581.2 | 628.4 | - | - | 0 | - |
| - | - | 565.1 | 630.5 | - | - | 0 | - |
| - | - | 623.6 | 650.1 | - | - | 0 | - |
| 3 | y | 807.8 | 654.4 | 0.008754 | 13.38 | +1 | 6 |
| 3 | y | 4344 | 655.4 | 0.005888 | 8.984 | +1 | 6 |
| - | - | 1761 | 656.4 | - | - | 0 | - |
| 3 | y | 1.1E+04 | 672.4 | 0.0006415 | 0.9541 | +1 | 6 |
| - | - | 3081 | 673.4 | - | - | 0 | - |
| - | - | 827 | 674.4 | - | - | 0 | - |
| - | - | 689.4 | 682.4 | - | - | 0 | - |
| 2 | y | 909.2 | 725.4 | 0.00241 | 3.323 | +1 | 7 |
| 2 | y | 4774 | 743.4 | 0.0005239 | 0.7047 | +1 | 7 |
| - | - | 1104 | 744.4 | - | - | 0 | - |
| - | - | 944.4 | 3081 | - | - | 0 | - |

m/z Charge Intensity FragmentType MassShift Position
120.08082580566406 0 2580.517
123.04452514648438 0 432.242
129.0183868408203 0 822.5594
129.10244750976562 0 1463.3381
130.08642578125 0 898.2298
136.07562255859375 0 1994.1184
140.081787109375 0 1079.7428
140.4561004638672 0 432.04175
141.1021728515625 0 1970.2804
142.10543823242188 0 433.4509
143.1123504638672 0 596.85333
143.1178741455078 0 6575.0454 a 1
145.09730529785156 0 554.86945
147.11273193359375 0 705.6716
148.95474243164062 0 778.00244
149.02328491210938 0 11299.311
150.02664184570312 0 5642.0435
150.2194366455078 0 379.0224
152.0439453125 0 459.18997
153.05413818359375 0 490.82968
155.1178741455078 0 656.8826
157.0125274658203 0 454.8623
157.13356018066406 0 4676.406
158.06072998046875 0 555.83496
158.0926055908203 0 715.3307 y Ammonia loss 7
158.13685607910156 0 1822.0554
159.9947509765625 0 458.70197
165.0545196533203 0 735.07227
167.0338897705078 0 637.6251
167.3312530517578 0 453.54062
168.03689575195312 0 524.78705
169.09706115722656 0 2433.8665
170.10043334960938 0 596.94214
171.11273193359375 0 5844.0825 b 1
171.13784790039062 0 513.2559
173.09181213378906 0 803.99585
173.12832641601562 0 6679.39
175.1189727783203 0 2379.5938 y 7
182.0811767578125 0 1819.0762
183.11276245117188 0 1696.3934
183.14895629882812 0 2169.5298
184.1531982421875 0 471.28525
184.17059326171875 0 445.75452
185.08151245117188 0 642.85443
185.12835693359375 0 1330.2878
187.1083984375 0 545.0538
193.0857696533203 0 642.4516
201.12318420410156 0 2595.9453
202.107666015625 0 597.84045
211.14398193359375 0 1450.809
212.13864135742188 0 550.80023
222.1234130859375 0 516.46014
226.15533447265625 0 973.65106 a Water loss 2
254.14996337890625 0 2164.5479 b Water loss 2
255.14491271972656 0 5456.0933 y Ammonia loss 6
256.1500549316406 0 625.05334
258.1446228027344 0 1744.6284
264.9909973144531 0 565.37744
270.1441650390625 0 1071.6019
272.15771484375 0 923.50946 b 2
272.17181396484375 0 2792.8994 y 6
276.1557312011719 0 1217.4706
277.1593933105469 0 637.0834
288.1552429199219 0 719.4142
293.09967041015625 0 1194.5968
302.9078369140625 0 828.7955
304.1080627441406 0 740.184
306.17974853515625 0 633.5838
306.2222900390625 0 535.59503
307.987060546875 0 700.0326
325.2236328125 0 1708.1008
327.6959228515625 0 2184.427 y Water loss 2
328.1976013183594 0 5045.233 y Ammonia loss 2
328.6983947753906 0 1598.8943
341.1871032714844 0 1159.4265
353.2177734375 0 879.6912 b Water loss 3
363.2180480957031 0 2308.2573 y Water loss 1
363.71990966796875 0 712.36176
368.2296142578125 0 655.0158 y Ammonia loss 5
385.2547607421875 0 1699.38 y 5
390.2440185546875 0 547.1565
391.2311096191406 0 647.1448
402.2085266113281 0 543.1356
403.81396484375 0 1374.4879
404.96795654296875 0 604.2552
412.75286865234375 0 2119.9907 Precursor Water loss
413.25457763671875 0 1293.0757
420.85809326171875 0 1370.9218
420.97100830078125 0 634.3693
421.9028015136719 0 762.3893
422.2408752441406 0 1144.4683
439.84442138671875 0 872.75085
472.28759765625 0 16318.209 y 4
473.290771484375 0 3516.4624
486.2914123535156 0 2592.5994
487.2944030761719 0 2958.0945
488.2960205078125 0 699.2449
558.3289184570312 0 716.4623
571.3560180664062 0 4857.6187 b 5
572.3561401367188 0 1018.99426
612.2794799804688 0 555.11523
628.3744506835938 0 581.2188
630.51171875 0 565.1264
650.1348876953125 0 623.6117
654.3845825195312 0 807.7789 y Water loss 2
655.3832397460938 0 4343.8867 y Ammonia loss 2
656.3866577148438 0 1761.3143
672.4032592773438 0 10999.635 y 2
673.4058837890625 0 3080.6946
674.4112548828125 0 826.9593
682.3880004882812 0 689.4422
725.4280395507812 0 909.232 y Water loss 1
743.4404907226562 0 4773.5493 y 1
744.4453125 0 1104.1891
3081.161376953125 0 944.3554

Spectrum Details

|  |  |
| --- | --- |
| Matched peaks? Matched peaks The total absolute number of peaks matched. Additionally in brackets the total fraction of peaks matched and the total number of peaks is shown. | 23 (20.00% of 115) |
| FDR? FDR The false discovery rate estimated for this peptide. It is calculated by matching all theoretical fragments with a non-integer shift with the raw peaks for this spectrum. This is done with 40 different shifts. The resulting percentage is the average number of annotated peaks over the number of annotated peaks with the correct spectrum. | 0.31% |
| Satellite FDR? Satellite FDR See the FDR for details on its calculation. This satellite ion specific FDR only contains the satellite ions (d/w) for I/L/J positions. | - |
| PSM Score? PSM Score The PSM Score as given by Hecklib to this annotated spectrum. It is shown with three significant figures. | 285 |

#### Spectrum 5920? Spectrum 5920 The raw spectrum of this peptide as annotated by Hecklib. The fragments are coloured according to ion type (see legend). Any peaks with a star '\*' as text can be hovered over to see the full details, first the ion type second the mass shift type. By hovering over the amino acids in the peptide or ions in the legend the corresponding peaks are highlighted. By toggling the 'Unassigned' label you can turn the background (unassigned) peaks on or off in the plot. By updating the slider in the Ion legend you can update the spectrum to only show the top X% of the peaks with labels. The top X% means any peak that is within X% of the highest intensity. By dragging in the spectrum you can zoom in to a specific part of the spectrum and use 'Zoom Out' to get back to the original zoom level. The annotation of the spectrum is based on the given sequence in the peptides file and is done with different software so inconsistencies are likely. The peaks are annotated based on the given sequence, with 20 ppm tolerance.

Copy Data

##### Spectrum 5920 (TSV)

###### Preview

```
Loading example...
```

*Click on the button to copy the data to your clipboard.*

Mz MinMz MaxIntensity Max

WidthHeightPeptide font sizePeptide stroke widthSpectrum font sizeSpectrum stroke widthCompact peptide

Ion legend

wxyz

abcd

OtherUnassignedIon^Charge^_Position_Show for top:%

VATVSJPR

01.17e+42.33e+43.50e+44.67e+4

Zoom Out

a^+1^_2_y^+1^_1_b^+1^_2_y^+1^_1_a^+1^_3_y^+2^_4_d^+1^_3_b^+1^_3_y^+1^_2_b^+1^_3_y^+1^_2_b^+2^_6_y^+2^_6_y^+2^_6_y^+2^_6_b^+1^_4_y^+2^_7_y^+1^_3_y^+1^_3_\*y^+1^_4_y^+1^_4_b^+1^_6_y^+1^_6_y^+1^_6_y^+1^_6_y^+1^_7_y^+1^_7_

0807161324203227

Fragment Matches Table

Show background peaks

| Position | Ion type | Intensity | mz Theoretical | mz Error (Th) | mz Error (ppm) | Charge | Series Number |
| --- | --- | --- | --- | --- | --- | --- | --- |
| - | - | 980.7 | 120.1 | - | - | 0 | - |
| - | - | 347.3 | 121.1 | - | - | 0 | - |
| - | - | 455.9 | 125.1 | - | - | 0 | - |
| - | - | 362 | 126.7 | - | - | 0 | - |
| - | - | 840.9 | 127.1 | - | - | 0 | - |
| - | - | 927.7 | 128.1 | - | - | 0 | - |
| - | - | 883.3 | 129 | - | - | 0 | - |
| - | - | 3827 | 129.1 | - | - | 0 | - |
| - | - | 313.2 | 129.3 | - | - | 0 | - |
| - | - | 898.1 | 130.1 | - | - | 0 | - |
| - | - | 387.3 | 130.1 | - | - | 0 | - |
| - | - | 391.1 | 133 | - | - | 0 | - |
| - | - | 584.4 | 136 | - | - | 0 | - |
| - | - | 394.5 | 138.1 | - | - | 0 | - |
| - | - | 399.1 | 138.7 | - | - | 0 | - |
| - | - | 540.6 | 139 | - | - | 0 | - |
| - | - | 3350 | 140.1 | - | - | 0 | - |
| - | - | 4865 | 141.1 | - | - | 0 | - |
| - | - | 1386 | 142.1 | - | - | 0 | - |
| 2 | a | 2.027E+04 | 143.1 | 0.0002135 | 1.492 | +1 | 2 |
| - | - | 1734 | 144.1 | - | - | 0 | - |
| - | - | 1223 | 145.1 | - | - | 0 | - |
| - | - | 1963 | 147.1 | - | - | 0 | - |
| - | - | 438.9 | 148.1 | - | - | 0 | - |
| - | - | 1.319E+04 | 149 | - | - | 0 | - |
| - | - | 6663 | 150 | - | - | 0 | - |
| - | - | 522.9 | 151 | - | - | 0 | - |
| - | - | 973.8 | 155.1 | - | - | 0 | - |
| - | - | 2587 | 155.1 | - | - | 0 | - |
| - | - | 610.4 | 157 | - | - | 0 | - |
| - | - | 713.7 | 157.1 | - | - | 0 | - |
| - | - | 1.052E+04 | 157.1 | - | - | 0 | - |
| 8 | y | 2539 | 158.1 | 3.468E-05 | 0.2194 | +1 | 1 |
| - | - | 3203 | 158.1 | - | - | 0 | - |
| - | - | 997.5 | 167 | - | - | 0 | - |
| - | - | 5295 | 169.1 | - | - | 0 | - |
| - | - | 1256 | 170.1 | - | - | 0 | - |
| 2 | b | 1.73E+04 | 171.1 | 0.0001719 | 1.005 | +1 | 2 |
| - | - | 1165 | 172.1 | - | - | 0 | - |
| - | - | 2319 | 173.1 | - | - | 0 | - |
| - | - | 1.986E+04 | 173.1 | - | - | 0 | - |
| - | - | 2151 | 173.4 | - | - | 0 | - |
| - | - | 1782 | 174.1 | - | - | 0 | - |
| 8 | y | 6587 | 175.1 | 0.0001732 | 0.989 | +1 | 1 |
| - | - | 4598 | 183.1 | - | - | 0 | - |
| - | - | 4672 | 183.1 | - | - | 0 | - |
| - | - | 913.7 | 184.2 | - | - | 0 | - |
| - | - | 1035 | 185.1 | - | - | 0 | - |
| - | - | 1680 | 185.1 | - | - | 0 | - |
| - | - | 1089 | 187.1 | - | - | 0 | - |
| - | - | 800.2 | 195.1 | - | - | 0 | - |
| - | - | 475.2 | 196.1 | - | - | 0 | - |
| - | - | 7564 | 201.1 | - | - | 0 | - |
| - | - | 659.7 | 202.1 | - | - | 0 | - |
| - | - | 559.8 | 210.1 | - | - | 0 | - |
| - | - | 683.3 | 211.1 | - | - | 0 | - |
| - | - | 2910 | 211.1 | - | - | 0 | - |
| - | - | 1702 | 212.1 | - | - | 0 | - |
| - | - | 659.5 | 213.1 | - | - | 0 | - |
| - | - | 570.8 | 215 | - | - | 0 | - |
| - | - | 629.2 | 215.1 | - | - | 0 | - |
| 3 | a | 3556 | 226.2 | 0.0001786 | 0.7896 | +1 | 3 |
| - | - | 1116 | 227.1 | - | - | 0 | - |
| 5 | y | 782.1 | 228.1 | 0.0005093 | 2.233 | +2 | 4 |
| 3 | d | 1470 | 230.1 | 0.0003811 | 1.656 | +1 | 3 |
| - | - | 527.1 | 237.1 | - | - | 0 | - |
| - | - | 1286 | 240.1 | - | - | 0 | - |
| - | - | 584.3 | 241.1 | - | - | 0 | - |
| 3 | b | 7405 | 254.1 | 3.018E-05 | 0.1187 | +1 | 3 |
| - | - | 855.8 | 254.2 | - | - | 0 | - |
| 7 | y | 1.892E+04 | 255.1 | 5.098E-05 | 0.1998 | +1 | 2 |
| - | - | 2203 | 256.1 | - | - | 0 | - |
| - | - | 3548 | 258.1 | - | - | 0 | - |
| - | - | 646.1 | 259.1 | - | - | 0 | - |
| - | - | 2722 | 270.1 | - | - | 0 | - |
| 3 | b | 3112 | 272.2 | 0.0006621 | 2.433 | +1 | 3 |
| 7 | y | 8275 | 272.2 | 0.0001285 | 0.472 | +1 | 2 |
| - | - | 2303 | 276.2 | - | - | 0 | - |
| - | - | 796.2 | 282.2 | - | - | 0 | - |
| 6 | b | 1017 | 286.2 | 0.005447 | 19.03 | +2 | 6 |
| - | - | 1498 | 288.2 | - | - | 0 | - |
| - | - | 893.9 | 291 | - | - | 0 | - |
| - | - | 806.8 | 291.1 | - | - | 0 | - |
| - | - | 1416 | 293.1 | - | - | 0 | - |
| - | - | 686.4 | 299.2 | - | - | 0 | - |
| - | - | 1313 | 306.2 | - | - | 0 | - |
| - | - | 703.1 | 312.2 | - | - | 0 | - |
| - | - | 607.3 | 324.9 | - | - | 0 | - |
| - | - | 3607 | 325.2 | - | - | 0 | - |
| 3 | y | 7647 | 327.7 | 0.001393 | 4.25 | +2 | 6 |
| 3 | y | 1.063E+04 | 328.2 | 0.004677 | 14.25 | +2 | 6 |
| - | - | 4199 | 328.7 | - | - | 0 | - |
| - | - | 763.8 | 329.2 | - | - | 0 | - |
| 3 | y | 629.7 | 336.7 | 0.0005271 | 1.565 | +2 | 6 |
| - | - | 2509 | 340.2 | - | - | 0 | - |
| - | - | 1326 | 341 | - | - | 0 | - |
| - | - | 1182 | 341.2 | - | - | 0 | - |
| - | - | 763.9 | 343.2 | - | - | 0 | - |
| - | - | 892 | 350.9 | - | - | 0 | - |
| 4 | b | 2360 | 353.2 | 0.0004364 | 1.235 | +1 | 4 |
| - | - | 1090 | 359 | - | - | 0 | - |
| 2 | y | 9492 | 363.2 | 0.0002352 | 0.6477 | +2 | 7 |
| - | - | 2775 | 363.7 | - | - | 0 | - |
| - | - | 844.9 | 364.2 | - | - | 0 | - |
| - | - | 1037 | 366.2 | - | - | 0 | - |
| 6 | y | 1285 | 368.2 | 0.0007763 | 2.108 | +1 | 3 |
| - | - | 670.9 | 380.2 | - | - | 0 | - |
| 6 | y | 5437 | 385.3 | 0.0001648 | 0.4277 | +1 | 3 |
| - | - | 643.4 | 386.2 | - | - | 0 | - |
| - | - | 836.3 | 390.2 | - | - | 0 | - |
| - | - | 2638 | 391.2 | - | - | 0 | - |
| - | - | 827.6 | 391.7 | - | - | 0 | - |
| - | - | 1453 | 402.2 | - | - | 0 | - |
| - | - | 1981 | 403.8 | - | - | 0 | - |
| 0 | Precursor | 5107 | 412.8 | 0.0004151 | 1.006 | +2 | -1 |
| - | - | 2723 | 413.3 | - | - | 0 | - |
| - | - | 751.2 | 413.8 | - | - | 0 | - |
| - | - | 1328 | 420.9 | - | - | 0 | - |
| - | - | 1237 | 422.2 | - | - | 0 | - |
| - | - | 1184 | 439.8 | - | - | 0 | - |
| - | - | 796.2 | 442.3 | - | - | 0 | - |
| 5 | y | 827.5 | 454.3 | 0.006675 | 14.69 | +1 | 4 |
| 5 | y | 4.623E+04 | 472.3 | 5.817E-05 | 0.1232 | +1 | 4 |
| - | - | 1.129E+04 | 473.3 | - | - | 0 | - |
| - | - | 1622 | 474.3 | - | - | 0 | - |
| - | - | 664.8 | 482.3 | - | - | 0 | - |
| - | - | 4855 | 486.3 | - | - | 0 | - |
| - | - | 7734 | 487.3 | - | - | 0 | - |
| - | - | 2235 | 488.3 | - | - | 0 | - |
| - | - | 637.1 | 508.3 | - | - | 0 | - |
| - | - | 659.1 | 509.3 | - | - | 0 | - |
| - | - | 965.5 | 557.3 | - | - | 0 | - |
| - | - | 1776 | 558.3 | - | - | 0 | - |
| - | - | 846.8 | 559.3 | - | - | 0 | - |
| 6 | b | 1.549E+04 | 571.3 | 0.0106 | 18.56 | +1 | 6 |
| - | - | 4424 | 572.4 | - | - | 0 | - |
| - | - | 701.9 | 573.4 | - | - | 0 | - |
| - | - | 704.2 | 615.6 | - | - | 0 | - |
| - | - | 1475 | 628.4 | - | - | 0 | - |
| 3 | y | 3472 | 654.4 | 0.00796 | 12.16 | +1 | 6 |
| 3 | y | 8047 | 655.4 | 0.007231 | 11.03 | +1 | 6 |
| - | - | 3376 | 656.4 | - | - | 0 | - |
| - | - | 729.9 | 657.4 | - | - | 0 | - |
| 3 | y | 3.386E+04 | 672.4 | 0.0009467 | 1.408 | +1 | 6 |
| - | - | 1.356E+04 | 673.4 | - | - | 0 | - |
| - | - | 2607 | 674.4 | - | - | 0 | - |
| - | - | 2957 | 682.4 | - | - | 0 | - |
| - | - | 1188 | 683.4 | - | - | 0 | - |
| 2 | y | 5046 | 725.4 | 0.001861 | 2.565 | +1 | 7 |
| - | - | 1539 | 726.4 | - | - | 0 | - |
| 2 | y | 1.315E+04 | 743.4 | 0.001256 | 1.69 | +1 | 7 |
| - | - | 5500 | 744.4 | - | - | 0 | - |
| - | - | 1415 | 745.4 | - | - | 0 | - |
| - | - | 626.6 | 767.5 | - | - | 0 | - |
| - | - | 766.3 | 1988 | - | - | 0 | - |
| - | - | 802.9 | 2590 | - | - | 0 | - |
| - | - | 744.6 | 2969 | - | - | 0 | - |
| - | - | 576.2 | 3079 | - | - | 0 | - |
| - | - | 740.9 | 3195 | - | - | 0 | - |

m/z Charge Intensity FragmentType MassShift Position
120.08118438720703 0 980.70056
121.08485412597656 0 347.30933
125.06694030761719 0 455.89566
126.67442321777344 0 362.0393
127.08682250976562 0 840.9438
128.10739135742188 0 927.7494
129.01844787597656 0 883.2822
129.10244750976562 0 3827.111
129.26385498046875 0 313.19635
130.08651733398438 0 898.1498
130.1056671142578 0 387.3426
132.99395751953125 0 391.13507
136.0217742919922 0 584.41125
138.1375274658203 0 394.51556
138.68051147460938 0 399.05322
139.00294494628906 0 540.59906
140.08206176757812 0 3350.037
141.10250854492188 0 4864.892
142.10572814941406 0 1385.8982
143.11810302734375 0 20265.48 a 1
144.12149047851562 0 1734.2268
145.0973358154297 0 1223.2618
147.11302185058594 0 1963.1946
148.11656188964844 0 438.86423
149.02349853515625 0 13193.568
150.0269012451172 0 6663.1025
151.0279998779297 0 522.8851
155.0817413330078 0 973.8337
155.11805725097656 0 2587.0952
157.01304626464844 0 610.434
157.10891723632812 0 713.71857
157.13377380371094 0 10522.432
158.09243774414062 0 2539.3613 y Ammonia loss 7
158.13706970214844 0 3202.645
167.0343017578125 0 997.5486
169.09735107421875 0 5295.208
170.100830078125 0 1256.3195
171.11297607421875 0 17295.104 b 1
172.11627197265625 0 1165.0472
173.09219360351562 0 2319.291
173.12867736816406 0 19860.303
173.43959045410156 0 2151.113
174.13189697265625 0 1781.6309
175.11912536621094 0 6587.479 y 7
183.11297607421875 0 4598.437
183.14942932128906 0 4671.8843
184.15318298339844 0 913.67236
185.08140563964844 0 1034.8727
185.12872314453125 0 1679.625
187.10760498046875 0 1088.7651
195.11280822753906 0 800.1556
196.0666961669922 0 475.2046
201.12351989746094 0 7564.289
202.12709045410156 0 659.7308
210.12399291992188 0 559.80634
211.10848999023438 0 683.2617
211.14422607421875 0 2910.3174
212.1394500732422 0 1701.9396
213.12315368652344 0 659.5341
215.03074645996094 0 570.8035
215.13858032226562 0 629.15643
226.15518188476562 0 3556.3843 a Water loss 2
227.13865661621094 0 1115.95
228.13375854492188 0 782.08826 y Ammonia loss 4
230.15029907226562 0 1469.9728 d 2
237.0908966064453 0 527.10144
240.13446044921875 0 1286.3372
241.1392822265625 0 584.2562
254.1499481201172 0 7405.3667 b Water loss 2
254.1625518798828 0 855.8315
255.1452178955078 0 18915.467 y Ammonia loss 6
256.1484680175781 0 2203.2065
258.1446838378906 0 3547.9946
259.1481628417969 0 646.1192
270.14483642578125 0 2722.218
272.1598205566406 0 3112.336 b 2
272.1718444824219 0 8275.314 y 6
276.15521240234375 0 2303.152
282.18084716796875 0 796.22003
286.18157958984375 0 1017.1244 b 5
288.1555480957031 0 1497.508
290.9600524902344 0 893.9185
291.0846252441406 0 806.8285
293.0992736816406 0 1415.513
299.1886291503906 0 686.35944
306.1791687011719 0 1312.6658
312.1921691894531 0 703.09094
324.8818359375 0 607.3186
325.2231140136719 0 3606.8035
327.69891357421875 0 7647.171 y Water loss 2
328.1969909667969 0 10634.724 y Ammonia loss 2
328.69818115234375 0 4199.423
329.198486328125 0 763.75287
336.70611572265625 0 629.72437 y 2
340.18695068359375 0 2509.2188
340.9647521972656 0 1325.6796
341.187744140625 0 1182.1798
343.23431396484375 0 763.8517
350.9267272949219 0 891.9791
353.2178955078125 0 2360.3337 b Water loss 3
358.9744873046875 0 1089.6144
363.2186279296875 0 9492.213 y Water loss 1
363.7206115722656 0 2774.9785
364.22296142578125 0 844.9227
366.1853942871094 0 1037.3014
368.22845458984375 0 1284.8977 y Ammonia loss 5
380.23626708984375 0 670.8987
385.255615234375 0 5436.6064 y 5
386.1517333984375 0 643.3686
390.2441101074219 0 836.3077
391.2315673828125 0 2637.6897
391.7322998046875 0 827.60333
402.2066650390625 0 1452.6997
403.8126220703125 0 1980.6859
412.7526550292969 0 5106.7124 Precursor Water loss
413.2548522949219 0 2722.5645
413.75592041015625 0 751.20685
420.8559875488281 0 1328.1957
422.2149353027344 0 1236.7839
439.8446960449219 0 1184.0999
442.2775573730469 0 796.1752
454.27056884765625 0 827.4951 y Water loss 4
472.2877502441406 0 46226.457 y 4
473.2904052734375 0 11293.197
474.2941589355469 0 1622.4089
482.2717590332031 0 664.8348
486.2920227050781 0 4854.7705
487.2951354980469 0 7734.466
488.2994079589844 0 2234.9294
508.27777099609375 0 637.0552
509.2802429199219 0 659.0808
557.3309936523438 0 965.4509
558.3313598632812 0 1775.845
559.3358764648438 0 846.842
571.3555908203125 0 15491.446 b 5
572.3590698242188 0 4424.2207
573.3594970703125 0 701.94653
615.5781860351562 0 704.16144
628.3770141601562 0 1474.5796
654.3853759765625 0 3471.53 y Water loss 2
655.3845825195312 0 8046.808 y Ammonia loss 2
656.3883056640625 0 3376.1003
657.3868408203125 0 729.92944
672.4029541015625 0 33855.527 y 2
673.4055786132812 0 13557.492
674.4097900390625 0 2607.177
682.38818359375 0 2956.644
683.392822265625 0 1187.9249
725.4285888671875 0 5046.0054 y Water loss 1
726.4300537109375 0 1538.8164
743.4397583007812 0 13149.34 y 1
744.4428100585938 0 5499.702
745.4442138671875 0 1414.6945
767.48681640625 0 626.6316
1987.9305419921875 0 766.3345
2589.752197265625 0 802.9103
2969.240966796875 0 744.5821
3079.377197265625 0 576.1708
3194.927001953125 0 740.91583

Spectrum Details

|  |  |
| --- | --- |
| Matched peaks? Matched peaks The total absolute number of peaks matched. Additionally in brackets the total fraction of peaks matched and the total number of peaks is shown. | 28 (17.61% of 159) |
| FDR? FDR The false discovery rate estimated for this peptide. It is calculated by matching all theoretical fragments with a non-integer shift with the raw peaks for this spectrum. This is done with 40 different shifts. The resulting percentage is the average number of annotated peaks over the number of annotated peaks with the correct spectrum. | 0.43% |
| Satellite FDR? Satellite FDR See the FDR for details on its calculation. This satellite ion specific FDR only contains the satellite ions (d/w) for I/L/J positions. | - |
| PSM Score? PSM Score The PSM Score as given by Hecklib to this annotated spectrum. It is shown with three significant figures. | 306 |

#### Spectrum 6159? Spectrum 6159 The raw spectrum of this peptide as annotated by Hecklib. The fragments are coloured according to ion type (see legend). Any peaks with a star '\*' as text can be hovered over to see the full details, first the ion type second the mass shift type. By hovering over the amino acids in the peptide or ions in the legend the corresponding peaks are highlighted. By toggling the 'Unassigned' label you can turn the background (unassigned) peaks on or off in the plot. By updating the slider in the Ion legend you can update the spectrum to only show the top X% of the peaks with labels. The top X% means any peak that is within X% of the highest intensity. By dragging in the spectrum you can zoom in to a specific part of the spectrum and use 'Zoom Out' to get back to the original zoom level. The annotation of the spectrum is based on the given sequence in the peptides file and is done with different software so inconsistencies are likely. The peaks are annotated based on the given sequence, with 20 ppm tolerance.

Copy Data

##### Spectrum 6159 (TSV)

###### Preview

```
Loading example...
```

*Click on the button to copy the data to your clipboard.*

Mz MinMz MaxIntensity Max

WidthHeightPeptide font sizePeptide stroke widthSpectrum font sizeSpectrum stroke widthCompact peptide

Ion legend

wxyz

abcd

OtherUnassignedIon^Charge^_Position_Show for top:%

VATVSJPR

08.48e+31.70e+42.54e+43.39e+4

Zoom Out

a^+1^_2_y^+1^_1_b^+1^_2_y^+1^_1_a^+1^_3_d^+1^_3_b^+1^_3_y^+1^_2_b^+1^_3_y^+1^_2_y^+2^_6_y^+2^_6_b^+1^_4_y^+2^_7_y^+1^_3_b^+1^_4_y^+1^_3_\*y^+1^_4_b^+1^_6_y^+1^_6_y^+1^_6_y^+1^_6_y^+1^_7_y^+1^_7_y^+1^_7_

0778155623343112

Fragment Matches Table

Show background peaks

| Position | Ion type | Intensity | mz Theoretical | mz Error (Th) | mz Error (ppm) | Charge | Series Number |
| --- | --- | --- | --- | --- | --- | --- | --- |
| - | - | 756.8 | 120.1 | - | - | 0 | - |
| - | - | 375.1 | 124.3 | - | - | 0 | - |
| - | - | 920.2 | 128.1 | - | - | 0 | - |
| - | - | 1506 | 129 | - | - | 0 | - |
| - | - | 2965 | 129.1 | - | - | 0 | - |
| - | - | 427.9 | 129.1 | - | - | 0 | - |
| - | - | 1007 | 130.1 | - | - | 0 | - |
| - | - | 1975 | 140.1 | - | - | 0 | - |
| - | - | 4231 | 141.1 | - | - | 0 | - |
| - | - | 718.5 | 142.1 | - | - | 0 | - |
| 2 | a | 1.397E+04 | 143.1 | 0.000183 | 1.278 | +1 | 2 |
| - | - | 470.2 | 144.1 | - | - | 0 | - |
| - | - | 1416 | 144.1 | - | - | 0 | - |
| - | - | 1130 | 145.1 | - | - | 0 | - |
| - | - | 1580 | 147.1 | - | - | 0 | - |
| - | - | 1.283E+04 | 149 | - | - | 0 | - |
| - | - | 5398 | 150 | - | - | 0 | - |
| - | - | 672.1 | 151 | - | - | 0 | - |
| - | - | 1716 | 155.1 | - | - | 0 | - |
| - | - | 559.8 | 157 | - | - | 0 | - |
| - | - | 9152 | 157.1 | - | - | 0 | - |
| 8 | y | 2146 | 158.1 | 3.468E-05 | 0.2194 | +1 | 1 |
| - | - | 528.2 | 158.1 | - | - | 0 | - |
| - | - | 2677 | 158.1 | - | - | 0 | - |
| - | - | 685 | 167 | - | - | 0 | - |
| - | - | 4261 | 169.1 | - | - | 0 | - |
| - | - | 1222 | 170.1 | - | - | 0 | - |
| 2 | b | 1.375E+04 | 171.1 | 9.562E-05 | 0.5588 | +1 | 2 |
| - | - | 1135 | 172.1 | - | - | 0 | - |
| - | - | 485 | 172.7 | - | - | 0 | - |
| - | - | 2242 | 173.1 | - | - | 0 | - |
| - | - | 1.631E+04 | 173.1 | - | - | 0 | - |
| - | - | 721.6 | 173.4 | - | - | 0 | - |
| - | - | 1256 | 174.1 | - | - | 0 | - |
| 8 | y | 3536 | 175.1 | 0.0001732 | 0.989 | +1 | 1 |
| - | - | 2958 | 183.1 | - | - | 0 | - |
| - | - | 3081 | 183.1 | - | - | 0 | - |
| - | - | 1624 | 184.2 | - | - | 0 | - |
| - | - | 671.6 | 185.1 | - | - | 0 | - |
| - | - | 2773 | 185.1 | - | - | 0 | - |
| - | - | 835.7 | 186.1 | - | - | 0 | - |
| - | - | 6633 | 201.1 | - | - | 0 | - |
| - | - | 699.9 | 202.1 | - | - | 0 | - |
| - | - | 1146 | 211.1 | - | - | 0 | - |
| - | - | 1237 | 212.1 | - | - | 0 | - |
| - | - | 743.3 | 212.1 | - | - | 0 | - |
| 3 | a | 1986 | 226.2 | 3.506E-05 | 0.155 | +1 | 3 |
| - | - | 1199 | 227.1 | - | - | 0 | - |
| 3 | d | 1165 | 230.1 | 0.0002445 | 1.062 | +1 | 3 |
| - | - | 456.8 | 234.5 | - | - | 0 | - |
| - | - | 718 | 237.1 | - | - | 0 | - |
| - | - | 628.1 | 240.1 | - | - | 0 | - |
| - | - | 516 | 245.9 | - | - | 0 | - |
| 3 | b | 5486 | 254.1 | 7.664E-05 | 0.3015 | +1 | 3 |
| - | - | 596.4 | 254.2 | - | - | 0 | - |
| 7 | y | 1.379E+04 | 255.1 | 1.006E-05 | 0.03942 | +1 | 2 |
| - | - | 884.3 | 256.1 | - | - | 0 | - |
| - | - | 628.9 | 257.1 | - | - | 0 | - |
| - | - | 2431 | 258.1 | - | - | 0 | - |
| - | - | 913.1 | 259.1 | - | - | 0 | - |
| - | - | 1896 | 270.1 | - | - | 0 | - |
| 3 | b | 2095 | 272.2 | 0.0007231 | 2.657 | +1 | 3 |
| 7 | y | 5937 | 272.2 | 0.000159 | 0.5841 | +1 | 2 |
| - | - | 1777 | 276.2 | - | - | 0 | - |
| - | - | 1014 | 277.2 | - | - | 0 | - |
| - | - | 569.1 | 286.1 | - | - | 0 | - |
| - | - | 2144 | 288.2 | - | - | 0 | - |
| - | - | 885.4 | 291 | - | - | 0 | - |
| - | - | 590.8 | 292 | - | - | 0 | - |
| - | - | 595.5 | 293.1 | - | - | 0 | - |
| - | - | 1014 | 306.2 | - | - | 0 | - |
| - | - | 667.4 | 312.2 | - | - | 0 | - |
| - | - | 1650 | 322.2 | - | - | 0 | - |
| - | - | 2303 | 325.2 | - | - | 0 | - |
| 3 | y | 5609 | 327.7 | 0.001759 | 5.367 | +2 | 6 |
| 3 | y | 7764 | 328.2 | 0.004921 | 14.99 | +2 | 6 |
| - | - | 3680 | 328.7 | - | - | 0 | - |
| - | - | 754.3 | 329.2 | - | - | 0 | - |
| - | - | 1601 | 340.2 | - | - | 0 | - |
| - | - | 1173 | 341.2 | - | - | 0 | - |
| - | - | 622.6 | 343.1 | - | - | 0 | - |
| - | - | 676.7 | 350.9 | - | - | 0 | - |
| - | - | 634.2 | 352.1 | - | - | 0 | - |
| 4 | b | 1749 | 353.2 | 0.0004364 | 1.235 | +1 | 4 |
| 2 | y | 6099 | 363.2 | 5.214E-05 | 0.1436 | +2 | 7 |
| - | - | 2087 | 363.7 | - | - | 0 | - |
| - | - | 939.2 | 364.2 | - | - | 0 | - |
| - | - | 1.192E+04 | 366.2 | - | - | 0 | - |
| - | - | 644.2 | 367.1 | - | - | 0 | - |
| - | - | 965.1 | 367.2 | - | - | 0 | - |
| - | - | 1150 | 368.2 | - | - | 0 | - |
| 6 | y | 878.2 | 368.2 | 0.0003491 | 0.948 | +1 | 3 |
| - | - | 754.2 | 368.9 | - | - | 0 | - |
| 4 | b | 700.6 | 371.2 | 0.001052 | 2.835 | +1 | 4 |
| - | - | 3756 | 384.2 | - | - | 0 | - |
| 6 | y | 3470 | 385.3 | 4.886E-05 | 0.1268 | +1 | 3 |
| - | - | 1231 | 391.2 | - | - | 0 | - |
| - | - | 816 | 391.7 | - | - | 0 | - |
| - | - | 8469 | 402.2 | - | - | 0 | - |
| - | - | 861.8 | 403.2 | - | - | 0 | - |
| - | - | 1448 | 403.8 | - | - | 0 | - |
| 0 | Precursor | 4302 | 412.8 | 0.0001647 | 0.3991 | +2 | -1 |
| - | - | 2282 | 413.3 | - | - | 0 | - |
| - | - | 1398 | 420.9 | - | - | 0 | - |
| - | - | 4345 | 422.2 | - | - | 0 | - |
| 5 | y | 3.358E+04 | 472.3 | 3.339E-05 | 0.07069 | +1 | 4 |
| - | - | 8410 | 473.3 | - | - | 0 | - |
| - | - | 916.4 | 474.3 | - | - | 0 | - |
| - | - | 5237 | 486.3 | - | - | 0 | - |
| - | - | 6052 | 487.3 | - | - | 0 | - |
| - | - | 1989 | 488.3 | - | - | 0 | - |
| - | - | 583.5 | 498.3 | - | - | 0 | - |
| - | - | 903.9 | 509.3 | - | - | 0 | - |
| - | - | 1085 | 558.3 | - | - | 0 | - |
| 6 | b | 1.261E+04 | 571.3 | 0.01103 | 19.3 | +1 | 6 |
| - | - | 2732 | 572.4 | - | - | 0 | - |
| - | - | 939.2 | 573.4 | - | - | 0 | - |
| - | - | 813.4 | 628.4 | - | - | 0 | - |
| 3 | y | 2743 | 654.4 | 0.009364 | 14.31 | +1 | 6 |
| 3 | y | 6086 | 655.4 | 0.007719 | 11.78 | +1 | 6 |
| - | - | 2728 | 656.4 | - | - | 0 | - |
| - | - | 682.5 | 657.4 | - | - | 0 | - |
| 3 | y | 2.644E+04 | 672.4 | 0.0007026 | 1.045 | +1 | 6 |
| - | - | 8421 | 673.4 | - | - | 0 | - |
| - | - | 2410 | 674.4 | - | - | 0 | - |
| - | - | 1816 | 682.4 | - | - | 0 | - |
| 2 | y | 2932 | 725.4 | 0.0008234 | 1.135 | +1 | 7 |
| 2 | y | 1374 | 726.4 | 0.01394 | 19.19 | +1 | 7 |
| 2 | y | 1.114E+04 | 743.4 | 0.000829 | 1.115 | +1 | 7 |
| - | - | 3185 | 744.4 | - | - | 0 | - |
| - | - | 715.8 | 745.4 | - | - | 0 | - |
| - | - | 545.9 | 830.6 | - | - | 0 | - |
| - | - | 628.2 | 1057 | - | - | 0 | - |
| - | - | 681.4 | 2341 | - | - | 0 | - |
| - | - | 714.4 | 3081 | - | - | 0 | - |

m/z Charge Intensity FragmentType MassShift Position
120.08092498779297 0 756.7933
124.34492492675781 0 375.07578
128.10740661621094 0 920.1918
129.01837158203125 0 1506.3865
129.1023712158203 0 2965.3032
129.1068115234375 0 427.93176
130.0863494873047 0 1006.889
140.08200073242188 0 1975.2689
141.1023406982422 0 4230.6763
142.1055908203125 0 718.5055
143.11807250976562 0 13965.812 a 1
144.09825134277344 0 470.1545
144.12156677246094 0 1415.6482
145.09732055664062 0 1130.3225
147.11309814453125 0 1580.0253
149.0234832763672 0 12833.123
150.02684020996094 0 5397.832
151.02886962890625 0 672.09424
155.11807250976562 0 1716.3463
157.0132598876953 0 559.76666
157.13368225097656 0 9152.311
158.09243774414062 0 2146.2903 y Ammonia loss 7
158.1307830810547 0 528.2152
158.1371307373047 0 2676.8926
167.0337371826172 0 684.96265
169.09730529785156 0 4260.7695
170.10079956054688 0 1221.5259
171.11289978027344 0 13745.822 b 1
172.1160430908203 0 1134.9407
172.67413330078125 0 485.01047
173.09213256835938 0 2241.8477
173.12860107421875 0 16309.277
173.43844604492188 0 721.6406
174.13204956054688 0 1255.6813
175.11912536621094 0 3535.931 y 7
183.1128387451172 0 2957.7744
183.1494598388672 0 3081.0881
184.15232849121094 0 1623.7407
185.0813751220703 0 671.64746
185.128662109375 0 2772.5486
186.1322784423828 0 835.7472
201.1234893798828 0 6632.736
202.12652587890625 0 699.86316
211.1435089111328 0 1145.9308
212.13841247558594 0 1236.9207
212.14857482910156 0 743.3363
226.15496826171875 0 1985.9329 a Water loss 2
227.1391143798828 0 1199.1837
230.14967346191406 0 1165.0094 d 2
234.48098754882812 0 456.82135
237.091796875 0 718.01056
240.1350860595703 0 628.11237
245.89173889160156 0 516.04083
254.14984130859375 0 5485.7007 b Water loss 2
254.16262817382812 0 596.42413
255.14515686035156 0 13791.987 y Ammonia loss 6
256.1484069824219 0 884.3072
257.0711975097656 0 628.88306
258.1448669433594 0 2431.145
259.1483459472656 0 913.0808
270.14471435546875 0 1896.4349
272.1597595214844 0 2094.617 b 2
272.171875 0 5937.027 y 6
276.1552429199219 0 1777.0267
277.158447265625 0 1014.26025
286.1246337890625 0 569.1209
288.1554870605469 0 2143.8787
290.9594421386719 0 885.42413
291.9598693847656 0 590.8227
293.1009521484375 0 595.47845
306.1788635253906 0 1013.6056
312.193603515625 0 667.35724
322.1953430175781 0 1650.3816
325.2236022949219 0 2302.7346
327.69854736328125 0 5608.8545 y Water loss 2
328.1972351074219 0 7764.444 y Ammonia loss 2
328.6974792480469 0 3679.695
329.2003479003906 0 754.30536
340.1864318847656 0 1601.4119
341.1872253417969 0 1172.9316
343.0699462890625 0 622.5866
350.9250793457031 0 676.67053
352.1141662597656 0 634.217
353.2178955078125 0 1749.2471 b Water loss 3
363.21881103515625 0 6098.627 y Water loss 1
363.7196044921875 0 2087.1045
364.21881103515625 0 939.23773
366.1853332519531 0 11917.86
367.11883544921875 0 644.17206
367.18988037109375 0 965.0723
368.18133544921875 0 1149.8577
368.2288818359375 0 878.2491 y Ammonia loss 5
368.93634033203125 0 754.2149
371.22784423828125 0 700.62573 b 3
384.19622802734375 0 3755.7102
385.2558288574219 0 3469.9707 y 5
391.2322998046875 0 1230.5941
391.7319030761719 0 815.99426
402.20654296875 0 8468.582
403.210693359375 0 861.76337
403.813232421875 0 1447.6571
412.75323486328125 0 4302.1206 Precursor Water loss
413.2545471191406 0 2282.3435
420.8580322265625 0 1397.7428
422.21270751953125 0 4345.173
472.287841796875 0 33581.168 y 4
473.2905578613281 0 8410.158
474.296875 0 916.38696
486.29193115234375 0 5236.533
487.2950744628906 0 6051.8813
488.2989196777344 0 1989.0779
498.2523193359375 0 583.50104
509.2789611816406 0 903.9326
558.3319091796875 0 1085.3015
571.3560180664062 0 12611.98 b 5
572.3575439453125 0 2731.6177
573.3618774414062 0 939.19226
628.3771362304688 0 813.42426
654.3839721679688 0 2742.8591 y Water loss 2
655.3850708007812 0 6085.8286 y Ammonia loss 2
656.3873901367188 0 2727.505
657.3894653320312 0 682.52686
672.4031982421875 0 26435.045 y 2
673.4059448242188 0 8420.849
674.4082641601562 0 2410.3647
682.3890380859375 0 1816.3665
725.4296264648438 0 2931.679 y Water loss 1
726.4284057617188 0 1374.1948 y Ammonia loss 1
743.440185546875 0 11139.965 y 1
744.44384765625 0 3184.5715
745.44384765625 0 715.8371
830.6194458007812 0 545.89703
1056.763916015625 0 628.21893
2341.186767578125 0 681.43396
3081.095703125 0 714.4476

Spectrum Details

|  |  |
| --- | --- |
| Matched peaks? Matched peaks The total absolute number of peaks matched. Additionally in brackets the total fraction of peaks matched and the total number of peaks is shown. | 26 (19.26% of 135) |
| FDR? FDR The false discovery rate estimated for this peptide. It is calculated by matching all theoretical fragments with a non-integer shift with the raw peaks for this spectrum. This is done with 40 different shifts. The resulting percentage is the average number of annotated peaks over the number of annotated peaks with the correct spectrum. | 0.73% |
| Satellite FDR? Satellite FDR See the FDR for details on its calculation. This satellite ion specific FDR only contains the satellite ions (d/w) for I/L/J positions. | - |
| PSM Score? PSM Score The PSM Score as given by Hecklib to this annotated spectrum. It is shown with three significant figures. | 327 |

#### Spectrum 5978? Spectrum 5978 The raw spectrum of this peptide as annotated by Hecklib. The fragments are coloured according to ion type (see legend). Any peaks with a star '\*' as text can be hovered over to see the full details, first the ion type second the mass shift type. By hovering over the amino acids in the peptide or ions in the legend the corresponding peaks are highlighted. By toggling the 'Unassigned' label you can turn the background (unassigned) peaks on or off in the plot. By updating the slider in the Ion legend you can update the spectrum to only show the top X% of the peaks with labels. The top X% means any peak that is within X% of the highest intensity. By dragging in the spectrum you can zoom in to a specific part of the spectrum and use 'Zoom Out' to get back to the original zoom level. The annotation of the spectrum is based on the given sequence in the peptides file and is done with different software so inconsistencies are likely. The peaks are annotated based on the given sequence, with 20 ppm tolerance.

Copy Data

##### Spectrum 5978 (TSV)

###### Preview

```
Loading example...
```

*Click on the button to copy the data to your clipboard.*

Mz MinMz MaxIntensity Max

WidthHeightPeptide font sizePeptide stroke widthSpectrum font sizeSpectrum stroke widthCompact peptide

Ion legend

wxyz

abcd

OtherUnassignedIon^Charge^_Position_Show for top:%

VATVSJPR

01.10e+42.21e+43.31e+44.42e+4

Zoom Out

a^+1^_2_y^+1^_1_b^+1^_2_y^+1^_1_a^+1^_3_y^+2^_4_d^+1^_3_b^+1^_3_y^+1^_2_b^+1^_3_y^+1^_2_y^+2^_5_y^+2^_6_y^+2^_6_y^+2^_6_b^+1^_4_y^+2^_7_y^+1^_3_y^+1^_3_\*y^+1^_4_y^+1^_4_b^+1^_6_y^+1^_6_y^+1^_6_y^+1^_6_y^+1^_7_y^+1^_7_

0727145421802907

Fragment Matches Table

Show background peaks

| Position | Ion type | Intensity | mz Theoretical | mz Error (Th) | mz Error (ppm) | Charge | Series Number |
| --- | --- | --- | --- | --- | --- | --- | --- |
| - | - | 2576 | 120.1 | - | - | 0 | - |
| - | - | 495.5 | 121.1 | - | - | 0 | - |
| - | - | 443.5 | 122.3 | - | - | 0 | - |
| - | - | 1130 | 127.1 | - | - | 0 | - |
| - | - | 1173 | 128.1 | - | - | 0 | - |
| - | - | 1222 | 129 | - | - | 0 | - |
| - | - | 367.3 | 129.1 | - | - | 0 | - |
| - | - | 4882 | 129.1 | - | - | 0 | - |
| - | - | 504.6 | 130 | - | - | 0 | - |
| - | - | 1297 | 130.1 | - | - | 0 | - |
| - | - | 748.8 | 130.1 | - | - | 0 | - |
| - | - | 415.2 | 133.1 | - | - | 0 | - |
| - | - | 407.9 | 134.9 | - | - | 0 | - |
| - | - | 410.6 | 135.2 | - | - | 0 | - |
| - | - | 483.3 | 136.1 | - | - | 0 | - |
| - | - | 3260 | 140.1 | - | - | 0 | - |
| - | - | 397.6 | 141.1 | - | - | 0 | - |
| - | - | 396.3 | 141.1 | - | - | 0 | - |
| - | - | 5776 | 141.1 | - | - | 0 | - |
| - | - | 986 | 142.1 | - | - | 0 | - |
| 2 | a | 1.821E+04 | 143.1 | 0.0001982 | 1.385 | +1 | 2 |
| - | - | 1355 | 144.1 | - | - | 0 | - |
| - | - | 1633 | 145.1 | - | - | 0 | - |
| - | - | 1919 | 147.1 | - | - | 0 | - |
| - | - | 1.322E+04 | 149 | - | - | 0 | - |
| - | - | 6062 | 150 | - | - | 0 | - |
| - | - | 557.9 | 153.1 | - | - | 0 | - |
| - | - | 1437 | 155.1 | - | - | 0 | - |
| - | - | 2840 | 155.1 | - | - | 0 | - |
| - | - | 663.5 | 157 | - | - | 0 | - |
| - | - | 940.2 | 157.1 | - | - | 0 | - |
| - | - | 1.099E+04 | 157.1 | - | - | 0 | - |
| 8 | y | 2507 | 158.1 | 0.0003551 | 2.246 | +1 | 1 |
| - | - | 2399 | 158.1 | - | - | 0 | - |
| - | - | 436 | 158.7 | - | - | 0 | - |
| - | - | 708.2 | 159.1 | - | - | 0 | - |
| - | - | 430.5 | 165.1 | - | - | 0 | - |
| - | - | 438.4 | 165.2 | - | - | 0 | - |
| - | - | 635 | 167 | - | - | 0 | - |
| - | - | 621 | 167.1 | - | - | 0 | - |
| - | - | 493.3 | 167.6 | - | - | 0 | - |
| - | - | 3773 | 169.1 | - | - | 0 | - |
| - | - | 1271 | 170.1 | - | - | 0 | - |
| 2 | b | 1.497E+04 | 171.1 | 0.0001719 | 1.005 | +1 | 2 |
| - | - | 1119 | 172.1 | - | - | 0 | - |
| - | - | 3328 | 173.1 | - | - | 0 | - |
| - | - | 2.03E+04 | 173.1 | - | - | 0 | - |
| - | - | 3878 | 173.5 | - | - | 0 | - |
| - | - | 1929 | 174.1 | - | - | 0 | - |
| 8 | y | 6730 | 175.1 | 0.0002037 | 1.163 | +1 | 1 |
| - | - | 495.6 | 181.9 | - | - | 0 | - |
| - | - | 3398 | 183.1 | - | - | 0 | - |
| - | - | 4221 | 183.1 | - | - | 0 | - |
| - | - | 1417 | 184.2 | - | - | 0 | - |
| - | - | 580.8 | 185.1 | - | - | 0 | - |
| - | - | 658.2 | 185.1 | - | - | 0 | - |
| - | - | 2856 | 185.1 | - | - | 0 | - |
| - | - | 682.1 | 186.1 | - | - | 0 | - |
| - | - | 974.6 | 187.1 | - | - | 0 | - |
| - | - | 6773 | 201.1 | - | - | 0 | - |
| - | - | 671.7 | 202.1 | - | - | 0 | - |
| - | - | 865.1 | 211.1 | - | - | 0 | - |
| - | - | 1835 | 211.1 | - | - | 0 | - |
| - | - | 2363 | 212.1 | - | - | 0 | - |
| - | - | 763.3 | 212.1 | - | - | 0 | - |
| - | - | 475.9 | 213.1 | - | - | 0 | - |
| - | - | 670.9 | 215.1 | - | - | 0 | - |
| - | - | 922.1 | 224.1 | - | - | 0 | - |
| 3 | a | 2736 | 226.2 | 0.0001023 | 0.4522 | +1 | 3 |
| - | - | 896.3 | 227.1 | - | - | 0 | - |
| 5 | y | 771.6 | 228.1 | 6.683E-05 | 0.2929 | +2 | 4 |
| 3 | d | 818.4 | 230.1 | 0.0001224 | 0.5319 | +1 | 3 |
| - | - | 480.5 | 234.2 | - | - | 0 | - |
| - | - | 725.2 | 239.1 | - | - | 0 | - |
| - | - | 1362 | 240.1 | - | - | 0 | - |
| - | - | 469.8 | 241.5 | - | - | 0 | - |
| - | - | 639.4 | 242.2 | - | - | 0 | - |
| - | - | 587.2 | 244.1 | - | - | 0 | - |
| 3 | b | 7036 | 254.1 | 0.000137 | 0.539 | +1 | 3 |
| - | - | 467.3 | 255.1 | - | - | 0 | - |
| 7 | y | 1.913E+04 | 255.1 | 0.0002188 | 0.8576 | +1 | 2 |
| - | - | 1649 | 256.1 | - | - | 0 | - |
| - | - | 1966 | 258.1 | - | - | 0 | - |
| - | - | 1080 | 259.1 | - | - | 0 | - |
| - | - | 533.6 | 266.2 | - | - | 0 | - |
| - | - | 2080 | 270.1 | - | - | 0 | - |
| 3 | b | 2052 | 272.2 | 0.0007841 | 2.881 | +1 | 3 |
| 7 | y | 7670 | 272.2 | 6.743E-05 | 0.2477 | +1 | 2 |
| - | - | 916 | 273.2 | - | - | 0 | - |
| - | - | 2031 | 276.2 | - | - | 0 | - |
| - | - | 755.6 | 277.2 | - | - | 0 | - |
| - | - | 510.3 | 281.2 | - | - | 0 | - |
| - | - | 813.4 | 282.2 | - | - | 0 | - |
| 4 | y | 947.6 | 286.2 | 0.001875 | 6.551 | +2 | 5 |
| - | - | 2003 | 288.2 | - | - | 0 | - |
| - | - | 1191 | 293.1 | - | - | 0 | - |
| - | - | 3279 | 303.2 | - | - | 0 | - |
| - | - | 1825 | 306.2 | - | - | 0 | - |
| - | - | 632.7 | 312.2 | - | - | 0 | - |
| - | - | 556.3 | 324.3 | - | - | 0 | - |
| - | - | 3237 | 325.2 | - | - | 0 | - |
| - | - | 848.9 | 326.2 | - | - | 0 | - |
| 3 | y | 7572 | 327.7 | 0.001576 | 4.809 | +2 | 6 |
| 3 | y | 1.009E+04 | 328.2 | 0.005135 | 15.65 | +2 | 6 |
| - | - | 3026 | 328.7 | - | - | 0 | - |
| 3 | y | 932.9 | 336.7 | 0.00048 | 1.426 | +2 | 6 |
| - | - | 677.8 | 337.2 | - | - | 0 | - |
| - | - | 1657 | 340.2 | - | - | 0 | - |
| - | - | 1013 | 341 | - | - | 0 | - |
| - | - | 1914 | 341.2 | - | - | 0 | - |
| - | - | 832.8 | 350.9 | - | - | 0 | - |
| 4 | b | 1651 | 353.2 | 0.0005889 | 1.667 | +1 | 4 |
| - | - | 555.1 | 359 | - | - | 0 | - |
| 2 | y | 6755 | 363.2 | 0.0001742 | 0.4796 | +2 | 7 |
| - | - | 2061 | 363.7 | - | - | 0 | - |
| 6 | y | 1054 | 368.2 | 0.0004406 | 1.197 | +1 | 3 |
| 6 | y | 3574 | 385.3 | 0.000232 | 0.6021 | +1 | 3 |
| - | - | 748.5 | 389.2 | - | - | 0 | - |
| - | - | 659.3 | 390.2 | - | - | 0 | - |
| - | - | 2331 | 391.2 | - | - | 0 | - |
| - | - | 718.7 | 397.8 | - | - | 0 | - |
| - | - | 621.7 | 403.7 | - | - | 0 | - |
| - | - | 1705 | 403.8 | - | - | 0 | - |
| - | - | 1100 | 410.2 | - | - | 0 | - |
| 0 | Precursor | 4712 | 412.8 | 0.0001647 | 0.3991 | +2 | -1 |
| - | - | 2076 | 413.3 | - | - | 0 | - |
| - | - | 1003 | 413.8 | - | - | 0 | - |
| - | - | 627.9 | 415 | - | - | 0 | - |
| - | - | 870.5 | 420.9 | - | - | 0 | - |
| - | - | 664.2 | 439.8 | - | - | 0 | - |
| 5 | y | 975.7 | 454.3 | 0.003501 | 7.707 | +1 | 4 |
| - | - | 2400 | 458.8 | - | - | 0 | - |
| - | - | 774.9 | 459.3 | - | - | 0 | - |
| - | - | 941.6 | 470.2 | - | - | 0 | - |
| 5 | y | 4.372E+04 | 472.3 | 0.0001555 | 0.3292 | +1 | 4 |
| - | - | 9701 | 473.3 | - | - | 0 | - |
| - | - | 1069 | 474.3 | - | - | 0 | - |
| - | - | 4226 | 486.3 | - | - | 0 | - |
| - | - | 5453 | 487.3 | - | - | 0 | - |
| - | - | 648.6 | 487.5 | - | - | 0 | - |
| - | - | 2092 | 488.3 | - | - | 0 | - |
| - | - | 737.9 | 557.3 | - | - | 0 | - |
| - | - | 2118 | 558.3 | - | - | 0 | - |
| - | - | 594.2 | 568 | - | - | 0 | - |
| 6 | b | 1.456E+04 | 571.3 | 0.01078 | 18.88 | +1 | 6 |
| - | - | 4916 | 572.4 | - | - | 0 | - |
| - | - | 732.8 | 585.3 | - | - | 0 | - |
| - | - | 850.1 | 611.4 | - | - | 0 | - |
| - | - | 1546 | 628.4 | - | - | 0 | - |
| 3 | y | 1992 | 654.4 | 0.007533 | 11.51 | +1 | 6 |
| 3 | y | 9222 | 655.4 | 0.00717 | 10.94 | +1 | 6 |
| - | - | 2641 | 656.4 | - | - | 0 | - |
| 3 | y | 3.284E+04 | 672.4 | 0.0007026 | 1.045 | +1 | 6 |
| - | - | 1.182E+04 | 673.4 | - | - | 0 | - |
| - | - | 2384 | 674.4 | - | - | 0 | - |
| - | - | 2156 | 682.4 | - | - | 0 | - |
| - | - | 1157 | 683.4 | - | - | 0 | - |
| - | - | 571.6 | 698.4 | - | - | 0 | - |
| 2 | y | 4022 | 725.4 | 0.001312 | 1.808 | +1 | 7 |
| - | - | 911.5 | 726.4 | - | - | 0 | - |
| 2 | y | 1.257E+04 | 743.4 | 0.000768 | 1.033 | +1 | 7 |
| - | - | 6149 | 744.4 | - | - | 0 | - |
| - | - | 1447 | 745.4 | - | - | 0 | - |
| - | - | 566.2 | 1012 | - | - | 0 | - |
| - | - | 643.6 | 1666 | - | - | 0 | - |
| - | - | 683.6 | 2878 | - | - | 0 | - |

m/z Charge Intensity FragmentType MassShift Position
120.08100128173828 0 2576.135
121.08440399169922 0 495.4804
122.34933471679688 0 443.4714
127.0868911743164 0 1130.3794
128.1071319580078 0 1173.4939
129.01856994628906 0 1222.0214
129.0668182373047 0 367.26572
129.1024932861328 0 4882.0986
130.04949951171875 0 504.59695
130.0867156982422 0 1296.6201
130.1058807373047 0 748.79816
133.06253051757812 0 415.20847
134.93756103515625 0 407.85446
135.2004852294922 0 410.61713
136.07601928710938 0 483.2885
140.08206176757812 0 3259.571
141.08624267578125 0 397.6246
141.09246826171875 0 396.3418
141.10247802734375 0 5776.192
142.1060333251953 0 986.009
143.1180877685547 0 18207.676 a 1
144.12158203125 0 1354.6566
145.09735107421875 0 1633.2738
147.11277770996094 0 1918.8965
149.02352905273438 0 13221.755
150.02694702148438 0 6062.338
153.10311889648438 0 557.89056
155.081787109375 0 1437.1952
155.11813354492188 0 2840.4497
157.01319885253906 0 663.48914
157.097412109375 0 940.18256
157.1337432861328 0 10994.002
158.09275817871094 0 2506.532 y Ammonia loss 7
158.13705444335938 0 2398.753
158.74169921875 0 436.02502
159.11279296875 0 708.2222
165.09246826171875 0 430.50812
165.20895385742188 0 438.4366
167.03427124023438 0 634.99896
167.08160400390625 0 621.0316
167.59103393554688 0 493.32822
169.09738159179688 0 3773.0715
170.10084533691406 0 1271.0908
171.11297607421875 0 14974.679 b 1
172.11634826660156 0 1118.9816
173.09228515625 0 3328.1274
173.12864685058594 0 20301.568
173.45045471191406 0 3877.6067
174.13218688964844 0 1928.8037
175.11915588378906 0 6729.569 y 7
181.933349609375 0 495.613
183.1130828857422 0 3398.1436
183.1494140625 0 4220.527
184.15264892578125 0 1417.0477
185.08103942871094 0 580.7799
185.0922088623047 0 658.15094
185.12863159179688 0 2855.976
186.1319580078125 0 682.1076
187.10794067382812 0 974.6172
201.12353515625 0 6772.743
202.12655639648438 0 671.71265
211.10791015625 0 865.0947
211.144287109375 0 1834.9639
212.13961791992188 0 2363.3547
212.1493682861328 0 763.30255
213.12400817871094 0 475.89624
215.1389923095703 0 670.8782
224.10328674316406 0 922.0771
226.1551055908203 0 2736.0874 a Water loss 2
227.13906860351562 0 896.34314
228.1342010498047 0 771.62256 y Ammonia loss 4
230.14979553222656 0 818.3642 d 2
234.1628875732422 0 480.49887
239.08868408203125 0 725.2084
240.13414001464844 0 1361.6505
241.47462463378906 0 469.77258
242.2221221923828 0 639.39844
244.1267852783203 0 587.1806
254.15005493164062 0 7036.3374 b Water loss 2
255.08392333984375 0 467.2854
255.1453857421875 0 19127.58 y Ammonia loss 6
256.1477966308594 0 1648.683
258.14501953125 0 1966.0387
259.1486511230469 0 1080.4763
266.1507873535156 0 533.62946
270.1446228027344 0 2080.0012
272.1596984863281 0 2051.9233 b 2
272.1717834472656 0 7669.511 y 6
273.1760559082031 0 916.0034
276.1552429199219 0 2030.815
277.1580810546875 0 755.6013
281.1683654785156 0 510.25772
282.1811218261719 0 813.4248
286.1836242675781 0 947.62024 y 3
288.1556396484375 0 2002.842
293.0993347167969 0 1190.6428
303.2142333984375 0 3278.652
306.1793518066406 0 1824.7998
312.1921081542969 0 632.7473
324.2769470214844 0 556.2901
325.223388671875 0 3236.6812
326.2267761230469 0 848.9493
327.69873046875 0 7571.5117 y Water loss 2
328.19744873046875 0 10089.364 y Ammonia loss 2
328.6981506347656 0 3025.7153
336.7051086425781 0 932.93884 y 2
337.20654296875 0 677.8326
340.18731689453125 0 1657.3011
340.9644470214844 0 1013.0819
341.18438720703125 0 1913.8826
350.9268798828125 0 832.82153
353.2177429199219 0 1651.1162 b Water loss 3
358.9718017578125 0 555.1073
363.21868896484375 0 6754.9795 y Water loss 1
363.71966552734375 0 2061.4167
368.2287902832031 0 1054.0385 y Ammonia loss 5
385.2560119628906 0 3573.6565 y 5
389.2392883300781 0 748.47095
390.2441711425781 0 659.3316
391.232666015625 0 2330.7043
397.80523681640625 0 718.6636
403.74725341796875 0 621.6703
403.8123474121094 0 1704.9707
410.2386474609375 0 1100.017
412.75323486328125 0 4711.594 Precursor Water loss
413.25457763671875 0 2075.8188
413.7561950683594 0 1003.022
414.97296142578125 0 627.861
420.8612060546875 0 870.5017
439.84423828125 0 664.1874
454.27374267578125 0 975.7475 y Water loss 4
458.7657165527344 0 2399.5
459.26678466796875 0 774.8836
470.2418212890625 0 941.6362
472.2879638671875 0 43718.047 y 4
473.29083251953125 0 9700.661
474.29400634765625 0 1068.5529
486.2925720214844 0 4225.731
487.29541015625 0 5453.1353
487.45745849609375 0 648.5794
488.2973937988281 0 2092.0552
557.3319091796875 0 737.92896
558.3317260742188 0 2117.871
567.9791259765625 0 594.2338
571.3557739257812 0 14555.862 b 5
572.35888671875 0 4916.3315
585.3027954101562 0 732.82184
611.3549194335938 0 850.08954
628.3753662109375 0 1545.8362
654.3858032226562 0 1991.9674 y Water loss 2
655.384521484375 0 9221.604 y Ammonia loss 2
656.3877563476562 0 2640.934
672.4031982421875 0 32842.81 y 2
673.4063720703125 0 11816.923
674.41015625 0 2384.0364
682.387451171875 0 2156.0867
683.390869140625 0 1157.06
698.388671875 0 571.6373
725.4291381835938 0 4021.783 y Water loss 1
726.4343872070312 0 911.46643
743.4402465820312 0 12567.88 y 1
744.442626953125 0 6148.8516
745.4473266601562 0 1447.1844
1011.5770263671875 0 566.2222
1665.5286865234375 0 643.558
2878.3193359375 0 683.6082

Spectrum Details

|  |  |
| --- | --- |
| Matched peaks? Matched peaks The total absolute number of peaks matched. Additionally in brackets the total fraction of peaks matched and the total number of peaks is shown. | 28 (16.87% of 166) |
| FDR? FDR The false discovery rate estimated for this peptide. It is calculated by matching all theoretical fragments with a non-integer shift with the raw peaks for this spectrum. This is done with 40 different shifts. The resulting percentage is the average number of annotated peaks over the number of annotated peaks with the correct spectrum. | 0.17% |
| Satellite FDR? Satellite FDR See the FDR for details on its calculation. This satellite ion specific FDR only contains the satellite ions (d/w) for I/L/J positions. | - |
| PSM Score? PSM Score The PSM Score as given by Hecklib to this annotated spectrum. It is shown with three significant figures. | 306 |

#### Spectrum 6362? Spectrum 6362 The raw spectrum of this peptide as annotated by Hecklib. The fragments are coloured according to ion type (see legend). Any peaks with a star '\*' as text can be hovered over to see the full details, first the ion type second the mass shift type. By hovering over the amino acids in the peptide or ions in the legend the corresponding peaks are highlighted. By toggling the 'Unassigned' label you can turn the background (unassigned) peaks on or off in the plot. By updating the slider in the Ion legend you can update the spectrum to only show the top X% of the peaks with labels. The top X% means any peak that is within X% of the highest intensity. By dragging in the spectrum you can zoom in to a specific part of the spectrum and use 'Zoom Out' to get back to the original zoom level. The annotation of the spectrum is based on the given sequence in the peptides file and is done with different software so inconsistencies are likely. The peaks are annotated based on the given sequence, with 20 ppm tolerance.

Copy Data

##### Spectrum 6362 (TSV)

###### Preview

```
Loading example...
```

*Click on the button to copy the data to your clipboard.*

Mz MinMz MaxIntensity Max

WidthHeightPeptide font sizePeptide stroke widthSpectrum font sizeSpectrum stroke widthCompact peptide

Ion legend

wxyz

abcd

OtherUnassignedIon^Charge^_Position_Show for top:%

VATVSJPR

07.09e+31.42e+42.13e+42.83e+4

Zoom Out

a^+1^_2_y^+1^_1_b^+1^_2_y^+1^_1_a^+1^_3_y^+2^_4_d^+1^_3_b^+1^_3_y^+1^_2_b^+1^_3_y^+1^_2_y^+2^_6_y^+2^_6_b^+1^_4_y^+2^_7_y^+1^_3_y^+1^_3_\*y^+1^_4_b^+1^_6_b^+1^_6_y^+1^_6_y^+1^_6_y^+1^_6_y^+1^_7_y^+1^_7_

0527105415822109

Fragment Matches Table

Show background peaks

| Position | Ion type | Intensity | mz Theoretical | mz Error (Th) | mz Error (ppm) | Charge | Series Number |
| --- | --- | --- | --- | --- | --- | --- | --- |
| - | - | 586.3 | 120.1 | - | - | 0 | - |
| - | - | 328.2 | 121.3 | - | - | 0 | - |
| - | - | 419.5 | 127.1 | - | - | 0 | - |
| - | - | 495.3 | 128.1 | - | - | 0 | - |
| - | - | 961.8 | 129 | - | - | 0 | - |
| - | - | 2853 | 129.1 | - | - | 0 | - |
| - | - | 687.9 | 130.1 | - | - | 0 | - |
| - | - | 865.7 | 136.1 | - | - | 0 | - |
| - | - | 1646 | 140.1 | - | - | 0 | - |
| - | - | 4120 | 141.1 | - | - | 0 | - |
| - | - | 1229 | 142.1 | - | - | 0 | - |
| 2 | a | 1.148E+04 | 143.1 | 0.0001985 | 1.387 | +1 | 2 |
| - | - | 451.4 | 144.1 | - | - | 0 | - |
| - | - | 962.9 | 145.1 | - | - | 0 | - |
| - | - | 1372 | 147.1 | - | - | 0 | - |
| - | - | 1.321E+04 | 149 | - | - | 0 | - |
| - | - | 583.4 | 150 | - | - | 0 | - |
| - | - | 6119 | 150 | - | - | 0 | - |
| - | - | 766.5 | 151 | - | - | 0 | - |
| - | - | 461.9 | 153.1 | - | - | 0 | - |
| - | - | 770.1 | 155.1 | - | - | 0 | - |
| - | - | 1364 | 155.1 | - | - | 0 | - |
| - | - | 713.1 | 157 | - | - | 0 | - |
| - | - | 8001 | 157.1 | - | - | 0 | - |
| 8 | y | 1912 | 158.1 | 0.0003163 | 2.001 | +1 | 1 |
| - | - | 2585 | 158.1 | - | - | 0 | - |
| - | - | 474.6 | 164.3 | - | - | 0 | - |
| - | - | 750.5 | 167 | - | - | 0 | - |
| - | - | 3303 | 169.1 | - | - | 0 | - |
| - | - | 849.9 | 170.1 | - | - | 0 | - |
| 2 | b | 1.022E+04 | 171.1 | 0.0003011 | 1.76 | +1 | 2 |
| - | - | 512.2 | 171.1 | - | - | 0 | - |
| - | - | 894.7 | 172.1 | - | - | 0 | - |
| - | - | 1552 | 173.1 | - | - | 0 | - |
| - | - | 738.7 | 173.1 | - | - | 0 | - |
| - | - | 1.222E+04 | 173.1 | - | - | 0 | - |
| - | - | 805.5 | 174.1 | - | - | 0 | - |
| 8 | y | 3427 | 175.1 | 0.0001778 | 1.015 | +1 | 1 |
| - | - | 467.7 | 178.6 | - | - | 0 | - |
| - | - | 2239 | 183.1 | - | - | 0 | - |
| - | - | 3530 | 183.1 | - | - | 0 | - |
| - | - | 983.2 | 184.2 | - | - | 0 | - |
| - | - | 938.9 | 185.1 | - | - | 0 | - |
| - | - | 515.6 | 185.1 | - | - | 0 | - |
| - | - | 1681 | 185.1 | - | - | 0 | - |
| - | - | 715.3 | 186.1 | - | - | 0 | - |
| - | - | 468.8 | 188.7 | - | - | 0 | - |
| - | - | 734.3 | 195.1 | - | - | 0 | - |
| - | - | 4997 | 201.1 | - | - | 0 | - |
| - | - | 457.3 | 202.1 | - | - | 0 | - |
| - | - | 537.4 | 205.1 | - | - | 0 | - |
| - | - | 1512 | 211.1 | - | - | 0 | - |
| - | - | 895.3 | 212.1 | - | - | 0 | - |
| - | - | 566.9 | 212.1 | - | - | 0 | - |
| 3 | a | 2024 | 226.2 | 0.0001419 | 0.6273 | +1 | 3 |
| - | - | 736.2 | 227.1 | - | - | 0 | - |
| - | - | 490.9 | 227.2 | - | - | 0 | - |
| 5 | y | 829.7 | 228.1 | 0.0006161 | 2.701 | +2 | 4 |
| 3 | d | 648.9 | 230.1 | 0.0002591 | 1.126 | +1 | 3 |
| - | - | 560.2 | 240.1 | - | - | 0 | - |
| - | - | 767.8 | 243.1 | - | - | 0 | - |
| - | - | 565.6 | 243.1 | - | - | 0 | - |
| - | - | 734.5 | 244.2 | - | - | 0 | - |
| 3 | b | 5186 | 254.1 | 0.0003971 | 1.562 | +1 | 3 |
| 7 | y | 1.053E+04 | 255.1 | 0.0004526 | 1.774 | +1 | 2 |
| - | - | 787.7 | 256.1 | - | - | 0 | - |
| - | - | 482.2 | 257.7 | - | - | 0 | - |
| - | - | 2497 | 258.1 | - | - | 0 | - |
| - | - | 497.3 | 259 | - | - | 0 | - |
| - | - | 1130 | 259.1 | - | - | 0 | - |
| - | - | 734.8 | 266.1 | - | - | 0 | - |
| - | - | 1527 | 270.1 | - | - | 0 | - |
| 3 | b | 1633 | 272.2 | 0.001456 | 5.348 | +1 | 3 |
| 7 | y | 5246 | 272.2 | 0.0003903 | 1.434 | +1 | 2 |
| - | - | 1971 | 276.2 | - | - | 0 | - |
| - | - | 1046 | 277.2 | - | - | 0 | - |
| - | - | 705.7 | 282.2 | - | - | 0 | - |
| - | - | 1577 | 288.2 | - | - | 0 | - |
| - | - | 613.4 | 289.2 | - | - | 0 | - |
| - | - | 1202 | 291 | - | - | 0 | - |
| - | - | 1759 | 293.1 | - | - | 0 | - |
| - | - | 572.6 | 322.2 | - | - | 0 | - |
| - | - | 533.8 | 325.1 | - | - | 0 | - |
| - | - | 1948 | 325.2 | - | - | 0 | - |
| 3 | y | 4719 | 327.7 | 0.002125 | 6.485 | +2 | 6 |
| 3 | y | 8566 | 328.2 | 0.004097 | 12.48 | +2 | 6 |
| - | - | 3132 | 328.7 | - | - | 0 | - |
| - | - | 1362 | 329.2 | - | - | 0 | - |
| - | - | 1813 | 340.2 | - | - | 0 | - |
| - | - | 1528 | 341.2 | - | - | 0 | - |
| - | - | 1343 | 350.9 | - | - | 0 | - |
| 4 | b | 840.9 | 353.2 | 0.001169 | 3.309 | +1 | 4 |
| - | - | 661.5 | 354.7 | - | - | 0 | - |
| 2 | y | 4904 | 363.2 | 0.001212 | 3.336 | +2 | 7 |
| - | - | 1988 | 363.7 | - | - | 0 | - |
| 6 | y | 945.3 | 368.2 | 0.0002308 | 0.6267 | +1 | 3 |
| 6 | y | 2617 | 385.3 | 0.0004699 | 1.22 | +1 | 3 |
| - | - | 753.8 | 386.3 | - | - | 0 | - |
| - | - | 1047 | 391.2 | - | - | 0 | - |
| - | - | 675.6 | 399.2 | - | - | 0 | - |
| - | - | 908.5 | 403 | - | - | 0 | - |
| - | - | 1108 | 403.8 | - | - | 0 | - |
| 0 | Precursor | 4401 | 412.8 | 0.0005372 | 1.301 | +2 | -1 |
| - | - | 1190 | 413.3 | - | - | 0 | - |
| - | - | 708.3 | 413.8 | - | - | 0 | - |
| - | - | 674.8 | 420.9 | - | - | 0 | - |
| - | - | 1064 | 439.8 | - | - | 0 | - |
| 5 | y | 2.806E+04 | 472.3 | 0.0001802 | 0.3816 | +1 | 4 |
| - | - | 6467 | 473.3 | - | - | 0 | - |
| - | - | 1115 | 474.3 | - | - | 0 | - |
| - | - | 3117 | 486.3 | - | - | 0 | - |
| - | - | 5204 | 487.3 | - | - | 0 | - |
| - | - | 1181 | 488.3 | - | - | 0 | - |
| 6 | b | 593.8 | 553.3 | 0.008105 | 14.65 | +1 | 6 |
| - | - | 1240 | 557.3 | - | - | 0 | - |
| - | - | 1164 | 558.3 | - | - | 0 | - |
| 6 | b | 9613 | 571.3 | 0.01078 | 18.88 | +1 | 6 |
| - | - | 2836 | 572.4 | - | - | 0 | - |
| - | - | 1129 | 644.4 | - | - | 0 | - |
| 3 | y | 2281 | 654.4 | 0.007594 | 11.6 | +1 | 6 |
| 3 | y | 6225 | 655.4 | 0.00717 | 10.94 | +1 | 6 |
| - | - | 1956 | 656.4 | - | - | 0 | - |
| - | - | 1003 | 657.4 | - | - | 0 | - |
| 3 | y | 1.867E+04 | 672.4 | 0.0006415 | 0.9541 | +1 | 6 |
| - | - | 1019 | 672.5 | - | - | 0 | - |
| - | - | 6125 | 673.4 | - | - | 0 | - |
| - | - | 1820 | 674.4 | - | - | 0 | - |
| - | - | 1677 | 682.4 | - | - | 0 | - |
| - | - | 804.9 | 683.4 | - | - | 0 | - |
| 2 | y | 2393 | 725.4 | 0.0007014 | 0.9668 | +1 | 7 |
| - | - | 1337 | 726.4 | - | - | 0 | - |
| 2 | y | 9384 | 743.4 | 0.0003408 | 0.4584 | +1 | 7 |
| - | - | 3601 | 744.4 | - | - | 0 | - |
| - | - | 990.3 | 745.5 | - | - | 0 | - |
| - | - | 621.3 | 1104 | - | - | 0 | - |
| - | - | 612 | 1310 | - | - | 0 | - |
| - | - | 637.5 | 1456 | - | - | 0 | - |
| - | - | 703.6 | 1578 | - | - | 0 | - |
| - | - | 762.7 | 2088 | - | - | 0 | - |

m/z Charge Intensity FragmentType MassShift Position
120.0809097290039 0 586.3193
121.33121490478516 0 328.19543
127.0865249633789 0 419.45898
128.10694885253906 0 495.31796
129.01824951171875 0 961.7541
129.10214233398438 0 2853.1345
130.08627319335938 0 687.9172
136.07577514648438 0 865.6908
140.08152770996094 0 1646.3662
141.10205078125 0 4119.5166
142.10569763183594 0 1229.1149
143.11769104003906 0 11481.27 a 1
144.12098693847656 0 451.39243
145.09689331054688 0 962.88794
147.1126708984375 0 1371.7241
149.0231170654297 0 13206.9375
150.02108764648438 0 583.40643
150.02651977539062 0 6119.2275
151.0283203125 0 766.4645
153.0543975830078 0 461.85437
155.08151245117188 0 770.1011
155.1181182861328 0 1363.5481
157.01312255859375 0 713.12885
157.13327026367188 0 8000.705
158.0920867919922 0 1912.1355 y Ammonia loss 7
158.1366424560547 0 2584.6182
164.26649475097656 0 474.60074
167.03379821777344 0 750.54144
169.0968780517578 0 3302.9207
170.10084533691406 0 849.94855
171.1125030517578 0 10219.284 b 1
171.13734436035156 0 512.22876
172.1160888671875 0 894.7185
173.09182739257812 0 1551.9844
173.12039184570312 0 738.73975
173.12820434570312 0 12216.909
174.13148498535156 0 805.4782
175.1187744140625 0 3426.5286 y 7
178.63522338867188 0 467.72556
183.11233520507812 0 2238.634
183.14892578125 0 3530.1724
184.15235900878906 0 983.2166
185.08009338378906 0 938.9337
185.1028594970703 0 515.5514
185.1280975341797 0 1681.3264
186.1320037841797 0 715.3387
188.7476348876953 0 468.8499
195.11264038085938 0 734.2684
201.1229705810547 0 4996.7104
202.1251678466797 0 457.25284
205.08297729492188 0 537.3662
211.14366149902344 0 1511.942
212.13819885253906 0 895.3025
212.14813232421875 0 566.88257
226.1548614501953 0 2023.5883 a Water loss 2
227.1388397216797 0 736.2455
227.16986083984375 0 490.8598
228.13365173339844 0 829.7179 y Ammonia loss 4
230.15017700195312 0 648.8843 d 2
240.133056640625 0 560.2268
243.0543670654297 0 767.8046
243.13421630859375 0 565.5646
244.1512451171875 0 734.495
254.14952087402344 0 5186.2383 b Water loss 2
255.14471435546875 0 10531.731 y Ammonia loss 6
256.1481628417969 0 787.69403
257.6512145996094 0 482.21185
258.1441345214844 0 2496.898
258.95233154296875 0 497.34262
259.1487731933594 0 1129.5043
266.1488037109375 0 734.7925
270.14410400390625 0 1526.9116
272.1590270996094 0 1632.9274 b 2
272.17132568359375 0 5246.009 y 6
276.15484619140625 0 1970.6387
277.1584777832031 0 1045.5288
282.1799621582031 0 705.7255
288.1553649902344 0 1576.8195
289.15765380859375 0 613.3625
290.9593811035156 0 1202.4839
293.0986022949219 0 1759.0522
322.1944274902344 0 572.63995
325.086181640625 0 533.77527
325.22296142578125 0 1947.549
327.69818115234375 0 4718.831 y Water loss 2
328.1964111328125 0 8566.158 y Ammonia loss 2
328.69781494140625 0 3132.2778
329.19879150390625 0 1361.9242
340.185791015625 0 1813.4288
341.18792724609375 0 1527.7181
350.9261474609375 0 1343.2312
353.2171630859375 0 840.8509 b Water loss 3
354.68585205078125 0 661.476
363.2176513671875 0 4904.129 y Water loss 1
363.72003173828125 0 1987.7524
368.2294616699219 0 945.255 y Ammonia loss 5
385.25531005859375 0 2616.6648 y 5
386.2564697265625 0 753.8235
391.23150634765625 0 1046.6321
399.19708251953125 0 675.62305
402.9612121582031 0 908.4809
403.8121643066406 0 1107.6656
412.7525329589844 0 4401.493 Precursor Water loss
413.25408935546875 0 1190.4246
413.757080078125 0 708.3372
420.8587951660156 0 674.7673
439.8431701660156 0 1063.9766
472.2876281738281 0 28062.84 y 4
473.290283203125 0 6467.2314
474.2923583984375 0 1115.2806
486.2925720214844 0 3116.8074
487.2947082519531 0 5204.2515
488.29736328125 0 1180.9668
553.342529296875 0 593.8098 b Water loss 5
557.3290405273438 0 1239.8835
558.3335571289062 0 1164.0404
571.3557739257812 0 9612.709 b 5
572.3580932617188 0 2836.3816
644.3706665039062 0 1129.1233
654.3857421875 0 2280.7942 y Water loss 2
655.384521484375 0 6225.448 y Ammonia loss 2
656.3893432617188 0 1955.8601
657.3938598632812 0 1003.43414
672.4032592773438 0 18668.406 y 2
672.4676513671875 0 1018.601
673.4061279296875 0 6125.214
674.4041748046875 0 1819.5093
682.3860473632812 0 1676.6892
683.389404296875 0 804.8877
725.4297485351562 0 2393.4893 y Water loss 1
726.4317016601562 0 1336.9404
743.440673828125 0 9383.928 y 1
744.4426879882812 0 3601.4243
745.451416015625 0 990.27576
1103.9742431640625 0 621.306
1310.2642822265625 0 611.98236
1456.40576171875 0 637.5493
1578.2901611328125 0 703.60504
2087.8310546875 0 762.65564

Spectrum Details

|  |  |
| --- | --- |
| Matched peaks? Matched peaks The total absolute number of peaks matched. Additionally in brackets the total fraction of peaks matched and the total number of peaks is shown. | 26 (18.71% of 139) |
| FDR? FDR The false discovery rate estimated for this peptide. It is calculated by matching all theoretical fragments with a non-integer shift with the raw peaks for this spectrum. This is done with 40 different shifts. The resulting percentage is the average number of annotated peaks over the number of annotated peaks with the correct spectrum. | 0.18% |
| Satellite FDR? Satellite FDR See the FDR for details on its calculation. This satellite ion specific FDR only contains the satellite ions (d/w) for I/L/J positions. | - |
| PSM Score? PSM Score The PSM Score as given by Hecklib to this annotated spectrum. It is shown with three significant figures. | 306 |

#### Spectrum 7447? Spectrum 7447 The raw spectrum of this peptide as annotated by Hecklib. The fragments are coloured according to ion type (see legend). Any peaks with a star '\*' as text can be hovered over to see the full details, first the ion type second the mass shift type. By hovering over the amino acids in the peptide or ions in the legend the corresponding peaks are highlighted. By toggling the 'Unassigned' label you can turn the background (unassigned) peaks on or off in the plot. By updating the slider in the Ion legend you can update the spectrum to only show the top X% of the peaks with labels. The top X% means any peak that is within X% of the highest intensity. By dragging in the spectrum you can zoom in to a specific part of the spectrum and use 'Zoom Out' to get back to the original zoom level. The annotation of the spectrum is based on the given sequence in the peptides file and is done with different software so inconsistencies are likely. The peaks are annotated based on the given sequence, with 20 ppm tolerance.

Copy Data

##### Spectrum 7447 (TSV)

###### Preview

```
Loading example...
```

*Click on the button to copy the data to your clipboard.*

Mz MinMz MaxIntensity Max

WidthHeightPeptide font sizePeptide stroke widthSpectrum font sizeSpectrum stroke widthCompact peptide

Ion legend

wxyz

abcd

OtherUnassignedIon^Charge^_Position_Show for top:%

VATVSJPR

03.57e+37.15e+31.07e+41.43e+4

Zoom Out

a^+1^_2_y^+1^_1_b^+1^_2_y^+1^_1_a^+1^_3_d^+1^_3_b^+1^_3_y^+1^_2_b^+1^_3_y^+1^_2_y^+2^_6_y^+2^_6_y^+2^_7_y^+1^_3_\*y^+1^_4_b^+1^_6_y^+1^_6_y^+1^_6_y^+1^_6_y^+1^_7_y^+1^_7_

0562112416862248

Fragment Matches Table

Show background peaks

| Position | Ion type | Intensity | mz Theoretical | mz Error (Th) | mz Error (ppm) | Charge | Series Number |
| --- | --- | --- | --- | --- | --- | --- | --- |
| - | - | 913.3 | 120.1 | - | - | 0 | - |
| - | - | 1156 | 129 | - | - | 0 | - |
| - | - | 1413 | 129.1 | - | - | 0 | - |
| - | - | 1208 | 130.1 | - | - | 0 | - |
| - | - | 421.2 | 132 | - | - | 0 | - |
| - | - | 447.6 | 133 | - | - | 0 | - |
| - | - | 495.5 | 136.1 | - | - | 0 | - |
| - | - | 408.6 | 139.6 | - | - | 0 | - |
| - | - | 1082 | 140.1 | - | - | 0 | - |
| - | - | 2583 | 141.1 | - | - | 0 | - |
| 2 | a | 5817 | 143.1 | 4.565E-05 | 0.3189 | +1 | 2 |
| - | - | 542 | 145 | - | - | 0 | - |
| - | - | 560.1 | 147 | - | - | 0 | - |
| - | - | 744 | 147.1 | - | - | 0 | - |
| - | - | 1.149E+04 | 149 | - | - | 0 | - |
| - | - | 5268 | 150 | - | - | 0 | - |
| - | - | 770.8 | 151 | - | - | 0 | - |
| - | - | 548.6 | 153.1 | - | - | 0 | - |
| - | - | 1053 | 155.1 | - | - | 0 | - |
| - | - | 562.9 | 157 | - | - | 0 | - |
| - | - | 4674 | 157.1 | - | - | 0 | - |
| 8 | y | 932.7 | 158.1 | 5.687E-05 | 0.3598 | +1 | 1 |
| - | - | 1369 | 158.1 | - | - | 0 | - |
| - | - | 806.9 | 167 | - | - | 0 | - |
| - | - | 2348 | 169.1 | - | - | 0 | - |
| 2 | b | 5225 | 171.1 | 4.071E-06 | 0.02379 | +1 | 2 |
| - | - | 965 | 173.1 | - | - | 0 | - |
| - | - | 6286 | 173.1 | - | - | 0 | - |
| - | - | 1393 | 173.4 | - | - | 0 | - |
| - | - | 584.7 | 174.1 | - | - | 0 | - |
| 8 | y | 1826 | 175.1 | 5.113E-05 | 0.292 | +1 | 1 |
| - | - | 496.3 | 179.1 | - | - | 0 | - |
| - | - | 572.5 | 183.1 | - | - | 0 | - |
| - | - | 1090 | 183.1 | - | - | 0 | - |
| - | - | 1525 | 183.1 | - | - | 0 | - |
| - | - | 658.3 | 184.2 | - | - | 0 | - |
| - | - | 614.2 | 185.1 | - | - | 0 | - |
| - | - | 1083 | 185.1 | - | - | 0 | - |
| - | - | 963.3 | 187.1 | - | - | 0 | - |
| - | - | 2212 | 201.1 | - | - | 0 | - |
| - | - | 1312 | 202.1 | - | - | 0 | - |
| - | - | 578.2 | 210.9 | - | - | 0 | - |
| - | - | 1213 | 211.1 | - | - | 0 | - |
| - | - | 634 | 212.1 | - | - | 0 | - |
| 3 | a | 1190 | 226.2 | 0.0004685 | 2.071 | +1 | 3 |
| - | - | 853.9 | 227.1 | - | - | 0 | - |
| 3 | d | 710.2 | 230.1 | 0.0007473 | 3.247 | +1 | 3 |
| - | - | 912.4 | 237.1 | - | - | 0 | - |
| - | - | 559.9 | 241.1 | - | - | 0 | - |
| - | - | 555 | 246.8 | - | - | 0 | - |
| - | - | 599.4 | 249.1 | - | - | 0 | - |
| 3 | b | 2625 | 254.1 | 0.0001072 | 0.4216 | +1 | 3 |
| - | - | 499.2 | 255 | - | - | 0 | - |
| 7 | y | 5190 | 255.1 | 0.0001779 | 0.6973 | +1 | 2 |
| - | - | 717.7 | 257.1 | - | - | 0 | - |
| - | - | 1338 | 258.1 | - | - | 0 | - |
| - | - | 706.1 | 259.1 | - | - | 0 | - |
| - | - | 583.2 | 269.1 | - | - | 0 | - |
| - | - | 603.2 | 269.9 | - | - | 0 | - |
| - | - | 579.8 | 271 | - | - | 0 | - |
| 3 | b | 1463 | 272.2 | 0.001547 | 5.684 | +1 | 3 |
| 7 | y | 2616 | 272.2 | 0.0001285 | 0.472 | +1 | 2 |
| - | - | 1697 | 276.2 | - | - | 0 | - |
| - | - | 2547 | 287.1 | - | - | 0 | - |
| - | - | 557.5 | 287.6 | - | - | 0 | - |
| - | - | 1052 | 288.2 | - | - | 0 | - |
| - | - | 759.5 | 291 | - | - | 0 | - |
| - | - | 1341 | 293.1 | - | - | 0 | - |
| - | - | 579.7 | 302.9 | - | - | 0 | - |
| - | - | 584.4 | 305.1 | - | - | 0 | - |
| - | - | 802.4 | 308 | - | - | 0 | - |
| - | - | 901.4 | 325.2 | - | - | 0 | - |
| 3 | y | 2187 | 327.7 | 0.0024 | 7.323 | +2 | 6 |
| 3 | y | 4004 | 328.2 | 0.00486 | 14.81 | +2 | 6 |
| - | - | 1517 | 328.7 | - | - | 0 | - |
| - | - | 825.9 | 329.2 | - | - | 0 | - |
| - | - | 856 | 341.2 | - | - | 0 | - |
| - | - | 628.6 | 350.9 | - | - | 0 | - |
| 2 | y | 1982 | 363.2 | 0.0007235 | 1.992 | +2 | 7 |
| - | - | 858.6 | 363.7 | - | - | 0 | - |
| - | - | 624.2 | 370.5 | - | - | 0 | - |
| - | - | 911.3 | 379 | - | - | 0 | - |
| 6 | y | 1187 | 385.3 | 0.0004699 | 1.22 | +1 | 3 |
| - | - | 562.1 | 390.2 | - | - | 0 | - |
| - | - | 776.1 | 403 | - | - | 0 | - |
| - | - | 868.3 | 403.8 | - | - | 0 | - |
| 0 | Precursor | 2080 | 412.8 | 0.0004761 | 1.154 | +2 | -1 |
| - | - | 1726 | 420.9 | - | - | 0 | - |
| - | - | 949.5 | 421 | - | - | 0 | - |
| - | - | 1012 | 439.8 | - | - | 0 | - |
| 5 | y | 1.416E+04 | 472.3 | 0.0001249 | 0.2645 | +1 | 4 |
| - | - | 2630 | 473.3 | - | - | 0 | - |
| - | - | 655.6 | 474.3 | - | - | 0 | - |
| - | - | 638.7 | 480.4 | - | - | 0 | - |
| - | - | 2558 | 486.3 | - | - | 0 | - |
| - | - | 3051 | 487.3 | - | - | 0 | - |
| - | - | 582.5 | 557.3 | - | - | 0 | - |
| - | - | 767.6 | 558.3 | - | - | 0 | - |
| 6 | b | 4906 | 571.3 | 0.01121 | 19.62 | +1 | 6 |
| - | - | 1258 | 572.4 | - | - | 0 | - |
| 3 | y | 1387 | 654.4 | 0.008387 | 12.82 | +1 | 6 |
| 3 | y | 2746 | 655.4 | 0.00717 | 10.94 | +1 | 6 |
| - | - | 1487 | 656.4 | - | - | 0 | - |
| 3 | y | 1.059E+04 | 672.4 | 0.0003974 | 0.591 | +1 | 6 |
| - | - | 2704 | 673.4 | - | - | 0 | - |
| - | - | 738.1 | 674.4 | - | - | 0 | - |
| 2 | y | 1283 | 725.4 | 0.001373 | 1.892 | +1 | 7 |
| 2 | y | 3703 | 743.4 | 0.001256 | 1.69 | +1 | 7 |
| - | - | 1624 | 744.4 | - | - | 0 | - |
| - | - | 568.5 | 976.8 | - | - | 0 | - |
| - | - | 552.3 | 992.4 | - | - | 0 | - |
| - | - | 734.5 | 1151 | - | - | 0 | - |
| - | - | 692.2 | 2174 | - | - | 0 | - |
| - | - | 738.2 | 2226 | - | - | 0 | - |

m/z Charge Intensity FragmentType MassShift Position
120.08076477050781 0 913.3193
129.018310546875 0 1156.1327
129.10235595703125 0 1413.3936
130.08628845214844 0 1207.5154
132.0215606689453 0 421.2267
132.95828247070312 0 447.64807
136.07615661621094 0 495.52652
139.576171875 0 408.62247
140.08200073242188 0 1081.7502
141.10227966308594 0 2582.8228
143.11793518066406 0 5816.7007 a 1
145.04922485351562 0 542.0426
147.04380798339844 0 560.06226
147.113037109375 0 743.99207
149.02337646484375 0 11492.48
150.02671813964844 0 5267.867
151.02833557128906 0 770.80145
153.05445861816406 0 548.62683
155.11795043945312 0 1053.4355
157.01312255859375 0 562.8952
157.1336212158203 0 4674.154
158.09234619140625 0 932.7413 y Ammonia loss 7
158.13705444335938 0 1369.3323
167.03399658203125 0 806.9268
169.09718322753906 0 2347.571
171.11280822753906 0 5224.5513 b 1
173.0924835205078 0 965.00085
173.1283721923828 0 6285.7285
173.4386444091797 0 1392.9701
174.1319580078125 0 584.65027
175.11900329589844 0 1826.0854 y 7
179.0704345703125 0 496.2615
183.06488037109375 0 572.4585
183.11276245117188 0 1090.4811
183.14935302734375 0 1524.6733
184.15277099609375 0 658.3242
185.0808868408203 0 614.2457
185.1287078857422 0 1082.6068
187.10794067382812 0 963.3006
201.12332153320312 0 2211.9817
202.1072235107422 0 1311.8473
210.90414428710938 0 578.1988
211.1445770263672 0 1212.6371
212.13938903808594 0 634.0211
226.1554718017578 0 1190.0637 a Water loss 2
227.13902282714844 0 853.8696
230.15066528320312 0 710.1791 d 2
237.09046936035156 0 912.40283
241.11997985839844 0 559.9064
246.84213256835938 0 554.9708
249.09117126464844 0 599.4311
254.14981079101562 0 2624.514 b Water loss 2
254.95260620117188 0 499.1534
255.14498901367188 0 5190.114 y Ammonia loss 6
257.096435546875 0 717.68695
258.1443176269531 0 1337.615
259.14837646484375 0 706.12054
269.0959777832031 0 583.1869
269.8777770996094 0 603.18524
271.0369873046875 0 579.8448
272.158935546875 0 1462.906 b 2
272.1718444824219 0 2615.5571 y 6
276.1549072265625 0 1696.9572
287.10687255859375 0 2547.338
287.5824890136719 0 557.51636
288.15557861328125 0 1051.813
290.9603271484375 0 759.51227
293.0997314453125 0 1340.5525
302.9089660644531 0 579.6771
305.098876953125 0 584.35095
307.98681640625 0 802.38763
325.2237548828125 0 901.3858
327.6979064941406 0 2187.2324 y Water loss 2
328.1971740722656 0 4004.0266 y Ammonia loss 2
328.6975402832031 0 1517.1205
329.1974792480469 0 825.93256
341.1880187988281 0 856.0211
350.9235534667969 0 628.6239
363.2181396484375 0 1982.4945 y Water loss 1
363.7184143066406 0 858.5814
370.5497131347656 0 624.2147
379.0335998535156 0 911.2752
385.25531005859375 0 1187.3993 y 5
390.2409362792969 0 562.1049
402.9623107910156 0 776.1485
403.8126525878906 0 868.2987
412.7525939941406 0 2080.1902 Precursor Water loss
420.8576965332031 0 1726.3691
420.9704284667969 0 949.51935
439.8415222167969 0 1011.62085
472.2879333496094 0 14156.011 y 4
473.2910461425781 0 2630.0286
474.2949523925781 0 655.6394
480.4432373046875 0 638.67645
486.2919921875 0 2558.378
487.2951354980469 0 3051.4521
557.3262939453125 0 582.5315
558.3319091796875 0 767.6286
571.356201171875 0 4906.1665 b 5
572.3590087890625 0 1257.6483
654.3849487304688 0 1386.8502 y Water loss 2
655.384521484375 0 2745.8694 y Ammonia loss 2
656.3881225585938 0 1486.5916
672.4035034179688 0 10587.497 y 2
673.4059448242188 0 2704.4094
674.4036865234375 0 738.12823
725.4290771484375 0 1283.367 y Water loss 1
743.4397583007812 0 3702.867 y 1
744.4404296875 0 1623.7075
976.7968139648438 0 568.51965
992.3721923828125 0 552.31995
1150.608642578125 0 734.4904
2174.26220703125 0 692.1885
2226.11767578125 0 738.2258

Spectrum Details

|  |  |
| --- | --- |
| Matched peaks? Matched peaks The total absolute number of peaks matched. Additionally in brackets the total fraction of peaks matched and the total number of peaks is shown. | 22 (19.30% of 114) |
| FDR? FDR The false discovery rate estimated for this peptide. It is calculated by matching all theoretical fragments with a non-integer shift with the raw peaks for this spectrum. This is done with 40 different shifts. The resulting percentage is the average number of annotated peaks over the number of annotated peaks with the correct spectrum. | 0.22% |
| Satellite FDR? Satellite FDR See the FDR for details on its calculation. This satellite ion specific FDR only contains the satellite ions (d/w) for I/L/J positions. | - |
| PSM Score? PSM Score The PSM Score as given by Hecklib to this annotated spectrum. It is shown with three significant figures. | 244 |

#### Spectrum 6104? Spectrum 6104 The raw spectrum of this peptide as annotated by Hecklib. The fragments are coloured according to ion type (see legend). Any peaks with a star '\*' as text can be hovered over to see the full details, first the ion type second the mass shift type. By hovering over the amino acids in the peptide or ions in the legend the corresponding peaks are highlighted. By toggling the 'Unassigned' label you can turn the background (unassigned) peaks on or off in the plot. By updating the slider in the Ion legend you can update the spectrum to only show the top X% of the peaks with labels. The top X% means any peak that is within X% of the highest intensity. By dragging in the spectrum you can zoom in to a specific part of the spectrum and use 'Zoom Out' to get back to the original zoom level. The annotation of the spectrum is based on the given sequence in the peptides file and is done with different software so inconsistencies are likely. The peaks are annotated based on the given sequence, with 20 ppm tolerance.

Copy Data

##### Spectrum 6104 (TSV)

###### Preview

```
Loading example...
```

*Click on the button to copy the data to your clipboard.*

Mz MinMz MaxIntensity Max

WidthHeightPeptide font sizePeptide stroke widthSpectrum font sizeSpectrum stroke widthCompact peptide

Ion legend

wxyz

abcd

OtherUnassignedIon^Charge^_Position_Show for top:%

VATVSJPR

09.88e+31.98e+42.96e+43.95e+4

Zoom Out

b^+2^_3_a^+1^_2_y^+1^_1_b^+1^_2_y^+1^_1_a^+1^_3_y^+2^_4_d^+1^_3_b^+1^_3_y^+1^_2_b^+1^_3_y^+1^_2_y^+2^_6_y^+2^_6_y^+2^_6_b^+1^_4_y^+2^_7_y^+1^_3_y^+1^_3_\*y^+1^_4_y^+1^_4_b^+1^_6_y^+1^_6_y^+1^_6_y^+1^_6_y^+1^_7_y^+1^_7_

0802160424063209

Fragment Matches Table

Show background peaks

| Position | Ion type | Intensity | mz Theoretical | mz Error (Th) | mz Error (ppm) | Charge | Series Number |
| --- | --- | --- | --- | --- | --- | --- | --- |
| - | - | 931.3 | 120.1 | - | - | 0 | - |
| - | - | 468.6 | 127.1 | - | - | 0 | - |
| - | - | 857.8 | 128.1 | - | - | 0 | - |
| - | - | 1418 | 129 | - | - | 0 | - |
| - | - | 3238 | 129.1 | - | - | 0 | - |
| - | - | 598.6 | 129.1 | - | - | 0 | - |
| - | - | 1542 | 130.1 | - | - | 0 | - |
| - | - | 1205 | 136.1 | - | - | 0 | - |
| 3 | b | 408.1 | 136.6 | 0.0005931 | 4.342 | +2 | 3 |
| - | - | 388.9 | 139.7 | - | - | 0 | - |
| - | - | 2740 | 140.1 | - | - | 0 | - |
| - | - | 366.6 | 141.1 | - | - | 0 | - |
| - | - | 4177 | 141.1 | - | - | 0 | - |
| - | - | 513.7 | 142.1 | - | - | 0 | - |
| 2 | a | 1.502E+04 | 143.1 | 0.000183 | 1.278 | +1 | 2 |
| - | - | 1191 | 144.1 | - | - | 0 | - |
| - | - | 1112 | 145.1 | - | - | 0 | - |
| - | - | 427.3 | 147.1 | - | - | 0 | - |
| - | - | 1250 | 147.1 | - | - | 0 | - |
| - | - | 1.258E+04 | 149 | - | - | 0 | - |
| - | - | 468.2 | 149.1 | - | - | 0 | - |
| - | - | 6507 | 150 | - | - | 0 | - |
| - | - | 649.7 | 151 | - | - | 0 | - |
| - | - | 449.6 | 151.1 | - | - | 0 | - |
| - | - | 993.6 | 155.1 | - | - | 0 | - |
| - | - | 1873 | 155.1 | - | - | 0 | - |
| - | - | 443.1 | 155.8 | - | - | 0 | - |
| - | - | 468.4 | 157 | - | - | 0 | - |
| - | - | 468.7 | 157.1 | - | - | 0 | - |
| - | - | 9631 | 157.1 | - | - | 0 | - |
| 8 | y | 2633 | 158.1 | 8.045E-05 | 0.5089 | +1 | 1 |
| - | - | 2578 | 158.1 | - | - | 0 | - |
| - | - | 395.9 | 159.5 | - | - | 0 | - |
| - | - | 574.7 | 162.7 | - | - | 0 | - |
| - | - | 4441 | 169.1 | - | - | 0 | - |
| - | - | 628.3 | 170.1 | - | - | 0 | - |
| 2 | b | 1.575E+04 | 171.1 | 0.0001414 | 0.8264 | +1 | 2 |
| - | - | 741.4 | 172.1 | - | - | 0 | - |
| - | - | 2204 | 173.1 | - | - | 0 | - |
| - | - | 1037 | 173.1 | - | - | 0 | - |
| - | - | 1.846E+04 | 173.1 | - | - | 0 | - |
| - | - | 1648 | 174.1 | - | - | 0 | - |
| 8 | y | 4069 | 175.1 | 0.0001885 | 1.076 | +1 | 1 |
| - | - | 3639 | 183.1 | - | - | 0 | - |
| - | - | 4033 | 183.1 | - | - | 0 | - |
| - | - | 435.9 | 183.5 | - | - | 0 | - |
| - | - | 916.1 | 184.2 | - | - | 0 | - |
| - | - | 1191 | 185.1 | - | - | 0 | - |
| - | - | 2105 | 185.1 | - | - | 0 | - |
| - | - | 1059 | 187.1 | - | - | 0 | - |
| - | - | 649.6 | 195.1 | - | - | 0 | - |
| - | - | 6754 | 201.1 | - | - | 0 | - |
| - | - | 673.7 | 202.1 | - | - | 0 | - |
| - | - | 1057 | 202.1 | - | - | 0 | - |
| - | - | 466.2 | 202.9 | - | - | 0 | - |
| - | - | 955.7 | 211.1 | - | - | 0 | - |
| - | - | 2120 | 211.1 | - | - | 0 | - |
| - | - | 1289 | 212.1 | - | - | 0 | - |
| - | - | 855.1 | 213.1 | - | - | 0 | - |
| - | - | 909.8 | 219.1 | - | - | 0 | - |
| - | - | 524.5 | 220.1 | - | - | 0 | - |
| 3 | a | 1887 | 226.2 | 0.0001938 | 0.857 | +1 | 3 |
| - | - | 1542 | 227.1 | - | - | 0 | - |
| 5 | y | 563.6 | 228.1 | 0.0004636 | 2.032 | +2 | 4 |
| 3 | d | 1246 | 230.1 | 0.0001529 | 0.6645 | +1 | 3 |
| - | - | 995.5 | 240.1 | - | - | 0 | - |
| - | - | 985 | 244.2 | - | - | 0 | - |
| 3 | b | 5496 | 254.1 | 3.086E-05 | 0.1214 | +1 | 3 |
| 7 | y | 1.59E+04 | 255.1 | 6.624E-05 | 0.2596 | +1 | 2 |
| - | - | 2159 | 256.1 | - | - | 0 | - |
| - | - | 1185 | 257.1 | - | - | 0 | - |
| - | - | 3187 | 258.1 | - | - | 0 | - |
| - | - | 739.2 | 259.1 | - | - | 0 | - |
| - | - | 861.8 | 266.1 | - | - | 0 | - |
| - | - | 2086 | 270.1 | - | - | 0 | - |
| - | - | 644.1 | 271 | - | - | 0 | - |
| 3 | b | 3048 | 272.2 | 0.0009367 | 3.442 | +1 | 3 |
| 7 | y | 5863 | 272.2 | 3.691E-05 | 0.1356 | +1 | 2 |
| - | - | 994.8 | 273.2 | - | - | 0 | - |
| - | - | 2131 | 276.2 | - | - | 0 | - |
| - | - | 1216 | 277.2 | - | - | 0 | - |
| - | - | 634.9 | 282.2 | - | - | 0 | - |
| - | - | 2100 | 288.2 | - | - | 0 | - |
| - | - | 1175 | 293.1 | - | - | 0 | - |
| - | - | 859.3 | 306.2 | - | - | 0 | - |
| - | - | 772.5 | 312.2 | - | - | 0 | - |
| - | - | 572.1 | 316.9 | - | - | 0 | - |
| - | - | 1661 | 322.2 | - | - | 0 | - |
| - | - | 3429 | 325.2 | - | - | 0 | - |
| 3 | y | 8320 | 327.7 | 0.001484 | 4.529 | +2 | 6 |
| 3 | y | 8979 | 328.2 | 0.005013 | 15.27 | +2 | 6 |
| - | - | 4577 | 328.7 | - | - | 0 | - |
| 3 | y | 791.9 | 336.7 | 0.0005105 | 1.516 | +2 | 6 |
| - | - | 830.4 | 340.2 | - | - | 0 | - |
| - | - | 1509 | 340.2 | - | - | 0 | - |
| - | - | 1081 | 341.2 | - | - | 0 | - |
| - | - | 1298 | 350.9 | - | - | 0 | - |
| - | - | 713.9 | 351.9 | - | - | 0 | - |
| 4 | b | 1727 | 353.2 | 0.000772 | 2.186 | +1 | 4 |
| 2 | y | 5815 | 363.2 | 0.0004794 | 1.32 | +2 | 7 |
| - | - | 3285 | 363.7 | - | - | 0 | - |
| - | - | 690.1 | 364.2 | - | - | 0 | - |
| - | - | 7127 | 366.2 | - | - | 0 | - |
| 6 | y | 735.8 | 368.2 | 0.0004406 | 1.197 | +1 | 3 |
| - | - | 625.8 | 368.9 | - | - | 0 | - |
| - | - | 613.2 | 376.8 | - | - | 0 | - |
| - | - | 1188 | 384.2 | - | - | 0 | - |
| 6 | y | 3535 | 385.3 | 7.321E-05 | 0.19 | +1 | 3 |
| - | - | 616.4 | 389.2 | - | - | 0 | - |
| - | - | 1219 | 391.2 | - | - | 0 | - |
| - | - | 4902 | 402.2 | - | - | 0 | - |
| - | - | 707.3 | 403 | - | - | 0 | - |
| - | - | 559.7 | 403.2 | - | - | 0 | - |
| - | - | 1626 | 403.8 | - | - | 0 | - |
| - | - | 564.4 | 406.9 | - | - | 0 | - |
| 0 | Precursor | 4088 | 412.8 | 1.838E-05 | 0.04453 | +2 | -1 |
| - | - | 2155 | 413.3 | - | - | 0 | - |
| - | - | 1149 | 413.8 | - | - | 0 | - |
| - | - | 1904 | 420.9 | - | - | 0 | - |
| - | - | 2408 | 422.2 | - | - | 0 | - |
| 5 | y | 667.7 | 454.3 | 0.001182 | 2.601 | +1 | 4 |
| - | - | 682.4 | 458.3 | - | - | 0 | - |
| 5 | y | 3.911E+04 | 472.3 | 0.0001497 | 0.317 | +1 | 4 |
| - | - | 9187 | 473.3 | - | - | 0 | - |
| - | - | 1464 | 474.3 | - | - | 0 | - |
| - | - | 4367 | 486.3 | - | - | 0 | - |
| - | - | 5804 | 487.3 | - | - | 0 | - |
| - | - | 1091 | 488.3 | - | - | 0 | - |
| - | - | 636.1 | 501.2 | - | - | 0 | - |
| - | - | 724.6 | 509.3 | - | - | 0 | - |
| - | - | 663.1 | 540.3 | - | - | 0 | - |
| - | - | 927.9 | 557.3 | - | - | 0 | - |
| - | - | 1591 | 558.3 | - | - | 0 | - |
| 6 | b | 1.349E+04 | 571.3 | 0.01024 | 17.92 | +1 | 6 |
| - | - | 3162 | 572.4 | - | - | 0 | - |
| - | - | 915 | 611.3 | - | - | 0 | - |
| - | - | 755.8 | 628.4 | - | - | 0 | - |
| 3 | y | 2468 | 654.4 | 0.009669 | 14.78 | +1 | 6 |
| 3 | y | 9093 | 655.4 | 0.006193 | 9.45 | +1 | 6 |
| - | - | 2782 | 656.4 | - | - | 0 | - |
| - | - | 991 | 657.4 | - | - | 0 | - |
| 3 | y | 2.882E+04 | 672.4 | 0.001496 | 2.225 | +1 | 6 |
| - | - | 1.093E+04 | 673.4 | - | - | 0 | - |
| - | - | 2155 | 674.4 | - | - | 0 | - |
| - | - | 1918 | 682.4 | - | - | 0 | - |
| 2 | y | 3467 | 725.4 | 0.001922 | 2.65 | +1 | 7 |
| - | - | 1065 | 726.4 | - | - | 0 | - |
| 2 | y | 1.26E+04 | 743.4 | 0.002355 | 3.168 | +1 | 7 |
| - | - | 4570 | 744.4 | - | - | 0 | - |
| - | - | 1569 | 745.4 | - | - | 0 | - |
| - | - | 652.6 | 1349 | - | - | 0 | - |
| - | - | 630.1 | 2828 | - | - | 0 | - |
| - | - | 747.5 | 3081 | - | - | 0 | - |
| - | - | 733.1 | 3177 | - | - | 0 | - |

m/z Charge Intensity FragmentType MassShift Position
120.0810546875 0 931.331
127.0867919921875 0 468.6316
128.10736083984375 0 857.7632
129.0184783935547 0 1418.1318
129.10244750976562 0 3237.7605
129.1073455810547 0 598.6321
130.08633422851562 0 1541.9479
136.07594299316406 0 1205.3828
136.58447265625 0 408.0848 b 2
139.6763916015625 0 388.90912
140.0820770263672 0 2739.916
141.09803771972656 0 366.5505
141.10240173339844 0 4176.807
142.10568237304688 0 513.6876
143.11807250976562 0 15024.617 a 1
144.12158203125 0 1190.765
145.097412109375 0 1112.3237
147.0752410888672 0 427.2861
147.11309814453125 0 1249.7445
149.02349853515625 0 12579.131
149.05987548828125 0 468.18326
150.02688598632812 0 6506.9272
151.02894592285156 0 649.7047
151.07571411132812 0 449.5807
155.0819549560547 0 993.5634
155.11795043945312 0 1873.0387
155.79615783691406 0 443.06262
157.01329040527344 0 468.37408
157.1088409423828 0 468.66913
157.13369750976562 0 9631.072
158.0924835205078 0 2633.1965 y Ammonia loss 7
158.1371612548828 0 2577.5415
159.49075317382812 0 395.85602
162.72848510742188 0 574.67944
169.09725952148438 0 4441.069
170.10122680664062 0 628.3387
171.11294555664062 0 15748.821 b 1
172.11634826660156 0 741.3836
173.09219360351562 0 2203.6772
173.12025451660156 0 1036.683
173.1286163330078 0 18456.549
174.1322784423828 0 1647.5596
175.119140625 0 4068.9363 y 7
183.11294555664062 0 3638.6248
183.1493682861328 0 4032.7544
183.5269317626953 0 435.85757
184.1529541015625 0 916.144
185.08091735839844 0 1190.7853
185.1285858154297 0 2105.4895
187.10768127441406 0 1058.9241
195.11279296875 0 649.6148
201.12347412109375 0 6754.1587
202.107421875 0 673.72314
202.1267852783203 0 1057.0852
202.85076904296875 0 466.23975
211.1081085205078 0 955.67755
211.1439208984375 0 2119.5332
212.1395721435547 0 1289.3275
213.123291015625 0 855.061
219.13365173339844 0 909.84784
220.13771057128906 0 524.52045
226.1551971435547 0 1887.4305 a Water loss 2
227.13919067382812 0 1542.3094
228.13380432128906 0 563.5986 y Ammonia loss 4
230.14976501464844 0 1246.1064 d 2
240.134033203125 0 995.46857
244.1507110595703 0 984.9971
254.14988708496094 0 5495.957 b Water loss 2
255.14523315429688 0 15902.671 y Ammonia loss 6
256.1484680175781 0 2159.124
257.0709533691406 0 1184.6582
258.14483642578125 0 3186.866
259.1484069824219 0 739.161
266.14990234375 0 861.7722
270.14453125 0 2085.9285
270.96978759765625 0 644.1183
272.1595458984375 0 3047.5454 b 2
272.1717529296875 0 5863.183 y 6
273.17535400390625 0 994.82367
276.15496826171875 0 2131.23
277.1591796875 0 1215.8759
282.1810302734375 0 634.8844
288.1557922363281 0 2100.1904
293.099853515625 0 1175.2947
306.1795654296875 0 859.3072
312.19012451171875 0 772.51086
316.8792419433594 0 572.1386
322.1956481933594 0 1660.9622
325.22314453125 0 3428.9448
327.6988220214844 0 8319.813 y Water loss 2
328.19732666015625 0 8978.97 y Ammonia loss 2
328.69781494140625 0 4577.039
336.705078125 0 791.923 y 2
340.1677551269531 0 830.39185
340.1867980957031 0 1508.7378
341.18798828125 0 1080.7812
350.9263000488281 0 1297.9698
351.9268493652344 0 713.9264
353.2175598144531 0 1726.8715 b Water loss 3
363.2183837890625 0 5815.117 y Water loss 1
363.72003173828125 0 3285.142
364.2193603515625 0 690.0956
366.1852111816406 0 7126.982
368.2287902832031 0 735.758 y Ammonia loss 5
368.9361572265625 0 625.8238
376.8458251953125 0 613.1521
384.1954650878906 0 1187.8668
385.2557067871094 0 3535.0183 y 5
389.2398986816406 0 616.3655
391.2319641113281 0 1219.0842
402.2062072753906 0 4902.01
402.96044921875 0 707.2995
403.2120666503906 0 559.71533
403.8121643066406 0 1626.0983
406.9405517578125 0 564.4379
412.7530517578125 0 4088.0962 Precursor Water loss
413.25445556640625 0 2154.6484
413.75567626953125 0 1148.5917
420.85711669921875 0 1903.9645
422.2116394042969 0 2407.7273
454.27606201171875 0 667.6796 y Water loss 4
458.2802429199219 0 682.3528
472.28765869140625 0 39113.863 y 4
473.2900390625 0 9187.41
474.29217529296875 0 1463.881
486.2916259765625 0 4367.0405
487.2947692871094 0 5804.484
488.2999572753906 0 1090.8164
501.2406005859375 0 636.0515
509.2831115722656 0 724.6071
540.325439453125 0 663.05005
557.3314819335938 0 927.9314
558.3309326171875 0 1590.6953
571.355224609375 0 13492.352 b 5
572.3584594726562 0 3162.2744
611.3497924804688 0 915.01465
628.3788452148438 0 755.76965
654.3836669921875 0 2468.3164 y Water loss 2
655.383544921875 0 9092.907 y Ammonia loss 2
656.3871459960938 0 2782.0498
657.3865356445312 0 990.9551
672.4024047851562 0 28819.932 y 2
673.4053344726562 0 10932.728
674.4078979492188 0 2154.8848
682.3870239257812 0 1917.9948
725.4285278320312 0 3467.1487 y Water loss 1
726.4317626953125 0 1065.4762
743.4386596679688 0 12597.604 y 1
744.4420166015625 0 4569.772
745.445068359375 0 1569.2657
1349.459716796875 0 652.62537
2828.4873046875 0 630.06744
3080.990966796875 0 747.50476
3176.867431640625 0 733.08014

Spectrum Details

|  |  |
| --- | --- |
| Matched peaks? Matched peaks The total absolute number of peaks matched. Additionally in brackets the total fraction of peaks matched and the total number of peaks is shown. | 28 (18.18% of 154) |
| FDR? FDR The false discovery rate estimated for this peptide. It is calculated by matching all theoretical fragments with a non-integer shift with the raw peaks for this spectrum. This is done with 40 different shifts. The resulting percentage is the average number of annotated peaks over the number of annotated peaks with the correct spectrum. | 0.34% |
| Satellite FDR? Satellite FDR See the FDR for details on its calculation. This satellite ion specific FDR only contains the satellite ions (d/w) for I/L/J positions. | - |
| PSM Score? PSM Score The PSM Score as given by Hecklib to this annotated spectrum. It is shown with three significant figures. | 306 |

#### Spectrum 4916? Spectrum 4916 The raw spectrum of this peptide as annotated by Hecklib. The fragments are coloured according to ion type (see legend). Any peaks with a star '\*' as text can be hovered over to see the full details, first the ion type second the mass shift type. By hovering over the amino acids in the peptide or ions in the legend the corresponding peaks are highlighted. By toggling the 'Unassigned' label you can turn the background (unassigned) peaks on or off in the plot. By updating the slider in the Ion legend you can update the spectrum to only show the top X% of the peaks with labels. The top X% means any peak that is within X% of the highest intensity. By dragging in the spectrum you can zoom in to a specific part of the spectrum and use 'Zoom Out' to get back to the original zoom level. The annotation of the spectrum is based on the given sequence in the peptides file and is done with different software so inconsistencies are likely. The peaks are annotated based on the given sequence, with 20 ppm tolerance.

Copy Data

##### Spectrum 4916 (TSV)

###### Preview

```
Loading example...
```

*Click on the button to copy the data to your clipboard.*

Mz MinMz MaxIntensity Max

WidthHeightPeptide font sizePeptide stroke widthSpectrum font sizeSpectrum stroke widthCompact peptide

Ion legend

wxyz

abcd

OtherUnassignedIon^Charge^_Position_Show for top:%

VATVSJPR

04.50e+48.99e+41.35e+51.80e+5

Zoom Out

a^+1^_2_y^+1^_1_b^+1^_2_y^+1^_1_a^+1^_3_y^+2^_4_d^+1^_3_a^+1^_3_b^+1^_3_y^+1^_2_b^+1^_3_y^+1^_2_b^+2^_6_b^+2^_6_y^+2^_6_y^+2^_6_y^+2^_6_b^+1^_4_y^+2^_7_y^+1^_3_b^+1^_4_y^+1^_3_\*y^+1^_4_y^+1^_4_y^+1^_4_b^+1^_6_y^+1^_5_b^+1^_6_y^+1^_6_y^+1^_6_y^+1^_6_y^+1^_7_y^+1^_7_

0874174826223495

Fragment Matches Table

Show background peaks

| Position | Ion type | Intensity | mz Theoretical | mz Error (Th) | mz Error (ppm) | Charge | Series Number |
| --- | --- | --- | --- | --- | --- | --- | --- |
| - | - | 1346 | 120.1 | - | - | 0 | - |
| - | - | 342.6 | 120.7 | - | - | 0 | - |
| - | - | 357.1 | 123.5 | - | - | 0 | - |
| - | - | 436.8 | 125.6 | - | - | 0 | - |
| - | - | 1561 | 127.1 | - | - | 0 | - |
| - | - | 1934 | 127.1 | - | - | 0 | - |
| - | - | 4175 | 128.1 | - | - | 0 | - |
| - | - | 820.8 | 129 | - | - | 0 | - |
| - | - | 1.283E+04 | 129.1 | - | - | 0 | - |
| - | - | 584.7 | 129.1 | - | - | 0 | - |
| - | - | 1049 | 130.1 | - | - | 0 | - |
| - | - | 5682 | 130.1 | - | - | 0 | - |
| - | - | 851.8 | 130.1 | - | - | 0 | - |
| - | - | 1614 | 130.1 | - | - | 0 | - |
| - | - | 772.8 | 131.1 | - | - | 0 | - |
| - | - | 556.7 | 133.1 | - | - | 0 | - |
| - | - | 865.8 | 136.1 | - | - | 0 | - |
| - | - | 2259 | 138.1 | - | - | 0 | - |
| - | - | 544.6 | 139.1 | - | - | 0 | - |
| - | - | 1.212E+04 | 140.1 | - | - | 0 | - |
| - | - | 726.3 | 141.1 | - | - | 0 | - |
| - | - | 2.121E+04 | 141.1 | - | - | 0 | - |
| - | - | 460.9 | 141.2 | - | - | 0 | - |
| - | - | 793.6 | 142.1 | - | - | 0 | - |
| - | - | 5109 | 142.1 | - | - | 0 | - |
| 2 | a | 7.548E+04 | 143.1 | 0.0003203 | 2.238 | +1 | 2 |
| - | - | 926.1 | 144.1 | - | - | 0 | - |
| - | - | 5730 | 144.1 | - | - | 0 | - |
| - | - | 4523 | 145.1 | - | - | 0 | - |
| - | - | 676.1 | 146.1 | - | - | 0 | - |
| - | - | 6754 | 147.1 | - | - | 0 | - |
| - | - | 1351 | 148.1 | - | - | 0 | - |
| - | - | 570.7 | 148.9 | - | - | 0 | - |
| - | - | 8220 | 149 | - | - | 0 | - |
| - | - | 5631 | 150 | - | - | 0 | - |
| - | - | 2622 | 155.1 | - | - | 0 | - |
| - | - | 9486 | 155.1 | - | - | 0 | - |
| - | - | 849.1 | 156.1 | - | - | 0 | - |
| - | - | 1451 | 157.1 | - | - | 0 | - |
| - | - | 2407 | 157.1 | - | - | 0 | - |
| - | - | 4.382E+04 | 157.1 | - | - | 0 | - |
| 8 | y | 9743 | 158.1 | 0.000233 | 1.474 | +1 | 1 |
| - | - | 539.4 | 158.1 | - | - | 0 | - |
| - | - | 1111 | 158.1 | - | - | 0 | - |
| - | - | 1.243E+04 | 158.1 | - | - | 0 | - |
| - | - | 467.8 | 159.1 | - | - | 0 | - |
| - | - | 1563 | 159.1 | - | - | 0 | - |
| - | - | 519.4 | 159.1 | - | - | 0 | - |
| - | - | 432.8 | 160.2 | - | - | 0 | - |
| - | - | 2.117E+04 | 169.1 | - | - | 0 | - |
| - | - | 7090 | 170.1 | - | - | 0 | - |
| 2 | b | 6.578E+04 | 171.1 | 0.000294 | 1.718 | +1 | 2 |
| - | - | 620.8 | 172.1 | - | - | 0 | - |
| - | - | 5821 | 172.1 | - | - | 0 | - |
| - | - | 1.036E+04 | 173.1 | - | - | 0 | - |
| - | - | 7.714E+04 | 173.1 | - | - | 0 | - |
| - | - | 596.5 | 174.1 | - | - | 0 | - |
| - | - | 626.3 | 174.1 | - | - | 0 | - |
| - | - | 6818 | 174.1 | - | - | 0 | - |
| 8 | y | 2.323E+04 | 175.1 | 0.00028 | 1.599 | +1 | 1 |
| - | - | 1526 | 176.1 | - | - | 0 | - |
| - | - | 541.6 | 181.1 | - | - | 0 | - |
| - | - | 1.624E+04 | 183.1 | - | - | 0 | - |
| - | - | 2.116E+04 | 183.1 | - | - | 0 | - |
| - | - | 1759 | 184.1 | - | - | 0 | - |
| - | - | 8494 | 184.2 | - | - | 0 | - |
| - | - | 1835 | 185.1 | - | - | 0 | - |
| - | - | 9349 | 185.1 | - | - | 0 | - |
| - | - | 658.6 | 185.2 | - | - | 0 | - |
| - | - | 2899 | 186.1 | - | - | 0 | - |
| - | - | 496.3 | 186.6 | - | - | 0 | - |
| - | - | 3934 | 187.1 | - | - | 0 | - |
| - | - | 773.7 | 194.1 | - | - | 0 | - |
| - | - | 3082 | 195.1 | - | - | 0 | - |
| - | - | 863.7 | 197.1 | - | - | 0 | - |
| - | - | 509.4 | 197.1 | - | - | 0 | - |
| - | - | 1184 | 199.1 | - | - | 0 | - |
| - | - | 2.745E+04 | 201.1 | - | - | 0 | - |
| - | - | 2680 | 202.1 | - | - | 0 | - |
| - | - | 1103 | 209.1 | - | - | 0 | - |
| - | - | 1643 | 210.1 | - | - | 0 | - |
| - | - | 3782 | 211.1 | - | - | 0 | - |
| - | - | 1.018E+04 | 211.1 | - | - | 0 | - |
| - | - | 879.4 | 211.2 | - | - | 0 | - |
| - | - | 6402 | 212.1 | - | - | 0 | - |
| - | - | 3363 | 212.1 | - | - | 0 | - |
| - | - | 1947 | 213.1 | - | - | 0 | - |
| - | - | 866.4 | 213.1 | - | - | 0 | - |
| - | - | 1736 | 215.1 | - | - | 0 | - |
| - | - | 727.7 | 226.1 | - | - | 0 | - |
| 3 | a | 1.016E+04 | 226.2 | 0.0003006 | 1.329 | +1 | 3 |
| - | - | 724.8 | 227.1 | - | - | 0 | - |
| - | - | 4846 | 227.1 | - | - | 0 | - |
| - | - | 986.2 | 227.2 | - | - | 0 | - |
| 5 | y | 2673 | 228.1 | 0.0003604 | 1.58 | +2 | 4 |
| - | - | 721.2 | 229.1 | - | - | 0 | - |
| 3 | d | 4273 | 230.1 | 0.0002743 | 1.192 | +1 | 3 |
| - | - | 638.2 | 238.1 | - | - | 0 | - |
| - | - | 4550 | 240.1 | - | - | 0 | - |
| - | - | 1574 | 241.1 | - | - | 0 | - |
| - | - | 1192 | 243.1 | - | - | 0 | - |
| - | - | 1968 | 243.7 | - | - | 0 | - |
| - | - | 1514 | 244.2 | - | - | 0 | - |
| 3 | a | 911.6 | 244.2 | 0.0008036 | 3.291 | +1 | 3 |
| - | - | 1365 | 252.1 | - | - | 0 | - |
| 3 | b | 2.851E+04 | 254.1 | 0.0001828 | 0.7191 | +1 | 3 |
| 7 | y | 6.985E+04 | 255.1 | 0.0002951 | 1.157 | +1 | 2 |
| - | - | 6735 | 256.1 | - | - | 0 | - |
| - | - | 580.9 | 257.1 | - | - | 0 | - |
| - | - | 1.383E+04 | 258.1 | - | - | 0 | - |
| - | - | 5130 | 259.1 | - | - | 0 | - |
| - | - | 524 | 260.1 | - | - | 0 | - |
| - | - | 928.9 | 264.2 | - | - | 0 | - |
| - | - | 853.9 | 265.2 | - | - | 0 | - |
| - | - | 3844 | 266.2 | - | - | 0 | - |
| - | - | 1560 | 267.2 | - | - | 0 | - |
| - | - | 1.069E+04 | 270.1 | - | - | 0 | - |
| - | - | 1299 | 271.1 | - | - | 0 | - |
| - | - | 828.6 | 271.2 | - | - | 0 | - |
| 3 | b | 1.161E+04 | 272.2 | 0.0005705 | 2.096 | +1 | 3 |
| 7 | y | 2.859E+04 | 272.2 | 0.0003421 | 1.257 | +1 | 2 |
| - | - | 1729 | 273.2 | - | - | 0 | - |
| - | - | 2943 | 273.2 | - | - | 0 | - |
| - | - | 1.125E+04 | 276.2 | - | - | 0 | - |
| - | - | 5172 | 277.2 | - | - | 0 | - |
| 6 | b | 1214 | 277.2 | 0.00487 | 17.57 | +2 | 6 |
| - | - | 1392 | 278.2 | - | - | 0 | - |
| - | - | 724.3 | 280.1 | - | - | 0 | - |
| - | - | 3056 | 282.2 | - | - | 0 | - |
| - | - | 695.7 | 283.2 | - | - | 0 | - |
| 6 | b | 2447 | 286.2 | 0.005691 | 19.89 | +2 | 6 |
| - | - | 9652 | 288.2 | - | - | 0 | - |
| - | - | 1599 | 293.1 | - | - | 0 | - |
| - | - | 638.9 | 297.2 | - | - | 0 | - |
| - | - | 1746 | 298.1 | - | - | 0 | - |
| - | - | 929.6 | 299.2 | - | - | 0 | - |
| - | - | 4927 | 306.2 | - | - | 0 | - |
| - | - | 1532 | 306.7 | - | - | 0 | - |
| - | - | 629.4 | 308 | - | - | 0 | - |
| - | - | 3243 | 312.2 | - | - | 0 | - |
| - | - | 539.9 | 313.2 | - | - | 0 | - |
| - | - | 646.2 | 316.9 | - | - | 0 | - |
| - | - | 621.5 | 318.7 | - | - | 0 | - |
| - | - | 764.4 | 319.7 | - | - | 0 | - |
| - | - | 1712 | 323.2 | - | - | 0 | - |
| - | - | 1.237E+04 | 325.2 | - | - | 0 | - |
| - | - | 2391 | 326.2 | - | - | 0 | - |
| 3 | y | 3.003E+04 | 327.7 | 0.001271 | 3.877 | +2 | 6 |
| 3 | y | 5.007E+04 | 328.2 | 0.005135 | 15.65 | +2 | 6 |
| - | - | 1.606E+04 | 328.7 | - | - | 0 | - |
| - | - | 4357 | 329.2 | - | - | 0 | - |
| - | - | 842.2 | 332.7 | - | - | 0 | - |
| 3 | y | 3129 | 336.7 | 9.985E-05 | 0.2965 | +2 | 6 |
| - | - | 1472 | 337.2 | - | - | 0 | - |
| - | - | 9296 | 340.2 | - | - | 0 | - |
| - | - | 7453 | 341.2 | - | - | 0 | - |
| - | - | 1014 | 341.7 | - | - | 0 | - |
| - | - | 1126 | 342.2 | - | - | 0 | - |
| - | - | 1977 | 343.2 | - | - | 0 | - |
| - | - | 687 | 349.2 | - | - | 0 | - |
| - | - | 868.9 | 350.9 | - | - | 0 | - |
| - | - | 1311 | 351.2 | - | - | 0 | - |
| - | - | 1732 | 352.2 | - | - | 0 | - |
| 4 | b | 8328 | 353.2 | 0.000174 | 0.4926 | +1 | 4 |
| - | - | 1355 | 354.2 | - | - | 0 | - |
| - | - | 1179 | 355.2 | - | - | 0 | - |
| - | - | 1824 | 358.2 | - | - | 0 | - |
| - | - | 2171 | 359.2 | - | - | 0 | - |
| 2 | y | 3.064E+04 | 363.2 | 6.993E-05 | 0.1925 | +2 | 7 |
| - | - | 1.213E+04 | 363.7 | - | - | 0 | - |
| - | - | 2984 | 364.2 | - | - | 0 | - |
| - | - | 576.5 | 365.2 | - | - | 0 | - |
| - | - | 610.4 | 367.2 | - | - | 0 | - |
| 6 | y | 4224 | 368.2 | 7.44E-05 | 0.2021 | +1 | 3 |
| - | - | 760.7 | 369.2 | - | - | 0 | - |
| 4 | b | 896.7 | 371.2 | 4.632E-05 | 0.1248 | +1 | 4 |
| - | - | 538.6 | 374.1 | - | - | 0 | - |
| - | - | 2019 | 379.2 | - | - | 0 | - |
| - | - | 566.4 | 379.7 | - | - | 0 | - |
| - | - | 2053 | 380.2 | - | - | 0 | - |
| - | - | 2358 | 383.2 | - | - | 0 | - |
| - | - | 1007 | 384.2 | - | - | 0 | - |
| - | - | 801.5 | 385.2 | - | - | 0 | - |
| 6 | y | 1.664E+04 | 385.3 | 0.0001709 | 0.4437 | +1 | 3 |
| - | - | 3579 | 386.3 | - | - | 0 | - |
| - | - | 3064 | 389.2 | - | - | 0 | - |
| - | - | 2919 | 390.2 | - | - | 0 | - |
| - | - | 6052 | 391.2 | - | - | 0 | - |
| - | - | 4176 | 391.7 | - | - | 0 | - |
| - | - | 681.6 | 401.2 | - | - | 0 | - |
| - | - | 983.7 | 403.7 | - | - | 0 | - |
| - | - | 1229 | 403.8 | - | - | 0 | - |
| - | - | 690.4 | 408.3 | - | - | 0 | - |
| - | - | 1719 | 411.2 | - | - | 0 | - |
| - | - | 1860 | 412.2 | - | - | 0 | - |
| 0 | Precursor | 2.379E+04 | 412.8 | 0.0003784 | 0.9167 | +2 | -1 |
| - | - | 8545 | 413.3 | - | - | 0 | - |
| - | - | 3484 | 413.8 | - | - | 0 | - |
| - | - | 1148 | 414.3 | - | - | 0 | - |
| - | - | 1147 | 420.9 | - | - | 0 | - |
| - | - | 1085 | 422.2 | - | - | 0 | - |
| - | - | 1280 | 426.3 | - | - | 0 | - |
| - | - | 2620 | 436.3 | - | - | 0 | - |
| - | - | 756.7 | 439.8 | - | - | 0 | - |
| - | - | 1525 | 442.3 | - | - | 0 | - |
| 5 | y | 2755 | 454.3 | 0.003165 | 6.968 | +1 | 4 |
| 5 | y | 796.1 | 455.3 | 0.004427 | 9.723 | +1 | 4 |
| - | - | 711.5 | 468.3 | - | - | 0 | - |
| - | - | 1436 | 469.3 | - | - | 0 | - |
| 5 | y | 1.78E+05 | 472.3 | 0.0004301 | 0.9107 | +1 | 4 |
| - | - | 4.245E+04 | 473.3 | - | - | 0 | - |
| - | - | 6672 | 474.3 | - | - | 0 | - |
| - | - | 596.6 | 480.2 | - | - | 0 | - |
| - | - | 1800 | 482.3 | - | - | 0 | - |
| - | - | 2.601E+04 | 486.3 | - | - | 0 | - |
| - | - | 3.15E+04 | 487.3 | - | - | 0 | - |
| - | - | 1.017E+04 | 488.3 | - | - | 0 | - |
| - | - | 1227 | 489.3 | - | - | 0 | - |
| - | - | 1950 | 508.3 | - | - | 0 | - |
| - | - | 3078 | 509.3 | - | - | 0 | - |
| - | - | 879.1 | 535.3 | - | - | 0 | - |
| - | - | 711.3 | 540.3 | - | - | 0 | - |
| 6 | b | 1710 | 553.3 | 0.00896 | 16.19 | +1 | 6 |
| 4 | y | 884.8 | 554.3 | 0.006875 | 12.4 | +1 | 5 |
| - | - | 4275 | 557.3 | - | - | 0 | - |
| - | - | 8559 | 558.3 | - | - | 0 | - |
| - | - | 2367 | 559.3 | - | - | 0 | - |
| 6 | b | 5.988E+04 | 571.3 | 0.01121 | 19.62 | +1 | 6 |
| - | - | 1.653E+04 | 572.4 | - | - | 0 | - |
| - | - | 3088 | 573.4 | - | - | 0 | - |
| - | - | 1271 | 581.3 | - | - | 0 | - |
| - | - | 716.9 | 592.2 | - | - | 0 | - |
| - | - | 649.8 | 597.3 | - | - | 0 | - |
| - | - | 2696 | 611.3 | - | - | 0 | - |
| - | - | 5353 | 628.4 | - | - | 0 | - |
| - | - | 1810 | 629.4 | - | - | 0 | - |
| - | - | 2254 | 637.4 | - | - | 0 | - |
| - | - | 1248 | 638.4 | - | - | 0 | - |
| 3 | y | 1.475E+04 | 654.4 | 0.008754 | 13.38 | +1 | 6 |
| 3 | y | 3.978E+04 | 655.4 | 0.00778 | 11.87 | +1 | 6 |
| - | - | 1.607E+04 | 656.4 | - | - | 0 | - |
| - | - | 3024 | 657.4 | - | - | 0 | - |
| - | - | 1045 | 664.4 | - | - | 0 | - |
| 3 | y | 1.428E+05 | 672.4 | 0.0002753 | 0.4094 | +1 | 6 |
| - | - | 5.089E+04 | 673.4 | - | - | 0 | - |
| - | - | 1.184E+04 | 674.4 | - | - | 0 | - |
| - | - | 1.007E+04 | 682.4 | - | - | 0 | - |
| - | - | 3062 | 683.4 | - | - | 0 | - |
| - | - | 876.7 | 684.4 | - | - | 0 | - |
| - | - | 924.9 | 699.4 | - | - | 0 | - |
| 2 | y | 1.714E+04 | 725.4 | 0.0006403 | 0.8827 | +1 | 7 |
| - | - | 6049 | 726.4 | - | - | 0 | - |
| - | - | 1951 | 727.4 | - | - | 0 | - |
| - | - | 731.2 | 735.4 | - | - | 0 | - |
| 2 | y | 5.524E+04 | 743.4 | 0.0005849 | 0.7868 | +1 | 7 |
| - | - | 2.432E+04 | 744.4 | - | - | 0 | - |
| - | - | 4486 | 745.4 | - | - | 0 | - |
| - | - | 1628 | 753.4 | - | - | 0 | - |
| - | - | 710 | 860.6 | - | - | 0 | - |
| - | - | 622 | 1260 | - | - | 0 | - |
| - | - | 577.3 | 1309 | - | - | 0 | - |
| - | - | 643 | 1941 | - | - | 0 | - |
| - | - | 608.5 | 2008 | - | - | 0 | - |
| - | - | 729.4 | 2545 | - | - | 0 | - |
| - | - | 700 | 2753 | - | - | 0 | - |
| - | - | 1115 | 3080 | - | - | 0 | - |
| - | - | 838.9 | 3461 | - | - | 0 | - |

m/z Charge Intensity FragmentType MassShift Position
120.08106231689453 0 1346.1273
120.67803955078125 0 342.6257
123.53107452392578 0 357.06934
125.63831329345703 0 436.77737
127.08674621582031 0 1561.1656
127.12326049804688 0 1934.0939
128.10726928710938 0 4174.6855
129.0187530517578 0 820.828
129.10252380371094 0 12834.452
129.1142578125 0 584.6635
130.05030822753906 0 1048.8984
130.0865478515625 0 5682.102
130.0980682373047 0 851.7973
130.10589599609375 0 1614.1987
131.08990478515625 0 772.8368
133.097412109375 0 556.7279
136.076171875 0 865.8467
138.09169006347656 0 2259.238
139.0867462158203 0 544.5909
140.08213806152344 0 12116.177
141.08563232421875 0 726.30457
141.1025390625 0 21209.867
141.18487548828125 0 460.90274
142.10008239746094 0 793.6287
142.10586547851562 0 5108.814
143.1182098388672 0 75480.59 a 1
144.11575317382812 0 926.1035
144.12158203125 0 5729.986
145.0974578857422 0 4523.265
146.10092163085938 0 676.12146
147.1130828857422 0 6753.6465
148.11647033691406 0 1350.7842
148.9473876953125 0 570.685
149.02365112304688 0 8220.259
150.02700805664062 0 5630.6406
155.08180236816406 0 2621.6118
155.11814880371094 0 9486.149
156.121337890625 0 849.1093
157.09722900390625 0 1450.5024
157.10877990722656 0 2406.5833
157.13381958007812 0 43815.14
158.09263610839844 0 9743.188 y Ammonia loss 7
158.1001739501953 0 539.38794
158.13052368164062 0 1111.452
158.13723754882812 0 12433.291
159.09642028808594 0 467.77988
159.1131134033203 0 1563.1111
159.1407928466797 0 519.3971
160.1926727294922 0 432.7667
169.09744262695312 0 21166.201
170.1008758544922 0 7090.1445
171.11309814453125 0 65783.02 b 1
172.10960388183594 0 620.82416
172.1165008544922 0 5821.3896
173.09231567382812 0 10357.174
173.12875366210938 0 77144.016
174.09661865234375 0 596.5308
174.1249542236328 0 626.3202
174.13214111328125 0 6818.027
175.11923217773438 0 23228.588 y 7
176.1225128173828 0 1526.2135
181.09683227539062 0 541.55804
183.1130828857422 0 16242.858
183.14950561523438 0 21157.66
184.1165771484375 0 1759.4955
184.15289306640625 0 8493.693
185.10369873046875 0 1834.5991
185.12872314453125 0 9348.789
185.15589904785156 0 658.59863
186.13209533691406 0 2899.1133
186.61700439453125 0 496.29376
187.1079559326172 0 3934.3618
194.12904357910156 0 773.65314
195.11325073242188 0 3082.4714
197.0923614501953 0 863.701
197.1278076171875 0 509.43152
199.14468383789062 0 1184.0283
201.12359619140625 0 27450.994
202.12704467773438 0 2680.4924
209.12860107421875 0 1102.6259
210.1238555908203 0 1642.5452
211.10797119140625 0 3782.184
211.14439392089844 0 10180.056
211.15478515625 0 879.4385
212.1395721435547 0 6401.62
212.14813232421875 0 3363.0908
213.12368774414062 0 1946.6952
213.142578125 0 866.3746
215.13931274414062 0 1735.6632
226.11953735351562 0 727.6621
226.15530395507812 0 10164.649 a Water loss 2
227.10284423828125 0 724.75336
227.1393280029297 0 4846.117
227.15887451171875 0 986.2494
228.13462829589844 0 2673.127 y Ammonia loss 4
229.1195831298828 0 721.16187
230.1501922607422 0 4272.538 d 2
238.1198272705078 0 638.24585
240.1343994140625 0 4549.723
241.13792419433594 0 1574.3297
243.1343231201172 0 1192.3711
243.6501922607422 0 1968.4387
244.15159606933594 0 1513.5013
244.16476440429688 0 911.5749 a 2
252.13478088378906 0 1364.9609
254.1501007080078 0 28512.156 b Water loss 2
255.1454620361328 0 69851.97 y Ammonia loss 6
256.1485595703125 0 6735.023
257.1493225097656 0 580.93146
258.14501953125 0 13830.552
259.1484375 0 5130.491
260.1495056152344 0 524.0437
264.1694030761719 0 928.861
265.17303466796875 0 853.9135
266.15008544921875 0 3844.4429
267.1526794433594 0 1560.17
270.1448669433594 0 10686.349
271.1482238769531 0 1298.8896
271.17681884765625 0 828.5903
272.159912109375 0 11611.368 b 2
272.17205810546875 0 28586.498 y 6
273.16229248046875 0 1729.3885
273.1750793457031 0 2942.991
276.1556396484375 0 11254.051
277.1587219238281 0 5172.339
277.17572021484375 0 1213.8613 b Water loss 5
278.16070556640625 0 1391.6353
280.1285705566406 0 724.3393
282.1812744140625 0 3056.4746
283.1847839355469 0 695.66516
286.18182373046875 0 2446.6343 b 5
288.15557861328125 0 9651.9
293.1001281738281 0 1599.0648
297.1741027832031 0 638.9224
298.140380859375 0 1745.6316
299.2086486816406 0 929.61505
306.1795349121094 0 4927.4775
306.68121337890625 0 1532.0886
307.9854736328125 0 629.4343
312.19244384765625 0 3242.9668
313.1924133300781 0 539.89056
316.8780822753906 0 646.23724
318.69488525390625 0 621.46967
319.68463134765625 0 764.3898
323.1719055175781 0 1711.9601
325.2237548828125 0 12368.369
326.22711181640625 0 2390.7234
327.69903564453125 0 30033.434 y Water loss 2
328.19744873046875 0 50071.566 y Ammonia loss 2
328.6983337402344 0 16055.118
329.19964599609375 0 4356.976
332.69305419921875 0 842.2313
336.7056884765625 0 3128.692 y 2
337.2079162597656 0 1472.3214
340.1870422363281 0 9296.414
341.18768310546875 0 7453.1294
341.6981201171875 0 1014.3136
342.1869812011719 0 1125.5171
343.23455810546875 0 1976.9741
349.2352294921875 0 686.9725
350.9280090332031 0 868.8852
351.2388916015625 0 1311.0051
352.2422180175781 0 1732.4615
353.218505859375 0 8328.354 b Water loss 3
354.2220764160156 0 1354.6301
355.2336730957031 0 1179.4158
358.197265625 0 1824.047
359.1947021484375 0 2170.654
363.21893310546875 0 30637.969 y Water loss 1
363.7202453613281 0 12125.926
364.22174072265625 0 2984.1196
365.21624755859375 0 576.5381
367.2468566894531 0 610.4277
368.2291564941406 0 4223.756 y Ammonia loss 5
369.2330322265625 0 760.7314
371.22894287109375 0 896.7481 b 3
374.0701599121094 0 538.6154
379.2341003417969 0 2018.8752
379.6829528808594 0 566.3865
380.236572265625 0 2053.1323
383.22906494140625 0 2357.838
384.2312316894531 0 1006.7671
385.21331787109375 0 801.47925
385.2559509277344 0 16637.031 y 5
386.2588195800781 0 3579.2588
389.2395935058594 0 3064.261
390.2435607910156 0 2918.586
391.2324523925781 0 6052.3926
391.7337341308594 0 4176.092
401.2401123046875 0 681.58844
403.749267578125 0 983.735
403.8113098144531 0 1228.7172
408.2607727050781 0 690.4451
411.22491455078125 0 1719.3191
412.2271728515625 0 1859.9021
412.7534484863281 0 23787.389 Precursor Water loss
413.25445556640625 0 8544.732
413.755859375 0 3484.3564
414.2596435546875 0 1148.3062
420.8567810058594 0 1147.1742
422.2396545410156 0 1085.4401
426.2713928222656 0 1279.9653
436.2569580078125 0 2619.7827
439.8407897949219 0 756.7151
442.2774963378906 0 1524.8723
454.2740783691406 0 2754.7507 y Water loss 4
455.26568603515625 0 796.12885 y Ammonia loss 4
468.28350830078125 0 711.4502
469.28631591796875 0 1435.6218
472.2882385253906 0 178029.38 y 4
473.2908935546875 0 42451.848
474.2937316894531 0 6671.6514
480.210693359375 0 596.63214
482.2736511230469 0 1800.0215
486.29254150390625 0 26006.61
487.2955322265625 0 31500.31
488.2981262207031 0 10173.558
489.2992858886719 0 1226.8253
508.27593994140625 0 1949.904
509.28021240234375 0 3077.8513
535.3252563476562 0 879.11115
540.3192749023438 0 711.3407
553.3433837890625 0 1709.8198 b Water loss 5
554.3365478515625 0 884.8238 y Ammonia loss 3
557.32861328125 0 4275.465
558.3318481445312 0 8558.56
559.3363647460938 0 2367.002
571.356201171875 0 59879.957 b 5
572.3590087890625 0 16529.531
573.3617553710938 0 3088.1924
581.34033203125 0 1271.3911
592.213134765625 0 716.9249
597.330810546875 0 649.7881
611.3499145507812 0 2695.9841
628.3770141601562 0 5353.2026
629.3805541992188 0 1809.9824
637.3733520507812 0 2254.4238
638.36279296875 0 1247.6497
654.3845825195312 0 14750.501 y Water loss 2
655.3851318359375 0 39780.555 y Ammonia loss 2
656.3875732421875 0 16072.583
657.390625 0 3023.7385
664.3770141601562 0 1045.2202
672.4036254882812 0 142793.62 y 2
673.4066772460938 0 50887.027
674.4090576171875 0 11835.711
682.3884887695312 0 10066.848
683.3888549804688 0 3061.5967
684.3911743164062 0 876.74426
699.4120483398438 0 924.9248
725.4298095703125 0 17139.307 y Water loss 1
726.432373046875 0 6048.558
727.4322509765625 0 1950.6215
735.4121704101562 0 731.1998
743.4404296875 0 55238.38 y 1
744.4434814453125 0 24323.564
745.4465942382812 0 4486.4595
753.4224243164062 0 1628.2039
860.6271362304688 0 709.9711
1260.06591796875 0 621.9833
1308.9144287109375 0 577.319
1940.978759765625 0 642.99664
2008.1300048828125 0 608.5299
2544.7314453125 0 729.3745
2753.360107421875 0 699.95465
3080.31298828125 0 1114.872
3460.8408203125 0 838.8961

Spectrum Details

|  |  |
| --- | --- |
| Matched peaks? Matched peaks The total absolute number of peaks matched. Additionally in brackets the total fraction of peaks matched and the total number of peaks is shown. | 34 (12.73% of 267) |
| FDR? FDR The false discovery rate estimated for this peptide. It is calculated by matching all theoretical fragments with a non-integer shift with the raw peaks for this spectrum. This is done with 40 different shifts. The resulting percentage is the average number of annotated peaks over the number of annotated peaks with the correct spectrum. | 0.14% |
| Satellite FDR? Satellite FDR See the FDR for details on its calculation. This satellite ion specific FDR only contains the satellite ions (d/w) for I/L/J positions. | - |
| PSM Score? PSM Score The PSM Score as given by Hecklib to this annotated spectrum. It is shown with three significant figures. | 418 |

#### Spectrum 6819? Spectrum 6819 The raw spectrum of this peptide as annotated by Hecklib. The fragments are coloured according to ion type (see legend). Any peaks with a star '\*' as text can be hovered over to see the full details, first the ion type second the mass shift type. By hovering over the amino acids in the peptide or ions in the legend the corresponding peaks are highlighted. By toggling the 'Unassigned' label you can turn the background (unassigned) peaks on or off in the plot. By updating the slider in the Ion legend you can update the spectrum to only show the top X% of the peaks with labels. The top X% means any peak that is within X% of the highest intensity. By dragging in the spectrum you can zoom in to a specific part of the spectrum and use 'Zoom Out' to get back to the original zoom level. The annotation of the spectrum is based on the given sequence in the peptides file and is done with different software so inconsistencies are likely. The peaks are annotated based on the given sequence, with 20 ppm tolerance.

Copy Data

##### Spectrum 6819 (TSV)

###### Preview

```
Loading example...
```

*Click on the button to copy the data to your clipboard.*

Mz MinMz MaxIntensity Max

WidthHeightPeptide font sizePeptide stroke widthSpectrum font sizeSpectrum stroke widthCompact peptide

Ion legend

wxyz

abcd

OtherUnassignedIon^Charge^_Position_Show for top:%

VATVSJPR

05.68e+31.14e+41.70e+42.27e+4

Zoom Out

a^+1^_2_y^+1^_1_b^+1^_2_y^+1^_1_a^+1^_3_b^+1^_3_y^+1^_2_b^+1^_3_y^+1^_2_y^+2^_6_y^+2^_6_b^+1^_4_y^+2^_7_y^+1^_3_\*y^+1^_4_b^+1^_6_y^+1^_6_y^+1^_6_y^+1^_7_y^+1^_7_

0777155523323110

Fragment Matches Table

Show background peaks

| Position | Ion type | Intensity | mz Theoretical | mz Error (Th) | mz Error (ppm) | Charge | Series Number |
| --- | --- | --- | --- | --- | --- | --- | --- |
| - | - | 727.8 | 120.1 | - | - | 0 | - |
| - | - | 344.5 | 120.1 | - | - | 0 | - |
| - | - | 335.4 | 120.9 | - | - | 0 | - |
| - | - | 371.1 | 123 | - | - | 0 | - |
| - | - | 439.6 | 126 | - | - | 0 | - |
| - | - | 559.3 | 127.1 | - | - | 0 | - |
| - | - | 355.9 | 128.7 | - | - | 0 | - |
| - | - | 1177 | 129 | - | - | 0 | - |
| - | - | 2591 | 129.1 | - | - | 0 | - |
| - | - | 1065 | 130.1 | - | - | 0 | - |
| - | - | 429.5 | 134.1 | - | - | 0 | - |
| - | - | 615.5 | 136 | - | - | 0 | - |
| - | - | 660.4 | 136.1 | - | - | 0 | - |
| - | - | 383.6 | 137.3 | - | - | 0 | - |
| - | - | 1151 | 140.1 | - | - | 0 | - |
| - | - | 1951 | 141.1 | - | - | 0 | - |
| - | - | 553.9 | 142.1 | - | - | 0 | - |
| - | - | 664.7 | 143.1 | - | - | 0 | - |
| 2 | a | 8616 | 143.1 | 3.065E-05 | 0.2141 | +1 | 2 |
| - | - | 726 | 144.1 | - | - | 0 | - |
| - | - | 491 | 145.1 | - | - | 0 | - |
| - | - | 864.5 | 147.1 | - | - | 0 | - |
| - | - | 1.111E+04 | 149 | - | - | 0 | - |
| - | - | 6135 | 150 | - | - | 0 | - |
| - | - | 769.5 | 151 | - | - | 0 | - |
| - | - | 392 | 151.2 | - | - | 0 | - |
| - | - | 908.7 | 155.1 | - | - | 0 | - |
| - | - | 601.8 | 157 | - | - | 0 | - |
| - | - | 442.7 | 157.1 | - | - | 0 | - |
| - | - | 635.6 | 157.1 | - | - | 0 | - |
| - | - | 6155 | 157.1 | - | - | 0 | - |
| 8 | y | 1488 | 158.1 | 0.00024 | 1.518 | +1 | 1 |
| - | - | 2073 | 158.1 | - | - | 0 | - |
| - | - | 442.7 | 159.1 | - | - | 0 | - |
| - | - | 493.4 | 159.1 | - | - | 0 | - |
| - | - | 644.1 | 167 | - | - | 0 | - |
| - | - | 2934 | 169.1 | - | - | 0 | - |
| 2 | b | 7758 | 171.1 | 0.0001027 | 0.6004 | +1 | 2 |
| - | - | 1004 | 173.1 | - | - | 0 | - |
| - | - | 9684 | 173.1 | - | - | 0 | - |
| - | - | 3078 | 173.5 | - | - | 0 | - |
| - | - | 585.4 | 174.1 | - | - | 0 | - |
| - | - | 432 | 175.1 | - | - | 0 | - |
| 8 | y | 3508 | 175.1 | 8.62E-05 | 0.4922 | +1 | 1 |
| - | - | 506.6 | 180.7 | - | - | 0 | - |
| - | - | 1314 | 183.1 | - | - | 0 | - |
| - | - | 2887 | 183.1 | - | - | 0 | - |
| - | - | 974.2 | 184.2 | - | - | 0 | - |
| - | - | 880.2 | 185.1 | - | - | 0 | - |
| - | - | 1335 | 185.1 | - | - | 0 | - |
| - | - | 412.7 | 186.1 | - | - | 0 | - |
| - | - | 490.2 | 199.1 | - | - | 0 | - |
| - | - | 2897 | 201.1 | - | - | 0 | - |
| - | - | 747.2 | 202.1 | - | - | 0 | - |
| - | - | 512.6 | 203 | - | - | 0 | - |
| - | - | 469.7 | 208.9 | - | - | 0 | - |
| - | - | 748.3 | 210.9 | - | - | 0 | - |
| - | - | 1307 | 211.1 | - | - | 0 | - |
| - | - | 499.9 | 212.6 | - | - | 0 | - |
| - | - | 678.3 | 223.1 | - | - | 0 | - |
| 3 | a | 1016 | 226.2 | 8.701E-05 | 0.3847 | +1 | 3 |
| - | - | 634.7 | 240.1 | - | - | 0 | - |
| - | - | 611.6 | 250.5 | - | - | 0 | - |
| 3 | b | 3208 | 254.1 | 0.000336 | 1.322 | +1 | 3 |
| 7 | y | 7398 | 255.1 | 0.0003 | 1.176 | +1 | 2 |
| - | - | 702.5 | 256.1 | - | - | 0 | - |
| - | - | 1961 | 258.1 | - | - | 0 | - |
| - | - | 1167 | 259.1 | - | - | 0 | - |
| - | - | 724.8 | 266.2 | - | - | 0 | - |
| - | - | 975.1 | 270.1 | - | - | 0 | - |
| 3 | b | 926.6 | 272.2 | 0.002646 | 9.721 | +1 | 3 |
| 7 | y | 3006 | 272.2 | 5.464E-05 | 0.2008 | +1 | 2 |
| - | - | 550.1 | 274.8 | - | - | 0 | - |
| - | - | 1777 | 276.2 | - | - | 0 | - |
| - | - | 618.6 | 277.2 | - | - | 0 | - |
| - | - | 1349 | 288.2 | - | - | 0 | - |
| - | - | 554.7 | 289.1 | - | - | 0 | - |
| - | - | 1699 | 293.1 | - | - | 0 | - |
| - | - | 1879 | 325.2 | - | - | 0 | - |
| 3 | y | 3088 | 327.7 | 0.002034 | 6.206 | +2 | 6 |
| 3 | y | 7720 | 328.2 | 0.004799 | 14.62 | +2 | 6 |
| - | - | 2810 | 328.7 | - | - | 0 | - |
| - | - | 1157 | 329.2 | - | - | 0 | - |
| - | - | 631.3 | 341.2 | - | - | 0 | - |
| 4 | b | 829.9 | 353.2 | 0.001761 | 4.985 | +1 | 4 |
| 2 | y | 3322 | 363.2 | 0.0009677 | 2.664 | +2 | 7 |
| - | - | 1029 | 363.7 | - | - | 0 | - |
| 6 | y | 2151 | 385.3 | 0.0001037 | 0.2693 | +1 | 3 |
| - | - | 1562 | 391.2 | - | - | 0 | - |
| - | - | 589.9 | 402.2 | - | - | 0 | - |
| - | - | 1299 | 403.8 | - | - | 0 | - |
| 0 | Precursor | 1874 | 412.8 | 0.0001037 | 0.2512 | +2 | -1 |
| - | - | 1423 | 413.3 | - | - | 0 | - |
| - | - | 1185 | 420.9 | - | - | 0 | - |
| - | - | 729.6 | 421.2 | - | - | 0 | - |
| - | - | 625.9 | 430.3 | - | - | 0 | - |
| - | - | 554.9 | 437.6 | - | - | 0 | - |
| - | - | 775.6 | 439.8 | - | - | 0 | - |
| 5 | y | 2.248E+04 | 472.3 | 5.817E-05 | 0.1232 | +1 | 4 |
| - | - | 4716 | 473.3 | - | - | 0 | - |
| - | - | 3593 | 486.3 | - | - | 0 | - |
| - | - | 4289 | 487.3 | - | - | 0 | - |
| - | - | 1697 | 488.3 | - | - | 0 | - |
| - | - | 740 | 509.3 | - | - | 0 | - |
| - | - | 1127 | 557.3 | - | - | 0 | - |
| - | - | 1093 | 558.3 | - | - | 0 | - |
| 6 | b | 7019 | 571.3 | 0.0106 | 18.56 | +1 | 6 |
| - | - | 2024 | 572.4 | - | - | 0 | - |
| - | - | 545.5 | 630.6 | - | - | 0 | - |
| - | - | 1055 | 654.4 | - | - | 0 | - |
| 3 | y | 4647 | 655.4 | 0.008085 | 12.34 | +1 | 6 |
| - | - | 1875 | 656.4 | - | - | 0 | - |
| 3 | y | 1.446E+04 | 672.4 | 0.0008246 | 1.226 | +1 | 6 |
| - | - | 5326 | 673.4 | - | - | 0 | - |
| - | - | 1572 | 674.4 | - | - | 0 | - |
| - | - | 1281 | 682.4 | - | - | 0 | - |
| 2 | y | 2035 | 725.4 | 0.002777 | 3.827 | +1 | 7 |
| - | - | 899 | 726.4 | - | - | 0 | - |
| 2 | y | 5943 | 743.4 | 0.0006459 | 0.8688 | +1 | 7 |
| - | - | 2908 | 744.4 | - | - | 0 | - |
| - | - | 690.7 | 882.7 | - | - | 0 | - |
| - | - | 596.8 | 3079 | - | - | 0 | - |

m/z Charge Intensity FragmentType MassShift Position
120.08086395263672 0 727.8393
120.1489486694336 0 344.46948
120.889892578125 0 335.39078
123.03369140625 0 371.1175
125.98970794677734 0 439.58426
127.0867919921875 0 559.2802
128.67970275878906 0 355.90915
129.01834106445312 0 1176.7406
129.1023406982422 0 2590.5015
130.08633422851562 0 1064.8951
134.14752197265625 0 429.4709
136.02178955078125 0 615.48505
136.07568359375 0 660.37225
137.26123046875 0 383.61172
140.0818634033203 0 1151.4272
141.10240173339844 0 1950.5574
142.1055145263672 0 553.8881
143.1125030517578 0 664.65497
143.11785888671875 0 8615.573 a 1
144.12127685546875 0 725.9689
145.06504821777344 0 490.9659
147.11288452148438 0 864.5432
149.02328491210938 0 11110.227
150.0266876220703 0 6134.7188
151.028076171875 0 769.5017
151.2295684814453 0 392.03702
155.1181182861328 0 908.74396
157.01275634765625 0 601.8158
157.10867309570312 0 442.68338
157.12767028808594 0 635.60583
157.1334991455078 0 6154.943
158.0921630859375 0 1488.4652 y Ammonia loss 7
158.13690185546875 0 2073.4934
159.05177307128906 0 442.72272
159.11257934570312 0 493.381
167.03407287597656 0 644.14453
169.09703063964844 0 2934.3167
171.11270141601562 0 7757.661 b 1
173.09181213378906 0 1003.71075
173.1283721923828 0 9683.561
173.45089721679688 0 3077.8022
174.13180541992188 0 585.38324
175.11195373535156 0 432.04056
175.11886596679688 0 3508.1504 y 7
180.6964874267578 0 506.61902
183.1129913330078 0 1314.127
183.1491241455078 0 2886.9302
184.1525421142578 0 974.1813
185.08087158203125 0 880.1933
185.12840270996094 0 1334.9855
186.0865936279297 0 412.68073
199.14381408691406 0 490.18164
201.12313842773438 0 2897.3884
202.1075439453125 0 747.15967
202.99783325195312 0 512.59796
208.92739868164062 0 469.7492
210.90440368652344 0 748.3196
211.14382934570312 0 1307.3354
212.56504821777344 0 499.8848
223.0575408935547 0 678.31146
226.15509033203125 0 1016.0346 a Water loss 2
240.1343231201172 0 634.66125
250.52955627441406 0 611.63916
254.1495819091797 0 3207.932 b Water loss 2
255.14486694335938 0 7397.7397 y Ammonia loss 6
256.1485900878906 0 702.5129
258.1444091796875 0 1961.2745
259.148193359375 0 1166.9518
266.1512756347656 0 724.7909
270.1443786621094 0 975.0841
272.1578369140625 0 926.5981 b 2
272.1716613769531 0 3006.2595 y 6
274.79339599609375 0 550.0661
276.1553955078125 0 1776.7599
277.1593017578125 0 618.60913
288.154296875 0 1349.0797
289.0934753417969 0 554.7214
293.0995178222656 0 1698.6965
325.2230529785156 0 1879.0747
327.6982727050781 0 3087.595 y Water loss 2
328.1971130371094 0 7719.954 y Ammonia loss 2
328.6969299316406 0 2810.0298
329.1986083984375 0 1157.1995
341.18780517578125 0 631.34375
353.2200927734375 0 829.8853 b Water loss 3
363.2178955078125 0 3321.6611 y Water loss 1
363.7203369140625 0 1029.3827
385.25567626953125 0 2150.7693 y 5
391.232177734375 0 1561.9857
402.2079772949219 0 589.855
403.8135986328125 0 1299.1455
412.753173828125 0 1873.9395 Precursor Water loss
413.25567626953125 0 1423.1753
420.85791015625 0 1184.803
421.1930847167969 0 729.6331
430.3356018066406 0 625.8742
437.5955505371094 0 554.8938
439.8426513671875 0 775.6463
472.2877502441406 0 22477.594 y 4
473.28955078125 0 4716.4155
486.292236328125 0 3593.282
487.2944641113281 0 4289.2607
488.29656982421875 0 1697.1996
509.2785949707031 0 740.0065
557.330322265625 0 1127.2909
558.3313598632812 0 1093.1368
571.3555908203125 0 7019.22 b 5
572.3583984375 0 2023.9984
630.6131591796875 0 545.4516
654.3800659179688 0 1055.1044
655.3854370117188 0 4646.5166 y Ammonia loss 2
656.3870239257812 0 1874.8503
672.403076171875 0 14457.462 y 2
673.40625 0 5326.2827
674.4069213867188 0 1572.43
682.3871459960938 0 1280.9995
725.4276733398438 0 2034.8551 y Water loss 1
726.436767578125 0 899.0249
743.4403686523438 0 5942.9897 y 1
744.4423828125 0 2908.4082
882.6829223632812 0 690.7093
3079.069091796875 0 596.80304

Spectrum Details

|  |  |
| --- | --- |
| Matched peaks? Matched peaks The total absolute number of peaks matched. Additionally in brackets the total fraction of peaks matched and the total number of peaks is shown. | 21 (17.21% of 122) |
| FDR? FDR The false discovery rate estimated for this peptide. It is calculated by matching all theoretical fragments with a non-integer shift with the raw peaks for this spectrum. This is done with 40 different shifts. The resulting percentage is the average number of annotated peaks over the number of annotated peaks with the correct spectrum. | 0.34% |
| Satellite FDR? Satellite FDR See the FDR for details on its calculation. This satellite ion specific FDR only contains the satellite ions (d/w) for I/L/J positions. | - |
| PSM Score? PSM Score The PSM Score as given by Hecklib to this annotated spectrum. It is shown with three significant figures. | 244 |

#### Spectrum 7964? Spectrum 7964 The raw spectrum of this peptide as annotated by Hecklib. The fragments are coloured according to ion type (see legend). Any peaks with a star '\*' as text can be hovered over to see the full details, first the ion type second the mass shift type. By hovering over the amino acids in the peptide or ions in the legend the corresponding peaks are highlighted. By toggling the 'Unassigned' label you can turn the background (unassigned) peaks on or off in the plot. By updating the slider in the Ion legend you can update the spectrum to only show the top X% of the peaks with labels. The top X% means any peak that is within X% of the highest intensity. By dragging in the spectrum you can zoom in to a specific part of the spectrum and use 'Zoom Out' to get back to the original zoom level. The annotation of the spectrum is based on the given sequence in the peptides file and is done with different software so inconsistencies are likely. The peaks are annotated based on the given sequence, with 20 ppm tolerance.

Copy Data

##### Spectrum 7964 (TSV)

###### Preview

```
Loading example...
```

*Click on the button to copy the data to your clipboard.*

Mz MinMz MaxIntensity Max

WidthHeightPeptide font sizePeptide stroke widthSpectrum font sizeSpectrum stroke widthCompact peptide

Ion legend

wxyz

abcd

OtherUnassignedIon^Charge^_Position_Show for top:%

VATVSJPR

03.15e+36.29e+39.44e+31.26e+4

Zoom Out

a^+1^_2_y^+1^_1_b^+1^_2_y^+1^_1_a^+1^_3_b^+1^_3_y^+1^_2_b^+1^_3_y^+1^_2_y^+2^_6_y^+2^_6_y^+2^_7_y^+1^_3_\*y^+1^_4_b^+1^_6_y^+1^_6_y^+1^_6_y^+1^_6_y^+1^_7_y^+1^_7_

0772154423163089

Fragment Matches Table

Show background peaks

| Position | Ion type | Intensity | mz Theoretical | mz Error (Th) | mz Error (ppm) | Charge | Series Number |
| --- | --- | --- | --- | --- | --- | --- | --- |
| - | - | 918.8 | 120 | - | - | 0 | - |
| - | - | 2833 | 120.1 | - | - | 0 | - |
| - | - | 455.1 | 123.1 | - | - | 0 | - |
| - | - | 438.7 | 124 | - | - | 0 | - |
| - | - | 357 | 127.1 | - | - | 0 | - |
| - | - | 641.7 | 128.1 | - | - | 0 | - |
| - | - | 1279 | 129 | - | - | 0 | - |
| - | - | 2237 | 129.1 | - | - | 0 | - |
| - | - | 459.4 | 130.1 | - | - | 0 | - |
| - | - | 1467 | 136.1 | - | - | 0 | - |
| - | - | 436.6 | 136.2 | - | - | 0 | - |
| - | - | 973.8 | 140.1 | - | - | 0 | - |
| - | - | 1910 | 141.1 | - | - | 0 | - |
| 2 | a | 4667 | 143.1 | 0.0001067 | 0.7454 | +1 | 2 |
| - | - | 9113 | 149 | - | - | 0 | - |
| - | - | 437.8 | 149.2 | - | - | 0 | - |
| - | - | 4457 | 150 | - | - | 0 | - |
| - | - | 455.9 | 151 | - | - | 0 | - |
| - | - | 440.2 | 152.1 | - | - | 0 | - |
| - | - | 502.5 | 155.1 | - | - | 0 | - |
| - | - | 581.4 | 155.1 | - | - | 0 | - |
| - | - | 519 | 157 | - | - | 0 | - |
| - | - | 3885 | 157.1 | - | - | 0 | - |
| 8 | y | 561.3 | 158.1 | 0.0001332 | 0.8423 | +1 | 1 |
| - | - | 972.6 | 158.1 | - | - | 0 | - |
| - | - | 420.9 | 160.1 | - | - | 0 | - |
| - | - | 477.1 | 165.1 | - | - | 0 | - |
| - | - | 2091 | 169.1 | - | - | 0 | - |
| 2 | b | 4338 | 171.1 | 0.0001109 | 0.648 | +1 | 2 |
| - | - | 554.5 | 172.1 | - | - | 0 | - |
| - | - | 439.1 | 172.4 | - | - | 0 | - |
| - | - | 837.5 | 173.1 | - | - | 0 | - |
| - | - | 5470 | 173.1 | - | - | 0 | - |
| - | - | 1805 | 173.5 | - | - | 0 | - |
| - | - | 615.3 | 174.1 | - | - | 0 | - |
| 8 | y | 1698 | 175.1 | 0.0002037 | 1.163 | +1 | 1 |
| - | - | 1112 | 183.1 | - | - | 0 | - |
| - | - | 2189 | 183.1 | - | - | 0 | - |
| - | - | 495.3 | 184.6 | - | - | 0 | - |
| - | - | 614.3 | 185.1 | - | - | 0 | - |
| - | - | 941.4 | 185.1 | - | - | 0 | - |
| - | - | 468.7 | 191.3 | - | - | 0 | - |
| - | - | 1866 | 201.1 | - | - | 0 | - |
| - | - | 1078 | 202.1 | - | - | 0 | - |
| - | - | 914.6 | 211.1 | - | - | 0 | - |
| - | - | 742.3 | 212.1 | - | - | 0 | - |
| - | - | 471 | 214 | - | - | 0 | - |
| 3 | a | 1022 | 226.2 | 1.072E-05 | 0.04738 | +1 | 3 |
| - | - | 652.3 | 237.2 | - | - | 0 | - |
| 3 | b | 2263 | 254.1 | 3.086E-05 | 0.1214 | +1 | 3 |
| 7 | y | 6071 | 255.1 | 0.000112 | 0.439 | +1 | 2 |
| - | - | 443.5 | 256.2 | - | - | 0 | - |
| - | - | 1495 | 258.1 | - | - | 0 | - |
| - | - | 1029 | 262.1 | - | - | 0 | - |
| - | - | 541.9 | 270.1 | - | - | 0 | - |
| 3 | b | 825.6 | 272.2 | 0.00289 | 10.62 | +1 | 3 |
| 7 | y | 2092 | 272.2 | 5.464E-05 | 0.2008 | +1 | 2 |
| - | - | 523.3 | 273.5 | - | - | 0 | - |
| - | - | 1482 | 276.2 | - | - | 0 | - |
| - | - | 616.3 | 284.7 | - | - | 0 | - |
| - | - | 679.1 | 288.2 | - | - | 0 | - |
| - | - | 522 | 298.2 | - | - | 0 | - |
| - | - | 526.5 | 308 | - | - | 0 | - |
| - | - | 514.4 | 309.3 | - | - | 0 | - |
| 3 | y | 2110 | 327.7 | 0.001515 | 4.622 | +2 | 6 |
| 3 | y | 4190 | 328.2 | 0.004585 | 13.97 | +2 | 6 |
| - | - | 1942 | 328.7 | - | - | 0 | - |
| - | - | 558.5 | 333.1 | - | - | 0 | - |
| - | - | 644.6 | 340.2 | - | - | 0 | - |
| 2 | y | 1578 | 363.2 | 0.0001132 | 0.3116 | +2 | 7 |
| - | - | 937.8 | 363.7 | - | - | 0 | - |
| 6 | y | 1162 | 385.3 | 0.0004151 | 1.077 | +1 | 3 |
| - | - | 2277 | 403.8 | - | - | 0 | - |
| 0 | Precursor | 1769 | 412.8 | 0.0002015 | 0.4881 | +2 | -1 |
| - | - | 1067 | 413.3 | - | - | 0 | - |
| - | - | 536.8 | 414.1 | - | - | 0 | - |
| - | - | 649.9 | 439.8 | - | - | 0 | - |
| - | - | 740.5 | 469.2 | - | - | 0 | - |
| 5 | y | 1.246E+04 | 472.3 | 0.000308 | 0.6522 | +1 | 4 |
| - | - | 2427 | 473.3 | - | - | 0 | - |
| - | - | 2188 | 486.3 | - | - | 0 | - |
| - | - | 2659 | 487.3 | - | - | 0 | - |
| - | - | 1153 | 488.3 | - | - | 0 | - |
| - | - | 635.6 | 514.3 | - | - | 0 | - |
| - | - | 564.4 | 559.3 | - | - | 0 | - |
| - | - | 611.2 | 568.3 | - | - | 0 | - |
| 6 | b | 3413 | 571.3 | 0.0103 | 18.02 | +1 | 6 |
| - | - | 879.5 | 572.4 | - | - | 0 | - |
| - | - | 596.6 | 640.1 | - | - | 0 | - |
| 3 | y | 1045 | 654.4 | 0.008082 | 12.35 | +1 | 6 |
| 3 | y | 2703 | 655.4 | 0.008513 | 12.99 | +1 | 6 |
| - | - | 1289 | 656.4 | - | - | 0 | - |
| 3 | y | 8447 | 672.4 | 0.0003974 | 0.591 | +1 | 6 |
| - | - | 2760 | 673.4 | - | - | 0 | - |
| - | - | 560.9 | 682.4 | - | - | 0 | - |
| - | - | 791.1 | 697.4 | - | - | 0 | - |
| 2 | y | 1045 | 725.4 | 0.00235 | 3.24 | +1 | 7 |
| - | - | 545.4 | 736.3 | - | - | 0 | - |
| 2 | y | 4222 | 743.4 | 0.0003306 | 0.4447 | +1 | 7 |
| - | - | 1538 | 744.4 | - | - | 0 | - |
| - | - | 656.5 | 3035 | - | - | 0 | - |
| - | - | 699.8 | 3058 | - | - | 0 | - |

m/z Charge Intensity FragmentType MassShift Position
120.04470825195312 0 918.83295
120.08094024658203 0 2832.63
123.11709594726562 0 455.07843
124.00328063964844 0 438.73364
127.07647705078125 0 357.01465
128.10723876953125 0 641.65137
129.01841735839844 0 1279.2621
129.1023712158203 0 2236.9888
130.08641052246094 0 459.3525
136.07594299316406 0 1467.1823
136.17108154296875 0 436.57498
140.08200073242188 0 973.75073
141.10220336914062 0 1910.0385
143.1179962158203 0 4667.142 a 1
149.02346801757812 0 9112.63
149.1987762451172 0 437.84586
150.02679443359375 0 4456.8364
151.02838134765625 0 455.94653
152.0889892578125 0 440.20786
155.0817108154297 0 502.49988
155.11770629882812 0 581.4178
157.01327514648438 0 518.9914
157.13368225097656 0 3885.189
158.09226989746094 0 561.2629 y Ammonia loss 7
158.13674926757812 0 972.59686
160.06858825683594 0 420.94214
165.09112548828125 0 477.061
169.0973663330078 0 2090.8096
171.1129150390625 0 4337.8965 b 1
172.11614990234375 0 554.46344
172.39598083496094 0 439.1224
173.09170532226562 0 837.4559
173.12860107421875 0 5470.289
173.45150756835938 0 1805.3721
174.0548553466797 0 615.28174
175.11915588378906 0 1697.6364 y 7
183.11270141601562 0 1111.748
183.14918518066406 0 2188.5745
184.58103942871094 0 495.3352
185.08114624023438 0 614.2585
185.12879943847656 0 941.3666
191.29983520507812 0 468.6583
201.12367248535156 0 1865.5359
202.10716247558594 0 1078.28
211.14414978027344 0 914.59906
212.13877868652344 0 742.32166
213.98338317871094 0 470.9715
226.15501403808594 0 1022.08307 a Water loss 2
237.16058349609375 0 652.28314
254.14988708496094 0 2263.0918 b Water loss 2
255.14527893066406 0 6070.8022 y Ammonia loss 6
256.15045166015625 0 443.48077
258.1446228027344 0 1495.4248
262.1184997558594 0 1028.516
270.146240234375 0 541.8504
272.1575927734375 0 825.6185 b 2
272.1716613769531 0 2092.0183 y 6
273.48809814453125 0 523.26526
276.15496826171875 0 1481.9634
284.68206787109375 0 616.29926
288.1564636230469 0 679.0897
298.2053527832031 0 521.98926
307.98504638671875 0 526.49646
309.3328857421875 0 514.35315
327.69879150390625 0 2110.2988 y Water loss 2
328.1968994140625 0 4189.5703 y Ammonia loss 2
328.697998046875 0 1941.5713
333.1142272949219 0 558.52594
340.1863708496094 0 644.5657
363.21875 0 1578.0192 y Water loss 1
363.7198181152344 0 937.78723
385.2561950683594 0 1161.7114 y 5
403.81329345703125 0 2277.2314
412.75286865234375 0 1769.2645 Precursor Water loss
413.2552795410156 0 1066.766
414.05194091796875 0 536.757
439.843994140625 0 649.87634
469.2478942871094 0 740.5424
472.2881164550781 0 12461.631 y 4
473.29046630859375 0 2426.5293
486.290771484375 0 2187.9587
487.2952880859375 0 2658.6956
488.2998962402344 0 1153.0043
514.2918090820312 0 635.6331
559.33203125 0 564.4023
568.3204345703125 0 611.2184
571.3552856445312 0 3413.22 b 5
572.3585205078125 0 879.52423
640.0850830078125 0 596.63947
654.38525390625 0 1045.489 y Water loss 2
655.3858642578125 0 2702.76 y Ammonia loss 2
656.38916015625 0 1289.4447
672.4035034179688 0 8447.353 y 2
673.4060668945312 0 2760.271
682.3871459960938 0 560.85474
697.3628540039062 0 791.0578
725.4328002929688 0 1044.9979 y Water loss 1
736.2877807617188 0 545.4291
743.4413452148438 0 4221.627 y 1
744.4439086914062 0 1538.4294
3035.363525390625 0 656.4573
3057.921875 0 699.84296

Spectrum Details

|  |  |
| --- | --- |
| Matched peaks? Matched peaks The total absolute number of peaks matched. Additionally in brackets the total fraction of peaks matched and the total number of peaks is shown. | 21 (20.59% of 102) |
| FDR? FDR The false discovery rate estimated for this peptide. It is calculated by matching all theoretical fragments with a non-integer shift with the raw peaks for this spectrum. This is done with 40 different shifts. The resulting percentage is the average number of annotated peaks over the number of annotated peaks with the correct spectrum. | 0.23% |
| Satellite FDR? Satellite FDR See the FDR for details on its calculation. This satellite ion specific FDR only contains the satellite ions (d/w) for I/L/J positions. | - |
| PSM Score? PSM Score The PSM Score as given by Hecklib to this annotated spectrum. It is shown with three significant figures. | 244 |

#### Spectrum 6296? Spectrum 6296 The raw spectrum of this peptide as annotated by Hecklib. The fragments are coloured according to ion type (see legend). Any peaks with a star '\*' as text can be hovered over to see the full details, first the ion type second the mass shift type. By hovering over the amino acids in the peptide or ions in the legend the corresponding peaks are highlighted. By toggling the 'Unassigned' label you can turn the background (unassigned) peaks on or off in the plot. By updating the slider in the Ion legend you can update the spectrum to only show the top X% of the peaks with labels. The top X% means any peak that is within X% of the highest intensity. By dragging in the spectrum you can zoom in to a specific part of the spectrum and use 'Zoom Out' to get back to the original zoom level. The annotation of the spectrum is based on the given sequence in the peptides file and is done with different software so inconsistencies are likely. The peaks are annotated based on the given sequence, with 20 ppm tolerance.

Copy Data

##### Spectrum 6296 (TSV)

###### Preview

```
Loading example...
```

*Click on the button to copy the data to your clipboard.*

Mz MinMz MaxIntensity Max

WidthHeightPeptide font sizePeptide stroke widthSpectrum font sizeSpectrum stroke widthCompact peptide

Ion legend

wxyz

abcd

OtherUnassignedIon^Charge^_Position_Show for top:%

VATVSJPR

07.52e+31.50e+42.26e+43.01e+4

Zoom Out

a^+1^_2_y^+1^_1_b^+1^_2_y^+1^_1_a^+1^_3_d^+1^_3_b^+1^_3_y^+1^_2_b^+1^_3_y^+1^_2_y^+2^_6_y^+2^_6_b^+1^_4_y^+2^_7_y^+1^_3_y^+1^_3_\*y^+1^_4_b^+1^_6_y^+1^_6_y^+1^_6_y^+1^_6_y^+1^_7_y^+1^_7_

0830166024903320

Fragment Matches Table

Show background peaks

| Position | Ion type | Intensity | mz Theoretical | mz Error (Th) | mz Error (ppm) | Charge | Series Number |
| --- | --- | --- | --- | --- | --- | --- | --- |
| - | - | 1053 | 120.1 | - | - | 0 | - |
| - | - | 455.5 | 125.1 | - | - | 0 | - |
| - | - | 592 | 127.1 | - | - | 0 | - |
| - | - | 699.6 | 128.1 | - | - | 0 | - |
| - | - | 1098 | 129 | - | - | 0 | - |
| - | - | 2921 | 129.1 | - | - | 0 | - |
| - | - | 1019 | 130.1 | - | - | 0 | - |
| - | - | 1973 | 140.1 | - | - | 0 | - |
| - | - | 422.9 | 141.1 | - | - | 0 | - |
| - | - | 4436 | 141.1 | - | - | 0 | - |
| - | - | 771.3 | 142.1 | - | - | 0 | - |
| 2 | a | 1.39E+04 | 143.1 | 0.0001982 | 1.385 | +1 | 2 |
| - | - | 1025 | 144.1 | - | - | 0 | - |
| - | - | 1026 | 145.1 | - | - | 0 | - |
| - | - | 720.4 | 147.1 | - | - | 0 | - |
| - | - | 1046 | 149 | - | - | 0 | - |
| - | - | 1.349E+04 | 149 | - | - | 0 | - |
| - | - | 6326 | 150 | - | - | 0 | - |
| - | - | 510.1 | 151 | - | - | 0 | - |
| - | - | 649.4 | 153.1 | - | - | 0 | - |
| - | - | 1707 | 155.1 | - | - | 0 | - |
| - | - | 9173 | 157.1 | - | - | 0 | - |
| 8 | y | 1720 | 158.1 | 0.000111 | 0.7019 | +1 | 1 |
| - | - | 2190 | 158.1 | - | - | 0 | - |
| - | - | 3248 | 169.1 | - | - | 0 | - |
| - | - | 1174 | 170.1 | - | - | 0 | - |
| - | - | 610.2 | 171.1 | - | - | 0 | - |
| 2 | b | 1.23E+04 | 171.1 | 0.0001567 | 0.9155 | +1 | 2 |
| - | - | 640.1 | 171.1 | - | - | 0 | - |
| - | - | 840.8 | 172.1 | - | - | 0 | - |
| - | - | 1692 | 173.1 | - | - | 0 | - |
| - | - | 1.328E+04 | 173.1 | - | - | 0 | - |
| - | - | 1045 | 174.1 | - | - | 0 | - |
| 8 | y | 4561 | 175.1 | 0.0001579 | 0.9019 | +1 | 1 |
| - | - | 1982 | 183.1 | - | - | 0 | - |
| - | - | 3865 | 183.1 | - | - | 0 | - |
| - | - | 862.8 | 184.2 | - | - | 0 | - |
| - | - | 1320 | 185.1 | - | - | 0 | - |
| - | - | 2388 | 185.1 | - | - | 0 | - |
| - | - | 410 | 185.8 | - | - | 0 | - |
| - | - | 718.4 | 186.1 | - | - | 0 | - |
| - | - | 537.9 | 186.8 | - | - | 0 | - |
| - | - | 739.6 | 187.1 | - | - | 0 | - |
| - | - | 425.1 | 187.8 | - | - | 0 | - |
| - | - | 527.9 | 195.1 | - | - | 0 | - |
| - | - | 5004 | 201.1 | - | - | 0 | - |
| - | - | 529.3 | 202.1 | - | - | 0 | - |
| - | - | 712 | 211.1 | - | - | 0 | - |
| - | - | 1475 | 211.1 | - | - | 0 | - |
| - | - | 1757 | 212.1 | - | - | 0 | - |
| 3 | a | 1810 | 226.2 | 8.084E-05 | 0.3574 | +1 | 3 |
| - | - | 699.5 | 227.1 | - | - | 0 | - |
| 3 | d | 738.1 | 230.1 | 0.0005337 | 2.319 | +1 | 3 |
| - | - | 516.8 | 230.5 | - | - | 0 | - |
| - | - | 671.5 | 237.1 | - | - | 0 | - |
| - | - | 895.8 | 240.1 | - | - | 0 | - |
| 3 | b | 4470 | 254.1 | 0.0001217 | 0.479 | +1 | 3 |
| - | - | 434.7 | 254.2 | - | - | 0 | - |
| 7 | y | 1.289E+04 | 255.1 | 0.000112 | 0.439 | +1 | 2 |
| - | - | 1437 | 256.1 | - | - | 0 | - |
| - | - | 2636 | 258.1 | - | - | 0 | - |
| - | - | 800.9 | 259.1 | - | - | 0 | - |
| - | - | 583.3 | 264.4 | - | - | 0 | - |
| - | - | 791.7 | 266.1 | - | - | 0 | - |
| - | - | 2500 | 270.1 | - | - | 0 | - |
| - | - | 1106 | 272.1 | - | - | 0 | - |
| 3 | b | 1021 | 272.2 | 0.002646 | 9.721 | +1 | 3 |
| 7 | y | 5968 | 272.2 | 6.743E-05 | 0.2477 | +1 | 2 |
| - | - | 690.6 | 273.2 | - | - | 0 | - |
| - | - | 596.9 | 275.1 | - | - | 0 | - |
| - | - | 1661 | 276.2 | - | - | 0 | - |
| - | - | 1032 | 277.2 | - | - | 0 | - |
| - | - | 860.4 | 282.2 | - | - | 0 | - |
| - | - | 1163 | 288.2 | - | - | 0 | - |
| - | - | 611.3 | 290 | - | - | 0 | - |
| - | - | 817.3 | 291 | - | - | 0 | - |
| - | - | 574.6 | 298.6 | - | - | 0 | - |
| - | - | 632.2 | 306.2 | - | - | 0 | - |
| - | - | 1280 | 312.2 | - | - | 0 | - |
| - | - | 538.8 | 314.9 | - | - | 0 | - |
| - | - | 697.5 | 316.9 | - | - | 0 | - |
| - | - | 1871 | 322.2 | - | - | 0 | - |
| - | - | 1706 | 325.2 | - | - | 0 | - |
| 3 | y | 4957 | 327.7 | 0.001515 | 4.622 | +2 | 6 |
| 3 | y | 8280 | 328.2 | 0.004646 | 14.16 | +2 | 6 |
| - | - | 3531 | 328.7 | - | - | 0 | - |
| - | - | 822.2 | 329.2 | - | - | 0 | - |
| - | - | 1410 | 340.2 | - | - | 0 | - |
| - | - | 1212 | 341.2 | - | - | 0 | - |
| - | - | 692.4 | 349.2 | - | - | 0 | - |
| - | - | 1091 | 350.9 | - | - | 0 | - |
| - | - | 682.3 | 352.9 | - | - | 0 | - |
| 4 | b | 1482 | 353.2 | 0.000772 | 2.186 | +1 | 4 |
| 2 | y | 5196 | 363.2 | 3.941E-05 | 0.1085 | +2 | 7 |
| - | - | 1359 | 363.7 | - | - | 0 | - |
| - | - | 7848 | 366.2 | - | - | 0 | - |
| - | - | 904.1 | 367.2 | - | - | 0 | - |
| - | - | 819.6 | 368.2 | - | - | 0 | - |
| 6 | y | 1021 | 368.2 | 0.001112 | 3.02 | +1 | 3 |
| - | - | 2427 | 384.2 | - | - | 0 | - |
| 6 | y | 2143 | 385.3 | 0.0003846 | 0.9982 | +1 | 3 |
| - | - | 646.2 | 386.3 | - | - | 0 | - |
| - | - | 607.6 | 386.9 | - | - | 0 | - |
| - | - | 519.7 | 390.2 | - | - | 0 | - |
| - | - | 1046 | 391.2 | - | - | 0 | - |
| - | - | 8412 | 402.2 | - | - | 0 | - |
| - | - | 2159 | 403.8 | - | - | 0 | - |
| 0 | Precursor | 3598 | 412.8 | 0.0005309 | 1.286 | +2 | -1 |
| - | - | 966.1 | 413.3 | - | - | 0 | - |
| - | - | 1227 | 420.9 | - | - | 0 | - |
| - | - | 741 | 421.2 | - | - | 0 | - |
| - | - | 629.6 | 421.3 | - | - | 0 | - |
| - | - | 713.7 | 422.2 | - | - | 0 | - |
| - | - | 3737 | 422.2 | - | - | 0 | - |
| - | - | 1410 | 439.8 | - | - | 0 | - |
| 5 | y | 2.977E+04 | 472.3 | 9.442E-05 | 0.1999 | +1 | 4 |
| - | - | 6776 | 473.3 | - | - | 0 | - |
| - | - | 1028 | 474.3 | - | - | 0 | - |
| - | - | 3751 | 486.3 | - | - | 0 | - |
| - | - | 6534 | 487.3 | - | - | 0 | - |
| - | - | 1540 | 488.3 | - | - | 0 | - |
| - | - | 1493 | 558.3 | - | - | 0 | - |
| 6 | b | 1.004E+04 | 571.3 | 0.01109 | 19.41 | +1 | 6 |
| - | - | 3063 | 572.4 | - | - | 0 | - |
| - | - | 1048 | 628.4 | - | - | 0 | - |
| - | - | 531.9 | 644.4 | - | - | 0 | - |
| 3 | y | 3037 | 654.4 | 0.008815 | 13.47 | +1 | 6 |
| 3 | y | 7103 | 655.4 | 0.00778 | 11.87 | +1 | 6 |
| - | - | 2614 | 656.4 | - | - | 0 | - |
| 3 | y | 2.372E+04 | 672.4 | 0.0004584 | 0.6818 | +1 | 6 |
| - | - | 8069 | 673.4 | - | - | 0 | - |
| - | - | 1987 | 674.4 | - | - | 0 | - |
| - | - | 2031 | 682.4 | - | - | 0 | - |
| - | - | 995.9 | 683.4 | - | - | 0 | - |
| - | - | 578.7 | 691 | - | - | 0 | - |
| - | - | 618.2 | 692.2 | - | - | 0 | - |
| 2 | y | 2976 | 725.4 | 0.0004583 | 0.6318 | +1 | 7 |
| 2 | y | 8807 | 743.4 | 0.0009511 | 1.279 | +1 | 7 |
| - | - | 4234 | 744.4 | - | - | 0 | - |
| - | - | 631.8 | 752.6 | - | - | 0 | - |
| - | - | 623.6 | 1544 | - | - | 0 | - |
| - | - | 642.3 | 1644 | - | - | 0 | - |
| - | - | 673 | 2095 | - | - | 0 | - |
| - | - | 665.6 | 3287 | - | - | 0 | - |

m/z Charge Intensity FragmentType MassShift Position
120.08094024658203 0 1053.069
125.05963134765625 0 455.47104
127.0870590209961 0 591.95276
128.10716247558594 0 699.6158
129.01829528808594 0 1097.765
129.10247802734375 0 2920.835
130.08641052246094 0 1018.79114
140.08197021484375 0 1973.4667
141.0655517578125 0 422.941
141.10238647460938 0 4435.6377
142.10572814941406 0 771.2547
143.1180877685547 0 13897.522 a 1
144.1213836669922 0 1024.776
145.0972442626953 0 1026.271
147.11279296875 0 720.3543
148.9546661376953 0 1046.3873
149.02352905273438 0 13486.3545
150.02688598632812 0 6325.56
151.02919006347656 0 510.1189
153.05477905273438 0 649.43695
155.1181640625 0 1706.6097
157.13368225097656 0 9173.348
158.09251403808594 0 1719.8549 y Ammonia loss 7
158.13719177246094 0 2190.1711
169.09725952148438 0 3247.6777
170.1006317138672 0 1174.2477
171.1045379638672 0 610.1953
171.1129608154297 0 12296.502 b 1
171.1378936767578 0 640.0659
172.11630249023438 0 840.7992
173.09225463867188 0 1691.5759
173.128662109375 0 13284.116
174.13182067871094 0 1044.6571
175.11911010742188 0 4560.56 y 7
183.11288452148438 0 1982.4451
183.1494140625 0 3865.0515
184.15306091308594 0 862.8464
185.08102416992188 0 1319.818
185.1285858154297 0 2387.953
185.7764434814453 0 410.02774
186.13247680664062 0 718.4296
186.78395080566406 0 537.9493
187.1077423095703 0 739.59845
187.84507751464844 0 425.06906
195.0771484375 0 527.88837
201.12350463867188 0 5003.592
202.1077117919922 0 529.3036
211.10848999023438 0 711.9607
211.1443634033203 0 1475.1396
212.13946533203125 0 1756.8668
226.15492248535156 0 1810.4069 a Water loss 2
227.13902282714844 0 699.49023
230.15045166015625 0 738.1005 d 2
230.45889282226562 0 516.7966
237.09201049804688 0 671.4515
240.13392639160156 0 895.7512
254.15003967285156 0 4470.429 b Water loss 2
254.1601104736328 0 434.67114
255.14527893066406 0 12886.367 y Ammonia loss 6
256.1486511230469 0 1437.3114
258.1448974609375 0 2636.265
259.1490783691406 0 800.9002
264.4088439941406 0 583.2897
266.1492614746094 0 791.6921
270.1449890136719 0 2500.3254
272.1078186035156 0 1106.2687
272.1578369140625 0 1021.4563 b 2
272.1717834472656 0 5968.466 y 6
273.1756286621094 0 690.57684
275.0902404785156 0 596.8659
276.15557861328125 0 1661.2004
277.1592102050781 0 1031.882
282.18121337890625 0 860.41614
288.1552429199219 0 1162.7643
289.9755554199219 0 611.338
290.96002197265625 0 817.3085
298.5677490234375 0 574.60864
306.17852783203125 0 632.192
312.1918640136719 0 1280.4355
314.86248779296875 0 538.8255
316.8773498535156 0 697.47125
322.1959228515625 0 1870.6179
325.223876953125 0 1706.1951
327.69879150390625 0 4957.478 y Water loss 2
328.19696044921875 0 8280.046 y Ammonia loss 2
328.6983947753906 0 3530.9026
329.1981201171875 0 822.2408
340.1868591308594 0 1410.3533
341.1853942871094 0 1212.373
349.1609802246094 0 692.39624
350.92645263671875 0 1090.6915
352.9344787597656 0 682.347
353.2175598144531 0 1482.1917 b Water loss 3
363.2189025878906 0 5196.321 y Water loss 1
363.71978759765625 0 1358.6326
366.1855773925781 0 7847.8716
367.1883850097656 0 904.1026
368.1814270019531 0 819.646
368.2281188964844 0 1021.0906 y Ammonia loss 5
384.1956481933594 0 2427.2163
385.25616455078125 0 2142.902 y 5
386.2590637207031 0 646.22797
386.8901672363281 0 607.61115
390.2405090332031 0 519.6801
391.2321472167969 0 1046.3286
402.2066345214844 0 8411.796
403.8128967285156 0 2158.8596
412.75360107421875 0 3597.5059 Precursor Water loss
413.2535705566406 0 966.1069
420.8575744628906 0 1227.468
421.2422180175781 0 741.02295
421.3285217285156 0 629.58307
422.164794921875 0 713.7179
422.2119140625 0 3736.9622
439.8416442871094 0 1409.844
472.28790283203125 0 29771.438 y 4
473.2904357910156 0 6775.8145
474.295166015625 0 1027.9528
486.29193115234375 0 3751.4497
487.2952880859375 0 6533.7954
488.2972717285156 0 1540.4459
558.3322143554688 0 1493.368
571.3560791015625 0 10035.155 b 5
572.3592529296875 0 3063.0293
628.3757934570312 0 1047.8264
644.3697509765625 0 531.93713
654.384521484375 0 3037.3142 y Water loss 2
655.3851318359375 0 7103.457 y Ammonia loss 2
656.388671875 0 2613.636
672.4034423828125 0 23724.844 y 2
673.4059448242188 0 8068.8965
674.40673828125 0 1987.439
682.38671875 0 2031.456
683.39111328125 0 995.87573
691.036376953125 0 578.70416
692.2479248046875 0 618.2112
725.430908203125 0 2976.4185 y Water loss 1
743.4400634765625 0 8807.296 y 1
744.4429321289062 0 4234.3857
752.640625 0 631.8273
1544.2723388671875 0 623.6013
1643.66064453125 0 642.33136
2094.998779296875 0 672.99927
3287.41796875 0 665.56946

Spectrum Details

|  |  |
| --- | --- |
| Matched peaks? Matched peaks The total absolute number of peaks matched. Additionally in brackets the total fraction of peaks matched and the total number of peaks is shown. | 24 (16.67% of 144) |
| FDR? FDR The false discovery rate estimated for this peptide. It is calculated by matching all theoretical fragments with a non-integer shift with the raw peaks for this spectrum. This is done with 40 different shifts. The resulting percentage is the average number of annotated peaks over the number of annotated peaks with the correct spectrum. | 0.10% |
| Satellite FDR? Satellite FDR See the FDR for details on its calculation. This satellite ion specific FDR only contains the satellite ions (d/w) for I/L/J positions. | - |
| PSM Score? PSM Score The PSM Score as given by Hecklib to this annotated spectrum. It is shown with three significant figures. | 285 |

#### Spectrum 6958? Spectrum 6958 The raw spectrum of this peptide as annotated by Hecklib. The fragments are coloured according to ion type (see legend). Any peaks with a star '\*' as text can be hovered over to see the full details, first the ion type second the mass shift type. By hovering over the amino acids in the peptide or ions in the legend the corresponding peaks are highlighted. By toggling the 'Unassigned' label you can turn the background (unassigned) peaks on or off in the plot. By updating the slider in the Ion legend you can update the spectrum to only show the top X% of the peaks with labels. The top X% means any peak that is within X% of the highest intensity. By dragging in the spectrum you can zoom in to a specific part of the spectrum and use 'Zoom Out' to get back to the original zoom level. The annotation of the spectrum is based on the given sequence in the peptides file and is done with different software so inconsistencies are likely. The peaks are annotated based on the given sequence, with 20 ppm tolerance.

Copy Data

##### Spectrum 6958 (TSV)

###### Preview

```
Loading example...
```

*Click on the button to copy the data to your clipboard.*

Mz MinMz MaxIntensity Max

WidthHeightPeptide font sizePeptide stroke widthSpectrum font sizeSpectrum stroke widthCompact peptide

Ion legend

wxyz

abcd

OtherUnassignedIon^Charge^_Position_Show for top:%

VATVSJPR

04.57e+39.13e+31.37e+41.83e+4

Zoom Out

a^+1^_2_y^+1^_1_b^+1^_2_y^+1^_1_a^+1^_3_b^+1^_3_y^+1^_2_b^+1^_3_y^+1^_2_b^+2^_6_y^+2^_6_y^+2^_6_y^+2^_6_y^+2^_7_y^+1^_3_\*y^+1^_4_b^+1^_6_y^+1^_6_y^+1^_6_y^+1^_6_y^+1^_7_y^+1^_7_

0549109816482197

Fragment Matches Table

Show background peaks

| Position | Ion type | Intensity | mz Theoretical | mz Error (Th) | mz Error (ppm) | Charge | Series Number |
| --- | --- | --- | --- | --- | --- | --- | --- |
| - | - | 845.9 | 120.1 | - | - | 0 | - |
| - | - | 372.6 | 120.5 | - | - | 0 | - |
| - | - | 536.2 | 128.1 | - | - | 0 | - |
| - | - | 381.3 | 128.9 | - | - | 0 | - |
| - | - | 703.7 | 129 | - | - | 0 | - |
| - | - | 1775 | 129.1 | - | - | 0 | - |
| - | - | 901 | 130.1 | - | - | 0 | - |
| - | - | 415.4 | 131.1 | - | - | 0 | - |
| - | - | 429.2 | 132.5 | - | - | 0 | - |
| - | - | 579 | 136 | - | - | 0 | - |
| - | - | 1152 | 140.1 | - | - | 0 | - |
| - | - | 2704 | 141.1 | - | - | 0 | - |
| - | - | 692.3 | 142.1 | - | - | 0 | - |
| - | - | 654.9 | 143.1 | - | - | 0 | - |
| 2 | a | 8199 | 143.1 | 6.117E-05 | 0.4274 | +1 | 2 |
| - | - | 787.8 | 144.1 | - | - | 0 | - |
| - | - | 403.8 | 144.4 | - | - | 0 | - |
| - | - | 425.5 | 144.4 | - | - | 0 | - |
| - | - | 515 | 145.1 | - | - | 0 | - |
| - | - | 744 | 147.1 | - | - | 0 | - |
| - | - | 1.137E+04 | 149 | - | - | 0 | - |
| - | - | 5914 | 150 | - | - | 0 | - |
| - | - | 403.1 | 150.5 | - | - | 0 | - |
| - | - | 917.9 | 151 | - | - | 0 | - |
| - | - | 572.5 | 152.1 | - | - | 0 | - |
| - | - | 853 | 155.1 | - | - | 0 | - |
| - | - | 777.7 | 157 | - | - | 0 | - |
| - | - | 6163 | 157.1 | - | - | 0 | - |
| 8 | y | 920.3 | 158.1 | 4.162E-05 | 0.2632 | +1 | 1 |
| - | - | 1600 | 158.1 | - | - | 0 | - |
| - | - | 484.2 | 159.1 | - | - | 0 | - |
| - | - | 633.2 | 167 | - | - | 0 | - |
| - | - | 2898 | 169.1 | - | - | 0 | - |
| - | - | 728.7 | 170.1 | - | - | 0 | - |
| 2 | b | 5938 | 171.1 | 0.000179 | 1.046 | +1 | 2 |
| - | - | 579.2 | 171.1 | - | - | 0 | - |
| - | - | 833 | 173.1 | - | - | 0 | - |
| - | - | 9526 | 173.1 | - | - | 0 | - |
| - | - | 3362 | 173.4 | - | - | 0 | - |
| - | - | 576.2 | 174.1 | - | - | 0 | - |
| - | - | 507 | 174.2 | - | - | 0 | - |
| 8 | y | 2608 | 175.1 | 0.000132 | 0.7536 | +1 | 1 |
| - | - | 1036 | 183.1 | - | - | 0 | - |
| - | - | 2804 | 183.1 | - | - | 0 | - |
| - | - | 678.4 | 184.2 | - | - | 0 | - |
| - | - | 742 | 185.1 | - | - | 0 | - |
| - | - | 1825 | 185.1 | - | - | 0 | - |
| - | - | 2579 | 201.1 | - | - | 0 | - |
| - | - | 638 | 202.1 | - | - | 0 | - |
| - | - | 1004 | 210.9 | - | - | 0 | - |
| - | - | 1528 | 211.1 | - | - | 0 | - |
| - | - | 1002 | 212.1 | - | - | 0 | - |
| - | - | 475.3 | 222 | - | - | 0 | - |
| 3 | a | 918.9 | 226.2 | 0.000447 | 1.977 | +1 | 3 |
| - | - | 501.5 | 226.6 | - | - | 0 | - |
| 3 | b | 2976 | 254.1 | 6.138E-05 | 0.2415 | +1 | 3 |
| 7 | y | 6385 | 255.1 | 0.0004373 | 1.714 | +1 | 2 |
| - | - | 1194 | 258.1 | - | - | 0 | - |
| - | - | 645.3 | 266.1 | - | - | 0 | - |
| - | - | 922.6 | 270.1 | - | - | 0 | - |
| 3 | b | 1798 | 272.2 | 0.001578 | 5.797 | +1 | 3 |
| 7 | y | 2801 | 272.2 | 5.464E-05 | 0.2008 | +1 | 2 |
| - | - | 618.2 | 276.2 | - | - | 0 | - |
| - | - | 834.4 | 277.2 | - | - | 0 | - |
| 6 | b | 664 | 286.2 | 0.005569 | 19.46 | +2 | 6 |
| - | - | 865.7 | 288.2 | - | - | 0 | - |
| - | - | 717.3 | 291 | - | - | 0 | - |
| - | - | 1350 | 293.1 | - | - | 0 | - |
| - | - | 1164 | 325.2 | - | - | 0 | - |
| 3 | y | 3011 | 327.7 | 0.002308 | 7.044 | +2 | 6 |
| 3 | y | 4730 | 328.2 | 0.004128 | 12.58 | +2 | 6 |
| - | - | 2433 | 328.7 | - | - | 0 | - |
| 3 | y | 698.5 | 336.7 | 0.0007241 | 2.151 | +2 | 6 |
| - | - | 859.6 | 340.2 | - | - | 0 | - |
| - | - | 1087 | 341.2 | - | - | 0 | - |
| - | - | 1021 | 350.9 | - | - | 0 | - |
| 2 | y | 3416 | 363.2 | 0.0006625 | 1.824 | +2 | 7 |
| - | - | 1615 | 363.7 | - | - | 0 | - |
| - | - | 696.9 | 366.2 | - | - | 0 | - |
| 6 | y | 1324 | 385.3 | 0.00163 | 4.23 | +1 | 3 |
| - | - | 873.8 | 391.2 | - | - | 0 | - |
| - | - | 761.9 | 403 | - | - | 0 | - |
| - | - | 763.3 | 403 | - | - | 0 | - |
| - | - | 1474 | 403.8 | - | - | 0 | - |
| 0 | Precursor | 2243 | 412.8 | 0.0006592 | 1.597 | +2 | -1 |
| - | - | 957.5 | 420.9 | - | - | 0 | - |
| - | - | 920.4 | 439.8 | - | - | 0 | - |
| - | - | 646.3 | 467.9 | - | - | 0 | - |
| 5 | y | 1.809E+04 | 472.3 | 0.0002718 | 0.5755 | +1 | 4 |
| - | - | 4795 | 473.3 | - | - | 0 | - |
| - | - | 927.3 | 474.3 | - | - | 0 | - |
| - | - | 2226 | 486.3 | - | - | 0 | - |
| - | - | 4819 | 487.3 | - | - | 0 | - |
| - | - | 1227 | 488.3 | - | - | 0 | - |
| - | - | 759.1 | 557.3 | - | - | 0 | - |
| - | - | 1134 | 558.3 | - | - | 0 | - |
| 6 | b | 6372 | 571.3 | 0.01085 | 18.98 | +1 | 6 |
| - | - | 2652 | 572.4 | - | - | 0 | - |
| 3 | y | 1427 | 654.4 | 0.008754 | 13.38 | +1 | 6 |
| 3 | y | 4227 | 655.4 | 0.006498 | 9.916 | +1 | 6 |
| - | - | 1609 | 656.4 | - | - | 0 | - |
| 3 | y | 1.443E+04 | 672.4 | 0.001069 | 1.589 | +1 | 6 |
| - | - | 3744 | 673.4 | - | - | 0 | - |
| - | - | 1089 | 674.4 | - | - | 0 | - |
| - | - | 1457 | 682.4 | - | - | 0 | - |
| 2 | y | 1830 | 725.4 | 0.001129 | 1.556 | +1 | 7 |
| - | - | 689.7 | 726.4 | - | - | 0 | - |
| 2 | y | 6313 | 743.4 | 0.001378 | 1.854 | +1 | 7 |
| - | - | 1729 | 744.4 | - | - | 0 | - |
| - | - | 633.4 | 1701 | - | - | 0 | - |
| - | - | 638.6 | 2175 | - | - | 0 | - |

m/z Charge Intensity FragmentType MassShift Position
120.08067321777344 0 845.9221
120.52798461914062 0 372.555
128.10694885253906 0 536.20526
128.85623168945312 0 381.32748
129.01824951171875 0 703.7271
129.1022186279297 0 1774.6758
130.08642578125 0 901.04346
131.1088104248047 0 415.3783
132.48007202148438 0 429.22818
136.0210723876953 0 579.0417
140.08209228515625 0 1151.59
141.10223388671875 0 2704.0806
142.10580444335938 0 692.2668
143.11241149902344 0 654.8678
143.11782836914062 0 8199.472 a 1
144.12132263183594 0 787.7693
144.3795928955078 0 403.82077
144.40701293945312 0 425.47614
145.06484985351562 0 515.0348
147.11285400390625 0 743.9789
149.02328491210938 0 11367.196
150.026611328125 0 5914.475
150.46890258789062 0 403.14032
151.0283203125 0 917.86786
152.1033477783203 0 572.5066
155.11790466308594 0 852.99255
157.0128936767578 0 777.68066
157.13343811035156 0 6163.1836
158.0923614501953 0 920.3025 y Ammonia loss 7
158.13682556152344 0 1599.8405
159.11331176757812 0 484.20718
167.0340118408203 0 633.18646
169.09698486328125 0 2897.9082
170.10037231445312 0 728.7345
171.1126251220703 0 5938.4507 b 1
171.13861083984375 0 579.2424
173.09214782714844 0 833.041
173.12831115722656 0 9526.118
173.4399871826172 0 3361.9712
174.13201904296875 0 576.2008
174.24171447753906 0 507.01584
175.1188201904297 0 2608.2148 y 7
183.11224365234375 0 1036.1339
183.14918518066406 0 2803.5706
184.1526641845703 0 678.37744
185.08029174804688 0 741.9529
185.12841796875 0 1824.7736
201.12327575683594 0 2579.236
202.1071319580078 0 638.00684
210.9049835205078 0 1004.15515
211.1439666748047 0 1528.2076
212.13900756835938 0 1002.3027
222.02529907226562 0 475.25735
226.15455627441406 0 918.9124 a Water loss 2
226.63003540039062 0 501.52924
254.1498565673828 0 2976.0322 b Water loss 2
255.1447296142578 0 6385.2676 y Ammonia loss 6
258.14447021484375 0 1194.126
266.1492614746094 0 645.32794
270.14471435546875 0 922.559
272.1589050292969 0 1798.392 b 2
272.1716613769531 0 2800.5369 y 6
276.15435791015625 0 618.21735
277.1584777832031 0 834.42676
286.18170166015625 0 664.02844 b 5
288.15582275390625 0 865.67896
290.9588317871094 0 717.27783
293.09918212890625 0 1350.4392
325.22344970703125 0 1164.489
327.697998046875 0 3011.341 y Water loss 2
328.1964416503906 0 4729.843 y Ammonia loss 2
328.6977233886719 0 2433.0063
336.7048645019531 0 698.45435 y 2
340.1856994628906 0 859.569
341.18695068359375 0 1086.5458
350.92626953125 0 1020.58026
363.21820068359375 0 3416.4766 y Water loss 1
363.7196960449219 0 1615.0964
366.1842346191406 0 696.8596
385.254150390625 0 1323.6176 y 5
391.2313232421875 0 873.8409
402.9642333984375 0 761.88135
402.9912109375 0 763.3097
403.812744140625 0 1473.6294
412.7524108886719 0 2242.9634 Precursor Water loss
420.85833740234375 0 957.5482
439.84356689453125 0 920.4079
467.8887939453125 0 646.27515
472.28753662109375 0 18088.17 y 4
473.2899475097656 0 4795.129
474.2914123535156 0 927.2717
486.29168701171875 0 2225.8477
487.2952880859375 0 4818.956
488.29522705078125 0 1226.9467
557.3302001953125 0 759.145
558.3310546875 0 1134.1265
571.3558349609375 0 6371.712 b 5
572.3582763671875 0 2652.2327
654.3845825195312 0 1427.4274 y Water loss 2
655.3838500976562 0 4226.635 y Ammonia loss 2
656.3885498046875 0 1609.0469
672.40283203125 0 14428.752 y 2
673.4066162109375 0 3743.8452
674.4083862304688 0 1089.4884
682.3906860351562 0 1457.3978
725.4293212890625 0 1829.9989 y Water loss 1
726.4306640625 0 689.65485
743.4396362304688 0 6313.0195 y 1
744.441650390625 0 1729.3833
1701.29443359375 0 633.4012
2175.0517578125 0 638.55023

Spectrum Details

|  |  |
| --- | --- |
| Matched peaks? Matched peaks The total absolute number of peaks matched. Additionally in brackets the total fraction of peaks matched and the total number of peaks is shown. | 23 (20.72% of 111) |
| FDR? FDR The false discovery rate estimated for this peptide. It is calculated by matching all theoretical fragments with a non-integer shift with the raw peaks for this spectrum. This is done with 40 different shifts. The resulting percentage is the average number of annotated peaks over the number of annotated peaks with the correct spectrum. | 0.21% |
| Satellite FDR? Satellite FDR See the FDR for details on its calculation. This satellite ion specific FDR only contains the satellite ions (d/w) for I/L/J positions. | - |
| PSM Score? PSM Score The PSM Score as given by Hecklib to this annotated spectrum. It is shown with three significant figures. | 244 |

#### Spectrum 6046? Spectrum 6046 The raw spectrum of this peptide as annotated by Hecklib. The fragments are coloured according to ion type (see legend). Any peaks with a star '\*' as text can be hovered over to see the full details, first the ion type second the mass shift type. By hovering over the amino acids in the peptide or ions in the legend the corresponding peaks are highlighted. By toggling the 'Unassigned' label you can turn the background (unassigned) peaks on or off in the plot. By updating the slider in the Ion legend you can update the spectrum to only show the top X% of the peaks with labels. The top X% means any peak that is within X% of the highest intensity. By dragging in the spectrum you can zoom in to a specific part of the spectrum and use 'Zoom Out' to get back to the original zoom level. The annotation of the spectrum is based on the given sequence in the peptides file and is done with different software so inconsistencies are likely. The peaks are annotated based on the given sequence, with 20 ppm tolerance.

Copy Data

##### Spectrum 6046 (TSV)

###### Preview

```
Loading example...
```

*Click on the button to copy the data to your clipboard.*

Mz MinMz MaxIntensity Max

WidthHeightPeptide font sizePeptide stroke widthSpectrum font sizeSpectrum stroke widthCompact peptide

Ion legend

wxyz

abcd

OtherUnassignedIon^Charge^_Position_Show for top:%

VATVSJPR

01.08e+42.15e+43.23e+44.31e+4

Zoom Out

a^+1^_2_y^+1^_1_b^+1^_2_y^+1^_1_a^+1^_3_y^+2^_4_b^+1^_3_y^+1^_2_b^+1^_3_y^+1^_2_b^+2^_6_y^+2^_6_y^+2^_6_y^+2^_6_b^+1^_4_y^+2^_7_y^+1^_3_y^+1^_3_\*y^+1^_4_y^+1^_4_b^+1^_6_b^+1^_6_y^+1^_6_y^+1^_6_y^+1^_6_y^+1^_7_y^+1^_7_

0869173726063475

Fragment Matches Table

Show background peaks

| Position | Ion type | Intensity | mz Theoretical | mz Error (Th) | mz Error (ppm) | Charge | Series Number |
| --- | --- | --- | --- | --- | --- | --- | --- |
| - | - | 999.3 | 120.1 | - | - | 0 | - |
| - | - | 397.1 | 121.6 | - | - | 0 | - |
| - | - | 392.5 | 123.8 | - | - | 0 | - |
| - | - | 382.7 | 125.4 | - | - | 0 | - |
| - | - | 463.5 | 126.2 | - | - | 0 | - |
| - | - | 384.4 | 127.1 | - | - | 0 | - |
| - | - | 505.4 | 127.1 | - | - | 0 | - |
| - | - | 381.5 | 127.1 | - | - | 0 | - |
| - | - | 703.8 | 128.1 | - | - | 0 | - |
| - | - | 896.8 | 129 | - | - | 0 | - |
| - | - | 3264 | 129.1 | - | - | 0 | - |
| - | - | 1318 | 130.1 | - | - | 0 | - |
| - | - | 390 | 130.1 | - | - | 0 | - |
| - | - | 472.7 | 130.9 | - | - | 0 | - |
| - | - | 707.5 | 136 | - | - | 0 | - |
| - | - | 562.2 | 136.1 | - | - | 0 | - |
| - | - | 495.1 | 139 | - | - | 0 | - |
| - | - | 2429 | 140.1 | - | - | 0 | - |
| - | - | 420.4 | 140.4 | - | - | 0 | - |
| - | - | 4373 | 141.1 | - | - | 0 | - |
| - | - | 1099 | 142.1 | - | - | 0 | - |
| 2 | a | 1.656E+04 | 143.1 | 0.000244 | 1.705 | +1 | 2 |
| - | - | 1339 | 144.1 | - | - | 0 | - |
| - | - | 1027 | 145.1 | - | - | 0 | - |
| - | - | 1496 | 147.1 | - | - | 0 | - |
| - | - | 553.2 | 148.1 | - | - | 0 | - |
| - | - | 1.108E+04 | 149 | - | - | 0 | - |
| - | - | 6545 | 150 | - | - | 0 | - |
| - | - | 387.3 | 151.6 | - | - | 0 | - |
| - | - | 552.8 | 155.1 | - | - | 0 | - |
| - | - | 1828 | 155.1 | - | - | 0 | - |
| - | - | 518.3 | 157 | - | - | 0 | - |
| - | - | 1.111E+04 | 157.1 | - | - | 0 | - |
| 8 | y | 3103 | 158.1 | 0.0002483 | 1.571 | +1 | 1 |
| - | - | 3072 | 158.1 | - | - | 0 | - |
| - | - | 473.3 | 167 | - | - | 0 | - |
| - | - | 5510 | 169.1 | - | - | 0 | - |
| - | - | 1422 | 170.1 | - | - | 0 | - |
| 2 | b | 1.45E+04 | 171.1 | 0.0002024 | 1.183 | +1 | 2 |
| - | - | 848.8 | 171.1 | - | - | 0 | - |
| - | - | 1318 | 172.1 | - | - | 0 | - |
| - | - | 3232 | 173.1 | - | - | 0 | - |
| - | - | 1.773E+04 | 173.1 | - | - | 0 | - |
| - | - | 1640 | 174.1 | - | - | 0 | - |
| 8 | y | 4912 | 175.1 | 0.0001274 | 0.7276 | +1 | 1 |
| - | - | 498.5 | 177.5 | - | - | 0 | - |
| - | - | 516.3 | 178.6 | - | - | 0 | - |
| - | - | 3439 | 183.1 | - | - | 0 | - |
| - | - | 5213 | 183.1 | - | - | 0 | - |
| - | - | 1917 | 184.2 | - | - | 0 | - |
| - | - | 849.3 | 185.1 | - | - | 0 | - |
| - | - | 460.4 | 185.1 | - | - | 0 | - |
| - | - | 454.9 | 185.1 | - | - | 0 | - |
| - | - | 2486 | 185.1 | - | - | 0 | - |
| - | - | 826.5 | 187.1 | - | - | 0 | - |
| - | - | 522.5 | 193.4 | - | - | 0 | - |
| - | - | 448.6 | 198 | - | - | 0 | - |
| - | - | 6118 | 201.1 | - | - | 0 | - |
| - | - | 554 | 202.1 | - | - | 0 | - |
| - | - | 1096 | 202.1 | - | - | 0 | - |
| - | - | 557.7 | 210.9 | - | - | 0 | - |
| - | - | 1247 | 211.1 | - | - | 0 | - |
| - | - | 2150 | 211.1 | - | - | 0 | - |
| - | - | 582.3 | 211.2 | - | - | 0 | - |
| - | - | 1577 | 212.1 | - | - | 0 | - |
| - | - | 925.2 | 212.1 | - | - | 0 | - |
| - | - | 570.5 | 213.1 | - | - | 0 | - |
| 3 | a | 1950 | 226.2 | 1.072E-05 | 0.04738 | +1 | 3 |
| - | - | 949.6 | 227.1 | - | - | 0 | - |
| 5 | y | 598.1 | 228.1 | 0.0002652 | 1.162 | +2 | 4 |
| - | - | 1057 | 240.1 | - | - | 0 | - |
| - | - | 575.4 | 243.1 | - | - | 0 | - |
| 3 | b | 6881 | 254.1 | 3.424E-07 | 0.001347 | +1 | 3 |
| - | - | 776.8 | 254.2 | - | - | 0 | - |
| 7 | y | 1.592E+04 | 255.1 | 0.0001273 | 0.4988 | +1 | 2 |
| - | - | 1498 | 256.1 | - | - | 0 | - |
| - | - | 843.3 | 257.1 | - | - | 0 | - |
| - | - | 3276 | 258.1 | - | - | 0 | - |
| - | - | 1228 | 259.1 | - | - | 0 | - |
| - | - | 680.4 | 266.9 | - | - | 0 | - |
| - | - | 563.9 | 267.2 | - | - | 0 | - |
| - | - | 1583 | 270.1 | - | - | 0 | - |
| 3 | b | 2869 | 272.2 | 0.0008147 | 2.993 | +1 | 3 |
| 7 | y | 6392 | 272.2 | 0.0001895 | 0.6962 | +1 | 2 |
| - | - | 1089 | 273.2 | - | - | 0 | - |
| - | - | 2062 | 276.2 | - | - | 0 | - |
| - | - | 946.4 | 277.2 | - | - | 0 | - |
| - | - | 558.7 | 280.1 | - | - | 0 | - |
| - | - | 860.2 | 282.2 | - | - | 0 | - |
| 6 | b | 714.1 | 286.2 | 0.004928 | 17.22 | +2 | 6 |
| - | - | 1598 | 288.2 | - | - | 0 | - |
| - | - | 1560 | 293.1 | - | - | 0 | - |
| - | - | 1166 | 306.2 | - | - | 0 | - |
| - | - | 2676 | 325.2 | - | - | 0 | - |
| 3 | y | 6488 | 327.7 | 0.001271 | 3.877 | +2 | 6 |
| 3 | y | 9721 | 328.2 | 0.004829 | 14.72 | +2 | 6 |
| - | - | 2709 | 328.7 | - | - | 0 | - |
| - | - | 1054 | 329.2 | - | - | 0 | - |
| - | - | 513 | 335 | - | - | 0 | - |
| 3 | y | 1016 | 336.7 | 0.0001914 | 0.5685 | +2 | 6 |
| - | - | 693.2 | 340.2 | - | - | 0 | - |
| - | - | 1681 | 340.2 | - | - | 0 | - |
| - | - | 1170 | 341.2 | - | - | 0 | - |
| - | - | 705 | 350.9 | - | - | 0 | - |
| 4 | b | 1791 | 353.2 | 3.963E-05 | 0.1122 | +1 | 4 |
| 2 | y | 7036 | 363.2 | 0.0003268 | 0.8997 | +2 | 7 |
| - | - | 2423 | 363.7 | - | - | 0 | - |
| 6 | y | 1168 | 368.2 | 0.0005359 | 1.455 | +1 | 3 |
| - | - | 639 | 383.2 | - | - | 0 | - |
| 6 | y | 3838 | 385.3 | 0.0003174 | 0.8237 | +1 | 3 |
| - | - | 551.7 | 385.8 | - | - | 0 | - |
| - | - | 660.5 | 391.2 | - | - | 0 | - |
| - | - | 606.3 | 393.1 | - | - | 0 | - |
| - | - | 735.8 | 403 | - | - | 0 | - |
| - | - | 593.6 | 403 | - | - | 0 | - |
| - | - | 1438 | 403.8 | - | - | 0 | - |
| 0 | Precursor | 4834 | 412.8 | 0.0001647 | 0.3991 | +2 | -1 |
| - | - | 2856 | 413.3 | - | - | 0 | - |
| - | - | 713 | 413.8 | - | - | 0 | - |
| - | - | 1488 | 420.9 | - | - | 0 | - |
| - | - | 790.1 | 421.9 | - | - | 0 | - |
| - | - | 718.7 | 440.8 | - | - | 0 | - |
| 5 | y | 909.4 | 454.3 | 0.002189 | 4.818 | +1 | 4 |
| 5 | y | 4.263E+04 | 472.3 | 9.442E-05 | 0.1999 | +1 | 4 |
| - | - | 8682 | 473.3 | - | - | 0 | - |
| - | - | 1826 | 474.3 | - | - | 0 | - |
| - | - | 625.6 | 480.3 | - | - | 0 | - |
| - | - | 4983 | 486.3 | - | - | 0 | - |
| - | - | 6994 | 487.3 | - | - | 0 | - |
| - | - | 1643 | 488.3 | - | - | 0 | - |
| - | - | 713 | 540 | - | - | 0 | - |
| 6 | b | 660.5 | 553.3 | 0.006396 | 11.56 | +1 | 6 |
| - | - | 936.7 | 557.3 | - | - | 0 | - |
| - | - | 1465 | 558.3 | - | - | 0 | - |
| 6 | b | 1.281E+04 | 571.3 | 0.01066 | 18.66 | +1 | 6 |
| - | - | 3697 | 572.4 | - | - | 0 | - |
| - | - | 1176 | 573.4 | - | - | 0 | - |
| - | - | 910.9 | 628.4 | - | - | 0 | - |
| - | - | 582.6 | 632.6 | - | - | 0 | - |
| 3 | y | 3145 | 654.4 | 0.008876 | 13.56 | +1 | 6 |
| 3 | y | 8517 | 655.4 | 0.007353 | 11.22 | +1 | 6 |
| - | - | 2169 | 656.4 | - | - | 0 | - |
| - | - | 830 | 657.4 | - | - | 0 | - |
| 3 | y | 3.227E+04 | 672.4 | 0.0009467 | 1.408 | +1 | 6 |
| - | - | 1.008E+04 | 673.4 | - | - | 0 | - |
| - | - | 1447 | 674.4 | - | - | 0 | - |
| - | - | 2153 | 682.4 | - | - | 0 | - |
| - | - | 1119 | 683.4 | - | - | 0 | - |
| 2 | y | 3629 | 725.4 | 0.0006403 | 0.8827 | +1 | 7 |
| - | - | 816.2 | 726.4 | - | - | 0 | - |
| 2 | y | 1.205E+04 | 743.4 | 0.001134 | 1.526 | +1 | 7 |
| - | - | 4270 | 744.4 | - | - | 0 | - |
| - | - | 1430 | 745.4 | - | - | 0 | - |
| - | - | 639.4 | 1483 | - | - | 0 | - |
| - | - | 670.3 | 2065 | - | - | 0 | - |
| - | - | 606.8 | 2364 | - | - | 0 | - |
| - | - | 863.7 | 2895 | - | - | 0 | - |
| - | - | 676 | 3440 | - | - | 0 | - |

m/z Charge Intensity FragmentType MassShift Position
120.0810546875 0 999.31433
121.62748718261719 0 397.06528
123.82511138916016 0 392.53384
125.44580078125 0 382.66028
126.1709976196289 0 463.4697
127.07575988769531 0 384.37152
127.08679962158203 0 505.35907
127.12360382080078 0 381.45264
128.1072998046875 0 703.78217
129.01849365234375 0 896.7626
129.10244750976562 0 3264.4055
130.08639526367188 0 1317.6528
130.1061248779297 0 389.9901
130.88267517089844 0 472.72345
136.02178955078125 0 707.51355
136.07614135742188 0 562.1787
139.00283813476562 0 495.13632
140.08212280273438 0 2428.8135
140.44615173339844 0 420.3539
141.10243225097656 0 4372.66
142.10589599609375 0 1099.3778
143.11813354492188 0 16556.025 a 1
144.12147521972656 0 1339.3428
145.0977020263672 0 1027.1625
147.11312866210938 0 1496.1619
148.1164093017578 0 553.1738
149.0235595703125 0 11081.38
150.0269775390625 0 6545.1704
151.5555419921875 0 387.27634
155.08108520507812 0 552.768
155.11830139160156 0 1828.1436
157.01280212402344 0 518.31665
157.1337432861328 0 11111.973
158.0926513671875 0 3102.8274 y Ammonia loss 7
158.1371612548828 0 3071.6978
167.0338592529297 0 473.28668
169.09732055664062 0 5510.367
170.10110473632812 0 1421.9828
171.11300659179688 0 14495.766 b 1
171.13796997070312 0 848.7579
172.11672973632812 0 1317.8225
173.09219360351562 0 3232.3276
173.128662109375 0 17726.738
174.13180541992188 0 1639.8682
175.11907958984375 0 4912.011 y 7
177.4850616455078 0 498.53162
178.5591583251953 0 516.2672
183.11289978027344 0 3438.7258
183.14947509765625 0 5213.468
184.1527862548828 0 1917.2808
185.0811767578125 0 849.3426
185.09130859375 0 460.44388
185.10292053222656 0 454.91687
185.12867736816406 0 2485.6472
187.10789489746094 0 826.5367
193.43997192382812 0 522.47565
198.036865234375 0 448.59454
201.12355041503906 0 6117.969
202.10763549804688 0 553.96936
202.12686157226562 0 1096.0469
210.92291259765625 0 557.71106
211.1077423095703 0 1247.0338
211.1441192626953 0 2150.0178
211.1545867919922 0 582.33203
212.13919067382812 0 1577.2968
212.14881896972656 0 925.2205
213.12332153320312 0 570.5313
226.15501403808594 0 1949.9606 a Water loss 2
227.13888549804688 0 949.5982
228.13400268554688 0 598.13074 y Ammonia loss 4
240.13458251953125 0 1056.5939
243.13441467285156 0 575.3614
254.14991760253906 0 6880.5854 b Water loss 2
254.16293334960938 0 776.8461
255.14529418945312 0 15919.376 y Ammonia loss 6
256.14837646484375 0 1498.3861
257.0714111328125 0 843.31085
258.14501953125 0 3276.2312
259.14837646484375 0 1227.9268
266.92657470703125 0 680.3502
267.15380859375 0 563.8647
270.1452941894531 0 1582.7743
272.15966796875 0 2868.6484 b 2
272.1719055175781 0 6391.667 y 6
273.1756286621094 0 1088.6515
276.1556701660156 0 2062.4097
277.16009521484375 0 946.4253
280.1300964355469 0 558.68365
282.1817321777344 0 860.16095
286.1810607910156 0 714.12305 b 5
288.1561279296875 0 1598.1593
293.1000061035156 0 1559.6302
306.179443359375 0 1165.7731
325.22369384765625 0 2676.2068
327.69903564453125 0 6488.139 y Water loss 2
328.1971435546875 0 9721.312 y Ammonia loss 2
328.6974182128906 0 2709.4133
329.1996765136719 0 1054.2743
335.0221862792969 0 512.96704
336.7057800292969 0 1016.4112 y 2
340.1675109863281 0 693.18427
340.18682861328125 0 1681.0623
341.18780517578125 0 1169.686
350.9273681640625 0 705.00397
353.2182922363281 0 1791.1519 b Water loss 3
363.2185363769531 0 7036.417 y Water loss 1
363.72039794921875 0 2423.359
368.2297668457031 0 1167.6724 y Ammonia loss 5
383.2304382324219 0 638.95776
385.2554626464844 0 3837.6772 y 5
385.845458984375 0 551.6937
391.2308349609375 0 660.53284
393.0856628417969 0 606.347
402.9665222167969 0 735.836
403.0064697265625 0 593.59796
403.8136901855469 0 1438.1675
412.75323486328125 0 4834.4507 Precursor Water loss
413.2547912597656 0 2855.606
413.7551574707031 0 712.9719
420.8577575683594 0 1487.5459
421.86236572265625 0 790.1141
440.8451843261719 0 718.6952
454.2750549316406 0 909.387 y Water loss 4
472.28790283203125 0 42629.832 y 4
473.2906188964844 0 8681.549
474.2923583984375 0 1826.2107
480.28662109375 0 625.6356
486.2919616699219 0 4983.0605
487.2950439453125 0 6994.028
488.2978210449219 0 1643.0533
540.0186157226562 0 713.04254
553.3408203125 0 660.51276 b Water loss 5
557.3308715820312 0 936.73444
558.3316650390625 0 1464.5317
571.3556518554688 0 12810.888 b 5
572.3590087890625 0 3696.806
573.3611450195312 0 1175.5857
628.3797607421875 0 910.87885
632.5987548828125 0 582.5686
654.3844604492188 0 3145.0132 y Water loss 2
655.3847045898438 0 8516.615 y Ammonia loss 2
656.3875732421875 0 2169.0537
657.3897094726562 0 830.01965
672.4029541015625 0 32273.773 y 2
673.40576171875 0 10084.913
674.4071655273438 0 1447.481
682.3867797851562 0 2153.3845
683.3907470703125 0 1119.0688
725.4298095703125 0 3628.6335 y Water loss 1
726.4332885742188 0 816.2224
743.4398803710938 0 12045.96 y 1
744.4432373046875 0 4269.565
745.4470825195312 0 1429.822
1482.9818115234375 0 639.4392
2065.41357421875 0 670.28577
2363.897705078125 0 606.79724
2895.206298828125 0 863.72174
3440.191162109375 0 676.0466

Spectrum Details

|  |  |
| --- | --- |
| Matched peaks? Matched peaks The total absolute number of peaks matched. Additionally in brackets the total fraction of peaks matched and the total number of peaks is shown. | 28 (17.72% of 158) |
| FDR? FDR The false discovery rate estimated for this peptide. It is calculated by matching all theoretical fragments with a non-integer shift with the raw peaks for this spectrum. This is done with 40 different shifts. The resulting percentage is the average number of annotated peaks over the number of annotated peaks with the correct spectrum. | 0.43% |
| Satellite FDR? Satellite FDR See the FDR for details on its calculation. This satellite ion specific FDR only contains the satellite ions (d/w) for I/L/J positions. | - |
| PSM Score? PSM Score The PSM Score as given by Hecklib to this annotated spectrum. It is shown with three significant figures. | 327 |

#### Spectrum 6665? Spectrum 6665 The raw spectrum of this peptide as annotated by Hecklib. The fragments are coloured according to ion type (see legend). Any peaks with a star '\*' as text can be hovered over to see the full details, first the ion type second the mass shift type. By hovering over the amino acids in the peptide or ions in the legend the corresponding peaks are highlighted. By toggling the 'Unassigned' label you can turn the background (unassigned) peaks on or off in the plot. By updating the slider in the Ion legend you can update the spectrum to only show the top X% of the peaks with labels. The top X% means any peak that is within X% of the highest intensity. By dragging in the spectrum you can zoom in to a specific part of the spectrum and use 'Zoom Out' to get back to the original zoom level. The annotation of the spectrum is based on the given sequence in the peptides file and is done with different software so inconsistencies are likely. The peaks are annotated based on the given sequence, with 20 ppm tolerance.

Copy Data

##### Spectrum 6665 (TSV)

###### Preview

```
Loading example...
```

*Click on the button to copy the data to your clipboard.*

Mz MinMz MaxIntensity Max

WidthHeightPeptide font sizePeptide stroke widthSpectrum font sizeSpectrum stroke widthCompact peptide

Ion legend

wxyz

abcd

OtherUnassignedIon^Charge^_Position_Show for top:%

VATVSJPR

06.04e+31.21e+41.81e+42.42e+4

Zoom Out

a^+1^_2_y^+1^_1_b^+1^_2_y^+1^_1_a^+1^_3_b^+1^_3_y^+1^_2_b^+1^_3_y^+1^_2_y^+2^_6_y^+2^_6_y^+2^_6_b^+1^_4_y^+2^_7_y^+1^_3_y^+1^_3_\*y^+1^_4_b^+1^_6_y^+1^_6_y^+1^_6_y^+1^_6_y^+1^_7_y^+1^_7_

043386613001733

Fragment Matches Table

Show background peaks

| Position | Ion type | Intensity | mz Theoretical | mz Error (Th) | mz Error (ppm) | Charge | Series Number |
| --- | --- | --- | --- | --- | --- | --- | --- |
| - | - | 364.2 | 125.2 | - | - | 0 | - |
| - | - | 406.4 | 126.8 | - | - | 0 | - |
| - | - | 638.3 | 129 | - | - | 0 | - |
| - | - | 2846 | 129.1 | - | - | 0 | - |
| - | - | 821.3 | 130.1 | - | - | 0 | - |
| - | - | 476 | 136.1 | - | - | 0 | - |
| - | - | 464.7 | 136.5 | - | - | 0 | - |
| - | - | 1629 | 140.1 | - | - | 0 | - |
| - | - | 3543 | 141.1 | - | - | 0 | - |
| 2 | a | 9544 | 143.1 | 7.616E-05 | 0.5322 | +1 | 2 |
| - | - | 431.1 | 144.1 | - | - | 0 | - |
| - | - | 959.6 | 147.1 | - | - | 0 | - |
| - | - | 696.7 | 149 | - | - | 0 | - |
| - | - | 1.115E+04 | 149 | - | - | 0 | - |
| - | - | 5872 | 150 | - | - | 0 | - |
| - | - | 447.2 | 152.8 | - | - | 0 | - |
| - | - | 1145 | 155.1 | - | - | 0 | - |
| - | - | 6859 | 157.1 | - | - | 0 | - |
| 8 | y | 1440 | 158.1 | 4.994E-05 | 0.3159 | +1 | 1 |
| - | - | 2167 | 158.1 | - | - | 0 | - |
| - | - | 668.3 | 160.2 | - | - | 0 | - |
| - | - | 422.5 | 162.3 | - | - | 0 | - |
| - | - | 738.9 | 167 | - | - | 0 | - |
| - | - | 642.7 | 168 | - | - | 0 | - |
| - | - | 3115 | 169.1 | - | - | 0 | - |
| - | - | 504.5 | 170.1 | - | - | 0 | - |
| - | - | 839 | 171.1 | - | - | 0 | - |
| 2 | b | 8491 | 171.1 | 1.933E-05 | 0.113 | +1 | 2 |
| - | - | 633.5 | 171.1 | - | - | 0 | - |
| - | - | 880.2 | 172.1 | - | - | 0 | - |
| - | - | 1345 | 173.1 | - | - | 0 | - |
| - | - | 1.064E+04 | 173.1 | - | - | 0 | - |
| - | - | 2307 | 173.5 | - | - | 0 | - |
| - | - | 595.6 | 174.1 | - | - | 0 | - |
| 8 | y | 3875 | 175.1 | 0.0001427 | 0.8148 | +1 | 1 |
| - | - | 571.3 | 181.5 | - | - | 0 | - |
| - | - | 2205 | 183.1 | - | - | 0 | - |
| - | - | 2900 | 183.1 | - | - | 0 | - |
| - | - | 831.8 | 184.2 | - | - | 0 | - |
| - | - | 1005 | 185.1 | - | - | 0 | - |
| - | - | 1321 | 185.1 | - | - | 0 | - |
| - | - | 583.1 | 186.1 | - | - | 0 | - |
| - | - | 450.6 | 196.4 | - | - | 0 | - |
| - | - | 2396 | 201.1 | - | - | 0 | - |
| - | - | 570.3 | 211.1 | - | - | 0 | - |
| - | - | 1352 | 211.1 | - | - | 0 | - |
| - | - | 1208 | 212.1 | - | - | 0 | - |
| - | - | 522.2 | 215.6 | - | - | 0 | - |
| - | - | 504.6 | 218.4 | - | - | 0 | - |
| - | - | 489.9 | 220.2 | - | - | 0 | - |
| - | - | 495.9 | 224.5 | - | - | 0 | - |
| 3 | a | 1225 | 226.2 | 0.0004928 | 2.179 | +1 | 3 |
| - | - | 682.8 | 227.1 | - | - | 0 | - |
| - | - | 751.3 | 240.1 | - | - | 0 | - |
| - | - | 488.5 | 247.3 | - | - | 0 | - |
| 3 | b | 3916 | 254.1 | 1.56E-05 | 0.06139 | +1 | 3 |
| 7 | y | 8867 | 255.1 | 4.058E-05 | 0.159 | +1 | 2 |
| - | - | 561.5 | 256 | - | - | 0 | - |
| - | - | 1648 | 258.1 | - | - | 0 | - |
| - | - | 555.3 | 259.1 | - | - | 0 | - |
| - | - | 717.1 | 266.2 | - | - | 0 | - |
| - | - | 1593 | 270.1 | - | - | 0 | - |
| 3 | b | 948.8 | 272.2 | 0.003012 | 11.07 | +1 | 3 |
| 7 | y | 3805 | 272.2 | 2.413E-05 | 0.08864 | +1 | 2 |
| - | - | 1393 | 276.2 | - | - | 0 | - |
| - | - | 585.1 | 277.2 | - | - | 0 | - |
| - | - | 1080 | 288.2 | - | - | 0 | - |
| - | - | 1700 | 293.1 | - | - | 0 | - |
| - | - | 558.9 | 313.1 | - | - | 0 | - |
| - | - | 1796 | 325.2 | - | - | 0 | - |
| 3 | y | 5388 | 327.7 | 0.001484 | 4.529 | +2 | 6 |
| 3 | y | 7089 | 328.2 | 0.004341 | 13.23 | +2 | 6 |
| - | - | 1484 | 328.7 | - | - | 0 | - |
| - | - | 681.5 | 329.2 | - | - | 0 | - |
| 3 | y | 665.5 | 336.7 | 0.002555 | 7.589 | +2 | 6 |
| - | - | 1397 | 340.2 | - | - | 0 | - |
| - | - | 729.3 | 341.2 | - | - | 0 | - |
| - | - | 563.6 | 349.9 | - | - | 0 | - |
| - | - | 1407 | 350.9 | - | - | 0 | - |
| 4 | b | 931.5 | 353.2 | 0.0005584 | 1.581 | +1 | 4 |
| - | - | 594.4 | 358.2 | - | - | 0 | - |
| - | - | 663.7 | 359.2 | - | - | 0 | - |
| 2 | y | 3781 | 363.2 | 0.0007235 | 1.992 | +2 | 7 |
| - | - | 1446 | 363.7 | - | - | 0 | - |
| - | - | 945.6 | 364.2 | - | - | 0 | - |
| 6 | y | 667.2 | 368.2 | 0.000658 | 1.787 | +1 | 3 |
| - | - | 618.5 | 368.9 | - | - | 0 | - |
| - | - | 613 | 383.2 | - | - | 0 | - |
| 6 | y | 2169 | 385.3 | 0.0004089 | 1.061 | +1 | 3 |
| - | - | 686.9 | 391.2 | - | - | 0 | - |
| - | - | 2263 | 403.8 | - | - | 0 | - |
| 0 | Precursor | 3421 | 412.8 | 1.214E-05 | 0.02941 | +2 | -1 |
| - | - | 1520 | 413.3 | - | - | 0 | - |
| - | - | 1107 | 420.9 | - | - | 0 | - |
| - | - | 619.5 | 421 | - | - | 0 | - |
| - | - | 675.7 | 439.8 | - | - | 0 | - |
| 5 | y | 2.393E+04 | 472.3 | 0.0001192 | 0.2524 | +1 | 4 |
| - | - | 5084 | 473.3 | - | - | 0 | - |
| - | - | 597.7 | 474.3 | - | - | 0 | - |
| - | - | 2635 | 486.3 | - | - | 0 | - |
| - | - | 4963 | 487.3 | - | - | 0 | - |
| - | - | 687.6 | 488.3 | - | - | 0 | - |
| - | - | 667.2 | 557.3 | - | - | 0 | - |
| - | - | 1082 | 558.3 | - | - | 0 | - |
| 6 | b | 6732 | 571.3 | 0.01066 | 18.66 | +1 | 6 |
| - | - | 2245 | 572.4 | - | - | 0 | - |
| - | - | 623.3 | 644.4 | - | - | 0 | - |
| 3 | y | 1633 | 654.4 | 0.0115 | 17.57 | +1 | 6 |
| 3 | y | 5384 | 655.4 | 0.00601 | 9.17 | +1 | 6 |
| - | - | 1941 | 656.4 | - | - | 0 | - |
| - | - | 765.6 | 657.4 | - | - | 0 | - |
| 3 | y | 1.657E+04 | 672.4 | 0.0008857 | 1.317 | +1 | 6 |
| - | - | 4477 | 673.4 | - | - | 0 | - |
| - | - | 748 | 674.4 | - | - | 0 | - |
| - | - | 1538 | 682.4 | - | - | 0 | - |
| 2 | y | 1581 | 725.4 | 0.0008845 | 1.219 | +1 | 7 |
| 2 | y | 7346 | 743.4 | 0.001439 | 1.936 | +1 | 7 |
| - | - | 2473 | 744.4 | - | - | 0 | - |
| - | - | 639.6 | 1348 | - | - | 0 | - |
| - | - | 628.1 | 1455 | - | - | 0 | - |
| - | - | 646.1 | 1486 | - | - | 0 | - |
| - | - | 694.7 | 1716 | - | - | 0 | - |

m/z Charge Intensity FragmentType MassShift Position
125.15078735351562 0 364.21143
126.79170989990234 0 406.43375
129.01820373535156 0 638.26385
129.10231018066406 0 2845.5017
130.08642578125 0 821.3123
136.07546997070312 0 476.0279
136.52743530273438 0 464.66702
140.0818634033203 0 1628.6599
141.10232543945312 0 3542.9844
143.1179656982422 0 9544.246 a 1
144.12127685546875 0 431.1177
147.11273193359375 0 959.6211
149.01698303222656 0 696.737
149.02342224121094 0 11153.812
150.02682495117188 0 5872.379
152.81871032714844 0 447.18835
155.1178741455078 0 1145.0573
157.13368225097656 0 6858.7734
158.0924530029297 0 1440.1715 y Ammonia loss 7
158.1369171142578 0 2166.9165
160.2301483154297 0 668.25507
162.30873107910156 0 422.5039
167.03387451171875 0 738.9207
168.03675842285156 0 642.7035
169.09707641601562 0 3114.7375
170.1007843017578 0 504.4848
171.10545349121094 0 838.9872
171.11282348632812 0 8490.926 b 1
171.13864135742188 0 633.49725
172.11688232421875 0 880.18243
173.0922088623047 0 1344.6581
173.12844848632812 0 10640.321
173.45050048828125 0 2306.9707
174.131591796875 0 595.6476
175.1190948486328 0 3875.271 y 7
181.48309326171875 0 571.335
183.1127166748047 0 2205.34
183.14923095703125 0 2900.1553
184.15269470214844 0 831.8054
185.0808563232422 0 1004.5743
185.1287078857422 0 1321.0548
186.13275146484375 0 583.1093
196.41152954101562 0 450.6338
201.12338256835938 0 2395.764
211.10614013671875 0 570.27277
211.143798828125 0 1352.3214
212.13856506347656 0 1207.6572
215.647705078125 0 522.23425
218.42686462402344 0 504.6216
220.2340087890625 0 489.92004
224.4701385498047 0 495.85413
226.15451049804688 0 1224.6473 a Water loss 2
227.13925170898438 0 682.7765
240.1337432861328 0 751.2981
247.33717346191406 0 488.49612
254.14990234375 0 3915.6047 b Water loss 2
255.14512634277344 0 8867.43 y Ammonia loss 6
256.0398864746094 0 561.51086
258.1444091796875 0 1648.228
259.1486511230469 0 555.3208
266.1512145996094 0 717.11914
270.1441345214844 0 1592.6476
272.157470703125 0 948.7862 b 2
272.17169189453125 0 3804.627 y 6
276.1554870605469 0 1393.0339
277.158203125 0 585.06726
288.1549072265625 0 1079.8276
293.09912109375 0 1700.2395
313.09686279296875 0 558.9182
325.22418212890625 0 1795.6204
327.6988220214844 0 5387.8784 y Water loss 2
328.1966552734375 0 7089.1216 y Ammonia loss 2
328.6971740722656 0 1483.7627
329.19610595703125 0 681.463
336.7030334472656 0 665.5178 y 2
340.1869201660156 0 1397.3201
341.1876525878906 0 729.2946
349.8991394042969 0 563.6458
350.92694091796875 0 1406.9773
353.2177734375 0 931.5461 b Water loss 3
358.1976623535156 0 594.42413
359.1947021484375 0 663.6779
363.2181396484375 0 3780.8125 y Water loss 1
363.7198486328125 0 1446.1968
364.22247314453125 0 945.5814
368.2298889160156 0 667.157 y Ammonia loss 5
368.935546875 0 618.5384
383.2315979003906 0 613.0072
385.25537109375 0 2168.9165 y 5
391.2314147949219 0 686.9001
403.8132019042969 0 2263.2197
412.7530822753906 0 3421.3442 Precursor Water loss
413.2554016113281 0 1519.5588
420.8578186035156 0 1106.6819
420.9998779296875 0 619.5218
439.843505859375 0 675.72614
472.2876892089844 0 23933.99 y 4
473.2904052734375 0 5084.3174
474.2913818359375 0 597.7149
486.2921447753906 0 2634.9739
487.29473876953125 0 4962.594
488.29833984375 0 687.59644
557.3243408203125 0 667.2073
558.3348388671875 0 1082.3049
571.3556518554688 0 6732.0767 b 5
572.3584594726562 0 2244.5107
644.3705444335938 0 623.277
654.3818359375 0 1632.6733 y Water loss 2
655.3833618164062 0 5384.479 y Ammonia loss 2
656.3857421875 0 1940.8951
657.3873291015625 0 765.55554
672.4030151367188 0 16568.88 y 2
673.4055786132812 0 4476.651
674.4093017578125 0 747.9571
682.3899536132812 0 1537.7274
725.4295654296875 0 1581.0911 y Water loss 1
743.4395751953125 0 7346.061 y 1
744.443359375 0 2473.1277
1347.7255859375 0 639.60864
1455.44482421875 0 628.05054
1485.658203125 0 646.1181
1715.6590576171875 0 694.73

Spectrum Details

|  |  |
| --- | --- |
| Matched peaks? Matched peaks The total absolute number of peaks matched. Additionally in brackets the total fraction of peaks matched and the total number of peaks is shown. | 24 (19.67% of 122) |
| FDR? FDR The false discovery rate estimated for this peptide. It is calculated by matching all theoretical fragments with a non-integer shift with the raw peaks for this spectrum. This is done with 40 different shifts. The resulting percentage is the average number of annotated peaks over the number of annotated peaks with the correct spectrum. | 0.10% |
| Satellite FDR? Satellite FDR See the FDR for details on its calculation. This satellite ion specific FDR only contains the satellite ions (d/w) for I/L/J positions. | - |
| PSM Score? PSM Score The PSM Score as given by Hecklib to this annotated spectrum. It is shown with three significant figures. | 285 |

#### Spectrum 3948? Spectrum 3948 The raw spectrum of this peptide as annotated by Hecklib. The fragments are coloured according to ion type (see legend). Any peaks with a star '\*' as text can be hovered over to see the full details, first the ion type second the mass shift type. By hovering over the amino acids in the peptide or ions in the legend the corresponding peaks are highlighted. By toggling the 'Unassigned' label you can turn the background (unassigned) peaks on or off in the plot. By updating the slider in the Ion legend you can update the spectrum to only show the top X% of the peaks with labels. The top X% means any peak that is within X% of the highest intensity. By dragging in the spectrum you can zoom in to a specific part of the spectrum and use 'Zoom Out' to get back to the original zoom level. The annotation of the spectrum is based on the given sequence in the peptides file and is done with different software so inconsistencies are likely. The peaks are annotated based on the given sequence, with 20 ppm tolerance.

Copy Data

##### Spectrum 3948 (TSV)

###### Preview

```
Loading example...
```

*Click on the button to copy the data to your clipboard.*

Mz MinMz MaxIntensity Max

WidthHeightPeptide font sizePeptide stroke widthSpectrum font sizeSpectrum stroke widthCompact peptide

Ion legend

wxyz

abcd

OtherUnassignedIon^Charge^_Position_Show for top:%

VATVSJPR

05.57e+31.11e+41.67e+42.23e+4

Zoom Out

a^+1^_2_y^+1^_1_b^+1^_2_y^+1^_1_a^+1^_3_d^+1^_3_b^+1^_3_y^+1^_2_b^+1^_3_y^+1^_2_y^+2^_6_y^+2^_6_b^+1^_4_y^+2^_7_y^+1^_3_\*y^+1^_4_b^+1^_6_y^+1^_6_y^+1^_6_y^+1^_7_y^+1^_7_

0778155623353113

Fragment Matches Table

Show background peaks

| Position | Ion type | Intensity | mz Theoretical | mz Error (Th) | mz Error (ppm) | Charge | Series Number |
| --- | --- | --- | --- | --- | --- | --- | --- |
| - | - | 1302 | 120.1 | - | - | 0 | - |
| - | - | 555 | 128.1 | - | - | 0 | - |
| - | - | 793.3 | 129 | - | - | 0 | - |
| - | - | 1829 | 129.1 | - | - | 0 | - |
| - | - | 374.3 | 129.6 | - | - | 0 | - |
| - | - | 510.4 | 130.1 | - | - | 0 | - |
| - | - | 406.1 | 134.1 | - | - | 0 | - |
| - | - | 1297 | 136.1 | - | - | 0 | - |
| - | - | 1897 | 140.1 | - | - | 0 | - |
| 2 | a | 1.202E+04 | 143.1 | 6.117E-05 | 0.4274 | +1 | 2 |
| - | - | 889.4 | 144.1 | - | - | 0 | - |
| - | - | 621 | 145.1 | - | - | 0 | - |
| - | - | 396.3 | 148.5 | - | - | 0 | - |
| - | - | 480.4 | 148.8 | - | - | 0 | - |
| - | - | 744.7 | 148.9 | - | - | 0 | - |
| - | - | 682 | 148.9 | - | - | 0 | - |
| - | - | 695.8 | 148.9 | - | - | 0 | - |
| - | - | 1083 | 148.9 | - | - | 0 | - |
| - | - | 1357 | 148.9 | - | - | 0 | - |
| - | - | 1376 | 148.9 | - | - | 0 | - |
| - | - | 3156 | 148.9 | - | - | 0 | - |
| - | - | 5543 | 148.9 | - | - | 0 | - |
| - | - | 3800 | 149 | - | - | 0 | - |
| - | - | 1850 | 149 | - | - | 0 | - |
| - | - | 1052 | 149 | - | - | 0 | - |
| - | - | 1292 | 149 | - | - | 0 | - |
| - | - | 585.1 | 149 | - | - | 0 | - |
| - | - | 448 | 149 | - | - | 0 | - |
| - | - | 655.5 | 149 | - | - | 0 | - |
| - | - | 595.4 | 149 | - | - | 0 | - |
| - | - | 9671 | 149 | - | - | 0 | - |
| - | - | 617.9 | 149 | - | - | 0 | - |
| - | - | 484.1 | 149.1 | - | - | 0 | - |
| - | - | 4756 | 150 | - | - | 0 | - |
| - | - | 956 | 155.1 | - | - | 0 | - |
| - | - | 596.6 | 157 | - | - | 0 | - |
| 8 | y | 2452 | 158.1 | 2.636E-05 | 0.1667 | +1 | 1 |
| - | - | 580.4 | 159.1 | - | - | 0 | - |
| - | - | 708.5 | 167 | - | - | 0 | - |
| - | - | 711.9 | 169.1 | - | - | 0 | - |
| 2 | b | 1.085E+04 | 171.1 | 0.0001027 | 0.6004 | +1 | 2 |
| - | - | 549.4 | 171.1 | - | - | 0 | - |
| - | - | 1611 | 173.1 | - | - | 0 | - |
| - | - | 8785 | 173.1 | - | - | 0 | - |
| - | - | 1067 | 174.1 | - | - | 0 | - |
| - | - | 446.6 | 174.2 | - | - | 0 | - |
| 8 | y | 4836 | 175.1 | 0.0001167 | 0.6665 | +1 | 1 |
| - | - | 1310 | 183.1 | - | - | 0 | - |
| - | - | 726.1 | 183.1 | - | - | 0 | - |
| - | - | 772 | 185.1 | - | - | 0 | - |
| - | - | 1261 | 187.1 | - | - | 0 | - |
| - | - | 496.7 | 188.8 | - | - | 0 | - |
| - | - | 560.5 | 195.1 | - | - | 0 | - |
| - | - | 570.5 | 199.1 | - | - | 0 | - |
| - | - | 3547 | 201.1 | - | - | 0 | - |
| - | - | 1280 | 212.1 | - | - | 0 | - |
| 3 | a | 1040 | 226.2 | 0.0002334 | 1.032 | +1 | 3 |
| - | - | 1046 | 227.1 | - | - | 0 | - |
| 3 | d | 898.4 | 230.1 | 7.595E-05 | 0.33 | +1 | 3 |
| - | - | 486.1 | 243.1 | - | - | 0 | - |
| - | - | 523.5 | 250.4 | - | - | 0 | - |
| 3 | b | 5642 | 254.1 | 0.0003666 | 1.442 | +1 | 3 |
| 7 | y | 1.021E+04 | 255.1 | 0.0002847 | 1.116 | +1 | 2 |
| - | - | 679.1 | 255.2 | - | - | 0 | - |
| - | - | 1469 | 270.1 | - | - | 0 | - |
| 3 | b | 1669 | 272.2 | 0.0009062 | 3.33 | +1 | 3 |
| 7 | y | 6357 | 272.2 | 0.0002988 | 1.098 | +1 | 2 |
| - | - | 636.7 | 275.1 | - | - | 0 | - |
| - | - | 1511 | 288.2 | - | - | 0 | - |
| - | - | 929.2 | 291 | - | - | 0 | - |
| - | - | 1726 | 293.1 | - | - | 0 | - |
| - | - | 536.3 | 307.7 | - | - | 0 | - |
| - | - | 838.2 | 308 | - | - | 0 | - |
| - | - | 710.7 | 323.2 | - | - | 0 | - |
| - | - | 1997 | 325.2 | - | - | 0 | - |
| 3 | y | 2987 | 327.7 | 0.001118 | 3.412 | +2 | 6 |
| - | - | 1111 | 328.2 | - | - | 0 | - |
| 3 | y | 631.9 | 336.7 | 0.0005881 | 1.747 | +2 | 6 |
| - | - | 1118 | 341.2 | - | - | 0 | - |
| - | - | 602.3 | 350.9 | - | - | 0 | - |
| 4 | b | 1089 | 353.2 | 0.00065 | 1.84 | +1 | 4 |
| 2 | y | 3691 | 363.2 | 0.0007846 | 2.16 | +2 | 7 |
| - | - | 1717 | 363.7 | - | - | 0 | - |
| - | - | 798.5 | 368.9 | - | - | 0 | - |
| 6 | y | 4178 | 385.3 | 0.0006225 | 1.616 | +1 | 3 |
| - | - | 557.2 | 385.8 | - | - | 0 | - |
| - | - | 1056 | 386.3 | - | - | 0 | - |
| - | - | 2180 | 403.8 | - | - | 0 | - |
| 0 | Precursor | 1705 | 412.8 | 0.0005982 | 1.449 | +2 | -1 |
| - | - | 831.4 | 413.3 | - | - | 0 | - |
| - | - | 1033 | 420.9 | - | - | 0 | - |
| - | - | 781.3 | 439.8 | - | - | 0 | - |
| 5 | y | 2.204E+04 | 472.3 | 0.0002413 | 0.5109 | +1 | 4 |
| - | - | 5749 | 473.3 | - | - | 0 | - |
| - | - | 812.2 | 474.3 | - | - | 0 | - |
| - | - | 1019 | 486.3 | - | - | 0 | - |
| - | - | 606.4 | 547.4 | - | - | 0 | - |
| 6 | b | 6982 | 571.3 | 0.0103 | 18.02 | +1 | 6 |
| - | - | 2506 | 572.4 | - | - | 0 | - |
| - | - | 821.5 | 573.4 | - | - | 0 | - |
| - | - | 717.5 | 611.3 | - | - | 0 | - |
| - | - | 834.7 | 628.4 | - | - | 0 | - |
| 3 | y | 1324 | 654.4 | 0.008632 | 13.19 | +1 | 6 |
| 3 | y | 1.899E+04 | 672.4 | 0.001008 | 1.499 | +1 | 6 |
| - | - | 6746 | 673.4 | - | - | 0 | - |
| - | - | 1283 | 674.4 | - | - | 0 | - |
| - | - | 994.2 | 682.4 | - | - | 0 | - |
| 2 | y | 2679 | 725.4 | 9.102E-05 | 0.1255 | +1 | 7 |
| - | - | 1003 | 726.4 | - | - | 0 | - |
| 2 | y | 8935 | 743.4 | 0.0009511 | 1.279 | +1 | 7 |
| - | - | 4120 | 744.4 | - | - | 0 | - |
| - | - | 705.8 | 745.4 | - | - | 0 | - |
| - | - | 661.1 | 951.4 | - | - | 0 | - |
| - | - | 566.7 | 1244 | - | - | 0 | - |
| - | - | 620.3 | 1284 | - | - | 0 | - |
| - | - | 765.1 | 1397 | - | - | 0 | - |
| - | - | 738 | 2436 | - | - | 0 | - |
| - | - | 757.9 | 3082 | - | - | 0 | - |

m/z Charge Intensity FragmentType MassShift Position
120.08075714111328 0 1302.0537
128.10708618164062 0 554.98535
129.01821899414062 0 793.2584
129.10220336914062 0 1828.9974
129.58944702148438 0 374.25363
130.08633422851562 0 510.37796
134.08953857421875 0 406.0817
136.0756378173828 0 1296.9304
140.08192443847656 0 1897.281
143.11782836914062 0 12016.331 a 1
144.121337890625 0 889.3598
145.09718322753906 0 620.96063
148.45298767089844 0 396.28546
148.83099365234375 0 480.36105
148.89491271972656 0 744.6604
148.90213012695312 0 681.9823
148.9095001220703 0 695.82745
148.9166717529297 0 1083.0332
148.92384338378906 0 1356.8047
148.930908203125 0 1375.614
148.9382781982422 0 3156.4915
148.94589233398438 0 5542.9897
148.96243286132812 0 3799.6067
148.9700469970703 0 1850.1594
148.97702026367188 0 1051.9451
148.98439025878906 0 1292.1731
148.99122619628906 0 585.0983
148.9986572265625 0 447.99606
149.0057830810547 0 655.49255
149.01303100585938 0 595.3965
149.02325439453125 0 9670.806
149.03372192382812 0 617.91486
149.0703125 0 484.1073
150.02670288085938 0 4756.1626
155.11778259277344 0 956.0117
157.0129852294922 0 596.61646
158.09237670898438 0 2452.4785 y Ammonia loss 7
159.11175537109375 0 580.4364
167.0337677001953 0 708.53326
169.09725952148438 0 711.87573
171.11270141601562 0 10846.574 b 1
171.1379852294922 0 549.4209
173.09201049804688 0 1610.8691
173.12831115722656 0 8784.919
174.1320037841797 0 1066.874
174.15298461914062 0 446.64178
175.11883544921875 0 4836.492 y 7
183.1125030517578 0 1310.4622
183.14918518066406 0 726.0938
185.0804443359375 0 772.0439
187.1076202392578 0 1261.1826
188.78387451171875 0 496.73978
195.11289978027344 0 560.4741
199.1437530517578 0 570.50867
201.12327575683594 0 3546.7927
212.13938903808594 0 1279.753
226.15476989746094 0 1040.1079 a Water loss 2
227.13917541503906 0 1046.1368
230.14999389648438 0 898.38324 d 2
243.14503479003906 0 486.08423
250.44589233398438 0 523.53284
254.14955139160156 0 5642.4185 b Water loss 2
255.14488220214844 0 10208.797 y Ammonia loss 6
255.15768432617188 0 679.06506
270.1456604003906 0 1468.7737
272.1595764160156 0 1669.0488 b 2
272.1714172363281 0 6356.5645 y 6
275.08856201171875 0 636.7371
288.1554870605469 0 1511.0139
290.9598083496094 0 929.1861
293.0987243652344 0 1725.7166
307.7248229980469 0 536.25586
307.985595703125 0 838.2042
323.1700134277344 0 710.6768
325.222900390625 0 1996.7618
327.6991882324219 0 2987.204 y Water loss 2
328.2010498046875 0 1111.0775
336.7061767578125 0 631.88727 y 2
341.1821594238281 0 1118.4287
350.9263610839844 0 602.34503
353.2176818847656 0 1088.8049 b Water loss 3
363.21807861328125 0 3690.6858 y Water loss 1
363.7186279296875 0 1717.2539
368.9373474121094 0 798.5017
385.2551574707031 0 4178.495 y 5
385.84521484375 0 557.2097
386.2570495605469 0 1056.1643
403.8125 0 2179.9907
412.7524719238281 0 1704.9833 Precursor Water loss
413.25408935546875 0 831.4125
420.86041259765625 0 1032.912
439.8431396484375 0 781.2726
472.2875671386719 0 22043.295 y 4
473.2900695800781 0 5749.1084
474.292236328125 0 812.2121
486.291015625 0 1018.929
547.3695068359375 0 606.3944
571.3552856445312 0 6982.383 b 5
572.357666015625 0 2506.221
573.3547973632812 0 821.54193
611.3499755859375 0 717.49164
628.3792114257812 0 834.6902
654.3847045898438 0 1324.0496 y Water loss 2
672.4028930664062 0 18986.305 y 2
673.405517578125 0 6746.044
674.4088745117188 0 1282.779
682.3831176757812 0 994.1806
725.4303588867188 0 2678.8618 y Water loss 1
726.4298706054688 0 1002.85486
743.4400634765625 0 8934.562 y 1
744.4427490234375 0 4119.603
745.4451293945312 0 705.80804
951.3501586914062 0 661.0757
1244.0338134765625 0 566.7274
1284.156494140625 0 620.2934
1397.4405517578125 0 765.0907
2435.613525390625 0 737.9749
3081.94140625 0 757.86273

Spectrum Details

|  |  |
| --- | --- |
| Matched peaks? Matched peaks The total absolute number of peaks matched. Additionally in brackets the total fraction of peaks matched and the total number of peaks is shown. | 22 (18.64% of 118) |
| FDR? FDR The false discovery rate estimated for this peptide. It is calculated by matching all theoretical fragments with a non-integer shift with the raw peaks for this spectrum. This is done with 40 different shifts. The resulting percentage is the average number of annotated peaks over the number of annotated peaks with the correct spectrum. | 0.43% |
| Satellite FDR? Satellite FDR See the FDR for details on its calculation. This satellite ion specific FDR only contains the satellite ions (d/w) for I/L/J positions. | - |
| PSM Score? PSM Score The PSM Score as given by Hecklib to this annotated spectrum. It is shown with three significant figures. | 244 |

#### Reverse Lookup? Reverse Lookup All places where this read could be placed.

| Group | Segment | Template | Template Part | Read Part | Score | Unique |
| --- | --- | --- | --- | --- | --- | --- |
| Decoy | Decoy | TRYP | [107..115] | [0..8] | 64 | True |

| Recombined | Template Part | Read Part | Score | Unique |
| --- | --- | --- | --- | --- |
| TRYP | [107..115] | [0..8] | 64 | True |

#### Meta Information from Multiple reads

##### Number of combined reads

24

##### Intensity

0.9151

##### TotalArea

5.177E+05

##### Changes to the peptide sequence

VATVSJPR

L→JNo support for either Leucine or Isoleucine based on side chain ions (Position: 6)

#### Positional Score

Copy Data

##### Positional Score (TSV)

###### Preview

```
Loading example...
```

*Click on the button to copy the data to your clipboard.*

1001234567

Label Value
"0" 0.878
"1" 0.888
"2" 0.878
"3" 0.865
"4" 0.873
"5" 0.883
"6" 0.884
"7" 0.908

#### Meta Information from PEAKS

##### Scan Identifier

F4:5082

##### Original sequence

V

A

T

V

S

L

P

R

##### Posttranslational Modifications

##### Source File

D:\separate\_stitch\_analyses\xle-disambiguation\raw\20210323\_F1\_UM1\_Peng0013\_SA\_F59\_ingel\_3ug\_tryp.raw

##### Fraction

4

##### Scan Feature

-

##### De Novo Score

99

##### ConfidenceScore

98

### m/z

421.7588

##### Mass

841.5021

##### Charge

2

##### Retention Time

27.17

##### Predicted Retention Time

27.27

##### Area

0

##### Parts Per Million

1

##### Fragmentation mode

HCD

##### Originating file

01 D:\separate\_stitch\_analyses\xle-disambiguation\20210325\_F59\_3ug\_DENOVO\_12.csv

#### Meta Information from PEAKS

##### Scan Identifier

F4:5133

##### Original sequence

V

A

T

V

S

L

P

R

##### Posttranslational Modifications

##### Source File

D:\separate\_stitch\_analyses\xle-disambiguation\raw\20210323\_F1\_UM1\_Peng0013\_SA\_F59\_ingel\_3ug\_tryp.raw

##### Fraction

4

##### Scan Feature

-

##### De Novo Score

99

##### ConfidenceScore

97

### m/z

421.7589

##### Mass

841.5021

##### Charge

2

##### Retention Time

27.45

##### Predicted Retention Time

27.27

##### Area

0

##### Parts Per Million

1.3

##### Fragmentation mode

HCD

##### Originating file

01 D:\separate\_stitch\_analyses\xle-disambiguation\20210325\_F59\_3ug\_DENOVO\_12.csv

#### Meta Information from PEAKS

##### Scan Identifier

F4:5189

##### Original sequence

V

A

T

V

S

L

P

R

##### Posttranslational Modifications

##### Source File

D:\separate\_stitch\_analyses\xle-disambiguation\raw\20210323\_F1\_UM1\_Peng0013\_SA\_F59\_ingel\_3ug\_tryp.raw

##### Fraction

4

##### Scan Feature

-

##### De Novo Score

99

##### ConfidenceScore

97

### m/z

421.7592

##### Mass

841.5021

##### Charge

2

##### Retention Time

27.75

##### Predicted Retention Time

27.27

##### Area

0

##### Parts Per Million

2.1

##### Fragmentation mode

HCD

##### Originating file

01 D:\separate\_stitch\_analyses\xle-disambiguation\20210325\_F59\_3ug\_DENOVO\_12.csv

#### Meta Information from PEAKS

##### Scan Identifier

F4:5298

##### Original sequence

V

A

T

V

S

L

P

R

##### Posttranslational Modifications

##### Source File

D:\separate\_stitch\_analyses\xle-disambiguation\raw\20210323\_F1\_UM1\_Peng0013\_SA\_F59\_ingel\_3ug\_tryp.raw

##### Fraction

4

##### Scan Feature

-

##### De Novo Score

98

##### ConfidenceScore

97

### m/z

421.7585

##### Mass

841.5021

##### Charge

2

##### Retention Time

28.34

##### Predicted Retention Time

27.27

##### Area

0

##### Parts Per Million

0.3

##### Fragmentation mode

HCD

##### Originating file

01 D:\separate\_stitch\_analyses\xle-disambiguation\20210325\_F59\_3ug\_DENOVO\_12.csv

#### Meta Information from PEAKS

##### Scan Identifier

F4:5587

##### Original sequence

V

A

T

V

S

L

P

R

##### Posttranslational Modifications

##### Source File

D:\separate\_stitch\_analyses\xle-disambiguation\raw\20210323\_F1\_UM1\_Peng0013\_SA\_F59\_ingel\_3ug\_tryp.raw

##### Fraction

4

##### Scan Feature

-

##### De Novo Score

98

##### ConfidenceScore

97

### m/z

421.7585

##### Mass

841.5021

##### Charge

2

##### Retention Time

29.93

##### Predicted Retention Time

27.27

##### Area

0

##### Parts Per Million

0.5

##### Fragmentation mode

HCD

##### Originating file

01 D:\separate\_stitch\_analyses\xle-disambiguation\20210325\_F59\_3ug\_DENOVO\_12.csv

#### Meta Information from PEAKS

##### Scan Identifier

F4:5424

##### Original sequence

V

A

T

V

S

L

P

R

##### Posttranslational Modifications

##### Source File

D:\separate\_stitch\_analyses\xle-disambiguation\raw\20210323\_F1\_UM1\_Peng0013\_SA\_F59\_ingel\_3ug\_tryp.raw

##### Fraction

4

##### Scan Feature

-

##### De Novo Score

98

##### ConfidenceScore

97

### m/z

421.7589

##### Mass

841.5021

##### Charge

2

##### Retention Time

29.04

##### Predicted Retention Time

27.27

##### Area

0

##### Parts Per Million

1.3

##### Fragmentation mode

HCD

##### Originating file

01 D:\separate\_stitch\_analyses\xle-disambiguation\20210325\_F59\_3ug\_DENOVO\_12.csv

#### Meta Information from PEAKS

##### Scan Identifier

F4:5365

##### Original sequence

V

A

T

V

S

L

P

R

##### Posttranslational Modifications

##### Source File

D:\separate\_stitch\_analyses\xle-disambiguation\raw\20210323\_F1\_UM1\_Peng0013\_SA\_F59\_ingel\_3ug\_tryp.raw

##### Fraction

4

##### Scan Feature

-

##### De Novo Score

98

##### ConfidenceScore

96

### m/z

421.7583

##### Mass

841.5021

##### Charge

2

##### Retention Time

28.7

##### Predicted Retention Time

27.27

##### Area

0

##### Fragmentation mode

HCD

##### Originating file

01 D:\separate\_stitch\_analyses\xle-disambiguation\20210325\_F59\_3ug\_DENOVO\_12.csv

#### Meta Information from PEAKS

##### Scan Identifier

F4:5646

##### Original sequence

V

A

T

V

S

L

P

R

##### Posttranslational Modifications

##### Source File

D:\separate\_stitch\_analyses\xle-disambiguation\raw\20210323\_F1\_UM1\_Peng0013\_SA\_F59\_ingel\_3ug\_tryp.raw

##### Fraction

4

##### Scan Feature

-

##### De Novo Score

98

##### ConfidenceScore

96

### m/z

421.7588

##### Mass

841.5021

##### Charge

2

##### Retention Time

30.23

##### Predicted Retention Time

27.27

##### Area

0

##### Parts Per Million

1.1

##### Fragmentation mode

HCD

##### Originating file

01 D:\separate\_stitch\_analyses\xle-disambiguation\20210325\_F59\_3ug\_DENOVO\_12.csv

#### Meta Information from PEAKS

##### Scan Identifier

F4:6234

##### Original sequence

V

A

T

V

S

L

P

R

##### Posttranslational Modifications

##### Source File

D:\separate\_stitch\_analyses\xle-disambiguation\raw\20210323\_F1\_UM1\_Peng0013\_SA\_F59\_ingel\_3ug\_tryp.raw

##### Fraction

4

##### Scan Feature

-

##### De Novo Score

98

##### ConfidenceScore

96

### m/z

421.7591

##### Mass

841.5021

##### Charge

2

##### Retention Time

33.39

##### Predicted Retention Time

27.27

##### Area

0

##### Parts Per Million

1.9

##### Fragmentation mode

HCD

##### Originating file

01 D:\separate\_stitch\_analyses\xle-disambiguation\20210325\_F59\_3ug\_DENOVO\_12.csv

#### Meta Information from PEAKS

##### Scan Identifier

F4:7239

##### Original sequence

V

A

T

V

S

L

P

R

##### Posttranslational Modifications

##### Source File

D:\separate\_stitch\_analyses\xle-disambiguation\raw\20210323\_F1\_UM1\_Peng0013\_SA\_F59\_ingel\_3ug\_tryp.raw

##### Fraction

4

##### Scan Feature

-

##### De Novo Score

98

##### ConfidenceScore

96

### m/z

421.7586

##### Mass

841.5021

##### Charge

2

##### Retention Time

39.09

##### Predicted Retention Time

27.27

##### Area

0

##### Parts Per Million

0.6

##### Fragmentation mode

HCD

##### Originating file

01 D:\separate\_stitch\_analyses\xle-disambiguation\20210325\_F59\_3ug\_DENOVO\_12.csv

#### Meta Information from PEAKS

##### Scan Identifier

F4:5920

##### Original sequence

V

A

T

V

S

L

P

R

##### Posttranslational Modifications

##### Source File

D:\separate\_stitch\_analyses\xle-disambiguation\raw\20210323\_F1\_UM1\_Peng0013\_SA\_F59\_ingel\_3ug\_tryp.raw

##### Fraction

4

##### Scan Feature

-

##### De Novo Score

98

##### ConfidenceScore

96

### m/z

421.759

##### Mass

841.5021

##### Charge

2

##### Retention Time

31.65

##### Predicted Retention Time

27.27

##### Area

0

##### Parts Per Million

1.6

##### Fragmentation mode

HCD

##### Originating file

01 D:\separate\_stitch\_analyses\xle-disambiguation\20210325\_F59\_3ug\_DENOVO\_12.csv

#### Meta Information from PEAKS

##### Scan Identifier

F4:6159

##### Original sequence

V

A

T

V

S

L

P

R

##### Posttranslational Modifications

##### Source File

D:\separate\_stitch\_analyses\xle-disambiguation\raw\20210323\_F1\_UM1\_Peng0013\_SA\_F59\_ingel\_3ug\_tryp.raw

##### Fraction

4

##### Scan Feature

-

##### De Novo Score

98

##### ConfidenceScore

96

### m/z

421.7588

##### Mass

841.5021

##### Charge

2

##### Retention Time

32.99

##### Predicted Retention Time

27.27

##### Area

0

##### Parts Per Million

1.1

##### Fragmentation mode

HCD

##### Originating file

01 D:\separate\_stitch\_analyses\xle-disambiguation\20210325\_F59\_3ug\_DENOVO\_12.csv

#### Meta Information from PEAKS

##### Scan Identifier

F4:5978

##### Original sequence

V

A

T

V

S

L

P

R

##### Posttranslational Modifications

##### Source File

D:\separate\_stitch\_analyses\xle-disambiguation\raw\20210323\_F1\_UM1\_Peng0013\_SA\_F59\_ingel\_3ug\_tryp.raw

##### Fraction

4

##### Scan Feature

F4:1151

##### De Novo Score

98

##### ConfidenceScore

96

### m/z

421.7586

##### Mass

841.5021

##### Charge

2

##### Retention Time

31.77

##### Predicted Retention Time

27.27

##### Area

3.202E+05

##### Parts Per Million

0.6

##### Fragmentation mode

HCD

##### Originating file

01 D:\separate\_stitch\_analyses\xle-disambiguation\20210325\_F59\_3ug\_DENOVO\_12.csv

#### Meta Information from PEAKS

##### Scan Identifier

F4:6362

##### Original sequence

V

A

T

V

S

L

P

R

##### Posttranslational Modifications

##### Source File

D:\separate\_stitch\_analyses\xle-disambiguation\raw\20210323\_F1\_UM1\_Peng0013\_SA\_F59\_ingel\_3ug\_tryp.raw

##### Fraction

4

##### Scan Feature

-

##### De Novo Score

97

##### ConfidenceScore

96

### m/z

421.7592

##### Mass

841.5021

##### Charge

2

##### Retention Time

34.15

##### Predicted Retention Time

27.27

##### Area

0

##### Parts Per Million

2.1

##### Fragmentation mode

HCD

##### Originating file

01 D:\separate\_stitch\_analyses\xle-disambiguation\20210325\_F59\_3ug\_DENOVO\_12.csv

#### Meta Information from PEAKS

##### Scan Identifier

F4:7447

##### Original sequence

V

A

T

V

S

L

P

R

##### Posttranslational Modifications

##### Source File

D:\separate\_stitch\_analyses\xle-disambiguation\raw\20210323\_F1\_UM1\_Peng0013\_SA\_F59\_ingel\_3ug\_tryp.raw

##### Fraction

4

##### Scan Feature

-

##### De Novo Score

97

##### ConfidenceScore

95

### m/z

421.7591

##### Mass

841.5021

##### Charge

2

##### Retention Time

40.22

##### Predicted Retention Time

27.27

##### Area

0

##### Parts Per Million

1.8

##### Fragmentation mode

HCD

##### Originating file

01 D:\separate\_stitch\_analyses\xle-disambiguation\20210325\_F59\_3ug\_DENOVO\_12.csv

#### Meta Information from PEAKS

##### Scan Identifier

F4:6104

##### Original sequence

V

A

T

V

S

L

P

R

##### Posttranslational Modifications

##### Source File

D:\separate\_stitch\_analyses\xle-disambiguation\raw\20210323\_F1\_UM1\_Peng0013\_SA\_F59\_ingel\_3ug\_tryp.raw

##### Fraction

4

##### Scan Feature

-

##### De Novo Score

97

##### ConfidenceScore

95

### m/z

421.7596

##### Mass

841.5021

##### Charge

2

##### Retention Time

32.68

##### Predicted Retention Time

27.27

##### Area

0

##### Parts Per Million

2.9

##### Fragmentation mode

HCD

##### Originating file

01 D:\separate\_stitch\_analyses\xle-disambiguation\20210325\_F59\_3ug\_DENOVO\_12.csv

#### Meta Information from PEAKS

##### Scan Identifier

F4:4916

##### Original sequence

V

A

T

V

S

L

P

R

##### Posttranslational Modifications

##### Source File

D:\separate\_stitch\_analyses\xle-disambiguation\raw\20210323\_F1\_UM1\_Peng0013\_SA\_F59\_ingel\_3ug\_tryp.raw

##### Fraction

4

##### Scan Feature

-

##### De Novo Score

97

##### ConfidenceScore

96

### m/z

421.759

##### Mass

841.5021

##### Charge

2

##### Retention Time

26.28

##### Predicted Retention Time

27.27

##### Area

0

##### Parts Per Million

1.6

##### Fragmentation mode

HCD

##### Originating file

01 D:\separate\_stitch\_analyses\xle-disambiguation\20210325\_F59\_3ug\_DENOVO\_12.csv

#### Meta Information from PEAKS

##### Scan Identifier

F4:6819

##### Original sequence

V

A

T

V

S

L

P

R

##### Posttranslational Modifications

##### Source File

D:\separate\_stitch\_analyses\xle-disambiguation\raw\20210323\_F1\_UM1\_Peng0013\_SA\_F59\_ingel\_3ug\_tryp.raw

##### Fraction

4

##### Scan Feature

-

##### De Novo Score

97

##### ConfidenceScore

96

### m/z

421.7591

##### Mass

841.5021

##### Charge

2

##### Retention Time

36.74

##### Predicted Retention Time

27.27

##### Area

0

##### Parts Per Million

1.8

##### Fragmentation mode

HCD

##### Originating file

01 D:\separate\_stitch\_analyses\xle-disambiguation\20210325\_F59\_3ug\_DENOVO\_12.csv

#### Meta Information from PEAKS

##### Scan Identifier

F4:7964

##### Original sequence

V

A

T

V

S

L

P

R

##### Posttranslational Modifications

##### Source File

D:\separate\_stitch\_analyses\xle-disambiguation\raw\20210323\_F1\_UM1\_Peng0013\_SA\_F59\_ingel\_3ug\_tryp.raw

##### Fraction

4

##### Scan Feature

F4:1147

##### De Novo Score

97

##### ConfidenceScore

95

### m/z

421.7582

##### Mass

841.5021

##### Charge

2

##### Retention Time

42.87

##### Predicted Retention Time

27.27

##### Area

9.603E+04

##### Fragmentation mode

HCD

##### Originating file

01 D:\separate\_stitch\_analyses\xle-disambiguation\20210325\_F59\_3ug\_DENOVO\_12.csv

#### Meta Information from PEAKS

##### Scan Identifier

F4:6296

##### Original sequence

V

A

T

V

S

L

P

R

##### Posttranslational Modifications

##### Source File

D:\separate\_stitch\_analyses\xle-disambiguation\raw\20210323\_F1\_UM1\_Peng0013\_SA\_F59\_ingel\_3ug\_tryp.raw

##### Fraction

4

##### Scan Feature

-

##### De Novo Score

97

##### ConfidenceScore

95

### m/z

421.7593

##### Mass

841.5021

##### Charge

2

##### Retention Time

33.76

##### Predicted Retention Time

27.27

##### Area

0

##### Parts Per Million

2.4

##### Fragmentation mode

HCD

##### Originating file

01 D:\separate\_stitch\_analyses\xle-disambiguation\20210325\_F59\_3ug\_DENOVO\_12.csv

#### Meta Information from PEAKS

##### Scan Identifier

F4:6958

##### Original sequence

V

A

T

V

S

L

P

R

##### Posttranslational Modifications

##### Source File

D:\separate\_stitch\_analyses\xle-disambiguation\raw\20210323\_F1\_UM1\_Peng0013\_SA\_F59\_ingel\_3ug\_tryp.raw

##### Fraction

4

##### Scan Feature

-

##### De Novo Score

97

##### ConfidenceScore

95

### m/z

421.7594

##### Mass

841.5021

##### Charge

2

##### Retention Time

37.49

##### Predicted Retention Time

27.27

##### Area

0

##### Parts Per Million

2.4

##### Fragmentation mode

HCD

##### Originating file

01 D:\separate\_stitch\_analyses\xle-disambiguation\20210325\_F59\_3ug\_DENOVO\_12.csv

#### Meta Information from PEAKS

##### Scan Identifier

F4:6046

##### Original sequence

V

A

T

V

S

L

P

R

##### Posttranslational Modifications

##### Source File

D:\separate\_stitch\_analyses\xle-disambiguation\raw\20210323\_F1\_UM1\_Peng0013\_SA\_F59\_ingel\_3ug\_tryp.raw

##### Fraction

4

##### Scan Feature

-

##### De Novo Score

97

##### ConfidenceScore

95

### m/z

421.7589

##### Mass

841.5021

##### Charge

2

##### Retention Time

32.32

##### Predicted Retention Time

27.27

##### Area

0

##### Parts Per Million

1.2

##### Fragmentation mode

HCD

##### Originating file

01 D:\separate\_stitch\_analyses\xle-disambiguation\20210325\_F59\_3ug\_DENOVO\_12.csv

#### Meta Information from PEAKS

##### Scan Identifier

F4:6665

##### Original sequence

V

A

T

V

S

L

P

R

##### Posttranslational Modifications

##### Source File

D:\separate\_stitch\_analyses\xle-disambiguation\raw\20210323\_F1\_UM1\_Peng0013\_SA\_F59\_ingel\_3ug\_tryp.raw

##### Fraction

4

##### Scan Feature

-

##### De Novo Score

97

##### ConfidenceScore

95

### m/z

421.7591

##### Mass

841.5021

##### Charge

2

##### Retention Time

35.82

##### Predicted Retention Time

27.27

##### Area

0

##### Parts Per Million

1.9

##### Fragmentation mode

HCD

##### Originating file

01 D:\separate\_stitch\_analyses\xle-disambiguation\20210325\_F59\_3ug\_DENOVO\_12.csv

#### Meta Information from PEAKS

##### Scan Identifier

F4:3948

##### Original sequence

V

A

T

V

S

L

P

R

##### Posttranslational Modifications

##### Source File

D:\separate\_stitch\_analyses\xle-disambiguation\raw\20210323\_F1\_UM1\_Peng0013\_SA\_F59\_ingel\_3ug\_tryp.raw

##### Fraction

4

##### Scan Feature

F4:1153

##### De Novo Score

92

##### ConfidenceScore

96

### m/z

421.7587

##### Mass

841.5021

##### Charge

2

##### Retention Time

21.18

##### Predicted Retention Time

27.27

##### Area

1.014E+05

##### Parts Per Million

0.8

##### Fragmentation mode

HCD

##### Originating file

01 D:\separate\_stitch\_analyses\xle-disambiguation\20210325\_F59\_3ug\_DENOVO\_12.csv
