## Supplementary material for "A handle on mass coincidence errors in *de novo* sequencing of antibodies by bottom-up proteomics": Combined_013.html

Details Combined\_013 | Stitch OverviewUndefined

### Read Combined\_013

#### Sequence (length=10)

JQAEDESMYF

#### Spectrum 6919? Spectrum 6919 The raw spectrum of this peptide as annotated by Hecklib. The fragments are coloured according to ion type (see legend). Any peaks with a star '\*' as text can be hovered over to see the full details, first the ion type second the mass shift type. By hovering over the amino acids in the peptide or ions in the legend the corresponding peaks are highlighted. By toggling the 'Unassigned' label you can turn the background (unassigned) peaks on or off in the plot. By updating the slider in the Ion legend you can update the spectrum to only show the top X% of the peaks with labels. The top X% means any peak that is within X% of the highest intensity. By dragging in the spectrum you can zoom in to a specific part of the spectrum and use 'Zoom Out' to get back to the original zoom level. The annotation of the spectrum is based on the given sequence in the peptides file and is done with different software so inconsistencies are likely. The peaks are annotated based on the given sequence, with 20 ppm tolerance.

Copy Data

##### Spectrum 6919 (TSV)

###### Preview

```
Loading example...
```

*Click on the button to copy the data to your clipboard.*

Mz MinMz MaxIntensity Max

WidthHeightPeptide font sizePeptide stroke widthSpectrum font sizeSpectrum stroke widthCompact peptide

Ion legend

wxyz

abcd

OtherUnassignedIon^Charge^_Position_Show for top:%

JQAEDESMYF

01.31e+52.63e+53.94e+55.26e+5

Zoom Out

y^+1^_1_c^+1^_2_c^+1^_2_c^+1^_3_y^+1^_2_c^+1^_3_c^+2^_6_c^+2^_7_c^+1^_4_c^+1^_4_c^+2^_8_y^+1^_3_z^+2^_8_c^+2^_9_y^+1^_4_c^+1^_5_y^+1^_4_c^+1^_5_y^+1^_5_z^+1^_5_c^+1^_6_y^+1^_5_c^+1^_6_z^+1^_6_c^+1^_7_y^+1^_6_c^+1^_7_z^+1^_6_y^+1^_6_w^+1^_7_c^+1^_8_y^+1^_7_c^+1^_8_z^+1^_8_y^+1^_8_z^+1^_8_y^+1^_8_w^+1^_9_c^+1^_9_c^+1^_9_z^+1^_9_y^+1^_9_

0658131519732630

Fragment Matches Table

Show background peaks

| Position | Ion type | Intensity | mz Theoretical | mz Error (Th) | mz Error (ppm) | Charge | Series Number |
| --- | --- | --- | --- | --- | --- | --- | --- |
| - | - | 4.38E+04 | 120.1 | - | - | 0 | - |
| - | - | 3666 | 121.1 | - | - | 0 | - |
| - | - | 457.8 | 127.2 | - | - | 0 | - |
| - | - | 418.1 | 128.1 | - | - | 0 | - |
| - | - | 1.283E+04 | 131.1 | - | - | 0 | - |
| - | - | 720.5 | 132.1 | - | - | 0 | - |
| - | - | 855.4 | 133.1 | - | - | 0 | - |
| - | - | 410.8 | 133.4 | - | - | 0 | - |
| - | - | 397.1 | 133.9 | - | - | 0 | - |
| - | - | 414 | 135.5 | - | - | 0 | - |
| - | - | 1.789E+05 | 136.1 | - | - | 0 | - |
| - | - | 1.338E+04 | 137.1 | - | - | 0 | - |
| - | - | 497.8 | 137.6 | - | - | 0 | - |
| - | - | 1470 | 149 | - | - | 0 | - |
| - | - | 563.6 | 149 | - | - | 0 | - |
| - | - | 943.5 | 149.1 | - | - | 0 | - |
| - | - | 4610 | 153.1 | - | - | 0 | - |
| - | - | 947.3 | 154.1 | - | - | 0 | - |
| - | - | 667.7 | 162.1 | - | - | 0 | - |
| - | - | 1582 | 166.1 | - | - | 0 | - |
| 10 | y | 1.122E+05 | 166.1 | 0.0004301 | 2.59 | +1 | 1 |
| - | - | 841.2 | 167.1 | - | - | 0 | - |
| - | - | 1.001E+04 | 167.1 | - | - | 0 | - |
| - | - | 558.6 | 168.1 | - | - | 0 | - |
| - | - | 762.6 | 171.1 | - | - | 0 | - |
| - | - | 497.7 | 182.1 | - | - | 0 | - |
| - | - | 840.7 | 197.1 | - | - | 0 | - |
| - | - | 1163 | 200 | - | - | 0 | - |
| - | - | 1153 | 200.1 | - | - | 0 | - |
| - | - | 1178 | 201.1 | - | - | 0 | - |
| - | - | 1058 | 206.1 | - | - | 0 | - |
| - | - | 459.5 | 212.6 | - | - | 0 | - |
| - | - | 3335 | 214.2 | - | - | 0 | - |
| - | - | 6264 | 217.1 | - | - | 0 | - |
| - | - | 939.5 | 218 | - | - | 0 | - |
| - | - | 617.4 | 218.1 | - | - | 0 | - |
| - | - | 3785 | 221.1 | - | - | 0 | - |
| - | - | 636.2 | 222.2 | - | - | 0 | - |
| - | - | 519.2 | 222.5 | - | - | 0 | - |
| - | - | 625.7 | 223.1 | - | - | 0 | - |
| - | - | 1375 | 225 | - | - | 0 | - |
| - | - | 1.286E+04 | 225.1 | - | - | 0 | - |
| - | - | 1436 | 226.1 | - | - | 0 | - |
| - | - | 555.7 | 229 | - | - | 0 | - |
| - | - | 555.4 | 232.8 | - | - | 0 | - |
| - | - | 1.755E+04 | 235.1 | - | - | 0 | - |
| - | - | 1150 | 236.1 | - | - | 0 | - |
| - | - | 4659 | 239.1 | - | - | 0 | - |
| - | - | 634.7 | 240.1 | - | - | 0 | - |
| - | - | 661.1 | 241.1 | - | - | 0 | - |
| 2 | c | 6.687E+04 | 242.1 | 0.0005642 | 2.33 | +1 | 2 |
| - | - | 486.1 | 243.1 | - | - | 0 | - |
| - | - | 599.1 | 243.1 | - | - | 0 | - |
| - | - | 7746 | 243.2 | - | - | 0 | - |
| - | - | 586.8 | 244.2 | - | - | 0 | - |
| - | - | 2907 | 245.1 | - | - | 0 | - |
| - | - | 532.1 | 248 | - | - | 0 | - |
| - | - | 598.1 | 257.1 | - | - | 0 | - |
| - | - | 475.9 | 259.1 | - | - | 0 | - |
| 2 | c | 907.6 | 259.2 | 0.0004434 | 1.711 | +1 | 2 |
| - | - | 671.7 | 268.2 | - | - | 0 | - |
| - | - | 5081 | 283.1 | - | - | 0 | - |
| - | - | 3952 | 283.1 | - | - | 0 | - |
| - | - | 6477 | 295.1 | - | - | 0 | - |
| - | - | 2549 | 296.1 | - | - | 0 | - |
| - | - | 642.5 | 296.2 | - | - | 0 | - |
| - | - | 1040 | 299.1 | - | - | 0 | - |
| - | - | 6197 | 311.1 | - | - | 0 | - |
| - | - | 1188 | 311.1 | - | - | 0 | - |
| - | - | 629.9 | 311.2 | - | - | 0 | - |
| 3 | c | 1.634E+04 | 313.2 | 0.0007124 | 2.275 | +1 | 3 |
| - | - | 1310 | 314.1 | - | - | 0 | - |
| - | - | 747.7 | 314.2 | - | - | 0 | - |
| - | - | 2508 | 314.2 | - | - | 0 | - |
| - | - | 1393 | 316.1 | - | - | 0 | - |
| - | - | 987.2 | 328.1 | - | - | 0 | - |
| 9 | y | 1.341E+05 | 329.1 | 0.000746 | 2.266 | +1 | 2 |
| - | - | 2.629E+04 | 330.2 | - | - | 0 | - |
| 3 | c | 4180 | 330.2 | 0.0005915 | 1.791 | +1 | 3 |
| - | - | 2882 | 331.2 | - | - | 0 | - |
| - | - | 835.4 | 332.1 | - | - | 0 | - |
| - | - | 721.1 | 334.1 | - | - | 0 | - |
| 6 | c | 1124 | 343.7 | 0.001261 | 3.67 | +2 | 6 |
| - | - | 4591 | 346.1 | - | - | 0 | - |
| - | - | 974.1 | 347.1 | - | - | 0 | - |
| - | - | 801.3 | 347.2 | - | - | 0 | - |
| - | - | 696.9 | 353.2 | - | - | 0 | - |
| - | - | 1195 | 355.1 | - | - | 0 | - |
| - | - | 876.8 | 356.1 | - | - | 0 | - |
| - | - | 1228 | 357.1 | - | - | 0 | - |
| - | - | 1067 | 358.2 | - | - | 0 | - |
| - | - | 510.1 | 364.1 | - | - | 0 | - |
| - | - | 2640 | 364.1 | - | - | 0 | - |
| - | - | 547.5 | 367.1 | - | - | 0 | - |
| - | - | 3900 | 369.1 | - | - | 0 | - |
| - | - | 1164 | 370.1 | - | - | 0 | - |
| - | - | 790.5 | 370.1 | - | - | 0 | - |
| - | - | 602.9 | 371.1 | - | - | 0 | - |
| - | - | 884.9 | 374.1 | - | - | 0 | - |
| - | - | 6923 | 378.2 | - | - | 0 | - |
| - | - | 2790 | 378.7 | - | - | 0 | - |
| - | - | 604 | 379.2 | - | - | 0 | - |
| - | - | 695.6 | 379.2 | - | - | 0 | - |
| - | - | 672.5 | 381.1 | - | - | 0 | - |
| - | - | 689.1 | 382.2 | - | - | 0 | - |
| - | - | 699.6 | 386.2 | - | - | 0 | - |
| 7 | c | 2493 | 387.2 | 0.001177 | 3.041 | +2 | 7 |
| - | - | 1536 | 389.2 | - | - | 0 | - |
| - | - | 761.9 | 397.1 | - | - | 0 | - |
| - | - | 1.015E+04 | 398.1 | - | - | 0 | - |
| - | - | 1084 | 399.1 | - | - | 0 | - |
| - | - | 1221 | 405.7 | - | - | 0 | - |
| - | - | 700 | 410.7 | - | - | 0 | - |
| - | - | 737.7 | 412.2 | - | - | 0 | - |
| - | - | 2557 | 414.2 | - | - | 0 | - |
| - | - | 615.8 | 415.2 | - | - | 0 | - |
| - | - | 1577 | 416.3 | - | - | 0 | - |
| - | - | 679.6 | 417.1 | - | - | 0 | - |
| - | - | 5330 | 419.7 | - | - | 0 | - |
| - | - | 3047 | 420.2 | - | - | 0 | - |
| - | - | 1191 | 424.2 | - | - | 0 | - |
| - | - | 760.6 | 425.1 | - | - | 0 | - |
| - | - | 2151 | 425.2 | - | - | 0 | - |
| - | - | 750.6 | 426.2 | - | - | 0 | - |
| - | - | 2349 | 427.1 | - | - | 0 | - |
| - | - | 867 | 441.2 | - | - | 0 | - |
| 4 | c | 1.545E+04 | 442.2 | 0.001241 | 2.805 | +1 | 4 |
| - | - | 4460 | 442.7 | - | - | 0 | - |
| - | - | 1918 | 443.2 | - | - | 0 | - |
| - | - | 3234 | 443.2 | - | - | 0 | - |
| - | - | 1238 | 443.7 | - | - | 0 | - |
| - | - | 3067 | 444.2 | - | - | 0 | - |
| - | - | 4436 | 445.2 | - | - | 0 | - |
| - | - | 832.3 | 446.2 | - | - | 0 | - |
| - | - | 5.744E+04 | 451.7 | - | - | 0 | - |
| - | - | 2.74E+04 | 452.2 | - | - | 0 | - |
| - | - | 1.024E+04 | 452.7 | - | - | 0 | - |
| - | - | 1884 | 453.2 | - | - | 0 | - |
| - | - | 1195 | 458.2 | - | - | 0 | - |
| - | - | 4589 | 458.2 | - | - | 0 | - |
| - | - | 1.211E+04 | 458.2 | - | - | 0 | - |
| 4 | c | 3.449E+04 | 459.3 | 0.0009061 | 1.973 | +1 | 4 |
| - | - | 7089 | 460.3 | - | - | 0 | - |
| 8 | c | 6864 | 460.7 | 0.003422 | 7.427 | +2 | 8 |
| - | - | 6108 | 461.1 | - | - | 0 | - |
| - | - | 3896 | 461.2 | - | - | 0 | - |
| - | - | 1021 | 461.3 | - | - | 0 | - |
| - | - | 1604 | 461.7 | - | - | 0 | - |
| - | - | 750 | 462.1 | - | - | 0 | - |
| 8 | y | 1.419E+04 | 476.2 | 0.005936 | 12.47 | +1 | 3 |
| - | - | 1841 | 476.2 | - | - | 0 | - |
| - | - | 3221 | 477.2 | - | - | 0 | - |
| - | - | 836.1 | 478.2 | - | - | 0 | - |
| - | - | 689 | 478.7 | - | - | 0 | - |
| - | - | 6120 | 479.1 | - | - | 0 | - |
| - | - | 872 | 480.1 | - | - | 0 | - |
| - | - | 1263 | 481.2 | - | - | 0 | - |
| - | - | 1222 | 481.2 | - | - | 0 | - |
| - | - | 1320 | 486.2 | - | - | 0 | - |
| - | - | 710.4 | 487.2 | - | - | 0 | - |
| - | - | 870.4 | 494.2 | - | - | 0 | - |
| 3 | z | 756.1 | 496.2 | 0.004292 | 8.65 | +2 | 8 |
| - | - | 1608 | 496.2 | - | - | 0 | - |
| - | - | 1353 | 496.7 | - | - | 0 | - |
| - | - | 647.1 | 497.2 | - | - | 0 | - |
| - | - | 1220 | 499.2 | - | - | 0 | - |
| - | - | 2082 | 509.2 | - | - | 0 | - |
| - | - | 1685 | 510.2 | - | - | 0 | - |
| - | - | 1021 | 510.7 | - | - | 0 | - |
| - | - | 1681 | 511.3 | - | - | 0 | - |
| - | - | 1111 | 514.2 | - | - | 0 | - |
| - | - | 1188 | 517.2 | - | - | 0 | - |
| - | - | 1098 | 519.2 | - | - | 0 | - |
| - | - | 600.7 | 519.7 | - | - | 0 | - |
| - | - | 1395 | 522.2 | - | - | 0 | - |
| - | - | 1514 | 527.2 | - | - | 0 | - |
| - | - | 915.2 | 527.2 | - | - | 0 | - |
| - | - | 2628 | 528.2 | - | - | 0 | - |
| - | - | 1811 | 528.7 | - | - | 0 | - |
| - | - | 3523 | 529.2 | - | - | 0 | - |
| - | - | 4782 | 530.3 | - | - | 0 | - |
| - | - | 7902 | 531.3 | - | - | 0 | - |
| - | - | 1777 | 532.2 | - | - | 0 | - |
| - | - | 1765 | 532.3 | - | - | 0 | - |
| - | - | 821.3 | 533.2 | - | - | 0 | - |
| - | - | 1539 | 536.2 | - | - | 0 | - |
| - | - | 684.5 | 536.7 | - | - | 0 | - |
| - | - | 4362 | 539.2 | - | - | 0 | - |
| - | - | 1947 | 540.2 | - | - | 0 | - |
| 9 | c | 1684 | 542.2 | 0.003953 | 7.291 | +2 | 9 |
| - | - | 842.2 | 542.7 | - | - | 0 | - |
| 7 | y | 891.4 | 545.2 | 0.01075 | 19.71 | +1 | 4 |
| - | - | 830.4 | 547.3 | - | - | 0 | - |
| - | - | 1106 | 549.2 | - | - | 0 | - |
| - | - | 591.5 | 554.2 | - | - | 0 | - |
| - | - | 1641 | 555.2 | - | - | 0 | - |
| - | - | 3327 | 556.2 | - | - | 0 | - |
| 5 | c | 1.188E+04 | 557.3 | 0.0009395 | 1.686 | +1 | 5 |
| - | - | 3147 | 558.3 | - | - | 0 | - |
| - | - | 735.5 | 559.3 | - | - | 0 | - |
| 7 | y | 1.715E+04 | 563.2 | 0.005799 | 10.3 | +1 | 4 |
| - | - | 4397 | 564.2 | - | - | 0 | - |
| - | - | 1011 | 565.2 | - | - | 0 | - |
| - | - | 2792 | 572.2 | - | - | 0 | - |
| - | - | 1211 | 573.2 | - | - | 0 | - |
| - | - | 3161 | 573.2 | - | - | 0 | - |
| - | - | 3.34E+04 | 573.3 | - | - | 0 | - |
| 5 | c | 1.617E+05 | 574.3 | 0.0009407 | 1.638 | +1 | 5 |
| - | - | 4162 | 575.2 | - | - | 0 | - |
| - | - | 4.447E+04 | 575.3 | - | - | 0 | - |
| - | - | 1407 | 576.2 | - | - | 0 | - |
| - | - | 7945 | 576.3 | - | - | 0 | - |
| - | - | 694.1 | 577.3 | - | - | 0 | - |
| - | - | 3319 | 583.8 | - | - | 0 | - |
| - | - | 3544 | 584.3 | - | - | 0 | - |
| - | - | 1681 | 584.8 | - | - | 0 | - |
| - | - | 611.6 | 585.3 | - | - | 0 | - |
| - | - | 892.8 | 587.2 | - | - | 0 | - |
| - | - | 1086 | 589.3 | - | - | 0 | - |
| - | - | 3036 | 590.2 | - | - | 0 | - |
| - | - | 2158 | 592.2 | - | - | 0 | - |
| - | - | 9954 | 592.8 | - | - | 0 | - |
| - | - | 6224 | 593.3 | - | - | 0 | - |
| - | - | 2064 | 593.8 | - | - | 0 | - |
| - | - | 870 | 595.3 | - | - | 0 | - |
| - | - | 833.8 | 597.2 | - | - | 0 | - |
| - | - | 4449 | 600.3 | - | - | 0 | - |
| - | - | 1483 | 601.3 | - | - | 0 | - |
| - | - | 846.8 | 605.3 | - | - | 0 | - |
| - | - | 583.1 | 606.3 | - | - | 0 | - |
| - | - | 1101 | 606.3 | - | - | 0 | - |
| - | - | 1840 | 608.2 | - | - | 0 | - |
| - | - | 760.6 | 609.2 | - | - | 0 | - |
| - | - | 686.8 | 609.2 | - | - | 0 | - |
| - | - | 865.5 | 610.2 | - | - | 0 | - |
| - | - | 1766 | 614.2 | - | - | 0 | - |
| - | - | 1150 | 615.2 | - | - | 0 | - |
| - | - | 1223 | 615.4 | - | - | 0 | - |
| - | - | 679.8 | 615.8 | - | - | 0 | - |
| - | - | 672.6 | 622.3 | - | - | 0 | - |
| - | - | 1100 | 623.3 | - | - | 0 | - |
| - | - | 1507 | 623.4 | - | - | 0 | - |
| - | - | 1349 | 624.2 | - | - | 0 | - |
| - | - | 683 | 624.4 | - | - | 0 | - |
| - | - | 1679 | 625.2 | - | - | 0 | - |
| - | - | 804.4 | 626.2 | - | - | 0 | - |
| - | - | 834.7 | 626.3 | - | - | 0 | - |
| - | - | 995 | 627.3 | - | - | 0 | - |
| - | - | 668.5 | 628.3 | - | - | 0 | - |
| - | - | 1687 | 630.3 | - | - | 0 | - |
| - | - | 1137 | 631.3 | - | - | 0 | - |
| - | - | 1553 | 639.3 | - | - | 0 | - |
| - | - | 1774 | 640.3 | - | - | 0 | - |
| - | - | 2158 | 641.3 | - | - | 0 | - |
| - | - | 5967 | 642.2 | - | - | 0 | - |
| - | - | 3197 | 643.2 | - | - | 0 | - |
| - | - | 978.3 | 644.2 | - | - | 0 | - |
| - | - | 507 | 645.3 | - | - | 0 | - |
| - | - | 835.3 | 650.3 | - | - | 0 | - |
| - | - | 936.8 | 651.3 | - | - | 0 | - |
| - | - | 4768 | 658.3 | - | - | 0 | - |
| - | - | 1.508E+04 | 659.3 | - | - | 0 | - |
| - | - | 1014 | 660.3 | - | - | 0 | - |
| - | - | 9798 | 660.3 | - | - | 0 | - |
| - | - | 8071 | 661.2 | - | - | 0 | - |
| - | - | 2432 | 661.3 | - | - | 0 | - |
| - | - | 1848 | 662.2 | - | - | 0 | - |
| - | - | 2053 | 663.2 | - | - | 0 | - |
| - | - | 662.6 | 664.2 | - | - | 0 | - |
| - | - | 7446 | 668.3 | - | - | 0 | - |
| - | - | 4101 | 669.3 | - | - | 0 | - |
| - | - | 1608 | 670.3 | - | - | 0 | - |
| - | - | 965.7 | 671.3 | - | - | 0 | - |
| 6 | y | 1132 | 674.2 | 0.002518 | 3.735 | +1 | 5 |
| 6 | z | 1271 | 676.2 | 0.005673 | 8.388 | +1 | 5 |
| - | - | 1966 | 676.3 | - | - | 0 | - |
| - | - | 630.9 | 678.2 | - | - | 0 | - |
| - | - | 8928 | 679.2 | - | - | 0 | - |
| - | - | 1751 | 680.2 | - | - | 0 | - |
| - | - | 5357 | 681.2 | - | - | 0 | - |
| - | - | 1418 | 682.2 | - | - | 0 | - |
| - | - | 756.4 | 685.3 | - | - | 0 | - |
| 6 | c | 2.815E+04 | 686.3 | 0.001254 | 1.827 | +1 | 6 |
| - | - | 8191 | 687.3 | - | - | 0 | - |
| - | - | 5009 | 688.3 | - | - | 0 | - |
| - | - | 1014 | 689.3 | - | - | 0 | - |
| - | - | 2945 | 691.3 | - | - | 0 | - |
| 6 | y | 9824 | 692.3 | 0.005869 | 8.479 | +1 | 5 |
| - | - | 2836 | 693.3 | - | - | 0 | - |
| - | - | 1650 | 694.3 | - | - | 0 | - |
| - | - | 786 | 695.2 | - | - | 0 | - |
| - | - | 1461 | 696.2 | - | - | 0 | - |
| - | - | 1.311E+05 | 702.3 | - | - | 0 | - |
| 6 | c | 7E+04 | 703.3 | 0.001308 | 1.86 | +1 | 6 |
| - | - | 1.763E+04 | 704.3 | - | - | 0 | - |
| - | - | 2470 | 705.3 | - | - | 0 | - |
| - | - | 785.2 | 710.3 | - | - | 0 | - |
| - | - | 1051 | 717.3 | - | - | 0 | - |
| - | - | 719.2 | 719.3 | - | - | 0 | - |
| - | - | 2727 | 720.3 | - | - | 0 | - |
| - | - | 894.2 | 721.3 | - | - | 0 | - |
| - | - | 1631 | 724.3 | - | - | 0 | - |
| - | - | 1632 | 728.3 | - | - | 0 | - |
| - | - | 2690 | 729.3 | - | - | 0 | - |
| - | - | 1814 | 730.3 | - | - | 0 | - |
| - | - | 3661 | 737.3 | - | - | 0 | - |
| - | - | 1560 | 738.3 | - | - | 0 | - |
| - | - | 957.8 | 739.3 | - | - | 0 | - |
| - | - | 2347 | 743.3 | - | - | 0 | - |
| - | - | 1062 | 744.3 | - | - | 0 | - |
| - | - | 1094 | 745.3 | - | - | 0 | - |
| - | - | 2265 | 746.3 | - | - | 0 | - |
| - | - | 2720 | 747.3 | - | - | 0 | - |
| - | - | 1358 | 747.4 | - | - | 0 | - |
| - | - | 2091 | 748.3 | - | - | 0 | - |
| - | - | 788.8 | 748.4 | - | - | 0 | - |
| - | - | 997.9 | 750.3 | - | - | 0 | - |
| - | - | 1798 | 753.2 | - | - | 0 | - |
| - | - | 1456 | 754.2 | - | - | 0 | - |
| - | - | 2.848E+04 | 755.3 | - | - | 0 | - |
| - | - | 1.122E+04 | 756.3 | - | - | 0 | - |
| - | - | 1892 | 757.3 | - | - | 0 | - |
| - | - | 951.3 | 760.3 | - | - | 0 | - |
| - | - | 918.3 | 768.3 | - | - | 0 | - |
| - | - | 2471 | 771.3 | - | - | 0 | - |
| - | - | 1625 | 771.3 | - | - | 0 | - |
| - | - | 4621 | 772.2 | - | - | 0 | - |
| 5 | z | 1389 | 773.3 | 0.000696 | 0.9001 | +1 | 6 |
| 7 | c | 2.086E+04 | 773.3 | 0.001294 | 1.674 | +1 | 7 |
| - | - | 6798 | 774.3 | - | - | 0 | - |
| - | - | 1973 | 775.3 | - | - | 0 | - |
| - | - | 1392 | 778.3 | - | - | 0 | - |
| - | - | 670.6 | 778.9 | - | - | 0 | - |
| 5 | y | 4158 | 789.3 | 0.005299 | 6.714 | +1 | 6 |
| - | - | 7.7E+04 | 789.4 | - | - | 0 | - |
| 7 | c | 1.543E+05 | 790.4 | 0.0009041 | 1.144 | +1 | 7 |
| 5 | z | 3170 | 791.3 | 0.01456 | 18.4 | +1 | 6 |
| - | - | 5.305E+04 | 791.4 | - | - | 0 | - |
| - | - | 3222 | 792.3 | - | - | 0 | - |
| - | - | 9583 | 792.4 | - | - | 0 | - |
| - | - | 1841 | 793.3 | - | - | 0 | - |
| - | - | 1107 | 803.4 | - | - | 0 | - |
| - | - | 1661 | 804.4 | - | - | 0 | - |
| - | - | 1128 | 805.4 | - | - | 0 | - |
| - | - | 4586 | 806.3 | - | - | 0 | - |
| 5 | y | 2.196E+04 | 807.3 | 0.005294 | 6.557 | +1 | 6 |
| - | - | 9625 | 808.3 | - | - | 0 | - |
| - | - | 1.002E+04 | 809.3 | - | - | 0 | - |
| - | - | 1989 | 810.3 | - | - | 0 | - |
| - | - | 826.9 | 811.4 | - | - | 0 | - |
| - | - | 2303 | 814.3 | - | - | 0 | - |
| - | - | 933.3 | 816.3 | - | - | 0 | - |
| - | - | 1782 | 816.4 | - | - | 0 | - |
| - | - | 1191 | 821.4 | - | - | 0 | - |
| - | - | 2438 | 822.3 | - | - | 0 | - |
| - | - | 620.9 | 823.4 | - | - | 0 | - |
| - | - | 2852 | 824.3 | - | - | 0 | - |
| - | - | 1031 | 825.3 | - | - | 0 | - |
| - | - | 3983 | 826.3 | - | - | 0 | - |
| - | - | 1781 | 827.3 | - | - | 0 | - |
| - | - | 867 | 828.3 | - | - | 0 | - |
| - | - | 3187 | 828.4 | - | - | 0 | - |
| - | - | 1553 | 829.4 | - | - | 0 | - |
| - | - | 1434 | 830.4 | - | - | 0 | - |
| - | - | 3773 | 831.4 | - | - | 0 | - |
| - | - | 1583 | 832.4 | - | - | 0 | - |
| - | - | 4218 | 838.4 | - | - | 0 | - |
| - | - | 4764 | 839.4 | - | - | 0 | - |
| - | - | 2014 | 840.4 | - | - | 0 | - |
| - | - | 8717 | 842.3 | - | - | 0 | - |
| - | - | 4718 | 843.3 | - | - | 0 | - |
| - | - | 1135 | 844.3 | - | - | 0 | - |
| - | - | 906.5 | 849.3 | - | - | 0 | - |
| - | - | 713.9 | 855.3 | - | - | 0 | - |
| - | - | 7280 | 856.4 | - | - | 0 | - |
| - | - | 3138 | 857.4 | - | - | 0 | - |
| - | - | 1436 | 858.4 | - | - | 0 | - |
| - | - | 883.9 | 859.4 | - | - | 0 | - |
| 4 | w | 1006 | 861.3 | 0.004006 | 4.651 | +1 | 7 |
| - | - | 1256 | 867.3 | - | - | 0 | - |
| - | - | 987.5 | 868.3 | - | - | 0 | - |
| - | - | 2404 | 872.3 | - | - | 0 | - |
| - | - | 849.7 | 873.3 | - | - | 0 | - |
| - | - | 2328 | 874.4 | - | - | 0 | - |
| - | - | 1925 | 875.4 | - | - | 0 | - |
| - | - | 1163 | 883.4 | - | - | 0 | - |
| - | - | 6875 | 884.3 | - | - | 0 | - |
| - | - | 5440 | 885.3 | - | - | 0 | - |
| - | - | 1956 | 886.3 | - | - | 0 | - |
| - | - | 781.5 | 887.3 | - | - | 0 | - |
| - | - | 860.3 | 889.4 | - | - | 0 | - |
| - | - | 902.4 | 890.3 | - | - | 0 | - |
| - | - | 823.8 | 891.4 | - | - | 0 | - |
| - | - | 5733 | 892.4 | - | - | 0 | - |
| - | - | 1.704E+04 | 893.4 | - | - | 0 | - |
| - | - | 1.047E+04 | 894.4 | - | - | 0 | - |
| - | - | 3203 | 895.4 | - | - | 0 | - |
| - | - | 753.6 | 901.4 | - | - | 0 | - |
| - | - | 4.052E+04 | 902.4 | - | - | 0 | - |
| - | - | 2.19E+04 | 903.4 | - | - | 0 | - |
| - | - | 680.3 | 903.9 | - | - | 0 | - |
| - | - | 1.516E+04 | 904.4 | - | - | 0 | - |
| - | - | 5672 | 905.4 | - | - | 0 | - |
| - | - | 2597 | 906.4 | - | - | 0 | - |
| - | - | 698.5 | 907.4 | - | - | 0 | - |
| - | - | 754.9 | 910.4 | - | - | 0 | - |
| - | - | 945.2 | 911.4 | - | - | 0 | - |
| - | - | 845.9 | 919.3 | - | - | 0 | - |
| 8 | c | 1.039E+05 | 920.4 | 0.006185 | 6.72 | +1 | 8 |
| - | - | 4.874E+04 | 921.4 | - | - | 0 | - |
| - | - | 3.93E+04 | 922.4 | - | - | 0 | - |
| - | - | 1.277E+04 | 923.4 | - | - | 0 | - |
| - | - | 3248 | 924.4 | - | - | 0 | - |
| - | - | 3128 | 925.4 | - | - | 0 | - |
| - | - | 1768 | 926.4 | - | - | 0 | - |
| - | - | 1329 | 927.4 | - | - | 0 | - |
| - | - | 1031 | 928.4 | - | - | 0 | - |
| - | - | 790.9 | 929.4 | - | - | 0 | - |
| - | - | 909.7 | 929.9 | - | - | 0 | - |
| - | - | 8257 | 935.3 | - | - | 0 | - |
| 4 | y | 1.876E+04 | 936.3 | 0.005913 | 6.315 | +1 | 7 |
| 8 | c | 2.369E+05 | 937.4 | 0.006613 | 7.055 | +1 | 8 |
| - | - | 1.134E+05 | 938.4 | - | - | 0 | - |
| - | - | 4.174E+04 | 939.4 | - | - | 0 | - |
| - | - | 835.2 | 939.5 | - | - | 0 | - |
| - | - | 5328 | 940.4 | - | - | 0 | - |
| - | - | 1.726E+04 | 943.4 | - | - | 0 | - |
| - | - | 7569 | 944.4 | - | - | 0 | - |
| - | - | 2966 | 945.4 | - | - | 0 | - |
| - | - | 1022 | 952.3 | - | - | 0 | - |
| - | - | 1318 | 953.3 | - | - | 0 | - |
| - | - | 5221 | 954.3 | - | - | 0 | - |
| - | - | 3171 | 955.4 | - | - | 0 | - |
| - | - | 2221 | 956.4 | - | - | 0 | - |
| - | - | 776.3 | 961.4 | - | - | 0 | - |
| - | - | 2771 | 970.3 | - | - | 0 | - |
| - | - | 2361 | 971.3 | - | - | 0 | - |
| - | - | 947 | 972.4 | - | - | 0 | - |
| 3 | z | 2457 | 973.3 | 0.0128 | 13.15 | +1 | 8 |
| - | - | 825.2 | 974.3 | - | - | 0 | - |
| - | - | 1003 | 984.4 | - | - | 0 | - |
| - | - | 2304 | 985.4 | - | - | 0 | - |
| - | - | 870.9 | 988.5 | - | - | 0 | - |
| 3 | y | 1895 | 989.4 | 0.004083 | 4.127 | +1 | 8 |
| - | - | 976.2 | 990.4 | - | - | 0 | - |
| 3 | z | 4.961E+04 | 991.3 | 0.007177 | 7.24 | +1 | 8 |
| - | - | 3.231E+04 | 992.4 | - | - | 0 | - |
| - | - | 1.109E+04 | 993.4 | - | - | 0 | - |
| - | - | 7931 | 993.5 | - | - | 0 | - |
| - | - | 2152 | 994.4 | - | - | 0 | - |
| - | - | 4624 | 994.5 | - | - | 0 | - |
| - | - | 1596 | 995.5 | - | - | 0 | - |
| - | - | 1876 | 1001 | - | - | 0 | - |
| - | - | 1553 | 1002 | - | - | 0 | - |
| - | - | 3184 | 1003 | - | - | 0 | - |
| - | - | 2229 | 1004 | - | - | 0 | - |
| - | - | 1844 | 1005 | - | - | 0 | - |
| - | - | 1.045E+04 | 1006 | - | - | 0 | - |
| - | - | 946 | 1006 | - | - | 0 | - |
| 3 | y | 4.922E+04 | 1007 | 0.006214 | 6.169 | +1 | 8 |
| - | - | 2.429E+04 | 1008 | - | - | 0 | - |
| - | - | 8125 | 1009 | - | - | 0 | - |
| - | - | 1483 | 1012 | - | - | 0 | - |
| - | - | 1360 | 1013 | - | - | 0 | - |
| - | - | 3924 | 1019 | - | - | 0 | - |
| - | - | 2321 | 1020 | - | - | 0 | - |
| - | - | 749.1 | 1021 | - | - | 0 | - |
| - | - | 3810 | 1024 | - | - | 0 | - |
| - | - | 1455 | 1025 | - | - | 0 | - |
| - | - | 1220 | 1037 | - | - | 0 | - |
| - | - | 2305 | 1038 | - | - | 0 | - |
| - | - | 1351 | 1039 | - | - | 0 | - |
| - | - | 1630 | 1041 | - | - | 0 | - |
| - | - | 1026 | 1042 | - | - | 0 | - |
| - | - | 1539 | 1047 | - | - | 0 | - |
| - | - | 1568 | 1048 | - | - | 0 | - |
| - | - | 2007 | 1049 | - | - | 0 | - |
| - | - | 903.7 | 1054 | - | - | 0 | - |
| - | - | 7092 | 1055 | - | - | 0 | - |
| - | - | 3.36E+04 | 1056 | - | - | 0 | - |
| - | - | 1.981E+04 | 1057 | - | - | 0 | - |
| - | - | 960.5 | 1058 | - | - | 0 | - |
| - | - | 6564 | 1058 | - | - | 0 | - |
| - | - | 1010 | 1059 | - | - | 0 | - |
| 2 | w | 3312 | 1061 | 0.008589 | 8.092 | +1 | 9 |
| - | - | 1333 | 1062 | - | - | 0 | - |
| - | - | 1.037E+04 | 1065 | - | - | 0 | - |
| - | - | 6687 | 1066 | - | - | 0 | - |
| - | - | 2.105E+04 | 1067 | - | - | 0 | - |
| - | - | 1.391E+04 | 1068 | - | - | 0 | - |
| - | - | 5317 | 1069 | - | - | 0 | - |
| - | - | 1094 | 1070 | - | - | 0 | - |
| - | - | 6998 | 1080 | - | - | 0 | - |
| - | - | 3872 | 1081 | - | - | 0 | - |
| - | - | 1654 | 1082 | - | - | 0 | - |
| 9 | c | 3.636E+04 | 1083 | 0.00676 | 6.24 | +1 | 9 |
| - | - | 2.136E+04 | 1084 | - | - | 0 | - |
| - | - | 1.068E+04 | 1085 | - | - | 0 | - |
| - | - | 3590 | 1086 | - | - | 0 | - |
| - | - | 1248 | 1087 | - | - | 0 | - |
| - | - | 1549 | 1097 | - | - | 0 | - |
| - | - | 1574 | 1098 | - | - | 0 | - |
| - | - | 958.1 | 1099 | - | - | 0 | - |
| 9 | c | 2.955E+05 | 1100 | 0.0067 | 6.089 | +1 | 9 |
| - | - | 1.696E+05 | 1101 | - | - | 0 | - |
| - | - | 6.653E+04 | 1102 | - | - | 0 | - |
| - | - | 9028 | 1103 | - | - | 0 | - |
| - | - | 797.7 | 1114 | - | - | 0 | - |
| - | - | 881.8 | 1115 | - | - | 0 | - |
| 2 | z | 7.668E+04 | 1119 | 0.006766 | 6.044 | +1 | 9 |
| - | - | 5.124E+04 | 1120 | - | - | 0 | - |
| - | - | 1.814E+04 | 1121 | - | - | 0 | - |
| - | - | 2785 | 1122 | - | - | 0 | - |
| - | - | 842.1 | 1126 | - | - | 0 | - |
| - | - | 2099 | 1134 | - | - | 0 | - |
| - | - | 2441 | 1134 | - | - | 0 | - |
| 2 | y | 1069 | 1135 | 0.0004835 | 0.4259 | +1 | 9 |
| - | - | 1790 | 1142 | - | - | 0 | - |
| - | - | 1202 | 1143 | - | - | 0 | - |
| - | - | 1087 | 1144 | - | - | 0 | - |
| - | - | 969.4 | 1145 | - | - | 0 | - |
| - | - | 1147 | 1148 | - | - | 0 | - |
| - | - | 946 | 1149 | - | - | 0 | - |
| - | - | 929.3 | 1149 | - | - | 0 | - |
| - | - | 1157 | 1151 | - | - | 0 | - |
| - | - | 863.3 | 1151 | - | - | 0 | - |
| - | - | 1.462E+04 | 1157 | - | - | 0 | - |
| - | - | 1.007E+04 | 1158 | - | - | 0 | - |
| - | - | 5519 | 1159 | - | - | 0 | - |
| - | - | 2391 | 1160 | - | - | 0 | - |
| - | - | 3880 | 1161 | - | - | 0 | - |
| - | - | 4182 | 1162 | - | - | 0 | - |
| - | - | 1770 | 1164 | - | - | 0 | - |
| - | - | 1219 | 1168 | - | - | 0 | - |
| - | - | 2141 | 1170 | - | - | 0 | - |
| - | - | 2227 | 1171 | - | - | 0 | - |
| - | - | 1219 | 1172 | - | - | 0 | - |
| - | - | 5831 | 1176 | - | - | 0 | - |
| - | - | 1.032E+04 | 1177 | - | - | 0 | - |
| - | - | 1.349E+04 | 1178 | - | - | 0 | - |
| - | - | 6711 | 1179 | - | - | 0 | - |
| - | - | 1767 | 1180 | - | - | 0 | - |
| - | - | 1.884E+04 | 1186 | - | - | 0 | - |
| - | - | 1.525E+04 | 1187 | - | - | 0 | - |
| - | - | 1.858E+04 | 1188 | - | - | 0 | - |
| - | - | 1.027E+04 | 1189 | - | - | 0 | - |
| - | - | 3.309E+04 | 1189 | - | - | 0 | - |
| - | - | 3.79E+04 | 1190 | - | - | 0 | - |
| - | - | 1.906E+04 | 1191 | - | - | 0 | - |
| - | - | 6385 | 1193 | - | - | 0 | - |
| - | - | 832.1 | 1194 | - | - | 0 | - |
| - | - | 1065 | 1202 | - | - | 0 | - |
| - | - | 8463 | 1204 | - | - | 0 | - |
| - | - | 4.775E+04 | 1204 | - | - | 0 | - |
| - | - | 7.016E+04 | 1206 | - | - | 0 | - |
| - | - | 4.167E+04 | 1207 | - | - | 0 | - |
| - | - | 1.5E+04 | 1208 | - | - | 0 | - |
| - | - | 2793 | 1209 | - | - | 0 | - |
| - | - | 1.004E+04 | 1214 | - | - | 0 | - |
| - | - | 1046 | 1215 | - | - | 0 | - |
| - | - | 6534 | 1215 | - | - | 0 | - |
| - | - | 4300 | 1216 | - | - | 0 | - |
| - | - | 2666 | 1217 | - | - | 0 | - |
| - | - | 1345 | 1218 | - | - | 0 | - |
| - | - | 1001 | 1220 | - | - | 0 | - |
| - | - | 2.923E+04 | 1222 | - | - | 0 | - |
| - | - | 1.824E+04 | 1223 | - | - | 0 | - |
| - | - | 7549 | 1224 | - | - | 0 | - |
| - | - | 1145 | 1225 | - | - | 0 | - |
| - | - | 969.2 | 1230 | - | - | 0 | - |
| - | - | 3549 | 1230 | - | - | 0 | - |
| - | - | 3.958E+04 | 1232 | - | - | 0 | - |
| - | - | 3.697E+05 | 1233 | - | - | 0 | - |
| - | - | 2.532E+05 | 1234 | - | - | 0 | - |
| - | - | 1.233E+05 | 1235 | - | - | 0 | - |
| - | - | 2.641E+04 | 1236 | - | - | 0 | - |
| - | - | 3730 | 1237 | - | - | 0 | - |
| - | - | 1166 | 1238 | - | - | 0 | - |
| - | - | 1725 | 1246 | - | - | 0 | - |
| - | - | 4570 | 1247 | - | - | 0 | - |
| - | - | 1.207E+05 | 1249 | - | - | 0 | - |
| - | - | 5.205E+05 | 1250 | - | - | 0 | - |
| - | - | 3.326E+05 | 1251 | - | - | 0 | - |
| - | - | 1.446E+05 | 1252 | - | - | 0 | - |
| - | - | 2.352E+04 | 1253 | - | - | 0 | - |
| - | - | 1184 | 1265 | - | - | 0 | - |
| - | - | 3166 | 1282 | - | - | 0 | - |
| - | - | 1945 | 1283 | - | - | 0 | - |
| - | - | 1120 | 1284 | - | - | 0 | - |
| - | - | 707.9 | 1799 | - | - | 0 | - |
| - | - | 811.4 | 1837 | - | - | 0 | - |
| - | - | 701.9 | 1838 | - | - | 0 | - |
| - | - | 1144 | 1876 | - | - | 0 | - |
| - | - | 655.9 | 2604 | - | - | 0 | - |

m/z Charge Intensity FragmentType MassShift Position
120.0811538696289 0 43798.484
121.08448791503906 0 3666.073
127.17645263671875 0 457.80228
128.10507202148438 0 418.14536
131.11827087402344 0 12830.791
132.12158203125 0 720.50055
133.0861053466797 0 855.4267
133.3738250732422 0 410.77115
133.9047088623047 0 397.11356
135.52880859375 0 414.0078
136.07611083984375 0 178899.55
137.07940673828125 0 13377.028
137.59886169433594 0 497.83414
148.95396423339844 0 1469.7875
149.0450897216797 0 563.6219
149.06033325195312 0 943.4731
153.0662841796875 0 4609.9077
154.05044555664062 0 947.32666
162.0558319091797 0 667.71313
166.05361938476562 0 1582.0696
166.08668518066406 0 112175.86 y 9
167.0829315185547 0 841.17865
167.08998107910156 0 10005.0625
168.09136962890625 0 558.5613
171.0767364501953 0 762.5606
182.0810546875 0 497.65363
197.12806701660156 0 840.67377
200.03793334960938 0 1162.8582
200.1031036376953 0 1152.912
201.0875244140625 0 1177.7883
206.08160400390625 0 1057.6665
212.58535766601562 0 459.49756
214.15562438964844 0 3334.962
217.0646209716797 0 6264.368
218.0480194091797 0 939.53253
218.06756591796875 0 617.37067
221.0850067138672 0 3784.8071
222.1925048828125 0 636.2323
222.52232360839844 0 519.2476
223.0805206298828 0 625.6727
225.04373168945312 0 1374.578
225.1238555908203 0 12856.695
226.1273956298828 0 1436.2587
229.0224609375 0 555.6724
232.7680206298828 0 555.35754
235.07525634765625 0 17547.342
236.0786895751953 0 1150.3329
239.0955352783203 0 4659.1445
240.09609985351562 0 634.6603
241.1428985595703 0 661.076
242.15048217773438 0 66872.414 c Ammonia loss 1
243.1352996826172 0 486.13605
243.13812255859375 0 599.0555
243.1536407470703 0 7746.001
244.15562438964844 0 586.82416
245.07720947265625 0 2906.827
248.02493286132812 0 532.0679
257.12445068359375 0 598.1293
259.0666809082031 0 475.867
259.1769104003906 0 907.5624 c 1
268.16571044921875 0 671.7312
283.11163330078125 0 5081.3306
283.1446533203125 0 3951.8745
295.1037292480469 0 6477.3506
296.1047668457031 0 2549.403
296.1611022949219 0 642.48175
299.06207275390625 0 1039.7717
311.106689453125 0 6197.227
311.1370544433594 0 1188.0187
311.17138671875 0 629.9456
313.187744140625 0 16336.846 c Ammonia loss 2
314.0992431640625 0 1310.0658
314.1725769042969 0 747.6895
314.19085693359375 0 2508.3071
316.114990234375 0 1393.1046
328.09783935546875 0 987.2071
329.15032958984375 0 134051.7 y 8
330.153564453125 0 26289.15
330.21417236328125 0 4179.639 c 2
331.156494140625 0 2881.5642
332.10986328125 0 835.39105
334.1412658691406 0 721.05927
343.65447998046875 0 1124.2098 c Ammonia loss 5
346.1075439453125 0 4590.8086
347.1097717285156 0 974.13873
347.1749572753906 0 801.2615
353.1810607910156 0 696.9152
355.0708923339844 0 1194.9368
356.1100769042969 0 876.82544
357.14105224609375 0 1228.0547
358.161376953125 0 1066.6836
364.0943908691406 0 510.09756
364.1175842285156 0 2640.3037
367.1285095214844 0 547.5262
369.12255859375 0 3900.2893
370.12237548828125 0 1164.081
370.1449279785156 0 790.5013
371.12139892578125 0 602.88513
374.1195068359375 0 884.86414
378.16497802734375 0 6922.5264
378.6662902832031 0 2790.227
379.1649169921875 0 603.95807
379.19818115234375 0 695.6243
381.100341796875 0 672.4637
382.2085876464844 0 689.1294
386.2274169921875 0 699.6436
387.17041015625 0 2492.5466 c Ammonia loss 6
389.1862487792969 0 1535.9963
397.1364440917969 0 761.91077
398.1390075683594 0 10145.266
399.14013671875 0 1083.7828
405.68524169921875 0 1220.5979
410.6777648925781 0 700.0297
412.1881408691406 0 737.6881
414.2364196777344 0 2557.346
415.2414245605469 0 615.81854
416.25140380859375 0 1576.5701
417.1434631347656 0 679.6127
419.6834411621094 0 5330.3174
420.1854248046875 0 3046.5261
424.2196960449219 0 1190.5718
425.12945556640625 0 760.5969
425.2038879394531 0 2150.8545
426.16290283203125 0 750.64746
427.14715576171875 0 2349.4377
441.1839294433594 0 867.0266
442.2308654785156 0 15453.343 c Ammonia loss 3
442.6778259277344 0 4460.262
443.1756896972656 0 1918.1387
443.2335510253906 0 3234.3906
443.6746520996094 0 1237.8147
444.17315673828125 0 3067.4395
445.15802001953125 0 4435.564
446.16259765625 0 832.3385
451.6827697753906 0 57436.242
452.18414306640625 0 27400.354
452.6839904785156 0 10237.449
453.1845397949219 0 1883.703
458.17413330078125 0 1194.9137
458.2084655761719 0 4588.858
458.2496643066406 0 12107.107
459.257080078125 0 34487.027 c 3
460.2596740722656 0 7088.966
460.6878967285156 0 6863.8696 c Ammonia loss 7
461.13531494140625 0 6107.5146
461.1893615722656 0 3895.774
461.263671875 0 1021.35486
461.6889343261719 0 1604.2081
462.1355895996094 0 749.9834
476.1860046386719 0 14185.228 y 7
476.21893310546875 0 1840.8319
477.18975830078125 0 3220.7312
478.1825256347656 0 836.07263
478.70709228515625 0 689.02814
479.14520263671875 0 6120.2603
480.1475830078125 0 871.9589
481.17584228515625 0 1262.9006
481.2116394042969 0 1221.6843
486.21966552734375 0 1319.695
487.2176208496094 0 710.42993
494.22686767578125 0 870.3637
496.170654296875 0 756.09973 z 2
496.2221374511719 0 1607.9521
496.7251892089844 0 1352.7535
497.2281799316406 0 647.0683
499.2210998535156 0 1220.2665
509.1705017089844 0 2081.6096
510.21954345703125 0 1685.1432
510.7113342285156 0 1021.2736
511.2533264160156 0 1681.4207
514.1787719726562 0 1110.6708
517.211669921875 0 1188.0597
519.2155151367188 0 1098.0209
519.7183227539062 0 600.67377
522.2211303710938 0 1395.4838
527.180419921875 0 1513.5184
527.2164916992188 0 915.1705
528.2218017578125 0 2628.4185
528.7243041992188 0 1810.609
529.24853515625 0 3522.6216
530.2694702148438 0 4781.9805
531.278076171875 0 7901.5337
532.1845703125 0 1777.2783
532.2801513671875 0 1765.034
533.1912231445312 0 821.3266
536.2177734375 0 1539.4529
536.7190551757812 0 684.47675
539.2474365234375 0 4361.8516
540.2357177734375 0 1947.1201
542.2200927734375 0 1684.0272 c Ammonia loss 8
542.7198486328125 0 842.18713
545.2122802734375 0 891.3605 y Water loss 6
547.260009765625 0 830.41223
549.2177734375 0 1105.7307
554.1533203125 0 591.45825
555.2057495117188 0 1640.5155
556.1904296875 0 3326.5679
557.2575073242188 0 11882.9 c Ammonia loss 4
558.2534790039062 0 3146.7424
559.2529296875 0 735.54364
563.2178955078125 0 17150.477 y 6
564.2214965820312 0 4396.643
565.224609375 0 1010.8809
572.166748046875 0 2791.7021
573.1690673828125 0 1211.0912
573.2186279296875 0 3160.7878
573.2765502929688 0 33404.566
574.2840576171875 0 161730.14 c 4
575.231689453125 0 4162.2466
575.2868041992188 0 44473.734
576.2347412109375 0 1406.8964
576.2888793945312 0 7945.3433
577.29296875 0 694.09326
583.7539672851562 0 3318.5137
584.2500610351562 0 3543.5234
584.7504272460938 0 1680.9552
585.2559814453125 0 611.6126
587.2474975585938 0 892.81354
589.2645263671875 0 1086.0143
590.177978515625 0 3035.9868
592.2294311523438 0 2158.017
592.760009765625 0 9953.57
593.2611694335938 0 6223.972
593.7623291015625 0 2063.8853
595.2718505859375 0 870.0376
597.2156982421875 0 833.7725
600.2994995117188 0 4448.5864
601.302490234375 0 1482.5778
605.2587890625 0 846.7579
606.2715454101562 0 583.1499
606.3395385742188 0 1100.6207
608.1890869140625 0 1839.9476
609.191162109375 0 760.5637
609.2462768554688 0 686.787
610.2041015625 0 865.5101
614.21337890625 0 1766.1748
615.2177734375 0 1149.8145
615.35009765625 0 1223.0618
615.7557373046875 0 679.7757
622.2634887695312 0 672.62024
623.2684326171875 0 1100.394
623.3683471679688 0 1507.0743
624.1975708007812 0 1348.9529
624.3804931640625 0 683.0431
625.2097778320312 0 1678.8124
626.2052612304688 0 804.4121
626.2820434570312 0 834.7126
627.2820434570312 0 994.97723
628.2658081054688 0 668.47174
630.2990112304688 0 1687.15
631.2868041992188 0 1137.3835
639.2817993164062 0 1553.0686
640.2857666015625 0 1774.42
641.275146484375 0 2158.1125
642.2091064453125 0 5966.8853
643.2107543945312 0 3197.1633
644.2057495117188 0 978.2501
645.2855224609375 0 506.95193
650.2770385742188 0 835.2549
651.2692260742188 0 936.7642
658.2943115234375 0 4768.2964
659.3126220703125 0 15081.359
660.252685546875 0 1013.6727
660.3187255859375 0 9797.545
661.214111328125 0 8071.201
661.3214721679688 0 2431.6309
662.2180786132812 0 1847.7311
663.22265625 0 2053.2593
664.2232666015625 0 662.5506
668.2896728515625 0 7446.196
669.2820434570312 0 4101.0503
670.2864990234375 0 1607.8384
671.2909545898438 0 965.74805
674.2466430664062 0 1131.6305 y Water loss 5
676.2416381835938 0 1270.737 z 5
676.2984008789062 0 1966.098
678.2379760742188 0 630.94885
679.2251586914062 0 8928.413
680.228759765625 0 1750.6584
681.2395629882812 0 5356.608
682.239990234375 0 1417.9269
685.2532348632812 0 756.38995
686.3004150390625 0 28146.24 c Ammonia loss 5
687.3038330078125 0 8190.586
688.3139038085938 0 5008.7275
689.3164672851562 0 1013.8377
691.2528076171875 0 2945.1865
692.2605590820312 0 9823.577 y 5
693.2645263671875 0 2836.168
694.2648315429688 0 1649.6852
695.2409057617188 0 786.0169
696.2484130859375 0 1460.5803
702.3193359375 0 131056.555
703.3244018554688 0 70000.84 c 5
704.3265991210938 0 17627.988
705.3335571289062 0 2469.9678
710.3143920898438 0 785.16406
717.3307495117188 0 1050.8058
719.3129272460938 0 719.15625
720.2879638671875 0 2726.5518
721.2857666015625 0 894.1812
724.3184204101562 0 1631.1703
728.3051147460938 0 1631.5333
729.3401489257812 0 2690.121
730.3422241210938 0 1813.9272
737.30908203125 0 3660.8018
738.302490234375 0 1560.2738
739.307373046875 0 957.764
743.2843627929688 0 2346.687
744.29296875 0 1062.3723
745.34033203125 0 1094.2968
746.34423828125 0 2264.7231
747.2813720703125 0 2719.8882
747.3543701171875 0 1357.7653
748.2842407226562 0 2091.372
748.3511962890625 0 788.83167
750.2933349609375 0 997.9214
753.2452392578125 0 1798.0658
754.2395629882812 0 1455.6189
755.321044921875 0 28484.504
756.32275390625 0 11216.52
757.3240356445312 0 1891.7709
760.2744750976562 0 951.3107
768.3215942382812 0 918.25354
771.2532958984375 0 2471.3022
771.3396606445312 0 1625.3447
772.2496337890625 0 4620.903
773.2516479492188 0 1389.4784 z Water loss 4
773.3298950195312 0 20856.75 c Ammonia loss 6
774.3283081054688 0 6797.937
775.3319091796875 0 1973.234
778.2886962890625 0 1391.6738
778.9306640625 0 670.62384
789.2763671875 0 4157.8975 y Water loss 4
789.3516235351562 0 76997.51
790.358642578125 0 154257.28 c 6
791.2774658203125 0 3170.0928 z 4
791.3612060546875 0 53054.35
792.2859497070312 0 3222.2852
792.3629760742188 0 9582.788
793.282958984375 0 1840.9346
803.354736328125 0 1107.3334
804.35400390625 0 1661.2444
805.3528442382812 0 1127.9371
806.279296875 0 4585.5303
807.2869262695312 0 21964.186 y 4
808.2894287109375 0 9624.63
809.29931640625 0 10020.62
810.3027954101562 0 1989.087
811.3511352539062 0 826.9155
814.295654296875 0 2303.498
816.2997436523438 0 933.26746
816.375 0 1782.3553
821.3501586914062 0 1191.194
822.341796875 0 2437.9912
823.3659057617188 0 620.92755
824.2783813476562 0 2851.5083
825.2763061523438 0 1031.3541
826.2720336914062 0 3982.7637
827.2747192382812 0 1780.8857
828.2870483398438 0 867.00824
828.373291015625 0 3186.5698
829.3807983398438 0 1552.6531
830.3839721679688 0 1433.9551
831.3972778320312 0 3772.9458
832.4013061523438 0 1582.5476
838.3582763671875 0 4217.5347
839.3594970703125 0 4764.48
840.3551025390625 0 2014.358
842.2886962890625 0 8716.882
843.2914428710938 0 4718.3936
844.2865600585938 0 1135.0311
849.3409423828125 0 906.5215
855.340576171875 0 713.9293
856.369140625 0 7280.2554
857.366943359375 0 3137.5454
858.3785400390625 0 1435.972
859.3895874023438 0 883.892
861.2962036132812 0 1005.72723 w 3
867.3257446289062 0 1256.1168
868.3139038085938 0 987.53394
872.3327026367188 0 2403.9695
873.3425903320312 0 849.73865
874.3681030273438 0 2328.4973
875.363525390625 0 1925.0846
883.3545532226562 0 1163.19
884.3471069335938 0 6875.403
885.3382568359375 0 5439.6
886.3388061523438 0 1956.1388
887.3431396484375 0 781.52454
889.3624877929688 0 860.26196
890.3251342773438 0 902.36884
891.364013671875 0 823.8348
892.3735961914062 0 5732.517
893.3809204101562 0 17042.88
894.3844604492188 0 10465.394
895.38525390625 0 3202.514
901.35498046875 0 753.55066
902.3574829101562 0 40523.42
903.3565673828125 0 21898.451
903.8758544921875 0 680.2781
904.3583374023438 0 15162.172
905.359130859375 0 5672.4927
906.3601684570312 0 2597.228
907.357666015625 0 698.462
910.3685913085938 0 754.9143
911.38427734375 0 945.2415
919.3467407226562 0 845.90845
920.3678588867188 0 103922.47 c Ammonia loss 7
921.3704223632812 0 48743.562
922.37890625 0 39297.97
923.3842163085938 0 12770.751
924.3861694335938 0 3247.6255
925.3611450195312 0 3128.4165
926.3645629882812 0 1767.9221
927.3759765625 0 1329.2684
928.3602905273438 0 1031.0984
929.3656005859375 0 790.8513
929.8751220703125 0 909.653
935.3226318359375 0 8256.951
936.3301391601562 0 18763.74 y 3
937.3948364257812 0 236866.62 c 7
938.3975219726562 0 113369.6
939.396728515625 0 41736.035
939.5121459960938 0 835.1515
940.3995971679688 0 5328.3857
943.369384765625 0 17256.273
944.3716430664062 0 7569.2207
945.3751831054688 0 2966.439
952.332763671875 0 1021.55646
953.3298950195312 0 1318.445
954.3309326171875 0 5220.5854
955.3706665039062 0 3170.7954
956.369384765625 0 2220.9595
961.3828735351562 0 776.2645
970.3491821289062 0 2770.909
971.3485717773438 0 2360.5674
972.3717041015625 0 947.00354
973.3448486328125 0 2456.735 z Water loss 2
974.340087890625 0 825.17004
984.4165649414062 0 1003.37726
985.4124145507812 0 2304.3806
988.4863891601562 0 870.85724
989.3548583984375 0 1895.4662 y Water loss 2
990.3534545898438 0 976.2336
991.3497924804688 0 49606.03 z 2
992.3524169921875 0 32312.602
993.3534545898438 0 11092.316
993.4573974609375 0 7931.4097
994.3533325195312 0 2152.2112
994.4580688476562 0 4623.9473
995.4593505859375 0 1595.5708
1001.4164428710938 0 1875.8333
1002.4183959960938 0 1553.3118
1003.4156494140625 0 3183.9536
1004.4251708984375 0 2229.4795
1005.4360961914062 0 1844.0836
1006.3603515625 0 10453.571
1006.4620971679688 0 945.9752
1007.3675537109375 0 49216.656 y 2
1008.3704833984375 0 24285.523
1009.3695678710938 0 8125.2173
1012.452392578125 0 1483.4996
1013.4551391601562 0 1360.3912
1019.4329833984375 0 3924.3186
1020.4353637695312 0 2321.413
1021.4305419921875 0 749.0569
1024.3564453125 0 3810.1924
1025.3548583984375 0 1454.6917
1037.4305419921875 0 1219.5403
1038.411376953125 0 2304.945
1039.4183349609375 0 1351.4896
1041.4300537109375 0 1629.5757
1042.4212646484375 0 1025.9656
1047.4049072265625 0 1538.6688
1048.406005859375 0 1567.8158
1049.4078369140625 0 2006.5148
1054.3936767578125 0 903.6802
1055.433349609375 0 7091.5474
1056.443359375 0 33598.27
1057.447265625 0 19811.215
1057.576171875 0 960.4511
1058.449951171875 0 6564.0693
1059.444091796875 0 1009.8132
1061.3804931640625 0 3312.4907 w 1
1062.3817138671875 0 1332.9856
1065.4217529296875 0 10367.337
1066.4180908203125 0 6686.8384
1067.4146728515625 0 21046.254
1068.4171142578125 0 13914.642
1069.4178466796875 0 5317.289
1070.422119140625 0 1094.1864
1080.467041015625 0 6997.878
1081.47021484375 0 3872.3801
1082.47216796875 0 1654.1313
1083.4317626953125 0 36363.08 c Ammonia loss 8
1084.434814453125 0 21358.633
1085.4376220703125 0 10682.398
1086.4442138671875 0 3590.4812
1087.4461669921875 0 1248.1796
1097.4521484375 0 1549.4236
1098.45263671875 0 1573.6382
1099.4468994140625 0 958.13947
1100.458251953125 0 295474.12 c 8
1101.4605712890625 0 169564.97
1102.460693359375 0 66529.17
1103.459228515625 0 9028.312
1114.4873046875 0 797.74274
1115.4869384765625 0 881.78076
1119.407958984375 0 76675.52 z 1
1120.4105224609375 0 51241.473
1121.4114990234375 0 18141.096
1122.41357421875 0 2784.5737
1126.4432373046875 0 842.1061
1133.5020751953125 0 2099.0952
1134.497314453125 0 2440.8306
1135.41943359375 0 1069.1467 y 1
1141.5267333984375 0 1790.2579
1142.502197265625 0 1202.0872
1143.515380859375 0 1087.4028
1144.5213623046875 0 969.3585
1148.428466796875 0 1146.889
1148.55810546875 0 946.03125
1149.49609375 0 929.33356
1150.503173828125 0 1157.0419
1151.3790283203125 0 863.29895
1157.4888916015625 0 14616.995
1158.4918212890625 0 10072.659
1159.49609375 0 5518.7256
1160.47509765625 0 2390.816
1161.4619140625 0 3879.7266
1162.4910888671875 0 4182.1055
1163.5147705078125 0 1769.8491
1168.4954833984375 0 1218.803
1169.5037841796875 0 2141.1519
1170.505126953125 0 2227.0342
1171.5003662109375 0 1218.5842
1176.467041015625 0 5830.775
1177.4893798828125 0 10315.436
1178.4874267578125 0 13492.474
1179.490234375 0 6710.976
1180.496337890625 0 1766.535
1185.520263671875 0 18839.586
1186.521484375 0 15247.244
1187.5198974609375 0 18583.729
1188.5196533203125 0 10266.447
1189.482666015625 0 33092.836
1190.4925537109375 0 37904.36
1191.498291015625 0 19057.525
1192.5029296875 0 6385.204
1193.521728515625 0 832.1333
1202.4998779296875 0 1065.0431
1203.51220703125 0 8463.075
1204.498779296875 0 47749.11
1205.51708984375 0 70156.14
1206.52392578125 0 41666.758
1207.5277099609375 0 15002.307
1208.5301513671875 0 2793.3691
1214.480224609375 0 10044.905
1214.6239013671875 0 1045.7885
1215.4830322265625 0 6534.111
1216.4852294921875 0 4300.4707
1217.486328125 0 2665.6572
1218.4937744140625 0 1345.1735
1219.5030517578125 0 1000.89685
1221.5220947265625 0 29226.61
1222.5260009765625 0 18240.273
1223.5289306640625 0 7548.9604
1224.5321044921875 0 1144.7936
1229.6109619140625 0 969.1698
1230.4913330078125 0 3549.3052
1231.5062255859375 0 39583.723
1232.5076904296875 0 369703.8
1233.51025390625 0 253219.55
1234.510498046875 0 123296.15
1235.508544921875 0 26413.19
1236.5015869140625 0 3729.875
1237.5084228515625 0 1166.3188
1246.490234375 0 1724.5146
1247.4986572265625 0 4570.4614
1248.509765625 0 120700.77
1249.5174560546875 0 520522
1250.519775390625 0 332587.7
1251.521240234375 0 144632.58
1252.522705078125 0 23523.701
1265.4912109375 0 1183.829
1281.5087890625 0 3165.9912
1282.512939453125 0 1944.7766
1283.5146484375 0 1120.299
1799.3681640625 0 707.8554
1836.747802734375 0 811.4168
1837.765625 0 701.9315
1875.84423828125 0 1144.4729
2604.28857421875 0 655.9122

Spectrum Details

|  |  |
| --- | --- |
| Matched peaks? Matched peaks The total absolute number of peaks matched. Additionally in brackets the total fraction of peaks matched and the total number of peaks is shown. | 42 (7.07% of 594) |
| FDR? FDR The false discovery rate estimated for this peptide. It is calculated by matching all theoretical fragments with a non-integer shift with the raw peaks for this spectrum. This is done with 40 different shifts. The resulting percentage is the average number of annotated peaks over the number of annotated peaks with the correct spectrum. | 5.27% |
| Satellite FDR? Satellite FDR See the FDR for details on its calculation. This satellite ion specific FDR only contains the satellite ions (d/w) for I/L/J positions. | - |
| PSM Score? PSM Score The PSM Score as given by Hecklib to this annotated spectrum. It is shown with three significant figures. | 558 |

#### Spectrum 6979? Spectrum 6979 The raw spectrum of this peptide as annotated by Hecklib. The fragments are coloured according to ion type (see legend). Any peaks with a star '\*' as text can be hovered over to see the full details, first the ion type second the mass shift type. By hovering over the amino acids in the peptide or ions in the legend the corresponding peaks are highlighted. By toggling the 'Unassigned' label you can turn the background (unassigned) peaks on or off in the plot. By updating the slider in the Ion legend you can update the spectrum to only show the top X% of the peaks with labels. The top X% means any peak that is within X% of the highest intensity. By dragging in the spectrum you can zoom in to a specific part of the spectrum and use 'Zoom Out' to get back to the original zoom level. The annotation of the spectrum is based on the given sequence in the peptides file and is done with different software so inconsistencies are likely. The peaks are annotated based on the given sequence, with 20 ppm tolerance.

Copy Data

##### Spectrum 6979 (TSV)

###### Preview

```
Loading example...
```

*Click on the button to copy the data to your clipboard.*

Mz MinMz MaxIntensity Max

WidthHeightPeptide font sizePeptide stroke widthSpectrum font sizeSpectrum stroke widthCompact peptide

Ion legend

wxyz

abcd

OtherUnassignedIon^Charge^_Position_Show for top:%

JQAEDESMYF

01.76e+43.52e+45.28e+47.04e+4

Zoom Out

y^+1^_1_c^+1^_2_c^+1^_3_y^+1^_2_c^+1^_4_c^+1^_4_c^+2^_8_y^+1^_3_c^+1^_5_y^+1^_4_c^+1^_5_c^+1^_6_y^+1^_5_c^+1^_6_c^+1^_7_c^+1^_7_y^+1^_6_c^+1^_8_y^+1^_7_c^+1^_8_z^+1^_8_y^+1^_8_c^+1^_9_c^+1^_9_z^+1^_9_

0775155123263102

Fragment Matches Table

Show background peaks

| Position | Ion type | Intensity | mz Theoretical | mz Error (Th) | mz Error (ppm) | Charge | Series Number |
| --- | --- | --- | --- | --- | --- | --- | --- |
| - | - | 3923 | 120.1 | - | - | 0 | - |
| - | - | 386.5 | 122.8 | - | - | 0 | - |
| - | - | 380.9 | 128.5 | - | - | 0 | - |
| - | - | 1038 | 131.1 | - | - | 0 | - |
| - | - | 599 | 133.1 | - | - | 0 | - |
| - | - | 374.8 | 135.2 | - | - | 0 | - |
| - | - | 939.6 | 136.1 | - | - | 0 | - |
| - | - | 1.648E+04 | 136.1 | - | - | 0 | - |
| - | - | 924 | 149 | - | - | 0 | - |
| - | - | 915 | 149 | - | - | 0 | - |
| - | - | 432.2 | 158.5 | - | - | 0 | - |
| - | - | 402.3 | 160.1 | - | - | 0 | - |
| - | - | 671.7 | 166.1 | - | - | 0 | - |
| 10 | y | 1.02E+04 | 166.1 | 9.443E-05 | 0.5686 | +1 | 1 |
| - | - | 834.7 | 167.1 | - | - | 0 | - |
| - | - | 5494 | 221.1 | - | - | 0 | - |
| - | - | 605.9 | 222.1 | - | - | 0 | - |
| - | - | 1103 | 225 | - | - | 0 | - |
| - | - | 902.2 | 225.1 | - | - | 0 | - |
| - | - | 1763 | 235.1 | - | - | 0 | - |
| - | - | 7679 | 239.1 | - | - | 0 | - |
| - | - | 1094 | 240.1 | - | - | 0 | - |
| 2 | c | 5820 | 242.1 | 3.086E-05 | 0.1274 | +1 | 2 |
| - | - | 848.7 | 283.1 | - | - | 0 | - |
| - | - | 8499 | 295.1 | - | - | 0 | - |
| - | - | 2266 | 296.1 | - | - | 0 | - |
| - | - | 1589 | 299.1 | - | - | 0 | - |
| - | - | 907.8 | 313.1 | - | - | 0 | - |
| 3 | c | 1600 | 313.2 | 0.0001631 | 0.5207 | +1 | 3 |
| 9 | y | 1.247E+04 | 329.1 | 1.357E-05 | 0.04124 | +1 | 2 |
| - | - | 1669 | 330.2 | - | - | 0 | - |
| - | - | 1434 | 355.1 | - | - | 0 | - |
| - | - | 5288 | 369.1 | - | - | 0 | - |
| - | - | 1049 | 370.1 | - | - | 0 | - |
| - | - | 777.7 | 378.2 | - | - | 0 | - |
| - | - | 890.3 | 419.7 | - | - | 0 | - |
| - | - | 594.9 | 421.2 | - | - | 0 | - |
| - | - | 653.6 | 429.1 | - | - | 0 | - |
| - | - | 892.9 | 439.2 | - | - | 0 | - |
| 4 | c | 1892 | 442.2 | 0.0005599 | 1.266 | +1 | 4 |
| - | - | 5985 | 451.7 | - | - | 0 | - |
| - | - | 2611 | 452.2 | - | - | 0 | - |
| - | - | 588.8 | 452.7 | - | - | 0 | - |
| - | - | 1425 | 458.2 | - | - | 0 | - |
| 4 | c | 5354 | 459.3 | 7.042E-05 | 0.1533 | +1 | 4 |
| - | - | 1487 | 460.3 | - | - | 0 | - |
| 8 | c | 1094 | 460.7 | 0.003269 | 7.096 | +2 | 8 |
| - | - | 564.4 | 475.2 | - | - | 0 | - |
| 8 | y | 1350 | 476.2 | 0.006364 | 13.36 | +1 | 3 |
| - | - | 559.7 | 490 | - | - | 0 | - |
| - | - | 940.2 | 529.2 | - | - | 0 | - |
| - | - | 1017 | 531.3 | - | - | 0 | - |
| - | - | 778 | 552.3 | - | - | 0 | - |
| 5 | c | 1163 | 557.3 | 0.000146 | 0.262 | +1 | 5 |
| 7 | y | 1220 | 563.2 | 0.004944 | 8.779 | +1 | 4 |
| - | - | 619.2 | 573.2 | - | - | 0 | - |
| - | - | 3607 | 573.3 | - | - | 0 | - |
| 5 | c | 2.102E+04 | 574.3 | 0.000219 | 0.3814 | +1 | 5 |
| - | - | 5422 | 575.3 | - | - | 0 | - |
| - | - | 894.1 | 576.3 | - | - | 0 | - |
| - | - | 694.3 | 592.8 | - | - | 0 | - |
| - | - | 825 | 600.3 | - | - | 0 | - |
| - | - | 1244 | 615.3 | - | - | 0 | - |
| - | - | 704 | 642.2 | - | - | 0 | - |
| - | - | 660.4 | 646.8 | - | - | 0 | - |
| - | - | 2624 | 659.3 | - | - | 0 | - |
| - | - | 1141 | 660.3 | - | - | 0 | - |
| - | - | 791.1 | 661.2 | - | - | 0 | - |
| - | - | 753.9 | 668.3 | - | - | 0 | - |
| - | - | 679.7 | 670.4 | - | - | 0 | - |
| - | - | 1023 | 679.2 | - | - | 0 | - |
| - | - | 747.3 | 681.2 | - | - | 0 | - |
| 6 | c | 2467 | 686.3 | 0.0003996 | 0.5822 | +1 | 6 |
| - | - | 888 | 687.3 | - | - | 0 | - |
| - | - | 1132 | 688.3 | - | - | 0 | - |
| 6 | y | 937.6 | 692.3 | 0.004466 | 6.451 | +1 | 5 |
| - | - | 2.038E+04 | 702.3 | - | - | 0 | - |
| 6 | c | 1.278E+04 | 703.3 | 0.002834 | 4.03 | +1 | 6 |
| - | - | 2474 | 704.3 | - | - | 0 | - |
| - | - | 567.5 | 705.3 | - | - | 0 | - |
| - | - | 844.2 | 750.3 | - | - | 0 | - |
| - | - | 2270 | 755.3 | - | - | 0 | - |
| - | - | 1032 | 756.3 | - | - | 0 | - |
| 7 | c | 3012 | 773.3 | 0.003675 | 4.752 | +1 | 7 |
| - | - | 1.022E+04 | 789.3 | - | - | 0 | - |
| 7 | c | 2.472E+04 | 790.4 | 0.0008659 | 1.096 | +1 | 7 |
| - | - | 8356 | 791.4 | - | - | 0 | - |
| - | - | 711.9 | 792.3 | - | - | 0 | - |
| - | - | 1758 | 792.4 | - | - | 0 | - |
| 5 | y | 2906 | 807.3 | 0.003768 | 4.667 | +1 | 6 |
| - | - | 782.3 | 808.3 | - | - | 0 | - |
| - | - | 1618 | 809.3 | - | - | 0 | - |
| - | - | 1007 | 842.3 | - | - | 0 | - |
| - | - | 823.3 | 885.3 | - | - | 0 | - |
| - | - | 2186 | 893.4 | - | - | 0 | - |
| - | - | 1471 | 894.4 | - | - | 0 | - |
| - | - | 3012 | 902.4 | - | - | 0 | - |
| - | - | 1941 | 903.4 | - | - | 0 | - |
| - | - | 1445 | 904.4 | - | - | 0 | - |
| - | - | 1087 | 905.4 | - | - | 0 | - |
| - | - | 1287 | 912.4 | - | - | 0 | - |
| 8 | c | 8928 | 920.4 | 0.003316 | 3.603 | +1 | 8 |
| - | - | 3894 | 921.4 | - | - | 0 | - |
| - | - | 4321 | 922.4 | - | - | 0 | - |
| - | - | 1673 | 923.4 | - | - | 0 | - |
| - | - | 672.6 | 935.3 | - | - | 0 | - |
| 4 | y | 2728 | 936.3 | 0.003472 | 3.708 | +1 | 7 |
| 8 | c | 3.749E+04 | 937.4 | 0.004538 | 4.841 | +1 | 8 |
| - | - | 1.647E+04 | 938.4 | - | - | 0 | - |
| - | - | 6343 | 939.4 | - | - | 0 | - |
| - | - | 1114 | 940.4 | - | - | 0 | - |
| - | - | 1349 | 943.4 | - | - | 0 | - |
| 3 | z | 6746 | 991.3 | 0.004674 | 4.715 | +1 | 8 |
| - | - | 3389 | 992.4 | - | - | 0 | - |
| - | - | 1235 | 993.4 | - | - | 0 | - |
| - | - | 1033 | 993.5 | - | - | 0 | - |
| - | - | 659.1 | 994.5 | - | - | 0 | - |
| - | - | 1293 | 1006 | - | - | 0 | - |
| 3 | y | 5283 | 1007 | 0.003712 | 3.685 | +1 | 8 |
| - | - | 2507 | 1008 | - | - | 0 | - |
| - | - | 1286 | 1009 | - | - | 0 | - |
| - | - | 4163 | 1056 | - | - | 0 | - |
| - | - | 2120 | 1057 | - | - | 0 | - |
| - | - | 641.3 | 1058 | - | - | 0 | - |
| - | - | 1101 | 1058 | - | - | 0 | - |
| - | - | 3586 | 1067 | - | - | 0 | - |
| - | - | 2332 | 1068 | - | - | 0 | - |
| - | - | 812.5 | 1069 | - | - | 0 | - |
| - | - | 952.9 | 1080 | - | - | 0 | - |
| 9 | c | 3146 | 1083 | 0.004685 | 4.324 | +1 | 9 |
| - | - | 1406 | 1084 | - | - | 0 | - |
| - | - | 1006 | 1085 | - | - | 0 | - |
| - | - | 797.8 | 1087 | - | - | 0 | - |
| 9 | c | 4.024E+04 | 1100 | 0.004869 | 4.425 | +1 | 9 |
| - | - | 2.392E+04 | 1101 | - | - | 0 | - |
| - | - | 8141 | 1102 | - | - | 0 | - |
| - | - | 1597 | 1103 | - | - | 0 | - |
| 2 | z | 8175 | 1119 | 0.003958 | 3.536 | +1 | 9 |
| - | - | 5160 | 1120 | - | - | 0 | - |
| - | - | 2773 | 1121 | - | - | 0 | - |
| - | - | 1441 | 1149 | - | - | 0 | - |
| - | - | 2751 | 1157 | - | - | 0 | - |
| - | - | 900.1 | 1158 | - | - | 0 | - |
| - | - | 873.3 | 1176 | - | - | 0 | - |
| - | - | 782.5 | 1177 | - | - | 0 | - |
| - | - | 2034 | 1178 | - | - | 0 | - |
| - | - | 2703 | 1186 | - | - | 0 | - |
| - | - | 2474 | 1187 | - | - | 0 | - |
| - | - | 2295 | 1188 | - | - | 0 | - |
| - | - | 1329 | 1189 | - | - | 0 | - |
| - | - | 4031 | 1189 | - | - | 0 | - |
| - | - | 4944 | 1190 | - | - | 0 | - |
| - | - | 2738 | 1191 | - | - | 0 | - |
| - | - | 715 | 1193 | - | - | 0 | - |
| - | - | 1496 | 1204 | - | - | 0 | - |
| - | - | 5547 | 1204 | - | - | 0 | - |
| - | - | 5973 | 1206 | - | - | 0 | - |
| - | - | 4621 | 1207 | - | - | 0 | - |
| - | - | 877.7 | 1208 | - | - | 0 | - |
| - | - | 1144 | 1214 | - | - | 0 | - |
| - | - | 859.3 | 1215 | - | - | 0 | - |
| - | - | 3257 | 1222 | - | - | 0 | - |
| - | - | 2107 | 1223 | - | - | 0 | - |
| - | - | 793.1 | 1224 | - | - | 0 | - |
| - | - | 4679 | 1232 | - | - | 0 | - |
| - | - | 5.168E+04 | 1233 | - | - | 0 | - |
| - | - | 3.279E+04 | 1234 | - | - | 0 | - |
| - | - | 1.53E+04 | 1235 | - | - | 0 | - |
| - | - | 2451 | 1236 | - | - | 0 | - |
| - | - | 858.9 | 1236 | - | - | 0 | - |
| - | - | 1.556E+04 | 1249 | - | - | 0 | - |
| - | - | 999.9 | 1249 | - | - | 0 | - |
| - | - | 6.966E+04 | 1250 | - | - | 0 | - |
| - | - | 4.33E+04 | 1251 | - | - | 0 | - |
| - | - | 1.872E+04 | 1252 | - | - | 0 | - |
| - | - | 3498 | 1253 | - | - | 0 | - |
| - | - | 706.7 | 1288 | - | - | 0 | - |
| - | - | 663.2 | 1720 | - | - | 0 | - |
| - | - | 769.2 | 1859 | - | - | 0 | - |
| - | - | 679.7 | 2453 | - | - | 0 | - |
| - | - | 725.6 | 2539 | - | - | 0 | - |
| - | - | 753.1 | 2651 | - | - | 0 | - |
| - | - | 695.3 | 3071 | - | - | 0 | - |

m/z Charge Intensity FragmentType MassShift Position
120.08090209960938 0 3922.604
122.81031799316406 0 386.5423
128.4912109375 0 380.85703
131.117919921875 0 1037.9407
133.08612060546875 0 598.98834
135.24908447265625 0 374.83862
136.07028198242188 0 939.59534
136.07579040527344 0 16484.229
148.95465087890625 0 924.0323
149.045166015625 0 915.0198
158.5304718017578 0 432.2001
160.11976623535156 0 402.33322
166.07891845703125 0 671.70197
166.0863494873047 0 10196.206 y 9
167.0892333984375 0 834.70703
221.0843048095703 0 5493.603
222.08477783203125 0 605.857
225.04296875 0 1102.6102
225.12319946289062 0 902.1965
235.0744171142578 0 1763.4791
239.09500122070312 0 7679.302
240.09481811523438 0 1093.831
242.14988708496094 0 5819.509 c Ammonia loss 1
283.11004638671875 0 848.72577
295.103271484375 0 8499.453
296.10406494140625 0 2266.4814
299.06182861328125 0 1588.5757
313.1138610839844 0 907.7868
313.18719482421875 0 1599.7185 c Ammonia loss 2
329.14959716796875 0 12472.553 y 8
330.15301513671875 0 1669.4802
355.06976318359375 0 1434.2842
369.1217956542969 0 5287.5293
370.12164306640625 0 1049.3954
378.1641845703125 0 777.6981
419.68218994140625 0 890.3436
421.2430114746094 0 594.91516
429.08856201171875 0 653.5987
439.1624755859375 0 892.913
442.22906494140625 0 1892.262 c Ammonia loss 3
451.6816711425781 0 5985.1987
452.18353271484375 0 2610.8306
452.68359375 0 588.78937
458.2483825683594 0 1424.5397
459.256103515625 0 5353.898 c 3
460.2593078613281 0 1486.5188
460.687744140625 0 1094.1161 c Ammonia loss 7
475.1723937988281 0 564.3514
476.1864318847656 0 1349.801 y 7
489.978759765625 0 559.68536
529.2433471679688 0 940.17456
531.2788696289062 0 1016.7135
552.266845703125 0 778.04956
557.2567138671875 0 1162.7147 c Ammonia loss 4
563.217041015625 0 1220.4431 y 6
573.228515625 0 619.20667
573.27490234375 0 3607.0842
574.2828979492188 0 21024.885 c 4
575.2860107421875 0 5421.8486
576.285888671875 0 894.1461
592.7570190429688 0 694.2665
600.2987060546875 0 825.00336
615.345947265625 0 1243.77
642.2097778320312 0 704.00775
646.8074340820312 0 660.36523
659.3124389648438 0 2624.0188
660.3155517578125 0 1141.2339
661.2110595703125 0 791.1397
668.2874755859375 0 753.92224
670.361328125 0 679.729
679.2251586914062 0 1022.6718
681.241943359375 0 747.3374
686.299560546875 0 2466.6165 c Ammonia loss 5
687.294189453125 0 887.9556
688.3131103515625 0 1131.5647
692.2591552734375 0 937.5587 y 5
702.3175659179688 0 20384.59
703.3228759765625 0 12778.578 c 5
704.327392578125 0 2474.2583
705.327392578125 0 567.5
750.3369750976562 0 844.1945
755.3177490234375 0 2269.6343
756.321533203125 0 1031.9578
773.3275146484375 0 3012.188 c Ammonia loss 6
789.3490600585938 0 10216.337
790.3568725585938 0 24723.18 c 6
791.359619140625 0 8356.079
792.2813110351562 0 711.91675
792.36279296875 0 1757.73
807.285400390625 0 2905.555 y 4
808.2854614257812 0 782.2742
809.29833984375 0 1618.4794
842.2908935546875 0 1006.9249
885.3396606445312 0 823.2515
893.3800659179688 0 2186.1616
894.3846435546875 0 1470.8319
902.3576049804688 0 3012.1782
903.3521728515625 0 1940.8608
904.355224609375 0 1445.2993
905.3678588867188 0 1086.9626
912.3688354492188 0 1287.1023
920.364990234375 0 8927.892 c Ammonia loss 7
921.3663940429688 0 3893.9685
922.375732421875 0 4320.9004
923.3814086914062 0 1672.7454
935.3131713867188 0 672.6018
936.3276977539062 0 2728.031 y 3
937.3927612304688 0 37488.395 c 7
938.3956909179688 0 16471.86
939.3958129882812 0 6342.5293
940.3932495117188 0 1114.4963
943.3685913085938 0 1349.2023
991.3472900390625 0 6745.599 z 2
992.3518676757812 0 3388.6306
993.350341796875 0 1235.3566
993.45703125 0 1033.2921
994.4580688476562 0 659.10614
1006.35791015625 0 1292.5748
1007.3650512695312 0 5283.1045 y 2
1008.3700561523438 0 2507.394
1009.3668823242188 0 1285.7274
1056.44189453125 0 4162.661
1057.443359375 0 2119.9395
1057.5672607421875 0 641.3314
1058.443115234375 0 1100.5774
1067.4130859375 0 3585.5688
1068.416259765625 0 2332.0713
1069.4134521484375 0 812.4846
1080.467529296875 0 952.9226
1083.4296875 0 3146.2925 c Ammonia loss 8
1084.42919921875 0 1406.1265
1085.4356689453125 0 1005.8681
1087.4901123046875 0 797.7772
1100.4564208984375 0 40243.625 c 8
1101.45849609375 0 23919.121
1102.458251953125 0 8141.325
1103.45947265625 0 1597.1794
1119.4051513671875 0 8174.6807 z 1
1120.409912109375 0 5159.8003
1121.408447265625 0 2773.42
1148.5472412109375 0 1440.8492
1157.4859619140625 0 2751.2656
1158.498291015625 0 900.0827
1176.4580078125 0 873.30963
1177.4888916015625 0 782.5244
1178.485107421875 0 2033.9761
1185.517333984375 0 2703.2063
1186.5205078125 0 2473.8955
1187.518798828125 0 2295.3237
1188.5201416015625 0 1328.771
1189.4716796875 0 4031.24
1190.4945068359375 0 4943.7744
1191.4990234375 0 2737.7407
1192.5076904296875 0 714.9534
1203.5081787109375 0 1495.8519
1204.496337890625 0 5547.035
1205.5206298828125 0 5972.5215
1206.5198974609375 0 4620.5444
1207.51708984375 0 877.68475
1214.4793701171875 0 1143.9392
1215.4713134765625 0 859.34314
1221.5177001953125 0 3257.3657
1222.523193359375 0 2106.6326
1223.527587890625 0 793.1081
1231.506103515625 0 4678.5337
1232.5050048828125 0 51679.594
1233.5062255859375 0 32786.76
1234.5101318359375 0 15301.775
1235.51220703125 0 2451.027
1236.48486328125 0 858.8983
1248.5069580078125 0 15561.967
1248.6531982421875 0 999.881
1249.5142822265625 0 69656.39
1250.517578125 0 43301.746
1251.5208740234375 0 18722.295
1252.5208740234375 0 3498.3523
1287.9517822265625 0 706.7484
1719.950439453125 0 663.2307
1858.7623291015625 0 769.1709
2453.44873046875 0 679.6946
2539.090087890625 0 725.62885
2651.076904296875 0 753.1405
3071.160888671875 0 695.26697

Spectrum Details

|  |  |
| --- | --- |
| Matched peaks? Matched peaks The total absolute number of peaks matched. Additionally in brackets the total fraction of peaks matched and the total number of peaks is shown. | 25 (13.66% of 183) |
| FDR? FDR The false discovery rate estimated for this peptide. It is calculated by matching all theoretical fragments with a non-integer shift with the raw peaks for this spectrum. This is done with 40 different shifts. The resulting percentage is the average number of annotated peaks over the number of annotated peaks with the correct spectrum. | 1.71% |
| Satellite FDR? Satellite FDR See the FDR for details on its calculation. This satellite ion specific FDR only contains the satellite ions (d/w) for I/L/J positions. | - |
| PSM Score? PSM Score The PSM Score as given by Hecklib to this annotated spectrum. It is shown with three significant figures. | 330 |

#### Spectrum 7038? Spectrum 7038 The raw spectrum of this peptide as annotated by Hecklib. The fragments are coloured according to ion type (see legend). Any peaks with a star '\*' as text can be hovered over to see the full details, first the ion type second the mass shift type. By hovering over the amino acids in the peptide or ions in the legend the corresponding peaks are highlighted. By toggling the 'Unassigned' label you can turn the background (unassigned) peaks on or off in the plot. By updating the slider in the Ion legend you can update the spectrum to only show the top X% of the peaks with labels. The top X% means any peak that is within X% of the highest intensity. By dragging in the spectrum you can zoom in to a specific part of the spectrum and use 'Zoom Out' to get back to the original zoom level. The annotation of the spectrum is based on the given sequence in the peptides file and is done with different software so inconsistencies are likely. The peaks are annotated based on the given sequence, with 20 ppm tolerance.

Copy Data

##### Spectrum 7038 (TSV)

###### Preview

```
Loading example...
```

*Click on the button to copy the data to your clipboard.*

Mz MinMz MaxIntensity Max

WidthHeightPeptide font sizePeptide stroke widthSpectrum font sizeSpectrum stroke widthCompact peptide

Ion legend

wxyz

abcd

OtherUnassignedIon^Charge^_Position_Show for top:%

JQAEDESMYF

07.02e+31.40e+42.11e+42.81e+4

Zoom Out

y^+1^_1_c^+1^_2_y^+1^_2_c^+1^_3_c^+1^_4_c^+1^_5_c^+1^_6_c^+1^_6_c^+1^_7_c^+1^_7_y^+1^_6_c^+1^_8_y^+1^_7_c^+1^_8_z^+1^_8_y^+1^_8_c^+1^_9_c^+1^_9_z^+1^_9_

0838167725153354

Fragment Matches Table

Show background peaks

| Position | Ion type | Intensity | mz Theoretical | mz Error (Th) | mz Error (ppm) | Charge | Series Number |
| --- | --- | --- | --- | --- | --- | --- | --- |
| - | - | 2121 | 120.1 | - | - | 0 | - |
| - | - | 337.9 | 122.4 | - | - | 0 | - |
| - | - | 443.1 | 131.5 | - | - | 0 | - |
| - | - | 619.5 | 133.1 | - | - | 0 | - |
| - | - | 6921 | 136.1 | - | - | 0 | - |
| - | - | 431.8 | 141.9 | - | - | 0 | - |
| - | - | 451.3 | 148.9 | - | - | 0 | - |
| - | - | 488.2 | 148.9 | - | - | 0 | - |
| - | - | 510.3 | 148.9 | - | - | 0 | - |
| - | - | 657.4 | 148.9 | - | - | 0 | - |
| - | - | 698.8 | 148.9 | - | - | 0 | - |
| - | - | 831.4 | 148.9 | - | - | 0 | - |
| - | - | 1217 | 148.9 | - | - | 0 | - |
| - | - | 879.4 | 148.9 | - | - | 0 | - |
| - | - | 2016 | 148.9 | - | - | 0 | - |
| - | - | 3813 | 148.9 | - | - | 0 | - |
| - | - | 4589 | 149 | - | - | 0 | - |
| - | - | 3013 | 149 | - | - | 0 | - |
| - | - | 1514 | 149 | - | - | 0 | - |
| - | - | 1444 | 149 | - | - | 0 | - |
| - | - | 891.5 | 149 | - | - | 0 | - |
| - | - | 685 | 149 | - | - | 0 | - |
| - | - | 713.2 | 149 | - | - | 0 | - |
| - | - | 551.7 | 149 | - | - | 0 | - |
| - | - | 1088 | 149 | - | - | 0 | - |
| 10 | y | 3576 | 166.1 | 0.0002012 | 1.212 | +1 | 1 |
| - | - | 443.4 | 166.8 | - | - | 0 | - |
| - | - | 484 | 169.3 | - | - | 0 | - |
| - | - | 976.4 | 173.4 | - | - | 0 | - |
| - | - | 433.1 | 183.8 | - | - | 0 | - |
| - | - | 513.8 | 211.3 | - | - | 0 | - |
| - | - | 4690 | 221.1 | - | - | 0 | - |
| - | - | 902.9 | 222.1 | - | - | 0 | - |
| - | - | 1476 | 225 | - | - | 0 | - |
| - | - | 703.5 | 235.1 | - | - | 0 | - |
| - | - | 7914 | 239.1 | - | - | 0 | - |
| - | - | 966.5 | 240.1 | - | - | 0 | - |
| 2 | c | 2185 | 242.1 | 0.0002133 | 0.8808 | +1 | 2 |
| - | - | 568.9 | 244.4 | - | - | 0 | - |
| - | - | 558.3 | 248.5 | - | - | 0 | - |
| - | - | 593.5 | 262.4 | - | - | 0 | - |
| - | - | 9312 | 295.1 | - | - | 0 | - |
| - | - | 2784 | 296.1 | - | - | 0 | - |
| - | - | 964.4 | 297.1 | - | - | 0 | - |
| - | - | 972.7 | 299.1 | - | - | 0 | - |
| - | - | 840.3 | 300.2 | - | - | 0 | - |
| - | - | 770.1 | 313.1 | - | - | 0 | - |
| - | - | 516.7 | 314.1 | - | - | 0 | - |
| - | - | 4691 | 326.1 | - | - | 0 | - |
| 9 | y | 3949 | 329.1 | 7.461E-05 | 0.2267 | +1 | 2 |
| - | - | 964.9 | 330.2 | - | - | 0 | - |
| 3 | c | 533.1 | 330.2 | 0.001446 | 4.379 | +1 | 3 |
| - | - | 545.3 | 353.6 | - | - | 0 | - |
| - | - | 1555 | 355.1 | - | - | 0 | - |
| - | - | 484.1 | 359 | - | - | 0 | - |
| - | - | 6870 | 369.1 | - | - | 0 | - |
| - | - | 2311 | 370.1 | - | - | 0 | - |
| - | - | 831.8 | 371.1 | - | - | 0 | - |
| - | - | 1122 | 383.2 | - | - | 0 | - |
| - | - | 2923 | 401.2 | - | - | 0 | - |
| - | - | 2830 | 451.7 | - | - | 0 | - |
| - | - | 942.5 | 452.2 | - | - | 0 | - |
| - | - | 522.2 | 452.7 | - | - | 0 | - |
| - | - | 781.9 | 458.2 | - | - | 0 | - |
| 4 | c | 2313 | 459.3 | 0.00112 | 2.438 | +1 | 4 |
| - | - | 584.5 | 476.9 | - | - | 0 | - |
| - | - | 556.6 | 520.2 | - | - | 0 | - |
| - | - | 803.3 | 552.3 | - | - | 0 | - |
| - | - | 1253 | 573.3 | - | - | 0 | - |
| 5 | c | 9035 | 574.3 | 9.694E-05 | 0.1688 | +1 | 5 |
| - | - | 2487 | 575.3 | - | - | 0 | - |
| - | - | 629.5 | 594.2 | - | - | 0 | - |
| - | - | 1684 | 607.3 | - | - | 0 | - |
| - | - | 2102 | 615.3 | - | - | 0 | - |
| - | - | 3338 | 625.3 | - | - | 0 | - |
| - | - | 645.4 | 642.3 | - | - | 0 | - |
| - | - | 1043 | 643.3 | - | - | 0 | - |
| 6 | c | 677.9 | 686.3 | 0.00137 | 1.997 | +1 | 6 |
| - | - | 627.6 | 687.3 | - | - | 0 | - |
| - | - | 636.5 | 687.4 | - | - | 0 | - |
| - | - | 8887 | 702.3 | - | - | 0 | - |
| 6 | c | 4886 | 703.3 | 0.003444 | 4.897 | +1 | 6 |
| - | - | 1635 | 704.3 | - | - | 0 | - |
| - | - | 1018 | 755.3 | - | - | 0 | - |
| 7 | c | 1101 | 773.3 | 0.000684 | 0.8845 | +1 | 7 |
| - | - | 4510 | 789.4 | - | - | 0 | - |
| 7 | c | 9655 | 790.4 | 0.0008659 | 1.096 | +1 | 7 |
| - | - | 2563 | 791.4 | - | - | 0 | - |
| - | - | 704.8 | 792.4 | - | - | 0 | - |
| 5 | y | 1162 | 807.3 | 0.003524 | 4.365 | +1 | 6 |
| - | - | 781.2 | 850.4 | - | - | 0 | - |
| - | - | 901.6 | 851.4 | - | - | 0 | - |
| - | - | 672.5 | 893.4 | - | - | 0 | - |
| - | - | 841.5 | 894.4 | - | - | 0 | - |
| - | - | 1605 | 902.4 | - | - | 0 | - |
| - | - | 809.2 | 903.4 | - | - | 0 | - |
| - | - | 1119 | 904.4 | - | - | 0 | - |
| - | - | 657.6 | 911.4 | - | - | 0 | - |
| 8 | c | 3536 | 920.4 | 0.001241 | 1.348 | +1 | 8 |
| - | - | 1940 | 921.4 | - | - | 0 | - |
| - | - | 2333 | 922.4 | - | - | 0 | - |
| - | - | 1045 | 935.3 | - | - | 0 | - |
| 4 | y | 1023 | 936.3 | 0.00251 | 2.68 | +1 | 7 |
| 8 | c | 1.625E+04 | 937.4 | 0.00466 | 4.971 | +1 | 8 |
| - | - | 6184 | 938.4 | - | - | 0 | - |
| - | - | 2307 | 939.4 | - | - | 0 | - |
| - | - | 852.3 | 988.5 | - | - | 0 | - |
| 3 | z | 2684 | 991.3 | 0.003576 | 3.607 | +1 | 8 |
| - | - | 1820 | 992.4 | - | - | 0 | - |
| 3 | y | 1911 | 1007 | 0.003834 | 3.806 | +1 | 8 |
| - | - | 1804 | 1008 | - | - | 0 | - |
| - | - | 2129 | 1056 | - | - | 0 | - |
| - | - | 933 | 1067 | - | - | 0 | - |
| - | - | 1111 | 1068 | - | - | 0 | - |
| 9 | c | 1542 | 1083 | 0.00908 | 8.38 | +1 | 9 |
| 9 | c | 1.628E+04 | 1100 | 0.004137 | 3.759 | +1 | 9 |
| - | - | 1.102E+04 | 1101 | - | - | 0 | - |
| - | - | 3188 | 1102 | - | - | 0 | - |
| 2 | z | 3802 | 1119 | 0.003958 | 3.536 | +1 | 9 |
| - | - | 2358 | 1120 | - | - | 0 | - |
| - | - | 743.4 | 1121 | - | - | 0 | - |
| - | - | 1623 | 1149 | - | - | 0 | - |
| - | - | 1284 | 1186 | - | - | 0 | - |
| - | - | 923.1 | 1187 | - | - | 0 | - |
| - | - | 772.4 | 1188 | - | - | 0 | - |
| - | - | 1117 | 1189 | - | - | 0 | - |
| - | - | 1677 | 1190 | - | - | 0 | - |
| - | - | 2656 | 1204 | - | - | 0 | - |
| - | - | 3423 | 1206 | - | - | 0 | - |
| - | - | 1011 | 1207 | - | - | 0 | - |
| - | - | 814.4 | 1208 | - | - | 0 | - |
| - | - | 1030 | 1222 | - | - | 0 | - |
| - | - | 1356 | 1223 | - | - | 0 | - |
| - | - | 2402 | 1232 | - | - | 0 | - |
| - | - | 2.001E+04 | 1233 | - | - | 0 | - |
| - | - | 1.2E+04 | 1234 | - | - | 0 | - |
| - | - | 6656 | 1235 | - | - | 0 | - |
| - | - | 1461 | 1236 | - | - | 0 | - |
| - | - | 5310 | 1249 | - | - | 0 | - |
| - | - | 2.779E+04 | 1250 | - | - | 0 | - |
| - | - | 1.712E+04 | 1251 | - | - | 0 | - |
| - | - | 7296 | 1252 | - | - | 0 | - |
| - | - | 684.8 | 1759 | - | - | 0 | - |
| - | - | 671.6 | 1843 | - | - | 0 | - |
| - | - | 722.2 | 1860 | - | - | 0 | - |
| - | - | 1299 | 1872 | - | - | 0 | - |
| - | - | 695.9 | 1873 | - | - | 0 | - |
| - | - | 621.3 | 1877 | - | - | 0 | - |
| - | - | 617.4 | 2010 | - | - | 0 | - |
| - | - | 602.9 | 2473 | - | - | 0 | - |
| - | - | 946.8 | 3071 | - | - | 0 | - |
| - | - | 885.2 | 3173 | - | - | 0 | - |
| - | - | 763.3 | 3299 | - | - | 0 | - |
| - | - | 666.4 | 3321 | - | - | 0 | - |

m/z Charge Intensity FragmentType MassShift Position
120.08100128173828 0 2121.3918
122.41539001464844 0 337.94864
131.5433349609375 0 443.0842
133.08616638183594 0 619.4584
136.07586669921875 0 6921.4297
141.91432189941406 0 431.82925
148.8697509765625 0 451.27283
148.87640380859375 0 488.15018
148.8911895751953 0 510.29965
148.89825439453125 0 657.40576
148.90538024902344 0 698.84735
148.9124755859375 0 831.37103
148.91969299316406 0 1216.6427
148.92758178710938 0 879.3552
148.93405151367188 0 2015.9725
148.94186401367188 0 3812.7542
148.95826721191406 0 4588.71
148.96603393554688 0 3012.6665
148.97341918945312 0 1514.3987
148.9805145263672 0 1443.6299
148.98765563964844 0 891.4577
148.9947052001953 0 684.96326
149.00228881835938 0 713.18494
149.0096435546875 0 551.7158
149.0452117919922 0 1087.9185
166.08645629882812 0 3575.744 y 9
166.81307983398438 0 443.44662
169.33981323242188 0 483.95422
173.44932556152344 0 976.4202
183.8438262939453 0 433.0922
211.33641052246094 0 513.82245
221.0845947265625 0 4689.8115
222.08522033691406 0 902.9349
225.0355224609375 0 1475.6523
235.07476806640625 0 703.478
239.09506225585938 0 7914.41
240.09474182128906 0 966.4814
242.15013122558594 0 2185.1006 c Ammonia loss 1
244.41864013671875 0 568.8692
248.50477600097656 0 558.309
262.44354248046875 0 593.48065
295.1033935546875 0 9312.007
296.1034851074219 0 2783.9807
297.1011657714844 0 964.4369
299.062255859375 0 972.73285
300.1915283203125 0 840.2902
313.11431884765625 0 770.06824
314.06903076171875 0 516.6926
326.084228515625 0 4691.318
329.149658203125 0 3948.8784 y 8
330.1534118652344 0 964.9442
330.21502685546875 0 533.07715 c 2
353.5710754394531 0 545.2997
355.0688171386719 0 1555.145
359.027587890625 0 484.07004
369.1221618652344 0 6870.451
370.12200927734375 0 2310.7297
371.1197814941406 0 831.8197
383.22906494140625 0 1122.2567
401.2399597167969 0 2923.3962
451.6819152832031 0 2830.2383
452.1839904785156 0 942.51245
452.6835632324219 0 522.23474
458.2467041015625 0 781.94745
459.2572937011719 0 2313.1628 c 3
476.8890075683594 0 584.5084
520.2471313476562 0 556.57996
552.2685546875 0 803.26263
573.2745361328125 0 1253.4749
574.2830200195312 0 9034.701 c 4
575.286376953125 0 2486.6575
594.2141723632812 0 629.5014
607.2584228515625 0 1684.0139
615.3443603515625 0 2101.6992
625.2687377929688 0 3338.424
642.3345336914062 0 645.43396
643.3445434570312 0 1043.4359
686.2977905273438 0 677.91187 c Ammonia loss 5
687.303955078125 0 627.6007
687.3721923828125 0 636.52563
702.3176879882812 0 8886.933
703.322265625 0 4886.0005 c 5
704.3206176757812 0 1634.584
755.320556640625 0 1017.57904
773.3305053710938 0 1100.813 c Ammonia loss 6
789.3507080078125 0 4510.1987
790.3568725585938 0 9655.491 c 6
791.3599243164062 0 2563.1956
792.3616333007812 0 704.7843
807.28515625 0 1161.7332 y 4
850.3645629882812 0 781.24146
851.38037109375 0 901.5704
893.3806762695312 0 672.4804
894.38671875 0 841.50775
902.3572998046875 0 1604.5645
903.35546875 0 809.1783
904.35693359375 0 1119.2473
911.3878784179688 0 657.619
920.3629150390625 0 3536.2612 c Ammonia loss 7
921.3698120117188 0 1940.3132
922.3792114257812 0 2333.1802
935.3173828125 0 1044.7926
936.3217163085938 0 1023.36523 y 3
937.3928833007812 0 16252.573 c 7
938.39599609375 0 6184.088
939.395751953125 0 2306.5103
988.489501953125 0 852.2519
991.34619140625 0 2684.229 z 2
992.351318359375 0 1819.7341
1007.3651733398438 0 1910.8915 y 2
1008.3677368164062 0 1803.9489
1056.4400634765625 0 2129.1606
1067.407958984375 0 933.0237
1068.4132080078125 0 1111.4634
1083.43408203125 0 1542.3585 c Ammonia loss 8
1100.4556884765625 0 16279.452 c 8
1101.458984375 0 11021.4
1102.4581298828125 0 3188.1492
1119.4051513671875 0 3802.2258 z 1
1120.4075927734375 0 2357.506
1120.540283203125 0 743.4034
1148.54833984375 0 1622.9274
1185.5050048828125 0 1283.5238
1186.5294189453125 0 923.086
1187.522216796875 0 772.43176
1189.47705078125 0 1116.9224
1190.48779296875 0 1676.7842
1204.4984130859375 0 2655.827
1205.517822265625 0 3422.613
1206.521484375 0 1011.18286
1207.5361328125 0 814.3895
1221.5247802734375 0 1030.386
1222.5213623046875 0 1356.1001
1231.506591796875 0 2401.8618
1232.50537109375 0 20013.846
1233.508544921875 0 11999.895
1234.5301513671875 0 6655.5938
1235.53076171875 0 1460.6365
1248.5062255859375 0 5310.4395
1249.5140380859375 0 27791.305
1250.5201416015625 0 17123.018
1251.525390625 0 7295.9326
1758.6683349609375 0 684.8017
1842.7222900390625 0 671.63916
1859.7388916015625 0 722.1893
1871.9093017578125 0 1298.9678
1872.9486083984375 0 695.8917
1876.730224609375 0 621.3424
2010.2535400390625 0 617.4463
2472.975341796875 0 602.87744
3071.369384765625 0 946.83716
3172.646240234375 0 885.18744
3298.70654296875 0 763.31366
3320.754638671875 0 666.4118

Spectrum Details

|  |  |
| --- | --- |
| Matched peaks? Matched peaks The total absolute number of peaks matched. Additionally in brackets the total fraction of peaks matched and the total number of peaks is shown. | 19 (12.34% of 154) |
| FDR? FDR The false discovery rate estimated for this peptide. It is calculated by matching all theoretical fragments with a non-integer shift with the raw peaks for this spectrum. This is done with 40 different shifts. The resulting percentage is the average number of annotated peaks over the number of annotated peaks with the correct spectrum. | 2.51% |
| Satellite FDR? Satellite FDR See the FDR for details on its calculation. This satellite ion specific FDR only contains the satellite ions (d/w) for I/L/J positions. | - |
| PSM Score? PSM Score The PSM Score as given by Hecklib to this annotated spectrum. It is shown with three significant figures. | 239 |

#### Reverse Lookup? Reverse Lookup All places where this read could be placed.

| Group | Segment | Template | Template Part | Read Part | Score | Unique |
| --- | --- | --- | --- | --- | --- | --- |
| Homo sapiens Light Chain | IGLV | IGLV2-14 | [79..89] | [0..10] | 53 | False |
| Homo sapiens Light Chain | IGLV | IGLV2-23 | [79..89] | [0..10] | 53 | False |
| Homo sapiens Light Chain | IGLV | IGLV2-18 | [79..89] | [0..10] | 53 | False |
| Homo sapiens Light Chain | IGLV | IGLV2-8 | [79..89] | [0..10] | 53 | False |
| Homo sapiens Light Chain | IGLV | IGLV2-11 | [79..89] | [0..10] | 53 | False |
| Homo sapiens Light Chain | IGLV | IGLV1-40 | [79..89] | [0..10] | 53 | False |
| Homo sapiens Light Chain | IGLV | IGLV8-61 | [80..89] | [0..10] | 48 | False |

| Recombined | Template Part | Read Part | Score | Unique |
| --- | --- | --- | --- | --- |
| REC-0-1\_002 | [79..89] | [0..10] | 80 | True |

#### Meta Information from Multiple reads

##### Number of combined reads

3

##### Intensity

0.6809

##### TotalArea

9.447E+07

##### Changes to the peptide sequence

JQAEDESMYF

L→JNo support for either Leucine or Isoleucine based on side chain ions (Position: 1)

#### Positional Score

Copy Data

##### Positional Score (TSV)

###### Preview

```
Loading example...
```

*Click on the button to copy the data to your clipboard.*

100123456789

Label Value
"0" 0.333
"1" 0.32
"2" 0.303
"3" 0.32
"4" 0.33
"5" 0.33
"6" 0.33
"7" 0.333
"8" 0.333
"9" 0.333

#### Meta Information from PEAKS

##### Scan Identifier

F2:6919

##### Original sequence

L

Q

A

E

D

E

S

M

+15.99

Y

F

##### Posttranslational Modifications

Oxidation (M)

##### Source File

D:\separate\_stitch\_analyses\xle-disambiguation\raw\20210323\_F1\_UM1\_Peng0013\_SA\_F59\_ingel\_3ug\_TL.raw

##### Fraction

2

##### Scan Feature

F2:9210

##### De Novo Score

99

##### ConfidenceScore

99

### m/z

624.7589

##### Mass

1247.5017

##### Charge

2

##### Retention Time

38.48

##### Predicted Retention Time

-

##### Area

3.149E+07

##### Parts Per Million

1.3

##### Fragmentation mode

ETHCD

##### Originating file

01 D:\separate\_stitch\_analyses\xle-disambiguation\20210325\_F59\_3ug\_DENOVO\_12.csv

#### Meta Information from PEAKS

##### Scan Identifier

F2:6979

##### Original sequence

L

Q

A

E

D

E

S

M

+15.99

Y

F

##### Posttranslational Modifications

Oxidation (M)

##### Source File

D:\separate\_stitch\_analyses\xle-disambiguation\raw\20210323\_F1\_UM1\_Peng0013\_SA\_F59\_ingel\_3ug\_TL.raw

##### Fraction

2

##### Scan Feature

F2:9210

##### De Novo Score

99

##### ConfidenceScore

99

### m/z

624.7589

##### Mass

1247.5017

##### Charge

2

##### Retention Time

38.48

##### Predicted Retention Time

-

##### Area

3.149E+07

##### Parts Per Million

1.3

##### Fragmentation mode

ETHCD

##### Originating file

01 D:\separate\_stitch\_analyses\xle-disambiguation\20210325\_F59\_3ug\_DENOVO\_12.csv

#### Meta Information from PEAKS

##### Scan Identifier

F2:7038

##### Original sequence

L

Q

A

E

D

E

S

M

+15.99

Y

F

##### Posttranslational Modifications

Oxidation (M)

##### Source File

D:\separate\_stitch\_analyses\xle-disambiguation\raw\20210323\_F1\_UM1\_Peng0013\_SA\_F59\_ingel\_3ug\_TL.raw

##### Fraction

2

##### Scan Feature

F2:9210

##### De Novo Score

97

##### ConfidenceScore

97

### m/z

624.7589

##### Mass

1247.5017

##### Charge

2

##### Retention Time

38.48

##### Predicted Retention Time

-

##### Area

3.149E+07

##### Parts Per Million

1.3

##### Fragmentation mode

ETHCD

##### Originating file

01 D:\separate\_stitch\_analyses\xle-disambiguation\20210325\_F59\_3ug\_DENOVO\_12.csv
