## Supplementary material for "A handle on mass coincidence errors in *de novo* sequencing of antibodies by bottom-up proteomics": Combined_014.html

Details Combined\_014 | Stitch OverviewUndefined

### Read Combined\_014

#### Sequence (length=14)

SWYQHHPGKAPKJJ

#### Spectrum 5528? Spectrum 5528 The raw spectrum of this peptide as annotated by Hecklib. The fragments are coloured according to ion type (see legend). Any peaks with a star '\*' as text can be hovered over to see the full details, first the ion type second the mass shift type. By hovering over the amino acids in the peptide or ions in the legend the corresponding peaks are highlighted. By toggling the 'Unassigned' label you can turn the background (unassigned) peaks on or off in the plot. By updating the slider in the Ion legend you can update the spectrum to only show the top X% of the peaks with labels. The top X% means any peak that is within X% of the highest intensity. By dragging in the spectrum you can zoom in to a specific part of the spectrum and use 'Zoom Out' to get back to the original zoom level. The annotation of the spectrum is based on the given sequence in the peptides file and is done with different software so inconsistencies are likely. The peaks are annotated based on the given sequence, with 20 ppm tolerance.

Copy Data

##### Spectrum 5528 (TSV)

###### Preview

```
Loading example...
```

*Click on the button to copy the data to your clipboard.*

Mz MinMz MaxIntensity Max

WidthHeightPeptide font sizePeptide stroke widthSpectrum font sizeSpectrum stroke widthCompact peptide

Ion legend

wxyz

abcd

OtherUnassignedIon^Charge^_Position_Show for top:%

SWYQHHPGKAPKJJ

01.06e+52.13e+53.19e+54.26e+5

Zoom Out

y^+1^_1_y^+2^_3_b^+3^_4_a^+2^_3_y^+2^_4_a^+1^_2_b^+3^_5_y^+1^_2_a^+1^_2_b^+1^_2_b^+1^_2_y^+2^_6_y^+2^_7_y^+1^_3_b^+2^_6_y^+2^_8_b^+1^_3_b^+2^_6_b^+1^_3_y^+3^_12_y^+1^_4_b^+3^_12_y^+2^_9_b^+2^_8_b^+3^_13_b^+3^_13_b^+3^_13_y^+3^_13_y^+3^_13_y^+2^_10_y^+1^_5_\*\*y^+2^_10_\*b^+2^_9_b^+1^_4_b^+2^_10_b^+2^_10_b^+2^_10_y^+2^_11_y^+2^_11_b^+2^_11_y^+1^_6_b^+1^_5_b^+1^_5_y^+2^_12_y^+2^_12_b^+2^_12_b^+1^_5_b^+2^_12_y^+1^_7_b^+2^_13_b^+2^_13_b^+2^_13_y^+2^_13_b^+1^_6_b^+1^_6_y^+1^_8_b^+1^_6_y^+1^_9_y^+1^_10_b^+1^_9_b^+1^_10_b^+1^_10_b^+1^_10_

0636127319092546

Fragment Matches Table

Show background peaks

| Position | Ion type | Intensity | mz Theoretical | mz Error (Th) | mz Error (ppm) | Charge | Series Number |
| --- | --- | --- | --- | --- | --- | --- | --- |
| - | - | 450.5 | 120.1 | - | - | 0 | - |
| - | - | 752.7 | 120.1 | - | - | 0 | - |
| - | - | 1250 | 120.1 | - | - | 0 | - |
| - | - | 986.9 | 122.1 | - | - | 0 | - |
| - | - | 1769 | 123.1 | - | - | 0 | - |
| - | - | 358.6 | 125.1 | - | - | 0 | - |
| - | - | 360.1 | 126 | - | - | 0 | - |
| - | - | 2329 | 127.1 | - | - | 0 | - |
| - | - | 3814 | 129.1 | - | - | 0 | - |
| - | - | 9.478E+04 | 129.1 | - | - | 0 | - |
| - | - | 2.505E+04 | 130.1 | - | - | 0 | - |
| - | - | 5536 | 130.1 | - | - | 0 | - |
| - | - | 388.5 | 130.5 | - | - | 0 | - |
| - | - | 2907 | 131.1 | - | - | 0 | - |
| - | - | 8.65E+04 | 132.1 | - | - | 0 | - |
| 14 | y | 1.877E+04 | 132.1 | 0.0004203 | 3.182 | +1 | 1 |
| - | - | 1058 | 133.1 | - | - | 0 | - |
| - | - | 8497 | 133.1 | - | - | 0 | - |
| - | - | 910 | 133.1 | - | - | 0 | - |
| - | - | 439.3 | 134.1 | - | - | 0 | - |
| - | - | 502.9 | 134.1 | - | - | 0 | - |
| - | - | 483.1 | 135.1 | - | - | 0 | - |
| - | - | 488.3 | 135.7 | - | - | 0 | - |
| - | - | 4.997E+04 | 136.1 | - | - | 0 | - |
| - | - | 761.3 | 137.1 | - | - | 0 | - |
| - | - | 4584 | 137.1 | - | - | 0 | - |
| - | - | 2541 | 138.1 | - | - | 0 | - |
| - | - | 1487 | 139.1 | - | - | 0 | - |
| - | - | 5957 | 141.1 | - | - | 0 | - |
| - | - | 645.9 | 142.1 | - | - | 0 | - |
| - | - | 579.3 | 143.1 | - | - | 0 | - |
| - | - | 1587 | 144.1 | - | - | 0 | - |
| - | - | 2479 | 145.1 | - | - | 0 | - |
| - | - | 1590 | 146.1 | - | - | 0 | - |
| - | - | 503.6 | 146.1 | - | - | 0 | - |
| - | - | 569.1 | 146.1 | - | - | 0 | - |
| - | - | 1092 | 147 | - | - | 0 | - |
| - | - | 638.9 | 149 | - | - | 0 | - |
| - | - | 1442 | 151.1 | - | - | 0 | - |
| - | - | 780.6 | 152.1 | - | - | 0 | - |
| - | - | 2306 | 152.1 | - | - | 0 | - |
| - | - | 2102 | 152.1 | - | - | 0 | - |
| - | - | 733.1 | 153.1 | - | - | 0 | - |
| - | - | 747.8 | 153.1 | - | - | 0 | - |
| - | - | 576.5 | 153.1 | - | - | 0 | - |
| - | - | 612.8 | 154.1 | - | - | 0 | - |
| - | - | 545.9 | 154.1 | - | - | 0 | - |
| - | - | 4506 | 155.1 | - | - | 0 | - |
| - | - | 3446 | 155.1 | - | - | 0 | - |
| - | - | 3688 | 155.1 | - | - | 0 | - |
| - | - | 4191 | 156.1 | - | - | 0 | - |
| - | - | 663.2 | 157.1 | - | - | 0 | - |
| - | - | 995.6 | 158.1 | - | - | 0 | - |
| - | - | 9116 | 158.1 | - | - | 0 | - |
| - | - | 4.055E+04 | 159.1 | - | - | 0 | - |
| - | - | 2167 | 160.1 | - | - | 0 | - |
| - | - | 4019 | 160.1 | - | - | 0 | - |
| - | - | 635.4 | 161.1 | - | - | 0 | - |
| - | - | 748.3 | 162.1 | - | - | 0 | - |
| - | - | 2086 | 163.1 | - | - | 0 | - |
| - | - | 545.3 | 164.1 | - | - | 0 | - |
| - | - | 762.2 | 165.1 | - | - | 0 | - |
| - | - | 721.6 | 165.1 | - | - | 0 | - |
| - | - | 3359 | 165.1 | - | - | 0 | - |
| - | - | 2499 | 166.1 | - | - | 0 | - |
| - | - | 700.5 | 167.1 | - | - | 0 | - |
| - | - | 795 | 167.1 | - | - | 0 | - |
| - | - | 865.2 | 168.1 | - | - | 0 | - |
| - | - | 4269 | 169.1 | - | - | 0 | - |
| - | - | 7627 | 169.1 | - | - | 0 | - |
| - | - | 1.476E+04 | 170.1 | - | - | 0 | - |
| - | - | 2150 | 171.1 | - | - | 0 | - |
| - | - | 837.7 | 171.1 | - | - | 0 | - |
| - | - | 1079 | 171.1 | - | - | 0 | - |
| - | - | 1335 | 171.1 | - | - | 0 | - |
| - | - | 5862 | 172.1 | - | - | 0 | - |
| - | - | 739.4 | 175.1 | - | - | 0 | - |
| - | - | 742.5 | 175.1 | - | - | 0 | - |
| - | - | 1413 | 176.1 | - | - | 0 | - |
| - | - | 663.1 | 178.1 | - | - | 0 | - |
| - | - | 521.8 | 179.1 | - | - | 0 | - |
| - | - | 917.1 | 180.1 | - | - | 0 | - |
| - | - | 1107 | 181.1 | - | - | 0 | - |
| - | - | 1.774E+04 | 181.1 | - | - | 0 | - |
| - | - | 2277 | 181.1 | - | - | 0 | - |
| - | - | 1592 | 182.1 | - | - | 0 | - |
| - | - | 3709 | 182.1 | - | - | 0 | - |
| - | - | 595.2 | 182.3 | - | - | 0 | - |
| - | - | 1.316E+04 | 183.1 | - | - | 0 | - |
| - | - | 1749 | 183.1 | - | - | 0 | - |
| - | - | 664.9 | 183.1 | - | - | 0 | - |
| - | - | 739 | 183.1 | - | - | 0 | - |
| - | - | 850.2 | 184.1 | - | - | 0 | - |
| - | - | 1048 | 184.1 | - | - | 0 | - |
| - | - | 2866 | 185.1 | - | - | 0 | - |
| - | - | 639.9 | 185.1 | - | - | 0 | - |
| - | - | 559.8 | 185.1 | - | - | 0 | - |
| - | - | 9998 | 186.1 | - | - | 0 | - |
| - | - | 8.048E+04 | 187.1 | - | - | 0 | - |
| - | - | 1714 | 187.1 | - | - | 0 | - |
| 12 | y | 1451 | 187.1 | 0.000427 | 2.281 | +2 | 3 |
| - | - | 8696 | 188.1 | - | - | 0 | - |
| 4 | b | 1895 | 189.1 | 0.002407 | 12.73 | +3 | 4 |
| - | - | 472.2 | 191.1 | - | - | 0 | - |
| - | - | 1063 | 195.1 | - | - | 0 | - |
| - | - | 1340 | 195.1 | - | - | 0 | - |
| - | - | 601.4 | 197.1 | - | - | 0 | - |
| - | - | 3751 | 197.2 | - | - | 0 | - |
| - | - | 641.2 | 198.1 | - | - | 0 | - |
| - | - | 490.1 | 198.2 | - | - | 0 | - |
| - | - | 2097 | 199.1 | - | - | 0 | - |
| - | - | 1817 | 199.1 | - | - | 0 | - |
| - | - | 919.4 | 199.1 | - | - | 0 | - |
| - | - | 3435 | 199.2 | - | - | 0 | - |
| - | - | 1162 | 200.1 | - | - | 0 | - |
| - | - | 607.1 | 201.1 | - | - | 0 | - |
| - | - | 6622 | 201.1 | - | - | 0 | - |
| - | - | 1196 | 201.1 | - | - | 0 | - |
| - | - | 1188 | 202.1 | - | - | 0 | - |
| - | - | 9106 | 202.1 | - | - | 0 | - |
| - | - | 530.3 | 203.1 | - | - | 0 | - |
| - | - | 695.9 | 203.1 | - | - | 0 | - |
| - | - | 1807 | 203.1 | - | - | 0 | - |
| - | - | 600.9 | 203.1 | - | - | 0 | - |
| - | - | 619.2 | 204.1 | - | - | 0 | - |
| - | - | 635.2 | 205.1 | - | - | 0 | - |
| 3 | a | 1070 | 205.1 | 0.0004716 | 2.3 | +2 | 3 |
| - | - | 1025 | 205.1 | - | - | 0 | - |
| - | - | 6483 | 207.1 | - | - | 0 | - |
| - | - | 559.7 | 207.1 | - | - | 0 | - |
| - | - | 695.7 | 207.1 | - | - | 0 | - |
| - | - | 3479 | 207.1 | - | - | 0 | - |
| - | - | 796.2 | 207.2 | - | - | 0 | - |
| - | - | 970.6 | 208.1 | - | - | 0 | - |
| - | - | 815.6 | 209.1 | - | - | 0 | - |
| - | - | 623.2 | 209.1 | - | - | 0 | - |
| - | - | 814.6 | 209.1 | - | - | 0 | - |
| - | - | 1605 | 211.1 | - | - | 0 | - |
| - | - | 687.8 | 211.1 | - | - | 0 | - |
| - | - | 1291 | 212.1 | - | - | 0 | - |
| - | - | 4326 | 214.1 | - | - | 0 | - |
| - | - | 2103 | 215.1 | - | - | 0 | - |
| - | - | 635.6 | 216.1 | - | - | 0 | - |
| - | - | 543.5 | 216.1 | - | - | 0 | - |
| - | - | 559 | 217 | - | - | 0 | - |
| - | - | 920.8 | 217.1 | - | - | 0 | - |
| - | - | 775.2 | 219.1 | - | - | 0 | - |
| - | - | 7066 | 220.1 | - | - | 0 | - |
| - | - | 9255 | 221.1 | - | - | 0 | - |
| - | - | 764.2 | 221.1 | - | - | 0 | - |
| - | - | 818.9 | 222.1 | - | - | 0 | - |
| - | - | 1089 | 222.1 | - | - | 0 | - |
| - | - | 758.7 | 223.1 | - | - | 0 | - |
| - | - | 1470 | 223.2 | - | - | 0 | - |
| - | - | 2068 | 224.1 | - | - | 0 | - |
| - | - | 4592 | 224.2 | - | - | 0 | - |
| - | - | 533.7 | 225 | - | - | 0 | - |
| - | - | 559.5 | 225.4 | - | - | 0 | - |
| - | - | 750.2 | 226.1 | - | - | 0 | - |
| - | - | 1.097E+05 | 226.2 | - | - | 0 | - |
| - | - | 2130 | 227.1 | - | - | 0 | - |
| - | - | 825.4 | 227.1 | - | - | 0 | - |
| 11 | y | 1.326E+04 | 227.2 | 0.001617 | 7.119 | +2 | 4 |
| - | - | 921.6 | 227.2 | - | - | 0 | - |
| 2 | a | 7124 | 228.1 | 0.0004937 | 2.164 | +1 | 2 |
| - | - | 946.8 | 228.1 | - | - | 0 | - |
| - | - | 1004 | 228.2 | - | - | 0 | - |
| 5 | b | 9273 | 229.1 | 0.001752 | 7.645 | +3 | 5 |
| - | - | 1507 | 229.1 | - | - | 0 | - |
| - | - | 943.9 | 229.2 | - | - | 0 | - |
| - | - | 775.1 | 229.2 | - | - | 0 | - |
| - | - | 2234 | 230.1 | - | - | 0 | - |
| - | - | 3172 | 231.1 | - | - | 0 | - |
| - | - | 1222 | 231.1 | - | - | 0 | - |
| - | - | 923.8 | 232.1 | - | - | 0 | - |
| - | - | 1701 | 233.1 | - | - | 0 | - |
| - | - | 735.7 | 234.1 | - | - | 0 | - |
| - | - | 1608 | 234.1 | - | - | 0 | - |
| - | - | 796.4 | 234.1 | - | - | 0 | - |
| - | - | 1.738E+04 | 235.1 | - | - | 0 | - |
| - | - | 2457 | 236.1 | - | - | 0 | - |
| - | - | 629.7 | 238.1 | - | - | 0 | - |
| - | - | 2023 | 238.1 | - | - | 0 | - |
| - | - | 1235 | 239.1 | - | - | 0 | - |
| - | - | 6210 | 239.2 | - | - | 0 | - |
| - | - | 897.2 | 240.1 | - | - | 0 | - |
| - | - | 637.7 | 240.2 | - | - | 0 | - |
| - | - | 1248 | 241.2 | - | - | 0 | - |
| - | - | 2528 | 242.2 | - | - | 0 | - |
| - | - | 786.9 | 243.2 | - | - | 0 | - |
| - | - | 862.1 | 244.1 | - | - | 0 | - |
| 13 | y | 3.787E+04 | 245.2 | 0.0006001 | 2.448 | +1 | 2 |
| 2 | a | 2.149E+05 | 246.1 | 0.0005644 | 2.293 | +1 | 2 |
| - | - | 4182 | 246.2 | - | - | 0 | - |
| - | - | 4.864E+04 | 247.1 | - | - | 0 | - |
| - | - | 1.112E+04 | 248.1 | - | - | 0 | - |
| - | - | 3513 | 248.1 | - | - | 0 | - |
| - | - | 1.572E+04 | 249.1 | - | - | 0 | - |
| - | - | 853 | 249.1 | - | - | 0 | - |
| - | - | 1060 | 249.1 | - | - | 0 | - |
| - | - | 1664 | 250.1 | - | - | 0 | - |
| - | - | 1281 | 250.1 | - | - | 0 | - |
| - | - | 5098 | 251.2 | - | - | 0 | - |
| - | - | 1486 | 252.2 | - | - | 0 | - |
| 2 | b | 5487 | 256.1 | 0.0005284 | 2.063 | +1 | 2 |
| - | - | 1591 | 257.1 | - | - | 0 | - |
| - | - | 1456 | 257.2 | - | - | 0 | - |
| - | - | 1095 | 258.1 | - | - | 0 | - |
| - | - | 2.556E+04 | 258.1 | - | - | 0 | - |
| - | - | 3956 | 259.1 | - | - | 0 | - |
| - | - | 747.7 | 259.1 | - | - | 0 | - |
| - | - | 1929 | 260.1 | - | - | 0 | - |
| - | - | 1459 | 260.2 | - | - | 0 | - |
| - | - | 972.6 | 263.1 | - | - | 0 | - |
| - | - | 1567 | 264.1 | - | - | 0 | - |
| - | - | 961.8 | 265.1 | - | - | 0 | - |
| - | - | 1241 | 265.2 | - | - | 0 | - |
| - | - | 1.38E+04 | 266.1 | - | - | 0 | - |
| - | - | 1836 | 267.1 | - | - | 0 | - |
| - | - | 608.9 | 267.1 | - | - | 0 | - |
| - | - | 6451 | 270.2 | - | - | 0 | - |
| 2 | b | 1.25E+05 | 274.1 | 0.0004923 | 1.796 | +1 | 2 |
| - | - | 3788 | 275.1 | - | - | 0 | - |
| - | - | 6.159E+04 | 275.1 | - | - | 0 | - |
| - | - | 2053 | 276.1 | - | - | 0 | - |
| - | - | 5810 | 276.1 | - | - | 0 | - |
| - | - | 698.3 | 276.2 | - | - | 0 | - |
| - | - | 1699 | 277.1 | - | - | 0 | - |
| - | - | 1081 | 279.2 | - | - | 0 | - |
| - | - | 662.5 | 280.2 | - | - | 0 | - |
| - | - | 1732 | 283.2 | - | - | 0 | - |
| - | - | 1.937E+04 | 283.2 | - | - | 0 | - |
| - | - | 2088 | 284.2 | - | - | 0 | - |
| - | - | 1989 | 285.1 | - | - | 0 | - |
| - | - | 5678 | 285.2 | - | - | 0 | - |
| - | - | 835.7 | 290.1 | - | - | 0 | - |
| - | - | 6597 | 290.2 | - | - | 0 | - |
| - | - | 1478 | 290.7 | - | - | 0 | - |
| - | - | 968.9 | 291.2 | - | - | 0 | - |
| - | - | 1.236E+04 | 292.1 | - | - | 0 | - |
| - | - | 586.9 | 292.2 | - | - | 0 | - |
| - | - | 1654 | 293.1 | - | - | 0 | - |
| - | - | 5983 | 294.1 | - | - | 0 | - |
| - | - | 736.9 | 295.1 | - | - | 0 | - |
| - | - | 689.1 | 297.2 | - | - | 0 | - |
| - | - | 1.699E+04 | 297.2 | - | - | 0 | - |
| - | - | 2865 | 298.2 | - | - | 0 | - |
| - | - | 916.3 | 299.1 | - | - | 0 | - |
| - | - | 800.3 | 300.1 | - | - | 0 | - |
| - | - | 788.5 | 301.1 | - | - | 0 | - |
| - | - | 3115 | 303.1 | - | - | 0 | - |
| - | - | 787.6 | 303.2 | - | - | 0 | - |
| - | - | 3150 | 305.1 | - | - | 0 | - |
| - | - | 1297 | 305.7 | - | - | 0 | - |
| - | - | 773.4 | 306.1 | - | - | 0 | - |
| - | - | 1012 | 306.1 | - | - | 0 | - |
| - | - | 2663 | 311.2 | - | - | 0 | - |
| - | - | 796.8 | 314.7 | - | - | 0 | - |
| - | - | 907.8 | 318.2 | - | - | 0 | - |
| - | - | 3497 | 320.1 | - | - | 0 | - |
| - | - | 1.653E+04 | 321.2 | - | - | 0 | - |
| - | - | 6158 | 322.2 | - | - | 0 | - |
| - | - | 3364 | 322.2 | - | - | 0 | - |
| - | - | 690.4 | 323.2 | - | - | 0 | - |
| - | - | 913.2 | 324.2 | - | - | 0 | - |
| - | - | 754.3 | 324.7 | - | - | 0 | - |
| - | - | 1048 | 325.2 | - | - | 0 | - |
| - | - | 3652 | 326.2 | - | - | 0 | - |
| - | - | 757.8 | 327.2 | - | - | 0 | - |
| - | - | 3530 | 332.7 | - | - | 0 | - |
| - | - | 1043 | 333.1 | - | - | 0 | - |
| - | - | 1326 | 333.2 | - | - | 0 | - |
| 9 | y | 721.3 | 335.2 | 0.001234 | 3.681 | +2 | 6 |
| - | - | 707.5 | 336.1 | - | - | 0 | - |
| - | - | 2.256E+04 | 336.2 | - | - | 0 | - |
| - | - | 731.1 | 337.2 | - | - | 0 | - |
| - | - | 4124 | 337.2 | - | - | 0 | - |
| - | - | 4647 | 337.7 | - | - | 0 | - |
| - | - | 739.3 | 338.1 | - | - | 0 | - |
| - | - | 887.6 | 339.2 | - | - | 0 | - |
| - | - | 1.675E+04 | 339.2 | - | - | 0 | - |
| - | - | 1444 | 340.2 | - | - | 0 | - |
| - | - | 992.4 | 340.2 | - | - | 0 | - |
| - | - | 3300 | 340.2 | - | - | 0 | - |
| - | - | 1183 | 341.2 | - | - | 0 | - |
| - | - | 1.657E+04 | 346.7 | - | - | 0 | - |
| - | - | 6946 | 347.2 | - | - | 0 | - |
| - | - | 1485 | 347.7 | - | - | 0 | - |
| - | - | 2978 | 350.2 | - | - | 0 | - |
| - | - | 831.1 | 351.2 | - | - | 0 | - |
| - | - | 1867 | 351.2 | - | - | 0 | - |
| - | - | 1062 | 353.2 | - | - | 0 | - |
| - | - | 2.993E+04 | 354.2 | - | - | 0 | - |
| - | - | 1271 | 355.1 | - | - | 0 | - |
| - | - | 5909 | 355.2 | - | - | 0 | - |
| - | - | 789.5 | 355.3 | - | - | 0 | - |
| - | - | 1366 | 355.7 | - | - | 0 | - |
| - | - | 725.8 | 356.2 | - | - | 0 | - |
| - | - | 973.1 | 357.2 | - | - | 0 | - |
| - | - | 3052 | 357.3 | - | - | 0 | - |
| - | - | 4709 | 358.2 | - | - | 0 | - |
| - | - | 2253 | 358.7 | - | - | 0 | - |
| - | - | 879.1 | 359.2 | - | - | 0 | - |
| - | - | 927 | 359.2 | - | - | 0 | - |
| - | - | 725.4 | 360.1 | - | - | 0 | - |
| - | - | 1227 | 360.7 | - | - | 0 | - |
| - | - | 754.1 | 361.7 | - | - | 0 | - |
| - | - | 1205 | 362.7 | - | - | 0 | - |
| 8 | y | 1088 | 363.7 | 0.001035 | 2.844 | +2 | 7 |
| - | - | 830.6 | 365.3 | - | - | 0 | - |
| - | - | 3377 | 366.2 | - | - | 0 | - |
| - | - | 1493 | 367.2 | - | - | 0 | - |
| - | - | 2008 | 368.1 | - | - | 0 | - |
| - | - | 1736 | 368.2 | - | - | 0 | - |
| - | - | 690.7 | 370.2 | - | - | 0 | - |
| 12 | y | 7964 | 373.3 | 0.00044 | 1.179 | +1 | 3 |
| - | - | 672.6 | 374.3 | - | - | 0 | - |
| - | - | 1106 | 374.3 | - | - | 0 | - |
| - | - | 1147 | 379.2 | - | - | 0 | - |
| - | - | 1800 | 380.2 | - | - | 0 | - |
| - | - | 752.9 | 381.2 | - | - | 0 | - |
| - | - | 1414 | 382.3 | - | - | 0 | - |
| - | - | 963.1 | 382.5 | - | - | 0 | - |
| - | - | 1531 | 383.3 | - | - | 0 | - |
| - | - | 4152 | 384.2 | - | - | 0 | - |
| - | - | 1.07E+04 | 385.2 | - | - | 0 | - |
| - | - | 3546 | 385.2 | - | - | 0 | - |
| - | - | 8128 | 386.2 | - | - | 0 | - |
| - | - | 888.4 | 386.2 | - | - | 0 | - |
| - | - | 1923 | 387.2 | - | - | 0 | - |
| - | - | 1081 | 389.8 | - | - | 0 | - |
| - | - | 919 | 392.2 | - | - | 0 | - |
| - | - | 3494 | 392.3 | - | - | 0 | - |
| - | - | 1554 | 393.2 | - | - | 0 | - |
| - | - | 1271 | 393.3 | - | - | 0 | - |
| - | - | 8028 | 394.2 | - | - | 0 | - |
| - | - | 710 | 394.3 | - | - | 0 | - |
| - | - | 2420 | 395.2 | - | - | 0 | - |
| - | - | 1534 | 396.3 | - | - | 0 | - |
| - | - | 906.7 | 397.2 | - | - | 0 | - |
| - | - | 974.4 | 398.2 | - | - | 0 | - |
| - | - | 1439 | 401.2 | - | - | 0 | - |
| - | - | 1666 | 401.3 | - | - | 0 | - |
| - | - | 4771 | 402.2 | - | - | 0 | - |
| - | - | 1.046E+04 | 403.2 | - | - | 0 | - |
| - | - | 780.9 | 403.2 | - | - | 0 | - |
| - | - | 2605 | 403.3 | - | - | 0 | - |
| - | - | 953.4 | 403.8 | - | - | 0 | - |
| - | - | 2300 | 404.2 | - | - | 0 | - |
| - | - | 5113 | 406.2 | - | - | 0 | - |
| - | - | 2709 | 406.3 | - | - | 0 | - |
| - | - | 2339 | 406.7 | - | - | 0 | - |
| - | - | 1638 | 406.8 | - | - | 0 | - |
| - | - | 1.005E+04 | 410.3 | - | - | 0 | - |
| 6 | b | 1.608E+04 | 411.2 | 0.0006014 | 1.463 | +2 | 6 |
| - | - | 1977 | 411.3 | - | - | 0 | - |
| - | - | 1998 | 412.2 | - | - | 0 | - |
| - | - | 2483 | 412.2 | - | - | 0 | - |
| 7 | y | 4.114E+04 | 412.3 | 0.0008672 | 2.103 | +2 | 8 |
| - | - | 2.171E+04 | 412.8 | - | - | 0 | - |
| - | - | 1583 | 413.2 | - | - | 0 | - |
| - | - | 6528 | 413.3 | - | - | 0 | - |
| - | - | 3639 | 415.3 | - | - | 0 | - |
| - | - | 1753 | 415.8 | - | - | 0 | - |
| 3 | b | 921.6 | 419.2 | 0.0002797 | 0.6673 | +1 | 3 |
| - | - | 621.5 | 419.2 | - | - | 0 | - |
| - | - | 1925 | 419.9 | - | - | 0 | - |
| 6 | b | 4673 | 420.2 | 0.000446 | 1.062 | +2 | 6 |
| - | - | 3612 | 420.2 | - | - | 0 | - |
| - | - | 3054 | 420.7 | - | - | 0 | - |
| - | - | 3734 | 422.3 | - | - | 0 | - |
| - | - | 1223 | 422.3 | - | - | 0 | - |
| - | - | 1834 | 423.3 | - | - | 0 | - |
| - | - | 1915 | 425.3 | - | - | 0 | - |
| - | - | 4676 | 427.2 | - | - | 0 | - |
| - | - | 2468 | 427.7 | - | - | 0 | - |
| - | - | 836.3 | 428.2 | - | - | 0 | - |
| - | - | 953.7 | 428.3 | - | - | 0 | - |
| - | - | 5.173E+04 | 429.2 | - | - | 0 | - |
| - | - | 1.287E+04 | 430.2 | - | - | 0 | - |
| - | - | 5557 | 431.2 | - | - | 0 | - |
| - | - | 1270 | 432.2 | - | - | 0 | - |
| - | - | 934.3 | 433.3 | - | - | 0 | - |
| - | - | 964.4 | 436.7 | - | - | 0 | - |
| 3 | b | 9560 | 437.2 | 0.0007014 | 1.604 | +1 | 3 |
| - | - | 3098 | 438.2 | - | - | 0 | - |
| - | - | 1234 | 438.6 | - | - | 0 | - |
| - | - | 1628 | 438.9 | - | - | 0 | - |
| - | - | 810.6 | 439.2 | - | - | 0 | - |
| - | - | 655.4 | 439.2 | - | - | 0 | - |
| - | - | 1276 | 439.3 | - | - | 0 | - |
| - | - | 1242 | 442.7 | - | - | 0 | - |
| - | - | 1417 | 444.2 | - | - | 0 | - |
| - | - | 1851 | 445.3 | - | - | 0 | - |
| - | - | 982.5 | 446.2 | - | - | 0 | - |
| - | - | 635.3 | 448.2 | - | - | 0 | - |
| - | - | 694.9 | 450.1 | - | - | 0 | - |
| - | - | 617.2 | 450.3 | - | - | 0 | - |
| - | - | 1638 | 451.2 | - | - | 0 | - |
| - | - | 3203 | 451.3 | - | - | 0 | - |
| - | - | 2167 | 451.7 | - | - | 0 | - |
| - | - | 922.9 | 451.7 | - | - | 0 | - |
| - | - | 766.7 | 452.2 | - | - | 0 | - |
| - | - | 795 | 452.3 | - | - | 0 | - |
| - | - | 1552 | 452.3 | - | - | 0 | - |
| - | - | 664.6 | 453.3 | - | - | 0 | - |
| - | - | 3288 | 454.2 | - | - | 0 | - |
| - | - | 1107 | 455.2 | - | - | 0 | - |
| - | - | 3624 | 455.3 | - | - | 0 | - |
| - | - | 924.3 | 456.3 | - | - | 0 | - |
| - | - | 2048 | 457.2 | - | - | 0 | - |
| - | - | 867.4 | 458.2 | - | - | 0 | - |
| - | - | 1998 | 460.2 | - | - | 0 | - |
| - | - | 696.4 | 460.7 | - | - | 0 | - |
| - | - | 1140 | 461.2 | - | - | 0 | - |
| 3 | y | 8465 | 463.6 | 0.0006605 | 1.425 | +3 | 12 |
| - | - | 8630 | 463.9 | - | - | 0 | - |
| - | - | 854.7 | 464.2 | - | - | 0 | - |
| - | - | 2649 | 464.3 | - | - | 0 | - |
| - | - | 1213 | 464.6 | - | - | 0 | - |
| - | - | 912.5 | 466.9 | - | - | 0 | - |
| - | - | 817.4 | 467.3 | - | - | 0 | - |
| - | - | 4585 | 467.3 | - | - | 0 | - |
| - | - | 2617 | 468.3 | - | - | 0 | - |
| - | - | 1251 | 469.8 | - | - | 0 | - |
| - | - | 1103 | 470.3 | - | - | 0 | - |
| 11 | y | 3.437E+04 | 470.3 | 0.0007157 | 1.522 | +1 | 4 |
| - | - | 1273 | 470.6 | - | - | 0 | - |
| - | - | 8478 | 471.3 | - | - | 0 | - |
| - | - | 2523 | 471.8 | - | - | 0 | - |
| - | - | 2664 | 472.3 | - | - | 0 | - |
| - | - | 1207 | 472.3 | - | - | 0 | - |
| - | - | 822.1 | 472.6 | - | - | 0 | - |
| 12 | b | 1.579E+04 | 473.2 | 0.001494 | 3.158 | +3 | 12 |
| - | - | 1.473E+04 | 473.6 | - | - | 0 | - |
| - | - | 1430 | 473.7 | - | - | 0 | - |
| - | - | 4820 | 473.9 | - | - | 0 | - |
| - | - | 1775 | 474.2 | - | - | 0 | - |
| - | - | 1566 | 474.3 | - | - | 0 | - |
| - | - | 2454 | 474.8 | - | - | 0 | - |
| - | - | 1906 | 475.3 | - | - | 0 | - |
| - | - | 787.5 | 475.8 | - | - | 0 | - |
| - | - | 5774 | 476.3 | - | - | 0 | - |
| - | - | 4418 | 476.6 | - | - | 0 | - |
| - | - | 2588 | 478.2 | - | - | 0 | - |
| 6 | y | 1.03E+04 | 480.8 | 0.0006166 | 1.282 | +2 | 9 |
| - | - | 5733 | 481.3 | - | - | 0 | - |
| - | - | 2002 | 481.9 | - | - | 0 | - |
| - | - | 3912 | 482.3 | - | - | 0 | - |
| - | - | 3728 | 482.3 | - | - | 0 | - |
| - | - | 2979 | 482.8 | - | - | 0 | - |
| - | - | 927.7 | 483.3 | - | - | 0 | - |
| - | - | 7099 | 483.8 | - | - | 0 | - |
| - | - | 3577 | 484.3 | - | - | 0 | - |
| - | - | 921.6 | 484.8 | - | - | 0 | - |
| - | - | 862.8 | 486.2 | - | - | 0 | - |
| - | - | 884.7 | 488.3 | - | - | 0 | - |
| - | - | 4760 | 491.3 | - | - | 0 | - |
| - | - | 1035 | 492.3 | - | - | 0 | - |
| - | - | 668.1 | 492.8 | - | - | 0 | - |
| - | - | 1495 | 494.2 | - | - | 0 | - |
| - | - | 1137 | 495.9 | - | - | 0 | - |
| - | - | 1625 | 496.3 | - | - | 0 | - |
| - | - | 867.6 | 496.8 | - | - | 0 | - |
| 8 | b | 1883 | 497.2 | 0.000289 | 0.5813 | +2 | 8 |
| - | - | 1948 | 497.7 | - | - | 0 | - |
| - | - | 7271 | 501.6 | - | - | 0 | - |
| - | - | 5848 | 501.9 | - | - | 0 | - |
| - | - | 3944 | 502.3 | - | - | 0 | - |
| - | - | 1583 | 502.6 | - | - | 0 | - |
| - | - | 2690 | 503.2 | - | - | 0 | - |
| 13 | b | 1.215E+04 | 504.9 | 0.0001473 | 0.2916 | +3 | 13 |
| 13 | b | 1.53E+04 | 505.3 | 0.005852 | 11.58 | +3 | 13 |
| - | - | 6882 | 505.6 | - | - | 0 | - |
| - | - | 3828 | 505.9 | - | - | 0 | - |
| - | - | 802.7 | 506.3 | - | - | 0 | - |
| - | - | 1851 | 508.8 | - | - | 0 | - |
| - | - | 1819 | 509.3 | - | - | 0 | - |
| 13 | b | 6.906E+04 | 510.9 | 0.000593 | 1.161 | +3 | 13 |
| - | - | 6.636E+04 | 511.3 | - | - | 0 | - |
| - | - | 2.819E+04 | 511.6 | - | - | 0 | - |
| - | - | 7660 | 511.9 | - | - | 0 | - |
| - | - | 2705 | 512.3 | - | - | 0 | - |
| - | - | 7593 | 516.9 | - | - | 0 | - |
| - | - | 9868 | 517.3 | - | - | 0 | - |
| - | - | 5356 | 517.6 | - | - | 0 | - |
| - | - | 1775 | 517.9 | - | - | 0 | - |
| - | - | 1254 | 517.9 | - | - | 0 | - |
| - | - | 847.9 | 519.6 | - | - | 0 | - |
| 2 | y | 2712 | 519.9 | 0.0001021 | 0.1963 | +3 | 13 |
| - | - | 872.1 | 520.2 | - | - | 0 | - |
| - | - | 2611 | 520.3 | - | - | 0 | - |
| - | - | 1791 | 520.4 | - | - | 0 | - |
| - | - | 1226 | 520.6 | - | - | 0 | - |
| - | - | 2490 | 521.2 | - | - | 0 | - |
| - | - | 930 | 523.4 | - | - | 0 | - |
| 2 | y | 1.171E+04 | 525.6 | 0.0001018 | 0.1937 | +3 | 13 |
| - | - | 9944 | 526 | - | - | 0 | - |
| - | - | 5653 | 526.3 | - | - | 0 | - |
| - | - | 2082 | 526.6 | - | - | 0 | - |
| - | - | 1962 | 528.2 | - | - | 0 | - |
| - | - | 907 | 529.2 | - | - | 0 | - |
| - | - | 1783 | 529.8 | - | - | 0 | - |
| - | - | 1723 | 530.3 | - | - | 0 | - |
| - | - | 1216 | 530.8 | - | - | 0 | - |
| - | - | 2329 | 531.2 | - | - | 0 | - |
| - | - | 1242 | 535.3 | - | - | 0 | - |
| - | - | 735.3 | 535.6 | - | - | 0 | - |
| - | - | 1047 | 537.3 | - | - | 0 | - |
| - | - | 823.7 | 537.6 | - | - | 0 | - |
| - | - | 2.747E+04 | 538.3 | - | - | 0 | - |
| - | - | 2192 | 538.4 | - | - | 0 | - |
| - | - | 4315 | 538.8 | - | - | 0 | - |
| - | - | 7805 | 539.3 | - | - | 0 | - |
| - | - | 7369 | 539.3 | - | - | 0 | - |
| - | - | 2044 | 539.6 | - | - | 0 | - |
| - | - | 2903 | 539.8 | - | - | 0 | - |
| - | - | 1016 | 540 | - | - | 0 | - |
| - | - | 1216 | 540.3 | - | - | 0 | - |
| - | - | 1022 | 540.3 | - | - | 0 | - |
| - | - | 804.8 | 540.6 | - | - | 0 | - |
| 5 | y | 781.6 | 540.8 | 0.006103 | 11.28 | +2 | 10 |
| - | - | 589.9 | 541.3 | - | - | 0 | - |
| 10 | y | 9739 | 541.4 | 0.0008333 | 1.539 | +1 | 5 |
| - | - | 879.1 | 542.3 | - | - | 0 | - |
| - | - | 3326 | 542.4 | - | - | 0 | - |
| - | - | 1852 | 543 | - | - | 0 | - |
| - | - | 1116 | 543.3 | - | - | 0 | - |
| - | - | 724 | 544.3 | - | - | 0 | - |
| - | - | 2640 | 547.8 | - | - | 0 | - |
| - | - | 1009 | 548.2 | - | - | 0 | - |
| - | - | 1948 | 548.3 | - | - | 0 | - |
| 0 | Precursor | 1.953E+04 | 548.6 | 0.0005156 | 0.9398 | +3 | -1 |
| 0 | Precursor | 2.31E+04 | 549 | 0.005579 | 10.16 | +3 | -1 |
| - | - | 1686 | 549.2 | - | - | 0 | - |
| - | - | 9260 | 549.3 | - | - | 0 | - |
| 5 | y | 1.671E+04 | 549.3 | 0.001068 | 1.944 | +2 | 10 |
| - | - | 7357 | 549.6 | - | - | 0 | - |
| - | - | 1.031E+04 | 549.8 | - | - | 0 | - |
| - | - | 1616 | 550 | - | - | 0 | - |
| - | - | 4840 | 550.3 | - | - | 0 | - |
| - | - | 1132 | 550.8 | - | - | 0 | - |
| - | - | 735.7 | 551.4 | - | - | 0 | - |
| - | - | 1456 | 553.3 | - | - | 0 | - |
| - | - | 3571 | 554 | - | - | 0 | - |
| - | - | 825.6 | 554.3 | - | - | 0 | - |
| - | - | 2627 | 554.3 | - | - | 0 | - |
| 0 | Precursor | 3.929E+05 | 554.6 | 0.0007172 | 1.293 | +3 | -1 |
| - | - | 4.214E+05 | 555 | - | - | 0 | - |
| - | - | 2.22E+05 | 555.3 | - | - | 0 | - |
| - | - | 5035 | 555.6 | - | - | 0 | - |
| - | - | 7.324E+04 | 555.6 | - | - | 0 | - |
| - | - | 2756 | 555.9 | - | - | 0 | - |
| - | - | 2.322E+04 | 556 | - | - | 0 | - |
| - | - | 1173 | 557.3 | - | - | 0 | - |
| 9 | b | 2376 | 561.3 | 0.0009032 | 1.609 | +2 | 9 |
| - | - | 919.8 | 561.3 | - | - | 0 | - |
| - | - | 3281 | 561.8 | - | - | 0 | - |
| - | - | 1480 | 562.3 | - | - | 0 | - |
| 4 | b | 2130 | 565.2 | 0.001114 | 1.971 | +1 | 4 |
| - | - | 3.405E+04 | 566.2 | - | - | 0 | - |
| - | - | 1.176E+04 | 567.2 | - | - | 0 | - |
| - | - | 1260 | 567.4 | - | - | 0 | - |
| - | - | 1713 | 568.3 | - | - | 0 | - |
| - | - | 899.3 | 568.4 | - | - | 0 | - |
| - | - | 4600 | 570.2 | - | - | 0 | - |
| - | - | 896.9 | 570.3 | - | - | 0 | - |
| - | - | 2124 | 571.2 | - | - | 0 | - |
| - | - | 1.82E+04 | 572.8 | - | - | 0 | - |
| - | - | 1.049E+04 | 573.3 | - | - | 0 | - |
| - | - | 5567 | 573.8 | - | - | 0 | - |
| - | - | 7046 | 577.4 | - | - | 0 | - |
| - | - | 2731 | 578.4 | - | - | 0 | - |
| - | - | 2.26E+04 | 579.4 | - | - | 0 | - |
| - | - | 6915 | 580.4 | - | - | 0 | - |
| - | - | 1218 | 581.4 | - | - | 0 | - |
| - | - | 1181 | 587.3 | - | - | 0 | - |
| 10 | b | 5180 | 587.8 | 0.0005681 | 0.9666 | +2 | 10 |
| 10 | b | 2873 | 588.3 | 0.009293 | 15.8 | +2 | 10 |
| - | - | 1384 | 588.8 | - | - | 0 | - |
| - | - | 7769 | 595.4 | - | - | 0 | - |
| - | - | 1916 | 595.8 | - | - | 0 | - |
| - | - | 848.1 | 596.3 | - | - | 0 | - |
| - | - | 2365 | 596.4 | - | - | 0 | - |
| 10 | b | 4899 | 596.8 | 7.554E-05 | 0.1266 | +2 | 10 |
| - | - | 1984 | 597.3 | - | - | 0 | - |
| - | - | 1604 | 597.8 | - | - | 0 | - |
| - | - | 2279 | 598.2 | - | - | 0 | - |
| - | - | 1067 | 602.3 | - | - | 0 | - |
| - | - | 9299 | 604.4 | - | - | 0 | - |
| 4 | y | 1.324E+04 | 604.8 | 0.004341 | 7.177 | +2 | 11 |
| - | - | 8385 | 605.4 | - | - | 0 | - |
| - | - | 3710 | 605.9 | - | - | 0 | - |
| - | - | 1442 | 606.8 | - | - | 0 | - |
| - | - | 4596 | 610.3 | - | - | 0 | - |
| - | - | 1201 | 611.3 | - | - | 0 | - |
| - | - | 2025 | 611.8 | - | - | 0 | - |
| - | - | 2697 | 612.3 | - | - | 0 | - |
| 4 | y | 1.291E+04 | 613.4 | 0.0007098 | 1.157 | +2 | 11 |
| - | - | 7769 | 613.9 | - | - | 0 | - |
| - | - | 3362 | 614.4 | - | - | 0 | - |
| - | - | 1765 | 615.3 | - | - | 0 | - |
| - | - | 8760 | 615.3 | - | - | 0 | - |
| - | - | 6706 | 615.9 | - | - | 0 | - |
| - | - | 2286 | 616.4 | - | - | 0 | - |
| - | - | 1.44E+04 | 620.3 | - | - | 0 | - |
| - | - | 1.279E+04 | 620.8 | - | - | 0 | - |
| - | - | 4196 | 621.3 | - | - | 0 | - |
| - | - | 1249 | 623.4 | - | - | 0 | - |
| - | - | 792.2 | 624.4 | - | - | 0 | - |
| - | - | 1021 | 628.3 | - | - | 0 | - |
| - | - | 3.717E+04 | 629.3 | - | - | 0 | - |
| - | - | 2.617E+04 | 629.8 | - | - | 0 | - |
| - | - | 1.053E+04 | 630.3 | - | - | 0 | - |
| - | - | 2519 | 630.8 | - | - | 0 | - |
| - | - | 982 | 632.3 | - | - | 0 | - |
| - | - | 4332 | 638.4 | - | - | 0 | - |
| - | - | 2539 | 638.9 | - | - | 0 | - |
| - | - | 970.8 | 643.3 | - | - | 0 | - |
| - | - | 756.8 | 644.8 | - | - | 0 | - |
| 11 | b | 2226 | 645.3 | 0.0001513 | 0.2345 | +2 | 11 |
| - | - | 1661 | 645.8 | - | - | 0 | - |
| - | - | 1420 | 647.4 | - | - | 0 | - |
| - | - | 2117 | 653.8 | - | - | 0 | - |
| - | - | 1631 | 654.3 | - | - | 0 | - |
| - | - | 1728 | 654.8 | - | - | 0 | - |
| - | - | 1679 | 656.3 | - | - | 0 | - |
| - | - | 2949 | 657.3 | - | - | 0 | - |
| - | - | 2403 | 658.3 | - | - | 0 | - |
| - | - | 943.9 | 659.3 | - | - | 0 | - |
| - | - | 6849 | 664.5 | - | - | 0 | - |
| - | - | 2754 | 665.5 | - | - | 0 | - |
| - | - | 1781 | 665.8 | - | - | 0 | - |
| - | - | 986.3 | 666.3 | - | - | 0 | - |
| - | - | 1004 | 666.5 | - | - | 0 | - |
| 9 | y | 3924 | 669.5 | 0.001207 | 1.803 | +1 | 6 |
| - | - | 1608 | 670.5 | - | - | 0 | - |
| - | - | 993.1 | 672.4 | - | - | 0 | - |
| - | - | 8872 | 674.3 | - | - | 0 | - |
| - | - | 2.604E+04 | 674.4 | - | - | 0 | - |
| - | - | 3178 | 675.3 | - | - | 0 | - |
| - | - | 1.139E+04 | 675.4 | - | - | 0 | - |
| - | - | 2025 | 676.4 | - | - | 0 | - |
| - | - | 4248 | 677.4 | - | - | 0 | - |
| - | - | 2655 | 677.9 | - | - | 0 | - |
| - | - | 2116 | 678.4 | - | - | 0 | - |
| - | - | 851.3 | 680.3 | - | - | 0 | - |
| 5 | b | 2119 | 684.3 | 0.0006797 | 0.9933 | +1 | 5 |
| 5 | b | 1802 | 685.3 | 0.01135 | 16.57 | +1 | 5 |
| - | - | 6949 | 685.9 | - | - | 0 | - |
| 3 | y | 7430 | 686.4 | 0.006734 | 9.811 | +2 | 12 |
| - | - | 3814 | 686.9 | - | - | 0 | - |
| - | - | 1410 | 687.4 | - | - | 0 | - |
| - | - | 2.448E+04 | 692.4 | - | - | 0 | - |
| - | - | 1.049E+04 | 693.4 | - | - | 0 | - |
| - | - | 2393 | 694.5 | - | - | 0 | - |
| 3 | y | 1.213E+05 | 694.9 | 0.0004176 | 0.601 | +2 | 12 |
| - | - | 1.019E+05 | 695.4 | - | - | 0 | - |
| - | - | 4.549E+04 | 695.9 | - | - | 0 | - |
| - | - | 1.422E+04 | 696.4 | - | - | 0 | - |
| - | - | 1941 | 696.9 | - | - | 0 | - |
| - | - | 1772 | 698.4 | - | - | 0 | - |
| 12 | b | 1498 | 700.8 | 0.00385 | 5.493 | +2 | 12 |
| 5 | b | 1.461E+04 | 702.3 | 0.00043 | 0.6123 | +1 | 5 |
| - | - | 6915 | 703.3 | - | - | 0 | - |
| - | - | 1190 | 704.3 | - | - | 0 | - |
| - | - | 3459 | 707.3 | - | - | 0 | - |
| - | - | 832.5 | 707.8 | - | - | 0 | - |
| - | - | 2098 | 708.3 | - | - | 0 | - |
| - | - | 1222 | 708.5 | - | - | 0 | - |
| 12 | b | 2.678E+04 | 709.4 | 0.0003408 | 0.4804 | +2 | 12 |
| - | - | 2.376E+04 | 709.9 | - | - | 0 | - |
| - | - | 666.2 | 710.3 | - | - | 0 | - |
| - | - | 1.045E+04 | 710.4 | - | - | 0 | - |
| - | - | 3730 | 710.5 | - | - | 0 | - |
| - | - | 3005 | 710.9 | - | - | 0 | - |
| - | - | 986.3 | 711.4 | - | - | 0 | - |
| - | - | 1032 | 711.5 | - | - | 0 | - |
| - | - | 1046 | 713.4 | - | - | 0 | - |
| - | - | 1260 | 713.9 | - | - | 0 | - |
| - | - | 1688 | 714.4 | - | - | 0 | - |
| - | - | 5204 | 716.4 | - | - | 0 | - |
| - | - | 1232 | 717.4 | - | - | 0 | - |
| - | - | 1131 | 720.3 | - | - | 0 | - |
| - | - | 972.7 | 720.4 | - | - | 0 | - |
| - | - | 2444 | 721.4 | - | - | 0 | - |
| - | - | 1094 | 722.4 | - | - | 0 | - |
| - | - | 833.5 | 722.9 | - | - | 0 | - |
| - | - | 2629 | 724.3 | - | - | 0 | - |
| - | - | 1090 | 725.3 | - | - | 0 | - |
| 8 | y | 9935 | 726.5 | 0.0004344 | 0.5979 | +1 | 7 |
| - | - | 3799 | 727.5 | - | - | 0 | - |
| - | - | 2666 | 735.3 | - | - | 0 | - |
| - | - | 836.7 | 736.3 | - | - | 0 | - |
| - | - | 1107 | 737.4 | - | - | 0 | - |
| - | - | 987.8 | 738.4 | - | - | 0 | - |
| - | - | 803 | 739.4 | - | - | 0 | - |
| - | - | 1582 | 740.3 | - | - | 0 | - |
| - | - | 2173 | 751.9 | - | - | 0 | - |
| - | - | 3452 | 752.3 | - | - | 0 | - |
| - | - | 2003 | 752.4 | - | - | 0 | - |
| - | - | 1067 | 753.3 | - | - | 0 | - |
| - | - | 949 | 756.4 | - | - | 0 | - |
| 13 | b | 2562 | 756.9 | 0.001741 | 2.3 | +2 | 13 |
| 13 | b | 2254 | 757.4 | 0.006679 | 8.818 | +2 | 13 |
| - | - | 1787 | 757.9 | - | - | 0 | - |
| - | - | 856 | 759 | - | - | 0 | - |
| 13 | b | 9393 | 765.9 | 4.171E-06 | 0.005445 | +2 | 13 |
| - | - | 6640 | 766.4 | - | - | 0 | - |
| - | - | 4914 | 766.9 | - | - | 0 | - |
| - | - | 1812 | 767.4 | - | - | 0 | - |
| - | - | 4087 | 774.9 | - | - | 0 | - |
| - | - | 3287 | 775.4 | - | - | 0 | - |
| - | - | 2111 | 775.9 | - | - | 0 | - |
| 2 | y | 3449 | 787.9 | 0.0001898 | 0.2409 | +2 | 13 |
| - | - | 2456 | 788.4 | - | - | 0 | - |
| - | - | 924.9 | 788.5 | - | - | 0 | - |
| - | - | 1784 | 788.9 | - | - | 0 | - |
| - | - | 2063 | 794.3 | - | - | 0 | - |
| - | - | 873.9 | 797.3 | - | - | 0 | - |
| - | - | 5493 | 805.5 | - | - | 0 | - |
| - | - | 2634 | 806.5 | - | - | 0 | - |
| - | - | 1.515E+04 | 811.4 | - | - | 0 | - |
| - | - | 4907 | 811.5 | - | - | 0 | - |
| - | - | 8035 | 812.4 | - | - | 0 | - |
| - | - | 2329 | 812.5 | - | - | 0 | - |
| - | - | 2195 | 813.4 | - | - | 0 | - |
| 6 | b | 2533 | 821.3 | 0.000676 | 0.823 | +1 | 6 |
| 6 | b | 1971 | 822.3 | 0.006092 | 7.408 | +1 | 6 |
| - | - | 1148 | 823.4 | - | - | 0 | - |
| 7 | y | 5.517E+04 | 823.5 | 3.871E-05 | 0.047 | +1 | 8 |
| - | - | 2.546E+04 | 824.5 | - | - | 0 | - |
| - | - | 1251 | 824.6 | - | - | 0 | - |
| - | - | 6520 | 825.5 | - | - | 0 | - |
| - | - | 1197 | 826.5 | - | - | 0 | - |
| - | - | 4368 | 829.5 | - | - | 0 | - |
| - | - | 2487 | 830.5 | - | - | 0 | - |
| 6 | b | 3.278E+04 | 839.4 | 1.019E-05 | 0.01214 | +1 | 6 |
| - | - | 1.623E+04 | 840.4 | - | - | 0 | - |
| - | - | 5366 | 841.4 | - | - | 0 | - |
| - | - | 4283 | 848.4 | - | - | 0 | - |
| - | - | 2473 | 849.4 | - | - | 0 | - |
| - | - | 3530 | 853.5 | - | - | 0 | - |
| - | - | 1454 | 854.5 | - | - | 0 | - |
| - | - | 795.7 | 855.5 | - | - | 0 | - |
| - | - | 2316 | 891.5 | - | - | 0 | - |
| - | - | 938.7 | 892.5 | - | - | 0 | - |
| - | - | 1.127E+04 | 901.4 | - | - | 0 | - |
| - | - | 6239 | 902.4 | - | - | 0 | - |
| - | - | 2002 | 903.4 | - | - | 0 | - |
| - | - | 6829 | 919.5 | - | - | 0 | - |
| - | - | 3325 | 920.5 | - | - | 0 | - |
| - | - | 4545 | 942.6 | - | - | 0 | - |
| - | - | 2756 | 943.6 | - | - | 0 | - |
| - | - | 1408 | 944.6 | - | - | 0 | - |
| - | - | 1009 | 948.5 | - | - | 0 | - |
| - | - | 743.6 | 949.6 | - | - | 0 | - |
| 6 | y | 1.082E+04 | 960.6 | 0.0002794 | 0.2909 | +1 | 9 |
| - | - | 698.8 | 961.5 | - | - | 0 | - |
| - | - | 6143 | 961.6 | - | - | 0 | - |
| - | - | 2274 | 962.6 | - | - | 0 | - |
| - | - | 2715 | 966.6 | - | - | 0 | - |
| - | - | 2126 | 967.6 | - | - | 0 | - |
| - | - | 750.4 | 1018 | - | - | 0 | - |
| - | - | 2329 | 1033 | - | - | 0 | - |
| - | - | 3303 | 1034 | - | - | 0 | - |
| - | - | 1104 | 1035 | - | - | 0 | - |
| - | - | 1152 | 1089 | - | - | 0 | - |
| 5 | y | 4453 | 1098 | 0.0006232 | 0.5677 | +1 | 10 |
| - | - | 3248 | 1099 | - | - | 0 | - |
| - | - | 1302 | 1100 | - | - | 0 | - |
| 9 | b | 3346 | 1122 | 0.002025 | 1.806 | +1 | 9 |
| - | - | 1560 | 1123 | - | - | 0 | - |
| - | - | 1566 | 1145 | - | - | 0 | - |
| 10 | b | 5477 | 1175 | 0.0009867 | 0.8401 | +1 | 10 |
| 10 | b | 4331 | 1176 | 0.01671 | 14.21 | +1 | 10 |
| - | - | 1446 | 1177 | - | - | 0 | - |
| 10 | b | 2267 | 1193 | 0.00496 | 4.159 | +1 | 10 |
| - | - | 1033 | 1194 | - | - | 0 | - |
| - | - | 856.3 | 1258 | - | - | 0 | - |
| - | - | 812.7 | 2521 | - | - | 0 | - |

m/z Charge Intensity FragmentType MassShift Position
120.0565185546875 0 450.45718
120.06600189208984 0 752.6646
120.08116912841797 0 1249.7837
122.07173919677734 0 986.8992
123.0558090209961 0 1769.2579
125.07100677490234 0 358.6497
126.03711700439453 0 360.0972
127.08700561523438 0 2328.773
129.0662384033203 0 3813.9534
129.1027069091797 0 94775.16
130.06556701660156 0 25048.703
130.10601806640625 0 5535.9824
130.531494140625 0 388.53265
131.06893920898438 0 2907.4116
132.08123779296875 0 86503.76
132.10232543945312 0 18769.807 y 13
133.06097412109375 0 1057.8383
133.08456420898438 0 8497.253
133.1058349609375 0 909.9638
134.07122802734375 0 439.25235
134.0882568359375 0 502.90842
135.0923614501953 0 483.10373
135.70071411132812 0 488.3128
136.07611083984375 0 49966.152
137.07382202148438 0 761.2919
137.07948303222656 0 4584.117
138.06668090820312 0 2540.881
139.0868682861328 0 1487.0582
141.10267639160156 0 5957.357
142.105712890625 0 645.9159
143.0821990966797 0 579.272
144.08108520507812 0 1586.9867
145.06121826171875 0 2478.9104
146.0605010986328 0 1590.0782
146.07168579101562 0 503.57452
146.12928771972656 0 569.11426
147.04473876953125 0 1091.5985
148.95370483398438 0 638.8678
151.08712768554688 0 1442.104
152.07098388671875 0 780.64044
152.082275390625 0 2305.6548
152.1437530517578 0 2101.7642
153.06642150878906 0 733.0885
153.10279846191406 0 747.7747
153.14781188964844 0 576.4647
154.0612335205078 0 612.83014
154.09796142578125 0 545.9254
155.0819091796875 0 4505.817
155.0931854248047 0 3446.2322
155.11830139160156 0 3687.556
156.07711791992188 0 4191.185
157.06167602539062 0 663.2315
158.06040954589844 0 995.56995
158.0843505859375 0 9115.543
159.09213256835938 0 40551.57
160.07614135742188 0 2167.3157
160.09548950195312 0 4019.4514
161.09237670898438 0 635.3711
162.05494689941406 0 748.30023
163.07162475585938 0 2086.318
164.07138061523438 0 545.31573
165.0662078857422 0 762.1528
165.07725524902344 0 721.5985
165.1027069091797 0 3359.4692
166.0614776611328 0 2499.1714
167.08163452148438 0 700.4559
167.1184844970703 0 795.011
168.11351013183594 0 865.23157
169.07647705078125 0 4269.276
169.09767150878906 0 7627.2607
170.0604705810547 0 14758.021
171.06405639648438 0 2149.6123
171.07730102539062 0 837.70807
171.1132354736328 0 1078.5592
171.14993286132812 0 1334.8263
172.1084747314453 0 5861.83
175.08714294433594 0 739.3643
175.09849548339844 0 742.5368
176.0822296142578 0 1412.8184
178.13438415527344 0 663.07294
179.08294677734375 0 521.78296
180.07752990722656 0 917.0762
181.0609893798828 0 1106.7305
181.09762573242188 0 17741.469
181.13392639160156 0 2277.1858
182.10110473632812 0 1591.8011
182.12928771972656 0 3709.4902
182.2926788330078 0 595.1994
183.0922088623047 0 13164.543
183.11294555664062 0 1749.451
183.1326446533203 0 664.9002
183.14898681640625 0 738.9846
184.07583618164062 0 850.21594
184.09580993652344 0 1047.9417
185.07130432128906 0 2865.8206
185.07980346679688 0 639.895
185.09226989746094 0 559.8101
186.12416076660156 0 9997.934
187.0870819091797 0 80476.055
187.10740661621094 0 1714.3689
187.14453125 0 1451.4321 y 11
188.09051513671875 0 8696.262
189.08743286132812 0 1895.2805 b 3
191.0816192626953 0 472.16507
195.11373901367188 0 1062.6682
195.1496124267578 0 1339.7548
197.12945556640625 0 601.3987
197.16529846191406 0 3751.281
198.08810424804688 0 641.22314
198.16851806640625 0 490.08954
199.08705139160156 0 2097.0151
199.10842895507812 0 1817.4072
199.14427185058594 0 919.4278
199.18081665039062 0 3434.918
200.139404296875 0 1161.5834
201.06739807128906 0 607.0745
201.10264587402344 0 6622.0483
201.12387084960938 0 1195.6694
202.083984375 0 1188.4794
202.10891723632812 0 9106.245
203.06607055664062 0 530.29675
203.09291076660156 0 695.871
203.10360717773438 0 1806.8868
203.11326599121094 0 600.9114
204.07777404785156 0 619.18427
205.08267211914062 0 635.21857
205.09762573242188 0 1070.2405 a 2
205.10838317871094 0 1025.3152
207.097900390625 0 6482.698
207.10972595214844 0 559.6531
207.114013671875 0 695.74365
207.14947509765625 0 3478.7979
207.15939331054688 0 796.20734
208.14462280273438 0 970.5968
209.0930633544922 0 815.61725
209.1033172607422 0 623.1864
209.12864685058594 0 814.5904
211.0869598388672 0 1604.5922
211.14419555664062 0 687.84375
212.13967895507812 0 1291.358
214.109130859375 0 4326.1973
215.13934326171875 0 2103.2717
216.09872436523438 0 635.5619
216.11257934570312 0 543.52167
217.04359436035156 0 559.0441
217.10841369628906 0 920.79425
219.076904296875 0 775.1659
220.11972045898438 0 7066.4893
221.10372924804688 0 9255.494
221.12301635742188 0 764.2453
222.10711669921875 0 818.87524
222.12420654296875 0 1088.5536
223.0638427734375 0 758.7145
223.15562438964844 0 1469.5099
224.11871337890625 0 2067.9727
224.1761932373047 0 4592.015
225.04339599609375 0 533.7187
225.42227172851562 0 559.52783
226.08314514160156 0 750.17725
226.15550231933594 0 109736.04
227.1029815673828 0 2129.5088
227.1394500732422 0 825.4359
227.15882873535156 0 13264.392 y Ammonia loss 10
227.17189025878906 0 921.5963
228.11363220214844 0 7123.5024 a Water loss 1
228.13455200195312 0 946.7726
228.16073608398438 0 1004.3101
229.09756469726562 0 9272.51 b Ammonia loss 4
229.1182861328125 0 1506.597
229.1546630859375 0 943.9004
229.166748046875 0 775.0593
230.10227966308594 0 2233.7644
231.08802795410156 0 3171.629
231.1353302001953 0 1222.4081
232.11927795410156 0 923.8188
233.10379028320312 0 1700.7045
234.10321044921875 0 735.73254
234.1241455078125 0 1608.4629
234.13601684570312 0 796.40826
235.11949157714844 0 17381.59
236.1224822998047 0 2457.3801
238.11807250976562 0 629.7453
238.1302490234375 0 2023.3734
239.08187866210938 0 1234.8708
239.15077209472656 0 6210.239
240.134765625 0 897.21954
240.15516662597656 0 637.67
241.155029296875 0 1248.0262
242.18695068359375 0 2528.3953
243.1819305419922 0 786.85236
244.09365844726562 0 862.13837
245.1865692138672 0 37871.996 y 12
246.124267578125 0 214894.94 a 1
246.1895751953125 0 4181.5234
247.12893676757812 0 48641.17
248.11460876464844 0 11121.167
248.13217163085938 0 3512.5535
249.0986785888672 0 15723.195
249.115234375 0 853.01636
249.11807250976562 0 1060.0356
250.1018524169922 0 1663.8105
250.12240600585938 0 1280.6588
251.1507568359375 0 5097.689
252.1708221435547 0 1486.1377
256.10858154296875 0 5487.0093 b Water loss 1
257.1138916015625 0 1590.6858
257.1607360839844 0 1455.9541
258.0760498046875 0 1095.342
258.0989685058594 0 25560.928
259.1024169921875 0 3956.1663
259.1297912597656 0 747.6749
260.10699462890625 0 1928.6362
260.19781494140625 0 1458.5767
263.1146545410156 0 972.5764
264.1343078613281 0 1567.3462
265.1410827636719 0 961.8413
265.166015625 0 1241.337
266.1251220703125 0 13801.409
267.12835693359375 0 1835.8499
267.14495849609375 0 608.87427
270.1816711425781 0 6451.3154
274.1191101074219 0 125047.9 b 1
275.10369873046875 0 3788.0845
275.1245422363281 0 61585.008
276.1091613769531 0 2052.6858
276.12786865234375 0 5809.585
276.1560363769531 0 698.2774
277.13336181640625 0 1698.8215
279.18121337890625 0 1080.6799
280.1665954589844 0 662.5218
283.1512451171875 0 1731.9713
283.177001953125 0 19365.838
284.1798095703125 0 2087.5952
285.1097412109375 0 1988.967
285.1562194824219 0 5677.5586
290.1147766113281 0 835.70026
290.18475341796875 0 6596.676
290.6861877441406 0 1477.8656
291.15814208984375 0 968.8777
292.1297912597656 0 12362.6045
292.1658935546875 0 586.9372
293.1332702636719 0 1653.8002
294.1202392578125 0 5982.77
295.1227111816406 0 736.85187
297.1565856933594 0 689.13086
297.19268798828125 0 16987.46
298.1959228515625 0 2865.3354
299.0631408691406 0 916.2675
300.0627136230469 0 800.315
301.1285095214844 0 788.546
303.12030029296875 0 3115.3623
303.21783447265625 0 787.6113
305.1287536621094 0 3150.493
305.6646728515625 0 1296.6882
306.1090087890625 0 773.35956
306.1325988769531 0 1011.7732
311.24517822265625 0 2662.9111
314.6693115234375 0 796.8298
318.193603515625 0 907.8465
320.1249694824219 0 3497.0728
321.2291564941406 0 16533.078
322.1556396484375 0 6157.9263
322.2322692871094 0 3363.908
323.1580810546875 0 690.4364
324.2168273925781 0 913.2492
324.71783447265625 0 754.32056
325.22406005859375 0 1047.9783
326.2195129394531 0 3651.7188
327.16815185546875 0 757.7804
332.7296142578125 0 3529.7349
333.1236572265625 0 1042.8165
333.231689453125 0 1326.177
335.23529052734375 0 721.2986 y 8
336.14593505859375 0 707.52576
336.2037048339844 0 22555.582
337.1509094238281 0 731.1097
337.2067565917969 0 4124.4404
337.7220153808594 0 4646.5005
338.1369323730469 0 739.2703
339.21636962890625 0 887.6239
339.23968505859375 0 16747.965
340.1516418457031 0 1444.3038
340.1876220703125 0 992.3884
340.2432861328125 0 3300.0684
341.2185363769531 0 1182.9708
346.72698974609375 0 16568.459
347.2283630371094 0 6945.6743
347.7295837402344 0 1484.7604
350.15020751953125 0 2978.3591
351.1520690917969 0 831.1227
351.23974609375 0 1866.5137
353.2190856933594 0 1062.4935
354.214111328125 0 29927.78
355.0696716308594 0 1271.0062
355.2168273925781 0 5908.7256
355.2707214355469 0 789.5367
355.7317199707031 0 1366.2035
356.23541259765625 0 725.8106
357.1786804199219 0 973.1107
357.25006103515625 0 3052.2961
358.1627197265625 0 4709.351
358.7140808105469 0 2253.1377
359.16680908203125 0 879.0555
359.21490478515625 0 926.9605
360.13604736328125 0 725.38165
360.689208984375 0 1227.054
361.6806640625 0 754.05963
362.6697692871094 0 1205.3159
363.748291015625 0 1088.1915 y 7
365.25482177734375 0 830.6
366.15618896484375 0 3376.9653
367.16162109375 0 1493.0847
368.14715576171875 0 2008.1107
368.1707763671875 0 1735.9988
370.2466125488281 0 690.65106
373.2813720703125 0 7963.9077 y 11
374.2576599121094 0 672.5519
374.28424072265625 0 1105.535
379.2080993652344 0 1147.2397
380.2296142578125 0 1799.7755
381.15203857421875 0 752.90533
382.2826232910156 0 1414.4648
382.54022216796875 0 963.14087
383.2657775878906 0 1531.3002
384.16705322265625 0 4151.5596
385.17303466796875 0 10701.049
385.1990966796875 0 3546.4058
386.1573486328125 0 8128.1143
386.2008361816406 0 888.43884
387.16015625 0 1923.3312
389.77520751953125 0 1080.8291
392.1624450683594 0 919.02325
392.26654052734375 0 3493.5576
393.1682434082031 0 1554.0375
393.26959228515625 0 1271.2758
394.15179443359375 0 8028.5
394.2809753417969 0 709.9993
395.15423583984375 0 2420.2312
396.26104736328125 0 1533.8588
397.2447814941406 0 906.74005
398.2430725097656 0 974.36597
401.1941833496094 0 1439.0667
401.259521484375 0 1666.4908
402.2253723144531 0 4771.2773
403.1842956542969 0 10460.554
403.22967529296875 0 780.9205
403.2692565917969 0 2605.071
403.7688293457031 0 953.4277
404.18719482421875 0 2300.4795
406.1861572265625 0 5112.9043
406.25164794921875 0 2709.2998
406.6869201660156 0 2339.2686
406.75201416015625 0 1638.1644
410.2768859863281 0 10048.322
411.1781311035156 0 16075.276 b Water loss 5
411.279541015625 0 1977.1611
412.1611328125 0 1998.166
412.1834716796875 0 2483.4158
412.2745056152344 0 41136.867 y 6
412.77587890625 0 21707.684
413.1687316894531 0 1582.6674
413.2778015136719 0 6527.6406
415.2564392089844 0 3639.4146
415.7587585449219 0 1753.4365
419.1716613769531 0 921.6499 b Water loss 2
419.21099853515625 0 621.4539
419.9006042480469 0 1924.5011
420.1832580566406 0 4673.124 b 5
420.2352294921875 0 3611.802
420.6845397949219 0 3053.528
422.2768249511719 0 3733.758
422.3092041015625 0 1222.7831
423.27410888671875 0 1834.4111
425.28790283203125 0 1915.1537
427.2441711425781 0 4676.213
427.74481201171875 0 2468.3162
428.249755859375 0 836.3319
428.2879638671875 0 953.6835
429.1888122558594 0 51725.746
430.192138671875 0 12865.838
431.1807556152344 0 5557.473
432.18145751953125 0 1270.4559
433.25653076171875 0 934.30054
436.6778869628906 0 964.4308
437.1826477050781 0 9560.015 b 2
438.1863708496094 0 3097.9207
438.5577087402344 0 1234.4707
438.8911437988281 0 1628.2212
439.19091796875 0 810.6022
439.2239074707031 0 655.368
439.30322265625 0 1275.9971
442.6789245605469 0 1242.3911
444.23138427734375 0 1416.8618
445.2689514160156 0 1850.8649
446.22283935546875 0 982.45966
448.2278137207031 0 635.3456
450.1085510253906 0 694.9254
450.2828063964844 0 617.2433
451.225341796875 0 1637.7008
451.26702880859375 0 3203.3616
451.6837158203125 0 2167.2764
451.72509765625 0 922.93616
452.2259826660156 0 766.6798
452.268798828125 0 794.9615
452.3242492675781 0 1552.2063
453.3251647949219 0 664.58563
454.2093811035156 0 3287.5684
455.2152404785156 0 1106.772
455.2525939941406 0 3623.808
456.2540283203125 0 924.3341
457.1828918457031 0 2048.3462
458.2347106933594 0 867.40424
460.2306213378906 0 1998.4507
460.7308044433594 0 696.36316
461.1816101074219 0 1139.5735
463.5987548828125 0 8465.268 y 2
463.9330139160156 0 8630.031
464.1665344238281 0 854.73975
464.26715087890625 0 2648.924
464.60223388671875 0 1212.9559
466.9178771972656 0 912.54333
467.25787353515625 0 817.3627
467.2983703613281 0 4584.6147
468.2968444824219 0 2617.4128
469.7869567871094 0 1250.76
470.2503356933594 0 1103.1603
470.33441162109375 0 34373.773 y 10
470.582275390625 0 1272.924
471.3377990722656 0 8477.502
471.7984924316406 0 2522.5774
472.301025390625 0 2664.2666
472.33721923828125 0 1207.0704
472.5941162109375 0 822.1406
473.2438049316406 0 15790.569 b 11
473.5773620605469 0 14727.41
473.6973876953125 0 1429.9797
473.9112243652344 0 4819.6455
474.2474365234375 0 1775.3962
474.2603454589844 0 1566.0276
474.78021240234375 0 2453.6082
475.28076171875 0 1905.81
475.7839660644531 0 787.49756
476.2527770996094 0 5773.5854
476.58746337890625 0 4417.578
478.21026611328125 0 2587.7336
480.8037109375 0 10304.003 y 5
481.305419921875 0 5732.552
481.9248046875 0 2002.0051
482.2631530761719 0 3911.676
482.3085632324219 0 3728.109
482.7629089355469 0 2979.2676
483.2637939453125 0 927.6903
483.78662109375 0 7098.626
484.2872314453125 0 3577.1616
484.7880554199219 0 921.616
486.1742248535156 0 862.75104
488.26513671875 0 884.6966
491.2731018066406 0 4759.865
492.27484130859375 0 1034.6089
492.79168701171875 0 668.0877
494.2340087890625 0 1494.9688
495.9336242675781 0 1137.216
496.262451171875 0 1624.644
496.75860595703125 0 867.6377
497.22021484375 0 1883.2834 b 7
497.7229919433594 0 1947.9014
501.6062927246094 0 7271.297
501.9404602050781 0 5848.139
502.2747497558594 0 3943.585
502.6073913574219 0 1583.1436
503.2160949707031 0 2690.0789
504.9336242675781 0 12151.53 b Water loss 12
505.267333984375 0 15300.683 b Ammonia loss 12
505.6015930175781 0 6882.415
505.9356689453125 0 3828.3936
506.26495361328125 0 802.65155
508.7563781738281 0 1850.6018
509.2605285644531 0 1818.928
510.9375915527344 0 69064.6 b 12
511.27191162109375 0 66357.37
511.6062927246094 0 28186.422
511.94024658203125 0 7660.295
512.2739868164062 0 2705.3184
516.9408569335938 0 7592.995
517.2748413085938 0 9868.379
517.6101684570312 0 5355.758
517.8829345703125 0 1775.0142
517.943603515625 0 1253.9347
519.6203002929688 0 847.93604
519.9489135742188 0 2712.0295 y Ammonia loss 1
520.246337890625 0 872.11053
520.2864990234375 0 2610.7915
520.3618774414062 0 1790.7996
520.6220092773438 0 1225.6769
521.2285766601562 0 2490.392
523.3628540039062 0 929.95197
525.6246337890625 0 11714.527 y 1
525.9592895507812 0 9943.726
526.2930297851562 0 5653.08
526.62841796875 0 2081.6956
528.2333984375 0 1961.5641
529.22265625 0 907.00397
529.8027954101562 0 1782.7307
530.2993774414062 0 1723.4545
530.7996826171875 0 1215.9053
531.2102661132812 0 2328.9368
535.293701171875 0 1241.5508
535.6129760742188 0 735.30835
537.2865600585938 0 1047.0834
537.62109375 0 823.7027
538.2523803710938 0 27466.426
538.37158203125 0 2192.2815
538.810791015625 0 4315.1675
539.2548217773438 0 7805.032
539.3048706054688 0 7369.3657
539.6322021484375 0 2044.0162
539.8051147460938 0 2902.7834
539.970947265625 0 1016.391
540.2568359375 0 1215.8821
540.2998046875 0 1022.4078
540.6224365234375 0 804.8151
540.8253784179688 0 781.6109 y Ammonia loss 4
541.32568359375 0 589.85803
541.3716430664062 0 9739.43 y 9
542.2510986328125 0 879.0648
542.3749389648438 0 3325.7236
542.9580688476562 0 1851.51
543.2928466796875 0 1116.1948
544.302001953125 0 724.0377
547.81494140625 0 2639.8372
548.2371215820312 0 1009.48566
548.3154907226562 0 1947.8187
548.6322021484375 0 19527.781 Precursor Water loss
548.9652709960938 0 23095.004 Precursor Ammonia loss
549.2238159179688 0 1686.1853
549.2974853515625 0 9259.645
549.3336181640625 0 16709.559 y 4
549.6326904296875 0 7357.077
549.8348388671875 0 10308.453
549.96484375 0 1616.0083
550.3359985351562 0 4840.373
550.840087890625 0 1131.678
551.375244140625 0 735.6548
553.3028564453125 0 1455.5933
553.9645385742188 0 3571.0823
554.2679443359375 0 825.58215
554.2982177734375 0 2626.5967
554.6359252929688 0 392883.84 Precursor
554.9700317382812 0 421384.03
555.3042602539062 0 222008.52
555.5818481445312 0 5035.367
555.6385498046875 0 73239.9
555.9149169921875 0 2755.8318
555.97265625 0 23224.922
557.2975463867188 0 1173.2229
561.268310546875 0 2376.016 b 8
561.349609375 0 919.7639
561.7692260742188 0 3280.9534
562.2703247070312 0 1480.27
565.2416381835938 0 2129.5928 b 3
566.2474975585938 0 34053.715
567.2498779296875 0 11762.273
567.3978881835938 0 1259.6799
568.2533569335938 0 1712.7361
568.4004516601562 0 899.2836
570.2460327148438 0 4600.322
570.3236694335938 0 896.9228
571.2484741210938 0 2124.362
572.8046875 0 18200.645
573.3056640625 0 10494.404
573.8067626953125 0 5567.213
577.3825073242188 0 7046.174
578.3845825195312 0 2730.7812
579.3619384765625 0 22603.66
580.3650512695312 0 6915.422
581.3702392578125 0 1218.2814
587.278564453125 0 1180.7612
587.78125 0 5179.915 b Water loss 9
588.281982421875 0 2872.7722 b Ammonia loss 9
588.7828369140625 0 1384.2516
595.3938598632812 0 7769.1553
595.8466186523438 0 1915.5405
596.3446655273438 0 848.11914
596.3951416015625 0 2364.5503
596.785888671875 0 4898.5225 b 9
597.2855224609375 0 1983.7014
597.7890014648438 0 1603.9176
598.241455078125 0 2279.2983
602.30224609375 0 1066.9891
604.3568115234375 0 9298.617
604.8529052734375 0 13236.628 y Ammonia loss 3
605.3526000976562 0 8385.451
605.8529663085938 0 3710.3745
606.83544921875 0 1441.5663
610.3209228515625 0 4596.133
611.3234252929688 0 1201.0475
611.8295288085938 0 2024.7878
612.3280029296875 0 2697.1567
613.362548828125 0 12907.66 y 3
613.8643188476562 0 7768.634
614.36474609375 0 3361.7236
615.2677001953125 0 1764.5215
615.3494262695312 0 8759.909
615.8505249023438 0 6706.457
616.3530883789062 0 2286.4358
620.3416137695312 0 14402.781
620.841796875 0 12792.505
621.3417358398438 0 4196.325
623.3877563476562 0 1249.2009
624.3877563476562 0 792.2381
628.3309936523438 0 1021.0147
629.3467407226562 0 37169.664
629.8480834960938 0 26170.562
630.349609375 0 10528.462
630.849853515625 0 2518.7559
632.29931640625 0 982.0021
638.3511962890625 0 4332.355
638.8541870117188 0 2538.9211
643.260986328125 0 970.8444
644.8289794921875 0 756.80396
645.3121948242188 0 2226.4192 b 10
645.8165283203125 0 1661.0149
647.4227905273438 0 1420.4349
653.827880859375 0 2117.3206
654.3289794921875 0 1631.3523
654.8256225585938 0 1728.4705
656.2931518554688 0 1679.1273
657.281005859375 0 2949.4275
658.307861328125 0 2403.49
659.3177490234375 0 943.90643
664.451171875 0 6848.9985
665.4529418945312 0 2753.688
665.8451538085938 0 1780.5361
666.3466186523438 0 986.33
666.4595336914062 0 1004.4538
669.4669799804688 0 3924.296 y 8
670.4689331054688 0 1608.2833
672.3914794921875 0 993.0914
674.3042602539062 0 8871.577
674.4351196289062 0 26044.682
675.3077392578125 0 3178.094
675.437255859375 0 11385.012
676.44140625 0 2024.6974
677.375244140625 0 4248.3677
677.8775024414062 0 2655.349
678.374267578125 0 2115.726
680.3147583007812 0 851.26996
684.28955078125 0 2119.2744 b Water loss 4
685.2842407226562 0 1802.1768 b Ammonia loss 4
685.8892211914062 0 6948.924
686.386962890625 0 7430.349 y Ammonia loss 2
686.8867797851562 0 3813.567
687.387939453125 0 1410.379
692.44580078125 0 24476.195
693.44873046875 0 10487.294
694.4510498046875 0 2392.7732
694.8939208984375 0 121263.766 y 2
695.3952026367188 0 101859.46
695.8966674804688 0 45493.645
696.398193359375 0 14216.433
696.8994750976562 0 1940.5979
698.4114990234375 0 1772.2881
700.8504028320312 0 1497.7838 b Ammonia loss 11
702.2998657226562 0 14612.0205 b 4
703.3021850585938 0 6914.571
704.3038330078125 0 1190.4498
707.3059692382812 0 3459.4377
707.794921875 0 832.4534
708.3099365234375 0 2098.1846
708.4794921875 0 1221.6395
709.3601684570312 0 26780.8 b 11
709.8614501953125 0 23764.951
710.297607421875 0 666.15576
710.3638916015625 0 10447.985
710.4548950195312 0 3730.354
710.8636474609375 0 3004.8162
711.3670654296875 0 986.2772
711.458740234375 0 1032.4366
713.378173828125 0 1046.2505
713.8829345703125 0 1259.8485
714.3800659179688 0 1687.5337
716.4205322265625 0 5203.98
717.4254150390625 0 1231.6758
720.317138671875 0 1131.081
720.3778076171875 0 972.731
721.3533325195312 0 2443.6838
722.3786010742188 0 1094.2709
722.8817138671875 0 833.5021
724.3335571289062 0 2628.8992
725.3333129882812 0 1089.5065
726.4876708984375 0 9935.101 y 7
727.4912109375 0 3799.4841
735.3020629882812 0 2665.5854
736.3004760742188 0 836.7209
737.35107421875 0 1106.8632
738.3740844726562 0 987.7514
739.36767578125 0 802.9957
740.2951049804688 0 1581.8499
751.9014282226562 0 2173.2568
752.3265991210938 0 3451.9314
752.4088134765625 0 2003.039
753.3297729492188 0 1067.1272
756.3894653320312 0 948.9949
756.8948364257812 0 2561.927 b Water loss 12
757.395263671875 0 2254.2117 b Ammonia loss 12
757.9032592773438 0 1786.5153
759.030029296875 0 856.0361
765.90185546875 0 9392.516 b 12
766.403076171875 0 6639.9663
766.9057006835938 0 4914.1323
767.4094848632812 0 1812.2142
774.9065551757812 0 4086.9705
775.4080810546875 0 3287.3591
775.909912109375 0 2110.7617
787.933349609375 0 3449.2983 y 1
788.4334106445312 0 2455.74
788.5131225585938 0 924.9032
788.9353637695312 0 1783.9329
794.3417358398438 0 2062.9878
797.3203735351562 0 873.9133
805.5292358398438 0 5493.0586
806.5335693359375 0 2634.1914
811.3635864257812 0 15153.042
811.4940185546875 0 4907.147
812.3656616210938 0 8035.0933
812.4990234375 0 2328.5176
813.3712158203125 0 2194.6675
821.3471069335938 0 2532.8 b Water loss 5
822.337890625 0 1970.8363 b Ammonia loss 5
823.351318359375 0 1148.2535
823.5400390625 0 55167.66 y 6
824.54296875 0 25459.72
824.6331176757812 0 1250.5078
825.544677734375 0 6519.9067
826.5498657226562 0 1197.4398
829.5038452148438 0 4368.289
830.5072021484375 0 2487.1792
839.3583374023438 0 32783.785 b 5
840.3610229492188 0 16234.791
841.3628540039062 0 5365.5024
848.4151611328125 0 4282.972
849.4205322265625 0 2473.4517
853.4778442382812 0 3529.846
854.4789428710938 0 1453.7216
855.4832763671875 0 795.73865
891.45751953125 0 2316.3345
892.4584350585938 0 938.6594
901.442138671875 0 11267.6045
902.4454956054688 0 6238.7056
903.4440307617188 0 2001.7255
919.4525146484375 0 6828.7104
920.4554443359375 0 3324.627
942.5863037109375 0 4544.667
943.592041015625 0 2755.8867
944.5927734375 0 1408.4716
948.5497436523438 0 1009.2304
949.5517578125 0 743.5562
960.5986328125 0 10819.561 y 5
961.5064697265625 0 698.84656
961.6012573242188 0 6143.423
962.605224609375 0 2273.7803
966.562255859375 0 2715.128
967.5645141601562 0 2126.4326
1017.5020141601562 0 750.4103
1032.5386962890625 0 2328.6743
1033.533203125 0 3302.9546
1034.5377197265625 0 1103.6049
1088.521484375 0 1152.0706
1097.658447265625 0 4452.955 y 4
1098.659423828125 0 3247.839
1099.659423828125 0 1301.7804
1121.5255126953125 0 3346.1326 b 8
1122.529052734375 0 1560.2318
1144.600830078125 0 1565.6581
1174.5531005859375 0 5477.4844 b Water loss 9
1175.5548095703125 0 4331.343 b Ammonia loss 9
1176.5599365234375 0 1446.0409
1192.5596923828125 0 2267.3677 b 9
1193.5689697265625 0 1032.934
1257.682861328125 0 856.3484
2520.533935546875 0 812.6525

Spectrum Details

|  |  |
| --- | --- |
| Matched peaks? Matched peaks The total absolute number of peaks matched. Additionally in brackets the total fraction of peaks matched and the total number of peaks is shown. | 66 (8.45% of 781) |
| FDR? FDR The false discovery rate estimated for this peptide. It is calculated by matching all theoretical fragments with a non-integer shift with the raw peaks for this spectrum. This is done with 40 different shifts. The resulting percentage is the average number of annotated peaks over the number of annotated peaks with the correct spectrum. | 3.46% |
| Satellite FDR? Satellite FDR See the FDR for details on its calculation. This satellite ion specific FDR only contains the satellite ions (d/w) for I/L/J positions. | - |
| PSM Score? PSM Score The PSM Score as given by Hecklib to this annotated spectrum. It is shown with three significant figures. | 360 |

#### Spectrum 5306? Spectrum 5306 The raw spectrum of this peptide as annotated by Hecklib. The fragments are coloured according to ion type (see legend). Any peaks with a star '\*' as text can be hovered over to see the full details, first the ion type second the mass shift type. By hovering over the amino acids in the peptide or ions in the legend the corresponding peaks are highlighted. By toggling the 'Unassigned' label you can turn the background (unassigned) peaks on or off in the plot. By updating the slider in the Ion legend you can update the spectrum to only show the top X% of the peaks with labels. The top X% means any peak that is within X% of the highest intensity. By dragging in the spectrum you can zoom in to a specific part of the spectrum and use 'Zoom Out' to get back to the original zoom level. The annotation of the spectrum is based on the given sequence in the peptides file and is done with different software so inconsistencies are likely. The peaks are annotated based on the given sequence, with 20 ppm tolerance.

Copy Data

##### Spectrum 5306 (TSV)

###### Preview

```
Loading example...
```

*Click on the button to copy the data to your clipboard.*

Mz MinMz MaxIntensity Max

WidthHeightPeptide font sizePeptide stroke widthSpectrum font sizeSpectrum stroke widthCompact peptide

Ion legend

wxyz

abcd

OtherUnassignedIon^Charge^_Position_Show for top:%

SWYQHHPGKAPKJJ

01.32e+62.63e+63.95e+65.26e+6

Zoom Out

y^+1^_1_a^+2^_3_y^+2^_4_a^+1^_2_b^+3^_5_y^+1^_2_a^+1^_2_b^+1^_2_y^+2^_5_b^+1^_2_y^+2^_6_y^+2^_7_y^+1^_3_b^+2^_6_y^+2^_8_b^+2^_6_b^+1^_3_y^+3^_12_y^+3^_12_b^+2^_7_y^+1^_4_b^+3^_12_y^+2^_9_b^+2^_8_b^+3^_13_b^+3^_13_b^+3^_13_y^+3^_13_y^+3^_13_y^+2^_10_y^+1^_5_\*\*y^+2^_10_\*b^+2^_9_b^+1^_4_b^+2^_10_b^+2^_10_b^+2^_10_y^+2^_11_y^+2^_11_b^+2^_11_y^+1^_6_b^+1^_5_b^+1^_5_y^+2^_12_y^+2^_12_b^+2^_12_b^+2^_12_b^+1^_5_b^+2^_12_y^+1^_7_b^+2^_13_b^+2^_13_b^+2^_13_y^+2^_13_y^+2^_13_b^+1^_6_b^+1^_6_y^+1^_8_b^+1^_6_y^+1^_9_b^+1^_8_y^+1^_10_b^+1^_9_b^+1^_10_b^+1^_10_b^+1^_10_

0833166624993332

Fragment Matches Table

Show background peaks

| Position | Ion type | Intensity | mz Theoretical | mz Error (Th) | mz Error (ppm) | Charge | Series Number |
| --- | --- | --- | --- | --- | --- | --- | --- |
| - | - | 9336 | 120.1 | - | - | 0 | - |
| - | - | 6660 | 122.1 | - | - | 0 | - |
| - | - | 1.908E+04 | 123.1 | - | - | 0 | - |
| - | - | 3437 | 124.7 | - | - | 0 | - |
| - | - | 3275 | 124.9 | - | - | 0 | - |
| - | - | 3.006E+04 | 127.1 | - | - | 0 | - |
| - | - | 8069 | 128.1 | - | - | 0 | - |
| - | - | 4.235E+04 | 129.1 | - | - | 0 | - |
| - | - | 1.118E+06 | 129.1 | - | - | 0 | - |
| - | - | 2.775E+05 | 130.1 | - | - | 0 | - |
| - | - | 9544 | 130.1 | - | - | 0 | - |
| - | - | 7.48E+04 | 130.1 | - | - | 0 | - |
| - | - | 3.304E+04 | 131.1 | - | - | 0 | - |
| - | - | 3758 | 131.1 | - | - | 0 | - |
| - | - | 1.143E+06 | 132.1 | - | - | 0 | - |
| 14 | y | 2.195E+05 | 132.1 | 0.0005119 | 3.875 | +1 | 1 |
| - | - | 3975 | 132.2 | - | - | 0 | - |
| - | - | 1.173E+05 | 133.1 | - | - | 0 | - |
| - | - | 5137 | 133.1 | - | - | 0 | - |
| - | - | 1.585E+04 | 133.1 | - | - | 0 | - |
| - | - | 8368 | 134.1 | - | - | 0 | - |
| - | - | 6.749E+05 | 136.1 | - | - | 0 | - |
| - | - | 6.088E+04 | 137.1 | - | - | 0 | - |
| - | - | 6767 | 137.1 | - | - | 0 | - |
| - | - | 2.552E+04 | 138.1 | - | - | 0 | - |
| - | - | 1.526E+04 | 139.1 | - | - | 0 | - |
| - | - | 8.456E+04 | 141.1 | - | - | 0 | - |
| - | - | 6308 | 142.1 | - | - | 0 | - |
| - | - | 5892 | 142.1 | - | - | 0 | - |
| - | - | 2.231E+04 | 144.1 | - | - | 0 | - |
| - | - | 6934 | 145.1 | - | - | 0 | - |
| - | - | 3908 | 146.1 | - | - | 0 | - |
| - | - | 1.218E+04 | 147 | - | - | 0 | - |
| - | - | 4032 | 149.1 | - | - | 0 | - |
| - | - | 8066 | 151.1 | - | - | 0 | - |
| - | - | 2.828E+04 | 152.1 | - | - | 0 | - |
| - | - | 2.413E+04 | 152.1 | - | - | 0 | - |
| - | - | 3983 | 152.2 | - | - | 0 | - |
| - | - | 4.693E+04 | 155.1 | - | - | 0 | - |
| - | - | 2.821E+04 | 155.1 | - | - | 0 | - |
| - | - | 4.02E+04 | 155.1 | - | - | 0 | - |
| - | - | 3.709E+04 | 156.1 | - | - | 0 | - |
| - | - | 5916 | 156.1 | - | - | 0 | - |
| - | - | 4379 | 156.6 | - | - | 0 | - |
| - | - | 5223 | 157.1 | - | - | 0 | - |
| - | - | 1.511E+04 | 158.1 | - | - | 0 | - |
| - | - | 1.135E+05 | 158.1 | - | - | 0 | - |
| - | - | 4796 | 158.1 | - | - | 0 | - |
| - | - | 5.233E+05 | 159.1 | - | - | 0 | - |
| - | - | 2.611E+04 | 160.1 | - | - | 0 | - |
| - | - | 9371 | 160.1 | - | - | 0 | - |
| - | - | 5.354E+04 | 160.1 | - | - | 0 | - |
| - | - | 3732 | 163 | - | - | 0 | - |
| - | - | 9646 | 164.1 | - | - | 0 | - |
| - | - | 3.165E+04 | 165.1 | - | - | 0 | - |
| - | - | 1.397E+04 | 166.1 | - | - | 0 | - |
| - | - | 5962 | 166.1 | - | - | 0 | - |
| - | - | 1.578E+04 | 168.1 | - | - | 0 | - |
| - | - | 5.779E+04 | 169.1 | - | - | 0 | - |
| - | - | 7119 | 169.1 | - | - | 0 | - |
| - | - | 1.042E+05 | 169.1 | - | - | 0 | - |
| - | - | 8155 | 169.1 | - | - | 0 | - |
| - | - | 6228 | 169.2 | - | - | 0 | - |
| - | - | 1.707E+05 | 170.1 | - | - | 0 | - |
| - | - | 7699 | 170.1 | - | - | 0 | - |
| - | - | 7505 | 170.1 | - | - | 0 | - |
| - | - | 2.509E+04 | 171.1 | - | - | 0 | - |
| - | - | 4834 | 171.1 | - | - | 0 | - |
| - | - | 7.416E+04 | 172.1 | - | - | 0 | - |
| - | - | 1.361E+04 | 173.5 | - | - | 0 | - |
| - | - | 7370 | 175.1 | - | - | 0 | - |
| - | - | 2.521E+04 | 176.1 | - | - | 0 | - |
| - | - | 5634 | 179.1 | - | - | 0 | - |
| - | - | 1.002E+04 | 180.1 | - | - | 0 | - |
| - | - | 6277 | 181.1 | - | - | 0 | - |
| - | - | 2.155E+05 | 181.1 | - | - | 0 | - |
| - | - | 2.112E+04 | 181.1 | - | - | 0 | - |
| - | - | 1.533E+04 | 182.1 | - | - | 0 | - |
| - | - | 3.763E+04 | 182.1 | - | - | 0 | - |
| - | - | 1.598E+05 | 183.1 | - | - | 0 | - |
| - | - | 7269 | 183.1 | - | - | 0 | - |
| - | - | 1.493E+04 | 184.1 | - | - | 0 | - |
| - | - | 2.081E+04 | 184.1 | - | - | 0 | - |
| - | - | 3.766E+04 | 185.1 | - | - | 0 | - |
| - | - | 5695 | 185.1 | - | - | 0 | - |
| - | - | 5268 | 186.1 | - | - | 0 | - |
| - | - | 1.181E+05 | 186.1 | - | - | 0 | - |
| - | - | 9.878E+05 | 187.1 | - | - | 0 | - |
| - | - | 8069 | 188.1 | - | - | 0 | - |
| - | - | 1.112E+05 | 188.1 | - | - | 0 | - |
| - | - | 5586 | 189.1 | - | - | 0 | - |
| - | - | 4645 | 193.1 | - | - | 0 | - |
| - | - | 1.392E+04 | 195.1 | - | - | 0 | - |
| - | - | 4200 | 195.5 | - | - | 0 | - |
| - | - | 5.755E+04 | 197.2 | - | - | 0 | - |
| - | - | 7548 | 198.1 | - | - | 0 | - |
| - | - | 7763 | 198.2 | - | - | 0 | - |
| - | - | 3.707E+04 | 199.1 | - | - | 0 | - |
| - | - | 7877 | 199.1 | - | - | 0 | - |
| - | - | 3.454E+04 | 199.2 | - | - | 0 | - |
| - | - | 5812 | 200.1 | - | - | 0 | - |
| - | - | 2.655E+04 | 200.1 | - | - | 0 | - |
| - | - | 8.836E+04 | 201.1 | - | - | 0 | - |
| - | - | 2.098E+04 | 202.1 | - | - | 0 | - |
| - | - | 1.122E+05 | 202.1 | - | - | 0 | - |
| - | - | 7746 | 203.1 | - | - | 0 | - |
| - | - | 1.205E+04 | 204.1 | - | - | 0 | - |
| 3 | a | 5666 | 205.1 | 0.0005632 | 2.746 | +2 | 3 |
| - | - | 5.135E+04 | 207.1 | - | - | 0 | - |
| - | - | 5430 | 208.2 | - | - | 0 | - |
| - | - | 8918 | 209.1 | - | - | 0 | - |
| - | - | 2.466E+04 | 211.1 | - | - | 0 | - |
| - | - | 7638 | 213.1 | - | - | 0 | - |
| - | - | 4.234E+04 | 214.1 | - | - | 0 | - |
| - | - | 1.127E+04 | 216.1 | - | - | 0 | - |
| - | - | 1.489E+04 | 217.1 | - | - | 0 | - |
| - | - | 7250 | 219.1 | - | - | 0 | - |
| - | - | 9.047E+04 | 220.1 | - | - | 0 | - |
| - | - | 1.157E+05 | 221.1 | - | - | 0 | - |
| - | - | 6808 | 221.1 | - | - | 0 | - |
| - | - | 9394 | 222.1 | - | - | 0 | - |
| - | - | 7204 | 222.1 | - | - | 0 | - |
| - | - | 2.124E+04 | 224.1 | - | - | 0 | - |
| - | - | 5149 | 224.1 | - | - | 0 | - |
| - | - | 6.215E+04 | 224.2 | - | - | 0 | - |
| - | - | 6525 | 225.1 | - | - | 0 | - |
| - | - | 1.109E+04 | 225.2 | - | - | 0 | - |
| - | - | 8316 | 225.2 | - | - | 0 | - |
| - | - | 1.407E+06 | 226.2 | - | - | 0 | - |
| - | - | 5360 | 227.1 | - | - | 0 | - |
| - | - | 7682 | 227.1 | - | - | 0 | - |
| 11 | y | 1.705E+05 | 227.2 | 0.001739 | 7.656 | +2 | 4 |
| - | - | 1.099E+04 | 227.2 | - | - | 0 | - |
| 2 | a | 1.034E+05 | 228.1 | 0.0006615 | 2.9 | +1 | 2 |
| - | - | 1.102E+04 | 228.1 | - | - | 0 | - |
| - | - | 1.437E+04 | 228.2 | - | - | 0 | - |
| 5 | b | 1.116E+05 | 229.1 | 0.002026 | 8.844 | +3 | 5 |
| - | - | 1.643E+04 | 229.1 | - | - | 0 | - |
| - | - | 3.062E+04 | 230.1 | - | - | 0 | - |
| - | - | 4.859E+04 | 231.1 | - | - | 0 | - |
| - | - | 2.24E+04 | 231.1 | - | - | 0 | - |
| - | - | 6240 | 232.1 | - | - | 0 | - |
| - | - | 1.428E+04 | 233.1 | - | - | 0 | - |
| - | - | 7601 | 234.1 | - | - | 0 | - |
| - | - | 2.017E+05 | 235.1 | - | - | 0 | - |
| - | - | 2.374E+04 | 236.1 | - | - | 0 | - |
| - | - | 9656 | 236.1 | - | - | 0 | - |
| - | - | 2.17E+04 | 238.1 | - | - | 0 | - |
| - | - | 1.083E+04 | 239.1 | - | - | 0 | - |
| - | - | 7.908E+04 | 239.2 | - | - | 0 | - |
| - | - | 1.032E+04 | 240.2 | - | - | 0 | - |
| - | - | 1.205E+04 | 241.2 | - | - | 0 | - |
| - | - | 2.549E+04 | 242.2 | - | - | 0 | - |
| - | - | 1.979E+04 | 243.2 | - | - | 0 | - |
| - | - | 7274 | 244.1 | - | - | 0 | - |
| - | - | 6257 | 244.2 | - | - | 0 | - |
| - | - | 6668 | 245.2 | - | - | 0 | - |
| - | - | 6263 | 245.2 | - | - | 0 | - |
| 13 | y | 4.462E+05 | 245.2 | 0.000768 | 3.132 | +1 | 2 |
| 2 | a | 2.716E+06 | 246.1 | 0.0007933 | 3.223 | +1 | 2 |
| - | - | 6.031E+04 | 246.2 | - | - | 0 | - |
| - | - | 6.202E+05 | 247.1 | - | - | 0 | - |
| - | - | 6480 | 247.2 | - | - | 0 | - |
| - | - | 1.549E+05 | 248.1 | - | - | 0 | - |
| - | - | 5.134E+04 | 248.1 | - | - | 0 | - |
| - | - | 1.972E+05 | 249.1 | - | - | 0 | - |
| - | - | 1.67E+04 | 249.1 | - | - | 0 | - |
| - | - | 2.139E+04 | 250.1 | - | - | 0 | - |
| - | - | 1.319E+04 | 250.1 | - | - | 0 | - |
| - | - | 7558 | 251.1 | - | - | 0 | - |
| - | - | 7001 | 251.2 | - | - | 0 | - |
| - | - | 2.616E+04 | 252.2 | - | - | 0 | - |
| - | - | 4742 | 253.2 | - | - | 0 | - |
| - | - | 6720 | 255.1 | - | - | 0 | - |
| 2 | b | 7.092E+04 | 256.1 | 0.000681 | 2.659 | +1 | 2 |
| - | - | 1.683E+04 | 257.1 | - | - | 0 | - |
| - | - | 2.265E+04 | 257.1 | - | - | 0 | - |
| - | - | 2.678E+04 | 257.2 | - | - | 0 | - |
| - | - | 1.449E+04 | 258.1 | - | - | 0 | - |
| - | - | 3.398E+05 | 258.1 | - | - | 0 | - |
| - | - | 5516 | 259.1 | - | - | 0 | - |
| - | - | 4.929E+04 | 259.1 | - | - | 0 | - |
| - | - | 1.25E+04 | 259.1 | - | - | 0 | - |
| - | - | 2.619E+04 | 260.1 | - | - | 0 | - |
| - | - | 2.04E+04 | 260.2 | - | - | 0 | - |
| - | - | 1.133E+04 | 262.2 | - | - | 0 | - |
| - | - | 1.244E+04 | 263.1 | - | - | 0 | - |
| - | - | 1.002E+04 | 264.1 | - | - | 0 | - |
| - | - | 2.336E+04 | 265.1 | - | - | 0 | - |
| - | - | 1.431E+04 | 265.2 | - | - | 0 | - |
| - | - | 1.774E+05 | 266.1 | - | - | 0 | - |
| - | - | 2.201E+04 | 267.1 | - | - | 0 | - |
| - | - | 7763 | 267.1 | - | - | 0 | - |
| - | - | 6.617E+04 | 270.2 | - | - | 0 | - |
| 10 | y | 1.613E+04 | 271.2 | 0.004351 | 16.04 | +2 | 5 |
| - | - | 1.276E+04 | 273.1 | - | - | 0 | - |
| - | - | 1.12E+04 | 273.1 | - | - | 0 | - |
| 2 | b | 1.647E+06 | 274.1 | 0.0007059 | 2.575 | +1 | 2 |
| - | - | 5.714E+04 | 275.1 | - | - | 0 | - |
| - | - | 7.047E+05 | 275.1 | - | - | 0 | - |
| - | - | 2.67E+04 | 276.1 | - | - | 0 | - |
| - | - | 8.422E+04 | 276.1 | - | - | 0 | - |
| - | - | 1.47E+04 | 277.1 | - | - | 0 | - |
| - | - | 1.956E+04 | 279.2 | - | - | 0 | - |
| - | - | 3.265E+04 | 283.2 | - | - | 0 | - |
| - | - | 2.255E+05 | 283.2 | - | - | 0 | - |
| - | - | 3.193E+04 | 284.2 | - | - | 0 | - |
| - | - | 1.463E+04 | 285.1 | - | - | 0 | - |
| - | - | 6.857E+04 | 285.2 | - | - | 0 | - |
| - | - | 6746 | 285.2 | - | - | 0 | - |
| - | - | 1.48E+04 | 286.2 | - | - | 0 | - |
| - | - | 7177 | 290.1 | - | - | 0 | - |
| - | - | 6.454E+04 | 290.2 | - | - | 0 | - |
| - | - | 2.212E+04 | 290.7 | - | - | 0 | - |
| - | - | 2.065E+04 | 291.2 | - | - | 0 | - |
| - | - | 1.84E+05 | 292.1 | - | - | 0 | - |
| - | - | 2.551E+04 | 293.1 | - | - | 0 | - |
| - | - | 8.47E+04 | 294.1 | - | - | 0 | - |
| - | - | 9712 | 295.1 | - | - | 0 | - |
| - | - | 1.902E+05 | 297.2 | - | - | 0 | - |
| - | - | 3.941E+04 | 298.2 | - | - | 0 | - |
| - | - | 8582 | 300.2 | - | - | 0 | - |
| - | - | 6115 | 301.1 | - | - | 0 | - |
| - | - | 4.453E+04 | 303.1 | - | - | 0 | - |
| - | - | 5624 | 303.2 | - | - | 0 | - |
| - | - | 4.751E+04 | 305.1 | - | - | 0 | - |
| - | - | 1.33E+04 | 305.7 | - | - | 0 | - |
| - | - | 9338 | 306.1 | - | - | 0 | - |
| - | - | 8133 | 308.1 | - | - | 0 | - |
| - | - | 8259 | 309.2 | - | - | 0 | - |
| - | - | 4.236E+04 | 311.2 | - | - | 0 | - |
| - | - | 7757 | 312.2 | - | - | 0 | - |
| - | - | 1.023E+04 | 313.2 | - | - | 0 | - |
| - | - | 6045 | 314.7 | - | - | 0 | - |
| - | - | 1.19E+04 | 318.2 | - | - | 0 | - |
| - | - | 6397 | 319.2 | - | - | 0 | - |
| - | - | 4.922E+04 | 320.1 | - | - | 0 | - |
| - | - | 7929 | 320.2 | - | - | 0 | - |
| - | - | 1.594E+04 | 321.1 | - | - | 0 | - |
| - | - | 2.324E+05 | 321.2 | - | - | 0 | - |
| - | - | 7.142E+04 | 322.2 | - | - | 0 | - |
| - | - | 4.558E+04 | 322.2 | - | - | 0 | - |
| - | - | 1.526E+04 | 323.2 | - | - | 0 | - |
| - | - | 6904 | 323.2 | - | - | 0 | - |
| - | - | 1.923E+04 | 324.2 | - | - | 0 | - |
| - | - | 5589 | 324.7 | - | - | 0 | - |
| - | - | 5249 | 325.2 | - | - | 0 | - |
| - | - | 4.758E+04 | 326.2 | - | - | 0 | - |
| - | - | 2.673E+04 | 332.7 | - | - | 0 | - |
| - | - | 1.443E+04 | 333.1 | - | - | 0 | - |
| - | - | 1.461E+04 | 333.2 | - | - | 0 | - |
| - | - | 7654 | 333.7 | - | - | 0 | - |
| 9 | y | 1.135E+04 | 335.2 | 0.001238 | 3.692 | +2 | 6 |
| - | - | 8532 | 336.1 | - | - | 0 | - |
| - | - | 2.898E+05 | 336.2 | - | - | 0 | - |
| - | - | 5.113E+04 | 337.2 | - | - | 0 | - |
| - | - | 5.685E+04 | 337.7 | - | - | 0 | - |
| - | - | 9334 | 338.2 | - | - | 0 | - |
| - | - | 2.202E+04 | 338.2 | - | - | 0 | - |
| - | - | 2.297E+05 | 339.2 | - | - | 0 | - |
| - | - | 2.1E+04 | 340.2 | - | - | 0 | - |
| - | - | 3.533E+04 | 340.2 | - | - | 0 | - |
| - | - | 6086 | 341.2 | - | - | 0 | - |
| - | - | 9682 | 342.2 | - | - | 0 | - |
| - | - | 6056 | 343.2 | - | - | 0 | - |
| - | - | 2.123E+05 | 346.7 | - | - | 0 | - |
| - | - | 8.97E+04 | 347.2 | - | - | 0 | - |
| - | - | 1.928E+04 | 347.7 | - | - | 0 | - |
| - | - | 7925 | 348.1 | - | - | 0 | - |
| - | - | 1.206E+04 | 349.7 | - | - | 0 | - |
| - | - | 2.411E+04 | 350.2 | - | - | 0 | - |
| - | - | 6491 | 350.2 | - | - | 0 | - |
| - | - | 2.947E+04 | 351.2 | - | - | 0 | - |
| - | - | 1.54E+04 | 353.2 | - | - | 0 | - |
| - | - | 1.228E+04 | 354.2 | - | - | 0 | - |
| - | - | 4.196E+05 | 354.2 | - | - | 0 | - |
| - | - | 8093 | 354.7 | - | - | 0 | - |
| - | - | 6.118E+04 | 355.2 | - | - | 0 | - |
| - | - | 7398 | 355.3 | - | - | 0 | - |
| - | - | 2.78E+04 | 355.7 | - | - | 0 | - |
| - | - | 7458 | 356.2 | - | - | 0 | - |
| - | - | 1.367E+04 | 357.2 | - | - | 0 | - |
| - | - | 4.489E+04 | 357.3 | - | - | 0 | - |
| - | - | 5.278E+04 | 358.2 | - | - | 0 | - |
| - | - | 8792 | 358.3 | - | - | 0 | - |
| - | - | 2.269E+04 | 358.7 | - | - | 0 | - |
| - | - | 9235 | 359.2 | - | - | 0 | - |
| - | - | 9228 | 359.2 | - | - | 0 | - |
| - | - | 9393 | 362.7 | - | - | 0 | - |
| 8 | y | 1.236E+04 | 363.7 | 0.000943 | 2.592 | +2 | 7 |
| - | - | 7155 | 365.3 | - | - | 0 | - |
| - | - | 2.812E+04 | 366.2 | - | - | 0 | - |
| - | - | 2.241E+04 | 367.2 | - | - | 0 | - |
| - | - | 3.413E+04 | 368.1 | - | - | 0 | - |
| - | - | 1.743E+04 | 368.2 | - | - | 0 | - |
| - | - | 9913 | 369.2 | - | - | 0 | - |
| - | - | 8174 | 369.2 | - | - | 0 | - |
| - | - | 1.165E+04 | 370.2 | - | - | 0 | - |
| 12 | y | 9.585E+04 | 373.3 | 0.0008977 | 2.405 | +1 | 3 |
| - | - | 7840 | 374.3 | - | - | 0 | - |
| - | - | 2.076E+04 | 374.3 | - | - | 0 | - |
| - | - | 8558 | 376.7 | - | - | 0 | - |
| - | - | 1.812E+04 | 380.2 | - | - | 0 | - |
| - | - | 1.518E+04 | 382.2 | - | - | 0 | - |
| - | - | 2.174E+04 | 382.3 | - | - | 0 | - |
| - | - | 1.818E+04 | 383.3 | - | - | 0 | - |
| - | - | 5.321E+04 | 384.2 | - | - | 0 | - |
| - | - | 1.52E+05 | 385.2 | - | - | 0 | - |
| - | - | 5.672E+04 | 385.2 | - | - | 0 | - |
| - | - | 8.485E+04 | 386.2 | - | - | 0 | - |
| - | - | 1.535E+04 | 386.2 | - | - | 0 | - |
| - | - | 2.026E+04 | 387.2 | - | - | 0 | - |
| - | - | 7292 | 388.2 | - | - | 0 | - |
| - | - | 6282 | 389.3 | - | - | 0 | - |
| - | - | 1.519E+04 | 392.2 | - | - | 0 | - |
| - | - | 5.973E+04 | 392.3 | - | - | 0 | - |
| - | - | 1.202E+04 | 393.2 | - | - | 0 | - |
| - | - | 1.442E+04 | 393.3 | - | - | 0 | - |
| - | - | 1.06E+05 | 394.2 | - | - | 0 | - |
| - | - | 1.101E+04 | 394.3 | - | - | 0 | - |
| - | - | 2.332E+04 | 395.2 | - | - | 0 | - |
| - | - | 2.531E+04 | 396.3 | - | - | 0 | - |
| - | - | 9456 | 397.2 | - | - | 0 | - |
| - | - | 9755 | 397.7 | - | - | 0 | - |
| - | - | 1.049E+04 | 398.2 | - | - | 0 | - |
| - | - | 1.981E+04 | 401.2 | - | - | 0 | - |
| - | - | 4.483E+04 | 402.2 | - | - | 0 | - |
| - | - | 1.415E+05 | 403.2 | - | - | 0 | - |
| - | - | 1.044E+04 | 403.2 | - | - | 0 | - |
| - | - | 2.803E+04 | 403.3 | - | - | 0 | - |
| - | - | 2.039E+04 | 403.8 | - | - | 0 | - |
| - | - | 3.103E+04 | 404.2 | - | - | 0 | - |
| - | - | 7759 | 404.3 | - | - | 0 | - |
| - | - | 6.808E+04 | 406.2 | - | - | 0 | - |
| - | - | 5.169E+04 | 406.3 | - | - | 0 | - |
| - | - | 3.117E+04 | 406.7 | - | - | 0 | - |
| - | - | 6070 | 406.7 | - | - | 0 | - |
| - | - | 3E+04 | 406.8 | - | - | 0 | - |
| - | - | 1.21E+04 | 407.2 | - | - | 0 | - |
| - | - | 6345 | 407.5 | - | - | 0 | - |
| - | - | 8628 | 408.2 | - | - | 0 | - |
| - | - | 1.223E+05 | 410.3 | - | - | 0 | - |
| 6 | b | 1.928E+05 | 411.2 | 0.00112 | 2.724 | +2 | 6 |
| - | - | 2.106E+04 | 411.3 | - | - | 0 | - |
| - | - | 1.858E+04 | 412.2 | - | - | 0 | - |
| - | - | 3.09E+04 | 412.2 | - | - | 0 | - |
| 7 | y | 5.548E+05 | 412.3 | 0.001203 | 2.918 | +2 | 8 |
| - | - | 2.432E+05 | 412.8 | - | - | 0 | - |
| - | - | 6892 | 413.2 | - | - | 0 | - |
| - | - | 6.579E+04 | 413.3 | - | - | 0 | - |
| - | - | 8732 | 413.8 | - | - | 0 | - |
| - | - | 5.103E+04 | 415.3 | - | - | 0 | - |
| - | - | 2.568E+04 | 415.8 | - | - | 0 | - |
| - | - | 2.143E+04 | 419.9 | - | - | 0 | - |
| 6 | b | 8.219E+04 | 420.2 | 0.0006291 | 1.497 | +2 | 6 |
| - | - | 3.55E+04 | 420.2 | - | - | 0 | - |
| - | - | 3.873E+04 | 420.7 | - | - | 0 | - |
| - | - | 8084 | 421.2 | - | - | 0 | - |
| - | - | 4.737E+04 | 422.3 | - | - | 0 | - |
| - | - | 1.071E+04 | 422.3 | - | - | 0 | - |
| - | - | 2.609E+04 | 423.3 | - | - | 0 | - |
| - | - | 1.455E+04 | 424.3 | - | - | 0 | - |
| - | - | 1.209E+04 | 424.7 | - | - | 0 | - |
| - | - | 1.816E+04 | 425.3 | - | - | 0 | - |
| - | - | 5873 | 425.7 | - | - | 0 | - |
| - | - | 3.198E+04 | 427.2 | - | - | 0 | - |
| - | - | 2.411E+04 | 427.7 | - | - | 0 | - |
| - | - | 6160 | 428.2 | - | - | 0 | - |
| - | - | 9592 | 428.3 | - | - | 0 | - |
| - | - | 6.662E+05 | 429.2 | - | - | 0 | - |
| - | - | 1.729E+05 | 430.2 | - | - | 0 | - |
| - | - | 8.7E+04 | 431.2 | - | - | 0 | - |
| - | - | 2.671E+04 | 432.2 | - | - | 0 | - |
| - | - | 1.162E+04 | 433.3 | - | - | 0 | - |
| 3 | b | 1.287E+05 | 437.2 | 0.00119 | 2.721 | +1 | 3 |
| - | - | 3.151E+04 | 438.2 | - | - | 0 | - |
| - | - | 1.801E+04 | 438.6 | - | - | 0 | - |
| - | - | 1.813E+04 | 438.9 | - | - | 0 | - |
| - | - | 1.785E+04 | 439.3 | - | - | 0 | - |
| - | - | 1.147E+04 | 444.2 | - | - | 0 | - |
| - | - | 1.073E+04 | 444.6 | - | - | 0 | - |
| - | - | 1.351E+04 | 445.3 | - | - | 0 | - |
| - | - | 1.148E+04 | 446.2 | - | - | 0 | - |
| - | - | 1.047E+04 | 450.3 | - | - | 0 | - |
| - | - | 2.665E+04 | 451.2 | - | - | 0 | - |
| - | - | 3.94E+04 | 451.3 | - | - | 0 | - |
| - | - | 1.424E+04 | 451.7 | - | - | 0 | - |
| - | - | 1.224E+04 | 451.9 | - | - | 0 | - |
| - | - | 1.514E+04 | 452.3 | - | - | 0 | - |
| - | - | 3.054E+04 | 454.2 | - | - | 0 | - |
| - | - | 1.067E+04 | 455.2 | - | - | 0 | - |
| - | - | 4.271E+04 | 455.3 | - | - | 0 | - |
| - | - | 1.38E+04 | 456.3 | - | - | 0 | - |
| - | - | 2.163E+04 | 457.2 | - | - | 0 | - |
| - | - | 1.208E+04 | 457.6 | - | - | 0 | - |
| 3 | y | 8695 | 457.9 | 0.005126 | 11.19 | +3 | 12 |
| - | - | 2.574E+04 | 460.2 | - | - | 0 | - |
| - | - | 1.212E+04 | 461.2 | - | - | 0 | - |
| - | - | 7658 | 461.2 | - | - | 0 | - |
| - | - | 1.281E+04 | 463.3 | - | - | 0 | - |
| 3 | y | 1.404E+05 | 463.6 | 0.001149 | 2.478 | +3 | 12 |
| - | - | 9.144E+04 | 463.9 | - | - | 0 | - |
| - | - | 3.14E+04 | 464.3 | - | - | 0 | - |
| - | - | 1.264E+04 | 464.6 | - | - | 0 | - |
| - | - | 8287 | 466.9 | - | - | 0 | - |
| - | - | 9579 | 467.3 | - | - | 0 | - |
| - | - | 6.135E+04 | 467.3 | - | - | 0 | - |
| - | - | 3.125E+04 | 468.3 | - | - | 0 | - |
| 7 | b | 8548 | 468.7 | 0.003147 | 6.715 | +2 | 7 |
| - | - | 1.12E+04 | 469.8 | - | - | 0 | - |
| - | - | 1.708E+04 | 470.3 | - | - | 0 | - |
| 11 | y | 4.137E+05 | 470.3 | 0.001234 | 2.625 | +1 | 4 |
| - | - | 2.259E+04 | 470.6 | - | - | 0 | - |
| - | - | 6507 | 470.9 | - | - | 0 | - |
| - | - | 1.137E+05 | 471.3 | - | - | 0 | - |
| - | - | 4.03E+04 | 471.8 | - | - | 0 | - |
| - | - | 2.037E+04 | 472.3 | - | - | 0 | - |
| - | - | 1.913E+04 | 472.3 | - | - | 0 | - |
| - | - | 7167 | 472.8 | - | - | 0 | - |
| 12 | b | 1.389E+05 | 473.2 | 0.001677 | 3.545 | +3 | 12 |
| - | - | 1.808E+05 | 473.6 | - | - | 0 | - |
| - | - | 8.836E+04 | 473.9 | - | - | 0 | - |
| - | - | 1.725E+04 | 474.3 | - | - | 0 | - |
| - | - | 1.136E+04 | 474.6 | - | - | 0 | - |
| - | - | 3.363E+04 | 474.8 | - | - | 0 | - |
| - | - | 1.478E+04 | 475.3 | - | - | 0 | - |
| - | - | 1.165E+04 | 475.8 | - | - | 0 | - |
| - | - | 1.809E+04 | 475.9 | - | - | 0 | - |
| - | - | 4.417E+04 | 476.3 | - | - | 0 | - |
| - | - | 4.409E+04 | 476.6 | - | - | 0 | - |
| - | - | 2.189E+04 | 476.9 | - | - | 0 | - |
| - | - | 3.157E+04 | 478.2 | - | - | 0 | - |
| - | - | 1.112E+04 | 479.2 | - | - | 0 | - |
| 6 | y | 1.333E+05 | 480.8 | 0.001196 | 2.488 | +2 | 9 |
| - | - | 6.539E+04 | 481.3 | - | - | 0 | - |
| - | - | 2.005E+04 | 481.8 | - | - | 0 | - |
| - | - | 3.377E+04 | 481.9 | - | - | 0 | - |
| - | - | 5.634E+04 | 482.3 | - | - | 0 | - |
| - | - | 4.285E+04 | 482.3 | - | - | 0 | - |
| - | - | 1.746E+04 | 482.6 | - | - | 0 | - |
| - | - | 3.077E+04 | 482.8 | - | - | 0 | - |
| - | - | 6826 | 482.9 | - | - | 0 | - |
| - | - | 1.298E+04 | 483.3 | - | - | 0 | - |
| - | - | 9.344E+04 | 483.8 | - | - | 0 | - |
| - | - | 3.604E+04 | 484.3 | - | - | 0 | - |
| - | - | 1.456E+04 | 484.8 | - | - | 0 | - |
| - | - | 5.651E+04 | 491.3 | - | - | 0 | - |
| - | - | 8424 | 491.8 | - | - | 0 | - |
| - | - | 1.111E+04 | 492.3 | - | - | 0 | - |
| - | - | 9358 | 492.8 | - | - | 0 | - |
| - | - | 6608 | 494.3 | - | - | 0 | - |
| - | - | 8744 | 495.9 | - | - | 0 | - |
| - | - | 1.354E+04 | 496.3 | - | - | 0 | - |
| 8 | b | 2.509E+04 | 497.2 | 0.0008994 | 1.809 | +2 | 8 |
| - | - | 1.132E+04 | 497.7 | - | - | 0 | - |
| - | - | 1.455E+04 | 499.3 | - | - | 0 | - |
| - | - | 9.883E+04 | 501.6 | - | - | 0 | - |
| - | - | 1.193E+05 | 501.9 | - | - | 0 | - |
| - | - | 4.923E+04 | 502.3 | - | - | 0 | - |
| - | - | 1.058E+04 | 502.6 | - | - | 0 | - |
| - | - | 7919 | 502.9 | - | - | 0 | - |
| - | - | 4.019E+04 | 503.2 | - | - | 0 | - |
| 13 | b | 1.55E+05 | 504.9 | 0.001337 | 2.649 | +3 | 13 |
| 13 | b | 1.536E+05 | 505.3 | 0.00631 | 12.49 | +3 | 13 |
| - | - | 8.892E+04 | 505.6 | - | - | 0 | - |
| - | - | 8554 | 505.7 | - | - | 0 | - |
| - | - | 3.782E+04 | 505.9 | - | - | 0 | - |
| - | - | 1.21E+04 | 506.3 | - | - | 0 | - |
| - | - | 2.267E+04 | 508.8 | - | - | 0 | - |
| - | - | 1.325E+04 | 509.3 | - | - | 0 | - |
| - | - | 7015 | 510.3 | - | - | 0 | - |
| - | - | 1.422E+04 | 510.4 | - | - | 0 | - |
| 13 | b | 7.69E+05 | 510.9 | 0.001203 | 2.355 | +3 | 13 |
| - | - | 7.766E+05 | 511.3 | - | - | 0 | - |
| - | - | 3.714E+05 | 511.6 | - | - | 0 | - |
| - | - | 1.262E+05 | 511.9 | - | - | 0 | - |
| - | - | 4.024E+04 | 512.3 | - | - | 0 | - |
| - | - | 6983 | 513.9 | - | - | 0 | - |
| - | - | 1.281E+04 | 514.3 | - | - | 0 | - |
| - | - | 1.082E+04 | 514.6 | - | - | 0 | - |
| - | - | 1.164E+05 | 516.9 | - | - | 0 | - |
| - | - | 1.506E+05 | 517.3 | - | - | 0 | - |
| - | - | 5.355E+04 | 517.6 | - | - | 0 | - |
| - | - | 1.88E+04 | 517.9 | - | - | 0 | - |
| - | - | 8721 | 519.6 | - | - | 0 | - |
| 2 | y | 3.27E+04 | 519.9 | 0.002706 | 5.204 | +3 | 13 |
| - | - | 1.353E+04 | 520.2 | - | - | 0 | - |
| - | - | 4.587E+04 | 520.3 | - | - | 0 | - |
| - | - | 1.961E+04 | 520.4 | - | - | 0 | - |
| - | - | 1.203E+04 | 520.6 | - | - | 0 | - |
| - | - | 1.169E+04 | 521 | - | - | 0 | - |
| - | - | 3.802E+04 | 521.2 | - | - | 0 | - |
| - | - | 9948 | 522.3 | - | - | 0 | - |
| - | - | 1.667E+04 | 523.4 | - | - | 0 | - |
| - | - | 8607 | 524.8 | - | - | 0 | - |
| - | - | 1.384E+04 | 525.3 | - | - | 0 | - |
| 2 | y | 1.524E+05 | 525.6 | 0.001139 | 2.168 | +3 | 13 |
| - | - | 1.231E+04 | 525.8 | - | - | 0 | - |
| - | - | 1.382E+05 | 526 | - | - | 0 | - |
| - | - | 6.464E+04 | 526.3 | - | - | 0 | - |
| - | - | 1.238E+04 | 526.6 | - | - | 0 | - |
| - | - | 2.247E+04 | 528.2 | - | - | 0 | - |
| - | - | 2.18E+04 | 529.2 | - | - | 0 | - |
| - | - | 2.009E+04 | 529.8 | - | - | 0 | - |
| - | - | 2.944E+04 | 530.3 | - | - | 0 | - |
| - | - | 1.88E+04 | 530.8 | - | - | 0 | - |
| - | - | 2.527E+04 | 531.2 | - | - | 0 | - |
| - | - | 9472 | 531.3 | - | - | 0 | - |
| - | - | 8071 | 535.3 | - | - | 0 | - |
| - | - | 8055 | 535.6 | - | - | 0 | - |
| - | - | 8404 | 536.3 | - | - | 0 | - |
| - | - | 8499 | 537.3 | - | - | 0 | - |
| - | - | 3.788E+05 | 538.3 | - | - | 0 | - |
| - | - | 2.689E+04 | 538.4 | - | - | 0 | - |
| - | - | 4.996E+04 | 538.8 | - | - | 0 | - |
| - | - | 8473 | 539 | - | - | 0 | - |
| - | - | 1.027E+05 | 539.3 | - | - | 0 | - |
| - | - | 1.067E+05 | 539.3 | - | - | 0 | - |
| - | - | 7952 | 539.4 | - | - | 0 | - |
| - | - | 2.794E+04 | 539.6 | - | - | 0 | - |
| - | - | 4.315E+04 | 539.8 | - | - | 0 | - |
| - | - | 2.265E+04 | 540 | - | - | 0 | - |
| - | - | 1.785E+04 | 540.3 | - | - | 0 | - |
| - | - | 1.591E+04 | 540.3 | - | - | 0 | - |
| - | - | 1.458E+04 | 540.3 | - | - | 0 | - |
| 5 | y | 1.041E+04 | 540.8 | 0.005736 | 10.61 | +2 | 10 |
| - | - | 9069 | 541.3 | - | - | 0 | - |
| 10 | y | 1.431E+05 | 541.4 | 0.001688 | 3.118 | +1 | 5 |
| - | - | 2.057E+04 | 542.3 | - | - | 0 | - |
| - | - | 4.282E+04 | 542.4 | - | - | 0 | - |
| - | - | 2.122E+04 | 543 | - | - | 0 | - |
| - | - | 8641 | 543.3 | - | - | 0 | - |
| - | - | 1.409E+04 | 544.6 | - | - | 0 | - |
| - | - | 1.065E+04 | 544.8 | - | - | 0 | - |
| - | - | 9553 | 545.3 | - | - | 0 | - |
| - | - | 5.767E+04 | 547.8 | - | - | 0 | - |
| - | - | 1.603E+04 | 548.2 | - | - | 0 | - |
| - | - | 3.28E+04 | 548.3 | - | - | 0 | - |
| 0 | Precursor | 2.247E+05 | 548.6 | 0.001431 | 2.608 | +3 | -1 |
| - | - | 1.343E+04 | 548.8 | - | - | 0 | - |
| 0 | Precursor | 2.798E+05 | 549 | 0.00619 | 11.28 | +3 | -1 |
| - | - | 2.167E+04 | 549.2 | - | - | 0 | - |
| - | - | 9.997E+04 | 549.3 | - | - | 0 | - |
| 5 | y | 2.152E+05 | 549.3 | 0.001556 | 2.833 | +2 | 10 |
| - | - | 1.019E+05 | 549.6 | - | - | 0 | - |
| - | - | 1.463E+05 | 549.8 | - | - | 0 | - |
| - | - | 3.011E+04 | 550 | - | - | 0 | - |
| - | - | 8883 | 550.2 | - | - | 0 | - |
| - | - | 1.027E+04 | 550.3 | - | - | 0 | - |
| - | - | 5.712E+04 | 550.3 | - | - | 0 | - |
| - | - | 9268 | 550.8 | - | - | 0 | - |
| - | - | 1.316E+04 | 553.2 | - | - | 0 | - |
| - | - | 1.012E+04 | 553.3 | - | - | 0 | - |
| - | - | 1.745E+04 | 554 | - | - | 0 | - |
| - | - | 5.19E+04 | 554.3 | - | - | 0 | - |
| 0 | Precursor | 5.189E+06 | 554.6 | 0.001266 | 2.283 | +3 | -1 |
| - | - | 5.211E+06 | 555 | - | - | 0 | - |
| - | - | 2.845E+06 | 555.3 | - | - | 0 | - |
| - | - | 1.007E+06 | 555.6 | - | - | 0 | - |
| - | - | 1.146E+04 | 555.8 | - | - | 0 | - |
| - | - | 2.984E+05 | 556 | - | - | 0 | - |
| - | - | 1.696E+04 | 557.3 | - | - | 0 | - |
| 9 | b | 4.451E+04 | 561.3 | 0.0007201 | 1.283 | +2 | 9 |
| - | - | 3.164E+04 | 561.8 | - | - | 0 | - |
| - | - | 1.741E+04 | 562.3 | - | - | 0 | - |
| 4 | b | 4.073E+04 | 565.2 | 0.0008091 | 1.431 | +1 | 4 |
| - | - | 7880 | 565.3 | - | - | 0 | - |
| - | - | 4.603E+05 | 566.2 | - | - | 0 | - |
| - | - | 1.386E+05 | 567.3 | - | - | 0 | - |
| - | - | 1.747E+04 | 567.4 | - | - | 0 | - |
| - | - | 2.916E+04 | 568.3 | - | - | 0 | - |
| - | - | 9577 | 568.4 | - | - | 0 | - |
| - | - | 8.017E+04 | 570.2 | - | - | 0 | - |
| - | - | 2.26E+04 | 571.2 | - | - | 0 | - |
| - | - | 2.05E+05 | 572.8 | - | - | 0 | - |
| - | - | 1.663E+05 | 573.3 | - | - | 0 | - |
| - | - | 6.107E+04 | 573.8 | - | - | 0 | - |
| - | - | 8.962E+04 | 577.4 | - | - | 0 | - |
| - | - | 3.04E+04 | 578.4 | - | - | 0 | - |
| - | - | 2.89E+05 | 579.4 | - | - | 0 | - |
| - | - | 7.716E+04 | 580.4 | - | - | 0 | - |
| - | - | 2.468E+04 | 581.4 | - | - | 0 | - |
| - | - | 1.214E+04 | 587.3 | - | - | 0 | - |
| 10 | b | 4.613E+04 | 587.8 | 0.001423 | 2.42 | +2 | 10 |
| 10 | b | 3.324E+04 | 588.3 | 0.009232 | 15.69 | +2 | 10 |
| - | - | 7819 | 588.3 | - | - | 0 | - |
| - | - | 1.163E+04 | 588.8 | - | - | 0 | - |
| - | - | 7333 | 589.3 | - | - | 0 | - |
| - | - | 6850 | 595.4 | - | - | 0 | - |
| - | - | 8.609E+04 | 595.4 | - | - | 0 | - |
| - | - | 1.836E+04 | 595.8 | - | - | 0 | - |
| - | - | 1.012E+04 | 596.3 | - | - | 0 | - |
| - | - | 2.911E+04 | 596.4 | - | - | 0 | - |
| 10 | b | 5.225E+04 | 596.8 | 0.0004738 | 0.7939 | +2 | 10 |
| - | - | 8374 | 596.8 | - | - | 0 | - |
| - | - | 1.735E+04 | 597.3 | - | - | 0 | - |
| - | - | 1.433E+04 | 597.8 | - | - | 0 | - |
| - | - | 3.298E+04 | 598.2 | - | - | 0 | - |
| - | - | 1.491E+04 | 599.2 | - | - | 0 | - |
| - | - | 2.174E+04 | 602.8 | - | - | 0 | - |
| - | - | 9925 | 603.3 | - | - | 0 | - |
| - | - | 1.137E+05 | 604.4 | - | - | 0 | - |
| 4 | y | 1.696E+05 | 604.8 | 0.005562 | 9.195 | +2 | 11 |
| - | - | 9.958E+04 | 605.4 | - | - | 0 | - |
| - | - | 3.652E+04 | 605.9 | - | - | 0 | - |
| - | - | 1.083E+04 | 606.4 | - | - | 0 | - |
| - | - | 1.084E+04 | 606.8 | - | - | 0 | - |
| - | - | 1.355E+04 | 607.3 | - | - | 0 | - |
| - | - | 4.753E+04 | 610.3 | - | - | 0 | - |
| - | - | 2.598E+04 | 611.3 | - | - | 0 | - |
| - | - | 2.778E+04 | 611.8 | - | - | 0 | - |
| - | - | 2.241E+04 | 612.3 | - | - | 0 | - |
| - | - | 1.084E+04 | 612.8 | - | - | 0 | - |
| 4 | y | 1.459E+05 | 613.4 | 0.001076 | 1.754 | +2 | 11 |
| - | - | 1.208E+05 | 613.9 | - | - | 0 | - |
| - | - | 4.962E+04 | 614.4 | - | - | 0 | - |
| - | - | 1.752E+04 | 615.3 | - | - | 0 | - |
| - | - | 9.504E+04 | 615.4 | - | - | 0 | - |
| - | - | 5.95E+04 | 615.9 | - | - | 0 | - |
| - | - | 2.524E+04 | 616.4 | - | - | 0 | - |
| - | - | 9211 | 616.9 | - | - | 0 | - |
| - | - | 8061 | 618.9 | - | - | 0 | - |
| - | - | 1.942E+05 | 620.3 | - | - | 0 | - |
| - | - | 1.117E+05 | 620.8 | - | - | 0 | - |
| - | - | 7.196E+04 | 621.3 | - | - | 0 | - |
| - | - | 1.15E+04 | 621.4 | - | - | 0 | - |
| - | - | 1.81E+04 | 621.8 | - | - | 0 | - |
| - | - | 2.157E+04 | 623.4 | - | - | 0 | - |
| - | - | 1.85E+04 | 628.3 | - | - | 0 | - |
| - | - | 4.447E+05 | 629.3 | - | - | 0 | - |
| - | - | 3.211E+05 | 629.8 | - | - | 0 | - |
| - | - | 1.228E+05 | 630.3 | - | - | 0 | - |
| - | - | 2.439E+04 | 630.9 | - | - | 0 | - |
| - | - | 1.425E+04 | 632.3 | - | - | 0 | - |
| - | - | 4.701E+04 | 638.4 | - | - | 0 | - |
| - | - | 2.743E+04 | 638.9 | - | - | 0 | - |
| - | - | 1.36E+04 | 639.4 | - | - | 0 | - |
| - | - | 8453 | 643.3 | - | - | 0 | - |
| 11 | b | 2.988E+04 | 645.3 | 3.179E-05 | 0.04926 | +2 | 11 |
| - | - | 1.783E+04 | 645.8 | - | - | 0 | - |
| - | - | 1.878E+04 | 647.4 | - | - | 0 | - |
| - | - | 2.811E+04 | 653.8 | - | - | 0 | - |
| - | - | 1.676E+04 | 654.3 | - | - | 0 | - |
| - | - | 1.552E+04 | 654.8 | - | - | 0 | - |
| - | - | 1.212E+04 | 656.3 | - | - | 0 | - |
| - | - | 3.188E+04 | 657.3 | - | - | 0 | - |
| - | - | 7241 | 657.3 | - | - | 0 | - |
| - | - | 9215 | 657.8 | - | - | 0 | - |
| - | - | 2.118E+04 | 658.3 | - | - | 0 | - |
| - | - | 1.614E+04 | 659.3 | - | - | 0 | - |
| - | - | 9.366E+04 | 664.5 | - | - | 0 | - |
| - | - | 3.479E+04 | 665.5 | - | - | 0 | - |
| - | - | 1.391E+04 | 665.8 | - | - | 0 | - |
| - | - | 8562 | 666.5 | - | - | 0 | - |
| - | - | 8398 | 667.4 | - | - | 0 | - |
| - | - | 1.468E+04 | 668.3 | - | - | 0 | - |
| 9 | y | 4.997E+04 | 669.5 | 0.001512 | 2.259 | +1 | 6 |
| - | - | 2.53E+04 | 670.5 | - | - | 0 | - |
| - | - | 1.354E+04 | 671.9 | - | - | 0 | - |
| - | - | 1.824E+04 | 672.4 | - | - | 0 | - |
| - | - | 7015 | 672.9 | - | - | 0 | - |
| - | - | 1.024E+05 | 674.3 | - | - | 0 | - |
| - | - | 3.096E+05 | 674.4 | - | - | 0 | - |
| - | - | 4.485E+04 | 675.3 | - | - | 0 | - |
| - | - | 1.346E+05 | 675.4 | - | - | 0 | - |
| - | - | 2.866E+04 | 676.4 | - | - | 0 | - |
| - | - | 5.597E+04 | 677.4 | - | - | 0 | - |
| - | - | 4.26E+04 | 677.9 | - | - | 0 | - |
| - | - | 2.535E+04 | 678.4 | - | - | 0 | - |
| 5 | b | 3.237E+04 | 684.3 | 0.00245 | 3.58 | +1 | 5 |
| 5 | b | 1.113E+04 | 685.3 | 0.01306 | 19.06 | +1 | 5 |
| - | - | 9.557E+04 | 685.9 | - | - | 0 | - |
| 3 | y | 8.985E+04 | 686.4 | 0.008138 | 11.86 | +2 | 12 |
| - | - | 4.306E+04 | 686.9 | - | - | 0 | - |
| - | - | 1.816E+04 | 687.4 | - | - | 0 | - |
| - | - | 3.284E+05 | 692.4 | - | - | 0 | - |
| - | - | 1.275E+05 | 693.4 | - | - | 0 | - |
| - | - | 8755 | 694.4 | - | - | 0 | - |
| - | - | 3.006E+04 | 694.5 | - | - | 0 | - |
| 3 | y | 1.608E+06 | 694.9 | 0.001333 | 1.918 | +2 | 12 |
| - | - | 1.235E+06 | 695.4 | - | - | 0 | - |
| - | - | 5.923E+05 | 695.9 | - | - | 0 | - |
| - | - | 1.617E+05 | 696.4 | - | - | 0 | - |
| - | - | 3.498E+04 | 696.9 | - | - | 0 | - |
| - | - | 3.149E+04 | 698.4 | - | - | 0 | - |
| 12 | b | 1.464E+04 | 700.4 | 0.01258 | 17.96 | +2 | 12 |
| 12 | b | 1.764E+04 | 700.8 | 0.005925 | 8.454 | +2 | 12 |
| - | - | 7202 | 701.4 | - | - | 0 | - |
| - | - | 8516 | 701.9 | - | - | 0 | - |
| 5 | b | 1.803E+05 | 702.3 | 0.001529 | 2.177 | +1 | 5 |
| - | - | 8.219E+04 | 703.3 | - | - | 0 | - |
| - | - | 2.179E+04 | 704.3 | - | - | 0 | - |
| - | - | 4.757E+04 | 707.3 | - | - | 0 | - |
| - | - | 2.438E+04 | 708.3 | - | - | 0 | - |
| - | - | 1.097E+04 | 708.5 | - | - | 0 | - |
| 12 | b | 3.542E+05 | 709.4 | 0.001562 | 2.201 | +2 | 12 |
| - | - | 2.816E+05 | 709.9 | - | - | 0 | - |
| - | - | 1.215E+05 | 710.4 | - | - | 0 | - |
| - | - | 4.644E+04 | 710.5 | - | - | 0 | - |
| - | - | 3.384E+04 | 710.9 | - | - | 0 | - |
| - | - | 1.672E+04 | 711.4 | - | - | 0 | - |
| - | - | 1.613E+04 | 711.5 | - | - | 0 | - |
| - | - | 8200 | 712.5 | - | - | 0 | - |
| - | - | 7646 | 713.4 | - | - | 0 | - |
| - | - | 1.313E+04 | 714.4 | - | - | 0 | - |
| - | - | 5.608E+04 | 716.4 | - | - | 0 | - |
| - | - | 2.424E+04 | 717.4 | - | - | 0 | - |
| - | - | 6861 | 718.4 | - | - | 0 | - |
| - | - | 1.107E+04 | 720.3 | - | - | 0 | - |
| - | - | 1.607E+04 | 720.4 | - | - | 0 | - |
| - | - | 2.52E+04 | 721.4 | - | - | 0 | - |
| - | - | 8852 | 722.4 | - | - | 0 | - |
| - | - | 1.607E+04 | 722.9 | - | - | 0 | - |
| - | - | 1.317E+04 | 723.4 | - | - | 0 | - |
| - | - | 2.841E+04 | 724.3 | - | - | 0 | - |
| - | - | 1.219E+04 | 725.3 | - | - | 0 | - |
| 8 | y | 1.241E+05 | 726.5 | 0.0009837 | 1.354 | +1 | 7 |
| - | - | 4.533E+04 | 727.5 | - | - | 0 | - |
| - | - | 4.587E+04 | 735.3 | - | - | 0 | - |
| - | - | 1.335E+04 | 736.3 | - | - | 0 | - |
| - | - | 1.352E+04 | 737.4 | - | - | 0 | - |
| - | - | 1.479E+04 | 738.4 | - | - | 0 | - |
| - | - | 1.147E+04 | 739.4 | - | - | 0 | - |
| - | - | 8379 | 740.4 | - | - | 0 | - |
| - | - | 1.574E+04 | 751.9 | - | - | 0 | - |
| - | - | 3.741E+04 | 752.3 | - | - | 0 | - |
| - | - | 2.184E+04 | 752.4 | - | - | 0 | - |
| - | - | 2.122E+04 | 753.3 | - | - | 0 | - |
| 13 | b | 3.511E+04 | 756.9 | 0.002104 | 2.78 | +2 | 13 |
| 13 | b | 4.248E+04 | 757.4 | 0.01077 | 14.22 | +2 | 13 |
| - | - | 3.792E+04 | 757.9 | - | - | 0 | - |
| - | - | 1.457E+04 | 758.4 | - | - | 0 | - |
| 13 | b | 1.155E+05 | 765.9 | 0.001949 | 2.545 | +2 | 13 |
| - | - | 9326 | 766.3 | - | - | 0 | - |
| - | - | 1.253E+05 | 766.4 | - | - | 0 | - |
| - | - | 5.288E+04 | 766.9 | - | - | 0 | - |
| - | - | 1.945E+04 | 767.4 | - | - | 0 | - |
| - | - | 5.696E+04 | 774.9 | - | - | 0 | - |
| - | - | 5.467E+04 | 775.4 | - | - | 0 | - |
| - | - | 1.802E+04 | 775.9 | - | - | 0 | - |
| - | - | 1.027E+04 | 776.4 | - | - | 0 | - |
| - | - | 1.147E+04 | 777.5 | - | - | 0 | - |
| 2 | y | 7881 | 779.4 | 0.006812 | 8.739 | +2 | 13 |
| - | - | 7160 | 780.3 | - | - | 0 | - |
| - | - | 9813 | 787.5 | - | - | 0 | - |
| 2 | y | 4.136E+04 | 787.9 | 0.002204 | 2.797 | +2 | 13 |
| - | - | 4.225E+04 | 788.4 | - | - | 0 | - |
| - | - | 1.707E+04 | 788.9 | - | - | 0 | - |
| - | - | 7528 | 789.4 | - | - | 0 | - |
| - | - | 9889 | 792.9 | - | - | 0 | - |
| - | - | 1.476E+04 | 793.4 | - | - | 0 | - |
| - | - | 2.456E+04 | 794.3 | - | - | 0 | - |
| - | - | 1.537E+04 | 795.4 | - | - | 0 | - |
| - | - | 1.869E+04 | 801.5 | - | - | 0 | - |
| - | - | 6.207E+04 | 805.5 | - | - | 0 | - |
| - | - | 3.124E+04 | 806.5 | - | - | 0 | - |
| - | - | 1.428E+04 | 807.5 | - | - | 0 | - |
| - | - | 1.94E+05 | 811.4 | - | - | 0 | - |
| - | - | 5.781E+04 | 811.5 | - | - | 0 | - |
| - | - | 8.429E+04 | 812.4 | - | - | 0 | - |
| - | - | 2.823E+04 | 812.5 | - | - | 0 | - |
| - | - | 2.217E+04 | 813.4 | - | - | 0 | - |
| - | - | 8896 | 814.4 | - | - | 0 | - |
| 6 | b | 2.843E+04 | 821.3 | 0.0003616 | 0.4403 | +1 | 6 |
| 6 | b | 1.934E+04 | 822.3 | 0.01268 | 15.42 | +1 | 6 |
| - | - | 8178 | 822.5 | - | - | 0 | - |
| - | - | 1.612E+04 | 823.3 | - | - | 0 | - |
| 7 | y | 7.55E+05 | 823.5 | 0.001259 | 1.529 | +1 | 8 |
| - | - | 3.434E+05 | 824.5 | - | - | 0 | - |
| - | - | 9.12E+04 | 825.5 | - | - | 0 | - |
| - | - | 1.205E+04 | 826.5 | - | - | 0 | - |
| - | - | 5.635E+04 | 829.5 | - | - | 0 | - |
| - | - | 2.715E+04 | 830.5 | - | - | 0 | - |
| - | - | 1.081E+04 | 831.5 | - | - | 0 | - |
| 6 | b | 4.082E+05 | 839.4 | 0.001027 | 1.224 | +1 | 6 |
| - | - | 2.089E+05 | 840.4 | - | - | 0 | - |
| - | - | 6.039E+04 | 841.4 | - | - | 0 | - |
| - | - | 8708 | 842.4 | - | - | 0 | - |
| - | - | 1.126E+04 | 847.5 | - | - | 0 | - |
| - | - | 6.824E+04 | 848.4 | - | - | 0 | - |
| - | - | 3.741E+04 | 849.4 | - | - | 0 | - |
| - | - | 7036 | 850.4 | - | - | 0 | - |
| - | - | 3.522E+04 | 853.5 | - | - | 0 | - |
| - | - | 2.209E+04 | 854.5 | - | - | 0 | - |
| - | - | 9240 | 866.4 | - | - | 0 | - |
| - | - | 2.363E+04 | 891.5 | - | - | 0 | - |
| - | - | 1.069E+04 | 892.5 | - | - | 0 | - |
| - | - | 1.39E+05 | 901.4 | - | - | 0 | - |
| - | - | 8.528E+04 | 902.4 | - | - | 0 | - |
| - | - | 1.874E+04 | 903.4 | - | - | 0 | - |
| - | - | 7.072E+04 | 919.5 | - | - | 0 | - |
| - | - | 5.287E+04 | 920.5 | - | - | 0 | - |
| - | - | 9001 | 921.5 | - | - | 0 | - |
| - | - | 4.106E+04 | 942.6 | - | - | 0 | - |
| - | - | 3.6E+04 | 943.6 | - | - | 0 | - |
| - | - | 7015 | 944.6 | - | - | 0 | - |
| - | - | 1.496E+04 | 948.6 | - | - | 0 | - |
| 6 | y | 1.381E+05 | 960.6 | 0.0007582 | 0.7893 | +1 | 9 |
| - | - | 9498 | 961.5 | - | - | 0 | - |
| - | - | 7.036E+04 | 961.6 | - | - | 0 | - |
| - | - | 1.848E+04 | 962.6 | - | - | 0 | - |
| - | - | 7141 | 963.6 | - | - | 0 | - |
| - | - | 1.157E+04 | 964.5 | - | - | 0 | - |
| - | - | 3.366E+04 | 966.6 | - | - | 0 | - |
| - | - | 1.855E+04 | 967.6 | - | - | 0 | - |
| 8 | b | 1.223E+04 | 993.4 | 0.00246 | 2.477 | +1 | 8 |
| - | - | 8758 | 994.4 | - | - | 0 | - |
| - | - | 1.618E+04 | 1010 | - | - | 0 | - |
| - | - | 1.013E+04 | 1011 | - | - | 0 | - |
| - | - | 7233 | 1017 | - | - | 0 | - |
| - | - | 2.248E+04 | 1033 | - | - | 0 | - |
| - | - | 3.18E+04 | 1034 | - | - | 0 | - |
| - | - | 1.132E+04 | 1035 | - | - | 0 | - |
| - | - | 1.346E+04 | 1088 | - | - | 0 | - |
| 5 | y | 4.842E+04 | 1098 | 0.0008673 | 0.7902 | +1 | 10 |
| - | - | 2.957E+04 | 1099 | - | - | 0 | - |
| - | - | 1.661E+04 | 1100 | - | - | 0 | - |
| 9 | b | 3.371E+04 | 1122 | 0.0004159 | 0.3708 | +1 | 9 |
| - | - | 2.669E+04 | 1123 | - | - | 0 | - |
| - | - | 1.856E+04 | 1145 | - | - | 0 | - |
| - | - | 1.174E+04 | 1146 | - | - | 0 | - |
| - | - | 1.094E+04 | 1147 | - | - | 0 | - |
| - | - | 8888 | 1165 | - | - | 0 | - |
| 10 | b | 7.48E+04 | 1175 | 0.0001322 | 0.1126 | +1 | 10 |
| 10 | b | 5.982E+04 | 1176 | 0.01781 | 15.15 | +1 | 10 |
| - | - | 1.398E+04 | 1177 | - | - | 0 | - |
| 10 | b | 2.849E+04 | 1193 | 0.0001674 | 0.1403 | +1 | 10 |
| - | - | 2.378E+04 | 1194 | - | - | 0 | - |
| - | - | 8384 | 1195 | - | - | 0 | - |
| - | - | 7175 | 2015 | - | - | 0 | - |
| - | - | 5725 | 2188 | - | - | 0 | - |
| - | - | 7062 | 2709 | - | - | 0 | - |
| - | - | 6881 | 2710 | - | - | 0 | - |
| - | - | 6924 | 2926 | - | - | 0 | - |
| - | - | 6994 | 3255 | - | - | 0 | - |
| - | - | 6799 | 3299 | - | - | 0 | - |

m/z Charge Intensity FragmentType MassShift Position
120.05616760253906 0 9335.62
122.07197570800781 0 6659.9927
123.0558090209961 0 19080.555
124.70914459228516 0 3436.7227
124.9344711303711 0 3274.5254
127.08711242675781 0 30055.32
128.07115173339844 0 8069.1245
129.0662841796875 0 42348.914
129.102783203125 0 1118315
130.06564331054688 0 277525.47
130.10052490234375 0 9543.871
130.1061248779297 0 74802.46
131.06900024414062 0 33037.586
131.07334899902344 0 3757.7832
132.08131408691406 0 1143495
132.1024169921875 0 219479.03 y 13
132.19078063964844 0 3974.8774
133.0846405029297 0 117340.555
133.100341796875 0 5136.6074
133.1058807373047 0 15852.439
134.08807373046875 0 8367.782
136.07620239257812 0 674938.4
137.0795135498047 0 60880.883
137.1079559326172 0 6766.6504
138.06661987304688 0 25519.441
139.08705139160156 0 15255.027
141.10279846191406 0 84560.4
142.06573486328125 0 6308.2915
142.10617065429688 0 5891.849
144.08135986328125 0 22313.717
145.06134033203125 0 6934.351
146.12893676757812 0 3907.8625
147.04469299316406 0 12179.828
149.07159423828125 0 4032.319
151.08688354492188 0 8066.132
152.08226013183594 0 28280.902
152.1438446044922 0 24125.053
152.1763916015625 0 3983.2795
155.0819549560547 0 46926.58
155.09329223632812 0 28209.715
155.11846923828125 0 40196.23
156.077392578125 0 37093.418
156.12193298339844 0 5915.679
156.56243896484375 0 4379.274
157.0765838623047 0 5222.6543
158.0603485107422 0 15114.788
158.0843963623047 0 113487.586
158.09616088867188 0 4796.414
159.0922088623047 0 523318.3
160.0760955810547 0 26112.61
160.08892822265625 0 9370.707
160.0955810546875 0 53544.457
163.0257568359375 0 3731.6477
164.07093811035156 0 9646.402
165.10275268554688 0 31646.193
166.0616455078125 0 13972.028
166.1056671142578 0 5961.577
168.11373901367188 0 15778.89
169.0766143798828 0 57790.215
169.08938598632812 0 7118.894
169.0977020263672 0 104202.33
169.1054229736328 0 8155.405
169.17025756835938 0 6227.7715
170.06057739257812 0 170683.12
170.07925415039062 0 7699.3335
170.10159301757812 0 7504.8706
171.06385803222656 0 25093.46
171.09307861328125 0 4834.1284
172.10861206054688 0 74161.164
173.45156860351562 0 13610.314
175.0980224609375 0 7369.801
176.0823974609375 0 25213.129
179.0823211669922 0 5633.8
180.0773162841797 0 10024.161
181.06118774414062 0 6276.806
181.09776306152344 0 215507.12
181.13426208496094 0 21115.016
182.10107421875 0 15332.58
182.1294403076172 0 37633.22
183.09225463867188 0 159801.31
183.1014404296875 0 7268.872
184.07595825195312 0 14925.828
184.09555053710938 0 20810.176
185.07151794433594 0 37661.85
185.10772705078125 0 5694.668
186.07937622070312 0 5268.1377
186.1243133544922 0 118114.24
187.0872039794922 0 987835.2
188.08184814453125 0 8069.2593
188.0905303955078 0 111190.48
189.09371948242188 0 5585.9883
193.1328887939453 0 4645.0835
195.14976501464844 0 13921.081
195.46792602539062 0 4199.9097
197.1653289794922 0 57549.9
198.10250854492188 0 7548.4243
198.16844177246094 0 7762.555
199.08706665039062 0 37072.043
199.10906982421875 0 7877.309
199.18084716796875 0 34543.895
200.08987426757812 0 5812.009
200.1400604248047 0 26552.207
201.10287475585938 0 88360.74
202.08676147460938 0 20983.51
202.10894775390625 0 112193.125
203.11375427246094 0 7745.506
204.07696533203125 0 12048.505
205.09771728515625 0 5666.3667 a 2
207.14979553222656 0 51353.83
208.1546630859375 0 5429.759
209.1288299560547 0 8918.301
211.08697509765625 0 24664.346
213.0769805908203 0 7637.709
214.109375 0 42344.957
216.11386108398438 0 11271.906
217.1086883544922 0 14891.007
219.0883026123047 0 7250.48
220.11990356445312 0 90473.81
221.10403442382812 0 115732.06
221.12278747558594 0 6808.0225
222.10667419433594 0 9394.17
222.1239471435547 0 7204.1943
224.11892700195312 0 21244.07
224.139892578125 0 5148.806
224.17640686035156 0 62148.953
225.0978546142578 0 6525.155
225.160400390625 0 11087.711
225.18038940429688 0 8315.584
226.1557159423828 0 1406759.4
227.08370971679688 0 5359.7646
227.1405029296875 0 7681.973
227.15895080566406 0 170510.66 y Ammonia loss 10
227.17684936523438 0 10991.877
228.11380004882812 0 103440.34 a Water loss 1
228.13424682617188 0 11024.911
228.16183471679688 0 14365.5205
229.09783935546875 0 111556.36 b Ammonia loss 4
229.11837768554688 0 16425.297
230.10284423828125 0 30623.05
231.08851623535156 0 48588.3
231.13583374023438 0 22396.572
232.11984252929688 0 6239.7407
233.1042022705078 0 14283.884
234.13497924804688 0 7601.397
235.11964416503906 0 201742.66
236.12283325195312 0 23739.8
236.13963317871094 0 9655.861
238.1304168701172 0 21697.193
239.08250427246094 0 10833.235
239.15101623535156 0 79080.57
240.15496826171875 0 10321.354
241.15573120117188 0 12052.871
242.1869659423828 0 25493.877
243.1824188232422 0 19785.414
244.10858154296875 0 7273.6934
244.1664276123047 0 6257.1113
245.17144775390625 0 6668.2812
245.1754608154297 0 6263.3955
245.18673706054688 0 446228.75 y 12
246.12449645996094 0 2715933.8 a 1
246.19009399414062 0 60310.336
247.1292266845703 0 620182.06
247.1925506591797 0 6479.733
248.11473083496094 0 154880.08
248.13165283203125 0 51341.92
249.0989227294922 0 197203.03
249.116943359375 0 16704.982
250.1021728515625 0 21389.295
250.12240600585938 0 13191.739
251.1294403076172 0 7558.378
251.1505126953125 0 7001.0874
252.1714630126953 0 26156.098
253.17433166503906 0 4741.88
255.07656860351562 0 6719.5337
256.1087341308594 0 70921.73 b Water loss 1
257.0926208496094 0 16829.86
257.1147155761719 0 22649.584
257.1615295410156 0 26784.984
258.07586669921875 0 14491.419
258.0992431640625 0 339805.75
259.0798645019531 0 5516.078
259.1025695800781 0 49287.285
259.13018798828125 0 12496.638
260.107177734375 0 26190.941
260.1977233886719 0 20399.387
262.15533447265625 0 11328.981
263.11383056640625 0 12441.438
264.1356506347656 0 10019.675
265.1410217285156 0 23355.547
265.16656494140625 0 14311.818
266.1253662109375 0 177414
267.12872314453125 0 22008.059
267.1448059082031 0 7762.5034
270.1818542480469 0 66167.695
271.1846923828125 0 16128.07 y 9
273.1098327636719 0 12760.724
273.1356201171875 0 11203.482
274.11932373046875 0 1647415.8 b 1
275.1036682128906 0 57140.742
275.12481689453125 0 704707.1
276.10833740234375 0 26697.016
276.1277160644531 0 84217.54
277.1334533691406 0 14701.645
279.1817626953125 0 19559.48
283.15191650390625 0 32652.26
283.1772766113281 0 225473.25
284.1805114746094 0 31925.945
285.1106872558594 0 14630.189
285.1564025878906 0 68566.94
285.1817321777344 0 6745.541
286.1598815917969 0 14801.721
290.11474609375 0 7176.827
290.1850280761719 0 64541.832
290.6862487792969 0 22121.176
291.15740966796875 0 20649.988
292.1299743652344 0 183955.62
293.133544921875 0 25510.998
294.1204833984375 0 84699.06
295.1250305175781 0 9712.438
297.1929931640625 0 190233.25
298.1966247558594 0 39406.207
300.2039794921875 0 8582.431
301.1283874511719 0 6115.1646
303.12103271484375 0 44529.656
303.21820068359375 0 5623.856
305.12921142578125 0 47512.19
305.6658630371094 0 13295.827
306.13250732421875 0 9337.615
308.1256408691406 0 8132.6724
309.1671142578125 0 8258.72
311.24505615234375 0 42363.547
312.24871826171875 0 7756.7427
313.2240295410156 0 10226.582
314.67071533203125 0 6045.4463
318.194091796875 0 11904.788
319.17791748046875 0 6397.0493
320.1252746582031 0 49220.637
320.1519470214844 0 7928.6553
321.1293640136719 0 15936.065
321.22943115234375 0 232441.69
322.1560363769531 0 71418.16
322.2326965332031 0 45583.86
323.15924072265625 0 15261.236
323.2347717285156 0 6903.809
324.2165222167969 0 19233.986
324.71795654296875 0 5589.138
325.2220458984375 0 5249.0967
326.2196350097656 0 47582.566
332.7294616699219 0 26728.617
333.1235656738281 0 14432.355
333.2313232421875 0 14610.018
333.7332763671875 0 7653.9946
335.2377624511719 0 11353.572 y 8
336.14605712890625 0 8532.301
336.2039794921875 0 289833.1
337.2067565917969 0 51131.3
337.72216796875 0 56847.055
338.1629943847656 0 9333.754
338.223388671875 0 22023.395
339.2400207519531 0 229689.67
340.15289306640625 0 21002.762
340.2432861328125 0 35333.688
341.2192687988281 0 6086.233
342.1678466796875 0 9682.417
343.21343994140625 0 6056.0127
346.7272033691406 0 212291.3
347.22845458984375 0 89695.195
347.7292175292969 0 19279.113
348.1461486816406 0 7925.216
349.7102966308594 0 12056.701
350.1512756347656 0 24106.982
350.2115173339844 0 6491.488
351.2400817871094 0 29466.773
353.2188720703125 0 15396.864
354.171875 0 12280.731
354.2144775390625 0 419646.88
354.6564636230469 0 8092.513
355.2177734375 0 61179.363
355.2704162597656 0 7398.4497
355.73291015625 0 27803.031
356.17333984375 0 7457.705
357.1798095703125 0 13665.792
357.25079345703125 0 44893.76
358.16326904296875 0 52778.547
358.2531433105469 0 8792.142
358.71514892578125 0 22694.41
359.16632080078125 0 9234.92
359.2159423828125 0 9228.004
362.6706237792969 0 9392.98
363.7481994628906 0 12359.7295 y 7
365.2541809082031 0 7155.263
366.1571350097656 0 28124.268
367.1623840332031 0 22410.143
368.1481628417969 0 34125.93
368.1725158691406 0 17430.443
369.1521911621094 0 9912.5205
369.1775817871094 0 8173.884
370.2459716796875 0 11646.16
373.2818298339844 0 95847.75 y 11
374.2586975097656 0 7839.5117
374.2848205566406 0 20762.229
376.66680908203125 0 8558.081
380.2291259765625 0 18118.654
382.2076416015625 0 15183.041
382.2816467285156 0 21739.969
383.2662658691406 0 18179.682
384.1678466796875 0 53207.24
385.17364501953125 0 151975.75
385.1997985839844 0 56716.3
386.1578063964844 0 84845.125
386.2010498046875 0 15352.845
387.16082763671875 0 20263.428
388.16351318359375 0 7292.1743
389.2701416015625 0 6281.5317
392.162109375 0 15188.744
392.2666015625 0 59727.902
393.1685485839844 0 12016.374
393.271728515625 0 14417.631
394.1519775390625 0 105970.32
394.2812194824219 0 11009.546
395.15460205078125 0 23317.27
396.2617492675781 0 25314.746
397.2431945800781 0 9455.714
397.6722412109375 0 9755.078
398.2414245605469 0 10493.849
401.1943664550781 0 19814.297
402.2259521484375 0 44825.773
403.18475341796875 0 141546.62
403.22723388671875 0 10435.997
403.2695617675781 0 28029.62
403.76861572265625 0 20392.943
404.1878356933594 0 31029.691
404.2680358886719 0 7758.9824
406.1861877441406 0 68078.18
406.2511901855469 0 51693.54
406.6879577636719 0 31167.176
406.72357177734375 0 6069.5327
406.7532958984375 0 29998.45
407.1920471191406 0 12102.561
407.51885986328125 0 6344.759
408.2220764160156 0 8628.369
410.2774658203125 0 122318.6
411.17864990234375 0 192810.92 b Water loss 5
411.2803039550781 0 21063.527
412.1603698730469 0 18577.014
412.1824035644531 0 30896.164
412.27484130859375 0 554760.8 y 6
412.77618408203125 0 243164.78
413.1649475097656 0 6892.3257
413.278076171875 0 65786.125
413.7796936035156 0 8732.272
415.2572021484375 0 51032.348
415.75921630859375 0 25679.229
419.8998718261719 0 21427.469
420.1834411621094 0 82191.53 b 5
420.2355041503906 0 35500.918
420.68560791015625 0 38731.918
421.18548583984375 0 8083.7734
422.2774353027344 0 47370.312
422.31011962890625 0 10708.876
423.27685546875 0 26093.96
424.3294372558594 0 14548.67
424.712890625 0 12093.402
425.2878723144531 0 18164.248
425.70782470703125 0 5872.761
427.2444152832031 0 31980.307
427.7464294433594 0 24108.012
428.2488098144531 0 6159.7964
428.2876281738281 0 9592.296
429.18939208984375 0 666157.9
430.1921081542969 0 172867.64
431.1805419921875 0 86999.875
432.1836853027344 0 26710.416
433.25567626953125 0 11618.637
437.1831359863281 0 128733.88 b 2
438.1865234375 0 31510.328
438.55767822265625 0 18013.38
438.8912658691406 0 18129.271
439.3043212890625 0 17853.748
444.23321533203125 0 11466.675
444.5667724609375 0 10729.029
445.2688903808594 0 13513.377
446.2198791503906 0 11475.445
450.2823181152344 0 10465.284
451.2268981933594 0 26646.562
451.26800537109375 0 39401.75
451.7270812988281 0 14244.078
451.9196472167969 0 12242.415
452.324951171875 0 15136.305
454.2099914550781 0 30542.754
455.2157897949219 0 10674.726
455.253173828125 0 42711.8
456.2566223144531 0 13795.997
457.183837890625 0 21628.625
457.5955810546875 0 12081.68
457.9277038574219 0 8694.713 y Ammonia loss 2
460.23193359375 0 25735.11
461.1826171875 0 12118.809
461.2418518066406 0 7658.0166
463.27801513671875 0 12810.578
463.5992431640625 0 140439.72 y 2
463.9337463378906 0 91437.46
464.2683410644531 0 31400.72
464.6015625 0 12638.36
466.9214782714844 0 8286.568
467.26031494140625 0 9578.594
467.2991027832031 0 61346.848
468.29754638671875 0 31254.605
468.71234130859375 0 8547.56 b 6
469.7890625 0 11200.057
470.25067138671875 0 17082.436
470.3349304199219 0 413731.84 y 10
470.58099365234375 0 22590.375
470.9086608886719 0 6506.874
471.3381042480469 0 113650.83
471.79949951171875 0 40300.55
472.2992858886719 0 20373.889
472.3403625488281 0 19130.426
472.80352783203125 0 7167.2866
473.2439880371094 0 138893.08 b 11
473.5778503417969 0 180771.47
473.9122009277344 0 88357.17
474.2660827636719 0 17248.443
474.5819396972656 0 11361.611
474.7809143066406 0 33626.98
475.280029296875 0 14776.197
475.7847900390625 0 11649.241
475.9239807128906 0 18085.26
476.2543640136719 0 44168.652
476.5879821777344 0 44088.97
476.9215087890625 0 21886.346
478.2090759277344 0 31570.867
479.2145080566406 0 11119.31
480.8042907714844 0 133343.08 y 5
481.30621337890625 0 65388.355
481.8053894042969 0 20047.326
481.9276428222656 0 33772.074
482.2629089355469 0 56340.824
482.3094482421875 0 42847.953
482.5958251953125 0 17464.348
482.7625732421875 0 30772.877
482.9305114746094 0 6825.8857
483.2604675292969 0 12977.503
483.7864074707031 0 93440.96
484.2870788574219 0 36042.438
484.7891540527344 0 14560.746
491.2732238769531 0 56511.215
491.77899169921875 0 8424.475
492.2769470214844 0 11108.826
492.7900695800781 0 9358.016
494.26556396484375 0 6607.859
495.93280029296875 0 8743.5205
496.265869140625 0 13540.742
497.2208251953125 0 25086.238 b 7
497.7239074707031 0 11323.206
499.25970458984375 0 14553.08
501.6067199707031 0 98825.87
501.94085693359375 0 119302.44
502.2747497558594 0 49225.74
502.6106872558594 0 10582.813
502.94207763671875 0 7919.054
503.2158508300781 0 40188.66
504.934814453125 0 154978.38 b Water loss 12
505.2677917480469 0 153604.84 b Ammonia loss 12
505.6026916503906 0 88921.43
505.7338562011719 0 8554.471
505.93609619140625 0 37820.62
506.26727294921875 0 12097.192
508.7579345703125 0 22672.393
509.2625427246094 0 13246.445
510.3001708984375 0 7014.767
510.37762451171875 0 14220.928
510.9382019042969 0 769010.06 b 12
511.27239990234375 0 776591.44
511.6067810058594 0 371375.94
511.9410095214844 0 126231.94
512.2753295898438 0 40242.703
513.9496459960938 0 6983.4536
514.2776489257812 0 12814.1875
514.6115112304688 0 10821.777
516.9414672851562 0 116424.086
517.2758178710938 0 150579.73
517.6099243164062 0 53545.152
517.9426879882812 0 18803.57
519.6194458007812 0 8721.36
519.9517211914062 0 32703.05 y Ammonia loss 1
520.2421264648438 0 13528.229
520.2855834960938 0 45870.504
520.3615112304688 0 19612.55
520.62060546875 0 12034.408
520.9528198242188 0 11685.277
521.2267456054688 0 38020.742
522.2622680664062 0 9948.439
523.3624877929688 0 16668.904
524.8131103515625 0 8606.761
525.3089599609375 0 13844.662
525.6256713867188 0 152426.69 y 1
525.8045043945312 0 12311.734
525.9596557617188 0 138174.84
526.2936401367188 0 64638.957
526.6278686523438 0 12384.66
528.2333374023438 0 22472.568
529.2237548828125 0 21795.92
529.8057861328125 0 20094.707
530.2991943359375 0 29436.701
530.7991943359375 0 18802.219
531.2116088867188 0 25269.398
531.3013305664062 0 9471.802
535.2919921875 0 8071.2036
535.613037109375 0 8054.752
536.2520141601562 0 8404.228
537.2905883789062 0 8499.489
538.2532958984375 0 378765.97
538.3727416992188 0 26886.879
538.8099365234375 0 49957.125
538.9566650390625 0 8472.692
539.2554321289062 0 102735.94
539.3036499023438 0 106694.94
539.3701171875 0 7952.215
539.633056640625 0 27936.771
539.8056640625 0 43154.414
539.9684448242188 0 22648.457
540.2586669921875 0 17846.023
540.2996215820312 0 15912.753
540.3289794921875 0 14582.265
540.8250122070312 0 10412.946 y Ammonia loss 4
541.3292236328125 0 9069.079
541.3724975585938 0 143052.06 y 9
542.2521362304688 0 20565.963
542.3756103515625 0 42819.65
542.9629516601562 0 21222.459
543.2950439453125 0 8641.029
544.6318359375 0 14093.142
544.75634765625 0 10647.947
545.30126953125 0 9552.956
547.816162109375 0 57665.32
548.2356567382812 0 16030.48
548.3155517578125 0 32796.484
548.6331176757812 0 224686.25 Precursor Water loss
548.8162231445312 0 13427.819
548.9658813476562 0 279822.84 Precursor Ammonia loss
549.2232055664062 0 21666.045
549.2965698242188 0 99974.76
549.3341064453125 0 215202.64 y 4
549.6321411132812 0 101863.12
549.8352661132812 0 146274.16
549.9641723632812 0 30106.924
550.2232666015625 0 8883.243
550.29443359375 0 10273.729
550.337158203125 0 57124.098
550.8380737304688 0 9267.578
553.2198486328125 0 13156.506
553.2649536132812 0 10118.671
553.9628295898438 0 17452.389
554.2999267578125 0 51903.438
554.636474609375 0 5188832.5 Precursor
554.970703125 0 5211137.5
555.3048706054688 0 2844816.5
555.6390991210938 0 1007323.75
555.7867431640625 0 11455.983
555.9730224609375 0 298427.53
557.296142578125 0 16956.984
561.2681274414062 0 44510.52 b 8
561.7697143554688 0 31639.406
562.2706298828125 0 17406.537
565.2413330078125 0 40734.516 b 3
565.3013305664062 0 7880.2285
566.2481079101562 0 460341.06
567.2508544921875 0 138557.02
567.3972778320312 0 17466.105
568.2539672851562 0 29160.809
568.4046630859375 0 9577.159
570.2470092773438 0 80165.59
571.2493286132812 0 22603.822
572.8052978515625 0 205006.2
573.3062744140625 0 166278.88
573.8082885742188 0 61065.27
577.3834228515625 0 89621.13
578.3846435546875 0 30400.09
579.362548828125 0 289011.2
580.3658447265625 0 77155.484
581.3677978515625 0 24679.145
587.2733154296875 0 12143.533
587.7821044921875 0 46128.69 b Water loss 9
588.2819213867188 0 33240.18 b Ammonia loss 9
588.3284301757812 0 7819.261
588.78466796875 0 11627.706
589.3261108398438 0 7333.1973
595.3538818359375 0 6849.5547
595.3941040039062 0 86090.54
595.8485107421875 0 18355.012
596.3461303710938 0 10115.279
596.3975219726562 0 29105.18
596.7864379882812 0 52245.383 b 9
596.8406982421875 0 8374.338
597.28857421875 0 17350.695
597.7889404296875 0 14333.792
598.2420654296875 0 32982.086
599.2451171875 0 14913.101
602.8228149414062 0 21735.152
603.3212280273438 0 9924.536
604.3582763671875 0 113661.35
604.8541259765625 0 169612.17 y Ammonia loss 3
605.3541259765625 0 99580.64
605.8549194335938 0 36515.004
606.350341796875 0 10831.089
606.8350830078125 0 10843.041
607.3406372070312 0 13552.047
610.3213500976562 0 47531.27
611.324951171875 0 25984.693
611.829833984375 0 27778.275
612.3312377929688 0 22414.871
612.8302001953125 0 10839.595
613.3629150390625 0 145948.61 y 3
613.8646240234375 0 120789.516
614.3652954101562 0 49616.383
615.2667236328125 0 17518.787
615.3501586914062 0 95035.2
615.8517456054688 0 59501.105
616.3523559570312 0 25244.889
616.85107421875 0 9211.406
618.8534545898438 0 8060.6797
620.3421630859375 0 194201.88
620.843017578125 0 111714.875
621.3432006835938 0 71957.35
621.4059448242188 0 11503.633
621.8419189453125 0 18097.955
623.3876953125 0 21567.426
628.3321533203125 0 18498.436
629.3472900390625 0 444729.9
629.8487548828125 0 321128.66
630.3499755859375 0 122822.67
630.8521728515625 0 24392.05
632.294189453125 0 14251.509
638.3528442382812 0 47010.41
638.8550415039062 0 27426.623
639.3544311523438 0 13602.242
643.2633056640625 0 8452.839
645.3123779296875 0 29877.223 b 10
645.8168334960938 0 17834.59
647.4287719726562 0 18779.059
653.827392578125 0 28112.904
654.3301391601562 0 16764.715
654.829345703125 0 15516.123
656.2946166992188 0 12120.752
657.2787475585938 0 31875.518
657.3306884765625 0 7241.3374
657.8353881835938 0 9215.283
658.309326171875 0 21175.812
659.3138427734375 0 16143.782
664.4518432617188 0 93656.8
665.4552612304688 0 34791.574
665.8462524414062 0 13914.127
666.4573974609375 0 8561.942
667.3516845703125 0 8397.533
668.3270263671875 0 14675.916
669.46728515625 0 49972.86 y 8
670.4694213867188 0 25304.402
671.8933715820312 0 13538.086
672.3947143554688 0 18244.207
672.8926391601562 0 7015.3823
674.3052368164062 0 102427.914
674.4364013671875 0 309593.94
675.30810546875 0 44854.688
675.4385986328125 0 134592.75
676.4420166015625 0 28661.389
677.3761596679688 0 55965.723
677.8779296875 0 42599.02
678.3755493164062 0 25352.832
684.2913208007812 0 32372.115 b Water loss 4
685.2859497070312 0 11130.45 b Ammonia loss 4
685.888916015625 0 95571.67
686.3883666992188 0 89847.12 y Ammonia loss 2
686.8873291015625 0 43059.76
687.3886108398438 0 18164.447
692.4465942382812 0 328385.94
693.4489135742188 0 127548.305
694.3912963867188 0 8754.574
694.4518432617188 0 30055.244
694.8948364257812 0 1607548 y 2
695.3963623046875 0 1235247.9
695.8973388671875 0 592262.6
696.3988647460938 0 161713.06
696.8988647460938 0 34979.93
698.4110107421875 0 31493.535
700.3671264648438 0 14635.796 b Water loss 11
700.8524780273438 0 17641.752 b Ammonia loss 11
701.3568115234375 0 7201.8193
701.8598022460938 0 8516.431
702.3009643554688 0 180251.84 b 4
703.3040771484375 0 82192.77
704.3037109375 0 21785.299
707.3063354492188 0 47573.438
708.3106079101562 0 24378.953
708.48193359375 0 10968.713
709.3613891601562 0 354174.62 b 11
709.8623657226562 0 281645.06
710.3638916015625 0 121548.15
710.458251953125 0 46436.06
710.8651123046875 0 33839.93
711.3672485351562 0 16716.002
711.4602661132812 0 16134.462
712.4742431640625 0 8199.533
713.3792114257812 0 7646.2065
714.382568359375 0 13134.001
716.421630859375 0 56076.598
717.4243774414062 0 24241.879
718.3670043945312 0 6861.422
720.3124389648438 0 11073.811
720.37109375 0 16066.729
721.3539428710938 0 25200.49
722.3877563476562 0 8851.758
722.88671875 0 16066.782
723.3818359375 0 13169.236
724.33251953125 0 28407.998
725.3347778320312 0 12194.5205
726.4882202148438 0 124112.125 y 7
727.4904174804688 0 45331
735.3007202148438 0 45867.145
736.3017578125 0 13354.381
737.3521728515625 0 13515.261
738.3805541992188 0 14785.625
739.371337890625 0 11471.859
740.3837890625 0 8378.789
751.9066162109375 0 15736.191
752.3280639648438 0 37412.61
752.4075927734375 0 21840.018
753.330810546875 0 21216.791
756.898681640625 0 35108.445 b Water loss 12
757.3993530273438 0 42478.215 b Ammonia loss 12
757.8993530273438 0 37916.36
758.408935546875 0 14570.23
765.90380859375 0 115548.91 b 12
766.3302001953125 0 9325.921
766.4047241210938 0 125282.266
766.9063720703125 0 52883.945
767.4124755859375 0 19452.518
774.90771484375 0 56959.582
775.4103393554688 0 54673.97
775.912353515625 0 18018.463
776.4122314453125 0 10272.81
777.5390014648438 0 11471.378
779.4266967773438 0 7881.037 y Ammonia loss 1
780.3218383789062 0 7160.2275
787.519287109375 0 9812.848
787.9353637695312 0 41356.527 y 1
788.4358520507812 0 42246.688
788.9366455078125 0 17071.73
789.4406127929688 0 7528.2925
792.9321899414062 0 9888.913
793.3544921875 0 14764.611
794.34130859375 0 24559.385
795.3526611328125 0 15368.612
801.5091552734375 0 18688.69
805.5320434570312 0 62068.574
806.5297241210938 0 31244.602
807.5337524414062 0 14284.98
811.364013671875 0 194022.9
811.494873046875 0 57813.867
812.3682250976562 0 84286.78
812.4966430664062 0 28232.75
813.3712158203125 0 22167.227
814.3839721679688 0 8896.208
821.34814453125 0 28428.12 b Water loss 5
822.344482421875 0 19339.172 b Ammonia loss 5
822.529296875 0 8178.104
823.3482666015625 0 16115.855
823.541259765625 0 754977.9 y 6
824.544189453125 0 343403.12
825.5470581054688 0 91199.336
826.5492553710938 0 12052.726
829.504150390625 0 56348.055
830.5095825195312 0 27150.928
831.513427734375 0 10805.503
839.359375 0 408180.38 b 5
840.362548828125 0 208921.62
841.3661499023438 0 60394.07
842.3718872070312 0 8707.856
847.5136108398438 0 11255.178
848.4170532226562 0 68237.8
849.4208984375 0 37410.03
850.4215087890625 0 7036.1323
853.4747924804688 0 35218.008
854.481201171875 0 22093.293
866.4033813476562 0 9240.018
891.4581298828125 0 23630.797
892.4611206054688 0 10688.214
901.4439697265625 0 139004.17
902.446044921875 0 85275.766
903.449951171875 0 18739.844
919.4539184570312 0 70718.05
920.4570922851562 0 52871.76
921.4625244140625 0 9000.983
942.59033203125 0 41057.195
943.5923461914062 0 35996.766
944.6041870117188 0 7015.4624
948.55712890625 0 14963.607
960.5996704101562 0 138073.6 y 5
961.4990844726562 0 9497.91
961.6018676757812 0 70358.09
962.6067504882812 0 18479.209
963.6104125976562 0 7141.216
964.5128173828125 0 11568.115
966.5637817382812 0 33662.8
967.5614624023438 0 18545.326
993.4301147460938 0 12225.598 b 7
994.4440307617188 0 8757.917
1010.4661865234375 0 16182.278
1011.4563598632812 0 10128.078
1016.5039672851562 0 7232.8823
1032.5382080078125 0 22480.008
1033.5374755859375 0 31798.658
1034.5482177734375 0 11320.731
1087.5211181640625 0 13459.846
1097.65869140625 0 48423.47 y 4
1098.6607666015625 0 29574.95
1099.6622314453125 0 16611.105
1121.5279541015625 0 33712.953 b 8
1122.5330810546875 0 26687.848
1144.6029052734375 0 18559.797
1145.60888671875 0 11743.084
1146.6087646484375 0 10935.851
1164.5703125 0 8887.645
1174.553955078125 0 74800.13 b Water loss 9
1175.555908203125 0 59817.574 b Ammonia loss 9
1176.5562744140625 0 13975.772
1192.5648193359375 0 28494.064 b 9
1193.566650390625 0 23783.994
1194.57861328125 0 8383.508
2015.07666015625 0 7175.4644
2187.9150390625 0 5724.7954
2709.309814453125 0 7062.3228
2709.870849609375 0 6881.481
2925.899169921875 0 6923.8784
3255.452880859375 0 6993.9653
3299.4013671875 0 6799.4385

Spectrum Details

|  |  |
| --- | --- |
| Matched peaks? Matched peaks The total absolute number of peaks matched. Additionally in brackets the total fraction of peaks matched and the total number of peaks is shown. | 69 (8.25% of 836) |
| FDR? FDR The false discovery rate estimated for this peptide. It is calculated by matching all theoretical fragments with a non-integer shift with the raw peaks for this spectrum. This is done with 40 different shifts. The resulting percentage is the average number of annotated peaks over the number of annotated peaks with the correct spectrum. | 3.73% |
| Satellite FDR? Satellite FDR See the FDR for details on its calculation. This satellite ion specific FDR only contains the satellite ions (d/w) for I/L/J positions. | - |
| PSM Score? PSM Score The PSM Score as given by Hecklib to this annotated spectrum. It is shown with three significant figures. | 360 |

#### Spectrum 5459? Spectrum 5459 The raw spectrum of this peptide as annotated by Hecklib. The fragments are coloured according to ion type (see legend). Any peaks with a star '\*' as text can be hovered over to see the full details, first the ion type second the mass shift type. By hovering over the amino acids in the peptide or ions in the legend the corresponding peaks are highlighted. By toggling the 'Unassigned' label you can turn the background (unassigned) peaks on or off in the plot. By updating the slider in the Ion legend you can update the spectrum to only show the top X% of the peaks with labels. The top X% means any peak that is within X% of the highest intensity. By dragging in the spectrum you can zoom in to a specific part of the spectrum and use 'Zoom Out' to get back to the original zoom level. The annotation of the spectrum is based on the given sequence in the peptides file and is done with different software so inconsistencies are likely. The peaks are annotated based on the given sequence, with 20 ppm tolerance.

Copy Data

##### Spectrum 5459 (TSV)

###### Preview

```
Loading example...
```

*Click on the button to copy the data to your clipboard.*

Mz MinMz MaxIntensity Max

WidthHeightPeptide font sizePeptide stroke widthSpectrum font sizeSpectrum stroke widthCompact peptide

Ion legend

wxyz

abcd

OtherUnassignedIon^Charge^_Position_Show for top:%

SWYQHHPGKAPKJJ

01.40e+52.80e+54.19e+55.59e+5

Zoom Out

y^+1^_1_y^+1^_2_z^+2^_7_z^+1^_3_y^+1^_3_y^+2^_8_c^+2^_6_w^+1^_4_c^+1^_3_y^+3^_12_y^+1^_4_c^+3^_12_y^+2^_9_c^+2^_8_c^+3^_13_z^+3^_13_z^+1^_5_y^+3^_13_z^+2^_10_y^+1^_5_y^+2^_10_c^+1^_4_c^+1^_4_w^+2^_11_c^+1^_4_w^+1^_6_c^+2^_10_y^+2^_11_z^+2^_11_y^+2^_11_z^+1^_6_c^+2^_11_y^+1^_6_y^+2^_12_z^+2^_12_y^+2^_12_c^+1^_5_c^+2^_12_z^+1^_7_c^+2^_12_c^+1^_5_y^+1^_7_c^+2^_13_c^+2^_13_y^+2^_13_z^+2^_13_w^+1^_8_y^+2^_13_y^+1^_8_c^+1^_6_z^+1^_9_c^+1^_7_y^+1^_9_c^+1^_8_y^+1^_10_z^+1^_10_y^+1^_10_c^+1^_9_c^+1^_9_y^+1^_11_z^+1^_11_y^+1^_11_c^+1^_11_z^+1^_12_y^+1^_12_c^+1^_12_c^+1^_13_z^+1^_13_

047194214131885

Fragment Matches Table

Show background peaks

| Position | Ion type | Intensity | mz Theoretical | mz Error (Th) | mz Error (ppm) | Charge | Series Number |
| --- | --- | --- | --- | --- | --- | --- | --- |
| - | - | 640.6 | 124.7 | - | - | 0 | - |
| - | - | 528.8 | 124.8 | - | - | 0 | - |
| - | - | 515.1 | 125.7 | - | - | 0 | - |
| - | - | 463.2 | 127.5 | - | - | 0 | - |
| - | - | 510.7 | 128.8 | - | - | 0 | - |
| - | - | 1918 | 129.1 | - | - | 0 | - |
| - | - | 1052 | 130.1 | - | - | 0 | - |
| - | - | 1845 | 132.1 | - | - | 0 | - |
| 14 | y | 3627 | 132.1 | 0.0002372 | 1.796 | +1 | 1 |
| - | - | 1253 | 136.1 | - | - | 0 | - |
| - | - | 527.4 | 141.5 | - | - | 0 | - |
| - | - | 1107 | 146.1 | - | - | 0 | - |
| - | - | 4361 | 153.1 | - | - | 0 | - |
| - | - | 622.3 | 164 | - | - | 0 | - |
| - | - | 633.2 | 168.1 | - | - | 0 | - |
| - | - | 616 | 172.3 | - | - | 0 | - |
| - | - | 1928 | 173.4 | - | - | 0 | - |
| - | - | 605.8 | 175.8 | - | - | 0 | - |
| - | - | 698.8 | 181.7 | - | - | 0 | - |
| - | - | 1639 | 186.1 | - | - | 0 | - |
| - | - | 3355 | 187.1 | - | - | 0 | - |
| - | - | 704.7 | 199.2 | - | - | 0 | - |
| - | - | 674.4 | 209.5 | - | - | 0 | - |
| - | - | 692.1 | 209.9 | - | - | 0 | - |
| - | - | 821.8 | 210.1 | - | - | 0 | - |
| - | - | 1073 | 214.2 | - | - | 0 | - |
| - | - | 1001 | 215.1 | - | - | 0 | - |
| - | - | 2237 | 223.1 | - | - | 0 | - |
| - | - | 3215 | 226.2 | - | - | 0 | - |
| - | - | 731.1 | 227.2 | - | - | 0 | - |
| - | - | 975 | 228.1 | - | - | 0 | - |
| - | - | 622.1 | 243.2 | - | - | 0 | - |
| - | - | 5270 | 245.1 | - | - | 0 | - |
| 13 | y | 6944 | 245.2 | 0.0002186 | 0.8917 | +1 | 2 |
| - | - | 2.101E+04 | 246.1 | - | - | 0 | - |
| - | - | 899.1 | 246.2 | - | - | 0 | - |
| - | - | 2814 | 247.1 | - | - | 0 | - |
| - | - | 1651 | 255.1 | - | - | 0 | - |
| - | - | 1729 | 258.1 | - | - | 0 | - |
| - | - | 830.5 | 267.1 | - | - | 0 | - |
| - | - | 1629 | 273.1 | - | - | 0 | - |
| - | - | 2.566E+04 | 274.1 | - | - | 0 | - |
| - | - | 4196 | 275.1 | - | - | 0 | - |
| - | - | 3090 | 281.2 | - | - | 0 | - |
| - | - | 849.2 | 282.2 | - | - | 0 | - |
| - | - | 3038 | 292.1 | - | - | 0 | - |
| - | - | 1332 | 297.2 | - | - | 0 | - |
| - | - | 1776 | 301.2 | - | - | 0 | - |
| - | - | 3342 | 309.2 | - | - | 0 | - |
| - | - | 2337 | 314.2 | - | - | 0 | - |
| - | - | 1719 | 338.1 | - | - | 0 | - |
| - | - | 902.6 | 338.2 | - | - | 0 | - |
| - | - | 3068 | 346.7 | - | - | 0 | - |
| 8 | z | 780 | 347.2 | 0.00475 | 13.68 | +2 | 7 |
| - | - | 2323 | 349.1 | - | - | 0 | - |
| - | - | 1026 | 350.3 | - | - | 0 | - |
| - | - | 788.1 | 355.1 | - | - | 0 | - |
| 12 | z | 1E+04 | 357.3 | 0.0001821 | 0.5097 | +1 | 3 |
| - | - | 3981 | 358.3 | - | - | 0 | - |
| - | - | 1257 | 371.1 | - | - | 0 | - |
| 12 | y | 2882 | 373.3 | 0.0001399 | 0.3747 | +1 | 3 |
| - | - | 1709 | 382.3 | - | - | 0 | - |
| - | - | 1783 | 394.3 | - | - | 0 | - |
| - | - | 1619 | 399.3 | - | - | 0 | - |
| 7 | y | 1.266E+04 | 412.3 | 0.000501 | 1.215 | +2 | 8 |
| - | - | 6346 | 412.8 | - | - | 0 | - |
| - | - | 3109 | 413.2 | - | - | 0 | - |
| - | - | 1280 | 413.3 | - | - | 0 | - |
| - | - | 980.6 | 414.2 | - | - | 0 | - |
| 6 | c | 1801 | 420.2 | 0.0008122 | 1.933 | +2 | 6 |
| - | - | 939.2 | 420.7 | - | - | 0 | - |
| - | - | 1569 | 423.2 | - | - | 0 | - |
| 11 | w | 5672 | 427.3 | 0.0004649 | 1.088 | +1 | 4 |
| - | - | 901.1 | 428.3 | - | - | 0 | - |
| - | - | 2249 | 428.3 | - | - | 0 | - |
| - | - | 1215 | 429.2 | - | - | 0 | - |
| - | - | 834.4 | 434.2 | - | - | 0 | - |
| - | - | 1591 | 435.2 | - | - | 0 | - |
| - | - | 1242 | 436.2 | - | - | 0 | - |
| - | - | 4531 | 437.2 | - | - | 0 | - |
| - | - | 1053 | 438.2 | - | - | 0 | - |
| - | - | 1240 | 439.2 | - | - | 0 | - |
| - | - | 1006 | 451.2 | - | - | 0 | - |
| - | - | 4933 | 453.2 | - | - | 0 | - |
| 3 | c | 9969 | 454.2 | 0.0001825 | 0.4017 | +1 | 3 |
| - | - | 2747 | 455.2 | - | - | 0 | - |
| 3 | y | 3674 | 463.6 | 0.0009047 | 1.951 | +3 | 12 |
| - | - | 2910 | 463.9 | - | - | 0 | - |
| - | - | 3579 | 465.2 | - | - | 0 | - |
| - | - | 1.055E+04 | 468.3 | - | - | 0 | - |
| - | - | 3329 | 469.3 | - | - | 0 | - |
| - | - | 2378 | 469.3 | - | - | 0 | - |
| - | - | 1628 | 470.3 | - | - | 0 | - |
| 11 | y | 4662 | 470.3 | 0.0001969 | 0.4185 | +1 | 4 |
| - | - | 1043 | 472.3 | - | - | 0 | - |
| 12 | c | 4621 | 473.2 | 0.0004568 | 0.9652 | +3 | 12 |
| - | - | 4526 | 473.6 | - | - | 0 | - |
| - | - | 1991 | 473.9 | - | - | 0 | - |
| - | - | 4072 | 477.2 | - | - | 0 | - |
| - | - | 1607 | 478.2 | - | - | 0 | - |
| - | - | 2054 | 480.2 | - | - | 0 | - |
| 6 | y | 1961 | 480.8 | 0.001624 | 3.377 | +2 | 9 |
| - | - | 1844 | 481.3 | - | - | 0 | - |
| - | - | 3145 | 482.3 | - | - | 0 | - |
| - | - | 1156 | 483.3 | - | - | 0 | - |
| 8 | c | 764.8 | 497.2 | 0.0007163 | 1.441 | +2 | 8 |
| - | - | 857.7 | 498.3 | - | - | 0 | - |
| - | - | 1269 | 501.6 | - | - | 0 | - |
| - | - | 999.7 | 501.9 | - | - | 0 | - |
| - | - | 6023 | 504.9 | - | - | 0 | - |
| - | - | 6193 | 505.3 | - | - | 0 | - |
| - | - | 1706 | 505.6 | - | - | 0 | - |
| - | - | 873.6 | 507.3 | - | - | 0 | - |
| 13 | c | 2.302E+04 | 510.9 | 0.0005319 | 1.041 | +3 | 13 |
| - | - | 1.822E+04 | 511.3 | - | - | 0 | - |
| - | - | 1.003E+04 | 511.6 | - | - | 0 | - |
| - | - | 3877 | 511.9 | - | - | 0 | - |
| - | - | 1122 | 512.3 | - | - | 0 | - |
| - | - | 5297 | 516.9 | - | - | 0 | - |
| - | - | 2998 | 517.3 | - | - | 0 | - |
| - | - | 1006 | 517.6 | - | - | 0 | - |
| - | - | 1169 | 517.9 | - | - | 0 | - |
| 2 | z | 1617 | 520.3 | 0.003768 | 7.243 | +3 | 13 |
| - | - | 2203 | 523.2 | - | - | 0 | - |
| 10 | z | 4.114E+04 | 525.4 | 0.0005144 | 0.9792 | +1 | 5 |
| 2 | y | 3434 | 525.6 | 0.0001423 | 0.2708 | +3 | 13 |
| - | - | 4485 | 526 | - | - | 0 | - |
| - | - | 1919 | 526.3 | - | - | 0 | - |
| - | - | 1.298E+04 | 526.4 | - | - | 0 | - |
| - | - | 2059 | 527.4 | - | - | 0 | - |
| - | - | 1008 | 536.2 | - | - | 0 | - |
| - | - | 1568 | 538.2 | - | - | 0 | - |
| - | - | 1770 | 538.3 | - | - | 0 | - |
| - | - | 2830 | 539.3 | - | - | 0 | - |
| - | - | 1172 | 540 | - | - | 0 | - |
| 5 | z | 1990 | 541.3 | 0.0007395 | 1.366 | +2 | 10 |
| 10 | y | 5638 | 541.4 | 0.001016 | 1.877 | +1 | 5 |
| - | - | 1166 | 541.8 | - | - | 0 | - |
| - | - | 1542 | 542.4 | - | - | 0 | - |
| - | - | 1988 | 546.2 | - | - | 0 | - |
| - | - | 765.8 | 547.2 | - | - | 0 | - |
| - | - | 5912 | 548.6 | - | - | 0 | - |
| - | - | 8449 | 549 | - | - | 0 | - |
| - | - | 4633 | 549.3 | - | - | 0 | - |
| 5 | y | 3692 | 549.3 | 0.002533 | 4.611 | +2 | 10 |
| - | - | 1520 | 549.6 | - | - | 0 | - |
| - | - | 3765 | 549.8 | - | - | 0 | - |
| - | - | 1084 | 550 | - | - | 0 | - |
| - | - | 690.9 | 553.3 | - | - | 0 | - |
| - | - | 1617 | 554 | - | - | 0 | - |
| - | - | 1944 | 554.3 | - | - | 0 | - |
| - | - | 1.936E+05 | 554.6 | - | - | 0 | - |
| - | - | 1.825E+05 | 555 | - | - | 0 | - |
| - | - | 9.881E+04 | 555.3 | - | - | 0 | - |
| - | - | 3.098E+04 | 555.6 | - | - | 0 | - |
| - | - | 1018 | 555.7 | - | - | 0 | - |
| - | - | 8261 | 556 | - | - | 0 | - |
| - | - | 5608 | 556.3 | - | - | 0 | - |
| - | - | 1116 | 561.8 | - | - | 0 | - |
| - | - | 1.199E+04 | 563.2 | - | - | 0 | - |
| 4 | c | 4267 | 564.3 | 0.006508 | 11.53 | +1 | 4 |
| 4 | c | 5120 | 565.2 | 0.001358 | 2.403 | +1 | 4 |
| - | - | 3924 | 566.2 | - | - | 0 | - |
| - | - | 3081 | 566.4 | - | - | 0 | - |
| - | - | 1334 | 567.3 | - | - | 0 | - |
| - | - | 979.9 | 569.8 | - | - | 0 | - |
| - | - | 2092 | 572.8 | - | - | 0 | - |
| - | - | 1394 | 573.3 | - | - | 0 | - |
| 4 | w | 1.136E+04 | 576.3 | 0.0007904 | 1.371 | +2 | 11 |
| - | - | 6942 | 576.8 | - | - | 0 | - |
| - | - | 2342 | 577.3 | - | - | 0 | - |
| - | - | 2020 | 577.8 | - | - | 0 | - |
| - | - | 1174 | 579.4 | - | - | 0 | - |
| - | - | 8264 | 581.3 | - | - | 0 | - |
| 4 | c | 4.522E+04 | 582.3 | 0.0005052 | 0.8676 | +1 | 4 |
| - | - | 3670 | 582.4 | - | - | 0 | - |
| - | - | 1.405E+04 | 583.3 | - | - | 0 | - |
| - | - | 2154 | 584.3 | - | - | 0 | - |
| 9 | w | 6817 | 595.4 | 0.0005836 | 0.9801 | +1 | 6 |
| - | - | 3418 | 596.4 | - | - | 0 | - |
| 10 | c | 1036 | 596.8 | 0.0003517 | 0.5893 | +2 | 10 |
| - | - | 1739 | 597.4 | - | - | 0 | - |
| - | - | 1523 | 604.4 | - | - | 0 | - |
| 4 | y | 3130 | 604.8 | 0.006355 | 10.51 | +2 | 11 |
| 4 | z | 3.266E+04 | 605.4 | 0.0005504 | 0.9092 | +2 | 11 |
| - | - | 3.432E+04 | 605.9 | - | - | 0 | - |
| - | - | 1.944E+04 | 606.4 | - | - | 0 | - |
| - | - | 6671 | 606.9 | - | - | 0 | - |
| - | - | 1037 | 607.4 | - | - | 0 | - |
| - | - | 1827 | 609.5 | - | - | 0 | - |
| - | - | 6650 | 610.4 | - | - | 0 | - |
| - | - | 2302 | 611.4 | - | - | 0 | - |
| - | - | 1206 | 612.9 | - | - | 0 | - |
| 4 | y | 1.095E+04 | 613.4 | 0.0002215 | 0.3612 | +2 | 11 |
| - | - | 6707 | 613.9 | - | - | 0 | - |
| - | - | 2146 | 614.4 | - | - | 0 | - |
| - | - | 949.5 | 617.3 | - | - | 0 | - |
| - | - | 1096 | 620.3 | - | - | 0 | - |
| - | - | 982.8 | 620.8 | - | - | 0 | - |
| - | - | 787 | 622.3 | - | - | 0 | - |
| - | - | 2093 | 624.4 | - | - | 0 | - |
| - | - | 5679 | 629.3 | - | - | 0 | - |
| - | - | 3597 | 629.8 | - | - | 0 | - |
| - | - | 1776 | 630.3 | - | - | 0 | - |
| - | - | 1200 | 637.4 | - | - | 0 | - |
| - | - | 1502 | 639.4 | - | - | 0 | - |
| - | - | 2508 | 645.8 | - | - | 0 | - |
| - | - | 1484 | 646.3 | - | - | 0 | - |
| - | - | 2502 | 651.4 | - | - | 0 | - |
| - | - | 2107 | 651.9 | - | - | 0 | - |
| 9 | z | 3.303E+04 | 653.4 | 0.0007052 | 1.079 | +1 | 6 |
| 11 | c | 2899 | 653.8 | 0.001955 | 2.99 | +2 | 11 |
| - | - | 1999 | 654.3 | - | - | 0 | - |
| - | - | 1.716E+04 | 654.5 | - | - | 0 | - |
| - | - | 3862 | 655.5 | - | - | 0 | - |
| - | - | 1241 | 657.3 | - | - | 0 | - |
| - | - | 4419 | 657.9 | - | - | 0 | - |
| - | - | 1303 | 658.4 | - | - | 0 | - |
| - | - | 1935 | 658.9 | - | - | 0 | - |
| - | - | 4740 | 659.3 | - | - | 0 | - |
| - | - | 1320 | 659.4 | - | - | 0 | - |
| - | - | 1444 | 660.3 | - | - | 0 | - |
| - | - | 1734 | 665.4 | - | - | 0 | - |
| - | - | 1956 | 665.9 | - | - | 0 | - |
| 9 | y | 1305 | 669.5 | 0.004591 | 6.858 | +1 | 6 |
| - | - | 3304 | 674.3 | - | - | 0 | - |
| - | - | 2349 | 675.3 | - | - | 0 | - |
| - | - | 1383 | 685.9 | - | - | 0 | - |
| 3 | y | 3514 | 686.4 | 0.005025 | 7.321 | +2 | 12 |
| 3 | z | 4.186E+04 | 686.9 | 0.0003802 | 0.5536 | +2 | 12 |
| - | - | 3.235E+04 | 687.4 | - | - | 0 | - |
| - | - | 1.629E+04 | 687.9 | - | - | 0 | - |
| - | - | 7667 | 688.4 | - | - | 0 | - |
| - | - | 1377 | 688.9 | - | - | 0 | - |
| - | - | 1410 | 692.4 | - | - | 0 | - |
| 3 | y | 4.183E+04 | 694.9 | 0.0006617 | 0.9523 | +2 | 12 |
| - | - | 3.373E+04 | 695.4 | - | - | 0 | - |
| - | - | 1.733E+04 | 695.9 | - | - | 0 | - |
| - | - | 4576 | 696.4 | - | - | 0 | - |
| - | - | 1567 | 696.9 | - | - | 0 | - |
| 5 | c | 3132 | 702.3 | 5.829E-05 | 0.083 | +1 | 5 |
| - | - | 1586 | 703.3 | - | - | 0 | - |
| - | - | 1510 | 708.4 | - | - | 0 | - |
| 12 | c | 8527 | 709.4 | 0.001134 | 1.599 | +2 | 12 |
| - | - | 6631 | 709.9 | - | - | 0 | - |
| - | - | 2875 | 710.4 | - | - | 0 | - |
| 8 | z | 5418 | 710.5 | 0.0001765 | 0.2485 | +1 | 7 |
| - | - | 4069 | 711.5 | - | - | 0 | - |
| - | - | 1211 | 712.5 | - | - | 0 | - |
| - | - | 2483 | 717.4 | - | - | 0 | - |
| 12 | c | 1.055E+05 | 717.9 | 0.0003719 | 0.5181 | +2 | 12 |
| - | - | 8.537E+04 | 718.4 | - | - | 0 | - |
| - | - | 4.025E+04 | 718.9 | - | - | 0 | - |
| 5 | c | 8.906E+04 | 719.3 | 0.0003701 | 0.5146 | +1 | 5 |
| - | - | 3165 | 719.9 | - | - | 0 | - |
| - | - | 4.106E+04 | 720.3 | - | - | 0 | - |
| - | - | 9045 | 721.3 | - | - | 0 | - |
| - | - | 1131 | 722.3 | - | - | 0 | - |
| 8 | y | 3321 | 726.5 | 6.819E-05 | 0.09386 | +1 | 7 |
| - | - | 1279 | 727.5 | - | - | 0 | - |
| - | - | 1561 | 730.9 | - | - | 0 | - |
| - | - | 1981 | 731.4 | - | - | 0 | - |
| - | - | 1029 | 731.9 | - | - | 0 | - |
| - | - | 1125 | 736.4 | - | - | 0 | - |
| - | - | 1159 | 744.9 | - | - | 0 | - |
| - | - | 1386 | 750.9 | - | - | 0 | - |
| - | - | 1964 | 751.4 | - | - | 0 | - |
| - | - | 3153 | 752.4 | - | - | 0 | - |
| - | - | 2263 | 752.9 | - | - | 0 | - |
| - | - | 1696 | 753.4 | - | - | 0 | - |
| - | - | 1334 | 756.9 | - | - | 0 | - |
| - | - | 1208 | 757.9 | - | - | 0 | - |
| - | - | 1544 | 758.4 | - | - | 0 | - |
| - | - | 4150 | 758.9 | - | - | 0 | - |
| - | - | 4225 | 759.4 | - | - | 0 | - |
| - | - | 1072 | 759.9 | - | - | 0 | - |
| 13 | c | 2915 | 765.9 | 0.002742 | 3.581 | +2 | 13 |
| - | - | 5658 | 766.4 | - | - | 0 | - |
| - | - | 4298 | 766.9 | - | - | 0 | - |
| - | - | 4999 | 767.4 | - | - | 0 | - |
| - | - | 1382 | 767.9 | - | - | 0 | - |
| - | - | 1150 | 768.4 | - | - | 0 | - |
| - | - | 1807 | 773.4 | - | - | 0 | - |
| - | - | 1369 | 773.9 | - | - | 0 | - |
| 13 | c | 6.673E+04 | 774.4 | 0.0007594 | 0.9806 | +2 | 13 |
| - | - | 6.683E+04 | 774.9 | - | - | 0 | - |
| - | - | 3.311E+04 | 775.4 | - | - | 0 | - |
| - | - | 9630 | 775.9 | - | - | 0 | - |
| - | - | 3204 | 776.4 | - | - | 0 | - |
| - | - | 2006 | 777.9 | - | - | 0 | - |
| - | - | 1575 | 778.4 | - | - | 0 | - |
| - | - | 2756 | 778.9 | - | - | 0 | - |
| 2 | y | 2298 | 779.4 | 0.002478 | 3.179 | +2 | 13 |
| 2 | z | 9584 | 779.9 | 0.0009459 | 1.213 | +2 | 13 |
| - | - | 1.337E+04 | 780.4 | - | - | 0 | - |
| 7 | w | 1541 | 780.5 | 0.0004898 | 0.6276 | +1 | 8 |
| - | - | 7264 | 780.9 | - | - | 0 | - |
| - | - | 2689 | 781.4 | - | - | 0 | - |
| - | - | 1417 | 781.5 | - | - | 0 | - |
| - | - | 1574 | 781.9 | - | - | 0 | - |
| - | - | 1716 | 782.4 | - | - | 0 | - |
| 2 | y | 6076 | 787.9 | 0.01043 | 13.24 | +2 | 13 |
| - | - | 5909 | 788.4 | - | - | 0 | - |
| - | - | 3718 | 788.9 | - | - | 0 | - |
| - | - | 1179 | 789.4 | - | - | 0 | - |
| - | - | 8495 | 790.9 | - | - | 0 | - |
| - | - | 7937 | 791.4 | - | - | 0 | - |
| - | - | 3493 | 791.9 | - | - | 0 | - |
| - | - | 3052 | 792.4 | - | - | 0 | - |
| - | - | 1518 | 792.9 | - | - | 0 | - |
| - | - | 1237 | 794.4 | - | - | 0 | - |
| - | - | 1443 | 794.4 | - | - | 0 | - |
| - | - | 1074 | 794.9 | - | - | 0 | - |
| - | - | 5683 | 795.4 | - | - | 0 | - |
| - | - | 7483 | 795.9 | - | - | 0 | - |
| - | - | 2398 | 796.4 | - | - | 0 | - |
| - | - | 6555 | 796.4 | - | - | 0 | - |
| - | - | 4874 | 796.9 | - | - | 0 | - |
| - | - | 2071 | 797.4 | - | - | 0 | - |
| - | - | 3217 | 797.4 | - | - | 0 | - |
| - | - | 1515 | 797.9 | - | - | 0 | - |
| - | - | 5869 | 800.4 | - | - | 0 | - |
| - | - | 7038 | 800.9 | - | - | 0 | - |
| - | - | 6868 | 801.4 | - | - | 0 | - |
| - | - | 4097 | 801.9 | - | - | 0 | - |
| - | - | 4390 | 802.4 | - | - | 0 | - |
| - | - | 4709 | 802.9 | - | - | 0 | - |
| - | - | 2643 | 803.4 | - | - | 0 | - |
| - | - | 3334 | 803.9 | - | - | 0 | - |
| - | - | 5210 | 804.4 | - | - | 0 | - |
| - | - | 2449 | 804.9 | - | - | 0 | - |
| - | - | 1863 | 809 | - | - | 0 | - |
| - | - | 1.382E+04 | 809.4 | - | - | 0 | - |
| - | - | 1.413E+04 | 809.9 | - | - | 0 | - |
| - | - | 8430 | 810.4 | - | - | 0 | - |
| - | - | 4828 | 810.9 | - | - | 0 | - |
| - | - | 1.061E+04 | 811.4 | - | - | 0 | - |
| - | - | 9401 | 812.4 | - | - | 0 | - |
| - | - | 2377 | 813.4 | - | - | 0 | - |
| - | - | 1066 | 814.4 | - | - | 0 | - |
| - | - | 3346 | 814.9 | - | - | 0 | - |
| - | - | 2039 | 815.4 | - | - | 0 | - |
| - | - | 1024 | 815.9 | - | - | 0 | - |
| - | - | 1837 | 822.4 | - | - | 0 | - |
| - | - | 2009 | 822.5 | - | - | 0 | - |
| - | - | 9069 | 822.9 | - | - | 0 | - |
| - | - | 5.319E+04 | 823.4 | - | - | 0 | - |
| 7 | y | 1.157E+04 | 823.5 | 3.871E-05 | 0.047 | +1 | 8 |
| - | - | 5.347E+04 | 823.9 | - | - | 0 | - |
| - | - | 2.698E+04 | 824.4 | - | - | 0 | - |
| - | - | 5990 | 824.5 | - | - | 0 | - |
| - | - | 9305 | 824.9 | - | - | 0 | - |
| - | - | 2745 | 825.4 | - | - | 0 | - |
| - | - | 1818 | 825.5 | - | - | 0 | - |
| - | - | 1297 | 830.4 | - | - | 0 | - |
| - | - | 7968 | 830.9 | - | - | 0 | - |
| - | - | 1.23E+05 | 831.5 | - | - | 0 | - |
| - | - | 5.537E+05 | 832 | - | - | 0 | - |
| - | - | 4.946E+05 | 832.5 | - | - | 0 | - |
| - | - | 2.61E+05 | 833 | - | - | 0 | - |
| - | - | 8.964E+04 | 833.5 | - | - | 0 | - |
| - | - | 2.426E+04 | 834 | - | - | 0 | - |
| 6 | c | 6989 | 839.4 | 0.0005391 | 0.6423 | +1 | 6 |
| - | - | 4187 | 840.4 | - | - | 0 | - |
| - | - | 1277 | 841.4 | - | - | 0 | - |
| - | - | 6522 | 850.5 | - | - | 0 | - |
| - | - | 3678 | 851.5 | - | - | 0 | - |
| - | - | 5924 | 852.5 | - | - | 0 | - |
| - | - | 2177 | 853.5 | - | - | 0 | - |
| - | - | 1793 | 854.5 | - | - | 0 | - |
| - | - | 2722 | 855.5 | - | - | 0 | - |
| - | - | 1419 | 856.5 | - | - | 0 | - |
| - | - | 1360 | 873.5 | - | - | 0 | - |
| - | - | 880.3 | 874.5 | - | - | 0 | - |
| - | - | 1868 | 883.4 | - | - | 0 | - |
| - | - | 984.8 | 884.4 | - | - | 0 | - |
| - | - | 1956 | 909.4 | - | - | 0 | - |
| - | - | 1366 | 910.4 | - | - | 0 | - |
| 6 | z | 8.995E+04 | 944.6 | 0.0003783 | 0.4005 | +1 | 9 |
| - | - | 5.354E+04 | 945.6 | - | - | 0 | - |
| - | - | 1.634E+04 | 946.6 | - | - | 0 | - |
| - | - | 4849 | 947.6 | - | - | 0 | - |
| 7 | c | 9581 | 953.4 | 0.001609 | 1.688 | +1 | 7 |
| - | - | 5432 | 954.4 | - | - | 0 | - |
| - | - | 1829 | 955.4 | - | - | 0 | - |
| 6 | y | 8268 | 960.6 | 0.0002089 | 0.2174 | +1 | 9 |
| - | - | 4787 | 961.6 | - | - | 0 | - |
| - | - | 1659 | 962.6 | - | - | 0 | - |
| - | - | 1854 | 968.6 | - | - | 0 | - |
| - | - | 4246 | 982.5 | - | - | 0 | - |
| - | - | 4269 | 983.6 | - | - | 0 | - |
| - | - | 1449 | 984.6 | - | - | 0 | - |
| - | - | 1329 | 988.6 | - | - | 0 | - |
| - | - | 3960 | 1009 | - | - | 0 | - |
| 8 | c | 7.73E+04 | 1010 | 0.0003484 | 0.3448 | +1 | 8 |
| - | - | 4.92E+04 | 1011 | - | - | 0 | - |
| - | - | 1.452E+04 | 1012 | - | - | 0 | - |
| - | - | 2970 | 1013 | - | - | 0 | - |
| 5 | y | 1584 | 1081 | 0.0001947 | 0.1802 | +1 | 10 |
| 5 | z | 4.574E+04 | 1082 | 0.0005484 | 0.507 | +1 | 10 |
| - | - | 1.004E+05 | 1083 | - | - | 0 | - |
| - | - | 5.747E+04 | 1084 | - | - | 0 | - |
| - | - | 1.989E+04 | 1085 | - | - | 0 | - |
| - | - | 3376 | 1086 | - | - | 0 | - |
| - | - | 3187 | 1095 | - | - | 0 | - |
| - | - | 2595 | 1096 | - | - | 0 | - |
| - | - | 2420 | 1096 | - | - | 0 | - |
| - | - | 3821 | 1097 | - | - | 0 | - |
| - | - | 759.4 | 1098 | - | - | 0 | - |
| 5 | y | 1.049E+04 | 1098 | 0.002062 | 1.879 | +1 | 10 |
| - | - | 4917 | 1099 | - | - | 0 | - |
| - | - | 1419 | 1100 | - | - | 0 | - |
| 9 | c | 1200 | 1122 | 0.0006601 | 0.5885 | +1 | 9 |
| - | - | 1295 | 1138 | - | - | 0 | - |
| 9 | c | 1.027E+05 | 1139 | 0.0007223 | 0.6344 | +1 | 9 |
| - | - | 6.936E+04 | 1140 | - | - | 0 | - |
| - | - | 2.694E+04 | 1141 | - | - | 0 | - |
| - | - | 5855 | 1142 | - | - | 0 | - |
| - | - | 2235 | 1146 | - | - | 0 | - |
| - | - | 2157 | 1147 | - | - | 0 | - |
| - | - | 989.1 | 1154 | - | - | 0 | - |
| - | - | 1845 | 1166 | - | - | 0 | - |
| - | - | 3700 | 1167 | - | - | 0 | - |
| - | - | 956.1 | 1168 | - | - | 0 | - |
| 4 | y | 926.3 | 1209 | 0.00162 | 1.34 | +1 | 11 |
| 4 | z | 1.155E+04 | 1210 | 0.0005647 | 0.4668 | +1 | 11 |
| - | - | 6.072E+04 | 1211 | - | - | 0 | - |
| - | - | 3.874E+04 | 1212 | - | - | 0 | - |
| - | - | 1.537E+04 | 1213 | - | - | 0 | - |
| - | - | 4427 | 1214 | - | - | 0 | - |
| 4 | y | 3461 | 1226 | 0.0007033 | 0.5738 | +1 | 11 |
| - | - | 2745 | 1227 | - | - | 0 | - |
| - | - | 941.7 | 1237 | - | - | 0 | - |
| - | - | 1817 | 1259 | - | - | 0 | - |
| - | - | 1684 | 1260 | - | - | 0 | - |
| - | - | 1229 | 1263 | - | - | 0 | - |
| - | - | 1954 | 1264 | - | - | 0 | - |
| - | - | 2793 | 1292 | - | - | 0 | - |
| - | - | 1493 | 1293 | - | - | 0 | - |
| - | - | 2650 | 1306 | - | - | 0 | - |
| 11 | c | 4.785E+04 | 1307 | 4.403E-05 | 0.0337 | +1 | 11 |
| - | - | 3.452E+04 | 1308 | - | - | 0 | - |
| - | - | 1.676E+04 | 1309 | - | - | 0 | - |
| - | - | 4606 | 1310 | - | - | 0 | - |
| 3 | z | 5787 | 1373 | 0.001363 | 0.9925 | +1 | 12 |
| - | - | 2.798E+04 | 1374 | - | - | 0 | - |
| - | - | 2.292E+04 | 1375 | - | - | 0 | - |
| - | - | 9859 | 1376 | - | - | 0 | - |
| - | - | 2838 | 1377 | - | - | 0 | - |
| 3 | y | 2116 | 1389 | 0.00506 | 3.643 | +1 | 12 |
| - | - | 2517 | 1390 | - | - | 0 | - |
| - | - | 1245 | 1402 | - | - | 0 | - |
| - | - | 2752 | 1419 | - | - | 0 | - |
| - | - | 4169 | 1420 | - | - | 0 | - |
| - | - | 1878 | 1421 | - | - | 0 | - |
| 12 | c | 1.57E+04 | 1435 | 0.001623 | 1.131 | +1 | 12 |
| - | - | 2.177E+04 | 1436 | - | - | 0 | - |
| - | - | 1.519E+04 | 1437 | - | - | 0 | - |
| - | - | 5321 | 1438 | - | - | 0 | - |
| - | - | 2061 | 1439 | - | - | 0 | - |
| - | - | 2642 | 1505 | - | - | 0 | - |
| - | - | 1330 | 1506 | - | - | 0 | - |
| - | - | 1985 | 1532 | - | - | 0 | - |
| - | - | 2783 | 1533 | - | - | 0 | - |
| - | - | 1497 | 1534 | - | - | 0 | - |
| 13 | c | 7720 | 1548 | 0.001093 | 0.7058 | +1 | 13 |
| - | - | 1.643E+04 | 1549 | - | - | 0 | - |
| - | - | 1.026E+04 | 1550 | - | - | 0 | - |
| - | - | 4307 | 1551 | - | - | 0 | - |
| - | - | 2011 | 1552 | - | - | 0 | - |
| 2 | z | 2992 | 1559 | 0.001574 | 1.01 | +1 | 13 |
| - | - | 1.026E+04 | 1560 | - | - | 0 | - |
| - | - | 7715 | 1561 | - | - | 0 | - |
| - | - | 5269 | 1562 | - | - | 0 | - |
| - | - | 1698 | 1563 | - | - | 0 | - |
| - | - | 3750 | 1581 | - | - | 0 | - |
| - | - | 2965 | 1582 | - | - | 0 | - |
| - | - | 1424 | 1583 | - | - | 0 | - |
| - | - | 1385 | 1584 | - | - | 0 | - |
| - | - | 1010 | 1591 | - | - | 0 | - |
| - | - | 1154 | 1601 | - | - | 0 | - |
| - | - | 1500 | 1602 | - | - | 0 | - |
| - | - | 1176 | 1603 | - | - | 0 | - |
| - | - | 4848 | 1618 | - | - | 0 | - |
| - | - | 1.271E+04 | 1619 | - | - | 0 | - |
| - | - | 9909 | 1620 | - | - | 0 | - |
| - | - | 5727 | 1621 | - | - | 0 | - |
| - | - | 1539 | 1622 | - | - | 0 | - |
| - | - | 6540 | 1629 | - | - | 0 | - |
| - | - | 5662 | 1630 | - | - | 0 | - |
| - | - | 2300 | 1631 | - | - | 0 | - |
| - | - | 5154 | 1636 | - | - | 0 | - |
| - | - | 3928 | 1637 | - | - | 0 | - |
| - | - | 2928 | 1638 | - | - | 0 | - |
| - | - | 1577 | 1645 | - | - | 0 | - |
| - | - | 1.229E+04 | 1646 | - | - | 0 | - |
| - | - | 5.273E+04 | 1647 | - | - | 0 | - |
| - | - | 4.824E+04 | 1648 | - | - | 0 | - |
| - | - | 2.278E+04 | 1649 | - | - | 0 | - |
| - | - | 7304 | 1650 | - | - | 0 | - |
| - | - | 1899 | 1651 | - | - | 0 | - |
| - | - | 1.117E+04 | 1662 | - | - | 0 | - |
| - | - | 6.117E+04 | 1663 | - | - | 0 | - |
| - | - | 2.693E+05 | 1664 | - | - | 0 | - |
| - | - | 2.393E+05 | 1665 | - | - | 0 | - |
| - | - | 1.207E+05 | 1666 | - | - | 0 | - |
| - | - | 3.684E+04 | 1667 | - | - | 0 | - |
| - | - | 1.213E+04 | 1668 | - | - | 0 | - |
| - | - | 1115 | 1866 | - | - | 0 | - |

m/z Charge Intensity FragmentType MassShift Position
124.71467590332031 0 640.5883
124.82473754882812 0 528.7971
125.71238708496094 0 515.0658
127.50818634033203 0 463.22693
128.7911834716797 0 510.7017
129.10238647460938 0 1917.6086
130.06536865234375 0 1051.7655
132.0810089111328 0 1844.5024
132.10214233398438 0 3627.4836 y 13
136.07589721679688 0 1253.3531
141.5152130126953 0 527.389
146.09278869628906 0 1106.6724
153.07858276367188 0 4361.2275
164.02366638183594 0 622.2674
168.10133361816406 0 633.17084
172.31005859375 0 616.0124
173.43862915039062 0 1928.4373
175.75599670410156 0 605.81714
181.7494659423828 0 698.8034
186.07847595214844 0 1638.8966
187.08673095703125 0 3355.1218
199.1806640625 0 704.7217
209.54022216796875 0 674.40967
209.9342803955078 0 692.0898
210.1494140625 0 821.80054
214.1551055908203 0 1073.3193
215.1389923095703 0 1001.3379
223.1191864013672 0 2236.5178
226.1551971435547 0 3214.6147
227.1754913330078 0 731.1262
228.11048889160156 0 974.9728
243.2021484375 0 622.0833
245.137451171875 0 5269.7373
245.18618774414062 0 6943.5303 y 12
246.1239471435547 0 21008.033
246.18959045410156 0 899.0804
247.12786865234375 0 2813.54
255.10096740722656 0 1650.6254
258.14532470703125 0 1729.3082
267.1064758300781 0 830.5342
273.1104736328125 0 1629.2363
274.1188049316406 0 25663.096
275.1229248046875 0 4195.8853
281.17352294921875 0 3089.8755
282.17950439453125 0 849.15686
292.12939453125 0 3038.0364
297.19146728515625 0 1332.4003
301.1998291015625 0 1775.5814
309.1558837890625 0 3342.1816
314.20770263671875 0 2336.9055
338.1348571777344 0 1719.4017
338.196533203125 0 902.5911
346.726806640625 0 3067.8975
347.2293701171875 0 779.9588 z Ammonia loss 7
349.1420593261719 0 2323.166
350.26812744140625 0 1026.183
355.06817626953125 0 788.13324
357.26239013671875 0 10004.924 z 11
358.267822265625 0 3981.185
371.09979248046875 0 1256.6587
373.2807922363281 0 2881.6611 y 11
382.28118896484375 0 1709.3297
394.2575988769531 0 1783.179
399.2613525390625 0 1618.704
412.2741394042969 0 12664.859 y 6
412.7757568359375 0 6345.9146
413.2281799316406 0 3109.486
413.27557373046875 0 1280.1488
414.23150634765625 0 980.6197
420.1836242675781 0 1801.2797 c Ammonia loss 5
420.6856994628906 0 939.19684
423.2243347167969 0 1568.8778
427.2919616699219 0 5671.5073 w 10
428.2640380859375 0 901.08734
428.2978210449219 0 2248.972
429.18597412109375 0 1214.9406
434.1932373046875 0 834.42505
435.19122314453125 0 1590.5074
436.17388916015625 0 1241.8254
437.18218994140625 0 4531.081
438.1857604980469 0 1052.8661
439.2200012207031 0 1239.7295
451.220947265625 0 1006.24615
453.2011413574219 0 4932.8667
454.20831298828125 0 9969.141 c 2
455.2120056152344 0 2747.4756
463.5989990234375 0 3673.713 y 2
463.9330749511719 0 2909.5579
465.2091064453125 0 3578.6475
468.3187561035156 0 10550.509
469.2897644042969 0 3329.2979
469.3233642578125 0 2377.7542
470.29486083984375 0 1627.9968
470.3338928222656 0 4662.2266 y 10
472.3034973144531 0 1042.5742
473.2427673339844 0 4620.9985 c Ammonia loss 11
473.5777587890625 0 4525.875
473.90997314453125 0 1990.9249
477.20166015625 0 4071.8867
478.20465087890625 0 1607.3646
480.22259521484375 0 2054.243
480.8047180175781 0 1960.7092 y 5
481.3049621582031 0 1844.0962
482.2983703613281 0 3145.0664
483.3006591796875 0 1156.1655
497.22064208984375 0 764.84235 c Ammonia loss 7
498.32867431640625 0 857.7322
501.60595703125 0 1268.5271
501.9413146972656 0 999.7493
504.93450927734375 0 6022.9766
505.26806640625 0 6192.5938
505.6025695800781 0 1706.4663
507.3424987792969 0 873.5628
510.9375305175781 0 23016.238 c Ammonia loss 12
511.2717590332031 0 18222.303
511.6063232421875 0 10032.104
511.9397277832031 0 3876.8906
512.2709350585938 0 1121.8223
516.941162109375 0 5297.3345
517.2754516601562 0 2998.0703
517.61328125 0 1005.6124
517.94873046875 0 1169.2219
520.2811889648438 0 1617.0875 z 1
523.2420043945312 0 2203.037
525.3526000976562 0 41141.473 z 9
525.6243896484375 0 3434.0613 y 1
525.9595947265625 0 4484.61
526.2941284179688 0 1919.1425
526.3563232421875 0 12976.219
527.3578491210938 0 2058.7314
536.1929321289062 0 1008.1378
538.2496337890625 0 1567.907
538.2889404296875 0 1770.2505
539.2613525390625 0 2830.309
539.9710693359375 0 1172.2559
541.3224487304688 0 1989.505 z 4
541.371826171875 0 5638.14 y 9
541.8252563476562 0 1166.1793
542.375 0 1541.6904
546.2218627929688 0 1988.3452
547.2241821289062 0 765.804
548.6322021484375 0 5912.004
548.96533203125 0 8449.321
549.2978515625 0 4633.356
549.3350830078125 0 3691.7559 y 4
549.6317138671875 0 1520.2092
549.8357543945312 0 3765.225
549.9630126953125 0 1084.3091
553.2877807617188 0 690.9115
553.9620361328125 0 1616.52
554.2923583984375 0 1944.189
554.6358032226562 0 193556.16
554.969970703125 0 182502.4
555.3041381835938 0 98810.46
555.638427734375 0 30978.799
555.7113037109375 0 1017.8056
555.9728393554688 0 8261.002
556.2979736328125 0 5608.114
561.7667236328125 0 1115.6196
563.24951171875 0 11990.408
564.25 0 4266.8013 c Water loss 3
565.2418823242188 0 5119.525 c Ammonia loss 3
566.2462158203125 0 3924.1584
566.4026489257812 0 3081.4392
567.2518920898438 0 1334.1322
569.8168334960938 0 979.8566
572.8050537109375 0 2091.673
573.3053588867188 0 1393.8931
576.338623046875 0 11361.1455 w 3
576.8399658203125 0 6941.5938
577.332275390625 0 2342.005
577.8230590820312 0 2019.9979
579.3635864257812 0 1174.4122
581.2601928710938 0 8263.537
582.267578125 0 45220.406 c 3
582.3737182617188 0 3669.6084
583.2706298828125 0 14049.167
584.2734985351562 0 2154.251
595.3819580078125 0 6816.912 w 8
596.38525390625 0 3417.5178
596.7863159179688 0 1035.6354 c Ammonia loss 9
597.3870849609375 0 1738.7267
604.3568725585938 0 1523.0048
604.8549194335938 0 3130.0952 y Ammonia loss 3
605.35302734375 0 32657.342 z 3
605.8553466796875 0 34320.61
606.357421875 0 19441.234
606.8590698242188 0 6671.055
607.3615112304688 0 1036.863
609.458251953125 0 1826.9169
610.39697265625 0 6649.7017
611.3995361328125 0 2302.1418
612.856689453125 0 1206.0596
613.362060546875 0 10945.606 y 3
613.8640747070312 0 6706.807
614.3648071289062 0 2145.7905
617.2804565429688 0 949.45404
620.3395385742188 0 1095.8892
620.8427734375 0 982.77496
622.32177734375 0 786.9802
624.4217529296875 0 2093.0818
629.3465576171875 0 5678.515
629.8486328125 0 3596.5986
630.3467407226562 0 1776.418
637.4302368164062 0 1199.8722
639.3961181640625 0 1501.7412
645.828125 0 2508.134
646.3255615234375 0 1484.0093
651.3571166992188 0 2502.3213
651.85888671875 0 2107.0422
653.44775390625 0 33025.42 z 8
653.8275756835938 0 2899.445 c 10
654.3236694335938 0 1998.8636
654.4515380859375 0 17159.04
655.455078125 0 3862.3586
657.304443359375 0 1241.3108
657.8662109375 0 4418.873
658.3728637695312 0 1302.8555
658.8568115234375 0 1935.4049
659.2955322265625 0 4739.7344
659.355224609375 0 1320.4569
660.2976684570312 0 1444.4414
665.3592529296875 0 1734.0854
665.8577880859375 0 1955.7449
669.461181640625 0 1305.4053 y 8
674.3057861328125 0 3304.3677
675.311279296875 0 2348.9329
685.8893432617188 0 1382.7634
686.38525390625 0 3513.8813 y Ammonia loss 2
686.884521484375 0 41860.863 z 2
687.38623046875 0 32351.145
687.8882446289062 0 16289.523
688.3895263671875 0 7667.447
688.893310546875 0 1376.7888
692.4469604492188 0 1409.6012
694.8941650390625 0 41826.516 y 2
695.3954467773438 0 33730.594
695.8965454101562 0 17334.504
696.3965454101562 0 4576.1685
696.900634765625 0 1566.6594
702.2993774414062 0 3131.7207 c Ammonia loss 4
703.2991333007812 0 1586.198
708.39013671875 0 1510.1346
709.3609619140625 0 8526.535 c Ammonia loss 11
709.8619995117188 0 6630.5557
710.3610229492188 0 2875.4712
710.4686889648438 0 5417.5044 z 7
711.4737548828125 0 4068.9429
712.4736328125 0 1211.0609
717.367431640625 0 2482.6812
717.8734741210938 0 105518.81 c 11
718.375 0 85365.34
718.876220703125 0 40247.367
719.3263549804688 0 89059.44 c 4
719.876708984375 0 3164.8096
720.32958984375 0 41064.285
721.3323974609375 0 9044.845
722.3372802734375 0 1130.9594
726.4873046875 0 3321.083 y 7
727.492431640625 0 1279.0773
730.8819580078125 0 1560.6257
731.3827514648438 0 1981.3398
731.8798828125 0 1029.3767
736.4064331054688 0 1125.0232
744.9028930664062 0 1159.3475
750.908935546875 0 1386.0026
751.411865234375 0 1963.6335
752.4059448242188 0 3152.9465
752.9094848632812 0 2262.5232
753.4118041992188 0 1696.3818
756.9041137695312 0 1333.6693
757.9078979492188 0 1208.2655
758.4102783203125 0 1544.3579
758.9132080078125 0 4150.0757
759.4137573242188 0 4225.165
759.9071044921875 0 1072.1198
765.9046020507812 0 2915.0227 c Ammonia loss 12
766.4111328125 0 5657.884
766.9154663085938 0 4298.481
767.421142578125 0 4998.5127
767.9329223632812 0 1381.6165
768.3718872070312 0 1149.7325
773.418212890625 0 1806.7052
773.9252319335938 0 1368.973
774.4158935546875 0 66730.83 c 12
774.9170532226562 0 66826.43
775.4183959960938 0 33113.945
775.9191284179688 0 9630.129
776.420166015625 0 3203.5244
777.9222412109375 0 2005.5104
778.4381713867188 0 1574.5787
778.9246826171875 0 2755.5198
779.42236328125 0 2297.7278 y Ammonia loss 1
779.9247436523438 0 9583.777 z 1
780.4248046875 0 13374.599
780.498291015625 0 1540.5652 w 6
780.92724609375 0 7263.8213
781.4232788085938 0 2689.172
781.50439453125 0 1416.8474
781.9232177734375 0 1574.296
782.4221801757812 0 1716.2446
787.9227294921875 0 6076.2764 y 1
788.424072265625 0 5908.6216
788.9288330078125 0 3717.7498
789.42626953125 0 1179.1816
790.9268798828125 0 8495.291
791.4290161132812 0 7936.637
791.9278564453125 0 3493.1284
792.4307861328125 0 3052.1934
792.9358520507812 0 1518.0884
794.3526000976562 0 1237.0023
794.4268798828125 0 1442.6404
794.8994140625 0 1073.598
795.4134521484375 0 5682.9985
795.9147338867188 0 7483.432
796.3530883789062 0 2398.446
796.4229125976562 0 6555.336
796.9218139648438 0 4874.3623
797.3594970703125 0 2070.9614
797.4285278320312 0 3217.4758
797.92919921875 0 1514.7122
800.4384765625 0 5869.3096
800.9418334960938 0 7038.347
801.444091796875 0 6868.261
801.9346923828125 0 4096.6416
802.4236450195312 0 4390.1685
802.9277954101562 0 4709.2383
803.426025390625 0 2642.6497
803.9207153320312 0 3333.6719
804.4236450195312 0 5210.117
804.9261474609375 0 2448.8801
808.9525146484375 0 1862.9606
809.44384765625 0 13819.534
809.9464111328125 0 14125.884
810.443359375 0 8429.679
810.944580078125 0 4828.013
811.3645629882812 0 10613.625
812.3690185546875 0 9401.038
813.3740234375 0 2377.235
814.4315185546875 0 1065.6099
814.9376220703125 0 3345.7517
815.4370727539062 0 2039.3654
815.9368896484375 0 1024.0663
822.4335327148438 0 1836.9218
822.5317993164062 0 2008.9421
822.9456176757812 0 9069.463
823.4418334960938 0 53191.547
823.5400390625 0 11571.972 y 6
823.9426879882812 0 53466.996
824.4434814453125 0 26978.473
824.5435180664062 0 5989.8687
824.9443969726562 0 9304.961
825.4453735351562 0 2744.556
825.5475463867188 0 1818.0363
830.4423217773438 0 1296.7317
830.9461059570312 0 7967.8267
831.4500732421875 0 122997.04
831.9535522460938 0 553663
832.4552001953125 0 494620.2
832.9564208984375 0 261011.86
833.4574584960938 0 89644.945
833.9593505859375 0 24255.748
839.35888671875 0 6988.7334 c Ammonia loss 5
840.3589477539062 0 4186.5435
841.3529052734375 0 1276.9333
850.5288696289062 0 6521.9526
851.5313110351562 0 3677.9563
852.5416870117188 0 5924.1606
853.542236328125 0 2177.089
854.4819946289062 0 1792.5436
855.4939575195312 0 2721.8552
856.4969482421875 0 1419.2634
873.5062866210938 0 1360.4526
874.5005493164062 0 880.30084
883.4102172851562 0 1867.9824
884.4158935546875 0 984.7586
909.4244384765625 0 1956.4403
910.424072265625 0 1365.6493
944.58056640625 0 89948.81 z 5
945.5844116210938 0 53543.56
946.5872192382812 0 16343.783
947.5929565429688 0 4848.56
953.4392700195312 0 9581.059 c 6
954.44189453125 0 5431.9395
955.44287109375 0 1829.0002
960.59912109375 0 8267.895 y 5
961.6024780273438 0 4786.882
962.6046142578125 0 1659.0905
968.5777587890625 0 1854.0248
982.546142578125 0 4246.3633
983.5531005859375 0 4268.5723
984.5596923828125 0 1449.1932
988.5908813476562 0 1329.1292
1009.451416015625 0 3959.734
1010.45947265625 0 77295.23 c 7
1011.4625244140625 0 49199.617
1012.4651489257812 0 14523.187
1013.47021484375 0 2970.2605
1080.6314697265625 0 1584.3158 y Ammonia loss 4
1081.6396484375 0 45737.47 z 4
1082.646240234375 0 100420.766
1083.649169921875 0 57474.36
1084.6533203125 0 19893.488
1085.6572265625 0 3376.4873
1094.5364990234375 0 3187.423
1095.5362548828125 0 2595.0818
1095.634033203125 0 2419.946
1096.6417236328125 0 3821.452
1097.5367431640625 0 759.35095
1097.65576171875 0 10493.479 y 4
1098.6605224609375 0 4917.364
1099.6640625 0 1418.7094
1121.5281982421875 0 1200.2357 c Ammonia loss 8
1137.538330078125 0 1294.9506
1138.5548095703125 0 102667.305 c 8
1139.55712890625 0 69357.35
1140.56103515625 0 26937.652
1141.5633544921875 0 5854.8057
1145.6103515625 0 2235.456
1146.61279296875 0 2157.3345
1153.646728515625 0 989.10223
1165.57763671875 0 1845.4886
1166.5838623046875 0 3700.1133
1167.5849609375 0 956.12384
1208.688232421875 0 926.34625 y Ammonia loss 3
1209.6982421875 0 11547.139 z 3
1210.7052001953125 0 60715.06
1211.708251953125 0 38744.727
1212.71142578125 0 15374.669
1213.7132568359375 0 4427.2495
1225.7156982421875 0 3460.9128 y 3
1226.72021484375 0 2744.947
1236.62060546875 0 941.6799
1258.6990966796875 0 1816.5231
1259.706298828125 0 1683.8561
1262.626708984375 0 1228.9135
1263.6390380859375 0 1954.2739
1291.6295166015625 0 2792.708
1292.6400146484375 0 1493.0145
1305.634033203125 0 2649.5027
1306.6439208984375 0 47845.742 c 10
1307.646484375 0 34521.58
1308.6492919921875 0 16759.58
1309.6522216796875 0 4605.6772
1372.7596435546875 0 5787.1904 z 2
1373.7679443359375 0 27975.28
1374.77099609375 0 22920.309
1375.7724609375 0 9858.637
1376.775634765625 0 2837.7085
1388.7847900390625 0 2116.0913 y 2
1389.781982421875 0 2516.7437
1401.7186279296875 0 1244.9264
1418.7225341796875 0 2752.1406
1419.722412109375 0 4169.163
1420.7225341796875 0 1877.9014
1434.7373046875 0 15703.218 c 11
1435.7430419921875 0 21765.49
1436.74658203125 0 15194.621
1437.752197265625 0 5320.9893
1438.750732421875 0 2061.3438
1504.8070068359375 0 2641.6748
1505.81103515625 0 1330.0642
1531.8040771484375 0 1984.5261
1532.8046875 0 2783.287
1533.80322265625 0 1496.86
1547.8218994140625 0 7719.8247 c 12
1548.8271484375 0 16434.426
1549.8299560546875 0 10264.35
1550.83203125 0 4306.884
1551.8388671875 0 2010.7697
1558.8387451171875 0 2991.902 z 1
1559.8463134765625 0 10262.709
1560.8492431640625 0 7715.373
1561.8497314453125 0 5268.817
1562.842041015625 0 1698.324
1580.8414306640625 0 3749.5283
1581.8509521484375 0 2965.4001
1582.8397216796875 0 1424.1637
1583.863037109375 0 1385.3141
1590.832275390625 0 1009.6437
1600.883056640625 0 1153.9957
1601.8773193359375 0 1499.7136
1602.862060546875 0 1176.0404
1617.882568359375 0 4848.3374
1618.8837890625 0 12711.949
1619.887451171875 0 9909.154
1620.88818359375 0 5726.889
1621.88671875 0 1538.9191
1628.8701171875 0 6540.3535
1629.869873046875 0 5661.98
1630.8795166015625 0 2300.3452
1635.9083251953125 0 5153.807
1636.9088134765625 0 3928.2778
1637.9189453125 0 2927.8975
1644.8753662109375 0 1576.5172
1645.8836669921875 0 12289.307
1646.8802490234375 0 52725.766
1647.8828125 0 48242.008
1648.8861083984375 0 22779.123
1649.888671875 0 7304.311
1650.8856201171875 0 1899.4266
1661.8876953125 0 11171.236
1662.8978271484375 0 61171
1663.90576171875 0 269288.3
1664.9083251953125 0 239314.78
1665.911865234375 0 120657.17
1666.91357421875 0 36843.89
1667.9161376953125 0 12128.67
1865.9827880859375 0 1115.4767

Spectrum Details

|  |  |
| --- | --- |
| Matched peaks? Matched peaks The total absolute number of peaks matched. Additionally in brackets the total fraction of peaks matched and the total number of peaks is shown. | 68 (13.36% of 509) |
| FDR? FDR The false discovery rate estimated for this peptide. It is calculated by matching all theoretical fragments with a non-integer shift with the raw peaks for this spectrum. This is done with 40 different shifts. The resulting percentage is the average number of annotated peaks over the number of annotated peaks with the correct spectrum. | 3.12% |
| Satellite FDR? Satellite FDR See the FDR for details on its calculation. This satellite ion specific FDR only contains the satellite ions (d/w) for I/L/J positions. | - |
| PSM Score? PSM Score The PSM Score as given by Hecklib to this annotated spectrum. It is shown with three significant figures. | 507 |

#### Spectrum 5375? Spectrum 5375 The raw spectrum of this peptide as annotated by Hecklib. The fragments are coloured according to ion type (see legend). Any peaks with a star '\*' as text can be hovered over to see the full details, first the ion type second the mass shift type. By hovering over the amino acids in the peptide or ions in the legend the corresponding peaks are highlighted. By toggling the 'Unassigned' label you can turn the background (unassigned) peaks on or off in the plot. By updating the slider in the Ion legend you can update the spectrum to only show the top X% of the peaks with labels. The top X% means any peak that is within X% of the highest intensity. By dragging in the spectrum you can zoom in to a specific part of the spectrum and use 'Zoom Out' to get back to the original zoom level. The annotation of the spectrum is based on the given sequence in the peptides file and is done with different software so inconsistencies are likely. The peaks are annotated based on the given sequence, with 20 ppm tolerance.

Copy Data

##### Spectrum 5375 (TSV)

###### Preview

```
Loading example...
```

*Click on the button to copy the data to your clipboard.*

Mz MinMz MaxIntensity Max

WidthHeightPeptide font sizePeptide stroke widthSpectrum font sizeSpectrum stroke widthCompact peptide

Ion legend

wxyz

abcd

OtherUnassignedIon^Charge^_Position_Show for top:%

SWYQHHPGKAPKJJ

04.15e+58.30e+51.25e+61.66e+6

Zoom Out

y^+1^_1_y^+1^_2_z^+1^_3_y^+1^_3_y^+2^_8_c^+2^_6_w^+1^_4_c^+1^_3_y^+3^_12_y^+1^_4_c^+3^_12_y^+2^_9_c^+3^_13_z^+1^_5_y^+3^_13_z^+2^_10_y^+1^_5_y^+2^_10_c^+2^_9_c^+1^_4_c^+1^_4_w^+2^_11_c^+1^_4_w^+1^_6_c^+2^_10_y^+2^_11_z^+2^_11_y^+2^_11_z^+1^_6_c^+2^_11_y^+1^_6_z^+2^_12_y^+2^_12_z^+2^_12_z^+1^_7_y^+2^_12_c^+1^_5_c^+2^_12_z^+1^_7_c^+2^_12_c^+1^_5_y^+1^_7_c^+2^_13_c^+2^_13_y^+2^_13_z^+2^_13_w^+1^_8_y^+2^_13_y^+1^_8_c^+1^_6_z^+1^_9_c^+1^_7_y^+1^_9_c^+1^_8_y^+1^_10_z^+1^_10_y^+1^_10_c^+1^_9_w^+1^_11_y^+1^_11_z^+1^_11_y^+1^_11_c^+1^_11_z^+1^_12_y^+1^_12_c^+1^_12_c^+1^_13_z^+1^_13_y^+1^_13_

042184212631685

Fragment Matches Table

Show background peaks

| Position | Ion type | Intensity | mz Theoretical | mz Error (Th) | mz Error (ppm) | Charge | Series Number |
| --- | --- | --- | --- | --- | --- | --- | --- |
| - | - | 1249 | 122.4 | - | - | 0 | - |
| - | - | 4828 | 129.1 | - | - | 0 | - |
| - | - | 2985 | 130.1 | - | - | 0 | - |
| - | - | 6440 | 132.1 | - | - | 0 | - |
| 14 | y | 8245 | 132.1 | 0.0001914 | 1.449 | +1 | 1 |
| - | - | 3833 | 136.1 | - | - | 0 | - |
| - | - | 1382 | 143.8 | - | - | 0 | - |
| - | - | 1590 | 146.1 | - | - | 0 | - |
| - | - | 1.035E+04 | 153.1 | - | - | 0 | - |
| - | - | 1469 | 168.1 | - | - | 0 | - |
| - | - | 1729 | 170.2 | - | - | 0 | - |
| - | - | 4463 | 186.1 | - | - | 0 | - |
| - | - | 8611 | 187.1 | - | - | 0 | - |
| - | - | 3241 | 199.2 | - | - | 0 | - |
| - | - | 2362 | 214.2 | - | - | 0 | - |
| - | - | 1758 | 216.8 | - | - | 0 | - |
| - | - | 4504 | 223.1 | - | - | 0 | - |
| - | - | 6103 | 226.2 | - | - | 0 | - |
| - | - | 2204 | 228.1 | - | - | 0 | - |
| - | - | 1.353E+04 | 245.1 | - | - | 0 | - |
| 13 | y | 1.764E+04 | 245.2 | 0.0001271 | 0.5183 | +1 | 2 |
| - | - | 5.268E+04 | 246.1 | - | - | 0 | - |
| - | - | 2943 | 246.1 | - | - | 0 | - |
| - | - | 2382 | 246.2 | - | - | 0 | - |
| - | - | 6505 | 247.1 | - | - | 0 | - |
| - | - | 3900 | 255.1 | - | - | 0 | - |
| - | - | 2743 | 255.2 | - | - | 0 | - |
| - | - | 1734 | 256.2 | - | - | 0 | - |
| - | - | 7359 | 258.1 | - | - | 0 | - |
| - | - | 1749 | 269 | - | - | 0 | - |
| - | - | 4372 | 273.1 | - | - | 0 | - |
| - | - | 6.865E+04 | 274.1 | - | - | 0 | - |
| - | - | 1.283E+04 | 275.1 | - | - | 0 | - |
| - | - | 8975 | 281.2 | - | - | 0 | - |
| - | - | 2602 | 282.2 | - | - | 0 | - |
| - | - | 9472 | 292.1 | - | - | 0 | - |
| - | - | 2497 | 299.2 | - | - | 0 | - |
| - | - | 5392 | 301.2 | - | - | 0 | - |
| - | - | 9426 | 309.2 | - | - | 0 | - |
| - | - | 2361 | 310.2 | - | - | 0 | - |
| - | - | 3925 | 314.2 | - | - | 0 | - |
| - | - | 2076 | 320.1 | - | - | 0 | - |
| - | - | 2684 | 323.2 | - | - | 0 | - |
| - | - | 5780 | 346.7 | - | - | 0 | - |
| - | - | 6037 | 349.1 | - | - | 0 | - |
| - | - | 2351 | 353.5 | - | - | 0 | - |
| 12 | z | 2.772E+04 | 357.3 | 0.0003652 | 1.022 | +1 | 3 |
| - | - | 1.217E+04 | 358.3 | - | - | 0 | - |
| - | - | 2795 | 359.3 | - | - | 0 | - |
| 12 | y | 6599 | 373.3 | 0.0002263 | 0.6063 | +1 | 3 |
| - | - | 5157 | 382.3 | - | - | 0 | - |
| - | - | 9072 | 394.3 | - | - | 0 | - |
| - | - | 1898 | 395.3 | - | - | 0 | - |
| - | - | 2172 | 399.3 | - | - | 0 | - |
| - | - | 2909 | 401.1 | - | - | 0 | - |
| - | - | 3468 | 406.2 | - | - | 0 | - |
| 7 | y | 3.741E+04 | 412.3 | 0.0007757 | 1.881 | +2 | 8 |
| - | - | 1.766E+04 | 412.8 | - | - | 0 | - |
| - | - | 6387 | 413.2 | - | - | 0 | - |
| - | - | 4602 | 413.3 | - | - | 0 | - |
| 6 | c | 2995 | 420.2 | 0.001972 | 4.693 | +2 | 6 |
| - | - | 3037 | 420.7 | - | - | 0 | - |
| - | - | 2305 | 426.2 | - | - | 0 | - |
| 11 | w | 1.511E+04 | 427.3 | 0.0007395 | 1.731 | +1 | 4 |
| - | - | 5877 | 428.3 | - | - | 0 | - |
| - | - | 2470 | 429.2 | - | - | 0 | - |
| - | - | 8335 | 435.2 | - | - | 0 | - |
| - | - | 2004 | 436.2 | - | - | 0 | - |
| - | - | 1.058E+04 | 437.2 | - | - | 0 | - |
| - | - | 2549 | 438.2 | - | - | 0 | - |
| - | - | 2385 | 439.2 | - | - | 0 | - |
| - | - | 1.49E+04 | 453.2 | - | - | 0 | - |
| 3 | c | 2.723E+04 | 454.2 | 0.0001532 | 0.3373 | +1 | 3 |
| - | - | 7865 | 455.2 | - | - | 0 | - |
| 3 | y | 1.167E+04 | 463.6 | 0.0008131 | 1.754 | +3 | 12 |
| - | - | 7311 | 463.9 | - | - | 0 | - |
| - | - | 9456 | 465.2 | - | - | 0 | - |
| - | - | 3.439E+04 | 468.3 | - | - | 0 | - |
| - | - | 8016 | 469.3 | - | - | 0 | - |
| - | - | 8695 | 469.3 | - | - | 0 | - |
| - | - | 2844 | 470.3 | - | - | 0 | - |
| 11 | y | 1.826E+04 | 470.3 | 0.0001694 | 0.3601 | +1 | 4 |
| - | - | 3744 | 471.3 | - | - | 0 | - |
| 12 | c | 1.852E+04 | 473.2 | 0.00122 | 2.577 | +3 | 12 |
| - | - | 1.008E+04 | 473.6 | - | - | 0 | - |
| - | - | 5249 | 473.9 | - | - | 0 | - |
| - | - | 2974 | 474.2 | - | - | 0 | - |
| - | - | 1.658E+04 | 477.2 | - | - | 0 | - |
| - | - | 4649 | 478.2 | - | - | 0 | - |
| - | - | 2261 | 479.2 | - | - | 0 | - |
| - | - | 3984 | 480.2 | - | - | 0 | - |
| 6 | y | 3070 | 480.8 | 0.000464 | 0.9651 | +2 | 9 |
| - | - | 4860 | 481.3 | - | - | 0 | - |
| - | - | 2435 | 482.3 | - | - | 0 | - |
| - | - | 9366 | 482.3 | - | - | 0 | - |
| - | - | 2620 | 483.8 | - | - | 0 | - |
| - | - | 3963 | 495.4 | - | - | 0 | - |
| - | - | 4099 | 498.3 | - | - | 0 | - |
| - | - | 4511 | 501.6 | - | - | 0 | - |
| - | - | 2694 | 501.9 | - | - | 0 | - |
| - | - | 1.635E+04 | 504.9 | - | - | 0 | - |
| - | - | 1.239E+04 | 505.3 | - | - | 0 | - |
| - | - | 6922 | 505.6 | - | - | 0 | - |
| - | - | 3207 | 507.3 | - | - | 0 | - |
| - | - | 2489 | 510.2 | - | - | 0 | - |
| 13 | c | 7.345E+04 | 510.9 | 0.0006845 | 1.34 | +3 | 13 |
| - | - | 7.02E+04 | 511.3 | - | - | 0 | - |
| - | - | 2.706E+04 | 511.6 | - | - | 0 | - |
| - | - | 9986 | 511.9 | - | - | 0 | - |
| - | - | 4112 | 512.3 | - | - | 0 | - |
| - | - | 8433 | 516.9 | - | - | 0 | - |
| - | - | 8331 | 517.3 | - | - | 0 | - |
| - | - | 3647 | 517.6 | - | - | 0 | - |
| - | - | 9106 | 523.2 | - | - | 0 | - |
| 10 | z | 1.1E+05 | 525.4 | 0.0006975 | 1.328 | +1 | 5 |
| 2 | y | 9103 | 525.6 | 0.000529 | 1.006 | +3 | 13 |
| - | - | 1.26E+04 | 526 | - | - | 0 | - |
| - | - | 8257 | 526.3 | - | - | 0 | - |
| - | - | 3.795E+04 | 526.4 | - | - | 0 | - |
| - | - | 7466 | 527.4 | - | - | 0 | - |
| - | - | 5996 | 538.3 | - | - | 0 | - |
| - | - | 4758 | 539.3 | - | - | 0 | - |
| - | - | 3610 | 539.3 | - | - | 0 | - |
| - | - | 2314 | 540.6 | - | - | 0 | - |
| 5 | z | 5943 | 541.3 | 0.002082 | 3.847 | +2 | 10 |
| 10 | y | 1.218E+04 | 541.4 | 0.001077 | 1.99 | +1 | 5 |
| - | - | 5613 | 541.8 | - | - | 0 | - |
| - | - | 4172 | 542.4 | - | - | 0 | - |
| - | - | 6143 | 546.2 | - | - | 0 | - |
| - | - | 2589 | 547.8 | - | - | 0 | - |
| - | - | 3592 | 548.6 | - | - | 0 | - |
| - | - | 1.901E+04 | 548.6 | - | - | 0 | - |
| - | - | 1.458E+04 | 549 | - | - | 0 | - |
| - | - | 6586 | 549.3 | - | - | 0 | - |
| 5 | y | 1.016E+04 | 549.3 | 0.00119 | 2.166 | +2 | 10 |
| - | - | 7886 | 549.6 | - | - | 0 | - |
| - | - | 1.06E+04 | 549.8 | - | - | 0 | - |
| - | - | 2519 | 550 | - | - | 0 | - |
| - | - | 2785 | 550.3 | - | - | 0 | - |
| - | - | 4489 | 553.4 | - | - | 0 | - |
| - | - | 6580 | 554 | - | - | 0 | - |
| - | - | 3949 | 554.3 | - | - | 0 | - |
| - | - | 5.548E+05 | 554.6 | - | - | 0 | - |
| - | - | 5.668E+05 | 555 | - | - | 0 | - |
| - | - | 3.046E+05 | 555.3 | - | - | 0 | - |
| - | - | 1.013E+05 | 555.6 | - | - | 0 | - |
| - | - | 2587 | 555.7 | - | - | 0 | - |
| - | - | 2.561E+04 | 556 | - | - | 0 | - |
| 9 | c | 3147 | 561.3 | 0.0009643 | 1.718 | +2 | 9 |
| - | - | 3.956E+04 | 563.2 | - | - | 0 | - |
| 4 | c | 1.178E+04 | 564.3 | 0.006874 | 12.18 | +1 | 4 |
| 4 | c | 1.678E+04 | 565.2 | 0.001908 | 3.375 | +1 | 4 |
| - | - | 1.289E+04 | 566.2 | - | - | 0 | - |
| - | - | 9462 | 566.4 | - | - | 0 | - |
| - | - | 3485 | 567.2 | - | - | 0 | - |
| - | - | 3096 | 567.4 | - | - | 0 | - |
| - | - | 2607 | 569.8 | - | - | 0 | - |
| - | - | 2682 | 572.3 | - | - | 0 | - |
| - | - | 5956 | 572.8 | - | - | 0 | - |
| - | - | 3077 | 573.3 | - | - | 0 | - |
| 4 | w | 3.223E+04 | 576.3 | 0.0007294 | 1.266 | +2 | 11 |
| - | - | 2.133E+04 | 576.8 | - | - | 0 | - |
| - | - | 9654 | 577.3 | - | - | 0 | - |
| - | - | 2429 | 579.4 | - | - | 0 | - |
| - | - | 2.385E+04 | 581.3 | - | - | 0 | - |
| 4 | c | 1.269E+05 | 582.3 | 0.0006272 | 1.077 | +1 | 4 |
| - | - | 1.168E+04 | 582.4 | - | - | 0 | - |
| - | - | 3.751E+04 | 583.3 | - | - | 0 | - |
| - | - | 3386 | 583.4 | - | - | 0 | - |
| - | - | 6991 | 584.3 | - | - | 0 | - |
| - | - | 2625 | 587.8 | - | - | 0 | - |
| 9 | w | 2.614E+04 | 595.4 | 0.001133 | 1.903 | +1 | 6 |
| - | - | 9695 | 596.4 | - | - | 0 | - |
| 10 | c | 2776 | 596.8 | 0.005601 | 9.385 | +2 | 10 |
| - | - | 4064 | 597.4 | - | - | 0 | - |
| - | - | 4992 | 604.4 | - | - | 0 | - |
| 4 | y | 7542 | 604.8 | 0.002083 | 3.443 | +2 | 11 |
| 4 | z | 9.228E+04 | 605.4 | 0.0006724 | 1.111 | +2 | 11 |
| - | - | 9.916E+04 | 605.9 | - | - | 0 | - |
| - | - | 4.995E+04 | 606.4 | - | - | 0 | - |
| - | - | 1.233E+04 | 606.9 | - | - | 0 | - |
| - | - | 4969 | 607.4 | - | - | 0 | - |
| - | - | 2355 | 609.4 | - | - | 0 | - |
| - | - | 6484 | 609.5 | - | - | 0 | - |
| - | - | 2.415E+04 | 610.4 | - | - | 0 | - |
| - | - | 7196 | 611.4 | - | - | 0 | - |
| 4 | y | 3.618E+04 | 613.4 | 0.0005877 | 0.9582 | +2 | 11 |
| - | - | 2.087E+04 | 613.9 | - | - | 0 | - |
| - | - | 6332 | 614.4 | - | - | 0 | - |
| - | - | 3601 | 620.3 | - | - | 0 | - |
| - | - | 3208 | 620.8 | - | - | 0 | - |
| - | - | 7468 | 624.4 | - | - | 0 | - |
| - | - | 1.484E+04 | 629.3 | - | - | 0 | - |
| - | - | 1.653E+04 | 629.8 | - | - | 0 | - |
| - | - | 5397 | 630.4 | - | - | 0 | - |
| - | - | 4372 | 637.4 | - | - | 0 | - |
| - | - | 2520 | 638.4 | - | - | 0 | - |
| - | - | 3475 | 645.8 | - | - | 0 | - |
| - | - | 1.041E+04 | 651.4 | - | - | 0 | - |
| - | - | 9722 | 651.9 | - | - | 0 | - |
| - | - | 2757 | 652.4 | - | - | 0 | - |
| 9 | z | 8.353E+04 | 653.4 | 0.001193 | 1.826 | +1 | 6 |
| 11 | c | 7844 | 653.8 | 0.000307 | 0.4696 | +2 | 11 |
| - | - | 8741 | 654.3 | - | - | 0 | - |
| - | - | 4.477E+04 | 654.5 | - | - | 0 | - |
| - | - | 3808 | 654.8 | - | - | 0 | - |
| - | - | 1.314E+04 | 655.5 | - | - | 0 | - |
| - | - | 2993 | 657.3 | - | - | 0 | - |
| - | - | 1.233E+04 | 657.9 | - | - | 0 | - |
| - | - | 8619 | 658.4 | - | - | 0 | - |
| - | - | 9792 | 658.9 | - | - | 0 | - |
| - | - | 1.41E+04 | 659.3 | - | - | 0 | - |
| - | - | 4580 | 659.4 | - | - | 0 | - |
| - | - | 3251 | 659.9 | - | - | 0 | - |
| - | - | 4465 | 660.3 | - | - | 0 | - |
| - | - | 2159 | 664.9 | - | - | 0 | - |
| - | - | 5175 | 665.4 | - | - | 0 | - |
| - | - | 3004 | 666.4 | - | - | 0 | - |
| 9 | y | 3469 | 669.5 | 0.00255 | 3.809 | +1 | 6 |
| - | - | 8097 | 674.3 | - | - | 0 | - |
| - | - | 9700 | 675.3 | - | - | 0 | - |
| - | - | 2795 | 676.3 | - | - | 0 | - |
| - | - | 3191 | 677.4 | - | - | 0 | - |
| - | - | 2707 | 677.9 | - | - | 0 | - |
| 3 | z | 2762 | 678.4 | 0.006331 | 9.332 | +2 | 12 |
| - | - | 7240 | 685.9 | - | - | 0 | - |
| 3 | y | 4684 | 686.4 | 0.008626 | 12.57 | +2 | 12 |
| 3 | z | 1.018E+05 | 686.9 | 0.0005633 | 0.8201 | +2 | 12 |
| - | - | 1.003E+05 | 687.4 | - | - | 0 | - |
| - | - | 5.311E+04 | 687.9 | - | - | 0 | - |
| - | - | 1.759E+04 | 688.4 | - | - | 0 | - |
| - | - | 4878 | 688.9 | - | - | 0 | - |
| - | - | 3507 | 692.4 | - | - | 0 | - |
| 8 | z | 2508 | 693.4 | 0.01025 | 14.78 | +1 | 7 |
| 3 | y | 1.17E+05 | 694.9 | 0.0009059 | 1.304 | +2 | 12 |
| - | - | 8.962E+04 | 695.4 | - | - | 0 | - |
| - | - | 4.765E+04 | 695.9 | - | - | 0 | - |
| - | - | 9127 | 696.4 | - | - | 0 | - |
| 5 | c | 6493 | 702.3 | 0.0009793 | 1.394 | +1 | 5 |
| - | - | 3506 | 703.3 | - | - | 0 | - |
| - | - | 4031 | 708.4 | - | - | 0 | - |
| 12 | c | 1.947E+04 | 709.4 | 0.002599 | 3.664 | +2 | 12 |
| - | - | 2.336E+04 | 709.9 | - | - | 0 | - |
| - | - | 9706 | 710.4 | - | - | 0 | - |
| 8 | z | 1.817E+04 | 710.5 | 0.001397 | 1.967 | +1 | 7 |
| - | - | 9380 | 711.5 | - | - | 0 | - |
| - | - | 2586 | 712.5 | - | - | 0 | - |
| - | - | 4059 | 714.4 | - | - | 0 | - |
| - | - | 4893 | 716.9 | - | - | 0 | - |
| - | - | 5156 | 717.4 | - | - | 0 | - |
| 12 | c | 3.194E+05 | 717.9 | 0.0006161 | 0.8582 | +2 | 12 |
| - | - | 2.664E+05 | 718.4 | - | - | 0 | - |
| - | - | 1.147E+05 | 718.9 | - | - | 0 | - |
| 5 | c | 2.62E+05 | 719.3 | 0.0006753 | 0.9388 | +1 | 5 |
| - | - | 7401 | 719.9 | - | - | 0 | - |
| - | - | 1.13E+05 | 720.3 | - | - | 0 | - |
| - | - | 2.791E+04 | 721.3 | - | - | 0 | - |
| - | - | 5149 | 722.3 | - | - | 0 | - |
| - | - | 2764 | 722.9 | - | - | 0 | - |
| 8 | y | 1.113E+04 | 726.5 | 0.0009084 | 1.25 | +1 | 7 |
| - | - | 2785 | 730.4 | - | - | 0 | - |
| - | - | 2792 | 731.4 | - | - | 0 | - |
| - | - | 3832 | 732.4 | - | - | 0 | - |
| - | - | 2628 | 735.9 | - | - | 0 | - |
| - | - | 3094 | 736.4 | - | - | 0 | - |
| - | - | 2262 | 737.4 | - | - | 0 | - |
| - | - | 3874 | 750.9 | - | - | 0 | - |
| - | - | 2409 | 751.3 | - | - | 0 | - |
| - | - | 4398 | 751.4 | - | - | 0 | - |
| - | - | 4790 | 751.9 | - | - | 0 | - |
| - | - | 8218 | 752.4 | - | - | 0 | - |
| - | - | 3824 | 752.9 | - | - | 0 | - |
| - | - | 2853 | 756.9 | - | - | 0 | - |
| - | - | 3667 | 757.4 | - | - | 0 | - |
| - | - | 4524 | 757.9 | - | - | 0 | - |
| - | - | 7851 | 758.4 | - | - | 0 | - |
| - | - | 1.457E+04 | 758.9 | - | - | 0 | - |
| - | - | 1.079E+04 | 759.4 | - | - | 0 | - |
| - | - | 6706 | 759.9 | - | - | 0 | - |
| 13 | c | 8685 | 765.9 | 0.003109 | 4.059 | +2 | 13 |
| - | - | 1.558E+04 | 766.4 | - | - | 0 | - |
| - | - | 1.209E+04 | 766.9 | - | - | 0 | - |
| - | - | 8321 | 767.4 | - | - | 0 | - |
| - | - | 7123 | 767.9 | - | - | 0 | - |
| - | - | 7253 | 773.4 | - | - | 0 | - |
| - | - | 7029 | 773.9 | - | - | 0 | - |
| 13 | c | 1.851E+05 | 774.4 | 0.001309 | 1.69 | +2 | 13 |
| - | - | 1.837E+05 | 774.9 | - | - | 0 | - |
| - | - | 9.109E+04 | 775.4 | - | - | 0 | - |
| - | - | 3.293E+04 | 775.9 | - | - | 0 | - |
| - | - | 7372 | 776.4 | - | - | 0 | - |
| - | - | 3470 | 778.4 | - | - | 0 | - |
| - | - | 4510 | 778.9 | - | - | 0 | - |
| 2 | y | 7150 | 779.4 | 0.00437 | 5.607 | +2 | 13 |
| 2 | z | 4.303E+04 | 779.9 | 9.143E-05 | 0.1172 | +2 | 13 |
| - | - | 3.411E+04 | 780.4 | - | - | 0 | - |
| 7 | w | 3828 | 780.5 | 0.001466 | 1.879 | +1 | 8 |
| - | - | 1.76E+04 | 780.9 | - | - | 0 | - |
| - | - | 8035 | 781.4 | - | - | 0 | - |
| - | - | 7379 | 781.9 | - | - | 0 | - |
| - | - | 4343 | 782.4 | - | - | 0 | - |
| - | - | 3368 | 787.4 | - | - | 0 | - |
| 2 | y | 2.019E+04 | 787.9 | 0.01202 | 15.25 | +2 | 13 |
| - | - | 1.098E+04 | 788.4 | - | - | 0 | - |
| - | - | 7157 | 788.9 | - | - | 0 | - |
| - | - | 2.513E+04 | 790.9 | - | - | 0 | - |
| - | - | 2.818E+04 | 791.4 | - | - | 0 | - |
| - | - | 1.728E+04 | 791.9 | - | - | 0 | - |
| - | - | 5290 | 792.4 | - | - | 0 | - |
| - | - | 4171 | 792.9 | - | - | 0 | - |
| - | - | 7542 | 794.9 | - | - | 0 | - |
| - | - | 1.793E+04 | 795.4 | - | - | 0 | - |
| - | - | 1.718E+04 | 795.9 | - | - | 0 | - |
| - | - | 9512 | 796.4 | - | - | 0 | - |
| - | - | 1.462E+04 | 796.4 | - | - | 0 | - |
| - | - | 1.63E+04 | 796.9 | - | - | 0 | - |
| - | - | 4595 | 797.4 | - | - | 0 | - |
| - | - | 6112 | 797.4 | - | - | 0 | - |
| - | - | 2878 | 797.9 | - | - | 0 | - |
| - | - | 1.819E+04 | 800.4 | - | - | 0 | - |
| - | - | 2.194E+04 | 800.9 | - | - | 0 | - |
| - | - | 2.34E+04 | 801.4 | - | - | 0 | - |
| - | - | 1.845E+04 | 801.9 | - | - | 0 | - |
| - | - | 9504 | 802.4 | - | - | 0 | - |
| - | - | 1.165E+04 | 802.9 | - | - | 0 | - |
| - | - | 9432 | 803.4 | - | - | 0 | - |
| - | - | 1.385E+04 | 803.9 | - | - | 0 | - |
| - | - | 8333 | 804.4 | - | - | 0 | - |
| - | - | 4140 | 804.9 | - | - | 0 | - |
| - | - | 5029 | 808.9 | - | - | 0 | - |
| - | - | 3.292E+04 | 809.4 | - | - | 0 | - |
| - | - | 3.498E+04 | 809.9 | - | - | 0 | - |
| - | - | 3.001E+04 | 810.4 | - | - | 0 | - |
| - | - | 1.276E+04 | 810.9 | - | - | 0 | - |
| - | - | 2.815E+04 | 811.4 | - | - | 0 | - |
| - | - | 5258 | 811.4 | - | - | 0 | - |
| - | - | 1.986E+04 | 812.4 | - | - | 0 | - |
| - | - | 7402 | 813.4 | - | - | 0 | - |
| - | - | 2980 | 813.9 | - | - | 0 | - |
| - | - | 4563 | 814.4 | - | - | 0 | - |
| - | - | 6001 | 814.9 | - | - | 0 | - |
| - | - | 2805 | 815.4 | - | - | 0 | - |
| - | - | 5109 | 822.5 | - | - | 0 | - |
| - | - | 2.273E+04 | 822.9 | - | - | 0 | - |
| - | - | 1.611E+05 | 823.4 | - | - | 0 | - |
| 7 | y | 3.394E+04 | 823.5 | 0.0002218 | 0.2693 | +1 | 8 |
| - | - | 1.522E+05 | 823.9 | - | - | 0 | - |
| - | - | 8.266E+04 | 824.4 | - | - | 0 | - |
| - | - | 1.584E+04 | 824.5 | - | - | 0 | - |
| - | - | 2.953E+04 | 824.9 | - | - | 0 | - |
| - | - | 9197 | 825.4 | - | - | 0 | - |
| - | - | 6227 | 825.5 | - | - | 0 | - |
| - | - | 4197 | 830.4 | - | - | 0 | - |
| - | - | 1.54E+04 | 830.9 | - | - | 0 | - |
| - | - | 3.654E+05 | 831.5 | - | - | 0 | - |
| - | - | 1.644E+06 | 832 | - | - | 0 | - |
| - | - | 1.475E+06 | 832.5 | - | - | 0 | - |
| - | - | 7.994E+05 | 833 | - | - | 0 | - |
| - | - | 2.498E+05 | 833.5 | - | - | 0 | - |
| - | - | 6.362E+04 | 834 | - | - | 0 | - |
| 6 | c | 2.511E+04 | 839.4 | 0.0006002 | 0.715 | +1 | 6 |
| - | - | 9449 | 840.4 | - | - | 0 | - |
| - | - | 2964 | 841.4 | - | - | 0 | - |
| - | - | 2.039E+04 | 850.5 | - | - | 0 | - |
| - | - | 1.119E+04 | 851.5 | - | - | 0 | - |
| - | - | 2.32E+04 | 852.5 | - | - | 0 | - |
| - | - | 8899 | 853.5 | - | - | 0 | - |
| - | - | 5336 | 854.5 | - | - | 0 | - |
| - | - | 7014 | 855.5 | - | - | 0 | - |
| - | - | 3241 | 856.5 | - | - | 0 | - |
| - | - | 2622 | 873.5 | - | - | 0 | - |
| - | - | 3539 | 883.4 | - | - | 0 | - |
| 6 | z | 2.508E+05 | 944.6 | 0.0009886 | 1.047 | +1 | 9 |
| - | - | 1.489E+05 | 945.6 | - | - | 0 | - |
| - | - | 5.26E+04 | 946.6 | - | - | 0 | - |
| - | - | 9259 | 947.6 | - | - | 0 | - |
| 7 | c | 2.879E+04 | 953.4 | 0.001182 | 1.24 | +1 | 7 |
| - | - | 1.777E+04 | 954.4 | - | - | 0 | - |
| - | - | 4772 | 955.4 | - | - | 0 | - |
| 6 | y | 2.599E+04 | 960.6 | 8.681E-05 | 0.09037 | +1 | 9 |
| - | - | 1.378E+04 | 961.6 | - | - | 0 | - |
| - | - | 3690 | 962.6 | - | - | 0 | - |
| - | - | 5023 | 968.6 | - | - | 0 | - |
| - | - | 3520 | 969.6 | - | - | 0 | - |
| - | - | 1.071E+04 | 982.5 | - | - | 0 | - |
| - | - | 1.401E+04 | 983.6 | - | - | 0 | - |
| - | - | 4762 | 984.6 | - | - | 0 | - |
| - | - | 3175 | 987.6 | - | - | 0 | - |
| - | - | 1.16E+04 | 1009 | - | - | 0 | - |
| 8 | c | 2.32E+05 | 1010 | 0.0008977 | 0.8884 | +1 | 8 |
| - | - | 1.301E+05 | 1011 | - | - | 0 | - |
| - | - | 4.566E+04 | 1012 | - | - | 0 | - |
| - | - | 1.303E+04 | 1013 | - | - | 0 | - |
| - | - | 3453 | 1026 | - | - | 0 | - |
| 5 | y | 4395 | 1081 | 0.003979 | 3.682 | +1 | 10 |
| 5 | z | 1.276E+05 | 1082 | 0.0007926 | 0.7327 | +1 | 10 |
| - | - | 2.848E+05 | 1083 | - | - | 0 | - |
| - | - | 1.662E+05 | 1084 | - | - | 0 | - |
| - | - | 5.727E+04 | 1085 | - | - | 0 | - |
| - | - | 1.151E+04 | 1086 | - | - | 0 | - |
| - | - | 3116 | 1087 | - | - | 0 | - |
| - | - | 7496 | 1095 | - | - | 0 | - |
| - | - | 7220 | 1096 | - | - | 0 | - |
| - | - | 5076 | 1096 | - | - | 0 | - |
| - | - | 1.189E+04 | 1097 | - | - | 0 | - |
| 5 | y | 2.552E+04 | 1098 | 0.0001092 | 0.09952 | +1 | 10 |
| - | - | 1.896E+04 | 1099 | - | - | 0 | - |
| - | - | 4664 | 1100 | - | - | 0 | - |
| - | - | 2815 | 1107 | - | - | 0 | - |
| - | - | 2295 | 1109 | - | - | 0 | - |
| - | - | 2414 | 1124 | - | - | 0 | - |
| - | - | 5503 | 1138 | - | - | 0 | - |
| 9 | c | 3.052E+05 | 1139 | 0.001455 | 1.278 | +1 | 9 |
| - | - | 2.06E+05 | 1140 | - | - | 0 | - |
| - | - | 7.398E+04 | 1141 | - | - | 0 | - |
| - | - | 1.8E+04 | 1142 | - | - | 0 | - |
| - | - | 4030 | 1143 | - | - | 0 | - |
| - | - | 6629 | 1146 | - | - | 0 | - |
| - | - | 4685 | 1147 | - | - | 0 | - |
| 4 | w | 2828 | 1152 | 0.0005421 | 0.4707 | +1 | 11 |
| - | - | 8262 | 1166 | - | - | 0 | - |
| - | - | 7551 | 1167 | - | - | 0 | - |
| - | - | 2714 | 1195 | - | - | 0 | - |
| 4 | y | 4195 | 1209 | 0.002841 | 2.35 | +1 | 11 |
| 4 | z | 3.829E+04 | 1210 | 0.001297 | 1.072 | +1 | 11 |
| - | - | 1.744E+05 | 1211 | - | - | 0 | - |
| - | - | 1.136E+05 | 1212 | - | - | 0 | - |
| - | - | 3.716E+04 | 1213 | - | - | 0 | - |
| - | - | 1.079E+04 | 1214 | - | - | 0 | - |
| 4 | y | 1.151E+04 | 1226 | 0.0001511 | 0.1233 | +1 | 11 |
| - | - | 6449 | 1227 | - | - | 0 | - |
| - | - | 4567 | 1237 | - | - | 0 | - |
| - | - | 5958 | 1259 | - | - | 0 | - |
| - | - | 3925 | 1260 | - | - | 0 | - |
| - | - | 3514 | 1263 | - | - | 0 | - |
| - | - | 6011 | 1264 | - | - | 0 | - |
| - | - | 4419 | 1265 | - | - | 0 | - |
| - | - | 8305 | 1306 | - | - | 0 | - |
| 11 | c | 1.344E+05 | 1307 | 0.0002001 | 0.1531 | +1 | 11 |
| - | - | 1.112E+05 | 1308 | - | - | 0 | - |
| - | - | 4.589E+04 | 1309 | - | - | 0 | - |
| - | - | 1.233E+04 | 1310 | - | - | 0 | - |
| - | - | 4934 | 1311 | - | - | 0 | - |
| 3 | z | 2.201E+04 | 1373 | 0.0002244 | 0.1635 | +1 | 12 |
| - | - | 9.256E+04 | 1374 | - | - | 0 | - |
| - | - | 6.026E+04 | 1375 | - | - | 0 | - |
| - | - | 2.752E+04 | 1376 | - | - | 0 | - |
| - | - | 8069 | 1377 | - | - | 0 | - |
| 3 | y | 4675 | 1389 | 0.001398 | 1.006 | +1 | 12 |
| - | - | 4336 | 1390 | - | - | 0 | - |
| - | - | 1.403E+04 | 1419 | - | - | 0 | - |
| - | - | 1.212E+04 | 1420 | - | - | 0 | - |
| - | - | 5970 | 1421 | - | - | 0 | - |
| 12 | c | 4.778E+04 | 1435 | 0.001379 | 0.9612 | +1 | 12 |
| - | - | 7.595E+04 | 1436 | - | - | 0 | - |
| - | - | 4.219E+04 | 1437 | - | - | 0 | - |
| - | - | 1.775E+04 | 1438 | - | - | 0 | - |
| - | - | 4968 | 1439 | - | - | 0 | - |
| - | - | 2618 | 1504 | - | - | 0 | - |
| - | - | 5266 | 1505 | - | - | 0 | - |
| - | - | 5355 | 1506 | - | - | 0 | - |
| - | - | 2471 | 1507 | - | - | 0 | - |
| - | - | 7264 | 1532 | - | - | 0 | - |
| - | - | 9160 | 1533 | - | - | 0 | - |
| - | - | 3635 | 1534 | - | - | 0 | - |
| 13 | c | 1.975E+04 | 1548 | 0.0001282 | 0.08282 | +1 | 13 |
| - | - | 4.461E+04 | 1549 | - | - | 0 | - |
| - | - | 3.223E+04 | 1550 | - | - | 0 | - |
| - | - | 1.156E+04 | 1551 | - | - | 0 | - |
| - | - | 4992 | 1552 | - | - | 0 | - |
| 2 | z | 5542 | 1559 | 0.00221 | 1.418 | +1 | 13 |
| - | - | 3.509E+04 | 1560 | - | - | 0 | - |
| - | - | 2.91E+04 | 1561 | - | - | 0 | - |
| - | - | 1.24E+04 | 1562 | - | - | 0 | - |
| - | - | 5820 | 1563 | - | - | 0 | - |
| 2 | y | 4192 | 1575 | 0.01078 | 6.843 | +1 | 13 |
| - | - | 2539 | 1576 | - | - | 0 | - |
| - | - | 5794 | 1581 | - | - | 0 | - |
| - | - | 7936 | 1582 | - | - | 0 | - |
| - | - | 3671 | 1583 | - | - | 0 | - |
| - | - | 3729 | 1591 | - | - | 0 | - |
| - | - | 5029 | 1601 | - | - | 0 | - |
| - | - | 4100 | 1602 | - | - | 0 | - |
| - | - | 5889 | 1618 | - | - | 0 | - |
| - | - | 3.77E+04 | 1619 | - | - | 0 | - |
| - | - | 3.174E+04 | 1620 | - | - | 0 | - |
| - | - | 1.64E+04 | 1621 | - | - | 0 | - |
| - | - | 4909 | 1622 | - | - | 0 | - |
| - | - | 2767 | 1628 | - | - | 0 | - |
| - | - | 2.053E+04 | 1629 | - | - | 0 | - |
| - | - | 1.675E+04 | 1630 | - | - | 0 | - |
| - | - | 8669 | 1631 | - | - | 0 | - |
| - | - | 3874 | 1635 | - | - | 0 | - |
| - | - | 1.521E+04 | 1636 | - | - | 0 | - |
| - | - | 1.077E+04 | 1637 | - | - | 0 | - |
| - | - | 8231 | 1638 | - | - | 0 | - |
| - | - | 2370 | 1639 | - | - | 0 | - |
| - | - | 4963 | 1645 | - | - | 0 | - |
| - | - | 4.153E+04 | 1646 | - | - | 0 | - |
| - | - | 1.533E+05 | 1647 | - | - | 0 | - |
| - | - | 1.35E+05 | 1648 | - | - | 0 | - |
| - | - | 7.493E+04 | 1649 | - | - | 0 | - |
| - | - | 2.317E+04 | 1650 | - | - | 0 | - |
| - | - | 5322 | 1651 | - | - | 0 | - |
| - | - | 2.987E+04 | 1662 | - | - | 0 | - |
| - | - | 1.733E+05 | 1663 | - | - | 0 | - |
| - | - | 7.792E+05 | 1664 | - | - | 0 | - |
| - | - | 6.916E+05 | 1665 | - | - | 0 | - |
| - | - | 3.6E+05 | 1666 | - | - | 0 | - |
| - | - | 1.173E+05 | 1667 | - | - | 0 | - |
| - | - | 3.336E+04 | 1668 | - | - | 0 | - |

m/z Charge Intensity FragmentType MassShift Position
122.43207550048828 0 1249.4977
129.10243225097656 0 4827.515
130.0655517578125 0 2984.52
132.0809783935547 0 6440.299
132.1020965576172 0 8245.447 y 13
136.0759735107422 0 3833.3726
143.7855682373047 0 1381.6019
146.09263610839844 0 1589.8337
153.07876586914062 0 10353.659
168.10198974609375 0 1468.5378
170.1539306640625 0 1729.2396
186.0789337158203 0 4462.808
187.0868682861328 0 8611.292
199.1807403564453 0 3240.7812
214.1556854248047 0 2362.4048
216.7711944580078 0 1757.8867
223.11924743652344 0 4504.3296
226.15525817871094 0 6103.2275
228.11184692382812 0 2204.3735
245.13729858398438 0 13528.748
245.18609619140625 0 17638.553 y 12
246.12396240234375 0 52684.34
246.13909912109375 0 2942.548
246.18991088867188 0 2381.674
247.12753295898438 0 6505.2275
255.10011291503906 0 3900.085
255.19451904296875 0 2743.0789
256.20074462890625 0 1733.9437
258.14520263671875 0 7358.9927
268.98162841796875 0 1749.1257
273.1115417480469 0 4372.2554
274.118896484375 0 68646.836
275.12255859375 0 12829.372
281.1737365722656 0 8975.436
282.1793212890625 0 2602.1294
292.12982177734375 0 9472.466
299.2088928222656 0 2496.9287
301.2001953125 0 5392.093
309.1564025878906 0 9425.667
310.1610107421875 0 2360.8784
314.2080078125 0 3924.8013
320.122802734375 0 2075.6526
323.185791015625 0 2684.3416
346.7266845703125 0 5779.834
349.1418762207031 0 6037.259
353.461669921875 0 2351.4072
357.2625732421875 0 27722.951 z 11
358.26800537109375 0 12173.412
359.27099609375 0 2794.5352
373.2811584472656 0 6598.901 y 11
382.2812194824219 0 5157.302
394.2582092285156 0 9072.2
395.2625732421875 0 1897.9485
399.26153564453125 0 2171.8816
401.13690185546875 0 2908.9495
406.18701171875 0 3467.818
412.2744140625 0 37411.62 y 6
412.7757568359375 0 17656.74
413.2276611328125 0 6386.9907
413.2772521972656 0 4601.618
420.1847839355469 0 2995.3442 c Ammonia loss 5
420.6858825683594 0 3037.3406
426.23944091796875 0 2304.6094
427.292236328125 0 15112.284 w 10
428.2980041503906 0 5876.708
429.1904296875 0 2470.3042
435.1915283203125 0 8335.249
436.17352294921875 0 2004.3734
437.1825256347656 0 10581.447
438.186279296875 0 2549.1116
439.1885070800781 0 2384.62
453.2014465332031 0 14896.41
454.2086486816406 0 27230.797 c 2
455.21209716796875 0 7865.043
463.5989074707031 0 11674.078 y 2
463.93365478515625 0 7310.754
465.2096252441406 0 9456.321
468.3187561035156 0 34387.594
469.2891845703125 0 8015.6416
469.3227233886719 0 8694.701
470.2937927246094 0 2843.5151
470.3335266113281 0 18261.09 y 10
471.3385925292969 0 3744.4612
473.2435302734375 0 18516.182 c Ammonia loss 11
473.5762939453125 0 10078.687
473.9121398925781 0 5248.5244
474.24493408203125 0 2973.537
477.2014465332031 0 16580.486
478.2055358886719 0 4649.397
479.2056884765625 0 2261.2122
480.2247009277344 0 3984.2976
480.8035583496094 0 3069.8635 y 5
481.3060607910156 0 4859.965
482.2643127441406 0 2435.062
482.2989501953125 0 9366.098
483.7891845703125 0 2619.8914
495.36529541015625 0 3962.539
498.327392578125 0 4099.111
501.6080017089844 0 4510.89
501.9422302246094 0 2694.0293
504.9329833984375 0 16352.533
505.26666259765625 0 12390.018
505.599365234375 0 6922.4326
507.3428039550781 0 3206.8286
510.2225646972656 0 2489.4775
510.93768310546875 0 73448.41 c Ammonia loss 12
511.27191162109375 0 70198.23
511.6062316894531 0 27057.674
511.940673828125 0 9985.823
512.2704467773438 0 4111.5083
516.940673828125 0 8432.708
517.2760620117188 0 8331.486
517.6087036132812 0 3646.966
523.2413330078125 0 9106.14
525.352783203125 0 110037.99 z 9
525.6250610351562 0 9102.734 y 1
525.9602661132812 0 12600.251
526.2945556640625 0 8257.48
526.3560180664062 0 37947.977
527.3585205078125 0 7465.8726
538.2537841796875 0 5996.472
539.258544921875 0 4757.7285
539.2984619140625 0 3609.6858
540.6119384765625 0 2314.163
541.3211059570312 0 5942.6772 z 4
541.3718872070312 0 12180.767 y 9
541.8262939453125 0 5612.5225
542.3726806640625 0 4172.027
546.2212524414062 0 6143.254
547.8137817382812 0 2588.605
548.5853881835938 0 3591.9333
548.6328735351562 0 19009.516
548.9658813476562 0 14577.756
549.298583984375 0 6586.121
549.333740234375 0 10164.823 y 4
549.6318969726562 0 7886.1772
549.834228515625 0 10603.67
549.9657592773438 0 2519.2056
550.3348388671875 0 2785.414
553.396484375 0 4489.0986
553.9655151367188 0 6579.732
554.3017578125 0 3949.4402
554.635986328125 0 554773.44
554.9700927734375 0 566830.06
555.304443359375 0 304554.66
555.6384887695312 0 101346.82
555.6881103515625 0 2586.7554
555.97265625 0 25607.203
561.2683715820312 0 3146.5222 c Ammonia loss 8
563.249267578125 0 39564.215
564.2496337890625 0 11779.334 c Water loss 3
565.242431640625 0 16780.031 c Ammonia loss 3
566.2451782226562 0 12894.201
566.4039916992188 0 9461.967
567.24951171875 0 3484.94
567.4049682617188 0 3096.2393
569.8164672851562 0 2606.749
572.2647705078125 0 2681.7905
572.8050537109375 0 5955.6055
573.3060302734375 0 3076.727
576.3385620117188 0 32233.705 w 3
576.8399047851562 0 21329.084
577.3355712890625 0 9653.8
579.3603515625 0 2429.169
581.26025390625 0 23847.096
582.2677001953125 0 126947.18 c 3
582.3748779296875 0 11682.403
583.270751953125 0 37512.598
583.37646484375 0 3385.572
584.2728881835938 0 6991.4956
587.7830200195312 0 2624.6555
595.3825073242188 0 26139.186 w 8
596.3860473632812 0 9695.353
596.7915649414062 0 2776.0513 c Ammonia loss 9
597.38916015625 0 4064.1514
604.3501586914062 0 4991.621
604.8506469726562 0 7541.8843 y Ammonia loss 3
605.3531494140625 0 92284.625 z 3
605.8556518554688 0 99156.89
606.357177734375 0 49953.1
606.8600463867188 0 12333.085
607.3580932617188 0 4968.6714
609.4007568359375 0 2354.8403
609.4574584960938 0 6483.727
610.3965454101562 0 24152.943
611.4015502929688 0 7195.6543
613.3624267578125 0 36184.02 y 3
613.8638305664062 0 20874.54
614.3641357421875 0 6331.802
620.3408813476562 0 3600.5435
620.8448486328125 0 3208.1514
624.4209594726562 0 7468.072
629.3464965820312 0 14841.712
629.8486938476562 0 16526.246
630.3504028320312 0 5396.7534
637.4267578125 0 4371.9453
638.4371337890625 0 2519.6284
645.8291625976562 0 3475.2295
651.3583374023438 0 10411.158
651.860107421875 0 9721.769
652.3580322265625 0 2756.8235
653.4482421875 0 83527.98 z 8
653.825927734375 0 7844.2534 c 10
654.3277587890625 0 8741.461
654.4520874023438 0 44767.184
654.8291625976562 0 3808.0662
655.455078125 0 13137.22
657.29931640625 0 2992.5125
657.8674926757812 0 12327.369
658.3692016601562 0 8619.499
658.8593139648438 0 9792.226
659.2944946289062 0 14103.604
659.3567504882812 0 4580.037
659.8544311523438 0 3250.7866
660.2973022460938 0 4464.8784
664.8843383789062 0 2158.6016
665.362060546875 0 5174.9683
666.3588256835938 0 3003.7903
669.4683227539062 0 3469.3293 y 8
674.3056640625 0 8096.9663
675.3126220703125 0 9700.456
676.3134155273438 0 2794.998
677.3758544921875 0 3190.5745
677.880859375 0 2707.0557
678.377197265625 0 2761.9783 z Ammonia loss 2
685.884765625 0 7239.8237
686.3888549804688 0 4684.2456 y Ammonia loss 2
686.8847045898438 0 101847.09 z 2
687.38623046875 0 100310.91
687.8881225585938 0 53112.43
688.3890991210938 0 17591.95
688.8906860351562 0 4877.922
692.4486083984375 0 3506.8618
693.4522094726562 0 2508.173 z Ammonia loss 7
694.8944091796875 0 116988.32 y 2
695.3955078125 0 89618.92
695.8966674804688 0 47646.195
696.396728515625 0 9126.745
702.3004150390625 0 6492.898 c Ammonia loss 4
703.3065795898438 0 3506.1636
708.3875122070312 0 4031.4937
709.3624267578125 0 19465.895 c Ammonia loss 11
709.8630981445312 0 23358.47
710.3619384765625 0 9705.576
710.4699096679688 0 18171.996 z 7
711.4718017578125 0 9379.784
712.4732666015625 0 2586.3738
714.3894653320312 0 4059.0593
716.86767578125 0 4892.984
717.3693237304688 0 5155.542
717.8737182617188 0 319442.4 c 11
718.375244140625 0 266358.12
718.8765258789062 0 114695.766
719.32666015625 0 262028.7 c 4
719.8799438476562 0 7400.528
720.329833984375 0 113008.37
721.33251953125 0 27910.354
722.3348999023438 0 5148.6406
722.8973388671875 0 2764.2874
726.486328125 0 11130.672 y 7
730.3890380859375 0 2785.0298
731.3817749023438 0 2792.3364
732.3899536132812 0 3831.7493
735.9143676757812 0 2627.8943
736.4043579101562 0 3094.3105
737.4027099609375 0 2262.3965
750.9096069335938 0 3874.0217
751.334228515625 0 2409.1802
751.410888671875 0 4398.0283
751.9071044921875 0 4790.3076
752.4119873046875 0 8217.637
752.9086303710938 0 3823.5127
756.9005737304688 0 2853.237
757.3974609375 0 3667.494
757.9223022460938 0 4523.5923
758.4152221679688 0 7850.538
758.9136352539062 0 14570.992
759.4136962890625 0 10786.553
759.9130859375 0 6706.4556
765.9049682617188 0 8685.055 c Ammonia loss 12
766.4124755859375 0 15578.088
766.9114379882812 0 12094.798
767.4194946289062 0 8320.646
767.9259643554688 0 7122.7974
773.4132690429688 0 7253.2085
773.918701171875 0 7029.288
774.4164428710938 0 185085.8 c 12
774.917236328125 0 183652.2
775.41845703125 0 91094.23
775.9198608398438 0 32930.73
776.419677734375 0 7371.9497
778.424072265625 0 3470.3025
778.9280395507812 0 4510.46
779.4242553710938 0 7149.87 y Ammonia loss 1
779.9238891601562 0 43033.152 z 1
780.4257202148438 0 34105.12
780.499267578125 0 3828.3943 w 6
780.9273071289062 0 17603.562
781.4276733398438 0 8034.596
781.919677734375 0 7378.8877
782.425537109375 0 4342.7407
787.4287719726562 0 3367.9263
787.921142578125 0 20193.412 y 1
788.4266967773438 0 10977.444
788.9268188476562 0 7157.32
790.9281005859375 0 25125.602
791.4285278320312 0 28183.865
791.92919921875 0 17279.477
792.4364013671875 0 5289.5073
792.9324340820312 0 4170.6094
794.9161987304688 0 7541.971
795.413818359375 0 17925.086
795.9125366210938 0 17178.54
796.3525390625 0 9511.785
796.4193725585938 0 14617.448
796.92431640625 0 16299.17
797.3541259765625 0 4594.599
797.4251708984375 0 6112.12
797.9280395507812 0 2877.8953
800.439208984375 0 18190.645
800.940185546875 0 21942.777
801.4439697265625 0 23403.87
801.9368286132812 0 18450.744
802.43017578125 0 9504.484
802.92529296875 0 11650.422
803.4262084960938 0 9432.384
803.9253540039062 0 13846.205
804.4251098632812 0 8332.692
804.9302978515625 0 4139.7485
808.9425659179688 0 5029.3765
809.4439697265625 0 32917.3
809.9463500976562 0 34981.668
810.4427490234375 0 30014.107
810.9385375976562 0 12758.009
811.3641967773438 0 28149.703
811.4386596679688 0 5257.708
812.3688354492188 0 19861.482
813.3704223632812 0 7401.8584
813.9224243164062 0 2979.6206
814.437255859375 0 4563.0103
814.9365844726562 0 6000.914
815.442626953125 0 2805.3691
822.5325317382812 0 5108.5083
822.9478149414062 0 22727.148
823.4422607421875 0 161117.67
823.5402221679688 0 33937.1 y 6
823.9425659179688 0 152160.86
824.4436645507812 0 82664.64
824.5433959960938 0 15843.895
824.9447021484375 0 29534.941
825.4465942382812 0 9197.242
825.5464477539062 0 6226.5293
830.4422607421875 0 4196.8496
830.9454956054688 0 15402.183
831.4505615234375 0 365386.9
831.9539794921875 0 1644090.1
832.4556274414062 0 1474952.5
832.9568481445312 0 799412.1
833.4578247070312 0 249797.75
833.9591674804688 0 63618.188
839.3589477539062 0 25106.707 c Ammonia loss 5
840.3640747070312 0 9448.764
841.3637084960938 0 2963.9226
850.52880859375 0 20392.004
851.53076171875 0 11186.624
852.5416870117188 0 23201.023
853.5464477539062 0 8899.493
854.485595703125 0 5336.4834
855.4956665039062 0 7014.3384
856.4948120117188 0 3241.3267
873.504638671875 0 2621.7358
883.40576171875 0 3539.0737
944.5811767578125 0 250821.14 z 5
945.5849609375 0 148921.44
946.5877075195312 0 52604.355
947.5928344726562 0 9258.889
953.4388427734375 0 28790.496 c 6
954.4424438476562 0 17773.154
955.4453125 0 4772.0977
960.5989990234375 0 25991.523 y 5
961.6041259765625 0 13779.491
962.6013793945312 0 3689.5737
968.5746459960938 0 5022.6655
969.5782470703125 0 3519.8535
982.5472412109375 0 10707.265
983.5532836914062 0 14011.256
984.5570068359375 0 4761.5737
987.5806274414062 0 3175.387
1009.4522705078125 0 11600.007
1010.4600219726562 0 231966.8 c 7
1011.462646484375 0 130094.375
1012.4652099609375 0 45662.875
1013.467041015625 0 13034.8
1025.582763671875 0 3453.3818
1080.63525390625 0 4394.6646 y Ammonia loss 4
1081.639892578125 0 127579.88 z 4
1082.6468505859375 0 284774.25
1083.6500244140625 0 166179
1084.6533203125 0 57273.33
1085.655517578125 0 11511.2
1086.668701171875 0 3115.5662
1094.5404052734375 0 7495.835
1095.5413818359375 0 7219.92
1095.6417236328125 0 5075.673
1096.640625 0 11889.191
1097.65771484375 0 25522.69 y 4
1098.6607666015625 0 18956.52
1099.6634521484375 0 4664.2773
1106.53369140625 0 2814.7363
1108.541748046875 0 2295.2212
1123.548583984375 0 2413.94
1137.544189453125 0 5502.5283
1138.5555419921875 0 305162.6 c 8
1139.557861328125 0 205968.55
1140.560546875 0 73977.39
1141.5634765625 0 18002.578
1142.5648193359375 0 4030.4153
1145.616455078125 0 6629.0327
1146.60986328125 0 4684.797
1151.6678466796875 0 2827.5918 w 3
1165.57666015625 0 8262.349
1166.5853271484375 0 7550.897
1194.575927734375 0 2714.1018
1208.68701171875 0 4195.0464 y Ammonia loss 3
1209.698974609375 0 38290.31 z 3
1210.7056884765625 0 174427.62
1211.7081298828125 0 113594.01
1212.7117919921875 0 37160.633
1213.71923828125 0 10792.3955
1225.716552734375 0 11512.668 y 3
1226.7181396484375 0 6448.596
1236.6258544921875 0 4567.0234
1258.694580078125 0 5957.973
1259.6986083984375 0 3925.2598
1262.6214599609375 0 3513.7856
1263.6351318359375 0 6010.517
1264.6427001953125 0 4418.5537
1305.638671875 0 8305.31
1306.6441650390625 0 134373.67 c 10
1307.64697265625 0 111209.63
1308.6502685546875 0 45892.223
1309.6536865234375 0 12329.886
1310.658447265625 0 4934.2026
1372.76123046875 0 22014.326 z 2
1373.7681884765625 0 92562.71
1374.771484375 0 60257.87
1375.7735595703125 0 27521.32
1376.7840576171875 0 8068.9795
1388.7811279296875 0 4675.0547 y 2
1389.7891845703125 0 4336.0845
1418.7196044921875 0 14030.027
1419.7205810546875 0 12124.85
1420.7183837890625 0 5969.6626
1434.737548828125 0 47779.24 c 11
1435.7432861328125 0 75950.86
1436.7469482421875 0 42189.215
1437.7510986328125 0 17750.857
1438.758544921875 0 4967.6494
1503.823486328125 0 2618.4565
1504.81298828125 0 5266.3257
1505.819091796875 0 5354.589
1506.818115234375 0 2470.6636
1531.80517578125 0 7263.957
1532.8077392578125 0 9159.73
1533.8079833984375 0 3635.3076
1547.8231201171875 0 19745.023 c 12
1548.8291015625 0 44614.293
1549.8321533203125 0 32228.443
1550.838623046875 0 11561.792
1551.844970703125 0 4991.51
1558.842529296875 0 5541.572 z 1
1559.8468017578125 0 35094.395
1560.8494873046875 0 29102.139
1561.853759765625 0 12398.014
1562.8453369140625 0 5820.2734
1574.8482666015625 0 4192.242 y 1
1575.8575439453125 0 2539.094
1580.8480224609375 0 5793.51
1581.8406982421875 0 7936.345
1582.8533935546875 0 3670.589
1590.8175048828125 0 3729.3223
1600.8714599609375 0 5028.568
1601.878173828125 0 4100.348
1617.877197265625 0 5889.2803
1618.886474609375 0 37698.508
1619.8870849609375 0 31739.402
1620.887451171875 0 16397.113
1621.8797607421875 0 4908.7705
1627.871826171875 0 2767.3618
1628.87255859375 0 20534.809
1629.8721923828125 0 16745.285
1630.8779296875 0 8668.644
1634.901123046875 0 3874.4192
1635.9107666015625 0 15207.048
1636.9107666015625 0 10771.299
1637.91796875 0 8231.446
1638.91796875 0 2369.6763
1644.870849609375 0 4962.8877
1645.87841796875 0 41531.76
1646.8809814453125 0 153254.12
1647.8839111328125 0 135033.16
1648.8870849609375 0 74929.75
1649.890380859375 0 23172.424
1650.8724365234375 0 5321.509
1661.891845703125 0 29873.715
1662.89892578125 0 173328.03
1663.9068603515625 0 779183.7
1664.9093017578125 0 691568.44
1665.912841796875 0 360012.38
1666.9149169921875 0 117335.94
1667.9158935546875 0 33357.41

Spectrum Details

|  |  |
| --- | --- |
| Matched peaks? Matched peaks The total absolute number of peaks matched. Additionally in brackets the total fraction of peaks matched and the total number of peaks is shown. | 69 (13.50% of 511) |
| FDR? FDR The false discovery rate estimated for this peptide. It is calculated by matching all theoretical fragments with a non-integer shift with the raw peaks for this spectrum. This is done with 40 different shifts. The resulting percentage is the average number of annotated peaks over the number of annotated peaks with the correct spectrum. | 3.07% |
| Satellite FDR? Satellite FDR See the FDR for details on its calculation. This satellite ion specific FDR only contains the satellite ions (d/w) for I/L/J positions. | - |
| PSM Score? PSM Score The PSM Score as given by Hecklib to this annotated spectrum. It is shown with three significant figures. | 527 |

#### Spectrum 4749? Spectrum 4749 The raw spectrum of this peptide as annotated by Hecklib. The fragments are coloured according to ion type (see legend). Any peaks with a star '\*' as text can be hovered over to see the full details, first the ion type second the mass shift type. By hovering over the amino acids in the peptide or ions in the legend the corresponding peaks are highlighted. By toggling the 'Unassigned' label you can turn the background (unassigned) peaks on or off in the plot. By updating the slider in the Ion legend you can update the spectrum to only show the top X% of the peaks with labels. The top X% means any peak that is within X% of the highest intensity. By dragging in the spectrum you can zoom in to a specific part of the spectrum and use 'Zoom Out' to get back to the original zoom level. The annotation of the spectrum is based on the given sequence in the peptides file and is done with different software so inconsistencies are likely. The peaks are annotated based on the given sequence, with 20 ppm tolerance.

Copy Data

##### Spectrum 4749 (TSV)

###### Preview

```
Loading example...
```

*Click on the button to copy the data to your clipboard.*

Mz MinMz MaxIntensity Max

WidthHeightPeptide font sizePeptide stroke widthSpectrum font sizeSpectrum stroke widthCompact peptide

Ion legend

wxyz

abcd

OtherUnassignedIon^Charge^_Position_Show for top:%

SWYQHHPGKAPKJJ

02.39e+44.78e+47.16e+49.55e+4

Zoom Out

y^+1^_1_y^+1^_2_c^+2^_4_z^+1^_3_y^+2^_8_w^+1^_4_c^+2^_6_c^+1^_3_y^+1^_4_c^+3^_12_c^+3^_13_z^+1^_5_y^+1^_5_c^+1^_4_w^+1^_6_c^+1^_4_z^+2^_11_y^+2^_11_z^+1^_6_z^+2^_12_y^+2^_12_z^+1^_7_c^+2^_12_c^+2^_12_y^+1^_7_c^+1^_5_c^+2^_13_w^+1^_8_c^+2^_13_y^+2^_13_z^+2^_13_y^+1^_8_c^+1^_6_z^+1^_9_y^+1^_9_c^+1^_7_c^+1^_8_z^+1^_10_y^+1^_10_c^+1^_9_z^+1^_11_c^+1^_11_z^+1^_12_y^+1^_12_c^+1^_12_c^+1^_13_

0769153723063074

Fragment Matches Table

Show background peaks

| Position | Ion type | Intensity | mz Theoretical | mz Error (Th) | mz Error (ppm) | Charge | Series Number |
| --- | --- | --- | --- | --- | --- | --- | --- |
| - | - | 346.8 | 127.6 | - | - | 0 | - |
| - | - | 1352 | 129.1 | - | - | 0 | - |
| - | - | 619.1 | 130.1 | - | - | 0 | - |
| 14 | y | 836.7 | 132.1 | 6.937E-05 | 0.5251 | +1 | 1 |
| - | - | 603.3 | 136.1 | - | - | 0 | - |
| - | - | 472.7 | 139.1 | - | - | 0 | - |
| - | - | 493.9 | 142.1 | - | - | 0 | - |
| - | - | 379.2 | 144.1 | - | - | 0 | - |
| - | - | 483.3 | 147.8 | - | - | 0 | - |
| - | - | 476.8 | 153.1 | - | - | 0 | - |
| - | - | 436.6 | 154.3 | - | - | 0 | - |
| - | - | 510.5 | 172.1 | - | - | 0 | - |
| - | - | 1339 | 173.5 | - | - | 0 | - |
| - | - | 471.3 | 198.2 | - | - | 0 | - |
| - | - | 465.2 | 211 | - | - | 0 | - |
| - | - | 589 | 226.2 | - | - | 0 | - |
| - | - | 543.5 | 236.8 | - | - | 0 | - |
| - | - | 600.3 | 241.2 | - | - | 0 | - |
| - | - | 487.2 | 244.1 | - | - | 0 | - |
| - | - | 621.6 | 245.1 | - | - | 0 | - |
| 13 | y | 1699 | 245.2 | 0.0002086 | 0.8508 | +1 | 2 |
| - | - | 684.4 | 255.1 | - | - | 0 | - |
| - | - | 1057 | 262.1 | - | - | 0 | - |
| - | - | 709.8 | 273.1 | - | - | 0 | - |
| - | - | 3735 | 290.1 | - | - | 0 | - |
| 4 | c | 945.7 | 291.1 | 0.001713 | 5.883 | +2 | 4 |
| - | - | 607.7 | 301.2 | - | - | 0 | - |
| - | - | 713.6 | 309.2 | - | - | 0 | - |
| - | - | 703.5 | 314.2 | - | - | 0 | - |
| 12 | z | 1383 | 357.3 | 0.0002757 | 0.7716 | +1 | 3 |
| - | - | 1000 | 358.3 | - | - | 0 | - |
| - | - | 830.8 | 369.2 | - | - | 0 | - |
| 7 | y | 2976 | 412.3 | 0.0002874 | 0.6971 | +2 | 8 |
| - | - | 805.3 | 412.8 | - | - | 0 | - |
| - | - | 616.2 | 413.3 | - | - | 0 | - |
| 11 | w | 1250 | 427.3 | 0.0008921 | 2.088 | +1 | 4 |
| 6 | c | 778 | 428.2 | 0.004073 | 9.512 | +2 | 6 |
| - | - | 565.9 | 432.4 | - | - | 0 | - |
| - | - | 959 | 451.2 | - | - | 0 | - |
| - | - | 661.2 | 453.2 | - | - | 0 | - |
| - | - | 2129 | 468.3 | - | - | 0 | - |
| - | - | 1195 | 469.2 | - | - | 0 | - |
| 3 | c | 3508 | 470.2 | 0.004416 | 9.392 | +1 | 3 |
| 11 | y | 1084 | 470.3 | 0.001817 | 3.864 | +1 | 4 |
| - | - | 810.4 | 471.2 | - | - | 0 | - |
| 12 | c | 789.7 | 478.6 | 0.001725 | 3.605 | +3 | 12 |
| - | - | 1557 | 478.9 | - | - | 0 | - |
| - | - | 594.6 | 479.2 | - | - | 0 | - |
| - | - | 665.9 | 506.9 | - | - | 0 | - |
| - | - | 679 | 507.3 | - | - | 0 | - |
| - | - | 950.6 | 510.3 | - | - | 0 | - |
| - | - | 824.3 | 510.6 | - | - | 0 | - |
| - | - | 764.6 | 510.9 | - | - | 0 | - |
| 13 | c | 5328 | 516.3 | 0.001861 | 3.605 | +3 | 13 |
| - | - | 4995 | 516.6 | - | - | 0 | - |
| - | - | 2692 | 516.9 | - | - | 0 | - |
| - | - | 1184 | 522.3 | - | - | 0 | - |
| 10 | z | 7730 | 525.4 | 0.0003313 | 0.6306 | +1 | 5 |
| - | - | 2330 | 526.4 | - | - | 0 | - |
| - | - | 755.6 | 531.3 | - | - | 0 | - |
| 10 | y | 835.4 | 541.4 | 0.001261 | 2.328 | +1 | 5 |
| - | - | 760.4 | 543.3 | - | - | 0 | - |
| - | - | 591.1 | 544.6 | - | - | 0 | - |
| - | - | 1464 | 554 | - | - | 0 | - |
| - | - | 1226 | 554.3 | - | - | 0 | - |
| - | - | 2065 | 554.6 | - | - | 0 | - |
| - | - | 1021 | 555 | - | - | 0 | - |
| - | - | 1535 | 559.3 | - | - | 0 | - |
| - | - | 3.21E+04 | 560 | - | - | 0 | - |
| - | - | 3.301E+04 | 560.3 | - | - | 0 | - |
| - | - | 1.617E+04 | 560.6 | - | - | 0 | - |
| - | - | 5472 | 561 | - | - | 0 | - |
| - | - | 3889 | 561.3 | - | - | 0 | - |
| - | - | 1960 | 579.2 | - | - | 0 | - |
| 4 | c | 1224 | 581.2 | 0.004217 | 7.256 | +1 | 4 |
| - | - | 891.2 | 582.4 | - | - | 0 | - |
| 9 | w | 2182 | 595.4 | 0.0008813 | 1.48 | +1 | 6 |
| - | - | 885.3 | 596.4 | - | - | 0 | - |
| - | - | 3853 | 597.3 | - | - | 0 | - |
| 4 | c | 8443 | 598.3 | 0.004829 | 8.072 | +1 | 4 |
| - | - | 3702 | 599.3 | - | - | 0 | - |
| 4 | z | 2567 | 605.4 | 0.0009166 | 1.514 | +2 | 11 |
| - | - | 3791 | 605.9 | - | - | 0 | - |
| - | - | 1893 | 606.4 | - | - | 0 | - |
| - | - | 1465 | 610.4 | - | - | 0 | - |
| 4 | y | 1452 | 613.4 | 0.001121 | 1.828 | +2 | 11 |
| - | - | 1301 | 613.9 | - | - | 0 | - |
| - | - | 724.6 | 629.3 | - | - | 0 | - |
| - | - | 737.1 | 629.8 | - | - | 0 | - |
| 9 | z | 4844 | 653.4 | 0.0002169 | 0.332 | +1 | 6 |
| - | - | 3203 | 654.5 | - | - | 0 | - |
| - | - | 947 | 655.5 | - | - | 0 | - |
| - | - | 929.8 | 658.4 | - | - | 0 | - |
| - | - | 694.2 | 667.4 | - | - | 0 | - |
| 3 | z | 5601 | 686.9 | 0.0005353 | 0.7793 | +2 | 12 |
| - | - | 8290 | 687.4 | - | - | 0 | - |
| - | - | 4021 | 687.9 | - | - | 0 | - |
| - | - | 1631 | 688.4 | - | - | 0 | - |
| - | - | 700.9 | 690.3 | - | - | 0 | - |
| 3 | y | 1.063E+04 | 694.9 | 0.000559 | 0.8044 | +2 | 12 |
| - | - | 8986 | 695.4 | - | - | 0 | - |
| - | - | 3673 | 695.9 | - | - | 0 | - |
| - | - | 651.6 | 696 | - | - | 0 | - |
| 8 | z | 2614 | 710.5 | 0.0001155 | 0.1626 | +1 | 7 |
| - | - | 1481 | 711.5 | - | - | 0 | - |
| - | - | 582.4 | 712.5 | - | - | 0 | - |
| 12 | c | 1531 | 717.4 | 0.004425 | 6.169 | +2 | 12 |
| - | - | 1.027E+04 | 721.9 | - | - | 0 | - |
| - | - | 8426 | 722.4 | - | - | 0 | - |
| - | - | 4923 | 722.9 | - | - | 0 | - |
| - | - | 1898 | 723.4 | - | - | 0 | - |
| - | - | 592.7 | 723.9 | - | - | 0 | - |
| 12 | c | 2.148E+04 | 725.9 | 0.00232 | 3.196 | +2 | 12 |
| - | - | 1.343E+04 | 726.4 | - | - | 0 | - |
| 8 | y | 667.8 | 726.5 | 0.002556 | 3.519 | +1 | 7 |
| - | - | 6351 | 726.9 | - | - | 0 | - |
| - | - | 2380 | 727.4 | - | - | 0 | - |
| - | - | 802.7 | 727.9 | - | - | 0 | - |
| 5 | c | 1.501E+04 | 735.3 | 0.00506 | 6.882 | +1 | 5 |
| - | - | 5465 | 736.3 | - | - | 0 | - |
| - | - | 1373 | 737.3 | - | - | 0 | - |
| - | - | 2250 | 758.9 | - | - | 0 | - |
| - | - | 2294 | 759.4 | - | - | 0 | - |
| - | - | 1683 | 765.4 | - | - | 0 | - |
| - | - | 1190 | 765.9 | - | - | 0 | - |
| - | - | 6799 | 766.4 | - | - | 0 | - |
| - | - | 4140 | 766.9 | - | - | 0 | - |
| - | - | 4876 | 767.4 | - | - | 0 | - |
| - | - | 2689 | 767.9 | - | - | 0 | - |
| - | - | 1534 | 768.4 | - | - | 0 | - |
| - | - | 1442 | 772.9 | - | - | 0 | - |
| 13 | c | 2462 | 773.4 | 0.006769 | 8.753 | +2 | 13 |
| - | - | 3145 | 773.9 | - | - | 0 | - |
| - | - | 4180 | 774.4 | - | - | 0 | - |
| 7 | w | 721.1 | 780.5 | 0.0008529 | 1.093 | +1 | 8 |
| 13 | c | 1.353E+04 | 782.4 | 0.002708 | 3.461 | +2 | 13 |
| - | - | 1.293E+04 | 782.9 | - | - | 0 | - |
| - | - | 6485 | 783.4 | - | - | 0 | - |
| - | - | 2246 | 783.9 | - | - | 0 | - |
| - | - | 1406 | 784.4 | - | - | 0 | - |
| 2 | y | 821.6 | 787.4 | 0.01071 | 13.61 | +2 | 13 |
| 2 | z | 900.4 | 787.9 | 0.006871 | 8.721 | +2 | 13 |
| - | - | 662.5 | 795.4 | - | - | 0 | - |
| - | - | 1300 | 798.9 | - | - | 0 | - |
| - | - | 1256 | 799.4 | - | - | 0 | - |
| - | - | 1177 | 799.9 | - | - | 0 | - |
| - | - | 1612 | 804.4 | - | - | 0 | - |
| - | - | 1108 | 804.9 | - | - | 0 | - |
| - | - | 1110 | 808.4 | - | - | 0 | - |
| - | - | 1023 | 808.9 | - | - | 0 | - |
| - | - | 931 | 809.4 | - | - | 0 | - |
| - | - | 915.3 | 809.9 | - | - | 0 | - |
| - | - | 1006 | 810.9 | - | - | 0 | - |
| - | - | 798.7 | 811.9 | - | - | 0 | - |
| - | - | 656.1 | 812.3 | - | - | 0 | - |
| - | - | 906 | 812.4 | - | - | 0 | - |
| - | - | 2080 | 817.4 | - | - | 0 | - |
| - | - | 3004 | 817.9 | - | - | 0 | - |
| - | - | 1466 | 818.4 | - | - | 0 | - |
| - | - | 1165 | 818.9 | - | - | 0 | - |
| - | - | 1330 | 821.9 | - | - | 0 | - |
| - | - | 3758 | 822.4 | - | - | 0 | - |
| - | - | 3377 | 822.9 | - | - | 0 | - |
| - | - | 1540 | 823.4 | - | - | 0 | - |
| 7 | y | 1909 | 823.5 | 0.0004049 | 0.4917 | +1 | 8 |
| - | - | 1076 | 824.5 | - | - | 0 | - |
| - | - | 1603 | 827.4 | - | - | 0 | - |
| - | - | 1422 | 828.4 | - | - | 0 | - |
| - | - | 809.7 | 829.4 | - | - | 0 | - |
| - | - | 823.8 | 829.9 | - | - | 0 | - |
| - | - | 2572 | 830.9 | - | - | 0 | - |
| - | - | 1.55E+04 | 831.4 | - | - | 0 | - |
| - | - | 1.344E+04 | 831.9 | - | - | 0 | - |
| - | - | 6336 | 832.4 | - | - | 0 | - |
| - | - | 3551 | 832.9 | - | - | 0 | - |
| - | - | 794.3 | 833.4 | - | - | 0 | - |
| - | - | 747.2 | 838.4 | - | - | 0 | - |
| - | - | 1139 | 838.9 | - | - | 0 | - |
| - | - | 2.186E+04 | 839.4 | - | - | 0 | - |
| - | - | 9.457E+04 | 840 | - | - | 0 | - |
| - | - | 8.02E+04 | 840.5 | - | - | 0 | - |
| - | - | 5.354E+04 | 841 | - | - | 0 | - |
| - | - | 2.213E+04 | 841.5 | - | - | 0 | - |
| - | - | 1.162E+04 | 841.9 | - | - | 0 | - |
| - | - | 724.5 | 850.5 | - | - | 0 | - |
| - | - | 891.2 | 852.5 | - | - | 0 | - |
| 6 | c | 1101 | 855.3 | 0.005168 | 6.042 | +1 | 6 |
| - | - | 677.1 | 899.4 | - | - | 0 | - |
| 6 | z | 1.369E+04 | 944.6 | 4.899E-05 | 0.05186 | +1 | 9 |
| - | - | 8730 | 945.6 | - | - | 0 | - |
| - | - | 2075 | 946.6 | - | - | 0 | - |
| - | - | 804 | 947.6 | - | - | 0 | - |
| 6 | y | 1030 | 960.6 | 0.001134 | 1.18 | +1 | 9 |
| - | - | 892.3 | 961.6 | - | - | 0 | - |
| 7 | c | 2829 | 969.4 | 0.007032 | 7.254 | +1 | 7 |
| - | - | 1171 | 970.4 | - | - | 0 | - |
| - | - | 898.7 | 983.6 | - | - | 0 | - |
| - | - | 1338 | 1025 | - | - | 0 | - |
| 8 | c | 1.505E+04 | 1026 | 0.004611 | 4.492 | +1 | 8 |
| - | - | 7575 | 1027 | - | - | 0 | - |
| - | - | 2505 | 1028 | - | - | 0 | - |
| 5 | z | 9212 | 1082 | 0.0001822 | 0.1685 | +1 | 10 |
| - | - | 2.149E+04 | 1083 | - | - | 0 | - |
| - | - | 1.259E+04 | 1084 | - | - | 0 | - |
| - | - | 4218 | 1085 | - | - | 0 | - |
| - | - | 580.5 | 1086 | - | - | 0 | - |
| - | - | 1287 | 1086 | - | - | 0 | - |
| 5 | y | 2111 | 1098 | 0.003161 | 2.88 | +1 | 10 |
| - | - | 1448 | 1099 | - | - | 0 | - |
| - | - | 765 | 1105 | - | - | 0 | - |
| - | - | 918.4 | 1109 | - | - | 0 | - |
| - | - | 923.4 | 1112 | - | - | 0 | - |
| - | - | 1576 | 1113 | - | - | 0 | - |
| - | - | 828.5 | 1119 | - | - | 0 | - |
| - | - | 835.1 | 1119 | - | - | 0 | - |
| - | - | 689.6 | 1120 | - | - | 0 | - |
| - | - | 914.5 | 1122 | - | - | 0 | - |
| - | - | 1111 | 1123 | - | - | 0 | - |
| 9 | c | 1.973E+04 | 1155 | 0.004863 | 4.212 | +1 | 9 |
| - | - | 1.195E+04 | 1156 | - | - | 0 | - |
| - | - | 4844 | 1157 | - | - | 0 | - |
| - | - | 1746 | 1158 | - | - | 0 | - |
| - | - | 1095 | 1159 | - | - | 0 | - |
| 4 | z | 3775 | 1210 | 0.002396 | 1.98 | +1 | 11 |
| - | - | 1.309E+04 | 1211 | - | - | 0 | - |
| - | - | 6818 | 1212 | - | - | 0 | - |
| - | - | 3162 | 1213 | - | - | 0 | - |
| - | - | 793.7 | 1214 | - | - | 0 | - |
| - | - | 1033 | 1322 | - | - | 0 | - |
| 11 | c | 8148 | 1323 | 0.003486 | 2.636 | +1 | 11 |
| - | - | 5172 | 1324 | - | - | 0 | - |
| - | - | 3087 | 1325 | - | - | 0 | - |
| - | - | 1063 | 1326 | - | - | 0 | - |
| 3 | z | 1077 | 1373 | 0.006978 | 5.083 | +1 | 12 |
| - | - | 4593 | 1374 | - | - | 0 | - |
| - | - | 5195 | 1375 | - | - | 0 | - |
| - | - | 1934 | 1376 | - | - | 0 | - |
| 3 | y | 2667 | 1389 | 0.003607 | 2.597 | +1 | 12 |
| - | - | 2354 | 1390 | - | - | 0 | - |
| - | - | 1193 | 1391 | - | - | 0 | - |
| - | - | 766.1 | 1443 | - | - | 0 | - |
| - | - | 2955 | 1444 | - | - | 0 | - |
| - | - | 2694 | 1445 | - | - | 0 | - |
| 12 | c | 3232 | 1451 | 0.001297 | 0.8938 | +1 | 12 |
| - | - | 5814 | 1452 | - | - | 0 | - |
| - | - | 4322 | 1453 | - | - | 0 | - |
| - | - | 1193 | 1454 | - | - | 0 | - |
| 13 | c | 1261 | 1564 | 0.004635 | 2.964 | +1 | 13 |
| - | - | 3700 | 1565 | - | - | 0 | - |
| - | - | 2595 | 1566 | - | - | 0 | - |
| - | - | 1510 | 1567 | - | - | 0 | - |
| - | - | 763.2 | 1568 | - | - | 0 | - |
| - | - | 720.5 | 1593 | - | - | 0 | - |
| - | - | 640.3 | 1599 | - | - | 0 | - |
| - | - | 1313 | 1634 | - | - | 0 | - |
| - | - | 2260 | 1635 | - | - | 0 | - |
| - | - | 1795 | 1636 | - | - | 0 | - |
| - | - | 1147 | 1637 | - | - | 0 | - |
| - | - | 1832 | 1645 | - | - | 0 | - |
| - | - | 2103 | 1646 | - | - | 0 | - |
| - | - | 1077 | 1647 | - | - | 0 | - |
| - | - | 1496 | 1652 | - | - | 0 | - |
| - | - | 791.9 | 1653 | - | - | 0 | - |
| - | - | 836.2 | 1654 | - | - | 0 | - |
| - | - | 5468 | 1662 | - | - | 0 | - |
| - | - | 1.408E+04 | 1663 | - | - | 0 | - |
| - | - | 1.219E+04 | 1664 | - | - | 0 | - |
| - | - | 6724 | 1665 | - | - | 0 | - |
| - | - | 3492 | 1666 | - | - | 0 | - |
| - | - | 1528 | 1667 | - | - | 0 | - |
| - | - | 2421 | 1678 | - | - | 0 | - |
| - | - | 1.183E+04 | 1679 | - | - | 0 | - |
| - | - | 5.063E+04 | 1680 | - | - | 0 | - |
| - | - | 4.71E+04 | 1681 | - | - | 0 | - |
| - | - | 3.06E+04 | 1682 | - | - | 0 | - |
| - | - | 1.228E+04 | 1683 | - | - | 0 | - |
| - | - | 9789 | 1684 | - | - | 0 | - |
| - | - | 690 | 2825 | - | - | 0 | - |
| - | - | 695 | 2952 | - | - | 0 | - |
| - | - | 812.9 | 3044 | - | - | 0 | - |

m/z Charge Intensity FragmentType MassShift Position
127.57176971435547 0 346.8379
129.10238647460938 0 1352.2083
130.06556701660156 0 619.09216
132.1019744873047 0 836.70984 y 13
136.0760955810547 0 603.3332
139.05718994140625 0 472.65103
142.09725952148438 0 493.91324
144.07852172851562 0 379.18918
147.820556640625 0 483.30408
153.07875061035156 0 476.84436
154.26792907714844 0 436.63046
172.0714569091797 0 510.46527
173.45132446289062 0 1339.2544
198.23947143554688 0 471.3028
210.9504852294922 0 465.19922
226.15423583984375 0 589.00543
236.8016815185547 0 543.48346
241.1549530029297 0 600.2743
244.09048461914062 0 487.22543
245.1365509033203 0 621.5664
245.18576049804688 0 1698.7666 y 12
255.0760955810547 0 684.3964
262.1186218261719 0 1056.7114
273.08697509765625 0 709.8057
290.113525390625 0 3735.358
291.1171875 0 945.71466 c Ammonia loss 3
301.19903564453125 0 607.7421
309.1553649902344 0 713.6311
314.2076721191406 0 703.4754
357.2619323730469 0 1382.5582 z 11
358.2684631347656 0 999.99164
369.2131652832031 0 830.7919
412.27392578125 0 2975.9104 y 6
412.77587890625 0 805.2595
413.27606201171875 0 616.1884
427.2923889160156 0 1250.33 w 10
428.181884765625 0 778.0131 c Ammonia loss 5
432.3693542480469 0 565.8935
451.1854553222656 0 958.99066
453.1808166503906 0 661.20526
468.3184509277344 0 2129.4463
469.19647216796875 0 1195.417
470.2029113769531 0 3508.345 c 2
470.3318786621094 0 1084.0232 y 10
471.20745849609375 0 810.4072
478.57403564453125 0 789.6961 c Ammonia loss 11
478.9090576171875 0 1556.6664
479.24737548828125 0 594.5946
506.93914794921875 0 665.9008
507.2711486816406 0 678.9867
510.26580810546875 0 950.63873
510.5977478027344 0 824.3108
510.93548583984375 0 764.5705
516.2688598632812 0 5327.8115 c Ammonia loss 12
516.6029663085938 0 4995.349
516.9367065429688 0 2692.0283
522.2732543945312 0 1183.8175
525.3524169921875 0 7729.6343 z 9
526.3560791015625 0 2329.6897
531.2904663085938 0 755.56116
541.3720703125 0 835.3821 y 9
543.2936401367188 0 760.3881
544.628173828125 0 591.0912
553.9638671875 0 1463.8802
554.2943115234375 0 1226.2606
554.6319580078125 0 2065.3418
554.9641723632812 0 1021.26605
559.3101196289062 0 1535.011
559.967041015625 0 32100.814
560.30126953125 0 33006.43
560.63525390625 0 16167.138
560.9680786132812 0 5472.402
561.3007202148438 0 3889.4165
579.2435302734375 0 1960.4489
581.2347412109375 0 1224.0504 c Ammonia loss 3
582.3731079101562 0 891.1929
595.3804931640625 0 2181.9197 w 8
596.3868408203125 0 885.27856
597.2545166015625 0 3852.8105
598.2619018554688 0 8442.835 c 3
599.2649536132812 0 3702.008
605.3533935546875 0 2566.6443 z 3
605.8562622070312 0 3790.6145
606.3573608398438 0 1893.3584
610.3958129882812 0 1465.3628
613.3607177734375 0 1452.3324 y 3
613.8623046875 0 1300.6127
629.3482055664062 0 724.5717
629.8484497070312 0 737.1432
653.447265625 0 4843.9683 z 8
654.4517211914062 0 3203.1345
655.4544677734375 0 947.0421
658.36083984375 0 929.78284
667.4063720703125 0 694.1824
686.8836059570312 0 5600.7563 z 2
687.3864135742188 0 8289.928
687.8881225585938 0 4021.333
688.3869018554688 0 1631.1047
690.2996215820312 0 700.92566
694.8929443359375 0 10630.335 y 2
695.3942260742188 0 8985.835
695.8967895507812 0 3673.2053
695.95751953125 0 651.61755
710.4686279296875 0 2613.86 z 7
711.47265625 0 1480.6473
712.4797973632812 0 582.4335
717.3592529296875 0 1531.2426 c Ammonia loss 11
721.8991088867188 0 10269.116
722.4000854492188 0 8425.558
722.9013671875 0 4922.9897
723.4033813476562 0 1898.2815
723.9154663085938 0 592.7497
725.8704223632812 0 21475.72 c 11
726.3717651367188 0 13431.157
726.4846801757812 0 667.82635 y 7
726.8734741210938 0 6350.967
727.3744506835938 0 2380.1702
727.8705444335938 0 802.6779
735.321044921875 0 15013.273 c 4
736.32373046875 0 5465.0933
737.3253173828125 0 1373.3882
758.9110107421875 0 2249.9253
759.4118041992188 0 2294.0654
765.4104614257812 0 1682.8535
765.9197998046875 0 1189.7692
766.41650390625 0 6798.6655
766.9188232421875 0 4139.888
767.4216918945312 0 4875.8076
767.9259033203125 0 2688.5981
768.4264526367188 0 1533.8558
772.9193115234375 0 1442.1062
773.41162109375 0 2461.781 c Water loss 12
773.9171752929688 0 3145.2776
774.4141845703125 0 4180.3506
780.4969482421875 0 721.08966 w 6
782.412841796875 0 13527.31 c 12
782.9143676757812 0 12933.002
783.415283203125 0 6484.5806
783.9147338867188 0 2245.6504
784.4132080078125 0 1405.5154
787.4255981445312 0 821.6098 y Ammonia loss 1
787.9119262695312 0 900.4297 z 1
795.4194946289062 0 662.4965
798.9220581054688 0 1300.2117
799.4247436523438 0 1256.3032
799.9212036132812 0 1177.0189
804.413330078125 0 1611.8716
804.9207763671875 0 1108.2281
808.4346313476562 0 1110.1937
808.9383544921875 0 1022.9709
809.4420776367188 0 930.9651
809.9083251953125 0 915.288
810.9203491210938 0 1006.27795
811.9129028320312 0 798.705
812.3430786132812 0 656.14703
812.4258422851562 0 905.9866
817.441162109375 0 2079.652
817.9429931640625 0 3004.3057
818.4349365234375 0 1466.1025
818.927734375 0 1165.1451
821.9409790039062 0 1329.6725
822.4338989257812 0 3757.5796
822.9334716796875 0 3376.9492
823.435546875 0 1539.8164
823.5404052734375 0 1908.5253 y 6
824.541748046875 0 1075.8765
827.356201171875 0 1602.9799
828.3583984375 0 1422.156
829.3750610351562 0 809.7358
829.9248657226562 0 823.77216
830.9434814453125 0 2572.0776
831.43994140625 0 15497.669
831.9404296875 0 13443.599
832.44091796875 0 6336.3228
832.9415893554688 0 3551.0928
833.4376831054688 0 794.31836
838.4263916015625 0 747.239
838.9466552734375 0 1138.8392
839.4472045898438 0 21855.508
839.9503784179688 0 94565.74
840.4512939453125 0 80197.23
840.9515380859375 0 53541.008
841.4511108398438 0 22125.557
841.9485473632812 0 11619.775
850.52978515625 0 724.5343
852.5421142578125 0 891.2276
855.353515625 0 1100.7804 c Ammonia loss 5
899.4166259765625 0 677.07275
944.5801391601562 0 13686.892 z 5
945.5833129882812 0 8729.501
946.5870361328125 0 2074.593
947.5894165039062 0 803.98883
960.5977783203125 0 1029.7671 y 5
961.6019287109375 0 892.3455
969.4346923828125 0 2828.5142 c 6
970.4349365234375 0 1170.6194
983.5519409179688 0 898.7015
1025.44921875 0 1338.0111
1026.4537353515625 0 15046.563 c 7
1027.457763671875 0 7575.234
1028.4619140625 0 2504.615
1081.6392822265625 0 9212.288 z 4
1082.6455078125 0 21492.045
1083.648681640625 0 12591.135
1084.65283203125 0 4217.855
1085.5235595703125 0 580.4688
1085.6510009765625 0 1287.1753
1097.6546630859375 0 2111.1882 y 4
1098.65625 0 1447.9132
1104.5162353515625 0 765.0269
1108.501953125 0 918.37177
1111.5404052734375 0 923.448
1112.5322265625 0 1576.3356
1118.531005859375 0 828.4741
1119.4541015625 0 835.07025
1120.0323486328125 0 689.6092
1121.5126953125 0 914.45233
1122.5157470703125 0 1111.0988
1154.5489501953125 0 19727.004 c 8
1155.5526123046875 0 11949.504
1156.5546875 0 4843.9443
1157.5638427734375 0 1745.9454
1158.54931640625 0 1094.9442
1209.7000732421875 0 3775.0312 z 3
1210.7042236328125 0 13085.309
1211.7076416015625 0 6817.5728
1212.7103271484375 0 3162.4285
1213.7196044921875 0 793.73975
1321.6192626953125 0 1033.3911
1322.637451171875 0 8148.4805 c 10
1323.6407470703125 0 5172.2734
1324.63916015625 0 3087.0044
1325.6473388671875 0 1063.2028
1372.7540283203125 0 1077.0286 z 2
1373.765625 0 4592.7944
1374.7708740234375 0 5194.9023
1375.769287109375 0 1934.2852
1388.776123046875 0 2667.0444 y 2
1389.78076171875 0 2353.669
1390.775634765625 0 1193.1385
1442.793701171875 0 766.09326
1443.798095703125 0 2955.1633
1444.797119140625 0 2693.8257
1450.730224609375 0 3231.7808 c 11
1451.7353515625 0 5813.884
1452.7425537109375 0 4322.1367
1453.734375 0 1193.2949
1563.817626953125 0 1261.4724 c 12
1564.8238525390625 0 3700.331
1565.83056640625 0 2594.8164
1566.8275146484375 0 1510.221
1567.8341064453125 0 763.21136
1592.84912109375 0 720.46155
1599.0814208984375 0 640.28217
1633.8807373046875 0 1313.0918
1634.87744140625 0 2260.012
1635.882568359375 0 1795.0111
1636.8974609375 0 1146.977
1644.8594970703125 0 1832.0201
1645.8629150390625 0 2102.7065
1646.879638671875 0 1076.7905
1651.906005859375 0 1496.149
1652.8994140625 0 791.8644
1653.893798828125 0 836.22614
1661.8824462890625 0 5467.752
1662.8787841796875 0 14080.976
1663.87939453125 0 12194.94
1664.8779296875 0 6723.572
1665.88037109375 0 3492.029
1666.861083984375 0 1527.507
1677.8790283203125 0 2420.5066
1678.8902587890625 0 11833.952
1679.900390625 0 50627.977
1680.9029541015625 0 47103
1681.9014892578125 0 30597.396
1682.898193359375 0 12278.841
1683.8916015625 0 9788.541
2825.337158203125 0 689.96
2951.9404296875 0 694.9821
3043.62890625 0 812.8615

Spectrum Details

|  |  |
| --- | --- |
| Matched peaks? Matched peaks The total absolute number of peaks matched. Additionally in brackets the total fraction of peaks matched and the total number of peaks is shown. | 46 (16.43% of 280) |
| FDR? FDR The false discovery rate estimated for this peptide. It is calculated by matching all theoretical fragments with a non-integer shift with the raw peaks for this spectrum. This is done with 40 different shifts. The resulting percentage is the average number of annotated peaks over the number of annotated peaks with the correct spectrum. | 2.90% |
| Satellite FDR? Satellite FDR See the FDR for details on its calculation. This satellite ion specific FDR only contains the satellite ions (d/w) for I/L/J positions. | - |
| PSM Score? PSM Score The PSM Score as given by Hecklib to this annotated spectrum. It is shown with three significant figures. | 344 |

#### Spectrum 5672? Spectrum 5672 The raw spectrum of this peptide as annotated by Hecklib. The fragments are coloured according to ion type (see legend). Any peaks with a star '\*' as text can be hovered over to see the full details, first the ion type second the mass shift type. By hovering over the amino acids in the peptide or ions in the legend the corresponding peaks are highlighted. By toggling the 'Unassigned' label you can turn the background (unassigned) peaks on or off in the plot. By updating the slider in the Ion legend you can update the spectrum to only show the top X% of the peaks with labels. The top X% means any peak that is within X% of the highest intensity. By dragging in the spectrum you can zoom in to a specific part of the spectrum and use 'Zoom Out' to get back to the original zoom level. The annotation of the spectrum is based on the given sequence in the peptides file and is done with different software so inconsistencies are likely. The peaks are annotated based on the given sequence, with 20 ppm tolerance.

Copy Data

##### Spectrum 5672 (TSV)

###### Preview

```
Loading example...
```

*Click on the button to copy the data to your clipboard.*

Mz MinMz MaxIntensity Max

WidthHeightPeptide font sizePeptide stroke widthSpectrum font sizeSpectrum stroke widthCompact peptide

Ion legend

wxyz

abcd

OtherUnassignedIon^Charge^_Position_Show for top:%

SWYQHHPGKAPKJJ

04.67e+49.35e+41.40e+51.87e+5

Zoom Out

y^+1^_1_y^+1^_2_z^+1^_3_y^+1^_3_y^+2^_8_w^+1^_4_c^+1^_3_y^+3^_12_y^+1^_4_c^+3^_12_c^+3^_13_z^+1^_5_y^+3^_13_y^+1^_5_y^+2^_10_c^+1^_4_c^+1^_4_w^+2^_11_c^+1^_4_w^+1^_6_y^+2^_11_z^+2^_11_y^+2^_11_z^+1^_6_c^+2^_11_y^+2^_12_z^+2^_12_z^+1^_7_y^+2^_12_c^+1^_5_c^+2^_12_z^+1^_7_c^+2^_12_c^+1^_5_y^+1^_7_c^+2^_13_z^+2^_13_y^+2^_13_y^+1^_8_c^+1^_6_z^+1^_9_c^+1^_7_y^+1^_9_c^+1^_8_z^+1^_10_y^+1^_10_c^+1^_9_y^+1^_11_z^+1^_11_y^+1^_11_c^+1^_11_z^+1^_12_y^+1^_12_c^+1^_12_c^+1^_13_z^+1^_13_

0689137920682757

Fragment Matches Table

Show background peaks

| Position | Ion type | Intensity | mz Theoretical | mz Error (Th) | mz Error (ppm) | Charge | Series Number |
| --- | --- | --- | --- | --- | --- | --- | --- |
| - | - | 363.9 | 120.1 | - | - | 0 | - |
| - | - | 1762 | 129.1 | - | - | 0 | - |
| - | - | 522.5 | 132.1 | - | - | 0 | - |
| 14 | y | 809.3 | 132.1 | 0.0001762 | 1.334 | +1 | 1 |
| - | - | 609.2 | 136.1 | - | - | 0 | - |
| - | - | 447.7 | 139.1 | - | - | 0 | - |
| - | - | 2199 | 153.1 | - | - | 0 | - |
| - | - | 4482 | 173.5 | - | - | 0 | - |
| - | - | 478.3 | 174.1 | - | - | 0 | - |
| - | - | 500.5 | 186.1 | - | - | 0 | - |
| - | - | 774.7 | 187.1 | - | - | 0 | - |
| - | - | 487.3 | 189.4 | - | - | 0 | - |
| - | - | 480.9 | 196.6 | - | - | 0 | - |
| - | - | 531.6 | 201.1 | - | - | 0 | - |
| - | - | 1147 | 211.1 | - | - | 0 | - |
| - | - | 2692 | 215.1 | - | - | 0 | - |
| - | - | 491.1 | 226.2 | - | - | 0 | - |
| - | - | 589.3 | 227.1 | - | - | 0 | - |
| - | - | 729.7 | 229.1 | - | - | 0 | - |
| - | - | 1.465E+04 | 229.2 | - | - | 0 | - |
| - | - | 1527 | 230.2 | - | - | 0 | - |
| - | - | 840.3 | 241.1 | - | - | 0 | - |
| - | - | 1440 | 245.1 | - | - | 0 | - |
| 13 | y | 2386 | 245.2 | 0.0002696 | 1.1 | +1 | 2 |
| - | - | 5670 | 246.1 | - | - | 0 | - |
| - | - | 836.6 | 246.1 | - | - | 0 | - |
| - | - | 518.9 | 246.2 | - | - | 0 | - |
| - | - | 983.2 | 247.1 | - | - | 0 | - |
| - | - | 570.5 | 247.3 | - | - | 0 | - |
| - | - | 642.4 | 258.1 | - | - | 0 | - |
| - | - | 469.8 | 265.1 | - | - | 0 | - |
| - | - | 7452 | 274.1 | - | - | 0 | - |
| - | - | 895.3 | 275.1 | - | - | 0 | - |
| - | - | 1154 | 281.2 | - | - | 0 | - |
| - | - | 1659 | 282.2 | - | - | 0 | - |
| - | - | 1511 | 292.1 | - | - | 0 | - |
| - | - | 934.1 | 299.1 | - | - | 0 | - |
| - | - | 556.3 | 299.2 | - | - | 0 | - |
| - | - | 573.5 | 300.1 | - | - | 0 | - |
| - | - | 1621 | 300.2 | - | - | 0 | - |
| - | - | 1155 | 301.2 | - | - | 0 | - |
| - | - | 1550 | 309.2 | - | - | 0 | - |
| - | - | 2736 | 314.2 | - | - | 0 | - |
| - | - | 646.5 | 315.2 | - | - | 0 | - |
| - | - | 624.3 | 322.3 | - | - | 0 | - |
| - | - | 560 | 331.2 | - | - | 0 | - |
| - | - | 4649 | 340.1 | - | - | 0 | - |
| - | - | 1023 | 341.1 | - | - | 0 | - |
| - | - | 594.2 | 343.2 | - | - | 0 | - |
| - | - | 797.4 | 355.1 | - | - | 0 | - |
| 12 | z | 2077 | 357.3 | 0.0002146 | 0.6008 | +1 | 3 |
| - | - | 1509 | 358.3 | - | - | 0 | - |
| - | - | 915.7 | 371.1 | - | - | 0 | - |
| 12 | y | 1245 | 373.3 | 0.0002314 | 0.62 | +1 | 3 |
| - | - | 760.8 | 382.3 | - | - | 0 | - |
| - | - | 917.6 | 394.3 | - | - | 0 | - |
| - | - | 2172 | 397.2 | - | - | 0 | - |
| - | - | 874 | 398.2 | - | - | 0 | - |
| - | - | 511.2 | 398.6 | - | - | 0 | - |
| - | - | 752.8 | 399.3 | - | - | 0 | - |
| 7 | y | 4300 | 412.3 | 0.0002925 | 0.7094 | +2 | 8 |
| - | - | 2205 | 412.8 | - | - | 0 | - |
| - | - | 567 | 418.2 | - | - | 0 | - |
| 11 | w | 1626 | 427.3 | 0.0002981 | 0.6976 | +1 | 4 |
| - | - | 1782 | 437.2 | - | - | 0 | - |
| - | - | 1176 | 453.2 | - | - | 0 | - |
| 3 | c | 3310 | 454.2 | 0.0006707 | 1.477 | +1 | 3 |
| - | - | 2074 | 455.8 | - | - | 0 | - |
| 3 | y | 1418 | 463.6 | 0.0004469 | 0.964 | +3 | 12 |
| - | - | 977.2 | 463.9 | - | - | 0 | - |
| - | - | 895.4 | 465.2 | - | - | 0 | - |
| - | - | 1760 | 467.8 | - | - | 0 | - |
| - | - | 923.4 | 468.3 | - | - | 0 | - |
| - | - | 3664 | 468.3 | - | - | 0 | - |
| - | - | 659.5 | 469.3 | - | - | 0 | - |
| - | - | 1182 | 469.3 | - | - | 0 | - |
| - | - | 811 | 470.3 | - | - | 0 | - |
| 11 | y | 1089 | 470.3 | 0.001756 | 3.734 | +1 | 4 |
| 12 | c | 1877 | 473.2 | 0.0005198 | 1.098 | +3 | 12 |
| - | - | 916.3 | 474.2 | - | - | 0 | - |
| - | - | 1903 | 477.2 | - | - | 0 | - |
| - | - | 1486 | 480.2 | - | - | 0 | - |
| - | - | 863.1 | 481.3 | - | - | 0 | - |
| - | - | 1250 | 482.3 | - | - | 0 | - |
| - | - | 818.5 | 495.4 | - | - | 0 | - |
| - | - | 990.9 | 501.6 | - | - | 0 | - |
| - | - | 665.4 | 501.9 | - | - | 0 | - |
| - | - | 658 | 504.9 | - | - | 0 | - |
| - | - | 1052 | 505.3 | - | - | 0 | - |
| - | - | 2.083E+04 | 510.3 | - | - | 0 | - |
| 13 | c | 7632 | 510.9 | 0.0005667 | 1.109 | +3 | 13 |
| - | - | 5795 | 511.3 | - | - | 0 | - |
| - | - | 5864 | 511.3 | - | - | 0 | - |
| - | - | 2178 | 511.6 | - | - | 0 | - |
| - | - | 775.9 | 511.9 | - | - | 0 | - |
| - | - | 645.5 | 513.7 | - | - | 0 | - |
| - | - | 2038 | 514.3 | - | - | 0 | - |
| - | - | 776.9 | 517.3 | - | - | 0 | - |
| - | - | 1264 | 523.2 | - | - | 0 | - |
| 10 | z | 1.313E+04 | 525.4 | 0.000279 | 0.5312 | +1 | 5 |
| 2 | y | 1695 | 525.6 | 0.0007121 | 1.355 | +3 | 13 |
| - | - | 1242 | 526 | - | - | 0 | - |
| - | - | 4523 | 526.4 | - | - | 0 | - |
| - | - | 648.3 | 528.2 | - | - | 0 | - |
| - | - | 952.1 | 539.3 | - | - | 0 | - |
| - | - | 659.7 | 539.3 | - | - | 0 | - |
| 10 | y | 1443 | 541.4 | 0.001322 | 2.441 | +1 | 5 |
| - | - | 653.4 | 541.8 | - | - | 0 | - |
| - | - | 2433 | 548.6 | - | - | 0 | - |
| - | - | 731.6 | 549 | - | - | 0 | - |
| - | - | 1359 | 549.3 | - | - | 0 | - |
| 5 | y | 957.5 | 549.3 | 0.005096 | 9.277 | +2 | 10 |
| - | - | 1271 | 549.6 | - | - | 0 | - |
| - | - | 876.9 | 550.3 | - | - | 0 | - |
| - | - | 2198 | 553.3 | - | - | 0 | - |
| - | - | 705.5 | 553.8 | - | - | 0 | - |
| - | - | 5611 | 554.2 | - | - | 0 | - |
| - | - | 734.9 | 554.3 | - | - | 0 | - |
| - | - | 5.642E+04 | 554.6 | - | - | 0 | - |
| - | - | 5.389E+04 | 555 | - | - | 0 | - |
| - | - | 1060 | 555.2 | - | - | 0 | - |
| - | - | 3.363E+04 | 555.3 | - | - | 0 | - |
| - | - | 9232 | 555.6 | - | - | 0 | - |
| - | - | 2535 | 556 | - | - | 0 | - |
| - | - | 3992 | 563.2 | - | - | 0 | - |
| 4 | c | 773.9 | 564.3 | 0.01072 | 19 | +1 | 4 |
| 4 | c | 2610 | 565.2 | 0.0001064 | 0.1882 | +1 | 4 |
| - | - | 912.9 | 566.2 | - | - | 0 | - |
| - | - | 1039 | 566.4 | - | - | 0 | - |
| - | - | 594.2 | 569.8 | - | - | 0 | - |
| - | - | 747.4 | 572.8 | - | - | 0 | - |
| 4 | w | 3152 | 576.3 | 0.0007965 | 1.382 | +2 | 11 |
| - | - | 2123 | 576.8 | - | - | 0 | - |
| - | - | 654 | 577.3 | - | - | 0 | - |
| - | - | 2055 | 581.3 | - | - | 0 | - |
| 4 | c | 1.167E+04 | 582.3 | 0.0007155 | 1.229 | +1 | 4 |
| - | - | 1912 | 582.4 | - | - | 0 | - |
| - | - | 4520 | 583.3 | - | - | 0 | - |
| - | - | 1211 | 584.3 | - | - | 0 | - |
| 9 | w | 3098 | 595.4 | 0.00198 | 3.325 | +1 | 6 |
| - | - | 1076 | 596.4 | - | - | 0 | - |
| - | - | 785.9 | 597.4 | - | - | 0 | - |
| 4 | y | 1178 | 604.8 | 0.001106 | 1.828 | +2 | 11 |
| 4 | z | 7713 | 605.4 | 0.0004262 | 0.7041 | +2 | 11 |
| - | - | 1.152E+04 | 605.9 | - | - | 0 | - |
| - | - | 5204 | 606.4 | - | - | 0 | - |
| - | - | 2111 | 606.9 | - | - | 0 | - |
| - | - | 949 | 609.5 | - | - | 0 | - |
| - | - | 1914 | 610.4 | - | - | 0 | - |
| - | - | 1105 | 611.4 | - | - | 0 | - |
| - | - | 883.8 | 612.9 | - | - | 0 | - |
| 4 | y | 4004 | 613.4 | 0.0002826 | 0.4607 | +2 | 11 |
| - | - | 2719 | 613.9 | - | - | 0 | - |
| - | - | 1195 | 614.4 | - | - | 0 | - |
| - | - | 925 | 620.3 | - | - | 0 | - |
| - | - | 962.1 | 624.4 | - | - | 0 | - |
| - | - | 956.7 | 628.4 | - | - | 0 | - |
| - | - | 1772 | 629.3 | - | - | 0 | - |
| - | - | 1103 | 629.8 | - | - | 0 | - |
| - | - | 764.7 | 637.4 | - | - | 0 | - |
| - | - | 921.2 | 645.8 | - | - | 0 | - |
| - | - | 789.8 | 651.4 | - | - | 0 | - |
| - | - | 931.6 | 651.9 | - | - | 0 | - |
| 9 | z | 9994 | 653.4 | 0.0003934 | 0.6021 | +1 | 6 |
| 11 | c | 1656 | 653.8 | 0.0006695 | 1.024 | +2 | 11 |
| - | - | 5514 | 654.5 | - | - | 0 | - |
| - | - | 1198 | 655.5 | - | - | 0 | - |
| - | - | 825.4 | 657.9 | - | - | 0 | - |
| - | - | 860.1 | 658.9 | - | - | 0 | - |
| - | - | 996.9 | 659.3 | - | - | 0 | - |
| - | - | 1032 | 660.3 | - | - | 0 | - |
| - | - | 667.4 | 660.4 | - | - | 0 | - |
| - | - | 1218 | 675.3 | - | - | 0 | - |
| - | - | 3289 | 682.4 | - | - | 0 | - |
| - | - | 1048 | 683.4 | - | - | 0 | - |
| 3 | y | 807.3 | 686.4 | 0.001261 | 1.838 | +2 | 12 |
| 3 | z | 1.012E+04 | 686.9 | 0.0009625 | 1.401 | +2 | 12 |
| - | - | 1.398E+04 | 687.4 | - | - | 0 | - |
| - | - | 6628 | 687.9 | - | - | 0 | - |
| - | - | 1072 | 688.4 | - | - | 0 | - |
| 8 | z | 1240 | 693.4 | 0.000435 | 0.6273 | +1 | 7 |
| 3 | y | 1.272E+04 | 694.9 | 0.0008031 | 1.156 | +2 | 12 |
| - | - | 8037 | 695.4 | - | - | 0 | - |
| - | - | 4023 | 695.9 | - | - | 0 | - |
| 5 | c | 1246 | 702.3 | 0.002133 | 3.038 | +1 | 5 |
| 12 | c | 1435 | 709.4 | 0.0009409 | 1.326 | +2 | 12 |
| - | - | 2310 | 709.9 | - | - | 0 | - |
| - | - | 915.2 | 710.4 | - | - | 0 | - |
| 8 | z | 1631 | 710.5 | 0.002936 | 4.133 | +1 | 7 |
| - | - | 1243 | 711.5 | - | - | 0 | - |
| 12 | c | 3.407E+04 | 717.9 | 0.0007877 | 1.097 | +2 | 12 |
| - | - | 2.797E+04 | 718.4 | - | - | 0 | - |
| - | - | 1.41E+04 | 718.9 | - | - | 0 | - |
| 5 | c | 2.91E+04 | 719.3 | 0.0006674 | 0.9279 | +1 | 5 |
| - | - | 1.407E+04 | 720.3 | - | - | 0 | - |
| - | - | 3353 | 721.3 | - | - | 0 | - |
| 8 | y | 1030 | 726.5 | 0.00219 | 3.015 | +1 | 7 |
| - | - | 655.1 | 740 | - | - | 0 | - |
| - | - | 1400 | 752.9 | - | - | 0 | - |
| - | - | 996.3 | 758.4 | - | - | 0 | - |
| - | - | 1475 | 758.9 | - | - | 0 | - |
| - | - | 927.8 | 759.4 | - | - | 0 | - |
| - | - | 1794 | 766.4 | - | - | 0 | - |
| - | - | 1063 | 766.9 | - | - | 0 | - |
| - | - | 1567 | 767.4 | - | - | 0 | - |
| - | - | 985.8 | 773.9 | - | - | 0 | - |
| 13 | c | 2.29E+04 | 774.4 | 0.0004613 | 0.5957 | +2 | 13 |
| - | - | 1.896E+04 | 774.9 | - | - | 0 | - |
| - | - | 1.116E+04 | 775.4 | - | - | 0 | - |
| - | - | 3212 | 775.9 | - | - | 0 | - |
| - | - | 1423 | 776.4 | - | - | 0 | - |
| 2 | z | 4735 | 779.9 | 0.001495 | 1.917 | +2 | 13 |
| - | - | 3441 | 780.4 | - | - | 0 | - |
| - | - | 2579 | 780.9 | - | - | 0 | - |
| - | - | 795 | 781.9 | - | - | 0 | - |
| 2 | y | 2003 | 787.9 | 0.01464 | 18.58 | +2 | 13 |
| - | - | 1037 | 788.4 | - | - | 0 | - |
| - | - | 1203 | 788.9 | - | - | 0 | - |
| - | - | 2929 | 790.9 | - | - | 0 | - |
| - | - | 3592 | 791.4 | - | - | 0 | - |
| - | - | 2530 | 791.9 | - | - | 0 | - |
| - | - | 827.7 | 792.4 | - | - | 0 | - |
| - | - | 1043 | 795.4 | - | - | 0 | - |
| - | - | 1485 | 795.9 | - | - | 0 | - |
| - | - | 1094 | 796.4 | - | - | 0 | - |
| - | - | 1805 | 796.9 | - | - | 0 | - |
| - | - | 1711 | 800.4 | - | - | 0 | - |
| - | - | 2111 | 800.9 | - | - | 0 | - |
| - | - | 2545 | 801.4 | - | - | 0 | - |
| - | - | 751.2 | 801.9 | - | - | 0 | - |
| - | - | 756.9 | 802.4 | - | - | 0 | - |
| - | - | 1396 | 802.9 | - | - | 0 | - |
| - | - | 1036 | 804.4 | - | - | 0 | - |
| - | - | 832.8 | 808.9 | - | - | 0 | - |
| - | - | 3304 | 809.4 | - | - | 0 | - |
| - | - | 5885 | 809.9 | - | - | 0 | - |
| - | - | 2578 | 810.4 | - | - | 0 | - |
| - | - | 1945 | 810.9 | - | - | 0 | - |
| - | - | 4468 | 811.4 | - | - | 0 | - |
| - | - | 807.5 | 811.4 | - | - | 0 | - |
| - | - | 3132 | 812.4 | - | - | 0 | - |
| - | - | 858.5 | 813.9 | - | - | 0 | - |
| - | - | 969.6 | 814.4 | - | - | 0 | - |
| - | - | 707.8 | 814.9 | - | - | 0 | - |
| - | - | 783.5 | 815.4 | - | - | 0 | - |
| - | - | 2499 | 822.9 | - | - | 0 | - |
| - | - | 1.716E+04 | 823.4 | - | - | 0 | - |
| 7 | y | 3639 | 823.5 | 0.005393 | 6.549 | +1 | 8 |
| - | - | 1.654E+04 | 823.9 | - | - | 0 | - |
| - | - | 6583 | 824.4 | - | - | 0 | - |
| - | - | 2086 | 824.5 | - | - | 0 | - |
| - | - | 4007 | 824.9 | - | - | 0 | - |
| - | - | 1250 | 825.4 | - | - | 0 | - |
| - | - | 789.2 | 830.4 | - | - | 0 | - |
| - | - | 2311 | 830.9 | - | - | 0 | - |
| - | - | 3.939E+04 | 831.4 | - | - | 0 | - |
| - | - | 1.851E+05 | 832 | - | - | 0 | - |
| - | - | 1.641E+05 | 832.5 | - | - | 0 | - |
| - | - | 8.84E+04 | 833 | - | - | 0 | - |
| - | - | 795.3 | 833.1 | - | - | 0 | - |
| - | - | 2.833E+04 | 833.5 | - | - | 0 | - |
| - | - | 7505 | 834 | - | - | 0 | - |
| - | - | 1501 | 838.5 | - | - | 0 | - |
| 6 | c | 2403 | 839.4 | 0.002146 | 2.557 | +1 | 6 |
| - | - | 856.7 | 839.4 | - | - | 0 | - |
| - | - | 1239 | 840.4 | - | - | 0 | - |
| - | - | 2132 | 850.5 | - | - | 0 | - |
| - | - | 989.6 | 851.5 | - | - | 0 | - |
| - | - | 2162 | 852.5 | - | - | 0 | - |
| - | - | 2101 | 855.5 | - | - | 0 | - |
| - | - | 739 | 855.9 | - | - | 0 | - |
| - | - | 1006 | 856.5 | - | - | 0 | - |
| - | - | 727.7 | 878.4 | - | - | 0 | - |
| - | - | 933.1 | 883.4 | - | - | 0 | - |
| - | - | 1550 | 894.5 | - | - | 0 | - |
| - | - | 762.2 | 903.3 | - | - | 0 | - |
| - | - | 7266 | 910.5 | - | - | 0 | - |
| - | - | 5956 | 911.5 | - | - | 0 | - |
| - | - | 2726 | 912.5 | - | - | 0 | - |
| - | - | 874 | 934.5 | - | - | 0 | - |
| - | - | 864.7 | 935.5 | - | - | 0 | - |
| 6 | z | 3.004E+04 | 944.6 | 0.0007814 | 0.8273 | +1 | 9 |
| - | - | 1.959E+04 | 945.6 | - | - | 0 | - |
| - | - | 5638 | 946.6 | - | - | 0 | - |
| - | - | 999.5 | 947.6 | - | - | 0 | - |
| 7 | c | 3540 | 953.4 | 0.001747 | 1.833 | +1 | 7 |
| - | - | 1792 | 954.4 | - | - | 0 | - |
| 6 | y | 3165 | 960.6 | 0.001439 | 1.498 | +1 | 9 |
| - | - | 997.2 | 961.6 | - | - | 0 | - |
| - | - | 769.9 | 964.4 | - | - | 0 | - |
| - | - | 701 | 968.6 | - | - | 0 | - |
| - | - | 1563 | 982.5 | - | - | 0 | - |
| - | - | 1391 | 983.6 | - | - | 0 | - |
| - | - | 1684 | 1009 | - | - | 0 | - |
| 8 | c | 2.722E+04 | 1010 | 0.0007502 | 0.7425 | +1 | 8 |
| - | - | 1.671E+04 | 1011 | - | - | 0 | - |
| - | - | 6403 | 1012 | - | - | 0 | - |
| - | - | 1157 | 1013 | - | - | 0 | - |
| - | - | 813 | 1081 | - | - | 0 | - |
| 5 | z | 1.37E+04 | 1082 | 0.001283 | 1.186 | +1 | 10 |
| - | - | 3.456E+04 | 1083 | - | - | 0 | - |
| - | - | 2.004E+04 | 1084 | - | - | 0 | - |
| - | - | 6493 | 1085 | - | - | 0 | - |
| - | - | 1994 | 1086 | - | - | 0 | - |
| - | - | 1094 | 1093 | - | - | 0 | - |
| - | - | 1024 | 1094 | - | - | 0 | - |
| - | - | 1827 | 1095 | - | - | 0 | - |
| - | - | 1044 | 1096 | - | - | 0 | - |
| - | - | 1577 | 1097 | - | - | 0 | - |
| 5 | y | 3750 | 1098 | 0.002917 | 2.657 | +1 | 10 |
| - | - | 1899 | 1099 | - | - | 0 | - |
| - | - | 909.4 | 1101 | - | - | 0 | - |
| - | - | 684.5 | 1102 | - | - | 0 | - |
| - | - | 891.2 | 1103 | - | - | 0 | - |
| - | - | 1144 | 1109 | - | - | 0 | - |
| - | - | 840.5 | 1109 | - | - | 0 | - |
| - | - | 1081 | 1110 | - | - | 0 | - |
| - | - | 1644 | 1111 | - | - | 0 | - |
| - | - | 1218 | 1112 | - | - | 0 | - |
| - | - | 661.3 | 1138 | - | - | 0 | - |
| 9 | c | 3.4E+04 | 1139 | 0.0006205 | 0.545 | +1 | 9 |
| - | - | 2.676E+04 | 1140 | - | - | 0 | - |
| - | - | 8013 | 1141 | - | - | 0 | - |
| - | - | 2227 | 1142 | - | - | 0 | - |
| - | - | 738.3 | 1166 | - | - | 0 | - |
| - | - | 728.6 | 1184 | - | - | 0 | - |
| 4 | y | 1056 | 1209 | 0.01016 | 8.41 | +1 | 11 |
| 4 | z | 4323 | 1210 | 0.0006867 | 0.5677 | +1 | 11 |
| - | - | 2.102E+04 | 1211 | - | - | 0 | - |
| - | - | 1.441E+04 | 1212 | - | - | 0 | - |
| - | - | 5128 | 1213 | - | - | 0 | - |
| - | - | 1217 | 1214 | - | - | 0 | - |
| 4 | y | 1605 | 1226 | 0.005464 | 4.458 | +1 | 11 |
| - | - | 903.3 | 1227 | - | - | 0 | - |
| - | - | 1270 | 1306 | - | - | 0 | - |
| 11 | c | 1.758E+04 | 1307 | 0.002119 | 1.622 | +1 | 11 |
| - | - | 1.259E+04 | 1308 | - | - | 0 | - |
| - | - | 5193 | 1309 | - | - | 0 | - |
| - | - | 1085 | 1310 | - | - | 0 | - |
| 3 | z | 2657 | 1373 | 0.002583 | 1.882 | +1 | 12 |
| - | - | 9237 | 1374 | - | - | 0 | - |
| - | - | 9321 | 1375 | - | - | 0 | - |
| - | - | 4108 | 1376 | - | - | 0 | - |
| - | - | 1736 | 1377 | - | - | 0 | - |
| 3 | y | 801.1 | 1389 | 0.003351 | 2.413 | +1 | 12 |
| - | - | 1689 | 1419 | - | - | 0 | - |
| - | - | 1375 | 1420 | - | - | 0 | - |
| 12 | c | 6075 | 1435 | 0.004919 | 3.429 | +1 | 12 |
| - | - | 8103 | 1436 | - | - | 0 | - |
| - | - | 7210 | 1437 | - | - | 0 | - |
| - | - | 1781 | 1438 | - | - | 0 | - |
| - | - | 997.5 | 1505 | - | - | 0 | - |
| - | - | 858.8 | 1532 | - | - | 0 | - |
| - | - | 1067 | 1533 | - | - | 0 | - |
| 13 | c | 1345 | 1548 | 0.005133 | 3.316 | +1 | 13 |
| - | - | 5293 | 1549 | - | - | 0 | - |
| - | - | 4899 | 1550 | - | - | 0 | - |
| - | - | 2406 | 1551 | - | - | 0 | - |
| - | - | 876.3 | 1552 | - | - | 0 | - |
| 2 | z | 845.7 | 1559 | 0.005384 | 3.454 | +1 | 13 |
| - | - | 3410 | 1560 | - | - | 0 | - |
| - | - | 3958 | 1561 | - | - | 0 | - |
| - | - | 1802 | 1562 | - | - | 0 | - |
| - | - | 1227 | 1581 | - | - | 0 | - |
| - | - | 1480 | 1582 | - | - | 0 | - |
| - | - | 911.3 | 1591 | - | - | 0 | - |
| - | - | 837.2 | 1602 | - | - | 0 | - |
| - | - | 1445 | 1618 | - | - | 0 | - |
| - | - | 3678 | 1619 | - | - | 0 | - |
| - | - | 4349 | 1620 | - | - | 0 | - |
| - | - | 1959 | 1621 | - | - | 0 | - |
| - | - | 1708 | 1629 | - | - | 0 | - |
| - | - | 2588 | 1630 | - | - | 0 | - |
| - | - | 1156 | 1631 | - | - | 0 | - |
| - | - | 1793 | 1636 | - | - | 0 | - |
| - | - | 1652 | 1637 | - | - | 0 | - |
| - | - | 783.4 | 1638 | - | - | 0 | - |
| - | - | 4887 | 1646 | - | - | 0 | - |
| - | - | 1.843E+04 | 1647 | - | - | 0 | - |
| - | - | 1.86E+04 | 1648 | - | - | 0 | - |
| - | - | 8040 | 1649 | - | - | 0 | - |
| - | - | 3663 | 1650 | - | - | 0 | - |
| - | - | 948.4 | 1651 | - | - | 0 | - |
| - | - | 4311 | 1662 | - | - | 0 | - |
| - | - | 2.453E+04 | 1663 | - | - | 0 | - |
| - | - | 9.213E+04 | 1664 | - | - | 0 | - |
| - | - | 8.881E+04 | 1665 | - | - | 0 | - |
| - | - | 1109 | 1665 | - | - | 0 | - |
| - | - | 4.865E+04 | 1666 | - | - | 0 | - |
| - | - | 1.658E+04 | 1667 | - | - | 0 | - |
| - | - | 3634 | 1668 | - | - | 0 | - |
| - | - | 622.6 | 1950 | - | - | 0 | - |
| - | - | 698.1 | 2730 | - | - | 0 | - |

m/z Charge Intensity FragmentType MassShift Position
120.06549072265625 0 363.87332
129.1020965576172 0 1762.232
132.0807647705078 0 522.45056
132.10208129882812 0 809.329 y 13
136.07559204101562 0 609.19574
139.07310485839844 0 447.66293
153.0783233642578 0 2198.8035
173.45123291015625 0 4482.144
174.0889434814453 0 478.2546
186.078369140625 0 500.45
187.0861358642578 0 774.6605
189.3968048095703 0 487.26755
196.61134338378906 0 480.9423
201.1235809326172 0 531.5845
211.14358520507812 0 1146.5282
215.13865661621094 0 2692.4426
226.15451049804688 0 491.0938
227.1390838623047 0 589.25494
229.1425018310547 0 729.742
229.15438842773438 0 14650.399
230.15774536132812 0 1526.6915
241.09178161621094 0 840.2639
245.13653564453125 0 1440.2024
245.18569946289062 0 2385.997 y 12
246.12339782714844 0 5670.117
246.1377410888672 0 836.64014
246.18792724609375 0 518.8598
247.1272430419922 0 983.18774
247.3211212158203 0 570.47626
258.1449279785156 0 642.36566
265.1280212402344 0 469.78247
274.11810302734375 0 7452.4097
275.1228942871094 0 895.2923
281.17352294921875 0 1154.2513
282.1807556152344 0 1659.1597
292.1290283203125 0 1510.5563
299.06158447265625 0 934.0572
299.207763671875 0 556.3352
300.06268310546875 0 573.47906
300.1914978027344 0 1621.3755
301.1985778808594 0 1154.9877
309.15545654296875 0 1550.3037
314.2072448730469 0 2736.0847
315.21014404296875 0 646.51825
322.3221130371094 0 624.33624
331.2091369628906 0 560.0445
340.0992431640625 0 4648.984
341.1037902832031 0 1022.8511
343.2439270019531 0 594.2177
355.0677490234375 0 797.36145
357.2619934082031 0 2076.654 z 11
358.26605224609375 0 1508.902
371.09991455078125 0 915.74286
373.28070068359375 0 1244.7853 y 11
382.2804870605469 0 760.7563
394.2567443847656 0 917.60144
397.24407958984375 0 2172.2095
398.2478332519531 0 873.9964
398.5660400390625 0 511.18787
399.25982666015625 0 752.7798
412.2733459472656 0 4300.058 y 6
412.7744445800781 0 2204.651
418.2406005859375 0 567.0105
427.29119873046875 0 1626.1702 w 10
437.1816711425781 0 1781.5381
453.20172119140625 0 1176.1031
454.20782470703125 0 3310.24 c 2
455.7709045410156 0 2073.5957
463.5985412597656 0 1417.7297 y 2
463.929931640625 0 977.1818
465.2079162597656 0 895.44037
467.7506103515625 0 1760.1935
468.2522888183594 0 923.44275
468.318115234375 0 3664.0088
469.28924560546875 0 659.45996
469.3222961425781 0 1181.9421
470.29437255859375 0 810.9928
470.3319396972656 0 1088.513 y 10
473.2417907714844 0 1876.5585 c Ammonia loss 11
474.2459716796875 0 916.2895
477.20013427734375 0 1902.7511
480.2259826660156 0 1485.6371
481.3087158203125 0 863.07355
482.29718017578125 0 1250.0802
495.3658142089844 0 818.4581
501.6058654785156 0 990.94147
501.937255859375 0 665.4354
504.9353942871094 0 657.9839
505.2698974609375 0 1052.3677
510.3283386230469 0 20829.705
510.9364318847656 0 7631.8057 c Ammonia loss 12
511.2706298828125 0 5794.758
511.33154296875 0 5863.9756
511.6044921875 0 2178.1929
511.93853759765625 0 775.85315
513.6951293945312 0 645.5405
514.2971801757812 0 2038.3545
517.2764282226562 0 776.8948
523.2399291992188 0 1263.8296
525.351806640625 0 13134.692 z 9
525.625244140625 0 1695.0101 y 1
525.9586791992188 0 1242.3995
526.354736328125 0 4523.4487
528.2423095703125 0 648.34753
539.2608032226562 0 952.093
539.3040771484375 0 659.7011
541.3721313476562 0 1442.7115 y 9
541.828857421875 0 653.4326
548.6319580078125 0 2432.8728
548.9625244140625 0 731.5532
549.297119140625 0 1359.3331
549.337646484375 0 957.4726 y 4
549.63037109375 0 1270.9879
550.33447265625 0 876.8975
553.3115844726562 0 2197.5164
553.7874145507812 0 705.52625
554.2304077148438 0 5610.514
554.2756958007812 0 734.94464
554.6347045898438 0 56415.91
554.9688720703125 0 53889.184
555.2297973632812 0 1060.2786
555.3031005859375 0 33629.4
555.6373291015625 0 9231.82
555.970703125 0 2535.4253
563.247802734375 0 3991.8752
564.2457885742188 0 773.92896 c Water loss 3
565.2404174804688 0 2609.692 c Ammonia loss 3
566.2440185546875 0 912.9472
566.4030151367188 0 1038.5848
569.8146362304688 0 594.2155
572.8004760742188 0 747.3925
576.3370361328125 0 3152.2668 w 3
576.8386840820312 0 2122.5752
577.3283081054688 0 653.95105
581.2581176757812 0 2054.8682
582.266357421875 0 11672.27 c 3
582.3737182617188 0 1912.1365
583.268310546875 0 4520.4106
584.2716064453125 0 1211.1141
595.37939453125 0 3098.1729 w 8
596.3837280273438 0 1076.0927
597.3859252929688 0 785.8905
604.8496704101562 0 1178.4614 y Ammonia loss 3
605.35205078125 0 7713.101 z 3
605.8536987304688 0 11517.347
606.3555297851562 0 5204.267
606.8572387695312 0 2111.3433
609.4559936523438 0 949.02234
610.3975219726562 0 1913.9965
611.3980102539062 0 1105.3522
612.8558349609375 0 883.7881
613.3621215820312 0 4003.6973 y 3
613.8622436523438 0 2718.8455
614.36279296875 0 1195.4514
620.3412475585938 0 924.9838
624.41748046875 0 962.1223
628.3750610351562 0 956.699
629.3455200195312 0 1771.8315
629.8445434570312 0 1102.7794
637.430419921875 0 764.70154
645.820556640625 0 921.1712
651.3624267578125 0 789.8407
651.8621826171875 0 931.59064
653.4466552734375 0 9994.468 z 8
653.824951171875 0 1656.249 c 10
654.4501342773438 0 5513.8965
655.4545288085938 0 1198.489
657.8656005859375 0 825.3696
658.8535766601562 0 860.106
659.2965698242188 0 996.8597
660.295166015625 0 1031.6653
660.3534545898438 0 667.38995
675.310546875 0 1218.2137
682.3873901367188 0 3289.0623
683.3897094726562 0 1048.0762
686.3789672851562 0 807.3461 y Ammonia loss 2
686.8831787109375 0 10117.445 z 2
687.385009765625 0 13983.05
687.886474609375 0 6628.4004
688.3900756835938 0 1071.5951
693.4415283203125 0 1239.9006 z Ammonia loss 7
694.8927001953125 0 12718.66 y 2
695.3932495117188 0 8037.382
695.8938598632812 0 4022.891
702.2973022460938 0 1245.5541 c Ammonia loss 4
709.35888671875 0 1435.306 c Ammonia loss 11
709.8618774414062 0 2309.5916
710.3637084960938 0 915.1768
710.465576171875 0 1630.7473 z 7
711.4700927734375 0 1243.1718
717.872314453125 0 34067.375 c 11
718.37353515625 0 27965.732
718.8748779296875 0 14101.073
719.3253173828125 0 29099.236 c 4
720.326904296875 0 14066.612
721.3302001953125 0 3352.8196
726.4850463867188 0 1029.8177 y 7
740.0234375 0 655.09216
752.91064453125 0 1399.7161
758.4188842773438 0 996.2716
758.9119262695312 0 1474.5627
759.4085693359375 0 927.79
766.4098510742188 0 1793.7478
766.9158325195312 0 1063.056
767.4080200195312 0 1567.4512
773.924560546875 0 985.8333
774.4146728515625 0 22898.043 c 12
774.91552734375 0 18959.396
775.4170532226562 0 11157.75
775.9174194335938 0 3212.2864
776.4186401367188 0 1422.7009
779.9223022460938 0 4734.697 z 1
780.4265747070312 0 3441.0425
780.9232177734375 0 2578.791
781.9112548828125 0 795.0204
787.9185180664062 0 2003.0654 y 1
788.4190673828125 0 1037.4958
788.9235229492188 0 1202.5133
790.9266967773438 0 2928.851
791.4260864257812 0 3591.9937
791.9273071289062 0 2530.3335
792.4403686523438 0 827.68915
795.4180908203125 0 1042.5347
795.90966796875 0 1485.4514
796.412353515625 0 1093.7395
796.9210815429688 0 1805.2917
800.4368286132812 0 1710.7722
800.9359130859375 0 2110.9114
801.4430541992188 0 2545.0107
801.94287109375 0 751.215
802.4134521484375 0 756.9184
802.9302368164062 0 1396.2778
804.4257202148438 0 1036.0736
808.9423828125 0 832.8243
809.4423828125 0 3304.087
809.9454956054688 0 5885.2896
810.4349365234375 0 2577.5518
810.9301147460938 0 1945.4702
811.3623657226562 0 4467.8057
811.4375610351562 0 807.5201
812.3684692382812 0 3132.3125
813.9107055664062 0 858.4595
814.4320068359375 0 969.6061
814.9439697265625 0 707.8284
815.4364624023438 0 783.51013
822.9458618164062 0 2499.1262
823.4400024414062 0 17157.762
823.5346069335938 0 3639.052 y 6
823.9407348632812 0 16536.777
824.4423828125 0 6582.8066
824.5391235351562 0 2085.736
824.94189453125 0 4007.465
825.4369506835938 0 1249.9875
830.4431762695312 0 789.16766
830.9432373046875 0 2311.2925
831.4488525390625 0 39386.375
831.9521484375 0 185105.62
832.4535522460938 0 164065.66
832.9542846679688 0 88403.305
833.142333984375 0 795.2832
833.4555053710938 0 28325.947
833.9560546875 0 7505.102
838.4521484375 0 1501.0068
839.356201171875 0 2402.626 c Ammonia loss 5
839.44970703125 0 856.65424
840.3558959960938 0 1238.8351
850.5271606445312 0 2132.3384
851.5308837890625 0 989.60834
852.5375366210938 0 2162.1736
855.4911499023438 0 2100.9805
855.9109497070312 0 738.9692
856.488525390625 0 1006.1152
878.4111938476562 0 727.68567
883.4086303710938 0 933.12976
894.51708984375 0 1549.6552
903.3319091796875 0 762.15137
910.5359497070312 0 7266.2725
911.5394897460938 0 5956.305
912.5433349609375 0 2726.1553
934.4880981445312 0 873.98065
935.5009155273438 0 864.7192
944.5794067382812 0 30040.398 z 5
945.5820922851562 0 19594.201
946.58544921875 0 5637.643
947.5807495117188 0 999.45087
953.4359130859375 0 3539.7764 c 6
954.4401245117188 0 1792.1396
960.5974731445312 0 3164.5273 y 5
961.6055297851562 0 997.1809
964.4495849609375 0 769.91986
968.5711669921875 0 701.0212
982.5469970703125 0 1563.1887
983.5552978515625 0 1391.2919
1009.4530639648438 0 1684.4597
1010.4583740234375 0 27218.014 c 7
1011.460693359375 0 16712.307
1012.4627685546875 0 6403.327
1013.4637451171875 0 1156.987
1081.0235595703125 0 812.9552
1081.6378173828125 0 13695.361 z 4
1082.64501953125 0 34555.266
1083.6474609375 0 20043.906
1084.6519775390625 0 6492.947
1085.654052734375 0 1993.5073
1092.5103759765625 0 1094.4537
1093.502197265625 0 1023.96265
1094.5362548828125 0 1827.185
1095.6395263671875 0 1044.3604
1096.639892578125 0 1577.317
1097.6549072265625 0 3749.7195 y 4
1098.6607666015625 0 1898.6729
1100.51318359375 0 909.37335
1102.028564453125 0 684.45435
1102.52294921875 0 891.15576
1108.5311279296875 0 1143.9673
1109.0433349609375 0 840.4865
1109.5189208984375 0 1081.1998
1110.5489501953125 0 1644.13
1111.57421875 0 1218.2007
1137.5472412109375 0 661.2727
1138.553466796875 0 33996.805 c 8
1139.5548095703125 0 26758.955
1140.5582275390625 0 8012.832
1141.5552978515625 0 2226.8276
1165.573486328125 0 738.2821
1183.678955078125 0 728.58075
1208.6796875 0 1056.0076 y Ammonia loss 3
1209.6983642578125 0 4322.9526 z 3
1210.7030029296875 0 21017.56
1211.706298828125 0 14412.792
1212.709716796875 0 5128.142
1213.7154541015625 0 1217.263
1225.7109375 0 1604.8579 y 3
1226.7091064453125 0 903.3484
1305.6387939453125 0 1269.8005
1306.641845703125 0 17582.14 c 10
1307.64453125 0 12586.857
1308.6446533203125 0 5193.0776
1309.647216796875 0 1085.4495
1372.7584228515625 0 2656.7366 z 2
1373.7642822265625 0 9236.557
1374.7696533203125 0 9320.732
1375.765625 0 4107.54
1376.771240234375 0 1735.8967
1388.7830810546875 0 801.1487 y 2
1418.71728515625 0 1688.7263
1419.7119140625 0 1374.7593
1434.7340087890625 0 6075.351 c 11
1435.740966796875 0 8102.6875
1436.74462890625 0 7209.7905
1437.74755859375 0 1781.1185
1504.810791015625 0 997.5412
1531.79541015625 0 858.8159
1532.80615234375 0 1067.458
1547.828125 0 1345.417 c 12
1548.8251953125 0 5293.0303
1549.82958984375 0 4899.1616
1550.8271484375 0 2405.9185
1551.8255615234375 0 876.29535
1558.845703125 0 845.7472 z 1
1559.8443603515625 0 3409.8872
1560.84716796875 0 3958.4316
1561.8485107421875 0 1802.4395
1580.8497314453125 0 1226.8027
1581.84521484375 0 1480.0914
1590.79931640625 0 911.3208
1601.885986328125 0 837.2047
1617.8760986328125 0 1444.5002
1618.880859375 0 3677.772
1619.87841796875 0 4349.316
1620.8856201171875 0 1958.7174
1628.8636474609375 0 1708.2147
1629.8603515625 0 2588.1675
1630.8623046875 0 1155.831
1635.9129638671875 0 1793.3148
1636.91357421875 0 1652.1682
1637.919921875 0 783.4158
1645.875732421875 0 4886.5776
1646.878173828125 0 18433.408
1647.8798828125 0 18596.125
1648.88232421875 0 8040.106
1649.8790283203125 0 3663.4185
1650.8779296875 0 948.4182
1661.88232421875 0 4311.471
1662.89599609375 0 24531.783
1663.904052734375 0 92133.69
1664.9063720703125 0 88806.85
1665.17724609375 0 1108.8507
1665.90966796875 0 48649.504
1666.9105224609375 0 16584.475
1667.9072265625 0 3633.7563
1950.453125 0 622.57025
2729.891357421875 0 698.065

Spectrum Details

|  |  |
| --- | --- |
| Matched peaks? Matched peaks The total absolute number of peaks matched. Additionally in brackets the total fraction of peaks matched and the total number of peaks is shown. | 56 (14.25% of 393) |
| FDR? FDR The false discovery rate estimated for this peptide. It is calculated by matching all theoretical fragments with a non-integer shift with the raw peaks for this spectrum. This is done with 40 different shifts. The resulting percentage is the average number of annotated peaks over the number of annotated peaks with the correct spectrum. | 2.68% |
| Satellite FDR? Satellite FDR See the FDR for details on its calculation. This satellite ion specific FDR only contains the satellite ions (d/w) for I/L/J positions. | - |
| PSM Score? PSM Score The PSM Score as given by Hecklib to this annotated spectrum. It is shown with three significant figures. | 469 |

#### Spectrum 5553? Spectrum 5553 The raw spectrum of this peptide as annotated by Hecklib. The fragments are coloured according to ion type (see legend). Any peaks with a star '\*' as text can be hovered over to see the full details, first the ion type second the mass shift type. By hovering over the amino acids in the peptide or ions in the legend the corresponding peaks are highlighted. By toggling the 'Unassigned' label you can turn the background (unassigned) peaks on or off in the plot. By updating the slider in the Ion legend you can update the spectrum to only show the top X% of the peaks with labels. The top X% means any peak that is within X% of the highest intensity. By dragging in the spectrum you can zoom in to a specific part of the spectrum and use 'Zoom Out' to get back to the original zoom level. The annotation of the spectrum is based on the given sequence in the peptides file and is done with different software so inconsistencies are likely. The peaks are annotated based on the given sequence, with 20 ppm tolerance.

Copy Data

##### Spectrum 5553 (TSV)

###### Preview

```
Loading example...
```

*Click on the button to copy the data to your clipboard.*

Mz MinMz MaxIntensity Max

WidthHeightPeptide font sizePeptide stroke widthSpectrum font sizeSpectrum stroke widthCompact peptide

Ion legend

wxyz

abcd

OtherUnassignedIon^Charge^_Position_Show for top:%

SWYQHHPGKAPKJJ

05.31e+41.06e+51.59e+52.12e+5

Zoom Out

y^+3^_3_y^+1^_1_b^+4^_4_b^+4^_5_y^+2^_3_a^+2^_3_y^+3^_6_y^+2^_4_a^+1^_2_b^+3^_5_y^+2^_4_y^+3^_7_y^+1^_2_a^+1^_2_b^+4^_8_b^+1^_2_y^+2^_5_b^+1^_2_y^+3^_8_y^+3^_9_b^+4^_11_y^+2^_6_y^+2^_7_y^+3^_10_y^+1^_3_b^+3^_9_y^+4^_13_b^+3^_10_b^+3^_10_y^+4^_13_b^+3^_10_y^+3^_11_y^+3^_11_b^+2^_6_\*\*y^+2^_8_\*b^+2^_6_b^+1^_3_y^+3^_12_y^+3^_12_y^+1^_4_b^+3^_12_y^+2^_9_b^+2^_8_b^+3^_13_b^+3^_13_y^+3^_13_y^+3^_13_y^+2^_10_y^+1^_5_y^+2^_10_b^+2^_9_b^+2^_9_b^+1^_4_b^+2^_10_b^+2^_10_b^+2^_10_y^+2^_11_y^+2^_11_y^+1^_6_b^+1^_5_y^+2^_12_b^+1^_5_y^+1^_7_y^+1^_8_b^+1^_6_y^+1^_9_y^+1^_9_

0757151422713027

Fragment Matches Table

Show background peaks

| Position | Ion type | Intensity | mz Theoretical | mz Error (Th) | mz Error (ppm) | Charge | Series Number |
| --- | --- | --- | --- | --- | --- | --- | --- |
| - | - | 478.3 | 120.1 | - | - | 0 | - |
| - | - | 728.8 | 120.1 | - | - | 0 | - |
| - | - | 1011 | 122.1 | - | - | 0 | - |
| - | - | 3355 | 123.1 | - | - | 0 | - |
| - | - | 748.1 | 123.1 | - | - | 0 | - |
| - | - | 435.6 | 125.1 | - | - | 0 | - |
| 12 | y | 424.1 | 125.1 | 0.002449 | 19.57 | +3 | 3 |
| - | - | 724.5 | 127.1 | - | - | 0 | - |
| - | - | 357.6 | 127.1 | - | - | 0 | - |
| - | - | 3831 | 127.1 | - | - | 0 | - |
| - | - | 490.5 | 128 | - | - | 0 | - |
| - | - | 491.8 | 128.1 | - | - | 0 | - |
| - | - | 716.9 | 128.1 | - | - | 0 | - |
| - | - | 5337 | 129.1 | - | - | 0 | - |
| - | - | 1.374E+05 | 129.1 | - | - | 0 | - |
| - | - | 3.143E+04 | 130.1 | - | - | 0 | - |
| - | - | 1205 | 130.1 | - | - | 0 | - |
| - | - | 8244 | 130.1 | - | - | 0 | - |
| - | - | 3917 | 131 | - | - | 0 | - |
| - | - | 3473 | 131.1 | - | - | 0 | - |
| - | - | 1124 | 131.1 | - | - | 0 | - |
| - | - | 1.193E+05 | 132.1 | - | - | 0 | - |
| 14 | y | 3.437E+04 | 132.1 | 0.0004966 | 3.759 | +1 | 1 |
| - | - | 435 | 133.1 | - | - | 0 | - |
| - | - | 1.09E+04 | 133.1 | - | - | 0 | - |
| - | - | 2277 | 133.1 | - | - | 0 | - |
| - | - | 720.6 | 134.1 | - | - | 0 | - |
| - | - | 804.8 | 134.1 | - | - | 0 | - |
| - | - | 1.609E+05 | 136.1 | - | - | 0 | - |
| - | - | 1203 | 137.1 | - | - | 0 | - |
| - | - | 1.434E+04 | 137.1 | - | - | 0 | - |
| - | - | 542.8 | 137.1 | - | - | 0 | - |
| - | - | 2503 | 138.1 | - | - | 0 | - |
| - | - | 466.1 | 138.1 | - | - | 0 | - |
| - | - | 2962 | 139.1 | - | - | 0 | - |
| - | - | 607.9 | 140.1 | - | - | 0 | - |
| - | - | 7518 | 141.1 | - | - | 0 | - |
| 4 | b | 1811 | 142.1 | 2.449E-05 | 0.1724 | +4 | 4 |
| - | - | 542.8 | 142.1 | - | - | 0 | - |
| - | - | 650.9 | 143.1 | - | - | 0 | - |
| - | - | 408.5 | 144.1 | - | - | 0 | - |
| - | - | 4700 | 144.1 | - | - | 0 | - |
| - | - | 1096 | 145.1 | - | - | 0 | - |
| - | - | 2038 | 146.1 | - | - | 0 | - |
| - | - | 558 | 146.1 | - | - | 0 | - |
| - | - | 6882 | 147 | - | - | 0 | - |
| - | - | 584.1 | 148 | - | - | 0 | - |
| - | - | 811.7 | 148.1 | - | - | 0 | - |
| - | - | 611.6 | 149 | - | - | 0 | - |
| - | - | 545.3 | 149 | - | - | 0 | - |
| - | - | 472.1 | 150.1 | - | - | 0 | - |
| - | - | 1986 | 151.1 | - | - | 0 | - |
| - | - | 1273 | 152.1 | - | - | 0 | - |
| - | - | 2979 | 152.1 | - | - | 0 | - |
| - | - | 1351 | 153.1 | - | - | 0 | - |
| - | - | 925.2 | 154.1 | - | - | 0 | - |
| - | - | 6837 | 155.1 | - | - | 0 | - |
| - | - | 2581 | 155.1 | - | - | 0 | - |
| - | - | 4810 | 155.1 | - | - | 0 | - |
| - | - | 2492 | 156.1 | - | - | 0 | - |
| - | - | 2944 | 156.1 | - | - | 0 | - |
| - | - | 680.8 | 156.1 | - | - | 0 | - |
| - | - | 1145 | 157.1 | - | - | 0 | - |
| - | - | 905.4 | 157.1 | - | - | 0 | - |
| - | - | 757.2 | 157.1 | - | - | 0 | - |
| - | - | 1880 | 158.1 | - | - | 0 | - |
| - | - | 1.504E+04 | 158.1 | - | - | 0 | - |
| - | - | 8.365E+04 | 159.1 | - | - | 0 | - |
| - | - | 1814 | 159.1 | - | - | 0 | - |
| - | - | 3841 | 160.1 | - | - | 0 | - |
| - | - | 9087 | 160.1 | - | - | 0 | - |
| - | - | 777 | 161.1 | - | - | 0 | - |
| - | - | 668.1 | 161.1 | - | - | 0 | - |
| - | - | 635.7 | 163.6 | - | - | 0 | - |
| - | - | 799.8 | 164 | - | - | 0 | - |
| - | - | 2112 | 164.1 | - | - | 0 | - |
| - | - | 1024 | 165.1 | - | - | 0 | - |
| - | - | 5462 | 165.1 | - | - | 0 | - |
| - | - | 2527 | 166.1 | - | - | 0 | - |
| - | - | 1196 | 168.1 | - | - | 0 | - |
| - | - | 8639 | 169.1 | - | - | 0 | - |
| - | - | 9766 | 169.1 | - | - | 0 | - |
| - | - | 839.1 | 169.2 | - | - | 0 | - |
| - | - | 3.958E+04 | 170.1 | - | - | 0 | - |
| - | - | 897.6 | 170.1 | - | - | 0 | - |
| - | - | 4350 | 171.1 | - | - | 0 | - |
| - | - | 1465 | 171.1 | - | - | 0 | - |
| 5 | b | 639.5 | 172.1 | 0.002432 | 14.13 | +4 | 5 |
| - | - | 533.1 | 172.1 | - | - | 0 | - |
| - | - | 8443 | 172.1 | - | - | 0 | - |
| - | - | 947.4 | 173.1 | - | - | 0 | - |
| - | - | 889 | 173.5 | - | - | 0 | - |
| - | - | 577.4 | 174.1 | - | - | 0 | - |
| - | - | 507.7 | 175.1 | - | - | 0 | - |
| - | - | 2129 | 175.1 | - | - | 0 | - |
| - | - | 3758 | 176.1 | - | - | 0 | - |
| - | - | 692.8 | 177.1 | - | - | 0 | - |
| - | - | 760.1 | 178.1 | - | - | 0 | - |
| - | - | 648.3 | 179.1 | - | - | 0 | - |
| - | - | 678.5 | 179.1 | - | - | 0 | - |
| - | - | 634.7 | 179.2 | - | - | 0 | - |
| - | - | 3188 | 181.1 | - | - | 0 | - |
| - | - | 4.097E+04 | 181.1 | - | - | 0 | - |
| - | - | 2413 | 181.1 | - | - | 0 | - |
| - | - | 595.8 | 182.1 | - | - | 0 | - |
| - | - | 537.5 | 182.1 | - | - | 0 | - |
| - | - | 3476 | 182.1 | - | - | 0 | - |
| - | - | 4789 | 182.1 | - | - | 0 | - |
| - | - | 1.688E+04 | 183.1 | - | - | 0 | - |
| - | - | 806.4 | 183.1 | - | - | 0 | - |
| - | - | 1652 | 183.1 | - | - | 0 | - |
| - | - | 601.9 | 183.1 | - | - | 0 | - |
| - | - | 1528 | 183.1 | - | - | 0 | - |
| - | - | 2050 | 184.1 | - | - | 0 | - |
| - | - | 2623 | 184.1 | - | - | 0 | - |
| - | - | 714.8 | 185.1 | - | - | 0 | - |
| - | - | 5237 | 185.1 | - | - | 0 | - |
| - | - | 665.5 | 185.1 | - | - | 0 | - |
| - | - | 1669 | 185.2 | - | - | 0 | - |
| - | - | 1.635E+04 | 186.1 | - | - | 0 | - |
| - | - | 8.867E+04 | 187.1 | - | - | 0 | - |
| - | - | 4179 | 187.1 | - | - | 0 | - |
| - | - | 1045 | 187.1 | - | - | 0 | - |
| 12 | y | 1313 | 187.1 | 0.0004575 | 2.445 | +2 | 3 |
| - | - | 709.8 | 188.1 | - | - | 0 | - |
| - | - | 9746 | 188.1 | - | - | 0 | - |
| - | - | 748.3 | 189.1 | - | - | 0 | - |
| - | - | 461.4 | 191.1 | - | - | 0 | - |
| - | - | 832.5 | 193.1 | - | - | 0 | - |
| - | - | 463.1 | 195.1 | - | - | 0 | - |
| - | - | 856.7 | 195.1 | - | - | 0 | - |
| - | - | 1205 | 195.1 | - | - | 0 | - |
| - | - | 1796 | 196.1 | - | - | 0 | - |
| - | - | 6359 | 197.2 | - | - | 0 | - |
| - | - | 1137 | 198.2 | - | - | 0 | - |
| - | - | 3374 | 199.1 | - | - | 0 | - |
| - | - | 1524 | 199.1 | - | - | 0 | - |
| - | - | 1984 | 199.2 | - | - | 0 | - |
| - | - | 4037 | 199.2 | - | - | 0 | - |
| - | - | 604.9 | 200.1 | - | - | 0 | - |
| - | - | 2307 | 200.1 | - | - | 0 | - |
| - | - | 703 | 201.1 | - | - | 0 | - |
| - | - | 8590 | 201.1 | - | - | 0 | - |
| - | - | 813.3 | 201.1 | - | - | 0 | - |
| - | - | 4651 | 202.1 | - | - | 0 | - |
| - | - | 1.433E+04 | 202.1 | - | - | 0 | - |
| - | - | 1259 | 203.1 | - | - | 0 | - |
| - | - | 1577 | 203.1 | - | - | 0 | - |
| - | - | 605.4 | 203.1 | - | - | 0 | - |
| - | - | 2148 | 204.1 | - | - | 0 | - |
| - | - | 2406 | 204.1 | - | - | 0 | - |
| 3 | a | 762.4 | 205.1 | 0.0005785 | 2.82 | +2 | 3 |
| - | - | 713.1 | 205.1 | - | - | 0 | - |
| - | - | 554.1 | 207.1 | - | - | 0 | - |
| - | - | 3403 | 207.1 | - | - | 0 | - |
| - | - | 897.3 | 208.1 | - | - | 0 | - |
| - | - | 567.8 | 209.1 | - | - | 0 | - |
| - | - | 1700 | 209.1 | - | - | 0 | - |
| - | - | 663.6 | 210.1 | - | - | 0 | - |
| - | - | 2679 | 211.1 | - | - | 0 | - |
| - | - | 803.9 | 212.1 | - | - | 0 | - |
| - | - | 2075 | 212.7 | - | - | 0 | - |
| - | - | 634.4 | 213.1 | - | - | 0 | - |
| - | - | 900.6 | 213.2 | - | - | 0 | - |
| - | - | 3961 | 214.1 | - | - | 0 | - |
| - | - | 1074 | 214.2 | - | - | 0 | - |
| - | - | 1136 | 215.1 | - | - | 0 | - |
| - | - | 1091 | 216.1 | - | - | 0 | - |
| - | - | 1234 | 216.1 | - | - | 0 | - |
| - | - | 585.2 | 216.1 | - | - | 0 | - |
| - | - | 1392 | 217.1 | - | - | 0 | - |
| - | - | 679.3 | 217.2 | - | - | 0 | - |
| 9 | y | 755 | 218.2 | 0.0009903 | 4.54 | +3 | 6 |
| - | - | 1.02E+04 | 220.1 | - | - | 0 | - |
| - | - | 1.469E+04 | 221.1 | - | - | 0 | - |
| - | - | 741.3 | 221.1 | - | - | 0 | - |
| - | - | 1678 | 222.1 | - | - | 0 | - |
| - | - | 1376 | 222.1 | - | - | 0 | - |
| - | - | 763.5 | 223.1 | - | - | 0 | - |
| - | - | 526.3 | 223.2 | - | - | 0 | - |
| - | - | 2369 | 224.1 | - | - | 0 | - |
| - | - | 6212 | 224.2 | - | - | 0 | - |
| - | - | 658.5 | 225.1 | - | - | 0 | - |
| - | - | 2100 | 225.2 | - | - | 0 | - |
| - | - | 822.2 | 225.2 | - | - | 0 | - |
| - | - | 1.134E+05 | 226.2 | - | - | 0 | - |
| - | - | 1049 | 227.1 | - | - | 0 | - |
| - | - | 818.7 | 227.1 | - | - | 0 | - |
| 11 | y | 1.407E+04 | 227.2 | 0.001709 | 7.522 | +2 | 4 |
| - | - | 755.9 | 227.2 | - | - | 0 | - |
| 2 | a | 7557 | 228.1 | 0.0006768 | 2.967 | +1 | 2 |
| - | - | 1466 | 228.1 | - | - | 0 | - |
| - | - | 910.4 | 228.2 | - | - | 0 | - |
| 5 | b | 9180 | 229.1 | 0.00195 | 8.511 | +3 | 5 |
| - | - | 1784 | 229.1 | - | - | 0 | - |
| - | - | 1197 | 229.2 | - | - | 0 | - |
| - | - | 1912 | 229.2 | - | - | 0 | - |
| - | - | 2505 | 230.1 | - | - | 0 | - |
| - | - | 4456 | 231.1 | - | - | 0 | - |
| - | - | 1403 | 231.1 | - | - | 0 | - |
| - | - | 2466 | 231.2 | - | - | 0 | - |
| - | - | 674.4 | 232.1 | - | - | 0 | - |
| - | - | 2636 | 233.1 | - | - | 0 | - |
| - | - | 897.9 | 234.1 | - | - | 0 | - |
| - | - | 992.5 | 234.1 | - | - | 0 | - |
| - | - | 1146 | 234.1 | - | - | 0 | - |
| - | - | 1.785E+04 | 235.1 | - | - | 0 | - |
| 11 | y | 1.691E+04 | 235.7 | 0.0005953 | 2.526 | +2 | 4 |
| - | - | 2586 | 236.1 | - | - | 0 | - |
| - | - | 789.7 | 236.1 | - | - | 0 | - |
| - | - | 5124 | 236.2 | - | - | 0 | - |
| - | - | 805.5 | 237.1 | - | - | 0 | - |
| 8 | y | 817.8 | 237.2 | 0.002094 | 8.829 | +3 | 7 |
| - | - | 1550 | 238.1 | - | - | 0 | - |
| - | - | 6777 | 239.2 | - | - | 0 | - |
| - | - | 1799 | 240.1 | - | - | 0 | - |
| - | - | 1071 | 240.2 | - | - | 0 | - |
| - | - | 1201 | 241.2 | - | - | 0 | - |
| - | - | 3921 | 242.2 | - | - | 0 | - |
| - | - | 1414 | 243.1 | - | - | 0 | - |
| - | - | 1334 | 243.2 | - | - | 0 | - |
| - | - | 644.3 | 244.1 | - | - | 0 | - |
| - | - | 619 | 244.2 | - | - | 0 | - |
| 13 | y | 2.714E+04 | 245.2 | 0.0006459 | 2.634 | +1 | 2 |
| 2 | a | 2.103E+05 | 246.1 | 0.0006559 | 2.665 | +1 | 2 |
| - | - | 3479 | 246.2 | - | - | 0 | - |
| - | - | 762.1 | 246.6 | - | - | 0 | - |
| - | - | 4.976E+04 | 247.1 | - | - | 0 | - |
| - | - | 1.594E+04 | 248.1 | - | - | 0 | - |
| - | - | 3511 | 248.1 | - | - | 0 | - |
| - | - | 2.228E+04 | 249.1 | - | - | 0 | - |
| 8 | b | 1435 | 249.1 | 0.0008093 | 3.249 | +4 | 8 |
| - | - | 914.3 | 249.1 | - | - | 0 | - |
| - | - | 2613 | 250.1 | - | - | 0 | - |
| - | - | 2946 | 250.1 | - | - | 0 | - |
| - | - | 1436 | 251.1 | - | - | 0 | - |
| - | - | 3270 | 251.2 | - | - | 0 | - |
| - | - | 516.4 | 252.2 | - | - | 0 | - |
| - | - | 2248 | 252.2 | - | - | 0 | - |
| - | - | 679.8 | 253.1 | - | - | 0 | - |
| - | - | 839.7 | 253.2 | - | - | 0 | - |
| - | - | 1243 | 255.1 | - | - | 0 | - |
| - | - | 1016 | 255.1 | - | - | 0 | - |
| 2 | b | 5655 | 256.1 | 0.0006505 | 2.54 | +1 | 2 |
| - | - | 1028 | 257.1 | - | - | 0 | - |
| - | - | 2574 | 257.1 | - | - | 0 | - |
| - | - | 2798 | 257.2 | - | - | 0 | - |
| - | - | 3555 | 258.1 | - | - | 0 | - |
| - | - | 3.817E+04 | 258.1 | - | - | 0 | - |
| - | - | 844.1 | 259.1 | - | - | 0 | - |
| - | - | 6307 | 259.1 | - | - | 0 | - |
| - | - | 1391 | 259.1 | - | - | 0 | - |
| - | - | 944.3 | 259.9 | - | - | 0 | - |
| - | - | 5128 | 260.1 | - | - | 0 | - |
| - | - | 3439 | 260.2 | - | - | 0 | - |
| - | - | 1028 | 261.1 | - | - | 0 | - |
| - | - | 1872 | 263.1 | - | - | 0 | - |
| - | - | 2095 | 264.1 | - | - | 0 | - |
| - | - | 1627 | 265.1 | - | - | 0 | - |
| - | - | 1615 | 265.2 | - | - | 0 | - |
| - | - | 928.1 | 266.1 | - | - | 0 | - |
| - | - | 1.391E+04 | 266.1 | - | - | 0 | - |
| - | - | 731.1 | 266.2 | - | - | 0 | - |
| - | - | 1665 | 267.1 | - | - | 0 | - |
| - | - | 1184 | 267.1 | - | - | 0 | - |
| - | - | 2789 | 268.2 | - | - | 0 | - |
| - | - | 2091 | 269.6 | - | - | 0 | - |
| - | - | 741.1 | 270.1 | - | - | 0 | - |
| - | - | 9391 | 270.2 | - | - | 0 | - |
| 10 | y | 5525 | 271.2 | 0.000414 | 1.526 | +2 | 5 |
| - | - | 1343 | 271.7 | - | - | 0 | - |
| - | - | 817.7 | 272.1 | - | - | 0 | - |
| - | - | 687.1 | 273.1 | - | - | 0 | - |
| 2 | b | 8.232E+04 | 274.1 | 0.0005228 | 1.907 | +1 | 2 |
| - | - | 1.055E+04 | 275.1 | - | - | 0 | - |
| - | - | 6.405E+04 | 275.1 | - | - | 0 | - |
| 7 | y | 2822 | 275.2 | 0.0005432 | 1.974 | +3 | 8 |
| - | - | 1423 | 275.5 | - | - | 0 | - |
| - | - | 2893 | 276.1 | - | - | 0 | - |
| - | - | 8850 | 276.1 | - | - | 0 | - |
| - | - | 884.1 | 277.1 | - | - | 0 | - |
| - | - | 553.9 | 277.1 | - | - | 0 | - |
| - | - | 3106 | 277.1 | - | - | 0 | - |
| - | - | 773.3 | 278.1 | - | - | 0 | - |
| - | - | 831.8 | 279.1 | - | - | 0 | - |
| - | - | 1979 | 279.2 | - | - | 0 | - |
| - | - | 1785 | 283.2 | - | - | 0 | - |
| - | - | 2.576E+04 | 283.2 | - | - | 0 | - |
| - | - | 2695 | 283.6 | - | - | 0 | - |
| - | - | 1554 | 284.1 | - | - | 0 | - |
| - | - | 3082 | 284.2 | - | - | 0 | - |
| - | - | 1563 | 285.1 | - | - | 0 | - |
| - | - | 5393 | 285.2 | - | - | 0 | - |
| - | - | 780.2 | 286.2 | - | - | 0 | - |
| - | - | 877 | 288.1 | - | - | 0 | - |
| - | - | 1029 | 290.1 | - | - | 0 | - |
| - | - | 9781 | 290.2 | - | - | 0 | - |
| - | - | 2332 | 290.7 | - | - | 0 | - |
| - | - | 1439 | 291.2 | - | - | 0 | - |
| - | - | 2.04E+04 | 292.1 | - | - | 0 | - |
| - | - | 744.9 | 292.2 | - | - | 0 | - |
| - | - | 3206 | 293.1 | - | - | 0 | - |
| - | - | 5434 | 294.1 | - | - | 0 | - |
| - | - | 2340 | 294.2 | - | - | 0 | - |
| - | - | 722.6 | 295.1 | - | - | 0 | - |
| - | - | 769.2 | 297.2 | - | - | 0 | - |
| - | - | 1.421E+04 | 297.2 | - | - | 0 | - |
| - | - | 2476 | 298.2 | - | - | 0 | - |
| - | - | 1884 | 300.1 | - | - | 0 | - |
| - | - | 1127 | 300.7 | - | - | 0 | - |
| - | - | 4107 | 303.1 | - | - | 0 | - |
| - | - | 677.1 | 303.2 | - | - | 0 | - |
| - | - | 7776 | 305.1 | - | - | 0 | - |
| - | - | 2389 | 305.7 | - | - | 0 | - |
| - | - | 1100 | 306.1 | - | - | 0 | - |
| - | - | 777.3 | 306.2 | - | - | 0 | - |
| - | - | 782.7 | 307.2 | - | - | 0 | - |
| - | - | 1171 | 309.2 | - | - | 0 | - |
| - | - | 746.1 | 309.2 | - | - | 0 | - |
| - | - | 711.2 | 311.2 | - | - | 0 | - |
| - | - | 3467 | 311.2 | - | - | 0 | - |
| - | - | 702.8 | 312.2 | - | - | 0 | - |
| - | - | 785.7 | 313.2 | - | - | 0 | - |
| - | - | 3692 | 314.7 | - | - | 0 | - |
| - | - | 1431 | 315.2 | - | - | 0 | - |
| - | - | 827.5 | 318.2 | - | - | 0 | - |
| - | - | 1018 | 318.2 | - | - | 0 | - |
| - | - | 885.1 | 320.1 | - | - | 0 | - |
| 6 | y | 3224 | 320.9 | 0.0007322 | 2.282 | +3 | 9 |
| - | - | 1118 | 321.2 | - | - | 0 | - |
| - | - | 1.628E+04 | 321.2 | - | - | 0 | - |
| - | - | 1.065E+04 | 322.2 | - | - | 0 | - |
| - | - | 543.9 | 322.2 | - | - | 0 | - |
| - | - | 3750 | 322.2 | - | - | 0 | - |
| - | - | 765 | 323 | - | - | 0 | - |
| 11 | b | 2668 | 323.2 | 0.00115 | 3.56 | +4 | 11 |
| - | - | 3625 | 324.2 | - | - | 0 | - |
| - | - | 1498 | 324.7 | - | - | 0 | - |
| - | - | 744.6 | 325.2 | - | - | 0 | - |
| - | - | 5052 | 326.2 | - | - | 0 | - |
| - | - | 711.1 | 326.7 | - | - | 0 | - |
| - | - | 770.3 | 327.2 | - | - | 0 | - |
| - | - | 1405 | 327.2 | - | - | 0 | - |
| - | - | 1493 | 327.3 | - | - | 0 | - |
| - | - | 5636 | 332.7 | - | - | 0 | - |
| - | - | 3233 | 333.1 | - | - | 0 | - |
| - | - | 2881 | 333.2 | - | - | 0 | - |
| - | - | 1013 | 334.2 | - | - | 0 | - |
| 9 | y | 2187 | 335.2 | 0.0009632 | 2.873 | +2 | 6 |
| - | - | 2.59E+04 | 336.2 | - | - | 0 | - |
| - | - | 4001 | 337.2 | - | - | 0 | - |
| - | - | 7119 | 337.7 | - | - | 0 | - |
| - | - | 627.8 | 338.2 | - | - | 0 | - |
| - | - | 3131 | 338.2 | - | - | 0 | - |
| - | - | 943.9 | 338.2 | - | - | 0 | - |
| - | - | 662 | 338.7 | - | - | 0 | - |
| - | - | 2.713E+04 | 339.2 | - | - | 0 | - |
| - | - | 674.6 | 339.8 | - | - | 0 | - |
| - | - | 2426 | 340.2 | - | - | 0 | - |
| - | - | 5354 | 340.2 | - | - | 0 | - |
| - | - | 718.9 | 341.2 | - | - | 0 | - |
| - | - | 770.6 | 341.2 | - | - | 0 | - |
| - | - | 1173 | 342.2 | - | - | 0 | - |
| - | - | 1235 | 345.6 | - | - | 0 | - |
| - | - | 931 | 346.1 | - | - | 0 | - |
| - | - | 1328 | 346.7 | - | - | 0 | - |
| - | - | 2.351E+04 | 346.7 | - | - | 0 | - |
| - | - | 8306 | 347.2 | - | - | 0 | - |
| - | - | 2170 | 347.7 | - | - | 0 | - |
| - | - | 2197 | 349.7 | - | - | 0 | - |
| - | - | 4128 | 350.2 | - | - | 0 | - |
| - | - | 2327 | 351.2 | - | - | 0 | - |
| - | - | 1429 | 353.2 | - | - | 0 | - |
| - | - | 2170 | 354.2 | - | - | 0 | - |
| - | - | 2376 | 354.2 | - | - | 0 | - |
| - | - | 3.361E+04 | 354.2 | - | - | 0 | - |
| - | - | 1888 | 354.5 | - | - | 0 | - |
| - | - | 1473 | 354.7 | - | - | 0 | - |
| - | - | 6204 | 355.2 | - | - | 0 | - |
| - | - | 5658 | 355.7 | - | - | 0 | - |
| - | - | 794.4 | 356.2 | - | - | 0 | - |
| - | - | 1513 | 356.2 | - | - | 0 | - |
| - | - | 664.8 | 356.7 | - | - | 0 | - |
| - | - | 901 | 357.2 | - | - | 0 | - |
| - | - | 7435 | 357.3 | - | - | 0 | - |
| - | - | 8458 | 358.2 | - | - | 0 | - |
| - | - | 1548 | 358.3 | - | - | 0 | - |
| - | - | 2055 | 358.7 | - | - | 0 | - |
| - | - | 1641 | 359.2 | - | - | 0 | - |
| - | - | 816.3 | 359.2 | - | - | 0 | - |
| - | - | 718.6 | 359.5 | - | - | 0 | - |
| - | - | 1607 | 359.9 | - | - | 0 | - |
| - | - | 911.6 | 360.1 | - | - | 0 | - |
| - | - | 1675 | 360.7 | - | - | 0 | - |
| - | - | 1274 | 360.7 | - | - | 0 | - |
| - | - | 858.1 | 361.2 | - | - | 0 | - |
| - | - | 903.6 | 362.2 | - | - | 0 | - |
| - | - | 2056 | 362.7 | - | - | 0 | - |
| - | - | 728.9 | 362.7 | - | - | 0 | - |
| 8 | y | 1670 | 363.7 | 0.0009735 | 2.676 | +2 | 7 |
| - | - | 724.4 | 365.3 | - | - | 0 | - |
| - | - | 3601 | 366.2 | - | - | 0 | - |
| 5 | y | 3312 | 366.6 | 0.0007991 | 2.18 | +3 | 10 |
| - | - | 2390 | 366.9 | - | - | 0 | - |
| - | - | 1030 | 367.1 | - | - | 0 | - |
| - | - | 2986 | 367.2 | - | - | 0 | - |
| - | - | 4760 | 368.1 | - | - | 0 | - |
| - | - | 892.9 | 368.2 | - | - | 0 | - |
| - | - | 2213 | 369.2 | - | - | 0 | - |
| - | - | 598.9 | 369.7 | - | - | 0 | - |
| - | - | 1071 | 370.5 | - | - | 0 | - |
| - | - | 686.2 | 370.7 | - | - | 0 | - |
| - | - | 1548 | 371.2 | - | - | 0 | - |
| 12 | y | 1.516E+04 | 373.3 | 0.0007146 | 1.914 | +1 | 3 |
| - | - | 785.5 | 374.3 | - | - | 0 | - |
| - | - | 2738 | 374.3 | - | - | 0 | - |
| 9 | b | 1003 | 374.5 | 0.0002519 | 0.6725 | +3 | 9 |
| - | - | 831.1 | 375.2 | - | - | 0 | - |
| - | - | 1542 | 376.7 | - | - | 0 | - |
| - | - | 832.6 | 377 | - | - | 0 | - |
| - | - | 1107 | 377.2 | - | - | 0 | - |
| - | - | 1358 | 378.7 | - | - | 0 | - |
| - | - | 2024 | 379 | - | - | 0 | - |
| - | - | 1142 | 379.2 | - | - | 0 | - |
| - | - | 1373 | 380.2 | - | - | 0 | - |
| - | - | 701.1 | 381.2 | - | - | 0 | - |
| - | - | 6741 | 382.2 | - | - | 0 | - |
| - | - | 1759 | 382.3 | - | - | 0 | - |
| - | - | 3337 | 382.5 | - | - | 0 | - |
| - | - | 754.5 | 382.7 | - | - | 0 | - |
| - | - | 1166 | 382.9 | - | - | 0 | - |
| - | - | 1826 | 383.3 | - | - | 0 | - |
| - | - | 3974 | 384.2 | - | - | 0 | - |
| - | - | 1.276E+04 | 385.2 | - | - | 0 | - |
| - | - | 3323 | 385.2 | - | - | 0 | - |
| - | - | 1.364E+04 | 386.2 | - | - | 0 | - |
| - | - | 1160 | 386.2 | - | - | 0 | - |
| - | - | 721.7 | 386.5 | - | - | 0 | - |
| - | - | 3122 | 387.2 | - | - | 0 | - |
| - | - | 958.6 | 388.2 | - | - | 0 | - |
| - | - | 2613 | 388.9 | - | - | 0 | - |
| - | - | 1486 | 389.2 | - | - | 0 | - |
| - | - | 1759 | 389.3 | - | - | 0 | - |
| - | - | 916.1 | 389.5 | - | - | 0 | - |
| - | - | 975.1 | 389.8 | - | - | 0 | - |
| 2 | y | 7197 | 390.2 | 0.001049 | 2.689 | +4 | 13 |
| - | - | 3916 | 390.5 | - | - | 0 | - |
| - | - | 4582 | 390.7 | - | - | 0 | - |
| 10 | b | 1765 | 392.2 | 0.0004862 | 1.24 | +3 | 10 |
| - | - | 5466 | 392.3 | - | - | 0 | - |
| 10 | b | 2016 | 392.5 | 0.005062 | 12.9 | +3 | 10 |
| - | - | 899.5 | 392.7 | - | - | 0 | - |
| - | - | 691.9 | 393 | - | - | 0 | - |
| - | - | 1237 | 393.2 | - | - | 0 | - |
| - | - | 1932 | 393.3 | - | - | 0 | - |
| - | - | 9941 | 394.2 | - | - | 0 | - |
| 2 | y | 3615 | 394.5 | 0.0003629 | 0.9201 | +4 | 13 |
| - | - | 1186 | 394.7 | - | - | 0 | - |
| - | - | 1399 | 395 | - | - | 0 | - |
| - | - | 2013 | 395.2 | - | - | 0 | - |
| - | - | 1457 | 396.3 | - | - | 0 | - |
| - | - | 655.4 | 397.2 | - | - | 0 | - |
| - | - | 1777 | 397.2 | - | - | 0 | - |
| - | - | 903.9 | 397.6 | - | - | 0 | - |
| 10 | b | 1676 | 398.2 | 0.0008099 | 2.034 | +3 | 10 |
| - | - | 986.9 | 398.2 | - | - | 0 | - |
| - | - | 1482 | 398.5 | - | - | 0 | - |
| - | - | 928.9 | 398.9 | - | - | 0 | - |
| - | - | 1609 | 399.2 | - | - | 0 | - |
| - | - | 762.5 | 400.2 | - | - | 0 | - |
| - | - | 981.1 | 400.2 | - | - | 0 | - |
| - | - | 1294 | 400.5 | - | - | 0 | - |
| - | - | 1095 | 401.2 | - | - | 0 | - |
| - | - | 809 | 401.3 | - | - | 0 | - |
| - | - | 4466 | 402.2 | - | - | 0 | - |
| - | - | 2216 | 402.5 | - | - | 0 | - |
| - | - | 1.174E+04 | 403.2 | - | - | 0 | - |
| - | - | 6311 | 403.2 | - | - | 0 | - |
| - | - | 3561 | 403.3 | - | - | 0 | - |
| 4 | y | 8966 | 403.6 | 0.00291 | 7.21 | +3 | 11 |
| - | - | 1141 | 403.8 | - | - | 0 | - |
| - | - | 5484 | 403.9 | - | - | 0 | - |
| - | - | 2111 | 404.2 | - | - | 0 | - |
| - | - | 2817 | 404.2 | - | - | 0 | - |
| - | - | 928.3 | 404.3 | - | - | 0 | - |
| - | - | 3432 | 404.5 | - | - | 0 | - |
| - | - | 3112 | 404.7 | - | - | 0 | - |
| - | - | 827.8 | 404.9 | - | - | 0 | - |
| - | - | 4729 | 405 | - | - | 0 | - |
| - | - | 3545 | 405.2 | - | - | 0 | - |
| - | - | 6124 | 406.2 | - | - | 0 | - |
| - | - | 3700 | 406.3 | - | - | 0 | - |
| - | - | 2607 | 406.7 | - | - | 0 | - |
| - | - | 2190 | 406.8 | - | - | 0 | - |
| - | - | 1537 | 407.2 | - | - | 0 | - |
| - | - | 2396 | 408.2 | - | - | 0 | - |
| - | - | 2292 | 408.6 | - | - | 0 | - |
| - | - | 1651 | 408.9 | - | - | 0 | - |
| 4 | y | 2885 | 409.2 | 0.001252 | 3.059 | +3 | 11 |
| - | - | 2891 | 409.6 | - | - | 0 | - |
| - | - | 1724 | 409.9 | - | - | 0 | - |
| - | - | 742.4 | 410.2 | - | - | 0 | - |
| - | - | 1.167E+04 | 410.3 | - | - | 0 | - |
| - | - | 4865 | 410.6 | - | - | 0 | - |
| - | - | 5026 | 410.9 | - | - | 0 | - |
| 6 | b | 1.132E+04 | 411.2 | 0.0007845 | 1.908 | +2 | 6 |
| - | - | 829.5 | 411.2 | - | - | 0 | - |
| - | - | 1596 | 411.2 | - | - | 0 | - |
| - | - | 2389 | 411.3 | - | - | 0 | - |
| 0 | Precursor | 1772 | 411.7 | 8.966E-05 | 0.2178 | +4 | -1 |
| 0 | Precursor | 2909 | 412 | 0.003113 | 7.556 | +4 | -1 |
| - | - | 2795 | 412.2 | - | - | 0 | - |
| - | - | 1998 | 412.2 | - | - | 0 | - |
| - | - | 2639 | 412.2 | - | - | 0 | - |
| 7 | y | 4.535E+04 | 412.3 | 0.0009893 | 2.4 | +2 | 8 |
| - | - | 864.5 | 412.5 | - | - | 0 | - |
| - | - | 2.158E+04 | 412.8 | - | - | 0 | - |
| - | - | 1728 | 413.2 | - | - | 0 | - |
| - | - | 6313 | 413.3 | - | - | 0 | - |
| - | - | 812.1 | 413.8 | - | - | 0 | - |
| - | - | 6678 | 413.9 | - | - | 0 | - |
| - | - | 753.5 | 414.2 | - | - | 0 | - |
| - | - | 6908 | 414.2 | - | - | 0 | - |
| - | - | 2821 | 414.6 | - | - | 0 | - |
| - | - | 701.3 | 414.9 | - | - | 0 | - |
| - | - | 837.3 | 415 | - | - | 0 | - |
| - | - | 5047 | 415.3 | - | - | 0 | - |
| - | - | 1672 | 415.8 | - | - | 0 | - |
| 0 | Precursor | 2.609E+04 | 416.2 | 0.0007482 | 1.798 | +4 | -1 |
| - | - | 3237 | 416.3 | - | - | 0 | - |
| - | - | 2.544E+04 | 416.5 | - | - | 0 | - |
| - | - | 1.297E+04 | 416.7 | - | - | 0 | - |
| - | - | 4009 | 417 | - | - | 0 | - |
| - | - | 2399 | 417.2 | - | - | 0 | - |
| - | - | 993.8 | 417.3 | - | - | 0 | - |
| - | - | 779.2 | 417.5 | - | - | 0 | - |
| - | - | 2.976E+04 | 419.9 | - | - | 0 | - |
| 6 | b | 2838 | 420.2 | 0.0006291 | 1.497 | +2 | 6 |
| - | - | 2.281E+04 | 420.2 | - | - | 0 | - |
| - | - | 8707 | 420.6 | - | - | 0 | - |
| - | - | 2460 | 420.7 | - | - | 0 | - |
| - | - | 2213 | 420.9 | - | - | 0 | - |
| - | - | 1277 | 421.2 | - | - | 0 | - |
| - | - | 3111 | 422.3 | - | - | 0 | - |
| - | - | 1645 | 423.3 | - | - | 0 | - |
| - | - | 3100 | 424.3 | - | - | 0 | - |
| - | - | 6618 | 424.7 | - | - | 0 | - |
| - | - | 3245 | 425.2 | - | - | 0 | - |
| - | - | 1772 | 425.3 | - | - | 0 | - |
| - | - | 845.2 | 425.3 | - | - | 0 | - |
| - | - | 6406 | 425.9 | - | - | 0 | - |
| - | - | 2399 | 426.2 | - | - | 0 | - |
| - | - | 930 | 426.6 | - | - | 0 | - |
| - | - | 5085 | 427.2 | - | - | 0 | - |
| - | - | 2075 | 427.7 | - | - | 0 | - |
| - | - | 1215 | 428.3 | - | - | 0 | - |
| - | - | 4.307E+04 | 429.2 | - | - | 0 | - |
| - | - | 1.171E+04 | 430.2 | - | - | 0 | - |
| - | - | 5478 | 431.2 | - | - | 0 | - |
| - | - | 1500 | 432.2 | - | - | 0 | - |
| - | - | 1906 | 433.7 | - | - | 0 | - |
| - | - | 682.8 | 434.2 | - | - | 0 | - |
| - | - | 913.5 | 436.6 | - | - | 0 | - |
| 3 | b | 2434 | 437.2 | 0.0009455 | 2.163 | +1 | 3 |
| - | - | 1630 | 438.2 | - | - | 0 | - |
| - | - | 778.4 | 438.2 | - | - | 0 | - |
| - | - | 2521 | 438.6 | - | - | 0 | - |
| - | - | 3287 | 438.9 | - | - | 0 | - |
| - | - | 2310 | 439.2 | - | - | 0 | - |
| - | - | 886.6 | 439.3 | - | - | 0 | - |
| - | - | 1972 | 442.7 | - | - | 0 | - |
| - | - | 1039 | 444.2 | - | - | 0 | - |
| - | - | 977.5 | 445.2 | - | - | 0 | - |
| - | - | 1719 | 445.3 | - | - | 0 | - |
| - | - | 765.7 | 445.9 | - | - | 0 | - |
| - | - | 4303 | 446.2 | - | - | 0 | - |
| - | - | 1864 | 446.7 | - | - | 0 | - |
| - | - | 760.8 | 447.2 | - | - | 0 | - |
| - | - | 7790 | 448.3 | - | - | 0 | - |
| - | - | 6623 | 448.6 | - | - | 0 | - |
| - | - | 2238 | 448.9 | - | - | 0 | - |
| - | - | 888.9 | 449.3 | - | - | 0 | - |
| - | - | 6348 | 451.2 | - | - | 0 | - |
| - | - | 3508 | 451.3 | - | - | 0 | - |
| - | - | 5089 | 451.7 | - | - | 0 | - |
| - | - | 1.055E+04 | 451.9 | - | - | 0 | - |
| - | - | 1082 | 452.2 | - | - | 0 | - |
| - | - | 8779 | 452.3 | - | - | 0 | - |
| - | - | 1537 | 452.3 | - | - | 0 | - |
| - | - | 5359 | 452.6 | - | - | 0 | - |
| - | - | 712.3 | 452.7 | - | - | 0 | - |
| - | - | 1356 | 452.9 | - | - | 0 | - |
| - | - | 1598 | 453.7 | - | - | 0 | - |
| - | - | 1909 | 454.2 | - | - | 0 | - |
| - | - | 4947 | 455.3 | - | - | 0 | - |
| - | - | 832.8 | 456.3 | - | - | 0 | - |
| - | - | 1807 | 457.2 | - | - | 0 | - |
| - | - | 6147 | 457.6 | - | - | 0 | - |
| 3 | y | 7096 | 457.9 | 0.004851 | 10.59 | +3 | 12 |
| - | - | 6550 | 458.3 | - | - | 0 | - |
| - | - | 1330 | 458.6 | - | - | 0 | - |
| - | - | 792.9 | 458.9 | - | - | 0 | - |
| - | - | 1019 | 459.2 | - | - | 0 | - |
| - | - | 8007 | 460.2 | - | - | 0 | - |
| - | - | 4922 | 460.7 | - | - | 0 | - |
| - | - | 1687 | 461.2 | - | - | 0 | - |
| - | - | 1523 | 462.9 | - | - | 0 | - |
| - | - | 1589 | 463.3 | - | - | 0 | - |
| 3 | y | 1.262E+05 | 463.6 | 0.0009657 | 2.083 | +3 | 12 |
| - | - | 1.025E+05 | 463.9 | - | - | 0 | - |
| - | - | 4.828E+04 | 464.3 | - | - | 0 | - |
| - | - | 1.089E+04 | 464.6 | - | - | 0 | - |
| - | - | 3937 | 464.9 | - | - | 0 | - |
| - | - | 801.4 | 465.3 | - | - | 0 | - |
| - | - | 1616 | 466.9 | - | - | 0 | - |
| - | - | 1670 | 467.3 | - | - | 0 | - |
| - | - | 3843 | 467.3 | - | - | 0 | - |
| - | - | 2370 | 468.3 | - | - | 0 | - |
| - | - | 2389 | 469.8 | - | - | 0 | - |
| - | - | 1825 | 470.2 | - | - | 0 | - |
| 11 | y | 5.232E+04 | 470.3 | 0.0008988 | 1.911 | +1 | 4 |
| - | - | 2882 | 470.6 | - | - | 0 | - |
| - | - | 1247 | 470.9 | - | - | 0 | - |
| - | - | 1.318E+04 | 471.3 | - | - | 0 | - |
| - | - | 4204 | 471.8 | - | - | 0 | - |
| - | - | 1962 | 472.3 | - | - | 0 | - |
| - | - | 2266 | 472.3 | - | - | 0 | - |
| - | - | 1726 | 472.6 | - | - | 0 | - |
| 12 | b | 873.6 | 473.2 | 0.007814 | 16.51 | +3 | 12 |
| - | - | 6296 | 473.3 | - | - | 0 | - |
| - | - | 2509 | 473.6 | - | - | 0 | - |
| - | - | 1535 | 473.9 | - | - | 0 | - |
| - | - | 2218 | 474.3 | - | - | 0 | - |
| - | - | 3174 | 474.8 | - | - | 0 | - |
| - | - | 2441 | 475.3 | - | - | 0 | - |
| - | - | 754.7 | 475.9 | - | - | 0 | - |
| - | - | 8384 | 476.3 | - | - | 0 | - |
| - | - | 1.049E+04 | 476.6 | - | - | 0 | - |
| - | - | 4240 | 476.9 | - | - | 0 | - |
| - | - | 1058 | 477.6 | - | - | 0 | - |
| - | - | 2638 | 478.2 | - | - | 0 | - |
| 6 | y | 1.127E+04 | 480.8 | 0.0009828 | 2.044 | +2 | 9 |
| - | - | 4694 | 481.3 | - | - | 0 | - |
| - | - | 1720 | 481.8 | - | - | 0 | - |
| - | - | 4797 | 481.9 | - | - | 0 | - |
| - | - | 4835 | 482.3 | - | - | 0 | - |
| - | - | 4129 | 482.3 | - | - | 0 | - |
| - | - | 3987 | 482.6 | - | - | 0 | - |
| - | - | 3151 | 482.8 | - | - | 0 | - |
| - | - | 2156 | 482.9 | - | - | 0 | - |
| - | - | 1518 | 483.3 | - | - | 0 | - |
| - | - | 7484 | 483.8 | - | - | 0 | - |
| - | - | 5355 | 484.3 | - | - | 0 | - |
| - | - | 1122 | 484.8 | - | - | 0 | - |
| - | - | 1178 | 487.9 | - | - | 0 | - |
| - | - | 1348 | 488.3 | - | - | 0 | - |
| - | - | 1143 | 488.6 | - | - | 0 | - |
| - | - | 1489 | 490.2 | - | - | 0 | - |
| - | - | 4946 | 491.3 | - | - | 0 | - |
| - | - | 885.2 | 492.3 | - | - | 0 | - |
| - | - | 1878 | 492.8 | - | - | 0 | - |
| - | - | 1764 | 493.3 | - | - | 0 | - |
| - | - | 4015 | 493.8 | - | - | 0 | - |
| - | - | 1571 | 494.2 | - | - | 0 | - |
| - | - | 1197 | 494.3 | - | - | 0 | - |
| 8 | b | 1349 | 497.2 | 0.0003213 | 0.6462 | +2 | 8 |
| - | - | 968.9 | 497.7 | - | - | 0 | - |
| - | - | 834.1 | 501.3 | - | - | 0 | - |
| - | - | 1520 | 501.6 | - | - | 0 | - |
| - | - | 925.1 | 501.9 | - | - | 0 | - |
| - | - | 879.6 | 502.3 | - | - | 0 | - |
| - | - | 3581 | 503.2 | - | - | 0 | - |
| - | - | 1095 | 504.2 | - | - | 0 | - |
| - | - | 1576 | 504.9 | - | - | 0 | - |
| 13 | b | 1660 | 505.3 | 0.008385 | 16.59 | +3 | 13 |
| - | - | 1116 | 505.6 | - | - | 0 | - |
| - | - | 1195 | 506.2 | - | - | 0 | - |
| - | - | 1069 | 508.8 | - | - | 0 | - |
| - | - | 3127 | 509.2 | - | - | 0 | - |
| - | - | 2073 | 509.7 | - | - | 0 | - |
| - | - | 1695 | 510.3 | - | - | 0 | - |
| - | - | 843.6 | 510.6 | - | - | 0 | - |
| 13 | b | 4153 | 510.9 | 0.0005319 | 1.041 | +3 | 13 |
| - | - | 4882 | 511.3 | - | - | 0 | - |
| - | - | 3945 | 511.6 | - | - | 0 | - |
| - | - | 1710 | 511.9 | - | - | 0 | - |
| - | - | 791.8 | 513.9 | - | - | 0 | - |
| - | - | 997 | 514.3 | - | - | 0 | - |
| - | - | 708.7 | 514.6 | - | - | 0 | - |
| - | - | 9579 | 516.9 | - | - | 0 | - |
| - | - | 1.286E+04 | 517.3 | - | - | 0 | - |
| - | - | 5414 | 517.6 | - | - | 0 | - |
| - | - | 1525 | 517.8 | - | - | 0 | - |
| - | - | 1232 | 517.9 | - | - | 0 | - |
| - | - | 1807 | 518.3 | - | - | 0 | - |
| 2 | y | 1.415E+04 | 519.9 | 0.001302 | 2.504 | +3 | 13 |
| - | - | 1170 | 520.2 | - | - | 0 | - |
| - | - | 1.03E+04 | 520.3 | - | - | 0 | - |
| - | - | 2347 | 520.4 | - | - | 0 | - |
| - | - | 5087 | 520.6 | - | - | 0 | - |
| - | - | 1422 | 521 | - | - | 0 | - |
| - | - | 4226 | 521.2 | - | - | 0 | - |
| - | - | 787.5 | 521.3 | - | - | 0 | - |
| - | - | 829.1 | 521.4 | - | - | 0 | - |
| - | - | 1179 | 522.2 | - | - | 0 | - |
| - | - | 1647 | 524.8 | - | - | 0 | - |
| - | - | 1956 | 525.3 | - | - | 0 | - |
| 2 | y | 1.52E+04 | 525.6 | 0.0002239 | 0.4259 | +3 | 13 |
| - | - | 1.808E+04 | 526 | - | - | 0 | - |
| - | - | 9693 | 526.3 | - | - | 0 | - |
| - | - | 916.4 | 526.3 | - | - | 0 | - |
| - | - | 2844 | 526.6 | - | - | 0 | - |
| - | - | 1618 | 527 | - | - | 0 | - |
| - | - | 762.8 | 529.2 | - | - | 0 | - |
| - | - | 1825 | 529.8 | - | - | 0 | - |
| - | - | 3483 | 530.3 | - | - | 0 | - |
| - | - | 1826 | 530.8 | - | - | 0 | - |
| - | - | 3265 | 531.2 | - | - | 0 | - |
| - | - | 1140 | 532.2 | - | - | 0 | - |
| - | - | 2651 | 535.8 | - | - | 0 | - |
| - | - | 1400 | 536.3 | - | - | 0 | - |
| - | - | 1160 | 536.8 | - | - | 0 | - |
| - | - | 819.9 | 537.8 | - | - | 0 | - |
| - | - | 2.62E+04 | 538.3 | - | - | 0 | - |
| - | - | 2585 | 538.4 | - | - | 0 | - |
| - | - | 861.7 | 538.8 | - | - | 0 | - |
| - | - | 4127 | 538.8 | - | - | 0 | - |
| - | - | 8008 | 539.3 | - | - | 0 | - |
| - | - | 8259 | 539.3 | - | - | 0 | - |
| - | - | 3114 | 539.8 | - | - | 0 | - |
| - | - | 1859 | 540.3 | - | - | 0 | - |
| - | - | 971.2 | 540.3 | - | - | 0 | - |
| 5 | y | 803.7 | 540.8 | 0.006103 | 11.28 | +2 | 10 |
| - | - | 864.8 | 541.3 | - | - | 0 | - |
| 10 | y | 1.112E+04 | 541.4 | 0.0007723 | 1.427 | +1 | 5 |
| - | - | 4817 | 542.3 | - | - | 0 | - |
| - | - | 3916 | 542.4 | - | - | 0 | - |
| - | - | 1532 | 543.3 | - | - | 0 | - |
| - | - | 1095 | 544.3 | - | - | 0 | - |
| - | - | 2548 | 544.8 | - | - | 0 | - |
| - | - | 1184 | 545.3 | - | - | 0 | - |
| - | - | 874.8 | 545.8 | - | - | 0 | - |
| - | - | 2851 | 547.8 | - | - | 0 | - |
| - | - | 1163 | 548.2 | - | - | 0 | - |
| - | - | 2766 | 548.3 | - | - | 0 | - |
| - | - | 1796 | 548.8 | - | - | 0 | - |
| - | - | 3587 | 549.2 | - | - | 0 | - |
| 5 | y | 2.908E+04 | 549.3 | 0.0008848 | 1.611 | +2 | 10 |
| - | - | 1.895E+04 | 549.8 | - | - | 0 | - |
| - | - | 7231 | 550.3 | - | - | 0 | - |
| - | - | 797.6 | 550.4 | - | - | 0 | - |
| - | - | 1516 | 550.8 | - | - | 0 | - |
| 9 | b | 718.8 | 552.8 | 0.007098 | 12.84 | +2 | 9 |
| - | - | 1526 | 553.2 | - | - | 0 | - |
| - | - | 1215 | 553.3 | - | - | 0 | - |
| - | - | 897.3 | 553.8 | - | - | 0 | - |
| - | - | 1659 | 555.3 | - | - | 0 | - |
| - | - | 1560 | 556.8 | - | - | 0 | - |
| - | - | 2803 | 557.3 | - | - | 0 | - |
| 9 | b | 7771 | 561.3 | 0.001025 | 1.827 | +2 | 9 |
| - | - | 4972 | 561.8 | - | - | 0 | - |
| - | - | 1420 | 562.3 | - | - | 0 | - |
| 4 | b | 752.2 | 565.2 | 0.001603 | 2.835 | +1 | 4 |
| - | - | 3.314E+04 | 566.2 | - | - | 0 | - |
| - | - | 8918 | 567.2 | - | - | 0 | - |
| - | - | 1453 | 567.4 | - | - | 0 | - |
| - | - | 1870 | 568.3 | - | - | 0 | - |
| - | - | 1.405E+04 | 570.2 | - | - | 0 | - |
| - | - | 4720 | 571.2 | - | - | 0 | - |
| - | - | 1088 | 572.3 | - | - | 0 | - |
| - | - | 1.651E+04 | 572.8 | - | - | 0 | - |
| - | - | 1.024E+04 | 573.3 | - | - | 0 | - |
| - | - | 3129 | 573.8 | - | - | 0 | - |
| - | - | 911.6 | 576.2 | - | - | 0 | - |
| - | - | 8804 | 577.4 | - | - | 0 | - |
| - | - | 1015 | 577.8 | - | - | 0 | - |
| - | - | 3323 | 578.4 | - | - | 0 | - |
| - | - | 1.89E+04 | 579.4 | - | - | 0 | - |
| - | - | 5078 | 580.4 | - | - | 0 | - |
| - | - | 1449 | 581.4 | - | - | 0 | - |
| - | - | 857.7 | 582.8 | - | - | 0 | - |
| - | - | 1516 | 587.3 | - | - | 0 | - |
| 10 | b | 1776 | 587.8 | 0.000385 | 0.655 | +2 | 10 |
| 10 | b | 2844 | 588.3 | 0.007279 | 12.37 | +2 | 10 |
| - | - | 761 | 594.2 | - | - | 0 | - |
| - | - | 787.5 | 595.3 | - | - | 0 | - |
| - | - | 6734 | 595.4 | - | - | 0 | - |
| - | - | 1618 | 595.8 | - | - | 0 | - |
| - | - | 1234 | 596.3 | - | - | 0 | - |
| - | - | 2489 | 596.4 | - | - | 0 | - |
| 10 | b | 1052 | 596.8 | 0.001206 | 2.021 | +2 | 10 |
| - | - | 2068 | 597.3 | - | - | 0 | - |
| - | - | 860.4 | 597.8 | - | - | 0 | - |
| - | - | 7317 | 598.2 | - | - | 0 | - |
| - | - | 2767 | 599.2 | - | - | 0 | - |
| - | - | 899.3 | 600.2 | - | - | 0 | - |
| - | - | 1007 | 600.3 | - | - | 0 | - |
| - | - | 5931 | 604.4 | - | - | 0 | - |
| 4 | y | 1.125E+04 | 604.8 | 0.00373 | 6.168 | +2 | 11 |
| - | - | 7554 | 605.4 | - | - | 0 | - |
| - | - | 2068 | 605.9 | - | - | 0 | - |
| - | - | 6006 | 610.3 | - | - | 0 | - |
| - | - | 2122 | 611.3 | - | - | 0 | - |
| - | - | 1256 | 611.8 | - | - | 0 | - |
| - | - | 1139 | 612.3 | - | - | 0 | - |
| 4 | y | 8720 | 613.4 | 0.0009539 | 1.555 | +2 | 11 |
| - | - | 698.3 | 613.4 | - | - | 0 | - |
| - | - | 4587 | 613.9 | - | - | 0 | - |
| - | - | 2241 | 614.4 | - | - | 0 | - |
| - | - | 743.1 | 614.9 | - | - | 0 | - |
| - | - | 2394 | 615.3 | - | - | 0 | - |
| - | - | 3788 | 615.3 | - | - | 0 | - |
| - | - | 2445 | 615.9 | - | - | 0 | - |
| - | - | 1072 | 616.4 | - | - | 0 | - |
| - | - | 805.7 | 618.9 | - | - | 0 | - |
| - | - | 5543 | 620.3 | - | - | 0 | - |
| - | - | 4312 | 620.8 | - | - | 0 | - |
| - | - | 2033 | 621.3 | - | - | 0 | - |
| - | - | 1164 | 623.4 | - | - | 0 | - |
| - | - | 957.8 | 628.3 | - | - | 0 | - |
| - | - | 1715 | 628.3 | - | - | 0 | - |
| - | - | 1.499E+04 | 629.3 | - | - | 0 | - |
| - | - | 9426 | 629.8 | - | - | 0 | - |
| - | - | 4687 | 630.3 | - | - | 0 | - |
| - | - | 933 | 630.8 | - | - | 0 | - |
| - | - | 8814 | 638.4 | - | - | 0 | - |
| - | - | 5145 | 638.9 | - | - | 0 | - |
| - | - | 4269 | 639.4 | - | - | 0 | - |
| - | - | 1245 | 647.4 | - | - | 0 | - |
| - | - | 773.3 | 648.8 | - | - | 0 | - |
| - | - | 1531 | 657.3 | - | - | 0 | - |
| - | - | 1245 | 657.3 | - | - | 0 | - |
| - | - | 2505 | 657.8 | - | - | 0 | - |
| - | - | 704.3 | 663.3 | - | - | 0 | - |
| - | - | 6514 | 664.5 | - | - | 0 | - |
| - | - | 2169 | 665.5 | - | - | 0 | - |
| - | - | 1620 | 665.8 | - | - | 0 | - |
| - | - | 1630 | 666.3 | - | - | 0 | - |
| - | - | 1307 | 667.3 | - | - | 0 | - |
| - | - | 1616 | 668.3 | - | - | 0 | - |
| 9 | y | 4300 | 669.5 | 0.0006579 | 0.9827 | +1 | 6 |
| - | - | 1629 | 670.5 | - | - | 0 | - |
| - | - | 2205 | 674.3 | - | - | 0 | - |
| - | - | 1.848E+04 | 674.4 | - | - | 0 | - |
| - | - | 933 | 675.3 | - | - | 0 | - |
| - | - | 8696 | 675.4 | - | - | 0 | - |
| - | - | 1752 | 676.4 | - | - | 0 | - |
| - | - | 760.4 | 679.3 | - | - | 0 | - |
| - | - | 1101 | 680.3 | - | - | 0 | - |
| 5 | b | 1448 | 684.3 | 0.0009239 | 1.35 | +1 | 5 |
| - | - | 1433 | 685.4 | - | - | 0 | - |
| - | - | 1677 | 690.3 | - | - | 0 | - |
| - | - | 1.91E+04 | 692.4 | - | - | 0 | - |
| - | - | 7840 | 693.4 | - | - | 0 | - |
| - | - | 1461 | 694.5 | - | - | 0 | - |
| 3 | y | 6299 | 694.9 | 0.0007838 | 1.128 | +2 | 12 |
| - | - | 4491 | 695.4 | - | - | 0 | - |
| - | - | 1265 | 695.9 | - | - | 0 | - |
| - | - | 1496 | 698.4 | - | - | 0 | - |
| 5 | b | 3041 | 702.3 | 0.001468 | 2.09 | +1 | 5 |
| - | - | 1234 | 703.3 | - | - | 0 | - |
| - | - | 8398 | 707.3 | - | - | 0 | - |
| - | - | 3840 | 708.3 | - | - | 0 | - |
| - | - | 938.5 | 708.5 | - | - | 0 | - |
| - | - | 6052 | 710.5 | - | - | 0 | - |
| - | - | 1738 | 711.5 | - | - | 0 | - |
| - | - | 1622 | 713.9 | - | - | 0 | - |
| - | - | 1445 | 714.4 | - | - | 0 | - |
| - | - | 902.4 | 714.9 | - | - | 0 | - |
| - | - | 3457 | 716.4 | - | - | 0 | - |
| - | - | 1765 | 717.4 | - | - | 0 | - |
| - | - | 906.5 | 718.4 | - | - | 0 | - |
| - | - | 1837 | 720.3 | - | - | 0 | - |
| - | - | 3049 | 721.4 | - | - | 0 | - |
| - | - | 1767 | 722.4 | - | - | 0 | - |
| - | - | 1792 | 722.9 | - | - | 0 | - |
| - | - | 2976 | 724.3 | - | - | 0 | - |
| - | - | 1046 | 725.3 | - | - | 0 | - |
| 8 | y | 9577 | 726.5 | 0.0003734 | 0.5139 | +1 | 7 |
| - | - | 4318 | 727.5 | - | - | 0 | - |
| - | - | 784 | 728.5 | - | - | 0 | - |
| - | - | 8619 | 735.3 | - | - | 0 | - |
| - | - | 3595 | 736.3 | - | - | 0 | - |
| - | - | 1048 | 737.4 | - | - | 0 | - |
| - | - | 1716 | 738.4 | - | - | 0 | - |
| - | - | 1542 | 739.4 | - | - | 0 | - |
| - | - | 5176 | 752.3 | - | - | 0 | - |
| - | - | 2068 | 753.3 | - | - | 0 | - |
| - | - | 1116 | 777.5 | - | - | 0 | - |
| - | - | 836.9 | 783.5 | - | - | 0 | - |
| - | - | 2405 | 805.5 | - | - | 0 | - |
| - | - | 1045 | 806.5 | - | - | 0 | - |
| - | - | 1994 | 811.4 | - | - | 0 | - |
| - | - | 2785 | 811.5 | - | - | 0 | - |
| - | - | 1398 | 812.4 | - | - | 0 | - |
| - | - | 1293 | 812.5 | - | - | 0 | - |
| 7 | y | 3.543E+04 | 823.5 | 0.000588 | 0.714 | +1 | 8 |
| - | - | 1.704E+04 | 824.5 | - | - | 0 | - |
| - | - | 4809 | 825.5 | - | - | 0 | - |
| - | - | 1229 | 826.6 | - | - | 0 | - |
| - | - | 2951 | 829.5 | - | - | 0 | - |
| - | - | 891.4 | 830.5 | - | - | 0 | - |
| 6 | b | 5466 | 839.4 | 0.0008443 | 1.006 | +1 | 6 |
| - | - | 2476 | 840.4 | - | - | 0 | - |
| - | - | 730.6 | 847.5 | - | - | 0 | - |
| - | - | 4627 | 848.4 | - | - | 0 | - |
| - | - | 2652 | 849.4 | - | - | 0 | - |
| - | - | 2348 | 853.5 | - | - | 0 | - |
| - | - | 1079 | 866.4 | - | - | 0 | - |
| - | - | 6422 | 901.4 | - | - | 0 | - |
| - | - | 3779 | 902.4 | - | - | 0 | - |
| - | - | 1174 | 903.4 | - | - | 0 | - |
| - | - | 787 | 906.4 | - | - | 0 | - |
| - | - | 729.5 | 907.4 | - | - | 0 | - |
| - | - | 3951 | 919.5 | - | - | 0 | - |
| - | - | 1379 | 920.5 | - | - | 0 | - |
| - | - | 1261 | 942.6 | - | - | 0 | - |
| 6 | y | 721.2 | 943.6 | 0.006067 | 6.43 | +1 | 9 |
| 6 | y | 4131 | 960.6 | 0.0003309 | 0.3445 | +1 | 9 |
| - | - | 1902 | 961.6 | - | - | 0 | - |
| - | - | 1214 | 966.6 | - | - | 0 | - |
| - | - | 908.5 | 1034 | - | - | 0 | - |
| - | - | 791.7 | 1035 | - | - | 0 | - |
| - | - | 937.3 | 1070 | - | - | 0 | - |
| - | - | 1316 | 1089 | - | - | 0 | - |
| - | - | 754.8 | 2997 | - | - | 0 | - |

m/z Charge Intensity FragmentType MassShift Position
120.06586456298828 0 478.2805
120.08129119873047 0 728.7647
122.07164764404297 0 1011.4923
123.05577850341797 0 3355.4436
123.11715698242188 0 748.1233
125.07156372070312 0 435.58987
125.0960464477539 0 424.07077 y 11
127.0504379272461 0 724.50726
127.08348846435547 0 357.60635
127.08705139160156 0 3831.4385
128.03472900390625 0 490.50363
128.07118225097656 0 491.77213
128.08230590820312 0 716.9423
129.0663604736328 0 5336.761
129.10276794433594 0 137426.23
130.0656280517578 0 31428.037
130.10037231445312 0 1205.4894
130.1060791015625 0 8244.339
131.04551696777344 0 3917.046
131.06903076171875 0 3473.3335
131.0818634033203 0 1124.3763
132.081298828125 0 119325.336
132.10240173339844 0 34366.676 y 13
133.06089782714844 0 434.9984
133.08482360839844 0 10895.505
133.105712890625 0 2277.4077
134.07192993164062 0 720.64417
134.08822631835938 0 804.81305
136.07620239257812 0 160920.33
137.07350158691406 0 1203.0891
137.07952880859375 0 14343.709
137.1080780029297 0 542.7981
138.06663513183594 0 2503.2593
138.1033477783203 0 466.0799
139.08706665039062 0 2962.1494
140.08236694335938 0 607.86633
141.10272216796875 0 7517.5894
142.06561279296875 0 1810.6866 b 3
142.10638427734375 0 542.83844
143.11856079101562 0 650.88715
144.0770263671875 0 408.54538
144.08120727539062 0 4699.9214
145.06130981445312 0 1096.1586
146.06065368652344 0 2038.2126
146.1292724609375 0 558.0367
147.04457092285156 0 6882.1997
148.0474395751953 0 584.05475
148.07615661621094 0 811.68134
148.9547882080078 0 611.625
149.02406311035156 0 545.32007
150.06614685058594 0 472.05478
151.08717346191406 0 1986.1028
152.08216857910156 0 1273.1146
152.1438446044922 0 2978.6628
153.10292053222656 0 1350.8275
154.098388671875 0 925.1538
155.0819854736328 0 6837.0767
155.09312438964844 0 2581.2231
155.11830139160156 0 4809.928
156.07667541503906 0 2492.1807
156.08164978027344 0 2944.2068
156.11395263671875 0 680.82965
157.07667541503906 0 1144.5494
157.0843505859375 0 905.36664
157.0972900390625 0 757.21246
158.0605010986328 0 1880.2104
158.08433532714844 0 15041.502
159.09217834472656 0 83646.04
159.1132354736328 0 1814.1372
160.07614135742188 0 3840.7437
160.09552001953125 0 9086.727
161.07937622070312 0 777.0305
161.09848022460938 0 668.11865
163.61387634277344 0 635.69025
164.03439331054688 0 799.842
164.0710906982422 0 2111.9167
165.07754516601562 0 1024.4872
165.10275268554688 0 5462.3896
166.06161499023438 0 2526.7275
168.11380004882812 0 1195.7145
169.0764923095703 0 8638.859
169.0976104736328 0 9765.549
169.17013549804688 0 839.1434
170.060546875 0 39581.82
170.10092163085938 0 897.6474
171.0639190673828 0 4349.9844
171.09210205078125 0 1465.2742
172.07611083984375 0 639.5452 b Ammonia loss 4
172.095947265625 0 533.08875
172.10855102539062 0 8443.239
173.07449340820312 0 947.44165
173.4521942138672 0 888.95496
174.05462646484375 0 577.4343
175.08892822265625 0 507.6931
175.09820556640625 0 2128.7153
176.08238220214844 0 3757.8044
177.11253356933594 0 692.76624
178.13424682617188 0 760.05707
179.08177185058594 0 648.3442
179.0933074951172 0 678.50275
179.1549530029297 0 634.7391
181.0613250732422 0 3188.1394
181.09771728515625 0 40970.254
181.13424682617188 0 2412.7297
182.06051635742188 0 595.77
182.0929718017578 0 537.48016
182.10101318359375 0 3476.2395
182.12930297851562 0 4789.354
183.09219360351562 0 16877.404
183.1016082763672 0 806.379
183.1131591796875 0 1651.7421
183.13272094726562 0 601.9376
183.14988708496094 0 1527.651
184.076171875 0 2049.7202
184.09552001953125 0 2623.151
185.0559539794922 0 714.7929
185.07151794433594 0 5237.0938
185.1077117919922 0 665.513
185.1653289794922 0 1669.0121
186.12420654296875 0 16353.415
187.087158203125 0 88669.234
187.10794067382812 0 4178.5454
187.12811279296875 0 1045.341
187.14456176757812 0 1312.5094 y 11
188.081787109375 0 709.75653
188.09056091308594 0 9746.041
189.1243896484375 0 748.31305
191.09298706054688 0 461.36606
193.09788513183594 0 832.49445
195.08816528320312 0 463.09415
195.11270141601562 0 856.7316
195.14964294433594 0 1204.8975
196.1085662841797 0 1796.305
197.1653289794922 0 6358.7803
198.1683349609375 0 1136.7758
199.08702087402344 0 3374.0576
199.10780334472656 0 1523.8497
199.1695556640625 0 1984.4026
199.18101501464844 0 4036.7422
200.09051513671875 0 604.9222
200.1398468017578 0 2306.9346
201.06918334960938 0 703.0004
201.10272216796875 0 8590.455
201.123779296875 0 813.28326
202.08657836914062 0 4651.185
202.10890197753906 0 14331.302
203.093017578125 0 1259.0276
203.1129608154297 0 1577.3965
203.1383819580078 0 605.408
204.07717895507812 0 2147.8696
204.13475036621094 0 2405.5059
205.0977325439453 0 762.4216 a 2
205.1083984375 0 713.0999
207.1238250732422 0 554.07367
207.14979553222656 0 3403.0676
208.14505004882812 0 897.29694
209.0933380126953 0 567.7719
209.12896728515625 0 1700.1293
210.12448120117188 0 663.6037
211.08702087402344 0 2678.6501
212.13958740234375 0 803.90967
212.66830444335938 0 2074.9956
213.12342834472656 0 634.37445
213.16957092285156 0 900.63995
214.1092529296875 0 3960.8052
214.1916046142578 0 1073.5156
215.13980102539062 0 1136.1351
216.11309814453125 0 1090.6078
216.12420654296875 0 1233.9465
216.135498046875 0 585.17816
217.10899353027344 0 1391.917
217.15560913085938 0 679.26587
218.1502685546875 0 754.95233 y Ammonia loss 8
220.1199188232422 0 10197.573
221.10386657714844 0 14689.4
221.12290954589844 0 741.3487
222.1062774658203 0 1678.108
222.1243896484375 0 1375.7515
223.1083984375 0 763.5498
223.1559295654297 0 526.313
224.11883544921875 0 2369.2847
224.1763153076172 0 6212.0166
225.12326049804688 0 658.5401
225.15997314453125 0 2099.884
225.1799774169922 0 822.16656
226.1555938720703 0 113350.5
227.0821990966797 0 1049.2467
227.1405487060547 0 818.74976
227.15892028808594 0 14071.483 y Ammonia loss 10
227.17379760742188 0 755.8508
228.1138153076172 0 7557.3657 a Water loss 1
228.13455200195312 0 1465.6088
228.16188049316406 0 910.4195
229.09776306152344 0 9179.932 b Ammonia loss 4
229.11805725097656 0 1783.6992
229.15447998046875 0 1196.554
229.1664581298828 0 1911.6348
230.10316467285156 0 2504.8315
231.0882568359375 0 4455.7646
231.1356658935547 0 1402.9724
231.17105102539062 0 2466.3193
232.11978149414062 0 674.4467
233.10414123535156 0 2635.737
234.10350036621094 0 897.8523
234.12356567382812 0 992.4653
234.13525390625 0 1145.9799
235.1195831298828 0 17850.361
235.67108154296875 0 16905.639 y 10
236.12277221679688 0 2586.493
236.1403045654297 0 789.66547
236.17279052734375 0 5124.2847
237.1351776123047 0 805.5322
237.16050720214844 0 817.75165 y Ammonia loss 7
238.13011169433594 0 1550.4283
239.15090942382812 0 6776.7227
240.13514709472656 0 1799.1636
240.15380859375 0 1071.2799
241.15621948242188 0 1200.6364
242.1869354248047 0 3921.3105
243.14756774902344 0 1413.6411
243.1814422607422 0 1333.8722
244.1082763671875 0 644.2555
244.16610717773438 0 619.0426
245.18661499023438 0 27141.67 y 12
246.12435913085938 0 210271.56 a 1
246.1896209716797 0 3478.6907
246.64111328125 0 762.1047
247.129150390625 0 49762.484
248.11465454101562 0 15944.805
248.1316680908203 0 3511.0059
249.09873962402344 0 22284.04
249.11441040039062 0 1435.3911 b 7
249.1326904296875 0 914.26276
250.10189819335938 0 2613.089
250.12313842773438 0 2945.816
251.1287841796875 0 1436.344
251.15101623535156 0 3269.8503
252.15621948242188 0 516.43994
252.17124938964844 0 2247.798
253.09658813476562 0 679.8184
253.1545867919922 0 839.73724
255.0765838623047 0 1243.343
255.1167755126953 0 1015.5749
256.10870361328125 0 5654.9995 b Water loss 1
257.09259033203125 0 1028.0612
257.1143798828125 0 2573.599
257.1614074707031 0 2798.4045
258.0765686035156 0 3554.5037
258.0990905761719 0 38165.188
259.080810546875 0 844.09216
259.10235595703125 0 6306.8184
259.1304626464844 0 1391.2186
259.8507385253906 0 944.2899
260.1072692871094 0 5128.2715
260.19744873046875 0 3438.7556
261.111083984375 0 1028.4609
263.1142883300781 0 1871.6348
264.1349792480469 0 2094.9512
265.1407775878906 0 1626.9808
265.16650390625 0 1614.8302
266.10931396484375 0 928.144
266.12530517578125 0 13905.358
266.1512451171875 0 731.0878
267.1287841796875 0 1665.0006
267.14501953125 0 1184.204
268.16632080078125 0 2788.9578
269.63018798828125 0 2091.3555
270.1264343261719 0 741.07086
270.1817626953125 0 9391.11
271.1886291503906 0 5524.9014 y 9
271.68994140625 0 1343.3794
272.1038513183594 0 817.7278
273.13519287109375 0 687.061
274.119140625 0 82315.055 b 1
275.10345458984375 0 10547.381
275.1250915527344 0 64052.395
275.1853942871094 0 2821.7312 y 6
275.519775390625 0 1422.5548
276.10821533203125 0 2893.2979
276.1285400390625 0 8850.289
277.09344482421875 0 884.09204
277.11590576171875 0 553.88306
277.1335144042969 0 3105.6765
278.1366271972656 0 773.3464
279.1472473144531 0 831.7749
279.18182373046875 0 1978.9163
283.15106201171875 0 1785.4048
283.1771240234375 0 25764.934
283.62738037109375 0 2694.5554
284.1289367675781 0 1553.7731
284.1803283691406 0 3081.7864
285.1101989746094 0 1563.1716
285.15625 0 5393.4634
286.1761169433594 0 780.21674
288.0996398925781 0 877.0204
290.1143798828125 0 1029.2722
290.18499755859375 0 9780.76
290.68682861328125 0 2331.5596
291.15740966796875 0 1438.9298
292.1297607421875 0 20400.158
292.18170166015625 0 744.9497
293.133056640625 0 3206.4744
294.120361328125 0 5433.8535
294.2178649902344 0 2339.7654
295.12322998046875 0 722.6099
297.15374755859375 0 769.1671
297.1927795410156 0 14210.68
298.1958923339844 0 2476.254
300.1372375488281 0 1884.0516
300.6728515625 0 1126.5056
303.12030029296875 0 4107.356
303.21826171875 0 677.12305
305.1293029785156 0 7775.5103
305.6649475097656 0 2389.261
306.1328125 0 1099.7083
306.18035888671875 0 777.32
307.1778564453125 0 782.6653
309.1669921875 0 1170.5894
309.19158935546875 0 746.109
311.173583984375 0 711.2378
311.24517822265625 0 3466.6797
312.2471008300781 0 702.79224
313.2237548828125 0 785.7349
314.6702880859375 0 3692.0674
315.1722106933594 0 1430.6539
318.15167236328125 0 827.4636
318.1923522949219 0 1018.18695
320.12384033203125 0 885.111
320.87188720703125 0 3224.3108 y 5
321.20794677734375 0 1118.4387
321.22930908203125 0 16279.199
322.1556396484375 0 10652.511
322.1754150390625 0 543.8933
322.232177734375 0 3750.2317
323.0200500488281 0 765.009
323.1586608886719 0 2667.594 b 10
324.21673583984375 0 3624.9683
324.7185363769531 0 1497.8057
325.2238464355469 0 744.60016
326.2191467285156 0 5052.1763
326.6630859375 0 711.10565
327.2025451660156 0 770.3383
327.2228698730469 0 1404.8394
327.2759704589844 0 1492.785
332.72979736328125 0 5636.1323
333.1241760253906 0 3233.3608
333.2310485839844 0 2880.717
334.17474365234375 0 1013.0246
335.23748779296875 0 2186.5208 y 8
336.2038269042969 0 25901.291
337.2070007324219 0 4000.9246
337.72198486328125 0 7118.721
338.1907958984375 0 627.773
338.2216796875 0 3130.5537
338.2452697753906 0 943.89014
338.71771240234375 0 661.9547
339.2398986816406 0 27130.508
339.8288879394531 0 674.6068
340.1528625488281 0 2426.2642
340.2431335449219 0 5353.5703
341.219970703125 0 718.93695
341.2442626953125 0 770.5983
342.1679992675781 0 1172.9312
345.6441955566406 0 1235.457
346.14434814453125 0 931.02704
346.66790771484375 0 1328.1757
346.72705078125 0 23506.295
347.2282409667969 0 8305.9375
347.7292175292969 0 2170.3662
349.7085876464844 0 2197.051
350.15045166015625 0 4128.4507
351.2403564453125 0 2327.07
353.2188720703125 0 1428.6982
354.1568298339844 0 2170.1746
354.1752014160156 0 2376.2397
354.2142333984375 0 33605.75
354.5103454589844 0 1887.926
354.6582336425781 0 1473.4225
355.21722412109375 0 6203.9414
355.7323913574219 0 5658.331
356.170654296875 0 794.38306
356.23345947265625 0 1512.7117
356.7328186035156 0 664.8174
357.1780700683594 0 900.9605
357.2503967285156 0 7434.975
358.16302490234375 0 8457.825
358.25286865234375 0 1547.948
358.7143249511719 0 2054.6797
359.1661376953125 0 1641.2067
359.2163391113281 0 816.29694
359.5417785644531 0 718.6086
359.87188720703125 0 1606.6519
360.13470458984375 0 911.58374
360.6648254394531 0 1674.9777
360.6888122558594 0 1273.6022
361.1853942871094 0 858.08636
362.15057373046875 0 903.6483
362.67022705078125 0 2056.2927
362.69677734375 0 728.8691
363.74822998046875 0 1669.902 y 7
365.2543029785156 0 724.38043
366.15692138671875 0 3601.2637
366.5582580566406 0 3312.4883 y 4
366.8927307128906 0 2389.61
367.1396789550781 0 1029.9824
367.162841796875 0 2985.9016
368.1484069824219 0 4760.3354
368.1683654785156 0 892.8579
369.1802062988281 0 2212.766
369.69329833984375 0 598.9474
370.527587890625 0 1070.5264
370.6891174316406 0 686.18207
371.22869873046875 0 1547.8363
373.2816467285156 0 15158.776 y 11
374.2597961425781 0 785.5197
374.2849426269531 0 2737.5876
374.5142822265625 0 1002.5985 b 8
375.185791015625 0 831.0862
376.669189453125 0 1542.2122
376.9589538574219 0 832.6275
377.1711120605469 0 1106.8848
378.7127685546875 0 1358.0746
378.96282958984375 0 2023.8695
379.2083740234375 0 1141.7968
380.230224609375 0 1372.9176
381.15008544921875 0 701.0955
382.2063293457031 0 6740.953
382.2825622558594 0 1759.3276
382.5399475097656 0 3336.9524
382.6721496582031 0 754.45105
382.8732604980469 0 1165.9366
383.2662048339844 0 1825.7911
384.1675720214844 0 3973.907
385.1734924316406 0 12755.176
385.19976806640625 0 3322.8909
386.1575927734375 0 13643.794
386.2018737792969 0 1160.1238
386.46160888671875 0 721.74194
387.1602478027344 0 3121.696
388.1634826660156 0 958.57025
388.8624267578125 0 2612.6099
389.19677734375 0 1485.9291
389.27117919921875 0 1758.5817
389.53216552734375 0 916.0874
389.7725830078125 0 975.09
390.2146301269531 0 7197.3667 y Ammonia loss 1
390.465087890625 0 3915.6377
390.71563720703125 0 4581.657
392.1900329589844 0 1765.4896 b Water loss 9
392.26678466796875 0 5466.0835
392.5226135253906 0 2015.5801 b Ammonia loss 9
392.7496337890625 0 899.5188
392.96893310546875 0 691.8534
393.1675109863281 0 1237.329
393.27032470703125 0 1931.6405
394.1518859863281 0 9941.298
394.4705810546875 0 3615.1711 y 1
394.7232666015625 0 1185.6547
394.972412109375 0 1398.8243
395.1556701660156 0 2013.4856
396.2611083984375 0 1457.4738
397.1779479980469 0 655.4429
397.24542236328125 0 1776.5956
397.5655517578125 0 903.9146
398.1938781738281 0 1676.2408 b 9
398.24432373046875 0 986.9216
398.528076171875 0 1481.8793
398.88970947265625 0 928.89905
399.22369384765625 0 1609.0022
400.174072265625 0 762.5311
400.22503662109375 0 981.129
400.4692077636719 0 1294.2036
401.1923828125 0 1094.7366
401.25921630859375 0 808.9601
402.22515869140625 0 4466.34
402.548828125 0 2216.4993
403.1845397949219 0 11738.4375
403.23992919921875 0 6311.0127
403.2693786621094 0 3561.044
403.571044921875 0 8966.269 y Ammonia loss 3
403.768798828125 0 1140.7913
403.904296875 0 5483.5234
404.187744140625 0 2110.7737
404.2383117675781 0 2816.727
404.26873779296875 0 928.2637
404.4703063964844 0 3432.3215
404.7243347167969 0 3112.307
404.89349365234375 0 827.7936
404.97589111328125 0 4729.1035
405.2279968261719 0 3544.7395
406.1860046386719 0 6123.719
406.2508239746094 0 3699.695
406.6873474121094 0 2606.9976
406.7523498535156 0 2190.3064
407.1918029785156 0 1537.247
408.22222900390625 0 2396.3455
408.5555725097656 0 2291.9363
408.88916015625 0 1651.061
409.2449035644531 0 2884.6348 y 3
409.5785217285156 0 2890.8062
409.912353515625 0 1724.4265
410.24554443359375 0 742.4259
410.2770690917969 0 11674.1
410.56939697265625 0 4865.388
410.9034423828125 0 5025.885
411.1783142089844 0 11318.826 b Water loss 5
411.2074890136719 0 829.46045
411.23785400390625 0 1595.739
411.2803649902344 0 2388.7295
411.7254943847656 0 1771.6936 Precursor Water loss
411.9747009277344 0 2909.2314 Precursor Ammonia loss
412.16131591796875 0 2794.7847
412.1844787597656 0 1998.4951
412.2248840332031 0 2639.437
412.2746276855469 0 45354.02 y 6
412.4781494140625 0 864.5461
412.7759704589844 0 21580.701
413.1673889160156 0 1727.5826
413.2774658203125 0 6312.977
413.7759704589844 0 812.0963
413.8973083496094 0 6678.0522
414.19512939453125 0 753.4658
414.2296142578125 0 6908.0894
414.5627746582031 0 2820.6414
414.8943786621094 0 701.26465
415.0377502441406 0 837.2714
415.25701904296875 0 5047.421
415.7579040527344 0 1671.6123
416.2289733886719 0 26092.908 Precursor
416.2868347167969 0 3236.618
416.4796447753906 0 25437.367
416.73040771484375 0 12972.685
416.9813232421875 0 4009.4883
417.23138427734375 0 2398.5518
417.2691650390625 0 993.77313
417.4835510253906 0 779.2108
419.9007263183594 0 29761.264
420.1834411621094 0 2837.738 b 5
420.2351379394531 0 22812.072
420.56890869140625 0 8707.44
420.6849060058594 0 2459.7688
420.9034729003906 0 2212.8784
421.23736572265625 0 1277.2372
422.27703857421875 0 3111.2053
423.2743225097656 0 1645.1565
424.32904052734375 0 3099.9666
424.7121887207031 0 6618.1367
425.2135009765625 0 3245.3901
425.288330078125 0 1771.9794
425.3338623046875 0 845.21576
425.90399169921875 0 6405.7944
426.2381286621094 0 2399.4907
426.5713806152344 0 930.00977
427.2442626953125 0 5084.9897
427.7448425292969 0 2075.1619
428.28802490234375 0 1214.6073
429.1891174316406 0 43071.285
430.1920471191406 0 11712.554
431.1805725097656 0 5478.4795
432.1818542480469 0 1499.6407
433.7040100097656 0 1906.0565
434.20062255859375 0 682.82214
436.5871276855469 0 913.4715
437.1828918457031 0 2433.6567 b 2
438.1845397949219 0 1630.458
438.22882080078125 0 778.41406
438.55645751953125 0 2521.468
438.8916931152344 0 3286.7227
439.2264404296875 0 2310.4797
439.3043212890625 0 886.61176
442.71295166015625 0 1971.7556
444.23150634765625 0 1039.4854
445.19091796875 0 977.48914
445.2688293457031 0 1719.4136
445.91680908203125 0 765.7239
446.2337646484375 0 4303.3887
446.7361145019531 0 1864.3279
447.2370910644531 0 760.7573
448.2638854980469 0 7789.656
448.5982971191406 0 6622.858
448.93304443359375 0 2237.9946
449.2699279785156 0 888.91895
451.2259216308594 0 6348.12
451.2672424316406 0 3508.4333
451.7244873046875 0 5089.432
451.920166015625 0 10553.876
452.2210998535156 0 1082.1035
452.254150390625 0 8778.786
452.3245544433594 0 1536.7867
452.5882873535156 0 5358.9766
452.7225646972656 0 712.34265
452.9203186035156 0 1355.9338
453.7059326171875 0 1598.0569
454.20855712890625 0 1909.2908
455.2523498535156 0 4946.5347
456.2577209472656 0 832.75757
457.1844482421875 0 1807.1676
457.5950927734375 0 6146.696
457.92742919921875 0 7095.8013 y Ammonia loss 2
458.26019287109375 0 6549.533
458.5927429199219 0 1329.5057
458.9273986816406 0 792.8523
459.2135009765625 0 1018.9614
460.2308654785156 0 8006.798
460.7322998046875 0 4921.975
461.242431640625 0 1687.4526
462.9256896972656 0 1523.064
463.27728271484375 0 1588.9844
463.59906005859375 0 126164 y 2
463.933349609375 0 102535.01
464.2675476074219 0 48283.12
464.6016540527344 0 10889.235
464.9361267089844 0 3936.69
465.2701721191406 0 801.4352
466.92041015625 0 1615.9926
467.2548522949219 0 1669.5879
467.2988586425781 0 3842.8286
468.29437255859375 0 2369.8835
469.7882080078125 0 2389.2808
470.2496032714844 0 1825.2098
470.3345947265625 0 52316.38 y 10
470.5814514160156 0 2882.0078
470.9169006347656 0 1247.0298
471.33770751953125 0 13177.758
471.7985534667969 0 4204.4497
472.2999267578125 0 1962.4484
472.3393859863281 0 2265.8508
472.5948486328125 0 1725.9066
473.2344970703125 0 873.5875 b 11
473.2625732421875 0 6295.865
473.57672119140625 0 2508.7253
473.9134521484375 0 1534.8903
474.2647705078125 0 2217.6438
474.7816162109375 0 3173.714
475.2809143066406 0 2441.1057
475.92425537109375 0 754.7083
476.2522277832031 0 8384.361
476.5868225097656 0 10488.152
476.9209289550781 0 4239.82
477.5929870605469 0 1058.1672
478.20947265625 0 2637.9849
480.8040771484375 0 11266.456 y 5
481.30596923828125 0 4693.9805
481.80596923828125 0 1719.6234
481.9271545410156 0 4797.44
482.2619323730469 0 4834.767
482.309326171875 0 4129.259
482.5924377441406 0 3986.9458
482.7619323730469 0 3151.3396
482.9231262207031 0 2155.522
483.2608947753906 0 1517.8431
483.78594970703125 0 7483.873
484.28765869140625 0 5354.8286
484.78875732421875 0 1121.9755
487.9300842285156 0 1177.6654
488.2648620605469 0 1347.9865
488.60003662109375 0 1143.0874
490.2352600097656 0 1489.3911
491.2732238769531 0 4945.7754
492.27435302734375 0 885.1896
492.79180908203125 0 1878.1367
493.2928771972656 0 1764.1328
493.7984924316406 0 4014.6482
494.2369689941406 0 1570.5985
494.2980651855469 0 1197.0868
497.2196044921875 0 1349.3058 b 7
497.7236633300781 0 968.8723
501.262939453125 0 834.1068
501.60528564453125 0 1519.637
501.9376525878906 0 925.1155
502.27313232421875 0 879.6082
503.2156982421875 0 3580.5586
504.21923828125 0 1094.7739
504.9471435546875 0 1575.9702
505.2698669433594 0 1660.2269 b Ammonia loss 12
505.6002502441406 0 1115.9622
506.2324523925781 0 1195.4846
508.75921630859375 0 1068.9619
509.2393798828125 0 3127.1243
509.7417907714844 0 2073.339
510.2907409667969 0 1695.3492
510.62591552734375 0 843.5817
510.9375305175781 0 4152.661 b 12
511.27178955078125 0 4882.4893
511.6067810058594 0 3944.968
511.94256591796875 0 1709.7119
513.9452514648438 0 791.8263
514.2998046875 0 996.9533
514.6096801757812 0 708.6558
516.9414672851562 0 9579.482
517.2750854492188 0 12862.245
517.610107421875 0 5414.326
517.7501831054688 0 1525.0826
517.9437255859375 0 1231.925
518.254638671875 0 1807.1666
519.9503173828125 0 14145.593 y Ammonia loss 1
520.2418212890625 0 1169.8694
520.2842407226562 0 10298.849
520.3602294921875 0 2347.0635
520.6188354492188 0 5086.9497
520.9500122070312 0 1422.1841
521.22705078125 0 4226.335
521.2847900390625 0 787.488
521.3605346679688 0 829.08997
522.224609375 0 1179.4227
524.8117065429688 0 1646.9825
525.3062744140625 0 1956.2665
525.624755859375 0 15200.531 y 1
525.9595336914062 0 18076.443
526.2937622070312 0 9693.33
526.3314819335938 0 916.43164
526.6285400390625 0 2843.677
526.9618530273438 0 1617.9537
529.2202758789062 0 762.82556
529.8038330078125 0 1825.044
530.2987060546875 0 3483.4697
530.7989501953125 0 1825.9385
531.2108764648438 0 3264.9
532.2132568359375 0 1140.3787
535.7518310546875 0 2650.9546
536.2511596679688 0 1400.023
536.7549438476562 0 1160.4177
537.75244140625 0 819.8763
538.2528076171875 0 26196.08
538.3719482421875 0 2585.0996
538.763671875 0 861.6617
538.8094482421875 0 4126.6865
539.2552490234375 0 8008.4565
539.3043823242188 0 8258.905
539.8038940429688 0 3113.559
540.255615234375 0 1859.2124
540.3018188476562 0 971.1688
540.8253784179688 0 803.69727 y Ammonia loss 4
541.32763671875 0 864.7838
541.37158203125 0 11124.784 y 9
542.251953125 0 4816.7783
542.3746948242188 0 3916.461
543.2574462890625 0 1531.8859
544.2687377929688 0 1094.5061
544.760009765625 0 2547.9841
545.2623291015625 0 1183.9728
545.7604370117188 0 874.7934
547.8146362304688 0 2851.1238
548.236328125 0 1163.0662
548.3123168945312 0 2766.2085
548.8109741210938 0 1795.9769
549.222900390625 0 3586.756
549.3334350585938 0 29076.45 y 4
549.8348388671875 0 18951.668
550.3360595703125 0 7230.795
550.3744506835938 0 797.64716
550.8377075195312 0 1516.4941
552.76123046875 0 718.8291 b Ammonia loss 8
553.2186889648438 0 1526.3779
553.2659301757812 0 1214.6996
553.7702026367188 0 897.286
555.28515625 0 1659.07
556.8201904296875 0 1559.5618
557.2958374023438 0 2803.4075
561.2684326171875 0 7771.0015 b 8
561.7693481445312 0 4972.451
562.2715454101562 0 1419.6116
565.2421264648438 0 752.23663 b 3
566.2476806640625 0 33137.67
567.2498168945312 0 8918.311
567.3997192382812 0 1452.8954
568.2526245117188 0 1870.0392
570.246826171875 0 14050.2
571.2499389648438 0 4720.15
572.2513427734375 0 1087.739
572.8050537109375 0 16513.4
573.3064575195312 0 10236.137
573.8074951171875 0 3129.1492
576.2357788085938 0 911.6224
577.3828735351562 0 8804.123
577.8414306640625 0 1014.7757
578.38427734375 0 3322.5059
579.3621826171875 0 18898.746
580.3649291992188 0 5078.2246
581.369140625 0 1448.6578
582.7914428710938 0 857.6677
587.2734985351562 0 1516.0586
587.7810668945312 0 1775.6959 b Water loss 9
588.2799682617188 0 2844.002 b Ammonia loss 9
594.2400512695312 0 761.0207
595.3477783203125 0 787.49066
595.3929443359375 0 6734.494
595.8462524414062 0 1618.0286
596.3486328125 0 1234.0723
596.3944702148438 0 2489.2893
596.7871704101562 0 1051.8776 b 9
597.2863159179688 0 2067.559
597.7880249023438 0 860.44025
598.242431640625 0 7316.959
599.244873046875 0 2767.25
600.2476806640625 0 899.3007
600.3341064453125 0 1006.9696
604.3562622070312 0 5931.2085
604.852294921875 0 11245.706 y Ammonia loss 3
605.3532104492188 0 7554.1064
605.852783203125 0 2067.7488
610.3209838867188 0 6006.117
611.322509765625 0 2121.87
611.8306274414062 0 1256.3807
612.32763671875 0 1139.0076
613.36279296875 0 8720.162 y 3
613.4064331054688 0 698.28186
613.8650512695312 0 4586.9297
614.3650512695312 0 2241.2798
614.865234375 0 743.1482
615.2673950195312 0 2394.1074
615.3483276367188 0 3787.652
615.8513793945312 0 2445.252
616.3521118164062 0 1072.17
618.8530883789062 0 805.65
620.342041015625 0 5542.959
620.8406372070312 0 4312.363
621.342041015625 0 2033.419
623.3883666992188 0 1164.1342
628.2755737304688 0 957.8485
628.3325805664062 0 1715.0889
629.3462524414062 0 14991.179
629.8482055664062 0 9426.403
630.34912109375 0 4686.822
630.8472900390625 0 932.9933
638.3521118164062 0 8813.692
638.8536987304688 0 5144.671
639.3522338867188 0 4269.0757
647.4268188476562 0 1245.0745
648.8363037109375 0 773.33215
657.2749633789062 0 1531.3921
657.3292846679688 0 1244.5532
657.8330688476562 0 2505.352
663.2986450195312 0 704.34753
664.45166015625 0 6514.2427
665.4541625976562 0 2168.5784
665.8457641601562 0 1619.5533
666.3460693359375 0 1629.5427
667.3446044921875 0 1306.553
668.3253173828125 0 1616.3574
669.4664306640625 0 4299.797 y 8
670.4678344726562 0 1629.2341
674.3026733398438 0 2205.268
674.435546875 0 18478.232
675.3076782226562 0 933.0043
675.4376220703125 0 8696.184
676.4384155273438 0 1751.9701
679.3114013671875 0 760.4277
680.3162841796875 0 1100.7562
684.289794921875 0 1448.4441 b Water loss 4
685.3525390625 0 1432.8654
690.2787475585938 0 1676.8901
692.445556640625 0 19102.676
693.4484252929688 0 7839.6016
694.4515380859375 0 1461.2635
694.894287109375 0 6298.5776 y 2
695.3956298828125 0 4490.897
695.8956298828125 0 1264.9873
698.4124145507812 0 1495.9235
702.3009033203125 0 3041.0576 b 4
703.2994384765625 0 1234.3473
707.3052978515625 0 8398.452
708.3084106445312 0 3840.4414
708.4765625 0 938.5104
710.456787109375 0 6051.834
711.4589233398438 0 1737.523
713.8800048828125 0 1622.3926
714.3788452148438 0 1444.6428
714.8810424804688 0 902.4323
716.4203491210938 0 3456.6174
717.4234619140625 0 1764.7249
718.3667602539062 0 906.528
720.3206176757812 0 1837.4054
721.3529052734375 0 3048.6404
722.3788452148438 0 1767.3489
722.8853759765625 0 1792.3607
724.3338012695312 0 2976.2349
725.3323974609375 0 1045.9764
726.4876098632812 0 9576.86 y 7
727.4898681640625 0 4318.469
728.4918823242188 0 784.03125
735.3005981445312 0 8619.387
736.3036499023438 0 3595.436
737.3541259765625 0 1047.949
738.376708984375 0 1716.3678
739.3688354492188 0 1542.3375
752.3287353515625 0 5175.827
753.3279418945312 0 2067.6235
777.53662109375 0 1115.8226
783.5028076171875 0 836.9417
805.53076171875 0 2404.7566
806.5355834960938 0 1044.7343
811.3657836914062 0 1994.3181
811.495849609375 0 2785.482
812.3681030273438 0 1397.8546
812.499267578125 0 1293.0173
823.5405883789062 0 35431.15 y 6
824.543212890625 0 17037.172
825.5465087890625 0 4808.7056
826.5519409179688 0 1229.3363
829.5040283203125 0 2951.1355
830.5070190429688 0 891.3503
839.3591918945312 0 5466.216 b 5
840.3615112304688 0 2476.1694
847.5133666992188 0 730.58014
848.4147338867188 0 4626.7397
849.4196166992188 0 2651.6074
853.4752197265625 0 2348.423
866.3933715820312 0 1078.7661
901.4420776367188 0 6421.8013
902.4450073242188 0 3778.9585
903.4485473632812 0 1173.6716
906.3983764648438 0 787.0142
907.396240234375 0 729.53174
919.4528198242188 0 3951.4846
920.4572143554688 0 1378.7917
942.5875244140625 0 1260.5247
943.5784301757812 0 721.24133 y Ammonia loss 5
960.5992431640625 0 4131.187 y 5
961.6027221679688 0 1901.5254
966.5628662109375 0 1214.0687
1033.5411376953125 0 908.5308
1034.5245361328125 0 791.7438
1070.498779296875 0 937.32623
1088.5179443359375 0 1316.1306
2997.419921875 0 754.8012

Spectrum Details

|  |  |
| --- | --- |
| Matched peaks? Matched peaks The total absolute number of peaks matched. Additionally in brackets the total fraction of peaks matched and the total number of peaks is shown. | 70 (7.56% of 926) |
| FDR? FDR The false discovery rate estimated for this peptide. It is calculated by matching all theoretical fragments with a non-integer shift with the raw peaks for this spectrum. This is done with 40 different shifts. The resulting percentage is the average number of annotated peaks over the number of annotated peaks with the correct spectrum. | 3.78% |
| Satellite FDR? Satellite FDR See the FDR for details on its calculation. This satellite ion specific FDR only contains the satellite ions (d/w) for I/L/J positions. | - |
| PSM Score? PSM Score The PSM Score as given by Hecklib to this annotated spectrum. It is shown with three significant figures. | 221 |

#### Spectrum 5359? Spectrum 5359 The raw spectrum of this peptide as annotated by Hecklib. The fragments are coloured according to ion type (see legend). Any peaks with a star '\*' as text can be hovered over to see the full details, first the ion type second the mass shift type. By hovering over the amino acids in the peptide or ions in the legend the corresponding peaks are highlighted. By toggling the 'Unassigned' label you can turn the background (unassigned) peaks on or off in the plot. By updating the slider in the Ion legend you can update the spectrum to only show the top X% of the peaks with labels. The top X% means any peak that is within X% of the highest intensity. By dragging in the spectrum you can zoom in to a specific part of the spectrum and use 'Zoom Out' to get back to the original zoom level. The annotation of the spectrum is based on the given sequence in the peptides file and is done with different software so inconsistencies are likely. The peaks are annotated based on the given sequence, with 20 ppm tolerance.

Copy Data

##### Spectrum 5359 (TSV)

###### Preview

```
Loading example...
```

*Click on the button to copy the data to your clipboard.*

Mz MinMz MaxIntensity Max

WidthHeightPeptide font sizePeptide stroke widthSpectrum font sizeSpectrum stroke widthCompact peptide

Ion legend

wxyz

abcd

OtherUnassignedIon^Charge^_Position_Show for top:%

SWYQHHPGKAPKJJ

03.56e+57.12e+51.07e+61.42e+6

Zoom Out

y^+1^_1_b^+4^_4_a^+2^_3_y^+2^_4_a^+1^_2_b^+3^_5_y^+2^_4_y^+1^_2_a^+1^_2_b^+4^_8_b^+1^_2_y^+2^_5_b^+1^_2_y^+3^_8_y^+3^_9_b^+4^_11_y^+2^_6_y^+2^_7_y^+3^_10_b^+3^_9_y^+1^_3_b^+3^_9_y^+4^_13_b^+3^_10_b^+3^_10_y^+4^_13_b^+3^_10_y^+3^_11_y^+3^_11_b^+2^_6_\*\*y^+2^_8_\*b^+2^_6_b^+1^_3_y^+3^_12_y^+3^_12_y^+1^_4_y^+2^_9_b^+2^_8_b^+3^_13_b^+3^_13_b^+3^_13_y^+3^_13_y^+3^_13_y^+2^_10_y^+1^_5_y^+2^_10_b^+2^_9_b^+1^_4_b^+2^_10_b^+2^_10_b^+2^_10_y^+2^_11_y^+2^_11_y^+1^_6_b^+1^_5_y^+2^_12_b^+1^_5_b^+2^_12_y^+1^_7_y^+1^_8_b^+1^_6_y^+1^_9_

02755498241098

Fragment Matches Table

Show background peaks

| Position | Ion type | Intensity | mz Theoretical | mz Error (Th) | mz Error (ppm) | Charge | Series Number |
| --- | --- | --- | --- | --- | --- | --- | --- |
| - | - | 4440 | 120.1 | - | - | 0 | - |
| - | - | 5189 | 120.1 | - | - | 0 | - |
| - | - | 3722 | 121 | - | - | 0 | - |
| - | - | 2397 | 122.1 | - | - | 0 | - |
| - | - | 5605 | 122.1 | - | - | 0 | - |
| - | - | 1.894E+04 | 123.1 | - | - | 0 | - |
| - | - | 2410 | 125.1 | - | - | 0 | - |
| - | - | 2300 | 125.1 | - | - | 0 | - |
| - | - | 2.429E+04 | 127.1 | - | - | 0 | - |
| - | - | 1910 | 127.1 | - | - | 0 | - |
| - | - | 2802 | 128.1 | - | - | 0 | - |
| - | - | 1894 | 128.1 | - | - | 0 | - |
| - | - | 2000 | 129 | - | - | 0 | - |
| - | - | 2.808E+04 | 129.1 | - | - | 0 | - |
| - | - | 8.622E+05 | 129.1 | - | - | 0 | - |
| - | - | 2.064E+05 | 130.1 | - | - | 0 | - |
| - | - | 3302 | 130.1 | - | - | 0 | - |
| - | - | 7462 | 130.1 | - | - | 0 | - |
| - | - | 5.296E+04 | 130.1 | - | - | 0 | - |
| - | - | 2534 | 131 | - | - | 0 | - |
| - | - | 2.157E+04 | 131.1 | - | - | 0 | - |
| - | - | 2691 | 131.1 | - | - | 0 | - |
| - | - | 8.282E+05 | 132.1 | - | - | 0 | - |
| 14 | y | 2.196E+05 | 132.1 | 0.0005424 | 4.106 | +1 | 1 |
| - | - | 8.572E+04 | 133.1 | - | - | 0 | - |
| - | - | 1.531E+04 | 133.1 | - | - | 0 | - |
| - | - | 3899 | 134.1 | - | - | 0 | - |
| - | - | 5178 | 134.1 | - | - | 0 | - |
| - | - | 1.093E+06 | 136.1 | - | - | 0 | - |
| - | - | 4826 | 137.1 | - | - | 0 | - |
| - | - | 9.543E+04 | 137.1 | - | - | 0 | - |
| - | - | 1.278E+04 | 138.1 | - | - | 0 | - |
| - | - | 2718 | 138.1 | - | - | 0 | - |
| - | - | 1.355E+04 | 139.1 | - | - | 0 | - |
| - | - | 4233 | 141.1 | - | - | 0 | - |
| - | - | 4.569E+04 | 141.1 | - | - | 0 | - |
| 4 | b | 1.951E+04 | 142.1 | 5.5E-05 | 0.3872 | +4 | 4 |
| - | - | 3.803E+04 | 144.1 | - | - | 0 | - |
| - | - | 3634 | 145.1 | - | - | 0 | - |
| - | - | 5377 | 146.1 | - | - | 0 | - |
| - | - | 2575 | 146.1 | - | - | 0 | - |
| - | - | 4.569E+04 | 147 | - | - | 0 | - |
| - | - | 2981 | 147.1 | - | - | 0 | - |
| - | - | 6079 | 148 | - | - | 0 | - |
| - | - | 2947 | 148.1 | - | - | 0 | - |
| - | - | 3756 | 149.1 | - | - | 0 | - |
| - | - | 2818 | 150.1 | - | - | 0 | - |
| - | - | 1.061E+04 | 151.1 | - | - | 0 | - |
| - | - | 4425 | 152.1 | - | - | 0 | - |
| - | - | 2.036E+04 | 152.1 | - | - | 0 | - |
| - | - | 2266 | 153.1 | - | - | 0 | - |
| - | - | 3324 | 153.1 | - | - | 0 | - |
| - | - | 6897 | 154.1 | - | - | 0 | - |
| - | - | 4.71E+04 | 155.1 | - | - | 0 | - |
| - | - | 1.468E+04 | 155.1 | - | - | 0 | - |
| - | - | 2.793E+04 | 155.1 | - | - | 0 | - |
| - | - | 8334 | 156.1 | - | - | 0 | - |
| - | - | 2.553E+04 | 156.1 | - | - | 0 | - |
| - | - | 4708 | 156.1 | - | - | 0 | - |
| - | - | 6541 | 157.1 | - | - | 0 | - |
| - | - | 3483 | 157.1 | - | - | 0 | - |
| - | - | 3014 | 157.1 | - | - | 0 | - |
| - | - | 1.185E+04 | 158.1 | - | - | 0 | - |
| - | - | 9.79E+04 | 158.1 | - | - | 0 | - |
| - | - | 5.728E+05 | 159.1 | - | - | 0 | - |
| - | - | 2.349E+04 | 160.1 | - | - | 0 | - |
| - | - | 4094 | 160.1 | - | - | 0 | - |
| - | - | 5.431E+04 | 160.1 | - | - | 0 | - |
| - | - | 2708 | 161.1 | - | - | 0 | - |
| - | - | 2530 | 162.1 | - | - | 0 | - |
| - | - | 2540 | 162.5 | - | - | 0 | - |
| - | - | 4688 | 163.6 | - | - | 0 | - |
| - | - | 3008 | 164 | - | - | 0 | - |
| - | - | 1.738E+04 | 164.1 | - | - | 0 | - |
| - | - | 2206 | 164.1 | - | - | 0 | - |
| - | - | 5006 | 165.1 | - | - | 0 | - |
| - | - | 3.367E+04 | 165.1 | - | - | 0 | - |
| - | - | 1.026E+04 | 166.1 | - | - | 0 | - |
| - | - | 3386 | 166.1 | - | - | 0 | - |
| - | - | 2505 | 166.2 | - | - | 0 | - |
| - | - | 2680 | 168.1 | - | - | 0 | - |
| - | - | 7618 | 168.1 | - | - | 0 | - |
| - | - | 6.1E+04 | 169.1 | - | - | 0 | - |
| - | - | 5.809E+04 | 169.1 | - | - | 0 | - |
| - | - | 3994 | 169.2 | - | - | 0 | - |
| - | - | 2.551E+05 | 170.1 | - | - | 0 | - |
| - | - | 2925 | 170.1 | - | - | 0 | - |
| - | - | 3.318E+04 | 171.1 | - | - | 0 | - |
| - | - | 1.117E+04 | 171.1 | - | - | 0 | - |
| - | - | 5.556E+04 | 172.1 | - | - | 0 | - |
| - | - | 3470 | 172.1 | - | - | 0 | - |
| - | - | 9142 | 173.4 | - | - | 0 | - |
| - | - | 2716 | 174.1 | - | - | 0 | - |
| - | - | 1.033E+04 | 175.1 | - | - | 0 | - |
| - | - | 2.341E+04 | 176.1 | - | - | 0 | - |
| - | - | 1.374E+04 | 177.1 | - | - | 0 | - |
| - | - | 3910 | 179.1 | - | - | 0 | - |
| - | - | 4346 | 179.1 | - | - | 0 | - |
| - | - | 2375 | 179.6 | - | - | 0 | - |
| - | - | 4788 | 180.1 | - | - | 0 | - |
| - | - | 1.242E+04 | 181.1 | - | - | 0 | - |
| - | - | 2.736E+05 | 181.1 | - | - | 0 | - |
| - | - | 1.501E+04 | 181.1 | - | - | 0 | - |
| - | - | 3400 | 181.2 | - | - | 0 | - |
| - | - | 4727 | 182.1 | - | - | 0 | - |
| - | - | 2527 | 182.1 | - | - | 0 | - |
| - | - | 2.227E+04 | 182.1 | - | - | 0 | - |
| - | - | 3.417E+04 | 182.1 | - | - | 0 | - |
| - | - | 1.232E+05 | 183.1 | - | - | 0 | - |
| - | - | 4434 | 183.1 | - | - | 0 | - |
| - | - | 4646 | 183.1 | - | - | 0 | - |
| - | - | 1.406E+04 | 184.1 | - | - | 0 | - |
| - | - | 1.51E+04 | 184.1 | - | - | 0 | - |
| - | - | 2.787E+04 | 185.1 | - | - | 0 | - |
| - | - | 2971 | 185.1 | - | - | 0 | - |
| - | - | 5858 | 185.1 | - | - | 0 | - |
| - | - | 6165 | 186.1 | - | - | 0 | - |
| - | - | 1.004E+05 | 186.1 | - | - | 0 | - |
| - | - | 5.796E+05 | 187.1 | - | - | 0 | - |
| - | - | 8509 | 187.1 | - | - | 0 | - |
| - | - | 7.166E+04 | 188.1 | - | - | 0 | - |
| - | - | 2486 | 189.1 | - | - | 0 | - |
| - | - | 4019 | 190.6 | - | - | 0 | - |
| - | - | 2918 | 191.1 | - | - | 0 | - |
| - | - | 2630 | 191.1 | - | - | 0 | - |
| - | - | 2313 | 191.1 | - | - | 0 | - |
| - | - | 3077 | 193.1 | - | - | 0 | - |
| - | - | 2295 | 195.1 | - | - | 0 | - |
| - | - | 4936 | 195.1 | - | - | 0 | - |
| - | - | 9612 | 195.1 | - | - | 0 | - |
| - | - | 6866 | 196.1 | - | - | 0 | - |
| - | - | 4.108E+04 | 197.2 | - | - | 0 | - |
| - | - | 3838 | 198.1 | - | - | 0 | - |
| - | - | 6408 | 198.2 | - | - | 0 | - |
| - | - | 2.665E+04 | 199.1 | - | - | 0 | - |
| - | - | 4317 | 199.1 | - | - | 0 | - |
| - | - | 1.999E+04 | 199.2 | - | - | 0 | - |
| - | - | 2824 | 200.1 | - | - | 0 | - |
| - | - | 1.432E+04 | 200.1 | - | - | 0 | - |
| - | - | 4221 | 201.1 | - | - | 0 | - |
| - | - | 6.169E+04 | 201.1 | - | - | 0 | - |
| - | - | 3867 | 201.1 | - | - | 0 | - |
| - | - | 3468 | 201.1 | - | - | 0 | - |
| - | - | 2.777E+04 | 202.1 | - | - | 0 | - |
| - | - | 9.236E+04 | 202.1 | - | - | 0 | - |
| - | - | 6696 | 203.1 | - | - | 0 | - |
| - | - | 6839 | 203.1 | - | - | 0 | - |
| - | - | 1.12E+04 | 204.1 | - | - | 0 | - |
| 3 | a | 4736 | 205.1 | 0.0007616 | 3.713 | +2 | 3 |
| - | - | 1.14E+04 | 205.1 | - | - | 0 | - |
| - | - | 2.985E+04 | 207.1 | - | - | 0 | - |
| - | - | 4523 | 208.1 | - | - | 0 | - |
| - | - | 2750 | 208.2 | - | - | 0 | - |
| - | - | 7569 | 209.1 | - | - | 0 | - |
| - | - | 4037 | 210.1 | - | - | 0 | - |
| - | - | 1.478E+04 | 211.1 | - | - | 0 | - |
| - | - | 3609 | 212.1 | - | - | 0 | - |
| - | - | 1.308E+04 | 212.1 | - | - | 0 | - |
| - | - | 1.873E+04 | 212.7 | - | - | 0 | - |
| - | - | 4526 | 213.1 | - | - | 0 | - |
| - | - | 2341 | 213.1 | - | - | 0 | - |
| - | - | 9780 | 213.2 | - | - | 0 | - |
| - | - | 3.113E+04 | 214.1 | - | - | 0 | - |
| - | - | 3349 | 214.1 | - | - | 0 | - |
| - | - | 7121 | 214.2 | - | - | 0 | - |
| - | - | 2686 | 215.1 | - | - | 0 | - |
| - | - | 3561 | 215.1 | - | - | 0 | - |
| - | - | 8134 | 216.1 | - | - | 0 | - |
| - | - | 7691 | 217.1 | - | - | 0 | - |
| - | - | 2885 | 219.1 | - | - | 0 | - |
| - | - | 7.102E+04 | 220.1 | - | - | 0 | - |
| - | - | 8.949E+04 | 221.1 | - | - | 0 | - |
| - | - | 8170 | 221.1 | - | - | 0 | - |
| - | - | 3619 | 221.1 | - | - | 0 | - |
| - | - | 8081 | 222.1 | - | - | 0 | - |
| - | - | 6837 | 222.1 | - | - | 0 | - |
| - | - | 2496 | 223.2 | - | - | 0 | - |
| - | - | 1.39E+04 | 224.1 | - | - | 0 | - |
| - | - | 4771 | 224.1 | - | - | 0 | - |
| - | - | 4.04E+04 | 224.2 | - | - | 0 | - |
| - | - | 2903 | 225.1 | - | - | 0 | - |
| - | - | 1.509E+04 | 225.2 | - | - | 0 | - |
| - | - | 4689 | 225.2 | - | - | 0 | - |
| - | - | 7.71E+05 | 226.2 | - | - | 0 | - |
| - | - | 2752 | 227.1 | - | - | 0 | - |
| - | - | 4198 | 227.1 | - | - | 0 | - |
| 11 | y | 8.416E+04 | 227.2 | 0.001754 | 7.723 | +2 | 4 |
| 2 | a | 5.011E+04 | 228.1 | 0.0007226 | 3.168 | +1 | 2 |
| - | - | 8199 | 228.1 | - | - | 0 | - |
| - | - | 6141 | 228.2 | - | - | 0 | - |
| 5 | b | 6.91E+04 | 229.1 | 0.001996 | 8.711 | +3 | 5 |
| - | - | 1.118E+04 | 229.1 | - | - | 0 | - |
| - | - | 9659 | 229.2 | - | - | 0 | - |
| - | - | 5933 | 230.1 | - | - | 0 | - |
| - | - | 2.261E+04 | 230.1 | - | - | 0 | - |
| - | - | 2.806E+04 | 231.1 | - | - | 0 | - |
| - | - | 1.172E+04 | 231.1 | - | - | 0 | - |
| - | - | 3870 | 232.1 | - | - | 0 | - |
| - | - | 6263 | 232.1 | - | - | 0 | - |
| - | - | 4712 | 232.1 | - | - | 0 | - |
| - | - | 2355 | 232.6 | - | - | 0 | - |
| - | - | 1.878E+04 | 233.1 | - | - | 0 | - |
| - | - | 2794 | 234.1 | - | - | 0 | - |
| - | - | 3006 | 234.1 | - | - | 0 | - |
| - | - | 1.427E+05 | 235.1 | - | - | 0 | - |
| 11 | y | 1.144E+05 | 235.7 | 0.0006716 | 2.85 | +2 | 4 |
| - | - | 1.156E+04 | 236.1 | - | - | 0 | - |
| - | - | 4559 | 236.1 | - | - | 0 | - |
| - | - | 3.318E+04 | 236.2 | - | - | 0 | - |
| - | - | 4821 | 237.1 | - | - | 0 | - |
| - | - | 8263 | 238.1 | - | - | 0 | - |
| - | - | 6550 | 239.1 | - | - | 0 | - |
| - | - | 5.553E+04 | 239.2 | - | - | 0 | - |
| - | - | 6265 | 240.1 | - | - | 0 | - |
| - | - | 7395 | 240.2 | - | - | 0 | - |
| - | - | 8523 | 241.2 | - | - | 0 | - |
| - | - | 2.606E+04 | 242.2 | - | - | 0 | - |
| - | - | 4226 | 243.2 | - | - | 0 | - |
| - | - | 7365 | 243.2 | - | - | 0 | - |
| - | - | 2792 | 244.1 | - | - | 0 | - |
| - | - | 6274 | 244.2 | - | - | 0 | - |
| - | - | 2666 | 245.2 | - | - | 0 | - |
| 13 | y | 1.807E+05 | 245.2 | 0.0007985 | 3.257 | +1 | 2 |
| 2 | a | 1.41E+06 | 246.1 | 0.0008085 | 3.285 | +1 | 2 |
| - | - | 2.071E+04 | 246.2 | - | - | 0 | - |
| - | - | 3.24E+05 | 247.1 | - | - | 0 | - |
| - | - | 1.055E+05 | 248.1 | - | - | 0 | - |
| - | - | 2.499E+04 | 248.1 | - | - | 0 | - |
| - | - | 1.472E+05 | 249.1 | - | - | 0 | - |
| 8 | b | 4875 | 249.1 | 0.0008398 | 3.371 | +4 | 8 |
| - | - | 4925 | 249.1 | - | - | 0 | - |
| - | - | 5111 | 249.1 | - | - | 0 | - |
| - | - | 1.596E+04 | 250.1 | - | - | 0 | - |
| - | - | 2.046E+04 | 250.1 | - | - | 0 | - |
| - | - | 7844 | 251.1 | - | - | 0 | - |
| - | - | 1.587E+04 | 252.2 | - | - | 0 | - |
| - | - | 5062 | 253.1 | - | - | 0 | - |
| - | - | 2517 | 253.2 | - | - | 0 | - |
| - | - | 2799 | 254.2 | - | - | 0 | - |
| - | - | 6014 | 254.2 | - | - | 0 | - |
| - | - | 2925 | 255.1 | - | - | 0 | - |
| 2 | b | 3.286E+04 | 256.1 | 0.00062 | 2.421 | +1 | 2 |
| - | - | 6094 | 257.1 | - | - | 0 | - |
| - | - | 1.795E+04 | 257.1 | - | - | 0 | - |
| - | - | 2.044E+04 | 257.2 | - | - | 0 | - |
| - | - | 2.587E+04 | 258.1 | - | - | 0 | - |
| - | - | 2.571E+05 | 258.1 | - | - | 0 | - |
| - | - | 4182 | 259.1 | - | - | 0 | - |
| - | - | 3.581E+04 | 259.1 | - | - | 0 | - |
| - | - | 9850 | 259.1 | - | - | 0 | - |
| - | - | 2917 | 259.8 | - | - | 0 | - |
| - | - | 3.262E+04 | 260.1 | - | - | 0 | - |
| - | - | 3084 | 260.1 | - | - | 0 | - |
| - | - | 1.65E+04 | 260.2 | - | - | 0 | - |
| - | - | 3435 | 261.1 | - | - | 0 | - |
| - | - | 3472 | 261.2 | - | - | 0 | - |
| - | - | 4013 | 262.2 | - | - | 0 | - |
| - | - | 7258 | 263.1 | - | - | 0 | - |
| - | - | 3172 | 264.1 | - | - | 0 | - |
| - | - | 1.677E+04 | 264.1 | - | - | 0 | - |
| - | - | 1.331E+04 | 265.1 | - | - | 0 | - |
| - | - | 1.04E+04 | 265.2 | - | - | 0 | - |
| - | - | 2410 | 266.1 | - | - | 0 | - |
| - | - | 5441 | 266.1 | - | - | 0 | - |
| - | - | 8.53E+04 | 266.1 | - | - | 0 | - |
| - | - | 1.344E+04 | 267.1 | - | - | 0 | - |
| - | - | 5656 | 267.1 | - | - | 0 | - |
| - | - | 1.656E+04 | 269.6 | - | - | 0 | - |
| - | - | 2844 | 270.1 | - | - | 0 | - |
| - | - | 5.5E+04 | 270.2 | - | - | 0 | - |
| 10 | y | 3.745E+04 | 271.2 | 1.329E-05 | 0.04901 | +2 | 5 |
| - | - | 8850 | 271.7 | - | - | 0 | - |
| - | - | 3320 | 272.2 | - | - | 0 | - |
| - | - | 4299 | 273.1 | - | - | 0 | - |
| 2 | b | 5.124E+05 | 274.1 | 0.0006754 | 2.464 | +1 | 2 |
| - | - | 7.063E+04 | 275.1 | - | - | 0 | - |
| - | - | 4.302E+05 | 275.1 | - | - | 0 | - |
| 7 | y | 1.696E+04 | 275.2 | 0.0006042 | 2.196 | +3 | 8 |
| - | - | 8046 | 275.5 | - | - | 0 | - |
| - | - | 4267 | 275.9 | - | - | 0 | - |
| - | - | 1.996E+04 | 276.1 | - | - | 0 | - |
| - | - | 5.457E+04 | 276.1 | - | - | 0 | - |
| - | - | 3583 | 277.1 | - | - | 0 | - |
| - | - | 2.347E+04 | 277.1 | - | - | 0 | - |
| - | - | 3757 | 278.1 | - | - | 0 | - |
| - | - | 5425 | 279.2 | - | - | 0 | - |
| - | - | 1.294E+04 | 279.2 | - | - | 0 | - |
| - | - | 3022 | 280.2 | - | - | 0 | - |
| - | - | 4512 | 282.2 | - | - | 0 | - |
| - | - | 1.698E+04 | 283.2 | - | - | 0 | - |
| - | - | 1.969E+05 | 283.2 | - | - | 0 | - |
| - | - | 1.927E+04 | 283.6 | - | - | 0 | - |
| - | - | 5480 | 284.1 | - | - | 0 | - |
| - | - | 2.748E+04 | 284.2 | - | - | 0 | - |
| - | - | 1.28E+04 | 285.1 | - | - | 0 | - |
| - | - | 3.775E+04 | 285.2 | - | - | 0 | - |
| - | - | 2649 | 286.1 | - | - | 0 | - |
| - | - | 6198 | 286.2 | - | - | 0 | - |
| - | - | 3506 | 288.1 | - | - | 0 | - |
| - | - | 2990 | 290.1 | - | - | 0 | - |
| - | - | 5.858E+04 | 290.2 | - | - | 0 | - |
| - | - | 2.349E+04 | 290.7 | - | - | 0 | - |
| - | - | 1.431E+04 | 291.2 | - | - | 0 | - |
| - | - | 3693 | 291.2 | - | - | 0 | - |
| - | - | 1.295E+05 | 292.1 | - | - | 0 | - |
| - | - | 1.882E+04 | 293.1 | - | - | 0 | - |
| - | - | 3439 | 293.2 | - | - | 0 | - |
| - | - | 3.808E+04 | 294.1 | - | - | 0 | - |
| - | - | 3883 | 294.1 | - | - | 0 | - |
| - | - | 1.328E+04 | 294.2 | - | - | 0 | - |
| - | - | 5317 | 295.1 | - | - | 0 | - |
| - | - | 1E+05 | 297.2 | - | - | 0 | - |
| - | - | 1.791E+04 | 298.2 | - | - | 0 | - |
| - | - | 6032 | 300.1 | - | - | 0 | - |
| - | - | 5880 | 300.2 | - | - | 0 | - |
| - | - | 6318 | 300.7 | - | - | 0 | - |
| - | - | 2.922E+04 | 303.1 | - | - | 0 | - |
| - | - | 2974 | 303.2 | - | - | 0 | - |
| - | - | 3029 | 304.1 | - | - | 0 | - |
| - | - | 5.652E+04 | 305.1 | - | - | 0 | - |
| - | - | 1.709E+04 | 305.7 | - | - | 0 | - |
| - | - | 1.786E+04 | 306.1 | - | - | 0 | - |
| - | - | 3072 | 306.2 | - | - | 0 | - |
| - | - | 3532 | 307.2 | - | - | 0 | - |
| - | - | 4523 | 309.2 | - | - | 0 | - |
| - | - | 4229 | 309.2 | - | - | 0 | - |
| - | - | 3970 | 311.2 | - | - | 0 | - |
| - | - | 2.205E+04 | 311.2 | - | - | 0 | - |
| - | - | 5015 | 312.2 | - | - | 0 | - |
| - | - | 3267 | 313.5 | - | - | 0 | - |
| - | - | 4166 | 314.2 | - | - | 0 | - |
| - | - | 2.457E+04 | 314.7 | - | - | 0 | - |
| - | - | 5386 | 315.2 | - | - | 0 | - |
| - | - | 2920 | 316.9 | - | - | 0 | - |
| - | - | 7802 | 318.2 | - | - | 0 | - |
| - | - | 3550 | 319.2 | - | - | 0 | - |
| - | - | 7209 | 320.1 | - | - | 0 | - |
| 6 | y | 1.61E+04 | 320.9 | 0.0007932 | 2.472 | +3 | 9 |
| - | - | 1.331E+04 | 321.2 | - | - | 0 | - |
| - | - | 1.183E+05 | 321.2 | - | - | 0 | - |
| - | - | 7.419E+04 | 322.2 | - | - | 0 | - |
| - | - | 4696 | 322.2 | - | - | 0 | - |
| - | - | 4942 | 322.2 | - | - | 0 | - |
| - | - | 2.395E+04 | 322.2 | - | - | 0 | - |
| - | - | 3913 | 323.1 | - | - | 0 | - |
| 11 | b | 1.167E+04 | 323.2 | 0.0005706 | 1.766 | +4 | 11 |
| - | - | 1.45E+04 | 324.2 | - | - | 0 | - |
| - | - | 5089 | 324.7 | - | - | 0 | - |
| - | - | 3893 | 325.2 | - | - | 0 | - |
| - | - | 4022 | 325.2 | - | - | 0 | - |
| - | - | 4.075E+04 | 326.2 | - | - | 0 | - |
| - | - | 7880 | 327.2 | - | - | 0 | - |
| - | - | 4407 | 327.2 | - | - | 0 | - |
| - | - | 7070 | 327.3 | - | - | 0 | - |
| - | - | 3261 | 327.7 | - | - | 0 | - |
| - | - | 5556 | 330.1 | - | - | 0 | - |
| - | - | 4135 | 330.2 | - | - | 0 | - |
| - | - | 2781 | 332.1 | - | - | 0 | - |
| - | - | 3268 | 332.7 | - | - | 0 | - |
| - | - | 4.339E+04 | 332.7 | - | - | 0 | - |
| - | - | 1.637E+04 | 333.1 | - | - | 0 | - |
| - | - | 1.476E+04 | 333.2 | - | - | 0 | - |
| - | - | 4139 | 334.1 | - | - | 0 | - |
| - | - | 6628 | 334.2 | - | - | 0 | - |
| - | - | 6826 | 334.7 | - | - | 0 | - |
| 9 | y | 8445 | 335.2 | 0.0005054 | 1.508 | +2 | 6 |
| - | - | 5458 | 335.7 | - | - | 0 | - |
| - | - | 1.638E+05 | 336.2 | - | - | 0 | - |
| - | - | 3232 | 337.2 | - | - | 0 | - |
| - | - | 3.314E+04 | 337.2 | - | - | 0 | - |
| - | - | 6.084E+04 | 337.7 | - | - | 0 | - |
| - | - | 3938 | 338.2 | - | - | 0 | - |
| - | - | 2.611E+04 | 338.2 | - | - | 0 | - |
| - | - | 3037 | 338.7 | - | - | 0 | - |
| - | - | 5249 | 339.2 | - | - | 0 | - |
| - | - | 1.726E+05 | 339.2 | - | - | 0 | - |
| - | - | 3386 | 339.5 | - | - | 0 | - |
| - | - | 2.211E+04 | 340.2 | - | - | 0 | - |
| - | - | 3807 | 340.2 | - | - | 0 | - |
| - | - | 3.092E+04 | 340.2 | - | - | 0 | - |
| - | - | 5545 | 341.2 | - | - | 0 | - |
| - | - | 6057 | 341.2 | - | - | 0 | - |
| - | - | 3319 | 341.2 | - | - | 0 | - |
| - | - | 9490 | 342.2 | - | - | 0 | - |
| - | - | 1.174E+04 | 345.6 | - | - | 0 | - |
| - | - | 5087 | 346.1 | - | - | 0 | - |
| - | - | 5402 | 346.7 | - | - | 0 | - |
| - | - | 1.476E+05 | 346.7 | - | - | 0 | - |
| - | - | 4785 | 347.2 | - | - | 0 | - |
| - | - | 5.357E+04 | 347.2 | - | - | 0 | - |
| - | - | 1.325E+04 | 347.7 | - | - | 0 | - |
| - | - | 9454 | 349.7 | - | - | 0 | - |
| - | - | 2.448E+04 | 350.2 | - | - | 0 | - |
| - | - | 7776 | 350.2 | - | - | 0 | - |
| - | - | 4505 | 350.5 | - | - | 0 | - |
| - | - | 4504 | 351.2 | - | - | 0 | - |
| - | - | 1.21E+04 | 351.2 | - | - | 0 | - |
| - | - | 4253 | 352.2 | - | - | 0 | - |
| - | - | 7130 | 353.2 | - | - | 0 | - |
| - | - | 4299 | 353.9 | - | - | 0 | - |
| - | - | 1.546E+04 | 354.2 | - | - | 0 | - |
| - | - | 9093 | 354.2 | - | - | 0 | - |
| - | - | 2.454E+05 | 354.2 | - | - | 0 | - |
| - | - | 8421 | 354.5 | - | - | 0 | - |
| - | - | 7114 | 354.7 | - | - | 0 | - |
| - | - | 4.008E+04 | 355.2 | - | - | 0 | - |
| - | - | 4784 | 355.3 | - | - | 0 | - |
| - | - | 2.22E+04 | 355.7 | - | - | 0 | - |
| - | - | 6092 | 356.2 | - | - | 0 | - |
| - | - | 3588 | 356.2 | - | - | 0 | - |
| - | - | 8331 | 357.2 | - | - | 0 | - |
| - | - | 5.047E+04 | 357.3 | - | - | 0 | - |
| - | - | 5692 | 357.5 | - | - | 0 | - |
| - | - | 3619 | 357.8 | - | - | 0 | - |
| - | - | 5.405E+04 | 358.2 | - | - | 0 | - |
| - | - | 1.104E+04 | 358.3 | - | - | 0 | - |
| - | - | 1.708E+04 | 358.7 | - | - | 0 | - |
| - | - | 7875 | 359.2 | - | - | 0 | - |
| - | - | 5191 | 359.2 | - | - | 0 | - |
| - | - | 7365 | 359.2 | - | - | 0 | - |
| - | - | 1.188E+04 | 359.5 | - | - | 0 | - |
| - | - | 1.525E+04 | 359.9 | - | - | 0 | - |
| - | - | 7304 | 360.2 | - | - | 0 | - |
| - | - | 1.461E+04 | 360.7 | - | - | 0 | - |
| - | - | 1.245E+04 | 360.7 | - | - | 0 | - |
| - | - | 8981 | 361.2 | - | - | 0 | - |
| - | - | 8165 | 361.7 | - | - | 0 | - |
| - | - | 6205 | 362.2 | - | - | 0 | - |
| - | - | 4861 | 362.7 | - | - | 0 | - |
| - | - | 6774 | 363.2 | - | - | 0 | - |
| - | - | 1.229E+04 | 363.5 | - | - | 0 | - |
| 8 | y | 6156 | 363.7 | 0.0006439 | 1.77 | +2 | 7 |
| - | - | 4648 | 364.2 | - | - | 0 | - |
| - | - | 5590 | 364.2 | - | - | 0 | - |
| - | - | 4160 | 365.5 | - | - | 0 | - |
| - | - | 2.911E+04 | 366.2 | - | - | 0 | - |
| 5 | y | 2.123E+04 | 366.6 | 0.0008906 | 2.43 | +3 | 10 |
| - | - | 2.033E+04 | 366.9 | - | - | 0 | - |
| - | - | 1.974E+04 | 367.2 | - | - | 0 | - |
| - | - | 4779 | 367.2 | - | - | 0 | - |
| - | - | 2.9E+04 | 368.1 | - | - | 0 | - |
| - | - | 7011 | 368.2 | - | - | 0 | - |
| - | - | 4208 | 368.7 | - | - | 0 | - |
| 9 | b | 3342 | 368.8 | 0.006579 | 17.84 | +3 | 9 |
| - | - | 3121 | 369.2 | - | - | 0 | - |
| - | - | 6705 | 369.2 | - | - | 0 | - |
| - | - | 3524 | 369.7 | - | - | 0 | - |
| - | - | 1.005E+04 | 370.2 | - | - | 0 | - |
| - | - | 3655 | 370.5 | - | - | 0 | - |
| - | - | 3082 | 370.7 | - | - | 0 | - |
| - | - | 3.485E+04 | 371.2 | - | - | 0 | - |
| - | - | 3521 | 372.2 | - | - | 0 | - |
| 12 | y | 9.197E+04 | 373.3 | 0.0008977 | 2.405 | +1 | 3 |
| - | - | 6150 | 374.3 | - | - | 0 | - |
| - | - | 1.256E+04 | 374.3 | - | - | 0 | - |
| 9 | b | 1.076E+04 | 374.5 | 0.001167 | 3.117 | +3 | 9 |
| - | - | 6172 | 375.2 | - | - | 0 | - |
| - | - | 5634 | 376.5 | - | - | 0 | - |
| - | - | 9322 | 376.7 | - | - | 0 | - |
| - | - | 3110 | 376.9 | - | - | 0 | - |
| - | - | 4002 | 377.2 | - | - | 0 | - |
| - | - | 1.008E+04 | 378.7 | - | - | 0 | - |
| - | - | 1.173E+04 | 379 | - | - | 0 | - |
| - | - | 7902 | 379.2 | - | - | 0 | - |
| - | - | 1.384E+04 | 380.2 | - | - | 0 | - |
| - | - | 3.436E+04 | 382.2 | - | - | 0 | - |
| - | - | 1.105E+04 | 382.3 | - | - | 0 | - |
| - | - | 5368 | 382.5 | - | - | 0 | - |
| - | - | 1.84E+04 | 382.5 | - | - | 0 | - |
| - | - | 4306 | 382.9 | - | - | 0 | - |
| - | - | 3470 | 383.2 | - | - | 0 | - |
| - | - | 4388 | 383.2 | - | - | 0 | - |
| - | - | 8710 | 383.3 | - | - | 0 | - |
| - | - | 3455 | 383.7 | - | - | 0 | - |
| - | - | 2.626E+04 | 384.2 | - | - | 0 | - |
| - | - | 9.717E+04 | 385.2 | - | - | 0 | - |
| - | - | 2.077E+04 | 385.2 | - | - | 0 | - |
| - | - | 9.587E+04 | 386.2 | - | - | 0 | - |
| - | - | 4300 | 386.2 | - | - | 0 | - |
| - | - | 7458 | 386.5 | - | - | 0 | - |
| - | - | 1.355E+04 | 387.2 | - | - | 0 | - |
| - | - | 4196 | 388.2 | - | - | 0 | - |
| - | - | 2.314E+04 | 388.9 | - | - | 0 | - |
| - | - | 1.477E+04 | 389.2 | - | - | 0 | - |
| - | - | 2.233E+04 | 389.3 | - | - | 0 | - |
| - | - | 3766 | 389.5 | - | - | 0 | - |
| - | - | 9065 | 389.8 | - | - | 0 | - |
| 2 | y | 5.712E+04 | 390.2 | 0.0009577 | 2.454 | +4 | 13 |
| - | - | 5.47E+04 | 390.5 | - | - | 0 | - |
| - | - | 1.853E+04 | 390.7 | - | - | 0 | - |
| - | - | 1.035E+04 | 391 | - | - | 0 | - |
| - | - | 3635 | 392.2 | - | - | 0 | - |
| 10 | b | 1.568E+04 | 392.2 | 0.0009745 | 2.485 | +3 | 10 |
| - | - | 3.664E+04 | 392.3 | - | - | 0 | - |
| 10 | b | 1.204E+04 | 392.5 | 0.003871 | 9.863 | +3 | 10 |
| - | - | 8311 | 392.9 | - | - | 0 | - |
| - | - | 5441 | 393.2 | - | - | 0 | - |
| - | - | 7178 | 393.3 | - | - | 0 | - |
| - | - | 5.852E+04 | 394.2 | - | - | 0 | - |
| - | - | 4066 | 394.3 | - | - | 0 | - |
| 2 | y | 3.78E+04 | 394.5 | 0.001156 | 2.932 | +4 | 13 |
| - | - | 2.33E+04 | 394.7 | - | - | 0 | - |
| - | - | 1.329E+04 | 395 | - | - | 0 | - |
| - | - | 1.371E+04 | 395.2 | - | - | 0 | - |
| - | - | 3998 | 395.2 | - | - | 0 | - |
| - | - | 7065 | 396.3 | - | - | 0 | - |
| - | - | 3705 | 397.2 | - | - | 0 | - |
| - | - | 7748 | 397.2 | - | - | 0 | - |
| - | - | 6927 | 397.6 | - | - | 0 | - |
| - | - | 7029 | 397.7 | - | - | 0 | - |
| - | - | 4408 | 397.9 | - | - | 0 | - |
| 10 | b | 8937 | 398.2 | 0.0005352 | 1.344 | +3 | 10 |
| - | - | 5546 | 398.2 | - | - | 0 | - |
| - | - | 1.469E+04 | 398.5 | - | - | 0 | - |
| - | - | 6930 | 398.9 | - | - | 0 | - |
| - | - | 3761 | 398.9 | - | - | 0 | - |
| - | - | 3.138E+04 | 399.2 | - | - | 0 | - |
| - | - | 9209 | 400.2 | - | - | 0 | - |
| - | - | 4219 | 400.5 | - | - | 0 | - |
| - | - | 7931 | 401.2 | - | - | 0 | - |
| - | - | 9517 | 401.3 | - | - | 0 | - |
| - | - | 3.587E+04 | 402.2 | - | - | 0 | - |
| - | - | 4974 | 402.6 | - | - | 0 | - |
| - | - | 8.748E+04 | 403.2 | - | - | 0 | - |
| - | - | 4.12E+04 | 403.2 | - | - | 0 | - |
| - | - | 1.591E+04 | 403.3 | - | - | 0 | - |
| 4 | y | 7.522E+04 | 403.6 | 0.003703 | 9.176 | +3 | 11 |
| - | - | 6235 | 403.8 | - | - | 0 | - |
| - | - | 2.519E+04 | 403.9 | - | - | 0 | - |
| - | - | 1.758E+04 | 404.2 | - | - | 0 | - |
| - | - | 1.339E+04 | 404.2 | - | - | 0 | - |
| - | - | 5089 | 404.3 | - | - | 0 | - |
| - | - | 1.016E+04 | 404.5 | - | - | 0 | - |
| - | - | 2.825E+04 | 404.7 | - | - | 0 | - |
| - | - | 1.039E+04 | 404.9 | - | - | 0 | - |
| - | - | 1.97E+04 | 405 | - | - | 0 | - |
| - | - | 2707 | 405.2 | - | - | 0 | - |
| - | - | 1.657E+04 | 405.2 | - | - | 0 | - |
| - | - | 3073 | 405.5 | - | - | 0 | - |
| - | - | 3.993E+04 | 406.2 | - | - | 0 | - |
| - | - | 3.505E+04 | 406.3 | - | - | 0 | - |
| - | - | 1.329E+04 | 406.7 | - | - | 0 | - |
| - | - | 1.056E+04 | 406.8 | - | - | 0 | - |
| - | - | 8976 | 407.2 | - | - | 0 | - |
| - | - | 5475 | 407.7 | - | - | 0 | - |
| - | - | 2.28E+04 | 408.2 | - | - | 0 | - |
| - | - | 1.477E+04 | 408.6 | - | - | 0 | - |
| - | - | 8647 | 408.9 | - | - | 0 | - |
| 4 | y | 2.053E+04 | 409.2 | 0.001252 | 3.059 | +3 | 11 |
| - | - | 1.172E+04 | 409.6 | - | - | 0 | - |
| - | - | 5256 | 409.9 | - | - | 0 | - |
| - | - | 8.245E+04 | 410.3 | - | - | 0 | - |
| - | - | 4.376E+04 | 410.6 | - | - | 0 | - |
| - | - | 3.219E+04 | 410.9 | - | - | 0 | - |
| 6 | b | 9.109E+04 | 411.2 | 0.00109 | 2.65 | +2 | 6 |
| - | - | 1.191E+04 | 411.2 | - | - | 0 | - |
| - | - | 1.72E+04 | 411.3 | - | - | 0 | - |
| - | - | 3705 | 411.6 | - | - | 0 | - |
| 0 | Precursor | 1.796E+04 | 411.7 | 0.00107 | 2.599 | +4 | -1 |
| 0 | Precursor | 2.1E+04 | 412 | 0.004029 | 9.779 | +4 | -1 |
| - | - | 1.887E+04 | 412.2 | - | - | 0 | - |
| - | - | 1.451E+04 | 412.2 | - | - | 0 | - |
| - | - | 1.228E+04 | 412.2 | - | - | 0 | - |
| 7 | y | 3.016E+05 | 412.3 | 0.001203 | 2.918 | +2 | 8 |
| - | - | 3986 | 412.5 | - | - | 0 | - |
| - | - | 1.407E+05 | 412.8 | - | - | 0 | - |
| - | - | 1.047E+04 | 413.2 | - | - | 0 | - |
| - | - | 4.078E+04 | 413.3 | - | - | 0 | - |
| - | - | 7430 | 413.8 | - | - | 0 | - |
| - | - | 4.685E+04 | 413.9 | - | - | 0 | - |
| - | - | 9733 | 414.2 | - | - | 0 | - |
| - | - | 4.299E+04 | 414.2 | - | - | 0 | - |
| - | - | 1.522E+04 | 414.6 | - | - | 0 | - |
| - | - | 5442 | 414.9 | - | - | 0 | - |
| - | - | 4214 | 415.2 | - | - | 0 | - |
| - | - | 5689 | 415.2 | - | - | 0 | - |
| - | - | 3.661E+04 | 415.3 | - | - | 0 | - |
| - | - | 8445 | 415.8 | - | - | 0 | - |
| 0 | Precursor | 1.645E+05 | 416.2 | 0.0008702 | 2.091 | +4 | -1 |
| - | - | 1.922E+05 | 416.5 | - | - | 0 | - |
| - | - | 6.357E+04 | 416.7 | - | - | 0 | - |
| - | - | 3.753E+04 | 417 | - | - | 0 | - |
| - | - | 1.645E+04 | 417.2 | - | - | 0 | - |
| - | - | 2.013E+05 | 419.9 | - | - | 0 | - |
| 6 | b | 2.269E+04 | 420.2 | 0.000385 | 0.9163 | +2 | 6 |
| - | - | 1.622E+05 | 420.2 | - | - | 0 | - |
| - | - | 5.307E+04 | 420.6 | - | - | 0 | - |
| - | - | 9145 | 420.7 | - | - | 0 | - |
| - | - | 1.406E+04 | 420.9 | - | - | 0 | - |
| - | - | 2.961E+04 | 422.3 | - | - | 0 | - |
| - | - | 1.13E+04 | 423.3 | - | - | 0 | - |
| - | - | 1.299E+04 | 424.3 | - | - | 0 | - |
| - | - | 2.394E+04 | 424.3 | - | - | 0 | - |
| - | - | 4.711E+04 | 424.7 | - | - | 0 | - |
| - | - | 3763 | 424.8 | - | - | 0 | - |
| - | - | 3532 | 424.9 | - | - | 0 | - |
| - | - | 1.421E+04 | 425.2 | - | - | 0 | - |
| - | - | 1.137E+04 | 425.3 | - | - | 0 | - |
| - | - | 4988 | 425.3 | - | - | 0 | - |
| - | - | 8093 | 425.7 | - | - | 0 | - |
| - | - | 2.486E+04 | 425.9 | - | - | 0 | - |
| - | - | 2.34E+04 | 426.2 | - | - | 0 | - |
| - | - | 8564 | 426.6 | - | - | 0 | - |
| - | - | 5790 | 426.9 | - | - | 0 | - |
| - | - | 2.549E+04 | 427.2 | - | - | 0 | - |
| - | - | 1.114E+04 | 427.7 | - | - | 0 | - |
| - | - | 4345 | 428.2 | - | - | 0 | - |
| - | - | 8853 | 428.3 | - | - | 0 | - |
| - | - | 2.88E+05 | 429.2 | - | - | 0 | - |
| - | - | 7.217E+04 | 430.2 | - | - | 0 | - |
| - | - | 3.959E+04 | 431.2 | - | - | 0 | - |
| - | - | 8719 | 432.2 | - | - | 0 | - |
| - | - | 3880 | 433.2 | - | - | 0 | - |
| - | - | 7269 | 433.7 | - | - | 0 | - |
| - | - | 6230 | 436.6 | - | - | 0 | - |
| 3 | b | 1.89E+04 | 437.2 | 0.001251 | 2.861 | +1 | 3 |
| - | - | 4968 | 438.2 | - | - | 0 | - |
| - | - | 4565 | 438.2 | - | - | 0 | - |
| - | - | 3.176E+04 | 438.6 | - | - | 0 | - |
| - | - | 2.106E+04 | 438.9 | - | - | 0 | - |
| - | - | 7413 | 439.2 | - | - | 0 | - |
| - | - | 5897 | 439.3 | - | - | 0 | - |
| - | - | 5436 | 442.2 | - | - | 0 | - |
| - | - | 1.06E+04 | 442.7 | - | - | 0 | - |
| - | - | 5735 | 443.2 | - | - | 0 | - |
| - | - | 7021 | 444.2 | - | - | 0 | - |
| - | - | 9385 | 444.6 | - | - | 0 | - |
| - | - | 8454 | 445.2 | - | - | 0 | - |
| - | - | 6898 | 445.3 | - | - | 0 | - |
| - | - | 6029 | 445.7 | - | - | 0 | - |
| - | - | 7815 | 445.9 | - | - | 0 | - |
| - | - | 3.403E+04 | 446.2 | - | - | 0 | - |
| - | - | 3472 | 446.6 | - | - | 0 | - |
| - | - | 2.073E+04 | 446.7 | - | - | 0 | - |
| - | - | 4638 | 447.2 | - | - | 0 | - |
| - | - | 4917 | 447.7 | - | - | 0 | - |
| - | - | 5.328E+04 | 448.3 | - | - | 0 | - |
| - | - | 3.517E+04 | 448.6 | - | - | 0 | - |
| - | - | 1.298E+04 | 448.9 | - | - | 0 | - |
| - | - | 5186 | 449.3 | - | - | 0 | - |
| - | - | 4.532E+04 | 451.2 | - | - | 0 | - |
| - | - | 1.453E+04 | 451.3 | - | - | 0 | - |
| - | - | 2.076E+04 | 451.7 | - | - | 0 | - |
| - | - | 7.112E+04 | 451.9 | - | - | 0 | - |
| - | - | 8713 | 452.2 | - | - | 0 | - |
| - | - | 5.808E+04 | 452.3 | - | - | 0 | - |
| - | - | 1.275E+04 | 452.3 | - | - | 0 | - |
| - | - | 3.071E+04 | 452.6 | - | - | 0 | - |
| - | - | 1.186E+04 | 452.9 | - | - | 0 | - |
| - | - | 1.001E+04 | 453.7 | - | - | 0 | - |
| - | - | 8861 | 454.2 | - | - | 0 | - |
| - | - | 3.045E+04 | 455.3 | - | - | 0 | - |
| - | - | 9164 | 456.3 | - | - | 0 | - |
| - | - | 8932 | 457.2 | - | - | 0 | - |
| - | - | 3.856E+04 | 457.6 | - | - | 0 | - |
| 3 | y | 5.281E+04 | 457.9 | 0.005004 | 10.93 | +3 | 12 |
| - | - | 3.183E+04 | 458.3 | - | - | 0 | - |
| - | - | 1.515E+04 | 458.6 | - | - | 0 | - |
| - | - | 7160 | 458.9 | - | - | 0 | - |
| - | - | 7175 | 459.2 | - | - | 0 | - |
| - | - | 6.215E+04 | 460.2 | - | - | 0 | - |
| - | - | 2.593E+04 | 460.7 | - | - | 0 | - |
| - | - | 6544 | 461.2 | - | - | 0 | - |
| - | - | 5605 | 461.2 | - | - | 0 | - |
| - | - | 4931 | 461.6 | - | - | 0 | - |
| - | - | 4471 | 461.9 | - | - | 0 | - |
| - | - | 4102 | 462.9 | - | - | 0 | - |
| - | - | 5120 | 463.3 | - | - | 0 | - |
| 3 | y | 8.344E+05 | 463.6 | 0.001179 | 2.544 | +3 | 12 |
| - | - | 6.962E+05 | 463.9 | - | - | 0 | - |
| - | - | 2.961E+05 | 464.3 | - | - | 0 | - |
| - | - | 8.326E+04 | 464.6 | - | - | 0 | - |
| - | - | 2.242E+04 | 464.9 | - | - | 0 | - |
| - | - | 2.05E+04 | 466.9 | - | - | 0 | - |
| - | - | 3447 | 467.2 | - | - | 0 | - |
| - | - | 1.276E+04 | 467.3 | - | - | 0 | - |
| - | - | 2.573E+04 | 467.3 | - | - | 0 | - |
| - | - | 9625 | 467.6 | - | - | 0 | - |
| - | - | 4032 | 468.3 | - | - | 0 | - |
| - | - | 1.118E+04 | 468.3 | - | - | 0 | - |
| - | - | 1.713E+04 | 469.8 | - | - | 0 | - |
| - | - | 1.606E+04 | 470.2 | - | - | 0 | - |
| 11 | y | 3.347E+05 | 470.3 | 0.001112 | 2.365 | +1 | 4 |
| - | - | 2.185E+04 | 470.6 | - | - | 0 | - |
| - | - | 8681 | 470.9 | - | - | 0 | - |
| - | - | 9.174E+04 | 471.3 | - | - | 0 | - |
| - | - | 2.151E+04 | 471.8 | - | - | 0 | - |
| - | - | 2.187E+04 | 472.3 | - | - | 0 | - |
| - | - | 1.476E+04 | 472.3 | - | - | 0 | - |
| - | - | 3821 | 472.6 | - | - | 0 | - |
| - | - | 5369 | 472.8 | - | - | 0 | - |
| - | - | 4430 | 472.9 | - | - | 0 | - |
| - | - | 3.774E+04 | 473.3 | - | - | 0 | - |
| - | - | 5206 | 473.3 | - | - | 0 | - |
| - | - | 1.47E+04 | 473.6 | - | - | 0 | - |
| - | - | 6914 | 473.9 | - | - | 0 | - |
| - | - | 1.404E+04 | 474.3 | - | - | 0 | - |
| - | - | 2.025E+04 | 474.8 | - | - | 0 | - |
| - | - | 1.578E+04 | 475.3 | - | - | 0 | - |
| - | - | 9194 | 475.9 | - | - | 0 | - |
| - | - | 5.497E+04 | 476.3 | - | - | 0 | - |
| - | - | 5.908E+04 | 476.6 | - | - | 0 | - |
| - | - | 1.965E+04 | 476.9 | - | - | 0 | - |
| - | - | 8969 | 477.3 | - | - | 0 | - |
| - | - | 1.101E+04 | 478.2 | - | - | 0 | - |
| - | - | 5671 | 479.2 | - | - | 0 | - |
| - | - | 5307 | 479.2 | - | - | 0 | - |
| - | - | 3533 | 479.6 | - | - | 0 | - |
| 6 | y | 7.48E+04 | 480.8 | 0.001196 | 2.488 | +2 | 9 |
| - | - | 3.993E+04 | 481.3 | - | - | 0 | - |
| - | - | 1.531E+04 | 481.8 | - | - | 0 | - |
| - | - | 2.877E+04 | 481.9 | - | - | 0 | - |
| - | - | 4.674E+04 | 482.3 | - | - | 0 | - |
| - | - | 2.347E+04 | 482.3 | - | - | 0 | - |
| - | - | 3.025E+04 | 482.6 | - | - | 0 | - |
| - | - | 2.951E+04 | 482.8 | - | - | 0 | - |
| - | - | 5810 | 482.9 | - | - | 0 | - |
| - | - | 4385 | 483.2 | - | - | 0 | - |
| - | - | 1.378E+04 | 483.3 | - | - | 0 | - |
| - | - | 5.165E+04 | 483.8 | - | - | 0 | - |
| - | - | 2.672E+04 | 484.3 | - | - | 0 | - |
| - | - | 1.217E+04 | 484.8 | - | - | 0 | - |
| - | - | 3644 | 485.3 | - | - | 0 | - |
| - | - | 3553 | 487.2 | - | - | 0 | - |
| - | - | 1.193E+04 | 487.9 | - | - | 0 | - |
| - | - | 5850 | 488.3 | - | - | 0 | - |
| - | - | 7038 | 488.6 | - | - | 0 | - |
| - | - | 2.916E+04 | 491.3 | - | - | 0 | - |
| - | - | 4485 | 491.8 | - | - | 0 | - |
| - | - | 7622 | 492.3 | - | - | 0 | - |
| - | - | 1.024E+04 | 492.8 | - | - | 0 | - |
| - | - | 5430 | 493.3 | - | - | 0 | - |
| - | - | 4458 | 494.8 | - | - | 0 | - |
| - | - | 7320 | 495.4 | - | - | 0 | - |
| - | - | 3248 | 495.6 | - | - | 0 | - |
| - | - | 4039 | 495.9 | - | - | 0 | - |
| 8 | b | 1.165E+04 | 497.2 | 0.001845 | 3.711 | +2 | 8 |
| - | - | 1.003E+04 | 497.7 | - | - | 0 | - |
| - | - | 3198 | 498.9 | - | - | 0 | - |
| - | - | 3483 | 499.9 | - | - | 0 | - |
| - | - | 1.567E+04 | 501.6 | - | - | 0 | - |
| - | - | 8209 | 501.9 | - | - | 0 | - |
| - | - | 2.709E+04 | 503.2 | - | - | 0 | - |
| - | - | 3784 | 504.6 | - | - | 0 | - |
| 13 | b | 1.24E+04 | 504.9 | 0.004633 | 9.176 | +3 | 13 |
| 13 | b | 1.889E+04 | 505.3 | 0.008965 | 17.74 | +3 | 13 |
| - | - | 7106 | 505.6 | - | - | 0 | - |
| - | - | 5915 | 505.7 | - | - | 0 | - |
| - | - | 4499 | 505.9 | - | - | 0 | - |
| - | - | 4970 | 506.2 | - | - | 0 | - |
| - | - | 1.419E+04 | 508.8 | - | - | 0 | - |
| - | - | 1.662E+04 | 509.2 | - | - | 0 | - |
| - | - | 9125 | 509.7 | - | - | 0 | - |
| - | - | 6192 | 510.2 | - | - | 0 | - |
| - | - | 6491 | 510.6 | - | - | 0 | - |
| 13 | b | 3.168E+04 | 510.9 | 0.0008371 | 1.638 | +3 | 13 |
| - | - | 4.23E+04 | 511.3 | - | - | 0 | - |
| - | - | 1.853E+04 | 511.6 | - | - | 0 | - |
| - | - | 8307 | 511.9 | - | - | 0 | - |
| - | - | 3339 | 512.3 | - | - | 0 | - |
| - | - | 5489 | 513.9 | - | - | 0 | - |
| - | - | 1.488E+04 | 514.3 | - | - | 0 | - |
| - | - | 1.163E+04 | 514.6 | - | - | 0 | - |
| - | - | 6.754E+04 | 516.9 | - | - | 0 | - |
| - | - | 7.828E+04 | 517.3 | - | - | 0 | - |
| - | - | 2.494E+04 | 517.6 | - | - | 0 | - |
| - | - | 1.088E+04 | 517.8 | - | - | 0 | - |
| - | - | 1.083E+04 | 517.9 | - | - | 0 | - |
| - | - | 1.313E+04 | 518.3 | - | - | 0 | - |
| - | - | 8644 | 519.6 | - | - | 0 | - |
| 2 | y | 7.371E+04 | 519.9 | 0.00118 | 2.269 | +3 | 13 |
| - | - | 4605 | 520.2 | - | - | 0 | - |
| - | - | 6.694E+04 | 520.3 | - | - | 0 | - |
| - | - | 2.171E+04 | 520.4 | - | - | 0 | - |
| - | - | 4.164E+04 | 520.6 | - | - | 0 | - |
| - | - | 1.436E+04 | 521 | - | - | 0 | - |
| - | - | 2.883E+04 | 521.2 | - | - | 0 | - |
| - | - | 4409 | 521.4 | - | - | 0 | - |
| - | - | 4254 | 522.3 | - | - | 0 | - |
| - | - | 9712 | 523.4 | - | - | 0 | - |
| - | - | 4523 | 525.2 | - | - | 0 | - |
| - | - | 1.026E+04 | 525.3 | - | - | 0 | - |
| 2 | y | 1.2E+05 | 525.6 | 0.0008342 | 1.587 | +3 | 13 |
| - | - | 8066 | 525.8 | - | - | 0 | - |
| - | - | 1.427E+05 | 526 | - | - | 0 | - |
| - | - | 6.154E+04 | 526.3 | - | - | 0 | - |
| - | - | 6259 | 526.3 | - | - | 0 | - |
| - | - | 2.185E+04 | 526.6 | - | - | 0 | - |
| - | - | 4046 | 527.2 | - | - | 0 | - |
| - | - | 5952 | 528.2 | - | - | 0 | - |
| - | - | 4838 | 529.2 | - | - | 0 | - |
| - | - | 6429 | 529.3 | - | - | 0 | - |
| - | - | 7758 | 529.8 | - | - | 0 | - |
| - | - | 2.065E+04 | 530.3 | - | - | 0 | - |
| - | - | 1.489E+04 | 530.8 | - | - | 0 | - |
| - | - | 2.047E+04 | 531.2 | - | - | 0 | - |
| - | - | 5367 | 532.2 | - | - | 0 | - |
| - | - | 4109 | 533.8 | - | - | 0 | - |
| - | - | 1.724E+04 | 535.8 | - | - | 0 | - |
| - | - | 1.404E+04 | 536.3 | - | - | 0 | - |
| - | - | 5421 | 536.8 | - | - | 0 | - |
| - | - | 1.873E+05 | 538.3 | - | - | 0 | - |
| - | - | 1.321E+04 | 538.4 | - | - | 0 | - |
| - | - | 2.626E+04 | 538.8 | - | - | 0 | - |
| - | - | 5.632E+04 | 539.3 | - | - | 0 | - |
| - | - | 5.513E+04 | 539.3 | - | - | 0 | - |
| - | - | 2.392E+04 | 539.8 | - | - | 0 | - |
| - | - | 5359 | 540.3 | - | - | 0 | - |
| - | - | 1.143E+04 | 540.3 | - | - | 0 | - |
| 5 | y | 9018 | 540.8 | 0.004211 | 7.786 | +2 | 10 |
| - | - | 5765 | 541.3 | - | - | 0 | - |
| 10 | y | 8.493E+04 | 541.4 | 0.001138 | 2.103 | +1 | 5 |
| - | - | 3.586E+04 | 542.3 | - | - | 0 | - |
| - | - | 2.503E+04 | 542.4 | - | - | 0 | - |
| - | - | 8927 | 543.3 | - | - | 0 | - |
| - | - | 4881 | 543.4 | - | - | 0 | - |
| - | - | 1.035E+04 | 544.3 | - | - | 0 | - |
| - | - | 2.238E+04 | 544.8 | - | - | 0 | - |
| - | - | 1.328E+04 | 545.3 | - | - | 0 | - |
| - | - | 4514 | 545.8 | - | - | 0 | - |
| - | - | 2.044E+04 | 547.8 | - | - | 0 | - |
| - | - | 5739 | 548.2 | - | - | 0 | - |
| - | - | 2.01E+04 | 548.3 | - | - | 0 | - |
| - | - | 4443 | 548.8 | - | - | 0 | - |
| - | - | 1.563E+04 | 549.2 | - | - | 0 | - |
| 5 | y | 2.22E+05 | 549.3 | 0.0008848 | 1.611 | +2 | 10 |
| - | - | 1.228E+05 | 549.8 | - | - | 0 | - |
| - | - | 5468 | 550.2 | - | - | 0 | - |
| - | - | 3.948E+04 | 550.3 | - | - | 0 | - |
| - | - | 3357 | 550.4 | - | - | 0 | - |
| - | - | 1.068E+04 | 550.8 | - | - | 0 | - |
| - | - | 1.508E+04 | 553.2 | - | - | 0 | - |
| - | - | 9219 | 553.3 | - | - | 0 | - |
| - | - | 7482 | 553.8 | - | - | 0 | - |
| - | - | 5303 | 555.3 | - | - | 0 | - |
| - | - | 5330 | 555.8 | - | - | 0 | - |
| - | - | 4194 | 556.8 | - | - | 0 | - |
| - | - | 2.079E+04 | 557.3 | - | - | 0 | - |
| - | - | 4497 | 558.3 | - | - | 0 | - |
| 9 | b | 5.098E+04 | 561.3 | 0.0008422 | 1.501 | +2 | 9 |
| - | - | 5302 | 561.3 | - | - | 0 | - |
| - | - | 3.493E+04 | 561.8 | - | - | 0 | - |
| - | - | 1.483E+04 | 562.3 | - | - | 0 | - |
| - | - | 4573 | 562.8 | - | - | 0 | - |
| - | - | 3408 | 564.3 | - | - | 0 | - |
| - | - | 4413 | 564.8 | - | - | 0 | - |
| 4 | b | 5619 | 565.2 | 0.002823 | 4.995 | +1 | 4 |
| - | - | 3998 | 565.3 | - | - | 0 | - |
| - | - | 2.28E+05 | 566.2 | - | - | 0 | - |
| - | - | 6.911E+04 | 567.3 | - | - | 0 | - |
| - | - | 1.708E+04 | 567.4 | - | - | 0 | - |
| - | - | 1.477E+04 | 568.3 | - | - | 0 | - |
| - | - | 5384 | 568.4 | - | - | 0 | - |
| - | - | 1.069E+05 | 570.2 | - | - | 0 | - |
| - | - | 2.807E+04 | 571.2 | - | - | 0 | - |
| - | - | 5228 | 572.3 | - | - | 0 | - |
| - | - | 9.255E+04 | 572.8 | - | - | 0 | - |
| - | - | 6.057E+04 | 573.3 | - | - | 0 | - |
| - | - | 1.981E+04 | 573.8 | - | - | 0 | - |
| - | - | 5832 | 574.3 | - | - | 0 | - |
| - | - | 6.863E+04 | 577.4 | - | - | 0 | - |
| - | - | 2.48E+04 | 578.4 | - | - | 0 | - |
| - | - | 1.28E+05 | 579.4 | - | - | 0 | - |
| - | - | 4.519E+04 | 580.4 | - | - | 0 | - |
| - | - | 1.114E+04 | 581.4 | - | - | 0 | - |
| - | - | 5622 | 581.8 | - | - | 0 | - |
| - | - | 8561 | 587.3 | - | - | 0 | - |
| 10 | b | 1.35E+04 | 587.8 | 0.001301 | 2.213 | +2 | 10 |
| 10 | b | 1.46E+04 | 588.3 | 0.007095 | 12.06 | +2 | 10 |
| - | - | 4030 | 588.3 | - | - | 0 | - |
| - | - | 8895 | 588.8 | - | - | 0 | - |
| - | - | 4049 | 591.4 | - | - | 0 | - |
| - | - | 4.209E+04 | 595.4 | - | - | 0 | - |
| - | - | 6923 | 595.8 | - | - | 0 | - |
| - | - | 6264 | 596.3 | - | - | 0 | - |
| - | - | 1.743E+04 | 596.4 | - | - | 0 | - |
| 10 | b | 1.41E+04 | 596.8 | 0.001084 | 1.817 | +2 | 10 |
| - | - | 4156 | 596.8 | - | - | 0 | - |
| - | - | 4088 | 597.3 | - | - | 0 | - |
| - | - | 4.511E+04 | 598.2 | - | - | 0 | - |
| - | - | 2.313E+04 | 599.2 | - | - | 0 | - |
| - | - | 7292 | 600.3 | - | - | 0 | - |
| - | - | 4.614E+04 | 604.4 | - | - | 0 | - |
| 4 | y | 7.726E+04 | 604.8 | 0.004341 | 7.177 | +2 | 11 |
| - | - | 5.316E+04 | 605.4 | - | - | 0 | - |
| - | - | 1.639E+04 | 605.9 | - | - | 0 | - |
| - | - | 8974 | 606.4 | - | - | 0 | - |
| - | - | 4.321E+04 | 610.3 | - | - | 0 | - |
| - | - | 1.523E+04 | 611.3 | - | - | 0 | - |
| - | - | 1.239E+04 | 611.8 | - | - | 0 | - |
| - | - | 9249 | 612.3 | - | - | 0 | - |
| 4 | y | 4.809E+04 | 613.4 | 0.0004657 | 0.7592 | +2 | 11 |
| - | - | 2907 | 613.4 | - | - | 0 | - |
| - | - | 3.885E+04 | 613.9 | - | - | 0 | - |
| - | - | 8831 | 614.4 | - | - | 0 | - |
| - | - | 1.072E+04 | 615.3 | - | - | 0 | - |
| - | - | 2.277E+04 | 615.4 | - | - | 0 | - |
| - | - | 1.511E+04 | 615.9 | - | - | 0 | - |
| - | - | 6898 | 616.4 | - | - | 0 | - |
| - | - | 4811 | 618.4 | - | - | 0 | - |
| - | - | 8540 | 618.8 | - | - | 0 | - |
| - | - | 4.299E+04 | 620.3 | - | - | 0 | - |
| - | - | 3.47E+04 | 620.8 | - | - | 0 | - |
| - | - | 1.105E+04 | 621.3 | - | - | 0 | - |
| - | - | 4464 | 621.8 | - | - | 0 | - |
| - | - | 7744 | 623.4 | - | - | 0 | - |
| - | - | 5590 | 624.4 | - | - | 0 | - |
| - | - | 1.252E+04 | 628.3 | - | - | 0 | - |
| - | - | 1.007E+05 | 629.3 | - | - | 0 | - |
| - | - | 7.124E+04 | 629.8 | - | - | 0 | - |
| - | - | 2.845E+04 | 630.3 | - | - | 0 | - |
| - | - | 9286 | 630.9 | - | - | 0 | - |
| - | - | 3803 | 631.3 | - | - | 0 | - |
| - | - | 6.093E+04 | 638.4 | - | - | 0 | - |
| - | - | 4.446E+04 | 638.9 | - | - | 0 | - |
| - | - | 1.532E+04 | 639.4 | - | - | 0 | - |
| - | - | 4400 | 639.9 | - | - | 0 | - |
| - | - | 3532 | 645.8 | - | - | 0 | - |
| - | - | 7803 | 647.4 | - | - | 0 | - |
| - | - | 5530 | 648.4 | - | - | 0 | - |
| - | - | 4976 | 653.8 | - | - | 0 | - |
| - | - | 5521 | 657.3 | - | - | 0 | - |
| - | - | 8150 | 657.3 | - | - | 0 | - |
| - | - | 7609 | 657.8 | - | - | 0 | - |
| - | - | 6017 | 663.3 | - | - | 0 | - |
| - | - | 3.928E+04 | 664.5 | - | - | 0 | - |
| - | - | 1.414E+04 | 665.5 | - | - | 0 | - |
| - | - | 1.485E+04 | 665.8 | - | - | 0 | - |
| - | - | 1.693E+04 | 666.3 | - | - | 0 | - |
| - | - | 6778 | 667.3 | - | - | 0 | - |
| - | - | 1.52E+04 | 668.3 | - | - | 0 | - |
| - | - | 6113 | 669.3 | - | - | 0 | - |
| 9 | y | 3.255E+04 | 669.5 | 0.0005358 | 0.8004 | +1 | 6 |
| - | - | 1.288E+04 | 670.5 | - | - | 0 | - |
| - | - | 1.934E+04 | 674.3 | - | - | 0 | - |
| - | - | 1.354E+05 | 674.4 | - | - | 0 | - |
| - | - | 6332 | 675.3 | - | - | 0 | - |
| - | - | 4.953E+04 | 675.4 | - | - | 0 | - |
| - | - | 1.236E+04 | 676.4 | - | - | 0 | - |
| - | - | 4622 | 677.4 | - | - | 0 | - |
| - | - | 6863 | 679.3 | - | - | 0 | - |
| - | - | 3948 | 680.3 | - | - | 0 | - |
| - | - | 3539 | 680.5 | - | - | 0 | - |
| 5 | b | 8264 | 684.3 | 0.001046 | 1.529 | +1 | 5 |
| - | - | 6084 | 685.3 | - | - | 0 | - |
| - | - | 1.197E+04 | 685.4 | - | - | 0 | - |
| - | - | 4602 | 686.4 | - | - | 0 | - |
| - | - | 9549 | 690.3 | - | - | 0 | - |
| - | - | 3719 | 691.3 | - | - | 0 | - |
| - | - | 1.179E+05 | 692.4 | - | - | 0 | - |
| - | - | 5.23E+04 | 693.4 | - | - | 0 | - |
| - | - | 1.133E+04 | 694.5 | - | - | 0 | - |
| 3 | y | 4.153E+04 | 694.9 | 0.001089 | 1.567 | +2 | 12 |
| - | - | 4.119E+04 | 695.4 | - | - | 0 | - |
| - | - | 1.755E+04 | 695.9 | - | - | 0 | - |
| - | - | 4080 | 696.4 | - | - | 0 | - |
| - | - | 1.122E+04 | 698.4 | - | - | 0 | - |
| - | - | 4484 | 699.4 | - | - | 0 | - |
| 5 | b | 2.56E+04 | 702.3 | 0.002444 | 3.48 | +1 | 5 |
| - | - | 9217 | 703.3 | - | - | 0 | - |
| - | - | 6.24E+04 | 707.3 | - | - | 0 | - |
| - | - | 2.51E+04 | 708.3 | - | - | 0 | - |
| - | - | 8614 | 708.5 | - | - | 0 | - |
| - | - | 6099 | 709.3 | - | - | 0 | - |
| 12 | b | 9674 | 709.4 | 0.003209 | 4.524 | +2 | 12 |
| - | - | 5709 | 709.9 | - | - | 0 | - |
| - | - | 4.382E+04 | 710.5 | - | - | 0 | - |
| - | - | 1.652E+04 | 711.5 | - | - | 0 | - |
| - | - | 4625 | 712.5 | - | - | 0 | - |
| - | - | 5835 | 713.4 | - | - | 0 | - |
| - | - | 8888 | 713.9 | - | - | 0 | - |
| - | - | 7947 | 714.4 | - | - | 0 | - |
| - | - | 4484 | 714.9 | - | - | 0 | - |
| - | - | 2.881E+04 | 716.4 | - | - | 0 | - |
| - | - | 1.026E+04 | 717.4 | - | - | 0 | - |
| - | - | 1.246E+04 | 720.3 | - | - | 0 | - |
| - | - | 7752 | 720.4 | - | - | 0 | - |
| - | - | 1.75E+04 | 721.4 | - | - | 0 | - |
| - | - | 4226 | 722.4 | - | - | 0 | - |
| - | - | 1.239E+04 | 722.9 | - | - | 0 | - |
| - | - | 3707 | 723.4 | - | - | 0 | - |
| - | - | 2.213E+04 | 724.3 | - | - | 0 | - |
| - | - | 7610 | 725.3 | - | - | 0 | - |
| 8 | y | 5.862E+04 | 726.5 | 0.0009227 | 1.27 | +1 | 7 |
| - | - | 2.524E+04 | 727.5 | - | - | 0 | - |
| - | - | 4238 | 728.4 | - | - | 0 | - |
| - | - | 4429 | 728.5 | - | - | 0 | - |
| - | - | 5882 | 731.4 | - | - | 0 | - |
| - | - | 4072 | 732.4 | - | - | 0 | - |
| - | - | 6.007E+04 | 735.3 | - | - | 0 | - |
| - | - | 2.72E+04 | 736.3 | - | - | 0 | - |
| - | - | 1.128E+04 | 737.3 | - | - | 0 | - |
| - | - | 7591 | 738.4 | - | - | 0 | - |
| - | - | 1.019E+04 | 739.4 | - | - | 0 | - |
| - | - | 5183 | 740.4 | - | - | 0 | - |
| - | - | 4677 | 749.3 | - | - | 0 | - |
| - | - | 3.085E+04 | 752.3 | - | - | 0 | - |
| - | - | 1.561E+04 | 753.3 | - | - | 0 | - |
| - | - | 3579 | 754.3 | - | - | 0 | - |
| - | - | 7135 | 756.4 | - | - | 0 | - |
| - | - | 3914 | 777.5 | - | - | 0 | - |
| - | - | 6075 | 793.5 | - | - | 0 | - |
| - | - | 4232 | 794.3 | - | - | 0 | - |
| - | - | 5213 | 801.5 | - | - | 0 | - |
| - | - | 2.063E+04 | 805.5 | - | - | 0 | - |
| - | - | 9636 | 806.5 | - | - | 0 | - |
| - | - | 3711 | 807.5 | - | - | 0 | - |
| - | - | 1.445E+04 | 811.4 | - | - | 0 | - |
| - | - | 1.69E+04 | 811.5 | - | - | 0 | - |
| - | - | 7730 | 812.4 | - | - | 0 | - |
| - | - | 7266 | 812.5 | - | - | 0 | - |
| 7 | y | 2.274E+05 | 823.5 | 0.0007711 | 0.9364 | +1 | 8 |
| - | - | 1.133E+05 | 824.5 | - | - | 0 | - |
| - | - | 2.926E+04 | 825.5 | - | - | 0 | - |
| - | - | 4301 | 826.5 | - | - | 0 | - |
| - | - | 1.441E+04 | 829.5 | - | - | 0 | - |
| - | - | 5224 | 830.5 | - | - | 0 | - |
| 6 | b | 3.517E+04 | 839.4 | 5.085E-05 | 0.06058 | +1 | 6 |
| - | - | 2.015E+04 | 840.4 | - | - | 0 | - |
| - | - | 7565 | 847.5 | - | - | 0 | - |
| - | - | 2.714E+04 | 848.4 | - | - | 0 | - |
| - | - | 1.256E+04 | 849.4 | - | - | 0 | - |
| - | - | 4884 | 850.4 | - | - | 0 | - |
| - | - | 5065 | 852.5 | - | - | 0 | - |
| - | - | 1.013E+04 | 853.5 | - | - | 0 | - |
| - | - | 8111 | 854.5 | - | - | 0 | - |
| - | - | 4312 | 866.4 | - | - | 0 | - |
| - | - | 5748 | 884.4 | - | - | 0 | - |
| - | - | 6342 | 891.5 | - | - | 0 | - |
| - | - | 4.381E+04 | 901.4 | - | - | 0 | - |
| - | - | 2.429E+04 | 902.4 | - | - | 0 | - |
| - | - | 4254 | 903.4 | - | - | 0 | - |
| - | - | 2.743E+04 | 919.5 | - | - | 0 | - |
| - | - | 9342 | 920.5 | - | - | 0 | - |
| - | - | 4675 | 921.5 | - | - | 0 | - |
| - | - | 1.125E+04 | 942.6 | - | - | 0 | - |
| 6 | y | 2.776E+04 | 960.6 | 0.0006361 | 0.6622 | +1 | 9 |
| - | - | 1.246E+04 | 961.6 | - | - | 0 | - |
| - | - | 3797 | 962.6 | - | - | 0 | - |
| - | - | 3968 | 964.5 | - | - | 0 | - |
| - | - | 1.054E+04 | 966.6 | - | - | 0 | - |
| - | - | 5170 | 967.6 | - | - | 0 | - |
| - | - | 5734 | 984.6 | - | - | 0 | - |
| - | - | 6840 | 1033 | - | - | 0 | - |
| - | - | 7009 | 1034 | - | - | 0 | - |
| - | - | 5277 | 1070 | - | - | 0 | - |
| - | - | 5556 | 1088 | - | - | 0 | - |

m/z Charge Intensity FragmentType MassShift Position
120.05255889892578 0 4440.105
120.05657196044922 0 5188.892
121.04007720947266 0 3721.8784
122.06037139892578 0 2396.8523
122.07203674316406 0 5605.316
123.05579376220703 0 18939.494
125.07168579101562 0 2410.2708
125.10835266113281 0 2300.2324
127.08711242675781 0 24292.947
127.12356567382812 0 1910.3345
128.0712127685547 0 2802.2808
128.08164978027344 0 1893.8969
129.03939819335938 0 2000.1936
129.0663299560547 0 28075.469
129.10281372070312 0 862213
130.065673828125 0 206395.05
130.08709716796875 0 3302.1257
130.09996032714844 0 7461.519
130.10610961914062 0 52964.54
131.04586791992188 0 2534.0432
131.0690155029297 0 21572.516
131.07369995117188 0 2690.8335
132.08135986328125 0 828188.5
132.10244750976562 0 219600.36 y 13
133.08514404296875 0 85720.766
133.10586547851562 0 15313.748
134.07179260253906 0 3898.5837
134.08824157714844 0 5178.491
136.07627868652344 0 1093121
137.07366943359375 0 4826.036
137.07958984375 0 95431.36
138.06658935546875 0 12780.958
138.08285522460938 0 2717.609
139.0870819091797 0 13549.642
141.09707641601562 0 4232.871
141.102783203125 0 45693.996
142.06564331054688 0 19511.725 b 3
144.08135986328125 0 38034.28
145.08480834960938 0 3634.0938
146.06063842773438 0 5377.473
146.07131958007812 0 2574.558
147.0446014404297 0 45686.53
147.11349487304688 0 2981.154
148.04782104492188 0 6079.395
148.07659912109375 0 2946.8896
149.08299255371094 0 3756.2507
150.06634521484375 0 2818.1309
151.08717346191406 0 10612.329
152.08248901367188 0 4425.1934
152.1439971923828 0 20363.16
153.0771026611328 0 2266.4004
153.10313415527344 0 3323.6282
154.09786987304688 0 6896.8164
155.08206176757812 0 47096.707
155.0933380126953 0 14684.081
155.1184539794922 0 27928.207
156.07627868652344 0 8333.88
156.08143615722656 0 25525.244
156.1217498779297 0 4707.871
157.0764617919922 0 6540.5347
157.08450317382812 0 3482.823
157.09774780273438 0 3014.0864
158.06053161621094 0 11851.107
158.0844268798828 0 97903.06
159.09225463867188 0 572843.9
160.0762939453125 0 23492.336
160.08848571777344 0 4093.8218
160.09564208984375 0 54314.934
161.08033752441406 0 2707.7654
162.10342407226562 0 2530.161
162.5108184814453 0 2539.87
163.61334228515625 0 4688.476
164.03489685058594 0 3008.265
164.0711212158203 0 17384.25
164.0824432373047 0 2205.5452
165.0776824951172 0 5006.3594
165.102783203125 0 33673.844
166.0616455078125 0 10258.404
166.10623168945312 0 3385.59
166.1678009033203 0 2504.9675
168.08143615722656 0 2679.776
168.11375427246094 0 7617.8564
169.07659912109375 0 60998.945
169.0977020263672 0 58086.297
169.1702880859375 0 3993.6602
170.06063842773438 0 255088
170.10137939453125 0 2924.7117
171.06407165527344 0 33183.375
171.09213256835938 0 11166.307
172.10858154296875 0 55564.953
172.14476013183594 0 3470.3748
173.43984985351562 0 9142.107
174.05612182617188 0 2715.628
175.09817504882812 0 10328.166
176.0824737548828 0 23406.066
177.11293029785156 0 13739.846
179.0824432373047 0 3909.775
179.09307861328125 0 4345.729
179.5849609375 0 2375.0298
180.07774353027344 0 4787.5776
181.0614013671875 0 12421.763
181.0977783203125 0 273610.97
181.1340789794922 0 15005.298
181.17034912109375 0 3400.43
182.06101989746094 0 4727.4604
182.0939178466797 0 2527.234
182.10125732421875 0 22271.477
182.12936401367188 0 34169.582
183.09230041503906 0 123221.734
183.11294555664062 0 4433.679
183.13279724121094 0 4646.417
184.0762939453125 0 14064.519
184.0955810546875 0 15099.1455
185.07151794433594 0 27874.586
185.0803680419922 0 2971.0156
185.10792541503906 0 5858.3174
186.07955932617188 0 6164.673
186.12432861328125 0 100377.68
187.0872344970703 0 579627.4
187.1275634765625 0 8508.955
188.09060668945312 0 71662.586
189.0933380126953 0 2485.747
190.61912536621094 0 4018.642
191.0821990966797 0 2917.5024
191.09335327148438 0 2629.7876
191.11949157714844 0 2312.5537
193.09854125976562 0 3077.1316
195.0886688232422 0 2294.8584
195.11314392089844 0 4935.527
195.14964294433594 0 9611.705
196.10873413085938 0 6865.6104
197.16549682617188 0 41080.625
198.10316467285156 0 3837.8457
198.16868591308594 0 6408.219
199.08714294433594 0 26649.037
199.10850524902344 0 4316.7495
199.18106079101562 0 19988.113
200.09088134765625 0 2823.862
200.14007568359375 0 14316.014
201.0928192138672 0 4220.713
201.10284423828125 0 61686.55
201.1239776611328 0 3866.6611
201.1430206298828 0 3467.795
202.0868377685547 0 27773.57
202.10911560058594 0 92364.6
203.09219360351562 0 6696.2056
203.11289978027344 0 6839.428
204.07728576660156 0 11197.305
205.09791564941406 0 4735.6567 a 2
205.10850524902344 0 11398.019
207.1498565673828 0 29852.314
208.1443634033203 0 4522.551
208.1543426513672 0 2749.7068
209.12918090820312 0 7569.264
210.12396240234375 0 4037.3267
211.0874786376953 0 14784.265
212.09153747558594 0 3609.0688
212.14022827148438 0 13084.619
212.668212890625 0 18733.338
213.0776824951172 0 4525.6055
213.1019744873047 0 2341.0552
213.1703643798828 0 9780.446
214.10926818847656 0 31128.725
214.11973571777344 0 3349.1353
214.1918487548828 0 7121.113
215.1119842529297 0 2686.3828
215.14004516601562 0 3561.023
216.11361694335938 0 8133.926
217.10873413085938 0 7691.1523
219.08937072753906 0 2884.9314
220.1199951171875 0 71016.99
221.10400390625 0 89493
221.1238250732422 0 8169.514
221.1380157470703 0 3619.286
222.10670471191406 0 8080.6235
222.12435913085938 0 6836.6006
223.18101501464844 0 2495.5186
224.11866760253906 0 13902.805
224.13992309570312 0 4770.843
224.1763916015625 0 40400.355
225.12258911132812 0 2902.9238
225.16064453125 0 15092.523
225.17994689941406 0 4688.5376
226.15574645996094 0 770983.75
227.0823211669922 0 2751.692
227.140380859375 0 4198.188
227.15896606445312 0 84159.586 y Ammonia loss 10
228.11386108398438 0 50106.715 a Water loss 1
228.1348114013672 0 8198.953
228.1597137451172 0 6141.459
229.09780883789062 0 69099.97 b Ammonia loss 4
229.11862182617188 0 11175.603
229.16656494140625 0 9658.609
230.0819854736328 0 5932.8154
230.1033477783203 0 22608.598
231.0883331298828 0 28057.533
231.13572692871094 0 11722.238
232.09153747558594 0 3870.0676
232.12008666992188 0 6262.8833
232.1431121826172 0 4712.447
232.64352416992188 0 2354.8977
233.10414123535156 0 18776.686
234.10284423828125 0 2793.511
234.13592529296875 0 3006.2407
235.1196746826172 0 142706.69
235.67115783691406 0 114353.17 y 10
236.12278747558594 0 11563.506
236.1393280029297 0 4559.0664
236.17276000976562 0 33181.617
237.1356964111328 0 4821.357
238.13027954101562 0 8263.258
239.0819091796875 0 6549.671
239.1509246826172 0 55534.24
240.13487243652344 0 6265.2656
240.15420532226562 0 7394.7476
241.15524291992188 0 8523.023
242.18714904785156 0 26058.473
243.16983032226562 0 4225.77
243.18243408203125 0 7364.7373
244.10806274414062 0 2792.4634
244.16592407226562 0 6273.807
245.17535400390625 0 2665.9941
245.186767578125 0 180724.48 y 12
246.12451171875 0 1410228.8 a 1
246.1902313232422 0 20712.162
247.12930297851562 0 324000.7
248.11477661132812 0 105523.27
248.1324920654297 0 24993.758
249.09893798828125 0 147234.44
249.11444091796875 0 4874.9717 b 7
249.12046813964844 0 4925.0864
249.13389587402344 0 5110.611
250.1019744873047 0 15960.9375
250.1232147216797 0 20461.943
251.12954711914062 0 7843.904
252.17129516601562 0 15870.509
253.09815979003906 0 5061.636
253.17648315429688 0 2516.6675
254.15078735351562 0 2798.9358
254.18679809570312 0 6013.5757
255.07742309570312 0 2924.8489
256.1086730957031 0 32856.445 b Water loss 1
257.0918884277344 0 6093.5913
257.1150207519531 0 17951.914
257.16162109375 0 20441.707
258.07623291015625 0 25871.865
258.0992736816406 0 257052.88
259.08111572265625 0 4181.862
259.1025695800781 0 35808.46
259.13043212890625 0 9849.7295
259.849365234375 0 2917.463
260.1075134277344 0 32615.264
260.1341247558594 0 3084.4575
260.1975402832031 0 16501.49
261.1119079589844 0 3435.3523
261.200927734375 0 3472.0457
262.15582275390625 0 4013.2866
263.1153869628906 0 7257.803
264.1212158203125 0 3172.04
264.1348876953125 0 16773.215
265.1408996582031 0 13310.365
265.1666564941406 0 10396.433
266.0782775878906 0 2409.8665
266.1091003417969 0 5441.3247
266.1253967285156 0 85296.18
267.12847900390625 0 13435.312
267.1448059082031 0 5656.3677
269.6302490234375 0 16564.697
270.1302795410156 0 2844.4917
270.1818542480469 0 55003.08
271.1890563964844 0 37449.938 y 9
271.6905212402344 0 8850.049
272.18994140625 0 3319.6807
273.13433837890625 0 4299.128
274.1192932128906 0 512369.4 b 1
275.1036071777344 0 70633.16
275.1253356933594 0 430219.7
275.1854553222656 0 16960.7 y 6
275.5197448730469 0 8046.406
275.85400390625 0 4267.207
276.1081848144531 0 19955.357
276.12841796875 0 54572.164
277.0943908691406 0 3582.991
277.1339111328125 0 23466.424
278.1371765136719 0 3756.5156
279.15203857421875 0 5424.627
279.1825256347656 0 12944.416
280.1658630371094 0 3022.4863
282.1833801269531 0 4512.1294
283.15130615234375 0 16981.799
283.1772766113281 0 196929.58
283.6282043457031 0 19268.572
284.1290588378906 0 5479.787
284.18060302734375 0 27481.27
285.1107177734375 0 12798.541
285.15655517578125 0 37748.89
286.115234375 0 2648.95
286.1599426269531 0 6198.2344
288.1075134277344 0 3505.8313
290.1132507324219 0 2990.376
290.1850891113281 0 58580.016
290.6864318847656 0 23492.008
291.1572265625 0 14314.326
291.18939208984375 0 3692.6992
292.1299133300781 0 129514.234
293.1330871582031 0 18819.719
293.1612548828125 0 3439.0264
294.12042236328125 0 38076.098
294.1367492675781 0 3883.3354
294.2182312011719 0 13276.04
295.1229553222656 0 5316.6626
297.1930236816406 0 100016.26
298.1967468261719 0 17909.078
300.13232421875 0 6031.6997
300.2030029296875 0 5880.4575
300.6723937988281 0 6317.667
303.12109375 0 29220.898
303.21923828125 0 2973.7002
304.12420654296875 0 3028.906
305.12933349609375 0 56520.23
305.6648254394531 0 17092.877
306.1328125 0 17857.9
306.1657409667969 0 3071.9514
307.176025390625 0 3532.3137
309.1676940917969 0 4523.224
309.1919250488281 0 4228.763
311.171630859375 0 3969.823
311.24517822265625 0 22045.979
312.2494201660156 0 5014.7563
313.52734375 0 3267.045
314.2192077636719 0 4166.141
314.67034912109375 0 24572.174
315.1703796386719 0 5386.4316
316.8572082519531 0 2919.7925
318.1925354003906 0 7801.6577
319.17596435546875 0 3549.756
320.12469482421875 0 7209.495
320.8719482421875 0 16103.373 y 5
321.2066650390625 0 13310.516
321.2294006347656 0 118319.95
322.1558837890625 0 74192.555
322.17535400390625 0 4696.2686
322.212890625 0 4941.9375
322.23284912109375 0 23953.889
323.12628173828125 0 3912.5696
323.15924072265625 0 11671.737 b 10
324.2164306640625 0 14502.749
324.71832275390625 0 5088.904
325.18804931640625 0 3892.5657
325.2240905761719 0 4022.2183
326.2197265625 0 40745.16
327.2012023925781 0 7880.245
327.221923828125 0 4406.514
327.27593994140625 0 7069.6694
327.6775207519531 0 3261.0642
330.1087951660156 0 5556.3315
330.176513671875 0 4135.281
332.1214294433594 0 2781.1533
332.6553955078125 0 3268.3494
332.7298583984375 0 43393.97
333.12457275390625 0 16370.226
333.2309265136719 0 14757.087
334.1272888183594 0 4139.2437
334.1756591796875 0 6627.5894
334.6671142578125 0 6825.7764
335.2370300292969 0 8444.782 y 8
335.73876953125 0 5458.3774
336.2039794921875 0 163822.45
337.18695068359375 0 3232.219
337.207275390625 0 33136.773
337.7220153808594 0 60838.113
338.16180419921875 0 3937.6018
338.22308349609375 0 26106.174
338.7249755859375 0 3036.7266
339.1673889160156 0 5248.5093
339.2400207519531 0 172560.69
339.5065612792969 0 3385.8406
340.15283203125 0 22108.137
340.22186279296875 0 3807.3975
340.2433776855469 0 30917.367
341.154052734375 0 5545.188
341.2191162109375 0 6056.5947
341.2445373535156 0 3318.5947
342.1684875488281 0 9490.312
345.64337158203125 0 11735.662
346.1469421386719 0 5086.9614
346.6692810058594 0 5401.735
346.72723388671875 0 147639.27
347.17047119140625 0 4785.1836
347.2287292480469 0 53568.438
347.72979736328125 0 13251.159
349.70916748046875 0 9453.849
350.1510314941406 0 24480.812
350.2098388671875 0 7775.87
350.541015625 0 4504.5103
351.15472412109375 0 4503.5303
351.2403869628906 0 12103.482
352.2418518066406 0 4253.209
353.21856689453125 0 7129.588
353.867431640625 0 4298.6606
354.1570739746094 0 15462.869
354.1757507324219 0 9092.839
354.21441650390625 0 245384.67
354.51043701171875 0 8421.296
354.65771484375 0 7113.9
355.2173767089844 0 40075.05
355.2706604003906 0 4784.1187
355.73248291015625 0 22199.268
356.17041015625 0 6091.9253
356.21875 0 3587.7603
357.17803955078125 0 8331.4795
357.25048828125 0 50468.473
357.50408935546875 0 5691.6704
357.832763671875 0 3618.7498
358.1631164550781 0 54054.15
358.25341796875 0 11038.81
358.7143249511719 0 17078.178
359.1660461425781 0 7875.2803
359.19195556640625 0 5190.761
359.2164611816406 0 7365.4927
359.5429382324219 0 11876.029
359.87249755859375 0 15247.13
360.20672607421875 0 7303.9336
360.66497802734375 0 14614.217
360.68865966796875 0 12448.303
361.1828308105469 0 8981.247
361.6836853027344 0 8165.1914
362.15008544921875 0 6205.4053
362.6702575683594 0 4861.272
363.19720458984375 0 6774.4854
363.5076904296875 0 12287.455
363.7466125488281 0 6156.4014 y 7
364.1766052246094 0 4647.6846
364.249755859375 0 5589.5493
365.5450439453125 0 4159.819
366.1572265625 0 29105.465
366.558349609375 0 21227.533 y 4
366.8925476074219 0 20328.498
367.16265869140625 0 19743.58
367.22857666015625 0 4778.777
368.1490478515625 0 28999.441
368.17193603515625 0 7011.416
368.6539611816406 0 4208.4033
368.8450927734375 0 3341.693 b Ammonia loss 8
369.15216064453125 0 3120.8176
369.1790771484375 0 6705.216
369.6960754394531 0 3523.667
370.1872253417969 0 10051.203
370.5270080566406 0 3655.409
370.6858215332031 0 3081.5251
371.228515625 0 34854.152
372.23187255859375 0 3521.4333
373.2818298339844 0 91970.016 y 11
374.2574768066406 0 6149.5405
374.2848815917969 0 12557.728
374.51519775390625 0 10763.262 b 8
375.18994140625 0 6171.514
376.45556640625 0 5634.3403
376.6676330566406 0 9322.1875
376.8702087402344 0 3110.0286
377.1687316894531 0 4002.045
378.7119445800781 0 10077.33
378.9624938964844 0 11733.722
379.212646484375 0 7902.3643
380.2301025390625 0 13839.593
382.2065734863281 0 34362.383
382.2825927734375 0 11046.868
382.5152893066406 0 5367.5684
382.5400390625 0 18404.611
382.87542724609375 0 4305.52
383.1852722167969 0 3470.1663
383.2140808105469 0 4387.864
383.26629638671875 0 8709.754
383.7059020996094 0 3455.4243
384.1676330566406 0 26256.102
385.1737060546875 0 97171.09
385.19976806640625 0 20770.477
386.15802001953125 0 95872.66
386.2012023925781 0 4299.7026
386.5249938964844 0 7457.773
387.1605224609375 0 13552.55
388.2371826171875 0 4196.1753
388.8625183105469 0 23135.832
389.196533203125 0 14772.616
389.2720642089844 0 22331.275
389.5328063964844 0 3765.9727
389.7737121582031 0 9065.46
390.21453857421875 0 57119.617 y Ammonia loss 1
390.4651794433594 0 54696.684
390.7165222167969 0 18526.305
390.967529296875 0 10349.713
392.1642761230469 0 3635.246
392.1905212402344 0 15680.695 b Water loss 9
392.266357421875 0 36641.63
392.52142333984375 0 12035.754 b Ammonia loss 9
392.8558654785156 0 8311.036
393.16558837890625 0 5440.859
393.26959228515625 0 7178.2593
394.15191650390625 0 58524.223
394.28302001953125 0 4066.3716
394.47137451171875 0 37804.883 y 1
394.7217102050781 0 23299.297
394.9726257324219 0 13286.892
395.1544189453125 0 13706.209
395.2235412597656 0 3997.5315
396.2616882324219 0 7064.6787
397.18218994140625 0 3704.713
397.24554443359375 0 7748.1704
397.5686950683594 0 6927.1777
397.7472229003906 0 7028.697
397.8948669433594 0 4407.762
398.193603515625 0 8936.874 b 9
398.2413635253906 0 5545.8765
398.52825927734375 0 14685.532
398.8631286621094 0 6930.0493
398.891357421875 0 3761.49
399.223876953125 0 31376.76
400.2264404296875 0 9208.636
400.4761657714844 0 4218.666
401.19488525390625 0 7931.041
401.2597351074219 0 9516.727
402.2250061035156 0 35869.99
402.55230712890625 0 4973.518
403.1846923828125 0 87480.305
403.240234375 0 41202.85
403.2703857421875 0 15914.011
403.57183837890625 0 75215.14 y Ammonia loss 3
403.7691955566406 0 6235.388
403.905029296875 0 25189.336
404.18701171875 0 17577.213
404.23779296875 0 13392.063
404.26702880859375 0 5089.3833
404.4713134765625 0 10157.092
404.7257385253906 0 28249.824
404.8947448730469 0 10392.89
404.9774169921875 0 19696.688
405.19146728515625 0 2706.563
405.22833251953125 0 16574.44
405.4809265136719 0 3073.207
406.1861267089844 0 39929.023
406.2508850097656 0 35048.97
406.6880798339844 0 13294.385
406.75213623046875 0 10564.352
407.1931457519531 0 8975.621
407.69281005859375 0 5474.7886
408.2225036621094 0 22801.445
408.5553283691406 0 14768.808
408.8872375488281 0 8646.693
409.2449035644531 0 20526.777 y 3
409.5788879394531 0 11723.912
409.9130859375 0 5256.1797
410.27728271484375 0 82454.78
410.5695495605469 0 43760.395
410.90362548828125 0 32188.523
411.1786193847656 0 91086.016 b Water loss 5
411.2384948730469 0 11913.83
411.2797546386719 0 17204.965
411.57330322265625 0 3705.109
411.7266540527344 0 17961.852 Precursor Water loss
411.9756164550781 0 20996.92 Precursor Ammonia loss
412.1612548828125 0 18865.271
412.1841735839844 0 14507.9375
412.2245178222656 0 12275.955
412.27484130859375 0 301598.4 y 6
412.47674560546875 0 3986.0498
412.7760314941406 0 140723
413.1665954589844 0 10474.657
413.27740478515625 0 40783.06
413.77886962890625 0 7430.0625
413.8974304199219 0 46852.51
414.1932067871094 0 9733.117
414.2294921875 0 42986.56
414.5627746582031 0 15222.325
414.89703369140625 0 5441.793
415.1996154785156 0 4213.878
415.22857666015625 0 5689.197
415.2568664550781 0 36610.438
415.7583312988281 0 8444.858
416.2290954589844 0 164515.55 Precursor
416.47991943359375 0 192199.66
416.7305603027344 0 63570.24
416.9811096191406 0 37527.688
417.2310485839844 0 16453.246
419.9009094238281 0 201318.88
420.1831970214844 0 22693.96 b 5
420.2351379394531 0 162173.47
420.56915283203125 0 53069.406
420.6863708496094 0 9145.457
420.9035949707031 0 14059.479
422.2773742675781 0 29607.209
423.2764892578125 0 11300.871
424.26251220703125 0 12992.376
424.32916259765625 0 23942.668
424.7127685546875 0 47112.766
424.7629089355469 0 3762.7075
424.88385009765625 0 3531.589
425.2140197753906 0 14209.748
425.2877502441406 0 11369.216
425.33184814453125 0 4987.8057
425.71502685546875 0 8093.3574
425.90435791015625 0 24860.605
426.2383728027344 0 23403.428
426.57379150390625 0 8564.206
426.9066467285156 0 5789.784
427.24420166015625 0 25485.084
427.7452697753906 0 11138.509
428.24993896484375 0 4344.964
428.28729248046875 0 8853.113
429.18939208984375 0 288010.28
430.1919250488281 0 72170.29
431.18072509765625 0 39588.31
432.1825256347656 0 8719.167
433.184814453125 0 3880.304
433.7082214355469 0 7268.7915
436.5856628417969 0 6230.429
437.1831970214844 0 18898.014 b 2
438.1869812011719 0 4968.1772
438.22784423828125 0 4564.9746
438.55731201171875 0 31763.752
438.892333984375 0 21059.846
439.22589111328125 0 7413.306
439.3026428222656 0 5896.532
442.1900939941406 0 5436.4487
442.7121276855469 0 10597.775
443.2157897949219 0 5734.517
444.23248291015625 0 7020.856
444.5665588378906 0 9385.261
445.1916809082031 0 8454.085
445.26824951171875 0 6897.562
445.6919860839844 0 6029.2856
445.9180908203125 0 7814.7095
446.2330017089844 0 34026.277
446.5763244628906 0 3472.081
446.735107421875 0 20732.438
447.2376403808594 0 4637.647
447.7398681640625 0 4917.2593
448.26416015625 0 53277.582
448.59844970703125 0 35168.418
448.9313659667969 0 12980.792
449.2647705078125 0 5186.2227
451.2259521484375 0 45320.383
451.2673645019531 0 14532.318
451.72430419921875 0 20759.744
451.9203796386719 0 71119.83
452.2203369140625 0 8712.969
452.2547302246094 0 58078.84
452.3245849609375 0 12745.679
452.5879821777344 0 30708.996
452.92218017578125 0 11862.611
453.7043762207031 0 10008.688
454.2088623046875 0 8860.945
455.25262451171875 0 30450.416
456.2547607421875 0 9163.685
457.18536376953125 0 8931.566
457.5956726074219 0 38560.086
457.9275817871094 0 52811.688 y Ammonia loss 2
458.2600402832031 0 31829.035
458.59375 0 15151.837
458.9283447265625 0 7160.417
459.2137451171875 0 7175.3975
460.2313232421875 0 62153.66
460.73333740234375 0 25927.83
461.18377685546875 0 6543.6807
461.23846435546875 0 5604.9375
461.5800476074219 0 4931.0005
461.912353515625 0 4471.13
462.9283752441406 0 4102.4556
463.2822570800781 0 5120.163
463.5992736816406 0 834353.44 y 2
463.93353271484375 0 696246.56
464.2677001953125 0 296128.7
464.6022644042969 0 83255.5
464.93634033203125 0 22420.85
466.9207763671875 0 20499.727
467.16851806640625 0 3446.846
467.25604248046875 0 12757.848
467.2988586425781 0 25732.746
467.5880432128906 0 9624.883
468.26165771484375 0 4031.7383
468.2977294921875 0 11175.164
469.7889709472656 0 17128.12
470.2494812011719 0 16056.157
470.3348083496094 0 334669.3 y 10
470.5814514160156 0 21853.254
470.91461181640625 0 8681.061
471.3378601074219 0 91740.836
471.7995300292969 0 21506.316
472.29949951171875 0 21868.102
472.3409729003906 0 14757.794
472.5960998535156 0 3821.4595
472.79876708984375 0 5368.6763
472.9299011230469 0 4429.6177
473.2626953125 0 37742.066
473.30450439453125 0 5206.246
473.5783996582031 0 14703.198
473.9124755859375 0 6913.622
474.2654724121094 0 14037.765
474.7808837890625 0 20251.541
475.2802734375 0 15775.653
475.9248352050781 0 9193.947
476.2530517578125 0 54968.684
476.5870361328125 0 59075.91
476.9202880859375 0 19648.12
477.2568359375 0 8968.508
478.2095031738281 0 11012.415
479.2151184082031 0 5671.3164
479.2498779296875 0 5307.3643
479.58154296875 0 3533.2598
480.8042907714844 0 74804.31 y 5
481.3057861328125 0 39926.38
481.8068542480469 0 15306.457
481.9284362792969 0 28768.32
482.2628479003906 0 46738.59
482.3092346191406 0 23465.645
482.5928649902344 0 30251.014
482.7630615234375 0 29507.986
482.9277648925781 0 5810.0396
483.21502685546875 0 4385.4224
483.2608642578125 0 13777.506
483.7862243652344 0 51651.805
484.2872314453125 0 26720.186
484.7880859375 0 12173.6875
485.2908020019531 0 3644.1865
487.220458984375 0 3553.266
487.9309387207031 0 11926.637
488.2643737792969 0 5849.928
488.5981750488281 0 7038.416
491.27349853515625 0 29156.514
491.7733154296875 0 4484.9814
492.2755432128906 0 7622.2324
492.792236328125 0 10244.112
493.2911376953125 0 5430.419
494.75860595703125 0 4458.192
495.3667297363281 0 7320.424
495.5988464355469 0 3247.668
495.9294738769531 0 4038.948
497.2217712402344 0 11654.03 b 7
497.7226867675781 0 10032.318
498.9390869140625 0 3197.8032
499.93896484375 0 3483.2278
501.6048889160156 0 15672.9
501.93951416015625 0 8208.813
503.2159729003906 0 27090.135
504.615234375 0 3783.6626
504.9381103515625 0 12403.653 b Water loss 12
505.27044677734375 0 18887.758 b Ammonia loss 12
505.6077880859375 0 7106.3066
505.7359924316406 0 5915.26
505.9339904785156 0 4499.0107
506.23504638671875 0 4970.2114
508.7574768066406 0 14185.966
509.2409362792969 0 16616.22
509.74267578125 0 9124.732
510.2414245605469 0 6192.2744
510.6250305175781 0 6491.2397
510.9378356933594 0 31684.572 b 12
511.27197265625 0 42301.973
511.6068420410156 0 18527.002
511.94293212890625 0 8306.575
512.2723388671875 0 3339.239
513.946044921875 0 5489.012
514.2777709960938 0 14884.866
514.6106567382812 0 11631.501
516.9418334960938 0 67538.33
517.2757568359375 0 78282.55
517.6099853515625 0 24935.855
517.75341796875 0 10882.858
517.9442749023438 0 10832.514
518.2545166015625 0 13134.98
519.6209716796875 0 8643.987
519.9501953125 0 73705.29 y Ammonia loss 1
520.2435302734375 0 4604.871
520.2847900390625 0 66942.74
520.361328125 0 21713.334
520.6195068359375 0 41641.688
520.9530639648438 0 14358.671
521.2266845703125 0 28830.854
521.3646850585938 0 4409.2236
522.2626953125 0 4253.9336
523.3616943359375 0 9712.095
525.2266235351562 0 4522.9014
525.3070678710938 0 10260.214
525.6253662109375 0 119966.375 y 1
525.8057250976562 0 8065.5513
525.9596557617188 0 142736.34
526.2939453125 0 61543.7
526.3316040039062 0 6259.27
526.6279296875 0 21850.723
527.2366943359375 0 4045.7412
528.2315673828125 0 5952.4233
529.2213134765625 0 4838.0176
529.2880249023438 0 6429.1914
529.8034057617188 0 7758.1885
530.300048828125 0 20648.69
530.7993774414062 0 14888.759
531.2113037109375 0 20472.635
532.2172241210938 0 5366.7505
533.8192138671875 0 4109.124
535.7520751953125 0 17235.691
536.2538452148438 0 14037.516
536.7550048828125 0 5420.574
538.2529296875 0 187259.4
538.3726196289062 0 13205.003
538.8097534179688 0 26255.498
539.2557983398438 0 56318.45
539.3046264648438 0 55128.95
539.80517578125 0 23921.248
540.301513671875 0 5358.6265
540.3265380859375 0 11428.378
540.823486328125 0 9018.277 y Ammonia loss 4
541.32373046875 0 5765.463
541.3719482421875 0 84929.13 y 9
542.2518310546875 0 35861.645
542.374755859375 0 25029.693
543.2562255859375 0 8926.628
543.3782348632812 0 4880.8545
544.2648315429688 0 10348.355
544.7601928710938 0 22381.984
545.2604370117188 0 13281.251
545.7633056640625 0 4513.8545
547.8154296875 0 20435.412
548.2400512695312 0 5738.9043
548.3148803710938 0 20097.234
548.8121337890625 0 4443.2495
549.2225952148438 0 15630.673
549.3334350585938 0 221962.9 y 4
549.8348999023438 0 122838.03
550.2249145507812 0 5468.1787
550.33642578125 0 39477.242
550.3746337890625 0 3356.6196
550.8363647460938 0 10677.335
553.2205810546875 0 15084.136
553.269775390625 0 9219.11
553.7695922851562 0 7482.09
555.2874755859375 0 5303.142
555.7868041992188 0 5329.7886
556.8223876953125 0 4193.892
557.2949829101562 0 20793.482
558.2615356445312 0 4497.048
561.2682495117188 0 50980.09 b 8
561.3470458984375 0 5301.9385
561.76953125 0 34933.06
562.26953125 0 14831.412
562.76953125 0 4572.5044
564.3475341796875 0 3408.06
564.7994384765625 0 4412.584
565.2433471679688 0 5618.6016 b 3
565.3012084960938 0 3998.2063
566.247802734375 0 228015.95
567.25048828125 0 69108.36
567.3989868164062 0 17083.518
568.2533569335938 0 14765.527
568.3967895507812 0 5384.2905
570.2469482421875 0 106945.19
571.249755859375 0 28066.785
572.2510375976562 0 5228.167
572.8050537109375 0 92553.414
573.3062744140625 0 60565.086
573.806640625 0 19812.74
574.3121948242188 0 5832.3965
577.3828125 0 68626.625
578.3843383789062 0 24795.451
579.3622436523438 0 127981.41
580.3654174804688 0 45191.203
581.3662109375 0 11139.4795
581.8070678710938 0 5622.1895
587.2730712890625 0 8561.249
587.781982421875 0 13504.057 b Water loss 9
588.27978515625 0 14598.581 b Ammonia loss 9
588.3267211914062 0 4030.0847
588.7814331054688 0 8895.094
591.3617553710938 0 4048.9915
595.3937377929688 0 42088.45
595.846923828125 0 6923.2017
596.34765625 0 6264.4165
596.396484375 0 17430.904
596.7870483398438 0 14095.577 b 9
596.8399047851562 0 4156.291
597.289306640625 0 4088.021
598.241943359375 0 45110.008
599.245361328125 0 23131.93
600.3365478515625 0 7292.1807
604.357666015625 0 46143.86
604.8529052734375 0 77262.805 y Ammonia loss 3
605.3529663085938 0 53157.207
605.8525390625 0 16390.818
606.3550415039062 0 8973.761
610.3212890625 0 43206.004
611.3245239257812 0 15234.312
611.82861328125 0 12388.097
612.3280639648438 0 9249.398
613.3623046875 0 48091.62 y 3
613.4081420898438 0 2907.1335
613.8646240234375 0 38848.363
614.3642578125 0 8830.517
615.2682495117188 0 10720.363
615.35009765625 0 22767.992
615.8511962890625 0 15107.165
616.3519287109375 0 6898.4634
618.351318359375 0 4810.814
618.849609375 0 8539.509
620.3419189453125 0 42990.434
620.8412475585938 0 34699.734
621.3409423828125 0 11050.995
621.83642578125 0 4464.17
623.3875732421875 0 7744.1704
624.3905639648438 0 5589.857
628.3307495117188 0 12519.094
629.3468017578125 0 100705.59
629.8477783203125 0 71236.1
630.3483276367188 0 28450.143
630.8511352539062 0 9286.077
631.3488159179688 0 3803.0984
638.3524169921875 0 60931.836
638.8536376953125 0 44455.363
639.3546752929688 0 15323.246
639.854248046875 0 4399.988
645.8224487304688 0 3532.4043
647.4257202148438 0 7802.8413
648.4269409179688 0 5529.8896
653.8285522460938 0 4976.233
657.2794799804688 0 5521.1113
657.3340454101562 0 8150.477
657.8346557617188 0 7609.1147
663.300537109375 0 6016.8213
664.4513549804688 0 39281.164
665.4544067382812 0 14142.632
665.84326171875 0 14851.4795
666.3447265625 0 16929.324
667.3436279296875 0 6777.964
668.32958984375 0 15198.853
669.3292846679688 0 6113.0723
669.46630859375 0 32552.834 y 8
670.4697875976562 0 12875.934
674.304443359375 0 19340.482
674.435791015625 0 135387.02
675.3071899414062 0 6331.696
675.4380493164062 0 49534.793
676.440185546875 0 12357.423
677.378173828125 0 4622.165
679.3109741210938 0 6863.016
680.3191528320312 0 3948.172
680.4876708984375 0 3538.7415
684.2899169921875 0 8263.982 b Water loss 4
685.2882690429688 0 6084.226
685.3531494140625 0 11966.622
686.3555297851562 0 4602.1753
690.2787475585938 0 9548.874
691.3073120117188 0 3719.4905
692.4462890625 0 117932.88
693.44873046875 0 52301.45
694.4523315429688 0 11328.956
694.8945922851562 0 41531.035 y 2
695.39599609375 0 41194.793
695.8955078125 0 17545.584
696.4022216796875 0 4080.0244
698.4098510742188 0 11219.408
699.4127197265625 0 4483.795
702.3018798828125 0 25603.03 b 4
703.3017578125 0 9217.393
707.3056030273438 0 62398.688
708.3093872070312 0 25100.54
708.4801635742188 0 8613.765
709.3031616210938 0 6099.4165
709.363037109375 0 9673.867 b 11
709.8613891601562 0 5708.853
710.4567260742188 0 43823.008
711.4601440429688 0 16517.99
712.4613647460938 0 4625.041
713.3781127929688 0 5834.6343
713.8788452148438 0 8887.716
714.3775634765625 0 7946.6685
714.8825073242188 0 4483.953
716.4212646484375 0 28807.447
717.423095703125 0 10255.486
720.3206787109375 0 12455.661
720.3760375976562 0 7752.237
721.3538818359375 0 17496.957
722.3621826171875 0 4226.0503
722.8861694335938 0 12391.148
723.3776245117188 0 3707.3533
724.33203125 0 22126.326
725.3329467773438 0 7609.919
726.4881591796875 0 58617.89 y 7
727.4907836914062 0 25244.69
728.3963012695312 0 4237.8984
728.4962768554688 0 4429.3247
731.3973388671875 0 5881.984
732.3930053710938 0 4071.6611
735.3009033203125 0 60073.293
736.3040161132812 0 27200.717
737.3482055664062 0 11282.87
738.376220703125 0 7590.938
739.365234375 0 10194.535
740.3707885742188 0 5182.774
749.34814453125 0 4677.4697
752.3270263671875 0 30845.953
753.3294067382812 0 15607.088
754.3343505859375 0 3579.161
756.3920288085938 0 7134.6704
777.531005859375 0 3914.1992
793.4815673828125 0 6075.222
794.3407592773438 0 4231.519
801.5072021484375 0 5213.2495
805.530029296875 0 20633.066
806.5299072265625 0 9636.388
807.5271606445312 0 3710.6357
811.3634643554688 0 14448.424
811.4937133789062 0 16899.16
812.3662719726562 0 7730.3457
812.4946899414062 0 7265.7803
823.540771484375 0 227439.17 y 6
824.5435791015625 0 113324.64
825.5462036132812 0 29257.795
826.5492553710938 0 4300.8403
829.5042724609375 0 14409.662
830.50732421875 0 5224.484
839.3583984375 0 35165.402 b 5
840.3615112304688 0 20148.178
847.517578125 0 7565.3022
848.41650390625 0 27136.928
849.4190063476562 0 12556.185
850.4205322265625 0 4883.9956
852.45751953125 0 5064.951
853.4759521484375 0 10130.47
854.47900390625 0 8110.6226
866.40673828125 0 4312.0806
884.41650390625 0 5747.6562
891.4595336914062 0 6341.562
901.443115234375 0 43814.63
902.4439086914062 0 24288.006
903.4458618164062 0 4253.567
919.4531860351562 0 27429.338
920.4573364257812 0 9341.816
921.4603881835938 0 4675.334
942.5826416015625 0 11247.698
960.5995483398438 0 27758.25 y 5
961.6033935546875 0 12457.802
962.6148071289062 0 3796.993
964.5136108398438 0 3968.1465
966.5626220703125 0 10542.179
967.5615844726562 0 5170.081
984.5675659179688 0 5733.8643
1032.5322265625 0 6840.2744
1033.535888671875 0 7009.0386
1070.4991455078125 0 5277.3955
1087.5162353515625 0 5556.287

Spectrum Details

|  |  |
| --- | --- |
| Matched peaks? Matched peaks The total absolute number of peaks matched. Additionally in brackets the total fraction of peaks matched and the total number of peaks is shown. | 65 (6.21% of 1047) |
| FDR? FDR The false discovery rate estimated for this peptide. It is calculated by matching all theoretical fragments with a non-integer shift with the raw peaks for this spectrum. This is done with 40 different shifts. The resulting percentage is the average number of annotated peaks over the number of annotated peaks with the correct spectrum. | 5.05% |
| Satellite FDR? Satellite FDR See the FDR for details on its calculation. This satellite ion specific FDR only contains the satellite ions (d/w) for I/L/J positions. | - |
| PSM Score? PSM Score The PSM Score as given by Hecklib to this annotated spectrum. It is shown with three significant figures. | 205 |

#### Spectrum 5748? Spectrum 5748 The raw spectrum of this peptide as annotated by Hecklib. The fragments are coloured according to ion type (see legend). Any peaks with a star '\*' as text can be hovered over to see the full details, first the ion type second the mass shift type. By hovering over the amino acids in the peptide or ions in the legend the corresponding peaks are highlighted. By toggling the 'Unassigned' label you can turn the background (unassigned) peaks on or off in the plot. By updating the slider in the Ion legend you can update the spectrum to only show the top X% of the peaks with labels. The top X% means any peak that is within X% of the highest intensity. By dragging in the spectrum you can zoom in to a specific part of the spectrum and use 'Zoom Out' to get back to the original zoom level. The annotation of the spectrum is based on the given sequence in the peptides file and is done with different software so inconsistencies are likely. The peaks are annotated based on the given sequence, with 20 ppm tolerance.

Copy Data

##### Spectrum 5748 (TSV)

###### Preview

```
Loading example...
```

*Click on the button to copy the data to your clipboard.*

Mz MinMz MaxIntensity Max

WidthHeightPeptide font sizePeptide stroke widthSpectrum font sizeSpectrum stroke widthCompact peptide

Ion legend

wxyz

abcd

OtherUnassignedIon^Charge^_Position_Show for top:%

SWYQHHPGKAPKJJ

03.30e+46.59e+49.89e+41.32e+5

Zoom Out

a^+3^_3_y^+1^_1_y^+2^_3_b^+3^_4_b^+2^_3_y^+2^_4_a^+1^_2_b^+3^_5_y^+1^_2_a^+1^_2_b^+1^_2_b^+1^_2_y^+1^_3_y^+3^_11_b^+2^_6_y^+2^_8_b^+2^_6_b^+1^_3_y^+3^_12_y^+1^_4_b^+3^_12_y^+2^_9_b^+2^_8_b^+3^_13_b^+3^_13_b^+3^_13_y^+3^_13_y^+3^_13_y^+1^_5_\*\*y^+2^_10_\*b^+2^_9_b^+2^_10_y^+2^_11_y^+2^_11_y^+1^_6_y^+2^_12_y^+2^_12_b^+1^_5_b^+2^_12_y^+1^_7_b^+2^_13_b^+2^_13_y^+2^_13_y^+1^_8_b^+1^_6_y^+1^_8_b^+1^_6_y^+1^_9_y^+1^_9_y^+1^_10_b^+1^_10_b^+1^_10_

0616123118472462

Fragment Matches Table

Show background peaks

| Position | Ion type | Intensity | mz Theoretical | mz Error (Th) | mz Error (ppm) | Charge | Series Number |
| --- | --- | --- | --- | --- | --- | --- | --- |
| - | - | 952.9 | 120.1 | - | - | 0 | - |
| - | - | 2014 | 120.1 | - | - | 0 | - |
| - | - | 376.8 | 122.1 | - | - | 0 | - |
| - | - | 351.1 | 122.9 | - | - | 0 | - |
| - | - | 543.4 | 123.1 | - | - | 0 | - |
| - | - | 897.1 | 127.1 | - | - | 0 | - |
| - | - | 368.2 | 127.7 | - | - | 0 | - |
| - | - | 496.7 | 128.1 | - | - | 0 | - |
| - | - | 480.4 | 128.1 | - | - | 0 | - |
| - | - | 393.7 | 128.2 | - | - | 0 | - |
| - | - | 1893 | 129.1 | - | - | 0 | - |
| - | - | 4.029E+04 | 129.1 | - | - | 0 | - |
| - | - | 477.7 | 130.1 | - | - | 0 | - |
| - | - | 588.5 | 130.1 | - | - | 0 | - |
| - | - | 8270 | 130.1 | - | - | 0 | - |
| - | - | 417.3 | 130.1 | - | - | 0 | - |
| - | - | 2857 | 130.1 | - | - | 0 | - |
| 3 | a | 364.5 | 131.1 | 0.002226 | 16.99 | +3 | 3 |
| - | - | 835.7 | 131.1 | - | - | 0 | - |
| - | - | 2.724E+04 | 132.1 | - | - | 0 | - |
| 14 | y | 6260 | 132.1 | 0.0002372 | 1.796 | +1 | 1 |
| - | - | 2605 | 133.1 | - | - | 0 | - |
| - | - | 1.971E+04 | 136.1 | - | - | 0 | - |
| - | - | 1861 | 137.1 | - | - | 0 | - |
| - | - | 1219 | 138.1 | - | - | 0 | - |
| - | - | 1078 | 139.1 | - | - | 0 | - |
| - | - | 525.1 | 140.1 | - | - | 0 | - |
| - | - | 527.3 | 141.1 | - | - | 0 | - |
| - | - | 3517 | 141.1 | - | - | 0 | - |
| - | - | 701.3 | 145.1 | - | - | 0 | - |
| - | - | 1089 | 146.1 | - | - | 0 | - |
| - | - | 510.4 | 146.1 | - | - | 0 | - |
| - | - | 560.7 | 147 | - | - | 0 | - |
| - | - | 1119 | 147.1 | - | - | 0 | - |
| - | - | 546 | 147.1 | - | - | 0 | - |
| - | - | 469.3 | 148.9 | - | - | 0 | - |
| - | - | 741.9 | 148.9 | - | - | 0 | - |
| - | - | 564 | 148.9 | - | - | 0 | - |
| - | - | 962.3 | 148.9 | - | - | 0 | - |
| - | - | 1162 | 148.9 | - | - | 0 | - |
| - | - | 2527 | 148.9 | - | - | 0 | - |
| - | - | 3858 | 148.9 | - | - | 0 | - |
| - | - | 3397 | 149 | - | - | 0 | - |
| - | - | 1689 | 149 | - | - | 0 | - |
| - | - | 1381 | 149 | - | - | 0 | - |
| - | - | 1246 | 149 | - | - | 0 | - |
| - | - | 689.2 | 149 | - | - | 0 | - |
| - | - | 761.5 | 149 | - | - | 0 | - |
| - | - | 567.2 | 149 | - | - | 0 | - |
| - | - | 537.1 | 149 | - | - | 0 | - |
| - | - | 651.7 | 149 | - | - | 0 | - |
| - | - | 409.2 | 150.1 | - | - | 0 | - |
| - | - | 1154 | 152.1 | - | - | 0 | - |
| - | - | 721.7 | 152.1 | - | - | 0 | - |
| - | - | 581.9 | 152.1 | - | - | 0 | - |
| - | - | 590.7 | 153.1 | - | - | 0 | - |
| - | - | 439.6 | 153.1 | - | - | 0 | - |
| - | - | 2148 | 155.1 | - | - | 0 | - |
| - | - | 1392 | 155.1 | - | - | 0 | - |
| - | - | 1326 | 155.1 | - | - | 0 | - |
| - | - | 1068 | 156.1 | - | - | 0 | - |
| - | - | 610.1 | 157.1 | - | - | 0 | - |
| - | - | 610 | 157.1 | - | - | 0 | - |
| - | - | 519.6 | 158.1 | - | - | 0 | - |
| - | - | 2306 | 158.1 | - | - | 0 | - |
| - | - | 700.3 | 159.1 | - | - | 0 | - |
| - | - | 1.35E+04 | 159.1 | - | - | 0 | - |
| - | - | 506.5 | 159.1 | - | - | 0 | - |
| - | - | 1110 | 160.1 | - | - | 0 | - |
| - | - | 1582 | 160.1 | - | - | 0 | - |
| - | - | 492.5 | 161.1 | - | - | 0 | - |
| - | - | 1156 | 165.1 | - | - | 0 | - |
| - | - | 1201 | 166.1 | - | - | 0 | - |
| - | - | 837.5 | 167.1 | - | - | 0 | - |
| - | - | 475.9 | 167.1 | - | - | 0 | - |
| - | - | 655.1 | 168.1 | - | - | 0 | - |
| - | - | 1714 | 169.1 | - | - | 0 | - |
| - | - | 4280 | 169.1 | - | - | 0 | - |
| - | - | 4037 | 170.1 | - | - | 0 | - |
| - | - | 1109 | 171.1 | - | - | 0 | - |
| - | - | 571.8 | 171.1 | - | - | 0 | - |
| - | - | 2579 | 172.1 | - | - | 0 | - |
| - | - | 465.3 | 172.1 | - | - | 0 | - |
| - | - | 1002 | 173.1 | - | - | 0 | - |
| - | - | 2614 | 173.1 | - | - | 0 | - |
| - | - | 811.1 | 175.1 | - | - | 0 | - |
| - | - | 611.4 | 175.1 | - | - | 0 | - |
| - | - | 970.5 | 181.1 | - | - | 0 | - |
| - | - | 5910 | 181.1 | - | - | 0 | - |
| - | - | 1158 | 181.1 | - | - | 0 | - |
| - | - | 542.2 | 182.1 | - | - | 0 | - |
| - | - | 816.4 | 182.1 | - | - | 0 | - |
| - | - | 3561 | 183.1 | - | - | 0 | - |
| - | - | 1415 | 183.1 | - | - | 0 | - |
| - | - | 918.2 | 183.1 | - | - | 0 | - |
| - | - | 522.4 | 184.1 | - | - | 0 | - |
| - | - | 1258 | 185.1 | - | - | 0 | - |
| - | - | 491.1 | 186.1 | - | - | 0 | - |
| - | - | 543.5 | 186.1 | - | - | 0 | - |
| - | - | 3537 | 186.1 | - | - | 0 | - |
| - | - | 2.535E+04 | 187.1 | - | - | 0 | - |
| - | - | 2056 | 187.1 | - | - | 0 | - |
| 12 | y | 3200 | 187.1 | 0.0002896 | 1.548 | +2 | 3 |
| - | - | 2679 | 188.1 | - | - | 0 | - |
| 4 | b | 566.5 | 189.1 | 0.001812 | 9.584 | +3 | 4 |
| - | - | 543.5 | 193.1 | - | - | 0 | - |
| - | - | 1133 | 195.1 | - | - | 0 | - |
| - | - | 615.5 | 195.2 | - | - | 0 | - |
| - | - | 623.4 | 196.1 | - | - | 0 | - |
| - | - | 476.7 | 196.3 | - | - | 0 | - |
| - | - | 842.4 | 197.1 | - | - | 0 | - |
| - | - | 797.1 | 197.1 | - | - | 0 | - |
| - | - | 1275 | 197.2 | - | - | 0 | - |
| - | - | 1004 | 199.1 | - | - | 0 | - |
| - | - | 1627 | 199.1 | - | - | 0 | - |
| - | - | 983.7 | 199.2 | - | - | 0 | - |
| - | - | 625.5 | 200.1 | - | - | 0 | - |
| - | - | 2055 | 201.1 | - | - | 0 | - |
| - | - | 926.7 | 201.1 | - | - | 0 | - |
| - | - | 668.7 | 202.1 | - | - | 0 | - |
| - | - | 2761 | 202.1 | - | - | 0 | - |
| - | - | 857.9 | 203.1 | - | - | 0 | - |
| - | - | 740.3 | 203.1 | - | - | 0 | - |
| - | - | 831.9 | 204.1 | - | - | 0 | - |
| - | - | 1186 | 205.1 | - | - | 0 | - |
| - | - | 555.4 | 207.1 | - | - | 0 | - |
| - | - | 738 | 207.1 | - | - | 0 | - |
| - | - | 1242 | 207.1 | - | - | 0 | - |
| - | - | 950.7 | 209.1 | - | - | 0 | - |
| - | - | 576.5 | 209.1 | - | - | 0 | - |
| - | - | 1191 | 211.1 | - | - | 0 | - |
| - | - | 575.7 | 213.1 | - | - | 0 | - |
| - | - | 1326 | 214.1 | - | - | 0 | - |
| - | - | 633.1 | 214.1 | - | - | 0 | - |
| - | - | 2544 | 215.1 | - | - | 0 | - |
| - | - | 838.2 | 216.1 | - | - | 0 | - |
| 3 | b | 701 | 219.1 | 0.00396 | 18.08 | +2 | 3 |
| - | - | 2353 | 220.1 | - | - | 0 | - |
| - | - | 2704 | 221.1 | - | - | 0 | - |
| - | - | 966.6 | 223.1 | - | - | 0 | - |
| - | - | 1316 | 223.2 | - | - | 0 | - |
| - | - | 530.7 | 224.2 | - | - | 0 | - |
| - | - | 764.3 | 225 | - | - | 0 | - |
| - | - | 635.4 | 225.2 | - | - | 0 | - |
| - | - | 697.2 | 226.1 | - | - | 0 | - |
| - | - | 946 | 226.1 | - | - | 0 | - |
| - | - | 3.597E+04 | 226.2 | - | - | 0 | - |
| - | - | 1366 | 227.1 | - | - | 0 | - |
| - | - | 964.4 | 227.1 | - | - | 0 | - |
| 11 | y | 3914 | 227.2 | 0.001083 | 4.768 | +2 | 4 |
| 2 | a | 2489 | 228.1 | 0.000219 | 0.9602 | +1 | 2 |
| - | - | 1134 | 228.1 | - | - | 0 | - |
| 5 | b | 2442 | 229.1 | 0.001233 | 5.381 | +3 | 5 |
| - | - | 1229 | 229.1 | - | - | 0 | - |
| - | - | 6713 | 229.2 | - | - | 0 | - |
| - | - | 692.4 | 230.2 | - | - | 0 | - |
| - | - | 808.4 | 231.1 | - | - | 0 | - |
| - | - | 936.6 | 231.1 | - | - | 0 | - |
| - | - | 1560 | 234.1 | - | - | 0 | - |
| - | - | 3647 | 235.1 | - | - | 0 | - |
| - | - | 2053 | 239.2 | - | - | 0 | - |
| - | - | 1365 | 240.1 | - | - | 0 | - |
| - | - | 698.9 | 241.2 | - | - | 0 | - |
| - | - | 2623 | 242.2 | - | - | 0 | - |
| - | - | 615.9 | 242.2 | - | - | 0 | - |
| - | - | 2279 | 243.1 | - | - | 0 | - |
| - | - | 587.8 | 244.1 | - | - | 0 | - |
| - | - | 550.4 | 244.2 | - | - | 0 | - |
| 13 | y | 1.184E+04 | 245.2 | 0.0002034 | 0.8295 | +1 | 2 |
| 2 | a | 6.887E+04 | 246.1 | 0.0001524 | 0.6192 | +1 | 2 |
| - | - | 1362 | 246.2 | - | - | 0 | - |
| - | - | 1.466E+04 | 247.1 | - | - | 0 | - |
| - | - | 3461 | 248.1 | - | - | 0 | - |
| - | - | 849.9 | 248.1 | - | - | 0 | - |
| - | - | 601.9 | 248.2 | - | - | 0 | - |
| - | - | 5223 | 249.1 | - | - | 0 | - |
| - | - | 624.6 | 249.1 | - | - | 0 | - |
| - | - | 1715 | 249.2 | - | - | 0 | - |
| - | - | 511.5 | 250.1 | - | - | 0 | - |
| - | - | 1002 | 250.1 | - | - | 0 | - |
| - | - | 4435 | 251.2 | - | - | 0 | - |
| - | - | 711.1 | 252.1 | - | - | 0 | - |
| - | - | 979.2 | 252.2 | - | - | 0 | - |
| 2 | b | 2376 | 256.1 | 0.0002232 | 0.8717 | +1 | 2 |
| - | - | 603.5 | 257.1 | - | - | 0 | - |
| - | - | 8607 | 258.1 | - | - | 0 | - |
| - | - | 659.9 | 258.1 | - | - | 0 | - |
| - | - | 1348 | 259.1 | - | - | 0 | - |
| - | - | 581.7 | 259.1 | - | - | 0 | - |
| - | - | 1581 | 260.1 | - | - | 0 | - |
| - | - | 2651 | 260.2 | - | - | 0 | - |
| - | - | 741.5 | 261.1 | - | - | 0 | - |
| - | - | 692.1 | 263.1 | - | - | 0 | - |
| - | - | 596.5 | 265.1 | - | - | 0 | - |
| - | - | 673.2 | 265.6 | - | - | 0 | - |
| - | - | 4348 | 266.1 | - | - | 0 | - |
| - | - | 1593 | 267.1 | - | - | 0 | - |
| - | - | 1648 | 270.2 | - | - | 0 | - |
| 2 | b | 4.098E+04 | 274.1 | 2.651E-05 | 0.0967 | +1 | 2 |
| - | - | 731.2 | 275.1 | - | - | 0 | - |
| - | - | 1.762E+04 | 275.1 | - | - | 0 | - |
| - | - | 714.9 | 276.1 | - | - | 0 | - |
| - | - | 2237 | 276.1 | - | - | 0 | - |
| - | - | 707.1 | 277.1 | - | - | 0 | - |
| - | - | 1506 | 278.1 | - | - | 0 | - |
| - | - | 523.8 | 282.2 | - | - | 0 | - |
| - | - | 636.7 | 283.2 | - | - | 0 | - |
| - | - | 5445 | 283.2 | - | - | 0 | - |
| - | - | 597.3 | 284.2 | - | - | 0 | - |
| - | - | 1148 | 285 | - | - | 0 | - |
| - | - | 1683 | 285.2 | - | - | 0 | - |
| - | - | 2632 | 290.2 | - | - | 0 | - |
| - | - | 730.7 | 290.7 | - | - | 0 | - |
| - | - | 4134 | 292.1 | - | - | 0 | - |
| - | - | 761.2 | 293.1 | - | - | 0 | - |
| - | - | 1601 | 294.1 | - | - | 0 | - |
| - | - | 970.8 | 297.2 | - | - | 0 | - |
| - | - | 4593 | 297.2 | - | - | 0 | - |
| - | - | 2690 | 299.1 | - | - | 0 | - |
| - | - | 1014 | 300.1 | - | - | 0 | - |
| - | - | 1175 | 301.1 | - | - | 0 | - |
| - | - | 615 | 303.1 | - | - | 0 | - |
| - | - | 1213 | 305.1 | - | - | 0 | - |
| - | - | 587.7 | 308.8 | - | - | 0 | - |
| - | - | 1301 | 311.2 | - | - | 0 | - |
| - | - | 834.5 | 314.2 | - | - | 0 | - |
| - | - | 1246 | 320.1 | - | - | 0 | - |
| - | - | 5322 | 321.2 | - | - | 0 | - |
| - | - | 1869 | 322.2 | - | - | 0 | - |
| - | - | 1552 | 322.2 | - | - | 0 | - |
| - | - | 949.9 | 326.2 | - | - | 0 | - |
| - | - | 800.6 | 326.7 | - | - | 0 | - |
| - | - | 656.3 | 327.2 | - | - | 0 | - |
| - | - | 1014 | 332.7 | - | - | 0 | - |
| - | - | 8384 | 336.2 | - | - | 0 | - |
| - | - | 1204 | 337.2 | - | - | 0 | - |
| - | - | 1320 | 337.7 | - | - | 0 | - |
| - | - | 1437 | 338.2 | - | - | 0 | - |
| - | - | 5598 | 339.2 | - | - | 0 | - |
| - | - | 1653 | 340.2 | - | - | 0 | - |
| - | - | 5648 | 346.7 | - | - | 0 | - |
| - | - | 2437 | 347.2 | - | - | 0 | - |
| - | - | 653.6 | 350.1 | - | - | 0 | - |
| - | - | 832.5 | 351.2 | - | - | 0 | - |
| - | - | 810.2 | 353.2 | - | - | 0 | - |
| - | - | 1.102E+04 | 354.2 | - | - | 0 | - |
| - | - | 1145 | 355.1 | - | - | 0 | - |
| - | - | 1465 | 355.2 | - | - | 0 | - |
| - | - | 1452 | 357.2 | - | - | 0 | - |
| - | - | 1458 | 358.2 | - | - | 0 | - |
| - | - | 1247 | 366.2 | - | - | 0 | - |
| - | - | 744 | 367.2 | - | - | 0 | - |
| - | - | 672.6 | 368.1 | - | - | 0 | - |
| - | - | 809.8 | 371.1 | - | - | 0 | - |
| 12 | y | 2296 | 373.3 | 4.322E-05 | 0.1158 | +1 | 3 |
| - | - | 1164 | 381.2 | - | - | 0 | - |
| - | - | 718.6 | 382.2 | - | - | 0 | - |
| - | - | 1422 | 384.2 | - | - | 0 | - |
| - | - | 3382 | 385.2 | - | - | 0 | - |
| - | - | 805.5 | 385.2 | - | - | 0 | - |
| - | - | 676.1 | 385.4 | - | - | 0 | - |
| - | - | 1851 | 386.2 | - | - | 0 | - |
| - | - | 1364 | 392.3 | - | - | 0 | - |
| - | - | 3469 | 394.2 | - | - | 0 | - |
| - | - | 807.1 | 395.2 | - | - | 0 | - |
| - | - | 650.8 | 396.3 | - | - | 0 | - |
| - | - | 693.7 | 397.2 | - | - | 0 | - |
| - | - | 1363 | 402.2 | - | - | 0 | - |
| - | - | 2983 | 403.2 | - | - | 0 | - |
| - | - | 739 | 403.3 | - | - | 0 | - |
| - | - | 958.2 | 404.2 | - | - | 0 | - |
| - | - | 1451 | 406.2 | - | - | 0 | - |
| - | - | 1244 | 406.3 | - | - | 0 | - |
| 4 | y | 591 | 409.2 | 0.002717 | 6.639 | +3 | 11 |
| - | - | 2554 | 410.3 | - | - | 0 | - |
| 6 | b | 5396 | 411.2 | 6.996E-05 | 0.1702 | +2 | 6 |
| - | - | 848.7 | 411.3 | - | - | 0 | - |
| - | - | 1521 | 412.2 | - | - | 0 | - |
| 7 | y | 1.41E+04 | 412.3 | 1.272E-05 | 0.03085 | +2 | 8 |
| - | - | 6299 | 412.8 | - | - | 0 | - |
| - | - | 682.5 | 413.2 | - | - | 0 | - |
| - | - | 1644 | 413.3 | - | - | 0 | - |
| - | - | 954.5 | 415.3 | - | - | 0 | - |
| 6 | b | 968.5 | 420.2 | 0.0005305 | 1.263 | +2 | 6 |
| - | - | 1283 | 420.2 | - | - | 0 | - |
| - | - | 953.1 | 420.7 | - | - | 0 | - |
| - | - | 1094 | 422.3 | - | - | 0 | - |
| - | - | 857 | 423.2 | - | - | 0 | - |
| - | - | 832.2 | 423.3 | - | - | 0 | - |
| - | - | 1.488E+04 | 429.2 | - | - | 0 | - |
| - | - | 4241 | 430.2 | - | - | 0 | - |
| - | - | 2132 | 431.2 | - | - | 0 | - |
| 3 | b | 2629 | 437.2 | 0.0002447 | 0.5597 | +1 | 3 |
| - | - | 1171 | 438.2 | - | - | 0 | - |
| - | - | 777.6 | 438.6 | - | - | 0 | - |
| - | - | 661.7 | 451.2 | - | - | 0 | - |
| - | - | 1204 | 454.2 | - | - | 0 | - |
| - | - | 1219 | 455.3 | - | - | 0 | - |
| - | - | 634.7 | 458.1 | - | - | 0 | - |
| 3 | y | 2738 | 463.6 | 0.0004076 | 0.8792 | +3 | 12 |
| - | - | 2981 | 463.9 | - | - | 0 | - |
| - | - | 674.5 | 464.3 | - | - | 0 | - |
| - | - | 1260 | 467.3 | - | - | 0 | - |
| - | - | 640.3 | 470.2 | - | - | 0 | - |
| 11 | y | 1.073E+04 | 470.3 | 7.478E-05 | 0.159 | +1 | 4 |
| - | - | 3155 | 471.3 | - | - | 0 | - |
| - | - | 1222 | 471.8 | - | - | 0 | - |
| - | - | 1168 | 472.3 | - | - | 0 | - |
| 12 | b | 3204 | 473.2 | 0.0005483 | 1.159 | +3 | 12 |
| - | - | 499.6 | 473.3 | - | - | 0 | - |
| - | - | 3566 | 473.6 | - | - | 0 | - |
| - | - | 2020 | 473.9 | - | - | 0 | - |
| - | - | 764.6 | 474.8 | - | - | 0 | - |
| - | - | 1007 | 475.3 | - | - | 0 | - |
| - | - | 1555 | 476.3 | - | - | 0 | - |
| - | - | 1973 | 476.6 | - | - | 0 | - |
| - | - | 845.5 | 478.2 | - | - | 0 | - |
| - | - | 1110 | 480.7 | - | - | 0 | - |
| 6 | y | 2790 | 480.8 | 6.252E-06 | 0.013 | +2 | 9 |
| - | - | 1594 | 481.3 | - | - | 0 | - |
| - | - | 758.8 | 481.7 | - | - | 0 | - |
| - | - | 595.6 | 481.9 | - | - | 0 | - |
| - | - | 1183 | 482.3 | - | - | 0 | - |
| - | - | 847.2 | 482.3 | - | - | 0 | - |
| - | - | 749.1 | 482.6 | - | - | 0 | - |
| - | - | 1731 | 483.8 | - | - | 0 | - |
| - | - | 852.1 | 484.3 | - | - | 0 | - |
| - | - | 1102 | 491.3 | - | - | 0 | - |
| - | - | 1589 | 494.2 | - | - | 0 | - |
| - | - | 705.4 | 496.3 | - | - | 0 | - |
| 8 | b | 847.1 | 497.2 | 0.0009604 | 1.932 | +2 | 8 |
| - | - | 2673 | 501.6 | - | - | 0 | - |
| - | - | 4606 | 501.9 | - | - | 0 | - |
| - | - | 1365 | 502.3 | - | - | 0 | - |
| - | - | 975.2 | 503.2 | - | - | 0 | - |
| 13 | b | 3777 | 504.9 | 0.0005852 | 1.159 | +3 | 13 |
| 13 | b | 5788 | 505.3 | 0.004753 | 9.407 | +3 | 13 |
| - | - | 1767 | 505.6 | - | - | 0 | - |
| - | - | 847.3 | 505.9 | - | - | 0 | - |
| - | - | 1667 | 510.3 | - | - | 0 | - |
| 13 | b | 2.327E+04 | 510.9 | 0.0003531 | 0.691 | +3 | 13 |
| - | - | 2.14E+04 | 511.3 | - | - | 0 | - |
| - | - | 9168 | 511.6 | - | - | 0 | - |
| - | - | 3796 | 511.9 | - | - | 0 | - |
| - | - | 778.8 | 512.3 | - | - | 0 | - |
| - | - | 3247 | 516.9 | - | - | 0 | - |
| - | - | 3284 | 517.3 | - | - | 0 | - |
| - | - | 983.4 | 517.6 | - | - | 0 | - |
| - | - | 812.7 | 519.3 | - | - | 0 | - |
| 2 | y | 1147 | 519.9 | 8.105E-05 | 0.1559 | +3 | 13 |
| - | - | 1036 | 520.3 | - | - | 0 | - |
| - | - | 1195 | 520.4 | - | - | 0 | - |
| - | - | 654.2 | 520.6 | - | - | 0 | - |
| - | - | 762.7 | 521.2 | - | - | 0 | - |
| 2 | y | 4072 | 525.6 | 0.0004475 | 0.8514 | +3 | 13 |
| - | - | 3883 | 526 | - | - | 0 | - |
| - | - | 723.7 | 526.3 | - | - | 0 | - |
| - | - | 1083 | 529.8 | - | - | 0 | - |
| - | - | 676.7 | 530.3 | - | - | 0 | - |
| - | - | 801.1 | 531.2 | - | - | 0 | - |
| - | - | 796.8 | 537 | - | - | 0 | - |
| - | - | 8752 | 538.3 | - | - | 0 | - |
| - | - | 615.4 | 538.4 | - | - | 0 | - |
| - | - | 856.6 | 538.8 | - | - | 0 | - |
| - | - | 2241 | 539.3 | - | - | 0 | - |
| - | - | 3027 | 539.3 | - | - | 0 | - |
| - | - | 770 | 539.6 | - | - | 0 | - |
| - | - | 1232 | 539.8 | - | - | 0 | - |
| - | - | 739.1 | 540.8 | - | - | 0 | - |
| 10 | y | 2251 | 541.4 | 0.00112 | 2.068 | +1 | 5 |
| - | - | 839 | 547.8 | - | - | 0 | - |
| - | - | 630.7 | 548.3 | - | - | 0 | - |
| 0 | Precursor | 4531 | 548.6 | 0.0008272 | 1.508 | +3 | -1 |
| 0 | Precursor | 6203 | 549 | 0.005274 | 9.607 | +3 | -1 |
| - | - | 2500 | 549.3 | - | - | 0 | - |
| 5 | y | 4548 | 549.3 | 3.073E-05 | 0.05595 | +2 | 10 |
| - | - | 2730 | 549.6 | - | - | 0 | - |
| - | - | 2949 | 549.8 | - | - | 0 | - |
| - | - | 727.5 | 550 | - | - | 0 | - |
| - | - | 1438 | 550.3 | - | - | 0 | - |
| - | - | 705.4 | 553.8 | - | - | 0 | - |
| - | - | 1507 | 553.9 | - | - | 0 | - |
| - | - | 1700 | 554 | - | - | 0 | - |
| - | - | 1409 | 554.3 | - | - | 0 | - |
| 0 | Precursor | 1.255E+05 | 554.6 | 0.0003815 | 0.6878 | +3 | -1 |
| - | - | 1.305E+05 | 555 | - | - | 0 | - |
| - | - | 6.372E+04 | 555.3 | - | - | 0 | - |
| - | - | 2.343E+04 | 555.6 | - | - | 0 | - |
| - | - | 8440 | 556 | - | - | 0 | - |
| 9 | b | 1045 | 561.3 | 0.001025 | 1.827 | +2 | 9 |
| - | - | 779.6 | 564.3 | - | - | 0 | - |
| - | - | 968.1 | 564.8 | - | - | 0 | - |
| - | - | 1.06E+04 | 566.2 | - | - | 0 | - |
| - | - | 2621 | 567.2 | - | - | 0 | - |
| - | - | 1277 | 570.2 | - | - | 0 | - |
| - | - | 4359 | 572.8 | - | - | 0 | - |
| - | - | 3677 | 573.3 | - | - | 0 | - |
| - | - | 1100 | 573.8 | - | - | 0 | - |
| - | - | 2256 | 577.4 | - | - | 0 | - |
| - | - | 925.7 | 578.4 | - | - | 0 | - |
| - | - | 682.6 | 579.3 | - | - | 0 | - |
| - | - | 6530 | 579.4 | - | - | 0 | - |
| - | - | 2037 | 580.4 | - | - | 0 | - |
| - | - | 796.1 | 587.3 | - | - | 0 | - |
| - | - | 855.2 | 589.3 | - | - | 0 | - |
| - | - | 1756 | 595.4 | - | - | 0 | - |
| 10 | b | 1608 | 596.8 | 0.001968 | 3.297 | +2 | 10 |
| - | - | 2226 | 604.4 | - | - | 0 | - |
| 4 | y | 3522 | 604.8 | 0.0007397 | 1.223 | +2 | 11 |
| - | - | 2762 | 605.4 | - | - | 0 | - |
| - | - | 1535 | 610.3 | - | - | 0 | - |
| - | - | 994.7 | 611.3 | - | - | 0 | - |
| 4 | y | 3959 | 613.4 | 0.0009992 | 1.629 | +2 | 11 |
| - | - | 2406 | 613.9 | - | - | 0 | - |
| - | - | 2680 | 615.3 | - | - | 0 | - |
| - | - | 1701 | 615.8 | - | - | 0 | - |
| - | - | 612.8 | 616.3 | - | - | 0 | - |
| - | - | 3318 | 620.3 | - | - | 0 | - |
| - | - | 3252 | 620.8 | - | - | 0 | - |
| - | - | 1252 | 621.3 | - | - | 0 | - |
| - | - | 840.8 | 628.3 | - | - | 0 | - |
| - | - | 9973 | 629.3 | - | - | 0 | - |
| - | - | 9222 | 629.8 | - | - | 0 | - |
| - | - | 3055 | 630.3 | - | - | 0 | - |
| - | - | 839.6 | 630.8 | - | - | 0 | - |
| - | - | 797.4 | 638.4 | - | - | 0 | - |
| - | - | 1668 | 638.9 | - | - | 0 | - |
| - | - | 1001 | 654.3 | - | - | 0 | - |
| - | - | 739.7 | 657.3 | - | - | 0 | - |
| - | - | 675.6 | 662 | - | - | 0 | - |
| - | - | 728.7 | 662.7 | - | - | 0 | - |
| - | - | 1819 | 664.4 | - | - | 0 | - |
| - | - | 1035 | 665.8 | - | - | 0 | - |
| 9 | y | 1887 | 669.5 | 0.0006579 | 0.9827 | +1 | 6 |
| - | - | 2359 | 674.3 | - | - | 0 | - |
| - | - | 8114 | 674.4 | - | - | 0 | - |
| - | - | 1098 | 675.3 | - | - | 0 | - |
| - | - | 3232 | 675.4 | - | - | 0 | - |
| - | - | 1203 | 677.4 | - | - | 0 | - |
| - | - | 1095 | 677.9 | - | - | 0 | - |
| - | - | 1551 | 680.3 | - | - | 0 | - |
| - | - | 2098 | 685.9 | - | - | 0 | - |
| 3 | y | 3078 | 686.4 | 0.005086 | 7.41 | +2 | 12 |
| - | - | 1341 | 686.9 | - | - | 0 | - |
| - | - | 7755 | 692.4 | - | - | 0 | - |
| - | - | 3734 | 693.4 | - | - | 0 | - |
| 3 | y | 3.28E+04 | 694.9 | 0.0009862 | 1.419 | +2 | 12 |
| - | - | 2.955E+04 | 695.4 | - | - | 0 | - |
| - | - | 1.541E+04 | 695.9 | - | - | 0 | - |
| - | - | 5564 | 696.4 | - | - | 0 | - |
| 5 | b | 4651 | 702.3 | 0.0003079 | 0.4384 | +1 | 5 |
| - | - | 1570 | 703.3 | - | - | 0 | - |
| - | - | 1730 | 707.3 | - | - | 0 | - |
| 12 | b | 9786 | 709.4 | 0.001307 | 1.843 | +2 | 12 |
| - | - | 6902 | 709.9 | - | - | 0 | - |
| - | - | 2709 | 710.4 | - | - | 0 | - |
| - | - | 686.9 | 710.5 | - | - | 0 | - |
| - | - | 794.6 | 713.9 | - | - | 0 | - |
| - | - | 1675 | 716.4 | - | - | 0 | - |
| - | - | 911.7 | 721.4 | - | - | 0 | - |
| - | - | 691.8 | 722.4 | - | - | 0 | - |
| - | - | 1151 | 724.3 | - | - | 0 | - |
| 8 | y | 2405 | 726.5 | 0.001702 | 2.343 | +1 | 7 |
| - | - | 1081 | 727.5 | - | - | 0 | - |
| - | - | 640.4 | 733.4 | - | - | 0 | - |
| - | - | 1029 | 735.3 | - | - | 0 | - |
| - | - | 886.2 | 739.4 | - | - | 0 | - |
| - | - | 703.5 | 752.3 | - | - | 0 | - |
| 13 | b | 1137 | 756.9 | 0.0003371 | 0.4453 | +2 | 13 |
| 13 | b | 2908 | 765.9 | 0.0007366 | 0.9617 | +2 | 13 |
| - | - | 2057 | 766.4 | - | - | 0 | - |
| - | - | 1601 | 766.9 | - | - | 0 | - |
| - | - | 1359 | 774.9 | - | - | 0 | - |
| - | - | 1332 | 775.4 | - | - | 0 | - |
| - | - | 758.2 | 775.9 | - | - | 0 | - |
| 2 | y | 1025 | 787.9 | 0.0001288 | 0.1635 | +2 | 13 |
| - | - | 664.5 | 788.4 | - | - | 0 | - |
| - | - | 682 | 795.3 | - | - | 0 | - |
| - | - | 1516 | 805.5 | - | - | 0 | - |
| 7 | y | 758.3 | 806.5 | 0.009132 | 11.32 | +1 | 8 |
| - | - | 944.3 | 809.4 | - | - | 0 | - |
| - | - | 4824 | 811.4 | - | - | 0 | - |
| - | - | 1005 | 811.5 | - | - | 0 | - |
| - | - | 1456 | 812.4 | - | - | 0 | - |
| 6 | b | 957.7 | 821.3 | 0.003474 | 4.23 | +1 | 6 |
| 7 | y | 1.597E+04 | 823.5 | 0.001914 | 2.325 | +1 | 8 |
| - | - | 8434 | 824.5 | - | - | 0 | - |
| - | - | 2305 | 825.5 | - | - | 0 | - |
| - | - | 1190 | 829.5 | - | - | 0 | - |
| - | - | 879 | 830.5 | - | - | 0 | - |
| 6 | b | 1.033E+04 | 839.4 | 0.001109 | 1.321 | +1 | 6 |
| - | - | 4333 | 840.4 | - | - | 0 | - |
| - | - | 1103 | 841.4 | - | - | 0 | - |
| - | - | 2015 | 848.4 | - | - | 0 | - |
| - | - | 702 | 849.4 | - | - | 0 | - |
| - | - | 813.3 | 853.5 | - | - | 0 | - |
| - | - | 3671 | 901.4 | - | - | 0 | - |
| - | - | 1959 | 902.4 | - | - | 0 | - |
| - | - | 1471 | 919.5 | - | - | 0 | - |
| - | - | 1527 | 942.6 | - | - | 0 | - |
| 6 | y | 1241 | 943.6 | 0.01577 | 16.71 | +1 | 9 |
| 6 | y | 4021 | 960.6 | 0.002355 | 2.451 | +1 | 9 |
| - | - | 1595 | 961.6 | - | - | 0 | - |
| - | - | 682.9 | 980.2 | - | - | 0 | - |
| 5 | y | 1039 | 1098 | 0.002184 | 1.99 | +1 | 10 |
| - | - | 883.8 | 1099 | - | - | 0 | - |
| - | - | 761.7 | 1145 | - | - | 0 | - |
| 10 | b | 1483 | 1175 | 0.001699 | 1.446 | +1 | 10 |
| 10 | b | 922 | 1176 | 0.01121 | 9.539 | +1 | 10 |
| - | - | 637 | 1685 | - | - | 0 | - |
| - | - | 717.2 | 2272 | - | - | 0 | - |
| - | - | 809.1 | 2438 | - | - | 0 | - |

m/z Charge Intensity FragmentType MassShift Position
120.06572723388672 0 952.9432
120.08094024658203 0 2014.3346
122.06057739257812 0 376.79172
122.90543365478516 0 351.14764
123.0556869506836 0 543.36725
127.08692932128906 0 897.1235
127.71723175048828 0 368.23715
128.07066345214844 0 496.73953
128.0822296142578 0 480.41922
128.2210693359375 0 393.68222
129.0660858154297 0 1893.0853
129.1024932861328 0 40290.406
130.05038452148438 0 477.73853
130.0605926513672 0 588.45215
130.06539916992188 0 8270.071
130.08660888671875 0 417.28568
130.10586547851562 0 2857.3735
131.0614471435547 0 364.4819 a Water loss 2
131.06884765625 0 835.7475
132.0810089111328 0 27241.166
132.10214233398438 0 6259.5347 y 13
133.08445739746094 0 2605.096
136.07594299316406 0 19707.242
137.07923889160156 0 1861.4321
138.06640625 0 1219.0282
139.08648681640625 0 1078.4891
140.0818328857422 0 525.12823
141.06565856933594 0 527.253
141.10244750976562 0 3516.528
145.0612335205078 0 701.2977
146.06031799316406 0 1088.8969
146.09254455566406 0 510.40228
147.04412841796875 0 560.6576
147.07675170898438 0 1118.7074
147.11306762695312 0 546.0384
148.8924560546875 0 469.25208
148.89952087402344 0 741.8759
148.91476440429688 0 563.9768
148.9218292236328 0 962.3262
148.92897033691406 0 1162.4663
148.9362030029297 0 2526.8564
148.94381713867188 0 3857.8657
148.96058654785156 0 3397.3508
148.9683074951172 0 1689.2134
148.97543334960938 0 1380.982
148.98304748535156 0 1245.8821
148.99012756347656 0 689.1802
148.9976806640625 0 761.537
149.0044403076172 0 567.214
149.0122528076172 0 537.12616
149.01910400390625 0 651.6724
150.0554962158203 0 409.20834
152.0707550048828 0 1153.5256
152.08200073242188 0 721.683
152.14352416992188 0 581.8573
153.06619262695312 0 590.6869
153.10263061523438 0 439.60474
155.08154296875 0 2147.9026
155.09291076660156 0 1392.2949
155.11817932128906 0 1326.485
156.07655334472656 0 1067.6521
157.06082153320312 0 610.09174
157.09744262695312 0 609.9988
158.06028747558594 0 519.57794
158.0840606689453 0 2305.5142
159.0852813720703 0 700.30585
159.09185791015625 0 13498.382
159.11251831054688 0 506.51755
160.07586669921875 0 1109.8279
160.09523010253906 0 1581.613
161.0924835205078 0 492.4805
165.1018829345703 0 1155.6707
166.06118774414062 0 1201.1971
167.0817413330078 0 837.4836
167.0931854248047 0 475.94568
168.06588745117188 0 655.06494
169.07601928710938 0 1714.0276
169.09732055664062 0 4279.611
170.06027221679688 0 4037.1929
171.07675170898438 0 1109.3938
171.1127471923828 0 571.7562
172.10836791992188 0 2579.3162
172.14596557617188 0 465.34048
173.09226989746094 0 1001.5995
173.12863159179688 0 2614.0305
175.08718872070312 0 811.10583
175.1075897216797 0 611.41925
181.06089782714844 0 970.4717
181.0973663330078 0 5910.446
181.1338653564453 0 1157.7573
182.10043334960938 0 542.2499
182.12869262695312 0 816.3849
183.0918731689453 0 3560.9312
183.11276245117188 0 1414.7904
183.14944458007812 0 918.15875
184.09585571289062 0 522.374
185.0711212158203 0 1257.703
186.0746612548828 0 491.05527
186.0880584716797 0 543.52264
186.1239471435547 0 3537.4395
187.08677673339844 0 25351.81
187.1078338623047 0 2056.059
187.14439392089844 0 3199.9668 y 11
188.0902557373047 0 2678.7876
189.0868377685547 0 566.49884 b 3
193.096923828125 0 543.5094
195.11300659179688 0 1133.4993
195.15037536621094 0 615.4765
196.1083526611328 0 623.4047
196.31845092773438 0 476.65463
197.0915985107422 0 842.3921
197.12855529785156 0 797.1461
197.1652374267578 0 1274.5905
199.08624267578125 0 1004.19
199.10757446289062 0 1627.1917
199.18008422851562 0 983.74023
200.1392364501953 0 625.4985
201.1022186279297 0 2054.6802
201.12362670898438 0 926.7166
202.0860595703125 0 668.7083
202.10865783691406 0 2760.5613
203.1028289794922 0 857.86993
203.11265563964844 0 740.3274
204.13409423828125 0 831.89386
205.1183319091797 0 1185.8563
207.09793090820312 0 555.3634
207.1129913330078 0 738.0381
207.1488037109375 0 1241.7181
209.093017578125 0 950.6865
209.12718200683594 0 576.51276
211.14476013183594 0 1191.4352
213.12283325195312 0 575.65094
214.1089324951172 0 1325.5177
214.11959838867188 0 633.1302
215.13917541503906 0 2543.8333
216.09820556640625 0 838.1537
219.09857177734375 0 701.0283 b 2
220.1193084716797 0 2352.51
221.1033477783203 0 2703.6318
223.0639190673828 0 966.5504
223.15505981445312 0 1315.5925
224.17544555664062 0 530.69183
225.0435333251953 0 764.2959
225.1588134765625 0 635.44293
226.0820770263672 0 697.1525
226.11907958984375 0 945.9797
226.15512084960938 0 35965.36
227.10289001464844 0 1365.8126
227.13914489746094 0 964.35236
227.15829467773438 0 3914.1296 y Ammonia loss 10
228.1133575439453 0 2489.1292 a Water loss 1
228.1342315673828 0 1134.0333
229.0970458984375 0 2442.0261 b Ammonia loss 4
229.1177978515625 0 1229.2177
229.15480041503906 0 6712.5444
230.15939331054688 0 692.42224
231.08837890625 0 808.42944
231.13497924804688 0 936.6329
234.1240234375 0 1560.0665
235.11903381347656 0 3647.1367
239.15013122558594 0 2053.2085
240.134521484375 0 1364.822
241.15499877929688 0 698.8828
242.1500701904297 0 2623.1948
242.1856689453125 0 615.9215
243.134033203125 0 2279.2188
244.09378051757812 0 587.84424
244.1661834716797 0 550.42334
245.18617248535156 0 11839.572 y 12
246.1238555908203 0 68866.41 a 1
246.19009399414062 0 1362.0916
247.12850952148438 0 14661.073
248.1142578125 0 3461.0386
248.128662109375 0 849.8907
248.15957641601562 0 601.8953
249.09837341308594 0 5223.326
249.12294006347656 0 624.62866
249.15994262695312 0 1715.0292
250.08433532714844 0 511.4871
250.1014404296875 0 1001.669
251.15042114257812 0 4434.9653
252.13427734375 0 711.0931
252.17068481445312 0 979.2302
256.1082763671875 0 2376.2136 b Water loss 1
257.1144714355469 0 603.49774
258.0984802246094 0 8607.197
258.1448669433594 0 659.8694
259.1023864746094 0 1347.8488
259.129150390625 0 581.7028
260.10614013671875 0 1581.4364
260.1606750488281 0 2650.6272
261.09033203125 0 741.4874
263.14007568359375 0 692.1135
265.1412658691406 0 596.5443
265.5841369628906 0 673.2004
266.12457275390625 0 4347.844
267.1337585449219 0 1593.4736
270.1809387207031 0 1648.4803
274.11859130859375 0 40979.88 b 1
275.1034240722656 0 731.23553
275.1240234375 0 17617.754
276.1088562011719 0 714.9424
276.1268005371094 0 2236.97
277.1326599121094 0 707.0739
278.1170349121094 0 1506.1487
282.18072509765625 0 523.80853
283.1528625488281 0 636.67413
283.1763610839844 0 5444.7837
284.17926025390625 0 597.3484
285.01007080078125 0 1147.9783
285.1559753417969 0 1683.2585
290.1843566894531 0 2632.0178
290.6855773925781 0 730.7037
292.12933349609375 0 4133.72
293.1338195800781 0 761.24414
294.12017822265625 0 1600.6946
297.1554260253906 0 970.8316
297.1925964355469 0 4592.9526
299.0616455078125 0 2690.2227
300.0625 0 1014.26263
301.05889892578125 0 1174.7318
303.1203308105469 0 614.95264
305.1281433105469 0 1213.4603
308.78509521484375 0 587.7147
311.24432373046875 0 1301.4204
314.1822814941406 0 834.4944
320.12518310546875 0 1246.132
321.22869873046875 0 5322.3726
322.155029296875 0 1868.7284
322.2321472167969 0 1551.5547
326.2191467285156 0 949.92664
326.6634216308594 0 800.6002
327.1672668457031 0 656.28394
332.72955322265625 0 1013.91974
336.2032470703125 0 8384.203
337.2060241699219 0 1204.1561
337.72119140625 0 1320.1449
338.2225036621094 0 1437.2008
339.2393493652344 0 5598.0195
340.2427062988281 0 1653.4364
346.7261962890625 0 5647.5815
347.2278747558594 0 2437.4395
350.1490478515625 0 653.5683
351.2392272949219 0 832.46014
353.2190246582031 0 810.23334
354.21368408203125 0 11020.199
355.07147216796875 0 1145.3325
355.2174072265625 0 1464.5424
357.24908447265625 0 1452.1138
358.1619873046875 0 1458.4484
366.1560974121094 0 1247.0696
367.1620788574219 0 743.9806
368.1474304199219 0 672.6231
371.099365234375 0 809.83545
373.2809753417969 0 2295.7832 y 11
381.1523742675781 0 1164.3503
382.20697021484375 0 718.587
384.1662902832031 0 1422.1328
385.1727600097656 0 3381.845
385.19757080078125 0 805.54553
385.39324951171875 0 676.117
386.1561584472656 0 1851.3657
392.26629638671875 0 1363.813
394.1510009765625 0 3468.917
395.1553955078125 0 807.11945
396.2591857910156 0 650.79767
397.2464904785156 0 693.6903
402.22406005859375 0 1362.5853
403.1835021972656 0 2982.702
403.26861572265625 0 739.0194
404.186767578125 0 958.2244
406.1847839355469 0 1451.1583
406.2511291503906 0 1243.962
409.2463684082031 0 590.9656 y 3
410.2757873535156 0 2554.244
411.1774597167969 0 5395.695 b Water loss 5
411.2776184082031 0 848.68335
412.1831359863281 0 1521.4767
412.2736511230469 0 14099.895 y 6
412.77532958984375 0 6299.4795
413.16375732421875 0 682.45447
413.27703857421875 0 1644.0916
415.2569580078125 0 954.4566
420.1822814941406 0 968.4781 b 5
420.2355041503906 0 1283.0042
420.6846008300781 0 953.05707
422.27734375 0 1094.3881
423.1885681152344 0 856.9694
423.27349853515625 0 832.16
429.1881103515625 0 14883.868
430.1917419433594 0 4241.2188
431.1791076660156 0 2131.8042
437.18170166015625 0 2629.2568 b 2
438.18670654296875 0 1171.0219
438.5582580566406 0 777.5782
451.22418212890625 0 661.7221
454.2102355957031 0 1204.1335
455.25250244140625 0 1219.1667
458.1286315917969 0 634.70935
463.5976867675781 0 2737.7024 y 2
463.932373046875 0 2981.251
464.26416015625 0 674.5265
467.297119140625 0 1260.0952
470.2467041015625 0 640.2721
470.3337707519531 0 10726.274 y 10
471.33648681640625 0 3154.7249
471.7976989746094 0 1222.1412
472.3017883300781 0 1167.724
473.24285888671875 0 3204.468 b 11
473.2686767578125 0 499.56042
473.57696533203125 0 3566.4033
473.9107971191406 0 2020.2887
474.77911376953125 0 764.55084
475.2635803222656 0 1007.1006
476.2520751953125 0 1555.0499
476.5850524902344 0 1972.7225
478.2068786621094 0 845.45233
480.7434997558594 0 1109.8237
480.8031005859375 0 2789.5635 y 5
481.3036804199219 0 1594.4491
481.740478515625 0 758.77856
481.92218017578125 0 595.6412
482.2627258300781 0 1182.7651
482.308349609375 0 847.1832
482.5945129394531 0 749.0783
483.7839050292969 0 1731.2708
484.2850646972656 0 852.0683
491.27105712890625 0 1102.0083
494.2352600097656 0 1589.3092
496.2631530761719 0 705.38605
497.22088623046875 0 847.08826 b 7
501.60455322265625 0 2673.1636
501.938720703125 0 4605.5513
502.2739562988281 0 1364.9971
503.2151794433594 0 975.1929
504.9328918457031 0 3776.5032 b Water loss 12
505.2662353515625 0 5787.811 b Ammonia loss 12
505.59918212890625 0 1767.2864
505.93145751953125 0 847.3127
510.32769775390625 0 1667.4521
510.9366455078125 0 23270.533 b 12
511.2709655761719 0 21396.58
511.60546875 0 9167.946
511.9392395019531 0 3795.8457
512.2716064453125 0 778.8076
516.9404296875 0 3247.0627
517.2745361328125 0 3284.081
517.6088256835938 0 983.3868
519.2799072265625 0 812.6554
519.9490966796875 0 1146.7964 y Ammonia loss 1
520.2863159179688 0 1036.0314
520.3597412109375 0 1195.3586
520.6153564453125 0 654.2386
521.2257690429688 0 762.6624
525.6240844726562 0 4071.5854 y 1
525.9575805664062 0 3882.617
526.289794921875 0 723.6735
529.804443359375 0 1083.1884
530.3010864257812 0 676.6555
531.2069702148438 0 801.13245
536.950927734375 0 796.8195
538.251708984375 0 8751.609
538.36376953125 0 615.4489
538.8063354492188 0 856.6218
539.251953125 0 2240.5503
539.301513671875 0 3027.388
539.6322021484375 0 770.0262
539.8042602539062 0 1232.4548
540.83056640625 0 739.1257
541.3696899414062 0 2251.4932 y 9
547.8139038085938 0 838.99084
548.3019409179688 0 630.7059
548.630859375 0 4531.4546 Precursor Water loss
548.9649658203125 0 6202.9385 Precursor Ammonia loss
549.2966918945312 0 2499.8247
549.33251953125 0 4547.588 y 4
549.6296997070312 0 2729.8613
549.8340454101562 0 2949.2434
549.9664916992188 0 727.468
550.3327026367188 0 1438.4136
553.7681274414062 0 705.43005
553.9224853515625 0 1506.579
553.9637451171875 0 1700.4199
554.3011474609375 0 1409.1136
554.6348266601562 0 125511.88 Precursor
554.9689331054688 0 130508.734
555.3029174804688 0 63717.316
555.6372680664062 0 23431.916
555.9718017578125 0 8440.456
561.2684326171875 0 1044.6356 b 8
564.2721557617188 0 779.5538
564.7769775390625 0 968.09875
566.2465209960938 0 10600.163
567.2483520507812 0 2620.8174
570.2454833984375 0 1276.6392
572.803466796875 0 4358.5776
573.3052978515625 0 3677.1365
573.803466796875 0 1099.5453
577.3826904296875 0 2256.4102
578.3858032226562 0 925.72876
579.2698974609375 0 682.5696
579.36083984375 0 6529.9634
580.3641357421875 0 2036.8302
587.3377075195312 0 796.0537
589.2734985351562 0 855.1545
595.39111328125 0 1755.859
596.7839965820312 0 1607.6133 b 9
604.35546875 0 2225.531
604.8493041992188 0 3522.494 y Ammonia loss 3
605.3504028320312 0 2762.045
610.3175048828125 0 1535.1512
611.322265625 0 994.7251
613.36083984375 0 3958.7505 y 3
613.8619384765625 0 2406.2715
615.3477172851562 0 2679.6445
615.8490600585938 0 1701.0425
616.3486938476562 0 612.818
620.3408813476562 0 3318.2156
620.8390502929688 0 3252.4253
621.3440551757812 0 1251.7764
628.3367309570312 0 840.7827
629.3451538085938 0 9973.145
629.8462524414062 0 9222.499
630.3485717773438 0 3055.198
630.8473510742188 0 839.5772
638.3505859375 0 797.41394
638.8538208007812 0 1668.2468
654.3242797851562 0 1000.7651
657.2699584960938 0 739.7023
661.9931640625 0 675.6156
662.6580810546875 0 728.7286
664.449951171875 0 1818.8282
665.84423828125 0 1034.7373
669.4664306640625 0 1886.8333 y 8
674.3045043945312 0 2359.2153
674.434326171875 0 8113.724
675.3093872070312 0 1098.162
675.4349365234375 0 3232.1736
677.3756713867188 0 1202.787
677.8755493164062 0 1094.534
680.3118896484375 0 1550.8983
685.8864135742188 0 2098.1968
686.3853149414062 0 3078.1401 y Ammonia loss 2
686.88916015625 0 1340.906
692.4445190429688 0 7755.4927
693.447509765625 0 3733.6187
694.8925170898438 0 32801.78 y 2
695.3938598632812 0 29553.262
695.8956298828125 0 15405.892
696.3969116210938 0 5563.578
702.2997436523438 0 4651.293 b 4
703.30224609375 0 1570.246
707.3067016601562 0 1730.4141
709.3585205078125 0 9786.165 b 11
709.8602905273438 0 6901.9746
710.361572265625 0 2709.2769
710.4602661132812 0 686.91113
713.8848266601562 0 794.61676
716.4175415039062 0 1674.8446
721.3560180664062 0 911.68646
722.38232421875 0 691.7794
724.331298828125 0 1150.9148
726.4855346679688 0 2405.4814 y 7
727.4866943359375 0 1080.702
733.3656616210938 0 640.3971
735.2979125976562 0 1029.3523
739.3590698242188 0 886.21313
752.3292846679688 0 703.49036
756.896240234375 0 1137.1782 b Water loss 12
765.901123046875 0 2907.6904 b 12
766.4036254882812 0 2057.0464
766.90283203125 0 1601.1783
774.9063110351562 0 1359.4594
775.4068603515625 0 1332.069
775.91552734375 0 758.2318
787.9332885742188 0 1025.1914 y 1
788.4344482421875 0 664.4599
795.3478393554688 0 681.9694
805.527099609375 0 1515.8519
806.5225830078125 0 758.3215 y Ammonia loss 6
809.4057006835938 0 944.25806
811.3616943359375 0 4823.59
811.489990234375 0 1004.5577
812.3638305664062 0 1455.8225
821.3512573242188 0 957.7071 b Water loss 5
823.5380859375 0 15968.044 y 6
824.5414428710938 0 8434.046
825.5454711914062 0 2305.4287
829.5049438476562 0 1189.7542
830.5021362304688 0 878.9549
839.3572387695312 0 10325.7295 b 5
840.360107421875 0 4333.1353
841.3648681640625 0 1103.4491
848.417236328125 0 2014.8975
849.4119262695312 0 701.95276
853.4806518554688 0 813.281
901.4391479492188 0 3671.2915
902.4422607421875 0 1958.8499
919.4520263671875 0 1470.5963
942.5825805664062 0 1526.7274
943.588134765625 0 1241.3785 y Ammonia loss 5
960.5965576171875 0 4020.707 y 5
961.5999145507812 0 1594.6741
980.1852416992188 0 682.9431
1097.6556396484375 0 1038.9814 y 4
1098.662353515625 0 883.8156
1144.5982666015625 0 761.7214
1174.5557861328125 0 1483.2632 b Water loss 9
1175.54931640625 0 921.9754 b Ammonia loss 9
1684.87744140625 0 636.9735
2271.670654296875 0 717.2212
2437.986328125 0 809.1443

Spectrum Details

|  |  |
| --- | --- |
| Matched peaks? Matched peaks The total absolute number of peaks matched. Additionally in brackets the total fraction of peaks matched and the total number of peaks is shown. | 55 (10.74% of 512) |
| FDR? FDR The false discovery rate estimated for this peptide. It is calculated by matching all theoretical fragments with a non-integer shift with the raw peaks for this spectrum. This is done with 40 different shifts. The resulting percentage is the average number of annotated peaks over the number of annotated peaks with the correct spectrum. | 1.90% |
| Satellite FDR? Satellite FDR See the FDR for details on its calculation. This satellite ion specific FDR only contains the satellite ions (d/w) for I/L/J positions. | - |
| PSM Score? PSM Score The PSM Score as given by Hecklib to this annotated spectrum. It is shown with three significant figures. | 271 |

#### Spectrum 5608? Spectrum 5608 The raw spectrum of this peptide as annotated by Hecklib. The fragments are coloured according to ion type (see legend). Any peaks with a star '\*' as text can be hovered over to see the full details, first the ion type second the mass shift type. By hovering over the amino acids in the peptide or ions in the legend the corresponding peaks are highlighted. By toggling the 'Unassigned' label you can turn the background (unassigned) peaks on or off in the plot. By updating the slider in the Ion legend you can update the spectrum to only show the top X% of the peaks with labels. The top X% means any peak that is within X% of the highest intensity. By dragging in the spectrum you can zoom in to a specific part of the spectrum and use 'Zoom Out' to get back to the original zoom level. The annotation of the spectrum is based on the given sequence in the peptides file and is done with different software so inconsistencies are likely. The peaks are annotated based on the given sequence, with 20 ppm tolerance.

Copy Data

##### Spectrum 5608 (TSV)

###### Preview

```
Loading example...
```

*Click on the button to copy the data to your clipboard.*

Mz MinMz MaxIntensity Max

WidthHeightPeptide font sizePeptide stroke widthSpectrum font sizeSpectrum stroke widthCompact peptide

Ion legend

wxyz

abcd

OtherUnassignedIon^Charge^_Position_Show for top:%

SWYQHHPGKAPKJJ

05.79e+41.16e+51.74e+52.31e+5

Zoom Out

y^+1^_1_c^+3^_4_y^+1^_2_z^+1^_3_y^+1^_3_y^+2^_8_w^+1^_4_c^+1^_3_y^+1^_4_y^+2^_9_c^+3^_12_y^+2^_9_c^+2^_8_c^+3^_13_z^+1^_5_y^+3^_13_y^+1^_5_c^+1^_4_c^+1^_4_w^+2^_11_c^+1^_4_y^+2^_11_z^+2^_11_y^+2^_11_z^+1^_6_c^+2^_11_y^+2^_12_z^+2^_12_z^+1^_7_y^+2^_12_c^+1^_5_c^+2^_12_z^+1^_7_c^+2^_12_c^+1^_5_y^+1^_7_c^+2^_13_c^+2^_13_z^+2^_13_w^+1^_8_y^+2^_13_c^+1^_6_z^+1^_9_c^+1^_7_y^+1^_9_c^+1^_8_y^+1^_10_z^+1^_10_y^+1^_10_c^+1^_9_z^+1^_11_y^+1^_11_c^+1^_11_z^+1^_12_y^+1^_12_c^+1^_12_c^+1^_13_z^+1^_13_

0882176426463528

Fragment Matches Table

Show background peaks

| Position | Ion type | Intensity | mz Theoretical | mz Error (Th) | mz Error (ppm) | Charge | Series Number |
| --- | --- | --- | --- | --- | --- | --- | --- |
| - | - | 332.2 | 123.5 | - | - | 0 | - |
| - | - | 2205 | 129.1 | - | - | 0 | - |
| 14 | y | 1042 | 132.1 | 0.0004661 | 3.528 | +1 | 1 |
| - | - | 388.6 | 136.1 | - | - | 0 | - |
| - | - | 767.9 | 146.1 | - | - | 0 | - |
| - | - | 516.4 | 148.8 | - | - | 0 | - |
| - | - | 523.4 | 148.8 | - | - | 0 | - |
| - | - | 532.9 | 148.9 | - | - | 0 | - |
| - | - | 588.5 | 148.9 | - | - | 0 | - |
| - | - | 562.6 | 148.9 | - | - | 0 | - |
| - | - | 855.9 | 148.9 | - | - | 0 | - |
| - | - | 1132 | 148.9 | - | - | 0 | - |
| - | - | 1155 | 148.9 | - | - | 0 | - |
| - | - | 2319 | 148.9 | - | - | 0 | - |
| - | - | 3949 | 148.9 | - | - | 0 | - |
| - | - | 3613 | 149 | - | - | 0 | - |
| - | - | 1734 | 149 | - | - | 0 | - |
| - | - | 1063 | 149 | - | - | 0 | - |
| - | - | 769.1 | 149 | - | - | 0 | - |
| - | - | 803.8 | 149 | - | - | 0 | - |
| - | - | 428.4 | 149 | - | - | 0 | - |
| - | - | 658 | 149 | - | - | 0 | - |
| - | - | 677.8 | 149 | - | - | 0 | - |
| - | - | 606.1 | 149 | - | - | 0 | - |
| - | - | 598 | 149 | - | - | 0 | - |
| - | - | 2153 | 153.1 | - | - | 0 | - |
| - | - | 422.4 | 160.8 | - | - | 0 | - |
| - | - | 698.5 | 173.4 | - | - | 0 | - |
| - | - | 896.7 | 175.1 | - | - | 0 | - |
| - | - | 438.5 | 178.1 | - | - | 0 | - |
| - | - | 461 | 184.2 | - | - | 0 | - |
| - | - | 816.2 | 186.1 | - | - | 0 | - |
| - | - | 1087 | 187.1 | - | - | 0 | - |
| 4 | c | 516 | 189.1 | 0.002621 | 13.86 | +3 | 4 |
| - | - | 924 | 215.1 | - | - | 0 | - |
| - | - | 986 | 223.1 | - | - | 0 | - |
| - | - | 852.3 | 226.2 | - | - | 0 | - |
| - | - | 1.648E+04 | 229.2 | - | - | 0 | - |
| - | - | 1815 | 230.2 | - | - | 0 | - |
| - | - | 1223 | 243.1 | - | - | 0 | - |
| - | - | 2236 | 245.1 | - | - | 0 | - |
| 13 | y | 3422 | 245.2 | 0.0005849 | 2.385 | +1 | 2 |
| - | - | 5785 | 246.1 | - | - | 0 | - |
| - | - | 775.3 | 246.2 | - | - | 0 | - |
| - | - | 1102 | 247.1 | - | - | 0 | - |
| - | - | 605.5 | 255.2 | - | - | 0 | - |
| - | - | 891.7 | 259.2 | - | - | 0 | - |
| - | - | 1123 | 260.2 | - | - | 0 | - |
| - | - | 1233 | 262.1 | - | - | 0 | - |
| - | - | 504.8 | 273.1 | - | - | 0 | - |
| - | - | 8200 | 274.1 | - | - | 0 | - |
| - | - | 1811 | 275.1 | - | - | 0 | - |
| - | - | 995.6 | 279.1 | - | - | 0 | - |
| - | - | 1605 | 281.2 | - | - | 0 | - |
| - | - | 1586 | 282.2 | - | - | 0 | - |
| - | - | 1846 | 292.1 | - | - | 0 | - |
| - | - | 974.2 | 300.2 | - | - | 0 | - |
| - | - | 1380 | 301.2 | - | - | 0 | - |
| - | - | 1523 | 309.2 | - | - | 0 | - |
| - | - | 549.4 | 310.7 | - | - | 0 | - |
| - | - | 1025 | 314.2 | - | - | 0 | - |
| - | - | 1115 | 314.2 | - | - | 0 | - |
| - | - | 3168 | 331.2 | - | - | 0 | - |
| - | - | 593.2 | 341 | - | - | 0 | - |
| - | - | 1267 | 346.7 | - | - | 0 | - |
| - | - | 1032 | 346.7 | - | - | 0 | - |
| - | - | 796.1 | 349.1 | - | - | 0 | - |
| - | - | 1176 | 355.1 | - | - | 0 | - |
| - | - | 574.9 | 357.2 | - | - | 0 | - |
| 12 | z | 3375 | 357.3 | 0.0003042 | 0.8514 | +1 | 3 |
| - | - | 1452 | 358.3 | - | - | 0 | - |
| - | - | 1555 | 363.2 | - | - | 0 | - |
| - | - | 1874 | 371.1 | - | - | 0 | - |
| - | - | 526.9 | 372.1 | - | - | 0 | - |
| 12 | y | 1197 | 373.3 | 0.0007756 | 2.078 | +1 | 3 |
| - | - | 1049 | 382.3 | - | - | 0 | - |
| - | - | 580.2 | 385.5 | - | - | 0 | - |
| - | - | 590.3 | 389.8 | - | - | 0 | - |
| - | - | 1431 | 394.3 | - | - | 0 | - |
| - | - | 2628 | 397.2 | - | - | 0 | - |
| - | - | 551.4 | 411.2 | - | - | 0 | - |
| 7 | y | 5490 | 412.3 | 0.0007757 | 1.881 | +2 | 8 |
| - | - | 2737 | 412.8 | - | - | 0 | - |
| - | - | 1327 | 413.2 | - | - | 0 | - |
| - | - | 997.5 | 413.3 | - | - | 0 | - |
| - | - | 599.4 | 413.8 | - | - | 0 | - |
| - | - | 1001 | 423.2 | - | - | 0 | - |
| - | - | 1363 | 424.2 | - | - | 0 | - |
| - | - | 3698 | 427.3 | - | - | 0 | - |
| 11 | w | 1767 | 427.3 | 0.001167 | 2.731 | +1 | 4 |
| - | - | 958.9 | 428.3 | - | - | 0 | - |
| - | - | 777.1 | 428.3 | - | - | 0 | - |
| - | - | 755.7 | 435.2 | - | - | 0 | - |
| - | - | 1856 | 437.2 | - | - | 0 | - |
| - | - | 606.8 | 439.2 | - | - | 0 | - |
| - | - | 598.6 | 441.2 | - | - | 0 | - |
| - | - | 568.4 | 442.2 | - | - | 0 | - |
| - | - | 641.5 | 443.2 | - | - | 0 | - |
| - | - | 673.3 | 451.2 | - | - | 0 | - |
| - | - | 1420 | 453.2 | - | - | 0 | - |
| 3 | c | 3626 | 454.2 | 0.0004279 | 0.942 | +1 | 3 |
| - | - | 1083 | 455.2 | - | - | 0 | - |
| - | - | 815 | 463.9 | - | - | 0 | - |
| - | - | 1181 | 465.2 | - | - | 0 | - |
| - | - | 4567 | 468.3 | - | - | 0 | - |
| - | - | 1316 | 469.3 | - | - | 0 | - |
| - | - | 1361 | 469.3 | - | - | 0 | - |
| - | - | 747.4 | 470.3 | - | - | 0 | - |
| 11 | y | 2010 | 470.3 | 1.677E-05 | 0.03566 | +1 | 4 |
| 6 | y | 696.9 | 472.3 | 0.009039 | 19.14 | +2 | 9 |
| 12 | c | 1080 | 473.2 | 0.0009145 | 1.932 | +3 | 12 |
| - | - | 815.8 | 473.6 | - | - | 0 | - |
| - | - | 1518 | 473.9 | - | - | 0 | - |
| - | - | 1627 | 477.2 | - | - | 0 | - |
| - | - | 699.5 | 480.2 | - | - | 0 | - |
| 6 | y | 1000 | 480.8 | 0.0007567 | 1.574 | +2 | 9 |
| - | - | 1066 | 482.3 | - | - | 0 | - |
| 8 | c | 789.5 | 497.2 | 0.0001975 | 0.3972 | +2 | 8 |
| - | - | 1064 | 501.6 | - | - | 0 | - |
| - | - | 2096 | 504.9 | - | - | 0 | - |
| - | - | 1174 | 505.3 | - | - | 0 | - |
| - | - | 815.1 | 505.6 | - | - | 0 | - |
| - | - | 689.4 | 510.2 | - | - | 0 | - |
| - | - | 1.41E+04 | 510.3 | - | - | 0 | - |
| 13 | c | 9369 | 510.9 | 0.0007761 | 1.519 | +3 | 13 |
| - | - | 8186 | 511.3 | - | - | 0 | - |
| - | - | 4544 | 511.3 | - | - | 0 | - |
| - | - | 5631 | 511.6 | - | - | 0 | - |
| - | - | 1438 | 511.9 | - | - | 0 | - |
| - | - | 1117 | 516.9 | - | - | 0 | - |
| - | - | 1438 | 517.3 | - | - | 0 | - |
| - | - | 1444 | 523.2 | - | - | 0 | - |
| 10 | z | 1.537E+04 | 525.4 | 0.0007585 | 1.444 | +1 | 5 |
| 2 | y | 1120 | 525.6 | 0.0001018 | 0.1937 | +3 | 13 |
| - | - | 1384 | 526 | - | - | 0 | - |
| - | - | 5300 | 526.4 | - | - | 0 | - |
| - | - | 936.6 | 527.4 | - | - | 0 | - |
| - | - | 1600 | 528.3 | - | - | 0 | - |
| - | - | 1099 | 538.3 | - | - | 0 | - |
| - | - | 1472 | 538.3 | - | - | 0 | - |
| - | - | 1022 | 539.3 | - | - | 0 | - |
| - | - | 864.6 | 539.3 | - | - | 0 | - |
| - | - | 786.2 | 541.3 | - | - | 0 | - |
| 10 | y | 1007 | 541.4 | 0.00474 | 8.755 | +1 | 5 |
| - | - | 810.3 | 541.8 | - | - | 0 | - |
| - | - | 751.8 | 547.8 | - | - | 0 | - |
| - | - | 2647 | 548.6 | - | - | 0 | - |
| - | - | 3749 | 549 | - | - | 0 | - |
| - | - | 3379 | 549.3 | - | - | 0 | - |
| - | - | 1107 | 549.8 | - | - | 0 | - |
| - | - | 1091 | 553.3 | - | - | 0 | - |
| - | - | 768.8 | 553.8 | - | - | 0 | - |
| - | - | 977 | 554.3 | - | - | 0 | - |
| - | - | 6.737E+04 | 554.6 | - | - | 0 | - |
| - | - | 6.472E+04 | 555 | - | - | 0 | - |
| - | - | 3.742E+04 | 555.3 | - | - | 0 | - |
| - | - | 1.082E+04 | 555.6 | - | - | 0 | - |
| - | - | 776.1 | 555.9 | - | - | 0 | - |
| - | - | 4170 | 556 | - | - | 0 | - |
| - | - | 2.617E+04 | 556.3 | - | - | 0 | - |
| - | - | 5968 | 563.2 | - | - | 0 | - |
| 4 | c | 2294 | 564.3 | 0.00541 | 9.587 | +1 | 4 |
| 4 | c | 2388 | 565.2 | 0.0003819 | 0.6756 | +1 | 4 |
| - | - | 2072 | 566.2 | - | - | 0 | - |
| - | - | 846.9 | 566.4 | - | - | 0 | - |
| 4 | w | 3588 | 576.3 | 0.0005463 | 0.9479 | +2 | 11 |
| - | - | 3002 | 576.8 | - | - | 0 | - |
| - | - | 1173 | 577.3 | - | - | 0 | - |
| - | - | 804.4 | 579.4 | - | - | 0 | - |
| - | - | 2982 | 581.3 | - | - | 0 | - |
| 4 | c | 1.75E+04 | 582.3 | 0.0003831 | 0.6579 | +1 | 4 |
| - | - | 1727 | 582.4 | - | - | 0 | - |
| - | - | 5614 | 583.3 | - | - | 0 | - |
| - | - | 1200 | 584.3 | - | - | 0 | - |
| - | - | 7168 | 595.4 | - | - | 0 | - |
| - | - | 2441 | 596.4 | - | - | 0 | - |
| - | - | 748 | 597.4 | - | - | 0 | - |
| - | - | 758.8 | 604.4 | - | - | 0 | - |
| 4 | y | 2181 | 604.8 | 0.00251 | 4.149 | +2 | 11 |
| 4 | z | 1.382E+04 | 605.4 | 0.0003062 | 0.5058 | +2 | 11 |
| - | - | 1.265E+04 | 605.9 | - | - | 0 | - |
| - | - | 6832 | 606.4 | - | - | 0 | - |
| - | - | 3040 | 606.9 | - | - | 0 | - |
| - | - | 709.1 | 609.5 | - | - | 0 | - |
| - | - | 3549 | 610.4 | - | - | 0 | - |
| - | - | 1176 | 612.9 | - | - | 0 | - |
| 4 | y | 4632 | 613.4 | 0.0009539 | 1.555 | +2 | 11 |
| - | - | 2213 | 613.9 | - | - | 0 | - |
| - | - | 1049 | 614.4 | - | - | 0 | - |
| - | - | 1343 | 624.4 | - | - | 0 | - |
| - | - | 1837 | 629.3 | - | - | 0 | - |
| - | - | 1252 | 629.8 | - | - | 0 | - |
| - | - | 1416 | 651.4 | - | - | 0 | - |
| - | - | 1526 | 651.9 | - | - | 0 | - |
| - | - | 1380 | 652.3 | - | - | 0 | - |
| 9 | z | 1.358E+04 | 653.4 | 0.0007662 | 1.173 | +1 | 6 |
| 11 | c | 1313 | 653.8 | 0.001284 | 1.963 | +2 | 11 |
| - | - | 1012 | 654.3 | - | - | 0 | - |
| - | - | 6723 | 654.5 | - | - | 0 | - |
| - | - | 876.8 | 655.5 | - | - | 0 | - |
| - | - | 1101 | 657.9 | - | - | 0 | - |
| - | - | 1493 | 658.4 | - | - | 0 | - |
| - | - | 1118 | 659.3 | - | - | 0 | - |
| - | - | 1020 | 660.3 | - | - | 0 | - |
| - | - | 809 | 665.4 | - | - | 0 | - |
| - | - | 1218 | 674.3 | - | - | 0 | - |
| - | - | 1056 | 675.3 | - | - | 0 | - |
| - | - | 1542 | 685.9 | - | - | 0 | - |
| 3 | y | 1183 | 686.4 | 0.007589 | 11.06 | +2 | 12 |
| 3 | z | 1.717E+04 | 686.9 | 0.0005023 | 0.7313 | +2 | 12 |
| - | - | 1.321E+04 | 687.4 | - | - | 0 | - |
| - | - | 7430 | 687.9 | - | - | 0 | - |
| - | - | 1956 | 688.4 | - | - | 0 | - |
| 8 | z | 1080 | 693.4 | 0.0007247 | 1.045 | +1 | 7 |
| 3 | y | 1.524E+04 | 694.9 | 0.0007228 | 1.04 | +2 | 12 |
| - | - | 1.135E+04 | 695.4 | - | - | 0 | - |
| - | - | 5622 | 695.9 | - | - | 0 | - |
| - | - | 1848 | 696.4 | - | - | 0 | - |
| 5 | c | 906.7 | 702.3 | 0.0001804 | 0.2568 | +1 | 5 |
| 12 | c | 4989 | 709.4 | 0.002172 | 3.062 | +2 | 12 |
| - | - | 4394 | 709.4 | - | - | 0 | - |
| - | - | 3817 | 709.9 | - | - | 0 | - |
| - | - | 1406 | 710.4 | - | - | 0 | - |
| 8 | z | 2334 | 710.5 | 0.005317 | 7.483 | +1 | 7 |
| - | - | 856.9 | 711.5 | - | - | 0 | - |
| - | - | 852.1 | 712.5 | - | - | 0 | - |
| - | - | 681.3 | 714.9 | - | - | 0 | - |
| - | - | 809.8 | 717.4 | - | - | 0 | - |
| 12 | c | 4.246E+04 | 717.9 | 0.0003109 | 0.4331 | +2 | 12 |
| - | - | 3.553E+04 | 718.4 | - | - | 0 | - |
| - | - | 1.542E+04 | 718.9 | - | - | 0 | - |
| 5 | c | 3.832E+04 | 719.3 | 0.0003091 | 0.4297 | +1 | 5 |
| - | - | 1260 | 719.9 | - | - | 0 | - |
| - | - | 1.943E+04 | 720.3 | - | - | 0 | - |
| - | - | 4200 | 721.3 | - | - | 0 | - |
| - | - | 842.7 | 722.3 | - | - | 0 | - |
| - | - | 684.3 | 723.3 | - | - | 0 | - |
| 8 | y | 1085 | 726.5 | 0.001472 | 2.026 | +1 | 7 |
| - | - | 804.8 | 731.4 | - | - | 0 | - |
| - | - | 911.8 | 736.4 | - | - | 0 | - |
| - | - | 1022 | 736.9 | - | - | 0 | - |
| - | - | 729.9 | 748.3 | - | - | 0 | - |
| - | - | 3983 | 751.4 | - | - | 0 | - |
| - | - | 725.2 | 751.9 | - | - | 0 | - |
| - | - | 1252 | 752.4 | - | - | 0 | - |
| - | - | 677.3 | 752.9 | - | - | 0 | - |
| - | - | 1178 | 757.4 | - | - | 0 | - |
| - | - | 3162 | 758.9 | - | - | 0 | - |
| - | - | 1524 | 759.4 | - | - | 0 | - |
| - | - | 795.6 | 759.9 | - | - | 0 | - |
| - | - | 983.6 | 765.4 | - | - | 0 | - |
| 13 | c | 1662 | 765.9 | 0.00201 | 2.624 | +2 | 13 |
| - | - | 1731 | 766.4 | - | - | 0 | - |
| - | - | 2768 | 766.9 | - | - | 0 | - |
| - | - | 776.5 | 767.4 | - | - | 0 | - |
| - | - | 873.7 | 773.4 | - | - | 0 | - |
| - | - | 999.7 | 773.9 | - | - | 0 | - |
| 13 | c | 2.856E+04 | 774.4 | 0.0006373 | 0.8229 | +2 | 13 |
| - | - | 2.466E+04 | 774.9 | - | - | 0 | - |
| - | - | 1.118E+04 | 775.4 | - | - | 0 | - |
| - | - | 6105 | 775.9 | - | - | 0 | - |
| - | - | 1174 | 776.4 | - | - | 0 | - |
| - | - | 667.7 | 778.4 | - | - | 0 | - |
| 2 | z | 5092 | 779.9 | 0.0003966 | 0.5085 | +2 | 13 |
| - | - | 4785 | 780.4 | - | - | 0 | - |
| 7 | w | 1977 | 780.5 | 0.008791 | 11.26 | +1 | 8 |
| - | - | 2491 | 780.9 | - | - | 0 | - |
| - | - | 814 | 781.4 | - | - | 0 | - |
| - | - | 1567 | 782.4 | - | - | 0 | - |
| 2 | y | 1709 | 787.9 | 0.01336 | 16.96 | +2 | 13 |
| - | - | 1708 | 788.4 | - | - | 0 | - |
| - | - | 908 | 788.9 | - | - | 0 | - |
| - | - | 912 | 789.4 | - | - | 0 | - |
| - | - | 4041 | 790.9 | - | - | 0 | - |
| - | - | 3477 | 791.4 | - | - | 0 | - |
| - | - | 2623 | 791.9 | - | - | 0 | - |
| - | - | 799.1 | 792.4 | - | - | 0 | - |
| - | - | 1513 | 795.4 | - | - | 0 | - |
| - | - | 2479 | 795.9 | - | - | 0 | - |
| - | - | 1393 | 796.3 | - | - | 0 | - |
| - | - | 3111 | 796.4 | - | - | 0 | - |
| - | - | 2567 | 796.9 | - | - | 0 | - |
| - | - | 981.2 | 797.4 | - | - | 0 | - |
| - | - | 2179 | 800.4 | - | - | 0 | - |
| - | - | 2281 | 800.9 | - | - | 0 | - |
| - | - | 2359 | 801.4 | - | - | 0 | - |
| - | - | 1648 | 801.9 | - | - | 0 | - |
| - | - | 701.6 | 802.4 | - | - | 0 | - |
| - | - | 1685 | 802.9 | - | - | 0 | - |
| - | - | 796.3 | 803.4 | - | - | 0 | - |
| - | - | 1425 | 803.9 | - | - | 0 | - |
| - | - | 1399 | 804.4 | - | - | 0 | - |
| - | - | 887.1 | 804.9 | - | - | 0 | - |
| - | - | 825.8 | 805.4 | - | - | 0 | - |
| - | - | 2742 | 807.5 | - | - | 0 | - |
| - | - | 1044 | 808.5 | - | - | 0 | - |
| - | - | 872.6 | 808.9 | - | - | 0 | - |
| - | - | 4961 | 809.4 | - | - | 0 | - |
| - | - | 4760 | 809.9 | - | - | 0 | - |
| - | - | 4166 | 810.4 | - | - | 0 | - |
| - | - | 2095 | 810.9 | - | - | 0 | - |
| - | - | 3117 | 811.4 | - | - | 0 | - |
| - | - | 910 | 811.4 | - | - | 0 | - |
| - | - | 2439 | 812.4 | - | - | 0 | - |
| - | - | 715.5 | 813.4 | - | - | 0 | - |
| - | - | 872.3 | 814.4 | - | - | 0 | - |
| - | - | 949.9 | 815.4 | - | - | 0 | - |
| - | - | 793.3 | 822.5 | - | - | 0 | - |
| - | - | 3077 | 822.9 | - | - | 0 | - |
| - | - | 1.962E+04 | 823.4 | - | - | 0 | - |
| - | - | 6342 | 823.5 | - | - | 0 | - |
| - | - | 1.913E+04 | 823.9 | - | - | 0 | - |
| - | - | 8589 | 824.4 | - | - | 0 | - |
| - | - | 9075 | 824.5 | - | - | 0 | - |
| - | - | 5209 | 824.9 | - | - | 0 | - |
| - | - | 1160 | 825.4 | - | - | 0 | - |
| - | - | 3499 | 825.5 | - | - | 0 | - |
| - | - | 748.3 | 826.5 | - | - | 0 | - |
| - | - | 2804 | 830.9 | - | - | 0 | - |
| - | - | 4.727E+04 | 831.5 | - | - | 0 | - |
| - | - | 2.292E+05 | 832 | - | - | 0 | - |
| - | - | 2.056E+05 | 832.5 | - | - | 0 | - |
| - | - | 1222 | 832.6 | - | - | 0 | - |
| - | - | 1.057E+05 | 833 | - | - | 0 | - |
| - | - | 3.845E+04 | 833.5 | - | - | 0 | - |
| - | - | 1.252E+04 | 834 | - | - | 0 | - |
| - | - | 2990 | 834.5 | - | - | 0 | - |
| 6 | c | 3488 | 839.4 | 0.0007833 | 0.9332 | +1 | 6 |
| - | - | 1801 | 840.4 | - | - | 0 | - |
| - | - | 2162 | 850.5 | - | - | 0 | - |
| - | - | 1458 | 851.5 | - | - | 0 | - |
| - | - | 2228 | 852.5 | - | - | 0 | - |
| - | - | 960.9 | 853.5 | - | - | 0 | - |
| - | - | 1929 | 855.5 | - | - | 0 | - |
| - | - | 1095 | 856.5 | - | - | 0 | - |
| - | - | 810.2 | 909.4 | - | - | 0 | - |
| 6 | z | 3.977E+04 | 944.6 | 0.0002562 | 0.2712 | +1 | 9 |
| - | - | 2.195E+04 | 945.6 | - | - | 0 | - |
| - | - | 7088 | 946.6 | - | - | 0 | - |
| - | - | 1718 | 947.6 | - | - | 0 | - |
| 7 | c | 4675 | 953.4 | 0.0007709 | 0.8085 | +1 | 7 |
| - | - | 2697 | 954.4 | - | - | 0 | - |
| 6 | y | 2360 | 960.6 | 0.002233 | 2.324 | +1 | 9 |
| - | - | 1881 | 961.6 | - | - | 0 | - |
| - | - | 890.7 | 962.6 | - | - | 0 | - |
| - | - | 1011 | 968.6 | - | - | 0 | - |
| - | - | 1826 | 982.5 | - | - | 0 | - |
| - | - | 2079 | 983.6 | - | - | 0 | - |
| - | - | 983 | 984.6 | - | - | 0 | - |
| - | - | 993.8 | 1009 | - | - | 0 | - |
| 8 | c | 3.462E+04 | 1010 | 0.0001399 | 0.1384 | +1 | 8 |
| - | - | 1.907E+04 | 1011 | - | - | 0 | - |
| - | - | 6968 | 1012 | - | - | 0 | - |
| - | - | 1245 | 1013 | - | - | 0 | - |
| - | - | 1213 | 1066 | - | - | 0 | - |
| 5 | y | 838.1 | 1081 | 0.00227 | 2.101 | +1 | 10 |
| - | - | 954.9 | 1082 | - | - | 0 | - |
| 5 | z | 1.884E+04 | 1082 | 0.0005484 | 0.507 | +1 | 10 |
| - | - | 4.36E+04 | 1083 | - | - | 0 | - |
| - | - | 2.53E+04 | 1084 | - | - | 0 | - |
| - | - | 8105 | 1085 | - | - | 0 | - |
| - | - | 1774 | 1086 | - | - | 0 | - |
| - | - | 839.9 | 1092 | - | - | 0 | - |
| - | - | 645.3 | 1092 | - | - | 0 | - |
| - | - | 4621 | 1094 | - | - | 0 | - |
| - | - | 4083 | 1095 | - | - | 0 | - |
| - | - | 1044 | 1096 | - | - | 0 | - |
| - | - | 1511 | 1097 | - | - | 0 | - |
| - | - | 817 | 1098 | - | - | 0 | - |
| 5 | y | 4760 | 1098 | 0.001574 | 1.434 | +1 | 10 |
| - | - | 2875 | 1099 | - | - | 0 | - |
| - | - | 838.3 | 1099 | - | - | 0 | - |
| - | - | 1447 | 1100 | - | - | 0 | - |
| - | - | 729.5 | 1102 | - | - | 0 | - |
| - | - | 703.7 | 1103 | - | - | 0 | - |
| - | - | 922.8 | 1111 | - | - | 0 | - |
| - | - | 2150 | 1112 | - | - | 0 | - |
| - | - | 804.2 | 1113 | - | - | 0 | - |
| - | - | 660.7 | 1138 | - | - | 0 | - |
| 9 | c | 4.539E+04 | 1139 | 0.0006002 | 0.5272 | +1 | 9 |
| - | - | 2.974E+04 | 1140 | - | - | 0 | - |
| - | - | 1.046E+04 | 1141 | - | - | 0 | - |
| - | - | 3100 | 1142 | - | - | 0 | - |
| - | - | 825 | 1146 | - | - | 0 | - |
| - | - | 1199 | 1166 | - | - | 0 | - |
| - | - | 998.3 | 1167 | - | - | 0 | - |
| 4 | z | 6464 | 1210 | 0.001877 | 1.551 | +1 | 11 |
| - | - | 2.465E+04 | 1211 | - | - | 0 | - |
| - | - | 1.769E+04 | 1212 | - | - | 0 | - |
| - | - | 6913 | 1213 | - | - | 0 | - |
| - | - | 1816 | 1214 | - | - | 0 | - |
| 4 | y | 1644 | 1226 | 0.001494 | 1.219 | +1 | 11 |
| - | - | 1557 | 1227 | - | - | 0 | - |
| - | - | 640.4 | 1237 | - | - | 0 | - |
| - | - | 1014 | 1259 | - | - | 0 | - |
| - | - | 979 | 1263 | - | - | 0 | - |
| - | - | 1418 | 1306 | - | - | 0 | - |
| 11 | c | 2.154E+04 | 1307 | 0.0005323 | 0.4074 | +1 | 11 |
| - | - | 1.525E+04 | 1308 | - | - | 0 | - |
| - | - | 6760 | 1309 | - | - | 0 | - |
| - | - | 1855 | 1310 | - | - | 0 | - |
| - | - | 738.7 | 1332 | - | - | 0 | - |
| 3 | z | 3826 | 1373 | 0.002461 | 1.793 | +1 | 12 |
| - | - | 1.315E+04 | 1374 | - | - | 0 | - |
| - | - | 1.034E+04 | 1375 | - | - | 0 | - |
| - | - | 3510 | 1376 | - | - | 0 | - |
| - | - | 1223 | 1377 | - | - | 0 | - |
| 3 | y | 1145 | 1389 | 0.003839 | 2.764 | +1 | 12 |
| - | - | 1308 | 1390 | - | - | 0 | - |
| - | - | 2281 | 1419 | - | - | 0 | - |
| - | - | 1859 | 1420 | - | - | 0 | - |
| 12 | c | 7656 | 1435 | 0.002234 | 1.557 | +1 | 12 |
| - | - | 1.017E+04 | 1436 | - | - | 0 | - |
| - | - | 5707 | 1437 | - | - | 0 | - |
| - | - | 2412 | 1438 | - | - | 0 | - |
| - | - | 994.6 | 1439 | - | - | 0 | - |
| - | - | 878.9 | 1505 | - | - | 0 | - |
| - | - | 1121 | 1533 | - | - | 0 | - |
| - | - | 820.8 | 1534 | - | - | 0 | - |
| 13 | c | 2757 | 1548 | 0.001093 | 0.7058 | +1 | 13 |
| - | - | 5502 | 1549 | - | - | 0 | - |
| - | - | 5047 | 1550 | - | - | 0 | - |
| - | - | 2742 | 1551 | - | - | 0 | - |
| 2 | z | 1894 | 1559 | 0.0003792 | 0.2433 | +1 | 13 |
| - | - | 4530 | 1560 | - | - | 0 | - |
| - | - | 4567 | 1561 | - | - | 0 | - |
| - | - | 1980 | 1562 | - | - | 0 | - |
| - | - | 893.4 | 1576 | - | - | 0 | - |
| - | - | 1346 | 1581 | - | - | 0 | - |
| - | - | 1231 | 1582 | - | - | 0 | - |
| - | - | 973.8 | 1591 | - | - | 0 | - |
| - | - | 1230 | 1592 | - | - | 0 | - |
| - | - | 1608 | 1618 | - | - | 0 | - |
| - | - | 6002 | 1619 | - | - | 0 | - |
| - | - | 5333 | 1620 | - | - | 0 | - |
| - | - | 2877 | 1621 | - | - | 0 | - |
| - | - | 1235 | 1622 | - | - | 0 | - |
| - | - | 3600 | 1629 | - | - | 0 | - |
| - | - | 2512 | 1630 | - | - | 0 | - |
| - | - | 1516 | 1631 | - | - | 0 | - |
| - | - | 2986 | 1636 | - | - | 0 | - |
| - | - | 1803 | 1637 | - | - | 0 | - |
| - | - | 5460 | 1646 | - | - | 0 | - |
| - | - | 2.532E+04 | 1647 | - | - | 0 | - |
| - | - | 2.221E+04 | 1648 | - | - | 0 | - |
| - | - | 1.08E+04 | 1649 | - | - | 0 | - |
| - | - | 4374 | 1650 | - | - | 0 | - |
| - | - | 948.6 | 1651 | - | - | 0 | - |
| - | - | 855.4 | 1661 | - | - | 0 | - |
| - | - | 6306 | 1662 | - | - | 0 | - |
| - | - | 2.701E+04 | 1663 | - | - | 0 | - |
| - | - | 1.236E+05 | 1664 | - | - | 0 | - |
| - | - | 1.056E+05 | 1665 | - | - | 0 | - |
| - | - | 5.479E+04 | 1666 | - | - | 0 | - |
| - | - | 1.996E+04 | 1667 | - | - | 0 | - |
| - | - | 6349 | 1668 | - | - | 0 | - |
| - | - | 1879 | 1669 | - | - | 0 | - |
| - | - | 775.3 | 2521 | - | - | 0 | - |
| - | - | 801.6 | 3219 | - | - | 0 | - |
| - | - | 699.6 | 3493 | - | - | 0 | - |

m/z Charge Intensity FragmentType MassShift Position
123.52214050292969 0 332.1582
129.1024169921875 0 2205.4543
132.1023712158203 0 1042.3175 y 13
136.07569885253906 0 388.56223
146.0929412841797 0 767.87994
148.8354949951172 0 516.41583
148.84226989746094 0 523.3579
148.89337158203125 0 532.93335
148.89964294433594 0 588.47046
148.90762329101562 0 562.6228
148.91461181640625 0 855.9345
148.92172241210938 0 1131.5883
148.92909240722656 0 1154.6843
148.9364013671875 0 2319.3184
148.94419860839844 0 3948.7036
148.96080017089844 0 3612.719
148.96868896484375 0 1733.8574
148.97581481933594 0 1062.9464
148.9832000732422 0 769.09375
148.990234375 0 803.8287
148.9970245361328 0 428.3853
149.0048065185547 0 658.0428
149.01217651367188 0 677.80817
149.01901245117188 0 606.1476
149.02671813964844 0 597.9797
153.0785369873047 0 2153.4258
160.8025665283203 0 422.37946
173.44178771972656 0 698.5333
175.108642578125 0 896.7428
178.08377075195312 0 438.49234
184.1870574951172 0 460.96872
186.07882690429688 0 816.20776
187.08694458007812 0 1086.9661
189.087646484375 0 515.9514 c Ammonia loss 3
215.13951110839844 0 924.02826
223.11932373046875 0 985.9802
226.15597534179688 0 852.3157
229.1549835205078 0 16481.596
230.1583251953125 0 1815.2279
243.14613342285156 0 1223.179
245.137451171875 0 2235.7288
245.18655395507812 0 3421.9304 y 12
246.12399291992188 0 5785.471
246.19056701660156 0 775.2927
247.1275634765625 0 1101.5834
255.1947021484375 0 605.50055
259.1636657714844 0 891.70056
260.1712951660156 0 1122.5958
262.1404113769531 0 1232.6583
273.1118469238281 0 504.81818
274.118896484375 0 8199.797
275.12249755859375 0 1811.2529
279.1346130371094 0 995.6044
281.17376708984375 0 1604.7888
282.1818542480469 0 1585.7332
292.1292724609375 0 1845.7422
300.1929931640625 0 974.2415
301.1999816894531 0 1380.4183
309.1556091308594 0 1522.5936
310.7340393066406 0 549.37115
314.1839904785156 0 1025.2443
314.2080078125 0 1114.9841
331.20916748046875 0 3167.8396
340.95391845703125 0 593.196
346.70745849609375 0 1267.1627
346.7279357910156 0 1031.894
349.1427307128906 0 796.09485
355.06982421875 0 1175.5825
357.17926025390625 0 574.94604
357.26251220703125 0 3374.933 z 11
358.26971435546875 0 1452.1476
363.18798828125 0 1555.3046
371.10162353515625 0 1874.0127
372.10162353515625 0 526.8543
373.2817077636719 0 1197.4507 y 11
382.28082275390625 0 1048.8922
385.5109558105469 0 580.2349
389.7866516113281 0 590.3134
394.25836181640625 0 1431.0774
397.2453308105469 0 2628.2195
411.23486328125 0 551.414
412.2744140625 0 5489.513 y 6
412.7760314941406 0 2736.597
413.22784423828125 0 1327.249
413.2767333984375 0 997.51984
413.848388671875 0 599.40686
423.2247619628906 0 1001.36945
424.2348937988281 0 1362.5984
427.2666015625 0 3698.077
427.29266357421875 0 1767.4122 w 10
428.2661437988281 0 958.9109
428.2972412109375 0 777.1167
435.1909484863281 0 755.74835
437.18194580078125 0 1856.4723
439.2197570800781 0 606.7996
441.2433776855469 0 598.60504
442.1958923339844 0 568.3736
443.2257995605469 0 641.5405
451.2188720703125 0 673.30695
453.2017822265625 0 1420.4408
454.20892333984375 0 3626.3828 c 2
455.21234130859375 0 1083.4967
463.9333190917969 0 815.0308
465.2103271484375 0 1181.455
468.3186340332031 0 4566.7886
469.2890625 0 1315.9717
469.32281494140625 0 1360.5432
470.2984619140625 0 747.38855
470.33367919921875 0 2010.1445 y 10
472.2988586425781 0 696.9288 y Ammonia loss 5
473.24322509765625 0 1080.3899 c Ammonia loss 11
473.5761413574219 0 815.8426
473.9110412597656 0 1517.7606
477.1993408203125 0 1626.8539
480.22650146484375 0 699.45386
480.8023376464844 0 1000.3968 y 5
482.2998352050781 0 1066.1838
497.2201232910156 0 789.4594 c Ammonia loss 7
501.6061096191406 0 1063.9569
504.9350280761719 0 2095.5994
505.2684020996094 0 1174.0642
505.59954833984375 0 815.0899
510.2268371582031 0 689.43274
510.3293151855469 0 14103.603
510.9377746582031 0 9369.406 c Ammonia loss 12
511.2718505859375 0 8186.0605
511.3314208984375 0 4544.065
511.6061706542969 0 5631.3403
511.9412536621094 0 1437.5303
516.9413452148438 0 1116.7439
517.2752685546875 0 1438.109
523.2421875 0 1444.4609
525.3528442382812 0 15367.276 z 9
525.6246337890625 0 1120.2183 y 1
525.9594116210938 0 1383.9241
526.35595703125 0 5300.0615
527.3599853515625 0 936.5984
528.3143310546875 0 1599.7695
538.2525634765625 0 1098.6829
538.2938842773438 0 1471.7416
539.2618408203125 0 1021.6521
539.2999267578125 0 864.60364
541.3411865234375 0 786.2133
541.3755493164062 0 1006.69214 y 9
541.8290405273438 0 810.2794
547.8139038085938 0 751.76587
548.6324462890625 0 2646.889
548.9658203125 0 3749.146
549.2985229492188 0 3378.7065
549.8358764648438 0 1107.2992
553.2951049804688 0 1091.3983
553.7589111328125 0 768.80493
554.28173828125 0 976.99036
554.6358032226562 0 67369.15
554.9698486328125 0 64720.504
555.3040771484375 0 37419.055
555.6385498046875 0 10817.552
555.9267578125 0 776.0789
555.972900390625 0 4170.367
556.308349609375 0 26169.258
563.2492065429688 0 5967.5547
564.2510986328125 0 2294.0432 c Water loss 3
565.2409057617188 0 2388.3245 c Ammonia loss 3
566.2467651367188 0 2072.1365
566.4067993164062 0 846.8802
576.33837890625 0 3588.4053 w 3
576.8387451171875 0 3002.391
577.3421630859375 0 1173.2794
579.3629760742188 0 804.3604
581.2592163085938 0 2982.3618
582.2674560546875 0 17495.76 c 3
582.3727416992188 0 1727.4952
583.270751953125 0 5614.1133
584.2753295898438 0 1200.1951
595.3593139648438 0 7167.544
596.3629760742188 0 2441.144
597.3779907226562 0 747.9929
604.3573608398438 0 758.7956
604.85107421875 0 2180.7668 y Ammonia loss 3
605.352783203125 0 13818.229 z 3
605.8554077148438 0 12647.631
606.3565673828125 0 6831.596
606.85791015625 0 3040.1653
609.4558715820312 0 709.1431
610.3977661132812 0 3548.7195
612.8578491210938 0 1176.2932
613.36279296875 0 4632.293 y 3
613.863525390625 0 2212.5469
614.3663330078125 0 1049.0277
624.4217529296875 0 1343.3611
629.3478393554688 0 1836.8551
629.8471069335938 0 1251.945
651.357177734375 0 1416.0624
651.8622436523438 0 1525.7217
652.2701416015625 0 1380.2153
653.4478149414062 0 13580.317 z 8
653.826904296875 0 1312.6226 c 10
654.3290405273438 0 1011.5448
654.4517822265625 0 6723.4966
655.45361328125 0 876.84937
657.8650512695312 0 1101.0388
658.36474609375 0 1492.8433
659.2946166992188 0 1118.0624
660.2965698242188 0 1019.67413
665.3585815429688 0 809.0359
674.3047485351562 0 1217.9187
675.3101196289062 0 1055.7616
685.8839111328125 0 1541.7095
686.3878173828125 0 1182.8884 y Ammonia loss 2
686.8846435546875 0 17167.197 z 2
687.386474609375 0 13205.851
687.8883666992188 0 7429.5283
688.392333984375 0 1955.8752
693.4426879882812 0 1080.05 z Ammonia loss 7
694.8942260742188 0 15238.998 y 2
695.3948974609375 0 11349.216
695.896240234375 0 5622.395
696.3939819335938 0 1847.5166
702.2992553710938 0 906.70734 c Ammonia loss 4
709.3619995117188 0 4988.699 c Ammonia loss 11
709.437255859375 0 4393.6055
709.8622436523438 0 3816.5999
710.3633422851562 0 1405.96
710.4631958007812 0 2333.6099 z 7
711.471923828125 0 856.86456
712.477294921875 0 852.13684
714.887939453125 0 681.2588
717.366943359375 0 809.753
717.8734130859375 0 42462.152 c 11
718.3748168945312 0 35526.79
718.876220703125 0 15422.752
719.3262939453125 0 38319.574 c 4
719.8777465820312 0 1260.333
720.3294067382812 0 19432.676
721.3324584960938 0 4200.009
722.334228515625 0 842.671
723.3419189453125 0 684.25684
726.4887084960938 0 1085.1678 y 7
731.3840942382812 0 804.829
736.400390625 0 911.7821
736.9080200195312 0 1021.5943
748.333984375 0 729.9095
751.4207763671875 0 3982.9553
751.9033813476562 0 725.20953
752.4170532226562 0 1251.8604
752.9116821289062 0 677.3093
757.3954467773438 0 1178.3966
758.9083251953125 0 3161.75
759.4155883789062 0 1523.6653
759.9127807617188 0 795.59503
765.3534545898438 0 983.58246
765.9038696289062 0 1661.5547 c Ammonia loss 12
766.4118041992188 0 1731.4312
766.91748046875 0 2767.952
767.4187622070312 0 776.5068
773.4107055664062 0 873.7083
773.91748046875 0 999.69653
774.415771484375 0 28562.105 c 12
774.916748046875 0 24655.887
775.4189453125 0 11182.147
775.919189453125 0 6104.6865
776.4210205078125 0 1174.2234
778.41552734375 0 667.7144
779.9241943359375 0 5091.8745 z 1
780.4249877929688 0 4784.6577
780.506591796875 0 1976.6615 w 6
780.927734375 0 2491.3765
781.4292602539062 0 813.97974
782.42578125 0 1566.9739
787.9197998046875 0 1709.161 y 1
788.42724609375 0 1708.2279
788.9210205078125 0 908.042
789.4356689453125 0 911.9836
790.9267578125 0 4040.8523
791.4287109375 0 3476.9954
791.9279174804688 0 2623.1697
792.432861328125 0 799.0556
795.4080810546875 0 1513.2606
795.9119873046875 0 2479.1526
796.3488159179688 0 1393.0931
796.4187622070312 0 3111.244
796.9187622070312 0 2566.6868
797.4258422851562 0 981.214
800.4369506835938 0 2179.4233
800.938720703125 0 2281.185
801.441162109375 0 2359.0288
801.9268798828125 0 1648.4886
802.4137573242188 0 701.63464
802.9222412109375 0 1685.1813
803.4332885742188 0 796.3402
803.9243774414062 0 1424.535
804.4244995117188 0 1399.0238
804.9260864257812 0 887.1335
805.4103393554688 0 825.76324
807.4855346679688 0 2742.1846
808.49267578125 0 1043.7461
808.9437255859375 0 872.6291
809.4451293945312 0 4960.7275
809.9465942382812 0 4760.2847
810.4400024414062 0 4165.855
810.9411010742188 0 2095.338
811.3632202148438 0 3116.5862
811.4381103515625 0 910.0109
812.3679809570312 0 2439.2107
813.37158203125 0 715.48773
814.4375 0 872.32495
815.4375610351562 0 949.9131
822.5349731445312 0 793.3303
822.9421997070312 0 3076.6677
823.44091796875 0 19616.877
823.5134887695312 0 6342.185
823.94287109375 0 19128.52
824.4410400390625 0 8588.55
824.515869140625 0 9075.476
824.9422607421875 0 5208.73
825.4400024414062 0 1159.8065
825.5160522460938 0 3499.1604
826.5115966796875 0 748.29144
830.9424438476562 0 2804.1956
831.4501953125 0 47274.395
831.953369140625 0 229198.64
832.455078125 0 205581.77
832.6387329101562 0 1221.9897
832.9561157226562 0 105725.836
833.4574584960938 0 38449.61
833.9578247070312 0 12523.847
834.4578247070312 0 2989.6133
839.359130859375 0 3487.5647 c Ammonia loss 5
840.3600463867188 0 1801.198
850.527587890625 0 2162.1543
851.5305786132812 0 1458.2719
852.544189453125 0 2228.033
853.5457153320312 0 960.9499
855.4939575195312 0 1928.9417
856.4912719726562 0 1095.3033
909.4361572265625 0 810.1615
944.5804443359375 0 39767.227 z 5
945.5839233398438 0 21953.086
946.5872802734375 0 7088.371
947.590576171875 0 1718.1643
953.4368896484375 0 4674.763 c 6
954.439453125 0 2697.4758
960.5966796875 0 2359.5522 y 5
961.6009521484375 0 1880.9152
962.6012573242188 0 890.70544
968.5811767578125 0 1011.022
982.5484619140625 0 1825.8904
983.55126953125 0 2078.865
984.5546875 0 982.96765
1009.4480590820312 0 993.8371
1010.458984375 0 34624.918 c 7
1011.4617309570312 0 19072.9
1012.4642944335938 0 6967.753
1013.4632568359375 0 1244.8778
1066.4942626953125 0 1212.6271
1080.633544921875 0 838.1021 y Ammonia loss 4
1081.508056640625 0 954.93774
1081.6396484375 0 18837.49 z 4
1082.6461181640625 0 43598.535
1083.6492919921875 0 25301.297
1084.6513671875 0 8105.467
1085.650390625 0 1774.3269
1091.5384521484375 0 839.8609
1091.980224609375 0 645.3388
1093.5042724609375 0 4620.5225
1094.5130615234375 0 4082.6733
1095.52783203125 0 1043.6665
1096.6451416015625 0 1511.1547
1097.5413818359375 0 817.0368
1097.65625 0 4759.5273 y 4
1098.6605224609375 0 2874.554
1099.034423828125 0 838.32104
1099.66455078125 0 1447.3528
1101.522705078125 0 729.4897
1102.5364990234375 0 703.718
1110.510498046875 0 922.7672
1111.513427734375 0 2150.204
1112.533935546875 0 804.1568
1137.55810546875 0 660.73004
1138.5546875 0 45392.188 c 8
1139.5567626953125 0 29738.21
1140.559814453125 0 10459.978
1141.5565185546875 0 3099.7488
1145.607666015625 0 824.9582
1165.57958984375 0 1199.145
1166.5789794921875 0 998.3274
1209.69580078125 0 6464.1636 z 3
1210.7056884765625 0 24652.262
1211.70751953125 0 17687.566
1212.7110595703125 0 6912.8213
1213.7137451171875 0 1816.1735
1225.7178955078125 0 1643.9504 y 3
1226.7139892578125 0 1557.1376
1236.5946044921875 0 640.38245
1258.6907958984375 0 1014.45557
1262.6199951171875 0 979.021
1305.63427734375 0 1418.3397
1306.6434326171875 0 21544.736 c 10
1307.645751953125 0 15252.036
1308.650146484375 0 6760.4243
1309.6539306640625 0 1855.2437
1331.692626953125 0 738.74426
1372.758544921875 0 3826.4272 z 2
1373.765625 0 13148.817
1374.7703857421875 0 10336.339
1375.7783203125 0 3510.2637
1376.78564453125 0 1223.4884
1388.7835693359375 0 1144.5609 y 2
1389.7852783203125 0 1308.3888
1418.7218017578125 0 2280.5005
1419.721923828125 0 1858.6166
1434.7366943359375 0 7655.5986 c 11
1435.742919921875 0 10170.507
1436.74755859375 0 5706.892
1437.746826171875 0 2412.4158
1438.755126953125 0 994.573
1504.8233642578125 0 878.9349
1532.800537109375 0 1121.4991
1533.8211669921875 0 820.8161
1547.8218994140625 0 2757.2441 c 12
1548.8291015625 0 5501.7085
1549.829345703125 0 5047.1167
1550.83349609375 0 2741.8376
1558.8406982421875 0 1893.6614 z 1
1559.846923828125 0 4529.689
1560.8441162109375 0 4567.2607
1561.85302734375 0 1980.2719
1575.853515625 0 893.3593
1580.84765625 0 1346.0145
1581.8216552734375 0 1231.0192
1590.8231201171875 0 973.75653
1591.83349609375 0 1230.4937
1617.8846435546875 0 1607.8235
1618.8814697265625 0 6002.0405
1619.888671875 0 5333.2744
1620.8846435546875 0 2877.0483
1621.8912353515625 0 1234.711
1628.87353515625 0 3599.5125
1629.86328125 0 2512.1812
1630.86328125 0 1516.4493
1635.905517578125 0 2985.504
1636.9114990234375 0 1803.3588
1645.8751220703125 0 5460.0933
1646.879150390625 0 25324.44
1647.8822021484375 0 22212.525
1648.8843994140625 0 10795.863
1649.880615234375 0 4374.0464
1650.8831787109375 0 948.5732
1660.8819580078125 0 855.3721
1661.8848876953125 0 6305.856
1662.89697265625 0 27009.805
1663.905029296875 0 123598.555
1664.9072265625 0 105639.555
1665.910888671875 0 54793.703
1666.9132080078125 0 19963.18
1667.9122314453125 0 6349.0815
1668.9107666015625 0 1878.5823
2521.2646484375 0 775.2649
3219.0263671875 0 801.6305
3492.976318359375 0 699.63727

Spectrum Details

|  |  |
| --- | --- |
| Matched peaks? Matched peaks The total absolute number of peaks matched. Additionally in brackets the total fraction of peaks matched and the total number of peaks is shown. | 58 (12.61% of 460) |
| FDR? FDR The false discovery rate estimated for this peptide. It is calculated by matching all theoretical fragments with a non-integer shift with the raw peaks for this spectrum. This is done with 40 different shifts. The resulting percentage is the average number of annotated peaks over the number of annotated peaks with the correct spectrum. | 3.00% |
| Satellite FDR? Satellite FDR See the FDR for details on its calculation. This satellite ion specific FDR only contains the satellite ions (d/w) for I/L/J positions. | - |
| PSM Score? PSM Score The PSM Score as given by Hecklib to this annotated spectrum. It is shown with three significant figures. | 451 |

#### Spectrum 4459? Spectrum 4459 The raw spectrum of this peptide as annotated by Hecklib. The fragments are coloured according to ion type (see legend). Any peaks with a star '\*' as text can be hovered over to see the full details, first the ion type second the mass shift type. By hovering over the amino acids in the peptide or ions in the legend the corresponding peaks are highlighted. By toggling the 'Unassigned' label you can turn the background (unassigned) peaks on or off in the plot. By updating the slider in the Ion legend you can update the spectrum to only show the top X% of the peaks with labels. The top X% means any peak that is within X% of the highest intensity. By dragging in the spectrum you can zoom in to a specific part of the spectrum and use 'Zoom Out' to get back to the original zoom level. The annotation of the spectrum is based on the given sequence in the peptides file and is done with different software so inconsistencies are likely. The peaks are annotated based on the given sequence, with 20 ppm tolerance.

Copy Data

##### Spectrum 4459 (TSV)

###### Preview

```
Loading example...
```

*Click on the button to copy the data to your clipboard.*

Mz MinMz MaxIntensity Max

WidthHeightPeptide font sizePeptide stroke widthSpectrum font sizeSpectrum stroke widthCompact peptide

Ion legend

wxyz

abcd

OtherUnassignedIon^Charge^_Position_Show for top:%

SWYQHHPGKAPKJJ

02.19e+44.38e+46.57e+48.76e+4

Zoom Out

y^+1^_1_b^+4^_4_y^+2^_3_y^+2^_4_y^+2^_4_y^+3^_7_y^+1^_2_a^+1^_2_y^+2^_5_b^+1^_2_y^+3^_8_b^+1^_2_b^+2^_4_y^+3^_9_y^+2^_6_b^+2^_5_y^+3^_10_y^+1^_3_y^+4^_13_b^+3^_10_b^+3^_10_y^+4^_13_b^+3^_10_y^+3^_11_y^+2^_8_y^+2^_8_\*\*b^+2^_6_\*b^+2^_6_b^+3^_11_b^+1^_3_b^+1^_3_y^+3^_12_y^+3^_12_y^+1^_4_y^+2^_9_b^+3^_12_y^+2^_9_b^+3^_13_b^+3^_13_b^+3^_13_y^+3^_13_y^+3^_13_y^+2^_10_y^+1^_5_y^+2^_10_b^+1^_4_b^+2^_9_b^+2^_10_b^+2^_10_b^+2^_10_y^+2^_11_y^+2^_11_b^+2^_11_y^+1^_6_y^+2^_12_b^+1^_5_b^+2^_12_b^+1^_5_y^+1^_7_y^+1^_8_y^+1^_8_b^+1^_6_y^+1^_9_

0625125118762501

Fragment Matches Table

Show background peaks

| Position | Ion type | Intensity | mz Theoretical | mz Error (Th) | mz Error (ppm) | Charge | Series Number |
| --- | --- | --- | --- | --- | --- | --- | --- |
| - | - | 364.6 | 120.1 | - | - | 0 | - |
| - | - | 3969 | 120.1 | - | - | 0 | - |
| - | - | 1112 | 122.1 | - | - | 0 | - |
| - | - | 1382 | 123.1 | - | - | 0 | - |
| - | - | 702.1 | 124 | - | - | 0 | - |
| - | - | 461.9 | 126.1 | - | - | 0 | - |
| - | - | 2898 | 127.1 | - | - | 0 | - |
| - | - | 415.7 | 127.1 | - | - | 0 | - |
| - | - | 740.8 | 128.1 | - | - | 0 | - |
| - | - | 2149 | 129 | - | - | 0 | - |
| - | - | 795.3 | 129.1 | - | - | 0 | - |
| - | - | 8.528E+04 | 129.1 | - | - | 0 | - |
| - | - | 8.676E+04 | 130.1 | - | - | 0 | - |
| - | - | 893 | 130.1 | - | - | 0 | - |
| - | - | 5589 | 130.1 | - | - | 0 | - |
| - | - | 8260 | 131.1 | - | - | 0 | - |
| - | - | 1283 | 131.1 | - | - | 0 | - |
| - | - | 4398 | 132 | - | - | 0 | - |
| - | - | 3131 | 132.1 | - | - | 0 | - |
| 14 | y | 2.059E+04 | 132.1 | 0.000344 | 2.604 | +1 | 1 |
| - | - | 465 | 133 | - | - | 0 | - |
| - | - | 1805 | 133.1 | - | - | 0 | - |
| - | - | 555 | 135.1 | - | - | 0 | - |
| - | - | 8.273E+04 | 136.1 | - | - | 0 | - |
| - | - | 6757 | 137.1 | - | - | 0 | - |
| - | - | 1619 | 138.1 | - | - | 0 | - |
| - | - | 604.3 | 139 | - | - | 0 | - |
| - | - | 1510 | 139.1 | - | - | 0 | - |
| - | - | 5006 | 141.1 | - | - | 0 | - |
| - | - | 492.4 | 144.1 | - | - | 0 | - |
| - | - | 817.2 | 145.1 | - | - | 0 | - |
| 4 | b | 3683 | 146.1 | 0.002786 | 19.07 | +4 | 4 |
| - | - | 4023 | 147 | - | - | 0 | - |
| - | - | 437 | 148 | - | - | 0 | - |
| - | - | 1049 | 148.1 | - | - | 0 | - |
| - | - | 6.028E+04 | 149 | - | - | 0 | - |
| - | - | 6225 | 150 | - | - | 0 | - |
| - | - | 1039 | 151.1 | - | - | 0 | - |
| - | - | 870.6 | 152.1 | - | - | 0 | - |
| - | - | 1145 | 152.1 | - | - | 0 | - |
| - | - | 1911 | 152.1 | - | - | 0 | - |
| - | - | 648 | 154.1 | - | - | 0 | - |
| - | - | 576.5 | 154.2 | - | - | 0 | - |
| - | - | 1310 | 155 | - | - | 0 | - |
| - | - | 1172 | 155.1 | - | - | 0 | - |
| - | - | 4123 | 155.1 | - | - | 0 | - |
| - | - | 1134 | 155.1 | - | - | 0 | - |
| - | - | 3494 | 155.1 | - | - | 0 | - |
| - | - | 2059 | 156.1 | - | - | 0 | - |
| - | - | 1635 | 157 | - | - | 0 | - |
| - | - | 2.095E+04 | 157.1 | - | - | 0 | - |
| - | - | 1367 | 157.1 | - | - | 0 | - |
| - | - | 1.541E+04 | 158.1 | - | - | 0 | - |
| - | - | 2323 | 158.1 | - | - | 0 | - |
| - | - | 1744 | 159.1 | - | - | 0 | - |
| - | - | 2758 | 159.1 | - | - | 0 | - |
| - | - | 518.9 | 159.1 | - | - | 0 | - |
| - | - | 623.3 | 159.1 | - | - | 0 | - |
| - | - | 492.9 | 160.1 | - | - | 0 | - |
| - | - | 437.9 | 160.1 | - | - | 0 | - |
| - | - | 512 | 162.1 | - | - | 0 | - |
| - | - | 1257 | 164.1 | - | - | 0 | - |
| - | - | 2980 | 165.1 | - | - | 0 | - |
| - | - | 1220 | 166.1 | - | - | 0 | - |
| - | - | 3687 | 167 | - | - | 0 | - |
| - | - | 1821 | 167.1 | - | - | 0 | - |
| - | - | 525.6 | 168 | - | - | 0 | - |
| - | - | 673.2 | 169.1 | - | - | 0 | - |
| - | - | 5081 | 169.1 | - | - | 0 | - |
| - | - | 479.6 | 170.1 | - | - | 0 | - |
| - | - | 1398 | 171.1 | - | - | 0 | - |
| - | - | 5113 | 172.1 | - | - | 0 | - |
| - | - | 539.5 | 172.1 | - | - | 0 | - |
| - | - | 3111 | 173.1 | - | - | 0 | - |
| - | - | 2579 | 173.1 | - | - | 0 | - |
| - | - | 1035 | 173.1 | - | - | 0 | - |
| - | - | 1076 | 174.1 | - | - | 0 | - |
| - | - | 2.434E+04 | 175.1 | - | - | 0 | - |
| - | - | 2115 | 176.1 | - | - | 0 | - |
| - | - | 3065 | 176.1 | - | - | 0 | - |
| - | - | 626.3 | 176.8 | - | - | 0 | - |
| - | - | 1451 | 181.1 | - | - | 0 | - |
| - | - | 3821 | 181.1 | - | - | 0 | - |
| - | - | 1.998E+04 | 181.1 | - | - | 0 | - |
| - | - | 1054 | 181.1 | - | - | 0 | - |
| - | - | 2218 | 182.1 | - | - | 0 | - |
| - | - | 2935 | 182.1 | - | - | 0 | - |
| - | - | 2.703E+04 | 183.1 | - | - | 0 | - |
| - | - | 819.4 | 183.1 | - | - | 0 | - |
| - | - | 915.9 | 183.1 | - | - | 0 | - |
| - | - | 852.7 | 183.2 | - | - | 0 | - |
| - | - | 2659 | 184.1 | - | - | 0 | - |
| - | - | 2.408E+04 | 185.1 | - | - | 0 | - |
| - | - | 2629 | 185.1 | - | - | 0 | - |
| - | - | 636.5 | 185.1 | - | - | 0 | - |
| - | - | 1400 | 186.1 | - | - | 0 | - |
| - | - | 1869 | 186.1 | - | - | 0 | - |
| - | - | 1.042E+04 | 186.1 | - | - | 0 | - |
| - | - | 3576 | 187.1 | - | - | 0 | - |
| - | - | 1.434E+04 | 187.1 | - | - | 0 | - |
| - | - | 938.6 | 187.1 | - | - | 0 | - |
| 12 | y | 1349 | 187.1 | 0.0003354 | 1.792 | +2 | 3 |
| - | - | 928.4 | 188.1 | - | - | 0 | - |
| - | - | 703.1 | 193.1 | - | - | 0 | - |
| - | - | 3323 | 195.1 | - | - | 0 | - |
| - | - | 769.1 | 195.1 | - | - | 0 | - |
| - | - | 1667 | 195.1 | - | - | 0 | - |
| - | - | 828.5 | 196.1 | - | - | 0 | - |
| - | - | 658.7 | 197.1 | - | - | 0 | - |
| - | - | 844.7 | 197.1 | - | - | 0 | - |
| - | - | 3492 | 197.2 | - | - | 0 | - |
| - | - | 2273 | 199.1 | - | - | 0 | - |
| - | - | 1040 | 199.1 | - | - | 0 | - |
| - | - | 1799 | 199.2 | - | - | 0 | - |
| - | - | 1023 | 200.1 | - | - | 0 | - |
| - | - | 830.1 | 201.1 | - | - | 0 | - |
| - | - | 1858 | 201.1 | - | - | 0 | - |
| - | - | 1814 | 202.1 | - | - | 0 | - |
| - | - | 7171 | 202.1 | - | - | 0 | - |
| - | - | 1281 | 203.1 | - | - | 0 | - |
| - | - | 836.8 | 203.1 | - | - | 0 | - |
| - | - | 743.6 | 204.1 | - | - | 0 | - |
| - | - | 2339 | 205.1 | - | - | 0 | - |
| - | - | 670.3 | 205.1 | - | - | 0 | - |
| - | - | 1102 | 205.1 | - | - | 0 | - |
| - | - | 659.5 | 207.1 | - | - | 0 | - |
| - | - | 2086 | 207.1 | - | - | 0 | - |
| - | - | 673.8 | 208.1 | - | - | 0 | - |
| - | - | 1918 | 209.1 | - | - | 0 | - |
| - | - | 903.7 | 209.1 | - | - | 0 | - |
| - | - | 556.9 | 210 | - | - | 0 | - |
| - | - | 744.3 | 210.1 | - | - | 0 | - |
| - | - | 1215 | 211.1 | - | - | 0 | - |
| - | - | 602.2 | 212.1 | - | - | 0 | - |
| - | - | 910.6 | 212.1 | - | - | 0 | - |
| - | - | 2933 | 212.7 | - | - | 0 | - |
| - | - | 1143 | 213.1 | - | - | 0 | - |
| - | - | 557.1 | 213.1 | - | - | 0 | - |
| - | - | 2725 | 214.1 | - | - | 0 | - |
| - | - | 1048 | 215.1 | - | - | 0 | - |
| - | - | 631.6 | 217 | - | - | 0 | - |
| - | - | 3897 | 217.1 | - | - | 0 | - |
| - | - | 738.5 | 217.1 | - | - | 0 | - |
| - | - | 7162 | 220.1 | - | - | 0 | - |
| - | - | 9395 | 221.1 | - | - | 0 | - |
| - | - | 1057 | 221.1 | - | - | 0 | - |
| - | - | 1079 | 222.1 | - | - | 0 | - |
| - | - | 1067 | 222.1 | - | - | 0 | - |
| - | - | 2345 | 223.1 | - | - | 0 | - |
| - | - | 511.5 | 224.1 | - | - | 0 | - |
| - | - | 3852 | 224.2 | - | - | 0 | - |
| - | - | 896.5 | 225.1 | - | - | 0 | - |
| - | - | 1588 | 225.2 | - | - | 0 | - |
| - | - | 833 | 225.2 | - | - | 0 | - |
| - | - | 2417 | 226.1 | - | - | 0 | - |
| - | - | 7.179E+04 | 226.2 | - | - | 0 | - |
| - | - | 3.135E+04 | 227.1 | - | - | 0 | - |
| - | - | 772.1 | 227.1 | - | - | 0 | - |
| 11 | y | 7470 | 227.2 | 0.001342 | 5.91 | +2 | 4 |
| - | - | 4853 | 228.1 | - | - | 0 | - |
| - | - | 3210 | 228.1 | - | - | 0 | - |
| - | - | 1257 | 228.1 | - | - | 0 | - |
| - | - | 717.3 | 228.2 | - | - | 0 | - |
| - | - | 669.7 | 229.1 | - | - | 0 | - |
| - | - | 726.3 | 229.1 | - | - | 0 | - |
| - | - | 637.3 | 229.2 | - | - | 0 | - |
| - | - | 764.9 | 230.1 | - | - | 0 | - |
| - | - | 3386 | 231.1 | - | - | 0 | - |
| - | - | 986.8 | 231.1 | - | - | 0 | - |
| - | - | 819.2 | 232.1 | - | - | 0 | - |
| - | - | 681.1 | 232.1 | - | - | 0 | - |
| - | - | 954.8 | 233.1 | - | - | 0 | - |
| - | - | 825.2 | 235.1 | - | - | 0 | - |
| - | - | 1.253E+04 | 235.1 | - | - | 0 | - |
| 11 | y | 1.184E+04 | 235.7 | 0.0002444 | 1.037 | +2 | 4 |
| - | - | 2046 | 236.1 | - | - | 0 | - |
| - | - | 2732 | 236.2 | - | - | 0 | - |
| - | - | 561 | 236.7 | - | - | 0 | - |
| - | - | 1342 | 237.1 | - | - | 0 | - |
| - | - | 1414 | 237.1 | - | - | 0 | - |
| 8 | y | 671.5 | 237.2 | 0.0008273 | 3.488 | +3 | 7 |
| - | - | 1386 | 238.1 | - | - | 0 | - |
| - | - | 576.4 | 239.1 | - | - | 0 | - |
| - | - | 4857 | 239.2 | - | - | 0 | - |
| - | - | 1390 | 240.1 | - | - | 0 | - |
| - | - | 649.9 | 241.1 | - | - | 0 | - |
| - | - | 3171 | 241.2 | - | - | 0 | - |
| - | - | 3195 | 242.2 | - | - | 0 | - |
| - | - | 737.5 | 243.2 | - | - | 0 | - |
| - | - | 4414 | 244.1 | - | - | 0 | - |
| - | - | 742.1 | 244.2 | - | - | 0 | - |
| - | - | 5391 | 245.1 | - | - | 0 | - |
| - | - | 619.5 | 245.2 | - | - | 0 | - |
| 13 | y | 1.39E+04 | 245.2 | 0.0003712 | 1.514 | +1 | 2 |
| - | - | 719.6 | 246.1 | - | - | 0 | - |
| - | - | 4516 | 246.1 | - | - | 0 | - |
| - | - | 1103 | 246.1 | - | - | 0 | - |
| - | - | 2023 | 246.2 | - | - | 0 | - |
| - | - | 883.5 | 247.1 | - | - | 0 | - |
| - | - | 1.578E+04 | 247.1 | - | - | 0 | - |
| - | - | 952.7 | 248.1 | - | - | 0 | - |
| - | - | 7902 | 248.1 | - | - | 0 | - |
| - | - | 1217 | 248.1 | - | - | 0 | - |
| - | - | 1.285E+04 | 249.1 | - | - | 0 | - |
| - | - | 648.4 | 249.1 | - | - | 0 | - |
| - | - | 1394 | 250.1 | - | - | 0 | - |
| - | - | 1030 | 251.2 | - | - | 0 | - |
| - | - | 1178 | 252.2 | - | - | 0 | - |
| - | - | 6057 | 254.1 | - | - | 0 | - |
| - | - | 4.091E+04 | 255.1 | - | - | 0 | - |
| - | - | 6573 | 256.1 | - | - | 0 | - |
| - | - | 1965 | 257.1 | - | - | 0 | - |
| - | - | 1495 | 257.2 | - | - | 0 | - |
| - | - | 2054 | 258.1 | - | - | 0 | - |
| - | - | 2.077E+04 | 258.1 | - | - | 0 | - |
| - | - | 2807 | 259.1 | - | - | 0 | - |
| - | - | 720.3 | 259.1 | - | - | 0 | - |
| - | - | 1322 | 259.2 | - | - | 0 | - |
| - | - | 1934 | 260.2 | - | - | 0 | - |
| 2 | a | 3.42E+04 | 262.1 | 0.005132 | 19.58 | +1 | 2 |
| - | - | 4635 | 263.1 | - | - | 0 | - |
| - | - | 669.5 | 265.1 | - | - | 0 | - |
| - | - | 1004 | 265.2 | - | - | 0 | - |
| - | - | 8668 | 266.1 | - | - | 0 | - |
| - | - | 1642 | 267.1 | - | - | 0 | - |
| - | - | 891.4 | 267.1 | - | - | 0 | - |
| - | - | 1237 | 269.6 | - | - | 0 | - |
| - | - | 5851 | 270.2 | - | - | 0 | - |
| 10 | y | 4454 | 271.2 | 0.0001698 | 0.6262 | +2 | 5 |
| - | - | 1644 | 271.7 | - | - | 0 | - |
| 2 | b | 5487 | 272.1 | 0.005096 | 18.73 | +1 | 2 |
| - | - | 1.115E+04 | 273.1 | - | - | 0 | - |
| - | - | 954.6 | 273.1 | - | - | 0 | - |
| - | - | 1580 | 274.1 | - | - | 0 | - |
| - | - | 9192 | 274.1 | - | - | 0 | - |
| - | - | 5492 | 275.1 | - | - | 0 | - |
| - | - | 2.96E+04 | 275.1 | - | - | 0 | - |
| 7 | y | 991.8 | 275.2 | 5.492E-05 | 0.1996 | +3 | 8 |
| - | - | 1805 | 276.1 | - | - | 0 | - |
| - | - | 3128 | 276.1 | - | - | 0 | - |
| - | - | 640.5 | 278.1 | - | - | 0 | - |
| - | - | 1246 | 279.2 | - | - | 0 | - |
| - | - | 964.4 | 283.2 | - | - | 0 | - |
| - | - | 1.818E+04 | 283.2 | - | - | 0 | - |
| - | - | 1563 | 283.6 | - | - | 0 | - |
| - | - | 814.2 | 284.1 | - | - | 0 | - |
| - | - | 2636 | 284.2 | - | - | 0 | - |
| - | - | 697.4 | 284.6 | - | - | 0 | - |
| - | - | 860.3 | 285.1 | - | - | 0 | - |
| - | - | 3428 | 285.2 | - | - | 0 | - |
| - | - | 962.4 | 287.1 | - | - | 0 | - |
| - | - | 1247 | 289.6 | - | - | 0 | - |
| 2 | b | 2.606E+04 | 290.1 | 0.005213 | 17.97 | +1 | 2 |
| - | - | 6336 | 290.2 | - | - | 0 | - |
| - | - | 749.6 | 290.6 | - | - | 0 | - |
| - | - | 1750 | 290.7 | - | - | 0 | - |
| 4 | b | 4158 | 291.1 | 0.001438 | 4.94 | +2 | 4 |
| - | - | 1686 | 291.2 | - | - | 0 | - |
| - | - | 9118 | 292.1 | - | - | 0 | - |
| - | - | 1623 | 293.1 | - | - | 0 | - |
| - | - | 7551 | 293.1 | - | - | 0 | - |
| - | - | 1496 | 293.2 | - | - | 0 | - |
| - | - | 545.1 | 293.6 | - | - | 0 | - |
| - | - | 842.4 | 294.1 | - | - | 0 | - |
| - | - | 1163 | 294.1 | - | - | 0 | - |
| - | - | 1388 | 294.2 | - | - | 0 | - |
| - | - | 9304 | 297.2 | - | - | 0 | - |
| - | - | 1113 | 298.2 | - | - | 0 | - |
| - | - | 2221 | 303.1 | - | - | 0 | - |
| - | - | 2064 | 305.7 | - | - | 0 | - |
| - | - | 1404 | 308.1 | - | - | 0 | - |
| - | - | 2703 | 311.2 | - | - | 0 | - |
| - | - | 1316 | 312.6 | - | - | 0 | - |
| - | - | 1450 | 314.7 | - | - | 0 | - |
| - | - | 778.1 | 315.2 | - | - | 0 | - |
| - | - | 1989 | 318.1 | - | - | 0 | - |
| - | - | 859.6 | 318.2 | - | - | 0 | - |
| - | - | 9517 | 320.1 | - | - | 0 | - |
| 6 | y | 2441 | 320.9 | 0.0002744 | 0.8552 | +3 | 9 |
| - | - | 1593 | 321.1 | - | - | 0 | - |
| - | - | 1414 | 321.2 | - | - | 0 | - |
| - | - | 1.024E+04 | 321.2 | - | - | 0 | - |
| - | - | 1688 | 322.2 | - | - | 0 | - |
| - | - | 1538 | 324.2 | - | - | 0 | - |
| - | - | 977.4 | 325.2 | - | - | 0 | - |
| - | - | 5034 | 326.2 | - | - | 0 | - |
| - | - | 644.3 | 327.2 | - | - | 0 | - |
| - | - | 1090 | 327.3 | - | - | 0 | - |
| - | - | 908.9 | 327.7 | - | - | 0 | - |
| - | - | 3549 | 332.7 | - | - | 0 | - |
| - | - | 1606 | 333.2 | - | - | 0 | - |
| 9 | y | 1511 | 335.2 | 7.817E-05 | 0.2332 | +2 | 6 |
| - | - | 1.583E+04 | 336.2 | - | - | 0 | - |
| - | - | 3226 | 337.2 | - | - | 0 | - |
| - | - | 4777 | 337.7 | - | - | 0 | - |
| - | - | 7873 | 338.2 | - | - | 0 | - |
| - | - | 2513 | 338.2 | - | - | 0 | - |
| - | - | 1949 | 339.2 | - | - | 0 | - |
| - | - | 1.739E+04 | 339.2 | - | - | 0 | - |
| - | - | 1758 | 340.2 | - | - | 0 | - |
| - | - | 3037 | 340.2 | - | - | 0 | - |
| - | - | 614.2 | 342.2 | - | - | 0 | - |
| - | - | 3952 | 345.1 | - | - | 0 | - |
| - | - | 2544 | 345.6 | - | - | 0 | - |
| - | - | 1374 | 346.1 | - | - | 0 | - |
| - | - | 1.494E+04 | 346.7 | - | - | 0 | - |
| - | - | 6374 | 347.2 | - | - | 0 | - |
| - | - | 1638 | 347.7 | - | - | 0 | - |
| - | - | 916.8 | 348.1 | - | - | 0 | - |
| - | - | 6668 | 348.1 | - | - | 0 | - |
| - | - | 1234 | 349.1 | - | - | 0 | - |
| - | - | 716.3 | 349.7 | - | - | 0 | - |
| - | - | 982.2 | 351.2 | - | - | 0 | - |
| - | - | 759.4 | 353.2 | - | - | 0 | - |
| - | - | 1017 | 354.2 | - | - | 0 | - |
| - | - | 2.316E+04 | 354.2 | - | - | 0 | - |
| - | - | 4187 | 355.2 | - | - | 0 | - |
| - | - | 2426 | 355.7 | - | - | 0 | - |
| - | - | 4042 | 356.1 | - | - | 0 | - |
| - | - | 930 | 356.6 | - | - | 0 | - |
| - | - | 4411 | 357.2 | - | - | 0 | - |
| - | - | 3771 | 358.2 | - | - | 0 | - |
| - | - | 1031 | 358.3 | - | - | 0 | - |
| - | - | 1395 | 358.7 | - | - | 0 | - |
| - | - | 3314 | 359.1 | - | - | 0 | - |
| - | - | 680.6 | 359.2 | - | - | 0 | - |
| - | - | 827.5 | 359.2 | - | - | 0 | - |
| 5 | b | 688 | 359.6 | 0.00471 | 13.1 | +2 | 5 |
| - | - | 1140 | 359.9 | - | - | 0 | - |
| - | - | 1716 | 360.7 | - | - | 0 | - |
| - | - | 1603 | 360.7 | - | - | 0 | - |
| - | - | 691 | 361.2 | - | - | 0 | - |
| - | - | 1856 | 361.7 | - | - | 0 | - |
| - | - | 703.9 | 361.8 | - | - | 0 | - |
| - | - | 1758 | 362.2 | - | - | 0 | - |
| - | - | 680 | 362.5 | - | - | 0 | - |
| - | - | 795.8 | 363.2 | - | - | 0 | - |
| - | - | 845 | 365.2 | - | - | 0 | - |
| - | - | 987.5 | 366.1 | - | - | 0 | - |
| 5 | y | 1664 | 366.6 | 0.000616 | 1.68 | +3 | 10 |
| - | - | 974.8 | 366.9 | - | - | 0 | - |
| - | - | 1074 | 367.2 | - | - | 0 | - |
| - | - | 8119 | 368.1 | - | - | 0 | - |
| - | - | 1096 | 368.2 | - | - | 0 | - |
| - | - | 2223 | 368.5 | - | - | 0 | - |
| - | - | 2321 | 368.6 | - | - | 0 | - |
| - | - | 804 | 369.2 | - | - | 0 | - |
| - | - | 1008 | 370.2 | - | - | 0 | - |
| 12 | y | 7312 | 373.3 | 0.0001958 | 0.5246 | +1 | 3 |
| - | - | 1243 | 374.3 | - | - | 0 | - |
| - | - | 1128 | 374.5 | - | - | 0 | - |
| - | - | 688.6 | 375 | - | - | 0 | - |
| - | - | 628.4 | 375.2 | - | - | 0 | - |
| - | - | 1350 | 375.5 | - | - | 0 | - |
| - | - | 1946 | 375.7 | - | - | 0 | - |
| - | - | 949.3 | 376.2 | - | - | 0 | - |
| - | - | 616.8 | 377.2 | - | - | 0 | - |
| - | - | 845.9 | 378.7 | - | - | 0 | - |
| - | - | 833.4 | 379.2 | - | - | 0 | - |
| - | - | 1101 | 380.2 | - | - | 0 | - |
| - | - | 1500 | 380.2 | - | - | 0 | - |
| - | - | 648.3 | 380.8 | - | - | 0 | - |
| - | - | 2255 | 382.2 | - | - | 0 | - |
| - | - | 1441 | 382.3 | - | - | 0 | - |
| - | - | 753.7 | 382.5 | - | - | 0 | - |
| - | - | 1718 | 382.5 | - | - | 0 | - |
| - | - | 764.2 | 383.3 | - | - | 0 | - |
| - | - | 2485 | 384.2 | - | - | 0 | - |
| - | - | 7662 | 385.2 | - | - | 0 | - |
| - | - | 2530 | 385.2 | - | - | 0 | - |
| - | - | 628.7 | 385.5 | - | - | 0 | - |
| - | - | 1011 | 385.8 | - | - | 0 | - |
| - | - | 7797 | 386.2 | - | - | 0 | - |
| - | - | 989.6 | 386.2 | - | - | 0 | - |
| - | - | 754.4 | 387 | - | - | 0 | - |
| - | - | 913.3 | 387.2 | - | - | 0 | - |
| - | - | 839.4 | 387.2 | - | - | 0 | - |
| - | - | 878.6 | 387.7 | - | - | 0 | - |
| - | - | 4269 | 389.2 | - | - | 0 | - |
| - | - | 1932 | 389.3 | - | - | 0 | - |
| - | - | 1172 | 389.7 | - | - | 0 | - |
| - | - | 1588 | 389.8 | - | - | 0 | - |
| - | - | 620 | 390.2 | - | - | 0 | - |
| - | - | 2101 | 391.5 | - | - | 0 | - |
| - | - | 1548 | 391.9 | - | - | 0 | - |
| - | - | 767.4 | 392.2 | - | - | 0 | - |
| - | - | 3346 | 392.3 | - | - | 0 | - |
| - | - | 664.6 | 393.3 | - | - | 0 | - |
| - | - | 1445 | 393.5 | - | - | 0 | - |
| - | - | 1020 | 393.7 | - | - | 0 | - |
| - | - | 2286 | 393.7 | - | - | 0 | - |
| - | - | 3973 | 394 | - | - | 0 | - |
| - | - | 5432 | 394.2 | - | - | 0 | - |
| - | - | 956.2 | 394.2 | - | - | 0 | - |
| 2 | y | 2474 | 394.2 | 0.006387 | 16.2 | +4 | 13 |
| - | - | 2016 | 394.5 | - | - | 0 | - |
| - | - | 1922 | 394.5 | - | - | 0 | - |
| - | - | 618.5 | 394.9 | - | - | 0 | - |
| - | - | 1400 | 395.2 | - | - | 0 | - |
| - | - | 653.3 | 396.7 | - | - | 0 | - |
| 10 | b | 2714 | 397.5 | 0.002121 | 5.335 | +3 | 10 |
| 10 | b | 1626 | 397.8 | 0.005903 | 14.84 | +3 | 10 |
| - | - | 1092 | 398.2 | - | - | 0 | - |
| - | - | 1002 | 398.2 | - | - | 0 | - |
| 2 | y | 5473 | 398.5 | 0.001459 | 3.662 | +4 | 13 |
| - | - | 3112 | 398.7 | - | - | 0 | - |
| - | - | 1242 | 399 | - | - | 0 | - |
| - | - | 806.9 | 399.2 | - | - | 0 | - |
| - | - | 652.3 | 399.5 | - | - | 0 | - |
| - | - | 1137 | 400 | - | - | 0 | - |
| - | - | 1061 | 401.2 | - | - | 0 | - |
| - | - | 3607 | 402.2 | - | - | 0 | - |
| - | - | 3441 | 402.7 | - | - | 0 | - |
| - | - | 1959 | 403 | - | - | 0 | - |
| - | - | 1129 | 403.2 | - | - | 0 | - |
| - | - | 7989 | 403.2 | - | - | 0 | - |
| - | - | 4677 | 403.2 | - | - | 0 | - |
| - | - | 1109 | 403.3 | - | - | 0 | - |
| 10 | b | 1847 | 403.5 | 0.002017 | 4.999 | +3 | 10 |
| 4 | y | 5491 | 403.6 | 0.002971 | 7.361 | +3 | 11 |
| 7 | y | 1467 | 403.8 | 0.006329 | 15.68 | +2 | 8 |
| - | - | 846.7 | 403.8 | - | - | 0 | - |
| - | - | 1210 | 403.9 | - | - | 0 | - |
| - | - | 4226 | 403.9 | - | - | 0 | - |
| - | - | 2066 | 404.2 | - | - | 0 | - |
| - | - | 865.7 | 404.2 | - | - | 0 | - |
| - | - | 1305 | 404.5 | - | - | 0 | - |
| - | - | 1236 | 404.7 | - | - | 0 | - |
| - | - | 727.6 | 404.9 | - | - | 0 | - |
| - | - | 1047 | 405.2 | - | - | 0 | - |
| - | - | 3897 | 406.3 | - | - | 0 | - |
| - | - | 1163 | 406.7 | - | - | 0 | - |
| - | - | 686.7 | 407 | - | - | 0 | - |
| - | - | 1855 | 407.2 | - | - | 0 | - |
| - | - | 785.6 | 407.5 | - | - | 0 | - |
| - | - | 3645 | 408.2 | - | - | 0 | - |
| - | - | 1560 | 408.5 | - | - | 0 | - |
| - | - | 3923 | 408.7 | - | - | 0 | - |
| - | - | 949.9 | 408.9 | - | - | 0 | - |
| - | - | 3521 | 409 | - | - | 0 | - |
| - | - | 855.3 | 409.2 | - | - | 0 | - |
| - | - | 951.2 | 409.5 | - | - | 0 | - |
| - | - | 1682 | 409.6 | - | - | 0 | - |
| - | - | 777.9 | 409.9 | - | - | 0 | - |
| - | - | 793.9 | 410.2 | - | - | 0 | - |
| - | - | 7749 | 410.3 | - | - | 0 | - |
| - | - | 3636 | 410.6 | - | - | 0 | - |
| - | - | 607.1 | 410.7 | - | - | 0 | - |
| - | - | 2341 | 410.9 | - | - | 0 | - |
| - | - | 7442 | 411.2 | - | - | 0 | - |
| - | - | 4517 | 411.2 | - | - | 0 | - |
| - | - | 1281 | 411.3 | - | - | 0 | - |
| - | - | 4730 | 411.5 | - | - | 0 | - |
| - | - | 2.446E+04 | 411.7 | - | - | 0 | - |
| - | - | 2285 | 411.7 | - | - | 0 | - |
| - | - | 970 | 412 | - | - | 0 | - |
| - | - | 1.159E+04 | 412.2 | - | - | 0 | - |
| 7 | y | 2.712E+04 | 412.3 | 0.0002569 | 0.623 | +2 | 8 |
| - | - | 3309 | 412.7 | - | - | 0 | - |
| - | - | 1.375E+04 | 412.8 | - | - | 0 | - |
| - | - | 723.7 | 413.2 | - | - | 0 | - |
| - | - | 2652 | 413.3 | - | - | 0 | - |
| - | - | 779.8 | 413.8 | - | - | 0 | - |
| - | - | 3651 | 413.9 | - | - | 0 | - |
| - | - | 3378 | 414.2 | - | - | 0 | - |
| - | - | 2189 | 414.2 | - | - | 0 | - |
| - | - | 2021 | 414.6 | - | - | 0 | - |
| - | - | 1372 | 414.7 | - | - | 0 | - |
| - | - | 744.5 | 414.9 | - | - | 0 | - |
| - | - | 2570 | 415.3 | - | - | 0 | - |
| 0 | Precursor | 3806 | 415.7 | 0.001556 | 3.743 | +4 | -1 |
| - | - | 1446 | 415.8 | - | - | 0 | - |
| 0 | Precursor | 4226 | 416 | 0.006254 | 15.03 | +4 | -1 |
| - | - | 2227 | 416.2 | - | - | 0 | - |
| - | - | 1923 | 417.2 | - | - | 0 | - |
| 6 | b | 1606 | 419.2 | 0.005876 | 14.02 | +2 | 6 |
| - | - | 1.18E+04 | 419.9 | - | - | 0 | - |
| 0 | Precursor | 2.206E+04 | 420.2 | 0.00395 | 9.4 | +4 | -1 |
| - | - | 2.001E+04 | 420.5 | - | - | 0 | - |
| - | - | 2225 | 420.6 | - | - | 0 | - |
| - | - | 4906 | 420.7 | - | - | 0 | - |
| - | - | 1.112E+04 | 420.7 | - | - | 0 | - |
| - | - | 1228 | 420.9 | - | - | 0 | - |
| - | - | 2750 | 421 | - | - | 0 | - |
| - | - | 1526 | 421.2 | - | - | 0 | - |
| - | - | 2552 | 421.2 | - | - | 0 | - |
| - | - | 1026 | 421.2 | - | - | 0 | - |
| - | - | 1404 | 421.3 | - | - | 0 | - |
| - | - | 898.9 | 421.5 | - | - | 0 | - |
| - | - | 1786 | 422.3 | - | - | 0 | - |
| - | - | 1885 | 423.3 | - | - | 0 | - |
| - | - | 2240 | 424.3 | - | - | 0 | - |
| - | - | 2988 | 424.7 | - | - | 0 | - |
| - | - | 1744 | 425.2 | - | - | 0 | - |
| - | - | 1437 | 425.9 | - | - | 0 | - |
| - | - | 1510 | 426.2 | - | - | 0 | - |
| - | - | 880.6 | 426.6 | - | - | 0 | - |
| - | - | 2952 | 427.2 | - | - | 0 | - |
| - | - | 1143 | 427.7 | - | - | 0 | - |
| 6 | b | 1643 | 428.2 | 0.002608 | 6.091 | +2 | 6 |
| - | - | 772.8 | 428.7 | - | - | 0 | - |
| - | - | 2.417E+04 | 429.2 | - | - | 0 | - |
| 11 | b | 6859 | 430.2 | 0.007799 | 18.13 | +3 | 11 |
| - | - | 695.3 | 430.5 | - | - | 0 | - |
| - | - | 3768 | 431.2 | - | - | 0 | - |
| 3 | b | 997.9 | 435.2 | 0.005672 | 13.03 | +1 | 3 |
| - | - | 737.4 | 439.3 | - | - | 0 | - |
| - | - | 961.7 | 439.8 | - | - | 0 | - |
| - | - | 975.5 | 443.6 | - | - | 0 | - |
| - | - | 1028 | 443.9 | - | - | 0 | - |
| - | - | 2688 | 446.2 | - | - | 0 | - |
| - | - | 983.6 | 446.7 | - | - | 0 | - |
| - | - | 4279 | 448.3 | - | - | 0 | - |
| - | - | 2932 | 448.6 | - | - | 0 | - |
| - | - | 1767 | 448.9 | - | - | 0 | - |
| - | - | 873.7 | 449.6 | - | - | 0 | - |
| - | - | 966.9 | 449.9 | - | - | 0 | - |
| - | - | 3807 | 451.2 | - | - | 0 | - |
| - | - | 1954 | 451.3 | - | - | 0 | - |
| - | - | 1772 | 451.7 | - | - | 0 | - |
| - | - | 3656 | 451.9 | - | - | 0 | - |
| - | - | 3473 | 452.3 | - | - | 0 | - |
| - | - | 1224 | 452.3 | - | - | 0 | - |
| - | - | 2553 | 452.6 | - | - | 0 | - |
| 3 | b | 1826 | 453.2 | 0.004384 | 9.674 | +1 | 3 |
| - | - | 2613 | 455.3 | - | - | 0 | - |
| - | - | 793.3 | 456.3 | - | - | 0 | - |
| - | - | 1907 | 457.2 | - | - | 0 | - |
| - | - | 2892 | 457.6 | - | - | 0 | - |
| 3 | y | 3359 | 457.9 | 0.003569 | 7.795 | +3 | 12 |
| - | - | 1268 | 458.2 | - | - | 0 | - |
| - | - | 2861 | 458.3 | - | - | 0 | - |
| - | - | 819.4 | 458.6 | - | - | 0 | - |
| - | - | 4326 | 460.2 | - | - | 0 | - |
| - | - | 3175 | 460.7 | - | - | 0 | - |
| - | - | 810.1 | 463.3 | - | - | 0 | - |
| 3 | y | 5.366E+04 | 463.6 | 0.0003248 | 0.7007 | +3 | 12 |
| - | - | 4.387E+04 | 463.9 | - | - | 0 | - |
| - | - | 2.2E+04 | 464.3 | - | - | 0 | - |
| - | - | 5799 | 464.6 | - | - | 0 | - |
| - | - | 1244 | 464.9 | - | - | 0 | - |
| - | - | 1465 | 466.9 | - | - | 0 | - |
| - | - | 2361 | 467.3 | - | - | 0 | - |
| - | - | 1246 | 468.3 | - | - | 0 | - |
| - | - | 1364 | 468.9 | - | - | 0 | - |
| - | - | 1412 | 469.8 | - | - | 0 | - |
| - | - | 1077 | 470.2 | - | - | 0 | - |
| 11 | y | 3.38E+04 | 470.3 | 0.0003494 | 0.743 | +1 | 4 |
| - | - | 9696 | 471.3 | - | - | 0 | - |
| - | - | 1852 | 471.8 | - | - | 0 | - |
| 6 | y | 1525 | 472.3 | 0.008581 | 18.17 | +2 | 9 |
| - | - | 1528 | 472.3 | - | - | 0 | - |
| - | - | 5993 | 473.3 | - | - | 0 | - |
| - | - | 1136 | 474.2 | - | - | 0 | - |
| - | - | 1161 | 474.3 | - | - | 0 | - |
| - | - | 1863 | 474.8 | - | - | 0 | - |
| - | - | 1117 | 475.3 | - | - | 0 | - |
| - | - | 1231 | 475.6 | - | - | 0 | - |
| - | - | 5724 | 476.2 | - | - | 0 | - |
| - | - | 1461 | 477.2 | - | - | 0 | - |
| 12 | b | 1629 | 478.6 | 0.003404 | 7.112 | +3 | 12 |
| - | - | 3176 | 478.9 | - | - | 0 | - |
| - | - | 1016 | 479.2 | - | - | 0 | - |
| 6 | y | 6855 | 480.8 | 0.0001283 | 0.2669 | +2 | 9 |
| - | - | 5322 | 481.3 | - | - | 0 | - |
| - | - | 3673 | 481.3 | - | - | 0 | - |
| - | - | 3251 | 481.6 | - | - | 0 | - |
| - | - | 1190 | 481.8 | - | - | 0 | - |
| - | - | 1609 | 481.9 | - | - | 0 | - |
| - | - | 1454 | 482.3 | - | - | 0 | - |
| - | - | 1820 | 482.3 | - | - | 0 | - |
| - | - | 2020 | 482.8 | - | - | 0 | - |
| - | - | 3709 | 483.8 | - | - | 0 | - |
| - | - | 2672 | 484.3 | - | - | 0 | - |
| - | - | 1037 | 486.3 | - | - | 0 | - |
| - | - | 2570 | 487.3 | - | - | 0 | - |
| - | - | 3779 | 487.6 | - | - | 0 | - |
| - | - | 1852 | 487.9 | - | - | 0 | - |
| - | - | 925.8 | 488.2 | - | - | 0 | - |
| - | - | 1160 | 490.2 | - | - | 0 | - |
| - | - | 2851 | 491.3 | - | - | 0 | - |
| - | - | 1947 | 494.2 | - | - | 0 | - |
| - | - | 935.5 | 495.4 | - | - | 0 | - |
| - | - | 760.8 | 498.9 | - | - | 0 | - |
| - | - | 2069 | 503.2 | - | - | 0 | - |
| - | - | 1220 | 504.9 | - | - | 0 | - |
| - | - | 1120 | 505.7 | - | - | 0 | - |
| - | - | 1616 | 506.9 | - | - | 0 | - |
| - | - | 1046 | 507.6 | - | - | 0 | - |
| - | - | 839.5 | 509.2 | - | - | 0 | - |
| - | - | 753.7 | 509.3 | - | - | 0 | - |
| 13 | b | 4084 | 510.3 | 0.001965 | 3.851 | +3 | 13 |
| 13 | b | 3983 | 510.6 | 0.007761 | 15.2 | +3 | 13 |
| - | - | 1184 | 510.9 | - | - | 0 | - |
| - | - | 726.5 | 513.7 | - | - | 0 | - |
| 13 | b | 4610 | 516.3 | 0.001983 | 3.842 | +3 | 13 |
| - | - | 5832 | 516.6 | - | - | 0 | - |
| - | - | 832.9 | 516.7 | - | - | 0 | - |
| - | - | 2290 | 516.9 | - | - | 0 | - |
| - | - | 1593 | 517.3 | - | - | 0 | - |
| - | - | 1175 | 519.3 | - | - | 0 | - |
| - | - | 1659 | 520.4 | - | - | 0 | - |
| - | - | 2400 | 521.2 | - | - | 0 | - |
| - | - | 801.6 | 521.4 | - | - | 0 | - |
| - | - | 7307 | 522.3 | - | - | 0 | - |
| - | - | 6099 | 522.6 | - | - | 0 | - |
| - | - | 1927 | 522.9 | - | - | 0 | - |
| - | - | 1046 | 523.3 | - | - | 0 | - |
| - | - | 3365 | 524.3 | - | - | 0 | - |
| - | - | 1801 | 524.6 | - | - | 0 | - |
| - | - | 1.076E+04 | 525 | - | - | 0 | - |
| 2 | y | 1.037E+04 | 525.3 | 0.008155 | 15.52 | +3 | 13 |
| - | - | 3415 | 525.6 | - | - | 0 | - |
| - | - | 1454 | 525.7 | - | - | 0 | - |
| - | - | 1211 | 526 | - | - | 0 | - |
| - | - | 835.4 | 526.2 | - | - | 0 | - |
| - | - | 1463 | 529.8 | - | - | 0 | - |
| - | - | 1529 | 530.3 | - | - | 0 | - |
| - | - | 864.6 | 530.8 | - | - | 0 | - |
| 2 | y | 6328 | 531 | 0.001218 | 2.293 | +3 | 13 |
| - | - | 1821 | 531.2 | - | - | 0 | - |
| - | - | 7571 | 531.3 | - | - | 0 | - |
| - | - | 3494 | 531.6 | - | - | 0 | - |
| - | - | 948.6 | 532 | - | - | 0 | - |
| - | - | 1.506E+04 | 538.3 | - | - | 0 | - |
| - | - | 1132 | 538.4 | - | - | 0 | - |
| - | - | 2478 | 538.8 | - | - | 0 | - |
| - | - | 3895 | 539.3 | - | - | 0 | - |
| - | - | 2800 | 539.3 | - | - | 0 | - |
| - | - | 2492 | 539.8 | - | - | 0 | - |
| - | - | 808.6 | 540.3 | - | - | 0 | - |
| 5 | y | 836.7 | 540.8 | 0.006347 | 11.74 | +2 | 10 |
| 10 | y | 7057 | 541.4 | 0.0003874 | 0.7156 | +1 | 5 |
| - | - | 1919 | 542.4 | - | - | 0 | - |
| - | - | 876.1 | 543.8 | - | - | 0 | - |
| - | - | 1125 | 545.2 | - | - | 0 | - |
| - | - | 1920 | 547.8 | - | - | 0 | - |
| - | - | 1094 | 548.3 | - | - | 0 | - |
| - | - | 1048 | 548.8 | - | - | 0 | - |
| - | - | 988.9 | 549.2 | - | - | 0 | - |
| 5 | y | 1.445E+04 | 549.3 | 0.0002134 | 0.3885 | +2 | 10 |
| - | - | 8092 | 549.8 | - | - | 0 | - |
| - | - | 2672 | 550.3 | - | - | 0 | - |
| - | - | 1045 | 550.8 | - | - | 0 | - |
| - | - | 1073 | 551.3 | - | - | 0 | - |
| - | - | 964.3 | 551.8 | - | - | 0 | - |
| - | - | 1588 | 552.3 | - | - | 0 | - |
| - | - | 993 | 552.8 | - | - | 0 | - |
| - | - | 1017 | 557.3 | - | - | 0 | - |
| - | - | 1081 | 561.3 | - | - | 0 | - |
| 4 | b | 906 | 563.2 | 0.00404 | 7.173 | +1 | 4 |
| - | - | 1.517E+04 | 566.2 | - | - | 0 | - |
| - | - | 5931 | 567.2 | - | - | 0 | - |
| - | - | 1320 | 567.4 | - | - | 0 | - |
| 9 | b | 5685 | 569.3 | 0.002363 | 4.151 | +2 | 9 |
| - | - | 2851 | 569.8 | - | - | 0 | - |
| - | - | 1617 | 570.3 | - | - | 0 | - |
| - | - | 8168 | 572.8 | - | - | 0 | - |
| - | - | 5151 | 573.3 | - | - | 0 | - |
| - | - | 2421 | 573.8 | - | - | 0 | - |
| - | - | 4564 | 577.4 | - | - | 0 | - |
| - | - | 2425 | 578.4 | - | - | 0 | - |
| - | - | 1.067E+04 | 579.4 | - | - | 0 | - |
| - | - | 3975 | 580.4 | - | - | 0 | - |
| - | - | 952.3 | 581.4 | - | - | 0 | - |
| - | - | 816 | 582.3 | - | - | 0 | - |
| - | - | 971.9 | 583.2 | - | - | 0 | - |
| - | - | 2897 | 585.3 | - | - | 0 | - |
| - | - | 889.1 | 587.3 | - | - | 0 | - |
| - | - | 3678 | 595.4 | - | - | 0 | - |
| 10 | b | 1689 | 595.8 | 0.00154 | 2.585 | +2 | 10 |
| 10 | b | 1566 | 596.3 | 0.01136 | 19.06 | +2 | 10 |
| - | - | 795 | 596.4 | - | - | 0 | - |
| - | - | 935.7 | 600.3 | - | - | 0 | - |
| - | - | 1121 | 603.3 | - | - | 0 | - |
| - | - | 3744 | 604.4 | - | - | 0 | - |
| 10 | b | 965.8 | 604.8 | 0.00822 | 13.59 | +2 | 10 |
| 4 | y | 6219 | 604.8 | 0.003425 | 5.663 | +2 | 11 |
| - | - | 996.9 | 605.3 | - | - | 0 | - |
| - | - | 3972 | 605.4 | - | - | 0 | - |
| - | - | 1821 | 605.9 | - | - | 0 | - |
| - | - | 3731 | 610.3 | - | - | 0 | - |
| - | - | 1613 | 611.2 | - | - | 0 | - |
| - | - | 992.7 | 611.3 | - | - | 0 | - |
| - | - | 984.1 | 611.8 | - | - | 0 | - |
| - | - | 4036 | 613.3 | - | - | 0 | - |
| 4 | y | 4986 | 613.4 | 0.0008771 | 1.43 | +2 | 11 |
| - | - | 1736 | 613.9 | - | - | 0 | - |
| - | - | 1880 | 614.3 | - | - | 0 | - |
| - | - | 1271 | 614.4 | - | - | 0 | - |
| - | - | 976.3 | 615.3 | - | - | 0 | - |
| - | - | 1479 | 615.8 | - | - | 0 | - |
| - | - | 3223 | 620.3 | - | - | 0 | - |
| - | - | 3412 | 620.8 | - | - | 0 | - |
| - | - | 1710 | 621.3 | - | - | 0 | - |
| - | - | 1514 | 628.3 | - | - | 0 | - |
| - | - | 7097 | 629.3 | - | - | 0 | - |
| - | - | 5316 | 629.8 | - | - | 0 | - |
| - | - | 2230 | 630.3 | - | - | 0 | - |
| - | - | 2327 | 631.3 | - | - | 0 | - |
| - | - | 3611 | 638.4 | - | - | 0 | - |
| - | - | 3967 | 638.9 | - | - | 0 | - |
| - | - | 1118 | 639.4 | - | - | 0 | - |
| - | - | 1278 | 647.4 | - | - | 0 | - |
| 11 | b | 1770 | 653.3 | 0.003086 | 4.723 | +2 | 11 |
| - | - | 904 | 654.4 | - | - | 0 | - |
| - | - | 4098 | 664.5 | - | - | 0 | - |
| - | - | 1427 | 664.8 | - | - | 0 | - |
| - | - | 1699 | 665.5 | - | - | 0 | - |
| - | - | 2001 | 668.3 | - | - | 0 | - |
| - | - | 746.9 | 669.3 | - | - | 0 | - |
| 9 | y | 2467 | 669.5 | 0.0001356 | 0.2025 | +1 | 6 |
| - | - | 1333 | 670.5 | - | - | 0 | - |
| - | - | 812.1 | 672.3 | - | - | 0 | - |
| - | - | 1120 | 673.3 | - | - | 0 | - |
| - | - | 1.198E+04 | 674.4 | - | - | 0 | - |
| - | - | 5285 | 675.4 | - | - | 0 | - |
| - | - | 1169 | 682.3 | - | - | 0 | - |
| - | - | 841.6 | 685.3 | - | - | 0 | - |
| - | - | 2823 | 690.3 | - | - | 0 | - |
| - | - | 845.8 | 691.3 | - | - | 0 | - |
| - | - | 1.117E+04 | 692.4 | - | - | 0 | - |
| - | - | 3907 | 693.4 | - | - | 0 | - |
| - | - | 1435 | 694.4 | - | - | 0 | - |
| 3 | y | 5752 | 694.9 | 0.0001927 | 0.2774 | +2 | 12 |
| - | - | 7252 | 695.4 | - | - | 0 | - |
| - | - | 2470 | 695.9 | - | - | 0 | - |
| - | - | 827 | 696.4 | - | - | 0 | - |
| - | - | 1009 | 698.4 | - | - | 0 | - |
| 5 | b | 1975 | 700.3 | 0.003966 | 5.663 | +1 | 5 |
| - | - | 746.1 | 701.3 | - | - | 0 | - |
| - | - | 3531 | 710.5 | - | - | 0 | - |
| - | - | 1117 | 711.5 | - | - | 0 | - |
| - | - | 2184 | 716.4 | - | - | 0 | - |
| 12 | b | 895.4 | 717.4 | 0.001496 | 2.085 | +2 | 12 |
| - | - | 1406 | 717.4 | - | - | 0 | - |
| 5 | b | 1843 | 718.3 | 0.003899 | 5.428 | +1 | 5 |
| - | - | 1125 | 719.3 | - | - | 0 | - |
| - | - | 1658 | 720.3 | - | - | 0 | - |
| - | - | 1559 | 721.4 | - | - | 0 | - |
| - | - | 1263 | 722.3 | - | - | 0 | - |
| - | - | 1205 | 726.4 | - | - | 0 | - |
| 8 | y | 4951 | 726.5 | 0.001336 | 1.838 | +1 | 7 |
| - | - | 687.1 | 726.9 | - | - | 0 | - |
| - | - | 1848 | 727.5 | - | - | 0 | - |
| - | - | 1029 | 730.4 | - | - | 0 | - |
| - | - | 1148 | 735.3 | - | - | 0 | - |
| - | - | 771.8 | 737.3 | - | - | 0 | - |
| - | - | 997.3 | 738.4 | - | - | 0 | - |
| - | - | 1371 | 740.3 | - | - | 0 | - |
| - | - | 1011 | 748.3 | - | - | 0 | - |
| - | - | 2545 | 750.3 | - | - | 0 | - |
| - | - | 2251 | 768.3 | - | - | 0 | - |
| - | - | 1128 | 769.3 | - | - | 0 | - |
| - | - | 1040 | 805.5 | - | - | 0 | - |
| 7 | y | 1020 | 806.5 | 0.01029 | 12.76 | +1 | 8 |
| - | - | 1561 | 811.5 | - | - | 0 | - |
| 7 | y | 1.881E+04 | 823.5 | 0.0008768 | 1.065 | +1 | 8 |
| - | - | 1.002E+04 | 824.5 | - | - | 0 | - |
| - | - | 2443 | 825.5 | - | - | 0 | - |
| - | - | 859.4 | 829.5 | - | - | 0 | - |
| - | - | 1185 | 838.3 | - | - | 0 | - |
| - | - | 671.7 | 847.5 | - | - | 0 | - |
| - | - | 2605 | 848.4 | - | - | 0 | - |
| - | - | 1275 | 849.4 | - | - | 0 | - |
| - | - | 816.7 | 853.5 | - | - | 0 | - |
| 6 | b | 3744 | 855.3 | 0.005168 | 6.042 | +1 | 6 |
| - | - | 1471 | 856.4 | - | - | 0 | - |
| - | - | 844 | 891.5 | - | - | 0 | - |
| - | - | 2942 | 901.4 | - | - | 0 | - |
| - | - | 2270 | 902.4 | - | - | 0 | - |
| - | - | 1844 | 919.4 | - | - | 0 | - |
| - | - | 820.2 | 920.5 | - | - | 0 | - |
| 6 | y | 2076 | 960.6 | 0.002233 | 2.324 | +1 | 9 |
| - | - | 1197 | 961.6 | - | - | 0 | - |
| - | - | 710.1 | 1034 | - | - | 0 | - |
| - | - | 650.4 | 1107 | - | - | 0 | - |
| - | - | 723 | 1123 | - | - | 0 | - |
| - | - | 647.4 | 1506 | - | - | 0 | - |
| - | - | 577.4 | 1861 | - | - | 0 | - |
| - | - | 638.3 | 2234 | - | - | 0 | - |
| - | - | 606.4 | 2476 | - | - | 0 | - |

m/z Charge Intensity FragmentType MassShift Position
120.05216979980469 0 364.6466
120.08110046386719 0 3969.0269
122.07166290283203 0 1111.6654
123.05573272705078 0 1381.8707
124.03952026367188 0 702.12195
126.05496978759766 0 461.86438
127.0868911743164 0 2898.343
127.12330627441406 0 415.73203
128.0708770751953 0 740.81396
129.0186004638672 0 2148.526
129.06626892089844 0 795.2512
129.1026153564453 0 85284.484
130.0654754638672 0 86761.35
130.1007537841797 0 892.9662
130.1059112548828 0 5588.624
131.06887817382812 0 8260.182
131.0815887451172 0 1282.9594
132.0447235107422 0 4398.1836
132.0811309814453 0 3130.794
132.1022491455078 0 20591.822 y 13
133.04783630371094 0 465.02365
133.10562133789062 0 1805.0123
135.06874084472656 0 555.03186
136.0760498046875 0 82734.53
137.0793914794922 0 6756.5107
138.06646728515625 0 1619.3785
139.0029754638672 0 604.2813
139.08702087402344 0 1510.4349
141.10263061523438 0 5005.5312
144.08096313476562 0 492.39963
145.07650756835938 0 817.1736
146.060302734375 0 3683.222 b 3
147.0443572998047 0 4023.3367
148.04867553710938 0 436.98654
148.076171875 0 1049.4672
149.023681640625 0 60279.375
150.0269775390625 0 6224.8833
151.0867462158203 0 1038.9191
152.07101440429688 0 870.57983
152.08233642578125 0 1144.645
152.14352416992188 0 1910.7184
154.0975799560547 0 648.0154
154.15940856933594 0 576.4859
155.04534912109375 0 1310.2998
155.06065368652344 0 1172.4962
155.0818328857422 0 4122.8887
155.09312438964844 0 1133.8225
155.1182098388672 0 3493.6328
156.07704162597656 0 2058.5417
157.013427734375 0 1635.4341
157.07638549804688 0 20952.408
157.1337127685547 0 1366.8693
158.06036376953125 0 15414.65
158.0797576904297 0 2323.4553
159.0636749267578 0 1743.6809
159.09185791015625 0 2758.1082
159.0984649658203 0 518.9066
159.11306762695312 0 623.2505
160.0762176513672 0 492.89664
160.09559631347656 0 437.8958
162.0550994873047 0 512.0308
164.07077026367188 0 1257.4148
165.10264587402344 0 2979.5986
166.0614471435547 0 1219.813
167.03424072265625 0 3687.223
167.08172607421875 0 1821.201
168.0368194580078 0 525.5646
169.0762481689453 0 673.1504
169.09750366210938 0 5080.551
170.0609588623047 0 479.64264
171.13858032226562 0 1397.7462
172.1083221435547 0 5113.2866
172.14511108398438 0 539.5353
173.07125854492188 0 3110.7544
173.0924072265625 0 2579.183
173.12840270996094 0 1034.5901
174.05538940429688 0 1076.0023
175.08688354492188 0 24338.719
176.08213806152344 0 2114.9426
176.09048461914062 0 3065.4968
176.7523193359375 0 626.34143
181.06129455566406 0 1451.0682
181.07644653320312 0 3820.7434
181.09750366210938 0 19979.676
181.13360595703125 0 1053.7804
182.1009063720703 0 2218.456
182.1291046142578 0 2934.793
183.0556182861328 0 27028.557
183.0916748046875 0 819.4066
183.11264038085938 0 915.91003
183.15003967285156 0 852.7098
184.05908203125 0 2659.1333
185.07127380371094 0 24084.549
185.0808563232422 0 2629.0671
185.12899780273438 0 636.5096
186.05519104003906 0 1399.9008
186.0746612548828 0 1868.6653
186.12400817871094 0 10416.392
187.08749389648438 0 3576.4595
187.10801696777344 0 14344.491
187.12747192382812 0 938.60693
187.14443969726562 0 1349.4159 y 11
188.111328125 0 928.37396
193.0976104736328 0 703.07916
195.07662963867188 0 3323.2546
195.11322021484375 0 769.05853
195.14950561523438 0 1666.5947
196.10841369628906 0 828.5079
197.07150268554688 0 658.7303
197.12852478027344 0 844.72943
197.1649627685547 0 3491.6003
199.0870361328125 0 2273.1182
199.10791015625 0 1040.2095
199.18087768554688 0 1798.7856
200.13951110839844 0 1022.9979
201.0664520263672 0 830.05023
201.12374877929688 0 1857.5469
202.08673095703125 0 1814.1602
202.10894775390625 0 7170.7173
203.08152770996094 0 1281.4729
203.1122283935547 0 836.7731
204.07662963867188 0 743.6067
205.06112670898438 0 2339.2483
205.09791564941406 0 670.2527
205.10867309570312 0 1102.216
207.12445068359375 0 659.48413
207.14935302734375 0 2086.078
208.14398193359375 0 673.77124
209.09231567382812 0 1917.9481
209.12876892089844 0 903.70996
209.9833526611328 0 556.9039
210.12350463867188 0 744.25256
211.0870361328125 0 1214.7443
212.10308837890625 0 602.1741
212.13966369628906 0 910.63666
212.66798400878906 0 2933.264
213.0763702392578 0 1142.7379
213.08743286132812 0 557.1481
214.10902404785156 0 2724.7622
215.13931274414062 0 1048.3801
217.03424072265625 0 631.62317
217.09742736816406 0 3897.0479
217.10809326171875 0 738.5066
220.11956787109375 0 7162.288
221.10357666015625 0 9394.525
221.12313842773438 0 1056.7216
222.1075897216797 0 1079.131
222.12413024902344 0 1066.7172
223.07162475585938 0 2345.2205
224.1395721435547 0 511.46317
224.17593383789062 0 3852.4568
225.06629943847656 0 896.49963
225.1602783203125 0 1587.8651
225.1798553466797 0 832.98
226.09791564941406 0 2417.0854
226.15530395507812 0 71792.695
227.08181762695312 0 31349.326
227.10232543945312 0 772.09314
227.15855407714844 0 7469.9775 y Ammonia loss 10
228.08526611328125 0 4853.129
228.11337280273438 0 3209.7932
228.13433837890625 0 1256.7886
228.16041564941406 0 717.2921
229.08665466308594 0 669.6997
229.11888122558594 0 726.3461
229.1656036376953 0 637.33716
230.114501953125 0 764.8741
231.08795166015625 0 3385.5913
231.1359405517578 0 986.8315
232.11953735351562 0 819.2492
232.1416778564453 0 681.11127
233.10403442382812 0 954.7648
235.10696411132812 0 825.18555
235.1193084716797 0 12529.341
235.6707305908203 0 11837.805 y 10
236.12228393554688 0 2045.5005
236.17257690429688 0 2731.9802
236.67311096191406 0 560.9706
237.06642150878906 0 1341.8425
237.13522338867188 0 1413.6136
237.15924072265625 0 671.51355 y Ammonia loss 7
238.1299591064453 0 1385.8492
239.11541748046875 0 576.43896
239.15037536621094 0 4856.8374
240.13467407226562 0 1389.7793
241.0831298828125 0 649.91626
241.1548309326172 0 3170.766
242.18655395507812 0 3195.2446
243.18182373046875 0 737.5225
244.108154296875 0 4414.2476
244.16671752929688 0 742.05493
245.09234619140625 0 5390.518
245.15011596679688 0 619.483
245.18634033203125 0 13900.77 y 12
246.09686279296875 0 719.6441
246.12405395507812 0 4516.2256
246.13795471191406 0 1103.4048
246.1896209716797 0 2023.3748
247.10801696777344 0 883.5298
247.13038635253906 0 15777
248.1028594970703 0 952.7321
248.11460876464844 0 7901.9194
248.13414001464844 0 1217.0328
249.0985107421875 0 12847.788
249.11422729492188 0 648.3673
250.1017608642578 0 1394.1764
251.15042114257812 0 1029.6332
252.1708984375 0 1177.7219
254.09254455566406 0 6057.4033
255.07672119140625 0 40906.434
256.08001708984375 0 6572.7554
257.11480712890625 0 1965.0244
257.16094970703125 0 1494.8213
258.0759582519531 0 2054.4766
258.0987854003906 0 20773.814
259.1020812988281 0 2807.4805
259.1305236816406 0 720.3048
259.1541442871094 0 1321.5514
260.1968994140625 0 1933.969
262.11883544921875 0 34201.023 a 1
263.1217041015625 0 4635.116
265.1402893066406 0 669.49554
265.166015625 0 1004.11755
266.12506103515625 0 8667.885
267.1255187988281 0 1641.7301
267.1449279785156 0 891.3588
269.6297607421875 0 1236.5616
270.1814880371094 0 5850.8745
271.1888732910156 0 4453.868 y 9
271.6904296875 0 1643.8925
272.1031494140625 0 5486.9883 b Water loss 1
273.0872497558594 0 11152.451
273.10443115234375 0 954.6259
274.0903625488281 0 1579.8447
274.1300964355469 0 9192.382
275.1029968261719 0 5492.325
275.1253356933594 0 29599.621
275.1849060058594 0 991.76886 y 6
276.1075439453125 0 1804.8446
276.1282653808594 0 3128.464
278.1128234863281 0 640.4626
279.1820373535156 0 1245.5215
283.1509704589844 0 964.36053
283.1767883300781 0 18184.824
283.6272277832031 0 1563.335
284.1296691894531 0 814.17505
284.1797180175781 0 2636.2556
284.6184387207031 0 697.4111
285.1106872558594 0 860.3021
285.1559753417969 0 3428.4944
287.11383056640625 0 962.4257
289.62896728515625 0 1246.9364
290.11383056640625 0 26058.543 b 1
290.1844177246094 0 6336.187
290.63714599609375 0 749.5868
290.6864318847656 0 1750.096
291.1174621582031 0 4157.7363 b 3
291.1565246582031 0 1685.6168
292.1292419433594 0 9117.841
293.09991455078125 0 1623.136
293.1296081542969 0 7550.697
293.17559814453125 0 1496.1415
293.6355285644531 0 545.0508
294.11907958984375 0 842.3822
294.1339416503906 0 1163.3104
294.2181701660156 0 1387.5367
297.19244384765625 0 9304.05
298.19525146484375 0 1112.6517
303.11993408203125 0 2221.1868
305.6643371582031 0 2063.854
308.11041259765625 0 1404.1517
311.2446594238281 0 2703.4827
312.6320495605469 0 1316.35
314.6697692871094 0 1449.9839
315.17071533203125 0 778.1496
318.12408447265625 0 1989.4786
318.1927795410156 0 859.55756
320.1397399902344 0 9517.062
320.8714294433594 0 2441.1694 y 5
321.1434326171875 0 1593.1959
321.2076416015625 0 1413.7145
321.2288818359375 0 10236.795
322.2318115234375 0 1687.9391
324.2160339355469 0 1537.6726
325.18896484375 0 977.3977
326.2191467285156 0 5033.998
327.2259826660156 0 644.3076
327.2751770019531 0 1090.0079
327.6952209472656 0 908.8943
332.7289733886719 0 3548.6597
333.2304992675781 0 1606.0829
335.2366027832031 0 1510.7906 y 8
336.2033386230469 0 15834.112
337.2062072753906 0 3226.391
337.7212829589844 0 4777.0854
338.15045166015625 0 7873.083
338.2223815917969 0 2512.8616
339.1530456542969 0 1948.736
339.2394104003906 0 17393.336
340.15185546875 0 1757.9755
340.24261474609375 0 3036.8918
342.1656188964844 0 614.2465
345.1448974609375 0 3951.7957
345.64666748046875 0 2543.699
346.1191711425781 0 1373.534
346.7265930175781 0 14942.576
347.2277526855469 0 6374.164
347.7277526855469 0 1637.739
348.0869140625 0 916.78876
348.1344909667969 0 6667.676
349.1385192871094 0 1233.9794
349.7083435058594 0 716.3208
351.2389221191406 0 982.2141
353.2182312011719 0 759.4205
354.1710510253906 0 1017.41364
354.2137451171875 0 23157.404
355.2169494628906 0 4187.302
355.7321472167969 0 2426.2705
356.1474609375 0 4041.9915
356.64996337890625 0 929.9765
357.24969482421875 0 4410.869
358.162353515625 0 3770.7986
358.25323486328125 0 1030.8896
358.7138366699219 0 1394.8047
359.1419372558594 0 3313.5735
359.16497802734375 0 680.6378
359.2152404785156 0 827.5263
359.6436462402344 0 687.96313 b 4
359.8728942871094 0 1140.2006
360.66290283203125 0 1715.6956
360.6894226074219 0 1602.8418
361.1844177246094 0 691.00146
361.662841796875 0 1856.0039
361.8353576660156 0 703.86414
362.1639709472656 0 1758.3411
362.5069274902344 0 679.9852
363.1949462890625 0 795.82166
365.16217041015625 0 845.019
366.14422607421875 0 987.53015
366.5580749511719 0 1663.9421 y 4
366.889892578125 0 974.8333
367.16217041015625 0 1074.4583
368.1470947265625 0 8119.316
368.1716613769531 0 1095.918
368.5099182128906 0 2222.6685
368.64697265625 0 2320.6113
369.1749572753906 0 803.9838
370.1871337890625 0 1007.56287
373.2811279296875 0 7312.0557 y 11
374.2841491699219 0 1242.987
374.5128173828125 0 1128.4752
374.9583740234375 0 688.5752
375.2115173339844 0 628.3654
375.4564208984375 0 1349.9358
375.6573486328125 0 1946.4565
376.1605224609375 0 949.32794
377.154296875 0 616.8362
378.6984558105469 0 845.87286
379.198974609375 0 833.3929
380.1803894042969 0 1101.3054
380.229248046875 0 1500.4463
380.7588195800781 0 648.25385
382.2061767578125 0 2254.651
382.28265380859375 0 1440.54
382.4660339355469 0 753.65155
382.5404052734375 0 1718.3317
383.26361083984375 0 764.24927
384.1671447753906 0 2485.3323
385.1726379394531 0 7662.006
385.1984558105469 0 2529.561
385.5164794921875 0 628.7028
385.8449401855469 0 1010.8058
386.1571960449219 0 7797.1216
386.18170166015625 0 989.5761
386.9698791503906 0 754.4114
387.1600341796875 0 913.3347
387.2166748046875 0 839.383
387.70501708984375 0 878.60767
389.16094970703125 0 4269.1357
389.27178955078125 0 1931.9424
389.66900634765625 0 1172.0447
389.7723388671875 0 1588.355
390.1659240722656 0 619.98804
391.5177307128906 0 2100.6587
391.8515930175781 0 1547.6005
392.1825866699219 0 767.4499
392.2658386230469 0 3345.5278
393.266845703125 0 664.63153
393.46319580078125 0 1445.3627
393.65252685546875 0 1019.88824
393.713623046875 0 2286.282
393.9649963378906 0 3972.853
394.15167236328125 0 5431.687
394.1910095214844 0 956.2101
394.21746826171875 0 2474.2852 y Ammonia loss 1
394.46673583984375 0 2016.2598
394.5281982421875 0 1922.466
394.8587951660156 0 618.5041
395.1551208496094 0 1399.5475
396.6722717285156 0 653.3043
397.52166748046875 0 2714.0337 b Water loss 9
397.85345458984375 0 1626.3828 b Ammonia loss 9
398.1879577636719 0 1091.736
398.24066162109375 0 1002.14044
398.46917724609375 0 5472.5825 y 1
398.71917724609375 0 3111.5078
398.9700622558594 0 1241.7161
399.2204895019531 0 806.87836
399.4732971191406 0 652.3497
399.9714660644531 0 1136.9758
401.1929016113281 0 1060.7094
402.224853515625 0 3606.5688
402.6581115722656 0 3441.4543
402.9641418457031 0 1958.9629
403.1576232910156 0 1128.9078
403.18414306640625 0 7988.7305
403.2400207519531 0 4676.89
403.26776123046875 0 1108.5614
403.52508544921875 0 1847.3871 b 9
403.57110595703125 0 5491.1206 y Ammonia loss 3
403.7666931152344 0 1467.0908 y Ammonia loss 6
403.8138122558594 0 846.7257
403.8583679199219 0 1210.1996
403.9045715332031 0 4225.5615
404.1895446777344 0 2065.8684
404.2424011230469 0 865.6635
404.470703125 0 1305.13
404.7227478027344 0 1236.2426
404.89642333984375 0 727.6226
405.17974853515625 0 1046.6074
406.25006103515625 0 3897.2822
406.7499694824219 0 1163.3138
406.9666748046875 0 686.6605
407.19329833984375 0 1854.5164
407.4653625488281 0 785.5832
408.22137451171875 0 3644.6157
408.47235107421875 0 1559.7004
408.7259521484375 0 3922.9224
408.8905029296875 0 949.9012
408.9767761230469 0 3521.4849
409.218994140625 0 855.33295
409.4786071777344 0 951.2078
409.5773620605469 0 1682.2184
409.91339111328125 0 777.8712
410.1692810058594 0 793.88617
410.276611328125 0 7748.639
410.56854248046875 0 3636.268
410.67120361328125 0 607.0839
410.903564453125 0 2340.9587
411.1778259277344 0 7441.9175
411.22259521484375 0 4517.145
411.2790222167969 0 1281.314
411.47186279296875 0 4730.146
411.66357421875 0 24459.393
411.72247314453125 0 2285.0393
411.9713439941406 0 969.95795
412.16400146484375 0 11591.801
412.2738952636719 0 27118.002 y 6
412.6641845703125 0 3308.7021
412.7755126953125 0 13745.794
413.1666564941406 0 723.7182
413.2766418457031 0 2652.072
413.77667236328125 0 779.80194
413.8973083496094 0 3651.1545
414.18304443359375 0 3377.6055
414.2281494140625 0 2188.8245
414.56304931640625 0 2021.4507
414.6839599609375 0 1371.5505
414.89471435546875 0 744.48004
415.256103515625 0 2570.0437
415.7246398925781 0 3806.0964 Precursor Water loss
415.7569580078125 0 1446.3795
415.975341796875 0 4226.1772 Precursor Ammonia loss
416.2240295410156 0 2226.9592
417.1551513671875 0 1923.2904
419.17840576171875 0 1605.9087 b Water loss 5
419.8999328613281 0 11799.603
420.22967529296875 0 22064.6 Precursor
420.4777526855469 0 20014.943
420.56756591796875 0 2225.239
420.6690368652344 0 4905.5107
420.72857666015625 0 11115.566
420.90191650390625 0 1228.0359
420.9800109863281 0 2749.8306
421.1684875488281 0 1526.3564
421.194091796875 0 2551.8052
421.23126220703125 0 1026.2787
421.3290100097656 0 1404.1226
421.48193359375 0 898.8527
422.2763366699219 0 1786.0822
423.2751770019531 0 1884.7078
424.32904052734375 0 2239.9846
424.71185302734375 0 2987.807
425.212646484375 0 1744.1532
425.90362548828125 0 1436.8357
426.2367248535156 0 1509.6183
426.5736083984375 0 880.6029
427.24346923828125 0 2952.0718
427.7450866699219 0 1143.4127
428.180419921875 0 1643.4075 b 5
428.6819763183594 0 772.77313
429.1882629394531 0 24167.092
430.190673828125 0 6858.5186 b Ammonia loss 10
430.5404357910156 0 695.29004
431.18084716796875 0 3767.6003
435.16705322265625 0 997.858 b Water loss 2
439.30389404296875 0 737.3657
439.8446960449219 0 961.6857
443.56201171875 0 975.48724
443.8931884765625 0 1028.3368
446.2331237792969 0 2688.3271
446.7361145019531 0 983.6431
448.263671875 0 4278.878
448.5978698730469 0 2932.4932
448.93170166015625 0 1766.7443
449.56402587890625 0 873.7163
449.8963928222656 0 966.9127
451.22509765625 0 3807.4824
451.2668762207031 0 1954.2126
451.72479248046875 0 1771.725
451.9195556640625 0 3655.755
452.2533264160156 0 3473.3245
452.3232421875 0 1224.3612
452.5859069824219 0 2553.2366
453.17633056640625 0 1825.6333 b 2
455.25146484375 0 2612.7603
456.2556457519531 0 793.29486
457.1839599609375 0 1907.1046
457.5946350097656 0 2892.2131
457.9261474609375 0 3359.0269 y Ammonia loss 2
458.1832580566406 0 1268.1111
458.2602844238281 0 2861.4336
458.590087890625 0 819.4424
460.2309265136719 0 4326.2637
460.73199462890625 0 3175.3376
463.2744445800781 0 810.1137
463.5984191894531 0 53663.844 y 2
463.9327697753906 0 43874.277
464.266845703125 0 21997.69
464.6012268066406 0 5798.895
464.9347839355469 0 1243.9735
466.9009094238281 0 1465.192
467.2976379394531 0 2360.657
468.29296875 0 1246.0321
468.9292907714844 0 1364.1338
469.7854309082031 0 1412.367
470.20672607421875 0 1077.4813
470.33404541015625 0 33802.58 y 10
471.3369445800781 0 9696.495
471.79833984375 0 1852.0262
472.29840087890625 0 1525.1417 y Ammonia loss 5
472.3417053222656 0 1527.6294
473.2621765136719 0 5992.9766
474.177978515625 0 1136.4269
474.2638854980469 0 1160.8448
474.7796325683594 0 1863.1692
475.2801818847656 0 1116.7671
475.5824279785156 0 1231.1953
476.193359375 0 5723.66
477.1939697265625 0 1460.7827
478.5757141113281 0 1628.5802 b 11
478.908935546875 0 3176.4902
479.2419738769531 0 1016.24274
480.80322265625 0 6855.239 y 5
481.254150390625 0 5322.228
481.30487060546875 0 3673.2095
481.5886535644531 0 3250.9973
481.8063049316406 0 1190.289
481.921875 0 1609.0441
482.2676086425781 0 1453.5414
482.3088073730469 0 1820.4443
482.7608947753906 0 2020.4519
483.78521728515625 0 3709.2483
484.287353515625 0 2671.61
486.29412841796875 0 1036.5675
487.25787353515625 0 2570.1296
487.5916748046875 0 3778.6177
487.9261779785156 0 1851.6194
488.2264709472656 0 925.82776
490.23046875 0 1159.8512
491.2718505859375 0 2851.4033
494.20330810546875 0 1947.003
495.3646545410156 0 935.48157
498.91644287109375 0 760.8015
503.2139892578125 0 2069.4626
504.93060302734375 0 1220.1089
505.71978759765625 0 1119.9006
506.9368591308594 0 1615.5782
507.60601806640625 0 1045.9197
509.1922607421875 0 839.4952
509.2549133300781 0 753.7388
510.26544189453125 0 4084.4934 b Water loss 12
510.5992431640625 0 3983.0828 b Ammonia loss 12
510.93548583984375 0 1184.2216
513.732177734375 0 726.47314
516.2689819335938 0 4610.4526 b 12
516.602294921875 0 5832.4375
516.7416381835938 0 832.92236
516.936767578125 0 2290.4314
517.2708740234375 0 1592.6624
519.2811889648438 0 1174.8851
520.3613891601562 0 1658.8552
521.2268676757812 0 2400.4478
521.3634643554688 0 801.5948
522.2721557617188 0 7307.323
522.6060180664062 0 6098.6523
522.9408569335938 0 1926.6968
523.2724609375 0 1046.3801
524.2806396484375 0 3365.156
524.6141357421875 0 1800.5406
524.952392578125 0 10759.839
525.2871704101562 0 10368.488 y Ammonia loss 1
525.6197509765625 0 3414.7075
525.747802734375 0 1454.268
525.9550170898438 0 1210.6676
526.2479858398438 0 835.4083
529.8026123046875 0 1462.8413
530.3007202148438 0 1528.9655
530.7979736328125 0 864.6442
530.9557495117188 0 6327.7607 y 1
531.2100830078125 0 1820.5009
531.2908325195312 0 7571.084
531.6241455078125 0 3493.5212
531.9590454101562 0 948.55927
538.252197265625 0 15062.139
538.37158203125 0 1131.6398
538.80908203125 0 2477.7373
539.2536010742188 0 3895.328
539.3048706054688 0 2800.0923
539.803466796875 0 2491.965
540.259033203125 0 808.62054
540.8256225585938 0 836.6667 y Ammonia loss 4
541.3704223632812 0 7057.041 y 9
542.37353515625 0 1918.9417
543.758544921875 0 876.1026
545.2130126953125 0 1124.6996
547.8132934570312 0 1919.6338
548.3118896484375 0 1093.646
548.8096313476562 0 1048.2085
549.22607421875 0 988.8539
549.332763671875 0 14445.664 y 4
549.8343505859375 0 8091.8057
550.3353881835938 0 2671.877
550.83740234375 0 1045.332
551.2554931640625 0 1072.7826
551.7561645507812 0 964.3454
552.2587890625 0 1587.7128
552.7625122070312 0 992.9859
557.2946166992188 0 1016.74365
561.2642211914062 0 1080.5072
563.2239990234375 0 905.99774 b Water loss 3
566.24658203125 0 15174.861
567.2496337890625 0 5930.786
567.3994140625 0 1319.5994
569.2647705078125 0 5685.0195 b 8
569.7662353515625 0 2850.578
570.2673950195312 0 1617.2985
572.8040161132812 0 8167.5176
573.3054809570312 0 5150.5635
573.8062744140625 0 2421.2964
577.3817138671875 0 4564.2617
578.3841552734375 0 2424.7893
579.3616333007812 0 10669.062
580.3643188476562 0 3974.8125
581.3641967773438 0 952.29486
582.3060913085938 0 815.95844
583.2421264648438 0 971.8903
585.2561645507812 0 2896.8062
587.2677612304688 0 889.0544
595.3917236328125 0 3678.4058
595.7772216796875 0 1689.328 b Water loss 9
596.279052734375 0 1566.3672 b Ammonia loss 9
596.3932495117188 0 794.96704
600.3345336914062 0 935.6782
603.2678833007812 0 1120.9601
604.356689453125 0 3743.7974
604.7891845703125 0 965.81213 b 9
604.8519897460938 0 6218.668 y Ammonia loss 3
605.2913208007812 0 996.88806
605.3521728515625 0 3971.953
605.8517456054688 0 1820.505
610.3207397460938 0 3731.091
611.23486328125 0 1612.8662
611.3225708007812 0 992.67426
611.8312377929688 0 984.0964
613.2503051757812 0 4036.3225
613.3609619140625 0 4986.2627 y 3
613.8642578125 0 1736.4608
614.2537231445312 0 1880.028
614.3639526367188 0 1270.8271
615.347900390625 0 976.2829
615.84912109375 0 1478.5648
620.3402709960938 0 3223.071
620.8405151367188 0 3412.3933
621.3419799804688 0 1710.4188
628.327392578125 0 1513.5264
629.3452758789062 0 7096.877
629.846435546875 0 5316.126
630.3468017578125 0 2230.3516
631.2608642578125 0 2326.9421
638.3506469726562 0 3611.3174
638.8529663085938 0 3967.302
639.3553466796875 0 1118.1926
647.4244384765625 0 1278.4994
653.3042602539062 0 1769.5135 b 10
654.3768920898438 0 903.9814
664.4506225585938 0 4098.1006
664.8350830078125 0 1427.3771
665.4520874023438 0 1698.7834
668.32763671875 0 2001.0542
669.3369750976562 0 746.90234
669.4656372070312 0 2467.3086 y 8
670.4718017578125 0 1332.9965
672.2897338867188 0 812.0752
673.28466796875 0 1119.9836
674.434326171875 0 11981.1
675.436279296875 0 5285.2373
682.2739868164062 0 1168.5349
685.3492431640625 0 841.5691
690.2992553710938 0 2822.8306
691.3035888671875 0 845.7534
692.4447021484375 0 11165.997
693.4475708007812 0 3907.0752
694.4493408203125 0 1434.7112
694.893310546875 0 5752.151 y 2
695.3937377929688 0 7252.3257
695.8955078125 0 2470.0535
696.399658203125 0 827.0405
698.4146118164062 0 1009.19507
700.2828369140625 0 1975.2957 b Water loss 4
701.2825927734375 0 746.12366
710.454345703125 0 3530.7246
711.45751953125 0 1116.9838
716.4199829101562 0 2183.5444
717.3563232421875 0 895.3859 b 11
717.423583984375 0 1406.1394
718.2933349609375 0 1842.9474 b 4
719.3004760742188 0 1124.9377
720.3084106445312 0 1657.9174
721.3582153320312 0 1559.1267
722.3123779296875 0 1262.661
726.3639526367188 0 1204.8129
726.4859008789062 0 4950.7173 y 7
726.86328125 0 687.1405
727.488525390625 0 1848.298
730.3878173828125 0 1029.4678
735.2828979492188 0 1148.2054
737.34619140625 0 771.8366
738.3771362304688 0 997.30396
740.3253784179688 0 1371.4474
748.2929077148438 0 1010.83606
750.3115234375 0 2545.3542
768.3177490234375 0 2250.7373
769.3232421875 0 1127.761
805.5324096679688 0 1040.1354
806.5237426757812 0 1019.7475 y Ammonia loss 6
811.49267578125 0 1560.7297
823.5391235351562 0 18808.889 y 6
824.54248046875 0 10023.306
825.5442504882812 0 2442.5898
829.5 0 859.413
838.3401489257812 0 1185.3068
847.5130615234375 0 671.7138
848.413818359375 0 2605.3306
849.4234008789062 0 1275.3784
853.47314453125 0 816.67737
855.353515625 0 3743.6438 b 5
856.3536987304688 0 1470.8679
891.4623413085938 0 844.006
901.4407958984375 0 2941.6943
902.443603515625 0 2270.489
919.4473266601562 0 1843.7687
920.45263671875 0 820.19006
960.5966796875 0 2076.024 y 5
961.5948486328125 0 1196.637
1033.54296875 0 710.0904
1107.31884765625 0 650.37006
1122.53857421875 0 722.965
1505.9610595703125 0 647.353
1860.939697265625 0 577.40186
2234.06591796875 0 638.28827
2476.438232421875 0 606.40173

Spectrum Details

|  |  |
| --- | --- |
| Matched peaks? Matched peaks The total absolute number of peaks matched. Additionally in brackets the total fraction of peaks matched and the total number of peaks is shown. | 66 (8.44% of 782) |
| FDR? FDR The false discovery rate estimated for this peptide. It is calculated by matching all theoretical fragments with a non-integer shift with the raw peaks for this spectrum. This is done with 40 different shifts. The resulting percentage is the average number of annotated peaks over the number of annotated peaks with the correct spectrum. | 4.83% |
| Satellite FDR? Satellite FDR See the FDR for details on its calculation. This satellite ion specific FDR only contains the satellite ions (d/w) for I/L/J positions. | - |
| PSM Score? PSM Score The PSM Score as given by Hecklib to this annotated spectrum. It is shown with three significant figures. | 221 |

#### Spectrum 5266? Spectrum 5266 The raw spectrum of this peptide as annotated by Hecklib. The fragments are coloured according to ion type (see legend). Any peaks with a star '\*' as text can be hovered over to see the full details, first the ion type second the mass shift type. By hovering over the amino acids in the peptide or ions in the legend the corresponding peaks are highlighted. By toggling the 'Unassigned' label you can turn the background (unassigned) peaks on or off in the plot. By updating the slider in the Ion legend you can update the spectrum to only show the top X% of the peaks with labels. The top X% means any peak that is within X% of the highest intensity. By dragging in the spectrum you can zoom in to a specific part of the spectrum and use 'Zoom Out' to get back to the original zoom level. The annotation of the spectrum is based on the given sequence in the peptides file and is done with different software so inconsistencies are likely. The peaks are annotated based on the given sequence, with 20 ppm tolerance.

Copy Data

##### Spectrum 5266 (TSV)

###### Preview

```
Loading example...
```

*Click on the button to copy the data to your clipboard.*

Mz MinMz MaxIntensity Max

WidthHeightPeptide font sizePeptide stroke widthSpectrum font sizeSpectrum stroke widthCompact peptide

Ion legend

wxyz

abcd

OtherUnassignedIon^Charge^_Position_Show for top:%

SWYQHHPGKAPKJJ

01.00e+52.00e+53.00e+54.00e+5

Zoom Out

y^+1^_1_b^+4^_4_b^+4^_5_a^+2^_3_b^+2^_3_y^+2^_4_y^+2^_4_y^+3^_7_y^+1^_2_y^+2^_5_y^+3^_8_b^+2^_4_y^+3^_9_y^+2^_6_y^+2^_6_y^+2^_7_y^+3^_10_y^+1^_3_b^+3^_9_b^+3^_9_y^+4^_13_b^+3^_10_b^+3^_10_y^+4^_13_b^+3^_10_y^+3^_11_y^+2^_8_y^+3^_11_y^+2^_8_\*\*b^+2^_6_\*b^+2^_6_b^+3^_11_b^+1^_3_y^+3^_12_y^+3^_12_y^+1^_4_b^+3^_12_y^+2^_9_b^+2^_8_b^+3^_13_b^+3^_13_b^+3^_13_y^+3^_13_y^+3^_13_y^+2^_10_y^+1^_5_y^+2^_10_b^+2^_9_b^+2^_9_b^+2^_10_b^+2^_10_y^+2^_11_y^+2^_11_y^+1^_6_y^+2^_12_y^+2^_12_b^+1^_5_b^+1^_5_y^+1^_7_y^+1^_8_b^+1^_6_b^+1^_6_y^+1^_9_y^+1^_10_b^+1^_10_

0552110516572209

Fragment Matches Table

Show background peaks

| Position | Ion type | Intensity | mz Theoretical | mz Error (Th) | mz Error (ppm) | Charge | Series Number |
| --- | --- | --- | --- | --- | --- | --- | --- |
| - | - | 2093 | 120 | - | - | 0 | - |
| - | - | 2877 | 120.1 | - | - | 0 | - |
| - | - | 654.3 | 120.1 | - | - | 0 | - |
| - | - | 3127 | 120.1 | - | - | 0 | - |
| - | - | 1063 | 121 | - | - | 0 | - |
| - | - | 812.4 | 121.1 | - | - | 0 | - |
| - | - | 1366 | 122.1 | - | - | 0 | - |
| - | - | 3495 | 122.1 | - | - | 0 | - |
| - | - | 3247 | 123 | - | - | 0 | - |
| - | - | 9624 | 123.1 | - | - | 0 | - |
| - | - | 917.9 | 123.1 | - | - | 0 | - |
| - | - | 920.8 | 124 | - | - | 0 | - |
| - | - | 941.5 | 124.1 | - | - | 0 | - |
| - | - | 2331 | 125.1 | - | - | 0 | - |
| - | - | 787.9 | 126.1 | - | - | 0 | - |
| - | - | 1214 | 126.1 | - | - | 0 | - |
| - | - | 705 | 126.1 | - | - | 0 | - |
| - | - | 1717 | 127.1 | - | - | 0 | - |
| - | - | 1.193E+04 | 127.1 | - | - | 0 | - |
| - | - | 916.7 | 127.1 | - | - | 0 | - |
| - | - | 1075 | 128 | - | - | 0 | - |
| - | - | 2.392E+04 | 128.1 | - | - | 0 | - |
| - | - | 1089 | 128.1 | - | - | 0 | - |
| - | - | 614.1 | 129.1 | - | - | 0 | - |
| - | - | 1.066E+04 | 129.1 | - | - | 0 | - |
| - | - | 3.964E+05 | 129.1 | - | - | 0 | - |
| - | - | 1.508E+05 | 130.1 | - | - | 0 | - |
| - | - | 2050 | 130.1 | - | - | 0 | - |
| - | - | 2719 | 130.1 | - | - | 0 | - |
| - | - | 2.401E+04 | 130.1 | - | - | 0 | - |
| - | - | 1162 | 131 | - | - | 0 | - |
| - | - | 1.639E+04 | 131.1 | - | - | 0 | - |
| - | - | 1432 | 131.1 | - | - | 0 | - |
| - | - | 842.6 | 131.1 | - | - | 0 | - |
| - | - | 5737 | 132 | - | - | 0 | - |
| - | - | 1.801E+05 | 132.1 | - | - | 0 | - |
| 14 | y | 1.001E+05 | 132.1 | 0.0006034 | 4.568 | +1 | 1 |
| - | - | 2846 | 133.1 | - | - | 0 | - |
| - | - | 1970 | 133.1 | - | - | 0 | - |
| - | - | 1492 | 133.1 | - | - | 0 | - |
| - | - | 1.746E+04 | 133.1 | - | - | 0 | - |
| - | - | 6743 | 133.1 | - | - | 0 | - |
| - | - | 856.8 | 134.1 | - | - | 0 | - |
| - | - | 1.12E+04 | 134.1 | - | - | 0 | - |
| - | - | 2015 | 134.1 | - | - | 0 | - |
| - | - | 697.8 | 134.1 | - | - | 0 | - |
| - | - | 1.027E+04 | 135 | - | - | 0 | - |
| - | - | 1077 | 135.1 | - | - | 0 | - |
| - | - | 1908 | 135.1 | - | - | 0 | - |
| - | - | 3.371E+05 | 136.1 | - | - | 0 | - |
| - | - | 2112 | 137.1 | - | - | 0 | - |
| - | - | 2.561E+04 | 137.1 | - | - | 0 | - |
| - | - | 1227 | 137.1 | - | - | 0 | - |
| - | - | 5946 | 138.1 | - | - | 0 | - |
| - | - | 1210 | 138.1 | - | - | 0 | - |
| - | - | 969.7 | 139.1 | - | - | 0 | - |
| - | - | 7369 | 139.1 | - | - | 0 | - |
| - | - | 958.8 | 140.1 | - | - | 0 | - |
| - | - | 2.38E+04 | 141.1 | - | - | 0 | - |
| - | - | 3722 | 142.1 | - | - | 0 | - |
| - | - | 1982 | 142.1 | - | - | 0 | - |
| - | - | 777.9 | 143.1 | - | - | 0 | - |
| - | - | 2654 | 144 | - | - | 0 | - |
| - | - | 1997 | 144.1 | - | - | 0 | - |
| - | - | 7992 | 144.1 | - | - | 0 | - |
| - | - | 4656 | 145.1 | - | - | 0 | - |
| - | - | 2585 | 145.1 | - | - | 0 | - |
| - | - | 951.5 | 145.1 | - | - | 0 | - |
| - | - | 1634 | 145.1 | - | - | 0 | - |
| 4 | b | 7.571E+04 | 146.1 | 0.002389 | 16.35 | +4 | 4 |
| - | - | 1416 | 146.1 | - | - | 0 | - |
| - | - | 2.338E+04 | 147 | - | - | 0 | - |
| - | - | 8092 | 147.1 | - | - | 0 | - |
| - | - | 1084 | 147.1 | - | - | 0 | - |
| - | - | 1662 | 148 | - | - | 0 | - |
| - | - | 8.718E+04 | 148.1 | - | - | 0 | - |
| - | - | 4426 | 148.1 | - | - | 0 | - |
| - | - | 2.425E+04 | 149 | - | - | 0 | - |
| - | - | 8244 | 149.1 | - | - | 0 | - |
| - | - | 1968 | 150 | - | - | 0 | - |
| - | - | 7739 | 150.1 | - | - | 0 | - |
| - | - | 892.4 | 150.1 | - | - | 0 | - |
| - | - | 7394 | 151.1 | - | - | 0 | - |
| - | - | 9.184E+04 | 152.1 | - | - | 0 | - |
| - | - | 3636 | 152.1 | - | - | 0 | - |
| - | - | 1.025E+04 | 152.1 | - | - | 0 | - |
| - | - | 1850 | 153.1 | - | - | 0 | - |
| - | - | 6643 | 153.1 | - | - | 0 | - |
| - | - | 4726 | 153.1 | - | - | 0 | - |
| - | - | 1159 | 153.1 | - | - | 0 | - |
| - | - | 895 | 154.1 | - | - | 0 | - |
| - | - | 1723 | 154.1 | - | - | 0 | - |
| - | - | 3179 | 155 | - | - | 0 | - |
| - | - | 1003 | 155.1 | - | - | 0 | - |
| - | - | 2.148E+04 | 155.1 | - | - | 0 | - |
| - | - | 8025 | 155.1 | - | - | 0 | - |
| - | - | 1520 | 155.1 | - | - | 0 | - |
| - | - | 1.583E+04 | 155.1 | - | - | 0 | - |
| - | - | 7386 | 156.1 | - | - | 0 | - |
| - | - | 2141 | 156.1 | - | - | 0 | - |
| - | - | 1167 | 156.1 | - | - | 0 | - |
| - | - | 1577 | 156.1 | - | - | 0 | - |
| - | - | 5122 | 157.1 | - | - | 0 | - |
| - | - | 1610 | 157.1 | - | - | 0 | - |
| - | - | 1.815E+04 | 157.1 | - | - | 0 | - |
| - | - | 1015 | 157.1 | - | - | 0 | - |
| - | - | 930.2 | 157.1 | - | - | 0 | - |
| - | - | 854.1 | 157.1 | - | - | 0 | - |
| - | - | 3.527E+04 | 158.1 | - | - | 0 | - |
| - | - | 1.786E+04 | 158.1 | - | - | 0 | - |
| - | - | 4298 | 159.1 | - | - | 0 | - |
| - | - | 1.249E+05 | 159.1 | - | - | 0 | - |
| - | - | 1.291E+04 | 160.1 | - | - | 0 | - |
| - | - | 1508 | 160.1 | - | - | 0 | - |
| - | - | 1.286E+04 | 160.1 | - | - | 0 | - |
| - | - | 2898 | 161.1 | - | - | 0 | - |
| - | - | 962 | 161.1 | - | - | 0 | - |
| - | - | 8216 | 162.1 | - | - | 0 | - |
| - | - | 924.2 | 162.1 | - | - | 0 | - |
| - | - | 1763 | 162.1 | - | - | 0 | - |
| - | - | 3112 | 163 | - | - | 0 | - |
| - | - | 1341 | 163.1 | - | - | 0 | - |
| - | - | 1798 | 163.6 | - | - | 0 | - |
| - | - | 978.8 | 164 | - | - | 0 | - |
| - | - | 8203 | 164.1 | - | - | 0 | - |
| - | - | 1226 | 164.1 | - | - | 0 | - |
| - | - | 1875 | 164.1 | - | - | 0 | - |
| - | - | 1808 | 165.1 | - | - | 0 | - |
| - | - | 4288 | 165.1 | - | - | 0 | - |
| - | - | 1.489E+04 | 165.1 | - | - | 0 | - |
| - | - | 1169 | 165.1 | - | - | 0 | - |
| - | - | 5442 | 166.1 | - | - | 0 | - |
| - | - | 1402 | 166.1 | - | - | 0 | - |
| - | - | 1805 | 167 | - | - | 0 | - |
| - | - | 1500 | 167.1 | - | - | 0 | - |
| - | - | 1037 | 167.1 | - | - | 0 | - |
| - | - | 2840 | 167.2 | - | - | 0 | - |
| - | - | 982 | 168.1 | - | - | 0 | - |
| - | - | 4087 | 168.1 | - | - | 0 | - |
| - | - | 921.2 | 169.1 | - | - | 0 | - |
| - | - | 1.631E+04 | 169.1 | - | - | 0 | - |
| - | - | 1398 | 169.1 | - | - | 0 | - |
| - | - | 2.767E+04 | 169.1 | - | - | 0 | - |
| - | - | 1111 | 169.1 | - | - | 0 | - |
| - | - | 1490 | 169.2 | - | - | 0 | - |
| - | - | 1.007E+05 | 170.1 | - | - | 0 | - |
| - | - | 1987 | 170.1 | - | - | 0 | - |
| - | - | 2471 | 170.1 | - | - | 0 | - |
| - | - | 1195 | 170.1 | - | - | 0 | - |
| - | - | 2619 | 171.1 | - | - | 0 | - |
| - | - | 1.156E+04 | 171.1 | - | - | 0 | - |
| - | - | 2580 | 171.1 | - | - | 0 | - |
| - | - | 968.6 | 172 | - | - | 0 | - |
| - | - | 939.2 | 172.1 | - | - | 0 | - |
| - | - | 4556 | 172.1 | - | - | 0 | - |
| - | - | 2.747E+04 | 172.1 | - | - | 0 | - |
| - | - | 2471 | 172.1 | - | - | 0 | - |
| - | - | 2.852E+04 | 173.1 | - | - | 0 | - |
| - | - | 5669 | 174.1 | - | - | 0 | - |
| - | - | 7465 | 174.1 | - | - | 0 | - |
| - | - | 7.243E+04 | 175.1 | - | - | 0 | - |
| - | - | 2454 | 175.1 | - | - | 0 | - |
| 5 | b | 2316 | 176.1 | 0.0006594 | 3.745 | +4 | 5 |
| - | - | 9773 | 176.1 | - | - | 0 | - |
| - | - | 7836 | 176.1 | - | - | 0 | - |
| - | - | 1044 | 177.1 | - | - | 0 | - |
| - | - | 1353 | 177.1 | - | - | 0 | - |
| - | - | 1022 | 177.1 | - | - | 0 | - |
| - | - | 913.6 | 177.1 | - | - | 0 | - |
| - | - | 4263 | 178.1 | - | - | 0 | - |
| - | - | 1318 | 178.1 | - | - | 0 | - |
| - | - | 2896 | 179.1 | - | - | 0 | - |
| - | - | 2136 | 179.1 | - | - | 0 | - |
| - | - | 2062 | 179.1 | - | - | 0 | - |
| - | - | 2024 | 179.2 | - | - | 0 | - |
| - | - | 2309 | 180.1 | - | - | 0 | - |
| - | - | 1775 | 180.1 | - | - | 0 | - |
| - | - | 7367 | 181.1 | - | - | 0 | - |
| - | - | 3137 | 181.1 | - | - | 0 | - |
| - | - | 8.38E+04 | 181.1 | - | - | 0 | - |
| - | - | 7329 | 181.1 | - | - | 0 | - |
| - | - | 1486 | 182.1 | - | - | 0 | - |
| - | - | 1397 | 182.1 | - | - | 0 | - |
| - | - | 7901 | 182.1 | - | - | 0 | - |
| - | - | 1.613E+04 | 182.1 | - | - | 0 | - |
| - | - | 1175 | 182.1 | - | - | 0 | - |
| - | - | 1124 | 183.1 | - | - | 0 | - |
| - | - | 772.1 | 183.1 | - | - | 0 | - |
| - | - | 2.895E+04 | 183.1 | - | - | 0 | - |
| - | - | 3157 | 183.1 | - | - | 0 | - |
| - | - | 1457 | 183.1 | - | - | 0 | - |
| - | - | 1397 | 183.2 | - | - | 0 | - |
| - | - | 2083 | 184 | - | - | 0 | - |
| - | - | 3648 | 184.1 | - | - | 0 | - |
| - | - | 3592 | 184.1 | - | - | 0 | - |
| - | - | 1.06E+04 | 185 | - | - | 0 | - |
| - | - | 3.264E+04 | 185.1 | - | - | 0 | - |
| - | - | 3000 | 185.1 | - | - | 0 | - |
| - | - | 980.7 | 185.1 | - | - | 0 | - |
| - | - | 5.159E+04 | 186.1 | - | - | 0 | - |
| - | - | 3479 | 186.1 | - | - | 0 | - |
| - | - | 3335 | 186.1 | - | - | 0 | - |
| - | - | 4.952E+04 | 186.1 | - | - | 0 | - |
| - | - | 1543 | 186.1 | - | - | 0 | - |
| - | - | 4962 | 187.1 | - | - | 0 | - |
| - | - | 1.507E+05 | 187.1 | - | - | 0 | - |
| - | - | 4273 | 187.1 | - | - | 0 | - |
| - | - | 3959 | 187.1 | - | - | 0 | - |
| - | - | 3165 | 188.1 | - | - | 0 | - |
| - | - | 1259 | 188.1 | - | - | 0 | - |
| - | - | 1.636E+04 | 188.1 | - | - | 0 | - |
| - | - | 1232 | 189.1 | - | - | 0 | - |
| - | - | 1181 | 190.1 | - | - | 0 | - |
| - | - | 1878 | 191.1 | - | - | 0 | - |
| - | - | 1701 | 191.1 | - | - | 0 | - |
| - | - | 1470 | 191.1 | - | - | 0 | - |
| - | - | 1888 | 192.1 | - | - | 0 | - |
| - | - | 2488 | 193.1 | - | - | 0 | - |
| - | - | 1169 | 193.1 | - | - | 0 | - |
| - | - | 2230 | 193.1 | - | - | 0 | - |
| - | - | 1055 | 193.6 | - | - | 0 | - |
| - | - | 1331 | 195.1 | - | - | 0 | - |
| - | - | 2791 | 195.1 | - | - | 0 | - |
| - | - | 1.634E+04 | 195.1 | - | - | 0 | - |
| - | - | 3606 | 196.1 | - | - | 0 | - |
| - | - | 2554 | 196.1 | - | - | 0 | - |
| - | - | 1870 | 196.2 | - | - | 0 | - |
| - | - | 5138 | 197.1 | - | - | 0 | - |
| - | - | 1088 | 197.1 | - | - | 0 | - |
| - | - | 2.088E+04 | 197.1 | - | - | 0 | - |
| - | - | 1.887E+04 | 197.2 | - | - | 0 | - |
| - | - | 1916 | 198.1 | - | - | 0 | - |
| - | - | 1081 | 198.1 | - | - | 0 | - |
| - | - | 1629 | 198.1 | - | - | 0 | - |
| - | - | 2516 | 198.2 | - | - | 0 | - |
| - | - | 1102 | 199.1 | - | - | 0 | - |
| - | - | 1.77E+04 | 199.1 | - | - | 0 | - |
| - | - | 4041 | 199.1 | - | - | 0 | - |
| - | - | 1031 | 199.1 | - | - | 0 | - |
| - | - | 1363 | 199.2 | - | - | 0 | - |
| - | - | 6707 | 199.2 | - | - | 0 | - |
| - | - | 1384 | 200.1 | - | - | 0 | - |
| - | - | 2025 | 200.1 | - | - | 0 | - |
| - | - | 6139 | 200.1 | - | - | 0 | - |
| - | - | 1089 | 200.2 | - | - | 0 | - |
| - | - | 4.184E+04 | 201.1 | - | - | 0 | - |
| - | - | 1.582E+04 | 201.1 | - | - | 0 | - |
| - | - | 2764 | 201.1 | - | - | 0 | - |
| - | - | 1342 | 202.1 | - | - | 0 | - |
| - | - | 4283 | 202.1 | - | - | 0 | - |
| - | - | 7485 | 202.1 | - | - | 0 | - |
| - | - | 3.659E+04 | 202.1 | - | - | 0 | - |
| - | - | 5.884E+04 | 203.1 | - | - | 0 | - |
| - | - | 4220 | 203.1 | - | - | 0 | - |
| - | - | 3709 | 203.1 | - | - | 0 | - |
| - | - | 2399 | 204.1 | - | - | 0 | - |
| 3 | a | 6041 | 204.1 | 0.001072 | 5.251 | +2 | 3 |
| - | - | 2066 | 204.1 | - | - | 0 | - |
| - | - | 6847 | 205.1 | - | - | 0 | - |
| - | - | 2936 | 205.1 | - | - | 0 | - |
| - | - | 872.8 | 205.1 | - | - | 0 | - |
| - | - | 1589 | 206.1 | - | - | 0 | - |
| - | - | 1855 | 207.1 | - | - | 0 | - |
| - | - | 890.4 | 207.1 | - | - | 0 | - |
| - | - | 1637 | 207.1 | - | - | 0 | - |
| - | - | 1453 | 207.1 | - | - | 0 | - |
| - | - | 1.244E+04 | 207.1 | - | - | 0 | - |
| - | - | 2531 | 208.1 | - | - | 0 | - |
| - | - | 960.4 | 208.2 | - | - | 0 | - |
| - | - | 2164 | 209.1 | - | - | 0 | - |
| - | - | 1107 | 209.1 | - | - | 0 | - |
| - | - | 3615 | 209.1 | - | - | 0 | - |
| - | - | 1096 | 209.2 | - | - | 0 | - |
| - | - | 2640 | 210.1 | - | - | 0 | - |
| - | - | 5124 | 211.1 | - | - | 0 | - |
| - | - | 4110 | 211.1 | - | - | 0 | - |
| - | - | 1344 | 211.7 | - | - | 0 | - |
| - | - | 791.6 | 212.1 | - | - | 0 | - |
| - | - | 4020 | 212.1 | - | - | 0 | - |
| - | - | 799.2 | 212.6 | - | - | 0 | - |
| - | - | 9061 | 212.7 | - | - | 0 | - |
| - | - | 2675 | 213.1 | - | - | 0 | - |
| - | - | 1712 | 213.1 | - | - | 0 | - |
| - | - | 1137 | 213.2 | - | - | 0 | - |
| - | - | 3907 | 214.1 | - | - | 0 | - |
| - | - | 1.288E+04 | 214.1 | - | - | 0 | - |
| - | - | 893.9 | 214.1 | - | - | 0 | - |
| - | - | 3287 | 214.2 | - | - | 0 | - |
| - | - | 1580 | 215.1 | - | - | 0 | - |
| - | - | 1.4E+04 | 215.1 | - | - | 0 | - |
| - | - | 1483 | 215.1 | - | - | 0 | - |
| - | - | 1.292E+04 | 215.1 | - | - | 0 | - |
| - | - | 1292 | 215.2 | - | - | 0 | - |
| - | - | 1005 | 216.1 | - | - | 0 | - |
| - | - | 2464 | 216.1 | - | - | 0 | - |
| - | - | 958.1 | 216.1 | - | - | 0 | - |
| - | - | 1079 | 217.1 | - | - | 0 | - |
| - | - | 6991 | 217.1 | - | - | 0 | - |
| - | - | 3515 | 217.1 | - | - | 0 | - |
| 3 | b | 4080 | 218.1 | 0.002557 | 11.73 | +2 | 3 |
| - | - | 1362 | 218.1 | - | - | 0 | - |
| - | - | 1874 | 219.1 | - | - | 0 | - |
| - | - | 1930 | 220.1 | - | - | 0 | - |
| - | - | 2.96E+04 | 220.1 | - | - | 0 | - |
| - | - | 1630 | 221.1 | - | - | 0 | - |
| - | - | 4.428E+04 | 221.1 | - | - | 0 | - |
| - | - | 3217 | 221.1 | - | - | 0 | - |
| - | - | 5041 | 222.1 | - | - | 0 | - |
| - | - | 3354 | 222.1 | - | - | 0 | - |
| - | - | 1418 | 222.1 | - | - | 0 | - |
| - | - | 912 | 222.2 | - | - | 0 | - |
| - | - | 1746 | 223.1 | - | - | 0 | - |
| - | - | 1563 | 223.1 | - | - | 0 | - |
| - | - | 2142 | 224.1 | - | - | 0 | - |
| - | - | 1.542E+04 | 224.1 | - | - | 0 | - |
| - | - | 2.013E+04 | 224.2 | - | - | 0 | - |
| - | - | 8228 | 225.1 | - | - | 0 | - |
| - | - | 5916 | 225.2 | - | - | 0 | - |
| - | - | 2280 | 225.2 | - | - | 0 | - |
| - | - | 1275 | 226.1 | - | - | 0 | - |
| - | - | 3083 | 226.1 | - | - | 0 | - |
| - | - | 3.392E+05 | 226.2 | - | - | 0 | - |
| - | - | 968.2 | 226.7 | - | - | 0 | - |
| - | - | 1148 | 227 | - | - | 0 | - |
| - | - | 2.9E+04 | 227.1 | - | - | 0 | - |
| - | - | 1250 | 227.1 | - | - | 0 | - |
| 11 | y | 4.001E+04 | 227.2 | 0.001953 | 8.597 | +2 | 4 |
| - | - | 4526 | 228.1 | - | - | 0 | - |
| - | - | 1.555E+04 | 228.1 | - | - | 0 | - |
| - | - | 3284 | 228.1 | - | - | 0 | - |
| - | - | 3494 | 228.2 | - | - | 0 | - |
| - | - | 3332 | 229.1 | - | - | 0 | - |
| - | - | 1.414E+04 | 229.1 | - | - | 0 | - |
| - | - | 3360 | 229.1 | - | - | 0 | - |
| - | - | 3670 | 229.2 | - | - | 0 | - |
| - | - | 1579 | 230.1 | - | - | 0 | - |
| - | - | 7378 | 230.1 | - | - | 0 | - |
| - | - | 2359 | 231.1 | - | - | 0 | - |
| - | - | 1.419E+04 | 231.1 | - | - | 0 | - |
| - | - | 987.9 | 231.1 | - | - | 0 | - |
| - | - | 5182 | 231.1 | - | - | 0 | - |
| - | - | 1375 | 232.1 | - | - | 0 | - |
| - | - | 3029 | 232.1 | - | - | 0 | - |
| - | - | 5496 | 232.1 | - | - | 0 | - |
| - | - | 1487 | 232.1 | - | - | 0 | - |
| - | - | 1527 | 233.1 | - | - | 0 | - |
| - | - | 7249 | 233.1 | - | - | 0 | - |
| - | - | 1524 | 233.1 | - | - | 0 | - |
| - | - | 1542 | 233.1 | - | - | 0 | - |
| - | - | 920.5 | 233.1 | - | - | 0 | - |
| - | - | 1277 | 233.2 | - | - | 0 | - |
| - | - | 1375 | 233.7 | - | - | 0 | - |
| - | - | 1022 | 234.1 | - | - | 0 | - |
| - | - | 1610 | 234.1 | - | - | 0 | - |
| - | - | 4529 | 234.1 | - | - | 0 | - |
| - | - | 824.8 | 234.2 | - | - | 0 | - |
| - | - | 5246 | 234.7 | - | - | 0 | - |
| - | - | 1192 | 235.1 | - | - | 0 | - |
| - | - | 5.902E+04 | 235.1 | - | - | 0 | - |
| 11 | y | 5.198E+04 | 235.7 | 0.00087 | 3.692 | +2 | 4 |
| - | - | 7264 | 236.1 | - | - | 0 | - |
| - | - | 2823 | 236.1 | - | - | 0 | - |
| - | - | 1.213E+04 | 236.2 | - | - | 0 | - |
| - | - | 1912 | 236.7 | - | - | 0 | - |
| - | - | 1161 | 237.1 | - | - | 0 | - |
| - | - | 2666 | 237.1 | - | - | 0 | - |
| 8 | y | 1501 | 237.2 | 0.002582 | 10.89 | +3 | 7 |
| - | - | 1625 | 238.1 | - | - | 0 | - |
| - | - | 5486 | 238.1 | - | - | 0 | - |
| - | - | 1491 | 239.1 | - | - | 0 | - |
| - | - | 2.074E+04 | 239.2 | - | - | 0 | - |
| - | - | 1081 | 240.1 | - | - | 0 | - |
| - | - | 9189 | 240.1 | - | - | 0 | - |
| - | - | 1823 | 240.2 | - | - | 0 | - |
| - | - | 904.7 | 240.2 | - | - | 0 | - |
| - | - | 1073 | 241.1 | - | - | 0 | - |
| - | - | 1161 | 241.1 | - | - | 0 | - |
| - | - | 2.158E+04 | 241.2 | - | - | 0 | - |
| - | - | 6649 | 242.1 | - | - | 0 | - |
| - | - | 2430 | 242.2 | - | - | 0 | - |
| - | - | 1.075E+04 | 242.2 | - | - | 0 | - |
| - | - | 7970 | 243.1 | - | - | 0 | - |
| - | - | 2723 | 243.2 | - | - | 0 | - |
| - | - | 2870 | 243.2 | - | - | 0 | - |
| - | - | 1265 | 243.7 | - | - | 0 | - |
| - | - | 1.123E+04 | 244.1 | - | - | 0 | - |
| - | - | 3240 | 244.2 | - | - | 0 | - |
| - | - | 1.167E+04 | 245.1 | - | - | 0 | - |
| - | - | 3238 | 245.1 | - | - | 0 | - |
| - | - | 966.4 | 245.2 | - | - | 0 | - |
| 13 | y | 7.132E+04 | 245.2 | 0.0009663 | 3.941 | +1 | 2 |
| - | - | 3.117E+05 | 246.1 | - | - | 0 | - |
| - | - | 8087 | 246.2 | - | - | 0 | - |
| - | - | 1181 | 246.6 | - | - | 0 | - |
| - | - | 2069 | 247.1 | - | - | 0 | - |
| - | - | 1.047E+05 | 247.1 | - | - | 0 | - |
| - | - | 4.528E+04 | 248.1 | - | - | 0 | - |
| - | - | 9758 | 248.1 | - | - | 0 | - |
| - | - | 6.18E+04 | 249.1 | - | - | 0 | - |
| - | - | 5558 | 249.1 | - | - | 0 | - |
| - | - | 1759 | 249.1 | - | - | 0 | - |
| - | - | 8756 | 250.1 | - | - | 0 | - |
| - | - | 1231 | 250.1 | - | - | 0 | - |
| - | - | 981.6 | 250.1 | - | - | 0 | - |
| - | - | 870.1 | 250.6 | - | - | 0 | - |
| - | - | 943.3 | 251.1 | - | - | 0 | - |
| - | - | 1500 | 251.1 | - | - | 0 | - |
| - | - | 3457 | 251.2 | - | - | 0 | - |
| - | - | 929 | 252.1 | - | - | 0 | - |
| - | - | 6128 | 252.2 | - | - | 0 | - |
| - | - | 3162 | 253.1 | - | - | 0 | - |
| - | - | 3818 | 254.1 | - | - | 0 | - |
| - | - | 2083 | 254.2 | - | - | 0 | - |
| - | - | 3.027E+04 | 255.1 | - | - | 0 | - |
| - | - | 4053 | 256.1 | - | - | 0 | - |
| - | - | 9925 | 256.1 | - | - | 0 | - |
| - | - | 1329 | 256.6 | - | - | 0 | - |
| - | - | 1653 | 257.1 | - | - | 0 | - |
| - | - | 8174 | 257.1 | - | - | 0 | - |
| - | - | 8893 | 257.2 | - | - | 0 | - |
| - | - | 688.5 | 257.7 | - | - | 0 | - |
| - | - | 6778 | 258.1 | - | - | 0 | - |
| - | - | 9.734E+04 | 258.1 | - | - | 0 | - |
| - | - | 1846 | 259.1 | - | - | 0 | - |
| - | - | 1.36E+04 | 259.1 | - | - | 0 | - |
| - | - | 5513 | 259.1 | - | - | 0 | - |
| - | - | 1.097E+04 | 260.1 | - | - | 0 | - |
| - | - | 7763 | 260.2 | - | - | 0 | - |
| - | - | 1568 | 261.1 | - | - | 0 | - |
| - | - | 2065 | 261.2 | - | - | 0 | - |
| - | - | 1374 | 261.2 | - | - | 0 | - |
| - | - | 1.495E+05 | 262.1 | - | - | 0 | - |
| - | - | 1733 | 263.1 | - | - | 0 | - |
| - | - | 2.201E+04 | 263.1 | - | - | 0 | - |
| - | - | 1974 | 264.1 | - | - | 0 | - |
| - | - | 4096 | 264.1 | - | - | 0 | - |
| - | - | 1442 | 265.1 | - | - | 0 | - |
| - | - | 3468 | 265.1 | - | - | 0 | - |
| - | - | 3946 | 265.2 | - | - | 0 | - |
| - | - | 3.833E+04 | 266.1 | - | - | 0 | - |
| - | - | 1487 | 266.2 | - | - | 0 | - |
| - | - | 5947 | 267.1 | - | - | 0 | - |
| - | - | 4702 | 267.1 | - | - | 0 | - |
| - | - | 967.9 | 267.2 | - | - | 0 | - |
| - | - | 1057 | 268.1 | - | - | 0 | - |
| - | - | 1566 | 268.2 | - | - | 0 | - |
| - | - | 1143 | 268.2 | - | - | 0 | - |
| - | - | 921.8 | 269.1 | - | - | 0 | - |
| - | - | 3237 | 269.6 | - | - | 0 | - |
| - | - | 5763 | 270.1 | - | - | 0 | - |
| - | - | 1221 | 270.1 | - | - | 0 | - |
| - | - | 2.832E+04 | 270.2 | - | - | 0 | - |
| - | - | 1060 | 270.6 | - | - | 0 | - |
| - | - | 2.234E+04 | 271.1 | - | - | 0 | - |
| - | - | 1855 | 271.1 | - | - | 0 | - |
| 10 | y | 1.449E+04 | 271.2 | 0.0001048 | 0.3866 | +2 | 5 |
| - | - | 4762 | 271.7 | - | - | 0 | - |
| - | - | 4094 | 272.1 | - | - | 0 | - |
| - | - | 1.02E+04 | 272.1 | - | - | 0 | - |
| - | - | 1723 | 272.2 | - | - | 0 | - |
| - | - | 964.9 | 272.2 | - | - | 0 | - |
| - | - | 9790 | 273.1 | - | - | 0 | - |
| - | - | 5063 | 273.1 | - | - | 0 | - |
| - | - | 1500 | 273.1 | - | - | 0 | - |
| - | - | 1246 | 273.1 | - | - | 0 | - |
| - | - | 4274 | 274.1 | - | - | 0 | - |
| - | - | 1.145E+05 | 274.1 | - | - | 0 | - |
| - | - | 2.358E+04 | 275.1 | - | - | 0 | - |
| - | - | 1.704E+05 | 275.1 | - | - | 0 | - |
| 7 | y | 4471 | 275.2 | 0.0008179 | 2.972 | +3 | 8 |
| - | - | 1391 | 275.5 | - | - | 0 | - |
| - | - | 8654 | 276.1 | - | - | 0 | - |
| - | - | 2.005E+04 | 276.1 | - | - | 0 | - |
| - | - | 1327 | 277.1 | - | - | 0 | - |
| - | - | 2391 | 277.1 | - | - | 0 | - |
| - | - | 3331 | 277.1 | - | - | 0 | - |
| - | - | 1146 | 277.6 | - | - | 0 | - |
| - | - | 1653 | 278.1 | - | - | 0 | - |
| - | - | 1190 | 279.2 | - | - | 0 | - |
| - | - | 5187 | 279.2 | - | - | 0 | - |
| - | - | 1687 | 280.1 | - | - | 0 | - |
| - | - | 1898 | 280.2 | - | - | 0 | - |
| - | - | 1367 | 281.2 | - | - | 0 | - |
| - | - | 1215 | 281.7 | - | - | 0 | - |
| - | - | 1198 | 282.2 | - | - | 0 | - |
| - | - | 4531 | 283.2 | - | - | 0 | - |
| - | - | 8.489E+04 | 283.2 | - | - | 0 | - |
| - | - | 4135 | 283.6 | - | - | 0 | - |
| - | - | 1.209E+04 | 284.2 | - | - | 0 | - |
| - | - | 5554 | 285.1 | - | - | 0 | - |
| - | - | 1.554E+04 | 285.2 | - | - | 0 | - |
| - | - | 1404 | 285.2 | - | - | 0 | - |
| - | - | 1084 | 285.7 | - | - | 0 | - |
| - | - | 1843 | 286.2 | - | - | 0 | - |
| - | - | 1.125E+04 | 288.1 | - | - | 0 | - |
| - | - | 1209 | 288.2 | - | - | 0 | - |
| - | - | 2377 | 289.1 | - | - | 0 | - |
| - | - | 1013 | 289.2 | - | - | 0 | - |
| - | - | 6.998E+04 | 290.1 | - | - | 0 | - |
| - | - | 2.243E+04 | 290.2 | - | - | 0 | - |
| - | - | 8129 | 290.7 | - | - | 0 | - |
| - | - | 5226 | 291.1 | - | - | 0 | - |
| 4 | b | 1.384E+04 | 291.1 | 0.0006445 | 2.214 | +2 | 4 |
| - | - | 7357 | 291.2 | - | - | 0 | - |
| - | - | 1131 | 291.2 | - | - | 0 | - |
| - | - | 1137 | 291.6 | - | - | 0 | - |
| - | - | 1610 | 292.1 | - | - | 0 | - |
| - | - | 3.81E+04 | 292.1 | - | - | 0 | - |
| - | - | 1103 | 292.2 | - | - | 0 | - |
| - | - | 1.368E+04 | 293.1 | - | - | 0 | - |
| - | - | 4517 | 293.2 | - | - | 0 | - |
| - | - | 1.651E+04 | 294.1 | - | - | 0 | - |
| - | - | 5334 | 294.2 | - | - | 0 | - |
| - | - | 1153 | 294.7 | - | - | 0 | - |
| - | - | 2005 | 295.1 | - | - | 0 | - |
| - | - | 1064 | 296.1 | - | - | 0 | - |
| - | - | 4.148E+04 | 297.2 | - | - | 0 | - |
| - | - | 6476 | 298.2 | - | - | 0 | - |
| - | - | 1839 | 299.2 | - | - | 0 | - |
| - | - | 1128 | 299.2 | - | - | 0 | - |
| - | - | 1972 | 300.2 | - | - | 0 | - |
| - | - | 2674 | 300.7 | - | - | 0 | - |
| - | - | 2213 | 302.1 | - | - | 0 | - |
| - | - | 1.218E+04 | 303.1 | - | - | 0 | - |
| - | - | 1271 | 303.2 | - | - | 0 | - |
| - | - | 1250 | 304.1 | - | - | 0 | - |
| - | - | 7419 | 305.1 | - | - | 0 | - |
| - | - | 6734 | 305.7 | - | - | 0 | - |
| - | - | 3328 | 306.1 | - | - | 0 | - |
| - | - | 1452 | 306.1 | - | - | 0 | - |
| - | - | 1767 | 306.2 | - | - | 0 | - |
| - | - | 1634 | 307.2 | - | - | 0 | - |
| - | - | 1.457E+04 | 308.1 | - | - | 0 | - |
| - | - | 1955 | 309.1 | - | - | 0 | - |
| - | - | 3361 | 309.2 | - | - | 0 | - |
| - | - | 3401 | 309.2 | - | - | 0 | - |
| - | - | 1358 | 310.7 | - | - | 0 | - |
| - | - | 4387 | 311.2 | - | - | 0 | - |
| - | - | 9638 | 311.2 | - | - | 0 | - |
| - | - | 1336 | 312.2 | - | - | 0 | - |
| - | - | 930.2 | 313.2 | - | - | 0 | - |
| - | - | 1200 | 314.1 | - | - | 0 | - |
| - | - | 8318 | 314.7 | - | - | 0 | - |
| - | - | 3919 | 315.2 | - | - | 0 | - |
| - | - | 2927 | 318.2 | - | - | 0 | - |
| - | - | 9107 | 319.1 | - | - | 0 | - |
| - | - | 4223 | 319.2 | - | - | 0 | - |
| - | - | 5432 | 320.1 | - | - | 0 | - |
| - | - | 1483 | 320.2 | - | - | 0 | - |
| 6 | y | 6694 | 320.9 | 0.001068 | 3.328 | +3 | 9 |
| - | - | 7712 | 321.1 | - | - | 0 | - |
| - | - | 4470 | 321.2 | - | - | 0 | - |
| - | - | 4.43E+04 | 321.2 | - | - | 0 | - |
| - | - | 1709 | 321.5 | - | - | 0 | - |
| - | - | 1244 | 322.1 | - | - | 0 | - |
| - | - | 6449 | 322.2 | - | - | 0 | - |
| - | - | 1212 | 322.2 | - | - | 0 | - |
| - | - | 8455 | 322.2 | - | - | 0 | - |
| - | - | 1576 | 323.1 | - | - | 0 | - |
| - | - | 1436 | 323.2 | - | - | 0 | - |
| - | - | 1026 | 323.2 | - | - | 0 | - |
| - | - | 6368 | 324.2 | - | - | 0 | - |
| - | - | 2186 | 324.7 | - | - | 0 | - |
| - | - | 941 | 325.2 | - | - | 0 | - |
| - | - | 3986 | 325.2 | - | - | 0 | - |
| - | - | 1.797E+04 | 326.2 | - | - | 0 | - |
| 9 | y | 1084 | 326.7 | 0.001726 | 5.281 | +2 | 6 |
| - | - | 3570 | 327.2 | - | - | 0 | - |
| - | - | 2716 | 327.3 | - | - | 0 | - |
| - | - | 1462 | 329.1 | - | - | 0 | - |
| - | - | 1040 | 330.1 | - | - | 0 | - |
| - | - | 1775 | 331.7 | - | - | 0 | - |
| - | - | 1050 | 332.1 | - | - | 0 | - |
| - | - | 1.259E+04 | 332.7 | - | - | 0 | - |
| - | - | 2272 | 333.1 | - | - | 0 | - |
| - | - | 5457 | 333.2 | - | - | 0 | - |
| - | - | 1147 | 333.7 | - | - | 0 | - |
| - | - | 908.3 | 334.1 | - | - | 0 | - |
| - | - | 1884 | 334.2 | - | - | 0 | - |
| - | - | 2497 | 334.7 | - | - | 0 | - |
| - | - | 1480 | 335.2 | - | - | 0 | - |
| - | - | 3425 | 335.2 | - | - | 0 | - |
| 9 | y | 3702 | 335.2 | 0.0009632 | 2.873 | +2 | 6 |
| - | - | 1349 | 335.7 | - | - | 0 | - |
| - | - | 4791 | 336.1 | - | - | 0 | - |
| - | - | 7.264E+04 | 336.2 | - | - | 0 | - |
| - | - | 2944 | 336.7 | - | - | 0 | - |
| - | - | 1385 | 337.2 | - | - | 0 | - |
| - | - | 1.389E+04 | 337.2 | - | - | 0 | - |
| - | - | 3478 | 337.2 | - | - | 0 | - |
| - | - | 2.049E+04 | 337.7 | - | - | 0 | - |
| - | - | 1.923E+04 | 338.2 | - | - | 0 | - |
| - | - | 7051 | 338.2 | - | - | 0 | - |
| - | - | 1450 | 338.7 | - | - | 0 | - |
| - | - | 2675 | 339.2 | - | - | 0 | - |
| - | - | 7.48E+04 | 339.2 | - | - | 0 | - |
| - | - | 2416 | 339.7 | - | - | 0 | - |
| - | - | 9726 | 340.2 | - | - | 0 | - |
| - | - | 1424 | 340.2 | - | - | 0 | - |
| - | - | 1.481E+04 | 340.2 | - | - | 0 | - |
| - | - | 1362 | 341.2 | - | - | 0 | - |
| - | - | 2628 | 341.2 | - | - | 0 | - |
| - | - | 1358 | 341.2 | - | - | 0 | - |
| - | - | 3309 | 342.2 | - | - | 0 | - |
| - | - | 1069 | 343.2 | - | - | 0 | - |
| - | - | 1305 | 344.7 | - | - | 0 | - |
| - | - | 3508 | 345.7 | - | - | 0 | - |
| - | - | 2370 | 346.2 | - | - | 0 | - |
| - | - | 1446 | 346.7 | - | - | 0 | - |
| - | - | 5.774E+04 | 346.7 | - | - | 0 | - |
| - | - | 5531 | 347.1 | - | - | 0 | - |
| - | - | 2.202E+04 | 347.2 | - | - | 0 | - |
| - | - | 6063 | 347.7 | - | - | 0 | - |
| - | - | 6418 | 348.1 | - | - | 0 | - |
| - | - | 1921 | 349.1 | - | - | 0 | - |
| - | - | 3739 | 349.7 | - | - | 0 | - |
| - | - | 2612 | 350.2 | - | - | 0 | - |
| - | - | 1537 | 350.2 | - | - | 0 | - |
| - | - | 2938 | 350.2 | - | - | 0 | - |
| - | - | 1783 | 350.5 | - | - | 0 | - |
| - | - | 1712 | 351.1 | - | - | 0 | - |
| - | - | 5570 | 351.2 | - | - | 0 | - |
| - | - | 1652 | 352.2 | - | - | 0 | - |
| - | - | 4660 | 352.2 | - | - | 0 | - |
| - | - | 6000 | 353.2 | - | - | 0 | - |
| - | - | 3392 | 353.7 | - | - | 0 | - |
| - | - | 2661 | 353.9 | - | - | 0 | - |
| - | - | 1.034E+05 | 354.2 | - | - | 0 | - |
| - | - | 1377 | 354.7 | - | - | 0 | - |
| - | - | 2801 | 354.7 | - | - | 0 | - |
| - | - | 1.944E+04 | 355.2 | - | - | 0 | - |
| - | - | 1753 | 355.3 | - | - | 0 | - |
| - | - | 1798 | 355.7 | - | - | 0 | - |
| - | - | 9151 | 355.7 | - | - | 0 | - |
| - | - | 1764 | 356.2 | - | - | 0 | - |
| - | - | 3843 | 356.2 | - | - | 0 | - |
| - | - | 2190 | 356.7 | - | - | 0 | - |
| - | - | 1473 | 356.7 | - | - | 0 | - |
| - | - | 3983 | 357.2 | - | - | 0 | - |
| - | - | 2.138E+04 | 357.3 | - | - | 0 | - |
| - | - | 2.079E+04 | 358.2 | - | - | 0 | - |
| - | - | 3314 | 358.3 | - | - | 0 | - |
| - | - | 8585 | 358.7 | - | - | 0 | - |
| - | - | 1317 | 358.8 | - | - | 0 | - |
| - | - | 4633 | 359.2 | - | - | 0 | - |
| - | - | 1804 | 359.2 | - | - | 0 | - |
| - | - | 2315 | 359.2 | - | - | 0 | - |
| - | - | 1951 | 359.5 | - | - | 0 | - |
| - | - | 3309 | 359.5 | - | - | 0 | - |
| - | - | 1539 | 359.7 | - | - | 0 | - |
| - | - | 5681 | 359.9 | - | - | 0 | - |
| - | - | 1366 | 360.1 | - | - | 0 | - |
| - | - | 1218 | 360.2 | - | - | 0 | - |
| - | - | 4072 | 360.7 | - | - | 0 | - |
| - | - | 5228 | 360.7 | - | - | 0 | - |
| - | - | 1559 | 361.1 | - | - | 0 | - |
| - | - | 4186 | 361.2 | - | - | 0 | - |
| - | - | 2644 | 361.7 | - | - | 0 | - |
| - | - | 4693 | 362.2 | - | - | 0 | - |
| - | - | 1366 | 362.7 | - | - | 0 | - |
| - | - | 1777 | 362.8 | - | - | 0 | - |
| - | - | 2270 | 363.2 | - | - | 0 | - |
| - | - | 1882 | 363.2 | - | - | 0 | - |
| - | - | 1520 | 363.2 | - | - | 0 | - |
| - | - | 1597 | 363.5 | - | - | 0 | - |
| 8 | y | 2607 | 363.7 | 0.001797 | 4.942 | +2 | 7 |
| - | - | 4603 | 364.1 | - | - | 0 | - |
| - | - | 1337 | 364.3 | - | - | 0 | - |
| - | - | 1583 | 364.5 | - | - | 0 | - |
| - | - | 1424 | 364.7 | - | - | 0 | - |
| - | - | 1733 | 365.3 | - | - | 0 | - |
| - | - | 1889 | 365.9 | - | - | 0 | - |
| - | - | 6824 | 366.2 | - | - | 0 | - |
| - | - | 2309 | 366.2 | - | - | 0 | - |
| 5 | y | 9829 | 366.6 | 0.001043 | 2.846 | +3 | 10 |
| - | - | 5125 | 366.9 | - | - | 0 | - |
| - | - | 6919 | 367.2 | - | - | 0 | - |
| - | - | 1762 | 367.2 | - | - | 0 | - |
| - | - | 1.214E+04 | 368.1 | - | - | 0 | - |
| - | - | 4623 | 368.2 | - | - | 0 | - |
| - | - | 1511 | 368.2 | - | - | 0 | - |
| - | - | 2505 | 368.5 | - | - | 0 | - |
| - | - | 2338 | 368.8 | - | - | 0 | - |
| - | - | 1985 | 369.2 | - | - | 0 | - |
| - | - | 2258 | 369.2 | - | - | 0 | - |
| - | - | 4406 | 369.7 | - | - | 0 | - |
| - | - | 4402 | 370.2 | - | - | 0 | - |
| - | - | 2331 | 370.2 | - | - | 0 | - |
| - | - | 1284 | 370.5 | - | - | 0 | - |
| - | - | 2165 | 370.7 | - | - | 0 | - |
| - | - | 1105 | 370.7 | - | - | 0 | - |
| - | - | 1146 | 371.2 | - | - | 0 | - |
| - | - | 1313 | 371.6 | - | - | 0 | - |
| - | - | 1498 | 371.9 | - | - | 0 | - |
| - | - | 1524 | 373.2 | - | - | 0 | - |
| 12 | y | 3.366E+04 | 373.3 | 0.001294 | 3.468 | +1 | 3 |
| 9 | b | 2001 | 373.8 | 0.002082 | 5.568 | +3 | 9 |
| - | - | 1151 | 374.2 | - | - | 0 | - |
| - | - | 6772 | 374.3 | - | - | 0 | - |
| - | - | 1974 | 374.8 | - | - | 0 | - |
| - | - | 3250 | 375.2 | - | - | 0 | - |
| - | - | 2340 | 375.3 | - | - | 0 | - |
| - | - | 1925 | 375.7 | - | - | 0 | - |
| - | - | 2422 | 376.1 | - | - | 0 | - |
| - | - | 3118 | 376.2 | - | - | 0 | - |
| - | - | 894.6 | 377.1 | - | - | 0 | - |
| - | - | 1291 | 377.2 | - | - | 0 | - |
| - | - | 1586 | 377.8 | - | - | 0 | - |
| - | - | 2546 | 378.1 | - | - | 0 | - |
| - | - | 1971 | 378.2 | - | - | 0 | - |
| - | - | 2503 | 378.5 | - | - | 0 | - |
| - | - | 1728 | 378.7 | - | - | 0 | - |
| - | - | 1473 | 379.2 | - | - | 0 | - |
| 9 | b | 1894 | 379.8 | 0.003107 | 8.18 | +3 | 9 |
| - | - | 1062 | 380.1 | - | - | 0 | - |
| - | - | 1282 | 380.2 | - | - | 0 | - |
| - | - | 6034 | 380.2 | - | - | 0 | - |
| - | - | 2720 | 382.2 | - | - | 0 | - |
| - | - | 9949 | 382.2 | - | - | 0 | - |
| - | - | 4375 | 382.3 | - | - | 0 | - |
| - | - | 5044 | 382.5 | - | - | 0 | - |
| - | - | 1743 | 382.7 | - | - | 0 | - |
| - | - | 2664 | 382.7 | - | - | 0 | - |
| - | - | 3405 | 383 | - | - | 0 | - |
| - | - | 1938 | 383.2 | - | - | 0 | - |
| - | - | 3918 | 383.3 | - | - | 0 | - |
| - | - | 1408 | 383.7 | - | - | 0 | - |
| - | - | 1676 | 384.1 | - | - | 0 | - |
| - | - | 1.04E+04 | 384.2 | - | - | 0 | - |
| - | - | 5406 | 384.6 | - | - | 0 | - |
| - | - | 1318 | 384.7 | - | - | 0 | - |
| - | - | 4274 | 384.9 | - | - | 0 | - |
| - | - | 4.239E+04 | 385.2 | - | - | 0 | - |
| - | - | 7754 | 385.2 | - | - | 0 | - |
| - | - | 4.24E+04 | 386.2 | - | - | 0 | - |
| - | - | 1306 | 386.2 | - | - | 0 | - |
| - | - | 1600 | 386.7 | - | - | 0 | - |
| - | - | 1275 | 387 | - | - | 0 | - |
| - | - | 6900 | 387.2 | - | - | 0 | - |
| - | - | 2834 | 387.5 | - | - | 0 | - |
| - | - | 1924 | 387.8 | - | - | 0 | - |
| - | - | 1844 | 388.1 | - | - | 0 | - |
| - | - | 2475 | 388.2 | - | - | 0 | - |
| - | - | 1166 | 388.5 | - | - | 0 | - |
| - | - | 2469 | 389.2 | - | - | 0 | - |
| - | - | 6833 | 389.3 | - | - | 0 | - |
| - | - | 2194 | 389.7 | - | - | 0 | - |
| - | - | 4313 | 389.8 | - | - | 0 | - |
| - | - | 2040 | 390 | - | - | 0 | - |
| - | - | 1845 | 390.7 | - | - | 0 | - |
| - | - | 1955 | 391.9 | - | - | 0 | - |
| - | - | 1786 | 392.2 | - | - | 0 | - |
| - | - | 1.536E+04 | 392.3 | - | - | 0 | - |
| - | - | 1926 | 393.2 | - | - | 0 | - |
| - | - | 1770 | 393.3 | - | - | 0 | - |
| - | - | 5198 | 393.7 | - | - | 0 | - |
| - | - | 8711 | 394 | - | - | 0 | - |
| - | - | 1.967E+04 | 394.2 | - | - | 0 | - |
| - | - | 1932 | 394.2 | - | - | 0 | - |
| 2 | y | 1.177E+04 | 394.2 | 0.003275 | 8.307 | +4 | 13 |
| - | - | 1293 | 394.3 | - | - | 0 | - |
| - | - | 1.233E+04 | 394.5 | - | - | 0 | - |
| - | - | 3611 | 394.5 | - | - | 0 | - |
| - | - | 6387 | 394.7 | - | - | 0 | - |
| - | - | 2020 | 394.9 | - | - | 0 | - |
| - | - | 1611 | 395 | - | - | 0 | - |
| - | - | 4917 | 395.2 | - | - | 0 | - |
| - | - | 2179 | 395.7 | - | - | 0 | - |
| - | - | 4712 | 396.3 | - | - | 0 | - |
| - | - | 1459 | 396.7 | - | - | 0 | - |
| - | - | 1166 | 396.9 | - | - | 0 | - |
| - | - | 2447 | 397 | - | - | 0 | - |
| - | - | 1743 | 397.2 | - | - | 0 | - |
| - | - | 2375 | 397.2 | - | - | 0 | - |
| - | - | 3666 | 397.2 | - | - | 0 | - |
| - | - | 2969 | 397.5 | - | - | 0 | - |
| 10 | b | 4319 | 397.5 | 0.002243 | 5.642 | +3 | 10 |
| - | - | 2502 | 397.6 | - | - | 0 | - |
| - | - | 1775 | 397.7 | - | - | 0 | - |
| 10 | b | 3606 | 397.8 | 0.005933 | 14.91 | +3 | 10 |
| - | - | 1495 | 397.9 | - | - | 0 | - |
| - | - | 3750 | 398 | - | - | 0 | - |
| - | - | 3775 | 398.2 | - | - | 0 | - |
| - | - | 2205 | 398.2 | - | - | 0 | - |
| - | - | 1833 | 398.2 | - | - | 0 | - |
| 2 | y | 9535 | 398.5 | 0.001947 | 4.887 | +4 | 13 |
| - | - | 1.043E+04 | 398.7 | - | - | 0 | - |
| - | - | 3905 | 399 | - | - | 0 | - |
| - | - | 1253 | 399.2 | - | - | 0 | - |
| - | - | 2687 | 399.2 | - | - | 0 | - |
| - | - | 1646 | 399.6 | - | - | 0 | - |
| - | - | 1579 | 400.2 | - | - | 0 | - |
| - | - | 2536 | 400.2 | - | - | 0 | - |
| - | - | 2217 | 401.2 | - | - | 0 | - |
| - | - | 5791 | 401.2 | - | - | 0 | - |
| - | - | 3115 | 401.3 | - | - | 0 | - |
| - | - | 3034 | 401.8 | - | - | 0 | - |
| - | - | 2021 | 402.2 | - | - | 0 | - |
| - | - | 1777 | 402.2 | - | - | 0 | - |
| - | - | 1.51E+04 | 402.2 | - | - | 0 | - |
| - | - | 1153 | 402.6 | - | - | 0 | - |
| - | - | 3.913E+04 | 403.2 | - | - | 0 | - |
| - | - | 1.536E+04 | 403.2 | - | - | 0 | - |
| - | - | 6698 | 403.3 | - | - | 0 | - |
| 10 | b | 3902 | 403.5 | 0.00217 | 5.377 | +3 | 10 |
| 4 | y | 2.589E+04 | 403.6 | 0.003795 | 9.403 | +3 | 11 |
| 7 | y | 3063 | 403.8 | 0.007825 | 19.38 | +2 | 8 |
| - | - | 5298 | 403.9 | - | - | 0 | - |
| - | - | 1.334E+04 | 403.9 | - | - | 0 | - |
| - | - | 2555 | 404 | - | - | 0 | - |
| - | - | 7221 | 404.2 | - | - | 0 | - |
| - | - | 3964 | 404.2 | - | - | 0 | - |
| - | - | 2216 | 404.2 | - | - | 0 | - |
| - | - | 4191 | 404.5 | - | - | 0 | - |
| - | - | 2895 | 404.7 | - | - | 0 | - |
| - | - | 1117 | 404.7 | - | - | 0 | - |
| - | - | 3353 | 405.2 | - | - | 0 | - |
| - | - | 1393 | 405.2 | - | - | 0 | - |
| - | - | 1328 | 405.2 | - | - | 0 | - |
| - | - | 1453 | 405.7 | - | - | 0 | - |
| - | - | 6345 | 406.2 | - | - | 0 | - |
| - | - | 1962 | 406.2 | - | - | 0 | - |
| - | - | 1.192E+04 | 406.3 | - | - | 0 | - |
| - | - | 2180 | 406.7 | - | - | 0 | - |
| - | - | 3921 | 406.8 | - | - | 0 | - |
| - | - | 1446 | 407 | - | - | 0 | - |
| - | - | 1357 | 407.2 | - | - | 0 | - |
| - | - | 1932 | 407.3 | - | - | 0 | - |
| - | - | 2982 | 407.9 | - | - | 0 | - |
| - | - | 2209 | 408 | - | - | 0 | - |
| - | - | 1.073E+04 | 408.2 | - | - | 0 | - |
| - | - | 1.252E+04 | 408.5 | - | - | 0 | - |
| - | - | 4970 | 408.6 | - | - | 0 | - |
| - | - | 1.224E+04 | 408.7 | - | - | 0 | - |
| - | - | 3033 | 408.9 | - | - | 0 | - |
| - | - | 1.376E+04 | 409 | - | - | 0 | - |
| - | - | 1353 | 409.2 | - | - | 0 | - |
| 4 | y | 6389 | 409.2 | 0.0005807 | 1.419 | +3 | 11 |
| - | - | 4847 | 409.6 | - | - | 0 | - |
| - | - | 7446 | 410.1 | - | - | 0 | - |
| - | - | 3.454E+04 | 410.3 | - | - | 0 | - |
| - | - | 9041 | 410.6 | - | - | 0 | - |
| - | - | 5076 | 410.9 | - | - | 0 | - |
| - | - | 3164 | 411 | - | - | 0 | - |
| - | - | 1196 | 411.1 | - | - | 0 | - |
| - | - | 2.842E+04 | 411.2 | - | - | 0 | - |
| - | - | 1808 | 411.2 | - | - | 0 | - |
| - | - | 1318 | 411.2 | - | - | 0 | - |
| - | - | 1.509E+04 | 411.3 | - | - | 0 | - |
| - | - | 6318 | 411.5 | - | - | 0 | - |
| - | - | 1246 | 411.6 | - | - | 0 | - |
| - | - | 3308 | 411.7 | - | - | 0 | - |
| - | - | 6298 | 411.8 | - | - | 0 | - |
| - | - | 7827 | 412.2 | - | - | 0 | - |
| - | - | 4089 | 412.2 | - | - | 0 | - |
| 7 | y | 9.955E+04 | 412.3 | 0.001508 | 3.658 | +2 | 8 |
| - | - | 4591 | 412.7 | - | - | 0 | - |
| - | - | 4.667E+04 | 412.8 | - | - | 0 | - |
| - | - | 1230 | 413 | - | - | 0 | - |
| - | - | 6288 | 413.2 | - | - | 0 | - |
| - | - | 1769 | 413.2 | - | - | 0 | - |
| - | - | 1.324E+04 | 413.3 | - | - | 0 | - |
| - | - | 3959 | 413.6 | - | - | 0 | - |
| - | - | 1074 | 413.7 | - | - | 0 | - |
| - | - | 2410 | 413.8 | - | - | 0 | - |
| - | - | 7741 | 413.9 | - | - | 0 | - |
| - | - | 8484 | 414.2 | - | - | 0 | - |
| - | - | 7454 | 414.2 | - | - | 0 | - |
| - | - | 5388 | 414.6 | - | - | 0 | - |
| - | - | 3714 | 414.7 | - | - | 0 | - |
| - | - | 1405 | 414.9 | - | - | 0 | - |
| - | - | 1679 | 415.2 | - | - | 0 | - |
| - | - | 5020 | 415.2 | - | - | 0 | - |
| - | - | 1.008E+04 | 415.3 | - | - | 0 | - |
| - | - | 4447 | 415.5 | - | - | 0 | - |
| - | - | 1254 | 415.6 | - | - | 0 | - |
| 0 | Precursor | 1.502E+04 | 415.7 | 0.001495 | 3.596 | +4 | -1 |
| - | - | 4730 | 415.8 | - | - | 0 | - |
| - | - | 3504 | 415.9 | - | - | 0 | - |
| 0 | Precursor | 1.596E+04 | 416 | 0.006376 | 15.33 | +4 | -1 |
| - | - | 1941 | 416.2 | - | - | 0 | - |
| - | - | 6761 | 416.2 | - | - | 0 | - |
| - | - | 1672 | 416.3 | - | - | 0 | - |
| - | - | 4485 | 416.5 | - | - | 0 | - |
| - | - | 2280 | 416.7 | - | - | 0 | - |
| - | - | 3220 | 418.6 | - | - | 0 | - |
| - | - | 1873 | 418.7 | - | - | 0 | - |
| - | - | 2034 | 418.9 | - | - | 0 | - |
| 6 | b | 2292 | 419.2 | 0.007158 | 17.08 | +2 | 6 |
| - | - | 8618 | 419.2 | - | - | 0 | - |
| - | - | 1.127E+04 | 419.3 | - | - | 0 | - |
| - | - | 6630 | 419.6 | - | - | 0 | - |
| - | - | 9760 | 419.7 | - | - | 0 | - |
| - | - | 9213 | 419.8 | - | - | 0 | - |
| - | - | 3.246E+04 | 419.9 | - | - | 0 | - |
| - | - | 9776 | 420 | - | - | 0 | - |
| - | - | 4290 | 420.2 | - | - | 0 | - |
| 0 | Precursor | 6.902E+04 | 420.2 | 0.004835 | 11.51 | +4 | -1 |
| - | - | 5736 | 420.3 | - | - | 0 | - |
| - | - | 5.686E+04 | 420.5 | - | - | 0 | - |
| - | - | 7502 | 420.6 | - | - | 0 | - |
| - | - | 1605 | 420.7 | - | - | 0 | - |
| - | - | 2.213E+04 | 420.7 | - | - | 0 | - |
| - | - | 2608 | 420.8 | - | - | 0 | - |
| - | - | 2472 | 420.9 | - | - | 0 | - |
| - | - | 1.198E+04 | 421 | - | - | 0 | - |
| - | - | 3237 | 421.2 | - | - | 0 | - |
| - | - | 5411 | 421.9 | - | - | 0 | - |
| - | - | 3457 | 422.2 | - | - | 0 | - |
| - | - | 1.058E+04 | 422.3 | - | - | 0 | - |
| - | - | 2349 | 422.3 | - | - | 0 | - |
| - | - | 1604 | 422.6 | - | - | 0 | - |
| - | - | 5274 | 423.3 | - | - | 0 | - |
| - | - | 2230 | 424.3 | - | - | 0 | - |
| - | - | 7617 | 424.3 | - | - | 0 | - |
| - | - | 5691 | 424.6 | - | - | 0 | - |
| - | - | 1.078E+04 | 424.7 | - | - | 0 | - |
| - | - | 1799 | 424.8 | - | - | 0 | - |
| - | - | 6026 | 424.9 | - | - | 0 | - |
| - | - | 1.413E+04 | 425.2 | - | - | 0 | - |
| - | - | 4741 | 425.3 | - | - | 0 | - |
| - | - | 1828 | 425.3 | - | - | 0 | - |
| - | - | 1.645E+04 | 425.6 | - | - | 0 | - |
| - | - | 1540 | 425.7 | - | - | 0 | - |
| - | - | 1.009E+04 | 425.9 | - | - | 0 | - |
| - | - | 4748 | 426.2 | - | - | 0 | - |
| - | - | 3322 | 426.6 | - | - | 0 | - |
| - | - | 9645 | 427.2 | - | - | 0 | - |
| - | - | 9065 | 427.2 | - | - | 0 | - |
| - | - | 4524 | 427.7 | - | - | 0 | - |
| - | - | 1324 | 427.9 | - | - | 0 | - |
| 6 | b | 8187 | 428.2 | 0.003218 | 7.516 | +2 | 6 |
| - | - | 1868 | 428.3 | - | - | 0 | - |
| - | - | 4191 | 428.3 | - | - | 0 | - |
| - | - | 2748 | 428.7 | - | - | 0 | - |
| - | - | 9.397E+04 | 429.2 | - | - | 0 | - |
| - | - | 1926 | 429.7 | - | - | 0 | - |
| 11 | b | 2.357E+04 | 430.2 | 0.005724 | 13.31 | +3 | 11 |
| - | - | 1698 | 430.9 | - | - | 0 | - |
| - | - | 1.626E+04 | 431.2 | - | - | 0 | - |
| - | - | 3354 | 431.2 | - | - | 0 | - |
| - | - | 3317 | 431.6 | - | - | 0 | - |
| - | - | 4468 | 432.2 | - | - | 0 | - |
| - | - | 7083 | 432.7 | - | - | 0 | - |
| - | - | 2542 | 433.2 | - | - | 0 | - |
| - | - | 2684 | 433.2 | - | - | 0 | - |
| - | - | 2063 | 433.3 | - | - | 0 | - |
| - | - | 3360 | 433.7 | - | - | 0 | - |
| - | - | 1730 | 435.3 | - | - | 0 | - |
| - | - | 3610 | 437.2 | - | - | 0 | - |
| - | - | 2003 | 438.2 | - | - | 0 | - |
| - | - | 1418 | 438.6 | - | - | 0 | - |
| - | - | 1518 | 439.2 | - | - | 0 | - |
| - | - | 2530 | 439.3 | - | - | 0 | - |
| - | - | 1533 | 442.6 | - | - | 0 | - |
| - | - | 4225 | 442.7 | - | - | 0 | - |
| - | - | 2802 | 443.2 | - | - | 0 | - |
| - | - | 2771 | 443.2 | - | - | 0 | - |
| - | - | 1832 | 443.6 | - | - | 0 | - |
| - | - | 1947 | 443.7 | - | - | 0 | - |
| - | - | 5571 | 443.9 | - | - | 0 | - |
| - | - | 3608 | 444.2 | - | - | 0 | - |
| - | - | 2.573E+04 | 445.2 | - | - | 0 | - |
| - | - | 3783 | 445.3 | - | - | 0 | - |
| - | - | 4496 | 446.2 | - | - | 0 | - |
| - | - | 6495 | 446.2 | - | - | 0 | - |
| - | - | 4450 | 446.7 | - | - | 0 | - |
| - | - | 1716 | 447.9 | - | - | 0 | - |
| - | - | 7385 | 448.3 | - | - | 0 | - |
| - | - | 3980 | 448.6 | - | - | 0 | - |
| - | - | 1508 | 448.9 | - | - | 0 | - |
| - | - | 1758 | 449.3 | - | - | 0 | - |
| - | - | 1362 | 449.3 | - | - | 0 | - |
| - | - | 3394 | 449.6 | - | - | 0 | - |
| - | - | 1373 | 449.9 | - | - | 0 | - |
| - | - | 1242 | 450.3 | - | - | 0 | - |
| - | - | 1381 | 451.2 | - | - | 0 | - |
| - | - | 9708 | 451.2 | - | - | 0 | - |
| - | - | 5655 | 451.3 | - | - | 0 | - |
| - | - | 7195 | 451.7 | - | - | 0 | - |
| - | - | 1.303E+04 | 451.9 | - | - | 0 | - |
| - | - | 9335 | 452.3 | - | - | 0 | - |
| - | - | 5031 | 452.3 | - | - | 0 | - |
| - | - | 6389 | 452.6 | - | - | 0 | - |
| - | - | 5917 | 452.9 | - | - | 0 | - |
| 3 | b | 4550 | 453.2 | 0.008718 | 19.24 | +1 | 3 |
| - | - | 1574 | 453.2 | - | - | 0 | - |
| - | - | 7626 | 453.6 | - | - | 0 | - |
| - | - | 5639 | 453.9 | - | - | 0 | - |
| - | - | 2370 | 454.2 | - | - | 0 | - |
| - | - | 3917 | 454.3 | - | - | 0 | - |
| - | - | 1932 | 454.7 | - | - | 0 | - |
| - | - | 1.312E+04 | 455.3 | - | - | 0 | - |
| - | - | 2775 | 456.3 | - | - | 0 | - |
| - | - | 4360 | 456.6 | - | - | 0 | - |
| - | - | 6349 | 456.9 | - | - | 0 | - |
| - | - | 4157 | 457.2 | - | - | 0 | - |
| - | - | 1.06E+04 | 457.3 | - | - | 0 | - |
| - | - | 1.37E+04 | 457.6 | - | - | 0 | - |
| 3 | y | 1.35E+04 | 457.9 | 0.002593 | 5.662 | +3 | 12 |
| - | - | 2199 | 458.2 | - | - | 0 | - |
| - | - | 1633 | 458.2 | - | - | 0 | - |
| - | - | 7644 | 458.3 | - | - | 0 | - |
| - | - | 2322 | 458.6 | - | - | 0 | - |
| - | - | 1441 | 458.9 | - | - | 0 | - |
| - | - | 1712 | 459.2 | - | - | 0 | - |
| - | - | 4998 | 459.2 | - | - | 0 | - |
| - | - | 3535 | 459.7 | - | - | 0 | - |
| - | - | 1.699E+04 | 460.2 | - | - | 0 | - |
| - | - | 9755 | 460.7 | - | - | 0 | - |
| - | - | 2002 | 461.2 | - | - | 0 | - |
| - | - | 2179 | 461.7 | - | - | 0 | - |
| - | - | 1664 | 462.2 | - | - | 0 | - |
| - | - | 3888 | 462.3 | - | - | 0 | - |
| - | - | 6878 | 462.6 | - | - | 0 | - |
| - | - | 2.842E+04 | 462.9 | - | - | 0 | - |
| - | - | 2.08E+04 | 463.3 | - | - | 0 | - |
| 3 | y | 1.452E+05 | 463.6 | 0.001332 | 2.873 | +3 | 12 |
| - | - | 1.187E+05 | 463.9 | - | - | 0 | - |
| - | - | 4.617E+04 | 464.3 | - | - | 0 | - |
| - | - | 1.715E+04 | 464.6 | - | - | 0 | - |
| - | - | 2997 | 464.9 | - | - | 0 | - |
| - | - | 2513 | 466.9 | - | - | 0 | - |
| - | - | 1579 | 467.3 | - | - | 0 | - |
| - | - | 1.015E+04 | 467.3 | - | - | 0 | - |
| - | - | 1372 | 467.6 | - | - | 0 | - |
| - | - | 1721 | 467.9 | - | - | 0 | - |
| - | - | 4937 | 468.2 | - | - | 0 | - |
| - | - | 3.607E+04 | 468.3 | - | - | 0 | - |
| - | - | 1343 | 468.3 | - | - | 0 | - |
| - | - | 1.083E+04 | 468.3 | - | - | 0 | - |
| - | - | 2.742E+04 | 468.6 | - | - | 0 | - |
| - | - | 3790 | 468.7 | - | - | 0 | - |
| - | - | 1.056E+05 | 468.9 | - | - | 0 | - |
| - | - | 3116 | 469.2 | - | - | 0 | - |
| - | - | 8.486E+04 | 469.3 | - | - | 0 | - |
| - | - | 3029 | 469.3 | - | - | 0 | - |
| - | - | 3.301E+04 | 469.6 | - | - | 0 | - |
| - | - | 5239 | 469.8 | - | - | 0 | - |
| - | - | 1.542E+04 | 469.9 | - | - | 0 | - |
| - | - | 2586 | 470.2 | - | - | 0 | - |
| 11 | y | 1.468E+05 | 470.3 | 0.001631 | 3.468 | +1 | 4 |
| - | - | 4.032E+04 | 471.3 | - | - | 0 | - |
| - | - | 2954 | 471.6 | - | - | 0 | - |
| - | - | 7724 | 471.8 | - | - | 0 | - |
| - | - | 1581 | 471.9 | - | - | 0 | - |
| - | - | 3032 | 472.3 | - | - | 0 | - |
| - | - | 6140 | 472.3 | - | - | 0 | - |
| - | - | 5781 | 472.3 | - | - | 0 | - |
| - | - | 3482 | 472.6 | - | - | 0 | - |
| - | - | 1918 | 472.8 | - | - | 0 | - |
| - | - | 1463 | 472.9 | - | - | 0 | - |
| - | - | 1.788E+04 | 473.3 | - | - | 0 | - |
| - | - | 5448 | 473.6 | - | - | 0 | - |
| - | - | 1890 | 473.9 | - | - | 0 | - |
| - | - | 5106 | 474.3 | - | - | 0 | - |
| - | - | 5890 | 474.8 | - | - | 0 | - |
| - | - | 1770 | 474.9 | - | - | 0 | - |
| - | - | 1739 | 475.2 | - | - | 0 | - |
| - | - | 4962 | 475.3 | - | - | 0 | - |
| - | - | 5277 | 475.6 | - | - | 0 | - |
| - | - | 3232 | 475.9 | - | - | 0 | - |
| - | - | 2625 | 476.2 | - | - | 0 | - |
| - | - | 9081 | 476.3 | - | - | 0 | - |
| - | - | 7841 | 476.6 | - | - | 0 | - |
| - | - | 4335 | 476.9 | - | - | 0 | - |
| - | - | 1342 | 477.6 | - | - | 0 | - |
| - | - | 1574 | 477.9 | - | - | 0 | - |
| 12 | b | 5318 | 478.6 | 0.002396 | 5.007 | +3 | 12 |
| - | - | 2666 | 478.8 | - | - | 0 | - |
| - | - | 8907 | 478.9 | - | - | 0 | - |
| - | - | 4381 | 479.2 | - | - | 0 | - |
| - | - | 1504 | 479.3 | - | - | 0 | - |
| - | - | 1317 | 479.6 | - | - | 0 | - |
| - | - | 2760 | 479.8 | - | - | 0 | - |
| 6 | y | 2.323E+04 | 480.8 | 0.001654 | 3.44 | +2 | 9 |
| - | - | 3724 | 480.9 | - | - | 0 | - |
| - | - | 6191 | 481.3 | - | - | 0 | - |
| - | - | 1.27E+04 | 481.3 | - | - | 0 | - |
| - | - | 1.53E+04 | 481.6 | - | - | 0 | - |
| - | - | 3829 | 481.8 | - | - | 0 | - |
| - | - | 1.334E+04 | 481.9 | - | - | 0 | - |
| - | - | 6332 | 482.3 | - | - | 0 | - |
| - | - | 1.131E+04 | 482.3 | - | - | 0 | - |
| - | - | 1929 | 482.6 | - | - | 0 | - |
| - | - | 8672 | 482.8 | - | - | 0 | - |
| - | - | 1454 | 483.2 | - | - | 0 | - |
| - | - | 4107 | 483.3 | - | - | 0 | - |
| - | - | 2780 | 483.3 | - | - | 0 | - |
| - | - | 1.565E+04 | 483.8 | - | - | 0 | - |
| - | - | 1847 | 484.2 | - | - | 0 | - |
| - | - | 8646 | 484.3 | - | - | 0 | - |
| - | - | 1424 | 484.3 | - | - | 0 | - |
| - | - | 2251 | 484.6 | - | - | 0 | - |
| - | - | 2547 | 484.8 | - | - | 0 | - |
| - | - | 3666 | 486.3 | - | - | 0 | - |
| - | - | 4096 | 486.6 | - | - | 0 | - |
| - | - | 4574 | 486.9 | - | - | 0 | - |
| - | - | 1.089E+04 | 487.3 | - | - | 0 | - |
| - | - | 1.101E+04 | 487.6 | - | - | 0 | - |
| - | - | 3190 | 487.8 | - | - | 0 | - |
| - | - | 5676 | 487.9 | - | - | 0 | - |
| - | - | 1849 | 488.3 | - | - | 0 | - |
| - | - | 1.295E+04 | 491.3 | - | - | 0 | - |
| - | - | 2293 | 491.6 | - | - | 0 | - |
| - | - | 2910 | 491.8 | - | - | 0 | - |
| - | - | 4281 | 492.3 | - | - | 0 | - |
| - | - | 2295 | 492.6 | - | - | 0 | - |
| - | - | 3623 | 492.8 | - | - | 0 | - |
| - | - | 2270 | 492.9 | - | - | 0 | - |
| - | - | 2036 | 493.3 | - | - | 0 | - |
| - | - | 2568 | 493.3 | - | - | 0 | - |
| - | - | 1599 | 493.4 | - | - | 0 | - |
| - | - | 3935 | 493.6 | - | - | 0 | - |
| - | - | 3490 | 494.2 | - | - | 0 | - |
| - | - | 1431 | 494.6 | - | - | 0 | - |
| - | - | 2127 | 494.9 | - | - | 0 | - |
| - | - | 1903 | 495.4 | - | - | 0 | - |
| - | - | 2395 | 495.6 | - | - | 0 | - |
| - | - | 1789 | 497.7 | - | - | 0 | - |
| - | - | 3663 | 500.6 | - | - | 0 | - |
| - | - | 2744 | 500.9 | - | - | 0 | - |
| - | - | 2572 | 501.3 | - | - | 0 | - |
| - | - | 7560 | 503.2 | - | - | 0 | - |
| - | - | 2501 | 504.2 | - | - | 0 | - |
| - | - | 1966 | 504.9 | - | - | 0 | - |
| 8 | b | 3231 | 505.2 | 0.003275 | 6.482 | +2 | 8 |
| - | - | 2790 | 505.3 | - | - | 0 | - |
| - | - | 2672 | 505.6 | - | - | 0 | - |
| - | - | 2466 | 505.7 | - | - | 0 | - |
| - | - | 1458 | 506.3 | - | - | 0 | - |
| - | - | 3383 | 506.6 | - | - | 0 | - |
| - | - | 5192 | 506.9 | - | - | 0 | - |
| - | - | 3399 | 507.3 | - | - | 0 | - |
| - | - | 2352 | 507.6 | - | - | 0 | - |
| - | - | 2550 | 508.2 | - | - | 0 | - |
| - | - | 1904 | 508.8 | - | - | 0 | - |
| - | - | 1493 | 509.6 | - | - | 0 | - |
| - | - | 1890 | 509.9 | - | - | 0 | - |
| 13 | b | 7967 | 510.3 | 0.003521 | 6.901 | +3 | 13 |
| 13 | b | 5627 | 510.6 | 0.008341 | 16.34 | +3 | 13 |
| - | - | 7537 | 510.9 | - | - | 0 | - |
| - | - | 8514 | 511.3 | - | - | 0 | - |
| - | - | 2829 | 511.6 | - | - | 0 | - |
| - | - | 1627 | 512.2 | - | - | 0 | - |
| - | - | 2408 | 513.7 | - | - | 0 | - |
| - | - | 2473 | 515.6 | - | - | 0 | - |
| - | - | 2906 | 515.9 | - | - | 0 | - |
| 13 | b | 1.75E+04 | 516.3 | 0.002655 | 5.142 | +3 | 13 |
| - | - | 1.397E+04 | 516.6 | - | - | 0 | - |
| - | - | 8311 | 516.9 | - | - | 0 | - |
| - | - | 1473 | 517.2 | - | - | 0 | - |
| - | - | 7593 | 517.3 | - | - | 0 | - |
| - | - | 2658 | 519.2 | - | - | 0 | - |
| - | - | 3990 | 519.3 | - | - | 0 | - |
| - | - | 4136 | 519.6 | - | - | 0 | - |
| - | - | 3124 | 519.9 | - | - | 0 | - |
| - | - | 2027 | 520.2 | - | - | 0 | - |
| - | - | 7815 | 520.4 | - | - | 0 | - |
| - | - | 9940 | 521.2 | - | - | 0 | - |
| - | - | 3150 | 521.6 | - | - | 0 | - |
| - | - | 4603 | 521.9 | - | - | 0 | - |
| - | - | 2857 | 522.2 | - | - | 0 | - |
| - | - | 2.505E+04 | 522.3 | - | - | 0 | - |
| - | - | 2.122E+04 | 522.6 | - | - | 0 | - |
| - | - | 8061 | 522.9 | - | - | 0 | - |
| - | - | 3936 | 523.3 | - | - | 0 | - |
| - | - | 1941 | 523.4 | - | - | 0 | - |
| - | - | 1940 | 523.6 | - | - | 0 | - |
| - | - | 1680 | 524.4 | - | - | 0 | - |
| - | - | 6292 | 524.6 | - | - | 0 | - |
| - | - | 1656 | 524.7 | - | - | 0 | - |
| - | - | 2222 | 524.8 | - | - | 0 | - |
| - | - | 1.474E+04 | 525 | - | - | 0 | - |
| 2 | y | 2.232E+04 | 525.3 | 0.005835 | 11.11 | +3 | 13 |
| - | - | 1.725E+04 | 525.6 | - | - | 0 | - |
| - | - | 2691 | 525.7 | - | - | 0 | - |
| - | - | 7247 | 526 | - | - | 0 | - |
| - | - | 1899 | 526.2 | - | - | 0 | - |
| - | - | 2916 | 526.3 | - | - | 0 | - |
| - | - | 2586 | 526.3 | - | - | 0 | - |
| - | - | 1699 | 526.8 | - | - | 0 | - |
| - | - | 2531 | 528.2 | - | - | 0 | - |
| - | - | 4177 | 529.8 | - | - | 0 | - |
| - | - | 1.9E+04 | 530.3 | - | - | 0 | - |
| - | - | 9860 | 530.6 | - | - | 0 | - |
| - | - | 3415 | 530.8 | - | - | 0 | - |
| 2 | y | 4.244E+04 | 531 | 0.002804 | 5.282 | +3 | 13 |
| - | - | 6790 | 531.2 | - | - | 0 | - |
| - | - | 3.61E+04 | 531.3 | - | - | 0 | - |
| - | - | 1.286E+04 | 531.6 | - | - | 0 | - |
| - | - | 5486 | 532 | - | - | 0 | - |
| - | - | 2743 | 533.8 | - | - | 0 | - |
| - | - | 1731 | 536.4 | - | - | 0 | - |
| - | - | 2664 | 537.2 | - | - | 0 | - |
| - | - | 2406 | 537.3 | - | - | 0 | - |
| - | - | 5.698E+04 | 538.3 | - | - | 0 | - |
| - | - | 4240 | 538.4 | - | - | 0 | - |
| - | - | 8735 | 538.8 | - | - | 0 | - |
| - | - | 1.632E+04 | 539.3 | - | - | 0 | - |
| - | - | 1.527E+04 | 539.3 | - | - | 0 | - |
| - | - | 2124 | 539.4 | - | - | 0 | - |
| - | - | 7829 | 539.8 | - | - | 0 | - |
| - | - | 2483 | 540.2 | - | - | 0 | - |
| - | - | 1970 | 540.3 | - | - | 0 | - |
| 5 | y | 1728 | 540.8 | 0.003539 | 6.544 | +2 | 10 |
| - | - | 1277 | 541.3 | - | - | 0 | - |
| 10 | y | 3.569E+04 | 541.4 | 0.001749 | 3.23 | +1 | 5 |
| - | - | 3144 | 542.3 | - | - | 0 | - |
| - | - | 1.019E+04 | 542.4 | - | - | 0 | - |
| - | - | 1888 | 543.3 | - | - | 0 | - |
| - | - | 1844 | 543.4 | - | - | 0 | - |
| - | - | 1955 | 543.6 | - | - | 0 | - |
| - | - | 3359 | 543.8 | - | - | 0 | - |
| - | - | 2805 | 544.3 | - | - | 0 | - |
| - | - | 2618 | 544.8 | - | - | 0 | - |
| - | - | 2116 | 545.7 | - | - | 0 | - |
| - | - | 1712 | 546.3 | - | - | 0 | - |
| - | - | 2059 | 547.2 | - | - | 0 | - |
| - | - | 1664 | 547.3 | - | - | 0 | - |
| - | - | 9691 | 547.8 | - | - | 0 | - |
| - | - | 1.08E+04 | 548.3 | - | - | 0 | - |
| - | - | 4953 | 548.8 | - | - | 0 | - |
| - | - | 6386 | 549.2 | - | - | 0 | - |
| 5 | y | 5.308E+04 | 549.3 | 0.001556 | 2.833 | +2 | 10 |
| - | - | 3.44E+04 | 549.8 | - | - | 0 | - |
| - | - | 1464 | 550.2 | - | - | 0 | - |
| - | - | 1.044E+04 | 550.3 | - | - | 0 | - |
| - | - | 2441 | 550.8 | - | - | 0 | - |
| - | - | 1567 | 551.3 | - | - | 0 | - |
| - | - | 2070 | 552.3 | - | - | 0 | - |
| - | - | 3694 | 552.8 | - | - | 0 | - |
| - | - | 1933 | 553.2 | - | - | 0 | - |
| - | - | 1597 | 553.3 | - | - | 0 | - |
| - | - | 1.626E+04 | 554.2 | - | - | 0 | - |
| - | - | 4391 | 555.3 | - | - | 0 | - |
| - | - | 8782 | 556.3 | - | - | 0 | - |
| - | - | 8448 | 556.8 | - | - | 0 | - |
| - | - | 1780 | 557.3 | - | - | 0 | - |
| - | - | 4029 | 557.3 | - | - | 0 | - |
| - | - | 5888 | 557.3 | - | - | 0 | - |
| - | - | 5800 | 557.8 | - | - | 0 | - |
| - | - | 4423 | 558.2 | - | - | 0 | - |
| - | - | 2597 | 558.3 | - | - | 0 | - |
| - | - | 1715 | 559.3 | - | - | 0 | - |
| - | - | 2454 | 559.8 | - | - | 0 | - |
| 9 | b | 1673 | 560.3 | 0.0009927 | 1.772 | +2 | 9 |
| - | - | 2392 | 560.8 | - | - | 0 | - |
| - | - | 9560 | 561.3 | - | - | 0 | - |
| - | - | 2343 | 561.4 | - | - | 0 | - |
| - | - | 4110 | 561.8 | - | - | 0 | - |
| - | - | 2230 | 562.3 | - | - | 0 | - |
| - | - | 1474 | 562.3 | - | - | 0 | - |
| - | - | 1704 | 563.3 | - | - | 0 | - |
| - | - | 2273 | 564.2 | - | - | 0 | - |
| - | - | 1743 | 564.3 | - | - | 0 | - |
| - | - | 1857 | 564.3 | - | - | 0 | - |
| - | - | 1559 | 565.2 | - | - | 0 | - |
| - | - | 5.959E+04 | 566.2 | - | - | 0 | - |
| - | - | 2.185E+04 | 567.3 | - | - | 0 | - |
| - | - | 3945 | 567.4 | - | - | 0 | - |
| - | - | 6004 | 568.3 | - | - | 0 | - |
| - | - | 1820 | 569.2 | - | - | 0 | - |
| 9 | b | 1.606E+04 | 569.3 | 0.003157 | 5.545 | +2 | 9 |
| - | - | 1.098E+04 | 569.8 | - | - | 0 | - |
| - | - | 9728 | 570.2 | - | - | 0 | - |
| - | - | 3491 | 571.3 | - | - | 0 | - |
| - | - | 1947 | 571.8 | - | - | 0 | - |
| - | - | 2.416E+04 | 572.8 | - | - | 0 | - |
| - | - | 1.734E+04 | 573.3 | - | - | 0 | - |
| - | - | 8639 | 573.8 | - | - | 0 | - |
| - | - | 1981 | 575.3 | - | - | 0 | - |
| - | - | 2375 | 576.2 | - | - | 0 | - |
| - | - | 2624 | 576.3 | - | - | 0 | - |
| - | - | 2485 | 576.8 | - | - | 0 | - |
| - | - | 2.095E+04 | 577.4 | - | - | 0 | - |
| - | - | 7054 | 578.4 | - | - | 0 | - |
| - | - | 5.264E+04 | 579.4 | - | - | 0 | - |
| - | - | 3722 | 579.8 | - | - | 0 | - |
| - | - | 1667 | 580.2 | - | - | 0 | - |
| - | - | 3207 | 580.3 | - | - | 0 | - |
| - | - | 1.683E+04 | 580.4 | - | - | 0 | - |
| - | - | 1.082E+04 | 580.8 | - | - | 0 | - |
| - | - | 7207 | 581.3 | - | - | 0 | - |
| - | - | 3626 | 581.4 | - | - | 0 | - |
| - | - | 3001 | 581.8 | - | - | 0 | - |
| - | - | 2.092E+04 | 582.2 | - | - | 0 | - |
| - | - | 5663 | 583.2 | - | - | 0 | - |
| - | - | 1961 | 585.3 | - | - | 0 | - |
| - | - | 1.574E+04 | 586.2 | - | - | 0 | - |
| - | - | 4134 | 587.2 | - | - | 0 | - |
| - | - | 1452 | 587.3 | - | - | 0 | - |
| - | - | 1781 | 591.4 | - | - | 0 | - |
| - | - | 2208 | 591.8 | - | - | 0 | - |
| - | - | 1713 | 593.3 | - | - | 0 | - |
| - | - | 4235 | 593.3 | - | - | 0 | - |
| - | - | 1988 | 594.2 | - | - | 0 | - |
| - | - | 1627 | 594.3 | - | - | 0 | - |
| - | - | 1359 | 595.4 | - | - | 0 | - |
| - | - | 1.813E+04 | 595.4 | - | - | 0 | - |
| 10 | b | 3870 | 595.8 | 0.003859 | 6.478 | +2 | 10 |
| - | - | 2114 | 595.8 | - | - | 0 | - |
| - | - | 3808 | 596.3 | - | - | 0 | - |
| - | - | 2214 | 596.3 | - | - | 0 | - |
| - | - | 7273 | 596.4 | - | - | 0 | - |
| - | - | 2188 | 596.8 | - | - | 0 | - |
| - | - | 5180 | 598.2 | - | - | 0 | - |
| - | - | 1764 | 599.2 | - | - | 0 | - |
| - | - | 2338 | 600.3 | - | - | 0 | - |
| - | - | 1762 | 601.3 | - | - | 0 | - |
| - | - | 2711 | 603.3 | - | - | 0 | - |
| - | - | 1900 | 603.8 | - | - | 0 | - |
| - | - | 1.404E+04 | 604.4 | - | - | 0 | - |
| 10 | b | 4911 | 604.8 | 0.005413 | 8.95 | +2 | 10 |
| 4 | y | 2.265E+04 | 604.8 | 0.005439 | 8.993 | +2 | 11 |
| - | - | 3909 | 605.3 | - | - | 0 | - |
| - | - | 1.589E+04 | 605.4 | - | - | 0 | - |
| - | - | 4333 | 605.9 | - | - | 0 | - |
| - | - | 1.709E+04 | 610.3 | - | - | 0 | - |
| - | - | 6179 | 611.3 | - | - | 0 | - |
| - | - | 4951 | 611.8 | - | - | 0 | - |
| - | - | 1924 | 612.2 | - | - | 0 | - |
| - | - | 4695 | 612.3 | - | - | 0 | - |
| - | - | 2568 | 612.9 | - | - | 0 | - |
| - | - | 4306 | 613.3 | - | - | 0 | - |
| 4 | y | 1.455E+04 | 613.4 | 0.001137 | 1.854 | +2 | 11 |
| - | - | 1.102E+04 | 613.9 | - | - | 0 | - |
| - | - | 7781 | 614.2 | - | - | 0 | - |
| - | - | 3035 | 614.4 | - | - | 0 | - |
| - | - | 1586 | 615.2 | - | - | 0 | - |
| - | - | 6332 | 615.4 | - | - | 0 | - |
| - | - | 3108 | 615.8 | - | - | 0 | - |
| - | - | 1651 | 616.4 | - | - | 0 | - |
| - | - | 1763 | 619.8 | - | - | 0 | - |
| - | - | 9028 | 620.3 | - | - | 0 | - |
| - | - | 6180 | 620.8 | - | - | 0 | - |
| - | - | 4741 | 621.3 | - | - | 0 | - |
| - | - | 2589 | 621.9 | - | - | 0 | - |
| - | - | 1431 | 622.8 | - | - | 0 | - |
| - | - | 2234 | 623.3 | - | - | 0 | - |
| - | - | 1401 | 626.3 | - | - | 0 | - |
| - | - | 2902 | 627.3 | - | - | 0 | - |
| - | - | 3339 | 627.8 | - | - | 0 | - |
| - | - | 9443 | 628.3 | - | - | 0 | - |
| - | - | 4091 | 628.8 | - | - | 0 | - |
| - | - | 2280 | 629.2 | - | - | 0 | - |
| - | - | 3.035E+04 | 629.3 | - | - | 0 | - |
| - | - | 1.817E+04 | 629.8 | - | - | 0 | - |
| - | - | 6759 | 630.4 | - | - | 0 | - |
| - | - | 3809 | 630.9 | - | - | 0 | - |
| - | - | 3228 | 631.3 | - | - | 0 | - |
| - | - | 1864 | 632.4 | - | - | 0 | - |
| - | - | 2228 | 632.9 | - | - | 0 | - |
| - | - | 4696 | 636.3 | - | - | 0 | - |
| - | - | 3831 | 636.8 | - | - | 0 | - |
| - | - | 1.142E+04 | 637.3 | - | - | 0 | - |
| - | - | 8406 | 637.8 | - | - | 0 | - |
| - | - | 1.282E+04 | 638.4 | - | - | 0 | - |
| - | - | 8718 | 638.9 | - | - | 0 | - |
| - | - | 2841 | 639.4 | - | - | 0 | - |
| - | - | 6406 | 641.4 | - | - | 0 | - |
| - | - | 4117 | 641.9 | - | - | 0 | - |
| - | - | 3458 | 645.3 | - | - | 0 | - |
| - | - | 1580 | 645.8 | - | - | 0 | - |
| - | - | 9289 | 646.3 | - | - | 0 | - |
| - | - | 4644 | 646.9 | - | - | 0 | - |
| - | - | 2913 | 647.4 | - | - | 0 | - |
| - | - | 1945 | 657.3 | - | - | 0 | - |
| - | - | 1.462E+04 | 664.5 | - | - | 0 | - |
| - | - | 1648 | 664.8 | - | - | 0 | - |
| - | - | 4078 | 665.3 | - | - | 0 | - |
| - | - | 6572 | 665.5 | - | - | 0 | - |
| - | - | 3520 | 665.8 | - | - | 0 | - |
| - | - | 1645 | 666.3 | - | - | 0 | - |
| - | - | 2718 | 667.3 | - | - | 0 | - |
| - | - | 4200 | 668.3 | - | - | 0 | - |
| - | - | 2171 | 669.3 | - | - | 0 | - |
| 9 | y | 9704 | 669.5 | 0.001573 | 2.35 | +1 | 6 |
| - | - | 2647 | 670.3 | - | - | 0 | - |
| - | - | 3310 | 670.5 | - | - | 0 | - |
| - | - | 3742 | 671.3 | - | - | 0 | - |
| - | - | 1591 | 672.4 | - | - | 0 | - |
| - | - | 2990 | 673.3 | - | - | 0 | - |
| - | - | 4738 | 673.8 | - | - | 0 | - |
| - | - | 1878 | 674.3 | - | - | 0 | - |
| - | - | 4.765E+04 | 674.4 | - | - | 0 | - |
| - | - | 2738 | 674.8 | - | - | 0 | - |
| - | - | 1.751E+04 | 675.4 | - | - | 0 | - |
| - | - | 4351 | 676.4 | - | - | 0 | - |
| - | - | 2473 | 678.4 | - | - | 0 | - |
| - | - | 1633 | 678.9 | - | - | 0 | - |
| - | - | 1380 | 685.3 | - | - | 0 | - |
| - | - | 4648 | 685.4 | - | - | 0 | - |
| 3 | y | 2601 | 686.4 | 0.004863 | 7.084 | +2 | 12 |
| - | - | 1637 | 686.9 | - | - | 0 | - |
| - | - | 1618 | 687.4 | - | - | 0 | - |
| - | - | 1658 | 688.3 | - | - | 0 | - |
| - | - | 4017 | 688.4 | - | - | 0 | - |
| - | - | 2012 | 689.4 | - | - | 0 | - |
| - | - | 6057 | 690.3 | - | - | 0 | - |
| - | - | 3643 | 690.4 | - | - | 0 | - |
| - | - | 2659 | 691.3 | - | - | 0 | - |
| - | - | 4.435E+04 | 692.4 | - | - | 0 | - |
| - | - | 1.84E+04 | 693.4 | - | - | 0 | - |
| - | - | 1628 | 693.9 | - | - | 0 | - |
| - | - | 2232 | 694.4 | - | - | 0 | - |
| - | - | 4412 | 694.5 | - | - | 0 | - |
| 3 | y | 1.269E+04 | 694.9 | 0.0002345 | 0.3375 | +2 | 12 |
| - | - | 1.051E+04 | 695.4 | - | - | 0 | - |
| - | - | 3594 | 695.9 | - | - | 0 | - |
| - | - | 1433 | 696.4 | - | - | 0 | - |
| - | - | 5515 | 698.4 | - | - | 0 | - |
| - | - | 1876 | 699.4 | - | - | 0 | - |
| 5 | b | 3035 | 700.3 | 0.004881 | 6.971 | +1 | 5 |
| - | - | 1620 | 701.3 | - | - | 0 | - |
| - | - | 4829 | 702.3 | - | - | 0 | - |
| - | - | 2312 | 702.4 | - | - | 0 | - |
| - | - | 5505 | 702.9 | - | - | 0 | - |
| - | - | 1832 | 703.3 | - | - | 0 | - |
| - | - | 5469 | 703.4 | - | - | 0 | - |
| - | - | 1509 | 706.3 | - | - | 0 | - |
| - | - | 4066 | 706.4 | - | - | 0 | - |
| - | - | 4063 | 707.3 | - | - | 0 | - |
| - | - | 1764 | 707.4 | - | - | 0 | - |
| - | - | 3142 | 708.3 | - | - | 0 | - |
| - | - | 1638 | 708.4 | - | - | 0 | - |
| - | - | 1466 | 709.5 | - | - | 0 | - |
| - | - | 1.352E+04 | 710.5 | - | - | 0 | - |
| - | - | 4598 | 711.5 | - | - | 0 | - |
| - | - | 1749 | 712.4 | - | - | 0 | - |
| - | - | 2159 | 713.9 | - | - | 0 | - |
| - | - | 1467 | 716.3 | - | - | 0 | - |
| - | - | 8099 | 716.4 | - | - | 0 | - |
| - | - | 3306 | 717.4 | - | - | 0 | - |
| - | - | 2654 | 717.9 | - | - | 0 | - |
| 5 | b | 7749 | 718.3 | 0.006341 | 8.827 | +1 | 5 |
| - | - | 1306 | 718.4 | - | - | 0 | - |
| - | - | 3355 | 719.3 | - | - | 0 | - |
| - | - | 3455 | 720.3 | - | - | 0 | - |
| - | - | 7215 | 721.4 | - | - | 0 | - |
| - | - | 2399 | 721.9 | - | - | 0 | - |
| - | - | 2815 | 722.4 | - | - | 0 | - |
| - | - | 9982 | 723.3 | - | - | 0 | - |
| - | - | 4295 | 724.3 | - | - | 0 | - |
| 8 | y | 2.25E+04 | 726.5 | 0.001838 | 2.53 | +1 | 7 |
| - | - | 9589 | 727.5 | - | - | 0 | - |
| - | - | 1951 | 728.5 | - | - | 0 | - |
| - | - | 2639 | 730.4 | - | - | 0 | - |
| - | - | 1677 | 730.9 | - | - | 0 | - |
| - | - | 2375 | 731.4 | - | - | 0 | - |
| - | - | 5486 | 735.3 | - | - | 0 | - |
| - | - | 3896 | 736.3 | - | - | 0 | - |
| - | - | 5305 | 737.3 | - | - | 0 | - |
| - | - | 3741 | 738.4 | - | - | 0 | - |
| - | - | 3057 | 739.4 | - | - | 0 | - |
| - | - | 1394 | 739.9 | - | - | 0 | - |
| - | - | 3858 | 740.3 | - | - | 0 | - |
| - | - | 1842 | 740.5 | - | - | 0 | - |
| - | - | 1863 | 741.3 | - | - | 0 | - |
| - | - | 1914 | 750.3 | - | - | 0 | - |
| - | - | 1.033E+04 | 751.3 | - | - | 0 | - |
| - | - | 4496 | 752.3 | - | - | 0 | - |
| - | - | 2571 | 753.3 | - | - | 0 | - |
| - | - | 2386 | 766.3 | - | - | 0 | - |
| - | - | 5562 | 768.3 | - | - | 0 | - |
| - | - | 3514 | 769.3 | - | - | 0 | - |
| - | - | 1435 | 777.5 | - | - | 0 | - |
| - | - | 1720 | 793.5 | - | - | 0 | - |
| - | - | 2336 | 801.5 | - | - | 0 | - |
| - | - | 6805 | 805.5 | - | - | 0 | - |
| - | - | 4007 | 806.5 | - | - | 0 | - |
| - | - | 2925 | 811.4 | - | - | 0 | - |
| - | - | 6172 | 811.5 | - | - | 0 | - |
| - | - | 2595 | 812.5 | - | - | 0 | - |
| - | - | 1372 | 820.5 | - | - | 0 | - |
| - | - | 6240 | 821.5 | - | - | 0 | - |
| - | - | 3801 | 822.5 | - | - | 0 | - |
| 7 | y | 8.023E+04 | 823.5 | 0.001687 | 2.048 | +1 | 8 |
| - | - | 3.637E+04 | 824.5 | - | - | 0 | - |
| - | - | 1428 | 825.3 | - | - | 0 | - |
| - | - | 7008 | 825.5 | - | - | 0 | - |
| - | - | 3587 | 827.4 | - | - | 0 | - |
| - | - | 1478 | 828.4 | - | - | 0 | - |
| - | - | 6651 | 829.5 | - | - | 0 | - |
| - | - | 3139 | 830.5 | - | - | 0 | - |
| 6 | b | 1916 | 837.3 | 0.003037 | 3.627 | +1 | 6 |
| - | - | 8434 | 837.5 | - | - | 0 | - |
| - | - | 3462 | 838.5 | - | - | 0 | - |
| - | - | 4785 | 839.4 | - | - | 0 | - |
| - | - | 3449 | 839.5 | - | - | 0 | - |
| - | - | 2887 | 840.4 | - | - | 0 | - |
| - | - | 1755 | 840.5 | - | - | 0 | - |
| - | - | 2668 | 847.5 | - | - | 0 | - |
| - | - | 9129 | 848.4 | - | - | 0 | - |
| - | - | 1317 | 848.5 | - | - | 0 | - |
| - | - | 4099 | 849.4 | - | - | 0 | - |
| - | - | 4388 | 853.5 | - | - | 0 | - |
| - | - | 2119 | 854.5 | - | - | 0 | - |
| 6 | b | 8740 | 855.3 | 0.006572 | 7.683 | +1 | 6 |
| - | - | 4537 | 856.4 | - | - | 0 | - |
| - | - | 1388 | 857.4 | - | - | 0 | - |
| - | - | 4615 | 864.4 | - | - | 0 | - |
| - | - | 2455 | 865.4 | - | - | 0 | - |
| - | - | 1767 | 866.4 | - | - | 0 | - |
| - | - | 1300 | 885.4 | - | - | 0 | - |
| - | - | 2411 | 891.5 | - | - | 0 | - |
| - | - | 1.14E+04 | 901.4 | - | - | 0 | - |
| - | - | 6726 | 902.4 | - | - | 0 | - |
| - | - | 2102 | 903.5 | - | - | 0 | - |
| - | - | 5151 | 917.4 | - | - | 0 | - |
| - | - | 2236 | 918.4 | - | - | 0 | - |
| - | - | 7390 | 919.5 | - | - | 0 | - |
| - | - | 3125 | 920.5 | - | - | 0 | - |
| - | - | 2112 | 935.5 | - | - | 0 | - |
| - | - | 4205 | 942.6 | - | - | 0 | - |
| - | - | 1841 | 943.6 | - | - | 0 | - |
| - | - | 1419 | 957.4 | - | - | 0 | - |
| 6 | y | 9080 | 960.6 | 0.0006361 | 0.6622 | +1 | 9 |
| - | - | 5471 | 961.6 | - | - | 0 | - |
| - | - | 3204 | 966.6 | - | - | 0 | - |
| - | - | 1769 | 967.6 | - | - | 0 | - |
| - | - | 1706 | 1034 | - | - | 0 | - |
| - | - | 1503 | 1086 | - | - | 0 | - |
| 5 | y | 2346 | 1098 | 0.002062 | 1.879 | +1 | 10 |
| - | - | 1890 | 1099 | - | - | 0 | - |
| - | - | 1181 | 1104 | - | - | 0 | - |
| - | - | 1791 | 1105 | - | - | 0 | - |
| - | - | 933.8 | 1176 | - | - | 0 | - |
| 10 | b | 1484 | 1191 | 0.006328 | 5.315 | +1 | 10 |
| - | - | 1081 | 2135 | - | - | 0 | - |
| - | - | 1138 | 2187 | - | - | 0 | - |

m/z Charge Intensity FragmentType MassShift Position
120.04502868652344 0 2092.8833
120.05640411376953 0 2877.0293
120.06645202636719 0 654.2718
120.08130645751953 0 3127.2458
121.0401611328125 0 1063.4185
121.06517791748047 0 812.3828
122.06072998046875 0 1366.0685
122.07186889648438 0 3494.964
123.04481506347656 0 3246.5735
123.05589294433594 0 9623.782
123.09223175048828 0 917.9494
124.03958129882812 0 920.829
124.07593536376953 0 941.4952
125.07159423828125 0 2330.6636
126.05511474609375 0 787.8989
126.06683349609375 0 1214.0419
126.0917739868164 0 704.98987
127.0508804321289 0 1717.2888
127.08721160888672 0 11925.55
127.12384796142578 0 916.7018
128.0349578857422 0 1074.9125
128.07122802734375 0 23922.469
128.0904083251953 0 1089.0302
129.05502319335938 0 614.1115
129.0665283203125 0 10663.19
129.10289001464844 0 396449.94
130.0657501220703 0 150846.58
130.08702087402344 0 2050.1387
130.10000610351562 0 2719.095
130.106201171875 0 24005.188
131.0498046875 0 1161.603
131.06907653808594 0 16390.39
131.08181762695312 0 1431.6227
131.11842346191406 0 842.638
132.04507446289062 0 5737.4
132.08140563964844 0 180145.4
132.10250854492188 0 100075.164 y 13
133.05279541015625 0 2845.7676
133.06155395507812 0 1970.0851
133.0789794921875 0 1492.299
133.0847625732422 0 17458.164
133.10586547851562 0 6743.4995
134.05526733398438 0 856.81024
134.06069946289062 0 11201
134.07199096679688 0 2014.5979
134.08775329589844 0 697.83777
135.04470825195312 0 10272.408
135.06349182128906 0 1077.2296
135.0688018798828 0 1907.7108
136.07635498046875 0 337092.03
137.0736083984375 0 2112.1912
137.0797119140625 0 25610.455
137.10813903808594 0 1227.1166
138.0667724609375 0 5946.012
138.1280517578125 0 1210.1625
139.05052185058594 0 969.7249
139.0872039794922 0 7369.091
140.10763549804688 0 958.789
141.10287475585938 0 23804.502
142.0657958984375 0 3721.9368
142.10604858398438 0 1982.216
143.07411193847656 0 777.93365
144.045166015625 0 2653.8047
144.0535888671875 0 1996.7466
144.08148193359375 0 7991.862
145.0613250732422 0 4655.7754
145.0768280029297 0 2585.4429
145.0847930908203 0 951.5136
145.0978546142578 0 1634.4825
146.06069946289062 0 75707.98 b 3
146.1294708251953 0 1415.5857
147.0447235107422 0 23382.406
147.06410217285156 0 8092.44
147.1133270263672 0 1083.7329
148.0477294921875 0 1661.6558
148.07635498046875 0 87176.74
148.0975341796875 0 4425.712
149.0239715576172 0 24252.514
149.07974243164062 0 8244.189
150.02731323242188 0 1968.4841
150.05564880371094 0 7738.6553
150.06658935546875 0 892.40393
151.08718872070312 0 7393.7256
152.0712890625 0 91841.88
152.08262634277344 0 3636.186
152.14401245117188 0 10247.361
153.06715393066406 0 1850.3975
153.07460021972656 0 6642.9404
153.1029815673828 0 4725.8228
153.14747619628906 0 1159.341
154.0614776611328 0 894.95667
154.09791564941406 0 1722.608
155.04563903808594 0 3179.4375
155.0611114501953 0 1003.3019
155.0821533203125 0 21484.719
155.09341430664062 0 8025.2896
155.1117401123047 0 1519.7563
155.11854553222656 0 15826.513
156.0775146484375 0 7386.4536
156.08274841308594 0 2141.2957
156.11431884765625 0 1167.321
156.1214141845703 0 1576.9824
157.0528106689453 0 5122.1567
157.06036376953125 0 1609.8466
157.07666015625 0 18146.008
157.08460998535156 0 1014.89844
157.09754943847656 0 930.1821
157.1343231201172 0 854.0943
158.06072998046875 0 35274.453
158.08447265625 0 17856.703
159.0640411376953 0 4298.138
159.0923309326172 0 124937.89
160.0763397216797 0 12905.353
160.0886688232422 0 1508.2627
160.0956573486328 0 12861.65
161.07162475585938 0 2898.4258
161.09927368164062 0 962.0257
162.0556182861328 0 8215.962
162.07992553710938 0 924.18286
162.10328674316406 0 1762.8221
163.0396270751953 0 3112.2366
163.0875244140625 0 1341.0172
163.6138153076172 0 1797.8901
164.03517150878906 0 978.8001
164.07131958007812 0 8203.216
164.0828094482422 0 1226.469
164.1079559326172 0 1874.8453
165.0670623779297 0 1807.7432
165.07754516601562 0 4288.3716
165.10293579101562 0 14893.768
165.1393280029297 0 1168.9359
166.0617218017578 0 5442.458
166.1062774658203 0 1402.4014
167.03445434570312 0 1804.7743
167.08192443847656 0 1500.4635
167.1188201904297 0 1037.2859
167.15484619140625 0 2840.4746
168.08126831054688 0 982.03033
168.1138916015625 0 4086.826
169.06292724609375 0 921.22107
169.07675170898438 0 16305.234
169.08950805664062 0 1397.8191
169.09786987304688 0 27672.318
169.13412475585938 0 1110.9446
169.17022705078125 0 1490.0444
170.06077575683594 0 100696.13
170.07888793945312 0 1987.2351
170.10113525390625 0 2471.0493
170.12940979003906 0 1194.9584
171.0561065673828 0 2619.026
171.064208984375 0 11555.587
171.0923614501953 0 2579.5088
172.0400390625 0 968.5794
172.06410217285156 0 939.21124
172.07643127441406 0 4555.6353
172.10873413085938 0 27471.89
172.14495849609375 0 2471.248
173.07167053222656 0 28516.008
174.05577087402344 0 5669.01
174.07933044433594 0 7465.153
175.08731079101562 0 72433.71
175.0978240966797 0 2453.527
176.07183837890625 0 2316.0903 b Ammonia loss 4
176.08255004882812 0 9773.283
176.0907745361328 0 7835.9756
177.07774353027344 0 1043.8546
177.0858154296875 0 1352.9785
177.10337829589844 0 1022.019
177.11351013183594 0 913.6357
178.05052185058594 0 4262.8486
178.13467407226562 0 1318.0093
179.08216857910156 0 2895.6174
179.0933380126953 0 2135.5515
179.11865234375 0 2061.933
179.155029296875 0 2024.3054
180.06655883789062 0 2309.2986
180.0774383544922 0 1775.435
181.0615234375 0 7366.5444
181.07704162597656 0 3136.6248
181.097900390625 0 83803.23
181.13417053222656 0 7328.9233
182.0602569580078 0 1485.8503
182.09385681152344 0 1396.9054
182.1012420654297 0 7901.0903
182.12960815429688 0 16131.731
182.1382598876953 0 1174.7045
183.05596923828125 0 1124.3688
183.06434631347656 0 772.0529
183.09242248535156 0 28954.533
183.11328125 0 3156.6301
183.13287353515625 0 1456.9125
183.15005493164062 0 1396.528
184.04006958007812 0 2082.9517
184.07630920410156 0 3647.696
184.09576416015625 0 3592.244
185.04788208007812 0 10602.512
185.0717010498047 0 32635.676
185.08067321777344 0 3000.1233
185.0909881591797 0 980.72015
186.0557098388672 0 51589.24
186.0764617919922 0 3479.3984
186.0781707763672 0 3334.5103
186.12448120117188 0 49518.477
186.13414001464844 0 1542.8796
187.05955505371094 0 4962.06
187.08734130859375 0 150744.08
187.10809326171875 0 4273.393
187.12799072265625 0 3959.0571
188.07107543945312 0 3164.9421
188.08226013183594 0 1258.9062
188.09085083007812 0 16361.529
189.06639099121094 0 1231.9542
190.0500030517578 0 1180.5669
191.08277893066406 0 1877.6018
191.09310913085938 0 1701.2129
191.1184539794922 0 1470.3785
192.077880859375 0 1888.1816
193.09805297851562 0 2487.772
193.10841369628906 0 1169.3593
193.1345672607422 0 2229.7078
193.5823516845703 0 1055.45
195.07687377929688 0 1330.8811
195.11338806152344 0 2790.842
195.14996337890625 0 16335.988
196.10879516601562 0 3606.2764
196.1341094970703 0 2553.7256
196.15357971191406 0 1869.7266
197.0717315673828 0 5137.6846
197.08180236816406 0 1087.9427
197.09283447265625 0 20877.582
197.1656036376953 0 18874.188
198.05577087402344 0 1916.1838
198.07786560058594 0 1080.7814
198.0960235595703 0 1629.1506
198.16912841796875 0 2515.5125
199.05108642578125 0 1102.0051
199.08726501464844 0 17701.482
199.10842895507812 0 4041.1907
199.14483642578125 0 1030.6191
199.17117309570312 0 1362.6111
199.18130493164062 0 6706.7715
200.0712890625 0 1383.8057
200.09130859375 0 2024.7073
200.14002990722656 0 6139.274
200.18434143066406 0 1089.3396
201.06658935546875 0 41844.266
201.1029510498047 0 15817.769
201.12399291992188 0 2764.4253
202.0508270263672 0 1342.2676
202.07342529296875 0 4282.605
202.08692932128906 0 7484.8564
202.10940551757812 0 36589.555
203.08221435546875 0 58837.312
203.09283447265625 0 4220.435
203.1128387451172 0 3709.43
204.07708740234375 0 2398.7407
204.08580017089844 0 6040.9893 a Water loss 2
204.11428833007812 0 2065.592
205.09788513183594 0 6846.519
205.1089324951172 0 2936.3967
205.11903381347656 0 872.78046
206.09315490722656 0 1589.3691
207.07711791992188 0 1855.4237
207.08807373046875 0 890.3732
207.1136932373047 0 1636.5558
207.12478637695312 0 1452.6041
207.14999389648438 0 12443.388
208.14500427246094 0 2530.6768
208.15457153320312 0 960.43713
209.0600128173828 0 2163.7537
209.07127380371094 0 1106.7308
209.12916564941406 0 3615.4285
209.16603088378906 0 1095.9623
210.1246795654297 0 2639.6768
211.08743286132812 0 5124.339
211.14480590820312 0 4109.9277
211.66065979003906 0 1344.1714
212.12950134277344 0 791.5754
212.14012145996094 0 4019.844
212.64215087890625 0 799.15674
212.6685333251953 0 9061.289
213.06642150878906 0 2674.9165
213.0773468017578 0 1712.124
213.1700439453125 0 1136.8381
214.09808349609375 0 3907.4453
214.1094512939453 0 12880.212
214.12026977539062 0 893.9107
214.19227600097656 0 3286.7483
215.0711669921875 0 1580.3339
215.082275390625 0 14001.086
215.10276794433594 0 1483.0225
215.1398468017578 0 12924.702
215.1757049560547 0 1292.4427
216.08676147460938 0 1004.72314
216.11419677734375 0 2463.6182
216.14346313476562 0 958.1241
217.06146240234375 0 1079.1042
217.0978546142578 0 6990.8267
217.1092529296875 0 3515.1924
218.08177185058594 0 4080.2808 b Water loss 2
218.09283447265625 0 1361.8955
219.08792114257812 0 1874.3394
220.10804748535156 0 1930.2416
220.1201629638672 0 29597.33
221.09368896484375 0 1630.4613
221.1042022705078 0 44275.363
221.12387084960938 0 3216.5444
222.1068115234375 0 5040.793
222.12477111816406 0 3353.8323
222.1359100341797 0 1418.1022
222.1604461669922 0 912.01166
223.11940002441406 0 1746.3457
223.14532470703125 0 1563.2574
224.1200408935547 0 2141.9302
224.1401824951172 0 15421.525
224.17654418945312 0 20132.223
225.0666961669922 0 8227.879
225.16053771972656 0 5915.8975
225.18052673339844 0 2280.0757
226.0706329345703 0 1274.6411
226.0985565185547 0 3083.1619
226.1559295654297 0 339154.06
226.66574096679688 0 968.1679
227.0458221435547 0 1147.8893
227.08236694335938 0 28995.084
227.101806640625 0 1250.4304
227.15916442871094 0 40006.11 y Ammonia loss 10
228.0858612060547 0 4525.719
228.11398315429688 0 15550.849
228.134765625 0 3284.1057
228.16184997558594 0 3494.339
229.06158447265625 0 3331.818
229.0980224609375 0 14139.932
229.11923217773438 0 3359.7976
229.16673278808594 0 3669.991
230.08230590820312 0 1579.0907
230.10374450683594 0 7378.4204
231.07687377929688 0 2359.142
231.08859252929688 0 14187.162
231.11383056640625 0 987.85333
231.13577270507812 0 5182.4624
232.09188842773438 0 1374.6144
232.1083984375 0 3028.9302
232.12033081054688 0 5496.191
232.14125061035156 0 1487.479
233.09194946289062 0 1527.0988
233.10414123535156 0 7249.031
233.11285400390625 0 1523.9099
233.11549377441406 0 1542.073
233.1261444091797 0 920.46484
233.1665496826172 0 1276.7946
233.65626525878906 0 1375.486
234.10687255859375 0 1022.05237
234.1239776611328 0 1609.9551
234.13587951660156 0 4529.0347
234.16128540039062 0 824.75024
234.6636199951172 0 5246.4204
235.0724334716797 0 1191.8704
235.11988830566406 0 59019.195
235.67135620117188 0 51981.55 y 10
236.12319946289062 0 7264.3047
236.1400909423828 0 2822.721
236.17303466796875 0 12127.897
236.67478942871094 0 1911.7537
237.06761169433594 0 1161.0226
237.13555908203125 0 2665.812
237.16099548339844 0 1500.7478 y Ammonia loss 7
238.11927795410156 0 1625.3975
238.1309356689453 0 5485.732
239.13739013671875 0 1490.8379
239.1511688232422 0 20740.348
240.1142120361328 0 1081.2562
240.13519287109375 0 9188.64
240.1551971435547 0 1823.0006
240.1714324951172 0 904.7482
241.06167602539062 0 1073.1235
241.141845703125 0 1161.2253
241.15570068359375 0 21580.541
242.0934295654297 0 6648.804
242.1503448486328 0 2430.2622
242.18728637695312 0 10752.817
243.07733154296875 0 7970.4
243.17100524902344 0 2722.6216
243.182861328125 0 2869.8484
243.6688690185547 0 1265.4512
244.10891723632812 0 11229.152
244.16680908203125 0 3240.29
245.093017578125 0 11667.314
245.11407470703125 0 3237.8936
245.1756591796875 0 966.40265
245.1869354248047 0 71323.9 y 12
246.12472534179688 0 311714.97
246.19027709960938 0 8087.326
246.6416778564453 0 1181.4716
247.08470153808594 0 2069.0022
247.12998962402344 0 104729.73
248.114990234375 0 45283.44
248.1324462890625 0 9757.557
249.09921264648438 0 61799.11
249.11805725097656 0 5557.7466
249.1353302001953 0 1759.3834
250.10205078125 0 8756.069
250.11697387695312 0 1231.2451
250.13206481933594 0 981.6218
250.6229248046875 0 870.08636
251.11524963378906 0 943.31177
251.1287078857422 0 1500.032
251.15113830566406 0 3456.755
252.11085510253906 0 929.03973
252.17144775390625 0 6128.307
253.0619659423828 0 3161.6907
254.09335327148438 0 3818.4236
254.1870880126953 0 2083.2263
255.07737731933594 0 30271.07
256.0810546875 0 4053.4875
256.1090087890625 0 9924.906
256.6231689453125 0 1328.671
257.092529296875 0 1652.8588
257.114990234375 0 8173.575
257.16168212890625 0 8892.891
257.7327880859375 0 688.50775
258.0762023925781 0 6777.5405
258.0994567871094 0 97343.24
259.0834045410156 0 1846.3376
259.10272216796875 0 13596.159
259.1312561035156 0 5513.2383
260.1051025390625 0 10968.954
260.1977844238281 0 7763.36
261.11944580078125 0 1568.3948
261.181884765625 0 2065.3884
261.2012023925781 0 1374.3633
262.1194763183594 0 149450.67
263.1015930175781 0 1733.2112
263.12274169921875 0 22010.074
264.1094970703125 0 1973.626
264.1349182128906 0 4095.9011
265.1084899902344 0 1441.5948
265.1415100097656 0 3467.6443
265.1669921875 0 3946.255
266.12542724609375 0 38326.91
266.1507873535156 0 1487.3632
267.12774658203125 0 5947.279
267.1461486816406 0 4702.376
267.18310546875 0 967.9024
268.1305847167969 0 1057.113
268.1667175292969 0 1565.596
268.2029724121094 0 1142.6838
269.1125183105469 0 921.7597
269.63055419921875 0 3236.922
270.0882263183594 0 5762.627
270.1297607421875 0 1220.6064
270.18218994140625 0 28323.28
270.62200927734375 0 1059.504
271.0723571777344 0 22337.363
271.0898132324219 0 1855.1443
271.18914794921875 0 14494.738 y 9
271.69122314453125 0 4761.9326
272.0753479003906 0 4093.7341
272.10406494140625 0 10203.333
272.161376953125 0 1722.6805
272.1910095214844 0 964.9187
273.0879821777344 0 9790.231
273.1092834472656 0 5063.346
273.13482666015625 0 1500.3593
273.1488342285156 0 1245.6581
274.0925598144531 0 4274.1406
274.1195068359375 0 114506.89
275.1038818359375 0 23583.07
275.1258239746094 0 170442.58
275.1856689453125 0 4470.73 y 6
275.5200500488281 0 1390.5447
276.1074523925781 0 8653.623
276.1290588378906 0 20046.158
277.09381103515625 0 1327.2136
277.1101379394531 0 2390.994
277.13385009765625 0 3331.0598
277.628662109375 0 1145.7286
278.1141357421875 0 1652.8798
279.1500244140625 0 1190.1912
279.1826477050781 0 5187.126
280.13018798828125 0 1686.6067
280.16595458984375 0 1897.6566
281.1619567871094 0 1367.1744
281.6712646484375 0 1215.2712
282.1825256347656 0 1198.4384
283.1512145996094 0 4531.296
283.17755126953125 0 84893.49
283.6280517578125 0 4134.864
284.18072509765625 0 12089.433
285.1106262207031 0 5554.3564
285.15673828125 0 15544.031
285.18377685546875 0 1403.5453
285.66119384765625 0 1084.0575
286.15936279296875 0 1842.9542
288.09912109375 0 11250.822
288.169921875 0 1209.0001
289.1033935546875 0 2376.5579
289.1964111328125 0 1013.179
290.11468505859375 0 69977.92
290.1853332519531 0 22433.57
290.68701171875 0 8129.454
291.0982360839844 0 5226.4146
291.1182556152344 0 13836.305 b 3
291.1574401855469 0 7357.1074
291.18609619140625 0 1130.9482
291.62615966796875 0 1136.9385
292.10272216796875 0 1609.8958
292.1302185058594 0 38100.285
292.2020568847656 0 1103.004
293.13116455078125 0 13676.377
293.1620788574219 0 4517.312
294.12066650390625 0 16509.82
294.21881103515625 0 5333.975
294.6667785644531 0 1153.3807
295.1236572265625 0 2004.586
296.1270446777344 0 1063.736
297.19329833984375 0 41479.734
298.1973876953125 0 6476.143
299.1728515625 0 1839.3033
299.19219970703125 0 1128.0626
300.20404052734375 0 1971.5748
300.6731872558594 0 2674.4844
302.11468505859375 0 2212.9866
303.12091064453125 0 12176.6045
303.2185974121094 0 1271.3821
304.12420654296875 0 1249.8508
305.1308288574219 0 7418.8384
305.6650695800781 0 6733.5825
306.1097106933594 0 3327.7026
306.1346740722656 0 1451.8003
306.1645202636719 0 1767.1313
307.17791748046875 0 1633.7457
308.1252136230469 0 14574.648
309.1282653808594 0 1955.0331
309.16790771484375 0 3361.4392
309.1935119628906 0 3400.5264
310.656494140625 0 1358.051
311.1722717285156 0 4386.7134
311.2454833984375 0 9638.208
312.2486877441406 0 1336.3695
313.22503662109375 0 930.2208
314.11456298828125 0 1200.0139
314.6705017089844 0 8317.842
315.1717834472656 0 3919.3552
318.19384765625 0 2926.843
319.1091613769531 0 9107.157
319.2140197753906 0 4222.872
320.1407470703125 0 5432.11
320.19854736328125 0 1483.3727
320.8722229003906 0 6694.376 y 5
321.1257629394531 0 7712.483
321.2073059082031 0 4469.663
321.22967529296875 0 44301.76
321.5406799316406 0 1709.4573
322.13812255859375 0 1243.6405
322.1565856933594 0 6448.5195
322.1746826171875 0 1211.9735
322.23309326171875 0 8454.899
323.12744140625 0 1575.5615
323.1588134765625 0 1436.3228
323.19122314453125 0 1025.9735
324.2167663574219 0 6368.29
324.7186279296875 0 2186.1401
325.1895446777344 0 940.99774
325.2239990234375 0 3986.2368
326.2199401855469 0 17966.94
326.7249755859375 0 1084.4518 y Ammonia loss 8
327.2237854003906 0 3570.0042
327.27679443359375 0 2716.3984
329.13641357421875 0 1461.5721
330.10931396484375 0 1039.885
331.72235107421875 0 1775.0193
332.1418762207031 0 1049.6678
332.7301330566406 0 12587.669
333.125244140625 0 2271.862
333.2314453125 0 5457.4497
333.7127685546875 0 1146.6796
334.12774658203125 0 908.27313
334.1905517578125 0 1883.7504
334.6688537597656 0 2497.3538
335.1700134277344 0 1480.1993
335.2093811035156 0 3425.122
335.23748779296875 0 3702.034 y 8
335.7386169433594 0 1349.0363
336.1357727050781 0 4790.8027
336.204345703125 0 72643.56
336.7148132324219 0 2944.187
337.1879577636719 0 1384.9769
337.2076416015625 0 13889.499
337.22509765625 0 3478.4333
337.7222595214844 0 20488.062
338.1514587402344 0 19228.297
338.2232666015625 0 7050.8823
338.7242126464844 0 1450.456
339.1540832519531 0 2674.6865
339.2402648925781 0 74798.6
339.72113037109375 0 2415.7708
340.1534729003906 0 9726.056
340.2212829589844 0 1423.9321
340.2436218261719 0 14812.745
341.155029296875 0 1362.1542
341.2195129394531 0 2627.9656
341.2442321777344 0 1358.2277
342.1683349609375 0 3308.8606
343.1770935058594 0 1069.4012
344.71234130859375 0 1304.7828
345.7193908691406 0 3507.7063
346.22052001953125 0 2370.4978
346.6689758300781 0 1446.4431
346.72760009765625 0 57738.918
347.10406494140625 0 5531.123
347.2291564941406 0 22023.715
347.7303771972656 0 6062.909
348.13555908203125 0 6417.774
349.11749267578125 0 1920.9064
349.70977783203125 0 3738.77
350.15228271484375 0 2611.8826
350.1849670410156 0 1536.5309
350.2104797363281 0 2937.8723
350.543701171875 0 1783.2336
351.1214294433594 0 1711.9835
351.2405700683594 0 5570.417
352.17633056640625 0 1652.2156
352.19976806640625 0 4659.712
353.2194519042969 0 6000.298
353.7175598144531 0 3391.7234
353.8689880371094 0 2661.1152
354.21484375 0 103381.266
354.6600341796875 0 1377.2938
354.7244567871094 0 2801.0835
355.2178955078125 0 19442.412
355.2705078125 0 1753.4087
355.69677734375 0 1798.333
355.7326354980469 0 9150.724
356.1706237792969 0 1763.9991
356.2347106933594 0 3843.2578
356.69921875 0 2189.544
356.7369689941406 0 1472.7983
357.17919921875 0 3983.4492
357.2509460449219 0 21382.67
358.1632995605469 0 20794.98
358.25396728515625 0 3314.3364
358.7145690917969 0 8585.07
358.8350524902344 0 1316.9377
359.1675720214844 0 4632.658
359.1949157714844 0 1803.7869
359.2166748046875 0 2314.9827
359.507568359375 0 1951.1958
359.54345703125 0 3309.3682
359.7169189453125 0 1538.6381
359.87481689453125 0 5680.863
360.13568115234375 0 1366.3445
360.205322265625 0 1218.1742
360.6647644042969 0 4072.0583
360.68963623046875 0 5227.689
361.1429748535156 0 1558.7216
361.18414306640625 0 4186.022
361.6639404296875 0 2644.3076
362.15350341796875 0 4692.7437
362.6546630859375 0 1365.8978
362.83624267578125 0 1776.6425
363.17169189453125 0 2270.246
363.196533203125 0 1882.0288
363.23486328125 0 1519.5063
363.5049743652344 0 1597.2736
363.7490539550781 0 2607.1504 y 7
364.1304931640625 0 4602.8213
364.25177001953125 0 1337.2792
364.510009765625 0 1582.89
364.7012939453125 0 1424.4386
365.255615234375 0 1733.3507
365.8829345703125 0 1889.3641
366.1554870605469 0 6823.8354
366.21697998046875 0 2309.0317
366.5585021972656 0 9829.471 y 4
366.8929138183594 0 5125.1787
367.1632995605469 0 6918.6577
367.2281188964844 0 1762.0521
368.1477355957031 0 12136.797
368.17181396484375 0 4623.242
368.1946105957031 0 1511.1927
368.5113525390625 0 2504.901
368.84130859375 0 2337.68
369.15087890625 0 1985.4149
369.1763916015625 0 2257.6245
369.6949157714844 0 4405.9346
370.1878662109375 0 4402.098
370.2447509765625 0 2330.981
370.5272521972656 0 1283.8096
370.6676025390625 0 2165.4849
370.691650390625 0 1105.2526
371.2164306640625 0 1145.8794
371.55181884765625 0 1312.9128
371.88836669921875 0 1498.0365
373.17462158203125 0 1523.594
373.2822265625 0 33661.812 y 11
373.84259033203125 0 2000.8269 b Water loss 8
374.15264892578125 0 1151.325
374.2852478027344 0 6772.1724
374.8497009277344 0 1973.5117
375.18939208984375 0 3249.8726
375.28765869140625 0 2340.2788
375.65972900390625 0 1925.124
376.1300354003906 0 2422.4807
376.15545654296875 0 3118.3345
377.1260681152344 0 894.64417
377.1635437011719 0 1290.9796
377.8428649902344 0 1586.2787
378.1468505859375 0 2545.6516
378.18133544921875 0 1970.5309
378.51593017578125 0 2502.8872
378.7013244628906 0 1728.2826
379.2077331542969 0 1473.43
379.8471374511719 0 1893.5684 b 8
380.1397399902344 0 1062.2865
380.17938232421875 0 1282.2788
380.23052978515625 0 6034.009
382.15313720703125 0 2720.0076
382.20745849609375 0 9948.949
382.28289794921875 0 4374.683
382.5401611328125 0 5043.794
382.6685791015625 0 1743.2711
382.7118225097656 0 2663.9446
382.9620666503906 0 3405.1177
383.2145080566406 0 1938.0437
383.2669372558594 0 3918.3079
383.711669921875 0 1408.4307
384.14215087890625 0 1676.2068
384.16796875 0 10403.46
384.56268310546875 0 5405.8516
384.6653747558594 0 1317.9429
384.8967590332031 0 4273.8213
385.1739196777344 0 42386.312
385.2001647949219 0 7753.9995
386.158447265625 0 42399.58
386.20477294921875 0 1305.7977
386.71246337890625 0 1599.5051
386.96282958984375 0 1275.4858
387.16131591796875 0 6899.7656
387.51953125 0 2833.569
387.84857177734375 0 1923.5747
388.1305847167969 0 1843.5881
388.23736572265625 0 2474.8396
388.5238342285156 0 1166.3379
389.16156005859375 0 2468.6836
389.2720642089844 0 6832.9126
389.71142578125 0 2194.095
389.7735290527344 0 4313.445
389.96246337890625 0 2040.2811
390.6666564941406 0 1844.5177
391.8518371582031 0 1954.8788
392.1907043457031 0 1785.7583
392.26666259765625 0 15362.361
393.168701171875 0 1925.892
393.2723693847656 0 1770.076
393.7095642089844 0 5197.9595
393.9630126953125 0 8710.984
394.1523132324219 0 19666.598
394.1902770996094 0 1932.4384
394.21435546875 0 11773.063 y Ammonia loss 1
394.2828063964844 0 1292.9955
394.46466064453125 0 12329.291
394.52923583984375 0 3610.6426
394.7162780761719 0 6387.0596
394.8625183105469 0 2019.9287
394.96527099609375 0 1610.7206
395.15631103515625 0 4917.0703
395.7458801269531 0 2178.7402
396.26226806640625 0 4712.352
396.7171325683594 0 1459.0715
396.85028076171875 0 1166.4535
396.9674377441406 0 2446.9463
397.1844177246094 0 1742.5032
397.21722412109375 0 2375.2869
397.2445373535156 0 3666.3997
397.46392822265625 0 2968.694
397.52178955078125 0 4319.0947 b Water loss 9
397.5687255859375 0 2501.7654
397.7484130859375 0 1775.1715
397.8534851074219 0 3606.4197 b Ammonia loss 9
397.900390625 0 1495.3893
397.9670715332031 0 3750.3005
398.1911315917969 0 3774.569
398.217041015625 0 2204.6086
398.2425537109375 0 1832.8358
398.46966552734375 0 9534.65 y 1
398.72052001953125 0 10425.23
398.9714660644531 0 3905.3994
399.1787109375 0 1252.7811
399.2232360839844 0 2687.2915
399.5533752441406 0 1645.5658
400.1678161621094 0 1578.8848
400.2181701660156 0 2536.2632
401.1684875488281 0 2217.2756
401.19476318359375 0 5791.294
401.25982666015625 0 3115.0935
401.7605285644531 0 3033.8652
402.1524963378906 0 2021.2628
402.1972351074219 0 1776.9753
402.22576904296875 0 15100.326
402.56719970703125 0 1152.9946
403.1852111816406 0 39134.46
403.2403869628906 0 15363.182
403.270263671875 0 6697.9644
403.5252380371094 0 3902.0918 b 9
403.5719299316406 0 25891.012 y Ammonia loss 3
403.7681884765625 0 3063.042 y Ammonia loss 6
403.8599853515625 0 5298.1396
403.90643310546875 0 13335.748
403.9668884277344 0 2554.6902
404.18817138671875 0 7221.0835
404.2205505371094 0 3963.5942
404.24383544921875 0 2216.437
404.47088623046875 0 4190.7163
404.66302490234375 0 2894.6682
404.72314453125 0 1117.4429
405.1580505371094 0 3352.9019
405.1867370605469 0 1392.5162
405.22296142578125 0 1328.1791
405.6709289550781 0 1452.7101
406.18670654296875 0 6344.733
406.2168884277344 0 1961.6508
406.25018310546875 0 11916.57
406.6866149902344 0 2180.0967
406.7530517578125 0 3921.2642
406.968505859375 0 1445.7642
407.217041015625 0 1357.0935
407.2527770996094 0 1932.0625
407.90106201171875 0 2981.687
407.9676513671875 0 2209.4773
408.2231750488281 0 10729.52
408.4710388183594 0 12518.943
408.5578918457031 0 4969.8105
408.7250671386719 0 12235.819
408.8974609375 0 3033.197
408.975341796875 0 13758.731
409.16290283203125 0 1353.422
409.2442321777344 0 6388.851 y 3
409.5791015625 0 4847.493
410.14727783203125 0 7445.7695
410.27777099609375 0 34535.92
410.56976318359375 0 9040.7295
410.9034118652344 0 5075.63
410.9655456542969 0 3164.0786
411.14990234375 0 1195.8918
411.17919921875 0 28422.83
411.2181091308594 0 1808.2585
411.2345886230469 0 1317.6655
411.2694396972656 0 15093.018
411.47100830078125 0 6317.8667
411.5718688964844 0 1246.2855
411.72015380859375 0 3307.7227
411.76922607421875 0 6298.2583
412.1632385253906 0 7826.7866
412.1863708496094 0 4088.8167
412.275146484375 0 99549.07 y 6
412.725341796875 0 4590.8438
412.7765197753906 0 46668.195
412.9765319824219 0 1230.1294
413.1720275878906 0 6288.0615
413.2442932128906 0 1768.572
413.2779846191406 0 13235.357
413.55462646484375 0 3958.571
413.7439880371094 0 1074.0962
413.7790832519531 0 2410.4763
413.8963623046875 0 7741.323
414.1838073730469 0 8484.216
414.2306823730469 0 7453.6343
414.5623779296875 0 5387.765
414.68560791015625 0 3714.261
414.8979797363281 0 1405.1637
415.1908874511719 0 1678.8007
415.22503662109375 0 5020.173
415.25732421875 0 10075.34
415.4720153808594 0 4447.068
415.562255859375 0 1253.7031
415.7245788574219 0 15015.82 Precursor Water loss
415.7580261230469 0 4730.335
415.90179443359375 0 3504.3242
415.9754638671875 0 15961.765 Precursor Ammonia loss
416.20111083984375 0 1940.797
416.2286071777344 0 6761.4434
416.2613830566406 0 1671.9551
416.4769287109375 0 4484.715
416.72344970703125 0 2280.3777
418.5569152832031 0 3219.9607
418.7147521972656 0 1872.8153
418.8918151855469 0 2033.6238
419.1796875 0 2291.8638 b Water loss 5
419.22894287109375 0 8617.61
419.2646484375 0 11270.421
419.56170654296875 0 6630.2173
419.7242736816406 0 9759.561
419.7666015625 0 9213.09
419.90093994140625 0 32457.537
419.9751281738281 0 9776.375
420.18133544921875 0 4290.2075
420.2305603027344 0 69015.08 Precursor
420.27001953125 0 5735.8047
420.4790954589844 0 56857.184
420.57000732421875 0 7501.5625
420.6856689453125 0 1604.8717
420.7298889160156 0 22132.635
420.7733154296875 0 2607.7058
420.9022521972656 0 2472.003
420.98065185546875 0 11975.6875
421.2342834472656 0 3237.4043
421.9051208496094 0 5410.603
422.24017333984375 0 3456.9822
422.2776794433594 0 10575.017
422.3135681152344 0 2349.4045
422.57452392578125 0 1604.1938
423.2755126953125 0 5274.2944
424.2608642578125 0 2230.3347
424.329833984375 0 7617.1094
424.5609436035156 0 5691.102
424.71258544921875 0 10781.053
424.7638854980469 0 1799.119
424.8954162597656 0 6025.983
425.23236083984375 0 14125.804
425.2879943847656 0 4740.681
425.3339538574219 0 1827.7786
425.5672912597656 0 16447.564
425.7176208496094 0 1540.1241
425.90325927734375 0 10093.763
426.23748779296875 0 4748.2563
426.57220458984375 0 3321.5415
427.1740417480469 0 9644.703
427.24420166015625 0 9064.7295
427.746337890625 0 4524.4355
427.9139404296875 0 1324.225
428.1810302734375 0 8186.9062 b 5
428.250732421875 0 1867.8274
428.2886657714844 0 4190.562
428.6827392578125 0 2747.931
429.18994140625 0 93968.7
429.7060852050781 0 1925.5228
430.1927490234375 0 23571.232 b Ammonia loss 10
430.8981628417969 0 1698.0812
431.18109130859375 0 16260.58
431.2379455566406 0 3354.334
431.5707702636719 0 3316.9907
432.1824951171875 0 4468.4473
432.7106628417969 0 7083.142
433.15325927734375 0 2541.9675
433.214111328125 0 2684.2031
433.257568359375 0 2063.259
433.7097473144531 0 3360.4836
435.2987365722656 0 1729.6588
437.183837890625 0 3609.964
438.2265319824219 0 2002.9564
438.5583190917969 0 1418.1252
439.1743469238281 0 1517.6207
439.3044738769531 0 2530.423
442.58984375 0 1532.976
442.7138366699219 0 4224.671
443.16876220703125 0 2802.0798
443.2145690917969 0 2771.4666
443.5550231933594 0 1831.8451
443.7037353515625 0 1947.2065
443.8896484375 0 5571.203
444.2252502441406 0 3607.542
445.1849060058594 0 25726.871
445.2688293457031 0 3783.2712
446.1873779296875 0 4496.1655
446.2334899902344 0 6495.1157
446.73577880859375 0 4450.474
447.92431640625 0 1715.7521
448.26458740234375 0 7384.629
448.5972900390625 0 3980.0122
448.9359130859375 0 1507.9807
449.25225830078125 0 1758.1923
449.2928771972656 0 1361.8923
449.56463623046875 0 3394.4456
449.89971923828125 0 1373.1243
450.2833557128906 0 1242.1847
451.1670837402344 0 1380.6023
451.2259826660156 0 9707.8125
451.267578125 0 5654.544
451.7250061035156 0 7194.8794
451.9208984375 0 13025.857
452.2551574707031 0 9335.488
452.32476806640625 0 5031.4243
452.58544921875 0 6388.9346
452.9203186035156 0 5916.9062
453.1806640625 0 4549.811 b 2
453.2220153808594 0 1574.4739
453.59521484375 0 7625.7603
453.93035888671875 0 5638.5967
454.23193359375 0 2370.3362
454.2625427246094 0 3916.8892
454.7352600097656 0 1932.194
455.2532653808594 0 13120.671
456.25640869140625 0 2775.3052
456.5788269042969 0 4360.473
456.9144287109375 0 6348.8423
457.1849365234375 0 4156.6216
457.2530517578125 0 10597.207
457.5918884277344 0 13696.113
457.9251708984375 0 13498.1875 y Ammonia loss 2
458.1859130859375 0 2199.1372
458.2185363769531 0 1633.0519
458.2585144042969 0 7643.878
458.5933532714844 0 2322.1633
458.92333984375 0 1440.6361
459.19061279296875 0 1711.5868
459.22412109375 0 4998.3325
459.7247619628906 0 3534.556
460.232177734375 0 16987.73
460.7333679199219 0 9754.6455
461.2341003417969 0 2001.9136
461.70343017578125 0 2178.7249
462.2062072753906 0 1664.4426
462.25390625 0 3888.2014
462.5862121582031 0 6878.3286
462.9272155761719 0 28422.338
463.2619934082031 0 20795.203
463.59942626953125 0 145213.34 y 2
463.9338684082031 0 118685.11
464.268310546875 0 46170.36
464.6023864746094 0 17145.441
464.935546875 0 2996.6558
466.9054260253906 0 2513.001
467.2569580078125 0 1578.509
467.29931640625 0 10151.646
467.57464599609375 0 1371.8527
467.91888427734375 0 1720.5682
468.2281188964844 0 4936.5366
468.25958251953125 0 36072.67
468.2915344238281 0 1342.765
468.3197021484375 0 10833.471
468.5936279296875 0 27419.316
468.7308654785156 0 3789.5098
468.93096923828125 0 105611.055
469.2366027832031 0 3115.9316
469.2657470703125 0 84862.06
469.3249816894531 0 3029.0002
469.5998229980469 0 33008.242
469.78900146484375 0 5239.256
469.93316650390625 0 15418.473
470.2057189941406 0 2585.6318
470.3353271484375 0 146808.56 y 10
471.33831787109375 0 40316.156
471.5798645019531 0 2953.6426
471.799560546875 0 7724.4297
471.911865234375 0 1580.6708
472.2537536621094 0 3031.5034
472.2994079589844 0 6140.4546
472.3409729003906 0 5781.4263
472.5853271484375 0 3482.3115
472.7978515625 0 1917.7756
472.9181213378906 0 1462.6821
473.2631530761719 0 17876.709
473.5789794921875 0 5447.9014
473.9153747558594 0 1890.2684
474.2650451660156 0 5106.2593
474.78106689453125 0 5890.3374
474.9091796875 0 1770.1136
475.2446594238281 0 1739.312
475.28118896484375 0 4962.4697
475.5799560546875 0 5276.687
475.9131164550781 0 3231.7393
476.19415283203125 0 2624.5405
476.2566223144531 0 9080.541
476.5927734375 0 7840.742
476.92657470703125 0 4335.122
477.58843994140625 0 1342.4752
477.9312438964844 0 1574.1039
478.57470703125 0 5317.816 b 11
478.78851318359375 0 2665.7644
478.9109802246094 0 8906.832
479.2455139160156 0 4380.58
479.2881164550781 0 1504.1344
479.5851135253906 0 1317.0178
479.79541015625 0 2760.1619
480.80474853515625 0 23234.111 y 5
480.9147644042969 0 3724.1377
481.254150390625 0 6190.7417
481.3061218261719 0 12699.32
481.5863342285156 0 15302.155
481.8074035644531 0 3828.5806
481.92041015625 0 13342.173
482.2645263671875 0 6332.453
482.31024169921875 0 11314.535
482.5890808105469 0 1928.6909
482.76519775390625 0 8672.225
483.2171936035156 0 1454.2567
483.26348876953125 0 4106.536
483.3138732910156 0 2779.578
483.78662109375 0 15648.289
484.19573974609375 0 1846.5385
484.2874450683594 0 8646.331
484.32110595703125 0 1423.6761
484.5804748535156 0 2251.015
484.7889709472656 0 2546.9978
486.3306884765625 0 3665.543
486.5887451171875 0 4096.125
486.9207763671875 0 4573.8887
487.2581787109375 0 10888.184
487.5916442871094 0 11007.166
487.7956848144531 0 3189.9768
487.92596435546875 0 5676.17
488.2611083984375 0 1848.9545
491.2742614746094 0 12952.005
491.59246826171875 0 2293.1794
491.7785949707031 0 2910.0422
492.2781982421875 0 4281.3643
492.5909118652344 0 2294.711
492.79205322265625 0 3622.789
492.9266052246094 0 2269.9292
493.260009765625 0 2036.0637
493.2938537597656 0 2568.2551
493.3502502441406 0 1598.7944
493.5968322753906 0 3935.274
494.20489501953125 0 3489.959
494.58941650390625 0 1430.6166
494.921630859375 0 2127.362
495.36578369140625 0 1902.6996
495.5957946777344 0 2395.2463
497.72186279296875 0 1789.165
500.5909729003906 0 3663.1416
500.9283752441406 0 2744.4033
501.2656555175781 0 2572.3862
503.21649169921875 0 7559.919
504.2173767089844 0 2500.917
504.932373046875 0 1965.923
505.21820068359375 0 3230.6008 b 7
505.26824951171875 0 2789.511
505.6007080078125 0 2672.062
505.7312927246094 0 2466.279
506.2701721191406 0 1458.1608
506.6008605957031 0 3383.211
506.93609619140625 0 5192.3677
507.2702331542969 0 3399.2327
507.6056213378906 0 2351.6155
508.1920471191406 0 2550.245
508.75732421875 0 1904.053
509.5950927734375 0 1492.8412
509.923095703125 0 1890.1729
510.2669982910156 0 7966.7754 b Water loss 12
510.5998229980469 0 5627.2876 b Ammonia loss 12
510.93603515625 0 7537.34
511.27252197265625 0 8514.286
511.6087951660156 0 2829.375
512.2374877929688 0 1627.184
513.7321166992188 0 2407.7673
515.5991821289062 0 2472.5737
515.931640625 0 2906.4573
516.2696533203125 0 17495.303 b 12
516.6046142578125 0 13970.528
516.93994140625 0 8311.088
517.2328491210938 0 1472.9901
517.2728881835938 0 7592.7275
519.2094116210938 0 2657.6604
519.27880859375 0 3990.2915
519.6116943359375 0 4135.943
519.9475708007812 0 3124.1716
520.2437744140625 0 2027.238
520.3623657226562 0 7815.051
521.2276000976562 0 9940.442
521.6030883789062 0 3149.9968
521.9362182617188 0 4602.585
522.2267456054688 0 2857.3286
522.2730102539062 0 25045.18
522.6076049804688 0 21218.254
522.9429321289062 0 8061.012
523.2747192382812 0 3936.4246
523.3609008789062 0 1940.8745
523.610595703125 0 1939.832
524.3600463867188 0 1679.6075
524.6107788085938 0 6292.1523
524.74365234375 0 1655.7643
524.81396484375 0 2221.8376
524.9512939453125 0 14736.19
525.2848510742188 0 22320.7 y Ammonia loss 1
525.6181030273438 0 17251.06
525.7498779296875 0 2691.1077
525.953125 0 7247.151
526.2468872070312 0 1899.3782
526.2869873046875 0 2916.1394
526.3334350585938 0 2586.4348
526.8330078125 0 1699.0995
528.2332763671875 0 2531.0625
529.8046875 0 4177.0923
530.2897338867188 0 18996.516
530.6205444335938 0 9860.04
530.7996826171875 0 3414.7888
530.9573364257812 0 42436.684 y 1
531.2119750976562 0 6790.191
531.2920532226562 0 36099.695
531.6265258789062 0 12859.772
531.9602661132812 0 5485.741
533.818603515625 0 2743.449
536.35888671875 0 1731.1436
537.2218627929688 0 2663.595
537.3421020507812 0 2405.5913
538.2537841796875 0 56982.758
538.372802734375 0 4239.6807
538.8106079101562 0 8734.607
539.2559814453125 0 16317.195
539.3056640625 0 15270.619
539.3570556640625 0 2123.99
539.8070678710938 0 7829.482
540.2320556640625 0 2482.6858
540.327392578125 0 1970.1917
540.8228149414062 0 1728.4303 y Ammonia loss 4
541.3272094726562 0 1277.0001
541.37255859375 0 35687.723 y 9
542.251708984375 0 3143.9966
542.3759765625 0 10186.465
543.2555541992188 0 1887.9456
543.3766479492188 0 1844.4103
543.6173095703125 0 1955.3644
543.75390625 0 3358.8557
544.2522583007812 0 2805.4744
544.7569580078125 0 2617.9463
545.749267578125 0 2115.9548
546.2979736328125 0 1712.4402
547.2066040039062 0 2058.7607
547.309814453125 0 1663.912
547.8156127929688 0 9690.908
548.3187866210938 0 10802.18
548.82080078125 0 4953.2144
549.2235717773438 0 6385.6665
549.3341064453125 0 53081.812 y 4
549.8357543945312 0 34404.35
550.225830078125 0 1464.3611
550.3367919921875 0 10442.674
550.838623046875 0 2441.432
551.25732421875 0 1566.9916
552.26806640625 0 2070.3196
552.761474609375 0 3694.123
553.2216796875 0 1933.2369
553.2660522460938 0 1597.3348
554.2489624023438 0 16255.182
555.2521362304688 0 4391.4946
556.323486328125 0 8782.341
556.8237915039062 0 8448.241
557.256103515625 0 1780.463
557.2946166992188 0 4028.8816
557.3309936523438 0 5888.05
557.8316040039062 0 5800.4863
558.2485961914062 0 4422.72
558.2991943359375 0 2597.4478
559.2511596679688 0 1714.6323
559.7503662109375 0 2453.547
560.2581176757812 0 1672.667 b Water loss 8
560.761962890625 0 2392.459
561.2679443359375 0 9560.37
561.3513793945312 0 2343.247
561.770263671875 0 4110.069
562.2698974609375 0 2230.3296
562.3302612304688 0 1473.8052
563.2870483398438 0 1703.6708
564.2308349609375 0 2273.381
564.2939453125 0 1742.8298
564.349609375 0 1857.0887
565.2156982421875 0 1558.5391
566.2485961914062 0 59586.824
567.251708984375 0 21846.723
567.3995361328125 0 3944.6072
568.2557373046875 0 6003.694
569.2150268554688 0 1819.777
569.2655639648438 0 16058.969 b 8
569.7676391601562 0 10981.997
570.249755859375 0 9727.569
571.2512817382812 0 3491.1702
571.796630859375 0 1946.6198
572.8060302734375 0 24157.402
573.3063354492188 0 17336.412
573.808349609375 0 8638.955
575.33056640625 0 1980.7509
576.2322998046875 0 2374.8525
576.3374633789062 0 2624.3784
576.8394165039062 0 2484.955
577.3836059570312 0 20954.734
578.3859252929688 0 7054.2056
579.3631591796875 0 52644.664
579.7948608398438 0 3722.1824
580.2265625 0 1666.974
580.2980346679688 0 3207.2073
580.365966796875 0 16827.838
580.8032836914062 0 10822.072
581.3070678710938 0 7207.282
581.3701782226562 0 3626.2014
581.811279296875 0 3001.3452
582.2437744140625 0 20917.938
583.2457275390625 0 5663.2744
585.2595825195312 0 1961.2594
586.2432250976562 0 15735.515
587.2470703125 0 4133.695
587.341064453125 0 1451.9141
591.3656616210938 0 1781.3801
591.79541015625 0 2207.5532
593.2628173828125 0 1713.4576
593.3417358398438 0 4235.0796
594.2457885742188 0 1987.9209
594.3450927734375 0 1627.0541
595.3546752929688 0 1358.531
595.3947143554688 0 18127.234
595.779541015625 0 3870.1826 b Water loss 9
595.845703125 0 2113.9653
596.28076171875 0 3808.09
596.3468627929688 0 2213.5737
596.396484375 0 7272.57
596.7849731445312 0 2188.0032
598.2431640625 0 5179.8945
599.245849609375 0 1763.7588
600.337890625 0 2337.971
601.252685546875 0 1762.0012
603.2673950195312 0 2710.708
603.8456420898438 0 1900.4097
604.3577270507812 0 14035.157
604.786376953125 0 4910.909 b 9
604.85400390625 0 22650.21 y Ammonia loss 3
605.285400390625 0 3909.0706
605.3549194335938 0 15887.101
605.8533935546875 0 4332.585
610.3228759765625 0 17087.709
611.32666015625 0 6178.9917
611.8380737304688 0 4951.405
612.2244873046875 0 1924.2234
612.3471069335938 0 4695.0527
612.851806640625 0 2567.701
613.2532958984375 0 4305.62
613.3629760742188 0 14550.549 y 3
613.865234375 0 11015.3125
614.2391967773438 0 7780.6323
614.3630981445312 0 3035.291
615.2340698242188 0 1586.1764
615.3504638671875 0 6331.5967
615.8497924804688 0 3107.935
616.3544311523438 0 1650.9432
619.8261108398438 0 1762.8285
620.3426513671875 0 9027.755
620.84423828125 0 6180.049
621.3485107421875 0 4740.83
621.8544921875 0 2589.4102
622.8400268554688 0 1430.9032
623.3458251953125 0 2234.427
626.3180541992188 0 1400.9567
627.3309326171875 0 2901.5835
627.831298828125 0 3338.916
628.3362426757812 0 9442.7
628.838134765625 0 4091.3525
629.2491455078125 0 2279.5469
629.3469848632812 0 30345.873
629.8489990234375 0 18168.213
630.3507690429688 0 6759.3833
630.8531494140625 0 3809.0222
631.26611328125 0 3228.4517
632.3529663085938 0 1863.5514
632.8533935546875 0 2228.092
636.3374633789062 0 4695.5615
636.8372192382812 0 3831.1936
637.3448486328125 0 11418.336
637.8477172851562 0 8405.994
638.3526611328125 0 12822.457
638.8545532226562 0 8718.261
639.3544311523438 0 2841.1328
641.3704223632812 0 6406.082
641.8713989257812 0 4117.064
645.3430786132812 0 3457.8872
645.8458251953125 0 1580.3711
646.3496704101562 0 9288.841
646.853271484375 0 4644.1997
647.4274291992188 0 2912.5864
657.3367919921875 0 1944.7021
664.452392578125 0 14615.635
664.8292236328125 0 1648.3724
665.33154296875 0 4077.9094
665.4544677734375 0 6572.302
665.836669921875 0 3520.2986
666.3460693359375 0 1645.1583
667.346435546875 0 2717.6504
668.330322265625 0 4199.837
669.3308715820312 0 2171.4995
669.4673461914062 0 9704.383 y 8
670.2763671875 0 2647.2495
670.4716796875 0 3309.9111
671.26220703125 0 3742.4167
672.4200439453125 0 1591.3224
673.2788696289062 0 2989.7756
673.84375 0 4738.1323
674.2993774414062 0 1878.3988
674.4368896484375 0 47648.812
674.84814453125 0 2737.562
675.4389038085938 0 17505.414
676.44091796875 0 4350.978
678.4317016601562 0 2473.4617
678.8678588867188 0 1633.024
685.289794921875 0 1379.992
685.3565063476562 0 4647.75
686.3753662109375 0 2601.3958 y Ammonia loss 2
686.8865356445312 0 1636.6697
687.379150390625 0 1617.6687
688.2842407226562 0 1657.6421
688.4163208007812 0 4016.6833
689.4202880859375 0 2011.924
690.298583984375 0 6057.277
690.4288940429688 0 3642.6536
691.3032836914062 0 2658.7222
692.4475708007812 0 44349.434
693.4495239257812 0 18404.76
693.8886108398438 0 1628.0219
694.3836059570312 0 2231.9497
694.4518432617188 0 4411.8696
694.8937377929688 0 12689.681 y 2
695.3961181640625 0 10507.926
695.8970336914062 0 3593.622
696.399169921875 0 1432.878
698.4114990234375 0 5515.003
699.4125366210938 0 1876.0643
700.2837524414062 0 3035.4238 b Water loss 4
701.282470703125 0 1620.0818
702.3046875 0 4829.038
702.383544921875 0 2311.5598
702.8927612304688 0 5505.2847
703.3056030273438 0 1831.585
703.3944702148438 0 5469.2856
706.2777099609375 0 1509.3977
706.428466796875 0 4066.0796
707.3064575195312 0 4062.575
707.4298706054688 0 1763.9392
708.3091430664062 0 3142.1873
708.4347534179688 0 1637.5724
709.4752807617188 0 1465.5289
710.458740234375 0 13516.414
711.461669921875 0 4598.347
712.3895263671875 0 1748.7872
713.8851928710938 0 2158.7852
716.2828979492188 0 1466.7953
716.4216918945312 0 8099.284
717.4251708984375 0 3306.4924
717.8618774414062 0 2653.866
718.2957763671875 0 7748.8276 b 4
718.4266967773438 0 1306.0026
719.2999877929688 0 3354.6968
720.3218383789062 0 3454.838
721.35791015625 0 7215.1553
721.874755859375 0 2399.2773
722.3698120117188 0 2815.0369
723.3016967773438 0 9982.442
724.308837890625 0 4294.6025
726.4890747070312 0 22501.986 y 7
727.4925537109375 0 9589.358
728.4932861328125 0 1950.6277
730.3905639648438 0 2639.4358
730.880859375 0 1676.6334
731.3882446289062 0 2375.2415
735.3020629882812 0 5486.3384
736.3101806640625 0 3895.613
737.3493041992188 0 5305.14
738.3793334960938 0 3740.6501
739.3721923828125 0 3057.4373
739.8914794921875 0 1394.2886
740.328857421875 0 3857.9521
740.4674682617188 0 1842.2058
741.3321533203125 0 1862.7377
750.3067626953125 0 1914.3407
751.2985229492188 0 10332.132
752.3137817382812 0 4495.503
753.3287353515625 0 2571.057
766.3085327148438 0 2386.0374
768.3216552734375 0 5562.001
769.3262329101562 0 3514.3687
777.5382690429688 0 1435.0001
793.4810180664062 0 1720.4923
801.510498046875 0 2336.0273
805.5304565429688 0 6804.8086
806.5316162109375 0 4006.7507
811.3635864257812 0 2925.1672
811.4959106445312 0 6171.773
812.4953002929688 0 2595.0317
820.50439453125 0 1372.3514
821.526611328125 0 6239.5674
822.5264282226562 0 3801.3738
823.5416870117188 0 80230.14 y 6
824.5447998046875 0 36374.984
825.3458251953125 0 1428.1045
825.5490112304688 0 7008.313
827.3609619140625 0 3586.78
828.3595581054688 0 1477.9796
829.5059204101562 0 6651.131
830.5079345703125 0 3139.4631
837.3408203125 0 1916.2449 b Water loss 5
837.5210571289062 0 8433.766
838.5218505859375 0 3462.4246
839.3596801757812 0 4784.927
839.534912109375 0 3449.2952
840.364013671875 0 2887.428
840.5411987304688 0 1755.4712
847.5139770507812 0 2668.2166
848.418212890625 0 9128.569
848.5113525390625 0 1317.1882
849.4210815429688 0 4098.784
853.4755249023438 0 4388.3564
854.486328125 0 2118.574
855.3549194335938 0 8740.139 b 5
856.357177734375 0 4536.624
857.3591918945312 0 1388.021
864.4132080078125 0 4614.523
865.4141845703125 0 2455.406
866.4100952148438 0 1766.651
885.4146728515625 0 1299.5087
891.4615478515625 0 2410.8896
901.4437255859375 0 11398.031
902.4428100585938 0 6725.807
903.4553833007812 0 2101.7126
917.4398803710938 0 5151.159
918.4378662109375 0 2236.3083
919.4532470703125 0 7390.168
920.457275390625 0 3125.2803
935.4509887695312 0 2111.857
942.5891723632812 0 4204.537
943.5923461914062 0 1840.7524
957.4335327148438 0 1418.734
960.5995483398438 0 9080.395 y 5
961.605224609375 0 5470.8564
966.564697265625 0 3203.7078
967.5709228515625 0 1768.8965
1033.54541015625 0 1706.0248
1086.49365234375 0 1502.836
1097.65576171875 0 2345.5764 y 4
1098.661865234375 0 1890.1628
1103.5223388671875 0 1180.6967
1104.514404296875 0 1791.0872
1175.5526123046875 0 933.7829
1190.5504150390625 0 1484.3167 b Water loss 9
2135.331298828125 0 1081.2963
2187.2958984375 0 1138.4489

Spectrum Details

|  |  |
| --- | --- |
| Matched peaks? Matched peaks The total absolute number of peaks matched. Additionally in brackets the total fraction of peaks matched and the total number of peaks is shown. | 68 (4.44% of 1531) |
| FDR? FDR The false discovery rate estimated for this peptide. It is calculated by matching all theoretical fragments with a non-integer shift with the raw peaks for this spectrum. This is done with 40 different shifts. The resulting percentage is the average number of annotated peaks over the number of annotated peaks with the correct spectrum. | 9.07% |
| Satellite FDR? Satellite FDR See the FDR for details on its calculation. This satellite ion specific FDR only contains the satellite ions (d/w) for I/L/J positions. | - |
| PSM Score? PSM Score The PSM Score as given by Hecklib to this annotated spectrum. It is shown with three significant figures. | 174 |

#### Spectrum 5331? Spectrum 5331 The raw spectrum of this peptide as annotated by Hecklib. The fragments are coloured according to ion type (see legend). Any peaks with a star '\*' as text can be hovered over to see the full details, first the ion type second the mass shift type. By hovering over the amino acids in the peptide or ions in the legend the corresponding peaks are highlighted. By toggling the 'Unassigned' label you can turn the background (unassigned) peaks on or off in the plot. By updating the slider in the Ion legend you can update the spectrum to only show the top X% of the peaks with labels. The top X% means any peak that is within X% of the highest intensity. By dragging in the spectrum you can zoom in to a specific part of the spectrum and use 'Zoom Out' to get back to the original zoom level. The annotation of the spectrum is based on the given sequence in the peptides file and is done with different software so inconsistencies are likely. The peaks are annotated based on the given sequence, with 20 ppm tolerance.

Copy Data

##### Spectrum 5331 (TSV)

###### Preview

```
Loading example...
```

*Click on the button to copy the data to your clipboard.*

Mz MinMz MaxIntensity Max

WidthHeightPeptide font sizePeptide stroke widthSpectrum font sizeSpectrum stroke widthCompact peptide

Ion legend

wxyz

abcd

OtherUnassignedIon^Charge^_Position_Show for top:%

SWYQHHPGKAPKJJ

05.34e+41.07e+51.60e+52.14e+5

Zoom Out

y^+1^_1_b^+4^_4_b^+4^_5_a^+2^_3_b^+2^_3_b^+2^_3_y^+2^_4_y^+2^_4_y^+3^_7_y^+1^_2_y^+2^_5_y^+3^_8_b^+2^_4_y^+3^_9_y^+2^_6_y^+2^_6_y^+2^_7_y^+1^_3_y^+2^_7_y^+3^_10_y^+1^_3_b^+4^_13_y^+4^_13_b^+3^_10_b^+3^_10_y^+4^_13_b^+3^_10_y^+3^_11_y^+3^_11_y^+2^_8_\*\*b^+2^_6_\*b^+2^_6_b^+3^_11_b^+1^_3_b^+1^_3_y^+3^_12_y^+3^_12_y^+1^_4_y^+2^_9_b^+3^_12_y^+2^_9_b^+2^_8_b^+3^_13_b^+3^_13_y^+3^_13_y^+3^_13_y^+2^_10_y^+1^_5_y^+2^_10_b^+2^_9_b^+2^_9_b^+2^_9_b^+1^_4_b^+2^_10_b^+2^_10_y^+2^_11_y^+2^_11_y^+1^_6_y^+2^_12_y^+2^_12_b^+1^_5_b^+1^_5_y^+1^_7_y^+1^_8_b^+1^_6_y^+1^_9_y^+1^_10_

02775558321110

Fragment Matches Table

Show background peaks

| Position | Ion type | Intensity | mz Theoretical | mz Error (Th) | mz Error (ppm) | Charge | Series Number |
| --- | --- | --- | --- | --- | --- | --- | --- |
| - | - | 1182 | 120 | - | - | 0 | - |
| - | - | 1261 | 120.1 | - | - | 0 | - |
| - | - | 693.8 | 120.1 | - | - | 0 | - |
| - | - | 621.6 | 120.1 | - | - | 0 | - |
| - | - | 2977 | 120.1 | - | - | 0 | - |
| - | - | 440.9 | 120.5 | - | - | 0 | - |
| - | - | 1272 | 121 | - | - | 0 | - |
| - | - | 869.6 | 122.1 | - | - | 0 | - |
| - | - | 1486 | 122.1 | - | - | 0 | - |
| - | - | 2206 | 123 | - | - | 0 | - |
| - | - | 4813 | 123.1 | - | - | 0 | - |
| - | - | 561.4 | 123.1 | - | - | 0 | - |
| - | - | 407 | 124.1 | - | - | 0 | - |
| - | - | 1659 | 125.1 | - | - | 0 | - |
| - | - | 652.7 | 125.1 | - | - | 0 | - |
| - | - | 464.3 | 126.1 | - | - | 0 | - |
| - | - | 666.2 | 126.1 | - | - | 0 | - |
| - | - | 511.6 | 126.5 | - | - | 0 | - |
| - | - | 1094 | 127.1 | - | - | 0 | - |
| - | - | 5540 | 127.1 | - | - | 0 | - |
| - | - | 474.5 | 127.1 | - | - | 0 | - |
| - | - | 466.2 | 128 | - | - | 0 | - |
| - | - | 748.9 | 128.1 | - | - | 0 | - |
| - | - | 1.223E+04 | 128.1 | - | - | 0 | - |
| - | - | 636 | 129.1 | - | - | 0 | - |
| - | - | 4928 | 129.1 | - | - | 0 | - |
| - | - | 2.115E+05 | 129.1 | - | - | 0 | - |
| - | - | 7.689E+04 | 130.1 | - | - | 0 | - |
| - | - | 955.5 | 130.1 | - | - | 0 | - |
| - | - | 1144 | 130.1 | - | - | 0 | - |
| - | - | 1.359E+04 | 130.1 | - | - | 0 | - |
| - | - | 7525 | 131.1 | - | - | 0 | - |
| - | - | 1138 | 131.1 | - | - | 0 | - |
| - | - | 741.2 | 131.1 | - | - | 0 | - |
| - | - | 3163 | 132 | - | - | 0 | - |
| - | - | 9.327E+04 | 132.1 | - | - | 0 | - |
| 14 | y | 5.219E+04 | 132.1 | 0.0006034 | 4.568 | +1 | 1 |
| - | - | 868.6 | 132.1 | - | - | 0 | - |
| - | - | 1746 | 133.1 | - | - | 0 | - |
| - | - | 1856 | 133.1 | - | - | 0 | - |
| - | - | 9533 | 133.1 | - | - | 0 | - |
| - | - | 3269 | 133.1 | - | - | 0 | - |
| - | - | 356.4 | 133.1 | - | - | 0 | - |
| - | - | 5667 | 134.1 | - | - | 0 | - |
| - | - | 1044 | 134.1 | - | - | 0 | - |
| - | - | 6553 | 135 | - | - | 0 | - |
| - | - | 1.755E+05 | 136.1 | - | - | 0 | - |
| - | - | 1281 | 137.1 | - | - | 0 | - |
| - | - | 1.435E+04 | 137.1 | - | - | 0 | - |
| - | - | 541.3 | 137.1 | - | - | 0 | - |
| - | - | 3271 | 138.1 | - | - | 0 | - |
| - | - | 570.6 | 138.1 | - | - | 0 | - |
| - | - | 535.3 | 139.1 | - | - | 0 | - |
| - | - | 4215 | 139.1 | - | - | 0 | - |
| - | - | 733.9 | 139.1 | - | - | 0 | - |
| - | - | 840.9 | 140.1 | - | - | 0 | - |
| - | - | 536.1 | 141.1 | - | - | 0 | - |
| - | - | 1.313E+04 | 141.1 | - | - | 0 | - |
| - | - | 2286 | 142.1 | - | - | 0 | - |
| - | - | 924.1 | 142.1 | - | - | 0 | - |
| - | - | 564.3 | 143 | - | - | 0 | - |
| - | - | 1386 | 144 | - | - | 0 | - |
| - | - | 623.2 | 144.1 | - | - | 0 | - |
| - | - | 4344 | 144.1 | - | - | 0 | - |
| - | - | 2453 | 145.1 | - | - | 0 | - |
| - | - | 1055 | 145.1 | - | - | 0 | - |
| - | - | 701.5 | 145.1 | - | - | 0 | - |
| 4 | b | 3.96E+04 | 146.1 | 0.00245 | 16.77 | +4 | 4 |
| - | - | 815.9 | 146.1 | - | - | 0 | - |
| - | - | 1.215E+04 | 147 | - | - | 0 | - |
| - | - | 3451 | 147.1 | - | - | 0 | - |
| - | - | 835.4 | 147.1 | - | - | 0 | - |
| - | - | 1115 | 148 | - | - | 0 | - |
| - | - | 4.542E+04 | 148.1 | - | - | 0 | - |
| - | - | 2022 | 148.1 | - | - | 0 | - |
| - | - | 3.06E+04 | 149 | - | - | 0 | - |
| - | - | 4221 | 149.1 | - | - | 0 | - |
| - | - | 2544 | 150 | - | - | 0 | - |
| - | - | 4069 | 150.1 | - | - | 0 | - |
| - | - | 1307 | 150.1 | - | - | 0 | - |
| - | - | 488 | 151.1 | - | - | 0 | - |
| - | - | 3799 | 151.1 | - | - | 0 | - |
| - | - | 4.549E+04 | 152.1 | - | - | 0 | - |
| - | - | 4963 | 152.1 | - | - | 0 | - |
| - | - | 1320 | 153.1 | - | - | 0 | - |
| - | - | 3751 | 153.1 | - | - | 0 | - |
| - | - | 1949 | 153.1 | - | - | 0 | - |
| - | - | 1540 | 154.1 | - | - | 0 | - |
| - | - | 1504 | 155 | - | - | 0 | - |
| - | - | 1.113E+04 | 155.1 | - | - | 0 | - |
| - | - | 4815 | 155.1 | - | - | 0 | - |
| - | - | 7423 | 155.1 | - | - | 0 | - |
| - | - | 4738 | 156.1 | - | - | 0 | - |
| - | - | 562 | 156.1 | - | - | 0 | - |
| - | - | 766.8 | 156.1 | - | - | 0 | - |
| - | - | 2501 | 157.1 | - | - | 0 | - |
| - | - | 1245 | 157.1 | - | - | 0 | - |
| - | - | 9232 | 157.1 | - | - | 0 | - |
| - | - | 1004 | 157.1 | - | - | 0 | - |
| - | - | 1.816E+04 | 158.1 | - | - | 0 | - |
| - | - | 1.043E+04 | 158.1 | - | - | 0 | - |
| - | - | 2139 | 159.1 | - | - | 0 | - |
| - | - | 7.051E+04 | 159.1 | - | - | 0 | - |
| - | - | 5929 | 160.1 | - | - | 0 | - |
| - | - | 863 | 160.1 | - | - | 0 | - |
| - | - | 7428 | 160.1 | - | - | 0 | - |
| - | - | 1360 | 161.1 | - | - | 0 | - |
| - | - | 581.3 | 161.1 | - | - | 0 | - |
| - | - | 4159 | 162.1 | - | - | 0 | - |
| - | - | 948.1 | 162.1 | - | - | 0 | - |
| - | - | 685.8 | 162.1 | - | - | 0 | - |
| - | - | 2029 | 163 | - | - | 0 | - |
| - | - | 900.2 | 163.1 | - | - | 0 | - |
| - | - | 1062 | 163.6 | - | - | 0 | - |
| - | - | 754.7 | 164 | - | - | 0 | - |
| - | - | 4595 | 164.1 | - | - | 0 | - |
| - | - | 1176 | 164.1 | - | - | 0 | - |
| - | - | 665.9 | 165.1 | - | - | 0 | - |
| - | - | 1093 | 165.1 | - | - | 0 | - |
| - | - | 7532 | 165.1 | - | - | 0 | - |
| - | - | 704.6 | 165.1 | - | - | 0 | - |
| - | - | 3498 | 166.1 | - | - | 0 | - |
| - | - | 812.4 | 166.1 | - | - | 0 | - |
| - | - | 695.3 | 166.1 | - | - | 0 | - |
| - | - | 1814 | 167 | - | - | 0 | - |
| - | - | 885.1 | 167.1 | - | - | 0 | - |
| - | - | 1494 | 167.2 | - | - | 0 | - |
| - | - | 1773 | 168.1 | - | - | 0 | - |
| - | - | 8178 | 169.1 | - | - | 0 | - |
| - | - | 1.494E+04 | 169.1 | - | - | 0 | - |
| - | - | 1055 | 169.2 | - | - | 0 | - |
| - | - | 5.183E+04 | 170.1 | - | - | 0 | - |
| - | - | 903.8 | 170.1 | - | - | 0 | - |
| - | - | 994 | 170.1 | - | - | 0 | - |
| - | - | 1522 | 171.1 | - | - | 0 | - |
| - | - | 6031 | 171.1 | - | - | 0 | - |
| - | - | 1784 | 171.1 | - | - | 0 | - |
| - | - | 553.5 | 171.1 | - | - | 0 | - |
| - | - | 664 | 171.2 | - | - | 0 | - |
| - | - | 574.2 | 172 | - | - | 0 | - |
| - | - | 693.3 | 172.1 | - | - | 0 | - |
| - | - | 1335 | 172.1 | - | - | 0 | - |
| - | - | 1.426E+04 | 172.1 | - | - | 0 | - |
| - | - | 1009 | 172.1 | - | - | 0 | - |
| - | - | 1.72E+04 | 173.1 | - | - | 0 | - |
| - | - | 4366 | 174.1 | - | - | 0 | - |
| - | - | 661.4 | 174.1 | - | - | 0 | - |
| - | - | 4559 | 174.1 | - | - | 0 | - |
| - | - | 3.887E+04 | 175.1 | - | - | 0 | - |
| - | - | 1275 | 175.1 | - | - | 0 | - |
| 5 | b | 1252 | 176.1 | 7.955E-05 | 0.4518 | +4 | 5 |
| - | - | 4806 | 176.1 | - | - | 0 | - |
| - | - | 3902 | 176.1 | - | - | 0 | - |
| - | - | 714.3 | 177.1 | - | - | 0 | - |
| - | - | 2940 | 178.1 | - | - | 0 | - |
| - | - | 664.6 | 178.1 | - | - | 0 | - |
| - | - | 2083 | 179.1 | - | - | 0 | - |
| - | - | 646.1 | 179.1 | - | - | 0 | - |
| - | - | 941.9 | 179.1 | - | - | 0 | - |
| - | - | 1775 | 180.1 | - | - | 0 | - |
| - | - | 865.6 | 180.1 | - | - | 0 | - |
| - | - | 3214 | 181.1 | - | - | 0 | - |
| - | - | 4.19E+04 | 181.1 | - | - | 0 | - |
| - | - | 3977 | 181.1 | - | - | 0 | - |
| - | - | 640.6 | 181.2 | - | - | 0 | - |
| - | - | 474.9 | 182.1 | - | - | 0 | - |
| - | - | 882.1 | 182.1 | - | - | 0 | - |
| - | - | 3847 | 182.1 | - | - | 0 | - |
| - | - | 7400 | 182.1 | - | - | 0 | - |
| - | - | 725.3 | 183.1 | - | - | 0 | - |
| - | - | 1.502E+04 | 183.1 | - | - | 0 | - |
| - | - | 1584 | 183.1 | - | - | 0 | - |
| - | - | 789.8 | 183.1 | - | - | 0 | - |
| - | - | 1380 | 184 | - | - | 0 | - |
| - | - | 2043 | 184.1 | - | - | 0 | - |
| - | - | 1581 | 184.1 | - | - | 0 | - |
| - | - | 5488 | 185 | - | - | 0 | - |
| - | - | 833.4 | 185.1 | - | - | 0 | - |
| - | - | 1.804E+04 | 185.1 | - | - | 0 | - |
| - | - | 1962 | 185.1 | - | - | 0 | - |
| - | - | 671.9 | 185.1 | - | - | 0 | - |
| - | - | 665.2 | 185.1 | - | - | 0 | - |
| - | - | 2.566E+04 | 186.1 | - | - | 0 | - |
| - | - | 966.4 | 186.1 | - | - | 0 | - |
| - | - | 2.66E+04 | 186.1 | - | - | 0 | - |
| - | - | 2130 | 187.1 | - | - | 0 | - |
| - | - | 7.604E+04 | 187.1 | - | - | 0 | - |
| - | - | 4116 | 187.1 | - | - | 0 | - |
| - | - | 2274 | 187.1 | - | - | 0 | - |
| - | - | 1586 | 188.1 | - | - | 0 | - |
| - | - | 789.1 | 188.1 | - | - | 0 | - |
| - | - | 8660 | 188.1 | - | - | 0 | - |
| - | - | 829.4 | 189.1 | - | - | 0 | - |
| - | - | 747.1 | 189.1 | - | - | 0 | - |
| - | - | 726.5 | 190.1 | - | - | 0 | - |
| - | - | 1393 | 191.1 | - | - | 0 | - |
| - | - | 974.5 | 191.1 | - | - | 0 | - |
| - | - | 759.1 | 191.1 | - | - | 0 | - |
| - | - | 1651 | 193.1 | - | - | 0 | - |
| - | - | 652.5 | 193.1 | - | - | 0 | - |
| - | - | 1260 | 193.1 | - | - | 0 | - |
| - | - | 1278 | 195.1 | - | - | 0 | - |
| - | - | 564.6 | 195.1 | - | - | 0 | - |
| - | - | 1417 | 195.1 | - | - | 0 | - |
| - | - | 8710 | 195.1 | - | - | 0 | - |
| - | - | 2162 | 196.1 | - | - | 0 | - |
| - | - | 1736 | 196.1 | - | - | 0 | - |
| - | - | 531.5 | 196.2 | - | - | 0 | - |
| - | - | 3764 | 197.1 | - | - | 0 | - |
| - | - | 1.004E+04 | 197.1 | - | - | 0 | - |
| - | - | 9490 | 197.2 | - | - | 0 | - |
| - | - | 1292 | 198.1 | - | - | 0 | - |
| - | - | 521.6 | 198.1 | - | - | 0 | - |
| - | - | 919.6 | 198.1 | - | - | 0 | - |
| - | - | 731.1 | 198.2 | - | - | 0 | - |
| - | - | 811.2 | 199.1 | - | - | 0 | - |
| - | - | 9258 | 199.1 | - | - | 0 | - |
| - | - | 1744 | 199.1 | - | - | 0 | - |
| - | - | 733.2 | 199.2 | - | - | 0 | - |
| - | - | 4745 | 199.2 | - | - | 0 | - |
| - | - | 966.1 | 200.1 | - | - | 0 | - |
| - | - | 1301 | 200.1 | - | - | 0 | - |
| - | - | 4145 | 200.1 | - | - | 0 | - |
| - | - | 1.889E+04 | 201.1 | - | - | 0 | - |
| - | - | 1149 | 201.1 | - | - | 0 | - |
| - | - | 6839 | 201.1 | - | - | 0 | - |
| - | - | 1393 | 201.1 | - | - | 0 | - |
| - | - | 720.4 | 202.1 | - | - | 0 | - |
| - | - | 3377 | 202.1 | - | - | 0 | - |
| - | - | 4669 | 202.1 | - | - | 0 | - |
| - | - | 1.943E+04 | 202.1 | - | - | 0 | - |
| - | - | 972.4 | 202.1 | - | - | 0 | - |
| - | - | 3.135E+04 | 203.1 | - | - | 0 | - |
| - | - | 2148 | 203.1 | - | - | 0 | - |
| - | - | 1259 | 203.1 | - | - | 0 | - |
| - | - | 1722 | 204.1 | - | - | 0 | - |
| 3 | a | 3665 | 204.1 | 0.000919 | 4.503 | +2 | 3 |
| - | - | 743.5 | 204.1 | - | - | 0 | - |
| - | - | 3933 | 205.1 | - | - | 0 | - |
| - | - | 1347 | 205.1 | - | - | 0 | - |
| - | - | 737.9 | 207.1 | - | - | 0 | - |
| - | - | 1160 | 207.1 | - | - | 0 | - |
| - | - | 1119 | 207.1 | - | - | 0 | - |
| - | - | 6420 | 207.1 | - | - | 0 | - |
| - | - | 1017 | 208.1 | - | - | 0 | - |
| - | - | 1064 | 209.1 | - | - | 0 | - |
| - | - | 2411 | 209.1 | - | - | 0 | - |
| - | - | 675.5 | 209.2 | - | - | 0 | - |
| - | - | 939.6 | 210.1 | - | - | 0 | - |
| - | - | 2937 | 211.1 | - | - | 0 | - |
| - | - | 1640 | 211.1 | - | - | 0 | - |
| - | - | 1826 | 212.1 | - | - | 0 | - |
| - | - | 6293 | 212.7 | - | - | 0 | - |
| - | - | 769.7 | 213.1 | - | - | 0 | - |
| - | - | 1025 | 213.1 | - | - | 0 | - |
| - | - | 845.5 | 213.2 | - | - | 0 | - |
| - | - | 2119 | 214.1 | - | - | 0 | - |
| - | - | 6979 | 214.1 | - | - | 0 | - |
| - | - | 1135 | 214.1 | - | - | 0 | - |
| - | - | 1562 | 214.2 | - | - | 0 | - |
| - | - | 7356 | 215.1 | - | - | 0 | - |
| - | - | 911.8 | 215.1 | - | - | 0 | - |
| - | - | 1227 | 215.1 | - | - | 0 | - |
| - | - | 677 | 215.2 | - | - | 0 | - |
| - | - | 884 | 216.1 | - | - | 0 | - |
| - | - | 1734 | 216.1 | - | - | 0 | - |
| - | - | 3188 | 217.1 | - | - | 0 | - |
| - | - | 1872 | 217.1 | - | - | 0 | - |
| - | - | 727.3 | 217.1 | - | - | 0 | - |
| 3 | b | 2120 | 218.1 | 0.002344 | 10.75 | +2 | 3 |
| - | - | 543.5 | 219.1 | - | - | 0 | - |
| - | - | 1016 | 219.1 | - | - | 0 | - |
| - | - | 984 | 220.1 | - | - | 0 | - |
| - | - | 1.61E+04 | 220.1 | - | - | 0 | - |
| - | - | 1085 | 220.1 | - | - | 0 | - |
| - | - | 2.406E+04 | 221.1 | - | - | 0 | - |
| - | - | 1333 | 221.1 | - | - | 0 | - |
| - | - | 2713 | 222.1 | - | - | 0 | - |
| - | - | 2181 | 222.1 | - | - | 0 | - |
| - | - | 610.5 | 222.2 | - | - | 0 | - |
| - | - | 955.8 | 223.1 | - | - | 0 | - |
| - | - | 724.8 | 223.1 | - | - | 0 | - |
| - | - | 1667 | 224.1 | - | - | 0 | - |
| - | - | 7998 | 224.1 | - | - | 0 | - |
| - | - | 9985 | 224.2 | - | - | 0 | - |
| - | - | 4279 | 225.1 | - | - | 0 | - |
| - | - | 1058 | 225.1 | - | - | 0 | - |
| - | - | 2801 | 225.2 | - | - | 0 | - |
| - | - | 849.8 | 225.2 | - | - | 0 | - |
| - | - | 1.764E+05 | 226.2 | - | - | 0 | - |
| - | - | 1.44E+04 | 227.1 | - | - | 0 | - |
| 3 | b | 1015 | 227.1 | 0.004276 | 18.83 | +2 | 3 |
| - | - | 1364 | 227.1 | - | - | 0 | - |
| 11 | y | 1.914E+04 | 227.2 | 0.001876 | 8.261 | +2 | 4 |
| - | - | 1153 | 227.2 | - | - | 0 | - |
| - | - | 1016 | 227.2 | - | - | 0 | - |
| - | - | 1924 | 228.1 | - | - | 0 | - |
| - | - | 8645 | 228.1 | - | - | 0 | - |
| - | - | 1977 | 228.1 | - | - | 0 | - |
| - | - | 906.6 | 228.2 | - | - | 0 | - |
| - | - | 1570 | 229.1 | - | - | 0 | - |
| - | - | 8343 | 229.1 | - | - | 0 | - |
| - | - | 2054 | 229.1 | - | - | 0 | - |
| - | - | 596.8 | 229.2 | - | - | 0 | - |
| - | - | 2002 | 229.2 | - | - | 0 | - |
| - | - | 775.5 | 230.1 | - | - | 0 | - |
| - | - | 4476 | 230.1 | - | - | 0 | - |
| - | - | 813.8 | 231.1 | - | - | 0 | - |
| - | - | 8195 | 231.1 | - | - | 0 | - |
| - | - | 2049 | 231.1 | - | - | 0 | - |
| - | - | 1426 | 232.1 | - | - | 0 | - |
| - | - | 1566 | 232.1 | - | - | 0 | - |
| - | - | 2605 | 232.1 | - | - | 0 | - |
| - | - | 2141 | 232.1 | - | - | 0 | - |
| - | - | 4913 | 233.1 | - | - | 0 | - |
| - | - | 1161 | 233.2 | - | - | 0 | - |
| - | - | 1994 | 234.1 | - | - | 0 | - |
| - | - | 770.1 | 234.7 | - | - | 0 | - |
| - | - | 2920 | 234.7 | - | - | 0 | - |
| - | - | 3.236E+04 | 235.1 | - | - | 0 | - |
| 11 | y | 2.797E+04 | 235.7 | 0.0007632 | 3.238 | +2 | 4 |
| - | - | 3188 | 236.1 | - | - | 0 | - |
| - | - | 1677 | 236.1 | - | - | 0 | - |
| - | - | 6587 | 236.2 | - | - | 0 | - |
| - | - | 816.5 | 236.7 | - | - | 0 | - |
| - | - | 1029 | 237.1 | - | - | 0 | - |
| 8 | y | 821 | 237.2 | 0.001667 | 7.027 | +3 | 7 |
| - | - | 774.8 | 238.1 | - | - | 0 | - |
| - | - | 985.7 | 238.1 | - | - | 0 | - |
| - | - | 2891 | 238.1 | - | - | 0 | - |
| - | - | 1398 | 239.1 | - | - | 0 | - |
| - | - | 519.9 | 239.1 | - | - | 0 | - |
| - | - | 9981 | 239.2 | - | - | 0 | - |
| - | - | 5223 | 240.1 | - | - | 0 | - |
| - | - | 1084 | 240.2 | - | - | 0 | - |
| - | - | 643.1 | 241.1 | - | - | 0 | - |
| - | - | 466.5 | 241.1 | - | - | 0 | - |
| - | - | 1.081E+04 | 241.2 | - | - | 0 | - |
| - | - | 2973 | 242.1 | - | - | 0 | - |
| - | - | 765.9 | 242.2 | - | - | 0 | - |
| - | - | 5497 | 242.2 | - | - | 0 | - |
| - | - | 3832 | 243.1 | - | - | 0 | - |
| - | - | 1380 | 243.2 | - | - | 0 | - |
| - | - | 1842 | 243.2 | - | - | 0 | - |
| - | - | 6357 | 244.1 | - | - | 0 | - |
| - | - | 709.8 | 244.1 | - | - | 0 | - |
| - | - | 1276 | 244.2 | - | - | 0 | - |
| - | - | 5339 | 245.1 | - | - | 0 | - |
| - | - | 1956 | 245.1 | - | - | 0 | - |
| - | - | 614 | 245.2 | - | - | 0 | - |
| 13 | y | 3.781E+04 | 245.2 | 0.0008595 | 3.506 | +1 | 2 |
| - | - | 1.751E+05 | 246.1 | - | - | 0 | - |
| - | - | 5126 | 246.2 | - | - | 0 | - |
| - | - | 1335 | 247.1 | - | - | 0 | - |
| - | - | 5.378E+04 | 247.1 | - | - | 0 | - |
| - | - | 2.455E+04 | 248.1 | - | - | 0 | - |
| - | - | 4690 | 248.1 | - | - | 0 | - |
| - | - | 983.8 | 248.2 | - | - | 0 | - |
| - | - | 3.532E+04 | 249.1 | - | - | 0 | - |
| - | - | 2216 | 249.1 | - | - | 0 | - |
| - | - | 4113 | 250.1 | - | - | 0 | - |
| - | - | 806.8 | 250.1 | - | - | 0 | - |
| - | - | 810.1 | 251.1 | - | - | 0 | - |
| - | - | 870.5 | 251.2 | - | - | 0 | - |
| - | - | 857.3 | 252.1 | - | - | 0 | - |
| - | - | 2711 | 252.2 | - | - | 0 | - |
| - | - | 1131 | 253.1 | - | - | 0 | - |
| - | - | 2003 | 254.1 | - | - | 0 | - |
| - | - | 1.696E+04 | 255.1 | - | - | 0 | - |
| - | - | 2122 | 256.1 | - | - | 0 | - |
| - | - | 4697 | 256.1 | - | - | 0 | - |
| - | - | 1032 | 257.1 | - | - | 0 | - |
| - | - | 4002 | 257.1 | - | - | 0 | - |
| - | - | 5039 | 257.2 | - | - | 0 | - |
| - | - | 3313 | 258.1 | - | - | 0 | - |
| - | - | 5.27E+04 | 258.1 | - | - | 0 | - |
| - | - | 947 | 259.1 | - | - | 0 | - |
| - | - | 6435 | 259.1 | - | - | 0 | - |
| - | - | 2539 | 259.1 | - | - | 0 | - |
| - | - | 6775 | 260.1 | - | - | 0 | - |
| - | - | 3657 | 260.2 | - | - | 0 | - |
| - | - | 635.2 | 260.7 | - | - | 0 | - |
| - | - | 711.7 | 261.1 | - | - | 0 | - |
| - | - | 792.1 | 261.2 | - | - | 0 | - |
| - | - | 7.921E+04 | 262.1 | - | - | 0 | - |
| - | - | 1520 | 262.2 | - | - | 0 | - |
| - | - | 1164 | 263.1 | - | - | 0 | - |
| - | - | 1.023E+04 | 263.1 | - | - | 0 | - |
| - | - | 718.7 | 264.1 | - | - | 0 | - |
| - | - | 2080 | 264.1 | - | - | 0 | - |
| - | - | 1825 | 265.1 | - | - | 0 | - |
| - | - | 2396 | 265.2 | - | - | 0 | - |
| - | - | 2.065E+04 | 266.1 | - | - | 0 | - |
| - | - | 2361 | 267.1 | - | - | 0 | - |
| - | - | 1823 | 267.1 | - | - | 0 | - |
| - | - | 756 | 268.7 | - | - | 0 | - |
| - | - | 1544 | 269.6 | - | - | 0 | - |
| - | - | 2998 | 270.1 | - | - | 0 | - |
| - | - | 678.6 | 270.1 | - | - | 0 | - |
| - | - | 1.586E+04 | 270.2 | - | - | 0 | - |
| - | - | 1.195E+04 | 271.1 | - | - | 0 | - |
| - | - | 1100 | 271.1 | - | - | 0 | - |
| 10 | y | 8227 | 271.2 | 1.723E-05 | 0.06352 | +2 | 5 |
| - | - | 1781 | 271.7 | - | - | 0 | - |
| - | - | 1960 | 272.1 | - | - | 0 | - |
| - | - | 5464 | 272.1 | - | - | 0 | - |
| - | - | 743.9 | 272.2 | - | - | 0 | - |
| - | - | 4757 | 273.1 | - | - | 0 | - |
| - | - | 2445 | 273.1 | - | - | 0 | - |
| - | - | 924.5 | 273.1 | - | - | 0 | - |
| - | - | 850 | 274.1 | - | - | 0 | - |
| - | - | 1276 | 274.1 | - | - | 0 | - |
| - | - | 6.29E+04 | 274.1 | - | - | 0 | - |
| - | - | 1.34E+04 | 275.1 | - | - | 0 | - |
| - | - | 8.688E+04 | 275.1 | - | - | 0 | - |
| 7 | y | 1938 | 275.2 | 0.0009094 | 3.305 | +3 | 8 |
| - | - | 898 | 275.5 | - | - | 0 | - |
| - | - | 4646 | 276.1 | - | - | 0 | - |
| - | - | 1.023E+04 | 276.1 | - | - | 0 | - |
| - | - | 791.1 | 277.1 | - | - | 0 | - |
| - | - | 1210 | 277.1 | - | - | 0 | - |
| - | - | 2507 | 277.1 | - | - | 0 | - |
| - | - | 643.7 | 278.1 | - | - | 0 | - |
| - | - | 586 | 279.1 | - | - | 0 | - |
| - | - | 3096 | 279.2 | - | - | 0 | - |
| - | - | 739.9 | 280.1 | - | - | 0 | - |
| - | - | 683.1 | 280.2 | - | - | 0 | - |
| - | - | 1440 | 281.2 | - | - | 0 | - |
| - | - | 722.7 | 282.2 | - | - | 0 | - |
| - | - | 2713 | 283.2 | - | - | 0 | - |
| - | - | 4.824E+04 | 283.2 | - | - | 0 | - |
| - | - | 2388 | 283.6 | - | - | 0 | - |
| - | - | 637.5 | 284.1 | - | - | 0 | - |
| - | - | 6631 | 284.2 | - | - | 0 | - |
| - | - | 2872 | 285.1 | - | - | 0 | - |
| - | - | 8221 | 285.2 | - | - | 0 | - |
| - | - | 1580 | 286.2 | - | - | 0 | - |
| - | - | 6059 | 288.1 | - | - | 0 | - |
| - | - | 1439 | 289.1 | - | - | 0 | - |
| - | - | 820.8 | 289.2 | - | - | 0 | - |
| - | - | 3.553E+04 | 290.1 | - | - | 0 | - |
| - | - | 1.242E+04 | 290.2 | - | - | 0 | - |
| - | - | 3611 | 290.7 | - | - | 0 | - |
| - | - | 2868 | 291.1 | - | - | 0 | - |
| 4 | b | 6815 | 291.1 | 0.0005225 | 1.795 | +2 | 4 |
| - | - | 3082 | 291.2 | - | - | 0 | - |
| - | - | 741.6 | 291.2 | - | - | 0 | - |
| - | - | 1113 | 291.6 | - | - | 0 | - |
| - | - | 993.4 | 292.1 | - | - | 0 | - |
| - | - | 2.08E+04 | 292.1 | - | - | 0 | - |
| - | - | 7724 | 293.1 | - | - | 0 | - |
| - | - | 2513 | 293.2 | - | - | 0 | - |
| - | - | 7660 | 294.1 | - | - | 0 | - |
| - | - | 2750 | 294.2 | - | - | 0 | - |
| - | - | 1172 | 295.1 | - | - | 0 | - |
| - | - | 677.5 | 295.2 | - | - | 0 | - |
| - | - | 821.4 | 297.2 | - | - | 0 | - |
| - | - | 2.058E+04 | 297.2 | - | - | 0 | - |
| - | - | 3838 | 298.2 | - | - | 0 | - |
| - | - | 650.6 | 300.1 | - | - | 0 | - |
| - | - | 843.3 | 300.2 | - | - | 0 | - |
| - | - | 957.7 | 300.7 | - | - | 0 | - |
| - | - | 791.4 | 302.1 | - | - | 0 | - |
| - | - | 5013 | 303.1 | - | - | 0 | - |
| - | - | 833.1 | 303.2 | - | - | 0 | - |
| - | - | 1010 | 304.1 | - | - | 0 | - |
| - | - | 3703 | 305.1 | - | - | 0 | - |
| - | - | 2678 | 305.7 | - | - | 0 | - |
| - | - | 1636 | 306.1 | - | - | 0 | - |
| - | - | 895.5 | 306.2 | - | - | 0 | - |
| - | - | 823 | 307.2 | - | - | 0 | - |
| - | - | 7155 | 308.1 | - | - | 0 | - |
| - | - | 1249 | 309.1 | - | - | 0 | - |
| - | - | 2023 | 309.2 | - | - | 0 | - |
| - | - | 1529 | 309.2 | - | - | 0 | - |
| - | - | 2775 | 311.2 | - | - | 0 | - |
| - | - | 5241 | 311.2 | - | - | 0 | - |
| - | - | 1315 | 312.2 | - | - | 0 | - |
| - | - | 1002 | 314.1 | - | - | 0 | - |
| - | - | 846.3 | 314.2 | - | - | 0 | - |
| - | - | 4980 | 314.7 | - | - | 0 | - |
| - | - | 1557 | 315.2 | - | - | 0 | - |
| - | - | 2296 | 318.2 | - | - | 0 | - |
| - | - | 4670 | 319.1 | - | - | 0 | - |
| - | - | 2788 | 319.2 | - | - | 0 | - |
| - | - | 3432 | 320.1 | - | - | 0 | - |
| 6 | y | 3218 | 320.9 | 0.0005796 | 1.806 | +3 | 9 |
| - | - | 4557 | 321.1 | - | - | 0 | - |
| - | - | 1470 | 321.2 | - | - | 0 | - |
| - | - | 2.346E+04 | 321.2 | - | - | 0 | - |
| - | - | 790.1 | 321.5 | - | - | 0 | - |
| - | - | 4673 | 322.2 | - | - | 0 | - |
| - | - | 5063 | 322.2 | - | - | 0 | - |
| - | - | 2931 | 324.2 | - | - | 0 | - |
| - | - | 1076 | 324.7 | - | - | 0 | - |
| - | - | 946.5 | 325.2 | - | - | 0 | - |
| - | - | 1240 | 325.2 | - | - | 0 | - |
| - | - | 1.033E+04 | 326.2 | - | - | 0 | - |
| 9 | y | 759.8 | 326.7 | 0.0004743 | 1.452 | +2 | 6 |
| - | - | 2066 | 327.2 | - | - | 0 | - |
| - | - | 863 | 329.1 | - | - | 0 | - |
| - | - | 834.2 | 330.1 | - | - | 0 | - |
| - | - | 7216 | 332.7 | - | - | 0 | - |
| - | - | 1626 | 333.1 | - | - | 0 | - |
| - | - | 2455 | 333.2 | - | - | 0 | - |
| - | - | 880.3 | 333.7 | - | - | 0 | - |
| - | - | 934.5 | 334.2 | - | - | 0 | - |
| - | - | 1315 | 334.7 | - | - | 0 | - |
| - | - | 2861 | 335.2 | - | - | 0 | - |
| 9 | y | 1608 | 335.2 | 0.0008411 | 2.509 | +2 | 6 |
| - | - | 834 | 335.7 | - | - | 0 | - |
| - | - | 2719 | 336.1 | - | - | 0 | - |
| - | - | 4.069E+04 | 336.2 | - | - | 0 | - |
| - | - | 1072 | 337.2 | - | - | 0 | - |
| - | - | 6135 | 337.2 | - | - | 0 | - |
| - | - | 2643 | 337.2 | - | - | 0 | - |
| - | - | 9549 | 337.7 | - | - | 0 | - |
| - | - | 1.098E+04 | 338.2 | - | - | 0 | - |
| - | - | 3342 | 338.2 | - | - | 0 | - |
| - | - | 697 | 338.2 | - | - | 0 | - |
| - | - | 708.5 | 338.7 | - | - | 0 | - |
| - | - | 2376 | 339.2 | - | - | 0 | - |
| - | - | 3.997E+04 | 339.2 | - | - | 0 | - |
| - | - | 760 | 339.7 | - | - | 0 | - |
| - | - | 3046 | 340.2 | - | - | 0 | - |
| - | - | 1224 | 340.2 | - | - | 0 | - |
| - | - | 7540 | 340.2 | - | - | 0 | - |
| - | - | 1124 | 341.2 | - | - | 0 | - |
| - | - | 1706 | 342.2 | - | - | 0 | - |
| - | - | 1058 | 342.2 | - | - | 0 | - |
| - | - | 1319 | 344.7 | - | - | 0 | - |
| - | - | 2087 | 345.7 | - | - | 0 | - |
| - | - | 953.3 | 346.2 | - | - | 0 | - |
| - | - | 3.108E+04 | 346.7 | - | - | 0 | - |
| - | - | 2691 | 347.1 | - | - | 0 | - |
| - | - | 9724 | 347.2 | - | - | 0 | - |
| - | - | 2336 | 347.7 | - | - | 0 | - |
| - | - | 1213 | 348.1 | - | - | 0 | - |
| - | - | 4693 | 348.1 | - | - | 0 | - |
| - | - | 1755 | 349.1 | - | - | 0 | - |
| - | - | 1612 | 349.7 | - | - | 0 | - |
| - | - | 1623 | 350.2 | - | - | 0 | - |
| - | - | 1146 | 350.2 | - | - | 0 | - |
| - | - | 832.9 | 350.2 | - | - | 0 | - |
| - | - | 775.6 | 350.5 | - | - | 0 | - |
| - | - | 2285 | 351.2 | - | - | 0 | - |
| - | - | 2742 | 352.2 | - | - | 0 | - |
| - | - | 3360 | 353.2 | - | - | 0 | - |
| - | - | 1115 | 353.6 | - | - | 0 | - |
| - | - | 2365 | 353.7 | - | - | 0 | - |
| - | - | 2041 | 353.9 | - | - | 0 | - |
| - | - | 1236 | 354.2 | - | - | 0 | - |
| - | - | 5.316E+04 | 354.2 | - | - | 0 | - |
| - | - | 729.5 | 354.7 | - | - | 0 | - |
| - | - | 1713 | 354.7 | - | - | 0 | - |
| - | - | 8794 | 355.2 | - | - | 0 | - |
| 8 | y | 650.9 | 355.2 | 0.005581 | 15.71 | +2 | 7 |
| - | - | 1000 | 355.3 | - | - | 0 | - |
| - | - | 4460 | 355.7 | - | - | 0 | - |
| - | - | 1792 | 356.2 | - | - | 0 | - |
| 12 | y | 722.7 | 356.3 | 0.001659 | 4.658 | +1 | 3 |
| - | - | 2430 | 357.2 | - | - | 0 | - |
| - | - | 9946 | 357.3 | - | - | 0 | - |
| - | - | 1.218E+04 | 358.2 | - | - | 0 | - |
| - | - | 1981 | 358.3 | - | - | 0 | - |
| - | - | 5398 | 358.7 | - | - | 0 | - |
| - | - | 2117 | 359.2 | - | - | 0 | - |
| - | - | 1039 | 359.2 | - | - | 0 | - |
| - | - | 1191 | 359.2 | - | - | 0 | - |
| - | - | 1431 | 359.5 | - | - | 0 | - |
| - | - | 1899 | 359.5 | - | - | 0 | - |
| - | - | 2185 | 359.9 | - | - | 0 | - |
| - | - | 1750 | 360.2 | - | - | 0 | - |
| - | - | 1973 | 360.7 | - | - | 0 | - |
| - | - | 2284 | 360.7 | - | - | 0 | - |
| - | - | 1366 | 361.2 | - | - | 0 | - |
| - | - | 2193 | 362.2 | - | - | 0 | - |
| - | - | 849.6 | 362.8 | - | - | 0 | - |
| - | - | 775.3 | 363.7 | - | - | 0 | - |
| 8 | y | 1640 | 363.7 | 0.001096 | 3.012 | +2 | 7 |
| - | - | 616.3 | 363.8 | - | - | 0 | - |
| - | - | 2355 | 364.1 | - | - | 0 | - |
| - | - | 766.7 | 364.2 | - | - | 0 | - |
| - | - | 747.5 | 364.2 | - | - | 0 | - |
| - | - | 779.6 | 365.3 | - | - | 0 | - |
| - | - | 567.2 | 365.5 | - | - | 0 | - |
| - | - | 3164 | 366.2 | - | - | 0 | - |
| - | - | 1266 | 366.2 | - | - | 0 | - |
| 5 | y | 3345 | 366.6 | 0.0009517 | 2.596 | +3 | 10 |
| - | - | 3351 | 366.9 | - | - | 0 | - |
| - | - | 2908 | 367.2 | - | - | 0 | - |
| - | - | 5283 | 368.1 | - | - | 0 | - |
| - | - | 891.4 | 368.2 | - | - | 0 | - |
| - | - | 1787 | 368.2 | - | - | 0 | - |
| - | - | 1228 | 368.7 | - | - | 0 | - |
| - | - | 1081 | 368.8 | - | - | 0 | - |
| - | - | 1035 | 369.2 | - | - | 0 | - |
| - | - | 2110 | 369.2 | - | - | 0 | - |
| - | - | 2050 | 369.7 | - | - | 0 | - |
| - | - | 2192 | 370.2 | - | - | 0 | - |
| - | - | 810.3 | 370.2 | - | - | 0 | - |
| - | - | 1365 | 370.7 | - | - | 0 | - |
| - | - | 906.6 | 371.2 | - | - | 0 | - |
| - | - | 732.7 | 371.3 | - | - | 0 | - |
| - | - | 835.2 | 371.6 | - | - | 0 | - |
| 12 | y | 1.723E+04 | 373.3 | 0.001142 | 3.059 | +1 | 3 |
| - | - | 1190 | 374.3 | - | - | 0 | - |
| - | - | 3910 | 374.3 | - | - | 0 | - |
| - | - | 1210 | 374.5 | - | - | 0 | - |
| - | - | 1238 | 375 | - | - | 0 | - |
| - | - | 1376 | 375.2 | - | - | 0 | - |
| - | - | 1070 | 375.3 | - | - | 0 | - |
| - | - | 1214 | 375.7 | - | - | 0 | - |
| - | - | 1713 | 376.1 | - | - | 0 | - |
| - | - | 725.3 | 376.2 | - | - | 0 | - |
| - | - | 746.2 | 376.7 | - | - | 0 | - |
| - | - | 1037 | 378.1 | - | - | 0 | - |
| - | - | 947.3 | 378.5 | - | - | 0 | - |
| - | - | 1168 | 378.7 | - | - | 0 | - |
| - | - | 2870 | 380.2 | - | - | 0 | - |
| - | - | 888.3 | 380.5 | - | - | 0 | - |
| - | - | 747.5 | 380.7 | - | - | 0 | - |
| - | - | 1125 | 381.8 | - | - | 0 | - |
| - | - | 2116 | 382.2 | - | - | 0 | - |
| - | - | 907.2 | 382.2 | - | - | 0 | - |
| - | - | 4267 | 382.2 | - | - | 0 | - |
| - | - | 3278 | 382.3 | - | - | 0 | - |
| - | - | 856.8 | 382.5 | - | - | 0 | - |
| - | - | 2461 | 382.5 | - | - | 0 | - |
| - | - | 1891 | 382.7 | - | - | 0 | - |
| 13 | b | 945.5 | 383.2 | 0.00198 | 5.167 | +4 | 13 |
| - | - | 1660 | 383.3 | - | - | 0 | - |
| - | - | 885 | 383.7 | - | - | 0 | - |
| - | - | 1092 | 384.1 | - | - | 0 | - |
| - | - | 6249 | 384.2 | - | - | 0 | - |
| - | - | 3881 | 384.6 | - | - | 0 | - |
| - | - | 1121 | 384.7 | - | - | 0 | - |
| - | - | 2194 | 384.9 | - | - | 0 | - |
| - | - | 2.205E+04 | 385.2 | - | - | 0 | - |
| - | - | 3368 | 385.2 | - | - | 0 | - |
| - | - | 2.041E+04 | 386.2 | - | - | 0 | - |
| - | - | 1239 | 386.7 | - | - | 0 | - |
| - | - | 1134 | 387 | - | - | 0 | - |
| - | - | 4578 | 387.2 | - | - | 0 | - |
| - | - | 1206 | 387.5 | - | - | 0 | - |
| - | - | 1160 | 387.5 | - | - | 0 | - |
| - | - | 1317 | 387.9 | - | - | 0 | - |
| - | - | 1044 | 388.1 | - | - | 0 | - |
| - | - | 1380 | 388.2 | - | - | 0 | - |
| - | - | 792.8 | 388.9 | - | - | 0 | - |
| - | - | 1773 | 389.2 | - | - | 0 | - |
| - | - | 5144 | 389.3 | - | - | 0 | - |
| - | - | 1183 | 389.8 | - | - | 0 | - |
| - | - | 794.1 | 390 | - | - | 0 | - |
| - | - | 931.3 | 390.7 | - | - | 0 | - |
| - | - | 1045 | 391.5 | - | - | 0 | - |
| - | - | 2037 | 391.8 | - | - | 0 | - |
| - | - | 1799 | 392.2 | - | - | 0 | - |
| - | - | 6830 | 392.3 | - | - | 0 | - |
| - | - | 1181 | 392.5 | - | - | 0 | - |
| - | - | 816.9 | 392.9 | - | - | 0 | - |
| - | - | 1651 | 393.3 | - | - | 0 | - |
| - | - | 896.8 | 393.5 | - | - | 0 | - |
| - | - | 2779 | 393.7 | - | - | 0 | - |
| - | - | 1204 | 393.9 | - | - | 0 | - |
| - | - | 977.8 | 393.9 | - | - | 0 | - |
| - | - | 5260 | 394 | - | - | 0 | - |
| - | - | 1.077E+04 | 394.2 | - | - | 0 | - |
| 2 | y | 7919 | 394.2 | 0.002603 | 6.604 | +4 | 13 |
| - | - | 1017 | 394.3 | - | - | 0 | - |
| - | - | 5864 | 394.5 | - | - | 0 | - |
| - | - | 2626 | 394.5 | - | - | 0 | - |
| - | - | 2942 | 394.7 | - | - | 0 | - |
| - | - | 1238 | 395 | - | - | 0 | - |
| - | - | 1942 | 395.2 | - | - | 0 | - |
| - | - | 2720 | 396.3 | - | - | 0 | - |
| - | - | 1318 | 396.9 | - | - | 0 | - |
| - | - | 882 | 397.2 | - | - | 0 | - |
| - | - | 1641 | 397.2 | - | - | 0 | - |
| - | - | 1927 | 397.2 | - | - | 0 | - |
| - | - | 795 | 397.3 | - | - | 0 | - |
| - | - | 1081 | 397.5 | - | - | 0 | - |
| 10 | b | 2975 | 397.5 | 0.002395 | 6.026 | +3 | 10 |
| - | - | 1116 | 397.7 | - | - | 0 | - |
| 10 | b | 1705 | 397.8 | 0.006757 | 16.98 | +3 | 10 |
| - | - | 811.1 | 397.9 | - | - | 0 | - |
| - | - | 1984 | 398 | - | - | 0 | - |
| - | - | 1878 | 398.2 | - | - | 0 | - |
| - | - | 808.6 | 398.2 | - | - | 0 | - |
| - | - | 1767 | 398.2 | - | - | 0 | - |
| 2 | y | 4384 | 398.5 | 0.001368 | 3.432 | +4 | 13 |
| - | - | 4201 | 398.7 | - | - | 0 | - |
| - | - | 838.5 | 398.9 | - | - | 0 | - |
| - | - | 1123 | 398.9 | - | - | 0 | - |
| - | - | 1697 | 399 | - | - | 0 | - |
| - | - | 1642 | 399.2 | - | - | 0 | - |
| - | - | 1185 | 400.2 | - | - | 0 | - |
| - | - | 1073 | 400.2 | - | - | 0 | - |
| - | - | 1231 | 401.2 | - | - | 0 | - |
| - | - | 3272 | 401.2 | - | - | 0 | - |
| - | - | 1752 | 401.3 | - | - | 0 | - |
| - | - | 1038 | 402.1 | - | - | 0 | - |
| - | - | 6958 | 402.2 | - | - | 0 | - |
| - | - | 921.4 | 402.6 | - | - | 0 | - |
| - | - | 1588 | 402.9 | - | - | 0 | - |
| - | - | 2.21E+04 | 403.2 | - | - | 0 | - |
| - | - | 7952 | 403.2 | - | - | 0 | - |
| - | - | 2478 | 403.3 | - | - | 0 | - |
| 10 | b | 2818 | 403.5 | 0.001865 | 4.621 | +3 | 10 |
| 4 | y | 1.16E+04 | 403.6 | 0.00294 | 7.286 | +3 | 11 |
| - | - | 1856 | 403.8 | - | - | 0 | - |
| - | - | 2754 | 403.9 | - | - | 0 | - |
| - | - | 5779 | 403.9 | - | - | 0 | - |
| - | - | 1579 | 404 | - | - | 0 | - |
| - | - | 3046 | 404.2 | - | - | 0 | - |
| - | - | 2832 | 404.2 | - | - | 0 | - |
| - | - | 981.4 | 404.3 | - | - | 0 | - |
| - | - | 1889 | 404.5 | - | - | 0 | - |
| - | - | 1595 | 404.7 | - | - | 0 | - |
| - | - | 800.1 | 404.7 | - | - | 0 | - |
| - | - | 1523 | 404.9 | - | - | 0 | - |
| - | - | 1465 | 405.2 | - | - | 0 | - |
| - | - | 1118 | 405.7 | - | - | 0 | - |
| - | - | 1299 | 406.1 | - | - | 0 | - |
| - | - | 2197 | 406.2 | - | - | 0 | - |
| - | - | 6901 | 406.2 | - | - | 0 | - |
| - | - | 2649 | 406.8 | - | - | 0 | - |
| - | - | 1285 | 407.2 | - | - | 0 | - |
| - | - | 1207 | 407.3 | - | - | 0 | - |
| - | - | 1636 | 407.5 | - | - | 0 | - |
| - | - | 847.5 | 407.9 | - | - | 0 | - |
| - | - | 4026 | 408.2 | - | - | 0 | - |
| - | - | 5952 | 408.5 | - | - | 0 | - |
| - | - | 826.5 | 408.6 | - | - | 0 | - |
| - | - | 1157 | 408.6 | - | - | 0 | - |
| - | - | 7594 | 408.7 | - | - | 0 | - |
| - | - | 1310 | 408.9 | - | - | 0 | - |
| - | - | 3873 | 409 | - | - | 0 | - |
| 4 | y | 4468 | 409.2 | 0.0003671 | 0.8969 | +3 | 11 |
| - | - | 2668 | 409.6 | - | - | 0 | - |
| - | - | 2093 | 409.9 | - | - | 0 | - |
| - | - | 2784 | 410.1 | - | - | 0 | - |
| - | - | 1.809E+04 | 410.3 | - | - | 0 | - |
| - | - | 4641 | 410.6 | - | - | 0 | - |
| - | - | 772.4 | 410.7 | - | - | 0 | - |
| - | - | 965.7 | 410.8 | - | - | 0 | - |
| - | - | 3621 | 410.9 | - | - | 0 | - |
| - | - | 1220 | 411 | - | - | 0 | - |
| - | - | 1035 | 411.1 | - | - | 0 | - |
| - | - | 1.569E+04 | 411.2 | - | - | 0 | - |
| - | - | 1821 | 411.2 | - | - | 0 | - |
| - | - | 7095 | 411.3 | - | - | 0 | - |
| - | - | 3813 | 411.5 | - | - | 0 | - |
| - | - | 1583 | 411.7 | - | - | 0 | - |
| - | - | 2968 | 411.8 | - | - | 0 | - |
| - | - | 3100 | 412.2 | - | - | 0 | - |
| - | - | 1569 | 412.2 | - | - | 0 | - |
| 7 | y | 5.556E+04 | 412.3 | 0.001294 | 3.14 | +2 | 8 |
| - | - | 3199 | 412.7 | - | - | 0 | - |
| - | - | 2.482E+04 | 412.8 | - | - | 0 | - |
| - | - | 1391 | 413 | - | - | 0 | - |
| - | - | 3256 | 413.2 | - | - | 0 | - |
| - | - | 1206 | 413.2 | - | - | 0 | - |
| - | - | 6422 | 413.3 | - | - | 0 | - |
| - | - | 1390 | 413.6 | - | - | 0 | - |
| - | - | 728 | 413.7 | - | - | 0 | - |
| - | - | 719.2 | 413.7 | - | - | 0 | - |
| - | - | 1979 | 413.8 | - | - | 0 | - |
| - | - | 4940 | 413.9 | - | - | 0 | - |
| - | - | 4943 | 414.2 | - | - | 0 | - |
| - | - | 4380 | 414.2 | - | - | 0 | - |
| - | - | 2244 | 414.6 | - | - | 0 | - |
| - | - | 2586 | 414.7 | - | - | 0 | - |
| - | - | 1091 | 415.2 | - | - | 0 | - |
| - | - | 5793 | 415.2 | - | - | 0 | - |
| - | - | 5546 | 415.3 | - | - | 0 | - |
| - | - | 2487 | 415.5 | - | - | 0 | - |
| 0 | Precursor | 6535 | 415.7 | 0.0004267 | 1.026 | +4 | -1 |
| - | - | 3480 | 415.8 | - | - | 0 | - |
| - | - | 1870 | 415.9 | - | - | 0 | - |
| 0 | Precursor | 8670 | 416 | 0.006803 | 16.36 | +4 | -1 |
| - | - | 4629 | 416.2 | - | - | 0 | - |
| - | - | 822.9 | 416.3 | - | - | 0 | - |
| - | - | 2409 | 416.5 | - | - | 0 | - |
| - | - | 2265 | 416.7 | - | - | 0 | - |
| - | - | 1047 | 418.6 | - | - | 0 | - |
| 6 | b | 1424 | 419.2 | 0.003679 | 8.776 | +2 | 6 |
| - | - | 3456 | 419.2 | - | - | 0 | - |
| - | - | 8200 | 419.3 | - | - | 0 | - |
| - | - | 3563 | 419.6 | - | - | 0 | - |
| - | - | 6196 | 419.7 | - | - | 0 | - |
| - | - | 3386 | 419.8 | - | - | 0 | - |
| - | - | 1.783E+04 | 419.9 | - | - | 0 | - |
| - | - | 3574 | 420 | - | - | 0 | - |
| 0 | Precursor | 4.006E+04 | 420.2 | 0.004835 | 11.51 | +4 | -1 |
| - | - | 3396 | 420.3 | - | - | 0 | - |
| - | - | 3.145E+04 | 420.5 | - | - | 0 | - |
| - | - | 5135 | 420.6 | - | - | 0 | - |
| - | - | 1081 | 420.7 | - | - | 0 | - |
| - | - | 1.255E+04 | 420.7 | - | - | 0 | - |
| - | - | 2243 | 420.9 | - | - | 0 | - |
| - | - | 7825 | 421 | - | - | 0 | - |
| - | - | 1459 | 421.2 | - | - | 0 | - |
| - | - | 3074 | 421.9 | - | - | 0 | - |
| - | - | 2219 | 422.2 | - | - | 0 | - |
| - | - | 5004 | 422.3 | - | - | 0 | - |
| - | - | 1210 | 422.3 | - | - | 0 | - |
| - | - | 917 | 422.6 | - | - | 0 | - |
| - | - | 1296 | 423.2 | - | - | 0 | - |
| - | - | 2409 | 423.3 | - | - | 0 | - |
| - | - | 1273 | 424.3 | - | - | 0 | - |
| - | - | 4759 | 424.3 | - | - | 0 | - |
| - | - | 3338 | 424.6 | - | - | 0 | - |
| - | - | 5650 | 424.7 | - | - | 0 | - |
| - | - | 2650 | 424.9 | - | - | 0 | - |
| - | - | 1.009E+04 | 425.2 | - | - | 0 | - |
| - | - | 1618 | 425.3 | - | - | 0 | - |
| - | - | 6538 | 425.6 | - | - | 0 | - |
| - | - | 3997 | 425.9 | - | - | 0 | - |
| - | - | 2673 | 426.2 | - | - | 0 | - |
| - | - | 4691 | 427.2 | - | - | 0 | - |
| - | - | 5284 | 427.2 | - | - | 0 | - |
| - | - | 4153 | 427.7 | - | - | 0 | - |
| 6 | b | 3567 | 428.2 | 0.002455 | 5.734 | +2 | 6 |
| - | - | 1101 | 428.2 | - | - | 0 | - |
| - | - | 1284 | 428.3 | - | - | 0 | - |
| - | - | 1609 | 428.7 | - | - | 0 | - |
| - | - | 4.857E+04 | 429.2 | - | - | 0 | - |
| 11 | b | 1.247E+04 | 430.2 | 0.006365 | 14.8 | +3 | 11 |
| - | - | 8566 | 431.2 | - | - | 0 | - |
| - | - | 1502 | 431.2 | - | - | 0 | - |
| - | - | 1013 | 431.6 | - | - | 0 | - |
| - | - | 1246 | 432.2 | - | - | 0 | - |
| - | - | 2375 | 432.7 | - | - | 0 | - |
| - | - | 1161 | 433.2 | - | - | 0 | - |
| - | - | 1086 | 433.3 | - | - | 0 | - |
| - | - | 1105 | 433.7 | - | - | 0 | - |
| - | - | 900.5 | 434.2 | - | - | 0 | - |
| 3 | b | 986.8 | 435.2 | 0.007075 | 16.26 | +1 | 3 |
| - | - | 1828 | 437.2 | - | - | 0 | - |
| - | - | 1224 | 439.2 | - | - | 0 | - |
| - | - | 1389 | 439.3 | - | - | 0 | - |
| - | - | 1707 | 442.7 | - | - | 0 | - |
| - | - | 914.8 | 443.2 | - | - | 0 | - |
| - | - | 1698 | 443.2 | - | - | 0 | - |
| - | - | 1029 | 443.6 | - | - | 0 | - |
| - | - | 3432 | 443.9 | - | - | 0 | - |
| - | - | 1718 | 444.2 | - | - | 0 | - |
| - | - | 1174 | 444.6 | - | - | 0 | - |
| - | - | 1.323E+04 | 445.2 | - | - | 0 | - |
| - | - | 1903 | 445.3 | - | - | 0 | - |
| - | - | 3451 | 446.2 | - | - | 0 | - |
| - | - | 3416 | 446.2 | - | - | 0 | - |
| - | - | 2063 | 446.7 | - | - | 0 | - |
| - | - | 959.4 | 447.2 | - | - | 0 | - |
| - | - | 4574 | 448.3 | - | - | 0 | - |
| - | - | 3367 | 448.6 | - | - | 0 | - |
| - | - | 1257 | 448.9 | - | - | 0 | - |
| - | - | 941.6 | 449.6 | - | - | 0 | - |
| - | - | 1671 | 449.9 | - | - | 0 | - |
| - | - | 3590 | 451.2 | - | - | 0 | - |
| - | - | 3328 | 451.3 | - | - | 0 | - |
| - | - | 744.8 | 451.6 | - | - | 0 | - |
| - | - | 4487 | 451.7 | - | - | 0 | - |
| - | - | 5782 | 451.9 | - | - | 0 | - |
| - | - | 915.3 | 452.2 | - | - | 0 | - |
| - | - | 6213 | 452.3 | - | - | 0 | - |
| - | - | 2927 | 452.3 | - | - | 0 | - |
| - | - | 2296 | 452.6 | - | - | 0 | - |
| - | - | 2245 | 452.9 | - | - | 0 | - |
| 3 | b | 3348 | 453.2 | 0.007314 | 16.14 | +1 | 3 |
| - | - | 1548 | 453.3 | - | - | 0 | - |
| - | - | 3432 | 453.6 | - | - | 0 | - |
| - | - | 3551 | 453.9 | - | - | 0 | - |
| - | - | 1472 | 454.2 | - | - | 0 | - |
| - | - | 1621 | 454.3 | - | - | 0 | - |
| - | - | 1964 | 454.7 | - | - | 0 | - |
| - | - | 930.3 | 455.2 | - | - | 0 | - |
| - | - | 5614 | 455.3 | - | - | 0 | - |
| - | - | 1186 | 456.3 | - | - | 0 | - |
| - | - | 2879 | 456.6 | - | - | 0 | - |
| - | - | 3190 | 456.9 | - | - | 0 | - |
| - | - | 1803 | 457.2 | - | - | 0 | - |
| - | - | 6138 | 457.3 | - | - | 0 | - |
| - | - | 4828 | 457.6 | - | - | 0 | - |
| 3 | y | 6412 | 457.9 | 0.002257 | 4.929 | +3 | 12 |
| - | - | 1279 | 458.2 | - | - | 0 | - |
| - | - | 999.2 | 458.2 | - | - | 0 | - |
| - | - | 3616 | 458.3 | - | - | 0 | - |
| - | - | 1758 | 458.6 | - | - | 0 | - |
| - | - | 3716 | 459.2 | - | - | 0 | - |
| - | - | 1759 | 459.7 | - | - | 0 | - |
| - | - | 9496 | 460.2 | - | - | 0 | - |
| - | - | 4437 | 460.7 | - | - | 0 | - |
| - | - | 1073 | 461.2 | - | - | 0 | - |
| - | - | 1057 | 461.7 | - | - | 0 | - |
| - | - | 2333 | 462.3 | - | - | 0 | - |
| - | - | 2486 | 462.6 | - | - | 0 | - |
| - | - | 1.511E+04 | 462.9 | - | - | 0 | - |
| - | - | 9541 | 463.3 | - | - | 0 | - |
| 3 | y | 8.558E+04 | 463.6 | 0.001118 | 2.412 | +3 | 12 |
| - | - | 6.696E+04 | 463.9 | - | - | 0 | - |
| - | - | 2.699E+04 | 464.3 | - | - | 0 | - |
| - | - | 7531 | 464.6 | - | - | 0 | - |
| - | - | 1121 | 464.9 | - | - | 0 | - |
| - | - | 1095 | 466.9 | - | - | 0 | - |
| - | - | 971.2 | 467.2 | - | - | 0 | - |
| - | - | 1208 | 467.3 | - | - | 0 | - |
| - | - | 6178 | 467.3 | - | - | 0 | - |
| - | - | 2556 | 468.2 | - | - | 0 | - |
| - | - | 2.357E+04 | 468.3 | - | - | 0 | - |
| - | - | 7373 | 468.3 | - | - | 0 | - |
| - | - | 1.617E+04 | 468.6 | - | - | 0 | - |
| - | - | 1683 | 468.7 | - | - | 0 | - |
| - | - | 5.637E+04 | 468.9 | - | - | 0 | - |
| - | - | 2424 | 469.2 | - | - | 0 | - |
| - | - | 4.165E+04 | 469.3 | - | - | 0 | - |
| - | - | 2718 | 469.3 | - | - | 0 | - |
| - | - | 1.511E+04 | 469.6 | - | - | 0 | - |
| - | - | 2082 | 469.8 | - | - | 0 | - |
| - | - | 5609 | 469.9 | - | - | 0 | - |
| 11 | y | 7.88E+04 | 470.3 | 0.001357 | 2.884 | +1 | 4 |
| - | - | 1.89E+04 | 471.3 | - | - | 0 | - |
| - | - | 1071 | 471.6 | - | - | 0 | - |
| - | - | 5198 | 471.8 | - | - | 0 | - |
| - | - | 1084 | 472.3 | - | - | 0 | - |
| 6 | y | 1777 | 472.3 | 0.009283 | 19.66 | +2 | 9 |
| - | - | 3144 | 472.3 | - | - | 0 | - |
| - | - | 1212 | 472.6 | - | - | 0 | - |
| - | - | 1173 | 472.8 | - | - | 0 | - |
| - | - | 1.042E+04 | 473.3 | - | - | 0 | - |
| - | - | 2918 | 473.6 | - | - | 0 | - |
| - | - | 1365 | 473.9 | - | - | 0 | - |
| - | - | 3107 | 474.3 | - | - | 0 | - |
| - | - | 3128 | 474.8 | - | - | 0 | - |
| - | - | 1189 | 474.9 | - | - | 0 | - |
| - | - | 966.6 | 475.2 | - | - | 0 | - |
| - | - | 1862 | 475.3 | - | - | 0 | - |
| - | - | 3372 | 475.6 | - | - | 0 | - |
| - | - | 2312 | 475.9 | - | - | 0 | - |
| - | - | 2410 | 476.2 | - | - | 0 | - |
| - | - | 6810 | 476.3 | - | - | 0 | - |
| - | - | 5250 | 476.6 | - | - | 0 | - |
| - | - | 2502 | 476.9 | - | - | 0 | - |
| 12 | b | 4967 | 478.6 | 0.003098 | 6.474 | +3 | 12 |
| - | - | 1588 | 478.8 | - | - | 0 | - |
| - | - | 4771 | 478.9 | - | - | 0 | - |
| - | - | 1727 | 479.2 | - | - | 0 | - |
| - | - | 1800 | 479.3 | - | - | 0 | - |
| - | - | 1777 | 479.6 | - | - | 0 | - |
| 6 | y | 1.45E+04 | 480.8 | 0.001288 | 2.679 | +2 | 9 |
| - | - | 2356 | 480.9 | - | - | 0 | - |
| - | - | 3723 | 481.3 | - | - | 0 | - |
| - | - | 7703 | 481.3 | - | - | 0 | - |
| - | - | 9919 | 481.6 | - | - | 0 | - |
| - | - | 2243 | 481.8 | - | - | 0 | - |
| - | - | 7259 | 481.9 | - | - | 0 | - |
| - | - | 4248 | 482.3 | - | - | 0 | - |
| - | - | 6054 | 482.3 | - | - | 0 | - |
| - | - | 1210 | 482.6 | - | - | 0 | - |
| - | - | 5599 | 482.8 | - | - | 0 | - |
| - | - | 973.1 | 483.2 | - | - | 0 | - |
| - | - | 2813 | 483.3 | - | - | 0 | - |
| - | - | 1843 | 483.3 | - | - | 0 | - |
| - | - | 8395 | 483.8 | - | - | 0 | - |
| - | - | 4941 | 484.3 | - | - | 0 | - |
| - | - | 1107 | 484.3 | - | - | 0 | - |
| - | - | 1263 | 484.6 | - | - | 0 | - |
| - | - | 2200 | 486.3 | - | - | 0 | - |
| - | - | 2222 | 486.6 | - | - | 0 | - |
| - | - | 3281 | 486.9 | - | - | 0 | - |
| - | - | 4910 | 487.3 | - | - | 0 | - |
| - | - | 8412 | 487.6 | - | - | 0 | - |
| - | - | 2130 | 487.8 | - | - | 0 | - |
| - | - | 2948 | 487.9 | - | - | 0 | - |
| - | - | 1114 | 488.3 | - | - | 0 | - |
| - | - | 1108 | 490.9 | - | - | 0 | - |
| - | - | 7340 | 491.3 | - | - | 0 | - |
| - | - | 2419 | 491.8 | - | - | 0 | - |
| - | - | 1395 | 492.2 | - | - | 0 | - |
| - | - | 1723 | 492.3 | - | - | 0 | - |
| - | - | 1051 | 492.6 | - | - | 0 | - |
| - | - | 2031 | 492.8 | - | - | 0 | - |
| - | - | 1223 | 492.9 | - | - | 0 | - |
| - | - | 1918 | 493.3 | - | - | 0 | - |
| - | - | 1809 | 494.2 | - | - | 0 | - |
| - | - | 1105 | 494.6 | - | - | 0 | - |
| - | - | 1123 | 494.9 | - | - | 0 | - |
| - | - | 1805 | 495.4 | - | - | 0 | - |
| - | - | 1949 | 500.9 | - | - | 0 | - |
| - | - | 1346 | 501.3 | - | - | 0 | - |
| - | - | 1686 | 501.6 | - | - | 0 | - |
| - | - | 920.8 | 501.9 | - | - | 0 | - |
| - | - | 4437 | 503.2 | - | - | 0 | - |
| - | - | 1011 | 504.6 | - | - | 0 | - |
| - | - | 2356 | 504.9 | - | - | 0 | - |
| 8 | b | 1705 | 505.2 | 0.003702 | 7.328 | +2 | 8 |
| - | - | 1083 | 505.3 | - | - | 0 | - |
| - | - | 1226 | 505.6 | - | - | 0 | - |
| - | - | 1181 | 505.7 | - | - | 0 | - |
| - | - | 1746 | 506.6 | - | - | 0 | - |
| - | - | 3626 | 506.9 | - | - | 0 | - |
| - | - | 1861 | 507.3 | - | - | 0 | - |
| - | - | 2028 | 508.8 | - | - | 0 | - |
| - | - | 853.1 | 509.3 | - | - | 0 | - |
| - | - | 1855 | 509.9 | - | - | 0 | - |
| 13 | b | 2481 | 510.3 | 0.002819 | 5.525 | +3 | 13 |
| - | - | 3041 | 510.6 | - | - | 0 | - |
| - | - | 4291 | 510.9 | - | - | 0 | - |
| - | - | 1893 | 511.3 | - | - | 0 | - |
| - | - | 977.5 | 511.6 | - | - | 0 | - |
| - | - | 1671 | 515.6 | - | - | 0 | - |
| 13 | b | 6364 | 516.3 | 0.002838 | 5.497 | +3 | 13 |
| - | - | 7990 | 516.6 | - | - | 0 | - |
| - | - | 1075 | 516.8 | - | - | 0 | - |
| - | - | 1125 | 516.8 | - | - | 0 | - |
| - | - | 4320 | 516.9 | - | - | 0 | - |
| - | - | 1379 | 517.2 | - | - | 0 | - |
| - | - | 2931 | 517.3 | - | - | 0 | - |
| - | - | 1064 | 517.7 | - | - | 0 | - |
| - | - | 1144 | 519.2 | - | - | 0 | - |
| - | - | 3904 | 519.3 | - | - | 0 | - |
| - | - | 2633 | 519.6 | - | - | 0 | - |
| - | - | 1156 | 519.9 | - | - | 0 | - |
| - | - | 3355 | 520.4 | - | - | 0 | - |
| - | - | 3926 | 521.2 | - | - | 0 | - |
| - | - | 2819 | 521.6 | - | - | 0 | - |
| - | - | 3216 | 521.9 | - | - | 0 | - |
| - | - | 1081 | 522.2 | - | - | 0 | - |
| - | - | 1.481E+04 | 522.3 | - | - | 0 | - |
| - | - | 1.123E+04 | 522.6 | - | - | 0 | - |
| - | - | 3156 | 522.9 | - | - | 0 | - |
| - | - | 1805 | 523.3 | - | - | 0 | - |
| - | - | 1526 | 523.4 | - | - | 0 | - |
| - | - | 3313 | 524.6 | - | - | 0 | - |
| - | - | 1000 | 524.8 | - | - | 0 | - |
| - | - | 8371 | 525 | - | - | 0 | - |
| 2 | y | 1.796E+04 | 525.3 | 0.004432 | 8.437 | +3 | 13 |
| - | - | 9534 | 525.6 | - | - | 0 | - |
| - | - | 1748 | 525.8 | - | - | 0 | - |
| - | - | 1267 | 525.8 | - | - | 0 | - |
| - | - | 5345 | 526 | - | - | 0 | - |
| - | - | 1449 | 526.2 | - | - | 0 | - |
| - | - | 1062 | 526.3 | - | - | 0 | - |
| - | - | 1385 | 526.8 | - | - | 0 | - |
| - | - | 1643 | 528.2 | - | - | 0 | - |
| - | - | 1743 | 529.8 | - | - | 0 | - |
| - | - | 1.017E+04 | 530.3 | - | - | 0 | - |
| - | - | 9291 | 530.6 | - | - | 0 | - |
| - | - | 2485 | 530.8 | - | - | 0 | - |
| 2 | y | 2.303E+04 | 531 | 0.002072 | 3.902 | +3 | 13 |
| - | - | 4196 | 531.2 | - | - | 0 | - |
| - | - | 1.757E+04 | 531.3 | - | - | 0 | - |
| - | - | 8271 | 531.6 | - | - | 0 | - |
| - | - | 3173 | 532 | - | - | 0 | - |
| - | - | 1720 | 532.3 | - | - | 0 | - |
| - | - | 951.1 | 533.8 | - | - | 0 | - |
| - | - | 945.2 | 534.3 | - | - | 0 | - |
| - | - | 1101 | 535.7 | - | - | 0 | - |
| - | - | 1208 | 537.2 | - | - | 0 | - |
| - | - | 1091 | 537.3 | - | - | 0 | - |
| - | - | 2.956E+04 | 538.3 | - | - | 0 | - |
| - | - | 2950 | 538.4 | - | - | 0 | - |
| - | - | 5360 | 538.8 | - | - | 0 | - |
| - | - | 9666 | 539.3 | - | - | 0 | - |
| - | - | 8572 | 539.3 | - | - | 0 | - |
| - | - | 4655 | 539.8 | - | - | 0 | - |
| 5 | y | 1761 | 540.8 | 0.006591 | 12.19 | +2 | 10 |
| - | - | 1078 | 541.3 | - | - | 0 | - |
| 10 | y | 1.708E+04 | 541.4 | 0.001383 | 2.554 | +1 | 5 |
| - | - | 1396 | 542.3 | - | - | 0 | - |
| - | - | 4752 | 542.4 | - | - | 0 | - |
| - | - | 1200 | 543.6 | - | - | 0 | - |
| - | - | 2628 | 543.8 | - | - | 0 | - |
| - | - | 1051 | 544.2 | - | - | 0 | - |
| - | - | 1240 | 545.3 | - | - | 0 | - |
| - | - | 1024 | 545.8 | - | - | 0 | - |
| - | - | 1131 | 546.8 | - | - | 0 | - |
| - | - | 1391 | 547.2 | - | - | 0 | - |
| - | - | 4605 | 547.8 | - | - | 0 | - |
| - | - | 4405 | 548.3 | - | - | 0 | - |
| - | - | 2634 | 548.8 | - | - | 0 | - |
| - | - | 3944 | 549.2 | - | - | 0 | - |
| 5 | y | 2.867E+04 | 549.3 | 0.001068 | 1.944 | +2 | 10 |
| - | - | 1.723E+04 | 549.8 | - | - | 0 | - |
| - | - | 5016 | 550.3 | - | - | 0 | - |
| - | - | 1121 | 550.8 | - | - | 0 | - |
| - | - | 2013 | 552.3 | - | - | 0 | - |
| - | - | 2395 | 552.8 | - | - | 0 | - |
| - | - | 1772 | 553.3 | - | - | 0 | - |
| - | - | 7735 | 554.2 | - | - | 0 | - |
| - | - | 2151 | 555.2 | - | - | 0 | - |
| - | - | 4523 | 556.3 | - | - | 0 | - |
| - | - | 4194 | 556.8 | - | - | 0 | - |
| - | - | 1218 | 557.3 | - | - | 0 | - |
| - | - | 1745 | 557.3 | - | - | 0 | - |
| - | - | 5230 | 557.3 | - | - | 0 | - |
| - | - | 2777 | 557.8 | - | - | 0 | - |
| - | - | 2052 | 558.2 | - | - | 0 | - |
| - | - | 1993 | 559.3 | - | - | 0 | - |
| - | - | 1241 | 559.8 | - | - | 0 | - |
| 9 | b | 1237 | 560.3 | 0.002824 | 5.04 | +2 | 9 |
| 9 | b | 1529 | 560.7 | 0.007032 | 12.54 | +2 | 9 |
| - | - | 3058 | 561.3 | - | - | 0 | - |
| - | - | 2549 | 561.8 | - | - | 0 | - |
| - | - | 1730 | 564.2 | - | - | 0 | - |
| - | - | 3.58E+04 | 566.2 | - | - | 0 | - |
| - | - | 9787 | 567.3 | - | - | 0 | - |
| - | - | 2540 | 567.4 | - | - | 0 | - |
| - | - | 2654 | 568.3 | - | - | 0 | - |
| - | - | 1302 | 568.8 | - | - | 0 | - |
| - | - | 1138 | 569.2 | - | - | 0 | - |
| 9 | b | 8125 | 569.3 | 0.003523 | 6.188 | +2 | 9 |
| - | - | 4924 | 569.8 | - | - | 0 | - |
| - | - | 4480 | 570.2 | - | - | 0 | - |
| - | - | 1044 | 571.2 | - | - | 0 | - |
| - | - | 1.483E+04 | 572.8 | - | - | 0 | - |
| - | - | 9749 | 573.3 | - | - | 0 | - |
| - | - | 2750 | 573.8 | - | - | 0 | - |
| - | - | 1064 | 574.8 | - | - | 0 | - |
| - | - | 2167 | 576.3 | - | - | 0 | - |
| - | - | 1271 | 576.8 | - | - | 0 | - |
| - | - | 1195 | 577.3 | - | - | 0 | - |
| - | - | 1.116E+04 | 577.4 | - | - | 0 | - |
| - | - | 3802 | 578.4 | - | - | 0 | - |
| - | - | 2.796E+04 | 579.4 | - | - | 0 | - |
| - | - | 2523 | 579.8 | - | - | 0 | - |
| - | - | 1139 | 580.3 | - | - | 0 | - |
| - | - | 8656 | 580.4 | - | - | 0 | - |
| - | - | 5706 | 580.8 | - | - | 0 | - |
| 4 | b | 1198 | 581.2 | 0.003729 | 6.416 | +1 | 4 |
| - | - | 3467 | 581.3 | - | - | 0 | - |
| - | - | 2605 | 581.4 | - | - | 0 | - |
| - | - | 1651 | 581.8 | - | - | 0 | - |
| - | - | 1.244E+04 | 582.2 | - | - | 0 | - |
| - | - | 4425 | 583.2 | - | - | 0 | - |
| - | - | 1613 | 585.3 | - | - | 0 | - |
| - | - | 6714 | 586.2 | - | - | 0 | - |
| - | - | 2368 | 587.2 | - | - | 0 | - |
| - | - | 2538 | 593.3 | - | - | 0 | - |
| - | - | 1076 | 595.3 | - | - | 0 | - |
| - | - | 1065 | 595.3 | - | - | 0 | - |
| - | - | 8694 | 595.4 | - | - | 0 | - |
| 10 | b | 2042 | 595.8 | 0.002272 | 3.814 | +2 | 10 |
| - | - | 2249 | 595.8 | - | - | 0 | - |
| - | - | 1246 | 596.3 | - | - | 0 | - |
| - | - | 1181 | 596.3 | - | - | 0 | - |
| - | - | 3217 | 596.4 | - | - | 0 | - |
| - | - | 1570 | 596.8 | - | - | 0 | - |
| - | - | 2194 | 598.2 | - | - | 0 | - |
| - | - | 1503 | 603.3 | - | - | 0 | - |
| - | - | 7709 | 604.4 | - | - | 0 | - |
| 10 | b | 2088 | 604.8 | 0.005047 | 8.344 | +2 | 10 |
| 4 | y | 1.407E+04 | 604.8 | 0.004646 | 7.681 | +2 | 11 |
| - | - | 1595 | 605.3 | - | - | 0 | - |
| - | - | 8095 | 605.4 | - | - | 0 | - |
| - | - | 3826 | 605.9 | - | - | 0 | - |
| - | - | 9488 | 610.3 | - | - | 0 | - |
| - | - | 2909 | 611.3 | - | - | 0 | - |
| - | - | 2658 | 611.8 | - | - | 0 | - |
| - | - | 2555 | 612.3 | - | - | 0 | - |
| - | - | 2811 | 612.9 | - | - | 0 | - |
| - | - | 1273 | 613.3 | - | - | 0 | - |
| 4 | y | 7984 | 613.4 | 0.0003888 | 0.6339 | +2 | 11 |
| - | - | 4468 | 613.9 | - | - | 0 | - |
| - | - | 4918 | 614.2 | - | - | 0 | - |
| - | - | 1958 | 614.4 | - | - | 0 | - |
| - | - | 1260 | 615.2 | - | - | 0 | - |
| - | - | 3169 | 615.4 | - | - | 0 | - |
| - | - | 2477 | 615.8 | - | - | 0 | - |
| - | - | 6213 | 620.3 | - | - | 0 | - |
| - | - | 5989 | 620.8 | - | - | 0 | - |
| - | - | 3139 | 621.3 | - | - | 0 | - |
| - | - | 1317 | 622.3 | - | - | 0 | - |
| - | - | 1212 | 623.3 | - | - | 0 | - |
| - | - | 1313 | 623.8 | - | - | 0 | - |
| - | - | 1287 | 625.3 | - | - | 0 | - |
| - | - | 1233 | 627.8 | - | - | 0 | - |
| - | - | 5064 | 628.3 | - | - | 0 | - |
| - | - | 3782 | 628.8 | - | - | 0 | - |
| - | - | 1288 | 629.3 | - | - | 0 | - |
| - | - | 1.524E+04 | 629.3 | - | - | 0 | - |
| - | - | 9931 | 629.8 | - | - | 0 | - |
| - | - | 4557 | 630.3 | - | - | 0 | - |
| - | - | 1352 | 630.9 | - | - | 0 | - |
| - | - | 1546 | 631.3 | - | - | 0 | - |
| - | - | 1439 | 632.4 | - | - | 0 | - |
| - | - | 1390 | 632.9 | - | - | 0 | - |
| - | - | 2281 | 636.3 | - | - | 0 | - |
| - | - | 2301 | 636.8 | - | - | 0 | - |
| - | - | 4982 | 637.3 | - | - | 0 | - |
| - | - | 5345 | 637.8 | - | - | 0 | - |
| - | - | 8942 | 638.4 | - | - | 0 | - |
| - | - | 4668 | 638.9 | - | - | 0 | - |
| - | - | 2238 | 639.4 | - | - | 0 | - |
| - | - | 2315 | 641.4 | - | - | 0 | - |
| - | - | 1210 | 641.9 | - | - | 0 | - |
| - | - | 1295 | 642.4 | - | - | 0 | - |
| - | - | 1617 | 645.8 | - | - | 0 | - |
| - | - | 3516 | 646.4 | - | - | 0 | - |
| - | - | 2345 | 646.9 | - | - | 0 | - |
| - | - | 1213 | 647.4 | - | - | 0 | - |
| - | - | 1919 | 647.4 | - | - | 0 | - |
| - | - | 918.9 | 650.4 | - | - | 0 | - |
| - | - | 7086 | 664.5 | - | - | 0 | - |
| - | - | 1310 | 664.8 | - | - | 0 | - |
| - | - | 2309 | 665.3 | - | - | 0 | - |
| - | - | 3424 | 665.5 | - | - | 0 | - |
| - | - | 1762 | 665.8 | - | - | 0 | - |
| - | - | 1202 | 666.3 | - | - | 0 | - |
| - | - | 1846 | 667.3 | - | - | 0 | - |
| - | - | 2623 | 668.3 | - | - | 0 | - |
| 9 | y | 5587 | 669.5 | 0.001085 | 1.621 | +1 | 6 |
| - | - | 1122 | 670.3 | - | - | 0 | - |
| - | - | 2480 | 670.5 | - | - | 0 | - |
| - | - | 1796 | 671.3 | - | - | 0 | - |
| - | - | 1170 | 673.3 | - | - | 0 | - |
| - | - | 2782 | 673.8 | - | - | 0 | - |
| - | - | 1564 | 674.3 | - | - | 0 | - |
| - | - | 2.357E+04 | 674.4 | - | - | 0 | - |
| - | - | 1002 | 675.3 | - | - | 0 | - |
| - | - | 9366 | 675.4 | - | - | 0 | - |
| - | - | 1351 | 676.4 | - | - | 0 | - |
| - | - | 1038 | 678.4 | - | - | 0 | - |
| - | - | 1224 | 678.4 | - | - | 0 | - |
| - | - | 2491 | 685.4 | - | - | 0 | - |
| - | - | 915.2 | 685.9 | - | - | 0 | - |
| 3 | y | 1063 | 686.4 | 0.006022 | 8.774 | +2 | 12 |
| - | - | 892.2 | 688.3 | - | - | 0 | - |
| - | - | 2529 | 688.4 | - | - | 0 | - |
| - | - | 3178 | 690.3 | - | - | 0 | - |
| - | - | 2123 | 690.4 | - | - | 0 | - |
| - | - | 1182 | 691.3 | - | - | 0 | - |
| - | - | 1111 | 691.4 | - | - | 0 | - |
| - | - | 2.565E+04 | 692.4 | - | - | 0 | - |
| - | - | 9026 | 693.4 | - | - | 0 | - |
| - | - | 1078 | 694.4 | - | - | 0 | - |
| - | - | 1841 | 694.5 | - | - | 0 | - |
| 3 | y | 7197 | 694.9 | 0.0008449 | 1.216 | +2 | 12 |
| - | - | 5454 | 695.4 | - | - | 0 | - |
| - | - | 2562 | 695.9 | - | - | 0 | - |
| - | - | 2017 | 698.4 | - | - | 0 | - |
| 5 | b | 1404 | 700.3 | 0.003417 | 4.879 | +1 | 5 |
| - | - | 2286 | 702.3 | - | - | 0 | - |
| - | - | 954.3 | 702.4 | - | - | 0 | - |
| - | - | 3417 | 702.9 | - | - | 0 | - |
| - | - | 1206 | 703.3 | - | - | 0 | - |
| - | - | 2977 | 703.4 | - | - | 0 | - |
| - | - | 1599 | 703.9 | - | - | 0 | - |
| - | - | 3123 | 706.4 | - | - | 0 | - |
| - | - | 3672 | 707.3 | - | - | 0 | - |
| - | - | 1357 | 708.3 | - | - | 0 | - |
| - | - | 1269 | 708.4 | - | - | 0 | - |
| - | - | 1109 | 709.4 | - | - | 0 | - |
| - | - | 966.4 | 710.4 | - | - | 0 | - |
| - | - | 8035 | 710.5 | - | - | 0 | - |
| - | - | 3003 | 711.5 | - | - | 0 | - |
| - | - | 1067 | 713.9 | - | - | 0 | - |
| - | - | 1160 | 714.4 | - | - | 0 | - |
| - | - | 5708 | 716.4 | - | - | 0 | - |
| - | - | 2757 | 717.4 | - | - | 0 | - |
| - | - | 1269 | 717.9 | - | - | 0 | - |
| 5 | b | 4340 | 718.3 | 0.00628 | 8.742 | +1 | 5 |
| - | - | 1173 | 718.4 | - | - | 0 | - |
| - | - | 1275 | 719.3 | - | - | 0 | - |
| - | - | 2807 | 720.3 | - | - | 0 | - |
| - | - | 4200 | 721.4 | - | - | 0 | - |
| - | - | 2331 | 721.9 | - | - | 0 | - |
| - | - | 1202 | 722.4 | - | - | 0 | - |
| - | - | 5966 | 723.3 | - | - | 0 | - |
| - | - | 2281 | 724.3 | - | - | 0 | - |
| - | - | 1081 | 724.4 | - | - | 0 | - |
| 8 | y | 1.198E+04 | 726.5 | 0.001228 | 1.69 | +1 | 7 |
| - | - | 3953 | 727.5 | - | - | 0 | - |
| - | - | 1173 | 728.5 | - | - | 0 | - |
| - | - | 1789 | 730.4 | - | - | 0 | - |
| - | - | 1893 | 730.9 | - | - | 0 | - |
| - | - | 2912 | 735.3 | - | - | 0 | - |
| - | - | 1639 | 736.3 | - | - | 0 | - |
| - | - | 2487 | 737.3 | - | - | 0 | - |
| - | - | 1697 | 738.4 | - | - | 0 | - |
| - | - | 2368 | 739.4 | - | - | 0 | - |
| - | - | 2881 | 740.3 | - | - | 0 | - |
| - | - | 1245 | 740.5 | - | - | 0 | - |
| - | - | 1513 | 741.3 | - | - | 0 | - |
| - | - | 957 | 742.4 | - | - | 0 | - |
| - | - | 819.9 | 750.3 | - | - | 0 | - |
| - | - | 5108 | 751.3 | - | - | 0 | - |
| - | - | 2190 | 752.3 | - | - | 0 | - |
| - | - | 929.8 | 756.4 | - | - | 0 | - |
| - | - | 1655 | 766.3 | - | - | 0 | - |
| - | - | 4675 | 768.3 | - | - | 0 | - |
| - | - | 1734 | 769.3 | - | - | 0 | - |
| - | - | 1260 | 777.5 | - | - | 0 | - |
| - | - | 1025 | 790.5 | - | - | 0 | - |
| - | - | 853 | 794.5 | - | - | 0 | - |
| - | - | 2586 | 805.5 | - | - | 0 | - |
| - | - | 1534 | 806.5 | - | - | 0 | - |
| - | - | 1009 | 808.3 | - | - | 0 | - |
| - | - | 1324 | 811.4 | - | - | 0 | - |
| - | - | 3317 | 811.5 | - | - | 0 | - |
| - | - | 967.1 | 812.4 | - | - | 0 | - |
| - | - | 897.1 | 819.5 | - | - | 0 | - |
| - | - | 3925 | 821.5 | - | - | 0 | - |
| - | - | 2160 | 822.5 | - | - | 0 | - |
| 7 | y | 4.401E+04 | 823.5 | 0.001076 | 1.307 | +1 | 8 |
| - | - | 1.874E+04 | 824.5 | - | - | 0 | - |
| - | - | 4178 | 825.5 | - | - | 0 | - |
| - | - | 962.6 | 826.6 | - | - | 0 | - |
| - | - | 2349 | 827.4 | - | - | 0 | - |
| - | - | 1063 | 829.4 | - | - | 0 | - |
| - | - | 2862 | 829.5 | - | - | 0 | - |
| - | - | 1734 | 830.5 | - | - | 0 | - |
| - | - | 5178 | 837.5 | - | - | 0 | - |
| - | - | 1549 | 838.5 | - | - | 0 | - |
| - | - | 2763 | 839.4 | - | - | 0 | - |
| - | - | 1920 | 839.5 | - | - | 0 | - |
| - | - | 926.9 | 840.4 | - | - | 0 | - |
| - | - | 1062 | 847.5 | - | - | 0 | - |
| - | - | 5923 | 848.4 | - | - | 0 | - |
| - | - | 3271 | 849.4 | - | - | 0 | - |
| - | - | 2184 | 853.5 | - | - | 0 | - |
| - | - | 932.5 | 854.5 | - | - | 0 | - |
| 6 | b | 5093 | 855.3 | 0.00645 | 7.541 | +1 | 6 |
| - | - | 2157 | 856.4 | - | - | 0 | - |
| - | - | 2071 | 864.4 | - | - | 0 | - |
| - | - | 896.9 | 866.4 | - | - | 0 | - |
| - | - | 852.6 | 891.5 | - | - | 0 | - |
| - | - | 6883 | 901.4 | - | - | 0 | - |
| - | - | 3443 | 902.4 | - | - | 0 | - |
| - | - | 1243 | 903.4 | - | - | 0 | - |
| - | - | 1911 | 917.4 | - | - | 0 | - |
| - | - | 1051 | 918.4 | - | - | 0 | - |
| - | - | 4161 | 919.5 | - | - | 0 | - |
| - | - | 1843 | 920.5 | - | - | 0 | - |
| - | - | 1346 | 935.4 | - | - | 0 | - |
| - | - | 981.9 | 936.4 | - | - | 0 | - |
| - | - | 2036 | 942.6 | - | - | 0 | - |
| - | - | 1423 | 943.6 | - | - | 0 | - |
| 6 | y | 5686 | 960.6 | 0.0005141 | 0.5351 | +1 | 9 |
| - | - | 2752 | 961.6 | - | - | 0 | - |
| - | - | 1041 | 962.6 | - | - | 0 | - |
| - | - | 1071 | 964.5 | - | - | 0 | - |
| - | - | 841.7 | 966.6 | - | - | 0 | - |
| - | - | 1104 | 1034 | - | - | 0 | - |
| - | - | 865.5 | 1086 | - | - | 0 | - |
| 5 | y | 1424 | 1098 | 0.001356 | 1.235 | +1 | 10 |
| - | - | 726.4 | 1099 | - | - | 0 | - |

m/z Charge Intensity FragmentType MassShift Position
120.04508972167969 0 1182.0309
120.052734375 0 1260.7505
120.05699920654297 0 693.81006
120.06645965576172 0 621.5933
120.08135986328125 0 2976.766
120.50309753417969 0 440.92883
121.0401611328125 0 1271.5809
122.06057739257812 0 869.63293
122.07190704345703 0 1485.9999
123.04464721679688 0 2206.1575
123.05585479736328 0 4813.489
123.09262084960938 0 561.3802
124.07610321044922 0 407.019
125.07154083251953 0 1658.8038
125.1081771850586 0 652.73645
126.05521392822266 0 464.3497
126.06670379638672 0 666.23193
126.48697662353516 0 511.57352
127.05087280273438 0 1093.8435
127.08711242675781 0 5539.5786
127.12398529052734 0 474.46753
128.03482055664062 0 466.16653
128.05014038085938 0 748.8901
128.07118225097656 0 12231.001
129.05531311035156 0 636.02747
129.06643676757812 0 4927.619
129.10284423828125 0 211486.08
130.0657196044922 0 76894.8
130.08712768554688 0 955.4768
130.10040283203125 0 1144.2052
130.1061553955078 0 13585.233
131.0690155029297 0 7524.5435
131.08206176757812 0 1138.1272
131.11856079101562 0 741.15344
132.04498291015625 0 3162.6924
132.0813751220703 0 93269.984
132.10250854492188 0 52191.81 y 13
132.11073303222656 0 868.6391
133.05271911621094 0 1746.0374
133.06141662597656 0 1856.08
133.08473205566406 0 9532.677
133.10581970214844 0 3268.658
133.10995483398438 0 356.37277
134.0605926513672 0 5667.187
134.0718231201172 0 1043.7953
135.04461669921875 0 6552.5283
136.07630920410156 0 175455.03
137.07342529296875 0 1281.4639
137.0796356201172 0 14349.988
137.10784912109375 0 541.2933
138.06689453125 0 3271.2217
138.10299682617188 0 570.6192
139.0507049560547 0 535.30865
139.08717346191406 0 4214.8286
139.12335205078125 0 733.87067
140.07106018066406 0 840.9109
141.0666961669922 0 536.06116
141.1028289794922 0 13132.015
142.06570434570312 0 2286.1125
142.10609436035156 0 924.0991
143.04571533203125 0 564.25214
144.04505920410156 0 1386.0267
144.0538330078125 0 623.24854
144.08140563964844 0 4343.943
145.06141662597656 0 2453.3684
145.07652282714844 0 1054.5353
145.09799194335938 0 701.49176
146.06063842773438 0 39602.977 b 3
146.1294708251953 0 815.8832
147.04464721679688 0 12149.983
147.06402587890625 0 3450.9692
147.1134490966797 0 835.41656
148.04808044433594 0 1115.1715
148.07630920410156 0 45421.492
148.09747314453125 0 2022.2554
149.02392578125 0 30599.057
149.07972717285156 0 4221.346
150.0272216796875 0 2543.869
150.0555419921875 0 4068.5835
150.0664825439453 0 1306.8137
151.0509490966797 0 488.02313
151.0871124267578 0 3799.0808
152.07122802734375 0 45492.99
152.14398193359375 0 4963.225
153.0670623779297 0 1319.5741
153.07460021972656 0 3750.9558
153.10296630859375 0 1948.9135
154.09799194335938 0 1539.743
155.04579162597656 0 1504.1572
155.08212280273438 0 11131.338
155.09329223632812 0 4814.5195
155.11842346191406 0 7423.242
156.07742309570312 0 4738.089
156.11395263671875 0 562.0321
156.12103271484375 0 766.7787
157.0526885986328 0 2500.6023
157.06068420410156 0 1245.1323
157.0766143798828 0 9231.543
157.08438110351562 0 1003.5684
158.0606689453125 0 18164.428
158.08445739746094 0 10432.104
159.06390380859375 0 2139.245
159.09228515625 0 70505.81
160.07626342773438 0 5928.517
160.08912658691406 0 863.0225
160.09568786621094 0 7428.219
161.07177734375 0 1360.2263
161.07864379882812 0 581.26495
162.05564880371094 0 4158.6074
162.0794677734375 0 948.1309
162.10293579101562 0 685.8234
163.03956604003906 0 2029.4023
163.08724975585938 0 900.2016
163.6136474609375 0 1061.5023
164.03428649902344 0 754.711
164.0711669921875 0 4594.581
164.08255004882812 0 1176.0287
165.0667266845703 0 665.8657
165.07733154296875 0 1093.0715
165.10284423828125 0 7531.526
165.13919067382812 0 704.60315
166.06175231933594 0 3497.6921
166.09779357910156 0 812.4077
166.1062469482422 0 695.3388
167.03440856933594 0 1813.8547
167.08261108398438 0 885.0871
167.15489196777344 0 1493.7887
168.1136474609375 0 1772.7637
169.0766143798828 0 8178.0234
169.0977783203125 0 14939.982
169.1705322265625 0 1054.7195
170.0607147216797 0 51826.543
170.09344482421875 0 903.841
170.10125732421875 0 994.0011
171.05593872070312 0 1522.0829
171.06407165527344 0 6031.184
171.09246826171875 0 1784.1565
171.1134490966797 0 553.45544
171.1500244140625 0 664.0381
172.04010009765625 0 574.1878
172.06387329101562 0 693.2759
172.0765838623047 0 1334.8873
172.10867309570312 0 14263.599
172.14500427246094 0 1008.978
173.07156372070312 0 17203.066
174.05572509765625 0 4366.4795
174.06678771972656 0 661.3813
174.07925415039062 0 4559.337
175.08724975585938 0 38866.11
175.09710693359375 0 1275.0504
176.07125854492188 0 1251.7214 b Ammonia loss 4
176.08251953125 0 4805.938
176.09083557128906 0 3901.8523
177.07723999023438 0 714.2612
178.05064392089844 0 2939.9814
178.13421630859375 0 664.6327
179.082275390625 0 2082.8506
179.0932159423828 0 646.0897
179.11842346191406 0 941.9202
180.0664825439453 0 1774.8065
180.077392578125 0 865.6175
181.06138610839844 0 3213.618
181.09783935546875 0 41900.19
181.1340789794922 0 3976.9917
181.17022705078125 0 640.63306
182.08477783203125 0 474.9347
182.09341430664062 0 882.12476
182.10133361816406 0 3847.122
182.12948608398438 0 7399.6133
183.0561981201172 0 725.2743
183.09234619140625 0 15020.161
183.11328125 0 1583.7455
183.14974975585938 0 789.80615
184.040283203125 0 1379.634
184.07608032226562 0 2043.2161
184.09576416015625 0 1581.0714
185.04779052734375 0 5487.6313
185.05592346191406 0 833.3527
185.0716094970703 0 18042.05
185.08058166503906 0 1962.2028
185.09182739257812 0 671.89624
185.10757446289062 0 665.2163
186.05563354492188 0 25657.076
186.0790557861328 0 966.4203
186.12440490722656 0 26604.283
187.05914306640625 0 2130.3533
187.08726501464844 0 76043.914
187.10813903808594 0 4116.158
187.12799072265625 0 2273.856
188.07119750976562 0 1585.7671
188.08230590820312 0 789.0957
188.09080505371094 0 8660.336
189.06658935546875 0 829.4332
189.093017578125 0 747.1028
190.05068969726562 0 726.543
191.0824737548828 0 1393.3292
191.09332275390625 0 974.4632
191.11837768554688 0 759.0984
193.0978546142578 0 1650.7797
193.1075439453125 0 652.5234
193.13442993164062 0 1259.5249
195.07797241210938 0 1277.8541
195.08740234375 0 564.60754
195.11349487304688 0 1417.0416
195.14981079101562 0 8710.487
196.1087188720703 0 2161.771
196.13389587402344 0 1736.003
196.1532440185547 0 531.4713
197.0715789794922 0 3763.74
197.09280395507812 0 10042.542
197.1655731201172 0 9489.855
198.05577087402344 0 1291.8982
198.07730102539062 0 521.5794
198.09579467773438 0 919.5904
198.1700439453125 0 731.143
199.05044555664062 0 811.21545
199.0872039794922 0 9258.456
199.1085205078125 0 1743.6307
199.17208862304688 0 733.23773
199.18116760253906 0 4744.8687
200.0711212158203 0 966.1357
200.09085083007812 0 1300.5046
200.1401824951172 0 4145.4966
201.06649780273438 0 18886
201.0770263671875 0 1148.5219
201.10293579101562 0 6838.5815
201.1240692138672 0 1392.5067
202.05076599121094 0 720.3859
202.07366943359375 0 3377.4724
202.08688354492188 0 4669.11
202.1092987060547 0 19430.584
202.12008666992188 0 972.3574
203.08213806152344 0 31347.707
203.09271240234375 0 2148.2542
203.11244201660156 0 1259.2759
204.0772247314453 0 1721.72
204.08595275878906 0 3665.4626 a Water loss 2
204.11349487304688 0 743.47455
205.0978240966797 0 3933.0183
205.1087646484375 0 1347.2363
207.0880889892578 0 737.9313
207.11428833007812 0 1159.8802
207.12486267089844 0 1119.1418
207.14991760253906 0 6420.451
208.1444549560547 0 1016.9676
209.0596923828125 0 1064.1001
209.12905883789062 0 2411.4832
209.16590881347656 0 675.45935
210.12423706054688 0 939.58624
211.08717346191406 0 2936.5786
211.14501953125 0 1639.7195
212.14007568359375 0 1825.8182
212.6684112548828 0 6292.9224
213.06680297851562 0 769.6887
213.07752990722656 0 1025.125
213.17018127441406 0 845.45123
214.0979766845703 0 2119.1414
214.109375 0 6979.059
214.1193389892578 0 1135.1488
214.19207763671875 0 1562.4291
215.0822296142578 0 7355.8003
215.11358642578125 0 911.8075
215.1393585205078 0 1226.7167
215.176513671875 0 677.0398
216.0863037109375 0 884.0149
216.1140594482422 0 1734.4269
217.09776306152344 0 3188.0728
217.10894775390625 0 1872.2705
217.11920166015625 0 727.29175
218.0819854736328 0 2120.0557 b Water loss 2
219.0766143798828 0 543.51227
219.08795166015625 0 1016.12823
220.1080780029297 0 983.99304
220.12002563476562 0 16097.214
220.13177490234375 0 1084.5782
221.1040802001953 0 24055.53
221.12258911132812 0 1333.139
222.1071014404297 0 2713.2993
222.12461853027344 0 2180.9832
222.15914916992188 0 610.51776
223.10838317871094 0 955.7726
223.1455841064453 0 724.8448
224.11883544921875 0 1667.2914
224.14012145996094 0 7997.983
224.17645263671875 0 9985.113
225.06658935546875 0 4279.206
225.14361572265625 0 1057.7501
225.16049194335938 0 2801.0554
225.18008422851562 0 849.84265
226.15582275390625 0 176396.84
227.08216857910156 0 14397.608
227.09388732910156 0 1015.1535 b 2
227.13961791992188 0 1363.5703
227.15908813476562 0 19139.725 y Ammonia loss 10
227.17413330078125 0 1152.8993
227.1763916015625 0 1015.7538
228.0862579345703 0 1923.7002
228.11387634277344 0 8645.051
228.1349334716797 0 1976.6387
228.16143798828125 0 906.62445
229.06143188476562 0 1570.4077
229.09780883789062 0 8343.168
229.11863708496094 0 2053.5735
229.1550750732422 0 596.80585
229.16656494140625 0 2001.8511
230.08248901367188 0 775.5252
230.10394287109375 0 4475.9307
231.07638549804688 0 813.8402
231.08843994140625 0 8194.638
231.13589477539062 0 2049.0415
232.09226989746094 0 1425.6575
232.1084747314453 0 1566.0352
232.12037658691406 0 2605.264
232.14215087890625 0 2141.1406
233.1041259765625 0 4912.8174
233.16542053222656 0 1160.5417
234.1359100341797 0 1993.8765
234.65121459960938 0 770.1356
234.6632537841797 0 2920.4714
235.11976623535156 0 32361.504
235.67124938964844 0 27970.818 y 10
236.12318420410156 0 3187.7043
236.13975524902344 0 1677.309
236.17286682128906 0 6587.387
236.67433166503906 0 816.49
237.1359405517578 0 1029.1135
237.1600799560547 0 820.9832 y Ammonia loss 7
238.0988006591797 0 774.77905
238.11865234375 0 985.70844
238.1306610107422 0 2891.3843
239.0821075439453 0 1397.8357
239.1386260986328 0 519.8809
239.151123046875 0 9981.361
240.13511657714844 0 5223.0806
240.15464782714844 0 1083.757
241.0618896484375 0 643.10114
241.14212036132812 0 466.47876
241.15562438964844 0 10814.779
242.0928955078125 0 2972.502
242.16000366210938 0 765.8676
242.18727111816406 0 5496.584
243.0771942138672 0 3831.8816
243.17079162597656 0 1379.5455
243.1825714111328 0 1841.5092
244.10894775390625 0 6356.589
244.12074279785156 0 709.84766
244.16566467285156 0 1276.3301
245.09280395507812 0 5338.8223
245.11453247070312 0 1956.435
245.17539978027344 0 614.0342
245.18682861328125 0 37805.953 y 12
246.1245880126953 0 175084.3
246.19007873535156 0 5126.3906
247.1069793701172 0 1335.1663
247.1299591064453 0 53775.445
248.1149139404297 0 24553.55
248.13278198242188 0 4689.5386
248.18682861328125 0 983.7686
249.09906005859375 0 35315.207
249.11788940429688 0 2215.6255
250.10208129882812 0 4113.008
250.12249755859375 0 806.78723
251.12954711914062 0 810.05743
251.15090942382812 0 870.5359
252.11083984375 0 857.2782
252.1712188720703 0 2710.5337
253.06130981445312 0 1130.6527
254.0933837890625 0 2003.0963
255.0772705078125 0 16956.379
256.080810546875 0 2121.9844
256.1087951660156 0 4697.278
257.0914611816406 0 1032.2415
257.114990234375 0 4002.0664
257.1615905761719 0 5039.017
258.076171875 0 3312.6384
258.0993347167969 0 52704.273
259.0826721191406 0 947.0404
259.10272216796875 0 6434.5913
259.1308288574219 0 2538.7344
260.1046142578125 0 6775.322
260.1976318359375 0 3657.064
260.6683044433594 0 635.19415
261.107421875 0 711.7271
261.1814270019531 0 792.1455
262.119384765625 0 79211.5
262.1546936035156 0 1520.2301
263.10345458984375 0 1164.0375
263.12255859375 0 10226.772
264.102783203125 0 718.7227
264.1350402832031 0 2080.0796
265.1412048339844 0 1825.4048
265.16656494140625 0 2395.7957
266.12530517578125 0 20649.621
267.1268615722656 0 2360.8555
267.1460266113281 0 1822.9799
268.6827697753906 0 755.9988
269.63018798828125 0 1544.0592
270.0881652832031 0 2998.4683
270.1298828125 0 678.5799
270.1820068359375 0 15855.271
271.0721740722656 0 11949.614
271.0871887207031 0 1100.3518
271.18902587890625 0 8227.425 y 9
271.6911315917969 0 1780.9792
272.07550048828125 0 1960.2395
272.1037292480469 0 5463.872
272.1921691894531 0 743.8508
273.08782958984375 0 4756.7427
273.1094055175781 0 2445.4985
273.1349792480469 0 924.5135
274.07232666015625 0 849.98627
274.0928649902344 0 1275.599
274.11932373046875 0 62899.875
275.1036376953125 0 13399.398
275.1256103515625 0 86880.04
275.1857604980469 0 1937.9182 y 6
275.5196228027344 0 898.0363
276.1064147949219 0 4646.1914
276.1287841796875 0 10234.584
277.0931091308594 0 791.11176
277.109375 0 1209.5491
277.1330871582031 0 2506.8596
278.11322021484375 0 643.6996
279.1491394042969 0 585.9805
279.1824035644531 0 3095.5535
280.13067626953125 0 739.92865
280.1647644042969 0 683.1243
281.16168212890625 0 1439.8777
282.1833801269531 0 722.65607
283.1515197753906 0 2712.8752
283.1773986816406 0 48243.977
283.6277770996094 0 2388.0159
284.103271484375 0 637.45544
284.1806640625 0 6631.462
285.1104736328125 0 2872.2507
285.1565856933594 0 8221.12
286.1597900390625 0 1580.276
288.098876953125 0 6058.5615
289.1025695800781 0 1439.0094
289.1784362792969 0 820.81055
290.1145324707031 0 35525.234
290.1851501464844 0 12415.69
290.68682861328125 0 3611.3591
291.0984802246094 0 2867.9883
291.1183776855469 0 6815.2983 b 3
291.1574401855469 0 3082.0024
291.1849670410156 0 741.5704
291.62554931640625 0 1113.196
292.10223388671875 0 993.3624
292.1300048828125 0 20797.814
293.1311950683594 0 7724.4062
293.1617126464844 0 2513.2134
294.1206970214844 0 7659.782
294.21844482421875 0 2749.5625
295.1223449707031 0 1171.9761
295.2226867675781 0 677.48816
297.1748352050781 0 821.4393
297.19317626953125 0 20576.893
298.1968078613281 0 3837.6523
300.1372375488281 0 650.58655
300.2037658691406 0 843.3019
300.67230224609375 0 957.73596
302.1145935058594 0 791.39246
303.1207275390625 0 5012.6055
303.2182312011719 0 833.08057
304.1243896484375 0 1009.8951
305.1304626464844 0 3703.2312
305.66522216796875 0 2678.3286
306.1095275878906 0 1635.8231
306.16595458984375 0 895.50604
307.1778869628906 0 822.987
308.12506103515625 0 7154.667
309.12823486328125 0 1249.3807
309.1675109863281 0 2023.1194
309.1921691894531 0 1529.1895
311.1722717285156 0 2774.8442
311.2450256347656 0 5241.3276
312.2481384277344 0 1315.1056
314.1145324707031 0 1002.405
314.2204895019531 0 846.2728
314.670166015625 0 4979.5854
315.170654296875 0 1557.2048
318.1929016113281 0 2295.7488
319.1090393066406 0 4670.154
319.2140808105469 0 2788.28
320.14068603515625 0 3431.6582
320.8717346191406 0 3217.5508 y 5
321.1260070800781 0 4556.6665
321.2082214355469 0 1470.2538
321.22955322265625 0 23461.123
321.5412292480469 0 790.06024
322.1563415527344 0 4672.5527
322.2331237792969 0 5062.519
324.2162170410156 0 2930.51
324.71759033203125 0 1076.398
325.18792724609375 0 946.5242
325.2237854003906 0 1239.7207
326.2196044921875 0 10334.13
326.7237243652344 0 759.8198 y Ammonia loss 8
327.2230224609375 0 2065.7385
329.13751220703125 0 862.98206
330.1097717285156 0 834.2265
332.72991943359375 0 7215.9478
333.125732421875 0 1626.293
333.23138427734375 0 2455.2358
333.7342529296875 0 880.3325
334.1904296875 0 934.5286
334.6675720214844 0 1315.4482
335.2088623046875 0 2860.6484
335.23736572265625 0 1608.2383 y 8
335.73883056640625 0 834.0428
336.13531494140625 0 2718.9346
336.2041931152344 0 40686.01
337.1860046386719 0 1071.6211
337.20684814453125 0 6135.4062
337.22540283203125 0 2643.4688
337.7220764160156 0 9549.466
338.1510925292969 0 10984.282
338.2234191894531 0 3342.2124
338.2440490722656 0 696.9885
338.716064453125 0 708.5331
339.15435791015625 0 2376.1865
339.2401123046875 0 39970.832
339.7197265625 0 760.0198
340.1528015136719 0 3046.4448
340.2215270996094 0 1223.5623
340.24334716796875 0 7540.495
341.2185363769531 0 1123.6877
342.1671447753906 0 1705.788
342.22662353515625 0 1058.3634
344.7115478515625 0 1319.4016
345.719970703125 0 2086.9707
346.22039794921875 0 953.33295
346.72735595703125 0 31080.264
347.1036376953125 0 2690.6042
347.2286376953125 0 9723.726
347.7301330566406 0 2336.1763
348.107421875 0 1213.358
348.13494873046875 0 4693.396
349.1191711425781 0 1755.1056
349.7103271484375 0 1611.6788
350.1517639160156 0 1623.0497
350.18402099609375 0 1146.3479
350.2110900878906 0 832.9306
350.54412841796875 0 775.6354
351.2397766113281 0 2285.0674
352.1988525390625 0 2741.8708
353.2196960449219 0 3360.2117
353.64111328125 0 1115.0271
353.7171630859375 0 2365.0269
353.8700866699219 0 2041.2937
354.1719055175781 0 1235.9705
354.2147216796875 0 53158.02
354.66162109375 0 729.4779
354.7238464355469 0 1712.5195
355.2174987792969 0 8793.734
355.23956298828125 0 650.9431 y Ammonia loss 7
355.2714538574219 0 1000.2091
355.7327575683594 0 4460.063
356.2333068847656 0 1791.7318
356.25604248046875 0 722.7044 y Ammonia loss 11
357.1788330078125 0 2430.2148
357.2507019042969 0 9945.976
358.16314697265625 0 12183.726
358.2533874511719 0 1981.3787
358.714599609375 0 5397.9785
359.1664733886719 0 2116.6667
359.1935729980469 0 1038.6376
359.21661376953125 0 1190.6943
359.50762939453125 0 1430.5587
359.5425720214844 0 1899.0486
359.87237548828125 0 2184.8362
360.2054138183594 0 1749.7635
360.6653747558594 0 1973.1533
360.689697265625 0 2284.454
361.1871337890625 0 1366.4393
362.1539001464844 0 2193.0652
362.8374938964844 0 849.6187
363.700439453125 0 775.26355
363.74835205078125 0 1640.2073 y 7
363.8405456542969 0 616.3264
364.13116455078125 0 2354.5054
364.1697082519531 0 766.65265
364.1984558105469 0 747.52954
365.25665283203125 0 779.55585
365.5459899902344 0 567.1504
366.15478515625 0 3163.519
366.2126770019531 0 1266.1266
366.55841064453125 0 3345.3154 y 4
366.8927917480469 0 3350.6719
367.162841796875 0 2908.478
368.1479187011719 0 5283.4
368.170166015625 0 891.43243
368.1929931640625 0 1787.1323
368.6617431640625 0 1228.247
368.84173583984375 0 1081.1292
369.15155029296875 0 1034.6082
369.17718505859375 0 2110.2832
369.6952209472656 0 2050.4224
370.1871337890625 0 2191.5176
370.2459411621094 0 810.33356
370.66754150390625 0 1365.1908
371.1678771972656 0 906.5601
371.2671203613281 0 732.72437
371.55364990234375 0 835.19995
373.2820739746094 0 17225.773 y 11
374.2588806152344 0 1190.4216
374.285400390625 0 3909.7556
374.5153503417969 0 1209.6602
374.958251953125 0 1238.0151
375.1895446777344 0 1376.2748
375.28668212890625 0 1069.5023
375.65966796875 0 1214.1128
376.13055419921875 0 1713.1045
376.15362548828125 0 725.2746
376.6513366699219 0 746.2326
378.1454162597656 0 1036.5367
378.51220703125 0 947.2792
378.70159912109375 0 1168.366
380.2307434082031 0 2870.0383
380.4563903808594 0 888.3446
380.7066345214844 0 747.48615
381.8434753417969 0 1125.3365
382.1523742675781 0 2115.7427
382.18218994140625 0 907.2397
382.2076416015625 0 4267.3047
382.2824401855469 0 3277.7266
382.51434326171875 0 856.8039
382.541259765625 0 2460.8171
382.7109680175781 0 1890.6881
383.1934509277344 0 945.49884 b Ammonia loss 12
383.2658386230469 0 1659.5884
383.6573486328125 0 884.9546
384.1419372558594 0 1092.1888
384.16754150390625 0 6248.5815
384.561767578125 0 3880.866
384.66351318359375 0 1120.5074
384.8966064453125 0 2194.3083
385.17401123046875 0 22052.951
385.19915771484375 0 3367.784
386.15826416015625 0 20414.934
386.71014404296875 0 1238.7241
386.9675598144531 0 1133.9883
387.1617736816406 0 4577.735
387.5152893066406 0 1206.4033
387.5406494140625 0 1160.4462
387.8507995605469 0 1316.577
388.1319580078125 0 1044.1328
388.23651123046875 0 1380.3645
388.8631286621094 0 792.7813
389.1631164550781 0 1773.2786
389.2723388671875 0 5143.66
389.7747802734375 0 1182.7343
389.9601135253906 0 794.06494
390.6664123535156 0 931.3209
391.5180358886719 0 1044.891
391.846435546875 0 2037.0449
392.19012451171875 0 1798.6342
392.2660827636719 0 6830.351
392.524169921875 0 1181.1995
392.8566589355469 0 816.8821
393.27142333984375 0 1650.9692
393.5214538574219 0 896.838
393.7101135253906 0 2779.3608
393.8566589355469 0 1203.9325
393.9068298339844 0 977.762
393.9624938964844 0 5259.7075
394.15216064453125 0 10771.714
394.21368408203125 0 7919.1167 y Ammonia loss 1
394.2806396484375 0 1017.4541
394.46435546875 0 5864.438
394.52801513671875 0 2625.8975
394.7158203125 0 2942.108
394.9650573730469 0 1238.3596
395.15521240234375 0 1941.7156
396.26141357421875 0 2719.903
396.8509216308594 0 1318.3999
397.18511962890625 0 882.0304
397.2155456542969 0 1641.1669
397.2448425292969 0 1926.81
397.2710266113281 0 794.958
397.4685363769531 0 1080.5472
397.5219421386719 0 2974.844 b Water loss 9
397.74554443359375 0 1116.3984
397.85430908203125 0 1705.0767 b Ammonia loss 9
397.9009094238281 0 811.10815
397.9654541015625 0 1983.8926
398.190185546875 0 1878.0856
398.2164611816406 0 808.6134
398.2420654296875 0 1767.3252
398.4690856933594 0 4384.0347 y 1
398.7207946777344 0 4201.235
398.8578186035156 0 838.4513
398.8892517089844 0 1122.555
398.9706726074219 0 1696.8094
399.2215270996094 0 1641.7947
400.1676940917969 0 1185.0303
400.220947265625 0 1073.08
401.1675720214844 0 1230.5485
401.19476318359375 0 3272.3232
401.2592468261719 0 1752.3499
402.14996337890625 0 1037.7645
402.2257080078125 0 6957.5293
402.57183837890625 0 921.3639
402.85430908203125 0 1588.0306
403.1849670410156 0 22103.564
403.24066162109375 0 7952.178
403.2701416015625 0 2477.9844
403.5249328613281 0 2817.6155 b 9
403.5710754394531 0 11597.036 y Ammonia loss 3
403.76873779296875 0 1855.8859
403.8604431152344 0 2753.5994
403.9051818847656 0 5779.072
403.9670104980469 0 1578.988
404.1869812011719 0 3045.766
404.2202453613281 0 2831.7715
404.2666015625 0 981.39264
404.47021484375 0 1888.6918
404.6647033691406 0 1595.402
404.7238464355469 0 800.14825
404.89398193359375 0 1523.3977
405.1563720703125 0 1465.2808
405.6749572753906 0 1117.615
406.1408386230469 0 1299.3198
406.18646240234375 0 2197.0444
406.2498779296875 0 6900.778
406.7519226074219 0 2649.1697
407.19512939453125 0 1284.8528
407.2540588378906 0 1206.5916
407.549560546875 0 1635.5325
407.90289306640625 0 847.4557
408.2232360839844 0 4025.8145
408.470458984375 0 5952.4976
408.5528869628906 0 826.54016
408.5740966796875 0 1157.4954
408.72393798828125 0 7593.9336
408.9014892578125 0 1309.9174
408.9734191894531 0 3873.1875
409.2440185546875 0 4468.3306 y 3
409.57855224609375 0 2667.5
409.9121398925781 0 2093.3943
410.1470031738281 0 2783.8245
410.2776184082031 0 18090.967
410.5688781738281 0 4640.602
410.7169189453125 0 772.3629
410.7599182128906 0 965.7297
410.9033203125 0 3620.6177
410.9632263183594 0 1220.0822
411.1495361328125 0 1034.9061
411.17864990234375 0 15690.731
411.2184753417969 0 1821.4167
411.2691650390625 0 7095.1606
411.4718322753906 0 3813.012
411.7207946777344 0 1583.4832
411.76934814453125 0 2967.7021
412.1614685058594 0 3099.5042
412.1860046386719 0 1569.3073
412.2749328613281 0 55561.668 y 6
412.72589111328125 0 3198.9932
412.7763977050781 0 24818.848
412.97601318359375 0 1391.2615
413.17083740234375 0 3256.2742
413.22540283203125 0 1205.5944
413.2777404785156 0 6422.042
413.5525817871094 0 1390.2272
413.6773681640625 0 728.0223
413.7457580566406 0 719.1584
413.778076171875 0 1978.7649
413.8974304199219 0 4939.68
414.1841735839844 0 4943.4287
414.2298583984375 0 4379.708
414.5628967285156 0 2244.1333
414.68560791015625 0 2585.972
415.1903991699219 0 1090.9883
415.2239685058594 0 5793.137
415.25726318359375 0 5545.7754
415.46856689453125 0 2486.79
415.7235107421875 0 6535.451 Precursor Water loss
415.75872802734375 0 3479.6594
415.90087890625 0 1870.2091
415.97589111328125 0 8670.427 Precursor Ammonia loss
416.2286682128906 0 4628.8013
416.2586669921875 0 822.8703
416.47772216796875 0 2408.8706
416.7254638671875 0 2265.4043
418.5574035644531 0 1046.888
419.17620849609375 0 1423.6433 b Water loss 5
419.2286682128906 0 3456.1184
419.2645263671875 0 8200.143
419.5624084472656 0 3562.5303
419.72430419921875 0 6195.8213
419.76568603515625 0 3386.347
419.9004211425781 0 17828.586
419.9739074707031 0 3573.593
420.2305603027344 0 40064.62 Precursor
420.2691345214844 0 3395.81
420.4786376953125 0 31446.908
420.5684814453125 0 5135.2266
420.6920471191406 0 1080.986
420.72998046875 0 12549.936
420.9036865234375 0 2243.0576
420.9803161621094 0 7825.1133
421.2320556640625 0 1458.8055
421.9047546386719 0 3073.6355
422.240478515625 0 2219.4832
422.2773132324219 0 5003.9136
422.31036376953125 0 1209.9342
422.57159423828125 0 916.95135
423.22747802734375 0 1296.4955
423.2761535644531 0 2408.6667
424.25909423828125 0 1273.1334
424.32928466796875 0 4759.296
424.56103515625 0 3338.463
424.71319580078125 0 5649.507
424.895751953125 0 2649.8455
425.2325439453125 0 10090.492
425.28790283203125 0 1617.8123
425.56695556640625 0 6537.8784
425.90380859375 0 3996.6345
426.23712158203125 0 2673.2393
427.173828125 0 4691.056
427.2445068359375 0 5284.0884
427.7458190917969 0 4153.159
428.1802673339844 0 3567.145 b 5
428.24761962890625 0 1100.5574
428.2879333496094 0 1284.4539
428.6822204589844 0 1609.3159
429.1894836425781 0 48569.8
430.1921081542969 0 12474.001 b Ammonia loss 10
431.18035888671875 0 8565.562
431.2356262207031 0 1502.2267
431.5677185058594 0 1013.3645
432.1816101074219 0 1246.3568
432.71014404296875 0 2375.119
433.2121887207031 0 1160.9612
433.2580261230469 0 1085.8015
433.70819091796875 0 1104.9238
434.2373352050781 0 900.5468
435.16845703125 0 986.80023 b Water loss 2
437.1833190917969 0 1828.085
439.1732482910156 0 1224.0365
439.3031311035156 0 1388.7666
442.7118225097656 0 1706.5187
443.1705627441406 0 914.82
443.2183837890625 0 1697.7037
443.5533447265625 0 1028.6055
443.89013671875 0 3431.9768
444.2237548828125 0 1717.6611
444.5628967285156 0 1174.466
445.18450927734375 0 13231.674
445.2677001953125 0 1903.3362
446.18670654296875 0 3451.3687
446.2334289550781 0 3415.9226
446.7353515625 0 2062.5322
447.23565673828125 0 959.3889
448.26446533203125 0 4573.793
448.5982971191406 0 3366.67
448.93560791015625 0 1257.3772
449.5646057128906 0 941.59125
449.90087890625 0 1670.617
451.22625732421875 0 3590.2275
451.2674560546875 0 3327.8386
451.58355712890625 0 744.82214
451.7264709472656 0 4487.151
451.9197998046875 0 5781.5884
452.2209777832031 0 915.28546
452.25421142578125 0 6212.881
452.32427978515625 0 2926.8215
452.5854187011719 0 2295.8674
452.9219055175781 0 2245.4595
453.17926025390625 0 3348.344 b 2
453.2560729980469 0 1548.2554
453.5947265625 0 3431.9998
453.9298400878906 0 3550.8784
454.23187255859375 0 1472.0808
454.263671875 0 1620.5394
454.7353820800781 0 1963.817
455.17156982421875 0 930.2776
455.2522888183594 0 5613.5234
456.2510070800781 0 1185.5873
456.58001708984375 0 2879.3752
456.91339111328125 0 3190.2778
457.1839294433594 0 1803.0198
457.2515869140625 0 6138.3945
457.5898742675781 0 4827.588
457.9248352050781 0 6412.366 y Ammonia loss 2
458.185302734375 0 1279.115
458.2206115722656 0 999.1977
458.2580261230469 0 3616.0806
458.5915832519531 0 1757.5574
459.22247314453125 0 3716.0347
459.7209167480469 0 1758.8854
460.23150634765625 0 9495.627
460.73199462890625 0 4437.4707
461.2366638183594 0 1072.6833
461.70269775390625 0 1057.2025
462.253173828125 0 2332.582
462.5873107910156 0 2486.1606
462.9268798828125 0 15113.935
463.2605895996094 0 9540.805
463.5992126464844 0 85575.5 y 2
463.9335632324219 0 66962.98
464.2680969238281 0 26992.18
464.6024169921875 0 7531.1494
464.9343566894531 0 1120.6473
466.9100341796875 0 1094.7161
467.168212890625 0 971.1552
467.2513732910156 0 1208.3695
467.2989807128906 0 6178.4336
468.22802734375 0 2555.6018
468.25927734375 0 23567.314
468.3192443847656 0 7373.008
468.59332275390625 0 16169.0625
468.7312316894531 0 1682.9962
468.9305419921875 0 56366.375
469.2339782714844 0 2423.9194
469.2653503417969 0 41649.055
469.32366943359375 0 2718.373
469.5997619628906 0 15108.526
469.7897644042969 0 2082.1472
469.933837890625 0 5608.583
470.3350524902344 0 78804.375 y 10
471.33831787109375 0 18901.36
471.5774230957031 0 1070.5918
471.7994079589844 0 5198.3877
472.2596740722656 0 1084.3248
472.2991027832031 0 1777.2275 y Ammonia loss 5
472.340576171875 0 3143.7131
472.5879211425781 0 1212.203
472.80047607421875 0 1172.736
473.2626037597656 0 10417.771
473.5774841308594 0 2917.8394
473.9091491699219 0 1364.7878
474.2651672363281 0 3106.946
474.7801818847656 0 3128.4568
474.90826416015625 0 1189.473
475.24676513671875 0 966.6172
475.28106689453125 0 1861.5305
475.5807800292969 0 3372.4976
475.914794921875 0 2312.296
476.19329833984375 0 2410.2588
476.2577819824219 0 6809.737
476.5928039550781 0 5249.948
476.92822265625 0 2501.7568
478.5754089355469 0 4967.48 b 11
478.78851318359375 0 1587.7745
478.9114990234375 0 4770.862
479.2493591308594 0 1726.7261
479.2909240722656 0 1799.6791
479.58233642578125 0 1776.8353
480.80438232421875 0 14495.16 y 5
480.9146423339844 0 2355.9058
481.25213623046875 0 3723.203
481.3054504394531 0 7703.085
481.58489990234375 0 9918.572
481.8074951171875 0 2242.699
481.9189758300781 0 7258.638
482.2609558105469 0 4247.8174
482.30950927734375 0 6054.4453
482.5917663574219 0 1209.9175
482.76324462890625 0 5599.006
483.21533203125 0 973.08057
483.26531982421875 0 2812.8167
483.31280517578125 0 1843.2407
483.7862548828125 0 8395.141
484.287353515625 0 4941.161
484.3219299316406 0 1106.5463
484.5777587890625 0 1263.1152
486.3304138183594 0 2199.6255
486.58795166015625 0 2222.3113
486.92047119140625 0 3280.5498
487.2582702636719 0 4909.595
487.5914306640625 0 8411.982
487.7940673828125 0 2129.581
487.9244079589844 0 2948.264
488.2594909667969 0 1114.0294
490.9275207519531 0 1108.4325
491.2734680175781 0 7340.17
491.7783508300781 0 2418.8699
492.1903991699219 0 1395.2032
492.2779235839844 0 1722.7002
492.5924377441406 0 1050.7775
492.7916259765625 0 2030.6501
492.9252014160156 0 1223.086
493.262939453125 0 1917.8293
494.2054138183594 0 1808.6418
494.59002685546875 0 1105.0304
494.9227294921875 0 1123.3053
495.36663818359375 0 1804.5228
500.9278564453125 0 1948.7173
501.25811767578125 0 1346.1421
501.60260009765625 0 1685.9629
501.939453125 0 920.8316
503.2164001464844 0 4436.8784
504.59625244140625 0 1011.2996
504.9320983886719 0 2356.3037
505.2186279296875 0 1704.5264 b 7
505.25823974609375 0 1082.968
505.6029968261719 0 1225.7563
505.72943115234375 0 1180.5657
506.598876953125 0 1745.8602
506.935791015625 0 3625.7444
507.2702331542969 0 1860.75
508.7569885253906 0 2027.6661
509.264892578125 0 853.08295
509.929443359375 0 1854.7183
510.26629638671875 0 2480.869 b Water loss 12
510.60211181640625 0 3040.8408
510.93792724609375 0 4290.6436
511.27325439453125 0 1892.8643
511.60919189453125 0 977.45844
515.5999755859375 0 1670.7808
516.2698364257812 0 6364.2974 b 12
516.6044921875 0 7989.612
516.752197265625 0 1074.6327
516.797607421875 0 1124.7623
516.9396362304688 0 4320.318
517.2335815429688 0 1378.8073
517.2734985351562 0 2931.289
517.7368774414062 0 1064.2219
519.2132568359375 0 1143.6714
519.2781982421875 0 3903.7695
519.6098022460938 0 2632.7625
519.9449462890625 0 1156.3524
520.3623657226562 0 3354.856
521.22705078125 0 3925.9915
521.6019897460938 0 2819.1335
521.9365844726562 0 3216.269
522.2272338867188 0 1080.9001
522.2726440429688 0 14813.235
522.607421875 0 11227.978
522.94189453125 0 3155.847
523.27294921875 0 1804.6958
523.3629760742188 0 1525.799
524.612548828125 0 3312.5276
524.8110961914062 0 1000.1671
524.9517211914062 0 8370.825
525.283447265625 0 17957.88 y Ammonia loss 1
525.6177368164062 0 9533.813
525.7509155273438 0 1748.2705
525.80810546875 0 1267.1594
525.9507446289062 0 5345.3403
526.2496337890625 0 1449.1443
526.3329467773438 0 1062.4629
526.8322143554688 0 1385.1107
528.2318725585938 0 1643.1396
529.8046875 0 1742.677
530.2875366210938 0 10174.224
530.6195068359375 0 9290.807
530.8004760742188 0 2485.0125
530.9566040039062 0 23028.857 y 1
531.2113647460938 0 4195.541
531.29150390625 0 17574.068
531.6246948242188 0 8271.447
531.9597778320312 0 3173.325
532.2940673828125 0 1719.8502
533.820556640625 0 951.1094
534.3223876953125 0 945.1949
535.74951171875 0 1101.036
537.221435546875 0 1207.7128
537.3419189453125 0 1090.6293
538.2533569335938 0 29562.682
538.3733520507812 0 2950.1765
538.8101196289062 0 5359.799
539.25537109375 0 9665.909
539.3058471679688 0 8572.001
539.8059692382812 0 4654.684
540.8258666992188 0 1761.1416 y Ammonia loss 4
541.327392578125 0 1077.5405
541.3721923828125 0 17083.867 y 9
542.2540893554688 0 1396.4927
542.3749389648438 0 4752.2524
543.6138916015625 0 1199.5004
543.7501220703125 0 2627.976
544.2487182617188 0 1050.9286
545.2576904296875 0 1239.888
545.7542724609375 0 1024.0615
546.8034057617188 0 1130.6207
547.20361328125 0 1390.5103
547.8153686523438 0 4604.6523
548.3189086914062 0 4404.5903
548.8206176757812 0 2634.0688
549.2215576171875 0 3943.738
549.3336181640625 0 28674.055 y 4
549.8353271484375 0 17226.898
550.3367309570312 0 5016.2124
550.83740234375 0 1121.449
552.2627563476562 0 2013.0985
552.7595825195312 0 2394.954
553.2637939453125 0 1771.9517
554.2484741210938 0 7735.3965
555.2492065429688 0 2151.4045
556.3226928710938 0 4522.7666
556.8250122070312 0 4194.4946
557.2546997070312 0 1218.105
557.2902221679688 0 1744.8674
557.3299560546875 0 5230.29
557.830322265625 0 2776.595
558.2481079101562 0 2051.6323
559.2504272460938 0 1993.4989
559.7523803710938 0 1240.829
560.2599487304688 0 1236.8545 b Water loss 8
560.7561645507812 0 1529.1458 b Ammonia loss 8
561.2657470703125 0 3057.7002
561.7700805664062 0 2548.9856
564.2320556640625 0 1730.0005
566.2483520507812 0 35799.793
567.2509765625 0 9786.811
567.3991088867188 0 2539.9077
568.2561645507812 0 2654.0435
568.7568359375 0 1302.188
569.2183227539062 0 1137.5017
569.2659301757812 0 8125.173 b 8
569.7669067382812 0 4924.1753
570.2496948242188 0 4480.007
571.2481689453125 0 1043.9855
572.8051147460938 0 14830.625
573.3062744140625 0 9748.732
573.809326171875 0 2749.7793
574.8081665039062 0 1064.0984
576.3397216796875 0 2167.097
576.8396606445312 0 1270.963
577.340087890625 0 1194.8264
577.3833618164062 0 11161.183
578.3846435546875 0 3802.0498
579.3626708984375 0 27956.785
579.7940673828125 0 2522.7861
580.2997436523438 0 1138.6285
580.364990234375 0 8656.393
580.8032836914062 0 5706.074
581.2342529296875 0 1198.0245 b 3
581.30615234375 0 3466.6387
581.3711547851562 0 2605.2332
581.8112182617188 0 1651.017
582.2434692382812 0 12436.995
583.245361328125 0 4424.603
585.2587280273438 0 1612.6699
586.2431030273438 0 6713.7812
587.24560546875 0 2368.1187
593.3408813476562 0 2538.2476
595.2785034179688 0 1076.009
595.3469848632812 0 1065.3887
595.3942260742188 0 8693.599
595.7779541015625 0 2041.8165 b Water loss 9
595.8447875976562 0 2249.2603
596.2827758789062 0 1245.6171
596.342529296875 0 1180.9634
596.3961791992188 0 3217.245
596.7835083007812 0 1570.2039
598.241943359375 0 2194.4592
603.2696533203125 0 1502.993
604.357177734375 0 7709.301
604.7860107421875 0 2087.5586 b 9
604.8532104492188 0 14067.397 y Ammonia loss 3
605.2876586914062 0 1595.0918
605.3543090820312 0 8095.4585
605.8533935546875 0 3825.7412
610.3217163085938 0 9487.854
611.3268432617188 0 2908.6606
611.8370361328125 0 2658.49
612.3458251953125 0 2554.64
612.850341796875 0 2811.2158
613.2538452148438 0 1273.4542
613.3614501953125 0 7983.943 y 3
613.864013671875 0 4467.9463
614.2378540039062 0 4918.3203
614.363037109375 0 1958.0609
615.2412109375 0 1260.0676
615.3511352539062 0 3168.7158
615.8492431640625 0 2476.7107
620.3427734375 0 6212.8755
620.8408813476562 0 5988.765
621.3499145507812 0 3138.9194
622.343017578125 0 1317.4792
623.3425903320312 0 1211.8036
623.8482666015625 0 1312.9631
625.25341796875 0 1287.4077
627.8302612304688 0 1233.3408
628.33447265625 0 5064.254
628.8396606445312 0 3781.7427
629.2501220703125 0 1287.9843
629.3466186523438 0 15237.562
629.8484497070312 0 9930.739
630.3496704101562 0 4556.713
630.8505249023438 0 1352.038
631.263671875 0 1545.8818
632.353515625 0 1438.863
632.8556518554688 0 1389.7784
636.3365478515625 0 2281.0842
636.8355712890625 0 2301.4092
637.3439331054688 0 4982.043
637.8472900390625 0 5344.8745
638.3515625 0 8942.227
638.854248046875 0 4667.8335
639.3560791015625 0 2237.5244
641.3702392578125 0 2315.3677
641.8717041015625 0 1209.6172
642.3765869140625 0 1294.5253
645.845458984375 0 1616.6605
646.3512573242188 0 3516.1248
646.8509521484375 0 2344.8381
647.3539428710938 0 1213.1766
647.4261474609375 0 1918.7318
650.428466796875 0 918.89264
664.4517822265625 0 7086.436
664.8310546875 0 1309.8345
665.3339233398438 0 2309.1653
665.4541625976562 0 3423.6003
665.836181640625 0 1761.81
666.3419189453125 0 1202.488
667.34326171875 0 1845.6128
668.33154296875 0 2622.9604
669.4668579101562 0 5587.407 y 8
670.2762451171875 0 1122.2223
670.470703125 0 2480.1895
671.264404296875 0 1795.8165
673.2748413085938 0 1170.485
673.841796875 0 2782.0518
674.2982788085938 0 1564.0728
674.4363403320312 0 23567.21
675.3077392578125 0 1001.97076
675.4376831054688 0 9365.536
676.4391479492188 0 1350.9634
678.3703002929688 0 1037.5282
678.4302978515625 0 1223.6725
685.3564453125 0 2490.927
685.8868408203125 0 915.1889
686.3742065429688 0 1063.3286 y Ammonia loss 2
688.2821655273438 0 892.1703
688.4136962890625 0 2528.869
690.2992553710938 0 3178.2947
690.4300537109375 0 2122.9546
691.303955078125 0 1181.6526
691.432373046875 0 1110.818
692.4464721679688 0 25645.844
693.4489135742188 0 9025.585
694.3886108398438 0 1077.9546
694.450927734375 0 1841.1467
694.8943481445312 0 7196.9756 y 2
695.3955078125 0 5454.442
695.8958129882812 0 2561.6653
698.4114379882812 0 2017.07
700.2822875976562 0 1404.3693 b Water loss 4
702.302490234375 0 2286.24
702.3771362304688 0 954.28125
702.8916015625 0 3417.1587
703.2993774414062 0 1206.4133
703.391845703125 0 2976.7466
703.8926391601562 0 1599.483
706.4266357421875 0 3123.2686
707.3042602539062 0 3671.5903
708.3117065429688 0 1356.9749
708.436279296875 0 1268.547
709.358154296875 0 1108.9751
710.38134765625 0 966.35455
710.4573974609375 0 8035.3574
711.4586791992188 0 3003.1284
713.881103515625 0 1067.311
714.3853149414062 0 1159.6178
716.4210205078125 0 5707.912
717.4247436523438 0 2756.5757
717.8609619140625 0 1268.8804
718.2957153320312 0 4340.3696 b 4
718.3629760742188 0 1172.9891
719.3001708984375 0 1274.8201
720.3208618164062 0 2807.342
721.3582153320312 0 4199.524
721.8767700195312 0 2331.4656
722.3762817382812 0 1201.5159
723.30224609375 0 5965.5903
724.3074340820312 0 2280.7808
724.4340209960938 0 1080.921
726.4884643554688 0 11979.525 y 7
727.4902954101562 0 3953.332
728.4949340820312 0 1173.0227
730.3897705078125 0 1788.6783
730.8866577148438 0 1893.0736
735.3016967773438 0 2912.407
736.31201171875 0 1638.8837
737.3468017578125 0 2487.236
738.3765258789062 0 1697.157
739.3707885742188 0 2367.8044
740.3277587890625 0 2880.8777
740.46728515625 0 1245.4253
741.3335571289062 0 1513.1869
742.41552734375 0 956.98285
750.3082885742188 0 819.9003
751.2974243164062 0 5107.7896
752.3081665039062 0 2190.377
756.390380859375 0 929.8046
766.3060913085938 0 1654.6324
768.3226318359375 0 4675.455
769.324951171875 0 1734.4346
777.5390014648438 0 1260.0895
790.4859619140625 0 1024.811
794.4805297851562 0 853.0135
805.5320434570312 0 2586.0276
806.5307006835938 0 1534.4072
808.3231811523438 0 1008.766
811.3616333007812 0 1323.6927
811.4954223632812 0 3317.1458
812.3668823242188 0 967.1061
819.5118408203125 0 897.083
821.524658203125 0 3925.434
822.5291137695312 0 2159.7605
823.5410766601562 0 44009.523 y 6
824.5440063476562 0 18743.906
825.5464477539062 0 4177.503
826.551025390625 0 962.6097
827.3589477539062 0 2349.1604
829.366455078125 0 1063.1433
829.5047607421875 0 2861.784
830.5066528320312 0 1734.0812
837.5206298828125 0 5178.337
838.52392578125 0 1548.6965
839.3571166992188 0 2763.1719
839.531005859375 0 1919.5096
840.3679809570312 0 926.89594
847.5187377929688 0 1062.4626
848.4171752929688 0 5922.799
849.4183959960938 0 3270.7825
853.473876953125 0 2183.9802
854.4823608398438 0 932.48376
855.3547973632812 0 5093.4297 b 5
856.3556518554688 0 2156.8855
864.4100952148438 0 2070.7036
866.4130249023438 0 896.86816
891.4632568359375 0 852.55536
901.44287109375 0 6883.0044
902.4448852539062 0 3443.1018
903.4415283203125 0 1243.3793
917.435546875 0 1911.2349
918.4374389648438 0 1050.7367
919.453125 0 4160.573
920.45703125 0 1842.8787
935.4439086914062 0 1346.4248
936.4481201171875 0 981.8824
942.58740234375 0 2036.0541
943.600341796875 0 1422.5974
960.5994262695312 0 5686.285 y 5
961.602294921875 0 2751.6335
962.6030883789062 0 1040.9307
964.5152587890625 0 1070.6223
966.5704956054688 0 841.68317
1033.5411376953125 0 1104.4949
1086.494384765625 0 865.471
1097.6591796875 0 1424.3438 y 4
1098.6607666015625 0 726.43005

Spectrum Details

|  |  |
| --- | --- |
| Matched peaks? Matched peaks The total absolute number of peaks matched. Additionally in brackets the total fraction of peaks matched and the total number of peaks is shown. | 70 (5.19% of 1348) |
| FDR? FDR The false discovery rate estimated for this peptide. It is calculated by matching all theoretical fragments with a non-integer shift with the raw peaks for this spectrum. This is done with 40 different shifts. The resulting percentage is the average number of annotated peaks over the number of annotated peaks with the correct spectrum. | 7.82% |
| Satellite FDR? Satellite FDR See the FDR for details on its calculation. This satellite ion specific FDR only contains the satellite ions (d/w) for I/L/J positions. | - |
| PSM Score? PSM Score The PSM Score as given by Hecklib to this annotated spectrum. It is shown with three significant figures. | 189 |

#### Spectrum 5226? Spectrum 5226 The raw spectrum of this peptide as annotated by Hecklib. The fragments are coloured according to ion type (see legend). Any peaks with a star '\*' as text can be hovered over to see the full details, first the ion type second the mass shift type. By hovering over the amino acids in the peptide or ions in the legend the corresponding peaks are highlighted. By toggling the 'Unassigned' label you can turn the background (unassigned) peaks on or off in the plot. By updating the slider in the Ion legend you can update the spectrum to only show the top X% of the peaks with labels. The top X% means any peak that is within X% of the highest intensity. By dragging in the spectrum you can zoom in to a specific part of the spectrum and use 'Zoom Out' to get back to the original zoom level. The annotation of the spectrum is based on the given sequence in the peptides file and is done with different software so inconsistencies are likely. The peaks are annotated based on the given sequence, with 20 ppm tolerance.

Copy Data

##### Spectrum 5226 (TSV)

###### Preview

```
Loading example...
```

*Click on the button to copy the data to your clipboard.*

Mz MinMz MaxIntensity Max

WidthHeightPeptide font sizePeptide stroke widthSpectrum font sizeSpectrum stroke widthCompact peptide

Ion legend

wxyz

abcd

OtherUnassignedIon^Charge^_Position_Show for top:%

SWYQHHPGKAPKJJ

01.87e+43.74e+45.62e+47.49e+4

Zoom Out

y^+1^_1_b^+4^_4_y^+2^_3_y^+2^_4_a^+1^_2_b^+3^_5_y^+2^_4_y^+1^_2_a^+1^_2_b^+1^_2_y^+2^_5_b^+1^_2_y^+3^_8_y^+3^_9_b^+4^_11_y^+3^_10_y^+1^_3_y^+4^_13_b^+3^_10_y^+4^_13_b^+3^_10_y^+3^_11_b^+2^_6_\*y^+2^_8_\*b^+2^_6_b^+1^_3_y^+3^_12_y^+3^_12_y^+1^_4_y^+2^_9_b^+3^_13_b^+3^_13_y^+3^_13_y^+3^_13_y^+1^_5_y^+2^_10_b^+2^_9_y^+2^_11_y^+2^_11_y^+1^_6_y^+2^_12_y^+1^_7_y^+1^_8_b^+1^_6_y^+1^_9_b^+1^_10_

0769153923083077

Fragment Matches Table

Show background peaks

| Position | Ion type | Intensity | mz Theoretical | mz Error (Th) | mz Error (ppm) | Charge | Series Number |
| --- | --- | --- | --- | --- | --- | --- | --- |
| - | - | 456.3 | 120.1 | - | - | 0 | - |
| - | - | 633.7 | 120.1 | - | - | 0 | - |
| - | - | 1185 | 120.1 | - | - | 0 | - |
| - | - | 803.8 | 122.1 | - | - | 0 | - |
| - | - | 1713 | 123.1 | - | - | 0 | - |
| - | - | 683 | 123.1 | - | - | 0 | - |
| - | - | 491.2 | 127.1 | - | - | 0 | - |
| - | - | 1905 | 127.1 | - | - | 0 | - |
| - | - | 501.5 | 128.1 | - | - | 0 | - |
| - | - | 759.8 | 128.1 | - | - | 0 | - |
| - | - | 481.9 | 129.1 | - | - | 0 | - |
| - | - | 3269 | 129.1 | - | - | 0 | - |
| - | - | 5.827E+04 | 129.1 | - | - | 0 | - |
| - | - | 1401 | 130 | - | - | 0 | - |
| - | - | 2570 | 130.1 | - | - | 0 | - |
| - | - | 1.343E+04 | 130.1 | - | - | 0 | - |
| - | - | 848.6 | 130.1 | - | - | 0 | - |
| - | - | 694.4 | 130.1 | - | - | 0 | - |
| - | - | 3413 | 130.1 | - | - | 0 | - |
| - | - | 4421 | 131 | - | - | 0 | - |
| - | - | 1549 | 131.1 | - | - | 0 | - |
| - | - | 1753 | 131.1 | - | - | 0 | - |
| - | - | 578.1 | 131.1 | - | - | 0 | - |
| - | - | 394.7 | 132 | - | - | 0 | - |
| - | - | 4.494E+04 | 132.1 | - | - | 0 | - |
| 14 | y | 1.28E+04 | 132.1 | 0.0001151 | 0.8716 | +1 | 1 |
| - | - | 665.6 | 133.1 | - | - | 0 | - |
| - | - | 5302 | 133.1 | - | - | 0 | - |
| - | - | 688.8 | 133.1 | - | - | 0 | - |
| - | - | 3884 | 134 | - | - | 0 | - |
| - | - | 6.254E+04 | 136.1 | - | - | 0 | - |
| - | - | 4707 | 137.1 | - | - | 0 | - |
| - | - | 1090 | 138.1 | - | - | 0 | - |
| - | - | 836 | 139.1 | - | - | 0 | - |
| - | - | 485.3 | 140.1 | - | - | 0 | - |
| - | - | 537.7 | 141.1 | - | - | 0 | - |
| - | - | 4090 | 141.1 | - | - | 0 | - |
| 4 | b | 853.7 | 142.1 | 0.0001739 | 1.224 | +4 | 4 |
| - | - | 519.2 | 142.1 | - | - | 0 | - |
| - | - | 1252 | 144.1 | - | - | 0 | - |
| - | - | 550.7 | 145.1 | - | - | 0 | - |
| - | - | 1543 | 146.1 | - | - | 0 | - |
| - | - | 3615 | 147 | - | - | 0 | - |
| - | - | 854.1 | 151.1 | - | - | 0 | - |
| - | - | 1828 | 152.1 | - | - | 0 | - |
| - | - | 484.2 | 152.1 | - | - | 0 | - |
| - | - | 1131 | 152.1 | - | - | 0 | - |
| - | - | 507.7 | 153.1 | - | - | 0 | - |
| - | - | 603 | 153.1 | - | - | 0 | - |
| - | - | 474.1 | 154.1 | - | - | 0 | - |
| - | - | 441 | 155.1 | - | - | 0 | - |
| - | - | 2555 | 155.1 | - | - | 0 | - |
| - | - | 1727 | 155.1 | - | - | 0 | - |
| - | - | 1553 | 155.1 | - | - | 0 | - |
| - | - | 640.6 | 156.1 | - | - | 0 | - |
| - | - | 786.7 | 156.1 | - | - | 0 | - |
| - | - | 646.1 | 157.1 | - | - | 0 | - |
| - | - | 972.3 | 158.1 | - | - | 0 | - |
| - | - | 4752 | 158.1 | - | - | 0 | - |
| - | - | 3.149E+04 | 159.1 | - | - | 0 | - |
| - | - | 1789 | 159.1 | - | - | 0 | - |
| - | - | 1695 | 160.1 | - | - | 0 | - |
| - | - | 3425 | 160.1 | - | - | 0 | - |
| - | - | 824.5 | 162.1 | - | - | 0 | - |
| - | - | 1086 | 164.1 | - | - | 0 | - |
| - | - | 2090 | 165.1 | - | - | 0 | - |
| - | - | 1333 | 166.1 | - | - | 0 | - |
| - | - | 936.4 | 168.1 | - | - | 0 | - |
| - | - | 3276 | 169.1 | - | - | 0 | - |
| - | - | 4059 | 169.1 | - | - | 0 | - |
| - | - | 1.383E+04 | 170.1 | - | - | 0 | - |
| - | - | 518.5 | 170.1 | - | - | 0 | - |
| - | - | 493.9 | 170.1 | - | - | 0 | - |
| - | - | 1986 | 171.1 | - | - | 0 | - |
| - | - | 689.8 | 171.1 | - | - | 0 | - |
| - | - | 695.1 | 171.1 | - | - | 0 | - |
| - | - | 773.3 | 171.1 | - | - | 0 | - |
| - | - | 3459 | 172.1 | - | - | 0 | - |
| - | - | 8406 | 173.1 | - | - | 0 | - |
| - | - | 1006 | 173.1 | - | - | 0 | - |
| - | - | 2644 | 173.4 | - | - | 0 | - |
| - | - | 1137 | 174.1 | - | - | 0 | - |
| - | - | 1008 | 175.1 | - | - | 0 | - |
| - | - | 643.3 | 175.1 | - | - | 0 | - |
| - | - | 913.3 | 176.1 | - | - | 0 | - |
| - | - | 1506 | 177.1 | - | - | 0 | - |
| - | - | 452.1 | 179.1 | - | - | 0 | - |
| - | - | 589.1 | 180.1 | - | - | 0 | - |
| - | - | 1612 | 181.1 | - | - | 0 | - |
| - | - | 1.611E+04 | 181.1 | - | - | 0 | - |
| - | - | 1257 | 182.1 | - | - | 0 | - |
| - | - | 1594 | 182.1 | - | - | 0 | - |
| - | - | 7521 | 183.1 | - | - | 0 | - |
| - | - | 882 | 183.1 | - | - | 0 | - |
| - | - | 868.1 | 184.1 | - | - | 0 | - |
| - | - | 848.1 | 184.1 | - | - | 0 | - |
| - | - | 1732 | 185.1 | - | - | 0 | - |
| - | - | 664.7 | 185.1 | - | - | 0 | - |
| - | - | 1179 | 185.2 | - | - | 0 | - |
| - | - | 5357 | 186.1 | - | - | 0 | - |
| - | - | 2.904E+04 | 187.1 | - | - | 0 | - |
| - | - | 6339 | 187.1 | - | - | 0 | - |
| 12 | y | 1652 | 187.1 | 0.0002591 | 1.385 | +2 | 3 |
| - | - | 3634 | 188.1 | - | - | 0 | - |
| - | - | 601.8 | 189.1 | - | - | 0 | - |
| - | - | 440 | 190.1 | - | - | 0 | - |
| - | - | 851.1 | 195.1 | - | - | 0 | - |
| - | - | 458.8 | 195.1 | - | - | 0 | - |
| - | - | 697.6 | 195.1 | - | - | 0 | - |
| - | - | 602.9 | 196.1 | - | - | 0 | - |
| - | - | 1363 | 197.1 | - | - | 0 | - |
| - | - | 2507 | 197.2 | - | - | 0 | - |
| - | - | 441.1 | 197.2 | - | - | 0 | - |
| - | - | 2878 | 199.1 | - | - | 0 | - |
| - | - | 1379 | 199.1 | - | - | 0 | - |
| - | - | 1543 | 199.1 | - | - | 0 | - |
| - | - | 2527 | 199.2 | - | - | 0 | - |
| - | - | 708.8 | 199.2 | - | - | 0 | - |
| - | - | 1158 | 200.1 | - | - | 0 | - |
| - | - | 3209 | 201.1 | - | - | 0 | - |
| - | - | 2040 | 201.1 | - | - | 0 | - |
| - | - | 1472 | 202.1 | - | - | 0 | - |
| - | - | 6139 | 202.1 | - | - | 0 | - |
| - | - | 825.9 | 204.1 | - | - | 0 | - |
| - | - | 629.1 | 207.1 | - | - | 0 | - |
| - | - | 1380 | 207.1 | - | - | 0 | - |
| - | - | 2795 | 209.1 | - | - | 0 | - |
| - | - | 829.5 | 211.1 | - | - | 0 | - |
| - | - | 790.4 | 212.7 | - | - | 0 | - |
| - | - | 2103 | 214.1 | - | - | 0 | - |
| - | - | 562.1 | 214.2 | - | - | 0 | - |
| - | - | 1871 | 215.1 | - | - | 0 | - |
| - | - | 1473 | 216.1 | - | - | 0 | - |
| - | - | 824.1 | 219.1 | - | - | 0 | - |
| - | - | 4519 | 220.1 | - | - | 0 | - |
| - | - | 5673 | 221.1 | - | - | 0 | - |
| - | - | 566.4 | 224.1 | - | - | 0 | - |
| - | - | 2469 | 224.2 | - | - | 0 | - |
| - | - | 1069 | 225.2 | - | - | 0 | - |
| - | - | 4.04E+04 | 226.2 | - | - | 0 | - |
| - | - | 2395 | 227.1 | - | - | 0 | - |
| - | - | 573.5 | 227.1 | - | - | 0 | - |
| 11 | y | 4959 | 227.2 | 0.001144 | 5.036 | +2 | 4 |
| 2 | a | 2460 | 228.1 | 0.0001624 | 0.7121 | +1 | 2 |
| 5 | b | 3753 | 229.1 | 0.00166 | 7.246 | +3 | 5 |
| - | - | 1708 | 229.1 | - | - | 0 | - |
| - | - | 819.4 | 229.2 | - | - | 0 | - |
| - | - | 940.6 | 230.1 | - | - | 0 | - |
| - | - | 1649 | 231.1 | - | - | 0 | - |
| - | - | 1380 | 233.1 | - | - | 0 | - |
| - | - | 733.7 | 234.1 | - | - | 0 | - |
| - | - | 7909 | 235.1 | - | - | 0 | - |
| 11 | y | 6891 | 235.7 | 0.0001528 | 0.6485 | +2 | 4 |
| - | - | 1081 | 236.1 | - | - | 0 | - |
| - | - | 1246 | 236.2 | - | - | 0 | - |
| - | - | 759.4 | 238.1 | - | - | 0 | - |
| - | - | 2437 | 239.2 | - | - | 0 | - |
| - | - | 670.8 | 240.1 | - | - | 0 | - |
| - | - | 1418 | 242.2 | - | - | 0 | - |
| - | - | 1382 | 243.1 | - | - | 0 | - |
| - | - | 8060 | 245.1 | - | - | 0 | - |
| 13 | y | 9584 | 245.2 | 9.657E-05 | 0.3939 | +1 | 2 |
| 2 | a | 7.413E+04 | 246.1 | 0.0001219 | 0.4952 | +1 | 2 |
| - | - | 1028 | 246.2 | - | - | 0 | - |
| - | - | 1.683E+04 | 247.1 | - | - | 0 | - |
| - | - | 6112 | 248.1 | - | - | 0 | - |
| - | - | 1623 | 248.1 | - | - | 0 | - |
| - | - | 9583 | 249.1 | - | - | 0 | - |
| - | - | 1755 | 249.2 | - | - | 0 | - |
| - | - | 617.5 | 250.1 | - | - | 0 | - |
| - | - | 972.5 | 250.1 | - | - | 0 | - |
| - | - | 885 | 251.1 | - | - | 0 | - |
| - | - | 995.4 | 252.2 | - | - | 0 | - |
| - | - | 819.7 | 253.2 | - | - | 0 | - |
| - | - | 806.6 | 255.2 | - | - | 0 | - |
| 2 | b | 1921 | 256.1 | 8.194E-05 | 0.3199 | +1 | 2 |
| - | - | 1128 | 257.1 | - | - | 0 | - |
| - | - | 875.1 | 257.2 | - | - | 0 | - |
| - | - | 941.3 | 258.1 | - | - | 0 | - |
| - | - | 1.354E+04 | 258.1 | - | - | 0 | - |
| - | - | 1701 | 259.1 | - | - | 0 | - |
| - | - | 905.6 | 259.1 | - | - | 0 | - |
| - | - | 1893 | 260.1 | - | - | 0 | - |
| - | - | 1637 | 260.2 | - | - | 0 | - |
| - | - | 1025 | 263.1 | - | - | 0 | - |
| - | - | 5482 | 266.1 | - | - | 0 | - |
| - | - | 1257 | 269.6 | - | - | 0 | - |
| - | - | 2735 | 270.2 | - | - | 0 | - |
| 10 | y | 2801 | 271.2 | 0.0006276 | 2.314 | +2 | 5 |
| 2 | b | 2.368E+04 | 274.1 | 8.754E-05 | 0.3194 | +1 | 2 |
| - | - | 2987 | 275.1 | - | - | 0 | - |
| - | - | 2.206E+04 | 275.1 | - | - | 0 | - |
| 7 | y | 695.8 | 275.2 | 0.0005859 | 2.129 | +3 | 8 |
| - | - | 911.7 | 276.1 | - | - | 0 | - |
| - | - | 2768 | 276.1 | - | - | 0 | - |
| - | - | 802.8 | 277.1 | - | - | 0 | - |
| - | - | 688.2 | 277.2 | - | - | 0 | - |
| - | - | 1071 | 279.2 | - | - | 0 | - |
| - | - | 683.8 | 282.2 | - | - | 0 | - |
| - | - | 602.6 | 283.2 | - | - | 0 | - |
| - | - | 686.2 | 283.2 | - | - | 0 | - |
| - | - | 8201 | 283.2 | - | - | 0 | - |
| - | - | 808.1 | 283.6 | - | - | 0 | - |
| - | - | 792.5 | 284.2 | - | - | 0 | - |
| - | - | 685.8 | 285.1 | - | - | 0 | - |
| - | - | 1355 | 285.2 | - | - | 0 | - |
| - | - | 1149 | 290 | - | - | 0 | - |
| - | - | 3028 | 290.2 | - | - | 0 | - |
| - | - | 1176 | 290.7 | - | - | 0 | - |
| - | - | 1387 | 291.2 | - | - | 0 | - |
| - | - | 6374 | 292.1 | - | - | 0 | - |
| - | - | 1198 | 293.1 | - | - | 0 | - |
| - | - | 2036 | 294.1 | - | - | 0 | - |
| - | - | 1066 | 294.2 | - | - | 0 | - |
| - | - | 4868 | 297.2 | - | - | 0 | - |
| - | - | 668.9 | 298.2 | - | - | 0 | - |
| - | - | 1253 | 303.1 | - | - | 0 | - |
| - | - | 2773 | 305.1 | - | - | 0 | - |
| - | - | 672.7 | 305.7 | - | - | 0 | - |
| - | - | 778.2 | 310.1 | - | - | 0 | - |
| - | - | 1095 | 311.2 | - | - | 0 | - |
| - | - | 2082 | 314.7 | - | - | 0 | - |
| - | - | 673.9 | 319.2 | - | - | 0 | - |
| - | - | 764.4 | 320.1 | - | - | 0 | - |
| 6 | y | 813.3 | 320.9 | 0.0007327 | 2.283 | +3 | 9 |
| - | - | 5742 | 321.2 | - | - | 0 | - |
| - | - | 3854 | 322.2 | - | - | 0 | - |
| - | - | 1246 | 322.2 | - | - | 0 | - |
| 11 | b | 948 | 323.2 | 0.002157 | 6.676 | +4 | 11 |
| - | - | 1794 | 326.2 | - | - | 0 | - |
| - | - | 821.4 | 327.2 | - | - | 0 | - |
| - | - | 1989 | 332.7 | - | - | 0 | - |
| - | - | 914.7 | 333.1 | - | - | 0 | - |
| - | - | 9500 | 336.2 | - | - | 0 | - |
| - | - | 1581 | 337.2 | - | - | 0 | - |
| - | - | 2747 | 337.7 | - | - | 0 | - |
| - | - | 8234 | 338.1 | - | - | 0 | - |
| - | - | 1232 | 338.2 | - | - | 0 | - |
| - | - | 573.3 | 338.7 | - | - | 0 | - |
| - | - | 1384 | 339.1 | - | - | 0 | - |
| - | - | 1.009E+04 | 339.2 | - | - | 0 | - |
| - | - | 708.3 | 340.2 | - | - | 0 | - |
| - | - | 1359 | 340.2 | - | - | 0 | - |
| - | - | 1685 | 340.2 | - | - | 0 | - |
| - | - | 607.2 | 342.2 | - | - | 0 | - |
| - | - | 1824 | 345 | - | - | 0 | - |
| - | - | 783.3 | 345.6 | - | - | 0 | - |
| - | - | 9410 | 346.7 | - | - | 0 | - |
| - | - | 677.9 | 347.1 | - | - | 0 | - |
| - | - | 3805 | 347.2 | - | - | 0 | - |
| - | - | 664.7 | 347.7 | - | - | 0 | - |
| - | - | 1242 | 350.1 | - | - | 0 | - |
| - | - | 1150 | 351.2 | - | - | 0 | - |
| - | - | 1699 | 354.1 | - | - | 0 | - |
| - | - | 694.4 | 354.2 | - | - | 0 | - |
| - | - | 1.173E+04 | 354.2 | - | - | 0 | - |
| - | - | 695.4 | 354.7 | - | - | 0 | - |
| - | - | 2682 | 355.2 | - | - | 0 | - |
| - | - | 1070 | 355.7 | - | - | 0 | - |
| - | - | 889.5 | 356.1 | - | - | 0 | - |
| - | - | 2823 | 357.2 | - | - | 0 | - |
| - | - | 3125 | 358.2 | - | - | 0 | - |
| - | - | 1465 | 358.7 | - | - | 0 | - |
| - | - | 1049 | 359.5 | - | - | 0 | - |
| - | - | 841.9 | 359.9 | - | - | 0 | - |
| - | - | 755.9 | 360.2 | - | - | 0 | - |
| - | - | 672 | 360.7 | - | - | 0 | - |
| - | - | 921.7 | 362.7 | - | - | 0 | - |
| - | - | 1404 | 366.2 | - | - | 0 | - |
| 5 | y | 1568 | 366.6 | 0.001001 | 2.732 | +3 | 10 |
| - | - | 1698 | 366.9 | - | - | 0 | - |
| - | - | 1499 | 367.2 | - | - | 0 | - |
| - | - | 1478 | 368.1 | - | - | 0 | - |
| - | - | 733.8 | 369.7 | - | - | 0 | - |
| - | - | 657.8 | 371.7 | - | - | 0 | - |
| 12 | y | 5058 | 373.3 | 7.885E-05 | 0.2112 | +1 | 3 |
| - | - | 694.4 | 381.2 | - | - | 0 | - |
| - | - | 1038 | 382.2 | - | - | 0 | - |
| - | - | 891.3 | 382.3 | - | - | 0 | - |
| - | - | 1623 | 382.5 | - | - | 0 | - |
| - | - | 1520 | 384.2 | - | - | 0 | - |
| - | - | 5847 | 385.2 | - | - | 0 | - |
| - | - | 1000 | 385.2 | - | - | 0 | - |
| - | - | 4092 | 386.2 | - | - | 0 | - |
| - | - | 1051 | 387.2 | - | - | 0 | - |
| - | - | 669.6 | 388 | - | - | 0 | - |
| - | - | 1700 | 388.9 | - | - | 0 | - |
| - | - | 984.6 | 389.2 | - | - | 0 | - |
| - | - | 1567 | 389.3 | - | - | 0 | - |
| 2 | y | 1535 | 390.2 | 0.0002935 | 0.7521 | +4 | 13 |
| - | - | 2655 | 390.5 | - | - | 0 | - |
| - | - | 1081 | 390.7 | - | - | 0 | - |
| - | - | 799.9 | 391 | - | - | 0 | - |
| - | - | 1887 | 392.3 | - | - | 0 | - |
| 10 | b | 1001 | 392.5 | 0.004726 | 12.04 | +3 | 10 |
| - | - | 814.7 | 392.9 | - | - | 0 | - |
| - | - | 4190 | 394.2 | - | - | 0 | - |
| 2 | y | 1659 | 394.5 | 0.0003935 | 0.9974 | +4 | 13 |
| - | - | 2449 | 394.7 | - | - | 0 | - |
| 10 | b | 2104 | 398.2 | 0.0004108 | 1.032 | +3 | 10 |
| - | - | 1063 | 399.2 | - | - | 0 | - |
| - | - | 800.1 | 400.1 | - | - | 0 | - |
| - | - | 873.1 | 400.5 | - | - | 0 | - |
| - | - | 1281 | 402.2 | - | - | 0 | - |
| - | - | 4801 | 403.2 | - | - | 0 | - |
| - | - | 2475 | 403.2 | - | - | 0 | - |
| - | - | 1246 | 403.3 | - | - | 0 | - |
| 4 | y | 3252 | 403.6 | 0.003581 | 8.874 | +3 | 11 |
| - | - | 2304 | 403.9 | - | - | 0 | - |
| - | - | 975.1 | 404.2 | - | - | 0 | - |
| - | - | 786.4 | 404.2 | - | - | 0 | - |
| - | - | 1070 | 404.7 | - | - | 0 | - |
| - | - | 677.5 | 404.9 | - | - | 0 | - |
| - | - | 1968 | 405 | - | - | 0 | - |
| - | - | 2219 | 406.2 | - | - | 0 | - |
| - | - | 1453 | 406.3 | - | - | 0 | - |
| - | - | 694.3 | 406.7 | - | - | 0 | - |
| - | - | 924.4 | 406.8 | - | - | 0 | - |
| - | - | 1648 | 408.6 | - | - | 0 | - |
| - | - | 636.7 | 409.2 | - | - | 0 | - |
| - | - | 3686 | 410.3 | - | - | 0 | - |
| - | - | 2109 | 410.6 | - | - | 0 | - |
| - | - | 1623 | 410.9 | - | - | 0 | - |
| 6 | b | 5134 | 411.2 | 0.0002657 | 0.6463 | +2 | 6 |
| - | - | 728.8 | 411.2 | - | - | 0 | - |
| 0 | Precursor | 683 | 411.7 | 0.0007 | 1.7 | +4 | -1 |
| - | - | 1205 | 412.2 | - | - | 0 | - |
| 7 | y | 1.426E+04 | 412.3 | 0.0003789 | 0.9191 | +2 | 8 |
| - | - | 7690 | 412.8 | - | - | 0 | - |
| - | - | 1722 | 413.3 | - | - | 0 | - |
| - | - | 850.4 | 413.8 | - | - | 0 | - |
| - | - | 1185 | 413.9 | - | - | 0 | - |
| - | - | 2634 | 414.2 | - | - | 0 | - |
| - | - | 1433 | 414.6 | - | - | 0 | - |
| - | - | 1284 | 415 | - | - | 0 | - |
| - | - | 1425 | 415.3 | - | - | 0 | - |
| - | - | 651.9 | 416.1 | - | - | 0 | - |
| 0 | Precursor | 1.173E+04 | 416.2 | 0.0002294 | 0.5511 | +4 | -1 |
| - | - | 948.2 | 416.3 | - | - | 0 | - |
| - | - | 1161 | 416.3 | - | - | 0 | - |
| - | - | 1.12E+04 | 416.5 | - | - | 0 | - |
| - | - | 3836 | 416.7 | - | - | 0 | - |
| - | - | 2130 | 417 | - | - | 0 | - |
| - | - | 855.8 | 417 | - | - | 0 | - |
| - | - | 6688 | 419.9 | - | - | 0 | - |
| 6 | b | 1139 | 420.2 | 0.0006597 | 1.57 | +2 | 6 |
| - | - | 7344 | 420.2 | - | - | 0 | - |
| - | - | 2164 | 420.6 | - | - | 0 | - |
| - | - | 710.1 | 422.3 | - | - | 0 | - |
| - | - | 4032 | 423.2 | - | - | 0 | - |
| - | - | 1113 | 423.3 | - | - | 0 | - |
| - | - | 926.9 | 424.3 | - | - | 0 | - |
| - | - | 2062 | 424.7 | - | - | 0 | - |
| - | - | 762.8 | 425.2 | - | - | 0 | - |
| - | - | 2227 | 425.9 | - | - | 0 | - |
| - | - | 1216 | 426.2 | - | - | 0 | - |
| - | - | 980.1 | 427.2 | - | - | 0 | - |
| - | - | 1.324E+04 | 429.2 | - | - | 0 | - |
| - | - | 3188 | 430.2 | - | - | 0 | - |
| - | - | 2354 | 431.2 | - | - | 0 | - |
| 3 | b | 721.6 | 437.2 | 0.001312 | 3 | +1 | 3 |
| - | - | 743 | 438.6 | - | - | 0 | - |
| - | - | 740.2 | 442.2 | - | - | 0 | - |
| - | - | 725.9 | 442.7 | - | - | 0 | - |
| - | - | 702.8 | 444.2 | - | - | 0 | - |
| - | - | 935 | 446.2 | - | - | 0 | - |
| - | - | 2867 | 448.3 | - | - | 0 | - |
| - | - | 1072 | 448.6 | - | - | 0 | - |
| - | - | 1029 | 448.9 | - | - | 0 | - |
| - | - | 4750 | 451.2 | - | - | 0 | - |
| - | - | 1400 | 451.3 | - | - | 0 | - |
| - | - | 1130 | 451.7 | - | - | 0 | - |
| - | - | 4031 | 451.9 | - | - | 0 | - |
| - | - | 1274 | 452.2 | - | - | 0 | - |
| - | - | 3371 | 452.3 | - | - | 0 | - |
| - | - | 1080 | 452.3 | - | - | 0 | - |
| - | - | 1710 | 452.6 | - | - | 0 | - |
| - | - | 652.1 | 455.2 | - | - | 0 | - |
| - | - | 1659 | 455.3 | - | - | 0 | - |
| - | - | 1851 | 457.6 | - | - | 0 | - |
| 3 | y | 3323 | 457.9 | 0.002929 | 6.395 | +3 | 12 |
| - | - | 1074 | 458.3 | - | - | 0 | - |
| - | - | 788.9 | 458.6 | - | - | 0 | - |
| - | - | 2026 | 460.2 | - | - | 0 | - |
| - | - | 1554 | 460.7 | - | - | 0 | - |
| - | - | 998.3 | 461.2 | - | - | 0 | - |
| 3 | y | 4.075E+04 | 463.6 | 0.0003859 | 0.8324 | +3 | 12 |
| - | - | 3.419E+04 | 463.9 | - | - | 0 | - |
| - | - | 1.448E+04 | 464.3 | - | - | 0 | - |
| - | - | 5080 | 464.6 | - | - | 0 | - |
| - | - | 1276 | 467.3 | - | - | 0 | - |
| - | - | 830.8 | 467.3 | - | - | 0 | - |
| - | - | 2585 | 469.2 | - | - | 0 | - |
| - | - | 628.6 | 470.2 | - | - | 0 | - |
| 11 | y | 1.682E+04 | 470.3 | 0.0003189 | 0.6781 | +1 | 4 |
| - | - | 1105 | 470.6 | - | - | 0 | - |
| - | - | 4649 | 471.3 | - | - | 0 | - |
| - | - | 990.9 | 471.8 | - | - | 0 | - |
| - | - | 707.1 | 472.3 | - | - | 0 | - |
| - | - | 1877 | 473.3 | - | - | 0 | - |
| - | - | 1465 | 473.6 | - | - | 0 | - |
| - | - | 929.2 | 474.8 | - | - | 0 | - |
| - | - | 3885 | 476.3 | - | - | 0 | - |
| - | - | 3316 | 476.6 | - | - | 0 | - |
| 6 | y | 3061 | 480.8 | 0.0003725 | 0.7747 | +2 | 9 |
| - | - | 1582 | 481.3 | - | - | 0 | - |
| - | - | 2877 | 481.9 | - | - | 0 | - |
| - | - | 2991 | 482.3 | - | - | 0 | - |
| - | - | 1348 | 482.3 | - | - | 0 | - |
| - | - | 1780 | 482.8 | - | - | 0 | - |
| - | - | 916 | 482.9 | - | - | 0 | - |
| - | - | 1176 | 483.3 | - | - | 0 | - |
| - | - | 2744 | 483.8 | - | - | 0 | - |
| - | - | 1910 | 484.3 | - | - | 0 | - |
| - | - | 1050 | 484.8 | - | - | 0 | - |
| - | - | 817.1 | 491.3 | - | - | 0 | - |
| - | - | 1374 | 494.2 | - | - | 0 | - |
| - | - | 974.9 | 501.9 | - | - | 0 | - |
| - | - | 1114 | 503.2 | - | - | 0 | - |
| - | - | 966.5 | 504.6 | - | - | 0 | - |
| 13 | b | 1092 | 505.3 | 0.004662 | 9.226 | +3 | 13 |
| - | - | 786.8 | 509.7 | - | - | 0 | - |
| 13 | b | 1708 | 510.9 | 0.00136 | 2.662 | +3 | 13 |
| - | - | 2000 | 511.3 | - | - | 0 | - |
| - | - | 1140 | 511.6 | - | - | 0 | - |
| - | - | 6666 | 516.9 | - | - | 0 | - |
| - | - | 3559 | 517.3 | - | - | 0 | - |
| - | - | 1775 | 517.6 | - | - | 0 | - |
| - | - | 1350 | 517.8 | - | - | 0 | - |
| - | - | 833 | 517.9 | - | - | 0 | - |
| - | - | 753.3 | 518.3 | - | - | 0 | - |
| 2 | y | 3914 | 519.9 | 0.0005083 | 0.9776 | +3 | 13 |
| - | - | 3707 | 520.3 | - | - | 0 | - |
| - | - | 825.1 | 520.4 | - | - | 0 | - |
| - | - | 1813 | 520.6 | - | - | 0 | - |
| - | - | 1609 | 521.2 | - | - | 0 | - |
| 2 | y | 6671 | 525.6 | 4.076E-05 | 0.07754 | +3 | 13 |
| - | - | 7342 | 526 | - | - | 0 | - |
| - | - | 1260 | 526.3 | - | - | 0 | - |
| - | - | 1374 | 526.6 | - | - | 0 | - |
| - | - | 679.2 | 529.8 | - | - | 0 | - |
| - | - | 820.5 | 530.3 | - | - | 0 | - |
| - | - | 697.7 | 535.8 | - | - | 0 | - |
| - | - | 749.1 | 536.3 | - | - | 0 | - |
| - | - | 690.5 | 536.8 | - | - | 0 | - |
| - | - | 8914 | 538.3 | - | - | 0 | - |
| - | - | 1011 | 538.8 | - | - | 0 | - |
| - | - | 2725 | 539.3 | - | - | 0 | - |
| - | - | 1512 | 539.3 | - | - | 0 | - |
| - | - | 1273 | 539.8 | - | - | 0 | - |
| - | - | 729.9 | 540.3 | - | - | 0 | - |
| - | - | 719.4 | 540.3 | - | - | 0 | - |
| 10 | y | 3949 | 541.4 | 0.0009554 | 1.765 | +1 | 5 |
| - | - | 2027 | 542.3 | - | - | 0 | - |
| - | - | 978.7 | 542.4 | - | - | 0 | - |
| - | - | 1322 | 547.8 | - | - | 0 | - |
| - | - | 997.8 | 549.2 | - | - | 0 | - |
| 5 | y | 8894 | 549.3 | 0.0002134 | 0.3885 | +2 | 10 |
| - | - | 6567 | 549.8 | - | - | 0 | - |
| - | - | 1611 | 550.3 | - | - | 0 | - |
| - | - | 1395 | 550.3 | - | - | 0 | - |
| 9 | b | 2979 | 561.3 | 0.0001954 | 0.3482 | +2 | 9 |
| - | - | 1976 | 561.8 | - | - | 0 | - |
| - | - | 1.026E+04 | 566.2 | - | - | 0 | - |
| - | - | 3033 | 567.2 | - | - | 0 | - |
| - | - | 1053 | 568.2 | - | - | 0 | - |
| - | - | 2587 | 568.3 | - | - | 0 | - |
| - | - | 4651 | 570.2 | - | - | 0 | - |
| - | - | 2017 | 571.2 | - | - | 0 | - |
| - | - | 5171 | 572.8 | - | - | 0 | - |
| - | - | 3464 | 573.3 | - | - | 0 | - |
| - | - | 872.3 | 573.8 | - | - | 0 | - |
| - | - | 2941 | 577.4 | - | - | 0 | - |
| - | - | 1029 | 578.4 | - | - | 0 | - |
| - | - | 7235 | 579.4 | - | - | 0 | - |
| - | - | 2552 | 580.4 | - | - | 0 | - |
| - | - | 2012 | 595.4 | - | - | 0 | - |
| - | - | 2473 | 598.2 | - | - | 0 | - |
| - | - | 2267 | 604.4 | - | - | 0 | - |
| 4 | y | 3586 | 604.8 | 0.003303 | 5.461 | +2 | 11 |
| - | - | 2629 | 605.4 | - | - | 0 | - |
| - | - | 2094 | 610.3 | - | - | 0 | - |
| 4 | y | 1982 | 613.4 | 0.0001447 | 0.2359 | +2 | 11 |
| - | - | 1164 | 613.9 | - | - | 0 | - |
| - | - | 1216 | 615.3 | - | - | 0 | - |
| - | - | 1035 | 615.9 | - | - | 0 | - |
| - | - | 1746 | 620.3 | - | - | 0 | - |
| - | - | 1213 | 620.8 | - | - | 0 | - |
| - | - | 1372 | 626.3 | - | - | 0 | - |
| - | - | 4537 | 629.3 | - | - | 0 | - |
| - | - | 3065 | 629.8 | - | - | 0 | - |
| - | - | 1140 | 630.3 | - | - | 0 | - |
| - | - | 3969 | 638.4 | - | - | 0 | - |
| - | - | 861.3 | 638.9 | - | - | 0 | - |
| - | - | 1411 | 639.4 | - | - | 0 | - |
| - | - | 918.7 | 657.3 | - | - | 0 | - |
| - | - | 1361 | 664.5 | - | - | 0 | - |
| - | - | 907.7 | 665.5 | - | - | 0 | - |
| - | - | 1106 | 665.8 | - | - | 0 | - |
| 9 | y | 1256 | 669.5 | 0.002028 | 3.029 | +1 | 6 |
| - | - | 6470 | 674.4 | - | - | 0 | - |
| - | - | 2236 | 675.4 | - | - | 0 | - |
| - | - | 6676 | 692.4 | - | - | 0 | - |
| - | - | 2105 | 693.4 | - | - | 0 | - |
| 3 | y | 2211 | 694.9 | 0.001028 | 1.479 | +2 | 12 |
| - | - | 1078 | 695.4 | - | - | 0 | - |
| - | - | 1015 | 695.9 | - | - | 0 | - |
| - | - | 3930 | 707.3 | - | - | 0 | - |
| - | - | 1058 | 708.3 | - | - | 0 | - |
| - | - | 620.3 | 710.4 | - | - | 0 | - |
| - | - | 1360 | 710.5 | - | - | 0 | - |
| - | - | 773 | 711.5 | - | - | 0 | - |
| - | - | 1613 | 716.4 | - | - | 0 | - |
| - | - | 827.5 | 720.3 | - | - | 0 | - |
| - | - | 1175 | 721.4 | - | - | 0 | - |
| - | - | 1013 | 724.3 | - | - | 0 | - |
| 8 | y | 3075 | 726.5 | 0.0003734 | 0.5139 | +1 | 7 |
| - | - | 984.3 | 727.5 | - | - | 0 | - |
| - | - | 2763 | 735.3 | - | - | 0 | - |
| - | - | 1030 | 736.3 | - | - | 0 | - |
| - | - | 779.5 | 737.3 | - | - | 0 | - |
| - | - | 1113 | 752.3 | - | - | 0 | - |
| - | - | 1375 | 805.5 | - | - | 0 | - |
| - | - | 1011 | 811.4 | - | - | 0 | - |
| 7 | y | 1.114E+04 | 823.5 | 9.974E-05 | 0.1211 | +1 | 8 |
| - | - | 6278 | 824.5 | - | - | 0 | - |
| - | - | 1372 | 825.5 | - | - | 0 | - |
| - | - | 933.1 | 829.5 | - | - | 0 | - |
| 6 | b | 1728 | 839.4 | 0.001475 | 1.757 | +1 | 6 |
| - | - | 937.6 | 840.4 | - | - | 0 | - |
| - | - | 1675 | 848.4 | - | - | 0 | - |
| - | - | 1422 | 901.4 | - | - | 0 | - |
| - | - | 1191 | 902.4 | - | - | 0 | - |
| 6 | y | 973.7 | 960.6 | 0.0004625 | 0.4815 | +1 | 9 |
| 10 | b | 763.6 | 1176 | 0.0211 | 17.95 | +1 | 10 |
| - | - | 742.3 | 1648 | - | - | 0 | - |
| - | - | 758 | 2596 | - | - | 0 | - |
| - | - | 687.2 | 2937 | - | - | 0 | - |
| - | - | 639.3 | 3047 | - | - | 0 | - |

m/z Charge Intensity FragmentType MassShift Position
120.05223846435547 0 456.31854
120.06588745117188 0 633.67395
120.08104705810547 0 1184.8188
122.07137298583984 0 803.76935
123.05535125732422 0 1713.2515
123.11723327636719 0 683.0293
127.05064392089844 0 491.18768
127.08663940429688 0 1904.7251
128.07077026367188 0 501.51685
128.08193969726562 0 759.83264
129.0545654296875 0 481.90344
129.06594848632812 0 3268.8865
129.10238647460938 0 58265.83
130.04998779296875 0 1401.4591
130.0611572265625 0 2570.3577
130.06532287597656 0 13430.928
130.08657836914062 0 848.6163
130.09786987304688 0 694.40015
130.10572814941406 0 3412.935
131.0452423095703 0 4421.052
131.06887817382812 0 1548.7618
131.0816650390625 0 1752.9131
131.11805725097656 0 578.05054
132.04481506347656 0 394.7219
132.08091735839844 0 44944.125
132.10202026367188 0 12803.296 y 13
133.0608367919922 0 665.64044
133.0848388671875 0 5302.011
133.10557556152344 0 688.75385
134.044921875 0 3883.59
136.07582092285156 0 62540.457
137.07920837402344 0 4706.768
138.0663604736328 0 1090.4381
139.0868682861328 0 835.9964
140.08197021484375 0 485.33057
141.06590270996094 0 537.6959
141.10238647460938 0 4089.736
142.06541442871094 0 853.7344 b 3
142.09765625 0 519.2156
144.08106994628906 0 1252.448
145.06082153320312 0 550.7491
146.06033325195312 0 1543.1282
147.04412841796875 0 3615.1052
151.086669921875 0 854.1211
152.07081604003906 0 1827.9863
152.08169555664062 0 484.22073
152.1436004638672 0 1131.1934
153.06643676757812 0 507.68085
153.0770721435547 0 603.03674
154.09706115722656 0 474.07822
155.0703125 0 441.0095
155.08152770996094 0 2555.0342
155.09295654296875 0 1727.4647
155.11798095703125 0 1553.0189
156.07582092285156 0 640.6435
156.08139038085938 0 786.7347
157.09695434570312 0 646.10864
158.0601806640625 0 972.28674
158.08395385742188 0 4751.633
159.0917510986328 0 31493.432
159.1129150390625 0 1788.6227
160.07569885253906 0 1695.3375
160.09510803222656 0 3425.0771
162.05508422851562 0 824.49994
164.07090759277344 0 1086.294
165.10226440429688 0 2090.0076
166.06106567382812 0 1333.1117
168.1133270263672 0 936.408
169.07620239257812 0 3275.621
169.09735107421875 0 4059.1206
170.06016540527344 0 13827.902
170.07989501953125 0 518.54517
170.0930938720703 0 493.8957
171.06353759765625 0 1985.5789
171.0771942138672 0 689.75397
171.09207153320312 0 695.08093
171.112548828125 0 773.31067
172.1082000732422 0 3459.4463
173.09219360351562 0 8405.913
173.128662109375 0 1006.00854
173.4402618408203 0 2644.2124
174.05517578125 0 1136.5969
175.08639526367188 0 1008.0802
175.09860229492188 0 643.3367
176.08229064941406 0 913.3169
177.11221313476562 0 1505.5146
179.09356689453125 0 452.06796
180.07679748535156 0 589.06494
181.06076049804688 0 1612.3121
181.0972137451172 0 16114.446
182.10093688964844 0 1257.3551
182.1287078857422 0 1594.4014
183.0917205810547 0 7521.3647
183.11264038085938 0 881.99207
184.0758819580078 0 868.096
184.09494018554688 0 848.1357
185.07083129882812 0 1732.4686
185.07962036132812 0 664.74493
185.16482543945312 0 1178.7844
186.12376403808594 0 5356.725
187.08670043945312 0 29039.434
187.10775756835938 0 6339.007
187.1443634033203 0 1651.6567 y 11
188.09014892578125 0 3633.7847
189.1240692138672 0 601.8272
190.0504913330078 0 440.04196
195.07656860351562 0 851.10815
195.087890625 0 458.7506
195.1494598388672 0 697.6386
196.10745239257812 0 602.90265
197.1285858154297 0 1362.7538
197.1648406982422 0 2507.3
197.1737518310547 0 441.13654
199.0714874267578 0 2878.4692
199.0863494873047 0 1379.0388
199.10760498046875 0 1543.2289
199.1695098876953 0 2526.677
199.1796417236328 0 708.8128
200.1395263671875 0 1157.9576
201.10220336914062 0 3209.426
201.12306213378906 0 2039.6108
202.08607482910156 0 1471.5114
202.1082305908203 0 6139.42
204.1346435546875 0 825.91846
207.0758514404297 0 629.127
207.149169921875 0 1379.9158
209.09217834472656 0 2795.0098
211.08660888671875 0 829.5241
212.66783142089844 0 790.3767
214.1085968017578 0 2102.9343
214.18991088867188 0 562.06165
215.13906860351562 0 1871.1469
216.1233367919922 0 1473.1394
219.13424682617188 0 824.0804
220.1193084716797 0 4518.653
221.1033477783203 0 5673.033
224.1184539794922 0 566.4376
224.17572021484375 0 2468.8616
225.1595458984375 0 1068.791
226.1550750732422 0 40404.68
227.10284423828125 0 2394.53
227.11517333984375 0 573.5224
227.15835571289062 0 4958.825 y Ammonia loss 10
228.11297607421875 0 2459.537 a Water loss 1
229.09747314453125 0 3753.4753 b Ammonia loss 4
229.11819458007812 0 1707.5026
229.15472412109375 0 819.396
230.10292053222656 0 940.5953
231.08749389648438 0 1648.9252
233.1038360595703 0 1380.2288
234.12393188476562 0 733.7158
235.11900329589844 0 7908.8486
235.67063903808594 0 6891.479 y 10
236.12283325195312 0 1081.088
236.1724090576172 0 1246.3245
238.13040161132812 0 759.3631
239.15037536621094 0 2436.7236
240.1343994140625 0 670.806
242.18634033203125 0 1418.2671
243.13400268554688 0 1381.5154
245.07687377929688 0 8059.9756
245.18606567382812 0 9583.855 y 12
246.1238250732422 0 74132.375 a 1
246.18905639648438 0 1027.6044
247.12863159179688 0 16831.977
248.1142120361328 0 6111.9766
248.13134765625 0 1622.502
249.0982666015625 0 9583.351
249.15945434570312 0 1755.456
250.10345458984375 0 617.5155
250.12191772460938 0 972.4653
251.14990234375 0 884.9548
252.17047119140625 0 995.3614
253.15467834472656 0 819.68274
255.1953125 0 806.6039
256.10797119140625 0 1921.0292 b Water loss 1
257.1146545410156 0 1127.9589
257.1609191894531 0 875.0589
258.07684326171875 0 941.2737
258.0986022949219 0 13540.763
259.1019592285156 0 1701.3544
259.1299133300781 0 905.64185
260.1059875488281 0 1892.6547
260.1968994140625 0 1636.9407
263.088134765625 0 1024.888
266.1247253417969 0 5482.188
269.6292419433594 0 1256.5928
270.18121337890625 0 2735.2573
271.18841552734375 0 2801.2522 y 9
274.1185302734375 0 23677.957 b 1
275.1031188964844 0 2987.3838
275.12469482421875 0 22064.627
275.18426513671875 0 695.8116 y 6
276.1082458496094 0 911.66797
276.1275329589844 0 2768.48
277.13385009765625 0 802.7894
277.1535339355469 0 688.17944
279.1813049316406 0 1070.6312
282.1823425292969 0 683.77106
283.15313720703125 0 602.58405
283.1564636230469 0 686.23425
283.17645263671875 0 8200.52
283.6259765625 0 808.1285
284.1815490722656 0 792.5433
285.1095275878906 0 685.84094
285.1561279296875 0 1355.3989
290.03826904296875 0 1148.7067
290.1842041015625 0 3027.9875
290.6849670410156 0 1175.9243
291.15631103515625 0 1386.7502
292.12933349609375 0 6373.5156
293.1339111328125 0 1197.7798
294.1195983886719 0 2036.3004
294.21649169921875 0 1066.3291
297.1922912597656 0 4868.471
298.1956787109375 0 668.8973
303.11962890625 0 1252.6346
305.1290588378906 0 2772.515
305.662841796875 0 672.72736
310.138671875 0 778.22375
311.2449645996094 0 1095.1006
314.6701354980469 0 2082.4216
319.15142822265625 0 673.903
320.1250305175781 0 764.3732
320.87042236328125 0 813.32697 y 5
321.22894287109375 0 5742.3447
322.1552429199219 0 3854.465
322.2315979003906 0 1245.5797
323.15765380859375 0 947.98737 b 10
326.2192077636719 0 1793.8514
327.20159912109375 0 821.3786
332.72900390625 0 1989.0819
333.122802734375 0 914.6701
336.2032165527344 0 9499.729
337.2066345214844 0 1581.4364
337.72125244140625 0 2746.5168
338.1347961425781 0 8233.687
338.22271728515625 0 1231.6449
338.72247314453125 0 573.317
339.1378173828125 0 1384.4951
339.23944091796875 0 10090.882
340.15374755859375 0 708.2979
340.1867370605469 0 1358.7373
340.2420654296875 0 1685.2552
342.1667785644531 0 607.18085
344.9761962890625 0 1823.9938
345.64276123046875 0 783.32886
346.7266845703125 0 9409.692
347.1295471191406 0 677.9122
347.22802734375 0 3804.811
347.7308044433594 0 664.6572
350.1495056152344 0 1241.9971
351.23980712890625 0 1150.167
354.1116943359375 0 1699.4755
354.1551513671875 0 694.4066
354.2137145996094 0 11726.59
354.6575622558594 0 695.4367
355.2165832519531 0 2682.0432
355.731689453125 0 1070.3109
356.1454162597656 0 889.4621
357.2498474121094 0 2822.534
358.1621398925781 0 3125.1238
358.7134704589844 0 1464.7655
359.5432434082031 0 1048.7062
359.875 0 841.8963
360.2083435058594 0 755.88086
360.66534423828125 0 671.96423
362.6637878417969 0 921.7202
366.1552734375 0 1404.0898
366.55645751953125 0 1568.4216 y 4
366.8912048339844 0 1698.2473
367.16265869140625 0 1499.1825
368.1475524902344 0 1477.6567
369.6947021484375 0 733.82446
371.66937255859375 0 657.83276
373.2808532714844 0 5057.715 y 11
381.15399169921875 0 694.42224
382.2064514160156 0 1038.0638
382.2790832519531 0 891.34784
382.5391540527344 0 1622.9669
384.16595458984375 0 1520.0903
385.173095703125 0 5846.5522
385.19873046875 0 999.9752
386.1568603515625 0 4092.3325
387.16021728515625 0 1051.4683
387.9622802734375 0 669.6022
388.8619384765625 0 1700.3387
389.19415283203125 0 984.60596
389.27081298828125 0 1566.5288
390.2132873535156 0 1535.0532 y Ammonia loss 1
390.4639892578125 0 2655.2036
390.7151184082031 0 1080.6957
390.9659118652344 0 799.90076
392.2654724121094 0 1886.5094
392.52227783203125 0 1000.91534 b Ammonia loss 9
392.8536376953125 0 814.6964
394.151123046875 0 4189.884
394.4706115722656 0 1658.6493 y 1
394.720703125 0 2448.5854
398.1926574707031 0 2103.9321 b 9
399.2223205566406 0 1063.271
400.1177673339844 0 800.13477
400.4716491699219 0 873.05005
402.2258605957031 0 1280.856
403.18389892578125 0 4801.3643
403.2405090332031 0 2475.2727
403.2699890136719 0 1245.9199
403.57171630859375 0 3252.4888 y Ammonia loss 3
403.903564453125 0 2304.1865
404.1865234375 0 975.1001
404.23846435546875 0 786.3623
404.73126220703125 0 1070.4718
404.89093017578125 0 677.4798
404.97625732421875 0 1968.3788
406.1858215332031 0 2219.1228
406.25140380859375 0 1452.8982
406.6878967285156 0 694.26245
406.7514953613281 0 924.44226
408.555908203125 0 1647.5082
409.21484375 0 636.70435
410.27593994140625 0 3686.3594
410.5684509277344 0 2109.2488
410.9030456542969 0 1622.9974
411.17779541015625 0 5134.33 b Water loss 5
411.2347412109375 0 728.7733
411.7248840332031 0 683.02783 Precursor Water loss
412.1611022949219 0 1204.8209
412.2740173339844 0 14259.053 y 6
412.77496337890625 0 7689.695
413.2759704589844 0 1721.7661
413.77923583984375 0 850.42267
413.89501953125 0 1184.9171
414.2301940917969 0 2634.1042
414.5601806640625 0 1433.1036
415.03515625 0 1284.0087
415.256103515625 0 1424.6143
416.1496887207031 0 651.85144
416.22845458984375 0 11729.079 Precursor
416.2572937011719 0 948.2235
416.2850036621094 0 1160.7388
416.47918701171875 0 11195.321
416.7290344238281 0 3835.7192
416.9798583984375 0 2130.07
417.0322570800781 0 855.83124
419.9006042480469 0 6688.4214
420.1834716796875 0 1139.4891 b 5
420.2345886230469 0 7344.289
420.5691833496094 0 2164.4539
422.277099609375 0 710.1033
423.224365234375 0 4031.685
423.27362060546875 0 1112.6965
424.327392578125 0 926.9312
424.7125549316406 0 2062.012
425.2140197753906 0 762.76373
425.9040222167969 0 2227.0195
426.23834228515625 0 1215.7543
427.2455749511719 0 980.0873
429.1884460449219 0 13241.541
430.1920166015625 0 3188.2725
431.1796875 0 2353.8079
437.1832580566406 0 721.5707 b 2
438.5562744140625 0 743.00946
442.18701171875 0 740.2455
442.71087646484375 0 725.86884
444.2330627441406 0 702.80383
446.2287292480469 0 935.0421
448.2631530761719 0 2867.363
448.59783935546875 0 1072.3263
448.93212890625 0 1029.3684
451.22198486328125 0 4750.4775
451.2652282714844 0 1399.7878
451.7259521484375 0 1129.639
451.9194030761719 0 4031.1548
452.22125244140625 0 1273.7754
452.2535400390625 0 3371.3591
452.3232727050781 0 1079.5575
452.587890625 0 1709.5635
455.2174072265625 0 652.13434
455.2519836425781 0 1659.2374
457.5947570800781 0 1851.3152
457.9255065917969 0 3322.785 y Ammonia loss 2
458.25958251953125 0 1073.7596
458.5913391113281 0 788.9492
460.2308044433594 0 2026.4507
460.7321472167969 0 1553.5988
461.181396484375 0 998.2789
463.5984802246094 0 40745.066 y 2
463.9325866699219 0 34190.883
464.26690673828125 0 14483.176
464.6007080078125 0 5080.1436
467.2554931640625 0 1276.4348
467.29620361328125 0 830.8101
469.23101806640625 0 2585.0527
470.2482604980469 0 628.6202
470.3340148925781 0 16820.104 y 10
470.58203125 0 1104.5348
471.33740234375 0 4649.224
471.7984313964844 0 990.856
472.30157470703125 0 707.0762
473.26263427734375 0 1876.5155
473.5751647949219 0 1464.6052
474.7794494628906 0 929.20935
476.2526550292969 0 3885.2078
476.58782958984375 0 3315.872
480.803466796875 0 3061.0784 y 5
481.3050537109375 0 1582.3864
481.92724609375 0 2876.6997
482.2598571777344 0 2991.1157
482.3084716796875 0 1348.36
482.7610168457031 0 1779.7854
482.9264221191406 0 915.9567
483.26116943359375 0 1175.6273
483.78515625 0 2744.3079
484.28631591796875 0 1910.2983
484.787353515625 0 1049.7135
491.27459716796875 0 817.06604
494.2324523925781 0 1374.2427
501.93975830078125 0 974.91785
503.2167663574219 0 1114.1605
504.61614990234375 0 966.5062
505.2661437988281 0 1091.7799 b Ammonia loss 12
509.7389221191406 0 786.7537
510.9356384277344 0 1708.0193 b 12
511.2684631347656 0 2000.0377
511.6077575683594 0 1140.1963
516.940673828125 0 6666.279
517.2737426757812 0 3558.5283
517.6109008789062 0 1774.8448
517.75341796875 0 1349.8848
517.9434204101562 0 832.98474
518.2760009765625 0 753.2918
519.9495239257812 0 3914.0623 y Ammonia loss 1
520.2838134765625 0 3707.1575
520.3589477539062 0 825.0508
520.617431640625 0 1812.8973
521.2268676757812 0 1608.8767
525.6245727539062 0 6670.797 y 1
525.959228515625 0 7342.3735
526.2939453125 0 1259.6853
526.6276245117188 0 1373.7911
529.8035278320312 0 679.1502
530.2945556640625 0 820.458
535.7528076171875 0 697.6828
536.2523193359375 0 749.12054
536.7527465820312 0 690.47546
538.252197265625 0 8913.99
538.8067016601562 0 1011.1144
539.2542724609375 0 2724.6343
539.3054809570312 0 1511.6198
539.801513671875 0 1272.6542
540.304443359375 0 729.9008
540.3290405273438 0 719.41376
541.3717651367188 0 3949.058 y 9
542.2501220703125 0 2027.0625
542.3720092773438 0 978.67236
547.8163452148438 0 1321.872
549.2232055664062 0 997.81287
549.332763671875 0 8893.731 y 4
549.834228515625 0 6566.8486
550.2882690429688 0 1610.9127
550.33544921875 0 1395.1942
561.2672119140625 0 2978.7957 b 8
561.767333984375 0 1976.0488
566.2470092773438 0 10264.914
567.2490234375 0 3033.4465
568.2474365234375 0 1053.3
568.2974853515625 0 2586.9058
570.2467651367188 0 4650.859
571.248779296875 0 2017.1255
572.80419921875 0 5171.385
573.3049926757812 0 3463.7046
573.8102416992188 0 872.33887
577.3811645507812 0 2940.9111
578.3832397460938 0 1028.5094
579.3616943359375 0 7235.314
580.3640747070312 0 2552.4182
595.39404296875 0 2011.9323
598.2413330078125 0 2472.76
604.354248046875 0 2266.7832
604.8518676757812 0 3586.4143 y Ammonia loss 3
605.351806640625 0 2628.6135
610.321533203125 0 2094.195
613.3616943359375 0 1981.6793 y 3
613.8658447265625 0 1163.8778
615.3482666015625 0 1216.3036
615.8507080078125 0 1035.4526
620.3409423828125 0 1745.6698
620.8382568359375 0 1213.3657
626.2573852539062 0 1372.1581
629.3460693359375 0 4536.896
629.8475952148438 0 3064.8574
630.3411865234375 0 1140.481
638.3522338867188 0 3969.1675
638.8514404296875 0 861.28125
639.35302734375 0 1411.4498
657.3320922851562 0 918.7074
664.4520263671875 0 1361.4149
665.453857421875 0 907.739
665.8424072265625 0 1105.748
669.4637451171875 0 1255.9305 y 8
674.4343872070312 0 6470.344
675.4378662109375 0 2235.869
692.4447631835938 0 6676.4043
693.4491577148438 0 2104.825
694.89453125 0 2211.0112 y 2
695.3934936523438 0 1077.5251
695.8973388671875 0 1014.75104
707.30517578125 0 3930.397
708.3082275390625 0 1057.6224
710.3790283203125 0 620.3247
710.455322265625 0 1360.4268
711.4627685546875 0 773.0212
716.421630859375 0 1613.2587
720.3169555664062 0 827.5096
721.3536376953125 0 1175.4652
724.330810546875 0 1013.3519
726.4876098632812 0 3074.6104 y 7
727.4896240234375 0 984.2573
735.3004150390625 0 2762.845
736.302001953125 0 1030.2844
737.348388671875 0 779.50183
752.32666015625 0 1112.6158
805.52880859375 0 1374.7556
811.3610229492188 0 1011.43494
823.5401000976562 0 11138.599 y 6
824.5421752929688 0 6277.715
825.5430297851562 0 1371.7733
829.5059204101562 0 933.05426
839.3568725585938 0 1727.558 b 5
840.360595703125 0 937.6113
848.4148559570312 0 1674.6921
901.438720703125 0 1422.0798
902.44580078125 0 1191.079
960.5984497070312 0 973.6766 y 5
1175.5592041015625 0 763.5761 b Ammonia loss 9
1648.355224609375 0 742.2921
2595.66259765625 0 758.00165
2936.9794921875 0 687.18915
3046.952880859375 0 639.31726

Spectrum Details

|  |  |
| --- | --- |
| Matched peaks? Matched peaks The total absolute number of peaks matched. Additionally in brackets the total fraction of peaks matched and the total number of peaks is shown. | 48 (8.91% of 539) |
| FDR? FDR The false discovery rate estimated for this peptide. It is calculated by matching all theoretical fragments with a non-integer shift with the raw peaks for this spectrum. This is done with 40 different shifts. The resulting percentage is the average number of annotated peaks over the number of annotated peaks with the correct spectrum. | 3.27% |
| Satellite FDR? Satellite FDR See the FDR for details on its calculation. This satellite ion specific FDR only contains the satellite ions (d/w) for I/L/J positions. | - |
| PSM Score? PSM Score The PSM Score as given by Hecklib to this annotated spectrum. It is shown with three significant figures. | 174 |

#### Spectrum 5622? Spectrum 5622 The raw spectrum of this peptide as annotated by Hecklib. The fragments are coloured according to ion type (see legend). Any peaks with a star '\*' as text can be hovered over to see the full details, first the ion type second the mass shift type. By hovering over the amino acids in the peptide or ions in the legend the corresponding peaks are highlighted. By toggling the 'Unassigned' label you can turn the background (unassigned) peaks on or off in the plot. By updating the slider in the Ion legend you can update the spectrum to only show the top X% of the peaks with labels. The top X% means any peak that is within X% of the highest intensity. By dragging in the spectrum you can zoom in to a specific part of the spectrum and use 'Zoom Out' to get back to the original zoom level. The annotation of the spectrum is based on the given sequence in the peptides file and is done with different software so inconsistencies are likely. The peaks are annotated based on the given sequence, with 20 ppm tolerance.

Copy Data

##### Spectrum 5622 (TSV)

###### Preview

```
Loading example...
```

*Click on the button to copy the data to your clipboard.*

Mz MinMz MaxIntensity Max

WidthHeightPeptide font sizePeptide stroke widthSpectrum font sizeSpectrum stroke widthCompact peptide

Ion legend

wxyz

abcd

OtherUnassignedIon^Charge^_Position_Show for top:%

SWYQHHPGKAPKJJ

07.12e+41.42e+52.14e+52.85e+5

Zoom Out

y^+1^_1_b^+4^_4_y^+2^_3_y^+2^_4_a^+1^_2_b^+3^_5_y^+2^_4_y^+1^_2_a^+1^_2_b^+4^_8_b^+1^_2_y^+2^_5_b^+1^_2_y^+3^_8_y^+3^_9_b^+4^_11_y^+2^_6_y^+2^_7_y^+3^_10_y^+1^_3_y^+4^_13_b^+3^_10_y^+4^_13_b^+3^_10_y^+3^_11_y^+2^_8_y^+3^_11_b^+2^_6_\*\*y^+2^_8_\*b^+2^_6_b^+1^_3_y^+3^_12_y^+3^_12_y^+1^_4_y^+2^_9_b^+2^_8_b^+3^_13_b^+3^_13_y^+3^_13_y^+3^_13_y^+2^_10_y^+1^_5_y^+2^_10_b^+2^_9_b^+2^_10_b^+2^_10_y^+2^_11_y^+2^_11_y^+1^_6_b^+1^_5_y^+2^_12_b^+1^_5_y^+1^_7_y^+1^_8_y^+1^_8_b^+1^_6_y^+1^_9_y^+1^_10_

0693138620792772

Fragment Matches Table

Show background peaks

| Position | Ion type | Intensity | mz Theoretical | mz Error (Th) | mz Error (ppm) | Charge | Series Number |
| --- | --- | --- | --- | --- | --- | --- | --- |
| - | - | 480.4 | 120.1 | - | - | 0 | - |
| - | - | 614.7 | 120.1 | - | - | 0 | - |
| - | - | 1216 | 120.1 | - | - | 0 | - |
| - | - | 546.8 | 122.1 | - | - | 0 | - |
| - | - | 1835 | 123.1 | - | - | 0 | - |
| - | - | 525.8 | 123.1 | - | - | 0 | - |
| - | - | 436.3 | 126.1 | - | - | 0 | - |
| - | - | 2944 | 127.1 | - | - | 0 | - |
| - | - | 4140 | 129.1 | - | - | 0 | - |
| - | - | 9.364E+04 | 129.1 | - | - | 0 | - |
| - | - | 470.8 | 129.5 | - | - | 0 | - |
| - | - | 2.102E+04 | 130.1 | - | - | 0 | - |
| - | - | 614.7 | 130.1 | - | - | 0 | - |
| - | - | 1073 | 130.1 | - | - | 0 | - |
| - | - | 6133 | 130.1 | - | - | 0 | - |
| - | - | 3971 | 131 | - | - | 0 | - |
| - | - | 4806 | 131.1 | - | - | 0 | - |
| - | - | 1215 | 131.1 | - | - | 0 | - |
| - | - | 8.194E+04 | 132.1 | - | - | 0 | - |
| 14 | y | 2.405E+04 | 132.1 | 0.0004508 | 3.413 | +1 | 1 |
| - | - | 792 | 132.1 | - | - | 0 | - |
| - | - | 2.232E+05 | 133.1 | - | - | 0 | - |
| - | - | 499.4 | 134.1 | - | - | 0 | - |
| - | - | 1.326E+04 | 134.1 | - | - | 0 | - |
| - | - | 1239 | 135.1 | - | - | 0 | - |
| - | - | 1.096E+05 | 136.1 | - | - | 0 | - |
| - | - | 721.9 | 137.1 | - | - | 0 | - |
| - | - | 9220 | 137.1 | - | - | 0 | - |
| - | - | 1211 | 138.1 | - | - | 0 | - |
| - | - | 378 | 138.1 | - | - | 0 | - |
| - | - | 1486 | 139.1 | - | - | 0 | - |
| - | - | 714.9 | 140.1 | - | - | 0 | - |
| - | - | 5597 | 141.1 | - | - | 0 | - |
| 4 | b | 1333 | 142.1 | 0.0003297 | 2.32 | +4 | 4 |
| - | - | 485.5 | 142.1 | - | - | 0 | - |
| - | - | 541.1 | 143.1 | - | - | 0 | - |
| - | - | 4000 | 144.1 | - | - | 0 | - |
| - | - | 808.8 | 145.1 | - | - | 0 | - |
| - | - | 1539 | 146.1 | - | - | 0 | - |
| - | - | 5352 | 147 | - | - | 0 | - |
| - | - | 456.6 | 147.1 | - | - | 0 | - |
| - | - | 1158 | 151.1 | - | - | 0 | - |
| - | - | 1717 | 151.1 | - | - | 0 | - |
| - | - | 605.3 | 152.1 | - | - | 0 | - |
| - | - | 748.8 | 152.1 | - | - | 0 | - |
| - | - | 1766 | 152.1 | - | - | 0 | - |
| - | - | 660.3 | 153.1 | - | - | 0 | - |
| - | - | 552.7 | 154.1 | - | - | 0 | - |
| - | - | 4890 | 155.1 | - | - | 0 | - |
| - | - | 1820 | 155.1 | - | - | 0 | - |
| - | - | 3539 | 155.1 | - | - | 0 | - |
| - | - | 1967 | 156.1 | - | - | 0 | - |
| - | - | 2035 | 156.1 | - | - | 0 | - |
| - | - | 573.9 | 157.1 | - | - | 0 | - |
| - | - | 874.4 | 157.1 | - | - | 0 | - |
| - | - | 531.1 | 157.1 | - | - | 0 | - |
| - | - | 441.6 | 157.8 | - | - | 0 | - |
| - | - | 1810 | 158.1 | - | - | 0 | - |
| - | - | 9201 | 158.1 | - | - | 0 | - |
| - | - | 5.532E+04 | 159.1 | - | - | 0 | - |
| - | - | 1141 | 159.1 | - | - | 0 | - |
| - | - | 3438 | 160.1 | - | - | 0 | - |
| - | - | 5882 | 160.1 | - | - | 0 | - |
| - | - | 8705 | 161.1 | - | - | 0 | - |
| - | - | 802.1 | 162.1 | - | - | 0 | - |
| - | - | 472.1 | 162.1 | - | - | 0 | - |
| - | - | 1451 | 164.1 | - | - | 0 | - |
| - | - | 657.3 | 165.1 | - | - | 0 | - |
| - | - | 3643 | 165.1 | - | - | 0 | - |
| - | - | 1386 | 166.1 | - | - | 0 | - |
| - | - | 587 | 168.1 | - | - | 0 | - |
| - | - | 1163 | 168.1 | - | - | 0 | - |
| - | - | 5631 | 169.1 | - | - | 0 | - |
| - | - | 5977 | 169.1 | - | - | 0 | - |
| - | - | 2.717E+04 | 170.1 | - | - | 0 | - |
| - | - | 759.4 | 170.1 | - | - | 0 | - |
| - | - | 544.7 | 170.1 | - | - | 0 | - |
| - | - | 3177 | 171.1 | - | - | 0 | - |
| - | - | 1083 | 171.1 | - | - | 0 | - |
| - | - | 540.8 | 171.1 | - | - | 0 | - |
| - | - | 4430 | 172.1 | - | - | 0 | - |
| - | - | 1099 | 173.1 | - | - | 0 | - |
| - | - | 3901 | 175.1 | - | - | 0 | - |
| - | - | 1789 | 176.1 | - | - | 0 | - |
| - | - | 9.565E+04 | 177.1 | - | - | 0 | - |
| - | - | 7713 | 178.1 | - | - | 0 | - |
| - | - | 533.6 | 178.1 | - | - | 0 | - |
| - | - | 952.4 | 179.1 | - | - | 0 | - |
| - | - | 2212 | 179.1 | - | - | 0 | - |
| - | - | 1078 | 179.1 | - | - | 0 | - |
| - | - | 1880 | 181.1 | - | - | 0 | - |
| - | - | 2.591E+04 | 181.1 | - | - | 0 | - |
| - | - | 1970 | 181.1 | - | - | 0 | - |
| - | - | 2884 | 182.1 | - | - | 0 | - |
| - | - | 3358 | 182.1 | - | - | 0 | - |
| - | - | 1.109E+04 | 183.1 | - | - | 0 | - |
| - | - | 1259 | 183.1 | - | - | 0 | - |
| - | - | 605.3 | 183.1 | - | - | 0 | - |
| - | - | 1399 | 184.1 | - | - | 0 | - |
| - | - | 1587 | 184.1 | - | - | 0 | - |
| - | - | 561.7 | 184.1 | - | - | 0 | - |
| - | - | 3069 | 185.1 | - | - | 0 | - |
| - | - | 959 | 185.1 | - | - | 0 | - |
| - | - | 1579 | 185.2 | - | - | 0 | - |
| - | - | 450.1 | 186.1 | - | - | 0 | - |
| - | - | 1.002E+04 | 186.1 | - | - | 0 | - |
| - | - | 6.013E+04 | 187.1 | - | - | 0 | - |
| - | - | 4411 | 187.1 | - | - | 0 | - |
| - | - | 724.5 | 187.1 | - | - | 0 | - |
| 12 | y | 1825 | 187.1 | 0.0007169 | 3.831 | +2 | 3 |
| - | - | 750.2 | 188.1 | - | - | 0 | - |
| - | - | 7547 | 188.1 | - | - | 0 | - |
| - | - | 547.7 | 193.1 | - | - | 0 | - |
| - | - | 4325 | 195.1 | - | - | 0 | - |
| - | - | 792.8 | 195.1 | - | - | 0 | - |
| - | - | 841.8 | 196.1 | - | - | 0 | - |
| - | - | 529 | 197.1 | - | - | 0 | - |
| - | - | 4126 | 197.2 | - | - | 0 | - |
| - | - | 549.8 | 198.1 | - | - | 0 | - |
| - | - | 2243 | 199.1 | - | - | 0 | - |
| - | - | 1340 | 199.1 | - | - | 0 | - |
| - | - | 2374 | 199.2 | - | - | 0 | - |
| - | - | 1611 | 199.2 | - | - | 0 | - |
| - | - | 1506 | 200.1 | - | - | 0 | - |
| - | - | 6655 | 201.1 | - | - | 0 | - |
| - | - | 1696 | 201.1 | - | - | 0 | - |
| - | - | 2946 | 202.1 | - | - | 0 | - |
| - | - | 1.105E+04 | 202.1 | - | - | 0 | - |
| - | - | 697.6 | 203.1 | - | - | 0 | - |
| - | - | 791.6 | 203.1 | - | - | 0 | - |
| - | - | 988.6 | 203.1 | - | - | 0 | - |
| - | - | 620.4 | 204.1 | - | - | 0 | - |
| - | - | 5225 | 204.1 | - | - | 0 | - |
| - | - | 5.376E+04 | 205.1 | - | - | 0 | - |
| - | - | 4916 | 206.1 | - | - | 0 | - |
| - | - | 2247 | 207.1 | - | - | 0 | - |
| - | - | 1994 | 209.1 | - | - | 0 | - |
| - | - | 1069 | 209.1 | - | - | 0 | - |
| - | - | 1771 | 211.1 | - | - | 0 | - |
| - | - | 454.3 | 211.1 | - | - | 0 | - |
| - | - | 679.9 | 212.1 | - | - | 0 | - |
| - | - | 1713 | 212.7 | - | - | 0 | - |
| - | - | 651.7 | 213.1 | - | - | 0 | - |
| - | - | 1461 | 213.2 | - | - | 0 | - |
| - | - | 779.9 | 213.2 | - | - | 0 | - |
| - | - | 2787 | 214.1 | - | - | 0 | - |
| - | - | 1584 | 215.1 | - | - | 0 | - |
| - | - | 838.8 | 216.1 | - | - | 0 | - |
| - | - | 1428 | 216.1 | - | - | 0 | - |
| - | - | 795.6 | 217.1 | - | - | 0 | - |
| - | - | 1422 | 219.1 | - | - | 0 | - |
| - | - | 6924 | 220.1 | - | - | 0 | - |
| - | - | 338.9 | 221.1 | - | - | 0 | - |
| - | - | 1.009E+04 | 221.1 | - | - | 0 | - |
| - | - | 991.8 | 221.1 | - | - | 0 | - |
| - | - | 2.033E+04 | 221.1 | - | - | 0 | - |
| - | - | 811.5 | 222.1 | - | - | 0 | - |
| - | - | 1946 | 222.1 | - | - | 0 | - |
| - | - | 731.8 | 223.2 | - | - | 0 | - |
| - | - | 1097 | 224.1 | - | - | 0 | - |
| - | - | 4071 | 224.2 | - | - | 0 | - |
| - | - | 853.1 | 225.1 | - | - | 0 | - |
| - | - | 1238 | 225.2 | - | - | 0 | - |
| - | - | 1180 | 226.1 | - | - | 0 | - |
| - | - | 7.75E+04 | 226.2 | - | - | 0 | - |
| - | - | 1151 | 227.1 | - | - | 0 | - |
| 11 | y | 9197 | 227.2 | 0.001587 | 6.984 | +2 | 4 |
| - | - | 846.1 | 227.2 | - | - | 0 | - |
| - | - | 648.9 | 227.2 | - | - | 0 | - |
| 2 | a | 5495 | 228.1 | 0.0004327 | 1.897 | +1 | 2 |
| - | - | 1160 | 228.1 | - | - | 0 | - |
| 5 | b | 6725 | 229.1 | 0.001813 | 7.912 | +3 | 5 |
| - | - | 1498 | 229.1 | - | - | 0 | - |
| - | - | 1917 | 229.2 | - | - | 0 | - |
| - | - | 676.2 | 229.2 | - | - | 0 | - |
| - | - | 1443 | 230.1 | - | - | 0 | - |
| - | - | 734.7 | 230.1 | - | - | 0 | - |
| - | - | 1083 | 230.2 | - | - | 0 | - |
| - | - | 3603 | 231.1 | - | - | 0 | - |
| - | - | 659.5 | 231.1 | - | - | 0 | - |
| - | - | 1014 | 231.2 | - | - | 0 | - |
| - | - | 674.4 | 232.1 | - | - | 0 | - |
| - | - | 1070 | 233.1 | - | - | 0 | - |
| - | - | 492.2 | 234.1 | - | - | 0 | - |
| - | - | 632.4 | 234.1 | - | - | 0 | - |
| - | - | 865 | 234.1 | - | - | 0 | - |
| - | - | 1.359E+04 | 235.1 | - | - | 0 | - |
| 11 | y | 1.297E+04 | 235.7 | 0.000519 | 2.202 | +2 | 4 |
| - | - | 1509 | 236.1 | - | - | 0 | - |
| - | - | 3949 | 236.2 | - | - | 0 | - |
| - | - | 878.9 | 236.7 | - | - | 0 | - |
| - | - | 802.3 | 237.1 | - | - | 0 | - |
| - | - | 772.1 | 238.1 | - | - | 0 | - |
| - | - | 755.9 | 239.1 | - | - | 0 | - |
| - | - | 1.284E+04 | 239.2 | - | - | 0 | - |
| - | - | 826.6 | 240.1 | - | - | 0 | - |
| - | - | 1503 | 240.2 | - | - | 0 | - |
| - | - | 802.1 | 241.2 | - | - | 0 | - |
| - | - | 2327 | 242.2 | - | - | 0 | - |
| - | - | 1196 | 243.1 | - | - | 0 | - |
| - | - | 677.1 | 243.2 | - | - | 0 | - |
| - | - | 576 | 244.1 | - | - | 0 | - |
| - | - | 615.7 | 244.2 | - | - | 0 | - |
| 13 | y | 1.701E+04 | 245.2 | 0.0006001 | 2.448 | +1 | 2 |
| 2 | a | 1.442E+05 | 246.1 | 0.0005949 | 2.417 | +1 | 2 |
| - | - | 2442 | 246.2 | - | - | 0 | - |
| - | - | 840.8 | 247.1 | - | - | 0 | - |
| - | - | 3.341E+04 | 247.1 | - | - | 0 | - |
| - | - | 1.195E+04 | 248.1 | - | - | 0 | - |
| - | - | 3545 | 248.1 | - | - | 0 | - |
| - | - | 1.461E+04 | 249.1 | - | - | 0 | - |
| 8 | b | 1419 | 249.1 | 0.004181 | 16.79 | +4 | 8 |
| - | - | 1.057E+04 | 249.1 | - | - | 0 | - |
| - | - | 1200 | 250.1 | - | - | 0 | - |
| - | - | 1575 | 250.1 | - | - | 0 | - |
| - | - | 1239 | 250.1 | - | - | 0 | - |
| - | - | 1008 | 251.2 | - | - | 0 | - |
| - | - | 2050 | 252.2 | - | - | 0 | - |
| - | - | 743.3 | 255.1 | - | - | 0 | - |
| 2 | b | 2977 | 256.1 | 0.0004674 | 1.825 | +1 | 2 |
| - | - | 2386 | 257.1 | - | - | 0 | - |
| - | - | 1936 | 257.2 | - | - | 0 | - |
| - | - | 2255 | 258.1 | - | - | 0 | - |
| - | - | 2.66E+04 | 258.1 | - | - | 0 | - |
| - | - | 688.7 | 259.1 | - | - | 0 | - |
| - | - | 2911 | 259.1 | - | - | 0 | - |
| - | - | 1148 | 259.1 | - | - | 0 | - |
| - | - | 3273 | 260.1 | - | - | 0 | - |
| - | - | 1815 | 260.2 | - | - | 0 | - |
| - | - | 777.7 | 261.1 | - | - | 0 | - |
| - | - | 535.5 | 262.2 | - | - | 0 | - |
| - | - | 1211 | 263.1 | - | - | 0 | - |
| - | - | 1597 | 264.1 | - | - | 0 | - |
| - | - | 1110 | 265.1 | - | - | 0 | - |
| - | - | 5626 | 265.2 | - | - | 0 | - |
| - | - | 9414 | 266.1 | - | - | 0 | - |
| - | - | 929.8 | 267.1 | - | - | 0 | - |
| - | - | 2192 | 267.1 | - | - | 0 | - |
| - | - | 855.6 | 267.2 | - | - | 0 | - |
| - | - | 626 | 268.1 | - | - | 0 | - |
| - | - | 680.6 | 268.1 | - | - | 0 | - |
| - | - | 972.5 | 268.2 | - | - | 0 | - |
| - | - | 1094 | 269.6 | - | - | 0 | - |
| - | - | 7019 | 270.2 | - | - | 0 | - |
| 10 | y | 3111 | 271.2 | 0.0007802 | 2.877 | +2 | 5 |
| - | - | 1159 | 271.7 | - | - | 0 | - |
| 2 | b | 5.504E+04 | 274.1 | 0.0004923 | 1.796 | +1 | 2 |
| - | - | 6670 | 275.1 | - | - | 0 | - |
| - | - | 4.62E+04 | 275.1 | - | - | 0 | - |
| 7 | y | 583.1 | 275.2 | 0.0008789 | 3.194 | +3 | 8 |
| - | - | 2132 | 276.1 | - | - | 0 | - |
| - | - | 5316 | 276.1 | - | - | 0 | - |
| - | - | 2253 | 277.1 | - | - | 0 | - |
| - | - | 1076 | 279.2 | - | - | 0 | - |
| - | - | 4.381E+04 | 283.2 | - | - | 0 | - |
| - | - | 2321 | 283.6 | - | - | 0 | - |
| - | - | 704.7 | 284.1 | - | - | 0 | - |
| - | - | 5329 | 284.2 | - | - | 0 | - |
| - | - | 1163 | 285.1 | - | - | 0 | - |
| - | - | 4572 | 285.2 | - | - | 0 | - |
| - | - | 5815 | 290.2 | - | - | 0 | - |
| - | - | 1927 | 290.7 | - | - | 0 | - |
| - | - | 1890 | 291.2 | - | - | 0 | - |
| - | - | 1.439E+04 | 292.1 | - | - | 0 | - |
| - | - | 2719 | 293.1 | - | - | 0 | - |
| - | - | 3412 | 293.2 | - | - | 0 | - |
| - | - | 3794 | 294.1 | - | - | 0 | - |
| - | - | 736.5 | 294.2 | - | - | 0 | - |
| - | - | 1249 | 294.2 | - | - | 0 | - |
| - | - | 660 | 295.1 | - | - | 0 | - |
| - | - | 1034 | 297.2 | - | - | 0 | - |
| - | - | 1.02E+04 | 297.2 | - | - | 0 | - |
| - | - | 1511 | 298.2 | - | - | 0 | - |
| - | - | 692.2 | 300.1 | - | - | 0 | - |
| - | - | 2762 | 303.1 | - | - | 0 | - |
| - | - | 5934 | 305.1 | - | - | 0 | - |
| - | - | 1508 | 305.7 | - | - | 0 | - |
| - | - | 3570 | 309.2 | - | - | 0 | - |
| - | - | 1825 | 311.2 | - | - | 0 | - |
| - | - | 2677 | 311.2 | - | - | 0 | - |
| - | - | 722.3 | 312.2 | - | - | 0 | - |
| - | - | 1631 | 314.2 | - | - | 0 | - |
| - | - | 2478 | 314.7 | - | - | 0 | - |
| - | - | 674.1 | 315.2 | - | - | 0 | - |
| - | - | 1076 | 318.2 | - | - | 0 | - |
| - | - | 638.4 | 319.7 | - | - | 0 | - |
| - | - | 1085 | 320.1 | - | - | 0 | - |
| 6 | y | 1761 | 320.9 | 0.0005185 | 1.616 | +3 | 9 |
| - | - | 1269 | 321.2 | - | - | 0 | - |
| - | - | 1.21E+04 | 321.2 | - | - | 0 | - |
| - | - | 6560 | 322.2 | - | - | 0 | - |
| - | - | 1022 | 322.2 | - | - | 0 | - |
| - | - | 2093 | 322.2 | - | - | 0 | - |
| - | - | 572.4 | 322.5 | - | - | 0 | - |
| - | - | 1202 | 323 | - | - | 0 | - |
| 11 | b | 1196 | 323.2 | 0.0009063 | 2.804 | +4 | 11 |
| - | - | 1162 | 324.2 | - | - | 0 | - |
| - | - | 976.4 | 324.7 | - | - | 0 | - |
| - | - | 3380 | 326.2 | - | - | 0 | - |
| - | - | 791.9 | 326.7 | - | - | 0 | - |
| - | - | 2.68E+04 | 327.2 | - | - | 0 | - |
| - | - | 1446 | 327.2 | - | - | 0 | - |
| - | - | 570.3 | 327.3 | - | - | 0 | - |
| - | - | 3928 | 328.2 | - | - | 0 | - |
| - | - | 3295 | 332.7 | - | - | 0 | - |
| - | - | 1509 | 333.1 | - | - | 0 | - |
| - | - | 1196 | 333.2 | - | - | 0 | - |
| 9 | y | 1547 | 335.2 | 0.00194 | 5.786 | +2 | 6 |
| - | - | 679.4 | 335.7 | - | - | 0 | - |
| - | - | 1.89E+04 | 336.2 | - | - | 0 | - |
| - | - | 2376 | 337.2 | - | - | 0 | - |
| - | - | 2642 | 337.2 | - | - | 0 | - |
| - | - | 5076 | 337.7 | - | - | 0 | - |
| - | - | 1820 | 338.2 | - | - | 0 | - |
| - | - | 724.1 | 338.2 | - | - | 0 | - |
| - | - | 1.857E+04 | 339.2 | - | - | 0 | - |
| - | - | 1502 | 340.2 | - | - | 0 | - |
| - | - | 2722 | 340.2 | - | - | 0 | - |
| - | - | 612.5 | 341.2 | - | - | 0 | - |
| - | - | 963.3 | 342.2 | - | - | 0 | - |
| - | - | 598.2 | 345 | - | - | 0 | - |
| - | - | 782.8 | 346.7 | - | - | 0 | - |
| - | - | 1.804E+04 | 346.7 | - | - | 0 | - |
| - | - | 6507 | 347.2 | - | - | 0 | - |
| - | - | 1405 | 347.7 | - | - | 0 | - |
| - | - | 1132 | 349.7 | - | - | 0 | - |
| - | - | 2678 | 350.2 | - | - | 0 | - |
| - | - | 1388 | 350.2 | - | - | 0 | - |
| - | - | 1476 | 351.2 | - | - | 0 | - |
| - | - | 4261 | 353.2 | - | - | 0 | - |
| - | - | 1990 | 354.2 | - | - | 0 | - |
| - | - | 1993 | 354.2 | - | - | 0 | - |
| - | - | 2.24E+04 | 354.2 | - | - | 0 | - |
| - | - | 806.8 | 354.5 | - | - | 0 | - |
| - | - | 614.8 | 354.7 | - | - | 0 | - |
| - | - | 3727 | 355.2 | - | - | 0 | - |
| - | - | 3010 | 355.2 | - | - | 0 | - |
| - | - | 638.4 | 355.3 | - | - | 0 | - |
| - | - | 2164 | 355.7 | - | - | 0 | - |
| - | - | 603 | 356.2 | - | - | 0 | - |
| - | - | 804.5 | 356.2 | - | - | 0 | - |
| - | - | 666.3 | 356.2 | - | - | 0 | - |
| - | - | 1010 | 357.2 | - | - | 0 | - |
| - | - | 631.3 | 357.2 | - | - | 0 | - |
| - | - | 4675 | 357.3 | - | - | 0 | - |
| - | - | 5222 | 358.2 | - | - | 0 | - |
| - | - | 1315 | 358.3 | - | - | 0 | - |
| - | - | 2389 | 358.7 | - | - | 0 | - |
| - | - | 1497 | 359.2 | - | - | 0 | - |
| - | - | 693.9 | 359.2 | - | - | 0 | - |
| - | - | 890.6 | 359.5 | - | - | 0 | - |
| - | - | 1678 | 359.9 | - | - | 0 | - |
| - | - | 622.4 | 360.2 | - | - | 0 | - |
| - | - | 1625 | 360.7 | - | - | 0 | - |
| - | - | 926.2 | 360.7 | - | - | 0 | - |
| - | - | 1141 | 361.2 | - | - | 0 | - |
| - | - | 805.2 | 362.7 | - | - | 0 | - |
| - | - | 553.2 | 363.2 | - | - | 0 | - |
| - | - | 1791 | 363.5 | - | - | 0 | - |
| 8 | y | 1529 | 363.7 | 0.0004547 | 1.25 | +2 | 7 |
| - | - | 639.7 | 364.2 | - | - | 0 | - |
| - | - | 3314 | 366.2 | - | - | 0 | - |
| 5 | y | 1960 | 366.6 | 0.001165 | 3.179 | +3 | 10 |
| - | - | 2362 | 366.9 | - | - | 0 | - |
| - | - | 1786 | 367.2 | - | - | 0 | - |
| - | - | 2646 | 368.1 | - | - | 0 | - |
| - | - | 855.7 | 369.2 | - | - | 0 | - |
| - | - | 1561 | 370.2 | - | - | 0 | - |
| - | - | 2.82E+05 | 371.2 | - | - | 0 | - |
| - | - | 4.815E+04 | 372.2 | - | - | 0 | - |
| 12 | y | 1.049E+04 | 373.3 | 0.0006841 | 1.833 | +1 | 3 |
| - | - | 1741 | 374.3 | - | - | 0 | - |
| - | - | 638.5 | 376.5 | - | - | 0 | - |
| - | - | 2149 | 379 | - | - | 0 | - |
| - | - | 891.1 | 380.2 | - | - | 0 | - |
| - | - | 8166 | 381.2 | - | - | 0 | - |
| - | - | 4435 | 382.2 | - | - | 0 | - |
| - | - | 1154 | 382.3 | - | - | 0 | - |
| - | - | 2444 | 382.5 | - | - | 0 | - |
| - | - | 1170 | 382.9 | - | - | 0 | - |
| - | - | 919.9 | 383.3 | - | - | 0 | - |
| - | - | 3568 | 384.2 | - | - | 0 | - |
| - | - | 9193 | 385.2 | - | - | 0 | - |
| - | - | 2086 | 385.2 | - | - | 0 | - |
| - | - | 1.214E+04 | 386.2 | - | - | 0 | - |
| - | - | 1558 | 387.2 | - | - | 0 | - |
| - | - | 745.2 | 388.2 | - | - | 0 | - |
| - | - | 2713 | 388.9 | - | - | 0 | - |
| - | - | 1495 | 389.2 | - | - | 0 | - |
| - | - | 1851 | 389.3 | - | - | 0 | - |
| - | - | 825.5 | 389.8 | - | - | 0 | - |
| 2 | y | 7004 | 390.2 | 0.0006831 | 1.751 | +4 | 13 |
| - | - | 3720 | 390.5 | - | - | 0 | - |
| - | - | 1506 | 390.7 | - | - | 0 | - |
| - | - | 1840 | 391 | - | - | 0 | - |
| - | - | 2674 | 392.3 | - | - | 0 | - |
| 10 | b | 989.3 | 392.5 | 0.00439 | 11.18 | +3 | 10 |
| - | - | 709.1 | 393 | - | - | 0 | - |
| - | - | 1206 | 393.2 | - | - | 0 | - |
| - | - | 913.8 | 393.3 | - | - | 0 | - |
| - | - | 720.9 | 393.5 | - | - | 0 | - |
| - | - | 6663 | 394.2 | - | - | 0 | - |
| 2 | y | 2883 | 394.5 | 0.0003019 | 0.7654 | +4 | 13 |
| - | - | 2932 | 394.7 | - | - | 0 | - |
| - | - | 1427 | 395 | - | - | 0 | - |
| - | - | 1388 | 395.2 | - | - | 0 | - |
| - | - | 1224 | 396.3 | - | - | 0 | - |
| - | - | 1261 | 397.2 | - | - | 0 | - |
| - | - | 781.7 | 397.9 | - | - | 0 | - |
| 10 | b | 1630 | 398.2 | 0.0006244 | 1.568 | +3 | 10 |
| - | - | 1006 | 398.2 | - | - | 0 | - |
| - | - | 1425 | 398.5 | - | - | 0 | - |
| - | - | 1.93E+05 | 399.2 | - | - | 0 | - |
| - | - | 3.535E+04 | 400.2 | - | - | 0 | - |
| - | - | 735.7 | 400.7 | - | - | 0 | - |
| - | - | 1171 | 401.8 | - | - | 0 | - |
| - | - | 3427 | 402.2 | - | - | 0 | - |
| - | - | 913.1 | 402.9 | - | - | 0 | - |
| - | - | 1.015E+04 | 403.2 | - | - | 0 | - |
| - | - | 4066 | 403.2 | - | - | 0 | - |
| - | - | 2049 | 403.3 | - | - | 0 | - |
| 4 | y | 7204 | 403.6 | 0.003032 | 7.513 | +3 | 11 |
| 7 | y | 1248 | 403.8 | 0.007733 | 19.15 | +2 | 8 |
| - | - | 4006 | 403.9 | - | - | 0 | - |
| - | - | 1930 | 404.2 | - | - | 0 | - |
| - | - | 2023 | 404.2 | - | - | 0 | - |
| - | - | 824.9 | 404.6 | - | - | 0 | - |
| - | - | 3010 | 404.7 | - | - | 0 | - |
| - | - | 1530 | 405 | - | - | 0 | - |
| - | - | 594 | 405.2 | - | - | 0 | - |
| - | - | 1011 | 405.2 | - | - | 0 | - |
| - | - | 651.2 | 405.7 | - | - | 0 | - |
| - | - | 3514 | 406.2 | - | - | 0 | - |
| - | - | 2575 | 406.3 | - | - | 0 | - |
| - | - | 2305 | 406.7 | - | - | 0 | - |
| - | - | 2311 | 406.8 | - | - | 0 | - |
| - | - | 738.5 | 407.7 | - | - | 0 | - |
| - | - | 3244 | 408.2 | - | - | 0 | - |
| - | - | 1942 | 408.6 | - | - | 0 | - |
| - | - | 1323 | 408.9 | - | - | 0 | - |
| 4 | y | 2316 | 409.2 | 0.0008553 | 2.09 | +3 | 11 |
| - | - | 877.8 | 409.6 | - | - | 0 | - |
| - | - | 702.8 | 409.9 | - | - | 0 | - |
| - | - | 7396 | 410.3 | - | - | 0 | - |
| - | - | 4232 | 410.6 | - | - | 0 | - |
| - | - | 3964 | 410.9 | - | - | 0 | - |
| 6 | b | 9830 | 411.2 | 0.000754 | 1.834 | +2 | 6 |
| - | - | 1371 | 411.2 | - | - | 0 | - |
| - | - | 1854 | 411.3 | - | - | 0 | - |
| 0 | Precursor | 1450 | 411.7 | 0.0004253 | 1.033 | +4 | -1 |
| 0 | Precursor | 1972 | 412 | 0.002075 | 5.038 | +4 | -1 |
| - | - | 1385 | 412.2 | - | - | 0 | - |
| - | - | 1591 | 412.2 | - | - | 0 | - |
| 7 | y | 3.231E+04 | 412.3 | 0.0008367 | 2.029 | +2 | 8 |
| - | - | 1.507E+04 | 412.8 | - | - | 0 | - |
| - | - | 1180 | 413.2 | - | - | 0 | - |
| - | - | 2640 | 413.3 | - | - | 0 | - |
| - | - | 997.2 | 413.8 | - | - | 0 | - |
| - | - | 4492 | 413.9 | - | - | 0 | - |
| - | - | 740.9 | 414.2 | - | - | 0 | - |
| - | - | 5038 | 414.2 | - | - | 0 | - |
| - | - | 2154 | 414.6 | - | - | 0 | - |
| - | - | 649.4 | 414.9 | - | - | 0 | - |
| - | - | 1220 | 415 | - | - | 0 | - |
| - | - | 3654 | 415.3 | - | - | 0 | - |
| - | - | 1384 | 415.8 | - | - | 0 | - |
| 0 | Precursor | 8390 | 416.2 | 0.001084 | 2.604 | +4 | -1 |
| - | - | 4.316E+04 | 416.3 | - | - | 0 | - |
| - | - | 4170 | 416.3 | - | - | 0 | - |
| - | - | 1.568E+04 | 416.5 | - | - | 0 | - |
| - | - | 8736 | 416.7 | - | - | 0 | - |
| - | - | 2120 | 417 | - | - | 0 | - |
| - | - | 1301 | 417.2 | - | - | 0 | - |
| - | - | 1.013E+04 | 417.3 | - | - | 0 | - |
| - | - | 866.9 | 417.3 | - | - | 0 | - |
| - | - | 735.5 | 418.7 | - | - | 0 | - |
| - | - | 1.534E+04 | 419.9 | - | - | 0 | - |
| 6 | b | 2509 | 420.2 | 0.0005681 | 1.352 | +2 | 6 |
| - | - | 1.709E+04 | 420.2 | - | - | 0 | - |
| - | - | 4226 | 420.6 | - | - | 0 | - |
| - | - | 1221 | 420.7 | - | - | 0 | - |
| - | - | 2525 | 420.9 | - | - | 0 | - |
| - | - | 2544 | 422.3 | - | - | 0 | - |
| - | - | 1765 | 423.3 | - | - | 0 | - |
| - | - | 715.7 | 424.2 | - | - | 0 | - |
| - | - | 1761 | 424.3 | - | - | 0 | - |
| - | - | 3991 | 424.7 | - | - | 0 | - |
| - | - | 1965 | 425.2 | - | - | 0 | - |
| - | - | 988.3 | 425.3 | - | - | 0 | - |
| - | - | 4707 | 425.9 | - | - | 0 | - |
| - | - | 2205 | 426.2 | - | - | 0 | - |
| - | - | 3679 | 427.2 | - | - | 0 | - |
| - | - | 1747 | 427.7 | - | - | 0 | - |
| - | - | 3.094E+04 | 429.2 | - | - | 0 | - |
| - | - | 6708 | 430.2 | - | - | 0 | - |
| - | - | 4533 | 431.2 | - | - | 0 | - |
| - | - | 1540 | 433.7 | - | - | 0 | - |
| 3 | b | 1541 | 437.2 | 0.0002447 | 0.5597 | +1 | 3 |
| - | - | 939.9 | 438.2 | - | - | 0 | - |
| - | - | 804.5 | 438.2 | - | - | 0 | - |
| - | - | 2627 | 438.6 | - | - | 0 | - |
| - | - | 1755 | 438.9 | - | - | 0 | - |
| - | - | 1275 | 439.2 | - | - | 0 | - |
| - | - | 923.6 | 442.2 | - | - | 0 | - |
| - | - | 943.7 | 442.6 | - | - | 0 | - |
| - | - | 902.1 | 444.2 | - | - | 0 | - |
| - | - | 739.3 | 444.9 | - | - | 0 | - |
| - | - | 873.6 | 445.7 | - | - | 0 | - |
| - | - | 2382 | 446.2 | - | - | 0 | - |
| - | - | 1755 | 446.7 | - | - | 0 | - |
| - | - | 4464 | 448.3 | - | - | 0 | - |
| - | - | 4192 | 448.6 | - | - | 0 | - |
| - | - | 3643 | 451.2 | - | - | 0 | - |
| - | - | 2040 | 451.3 | - | - | 0 | - |
| - | - | 3638 | 451.7 | - | - | 0 | - |
| - | - | 7767 | 451.9 | - | - | 0 | - |
| - | - | 820 | 452.2 | - | - | 0 | - |
| - | - | 4052 | 452.3 | - | - | 0 | - |
| - | - | 1160 | 452.3 | - | - | 0 | - |
| - | - | 2430 | 452.6 | - | - | 0 | - |
| - | - | 1609 | 452.9 | - | - | 0 | - |
| - | - | 1388 | 454.2 | - | - | 0 | - |
| - | - | 595 | 455.2 | - | - | 0 | - |
| - | - | 3305 | 455.3 | - | - | 0 | - |
| - | - | 995.9 | 457.2 | - | - | 0 | - |
| - | - | 4060 | 457.6 | - | - | 0 | - |
| 3 | y | 5396 | 457.9 | 0.003844 | 8.395 | +3 | 12 |
| - | - | 4827 | 458.3 | - | - | 0 | - |
| - | - | 991 | 458.6 | - | - | 0 | - |
| - | - | 1512 | 458.9 | - | - | 0 | - |
| - | - | 5217 | 460.2 | - | - | 0 | - |
| - | - | 4115 | 460.7 | - | - | 0 | - |
| - | - | 1481 | 461.2 | - | - | 0 | - |
| - | - | 687.3 | 463.3 | - | - | 0 | - |
| 3 | y | 8.732E+04 | 463.6 | 0.0009047 | 1.951 | +3 | 12 |
| - | - | 7.404E+04 | 463.9 | - | - | 0 | - |
| - | - | 2.706E+04 | 464.3 | - | - | 0 | - |
| - | - | 8965 | 464.6 | - | - | 0 | - |
| - | - | 1082 | 464.9 | - | - | 0 | - |
| - | - | 2658 | 466.9 | - | - | 0 | - |
| - | - | 1046 | 467.3 | - | - | 0 | - |
| - | - | 2592 | 467.3 | - | - | 0 | - |
| - | - | 1798 | 468.3 | - | - | 0 | - |
| - | - | 1901 | 469.8 | - | - | 0 | - |
| - | - | 1749 | 470.2 | - | - | 0 | - |
| 11 | y | 3.635E+04 | 470.3 | 0.0008377 | 1.781 | +1 | 4 |
| - | - | 821.1 | 470.6 | - | - | 0 | - |
| - | - | 1123 | 470.9 | - | - | 0 | - |
| - | - | 694.1 | 471.2 | - | - | 0 | - |
| - | - | 1.002E+04 | 471.3 | - | - | 0 | - |
| - | - | 2630 | 471.8 | - | - | 0 | - |
| - | - | 1034 | 472.3 | - | - | 0 | - |
| - | - | 1816 | 472.3 | - | - | 0 | - |
| - | - | 4875 | 473.3 | - | - | 0 | - |
| - | - | 1639 | 473.6 | - | - | 0 | - |
| - | - | 1159 | 474.3 | - | - | 0 | - |
| - | - | 1873 | 474.8 | - | - | 0 | - |
| - | - | 1877 | 475.3 | - | - | 0 | - |
| - | - | 1219 | 475.9 | - | - | 0 | - |
| - | - | 7501 | 476.3 | - | - | 0 | - |
| - | - | 7502 | 476.6 | - | - | 0 | - |
| - | - | 2224 | 476.9 | - | - | 0 | - |
| - | - | 1041 | 478.2 | - | - | 0 | - |
| 6 | y | 7149 | 480.8 | 0.0009218 | 1.917 | +2 | 9 |
| - | - | 3954 | 481.3 | - | - | 0 | - |
| - | - | 4276 | 481.9 | - | - | 0 | - |
| - | - | 4872 | 482.3 | - | - | 0 | - |
| - | - | 2980 | 482.3 | - | - | 0 | - |
| - | - | 1810 | 482.6 | - | - | 0 | - |
| - | - | 2838 | 482.8 | - | - | 0 | - |
| - | - | 2164 | 483.3 | - | - | 0 | - |
| - | - | 745.2 | 483.3 | - | - | 0 | - |
| - | - | 5283 | 483.8 | - | - | 0 | - |
| - | - | 3387 | 484.3 | - | - | 0 | - |
| - | - | 1560 | 484.8 | - | - | 0 | - |
| - | - | 1451 | 487.9 | - | - | 0 | - |
| - | - | 3578 | 491.3 | - | - | 0 | - |
| - | - | 1793 | 492.8 | - | - | 0 | - |
| - | - | 1184 | 493.8 | - | - | 0 | - |
| - | - | 929.2 | 495.9 | - | - | 0 | - |
| 8 | b | 1498 | 497.2 | 0.0005637 | 1.134 | +2 | 8 |
| - | - | 830.7 | 501.6 | - | - | 0 | - |
| - | - | 964.7 | 502.3 | - | - | 0 | - |
| - | - | 2936 | 503.2 | - | - | 0 | - |
| - | - | 798.8 | 504.2 | - | - | 0 | - |
| - | - | 716.7 | 504.9 | - | - | 0 | - |
| 13 | b | 1468 | 505.3 | 0.004692 | 9.286 | +3 | 13 |
| - | - | 932.3 | 508.8 | - | - | 0 | - |
| - | - | 1913 | 509.2 | - | - | 0 | - |
| - | - | 1129 | 509.7 | - | - | 0 | - |
| - | - | 1370 | 510.6 | - | - | 0 | - |
| 13 | b | 4267 | 510.9 | 7.418E-05 | 0.1452 | +3 | 13 |
| - | - | 3000 | 511.3 | - | - | 0 | - |
| - | - | 3088 | 511.6 | - | - | 0 | - |
| - | - | 1559 | 514.3 | - | - | 0 | - |
| - | - | 7742 | 516.9 | - | - | 0 | - |
| - | - | 7097 | 517.3 | - | - | 0 | - |
| - | - | 3241 | 517.6 | - | - | 0 | - |
| - | - | 1750 | 517.8 | - | - | 0 | - |
| - | - | 1322 | 517.9 | - | - | 0 | - |
| - | - | 1050 | 518.3 | - | - | 0 | - |
| - | - | 704.5 | 519.6 | - | - | 0 | - |
| 2 | y | 7691 | 519.9 | 0.0009355 | 1.799 | +3 | 13 |
| - | - | 6564 | 520.3 | - | - | 0 | - |
| - | - | 2088 | 520.4 | - | - | 0 | - |
| - | - | 2098 | 520.6 | - | - | 0 | - |
| - | - | 905.5 | 520.9 | - | - | 0 | - |
| - | - | 2200 | 521.2 | - | - | 0 | - |
| 2 | y | 1.191E+04 | 525.6 | 0.000529 | 1.006 | +3 | 13 |
| - | - | 842 | 525.8 | - | - | 0 | - |
| - | - | 1.188E+04 | 526 | - | - | 0 | - |
| - | - | 6054 | 526.3 | - | - | 0 | - |
| - | - | 765.3 | 526.3 | - | - | 0 | - |
| - | - | 3473 | 526.6 | - | - | 0 | - |
| - | - | 1779 | 530.3 | - | - | 0 | - |
| - | - | 844.1 | 530.8 | - | - | 0 | - |
| - | - | 1681 | 530.8 | - | - | 0 | - |
| - | - | 2408 | 531.2 | - | - | 0 | - |
| - | - | 1098 | 532.2 | - | - | 0 | - |
| - | - | 1001 | 533.8 | - | - | 0 | - |
| - | - | 1502 | 535.8 | - | - | 0 | - |
| - | - | 2147 | 536.3 | - | - | 0 | - |
| - | - | 1.844E+04 | 538.3 | - | - | 0 | - |
| - | - | 1449 | 538.4 | - | - | 0 | - |
| - | - | 3448 | 538.8 | - | - | 0 | - |
| - | - | 5840 | 539.3 | - | - | 0 | - |
| - | - | 4542 | 539.3 | - | - | 0 | - |
| - | - | 2458 | 539.8 | - | - | 0 | - |
| 5 | y | 877.2 | 540.8 | 0.009948 | 18.39 | +2 | 10 |
| 10 | y | 7029 | 541.4 | 0.0008943 | 1.652 | +1 | 5 |
| - | - | 3272 | 542.3 | - | - | 0 | - |
| - | - | 2409 | 542.4 | - | - | 0 | - |
| - | - | 907.5 | 543.3 | - | - | 0 | - |
| - | - | 1234 | 544.3 | - | - | 0 | - |
| - | - | 2410 | 544.8 | - | - | 0 | - |
| - | - | 1688 | 545.3 | - | - | 0 | - |
| - | - | 812.4 | 545.8 | - | - | 0 | - |
| - | - | 748.6 | 546.3 | - | - | 0 | - |
| - | - | 2183 | 547.8 | - | - | 0 | - |
| - | - | 707.3 | 548.2 | - | - | 0 | - |
| - | - | 2045 | 548.3 | - | - | 0 | - |
| - | - | 2156 | 549.2 | - | - | 0 | - |
| 5 | y | 2.016E+04 | 549.3 | 0.0005186 | 0.944 | +2 | 10 |
| - | - | 1.073E+04 | 549.8 | - | - | 0 | - |
| - | - | 939.7 | 550.2 | - | - | 0 | - |
| - | - | 3755 | 550.3 | - | - | 0 | - |
| - | - | 1056 | 550.8 | - | - | 0 | - |
| - | - | 2151 | 553.2 | - | - | 0 | - |
| - | - | 1100 | 553.3 | - | - | 0 | - |
| - | - | 1150 | 555.3 | - | - | 0 | - |
| - | - | 2677 | 557.3 | - | - | 0 | - |
| 9 | b | 6250 | 561.3 | 0.0005981 | 1.066 | +2 | 9 |
| - | - | 3116 | 561.8 | - | - | 0 | - |
| - | - | 1135 | 562.3 | - | - | 0 | - |
| - | - | 2.337E+04 | 566.2 | - | - | 0 | - |
| - | - | 7667 | 567.2 | - | - | 0 | - |
| - | - | 925.7 | 567.4 | - | - | 0 | - |
| - | - | 1410 | 568.3 | - | - | 0 | - |
| - | - | 9045 | 570.2 | - | - | 0 | - |
| - | - | 4512 | 571.2 | - | - | 0 | - |
| - | - | 1.143E+04 | 572.8 | - | - | 0 | - |
| - | - | 6753 | 573.3 | - | - | 0 | - |
| - | - | 1301 | 573.8 | - | - | 0 | - |
| - | - | 7230 | 577.4 | - | - | 0 | - |
| - | - | 2490 | 578.4 | - | - | 0 | - |
| - | - | 1.452E+04 | 579.4 | - | - | 0 | - |
| - | - | 3770 | 580.4 | - | - | 0 | - |
| - | - | 695.1 | 581.2 | - | - | 0 | - |
| - | - | 853.9 | 582.8 | - | - | 0 | - |
| - | - | 729 | 587.3 | - | - | 0 | - |
| 10 | b | 1538 | 588.3 | 0.005142 | 8.741 | +2 | 10 |
| - | - | 4357 | 595.4 | - | - | 0 | - |
| - | - | 958 | 596.3 | - | - | 0 | - |
| - | - | 1696 | 596.4 | - | - | 0 | - |
| 10 | b | 1147 | 596.8 | 0.001235 | 2.07 | +2 | 10 |
| - | - | 863.6 | 597.4 | - | - | 0 | - |
| - | - | 5379 | 598.2 | - | - | 0 | - |
| - | - | 2178 | 599.2 | - | - | 0 | - |
| - | - | 819.1 | 600.3 | - | - | 0 | - |
| - | - | 897.6 | 601.8 | - | - | 0 | - |
| - | - | 4641 | 604.4 | - | - | 0 | - |
| 4 | y | 7076 | 604.8 | 0.003975 | 6.571 | +2 | 11 |
| - | - | 5190 | 605.4 | - | - | 0 | - |
| - | - | 1782 | 605.9 | - | - | 0 | - |
| - | - | 852.3 | 606.4 | - | - | 0 | - |
| - | - | 815.7 | 606.8 | - | - | 0 | - |
| - | - | 3842 | 610.3 | - | - | 0 | - |
| - | - | 675.5 | 611.3 | - | - | 0 | - |
| 4 | y | 3893 | 613.4 | 0.0002215 | 0.3612 | +2 | 11 |
| - | - | 637.9 | 613.4 | - | - | 0 | - |
| - | - | 3743 | 613.9 | - | - | 0 | - |
| - | - | 1340 | 614.4 | - | - | 0 | - |
| - | - | 1332 | 615.3 | - | - | 0 | - |
| - | - | 2683 | 615.4 | - | - | 0 | - |
| - | - | 1554 | 615.8 | - | - | 0 | - |
| - | - | 894.7 | 616.4 | - | - | 0 | - |
| - | - | 3776 | 620.3 | - | - | 0 | - |
| - | - | 4243 | 620.8 | - | - | 0 | - |
| - | - | 1827 | 621.3 | - | - | 0 | - |
| - | - | 706.2 | 621.4 | - | - | 0 | - |
| - | - | 1155 | 623.4 | - | - | 0 | - |
| - | - | 1899 | 628.3 | - | - | 0 | - |
| - | - | 9540 | 629.3 | - | - | 0 | - |
| - | - | 6243 | 629.8 | - | - | 0 | - |
| - | - | 2444 | 630.3 | - | - | 0 | - |
| - | - | 977.8 | 630.8 | - | - | 0 | - |
| - | - | 4992 | 638.4 | - | - | 0 | - |
| - | - | 5566 | 638.9 | - | - | 0 | - |
| - | - | 1904 | 639.4 | - | - | 0 | - |
| - | - | 1289 | 639.9 | - | - | 0 | - |
| - | - | 754.2 | 657.3 | - | - | 0 | - |
| - | - | 1726 | 657.3 | - | - | 0 | - |
| - | - | 1061 | 657.8 | - | - | 0 | - |
| - | - | 3861 | 664.5 | - | - | 0 | - |
| - | - | 1270 | 665.5 | - | - | 0 | - |
| - | - | 1927 | 665.8 | - | - | 0 | - |
| - | - | 1226 | 666.3 | - | - | 0 | - |
| - | - | 1858 | 668.3 | - | - | 0 | - |
| 9 | y | 3553 | 669.5 | 0.0005969 | 0.8915 | +1 | 6 |
| - | - | 1735 | 670.5 | - | - | 0 | - |
| - | - | 1467 | 674.3 | - | - | 0 | - |
| - | - | 1.537E+04 | 674.4 | - | - | 0 | - |
| - | - | 1319 | 675.3 | - | - | 0 | - |
| - | - | 5940 | 675.4 | - | - | 0 | - |
| - | - | 1320 | 676.4 | - | - | 0 | - |
| - | - | 780.3 | 680.3 | - | - | 0 | - |
| 5 | b | 924.6 | 684.3 | 0.002511 | 3.669 | +1 | 5 |
| - | - | 863.5 | 690.3 | - | - | 0 | - |
| - | - | 1.37E+04 | 692.4 | - | - | 0 | - |
| - | - | 5547 | 693.4 | - | - | 0 | - |
| 3 | y | 3783 | 694.9 | 0.0003758 | 0.5409 | +2 | 12 |
| - | - | 4203 | 695.4 | - | - | 0 | - |
| - | - | 1511 | 695.9 | - | - | 0 | - |
| - | - | 1473 | 698.4 | - | - | 0 | - |
| 5 | b | 2894 | 702.3 | 0.001651 | 2.35 | +1 | 5 |
| - | - | 1570 | 703.3 | - | - | 0 | - |
| - | - | 4357 | 707.3 | - | - | 0 | - |
| - | - | 2831 | 708.3 | - | - | 0 | - |
| - | - | 699.8 | 708.5 | - | - | 0 | - |
| - | - | 718.6 | 709.5 | - | - | 0 | - |
| - | - | 1013 | 709.9 | - | - | 0 | - |
| - | - | 638.9 | 710.3 | - | - | 0 | - |
| - | - | 3053 | 710.5 | - | - | 0 | - |
| - | - | 1895 | 711.5 | - | - | 0 | - |
| - | - | 1620 | 713.9 | - | - | 0 | - |
| - | - | 1347 | 714.4 | - | - | 0 | - |
| - | - | 706.7 | 714.9 | - | - | 0 | - |
| - | - | 2737 | 716.4 | - | - | 0 | - |
| - | - | 1354 | 717.4 | - | - | 0 | - |
| - | - | 1606 | 720.3 | - | - | 0 | - |
| - | - | 1047 | 720.4 | - | - | 0 | - |
| - | - | 2164 | 721.4 | - | - | 0 | - |
| - | - | 1058 | 722.9 | - | - | 0 | - |
| - | - | 2771 | 724.3 | - | - | 0 | - |
| - | - | 1149 | 725.3 | - | - | 0 | - |
| 8 | y | 6539 | 726.5 | 0.0004811 | 0.6623 | +1 | 7 |
| - | - | 3012 | 727.5 | - | - | 0 | - |
| - | - | 5802 | 735.3 | - | - | 0 | - |
| - | - | 2238 | 736.3 | - | - | 0 | - |
| - | - | 743.7 | 738.4 | - | - | 0 | - |
| - | - | 804.2 | 739.4 | - | - | 0 | - |
| - | - | 2278 | 752.3 | - | - | 0 | - |
| - | - | 1627 | 753.3 | - | - | 0 | - |
| - | - | 870.2 | 777.5 | - | - | 0 | - |
| - | - | 2126 | 805.5 | - | - | 0 | - |
| 7 | y | 1120 | 806.5 | 0.009803 | 12.15 | +1 | 8 |
| - | - | 1741 | 811.4 | - | - | 0 | - |
| - | - | 1665 | 811.5 | - | - | 0 | - |
| - | - | 889.4 | 812.5 | - | - | 0 | - |
| 7 | y | 2.532E+04 | 823.5 | 0.0001608 | 0.1952 | +1 | 8 |
| - | - | 1.09E+04 | 824.5 | - | - | 0 | - |
| - | - | 2407 | 825.5 | - | - | 0 | - |
| - | - | 1244 | 829.5 | - | - | 0 | - |
| 6 | b | 2394 | 839.4 | 0.00117 | 1.394 | +1 | 6 |
| - | - | 1555 | 840.4 | - | - | 0 | - |
| - | - | 3523 | 848.4 | - | - | 0 | - |
| - | - | 1224 | 849.4 | - | - | 0 | - |
| - | - | 1198 | 853.5 | - | - | 0 | - |
| - | - | 677.5 | 889.4 | - | - | 0 | - |
| - | - | 734.2 | 892.5 | - | - | 0 | - |
| - | - | 3654 | 901.4 | - | - | 0 | - |
| - | - | 3298 | 902.4 | - | - | 0 | - |
| - | - | 1802 | 919.5 | - | - | 0 | - |
| - | - | 1158 | 920.5 | - | - | 0 | - |
| - | - | 682 | 943.6 | - | - | 0 | - |
| 6 | y | 2162 | 960.6 | 0.0001478 | 0.1539 | +1 | 9 |
| - | - | 1614 | 961.6 | - | - | 0 | - |
| - | - | 1113 | 1034 | - | - | 0 | - |
| - | - | 818.4 | 1071 | - | - | 0 | - |
| - | - | 608.6 | 1082 | - | - | 0 | - |
| - | - | 624.5 | 1088 | - | - | 0 | - |
| 5 | y | 695 | 1098 | 0.00514 | 4.683 | +1 | 10 |
| - | - | 786.4 | 2745 | - | - | 0 | - |

m/z Charge Intensity FragmentType MassShift Position
120.0565414428711 0 480.40823
120.06602478027344 0 614.7225
120.08130645751953 0 1216.486
122.07160949707031 0 546.7702
123.0556640625 0 1835.1492
123.11750030517578 0 525.81146
126.10301208496094 0 436.29544
127.08711242675781 0 2943.5576
129.06634521484375 0 4139.621
129.10272216796875 0 93638.7
129.52960205078125 0 470.84543
130.0656280517578 0 21016.395
130.08656311035156 0 614.6619
130.10011291503906 0 1072.9133
130.10604858398438 0 6132.6577
131.04556274414062 0 3971.39
131.0704345703125 0 4806.166
131.08192443847656 0 1214.8688
132.08126831054688 0 81939.21
132.10235595703125 0 24049.17 y 13
132.10824584960938 0 791.9551
133.08639526367188 0 223187.36
134.07196044921875 0 499.40872
134.08973693847656 0 13262.066
135.10206604003906 0 1238.8623
136.07615661621094 0 109610.22
137.07322692871094 0 721.8521
137.07952880859375 0 9220.319
138.06655883789062 0 1211.3667
138.0807342529297 0 377.99808
139.0872344970703 0 1486.4816
140.08245849609375 0 714.941
141.1026611328125 0 5597.4756
142.06591796875 0 1332.7709 b 3
142.1056365966797 0 485.5298
143.08181762695312 0 541.0984
144.08135986328125 0 3999.911
145.0613250732422 0 808.82587
146.0605010986328 0 1539.2177
147.04449462890625 0 5352.3066
147.06500244140625 0 456.6437
151.0872344970703 0 1157.5001
151.0970458984375 0 1716.6251
152.0711669921875 0 605.27515
152.08187866210938 0 748.79803
152.1437530517578 0 1765.6797
153.10267639160156 0 660.2567
154.09805297851562 0 552.73584
155.0819549560547 0 4889.9795
155.09307861328125 0 1819.5137
155.11831665039062 0 3539.4126
156.07672119140625 0 1966.9788
156.08200073242188 0 2034.6641
157.0621795654297 0 573.9312
157.07647705078125 0 874.3823
157.08547973632812 0 531.06604
157.81053161621094 0 441.58588
158.06028747558594 0 1809.819
158.08433532714844 0 9200.637
159.09214782714844 0 55320.99
159.1128692626953 0 1140.6599
160.07620239257812 0 3437.6108
160.0955352783203 0 5881.832
161.08128356933594 0 8704.9
162.08494567871094 0 802.0774
162.10333251953125 0 472.05566
164.071044921875 0 1450.79
165.07765197753906 0 657.3367
165.10264587402344 0 3642.583
166.06141662597656 0 1385.5623
168.08241271972656 0 587.0223
168.11399841308594 0 1162.6056
169.0764923095703 0 5630.671
169.0976104736328 0 5976.813
170.0605010986328 0 27171.982
170.07923889160156 0 759.4148
170.10150146484375 0 544.7079
171.06390380859375 0 3177.1548
171.0921630859375 0 1083.1445
171.1128387451172 0 540.8088
172.10853576660156 0 4430.478
173.09249877929688 0 1099.2021
175.09719848632812 0 3900.7307
176.08230590820312 0 1788.555
177.1126708984375 0 95645.62
178.11593627929688 0 7713.218
178.13462829589844 0 533.61176
179.0829315185547 0 952.4488
179.092041015625 0 2212.0942
179.1284637451172 0 1077.8516
181.0611114501953 0 1880.263
181.09767150878906 0 25908.758
181.13389587402344 0 1970.1599
182.1010284423828 0 2883.9333
182.12928771972656 0 3358.4302
183.09217834472656 0 11091.711
183.11294555664062 0 1258.9675
183.14964294433594 0 605.3041
184.0762939453125 0 1398.7682
184.09571838378906 0 1587.1097
184.10853576660156 0 561.67725
185.0712432861328 0 3069.078
185.08041381835938 0 959.0034
185.1653289794922 0 1578.9242
186.07594299316406 0 450.1318
186.12423706054688 0 10023.43
187.08709716796875 0 60126.496
187.10794067382812 0 4410.782
187.12754821777344 0 724.53973
187.1448211669922 0 1824.6505 y 11
188.0820770263672 0 750.1665
188.09046936035156 0 7546.889
193.10794067382812 0 547.7029
195.12330627441406 0 4325.342
195.1494140625 0 792.7815
196.10877990722656 0 841.8066
197.10354614257812 0 529.0251
197.1653594970703 0 4125.5156
198.1248779296875 0 549.82355
199.08673095703125 0 2242.6265
199.10792541503906 0 1339.6663
199.1695556640625 0 2374.24
199.18096923828125 0 1611.1417
200.14002990722656 0 1505.7117
201.1026611328125 0 6654.9023
201.1238250732422 0 1696.3273
202.08660888671875 0 2945.8757
202.10885620117188 0 11052.314
203.09230041503906 0 697.5687
203.11326599121094 0 791.624
203.1399383544922 0 988.60004
204.07737731933594 0 620.4282
204.13473510742188 0 5224.7134
205.10751342773438 0 53758.246
206.11093139648438 0 4915.8823
207.1498565673828 0 2246.5554
209.09246826171875 0 1994.1833
209.12921142578125 0 1068.5146
211.08738708496094 0 1770.682
211.10696411132812 0 454.34622
212.1398162841797 0 679.8929
212.66815185546875 0 1712.899
213.0778350830078 0 651.74396
213.1600341796875 0 1460.7961
213.1703643798828 0 779.9441
214.10919189453125 0 2787.0513
215.1393585205078 0 1584.2094
216.11318969726562 0 838.75543
216.12376403808594 0 1428.4391
217.1085205078125 0 795.5712
219.12330627441406 0 1421.675
220.11981201171875 0 6924.0215
221.09225463867188 0 338.85986
221.10389709472656 0 10093.894
221.12570190429688 0 991.84076
221.1388702392578 0 20330.082
222.10682678222656 0 811.529
222.14231872558594 0 1945.6857
223.15554809570312 0 731.7871
224.11911010742188 0 1097.0801
224.1762237548828 0 4070.7197
225.0989990234375 0 853.0942
225.16009521484375 0 1237.6211
226.1201934814453 0 1180.3179
226.15553283691406 0 77495.13
227.1398468017578 0 1150.6499
227.15879821777344 0 9197.397 y Ammonia loss 10
227.1719970703125 0 846.1436
227.17652893066406 0 648.8549
228.1135711669922 0 5494.567 a Water loss 1
228.13427734375 0 1160.4076
229.09762573242188 0 6724.883 b Ammonia loss 4
229.11830139160156 0 1497.8219
229.15499877929688 0 1916.6167
229.1666717529297 0 676.1859
230.10281372070312 0 1443.4734
230.11460876464844 0 734.7232
230.15872192382812 0 1082.7225
231.08824157714844 0 3603.3306
231.13558959960938 0 659.52954
231.1714630126953 0 1014.3548
232.09182739257812 0 674.441
233.10389709472656 0 1070.0035
234.10572814941406 0 492.22812
234.12315368652344 0 632.41
234.13528442382812 0 864.9507
235.11952209472656 0 13594.857
235.67100524902344 0 12969.9795 y 10
236.12271118164062 0 1508.6322
236.17279052734375 0 3949.4373
236.67393493652344 0 878.91595
237.13516235351562 0 802.32074
238.1300811767578 0 772.10376
239.08184814453125 0 755.8744
239.15000915527344 0 12844.935
240.13453674316406 0 826.6016
240.15371704101562 0 1503.4286
241.15521240234375 0 802.0991
242.18670654296875 0 2326.8782
243.14633178710938 0 1196.3374
243.18153381347656 0 677.0667
244.05148315429688 0 576.01544
244.16598510742188 0 615.70276
245.1865692138672 0 17008.07 y 12
246.12429809570312 0 144208.19 a 1
246.189697265625 0 2441.8103
247.10691833496094 0 840.79193
247.1291046142578 0 33410.37
248.11459350585938 0 11949.349
248.132080078125 0 3545.3552
249.09873962402344 0 14613.48
249.11778259277344 0 1419.1793 b 7
249.1338348388672 0 10566.057
250.10311889648438 0 1199.6501
250.1229248046875 0 1574.5636
250.13697814941406 0 1238.9076
251.15057373046875 0 1007.91547
252.17115783691406 0 2049.6057
255.06539916992188 0 743.2587
256.1085205078125 0 2976.9604 b Water loss 1
257.1143798828125 0 2385.5325
257.16107177734375 0 1935.7104
258.07598876953125 0 2255.3252
258.0990295410156 0 26595.752
259.0799255371094 0 688.65936
259.1022033691406 0 2910.6743
259.130126953125 0 1148.263
260.10699462890625 0 3273.0957
260.1968688964844 0 1815.4001
261.1109313964844 0 777.71643
262.15411376953125 0 535.5149
263.1153869628906 0 1211.4285
264.13458251953125 0 1596.7191
265.1412353515625 0 1109.863
265.1652526855469 0 5625.815
266.1250915527344 0 9413.559
267.130126953125 0 929.81757
267.1454162597656 0 2191.824
267.1819152832031 0 855.62506
268.1303405761719 0 626.0327
268.1485900878906 0 680.6237
268.1654968261719 0 972.5039
269.6300354003906 0 1093.7177
270.1816101074219 0 7019.049
271.1882629394531 0 3110.708 y 9
271.69085693359375 0 1158.617
274.1191101074219 0 55041.113 b 1
275.1033020019531 0 6669.689
275.1251525878906 0 46198.94
275.18572998046875 0 583.0802 y 6
276.1074523925781 0 2132.0361
276.1282653808594 0 5315.705
277.133056640625 0 2252.612
279.1822509765625 0 1076.4828
283.1762390136719 0 43807.438
283.6279296875 0 2320.665
284.1297912597656 0 704.669
284.1795959472656 0 5329.0034
285.11004638671875 0 1163.1417
285.1562805175781 0 4571.5273
290.1852111816406 0 5815.1616
290.686767578125 0 1926.7555
291.1573181152344 0 1890.2377
292.1297912597656 0 14385.126
293.13348388671875 0 2719.0176
293.16033935546875 0 3412.289
294.1199035644531 0 3794.0103
294.1617431640625 0 736.5376
294.2181396484375 0 1249.39
295.12176513671875 0 659.9846
297.1556091308594 0 1033.5562
297.19281005859375 0 10200.423
298.1974182128906 0 1511.3673
300.1379699707031 0 692.1774
303.12054443359375 0 2761.5242
305.1291809082031 0 5933.652
305.6647644042969 0 1508.3298
309.1918640136719 0 3570.1462
311.1707458496094 0 1825.3053
311.2450866699219 0 2677.2534
312.2462158203125 0 722.25287
314.1830139160156 0 1631.2236
314.6695251464844 0 2478.3828
315.1717834472656 0 674.0586
318.1923828125 0 1076.4099
319.6802673339844 0 638.37146
320.1261291503906 0 1085.1229
320.8716735839844 0 1761.1362 y 5
321.207763671875 0 1269.3176
321.2291259765625 0 12095.351
322.15557861328125 0 6560.0825
322.1763916015625 0 1022.49744
322.2325134277344 0 2093.3447
322.4861145019531 0 572.3992
323.01910400390625 0 1202.3218
323.1589050292969 0 1195.863 b 10
324.21685791015625 0 1161.6104
324.71771240234375 0 976.43207
326.2195129394531 0 3379.9429
326.66363525390625 0 791.94946
327.2020568847656 0 26803.738
327.2225036621094 0 1446.1133
327.2742614746094 0 570.31055
328.20556640625 0 3927.6182
332.7298889160156 0 3294.8486
333.1250915527344 0 1508.5413
333.23052978515625 0 1195.7803
335.23846435546875 0 1547.1638 y 8
335.736328125 0 679.4483
336.20367431640625 0 18901.707
337.18621826171875 0 2376.2515
337.2078857421875 0 2642.2092
337.7216491699219 0 5075.6523
338.22113037109375 0 1820.4597
338.24407958984375 0 724.0511
339.2397155761719 0 18574.041
340.1531982421875 0 1502.248
340.24334716796875 0 2721.6133
341.2198791503906 0 612.5266
342.167724609375 0 963.27594
344.9772644042969 0 598.1762
346.6671142578125 0 782.78656
346.72686767578125 0 18038.77
347.2279357910156 0 6507.139
347.7300720214844 0 1405.1421
349.70989990234375 0 1132.2946
350.1506042480469 0 2678.14
350.2108459472656 0 1388.1432
351.2408142089844 0 1476.2368
353.21783447265625 0 4260.814
354.15679931640625 0 1990.4755
354.17474365234375 0 1992.5721
354.2143249511719 0 22398.303
354.5110778808594 0 806.8025
354.6587829589844 0 614.79816
355.1966552734375 0 3726.763
355.2172546386719 0 3009.516
355.2694091796875 0 638.35547
355.7314453125 0 2163.9229
356.16986083984375 0 602.9792
356.1975402832031 0 804.4909
356.23406982421875 0 666.2618
357.1788024902344 0 1010.1601
357.22991943359375 0 631.2583
357.25030517578125 0 4675.4634
358.1631774902344 0 5222.367
358.25335693359375 0 1315.4928
358.7138977050781 0 2388.6726
359.1663513183594 0 1496.67
359.1910705566406 0 693.8614
359.5439147949219 0 890.6221
359.8721008300781 0 1678.4215
360.2069091796875 0 622.446
360.6644287109375 0 1625.4934
360.68841552734375 0 926.24603
361.1849060058594 0 1140.9585
362.67120361328125 0 805.1924
363.1969909667969 0 553.161
363.5087890625 0 1791.1372
363.7477111816406 0 1529.4791 y 7
364.17498779296875 0 639.65027
366.15679931640625 0 3313.691
366.5586242675781 0 1960.0665 y 4
366.8921813964844 0 2362.4192
367.1627197265625 0 1786.4148
368.14825439453125 0 2646.3542
369.17706298828125 0 855.7382
370.1867370605469 0 1561.0697
371.22821044921875 0 282009.97
372.2315368652344 0 48146.84
373.2816162109375 0 10485.265 y 11
374.2841796875 0 1740.6704
376.5292053222656 0 638.4847
378.9637451171875 0 2148.6736
380.22821044921875 0 891.10333
381.2124328613281 0 8165.72
382.20709228515625 0 4435.0327
382.2817077636719 0 1154.1742
382.539794921875 0 2443.7546
382.87335205078125 0 1170.3564
383.26397705078125 0 919.8906
384.1673583984375 0 3568.4167
385.17352294921875 0 9193.437
385.198974609375 0 2086.1619
386.1577453613281 0 12138.841
387.16058349609375 0 1558.4484
388.2336120605469 0 745.2372
388.8616943359375 0 2713.2173
389.1963806152344 0 1494.5359
389.27093505859375 0 1850.6117
389.7725524902344 0 825.4908
390.2142639160156 0 7004.0723 y Ammonia loss 1
390.46514892578125 0 3720.262
390.7157897949219 0 1505.7338
390.9669189453125 0 1839.8357
392.2653503417969 0 2674.24
392.5219421386719 0 989.2998 b Ammonia loss 9
392.9688720703125 0 709.10406
393.16748046875 0 1206.4431
393.2699279785156 0 913.8216
393.46490478515625 0 720.8604
394.15167236328125 0 6663.096
394.47052001953125 0 2883.468 y 1
394.7222595214844 0 2931.7542
394.9729919433594 0 1426.8422
395.15582275390625 0 1388.06
396.2614440917969 0 1223.5057
397.24456787109375 0 1260.5074
397.8970947265625 0 781.74524
398.19244384765625 0 1630.2936 b 9
398.2411804199219 0 1005.92
398.52874755859375 0 1424.9325
399.2232971191406 0 192950.17
400.2266540527344 0 35348.773
400.72418212890625 0 735.6899
401.7604675292969 0 1171.014
402.2238464355469 0 3426.9133
402.88653564453125 0 913.05115
403.184326171875 0 10152.582
403.24072265625 0 4066.1108
403.2701721191406 0 2049.4126
403.5711669921875 0 7204.1772 y Ammonia loss 3
403.7680969238281 0 1247.7969 y Ammonia loss 6
403.9046630859375 0 4005.773
404.1866760253906 0 1930.488
404.2378234863281 0 2022.6279
404.5733947753906 0 824.89557
404.7255859375 0 3009.8828
404.9786376953125 0 1529.854
405.1920471191406 0 593.98157
405.2248840332031 0 1011.10535
405.7400817871094 0 651.2157
406.1861877441406 0 3513.9006
406.2511291503906 0 2575.0122
406.68792724609375 0 2305.0176
406.751953125 0 2311.0088
407.6916809082031 0 738.49774
408.2215270996094 0 3244.4272
408.5548400878906 0 1942.4651
408.8902282714844 0 1322.6455
409.2445068359375 0 2315.6626 y 3
409.57958984375 0 877.81573
409.9125671386719 0 702.8482
410.27691650390625 0 7395.92
410.56890869140625 0 4231.6343
410.9029541015625 0 3964.408
411.17828369140625 0 9830.162 b Water loss 5
411.23797607421875 0 1371.2258
411.2793273925781 0 1853.803
411.72515869140625 0 1449.8043 Precursor Water loss
411.9736633300781 0 1971.602 Precursor Ammonia loss
412.15875244140625 0 1385.195
412.1834716796875 0 1591.2063
412.27447509765625 0 32306.014 y 6
412.77587890625 0 15068.056
413.1656494140625 0 1180.3479
413.2767333984375 0 2639.5576
413.7775573730469 0 997.22626
413.8968505859375 0 4492.3877
414.1954650878906 0 740.9215
414.2289123535156 0 5037.9995
414.5609130859375 0 2154.13
414.8989562988281 0 649.4405
415.03753662109375 0 1220.0203
415.2592468261719 0 3653.8274
415.75897216796875 0 1384.3453
416.22930908203125 0 8389.555 Precursor
416.2500305175781 0 43162.836
416.2870178222656 0 4170.3228
416.4795227050781 0 15675.15
416.7299499511719 0 8736.127
416.9806823730469 0 2119.5505
417.2269592285156 0 1300.6006
417.25347900390625 0 10131.391
417.2880859375 0 866.8884
418.7111511230469 0 735.47943
419.9004211425781 0 15337.752
420.1833801269531 0 2508.5347 b 5
420.2349853515625 0 17085.088
420.5689392089844 0 4225.798
420.6858215332031 0 1220.9177
420.9042053222656 0 2524.6099
422.2772521972656 0 2544.0273
423.2744140625 0 1764.6166
424.22015380859375 0 715.6974
424.32879638671875 0 1760.7632
424.7123718261719 0 3991.0996
425.2134094238281 0 1965.1512
425.2860107421875 0 988.2659
425.9039611816406 0 4706.582
426.2379150390625 0 2204.519
427.2435607910156 0 3679.2764
427.7455749511719 0 1747.2653
429.1890563964844 0 30938.396
430.1921081542969 0 6707.5483
431.17987060546875 0 4532.7305
433.70703125 0 1539.9053
437.18170166015625 0 1541.3158 b 2
438.1872253417969 0 939.8921
438.2298889160156 0 804.4851
438.55718994140625 0 2626.7065
438.8913879394531 0 1755.0782
439.2237548828125 0 1275.288
442.187255859375 0 923.5798
442.58941650390625 0 943.7196
444.23406982421875 0 902.0683
444.90362548828125 0 739.29333
445.6941833496094 0 873.647
446.2323303222656 0 2382.4253
446.7350769042969 0 1754.7871
448.2634582519531 0 4463.5845
448.5990295410156 0 4191.8374
451.2257385253906 0 3642.962
451.26605224609375 0 2039.9769
451.72412109375 0 3637.8994
451.9199523925781 0 7766.647
452.22088623046875 0 819.9946
452.25341796875 0 4051.872
452.3255920410156 0 1160.4531
452.5874938964844 0 2429.5928
452.9220886230469 0 1608.5424
454.209716796875 0 1387.8102
455.2176208496094 0 595.0026
455.25213623046875 0 3305.0703
457.1848449707031 0 995.90375
457.5947570800781 0 4060.1626
457.9264221191406 0 5396.2197 y Ammonia loss 2
458.259521484375 0 4826.838
458.59393310546875 0 991.02515
458.92755126953125 0 1512.0359
460.23065185546875 0 5217.234
460.73199462890625 0 4115.444
461.240234375 0 1481.1416
463.28033447265625 0 687.2642
463.5989990234375 0 87315.43 y 2
463.9331970214844 0 74037.695
464.267578125 0 27056.201
464.60186767578125 0 8964.556
464.9367370605469 0 1081.8799
466.9195556640625 0 2657.824
467.2587585449219 0 1045.6456
467.2987365722656 0 2591.6
468.294677734375 0 1798.1736
469.78863525390625 0 1900.6566
470.2493591308594 0 1748.8048
470.33453369140625 0 36347.09 y 10
470.5832214355469 0 821.09564
470.9164123535156 0 1122.7078
471.24755859375 0 694.0678
471.337646484375 0 10020.93
471.7981872558594 0 2630.397
472.3013916015625 0 1034.4207
472.3400573730469 0 1816.37
473.2622375488281 0 4875.4624
473.5775146484375 0 1638.6373
474.2663269042969 0 1159.1328
474.7814025878906 0 1873.2023
475.28021240234375 0 1877.322
475.9248962402344 0 1219.4254
476.25390625 0 7500.6772
476.58782958984375 0 7502.1514
476.92059326171875 0 2224.424
478.2075500488281 0 1041.4028
480.80401611328125 0 7148.673 y 5
481.3052062988281 0 3954.4043
481.9273986816406 0 4275.8306
482.2614440917969 0 4871.6035
482.30908203125 0 2979.8647
482.5944519042969 0 1810.0568
482.76239013671875 0 2837.8499
483.26220703125 0 2163.9143
483.3117370605469 0 745.1765
483.7860412597656 0 5282.557
484.28680419921875 0 3387.237
484.78790283203125 0 1560.1831
487.92987060546875 0 1450.6577
491.2734375 0 3577.8643
492.7911376953125 0 1793.4336
493.7992248535156 0 1183.6552
495.9276428222656 0 929.2206
497.2204895019531 0 1497.5328 b 7
501.6069030761719 0 830.6979
502.2713623046875 0 964.72284
503.21588134765625 0 2935.7708
504.2150573730469 0 798.81177
504.9436950683594 0 716.73267
505.26617431640625 0 1468.304 b Ammonia loss 12
508.7578430175781 0 932.32684
509.23858642578125 0 1913.1779
509.7398986816406 0 1129.1288
510.6248779296875 0 1369.8312
510.93707275390625 0 4267.375 b 12
511.2714538574219 0 2999.507
511.6071472167969 0 3087.7375
514.2809448242188 0 1558.8617
516.9408569335938 0 7742.298
517.2758178710938 0 7097.204
517.6085205078125 0 3241.2534
517.7518310546875 0 1750.0658
517.94140625 0 1322.0815
518.2756958007812 0 1049.9609
519.6221923828125 0 704.5326
519.949951171875 0 7690.8813 y Ammonia loss 1
520.2843627929688 0 6564.0737
520.3609619140625 0 2088.4863
520.6187133789062 0 2097.9922
520.9497680664062 0 905.5444
521.225830078125 0 2200.2202
525.6250610351562 0 11911.833 y 1
525.80810546875 0 841.9654
525.9595336914062 0 11880.129
526.2935791015625 0 6054.3677
526.3316040039062 0 765.26953
526.627685546875 0 3473.4675
530.2987060546875 0 1779.4236
530.7564697265625 0 844.1065
530.8018188476562 0 1681.1464
531.210693359375 0 2408.185
532.2122802734375 0 1097.749
533.8189086914062 0 1000.5999
535.75341796875 0 1502.1134
536.252197265625 0 2146.8372
538.2525024414062 0 18441.83
538.3724365234375 0 1449.0161
538.8099975585938 0 3448.238
539.2545776367188 0 5840.4365
539.3032836914062 0 4541.5264
539.8035278320312 0 2457.6323
540.8292236328125 0 877.2276 y Ammonia loss 4
541.3717041015625 0 7029.3726 y 9
542.2518310546875 0 3272.0537
542.3733520507812 0 2409.2441
543.2555541992188 0 907.48486
544.2647705078125 0 1233.7076
544.7598876953125 0 2409.977
545.2598876953125 0 1687.7085
545.7588500976562 0 812.4102
546.255126953125 0 748.628
547.8133544921875 0 2183.0305
548.2341918945312 0 707.33936
548.3135375976562 0 2045.1105
549.2214965820312 0 2155.6948
549.3330688476562 0 20156.09 y 4
549.83447265625 0 10725.729
550.2225952148438 0 939.7108
550.3352661132812 0 3754.5984
550.8372192382812 0 1056.2554
553.2216186523438 0 2150.8535
553.2682495117188 0 1100.1741
555.2866821289062 0 1149.5409
557.2952270507812 0 2677.4941
561.2680053710938 0 6249.6226 b 8
561.76953125 0 3115.5833
562.2695922851562 0 1134.8711
566.2473754882812 0 23372.717
567.2499389648438 0 7667.197
567.3958740234375 0 925.6697
568.2516479492188 0 1410.2806
570.2464599609375 0 9045.477
571.249755859375 0 4512.1875
572.8045043945312 0 11433.276
573.3062133789062 0 6753.4727
573.8069458007812 0 1300.6361
577.382568359375 0 7230.493
578.3855590820312 0 2490.179
579.3619384765625 0 14520.474
580.3652954101562 0 3770.453
581.21533203125 0 695.05237
582.7874145507812 0 853.86053
587.2724609375 0 729.0014
588.27783203125 0 1537.7068 b Ammonia loss 9
595.3931884765625 0 4357.452
596.345458984375 0 957.9698
596.3949584960938 0 1695.9059
596.7847290039062 0 1146.9148 b 9
597.3668212890625 0 863.64276
598.2411499023438 0 5379.406
599.2448120117188 0 2177.7559
600.3352661132812 0 819.09143
601.8014526367188 0 897.587
604.355712890625 0 4640.8
604.8525390625 0 7076.466 y Ammonia loss 3
605.3524169921875 0 5189.976
605.8531494140625 0 1782.3752
606.3558349609375 0 852.26245
606.8362426757812 0 815.6999
610.32177734375 0 3841.8281
611.3284301757812 0 675.52
613.362060546875 0 3892.5562 y 3
613.4083251953125 0 637.92737
613.86328125 0 3743.0454
614.3632202148438 0 1340.2169
615.2670288085938 0 1332.2383
615.3507080078125 0 2683.2588
615.8489379882812 0 1553.619
616.3521728515625 0 894.6849
620.341552734375 0 3776.4062
620.8394775390625 0 4242.7266
621.3429565429688 0 1827.3428
621.4041137695312 0 706.1971
623.38818359375 0 1155.0776
628.3309936523438 0 1899.3992
629.3466186523438 0 9540.183
629.8473510742188 0 6243.4907
630.3477172851562 0 2443.8137
630.8479614257812 0 977.8337
638.3514404296875 0 4991.626
638.8536376953125 0 5565.777
639.356201171875 0 1904.3346
639.85888671875 0 1288.7217
657.2776489257812 0 754.22565
657.3335571289062 0 1725.5347
657.8328857421875 0 1060.7798
664.450439453125 0 3860.9255
665.452880859375 0 1270.3062
665.8443603515625 0 1927.215
666.3447875976562 0 1226.483
668.3271484375 0 1857.5402
669.4663696289062 0 3553.175 y 8
670.470703125 0 1735.0414
674.3016967773438 0 1467.1134
674.4356079101562 0 15372.095
675.30712890625 0 1318.7589
675.4359130859375 0 5939.7944
676.43994140625 0 1319.993
680.3187255859375 0 780.3122
684.2913818359375 0 924.5672 b Water loss 4
690.2798461914062 0 863.52765
692.4458618164062 0 13704.533
693.448486328125 0 5547.2104
694.8931274414062 0 3782.7202 y 2
695.394287109375 0 4202.8677
695.897216796875 0 1510.8202
698.4098510742188 0 1473.063
702.3010864257812 0 2894.3425 b 4
703.30419921875 0 1570.1067
707.3045043945312 0 4357.186
708.3082885742188 0 2830.8298
708.47998046875 0 699.8102
709.4776611328125 0 718.5925
709.8595581054688 0 1012.8565
710.3095092773438 0 638.89886
710.4577026367188 0 3053.4565
711.461669921875 0 1895.3992
713.8773193359375 0 1619.7848
714.381591796875 0 1347.1392
714.8719482421875 0 706.7322
716.421630859375 0 2736.7385
717.4231567382812 0 1353.9059
720.3191528320312 0 1605.91
720.3786010742188 0 1047.0637
721.3531494140625 0 2163.5063
722.8860473632812 0 1057.7681
724.3317260742188 0 2770.8662
725.3329467773438 0 1148.5365
726.4867553710938 0 6539.428 y 7
727.4918212890625 0 3011.62
735.3009033203125 0 5801.9785
736.3043212890625 0 2238.2507
738.3767700195312 0 743.6622
739.3721313476562 0 804.1697
752.3248901367188 0 2278.2505
753.3299560546875 0 1627.3119
777.5303344726562 0 870.1811
805.5289916992188 0 2125.9224
806.5232543945312 0 1120.034 y Ammonia loss 6
811.3619995117188 0 1740.6445
811.495361328125 0 1664.7903
812.4891967773438 0 889.40204
823.5401611328125 0 25316.432 y 6
824.543212890625 0 10896.506
825.5477294921875 0 2407.3232
829.502685546875 0 1244.113
839.357177734375 0 2394.2507 b 5
840.3631591796875 0 1555.0754
848.41552734375 0 3522.6145
849.4183959960938 0 1223.6924
853.4747924804688 0 1197.6139
889.3732299804688 0 677.47943
892.4641723632812 0 734.15753
901.44189453125 0 3653.6921
902.4444580078125 0 3298.1177
919.4537353515625 0 1801.7046
920.4508666992188 0 1157.6504
943.5943603515625 0 681.9963
960.5990600585938 0 2161.849 y 5
961.6031494140625 0 1613.6115
1033.531005859375 0 1112.9849
1071.48583984375 0 818.39166
1082.322021484375 0 608.62946
1087.526123046875 0 624.5279
1097.6629638671875 0 694.95746 y 4
2744.77734375 0 786.3673

Spectrum Details

|  |  |
| --- | --- |
| Matched peaks? Matched peaks The total absolute number of peaks matched. Additionally in brackets the total fraction of peaks matched and the total number of peaks is shown. | 61 (7.69% of 793) |
| FDR? FDR The false discovery rate estimated for this peptide. It is calculated by matching all theoretical fragments with a non-integer shift with the raw peaks for this spectrum. This is done with 40 different shifts. The resulting percentage is the average number of annotated peaks over the number of annotated peaks with the correct spectrum. | 4.02% |
| Satellite FDR? Satellite FDR See the FDR for details on its calculation. This satellite ion specific FDR only contains the satellite ions (d/w) for I/L/J positions. | - |
| PSM Score? PSM Score The PSM Score as given by Hecklib to this annotated spectrum. It is shown with three significant figures. | 221 |

#### Spectrum 5292? Spectrum 5292 The raw spectrum of this peptide as annotated by Hecklib. The fragments are coloured according to ion type (see legend). Any peaks with a star '\*' as text can be hovered over to see the full details, first the ion type second the mass shift type. By hovering over the amino acids in the peptide or ions in the legend the corresponding peaks are highlighted. By toggling the 'Unassigned' label you can turn the background (unassigned) peaks on or off in the plot. By updating the slider in the Ion legend you can update the spectrum to only show the top X% of the peaks with labels. The top X% means any peak that is within X% of the highest intensity. By dragging in the spectrum you can zoom in to a specific part of the spectrum and use 'Zoom Out' to get back to the original zoom level. The annotation of the spectrum is based on the given sequence in the peptides file and is done with different software so inconsistencies are likely. The peaks are annotated based on the given sequence, with 20 ppm tolerance.

Copy Data

##### Spectrum 5292 (TSV)

###### Preview

```
Loading example...
```

*Click on the button to copy the data to your clipboard.*

Mz MinMz MaxIntensity Max

WidthHeightPeptide font sizePeptide stroke widthSpectrum font sizeSpectrum stroke widthCompact peptide

Ion legend

wxyz

abcd

OtherUnassignedIon^Charge^_Position_Show for top:%

SWYQHHPGKAPKJJ

08.14e+51.63e+62.44e+63.26e+6

Zoom Out

y^+1^_1_b^+4^_4_b^+4^_5_a^+2^_3_y^+2^_4_a^+1^_2_b^+3^_5_y^+2^_4_y^+3^_7_y^+1^_2_a^+1^_2_b^+4^_8_b^+1^_2_y^+2^_5_b^+1^_2_y^+3^_8_y^+3^_9_y^+3^_9_b^+4^_11_y^+2^_6_b^+3^_8_y^+2^_6_y^+4^_12_b^+2^_5_y^+3^_10_y^+2^_7_y^+3^_10_y^+1^_3_b^+3^_9_b^+4^_13_y^+4^_13_b^+3^_10_b^+3^_10_y^+4^_13_b^+3^_10_y^+3^_11_y^+2^_8_y^+3^_11_b^+2^_6_\*\*y^+2^_8_\*b^+2^_6_b^+3^_11_b^+1^_3_y^+3^_12_y^+3^_12_y^+1^_4_y^+2^_9_b^+2^_8_b^+3^_13_b^+3^_13_b^+3^_13_y^+3^_13_y^+3^_13_y^+2^_10_y^+1^_5_y^+2^_10_b^+2^_9_b^+2^_9_b^+1^_4_b^+2^_10_b^+2^_10_b^+2^_10_y^+2^_11_y^+2^_11_y^+1^_6_b^+1^_5_b^+1^_5_y^+2^_12_y^+2^_12_b^+1^_5_b^+2^_12_y^+1^_7_y^+1^_8_b^+1^_6_y^+1^_8_b^+1^_6_y^+1^_9_b^+1^_8_y^+1^_10_

0871174126123482

Fragment Matches Table

Show background peaks

| Position | Ion type | Intensity | mz Theoretical | mz Error (Th) | mz Error (ppm) | Charge | Series Number |
| --- | --- | --- | --- | --- | --- | --- | --- |
| - | - | 8321 | 120.1 | - | - | 0 | - |
| - | - | 1.544E+04 | 120.1 | - | - | 0 | - |
| - | - | 3693 | 121 | - | - | 0 | - |
| - | - | 5925 | 122.1 | - | - | 0 | - |
| - | - | 1.422E+04 | 122.1 | - | - | 0 | - |
| - | - | 4.594E+04 | 123.1 | - | - | 0 | - |
| - | - | 3298 | 123.1 | - | - | 0 | - |
| - | - | 5.137E+04 | 127.1 | - | - | 0 | - |
| - | - | 3315 | 128 | - | - | 0 | - |
| - | - | 6128 | 128.1 | - | - | 0 | - |
| - | - | 4615 | 128.1 | - | - | 0 | - |
| - | - | 6.296E+04 | 129.1 | - | - | 0 | - |
| - | - | 1.896E+06 | 129.1 | - | - | 0 | - |
| - | - | 4.374E+05 | 130.1 | - | - | 0 | - |
| - | - | 1.482E+04 | 130.1 | - | - | 0 | - |
| - | - | 1.228E+05 | 130.1 | - | - | 0 | - |
| - | - | 3.691E+04 | 131.1 | - | - | 0 | - |
| - | - | 6397 | 131.1 | - | - | 0 | - |
| - | - | 5116 | 131.1 | - | - | 0 | - |
| - | - | 3543 | 132 | - | - | 0 | - |
| - | - | 1.808E+06 | 132.1 | - | - | 0 | - |
| 14 | y | 4.837E+05 | 132.1 | 0.0005119 | 3.875 | +1 | 1 |
| - | - | 1.662E+05 | 133.1 | - | - | 0 | - |
| - | - | 2.896E+04 | 133.1 | - | - | 0 | - |
| - | - | 1.236E+04 | 134.1 | - | - | 0 | - |
| - | - | 7837 | 134.1 | - | - | 0 | - |
| - | - | 2955 | 134.1 | - | - | 0 | - |
| - | - | 2.442E+06 | 136.1 | - | - | 0 | - |
| - | - | 3457 | 136.5 | - | - | 0 | - |
| - | - | 1.004E+04 | 137.1 | - | - | 0 | - |
| - | - | 2.028E+05 | 137.1 | - | - | 0 | - |
| - | - | 3.209E+04 | 138.1 | - | - | 0 | - |
| - | - | 6320 | 138.1 | - | - | 0 | - |
| - | - | 4724 | 138.1 | - | - | 0 | - |
| - | - | 2859 | 138.4 | - | - | 0 | - |
| - | - | 2.673E+04 | 139.1 | - | - | 0 | - |
| - | - | 1.202E+05 | 141.1 | - | - | 0 | - |
| 4 | b | 3.819E+04 | 142.1 | 0.000116 | 0.8168 | +4 | 4 |
| - | - | 8990 | 142.1 | - | - | 0 | - |
| - | - | 8.592E+04 | 144.1 | - | - | 0 | - |
| - | - | 7723 | 145.1 | - | - | 0 | - |
| - | - | 1.209E+04 | 145.1 | - | - | 0 | - |
| - | - | 9778 | 146.1 | - | - | 0 | - |
| - | - | 3564 | 146.1 | - | - | 0 | - |
| - | - | 1.157E+05 | 147 | - | - | 0 | - |
| - | - | 5438 | 147.1 | - | - | 0 | - |
| - | - | 8039 | 148 | - | - | 0 | - |
| - | - | 7958 | 148.1 | - | - | 0 | - |
| - | - | 5830 | 149.1 | - | - | 0 | - |
| - | - | 3625 | 150.1 | - | - | 0 | - |
| - | - | 1.962E+04 | 151.1 | - | - | 0 | - |
| - | - | 4242 | 151.9 | - | - | 0 | - |
| - | - | 4179 | 152.1 | - | - | 0 | - |
| - | - | 1.433E+04 | 152.1 | - | - | 0 | - |
| - | - | 5.015E+04 | 152.1 | - | - | 0 | - |
| - | - | 5862 | 153.1 | - | - | 0 | - |
| - | - | 9353 | 153.1 | - | - | 0 | - |
| - | - | 6655 | 154.1 | - | - | 0 | - |
| - | - | 9349 | 154.1 | - | - | 0 | - |
| - | - | 3572 | 154.2 | - | - | 0 | - |
| - | - | 1.025E+05 | 155.1 | - | - | 0 | - |
| - | - | 2.689E+04 | 155.1 | - | - | 0 | - |
| - | - | 6.173E+04 | 155.1 | - | - | 0 | - |
| - | - | 3.617E+04 | 156.1 | - | - | 0 | - |
| - | - | 4.9E+04 | 156.1 | - | - | 0 | - |
| - | - | 1.724E+04 | 157.1 | - | - | 0 | - |
| - | - | 6454 | 157.1 | - | - | 0 | - |
| - | - | 7128 | 157.1 | - | - | 0 | - |
| - | - | 3533 | 157.1 | - | - | 0 | - |
| - | - | 2.635E+04 | 158.1 | - | - | 0 | - |
| - | - | 2.135E+05 | 158.1 | - | - | 0 | - |
| - | - | 1.256E+06 | 159.1 | - | - | 0 | - |
| - | - | 6.292E+04 | 160.1 | - | - | 0 | - |
| - | - | 6379 | 160.1 | - | - | 0 | - |
| - | - | 1.279E+05 | 160.1 | - | - | 0 | - |
| - | - | 8853 | 161.1 | - | - | 0 | - |
| - | - | 5722 | 161.1 | - | - | 0 | - |
| - | - | 3018 | 161.4 | - | - | 0 | - |
| - | - | 4031 | 162.1 | - | - | 0 | - |
| - | - | 7464 | 162.1 | - | - | 0 | - |
| - | - | 3960 | 163.1 | - | - | 0 | - |
| - | - | 1.119E+04 | 163.6 | - | - | 0 | - |
| - | - | 8069 | 164 | - | - | 0 | - |
| - | - | 3.217E+04 | 164.1 | - | - | 0 | - |
| - | - | 1.084E+04 | 165.1 | - | - | 0 | - |
| - | - | 7.98E+04 | 165.1 | - | - | 0 | - |
| - | - | 2.648E+04 | 166.1 | - | - | 0 | - |
| - | - | 9036 | 166.1 | - | - | 0 | - |
| - | - | 3776 | 166.9 | - | - | 0 | - |
| - | - | 6801 | 167.2 | - | - | 0 | - |
| - | - | 7339 | 168.1 | - | - | 0 | - |
| - | - | 1.778E+04 | 168.1 | - | - | 0 | - |
| - | - | 3951 | 169 | - | - | 0 | - |
| - | - | 1.31E+05 | 169.1 | - | - | 0 | - |
| - | - | 1.294E+05 | 169.1 | - | - | 0 | - |
| - | - | 6752 | 169.2 | - | - | 0 | - |
| - | - | 6.147E+05 | 170.1 | - | - | 0 | - |
| - | - | 1.388E+04 | 170.1 | - | - | 0 | - |
| - | - | 9376 | 170.1 | - | - | 0 | - |
| - | - | 7.33E+04 | 171.1 | - | - | 0 | - |
| - | - | 2.182E+04 | 171.1 | - | - | 0 | - |
| - | - | 3769 | 172.1 | - | - | 0 | - |
| 5 | b | 7244 | 172.1 | 0.002325 | 13.51 | +4 | 5 |
| - | - | 1.207E+05 | 172.1 | - | - | 0 | - |
| - | - | 9067 | 172.1 | - | - | 0 | - |
| - | - | 4689 | 174.1 | - | - | 0 | - |
| - | - | 1.869E+04 | 175.1 | - | - | 0 | - |
| - | - | 4.612E+04 | 176.1 | - | - | 0 | - |
| - | - | 7772 | 179.1 | - | - | 0 | - |
| - | - | 7233 | 179.2 | - | - | 0 | - |
| - | - | 4393 | 179.6 | - | - | 0 | - |
| - | - | 7142 | 180.1 | - | - | 0 | - |
| - | - | 3.323E+04 | 181.1 | - | - | 0 | - |
| - | - | 5.821E+05 | 181.1 | - | - | 0 | - |
| - | - | 3.265E+04 | 181.1 | - | - | 0 | - |
| - | - | 6696 | 181.2 | - | - | 0 | - |
| - | - | 1.313E+04 | 182.1 | - | - | 0 | - |
| - | - | 6.016E+04 | 182.1 | - | - | 0 | - |
| - | - | 6.921E+04 | 182.1 | - | - | 0 | - |
| - | - | 2.599E+05 | 183.1 | - | - | 0 | - |
| - | - | 1.633E+04 | 183.1 | - | - | 0 | - |
| - | - | 7098 | 183.1 | - | - | 0 | - |
| - | - | 5550 | 183.1 | - | - | 0 | - |
| - | - | 3.388E+04 | 184.1 | - | - | 0 | - |
| - | - | 3.184E+04 | 184.1 | - | - | 0 | - |
| - | - | 6.591E+04 | 185.1 | - | - | 0 | - |
| - | - | 8721 | 185.1 | - | - | 0 | - |
| - | - | 1.1E+04 | 185.1 | - | - | 0 | - |
| - | - | 1.35E+04 | 186.1 | - | - | 0 | - |
| - | - | 2.482E+05 | 186.1 | - | - | 0 | - |
| - | - | 1.322E+06 | 187.1 | - | - | 0 | - |
| - | - | 1.941E+04 | 187.1 | - | - | 0 | - |
| - | - | 9333 | 188.1 | - | - | 0 | - |
| - | - | 1.672E+05 | 188.1 | - | - | 0 | - |
| - | - | 7904 | 189.1 | - | - | 0 | - |
| - | - | 4619 | 190.6 | - | - | 0 | - |
| - | - | 8641 | 191.1 | - | - | 0 | - |
| - | - | 9876 | 191.1 | - | - | 0 | - |
| - | - | 6131 | 191.1 | - | - | 0 | - |
| - | - | 6968 | 193.1 | - | - | 0 | - |
| - | - | 4069 | 193.1 | - | - | 0 | - |
| - | - | 9708 | 195.1 | - | - | 0 | - |
| - | - | 2.378E+04 | 195.1 | - | - | 0 | - |
| - | - | 1.05E+04 | 196.1 | - | - | 0 | - |
| - | - | 6438 | 197.1 | - | - | 0 | - |
| - | - | 4170 | 197.1 | - | - | 0 | - |
| - | - | 9.456E+04 | 197.2 | - | - | 0 | - |
| - | - | 4550 | 197.6 | - | - | 0 | - |
| - | - | 1.032E+04 | 198.1 | - | - | 0 | - |
| - | - | 8099 | 198.2 | - | - | 0 | - |
| - | - | 5.637E+04 | 199.1 | - | - | 0 | - |
| - | - | 9803 | 199.1 | - | - | 0 | - |
| - | - | 5761 | 199.2 | - | - | 0 | - |
| - | - | 3.939E+04 | 199.2 | - | - | 0 | - |
| - | - | 6605 | 200.1 | - | - | 0 | - |
| - | - | 3.179E+04 | 200.1 | - | - | 0 | - |
| - | - | 4922 | 200.2 | - | - | 0 | - |
| - | - | 1.482E+05 | 201.1 | - | - | 0 | - |
| - | - | 6690 | 201.1 | - | - | 0 | - |
| - | - | 5.687E+04 | 202.1 | - | - | 0 | - |
| - | - | 2.079E+05 | 202.1 | - | - | 0 | - |
| - | - | 1.726E+04 | 203.1 | - | - | 0 | - |
| - | - | 2.141E+04 | 203.1 | - | - | 0 | - |
| - | - | 2.352E+04 | 204.1 | - | - | 0 | - |
| 3 | a | 7014 | 205.1 | 0.001036 | 5.052 | +2 | 3 |
| - | - | 1.244E+04 | 205.1 | - | - | 0 | - |
| - | - | 9024 | 207.1 | - | - | 0 | - |
| - | - | 6.737E+04 | 207.1 | - | - | 0 | - |
| - | - | 1.302E+04 | 208.1 | - | - | 0 | - |
| - | - | 2.539E+04 | 209.1 | - | - | 0 | - |
| - | - | 9277 | 209.2 | - | - | 0 | - |
| - | - | 5885 | 210.1 | - | - | 0 | - |
| - | - | 3.341E+04 | 211.1 | - | - | 0 | - |
| - | - | 7700 | 212.1 | - | - | 0 | - |
| - | - | 1.833E+04 | 212.1 | - | - | 0 | - |
| - | - | 3970 | 212.2 | - | - | 0 | - |
| - | - | 4.701E+04 | 212.7 | - | - | 0 | - |
| - | - | 1.533E+04 | 213.1 | - | - | 0 | - |
| - | - | 1.192E+04 | 213.2 | - | - | 0 | - |
| - | - | 6.783E+04 | 214.1 | - | - | 0 | - |
| - | - | 4655 | 214.1 | - | - | 0 | - |
| - | - | 1.888E+04 | 214.2 | - | - | 0 | - |
| - | - | 6851 | 215.1 | - | - | 0 | - |
| - | - | 2.19E+04 | 216.1 | - | - | 0 | - |
| - | - | 1.959E+04 | 217.1 | - | - | 0 | - |
| - | - | 1.557E+05 | 220.1 | - | - | 0 | - |
| - | - | 2.223E+05 | 221.1 | - | - | 0 | - |
| - | - | 1.193E+04 | 221.1 | - | - | 0 | - |
| - | - | 2.424E+04 | 222.1 | - | - | 0 | - |
| - | - | 1.208E+04 | 222.1 | - | - | 0 | - |
| - | - | 5101 | 223.2 | - | - | 0 | - |
| - | - | 3.447E+04 | 224.1 | - | - | 0 | - |
| - | - | 8251 | 224.1 | - | - | 0 | - |
| - | - | 6048 | 224.2 | - | - | 0 | - |
| - | - | 1.017E+05 | 224.2 | - | - | 0 | - |
| - | - | 4501 | 225.1 | - | - | 0 | - |
| - | - | 4587 | 225.1 | - | - | 0 | - |
| - | - | 3.146E+04 | 225.2 | - | - | 0 | - |
| - | - | 1.083E+04 | 225.2 | - | - | 0 | - |
| - | - | 1.778E+06 | 226.2 | - | - | 0 | - |
| - | - | 7912 | 227.1 | - | - | 0 | - |
| - | - | 8408 | 227.1 | - | - | 0 | - |
| 11 | y | 2.136E+05 | 227.2 | 0.001846 | 8.126 | +2 | 4 |
| - | - | 8808 | 227.2 | - | - | 0 | - |
| 2 | a | 1.207E+05 | 228.1 | 0.0006158 | 2.699 | +1 | 2 |
| - | - | 1.171E+04 | 228.1 | - | - | 0 | - |
| - | - | 1.785E+04 | 228.2 | - | - | 0 | - |
| 5 | b | 1.432E+05 | 229.1 | 0.00195 | 8.511 | +3 | 5 |
| - | - | 1.895E+04 | 229.1 | - | - | 0 | - |
| - | - | 3959 | 229.1 | - | - | 0 | - |
| - | - | 2.292E+04 | 229.2 | - | - | 0 | - |
| - | - | 8097 | 230.1 | - | - | 0 | - |
| - | - | 3.928E+04 | 230.1 | - | - | 0 | - |
| - | - | 7741 | 231.1 | - | - | 0 | - |
| - | - | 8.173E+04 | 231.1 | - | - | 0 | - |
| - | - | 2.023E+04 | 231.1 | - | - | 0 | - |
| - | - | 1.114E+04 | 232.1 | - | - | 0 | - |
| - | - | 1.588E+04 | 232.1 | - | - | 0 | - |
| - | - | 6452 | 232.1 | - | - | 0 | - |
| - | - | 4.768E+04 | 233.1 | - | - | 0 | - |
| - | - | 1.262E+04 | 234.1 | - | - | 0 | - |
| - | - | 9854 | 234.1 | - | - | 0 | - |
| - | - | 2.988E+05 | 235.1 | - | - | 0 | - |
| 11 | y | 2.642E+05 | 235.7 | 0.0007174 | 3.044 | +2 | 4 |
| - | - | 3.237E+04 | 236.1 | - | - | 0 | - |
| - | - | 8017 | 236.1 | - | - | 0 | - |
| - | - | 6.569E+04 | 236.2 | - | - | 0 | - |
| - | - | 7457 | 236.7 | - | - | 0 | - |
| - | - | 8044 | 237.1 | - | - | 0 | - |
| 8 | y | 1.145E+04 | 237.2 | 0.00156 | 6.577 | +3 | 7 |
| - | - | 6305 | 237.6 | - | - | 0 | - |
| - | - | 2.336E+04 | 238.1 | - | - | 0 | - |
| - | - | 1.498E+04 | 239.1 | - | - | 0 | - |
| - | - | 1.174E+05 | 239.2 | - | - | 0 | - |
| - | - | 1.292E+04 | 240.1 | - | - | 0 | - |
| - | - | 1.396E+04 | 240.2 | - | - | 0 | - |
| - | - | 2.255E+04 | 241.2 | - | - | 0 | - |
| - | - | 6.408E+04 | 242.2 | - | - | 0 | - |
| - | - | 7489 | 243.2 | - | - | 0 | - |
| - | - | 2.018E+04 | 243.2 | - | - | 0 | - |
| - | - | 6115 | 244.1 | - | - | 0 | - |
| - | - | 1.468E+04 | 244.2 | - | - | 0 | - |
| - | - | 5104 | 245.2 | - | - | 0 | - |
| 13 | y | 4.03E+05 | 245.2 | 0.0007985 | 3.257 | +1 | 2 |
| 2 | a | 3.225E+06 | 246.1 | 0.0008085 | 3.285 | +1 | 2 |
| - | - | 5.346E+04 | 246.2 | - | - | 0 | - |
| - | - | 4061 | 246.6 | - | - | 0 | - |
| - | - | 7.394E+05 | 247.1 | - | - | 0 | - |
| - | - | 2.411E+05 | 248.1 | - | - | 0 | - |
| - | - | 6.61E+04 | 248.1 | - | - | 0 | - |
| - | - | 5373 | 248.2 | - | - | 0 | - |
| - | - | 3.238E+05 | 249.1 | - | - | 0 | - |
| 8 | b | 2.034E+04 | 249.1 | 0.003647 | 14.64 | +4 | 8 |
| - | - | 8693 | 249.1 | - | - | 0 | - |
| - | - | 3.508E+04 | 250.1 | - | - | 0 | - |
| - | - | 4.606E+04 | 250.1 | - | - | 0 | - |
| - | - | 2.327E+04 | 251.1 | - | - | 0 | - |
| - | - | 4262 | 251.9 | - | - | 0 | - |
| - | - | 5497 | 252.1 | - | - | 0 | - |
| - | - | 4.414E+04 | 252.2 | - | - | 0 | - |
| - | - | 4757 | 253.1 | - | - | 0 | - |
| - | - | 4106 | 253.2 | - | - | 0 | - |
| - | - | 5452 | 254.2 | - | - | 0 | - |
| - | - | 7487 | 254.2 | - | - | 0 | - |
| - | - | 1.012E+04 | 255.1 | - | - | 0 | - |
| - | - | 4755 | 255.2 | - | - | 0 | - |
| 2 | b | 7.086E+04 | 256.1 | 0.000681 | 2.659 | +1 | 2 |
| - | - | 9547 | 257.1 | - | - | 0 | - |
| - | - | 3.273E+04 | 257.1 | - | - | 0 | - |
| - | - | 4.044E+04 | 257.2 | - | - | 0 | - |
| - | - | 5.683E+04 | 258.1 | - | - | 0 | - |
| - | - | 5.884E+05 | 258.1 | - | - | 0 | - |
| - | - | 7503 | 259.1 | - | - | 0 | - |
| - | - | 7.827E+04 | 259.1 | - | - | 0 | - |
| - | - | 2.132E+04 | 259.1 | - | - | 0 | - |
| - | - | 9224 | 259.9 | - | - | 0 | - |
| - | - | 7.775E+04 | 260.1 | - | - | 0 | - |
| - | - | 4.004E+04 | 260.2 | - | - | 0 | - |
| - | - | 1.339E+04 | 261.1 | - | - | 0 | - |
| - | - | 6495 | 261.2 | - | - | 0 | - |
| - | - | 5397 | 261.2 | - | - | 0 | - |
| - | - | 5325 | 261.6 | - | - | 0 | - |
| - | - | 6288 | 262.1 | - | - | 0 | - |
| - | - | 1.38E+04 | 262.2 | - | - | 0 | - |
| - | - | 3949 | 263 | - | - | 0 | - |
| - | - | 1.486E+04 | 263.1 | - | - | 0 | - |
| - | - | 4847 | 263.1 | - | - | 0 | - |
| - | - | 7490 | 264.1 | - | - | 0 | - |
| - | - | 3.11E+04 | 264.1 | - | - | 0 | - |
| - | - | 2.098E+04 | 265.1 | - | - | 0 | - |
| - | - | 2.185E+04 | 265.2 | - | - | 0 | - |
| - | - | 6991 | 266.1 | - | - | 0 | - |
| - | - | 1.905E+05 | 266.1 | - | - | 0 | - |
| - | - | 2.459E+04 | 267.1 | - | - | 0 | - |
| - | - | 1.037E+04 | 267.1 | - | - | 0 | - |
| - | - | 4505 | 267.2 | - | - | 0 | - |
| - | - | 4858 | 268.1 | - | - | 0 | - |
| - | - | 4901 | 269.2 | - | - | 0 | - |
| - | - | 4.013E+04 | 269.6 | - | - | 0 | - |
| - | - | 1.572E+04 | 270.1 | - | - | 0 | - |
| - | - | 1.341E+05 | 270.2 | - | - | 0 | - |
| 10 | y | 8.984E+04 | 271.2 | 1.329E-05 | 0.04901 | +2 | 5 |
| - | - | 2.124E+04 | 271.7 | - | - | 0 | - |
| - | - | 6363 | 272.2 | - | - | 0 | - |
| - | - | 1.137E+04 | 273.1 | - | - | 0 | - |
| 2 | b | 1.176E+06 | 274.1 | 0.0007364 | 2.687 | +1 | 2 |
| - | - | 1.68E+05 | 275.1 | - | - | 0 | - |
| - | - | 9.938E+05 | 275.1 | - | - | 0 | - |
| 7 | y | 2.678E+04 | 275.2 | 0.0005127 | 1.863 | +3 | 8 |
| - | - | 1.914E+04 | 275.5 | - | - | 0 | - |
| - | - | 4.462E+04 | 276.1 | - | - | 0 | - |
| - | - | 1.084E+05 | 276.1 | - | - | 0 | - |
| - | - | 1.16E+04 | 277.1 | - | - | 0 | - |
| - | - | 5538 | 277.1 | - | - | 0 | - |
| - | - | 4536 | 277.1 | - | - | 0 | - |
| - | - | 4.593E+04 | 277.1 | - | - | 0 | - |
| - | - | 8056 | 278.1 | - | - | 0 | - |
| - | - | 6929 | 279.2 | - | - | 0 | - |
| - | - | 2.68E+04 | 279.2 | - | - | 0 | - |
| - | - | 4828 | 281.7 | - | - | 0 | - |
| - | - | 4375 | 282.2 | - | - | 0 | - |
| - | - | 3.086E+04 | 283.2 | - | - | 0 | - |
| - | - | 4.359E+05 | 283.2 | - | - | 0 | - |
| - | - | 3.637E+04 | 283.6 | - | - | 0 | - |
| - | - | 1.079E+04 | 284.1 | - | - | 0 | - |
| - | - | 7486 | 284.2 | - | - | 0 | - |
| - | - | 5.925E+04 | 284.2 | - | - | 0 | - |
| - | - | 2.399E+04 | 285.1 | - | - | 0 | - |
| - | - | 8.299E+04 | 285.2 | - | - | 0 | - |
| - | - | 5358 | 285.2 | - | - | 0 | - |
| - | - | 7909 | 286.1 | - | - | 0 | - |
| - | - | 8899 | 286.2 | - | - | 0 | - |
| - | - | 5807 | 288.1 | - | - | 0 | - |
| - | - | 8957 | 290.1 | - | - | 0 | - |
| - | - | 1.408E+05 | 290.2 | - | - | 0 | - |
| - | - | 3.322E+04 | 290.7 | - | - | 0 | - |
| - | - | 4.028E+04 | 291.2 | - | - | 0 | - |
| - | - | 6694 | 291.2 | - | - | 0 | - |
| - | - | 3.03E+05 | 292.1 | - | - | 0 | - |
| - | - | 4.53E+04 | 293.1 | - | - | 0 | - |
| - | - | 8199 | 293.2 | - | - | 0 | - |
| - | - | 7.971E+04 | 294.1 | - | - | 0 | - |
| - | - | 7918 | 294.1 | - | - | 0 | - |
| - | - | 3.301E+04 | 294.2 | - | - | 0 | - |
| - | - | 1.146E+04 | 295.1 | - | - | 0 | - |
| - | - | 6446 | 295.2 | - | - | 0 | - |
| - | - | 6749 | 296.2 | - | - | 0 | - |
| - | - | 2.264E+05 | 297.2 | - | - | 0 | - |
| - | - | 3.281E+04 | 298.2 | - | - | 0 | - |
| - | - | 9597 | 300.2 | - | - | 0 | - |
| - | - | 2.029E+04 | 300.7 | - | - | 0 | - |
| - | - | 7931 | 302.1 | - | - | 0 | - |
| - | - | 5.924E+04 | 303.1 | - | - | 0 | - |
| - | - | 8216 | 303.2 | - | - | 0 | - |
| - | - | 8878 | 304.1 | - | - | 0 | - |
| - | - | 4355 | 304.2 | - | - | 0 | - |
| - | - | 1.186E+05 | 305.1 | - | - | 0 | - |
| - | - | 4.729E+04 | 305.7 | - | - | 0 | - |
| - | - | 6551 | 305.9 | - | - | 0 | - |
| - | - | 2.742E+04 | 306.1 | - | - | 0 | - |
| - | - | 1.095E+04 | 306.2 | - | - | 0 | - |
| - | - | 8146 | 307.2 | - | - | 0 | - |
| - | - | 5055 | 308.1 | - | - | 0 | - |
| - | - | 9358 | 309.2 | - | - | 0 | - |
| - | - | 1.728E+04 | 309.2 | - | - | 0 | - |
| - | - | 5480 | 311.2 | - | - | 0 | - |
| - | - | 5164 | 311.2 | - | - | 0 | - |
| - | - | 7.075E+04 | 311.2 | - | - | 0 | - |
| - | - | 1.285E+04 | 312.2 | - | - | 0 | - |
| - | - | 4742 | 313.2 | - | - | 0 | - |
| - | - | 5380 | 314.2 | - | - | 0 | - |
| - | - | 4.614E+04 | 314.7 | - | - | 0 | - |
| - | - | 1.969E+04 | 315.2 | - | - | 0 | - |
| 6 | y | 5520 | 315.2 | 0.002207 | 7.001 | +3 | 9 |
| - | - | 4640 | 315.7 | - | - | 0 | - |
| - | - | 1.549E+04 | 318.2 | - | - | 0 | - |
| - | - | 2.184E+04 | 320.1 | - | - | 0 | - |
| 6 | y | 5.882E+04 | 320.9 | 0.0009763 | 3.043 | +3 | 9 |
| - | - | 2.293E+04 | 321.2 | - | - | 0 | - |
| - | - | 2.755E+05 | 321.2 | - | - | 0 | - |
| - | - | 8447 | 321.5 | - | - | 0 | - |
| - | - | 1.03E+04 | 321.8 | - | - | 0 | - |
| - | - | 1.93E+05 | 322.2 | - | - | 0 | - |
| - | - | 8276 | 322.2 | - | - | 0 | - |
| - | - | 4.852E+04 | 322.2 | - | - | 0 | - |
| - | - | 7167 | 322.9 | - | - | 0 | - |
| - | - | 5315 | 323.1 | - | - | 0 | - |
| 11 | b | 3.593E+04 | 323.2 | 0.0006621 | 2.049 | +4 | 11 |
| - | - | 8038 | 323.2 | - | - | 0 | - |
| - | - | 5965 | 323.2 | - | - | 0 | - |
| - | - | 5721 | 324.2 | - | - | 0 | - |
| - | - | 4.015E+04 | 324.2 | - | - | 0 | - |
| - | - | 1.313E+04 | 324.7 | - | - | 0 | - |
| - | - | 4614 | 325.1 | - | - | 0 | - |
| - | - | 6786 | 325.2 | - | - | 0 | - |
| - | - | 1.003E+04 | 325.2 | - | - | 0 | - |
| - | - | 1.053E+05 | 326.2 | - | - | 0 | - |
| 9 | y | 7156 | 326.7 | 0.001176 | 3.6 | +2 | 6 |
| - | - | 2.322E+04 | 327.2 | - | - | 0 | - |
| - | - | 2.401E+04 | 327.3 | - | - | 0 | - |
| - | - | 6905 | 330.1 | - | - | 0 | - |
| 8 | b | 5626 | 331.8 | 0.002009 | 6.055 | +3 | 8 |
| - | - | 1.164E+05 | 332.7 | - | - | 0 | - |
| - | - | 3.557E+04 | 333.1 | - | - | 0 | - |
| - | - | 2.652E+04 | 333.2 | - | - | 0 | - |
| - | - | 8871 | 334.1 | - | - | 0 | - |
| - | - | 1.084E+04 | 334.2 | - | - | 0 | - |
| - | - | 1.189E+04 | 334.7 | - | - | 0 | - |
| - | - | 6600 | 335.2 | - | - | 0 | - |
| 9 | y | 2.76E+04 | 335.2 | 0.0008106 | 2.418 | +2 | 6 |
| - | - | 9523 | 335.7 | - | - | 0 | - |
| - | - | 4.089E+05 | 336.2 | - | - | 0 | - |
| - | - | 8923 | 337.2 | - | - | 0 | - |
| - | - | 7.233E+04 | 337.2 | - | - | 0 | - |
| - | - | 5993 | 337.3 | - | - | 0 | - |
| - | - | 1.231E+05 | 337.7 | - | - | 0 | - |
| - | - | 6768 | 338.2 | - | - | 0 | - |
| - | - | 3.25E+04 | 338.2 | - | - | 0 | - |
| - | - | 9678 | 338.7 | - | - | 0 | - |
| - | - | 8559 | 339.2 | - | - | 0 | - |
| - | - | 3.959E+05 | 339.2 | - | - | 0 | - |
| - | - | 4348 | 339.8 | - | - | 0 | - |
| - | - | 3.674E+04 | 340.2 | - | - | 0 | - |
| - | - | 7.126E+04 | 340.2 | - | - | 0 | - |
| - | - | 7759 | 341.2 | - | - | 0 | - |
| - | - | 9007 | 341.2 | - | - | 0 | - |
| - | - | 1.114E+04 | 341.2 | - | - | 0 | - |
| - | - | 1.527E+04 | 342.2 | - | - | 0 | - |
| - | - | 6016 | 342.3 | - | - | 0 | - |
| - | - | 1.754E+04 | 345.6 | - | - | 0 | - |
| - | - | 1.15E+04 | 346.1 | - | - | 0 | - |
| - | - | 1.428E+04 | 346.7 | - | - | 0 | - |
| - | - | 3.43E+05 | 346.7 | - | - | 0 | - |
| - | - | 1.363E+05 | 347.2 | - | - | 0 | - |
| - | - | 4.339E+04 | 347.7 | - | - | 0 | - |
| 3 | y | 1.001E+04 | 348 | 0.001667 | 4.791 | +4 | 12 |
| - | - | 6789 | 348.1 | - | - | 0 | - |
| - | - | 5893 | 348.2 | - | - | 0 | - |
| - | - | 7626 | 349.1 | - | - | 0 | - |
| - | - | 5033 | 349.2 | - | - | 0 | - |
| - | - | 1.919E+04 | 349.7 | - | - | 0 | - |
| - | - | 4.449E+04 | 350.2 | - | - | 0 | - |
| - | - | 1.719E+04 | 350.2 | - | - | 0 | - |
| - | - | 6990 | 350.5 | - | - | 0 | - |
| - | - | 4434 | 350.7 | - | - | 0 | - |
| - | - | 7764 | 350.9 | - | - | 0 | - |
| - | - | 4956 | 351.1 | - | - | 0 | - |
| - | - | 8931 | 351.2 | - | - | 0 | - |
| - | - | 2.959E+04 | 351.2 | - | - | 0 | - |
| 5 | b | 5181 | 351.7 | 0.003596 | 10.23 | +2 | 5 |
| - | - | 5711 | 351.8 | - | - | 0 | - |
| - | - | 6413 | 352.2 | - | - | 0 | - |
| - | - | 5935 | 352.2 | - | - | 0 | - |
| - | - | 1.686E+04 | 353.2 | - | - | 0 | - |
| - | - | 6631 | 353.9 | - | - | 0 | - |
| - | - | 5.174E+04 | 354.2 | - | - | 0 | - |
| - | - | 2.405E+04 | 354.2 | - | - | 0 | - |
| - | - | 5.14E+05 | 354.2 | - | - | 0 | - |
| - | - | 1.08E+04 | 354.5 | - | - | 0 | - |
| - | - | 4998 | 354.5 | - | - | 0 | - |
| - | - | 2.687E+04 | 354.7 | - | - | 0 | - |
| - | - | 5155 | 354.8 | - | - | 0 | - |
| - | - | 4507 | 355.2 | - | - | 0 | - |
| - | - | 9.471E+04 | 355.2 | - | - | 0 | - |
| - | - | 1.07E+04 | 355.3 | - | - | 0 | - |
| - | - | 5.536E+04 | 355.7 | - | - | 0 | - |
| - | - | 1.255E+04 | 356.2 | - | - | 0 | - |
| - | - | 2.081E+04 | 356.2 | - | - | 0 | - |
| - | - | 8691 | 356.7 | - | - | 0 | - |
| - | - | 1.868E+04 | 357.2 | - | - | 0 | - |
| - | - | 1.179E+05 | 357.3 | - | - | 0 | - |
| - | - | 1.167E+04 | 357.5 | - | - | 0 | - |
| - | - | 1.976E+04 | 357.8 | - | - | 0 | - |
| - | - | 1.076E+05 | 358.2 | - | - | 0 | - |
| - | - | 1.917E+04 | 358.3 | - | - | 0 | - |
| - | - | 3.856E+04 | 358.7 | - | - | 0 | - |
| - | - | 5198 | 359.1 | - | - | 0 | - |
| - | - | 2.481E+04 | 359.2 | - | - | 0 | - |
| - | - | 9303 | 359.2 | - | - | 0 | - |
| - | - | 1.751E+04 | 359.2 | - | - | 0 | - |
| - | - | 2.622E+04 | 359.5 | - | - | 0 | - |
| - | - | 5800 | 359.7 | - | - | 0 | - |
| - | - | 5690 | 359.9 | - | - | 0 | - |
| - | - | 2.664E+04 | 359.9 | - | - | 0 | - |
| - | - | 1.106E+04 | 360.1 | - | - | 0 | - |
| - | - | 7161 | 360.2 | - | - | 0 | - |
| - | - | 3.394E+04 | 360.7 | - | - | 0 | - |
| - | - | 2.937E+04 | 360.7 | - | - | 0 | - |
| 5 | y | 6394 | 360.9 | 0.0009612 | 2.663 | +3 | 10 |
| - | - | 1.503E+04 | 361.2 | - | - | 0 | - |
| - | - | 1.051E+04 | 362.2 | - | - | 0 | - |
| - | - | 8425 | 362.7 | - | - | 0 | - |
| - | - | 7216 | 363.2 | - | - | 0 | - |
| - | - | 9661 | 363.2 | - | - | 0 | - |
| - | - | 3.201E+04 | 363.5 | - | - | 0 | - |
| 8 | y | 2.355E+04 | 363.7 | 0.0006988 | 1.921 | +2 | 7 |
| - | - | 1.493E+04 | 363.8 | - | - | 0 | - |
| - | - | 6568 | 364.2 | - | - | 0 | - |
| - | - | 7944 | 364.3 | - | - | 0 | - |
| - | - | 1.125E+04 | 365.3 | - | - | 0 | - |
| - | - | 1.347E+04 | 365.5 | - | - | 0 | - |
| - | - | 5638 | 365.9 | - | - | 0 | - |
| - | - | 5.302E+04 | 366.2 | - | - | 0 | - |
| - | - | 5269 | 366.3 | - | - | 0 | - |
| 5 | y | 6.438E+04 | 366.6 | 0.000616 | 1.68 | +3 | 10 |
| - | - | 2.886E+04 | 366.9 | - | - | 0 | - |
| - | - | 8381 | 367.1 | - | - | 0 | - |
| - | - | 4.457E+04 | 367.2 | - | - | 0 | - |
| - | - | 1.697E+04 | 367.2 | - | - | 0 | - |
| - | - | 5.921E+04 | 368.1 | - | - | 0 | - |
| - | - | 1.462E+04 | 368.2 | - | - | 0 | - |
| - | - | 1.753E+04 | 368.7 | - | - | 0 | - |
| - | - | 1.523E+04 | 369.2 | - | - | 0 | - |
| - | - | 1.918E+04 | 369.2 | - | - | 0 | - |
| - | - | 1.824E+04 | 369.7 | - | - | 0 | - |
| - | - | 2.518E+04 | 370.2 | - | - | 0 | - |
| - | - | 1.154E+04 | 370.2 | - | - | 0 | - |
| - | - | 4407 | 370.5 | - | - | 0 | - |
| - | - | 7577 | 370.7 | - | - | 0 | - |
| - | - | 6154 | 370.9 | - | - | 0 | - |
| - | - | 5576 | 373.2 | - | - | 0 | - |
| 12 | y | 2.442E+05 | 373.3 | 0.0009893 | 2.65 | +1 | 3 |
| - | - | 9796 | 374.2 | - | - | 0 | - |
| - | - | 8400 | 374.3 | - | - | 0 | - |
| - | - | 3.568E+04 | 374.3 | - | - | 0 | - |
| 9 | b | 9401 | 374.5 | 3.824E-05 | 0.1021 | +3 | 9 |
| - | - | 1.771E+04 | 374.8 | - | - | 0 | - |
| - | - | 1.29E+04 | 375.2 | - | - | 0 | - |
| - | - | 5543 | 375.3 | - | - | 0 | - |
| - | - | 1.067E+04 | 376.5 | - | - | 0 | - |
| - | - | 8147 | 376.5 | - | - | 0 | - |
| - | - | 1.64E+04 | 376.7 | - | - | 0 | - |
| - | - | 6011 | 376.9 | - | - | 0 | - |
| - | - | 5106 | 377 | - | - | 0 | - |
| - | - | 4991 | 377.1 | - | - | 0 | - |
| - | - | 8223 | 377.2 | - | - | 0 | - |
| - | - | 4731 | 377.5 | - | - | 0 | - |
| - | - | 5188 | 378.2 | - | - | 0 | - |
| - | - | 2.087E+04 | 378.7 | - | - | 0 | - |
| - | - | 2.657E+04 | 379 | - | - | 0 | - |
| - | - | 1.282E+04 | 379.2 | - | - | 0 | - |
| - | - | 7197 | 379.5 | - | - | 0 | - |
| - | - | 2.71E+04 | 380.2 | - | - | 0 | - |
| - | - | 5201 | 381.2 | - | - | 0 | - |
| - | - | 1.073E+05 | 382.2 | - | - | 0 | - |
| - | - | 2.619E+04 | 382.3 | - | - | 0 | - |
| - | - | 5.491E+04 | 382.5 | - | - | 0 | - |
| - | - | 6877 | 382.7 | - | - | 0 | - |
| - | - | 9636 | 382.9 | - | - | 0 | - |
| - | - | 7209 | 383.2 | - | - | 0 | - |
| - | - | 2.123E+04 | 383.3 | - | - | 0 | - |
| 13 | b | 7656 | 383.5 | 0.0001439 | 0.3752 | +4 | 13 |
| - | - | 7734 | 383.7 | - | - | 0 | - |
| - | - | 7.181E+04 | 384.2 | - | - | 0 | - |
| - | - | 2.15E+05 | 385.2 | - | - | 0 | - |
| - | - | 6.191E+04 | 385.2 | - | - | 0 | - |
| - | - | 6351 | 386 | - | - | 0 | - |
| - | - | 2.115E+05 | 386.2 | - | - | 0 | - |
| - | - | 1.691E+04 | 386.2 | - | - | 0 | - |
| - | - | 8932 | 386.5 | - | - | 0 | - |
| - | - | 8389 | 386.9 | - | - | 0 | - |
| - | - | 5.266E+04 | 387.2 | - | - | 0 | - |
| - | - | 7357 | 388.2 | - | - | 0 | - |
| - | - | 1.017E+04 | 388.2 | - | - | 0 | - |
| - | - | 6354 | 388.6 | - | - | 0 | - |
| - | - | 5.617E+04 | 388.9 | - | - | 0 | - |
| - | - | 4.1E+04 | 389.2 | - | - | 0 | - |
| - | - | 5.132E+04 | 389.3 | - | - | 0 | - |
| - | - | 6256 | 389.5 | - | - | 0 | - |
| - | - | 2.157E+04 | 389.8 | - | - | 0 | - |
| 2 | y | 1.592E+05 | 390.2 | 0.001049 | 2.689 | +4 | 13 |
| - | - | 5973 | 390.3 | - | - | 0 | - |
| - | - | 1.13E+05 | 390.5 | - | - | 0 | - |
| - | - | 3.776E+04 | 390.7 | - | - | 0 | - |
| - | - | 1.425E+04 | 391 | - | - | 0 | - |
| - | - | 6870 | 392.2 | - | - | 0 | - |
| 10 | b | 2.687E+04 | 392.2 | 0.0007304 | 1.862 | +3 | 10 |
| - | - | 6.903E+04 | 392.3 | - | - | 0 | - |
| 10 | b | 2.992E+04 | 392.5 | 0.005428 | 13.83 | +3 | 10 |
| - | - | 8660 | 392.8 | - | - | 0 | - |
| - | - | 1.58E+04 | 392.9 | - | - | 0 | - |
| - | - | 1.742E+04 | 393.2 | - | - | 0 | - |
| - | - | 9309 | 393.2 | - | - | 0 | - |
| - | - | 1.92E+04 | 393.3 | - | - | 0 | - |
| - | - | 1.692E+05 | 394.2 | - | - | 0 | - |
| 2 | y | 6.335E+04 | 394.5 | 0.0007902 | 2.003 | +4 | 13 |
| - | - | 6.022E+04 | 394.7 | - | - | 0 | - |
| - | - | 3.531E+04 | 395 | - | - | 0 | - |
| - | - | 3.066E+04 | 395.2 | - | - | 0 | - |
| - | - | 1.172E+04 | 395.2 | - | - | 0 | - |
| - | - | 5305 | 395.5 | - | - | 0 | - |
| - | - | 5498 | 396 | - | - | 0 | - |
| - | - | 5352 | 396.2 | - | - | 0 | - |
| - | - | 2.083E+04 | 396.3 | - | - | 0 | - |
| - | - | 9602 | 397.2 | - | - | 0 | - |
| - | - | 1.872E+04 | 397.2 | - | - | 0 | - |
| - | - | 1.438E+04 | 397.6 | - | - | 0 | - |
| - | - | 5987 | 397.7 | - | - | 0 | - |
| - | - | 9528 | 397.7 | - | - | 0 | - |
| 10 | b | 4.137E+04 | 398.2 | 0.0008404 | 2.111 | +3 | 10 |
| - | - | 1.035E+04 | 398.2 | - | - | 0 | - |
| - | - | 2.847E+04 | 398.5 | - | - | 0 | - |
| - | - | 8863 | 398.9 | - | - | 0 | - |
| - | - | 1.808E+04 | 398.9 | - | - | 0 | - |
| - | - | 7638 | 399.2 | - | - | 0 | - |
| - | - | 4483 | 399.6 | - | - | 0 | - |
| - | - | 1.049E+04 | 400.2 | - | - | 0 | - |
| - | - | 5826 | 400.2 | - | - | 0 | - |
| - | - | 7641 | 400.5 | - | - | 0 | - |
| - | - | 1.455E+04 | 400.7 | - | - | 0 | - |
| - | - | 1.856E+04 | 401.2 | - | - | 0 | - |
| - | - | 2.505E+04 | 401.3 | - | - | 0 | - |
| - | - | 9741 | 401.8 | - | - | 0 | - |
| - | - | 7.982E+04 | 402.2 | - | - | 0 | - |
| - | - | 1.26E+04 | 402.6 | - | - | 0 | - |
| - | - | 8789 | 402.9 | - | - | 0 | - |
| - | - | 2.087E+05 | 403.2 | - | - | 0 | - |
| - | - | 1.059E+05 | 403.2 | - | - | 0 | - |
| - | - | 5.698E+04 | 403.3 | - | - | 0 | - |
| 4 | y | 1.715E+05 | 403.6 | 0.003612 | 8.949 | +3 | 11 |
| 7 | y | 3.264E+04 | 403.8 | 0.006299 | 15.6 | +2 | 8 |
| - | - | 7.326E+04 | 403.9 | - | - | 0 | - |
| - | - | 3.867E+04 | 404.2 | - | - | 0 | - |
| - | - | 3.245E+04 | 404.2 | - | - | 0 | - |
| - | - | 7705 | 404.3 | - | - | 0 | - |
| - | - | 2.473E+04 | 404.5 | - | - | 0 | - |
| - | - | 8419 | 404.6 | - | - | 0 | - |
| - | - | 6.661E+04 | 404.7 | - | - | 0 | - |
| - | - | 2.071E+04 | 404.9 | - | - | 0 | - |
| - | - | 6.836E+04 | 405 | - | - | 0 | - |
| - | - | 4.567E+04 | 405.2 | - | - | 0 | - |
| - | - | 1.322E+04 | 405.5 | - | - | 0 | - |
| - | - | 7.26E+04 | 406.2 | - | - | 0 | - |
| - | - | 7.037E+04 | 406.3 | - | - | 0 | - |
| - | - | 3.202E+04 | 406.7 | - | - | 0 | - |
| - | - | 2.73E+04 | 406.8 | - | - | 0 | - |
| - | - | 3.186E+04 | 407.2 | - | - | 0 | - |
| - | - | 1.296E+04 | 407.3 | - | - | 0 | - |
| - | - | 5423 | 407.5 | - | - | 0 | - |
| - | - | 6026 | 407.7 | - | - | 0 | - |
| - | - | 5.847E+04 | 408.2 | - | - | 0 | - |
| - | - | 3.714E+04 | 408.6 | - | - | 0 | - |
| - | - | 1.931E+04 | 408.9 | - | - | 0 | - |
| - | - | 7669 | 409.2 | - | - | 0 | - |
| 4 | y | 4.283E+04 | 409.2 | 0.001161 | 2.836 | +3 | 11 |
| - | - | 3.493E+04 | 409.6 | - | - | 0 | - |
| - | - | 8147 | 409.9 | - | - | 0 | - |
| - | - | 5020 | 410.3 | - | - | 0 | - |
| - | - | 1.885E+05 | 410.3 | - | - | 0 | - |
| - | - | 9.414E+04 | 410.6 | - | - | 0 | - |
| - | - | 7.852E+04 | 410.9 | - | - | 0 | - |
| 6 | b | 1.974E+05 | 411.2 | 0.001059 | 2.576 | +2 | 6 |
| - | - | 2.966E+04 | 411.2 | - | - | 0 | - |
| - | - | 3.791E+04 | 411.3 | - | - | 0 | - |
| - | - | 6209 | 411.6 | - | - | 0 | - |
| 0 | Precursor | 4.673E+04 | 411.7 | 0.0007343 | 1.784 | +4 | -1 |
| 0 | Precursor | 5.228E+04 | 412 | 0.003266 | 7.927 | +4 | -1 |
| - | - | 4.581E+04 | 412.2 | - | - | 0 | - |
| - | - | 2.452E+04 | 412.2 | - | - | 0 | - |
| - | - | 3.918E+04 | 412.2 | - | - | 0 | - |
| 7 | y | 6.97E+05 | 412.3 | 0.001203 | 2.918 | +2 | 8 |
| - | - | 1.526E+04 | 412.5 | - | - | 0 | - |
| - | - | 3.509E+05 | 412.8 | - | - | 0 | - |
| - | - | 2.116E+04 | 413.2 | - | - | 0 | - |
| - | - | 7.911E+04 | 413.3 | - | - | 0 | - |
| - | - | 1.74E+04 | 413.8 | - | - | 0 | - |
| - | - | 1.103E+05 | 413.9 | - | - | 0 | - |
| - | - | 1.63E+04 | 414.2 | - | - | 0 | - |
| - | - | 1.118E+05 | 414.2 | - | - | 0 | - |
| - | - | 5605 | 414.3 | - | - | 0 | - |
| - | - | 4.957E+04 | 414.6 | - | - | 0 | - |
| - | - | 1.559E+04 | 414.9 | - | - | 0 | - |
| - | - | 5075 | 415.2 | - | - | 0 | - |
| - | - | 6.752E+04 | 415.3 | - | - | 0 | - |
| - | - | 3.299E+04 | 415.8 | - | - | 0 | - |
| 0 | Precursor | 4.028E+05 | 416.2 | 0.001084 | 2.604 | +4 | -1 |
| - | - | 3.871E+05 | 416.5 | - | - | 0 | - |
| - | - | 1.804E+05 | 416.7 | - | - | 0 | - |
| - | - | 9.583E+04 | 417 | - | - | 0 | - |
| - | - | 1.787E+04 | 417.2 | - | - | 0 | - |
| - | - | 1.088E+04 | 418.7 | - | - | 0 | - |
| - | - | 6516 | 419.6 | - | - | 0 | - |
| - | - | 3.636E+05 | 419.9 | - | - | 0 | - |
| 6 | b | 5.03E+04 | 420.2 | 0.0006597 | 1.57 | +2 | 6 |
| - | - | 3.327E+05 | 420.2 | - | - | 0 | - |
| - | - | 1.166E+05 | 420.6 | - | - | 0 | - |
| - | - | 1.773E+04 | 420.7 | - | - | 0 | - |
| - | - | 4.205E+04 | 420.9 | - | - | 0 | - |
| - | - | 7852 | 421.2 | - | - | 0 | - |
| - | - | 1.878E+04 | 421.2 | - | - | 0 | - |
| - | - | 5.35E+04 | 422.3 | - | - | 0 | - |
| - | - | 2.664E+04 | 423.3 | - | - | 0 | - |
| - | - | 9715 | 424.3 | - | - | 0 | - |
| - | - | 5.531E+04 | 424.3 | - | - | 0 | - |
| - | - | 1.168E+05 | 424.7 | - | - | 0 | - |
| - | - | 6892 | 424.8 | - | - | 0 | - |
| - | - | 4.96E+04 | 425.2 | - | - | 0 | - |
| - | - | 2.567E+04 | 425.3 | - | - | 0 | - |
| - | - | 1.674E+04 | 425.3 | - | - | 0 | - |
| - | - | 1.545E+04 | 425.7 | - | - | 0 | - |
| - | - | 7.931E+04 | 425.9 | - | - | 0 | - |
| - | - | 5.219E+04 | 426.2 | - | - | 0 | - |
| - | - | 6618 | 426.3 | - | - | 0 | - |
| - | - | 1.009E+04 | 426.6 | - | - | 0 | - |
| - | - | 6538 | 426.9 | - | - | 0 | - |
| - | - | 6.908E+04 | 427.2 | - | - | 0 | - |
| - | - | 4.338E+04 | 427.7 | - | - | 0 | - |
| - | - | 1.169E+04 | 428.2 | - | - | 0 | - |
| - | - | 1.39E+04 | 428.3 | - | - | 0 | - |
| - | - | 5960 | 428.7 | - | - | 0 | - |
| - | - | 6.901E+05 | 429.2 | - | - | 0 | - |
| - | - | 1.744E+05 | 430.2 | - | - | 0 | - |
| 11 | b | 6845 | 430.5 | 0.00435 | 10.1 | +3 | 11 |
| - | - | 9.287E+04 | 431.2 | - | - | 0 | - |
| - | - | 2.126E+04 | 432.2 | - | - | 0 | - |
| - | - | 6632 | 432.2 | - | - | 0 | - |
| - | - | 1.121E+04 | 433.3 | - | - | 0 | - |
| - | - | 1.531E+04 | 433.7 | - | - | 0 | - |
| - | - | 1.368E+04 | 434.2 | - | - | 0 | - |
| - | - | 9723 | 436.6 | - | - | 0 | - |
| - | - | 7743 | 436.9 | - | - | 0 | - |
| 3 | b | 4.44E+04 | 437.2 | 0.001556 | 3.559 | +1 | 3 |
| - | - | 5488 | 437.3 | - | - | 0 | - |
| - | - | 7495 | 437.3 | - | - | 0 | - |
| - | - | 8413 | 438.2 | - | - | 0 | - |
| - | - | 8174 | 438.2 | - | - | 0 | - |
| - | - | 6.548E+04 | 438.6 | - | - | 0 | - |
| - | - | 4.979E+04 | 438.9 | - | - | 0 | - |
| - | - | 1.18E+04 | 439.2 | - | - | 0 | - |
| - | - | 8650 | 439.2 | - | - | 0 | - |
| - | - | 8965 | 439.3 | - | - | 0 | - |
| - | - | 1.084E+04 | 439.6 | - | - | 0 | - |
| - | - | 1.426E+04 | 442.2 | - | - | 0 | - |
| - | - | 2.329E+04 | 442.7 | - | - | 0 | - |
| - | - | 1.374E+04 | 443.2 | - | - | 0 | - |
| - | - | 9849 | 443.7 | - | - | 0 | - |
| - | - | 1.507E+04 | 444.2 | - | - | 0 | - |
| - | - | 1.53E+04 | 444.6 | - | - | 0 | - |
| - | - | 7493 | 444.9 | - | - | 0 | - |
| - | - | 1.03E+04 | 445.2 | - | - | 0 | - |
| - | - | 2.474E+04 | 445.3 | - | - | 0 | - |
| - | - | 1.142E+04 | 445.9 | - | - | 0 | - |
| - | - | 5.955E+04 | 446.2 | - | - | 0 | - |
| - | - | 1.464E+04 | 446.6 | - | - | 0 | - |
| - | - | 4.344E+04 | 446.7 | - | - | 0 | - |
| - | - | 1.094E+04 | 447.2 | - | - | 0 | - |
| - | - | 1.208E+05 | 448.3 | - | - | 0 | - |
| - | - | 7.482E+04 | 448.6 | - | - | 0 | - |
| - | - | 3.019E+04 | 448.9 | - | - | 0 | - |
| - | - | 8450 | 449.3 | - | - | 0 | - |
| - | - | 8549 | 450.3 | - | - | 0 | - |
| - | - | 9.171E+04 | 451.2 | - | - | 0 | - |
| - | - | 5.451E+04 | 451.3 | - | - | 0 | - |
| - | - | 5774 | 451.6 | - | - | 0 | - |
| - | - | 6.23E+04 | 451.7 | - | - | 0 | - |
| - | - | 1.663E+05 | 451.9 | - | - | 0 | - |
| - | - | 1.277E+04 | 452.2 | - | - | 0 | - |
| - | - | 1.295E+05 | 452.3 | - | - | 0 | - |
| - | - | 2.65E+04 | 452.3 | - | - | 0 | - |
| - | - | 5.014E+04 | 452.6 | - | - | 0 | - |
| - | - | 2.665E+04 | 452.9 | - | - | 0 | - |
| - | - | 1.055E+04 | 453.3 | - | - | 0 | - |
| - | - | 1.441E+04 | 453.7 | - | - | 0 | - |
| - | - | 3.213E+04 | 454.2 | - | - | 0 | - |
| - | - | 6275 | 455.2 | - | - | 0 | - |
| - | - | 6.833E+04 | 455.3 | - | - | 0 | - |
| - | - | 1.336E+04 | 456.3 | - | - | 0 | - |
| - | - | 1.923E+04 | 457.2 | - | - | 0 | - |
| - | - | 7189 | 457.6 | - | - | 0 | - |
| - | - | 7.918E+04 | 457.6 | - | - | 0 | - |
| - | - | 9407 | 457.8 | - | - | 0 | - |
| 3 | y | 1.346E+05 | 457.9 | 0.004821 | 10.53 | +3 | 12 |
| - | - | 7.845E+04 | 458.3 | - | - | 0 | - |
| - | - | 2.923E+04 | 458.6 | - | - | 0 | - |
| - | - | 1.102E+04 | 458.9 | - | - | 0 | - |
| - | - | 1.763E+04 | 459.2 | - | - | 0 | - |
| - | - | 1.468E+05 | 460.2 | - | - | 0 | - |
| - | - | 6.631E+04 | 460.7 | - | - | 0 | - |
| - | - | 1.517E+04 | 461.2 | - | - | 0 | - |
| - | - | 2.155E+04 | 461.2 | - | - | 0 | - |
| - | - | 7919 | 461.9 | - | - | 0 | - |
| - | - | 7771 | 462.8 | - | - | 0 | - |
| - | - | 1.255E+04 | 462.9 | - | - | 0 | - |
| - | - | 1.874E+04 | 463.3 | - | - | 0 | - |
| 3 | y | 1.913E+06 | 463.6 | 0.001271 | 2.741 | +3 | 12 |
| - | - | 1.611E+06 | 463.9 | - | - | 0 | - |
| - | - | 6.144E+05 | 464.3 | - | - | 0 | - |
| - | - | 2.2E+05 | 464.6 | - | - | 0 | - |
| - | - | 4.635E+04 | 464.9 | - | - | 0 | - |
| - | - | 4.101E+04 | 466.9 | - | - | 0 | - |
| - | - | 2.98E+04 | 467.3 | - | - | 0 | - |
| - | - | 5.633E+04 | 467.3 | - | - | 0 | - |
| - | - | 1.894E+04 | 467.6 | - | - | 0 | - |
| - | - | 9754 | 467.9 | - | - | 0 | - |
| - | - | 2.689E+04 | 468.3 | - | - | 0 | - |
| - | - | 3.804E+04 | 469.8 | - | - | 0 | - |
| - | - | 3.329E+04 | 470.3 | - | - | 0 | - |
| 11 | y | 7.934E+05 | 470.3 | 0.001234 | 2.625 | +1 | 4 |
| - | - | 5.916E+04 | 470.6 | - | - | 0 | - |
| - | - | 1.773E+04 | 470.9 | - | - | 0 | - |
| - | - | 1.28E+04 | 471.3 | - | - | 0 | - |
| - | - | 2.081E+05 | 471.3 | - | - | 0 | - |
| - | - | 5.269E+04 | 471.8 | - | - | 0 | - |
| - | - | 3.371E+04 | 472.3 | - | - | 0 | - |
| - | - | 3.565E+04 | 472.3 | - | - | 0 | - |
| - | - | 1.823E+04 | 472.6 | - | - | 0 | - |
| - | - | 9569 | 472.8 | - | - | 0 | - |
| - | - | 2.086E+04 | 472.9 | - | - | 0 | - |
| - | - | 9.352E+04 | 473.3 | - | - | 0 | - |
| - | - | 4.487E+04 | 473.6 | - | - | 0 | - |
| - | - | 9004 | 473.9 | - | - | 0 | - |
| - | - | 2.327E+04 | 474.3 | - | - | 0 | - |
| - | - | 5.417E+04 | 474.8 | - | - | 0 | - |
| - | - | 2.462E+04 | 475.3 | - | - | 0 | - |
| - | - | 1.332E+04 | 475.8 | - | - | 0 | - |
| - | - | 2.148E+04 | 475.9 | - | - | 0 | - |
| - | - | 1.382E+05 | 476.3 | - | - | 0 | - |
| - | - | 1.363E+05 | 476.6 | - | - | 0 | - |
| - | - | 4.031E+04 | 476.9 | - | - | 0 | - |
| - | - | 1.821E+04 | 477.3 | - | - | 0 | - |
| - | - | 2.585E+04 | 478.2 | - | - | 0 | - |
| - | - | 1.401E+04 | 479.2 | - | - | 0 | - |
| - | - | 7892 | 479.2 | - | - | 0 | - |
| - | - | 1.024E+04 | 479.6 | - | - | 0 | - |
| 6 | y | 1.684E+05 | 480.8 | 0.001196 | 2.488 | +2 | 9 |
| - | - | 9.829E+04 | 481.3 | - | - | 0 | - |
| - | - | 1.771E+04 | 481.8 | - | - | 0 | - |
| - | - | 7.469E+04 | 481.9 | - | - | 0 | - |
| - | - | 9.292E+04 | 482.3 | - | - | 0 | - |
| - | - | 5.723E+04 | 482.3 | - | - | 0 | - |
| - | - | 4.939E+04 | 482.6 | - | - | 0 | - |
| - | - | 4.906E+04 | 482.8 | - | - | 0 | - |
| - | - | 2.097E+04 | 482.9 | - | - | 0 | - |
| - | - | 9978 | 483.2 | - | - | 0 | - |
| - | - | 3.48E+04 | 483.3 | - | - | 0 | - |
| - | - | 1.714E+04 | 483.3 | - | - | 0 | - |
| - | - | 1.157E+05 | 483.8 | - | - | 0 | - |
| - | - | 1.068E+04 | 484.2 | - | - | 0 | - |
| - | - | 7.516E+04 | 484.3 | - | - | 0 | - |
| - | - | 2.103E+04 | 484.8 | - | - | 0 | - |
| - | - | 8385 | 485.3 | - | - | 0 | - |
| - | - | 3.301E+04 | 487.9 | - | - | 0 | - |
| - | - | 1.654E+04 | 488.3 | - | - | 0 | - |
| - | - | 8639 | 488.6 | - | - | 0 | - |
| - | - | 7604 | 489.9 | - | - | 0 | - |
| - | - | 7568 | 491.2 | - | - | 0 | - |
| - | - | 6.95E+04 | 491.3 | - | - | 0 | - |
| - | - | 1.366E+04 | 491.8 | - | - | 0 | - |
| - | - | 1.436E+04 | 492.3 | - | - | 0 | - |
| - | - | 4.058E+04 | 492.8 | - | - | 0 | - |
| - | - | 1.356E+04 | 493.2 | - | - | 0 | - |
| - | - | 2.061E+04 | 493.3 | - | - | 0 | - |
| - | - | 9203 | 493.4 | - | - | 0 | - |
| - | - | 9373 | 494.8 | - | - | 0 | - |
| - | - | 1.113E+04 | 495.3 | - | - | 0 | - |
| - | - | 1.829E+04 | 495.4 | - | - | 0 | - |
| - | - | 6267 | 495.6 | - | - | 0 | - |
| - | - | 1.197E+04 | 495.9 | - | - | 0 | - |
| - | - | 7591 | 496.3 | - | - | 0 | - |
| 8 | b | 2.614E+04 | 497.2 | 0.001723 | 3.466 | +2 | 8 |
| - | - | 1.542E+04 | 497.7 | - | - | 0 | - |
| - | - | 9602 | 500.2 | - | - | 0 | - |
| - | - | 9651 | 501.3 | - | - | 0 | - |
| - | - | 2.073E+04 | 501.6 | - | - | 0 | - |
| - | - | 3.405E+04 | 501.9 | - | - | 0 | - |
| - | - | 6319 | 502.3 | - | - | 0 | - |
| - | - | 5.894E+04 | 503.2 | - | - | 0 | - |
| - | - | 1.408E+04 | 504.2 | - | - | 0 | - |
| - | - | 1.023E+04 | 504.6 | - | - | 0 | - |
| 13 | b | 3.268E+04 | 504.9 | 0.002345 | 4.643 | +3 | 13 |
| 13 | b | 3.088E+04 | 505.3 | 0.008751 | 17.32 | +3 | 13 |
| - | - | 2.098E+04 | 505.6 | - | - | 0 | - |
| - | - | 1.408E+04 | 505.7 | - | - | 0 | - |
| - | - | 6292 | 506.2 | - | - | 0 | - |
| - | - | 3.275E+04 | 508.8 | - | - | 0 | - |
| - | - | 3.688E+04 | 509.2 | - | - | 0 | - |
| - | - | 2.102E+04 | 509.7 | - | - | 0 | - |
| - | - | 8863 | 510.2 | - | - | 0 | - |
| - | - | 1.17E+04 | 510.3 | - | - | 0 | - |
| - | - | 8693 | 510.4 | - | - | 0 | - |
| - | - | 2.382E+04 | 510.6 | - | - | 0 | - |
| 13 | b | 7.237E+04 | 510.9 | 0.001417 | 2.773 | +3 | 13 |
| - | - | 9.581E+04 | 511.3 | - | - | 0 | - |
| - | - | 4.429E+04 | 511.6 | - | - | 0 | - |
| - | - | 1.906E+04 | 511.9 | - | - | 0 | - |
| - | - | 1.762E+04 | 513.9 | - | - | 0 | - |
| - | - | 1.541E+04 | 514.3 | - | - | 0 | - |
| - | - | 2.917E+04 | 514.6 | - | - | 0 | - |
| - | - | 1.923E+05 | 516.9 | - | - | 0 | - |
| - | - | 1.705E+05 | 517.3 | - | - | 0 | - |
| - | - | 6.805E+04 | 517.6 | - | - | 0 | - |
| - | - | 2.58E+04 | 517.8 | - | - | 0 | - |
| - | - | 2.766E+04 | 517.9 | - | - | 0 | - |
| - | - | 2.522E+04 | 518.3 | - | - | 0 | - |
| - | - | 1.286E+04 | 519.6 | - | - | 0 | - |
| 2 | y | 1.909E+05 | 519.9 | 0.00118 | 2.269 | +3 | 13 |
| - | - | 9153 | 520.2 | - | - | 0 | - |
| - | - | 1.289E+05 | 520.3 | - | - | 0 | - |
| - | - | 4.67E+04 | 520.4 | - | - | 0 | - |
| - | - | 9.068E+04 | 520.6 | - | - | 0 | - |
| - | - | 3.163E+04 | 521 | - | - | 0 | - |
| - | - | 6.908E+04 | 521.2 | - | - | 0 | - |
| - | - | 1.002E+04 | 521.3 | - | - | 0 | - |
| - | - | 1.176E+04 | 521.4 | - | - | 0 | - |
| - | - | 2.133E+04 | 522.2 | - | - | 0 | - |
| - | - | 1.469E+04 | 523.4 | - | - | 0 | - |
| - | - | 8106 | 524.4 | - | - | 0 | - |
| - | - | 1.21E+04 | 524.8 | - | - | 0 | - |
| - | - | 1.308E+04 | 525.2 | - | - | 0 | - |
| - | - | 2.939E+04 | 525.3 | - | - | 0 | - |
| 2 | y | 2.807E+05 | 525.6 | 0.001139 | 2.168 | +3 | 13 |
| - | - | 1.496E+04 | 525.8 | - | - | 0 | - |
| - | - | 3.141E+05 | 526 | - | - | 0 | - |
| - | - | 1.175E+05 | 526.3 | - | - | 0 | - |
| - | - | 9304 | 526.3 | - | - | 0 | - |
| - | - | 5.44E+04 | 526.6 | - | - | 0 | - |
| - | - | 9196 | 526.8 | - | - | 0 | - |
| - | - | 6437 | 527 | - | - | 0 | - |
| - | - | 8840 | 527.2 | - | - | 0 | - |
| - | - | 2.138E+04 | 528.2 | - | - | 0 | - |
| - | - | 7449 | 529 | - | - | 0 | - |
| - | - | 9444 | 529.2 | - | - | 0 | - |
| - | - | 7779 | 529.3 | - | - | 0 | - |
| - | - | 2.756E+04 | 529.8 | - | - | 0 | - |
| - | - | 4.449E+04 | 530.3 | - | - | 0 | - |
| - | - | 8204 | 530.8 | - | - | 0 | - |
| - | - | 2.534E+04 | 530.8 | - | - | 0 | - |
| - | - | 5.682E+04 | 531.2 | - | - | 0 | - |
| - | - | 1.124E+04 | 531.3 | - | - | 0 | - |
| - | - | 1.764E+04 | 532.2 | - | - | 0 | - |
| - | - | 9644 | 533.8 | - | - | 0 | - |
| - | - | 5.086E+04 | 535.8 | - | - | 0 | - |
| - | - | 3.396E+04 | 536.3 | - | - | 0 | - |
| - | - | 7907 | 536.8 | - | - | 0 | - |
| - | - | 4.224E+05 | 538.3 | - | - | 0 | - |
| - | - | 3.278E+04 | 538.4 | - | - | 0 | - |
| - | - | 5.656E+04 | 538.8 | - | - | 0 | - |
| - | - | 1.162E+05 | 539.3 | - | - | 0 | - |
| - | - | 8.605E+04 | 539.3 | - | - | 0 | - |
| - | - | 7350 | 539.4 | - | - | 0 | - |
| - | - | 6.022E+04 | 539.8 | - | - | 0 | - |
| - | - | 1.81E+04 | 540.3 | - | - | 0 | - |
| - | - | 5925 | 540.3 | - | - | 0 | - |
| - | - | 2.23E+04 | 540.3 | - | - | 0 | - |
| 5 | y | 2.687E+04 | 540.8 | 0.009521 | 17.6 | +2 | 10 |
| - | - | 9766 | 541.3 | - | - | 0 | - |
| 10 | y | 1.902E+05 | 541.4 | 0.001261 | 2.328 | +1 | 5 |
| - | - | 7.563E+04 | 542.3 | - | - | 0 | - |
| - | - | 6.741E+04 | 542.4 | - | - | 0 | - |
| - | - | 2.779E+04 | 543.3 | - | - | 0 | - |
| - | - | 7722 | 543.4 | - | - | 0 | - |
| - | - | 1.992E+04 | 544.3 | - | - | 0 | - |
| - | - | 4.214E+04 | 544.8 | - | - | 0 | - |
| - | - | 3.173E+04 | 545.3 | - | - | 0 | - |
| - | - | 1.393E+04 | 545.8 | - | - | 0 | - |
| - | - | 5.412E+04 | 547.8 | - | - | 0 | - |
| - | - | 1.69E+04 | 548.2 | - | - | 0 | - |
| - | - | 3.894E+04 | 548.3 | - | - | 0 | - |
| - | - | 2.578E+04 | 548.8 | - | - | 0 | - |
| - | - | 4.06E+04 | 549.2 | - | - | 0 | - |
| 5 | y | 4.782E+05 | 549.3 | 0.001068 | 1.944 | +2 | 10 |
| - | - | 3.209E+05 | 549.8 | - | - | 0 | - |
| - | - | 1.175E+04 | 550.2 | - | - | 0 | - |
| - | - | 9.19E+04 | 550.3 | - | - | 0 | - |
| - | - | 3.568E+04 | 550.8 | - | - | 0 | - |
| 9 | b | 7280 | 552.3 | 0.003805 | 6.89 | +2 | 9 |
| - | - | 2.911E+04 | 553.2 | - | - | 0 | - |
| - | - | 2.099E+04 | 553.3 | - | - | 0 | - |
| - | - | 1.334E+04 | 553.8 | - | - | 0 | - |
| - | - | 8126 | 554.2 | - | - | 0 | - |
| - | - | 7196 | 554.3 | - | - | 0 | - |
| - | - | 1.871E+04 | 555.3 | - | - | 0 | - |
| - | - | 1.451E+04 | 555.8 | - | - | 0 | - |
| - | - | 8143 | 556.3 | - | - | 0 | - |
| - | - | 1.462E+04 | 556.8 | - | - | 0 | - |
| - | - | 7089 | 557.3 | - | - | 0 | - |
| - | - | 4.321E+04 | 557.3 | - | - | 0 | - |
| - | - | 8604 | 558.3 | - | - | 0 | - |
| 9 | b | 1.334E+05 | 561.3 | 0.0009032 | 1.609 | +2 | 9 |
| - | - | 1.103E+04 | 561.4 | - | - | 0 | - |
| - | - | 9.14E+04 | 561.8 | - | - | 0 | - |
| - | - | 2.613E+04 | 562.3 | - | - | 0 | - |
| - | - | 7070 | 562.3 | - | - | 0 | - |
| - | - | 1.484E+04 | 562.8 | - | - | 0 | - |
| - | - | 1.325E+04 | 564.3 | - | - | 0 | - |
| - | - | 7523 | 564.8 | - | - | 0 | - |
| 4 | b | 1.697E+04 | 565.2 | 0.001725 | 3.051 | +1 | 4 |
| - | - | 5.04E+05 | 566.2 | - | - | 0 | - |
| - | - | 1.496E+05 | 567.3 | - | - | 0 | - |
| - | - | 2.898E+04 | 567.4 | - | - | 0 | - |
| - | - | 3.134E+04 | 568.3 | - | - | 0 | - |
| - | - | 2.332E+05 | 570.2 | - | - | 0 | - |
| - | - | 7.812E+04 | 571.3 | - | - | 0 | - |
| - | - | 1.251E+04 | 572.3 | - | - | 0 | - |
| - | - | 2.365E+05 | 572.8 | - | - | 0 | - |
| - | - | 1.466E+05 | 573.3 | - | - | 0 | - |
| - | - | 5.812E+04 | 573.8 | - | - | 0 | - |
| - | - | 1.276E+04 | 574.3 | - | - | 0 | - |
| - | - | 1.512E+05 | 577.4 | - | - | 0 | - |
| - | - | 5.173E+04 | 578.4 | - | - | 0 | - |
| - | - | 9000 | 579.2 | - | - | 0 | - |
| - | - | 2.958E+05 | 579.4 | - | - | 0 | - |
| - | - | 9.708E+04 | 580.4 | - | - | 0 | - |
| - | - | 1.419E+04 | 581.2 | - | - | 0 | - |
| - | - | 2.495E+04 | 581.4 | - | - | 0 | - |
| - | - | 1.281E+04 | 581.8 | - | - | 0 | - |
| - | - | 9038 | 582.8 | - | - | 0 | - |
| - | - | 7102 | 583.3 | - | - | 0 | - |
| - | - | 1.697E+04 | 587.3 | - | - | 0 | - |
| 10 | b | 2.844E+04 | 587.8 | 0.001728 | 2.94 | +2 | 10 |
| 10 | b | 3.154E+04 | 588.3 | 0.008255 | 14.03 | +2 | 10 |
| - | - | 1.075E+04 | 588.8 | - | - | 0 | - |
| - | - | 1.189E+04 | 594.2 | - | - | 0 | - |
| - | - | 1.053E+05 | 595.4 | - | - | 0 | - |
| - | - | 1.611E+04 | 595.8 | - | - | 0 | - |
| - | - | 1.719E+04 | 596.3 | - | - | 0 | - |
| - | - | 3.663E+04 | 596.4 | - | - | 0 | - |
| 10 | b | 3.505E+04 | 596.8 | 0.0005348 | 0.8962 | +2 | 10 |
| - | - | 8091 | 596.8 | - | - | 0 | - |
| - | - | 2.602E+04 | 597.3 | - | - | 0 | - |
| - | - | 1.096E+05 | 598.2 | - | - | 0 | - |
| - | - | 3.465E+04 | 599.2 | - | - | 0 | - |
| - | - | 9523 | 600.2 | - | - | 0 | - |
| - | - | 1.199E+04 | 600.3 | - | - | 0 | - |
| - | - | 1.491E+04 | 601.8 | - | - | 0 | - |
| - | - | 1.111E+05 | 604.4 | - | - | 0 | - |
| 4 | y | 1.898E+05 | 604.8 | 0.003975 | 6.571 | +2 | 11 |
| - | - | 1.248E+05 | 605.4 | - | - | 0 | - |
| - | - | 4.324E+04 | 605.9 | - | - | 0 | - |
| - | - | 1.934E+04 | 606.4 | - | - | 0 | - |
| - | - | 1.307E+04 | 606.8 | - | - | 0 | - |
| - | - | 6299 | 607.3 | - | - | 0 | - |
| - | - | 1.048E+05 | 610.3 | - | - | 0 | - |
| - | - | 3.786E+04 | 611.3 | - | - | 0 | - |
| - | - | 2.29E+04 | 611.8 | - | - | 0 | - |
| - | - | 1.57E+04 | 612.3 | - | - | 0 | - |
| - | - | 7358 | 612.8 | - | - | 0 | - |
| 4 | y | 1.015E+05 | 613.4 | 0.0008319 | 1.356 | +2 | 11 |
| - | - | 1.157E+04 | 613.4 | - | - | 0 | - |
| - | - | 6.932E+04 | 613.9 | - | - | 0 | - |
| - | - | 2.217E+04 | 614.4 | - | - | 0 | - |
| - | - | 7678 | 614.9 | - | - | 0 | - |
| - | - | 2.631E+04 | 615.3 | - | - | 0 | - |
| - | - | 6.069E+04 | 615.3 | - | - | 0 | - |
| - | - | 4.159E+04 | 615.9 | - | - | 0 | - |
| - | - | 1.141E+04 | 616.3 | - | - | 0 | - |
| - | - | 1.372E+04 | 616.4 | - | - | 0 | - |
| - | - | 1.904E+04 | 618.8 | - | - | 0 | - |
| - | - | 8385 | 619.4 | - | - | 0 | - |
| - | - | 8.504E+04 | 620.3 | - | - | 0 | - |
| - | - | 7.787E+04 | 620.8 | - | - | 0 | - |
| - | - | 4.377E+04 | 621.3 | - | - | 0 | - |
| - | - | 1.164E+04 | 621.8 | - | - | 0 | - |
| - | - | 1.574E+04 | 623.4 | - | - | 0 | - |
| - | - | 3.813E+04 | 628.3 | - | - | 0 | - |
| - | - | 2.149E+05 | 629.3 | - | - | 0 | - |
| - | - | 1.435E+05 | 629.8 | - | - | 0 | - |
| - | - | 6.869E+04 | 630.3 | - | - | 0 | - |
| - | - | 2.215E+04 | 630.9 | - | - | 0 | - |
| - | - | 6316 | 631.4 | - | - | 0 | - |
| - | - | 1.441E+05 | 638.4 | - | - | 0 | - |
| - | - | 1.094E+05 | 638.9 | - | - | 0 | - |
| - | - | 3.184E+04 | 639.4 | - | - | 0 | - |
| - | - | 7747 | 639.9 | - | - | 0 | - |
| - | - | 1.986E+04 | 647.4 | - | - | 0 | - |
| - | - | 7570 | 648.3 | - | - | 0 | - |
| - | - | 7685 | 648.4 | - | - | 0 | - |
| - | - | 9734 | 648.8 | - | - | 0 | - |
| - | - | 1.828E+04 | 657.3 | - | - | 0 | - |
| - | - | 2.755E+04 | 657.3 | - | - | 0 | - |
| - | - | 2.092E+04 | 657.8 | - | - | 0 | - |
| - | - | 9116 | 658.3 | - | - | 0 | - |
| - | - | 7423 | 663.3 | - | - | 0 | - |
| - | - | 8.112E+04 | 664.5 | - | - | 0 | - |
| - | - | 2.993E+04 | 665.5 | - | - | 0 | - |
| - | - | 3.154E+04 | 665.8 | - | - | 0 | - |
| - | - | 2.904E+04 | 666.3 | - | - | 0 | - |
| - | - | 9599 | 666.5 | - | - | 0 | - |
| - | - | 2.028E+04 | 667.3 | - | - | 0 | - |
| - | - | 2.645E+04 | 668.3 | - | - | 0 | - |
| - | - | 1.252E+04 | 669.3 | - | - | 0 | - |
| 9 | y | 6.859E+04 | 669.5 | 0.001268 | 1.894 | +1 | 6 |
| - | - | 2.299E+04 | 670.5 | - | - | 0 | - |
| - | - | 4.86E+04 | 674.3 | - | - | 0 | - |
| - | - | 3.144E+05 | 674.4 | - | - | 0 | - |
| - | - | 1.584E+04 | 675.3 | - | - | 0 | - |
| - | - | 1.222E+05 | 675.4 | - | - | 0 | - |
| - | - | 2.151E+04 | 676.4 | - | - | 0 | - |
| - | - | 9357 | 679.3 | - | - | 0 | - |
| - | - | 7431 | 680.3 | - | - | 0 | - |
| 5 | b | 1.337E+04 | 684.3 | 0.003854 | 5.631 | +1 | 5 |
| 5 | b | 1.05E+04 | 685.3 | 0.01196 | 17.46 | +1 | 5 |
| - | - | 2.268E+04 | 685.4 | - | - | 0 | - |
| 3 | y | 1.129E+04 | 686.4 | 0.009053 | 13.19 | +2 | 12 |
| - | - | 2.137E+04 | 690.3 | - | - | 0 | - |
| - | - | 2.894E+05 | 692.4 | - | - | 0 | - |
| - | - | 1.182E+05 | 693.4 | - | - | 0 | - |
| - | - | 2.039E+04 | 694.5 | - | - | 0 | - |
| 3 | y | 1.049E+05 | 694.9 | 0.001272 | 1.831 | +2 | 12 |
| - | - | 7.333E+04 | 695.4 | - | - | 0 | - |
| - | - | 3.074E+04 | 695.9 | - | - | 0 | - |
| - | - | 1.09E+04 | 696.4 | - | - | 0 | - |
| - | - | 2.458E+04 | 698.4 | - | - | 0 | - |
| - | - | 1.331E+04 | 699.4 | - | - | 0 | - |
| - | - | 7848 | 699.9 | - | - | 0 | - |
| 5 | b | 6.442E+04 | 702.3 | 0.001895 | 2.698 | +1 | 5 |
| - | - | 2.485E+04 | 703.3 | - | - | 0 | - |
| - | - | 1.383E+05 | 707.3 | - | - | 0 | - |
| - | - | 6.266E+04 | 708.3 | - | - | 0 | - |
| - | - | 1.856E+04 | 708.5 | - | - | 0 | - |
| - | - | 1.272E+04 | 709.3 | - | - | 0 | - |
| 12 | b | 9330 | 709.4 | 0.009435 | 13.3 | +2 | 12 |
| - | - | 1.08E+04 | 709.5 | - | - | 0 | - |
| - | - | 1.448E+04 | 709.9 | - | - | 0 | - |
| - | - | 7.49E+04 | 710.5 | - | - | 0 | - |
| - | - | 3.762E+04 | 711.5 | - | - | 0 | - |
| - | - | 7600 | 712.5 | - | - | 0 | - |
| - | - | 1.111E+04 | 713.4 | - | - | 0 | - |
| - | - | 3.735E+04 | 713.9 | - | - | 0 | - |
| - | - | 2.245E+04 | 714.4 | - | - | 0 | - |
| - | - | 1.456E+04 | 714.9 | - | - | 0 | - |
| - | - | 6.429E+04 | 716.4 | - | - | 0 | - |
| - | - | 2.182E+04 | 717.4 | - | - | 0 | - |
| - | - | 7223 | 718.9 | - | - | 0 | - |
| - | - | 3.287E+04 | 720.3 | - | - | 0 | - |
| - | - | 3.278E+04 | 721.4 | - | - | 0 | - |
| - | - | 2.169E+04 | 722.4 | - | - | 0 | - |
| - | - | 1.974E+04 | 722.9 | - | - | 0 | - |
| - | - | 1.585E+04 | 723.4 | - | - | 0 | - |
| - | - | 4.117E+04 | 724.3 | - | - | 0 | - |
| - | - | 1.666E+04 | 725.3 | - | - | 0 | - |
| 8 | y | 1.239E+05 | 726.5 | 0.001045 | 1.438 | +1 | 7 |
| - | - | 6.402E+04 | 727.5 | - | - | 0 | - |
| - | - | 1.033E+04 | 728.4 | - | - | 0 | - |
| - | - | 1.285E+04 | 728.5 | - | - | 0 | - |
| - | - | 9390 | 731.4 | - | - | 0 | - |
| - | - | 7638 | 732.4 | - | - | 0 | - |
| - | - | 1.344E+05 | 735.3 | - | - | 0 | - |
| - | - | 5.779E+04 | 736.3 | - | - | 0 | - |
| - | - | 6617 | 736.5 | - | - | 0 | - |
| - | - | 2.102E+04 | 737.3 | - | - | 0 | - |
| - | - | 1.766E+04 | 738.4 | - | - | 0 | - |
| - | - | 1.852E+04 | 739.4 | - | - | 0 | - |
| - | - | 8468 | 740.4 | - | - | 0 | - |
| - | - | 6.833E+04 | 752.3 | - | - | 0 | - |
| - | - | 2.934E+04 | 753.3 | - | - | 0 | - |
| - | - | 7495 | 754.3 | - | - | 0 | - |
| - | - | 1.335E+04 | 756.4 | - | - | 0 | - |
| - | - | 6915 | 760.5 | - | - | 0 | - |
| - | - | 1.848E+04 | 777.5 | - | - | 0 | - |
| - | - | 5721 | 780.3 | - | - | 0 | - |
| - | - | 5998 | 783.5 | - | - | 0 | - |
| - | - | 6309 | 793.3 | - | - | 0 | - |
| - | - | 6631 | 793.5 | - | - | 0 | - |
| - | - | 6741 | 794.3 | - | - | 0 | - |
| - | - | 5539 | 794.5 | - | - | 0 | - |
| - | - | 1.019E+04 | 801.5 | - | - | 0 | - |
| - | - | 7738 | 802.5 | - | - | 0 | - |
| - | - | 4.786E+04 | 805.5 | - | - | 0 | - |
| 7 | y | 2.033E+04 | 806.5 | 0.0134 | 16.62 | +1 | 8 |
| - | - | 6787 | 807.5 | - | - | 0 | - |
| - | - | 3.975E+04 | 811.4 | - | - | 0 | - |
| - | - | 3.862E+04 | 811.5 | - | - | 0 | - |
| - | - | 1.834E+04 | 812.4 | - | - | 0 | - |
| - | - | 1.902E+04 | 812.5 | - | - | 0 | - |
| - | - | 7216 | 813.4 | - | - | 0 | - |
| - | - | 6137 | 813.5 | - | - | 0 | - |
| 6 | b | 8464 | 821.3 | 0.001765 | 2.149 | +1 | 6 |
| - | - | 8721 | 822.3 | - | - | 0 | - |
| 7 | y | 5.475E+05 | 823.5 | 0.0008932 | 1.085 | +1 | 8 |
| - | - | 2.589E+05 | 824.5 | - | - | 0 | - |
| - | - | 6.304E+04 | 825.5 | - | - | 0 | - |
| - | - | 3.686E+04 | 829.5 | - | - | 0 | - |
| - | - | 1.965E+04 | 830.5 | - | - | 0 | - |
| - | - | 6147 | 831.5 | - | - | 0 | - |
| 6 | b | 7.481E+04 | 839.4 | 0.0001119 | 0.1333 | +1 | 6 |
| - | - | 3.438E+04 | 840.4 | - | - | 0 | - |
| - | - | 1.082E+04 | 841.4 | - | - | 0 | - |
| - | - | 2.148E+04 | 847.5 | - | - | 0 | - |
| - | - | 7.938E+04 | 848.4 | - | - | 0 | - |
| - | - | 6921 | 848.5 | - | - | 0 | - |
| - | - | 3.851E+04 | 849.4 | - | - | 0 | - |
| - | - | 1.424E+04 | 850.4 | - | - | 0 | - |
| - | - | 7479 | 852.5 | - | - | 0 | - |
| - | - | 2.851E+04 | 853.5 | - | - | 0 | - |
| - | - | 1.696E+04 | 854.5 | - | - | 0 | - |
| - | - | 5667 | 866.4 | - | - | 0 | - |
| - | - | 6085 | 874.4 | - | - | 0 | - |
| - | - | 1.102E+04 | 884.4 | - | - | 0 | - |
| - | - | 6773 | 885.4 | - | - | 0 | - |
| - | - | 1.032E+04 | 889.4 | - | - | 0 | - |
| - | - | 8751 | 890.4 | - | - | 0 | - |
| - | - | 1.329E+04 | 891.5 | - | - | 0 | - |
| - | - | 7023 | 892.5 | - | - | 0 | - |
| - | - | 9.447E+04 | 901.4 | - | - | 0 | - |
| - | - | 5.306E+04 | 902.4 | - | - | 0 | - |
| - | - | 1.175E+04 | 903.4 | - | - | 0 | - |
| - | - | 1.181E+04 | 906.4 | - | - | 0 | - |
| - | - | 5.245E+04 | 919.5 | - | - | 0 | - |
| - | - | 3.452E+04 | 920.5 | - | - | 0 | - |
| - | - | 9520 | 921.5 | - | - | 0 | - |
| - | - | 2.231E+04 | 942.6 | - | - | 0 | - |
| - | - | 9744 | 943.6 | - | - | 0 | - |
| - | - | 6250 | 948.6 | - | - | 0 | - |
| 6 | y | 5.688E+04 | 960.6 | 0.0006972 | 0.7258 | +1 | 9 |
| - | - | 3.411E+04 | 961.6 | - | - | 0 | - |
| - | - | 9754 | 962.6 | - | - | 0 | - |
| - | - | 5452 | 964.5 | - | - | 0 | - |
| - | - | 1.331E+04 | 966.6 | - | - | 0 | - |
| - | - | 9092 | 967.6 | - | - | 0 | - |
| - | - | 4970 | 984.6 | - | - | 0 | - |
| 8 | b | 8109 | 993.4 | 0.0008735 | 0.8793 | +1 | 8 |
| - | - | 6402 | 1005 | - | - | 0 | - |
| - | - | 7759 | 1017 | - | - | 0 | - |
| - | - | 7327 | 1018 | - | - | 0 | - |
| - | - | 1.079E+04 | 1033 | - | - | 0 | - |
| - | - | 2.099E+04 | 1034 | - | - | 0 | - |
| - | - | 1.061E+04 | 1035 | - | - | 0 | - |
| - | - | 1.964E+04 | 1070 | - | - | 0 | - |
| - | - | 1.147E+04 | 1071 | - | - | 0 | - |
| - | - | 4901 | 1072 | - | - | 0 | - |
| - | - | 1.228E+04 | 1088 | - | - | 0 | - |
| - | - | 6116 | 1089 | - | - | 0 | - |
| 5 | y | 6705 | 1098 | 0.00426 | 3.881 | +1 | 10 |
| - | - | 5570 | 2341 | - | - | 0 | - |
| - | - | 5718 | 3448 | - | - | 0 | - |

m/z Charge Intensity FragmentType MassShift Position
120.05243682861328 0 8321.041
120.05620574951172 0 15439.75
121.03978729248047 0 3693.2527
122.06065368652344 0 5924.8257
122.07179260253906 0 14218.842
123.05582427978516 0 45942.52
123.09215545654297 0 3297.9033
127.08710479736328 0 51365.66
128.03480529785156 0 3314.9348
128.07115173339844 0 6127.956
128.090576171875 0 4614.763
129.06642150878906 0 62958.58
129.10279846191406 0 1895711.2
130.065673828125 0 437380.06
130.09999084472656 0 14824.856
130.10610961914062 0 122815.45
131.06898498535156 0 36910.25
131.07373046875 0 6396.8447
131.1182861328125 0 5116.452
131.99795532226562 0 3543.0884
132.0813446044922 0 1807736.1
132.1024169921875 0 483734.72 y 13
133.08465576171875 0 166246
133.10585021972656 0 28962.406
134.0716552734375 0 12358.851
134.08787536621094 0 7836.753
134.1443328857422 0 2955.0486
136.0762481689453 0 2441922.5
136.4864959716797 0 3456.5947
137.07357788085938 0 10036.283
137.07957458496094 0 202772.3
138.06661987304688 0 32090.383
138.08216857910156 0 6319.962
138.1282958984375 0 4723.7803
138.39471435546875 0 2858.829
139.08709716796875 0 26732.174
141.102783203125 0 120150.625
142.06570434570312 0 38186.7 b 3
142.10606384277344 0 8989.514
144.081298828125 0 85918.87
145.07666015625 0 7722.8335
145.08477783203125 0 12087.292
146.0607452392578 0 9777.613
146.1295928955078 0 3564.4844
147.0446014404297 0 115708.66
147.11329650878906 0 5438.488
148.04808044433594 0 8039.1904
148.07618713378906 0 7957.535
149.08294677734375 0 5830.209
150.0663604736328 0 3625.2097
151.0871124267578 0 19624.975
151.92294311523438 0 4242.0522
152.0712890625 0 4178.6855
152.0823516845703 0 14329.473
152.1439971923828 0 50153.055
153.06643676757812 0 5862.2705
153.10287475585938 0 9352.977
154.065673828125 0 6654.7427
154.09803771972656 0 9349.486
154.1588897705078 0 3572.3496
155.08206176757812 0 102485.74
155.09332275390625 0 26891.605
155.1184539794922 0 61731.707
156.07664489746094 0 36174.15
156.08163452148438 0 48998.27
157.0764923095703 0 17237.7
157.08473205566406 0 6454.289
157.0973358154297 0 7127.6606
157.1350860595703 0 3532.8938
158.0606231689453 0 26349.676
158.0843963623047 0 213536.39
159.09222412109375 0 1256059.5
160.07620239257812 0 62918.273
160.08932495117188 0 6378.5938
160.0955810546875 0 127895.555
161.07958984375 0 8852.834
161.09889221191406 0 5721.782
161.36233520507812 0 3018.0266
162.05528259277344 0 4030.9756
162.1031951904297 0 7463.928
163.08641052246094 0 3960.398
163.6134490966797 0 11190.112
164.03431701660156 0 8069.394
164.0711669921875 0 32171.06
165.07711791992188 0 10842.665
165.10284423828125 0 79803.266
166.0615997314453 0 26482.76
166.1062469482422 0 9036.297
166.9498748779297 0 3775.6025
167.15487670898438 0 6800.762
168.08157348632812 0 7338.8716
168.11380004882812 0 17782.408
168.97137451171875 0 3950.8582
169.07659912109375 0 130954.75
169.0977325439453 0 129357.445
169.17022705078125 0 6752.445
170.06065368652344 0 614657.9
170.07936096191406 0 13883.438
170.1014862060547 0 9375.791
171.06402587890625 0 73303.6
171.0921630859375 0 21816.072
172.06781005859375 0 3769.4126
172.0760040283203 0 7243.6304 b Ammonia loss 4
172.10861206054688 0 120728.89
172.1454315185547 0 9067.126
174.05482482910156 0 4688.5713
175.09825134277344 0 18692.922
176.08245849609375 0 46117.652
179.0818328857422 0 7772.0923
179.15447998046875 0 7232.806
179.58547973632812 0 4392.605
180.0779571533203 0 7141.9
181.06146240234375 0 33230.496
181.0977783203125 0 582098.2
181.13409423828125 0 32653.244
181.1698455810547 0 6696.0234
182.06051635742188 0 13126.538
182.10113525390625 0 60161.863
182.12939453125 0 69210.88
183.09228515625 0 259886.23
183.11294555664062 0 16331.678
183.13282775878906 0 7098.402
183.14923095703125 0 5549.9453
184.07626342773438 0 33877.285
184.0956268310547 0 31837.514
185.07156372070312 0 65906.43
185.08074951171875 0 8720.51
185.1079864501953 0 11000.425
186.07916259765625 0 13497.287
186.1243438720703 0 248212.56
187.0872344970703 0 1322461.1
187.1276397705078 0 19413.777
188.08189392089844 0 9333.019
188.09059143066406 0 167188.2
189.09393310546875 0 7904.2144
190.61854553222656 0 4619.2217
191.08233642578125 0 8640.701
191.0935516357422 0 9876.262
191.11895751953125 0 6131.1704
193.0976104736328 0 6968.3867
193.1088104248047 0 4069.3362
195.1130828857422 0 9707.943
195.149658203125 0 23780.47
196.10877990722656 0 10496.401
197.07147216796875 0 6437.7554
197.08145141601562 0 4170.1904
197.1654510498047 0 94558.24
197.64532470703125 0 4549.6787
198.10313415527344 0 10320.706
198.1691436767578 0 8099.3584
199.0870819091797 0 56365.43
199.10841369628906 0 9802.794
199.1712188720703 0 5761.347
199.18109130859375 0 39389.87
200.09071350097656 0 6605.061
200.13998413085938 0 31794.322
200.18434143066406 0 4921.63
201.1028594970703 0 148173.06
201.12437438964844 0 6689.989
202.08676147460938 0 56869.18
202.10914611816406 0 207910.48
203.09246826171875 0 17261.963
203.1124725341797 0 21408.672
204.07733154296875 0 23518.746
205.0981903076172 0 7014.164 a 2
205.10916137695312 0 12435.825
207.12452697753906 0 9023.567
207.1497039794922 0 67373.64
208.14488220214844 0 13024.026
209.1287841796875 0 25393.533
209.16525268554688 0 9277.28
210.12452697753906 0 5884.8633
211.0872802734375 0 33411.797
212.09263610839844 0 7700.2974
212.13980102539062 0 18328.021
212.17608642578125 0 3969.7458
212.6683349609375 0 47007.285
213.07772827148438 0 15329.643
213.17005920410156 0 11919.705
214.10928344726562 0 67833.055
214.1198272705078 0 4654.592
214.1922149658203 0 18879.072
215.11309814453125 0 6851.485
216.1139373779297 0 21899.035
217.1090850830078 0 19588.295
220.11996459960938 0 155715.12
221.10401916503906 0 222258.89
221.12364196777344 0 11928.102
222.10633850097656 0 24237.367
222.12449645996094 0 12077.34
223.18145751953125 0 5101.4565
224.11880493164062 0 34465.137
224.13954162597656 0 8251.187
224.1636505126953 0 6047.8896
224.1763916015625 0 101659.31
225.10186767578125 0 4500.6963
225.12269592285156 0 4587.016
225.1603546142578 0 31455.773
225.1798858642578 0 10829.56
226.15574645996094 0 1778059.9
227.08236694335938 0 7911.618
227.13829040527344 0 8408.1455
227.1590576171875 0 213579.89 y Ammonia loss 10
227.1760711669922 0 8808.146
228.11375427246094 0 120669.91 a Water loss 1
228.13490295410156 0 11709.776
228.16128540039062 0 17852.525
229.09776306152344 0 143169.5 b Ammonia loss 4
229.11875915527344 0 18954.205
229.13778686523438 0 3958.7908
229.1665496826172 0 22920.922
230.08270263671875 0 8097.304
230.1029510498047 0 39281.973
231.07594299316406 0 7741.1636
231.0883331298828 0 81732.164
231.1359100341797 0 20229.367
232.0921173095703 0 11142.143
232.1200408935547 0 15884.338
232.14337158203125 0 6451.886
233.1040496826172 0 47680.19
234.10470581054688 0 12622.859
234.1353759765625 0 9853.759
235.1197052001953 0 298839.25
235.67120361328125 0 264175.03 y 10
236.12258911132812 0 32368.758
236.13827514648438 0 8017.157
236.1728515625 0 65694.305
236.6754913330078 0 7457.268
237.1356658935547 0 8043.841
237.15997314453125 0 11453.6045 y Ammonia loss 7
237.6268310546875 0 6304.72
238.13052368164062 0 23356.838
239.08229064941406 0 14976.385
239.15097045898438 0 117380.08
240.13497924804688 0 12923.316
240.15452575683594 0 13964.863
241.15579223632812 0 22546.727
242.18704223632812 0 64076.43
243.17071533203125 0 7488.6196
243.18246459960938 0 20176.404
244.10975646972656 0 6114.6333
244.16641235351562 0 14682.692
245.17152404785156 0 5103.731
245.186767578125 0 403032.5 y 12
246.12451171875 0 3225113.8 a 1
246.1898651123047 0 53455.508
246.64268493652344 0 4060.8042
247.1292724609375 0 739434.56
248.11474609375 0 241079.06
248.13259887695312 0 66104.96
248.18560791015625 0 5373.17
249.0989227294922 0 323765.66
249.11724853515625 0 20339.59 b 7
249.13351440429688 0 8693.188
250.10194396972656 0 35079.004
250.12344360351562 0 46064.406
251.12899780273438 0 23267.762
251.9132080078125 0 4262.1157
252.11074829101562 0 5496.5723
252.17124938964844 0 44144.465
253.0983428955078 0 4756.6333
253.17327880859375 0 4105.539
254.15084838867188 0 5451.847
254.186767578125 0 7487.219
255.07679748535156 0 10120.499
255.17201232910156 0 4755.372
256.1087341308594 0 70858.95 b Water loss 1
257.0924072265625 0 9546.729
257.1148681640625 0 32734.371
257.1614074707031 0 40435.984
258.0763244628906 0 56833.312
258.0992736816406 0 588415.44
259.0816345214844 0 7503.111
259.1025085449219 0 78273.56
259.130615234375 0 21321.371
259.85003662109375 0 9224.394
260.107421875 0 77750
260.1976623535156 0 40042.742
261.11260986328125 0 13385.767
261.1764221191406 0 6495.114
261.2000427246094 0 5397.489
261.6195983886719 0 5324.533
262.1197814941406 0 6288.344
262.1554260253906 0 13799.974
263.02850341796875 0 3949.3765
263.114990234375 0 14861.874
263.1299743652344 0 4846.999
264.1194763183594 0 7489.7627
264.13494873046875 0 31095.713
265.14044189453125 0 20975.031
265.16656494140625 0 21851.006
266.1101989746094 0 6991.358
266.1253967285156 0 190532.42
267.1282958984375 0 24585.53
267.14556884765625 0 10369.171
267.1820983886719 0 4505.2993
268.13006591796875 0 4857.665
269.1968994140625 0 4901.0825
269.6302490234375 0 40127.336
270.1295166015625 0 15718.685
270.18194580078125 0 134127.5
271.1890563964844 0 89843.83 y 9
271.69110107421875 0 21240.324
272.19146728515625 0 6362.5425
273.1359558105469 0 11371.398
274.1193542480469 0 1175764 b 1
275.1036682128906 0 167965.52
275.1253356933594 0 993777.7
275.18536376953125 0 26775.545 y 6
275.5198669433594 0 19142.398
276.10821533203125 0 44616.25
276.12847900390625 0 108404.18
277.0937194824219 0 11595.063
277.1092224121094 0 5538.1113
277.1157531738281 0 4535.5576
277.1340026855469 0 45930.92
278.13775634765625 0 8056.4224
279.1523132324219 0 6929.344
279.18194580078125 0 26801.012
281.6723327636719 0 4827.686
282.1826477050781 0 4374.932
283.1512451171875 0 30857.893
283.1773376464844 0 435930.88
283.62799072265625 0 36370.39
284.12982177734375 0 10790.776
284.1544494628906 0 7485.845
284.180419921875 0 59247.727
285.11016845703125 0 23986.672
285.1564636230469 0 82994.414
285.229248046875 0 5357.9062
286.11383056640625 0 7909.282
286.15972900390625 0 8898.839
288.1084899902344 0 5806.89
290.1141357421875 0 8956.904
290.1850891113281 0 140764.45
290.6866149902344 0 33220.33
291.1572265625 0 40279.62
291.1864318847656 0 6694.105
292.1299743652344 0 302981.06
293.13336181640625 0 45304.645
293.1617736816406 0 8198.556
294.12042236328125 0 79705.88
294.1368408203125 0 7918.071
294.2183837890625 0 33014.535
295.12286376953125 0 11463.081
295.2217102050781 0 6445.8696
296.1974182128906 0 6749.13
297.19305419921875 0 226379.62
298.1966247558594 0 32812.203
300.2032470703125 0 9596.621
300.6728820800781 0 20291.264
302.11431884765625 0 7931.0117
303.1206359863281 0 59241.195
303.21820068359375 0 8215.518
304.1241760253906 0 8878.154
304.2214050292969 0 4354.9097
305.1292724609375 0 118568.086
305.66497802734375 0 47293.523
305.86956787109375 0 6551.2246
306.1328430175781 0 27417.887
306.1658630371094 0 10953.699
307.1772155761719 0 8145.765
308.1253662109375 0 5055.3486
309.16937255859375 0 9357.855
309.1924133300781 0 17282.512
311.17254638671875 0 5480.1284
311.21173095703125 0 5163.8745
311.24505615234375 0 70749.7
312.24847412109375 0 12848.806
313.22369384765625 0 4741.991
314.21954345703125 0 5380.1465
314.6700439453125 0 46142.93
315.17138671875 0 19690.883
315.1978454589844 0 5520.4287 y Ammonia loss 5
315.67376708984375 0 4640.119
318.1932067871094 0 15490.547
320.1246643066406 0 21835.318
320.87213134765625 0 58824.5 y 5
321.20745849609375 0 22931.635
321.22943115234375 0 275500.6
321.5401916503906 0 8446.886
321.8478088378906 0 10304.94
322.1559143066406 0 193002
322.2131652832031 0 8275.55
322.23260498046875 0 48521.47
322.85919189453125 0 7167.326
323.12677001953125 0 5315.3237
323.1591491699219 0 35934.863 b 10
323.1938171386719 0 8038.2373
323.236328125 0 5965.4478
324.1631774902344 0 5721.472
324.2167663574219 0 40151.81
324.7177734375 0 13134.371
325.1423645019531 0 4614.3936
325.18780517578125 0 6786.1724
325.22296142578125 0 10032.203
326.2197265625 0 105296.66
326.72442626953125 0 7156.436 y Ammonia loss 8
327.22344970703125 0 23221.408
327.2768859863281 0 24008.666
330.10906982421875 0 6904.605
331.8177185058594 0 5626.195 b 7
332.72991943359375 0 116400.625
333.1241760253906 0 35570.58
333.23089599609375 0 26520.656
334.1272277832031 0 8870.91
334.175048828125 0 10837.2705
334.6683044433594 0 11891.145
335.16973876953125 0 6600.3555
335.2373352050781 0 27599.957 y 8
335.7389221191406 0 9522.528
336.20404052734375 0 408907.38
337.1864013671875 0 8922.713
337.2069396972656 0 72330
337.26055908203125 0 5993.179
337.7220153808594 0 123120.06
338.16064453125 0 6767.8325
338.2221984863281 0 32504.584
338.7240905761719 0 9678.402
339.1692199707031 0 8559.464
339.2400207519531 0 395881.78
339.83013916015625 0 4348.4106
340.15301513671875 0 36744.848
340.2432861328125 0 71256.04
341.15625 0 7758.9736
341.21978759765625 0 9006.607
341.2464294433594 0 11138.602
342.1686706542969 0 15266.286
342.25006103515625 0 6015.859
345.6446228027344 0 17541.723
346.1447448730469 0 11500.448
346.66827392578125 0 14276.794
346.727294921875 0 343035.84
347.22857666015625 0 136327.92
347.7300109863281 0 43391.957
347.9520568847656 0 10006.621 y 2
348.14691162109375 0 6789.443
348.2301025390625 0 5893.3335
349.13079833984375 0 7625.6787
349.1986083984375 0 5033.3057
349.7095642089844 0 19194.342
350.151123046875 0 44485.08
350.2103271484375 0 17190.312
350.5416564941406 0 6990.244
350.69189453125 0 4434.331
350.8775634765625 0 7763.975
351.1207275390625 0 4955.649
351.1534423828125 0 8930.923
351.2400817871094 0 29593.26
351.6569519042969 0 5181.021 b 4
351.8283386230469 0 5711.4116
352.15594482421875 0 6413.4727
352.24114990234375 0 5934.7583
353.2188415527344 0 16864.406
353.87103271484375 0 6631.356
354.1575622558594 0 51743.05
354.17510986328125 0 24053.72
354.2144775390625 0 514049.03
354.5099182128906 0 10801.118
354.5357971191406 0 4998.4414
354.6582946777344 0 26874.773
354.84130859375 0 5154.78
355.1586608886719 0 4507.0117
355.21771240234375 0 94710.28
355.2706298828125 0 10700.859
355.7325744628906 0 55362.414
356.1722412109375 0 12550.324
356.2333679199219 0 20809.447
356.7356262207031 0 8690.529
357.1788635253906 0 18677.182
357.2505798339844 0 117883.79
357.504150390625 0 11670.585
357.8336181640625 0 19761.02
358.16326904296875 0 107592.38
358.25421142578125 0 19165.123
358.71478271484375 0 38555.34
359.1415100097656 0 5197.9194
359.1663818359375 0 24808.979
359.1941833496094 0 9302.507
359.2162780761719 0 17511.434
359.5431823730469 0 26217.73
359.7177429199219 0 5799.5483
359.85052490234375 0 5690.1313
359.8739318847656 0 26641.508
360.1353759765625 0 11064.293
360.203369140625 0 7160.95
360.6651916503906 0 33935.918
360.68975830078125 0 29366.658
360.8809814453125 0 6394.062 y Ammonia loss 4
361.18499755859375 0 15025.57
362.151123046875 0 10510.463
362.66961669921875 0 8425.471
363.1754455566406 0 7215.7104
363.1968688964844 0 9661.319
363.50775146484375 0 32010.588
363.7479553222656 0 23551.812 y 7
363.8428955078125 0 14928.705
364.19915771484375 0 6568.112
364.25006103515625 0 7943.798
365.2555236816406 0 11250.171
365.5465393066406 0 13469.304
365.8770446777344 0 5638.2876
366.15704345703125 0 53016.375
366.2596740722656 0 5269.4985
366.5580749511719 0 64378.895 y 4
366.8926086425781 0 28864.17
367.1393127441406 0 8381.239
367.162841796875 0 44568.703
367.2275390625 0 16965.29
368.1486511230469 0 59205.656
368.1722106933594 0 14623.4
368.65625 0 17529.719
369.1527099609375 0 15229.061
369.1812438964844 0 19179.633
369.6938171386719 0 18237.47
370.1869201660156 0 25178.22
370.2459716796875 0 11540.948
370.5272521972656 0 4406.833
370.6885986328125 0 7576.907
370.8599853515625 0 6153.7646
373.1849670410156 0 5575.9683
373.28192138671875 0 244248.33 y 11
374.1512756347656 0 9795.886
374.25714111328125 0 8399.92
374.2848815917969 0 35680.785
374.5140686035156 0 9400.6 b 8
374.8489990234375 0 17708.846
375.1885681152344 0 12898.816
375.28814697265625 0 5542.5757
376.45654296875 0 10665.936
376.5313720703125 0 8147.0103
376.6676025390625 0 16395.44
376.86920166015625 0 6010.5195
376.9572448730469 0 5106.274
377.1254577636719 0 4991.451
377.1702575683594 0 8222.659
377.5386047363281 0 4730.5625
378.2493896484375 0 5187.893
378.7124938964844 0 20869.023
378.9623107910156 0 26565.26
379.2123718261719 0 12818.746
379.4668884277344 0 7197.068
380.2302551269531 0 27096.734
381.2319641113281 0 5201.467
382.20660400390625 0 107309.125
382.2823181152344 0 26191.17
382.5402526855469 0 54908.8
382.6678466796875 0 6877.085
382.8764953613281 0 9635.899
383.1887512207031 0 7208.933
383.2663879394531 0 21231.309
383.4547119140625 0 7655.6846 b 12
383.7410888671875 0 7734.1196
384.1676025390625 0 71814.125
385.1736755371094 0 214965.12
385.19970703125 0 61909.586
385.9567565917969 0 6350.7495
386.1579895019531 0 211453.86
386.2026062011719 0 16909.049
386.52642822265625 0 8932.327
386.8614196777344 0 8389.299
387.16094970703125 0 52664.043
388.1629943847656 0 7356.5366
388.2374267578125 0 10170.939
388.5688781738281 0 6354.4556
388.8624267578125 0 56168.707
389.1968994140625 0 40998.08
389.2716979980469 0 51322.93
389.5311584472656 0 6255.6245
389.7742919921875 0 21573.104
390.2146301269531 0 159214.86 y Ammonia loss 1
390.27276611328125 0 5973.1714
390.4652099609375 0 113045.09
390.7159423828125 0 37758.37
390.9659423828125 0 14254.607
392.162109375 0 6870.278
392.1902770996094 0 26867.104 b Water loss 9
392.2663269042969 0 69026.42
392.5229797363281 0 29920.303 b Ammonia loss 9
392.7555847167969 0 8659.608
392.8548889160156 0 15801.75
393.167724609375 0 17421.924
393.2174377441406 0 9309.076
393.269775390625 0 19200.258
394.1521301269531 0 169230.3
394.47100830078125 0 63348.64 y 1
394.7216796875 0 60224.242
394.9727783203125 0 35311.996
395.15472412109375 0 30656.605
395.22430419921875 0 11724.634
395.4739685058594 0 5304.628
395.9692077636719 0 5498.139
396.1617126464844 0 5351.638
396.2618103027344 0 20830.105
397.17987060546875 0 9601.549
397.245849609375 0 18723.8
397.5676574707031 0 14381.106
397.6741027832031 0 5987.4805
397.7474365234375 0 9528.406
398.19390869140625 0 41368.516 b 9
398.2408447265625 0 10347.483
398.52777099609375 0 28471.393
398.86212158203125 0 8863.239
398.8915100097656 0 18077.213
399.22125244140625 0 7638.203
399.56280517578125 0 4483.028
400.17340087890625 0 10489.17
400.2218017578125 0 5825.6274
400.4747619628906 0 7641.055
400.7220764160156 0 14551.314
401.195556640625 0 18560.25
401.25927734375 0 25053.652
401.7602233886719 0 9741.081
402.2249755859375 0 79824.9
402.55194091796875 0 12599.341
402.88232421875 0 8789.2
403.1847229003906 0 208668.31
403.2403869628906 0 105942.93
403.2700500488281 0 56980.027
403.5717468261719 0 171494.19 y Ammonia loss 3
403.76666259765625 0 32643.059 y Ammonia loss 6
403.9046325683594 0 73257.35
404.1876220703125 0 38666.49
404.2386169433594 0 32450.805
404.2660217285156 0 7705.2896
404.47039794921875 0 24732.217
404.5727233886719 0 8418.749
404.7251892089844 0 66614.56
404.89447021484375 0 20709.268
404.97760009765625 0 68355.92
405.2284851074219 0 45671.117
405.48028564453125 0 13218.472
406.18646240234375 0 72599.69
406.2514343261719 0 70374.57
406.6882629394531 0 32015.031
406.75274658203125 0 27298.322
407.1928405761719 0 31855.08
407.2538146972656 0 12961.727
407.4700012207031 0 5422.505
407.6930236816406 0 6025.842
408.2221984863281 0 58469.574
408.5551452636719 0 37143.168
408.888671875 0 19305.648
409.21685791015625 0 7669.022
409.24481201171875 0 42833.184 y 3
409.5787658691406 0 34927.074
409.9140930175781 0 8146.513
410.2513427734375 0 5020.3643
410.2774658203125 0 188540.28
410.5693664550781 0 94140.84
410.9035339355469 0 78524.86
411.1785888671875 0 197440.16 b Water loss 5
411.23760986328125 0 29655.736
411.2798156738281 0 37908.504
411.57049560546875 0 6209.0425
411.726318359375 0 46729.29 Precursor Water loss
411.974853515625 0 52282.85 Precursor Ammonia loss
412.1623840332031 0 45806.418
412.1853332519531 0 24523.6
412.22552490234375 0 39182.312
412.27484130859375 0 697009.7 y 6
412.4749450683594 0 15258.341
412.7762451171875 0 350922.94
413.1676025390625 0 21155.053
413.27789306640625 0 79111.16
413.7789001464844 0 17397.498
413.8975524902344 0 110281.664
414.1939392089844 0 16304.841
414.2300720214844 0 111754.44
414.2799987792969 0 5604.8823
414.5634765625 0 49570.492
414.8970642089844 0 15594.369
415.1546630859375 0 5075.24
415.25665283203125 0 67521.234
415.7586975097656 0 32989.934
416.22930908203125 0 402776.22 Precursor
416.47998046875 0 387122.88
416.73077392578125 0 180428.55
416.9814147949219 0 95832.41
417.2303466796875 0 17869.36
418.7143249511719 0 10879.555
419.5641784667969 0 6516.1226
419.9010314941406 0 363560.6
420.1834716796875 0 50302.363 b 5
420.23541259765625 0 332678.62
420.56964111328125 0 116593.695
420.685791015625 0 17725.455
420.9040222167969 0 42046.027
421.189208984375 0 7851.9844
421.23980712890625 0 18776.434
422.27734375 0 53495.6
423.2750549316406 0 26638.031
424.2620544433594 0 9715.086
424.3296203613281 0 55307.707
424.7127380371094 0 116805.445
424.7625732421875 0 6892.052
425.2142639160156 0 49604.758
425.28839111328125 0 25671.271
425.3316955566406 0 16741.316
425.71539306640625 0 15452.488
425.9045715332031 0 79305.414
426.23876953125 0 52194.55
426.2900390625 0 6618.25
426.5730285644531 0 10089.811
426.90869140625 0 6538.36
427.24383544921875 0 69081.51
427.7458190917969 0 43375.555
428.2453918457031 0 11693.232
428.2874450683594 0 13904.944
428.7174377441406 0 5960.2417
429.1894226074219 0 690128.44
430.192138671875 0 174370.75
430.54833984375 0 6844.876 b 10
431.180419921875 0 92871.95
432.1835632324219 0 21263.371
432.2343444824219 0 6631.7983
433.2570495605469 0 11207.096
433.7069091796875 0 15307.8125
434.2083740234375 0 13682.824
436.5844421386719 0 9723.421
436.9169921875 0 7742.9844
437.1835021972656 0 44403.895 b 2
437.2600402832031 0 5487.759
437.29058837890625 0 7495.1
438.1893615722656 0 8412.509
438.2220764160156 0 8173.8555
438.55712890625 0 65477.824
438.891845703125 0 49785.062
439.1741943359375 0 11800.249
439.2270812988281 0 8650.273
439.30255126953125 0 8965.471
439.5595703125 0 10837.891
442.18804931640625 0 14256
442.71246337890625 0 23289.562
443.2144775390625 0 13743.635
443.70623779296875 0 9849.396
444.234130859375 0 15066.053
444.56658935546875 0 15298.192
444.9017639160156 0 7493.2183
445.19256591796875 0 10302.374
445.2682189941406 0 24737.178
445.9166259765625 0 11418.401
446.2337646484375 0 59549.746
446.5782775878906 0 14636.261
446.7355041503906 0 43439.51
447.2360534667969 0 10938.365
448.2641906738281 0 120779.09
448.5986633300781 0 74824.5
448.9325256347656 0 30191.686
449.26171875 0 8449.93
450.2816162109375 0 8548.773
451.2264709472656 0 91711.33
451.2674255371094 0 54506.957
451.5931701660156 0 5774.041
451.7254333496094 0 62300.5
451.9203796386719 0 166260.23
452.2226257324219 0 12772.945
452.25445556640625 0 129533.36
452.3246154785156 0 26498.506
452.5881652832031 0 50144.95
452.9219970703125 0 26647.447
453.25714111328125 0 10552.47
453.7038879394531 0 14408.737
454.2082214355469 0 32132.992
455.2162780761719 0 6275.062
455.2526550292969 0 68325.766
456.2541198730469 0 13361.745
457.1838684082031 0 19231.621
457.5623779296875 0 7189.49
457.5959167480469 0 79175.56
457.80072021484375 0 9407.3125
457.9273986816406 0 134635 y Ammonia loss 2
458.25982666015625 0 78453.53
458.59405517578125 0 29229.44
458.9267883300781 0 11023.167
459.21502685546875 0 17631.99
460.2312316894531 0 146776.45
460.73284912109375 0 66309.84
461.1822509765625 0 15172.955
461.2410888671875 0 21547.342
461.91387939453125 0 7918.983
462.7922058105469 0 7770.6616
462.9281921386719 0 12554.429
463.2765808105469 0 18735.768
463.599365234375 0 1913260.5 y 2
463.9336242675781 0 1610667.6
464.2679138183594 0 614425
464.6021728515625 0 219969.88
464.936279296875 0 46352.95
466.92022705078125 0 41007.23
467.25469970703125 0 29798.146
467.298828125 0 56326.08
467.5870056152344 0 18943.4
467.9124755859375 0 9753.897
468.29693603515625 0 26889.469
469.7892761230469 0 38044.04
470.250244140625 0 33288.6
470.3349304199219 0 793430.44 y 10
470.5814208984375 0 59161.816
470.91705322265625 0 17730.541
471.2503356933594 0 12803.331
471.3380432128906 0 208108.17
471.7994079589844 0 52687.426
472.2994079589844 0 33713.83
472.3408203125 0 35647.707
472.5958251953125 0 18231.102
472.79974365234375 0 9568.536
472.930908203125 0 20863.877
473.2621765136719 0 93523.35
473.5771789550781 0 44873.598
473.9158020019531 0 9003.929
474.26580810546875 0 23273.896
474.78057861328125 0 54168.656
475.280517578125 0 24616.752
475.7820739746094 0 13320.655
475.9235534667969 0 21478.83
476.25384521484375 0 138171.84
476.5874938964844 0 136302.48
476.92169189453125 0 40311.062
477.2548828125 0 18211.771
478.2096862792969 0 25854.943
479.21575927734375 0 14005.059
479.2498779296875 0 7892.029
479.58154296875 0 10243.217
480.8042907714844 0 168395.02 y 5
481.30596923828125 0 98294.06
481.8063659667969 0 17710.229
481.9272766113281 0 74692.01
482.2611389160156 0 92921.836
482.3094482421875 0 57232.793
482.5926818847656 0 49391.34
482.7616882324219 0 49060.246
482.92889404296875 0 20970.885
483.2208251953125 0 9977.581
483.26263427734375 0 34802.434
483.311767578125 0 17142.725
483.78631591796875 0 115654.5
484.19580078125 0 10681.162
484.2879333496094 0 75161.19
484.7892761230469 0 21028.863
485.2900390625 0 8384.983
487.93035888671875 0 33005.742
488.2653503417969 0 16544.14
488.5979919433594 0 8638.964
489.93414306640625 0 7604.106
491.2358703613281 0 7568.2554
491.2735900878906 0 69500.125
491.7756652832031 0 13659.004
492.277099609375 0 14356.617
492.7919006347656 0 40579.727
493.2328186035156 0 13560.603
493.2931823730469 0 20609.45
493.3505859375 0 9203.232
494.76361083984375 0 9372.983
495.26312255859375 0 11134.756
495.36578369140625 0 18294.246
495.5986633300781 0 6267.1567
495.9293212890625 0 11965.13
496.26763916015625 0 7591.195
497.2216491699219 0 26141.799 b 7
497.72296142578125 0 15418.796
500.23760986328125 0 9602.011
501.262939453125 0 9650.974
501.60418701171875 0 20728.22
501.93963623046875 0 34054.613
502.2769775390625 0 6319.274
503.2162780761719 0 58939.81
504.2169189453125 0 14078.909
504.6152038574219 0 10225.111
504.9358215332031 0 32680.71 b Water loss 12
505.2702331542969 0 30875.865 b Ammonia loss 12
505.6053771972656 0 20984.967
505.73358154296875 0 14081.017
506.23321533203125 0 6292.225
508.7576904296875 0 32749.713
509.2412109375 0 36881.645
509.7417907714844 0 21016.113
510.2435607910156 0 8863.203
510.2925109863281 0 11704.473
510.3774719238281 0 8692.701
510.62457275390625 0 23821.443
510.93841552734375 0 72367.32 b 12
511.27227783203125 0 95805.57
511.607177734375 0 44290.234
511.9449462890625 0 19061.441
513.9449462890625 0 17619.316
514.2786865234375 0 15409.928
514.6124267578125 0 29171.654
516.9417114257812 0 192321.7
517.275390625 0 170537.61
517.6103515625 0 68054.57
517.7536010742188 0 25799.43
517.94482421875 0 27659.86
518.2536010742188 0 25221.857
519.6217651367188 0 12857.057
519.9501953125 0 190943.95 y Ammonia loss 1
520.2429809570312 0 9153.475
520.2843627929688 0 128913.2
520.3616943359375 0 46704.04
520.6182861328125 0 90681.96
520.9527587890625 0 31630.328
521.2269897460938 0 69078.4
521.2852172851562 0 10024.614
521.361083984375 0 11756.11
522.226806640625 0 21333.549
523.3612060546875 0 14692.8955
524.3574829101562 0 8106.01
524.8134155273438 0 12103.335
525.2246704101562 0 13080.173
525.3060913085938 0 29388.426
525.6256713867188 0 280659.22 y 1
525.8070068359375 0 14963.544
525.9600219726562 0 314069.3
526.29443359375 0 117543.37
526.33349609375 0 9303.814
526.6281127929688 0 54397.58
526.8319091796875 0 9196.138
526.9622802734375 0 6437.4756
527.2368774414062 0 8840.5
528.2337646484375 0 21380.576
528.9528198242188 0 7449.443
529.2244262695312 0 9443.65
529.2901611328125 0 7779.0703
529.8035278320312 0 27557.002
530.2993774414062 0 44493.44
530.7579956054688 0 8204.156
530.7988891601562 0 25337.547
531.2116088867188 0 56818.082
531.299560546875 0 11236.637
532.21142578125 0 17640.086
533.8193359375 0 9643.595
535.7523803710938 0 50857.527
536.2535400390625 0 33958.11
536.7540283203125 0 7906.726
538.2530517578125 0 422435.03
538.3729858398438 0 32780.844
538.8109741210938 0 56562.523
539.2554321289062 0 116161.85
539.3046875 0 86050.63
539.3735961914062 0 7349.846
539.804931640625 0 60217.96
540.2579956054688 0 18102.66
540.302978515625 0 5924.6836
540.3264770507812 0 22301.074
540.8287963867188 0 26871.953 y Ammonia loss 4
541.3271484375 0 9766.014
541.3720703125 0 190158.62 y 9
542.2521362304688 0 75626.016
542.375 0 67410.99
543.2555541992188 0 27789.793
543.3775024414062 0 7721.6484
544.2647094726562 0 19921.271
544.7609252929688 0 42138.816
545.2609252929688 0 31727.883
545.759033203125 0 13933.325
547.8157958984375 0 54119.027
548.2367553710938 0 16902.066
548.3128662109375 0 38943.215
548.8125610351562 0 25783.855
549.2221069335938 0 40596.805
549.3336181640625 0 478154.03 y 4
549.8350830078125 0 320914.16
550.2229614257812 0 11746.347
550.3364868164062 0 91901.09
550.8373413085938 0 35677.75
552.2659301757812 0 7280.1333 b Water loss 8
553.2200927734375 0 29111.346
553.2680053710938 0 20991.154
553.7700805664062 0 13343.572
554.2227172851562 0 8125.6113
554.3241577148438 0 7196.251
555.2850952148438 0 18714.88
555.7872314453125 0 14508.633
556.2811889648438 0 8142.932
556.820068359375 0 14623.769
557.26171875 0 7088.5757
557.2958984375 0 43207.5
558.3015747070312 0 8604.206
561.268310546875 0 133371.06 b 8
561.3504028320312 0 11029.858
561.7694702148438 0 91397.35
562.2699584960938 0 26126.748
562.34716796875 0 7070.446
562.7736206054688 0 14840.869
564.2942504882812 0 13251.52
564.7982177734375 0 7523.321
565.2422485351562 0 16970.06 b 3
566.2479858398438 0 504002.4
567.2506713867188 0 149584.94
567.3986206054688 0 28978.203
568.2530517578125 0 31342.354
570.2471923828125 0 233212.97
571.2501831054688 0 78120.8
572.2525024414062 0 12507.491
572.8051147460938 0 236536.06
573.3065795898438 0 146606.39
573.8084716796875 0 58120.84
574.3124389648438 0 12756.533
577.3832397460938 0 151178.42
578.3839111328125 0 51728.78
579.2487182617188 0 9000.483
579.3624877929688 0 295816.34
580.3656005859375 0 97078.266
581.215576171875 0 14189.124
581.3653564453125 0 24945.84
581.8102416992188 0 12805.68
582.7890625 0 9038.3955
583.2892456054688 0 7101.6704
587.2732543945312 0 16971.34
587.7824096679688 0 28442.496 b Water loss 9
588.2809448242188 0 31538.816 b Ammonia loss 9
588.7841796875 0 10753.017
594.241943359375 0 11885.485
595.393798828125 0 105328.06
595.8460693359375 0 16114.257
596.3447265625 0 17194.426
596.39697265625 0 36632.094
596.7864990234375 0 35052.535 b 9
596.8389892578125 0 8091.277
597.28759765625 0 26020.29
598.2420043945312 0 109576.7
599.24462890625 0 34647.895
600.2490844726562 0 9523.473
600.3363647460938 0 11993.029
601.7968139648438 0 14907.627
604.3578491210938 0 111118.48
604.8525390625 0 189812.3 y Ammonia loss 3
605.353271484375 0 124849.65
605.8525390625 0 43239.3
606.3563232421875 0 19336.178
606.8390502929688 0 13072.819
607.34521484375 0 6298.814
610.3212890625 0 104775.41
611.3237915039062 0 37856.39
611.8292236328125 0 22896.846
612.3273315429688 0 15698.173
612.8329467773438 0 7357.893
613.3626708984375 0 101473.25 y 3
613.4073486328125 0 11569.128
613.86474609375 0 69318.08
614.3644409179688 0 22170.39
614.8675537109375 0 7678.4116
615.2686157226562 0 26307.465
615.3497924804688 0 60687.94
615.8512573242188 0 41594.87
616.2725219726562 0 11405.46
616.3527221679688 0 13722.8955
618.849365234375 0 19039.648
619.3504638671875 0 8384.831
620.342041015625 0 85041.91
620.8407592773438 0 77874.69
621.3408203125 0 43768.094
621.8424072265625 0 11642.095
623.3890991210938 0 15736.714
628.3323364257812 0 38134.418
629.3469848632812 0 214921.75
629.8482666015625 0 143456.27
630.3489990234375 0 68694.055
630.850341796875 0 22154.615
631.3530883789062 0 6316.48
638.352294921875 0 144148.45
638.8540649414062 0 109430.586
639.3551025390625 0 31840.11
639.8546752929688 0 7746.557
647.4254150390625 0 19863.459
648.3359375 0 7569.857
648.4280395507812 0 7684.8696
648.8358154296875 0 9734.242
657.278076171875 0 18275.572
657.3334350585938 0 27553.027
657.8344116210938 0 20917.133
658.3286743164062 0 9115.912
663.3038330078125 0 7423.3994
664.45166015625 0 81117.03
665.4539184570312 0 29933.904
665.8455810546875 0 31536.58
666.344482421875 0 29041.578
666.4553833007812 0 9599.186
667.3479614257812 0 20276.32
668.3294677734375 0 26453.926
669.3338623046875 0 12522.485
669.467041015625 0 68585.95 y 8
670.4701538085938 0 22985.2
674.3050537109375 0 48597.88
674.4362182617188 0 314350.25
675.3074340820312 0 15842.346
675.43798828125 0 122221.62
676.4422607421875 0 21510.314
679.3096923828125 0 9357.02
680.3157348632812 0 7431.071
684.292724609375 0 13368.609 b Water loss 4
685.2848510742188 0 10496.37 b Ammonia loss 4
685.3524169921875 0 22676.258
686.3892822265625 0 11289.9795 y Ammonia loss 2
690.28076171875 0 21365.463
692.4464111328125 0 289355.16
693.4490966796875 0 118150.48
694.4515991210938 0 20393.988
694.894775390625 0 104944.81 y 2
695.3965454101562 0 73326.47
695.8971557617188 0 30740.05
696.4005126953125 0 10896.079
698.4103393554688 0 24575.934
699.4118041992188 0 13305.102
699.882568359375 0 7848.177
702.3013305664062 0 64420.953 b 4
703.3043212890625 0 24846.912
707.3062744140625 0 138265.44
708.309326171875 0 62656.793
708.4771728515625 0 18562.623
709.3111572265625 0 12720.632
709.3692626953125 0 9330.479 b 11
709.4765014648438 0 10797.807
709.8622436523438 0 14476.65
710.4570922851562 0 74898.836
711.46044921875 0 37621.742
712.4642333984375 0 7600.449
713.3796997070312 0 11114.856
713.8800048828125 0 37347.457
714.3798828125 0 22454.602
714.88232421875 0 14555.734
716.4212646484375 0 64291.184
717.4236450195312 0 21818.281
718.8677978515625 0 7222.768
720.3221435546875 0 32873.117
721.35205078125 0 32777.223
722.3678588867188 0 21687.25
722.884033203125 0 19737.883
723.3832397460938 0 15850.413
724.3333129882812 0 41170.395
725.3348388671875 0 16662.87
726.48828125 0 123865.12 y 7
727.4920043945312 0 64016.156
728.39990234375 0 10326.756
728.4932250976562 0 12849.66
731.3920288085938 0 9389.674
732.3941650390625 0 7637.9614
735.30078125 0 134369.12
736.303466796875 0 57791.188
736.470947265625 0 6616.5815
737.3489379882812 0 21016.875
738.3743896484375 0 17658.023
739.369384765625 0 18520.676
740.37353515625 0 8468.161
752.3280639648438 0 68325.73
753.3296508789062 0 29339.3
754.3348999023438 0 7495.017
756.39208984375 0 13350.134
760.5165405273438 0 6915.457
777.5347290039062 0 18483.035
780.3225708007812 0 5720.512
783.5018920898438 0 5997.8457
793.340576171875 0 6308.762
793.4862670898438 0 6630.6743
794.3425903320312 0 6741.1353
794.482177734375 0 5538.69
801.509033203125 0 10189.608
802.5117797851562 0 7737.5093
805.530517578125 0 47862.094
806.52685546875 0 20325.367 y Ammonia loss 6
807.52734375 0 6787.372
811.3639526367188 0 39747.297
811.4949340820312 0 38622.46
812.3670043945312 0 18341.254
812.4959106445312 0 19024.596
813.3803100585938 0 7215.724
813.5004272460938 0 6136.63
821.3495483398438 0 8464.448 b Water loss 5
822.34912109375 0 8720.872
823.5408935546875 0 547495.5 y 6
824.5438842773438 0 258897.38
825.5465698242188 0 63035.895
829.5053100585938 0 36864.594
830.5059814453125 0 19654.549
831.512451171875 0 6146.9404
839.3584594726562 0 74813.77 b 5
840.3623046875 0 34381.62
841.3668823242188 0 10822.938
847.5172729492188 0 21479.984
848.4166259765625 0 79379.055
848.5106811523438 0 6920.6396
849.4204711914062 0 38509.715
850.419189453125 0 14243.999
852.451171875 0 7479.0273
853.4763793945312 0 28513.195
854.4747314453125 0 16959.59
866.4109497070312 0 5666.6016
874.4365234375 0 6084.6196
884.4208984375 0 11020.697
885.4114990234375 0 6772.87
889.3724975585938 0 10317.656
890.3790893554688 0 8750.601
891.458740234375 0 13287.936
892.4576416015625 0 7022.7812
901.4424438476562 0 94466.19
902.4435424804688 0 53056.582
903.442138671875 0 11753.452
906.39794921875 0 11811.718
919.4535522460938 0 52452.453
920.4569091796875 0 34521.36
921.4613647460938 0 9519.788
942.5889892578125 0 22311.584
943.5934448242188 0 9743.558
948.5516357421875 0 6249.844
960.599609375 0 56881.516 y 5
961.6024169921875 0 34109.26
962.6112060546875 0 9754.16
964.5060424804688 0 5452.027
966.5615844726562 0 13307.381
967.5648803710938 0 9092.271
984.57763671875 0 4969.9683
993.4317016601562 0 8109.2754 b 7
1004.542236328125 0 6402.35
1017.4722290039062 0 7758.6094
1018.4794921875 0 7326.6235
1032.53955078125 0 10793.023
1033.5352783203125 0 20987.947
1034.5369873046875 0 10614.67
1070.4969482421875 0 19636.754
1071.496826171875 0 11465.316
1072.4869384765625 0 4900.7227
1087.524658203125 0 12277.562
1088.51806640625 0 6116.1562
1097.653564453125 0 6704.9106 y 4
2340.854736328125 0 5569.943
3447.593994140625 0 5717.509

Spectrum Details

|  |  |
| --- | --- |
| Matched peaks? Matched peaks The total absolute number of peaks matched. Additionally in brackets the total fraction of peaks matched and the total number of peaks is shown. | 82 (6.69% of 1226) |
| FDR? FDR The false discovery rate estimated for this peptide. It is calculated by matching all theoretical fragments with a non-integer shift with the raw peaks for this spectrum. This is done with 40 different shifts. The resulting percentage is the average number of annotated peaks over the number of annotated peaks with the correct spectrum. | 5.23% |
| Satellite FDR? Satellite FDR See the FDR for details on its calculation. This satellite ion specific FDR only contains the satellite ions (d/w) for I/L/J positions. | - |
| PSM Score? PSM Score The PSM Score as given by Hecklib to this annotated spectrum. It is shown with three significant figures. | 288 |

#### Spectrum 4737? Spectrum 4737 The raw spectrum of this peptide as annotated by Hecklib. The fragments are coloured according to ion type (see legend). Any peaks with a star '\*' as text can be hovered over to see the full details, first the ion type second the mass shift type. By hovering over the amino acids in the peptide or ions in the legend the corresponding peaks are highlighted. By toggling the 'Unassigned' label you can turn the background (unassigned) peaks on or off in the plot. By updating the slider in the Ion legend you can update the spectrum to only show the top X% of the peaks with labels. The top X% means any peak that is within X% of the highest intensity. By dragging in the spectrum you can zoom in to a specific part of the spectrum and use 'Zoom Out' to get back to the original zoom level. The annotation of the spectrum is based on the given sequence in the peptides file and is done with different software so inconsistencies are likely. The peaks are annotated based on the given sequence, with 20 ppm tolerance.

Copy Data

##### Spectrum 4737 (TSV)

###### Preview

```
Loading example...
```

*Click on the button to copy the data to your clipboard.*

Mz MinMz MaxIntensity Max

WidthHeightPeptide font sizePeptide stroke widthSpectrum font sizeSpectrum stroke widthCompact peptide

Ion legend

wxyz

abcd

OtherUnassignedIon^Charge^_Position_Show for top:%

SWYQHHPGKAPKJJ

01.40e+42.79e+44.19e+45.58e+4

Zoom Out

y^+1^_1_b^+4^_4_y^+2^_3_y^+3^_6_y^+2^_4_y^+2^_4_y^+1^_2_a^+1^_2_y^+2^_5_b^+1^_2_b^+1^_2_b^+2^_4_y^+3^_9_y^+3^_10_y^+1^_3_y^+4^_13_b^+3^_10_y^+4^_13_b^+3^_10_y^+3^_11_y^+2^_8_y^+2^_8_\*b^+2^_6_\*b^+2^_6_b^+3^_11_b^+1^_3_b^+1^_3_y^+3^_12_y^+3^_12_y^+1^_4_y^+2^_9_b^+3^_12_y^+2^_9_b^+3^_13_b^+3^_13_b^+3^_13_y^+3^_13_y^+3^_13_y^+1^_5_y^+2^_10_b^+2^_9_b^+1^_4_y^+2^_11_y^+2^_11_y^+1^_6_y^+2^_12_b^+1^_5_b^+1^_5_y^+1^_7_y^+1^_8_b^+1^_6_y^+1^_9_

0846169225383384

Fragment Matches Table

Show background peaks

| Position | Ion type | Intensity | mz Theoretical | mz Error (Th) | mz Error (ppm) | Charge | Series Number |
| --- | --- | --- | --- | --- | --- | --- | --- |
| - | - | 1864 | 120 | - | - | 0 | - |
| - | - | 696.1 | 120.1 | - | - | 0 | - |
| - | - | 3025 | 120.1 | - | - | 0 | - |
| - | - | 626.2 | 122.1 | - | - | 0 | - |
| - | - | 395.9 | 123 | - | - | 0 | - |
| - | - | 603 | 123.1 | - | - | 0 | - |
| - | - | 564.5 | 123.1 | - | - | 0 | - |
| - | - | 677.9 | 124 | - | - | 0 | - |
| - | - | 418.2 | 126.1 | - | - | 0 | - |
| - | - | 497.9 | 127.1 | - | - | 0 | - |
| - | - | 417 | 127.1 | - | - | 0 | - |
| - | - | 1627 | 127.1 | - | - | 0 | - |
| - | - | 427.9 | 127.1 | - | - | 0 | - |
| - | - | 347.3 | 127.9 | - | - | 0 | - |
| - | - | 899.8 | 128.1 | - | - | 0 | - |
| - | - | 834.2 | 129 | - | - | 0 | - |
| - | - | 559.7 | 129.1 | - | - | 0 | - |
| - | - | 5.526E+04 | 129.1 | - | - | 0 | - |
| - | - | 4.408E+04 | 130.1 | - | - | 0 | - |
| - | - | 632 | 130.1 | - | - | 0 | - |
| - | - | 4013 | 130.1 | - | - | 0 | - |
| - | - | 381.7 | 130.4 | - | - | 0 | - |
| - | - | 512.7 | 131 | - | - | 0 | - |
| - | - | 4636 | 131.1 | - | - | 0 | - |
| - | - | 1167 | 131.1 | - | - | 0 | - |
| - | - | 2732 | 132 | - | - | 0 | - |
| - | - | 1296 | 132.1 | - | - | 0 | - |
| 14 | y | 1.212E+04 | 132.1 | 0.0003135 | 2.373 | +1 | 1 |
| - | - | 3421 | 133.1 | - | - | 0 | - |
| - | - | 723.9 | 133.1 | - | - | 0 | - |
| - | - | 445.1 | 134 | - | - | 0 | - |
| - | - | 463.1 | 134.1 | - | - | 0 | - |
| - | - | 415.6 | 134.5 | - | - | 0 | - |
| - | - | 445.4 | 135.1 | - | - | 0 | - |
| - | - | 482.6 | 135.1 | - | - | 0 | - |
| - | - | 646.5 | 136 | - | - | 0 | - |
| - | - | 4.546E+04 | 136.1 | - | - | 0 | - |
| - | - | 3871 | 137.1 | - | - | 0 | - |
| - | - | 1024 | 138.1 | - | - | 0 | - |
| - | - | 419 | 139.1 | - | - | 0 | - |
| - | - | 1231 | 139.1 | - | - | 0 | - |
| - | - | 3749 | 141.1 | - | - | 0 | - |
| - | - | 533.6 | 142.1 | - | - | 0 | - |
| - | - | 389.5 | 142.6 | - | - | 0 | - |
| - | - | 773.8 | 143 | - | - | 0 | - |
| - | - | 1295 | 145.1 | - | - | 0 | - |
| 4 | b | 7628 | 146.1 | 0.00274 | 18.76 | +4 | 4 |
| - | - | 2180 | 147 | - | - | 0 | - |
| - | - | 900.8 | 147.1 | - | - | 0 | - |
| - | - | 445.7 | 147.1 | - | - | 0 | - |
| - | - | 507.3 | 147.1 | - | - | 0 | - |
| - | - | 447.3 | 148.1 | - | - | 0 | - |
| - | - | 986 | 149 | - | - | 0 | - |
| - | - | 4.651E+04 | 149 | - | - | 0 | - |
| - | - | 401.8 | 149.2 | - | - | 0 | - |
| - | - | 4621 | 150 | - | - | 0 | - |
| - | - | 932.6 | 151.1 | - | - | 0 | - |
| - | - | 3310 | 152.1 | - | - | 0 | - |
| - | - | 1644 | 152.1 | - | - | 0 | - |
| - | - | 428.6 | 153.1 | - | - | 0 | - |
| - | - | 438.2 | 153.6 | - | - | 0 | - |
| - | - | 496.4 | 155 | - | - | 0 | - |
| - | - | 2809 | 155.1 | - | - | 0 | - |
| - | - | 1213 | 155.1 | - | - | 0 | - |
| - | - | 2005 | 155.1 | - | - | 0 | - |
| - | - | 1908 | 156.1 | - | - | 0 | - |
| - | - | 642 | 157 | - | - | 0 | - |
| - | - | 666.2 | 157.1 | - | - | 0 | - |
| - | - | 8427 | 157.1 | - | - | 0 | - |
| - | - | 783.8 | 157.1 | - | - | 0 | - |
| - | - | 9864 | 158.1 | - | - | 0 | - |
| - | - | 723.6 | 158.1 | - | - | 0 | - |
| - | - | 668.2 | 159.1 | - | - | 0 | - |
| - | - | 1223 | 159.1 | - | - | 0 | - |
| - | - | 2996 | 162.1 | - | - | 0 | - |
| - | - | 751.3 | 163.1 | - | - | 0 | - |
| - | - | 573.8 | 164.1 | - | - | 0 | - |
| - | - | 1977 | 165.1 | - | - | 0 | - |
| - | - | 1421 | 166.1 | - | - | 0 | - |
| - | - | 3388 | 167 | - | - | 0 | - |
| - | - | 605.8 | 168 | - | - | 0 | - |
| - | - | 582.1 | 168.1 | - | - | 0 | - |
| - | - | 483.3 | 169.1 | - | - | 0 | - |
| - | - | 4212 | 169.1 | - | - | 0 | - |
| - | - | 630.8 | 170.1 | - | - | 0 | - |
| - | - | 783.7 | 171.1 | - | - | 0 | - |
| - | - | 3359 | 172.1 | - | - | 0 | - |
| - | - | 1422 | 173.1 | - | - | 0 | - |
| - | - | 678.2 | 173.1 | - | - | 0 | - |
| - | - | 715.5 | 173.1 | - | - | 0 | - |
| - | - | 1483 | 173.5 | - | - | 0 | - |
| - | - | 8941 | 174.1 | - | - | 0 | - |
| - | - | 620 | 175.1 | - | - | 0 | - |
| - | - | 1.335E+04 | 175.1 | - | - | 0 | - |
| - | - | 949 | 176.1 | - | - | 0 | - |
| - | - | 1222 | 176.1 | - | - | 0 | - |
| - | - | 3354 | 177.1 | - | - | 0 | - |
| - | - | 602.6 | 177.1 | - | - | 0 | - |
| - | - | 547.5 | 177.1 | - | - | 0 | - |
| - | - | 470.9 | 178.1 | - | - | 0 | - |
| - | - | 621.1 | 181.1 | - | - | 0 | - |
| - | - | 571.3 | 181.1 | - | - | 0 | - |
| - | - | 9672 | 181.1 | - | - | 0 | - |
| - | - | 771.3 | 181.1 | - | - | 0 | - |
| - | - | 477.6 | 181.6 | - | - | 0 | - |
| - | - | 782.3 | 182.1 | - | - | 0 | - |
| - | - | 1623 | 182.1 | - | - | 0 | - |
| - | - | 788.5 | 183.1 | - | - | 0 | - |
| - | - | 596.9 | 183.1 | - | - | 0 | - |
| - | - | 3372 | 185.1 | - | - | 0 | - |
| - | - | 5431 | 185.1 | - | - | 0 | - |
| - | - | 1617 | 185.1 | - | - | 0 | - |
| - | - | 930 | 185.1 | - | - | 0 | - |
| - | - | 636.4 | 185.1 | - | - | 0 | - |
| - | - | 1093 | 186.1 | - | - | 0 | - |
| - | - | 481.6 | 186.1 | - | - | 0 | - |
| - | - | 7456 | 186.1 | - | - | 0 | - |
| - | - | 905.7 | 187.1 | - | - | 0 | - |
| - | - | 4661 | 187.1 | - | - | 0 | - |
| 12 | y | 1458 | 187.1 | 0.000427 | 2.281 | +2 | 3 |
| - | - | 763.5 | 189.1 | - | - | 0 | - |
| - | - | 500.3 | 189.1 | - | - | 0 | - |
| - | - | 4484 | 190.1 | - | - | 0 | - |
| - | - | 1298 | 191.1 | - | - | 0 | - |
| - | - | 649.2 | 193.1 | - | - | 0 | - |
| - | - | 558.9 | 195.1 | - | - | 0 | - |
| - | - | 509.3 | 195.1 | - | - | 0 | - |
| - | - | 538.8 | 195.1 | - | - | 0 | - |
| - | - | 1107 | 195.1 | - | - | 0 | - |
| - | - | 519 | 197.1 | - | - | 0 | - |
| - | - | 2192 | 197.2 | - | - | 0 | - |
| - | - | 1865 | 199.1 | - | - | 0 | - |
| - | - | 798.8 | 199.2 | - | - | 0 | - |
| - | - | 530.7 | 199.2 | - | - | 0 | - |
| - | - | 777.3 | 200.1 | - | - | 0 | - |
| - | - | 618.1 | 201.1 | - | - | 0 | - |
| - | - | 639.3 | 202.1 | - | - | 0 | - |
| - | - | 5185 | 202.1 | - | - | 0 | - |
| - | - | 624.7 | 203.1 | - | - | 0 | - |
| - | - | 868.3 | 203.1 | - | - | 0 | - |
| - | - | 547.1 | 203.1 | - | - | 0 | - |
| - | - | 767.4 | 205.1 | - | - | 0 | - |
| - | - | 1087 | 205.1 | - | - | 0 | - |
| - | - | 588.4 | 206.1 | - | - | 0 | - |
| - | - | 861.9 | 207.1 | - | - | 0 | - |
| - | - | 1447 | 207.1 | - | - | 0 | - |
| - | - | 687.6 | 209.1 | - | - | 0 | - |
| - | - | 550.9 | 211.1 | - | - | 0 | - |
| - | - | 769.8 | 212.1 | - | - | 0 | - |
| - | - | 1512 | 212.7 | - | - | 0 | - |
| - | - | 1462 | 214.1 | - | - | 0 | - |
| - | - | 541.1 | 214.2 | - | - | 0 | - |
| - | - | 1131 | 215.1 | - | - | 0 | - |
| - | - | 1462 | 217.1 | - | - | 0 | - |
| 9 | y | 749.7 | 218.2 | 0.001784 | 8.177 | +3 | 6 |
| - | - | 2890 | 220.1 | - | - | 0 | - |
| - | - | 6363 | 221.1 | - | - | 0 | - |
| - | - | 895.1 | 224.1 | - | - | 0 | - |
| - | - | 2494 | 224.2 | - | - | 0 | - |
| - | - | 856.1 | 225.1 | - | - | 0 | - |
| - | - | 1173 | 225.1 | - | - | 0 | - |
| - | - | 685.1 | 225.2 | - | - | 0 | - |
| - | - | 625.4 | 226.1 | - | - | 0 | - |
| - | - | 1265 | 226.1 | - | - | 0 | - |
| - | - | 4.394E+04 | 226.2 | - | - | 0 | - |
| - | - | 1.388E+04 | 227.1 | - | - | 0 | - |
| 11 | y | 4398 | 227.2 | 0.001358 | 5.977 | +2 | 4 |
| - | - | 1918 | 228.1 | - | - | 0 | - |
| - | - | 1625 | 228.1 | - | - | 0 | - |
| - | - | 2125 | 231.1 | - | - | 0 | - |
| - | - | 640.5 | 231.1 | - | - | 0 | - |
| - | - | 934.1 | 233.1 | - | - | 0 | - |
| - | - | 831.6 | 234.1 | - | - | 0 | - |
| - | - | 6555 | 235.1 | - | - | 0 | - |
| 11 | y | 7969 | 235.7 | 0.0001986 | 0.8427 | +2 | 4 |
| - | - | 1702 | 236.2 | - | - | 0 | - |
| - | - | 689.9 | 237.1 | - | - | 0 | - |
| - | - | 657 | 238.1 | - | - | 0 | - |
| - | - | 524.5 | 239.1 | - | - | 0 | - |
| - | - | 2988 | 239.2 | - | - | 0 | - |
| - | - | 1420 | 240.1 | - | - | 0 | - |
| - | - | 1441 | 241.2 | - | - | 0 | - |
| - | - | 1137 | 242.2 | - | - | 0 | - |
| - | - | 546.8 | 243.1 | - | - | 0 | - |
| - | - | 1735 | 244.1 | - | - | 0 | - |
| - | - | 590.9 | 244.2 | - | - | 0 | - |
| - | - | 2470 | 245.1 | - | - | 0 | - |
| 13 | y | 9413 | 245.2 | 0.0002492 | 1.016 | +1 | 2 |
| - | - | 1130 | 246.1 | - | - | 0 | - |
| - | - | 689.7 | 246.2 | - | - | 0 | - |
| - | - | 9597 | 247.1 | - | - | 0 | - |
| - | - | 5837 | 248.1 | - | - | 0 | - |
| - | - | 1110 | 248.1 | - | - | 0 | - |
| - | - | 6815 | 249.1 | - | - | 0 | - |
| - | - | 775.7 | 249.2 | - | - | 0 | - |
| - | - | 1012 | 250.1 | - | - | 0 | - |
| - | - | 753.3 | 251.2 | - | - | 0 | - |
| - | - | 981.6 | 252.2 | - | - | 0 | - |
| - | - | 1344 | 254.1 | - | - | 0 | - |
| - | - | 1.564E+04 | 255.1 | - | - | 0 | - |
| - | - | 2038 | 256.1 | - | - | 0 | - |
| - | - | 698.3 | 256.1 | - | - | 0 | - |
| - | - | 1296 | 257.2 | - | - | 0 | - |
| - | - | 1051 | 258.1 | - | - | 0 | - |
| - | - | 1.058E+04 | 258.1 | - | - | 0 | - |
| - | - | 1581 | 259.1 | - | - | 0 | - |
| - | - | 735.8 | 260.2 | - | - | 0 | - |
| - | - | 769.9 | 261.1 | - | - | 0 | - |
| 2 | a | 1.789E+04 | 262.1 | 0.00498 | 19 | +1 | 2 |
| - | - | 2815 | 263.1 | - | - | 0 | - |
| - | - | 5045 | 266.1 | - | - | 0 | - |
| - | - | 677.7 | 267.1 | - | - | 0 | - |
| - | - | 1190 | 267.1 | - | - | 0 | - |
| - | - | 3146 | 270.2 | - | - | 0 | - |
| 10 | y | 1467 | 271.2 | 1.329E-05 | 0.04901 | +2 | 5 |
| - | - | 859.2 | 271.7 | - | - | 0 | - |
| 2 | b | 2618 | 272.1 | 0.004822 | 17.72 | +1 | 2 |
| - | - | 4748 | 273.1 | - | - | 0 | - |
| - | - | 4960 | 274.1 | - | - | 0 | - |
| - | - | 1700 | 275.1 | - | - | 0 | - |
| - | - | 1.684E+04 | 275.1 | - | - | 0 | - |
| - | - | 1014 | 276.1 | - | - | 0 | - |
| - | - | 2655 | 276.1 | - | - | 0 | - |
| - | - | 2218 | 278.1 | - | - | 0 | - |
| - | - | 1.084E+04 | 283.2 | - | - | 0 | - |
| - | - | 1061 | 283.6 | - | - | 0 | - |
| - | - | 1533 | 284.2 | - | - | 0 | - |
| - | - | 1416 | 285.1 | - | - | 0 | - |
| - | - | 1779 | 285.2 | - | - | 0 | - |
| 2 | b | 1.268E+04 | 290.1 | 0.005182 | 17.86 | +1 | 2 |
| - | - | 3091 | 290.2 | - | - | 0 | - |
| - | - | 2357 | 290.6 | - | - | 0 | - |
| - | - | 1301 | 290.7 | - | - | 0 | - |
| 4 | b | 2157 | 291.1 | 0.001835 | 6.302 | +2 | 4 |
| - | - | 4497 | 292.1 | - | - | 0 | - |
| - | - | 842 | 293.1 | - | - | 0 | - |
| - | - | 3484 | 293.1 | - | - | 0 | - |
| - | - | 1044 | 293.2 | - | - | 0 | - |
| - | - | 850.2 | 294.1 | - | - | 0 | - |
| - | - | 562 | 294.2 | - | - | 0 | - |
| - | - | 776.6 | 294.2 | - | - | 0 | - |
| - | - | 5020 | 297.2 | - | - | 0 | - |
| - | - | 730.1 | 300.1 | - | - | 0 | - |
| - | - | 782.7 | 302.1 | - | - | 0 | - |
| - | - | 986.2 | 303.1 | - | - | 0 | - |
| - | - | 695.8 | 305.7 | - | - | 0 | - |
| - | - | 777.1 | 307.2 | - | - | 0 | - |
| - | - | 627.9 | 309.2 | - | - | 0 | - |
| - | - | 649.5 | 311.2 | - | - | 0 | - |
| - | - | 2809 | 320.1 | - | - | 0 | - |
| 6 | y | 1919 | 320.9 | 0.0007932 | 2.472 | +3 | 9 |
| - | - | 5339 | 321.2 | - | - | 0 | - |
| - | - | 775.3 | 322.2 | - | - | 0 | - |
| - | - | 1057 | 324.2 | - | - | 0 | - |
| - | - | 704.2 | 325.6 | - | - | 0 | - |
| - | - | 2879 | 326.2 | - | - | 0 | - |
| - | - | 875.6 | 327.7 | - | - | 0 | - |
| - | - | 2118 | 332.7 | - | - | 0 | - |
| - | - | 669.4 | 333.2 | - | - | 0 | - |
| - | - | 8937 | 336.2 | - | - | 0 | - |
| - | - | 444.8 | 336.2 | - | - | 0 | - |
| - | - | 1301 | 337.2 | - | - | 0 | - |
| - | - | 2595 | 337.7 | - | - | 0 | - |
| - | - | 4230 | 338.1 | - | - | 0 | - |
| - | - | 9708 | 339.2 | - | - | 0 | - |
| - | - | 1003 | 340.2 | - | - | 0 | - |
| - | - | 2212 | 340.2 | - | - | 0 | - |
| - | - | 7613 | 346.7 | - | - | 0 | - |
| - | - | 2169 | 347.2 | - | - | 0 | - |
| - | - | 933.3 | 347.7 | - | - | 0 | - |
| - | - | 2447 | 348.1 | - | - | 0 | - |
| - | - | 625.2 | 349.7 | - | - | 0 | - |
| - | - | 638.5 | 351.2 | - | - | 0 | - |
| - | - | 628.4 | 352.2 | - | - | 0 | - |
| - | - | 932.3 | 353.2 | - | - | 0 | - |
| - | - | 1.127E+04 | 354.2 | - | - | 0 | - |
| - | - | 2077 | 355.2 | - | - | 0 | - |
| - | - | 1068 | 355.7 | - | - | 0 | - |
| - | - | 2656 | 357.3 | - | - | 0 | - |
| - | - | 2963 | 358.2 | - | - | 0 | - |
| - | - | 1148 | 358.7 | - | - | 0 | - |
| - | - | 841.2 | 359.5 | - | - | 0 | - |
| - | - | 787.5 | 360.7 | - | - | 0 | - |
| - | - | 646.7 | 360.7 | - | - | 0 | - |
| - | - | 1246 | 361.7 | - | - | 0 | - |
| - | - | 670.6 | 362.2 | - | - | 0 | - |
| 5 | y | 1683 | 366.6 | 0.0004522 | 1.234 | +3 | 10 |
| - | - | 1582 | 366.7 | - | - | 0 | - |
| - | - | 621.2 | 366.9 | - | - | 0 | - |
| - | - | 1289 | 368.1 | - | - | 0 | - |
| - | - | 1118 | 368.8 | - | - | 0 | - |
| - | - | 675.8 | 369.1 | - | - | 0 | - |
| - | - | 882.7 | 370.7 | - | - | 0 | - |
| - | - | 893.3 | 371.7 | - | - | 0 | - |
| 12 | y | 4983 | 373.3 | 0.0002314 | 0.62 | +1 | 3 |
| - | - | 740.2 | 374.3 | - | - | 0 | - |
| - | - | 729.1 | 374.7 | - | - | 0 | - |
| - | - | 737.1 | 375.2 | - | - | 0 | - |
| - | - | 1077 | 375.7 | - | - | 0 | - |
| - | - | 1438 | 380.2 | - | - | 0 | - |
| - | - | 831.1 | 380.7 | - | - | 0 | - |
| - | - | 949.1 | 382.2 | - | - | 0 | - |
| - | - | 849 | 382.2 | - | - | 0 | - |
| - | - | 870.9 | 382.3 | - | - | 0 | - |
| - | - | 2419 | 383.2 | - | - | 0 | - |
| - | - | 1502 | 383.7 | - | - | 0 | - |
| - | - | 1644 | 384.2 | - | - | 0 | - |
| - | - | 4618 | 385.2 | - | - | 0 | - |
| - | - | 750.3 | 385.2 | - | - | 0 | - |
| - | - | 873.9 | 385.8 | - | - | 0 | - |
| - | - | 5196 | 386.2 | - | - | 0 | - |
| - | - | 991.4 | 386.7 | - | - | 0 | - |
| - | - | 817.6 | 388.2 | - | - | 0 | - |
| - | - | 1329 | 389.3 | - | - | 0 | - |
| - | - | 2729 | 389.7 | - | - | 0 | - |
| - | - | 817.8 | 390.7 | - | - | 0 | - |
| - | - | 1641 | 392.3 | - | - | 0 | - |
| - | - | 2869 | 394.2 | - | - | 0 | - |
| 2 | y | 2188 | 394.2 | 0.006296 | 15.97 | +4 | 13 |
| - | - | 758.5 | 394.5 | - | - | 0 | - |
| - | - | 1290 | 394.5 | - | - | 0 | - |
| - | - | 892.2 | 397.2 | - | - | 0 | - |
| 10 | b | 1143 | 397.8 | 0.007368 | 18.52 | +3 | 10 |
| - | - | 642.9 | 398 | - | - | 0 | - |
| 2 | y | 1629 | 398.5 | 0.001398 | 3.509 | +4 | 13 |
| - | - | 740.9 | 398.7 | - | - | 0 | - |
| - | - | 1060 | 398.7 | - | - | 0 | - |
| - | - | 699.4 | 400.1 | - | - | 0 | - |
| - | - | 699.9 | 401.5 | - | - | 0 | - |
| - | - | 1490 | 402.2 | - | - | 0 | - |
| - | - | 1976 | 402.7 | - | - | 0 | - |
| - | - | 3961 | 403.2 | - | - | 0 | - |
| - | - | 2745 | 403.2 | - | - | 0 | - |
| - | - | 1136 | 403.3 | - | - | 0 | - |
| 10 | b | 907.8 | 403.5 | 0.002841 | 7.041 | +3 | 10 |
| 4 | y | 3251 | 403.6 | 0.00175 | 4.337 | +3 | 11 |
| 7 | y | 876.6 | 403.8 | 0.007458 | 18.47 | +2 | 8 |
| - | - | 928.3 | 403.9 | - | - | 0 | - |
| - | - | 1487 | 403.9 | - | - | 0 | - |
| - | - | 1247 | 404.5 | - | - | 0 | - |
| - | - | 1677 | 406.3 | - | - | 0 | - |
| - | - | 1923 | 408.2 | - | - | 0 | - |
| - | - | 648.1 | 408.6 | - | - | 0 | - |
| - | - | 2718 | 408.7 | - | - | 0 | - |
| - | - | 825.1 | 409 | - | - | 0 | - |
| - | - | 4250 | 410.3 | - | - | 0 | - |
| - | - | 2149 | 410.6 | - | - | 0 | - |
| - | - | 704 | 410.9 | - | - | 0 | - |
| - | - | 6424 | 411.2 | - | - | 0 | - |
| - | - | 1916 | 411.2 | - | - | 0 | - |
| - | - | 814.3 | 411.3 | - | - | 0 | - |
| - | - | 1461 | 411.5 | - | - | 0 | - |
| - | - | 1196 | 411.7 | - | - | 0 | - |
| - | - | 1090 | 411.7 | - | - | 0 | - |
| 7 | y | 1.293E+04 | 412.3 | 0.0001653 | 0.401 | +2 | 8 |
| - | - | 7133 | 412.8 | - | - | 0 | - |
| - | - | 1180 | 413.3 | - | - | 0 | - |
| - | - | 854.6 | 413.9 | - | - | 0 | - |
| - | - | 1357 | 414.2 | - | - | 0 | - |
| - | - | 1057 | 414.2 | - | - | 0 | - |
| - | - | 760.2 | 414.7 | - | - | 0 | - |
| - | - | 2176 | 415.3 | - | - | 0 | - |
| 0 | Precursor | 1771 | 416 | 0.004575 | 11 | +4 | -1 |
| - | - | 1479 | 416.2 | - | - | 0 | - |
| - | - | 1136 | 416.5 | - | - | 0 | - |
| - | - | 693.8 | 417.2 | - | - | 0 | - |
| - | - | 5954 | 418.7 | - | - | 0 | - |
| 6 | b | 4626 | 419.2 | 0.00612 | 14.6 | +2 | 6 |
| - | - | 1041 | 419.7 | - | - | 0 | - |
| - | - | 3437 | 419.9 | - | - | 0 | - |
| 0 | Precursor | 1.104E+04 | 420.2 | 0.004621 | 11 | +4 | -1 |
| - | - | 6917 | 420.5 | - | - | 0 | - |
| - | - | 2756 | 420.6 | - | - | 0 | - |
| - | - | 3004 | 420.7 | - | - | 0 | - |
| - | - | 1882 | 421 | - | - | 0 | - |
| - | - | 1613 | 421.2 | - | - | 0 | - |
| - | - | 947.4 | 422.3 | - | - | 0 | - |
| - | - | 721.7 | 423.3 | - | - | 0 | - |
| - | - | 1195 | 424.3 | - | - | 0 | - |
| - | - | 995.8 | 424.7 | - | - | 0 | - |
| - | - | 972 | 425.2 | - | - | 0 | - |
| - | - | 1352 | 425.9 | - | - | 0 | - |
| - | - | 822.7 | 426.2 | - | - | 0 | - |
| - | - | 1603 | 427.2 | - | - | 0 | - |
| 6 | b | 1544 | 428.2 | 0.002516 | 5.877 | +2 | 6 |
| - | - | 729.4 | 428.3 | - | - | 0 | - |
| - | - | 1.156E+04 | 429.2 | - | - | 0 | - |
| 11 | b | 2704 | 430.2 | 0.00664 | 15.43 | +3 | 11 |
| - | - | 1577 | 431.2 | - | - | 0 | - |
| 3 | b | 1048 | 435.2 | 0.004359 | 10.02 | +1 | 3 |
| - | - | 944.9 | 439.2 | - | - | 0 | - |
| - | - | 681.2 | 441.2 | - | - | 0 | - |
| - | - | 1624 | 446.2 | - | - | 0 | - |
| - | - | 874.6 | 446.7 | - | - | 0 | - |
| - | - | 832.1 | 448.3 | - | - | 0 | - |
| - | - | 1247 | 448.6 | - | - | 0 | - |
| - | - | 1688 | 451.2 | - | - | 0 | - |
| - | - | 1099 | 451.3 | - | - | 0 | - |
| - | - | 1020 | 451.7 | - | - | 0 | - |
| - | - | 2828 | 451.9 | - | - | 0 | - |
| - | - | 1856 | 452.3 | - | - | 0 | - |
| - | - | 758 | 452.6 | - | - | 0 | - |
| 3 | b | 1631 | 453.2 | 0.0053 | 11.69 | +1 | 3 |
| - | - | 1491 | 455.3 | - | - | 0 | - |
| - | - | 2156 | 457.6 | - | - | 0 | - |
| 3 | y | 757.4 | 457.9 | 0.002715 | 5.929 | +3 | 12 |
| - | - | 679.9 | 458.2 | - | - | 0 | - |
| - | - | 1084 | 458.3 | - | - | 0 | - |
| - | - | 2299 | 460.2 | - | - | 0 | - |
| 3 | y | 2.123E+04 | 463.6 | 0.0003248 | 0.7007 | +3 | 12 |
| - | - | 1.891E+04 | 463.9 | - | - | 0 | - |
| - | - | 9490 | 464.3 | - | - | 0 | - |
| - | - | 1892 | 464.6 | - | - | 0 | - |
| - | - | 1080 | 467.3 | - | - | 0 | - |
| - | - | 3363 | 468.9 | - | - | 0 | - |
| - | - | 852.7 | 469.3 | - | - | 0 | - |
| 11 | y | 1.869E+04 | 470.3 | 0.0003189 | 0.6781 | +1 | 4 |
| - | - | 5385 | 471.3 | - | - | 0 | - |
| - | - | 1272 | 471.8 | - | - | 0 | - |
| - | - | 864.5 | 471.9 | - | - | 0 | - |
| - | - | 746.5 | 472.2 | - | - | 0 | - |
| 6 | y | 920.5 | 472.3 | 0.008734 | 18.49 | +2 | 9 |
| - | - | 1011 | 472.3 | - | - | 0 | - |
| - | - | 2060 | 473.3 | - | - | 0 | - |
| - | - | 898.3 | 474.8 | - | - | 0 | - |
| - | - | 2411 | 476.2 | - | - | 0 | - |
| - | - | 836.1 | 477.2 | - | - | 0 | - |
| 12 | b | 1756 | 478.6 | 0.001817 | 3.796 | +3 | 12 |
| 6 | y | 3257 | 480.8 | 0.0001283 | 0.2669 | +2 | 9 |
| - | - | 2606 | 481.3 | - | - | 0 | - |
| - | - | 977 | 481.3 | - | - | 0 | - |
| - | - | 1178 | 481.6 | - | - | 0 | - |
| - | - | 879.5 | 481.8 | - | - | 0 | - |
| - | - | 824.8 | 481.9 | - | - | 0 | - |
| - | - | 922.3 | 482.3 | - | - | 0 | - |
| - | - | 1492 | 482.3 | - | - | 0 | - |
| - | - | 1115 | 482.8 | - | - | 0 | - |
| - | - | 723.4 | 483.2 | - | - | 0 | - |
| - | - | 1132 | 483.2 | - | - | 0 | - |
| - | - | 1402 | 483.8 | - | - | 0 | - |
| - | - | 1268 | 487.3 | - | - | 0 | - |
| - | - | 1270 | 487.6 | - | - | 0 | - |
| - | - | 931.6 | 487.9 | - | - | 0 | - |
| - | - | 1440 | 491.3 | - | - | 0 | - |
| - | - | 730.2 | 492.8 | - | - | 0 | - |
| - | - | 1193 | 494.2 | - | - | 0 | - |
| - | - | 1125 | 503.2 | - | - | 0 | - |
| - | - | 714.3 | 506.9 | - | - | 0 | - |
| - | - | 917.7 | 508.8 | - | - | 0 | - |
| 13 | b | 1805 | 510.3 | 0.002514 | 4.927 | +3 | 13 |
| 13 | b | 1775 | 510.6 | 0.007456 | 14.6 | +3 | 13 |
| - | - | 1024 | 510.9 | - | - | 0 | - |
| 13 | b | 2558 | 516.3 | 0.0005795 | 1.123 | +3 | 13 |
| - | - | 3347 | 516.6 | - | - | 0 | - |
| - | - | 902.1 | 520.4 | - | - | 0 | - |
| - | - | 833.2 | 520.6 | - | - | 0 | - |
| - | - | 923.3 | 521.2 | - | - | 0 | - |
| - | - | 5528 | 522.3 | - | - | 0 | - |
| - | - | 3064 | 522.6 | - | - | 0 | - |
| - | - | 1373 | 522.9 | - | - | 0 | - |
| - | - | 2628 | 525 | - | - | 0 | - |
| 2 | y | 2202 | 525.3 | 0.007605 | 14.48 | +3 | 13 |
| - | - | 1554 | 525.6 | - | - | 0 | - |
| - | - | 804.6 | 525.7 | - | - | 0 | - |
| - | - | 1462 | 526.6 | - | - | 0 | - |
| - | - | 845.2 | 526.9 | - | - | 0 | - |
| 2 | y | 3723 | 531 | 0.001156 | 2.178 | +3 | 13 |
| - | - | 2456 | 531.3 | - | - | 0 | - |
| - | - | 1742 | 531.6 | - | - | 0 | - |
| - | - | 1671 | 532 | - | - | 0 | - |
| - | - | 874.9 | 532.3 | - | - | 0 | - |
| - | - | 7094 | 538.3 | - | - | 0 | - |
| - | - | 1427 | 539.3 | - | - | 0 | - |
| - | - | 2148 | 539.3 | - | - | 0 | - |
| - | - | 812.8 | 539.8 | - | - | 0 | - |
| 10 | y | 4103 | 541.4 | 3.985E-05 | 0.07361 | +1 | 5 |
| - | - | 852.2 | 547.8 | - | - | 0 | - |
| - | - | 1248 | 548.3 | - | - | 0 | - |
| 5 | y | 7856 | 549.3 | 0.000458 | 0.8337 | +2 | 10 |
| - | - | 4752 | 549.8 | - | - | 0 | - |
| - | - | 657.1 | 552.3 | - | - | 0 | - |
| - | - | 716.8 | 561.3 | - | - | 0 | - |
| - | - | 7422 | 566.2 | - | - | 0 | - |
| - | - | 3006 | 567.3 | - | - | 0 | - |
| - | - | 1027 | 567.4 | - | - | 0 | - |
| 9 | b | 2587 | 569.3 | 0.00157 | 2.757 | +2 | 9 |
| - | - | 1932 | 569.8 | - | - | 0 | - |
| - | - | 824.6 | 570.3 | - | - | 0 | - |
| - | - | 2989 | 572.8 | - | - | 0 | - |
| - | - | 2057 | 573.3 | - | - | 0 | - |
| - | - | 953.1 | 573.8 | - | - | 0 | - |
| - | - | 1967 | 577.4 | - | - | 0 | - |
| - | - | 1253 | 578.4 | - | - | 0 | - |
| - | - | 5512 | 579.4 | - | - | 0 | - |
| - | - | 1240 | 580.4 | - | - | 0 | - |
| 4 | b | 914.8 | 581.2 | 0.0009825 | 1.69 | +1 | 4 |
| - | - | 1785 | 585.3 | - | - | 0 | - |
| - | - | 2124 | 595.4 | - | - | 0 | - |
| - | - | 1112 | 596.4 | - | - | 0 | - |
| - | - | 945.2 | 603.3 | - | - | 0 | - |
| - | - | 1713 | 604.4 | - | - | 0 | - |
| 4 | y | 3110 | 604.8 | 0.003181 | 5.259 | +2 | 11 |
| - | - | 1382 | 605.4 | - | - | 0 | - |
| - | - | 1702 | 610.3 | - | - | 0 | - |
| - | - | 2574 | 613.3 | - | - | 0 | - |
| 4 | y | 1058 | 613.4 | 0.00222 | 3.619 | +2 | 11 |
| - | - | 2515 | 620.3 | - | - | 0 | - |
| - | - | 1012 | 620.8 | - | - | 0 | - |
| - | - | 722.4 | 628.3 | - | - | 0 | - |
| - | - | 2993 | 629.3 | - | - | 0 | - |
| - | - | 2025 | 629.8 | - | - | 0 | - |
| - | - | 748 | 630.3 | - | - | 0 | - |
| - | - | 1779 | 638.4 | - | - | 0 | - |
| - | - | 1086 | 638.9 | - | - | 0 | - |
| - | - | 893.2 | 639.4 | - | - | 0 | - |
| - | - | 823 | 649.3 | - | - | 0 | - |
| - | - | 663 | 650.3 | - | - | 0 | - |
| - | - | 860.2 | 661.3 | - | - | 0 | - |
| - | - | 1692 | 664.5 | - | - | 0 | - |
| - | - | 800.6 | 665.5 | - | - | 0 | - |
| 9 | y | 1538 | 669.5 | 0.0005358 | 0.8004 | +1 | 6 |
| - | - | 5329 | 674.4 | - | - | 0 | - |
| - | - | 2892 | 675.4 | - | - | 0 | - |
| - | - | 951.5 | 690.3 | - | - | 0 | - |
| - | - | 6306 | 692.4 | - | - | 0 | - |
| - | - | 2710 | 693.4 | - | - | 0 | - |
| - | - | 790.6 | 694.4 | - | - | 0 | - |
| 3 | y | 1403 | 694.9 | 0.001902 | 2.737 | +2 | 12 |
| - | - | 1793 | 695.4 | - | - | 0 | - |
| 5 | b | 1008 | 700.3 | 0.001524 | 2.177 | +1 | 5 |
| - | - | 1601 | 710.5 | - | - | 0 | - |
| - | - | 1059 | 716.4 | - | - | 0 | - |
| 5 | b | 1293 | 718.3 | 0.005486 | 7.638 | +1 | 5 |
| - | - | 740.9 | 719.3 | - | - | 0 | - |
| - | - | 1166 | 719.4 | - | - | 0 | - |
| - | - | 848.6 | 720.3 | - | - | 0 | - |
| - | - | 823.3 | 721.4 | - | - | 0 | - |
| 8 | y | 2448 | 726.5 | 0.003106 | 4.275 | +1 | 7 |
| - | - | 1003 | 727.5 | - | - | 0 | - |
| - | - | 767.7 | 736.4 | - | - | 0 | - |
| - | - | 868.1 | 738.4 | - | - | 0 | - |
| - | - | 944.7 | 740.3 | - | - | 0 | - |
| - | - | 983.6 | 742.3 | - | - | 0 | - |
| - | - | 815.2 | 750.3 | - | - | 0 | - |
| - | - | 1272 | 768.3 | - | - | 0 | - |
| - | - | 1162 | 805.5 | - | - | 0 | - |
| - | - | 948 | 811.5 | - | - | 0 | - |
| 7 | y | 9040 | 823.5 | 0.0005716 | 0.6941 | +1 | 8 |
| - | - | 4744 | 824.5 | - | - | 0 | - |
| - | - | 910 | 825.5 | - | - | 0 | - |
| - | - | 1675 | 848.4 | - | - | 0 | - |
| - | - | 692.2 | 852.2 | - | - | 0 | - |
| 6 | b | 2002 | 855.3 | 0.003215 | 3.759 | +1 | 6 |
| - | - | 1895 | 901.4 | - | - | 0 | - |
| - | - | 959.1 | 902.4 | - | - | 0 | - |
| - | - | 1059 | 919.5 | - | - | 0 | - |
| 6 | y | 808.3 | 960.6 | 0.001256 | 1.307 | +1 | 9 |
| - | - | 627.1 | 1666 | - | - | 0 | - |
| - | - | 733.3 | 1686 | - | - | 0 | - |
| - | - | 704.6 | 2271 | - | - | 0 | - |
| - | - | 762.3 | 2695 | - | - | 0 | - |
| - | - | 670.8 | 3019 | - | - | 0 | - |
| - | - | 936.9 | 3044 | - | - | 0 | - |
| - | - | 841 | 3045 | - | - | 0 | - |
| - | - | 793.2 | 3350 | - | - | 0 | - |

m/z Charge Intensity FragmentType MassShift Position
120.04458618164062 0 1863.7544
120.06582641601562 0 696.1044
120.08106231689453 0 3024.7312
122.07145690917969 0 626.2094
123.04464721679688 0 395.93423
123.05561828613281 0 602.9568
123.11734771728516 0 564.45953
124.03961181640625 0 677.8872
126.05552673339844 0 418.21686
127.05073547363281 0 497.87808
127.07520294189453 0 417.0187
127.08683013916016 0 1626.5768
127.11205291748047 0 427.93472
127.9197769165039 0 347.343
128.07093811035156 0 899.8343
129.01873779296875 0 834.1903
129.06600952148438 0 559.6915
129.10255432128906 0 55255.914
130.0654296875 0 44077.156
130.0868682861328 0 631.9586
130.1059112548828 0 4012.5994
130.37477111816406 0 381.6974
131.04537963867188 0 512.71747
131.06884765625 0 4636.153
131.08180236816406 0 1167.3278
132.0447998046875 0 2731.9365
132.0813446044922 0 1295.6666
132.1022186279297 0 12120.2705 y 13
133.0609893798828 0 3420.7454
133.10557556152344 0 723.8798
134.0453643798828 0 445.0923
134.0600128173828 0 463.14127
134.52549743652344 0 415.59097
135.06796264648438 0 445.35837
135.0764923095703 0 482.61743
136.0399627685547 0 646.4529
136.0760040283203 0 45459.168
137.0793914794922 0 3871.0798
138.06661987304688 0 1024.112
139.05091857910156 0 418.99658
139.0869598388672 0 1231.2777
141.10252380371094 0 3748.718
142.0613555908203 0 533.5851
142.5559539794922 0 389.47498
143.04563903808594 0 773.75757
145.06134033203125 0 1294.7866
146.0603485107422 0 7627.5664 b 3
147.04441833496094 0 2179.8147
147.0647430419922 0 900.80475
147.07696533203125 0 445.67084
147.11343383789062 0 507.34006
148.0757598876953 0 447.29742
148.9554901123047 0 986.0283
149.02365112304688 0 46513.1
149.21324157714844 0 401.7781
150.02696228027344 0 4620.5625
151.086669921875 0 932.5783
152.07089233398438 0 3310.2573
152.14352416992188 0 1643.6758
153.06617736816406 0 428.60355
153.63311767578125 0 438.23477
154.97097778320312 0 496.41083
155.0817413330078 0 2808.6277
155.09315490722656 0 1212.9658
155.1180877685547 0 2004.9006
156.0769805908203 0 1908.4059
157.01368713378906 0 642.0061
157.06106567382812 0 666.15814
157.07627868652344 0 8426.694
157.1340789794922 0 783.7649
158.060302734375 0 9863.735
158.079833984375 0 723.6431
159.06361389160156 0 668.1839
159.0919647216797 0 1222.8964
162.0552978515625 0 2995.7795
163.08729553222656 0 751.25134
164.07147216796875 0 573.78613
165.1024169921875 0 1976.9708
166.06109619140625 0 1420.5759
167.0341339111328 0 3387.7222
168.03765869140625 0 605.7822
168.11305236816406 0 582.12384
169.07623291015625 0 483.34476
169.09754943847656 0 4212.206
170.0597381591797 0 630.8371
171.13809204101562 0 783.68414
172.10826110839844 0 3359.0774
173.07093811035156 0 1421.5609
173.09228515625 0 678.1559
173.12889099121094 0 715.5293
173.4515380859375 0 1483.0936
174.05523681640625 0 8940.826
175.05825805664062 0 620.0101
175.08689880371094 0 13351.779
176.082275390625 0 948.9519
176.08999633789062 0 1222.196
177.06614685058594 0 3354.013
177.07704162597656 0 602.634
177.1024169921875 0 547.47327
178.1345672607422 0 470.88974
181.06163024902344 0 621.14014
181.07586669921875 0 571.29517
181.09751892089844 0 9671.651
181.13380432128906 0 771.31195
181.59622192382812 0 477.62027
182.10031127929688 0 782.29626
182.12933349609375 0 1623.3256
183.05592346191406 0 788.4683
183.11354064941406 0 596.85254
185.0559844970703 0 3372.211
185.0712890625 0 5431.244
185.08139038085938 0 1616.7023
185.0922088623047 0 929.97125
185.1288604736328 0 636.363
186.05490112304688 0 1092.6393
186.0742645263672 0 481.60175
186.1239471435547 0 7456.281
187.0868682861328 0 905.7156
187.1079864501953 0 4660.696
187.14453125 0 1458.0153 y 11
189.08767700195312 0 763.4819
189.13555908203125 0 500.3461
190.0501708984375 0 4484.075
191.08180236816406 0 1298.0808
193.1088104248047 0 649.2313
195.07627868652344 0 558.928
195.08714294433594 0 509.2647
195.11331176757812 0 538.79083
195.14947509765625 0 1107.0378
197.07147216796875 0 518.9693
197.16488647460938 0 2192.3171
199.08685302734375 0 1865.0374
199.16946411132812 0 798.80994
199.1806182861328 0 530.7335
200.13987731933594 0 777.31573
201.12351989746094 0 618.061
202.086181640625 0 639.31305
202.1087646484375 0 5184.873
203.08201599121094 0 624.7386
203.0924072265625 0 868.30176
203.11288452148438 0 547.09174
205.06040954589844 0 767.4129
205.09722900390625 0 1086.589
206.0598602294922 0 588.35706
207.07666015625 0 861.92865
207.149169921875 0 1446.7916
209.09237670898438 0 687.64685
211.1076202392578 0 550.9003
212.13917541503906 0 769.82654
212.66798400878906 0 1511.8037
214.1088409423828 0 1461.6194
214.1916961669922 0 541.1457
215.1391143798828 0 1131.0835
217.0975799560547 0 1461.8854
218.14947509765625 0 749.7087 y Ammonia loss 8
220.1194610595703 0 2890.418
221.10350036621094 0 6362.903
224.139892578125 0 895.06616
224.1759033203125 0 2493.9624
225.06578063964844 0 856.10815
225.09848022460938 0 1173.3794
225.1597442626953 0 685.05316
226.09779357910156 0 625.40265
226.11924743652344 0 1264.9171
226.15525817871094 0 43943.09
227.0817413330078 0 13882.31
227.1585693359375 0 4398.1406 y Ammonia loss 10
228.0848846435547 0 1917.8185
228.1131134033203 0 1625.2943
231.08822631835938 0 2124.5144
231.13389587402344 0 640.51715
233.10386657714844 0 934.1107
234.12432861328125 0 831.5829
235.11929321289062 0 6555.203
235.67068481445312 0 7969.386 y 10
236.17221069335938 0 1701.7302
237.13436889648438 0 689.85876
238.1302032470703 0 657.04565
239.1383056640625 0 524.525
239.1507110595703 0 2988.4766
240.1349639892578 0 1419.9635
241.15484619140625 0 1440.9105
242.18663024902344 0 1136.591
243.14552307128906 0 546.79663
244.10821533203125 0 1735.4283
244.1658477783203 0 590.9165
245.09263610839844 0 2469.7751
245.18621826171875 0 9413.424 y 12
246.13967895507812 0 1130.1409
246.18939208984375 0 689.69507
247.13046264648438 0 9597.007
248.1145477294922 0 5837.275
248.13442993164062 0 1109.6606
249.0985565185547 0 6814.8467
249.1607666015625 0 775.7045
250.10252380371094 0 1011.802
251.15089416503906 0 753.3237
252.17108154296875 0 981.61615
254.09222412109375 0 1344.4415
255.07666015625 0 15644.65
256.07958984375 0 2038.3134
256.10845947265625 0 698.3386
257.1609802246094 0 1295.6666
258.0763244628906 0 1050.8329
258.0987548828125 0 10583.004
259.1023864746094 0 1581.0598
260.1969299316406 0 735.82825
261.08758544921875 0 769.8657
262.1186828613281 0 17888.432 a 1
263.1212463378906 0 2814.6833
266.1249084472656 0 5045.233
267.1093444824219 0 677.7143
267.1246643066406 0 1190.4609
270.1813659667969 0 3145.508
271.1890563964844 0 1466.8818 y 9
271.6910400390625 0 859.18207
272.1028747558594 0 2618.2397 b Water loss 1
273.0870056152344 0 4748.2334
274.1302490234375 0 4960.0366
275.103271484375 0 1699.9209
275.12530517578125 0 16844.998
276.10833740234375 0 1014.02563
276.128662109375 0 2655.054
278.11370849609375 0 2218.1985
283.1766662597656 0 10843.531
283.6275329589844 0 1061.4205
284.1800231933594 0 1532.9484
285.1097412109375 0 1416.2388
285.1561279296875 0 1779.0215
290.1138000488281 0 12683.786 b 1
290.1844177246094 0 3090.5466
290.6372985839844 0 2357.0464
290.6868591308594 0 1300.5667
291.1170654296875 0 2157.4392 b 3
292.1293029785156 0 4496.958
293.1005859375 0 842.0099
293.1292724609375 0 3483.8054
293.1758117675781 0 1044.4285
294.11907958984375 0 850.16907
294.1574401855469 0 562.02747
294.218505859375 0 776.6352
297.19244384765625 0 5019.7627
300.0644226074219 0 730.0541
302.1146240234375 0 782.7155
303.1205749511719 0 986.19824
305.6644592285156 0 695.8235
307.17742919921875 0 777.078
309.1689147949219 0 627.8685
311.24371337890625 0 649.5146
320.13995361328125 0 2808.843
320.8719482421875 0 1919.2498 y 5
321.2287292480469 0 5339.3164
322.2322692871094 0 775.2642
324.2154541015625 0 1057.3385
325.6261291503906 0 704.1916
326.2189025878906 0 2879.2878
327.69427490234375 0 875.5872
332.7290344238281 0 2118.394
333.2315673828125 0 669.41907
336.2032470703125 0 8937.142
336.2245178222656 0 444.80908
337.20574951171875 0 1301.3264
337.7214050292969 0 2594.8813
338.14984130859375 0 4230.1943
339.2394104003906 0 9707.714
340.1531066894531 0 1003.35297
340.242431640625 0 2212.1816
346.7265625 0 7613.1753
347.228271484375 0 2168.6655
347.7305908203125 0 933.3241
348.1342468261719 0 2447.1543
349.70660400390625 0 625.22906
351.2383728027344 0 638.47705
352.177490234375 0 628.374
353.2195739746094 0 932.27527
354.21392822265625 0 11266.796
355.21697998046875 0 2076.9988
355.73162841796875 0 1068.0697
357.2500305175781 0 2655.903
358.16162109375 0 2962.9968
358.7137756347656 0 1147.6023
359.5431213378906 0 841.16064
360.66363525390625 0 787.4981
360.6878662109375 0 646.70355
361.6632995605469 0 1245.7689
362.1625061035156 0 670.5686
366.5570068359375 0 1682.6968 y 4
366.6693420410156 0 1581.7754
366.89202880859375 0 621.1683
368.14739990234375 0 1289.007
368.84429931640625 0 1118.186
369.1489562988281 0 675.7585
370.6657409667969 0 882.68225
371.6551818847656 0 893.32965
373.28070068359375 0 4982.6694 y 11
374.2835693359375 0 740.182
374.6634216308594 0 729.1417
375.2091064453125 0 737.0769
375.65850830078125 0 1077.1669
380.229248046875 0 1438.3219
380.6615905761719 0 831.10785
382.1763000488281 0 949.0763
382.2054443359375 0 849.0259
382.2821350097656 0 870.884
383.1580810546875 0 2419.3289
383.6596984863281 0 1501.9525
384.16534423828125 0 1644.2908
385.1724548339844 0 4617.818
385.1971130371094 0 750.2676
385.8437805175781 0 873.85455
386.1568908691406 0 5196.4004
386.6737976074219 0 991.44476
388.189697265625 0 817.62714
389.27130126953125 0 1328.8231
389.6689147949219 0 2728.9834
390.7067565917969 0 817.805
392.265625 0 1641.2969
394.15045166015625 0 2869.426
394.2173767089844 0 2187.7607 y Ammonia loss 1
394.4663391113281 0 758.4957
394.52752685546875 0 1290.4446
397.24591064453125 0 892.2354
397.85491943359375 0 1143.417 b Ammonia loss 9
397.95989990234375 0 642.90643
398.4691162109375 0 1629.1544 y 1
398.6607360839844 0 740.8826
398.72198486328125 0 1059.6404
400.1304016113281 0 699.3579
401.4600524902344 0 699.9407
402.22442626953125 0 1490.0786
402.66265869140625 0 1975.6245
403.1837158203125 0 3961.438
403.2402038574219 0 2744.7705
403.2684631347656 0 1136.1575
403.5259094238281 0 907.8332 b 9
403.56988525390625 0 3250.5583 y Ammonia loss 3
403.767822265625 0 876.5914 y Ammonia loss 6
403.8576354980469 0 928.2555
403.9038391113281 0 1486.8088
404.4700927734375 0 1246.6371
406.2515869140625 0 1677.0054
408.2218017578125 0 1923.3073
408.5543518066406 0 648.0606
408.7258605957031 0 2717.543
408.97674560546875 0 825.1098
410.2767333984375 0 4249.6865
410.5690002441406 0 2149.405
410.904052734375 0 703.9513
411.1767578125 0 6423.526
411.222412109375 0 1916.0875
411.2652282714844 0 814.2733
411.4707946777344 0 1461.2579
411.6803894042969 0 1196.4017
411.72296142578125 0 1090.1287
412.2738037109375 0 12930.282 y 6
412.77508544921875 0 7133.4907
413.2760925292969 0 1179.9795
413.8950500488281 0 854.623
414.18316650390625 0 1357.2767
414.2290954589844 0 1056.7694
414.6876220703125 0 760.219
415.2552490234375 0 2176.3699
415.9736633300781 0 1771.3723 Precursor Ammonia loss
416.2245788574219 0 1478.918
416.471923828125 0 1136.4285
417.1579895019531 0 693.81134
418.6767272949219 0 5954.2026
419.17864990234375 0 4626.0835 b Water loss 5
419.6780700683594 0 1040.7646
419.89971923828125 0 3437.4958
420.2303466796875 0 11037.443 Precursor
420.47802734375 0 6917.08
420.5680847167969 0 2756.426
420.7282409667969 0 3004.233
420.9760437011719 0 1882.4044
421.2260437011719 0 1612.992
422.2760009765625 0 947.44055
423.2739562988281 0 721.6721
424.3279724121094 0 1194.5696
424.71221923828125 0 995.8043
425.21533203125 0 971.9565
425.9046325683594 0 1352.2745
426.23834228515625 0 822.67365
427.24395751953125 0 1603.1882
428.1803283691406 0 1544.3246 b 5
428.28607177734375 0 729.4333
429.1884765625 0 11555.884
430.19183349609375 0 2704.282 b Ammonia loss 10
431.177978515625 0 1577.4059
435.1657409667969 0 1048.3398 b Water loss 2
439.16229248046875 0 944.91626
441.18438720703125 0 681.20557
446.2338562011719 0 1623.7246
446.732421875 0 874.5801
448.2640075683594 0 832.0521
448.5953063964844 0 1246.8486
451.2250671386719 0 1687.6443
451.2659912109375 0 1098.7123
451.7261962890625 0 1020.21185
451.91937255859375 0 2827.855
452.2539978027344 0 1856.1119
452.5899658203125 0 758.0343
453.17724609375 0 1631.39 b 2
455.251953125 0 1490.7029
457.5944519042969 0 2155.9219
457.92529296875 0 757.4449 y Ammonia loss 2
458.1829528808594 0 679.88654
458.26202392578125 0 1083.7039
460.22967529296875 0 2298.8345
463.5984191894531 0 21232.543 y 2
463.93231201171875 0 18906.516
464.2668151855469 0 9490.096
464.60150146484375 0 1891.6123
467.2978820800781 0 1079.6853
468.9285888671875 0 3362.8997
469.2652282714844 0 852.7423
470.3340148925781 0 18686.465 y 10
471.3369445800781 0 5384.588
471.7977600097656 0 1271.5205
471.9245300292969 0 864.51794
472.2333068847656 0 746.48975
472.2985534667969 0 920.50946 y Ammonia loss 5
472.3374938964844 0 1011.19946
473.26104736328125 0 2060.2432
474.7813415527344 0 898.2871
476.19219970703125 0 2411.3796
477.2000427246094 0 836.0885
478.5741271972656 0 1755.799 b 11
480.80322265625 0 3256.8948 y 5
481.2540588378906 0 2606.0857
481.3045349121094 0 977.03
481.590087890625 0 1178.066
481.8077087402344 0 879.49274
481.9248046875 0 824.8319
482.26763916015625 0 922.31824
482.3093566894531 0 1492.1162
482.76165771484375 0 1114.6381
483.208984375 0 723.3892
483.248291015625 0 1132.3035
483.785888671875 0 1401.6213
487.2585144042969 0 1267.878
487.59228515625 0 1269.592
487.9207458496094 0 931.5802
491.27197265625 0 1439.6057
492.7922668457031 0 730.15314
494.2038269042969 0 1192.6796
503.2137756347656 0 1125.1532
506.9377136230469 0 714.30084
508.75567626953125 0 917.6732
510.2659912109375 0 1804.8204 b Water loss 12
510.59893798828125 0 1775.3334 b Ammonia loss 12
510.93255615234375 0 1023.50385
516.267578125 0 2557.541 b 12
516.6027221679688 0 3347.3818
520.3612670898438 0 902.0617
520.6087036132812 0 833.1943
521.2271728515625 0 923.3362
522.2717895507812 0 5527.5415
522.60693359375 0 3064.3894
522.9403076171875 0 1373.4496
524.9522705078125 0 2627.8115
525.28662109375 0 2202.2754 y Ammonia loss 1
525.6181030273438 0 1553.7446
525.7484741210938 0 804.5614
526.6116943359375 0 1461.5385
526.9445190429688 0 845.16736
530.9556884765625 0 3722.7407 y 1
531.290771484375 0 2455.5527
531.619140625 0 1741.8887
531.95166015625 0 1670.6559
532.2874755859375 0 874.87787
538.2523193359375 0 7094.1836
539.2538452148438 0 1426.538
539.302734375 0 2147.7568
539.802001953125 0 812.80505
541.370849609375 0 4102.524 y 9
547.8137817382812 0 852.1802
548.3121948242188 0 1248.3816
549.3320922851562 0 7856.2837 y 4
549.8335571289062 0 4751.892
552.259521484375 0 657.0526
561.2657470703125 0 716.77234
566.2467651367188 0 7421.5757
567.2501220703125 0 3005.9373
567.3970336914062 0 1027.1975
569.2639770507812 0 2587.3713 b 8
569.7653198242188 0 1932.4756
570.2645874023438 0 824.57434
572.8037719726562 0 2988.5347
573.3038330078125 0 2057.4443
573.8104858398438 0 953.1389
577.3832397460938 0 1967.1049
578.3837890625 0 1252.8079
579.3616943359375 0 5511.611
580.3623046875 0 1240.1163
581.2315063476562 0 914.7684 b 3
585.2567749023438 0 1784.744
595.3922729492188 0 2124.299
596.39306640625 0 1112.2645
603.2669067382812 0 945.22217
604.35400390625 0 1712.9019
604.8517456054688 0 3110.1572 y Ammonia loss 3
605.3500366210938 0 1381.6711
610.3201904296875 0 1702.4734
613.2506103515625 0 2573.976
613.359619140625 0 1057.7399 y 3
620.340576171875 0 2514.846
620.8414916992188 0 1011.87213
628.3316040039062 0 722.35394
629.3451538085938 0 2993.479
629.8462524414062 0 2024.5801
630.3477172851562 0 748.0412
638.3510131835938 0 1778.7152
638.8558959960938 0 1085.8629
639.3533935546875 0 893.2419
649.329833984375 0 822.96967
650.3394775390625 0 663.03467
661.3267822265625 0 860.1924
664.4509887695312 0 1691.6239
665.4580688476562 0 800.63135
669.46630859375 0 1537.8975 y 8
674.43359375 0 5329.3667
675.4373779296875 0 2891.5671
690.2982788085938 0 951.50104
692.4442138671875 0 6305.8213
693.447021484375 0 2709.7954
694.448486328125 0 790.56036
694.8916015625 0 1402.5652 y 2
695.3946533203125 0 1792.6212
700.2803955078125 0 1007.80286 b Water loss 4
710.455322265625 0 1601.3966
716.4220581054688 0 1058.564
718.294921875 0 1293.1919 b 4
719.298583984375 0 740.8531
719.3690795898438 0 1165.8135
720.316650390625 0 848.6466
721.3597412109375 0 823.2981
726.484130859375 0 2448.4172 y 7
727.48876953125 0 1002.52734
736.3641967773438 0 767.7083
738.3777465820312 0 868.13245
740.3199462890625 0 944.671
742.3065795898438 0 983.6485
750.3129272460938 0 815.1768
768.3185424804688 0 1271.71
805.5300903320312 0 1162.2277
811.4948120117188 0 947.9681
823.5394287109375 0 9039.835 y 6
824.5408935546875 0 4743.8003
825.5440673828125 0 909.9552
848.4136962890625 0 1674.944
852.1517333984375 0 692.2087
855.3515625 0 2002.4491 b 5
901.44189453125 0 1894.74
902.4390869140625 0 959.11224
919.4508666992188 0 1058.7429
960.59765625 0 808.2955 y 5
1666.249267578125 0 627.1165
1685.992431640625 0 733.3311
2270.565185546875 0 704.6425
2695.34765625 0 762.2665
3019.1728515625 0 670.8262
3044.292236328125 0 936.91425
3044.8369140625 0 841.0093
3350.053466796875 0 793.2113

Spectrum Details

|  |  |
| --- | --- |
| Matched peaks? Matched peaks The total absolute number of peaks matched. Additionally in brackets the total fraction of peaks matched and the total number of peaks is shown. | 54 (9.56% of 565) |
| FDR? FDR The false discovery rate estimated for this peptide. It is calculated by matching all theoretical fragments with a non-integer shift with the raw peaks for this spectrum. This is done with 40 different shifts. The resulting percentage is the average number of annotated peaks over the number of annotated peaks with the correct spectrum. | 4.54% |
| Satellite FDR? Satellite FDR See the FDR for details on its calculation. This satellite ion specific FDR only contains the satellite ions (d/w) for I/L/J positions. | - |
| PSM Score? PSM Score The PSM Score as given by Hecklib to this annotated spectrum. It is shown with three significant figures. | 205 |

#### Spectrum 5686? Spectrum 5686 The raw spectrum of this peptide as annotated by Hecklib. The fragments are coloured according to ion type (see legend). Any peaks with a star '\*' as text can be hovered over to see the full details, first the ion type second the mass shift type. By hovering over the amino acids in the peptide or ions in the legend the corresponding peaks are highlighted. By toggling the 'Unassigned' label you can turn the background (unassigned) peaks on or off in the plot. By updating the slider in the Ion legend you can update the spectrum to only show the top X% of the peaks with labels. The top X% means any peak that is within X% of the highest intensity. By dragging in the spectrum you can zoom in to a specific part of the spectrum and use 'Zoom Out' to get back to the original zoom level. The annotation of the spectrum is based on the given sequence in the peptides file and is done with different software so inconsistencies are likely. The peaks are annotated based on the given sequence, with 20 ppm tolerance.

Copy Data

##### Spectrum 5686 (TSV)

###### Preview

```
Loading example...
```

*Click on the button to copy the data to your clipboard.*

Mz MinMz MaxIntensity Max

WidthHeightPeptide font sizePeptide stroke widthSpectrum font sizeSpectrum stroke widthCompact peptide

Ion legend

wxyz

abcd

OtherUnassignedIon^Charge^_Position_Show for top:%

SWYQHHPGKAPKJJ

04.00e+47.99e+41.20e+51.60e+5

Zoom Out

y^+1^_1_b^+4^_4_y^+2^_3_y^+2^_4_a^+1^_2_b^+3^_5_y^+2^_4_y^+1^_2_a^+1^_2_b^+1^_2_y^+2^_5_b^+1^_2_y^+3^_8_y^+3^_9_b^+4^_11_y^+2^_6_y^+2^_7_y^+3^_10_y^+1^_3_b^+4^_13_y^+4^_13_b^+3^_10_y^+4^_13_b^+3^_10_y^+3^_11_y^+2^_8_y^+3^_11_b^+2^_6_\*\*y^+2^_8_\*b^+2^_6_b^+1^_3_y^+3^_12_y^+3^_12_y^+1^_4_y^+2^_9_b^+2^_8_b^+3^_13_b^+3^_13_b^+3^_13_y^+3^_13_y^+3^_13_y^+1^_5_y^+2^_10_b^+2^_9_b^+2^_10_b^+2^_10_b^+2^_10_y^+2^_11_y^+2^_11_y^+1^_6_y^+2^_12_b^+1^_5_y^+1^_7_b^+1^_6_y^+1^_8_b^+1^_6_y^+1^_9_

0796159323893186

Fragment Matches Table

Show background peaks

| Position | Ion type | Intensity | mz Theoretical | mz Error (Th) | mz Error (ppm) | Charge | Series Number |
| --- | --- | --- | --- | --- | --- | --- | --- |
| - | - | 522.7 | 120.1 | - | - | 0 | - |
| - | - | 867.1 | 120.1 | - | - | 0 | - |
| - | - | 1545 | 120.1 | - | - | 0 | - |
| - | - | 863.4 | 122 | - | - | 0 | - |
| - | - | 561.9 | 122.1 | - | - | 0 | - |
| - | - | 1577 | 123.1 | - | - | 0 | - |
| - | - | 575.4 | 123.1 | - | - | 0 | - |
| - | - | 527.7 | 125.1 | - | - | 0 | - |
| - | - | 418.1 | 126.1 | - | - | 0 | - |
| - | - | 954.4 | 126.1 | - | - | 0 | - |
| - | - | 994.1 | 126.1 | - | - | 0 | - |
| - | - | 458.3 | 127.1 | - | - | 0 | - |
| - | - | 3235 | 127.1 | - | - | 0 | - |
| - | - | 678.5 | 128.1 | - | - | 0 | - |
| - | - | 946.1 | 128.1 | - | - | 0 | - |
| - | - | 378.5 | 128.6 | - | - | 0 | - |
| - | - | 692 | 129.1 | - | - | 0 | - |
| - | - | 572.6 | 129.1 | - | - | 0 | - |
| - | - | 4231 | 129.1 | - | - | 0 | - |
| - | - | 8.309E+04 | 129.1 | - | - | 0 | - |
| - | - | 2058 | 130.1 | - | - | 0 | - |
| - | - | 2.068E+04 | 130.1 | - | - | 0 | - |
| - | - | 745.3 | 130.1 | - | - | 0 | - |
| - | - | 866.9 | 130.1 | - | - | 0 | - |
| - | - | 6123 | 130.1 | - | - | 0 | - |
| - | - | 3860 | 131 | - | - | 0 | - |
| - | - | 1792 | 131.1 | - | - | 0 | - |
| - | - | 2560 | 131.1 | - | - | 0 | - |
| - | - | 7.264E+04 | 132.1 | - | - | 0 | - |
| 14 | y | 1.999E+04 | 132.1 | 0.0004051 | 3.066 | +1 | 1 |
| - | - | 424.7 | 133.1 | - | - | 0 | - |
| - | - | 2.624E+04 | 133.1 | - | - | 0 | - |
| - | - | 1557 | 133.1 | - | - | 0 | - |
| - | - | 429.7 | 134.1 | - | - | 0 | - |
| - | - | 555.9 | 134.1 | - | - | 0 | - |
| - | - | 1708 | 134.1 | - | - | 0 | - |
| - | - | 473.9 | 136 | - | - | 0 | - |
| - | - | 1.583E+05 | 136.1 | - | - | 0 | - |
| - | - | 454.7 | 137.1 | - | - | 0 | - |
| - | - | 1167 | 137.1 | - | - | 0 | - |
| - | - | 1.482E+04 | 137.1 | - | - | 0 | - |
| - | - | 1595 | 138.1 | - | - | 0 | - |
| - | - | 1711 | 139.1 | - | - | 0 | - |
| - | - | 703.1 | 140.1 | - | - | 0 | - |
| - | - | 790.2 | 141.1 | - | - | 0 | - |
| - | - | 5811 | 141.1 | - | - | 0 | - |
| 4 | b | 1480 | 142.1 | 0.0001466 | 1.032 | +4 | 4 |
| - | - | 3736 | 142.1 | - | - | 0 | - |
| - | - | 4663 | 143.1 | - | - | 0 | - |
| - | - | 2872 | 144.1 | - | - | 0 | - |
| - | - | 445.8 | 144.1 | - | - | 0 | - |
| - | - | 1270 | 146.1 | - | - | 0 | - |
| - | - | 5276 | 147 | - | - | 0 | - |
| - | - | 607.1 | 147.1 | - | - | 0 | - |
| - | - | 582.1 | 148 | - | - | 0 | - |
| - | - | 496 | 148.1 | - | - | 0 | - |
| - | - | 1504 | 151.1 | - | - | 0 | - |
| - | - | 3733 | 152 | - | - | 0 | - |
| - | - | 554.4 | 152 | - | - | 0 | - |
| - | - | 540.4 | 152.1 | - | - | 0 | - |
| - | - | 832.3 | 152.1 | - | - | 0 | - |
| - | - | 609.9 | 152.1 | - | - | 0 | - |
| - | - | 1654 | 152.1 | - | - | 0 | - |
| - | - | 435.7 | 153.1 | - | - | 0 | - |
| - | - | 457.6 | 153.1 | - | - | 0 | - |
| - | - | 747.6 | 153.1 | - | - | 0 | - |
| - | - | 436.6 | 154 | - | - | 0 | - |
| - | - | 929 | 154.1 | - | - | 0 | - |
| - | - | 3848 | 155.1 | - | - | 0 | - |
| - | - | 1435 | 155.1 | - | - | 0 | - |
| - | - | 3558 | 155.1 | - | - | 0 | - |
| - | - | 1469 | 156.1 | - | - | 0 | - |
| - | - | 1823 | 156.1 | - | - | 0 | - |
| - | - | 485 | 157.1 | - | - | 0 | - |
| - | - | 1445 | 158.1 | - | - | 0 | - |
| - | - | 9067 | 158.1 | - | - | 0 | - |
| - | - | 1694 | 159.1 | - | - | 0 | - |
| - | - | 5.116E+04 | 159.1 | - | - | 0 | - |
| - | - | 2468 | 160.1 | - | - | 0 | - |
| - | - | 547.2 | 160.1 | - | - | 0 | - |
| - | - | 4847 | 160.1 | - | - | 0 | - |
| - | - | 1045 | 161.1 | - | - | 0 | - |
| - | - | 446.1 | 162.1 | - | - | 0 | - |
| - | - | 492.6 | 162.1 | - | - | 0 | - |
| - | - | 478.1 | 162.3 | - | - | 0 | - |
| - | - | 652.1 | 163.6 | - | - | 0 | - |
| - | - | 1448 | 164.1 | - | - | 0 | - |
| - | - | 475.1 | 165.1 | - | - | 0 | - |
| - | - | 2971 | 165.1 | - | - | 0 | - |
| - | - | 985.8 | 166.1 | - | - | 0 | - |
| - | - | 710.3 | 166.1 | - | - | 0 | - |
| - | - | 528.2 | 167.1 | - | - | 0 | - |
| - | - | 5121 | 169.1 | - | - | 0 | - |
| - | - | 5695 | 169.1 | - | - | 0 | - |
| - | - | 1746 | 169.1 | - | - | 0 | - |
| - | - | 2.475E+04 | 170.1 | - | - | 0 | - |
| - | - | 3018 | 171.1 | - | - | 0 | - |
| - | - | 776.1 | 171.1 | - | - | 0 | - |
| - | - | 387.9 | 172.1 | - | - | 0 | - |
| - | - | 598.9 | 172.1 | - | - | 0 | - |
| - | - | 4935 | 172.1 | - | - | 0 | - |
| - | - | 544.6 | 172.1 | - | - | 0 | - |
| - | - | 668.2 | 173.1 | - | - | 0 | - |
| - | - | 818.8 | 173.5 | - | - | 0 | - |
| - | - | 643.5 | 174.1 | - | - | 0 | - |
| - | - | 853 | 175.1 | - | - | 0 | - |
| - | - | 1291 | 175.1 | - | - | 0 | - |
| - | - | 478.5 | 175.1 | - | - | 0 | - |
| - | - | 1477 | 176.1 | - | - | 0 | - |
| - | - | 707 | 177 | - | - | 0 | - |
| - | - | 9903 | 177.1 | - | - | 0 | - |
| - | - | 521.2 | 179.1 | - | - | 0 | - |
| - | - | 1398 | 181.1 | - | - | 0 | - |
| - | - | 2.363E+04 | 181.1 | - | - | 0 | - |
| - | - | 1805 | 181.1 | - | - | 0 | - |
| - | - | 2020 | 182.1 | - | - | 0 | - |
| - | - | 2965 | 182.1 | - | - | 0 | - |
| - | - | 1.046E+04 | 183.1 | - | - | 0 | - |
| - | - | 696.1 | 183.1 | - | - | 0 | - |
| - | - | 1233 | 183.1 | - | - | 0 | - |
| - | - | 730.6 | 183.1 | - | - | 0 | - |
| - | - | 1013 | 184.1 | - | - | 0 | - |
| - | - | 1187 | 184.1 | - | - | 0 | - |
| - | - | 3436 | 185.1 | - | - | 0 | - |
| - | - | 574.7 | 185.1 | - | - | 0 | - |
| - | - | 656.5 | 185.1 | - | - | 0 | - |
| - | - | 3704 | 185.2 | - | - | 0 | - |
| - | - | 8979 | 186.1 | - | - | 0 | - |
| - | - | 5.222E+04 | 187.1 | - | - | 0 | - |
| - | - | 5714 | 187.1 | - | - | 0 | - |
| 12 | y | 1.171E+05 | 187.1 | 0.0004575 | 2.445 | +2 | 3 |
| - | - | 6103 | 188.1 | - | - | 0 | - |
| - | - | 1.082E+04 | 188.1 | - | - | 0 | - |
| - | - | 525.1 | 189.1 | - | - | 0 | - |
| - | - | 549.6 | 190.5 | - | - | 0 | - |
| - | - | 502 | 191.1 | - | - | 0 | - |
| - | - | 589.3 | 193.1 | - | - | 0 | - |
| - | - | 1313 | 195 | - | - | 0 | - |
| - | - | 1014 | 195.1 | - | - | 0 | - |
| - | - | 1244 | 197.1 | - | - | 0 | - |
| - | - | 3150 | 197.2 | - | - | 0 | - |
| - | - | 671.5 | 198.2 | - | - | 0 | - |
| - | - | 2514 | 199.1 | - | - | 0 | - |
| - | - | 1160 | 199.1 | - | - | 0 | - |
| - | - | 1437 | 199.2 | - | - | 0 | - |
| - | - | 1742 | 199.2 | - | - | 0 | - |
| - | - | 1556 | 200.1 | - | - | 0 | - |
| - | - | 5282 | 201.1 | - | - | 0 | - |
| - | - | 1414 | 201.1 | - | - | 0 | - |
| - | - | 2421 | 202.1 | - | - | 0 | - |
| - | - | 1.036E+04 | 202.1 | - | - | 0 | - |
| - | - | 636.5 | 203.1 | - | - | 0 | - |
| - | - | 844.1 | 203.1 | - | - | 0 | - |
| - | - | 757.4 | 204.1 | - | - | 0 | - |
| - | - | 3692 | 204.1 | - | - | 0 | - |
| - | - | 6729 | 205.1 | - | - | 0 | - |
| - | - | 2634 | 207.1 | - | - | 0 | - |
| - | - | 697.7 | 209.1 | - | - | 0 | - |
| - | - | 1933 | 211.1 | - | - | 0 | - |
| - | - | 548.4 | 211.1 | - | - | 0 | - |
| - | - | 728.2 | 212.1 | - | - | 0 | - |
| - | - | 2142 | 212.7 | - | - | 0 | - |
| - | - | 1416 | 213.2 | - | - | 0 | - |
| - | - | 2388 | 214.1 | - | - | 0 | - |
| - | - | 715.5 | 214.2 | - | - | 0 | - |
| - | - | 4.498E+04 | 215.1 | - | - | 0 | - |
| - | - | 616.9 | 216.1 | - | - | 0 | - |
| - | - | 993.5 | 216.1 | - | - | 0 | - |
| - | - | 5424 | 216.1 | - | - | 0 | - |
| - | - | 982.1 | 217.1 | - | - | 0 | - |
| - | - | 576.8 | 217.1 | - | - | 0 | - |
| - | - | 9410 | 219.1 | - | - | 0 | - |
| - | - | 1463 | 220.1 | - | - | 0 | - |
| - | - | 6045 | 220.1 | - | - | 0 | - |
| - | - | 6909 | 221.1 | - | - | 0 | - |
| - | - | 2469 | 221.1 | - | - | 0 | - |
| - | - | 602.6 | 224.1 | - | - | 0 | - |
| - | - | 948 | 224.1 | - | - | 0 | - |
| - | - | 3472 | 224.2 | - | - | 0 | - |
| - | - | 1572 | 225.2 | - | - | 0 | - |
| - | - | 6.529E+04 | 226.2 | - | - | 0 | - |
| - | - | 745.5 | 227.1 | - | - | 0 | - |
| 11 | y | 8196 | 227.2 | 0.001632 | 7.186 | +2 | 4 |
| - | - | 887.9 | 227.2 | - | - | 0 | - |
| 2 | a | 5187 | 228.1 | 0.0004021 | 1.763 | +1 | 2 |
| - | - | 709 | 228.1 | - | - | 0 | - |
| 5 | b | 6274 | 229.1 | 0.001782 | 7.779 | +3 | 5 |
| - | - | 988.9 | 229.1 | - | - | 0 | - |
| - | - | 753.1 | 229.2 | - | - | 0 | - |
| - | - | 2009 | 230.1 | - | - | 0 | - |
| - | - | 2485 | 231.1 | - | - | 0 | - |
| - | - | 1218 | 231.1 | - | - | 0 | - |
| - | - | 956 | 232.1 | - | - | 0 | - |
| - | - | 823.9 | 233.1 | - | - | 0 | - |
| - | - | 1123 | 233.1 | - | - | 0 | - |
| - | - | 617.2 | 235.1 | - | - | 0 | - |
| - | - | 1.23E+04 | 235.1 | - | - | 0 | - |
| 11 | y | 9716 | 235.7 | 0.0003817 | 1.62 | +2 | 4 |
| - | - | 1639 | 236.1 | - | - | 0 | - |
| - | - | 2076 | 236.2 | - | - | 0 | - |
| - | - | 595.7 | 237.1 | - | - | 0 | - |
| - | - | 5496 | 239.2 | - | - | 0 | - |
| - | - | 669.4 | 240.2 | - | - | 0 | - |
| - | - | 3.57E+04 | 241 | - | - | 0 | - |
| - | - | 1247 | 241.2 | - | - | 0 | - |
| - | - | 2428 | 242 | - | - | 0 | - |
| - | - | 3477 | 242.2 | - | - | 0 | - |
| - | - | 3580 | 243 | - | - | 0 | - |
| 13 | y | 1.569E+04 | 245.2 | 0.0005086 | 2.074 | +1 | 2 |
| 2 | a | 1.199E+05 | 246.1 | 0.0004881 | 1.983 | +1 | 2 |
| - | - | 2341 | 246.2 | - | - | 0 | - |
| - | - | 2.723E+04 | 247.1 | - | - | 0 | - |
| - | - | 8523 | 248.1 | - | - | 0 | - |
| - | - | 2467 | 248.1 | - | - | 0 | - |
| - | - | 535.8 | 248.2 | - | - | 0 | - |
| - | - | 1.214E+04 | 249.1 | - | - | 0 | - |
| - | - | 873.7 | 249.1 | - | - | 0 | - |
| - | - | 1720 | 249.1 | - | - | 0 | - |
| - | - | 5.092E+04 | 249.2 | - | - | 0 | - |
| - | - | 1941 | 250.1 | - | - | 0 | - |
| - | - | 1797 | 250.1 | - | - | 0 | - |
| - | - | 634.5 | 250.1 | - | - | 0 | - |
| - | - | 9901 | 250.2 | - | - | 0 | - |
| - | - | 696.8 | 251.1 | - | - | 0 | - |
| - | - | 1121 | 251.2 | - | - | 0 | - |
| - | - | 615.7 | 251.2 | - | - | 0 | - |
| - | - | 1944 | 252.2 | - | - | 0 | - |
| - | - | 917.5 | 255.1 | - | - | 0 | - |
| 2 | b | 2469 | 256.1 | 0.0005894 | 2.302 | +1 | 2 |
| - | - | 1781 | 257.1 | - | - | 0 | - |
| - | - | 1371 | 257.2 | - | - | 0 | - |
| - | - | 2154 | 258.1 | - | - | 0 | - |
| - | - | 2.1E+04 | 258.1 | - | - | 0 | - |
| - | - | 3039 | 259.1 | - | - | 0 | - |
| - | - | 631.8 | 259.1 | - | - | 0 | - |
| - | - | 2789 | 260.1 | - | - | 0 | - |
| - | - | 1516 | 260.2 | - | - | 0 | - |
| - | - | 769.1 | 262.2 | - | - | 0 | - |
| - | - | 1276 | 264.1 | - | - | 0 | - |
| - | - | 1307 | 265.2 | - | - | 0 | - |
| - | - | 8191 | 266.1 | - | - | 0 | - |
| - | - | 788.5 | 266.1 | - | - | 0 | - |
| - | - | 1106 | 267.1 | - | - | 0 | - |
| - | - | 809.8 | 267.1 | - | - | 0 | - |
| - | - | 681.7 | 268.2 | - | - | 0 | - |
| - | - | 1003 | 269.2 | - | - | 0 | - |
| - | - | 1805 | 269.6 | - | - | 0 | - |
| - | - | 5089 | 270.2 | - | - | 0 | - |
| 10 | y | 4391 | 271.2 | 4.381E-05 | 0.1615 | +2 | 5 |
| 2 | b | 4.467E+04 | 274.1 | 0.0003702 | 1.351 | +1 | 2 |
| - | - | 4700 | 275.1 | - | - | 0 | - |
| - | - | 3.94E+04 | 275.1 | - | - | 0 | - |
| 7 | y | 612.6 | 275.2 | 0.0008179 | 2.972 | +3 | 8 |
| - | - | 1843 | 276.1 | - | - | 0 | - |
| - | - | 5066 | 276.1 | - | - | 0 | - |
| - | - | 1452 | 277.1 | - | - | 0 | - |
| - | - | 1.156E+04 | 277.2 | - | - | 0 | - |
| - | - | 2942 | 278.2 | - | - | 0 | - |
| - | - | 568.7 | 279.2 | - | - | 0 | - |
| - | - | 1425 | 279.2 | - | - | 0 | - |
| - | - | 1773 | 283.2 | - | - | 0 | - |
| - | - | 1.769E+04 | 283.2 | - | - | 0 | - |
| - | - | 1213 | 283.6 | - | - | 0 | - |
| - | - | 3423 | 284.2 | - | - | 0 | - |
| - | - | 1180 | 285.1 | - | - | 0 | - |
| - | - | 3366 | 285.2 | - | - | 0 | - |
| - | - | 6025 | 290.2 | - | - | 0 | - |
| - | - | 1766 | 290.7 | - | - | 0 | - |
| - | - | 1263 | 291.2 | - | - | 0 | - |
| - | - | 1.226E+04 | 292.1 | - | - | 0 | - |
| - | - | 1882 | 293.1 | - | - | 0 | - |
| - | - | 3540 | 294.1 | - | - | 0 | - |
| - | - | 821.3 | 294.2 | - | - | 0 | - |
| - | - | 681.9 | 294.7 | - | - | 0 | - |
| - | - | 701.3 | 295.1 | - | - | 0 | - |
| - | - | 7733 | 297.2 | - | - | 0 | - |
| - | - | 1126 | 298.2 | - | - | 0 | - |
| - | - | 2658 | 303.1 | - | - | 0 | - |
| - | - | 4451 | 305.1 | - | - | 0 | - |
| - | - | 2044 | 305.7 | - | - | 0 | - |
| - | - | 1298 | 306.1 | - | - | 0 | - |
| - | - | 707.6 | 309.2 | - | - | 0 | - |
| - | - | 2994 | 311.2 | - | - | 0 | - |
| - | - | 708.5 | 313.2 | - | - | 0 | - |
| - | - | 4952 | 314.2 | - | - | 0 | - |
| - | - | 1212 | 314.7 | - | - | 0 | - |
| - | - | 899.5 | 315.2 | - | - | 0 | - |
| - | - | 514.2 | 318.2 | - | - | 0 | - |
| - | - | 989.7 | 320.1 | - | - | 0 | - |
| 6 | y | 2018 | 320.9 | 9.13E-05 | 0.2845 | +3 | 9 |
| - | - | 911.6 | 321.2 | - | - | 0 | - |
| - | - | 1.086E+04 | 321.2 | - | - | 0 | - |
| - | - | 688 | 321.5 | - | - | 0 | - |
| - | - | 771.1 | 322.1 | - | - | 0 | - |
| - | - | 5595 | 322.2 | - | - | 0 | - |
| - | - | 1903 | 322.2 | - | - | 0 | - |
| 11 | b | 1250 | 323.2 | 0.0002229 | 0.6897 | +4 | 11 |
| - | - | 2036 | 324.2 | - | - | 0 | - |
| - | - | 805.3 | 325.2 | - | - | 0 | - |
| - | - | 3815 | 326.2 | - | - | 0 | - |
| - | - | 586.1 | 326.7 | - | - | 0 | - |
| - | - | 3479 | 327.2 | - | - | 0 | - |
| - | - | 855 | 327.2 | - | - | 0 | - |
| - | - | 650.6 | 328.2 | - | - | 0 | - |
| - | - | 2698 | 332.7 | - | - | 0 | - |
| - | - | 1311 | 333.1 | - | - | 0 | - |
| - | - | 931.7 | 333.2 | - | - | 0 | - |
| - | - | 656.7 | 334.1 | - | - | 0 | - |
| - | - | 1465 | 334.7 | - | - | 0 | - |
| 9 | y | 1131 | 335.2 | 0.001055 | 3.146 | +2 | 6 |
| - | - | 1.526E+04 | 336.2 | - | - | 0 | - |
| - | - | 2503 | 337.2 | - | - | 0 | - |
| - | - | 5058 | 337.7 | - | - | 0 | - |
| - | - | 2094 | 338.2 | - | - | 0 | - |
| - | - | 634.5 | 338.7 | - | - | 0 | - |
| - | - | 1.596E+04 | 339.2 | - | - | 0 | - |
| - | - | 5.332E+04 | 340.1 | - | - | 0 | - |
| - | - | 2987 | 340.2 | - | - | 0 | - |
| - | - | 8345 | 341.1 | - | - | 0 | - |
| - | - | 5233 | 342.1 | - | - | 0 | - |
| - | - | 767.6 | 342.2 | - | - | 0 | - |
| - | - | 596.4 | 343.1 | - | - | 0 | - |
| - | - | 558.3 | 344.9 | - | - | 0 | - |
| - | - | 1524 | 345.6 | - | - | 0 | - |
| - | - | 713.7 | 346.7 | - | - | 0 | - |
| - | - | 1.385E+04 | 346.7 | - | - | 0 | - |
| - | - | 740.8 | 347.2 | - | - | 0 | - |
| - | - | 5530 | 347.2 | - | - | 0 | - |
| - | - | 1092 | 347.7 | - | - | 0 | - |
| - | - | 1641 | 350.1 | - | - | 0 | - |
| - | - | 1098 | 351.2 | - | - | 0 | - |
| - | - | 673.6 | 352.2 | - | - | 0 | - |
| - | - | 991.8 | 353.2 | - | - | 0 | - |
| - | - | 6487 | 354.1 | - | - | 0 | - |
| - | - | 1782 | 354.2 | - | - | 0 | - |
| - | - | 1.995E+04 | 354.2 | - | - | 0 | - |
| - | - | 1070 | 354.7 | - | - | 0 | - |
| - | - | 1051 | 355.1 | - | - | 0 | - |
| - | - | 4100 | 355.2 | - | - | 0 | - |
| - | - | 1484 | 355.7 | - | - | 0 | - |
| - | - | 525.7 | 356.2 | - | - | 0 | - |
| - | - | 1269 | 356.2 | - | - | 0 | - |
| - | - | 4775 | 357.2 | - | - | 0 | - |
| - | - | 1337 | 357.8 | - | - | 0 | - |
| - | - | 4616 | 358.2 | - | - | 0 | - |
| - | - | 2141 | 358.7 | - | - | 0 | - |
| - | - | 785.7 | 359.2 | - | - | 0 | - |
| - | - | 1344 | 359.2 | - | - | 0 | - |
| - | - | 826.1 | 359.5 | - | - | 0 | - |
| - | - | 1384 | 359.9 | - | - | 0 | - |
| - | - | 900 | 360.2 | - | - | 0 | - |
| - | - | 922.8 | 360.7 | - | - | 0 | - |
| - | - | 1164 | 363.5 | - | - | 0 | - |
| 8 | y | 859.5 | 363.7 | 0.0009735 | 2.676 | +2 | 7 |
| - | - | 1163 | 363.8 | - | - | 0 | - |
| - | - | 546.6 | 365.7 | - | - | 0 | - |
| - | - | 1557 | 366.2 | - | - | 0 | - |
| 5 | y | 2477 | 366.6 | 0.0001165 | 0.3177 | +3 | 10 |
| - | - | 860 | 366.9 | - | - | 0 | - |
| - | - | 1043 | 367.2 | - | - | 0 | - |
| - | - | 2570 | 368.1 | - | - | 0 | - |
| - | - | 693.1 | 370.2 | - | - | 0 | - |
| - | - | 753.2 | 370.5 | - | - | 0 | - |
| - | - | 3.036E+04 | 371.2 | - | - | 0 | - |
| - | - | 6577 | 372.2 | - | - | 0 | - |
| 12 | y | 8388 | 373.3 | 0.0001348 | 0.3611 | +1 | 3 |
| - | - | 1663 | 374.3 | - | - | 0 | - |
| - | - | 947.8 | 374.8 | - | - | 0 | - |
| - | - | 677.2 | 376.5 | - | - | 0 | - |
| - | - | 935.5 | 376.7 | - | - | 0 | - |
| - | - | 544.7 | 376.7 | - | - | 0 | - |
| 13 | b | 832.7 | 379 | 0.004189 | 11.05 | +4 | 13 |
| - | - | 591.5 | 380.2 | - | - | 0 | - |
| - | - | 1271 | 380.2 | - | - | 0 | - |
| - | - | 4056 | 382.2 | - | - | 0 | - |
| - | - | 615.3 | 382.3 | - | - | 0 | - |
| - | - | 1691 | 382.5 | - | - | 0 | - |
| - | - | 1247 | 382.8 | - | - | 0 | - |
| - | - | 1149 | 382.9 | - | - | 0 | - |
| - | - | 1206 | 383.3 | - | - | 0 | - |
| - | - | 2280 | 384.2 | - | - | 0 | - |
| - | - | 9999 | 385.2 | - | - | 0 | - |
| - | - | 2070 | 385.2 | - | - | 0 | - |
| - | - | 8848 | 386.2 | - | - | 0 | - |
| - | - | 1262 | 387.2 | - | - | 0 | - |
| - | - | 656.5 | 388.2 | - | - | 0 | - |
| - | - | 2041 | 388.9 | - | - | 0 | - |
| - | - | 1993 | 389.2 | - | - | 0 | - |
| - | - | 2352 | 389.3 | - | - | 0 | - |
| - | - | 1042 | 389.8 | - | - | 0 | - |
| 2 | y | 5716 | 390.2 | 1.169E-05 | 0.02997 | +4 | 13 |
| - | - | 5125 | 390.5 | - | - | 0 | - |
| - | - | 2084 | 390.7 | - | - | 0 | - |
| - | - | 3685 | 392.3 | - | - | 0 | - |
| 10 | b | 890.6 | 392.5 | 0.006954 | 17.72 | +3 | 10 |
| - | - | 5945 | 394.2 | - | - | 0 | - |
| - | - | 787.4 | 394.2 | - | - | 0 | - |
| 2 | y | 2591 | 394.5 | 0.0009123 | 2.313 | +4 | 13 |
| - | - | 2514 | 394.7 | - | - | 0 | - |
| - | - | 1619 | 395 | - | - | 0 | - |
| - | - | 1439 | 396.3 | - | - | 0 | - |
| - | - | 1513 | 397.2 | - | - | 0 | - |
| - | - | 801.6 | 397.6 | - | - | 0 | - |
| 10 | b | 1621 | 398.2 | 0.0007794 | 1.957 | +3 | 10 |
| - | - | 802.7 | 398.2 | - | - | 0 | - |
| - | - | 804.6 | 398.9 | - | - | 0 | - |
| - | - | 2.112E+04 | 399.2 | - | - | 0 | - |
| - | - | 3882 | 400.2 | - | - | 0 | - |
| - | - | 1029 | 401.2 | - | - | 0 | - |
| - | - | 916.9 | 401.3 | - | - | 0 | - |
| - | - | 3373 | 402.2 | - | - | 0 | - |
| - | - | 944.7 | 402.5 | - | - | 0 | - |
| - | - | 683 | 402.9 | - | - | 0 | - |
| - | - | 6445 | 403.2 | - | - | 0 | - |
| - | - | 3120 | 403.2 | - | - | 0 | - |
| - | - | 2215 | 403.3 | - | - | 0 | - |
| 4 | y | 5302 | 403.6 | 0.00291 | 7.21 | +3 | 11 |
| 7 | y | 785.6 | 403.8 | 0.004559 | 11.29 | +2 | 8 |
| - | - | 2528 | 403.9 | - | - | 0 | - |
| - | - | 1495 | 404.2 | - | - | 0 | - |
| - | - | 1401 | 404.2 | - | - | 0 | - |
| - | - | 998.4 | 404.5 | - | - | 0 | - |
| - | - | 3443 | 404.7 | - | - | 0 | - |
| - | - | 1401 | 404.9 | - | - | 0 | - |
| - | - | 3314 | 405 | - | - | 0 | - |
| - | - | 1068 | 405.2 | - | - | 0 | - |
| - | - | 2507 | 406.2 | - | - | 0 | - |
| - | - | 3207 | 406.3 | - | - | 0 | - |
| - | - | 1082 | 406.7 | - | - | 0 | - |
| - | - | 1851 | 406.8 | - | - | 0 | - |
| - | - | 1851 | 408.2 | - | - | 0 | - |
| - | - | 2017 | 408.6 | - | - | 0 | - |
| 4 | y | 1041 | 409.2 | 0.0002145 | 0.5241 | +3 | 11 |
| - | - | 866.2 | 409.6 | - | - | 0 | - |
| - | - | 6361 | 410.3 | - | - | 0 | - |
| - | - | 2599 | 410.6 | - | - | 0 | - |
| - | - | 3957 | 410.9 | - | - | 0 | - |
| 6 | b | 8738 | 411.2 | 0.0005404 | 1.314 | +2 | 6 |
| - | - | 740.5 | 411.2 | - | - | 0 | - |
| - | - | 1535 | 411.3 | - | - | 0 | - |
| 0 | Precursor | 1511 | 411.7 | 0.001341 | 3.257 | +4 | -1 |
| 0 | Precursor | 2014 | 412 | 0.003388 | 8.223 | +4 | -1 |
| - | - | 2087 | 412.2 | - | - | 0 | - |
| - | - | 1693 | 412.2 | - | - | 0 | - |
| 7 | y | 2.471E+04 | 412.3 | 0.0007451 | 1.807 | +2 | 8 |
| - | - | 1.299E+04 | 412.8 | - | - | 0 | - |
| - | - | 3732 | 413.3 | - | - | 0 | - |
| - | - | 2794 | 413.9 | - | - | 0 | - |
| - | - | 680 | 414.2 | - | - | 0 | - |
| - | - | 3907 | 414.2 | - | - | 0 | - |
| - | - | 2122 | 414.6 | - | - | 0 | - |
| - | - | 964.5 | 414.9 | - | - | 0 | - |
| - | - | 3527 | 415.3 | - | - | 0 | - |
| - | - | 1656 | 415.8 | - | - | 0 | - |
| 0 | Precursor | 1.498E+04 | 416.2 | 0.0002904 | 0.6977 | +4 | -1 |
| - | - | 2768 | 416.3 | - | - | 0 | - |
| - | - | 2457 | 416.3 | - | - | 0 | - |
| - | - | 1.155E+04 | 416.5 | - | - | 0 | - |
| - | - | 6591 | 416.7 | - | - | 0 | - |
| - | - | 2364 | 417 | - | - | 0 | - |
| - | - | 1586 | 417.2 | - | - | 0 | - |
| - | - | 1.362E+04 | 419.9 | - | - | 0 | - |
| 6 | b | 1041 | 420.2 | 0.001026 | 2.441 | +2 | 6 |
| - | - | 1.322E+04 | 420.2 | - | - | 0 | - |
| - | - | 5590 | 420.6 | - | - | 0 | - |
| - | - | 1233 | 420.7 | - | - | 0 | - |
| - | - | 1148 | 420.9 | - | - | 0 | - |
| - | - | 2289 | 422.3 | - | - | 0 | - |
| - | - | 911.3 | 423.3 | - | - | 0 | - |
| - | - | 1619 | 424.3 | - | - | 0 | - |
| - | - | 4429 | 424.7 | - | - | 0 | - |
| - | - | 2036 | 425.2 | - | - | 0 | - |
| - | - | 723.8 | 425.7 | - | - | 0 | - |
| - | - | 3065 | 425.9 | - | - | 0 | - |
| - | - | 2255 | 426.2 | - | - | 0 | - |
| - | - | 1696 | 427.2 | - | - | 0 | - |
| - | - | 1476 | 427.7 | - | - | 0 | - |
| - | - | 2.431E+04 | 429.2 | - | - | 0 | - |
| - | - | 6491 | 430.2 | - | - | 0 | - |
| - | - | 2931 | 431.2 | - | - | 0 | - |
| - | - | 671.7 | 432.2 | - | - | 0 | - |
| 3 | b | 1869 | 437.2 | 5.543E-07 | 0.001268 | +1 | 3 |
| - | - | 927.9 | 438.2 | - | - | 0 | - |
| - | - | 987.8 | 438.2 | - | - | 0 | - |
| - | - | 2134 | 438.6 | - | - | 0 | - |
| - | - | 1265 | 438.9 | - | - | 0 | - |
| - | - | 1456 | 439.2 | - | - | 0 | - |
| - | - | 660 | 442.7 | - | - | 0 | - |
| - | - | 832.6 | 444.2 | - | - | 0 | - |
| - | - | 974.2 | 444.6 | - | - | 0 | - |
| - | - | 794 | 444.9 | - | - | 0 | - |
| - | - | 1904 | 446.2 | - | - | 0 | - |
| - | - | 1244 | 446.7 | - | - | 0 | - |
| - | - | 4302 | 448.3 | - | - | 0 | - |
| - | - | 2750 | 448.6 | - | - | 0 | - |
| - | - | 1559 | 448.9 | - | - | 0 | - |
| - | - | 3780 | 451.2 | - | - | 0 | - |
| - | - | 2282 | 451.3 | - | - | 0 | - |
| - | - | 3290 | 451.7 | - | - | 0 | - |
| - | - | 6263 | 451.9 | - | - | 0 | - |
| - | - | 4661 | 452.3 | - | - | 0 | - |
| - | - | 1478 | 452.3 | - | - | 0 | - |
| - | - | 3261 | 452.6 | - | - | 0 | - |
| - | - | 786.6 | 452.9 | - | - | 0 | - |
| - | - | 1.028E+04 | 453.2 | - | - | 0 | - |
| - | - | 1660 | 454.2 | - | - | 0 | - |
| - | - | 1350 | 455.2 | - | - | 0 | - |
| - | - | 2431 | 455.3 | - | - | 0 | - |
| - | - | 3451 | 457.6 | - | - | 0 | - |
| 3 | y | 5375 | 457.9 | 0.0036 | 7.862 | +3 | 12 |
| - | - | 3546 | 458.3 | - | - | 0 | - |
| - | - | 665.9 | 458.6 | - | - | 0 | - |
| - | - | 5380 | 460.2 | - | - | 0 | - |
| - | - | 2644 | 460.7 | - | - | 0 | - |
| - | - | 593.6 | 461.6 | - | - | 0 | - |
| - | - | 678.9 | 462.2 | - | - | 0 | - |
| 3 | y | 7.33E+04 | 463.6 | 0.0006605 | 1.425 | +3 | 12 |
| - | - | 6.028E+04 | 463.9 | - | - | 0 | - |
| - | - | 2.345E+04 | 464.3 | - | - | 0 | - |
| - | - | 6510 | 464.6 | - | - | 0 | - |
| - | - | 1521 | 464.9 | - | - | 0 | - |
| - | - | 1390 | 466.9 | - | - | 0 | - |
| - | - | 1526 | 467.3 | - | - | 0 | - |
| - | - | 2399 | 467.3 | - | - | 0 | - |
| - | - | 1717 | 468.3 | - | - | 0 | - |
| - | - | 942.2 | 469.8 | - | - | 0 | - |
| - | - | 1084 | 470.3 | - | - | 0 | - |
| 11 | y | 2.783E+04 | 470.3 | 0.0006546 | 1.392 | +1 | 4 |
| - | - | 2427 | 470.6 | - | - | 0 | - |
| - | - | 723.2 | 470.8 | - | - | 0 | - |
| - | - | 937.4 | 470.9 | - | - | 0 | - |
| - | - | 745.9 | 471.2 | - | - | 0 | - |
| - | - | 8231 | 471.3 | - | - | 0 | - |
| - | - | 3498 | 471.8 | - | - | 0 | - |
| - | - | 1087 | 472.3 | - | - | 0 | - |
| - | - | 1513 | 472.3 | - | - | 0 | - |
| - | - | 867.7 | 472.6 | - | - | 0 | - |
| - | - | 3825 | 473.3 | - | - | 0 | - |
| - | - | 1810 | 473.6 | - | - | 0 | - |
| - | - | 792.4 | 474.3 | - | - | 0 | - |
| - | - | 2049 | 474.8 | - | - | 0 | - |
| - | - | 5701 | 476.3 | - | - | 0 | - |
| - | - | 4224 | 476.6 | - | - | 0 | - |
| - | - | 1749 | 476.9 | - | - | 0 | - |
| 6 | y | 6790 | 480.8 | 0.0005251 | 1.092 | +2 | 9 |
| - | - | 4595 | 481.3 | - | - | 0 | - |
| - | - | 1054 | 481.8 | - | - | 0 | - |
| - | - | 2313 | 481.9 | - | - | 0 | - |
| - | - | 5019 | 482.3 | - | - | 0 | - |
| - | - | 2667 | 482.3 | - | - | 0 | - |
| - | - | 1479 | 482.6 | - | - | 0 | - |
| - | - | 2039 | 482.8 | - | - | 0 | - |
| - | - | 666.7 | 482.9 | - | - | 0 | - |
| - | - | 1101 | 483.3 | - | - | 0 | - |
| - | - | 4742 | 483.8 | - | - | 0 | - |
| - | - | 2125 | 484.3 | - | - | 0 | - |
| - | - | 833.1 | 488.3 | - | - | 0 | - |
| - | - | 2047 | 491.3 | - | - | 0 | - |
| - | - | 1184 | 492.8 | - | - | 0 | - |
| - | - | 917.6 | 493.3 | - | - | 0 | - |
| - | - | 2074 | 493.8 | - | - | 0 | - |
| 8 | b | 1061 | 497.2 | 0.0001077 | 0.2166 | +2 | 8 |
| - | - | 1032 | 501.9 | - | - | 0 | - |
| - | - | 1719 | 503.2 | - | - | 0 | - |
| - | - | 981.1 | 504.2 | - | - | 0 | - |
| 13 | b | 1434 | 504.9 | 0.006617 | 13.1 | +3 | 13 |
| 13 | b | 1462 | 505.3 | 0.006737 | 13.33 | +3 | 13 |
| - | - | 1041 | 505.7 | - | - | 0 | - |
| - | - | 1492 | 508.8 | - | - | 0 | - |
| - | - | 726.3 | 509.2 | - | - | 0 | - |
| - | - | 801.1 | 509.7 | - | - | 0 | - |
| 13 | b | 4612 | 510.9 | 0.0003794 | 0.7425 | +3 | 13 |
| - | - | 3845 | 511.3 | - | - | 0 | - |
| - | - | 1696 | 511.6 | - | - | 0 | - |
| - | - | 965.8 | 513.9 | - | - | 0 | - |
| - | - | 637.3 | 514.3 | - | - | 0 | - |
| - | - | 710.8 | 514.6 | - | - | 0 | - |
| - | - | 6664 | 516.9 | - | - | 0 | - |
| - | - | 7621 | 517.2 | - | - | 0 | - |
| - | - | 6470 | 517.3 | - | - | 0 | - |
| - | - | 3298 | 517.6 | - | - | 0 | - |
| - | - | 1231 | 517.8 | - | - | 0 | - |
| - | - | 1270 | 517.9 | - | - | 0 | - |
| - | - | 1988 | 518.2 | - | - | 0 | - |
| - | - | 842.5 | 519.2 | - | - | 0 | - |
| 2 | y | 4869 | 519.9 | 0.0002642 | 0.508 | +3 | 13 |
| - | - | 4873 | 520.3 | - | - | 0 | - |
| - | - | 1789 | 520.4 | - | - | 0 | - |
| - | - | 2698 | 520.6 | - | - | 0 | - |
| - | - | 1480 | 521 | - | - | 0 | - |
| - | - | 1730 | 521.2 | - | - | 0 | - |
| 2 | y | 1.359E+04 | 525.6 | 0.0001628 | 0.3098 | +3 | 13 |
| - | - | 1.045E+04 | 526 | - | - | 0 | - |
| - | - | 6220 | 526.3 | - | - | 0 | - |
| - | - | 680.7 | 526.3 | - | - | 0 | - |
| - | - | 1620 | 526.6 | - | - | 0 | - |
| - | - | 785.7 | 528.2 | - | - | 0 | - |
| - | - | 1557 | 530.3 | - | - | 0 | - |
| - | - | 1535 | 530.8 | - | - | 0 | - |
| - | - | 1951 | 531.2 | - | - | 0 | - |
| - | - | 2244 | 535.8 | - | - | 0 | - |
| - | - | 1453 | 536.3 | - | - | 0 | - |
| - | - | 1.584E+04 | 538.3 | - | - | 0 | - |
| - | - | 1045 | 538.4 | - | - | 0 | - |
| - | - | 1771 | 538.8 | - | - | 0 | - |
| - | - | 4579 | 539.3 | - | - | 0 | - |
| - | - | 3475 | 539.3 | - | - | 0 | - |
| - | - | 2373 | 539.8 | - | - | 0 | - |
| - | - | 850.4 | 540.3 | - | - | 0 | - |
| 10 | y | 6505 | 541.4 | 0.0006502 | 1.201 | +1 | 5 |
| - | - | 2264 | 542.3 | - | - | 0 | - |
| - | - | 1803 | 542.4 | - | - | 0 | - |
| - | - | 1503 | 543.3 | - | - | 0 | - |
| - | - | 1887 | 544.8 | - | - | 0 | - |
| - | - | 668.3 | 545.3 | - | - | 0 | - |
| - | - | 998.6 | 547.8 | - | - | 0 | - |
| - | - | 1223 | 548.3 | - | - | 0 | - |
| - | - | 1910 | 549.2 | - | - | 0 | - |
| 5 | y | 1.843E+04 | 549.3 | 0.0004575 | 0.8329 | +2 | 10 |
| - | - | 1.07E+04 | 549.8 | - | - | 0 | - |
| - | - | 3575 | 550.3 | - | - | 0 | - |
| - | - | 1555 | 553.2 | - | - | 0 | - |
| - | - | 7134 | 554.2 | - | - | 0 | - |
| - | - | 2485 | 555.2 | - | - | 0 | - |
| - | - | 896.7 | 556.8 | - | - | 0 | - |
| - | - | 1964 | 557.3 | - | - | 0 | - |
| 9 | b | 5424 | 561.3 | 0.000537 | 0.9568 | +2 | 9 |
| - | - | 3635 | 561.8 | - | - | 0 | - |
| - | - | 1410 | 562.3 | - | - | 0 | - |
| - | - | 1.885E+04 | 566.2 | - | - | 0 | - |
| - | - | 5401 | 567.2 | - | - | 0 | - |
| - | - | 1134 | 567.4 | - | - | 0 | - |
| - | - | 904.3 | 568.3 | - | - | 0 | - |
| - | - | 8995 | 570.2 | - | - | 0 | - |
| - | - | 2560 | 571.2 | - | - | 0 | - |
| - | - | 787.4 | 572.2 | - | - | 0 | - |
| - | - | 6755 | 572.8 | - | - | 0 | - |
| - | - | 5024 | 573.3 | - | - | 0 | - |
| - | - | 2222 | 573.8 | - | - | 0 | - |
| - | - | 6015 | 577.4 | - | - | 0 | - |
| - | - | 2138 | 578.4 | - | - | 0 | - |
| - | - | 1.109E+04 | 579.4 | - | - | 0 | - |
| - | - | 4411 | 580.4 | - | - | 0 | - |
| - | - | 1273 | 581.2 | - | - | 0 | - |
| - | - | 697 | 581.4 | - | - | 0 | - |
| 10 | b | 1510 | 587.8 | 0.0002253 | 0.3834 | +2 | 10 |
| 10 | b | 1359 | 588.3 | 0.01051 | 17.87 | +2 | 10 |
| - | - | 3966 | 595.4 | - | - | 0 | - |
| - | - | 1834 | 596.4 | - | - | 0 | - |
| 10 | b | 1818 | 596.8 | 0.0004738 | 0.7939 | +2 | 10 |
| - | - | 1130 | 597.3 | - | - | 0 | - |
| - | - | 4344 | 598.2 | - | - | 0 | - |
| - | - | 1069 | 599.2 | - | - | 0 | - |
| - | - | 3299 | 604.4 | - | - | 0 | - |
| 4 | y | 5929 | 604.8 | 0.004402 | 7.278 | +2 | 11 |
| - | - | 4706 | 605.4 | - | - | 0 | - |
| - | - | 797.5 | 605.9 | - | - | 0 | - |
| - | - | 3768 | 610.3 | - | - | 0 | - |
| - | - | 1205 | 611.3 | - | - | 0 | - |
| - | - | 884.2 | 611.8 | - | - | 0 | - |
| - | - | 686 | 612.3 | - | - | 0 | - |
| 4 | y | 4560 | 613.4 | 0.0004499 | 0.7334 | +2 | 11 |
| - | - | 3115 | 613.9 | - | - | 0 | - |
| - | - | 784.3 | 614.4 | - | - | 0 | - |
| - | - | 715 | 615.3 | - | - | 0 | - |
| - | - | 1135 | 615.4 | - | - | 0 | - |
| - | - | 1649 | 615.8 | - | - | 0 | - |
| - | - | 2.056E+04 | 616.2 | - | - | 0 | - |
| - | - | 6777 | 617.3 | - | - | 0 | - |
| - | - | 3459 | 618.2 | - | - | 0 | - |
| - | - | 921.1 | 619.2 | - | - | 0 | - |
| - | - | 2714 | 620.3 | - | - | 0 | - |
| - | - | 2386 | 620.8 | - | - | 0 | - |
| - | - | 757.9 | 621.3 | - | - | 0 | - |
| - | - | 7602 | 629.3 | - | - | 0 | - |
| - | - | 8757 | 629.8 | - | - | 0 | - |
| - | - | 2741 | 630.3 | - | - | 0 | - |
| - | - | 1035 | 630.8 | - | - | 0 | - |
| - | - | 760.1 | 631.4 | - | - | 0 | - |
| - | - | 5329 | 638.4 | - | - | 0 | - |
| - | - | 2986 | 638.9 | - | - | 0 | - |
| - | - | 1660 | 639.4 | - | - | 0 | - |
| - | - | 1097 | 647.4 | - | - | 0 | - |
| - | - | 1748 | 656.3 | - | - | 0 | - |
| - | - | 772.4 | 657.3 | - | - | 0 | - |
| - | - | 2238 | 657.3 | - | - | 0 | - |
| - | - | 1122 | 657.8 | - | - | 0 | - |
| - | - | 3312 | 664.5 | - | - | 0 | - |
| - | - | 1553 | 665.5 | - | - | 0 | - |
| - | - | 1864 | 665.8 | - | - | 0 | - |
| - | - | 1171 | 666.8 | - | - | 0 | - |
| - | - | 1917 | 667.3 | - | - | 0 | - |
| - | - | 1739 | 668.3 | - | - | 0 | - |
| 9 | y | 3052 | 669.5 | 0.001573 | 2.35 | +1 | 6 |
| - | - | 967.9 | 670.5 | - | - | 0 | - |
| - | - | 1969 | 674.3 | - | - | 0 | - |
| - | - | 9270 | 674.4 | - | - | 0 | - |
| - | - | 784 | 675.3 | - | - | 0 | - |
| - | - | 3934 | 675.4 | - | - | 0 | - |
| - | - | 793.3 | 676.4 | - | - | 0 | - |
| - | - | 758.8 | 686.4 | - | - | 0 | - |
| - | - | 1.241E+04 | 692.4 | - | - | 0 | - |
| - | - | 3996 | 693.4 | - | - | 0 | - |
| - | - | 1456 | 694.5 | - | - | 0 | - |
| 3 | y | 4892 | 694.9 | 7.067E-05 | 0.1017 | +2 | 12 |
| - | - | 1847 | 695.4 | - | - | 0 | - |
| - | - | 1438 | 695.9 | - | - | 0 | - |
| - | - | 1068 | 698.4 | - | - | 0 | - |
| 5 | b | 2151 | 702.3 | 0.0007962 | 1.134 | +1 | 5 |
| - | - | 984.8 | 703.3 | - | - | 0 | - |
| - | - | 4997 | 707.3 | - | - | 0 | - |
| - | - | 2166 | 708.3 | - | - | 0 | - |
| - | - | 3098 | 710.5 | - | - | 0 | - |
| - | - | 1442 | 711.5 | - | - | 0 | - |
| - | - | 743.8 | 713.9 | - | - | 0 | - |
| - | - | 1736 | 716.4 | - | - | 0 | - |
| - | - | 1017 | 717.4 | - | - | 0 | - |
| - | - | 1131 | 720.3 | - | - | 0 | - |
| - | - | 1147 | 721.4 | - | - | 0 | - |
| - | - | 827.6 | 723.4 | - | - | 0 | - |
| - | - | 2203 | 724.3 | - | - | 0 | - |
| 8 | y | 5867 | 726.5 | 0.000237 | 0.3262 | +1 | 7 |
| - | - | 2333 | 727.5 | - | - | 0 | - |
| - | - | 714.7 | 729.3 | - | - | 0 | - |
| - | - | 789.1 | 731.4 | - | - | 0 | - |
| - | - | 5594 | 735.3 | - | - | 0 | - |
| - | - | 2631 | 736.3 | - | - | 0 | - |
| - | - | 1084 | 737.4 | - | - | 0 | - |
| - | - | 837.6 | 738.4 | - | - | 0 | - |
| - | - | 2508 | 752.3 | - | - | 0 | - |
| - | - | 1372 | 753.3 | - | - | 0 | - |
| - | - | 893.3 | 777.5 | - | - | 0 | - |
| - | - | 2484 | 805.5 | - | - | 0 | - |
| - | - | 1245 | 811.4 | - | - | 0 | - |
| - | - | 1215 | 811.5 | - | - | 0 | - |
| - | - | 902.6 | 812.5 | - | - | 0 | - |
| 6 | b | 781.5 | 821.3 | 0.005305 | 6.459 | +1 | 6 |
| 7 | y | 2.067E+04 | 823.5 | 0.0002665 | 0.3236 | +1 | 8 |
| - | - | 8809 | 824.5 | - | - | 0 | - |
| - | - | 2189 | 825.5 | - | - | 0 | - |
| - | - | 1751 | 829.5 | - | - | 0 | - |
| 6 | b | 2588 | 839.4 | 0.0001323 | 0.1576 | +1 | 6 |
| - | - | 1828 | 840.4 | - | - | 0 | - |
| - | - | 865.4 | 847.5 | - | - | 0 | - |
| - | - | 3161 | 848.4 | - | - | 0 | - |
| - | - | 1110 | 849.4 | - | - | 0 | - |
| - | - | 1104 | 853.5 | - | - | 0 | - |
| - | - | 802.9 | 891.5 | - | - | 0 | - |
| - | - | 3925 | 901.4 | - | - | 0 | - |
| - | - | 1398 | 902.4 | - | - | 0 | - |
| - | - | 1953 | 919.5 | - | - | 0 | - |
| - | - | 904.3 | 920.5 | - | - | 0 | - |
| 6 | y | 1490 | 960.6 | 0.0005751 | 0.5987 | +1 | 9 |
| - | - | 1078 | 961.6 | - | - | 0 | - |
| - | - | 876.4 | 1070 | - | - | 0 | - |
| - | - | 647.9 | 1371 | - | - | 0 | - |
| - | - | 736.3 | 2953 | - | - | 0 | - |
| - | - | 680.8 | 3154 | - | - | 0 | - |

m/z Charge Intensity FragmentType MassShift Position
120.05241394042969 0 522.70465
120.0659408569336 0 867.08936
120.08114624023438 0 1544.5773
122.02733612060547 0 863.4428
122.07195281982422 0 561.8985
123.05567932128906 0 1577.4445
123.1170654296875 0 575.4189
125.07111358642578 0 527.67084
126.05535125732422 0 418.07718
126.09165954589844 0 954.4382
126.1281509399414 0 994.1048
127.05084228515625 0 458.27948
127.0870361328125 0 3234.561
128.07081604003906 0 678.4901
128.0822296142578 0 946.1428
128.5526580810547 0 378.51987
129.05519104003906 0 691.9675
129.06173706054688 0 572.5529
129.0662384033203 0 4231.4517
129.10269165039062 0 83086.91
130.0614471435547 0 2058.3809
130.06558227539062 0 20675.277
130.08670043945312 0 745.3272
130.1003875732422 0 866.9065
130.1060028076172 0 6123.046
131.04559326171875 0 3860.4204
131.0697021484375 0 1791.8339
131.08192443847656 0 2559.807
132.0812225341797 0 72642.57
132.10231018066406 0 19989.69 y 13
133.0609130859375 0 424.7323
133.0861053466797 0 26240.803
133.10580444335938 0 1556.7456
134.0608367919922 0 429.71487
134.07200622558594 0 555.90125
134.08966064453125 0 1707.8014
136.01864624023438 0 473.87897
136.07614135742188 0 158301.62
137.0602264404297 0 454.72684
137.0733184814453 0 1167.0634
137.07945251464844 0 14820.394
138.066650390625 0 1595.2592
139.0871124267578 0 1711.3662
140.08255004882812 0 703.1203
141.0662841796875 0 790.23676
141.10267639160156 0 5810.8853
142.06573486328125 0 1480.2667 b 3
142.12301635742188 0 3736.1501
143.1183319091797 0 4662.6245
144.08123779296875 0 2872.2473
144.1220245361328 0 445.77087
146.06045532226562 0 1269.7719
147.04444885253906 0 5275.947
147.1130828857422 0 607.0546
148.0478973388672 0 582.1101
148.07611083984375 0 495.97714
151.0868377685547 0 1504.0763
151.98390197753906 0 3733.4788
151.98968505859375 0 554.43616
152.07127380371094 0 540.38226
152.082275390625 0 832.3064
152.1072540283203 0 609.8963
152.1439208984375 0 1654.101
153.06617736816406 0 435.6827
153.07618713378906 0 457.6391
153.1024932861328 0 747.6111
153.99948120117188 0 436.57385
154.0979461669922 0 929.01953
155.08192443847656 0 3848.0024
155.09320068359375 0 1435.408
155.1182098388672 0 3557.8157
156.07643127441406 0 1468.9264
156.08148193359375 0 1822.949
157.06121826171875 0 484.97684
158.0605010986328 0 1444.7255
158.08425903320312 0 9066.536
159.07669067382812 0 1694.3894
159.0921173095703 0 51155.426
160.0760498046875 0 2468.3296
160.089111328125 0 547.1726
160.09544372558594 0 4847.1733
161.0810546875 0 1044.7351
162.05496215820312 0 446.05005
162.1030731201172 0 492.58035
162.25662231445312 0 478.05936
163.61337280273438 0 652.06116
164.07083129882812 0 1447.9497
165.07789611816406 0 475.08917
165.1027374267578 0 2971.232
166.06126403808594 0 985.77875
166.1057586669922 0 710.27936
167.08189392089844 0 528.20886
169.0763702392578 0 5121.471
169.09756469726562 0 5695.0605
169.1338348388672 0 1746.1525
170.06044006347656 0 24748.8
171.06375122070312 0 3018.1982
171.0924835205078 0 776.1216
172.09756469726562 0 387.9442
172.1005096435547 0 598.92346
172.10853576660156 0 4934.6074
172.14540100097656 0 544.55347
173.07154846191406 0 668.249
173.45196533203125 0 818.80475
174.05540466308594 0 643.5289
175.08743286132812 0 853.04767
175.0974578857422 0 1290.6615
175.11947631835938 0 478.49142
176.0823211669922 0 1477.3553
177.04571533203125 0 706.981
177.1125946044922 0 9902.728
179.0918731689453 0 521.1928
181.06134033203125 0 1397.9078
181.0976104736328 0 23634.76
181.13414001464844 0 1804.5043
182.10092163085938 0 2020.4924
182.12936401367188 0 2964.7104
183.0921630859375 0 10459.083
183.1014862060547 0 696.07227
183.11285400390625 0 1233.4595
183.1492156982422 0 730.6119
184.0758056640625 0 1012.7041
184.09555053710938 0 1186.5237
185.07147216796875 0 3435.5857
185.0795135498047 0 574.7483
185.1291046142578 0 656.47845
185.1652069091797 0 3703.989
186.12411499023438 0 8978.96
187.0870361328125 0 52222.227
187.10801696777344 0 5713.882
187.14456176757812 0 117094.14 y 11
188.09046936035156 0 6103.15
188.14794921875 0 10824.703
189.0780487060547 0 525.1211
190.53692626953125 0 549.6365
191.09291076660156 0 501.96582
193.09689331054688 0 589.34546
195.0256805419922 0 1312.6655
195.1495361328125 0 1013.6444
197.1287841796875 0 1243.7782
197.16525268554688 0 3149.7383
198.16793823242188 0 671.4917
199.08668518066406 0 2514.4456
199.10830688476562 0 1160.3082
199.16943359375 0 1436.9888
199.1806640625 0 1742.3782
200.14007568359375 0 1555.5676
201.1027069091797 0 5282.3574
201.1236114501953 0 1413.8804
202.08627319335938 0 2420.7607
202.10870361328125 0 10359.168
203.08319091796875 0 636.5315
203.0932159423828 0 844.0993
204.07748413085938 0 757.38403
204.13551330566406 0 3691.8367
205.10757446289062 0 6729.261
207.149658203125 0 2633.9663
209.1287078857422 0 697.68225
211.0867156982422 0 1932.7416
211.10833740234375 0 548.37994
212.1395263671875 0 728.21344
212.66810607910156 0 2141.8489
213.16026306152344 0 1416.2412
214.1089324951172 0 2387.6528
214.19146728515625 0 715.455
215.13943481445312 0 44984.527
216.1134033203125 0 616.9012
216.1243896484375 0 993.45087
216.14303588867188 0 5423.735
217.10816955566406 0 982.1475
217.14491271972656 0 576.75836
219.0802001953125 0 9409.87
220.0831756591797 0 1463.0613
220.11985778808594 0 6045.453
221.1038360595703 0 6909.346
221.13868713378906 0 2468.9712
224.0675048828125 0 602.57715
224.118896484375 0 948.0118
224.1760711669922 0 3471.8018
225.16006469726562 0 1572.0793
226.1554718017578 0 65292.72
227.082763671875 0 745.5202
227.15884399414062 0 8195.51 y Ammonia loss 10
227.17213439941406 0 887.919
228.11354064941406 0 5186.62 a Water loss 1
228.13360595703125 0 708.9609
229.09759521484375 0 6273.7764 b Ammonia loss 4
229.11837768554688 0 988.9268
229.16635131835938 0 753.1322
230.10301208496094 0 2008.8933
231.08822631835938 0 2485.104
231.13548278808594 0 1218.3938
232.1205596923828 0 955.9955
233.0939483642578 0 823.86127
233.10525512695312 0 1122.6812
235.10726928710938 0 617.1963
235.1194610595703 0 12297.315
235.67086791992188 0 9715.67 y 10
236.12265014648438 0 1639.0768
236.17222595214844 0 2076.0325
237.13473510742188 0 595.65643
239.15049743652344 0 5495.7114
240.15391540527344 0 669.40857
241.03164672851562 0 35696.727
241.15509033203125 0 1246.6482
242.0342254638672 0 2427.8623
242.18679809570312 0 3477.0728
243.02740478515625 0 3580.143
245.1864776611328 0 15690.121 y 12
246.1241912841797 0 119896.85 a 1
246.18975830078125 0 2340.5254
247.1291046142578 0 27230.996
248.11453247070312 0 8522.613
248.13238525390625 0 2466.9749
248.18667602539062 0 535.76306
249.0986328125 0 12143.76
249.12001037597656 0 873.742
249.1352081298828 0 1720.1721
249.16029357910156 0 50921.008
250.1016387939453 0 1940.6118
250.1239471435547 0 1797.2185
250.1422576904297 0 634.45593
250.16348266601562 0 9901.122
251.12783813476562 0 696.8305
251.15090942382812 0 1120.747
251.16558837890625 0 615.7172
252.17141723632812 0 1944.1289
255.06558227539062 0 917.4703
256.108642578125 0 2469.0183 b Water loss 1
257.1142883300781 0 1781.4862
257.1612548828125 0 1371.4948
258.0760192871094 0 2153.5432
258.09893798828125 0 21000.758
259.1021423339844 0 3038.9182
259.1294860839844 0 631.76685
260.1072998046875 0 2789.1956
260.1972351074219 0 1516.028
262.1549987792969 0 769.10657
264.134765625 0 1275.6716
265.1658630371094 0 1307.2937
266.1251525878906 0 8190.729
266.1499938964844 0 788.5216
267.1282653808594 0 1105.998
267.1441650390625 0 809.78076
268.16546630859375 0 681.7483
269.1861572265625 0 1002.5427
269.6302185058594 0 1804.5099
270.1814270019531 0 5088.516
271.1890869140625 0 4390.5225 y 9
274.1189880371094 0 44665.53 b 1
275.10345458984375 0 4699.926
275.125 0 39396.754
275.1856689453125 0 612.6399 y 6
276.1077575683594 0 1843.3239
276.1280822753906 0 5066.4644
277.1346740722656 0 1451.9065
277.1551208496094 0 11556.255
278.15863037109375 0 2942.2576
279.1612243652344 0 568.7487
279.1819152832031 0 1425.2551
283.15179443359375 0 1773.3975
283.1767578125 0 17694.084
283.62750244140625 0 1212.5657
284.1803894042969 0 3422.728
285.1094665527344 0 1180.3705
285.1561584472656 0 3366.153
290.18463134765625 0 6025.3438
290.686279296875 0 1765.7352
291.15826416015625 0 1262.7856
292.12957763671875 0 12255.644
293.1327819824219 0 1881.7196
294.12054443359375 0 3540.365
294.218017578125 0 821.3149
294.6681213378906 0 681.9184
295.1239929199219 0 701.29456
297.1926574707031 0 7732.6465
298.1963195800781 0 1126.0778
303.12017822265625 0 2657.8945
305.12872314453125 0 4451.236
305.66455078125 0 2043.5083
306.1324462890625 0 1297.7048
309.1937255859375 0 707.63586
311.2447204589844 0 2993.887
313.2235412597656 0 708.49194
314.20806884765625 0 4951.5425
314.66943359375 0 1211.945
315.2104187011719 0 899.5127
318.15081787109375 0 514.24054
320.12481689453125 0 989.70526
320.8712463378906 0 2018.199 y 5
321.2076110839844 0 911.55896
321.2291259765625 0 10863.576
321.5400695800781 0 688.0173
322.0899963378906 0 771.1184
322.1554870605469 0 5594.9136
322.2328796386719 0 1903.0074
323.1600341796875 0 1250.3367 b 10
324.216064453125 0 2035.7544
325.2220764160156 0 805.27545
326.2193603515625 0 3814.6921
326.664794921875 0 586.0784
327.2016296386719 0 3478.6064
327.2217102050781 0 854.96344
328.20465087890625 0 650.5521
332.7290954589844 0 2698.0813
333.12396240234375 0 1311.0386
333.229248046875 0 931.66473
334.1296081542969 0 656.71466
334.6676330566406 0 1464.9585
335.2375793457031 0 1130.8774 y 8
336.20361328125 0 15263.127
337.206787109375 0 2503.4534
337.72137451171875 0 5057.8496
338.2221984863281 0 2093.842
338.7212829589844 0 634.45874
339.23968505859375 0 15959.827
340.1001281738281 0 53319.656
340.2431335449219 0 2986.8157
341.1027526855469 0 8344.68
342.0961608886719 0 5232.776
342.1670227050781 0 767.62994
343.10015869140625 0 596.4082
344.9319152832031 0 558.27905
345.6438293457031 0 1524.2958
346.6698303222656 0 713.73956
346.7268371582031 0 13845.703
347.16949462890625 0 740.82056
347.2282409667969 0 5530.139
347.7292785644531 0 1092.4075
350.1499328613281 0 1640.8035
351.23895263671875 0 1097.9784
352.2369689941406 0 673.5557
353.21905517578125 0 991.7526
354.11566162109375 0 6487.0903
354.1573486328125 0 1782.2476
354.2140808105469 0 19950.057
354.6605529785156 0 1069.8124
355.11944580078125 0 1051.4377
355.21685791015625 0 4099.8574
355.7328796386719 0 1483.5327
356.19140625 0 525.6735
356.23480224609375 0 1268.5381
357.2498779296875 0 4775.4067
357.8319396972656 0 1336.5676
358.162841796875 0 4616.295
358.7136535644531 0 2141.3118
359.16400146484375 0 785.731
359.2145080566406 0 1343.9447
359.5416259765625 0 826.1366
359.87176513671875 0 1383.8734
360.20452880859375 0 900
360.665283203125 0 922.7585
363.50677490234375 0 1164.4897
363.74822998046875 0 859.5333 y 7
363.8405456542969 0 1162.7827
365.67425537109375 0 546.60596
366.156005859375 0 1556.7473
366.5573425292969 0 2477.4548 y 4
366.8938903808594 0 859.9757
367.162109375 0 1042.938
368.1494140625 0 2569.799
370.1869201660156 0 693.06866
370.5256042480469 0 753.1645
371.22796630859375 0 30358.723
372.2312927246094 0 6576.521
373.28106689453125 0 8387.57 y 11
374.28521728515625 0 1663.2811
374.8486022949219 0 947.79645
376.4595947265625 0 677.171
376.6684265136719 0 935.50244
376.7022399902344 0 544.69525
378.95611572265625 0 832.714 b Water loss 12
380.1595458984375 0 591.5374
380.22882080078125 0 1270.8173
382.20623779296875 0 4055.7366
382.28271484375 0 615.3156
382.5401611328125 0 1691.2234
382.8486328125 0 1247.2678
382.87408447265625 0 1149.1726
383.26483154296875 0 1205.7166
384.1668701171875 0 2280.2039
385.1733093261719 0 9998.571
385.1995544433594 0 2069.5369
386.1573791503906 0 8847.627
387.1603088378906 0 1262.0586
388.2328186035156 0 656.4808
388.86163330078125 0 2041.1116
389.1963195800781 0 1993.0156
389.27178955078125 0 2351.7234
389.7722473144531 0 1041.8663
390.2135925292969 0 5715.922 y Ammonia loss 1
390.46435546875 0 5124.613
390.7157287597656 0 2084.4563
392.2660217285156 0 3685.1897
392.5245056152344 0 890.579 b Ammonia loss 9
394.1513671875 0 5945.0522
394.24566650390625 0 787.36835
394.47113037109375 0 2590.8628 y 1
394.7212829589844 0 2514.2317
394.972900390625 0 1618.6931
396.2615051269531 0 1438.5286
397.2451171875 0 1512.5834
397.56561279296875 0 801.55084
398.19384765625 0 1621.1617 b 9
398.23858642578125 0 802.73315
398.89105224609375 0 804.58057
399.2231750488281 0 21120.396
400.2259521484375 0 3882.309
401.1925354003906 0 1029.3439
401.2588806152344 0 916.85565
402.2234191894531 0 3373.1597
402.548583984375 0 944.73254
402.880615234375 0 682.99426
403.1845703125 0 6445.3154
403.2403259277344 0 3119.6086
403.26885986328125 0 2214.5635
403.571044921875 0 5302.2734 y Ammonia loss 3
403.7649230957031 0 785.5978 y Ammonia loss 6
403.9050598144531 0 2527.6282
404.185791015625 0 1495.2332
404.2355041503906 0 1401.494
404.47088623046875 0 998.43695
404.72442626953125 0 3442.523
404.8937072753906 0 1401.0995
404.9768981933594 0 3314.3699
405.2300109863281 0 1067.5082
406.1854553222656 0 2507.005
406.2513122558594 0 3207.2144
406.68902587890625 0 1082.4608
406.75140380859375 0 1850.7838
408.2216491699219 0 1850.7172
408.5567321777344 0 2017.363
409.2438659667969 0 1041.3705 y 3
409.577880859375 0 866.1973
410.2764892578125 0 6360.787
410.56884765625 0 2599.2327
410.9029235839844 0 3957.4343
411.1780700683594 0 8737.731 b Water loss 5
411.23590087890625 0 740.47833
411.2798767089844 0 1534.7831
411.7242431640625 0 1510.9813 Precursor Water loss
411.9749755859375 0 2013.794 Precursor Ammonia loss
412.1625061035156 0 2086.6523
412.2248229980469 0 1693.4252
412.2743835449219 0 24710.479 y 6
412.7757873535156 0 12993.93
413.2772521972656 0 3732.49
413.8972473144531 0 2794.204
414.1973876953125 0 679.96387
414.22869873046875 0 3907.4119
414.5625 0 2121.5103
414.8952331542969 0 964.5049
415.2566833496094 0 3526.7344
415.75732421875 0 1655.8002
416.228515625 0 14983.258 Precursor
416.2520446777344 0 2768.006
416.2886047363281 0 2457.212
416.47955322265625 0 11545.662
416.7301025390625 0 6590.991
416.9810791015625 0 2364.446
417.2319030761719 0 1586.3983
419.90032958984375 0 13616.238
420.183837890625 0 1040.5297 b 5
420.2348327636719 0 13223.853
420.5687255859375 0 5589.531
420.685302734375 0 1232.5516
420.9024353027344 0 1148.1117
422.27630615234375 0 2288.7625
423.2719421386719 0 911.3467
424.3290100097656 0 1619.1914
424.7117919921875 0 4428.862
425.2134094238281 0 2036.2358
425.7148132324219 0 723.7717
425.9034118652344 0 3065.3945
426.2391357421875 0 2255.4536
427.2431640625 0 1695.7332
427.74462890625 0 1475.614
429.1888427734375 0 24312.857
430.191650390625 0 6491.113
431.17950439453125 0 2931.4712
432.2326965332031 0 671.6929
437.18194580078125 0 1869.1758 b 2
438.187255859375 0 927.8678
438.2268371582031 0 987.80334
438.55792236328125 0 2134.2844
438.8916320800781 0 1264.652
439.2251892089844 0 1456.0923
442.7115783691406 0 660.04407
444.230224609375 0 832.6267
444.568115234375 0 974.19257
444.9036560058594 0 793.98773
446.2344970703125 0 1903.9302
446.73614501953125 0 1243.7257
448.26348876953125 0 4301.8047
448.59796142578125 0 2750.369
448.9333801269531 0 1559.4407
451.2256164550781 0 3780.123
451.2675476074219 0 2281.9695
451.72698974609375 0 3290.0527
451.9198913574219 0 6262.817
452.2536926269531 0 4660.597
452.32330322265625 0 1477.7189
452.5859069824219 0 3260.5803
452.921875 0 786.6068
453.1841125488281 0 10278.057
454.18792724609375 0 1659.5406
455.18023681640625 0 1349.5001
455.2522888183594 0 2431.3918
457.5950927734375 0 3451.078
457.9261779785156 0 5374.76 y Ammonia loss 2
458.2605285644531 0 3546.4978
458.59356689453125 0 665.8874
460.2308044433594 0 5380.199
460.732666015625 0 2643.8237
461.5814514160156 0 593.6474
462.2471923828125 0 678.9318
463.5987548828125 0 73298.88 y 2
463.9329528808594 0 60283.684
464.2671203613281 0 23447
464.6014099121094 0 6509.8574
464.9358215332031 0 1520.5172
466.92041015625 0 1390.0388
467.2532958984375 0 1526.3905
467.2987365722656 0 2398.563
468.2980041503906 0 1716.9281
469.7885437011719 0 942.1754
470.25543212890625 0 1084.1295
470.3343505859375 0 27831.918 y 10
470.581298828125 0 2427.209
470.7922668457031 0 723.18933
470.9166259765625 0 937.3776
471.24755859375 0 745.9094
471.3373718261719 0 8230.504
471.79833984375 0 3498.215
472.30108642578125 0 1087.1794
472.3398132324219 0 1513.3949
472.5914001464844 0 867.69775
473.2621765136719 0 3824.7893
473.57647705078125 0 1809.6896
474.2649230957031 0 792.3882
474.7793884277344 0 2049.404
476.2522277832031 0 5701.3003
476.58697509765625 0 4224.2446
476.91961669921875 0 1749.2988
480.8036193847656 0 6790.18 y 5
481.30548095703125 0 4595.041
481.8044128417969 0 1054.1221
481.9277648925781 0 2313.0325
482.26068115234375 0 5019.1963
482.30859375 0 2667.1726
482.5912780761719 0 1478.9299
482.76202392578125 0 2039.3757
482.9241638183594 0 666.69916
483.2643737792969 0 1101.1367
483.7857971191406 0 4741.869
484.2877197265625 0 2125.4348
488.2642822265625 0 833.11237
491.2734680175781 0 2047.3896
492.79180908203125 0 1184.198
493.2930603027344 0 917.5739
493.79644775390625 0 2074.2517
497.2198181152344 0 1060.6956 b 7
501.9415283203125 0 1031.6337
503.21563720703125 0 1719.4838
504.21783447265625 0 981.0891
504.9400939941406 0 1434.1378 b Water loss 12
505.2682189941406 0 1461.5571 b Ammonia loss 12
505.73382568359375 0 1040.8177
508.7578125 0 1492.1241
509.24005126953125 0 726.34845
509.7383117675781 0 801.06287
510.9373779296875 0 4611.994 b 12
511.2716979980469 0 3845.4985
511.61041259765625 0 1695.925
513.9470825195312 0 965.83826
514.2783203125 0 637.2683
514.6087036132812 0 710.8301
516.940673828125 0 6664.0317
517.1787719726562 0 7621.351
517.2748413085938 0 6469.762
517.6089477539062 0 3297.6702
517.7534790039062 0 1230.9458
517.9429321289062 0 1269.6282
518.1802368164062 0 1988.4211
519.1741943359375 0 842.472
519.9492797851562 0 4869.272 y Ammonia loss 1
520.2841796875 0 4872.798
520.3604125976562 0 1789.2759
520.61865234375 0 2698.1973
520.95068359375 0 1479.9106
521.226318359375 0 1729.9036
525.6246948242188 0 13585.397 y 1
525.9593505859375 0 10446.445
526.2933349609375 0 6220.3647
526.3239135742188 0 680.7462
526.6279296875 0 1619.6199
528.2337036132812 0 785.67004
530.2976684570312 0 1556.9055
530.801513671875 0 1534.8391
531.2102661132812 0 1951.3191
535.7509765625 0 2243.5884
536.2530517578125 0 1453.1592
538.2523803710938 0 15837.143
538.3720703125 0 1044.8312
538.8097534179688 0 1771.092
539.25537109375 0 4579.0415
539.3048095703125 0 3474.6375
539.8035888671875 0 2373.1584
540.3157348632812 0 850.4313
541.3714599609375 0 6505.3823 y 9
542.2501831054688 0 2263.9924
542.374267578125 0 1802.9268
543.25439453125 0 1502.6685
544.7582397460938 0 1886.949
545.2637939453125 0 668.26416
547.8146362304688 0 998.5915
548.3135986328125 0 1222.8502
549.2214965820312 0 1910.3928
549.3330078125 0 18433.756 y 4
549.8344116210938 0 10699.392
550.3363647460938 0 3574.805
553.2224731445312 0 1554.8691
554.23046875 0 7134.1763
555.2349243164062 0 2484.5525
556.8209838867188 0 896.6937
557.2939453125 0 1963.9536
561.2679443359375 0 5424.3857 b 8
561.7688598632812 0 3634.7
562.2730102539062 0 1410.1096
566.2476196289062 0 18846.646
567.2494506835938 0 5401.3457
567.3983764648438 0 1133.6354
568.2515869140625 0 904.2561
570.2462158203125 0 8994.546
571.248046875 0 2560.368
572.2461547851562 0 787.4399
572.8038940429688 0 6755.445
573.3057861328125 0 5024.4634
573.8070068359375 0 2222.0466
577.3819580078125 0 6015.4346
578.3831176757812 0 2137.819
579.3616333007812 0 11093.357
580.3648681640625 0 4410.683
581.2134399414062 0 1273.3794
581.3635864257812 0 697.02814
587.7804565429688 0 1509.6028 b Water loss 9
588.283203125 0 1359.4386 b Ammonia loss 9
595.3925170898438 0 3965.5479
596.3949584960938 0 1834.3662
596.7864379882812 0 1817.7401 b 9
597.2865600585938 0 1130.3849
598.240478515625 0 4343.8887
599.2452392578125 0 1068.9121
604.3545532226562 0 3298.67
604.8529663085938 0 5929.159 y Ammonia loss 3
605.3513793945312 0 4706.2646
605.8511962890625 0 797.5023
610.3206176757812 0 3768.2407
611.3226928710938 0 1205.0176
611.8262939453125 0 884.19653
612.3262329101562 0 685.9611
613.3613891601562 0 4559.8765 y 3
613.8633422851562 0 3114.9434
614.3631591796875 0 784.31244
615.2664794921875 0 714.96075
615.3518676757812 0 1134.5812
615.8499755859375 0 1648.5474
616.2473754882812 0 20560.78
617.2503662109375 0 6776.9106
618.2485961914062 0 3459.1357
619.2474975585938 0 921.07983
620.3411254882812 0 2714.4976
620.83837890625 0 2385.8887
621.343017578125 0 757.8568
629.3467407226562 0 7602.1377
629.8469848632812 0 8756.55
630.3486328125 0 2741.2368
630.849609375 0 1035.4838
631.352294921875 0 760.0668
638.35107421875 0 5328.9062
638.85302734375 0 2985.6821
639.353271484375 0 1660.1323
647.424560546875 0 1097.3932
656.337646484375 0 1747.825
657.2764892578125 0 772.4203
657.334228515625 0 2237.605
657.8316040039062 0 1122.0406
664.4514770507812 0 3311.567
665.45458984375 0 1552.5275
665.8455200195312 0 1864.2236
666.8463134765625 0 1170.5192
667.3185424804688 0 1917.0394
668.3280029296875 0 1738.96
669.4673461914062 0 3051.5024 y 8
670.4667358398438 0 967.87
674.3031005859375 0 1968.6547
674.4344482421875 0 9269.601
675.307373046875 0 783.99194
675.4381713867188 0 3934.3003
676.4429931640625 0 793.3021
686.3557739257812 0 758.83057
692.4454345703125 0 12414.93
693.4486083984375 0 3996.0728
694.452392578125 0 1455.972
694.8934326171875 0 4892.316 y 2
695.3952026367188 0 1847.3114
695.8959350585938 0 1438.0936
698.4114379882812 0 1068.0839
702.3002319335938 0 2150.8784 b 4
703.297607421875 0 984.7977
707.3042602539062 0 4996.96
708.3068237304688 0 2165.9087
710.4564208984375 0 3098.1992
711.461669921875 0 1441.627
713.8782958984375 0 743.78107
716.4219360351562 0 1735.5851
717.4254150390625 0 1016.76575
720.3167724609375 0 1131.1127
721.353515625 0 1147.333
723.3853759765625 0 827.63055
724.331298828125 0 2203.3022
726.4869995117188 0 5867.3813 y 7
727.489013671875 0 2333.1392
729.333984375 0 714.7427
731.373779296875 0 789.05273
735.2996215820312 0 5594.2246
736.3028564453125 0 2631.0454
737.3504028320312 0 1083.7491
738.3768310546875 0 837.60516
752.3280639648438 0 2507.886
753.3270874023438 0 1372.033
777.5433959960938 0 893.263
805.5304565429688 0 2483.9805
811.3607177734375 0 1244.62
811.498046875 0 1214.8278
812.498291015625 0 902.5694
821.3530883789062 0 781.53864 b Water loss 5
823.5397338867188 0 20665.814 y 6
824.5426025390625 0 8809.216
825.5437622070312 0 2188.8047
829.5040283203125 0 1751.1329
839.3582153320312 0 2587.998 b 5
840.3613891601562 0 1828.1708
847.51513671875 0 865.3752
848.4150390625 0 3160.7615
849.4138793945312 0 1110.3254
853.4774169921875 0 1103.8628
891.4589233398438 0 802.8919
901.4417724609375 0 3924.9873
902.4389038085938 0 1397.5853
919.45458984375 0 1953.0856
920.4578857421875 0 904.317
960.5994873046875 0 1489.5878 y 5
961.60302734375 0 1077.9642
1070.491455078125 0 876.38403
1370.79150390625 0 647.8803
2952.703857421875 0 736.3475
3154.11669921875 0 680.75464

Spectrum Details

|  |  |
| --- | --- |
| Matched peaks? Matched peaks The total absolute number of peaks matched. Additionally in brackets the total fraction of peaks matched and the total number of peaks is shown. | 60 (7.92% of 758) |
| FDR? FDR The false discovery rate estimated for this peptide. It is calculated by matching all theoretical fragments with a non-integer shift with the raw peaks for this spectrum. This is done with 40 different shifts. The resulting percentage is the average number of annotated peaks over the number of annotated peaks with the correct spectrum. | 3.49% |
| Satellite FDR? Satellite FDR See the FDR for details on its calculation. This satellite ion specific FDR only contains the satellite ions (d/w) for I/L/J positions. | - |
| PSM Score? PSM Score The PSM Score as given by Hecklib to this annotated spectrum. It is shown with three significant figures. | 189 |

#### Spectrum 4404? Spectrum 4404 The raw spectrum of this peptide as annotated by Hecklib. The fragments are coloured according to ion type (see legend). Any peaks with a star '\*' as text can be hovered over to see the full details, first the ion type second the mass shift type. By hovering over the amino acids in the peptide or ions in the legend the corresponding peaks are highlighted. By toggling the 'Unassigned' label you can turn the background (unassigned) peaks on or off in the plot. By updating the slider in the Ion legend you can update the spectrum to only show the top X% of the peaks with labels. The top X% means any peak that is within X% of the highest intensity. By dragging in the spectrum you can zoom in to a specific part of the spectrum and use 'Zoom Out' to get back to the original zoom level. The annotation of the spectrum is based on the given sequence in the peptides file and is done with different software so inconsistencies are likely. The peaks are annotated based on the given sequence, with 20 ppm tolerance.

Copy Data

##### Spectrum 4404 (TSV)

###### Preview

```
Loading example...
```

*Click on the button to copy the data to your clipboard.*

Mz MinMz MaxIntensity Max

WidthHeightPeptide font sizePeptide stroke widthSpectrum font sizeSpectrum stroke widthCompact peptide

Ion legend

wxyz

abcd

OtherUnassignedIon^Charge^_Position_Show for top:%

SWYQHHPGKAPKJJ

01.88e+43.76e+45.64e+47.52e+4

Zoom Out

y^+1^_1_b^+4^_4_y^+2^_3_y^+3^_6_y^+2^_4_y^+2^_4_y^+1^_2_a^+1^_2_y^+2^_5_b^+1^_2_b^+1^_2_b^+2^_4_y^+3^_9_y^+2^_6_y^+3^_10_y^+1^_3_y^+4^_13_b^+3^_10_y^+4^_13_b^+3^_10_y^+3^_11_y^+2^_8_\*\*\*b^+2^_6_b^+3^_11_b^+1^_3_y^+3^_12_y^+3^_12_y^+1^_4_b^+3^_12_y^+2^_9_b^+3^_13_b^+3^_13_b^+3^_13_y^+3^_13_y^+3^_13_y^+1^_5_y^+2^_10_b^+2^_9_y^+2^_11_y^+2^_11_b^+2^_11_y^+1^_6_y^+2^_12_b^+1^_5_y^+1^_7_y^+1^_8_b^+1^_6_y^+1^_9_

048196114421922

Fragment Matches Table

Show background peaks

| Position | Ion type | Intensity | mz Theoretical | mz Error (Th) | mz Error (ppm) | Charge | Series Number |
| --- | --- | --- | --- | --- | --- | --- | --- |
| - | - | 644.2 | 120.1 | - | - | 0 | - |
| - | - | 3718 | 120.1 | - | - | 0 | - |
| - | - | 364.9 | 121 | - | - | 0 | - |
| - | - | 675.7 | 122.1 | - | - | 0 | - |
| - | - | 764.8 | 123.1 | - | - | 0 | - |
| - | - | 443.4 | 123.1 | - | - | 0 | - |
| - | - | 357 | 124.5 | - | - | 0 | - |
| - | - | 474.3 | 125.1 | - | - | 0 | - |
| - | - | 551.7 | 126.1 | - | - | 0 | - |
| - | - | 1829 | 127.1 | - | - | 0 | - |
| - | - | 2562 | 129 | - | - | 0 | - |
| - | - | 455.2 | 129.1 | - | - | 0 | - |
| - | - | 5.666E+04 | 129.1 | - | - | 0 | - |
| - | - | 3.493E+04 | 130.1 | - | - | 0 | - |
| - | - | 582.2 | 130.1 | - | - | 0 | - |
| - | - | 3638 | 130.1 | - | - | 0 | - |
| - | - | 467.2 | 131.1 | - | - | 0 | - |
| - | - | 3297 | 131.1 | - | - | 0 | - |
| - | - | 1745 | 131.1 | - | - | 0 | - |
| - | - | 3398 | 132 | - | - | 0 | - |
| - | - | 2132 | 132.1 | - | - | 0 | - |
| 14 | y | 1.55E+04 | 132.1 | 0.0001914 | 1.449 | +1 | 1 |
| - | - | 1815 | 133.1 | - | - | 0 | - |
| - | - | 942.5 | 133.1 | - | - | 0 | - |
| - | - | 525.2 | 135.5 | - | - | 0 | - |
| - | - | 4.74E+04 | 136.1 | - | - | 0 | - |
| - | - | 2305 | 137.1 | - | - | 0 | - |
| - | - | 999.9 | 138.1 | - | - | 0 | - |
| - | - | 958.2 | 139 | - | - | 0 | - |
| - | - | 1142 | 139.1 | - | - | 0 | - |
| - | - | 3575 | 141.1 | - | - | 0 | - |
| - | - | 521.9 | 143 | - | - | 0 | - |
| - | - | 593.8 | 143.1 | - | - | 0 | - |
| - | - | 652.7 | 144.1 | - | - | 0 | - |
| - | - | 580.8 | 145.1 | - | - | 0 | - |
| 4 | b | 4179 | 146.1 | 0.002892 | 19.8 | +4 | 4 |
| - | - | 2320 | 147 | - | - | 0 | - |
| - | - | 1171 | 148.1 | - | - | 0 | - |
| - | - | 739.6 | 148.1 | - | - | 0 | - |
| - | - | 726.6 | 149 | - | - | 0 | - |
| - | - | 7.446E+04 | 149 | - | - | 0 | - |
| - | - | 6987 | 150 | - | - | 0 | - |
| - | - | 505.2 | 150.1 | - | - | 0 | - |
| - | - | 582.7 | 151.1 | - | - | 0 | - |
| - | - | 1182 | 152.1 | - | - | 0 | - |
| - | - | 641.8 | 153.1 | - | - | 0 | - |
| - | - | 3142 | 155.1 | - | - | 0 | - |
| - | - | 754 | 155.1 | - | - | 0 | - |
| - | - | 2032 | 155.1 | - | - | 0 | - |
| - | - | 510.4 | 156.1 | - | - | 0 | - |
| - | - | 1421 | 156.1 | - | - | 0 | - |
| - | - | 2169 | 157 | - | - | 0 | - |
| - | - | 1552 | 157.1 | - | - | 0 | - |
| - | - | 1.213E+04 | 157.1 | - | - | 0 | - |
| - | - | 1432 | 157.1 | - | - | 0 | - |
| - | - | 9175 | 158.1 | - | - | 0 | - |
| - | - | 982.9 | 158.1 | - | - | 0 | - |
| - | - | 417.2 | 158.9 | - | - | 0 | - |
| - | - | 948.2 | 159.1 | - | - | 0 | - |
| - | - | 3043 | 159.1 | - | - | 0 | - |
| - | - | 992.8 | 159.1 | - | - | 0 | - |
| - | - | 508.8 | 160.1 | - | - | 0 | - |
| - | - | 478.2 | 163.6 | - | - | 0 | - |
| - | - | 501.1 | 164.1 | - | - | 0 | - |
| - | - | 2249 | 165.1 | - | - | 0 | - |
| - | - | 1221 | 166.1 | - | - | 0 | - |
| - | - | 666.8 | 166.1 | - | - | 0 | - |
| - | - | 5075 | 167 | - | - | 0 | - |
| - | - | 1300 | 167.1 | - | - | 0 | - |
| - | - | 709.7 | 168 | - | - | 0 | - |
| - | - | 1243 | 169.1 | - | - | 0 | - |
| - | - | 4161 | 169.1 | - | - | 0 | - |
| - | - | 1408 | 170.1 | - | - | 0 | - |
| - | - | 444.3 | 171.1 | - | - | 0 | - |
| - | - | 536.8 | 171.1 | - | - | 0 | - |
| - | - | 1278 | 171.1 | - | - | 0 | - |
| - | - | 1154 | 171.1 | - | - | 0 | - |
| - | - | 641.4 | 172 | - | - | 0 | - |
| - | - | 3565 | 172.1 | - | - | 0 | - |
| - | - | 1307 | 173.1 | - | - | 0 | - |
| - | - | 3093 | 173.1 | - | - | 0 | - |
| - | - | 1163 | 173.1 | - | - | 0 | - |
| - | - | 1355 | 174.1 | - | - | 0 | - |
| - | - | 597.3 | 174.1 | - | - | 0 | - |
| - | - | 874.2 | 175.1 | - | - | 0 | - |
| - | - | 9677 | 175.1 | - | - | 0 | - |
| - | - | 936.5 | 175.1 | - | - | 0 | - |
| - | - | 1315 | 176.1 | - | - | 0 | - |
| - | - | 1043 | 176.1 | - | - | 0 | - |
| - | - | 645.2 | 177.1 | - | - | 0 | - |
| - | - | 579 | 181.1 | - | - | 0 | - |
| - | - | 880.4 | 181.1 | - | - | 0 | - |
| - | - | 8633 | 181.1 | - | - | 0 | - |
| - | - | 1071 | 181.1 | - | - | 0 | - |
| - | - | 2092 | 182.1 | - | - | 0 | - |
| - | - | 4183 | 183.1 | - | - | 0 | - |
| - | - | 1336 | 183.1 | - | - | 0 | - |
| - | - | 720.6 | 183.1 | - | - | 0 | - |
| - | - | 608.2 | 184.1 | - | - | 0 | - |
| - | - | 873.7 | 185.1 | - | - | 0 | - |
| - | - | 2.342E+04 | 185.1 | - | - | 0 | - |
| - | - | 3144 | 185.1 | - | - | 0 | - |
| - | - | 1136 | 186.1 | - | - | 0 | - |
| - | - | 2979 | 186.1 | - | - | 0 | - |
| - | - | 7569 | 186.1 | - | - | 0 | - |
| - | - | 1284 | 187.1 | - | - | 0 | - |
| - | - | 1.862E+04 | 187.1 | - | - | 0 | - |
| 12 | y | 1286 | 187.1 | 0.0001828 | 0.9769 | +2 | 3 |
| - | - | 564.4 | 188.1 | - | - | 0 | - |
| - | - | 498.4 | 188.1 | - | - | 0 | - |
| - | - | 664.1 | 188.1 | - | - | 0 | - |
| - | - | 1602 | 190.1 | - | - | 0 | - |
| - | - | 820.3 | 193.1 | - | - | 0 | - |
| - | - | 761.2 | 193.1 | - | - | 0 | - |
| - | - | 598.9 | 195 | - | - | 0 | - |
| - | - | 2455 | 195.1 | - | - | 0 | - |
| - | - | 578.6 | 197.1 | - | - | 0 | - |
| - | - | 2080 | 197.2 | - | - | 0 | - |
| - | - | 556.1 | 199.1 | - | - | 0 | - |
| - | - | 1013 | 199.1 | - | - | 0 | - |
| - | - | 675.8 | 199.2 | - | - | 0 | - |
| - | - | 1171 | 200.1 | - | - | 0 | - |
| - | - | 1757 | 201.1 | - | - | 0 | - |
| - | - | 4519 | 202.1 | - | - | 0 | - |
| - | - | 931 | 203.1 | - | - | 0 | - |
| - | - | 685.6 | 203.1 | - | - | 0 | - |
| - | - | 637.8 | 203.1 | - | - | 0 | - |
| - | - | 592.6 | 204.1 | - | - | 0 | - |
| - | - | 2016 | 205.1 | - | - | 0 | - |
| - | - | 798 | 205.1 | - | - | 0 | - |
| - | - | 670.4 | 205.1 | - | - | 0 | - |
| - | - | 1766 | 207.1 | - | - | 0 | - |
| - | - | 780 | 209.1 | - | - | 0 | - |
| - | - | 1217 | 209.1 | - | - | 0 | - |
| - | - | 881.7 | 210.9 | - | - | 0 | - |
| - | - | 487.9 | 211.1 | - | - | 0 | - |
| - | - | 2157 | 212 | - | - | 0 | - |
| - | - | 890.3 | 212.1 | - | - | 0 | - |
| - | - | 1418 | 212.7 | - | - | 0 | - |
| - | - | 844.6 | 213.1 | - | - | 0 | - |
| - | - | 570.2 | 213.1 | - | - | 0 | - |
| - | - | 1894 | 214.1 | - | - | 0 | - |
| - | - | 868.7 | 215.1 | - | - | 0 | - |
| - | - | 1543 | 215.1 | - | - | 0 | - |
| - | - | 608.5 | 216.1 | - | - | 0 | - |
| - | - | 639.6 | 216.3 | - | - | 0 | - |
| - | - | 1062 | 217 | - | - | 0 | - |
| - | - | 1520 | 217.1 | - | - | 0 | - |
| - | - | 2093 | 218.1 | - | - | 0 | - |
| 9 | y | 603.8 | 218.2 | 0.001509 | 6.918 | +3 | 6 |
| - | - | 561.3 | 219.1 | - | - | 0 | - |
| - | - | 716.1 | 220.1 | - | - | 0 | - |
| - | - | 3523 | 220.1 | - | - | 0 | - |
| - | - | 984.3 | 221.1 | - | - | 0 | - |
| - | - | 6827 | 221.1 | - | - | 0 | - |
| - | - | 678.9 | 222.1 | - | - | 0 | - |
| - | - | 1721 | 223.1 | - | - | 0 | - |
| - | - | 2503 | 224.2 | - | - | 0 | - |
| - | - | 1486 | 225.1 | - | - | 0 | - |
| - | - | 974 | 225.2 | - | - | 0 | - |
| - | - | 3039 | 226.1 | - | - | 0 | - |
| - | - | 4.531E+04 | 226.2 | - | - | 0 | - |
| - | - | 1.147E+04 | 227.1 | - | - | 0 | - |
| - | - | 774 | 227.1 | - | - | 0 | - |
| - | - | 590.4 | 227.1 | - | - | 0 | - |
| 11 | y | 4948 | 227.2 | 0.001266 | 5.574 | +2 | 4 |
| - | - | 1977 | 228.1 | - | - | 0 | - |
| - | - | 1820 | 228.1 | - | - | 0 | - |
| - | - | 788.5 | 228.1 | - | - | 0 | - |
| - | - | 501.7 | 228.3 | - | - | 0 | - |
| - | - | 743.1 | 229.1 | - | - | 0 | - |
| - | - | 1092 | 230.1 | - | - | 0 | - |
| - | - | 746.6 | 231.1 | - | - | 0 | - |
| - | - | 1746 | 231.1 | - | - | 0 | - |
| - | - | 772.4 | 232.1 | - | - | 0 | - |
| - | - | 788.6 | 233.1 | - | - | 0 | - |
| - | - | 942.9 | 233.1 | - | - | 0 | - |
| - | - | 867.8 | 235.1 | - | - | 0 | - |
| - | - | 6623 | 235.1 | - | - | 0 | - |
| 11 | y | 8872 | 235.7 | 0.0001376 | 0.5837 | +2 | 4 |
| - | - | 825.2 | 236.1 | - | - | 0 | - |
| - | - | 2683 | 236.2 | - | - | 0 | - |
| - | - | 950.5 | 237.1 | - | - | 0 | - |
| - | - | 1014 | 237.1 | - | - | 0 | - |
| - | - | 922.9 | 238.1 | - | - | 0 | - |
| - | - | 1056 | 239.1 | - | - | 0 | - |
| - | - | 3327 | 239.2 | - | - | 0 | - |
| - | - | 949.5 | 240.1 | - | - | 0 | - |
| - | - | 582.7 | 241.1 | - | - | 0 | - |
| - | - | 901.6 | 241.2 | - | - | 0 | - |
| - | - | 1740 | 242.2 | - | - | 0 | - |
| - | - | 929.5 | 243.1 | - | - | 0 | - |
| - | - | 1970 | 244.1 | - | - | 0 | - |
| - | - | 629.9 | 244.2 | - | - | 0 | - |
| - | - | 1760 | 245.1 | - | - | 0 | - |
| 13 | y | 8237 | 245.2 | 0.0002492 | 1.016 | +1 | 2 |
| - | - | 1811 | 246.1 | - | - | 0 | - |
| - | - | 565.9 | 246.1 | - | - | 0 | - |
| - | - | 894.6 | 246.2 | - | - | 0 | - |
| - | - | 8159 | 247.1 | - | - | 0 | - |
| - | - | 5919 | 248.1 | - | - | 0 | - |
| - | - | 857 | 248.1 | - | - | 0 | - |
| - | - | 614.8 | 248.1 | - | - | 0 | - |
| - | - | 7445 | 249.1 | - | - | 0 | - |
| - | - | 822.8 | 250.1 | - | - | 0 | - |
| - | - | 1135 | 251.2 | - | - | 0 | - |
| - | - | 788.4 | 253.1 | - | - | 0 | - |
| - | - | 5510 | 254.1 | - | - | 0 | - |
| - | - | 1.195E+04 | 255.1 | - | - | 0 | - |
| - | - | 854 | 255.1 | - | - | 0 | - |
| - | - | 646.6 | 255.1 | - | - | 0 | - |
| - | - | 1703 | 256.1 | - | - | 0 | - |
| - | - | 1151 | 257.1 | - | - | 0 | - |
| - | - | 1243 | 257.2 | - | - | 0 | - |
| - | - | 1259 | 258.1 | - | - | 0 | - |
| - | - | 1.096E+04 | 258.1 | - | - | 0 | - |
| - | - | 962.5 | 258.1 | - | - | 0 | - |
| - | - | 1177 | 259.1 | - | - | 0 | - |
| - | - | 844.2 | 259.1 | - | - | 0 | - |
| - | - | 1411 | 259.2 | - | - | 0 | - |
| - | - | 1436 | 260.2 | - | - | 0 | - |
| - | - | 1207 | 261.1 | - | - | 0 | - |
| 2 | a | 1.164E+04 | 262.1 | 0.00501 | 19.11 | +1 | 2 |
| - | - | 1266 | 263.1 | - | - | 0 | - |
| - | - | 731.1 | 264.1 | - | - | 0 | - |
| - | - | 4770 | 266.1 | - | - | 0 | - |
| - | - | 1106 | 267.1 | - | - | 0 | - |
| - | - | 705.2 | 269.6 | - | - | 0 | - |
| - | - | 3895 | 270.2 | - | - | 0 | - |
| - | - | 1567 | 271.1 | - | - | 0 | - |
| 10 | y | 2631 | 271.2 | 0.0001088 | 0.4011 | +2 | 5 |
| 2 | b | 2004 | 272.1 | 0.0047 | 17.27 | +1 | 2 |
| - | - | 2802 | 273.1 | - | - | 0 | - |
| - | - | 750.1 | 274.1 | - | - | 0 | - |
| - | - | 5037 | 274.1 | - | - | 0 | - |
| - | - | 839.6 | 275.1 | - | - | 0 | - |
| - | - | 2546 | 275.1 | - | - | 0 | - |
| - | - | 1.594E+04 | 275.1 | - | - | 0 | - |
| - | - | 619.4 | 275.5 | - | - | 0 | - |
| - | - | 885.7 | 276.1 | - | - | 0 | - |
| - | - | 1647 | 276.1 | - | - | 0 | - |
| - | - | 741.3 | 279.2 | - | - | 0 | - |
| - | - | 628.1 | 280.2 | - | - | 0 | - |
| - | - | 681.1 | 283.2 | - | - | 0 | - |
| - | - | 1.135E+04 | 283.2 | - | - | 0 | - |
| - | - | 1587 | 284.2 | - | - | 0 | - |
| - | - | 1685 | 285.1 | - | - | 0 | - |
| - | - | 3198 | 285.2 | - | - | 0 | - |
| - | - | 1559 | 288.1 | - | - | 0 | - |
| 2 | b | 9440 | 290.1 | 0.005091 | 17.55 | +1 | 2 |
| - | - | 4841 | 290.2 | - | - | 0 | - |
| - | - | 1876 | 290.7 | - | - | 0 | - |
| 4 | b | 2054 | 291.1 | 0.001377 | 4.73 | +2 | 4 |
| - | - | 3913 | 292.1 | - | - | 0 | - |
| - | - | 2199 | 293.1 | - | - | 0 | - |
| - | - | 3907 | 293.1 | - | - | 0 | - |
| - | - | 2139 | 293.2 | - | - | 0 | - |
| - | - | 976.7 | 294.1 | - | - | 0 | - |
| - | - | 509.8 | 297.2 | - | - | 0 | - |
| - | - | 4587 | 297.2 | - | - | 0 | - |
| - | - | 902.8 | 298.1 | - | - | 0 | - |
| - | - | 530.2 | 298.2 | - | - | 0 | - |
| - | - | 693.8 | 300.7 | - | - | 0 | - |
| - | - | 544.9 | 303.1 | - | - | 0 | - |
| - | - | 993.5 | 305.7 | - | - | 0 | - |
| - | - | 915.8 | 306.1 | - | - | 0 | - |
| - | - | 539.2 | 309.4 | - | - | 0 | - |
| - | - | 1735 | 311.2 | - | - | 0 | - |
| - | - | 645 | 315.2 | - | - | 0 | - |
| - | - | 795.5 | 320.1 | - | - | 0 | - |
| - | - | 8878 | 320.1 | - | - | 0 | - |
| 6 | y | 648.3 | 320.9 | 0.001007 | 3.138 | +3 | 9 |
| - | - | 1421 | 321.1 | - | - | 0 | - |
| - | - | 1023 | 321.2 | - | - | 0 | - |
| - | - | 5674 | 321.2 | - | - | 0 | - |
| - | - | 1105 | 322.2 | - | - | 0 | - |
| - | - | 1128 | 324.2 | - | - | 0 | - |
| - | - | 2800 | 326.2 | - | - | 0 | - |
| - | - | 593.4 | 328.1 | - | - | 0 | - |
| - | - | 2664 | 332.7 | - | - | 0 | - |
| - | - | 1035 | 333.2 | - | - | 0 | - |
| 9 | y | 580.9 | 335.2 | 0.0003491 | 1.041 | +2 | 6 |
| - | - | 1.063E+04 | 336.2 | - | - | 0 | - |
| - | - | 1461 | 337.2 | - | - | 0 | - |
| - | - | 3769 | 337.7 | - | - | 0 | - |
| - | - | 2871 | 338.2 | - | - | 0 | - |
| - | - | 823.4 | 339.2 | - | - | 0 | - |
| - | - | 1.174E+04 | 339.2 | - | - | 0 | - |
| - | - | 657 | 340.2 | - | - | 0 | - |
| - | - | 2147 | 340.2 | - | - | 0 | - |
| - | - | 9370 | 346.7 | - | - | 0 | - |
| - | - | 3757 | 347.2 | - | - | 0 | - |
| - | - | 4578 | 348.1 | - | - | 0 | - |
| - | - | 1125 | 349.1 | - | - | 0 | - |
| - | - | 1154 | 351.2 | - | - | 0 | - |
| - | - | 1083 | 353.7 | - | - | 0 | - |
| - | - | 698.7 | 354.2 | - | - | 0 | - |
| - | - | 1.569E+04 | 354.2 | - | - | 0 | - |
| - | - | 1734 | 355.2 | - | - | 0 | - |
| - | - | 1578 | 355.7 | - | - | 0 | - |
| - | - | 855.5 | 356.1 | - | - | 0 | - |
| - | - | 733.5 | 356.2 | - | - | 0 | - |
| - | - | 2590 | 357.2 | - | - | 0 | - |
| - | - | 2931 | 358.2 | - | - | 0 | - |
| - | - | 1268 | 358.7 | - | - | 0 | - |
| - | - | 1439 | 359.9 | - | - | 0 | - |
| - | - | 610.2 | 360.7 | - | - | 0 | - |
| - | - | 1500 | 361.7 | - | - | 0 | - |
| - | - | 1018 | 362.2 | - | - | 0 | - |
| - | - | 1358 | 362.5 | - | - | 0 | - |
| 5 | y | 1901 | 366.6 | 0.0009211 | 2.513 | +3 | 10 |
| - | - | 931.8 | 366.9 | - | - | 0 | - |
| - | - | 1004 | 367.2 | - | - | 0 | - |
| - | - | 1783 | 368.1 | - | - | 0 | - |
| - | - | 1426 | 368.2 | - | - | 0 | - |
| - | - | 1210 | 369.7 | - | - | 0 | - |
| - | - | 857.7 | 371.6 | - | - | 0 | - |
| 12 | y | 5126 | 373.3 | 0.0007146 | 1.914 | +1 | 3 |
| - | - | 861.3 | 375 | - | - | 0 | - |
| - | - | 1067 | 375.7 | - | - | 0 | - |
| - | - | 631.7 | 378.5 | - | - | 0 | - |
| - | - | 872.8 | 379.2 | - | - | 0 | - |
| - | - | 662.7 | 380.2 | - | - | 0 | - |
| - | - | 1014 | 382.2 | - | - | 0 | - |
| - | - | 621.1 | 382.5 | - | - | 0 | - |
| - | - | 1162 | 384.2 | - | - | 0 | - |
| - | - | 4723 | 385.2 | - | - | 0 | - |
| - | - | 622.4 | 385.2 | - | - | 0 | - |
| - | - | 789.7 | 385.5 | - | - | 0 | - |
| - | - | 965.2 | 385.8 | - | - | 0 | - |
| - | - | 4156 | 386.2 | - | - | 0 | - |
| - | - | 911.7 | 386.2 | - | - | 0 | - |
| - | - | 843.5 | 387.2 | - | - | 0 | - |
| - | - | 714.4 | 387.5 | - | - | 0 | - |
| - | - | 4084 | 389.2 | - | - | 0 | - |
| - | - | 657.2 | 389.7 | - | - | 0 | - |
| - | - | 1942 | 391.5 | - | - | 0 | - |
| - | - | 761.8 | 391.9 | - | - | 0 | - |
| - | - | 793.2 | 392.2 | - | - | 0 | - |
| - | - | 1780 | 392.3 | - | - | 0 | - |
| - | - | 1780 | 393.7 | - | - | 0 | - |
| - | - | 6856 | 394 | - | - | 0 | - |
| - | - | 1961 | 394.2 | - | - | 0 | - |
| 2 | y | 3962 | 394.2 | 0.003916 | 9.933 | +4 | 13 |
| - | - | 1031 | 394.5 | - | - | 0 | - |
| - | - | 657.6 | 394.5 | - | - | 0 | - |
| - | - | 703.2 | 395.2 | - | - | 0 | - |
| - | - | 1642 | 396.2 | - | - | 0 | - |
| - | - | 700 | 397.2 | - | - | 0 | - |
| 10 | b | 1246 | 397.8 | 0.002332 | 5.862 | +3 | 10 |
| 2 | y | 1114 | 398.5 | 0.002131 | 5.347 | +4 | 13 |
| - | - | 690.4 | 398.7 | - | - | 0 | - |
| - | - | 1200 | 399.7 | - | - | 0 | - |
| - | - | 775 | 401 | - | - | 0 | - |
| - | - | 883.6 | 401.3 | - | - | 0 | - |
| - | - | 2135 | 402.2 | - | - | 0 | - |
| - | - | 1145 | 402.7 | - | - | 0 | - |
| - | - | 4322 | 403.2 | - | - | 0 | - |
| - | - | 1533 | 403.2 | - | - | 0 | - |
| - | - | 983.4 | 403.3 | - | - | 0 | - |
| 10 | b | 783.9 | 403.5 | 0.001315 | 3.259 | +3 | 10 |
| 4 | y | 3438 | 403.6 | 0.004802 | 11.9 | +3 | 11 |
| - | - | 661.4 | 403.9 | - | - | 0 | - |
| - | - | 1440 | 403.9 | - | - | 0 | - |
| - | - | 1228 | 404.2 | - | - | 0 | - |
| - | - | 962.6 | 404.2 | - | - | 0 | - |
| - | - | 2136 | 405.2 | - | - | 0 | - |
| - | - | 659 | 405.7 | - | - | 0 | - |
| - | - | 1170 | 406.3 | - | - | 0 | - |
| - | - | 1212 | 408.2 | - | - | 0 | - |
| - | - | 969.4 | 408.5 | - | - | 0 | - |
| - | - | 1416 | 408.7 | - | - | 0 | - |
| - | - | 667.1 | 409.9 | - | - | 0 | - |
| - | - | 4920 | 410.3 | - | - | 0 | - |
| - | - | 742.7 | 410.6 | - | - | 0 | - |
| - | - | 701.3 | 410.7 | - | - | 0 | - |
| - | - | 752.2 | 410.9 | - | - | 0 | - |
| - | - | 2959 | 411.2 | - | - | 0 | - |
| - | - | 3263 | 411.2 | - | - | 0 | - |
| - | - | 1369 | 411.3 | - | - | 0 | - |
| - | - | 6057 | 411.5 | - | - | 0 | - |
| - | - | 3902 | 411.7 | - | - | 0 | - |
| - | - | 2404 | 411.7 | - | - | 0 | - |
| - | - | 1662 | 412 | - | - | 0 | - |
| - | - | 2060 | 412.2 | - | - | 0 | - |
| - | - | 894.9 | 412.2 | - | - | 0 | - |
| 7 | y | 1.482E+04 | 412.3 | 0.0006231 | 1.511 | +2 | 8 |
| - | - | 6285 | 412.8 | - | - | 0 | - |
| - | - | 820.1 | 413.2 | - | - | 0 | - |
| - | - | 1197 | 413.3 | - | - | 0 | - |
| - | - | 785.8 | 413.9 | - | - | 0 | - |
| - | - | 931.4 | 414.2 | - | - | 0 | - |
| - | - | 1845 | 414.2 | - | - | 0 | - |
| - | - | 898.4 | 414.7 | - | - | 0 | - |
| - | - | 1712 | 415.3 | - | - | 0 | - |
| 0 | Precursor | 2042 | 415.7 | 0.001586 | 3.816 | +4 | -1 |
| - | - | 937.8 | 415.8 | - | - | 0 | - |
| 0 | Precursor | 2537 | 416 | 0.006254 | 15.03 | +4 | -1 |
| - | - | 968.4 | 417.2 | - | - | 0 | - |
| - | - | 5064 | 419.9 | - | - | 0 | - |
| 0 | Precursor | 6084 | 420.2 | 0.0063 | 14.99 | +4 | -1 |
| - | - | 5440 | 420.5 | - | - | 0 | - |
| - | - | 1501 | 420.6 | - | - | 0 | - |
| - | - | 2383 | 420.7 | - | - | 0 | - |
| - | - | 1227 | 421.2 | - | - | 0 | - |
| - | - | 1319 | 421.3 | - | - | 0 | - |
| - | - | 1137 | 422.3 | - | - | 0 | - |
| - | - | 805.6 | 423.3 | - | - | 0 | - |
| - | - | 1134 | 424.3 | - | - | 0 | - |
| - | - | 1490 | 424.7 | - | - | 0 | - |
| - | - | 818.6 | 425.2 | - | - | 0 | - |
| - | - | 1153 | 427.2 | - | - | 0 | - |
| 6 | b | 803.4 | 428.2 | 0.004958 | 11.58 | +2 | 6 |
| - | - | 1.016E+04 | 429.2 | - | - | 0 | - |
| 11 | b | 2996 | 430.2 | 0.006945 | 16.14 | +3 | 11 |
| - | - | 1646 | 431.2 | - | - | 0 | - |
| - | - | 1014 | 443.6 | - | - | 0 | - |
| - | - | 1237 | 443.9 | - | - | 0 | - |
| - | - | 662.4 | 449.6 | - | - | 0 | - |
| - | - | 1583 | 451.2 | - | - | 0 | - |
| - | - | 1414 | 451.3 | - | - | 0 | - |
| - | - | 1629 | 451.7 | - | - | 0 | - |
| - | - | 2530 | 451.9 | - | - | 0 | - |
| - | - | 1720 | 452.3 | - | - | 0 | - |
| - | - | 934.8 | 452.3 | - | - | 0 | - |
| - | - | 1248 | 452.6 | - | - | 0 | - |
| 3 | b | 1062 | 453.2 | 0.004079 | 9.001 | +1 | 3 |
| - | - | 1557 | 455.3 | - | - | 0 | - |
| - | - | 820.6 | 456.3 | - | - | 0 | - |
| - | - | 1078 | 457.6 | - | - | 0 | - |
| 3 | y | 2375 | 457.9 | 0.004088 | 8.928 | +3 | 12 |
| - | - | 838.4 | 458.3 | - | - | 0 | - |
| - | - | 2443 | 460.2 | - | - | 0 | - |
| - | - | 1157 | 460.7 | - | - | 0 | - |
| - | - | 677.3 | 461.6 | - | - | 0 | - |
| - | - | 869.5 | 462.2 | - | - | 0 | - |
| 3 | y | 2.403E+04 | 463.6 | 0.0006605 | 1.425 | +3 | 12 |
| - | - | 1.829E+04 | 463.9 | - | - | 0 | - |
| - | - | 5708 | 464.3 | - | - | 0 | - |
| - | - | 1184 | 464.6 | - | - | 0 | - |
| - | - | 916.1 | 464.9 | - | - | 0 | - |
| - | - | 1685 | 466.9 | - | - | 0 | - |
| - | - | 1184 | 467.3 | - | - | 0 | - |
| - | - | 896.7 | 468.3 | - | - | 0 | - |
| 11 | y | 2.122E+04 | 470.3 | 0.0006241 | 1.327 | +1 | 4 |
| - | - | 5100 | 471.3 | - | - | 0 | - |
| - | - | 830.1 | 471.8 | - | - | 0 | - |
| - | - | 651.8 | 471.9 | - | - | 0 | - |
| - | - | 753.2 | 472.3 | - | - | 0 | - |
| - | - | 957.8 | 472.6 | - | - | 0 | - |
| - | - | 2559 | 473.3 | - | - | 0 | - |
| - | - | 813.7 | 474.8 | - | - | 0 | - |
| - | - | 1225 | 475.6 | - | - | 0 | - |
| - | - | 4505 | 476.2 | - | - | 0 | - |
| - | - | 711.9 | 476.9 | - | - | 0 | - |
| - | - | 879.3 | 477.2 | - | - | 0 | - |
| 12 | b | 1200 | 478.6 | 0.001237 | 2.584 | +3 | 12 |
| - | - | 696.2 | 478.9 | - | - | 0 | - |
| 6 | y | 2532 | 480.8 | 0.001074 | 2.235 | +2 | 9 |
| - | - | 3970 | 481.3 | - | - | 0 | - |
| - | - | 2340 | 481.3 | - | - | 0 | - |
| - | - | 3264 | 481.6 | - | - | 0 | - |
| - | - | 1082 | 481.9 | - | - | 0 | - |
| - | - | 1264 | 482.3 | - | - | 0 | - |
| - | - | 1423 | 482.3 | - | - | 0 | - |
| - | - | 736.6 | 482.8 | - | - | 0 | - |
| - | - | 928.2 | 483.3 | - | - | 0 | - |
| - | - | 2173 | 483.8 | - | - | 0 | - |
| - | - | 1136 | 484.3 | - | - | 0 | - |
| - | - | 2293 | 487.3 | - | - | 0 | - |
| - | - | 1587 | 487.6 | - | - | 0 | - |
| - | - | 2252 | 488.2 | - | - | 0 | - |
| - | - | 1204 | 490.3 | - | - | 0 | - |
| - | - | 1128 | 490.6 | - | - | 0 | - |
| - | - | 1477 | 491.3 | - | - | 0 | - |
| - | - | 1158 | 494.2 | - | - | 0 | - |
| - | - | 933.1 | 495.9 | - | - | 0 | - |
| - | - | 694.6 | 500.3 | - | - | 0 | - |
| - | - | 908.8 | 504.3 | - | - | 0 | - |
| - | - | 721.5 | 504.6 | - | - | 0 | - |
| - | - | 2806 | 505.3 | - | - | 0 | - |
| - | - | 2162 | 505.6 | - | - | 0 | - |
| - | - | 2072 | 506.2 | - | - | 0 | - |
| - | - | 1150 | 508.2 | - | - | 0 | - |
| - | - | 1025 | 509.6 | - | - | 0 | - |
| 13 | b | 1620 | 510.3 | 0.001202 | 2.356 | +3 | 13 |
| 13 | b | 2038 | 510.6 | 0.008402 | 16.46 | +3 | 13 |
| - | - | 1105 | 510.9 | - | - | 0 | - |
| 13 | b | 2305 | 516.3 | 0.001678 | 3.251 | +3 | 13 |
| - | - | 1945 | 516.6 | - | - | 0 | - |
| - | - | 1357 | 516.7 | - | - | 0 | - |
| - | - | 1089 | 516.9 | - | - | 0 | - |
| - | - | 1424 | 517.3 | - | - | 0 | - |
| - | - | 1019 | 518.9 | - | - | 0 | - |
| - | - | 1689 | 519.3 | - | - | 0 | - |
| - | - | 828.5 | 520.4 | - | - | 0 | - |
| - | - | 897.9 | 521.2 | - | - | 0 | - |
| - | - | 804.1 | 521.9 | - | - | 0 | - |
| - | - | 2773 | 522.3 | - | - | 0 | - |
| - | - | 2043 | 522.6 | - | - | 0 | - |
| - | - | 736.8 | 524.6 | - | - | 0 | - |
| - | - | 1.311E+04 | 525 | - | - | 0 | - |
| 2 | y | 9444 | 525.3 | 0.008033 | 15.29 | +3 | 13 |
| - | - | 3913 | 525.6 | - | - | 0 | - |
| - | - | 1786 | 526 | - | - | 0 | - |
| - | - | 1228 | 526.2 | - | - | 0 | - |
| - | - | 1154 | 528.3 | - | - | 0 | - |
| 2 | y | 2016 | 531 | 0.001401 | 2.638 | +3 | 13 |
| - | - | 946.3 | 531.2 | - | - | 0 | - |
| - | - | 2975 | 531.3 | - | - | 0 | - |
| - | - | 6272 | 538.3 | - | - | 0 | - |
| - | - | 963.5 | 538.4 | - | - | 0 | - |
| - | - | 2757 | 539.3 | - | - | 0 | - |
| - | - | 2299 | 539.3 | - | - | 0 | - |
| - | - | 1120 | 539.8 | - | - | 0 | - |
| - | - | 766.8 | 540.3 | - | - | 0 | - |
| 10 | y | 4515 | 541.4 | 0.00181 | 3.343 | +1 | 5 |
| - | - | 1388 | 542.4 | - | - | 0 | - |
| - | - | 1075 | 543.3 | - | - | 0 | - |
| - | - | 813.4 | 543.8 | - | - | 0 | - |
| - | - | 1081 | 547.8 | - | - | 0 | - |
| 5 | y | 7307 | 549.3 | 0.0005186 | 0.944 | +2 | 10 |
| - | - | 4309 | 549.8 | - | - | 0 | - |
| - | - | 2460 | 550.3 | - | - | 0 | - |
| - | - | 849.1 | 551.3 | - | - | 0 | - |
| - | - | 1878 | 552.3 | - | - | 0 | - |
| - | - | 744.5 | 554.2 | - | - | 0 | - |
| - | - | 1094 | 557.3 | - | - | 0 | - |
| - | - | 7757 | 566.2 | - | - | 0 | - |
| - | - | 2601 | 567.2 | - | - | 0 | - |
| 9 | b | 1573 | 569.3 | 0.001997 | 3.508 | +2 | 9 |
| - | - | 1062 | 569.8 | - | - | 0 | - |
| - | - | 976.2 | 570.3 | - | - | 0 | - |
| - | - | 3171 | 572.8 | - | - | 0 | - |
| - | - | 1596 | 573.3 | - | - | 0 | - |
| - | - | 3092 | 577.4 | - | - | 0 | - |
| - | - | 1477 | 578.4 | - | - | 0 | - |
| - | - | 7028 | 579.4 | - | - | 0 | - |
| - | - | 1716 | 580.4 | - | - | 0 | - |
| - | - | 1758 | 585.3 | - | - | 0 | - |
| - | - | 744.3 | 590.3 | - | - | 0 | - |
| - | - | 2049 | 595.4 | - | - | 0 | - |
| - | - | 937 | 596.3 | - | - | 0 | - |
| - | - | 710.2 | 600.3 | - | - | 0 | - |
| 4 | y | 3535 | 604.8 | 0.003669 | 6.067 | +2 | 11 |
| - | - | 2034 | 605.4 | - | - | 0 | - |
| - | - | 1363 | 605.9 | - | - | 0 | - |
| - | - | 1141 | 610.3 | - | - | 0 | - |
| - | - | 770.3 | 611.3 | - | - | 0 | - |
| - | - | 3182 | 613.2 | - | - | 0 | - |
| 4 | y | 1502 | 613.4 | 0.002891 | 4.714 | +2 | 11 |
| - | - | 1246 | 613.9 | - | - | 0 | - |
| - | - | 1454 | 614.3 | - | - | 0 | - |
| - | - | 1006 | 614.4 | - | - | 0 | - |
| - | - | 932.8 | 615.3 | - | - | 0 | - |
| - | - | 951.2 | 618.4 | - | - | 0 | - |
| - | - | 1258 | 620.3 | - | - | 0 | - |
| - | - | 1021 | 620.8 | - | - | 0 | - |
| - | - | 767.1 | 621.3 | - | - | 0 | - |
| - | - | 826 | 627.4 | - | - | 0 | - |
| - | - | 2606 | 629.3 | - | - | 0 | - |
| - | - | 3432 | 629.8 | - | - | 0 | - |
| - | - | 1228 | 630.4 | - | - | 0 | - |
| - | - | 801.9 | 633.3 | - | - | 0 | - |
| - | - | 3616 | 635.3 | - | - | 0 | - |
| - | - | 1590 | 636.3 | - | - | 0 | - |
| - | - | 840.5 | 637.3 | - | - | 0 | - |
| - | - | 1541 | 638.4 | - | - | 0 | - |
| - | - | 1134 | 638.9 | - | - | 0 | - |
| 11 | b | 4623 | 653.3 | 0.001987 | 3.042 | +2 | 11 |
| - | - | 1268 | 654.4 | - | - | 0 | - |
| - | - | 1985 | 664.5 | - | - | 0 | - |
| - | - | 752.8 | 664.8 | - | - | 0 | - |
| 9 | y | 947.2 | 669.5 | 0.001451 | 2.168 | +1 | 6 |
| - | - | 6072 | 674.4 | - | - | 0 | - |
| - | - | 3127 | 675.4 | - | - | 0 | - |
| - | - | 1529 | 682.3 | - | - | 0 | - |
| - | - | 7365 | 692.4 | - | - | 0 | - |
| - | - | 2323 | 693.4 | - | - | 0 | - |
| 3 | y | 3750 | 694.9 | 9.638E-06 | 0.01387 | +2 | 12 |
| - | - | 4028 | 695.4 | - | - | 0 | - |
| - | - | 1242 | 695.9 | - | - | 0 | - |
| - | - | 2336 | 710.5 | - | - | 0 | - |
| - | - | 825.6 | 711.5 | - | - | 0 | - |
| - | - | 1286 | 716.4 | - | - | 0 | - |
| 5 | b | 914.9 | 718.3 | 0.008599 | 11.97 | +1 | 5 |
| - | - | 1678 | 721.4 | - | - | 0 | - |
| - | - | 1000 | 722.3 | - | - | 0 | - |
| 8 | y | 2317 | 726.5 | 0.0003734 | 0.5139 | +1 | 7 |
| - | - | 975.8 | 727.5 | - | - | 0 | - |
| - | - | 1868 | 750.3 | - | - | 0 | - |
| - | - | 911.5 | 751.3 | - | - | 0 | - |
| - | - | 1423 | 757.4 | - | - | 0 | - |
| - | - | 1161 | 768.3 | - | - | 0 | - |
| - | - | 1041 | 805.5 | - | - | 0 | - |
| - | - | 755.6 | 811.5 | - | - | 0 | - |
| - | - | 1085 | 819.3 | - | - | 0 | - |
| 7 | y | 1.102E+04 | 823.5 | 0.001076 | 1.307 | +1 | 8 |
| - | - | 4267 | 824.5 | - | - | 0 | - |
| - | - | 1486 | 825.5 | - | - | 0 | - |
| - | - | 852.9 | 827.4 | - | - | 0 | - |
| - | - | 803.8 | 829.5 | - | - | 0 | - |
| - | - | 657.9 | 831.9 | - | - | 0 | - |
| - | - | 872 | 848.4 | - | - | 0 | - |
| 6 | b | 1103 | 855.3 | 0.008159 | 9.539 | +1 | 6 |
| - | - | 640.9 | 856.4 | - | - | 0 | - |
| - | - | 1547 | 901.4 | - | - | 0 | - |
| - | - | 1390 | 919.5 | - | - | 0 | - |
| - | - | 805.9 | 942.6 | - | - | 0 | - |
| - | - | 905.2 | 943.6 | - | - | 0 | - |
| 6 | y | 1266 | 960.6 | 0.001002 | 1.043 | +1 | 9 |
| - | - | 682.6 | 1903 | - | - | 0 | - |

m/z Charge Intensity FragmentType MassShift Position
120.06565856933594 0 644.1935
120.08090209960938 0 3718.3203
121.02838897705078 0 364.94617
122.07172393798828 0 675.68646
123.05537414550781 0 764.82697
123.11678314208984 0 443.40866
124.48589324951172 0 356.9859
125.10750579833984 0 474.2679
126.05519104003906 0 551.66754
127.08675384521484 0 1828.6892
129.0185089111328 0 2561.5198
129.06634521484375 0 455.20035
129.10243225097656 0 56660.94
130.06533813476562 0 34928.266
130.10061645507812 0 582.2183
130.1057891845703 0 3637.7883
131.06419372558594 0 467.18665
131.06874084472656 0 3296.8638
131.08168029785156 0 1744.8905
132.04458618164062 0 3397.7524
132.0809783935547 0 2131.814
132.1020965576172 0 15499.676 y 13
133.05252075195312 0 1815.0836
133.10528564453125 0 942.5168
135.54837036132812 0 525.1714
136.07591247558594 0 47401.973
137.0792694091797 0 2304.5286
138.0665283203125 0 999.91864
139.00282287597656 0 958.2348
139.0869598388672 0 1142.4873
141.10247802734375 0 3575.1777
143.04541015625 0 521.8618
143.11851501464844 0 593.7949
144.08108520507812 0 652.66583
145.07647705078125 0 580.79474
146.06019592285156 0 4178.553 b 3
147.044189453125 0 2319.676
148.0633087158203 0 1170.5082
148.07598876953125 0 739.5657
148.95428466796875 0 726.64514
149.02354431152344 0 74456.55
150.0269317626953 0 6986.7607
150.06661987304688 0 505.2201
151.0867156982422 0 582.70215
152.14364624023438 0 1182.0599
153.10260009765625 0 641.7961
155.0817108154297 0 3142.2222
155.09295654296875 0 754.03735
155.11810302734375 0 2032.2714
156.06814575195312 0 510.36722
156.07687377929688 0 1420.9264
157.01329040527344 0 2168.8848
157.0610809326172 0 1552.3817
157.07618713378906 0 12127.193
157.13348388671875 0 1432.4402
158.0602264404297 0 9174.891
158.07960510253906 0 982.86945
158.92115783691406 0 417.16415
159.06382751464844 0 948.24646
159.09181213378906 0 3043.0647
159.11260986328125 0 992.8092
160.0757598876953 0 508.7805
163.61322021484375 0 478.15082
164.07028198242188 0 501.13074
165.1024932861328 0 2249.1099
166.06092834472656 0 1220.599
166.0861358642578 0 666.8423
167.03408813476562 0 5074.9175
167.08177185058594 0 1299.6428
168.03726196289062 0 709.67993
169.0612030029297 0 1243.0487
169.09739685058594 0 4160.8247
170.06027221679688 0 1407.533
171.06448364257812 0 444.26733
171.0770721435547 0 536.8288
171.11309814453125 0 1277.5051
171.13833618164062 0 1153.9261
172.03936767578125 0 641.3882
172.10824584960938 0 3564.5037
173.07101440429688 0 1306.5945
173.09219360351562 0 3093.4998
173.12860107421875 0 1162.9438
174.05528259277344 0 1355.0642
174.0662078857422 0 597.27563
175.07176208496094 0 874.1597
175.08682250976562 0 9677.179
175.09605407714844 0 936.51404
176.08212280273438 0 1314.8591
176.0901336669922 0 1042.9438
177.1029052734375 0 645.2183
181.0606689453125 0 579.03156
181.076171875 0 880.36865
181.0972900390625 0 8632.694
181.1337432861328 0 1070.7749
182.1289825439453 0 2092.4734
183.05548095703125 0 4183.2246
183.11302185058594 0 1336.4265
183.14926147460938 0 720.6239
184.06361389160156 0 608.2375
185.05628967285156 0 873.67523
185.07110595703125 0 23423.443
185.0808868408203 0 3144.0603
186.05471801757812 0 1136.373
186.07444763183594 0 2978.872
186.12393188476562 0 7568.664
187.08718872070312 0 1283.9288
187.10794067382812 0 18615.33
187.144287109375 0 1285.8804 y 11
188.07037353515625 0 564.4076
188.08123779296875 0 498.38562
188.1114959716797 0 664.05615
190.0862274169922 0 1601.5287
193.09744262695312 0 820.32007
193.1085205078125 0 761.18494
194.97959899902344 0 598.86224
195.0767364501953 0 2454.6377
197.1288604736328 0 578.577
197.1649627685547 0 2079.823
199.07135009765625 0 556.1376
199.08665466308594 0 1012.60474
199.1805877685547 0 675.78534
200.13958740234375 0 1171.0203
201.12368774414062 0 1756.828
202.10870361328125 0 4518.9805
203.08177185058594 0 931.0469
203.09251403808594 0 685.62555
203.11297607421875 0 637.7552
204.0773468017578 0 592.6256
205.06100463867188 0 2015.9521
205.09671020507812 0 797.9983
205.10800170898438 0 670.42096
207.14935302734375 0 1765.9423
209.0706329345703 0 780.02734
209.09228515625 0 1216.7391
210.9053497314453 0 881.70557
211.14508056640625 0 487.85907
212.00462341308594 0 2156.6082
212.1389923095703 0 890.2593
212.6678466796875 0 1417.6866
213.07598876953125 0 844.5741
213.086669921875 0 570.2322
214.1092529296875 0 1893.8744
215.08213806152344 0 868.70575
215.13966369628906 0 1543.1721
216.0665740966797 0 608.534
216.25216674804688 0 639.6126
217.03497314453125 0 1061.632
217.09754943847656 0 1520.1755
218.06875610351562 0 2093.1626
218.14974975585938 0 603.8275 y Ammonia loss 8
219.13597106933594 0 561.2801
220.1087646484375 0 716.0853
220.1195526123047 0 3522.5422
221.09268188476562 0 984.26904
221.10353088378906 0 6827.009
222.10723876953125 0 678.8886
223.0713348388672 0 1720.5867
224.1758575439453 0 2503.4153
225.06613159179688 0 1485.6418
225.15975952148438 0 973.9775
226.09776306152344 0 3038.938
226.15518188476562 0 45311.445
227.08177185058594 0 11472.105
227.1015167236328 0 773.9732
227.13844299316406 0 590.39923
227.15847778320312 0 4947.8193 y Ammonia loss 10
228.08514404296875 0 1977.0396
228.11325073242188 0 1819.7668
228.1333465576172 0 788.51105
228.34519958496094 0 501.68948
229.11752319335938 0 743.1341
230.11419677734375 0 1092.191
231.076416015625 0 746.5958
231.088134765625 0 1745.8508
232.142822265625 0 772.39526
233.0919952392578 0 788.60535
233.103759765625 0 942.91345
235.0959930419922 0 867.76556
235.1190948486328 0 6623.443
235.67062377929688 0 8872.329 y 10
236.1232452392578 0 825.20593
236.17237854003906 0 2683.367
237.0912322998047 0 950.47034
237.135498046875 0 1013.5461
238.13067626953125 0 922.94934
239.1138916015625 0 1055.7301
239.150390625 0 3326.8977
240.13409423828125 0 949.5007
241.1420135498047 0 582.67755
241.15518188476562 0 901.58154
242.1865997314453 0 1739.8182
243.14578247070312 0 929.47766
244.1083221435547 0 1969.9979
244.1649627685547 0 629.88586
245.0930633544922 0 1760.3818
245.18621826171875 0 8236.838 y 12
246.123779296875 0 1811.3063
246.13746643066406 0 565.8818
246.1900634765625 0 894.6146
247.13027954101562 0 8159.0664
248.11439514160156 0 5919.4644
248.1282196044922 0 857.0431
248.13502502441406 0 614.8317
249.09840393066406 0 7444.8677
250.10107421875 0 822.7956
251.15008544921875 0 1135.0634
253.06161499023438 0 788.3977
254.09268188476562 0 5510.0415
255.0767059326172 0 11945.3125
255.09317016601562 0 854.0201
255.09654235839844 0 646.56195
256.0796203613281 0 1703.2689
257.11456298828125 0 1150.9415
257.1612548828125 0 1243.15
258.0752868652344 0 1258.5312
258.0987243652344 0 10958.202
258.1129455566406 0 962.45337
259.1031799316406 0 1176.8816
259.1302490234375 0 844.1823
259.15411376953125 0 1411.176
260.19720458984375 0 1435.5082
261.08612060546875 0 1206.7517
262.11871337890625 0 11641.484 a 1
263.12298583984375 0 1265.9436
264.135009765625 0 731.1411
266.1248474121094 0 4770.249
267.1242370605469 0 1106.323
269.6304931640625 0 705.2171
270.1812744140625 0 3894.998
271.0710754394531 0 1566.7198
271.1889343261719 0 2630.8413 y 9
272.1027526855469 0 2004.128 b Water loss 1
273.08709716796875 0 2801.5264
274.09100341796875 0 750.0757
274.13006591796875 0 5036.9785
275.0887145996094 0 839.58093
275.10321044921875 0 2546.3955
275.1253662109375 0 15943.228
275.517333984375 0 619.3545
276.1105041503906 0 885.6873
276.1282653808594 0 1646.5106
279.1821594238281 0 741.2955
280.1664123535156 0 628.14374
283.1514892578125 0 681.14
283.17669677734375 0 11345.679
284.1794128417969 0 1586.7035
285.109619140625 0 1685.4471
285.1561584472656 0 3197.9004
288.0981140136719 0 1559.0863
290.11370849609375 0 9439.937 b 1
290.1847839355469 0 4841.2905
290.6863098144531 0 1876.0687
291.1175231933594 0 2053.8677 b 3
292.1293029785156 0 3913.33
293.09979248046875 0 2199.1494
293.1300048828125 0 3906.7559
293.17529296875 0 2138.719
294.121337890625 0 976.69415
297.1571044921875 0 509.81833
297.1925048828125 0 4586.6445
298.1401672363281 0 902.8344
298.1961669921875 0 530.19574
300.6712646484375 0 693.75244
303.119140625 0 544.8653
305.6643981933594 0 993.4678
306.10791015625 0 915.7898
309.38848876953125 0 539.19196
311.2445983886719 0 1735.4987
315.1688537597656 0 644.9958
320.12237548828125 0 795.47284
320.1397399902344 0 8878.179
320.8721618652344 0 648.2989 y 5
321.14312744140625 0 1421.1339
321.20703125 0 1022.9173
321.22894287109375 0 5673.817
322.2328796386719 0 1104.9224
324.2168273925781 0 1127.6045
326.2195739746094 0 2799.6062
328.1351623535156 0 593.4142
332.7299499511719 0 2664.4087
333.2313537597656 0 1035.468
335.2361755371094 0 580.86346 y 8
336.2033996582031 0 10632.299
337.2060241699219 0 1460.8635
337.7211608886719 0 3769.1692
338.150390625 0 2871.2236
339.1532287597656 0 823.3594
339.2395324707031 0 11743.647
340.1520690917969 0 657.03534
340.2425231933594 0 2146.9146
346.72662353515625 0 9370.398
347.2276611328125 0 3757.1846
348.1344909667969 0 4578.332
349.1369934082031 0 1124.5723
351.2400207519531 0 1154.2831
353.6656494140625 0 1082.8505
354.1911315917969 0 698.65094
354.2138977050781 0 15687.018
355.21746826171875 0 1733.978
355.7315979003906 0 1578.1799
356.1475524902344 0 855.49097
356.2323913574219 0 733.54315
357.2499084472656 0 2590.1084
358.16204833984375 0 2930.7803
358.71343994140625 0 1267.7102
359.8710021972656 0 1438.551
360.6867370605469 0 610.17584
361.6593322753906 0 1499.5166
362.16168212890625 0 1018.16504
362.50653076171875 0 1358.2463
366.5583801269531 0 1901.0547 y 4
366.89337158203125 0 931.8378
367.1640930175781 0 1003.69354
368.1473083496094 0 1782.5254
368.1732482910156 0 1426.1587
369.69390869140625 0 1209.9042
371.64697265625 0 857.67786
373.2816467285156 0 5125.9204 y 11
374.9593505859375 0 861.3234
375.6560974121094 0 1067.2842
378.5097351074219 0 631.6574
379.2021789550781 0 872.8147
380.228515625 0 662.6957
382.2059326171875 0 1013.5374
382.4574279785156 0 621.1344
384.16619873046875 0 1161.524
385.1732482910156 0 4722.539
385.1973571777344 0 622.42615
385.513427734375 0 789.7437
385.8433837890625 0 965.1566
386.15679931640625 0 4156.009
386.1756896972656 0 911.6823
387.1599426269531 0 843.45
387.4681091308594 0 714.4062
389.1610412597656 0 4084.0415
389.71185302734375 0 657.2298
391.5177917480469 0 1942.4667
391.85137939453125 0 761.80786
392.18524169921875 0 793.22266
392.26593017578125 0 1779.688
393.7156066894531 0 1779.9149
393.9645080566406 0 6856.405
394.1510314941406 0 1961.2753
394.2149963378906 0 3961.59 y Ammonia loss 1
394.4664001464844 0 1030.8702
394.5267639160156 0 657.59174
395.1570129394531 0 703.1698
396.1716003417969 0 1641.5516
397.17388916015625 0 699.95953
397.8498840332031 0 1245.6478 b Ammonia loss 9
398.4698486328125 0 1113.624 y 1
398.72119140625 0 690.35406
399.7203063964844 0 1199.7178
400.96527099609375 0 775.0152
401.2590637207031 0 883.5532
402.22528076171875 0 2135.251
402.65826416015625 0 1145.4545
403.1846923828125 0 4321.905
403.23907470703125 0 1533.254
403.268798828125 0 983.3538
403.5243835449219 0 783.89185 b 9
403.57293701171875 0 3438.2769 y Ammonia loss 3
403.85870361328125 0 661.4034
403.9039611816406 0 1440.3134
404.1903381347656 0 1228.0181
404.23388671875 0 962.5885
405.1873779296875 0 2135.7544
405.6788330078125 0 658.98126
406.251220703125 0 1169.6171
408.2218017578125 0 1212.2876
408.471923828125 0 969.37537
408.72509765625 0 1415.7375
409.9124755859375 0 667.1464
410.27667236328125 0 4920.4287
410.56976318359375 0 742.71405
410.6656799316406 0 701.29767
410.9040222167969 0 752.1743
411.1780090332031 0 2959.1667
411.22283935546875 0 3263.0718
411.28021240234375 0 1368.9998
411.4717102050781 0 6056.9414
411.6640319824219 0 3902.3645
411.7218017578125 0 2403.8599
411.97369384765625 0 1661.6926
412.163818359375 0 2059.6558
412.2226257324219 0 894.89545
412.2742614746094 0 14824.142 y 6
412.7756042480469 0 6284.622
413.1658935546875 0 820.11554
413.2792663574219 0 1196.5873
413.8971862792969 0 785.75806
414.1825256347656 0 931.37103
414.2292175292969 0 1844.8657
414.6855773925781 0 898.42975
415.2559814453125 0 1712.4833
415.72467041015625 0 2041.9742 Precursor Water loss
415.7558288574219 0 937.75586
415.975341796875 0 2537.1064 Precursor Ammonia loss
417.1561279296875 0 968.3726
419.90081787109375 0 5063.742
420.2320251464844 0 6083.6216 Precursor
420.47821044921875 0 5440.2134
420.5687255859375 0 1501.271
420.7292175292969 0 2382.9502
421.1693115234375 0 1226.7123
421.32904052734375 0 1318.8944
422.27716064453125 0 1136.7345
423.2716979980469 0 805.57465
424.32806396484375 0 1133.772
424.7118225097656 0 1490.4443
425.2118835449219 0 818.5561
427.2432556152344 0 1152.776
428.1827697753906 0 803.3937 b 5
429.1884460449219 0 10157.09
430.1915283203125 0 2996.1667 b Ammonia loss 10
431.1805114746094 0 1645.6614
443.5593566894531 0 1014.15576
443.8937683105469 0 1236.6545
449.56634521484375 0 662.4264
451.22314453125 0 1582.9979
451.26611328125 0 1413.6661
451.7255859375 0 1629.1716
451.9200439453125 0 2529.691
452.2531433105469 0 1720.4397
452.3248596191406 0 934.7959
452.5876770019531 0 1247.6036
453.176025390625 0 1062.1989 b 2
455.2500915527344 0 1556.8014
456.2526550292969 0 820.6383
457.5943603515625 0 1078.1483
457.9266662597656 0 2374.6348 y Ammonia loss 2
458.25799560546875 0 838.42017
460.2305908203125 0 2442.7812
460.73284912109375 0 1157.3265
461.57037353515625 0 677.2698
462.2367858886719 0 869.5372
463.5987548828125 0 24030.324 y 2
463.93304443359375 0 18287.25
464.26727294921875 0 5708.248
464.6025390625 0 1184.4226
464.9342956542969 0 916.11
466.92510986328125 0 1685.0695
467.2986145019531 0 1183.6387
468.2941589355469 0 896.68695
470.3343200683594 0 21221.37 y 10
471.3372802734375 0 5100.0273
471.7975769042969 0 830.0806
471.91998291015625 0 651.836
472.25408935546875 0 753.2255
472.5930480957031 0 957.80334
473.26275634765625 0 2559.093
474.77972412109375 0 813.6746
475.5768737792969 0 1225.4723
476.1931457519531 0 4504.642
476.91168212890625 0 711.88495
477.1968688964844 0 879.32074
478.57354736328125 0 1199.5731 b 11
478.9062194824219 0 696.2311
480.8041687011719 0 2532.153 y 5
481.2540588378906 0 3970.165
481.3043518066406 0 2340.1968
481.5891418457031 0 3263.6836
481.9159851074219 0 1082.2559
482.2713623046875 0 1264.4954
482.30902099609375 0 1423.2332
482.7639465332031 0 736.6047
483.2590026855469 0 928.2092
483.7862548828125 0 2173.492
484.286865234375 0 1136.3538
487.2579650878906 0 2292.5518
487.5915832519531 0 1586.9392
488.22576904296875 0 2251.7883
490.2640075683594 0 1203.5099
490.6021728515625 0 1128.02
491.272216796875 0 1476.9928
494.2040710449219 0 1157.7463
495.9351806640625 0 933.1436
500.26531982421875 0 694.59296
504.2641296386719 0 908.7641
504.5934753417969 0 721.4853
505.26824951171875 0 2806.3347
505.603271484375 0 2162.3972
506.2359924316406 0 2072.1782
508.237060546875 0 1149.9501
509.6157531738281 0 1024.5206
510.2646789550781 0 1619.9259 b Water loss 12
510.5998840332031 0 2038.0385 b Ammonia loss 12
510.9300231933594 0 1104.6449
516.2686767578125 0 2305.2493 b 12
516.6016845703125 0 1944.7144
516.7411499023438 0 1356.6238
516.9381103515625 0 1088.737
517.2724609375 0 1424.1111
518.9487915039062 0 1018.92236
519.2843017578125 0 1689.4038
520.3629150390625 0 828.49524
521.2267456054688 0 897.9277
521.932373046875 0 804.0539
522.2722778320312 0 2772.7063
522.6061401367188 0 2042.573
524.6168212890625 0 736.825
524.95263671875 0 13110.84
525.2870483398438 0 9443.734 y Ammonia loss 1
525.6203002929688 0 3913.1367
525.9560546875 0 1785.9369
526.2050170898438 0 1227.7015
528.3137817382812 0 1154.2554
530.9559326171875 0 2016.1235 y 1
531.2108764648438 0 946.27386
531.291259765625 0 2974.552
538.2534790039062 0 6272.168
538.3734130859375 0 963.50146
539.2556762695312 0 2756.9934
539.3045043945312 0 2299.4956
539.8043823242188 0 1120.0013
540.326416015625 0 766.77875
541.3726196289062 0 4515.3154 y 9
542.3738403320312 0 1387.7461
543.2537231445312 0 1074.9928
543.753662109375 0 813.35565
547.8143310546875 0 1080.8009
549.3330688476562 0 7307.4697 y 4
549.8348999023438 0 4308.9517
550.3358154296875 0 2460.0813
551.2525024414062 0 849.12396
552.2607421875 0 1878.0226
554.2010498046875 0 744.4755
557.29736328125 0 1093.7592
566.2473754882812 0 7757.2666
567.249267578125 0 2601.269
569.264404296875 0 1573.0646 b 8
569.765869140625 0 1061.8159
570.2664184570312 0 976.17334
572.804931640625 0 3171.1775
573.3053588867188 0 1595.6196
577.3829956054688 0 3091.942
578.3837890625 0 1477.094
579.3621826171875 0 7027.6235
580.3638305664062 0 1716.2208
585.2561645507812 0 1758.4977
590.2949829101562 0 744.27686
595.3930053710938 0 2048.8887
596.3477172851562 0 937.03485
600.3355712890625 0 710.18787
604.8522338867188 0 3534.8804 y Ammonia loss 3
605.3524169921875 0 2034.3638
605.85107421875 0 1362.51
610.3226318359375 0 1140.9443
611.3261108398438 0 770.294
613.2492065429688 0 3182.4915
613.3589477539062 0 1501.9187 y 3
613.8650512695312 0 1246.3573
614.25048828125 0 1454.3577
614.3646240234375 0 1005.75745
615.3497314453125 0 932.7829
618.3526611328125 0 951.15485
620.341796875 0 1257.5786
620.841064453125 0 1021.0228
621.3431396484375 0 767.1117
627.3851928710938 0 825.9944
629.3478393554688 0 2605.7063
629.8472900390625 0 3431.6516
630.3516845703125 0 1227.8927
633.2947387695312 0 801.87164
635.298828125 0 3616.126
636.298583984375 0 1590.4971
637.3055419921875 0 840.46045
638.3534545898438 0 1540.804
638.8546142578125 0 1133.5338
653.3053588867188 0 4622.9434 b 10
654.3823852539062 0 1267.5042
664.4525756835938 0 1984.9126
664.8321533203125 0 752.8457
669.4672241210938 0 947.2325 y 8
674.436279296875 0 6072.457
675.4368896484375 0 3126.7136
682.2734375 0 1529.4746
692.4451904296875 0 7365.093
693.4487915039062 0 2323.017
694.8934936523438 0 3749.8386 y 2
695.3944702148438 0 4027.816
695.8976440429688 0 1242.367
710.4564819335938 0 2335.9988
711.465087890625 0 825.6002
716.4205932617188 0 1285.9795
718.2980346679688 0 914.85803 b 4
721.3601684570312 0 1678.3848
722.3128051757812 0 1000.0092
726.4876098632812 0 2316.6196 y 7
727.4903564453125 0 975.765
750.3106689453125 0 1867.5409
751.3065795898438 0 911.5128
757.3956298828125 0 1422.7639
768.3211669921875 0 1161.4341
805.5323486328125 0 1041.2013
811.493408203125 0 755.5813
819.3316650390625 0 1085.3439
823.5410766601562 0 11024.146 y 6
824.5433959960938 0 4266.674
825.5450439453125 0 1485.6576
827.35302734375 0 852.925
829.5042114257812 0 803.8439
831.94189453125 0 657.9022
848.4182739257812 0 872.0464
855.3565063476562 0 1102.8019 b 5
856.3656005859375 0 640.9196
901.4425048828125 0 1546.5845
919.4532470703125 0 1390.0732
942.5865478515625 0 805.92084
943.5919189453125 0 905.21606
960.5999145507812 0 1265.7496 y 5
1903.3800048828125 0 682.61865

Spectrum Details

|  |  |
| --- | --- |
| Matched peaks? Matched peaks The total absolute number of peaks matched. Additionally in brackets the total fraction of peaks matched and the total number of peaks is shown. | 51 (8.33% of 612) |
| FDR? FDR The false discovery rate estimated for this peptide. It is calculated by matching all theoretical fragments with a non-integer shift with the raw peaks for this spectrum. This is done with 40 different shifts. The resulting percentage is the average number of annotated peaks over the number of annotated peaks with the correct spectrum. | 4.48% |
| Satellite FDR? Satellite FDR See the FDR for details on its calculation. This satellite ion specific FDR only contains the satellite ions (d/w) for I/L/J positions. | - |
| PSM Score? PSM Score The PSM Score as given by Hecklib to this annotated spectrum. It is shown with three significant figures. | 159 |

#### Reverse Lookup? Reverse Lookup All places where this read could be placed.

| Group | Segment | Template | Template Part | Read Part | Score | Unique |
| --- | --- | --- | --- | --- | --- | --- |
| Homo sapiens Light Chain | IGLV | IGLV2-14 | [35..49] | [0..14] | 94 | False |
| Homo sapiens Light Chain | IGLV | IGLV2-23 | [35..49] | [0..14] | 94 | False |
| Homo sapiens Light Chain | IGLV | IGLV2-8 | [35..49] | [0..14] | 94 | False |
| Homo sapiens Light Chain | IGLV | IGLV2-11 | [35..49] | [0..14] | 94 | False |
| Homo sapiens Light Chain | IGLV | IGLV1-36 | [35..48] | [0..14] | 85 | False |
| Homo sapiens Light Chain | IGLV | IGLV1-51 | [34..48] | [0..14] | 85 | False |
| Homo sapiens Light Chain | IGLV | IGKV1-9 | [34..47] | [0..14] | 85 | False |
| Homo sapiens Light Chain | IGLV | IGKV1-33 | [34..47] | [0..14] | 85 | False |
| Homo sapiens Light Chain | IGLV | IGKV1-39 | [34..47] | [0..14] | 85 | False |
| Homo sapiens Light Chain | IGLV | IGKV1-8 | [34..47] | [0..14] | 85 | False |
| Homo sapiens Light Chain | IGLV | IGKV1D-13 | [34..47] | [0..14] | 85 | False |
| Homo sapiens Light Chain | IGLV | IGKV1-5 | [34..47] | [0..14] | 85 | False |
| Homo sapiens Light Chain | IGLV | IGKV1-6 | [34..47] | [0..14] | 85 | False |
| Homo sapiens Light Chain | IGLV | IGKV1-NL1 | [34..47] | [0..14] | 85 | False |
| Homo sapiens Light Chain | IGLV | IGKV1-12 | [34..47] | [0..14] | 85 | False |

| Recombined | Template Part | Read Part | Score | Unique |
| --- | --- | --- | --- | --- |
| REC-0-1\_002 | [35..49] | [0..14] | 112 | True |

#### Meta Information from Multiple reads

##### Number of combined reads

19

##### Intensity

0.8166

##### TotalArea

4.162E+09

##### Changes to the peptide sequence

SWYQHHPGKAPKJJ

L→JNo support for either Leucine or Isoleucine based on side chain ions (Position: 14)

L→JNo support for either Leucine or Isoleucine based on side chain ions (Position: 13)

#### Positional Score

Copy Data

##### Positional Score (TSV)

###### Preview

```
Loading example...
```

*Click on the button to copy the data to your clipboard.*

10012345678910111213

Label Value
"0" 0.879
"1" 0.876
"2" 0.879
"3" 0.868
"4" 0.883
"5" 0.886
"6" 0.882
"7" 0.877
"8" 0.815
"9" 0.83
"10" 0.859
"11" 0.878
"12" 0.892
"13" 0.894

#### Meta Information from PEAKS

##### Scan Identifier

F1:5528

##### Original sequence

S

W

Y

Q

H

H

P

G

K

A

P

K

L

L

##### Posttranslational Modifications

##### Source File

D:\separate\_stitch\_analyses\xle-disambiguation\raw\20210323\_F1\_UM1\_Peng0013\_SA\_F59\_ingel\_3ug\_ELA.raw

##### Fraction

1

##### Scan Feature

F1:6395

##### De Novo Score

99

##### ConfidenceScore

99

### m/z

554.636

##### Mass

1660.8838

##### Charge

3

##### Retention Time

28.16

##### Predicted Retention Time

-

##### Area

3.634E+08

##### Parts Per Million

1.5

##### Fragmentation mode

HCD

##### Originating file

01 D:\separate\_stitch\_analyses\xle-disambiguation\20210325\_F59\_3ug\_DENOVO\_12.csv

#### Meta Information from PEAKS

##### Scan Identifier

F1:5306

##### Original sequence

S

W

Y

Q

H

H

P

G

K

A

P

K

L

L

##### Posttranslational Modifications

##### Source File

D:\separate\_stitch\_analyses\xle-disambiguation\raw\20210323\_F1\_UM1\_Peng0013\_SA\_F59\_ingel\_3ug\_ELA.raw

##### Fraction

1

##### Scan Feature

F1:6395

##### De Novo Score

99

##### ConfidenceScore

99

### m/z

554.636

##### Mass

1660.8838

##### Charge

3

##### Retention Time

28.16

##### Predicted Retention Time

-

##### Area

3.634E+08

##### Parts Per Million

1.5

##### Fragmentation mode

HCD

##### Originating file

01 D:\separate\_stitch\_analyses\xle-disambiguation\20210325\_F59\_3ug\_DENOVO\_12.csv

#### Meta Information from PEAKS

##### Scan Identifier

F1:5459

##### Original sequence

S

W

Y

Q

H

H

P

G

K

A

P

K

L

L

##### Posttranslational Modifications

##### Source File

D:\separate\_stitch\_analyses\xle-disambiguation\raw\20210323\_F1\_UM1\_Peng0013\_SA\_F59\_ingel\_3ug\_ELA.raw

##### Fraction

1

##### Scan Feature

F1:6395

##### De Novo Score

99

##### ConfidenceScore

99

### m/z

554.636

##### Mass

1660.8838

##### Charge

3

##### Retention Time

28.16

##### Predicted Retention Time

-

##### Area

3.634E+08

##### Parts Per Million

1.5

##### Fragmentation mode

ETHCD

##### Originating file

01 D:\separate\_stitch\_analyses\xle-disambiguation\20210325\_F59\_3ug\_DENOVO\_12.csv

#### Meta Information from PEAKS

##### Scan Identifier

F1:5375

##### Original sequence

S

W

Y

Q

H

H

P

G

K

A

P

K

L

L

##### Posttranslational Modifications

##### Source File

D:\separate\_stitch\_analyses\xle-disambiguation\raw\20210323\_F1\_UM1\_Peng0013\_SA\_F59\_ingel\_3ug\_ELA.raw

##### Fraction

1

##### Scan Feature

F1:6395

##### De Novo Score

99

##### ConfidenceScore

99

### m/z

554.636

##### Mass

1660.8838

##### Charge

3

##### Retention Time

28.16

##### Predicted Retention Time

-

##### Area

3.634E+08

##### Parts Per Million

1.5

##### Fragmentation mode

ETHCD

##### Originating file

01 D:\separate\_stitch\_analyses\xle-disambiguation\20210325\_F59\_3ug\_DENOVO\_12.csv

#### Meta Information from PEAKS

##### Scan Identifier

F1:4749

##### Original sequence

S

W

+15.99

Y

Q

H

H

P

G

K

A

P

K

L

L

##### Posttranslational Modifications

Oxidation (HW)

##### Source File

D:\separate\_stitch\_analyses\xle-disambiguation\raw\20210323\_F1\_UM1\_Peng0013\_SA\_F59\_ingel\_3ug\_ELA.raw

##### Fraction

1

##### Scan Feature

F1:6624

##### De Novo Score

98

##### ConfidenceScore

98

### m/z

559.9669

##### Mass

1676.8787

##### Charge

3

##### Retention Time

25.33

##### Predicted Retention Time

-

##### Area

3.862E+06

##### Parts Per Million

0.2

##### Fragmentation mode

ETHCD

##### Originating file

01 D:\separate\_stitch\_analyses\xle-disambiguation\20210325\_F59\_3ug\_DENOVO\_12.csv

#### Meta Information from PEAKS

##### Scan Identifier

F1:5672

##### Original sequence

S

W

Y

Q

H

H

P

G

K

A

P

K

L

L

##### Posttranslational Modifications

##### Source File

D:\separate\_stitch\_analyses\xle-disambiguation\raw\20210323\_F1\_UM1\_Peng0013\_SA\_F59\_ingel\_3ug\_ELA.raw

##### Fraction

1

##### Scan Feature

F1:6395

##### De Novo Score

98

##### ConfidenceScore

98

### m/z

554.636

##### Mass

1660.8838

##### Charge

3

##### Retention Time

28.16

##### Predicted Retention Time

-

##### Area

3.634E+08

##### Parts Per Million

1.5

##### Fragmentation mode

ETHCD

##### Originating file

01 D:\separate\_stitch\_analyses\xle-disambiguation\20210325\_F59\_3ug\_DENOVO\_12.csv

#### Meta Information from PEAKS

##### Scan Identifier

F1:5553

##### Original sequence

S

W

Y

Q

H

H

P

G

K

A

P

K

L

L

##### Posttranslational Modifications

##### Source File

D:\separate\_stitch\_analyses\xle-disambiguation\raw\20210323\_F1\_UM1\_Peng0013\_SA\_F59\_ingel\_3ug\_ELA.raw

##### Fraction

1

##### Scan Feature

F1:1150

##### De Novo Score

98

##### ConfidenceScore

98

### m/z

416.2288

##### Mass

1660.8838

##### Charge

4

##### Retention Time

28.16

##### Predicted Retention Time

-

##### Area

3.929E+08

##### Parts Per Million

1.4

##### Fragmentation mode

HCD

##### Originating file

01 D:\separate\_stitch\_analyses\xle-disambiguation\20210325\_F59\_3ug\_DENOVO\_12.csv

#### Meta Information from PEAKS

##### Scan Identifier

F1:5359

##### Original sequence

S

W

Y

Q

H

H

P

G

K

A

P

K

L

L

##### Posttranslational Modifications

##### Source File

D:\separate\_stitch\_analyses\xle-disambiguation\raw\20210323\_F1\_UM1\_Peng0013\_SA\_F59\_ingel\_3ug\_ELA.raw

##### Fraction

1

##### Scan Feature

F1:1150

##### De Novo Score

97

##### ConfidenceScore

97

### m/z

416.2288

##### Mass

1660.8838

##### Charge

4

##### Retention Time

28.16

##### Predicted Retention Time

-

##### Area

3.929E+08

##### Parts Per Million

1.4

##### Fragmentation mode

HCD

##### Originating file

01 D:\separate\_stitch\_analyses\xle-disambiguation\20210325\_F59\_3ug\_DENOVO\_12.csv

#### Meta Information from PEAKS

##### Scan Identifier

F1:5748

##### Original sequence

S

W

Y

Q

H

H

P

G

K

A

P

K

L

L

##### Posttranslational Modifications

##### Source File

D:\separate\_stitch\_analyses\xle-disambiguation\raw\20210323\_F1\_UM1\_Peng0013\_SA\_F59\_ingel\_3ug\_ELA.raw

##### Fraction

1

##### Scan Feature

-

##### De Novo Score

97

##### ConfidenceScore

97

### m/z

554.6357

##### Mass

1660.8838

##### Charge

3

##### Retention Time

30.76

##### Predicted Retention Time

-

##### Area

0

##### Parts Per Million

1

##### Fragmentation mode

HCD

##### Originating file

01 D:\separate\_stitch\_analyses\xle-disambiguation\20210325\_F59\_3ug\_DENOVO\_12.csv

#### Meta Information from PEAKS

##### Scan Identifier

F1:5608

##### Original sequence

S

W

Y

Q

H

H

P

G

K

A

P

K

L

L

##### Posttranslational Modifications

##### Source File

D:\separate\_stitch\_analyses\xle-disambiguation\raw\20210323\_F1\_UM1\_Peng0013\_SA\_F59\_ingel\_3ug\_ELA.raw

##### Fraction

1

##### Scan Feature

F1:6395

##### De Novo Score

97

##### ConfidenceScore

97

### m/z

554.636

##### Mass

1660.8838

##### Charge

3

##### Retention Time

28.16

##### Predicted Retention Time

-

##### Area

3.634E+08

##### Parts Per Million

1.5

##### Fragmentation mode

ETHCD

##### Originating file

01 D:\separate\_stitch\_analyses\xle-disambiguation\20210325\_F59\_3ug\_DENOVO\_12.csv

#### Meta Information from PEAKS

##### Scan Identifier

F1:4459

##### Original sequence

S

W

+15.99

Y

Q

H

H

P

G

K

A

P

K

L

L

##### Posttranslational Modifications

Oxidation (HW)

##### Source File

D:\separate\_stitch\_analyses\xle-disambiguation\raw\20210323\_F1\_UM1\_Peng0013\_SA\_F59\_ingel\_3ug\_ELA.raw

##### Fraction

1

##### Scan Feature

F1:1277

##### De Novo Score

97

##### ConfidenceScore

97

### m/z

420.2282

##### Mass

1676.8787

##### Charge

4

##### Retention Time

23.6

##### Predicted Retention Time

-

##### Area

4.922E+06

##### Parts Per Million

3

##### Fragmentation mode

HCD

##### Originating file

01 D:\separate\_stitch\_analyses\xle-disambiguation\20210325\_F59\_3ug\_DENOVO\_12.csv

#### Meta Information from PEAKS

##### Scan Identifier

F1:5266

##### Original sequence

S

W

+15.99

Y

Q

H

H

P

G

K

A

P

K

L

L

##### Posttranslational Modifications

Oxidation (HW)

##### Source File

D:\separate\_stitch\_analyses\xle-disambiguation\raw\20210323\_F1\_UM1\_Peng0013\_SA\_F59\_ingel\_3ug\_ELA.raw

##### Fraction

1

##### Scan Feature

-

##### De Novo Score

97

##### ConfidenceScore

97

### m/z

420.2263

##### Mass

1676.8787

##### Charge

4

##### Retention Time

28.13

##### Predicted Retention Time

-

##### Area

0

##### Fragmentation mode

HCD

##### Originating file

01 D:\separate\_stitch\_analyses\xle-disambiguation\20210325\_F59\_3ug\_DENOVO\_12.csv

#### Meta Information from PEAKS

##### Scan Identifier

F1:5331

##### Original sequence

S

W

+15.99

Y

Q

H

H

P

G

K

A

P

K

L

L

##### Posttranslational Modifications

Oxidation (HW)

##### Source File

D:\separate\_stitch\_analyses\xle-disambiguation\raw\20210323\_F1\_UM1\_Peng0013\_SA\_F59\_ingel\_3ug\_ELA.raw

##### Fraction

1

##### Scan Feature

-

##### De Novo Score

97

##### ConfidenceScore

97

### m/z

420.227

##### Mass

1676.8787

##### Charge

4

##### Retention Time

28.49

##### Predicted Retention Time

-

##### Area

0

##### Parts Per Million

0

##### Fragmentation mode

HCD

##### Originating file

01 D:\separate\_stitch\_analyses\xle-disambiguation\20210325\_F59\_3ug\_DENOVO\_12.csv

#### Meta Information from PEAKS

##### Scan Identifier

F1:5226

##### Original sequence

S

W

Y

Q

H

H

P

G

K

A

P

K

L

L

##### Posttranslational Modifications

##### Source File

D:\separate\_stitch\_analyses\xle-disambiguation\raw\20210323\_F1\_UM1\_Peng0013\_SA\_F59\_ingel\_3ug\_ELA.raw

##### Fraction

1

##### Scan Feature

-

##### De Novo Score

97

##### ConfidenceScore

97

### m/z

416.2287

##### Mass

1660.8838

##### Charge

4

##### Retention Time

27.91

##### Predicted Retention Time

-

##### Area

0

##### Parts Per Million

1.1

##### Fragmentation mode

HCD

##### Originating file

01 D:\separate\_stitch\_analyses\xle-disambiguation\20210325\_F59\_3ug\_DENOVO\_12.csv

#### Meta Information from PEAKS

##### Scan Identifier

F1:5622

##### Original sequence

S

W

Y

Q

H

H

P

G

K

A

P

K

L

L

##### Posttranslational Modifications

##### Source File

D:\separate\_stitch\_analyses\xle-disambiguation\raw\20210323\_F1\_UM1\_Peng0013\_SA\_F59\_ingel\_3ug\_ELA.raw

##### Fraction

1

##### Scan Feature

F1:1150

##### De Novo Score

96

##### ConfidenceScore

96

### m/z

416.2288

##### Mass

1660.8838

##### Charge

4

##### Retention Time

28.16

##### Predicted Retention Time

-

##### Area

3.929E+08

##### Parts Per Million

1.4

##### Fragmentation mode

HCD

##### Originating file

01 D:\separate\_stitch\_analyses\xle-disambiguation\20210325\_F59\_3ug\_DENOVO\_12.csv

#### Meta Information from PEAKS

##### Scan Identifier

F1:5292

##### Original sequence

S

W

Y

Q

H

H

P

G

K

A

P

K

L

L

##### Posttranslational Modifications

##### Source File

D:\separate\_stitch\_analyses\xle-disambiguation\raw\20210323\_F1\_UM1\_Peng0013\_SA\_F59\_ingel\_3ug\_ELA.raw

##### Fraction

1

##### Scan Feature

F1:1150

##### De Novo Score

96

##### ConfidenceScore

96

### m/z

416.2288

##### Mass

1660.8838

##### Charge

4

##### Retention Time

28.16

##### Predicted Retention Time

-

##### Area

3.929E+08

##### Parts Per Million

1.4

##### Fragmentation mode

HCD

##### Originating file

01 D:\separate\_stitch\_analyses\xle-disambiguation\20210325\_F59\_3ug\_DENOVO\_12.csv

#### Meta Information from PEAKS

##### Scan Identifier

F1:4737

##### Original sequence

S

W

+15.99

Y

Q

H

H

P

G

K

A

P

K

L

L

##### Posttranslational Modifications

Oxidation (HW)

##### Source File

D:\separate\_stitch\_analyses\xle-disambiguation\raw\20210323\_F1\_UM1\_Peng0013\_SA\_F59\_ingel\_3ug\_ELA.raw

##### Fraction

1

##### Scan Feature

F1:1276

##### De Novo Score

96

##### ConfidenceScore

96

### m/z

420.2271

##### Mass

1676.8787

##### Charge

4

##### Retention Time

25.36

##### Predicted Retention Time

-

##### Area

4.079E+06

##### Parts Per Million

0.5

##### Fragmentation mode

HCD

##### Originating file

01 D:\separate\_stitch\_analyses\xle-disambiguation\20210325\_F59\_3ug\_DENOVO\_12.csv

#### Meta Information from PEAKS

##### Scan Identifier

F1:5686

##### Original sequence

S

W

Y

Q

H

H

P

G

K

A

P

K

L

L

##### Posttranslational Modifications

##### Source File

D:\separate\_stitch\_analyses\xle-disambiguation\raw\20210323\_F1\_UM1\_Peng0013\_SA\_F59\_ingel\_3ug\_ELA.raw

##### Fraction

1

##### Scan Feature

F1:1150

##### De Novo Score

95

##### ConfidenceScore

95

### m/z

416.2288

##### Mass

1660.8838

##### Charge

4

##### Retention Time

28.16

##### Predicted Retention Time

-

##### Area

3.929E+08

##### Parts Per Million

1.4

##### Fragmentation mode

HCD

##### Originating file

01 D:\separate\_stitch\_analyses\xle-disambiguation\20210325\_F59\_3ug\_DENOVO\_12.csv

#### Meta Information from PEAKS

##### Scan Identifier

F1:4404

##### Original sequence

S

W

+15.99

Y

Q

H

H

P

G

K

A

P

K

L

L

##### Posttranslational Modifications

Oxidation (HW)

##### Source File

D:\separate\_stitch\_analyses\xle-disambiguation\raw\20210323\_F1\_UM1\_Peng0013\_SA\_F59\_ingel\_3ug\_ELA.raw

##### Fraction

1

##### Scan Feature

F1:1277

##### De Novo Score

95

##### ConfidenceScore

95

### m/z

420.2282

##### Mass

1676.8787

##### Charge

4

##### Retention Time

23.6

##### Predicted Retention Time

-

##### Area

4.922E+06

##### Parts Per Million

3

##### Fragmentation mode

HCD

##### Originating file

01 D:\separate\_stitch\_analyses\xle-disambiguation\20210325\_F59\_3ug\_DENOVO\_12.csv
