## Supplementary material for "A handle on mass coincidence errors in *de novo* sequencing of antibodies by bottom-up proteomics": Combined_015.html

Details Combined\_015 | Stitch OverviewUndefined

### Read Combined\_015

#### Sequence (length=11)

FVVFGGGTKJT

#### Spectrum 8344? Spectrum 8344 The raw spectrum of this peptide as annotated by Hecklib. The fragments are coloured according to ion type (see legend). Any peaks with a star '\*' as text can be hovered over to see the full details, first the ion type second the mass shift type. By hovering over the amino acids in the peptide or ions in the legend the corresponding peaks are highlighted. By toggling the 'Unassigned' label you can turn the background (unassigned) peaks on or off in the plot. By updating the slider in the Ion legend you can update the spectrum to only show the top X% of the peaks with labels. The top X% means any peak that is within X% of the highest intensity. By dragging in the spectrum you can zoom in to a specific part of the spectrum and use 'Zoom Out' to get back to the original zoom level. The annotation of the spectrum is based on the given sequence in the peptides file and is done with different software so inconsistencies are likely. The peaks are annotated based on the given sequence, with 20 ppm tolerance.

Copy Data

##### Spectrum 8344 (TSV)

###### Preview

```
Loading example...
```

*Click on the button to copy the data to your clipboard.*

Mz MinMz MaxIntensity Max

WidthHeightPeptide font sizePeptide stroke widthSpectrum font sizeSpectrum stroke widthCompact peptide

Ion legend

wxyz

abcd

OtherUnassignedIon^Charge^_Position_Show for top:%

FVVFGGGTKJT

02.82e+45.63e+48.45e+41.13e+5

Zoom Out

y^+1^_1_y^+1^_2_y^+1^_2_y^+1^_3_y^+2^_8_y^+2^_9_y^+1^_4_c^+2^_9_y^+1^_4_y^+2^_10_c^+2^_10_c^+2^_10_y^+1^_5_z^+1^_6_c^+1^_5_y^+1^_6_y^+1^_7_z^+1^_7_c^+1^_6_y^+1^_7_c^+1^_7_z^+1^_8_y^+1^_8_c^+1^_8_y^+1^_8_c^+1^_8_z^+1^_9_y^+1^_9_c^+1^_9_c^+1^_9_z^+1^_10_y^+1^_10_c^+1^_10_c^+1^_10_

0851170225523403

Fragment Matches Table

Show background peaks

| Position | Ion type | Intensity | mz Theoretical | mz Error (Th) | mz Error (ppm) | Charge | Series Number |
| --- | --- | --- | --- | --- | --- | --- | --- |
| 11 | y | 3304 | 120.1 | 0.000139 | 1.157 | +1 | 1 |
| - | - | 5498 | 120.1 | - | - | 0 | - |
| - | - | 472.5 | 121.1 | - | - | 0 | - |
| - | - | 430.1 | 125.5 | - | - | 0 | - |
| - | - | 357.4 | 127.4 | - | - | 0 | - |
| - | - | 1412 | 129.1 | - | - | 0 | - |
| - | - | 392.1 | 132.2 | - | - | 0 | - |
| - | - | 1002 | 133.1 | - | - | 0 | - |
| - | - | 462.6 | 169.8 | - | - | 0 | - |
| - | - | 1658 | 173.4 | - | - | 0 | - |
| - | - | 1165 | 177.1 | - | - | 0 | - |
| - | - | 511.6 | 202.1 | - | - | 0 | - |
| - | - | 817.9 | 207.1 | - | - | 0 | - |
| 10 | y | 1179 | 215.1 | 0.0001487 | 0.691 | +1 | 2 |
| - | - | 1405 | 219.1 | - | - | 0 | - |
| - | - | 2.495E+04 | 219.1 | - | - | 0 | - |
| - | - | 3127 | 220.2 | - | - | 0 | - |
| 10 | y | 2668 | 233.1 | 0.0001238 | 0.5308 | +1 | 2 |
| - | - | 3.555E+04 | 247.1 | - | - | 0 | - |
| - | - | 5594 | 248.1 | - | - | 0 | - |
| - | - | 2042 | 300.2 | - | - | 0 | - |
| - | - | 1626 | 301.2 | - | - | 0 | - |
| - | - | 1362 | 332.2 | - | - | 0 | - |
| - | - | 1184 | 344.2 | - | - | 0 | - |
| - | - | 532.2 | 344.9 | - | - | 0 | - |
| - | - | 1.499E+04 | 346.2 | - | - | 0 | - |
| - | - | 526.1 | 346.2 | - | - | 0 | - |
| - | - | 3132 | 347.2 | - | - | 0 | - |
| - | - | 970 | 358.2 | - | - | 0 | - |
| 9 | y | 1256 | 361.2 | 0.0006011 | 1.664 | +1 | 3 |
| - | - | 608.9 | 383.2 | - | - | 0 | - |
| 4 | y | 1413 | 390.7 | 2.783E-05 | 0.07122 | +2 | 8 |
| - | - | 713.4 | 391.2 | - | - | 0 | - |
| - | - | 992.4 | 394.2 | - | - | 0 | - |
| - | - | 1310 | 401.2 | - | - | 0 | - |
| - | - | 1197 | 401.2 | - | - | 0 | - |
| - | - | 566.3 | 410.2 | - | - | 0 | - |
| - | - | 948.6 | 433.2 | - | - | 0 | - |
| - | - | 575.4 | 436.9 | - | - | 0 | - |
| 3 | y | 4205 | 440.3 | 0.0003111 | 0.7067 | +2 | 9 |
| - | - | 1486 | 440.8 | - | - | 0 | - |
| - | - | 1151 | 444.3 | - | - | 0 | - |
| - | - | 771.3 | 445.2 | - | - | 0 | - |
| 8 | y | 3862 | 445.3 | 8.15E-05 | 0.183 | +1 | 4 |
| 9 | c | 1302 | 447.2 | 2.953E-05 | 0.06602 | +2 | 9 |
| - | - | 1846 | 447.3 | - | - | 0 | - |
| - | - | 641.6 | 447.7 | - | - | 0 | - |
| - | - | 700.6 | 451.2 | - | - | 0 | - |
| - | - | 636.7 | 457.3 | - | - | 0 | - |
| - | - | 926.5 | 459.3 | - | - | 0 | - |
| 8 | y | 1038 | 462.3 | 0.002117 | 4.579 | +1 | 4 |
| - | - | 1623 | 465.3 | - | - | 0 | - |
| - | - | 652 | 466.3 | - | - | 0 | - |
| - | - | 1123 | 470.3 | - | - | 0 | - |
| 2 | y | 1003 | 489.8 | 0.006509 | 13.29 | +2 | 10 |
| - | - | 902.7 | 490.3 | - | - | 0 | - |
| - | - | 1793 | 493.3 | - | - | 0 | - |
| - | - | 936.4 | 494.3 | - | - | 0 | - |
| - | - | 564.6 | 495.3 | - | - | 0 | - |
| - | - | 1677 | 500.3 | - | - | 0 | - |
| 10 | c | 703.6 | 503.3 | 0.001214 | 2.412 | +2 | 10 |
| 10 | c | 1665 | 503.8 | 0.000407 | 0.8079 | +2 | 10 |
| - | - | 1.15E+04 | 504.3 | - | - | 0 | - |
| - | - | 9973 | 504.3 | - | - | 0 | - |
| - | - | 3225 | 505.3 | - | - | 0 | - |
| - | - | 2323 | 505.3 | - | - | 0 | - |
| - | - | 2316 | 514.3 | - | - | 0 | - |
| - | - | 913.8 | 516.3 | - | - | 0 | - |
| - | - | 922.5 | 518.3 | - | - | 0 | - |
| 7 | y | 2958 | 519.3 | 0.0003343 | 0.6438 | +1 | 5 |
| - | - | 1075 | 520.3 | - | - | 0 | - |
| - | - | 625.1 | 522.3 | - | - | 0 | - |
| - | - | 3461 | 536.3 | - | - | 0 | - |
| - | - | 1726 | 546.3 | - | - | 0 | - |
| - | - | 1545 | 547.3 | - | - | 0 | - |
| - | - | 1630 | 548.3 | - | - | 0 | - |
| - | - | 825.9 | 550.3 | - | - | 0 | - |
| - | - | 784.8 | 554.3 | - | - | 0 | - |
| - | - | 1057 | 554.8 | - | - | 0 | - |
| 6 | z | 1.144E+04 | 560.3 | 8.325E-05 | 0.1486 | +1 | 6 |
| - | - | 3.573E+04 | 561.3 | - | - | 0 | - |
| - | - | 570.6 | 562.2 | - | - | 0 | - |
| - | - | 8083 | 562.3 | - | - | 0 | - |
| - | - | 3625 | 563.3 | - | - | 0 | - |
| - | - | 1656 | 563.8 | - | - | 0 | - |
| - | - | 4174 | 564.3 | - | - | 0 | - |
| - | - | 652.4 | 564.3 | - | - | 0 | - |
| 5 | c | 4543 | 567.3 | 0.0001566 | 0.2761 | +1 | 5 |
| - | - | 1192 | 568.3 | - | - | 0 | - |
| - | - | 2574 | 575.3 | - | - | 0 | - |
| 6 | y | 1.741E+04 | 576.3 | 0.0005578 | 0.9679 | +1 | 6 |
| - | - | 3681 | 577.3 | - | - | 0 | - |
| - | - | 591.8 | 578.3 | - | - | 0 | - |
| - | - | 846.7 | 607.3 | - | - | 0 | - |
| 5 | y | 1209 | 615.3 | 0.0006892 | 1.12 | +1 | 7 |
| 5 | z | 8655 | 617.3 | 6.26E-05 | 0.1014 | +1 | 7 |
| - | - | 1.96E+04 | 618.3 | - | - | 0 | - |
| - | - | 6591 | 619.3 | - | - | 0 | - |
| - | - | 1236 | 620.4 | - | - | 0 | - |
| - | - | 4707 | 623.3 | - | - | 0 | - |
| 6 | c | 4996 | 624.4 | 0.001715 | 2.747 | +1 | 6 |
| - | - | 1836 | 625.4 | - | - | 0 | - |
| - | - | 1.875E+04 | 632.3 | - | - | 0 | - |
| 5 | y | 4.087E+04 | 633.4 | 0.001453 | 2.294 | +1 | 7 |
| - | - | 1.118E+04 | 634.4 | - | - | 0 | - |
| - | - | 2181 | 635.4 | - | - | 0 | - |
| - | - | 705.6 | 643.4 | - | - | 0 | - |
| - | - | 1330 | 647.3 | - | - | 0 | - |
| - | - | 635.2 | 657.5 | - | - | 0 | - |
| - | - | 2848 | 661.4 | - | - | 0 | - |
| - | - | 1602 | 663.4 | - | - | 0 | - |
| - | - | 5333 | 680.4 | - | - | 0 | - |
| 7 | c | 2682 | 681.4 | 0.003098 | 4.547 | +1 | 7 |
| 4 | z | 777.7 | 746.4 | 0.001475 | 1.977 | +1 | 8 |
| - | - | 2972 | 760.4 | - | - | 0 | - |
| - | - | 1575 | 761.4 | - | - | 0 | - |
| 4 | y | 1061 | 763.4 | 0.00429 | 5.62 | +1 | 8 |
| 8 | c | 1.883E+04 | 764.4 | 0.003347 | 4.378 | +1 | 8 |
| - | - | 1.799E+04 | 765.4 | - | - | 0 | - |
| - | - | 4899 | 766.4 | - | - | 0 | - |
| - | - | 756.8 | 767.4 | - | - | 0 | - |
| - | - | 4320 | 779.4 | - | - | 0 | - |
| 4 | y | 3.106E+04 | 780.4 | 0.0009579 | 1.227 | +1 | 8 |
| - | - | 1.588E+04 | 781.4 | - | - | 0 | - |
| 8 | c | 5.417E+04 | 782.4 | 0.0003614 | 0.4619 | +1 | 8 |
| - | - | 2.167E+04 | 783.4 | - | - | 0 | - |
| - | - | 4638 | 784.4 | - | - | 0 | - |
| - | - | 1715 | 785.4 | - | - | 0 | - |
| - | - | 633.7 | 807.4 | - | - | 0 | - |
| - | - | 772.7 | 817.5 | - | - | 0 | - |
| 3 | z | 1.282E+04 | 863.5 | 0.0001717 | 0.1988 | +1 | 9 |
| - | - | 5733 | 864.5 | - | - | 0 | - |
| - | - | 2397 | 865.5 | - | - | 0 | - |
| - | - | 1189 | 866.5 | - | - | 0 | - |
| - | - | 679.7 | 875.5 | - | - | 0 | - |
| - | - | 935.9 | 876.5 | - | - | 0 | - |
| 3 | y | 2.705E+04 | 879.5 | 0.001073 | 1.221 | +1 | 9 |
| - | - | 1.367E+04 | 880.5 | - | - | 0 | - |
| - | - | 3785 | 881.5 | - | - | 0 | - |
| 9 | c | 4927 | 893.5 | 0.001087 | 1.217 | +1 | 9 |
| - | - | 3135 | 894.5 | - | - | 0 | - |
| 9 | c | 3.339E+04 | 910.5 | 0.001025 | 1.126 | +1 | 9 |
| - | - | 1.988E+04 | 911.5 | - | - | 0 | - |
| - | - | 357.8 | 911.6 | - | - | 0 | - |
| - | - | 4983 | 912.5 | - | - | 0 | - |
| - | - | 1104 | 936.5 | - | - | 0 | - |
| 2 | z | 9416 | 962.5 | 0.001081 | 1.123 | +1 | 10 |
| - | - | 4523 | 963.5 | - | - | 0 | - |
| - | - | 1514 | 964.6 | - | - | 0 | - |
| 2 | y | 1495 | 978.6 | 0.002656 | 2.714 | +1 | 10 |
| - | - | 1.204E+04 | 979.6 | - | - | 0 | - |
| - | - | 7121 | 980.6 | - | - | 0 | - |
| - | - | 1542 | 981.6 | - | - | 0 | - |
| 10 | c | 3976 | 1007 | 0.001594 | 1.584 | +1 | 10 |
| - | - | 2042 | 1008 | - | - | 0 | - |
| - | - | 3683 | 1009 | - | - | 0 | - |
| - | - | 1866 | 1010 | - | - | 0 | - |
| 10 | c | 5.19E+04 | 1024 | 0.001349 | 1.318 | +1 | 10 |
| - | - | 3.167E+04 | 1025 | - | - | 0 | - |
| - | - | 1.001E+04 | 1026 | - | - | 0 | - |
| - | - | 1662 | 1027 | - | - | 0 | - |
| - | - | 1586 | 1049 | - | - | 0 | - |
| - | - | 1182 | 1050 | - | - | 0 | - |
| - | - | 669.4 | 1053 | - | - | 0 | - |
| - | - | 1164 | 1054 | - | - | 0 | - |
| - | - | 675.6 | 1055 | - | - | 0 | - |
| - | - | 2235 | 1067 | - | - | 0 | - |
| - | - | 868.2 | 1068 | - | - | 0 | - |
| - | - | 8047 | 1071 | - | - | 0 | - |
| - | - | 5789 | 1072 | - | - | 0 | - |
| - | - | 1476 | 1073 | - | - | 0 | - |
| - | - | 894.4 | 1081 | - | - | 0 | - |
| - | - | 2424 | 1099 | - | - | 0 | - |
| - | - | 2062 | 1100 | - | - | 0 | - |
| - | - | 1235 | 1109 | - | - | 0 | - |
| - | - | 1.115E+05 | 1110 | - | - | 0 | - |
| - | - | 7.673E+04 | 1111 | - | - | 0 | - |
| - | - | 2.71E+04 | 1112 | - | - | 0 | - |
| - | - | 3238 | 1113 | - | - | 0 | - |
| - | - | 663.3 | 1124 | - | - | 0 | - |
| - | - | 1714 | 1125 | - | - | 0 | - |
| - | - | 4.751E+04 | 1126 | - | - | 0 | - |
| - | - | 9.181E+04 | 1127 | - | - | 0 | - |
| - | - | 4.963E+04 | 1128 | - | - | 0 | - |
| - | - | 4828 | 1129 | - | - | 0 | - |
| - | - | 1.305E+04 | 1129 | - | - | 0 | - |
| - | - | 3995 | 1130 | - | - | 0 | - |
| - | - | 635.8 | 1142 | - | - | 0 | - |
| - | - | 806.9 | 1480 | - | - | 0 | - |
| - | - | 657.1 | 1566 | - | - | 0 | - |
| - | - | 624.6 | 2007 | - | - | 0 | - |
| - | - | 612.7 | 3239 | - | - | 0 | - |
| - | - | 689.7 | 3369 | - | - | 0 | - |

m/z Charge Intensity FragmentType MassShift Position
120.06565856933594 0 3303.7593 y 10
120.0809326171875 0 5498.086
121.08419799804688 0 472.49213
125.4717025756836 0 430.12363
127.35767364501953 0 357.35168
129.10231018066406 0 1412.1506
132.1802978515625 0 392.0715
133.08599853515625 0 1002.26917
169.83399963378906 0 462.5703
173.43905639648438 0 1658.4547
177.11270141601562 0 1164.7604
202.05116271972656 0 511.63663
207.076171875 0 817.87494
215.1388702392578 0 1179.2496 y Water loss 9
219.135009765625 0 1405.245
219.1492156982422 0 24954.56
220.1526336669922 0 3127.1714
233.1494598388672 0 2668.4038 y 9
247.1441192626953 0 35550.953
248.1473846435547 0 5593.625
300.19171142578125 0 2042.266
301.19140625 0 1626.2378
332.21856689453125 0 1362.098
344.18115234375 0 1183.6216
344.89166259765625 0 532.20764
346.21234130859375 0 14993.988
346.2310485839844 0 526.072
347.21563720703125 0 3131.5837
358.19586181640625 0 970.00146
361.2451477050781 0 1255.7993 y 8
383.22698974609375 0 608.86115
390.71612548828125 0 1412.8883 y 3
391.2184753417969 0 713.3615
394.21380615234375 0 992.36035
401.2140808105469 0 1309.9244
401.2400817871094 0 1196.5658
410.2216796875 0 566.2668
433.1740417480469 0 948.6175
436.8644714355469 0 575.37024
440.25067138671875 0 4205.2295 y 2
440.7524108886719 0 1486.3091
444.25634765625 0 1151.1986
445.2311706542969 0 771.31024
445.2655944824219 0 3861.9543 y Ammonia loss 7
447.2476501464844 0 1302.3843 c Ammonia loss 8
447.2816162109375 0 1846.006
447.7486877441406 0 641.64197
451.2338562011719 0 700.612
457.2776794433594 0 636.6587
459.28778076171875 0 926.5366
462.2943420410156 0 1037.9745 y 7
465.2857360839844 0 1622.9668
466.289306640625 0 652.02716
470.26251220703125 0 1123.1611
489.79107666015625 0 1002.94586 y 1
490.2920837402344 0 902.74493
493.279541015625 0 1792.822
494.28533935546875 0 936.36505
495.280029296875 0 564.57874
500.308837890625 0 1677.124
503.2988586425781 0 703.64246 c Water loss 9
503.78924560546875 0 1664.9423 c Ammonia loss 9
504.253662109375 0 11499.195
504.3025207519531 0 9972.819
505.25823974609375 0 3224.7095
505.3062438964844 0 2323.3306
514.2982788085938 0 2315.5266
516.3131103515625 0 913.8203
518.3115234375 0 922.54
519.3133544921875 0 2958.2576 y 6
520.3175048828125 0 1075.2737
522.30859375 0 625.12854
536.271484375 0 3461.2622
546.2552490234375 0 1726.2384
547.33154296875 0 1545.374
548.2824096679688 0 1630.1055
550.3010864257812 0 825.9447
554.314208984375 0 784.8295
554.8165893554688 0 1057.3423
560.3163452148438 0 11442.894 z 5
561.3235473632812 0 35727.3
562.2410278320312 0 570.63806
562.326904296875 0 8082.59
563.3207397460938 0 3624.9858
563.8192749023438 0 1655.9286
564.2657470703125 0 4174.12
564.310302734375 0 652.354
567.3291015625 0 4542.6973 c 4
568.330810546875 0 1191.5515
575.3274536132812 0 2574.0564
576.3345947265625 0 17410.674 y 5
577.3380737304688 0 3680.635
578.3375854492188 0 591.77875
607.321533203125 0 846.718
615.3467407226562 0 1208.7925 y Water loss 4
617.3378295898438 0 8654.793 z 4
618.3446655273438 0 19595.566
619.3484497070312 0 6590.7817
620.3521118164062 0 1236.4762
623.3424682617188 0 4706.872
624.3486938476562 0 4996.293 c 5
625.35302734375 0 1835.9155
632.3482666015625 0 18745.172
633.3551635742188 0 40870.906 y 4
634.359375 0 11182.44
635.3601684570312 0 2181.0251
643.357666015625 0 705.5799
647.3497924804688 0 1330.3171
657.5451049804688 0 635.19806
661.3663330078125 0 2848.2485
663.3820190429688 0 1602.4755
680.3637084960938 0 5332.752
681.3687744140625 0 2681.6978 c 6
746.397216796875 0 777.67865 z Water loss 3
760.4342651367188 0 2971.5515
761.4390869140625 0 1574.5636
763.4027709960938 0 1060.7739 y Ammonia loss 3
764.4056396484375 0 18830.418 c Water loss 7
765.4089965820312 0 17993.68
766.4133911132812 0 4898.773
767.4190673828125 0 756.78107
779.4171752929688 0 4320.373
780.424072265625 0 31064.293 y 3
781.4224243164062 0 15881.989
782.419189453125 0 54165.95 c 7
783.4216918945312 0 21672.926
784.4246826171875 0 4637.681
785.4197998046875 0 1714.9076
807.4036254882812 0 633.6818
817.4976806640625 0 772.66956
863.4745483398438 0 12818.224 z 2
864.4774780273438 0 5733.4976
865.4832763671875 0 2396.6865
866.4942626953125 0 1188.875
875.485107421875 0 679.7049
876.48193359375 0 935.857
879.4923706054688 0 27050.092 y 2
880.4957885742188 0 13670.276
881.497314453125 0 3784.8625
893.4868774414062 0 4926.599 c Ammonia loss 8
894.4917602539062 0 3134.6062
910.5134887695312 0 33387.305 c 8
911.5162353515625 0 19882.65
911.5933227539062 0 357.81995
912.5194702148438 0 4982.744
936.5350341796875 0 1103.6943
962.5420532226562 0 9415.528 z 1
963.5458374023438 0 4523.091
964.5531005859375 0 1513.508
978.5645141601562 0 1495.0754 y 1
979.5823364257812 0 12036.536
980.5859375 0 7120.9233
981.58837890625 0 1541.6982
1006.5704345703125 0 3976.246 c Ammonia loss 9
1007.5762329101562 0 2041.8013
1008.582763671875 0 3683.4238
1009.5906982421875 0 1866.2406
1023.5972290039062 0 51896.973 c 9
1024.6002197265625 0 31669.348
1025.60302734375 0 10006.091
1026.6004638671875 0 1661.6859
1048.5928955078125 0 1585.5881
1049.596923828125 0 1182.3722
1053.4210205078125 0 669.3832
1053.54638671875 0 1163.8081
1054.54541015625 0 675.6457
1066.5980224609375 0 2235.3
1067.6007080078125 0 868.1642
1070.5733642578125 0 8047.4062
1071.5770263671875 0 5789.104
1072.574462890625 0 1475.8013
1080.6265869140625 0 894.405
1098.6431884765625 0 2423.6074
1099.6419677734375 0 2061.6707
1108.6168212890625 0 1235.2383
1109.6099853515625 0 111497.16
1110.61279296875 0 76729.47
1111.6156005859375 0 27099.389
1112.6160888671875 0 3237.9346
1123.6036376953125 0 663.33026
1124.6123046875 0 1713.5984
1125.6285400390625 0 47506.87
1126.634765625 0 91812.27
1127.6383056640625 0 49632.566
1128.534912109375 0 4828.492
1128.6441650390625 0 13053.586
1129.545166015625 0 3994.8079
1141.5966796875 0 635.80493
1479.7281494140625 0 806.8507
1565.861328125 0 657.09296
2007.2567138671875 0 624.6285
3239.284423828125 0 612.71136
3369.461669921875 0 689.72723

Spectrum Details

|  |  |
| --- | --- |
| Matched peaks? Matched peaks The total absolute number of peaks matched. Additionally in brackets the total fraction of peaks matched and the total number of peaks is shown. | 34 (17.62% of 193) |
| FDR? FDR The false discovery rate estimated for this peptide. It is calculated by matching all theoretical fragments with a non-integer shift with the raw peaks for this spectrum. This is done with 40 different shifts. The resulting percentage is the average number of annotated peaks over the number of annotated peaks with the correct spectrum. | 0.42% |
| Satellite FDR? Satellite FDR See the FDR for details on its calculation. This satellite ion specific FDR only contains the satellite ions (d/w) for I/L/J positions. | - |
| PSM Score? PSM Score The PSM Score as given by Hecklib to this annotated spectrum. It is shown with three significant figures. | 386 |

#### Spectrum 8277? Spectrum 8277 The raw spectrum of this peptide as annotated by Hecklib. The fragments are coloured according to ion type (see legend). Any peaks with a star '\*' as text can be hovered over to see the full details, first the ion type second the mass shift type. By hovering over the amino acids in the peptide or ions in the legend the corresponding peaks are highlighted. By toggling the 'Unassigned' label you can turn the background (unassigned) peaks on or off in the plot. By updating the slider in the Ion legend you can update the spectrum to only show the top X% of the peaks with labels. The top X% means any peak that is within X% of the highest intensity. By dragging in the spectrum you can zoom in to a specific part of the spectrum and use 'Zoom Out' to get back to the original zoom level. The annotation of the spectrum is based on the given sequence in the peptides file and is done with different software so inconsistencies are likely. The peaks are annotated based on the given sequence, with 20 ppm tolerance.

Copy Data

##### Spectrum 8277 (TSV)

###### Preview

```
Loading example...
```

*Click on the button to copy the data to your clipboard.*

Mz MinMz MaxIntensity Max

WidthHeightPeptide font sizePeptide stroke widthSpectrum font sizeSpectrum stroke widthCompact peptide

Ion legend

wxyz

abcd

OtherUnassignedIon^Charge^_Position_Show for top:%

FVVFGGGTKJT

01.41e+52.81e+54.22e+55.62e+5

Zoom Out

y^+1^_1_y^+1^_2_a^+1^_2_y^+1^_2_b^+1^_2_y^+2^_5_b^+2^_6_y^+2^_7_y^+1^_3_b^+1^_3_y^+1^_3_y^+2^_8_b^+2^_8_y^+2^_8_y^+2^_9_b^+2^_9_y^+2^_9_y^+1^_4_b^+2^_9_y^+1^_4_y^+2^_10_b^+1^_4_b^+2^_10_b^+2^_10_y^+1^_5_b^+2^_10_y^+1^_5_b^+1^_5_\*\*y^+1^_6_\*y^+1^_6_b^+1^_6_y^+1^_7_y^+1^_7_b^+1^_7_b^+1^_8_y^+1^_8_b^+1^_8_y^+1^_8_y^+1^_9_b^+1^_9_y^+1^_9_b^+1^_9_y^+1^_10_b^+1^_10_b^+1^_10_b^+1^_10_

0627125518822510

Fragment Matches Table

Show background peaks

| Position | Ion type | Intensity | mz Theoretical | mz Error (Th) | mz Error (ppm) | Charge | Series Number |
| --- | --- | --- | --- | --- | --- | --- | --- |
| 11 | y | 2.954E+04 | 120.1 | 0.0003831 | 3.191 | +1 | 1 |
| - | - | 4.17E+05 | 120.1 | - | - | 0 | - |
| - | - | 1245 | 121.1 | - | - | 0 | - |
| - | - | 3.461E+04 | 121.1 | - | - | 0 | - |
| - | - | 857.5 | 122.1 | - | - | 0 | - |
| - | - | 412.4 | 126.1 | - | - | 0 | - |
| - | - | 1899 | 127.1 | - | - | 0 | - |
| - | - | 623.5 | 127.1 | - | - | 0 | - |
| - | - | 2346 | 128.1 | - | - | 0 | - |
| - | - | 1.847E+05 | 129.1 | - | - | 0 | - |
| - | - | 561.6 | 130.1 | - | - | 0 | - |
| - | - | 876 | 130.1 | - | - | 0 | - |
| - | - | 1.106E+04 | 130.1 | - | - | 0 | - |
| - | - | 485.8 | 131 | - | - | 0 | - |
| - | - | 583.8 | 131.1 | - | - | 0 | - |
| - | - | 6410 | 131.1 | - | - | 0 | - |
| - | - | 935.9 | 132.1 | - | - | 0 | - |
| - | - | 502 | 132.1 | - | - | 0 | - |
| - | - | 6556 | 133.1 | - | - | 0 | - |
| - | - | 1562 | 136.1 | - | - | 0 | - |
| - | - | 425.9 | 138.1 | - | - | 0 | - |
| - | - | 396.2 | 138.7 | - | - | 0 | - |
| - | - | 847 | 139.1 | - | - | 0 | - |
| - | - | 476.2 | 140.1 | - | - | 0 | - |
| - | - | 3205 | 141.1 | - | - | 0 | - |
| - | - | 680.6 | 141.1 | - | - | 0 | - |
| - | - | 389.8 | 141.6 | - | - | 0 | - |
| - | - | 572.2 | 142.1 | - | - | 0 | - |
| - | - | 512.6 | 146.1 | - | - | 0 | - |
| - | - | 443 | 147.1 | - | - | 0 | - |
| - | - | 733.1 | 149 | - | - | 0 | - |
| - | - | 435.6 | 149.1 | - | - | 0 | - |
| - | - | 457.7 | 151.1 | - | - | 0 | - |
| - | - | 769 | 152.1 | - | - | 0 | - |
| - | - | 1529 | 152.1 | - | - | 0 | - |
| - | - | 758.2 | 153.1 | - | - | 0 | - |
| - | - | 536.7 | 153.1 | - | - | 0 | - |
| - | - | 762.8 | 154.1 | - | - | 0 | - |
| - | - | 1087 | 155.1 | - | - | 0 | - |
| - | - | 4297 | 155.1 | - | - | 0 | - |
| - | - | 612.2 | 156.1 | - | - | 0 | - |
| - | - | 2555 | 158.1 | - | - | 0 | - |
| - | - | 5597 | 159.1 | - | - | 0 | - |
| - | - | 677.4 | 160.1 | - | - | 0 | - |
| - | - | 488.4 | 166.1 | - | - | 0 | - |
| - | - | 800.1 | 167.1 | - | - | 0 | - |
| - | - | 620.2 | 169.1 | - | - | 0 | - |
| - | - | 2932 | 170.1 | - | - | 0 | - |
| - | - | 1470 | 171.1 | - | - | 0 | - |
| - | - | 1.05E+04 | 171.1 | - | - | 0 | - |
| - | - | 6489 | 172.1 | - | - | 0 | - |
| - | - | 735.1 | 172.2 | - | - | 0 | - |
| - | - | 889.6 | 173.1 | - | - | 0 | - |
| - | - | 2849 | 173.5 | - | - | 0 | - |
| - | - | 3054 | 174.1 | - | - | 0 | - |
| - | - | 6556 | 176.1 | - | - | 0 | - |
| - | - | 7598 | 177.1 | - | - | 0 | - |
| - | - | 3132 | 177.1 | - | - | 0 | - |
| - | - | 613 | 178.1 | - | - | 0 | - |
| - | - | 583.8 | 181.1 | - | - | 0 | - |
| - | - | 892.1 | 182.1 | - | - | 0 | - |
| - | - | 981.1 | 183.1 | - | - | 0 | - |
| - | - | 581.8 | 183.1 | - | - | 0 | - |
| - | - | 3094 | 185.1 | - | - | 0 | - |
| - | - | 2022 | 186.1 | - | - | 0 | - |
| - | - | 1880 | 187.1 | - | - | 0 | - |
| - | - | 681.5 | 187.1 | - | - | 0 | - |
| - | - | 1501 | 188.1 | - | - | 0 | - |
| - | - | 708.1 | 188.1 | - | - | 0 | - |
| - | - | 741.7 | 194.1 | - | - | 0 | - |
| - | - | 630.3 | 195.1 | - | - | 0 | - |
| - | - | 716.1 | 197.1 | - | - | 0 | - |
| - | - | 7621 | 197.2 | - | - | 0 | - |
| - | - | 6736 | 198.1 | - | - | 0 | - |
| - | - | 518.8 | 198.2 | - | - | 0 | - |
| - | - | 7856 | 199.1 | - | - | 0 | - |
| - | - | 720 | 200.1 | - | - | 0 | - |
| - | - | 539.8 | 201.1 | - | - | 0 | - |
| - | - | 1102 | 201.1 | - | - | 0 | - |
| - | - | 2072 | 203.1 | - | - | 0 | - |
| - | - | 757 | 203.2 | - | - | 0 | - |
| - | - | 1.695E+04 | 205.1 | - | - | 0 | - |
| - | - | 868.5 | 205.1 | - | - | 0 | - |
| - | - | 1824 | 206.1 | - | - | 0 | - |
| - | - | 3086 | 207.1 | - | - | 0 | - |
| - | - | 2742 | 208.1 | - | - | 0 | - |
| - | - | 1152 | 209.1 | - | - | 0 | - |
| - | - | 794.3 | 209.1 | - | - | 0 | - |
| - | - | 1411 | 210.1 | - | - | 0 | - |
| - | - | 736.7 | 211.1 | - | - | 0 | - |
| - | - | 2.11E+04 | 212.1 | - | - | 0 | - |
| - | - | 661.5 | 213.1 | - | - | 0 | - |
| - | - | 2637 | 213.1 | - | - | 0 | - |
| - | - | 750.1 | 214.1 | - | - | 0 | - |
| - | - | 2004 | 214.2 | - | - | 0 | - |
| - | - | 3195 | 215.1 | - | - | 0 | - |
| 10 | y | 5542 | 215.1 | 0.0002938 | 1.366 | +1 | 2 |
| - | - | 5423 | 216.1 | - | - | 0 | - |
| - | - | 586.3 | 216.1 | - | - | 0 | - |
| - | - | 3218 | 217.1 | - | - | 0 | - |
| - | - | 595.1 | 217.1 | - | - | 0 | - |
| 2 | a | 5.567E+05 | 219.1 | 0.0005296 | 2.417 | +1 | 2 |
| - | - | 8.074E+04 | 220.2 | - | - | 0 | - |
| - | - | 1276 | 221.1 | - | - | 0 | - |
| - | - | 761.3 | 221.1 | - | - | 0 | - |
| - | - | 4321 | 221.2 | - | - | 0 | - |
| - | - | 981 | 222.1 | - | - | 0 | - |
| - | - | 6090 | 224.2 | - | - | 0 | - |
| - | - | 3115 | 225.1 | - | - | 0 | - |
| - | - | 1409 | 225.2 | - | - | 0 | - |
| - | - | 898.7 | 225.2 | - | - | 0 | - |
| - | - | 9875 | 227.1 | - | - | 0 | - |
| - | - | 4328 | 228.1 | - | - | 0 | - |
| - | - | 641.7 | 228.1 | - | - | 0 | - |
| - | - | 614.7 | 228.1 | - | - | 0 | - |
| - | - | 1491 | 229.1 | - | - | 0 | - |
| - | - | 701.6 | 229.1 | - | - | 0 | - |
| - | - | 3.947E+04 | 230.2 | - | - | 0 | - |
| - | - | 1895 | 231.1 | - | - | 0 | - |
| - | - | 5446 | 231.2 | - | - | 0 | - |
| - | - | 644.8 | 232.2 | - | - | 0 | - |
| - | - | 892.8 | 233.1 | - | - | 0 | - |
| 10 | y | 2.087E+04 | 233.1 | 0.0004866 | 2.087 | +1 | 2 |
| - | - | 1172 | 233.2 | - | - | 0 | - |
| - | - | 1171 | 234.1 | - | - | 0 | - |
| - | - | 2082 | 234.2 | - | - | 0 | - |
| - | - | 1958 | 237.1 | - | - | 0 | - |
| - | - | 1667 | 240.1 | - | - | 0 | - |
| - | - | 1561 | 241.2 | - | - | 0 | - |
| - | - | 7997 | 242.2 | - | - | 0 | - |
| - | - | 3023 | 243.1 | - | - | 0 | - |
| - | - | 1765 | 243.1 | - | - | 0 | - |
| - | - | 1587 | 243.2 | - | - | 0 | - |
| - | - | 1125 | 244.1 | - | - | 0 | - |
| - | - | 8282 | 245.1 | - | - | 0 | - |
| 2 | b | 4.441E+05 | 247.1 | 0.0006101 | 2.468 | +1 | 2 |
| - | - | 6.972E+04 | 248.1 | - | - | 0 | - |
| - | - | 735.7 | 249.1 | - | - | 0 | - |
| - | - | 4428 | 249.2 | - | - | 0 | - |
| 7 | y | 1598 | 251.2 | 0.004825 | 19.21 | +2 | 5 |
| - | - | 722.4 | 252.2 | - | - | 0 | - |
| - | - | 1103 | 253.2 | - | - | 0 | - |
| - | - | 2.094E+04 | 255.1 | - | - | 0 | - |
| - | - | 2694 | 256.1 | - | - | 0 | - |
| - | - | 657.7 | 257.2 | - | - | 0 | - |
| - | - | 5094 | 259.1 | - | - | 0 | - |
| - | - | 828.6 | 260.2 | - | - | 0 | - |
| - | - | 2563 | 261.1 | - | - | 0 | - |
| - | - | 9274 | 262.1 | - | - | 0 | - |
| - | - | 949.7 | 263.1 | - | - | 0 | - |
| - | - | 781 | 264.2 | - | - | 0 | - |
| - | - | 3068 | 267.1 | - | - | 0 | - |
| - | - | 2.307E+04 | 269.2 | - | - | 0 | - |
| - | - | 3354 | 270.2 | - | - | 0 | - |
| - | - | 2889 | 270.2 | - | - | 0 | - |
| - | - | 1263 | 271.1 | - | - | 0 | - |
| - | - | 2018 | 271.1 | - | - | 0 | - |
| - | - | 2425 | 272.1 | - | - | 0 | - |
| - | - | 2.886E+04 | 273.1 | - | - | 0 | - |
| - | - | 2832 | 273.2 | - | - | 0 | - |
| - | - | 5577 | 274.1 | - | - | 0 | - |
| - | - | 1066 | 275.1 | - | - | 0 | - |
| - | - | 2062 | 275.2 | - | - | 0 | - |
| - | - | 991.4 | 279.1 | - | - | 0 | - |
| - | - | 572.9 | 281.1 | - | - | 0 | - |
| - | - | 946.1 | 281.2 | - | - | 0 | - |
| - | - | 2004 | 282.2 | - | - | 0 | - |
| - | - | 939.2 | 285.2 | - | - | 0 | - |
| - | - | 7633 | 287.2 | - | - | 0 | - |
| - | - | 1769 | 287.2 | - | - | 0 | - |
| - | - | 1253 | 288.1 | - | - | 0 | - |
| - | - | 1096 | 288.2 | - | - | 0 | - |
| - | - | 583.3 | 290.2 | - | - | 0 | - |
| - | - | 5415 | 291.1 | - | - | 0 | - |
| - | - | 885.8 | 292.1 | - | - | 0 | - |
| - | - | 5151 | 295.1 | - | - | 0 | - |
| - | - | 581.2 | 295.2 | - | - | 0 | - |
| - | - | 1397 | 296.1 | - | - | 0 | - |
| - | - | 688.9 | 296.2 | - | - | 0 | - |
| - | - | 641.8 | 297.2 | - | - | 0 | - |
| - | - | 866.8 | 299.1 | - | - | 0 | - |
| - | - | 1196 | 300.1 | - | - | 0 | - |
| - | - | 1.885E+04 | 301.2 | - | - | 0 | - |
| - | - | 3532 | 302.2 | - | - | 0 | - |
| 6 | b | 1.019E+04 | 304.2 | 0.0006307 | 2.074 | +2 | 6 |
| - | - | 2010 | 305.2 | - | - | 0 | - |
| - | - | 636.8 | 306.1 | - | - | 0 | - |
| - | - | 1096 | 307.2 | - | - | 0 | - |
| 5 | y | 1182 | 308.2 | 0.003782 | 12.27 | +2 | 7 |
| - | - | 1237 | 308.2 | - | - | 0 | - |
| - | - | 1461 | 309.2 | - | - | 0 | - |
| - | - | 822.3 | 315.2 | - | - | 0 | - |
| - | - | 1610 | 316.2 | - | - | 0 | - |
| - | - | 629.1 | 317.4 | - | - | 0 | - |
| - | - | 2256 | 318.1 | - | - | 0 | - |
| - | - | 9279 | 319.1 | - | - | 0 | - |
| - | - | 1470 | 320.1 | - | - | 0 | - |
| - | - | 645.3 | 321.2 | - | - | 0 | - |
| - | - | 2895 | 321.2 | - | - | 0 | - |
| - | - | 4541 | 325.2 | - | - | 0 | - |
| - | - | 2.261E+04 | 326.2 | - | - | 0 | - |
| - | - | 1110 | 326.2 | - | - | 0 | - |
| - | - | 3533 | 327.2 | - | - | 0 | - |
| - | - | 1308 | 328.1 | - | - | 0 | - |
| - | - | 730.4 | 328.2 | - | - | 0 | - |
| - | - | 2414 | 328.2 | - | - | 0 | - |
| - | - | 1429 | 333.2 | - | - | 0 | - |
| - | - | 1161 | 336.2 | - | - | 0 | - |
| - | - | 2643 | 339.2 | - | - | 0 | - |
| - | - | 911.3 | 340.2 | - | - | 0 | - |
| - | - | 1095 | 341.2 | - | - | 0 | - |
| 9 | y | 7597 | 343.2 | 0.0006372 | 1.857 | +1 | 3 |
| - | - | 5646 | 344.2 | - | - | 0 | - |
| - | - | 1830 | 344.2 | - | - | 0 | - |
| - | - | 963.5 | 345.2 | - | - | 0 | - |
| - | - | 1586 | 345.2 | - | - | 0 | - |
| 3 | b | 1.122E+05 | 346.2 | 0.0006166 | 1.781 | +1 | 3 |
| - | - | 2.433E+04 | 347.2 | - | - | 0 | - |
| - | - | 4172 | 348.2 | - | - | 0 | - |
| - | - | 1997 | 348.2 | - | - | 0 | - |
| - | - | 823.6 | 349.2 | - | - | 0 | - |
| - | - | 4641 | 349.2 | - | - | 0 | - |
| - | - | 3953 | 352.2 | - | - | 0 | - |
| - | - | 880.5 | 353.2 | - | - | 0 | - |
| - | - | 3803 | 354.2 | - | - | 0 | - |
| - | - | 838 | 356.2 | - | - | 0 | - |
| - | - | 1120 | 356.2 | - | - | 0 | - |
| - | - | 6892 | 357.2 | - | - | 0 | - |
| - | - | 1253 | 358.2 | - | - | 0 | - |
| - | - | 1777 | 358.2 | - | - | 0 | - |
| - | - | 1168 | 360.2 | - | - | 0 | - |
| - | - | 2923 | 361.2 | - | - | 0 | - |
| 9 | y | 3.216E+04 | 361.2 | 0.000418 | 1.157 | +1 | 3 |
| - | - | 613.8 | 362.2 | - | - | 0 | - |
| - | - | 5953 | 362.2 | - | - | 0 | - |
| - | - | 937.7 | 364.2 | - | - | 0 | - |
| - | - | 4051 | 365.2 | - | - | 0 | - |
| - | - | 1218 | 366.2 | - | - | 0 | - |
| - | - | 1083 | 366.2 | - | - | 0 | - |
| - | - | 940.3 | 366.7 | - | - | 0 | - |
| - | - | 766.4 | 367.2 | - | - | 0 | - |
| - | - | 1637 | 368.2 | - | - | 0 | - |
| - | - | 1162 | 372.2 | - | - | 0 | - |
| - | - | 1251 | 372.3 | - | - | 0 | - |
| - | - | 1285 | 373.2 | - | - | 0 | - |
| - | - | 6989 | 374.2 | - | - | 0 | - |
| - | - | 593.5 | 374.2 | - | - | 0 | - |
| - | - | 3527 | 375.2 | - | - | 0 | - |
| - | - | 890.5 | 375.2 | - | - | 0 | - |
| - | - | 1181 | 376.2 | - | - | 0 | - |
| - | - | 632.5 | 376.2 | - | - | 0 | - |
| - | - | 766.1 | 381.2 | - | - | 0 | - |
| 4 | y | 1349 | 381.7 | 0.0004327 | 1.134 | +2 | 8 |
| - | - | 5419 | 382.2 | - | - | 0 | - |
| 8 | b | 2.647E+04 | 383.2 | 0.004146 | 10.82 | +2 | 8 |
| - | - | 1077 | 383.2 | - | - | 0 | - |
| - | - | 1328 | 384.2 | - | - | 0 | - |
| - | - | 5350 | 384.2 | - | - | 0 | - |
| - | - | 755.2 | 385.2 | - | - | 0 | - |
| - | - | 956.4 | 386.2 | - | - | 0 | - |
| - | - | 1504 | 387.2 | - | - | 0 | - |
| - | - | 4526 | 390.2 | - | - | 0 | - |
| 4 | y | 1.188E+04 | 390.7 | 0.000613 | 1.569 | +2 | 8 |
| - | - | 6317 | 391.2 | - | - | 0 | - |
| - | - | 1508 | 391.7 | - | - | 0 | - |
| - | - | 8430 | 392.2 | - | - | 0 | - |
| - | - | 1198 | 393.2 | - | - | 0 | - |
| - | - | 8611 | 394.2 | - | - | 0 | - |
| - | - | 2489 | 395.2 | - | - | 0 | - |
| - | - | 8437 | 400.3 | - | - | 0 | - |
| - | - | 827 | 401.2 | - | - | 0 | - |
| - | - | 2.272E+04 | 401.2 | - | - | 0 | - |
| - | - | 2037 | 401.3 | - | - | 0 | - |
| - | - | 2.741E+04 | 402.2 | - | - | 0 | - |
| - | - | 4440 | 402.2 | - | - | 0 | - |
| - | - | 6807 | 403.2 | - | - | 0 | - |
| - | - | 3536 | 403.2 | - | - | 0 | - |
| - | - | 758.3 | 404.2 | - | - | 0 | - |
| - | - | 1020 | 404.2 | - | - | 0 | - |
| - | - | 2267 | 406.2 | - | - | 0 | - |
| - | - | 612.8 | 407.2 | - | - | 0 | - |
| - | - | 1745 | 413.3 | - | - | 0 | - |
| - | - | 3502 | 418.2 | - | - | 0 | - |
| - | - | 2069 | 419.2 | - | - | 0 | - |
| - | - | 1.493E+04 | 420.2 | - | - | 0 | - |
| - | - | 3475 | 421.2 | - | - | 0 | - |
| - | - | 1211 | 421.3 | - | - | 0 | - |
| - | - | 4845 | 423.2 | - | - | 0 | - |
| - | - | 1093 | 424.2 | - | - | 0 | - |
| - | - | 3108 | 429.3 | - | - | 0 | - |
| - | - | 1181 | 430.3 | - | - | 0 | - |
| 3 | y | 3130 | 431.2 | 0.0001003 | 0.2326 | +2 | 9 |
| - | - | 2047 | 431.7 | - | - | 0 | - |
| - | - | 1160 | 437.2 | - | - | 0 | - |
| 9 | b | 2658 | 438.2 | 0.001345 | 3.068 | +2 | 9 |
| - | - | 1145 | 438.7 | - | - | 0 | - |
| - | - | 8479 | 439.3 | - | - | 0 | - |
| 3 | y | 1.862E+04 | 440.3 | 0.0009215 | 2.093 | +2 | 9 |
| - | - | 1.05E+04 | 440.8 | - | - | 0 | - |
| - | - | 2608 | 441.3 | - | - | 0 | - |
| 8 | y | 1595 | 444.3 | 0.0005051 | 1.137 | +1 | 4 |
| 9 | b | 6301 | 447.2 | 0.0001536 | 0.3434 | +2 | 9 |
| - | - | 4236 | 447.7 | - | - | 0 | - |
| - | - | 3875 | 448.3 | - | - | 0 | - |
| - | - | 1715 | 449.3 | - | - | 0 | - |
| - | - | 6987 | 451.2 | - | - | 0 | - |
| - | - | 1196 | 451.3 | - | - | 0 | - |
| - | - | 2046 | 452.2 | - | - | 0 | - |
| - | - | 1031 | 452.3 | - | - | 0 | - |
| - | - | 2452 | 456.2 | - | - | 0 | - |
| - | - | 1.906E+04 | 457.3 | - | - | 0 | - |
| - | - | 4465 | 458.3 | - | - | 0 | - |
| - | - | 687.8 | 459.3 | - | - | 0 | - |
| - | - | 1792 | 460.3 | - | - | 0 | - |
| - | - | 694.1 | 461.3 | - | - | 0 | - |
| 8 | y | 7725 | 462.3 | 0.0009268 | 2.005 | +1 | 4 |
| - | - | 712.2 | 463.2 | - | - | 0 | - |
| - | - | 1673 | 463.3 | - | - | 0 | - |
| - | - | 1.342E+04 | 465.3 | - | - | 0 | - |
| - | - | 5387 | 466.3 | - | - | 0 | - |
| - | - | 707.8 | 470.2 | - | - | 0 | - |
| - | - | 711.3 | 470.3 | - | - | 0 | - |
| - | - | 2935 | 473.3 | - | - | 0 | - |
| - | - | 991.7 | 474.2 | - | - | 0 | - |
| - | - | 1022 | 475.2 | - | - | 0 | - |
| - | - | 2893 | 478.3 | - | - | 0 | - |
| - | - | 947.8 | 480.3 | - | - | 0 | - |
| - | - | 1100 | 483.2 | - | - | 0 | - |
| - | - | 1.207E+04 | 486.3 | - | - | 0 | - |
| - | - | 2967 | 487.3 | - | - | 0 | - |
| - | - | 1235 | 488.3 | - | - | 0 | - |
| 2 | y | 5376 | 489.8 | 0.007761 | 15.85 | +2 | 10 |
| - | - | 5002 | 490.3 | - | - | 0 | - |
| - | - | 2443 | 490.8 | - | - | 0 | - |
| - | - | 4858 | 491.3 | - | - | 0 | - |
| - | - | 746.6 | 492.3 | - | - | 0 | - |
| 4 | b | 1.28E+04 | 493.3 | 0.0008367 | 1.696 | +1 | 4 |
| - | - | 4003 | 494.3 | - | - | 0 | - |
| 10 | b | 1847 | 494.8 | 0.004249 | 8.588 | +2 | 10 |
| - | - | 896.5 | 495.2 | - | - | 0 | - |
| 10 | b | 1516 | 495.3 | 0.007893 | 15.94 | +2 | 10 |
| - | - | 2.387E+04 | 496.3 | - | - | 0 | - |
| - | - | 6351 | 497.3 | - | - | 0 | - |
| - | - | 1745 | 498.3 | - | - | 0 | - |
| - | - | 1.141E+04 | 501.2 | - | - | 0 | - |
| 7 | y | 1339 | 501.3 | 3.75E-05 | 0.0748 | +1 | 5 |
| - | - | 4194 | 502.2 | - | - | 0 | - |
| - | - | 927.6 | 503.3 | - | - | 0 | - |
| - | - | 812.9 | 503.3 | - | - | 0 | - |
| 10 | b | 4140 | 503.8 | 0.0005696 | 1.131 | +2 | 10 |
| - | - | 2692 | 504.3 | - | - | 0 | - |
| - | - | 2294 | 505.3 | - | - | 0 | - |
| - | - | 730.4 | 506.3 | - | - | 0 | - |
| - | - | 1960 | 508.3 | - | - | 0 | - |
| - | - | 1103 | 509.3 | - | - | 0 | - |
| - | - | 4288 | 512.3 | - | - | 0 | - |
| - | - | 1207 | 513.3 | - | - | 0 | - |
| - | - | 5.555E+04 | 514.3 | - | - | 0 | - |
| - | - | 1.57E+04 | 515.3 | - | - | 0 | - |
| - | - | 2519 | 516.3 | - | - | 0 | - |
| - | - | 623.6 | 517.3 | - | - | 0 | - |
| - | - | 5738 | 519.3 | - | - | 0 | - |
| 7 | y | 1.279E+04 | 519.3 | 0.0007033 | 1.354 | +1 | 5 |
| - | - | 2831 | 520.3 | - | - | 0 | - |
| - | - | 2632 | 520.3 | - | - | 0 | - |
| - | - | 777.8 | 521.3 | - | - | 0 | - |
| - | - | 4721 | 522.3 | - | - | 0 | - |
| - | - | 1758 | 523.3 | - | - | 0 | - |
| - | - | 1.605E+04 | 530.3 | - | - | 0 | - |
| - | - | 3270 | 531.3 | - | - | 0 | - |
| - | - | 1235 | 532.3 | - | - | 0 | - |
| - | - | 742.1 | 536.3 | - | - | 0 | - |
| - | - | 1123 | 540.3 | - | - | 0 | - |
| - | - | 4202 | 547.3 | - | - | 0 | - |
| - | - | 2.927E+04 | 548.3 | - | - | 0 | - |
| - | - | 8815 | 549.3 | - | - | 0 | - |
| 5 | b | 5350 | 550.3 | 0.001431 | 2.601 | +1 | 5 |
| - | - | 2145 | 551.3 | - | - | 0 | - |
| 0 | Precursor | 9477 | 554.3 | 0.0009002 | 1.624 | +2 | -1 |
| 0 | Precursor | 6541 | 554.8 | 0.009686 | 17.46 | +2 | -1 |
| - | - | 1833 | 555.3 | - | - | 0 | - |
| - | - | 902.4 | 558.3 | - | - | 0 | - |
| 6 | y | 3873 | 558.3 | 5.815E-05 | 0.1041 | +1 | 6 |
| - | - | 1452 | 559.3 | - | - | 0 | - |
| - | - | 741.7 | 560.3 | - | - | 0 | - |
| - | - | 919.1 | 562.3 | - | - | 0 | - |
| 0 | Precursor | 1.316E+04 | 563.3 | 0.0003786 | 0.6722 | +2 | -1 |
| - | - | 8978 | 563.8 | - | - | 0 | - |
| - | - | 2469 | 564.3 | - | - | 0 | - |
| - | - | 1340 | 565.3 | - | - | 0 | - |
| - | - | 1984 | 571.4 | - | - | 0 | - |
| - | - | 722.9 | 572.4 | - | - | 0 | - |
| 6 | y | 3.881E+04 | 576.3 | 0.0004798 | 0.8325 | +1 | 6 |
| - | - | 1.208E+04 | 577.3 | - | - | 0 | - |
| - | - | 1956 | 578.3 | - | - | 0 | - |
| - | - | 2089 | 579.3 | - | - | 0 | - |
| - | - | 1161 | 587.4 | - | - | 0 | - |
| - | - | 2064 | 597.3 | - | - | 0 | - |
| - | - | 1010 | 598.3 | - | - | 0 | - |
| - | - | 1512 | 599.3 | - | - | 0 | - |
| - | - | 1029 | 600.3 | - | - | 0 | - |
| 6 | b | 3365 | 607.3 | 0.002068 | 3.405 | +1 | 6 |
| - | - | 1095 | 608.3 | - | - | 0 | - |
| - | - | 1558 | 611.3 | - | - | 0 | - |
| 5 | y | 1.2E+04 | 615.3 | 0.001238 | 2.013 | +1 | 7 |
| - | - | 4267 | 616.3 | - | - | 0 | - |
| - | - | 1802 | 617.3 | - | - | 0 | - |
| - | - | 1511 | 618.3 | - | - | 0 | - |
| - | - | 1192 | 619.3 | - | - | 0 | - |
| - | - | 648.1 | 620.3 | - | - | 0 | - |
| - | - | 2765 | 625.3 | - | - | 0 | - |
| - | - | 1239 | 626.3 | - | - | 0 | - |
| - | - | 1.013E+04 | 629.3 | - | - | 0 | - |
| - | - | 4530 | 630.3 | - | - | 0 | - |
| - | - | 924.6 | 631.3 | - | - | 0 | - |
| 5 | y | 1.951E+05 | 633.4 | 0.001355 | 2.139 | +1 | 7 |
| - | - | 6.503E+04 | 634.4 | - | - | 0 | - |
| - | - | 1.173E+04 | 635.4 | - | - | 0 | - |
| - | - | 950.2 | 636.4 | - | - | 0 | - |
| - | - | 2.848E+04 | 643.4 | - | - | 0 | - |
| - | - | 1.088E+04 | 644.4 | - | - | 0 | - |
| - | - | 1903 | 645.4 | - | - | 0 | - |
| - | - | 2.014E+04 | 647.4 | - | - | 0 | - |
| - | - | 655.7 | 648.3 | - | - | 0 | - |
| - | - | 8075 | 648.4 | - | - | 0 | - |
| - | - | 1607 | 649.4 | - | - | 0 | - |
| - | - | 764.1 | 653.3 | - | - | 0 | - |
| - | - | 5.505E+04 | 661.4 | - | - | 0 | - |
| - | - | 2.24E+04 | 662.4 | - | - | 0 | - |
| - | - | 4355 | 663.4 | - | - | 0 | - |
| 7 | b | 2739 | 664.3 | 0.001417 | 2.134 | +1 | 7 |
| - | - | 735.2 | 665.3 | - | - | 0 | - |
| - | - | 1408 | 666.3 | - | - | 0 | - |
| - | - | 645.9 | 667.3 | - | - | 0 | - |
| - | - | 1127 | 679.4 | - | - | 0 | - |
| - | - | 700.1 | 696.4 | - | - | 0 | - |
| - | - | 1025 | 713.4 | - | - | 0 | - |
| - | - | 2405 | 718.4 | - | - | 0 | - |
| - | - | 1697 | 720.4 | - | - | 0 | - |
| - | - | 1656 | 724.4 | - | - | 0 | - |
| - | - | 903.8 | 726.4 | - | - | 0 | - |
| - | - | 7176 | 732.4 | - | - | 0 | - |
| - | - | 2781 | 733.4 | - | - | 0 | - |
| - | - | 752.5 | 734.4 | - | - | 0 | - |
| - | - | 1868 | 737.4 | - | - | 0 | - |
| - | - | 1.587E+04 | 742.4 | - | - | 0 | - |
| - | - | 7492 | 743.4 | - | - | 0 | - |
| - | - | 2772 | 744.4 | - | - | 0 | - |
| - | - | 786.4 | 745.4 | - | - | 0 | - |
| - | - | 709.9 | 746.4 | - | - | 0 | - |
| 8 | b | 4345 | 747.4 | 0.0007842 | 1.049 | +1 | 8 |
| - | - | 1962 | 748.4 | - | - | 0 | - |
| - | - | 1766 | 757.4 | - | - | 0 | - |
| - | - | 4.645E+04 | 760.4 | - | - | 0 | - |
| - | - | 2.254E+04 | 761.4 | - | - | 0 | - |
| 4 | y | 1.126E+04 | 762.4 | 0.006982 | 9.158 | +1 | 8 |
| - | - | 5559 | 763.4 | - | - | 0 | - |
| - | - | 1472 | 764.4 | - | - | 0 | - |
| 8 | b | 1.067E+04 | 765.4 | 0.000736 | 0.9616 | +1 | 8 |
| - | - | 5610 | 766.4 | - | - | 0 | - |
| - | - | 1129 | 767.4 | - | - | 0 | - |
| - | - | 984.9 | 778.4 | - | - | 0 | - |
| 4 | y | 2.169E+05 | 780.4 | 0.0005069 | 0.6496 | +1 | 8 |
| - | - | 1.032E+05 | 781.4 | - | - | 0 | - |
| - | - | 2.544E+04 | 782.4 | - | - | 0 | - |
| - | - | 2605 | 783.4 | - | - | 0 | - |
| - | - | 3995 | 785.4 | - | - | 0 | - |
| - | - | 1180 | 786.4 | - | - | 0 | - |
| - | - | 913.1 | 799.4 | - | - | 0 | - |
| - | - | 1626 | 817.5 | - | - | 0 | - |
| - | - | 835.2 | 818.5 | - | - | 0 | - |
| - | - | 625.5 | 831.5 | - | - | 0 | - |
| - | - | 906.1 | 833.5 | - | - | 0 | - |
| - | - | 600.9 | 841.5 | - | - | 0 | - |
| - | - | 2036 | 843.5 | - | - | 0 | - |
| - | - | 1690 | 844.5 | - | - | 0 | - |
| - | - | 2938 | 859.5 | - | - | 0 | - |
| - | - | 1977 | 860.5 | - | - | 0 | - |
| 3 | y | 9177 | 861.5 | 0.0002749 | 0.3191 | +1 | 9 |
| - | - | 4268 | 862.5 | - | - | 0 | - |
| - | - | 1010 | 863.5 | - | - | 0 | - |
| 9 | b | 5855 | 875.5 | 0.0003221 | 0.3679 | +1 | 9 |
| - | - | 2412 | 876.5 | - | - | 0 | - |
| - | - | 2005 | 877.5 | - | - | 0 | - |
| 3 | y | 2.005E+05 | 879.5 | 0.0003914 | 0.445 | +1 | 9 |
| - | - | 1.049E+05 | 880.5 | - | - | 0 | - |
| - | - | 2.744E+04 | 881.5 | - | - | 0 | - |
| - | - | 3017 | 882.5 | - | - | 0 | - |
| 9 | b | 2.522E+04 | 893.5 | 0.0002554 | 0.2859 | +1 | 9 |
| - | - | 1.533E+04 | 894.5 | - | - | 0 | - |
| - | - | 3671 | 895.5 | - | - | 0 | - |
| - | - | 2075 | 907.5 | - | - | 0 | - |
| - | - | 1139 | 908.5 | - | - | 0 | - |
| 2 | y | 1.306E+04 | 978.6 | 0.001619 | 1.654 | +1 | 10 |
| - | - | 7233 | 979.6 | - | - | 0 | - |
| - | - | 2496 | 980.6 | - | - | 0 | - |
| 10 | b | 4297 | 988.6 | 0.00287 | 2.904 | +1 | 10 |
| 10 | b | 2568 | 989.5 | 0.01769 | 17.88 | +1 | 10 |
| 10 | b | 2.649E+04 | 1007 | 0.0001293 | 0.1285 | +1 | 10 |
| - | - | 1.838E+04 | 1008 | - | - | 0 | - |
| - | - | 5038 | 1009 | - | - | 0 | - |
| - | - | 958.5 | 1010 | - | - | 0 | - |
| - | - | 614.1 | 2485 | - | - | 0 | - |

m/z Charge Intensity FragmentType MassShift Position
120.06590270996094 0 29540.938 y 10
120.08122253417969 0 417039.88
121.06938934326172 0 1244.8359
121.0844955444336 0 34605.04
122.08755493164062 0 857.46515
126.12836456298828 0 412.37082
127.05069732666016 0 1898.8956
127.08733367919922 0 623.4604
128.1073455810547 0 2345.5981
129.10264587402344 0 184720.39
130.05007934570312 0 561.6482
130.10031127929688 0 875.9522
130.10594177246094 0 11061.099
131.04920959472656 0 485.78372
131.07083129882812 0 583.77484
131.08180236816406 0 6409.508
132.0767822265625 0 935.92786
132.08157348632812 0 502.0353
133.0862579345703 0 6555.9155
136.0759735107422 0 1561.664
138.06524658203125 0 425.86072
138.7152862548828 0 396.19965
139.087158203125 0 847.0347
140.14332580566406 0 476.24576
141.0662078857422 0 3204.9387
141.10260009765625 0 680.6067
141.64199829101562 0 389.79114
142.1228790283203 0 572.2319
146.1293487548828 0 512.6133
147.1451873779297 0 442.98508
148.953857421875 0 733.124
149.07958984375 0 435.64276
151.0870361328125 0 457.6942
152.07095336914062 0 768.9635
152.14361572265625 0 1529.0896
153.0664825439453 0 758.22595
153.1464385986328 0 536.6838
154.06137084960938 0 762.7737
155.081787109375 0 1087.3335
155.11825561523438 0 4296.5005
156.0768280029297 0 612.194
158.0928497314453 0 2554.9995
159.0768585205078 0 5596.9526
160.08079528808594 0 677.4281
166.0865020751953 0 488.40143
167.118408203125 0 800.11816
169.09762573242188 0 620.16235
170.09280395507812 0 2932.4678
171.07676696777344 0 1470.3673
171.1495819091797 0 10496.38
172.0720672607422 0 6489.1396
172.15325927734375 0 735.1174
173.1288299560547 0 889.62177
173.45132446289062 0 2849.0112
174.12814331054688 0 3053.6528
176.10733032226562 0 6556.2944
177.10263061523438 0 7598.1426
177.1121826171875 0 3131.924
178.10614013671875 0 612.97894
181.13316345214844 0 583.77875
182.0818634033203 0 892.08527
183.11302185058594 0 981.0704
183.1499481201172 0 581.8469
185.1287841796875 0 3094.2502
186.1243896484375 0 2021.6327
187.10797119140625 0 1880.1035
187.14468383789062 0 681.5056
188.10330200195312 0 1501.1888
188.1436767578125 0 708.0949
194.1292266845703 0 741.67596
195.1497039794922 0 630.2572
197.12844848632812 0 716.1384
197.16513061523438 0 7620.7295
198.0876922607422 0 6736.1074
198.168212890625 0 518.8134
199.14450073242188 0 7856.0903
200.14759826660156 0 719.96173
201.09844970703125 0 539.75446
201.12396240234375 0 1101.9015
203.1182098388672 0 2071.7874
203.1543731689453 0 757.02875
205.09751892089844 0 16950.828
205.107177734375 0 868.50745
206.10098266601562 0 1823.6719
207.1494140625 0 3085.9922
208.10841369628906 0 2742.2522
209.10302734375 0 1152.3607
209.1286163330078 0 794.27985
210.08729553222656 0 1410.8452
211.10850524902344 0 736.6798
212.13973999023438 0 21097.947
213.12451171875 0 661.47626
213.1431427001953 0 2636.944
214.1188201904297 0 750.0762
214.19183349609375 0 2004.2781
215.1148681640625 0 3195.231
215.13931274414062 0 5541.758 y Water loss 9
216.09837341308594 0 5422.8584
216.14244079589844 0 586.2555
217.0977325439453 0 3218.0745
217.13436889648438 0 595.0945
219.14971923828125 0 556705.1 a 1
220.15293884277344 0 80743.22
221.12831115722656 0 1275.819
221.139404296875 0 761.2941
221.15599060058594 0 4320.9854
222.12664794921875 0 980.969
224.17608642578125 0 6089.6846
225.13525390625 0 3115.391
225.1599884033203 0 1408.6769
225.1798858642578 0 898.6829
227.11436462402344 0 9875.008
228.09828186035156 0 4327.587
228.10916137695312 0 641.7352
228.133544921875 0 614.6585
229.09315490722656 0 1490.8671
229.1296844482422 0 701.56647
230.15037536621094 0 39469.2
231.1132049560547 0 1894.9093
231.152099609375 0 5445.535
232.1540069580078 0 644.79425
233.12908935546875 0 892.76807
233.1500701904297 0 20874.22 y 9
233.16366577148438 0 1171.9266
234.1243438720703 0 1171.1332
234.15341186523438 0 2082.054
237.09849548339844 0 1957.7123
240.1349639892578 0 1666.7024
241.1554412841797 0 1560.9789
242.18679809570312 0 7996.834
243.1130828857422 0 3022.5354
243.1459503173828 0 1765.3125
243.1902618408203 0 1586.6301
244.10757446289062 0 1124.7218
245.1251678466797 0 8281.832
247.14471435546875 0 444076.44 b 1
248.14796447753906 0 69722.16
249.1239013671875 0 735.73615
249.1507110595703 0 4427.732
251.15037536621094 0 1598.0602 y Water loss 6
252.17022705078125 0 722.36786
253.1664581298828 0 1102.6564
255.10922241210938 0 20943.01
256.11236572265625 0 2694.41
257.1611328125 0 657.73615
259.1444091796875 0 5094.0986
260.1978759765625 0 828.6079
261.1234130859375 0 2563.4995
262.1190490722656 0 9273.597
263.122314453125 0 949.7414
264.1714172363281 0 781.04285
267.14935302734375 0 3068.0984
269.1612854003906 0 23074.137
270.1641540527344 0 3353.7358
270.1817321777344 0 2888.6172
271.1077880859375 0 1262.5914
271.14068603515625 0 2017.6342
272.13604736328125 0 2425.279
273.1197509765625 0 28857.89
273.19635009765625 0 2831.8018
274.1204528808594 0 5576.851
275.1236267089844 0 1066.2761
275.17547607421875 0 2062.2651
279.1474914550781 0 991.43176
281.0897521972656 0 572.9454
281.19891357421875 0 946.0989
282.15631103515625 0 2003.964
285.15777587890625 0 939.2312
287.171875 0 7633.4463
287.212158203125 0 1769.2544
288.1348571777344 0 1253.3353
288.1752624511719 0 1096.256
290.1844787597656 0 583.31683
291.1455993652344 0 5414.567
292.1490173339844 0 885.8046
295.1446228027344 0 5150.8896
295.17645263671875 0 581.1546
296.1479797363281 0 1396.6252
296.1965026855469 0 688.8654
297.1549987792969 0 641.80255
299.06304931640625 0 866.8329
300.1349182128906 0 1196.3285
301.1916198730469 0 18847.43
302.1950988769531 0 3531.5776
304.16619873046875 0 10193.002 b 5
305.1690673828125 0 2010.34
306.1452941894531 0 636.8353
307.2135009765625 0 1096.0098
308.1728820800781 0 1181.8044 y Water loss 4
308.19683837890625 0 1237.055
309.1561279296875 0 1461.151
315.2398986816406 0 822.3205
316.1663513183594 0 1610.0115
317.4058532714844 0 629.0688
318.1445617675781 0 2256.4722
319.1408996582031 0 9278.952
320.1441345214844 0 1470.0549
321.1670227050781 0 645.31116
321.19635009765625 0 2895.274
325.2242126464844 0 4540.9565
326.1829528808594 0 22610.555
326.20556640625 0 1110.011
327.18603515625 0 3532.8093
328.1287841796875 0 1308.2927
328.16241455078125 0 730.4154
328.2029113769531 0 2414.4343
333.1933898925781 0 1428.7249
336.1679382324219 0 1160.6053
339.1782531738281 0 2642.7493
340.19891357421875 0 911.26086
341.18194580078125 0 1094.9592
343.234619140625 0 7596.9893 y Water loss 8
344.1932678222656 0 5646.118
344.23785400390625 0 1830.2776
345.1567687988281 0 963.4785
345.1953125 0 1586.0474
346.213134765625 0 112214.37 b 2
347.2163391113281 0 24334.658
348.1673583984375 0 4172.2227
348.218994140625 0 1996.9198
349.1702880859375 0 823.5985
349.1916198730469 0 4641.412
352.165771484375 0 3952.5312
353.169677734375 0 880.52856
354.177490234375 0 3802.8179
356.17218017578125 0 838.03284
356.22845458984375 0 1119.7709
357.1565856933594 0 6891.941
358.1588134765625 0 1252.842
358.21173095703125 0 1777.3945
360.19244384765625 0 1167.8917
361.1875915527344 0 2922.8965
361.2449645996094 0 32158.41 y 8
362.1914978027344 0 613.8366
362.2475891113281 0 5952.972
364.23382568359375 0 937.73083
365.193359375 0 4051.4082
366.1759948730469 0 1218.4163
366.19842529296875 0 1082.8291
366.7248229980469 0 940.2997
367.2243957519531 0 766.3679
368.1940002441406 0 1637.0547
372.18853759765625 0 1161.5402
372.25921630859375 0 1250.7864
373.1886901855469 0 1284.6672
374.1830139160156 0 6989.4307
374.2084045410156 0 593.53845
375.16693115234375 0 3526.899
375.23968505859375 0 890.4823
376.1634216308594 0 1181.0969
376.1998596191406 0 632.54065
381.192138671875 0 766.1063
381.7113037109375 0 1348.7659 y Water loss 3
382.2454833984375 0 5418.83
383.20428466796875 0 26465.146 b 7
383.2469482421875 0 1076.5347
384.1687316894531 0 1328.2231
384.20684814453125 0 5349.893
385.1507263183594 0 755.15497
386.2021484375 0 956.4272
387.2027893066406 0 1503.5712
390.2141418457031 0 4525.573
390.7167663574219 0 11878.666 y 3
391.2180480957031 0 6317.406
391.719482421875 0 1507.6085
392.1934814453125 0 8429.79
393.1942443847656 0 1197.7173
394.2132263183594 0 8610.624
395.2161560058594 0 2489.1672
400.25604248046875 0 8437.305
401.1859130859375 0 826.9865
401.2149353027344 0 22718.373
401.2604064941406 0 2036.5853
402.1778869628906 0 27412.264
402.2169189453125 0 4440.1377
403.1811828613281 0 6806.958
403.2343444824219 0 3535.9072
404.184814453125 0 758.30054
404.2359924316406 0 1019.5042
406.2129821777344 0 2266.533
407.2138977050781 0 612.8444
413.25213623046875 0 1744.7281
418.2092590332031 0 3501.9934
419.2055969238281 0 2068.8535
420.1885070800781 0 14928.881
421.1908874511719 0 3474.7153
421.2552185058594 0 1211.0916
423.2392883300781 0 4844.9517
424.2425842285156 0 1092.5951
429.2834167480469 0 3108.1357
430.2852783203125 0 1180.8931
431.24517822265625 0 3129.5872 y Water loss 2
431.74652099609375 0 2047.4559
437.2125244140625 0 1160.133
438.2436828613281 0 2658.0808 b Water loss 8
438.7431945800781 0 1144.7722
439.2669982910156 0 8478.829
440.25128173828125 0 18617.871 y 2
440.75262451171875 0 10500.552
441.2525634765625 0 2608.3577
444.28216552734375 0 1594.6918 y Water loss 7
447.2474670410156 0 6300.7866 b 8
447.74945068359375 0 4236.0566
448.25860595703125 0 3875.0354
449.2635803222656 0 1714.9276
451.2348327636719 0 6986.7114
451.267333984375 0 1195.7897
452.2366027832031 0 2045.8685
452.2674255371094 0 1030.8352
456.2249755859375 0 2451.641
457.2778015136719 0 19064.15
458.28118896484375 0 4465.018
459.2806091308594 0 687.8062
460.2566223144531 0 1792.0826
461.2550048828125 0 694.07465
462.29315185546875 0 7724.904 y 7
463.2325134277344 0 712.1715
463.2965393066406 0 1672.509
465.286865234375 0 13415.029
466.2906494140625 0 5387.148
470.2396240234375 0 707.82153
470.27386474609375 0 711.3271
473.2512512207031 0 2934.8523
474.2393798828125 0 991.65875
475.231201171875 0 1021.775
478.2788391113281 0 2892.8167
480.26092529296875 0 947.78406
483.23492431640625 0 1099.9441
486.30438232421875 0 12066.722
487.3079833984375 0 2966.547
488.2501220703125 0 1235.2743
489.7923278808594 0 5376.1577 y 1
490.2934875488281 0 5002.0923
490.79388427734375 0 2443.2915
491.2618713378906 0 4858.0938
492.2649841308594 0 746.6228
493.2817687988281 0 12796.135 b 3
494.28485107421875 0 4002.9192
494.7801208496094 0 1846.7415 b Water loss 9
495.2370300292969 0 896.45886
495.2842712402344 0 1515.9847 b Ammonia loss 9
496.2886657714844 0 23869.227
497.2911376953125 0 6350.996
498.295654296875 0 1745.219
501.24627685546875 0 11412.825
501.30316162109375 0 1339.0657 y Water loss 6
502.2496032714844 0 4193.7163
503.2502746582031 0 927.60114
503.30828857421875 0 812.8792
503.79022216796875 0 4140.2993 b 9
504.2928466796875 0 2691.6917
505.28204345703125 0 2293.9604
506.2845153808594 0 730.3958
508.25592041015625 0 1960.3871
509.2590637207031 0 1103.1489
512.26220703125 0 4288.1494
513.2630004882812 0 1206.6743
514.2989501953125 0 55550.73
515.3020629882812 0 15703.991
516.3037109375 0 2519.3406
517.2717895507812 0 623.636
519.2574462890625 0 5738.2896
519.3143920898438 0 12794.481 y 6
520.2594604492188 0 2830.6516
520.3175048828125 0 2632.1697
521.2520751953125 0 777.7882
522.3075561523438 0 4720.9995
523.3115844726562 0 1757.9163
530.2728881835938 0 16045.136
531.2752075195312 0 3270.45
532.3091430664062 0 1234.6523
536.285888671875 0 742.09247
540.3165283203125 0 1122.5032
547.3324584960938 0 4201.623
548.2833862304688 0 29271.812
549.286376953125 0 8814.899
550.3009643554688 0 5350.4766 b 4
551.3056030273438 0 2144.7795
554.3143920898438 0 9477.098 Precursor Water loss
554.815185546875 0 6541.296 Precursor Ammonia loss
555.3171997070312 0 1833.2468
558.27587890625 0 902.39056
558.3246459960938 0 3873.317 y Water loss 5
559.3231201171875 0 1452.1768
560.3206176757812 0 741.71564
562.302978515625 0 919.11816
563.3191528320312 0 13163.027 Precursor
563.8203735351562 0 8978.16
564.3211669921875 0 2468.8901
565.2764892578125 0 1340.353
571.3551025390625 0 1983.5798
572.366455078125 0 722.9341
576.3356323242188 0 38806.76 y 5
577.3385620117188 0 12083.2
578.3400268554688 0 1956.1157
579.3302612304688 0 2089.477
587.3501586914062 0 1161.4927
597.3380737304688 0 2063.5146
598.333251953125 0 1009.82166
599.3316650390625 0 1512.0651
600.3213500976562 0 1029.4976
607.325927734375 0 3365.4524 b 5
608.3258056640625 0 1095.4843
611.3308715820312 0 1557.6948
615.3472900390625 0 11998.199 y Water loss 4
616.3483276367188 0 4266.596
617.3455810546875 0 1801.8722
618.3274536132812 0 1511.4421
619.3270874023438 0 1191.8004
620.3285522460938 0 648.129
625.345947265625 0 2764.7612
626.3452758789062 0 1238.8873
629.3408813476562 0 10128.336
630.3446655273438 0 4529.817
631.3441772460938 0 924.6013
633.3579711914062 0 195060.95 y 4
634.3609619140625 0 65028.78
635.3629150390625 0 11730.865
636.358642578125 0 950.19403
643.3568725585938 0 28478.63
644.3600463867188 0 10878.361
645.3645629882812 0 1902.7993
647.3515625 0 20144.68
648.308349609375 0 655.703
648.3550415039062 0 8075.167
649.35498046875 0 1607.0309
653.3383178710938 0 764.05945
661.3673095703125 0 55053.043
662.3704833984375 0 22398.908
663.3731079101562 0 4354.5737
664.3467407226562 0 2738.7756 b 6
665.348876953125 0 735.2016
666.3226318359375 0 1407.5278
667.3250122070312 0 645.8755
679.37744140625 0 1127.0099
696.36865234375 0 700.1451
713.3952026367188 0 1025.0868
718.4229736328125 0 2404.6267
720.37646484375 0 1696.5853
724.4163208007812 0 1655.5417
726.4025268554688 0 903.8077
732.4406127929688 0 7176.2246
733.44384765625 0 2780.7803
734.4292602539062 0 752.4989
737.3984985351562 0 1868.4851
742.42529296875 0 15872.255
743.4277954101562 0 7492.427
744.417236328125 0 2772.357
745.4022216796875 0 786.4087
746.4185180664062 0 709.94135
747.3816528320312 0 4344.543 b Water loss 7
748.3859252929688 0 1961.6523
757.4188232421875 0 1765.9266
760.4356689453125 0 46449.14
761.438720703125 0 22537.117
762.4214477539062 0 11258.078 y Water loss 3
763.417236328125 0 5559.1724
764.4157104492188 0 1472.3633
765.3937377929688 0 10671.273 b 7
766.3965454101562 0 5610.058
767.3977661132812 0 1129.0298
778.429443359375 0 984.92914
780.425537109375 0 216903.44 y 3
781.4285888671875 0 103203.625
782.4310302734375 0 25443.807
783.4299926757812 0 2604.8838
785.4160766601562 0 3994.9526
786.4241943359375 0 1180.4703
799.3841552734375 0 913.1268
817.4933471679688 0 1625.7023
818.4966430664062 0 835.19507
831.5206909179688 0 625.45013
833.4871826171875 0 906.12524
841.4926147460938 0 600.87756
843.47216796875 0 2036.2051
844.4700927734375 0 1689.589
859.504638671875 0 2937.5903
860.5090942382812 0 1976.8192
861.483154296875 0 9177.219 y Water loss 2
862.484619140625 0 4268.3413
863.4901733398438 0 1010.43835
875.4777221679688 0 5854.9604 b Water loss 8
876.4808959960938 0 2412.12
877.4795532226562 0 2005.3712
879.4938354492188 0 200485.8 y 2
880.4968872070312 0 104941.21
881.4996948242188 0 27441.8
882.5009155273438 0 3016.6729
893.4882202148438 0 25215.213 b 8
894.4911499023438 0 15327.782
895.4940185546875 0 3670.937
907.5056762695312 0 2075.1323
908.51171875 0 1139.3267
978.5634765625 0 13055.225 y 1
979.5654907226562 0 7232.934
980.569091796875 0 2495.628
988.55859375 0 4297.392 b Water loss 9
989.5631713867188 0 2567.7012 b Ammonia loss 9
1006.5718994140625 0 26493.818 b 9
1007.5755615234375 0 18377.664
1008.5768432617188 0 5037.871
1009.5681762695312 0 958.4889
2484.978515625 0 614.0941

Spectrum Details

|  |  |
| --- | --- |
| Matched peaks? Matched peaks The total absolute number of peaks matched. Additionally in brackets the total fraction of peaks matched and the total number of peaks is shown. | 49 (9.74% of 503) |
| FDR? FDR The false discovery rate estimated for this peptide. It is calculated by matching all theoretical fragments with a non-integer shift with the raw peaks for this spectrum. This is done with 40 different shifts. The resulting percentage is the average number of annotated peaks over the number of annotated peaks with the correct spectrum. | 0.10% |
| Satellite FDR? Satellite FDR See the FDR for details on its calculation. This satellite ion specific FDR only contains the satellite ions (d/w) for I/L/J positions. | - |
| PSM Score? PSM Score The PSM Score as given by Hecklib to this annotated spectrum. It is shown with three significant figures. | 522 |

#### Spectrum 8401? Spectrum 8401 The raw spectrum of this peptide as annotated by Hecklib. The fragments are coloured according to ion type (see legend). Any peaks with a star '\*' as text can be hovered over to see the full details, first the ion type second the mass shift type. By hovering over the amino acids in the peptide or ions in the legend the corresponding peaks are highlighted. By toggling the 'Unassigned' label you can turn the background (unassigned) peaks on or off in the plot. By updating the slider in the Ion legend you can update the spectrum to only show the top X% of the peaks with labels. The top X% means any peak that is within X% of the highest intensity. By dragging in the spectrum you can zoom in to a specific part of the spectrum and use 'Zoom Out' to get back to the original zoom level. The annotation of the spectrum is based on the given sequence in the peptides file and is done with different software so inconsistencies are likely. The peaks are annotated based on the given sequence, with 20 ppm tolerance.

Copy Data

##### Spectrum 8401 (TSV)

###### Preview

```
Loading example...
```

*Click on the button to copy the data to your clipboard.*

Mz MinMz MaxIntensity Max

WidthHeightPeptide font sizePeptide stroke widthSpectrum font sizeSpectrum stroke widthCompact peptide

Ion legend

wxyz

abcd

OtherUnassignedIon^Charge^_Position_Show for top:%

FVVFGGGTKJT

03.45e+46.90e+41.04e+51.38e+5

Zoom Out

y^+1^_1_y^+1^_2_a^+1^_2_y^+1^_2_b^+1^_2_y^+2^_5_b^+2^_6_y^+1^_3_b^+1^_3_y^+1^_3_y^+2^_8_b^+2^_8_y^+2^_8_y^+2^_9_b^+2^_9_y^+2^_9_b^+2^_9_y^+1^_4_y^+2^_10_b^+1^_4_b^+2^_10_y^+1^_5_b^+1^_5_\*y^+1^_6_\*y^+1^_6_y^+1^_7_y^+1^_7_b^+1^_7_b^+1^_8_y^+1^_8_y^+1^_8_b^+1^_8_y^+1^_8_y^+1^_9_b^+1^_9_y^+1^_9_b^+1^_9_y^+1^_10_b^+1^_10_b^+1^_10_

0776155323293105

Fragment Matches Table

Show background peaks

| Position | Ion type | Intensity | mz Theoretical | mz Error (Th) | mz Error (ppm) | Charge | Series Number |
| --- | --- | --- | --- | --- | --- | --- | --- |
| 11 | y | 7876 | 120.1 | 0.0002687 | 2.238 | +1 | 1 |
| - | - | 1.049E+05 | 120.1 | - | - | 0 | - |
| - | - | 490.6 | 121.1 | - | - | 0 | - |
| - | - | 8484 | 121.1 | - | - | 0 | - |
| - | - | 579.6 | 127.1 | - | - | 0 | - |
| - | - | 757.1 | 127.1 | - | - | 0 | - |
| - | - | 742.7 | 128.1 | - | - | 0 | - |
| - | - | 4.773E+04 | 129.1 | - | - | 0 | - |
| - | - | 2532 | 130.1 | - | - | 0 | - |
| - | - | 506.8 | 131.1 | - | - | 0 | - |
| - | - | 1504 | 131.1 | - | - | 0 | - |
| - | - | 427.6 | 132.1 | - | - | 0 | - |
| - | - | 448.9 | 133.1 | - | - | 0 | - |
| - | - | 4280 | 133.1 | - | - | 0 | - |
| - | - | 1624 | 136.1 | - | - | 0 | - |
| - | - | 393.4 | 137.2 | - | - | 0 | - |
| - | - | 646.6 | 139.1 | - | - | 0 | - |
| - | - | 622.8 | 140.1 | - | - | 0 | - |
| - | - | 846.3 | 141.1 | - | - | 0 | - |
| - | - | 573 | 141.1 | - | - | 0 | - |
| - | - | 481.9 | 142.1 | - | - | 0 | - |
| - | - | 637.6 | 144.1 | - | - | 0 | - |
| - | - | 839.8 | 146.1 | - | - | 0 | - |
| - | - | 576.8 | 147.1 | - | - | 0 | - |
| - | - | 530.6 | 153.2 | - | - | 0 | - |
| - | - | 468.1 | 153.7 | - | - | 0 | - |
| - | - | 961.8 | 155.1 | - | - | 0 | - |
| - | - | 729.5 | 155.1 | - | - | 0 | - |
| - | - | 826.1 | 158.1 | - | - | 0 | - |
| - | - | 1397 | 159.1 | - | - | 0 | - |
| - | - | 498.5 | 166.1 | - | - | 0 | - |
| - | - | 442.3 | 166.1 | - | - | 0 | - |
| - | - | 809.5 | 170.1 | - | - | 0 | - |
| - | - | 1002 | 171.1 | - | - | 0 | - |
| - | - | 2331 | 171.1 | - | - | 0 | - |
| - | - | 2159 | 172.1 | - | - | 0 | - |
| - | - | 457.3 | 172.1 | - | - | 0 | - |
| - | - | 919.6 | 173.1 | - | - | 0 | - |
| - | - | 2162 | 173.4 | - | - | 0 | - |
| - | - | 678.1 | 174.1 | - | - | 0 | - |
| - | - | 1070 | 175.1 | - | - | 0 | - |
| - | - | 429.1 | 175.6 | - | - | 0 | - |
| - | - | 1401 | 176.1 | - | - | 0 | - |
| - | - | 1127 | 177.1 | - | - | 0 | - |
| - | - | 2289 | 177.1 | - | - | 0 | - |
| - | - | 448 | 181.8 | - | - | 0 | - |
| - | - | 565.1 | 182.1 | - | - | 0 | - |
| - | - | 880 | 183.1 | - | - | 0 | - |
| - | - | 532.8 | 185.1 | - | - | 0 | - |
| - | - | 798.4 | 185.1 | - | - | 0 | - |
| - | - | 2784 | 185.2 | - | - | 0 | - |
| - | - | 534.5 | 186.1 | - | - | 0 | - |
| - | - | 452.4 | 186.1 | - | - | 0 | - |
| - | - | 719.9 | 187.1 | - | - | 0 | - |
| - | - | 810.6 | 187.1 | - | - | 0 | - |
| - | - | 695.3 | 188.1 | - | - | 0 | - |
| - | - | 641.9 | 191.1 | - | - | 0 | - |
| - | - | 464.5 | 193.7 | - | - | 0 | - |
| - | - | 1614 | 197.2 | - | - | 0 | - |
| - | - | 1038 | 198.1 | - | - | 0 | - |
| - | - | 1986 | 199.1 | - | - | 0 | - |
| - | - | 1147 | 201.1 | - | - | 0 | - |
| - | - | 3346 | 205.1 | - | - | 0 | - |
| - | - | 631.3 | 206.1 | - | - | 0 | - |
| - | - | 763.1 | 208.1 | - | - | 0 | - |
| - | - | 4884 | 212.1 | - | - | 0 | - |
| - | - | 458.7 | 213.1 | - | - | 0 | - |
| - | - | 1438 | 213.2 | - | - | 0 | - |
| - | - | 535.1 | 214.2 | - | - | 0 | - |
| 10 | y | 2267 | 215.1 | 0.0004617 | 2.146 | +1 | 2 |
| - | - | 1371 | 216.1 | - | - | 0 | - |
| - | - | 754.8 | 217.1 | - | - | 0 | - |
| 2 | a | 1.367E+05 | 219.1 | 0.000377 | 1.72 | +1 | 2 |
| - | - | 1.748E+04 | 220.2 | - | - | 0 | - |
| - | - | 1026 | 221.2 | - | - | 0 | - |
| - | - | 1673 | 224.2 | - | - | 0 | - |
| - | - | 960.4 | 225.1 | - | - | 0 | - |
| - | - | 1915 | 227.1 | - | - | 0 | - |
| - | - | 1040 | 228.1 | - | - | 0 | - |
| - | - | 1.003E+04 | 230.2 | - | - | 0 | - |
| - | - | 695.6 | 231.2 | - | - | 0 | - |
| - | - | 596.6 | 233.1 | - | - | 0 | - |
| 10 | y | 6228 | 233.1 | 0.0003493 | 1.498 | +1 | 2 |
| - | - | 856.8 | 234.2 | - | - | 0 | - |
| - | - | 1181 | 239.1 | - | - | 0 | - |
| - | - | 2157 | 242.2 | - | - | 0 | - |
| - | - | 1185 | 243.1 | - | - | 0 | - |
| - | - | 2069 | 245.1 | - | - | 0 | - |
| 2 | b | 1.075E+05 | 247.1 | 0.0003507 | 1.419 | +1 | 2 |
| - | - | 1.691E+04 | 248.1 | - | - | 0 | - |
| - | - | 1191 | 249.2 | - | - | 0 | - |
| 7 | y | 1001 | 251.2 | 0.004077 | 16.23 | +2 | 5 |
| - | - | 675.6 | 253.2 | - | - | 0 | - |
| - | - | 6271 | 255.1 | - | - | 0 | - |
| - | - | 874.8 | 259.1 | - | - | 0 | - |
| - | - | 735.7 | 261.1 | - | - | 0 | - |
| - | - | 2410 | 262.1 | - | - | 0 | - |
| - | - | 739.2 | 267.1 | - | - | 0 | - |
| - | - | 5378 | 269.2 | - | - | 0 | - |
| - | - | 1126 | 270.2 | - | - | 0 | - |
| - | - | 576.8 | 271.1 | - | - | 0 | - |
| - | - | 5971 | 273.1 | - | - | 0 | - |
| - | - | 750.3 | 273.2 | - | - | 0 | - |
| - | - | 1018 | 274.1 | - | - | 0 | - |
| - | - | 555.9 | 274.5 | - | - | 0 | - |
| - | - | 541.1 | 279.3 | - | - | 0 | - |
| - | - | 796.4 | 283.2 | - | - | 0 | - |
| - | - | 1938 | 287.2 | - | - | 0 | - |
| - | - | 915.2 | 291.1 | - | - | 0 | - |
| - | - | 1047 | 295.1 | - | - | 0 | - |
| - | - | 1020 | 296.2 | - | - | 0 | - |
| - | - | 672.9 | 297.9 | - | - | 0 | - |
| - | - | 836.6 | 299.1 | - | - | 0 | - |
| - | - | 494.5 | 300.1 | - | - | 0 | - |
| - | - | 4446 | 301.2 | - | - | 0 | - |
| - | - | 942.4 | 302.2 | - | - | 0 | - |
| 6 | b | 2928 | 304.2 | 0.0003561 | 1.171 | +2 | 6 |
| - | - | 2604 | 319.1 | - | - | 0 | - |
| - | - | 621.5 | 321.2 | - | - | 0 | - |
| - | - | 866.6 | 325.2 | - | - | 0 | - |
| - | - | 4765 | 326.2 | - | - | 0 | - |
| - | - | 646.5 | 336.1 | - | - | 0 | - |
| - | - | 876.3 | 339.2 | - | - | 0 | - |
| 9 | y | 1854 | 343.2 | 0.0007288 | 2.123 | +1 | 3 |
| - | - | 702.7 | 344.2 | - | - | 0 | - |
| 3 | b | 2.641E+04 | 346.2 | 0.0003114 | 0.8994 | +1 | 3 |
| - | - | 5205 | 347.2 | - | - | 0 | - |
| - | - | 1453 | 348.2 | - | - | 0 | - |
| - | - | 1542 | 349.2 | - | - | 0 | - |
| - | - | 913.4 | 354.2 | - | - | 0 | - |
| - | - | 836.3 | 357.2 | - | - | 0 | - |
| - | - | 992.3 | 361.2 | - | - | 0 | - |
| 9 | y | 7136 | 361.2 | 0.0002349 | 0.6502 | +1 | 3 |
| - | - | 1464 | 362.2 | - | - | 0 | - |
| - | - | 1331 | 365.2 | - | - | 0 | - |
| - | - | 2348 | 374.2 | - | - | 0 | - |
| - | - | 700.5 | 379.2 | - | - | 0 | - |
| 4 | y | 688.2 | 381.7 | 0.0005438 | 1.425 | +2 | 8 |
| - | - | 1446 | 382.2 | - | - | 0 | - |
| 8 | b | 6450 | 383.2 | 0.003779 | 9.863 | +2 | 8 |
| - | - | 940 | 384.2 | - | - | 0 | - |
| - | - | 1139 | 390.2 | - | - | 0 | - |
| 4 | y | 3075 | 390.7 | 0.0003079 | 0.788 | +2 | 8 |
| - | - | 1978 | 392.2 | - | - | 0 | - |
| - | - | 645.6 | 393.2 | - | - | 0 | - |
| - | - | 2642 | 394.2 | - | - | 0 | - |
| - | - | 2488 | 400.3 | - | - | 0 | - |
| - | - | 5392 | 401.2 | - | - | 0 | - |
| - | - | 6784 | 402.2 | - | - | 0 | - |
| - | - | 756.3 | 402.2 | - | - | 0 | - |
| - | - | 1402 | 403.2 | - | - | 0 | - |
| - | - | 1042 | 403.2 | - | - | 0 | - |
| - | - | 1049 | 418.2 | - | - | 0 | - |
| - | - | 3488 | 420.2 | - | - | 0 | - |
| - | - | 797.9 | 421.2 | - | - | 0 | - |
| - | - | 649.7 | 423.2 | - | - | 0 | - |
| 3 | y | 775.9 | 431.2 | 0.0008938 | 2.072 | +2 | 9 |
| 9 | b | 679.5 | 438.7 | 0.008574 | 19.54 | +2 | 9 |
| - | - | 1861 | 439.3 | - | - | 0 | - |
| 3 | y | 5832 | 440.3 | 0.0006468 | 1.469 | +2 | 9 |
| - | - | 2688 | 440.8 | - | - | 0 | - |
| 9 | b | 2594 | 447.2 | 0.0006704 | 1.499 | +2 | 9 |
| - | - | 1130 | 448.3 | - | - | 0 | - |
| - | - | 2132 | 451.2 | - | - | 0 | - |
| - | - | 2921 | 457.3 | - | - | 0 | - |
| - | - | 843.8 | 458.3 | - | - | 0 | - |
| 8 | y | 1688 | 462.3 | 4.176E-05 | 0.09033 | +1 | 4 |
| - | - | 2960 | 465.3 | - | - | 0 | - |
| - | - | 1181 | 466.3 | - | - | 0 | - |
| - | - | 583.9 | 474.2 | - | - | 0 | - |
| - | - | 1223 | 478.3 | - | - | 0 | - |
| - | - | 2397 | 486.3 | - | - | 0 | - |
| - | - | 789.3 | 487.3 | - | - | 0 | - |
| 2 | y | 2092 | 489.8 | 0.006479 | 13.23 | +2 | 10 |
| - | - | 1722 | 490.3 | - | - | 0 | - |
| - | - | 1066 | 491.3 | - | - | 0 | - |
| 4 | b | 2808 | 493.3 | 0.0001348 | 0.2732 | +1 | 4 |
| 10 | b | 782.4 | 495.3 | 0.007222 | 14.58 | +2 | 10 |
| - | - | 6170 | 496.3 | - | - | 0 | - |
| - | - | 1212 | 497.3 | - | - | 0 | - |
| - | - | 2209 | 501.2 | - | - | 0 | - |
| - | - | 836.9 | 502.2 | - | - | 0 | - |
| - | - | 645.9 | 508.3 | - | - | 0 | - |
| - | - | 996.4 | 512.3 | - | - | 0 | - |
| - | - | 1.489E+04 | 514.3 | - | - | 0 | - |
| - | - | 4188 | 515.3 | - | - | 0 | - |
| - | - | 903.8 | 516.3 | - | - | 0 | - |
| - | - | 1884 | 519.3 | - | - | 0 | - |
| 7 | y | 2616 | 519.3 | 3.189E-05 | 0.06141 | +1 | 5 |
| - | - | 787.6 | 520.3 | - | - | 0 | - |
| - | - | 822.1 | 520.3 | - | - | 0 | - |
| - | - | 1171 | 522.3 | - | - | 0 | - |
| - | - | 973.9 | 527.3 | - | - | 0 | - |
| - | - | 3616 | 530.3 | - | - | 0 | - |
| - | - | 773.6 | 531.3 | - | - | 0 | - |
| - | - | 2051 | 531.3 | - | - | 0 | - |
| - | - | 678 | 545.3 | - | - | 0 | - |
| - | - | 667.4 | 547.3 | - | - | 0 | - |
| - | - | 7546 | 548.3 | - | - | 0 | - |
| - | - | 2569 | 549.3 | - | - | 0 | - |
| 5 | b | 1162 | 550.3 | 0.0006437 | 1.17 | +1 | 5 |
| 0 | Precursor | 1163 | 554.3 | 0.0004425 | 0.7983 | +2 | -1 |
| 6 | y | 884.2 | 558.3 | 0.003842 | 6.882 | +1 | 6 |
| - | - | 1557 | 562.3 | - | - | 0 | - |
| 0 | Precursor | 2219 | 563.3 | 0.001269 | 2.253 | +2 | -1 |
| - | - | 2511 | 563.8 | - | - | 0 | - |
| - | - | 695.9 | 564.8 | - | - | 0 | - |
| 6 | y | 8652 | 576.3 | 6.953E-05 | 0.1206 | +1 | 6 |
| - | - | 2994 | 577.3 | - | - | 0 | - |
| - | - | 835 | 578.3 | - | - | 0 | - |
| - | - | 1449 | 613.3 | - | - | 0 | - |
| 5 | y | 2533 | 615.3 | 0.0007502 | 1.219 | +1 | 7 |
| - | - | 1079 | 616.3 | - | - | 0 | - |
| - | - | 1680 | 629.3 | - | - | 0 | - |
| 5 | y | 4.181E+04 | 633.4 | 0.0005615 | 0.8865 | +1 | 7 |
| - | - | 1.392E+04 | 634.4 | - | - | 0 | - |
| - | - | 2564 | 635.4 | - | - | 0 | - |
| - | - | 6556 | 643.4 | - | - | 0 | - |
| - | - | 2684 | 644.4 | - | - | 0 | - |
| - | - | 676.4 | 645.4 | - | - | 0 | - |
| - | - | 4280 | 647.4 | - | - | 0 | - |
| - | - | 1796 | 648.4 | - | - | 0 | - |
| - | - | 1.344E+04 | 661.4 | - | - | 0 | - |
| - | - | 5157 | 662.4 | - | - | 0 | - |
| - | - | 716.7 | 663.4 | - | - | 0 | - |
| 7 | b | 1209 | 664.3 | 0.0006578 | 0.9901 | +1 | 7 |
| - | - | 837.8 | 665.3 | - | - | 0 | - |
| - | - | 1880 | 732.4 | - | - | 0 | - |
| - | - | 807 | 733.4 | - | - | 0 | - |
| - | - | 3678 | 742.4 | - | - | 0 | - |
| - | - | 1723 | 743.4 | - | - | 0 | - |
| 8 | b | 1225 | 747.4 | 0.0009247 | 1.237 | +1 | 8 |
| - | - | 1.174E+04 | 760.4 | - | - | 0 | - |
| - | - | 4000 | 761.4 | - | - | 0 | - |
| 4 | y | 2608 | 762.4 | 0.005456 | 7.157 | +1 | 8 |
| 4 | y | 766.3 | 763.4 | 0.01454 | 19.05 | +1 | 8 |
| 8 | b | 2713 | 765.4 | 0.0007899 | 1.032 | +1 | 8 |
| - | - | 1312 | 766.4 | - | - | 0 | - |
| - | - | 662 | 778.2 | - | - | 0 | - |
| 4 | y | 5.242E+04 | 780.4 | 0.0001645 | 0.2107 | +1 | 8 |
| - | - | 2.27E+04 | 781.4 | - | - | 0 | - |
| - | - | 5935 | 782.4 | - | - | 0 | - |
| - | - | 2459 | 785.4 | - | - | 0 | - |
| 3 | y | 1926 | 861.5 | 0.004059 | 4.711 | +1 | 9 |
| - | - | 1196 | 862.5 | - | - | 0 | - |
| 9 | b | 1348 | 875.5 | 4.414E-05 | 0.05042 | +1 | 9 |
| - | - | 920.7 | 876.5 | - | - | 0 | - |
| 3 | y | 4.466E+04 | 879.5 | 0.0005242 | 0.596 | +1 | 9 |
| - | - | 2.212E+04 | 880.5 | - | - | 0 | - |
| - | - | 5462 | 881.5 | - | - | 0 | - |
| 9 | b | 5658 | 893.5 | 0.001148 | 1.285 | +1 | 9 |
| - | - | 2578 | 894.5 | - | - | 0 | - |
| - | - | 1300 | 895.5 | - | - | 0 | - |
| - | - | 833 | 898.4 | - | - | 0 | - |
| - | - | 1480 | 914.4 | - | - | 0 | - |
| - | - | 621.8 | 915.4 | - | - | 0 | - |
| 2 | y | 2584 | 978.6 | 0.0002147 | 0.2195 | +1 | 10 |
| - | - | 1474 | 979.6 | - | - | 0 | - |
| 10 | b | 1020 | 988.6 | 0.00049 | 0.4956 | +1 | 10 |
| 10 | b | 5899 | 1007 | 0.001472 | 1.463 | +1 | 10 |
| - | - | 3692 | 1008 | - | - | 0 | - |
| - | - | 1064 | 1009 | - | - | 0 | - |
| - | - | 648.1 | 1171 | - | - | 0 | - |
| - | - | 603.8 | 2374 | - | - | 0 | - |
| - | - | 774.1 | 3026 | - | - | 0 | - |
| - | - | 662.3 | 3074 | - | - | 0 | - |
| - | - | 934.7 | 3074 | - | - | 0 | - |

m/z Charge Intensity FragmentType MassShift Position
120.06578826904297 0 7876.2617 y 10
120.08112335205078 0 104916.39
121.07891082763672 0 490.55603
121.08441925048828 0 8484.112
127.05045318603516 0 579.55054
127.07573699951172 0 757.0655
128.10736083984375 0 742.6664
129.1025390625 0 47728.324
130.10585021972656 0 2532.3057
131.07061767578125 0 506.78522
131.0817108154297 0 1503.7627
132.10214233398438 0 427.56537
133.06150817871094 0 448.86737
133.08621215820312 0 4280.317
136.07603454589844 0 1623.671
137.2362518310547 0 393.3744
139.0873260498047 0 646.62396
140.08216857910156 0 622.80664
141.06617736816406 0 846.2707
141.10263061523438 0 573.02026
142.06153869628906 0 481.9299
144.07728576660156 0 637.5955
146.06036376953125 0 839.7935
147.0647430419922 0 576.7921
153.19931030273438 0 530.64984
153.67466735839844 0 468.08716
155.0814666748047 0 961.8176
155.11814880371094 0 729.5063
158.09275817871094 0 826.06915
159.0765380859375 0 1397.4185
166.0539093017578 0 498.46686
166.08697509765625 0 442.25534
170.0927276611328 0 809.5459
171.07662963867188 0 1002.2456
171.1494598388672 0 2330.5762
172.0718231201172 0 2159.088
172.07925415039062 0 457.2779
173.12904357910156 0 919.55237
173.43942260742188 0 2161.9941
174.12818908691406 0 678.08307
175.09674072265625 0 1069.8329
175.56813049316406 0 429.09393
176.10702514648438 0 1400.8124
177.10301208496094 0 1127.453
177.11239624023438 0 2289.4514
181.8208465576172 0 447.95444
182.08175659179688 0 565.08203
183.1134033203125 0 879.9732
185.09274291992188 0 532.8116
185.12879943847656 0 798.3974
185.16514587402344 0 2783.8142
186.1232147216797 0 534.5466
186.1317901611328 0 452.35764
187.107177734375 0 719.88605
187.144287109375 0 810.5911
188.10321044921875 0 695.2698
191.0914764404297 0 641.8981
193.72377014160156 0 464.45453
197.1649932861328 0 1614.396
198.08737182617188 0 1038.182
199.14431762695312 0 1986.205
201.12380981445312 0 1147.1044
205.09742736816406 0 3345.6133
206.1019287109375 0 631.2694
208.10772705078125 0 763.07324
212.13966369628906 0 4883.5415
213.14251708984375 0 458.7142
213.16012573242188 0 1437.57
214.19149780273438 0 535.1048
215.1394805908203 0 2266.5413 y Water loss 9
216.09823608398438 0 1371.2874
217.0972137451172 0 754.77856
219.14956665039062 0 136706.03 a 1
220.15283203125 0 17484.752
221.15615844726562 0 1026.2437
224.1763153076172 0 1673.3982
225.1349334716797 0 960.4391
227.1142120361328 0 1914.8531
228.09835815429688 0 1040.365
230.1501922607422 0 10030.044
231.15220642089844 0 695.5591
233.1295166015625 0 596.6039
233.14993286132812 0 6228.362 y 9
234.15341186523438 0 856.7584
239.14920043945312 0 1181.1505
242.18637084960938 0 2157.2625
243.1133270263672 0 1184.906
245.12478637695312 0 2069.358
247.1444549560547 0 107492.445 b 1
248.14772033691406 0 16914.842
249.1502227783203 0 1191.0236
251.151123046875 0 1000.59033 y Water loss 6
253.16661071777344 0 675.63837
255.10903930664062 0 6271.4614
259.1446228027344 0 874.8282
261.1233825683594 0 735.7149
262.1191711425781 0 2410.2932
267.148681640625 0 739.20917
269.1609802246094 0 5378.4746
270.1649475097656 0 1126.034
271.14013671875 0 576.78827
273.1194763183594 0 5971.283
273.19683837890625 0 750.29877
274.12164306640625 0 1017.5504
274.50042724609375 0 555.9097
279.2686767578125 0 541.1283
283.1758117675781 0 796.3549
287.1722412109375 0 1938.1738
291.14508056640625 0 915.2085
295.14501953125 0 1047.469
296.1973876953125 0 1020.2767
297.8651123046875 0 672.8867
299.0623779296875 0 836.64557
300.0629577636719 0 494.54153
301.19146728515625 0 4446.3916
302.1941223144531 0 942.4298
304.1659240722656 0 2927.9512 b 5
319.1407775878906 0 2604.1362
321.1973571777344 0 621.46643
325.2232666015625 0 866.64404
326.1824645996094 0 4764.9834
336.1443176269531 0 646.52496
339.1796875 0 876.26733
343.2347106933594 0 1854.3351 y Water loss 8
344.19439697265625 0 702.6773
346.21282958984375 0 26407.87 b 2
347.2159118652344 0 5205.496
348.16778564453125 0 1452.897
349.19085693359375 0 1542.1168
354.17645263671875 0 913.3851
357.15606689453125 0 836.3373
361.1871337890625 0 992.2988
361.2447814941406 0 7135.7026 y 8
362.24835205078125 0 1463.9545
365.1928405761719 0 1331.1519
374.18243408203125 0 2348.3018
379.21075439453125 0 700.4901
381.7103271484375 0 688.1868 y Water loss 3
382.2466735839844 0 1446.2856
383.20391845703125 0 6449.76 b 7
384.2071533203125 0 940.04767
390.2144775390625 0 1138.8027
390.7164611816406 0 3074.5334 y 3
392.193359375 0 1977.6101
393.1969909667969 0 645.57
394.2129211425781 0 2641.851
400.2552795410156 0 2487.571
401.2147216796875 0 5392.395
402.17767333984375 0 6784.4146
402.2184143066406 0 756.3284
403.1794738769531 0 1402.4583
403.2342834472656 0 1041.756
418.20794677734375 0 1049.4698
420.1883239746094 0 3487.7185
421.1919250488281 0 797.9058
423.24090576171875 0 649.7478
431.2459716796875 0 775.8939 y Water loss 2
438.742919921875 0 679.46875 b Ammonia loss 8
439.2671203613281 0 1860.9502
440.2510070800781 0 5832.151 y 2
440.7520751953125 0 2688.3237
447.248291015625 0 2593.7876 b 8
448.2572326660156 0 1129.5939
451.2348327636719 0 2131.8062
457.27752685546875 0 2920.7942
458.28143310546875 0 843.75977
462.2922668457031 0 1687.545 y 7
465.2862854003906 0 2960.4363
466.2907409667969 0 1181.4834
474.2400817871094 0 583.9407
478.2775573730469 0 1223.479
486.30401611328125 0 2397.1235
487.3056945800781 0 789.27466
489.7910461425781 0 2092.1008 y 1
490.29278564453125 0 1722.2457
491.2626037597656 0 1065.9952
493.28106689453125 0 2807.515 b 3
495.2835998535156 0 782.4424 b Ammonia loss 9
496.2878112792969 0 6170.426
497.2921142578125 0 1212.2476
501.24664306640625 0 2208.6338
502.24993896484375 0 836.92206
508.2586975097656 0 645.90564
512.2625732421875 0 996.37683
514.298583984375 0 14892.674
515.3018798828125 0 4187.97
516.3043823242188 0 903.81915
519.2568969726562 0 1884.3153
519.313720703125 0 2616.2004 y 6
520.257080078125 0 787.6265
520.31640625 0 822.06934
522.305908203125 0 1170.7649
527.306640625 0 973.85956
530.2726440429688 0 3616.1113
531.270263671875 0 773.6215
531.3372802734375 0 2051.4495
545.31787109375 0 678.00867
547.3321533203125 0 667.3509
548.282470703125 0 7546.441
549.2869873046875 0 2568.6404
550.3030395507812 0 1161.6227 b 4
554.3130493164062 0 1162.8052 Precursor Water loss
558.3284301757812 0 884.1667 y Water loss 5
562.343017578125 0 1557.3239
563.3175048828125 0 2218.6804 Precursor
563.8197631835938 0 2511.4297
564.7581787109375 0 695.9308
576.3350830078125 0 8652.064 y 5
577.3392944335938 0 2993.702
578.3399047851562 0 834.99536
613.3286743164062 0 1449.2053
615.3468017578125 0 2533.4382 y Water loss 4
616.3439331054688 0 1078.7129
629.339599609375 0 1680.4663
633.357177734375 0 41814.656 y 4
634.3606567382812 0 13919.315
635.3630981445312 0 2564.044
643.356201171875 0 6555.716
644.3587036132812 0 2683.882
645.3644409179688 0 676.44
647.3523559570312 0 4280.15
648.3530883789062 0 1795.8734
661.366943359375 0 13443.965
662.369140625 0 5156.7183
663.3702392578125 0 716.6928
664.3446655273438 0 1209.1594 b 6
665.3494262695312 0 837.7972
732.4422607421875 0 1880.2908
733.443359375 0 806.9928
742.4251098632812 0 3678.1167
743.4262084960938 0 1722.8322
747.3833618164062 0 1224.7572 b Water loss 7
760.43505859375 0 11735.72
761.4393310546875 0 4000.04
762.419921875 0 2607.9968 y Water loss 3
763.4130249023438 0 766.3156 y Ammonia loss 3
765.3922119140625 0 2713.2524 b 7
766.3944091796875 0 1312.2194
778.1591796875 0 661.9979
780.4248657226562 0 52422.37 y 3
781.4278564453125 0 22698.977
782.4296264648438 0 5935.2666
785.413330078125 0 2458.8652
861.4788208007812 0 1926.3092 y Water loss 2
862.4849243164062 0 1196.2983
875.4773559570312 0 1348.3646 b Water loss 8
876.4810791015625 0 920.7095
879.492919921875 0 44656.48 y 2
880.4961547851562 0 22118.625
881.4974975585938 0 5462.071
893.48681640625 0 5657.874 b 8
894.49072265625 0 2577.6792
895.4905395507812 0 1299.7368
898.4430541992188 0 833.00946
914.43505859375 0 1479.7014
915.4414672851562 0 621.7996
978.5620727539062 0 2584.0923 y 1
979.5665893554688 0 1473.5024
988.5609741210938 0 1020.2086 b Water loss 9
1006.570556640625 0 5898.902 b 9
1007.572509765625 0 3692.392
1008.5792236328125 0 1063.5049
1170.5736083984375 0 648.05835
2374.168701171875 0 603.8232
3025.90234375 0 774.1399
3073.71923828125 0 662.2555
3074.403076171875 0 934.6561

Spectrum Details

|  |  |
| --- | --- |
| Matched peaks? Matched peaks The total absolute number of peaks matched. Additionally in brackets the total fraction of peaks matched and the total number of peaks is shown. | 42 (15.73% of 267) |
| FDR? FDR The false discovery rate estimated for this peptide. It is calculated by matching all theoretical fragments with a non-integer shift with the raw peaks for this spectrum. This is done with 40 different shifts. The resulting percentage is the average number of annotated peaks over the number of annotated peaks with the correct spectrum. | 0.17% |
| Satellite FDR? Satellite FDR See the FDR for details on its calculation. This satellite ion specific FDR only contains the satellite ions (d/w) for I/L/J positions. | - |
| PSM Score? PSM Score The PSM Score as given by Hecklib to this annotated spectrum. It is shown with three significant figures. | 454 |

#### Spectrum 8454? Spectrum 8454 The raw spectrum of this peptide as annotated by Hecklib. The fragments are coloured according to ion type (see legend). Any peaks with a star '\*' as text can be hovered over to see the full details, first the ion type second the mass shift type. By hovering over the amino acids in the peptide or ions in the legend the corresponding peaks are highlighted. By toggling the 'Unassigned' label you can turn the background (unassigned) peaks on or off in the plot. By updating the slider in the Ion legend you can update the spectrum to only show the top X% of the peaks with labels. The top X% means any peak that is within X% of the highest intensity. By dragging in the spectrum you can zoom in to a specific part of the spectrum and use 'Zoom Out' to get back to the original zoom level. The annotation of the spectrum is based on the given sequence in the peptides file and is done with different software so inconsistencies are likely. The peaks are annotated based on the given sequence, with 20 ppm tolerance.

Copy Data

##### Spectrum 8454 (TSV)

###### Preview

```
Loading example...
```

*Click on the button to copy the data to your clipboard.*

Mz MinMz MaxIntensity Max

WidthHeightPeptide font sizePeptide stroke widthSpectrum font sizeSpectrum stroke widthCompact peptide

Ion legend

wxyz

abcd

OtherUnassignedIon^Charge^_Position_Show for top:%

FVVFGGGTKJT

02.51e+45.03e+47.54e+41.01e+5

Zoom Out

y^+1^_1_y^+1^_2_a^+1^_2_y^+1^_2_b^+1^_2_b^+2^_6_y^+1^_3_b^+1^_3_y^+1^_3_b^+2^_8_y^+2^_8_y^+2^_9_b^+2^_9_y^+1^_4_y^+2^_10_b^+1^_4_b^+2^_10_y^+1^_5_b^+1^_5_\*\*\*y^+1^_6_y^+1^_7_y^+1^_7_y^+1^_8_b^+1^_8_y^+1^_8_y^+1^_9_b^+1^_9_y^+1^_9_b^+1^_9_y^+1^_10_b^+1^_10_

0776155323293106

Fragment Matches Table

Show background peaks

| Position | Ion type | Intensity | mz Theoretical | mz Error (Th) | mz Error (ppm) | Charge | Series Number |
| --- | --- | --- | --- | --- | --- | --- | --- |
| 11 | y | 5225 | 120.1 | 0.0002305 | 1.92 | +1 | 1 |
| - | - | 7.376E+04 | 120.1 | - | - | 0 | - |
| - | - | 6648 | 121.1 | - | - | 0 | - |
| - | - | 470.2 | 123.1 | - | - | 0 | - |
| - | - | 853.9 | 128.1 | - | - | 0 | - |
| - | - | 3.374E+04 | 129.1 | - | - | 0 | - |
| - | - | 471.8 | 130.1 | - | - | 0 | - |
| - | - | 382.8 | 130.1 | - | - | 0 | - |
| - | - | 2080 | 130.1 | - | - | 0 | - |
| - | - | 448.6 | 131.1 | - | - | 0 | - |
| - | - | 1010 | 131.1 | - | - | 0 | - |
| - | - | 6254 | 133.1 | - | - | 0 | - |
| - | - | 389.2 | 135.2 | - | - | 0 | - |
| - | - | 2218 | 136.1 | - | - | 0 | - |
| - | - | 559.8 | 141.1 | - | - | 0 | - |
| - | - | 783.3 | 144.1 | - | - | 0 | - |
| - | - | 468.1 | 144.6 | - | - | 0 | - |
| - | - | 439.5 | 147.1 | - | - | 0 | - |
| - | - | 743.2 | 147.1 | - | - | 0 | - |
| - | - | 511.5 | 152.1 | - | - | 0 | - |
| - | - | 485.2 | 155.1 | - | - | 0 | - |
| - | - | 609.5 | 155.1 | - | - | 0 | - |
| - | - | 816.6 | 158.1 | - | - | 0 | - |
| - | - | 1170 | 159.1 | - | - | 0 | - |
| - | - | 423.7 | 163.1 | - | - | 0 | - |
| - | - | 469.9 | 166.1 | - | - | 0 | - |
| - | - | 615.7 | 167.1 | - | - | 0 | - |
| - | - | 863.4 | 170.1 | - | - | 0 | - |
| - | - | 2340 | 171.1 | - | - | 0 | - |
| - | - | 1602 | 172.1 | - | - | 0 | - |
| - | - | 2746 | 173.5 | - | - | 0 | - |
| - | - | 745.2 | 175.1 | - | - | 0 | - |
| - | - | 1238 | 176.1 | - | - | 0 | - |
| - | - | 1331 | 177.1 | - | - | 0 | - |
| - | - | 1952 | 177.1 | - | - | 0 | - |
| - | - | 945.9 | 182.1 | - | - | 0 | - |
| - | - | 564.3 | 183.1 | - | - | 0 | - |
| - | - | 602.4 | 185.1 | - | - | 0 | - |
| - | - | 732 | 186.1 | - | - | 0 | - |
| - | - | 571.7 | 187.1 | - | - | 0 | - |
| - | - | 824.7 | 197.2 | - | - | 0 | - |
| - | - | 1037 | 198.1 | - | - | 0 | - |
| - | - | 1203 | 199.1 | - | - | 0 | - |
| - | - | 2942 | 205.1 | - | - | 0 | - |
| - | - | 650.7 | 208.1 | - | - | 0 | - |
| - | - | 555.5 | 210.1 | - | - | 0 | - |
| - | - | 3735 | 212.1 | - | - | 0 | - |
| - | - | 876.9 | 215.1 | - | - | 0 | - |
| 10 | y | 969.2 | 215.1 | 0.0003701 | 1.72 | +1 | 2 |
| - | - | 803 | 216.1 | - | - | 0 | - |
| 2 | a | 9.96E+04 | 219.1 | 0.0002091 | 0.9543 | +1 | 2 |
| - | - | 1.356E+04 | 220.2 | - | - | 0 | - |
| - | - | 628.6 | 221.1 | - | - | 0 | - |
| - | - | 643.5 | 221.2 | - | - | 0 | - |
| - | - | 1163 | 224.2 | - | - | 0 | - |
| - | - | 610.6 | 227.1 | - | - | 0 | - |
| - | - | 2084 | 227.1 | - | - | 0 | - |
| - | - | 763.9 | 228.1 | - | - | 0 | - |
| - | - | 6070 | 230.1 | - | - | 0 | - |
| - | - | 682 | 231.2 | - | - | 0 | - |
| 10 | y | 4028 | 233.1 | 0.0001814 | 0.7781 | +1 | 2 |
| - | - | 735.9 | 239.1 | - | - | 0 | - |
| - | - | 1633 | 242.2 | - | - | 0 | - |
| - | - | 618.4 | 243.1 | - | - | 0 | - |
| - | - | 586.9 | 243.1 | - | - | 0 | - |
| - | - | 1280 | 245.1 | - | - | 0 | - |
| 2 | b | 7.481E+04 | 247.1 | 0.0002133 | 0.8632 | +1 | 2 |
| - | - | 1.049E+04 | 248.1 | - | - | 0 | - |
| - | - | 750.7 | 253.2 | - | - | 0 | - |
| - | - | 3262 | 255.1 | - | - | 0 | - |
| - | - | 662.4 | 256.1 | - | - | 0 | - |
| - | - | 890.6 | 259.1 | - | - | 0 | - |
| - | - | 531.1 | 261.1 | - | - | 0 | - |
| - | - | 1764 | 262.1 | - | - | 0 | - |
| - | - | 4864 | 269.2 | - | - | 0 | - |
| - | - | 5254 | 273.1 | - | - | 0 | - |
| - | - | 838.1 | 274.1 | - | - | 0 | - |
| - | - | 526.3 | 286.2 | - | - | 0 | - |
| - | - | 1254 | 287.2 | - | - | 0 | - |
| - | - | 1305 | 291.1 | - | - | 0 | - |
| - | - | 1102 | 295.1 | - | - | 0 | - |
| - | - | 901.5 | 299.1 | - | - | 0 | - |
| - | - | 634 | 299.1 | - | - | 0 | - |
| - | - | 2109 | 301.2 | - | - | 0 | - |
| 6 | b | 1674 | 304.2 | 0.0003458 | 1.137 | +2 | 6 |
| - | - | 1258 | 319.1 | - | - | 0 | - |
| - | - | 617.4 | 321.2 | - | - | 0 | - |
| - | - | 4045 | 326.2 | - | - | 0 | - |
| - | - | 642.5 | 327.2 | - | - | 0 | - |
| - | - | 612.2 | 328.2 | - | - | 0 | - |
| - | - | 1049 | 339.2 | - | - | 0 | - |
| 9 | y | 1601 | 343.2 | 0.0003626 | 1.056 | +1 | 3 |
| - | - | 907.9 | 344.2 | - | - | 0 | - |
| 3 | b | 1.706E+04 | 346.2 | 0.0001283 | 0.3705 | +1 | 3 |
| - | - | 4543 | 347.2 | - | - | 0 | - |
| - | - | 532.8 | 355.2 | - | - | 0 | - |
| - | - | 1179 | 357.2 | - | - | 0 | - |
| - | - | 600 | 361.2 | - | - | 0 | - |
| 9 | y | 4296 | 361.2 | 2.126E-05 | 0.05885 | +1 | 3 |
| - | - | 1039 | 362.2 | - | - | 0 | - |
| - | - | 1149 | 365.2 | - | - | 0 | - |
| - | - | 538.3 | 371.2 | - | - | 0 | - |
| - | - | 1572 | 374.2 | - | - | 0 | - |
| - | - | 809.8 | 375.2 | - | - | 0 | - |
| 8 | b | 3836 | 383.2 | 0.003627 | 9.464 | +2 | 8 |
| - | - | 996.4 | 384.2 | - | - | 0 | - |
| 4 | y | 1648 | 390.7 | 0.0007351 | 1.881 | +2 | 8 |
| - | - | 713.7 | 391.2 | - | - | 0 | - |
| - | - | 1387 | 392.2 | - | - | 0 | - |
| - | - | 572.6 | 392.8 | - | - | 0 | - |
| - | - | 1525 | 394.2 | - | - | 0 | - |
| - | - | 1745 | 400.3 | - | - | 0 | - |
| - | - | 3807 | 401.2 | - | - | 0 | - |
| - | - | 4021 | 402.2 | - | - | 0 | - |
| - | - | 1078 | 402.2 | - | - | 0 | - |
| - | - | 646.4 | 403.2 | - | - | 0 | - |
| - | - | 1431 | 420.2 | - | - | 0 | - |
| - | - | 697.7 | 421.2 | - | - | 0 | - |
| - | - | 1552 | 439.3 | - | - | 0 | - |
| 3 | y | 2958 | 440.3 | 0.001318 | 2.994 | +2 | 9 |
| - | - | 1694 | 440.8 | - | - | 0 | - |
| 9 | b | 1644 | 447.2 | 0.0001211 | 0.2707 | +2 | 9 |
| - | - | 654.8 | 447.7 | - | - | 0 | - |
| - | - | 1172 | 451.2 | - | - | 0 | - |
| - | - | 715.4 | 456.2 | - | - | 0 | - |
| - | - | 3553 | 457.3 | - | - | 0 | - |
| 8 | y | 1334 | 462.3 | 0.0002634 | 0.5698 | +1 | 4 |
| - | - | 1841 | 465.3 | - | - | 0 | - |
| - | - | 675.7 | 466.3 | - | - | 0 | - |
| - | - | 1401 | 486.3 | - | - | 0 | - |
| - | - | 751.2 | 487.3 | - | - | 0 | - |
| 2 | y | 1198 | 489.8 | 0.008249 | 16.84 | +2 | 10 |
| - | - | 1124 | 490.3 | - | - | 0 | - |
| - | - | 659.5 | 491.3 | - | - | 0 | - |
| 4 | b | 2260 | 493.3 | 0.0009893 | 2.005 | +1 | 4 |
| - | - | 3402 | 496.3 | - | - | 0 | - |
| - | - | 647.8 | 497.3 | - | - | 0 | - |
| - | - | 2029 | 501.2 | - | - | 0 | - |
| 10 | b | 589.4 | 503.8 | 0.001058 | 2.1 | +2 | 10 |
| - | - | 599.1 | 508.3 | - | - | 0 | - |
| - | - | 9398 | 514.3 | - | - | 0 | - |
| - | - | 2636 | 515.3 | - | - | 0 | - |
| - | - | 650.9 | 519.3 | - | - | 0 | - |
| 7 | y | 1662 | 519.3 | 0.001189 | 2.289 | +1 | 5 |
| - | - | 662.9 | 520.3 | - | - | 0 | - |
| - | - | 774.4 | 522.3 | - | - | 0 | - |
| - | - | 636.3 | 527.3 | - | - | 0 | - |
| - | - | 2858 | 530.3 | - | - | 0 | - |
| - | - | 827.8 | 531.3 | - | - | 0 | - |
| - | - | 4479 | 545.3 | - | - | 0 | - |
| - | - | 2090 | 546.3 | - | - | 0 | - |
| - | - | 1751 | 547.3 | - | - | 0 | - |
| - | - | 4545 | 548.3 | - | - | 0 | - |
| - | - | 1218 | 549.3 | - | - | 0 | - |
| 5 | b | 1011 | 550.3 | 0.003446 | 6.261 | +1 | 5 |
| 0 | Precursor | 1293 | 554.3 | 0.001144 | 2.065 | +2 | -1 |
| 0 | Precursor | 670.2 | 554.8 | 0.008892 | 16.03 | +2 | -1 |
| - | - | 1916 | 562.3 | - | - | 0 | - |
| 0 | Precursor | 1715 | 563.3 | 0.002185 | 3.879 | +2 | -1 |
| - | - | 1493 | 563.8 | - | - | 0 | - |
| 6 | y | 5665 | 576.3 | 0.0002526 | 0.4383 | +1 | 6 |
| - | - | 1853 | 577.3 | - | - | 0 | - |
| - | - | 838 | 613.3 | - | - | 0 | - |
| 5 | y | 1948 | 615.3 | 7.88E-05 | 0.1281 | +1 | 7 |
| - | - | 670 | 616.3 | - | - | 0 | - |
| 5 | y | 3.033E+04 | 633.4 | 0.0003784 | 0.5974 | +1 | 7 |
| - | - | 9492 | 634.4 | - | - | 0 | - |
| - | - | 1935 | 635.4 | - | - | 0 | - |
| - | - | 4623 | 643.4 | - | - | 0 | - |
| - | - | 1368 | 644.4 | - | - | 0 | - |
| - | - | 3237 | 647.4 | - | - | 0 | - |
| - | - | 800.3 | 648.4 | - | - | 0 | - |
| - | - | 8833 | 661.4 | - | - | 0 | - |
| - | - | 3676 | 662.4 | - | - | 0 | - |
| - | - | 725.8 | 663.4 | - | - | 0 | - |
| - | - | 1016 | 732.4 | - | - | 0 | - |
| - | - | 2147 | 742.4 | - | - | 0 | - |
| - | - | 761.4 | 743.4 | - | - | 0 | - |
| - | - | 679.2 | 757.4 | - | - | 0 | - |
| - | - | 7593 | 760.4 | - | - | 0 | - |
| - | - | 3318 | 761.4 | - | - | 0 | - |
| 4 | y | 2040 | 762.4 | 0.003442 | 4.515 | +1 | 8 |
| - | - | 1059 | 763.4 | - | - | 0 | - |
| 8 | b | 1993 | 765.4 | 0.0002477 | 0.3237 | +1 | 8 |
| 4 | y | 3.297E+04 | 780.4 | 0.0005917 | 0.7582 | +1 | 8 |
| - | - | 1.555E+04 | 781.4 | - | - | 0 | - |
| - | - | 3578 | 782.4 | - | - | 0 | - |
| - | - | 2136 | 785.4 | - | - | 0 | - |
| 3 | y | 1702 | 861.5 | 0.001434 | 1.665 | +1 | 9 |
| 9 | b | 751.7 | 875.5 | 0.004045 | 4.621 | +1 | 9 |
| 3 | y | 3.105E+04 | 879.5 | 0.001196 | 1.359 | +1 | 9 |
| - | - | 1.429E+04 | 880.5 | - | - | 0 | - |
| - | - | 4304 | 881.5 | - | - | 0 | - |
| 9 | b | 4588 | 893.5 | 0.001515 | 1.695 | +1 | 9 |
| - | - | 2105 | 894.5 | - | - | 0 | - |
| - | - | 792 | 895.5 | - | - | 0 | - |
| 2 | y | 1130 | 978.6 | 0.006807 | 6.956 | +1 | 10 |
| - | - | 1132 | 979.6 | - | - | 0 | - |
| 10 | b | 4030 | 1007 | 0.002082 | 2.069 | +1 | 10 |
| - | - | 3418 | 1008 | - | - | 0 | - |
| - | - | 674 | 1014 | - | - | 0 | - |
| - | - | 631.7 | 1095 | - | - | 0 | - |
| - | - | 673.9 | 1280 | - | - | 0 | - |
| - | - | 591.3 | 1361 | - | - | 0 | - |
| - | - | 582.3 | 1487 | - | - | 0 | - |
| - | - | 763.7 | 2002 | - | - | 0 | - |
| - | - | 761.1 | 2273 | - | - | 0 | - |
| - | - | 667.7 | 2583 | - | - | 0 | - |
| - | - | 1093 | 3075 | - | - | 0 | - |

m/z Charge Intensity FragmentType MassShift Position
120.06575012207031 0 5224.811 y 10
120.08103942871094 0 73758.83
121.0843505859375 0 6648.1143
123.08038330078125 0 470.22665
128.10726928710938 0 853.9482
129.1024627685547 0 33744.824
130.06101989746094 0 471.84036
130.099365234375 0 382.7595
130.10568237304688 0 2080.0288
131.07029724121094 0 448.60657
131.0817108154297 0 1010.03876
133.08619689941406 0 6254.1387
135.18118286132812 0 389.2341
136.07589721679688 0 2218.282
141.06594848632812 0 559.8024
144.0764617919922 0 783.29755
144.5637664794922 0 468.0649
147.06419372558594 0 439.45908
147.10169982910156 0 743.24744
152.1437530517578 0 511.46234
155.07020568847656 0 485.16922
155.11854553222656 0 609.5447
158.0924530029297 0 816.64795
159.07676696777344 0 1170.2742
163.05308532714844 0 423.71994
166.08648681640625 0 469.86823
167.1180419921875 0 615.7259
170.09263610839844 0 863.4135
171.14932250976562 0 2340.0312
172.07211303710938 0 1601.9667
173.45140075683594 0 2745.5637
175.09661865234375 0 745.20123
176.10755920410156 0 1238.0386
177.102294921875 0 1331.2802
177.11219787597656 0 1951.6377
182.0812530517578 0 945.9104
183.1132049560547 0 564.27246
185.09190368652344 0 602.42346
186.12364196777344 0 731.95325
187.10787963867188 0 571.6864
197.1652069091797 0 824.66034
198.0876922607422 0 1036.89
199.14405822753906 0 1203.0321
205.09732055664062 0 2941.5164
208.1078643798828 0 650.7055
210.0884552001953 0 555.4565
212.1396484375 0 3734.859
215.11436462402344 0 876.924
215.13938903808594 0 969.2113 y Water loss 9
216.09762573242188 0 803.00916
219.14939880371094 0 99603.32 a 1
220.1527862548828 0 13557.564
221.13902282714844 0 628.6093
221.15589904785156 0 643.5159
224.17544555664062 0 1163.1564
227.10240173339844 0 610.5977
227.11416625976562 0 2083.5283
228.0979461669922 0 763.8888
230.14999389648438 0 6070.244
231.15090942382812 0 681.9544
233.14976501464844 0 4028.155 y 9
239.1494140625 0 735.9134
242.18621826171875 0 1633.1073
243.11334228515625 0 618.4264
243.14576721191406 0 586.94916
245.1249237060547 0 1279.958
247.14431762695312 0 74813.95 b 1
248.147705078125 0 10491.901
253.1667022705078 0 750.65857
255.1090850830078 0 3261.6743
256.1116943359375 0 662.3794
259.1440124511719 0 890.555
261.1226806640625 0 531.126
262.1188659667969 0 1763.9489
269.16082763671875 0 4864.0283
273.1194763183594 0 5254.362
274.12060546875 0 838.1222
286.2140808105469 0 526.3382
287.1719055175781 0 1253.9198
291.1454772949219 0 1304.618
295.1444396972656 0 1102.25
299.0621032714844 0 901.4685
299.139404296875 0 634
301.19134521484375 0 2109.024
304.16522216796875 0 1673.8646 b 5
319.14019775390625 0 1257.7139
321.1954040527344 0 617.43713
326.1824645996094 0 4045.4475
327.2013854980469 0 642.53613
328.1671142578125 0 612.21576
339.17730712890625 0 1049.2522
343.2343444824219 0 1601.4377 y Water loss 8
344.193603515625 0 907.8795
346.212646484375 0 17055.576 b 2
347.2160339355469 0 4542.835
355.2232666015625 0 532.8102
357.1552429199219 0 1179.3385
361.185302734375 0 599.9859
361.24456787109375 0 4296.394 y 8
362.2455749511719 0 1038.556
365.1932678222656 0 1148.5966
371.2288513183594 0 538.3028
374.1828308105469 0 1572.4004
375.1659851074219 0 809.75616
383.2037658691406 0 3836.4514 b 7
384.2063293457031 0 996.43823
390.7168884277344 0 1647.5743 y 3
391.2159729003906 0 713.67725
392.1924133300781 0 1387.1632
392.77850341796875 0 572.6493
394.2121887207031 0 1525.2048
400.2557678222656 0 1745.3502
401.21435546875 0 3806.592
402.17742919921875 0 4020.7915
402.2166442871094 0 1077.6553
403.1770935058594 0 646.39984
420.18743896484375 0 1431.0001
421.1888122558594 0 697.7213
439.265380859375 0 1552.3812
440.2516784667969 0 2958.2769 y 2
440.7528991699219 0 1693.6432
447.24774169921875 0 1644.4879 b 8
447.74859619140625 0 654.8439
451.2360534667969 0 1172.1233
456.22418212890625 0 715.3526
457.2768859863281 0 3552.8416
462.2919616699219 0 1334.0803 y 7
465.2867736816406 0 1840.8182
466.2902526855469 0 675.7479
486.30462646484375 0 1400.742
487.3055419921875 0 751.20764
489.7928161621094 0 1198.4071 y 1
490.2942199707031 0 1124.4929
491.2630920410156 0 659.5068
493.28192138671875 0 2259.8108 b 3
496.2873229980469 0 3402.0789
497.2962646484375 0 647.8065
501.2461242675781 0 2028.992
503.79071044921875 0 589.3989 b 9
508.2566223144531 0 599.1224
514.2981567382812 0 9398.274
515.3014526367188 0 2635.705
519.2552490234375 0 650.8693
519.3125 0 1662.3805 y 6
520.3181762695312 0 662.9332
522.3053588867188 0 774.36993
527.3095092773438 0 636.2765
530.2711791992188 0 2857.6904
531.2779541015625 0 827.8179
545.3163452148438 0 4479.0317
546.3202514648438 0 2089.6965
547.3296508789062 0 1751.4556
548.2823486328125 0 4545.202
549.2863159179688 0 1218.2673
550.2989501953125 0 1011.40344 b 4
554.3146362304688 0 1292.5125 Precursor Water loss
554.8143920898438 0 670.1724 Precursor Ammonia loss
562.343505859375 0 1916.4993
563.3165893554688 0 1715.4354 Precursor
563.8186645507812 0 1492.6141
576.3348999023438 0 5665.1196 y 5
577.33837890625 0 1852.7769
613.3303833007812 0 838.02545
615.3461303710938 0 1947.6415 y Water loss 4
616.347900390625 0 669.9853
633.3569946289062 0 30330.33 y 4
634.3599243164062 0 9492.1
635.3613891601562 0 1935.3107
643.355712890625 0 4622.917
644.35693359375 0 1368.2228
647.3502807617188 0 3237.1726
648.3518676757812 0 800.34125
661.3668212890625 0 8832.747
662.3692016601562 0 3676.4343
663.3663940429688 0 725.7741
732.4404296875 0 1015.7139
742.4240112304688 0 2146.829
743.4320678710938 0 761.3609
757.4234619140625 0 679.192
760.4337768554688 0 7592.5547
761.4371337890625 0 3317.7283
762.4179077148438 0 2040.2981 y Water loss 3
763.4191284179688 0 1058.7815
765.3932495117188 0 1992.8308 b 7
780.4244384765625 0 32974.152 y 3
781.4271850585938 0 15548.909
782.4296264648438 0 3577.774
785.4158325195312 0 2136.4963
861.4814453125 0 1702.3191 y Water loss 2
875.4814453125 0 751.6976 b Water loss 8
879.4922485351562 0 31052.49 y 2
880.4954223632812 0 14293.788
881.4972534179688 0 4303.651
893.4864501953125 0 4588.4004 b 8
894.4885864257812 0 2105.0588
895.4951782226562 0 791.97766
978.5686645507812 0 1129.8003 y 1
979.5633544921875 0 1131.6635
1006.5699462890625 0 4030.392 b 9
1007.5755615234375 0 3417.8606
1013.8834838867188 0 674.01385
1094.5284423828125 0 631.7493
1279.724853515625 0 673.85333
1360.8494873046875 0 591.3187
1487.283447265625 0 582.3247
2002.329345703125 0 763.68286
2273.465087890625 0 761.13776
2583.433349609375 0 667.6547
3074.9033203125 0 1093.4365

Spectrum Details

|  |  |
| --- | --- |
| Matched peaks? Matched peaks The total absolute number of peaks matched. Additionally in brackets the total fraction of peaks matched and the total number of peaks is shown. | 34 (16.27% of 209) |
| FDR? FDR The false discovery rate estimated for this peptide. It is calculated by matching all theoretical fragments with a non-integer shift with the raw peaks for this spectrum. This is done with 40 different shifts. The resulting percentage is the average number of annotated peaks over the number of annotated peaks with the correct spectrum. | 0.35% |
| Satellite FDR? Satellite FDR See the FDR for details on its calculation. This satellite ion specific FDR only contains the satellite ions (d/w) for I/L/J positions. | - |
| PSM Score? PSM Score The PSM Score as given by Hecklib to this annotated spectrum. It is shown with three significant figures. | 349 |

#### Spectrum 8568? Spectrum 8568 The raw spectrum of this peptide as annotated by Hecklib. The fragments are coloured according to ion type (see legend). Any peaks with a star '\*' as text can be hovered over to see the full details, first the ion type second the mass shift type. By hovering over the amino acids in the peptide or ions in the legend the corresponding peaks are highlighted. By toggling the 'Unassigned' label you can turn the background (unassigned) peaks on or off in the plot. By updating the slider in the Ion legend you can update the spectrum to only show the top X% of the peaks with labels. The top X% means any peak that is within X% of the highest intensity. By dragging in the spectrum you can zoom in to a specific part of the spectrum and use 'Zoom Out' to get back to the original zoom level. The annotation of the spectrum is based on the given sequence in the peptides file and is done with different software so inconsistencies are likely. The peaks are annotated based on the given sequence, with 20 ppm tolerance.

Copy Data

##### Spectrum 8568 (TSV)

###### Preview

```
Loading example...
```

*Click on the button to copy the data to your clipboard.*

Mz MinMz MaxIntensity Max

WidthHeightPeptide font sizePeptide stroke widthSpectrum font sizeSpectrum stroke widthCompact peptide

Ion legend

wxyz

abcd

OtherUnassignedIon^Charge^_Position_Show for top:%

FVVFGGGTKJT

01.38e+42.76e+44.15e+45.53e+4

Zoom Out

y^+1^_1_y^+1^_2_a^+1^_2_y^+1^_2_b^+1^_2_b^+2^_6_b^+1^_3_y^+1^_3_b^+2^_8_y^+2^_8_b^+2^_9_y^+2^_9_b^+2^_9_y^+1^_4_b^+1^_4_b^+2^_10_y^+1^_5_y^+1^_6_y^+1^_7_y^+1^_7_b^+1^_8_y^+1^_8_y^+1^_9_y^+1^_9_b^+1^_9_y^+1^_10_b^+1^_10_

043286312951726

Fragment Matches Table

Show background peaks

| Position | Ion type | Intensity | mz Theoretical | mz Error (Th) | mz Error (ppm) | Charge | Series Number |
| --- | --- | --- | --- | --- | --- | --- | --- |
| 11 | y | 2953 | 120.1 | 0.0001237 | 1.03 | +1 | 1 |
| - | - | 4.58E+04 | 120.1 | - | - | 0 | - |
| - | - | 3646 | 121.1 | - | - | 0 | - |
| - | - | 352.5 | 121.2 | - | - | 0 | - |
| - | - | 396.2 | 127.1 | - | - | 0 | - |
| - | - | 392.4 | 127.1 | - | - | 0 | - |
| - | - | 637.1 | 128.1 | - | - | 0 | - |
| - | - | 446.8 | 128.1 | - | - | 0 | - |
| - | - | 1.943E+04 | 129.1 | - | - | 0 | - |
| - | - | 436.6 | 130.1 | - | - | 0 | - |
| - | - | 1369 | 130.1 | - | - | 0 | - |
| - | - | 1215 | 131.1 | - | - | 0 | - |
| - | - | 832.1 | 131.1 | - | - | 0 | - |
| - | - | 639.2 | 131.1 | - | - | 0 | - |
| - | - | 457.2 | 131.3 | - | - | 0 | - |
| - | - | 588.7 | 133.1 | - | - | 0 | - |
| - | - | 9468 | 133.1 | - | - | 0 | - |
| - | - | 496.9 | 133.1 | - | - | 0 | - |
| - | - | 694.3 | 134.1 | - | - | 0 | - |
| - | - | 415.4 | 134.3 | - | - | 0 | - |
| - | - | 1863 | 136.1 | - | - | 0 | - |
| - | - | 854.5 | 140.1 | - | - | 0 | - |
| - | - | 569.7 | 141.1 | - | - | 0 | - |
| - | - | 535.1 | 147.1 | - | - | 0 | - |
| - | - | 700.7 | 147.1 | - | - | 0 | - |
| - | - | 456.2 | 147.1 | - | - | 0 | - |
| - | - | 416.5 | 151.1 | - | - | 0 | - |
| - | - | 632.1 | 155.1 | - | - | 0 | - |
| - | - | 564.7 | 155.1 | - | - | 0 | - |
| - | - | 977.6 | 155.1 | - | - | 0 | - |
| - | - | 694.8 | 156.1 | - | - | 0 | - |
| - | - | 544.6 | 157.1 | - | - | 0 | - |
| - | - | 1159 | 158.1 | - | - | 0 | - |
| - | - | 1068 | 159.1 | - | - | 0 | - |
| - | - | 688.5 | 165.1 | - | - | 0 | - |
| - | - | 465.3 | 169.1 | - | - | 0 | - |
| - | - | 532.2 | 169.1 | - | - | 0 | - |
| - | - | 487.8 | 170.1 | - | - | 0 | - |
| - | - | 1457 | 171.1 | - | - | 0 | - |
| - | - | 1333 | 172.1 | - | - | 0 | - |
| - | - | 553.7 | 172.1 | - | - | 0 | - |
| - | - | 651.4 | 173.1 | - | - | 0 | - |
| - | - | 1281 | 173.4 | - | - | 0 | - |
| - | - | 681.4 | 175.1 | - | - | 0 | - |
| - | - | 490.6 | 175.2 | - | - | 0 | - |
| - | - | 621.7 | 176.1 | - | - | 0 | - |
| - | - | 517.9 | 177.1 | - | - | 0 | - |
| - | - | 3153 | 177.1 | - | - | 0 | - |
| - | - | 439.8 | 181.7 | - | - | 0 | - |
| - | - | 735.8 | 182.1 | - | - | 0 | - |
| - | - | 491.7 | 187.1 | - | - | 0 | - |
| - | - | 715 | 187.1 | - | - | 0 | - |
| - | - | 531.2 | 188.1 | - | - | 0 | - |
| - | - | 736.5 | 191.1 | - | - | 0 | - |
| - | - | 1378 | 195.1 | - | - | 0 | - |
| - | - | 813.1 | 197.2 | - | - | 0 | - |
| - | - | 878.1 | 198.1 | - | - | 0 | - |
| - | - | 821.2 | 199.1 | - | - | 0 | - |
| - | - | 1461 | 205.1 | - | - | 0 | - |
| - | - | 533.5 | 207.1 | - | - | 0 | - |
| - | - | 2370 | 212.1 | - | - | 0 | - |
| - | - | 780.6 | 213.2 | - | - | 0 | - |
| 10 | y | 717.7 | 215.1 | 0.0001182 | 0.5492 | +1 | 2 |
| - | - | 792.4 | 216.1 | - | - | 0 | - |
| 2 | a | 5.474E+04 | 219.1 | 0.0001481 | 0.6758 | +1 | 2 |
| - | - | 8034 | 220.2 | - | - | 0 | - |
| - | - | 1330 | 221.1 | - | - | 0 | - |
| - | - | 1493 | 227.1 | - | - | 0 | - |
| - | - | 642.9 | 229.2 | - | - | 0 | - |
| - | - | 3791 | 230.1 | - | - | 0 | - |
| - | - | 655.4 | 231.1 | - | - | 0 | - |
| 10 | y | 2350 | 233.1 | 0.0003187 | 1.367 | +1 | 2 |
| - | - | 709.2 | 235.2 | - | - | 0 | - |
| - | - | 547.4 | 236.5 | - | - | 0 | - |
| - | - | 1413 | 239.1 | - | - | 0 | - |
| 2 | b | 4.164E+04 | 247.1 | 0.0001065 | 0.431 | +1 | 2 |
| - | - | 7244 | 248.1 | - | - | 0 | - |
| - | - | 647.7 | 253.2 | - | - | 0 | - |
| - | - | 2222 | 255.1 | - | - | 0 | - |
| - | - | 615.9 | 261.1 | - | - | 0 | - |
| - | - | 703.6 | 262.1 | - | - | 0 | - |
| - | - | 2483 | 269.2 | - | - | 0 | - |
| - | - | 2572 | 273.1 | - | - | 0 | - |
| - | - | 747.4 | 295.1 | - | - | 0 | - |
| - | - | 895.4 | 299.1 | - | - | 0 | - |
| - | - | 580.8 | 300.1 | - | - | 0 | - |
| - | - | 2616 | 301.2 | - | - | 0 | - |
| 6 | b | 1056 | 304.2 | 0.0009359 | 3.077 | +2 | 6 |
| - | - | 984.9 | 319.1 | - | - | 0 | - |
| - | - | 2166 | 326.2 | - | - | 0 | - |
| 3 | b | 1.062E+04 | 346.2 | 6.212E-06 | 0.01794 | +1 | 3 |
| - | - | 2777 | 347.2 | - | - | 0 | - |
| 9 | y | 2480 | 361.2 | 0.000345 | 0.9549 | +1 | 3 |
| - | - | 823.7 | 362.2 | - | - | 0 | - |
| - | - | 721.8 | 374.2 | - | - | 0 | - |
| 8 | b | 2208 | 383.2 | 0.00381 | 9.942 | +2 | 8 |
| - | - | 805.5 | 384.2 | - | - | 0 | - |
| 4 | y | 1007 | 390.7 | 0.0008213 | 2.102 | +2 | 8 |
| - | - | 816.3 | 391.2 | - | - | 0 | - |
| - | - | 580 | 391.3 | - | - | 0 | - |
| - | - | 616.2 | 392.2 | - | - | 0 | - |
| - | - | 676.7 | 400.3 | - | - | 0 | - |
| - | - | 1875 | 401.2 | - | - | 0 | - |
| - | - | 2306 | 402.2 | - | - | 0 | - |
| - | - | 1766 | 420.2 | - | - | 0 | - |
| 9 | b | 640.5 | 438.2 | 0.001585 | 3.617 | +2 | 9 |
| - | - | 614 | 439.3 | - | - | 0 | - |
| 3 | y | 1775 | 440.3 | 0.001379 | 3.133 | +2 | 9 |
| 9 | b | 683.7 | 447.2 | 0.0001211 | 0.2707 | +2 | 9 |
| - | - | 1104 | 451.2 | - | - | 0 | - |
| - | - | 1747 | 457.3 | - | - | 0 | - |
| 8 | y | 649.1 | 462.3 | 0.002483 | 5.371 | +1 | 4 |
| - | - | 1650 | 465.3 | - | - | 0 | - |
| - | - | 625.3 | 473.3 | - | - | 0 | - |
| - | - | 634.3 | 479.2 | - | - | 0 | - |
| - | - | 987.4 | 486.3 | - | - | 0 | - |
| - | - | 591.6 | 487.3 | - | - | 0 | - |
| - | - | 1006 | 491.3 | - | - | 0 | - |
| 4 | b | 1029 | 493.3 | 0.001269 | 2.573 | +1 | 4 |
| - | - | 2342 | 496.3 | - | - | 0 | - |
| - | - | 686.5 | 497.3 | - | - | 0 | - |
| - | - | 811.8 | 501.2 | - | - | 0 | - |
| 10 | b | 650.6 | 503.8 | 0.0003254 | 0.6459 | +2 | 10 |
| - | - | 5252 | 514.3 | - | - | 0 | - |
| - | - | 1483 | 515.3 | - | - | 0 | - |
| 7 | y | 1132 | 519.3 | 0.0009474 | 1.824 | +1 | 5 |
| - | - | 1297 | 530.3 | - | - | 0 | - |
| - | - | 627.4 | 546.6 | - | - | 0 | - |
| - | - | 883.1 | 547.3 | - | - | 0 | - |
| - | - | 2831 | 548.3 | - | - | 0 | - |
| - | - | 1206 | 549.3 | - | - | 0 | - |
| - | - | 654.6 | 559 | - | - | 0 | - |
| - | - | 1153 | 562.4 | - | - | 0 | - |
| - | - | 1122 | 563.8 | - | - | 0 | - |
| - | - | 556.6 | 574.9 | - | - | 0 | - |
| 6 | y | 3208 | 576.3 | 0.0004357 | 0.756 | +1 | 6 |
| - | - | 687.2 | 577.3 | - | - | 0 | - |
| - | - | 917.7 | 611.3 | - | - | 0 | - |
| 5 | y | 1146 | 615.3 | 0.0006536 | 1.062 | +1 | 7 |
| - | - | 866 | 616.3 | - | - | 0 | - |
| - | - | 713.5 | 629.3 | - | - | 0 | - |
| 5 | y | 1.674E+04 | 633.4 | 0.0003784 | 0.5974 | +1 | 7 |
| - | - | 6758 | 634.4 | - | - | 0 | - |
| - | - | 1213 | 635.4 | - | - | 0 | - |
| - | - | 2245 | 643.4 | - | - | 0 | - |
| - | - | 896.4 | 644.4 | - | - | 0 | - |
| - | - | 1494 | 647.4 | - | - | 0 | - |
| - | - | 5242 | 661.4 | - | - | 0 | - |
| - | - | 951.7 | 662.4 | - | - | 0 | - |
| - | - | 595.4 | 739.4 | - | - | 0 | - |
| - | - | 1621 | 742.4 | - | - | 0 | - |
| - | - | 839.2 | 746.4 | - | - | 0 | - |
| - | - | 681.8 | 757.4 | - | - | 0 | - |
| - | - | 3780 | 760.4 | - | - | 0 | - |
| - | - | 1978 | 761.4 | - | - | 0 | - |
| 8 | b | 1386 | 765.4 | 0.002621 | 3.424 | +1 | 8 |
| - | - | 759.6 | 766.4 | - | - | 0 | - |
| 4 | y | 1.879E+04 | 780.4 | 0.00108 | 1.384 | +1 | 8 |
| - | - | 8117 | 781.4 | - | - | 0 | - |
| - | - | 2144 | 782.4 | - | - | 0 | - |
| - | - | 2679 | 785.4 | - | - | 0 | - |
| 3 | y | 870.2 | 861.5 | 3.028E-05 | 0.03515 | +1 | 9 |
| 3 | y | 1.526E+04 | 879.5 | 0.00144 | 1.637 | +1 | 9 |
| - | - | 8380 | 880.5 | - | - | 0 | - |
| - | - | 2066 | 881.5 | - | - | 0 | - |
| 9 | b | 1902 | 893.5 | 0.0001334 | 0.1493 | +1 | 9 |
| - | - | 1138 | 894.5 | - | - | 0 | - |
| 2 | y | 948.6 | 978.6 | 0.00186 | 1.901 | +1 | 10 |
| 10 | b | 2087 | 1007 | 0.001946 | 1.933 | +1 | 10 |
| - | - | 1309 | 1008 | - | - | 0 | - |
| - | - | 739 | 1567 | - | - | 0 | - |
| - | - | 655.7 | 1709 | - | - | 0 | - |

m/z Charge Intensity FragmentType MassShift Position
120.06564331054688 0 2952.921 y 10
120.08097839355469 0 45798.91
121.08428955078125 0 3646.2534
121.18601989746094 0 352.46152
127.07585906982422 0 396.15793
127.08684539794922 0 392.41238
128.10728454589844 0 637.06244
128.14451599121094 0 446.75772
129.1024169921875 0 19432.4
130.08697509765625 0 436.55316
130.1058349609375 0 1368.5503
131.0706329345703 0 1215.479
131.08175659179688 0 832.112
131.10665893554688 0 639.24634
131.34259033203125 0 457.1678
133.06097412109375 0 588.6995
133.0861053466797 0 9467.893
133.09190368652344 0 496.89697
134.0895538330078 0 694.3191
134.2777099609375 0 415.37756
136.0758514404297 0 1862.6505
140.08197021484375 0 854.513
141.06600952148438 0 569.72235
147.0765838623047 0 535.0509
147.10208129882812 0 700.6777
147.11233520507812 0 456.16757
151.07301330566406 0 416.5041
155.0706787109375 0 632.05505
155.08213806152344 0 564.6633
155.11842346191406 0 977.56604
156.07681274414062 0 694.77374
157.08633422851562 0 544.62537
158.09268188476562 0 1158.6737
159.07659912109375 0 1067.887
165.1027069091797 0 688.5108
169.07286071777344 0 465.28598
169.0977783203125 0 532.1751
170.09324645996094 0 487.7968
171.14964294433594 0 1456.9376
172.07168579101562 0 1333.2769
172.10800170898438 0 553.72614
173.12832641601562 0 651.4323
173.43887329101562 0 1281.15
175.09674072265625 0 681.3984
175.21055603027344 0 490.6338
176.10728454589844 0 621.72485
177.10400390625 0 517.8509
177.11221313476562 0 3153.1792
181.69757080078125 0 439.79956
182.0817108154297 0 735.8484
187.07203674316406 0 491.71655
187.1080780029297 0 714.976
188.06597900390625 0 531.17645
191.12750244140625 0 736.4767
195.1229248046875 0 1377.7991
197.1649932861328 0 813.1377
198.08743286132812 0 878.14105
199.1444854736328 0 821.1718
205.09713745117188 0 1461.2152
207.14903259277344 0 533.4603
212.13926696777344 0 2369.7021
213.16001892089844 0 780.5966
215.13890075683594 0 717.66235 y Water loss 9
216.0980987548828 0 792.3539
219.1493377685547 0 54742.695 a 1
220.1527099609375 0 8034.0264
221.13783264160156 0 1330.4924
227.11322021484375 0 1492.6527
229.1548309326172 0 642.9399
230.1497802734375 0 3791.3083
231.11309814453125 0 655.3775
233.14990234375 0 2350.2302 y 9
235.15614318847656 0 709.16455
236.4979248046875 0 547.4197
239.14907836914062 0 1412.9767
247.1442108154297 0 41638.766 b 1
248.14759826660156 0 7243.933
253.16661071777344 0 647.7341
255.10906982421875 0 2221.8962
261.1246032714844 0 615.9198
262.1192626953125 0 703.5853
269.1610107421875 0 2482.6885
273.1192626953125 0 2572.4883
295.14404296875 0 747.4153
299.0629577636719 0 895.385
300.0617370605469 0 580.77356
301.1912841796875 0 2616.267
304.16650390625 0 1055.8785 b 5
319.1407775878906 0 984.941
326.182861328125 0 2165.6855
346.2125244140625 0 10618.97 b 2
347.21563720703125 0 2776.537
361.24420166015625 0 2480.317 y 8
362.2463684082031 0 823.6555
374.1842956542969 0 721.84503
383.2039489746094 0 2208.1147 b 7
384.2062072753906 0 805.53015
390.71533203125 0 1006.7907 y 3
391.21807861328125 0 816.3171
391.282470703125 0 579.9861
392.1922912597656 0 616.2397
400.2582702636719 0 676.69714
401.2142639160156 0 1874.7522
402.17742919921875 0 2305.6943
420.1868896484375 0 1766.4021
438.2407531738281 0 640.51764 b Water loss 8
439.26702880859375 0 613.9848
440.2517395019531 0 1775.4055 y 2
447.24774169921875 0 683.6962 b 8
451.2318420410156 0 1104.0853
457.2773132324219 0 1747.0353
462.2947082519531 0 649.0843 y 7
465.2864990234375 0 1649.5796
473.25048828125 0 625.31683
479.24871826171875 0 634.2906
486.3035888671875 0 987.40796
487.30487060546875 0 591.56537
491.261962890625 0 1005.6863
493.2796630859375 0 1028.5906 b 3
496.2879943847656 0 2341.9775
497.291748046875 0 686.52405
501.2485046386719 0 811.8458
503.78997802734375 0 650.6153 b 9
514.2985229492188 0 5251.942
515.30224609375 0 1482.8287
519.3146362304688 0 1132.0681 y 6
530.2733764648438 0 1297.2906
546.638916015625 0 627.39514
547.3346557617188 0 883.13696
548.2822875976562 0 2830.6443
549.2857055664062 0 1205.9015
559.01611328125 0 654.6328
562.3800659179688 0 1152.9443
563.8197021484375 0 1121.6617
574.9459838867188 0 556.5737
576.334716796875 0 3208.2595 y 5
577.3357543945312 0 687.15173
611.3263549804688 0 917.6787
615.3453979492188 0 1146.0101 y Water loss 4
616.3466796875 0 866.0307
629.3430786132812 0 713.5197
633.3569946289062 0 16742.762 y 4
634.3597412109375 0 6757.9717
635.35791015625 0 1213.0764
643.355224609375 0 2245.117
644.3626098632812 0 896.3574
647.3518676757812 0 1493.6979
661.3665161132812 0 5241.652
662.3681030273438 0 951.73926
739.4097900390625 0 595.3735
742.4248046875 0 1621.4886
746.3671875 0 839.1712
757.4194946289062 0 681.8337
760.4337158203125 0 3780.3523
761.4359741210938 0 1978.3904
765.390380859375 0 1385.5164 b 7
766.3970947265625 0 759.6401
780.4239501953125 0 18791.059 y 3
781.427734375 0 8116.675
782.431640625 0 2143.6987
785.4149169921875 0 2678.6533
861.4828491210938 0 870.2006 y Water loss 2
879.4920043945312 0 15264.286 y 2
880.4956665039062 0 8380.021
881.4991455078125 0 2065.742
893.4880981445312 0 1902.093 b 8
894.4935913085938 0 1137.6382
978.5599975585938 0 948.6193 y 1
1006.573974609375 0 2086.6843 b 9
1007.5706176757812 0 1309.2473
1567.2696533203125 0 739.0318
1708.9984130859375 0 655.72705

Spectrum Details

|  |  |
| --- | --- |
| Matched peaks? Matched peaks The total absolute number of peaks matched. Additionally in brackets the total fraction of peaks matched and the total number of peaks is shown. | 27 (15.70% of 172) |
| FDR? FDR The false discovery rate estimated for this peptide. It is calculated by matching all theoretical fragments with a non-integer shift with the raw peaks for this spectrum. This is done with 40 different shifts. The resulting percentage is the average number of annotated peaks over the number of annotated peaks with the correct spectrum. | 0.09% |
| Satellite FDR? Satellite FDR See the FDR for details on its calculation. This satellite ion specific FDR only contains the satellite ions (d/w) for I/L/J positions. | - |
| PSM Score? PSM Score The PSM Score as given by Hecklib to this annotated spectrum. It is shown with three significant figures. | 271 |

#### Spectrum 8732? Spectrum 8732 The raw spectrum of this peptide as annotated by Hecklib. The fragments are coloured according to ion type (see legend). Any peaks with a star '\*' as text can be hovered over to see the full details, first the ion type second the mass shift type. By hovering over the amino acids in the peptide or ions in the legend the corresponding peaks are highlighted. By toggling the 'Unassigned' label you can turn the background (unassigned) peaks on or off in the plot. By updating the slider in the Ion legend you can update the spectrum to only show the top X% of the peaks with labels. The top X% means any peak that is within X% of the highest intensity. By dragging in the spectrum you can zoom in to a specific part of the spectrum and use 'Zoom Out' to get back to the original zoom level. The annotation of the spectrum is based on the given sequence in the peptides file and is done with different software so inconsistencies are likely. The peaks are annotated based on the given sequence, with 20 ppm tolerance.

Copy Data

##### Spectrum 8732 (TSV)

###### Preview

```
Loading example...
```

*Click on the button to copy the data to your clipboard.*

Mz MinMz MaxIntensity Max

WidthHeightPeptide font sizePeptide stroke widthSpectrum font sizeSpectrum stroke widthCompact peptide

Ion legend

wxyz

abcd

OtherUnassignedIon^Charge^_Position_Show for top:%

FVVFGGGTKJT

08.51e+31.70e+42.55e+43.41e+4

Zoom Out

y^+1^_1_a^+1^_2_y^+1^_2_b^+1^_2_b^+2^_6_b^+1^_3_y^+1^_3_b^+2^_8_y^+2^_9_y^+1^_4_y^+2^_10_y^+1^_5_b^+1^_5_y^+1^_6_y^+1^_7_y^+1^_8_b^+1^_8_y^+1^_8_y^+1^_9_b^+1^_9_y^+1^_10_b^+1^_10_

0623124718702494

Fragment Matches Table

Show background peaks

| Position | Ion type | Intensity | mz Theoretical | mz Error (Th) | mz Error (ppm) | Charge | Series Number |
| --- | --- | --- | --- | --- | --- | --- | --- |
| 11 | y | 1817 | 120.1 | 4.74E-05 | 0.3948 | +1 | 1 |
| - | - | 3.22E+04 | 120.1 | - | - | 0 | - |
| - | - | 2536 | 121.1 | - | - | 0 | - |
| - | - | 452.4 | 121.1 | - | - | 0 | - |
| - | - | 416.3 | 127.1 | - | - | 0 | - |
| - | - | 593.1 | 128.1 | - | - | 0 | - |
| - | - | 1.326E+04 | 129.1 | - | - | 0 | - |
| - | - | 458.6 | 130.1 | - | - | 0 | - |
| - | - | 391.3 | 130.1 | - | - | 0 | - |
| - | - | 1345 | 131.1 | - | - | 0 | - |
| - | - | 503 | 131.1 | - | - | 0 | - |
| - | - | 443.9 | 131.4 | - | - | 0 | - |
| - | - | 919.3 | 133.1 | - | - | 0 | - |
| - | - | 1.139E+04 | 133.1 | - | - | 0 | - |
| - | - | 773.1 | 134.1 | - | - | 0 | - |
| - | - | 1981 | 136.1 | - | - | 0 | - |
| - | - | 457.9 | 140.1 | - | - | 0 | - |
| - | - | 583.4 | 141.1 | - | - | 0 | - |
| - | - | 397 | 141.9 | - | - | 0 | - |
| - | - | 1552 | 147.1 | - | - | 0 | - |
| - | - | 1046 | 149 | - | - | 0 | - |
| - | - | 411.4 | 157.5 | - | - | 0 | - |
| - | - | 566.2 | 159.1 | - | - | 0 | - |
| - | - | 666.9 | 166.1 | - | - | 0 | - |
| - | - | 487.9 | 167.1 | - | - | 0 | - |
| - | - | 512.8 | 170.6 | - | - | 0 | - |
| - | - | 827.3 | 171.1 | - | - | 0 | - |
| - | - | 843.5 | 172.1 | - | - | 0 | - |
| - | - | 456.9 | 172.5 | - | - | 0 | - |
| - | - | 615.8 | 173.1 | - | - | 0 | - |
| - | - | 2749 | 173.5 | - | - | 0 | - |
| - | - | 3790 | 177.1 | - | - | 0 | - |
| - | - | 639.6 | 182.1 | - | - | 0 | - |
| - | - | 851.8 | 187.1 | - | - | 0 | - |
| - | - | 1458 | 191.1 | - | - | 0 | - |
| - | - | 524.2 | 192.1 | - | - | 0 | - |
| - | - | 758.3 | 195.1 | - | - | 0 | - |
| - | - | 536.1 | 198.6 | - | - | 0 | - |
| - | - | 545.5 | 199.1 | - | - | 0 | - |
| - | - | 501.6 | 199.1 | - | - | 0 | - |
| - | - | 1218 | 205.1 | - | - | 0 | - |
| - | - | 492.1 | 206.1 | - | - | 0 | - |
| - | - | 550.8 | 207.4 | - | - | 0 | - |
| - | - | 485.2 | 211.1 | - | - | 0 | - |
| - | - | 1274 | 212.1 | - | - | 0 | - |
| - | - | 626.3 | 219.1 | - | - | 0 | - |
| 2 | a | 3.371E+04 | 219.1 | 4.129E-05 | 0.1884 | +1 | 2 |
| - | - | 485.2 | 219.5 | - | - | 0 | - |
| - | - | 4096 | 220.2 | - | - | 0 | - |
| - | - | 621.3 | 221.1 | - | - | 0 | - |
| - | - | 629.8 | 227.1 | - | - | 0 | - |
| - | - | 1396 | 230.1 | - | - | 0 | - |
| 10 | y | 1068 | 233.1 | 0.0003187 | 1.367 | +1 | 2 |
| - | - | 790.3 | 235.2 | - | - | 0 | - |
| - | - | 572.5 | 238.1 | - | - | 0 | - |
| 2 | b | 2.475E+04 | 247.1 | 0.0001218 | 0.4928 | +1 | 2 |
| - | - | 4126 | 248.1 | - | - | 0 | - |
| - | - | 532.5 | 249.3 | - | - | 0 | - |
| - | - | 727.1 | 253.2 | - | - | 0 | - |
| - | - | 1186 | 255.1 | - | - | 0 | - |
| - | - | 1472 | 269.2 | - | - | 0 | - |
| - | - | 2018 | 273.1 | - | - | 0 | - |
| - | - | 548.2 | 274.1 | - | - | 0 | - |
| - | - | 1213 | 283.2 | - | - | 0 | - |
| - | - | 1080 | 299.1 | - | - | 0 | - |
| - | - | 1060 | 301.2 | - | - | 0 | - |
| 6 | b | 901.9 | 304.2 | 0.001119 | 3.679 | +2 | 6 |
| - | - | 815.3 | 326.2 | - | - | 0 | - |
| - | - | 1018 | 343.1 | - | - | 0 | - |
| 3 | b | 6599 | 346.2 | 3.673E-05 | 0.1061 | +1 | 3 |
| - | - | 1245 | 347.2 | - | - | 0 | - |
| 9 | y | 1279 | 361.2 | 0.0002839 | 0.7859 | +1 | 3 |
| - | - | 631 | 363.6 | - | - | 0 | - |
| - | - | 585 | 373.2 | - | - | 0 | - |
| 8 | b | 1683 | 383.2 | 0.003535 | 9.225 | +2 | 8 |
| - | - | 640.5 | 392.2 | - | - | 0 | - |
| - | - | 680.2 | 394.2 | - | - | 0 | - |
| - | - | 1080 | 401.2 | - | - | 0 | - |
| - | - | 1859 | 402.2 | - | - | 0 | - |
| - | - | 871 | 420.2 | - | - | 0 | - |
| 3 | y | 1067 | 440.3 | 0.001196 | 2.717 | +2 | 9 |
| - | - | 670.7 | 440.8 | - | - | 0 | - |
| - | - | 787.5 | 451.2 | - | - | 0 | - |
| 8 | y | 802.7 | 462.3 | 0.002308 | 4.993 | +1 | 4 |
| - | - | 533.4 | 481.2 | - | - | 0 | - |
| - | - | 688.1 | 484.6 | - | - | 0 | - |
| - | - | 724.6 | 486.3 | - | - | 0 | - |
| 2 | y | 788 | 489.8 | 0.004953 | 10.11 | +2 | 10 |
| - | - | 1056 | 496.3 | - | - | 0 | - |
| - | - | 733.5 | 498.3 | - | - | 0 | - |
| - | - | 2625 | 514.3 | - | - | 0 | - |
| - | - | 717.3 | 515.3 | - | - | 0 | - |
| 7 | y | 887 | 519.3 | 0.0009447 | 1.819 | +1 | 5 |
| - | - | 887.7 | 530.3 | - | - | 0 | - |
| - | - | 568.7 | 544.2 | - | - | 0 | - |
| - | - | 1658 | 545.3 | - | - | 0 | - |
| - | - | 1722 | 545.4 | - | - | 0 | - |
| - | - | 737.5 | 546.3 | - | - | 0 | - |
| - | - | 820.9 | 547.3 | - | - | 0 | - |
| - | - | 1728 | 548.3 | - | - | 0 | - |
| - | - | 1125 | 549.3 | - | - | 0 | - |
| 5 | b | 660.2 | 550.3 | 0.002164 | 3.932 | +1 | 5 |
| - | - | 825.8 | 562.4 | - | - | 0 | - |
| - | - | 799.3 | 563.8 | - | - | 0 | - |
| 6 | y | 1406 | 576.3 | 0.0009681 | 1.68 | +1 | 6 |
| - | - | 1744 | 613.3 | - | - | 0 | - |
| 5 | y | 1.14E+04 | 633.4 | 0.0005004 | 0.7901 | +1 | 7 |
| - | - | 2897 | 634.4 | - | - | 0 | - |
| - | - | 712.3 | 635.4 | - | - | 0 | - |
| - | - | 1109 | 643.4 | - | - | 0 | - |
| - | - | 1121 | 647.3 | - | - | 0 | - |
| - | - | 2747 | 661.4 | - | - | 0 | - |
| - | - | 938.1 | 662.4 | - | - | 0 | - |
| - | - | 843.9 | 714.4 | - | - | 0 | - |
| - | - | 721.4 | 742.4 | - | - | 0 | - |
| - | - | 1175 | 757.4 | - | - | 0 | - |
| - | - | 3627 | 760.4 | - | - | 0 | - |
| 4 | y | 844.3 | 762.4 | 3.678E-05 | 0.04824 | +1 | 8 |
| 8 | b | 597.8 | 765.4 | 5.743E-05 | 0.07503 | +1 | 8 |
| 4 | y | 1.188E+04 | 780.4 | 0.0002255 | 0.2889 | +1 | 8 |
| - | - | 4499 | 781.4 | - | - | 0 | - |
| - | - | 985.9 | 782.4 | - | - | 0 | - |
| - | - | 4591 | 785.4 | - | - | 0 | - |
| - | - | 684.8 | 786.4 | - | - | 0 | - |
| 3 | y | 1.091E+04 | 879.5 | 8.618E-05 | 0.09799 | +1 | 9 |
| - | - | 4166 | 880.5 | - | - | 0 | - |
| - | - | 1196 | 881.5 | - | - | 0 | - |
| 9 | b | 1287 | 893.5 | 0.001087 | 1.217 | +1 | 9 |
| - | - | 607.2 | 953.7 | - | - | 0 | - |
| 2 | y | 865.1 | 978.6 | 0.002227 | 2.275 | +1 | 10 |
| 10 | b | 1677 | 1007 | 0.0009838 | 0.9774 | +1 | 10 |
| - | - | 620.1 | 1079 | - | - | 0 | - |
| - | - | 578.3 | 1751 | - | - | 0 | - |
| - | - | 690.9 | 2469 | - | - | 0 | - |

m/z Charge Intensity FragmentType MassShift Position
120.06556701660156 0 1816.6678 y 10
120.0809326171875 0 32203.031
121.08428192138672 0 2536.381
121.08834838867188 0 452.37628
127.0757064819336 0 416.2979
128.10708618164062 0 593.1122
129.10240173339844 0 13255.053
130.0610809326172 0 458.6409
130.10597229003906 0 391.281
131.0704345703125 0 1345.1511
131.08160400390625 0 502.99765
131.402587890625 0 443.91443
133.08079528808594 0 919.3087
133.0860595703125 0 11385.341
134.0895233154297 0 773.14386
136.0757293701172 0 1981.3062
140.0817413330078 0 457.89365
141.0659637451172 0 583.3989
141.92034912109375 0 397.0361
147.10166931152344 0 1551.8983
148.9547119140625 0 1045.6752
157.49508666992188 0 411.41574
159.09165954589844 0 566.23987
166.08631896972656 0 666.8952
167.05551147460938 0 487.91165
170.64236450195312 0 512.8128
171.14932250976562 0 827.2803
172.0718231201172 0 843.4875
172.48121643066406 0 456.8956
173.1285858154297 0 615.8014
173.45167541503906 0 2749.3645
177.11224365234375 0 3789.9265
182.08102416992188 0 639.6258
187.10801696777344 0 851.8069
191.1278839111328 0 1458.3403
192.08689880371094 0 524.24536
195.1229705810547 0 758.27496
198.58819580078125 0 536.07635
199.0825653076172 0 545.5159
199.14405822753906 0 501.58627
205.09686279296875 0 1217.6768
206.1011962890625 0 492.1155
207.38914489746094 0 550.8029
211.1083526611328 0 485.21582
212.13922119140625 0 1273.553
219.06517028808594 0 626.3313
219.14923095703125 0 33714.574 a 1
219.48851013183594 0 485.15714
220.1526641845703 0 4095.6035
221.1389617919922 0 621.296
227.11412048339844 0 629.84454
230.14971923828125 0 1395.7869
233.14990234375 0 1067.8276 y 9
235.15447998046875 0 790.296
238.1416015625 0 572.5021
247.14422607421875 0 24751.488 b 1
248.1475372314453 0 4126.277
249.3133544921875 0 532.48047
253.16673278808594 0 727.13684
255.10891723632812 0 1185.8711
269.1606750488281 0 1471.8037
273.11932373046875 0 2017.9979
274.1207580566406 0 548.1736
283.17474365234375 0 1212.7488
299.0613098144531 0 1080.4702
301.19140625 0 1059.6238
304.16668701171875 0 901.9098 b 5
326.1828308105469 0 815.2809
343.1293029785156 0 1018.2268
346.2125549316406 0 6598.6943 b 2
347.2171325683594 0 1244.8944
361.2442626953125 0 1279.4537 y 8
363.6388244628906 0 631.0227
373.19110107421875 0 585.0259
383.20367431640625 0 1682.8425 b 7
392.19580078125 0 640.4597
394.2137756347656 0 680.2049
401.2158203125 0 1080.1594
402.1771545410156 0 1859.2292
420.18707275390625 0 870.9841
440.2515563964844 0 1066.5862 y 2
440.75634765625 0 670.6781
451.2337951660156 0 787.5116
462.2899169921875 0 802.6772 y 7
481.16229248046875 0 533.3899
484.576904296875 0 688.13074
486.30511474609375 0 724.62366
489.7895202636719 0 788.0492 y 1
496.2872314453125 0 1055.7617
498.3031311035156 0 733.49414
514.2984619140625 0 2625.4592
515.30078125 0 717.3198
519.312744140625 0 887.02136 y 6
530.2726440429688 0 887.7364
544.2271728515625 0 568.673
545.3150024414062 0 1658.2222
545.35498046875 0 1721.9753
546.3174438476562 0 737.5039
547.330810546875 0 820.8512
548.2839965820312 0 1727.8108
549.288330078125 0 1124.9003
550.3002319335938 0 660.1812 b 4
562.3816528320312 0 825.76996
563.8159790039062 0 799.33057
576.3361206054688 0 1405.6819 y 5
613.3297729492188 0 1743.7681
633.3571166992188 0 11404.764 y 4
634.35986328125 0 2897.2583
635.3653564453125 0 712.30005
643.3582763671875 0 1109.3427
647.3486938476562 0 1121.0231
661.3670043945312 0 2746.676
662.3677978515625 0 938.1099
714.3815307617188 0 843.87524
742.4212646484375 0 721.3873
757.4224243164062 0 1175.4615
760.43505859375 0 3626.5466
762.4144287109375 0 844.32446 y Water loss 3
765.3929443359375 0 597.8057 b 7
780.4248046875 0 11884.726 y 3
781.4261474609375 0 4499.0923
782.4332275390625 0 985.85974
785.4148559570312 0 4590.9487
786.4210205078125 0 684.7516
879.4935302734375 0 10906.124 y 2
880.495849609375 0 4166.2456
881.5018920898438 0 1196.2273
893.4868774414062 0 1286.8638 b 8
953.6849975585938 0 607.2395
978.5596313476562 0 865.0951 y 1
1006.571044921875 0 1677.3374 b 9
1078.6455078125 0 620.09875
1751.22900390625 0 578.26324
2469.244873046875 0 690.8816

Spectrum Details

|  |  |
| --- | --- |
| Matched peaks? Matched peaks The total absolute number of peaks matched. Additionally in brackets the total fraction of peaks matched and the total number of peaks is shown. | 22 (16.42% of 134) |
| FDR? FDR The false discovery rate estimated for this peptide. It is calculated by matching all theoretical fragments with a non-integer shift with the raw peaks for this spectrum. This is done with 40 different shifts. The resulting percentage is the average number of annotated peaks over the number of annotated peaks with the correct spectrum. | 0.22% |
| Satellite FDR? Satellite FDR See the FDR for details on its calculation. This satellite ion specific FDR only contains the satellite ions (d/w) for I/L/J positions. | - |
| PSM Score? PSM Score The PSM Score as given by Hecklib to this annotated spectrum. It is shown with three significant figures. | 235 |

#### Reverse Lookup? Reverse Lookup All places where this read could be placed.

| Group | Segment | Template | Template Part | Read Part | Score | Unique |
| --- | --- | --- | --- | --- | --- | --- |
| Homo sapiens Light Chain | IGLJ | IGLJ2 | [0..10] | [1..11] | 80 | True |

| Recombined | Template Part | Read Part | Score | Unique |
| --- | --- | --- | --- | --- |
| REC-0-1\_002 | [98..109] | [0..11] | 88 | True |

#### Meta Information from Multiple reads

##### Number of combined reads

6

##### Intensity

0.7907

##### TotalArea

1.36E+08

##### Changes to the peptide sequence

FVVFGGGTKJT

L→JNo support for either Leucine or Isoleucine based on side chain ions (Position: 10)

#### Positional Score

Copy Data

##### Positional Score (TSV)

###### Preview

```
Loading example...
```

*Click on the button to copy the data to your clipboard.*

10012345678910

Label Value
"0" 0.64
"1" 0.658
"2" 0.667
"3" 0.667
"4" 0.662
"5" 0.64
"6" 0.607
"7" 0.63
"8" 0.657
"9" 0.648
"10" 0.657

#### Meta Information from PEAKS

##### Scan Identifier

F2:8344

##### Original sequence

F

V

V

F

G

G

G

T

K

L

T

##### Posttranslational Modifications

##### Source File

D:\separate\_stitch\_analyses\xle-disambiguation\raw\20210323\_F1\_UM1\_Peng0013\_SA\_F59\_ingel\_3ug\_TL.raw

##### Fraction

2

##### Scan Feature

F2:6855

##### De Novo Score

99

##### ConfidenceScore

99

### m/z

563.3195

##### Mass

1124.623

##### Charge

2

##### Retention Time

46.76

##### Predicted Retention Time

-

##### Area

3.401E+07

##### Parts Per Million

1.2

##### Fragmentation mode

ETHCD

##### Originating file

01 D:\separate\_stitch\_analyses\xle-disambiguation\20210325\_F59\_3ug\_DENOVO\_12.csv

#### Meta Information from PEAKS

##### Scan Identifier

F2:8277

##### Original sequence

F

V

V

F

G

G

G

T

K

L

T

##### Posttranslational Modifications

##### Source File

D:\separate\_stitch\_analyses\xle-disambiguation\raw\20210323\_F1\_UM1\_Peng0013\_SA\_F59\_ingel\_3ug\_TL.raw

##### Fraction

2

##### Scan Feature

F2:6855

##### De Novo Score

99

##### ConfidenceScore

99

### m/z

563.3195

##### Mass

1124.623

##### Charge

2

##### Retention Time

46.76

##### Predicted Retention Time

-

##### Area

3.401E+07

##### Parts Per Million

1.2

##### Fragmentation mode

HCD

##### Originating file

01 D:\separate\_stitch\_analyses\xle-disambiguation\20210325\_F59\_3ug\_DENOVO\_12.csv

#### Meta Information from PEAKS

##### Scan Identifier

F2:8401

##### Original sequence

F

V

V

F

G

G

G

T

K

L

T

##### Posttranslational Modifications

##### Source File

D:\separate\_stitch\_analyses\xle-disambiguation\raw\20210323\_F1\_UM1\_Peng0013\_SA\_F59\_ingel\_3ug\_TL.raw

##### Fraction

2

##### Scan Feature

F2:6855

##### De Novo Score

99

##### ConfidenceScore

99

### m/z

563.3195

##### Mass

1124.623

##### Charge

2

##### Retention Time

46.76

##### Predicted Retention Time

-

##### Area

3.401E+07

##### Parts Per Million

1.2

##### Fragmentation mode

HCD

##### Originating file

01 D:\separate\_stitch\_analyses\xle-disambiguation\20210325\_F59\_3ug\_DENOVO\_12.csv

#### Meta Information from PEAKS

##### Scan Identifier

F2:8454

##### Original sequence

F

V

V

F

G

G

G

T

K

L

T

##### Posttranslational Modifications

##### Source File

D:\separate\_stitch\_analyses\xle-disambiguation\raw\20210323\_F1\_UM1\_Peng0013\_SA\_F59\_ingel\_3ug\_TL.raw

##### Fraction

2

##### Scan Feature

F2:6855

##### De Novo Score

98

##### ConfidenceScore

98

### m/z

563.3195

##### Mass

1124.623

##### Charge

2

##### Retention Time

46.76

##### Predicted Retention Time

-

##### Area

3.401E+07

##### Parts Per Million

1.2

##### Fragmentation mode

HCD

##### Originating file

01 D:\separate\_stitch\_analyses\xle-disambiguation\20210325\_F59\_3ug\_DENOVO\_12.csv

#### Meta Information from PEAKS

##### Scan Identifier

F2:8568

##### Original sequence

F

V

V

F

G

G

G

T

K

L

T

##### Posttranslational Modifications

##### Source File

D:\separate\_stitch\_analyses\xle-disambiguation\raw\20210323\_F1\_UM1\_Peng0013\_SA\_F59\_ingel\_3ug\_TL.raw

##### Fraction

2

##### Scan Feature

-

##### De Novo Score

96

##### ConfidenceScore

96

### m/z

563.3193

##### Mass

1124.623

##### Charge

2

##### Retention Time

48.25

##### Predicted Retention Time

-

##### Area

0

##### Parts Per Million

1

##### Fragmentation mode

HCD

##### Originating file

01 D:\separate\_stitch\_analyses\xle-disambiguation\20210325\_F59\_3ug\_DENOVO\_12.csv

#### Meta Information from PEAKS

##### Scan Identifier

F2:8732

##### Original sequence

F

V

V

F

G

G

G

T

K

L

T

##### Posttranslational Modifications

##### Source File

D:\separate\_stitch\_analyses\xle-disambiguation\raw\20210323\_F1\_UM1\_Peng0013\_SA\_F59\_ingel\_3ug\_TL.raw

##### Fraction

2

##### Scan Feature

-

##### De Novo Score

95

##### ConfidenceScore

95

### m/z

563.319

##### Mass

1124.623

##### Charge

2

##### Retention Time

49.18

##### Predicted Retention Time

-

##### Area

0

##### Parts Per Million

0.4

##### Fragmentation mode

HCD

##### Originating file

01 D:\separate\_stitch\_analyses\xle-disambiguation\20210325\_F59\_3ug\_DENOVO\_12.csv
