## Supplementary material for "A handle on mass coincidence errors in *de novo* sequencing of antibodies by bottom-up proteomics": Combined_016.html

Details Combined\_016 | Stitch OverviewUndefined

### Read Combined\_016

#### Sequence (length=12)

TISRDNAKNSJY

#### Spectrum 3437? Spectrum 3437 The raw spectrum of this peptide as annotated by Hecklib. The fragments are coloured according to ion type (see legend). Any peaks with a star '\*' as text can be hovered over to see the full details, first the ion type second the mass shift type. By hovering over the amino acids in the peptide or ions in the legend the corresponding peaks are highlighted. By toggling the 'Unassigned' label you can turn the background (unassigned) peaks on or off in the plot. By updating the slider in the Ion legend you can update the spectrum to only show the top X% of the peaks with labels. The top X% means any peak that is within X% of the highest intensity. By dragging in the spectrum you can zoom in to a specific part of the spectrum and use 'Zoom Out' to get back to the original zoom level. The annotation of the spectrum is based on the given sequence in the peptides file and is done with different software so inconsistencies are likely. The peaks are annotated based on the given sequence, with 20 ppm tolerance.

Copy Data

##### Spectrum 3437 (TSV)

###### Preview

```
Loading example...
```

*Click on the button to copy the data to your clipboard.*

Mz MinMz MaxIntensity Max

WidthHeightPeptide font sizePeptide stroke widthSpectrum font sizeSpectrum stroke widthCompact peptide

Ion legend

wxyz

abcd

OtherUnassignedIon^Charge^_Position_Show for top:%

TISRDNAKNSJY

02.85e+55.69e+58.54e+51.14e+6

Zoom Out

y^+1^_1_w^+3^_5_c^+1^_2_y^+1^_2_c^+3^_8_c^+1^_3_c^+2^_6_c^+3^_10_y^+1^_3_c^+2^_8_c^+2^_8_c^+2^_8_c^+1^_4_z^+1^_4_c^+1^_4_z^+1^_4_y^+1^_4_c^+2^_9_c^+2^_9_c^+2^_9_z^+2^_9_z^+2^_9_y^+2^_9_c^+2^_10_c^+2^_10_c^+2^_10_w^+2^_10_c^+1^_5_y^+2^_10_y^+2^_10_z^+2^_10_y^+2^_10_c^+1^_5_c^+2^_11_z^+1^_5_c^+2^_11_w^+2^_11_y^+1^_5_y^+2^_11_y^+2^_11_z^+2^_11_y^+2^_11_z^+1^_6_z^+1^_6_c^+1^_6_y^+1^_6_c^+1^_6_w^+1^_7_c^+1^_7_c^+1^_7_z^+1^_7_y^+1^_7_c^+1^_8_c^+1^_8_c^+1^_8_z^+1^_8_y^+1^_8_w^+1^_9_c^+1^_9_c^+1^_9_c^+1^_9_y^+1^_9_z^+1^_9_y^+1^_9_c^+1^_10_c^+1^_10_z^+1^_10_z^+1^_10_y^+1^_10_c^+1^_11_z^+1^_11_y^+1^_11_

0551110216532204

Fragment Matches Table

Show background peaks

| Position | Ion type | Intensity | mz Theoretical | mz Error (Th) | mz Error (ppm) | Charge | Series Number |
| --- | --- | --- | --- | --- | --- | --- | --- |
| - | - | 1048 | 122.4 | - | - | 0 | - |
| - | - | 1092 | 122.4 | - | - | 0 | - |
| - | - | 1907 | 123 | - | - | 0 | - |
| - | - | 1381 | 123.1 | - | - | 0 | - |
| - | - | 1.448E+04 | 129.1 | - | - | 0 | - |
| - | - | 1148 | 134.5 | - | - | 0 | - |
| - | - | 1241 | 135 | - | - | 0 | - |
| - | - | 5.716E+04 | 136.1 | - | - | 0 | - |
| - | - | 3707 | 137.1 | - | - | 0 | - |
| - | - | 1171 | 140.1 | - | - | 0 | - |
| - | - | 1179 | 140.7 | - | - | 0 | - |
| - | - | 4665 | 142.1 | - | - | 0 | - |
| - | - | 3628 | 143.1 | - | - | 0 | - |
| - | - | 2223 | 147 | - | - | 0 | - |
| - | - | 4617 | 149 | - | - | 0 | - |
| - | - | 1262 | 154.8 | - | - | 0 | - |
| - | - | 1694 | 155.1 | - | - | 0 | - |
| - | - | 1376 | 156.9 | - | - | 0 | - |
| - | - | 1691 | 157.1 | - | - | 0 | - |
| - | - | 1296 | 157.1 | - | - | 0 | - |
| - | - | 3.797E+04 | 165.1 | - | - | 0 | - |
| - | - | 3672 | 166.1 | - | - | 0 | - |
| - | - | 1808 | 167.1 | - | - | 0 | - |
| - | - | 3505 | 169.1 | - | - | 0 | - |
| - | - | 5988 | 173.1 | - | - | 0 | - |
| - | - | 2762 | 173.5 | - | - | 0 | - |
| - | - | 1578 | 175.1 | - | - | 0 | - |
| - | - | 2057 | 181.1 | - | - | 0 | - |
| 12 | y | 1.624E+05 | 182.1 | 0.000236 | 1.296 | +1 | 1 |
| - | - | 1.402E+04 | 183.1 | - | - | 0 | - |
| 8 | w | 2598 | 184.1 | 0.001964 | 10.67 | +3 | 5 |
| - | - | 2363 | 186.1 | - | - | 0 | - |
| - | - | 2.952E+05 | 187.1 | - | - | 0 | - |
| - | - | 1874 | 188.1 | - | - | 0 | - |
| - | - | 2.488E+04 | 188.1 | - | - | 0 | - |
| - | - | 1635 | 197.1 | - | - | 0 | - |
| - | - | 2812 | 200.1 | - | - | 0 | - |
| - | - | 9390 | 201.1 | - | - | 0 | - |
| - | - | 3675 | 201.1 | - | - | 0 | - |
| - | - | 2766 | 201.1 | - | - | 0 | - |
| - | - | 4485 | 202.1 | - | - | 0 | - |
| - | - | 6.653E+04 | 215.1 | - | - | 0 | - |
| - | - | 4976 | 216.1 | - | - | 0 | - |
| - | - | 1706 | 217.1 | - | - | 0 | - |
| - | - | 1907 | 225 | - | - | 0 | - |
| - | - | 5861 | 229.1 | - | - | 0 | - |
| 2 | c | 4411 | 232.2 | 9.666E-05 | 0.4164 | +1 | 2 |
| - | - | 1496 | 238.1 | - | - | 0 | - |
| - | - | 3969 | 239.1 | - | - | 0 | - |
| - | - | 1782 | 243 | - | - | 0 | - |
| - | - | 3511 | 244.1 | - | - | 0 | - |
| - | - | 2519 | 245.2 | - | - | 0 | - |
| - | - | 3810 | 259.1 | - | - | 0 | - |
| - | - | 1594 | 267.7 | - | - | 0 | - |
| - | - | 3689 | 271.1 | - | - | 0 | - |
| - | - | 2839 | 279.1 | - | - | 0 | - |
| - | - | 1728 | 281.1 | - | - | 0 | - |
| - | - | 1.48E+04 | 282.1 | - | - | 0 | - |
| - | - | 1988 | 283.1 | - | - | 0 | - |
| - | - | 3678 | 287.2 | - | - | 0 | - |
| - | - | 1534 | 290.9 | - | - | 0 | - |
| - | - | 2599 | 292.1 | - | - | 0 | - |
| 11 | y | 1.419E+04 | 295.2 | 0.0002632 | 0.8916 | +1 | 2 |
| - | - | 1626 | 296.1 | - | - | 0 | - |
| 8 | c | 1914 | 296.2 | 0.005266 | 17.78 | +3 | 8 |
| - | - | 5081 | 297.2 | - | - | 0 | - |
| - | - | 2009 | 298.2 | - | - | 0 | - |
| - | - | 2005 | 299.1 | - | - | 0 | - |
| - | - | 4487 | 299.1 | - | - | 0 | - |
| - | - | 3663 | 299.2 | - | - | 0 | - |
| - | - | 4.297E+04 | 300.2 | - | - | 0 | - |
| - | - | 6484 | 301.2 | - | - | 0 | - |
| - | - | 8966 | 315.2 | - | - | 0 | - |
| - | - | 2112 | 315.2 | - | - | 0 | - |
| - | - | 1979 | 316.2 | - | - | 0 | - |
| - | - | 2.084E+04 | 317.2 | - | - | 0 | - |
| - | - | 1784 | 317.4 | - | - | 0 | - |
| - | - | 2921 | 318.2 | - | - | 0 | - |
| 3 | c | 1.818E+04 | 319.2 | 0.0004321 | 1.354 | +1 | 3 |
| - | - | 1862 | 320.2 | - | - | 0 | - |
| - | - | 1.546E+04 | 341 | - | - | 0 | - |
| 6 | c | 2001 | 344.2 | 0.0005499 | 1.598 | +2 | 6 |
| - | - | 4.259E+04 | 359 | - | - | 0 | - |
| - | - | 2326 | 359.2 | - | - | 0 | - |
| 10 | c | 3016 | 363.2 | 0.003443 | 9.481 | +3 | 10 |
| - | - | 2473 | 367.2 | - | - | 0 | - |
| - | - | 1669 | 368.4 | - | - | 0 | - |
| - | - | 2678 | 370.1 | - | - | 0 | - |
| - | - | 4901 | 372.7 | - | - | 0 | - |
| - | - | 3366 | 373.2 | - | - | 0 | - |
| - | - | 3383 | 374.2 | - | - | 0 | - |
| - | - | 3400 | 375.9 | - | - | 0 | - |
| 10 | y | 9313 | 382.2 | 0.0005152 | 1.348 | +1 | 3 |
| - | - | 3.297E+04 | 385.2 | - | - | 0 | - |
| - | - | 8207 | 385.5 | - | - | 0 | - |
| - | - | 9512 | 385.9 | - | - | 0 | - |
| - | - | 2730 | 386.2 | - | - | 0 | - |
| - | - | 1.469E+04 | 386.2 | - | - | 0 | - |
| - | - | 2127 | 387.2 | - | - | 0 | - |
| - | - | 3.584E+04 | 391.6 | - | - | 0 | - |
| - | - | 2.131E+04 | 391.9 | - | - | 0 | - |
| - | - | 7488 | 392.2 | - | - | 0 | - |
| - | - | 2034 | 392.6 | - | - | 0 | - |
| - | - | 4404 | 393.7 | - | - | 0 | - |
| - | - | 1920 | 394.2 | - | - | 0 | - |
| - | - | 2640 | 394.7 | - | - | 0 | - |
| - | - | 2590 | 395.2 | - | - | 0 | - |
| - | - | 1708 | 403.5 | - | - | 0 | - |
| - | - | 2180 | 407.2 | - | - | 0 | - |
| - | - | 8910 | 415.3 | - | - | 0 | - |
| - | - | 4963 | 415.7 | - | - | 0 | - |
| - | - | 1.728E+04 | 419.2 | - | - | 0 | - |
| - | - | 4384 | 420.2 | - | - | 0 | - |
| - | - | 3743 | 424.7 | - | - | 0 | - |
| - | - | 4007 | 425.2 | - | - | 0 | - |
| - | - | 3121 | 425.3 | - | - | 0 | - |
| - | - | 5066 | 426.2 | - | - | 0 | - |
| - | - | 1.172E+05 | 429.1 | - | - | 0 | - |
| - | - | 2163 | 429.2 | - | - | 0 | - |
| - | - | 9298 | 429.8 | - | - | 0 | - |
| - | - | 2887 | 430.2 | - | - | 0 | - |
| - | - | 1.628E+04 | 430.2 | - | - | 0 | - |
| - | - | 3883 | 430.3 | - | - | 0 | - |
| - | - | 3091 | 431.2 | - | - | 0 | - |
| - | - | 3046 | 431.3 | - | - | 0 | - |
| - | - | 1893 | 434.2 | - | - | 0 | - |
| - | - | 4981 | 435.2 | - | - | 0 | - |
| - | - | 2697 | 436.2 | - | - | 0 | - |
| - | - | 3966 | 437.2 | - | - | 0 | - |
| - | - | 1966 | 437.7 | - | - | 0 | - |
| - | - | 6408 | 438.7 | - | - | 0 | - |
| - | - | 4241 | 439.2 | - | - | 0 | - |
| - | - | 2.251E+04 | 442.7 | - | - | 0 | - |
| 8 | c | 9822 | 443.2 | 0.008092 | 18.26 | +2 | 8 |
| 8 | c | 1.328E+04 | 443.7 | 0.002351 | 5.298 | +2 | 8 |
| - | - | 3612 | 444.2 | - | - | 0 | - |
| - | - | 2223 | 444.3 | - | - | 0 | - |
| - | - | 6.869E+04 | 445.1 | - | - | 0 | - |
| - | - | 1.45E+04 | 446.2 | - | - | 0 | - |
| - | - | 2765 | 447.2 | - | - | 0 | - |
| - | - | 6.874E+04 | 451.8 | - | - | 0 | - |
| 8 | c | 2.209E+05 | 452.3 | 6.273E-05 | 0.1387 | +2 | 8 |
| - | - | 1.014E+05 | 452.8 | - | - | 0 | - |
| - | - | 2.377E+04 | 453.3 | - | - | 0 | - |
| - | - | 3454 | 453.8 | - | - | 0 | - |
| - | - | 3012 | 454.3 | - | - | 0 | - |
| - | - | 2634 | 454.6 | - | - | 0 | - |
| - | - | 1840 | 454.9 | - | - | 0 | - |
| - | - | 3317 | 455.2 | - | - | 0 | - |
| - | - | 4255 | 455.6 | - | - | 0 | - |
| - | - | 2330 | 455.9 | - | - | 0 | - |
| - | - | 3074 | 456.3 | - | - | 0 | - |
| 4 | c | 9910 | 458.3 | 0.0005214 | 1.138 | +1 | 4 |
| - | - | 2190 | 458.8 | - | - | 0 | - |
| - | - | 2517 | 459.3 | - | - | 0 | - |
| - | - | 7.907E+04 | 460.2 | - | - | 0 | - |
| - | - | 3965 | 460.3 | - | - | 0 | - |
| - | - | 4.988E+04 | 460.7 | - | - | 0 | - |
| - | - | 2.321E+04 | 461.2 | - | - | 0 | - |
| - | - | 5870 | 461.6 | - | - | 0 | - |
| - | - | 3197 | 461.7 | - | - | 0 | - |
| 9 | z | 3165 | 462.2 | 0.002191 | 4.74 | +1 | 4 |
| - | - | 3273 | 462.2 | - | - | 0 | - |
| - | - | 3452 | 463.2 | - | - | 0 | - |
| - | - | 1974 | 465.3 | - | - | 0 | - |
| - | - | 3386 | 465.8 | - | - | 0 | - |
| - | - | 2534 | 466.7 | - | - | 0 | - |
| - | - | 3269 | 467.2 | - | - | 0 | - |
| - | - | 5315 | 470.8 | - | - | 0 | - |
| - | - | 7814 | 471.2 | - | - | 0 | - |
| - | - | 1878 | 471.7 | - | - | 0 | - |
| - | - | 9221 | 472.3 | - | - | 0 | - |
| - | - | 2742 | 473.3 | - | - | 0 | - |
| - | - | 3738 | 474.3 | - | - | 0 | - |
| 4 | c | 5.882E+05 | 475.3 | 0.0006395 | 1.345 | +1 | 4 |
| - | - | 4228 | 476.2 | - | - | 0 | - |
| - | - | 1.449E+05 | 476.3 | - | - | 0 | - |
| - | - | 2491 | 476.7 | - | - | 0 | - |
| - | - | 2576 | 477.2 | - | - | 0 | - |
| - | - | 2.628E+04 | 477.3 | - | - | 0 | - |
| - | - | 5187 | 479.3 | - | - | 0 | - |
| - | - | 7.892E+04 | 479.8 | - | - | 0 | - |
| 9 | z | 8684 | 480.2 | 0.0008234 | 1.715 | +1 | 4 |
| - | - | 3.832E+04 | 480.3 | - | - | 0 | - |
| - | - | 1.342E+04 | 480.8 | - | - | 0 | - |
| - | - | 4.478E+04 | 481.2 | - | - | 0 | - |
| - | - | 1.058E+04 | 482.2 | - | - | 0 | - |
| - | - | 8657 | 484.7 | - | - | 0 | - |
| - | - | 1.147E+04 | 485.2 | - | - | 0 | - |
| - | - | 2182 | 485.7 | - | - | 0 | - |
| - | - | 2768 | 486.2 | - | - | 0 | - |
| - | - | 1.113E+04 | 486.8 | - | - | 0 | - |
| - | - | 2.128E+04 | 487.3 | - | - | 0 | - |
| - | - | 8745 | 487.8 | - | - | 0 | - |
| - | - | 3354 | 488.3 | - | - | 0 | - |
| - | - | 2426 | 488.8 | - | - | 0 | - |
| - | - | 5364 | 491.8 | - | - | 0 | - |
| - | - | 5685 | 492.3 | - | - | 0 | - |
| - | - | 1.846E+04 | 493.8 | - | - | 0 | - |
| - | - | 7646 | 494.3 | - | - | 0 | - |
| - | - | 3652 | 494.8 | - | - | 0 | - |
| - | - | 5825 | 495.2 | - | - | 0 | - |
| 9 | y | 2.261E+04 | 496.2 | 0.0002585 | 0.5209 | +1 | 4 |
| - | - | 3814 | 497.2 | - | - | 0 | - |
| - | - | 9.547E+04 | 499.8 | - | - | 0 | - |
| 9 | c | 4.548E+04 | 500.3 | 0.002782 | 5.562 | +2 | 9 |
| 9 | c | 3.931E+04 | 500.8 | 0.002188 | 4.37 | +2 | 9 |
| - | - | 1.583E+04 | 501.3 | - | - | 0 | - |
| - | - | 1.566E+04 | 501.8 | - | - | 0 | - |
| - | - | 6.622E+04 | 502.3 | - | - | 0 | - |
| - | - | 2783 | 502.8 | - | - | 0 | - |
| - | - | 1.656E+04 | 503.3 | - | - | 0 | - |
| - | - | 3230 | 504.3 | - | - | 0 | - |
| - | - | 4217 | 508.3 | - | - | 0 | - |
| - | - | 1.807E+05 | 508.8 | - | - | 0 | - |
| 9 | c | 3.061E+05 | 509.3 | 0.0008627 | 1.694 | +2 | 9 |
| - | - | 1.405E+05 | 509.8 | - | - | 0 | - |
| - | - | 3.972E+04 | 510.3 | - | - | 0 | - |
| - | - | 1.33E+04 | 510.8 | - | - | 0 | - |
| - | - | 3888 | 511.3 | - | - | 0 | - |
| - | - | 1972 | 514.3 | - | - | 0 | - |
| - | - | 3474 | 521.3 | - | - | 0 | - |
| - | - | 3964 | 521.8 | - | - | 0 | - |
| - | - | 2271 | 522.3 | - | - | 0 | - |
| - | - | 4831 | 522.8 | - | - | 0 | - |
| - | - | 4158 | 523.3 | - | - | 0 | - |
| 4 | z | 2363 | 523.8 | 0.002139 | 4.083 | +2 | 9 |
| - | - | 7457 | 525.4 | - | - | 0 | - |
| - | - | 2247 | 525.7 | - | - | 0 | - |
| - | - | 1.5E+04 | 526.3 | - | - | 0 | - |
| - | - | 2325 | 526.4 | - | - | 0 | - |
| - | - | 7041 | 526.8 | - | - | 0 | - |
| - | - | 3421 | 527.3 | - | - | 0 | - |
| - | - | 2113 | 529.3 | - | - | 0 | - |
| - | - | 1.105E+04 | 530.3 | - | - | 0 | - |
| - | - | 2960 | 531.3 | - | - | 0 | - |
| 4 | z | 2.687E+04 | 532.8 | 0.0001521 | 0.2856 | +2 | 9 |
| - | - | 1.534E+04 | 533.3 | - | - | 0 | - |
| - | - | 2177 | 533.8 | - | - | 0 | - |
| - | - | 6589 | 534.3 | - | - | 0 | - |
| - | - | 9324 | 535.3 | - | - | 0 | - |
| - | - | 7628 | 535.3 | - | - | 0 | - |
| - | - | 7164 | 535.8 | - | - | 0 | - |
| - | - | 2654 | 535.8 | - | - | 0 | - |
| - | - | 6476 | 536.3 | - | - | 0 | - |
| - | - | 2433 | 540.3 | - | - | 0 | - |
| 4 | y | 6451 | 540.8 | 0.002923 | 5.406 | +2 | 9 |
| - | - | 3550 | 541.3 | - | - | 0 | - |
| - | - | 3194 | 541.8 | - | - | 0 | - |
| - | - | 2781 | 543.3 | - | - | 0 | - |
| 10 | c | 3412 | 543.8 | 0.009371 | 17.23 | +2 | 10 |
| 10 | c | 3.718E+04 | 544.3 | 0.0006396 | 1.175 | +2 | 10 |
| - | - | 1.679E+04 | 544.8 | - | - | 0 | - |
| - | - | 2.065E+04 | 545.3 | - | - | 0 | - |
| - | - | 2626 | 545.8 | - | - | 0 | - |
| - | - | 8550 | 546.3 | - | - | 0 | - |
| - | - | 3.046E+04 | 548.3 | - | - | 0 | - |
| - | - | 1.815E+04 | 548.8 | - | - | 0 | - |
| - | - | 7497 | 549.3 | - | - | 0 | - |
| - | - | 4601 | 550.3 | - | - | 0 | - |
| - | - | 3665 | 550.8 | - | - | 0 | - |
| - | - | 4523 | 552.3 | - | - | 0 | - |
| 10 | c | 2.212E+05 | 552.8 | 0.0002435 | 0.4405 | +2 | 10 |
| - | - | 1.374E+05 | 553.3 | - | - | 0 | - |
| - | - | 5.535E+04 | 553.8 | - | - | 0 | - |
| - | - | 1.511E+04 | 554.3 | - | - | 0 | - |
| - | - | 8510 | 554.8 | - | - | 0 | - |
| - | - | 3066 | 555.3 | - | - | 0 | - |
| - | - | 2235 | 557.3 | - | - | 0 | - |
| - | - | 2265 | 557.8 | - | - | 0 | - |
| - | - | 4529 | 561.3 | - | - | 0 | - |
| - | - | 2306 | 563.3 | - | - | 0 | - |
| - | - | 2.041E+04 | 564.3 | - | - | 0 | - |
| - | - | 3523 | 565.3 | - | - | 0 | - |
| - | - | 1.033E+04 | 565.3 | - | - | 0 | - |
| - | - | 1.832E+04 | 565.8 | - | - | 0 | - |
| - | - | 1.306E+04 | 566.3 | - | - | 0 | - |
| - | - | 4804 | 566.8 | - | - | 0 | - |
| - | - | 1.03E+04 | 567.3 | - | - | 0 | - |
| 3 | w | 1.005E+04 | 567.8 | 0.0007666 | 1.35 | +2 | 10 |
| - | - | 9352 | 568.3 | - | - | 0 | - |
| - | - | 7069 | 568.8 | - | - | 0 | - |
| - | - | 3502 | 569.8 | - | - | 0 | - |
| - | - | 2287 | 570.3 | - | - | 0 | - |
| - | - | 1931 | 570.8 | - | - | 0 | - |
| 5 | c | 1.008E+04 | 573.3 | 0.004854 | 8.466 | +1 | 5 |
| - | - | 6674 | 574.3 | - | - | 0 | - |
| - | - | 5064 | 574.8 | - | - | 0 | - |
| 3 | y | 8807 | 575.3 | 0.0005326 | 0.9258 | +2 | 10 |
| 3 | y | 1.631E+04 | 575.8 | 0.002272 | 3.945 | +2 | 10 |
| 3 | z | 8.671E+04 | 576.3 | 0.0002512 | 0.4359 | +2 | 10 |
| - | - | 6.075E+04 | 576.8 | - | - | 0 | - |
| - | - | 2.169E+04 | 577.3 | - | - | 0 | - |
| - | - | 5537 | 577.8 | - | - | 0 | - |
| - | - | 1.309E+04 | 577.8 | - | - | 0 | - |
| - | - | 9758 | 578.3 | - | - | 0 | - |
| - | - | 1E+04 | 578.8 | - | - | 0 | - |
| - | - | 1.309E+04 | 579.3 | - | - | 0 | - |
| - | - | 5040 | 579.8 | - | - | 0 | - |
| - | - | 1.026E+04 | 583.3 | - | - | 0 | - |
| - | - | 7541 | 583.8 | - | - | 0 | - |
| 3 | y | 2.828E+05 | 584.3 | 0.0002276 | 0.3895 | +2 | 10 |
| - | - | 1.742E+05 | 584.8 | - | - | 0 | - |
| - | - | 6.491E+04 | 585.3 | - | - | 0 | - |
| - | - | 1.704E+04 | 585.8 | - | - | 0 | - |
| - | - | 3.151E+04 | 586.8 | - | - | 0 | - |
| - | - | 3.12E+04 | 587.3 | - | - | 0 | - |
| - | - | 1.923E+04 | 587.8 | - | - | 0 | - |
| - | - | 1.573E+04 | 588.3 | - | - | 0 | - |
| - | - | 5704 | 588.3 | - | - | 0 | - |
| - | - | 1.07E+04 | 588.8 | - | - | 0 | - |
| - | - | 2176 | 589.3 | - | - | 0 | - |
| - | - | 1.238E+04 | 589.3 | - | - | 0 | - |
| 5 | c | 1.127E+06 | 590.3 | 0.0003993 | 0.6764 | +1 | 5 |
| - | - | 3.114E+05 | 591.3 | - | - | 0 | - |
| - | - | 2.723E+04 | 591.8 | - | - | 0 | - |
| - | - | 6.885E+04 | 592.3 | - | - | 0 | - |
| - | - | 1.684E+04 | 592.8 | - | - | 0 | - |
| - | - | 8785 | 593.3 | - | - | 0 | - |
| - | - | 2973 | 593.8 | - | - | 0 | - |
| - | - | 5201 | 597.8 | - | - | 0 | - |
| - | - | 3780 | 598.3 | - | - | 0 | - |
| - | - | 2369 | 599.8 | - | - | 0 | - |
| 11 | c | 5.099E+04 | 600.8 | 0.0002336 | 0.3888 | +2 | 11 |
| - | - | 3.333E+04 | 601.3 | - | - | 0 | - |
| - | - | 2.327E+04 | 601.8 | - | - | 0 | - |
| - | - | 6340 | 602.3 | - | - | 0 | - |
| - | - | 2249 | 602.8 | - | - | 0 | - |
| - | - | 1.255E+04 | 607.3 | - | - | 0 | - |
| - | - | 6132 | 607.8 | - | - | 0 | - |
| 8 | z | 6.069E+04 | 608.3 | 0.0009543 | 1.569 | +1 | 5 |
| - | - | 3387 | 608.8 | - | - | 0 | - |
| 11 | c | 5.919E+05 | 609.3 | 4.045E-05 | 0.06639 | +2 | 11 |
| - | - | 3.852E+05 | 609.8 | - | - | 0 | - |
| - | - | 1.547E+05 | 610.3 | - | - | 0 | - |
| - | - | 4.238E+04 | 610.8 | - | - | 0 | - |
| - | - | 6635 | 611.3 | - | - | 0 | - |
| - | - | 1.098E+04 | 616.3 | - | - | 0 | - |
| - | - | 2610 | 616.8 | - | - | 0 | - |
| 2 | w | 3.686E+04 | 618.3 | 0.0001235 | 0.1997 | +2 | 11 |
| - | - | 2.475E+04 | 618.8 | - | - | 0 | - |
| - | - | 1.101E+04 | 619.3 | - | - | 0 | - |
| - | - | 3812 | 619.8 | - | - | 0 | - |
| - | - | 4635 | 620.8 | - | - | 0 | - |
| - | - | 2512 | 621.3 | - | - | 0 | - |
| - | - | 2868 | 621.8 | - | - | 0 | - |
| - | - | 5767 | 623.8 | - | - | 0 | - |
| 8 | y | 9.69E+04 | 624.3 | 0.0001136 | 0.1819 | +1 | 5 |
| - | - | 3.112E+04 | 625.3 | - | - | 0 | - |
| - | - | 2314 | 625.8 | - | - | 0 | - |
| - | - | 7252 | 626.3 | - | - | 0 | - |
| - | - | 2284 | 626.8 | - | - | 0 | - |
| - | - | 1.84E+04 | 630.3 | - | - | 0 | - |
| - | - | 1.421E+04 | 630.8 | - | - | 0 | - |
| - | - | 6446 | 631.3 | - | - | 0 | - |
| 2 | y | 4569 | 631.8 | 0.008217 | 13 | +2 | 11 |
| 2 | y | 5479 | 632.3 | 0.0002096 | 0.3314 | +2 | 11 |
| 2 | z | 1.193E+04 | 632.8 | 0.001127 | 1.781 | +2 | 11 |
| - | - | 2.475E+04 | 633.3 | - | - | 0 | - |
| - | - | 1.542E+04 | 633.8 | - | - | 0 | - |
| - | - | 1.13E+04 | 634.3 | - | - | 0 | - |
| - | - | 5207 | 635.4 | - | - | 0 | - |
| - | - | 1.423E+04 | 636.3 | - | - | 0 | - |
| - | - | 4145 | 637.4 | - | - | 0 | - |
| - | - | 8676 | 637.8 | - | - | 0 | - |
| - | - | 1.019E+04 | 638.3 | - | - | 0 | - |
| - | - | 6784 | 638.8 | - | - | 0 | - |
| - | - | 7185 | 639.3 | - | - | 0 | - |
| - | - | 6220 | 639.8 | - | - | 0 | - |
| - | - | 7855 | 640.3 | - | - | 0 | - |
| 2 | y | 3.26E+04 | 640.8 | 0.000676 | 1.055 | +2 | 11 |
| - | - | 2.94E+04 | 641.3 | - | - | 0 | - |
| - | - | 1.017E+04 | 641.8 | - | - | 0 | - |
| - | - | 7219 | 642.3 | - | - | 0 | - |
| - | - | 7744 | 644.3 | - | - | 0 | - |
| - | - | 2553 | 645.8 | - | - | 0 | - |
| - | - | 3216 | 646.3 | - | - | 0 | - |
| - | - | 2435 | 646.8 | - | - | 0 | - |
| - | - | 3.391E+04 | 647.4 | - | - | 0 | - |
| - | - | 2.739E+04 | 647.9 | - | - | 0 | - |
| - | - | 1.35E+04 | 648.4 | - | - | 0 | - |
| - | - | 3347 | 648.9 | - | - | 0 | - |
| - | - | 1.834E+04 | 649.3 | - | - | 0 | - |
| - | - | 1.017E+04 | 650.3 | - | - | 0 | - |
| - | - | 2702 | 651.3 | - | - | 0 | - |
| - | - | 2901 | 652.3 | - | - | 0 | - |
| - | - | 5197 | 652.8 | - | - | 0 | - |
| - | - | 1.396E+04 | 653.3 | - | - | 0 | - |
| - | - | 8390 | 653.4 | - | - | 0 | - |
| - | - | 7818 | 653.8 | - | - | 0 | - |
| - | - | 6406 | 654.3 | - | - | 0 | - |
| - | - | 3281 | 654.5 | - | - | 0 | - |
| - | - | 4401 | 654.8 | - | - | 0 | - |
| - | - | 4.407E+04 | 655.3 | - | - | 0 | - |
| - | - | 2.846E+04 | 655.8 | - | - | 0 | - |
| - | - | 1.279E+04 | 656.3 | - | - | 0 | - |
| - | - | 3753 | 656.8 | - | - | 0 | - |
| - | - | 5.718E+04 | 659.3 | - | - | 0 | - |
| - | - | 2701 | 659.9 | - | - | 0 | - |
| - | - | 9.288E+04 | 660.3 | - | - | 0 | - |
| - | - | 6.066E+04 | 660.9 | - | - | 0 | - |
| 7 | z | 1.023E+05 | 661.3 | 0.009195 | 13.9 | +1 | 6 |
| - | - | 7.027E+04 | 661.8 | - | - | 0 | - |
| - | - | 9.258E+04 | 662.3 | - | - | 0 | - |
| - | - | 5.159E+04 | 662.8 | - | - | 0 | - |
| - | - | 1.826E+04 | 663.3 | - | - | 0 | - |
| - | - | 6491 | 663.8 | - | - | 0 | - |
| - | - | 2.836E+04 | 668.8 | - | - | 0 | - |
| - | - | 1.173E+05 | 669.3 | - | - | 0 | - |
| - | - | 7.128E+04 | 669.8 | - | - | 0 | - |
| - | - | 9.318E+04 | 670.3 | - | - | 0 | - |
| - | - | 4.993E+04 | 670.8 | - | - | 0 | - |
| - | - | 2.388E+04 | 671.3 | - | - | 0 | - |
| - | - | 5343 | 671.8 | - | - | 0 | - |
| - | - | 1.109E+04 | 672.3 | - | - | 0 | - |
| - | - | 1.039E+04 | 672.9 | - | - | 0 | - |
| - | - | 6216 | 673.4 | - | - | 0 | - |
| - | - | 1.309E+04 | 673.9 | - | - | 0 | - |
| - | - | 2.281E+04 | 674.3 | - | - | 0 | - |
| - | - | 5.338E+04 | 674.8 | - | - | 0 | - |
| - | - | 3.91E+04 | 675.3 | - | - | 0 | - |
| - | - | 5.223E+04 | 675.8 | - | - | 0 | - |
| - | - | 3.031E+04 | 676.3 | - | - | 0 | - |
| - | - | 1.663E+04 | 676.8 | - | - | 0 | - |
| - | - | 4595 | 677.3 | - | - | 0 | - |
| 7 | z | 2.324E+05 | 679.4 | 0.0003396 | 0.4999 | +1 | 6 |
| - | - | 8.702E+04 | 680.4 | - | - | 0 | - |
| - | - | 8875 | 680.9 | - | - | 0 | - |
| - | - | 2.686E+04 | 681.4 | - | - | 0 | - |
| - | - | 1.395E+04 | 681.9 | - | - | 0 | - |
| - | - | 7897 | 682.4 | - | - | 0 | - |
| - | - | 2.386E+04 | 682.9 | - | - | 0 | - |
| - | - | 3.713E+05 | 683.3 | - | - | 0 | - |
| - | - | 2.721E+05 | 683.8 | - | - | 0 | - |
| - | - | 1.329E+05 | 684.3 | - | - | 0 | - |
| - | - | 4.704E+04 | 684.8 | - | - | 0 | - |
| - | - | 6662 | 685.4 | - | - | 0 | - |
| 6 | c | 1.689E+04 | 687.3 | 0.00752 | 10.94 | +1 | 6 |
| - | - | 5419 | 688.4 | - | - | 0 | - |
| - | - | 7904 | 689.9 | - | - | 0 | - |
| - | - | 1.926E+04 | 690.4 | - | - | 0 | - |
| - | - | 1.444E+04 | 690.9 | - | - | 0 | - |
| - | - | 3.26E+05 | 691.4 | - | - | 0 | - |
| - | - | 2.576E+05 | 691.9 | - | - | 0 | - |
| - | - | 1.256E+05 | 692.4 | - | - | 0 | - |
| - | - | 4.124E+04 | 692.9 | - | - | 0 | - |
| - | - | 1.472E+04 | 693.4 | - | - | 0 | - |
| 7 | y | 5.318E+04 | 695.4 | 0.0003533 | 0.5081 | +1 | 6 |
| - | - | 2.372E+04 | 696.4 | - | - | 0 | - |
| - | - | 5789 | 697.4 | - | - | 0 | - |
| 6 | c | 3.151E+05 | 704.4 | 0.0002575 | 0.3656 | +1 | 6 |
| - | - | 1.085E+05 | 705.4 | - | - | 0 | - |
| - | - | 3348 | 705.4 | - | - | 0 | - |
| - | - | 2.372E+04 | 706.4 | - | - | 0 | - |
| - | - | 2782 | 707.4 | - | - | 0 | - |
| - | - | 1.184E+04 | 709.3 | - | - | 0 | - |
| - | - | 5884 | 710.3 | - | - | 0 | - |
| - | - | 3437 | 711.3 | - | - | 0 | - |
| - | - | 3366 | 730.4 | - | - | 0 | - |
| - | - | 4.983E+04 | 731.4 | - | - | 0 | - |
| - | - | 1.628E+04 | 732.4 | - | - | 0 | - |
| - | - | 5302 | 733.4 | - | - | 0 | - |
| - | - | 1.353E+04 | 737.3 | - | - | 0 | - |
| - | - | 3761 | 738.3 | - | - | 0 | - |
| - | - | 2830 | 740.4 | - | - | 0 | - |
| - | - | 3470 | 741.4 | - | - | 0 | - |
| - | - | 2554 | 743.4 | - | - | 0 | - |
| 6 | w | 1.577E+05 | 749.4 | 0.001385 | 1.849 | +1 | 7 |
| - | - | 6.512E+04 | 750.4 | - | - | 0 | - |
| - | - | 2.085E+04 | 751.4 | - | - | 0 | - |
| - | - | 4477 | 752.4 | - | - | 0 | - |
| - | - | 7724 | 755.4 | - | - | 0 | - |
| - | - | 3646 | 756.4 | - | - | 0 | - |
| - | - | 2333 | 757.4 | - | - | 0 | - |
| 7 | c | 3.345E+04 | 758.4 | 0.0008623 | 1.137 | +1 | 7 |
| - | - | 1.036E+04 | 759.4 | - | - | 0 | - |
| - | - | 3374 | 760.4 | - | - | 0 | - |
| - | - | 3182 | 761.4 | - | - | 0 | - |
| - | - | 2436 | 762.4 | - | - | 0 | - |
| - | - | 2637 | 774.4 | - | - | 0 | - |
| 7 | c | 2.919E+05 | 775.4 | 0.0002352 | 0.3033 | +1 | 7 |
| - | - | 1.125E+05 | 776.4 | - | - | 0 | - |
| - | - | 2.345E+04 | 777.4 | - | - | 0 | - |
| - | - | 4779 | 778.4 | - | - | 0 | - |
| - | - | 2479 | 785.4 | - | - | 0 | - |
| - | - | 6366 | 786.4 | - | - | 0 | - |
| - | - | 8488 | 787.4 | - | - | 0 | - |
| - | - | 6563 | 788.4 | - | - | 0 | - |
| - | - | 2862 | 789.4 | - | - | 0 | - |
| 6 | z | 7.174E+05 | 793.4 | 0.0002588 | 0.3262 | +1 | 7 |
| - | - | 3.192E+05 | 794.4 | - | - | 0 | - |
| - | - | 8.762E+04 | 795.4 | - | - | 0 | - |
| - | - | 1.77E+04 | 796.4 | - | - | 0 | - |
| - | - | 2338 | 797.4 | - | - | 0 | - |
| - | - | 3711 | 803.4 | - | - | 0 | - |
| - | - | 5942 | 808.4 | - | - | 0 | - |
| 6 | y | 8257 | 809.4 | 0.0009482 | 1.171 | +1 | 7 |
| - | - | 5477 | 810.4 | - | - | 0 | - |
| - | - | 2948 | 811.4 | - | - | 0 | - |
| - | - | 2401 | 815.5 | - | - | 0 | - |
| - | - | 3215 | 820.4 | - | - | 0 | - |
| - | - | 2997 | 821.4 | - | - | 0 | - |
| - | - | 2722 | 823.5 | - | - | 0 | - |
| - | - | 4.133E+04 | 832.4 | - | - | 0 | - |
| - | - | 2.482E+04 | 833.4 | - | - | 0 | - |
| - | - | 5583 | 834.5 | - | - | 0 | - |
| - | - | 5256 | 843.5 | - | - | 0 | - |
| - | - | 2949 | 844.5 | - | - | 0 | - |
| - | - | 3147 | 848.5 | - | - | 0 | - |
| - | - | 2833 | 849.5 | - | - | 0 | - |
| - | - | 4613 | 852.4 | - | - | 0 | - |
| - | - | 1.7E+04 | 855.4 | - | - | 0 | - |
| - | - | 9299 | 856.4 | - | - | 0 | - |
| - | - | 8321 | 858.5 | - | - | 0 | - |
| - | - | 2.588E+04 | 859.5 | - | - | 0 | - |
| - | - | 1.974E+04 | 860.5 | - | - | 0 | - |
| - | - | 6448 | 861.5 | - | - | 0 | - |
| - | - | 2.3E+05 | 864.4 | - | - | 0 | - |
| - | - | 1.109E+05 | 865.4 | - | - | 0 | - |
| - | - | 3.56E+04 | 866.4 | - | - | 0 | - |
| - | - | 8002 | 867.4 | - | - | 0 | - |
| - | - | 2178 | 871.5 | - | - | 0 | - |
| - | - | 7040 | 873.4 | - | - | 0 | - |
| - | - | 7815 | 874.4 | - | - | 0 | - |
| - | - | 1.28E+04 | 875.4 | - | - | 0 | - |
| - | - | 7590 | 876.5 | - | - | 0 | - |
| - | - | 3070 | 877.5 | - | - | 0 | - |
| - | - | 1.48E+04 | 884.5 | - | - | 0 | - |
| 8 | c | 1.162E+04 | 885.5 | 0.006264 | 7.074 | +1 | 8 |
| 8 | c | 3.382E+04 | 886.5 | 0.0006868 | 0.7748 | +1 | 8 |
| - | - | 1.468E+04 | 887.5 | - | - | 0 | - |
| - | - | 7165 | 888.5 | - | - | 0 | - |
| - | - | 2.579E+04 | 900.5 | - | - | 0 | - |
| - | - | 1.401E+04 | 901.5 | - | - | 0 | - |
| - | - | 9.929E+04 | 902.5 | - | - | 0 | - |
| 8 | c | 3.665E+05 | 903.5 | 0.0008378 | 0.9273 | +1 | 8 |
| - | - | 1.508E+05 | 904.5 | - | - | 0 | - |
| - | - | 4.379E+04 | 905.5 | - | - | 0 | - |
| - | - | 7815 | 906.5 | - | - | 0 | - |
| 5 | z | 2.364E+05 | 908.4 | 0.0004391 | 0.4834 | +1 | 8 |
| - | - | 1.105E+05 | 909.4 | - | - | 0 | - |
| - | - | 3.787E+04 | 910.4 | - | - | 0 | - |
| - | - | 7113 | 911.4 | - | - | 0 | - |
| - | - | 1.004E+04 | 919.5 | - | - | 0 | - |
| - | - | 3.601E+04 | 920.5 | - | - | 0 | - |
| - | - | 2.143E+04 | 921.5 | - | - | 0 | - |
| - | - | 9083 | 922.5 | - | - | 0 | - |
| - | - | 2694 | 923.5 | - | - | 0 | - |
| 5 | y | 1.915E+04 | 924.4 | 0.0001239 | 0.1341 | +1 | 8 |
| - | - | 1.061E+04 | 925.4 | - | - | 0 | - |
| - | - | 3039 | 926.4 | - | - | 0 | - |
| - | - | 2318 | 943.5 | - | - | 0 | - |
| - | - | 2404 | 957.5 | - | - | 0 | - |
| - | - | 2405 | 960.5 | - | - | 0 | - |
| - | - | 3867 | 961.5 | - | - | 0 | - |
| - | - | 5211 | 968.5 | - | - | 0 | - |
| - | - | 1.089E+04 | 969.5 | - | - | 0 | - |
| - | - | 3784 | 970.5 | - | - | 0 | - |
| - | - | 2387 | 972.5 | - | - | 0 | - |
| - | - | 4427 | 973.5 | - | - | 0 | - |
| - | - | 6264 | 974.5 | - | - | 0 | - |
| 4 | w | 3105 | 978.5 | 0.001224 | 1.251 | +1 | 9 |
| - | - | 2186 | 979.5 | - | - | 0 | - |
| - | - | 2052 | 981.8 | - | - | 0 | - |
| - | - | 2.004E+04 | 986.5 | - | - | 0 | - |
| - | - | 2.813E+04 | 987.5 | - | - | 0 | - |
| - | - | 1.466E+04 | 988.5 | - | - | 0 | - |
| - | - | 5380 | 989.5 | - | - | 0 | - |
| 9 | c | 1.68E+04 | 999.5 | 0.000852 | 0.8524 | +1 | 9 |
| 9 | c | 2.438E+04 | 1001 | 0.005001 | 4.998 | +1 | 9 |
| - | - | 1.395E+04 | 1002 | - | - | 0 | - |
| - | - | 1.332E+04 | 1003 | - | - | 0 | - |
| - | - | 6401 | 1004 | - | - | 0 | - |
| - | - | 2628 | 1005 | - | - | 0 | - |
| - | - | 2.83E+04 | 1017 | - | - | 0 | - |
| 9 | c | 2.17E+05 | 1018 | 0.0007965 | 0.7828 | +1 | 9 |
| - | - | 1.11E+05 | 1019 | - | - | 0 | - |
| - | - | 3.489E+04 | 1020 | - | - | 0 | - |
| - | - | 1.499E+04 | 1021 | - | - | 0 | - |
| - | - | 3564 | 1022 | - | - | 0 | - |
| - | - | 2782 | 1023 | - | - | 0 | - |
| - | - | 3782 | 1024 | - | - | 0 | - |
| - | - | 3202 | 1025 | - | - | 0 | - |
| - | - | 2705 | 1052 | - | - | 0 | - |
| - | - | 5937 | 1061 | - | - | 0 | - |
| - | - | 2435 | 1062 | - | - | 0 | - |
| - | - | 5795 | 1063 | - | - | 0 | - |
| 4 | y | 3132 | 1064 | 0.008936 | 8.402 | +1 | 9 |
| 4 | z | 9.299E+04 | 1065 | 0.0003538 | 0.3324 | +1 | 9 |
| - | - | 8.927E+04 | 1066 | - | - | 0 | - |
| - | - | 4.781E+04 | 1067 | - | - | 0 | - |
| - | - | 1.471E+04 | 1068 | - | - | 0 | - |
| - | - | 3778 | 1069 | - | - | 0 | - |
| - | - | 8039 | 1072 | - | - | 0 | - |
| - | - | 4888 | 1073 | - | - | 0 | - |
| - | - | 2951 | 1080 | - | - | 0 | - |
| 4 | y | 9275 | 1081 | 0.004552 | 4.212 | +1 | 9 |
| - | - | 3793 | 1082 | - | - | 0 | - |
| - | - | 3373 | 1083 | - | - | 0 | - |
| 10 | c | 2289 | 1087 | 0.01467 | 13.5 | +1 | 10 |
| - | - | 1.317E+04 | 1089 | - | - | 0 | - |
| - | - | 1.081E+04 | 1090 | - | - | 0 | - |
| - | - | 6089 | 1091 | - | - | 0 | - |
| - | - | 4549 | 1095 | - | - | 0 | - |
| - | - | 7378 | 1096 | - | - | 0 | - |
| 10 | c | 6.473E+04 | 1105 | 0.001148 | 1.039 | +1 | 10 |
| - | - | 3.955E+04 | 1106 | - | - | 0 | - |
| - | - | 1.83E+04 | 1107 | - | - | 0 | - |
| - | - | 5233 | 1108 | - | - | 0 | - |
| - | - | 2874 | 1111 | - | - | 0 | - |
| 3 | z | 7248 | 1135 | 0.02237 | 19.71 | +1 | 10 |
| - | - | 7234 | 1136 | - | - | 0 | - |
| - | - | 4101 | 1137 | - | - | 0 | - |
| 3 | z | 5.641E+04 | 1152 | 3.362E-05 | 0.02919 | +1 | 10 |
| - | - | 1.124E+05 | 1153 | - | - | 0 | - |
| - | - | 5.712E+04 | 1154 | - | - | 0 | - |
| - | - | 2.212E+04 | 1155 | - | - | 0 | - |
| - | - | 5482 | 1156 | - | - | 0 | - |
| - | - | 6297 | 1158 | - | - | 0 | - |
| - | - | 6001 | 1159 | - | - | 0 | - |
| - | - | 5654 | 1160 | - | - | 0 | - |
| - | - | 4715 | 1161 | - | - | 0 | - |
| 3 | y | 4560 | 1168 | 0.004924 | 4.217 | +1 | 10 |
| - | - | 5971 | 1169 | - | - | 0 | - |
| - | - | 3720 | 1170 | - | - | 0 | - |
| - | - | 1.418E+04 | 1174 | - | - | 0 | - |
| - | - | 3.404E+04 | 1175 | - | - | 0 | - |
| - | - | 2.203E+04 | 1176 | - | - | 0 | - |
| - | - | 9094 | 1177 | - | - | 0 | - |
| - | - | 3123 | 1178 | - | - | 0 | - |
| - | - | 4006 | 1185 | - | - | 0 | - |
| - | - | 2991 | 1186 | - | - | 0 | - |
| - | - | 3742 | 1201 | - | - | 0 | - |
| - | - | 4.71E+04 | 1202 | - | - | 0 | - |
| - | - | 2.713E+04 | 1203 | - | - | 0 | - |
| - | - | 1.011E+04 | 1204 | - | - | 0 | - |
| - | - | 5523 | 1205 | - | - | 0 | - |
| 11 | c | 3.7E+04 | 1218 | 0.001105 | 0.9077 | +1 | 11 |
| - | - | 8.035E+04 | 1219 | - | - | 0 | - |
| - | - | 4.05E+04 | 1220 | - | - | 0 | - |
| - | - | 1.758E+04 | 1221 | - | - | 0 | - |
| - | - | 3653 | 1222 | - | - | 0 | - |
| 2 | z | 3015 | 1265 | 0.007089 | 5.605 | +1 | 11 |
| - | - | 2.248E+04 | 1266 | - | - | 0 | - |
| - | - | 1.477E+04 | 1267 | - | - | 0 | - |
| - | - | 7315 | 1268 | - | - | 0 | - |
| 2 | y | 4446 | 1281 | 0.01261 | 9.848 | +1 | 11 |
| - | - | 2674 | 1282 | - | - | 0 | - |
| - | - | 4085 | 1295 | - | - | 0 | - |
| - | - | 3521 | 1296 | - | - | 0 | - |
| - | - | 3819 | 1305 | - | - | 0 | - |
| - | - | 1.763E+04 | 1306 | - | - | 0 | - |
| - | - | 1.751E+04 | 1307 | - | - | 0 | - |
| - | - | 1.188E+04 | 1308 | - | - | 0 | - |
| - | - | 6883 | 1309 | - | - | 0 | - |
| - | - | 4536 | 1310 | - | - | 0 | - |
| - | - | 3206 | 1320 | - | - | 0 | - |
| - | - | 1.058E+04 | 1321 | - | - | 0 | - |
| - | - | 2.262E+04 | 1322 | - | - | 0 | - |
| - | - | 1.595E+04 | 1323 | - | - | 0 | - |
| - | - | 4.022E+04 | 1324 | - | - | 0 | - |
| - | - | 7.073E+04 | 1325 | - | - | 0 | - |
| - | - | 4.004E+04 | 1326 | - | - | 0 | - |
| - | - | 1.982E+04 | 1327 | - | - | 0 | - |
| - | - | 5452 | 1328 | - | - | 0 | - |
| - | - | 1.989E+04 | 1338 | - | - | 0 | - |
| - | - | 1.107E+05 | 1339 | - | - | 0 | - |
| - | - | 7.342E+04 | 1340 | - | - | 0 | - |
| - | - | 3.514E+04 | 1341 | - | - | 0 | - |
| - | - | 9829 | 1342 | - | - | 0 | - |
| - | - | 3307 | 1343 | - | - | 0 | - |
| - | - | 2.42E+04 | 1349 | - | - | 0 | - |
| - | - | 4.766E+04 | 1350 | - | - | 0 | - |
| - | - | 2.733E+04 | 1351 | - | - | 0 | - |
| - | - | 1.204E+04 | 1352 | - | - | 0 | - |
| - | - | 2565 | 1353 | - | - | 0 | - |
| - | - | 2409 | 1355 | - | - | 0 | - |
| - | - | 5179 | 1356 | - | - | 0 | - |
| - | - | 5027 | 1357 | - | - | 0 | - |
| - | - | 2933 | 1363 | - | - | 0 | - |
| - | - | 7998 | 1364 | - | - | 0 | - |
| - | - | 5571 | 1365 | - | - | 0 | - |
| - | - | 5.986E+04 | 1366 | - | - | 0 | - |
| - | - | 3.541E+05 | 1367 | - | - | 0 | - |
| - | - | 2.497E+05 | 1368 | - | - | 0 | - |
| - | - | 1.204E+05 | 1369 | - | - | 0 | - |
| - | - | 3.355E+04 | 1370 | - | - | 0 | - |
| - | - | 5096 | 1371 | - | - | 0 | - |
| - | - | 4537 | 1380 | - | - | 0 | - |
| - | - | 1.089E+04 | 1381 | - | - | 0 | - |
| - | - | 1.76E+04 | 1382 | - | - | 0 | - |
| - | - | 4.018E+04 | 1383 | - | - | 0 | - |
| - | - | 7.006E+04 | 1384 | - | - | 0 | - |
| - | - | 5.139E+04 | 1385 | - | - | 0 | - |
| - | - | 2.286E+04 | 1386 | - | - | 0 | - |
| - | - | 7102 | 1387 | - | - | 0 | - |
| - | - | 2012 | 2183 | - | - | 0 | - |

m/z Charge Intensity FragmentType MassShift Position
122.41844940185547 0 1047.7313
122.42354583740234 0 1092.2059
123.04405975341797 0 1907.4362
123.10458374023438 0 1380.5104
129.10247802734375 0 14482.941
134.47012329101562 0 1147.798
134.97645568847656 0 1240.8873
136.075927734375 0 57157.773
137.07933044433594 0 3707.3513
140.06455993652344 0 1170.8918
140.69766235351562 0 1179.3236
142.1229705810547 0 4664.6006
143.11817932128906 0 3627.8123
147.04432678222656 0 2223.065
149.04502868652344 0 4617.0015
154.7826690673828 0 1261.8231
155.1176300048828 0 1693.9437
156.8673858642578 0 1376.0559
157.06077575683594 0 1690.7605
157.13409423828125 0 1296.0868
165.05484008789062 0 37974.445
166.05795288085938 0 3672.1665
167.05601501464844 0 1808.3086
169.1339111328125 0 3504.5222
173.12847900390625 0 5987.5176
173.4519500732422 0 2762.0242
175.07183837890625 0 1578.4751
181.09739685058594 0 2057.3923
182.08140563964844 0 162440.2 y 11
183.08482360839844 0 14020.469
184.0864715576172 0 2598.218 w 7
186.11224365234375 0 2362.8743
187.14437866210938 0 295162.5
188.14031982421875 0 1873.9855
188.147705078125 0 24878.043
197.12867736816406 0 1634.7573
200.13929748535156 0 2811.522
201.12355041503906 0 9389.519
201.13392639160156 0 3674.8196
201.14697265625 0 2765.558
202.08255004882812 0 4485.145
215.13925170898438 0 66528.55
216.14231872558594 0 4975.7495
217.14598083496094 0 1706.3054
225.04273986816406 0 1907.3138
229.12965393066406 0 5861.031
232.16566467285156 0 4411.3247 c 1
238.13018798828125 0 1495.5281
239.09481811523438 0 3968.5208
243.0443115234375 0 1781.5952
244.14105224609375 0 3511.2651
245.16015625 0 2519.3403
259.1174621582031 0 3810.252
267.6749267578125 0 1593.6473
271.1400451660156 0 3689.4236
279.1346740722656 0 2838.7327
281.1371154785156 0 1728.29
282.14501953125 0 14800.75
283.1290283203125 0 1988.3658
287.1729431152344 0 3677.9802
290.86517333984375 0 1533.8689
292.1299743652344 0 2598.5803
295.1654968261719 0 14188.506 y 10
296.1046142578125 0 1625.8147
296.16815185546875 0 1914.1879 c Ammonia loss 7
297.15625 0 5081.3105
298.1617431640625 0 2008.848
299.0624694824219 0 2005.3008
299.1476135253906 0 4486.6714
299.1712341308594 0 3662.8777
300.1556701660156 0 42972.168
301.15911865234375 0 6484.374
315.1665344238281 0 8966.076
315.1868896484375 0 2111.8928
316.1708984375 0 1978.962
317.1823425292969 0 20844.074
317.4248352050781 0 1784.4186
318.185791015625 0 2921.3433
319.1980285644531 0 18182.926 c 2
320.2013244628906 0 1862.4448
341.0181579589844 0 15455.715
344.1741027832031 0 2001.1327 c Ammonia loss 5
359.028564453125 0 42586.32
359.1662902832031 0 2326.2593
363.1913146972656 0 3015.8513 c Ammonia loss 9
367.1846008300781 0 2473.1096
368.40625 0 1668.7644
370.1234130859375 0 2677.7913
372.69671630859375 0 4901.3716
373.198974609375 0 3366.2686
374.17877197265625 0 3383.173
375.8616943359375 0 3400.4512
382.1967468261719 0 9313.38 y 9
385.2196350097656 0 32969.402
385.5469970703125 0 8207.446
385.88214111328125 0 9511.765
386.1665344238281 0 2730.3594
386.2242126464844 0 14690.204
387.2313537597656 0 2127.4414
391.5510559082031 0 35837.375
391.8846740722656 0 21308.46
392.2178039550781 0 7487.769
392.55230712890625 0 2034.0315
393.6955871582031 0 4403.628
394.1969909667969 0 1920.2194
394.70489501953125 0 2639.74
395.20941162109375 0 2590.352
403.524169921875 0 1707.5039
407.1930236816406 0 2179.7197
415.2674560546875 0 8909.783
415.7094421386719 0 4962.5547
419.21624755859375 0 17278.625
420.221923828125 0 4383.6187
424.7363586425781 0 3743.238
425.2016906738281 0 4007.4211
425.2523498535156 0 3120.612
426.2342834472656 0 5065.958
429.0892028808594 0 117216.586
429.2358093261719 0 2162.5425
429.7547607421875 0 9298.286
430.2060241699219 0 2886.654
430.24212646484375 0 16278.157
430.2777099609375 0 3883.0107
431.246337890625 0 3090.667
431.2782287597656 0 3046.2969
434.22412109375 0 1892.7222
435.2347106933594 0 4980.8613
436.23419189453125 0 2697.2366
437.21148681640625 0 3965.9036
437.7122802734375 0 1966.2797
438.7327880859375 0 6408.4707
439.2333984375 0 4241.4873
442.7452697753906 0 22509.871
443.25677490234375 0 9821.917 c Water loss 7
443.7430419921875 0 13277.935 c Ammonia loss 7
444.2404479980469 0 3612.4028
444.2700500488281 0 2222.7087
445.1204833984375 0 68692.59
446.2126770019531 0 14504.244
447.2179870605469 0 2765.2295
451.75054931640625 0 68740.08
452.2540283203125 0 220922.3 c 7
452.7554626464844 0 101409.65
453.2569580078125 0 23774.553
453.7588195800781 0 3453.7463
454.25445556640625 0 3011.972
454.5861511230469 0 2634.197
454.9188232421875 0 1839.6984
455.23931884765625 0 3316.5405
455.57220458984375 0 4255.319
455.9051208496094 0 2330.2915
456.25823974609375 0 3074.321
458.2716369628906 0 9909.932 c Ammonia loss 3
458.75946044921875 0 2189.5396
459.2745361328125 0 2517.434
460.23577880859375 0 79066.65
460.2728576660156 0 3965.2075
460.73687744140625 0 49879.53
461.23944091796875 0 23211.72
461.5754089355469 0 5870.038
461.7391662597656 0 3196.8142
462.2087097167969 0 3164.5947 z Water loss 8
462.2426452636719 0 3272.955
463.2206726074219 0 3452.0596
465.2651062011719 0 1973.6378
465.7648010253906 0 3385.7122
466.7328186035156 0 2534.2327
467.2333679199219 0 3268.9143
470.75091552734375 0 5314.9077
471.24713134765625 0 7814.3105
471.7441101074219 0 1878.4509
472.2508239746094 0 9221.137
473.2566833496094 0 2741.6423
474.29229736328125 0 3738.4011
475.2993469238281 0 588248.6 c 3
476.2379150390625 0 4227.6997
476.302001953125 0 144939.12
476.7381286621094 0 2491.361
477.234375 0 2576.1882
477.3039245605469 0 26276.799
479.2582092285156 0 5186.8623
479.7583923339844 0 78917.94
480.22064208984375 0 8683.861 z 8
480.2591857910156 0 38319.793
480.75689697265625 0 13422.935
481.22930908203125 0 44776.94
482.23175048828125 0 10580.34
484.7496643066406 0 8656.728
485.24237060546875 0 11474.803
485.7484130859375 0 2182.0352
486.2435302734375 0 2768.4849
486.7682800292969 0 11126.509
487.2690124511719 0 21276.734
487.7701721191406 0 8744.906
488.27337646484375 0 3354.4019
488.7713317871094 0 2426.199
491.7567443847656 0 5363.7393
492.2554931640625 0 5685.4106
493.7547302246094 0 18459.96
494.25445556640625 0 7646.227
494.7590026855469 0 3651.5603
495.230712890625 0 5824.5156
496.2404479980469 0 22609.828 y 8
497.2413330078125 0 3814.4075
499.7666320800781 0 95474.38
500.2673645019531 0 45481.38 c Water loss 8
500.76434326171875 0 39312.28 c Ammonia loss 8
501.26239013671875 0 15827.862
501.7681579589844 0 15664.549
502.2745361328125 0 66223.68
502.7656555175781 0 2782.5188
503.27490234375 0 16561
504.2799072265625 0 3230.2334
508.26422119140625 0 4217.0293
508.7716979980469 0 180692.42
509.2745666503906 0 306121.1 c 8
509.7749938964844 0 140513.1
510.2762451171875 0 39723.863
510.77398681640625 0 13295.358
511.27008056640625 0 3887.7834
514.2981567382812 0 1972.0322
521.2732543945312 0 3474.3088
521.7777709960938 0 3963.9148
522.2696533203125 0 2270.8865
522.7764892578125 0 4830.8926
523.2661743164062 0 4158.3027
523.7627563476562 0 2362.7368 z Water loss 3
525.35302734375 0 7456.8174
525.7474975585938 0 2246.8901
526.2545776367188 0 15004.072
526.3554077148438 0 2325.15
526.7579956054688 0 7041.246
527.2969970703125 0 3421.4177
529.311279296875 0 2112.504
530.2882080078125 0 11049.037
531.2942504882812 0 2960.3828
532.7660522460938 0 26867.207 z 3
533.2669067382812 0 15343.246
533.766357421875 0 2176.5688
534.3047485351562 0 6589.261
535.2734375 0 9324.168
535.3096313476562 0 7627.793
535.7713623046875 0 7164.115
535.81591796875 0 2654.3718
536.30078125 0 6476.256
540.2734985351562 0 2433.1484
540.7723388671875 0 6451.1553 y 3
541.2748413085938 0 3550.4429
541.7830810546875 0 3193.8337
543.2882690429688 0 2780.8853
543.7955322265625 0 3412.4902 c Water loss 9
544.27880859375 0 37182.84 c Ammonia loss 9
544.7791748046875 0 16788.924
545.303466796875 0 20651.512
545.7775268554688 0 2626.21
546.30908203125 0 8550.064
548.2509765625 0 30458.97
548.75244140625 0 18149.451
549.2514038085938 0 7496.8013
550.2955322265625 0 4600.64
550.7963256835938 0 3664.5247
552.2864990234375 0 4523.154
552.7916870117188 0 221174.03 c 9
553.291748046875 0 137427.66
553.7920532226562 0 55345.36
554.2868041992188 0 15112.879
554.7807006835938 0 8510.073
555.2885131835938 0 3065.9763
557.3082885742188 0 2234.8804
557.8092041015625 0 2264.7434
561.2866821289062 0 4528.5645
563.3173217773438 0 2305.7124
564.3037719726562 0 20406.633
565.26025390625 0 3523.324
565.308837890625 0 10326.986
565.7991943359375 0 18317.354
566.2987060546875 0 13055.223
566.7734985351562 0 4803.597
567.2943725585938 0 10299.157
567.7813110351562 0 10051.394 w 2
568.2799072265625 0 9352.339
568.7809448242188 0 7069.4434
569.8035278320312 0 3501.8833
570.2974243164062 0 2286.6194
570.809326171875 0 1930.8049
573.303955078125 0 10077.324 c Ammonia loss 4
574.3003540039062 0 6674.289
574.7924194335938 0 5063.943
575.2854614257812 0 8806.852 y Water loss 2
575.7802734375 0 16312.683 y Ammonia loss 2
576.2821655273438 0 86713.016 z 2
576.7825927734375 0 60752.32
577.2841796875 0 21690.932
577.7764282226562 0 5536.738
577.817626953125 0 13093.764
578.3156127929688 0 9758.1875
578.8172607421875 0 10000.625
579.31640625 0 13091.033
579.8125 0 5039.8325
583.3021240234375 0 10258.723
583.8011474609375 0 7541.2266
584.29150390625 0 282773.47 y 2
584.7918090820312 0 174227.31
585.29248046875 0 64907.21
585.7933959960938 0 17037.398
586.8228149414062 0 31514.408
587.3239135742188 0 31201.678
587.8248291015625 0 19231.422
588.275634765625 0 15731.859
588.3271484375 0 5704.1895
588.7767333984375 0 10695.816
589.2767333984375 0 2176.2986
589.3212890625 0 12378.747
590.3260498046875 0 1127278 c 4
591.32861328125 0 311403.12
591.8154907226562 0 27234.758
592.3280639648438 0 68853.62
592.8108520507812 0 16836.809
593.328857421875 0 8785.111
593.8156127929688 0 2972.921
597.7835083007812 0 5200.8833
598.2882690429688 0 3779.918
599.8433227539062 0 2368.8806
600.8204345703125 0 50989.836 c Ammonia loss 10
601.3208618164062 0 33326.566
601.8244018554688 0 23272.055
602.3228149414062 0 6340.141
602.8311767578125 0 2249.156
607.3272705078125 0 12553.2
607.830078125 0 6132.0464
608.3173828125 0 60687.516 z 7
608.82763671875 0 3386.7646
609.3334350585938 0 591861.1 c 10
609.8339233398438 0 385217.56
610.3341064453125 0 154664.22
610.8347778320312 0 42383.45
611.3289184570312 0 6635.142
616.3427124023438 0 10981.7705
616.8279418945312 0 2610.4573
618.3042602539062 0 36863.336 w 1
618.8048706054688 0 24750.932
619.3060302734375 0 11011.375
619.8126220703125 0 3811.5107
620.8287353515625 0 4635.398
621.32763671875 0 2511.7502
621.832275390625 0 2867.5361
623.8350219726562 0 5767.432
624.3352661132812 0 96904.56 y 7
625.3357543945312 0 31124.195
625.8126831054688 0 2314.2527
626.328857421875 0 7252.1724
626.804443359375 0 2284.14
630.32763671875 0 18397.424
630.8280029296875 0 14212.028
631.3324584960938 0 6446.4067
631.8362426757812 0 4569.271 y Water loss 1
632.31982421875 0 5479.4053 y Ammonia loss 1
632.8250732421875 0 11932.365 z 1
633.3212890625 0 24751.887
633.8224487304688 0 15418.48
634.3195190429688 0 11296.329
635.3599243164062 0 5207.1533
636.3493041992188 0 14234.053
637.3532104492188 0 4145.1772
637.8334350585938 0 8675.661
638.3469848632812 0 10189.02
638.8457641601562 0 6783.5527
639.3447265625 0 7184.985
639.8363647460938 0 6219.5234
640.334716796875 0 7854.5483
640.833984375 0 32595.615 y 1
641.333251953125 0 29401.883
641.83251953125 0 10171.504
642.3397216796875 0 7218.9233
644.3370971679688 0 7744.036
645.8290405273438 0 2553.094
646.3148193359375 0 3216.2983
646.8058471679688 0 2434.7356
647.3523559570312 0 33911.67
647.8511962890625 0 27386.64
648.3555297851562 0 13501.364
648.8613891601562 0 3347.396
649.3311767578125 0 18339.057
650.332763671875 0 10170.316
651.3292846679688 0 2702.4668
652.3450317382812 0 2900.9985
652.8438110351562 0 5196.915
653.33935546875 0 13959.412
653.44775390625 0 8389.874
653.8363037109375 0 7817.8286
654.3338012695312 0 6406.0493
654.4519653320312 0 3280.5635
654.8298950195312 0 4401.34
655.3172607421875 0 44065.062
655.8174438476562 0 28456.164
656.31787109375 0 12786.771
656.8202514648438 0 3753.2463
659.3474731445312 0 57175.496
659.85498046875 0 2700.881
660.348388671875 0 92880.71
660.8518676757812 0 60658.758
661.3521728515625 0 102251.45 z Water loss 6
661.8492431640625 0 70268.13
662.3456420898438 0 92583.11
662.8447265625 0 51591.836
663.3441772460938 0 18263.15
663.8460083007812 0 6491.098
668.8295288085938 0 28355.908
669.3480224609375 0 117312.11
669.8499145507812 0 71282.625
670.3485107421875 0 93177.56
670.8475952148438 0 49933.35
671.3473510742188 0 23879.01
671.8472900390625 0 5342.5215
672.3462524414062 0 11089.982
672.8641357421875 0 10385.984
673.3627319335938 0 6215.7397
673.8517456054688 0 13085.093
674.347412109375 0 22808.791
674.845703125 0 53375.438
675.3457641601562 0 39099.09
675.8394165039062 0 52229.082
676.3380126953125 0 30310.168
676.83642578125 0 16634.309
677.3377075195312 0 4595.3306
679.3538818359375 0 232415.36 z 6
680.3540649414062 0 87018.99
680.8724975585938 0 8874.819
681.3607177734375 0 26859.482
681.87548828125 0 13950.7295
682.3659057617188 0 7896.5933
682.8552856445312 0 23862.379
683.3483276367188 0 371346.34
683.8482666015625 0 272102.1
684.3486938476562 0 132923.31
684.849609375 0 47039.117
685.3505249023438 0 6661.932
687.3495483398438 0 16894.846 c Ammonia loss 5
688.3523559570312 0 5418.687
689.8764038085938 0 7903.8184
690.380615234375 0 19263.172
690.8818359375 0 14438.673
691.3572998046875 0 325968.2
691.8573608398438 0 257579
692.35791015625 0 125593.34
692.8589477539062 0 41237.914
693.3552856445312 0 14718.743
695.3726196289062 0 53176.57 y 6
696.3722534179688 0 23720.348
697.369384765625 0 5789.339
704.3688354492188 0 315070.3 c 5
705.3713989257812 0 108461.81
705.444580078125 0 3348.0464
706.373046875 0 23717.584
707.3800048828125 0 2781.7595
709.3049926757812 0 11838.674
710.3085327148438 0 5883.989
711.3402099609375 0 3436.8735
730.388916015625 0 3366.04
731.3919677734375 0 49833.562
732.394775390625 0 16277.349
733.3958129882812 0 5302.269
737.3341674804688 0 13534.465
738.3328857421875 0 3760.803
740.3721313476562 0 2829.8252
741.3558349609375 0 3469.7239
743.3782348632812 0 2554.0994
749.3842163085938 0 157722.28 w 5
750.3861083984375 0 65116.81
751.3887329101562 0 20852.643
752.3910522460938 0 4476.8994
755.414794921875 0 7724.4595
756.4215087890625 0 3645.5679
757.4110107421875 0 2333.3975
758.3800048828125 0 33454.395 c Ammonia loss 6
759.3825073242188 0 10356.791
760.38671875 0 3374.0864
761.4046020507812 0 3182.1165
762.41064453125 0 2435.8914
774.3975219726562 0 2636.9775
775.4054565429688 0 291912.94 c 6
776.407470703125 0 112521.64
777.4088745117188 0 23446.19
778.4126586914062 0 4779.1006
785.4290161132812 0 2479.253
786.383544921875 0 6365.8154
787.39111328125 0 8488.11
788.3972778320312 0 6562.9053
789.3922729492188 0 2861.946
793.396728515625 0 717422.4 z 5
794.3973388671875 0 319228.62
795.3989868164062 0 87623.555
796.3995361328125 0 17702.934
797.4036254882812 0 2338.421
803.4118041992188 0 3710.5508
808.3731689453125 0 5942.094
809.4142456054688 0 8257.008 y 5
810.4099731445312 0 5477.314
811.414794921875 0 2948.1663
815.4882202148438 0 2400.5671
820.4273681640625 0 3215.4263
821.434814453125 0 2996.8547
823.5386962890625 0 2721.7896
832.4437255859375 0 41331.914
833.4461059570312 0 24824.904
834.4501342773438 0 5582.9478
843.4641723632812 0 5255.5806
844.466064453125 0 2949.4868
848.4647216796875 0 3146.5684
849.4650268554688 0 2832.6272
852.35888671875 0 4613.223
855.410888671875 0 17001.719
856.412109375 0 9299.433
858.4918823242188 0 8320.979
859.4871215820312 0 25878.346
860.4915161132812 0 19743.525
861.4981689453125 0 6447.9897
864.4330444335938 0 230038.89
865.4331665039062 0 110854.234
866.43359375 0 35597.113
867.43505859375 0 8001.6406
871.470947265625 0 2178.0457
873.420166015625 0 7039.9214
874.421875 0 7814.857
875.4484252929688 0 12797.428
876.4523315429688 0 7589.5156
877.4561157226562 0 3069.9539
884.4812622070312 0 14797.795
885.4838256835938 0 11623.989 c Water loss 7
886.4747924804688 0 33821.26 c Ammonia loss 7
887.4765625 0 14676.251
888.4822998046875 0 7164.738
900.4761352539062 0 25785.639
901.4778442382812 0 14013.205
902.4917602539062 0 99294.55
903.4998168945312 0 366452.97 c 7
904.5025634765625 0 150796.6
905.5047607421875 0 43793.812
906.5076904296875 0 7814.8867
908.4229736328125 0 236356.42 z 4
909.42333984375 0 110501.34
910.4242553710938 0 37872.04
911.426513671875 0 7112.9233
919.4639892578125 0 10038.082
920.4688110351562 0 36010.56
921.4713745117188 0 21426.875
922.4677124023438 0 9082.99
923.4720458984375 0 2694.2603
924.4422607421875 0 19151.268 y 4
925.4390258789062 0 10610.848
926.4466552734375 0 3039.1387
943.4888305664062 0 2317.7107
957.5086059570312 0 2404.0408
960.4954223632812 0 2404.8755
961.5086669921875 0 3866.9902
968.4944458007812 0 5210.8057
969.4769287109375 0 10885.716
970.4865112304688 0 3784.2034
972.5184936523438 0 2387.0417
973.5260009765625 0 4427.041
974.5341186523438 0 6264.393
978.4514770507812 0 3105.2024 w 3
979.4597778320312 0 2186.4485
981.8323364257812 0 2051.6216
986.5010375976562 0 20038.871
987.5036010742188 0 28133.256
988.5067749023438 0 14657.516
989.5082397460938 0 5379.5757
999.5321655273438 0 16800.436 c Water loss 8
1000.5220336914062 0 24380.412 c Ammonia loss 8
1001.5210571289062 0 13950.395
1002.5291748046875 0 13316.293
1003.5279541015625 0 6401.106
1004.5253295898438 0 2627.9382
1016.5347900390625 0 28300.434
1017.5427856445312 0 217015.64 c 8
1018.5428466796875 0 110998.8
1019.5447387695312 0 34890.188
1020.5405883789062 0 14991.727
1021.5347290039062 0 3564.4695
1022.515869140625 0 2781.5093
1023.508056640625 0 3782.2412
1024.500244140625 0 3201.612
1052.470947265625 0 2704.539
1060.5621337890625 0 5936.825
1061.5654296875 0 2434.7266
1062.5572509765625 0 5795.483
1063.525634765625 0 3131.6162 y Ammonia loss 3
1064.524169921875 0 92990.67 z 3
1065.5267333984375 0 89271.08
1066.528076171875 0 47812.844
1067.529541015625 0 14713.119
1068.5372314453125 0 3778.4243
1071.607666015625 0 8038.7847
1072.6092529296875 0 4888.164
1079.536865234375 0 2950.7932
1080.5386962890625 0 9274.629 y 3
1081.54248046875 0 3792.5898
1082.55224609375 0 3372.6135
1086.5797119140625 0 2288.7415 c Water loss 9
1088.5565185546875 0 13169.719
1089.5570068359375 0 10810.561
1090.5562744140625 0 6088.5684
1095.4984130859375 0 4549.033
1096.4971923828125 0 7377.8423
1104.574462890625 0 64726.043 c 9
1105.575927734375 0 39550.164
1106.5771484375 0 18300.754
1107.568603515625 0 5232.6875
1110.5379638671875 0 2873.917
1134.5523681640625 0 7247.5273 z Ammonia loss 2
1135.544189453125 0 7234.049
1136.543212890625 0 4101.044
1151.5565185546875 0 56414.723 z 2
1152.5623779296875 0 112379.35
1153.5634765625 0 57124.266
1154.5638427734375 0 22117.002
1155.572998046875 0 5481.9785
1157.6317138671875 0 6297.0156
1158.6300048828125 0 6000.954
1159.6295166015625 0 5654.4395
1160.6328125 0 4715.1226
1167.5802001953125 0 4559.9243 y 2
1168.581298828125 0 5971.3647
1169.5810546875 0 3719.8682
1173.6458740234375 0 14180.133
1174.651123046875 0 34039.766
1175.64794921875 0 22028.182
1176.6502685546875 0 9094.25
1177.6541748046875 0 3123.0562
1184.6324462890625 0 4006.1672
1185.628173828125 0 2990.7415
1200.661376953125 0 3742.3542
1201.639892578125 0 47099.363
1202.64208984375 0 27132.09
1203.6434326171875 0 10113.726
1204.6475830078125 0 5522.856
1217.6585693359375 0 36995.562 c 10
1218.66259765625 0 80352.805
1219.6644287109375 0 40502.14
1220.6661376953125 0 17582.889
1221.6588134765625 0 3653.4204
1264.647705078125 0 3014.5117 z 1
1265.6484375 0 22479.182
1266.6400146484375 0 14772.605
1267.64453125 0 7314.67
1280.646728515625 0 4446.482 y 1
1281.630126953125 0 2674.2415
1294.6390380859375 0 4085.287
1295.638671875 0 3521.4094
1304.69287109375 0 3819.3442
1305.6591796875 0 17625.908
1306.655029296875 0 17514.125
1307.655517578125 0 11882.314
1308.6578369140625 0 6882.635
1309.65234375 0 4535.696
1319.6961669921875 0 3206.0732
1320.6934814453125 0 10584.956
1321.694091796875 0 22624.574
1322.6959228515625 0 15950.641
1323.6795654296875 0 40221.055
1324.6796875 0 70727.945
1325.6798095703125 0 40042.637
1326.6810302734375 0 19819.121
1327.677490234375 0 5451.6226
1337.6943359375 0 19890.164
1338.698486328125 0 110679.53
1339.7003173828125 0 73417.26
1340.6995849609375 0 35139.38
1341.701416015625 0 9828.91
1342.707763671875 0 3306.9722
1348.68505859375 0 24198.15
1349.673828125 0 47657.07
1350.6702880859375 0 27330.47
1351.67236328125 0 12041.921
1352.6590576171875 0 2565.112
1354.715576171875 0 2408.6333
1355.725830078125 0 5179.3716
1356.7161865234375 0 5027.1035
1362.7442626953125 0 2933.384
1363.7333984375 0 7998.451
1364.7236328125 0 5570.6494
1365.6922607421875 0 59857.34
1366.6932373046875 0 354074.62
1367.6944580078125 0 249702.56
1368.6953125 0 120393.31
1369.696044921875 0 33549.832
1370.69580078125 0 5096.237
1379.742919921875 0 4536.782
1380.7569580078125 0 10885.0205
1381.722412109375 0 17596.38
1382.71484375 0 40183.137
1383.717041015625 0 70061.266
1384.717529296875 0 51389.06
1385.71875 0 22858.22
1386.72265625 0 7101.721
2182.65087890625 0 2012.172

Spectrum Details

|  |  |
| --- | --- |
| Matched peaks? Matched peaks The total absolute number of peaks matched. Additionally in brackets the total fraction of peaks matched and the total number of peaks is shown. | 72 (10.33% of 697) |
| FDR? FDR The false discovery rate estimated for this peptide. It is calculated by matching all theoretical fragments with a non-integer shift with the raw peaks for this spectrum. This is done with 40 different shifts. The resulting percentage is the average number of annotated peaks over the number of annotated peaks with the correct spectrum. | 1.36% |
| Satellite FDR? Satellite FDR See the FDR for details on its calculation. This satellite ion specific FDR only contains the satellite ions (d/w) for I/L/J positions. | 0.00% |
| PSM Score? PSM Score The PSM Score as given by Hecklib to this annotated spectrum. It is shown with three significant figures. | 600 |

#### Spectrum 3360? Spectrum 3360 The raw spectrum of this peptide as annotated by Hecklib. The fragments are coloured according to ion type (see legend). Any peaks with a star '\*' as text can be hovered over to see the full details, first the ion type second the mass shift type. By hovering over the amino acids in the peptide or ions in the legend the corresponding peaks are highlighted. By toggling the 'Unassigned' label you can turn the background (unassigned) peaks on or off in the plot. By updating the slider in the Ion legend you can update the spectrum to only show the top X% of the peaks with labels. The top X% means any peak that is within X% of the highest intensity. By dragging in the spectrum you can zoom in to a specific part of the spectrum and use 'Zoom Out' to get back to the original zoom level. The annotation of the spectrum is based on the given sequence in the peptides file and is done with different software so inconsistencies are likely. The peaks are annotated based on the given sequence, with 20 ppm tolerance.

Copy Data

##### Spectrum 3360 (TSV)

###### Preview

```
Loading example...
```

*Click on the button to copy the data to your clipboard.*

Mz MinMz MaxIntensity Max

WidthHeightPeptide font sizePeptide stroke widthSpectrum font sizeSpectrum stroke widthCompact peptide

Ion legend

wxyz

abcd

OtherUnassignedIon^Charge^_Position_Show for top:%

TISRDNAKNSJY

09.46e+51.89e+62.84e+63.78e+6

Zoom Out

y^+1^_1_c^+1^_2_y^+1^_2_c^+1^_3_z^+1^_3_y^+1^_3_c^+2^_8_c^+2^_8_c^+2^_8_c^+1^_4_c^+1^_4_z^+1^_4_y^+1^_4_c^+2^_9_c^+2^_9_c^+2^_9_z^+2^_9_z^+2^_9_y^+2^_9_c^+2^_10_c^+2^_10_c^+2^_10_z^+2^_10_z^+2^_10_c^+1^_5_y^+2^_10_y^+2^_10_z^+2^_10_y^+2^_10_c^+1^_5_c^+2^_11_z^+1^_5_c^+2^_11_w^+2^_11_y^+1^_5_y^+2^_11_y^+2^_11_z^+2^_11_y^+2^_11_z^+1^_6_z^+1^_6_c^+1^_6_y^+1^_6_c^+1^_6_w^+1^_7_c^+1^_7_c^+1^_7_z^+1^_7_y^+1^_7_c^+1^_8_c^+1^_8_c^+1^_8_z^+1^_8_y^+1^_8_c^+1^_9_c^+1^_9_c^+1^_9_y^+1^_9_z^+1^_9_y^+1^_9_c^+1^_10_z^+1^_10_z^+1^_10_y^+1^_10_c^+1^_11_z^+1^_11_y^+1^_11_

0844168825323376

Fragment Matches Table

Show background peaks

| Position | Ion type | Intensity | mz Theoretical | mz Error (Th) | mz Error (ppm) | Charge | Series Number |
| --- | --- | --- | --- | --- | --- | --- | --- |
| - | - | 6709 | 123 | - | - | 0 | - |
| - | - | 8396 | 123.1 | - | - | 0 | - |
| - | - | 7239 | 129.1 | - | - | 0 | - |
| - | - | 1.352E+05 | 136.1 | - | - | 0 | - |
| - | - | 5018 | 136.3 | - | - | 0 | - |
| - | - | 9681 | 137.1 | - | - | 0 | - |
| - | - | 9372 | 142.1 | - | - | 0 | - |
| - | - | 9787 | 143.1 | - | - | 0 | - |
| - | - | 4896 | 143.9 | - | - | 0 | - |
| - | - | 5390 | 145.1 | - | - | 0 | - |
| - | - | 6756 | 149 | - | - | 0 | - |
| - | - | 6584 | 155.1 | - | - | 0 | - |
| - | - | 5674 | 157.1 | - | - | 0 | - |
| - | - | 1.272E+05 | 165.1 | - | - | 0 | - |
| - | - | 1.146E+04 | 166.1 | - | - | 0 | - |
| - | - | 8765 | 169.1 | - | - | 0 | - |
| - | - | 5373 | 171.8 | - | - | 0 | - |
| - | - | 1.305E+04 | 173.1 | - | - | 0 | - |
| - | - | 9170 | 173.5 | - | - | 0 | - |
| 12 | y | 4.116E+05 | 182.1 | 0.0003428 | 1.883 | +1 | 1 |
| - | - | 4.625E+04 | 183.1 | - | - | 0 | - |
| - | - | 7.576E+05 | 187.1 | - | - | 0 | - |
| - | - | 9.209E+04 | 188.1 | - | - | 0 | - |
| - | - | 9298 | 200.1 | - | - | 0 | - |
| - | - | 1.972E+04 | 201.1 | - | - | 0 | - |
| - | - | 7932 | 202.1 | - | - | 0 | - |
| - | - | 2.054E+05 | 215.1 | - | - | 0 | - |
| - | - | 2.309E+04 | 216.1 | - | - | 0 | - |
| - | - | 6632 | 228.7 | - | - | 0 | - |
| - | - | 1.578E+04 | 229.1 | - | - | 0 | - |
| - | - | 5702 | 231.2 | - | - | 0 | - |
| 2 | c | 1.006E+04 | 232.2 | 0.0002035 | 0.8764 | +1 | 2 |
| - | - | 7782 | 239.1 | - | - | 0 | - |
| - | - | 1.064E+04 | 245.2 | - | - | 0 | - |
| - | - | 5457 | 274.2 | - | - | 0 | - |
| - | - | 6173 | 281.1 | - | - | 0 | - |
| - | - | 4.788E+04 | 282.1 | - | - | 0 | - |
| - | - | 9934 | 287.2 | - | - | 0 | - |
| 11 | y | 3.317E+04 | 295.2 | 1.903E-05 | 0.06446 | +1 | 2 |
| - | - | 7830 | 297.2 | - | - | 0 | - |
| - | - | 6813 | 298.1 | - | - | 0 | - |
| - | - | 8219 | 299.1 | - | - | 0 | - |
| - | - | 1.631E+04 | 299.2 | - | - | 0 | - |
| - | - | 1.438E+05 | 300.2 | - | - | 0 | - |
| - | - | 1.829E+04 | 301.2 | - | - | 0 | - |
| - | - | 1.422E+04 | 302.2 | - | - | 0 | - |
| - | - | 7076 | 314.2 | - | - | 0 | - |
| - | - | 2.716E+04 | 315.2 | - | - | 0 | - |
| - | - | 7.556E+04 | 317.2 | - | - | 0 | - |
| - | - | 8115 | 318.2 | - | - | 0 | - |
| 3 | c | 6.07E+04 | 319.2 | 0.0003711 | 1.163 | +1 | 3 |
| - | - | 1.443E+04 | 320.2 | - | - | 0 | - |
| - | - | 8091 | 341 | - | - | 0 | - |
| - | - | 9426 | 357.5 | - | - | 0 | - |
| - | - | 4.069E+04 | 359 | - | - | 0 | - |
| 10 | z | 6672 | 366.2 | 0.001658 | 4.528 | +1 | 3 |
| - | - | 6128 | 367.2 | - | - | 0 | - |
| - | - | 1.137E+04 | 372.7 | - | - | 0 | - |
| - | - | 7605 | 373.2 | - | - | 0 | - |
| - | - | 9309 | 374.2 | - | - | 0 | - |
| 10 | y | 3.143E+04 | 382.2 | 0.0002172 | 0.5682 | +1 | 3 |
| - | - | 1.057E+05 | 385.2 | - | - | 0 | - |
| - | - | 3.658E+04 | 385.5 | - | - | 0 | - |
| - | - | 2.031E+04 | 385.9 | - | - | 0 | - |
| - | - | 2.818E+04 | 386.2 | - | - | 0 | - |
| - | - | 8.84E+04 | 391.6 | - | - | 0 | - |
| - | - | 5.068E+04 | 391.9 | - | - | 0 | - |
| - | - | 2.076E+04 | 392.2 | - | - | 0 | - |
| - | - | 1.486E+04 | 394.7 | - | - | 0 | - |
| - | - | 1.6E+04 | 400.3 | - | - | 0 | - |
| - | - | 6984 | 408.2 | - | - | 0 | - |
| - | - | 1.836E+04 | 415.3 | - | - | 0 | - |
| - | - | 9830 | 425.2 | - | - | 0 | - |
| - | - | 6979 | 428.7 | - | - | 0 | - |
| - | - | 5570 | 429.1 | - | - | 0 | - |
| - | - | 1.172E+05 | 429.1 | - | - | 0 | - |
| - | - | 3.519E+04 | 429.8 | - | - | 0 | - |
| - | - | 8822 | 430.2 | - | - | 0 | - |
| - | - | 3.74E+04 | 430.2 | - | - | 0 | - |
| - | - | 1.17E+04 | 430.3 | - | - | 0 | - |
| - | - | 9640 | 431.3 | - | - | 0 | - |
| - | - | 2.136E+04 | 437.2 | - | - | 0 | - |
| - | - | 8002 | 439.9 | - | - | 0 | - |
| - | - | 5.672E+04 | 442.7 | - | - | 0 | - |
| 8 | c | 3.571E+04 | 443.2 | 0.0004928 | 1.112 | +2 | 8 |
| 8 | c | 3.773E+04 | 443.7 | 0.001252 | 2.822 | +2 | 8 |
| - | - | 2.117E+04 | 444.2 | - | - | 0 | - |
| - | - | 6.153E+04 | 445.1 | - | - | 0 | - |
| - | - | 5.806E+04 | 446.2 | - | - | 0 | - |
| - | - | 2.322E+05 | 451.8 | - | - | 0 | - |
| 8 | c | 7.013E+05 | 452.3 | 0.0002458 | 0.5436 | +2 | 8 |
| - | - | 2.729E+05 | 452.8 | - | - | 0 | - |
| - | - | 6.537E+04 | 453.3 | - | - | 0 | - |
| - | - | 1.094E+04 | 453.8 | - | - | 0 | - |
| - | - | 1.172E+04 | 455.2 | - | - | 0 | - |
| - | - | 8488 | 456.3 | - | - | 0 | - |
| - | - | 6965 | 457.3 | - | - | 0 | - |
| 4 | c | 2.291E+04 | 458.3 | 0.000674 | 1.471 | +1 | 4 |
| - | - | 1.511E+04 | 461.2 | - | - | 0 | - |
| - | - | 8408 | 461.6 | - | - | 0 | - |
| - | - | 1.406E+04 | 463.2 | - | - | 0 | - |
| - | - | 2.52E+04 | 466.7 | - | - | 0 | - |
| - | - | 1.214E+04 | 467.2 | - | - | 0 | - |
| - | - | 1.351E+04 | 470.8 | - | - | 0 | - |
| - | - | 1.496E+04 | 471.2 | - | - | 0 | - |
| - | - | 3.478E+04 | 472.3 | - | - | 0 | - |
| - | - | 9145 | 473.3 | - | - | 0 | - |
| - | - | 8009 | 474.3 | - | - | 0 | - |
| 4 | c | 1.94E+06 | 475.3 | 0.0008226 | 1.731 | +1 | 4 |
| - | - | 4.367E+05 | 476.3 | - | - | 0 | - |
| - | - | 6.227E+04 | 477.3 | - | - | 0 | - |
| - | - | 1.84E+04 | 479.3 | - | - | 0 | - |
| - | - | 2.046E+05 | 479.8 | - | - | 0 | - |
| 9 | z | 2.334E+04 | 480.2 | 3.114E-05 | 0.06484 | +1 | 4 |
| - | - | 1.001E+05 | 480.3 | - | - | 0 | - |
| - | - | 5.617E+04 | 480.8 | - | - | 0 | - |
| - | - | 1.489E+05 | 481.2 | - | - | 0 | - |
| - | - | 3.077E+04 | 482.2 | - | - | 0 | - |
| - | - | 9383 | 483.2 | - | - | 0 | - |
| - | - | 2.065E+04 | 484.7 | - | - | 0 | - |
| - | - | 3.491E+04 | 485.2 | - | - | 0 | - |
| - | - | 1.366E+04 | 485.7 | - | - | 0 | - |
| - | - | 8144 | 486.3 | - | - | 0 | - |
| - | - | 4.199E+04 | 486.8 | - | - | 0 | - |
| - | - | 5.774E+04 | 487.3 | - | - | 0 | - |
| - | - | 2.032E+04 | 487.8 | - | - | 0 | - |
| - | - | 1.19E+04 | 488.3 | - | - | 0 | - |
| - | - | 1.072E+04 | 491.8 | - | - | 0 | - |
| - | - | 4.484E+04 | 493.8 | - | - | 0 | - |
| - | - | 1.902E+04 | 494.3 | - | - | 0 | - |
| - | - | 1.44E+04 | 494.8 | - | - | 0 | - |
| - | - | 1.427E+04 | 495.2 | - | - | 0 | - |
| 9 | y | 7.457E+04 | 496.2 | 0.0005637 | 1.136 | +1 | 4 |
| - | - | 1.315E+04 | 497.2 | - | - | 0 | - |
| - | - | 2.581E+05 | 499.8 | - | - | 0 | - |
| 9 | c | 1.244E+05 | 500.3 | 0.002691 | 5.379 | +2 | 9 |
| 9 | c | 8.605E+04 | 500.8 | 0.002921 | 5.833 | +2 | 9 |
| - | - | 5.622E+04 | 501.3 | - | - | 0 | - |
| - | - | 3.983E+04 | 501.8 | - | - | 0 | - |
| - | - | 2.244E+05 | 502.3 | - | - | 0 | - |
| - | - | 1.092E+04 | 502.8 | - | - | 0 | - |
| - | - | 5.247E+04 | 503.3 | - | - | 0 | - |
| - | - | 1.892E+04 | 508.3 | - | - | 0 | - |
| - | - | 5.588E+05 | 508.8 | - | - | 0 | - |
| 9 | c | 9.915E+05 | 509.3 | 0.0006185 | 1.215 | +2 | 9 |
| - | - | 4.433E+05 | 509.8 | - | - | 0 | - |
| - | - | 1.355E+05 | 510.3 | - | - | 0 | - |
| - | - | 4.057E+04 | 510.8 | - | - | 0 | - |
| - | - | 8541 | 514.3 | - | - | 0 | - |
| - | - | 1.158E+04 | 521.3 | - | - | 0 | - |
| - | - | 8975 | 521.8 | - | - | 0 | - |
| - | - | 1.279E+04 | 522.8 | - | - | 0 | - |
| - | - | 1.4E+04 | 523.3 | - | - | 0 | - |
| 4 | z | 1.017E+04 | 523.8 | 0.003115 | 5.948 | +2 | 9 |
| - | - | 1.005E+04 | 525.8 | - | - | 0 | - |
| - | - | 4.399E+04 | 526.3 | - | - | 0 | - |
| - | - | 3.661E+04 | 526.8 | - | - | 0 | - |
| - | - | 4.146E+04 | 530.3 | - | - | 0 | - |
| - | - | 1.089E+04 | 530.8 | - | - | 0 | - |
| - | - | 1.043E+04 | 531.3 | - | - | 0 | - |
| 4 | z | 9.057E+04 | 532.8 | 0.00119 | 2.233 | +2 | 9 |
| - | - | 5.173E+04 | 533.3 | - | - | 0 | - |
| - | - | 1.726E+04 | 533.8 | - | - | 0 | - |
| - | - | 9347 | 534.3 | - | - | 0 | - |
| - | - | 3.645E+04 | 535.3 | - | - | 0 | - |
| - | - | 2.043E+04 | 535.8 | - | - | 0 | - |
| - | - | 7022 | 536.6 | - | - | 0 | - |
| - | - | 1.321E+04 | 540.3 | - | - | 0 | - |
| 4 | y | 2.361E+04 | 540.8 | 0.0002377 | 0.4396 | +2 | 9 |
| - | - | 1.205E+04 | 541.3 | - | - | 0 | - |
| - | - | 1.031E+04 | 541.8 | - | - | 0 | - |
| 10 | c | 6754 | 543.8 | 0.001737 | 3.195 | +2 | 10 |
| 10 | c | 1.063E+05 | 544.3 | 0.0007007 | 1.287 | +2 | 10 |
| - | - | 6.408E+04 | 544.8 | - | - | 0 | - |
| - | - | 6.393E+04 | 545.3 | - | - | 0 | - |
| - | - | 7708 | 546.3 | - | - | 0 | - |
| - | - | 2.802E+04 | 546.3 | - | - | 0 | - |
| - | - | 1.026E+05 | 548.3 | - | - | 0 | - |
| - | - | 5.915E+04 | 548.8 | - | - | 0 | - |
| - | - | 1.649E+04 | 549.3 | - | - | 0 | - |
| - | - | 9596 | 550.3 | - | - | 0 | - |
| - | - | 9315 | 552.3 | - | - | 0 | - |
| 10 | c | 7.595E+05 | 552.8 | 0.0006097 | 1.103 | +2 | 10 |
| - | - | 4.05E+05 | 553.3 | - | - | 0 | - |
| - | - | 1.409E+05 | 553.8 | - | - | 0 | - |
| - | - | 6.165E+04 | 554.3 | - | - | 0 | - |
| - | - | 3.157E+04 | 554.8 | - | - | 0 | - |
| - | - | 9938 | 558.3 | - | - | 0 | - |
| - | - | 1.087E+04 | 561.3 | - | - | 0 | - |
| - | - | 1.291E+04 | 561.8 | - | - | 0 | - |
| - | - | 9199 | 562.3 | - | - | 0 | - |
| - | - | 1.098E+04 | 563.3 | - | - | 0 | - |
| - | - | 5.818E+04 | 564.3 | - | - | 0 | - |
| - | - | 1.146E+04 | 565.3 | - | - | 0 | - |
| - | - | 2.796E+04 | 565.3 | - | - | 0 | - |
| - | - | 6.412E+04 | 565.8 | - | - | 0 | - |
| - | - | 3.445E+04 | 566.3 | - | - | 0 | - |
| - | - | 2.122E+04 | 566.8 | - | - | 0 | - |
| 3 | z | 1.761E+04 | 567.3 | 0.0003258 | 0.5744 | +2 | 10 |
| 3 | z | 1.513E+04 | 567.8 | 0.008338 | 14.69 | +2 | 10 |
| - | - | 2.305E+04 | 568.3 | - | - | 0 | - |
| - | - | 2.817E+04 | 568.8 | - | - | 0 | - |
| 5 | c | 3.242E+04 | 573.3 | 0.007417 | 12.94 | +1 | 5 |
| - | - | 1.369E+04 | 574.3 | - | - | 0 | - |
| - | - | 8529 | 574.8 | - | - | 0 | - |
| 3 | y | 2.37E+04 | 575.3 | 0.003801 | 6.607 | +2 | 10 |
| 3 | y | 5.25E+04 | 575.8 | 0.003248 | 5.641 | +2 | 10 |
| 3 | z | 2.695E+05 | 576.3 | 0.0004343 | 0.7537 | +2 | 10 |
| - | - | 1.827E+05 | 576.8 | - | - | 0 | - |
| - | - | 7.508E+04 | 577.3 | - | - | 0 | - |
| - | - | 1.334E+04 | 577.8 | - | - | 0 | - |
| - | - | 5.239E+04 | 578.3 | - | - | 0 | - |
| - | - | 2.317E+04 | 578.8 | - | - | 0 | - |
| - | - | 2.845E+04 | 579.3 | - | - | 0 | - |
| - | - | 1.14E+04 | 579.8 | - | - | 0 | - |
| - | - | 8485 | 582.8 | - | - | 0 | - |
| - | - | 3.251E+04 | 583.3 | - | - | 0 | - |
| - | - | 1.813E+04 | 583.8 | - | - | 0 | - |
| 3 | y | 8.241E+05 | 584.3 | 0.0004107 | 0.7028 | +2 | 10 |
| - | - | 5.1E+05 | 584.8 | - | - | 0 | - |
| - | - | 1.942E+05 | 585.3 | - | - | 0 | - |
| - | - | 4.713E+04 | 585.8 | - | - | 0 | - |
| - | - | 9.212E+04 | 586.8 | - | - | 0 | - |
| - | - | 1.045E+05 | 587.3 | - | - | 0 | - |
| - | - | 4.626E+04 | 587.8 | - | - | 0 | - |
| - | - | 3.285E+04 | 588.3 | - | - | 0 | - |
| - | - | 1.282E+04 | 588.3 | - | - | 0 | - |
| - | - | 2.788E+04 | 588.8 | - | - | 0 | - |
| - | - | 1.72E+04 | 589.3 | - | - | 0 | - |
| - | - | 1.472E+04 | 589.3 | - | - | 0 | - |
| 5 | c | 3.745E+06 | 590.3 | 0.0006435 | 1.09 | +1 | 5 |
| - | - | 1.059E+06 | 591.3 | - | - | 0 | - |
| - | - | 9923 | 591.4 | - | - | 0 | - |
| - | - | 7.559E+04 | 591.8 | - | - | 0 | - |
| - | - | 1.921E+05 | 592.3 | - | - | 0 | - |
| - | - | 4.336E+04 | 592.8 | - | - | 0 | - |
| - | - | 2.344E+04 | 593.3 | - | - | 0 | - |
| - | - | 1.579E+04 | 597.8 | - | - | 0 | - |
| - | - | 8732 | 600.3 | - | - | 0 | - |
| 11 | c | 1.446E+05 | 600.8 | 0.0006609 | 1.1 | +2 | 11 |
| - | - | 1.035E+05 | 601.3 | - | - | 0 | - |
| - | - | 5.979E+04 | 601.8 | - | - | 0 | - |
| - | - | 1.77E+04 | 602.3 | - | - | 0 | - |
| - | - | 3.667E+04 | 607.3 | - | - | 0 | - |
| - | - | 2.513E+04 | 607.8 | - | - | 0 | - |
| 8 | z | 2.068E+05 | 608.3 | 0.001076 | 1.77 | +1 | 5 |
| - | - | 1.563E+04 | 608.8 | - | - | 0 | - |
| 11 | c | 1.852E+06 | 609.3 | 0.0002037 | 0.3343 | +2 | 11 |
| - | - | 1.103E+06 | 609.8 | - | - | 0 | - |
| - | - | 4.707E+05 | 610.3 | - | - | 0 | - |
| - | - | 1.392E+05 | 610.8 | - | - | 0 | - |
| - | - | 1.828E+04 | 611.3 | - | - | 0 | - |
| - | - | 1.093E+04 | 612.3 | - | - | 0 | - |
| - | - | 2.633E+04 | 616.3 | - | - | 0 | - |
| 2 | w | 9.598E+04 | 618.3 | 0.0009752 | 1.577 | +2 | 11 |
| - | - | 6.817E+04 | 618.8 | - | - | 0 | - |
| - | - | 3.121E+04 | 619.3 | - | - | 0 | - |
| - | - | 1.214E+04 | 620.8 | - | - | 0 | - |
| - | - | 1.797E+04 | 621.3 | - | - | 0 | - |
| 8 | y | 2.915E+05 | 624.3 | 0.0002967 | 0.4752 | +1 | 5 |
| - | - | 8.464E+04 | 625.3 | - | - | 0 | - |
| - | - | 2.517E+04 | 626.3 | - | - | 0 | - |
| - | - | 6.272E+04 | 630.3 | - | - | 0 | - |
| - | - | 4.117E+04 | 630.8 | - | - | 0 | - |
| - | - | 2.266E+04 | 631.3 | - | - | 0 | - |
| 2 | y | 1.129E+04 | 631.8 | 0.003014 | 4.77 | +2 | 11 |
| 2 | y | 1.172E+04 | 632.3 | 0.001927 | 3.047 | +2 | 11 |
| 2 | z | 4.539E+04 | 632.8 | 0.002169 | 3.427 | +2 | 11 |
| - | - | 8.973E+04 | 633.3 | - | - | 0 | - |
| - | - | 3.473E+04 | 633.8 | - | - | 0 | - |
| - | - | 1.29E+04 | 634.3 | - | - | 0 | - |
| - | - | 9163 | 634.8 | - | - | 0 | - |
| - | - | 1.394E+04 | 635.3 | - | - | 0 | - |
| - | - | 3.77E+04 | 636.3 | - | - | 0 | - |
| - | - | 9287 | 637.3 | - | - | 0 | - |
| - | - | 3.252E+04 | 637.8 | - | - | 0 | - |
| - | - | 3.714E+04 | 638.3 | - | - | 0 | - |
| - | - | 2.48E+04 | 638.8 | - | - | 0 | - |
| - | - | 1.158E+04 | 639.3 | - | - | 0 | - |
| - | - | 2.468E+04 | 639.8 | - | - | 0 | - |
| - | - | 2.586E+04 | 640.3 | - | - | 0 | - |
| 2 | y | 1.003E+05 | 640.8 | 0.0004226 | 0.6594 | +2 | 11 |
| - | - | 8.278E+04 | 641.3 | - | - | 0 | - |
| - | - | 2.361E+04 | 641.8 | - | - | 0 | - |
| - | - | 1.418E+04 | 642.3 | - | - | 0 | - |
| - | - | 1.956E+04 | 644.3 | - | - | 0 | - |
| - | - | 1.359E+04 | 646.8 | - | - | 0 | - |
| - | - | 1.255E+05 | 647.4 | - | - | 0 | - |
| - | - | 7.503E+04 | 647.9 | - | - | 0 | - |
| - | - | 1.891E+04 | 648.4 | - | - | 0 | - |
| - | - | 2.097E+04 | 652.3 | - | - | 0 | - |
| - | - | 2.02E+04 | 652.8 | - | - | 0 | - |
| - | - | 4.687E+04 | 653.3 | - | - | 0 | - |
| - | - | 2.011E+04 | 653.8 | - | - | 0 | - |
| - | - | 1.24E+04 | 654.3 | - | - | 0 | - |
| - | - | 1.945E+04 | 654.8 | - | - | 0 | - |
| - | - | 1.271E+05 | 655.3 | - | - | 0 | - |
| - | - | 8.131E+04 | 655.8 | - | - | 0 | - |
| - | - | 2.811E+04 | 656.3 | - | - | 0 | - |
| - | - | 1.631E+04 | 656.8 | - | - | 0 | - |
| - | - | 1.891E+05 | 659.3 | - | - | 0 | - |
| - | - | 2.35E+05 | 660.3 | - | - | 0 | - |
| - | - | 1.648E+05 | 660.9 | - | - | 0 | - |
| 7 | z | 3.531E+05 | 661.3 | 0.009134 | 13.81 | +1 | 6 |
| - | - | 2.367E+05 | 661.9 | - | - | 0 | - |
| - | - | 2.953E+05 | 662.3 | - | - | 0 | - |
| - | - | 1.646E+05 | 662.8 | - | - | 0 | - |
| - | - | 8.462E+04 | 663.3 | - | - | 0 | - |
| - | - | 1.682E+04 | 663.9 | - | - | 0 | - |
| - | - | 1.105E+05 | 668.8 | - | - | 0 | - |
| - | - | 3.702E+05 | 669.3 | - | - | 0 | - |
| - | - | 2.313E+05 | 669.9 | - | - | 0 | - |
| - | - | 2.701E+05 | 670.3 | - | - | 0 | - |
| - | - | 1.67E+05 | 670.8 | - | - | 0 | - |
| - | - | 6.746E+04 | 671.3 | - | - | 0 | - |
| - | - | 1.697E+04 | 671.9 | - | - | 0 | - |
| - | - | 3.428E+04 | 672.3 | - | - | 0 | - |
| - | - | 1.117E+04 | 673.3 | - | - | 0 | - |
| - | - | 4.179E+04 | 673.9 | - | - | 0 | - |
| - | - | 8.312E+04 | 674.3 | - | - | 0 | - |
| - | - | 1.938E+05 | 674.8 | - | - | 0 | - |
| - | - | 1.361E+05 | 675.3 | - | - | 0 | - |
| - | - | 1.705E+05 | 675.8 | - | - | 0 | - |
| - | - | 1.08E+05 | 676.3 | - | - | 0 | - |
| - | - | 4.268E+04 | 676.8 | - | - | 0 | - |
| - | - | 1.695E+04 | 677.3 | - | - | 0 | - |
| 7 | z | 8.048E+05 | 679.4 | 0.0005227 | 0.7694 | +1 | 6 |
| - | - | 3.013E+05 | 680.4 | - | - | 0 | - |
| - | - | 6.801E+04 | 681.4 | - | - | 0 | - |
| - | - | 8964 | 682.4 | - | - | 0 | - |
| - | - | 6.758E+04 | 682.9 | - | - | 0 | - |
| - | - | 1.134E+06 | 683.3 | - | - | 0 | - |
| - | - | 7.846E+05 | 683.8 | - | - | 0 | - |
| - | - | 3.866E+05 | 684.3 | - | - | 0 | - |
| - | - | 1.191E+05 | 684.8 | - | - | 0 | - |
| - | - | 3.47E+04 | 685.4 | - | - | 0 | - |
| 6 | c | 4.96E+04 | 687.3 | 0.008618 | 12.54 | +1 | 6 |
| - | - | 1.55E+04 | 688.4 | - | - | 0 | - |
| - | - | 1.107E+04 | 689.4 | - | - | 0 | - |
| - | - | 1.044E+06 | 691.4 | - | - | 0 | - |
| - | - | 7.506E+05 | 691.9 | - | - | 0 | - |
| - | - | 3.664E+05 | 692.4 | - | - | 0 | - |
| - | - | 9.291E+04 | 692.9 | - | - | 0 | - |
| - | - | 3.747E+04 | 693.4 | - | - | 0 | - |
| 7 | y | 1.918E+05 | 695.4 | 0.0005364 | 0.7714 | +1 | 6 |
| - | - | 7.243E+04 | 696.4 | - | - | 0 | - |
| - | - | 9520 | 703.4 | - | - | 0 | - |
| 6 | c | 1.009E+06 | 704.4 | 0.0005627 | 0.7989 | +1 | 6 |
| - | - | 3.401E+05 | 705.4 | - | - | 0 | - |
| - | - | 8.209E+04 | 706.4 | - | - | 0 | - |
| - | - | 1.011E+04 | 707.4 | - | - | 0 | - |
| - | - | 1.407E+04 | 711.3 | - | - | 0 | - |
| - | - | 1.003E+04 | 712.3 | - | - | 0 | - |
| - | - | 1.05E+04 | 722.4 | - | - | 0 | - |
| - | - | 1.09E+04 | 730.4 | - | - | 0 | - |
| - | - | 1.413E+05 | 731.4 | - | - | 0 | - |
| - | - | 5.073E+04 | 732.4 | - | - | 0 | - |
| - | - | 1.177E+04 | 733.4 | - | - | 0 | - |
| - | - | 4.633E+04 | 737.3 | - | - | 0 | - |
| - | - | 1.732E+04 | 738.3 | - | - | 0 | - |
| - | - | 8348 | 742.4 | - | - | 0 | - |
| 6 | w | 4.643E+05 | 749.4 | 0.00169 | 2.256 | +1 | 7 |
| - | - | 2.19E+05 | 750.4 | - | - | 0 | - |
| - | - | 5.902E+04 | 751.4 | - | - | 0 | - |
| - | - | 1.892E+04 | 752.4 | - | - | 0 | - |
| 7 | c | 8.578E+04 | 758.4 | 0.001534 | 2.022 | +1 | 7 |
| - | - | 3.749E+04 | 759.4 | - | - | 0 | - |
| - | - | 1.432E+04 | 760.4 | - | - | 0 | - |
| 7 | c | 9.36E+05 | 775.4 | 0.0003141 | 0.4051 | +1 | 7 |
| - | - | 3.52E+05 | 776.4 | - | - | 0 | - |
| - | - | 9.132E+04 | 777.4 | - | - | 0 | - |
| - | - | 1.163E+04 | 778.4 | - | - | 0 | - |
| - | - | 1.057E+04 | 786.4 | - | - | 0 | - |
| - | - | 3.81E+04 | 787.4 | - | - | 0 | - |
| - | - | 2.456E+04 | 788.4 | - | - | 0 | - |
| 6 | z | 2.402E+06 | 793.4 | 0.000503 | 0.6339 | +1 | 7 |
| - | - | 1.017E+06 | 794.4 | - | - | 0 | - |
| - | - | 2.891E+05 | 795.4 | - | - | 0 | - |
| - | - | 5.835E+04 | 796.4 | - | - | 0 | - |
| - | - | 1.058E+04 | 803.4 | - | - | 0 | - |
| - | - | 2.463E+04 | 808.4 | - | - | 0 | - |
| 6 | y | 2.292E+04 | 809.4 | 0.003268 | 4.037 | +1 | 7 |
| - | - | 1.507E+04 | 810.4 | - | - | 0 | - |
| - | - | 9533 | 815.5 | - | - | 0 | - |
| - | - | 1.804E+04 | 820.4 | - | - | 0 | - |
| - | - | 9638 | 825.4 | - | - | 0 | - |
| - | - | 2.042E+04 | 852.4 | - | - | 0 | - |
| - | - | 3.048E+04 | 858.5 | - | - | 0 | - |
| - | - | 8.081E+04 | 859.5 | - | - | 0 | - |
| - | - | 8.056E+04 | 860.5 | - | - | 0 | - |
| - | - | 3.194E+04 | 861.5 | - | - | 0 | - |
| - | - | 7.644E+05 | 864.4 | - | - | 0 | - |
| - | - | 3.69E+05 | 865.4 | - | - | 0 | - |
| - | - | 1.208E+05 | 866.4 | - | - | 0 | - |
| - | - | 3.212E+04 | 867.4 | - | - | 0 | - |
| - | - | 1.355E+04 | 869.5 | - | - | 0 | - |
| - | - | 1.52E+04 | 873.4 | - | - | 0 | - |
| - | - | 2.203E+04 | 874.4 | - | - | 0 | - |
| - | - | 1.187E+04 | 875.4 | - | - | 0 | - |
| - | - | 4.836E+04 | 884.5 | - | - | 0 | - |
| 8 | c | 2.761E+04 | 885.5 | 0.006386 | 7.212 | +1 | 8 |
| 8 | c | 9.852E+04 | 886.5 | 0.0038 | 4.286 | +1 | 8 |
| - | - | 5.541E+04 | 887.5 | - | - | 0 | - |
| - | - | 2.542E+04 | 888.5 | - | - | 0 | - |
| - | - | 1.033E+04 | 889.5 | - | - | 0 | - |
| - | - | 1.745E+04 | 896.4 | - | - | 0 | - |
| - | - | 8.205E+04 | 900.5 | - | - | 0 | - |
| - | - | 6.451E+04 | 901.5 | - | - | 0 | - |
| - | - | 3.148E+05 | 902.5 | - | - | 0 | - |
| 8 | c | 1.249E+06 | 903.5 | 0.0004106 | 0.4544 | +1 | 8 |
| - | - | 5.131E+05 | 904.5 | - | - | 0 | - |
| - | - | 1.569E+05 | 905.5 | - | - | 0 | - |
| - | - | 2.796E+04 | 906.5 | - | - | 0 | - |
| 5 | z | 7.958E+05 | 908.4 | 0.0001339 | 0.1474 | +1 | 8 |
| - | - | 3.737E+05 | 909.4 | - | - | 0 | - |
| - | - | 1.153E+05 | 910.4 | - | - | 0 | - |
| - | - | 2.92E+04 | 911.4 | - | - | 0 | - |
| 5 | y | 6.597E+04 | 924.4 | 0.0008526 | 0.9223 | +1 | 8 |
| - | - | 4E+04 | 925.4 | - | - | 0 | - |
| - | - | 9918 | 926.4 | - | - | 0 | - |
| - | - | 1.073E+04 | 943.5 | - | - | 0 | - |
| - | - | 1.143E+04 | 951.5 | - | - | 0 | - |
| - | - | 9418 | 959.5 | - | - | 0 | - |
| - | - | 1.305E+04 | 961.5 | - | - | 0 | - |
| - | - | 2.167E+04 | 968.5 | - | - | 0 | - |
| - | - | 1.686E+04 | 969.5 | - | - | 0 | - |
| - | - | 1.896E+04 | 973.5 | - | - | 0 | - |
| - | - | 1.974E+04 | 974.5 | - | - | 0 | - |
| - | - | 1.495E+04 | 975.5 | - | - | 0 | - |
| - | - | 6.403E+04 | 986.5 | - | - | 0 | - |
| - | - | 9.307E+04 | 987.5 | - | - | 0 | - |
| - | - | 4.057E+04 | 988.5 | - | - | 0 | - |
| - | - | 1.806E+04 | 989.5 | - | - | 0 | - |
| - | - | 9165 | 998.5 | - | - | 0 | - |
| 9 | c | 5.821E+04 | 999.5 | 0.003115 | 3.117 | +1 | 9 |
| 9 | c | 7.451E+04 | 1001 | 0.00671 | 6.706 | +1 | 9 |
| - | - | 4.678E+04 | 1002 | - | - | 0 | - |
| - | - | 3.653E+04 | 1003 | - | - | 0 | - |
| - | - | 2.354E+04 | 1004 | - | - | 0 | - |
| - | - | 9.875E+04 | 1017 | - | - | 0 | - |
| 9 | c | 7.743E+05 | 1018 | 0.0005524 | 0.5429 | +1 | 9 |
| - | - | 3.894E+05 | 1019 | - | - | 0 | - |
| - | - | 1.402E+05 | 1020 | - | - | 0 | - |
| - | - | 5.163E+04 | 1021 | - | - | 0 | - |
| - | - | 1.388E+04 | 1022 | - | - | 0 | - |
| - | - | 1.699E+04 | 1023 | - | - | 0 | - |
| - | - | 1.246E+04 | 1024 | - | - | 0 | - |
| - | - | 1.137E+04 | 1047 | - | - | 0 | - |
| - | - | 1.185E+04 | 1049 | - | - | 0 | - |
| - | - | 2.359E+04 | 1061 | - | - | 0 | - |
| - | - | 2.445E+04 | 1062 | - | - | 0 | - |
| 4 | y | 2.044E+04 | 1063 | 0.02091 | 19.68 | +1 | 9 |
| 4 | z | 2.846E+05 | 1065 | 0.000598 | 0.5617 | +1 | 9 |
| - | - | 3.46E+05 | 1066 | - | - | 0 | - |
| - | - | 1.712E+05 | 1067 | - | - | 0 | - |
| - | - | 5.247E+04 | 1068 | - | - | 0 | - |
| - | - | 1.144E+04 | 1080 | - | - | 0 | - |
| 4 | y | 2.723E+04 | 1081 | 0.003697 | 3.421 | +1 | 9 |
| - | - | 1.176E+04 | 1082 | - | - | 0 | - |
| - | - | 5.656E+04 | 1089 | - | - | 0 | - |
| - | - | 3.498E+04 | 1090 | - | - | 0 | - |
| - | - | 1.803E+04 | 1091 | - | - | 0 | - |
| - | - | 9766 | 1095 | - | - | 0 | - |
| - | - | 1.32E+04 | 1096 | - | - | 0 | - |
| - | - | 1.472E+04 | 1097 | - | - | 0 | - |
| 10 | c | 2.367E+05 | 1105 | 0.0007815 | 0.7075 | +1 | 10 |
| - | - | 1.588E+05 | 1106 | - | - | 0 | - |
| - | - | 6.776E+04 | 1107 | - | - | 0 | - |
| - | - | 2.285E+04 | 1108 | - | - | 0 | - |
| - | - | 1.022E+04 | 1109 | - | - | 0 | - |
| - | - | 1.623E+04 | 1110 | - | - | 0 | - |
| 3 | z | 2.496E+04 | 1135 | 0.02017 | 17.78 | +1 | 10 |
| - | - | 1.798E+04 | 1136 | - | - | 0 | - |
| - | - | 9620 | 1137 | - | - | 0 | - |
| 3 | z | 2.07E+05 | 1152 | 0.0005219 | 0.4532 | +1 | 10 |
| - | - | 3.805E+05 | 1153 | - | - | 0 | - |
| - | - | 1.958E+05 | 1154 | - | - | 0 | - |
| - | - | 7.903E+04 | 1155 | - | - | 0 | - |
| - | - | 2.375E+04 | 1156 | - | - | 0 | - |
| - | - | 2.664E+04 | 1158 | - | - | 0 | - |
| - | - | 1.965E+04 | 1159 | - | - | 0 | - |
| - | - | 1.927E+04 | 1160 | - | - | 0 | - |
| - | - | 1.604E+04 | 1161 | - | - | 0 | - |
| 3 | y | 1.881E+04 | 1168 | 0.001628 | 1.394 | +1 | 10 |
| - | - | 2.759E+04 | 1169 | - | - | 0 | - |
| - | - | 1.288E+04 | 1170 | - | - | 0 | - |
| - | - | 5.949E+04 | 1174 | - | - | 0 | - |
| - | - | 1.062E+05 | 1175 | - | - | 0 | - |
| - | - | 6.603E+04 | 1176 | - | - | 0 | - |
| - | - | 2.693E+04 | 1177 | - | - | 0 | - |
| - | - | 9053 | 1185 | - | - | 0 | - |
| - | - | 8345 | 1188 | - | - | 0 | - |
| - | - | 1.652E+05 | 1202 | - | - | 0 | - |
| - | - | 9.341E+04 | 1203 | - | - | 0 | - |
| - | - | 5.421E+04 | 1204 | - | - | 0 | - |
| - | - | 1.138E+04 | 1205 | - | - | 0 | - |
| 11 | c | 1.314E+05 | 1218 | 0.001471 | 1.208 | +1 | 11 |
| - | - | 2.888E+05 | 1219 | - | - | 0 | - |
| - | - | 1.488E+05 | 1220 | - | - | 0 | - |
| - | - | 5.83E+04 | 1221 | - | - | 0 | - |
| - | - | 1.122E+04 | 1222 | - | - | 0 | - |
| 2 | z | 1.038E+04 | 1265 | 0.005851 | 4.626 | +1 | 11 |
| - | - | 7.645E+04 | 1266 | - | - | 0 | - |
| - | - | 4.922E+04 | 1267 | - | - | 0 | - |
| - | - | 2.716E+04 | 1268 | - | - | 0 | - |
| 2 | y | 1.838E+04 | 1281 | 0.01566 | 12.23 | +1 | 11 |
| - | - | 9138 | 1295 | - | - | 0 | - |
| - | - | 1.656E+04 | 1305 | - | - | 0 | - |
| - | - | 6.567E+04 | 1306 | - | - | 0 | - |
| - | - | 4.205E+04 | 1307 | - | - | 0 | - |
| - | - | 6.103E+04 | 1308 | - | - | 0 | - |
| - | - | 2.889E+04 | 1309 | - | - | 0 | - |
| - | - | 1.312E+04 | 1310 | - | - | 0 | - |
| - | - | 1.284E+04 | 1320 | - | - | 0 | - |
| - | - | 3.667E+04 | 1321 | - | - | 0 | - |
| - | - | 7.869E+04 | 1322 | - | - | 0 | - |
| - | - | 5.22E+04 | 1323 | - | - | 0 | - |
| - | - | 1.399E+05 | 1324 | - | - | 0 | - |
| - | - | 2.627E+05 | 1325 | - | - | 0 | - |
| - | - | 1.536E+05 | 1326 | - | - | 0 | - |
| - | - | 7.65E+04 | 1327 | - | - | 0 | - |
| - | - | 1.784E+04 | 1328 | - | - | 0 | - |
| - | - | 6.708E+04 | 1338 | - | - | 0 | - |
| - | - | 4.238E+05 | 1339 | - | - | 0 | - |
| - | - | 2.891E+05 | 1340 | - | - | 0 | - |
| - | - | 1.454E+05 | 1341 | - | - | 0 | - |
| - | - | 5.596E+04 | 1342 | - | - | 0 | - |
| - | - | 1.103E+04 | 1343 | - | - | 0 | - |
| - | - | 1.546E+04 | 1348 | - | - | 0 | - |
| - | - | 8.3E+04 | 1349 | - | - | 0 | - |
| - | - | 1.66E+05 | 1350 | - | - | 0 | - |
| - | - | 9.762E+04 | 1351 | - | - | 0 | - |
| - | - | 4.088E+04 | 1352 | - | - | 0 | - |
| - | - | 8831 | 1353 | - | - | 0 | - |
| - | - | 1.406E+04 | 1355 | - | - | 0 | - |
| - | - | 2.653E+04 | 1356 | - | - | 0 | - |
| - | - | 1.144E+04 | 1357 | - | - | 0 | - |
| - | - | 9988 | 1365 | - | - | 0 | - |
| - | - | 2.112E+05 | 1366 | - | - | 0 | - |
| - | - | 1.317E+06 | 1367 | - | - | 0 | - |
| - | - | 9.123E+05 | 1368 | - | - | 0 | - |
| - | - | 4.206E+05 | 1369 | - | - | 0 | - |
| - | - | 1.383E+05 | 1370 | - | - | 0 | - |
| - | - | 2.609E+04 | 1371 | - | - | 0 | - |
| - | - | 5.702E+04 | 1382 | - | - | 0 | - |
| - | - | 1.687E+05 | 1383 | - | - | 0 | - |
| - | - | 2.791E+05 | 1384 | - | - | 0 | - |
| - | - | 1.715E+05 | 1385 | - | - | 0 | - |
| - | - | 7.671E+04 | 1386 | - | - | 0 | - |
| - | - | 2.109E+04 | 1387 | - | - | 0 | - |
| - | - | 8534 | 1897 | - | - | 0 | - |
| - | - | 1.298E+04 | 3084 | - | - | 0 | - |
| - | - | 9327 | 3085 | - | - | 0 | - |
| - | - | 8611 | 3343 | - | - | 0 | - |

m/z Charge Intensity FragmentType MassShift Position
123.0443115234375 0 6709.2065
123.10479736328125 0 8395.733
129.10256958007812 0 7238.8867
136.07598876953125 0 135160.12
136.3201904296875 0 5018.1333
137.07940673828125 0 9681.107
142.1228790283203 0 9371.791
143.11842346191406 0 9787.361
143.94691467285156 0 4895.96
145.1215362548828 0 5389.8633
149.04556274414062 0 6755.806
155.1183624267578 0 6583.871
157.1338348388672 0 5673.6865
165.05487060546875 0 127190.71
166.05807495117188 0 11461.008
169.1338653564453 0 8764.781
171.79342651367188 0 5373.1616
173.12881469726562 0 13054.6
173.45211791992188 0 9169.78
182.08151245117188 0 411558.34 y 11
183.08486938476562 0 46250.03
187.14447021484375 0 757644.56
188.14779663085938 0 92093.695
200.1393585205078 0 9298.326
201.1239471435547 0 19724.664
202.0828094482422 0 7932.2188
215.1393280029297 0 205360.95
216.14268493652344 0 23087.838
228.69961547851562 0 6631.8066
229.12957763671875 0 15776.961
231.15907287597656 0 5702.313
232.165771484375 0 10056.311 c 1
239.09564208984375 0 7782.1587
245.16091918945312 0 10640.018
274.2394104003906 0 5456.8096
281.1372985839844 0 6172.7627
282.1449279785156 0 47876.82
287.1726379394531 0 9933.733
295.1652526855469 0 33168.25 y 10
297.1563720703125 0 7829.9927
298.14111328125 0 6813.2764
299.14837646484375 0 8219.005
299.1712646484375 0 16307.719
300.1557312011719 0 143835.42
301.15960693359375 0 18286.361
302.1717529296875 0 14219.322
314.1835021972656 0 7076.0024
315.1669616699219 0 27157.984
317.18243408203125 0 75562.74
318.18524169921875 0 8114.525
319.1979675292969 0 60699.215 c 2
320.2012023925781 0 14431.425
341.01800537109375 0 8091.161
357.51483154296875 0 9426.15
359.02899169921875 0 40690.87
366.1768798828125 0 6671.529 z 9
367.1833190917969 0 6127.7485
372.6985778808594 0 11365.028
373.1993103027344 0 7604.846
374.1814880371094 0 9308.944
382.1974792480469 0 31433.652 y 9
385.21990966796875 0 105737.77
385.54803466796875 0 36581.4
385.88043212890625 0 20312.744
386.2250671386719 0 28180.629
391.55120849609375 0 88396.445
391.8852233886719 0 50682.69
392.21875 0 20761.697
394.70428466796875 0 14859.325
400.25640869140625 0 16003.694
408.2216491699219 0 6983.707
415.2686462402344 0 18357.48
425.24957275390625 0 9829.519
428.7001037597656 0 6978.919
429.05712890625 0 5570.1
429.0894775390625 0 117197.766
429.75531005859375 0 35186.74
430.2106018066406 0 8822.468
430.2447204589844 0 37401.188
430.28021240234375 0 11695.386
431.28021240234375 0 9640.231
437.212890625 0 21364.395
439.90533447265625 0 8001.9673
442.74481201171875 0 56722.848
443.2491760253906 0 35707.13 c Water loss 7
443.741943359375 0 37729.99 c Ammonia loss 7
444.2444763183594 0 21166.15
445.1210021972656 0 61533.887
446.21295166015625 0 58059.72
451.7507629394531 0 232189.58
452.25421142578125 0 701286.94 c 7
452.75579833984375 0 272926.03
453.2565002441406 0 65374.574
453.7582702636719 0 10937.489
455.2386169433594 0 11717.427
456.2586975097656 0 8488.335
457.261474609375 0 6965.103
458.271484375 0 22907.65 c Ammonia loss 3
461.2400817871094 0 15105.774
461.5748291015625 0 8408.033
463.2195739746094 0 14058.491
466.7313537597656 0 25203.477
467.2314453125 0 12136.458
470.75140380859375 0 13512.843
471.2469787597656 0 14958.779
472.252197265625 0 34783.754
473.25897216796875 0 9144.93
474.2920227050781 0 8009.409
475.2995300292969 0 1940294.4 c 3
476.3022766113281 0 436746.03
477.3045959472656 0 62267.684
479.2626953125 0 18401.355
479.7585754394531 0 204594.08
480.22149658203125 0 23336.45 z 8
480.2601013183594 0 100113.85
480.7569274902344 0 56172.734
481.2297058105469 0 148915.58
482.2320861816406 0 30766.336
483.237060546875 0 9382.7295
484.7484130859375 0 20645.79
485.24456787109375 0 34908.69
485.7430419921875 0 13661.66
486.25213623046875 0 8143.6313
486.76861572265625 0 41992.105
487.2691650390625 0 57737.082
487.7698974609375 0 20324.084
488.27349853515625 0 11898.984
491.7560729980469 0 10719.054
493.7556457519531 0 44838.11
494.2571105957031 0 19024.07
494.7578430175781 0 14398.099
495.2282409667969 0 14271.631
496.2407531738281 0 74565.92 y 8
497.2401123046875 0 13150.068
499.76690673828125 0 258073.36
500.2674560546875 0 124392.195 c Water loss 8
500.76507568359375 0 86045.086 c Ammonia loss 8
501.26409912109375 0 56224.72
501.7693786621094 0 39829.375
502.2750244140625 0 224393.61
502.7672424316406 0 10921.861
503.27630615234375 0 52471.31
508.2658996582031 0 18915.361
508.7720031738281 0 558751.06
509.2748107910156 0 991524.8 c 8
509.77593994140625 0 443299.94
510.27685546875 0 135516.38
510.77581787109375 0 40572.426
514.25927734375 0 8541.38
521.2744750976562 0 11577.963
521.7769775390625 0 8975.028
522.7750244140625 0 12789.339
523.2720947265625 0 13996.755
523.7637329101562 0 10174.681 z Water loss 3
525.7522583007812 0 10050.743
526.255126953125 0 43989.426
526.75732421875 0 36611.773
530.2885131835938 0 41463.15
530.7815551757812 0 10885.795
531.283203125 0 10430.862
532.76708984375 0 90568.6 z 3
533.2673950195312 0 51734.28
533.7694091796875 0 17263.24
534.2685546875 0 9346.67
535.2741088867188 0 36445.867
535.7699584960938 0 20433.498
536.6134033203125 0 7021.5938
540.2718505859375 0 13205.26
540.7750244140625 0 23609.31 y 3
541.2770385742188 0 12054.131
541.780029296875 0 10305.445
543.784423828125 0 6754.286 c Water loss 9
544.2788696289062 0 106316.836 c Ammonia loss 9
544.7792358398438 0 64077.21
545.3046264648438 0 63929.81
546.2636108398438 0 7707.6514
546.308837890625 0 28020.58
548.2510375976562 0 102567.84
548.7531127929688 0 59148.375
549.25146484375 0 16493.387
550.2962646484375 0 9596.415
552.2511596679688 0 9315.484
552.7920532226562 0 759482.3 c 9
553.2925415039062 0 405001.66
553.7929077148438 0 140913.6
554.2908935546875 0 61646.33
554.7767944335938 0 31572.68
558.29443359375 0 9938.133
561.2797241210938 0 10865.013
561.7911376953125 0 12906.796
562.2843017578125 0 9199.43
563.321044921875 0 10975.09
564.3049926757812 0 58178.355
565.2611694335938 0 11455.195
565.3095092773438 0 27962.35
565.80078125 0 64119.234
566.300537109375 0 34453.01
566.7736206054688 0 21224.56
567.2763061523438 0 17613.8 z Water loss 2
567.7769775390625 0 15126.636 z Ammonia loss 2
568.279052734375 0 23048.338
568.7797241210938 0 28169.2
573.3065185546875 0 32416.992 c Ammonia loss 4
574.3049926757812 0 13690.387
574.789306640625 0 8528.833
575.289794921875 0 23701.658 y Water loss 2
575.78125 0 52499.336 y Ammonia loss 2
576.2823486328125 0 269469.6 z 2
576.7822875976562 0 182728.34
577.2844848632812 0 75083.64
577.823974609375 0 13341.809
578.3148803710938 0 52389.957
578.8148803710938 0 23167.83
579.3139038085938 0 28446.81
579.81591796875 0 11402.055
582.8080444335938 0 8484.654
583.3032836914062 0 32512.486
583.8003540039062 0 18134.8
584.2916870117188 0 824133.75 y 2
584.7924194335938 0 509967.94
585.292724609375 0 194242.78
585.794921875 0 47129.883
586.822998046875 0 92123.305
587.32470703125 0 104521
587.8250732421875 0 46256.023
588.2755126953125 0 32850.26
588.3238525390625 0 12823.962
588.7777709960938 0 27882.545
589.2745971679688 0 17203.148
589.3192749023438 0 14723.623
590.3262939453125 0 3744706 c 4
591.3287963867188 0 1058628.6
591.3854370117188 0 9922.719
591.8148193359375 0 75586.28
592.3284301757812 0 192133.58
592.8096313476562 0 43356.086
593.3251953125 0 23444.822
597.78466796875 0 15791.372
600.3409423828125 0 8731.712
600.8208618164062 0 144619.8 c Ammonia loss 10
601.3216552734375 0 103488.54
601.8229370117188 0 59794.812
602.3330688476562 0 17696.273
607.3296508789062 0 36667.445
607.8284912109375 0 25130.412
608.3175048828125 0 206828 z 7
608.8312377929688 0 15632.152
609.3336791992188 0 1852352.4 c 10
609.83447265625 0 1103096
610.3347778320312 0 470712.38
610.8358764648438 0 139206.72
611.3342895507812 0 18283.156
612.3212280273438 0 10931.51
616.34326171875 0 26334.54
618.3053588867188 0 95975.02 w 1
618.804931640625 0 68173.805
619.3096923828125 0 31207.898
620.8263549804688 0 12135.49
621.328857421875 0 17966.92
624.33544921875 0 291456.3 y 7
625.334228515625 0 84643.85
626.3367919921875 0 25171.102
630.3274536132812 0 62723.895
630.8281860351562 0 41171.055
631.3326416015625 0 22663.479
631.8250122070312 0 11291.005 y Water loss 1
632.3219604492188 0 11716.3955 y Ammonia loss 1
632.82177734375 0 45393.82 z 1
633.32177734375 0 89733.63
633.8203735351562 0 34727.152
634.3191528320312 0 12896.465
634.8259887695312 0 9162.967
635.3472290039062 0 13935.502
636.3488159179688 0 37702.65
637.348388671875 0 9286.561
637.8330078125 0 32518.107
638.3455200195312 0 37141.11
638.8453369140625 0 24800.072
639.3449096679688 0 11577.362
639.8291015625 0 24679.889
640.3385009765625 0 25860.812
640.8328857421875 0 100257.164 y 1
641.33154296875 0 82776.15
641.8336791992188 0 23610.574
642.34423828125 0 14181.239
644.3366088867188 0 19559.842
646.8137817382812 0 13592.811
647.3515625 0 125509.26
647.8541259765625 0 75025.71
648.3536376953125 0 18907.846
652.3460083007812 0 20969.438
652.8414916992188 0 20198.797
653.339111328125 0 46869.543
653.8395385742188 0 20109.457
654.3328857421875 0 12401.387
654.8309936523438 0 19445.432
655.3181762695312 0 127104.945
655.81787109375 0 81309.87
656.3203735351562 0 28113.605
656.818603515625 0 16308.176
659.34765625 0 189120.36
660.349365234375 0 234993.12
660.8506469726562 0 164843.77
661.3521118164062 0 353114 z Water loss 6
661.8504028320312 0 236714.3
662.345947265625 0 295325.03
662.8464965820312 0 164568.16
663.3450317382812 0 84618.8
663.8513793945312 0 16820.955
668.8297729492188 0 110493.445
669.3486328125 0 370233.75
669.8511962890625 0 231315.25
670.3484497070312 0 270130.03
670.8485107421875 0 167043.36
671.3472900390625 0 67459.47
671.8507080078125 0 16971.701
672.3429565429688 0 34278.29
673.3482666015625 0 11166.808
673.8511962890625 0 41787.14
674.3455200195312 0 83124.25
674.8457641601562 0 193775.28
675.3457641601562 0 136066.33
675.8406372070312 0 170496.48
676.3388671875 0 108040.66
676.837890625 0 42680.77
677.3434448242188 0 16946.562
679.3540649414062 0 804763.3 z 6
680.3546142578125 0 301250.22
681.3574829101562 0 68007.6
682.3532104492188 0 8963.542
682.8551635742188 0 67575.37
683.3485717773438 0 1133608
683.8486328125 0 784636
684.3494873046875 0 386608.66
684.849853515625 0 119076.516
685.351318359375 0 34696.18
687.3506469726562 0 49604.19 c Ammonia loss 5
688.3565673828125 0 15498.037
689.3612670898438 0 11069.621
691.3575439453125 0 1043738.94
691.8578491210938 0 750620.6
692.358642578125 0 366379.94
692.8594360351562 0 92914.98
693.3555297851562 0 37473.152
695.372802734375 0 191810.78 y 6
696.373046875 0 72425.805
703.3638305664062 0 9520.446
704.369140625 0 1009072.1 c 5
705.3716430664062 0 340060.47
706.3745727539062 0 82085.15
707.3756713867188 0 10107.578
711.3410034179688 0 14073.711
712.349853515625 0 10034.413
722.364990234375 0 10496.245
730.3903198242188 0 10896.063
731.3919677734375 0 141320.23
732.3953247070312 0 50734.17
733.3987426757812 0 11767.972
737.3341674804688 0 46333.203
738.3331909179688 0 17316.674
742.3528442382812 0 8347.689
749.384521484375 0 464278.4 w 5
750.3865356445312 0 218974.81
751.3898315429688 0 59015.555
752.389892578125 0 18919.36
758.3806762695312 0 85784.28 c Ammonia loss 6
759.38330078125 0 37488.3
760.3876953125 0 14316.897
775.406005859375 0 936046.7 c 6
776.40771484375 0 352028.03
777.41064453125 0 91318.79
778.410400390625 0 11629.606
786.382080078125 0 10570.382
787.3893432617188 0 38098.5
788.3961181640625 0 24557.549
793.39697265625 0 2402150.8 z 5
794.3978271484375 0 1016600.44
795.4000854492188 0 289133.53
796.4007568359375 0 58353.17
803.4139404296875 0 10576.797
808.3717041015625 0 24629.875
809.4119262695312 0 22922.676 y 5
810.4141235351562 0 15073.386
815.4967651367188 0 9533.275
820.4354248046875 0 18042.5
825.38623046875 0 9638.1
852.3577270507812 0 20416.436
858.492919921875 0 30479.625
859.488037109375 0 80805.39
860.49267578125 0 80555.1
861.4953002929688 0 31940.254
864.4334716796875 0 764439.5
865.4345703125 0 369008.5
866.43408203125 0 120834.27
867.4376831054688 0 32119.217
869.4548950195312 0 13547.664
873.4195556640625 0 15198.223
874.4243774414062 0 22026.398
875.4230346679688 0 11871.644
884.4819946289062 0 48359.137
885.4837036132812 0 27605.24 c Water loss 7
886.4779052734375 0 98517.086 c Ammonia loss 7
887.4791259765625 0 55405.863
888.482666015625 0 25424.035
889.491943359375 0 10332.157
896.4268188476562 0 17448.314
900.4769897460938 0 82049.91
901.4794921875 0 64513.59
902.4923095703125 0 314818.38
903.500244140625 0 1248626.1 c 7
904.5025024414062 0 513056.38
905.5047607421875 0 156859.23
906.5065307617188 0 27962.707
908.4232788085938 0 795844 z 4
909.4256591796875 0 373731.2
910.4262084960938 0 115339.99
911.427001953125 0 29197.732
924.4412841796875 0 65965.33 y 4
925.4417724609375 0 39999.67
926.4379272460938 0 9918.008
943.4992065429688 0 10734.232
951.4609985351562 0 11430.335
959.5078735351562 0 9418.441
961.51025390625 0 13046.804
968.4931030273438 0 21670.66
969.4822998046875 0 16856.883
973.5333862304688 0 18964.396
974.534423828125 0 19743.594
975.5343017578125 0 14951.432
986.5001831054688 0 64034.754
987.5055541992188 0 93069.734
988.5066528320312 0 40571.535
989.5031127929688 0 18057.145
998.5279541015625 0 9165.313
999.5361328125 0 58210.297 c Water loss 8
1000.5237426757812 0 74512.14 c Ammonia loss 8
1001.521484375 0 46784.64
1002.5278930664062 0 36528.688
1003.5307006835938 0 23537.176
1016.535888671875 0 98754.27
1017.5430297851562 0 774305.8 c 8
1018.5438842773438 0 389386.28
1019.5458374023438 0 140206.28
1020.5427856445312 0 51634.035
1021.5388793945312 0 13879.318
1022.5149536132812 0 16985.889
1023.5125122070312 0 12461.908
1046.53564453125 0 11371.263
1048.509521484375 0 11854.086
1060.5611572265625 0 23585.352
1061.56884765625 0 24445.555
1062.5535888671875 0 20438.271 y Water loss 3
1064.52392578125 0 284571.8 z 3
1065.5277099609375 0 346044.62
1066.529052734375 0 171192.72
1067.5302734375 0 52472.004
1079.5296630859375 0 11443.498
1080.53955078125 0 27225.906 y 3
1081.545654296875 0 11762.584
1088.5576171875 0 56555.42
1089.5574951171875 0 34977.066
1090.5625 0 18025.309
1094.5203857421875 0 9765.699
1095.5003662109375 0 13197.385
1096.5103759765625 0 14722.775
1104.5748291015625 0 236735.5 c 9
1105.577880859375 0 158780.84
1106.579345703125 0 67756.35
1107.572509765625 0 22849.223
1108.563720703125 0 10216.924
1109.5428466796875 0 16226.297
1134.5501708984375 0 24958.074 z Ammonia loss 2
1135.54541015625 0 17983.602
1136.54931640625 0 9619.533
1151.5560302734375 0 206984.05 z 2
1152.56201171875 0 380503.56
1153.5626220703125 0 195845.61
1154.5655517578125 0 79028.64
1155.57373046875 0 23746.893
1157.6268310546875 0 26635.63
1158.63623046875 0 19648.275
1159.63037109375 0 19266.56
1160.6243896484375 0 16035.171
1167.576904296875 0 18808.273 y 2
1168.5792236328125 0 27587.533
1169.5892333984375 0 12884.388
1173.64599609375 0 59486.953
1174.6544189453125 0 106246.86
1175.6527099609375 0 66032.484
1176.6519775390625 0 26931.514
1184.625 0 9052.678
1187.898193359375 0 8344.854
1201.6402587890625 0 165247.7
1202.6422119140625 0 93411.484
1203.6424560546875 0 54210.48
1204.6505126953125 0 11377.21
1217.658203125 0 131359.86 c 10
1218.6629638671875 0 288804.84
1219.6654052734375 0 148758.48
1220.66552734375 0 58303.098
1221.66259765625 0 11219.163
1264.634765625 0 10375.085 z 1
1265.646728515625 0 76450.04
1266.6431884765625 0 49221.617
1267.640380859375 0 27162.752
1280.6436767578125 0 18382.01 y 1
1294.6414794921875 0 9137.934
1304.7012939453125 0 16556.803
1305.6585693359375 0 65674.1
1306.657470703125 0 42046.195
1307.6571044921875 0 61028.137
1308.65966796875 0 28893.1
1309.6563720703125 0 13121.055
1319.68017578125 0 12838.354
1320.6915283203125 0 36670.707
1321.692626953125 0 78685.57
1322.6934814453125 0 52199.3
1323.6827392578125 0 139949.48
1324.679931640625 0 262717.38
1325.68359375 0 153616.2
1326.6832275390625 0 76503.02
1327.672607421875 0 17840.627
1337.691650390625 0 67084.305
1338.699462890625 0 423770.1
1339.7012939453125 0 289077.53
1340.7008056640625 0 145418.95
1341.7003173828125 0 55963.74
1342.707275390625 0 11025.25
1347.7003173828125 0 15459.694
1348.684814453125 0 83004.76
1349.6744384765625 0 166040.81
1350.6773681640625 0 97620.15
1351.674072265625 0 40878.613
1352.6605224609375 0 8830.693
1354.7227783203125 0 14062.278
1355.7213134765625 0 26532.27
1356.722900390625 0 11440.755
1364.70458984375 0 9987.851
1365.690673828125 0 211150
1366.693359375 0 1317233.6
1367.6949462890625 0 912344.2
1368.6961669921875 0 420558.4
1369.69921875 0 138320.95
1370.7008056640625 0 26085.719
1381.7047119140625 0 57022.93
1382.7119140625 0 168678.25
1383.717041015625 0 279068.97
1384.7178955078125 0 171505.44
1385.720947265625 0 76713.76
1386.72119140625 0 21087.672
1897.2724609375 0 8533.529
3083.61962890625 0 12980.386
3084.55224609375 0 9327.01
3342.85693359375 0 8611.294

Spectrum Details

|  |  |
| --- | --- |
| Matched peaks? Matched peaks The total absolute number of peaks matched. Additionally in brackets the total fraction of peaks matched and the total number of peaks is shown. | 67 (12.09% of 554) |
| FDR? FDR The false discovery rate estimated for this peptide. It is calculated by matching all theoretical fragments with a non-integer shift with the raw peaks for this spectrum. This is done with 40 different shifts. The resulting percentage is the average number of annotated peaks over the number of annotated peaks with the correct spectrum. | 1.39% |
| Satellite FDR? Satellite FDR See the FDR for details on its calculation. This satellite ion specific FDR only contains the satellite ions (d/w) for I/L/J positions. | 0.00% |
| PSM Score? PSM Score The PSM Score as given by Hecklib to this annotated spectrum. It is shown with three significant figures. | 580 |

#### Spectrum 3614? Spectrum 3614 The raw spectrum of this peptide as annotated by Hecklib. The fragments are coloured according to ion type (see legend). Any peaks with a star '\*' as text can be hovered over to see the full details, first the ion type second the mass shift type. By hovering over the amino acids in the peptide or ions in the legend the corresponding peaks are highlighted. By toggling the 'Unassigned' label you can turn the background (unassigned) peaks on or off in the plot. By updating the slider in the Ion legend you can update the spectrum to only show the top X% of the peaks with labels. The top X% means any peak that is within X% of the highest intensity. By dragging in the spectrum you can zoom in to a specific part of the spectrum and use 'Zoom Out' to get back to the original zoom level. The annotation of the spectrum is based on the given sequence in the peptides file and is done with different software so inconsistencies are likely. The peaks are annotated based on the given sequence, with 20 ppm tolerance.

Copy Data

##### Spectrum 3614 (TSV)

###### Preview

```
Loading example...
```

*Click on the button to copy the data to your clipboard.*

Mz MinMz MaxIntensity Max

WidthHeightPeptide font sizePeptide stroke widthSpectrum font sizeSpectrum stroke widthCompact peptide

Ion legend

wxyz

abcd

OtherUnassignedIon^Charge^_Position_Show for top:%

TISRDNAKNSJY

09.39e+41.88e+52.82e+53.75e+5

Zoom Out

y^+1^_1_c^+1^_2_y^+1^_2_c^+1^_3_z^+1^_3_y^+1^_3_y^+3^_10_c^+3^_11_c^+2^_8_c^+2^_8_c^+2^_8_c^+1^_4_c^+1^_4_z^+1^_4_y^+1^_4_c^+2^_9_c^+2^_9_c^+2^_9_z^+2^_9_z^+2^_9_y^+2^_9_c^+2^_10_c^+2^_10_z^+2^_10_z^+2^_10_c^+1^_5_y^+2^_10_y^+2^_10_z^+2^_10_y^+2^_10_c^+1^_5_c^+2^_11_c^+2^_11_z^+1^_5_c^+2^_11_w^+2^_11_y^+1^_5_y^+2^_11_z^+2^_11_y^+2^_11_z^+1^_6_z^+1^_6_c^+1^_6_y^+1^_6_c^+1^_6_w^+1^_7_c^+1^_7_c^+1^_7_c^+1^_7_z^+1^_7_y^+1^_7_c^+1^_8_c^+1^_8_c^+1^_8_z^+1^_8_y^+1^_8_c^+1^_9_c^+1^_9_c^+1^_9_y^+1^_9_z^+1^_9_y^+1^_9_c^+1^_10_w^+1^_10_z^+1^_10_c^+1^_11_z^+1^_11_

0789157823663155

Fragment Matches Table

Show background peaks

| Position | Ion type | Intensity | mz Theoretical | mz Error (Th) | mz Error (ppm) | Charge | Series Number |
| --- | --- | --- | --- | --- | --- | --- | --- |
| - | - | 426.7 | 122.4 | - | - | 0 | - |
| - | - | 901 | 123.1 | - | - | 0 | - |
| - | - | 1521 | 129.1 | - | - | 0 | - |
| - | - | 476.3 | 131.1 | - | - | 0 | - |
| - | - | 512.8 | 132.1 | - | - | 0 | - |
| - | - | 4.623E+04 | 136.1 | - | - | 0 | - |
| - | - | 822.3 | 137.1 | - | - | 0 | - |
| - | - | 8227 | 137.1 | - | - | 0 | - |
| - | - | 2815 | 140.1 | - | - | 0 | - |
| - | - | 2077 | 141.1 | - | - | 0 | - |
| - | - | 1259 | 142.1 | - | - | 0 | - |
| - | - | 1531 | 143.1 | - | - | 0 | - |
| - | - | 4435 | 149 | - | - | 0 | - |
| - | - | 1.319E+04 | 156.1 | - | - | 0 | - |
| - | - | 2435 | 157.1 | - | - | 0 | - |
| - | - | 516.5 | 159.5 | - | - | 0 | - |
| - | - | 1.222E+04 | 165.1 | - | - | 0 | - |
| - | - | 984.2 | 166.1 | - | - | 0 | - |
| - | - | 1210 | 167.1 | - | - | 0 | - |
| - | - | 678 | 169.1 | - | - | 0 | - |
| - | - | 570.2 | 171.1 | - | - | 0 | - |
| - | - | 1849 | 173.1 | - | - | 0 | - |
| - | - | 2705 | 173.5 | - | - | 0 | - |
| 12 | y | 4.467E+04 | 182.1 | 0.0003886 | 2.134 | +1 | 1 |
| - | - | 4902 | 183.1 | - | - | 0 | - |
| - | - | 1128 | 186.1 | - | - | 0 | - |
| - | - | 9.111E+04 | 187.1 | - | - | 0 | - |
| - | - | 609.9 | 188.1 | - | - | 0 | - |
| - | - | 8512 | 188.1 | - | - | 0 | - |
| - | - | 948.8 | 190.1 | - | - | 0 | - |
| - | - | 596 | 193.8 | - | - | 0 | - |
| - | - | 636.4 | 199.2 | - | - | 0 | - |
| - | - | 1171 | 200.1 | - | - | 0 | - |
| - | - | 3980 | 201.1 | - | - | 0 | - |
| - | - | 1340 | 201.1 | - | - | 0 | - |
| - | - | 1459 | 202.1 | - | - | 0 | - |
| - | - | 773.5 | 203.1 | - | - | 0 | - |
| - | - | 632.1 | 213.1 | - | - | 0 | - |
| - | - | 2.538E+04 | 215.1 | - | - | 0 | - |
| - | - | 2611 | 216.1 | - | - | 0 | - |
| - | - | 652.3 | 219.1 | - | - | 0 | - |
| - | - | 2459 | 221.1 | - | - | 0 | - |
| - | - | 1039 | 222.1 | - | - | 0 | - |
| - | - | 1507 | 223.1 | - | - | 0 | - |
| - | - | 1720 | 225 | - | - | 0 | - |
| - | - | 1627 | 229.1 | - | - | 0 | - |
| 2 | c | 703 | 232.2 | 0.0001882 | 0.8107 | +1 | 2 |
| - | - | 6.544E+04 | 235.1 | - | - | 0 | - |
| - | - | 3.141E+04 | 236.1 | - | - | 0 | - |
| - | - | 3114 | 237.2 | - | - | 0 | - |
| - | - | 1514 | 238.1 | - | - | 0 | - |
| - | - | 2036 | 239.1 | - | - | 0 | - |
| - | - | 1796 | 240.1 | - | - | 0 | - |
| - | - | 1152 | 241.1 | - | - | 0 | - |
| - | - | 1011 | 244.1 | - | - | 0 | - |
| - | - | 655.6 | 245.2 | - | - | 0 | - |
| - | - | 564.7 | 249.4 | - | - | 0 | - |
| - | - | 7021 | 255.1 | - | - | 0 | - |
| - | - | 2041 | 256.1 | - | - | 0 | - |
| - | - | 686.9 | 257.1 | - | - | 0 | - |
| - | - | 1354 | 260.1 | - | - | 0 | - |
| - | - | 2.389E+04 | 263.1 | - | - | 0 | - |
| - | - | 639.5 | 264 | - | - | 0 | - |
| - | - | 623.7 | 264 | - | - | 0 | - |
| - | - | 1.231E+04 | 264.1 | - | - | 0 | - |
| - | - | 1526 | 265.1 | - | - | 0 | - |
| - | - | 625.1 | 266.2 | - | - | 0 | - |
| - | - | 645.9 | 269.2 | - | - | 0 | - |
| - | - | 1451 | 272.1 | - | - | 0 | - |
| - | - | 646.7 | 273.2 | - | - | 0 | - |
| - | - | 3760 | 282.1 | - | - | 0 | - |
| - | - | 971 | 287.2 | - | - | 0 | - |
| - | - | 1744 | 295.1 | - | - | 0 | - |
| 11 | y | 3399 | 295.2 | 0.0006599 | 2.236 | +1 | 2 |
| - | - | 2584 | 296.1 | - | - | 0 | - |
| - | - | 1700 | 297.1 | - | - | 0 | - |
| - | - | 1067 | 297.2 | - | - | 0 | - |
| - | - | 1065 | 298.1 | - | - | 0 | - |
| - | - | 1947 | 299.1 | - | - | 0 | - |
| - | - | 1343 | 299.1 | - | - | 0 | - |
| - | - | 1338 | 299.2 | - | - | 0 | - |
| - | - | 1756 | 300.1 | - | - | 0 | - |
| - | - | 1.185E+04 | 300.2 | - | - | 0 | - |
| - | - | 750.2 | 300.2 | - | - | 0 | - |
| - | - | 1391 | 301.1 | - | - | 0 | - |
| - | - | 651.4 | 301.1 | - | - | 0 | - |
| - | - | 1895 | 301.2 | - | - | 0 | - |
| - | - | 1064 | 302.2 | - | - | 0 | - |
| - | - | 3261 | 309.2 | - | - | 0 | - |
| - | - | 1835 | 310.2 | - | - | 0 | - |
| - | - | 3913 | 315.2 | - | - | 0 | - |
| - | - | 7710 | 317.2 | - | - | 0 | - |
| 3 | c | 7330 | 319.2 | 0.0005237 | 1.641 | +1 | 3 |
| - | - | 1.388E+04 | 341 | - | - | 0 | - |
| - | - | 1654 | 355.1 | - | - | 0 | - |
| - | - | 4.231E+04 | 359 | - | - | 0 | - |
| 10 | z | 964.4 | 366.2 | 0.0004476 | 1.222 | +1 | 3 |
| - | - | 2009 | 367.2 | - | - | 0 | - |
| - | - | 8387 | 368.2 | - | - | 0 | - |
| - | - | 1046 | 369.1 | - | - | 0 | - |
| - | - | 9600 | 369.2 | - | - | 0 | - |
| - | - | 3490 | 370.1 | - | - | 0 | - |
| - | - | 5470 | 370.2 | - | - | 0 | - |
| - | - | 2918 | 371.1 | - | - | 0 | - |
| - | - | 727.5 | 371.2 | - | - | 0 | - |
| - | - | 2033 | 372.7 | - | - | 0 | - |
| - | - | 1161 | 374.2 | - | - | 0 | - |
| - | - | 787 | 375.9 | - | - | 0 | - |
| - | - | 2634 | 378.2 | - | - | 0 | - |
| - | - | 1272 | 378.2 | - | - | 0 | - |
| - | - | 2494 | 379.2 | - | - | 0 | - |
| 10 | y | 1906 | 382.2 | 0.001652 | 4.321 | +1 | 3 |
| 3 | y | 3948 | 384.2 | 0.0001977 | 0.5147 | +3 | 10 |
| - | - | 2421 | 385.2 | - | - | 0 | - |
| - | - | 8831 | 385.2 | - | - | 0 | - |
| - | - | 3246 | 385.5 | - | - | 0 | - |
| - | - | 2101 | 385.9 | - | - | 0 | - |
| - | - | 2554 | 386.2 | - | - | 0 | - |
| - | - | 1024 | 386.5 | - | - | 0 | - |
| - | - | 723 | 387.2 | - | - | 0 | - |
| - | - | 7763 | 391.6 | - | - | 0 | - |
| - | - | 6553 | 391.9 | - | - | 0 | - |
| - | - | 2485 | 392.2 | - | - | 0 | - |
| - | - | 1302 | 394.7 | - | - | 0 | - |
| - | - | 921.4 | 400.3 | - | - | 0 | - |
| 11 | c | 821.2 | 400.9 | 0.0003446 | 0.8596 | +3 | 11 |
| - | - | 3631 | 401.2 | - | - | 0 | - |
| - | - | 2337 | 402.2 | - | - | 0 | - |
| - | - | 722.1 | 403.2 | - | - | 0 | - |
| - | - | 835.8 | 406.2 | - | - | 0 | - |
| - | - | 2192 | 408.2 | - | - | 0 | - |
| - | - | 814.4 | 409.2 | - | - | 0 | - |
| - | - | 1356 | 413.2 | - | - | 0 | - |
| - | - | 827.8 | 414.2 | - | - | 0 | - |
| - | - | 2020 | 415.3 | - | - | 0 | - |
| - | - | 945.7 | 416.2 | - | - | 0 | - |
| - | - | 1231 | 425.3 | - | - | 0 | - |
| - | - | 928.7 | 426.2 | - | - | 0 | - |
| - | - | 1.14E+05 | 429.1 | - | - | 0 | - |
| - | - | 2445 | 429.8 | - | - | 0 | - |
| - | - | 4196 | 430.2 | - | - | 0 | - |
| - | - | 1722 | 430.3 | - | - | 0 | - |
| - | - | 911.4 | 431.3 | - | - | 0 | - |
| - | - | 919.3 | 435.2 | - | - | 0 | - |
| - | - | 721.3 | 435.2 | - | - | 0 | - |
| - | - | 828.1 | 436.2 | - | - | 0 | - |
| - | - | 2398 | 437.2 | - | - | 0 | - |
| - | - | 1745 | 438.7 | - | - | 0 | - |
| - | - | 1432 | 439.2 | - | - | 0 | - |
| - | - | 4545 | 442.7 | - | - | 0 | - |
| 8 | c | 3431 | 443.2 | 0.007167 | 16.17 | +2 | 8 |
| - | - | 1896 | 443.3 | - | - | 0 | - |
| 8 | c | 4145 | 443.7 | 0.001313 | 2.96 | +2 | 8 |
| - | - | 2233 | 444.2 | - | - | 0 | - |
| - | - | 6.323E+04 | 445.1 | - | - | 0 | - |
| - | - | 5449 | 446.2 | - | - | 0 | - |
| - | - | 744.1 | 447.1 | - | - | 0 | - |
| - | - | 962.7 | 450.7 | - | - | 0 | - |
| - | - | 1304 | 451.2 | - | - | 0 | - |
| - | - | 1237 | 451.7 | - | - | 0 | - |
| - | - | 1.814E+04 | 451.8 | - | - | 0 | - |
| 8 | c | 6.273E+04 | 452.3 | 0.0003679 | 0.8135 | +2 | 8 |
| - | - | 2.928E+04 | 452.8 | - | - | 0 | - |
| - | - | 7119 | 453.3 | - | - | 0 | - |
| - | - | 2555 | 453.8 | - | - | 0 | - |
| - | - | 893.2 | 454.3 | - | - | 0 | - |
| - | - | 831.4 | 455.2 | - | - | 0 | - |
| - | - | 1053 | 455.6 | - | - | 0 | - |
| - | - | 1290 | 456.2 | - | - | 0 | - |
| - | - | 1140 | 457.2 | - | - | 0 | - |
| - | - | 944.7 | 457.8 | - | - | 0 | - |
| - | - | 889.5 | 458.2 | - | - | 0 | - |
| 4 | c | 3050 | 458.3 | 0.000974 | 2.125 | +1 | 4 |
| - | - | 918.9 | 459.3 | - | - | 0 | - |
| - | - | 1.116E+04 | 459.7 | - | - | 0 | - |
| - | - | 912.1 | 459.8 | - | - | 0 | - |
| - | - | 5479 | 460.2 | - | - | 0 | - |
| - | - | 2006 | 460.7 | - | - | 0 | - |
| - | - | 1782 | 461.2 | - | - | 0 | - |
| - | - | 1926 | 461.6 | - | - | 0 | - |
| - | - | 895.3 | 461.9 | - | - | 0 | - |
| - | - | 780.3 | 465.3 | - | - | 0 | - |
| - | - | 1368 | 465.8 | - | - | 0 | - |
| - | - | 1353 | 466.3 | - | - | 0 | - |
| - | - | 1167 | 466.7 | - | - | 0 | - |
| - | - | 4182 | 467.2 | - | - | 0 | - |
| - | - | 5825 | 468.2 | - | - | 0 | - |
| - | - | 2446 | 469.2 | - | - | 0 | - |
| - | - | 1539 | 470.8 | - | - | 0 | - |
| - | - | 1807 | 471.2 | - | - | 0 | - |
| - | - | 1312 | 471.7 | - | - | 0 | - |
| - | - | 2193 | 472.2 | - | - | 0 | - |
| - | - | 1181 | 473.2 | - | - | 0 | - |
| 4 | c | 1.961E+05 | 475.3 | 0.001006 | 2.116 | +1 | 4 |
| - | - | 4.39E+04 | 476.3 | - | - | 0 | - |
| - | - | 882.1 | 477.2 | - | - | 0 | - |
| - | - | 7042 | 477.3 | - | - | 0 | - |
| - | - | 899.6 | 478.3 | - | - | 0 | - |
| - | - | 1353 | 479.3 | - | - | 0 | - |
| - | - | 1.92E+04 | 479.8 | - | - | 0 | - |
| 9 | z | 1913 | 480.2 | 0.0008234 | 1.715 | +1 | 4 |
| - | - | 1.068E+04 | 480.3 | - | - | 0 | - |
| - | - | 4837 | 480.8 | - | - | 0 | - |
| - | - | 1.378E+04 | 481.2 | - | - | 0 | - |
| - | - | 725.8 | 481.8 | - | - | 0 | - |
| - | - | 3377 | 482.2 | - | - | 0 | - |
| - | - | 1107 | 483.3 | - | - | 0 | - |
| - | - | 3148 | 484.3 | - | - | 0 | - |
| - | - | 1821 | 484.7 | - | - | 0 | - |
| - | - | 2227 | 485.2 | - | - | 0 | - |
| - | - | 1236 | 485.7 | - | - | 0 | - |
| - | - | 2606 | 486.8 | - | - | 0 | - |
| - | - | 5832 | 487.3 | - | - | 0 | - |
| - | - | 3217 | 487.8 | - | - | 0 | - |
| - | - | 1085 | 488.3 | - | - | 0 | - |
| - | - | 888 | 492.3 | - | - | 0 | - |
| - | - | 4183 | 493.8 | - | - | 0 | - |
| - | - | 3167 | 494.3 | - | - | 0 | - |
| - | - | 1136 | 494.8 | - | - | 0 | - |
| - | - | 1207 | 495.2 | - | - | 0 | - |
| - | - | 888.6 | 495.7 | - | - | 0 | - |
| 9 | y | 6613 | 496.2 | 0.0008994 | 1.812 | +1 | 4 |
| - | - | 977.5 | 497.2 | - | - | 0 | - |
| - | - | 2.08E+04 | 499.8 | - | - | 0 | - |
| 9 | c | 1.368E+04 | 500.3 | 0.003912 | 7.819 | +2 | 9 |
| 9 | c | 1.037E+04 | 500.8 | 0.001731 | 3.456 | +2 | 9 |
| - | - | 6468 | 501.3 | - | - | 0 | - |
| - | - | 3795 | 501.8 | - | - | 0 | - |
| - | - | 1314 | 502.2 | - | - | 0 | - |
| - | - | 2.15E+04 | 502.3 | - | - | 0 | - |
| - | - | 1286 | 502.8 | - | - | 0 | - |
| - | - | 3658 | 503.3 | - | - | 0 | - |
| - | - | 1467 | 508.3 | - | - | 0 | - |
| - | - | 4.817E+04 | 508.8 | - | - | 0 | - |
| 9 | c | 8.231E+04 | 509.3 | 0.001412 | 2.773 | +2 | 9 |
| - | - | 5.83E+04 | 509.8 | - | - | 0 | - |
| - | - | 790.6 | 509.8 | - | - | 0 | - |
| - | - | 1.935E+04 | 510.3 | - | - | 0 | - |
| - | - | 7912 | 510.8 | - | - | 0 | - |
| - | - | 2680 | 511.3 | - | - | 0 | - |
| - | - | 1850 | 521.3 | - | - | 0 | - |
| - | - | 926.2 | 521.8 | - | - | 0 | - |
| - | - | 1389 | 522.8 | - | - | 0 | - |
| - | - | 1819 | 523.3 | - | - | 0 | - |
| 4 | z | 2625 | 524.3 | 0.006102 | 11.64 | +2 | 9 |
| - | - | 7067 | 525.3 | - | - | 0 | - |
| - | - | 7225 | 526.3 | - | - | 0 | - |
| - | - | 2505 | 526.8 | - | - | 0 | - |
| - | - | 788.3 | 527.3 | - | - | 0 | - |
| - | - | 3988 | 530.3 | - | - | 0 | - |
| - | - | 1481 | 530.8 | - | - | 0 | - |
| - | - | 1016 | 531.3 | - | - | 0 | - |
| 4 | z | 7239 | 532.8 | 0.0008846 | 1.66 | +2 | 9 |
| - | - | 4718 | 533.3 | - | - | 0 | - |
| - | - | 2245 | 533.8 | - | - | 0 | - |
| - | - | 2247 | 534.3 | - | - | 0 | - |
| - | - | 4469 | 535.3 | - | - | 0 | - |
| - | - | 4956 | 535.8 | - | - | 0 | - |
| - | - | 1365 | 536.3 | - | - | 0 | - |
| - | - | 969.6 | 536.8 | - | - | 0 | - |
| - | - | 1.34E+04 | 540.3 | - | - | 0 | - |
| 4 | y | 2967 | 540.8 | 0.0005429 | 1.004 | +2 | 9 |
| - | - | 1.875E+04 | 541.3 | - | - | 0 | - |
| - | - | 876.3 | 541.8 | - | - | 0 | - |
| - | - | 5189 | 542.3 | - | - | 0 | - |
| - | - | 1152 | 543.3 | - | - | 0 | - |
| 10 | c | 8985 | 544.3 | 0.001006 | 1.848 | +2 | 10 |
| - | - | 5932 | 544.8 | - | - | 0 | - |
| - | - | 1238 | 545.3 | - | - | 0 | - |
| - | - | 6463 | 545.3 | - | - | 0 | - |
| - | - | 919.3 | 545.8 | - | - | 0 | - |
| - | - | 5843 | 546.3 | - | - | 0 | - |
| - | - | 1074 | 547.3 | - | - | 0 | - |
| - | - | 8265 | 548.3 | - | - | 0 | - |
| - | - | 4575 | 548.8 | - | - | 0 | - |
| - | - | 3512 | 549.3 | - | - | 0 | - |
| - | - | 1040 | 549.8 | - | - | 0 | - |
| - | - | 947.7 | 550.3 | - | - | 0 | - |
| - | - | 1782 | 551.3 | - | - | 0 | - |
| - | - | 2122 | 552.3 | - | - | 0 | - |
| 10 | c | 5.939E+04 | 552.8 | 0.0006707 | 1.213 | +2 | 10 |
| - | - | 5.13E+04 | 553.3 | - | - | 0 | - |
| - | - | 2.576E+04 | 553.8 | - | - | 0 | - |
| - | - | 1.111E+04 | 554.3 | - | - | 0 | - |
| - | - | 2347 | 554.8 | - | - | 0 | - |
| - | - | 877.4 | 555.3 | - | - | 0 | - |
| - | - | 717.8 | 558.8 | - | - | 0 | - |
| - | - | 749.1 | 559.3 | - | - | 0 | - |
| - | - | 1072 | 561.3 | - | - | 0 | - |
| - | - | 935.6 | 561.8 | - | - | 0 | - |
| - | - | 6231 | 564.3 | - | - | 0 | - |
| - | - | 3022 | 565.3 | - | - | 0 | - |
| - | - | 3458 | 565.8 | - | - | 0 | - |
| - | - | 3446 | 566.3 | - | - | 0 | - |
| - | - | 2068 | 566.8 | - | - | 0 | - |
| 3 | z | 1201 | 567.3 | 0.009969 | 17.57 | +2 | 10 |
| 3 | z | 2155 | 567.8 | 0.009986 | 17.59 | +2 | 10 |
| - | - | 2181 | 568.3 | - | - | 0 | - |
| - | - | 918.9 | 568.8 | - | - | 0 | - |
| - | - | 844 | 569.3 | - | - | 0 | - |
| - | - | 1079 | 569.8 | - | - | 0 | - |
| 5 | c | 2770 | 573.3 | 0.006013 | 10.49 | +1 | 5 |
| - | - | 1181 | 574.3 | - | - | 0 | - |
| 3 | y | 1861 | 575.3 | 0.005632 | 9.79 | +2 | 10 |
| 3 | y | 4963 | 575.8 | 0.001234 | 2.143 | +2 | 10 |
| 3 | z | 2.236E+04 | 576.3 | 0.0006174 | 1.071 | +2 | 10 |
| - | - | 1.78E+04 | 576.8 | - | - | 0 | - |
| - | - | 7834 | 577.3 | - | - | 0 | - |
| - | - | 1952 | 577.8 | - | - | 0 | - |
| - | - | 1323 | 577.8 | - | - | 0 | - |
| - | - | 3846 | 578.3 | - | - | 0 | - |
| - | - | 2583 | 578.8 | - | - | 0 | - |
| - | - | 2396 | 579.3 | - | - | 0 | - |
| - | - | 2192 | 579.8 | - | - | 0 | - |
| - | - | 3519 | 583.3 | - | - | 0 | - |
| - | - | 2283 | 583.8 | - | - | 0 | - |
| 3 | y | 6.901E+04 | 584.3 | 0.0007769 | 1.33 | +2 | 10 |
| - | - | 6.004E+04 | 584.8 | - | - | 0 | - |
| - | - | 3.04E+04 | 585.3 | - | - | 0 | - |
| - | - | 1.153E+04 | 585.8 | - | - | 0 | - |
| - | - | 1918 | 586.3 | - | - | 0 | - |
| - | - | 1.073E+04 | 586.8 | - | - | 0 | - |
| - | - | 9610 | 587.3 | - | - | 0 | - |
| - | - | 6987 | 587.8 | - | - | 0 | - |
| - | - | 4468 | 588.3 | - | - | 0 | - |
| - | - | 2526 | 588.3 | - | - | 0 | - |
| - | - | 4103 | 588.8 | - | - | 0 | - |
| - | - | 918.8 | 588.8 | - | - | 0 | - |
| - | - | 2441 | 589.3 | - | - | 0 | - |
| - | - | 1498 | 589.3 | - | - | 0 | - |
| 5 | c | 3.717E+05 | 590.3 | 0.0008266 | 1.4 | +1 | 5 |
| - | - | 1.014E+05 | 591.3 | - | - | 0 | - |
| - | - | 6604 | 591.8 | - | - | 0 | - |
| - | - | 2.277E+04 | 592.3 | - | - | 0 | - |
| - | - | 4798 | 592.8 | - | - | 0 | - |
| - | - | 2212 | 593.3 | - | - | 0 | - |
| - | - | 2131 | 595.3 | - | - | 0 | - |
| - | - | 4554 | 596.3 | - | - | 0 | - |
| - | - | 1147 | 597.3 | - | - | 0 | - |
| 11 | c | 707.5 | 600.3 | 0.002068 | 3.445 | +2 | 11 |
| 11 | c | 1.313E+04 | 600.8 | 0.0003557 | 0.592 | +2 | 11 |
| - | - | 9176 | 601.3 | - | - | 0 | - |
| - | - | 6992 | 601.8 | - | - | 0 | - |
| - | - | 2797 | 602.3 | - | - | 0 | - |
| - | - | 1666 | 602.8 | - | - | 0 | - |
| - | - | 1023 | 603.3 | - | - | 0 | - |
| - | - | 1762 | 605.3 | - | - | 0 | - |
| - | - | 1821 | 607.3 | - | - | 0 | - |
| - | - | 2481 | 607.8 | - | - | 0 | - |
| 8 | z | 2.065E+04 | 608.3 | 0.001382 | 2.271 | +1 | 5 |
| 11 | c | 1.518E+05 | 609.3 | 0.0004478 | 0.7349 | +2 | 11 |
| - | - | 1.323E+05 | 609.8 | - | - | 0 | - |
| - | - | 6.082E+04 | 610.3 | - | - | 0 | - |
| - | - | 2.38E+04 | 610.8 | - | - | 0 | - |
| - | - | 5232 | 611.3 | - | - | 0 | - |
| - | - | 2757 | 616.3 | - | - | 0 | - |
| - | - | 1060 | 617.3 | - | - | 0 | - |
| 2 | w | 7744 | 618.3 | 0.0001207 | 0.1952 | +2 | 11 |
| - | - | 6611 | 618.8 | - | - | 0 | - |
| - | - | 4613 | 619.3 | - | - | 0 | - |
| - | - | 1416 | 621.3 | - | - | 0 | - |
| 8 | y | 2.85E+04 | 624.3 | 0.0006629 | 1.062 | +1 | 5 |
| - | - | 9255 | 625.3 | - | - | 0 | - |
| - | - | 798.6 | 625.8 | - | - | 0 | - |
| - | - | 2553 | 626.3 | - | - | 0 | - |
| - | - | 814.7 | 626.8 | - | - | 0 | - |
| - | - | 765.2 | 628.8 | - | - | 0 | - |
| - | - | 906.2 | 629.3 | - | - | 0 | - |
| - | - | 5254 | 630.3 | - | - | 0 | - |
| - | - | 5171 | 630.8 | - | - | 0 | - |
| - | - | 2547 | 631.3 | - | - | 0 | - |
| 2 | y | 1657 | 632.3 | 0.004116 | 6.509 | +2 | 11 |
| 2 | z | 2862 | 632.8 | 0.00339 | 5.356 | +2 | 11 |
| - | - | 5727 | 633.3 | - | - | 0 | - |
| - | - | 4170 | 633.8 | - | - | 0 | - |
| - | - | 3584 | 634.3 | - | - | 0 | - |
| - | - | 1002 | 634.8 | - | - | 0 | - |
| - | - | 861.2 | 635.4 | - | - | 0 | - |
| - | - | 4368 | 636.3 | - | - | 0 | - |
| - | - | 2318 | 637.3 | - | - | 0 | - |
| - | - | 2223 | 637.8 | - | - | 0 | - |
| - | - | 2080 | 638.3 | - | - | 0 | - |
| - | - | 2405 | 638.3 | - | - | 0 | - |
| - | - | 2565 | 638.8 | - | - | 0 | - |
| - | - | 2290 | 639.3 | - | - | 0 | - |
| - | - | 1089 | 639.3 | - | - | 0 | - |
| - | - | 1289 | 639.8 | - | - | 0 | - |
| - | - | 3351 | 640.3 | - | - | 0 | - |
| - | - | 1076 | 640.3 | - | - | 0 | - |
| 2 | y | 9269 | 640.8 | 6.569E-05 | 0.1025 | +2 | 11 |
| - | - | 7887 | 641.3 | - | - | 0 | - |
| - | - | 3755 | 641.8 | - | - | 0 | - |
| - | - | 1899 | 642.3 | - | - | 0 | - |
| - | - | 2296 | 644.3 | - | - | 0 | - |
| - | - | 7886 | 647.4 | - | - | 0 | - |
| - | - | 1.016E+04 | 647.9 | - | - | 0 | - |
| - | - | 5615 | 648.3 | - | - | 0 | - |
| - | - | 1956 | 648.9 | - | - | 0 | - |
| - | - | 2009 | 652.8 | - | - | 0 | - |
| - | - | 5988 | 653.3 | - | - | 0 | - |
| - | - | 2703 | 653.8 | - | - | 0 | - |
| - | - | 2602 | 654.3 | - | - | 0 | - |
| - | - | 2080 | 654.8 | - | - | 0 | - |
| - | - | 4.721E+04 | 655.3 | - | - | 0 | - |
| - | - | 9994 | 655.8 | - | - | 0 | - |
| - | - | 9.378E+04 | 656.3 | - | - | 0 | - |
| - | - | 2301 | 656.8 | - | - | 0 | - |
| - | - | 2.802E+04 | 657.3 | - | - | 0 | - |
| - | - | 5355 | 658.3 | - | - | 0 | - |
| - | - | 1.377E+04 | 659.3 | - | - | 0 | - |
| - | - | 2.15E+04 | 660.3 | - | - | 0 | - |
| - | - | 1.768E+04 | 660.9 | - | - | 0 | - |
| 7 | z | 2.906E+04 | 661.3 | 0.007975 | 12.06 | +1 | 6 |
| - | - | 2.494E+04 | 661.8 | - | - | 0 | - |
| - | - | 3.261E+04 | 662.3 | - | - | 0 | - |
| - | - | 2.153E+04 | 662.8 | - | - | 0 | - |
| - | - | 8182 | 663.3 | - | - | 0 | - |
| - | - | 1935 | 663.8 | - | - | 0 | - |
| - | - | 8105 | 668.8 | - | - | 0 | - |
| - | - | 2.835E+04 | 669.3 | - | - | 0 | - |
| - | - | 2.549E+04 | 669.8 | - | - | 0 | - |
| - | - | 2.276E+04 | 670.3 | - | - | 0 | - |
| - | - | 1.953E+04 | 670.8 | - | - | 0 | - |
| - | - | 8829 | 671.3 | - | - | 0 | - |
| - | - | 3023 | 671.8 | - | - | 0 | - |
| - | - | 3701 | 672.3 | - | - | 0 | - |
| - | - | 1443 | 673.3 | - | - | 0 | - |
| - | - | 2923 | 673.9 | - | - | 0 | - |
| - | - | 1.034E+04 | 674.3 | - | - | 0 | - |
| - | - | 1.73E+04 | 674.8 | - | - | 0 | - |
| - | - | 1.614E+04 | 675.3 | - | - | 0 | - |
| - | - | 1.706E+04 | 675.8 | - | - | 0 | - |
| - | - | 1.035E+04 | 676.3 | - | - | 0 | - |
| - | - | 5559 | 676.8 | - | - | 0 | - |
| - | - | 1986 | 677.3 | - | - | 0 | - |
| 7 | z | 7.075E+04 | 679.4 | 0.0008889 | 1.308 | +1 | 6 |
| - | - | 3.102E+04 | 680.4 | - | - | 0 | - |
| - | - | 1.234E+04 | 681.3 | - | - | 0 | - |
| - | - | 2252 | 681.9 | - | - | 0 | - |
| - | - | 3547 | 682.3 | - | - | 0 | - |
| - | - | 5323 | 682.9 | - | - | 0 | - |
| - | - | 1.012E+05 | 683.3 | - | - | 0 | - |
| - | - | 9.911E+04 | 683.8 | - | - | 0 | - |
| - | - | 5.548E+04 | 684.3 | - | - | 0 | - |
| - | - | 1.666E+04 | 684.8 | - | - | 0 | - |
| - | - | 4571 | 685.3 | - | - | 0 | - |
| 6 | c | 3305 | 687.3 | 0.009045 | 13.16 | +1 | 6 |
| - | - | 1249 | 688.3 | - | - | 0 | - |
| - | - | 3563 | 690.4 | - | - | 0 | - |
| - | - | 2076 | 690.9 | - | - | 0 | - |
| - | - | 8.502E+04 | 691.4 | - | - | 0 | - |
| - | - | 9.02E+04 | 691.9 | - | - | 0 | - |
| - | - | 5.424E+04 | 692.4 | - | - | 0 | - |
| - | - | 2.154E+04 | 692.9 | - | - | 0 | - |
| - | - | 4680 | 693.4 | - | - | 0 | - |
| - | - | 1115 | 694.3 | - | - | 0 | - |
| 7 | y | 1.489E+04 | 695.4 | 0.0005364 | 0.7714 | +1 | 6 |
| - | - | 6607 | 696.4 | - | - | 0 | - |
| - | - | 1485 | 697.4 | - | - | 0 | - |
| - | - | 1037 | 703.4 | - | - | 0 | - |
| 6 | c | 8.691E+04 | 704.4 | 0.0008679 | 1.232 | +1 | 6 |
| - | - | 3.626E+04 | 705.4 | - | - | 0 | - |
| - | - | 9132 | 706.4 | - | - | 0 | - |
| - | - | 2248 | 707.4 | - | - | 0 | - |
| - | - | 1341 | 711.3 | - | - | 0 | - |
| - | - | 1021 | 719.4 | - | - | 0 | - |
| - | - | 1071 | 723.3 | - | - | 0 | - |
| - | - | 1150 | 730.4 | - | - | 0 | - |
| - | - | 1.171E+04 | 731.4 | - | - | 0 | - |
| - | - | 5273 | 732.4 | - | - | 0 | - |
| - | - | 2559 | 733.4 | - | - | 0 | - |
| - | - | 2206 | 737.3 | - | - | 0 | - |
| - | - | 1230 | 738.3 | - | - | 0 | - |
| - | - | 1125 | 739.4 | - | - | 0 | - |
| - | - | 867.8 | 740.4 | - | - | 0 | - |
| - | - | 834.1 | 741.4 | - | - | 0 | - |
| 6 | w | 3.807E+04 | 749.4 | 0.002362 | 3.152 | +1 | 7 |
| - | - | 5.295E+04 | 750.4 | - | - | 0 | - |
| - | - | 1.906E+04 | 751.4 | - | - | 0 | - |
| - | - | 3404 | 752.4 | - | - | 0 | - |
| - | - | 2807 | 754.4 | - | - | 0 | - |
| - | - | 1.087E+04 | 755.4 | - | - | 0 | - |
| - | - | 4147 | 756.4 | - | - | 0 | - |
| 7 | c | 1772 | 757.4 | 0.01476 | 19.48 | +1 | 7 |
| 7 | c | 7721 | 758.4 | 0.001717 | 2.264 | +1 | 7 |
| - | - | 3778 | 759.4 | - | - | 0 | - |
| - | - | 1217 | 760.4 | - | - | 0 | - |
| - | - | 2050 | 763.4 | - | - | 0 | - |
| - | - | 1182 | 764.4 | - | - | 0 | - |
| - | - | 1394 | 774.4 | - | - | 0 | - |
| 7 | c | 7.934E+04 | 775.4 | 0.0006804 | 0.8774 | +1 | 7 |
| - | - | 3.654E+04 | 776.4 | - | - | 0 | - |
| - | - | 1.043E+04 | 777.4 | - | - | 0 | - |
| - | - | 1847 | 778.4 | - | - | 0 | - |
| - | - | 6979 | 779.4 | - | - | 0 | - |
| - | - | 3.886E+04 | 780.4 | - | - | 0 | - |
| - | - | 1.476E+04 | 781.4 | - | - | 0 | - |
| - | - | 3116 | 782.4 | - | - | 0 | - |
| - | - | 1014 | 786.4 | - | - | 0 | - |
| - | - | 3098 | 787.4 | - | - | 0 | - |
| - | - | 1943 | 788.4 | - | - | 0 | - |
| - | - | 1068 | 789.4 | - | - | 0 | - |
| 6 | z | 1.91E+05 | 793.4 | 0.0008692 | 1.096 | +1 | 7 |
| - | - | 9.496E+04 | 794.4 | - | - | 0 | - |
| - | - | 2.965E+04 | 795.4 | - | - | 0 | - |
| - | - | 6630 | 796.4 | - | - | 0 | - |
| - | - | 1274 | 797.4 | - | - | 0 | - |
| - | - | 1318 | 808.4 | - | - | 0 | - |
| 6 | y | 2817 | 809.4 | 0.001131 | 1.398 | +1 | 7 |
| - | - | 4569 | 810.4 | - | - | 0 | - |
| - | - | 5883 | 811.4 | - | - | 0 | - |
| - | - | 1047 | 812.4 | - | - | 0 | - |
| - | - | 1913 | 821.4 | - | - | 0 | - |
| - | - | 1152 | 822.4 | - | - | 0 | - |
| - | - | 2194 | 830.4 | - | - | 0 | - |
| - | - | 1256 | 831.4 | - | - | 0 | - |
| - | - | 3017 | 832.4 | - | - | 0 | - |
| - | - | 2454 | 833.4 | - | - | 0 | - |
| - | - | 1738 | 838.4 | - | - | 0 | - |
| - | - | 1442 | 852.4 | - | - | 0 | - |
| - | - | 1220 | 855.4 | - | - | 0 | - |
| - | - | 2766 | 858.4 | - | - | 0 | - |
| - | - | 1123 | 858.5 | - | - | 0 | - |
| - | - | 8119 | 859.4 | - | - | 0 | - |
| - | - | 5758 | 859.5 | - | - | 0 | - |
| - | - | 2759 | 860.4 | - | - | 0 | - |
| - | - | 5516 | 860.5 | - | - | 0 | - |
| - | - | 3082 | 861.5 | - | - | 0 | - |
| - | - | 1079 | 862.5 | - | - | 0 | - |
| - | - | 5.82E+04 | 864.4 | - | - | 0 | - |
| - | - | 4.113E+04 | 865.4 | - | - | 0 | - |
| - | - | 1.616E+04 | 866.4 | - | - | 0 | - |
| - | - | 4151 | 867.4 | - | - | 0 | - |
| - | - | 1295 | 873.4 | - | - | 0 | - |
| - | - | 2833 | 874.4 | - | - | 0 | - |
| - | - | 1202 | 875.4 | - | - | 0 | - |
| - | - | 3632 | 884.5 | - | - | 0 | - |
| 8 | c | 4367 | 885.5 | 0.003518 | 3.973 | +1 | 8 |
| 8 | c | 9843 | 886.5 | 0.001114 | 1.257 | +1 | 8 |
| - | - | 5075 | 887.5 | - | - | 0 | - |
| - | - | 3160 | 888.5 | - | - | 0 | - |
| - | - | 909 | 889.5 | - | - | 0 | - |
| - | - | 1052 | 899.5 | - | - | 0 | - |
| - | - | 6069 | 900.5 | - | - | 0 | - |
| - | - | 3908 | 901.5 | - | - | 0 | - |
| - | - | 6.105E+04 | 902.4 | - | - | 0 | - |
| - | - | 1.461E+04 | 902.5 | - | - | 0 | - |
| - | - | 1.823E+04 | 903.4 | - | - | 0 | - |
| 8 | c | 9.037E+04 | 903.5 | 0.0001387 | 0.1535 | +1 | 8 |
| - | - | 5.182E+04 | 904.5 | - | - | 0 | - |
| - | - | 1.895E+04 | 905.5 | - | - | 0 | - |
| - | - | 4532 | 906.5 | - | - | 0 | - |
| 5 | z | 5.977E+04 | 908.4 | 0.0004764 | 0.5245 | +1 | 8 |
| - | - | 4.189E+04 | 909.4 | - | - | 0 | - |
| - | - | 1.688E+04 | 910.4 | - | - | 0 | - |
| - | - | 5126 | 911.4 | - | - | 0 | - |
| - | - | 1713 | 912.4 | - | - | 0 | - |
| - | - | 3.377E+04 | 918.4 | - | - | 0 | - |
| - | - | 1.116E+05 | 919.4 | - | - | 0 | - |
| - | - | 4.949E+04 | 920.5 | - | - | 0 | - |
| - | - | 1.485E+04 | 921.5 | - | - | 0 | - |
| - | - | 3948 | 922.5 | - | - | 0 | - |
| - | - | 1207 | 923.5 | - | - | 0 | - |
| 5 | y | 4379 | 924.4 | 0.003481 | 3.765 | +1 | 8 |
| - | - | 3244 | 925.4 | - | - | 0 | - |
| - | - | 1914 | 926.4 | - | - | 0 | - |
| - | - | 1412 | 957.5 | - | - | 0 | - |
| - | - | 1276 | 959.5 | - | - | 0 | - |
| - | - | 1109 | 960.5 | - | - | 0 | - |
| - | - | 1038 | 961.5 | - | - | 0 | - |
| - | - | 1131 | 968.5 | - | - | 0 | - |
| - | - | 3026 | 969.5 | - | - | 0 | - |
| - | - | 1430 | 970.5 | - | - | 0 | - |
| - | - | 1917 | 974.5 | - | - | 0 | - |
| - | - | 1670 | 975.5 | - | - | 0 | - |
| - | - | 4720 | 986.5 | - | - | 0 | - |
| - | - | 6502 | 987.5 | - | - | 0 | - |
| - | - | 5708 | 988.5 | - | - | 0 | - |
| - | - | 1831 | 989.5 | - | - | 0 | - |
| 9 | c | 3942 | 999.5 | 0.0004298 | 0.43 | +1 | 9 |
| 9 | c | 6379 | 1001 | 0.006038 | 6.035 | +1 | 9 |
| - | - | 6044 | 1002 | - | - | 0 | - |
| - | - | 3583 | 1003 | - | - | 0 | - |
| - | - | 3365 | 1004 | - | - | 0 | - |
| - | - | 2053 | 1005 | - | - | 0 | - |
| - | - | 7845 | 1017 | - | - | 0 | - |
| 9 | c | 5.578E+04 | 1018 | 0.0006745 | 0.6628 | +1 | 9 |
| - | - | 4.056E+04 | 1019 | - | - | 0 | - |
| - | - | 2.138E+04 | 1020 | - | - | 0 | - |
| - | - | 7785 | 1021 | - | - | 0 | - |
| - | - | 2655 | 1022 | - | - | 0 | - |
| - | - | 1472 | 1023 | - | - | 0 | - |
| - | - | 1210 | 1024 | - | - | 0 | - |
| - | - | 870.6 | 1048 | - | - | 0 | - |
| - | - | 1507 | 1049 | - | - | 0 | - |
| - | - | 1735 | 1061 | - | - | 0 | - |
| - | - | 2461 | 1062 | - | - | 0 | - |
| - | - | 1679 | 1063 | - | - | 0 | - |
| 4 | y | 1402 | 1064 | 0.01394 | 13.11 | +1 | 9 |
| 4 | z | 2.272E+04 | 1065 | 0.0002565 | 0.241 | +1 | 9 |
| - | - | 3.359E+04 | 1066 | - | - | 0 | - |
| - | - | 2.263E+04 | 1067 | - | - | 0 | - |
| - | - | 6947 | 1068 | - | - | 0 | - |
| - | - | 2286 | 1069 | - | - | 0 | - |
| 4 | y | 2044 | 1081 | 0.004674 | 4.325 | +1 | 9 |
| - | - | 2073 | 1082 | - | - | 0 | - |
| - | - | 1330 | 1083 | - | - | 0 | - |
| - | - | 3720 | 1089 | - | - | 0 | - |
| - | - | 3829 | 1090 | - | - | 0 | - |
| - | - | 1682 | 1091 | - | - | 0 | - |
| - | - | 1268 | 1092 | - | - | 0 | - |
| - | - | 1096 | 1094 | - | - | 0 | - |
| - | - | 1682 | 1095 | - | - | 0 | - |
| - | - | 1423 | 1097 | - | - | 0 | - |
| 10 | c | 1.751E+04 | 1105 | 4.907E-05 | 0.04442 | +1 | 10 |
| - | - | 1.399E+04 | 1106 | - | - | 0 | - |
| - | - | 9146 | 1107 | - | - | 0 | - |
| - | - | 3880 | 1108 | - | - | 0 | - |
| - | - | 903.7 | 1109 | - | - | 0 | - |
| 3 | w | 1822 | 1135 | 0.001241 | 1.094 | +1 | 10 |
| - | - | 1953 | 1136 | - | - | 0 | - |
| - | - | 1286 | 1137 | - | - | 0 | - |
| 3 | z | 1.596E+04 | 1152 | 8.845E-05 | 0.07681 | +1 | 10 |
| - | - | 3.507E+04 | 1153 | - | - | 0 | - |
| - | - | 2.291E+04 | 1154 | - | - | 0 | - |
| - | - | 9510 | 1155 | - | - | 0 | - |
| - | - | 3249 | 1156 | - | - | 0 | - |
| - | - | 1314 | 1158 | - | - | 0 | - |
| - | - | 1905 | 1159 | - | - | 0 | - |
| - | - | 3085 | 1160 | - | - | 0 | - |
| - | - | 1159 | 1161 | - | - | 0 | - |
| - | - | 1904 | 1169 | - | - | 0 | - |
| - | - | 1868 | 1170 | - | - | 0 | - |
| - | - | 3695 | 1174 | - | - | 0 | - |
| - | - | 7734 | 1175 | - | - | 0 | - |
| - | - | 8128 | 1176 | - | - | 0 | - |
| - | - | 4187 | 1177 | - | - | 0 | - |
| - | - | 1752 | 1178 | - | - | 0 | - |
| - | - | 959.7 | 1200 | - | - | 0 | - |
| - | - | 1.188E+04 | 1202 | - | - | 0 | - |
| - | - | 9079 | 1203 | - | - | 0 | - |
| - | - | 6315 | 1204 | - | - | 0 | - |
| - | - | 2276 | 1205 | - | - | 0 | - |
| 11 | c | 1.051E+04 | 1218 | 0.001349 | 1.108 | +1 | 11 |
| - | - | 2.509E+04 | 1219 | - | - | 0 | - |
| - | - | 1.587E+04 | 1220 | - | - | 0 | - |
| - | - | 8300 | 1221 | - | - | 0 | - |
| - | - | 1960 | 1222 | - | - | 0 | - |
| 2 | z | 1020 | 1265 | 0.00878 | 6.943 | +1 | 11 |
| - | - | 4420 | 1266 | - | - | 0 | - |
| - | - | 4845 | 1267 | - | - | 0 | - |
| - | - | 3378 | 1268 | - | - | 0 | - |
| - | - | 923.1 | 1280 | - | - | 0 | - |
| - | - | 1069 | 1295 | - | - | 0 | - |
| - | - | 1046 | 1297 | - | - | 0 | - |
| - | - | 4880 | 1306 | - | - | 0 | - |
| - | - | 5678 | 1307 | - | - | 0 | - |
| - | - | 6077 | 1308 | - | - | 0 | - |
| - | - | 3627 | 1309 | - | - | 0 | - |
| - | - | 1523 | 1310 | - | - | 0 | - |
| - | - | 1101 | 1311 | - | - | 0 | - |
| - | - | 3371 | 1321 | - | - | 0 | - |
| - | - | 6819 | 1322 | - | - | 0 | - |
| - | - | 7486 | 1323 | - | - | 0 | - |
| - | - | 1.41E+04 | 1324 | - | - | 0 | - |
| - | - | 2.429E+04 | 1325 | - | - | 0 | - |
| - | - | 1.796E+04 | 1326 | - | - | 0 | - |
| - | - | 9162 | 1327 | - | - | 0 | - |
| - | - | 2047 | 1328 | - | - | 0 | - |
| - | - | 4947 | 1338 | - | - | 0 | - |
| - | - | 3.006E+04 | 1339 | - | - | 0 | - |
| - | - | 2.686E+04 | 1340 | - | - | 0 | - |
| - | - | 1.759E+04 | 1341 | - | - | 0 | - |
| - | - | 5934 | 1342 | - | - | 0 | - |
| - | - | 1999 | 1343 | - | - | 0 | - |
| - | - | 8481 | 1349 | - | - | 0 | - |
| - | - | 1.493E+04 | 1350 | - | - | 0 | - |
| - | - | 1.148E+04 | 1351 | - | - | 0 | - |
| - | - | 7441 | 1352 | - | - | 0 | - |
| - | - | 1903 | 1353 | - | - | 0 | - |
| - | - | 1454 | 1356 | - | - | 0 | - |
| - | - | 1954 | 1357 | - | - | 0 | - |
| - | - | 1255 | 1358 | - | - | 0 | - |
| - | - | 1150 | 1364 | - | - | 0 | - |
| - | - | 1169 | 1365 | - | - | 0 | - |
| - | - | 1.763E+04 | 1366 | - | - | 0 | - |
| - | - | 1.013E+05 | 1367 | - | - | 0 | - |
| - | - | 9.609E+04 | 1368 | - | - | 0 | - |
| - | - | 5.475E+04 | 1369 | - | - | 0 | - |
| - | - | 2.065E+04 | 1370 | - | - | 0 | - |
| - | - | 3740 | 1371 | - | - | 0 | - |
| - | - | 1665 | 1381 | - | - | 0 | - |
| - | - | 4712 | 1382 | - | - | 0 | - |
| - | - | 1.464E+04 | 1383 | - | - | 0 | - |
| - | - | 2.554E+04 | 1384 | - | - | 0 | - |
| - | - | 1.98E+04 | 1385 | - | - | 0 | - |
| - | - | 1.089E+04 | 1386 | - | - | 0 | - |
| - | - | 3030 | 1387 | - | - | 0 | - |
| - | - | 1141 | 1388 | - | - | 0 | - |
| - | - | 860.1 | 2680 | - | - | 0 | - |
| - | - | 841.5 | 3124 | - | - | 0 | - |

m/z Charge Intensity FragmentType MassShift Position
122.37100219726562 0 426.6859
123.10454559326172 0 901.0238
129.10272216796875 0 1520.5752
131.0865020751953 0 476.2706
132.07696533203125 0 512.7776
136.0760498046875 0 46225.684
137.0735321044922 0 822.3185
137.07940673828125 0 8227.186
140.05838012695312 0 2814.6243
141.0663299560547 0 2077.2393
142.12310791015625 0 1258.822
143.1182403564453 0 1530.6038
149.04527282714844 0 4434.574
156.0771026611328 0 13185.071
157.0804443359375 0 2434.9219
159.46250915527344 0 516.50256
165.05496215820312 0 12221.281
166.0583038330078 0 984.1619
167.055419921875 0 1209.8068
169.1343231201172 0 678
171.1250457763672 0 570.2091
173.1289520263672 0 1849.3124
173.4510040283203 0 2704.8984
182.08155822753906 0 44666.6 y 11
183.08499145507812 0 4901.873
186.1129913330078 0 1128.4471
187.14450073242188 0 91107.44
188.14036560058594 0 609.93414
188.14788818359375 0 8511.663
190.12289428710938 0 948.8209
193.75306701660156 0 595.9772
199.16973876953125 0 636.3821
200.13987731933594 0 1171.0002
201.12384033203125 0 3979.8606
201.13429260253906 0 1340.2354
202.08291625976562 0 1459.4985
203.13914489746094 0 773.47253
213.14862060546875 0 632.1375
215.13938903808594 0 25375.65
216.14283752441406 0 2610.939
219.1346435546875 0 652.29663
221.0849151611328 0 2458.6714
222.0853271484375 0 1038.9491
223.06446838378906 0 1507.3146
225.04318237304688 0 1720.1067
229.12991333007812 0 1627.3054
232.16575622558594 0 703.03705 c 1
235.14459228515625 0 65439.85
236.1479034423828 0 31414.295
237.15032958984375 0 3113.9167
238.11935424804688 0 1514.1556
239.09530639648438 0 2035.5726
240.0960235595703 0 1795.646
241.09278869628906 0 1152.4601
244.14102172851562 0 1011.23615
245.1617431640625 0 655.5825
249.42337036132812 0 564.73773
255.14547729492188 0 7020.5356
256.1488342285156 0 2041.44
257.1495666503906 0 686.8702
260.1238098144531 0 1354.2872
263.1393737792969 0 23892.264
263.966552734375 0 639.46356
263.9815673828125 0 623.70215
264.14276123046875 0 12307.29
265.1454772949219 0 1525.7229
266.1877746582031 0 625.12506
269.1608581542969 0 645.916
272.12518310546875 0 1451.4312
273.16497802734375 0 646.7121
282.1449279785156 0 3760.0383
287.1713562011719 0 971.0472
295.1036682128906 0 1743.6199
295.1658935546875 0 3399.3943 y 10
296.1045227050781 0 2583.7876
297.1012268066406 0 1699.9628
297.15594482421875 0 1067.1388
298.14080810546875 0 1065.0701
299.0625 0 1946.851
299.1481628417969 0 1343.1924
299.1717529296875 0 1337.6622
300.0627136230469 0 1756.3838
300.1558837890625 0 11851.777
300.17431640625 0 750.2382
301.0600891113281 0 1391.0287
301.1401062011719 0 651.41034
301.1590270996094 0 1894.7152
302.1718444824219 0 1064.4661
309.15594482421875 0 3261.1775
310.15972900390625 0 1834.9019
315.167236328125 0 3913.1426
317.1824645996094 0 7709.8906
319.1981201171875 0 7330.2393 c 2
341.0185546875 0 13877.755
355.0699157714844 0 1653.7352
359.0289611816406 0 42306.223
366.1789855957031 0 964.3701 z 9
367.18426513671875 0 2008.7891
368.1695556640625 0 8386.606
369.1230773925781 0 1045.9706
369.17498779296875 0 9599.669
370.1236267089844 0 3489.7146
370.178955078125 0 5469.979
371.1207580566406 0 2917.5286
371.18304443359375 0 727.4989
372.69732666015625 0 2033.1953
374.1811828613281 0 1160.7903
375.8612060546875 0 787.0159
378.1660461425781 0 2634.3406
378.1909484863281 0 1271.8677
379.1708679199219 0 2494.1758
382.19891357421875 0 1905.9312 y 9
384.1879577636719 0 3948.477 y Ammonia loss 2
385.19073486328125 0 2421.2449
385.2198791503906 0 8830.998
385.5480041503906 0 3246.4524
385.8796081542969 0 2101.459
386.22369384765625 0 2554.076
386.5454406738281 0 1024.2764
387.2324523925781 0 722.9704
391.55133056640625 0 7763.3975
391.8837890625 0 6553.04
392.21539306640625 0 2484.6707
394.7027893066406 0 1301.6588
400.2563171386719 0 921.38403
400.8829040527344 0 821.19507 c Ammonia loss 10
401.166748046875 0 3631.4314
402.1705017089844 0 2336.8452
403.1727600097656 0 722.10065
406.1954650878906 0 835.76807
408.2250061035156 0 2191.8374
409.2291564941406 0 814.4145
413.23968505859375 0 1355.5178
414.24261474609375 0 827.7559
415.2669982910156 0 2019.8846
416.2022705078125 0 945.7011
425.2510070800781 0 1231.4965
426.2342834472656 0 928.6758
429.089599609375 0 113984.35
429.7541198730469 0 2444.7979
430.242919921875 0 4196.2617
430.2794189453125 0 1722.4077
431.2834167480469 0 911.4051
435.1868896484375 0 919.32745
435.23193359375 0 721.294
436.1878356933594 0 828.0988
437.2139587402344 0 2397.9507
438.7337951660156 0 1745.3318
439.2347106933594 0 1432.3617
442.74578857421875 0 4545.3545
443.24151611328125 0 3430.6519 c Water loss 7
443.2635803222656 0 1895.76
443.74200439453125 0 4144.881 c Ammonia loss 7
444.2420959472656 0 2233.0261
445.1208801269531 0 63225.047
446.2132873535156 0 5449.1357
447.1022033691406 0 744.0507
450.71954345703125 0 962.74347
451.21075439453125 0 1303.7721
451.7148132324219 0 1237.46
451.7508544921875 0 18144.018
452.25433349609375 0 62726.07 c 7
452.7538757324219 0 29280.426
453.25286865234375 0 7119.407
453.756103515625 0 2555.3723
454.2529296875 0 893.16425
455.23663330078125 0 831.3528
455.5682373046875 0 1053.043
456.2255554199219 0 1289.7312
457.2275695800781 0 1139.9386
457.7537841796875 0 944.6593
458.2376708984375 0 889.51263
458.27313232421875 0 3050.148 c Ammonia loss 3
459.27130126953125 0 918.85236
459.72412109375 0 11164.341
459.75439453125 0 912.0996
460.2270202636719 0 5479.365
460.7257385253906 0 2005.9735
461.2390441894531 0 1781.8561
461.57440185546875 0 1925.9453
461.90496826171875 0 895.3025
465.2537841796875 0 780.3465
465.7648010253906 0 1367.5667
466.253173828125 0 1353.154
466.7316589355469 0 1167.1774
467.23681640625 0 4181.949
468.2426452636719 0 5824.793
469.2479248046875 0 2446.263
470.75091552734375 0 1539.1866
471.2477722167969 0 1806.747
471.7432556152344 0 1311.7457
472.2489013671875 0 2192.5405
473.24603271484375 0 1180.5276
475.2997131347656 0 196135.02 c 3
476.30242919921875 0 43899.42
477.23443603515625 0 882.1146
477.30517578125 0 7042.0728
478.3097229003906 0 899.6214
479.2601623535156 0 1353.3427
479.7584533691406 0 19195.404
480.22064208984375 0 1912.9519 z 8
480.25665283203125 0 10681.486
480.7555847167969 0 4836.955
481.23004150390625 0 13776.938
481.7521057128906 0 725.79987
482.2314758300781 0 3377.4175
483.2569580078125 0 1106.5482
484.260498046875 0 3147.7825
484.74945068359375 0 1820.9136
485.24517822265625 0 2226.5774
485.7414855957031 0 1235.5935
486.7711181640625 0 2605.6624
487.269775390625 0 5831.5728
487.7666320800781 0 3217.2607
488.2655334472656 0 1084.9886
492.25347900390625 0 887.96356
493.7552490234375 0 4182.699
494.2527770996094 0 3166.669
494.7547607421875 0 1135.8853
495.2322692871094 0 1206.7864
495.7285461425781 0 888.5629
496.2410888671875 0 6612.7207 y 8
497.2267761230469 0 977.4834
499.7671813964844 0 20802.023
500.2662353515625 0 13681.645 c Water loss 8
500.7638854980469 0 10366.479 c Ammonia loss 8
501.2618713378906 0 6467.7617
501.7672119140625 0 3795.0137
502.2371520996094 0 1314.3049
502.27490234375 0 21498.783
502.76556396484375 0 1285.6051
503.2752380371094 0 3657.9458
508.2644348144531 0 1466.8667
508.7724304199219 0 48165.504
509.2740173339844 0 82314.37 c 8
509.7723083496094 0 58299.027
509.83441162109375 0 790.62823
510.2711181640625 0 19347.068
510.7719421386719 0 7912.0024
511.2701110839844 0 2679.7788
521.2713623046875 0 1849.5048
521.7780151367188 0 926.238
522.7767944335938 0 1389.3794
523.268310546875 0 1819.1771
524.2587280273438 0 2624.5696 z Ammonia loss 3
525.2649536132812 0 7067.3613
526.267333984375 0 7225.177
526.7570190429688 0 2504.8054
527.273193359375 0 788.2534
530.288330078125 0 3988.06
530.7789306640625 0 1481.3524
531.2984619140625 0 1015.78796
532.7667846679688 0 7238.591 z 3
533.2634887695312 0 4717.6104
533.7636108398438 0 2244.5776
534.25830078125 0 2246.7976
535.271240234375 0 4468.8945
535.7677001953125 0 4956.35
536.2642822265625 0 1364.8188
536.8028564453125 0 969.6409
540.2781982421875 0 13402.555
540.7747192382812 0 2967.2263 y 3
541.2805786132812 0 18752.555
541.774658203125 0 876.33484
542.2841796875 0 5189.264
543.2821044921875 0 1151.5896
544.2791748046875 0 8984.5205 c Ammonia loss 9
544.7778930664062 0 5931.9307
545.2643432617188 0 1237.8011
545.3043212890625 0 6463.374
545.7725219726562 0 919.2794
546.3106689453125 0 5843.309
547.3115844726562 0 1074.3931
548.25146484375 0 8265.446
548.7505493164062 0 4575.2524
549.251708984375 0 3512.2458
549.7527465820312 0 1039.746
550.2938232421875 0 947.672
551.2825927734375 0 1782.3097
552.2881469726562 0 2121.7747
552.7921142578125 0 59386.04 c 9
553.2890014648438 0 51304.418
553.7879638671875 0 25758.744
554.2859497070312 0 11109.163
554.784423828125 0 2346.6426
555.2861328125 0 877.3503
558.7989501953125 0 717.8205
559.3005981445312 0 749.08545
561.2791748046875 0 1071.568
561.7858276367188 0 935.63806
564.304443359375 0 6231.334
565.31005859375 0 3021.919
565.8002319335938 0 3457.842
566.301513671875 0 3445.5857
566.7774047851562 0 2068.2766
567.2666625976562 0 1201.4663 z Water loss 2
567.7786254882812 0 2154.9517 z Ammonia loss 2
568.2755737304688 0 2180.7236
568.7656860351562 0 918.9494
569.2706298828125 0 843.9559
569.8065795898438 0 1078.7021
573.3051147460938 0 2770.4302 c Ammonia loss 4
574.3013305664062 0 1181.3568
575.2916259765625 0 1860.832 y Water loss 2
575.7792358398438 0 4963.3135 y Ammonia loss 2
576.2825317382812 0 22358.148 z 2
576.78076171875 0 17803.15
577.2808227539062 0 7834.0347
577.7799682617188 0 1952.3308
577.822998046875 0 1322.7926
578.31103515625 0 3846.3489
578.814697265625 0 2583.1719
579.3065185546875 0 2396.3423
579.8097534179688 0 2191.601
583.30419921875 0 3518.9885
583.79833984375 0 2283.1858
584.2920532226562 0 69012.99 y 2
584.7896728515625 0 60035.02
585.2888793945312 0 30404.662
585.78857421875 0 11531.045
586.2891235351562 0 1918.0571
586.8232421875 0 10731.814
587.3233642578125 0 9610.449
587.8223876953125 0 6987.396
588.275146484375 0 4467.71
588.3221435546875 0 2525.5576
588.7736206054688 0 4102.7603
588.8221435546875 0 918.7819
589.2744140625 0 2441.0376
589.3203735351562 0 1497.6685
590.3264770507812 0 371708.44 c 4
591.3291015625 0 101440.414
591.815185546875 0 6603.549
592.3287963867188 0 22766.04
592.8093872070312 0 4798.4785
593.3286743164062 0 2211.6924
595.2985229492188 0 2130.5464
596.300048828125 0 4553.531
597.3016357421875 0 1147.2959
600.3302612304688 0 707.5281 c Water loss 10
600.820556640625 0 13128.724 c Ammonia loss 10
601.3201293945312 0 9176.315
601.8216552734375 0 6991.7397
602.3070068359375 0 2796.737
602.7922973632812 0 1665.6594
603.3113403320312 0 1023.30035
605.296630859375 0 1762.118
607.3273315429688 0 1821.085
607.8264770507812 0 2480.6904
608.3178100585938 0 20648.01 z 7
609.3339233398438 0 151755.36 c 10
609.8316650390625 0 132334.08
610.3308715820312 0 60816.9
610.830810546875 0 23797.145
611.3280639648438 0 5231.5596
616.3414916992188 0 2756.6226
617.3414306640625 0 1060.4949
618.3045043945312 0 7743.954 w 1
618.802734375 0 6610.557
619.3012084960938 0 4612.9473
621.3269653320312 0 1416.419
624.3358154296875 0 28498.545 y 7
625.332275390625 0 9254.92
625.8113403320312 0 798.64264
626.33251953125 0 2552.6255
626.80615234375 0 814.72546
628.7562255859375 0 765.2333
629.3310546875 0 906.2474
630.3292846679688 0 5254.1543
630.824462890625 0 5171.233
631.3296508789062 0 2546.8188
632.31591796875 0 1656.9458 y Ammonia loss 1
632.820556640625 0 2861.9094 z 1
633.3209838867188 0 5726.77
633.8201904296875 0 4170.252
634.3185424804688 0 3583.9258
634.8192749023438 0 1002.3309
635.357421875 0 861.158
636.3497314453125 0 4368.122
637.303955078125 0 2318.2922
637.8333129882812 0 2222.846
638.2966918945312 0 2079.964
638.3451538085938 0 2405.4832
638.8355712890625 0 2564.981
639.2910766601562 0 2289.8506
639.34716796875 0 1088.7864
639.8348388671875 0 1289.2146
640.291015625 0 3350.6108
640.3444213867188 0 1076.1644
640.8333740234375 0 9268.843 y 1
641.330810546875 0 7886.742
641.8291015625 0 3754.5466
642.3299560546875 0 1898.8644
644.3414916992188 0 2296.2974
647.3514404296875 0 7886.169
647.85107421875 0 10160.586
648.3499145507812 0 5615.4736
648.8547973632812 0 1956.033
652.8482666015625 0 2008.6918
653.339111328125 0 5987.5376
653.8366088867188 0 2703.3552
654.3364868164062 0 2601.7117
654.8298950195312 0 2079.7192
655.3075561523438 0 47213.445
655.8145751953125 0 9994.188
656.3084716796875 0 93777.71
656.8150024414062 0 2301.1235
657.3111572265625 0 28017.799
658.3134155273438 0 5355.2915
659.3473510742188 0 13767.611
660.3465576171875 0 21498.559
660.8514404296875 0 17680.557
661.3509521484375 0 29055.703 z Water loss 6
661.8491821289062 0 24941.879
662.3455810546875 0 32608.35
662.8440551757812 0 21533.178
663.3434448242188 0 8181.849
663.8419799804688 0 1934.8835
668.8303833007812 0 8105.392
669.3482666015625 0 28350.656
669.8489990234375 0 25491.377
670.3473510742188 0 22758.857
670.8455200195312 0 19534.332
671.34326171875 0 8829.375
671.8399658203125 0 3023.2427
672.3453369140625 0 3701.376
673.34326171875 0 1442.961
673.8519287109375 0 2922.5771
674.3458862304688 0 10344.743
674.8450317382812 0 17298.576
675.3427124023438 0 16137.548
675.8406372070312 0 17060.137
676.3359985351562 0 10350.871
676.8336791992188 0 5558.9595
677.3348388671875 0 1986.2571
679.3544311523438 0 70749.66 z 6
680.3508911132812 0 31018.07
681.3340454101562 0 12340.38
681.874755859375 0 2251.8975
682.3342895507812 0 3546.6223
682.8555297851562 0 5323.3335
683.3486938476562 0 101184.26
683.8472900390625 0 99110.664
684.3463134765625 0 55481.895
684.8451538085938 0 16663.514
685.3447875976562 0 4571.261
687.35107421875 0 3304.629 c Ammonia loss 5
688.3356323242188 0 1249.4362
690.3812255859375 0 3562.9973
690.8816528320312 0 2076.347
691.3579711914062 0 85015.08
691.8562622070312 0 90204.25
692.3557739257812 0 54244.67
692.854736328125 0 21541.273
693.352783203125 0 4679.9526
694.3341064453125 0 1115.2487
695.372802734375 0 14892.593 y 6
696.367431640625 0 6607.099
697.36669921875 0 1484.7576
703.3575439453125 0 1036.6034
704.3694458007812 0 86909.9 c 5
705.3655395507812 0 36256.156
706.36767578125 0 9131.871
707.368408203125 0 2248.3745
711.341796875 0 1341.2242
719.3655395507812 0 1020.91907
723.3367919921875 0 1070.9943
730.3861083984375 0 1150.3137
731.3921508789062 0 11710.161
732.3861694335938 0 5273.0044
733.3840942382812 0 2558.9026
737.3336791992188 0 2205.5447
738.334716796875 0 1230.1165
739.3576049804688 0 1124.8516
740.3626098632812 0 867.8096
741.3561401367188 0 834.05505
749.3851928710938 0 38070.95 w 5
750.3892211914062 0 52954.11
751.3909912109375 0 19057.8
752.3922119140625 0 3404.2153
754.3763427734375 0 2806.8586
755.3765869140625 0 10870.335
756.3819580078125 0 4147.3574
757.38037109375 0 1771.7242 c Water loss 6
758.380859375 0 7720.5645 c Ammonia loss 6
759.3819580078125 0 3778.1255
760.3809204101562 0 1217.2363
763.3702392578125 0 2049.9763
764.3707885742188 0 1181.9685
774.3970336914062 0 1393.5347
775.4063720703125 0 79342.34 c 6
776.4022216796875 0 36544.598
777.4021606445312 0 10426.81
778.3953857421875 0 1846.8955
779.3948974609375 0 6978.7847
780.397216796875 0 38858.1
781.3995971679688 0 14758.606
782.3988037109375 0 3116.112
786.3843383789062 0 1013.6241
787.3909301757812 0 3097.5479
788.3912963867188 0 1942.6079
789.3895263671875 0 1068.0963
793.3973388671875 0 191046.48 z 5
794.3948974609375 0 94963.73
795.3930053710938 0 29653.473
796.3915405273438 0 6629.626
797.3886108398438 0 1274.4154
808.3768920898438 0 1317.8193
809.4140625 0 2817.4563 y 5
810.4041137695312 0 4568.7866
811.3912963867188 0 5883.326
812.3964233398438 0 1047.3418
821.4237670898438 0 1912.5682
822.432373046875 0 1152.3336
830.3995971679688 0 2194.155
831.408203125 0 1255.9155
832.44287109375 0 3016.76
833.4459838867188 0 2453.6265
838.4051513671875 0 1737.8822
852.3599243164062 0 1441.7434
855.4157104492188 0 1220.4158
858.430908203125 0 2765.5247
858.50927734375 0 1122.9012
859.4261474609375 0 8118.5234
859.4938354492188 0 5758.4805
860.4217529296875 0 2759.187
860.490966796875 0 5516.2783
861.486083984375 0 3082.2876
862.4891967773438 0 1078.9171
864.43408203125 0 58203
865.4279174804688 0 41133.336
866.4264526367188 0 16160.791
867.427490234375 0 4150.6597
873.434326171875 0 1295.1204
874.4304809570312 0 2832.8206
875.4234619140625 0 1202.2174
884.4835205078125 0 3631.7815
885.486572265625 0 4366.6973 c Water loss 7
886.4752197265625 0 9843.194 c Ammonia loss 7
887.4739379882812 0 5074.7925
888.4799194335938 0 3159.982
889.4923706054688 0 908.9799
899.4694213867188 0 1051.5691
900.4771118164062 0 6069.392
901.4703979492188 0 3908.2766
902.4205322265625 0 61045.3
902.4971923828125 0 14613.976
903.4208374023438 0 18232.377
903.5007934570312 0 90371.26 c 7
904.4984130859375 0 51820.492
905.4978637695312 0 18953.447
906.4976196289062 0 4532.4473
908.4238891601562 0 59772.055 z 4
909.41845703125 0 41886.555
910.41748046875 0 16878.197
911.4190673828125 0 5126.1675
912.4291381835938 0 1712.546
918.4398193359375 0 33772.355
919.4470825195312 0 111560
920.4500732421875 0 49494.44
921.452392578125 0 14849.847
922.4570922851562 0 3947.8086
923.464111328125 0 1206.6011
924.4456176757812 0 4378.906 y 4
925.4376831054688 0 3244.0132
926.434326171875 0 1914.0442
957.5150146484375 0 1411.5303
959.501220703125 0 1275.7196
960.5086669921875 0 1108.5609
961.502685546875 0 1038.2177
968.4937133789062 0 1130.5521
969.4783325195312 0 3026.3848
970.4771728515625 0 1430.2633
974.533447265625 0 1916.7915
975.528076171875 0 1669.7599
986.5015869140625 0 4719.7476
987.5033569335938 0 6501.987
988.501220703125 0 5707.9272
989.5028686523438 0 1830.931
999.533447265625 0 3942.0576 c Water loss 8
1000.5230712890625 0 6379.0215 c Ammonia loss 8
1001.5186157226562 0 6043.7476
1002.5213623046875 0 3582.5574
1003.5271606445312 0 3364.7615
1004.5181884765625 0 2052.7744
1016.5349731445312 0 7844.57
1017.5429077148438 0 55778.867 c 8
1018.5387573242188 0 40561.562
1019.5364379882812 0 21381.412
1020.5349731445312 0 7784.924
1021.5288696289062 0 2654.5754
1022.5186157226562 0 1471.8469
1023.5001831054688 0 1210.2651
1047.53515625 0 870.58215
1048.503173828125 0 1506.6466
1060.5574951171875 0 1735.1418
1061.5638427734375 0 2461.0215
1062.5650634765625 0 1678.7128
1063.5306396484375 0 1401.5216 y Ammonia loss 3
1064.5247802734375 0 22724.328 z 3
1065.5224609375 0 33592.492
1066.5189208984375 0 22632.438
1067.5206298828125 0 6947.106
1068.5263671875 0 2285.928
1080.53857421875 0 2044.3221 y 3
1081.529296875 0 2072.6882
1082.5406494140625 0 1330.2849
1088.5552978515625 0 3720.4644
1089.5546875 0 3829.0977
1090.55078125 0 1682.1571
1091.555908203125 0 1267.6406
1093.5103759765625 0 1095.7837
1095.493408203125 0 1681.6278
1096.5030517578125 0 1423.227
1104.5755615234375 0 17508.451 c 9
1105.5711669921875 0 13989.089
1106.570556640625 0 9145.992
1107.56396484375 0 3879.8105
1108.55712890625 0 903.7306
1134.5550537109375 0 1822.4565 w 2
1135.5469970703125 0 1952.8759
1136.53515625 0 1286.2164
1151.556640625 0 15960.519 z 2
1152.558349609375 0 35067.617
1153.5562744140625 0 22911.635
1154.56005859375 0 9509.733
1155.563232421875 0 3248.935
1157.6341552734375 0 1314.2286
1158.62939453125 0 1905.291
1159.62744140625 0 3085.4885
1160.628173828125 0 1158.6852
1168.579833984375 0 1904.4126
1169.5775146484375 0 1868.4496
1173.644775390625 0 3694.672
1174.6519775390625 0 7733.8853
1175.6478271484375 0 8128.1357
1176.6473388671875 0 4187.0596
1177.647216796875 0 1752.0419
1199.6119384765625 0 959.70087
1201.639892578125 0 11880.1
1202.63525390625 0 9079.031
1203.63525390625 0 6314.6577
1204.6395263671875 0 2276.4243
1217.6583251953125 0 10507.966 c 10
1218.66064453125 0 25093.998
1219.6590576171875 0 15872.6
1220.660888671875 0 8299.614
1221.6556396484375 0 1960.4609
1264.6318359375 0 1019.9369 z 1
1265.643310546875 0 4420.4355
1266.64111328125 0 4845.494
1267.639892578125 0 3377.7446
1279.647705078125 0 923.0982
1294.6436767578125 0 1069.2432
1296.649658203125 0 1046.4661
1305.6593017578125 0 4880.241
1306.65478515625 0 5678.1274
1307.6512451171875 0 6077.281
1308.6529541015625 0 3627.022
1309.647216796875 0 1522.6877
1310.6641845703125 0 1100.9866
1320.69580078125 0 3370.7046
1321.6893310546875 0 6818.653
1322.691650390625 0 7486.3076
1323.6798095703125 0 14104.629
1324.681396484375 0 24294.51
1325.6751708984375 0 17957.291
1326.6776123046875 0 9161.539
1327.6778564453125 0 2046.7015
1337.6949462890625 0 4946.767
1338.7003173828125 0 30063.113
1339.6971435546875 0 26857.08
1340.69775390625 0 17588.32
1341.692626953125 0 5933.6514
1342.6983642578125 0 1998.715
1348.68310546875 0 8481.128
1349.673583984375 0 14926.25
1350.6697998046875 0 11476.911
1351.66845703125 0 7441.053
1352.665771484375 0 1902.8685
1355.72021484375 0 1453.7794
1356.71484375 0 1953.8975
1357.710205078125 0 1254.7756
1363.7344970703125 0 1149.7323
1364.710205078125 0 1169.0399
1365.6925048828125 0 17633.555
1366.693359375 0 101264.12
1367.68994140625 0 96092.62
1368.6888427734375 0 54752.363
1369.687744140625 0 20653.412
1370.688720703125 0 3740.334
1380.7535400390625 0 1664.6694
1381.719482421875 0 4711.964
1382.7119140625 0 14643.23
1383.7137451171875 0 25535.639
1384.712890625 0 19797.83
1385.7113037109375 0 10893.658
1386.713134765625 0 3030.2983
1387.716552734375 0 1141.2915
2679.982177734375 0 860.05536
3124.057861328125 0 841.4896

Spectrum Details

|  |  |
| --- | --- |
| Matched peaks? Matched peaks The total absolute number of peaks matched. Additionally in brackets the total fraction of peaks matched and the total number of peaks is shown. | 67 (9.56% of 701) |
| FDR? FDR The false discovery rate estimated for this peptide. It is calculated by matching all theoretical fragments with a non-integer shift with the raw peaks for this spectrum. This is done with 40 different shifts. The resulting percentage is the average number of annotated peaks over the number of annotated peaks with the correct spectrum. | 1.21% |
| Satellite FDR? Satellite FDR See the FDR for details on its calculation. This satellite ion specific FDR only contains the satellite ions (d/w) for I/L/J positions. | 0.00% |
| PSM Score? PSM Score The PSM Score as given by Hecklib to this annotated spectrum. It is shown with three significant figures. | 541 |

#### Spectrum 4261? Spectrum 4261 The raw spectrum of this peptide as annotated by Hecklib. The fragments are coloured according to ion type (see legend). Any peaks with a star '\*' as text can be hovered over to see the full details, first the ion type second the mass shift type. By hovering over the amino acids in the peptide or ions in the legend the corresponding peaks are highlighted. By toggling the 'Unassigned' label you can turn the background (unassigned) peaks on or off in the plot. By updating the slider in the Ion legend you can update the spectrum to only show the top X% of the peaks with labels. The top X% means any peak that is within X% of the highest intensity. By dragging in the spectrum you can zoom in to a specific part of the spectrum and use 'Zoom Out' to get back to the original zoom level. The annotation of the spectrum is based on the given sequence in the peptides file and is done with different software so inconsistencies are likely. The peaks are annotated based on the given sequence, with 20 ppm tolerance.

Copy Data

##### Spectrum 4261 (TSV)

###### Preview

```
Loading example...
```

*Click on the button to copy the data to your clipboard.*

Mz MinMz MaxIntensity Max

WidthHeightPeptide font sizePeptide stroke widthSpectrum font sizeSpectrum stroke widthCompact peptide

Ion legend

wxyz

abcd

OtherUnassignedIon^Charge^_Position_Show for top:%

TISRDNAKNSJY

03.08e+46.16e+49.24e+41.23e+5

Zoom Out

y^+1^_1_y^+1^_2_c^+1^_3_c^+2^_7_y^+1^_3_c^+2^_8_c^+2^_8_c^+2^_8_c^+1^_4_c^+1^_4_z^+1^_4_y^+1^_4_c^+2^_9_c^+2^_9_c^+2^_9_z^+2^_9_c^+2^_10_c^+2^_10_c^+1^_5_y^+2^_10_z^+2^_10_y^+2^_10_c^+1^_5_c^+2^_11_z^+1^_5_c^+2^_11_w^+2^_11_y^+1^_5_z^+2^_11_y^+2^_11_z^+1^_6_z^+1^_6_c^+1^_6_y^+1^_6_c^+1^_6_w^+1^_7_c^+1^_7_c^+1^_7_z^+1^_7_y^+1^_7_c^+1^_8_c^+1^_8_c^+1^_8_z^+1^_8_y^+1^_8_c^+1^_9_c^+1^_9_c^+1^_9_y^+1^_9_z^+1^_9_c^+1^_10_z^+1^_10_y^+1^_10_c^+1^_11_

035070010501400

Fragment Matches Table

Show background peaks

| Position | Ion type | Intensity | mz Theoretical | mz Error (Th) | mz Error (ppm) | Charge | Series Number |
| --- | --- | --- | --- | --- | --- | --- | --- |
| - | - | 422.9 | 129.1 | - | - | 0 | - |
| - | - | 512.4 | 133.1 | - | - | 0 | - |
| - | - | 3457 | 136.1 | - | - | 0 | - |
| - | - | 950.5 | 142.1 | - | - | 0 | - |
| - | - | 654.5 | 143.1 | - | - | 0 | - |
| - | - | 418.3 | 145 | - | - | 0 | - |
| - | - | 502.6 | 148.9 | - | - | 0 | - |
| - | - | 4698 | 149 | - | - | 0 | - |
| - | - | 446.8 | 160.8 | - | - | 0 | - |
| - | - | 2831 | 165.1 | - | - | 0 | - |
| - | - | 1793 | 167.1 | - | - | 0 | - |
| - | - | 2871 | 173.1 | - | - | 0 | - |
| - | - | 531.6 | 173.1 | - | - | 0 | - |
| - | - | 547.1 | 173.5 | - | - | 0 | - |
| 12 | y | 1.133E+04 | 182.1 | 0.0001139 | 0.6255 | +1 | 1 |
| - | - | 485.1 | 183.1 | - | - | 0 | - |
| - | - | 475.9 | 183.2 | - | - | 0 | - |
| - | - | 2.054E+04 | 187.1 | - | - | 0 | - |
| - | - | 1839 | 188.1 | - | - | 0 | - |
| - | - | 574.4 | 190 | - | - | 0 | - |
| - | - | 973.2 | 200.1 | - | - | 0 | - |
| - | - | 678.6 | 201.1 | - | - | 0 | - |
| - | - | 651.3 | 202.1 | - | - | 0 | - |
| - | - | 546.9 | 207.7 | - | - | 0 | - |
| - | - | 5184 | 215.1 | - | - | 0 | - |
| - | - | 858.6 | 216.1 | - | - | 0 | - |
| - | - | 2774 | 221.1 | - | - | 0 | - |
| - | - | 1398 | 222.1 | - | - | 0 | - |
| - | - | 1313 | 223.1 | - | - | 0 | - |
| - | - | 859.3 | 223.1 | - | - | 0 | - |
| - | - | 2458 | 225 | - | - | 0 | - |
| - | - | 659.7 | 226 | - | - | 0 | - |
| - | - | 615.3 | 229.1 | - | - | 0 | - |
| - | - | 555.1 | 235.1 | - | - | 0 | - |
| - | - | 3322 | 239.1 | - | - | 0 | - |
| - | - | 731.4 | 239.1 | - | - | 0 | - |
| - | - | 2220 | 240.1 | - | - | 0 | - |
| - | - | 708.9 | 241.1 | - | - | 0 | - |
| - | - | 1712 | 260.1 | - | - | 0 | - |
| - | - | 1213 | 282.1 | - | - | 0 | - |
| - | - | 616.3 | 287.2 | - | - | 0 | - |
| - | - | 2284 | 295.1 | - | - | 0 | - |
| 11 | y | 760.9 | 295.2 | 0.0008049 | 2.727 | +1 | 2 |
| - | - | 3000 | 296.1 | - | - | 0 | - |
| - | - | 1650 | 297.1 | - | - | 0 | - |
| - | - | 1948 | 299.1 | - | - | 0 | - |
| - | - | 1772 | 300.1 | - | - | 0 | - |
| - | - | 3677 | 300.2 | - | - | 0 | - |
| - | - | 1553 | 301.1 | - | - | 0 | - |
| - | - | 671.6 | 301.2 | - | - | 0 | - |
| - | - | 509.8 | 303.3 | - | - | 0 | - |
| - | - | 745.3 | 313.1 | - | - | 0 | - |
| - | - | 1046 | 315.2 | - | - | 0 | - |
| - | - | 1750 | 317.2 | - | - | 0 | - |
| 3 | c | 1443 | 319.2 | 0.0008289 | 2.597 | +1 | 3 |
| - | - | 1.23E+04 | 341 | - | - | 0 | - |
| - | - | 803.5 | 355.1 | - | - | 0 | - |
| - | - | 4.842E+04 | 359 | - | - | 0 | - |
| - | - | 790.1 | 369.1 | - | - | 0 | - |
| - | - | 2680 | 370.1 | - | - | 0 | - |
| - | - | 1749 | 371.1 | - | - | 0 | - |
| - | - | 933 | 372.1 | - | - | 0 | - |
| - | - | 845.4 | 375.9 | - | - | 0 | - |
| 7 | c | 868.6 | 379.2 | 0.005891 | 15.53 | +2 | 7 |
| 10 | y | 903.4 | 382.2 | 0.0003087 | 0.8078 | +1 | 3 |
| - | - | 1901 | 385.2 | - | - | 0 | - |
| - | - | 735.9 | 385.9 | - | - | 0 | - |
| - | - | 1018 | 386.2 | - | - | 0 | - |
| - | - | 1476 | 391.6 | - | - | 0 | - |
| - | - | 1157 | 391.9 | - | - | 0 | - |
| - | - | 605.9 | 415.3 | - | - | 0 | - |
| - | - | 639.9 | 421.2 | - | - | 0 | - |
| - | - | 1375 | 427.2 | - | - | 0 | - |
| - | - | 1.22E+05 | 429.1 | - | - | 0 | - |
| - | - | 1535 | 429.8 | - | - | 0 | - |
| - | - | 788.9 | 430.2 | - | - | 0 | - |
| - | - | 735.2 | 430.3 | - | - | 0 | - |
| - | - | 576.5 | 437.2 | - | - | 0 | - |
| - | - | 1519 | 442.7 | - | - | 0 | - |
| 8 | c | 1471 | 443.2 | 0.005702 | 12.86 | +2 | 8 |
| 8 | c | 861.3 | 443.7 | 0.0005809 | 1.309 | +2 | 8 |
| - | - | 2004 | 444.2 | - | - | 0 | - |
| - | - | 6.286E+04 | 445.1 | - | - | 0 | - |
| - | - | 4366 | 445.3 | - | - | 0 | - |
| - | - | 1171 | 446.2 | - | - | 0 | - |
| - | - | 768.4 | 447.1 | - | - | 0 | - |
| - | - | 5917 | 451.8 | - | - | 0 | - |
| 8 | c | 1.84E+04 | 452.3 | 0.0002119 | 0.4686 | +2 | 8 |
| - | - | 6677 | 452.8 | - | - | 0 | - |
| - | - | 1856 | 453.3 | - | - | 0 | - |
| 4 | c | 734.3 | 458.3 | 9.414E-05 | 0.2054 | +1 | 4 |
| - | - | 1234 | 459.7 | - | - | 0 | - |
| - | - | 1162 | 460.2 | - | - | 0 | - |
| - | - | 1084 | 460.2 | - | - | 0 | - |
| - | - | 670.8 | 462.1 | - | - | 0 | - |
| - | - | 700.1 | 472.3 | - | - | 0 | - |
| 4 | c | 5.656E+04 | 475.3 | 0.0002122 | 0.4465 | +1 | 4 |
| - | - | 1.192E+04 | 476.3 | - | - | 0 | - |
| - | - | 2063 | 477.3 | - | - | 0 | - |
| - | - | 5649 | 479.8 | - | - | 0 | - |
| 9 | z | 717.6 | 480.2 | 0.0007025 | 1.463 | +1 | 4 |
| - | - | 2564 | 480.3 | - | - | 0 | - |
| - | - | 987.9 | 480.8 | - | - | 0 | - |
| - | - | 3325 | 481.2 | - | - | 0 | - |
| - | - | 1009 | 482.2 | - | - | 0 | - |
| - | - | 703.9 | 485.2 | - | - | 0 | - |
| - | - | 1501 | 486.8 | - | - | 0 | - |
| - | - | 1760 | 487.3 | - | - | 0 | - |
| - | - | 614.5 | 487.8 | - | - | 0 | - |
| - | - | 695.8 | 491.8 | - | - | 0 | - |
| - | - | 1407 | 493.8 | - | - | 0 | - |
| 9 | y | 1938 | 496.2 | 0.0005045 | 1.017 | +1 | 4 |
| - | - | 6304 | 499.8 | - | - | 0 | - |
| 9 | c | 3302 | 500.3 | 0.00794 | 15.87 | +2 | 9 |
| 9 | c | 3637 | 500.8 | 0.0009373 | 1.872 | +2 | 9 |
| - | - | 1537 | 501.3 | - | - | 0 | - |
| - | - | 1329 | 501.8 | - | - | 0 | - |
| - | - | 5365 | 502.3 | - | - | 0 | - |
| - | - | 949.7 | 503.3 | - | - | 0 | - |
| - | - | 1387 | 508.3 | - | - | 0 | - |
| - | - | 1.221E+04 | 508.8 | - | - | 0 | - |
| 9 | c | 2.04E+04 | 509.3 | 0.001961 | 3.851 | +2 | 9 |
| - | - | 1.307E+04 | 509.8 | - | - | 0 | - |
| - | - | 5192 | 510.3 | - | - | 0 | - |
| - | - | 996.5 | 510.8 | - | - | 0 | - |
| - | - | 845.1 | 526.3 | - | - | 0 | - |
| - | - | 794.1 | 526.8 | - | - | 0 | - |
| - | - | 879.7 | 530.3 | - | - | 0 | - |
| 4 | z | 2128 | 532.8 | 0.0006404 | 1.202 | +2 | 9 |
| - | - | 650.6 | 533.3 | - | - | 0 | - |
| - | - | 946.7 | 535.3 | - | - | 0 | - |
| - | - | 1079 | 535.8 | - | - | 0 | - |
| - | - | 854.6 | 541.3 | - | - | 0 | - |
| 10 | c | 1131 | 544.3 | 0.002898 | 5.324 | +2 | 10 |
| - | - | 1662 | 544.8 | - | - | 0 | - |
| - | - | 2139 | 545.3 | - | - | 0 | - |
| - | - | 1075 | 546.3 | - | - | 0 | - |
| - | - | 2319 | 548.3 | - | - | 0 | - |
| - | - | 1620 | 548.7 | - | - | 0 | - |
| - | - | 611.1 | 551.3 | - | - | 0 | - |
| 10 | c | 1.469E+04 | 552.8 | 0.0001227 | 0.222 | +2 | 10 |
| - | - | 1.357E+04 | 553.3 | - | - | 0 | - |
| - | - | 4386 | 553.8 | - | - | 0 | - |
| - | - | 3401 | 554.3 | - | - | 0 | - |
| - | - | 899.5 | 554.8 | - | - | 0 | - |
| - | - | 547 | 562.3 | - | - | 0 | - |
| - | - | 2471 | 564.3 | - | - | 0 | - |
| - | - | 790.6 | 565.3 | - | - | 0 | - |
| - | - | 648.8 | 566.8 | - | - | 0 | - |
| - | - | 563 | 568.8 | - | - | 0 | - |
| 5 | c | 790.3 | 573.3 | 0.00638 | 11.13 | +1 | 5 |
| - | - | 596 | 573.8 | - | - | 0 | - |
| - | - | 676.7 | 574.8 | - | - | 0 | - |
| 3 | y | 627.2 | 575.8 | 0.003187 | 5.535 | +2 | 10 |
| 3 | z | 6408 | 576.3 | 0.001092 | 1.894 | +2 | 10 |
| - | - | 4876 | 576.8 | - | - | 0 | - |
| - | - | 2487 | 577.3 | - | - | 0 | - |
| - | - | 1663 | 578.3 | - | - | 0 | - |
| - | - | 1156 | 579.3 | - | - | 0 | - |
| - | - | 1055 | 583.8 | - | - | 0 | - |
| 3 | y | 1.703E+04 | 584.3 | 0.0005049 | 0.8641 | +2 | 10 |
| - | - | 1.417E+04 | 584.8 | - | - | 0 | - |
| - | - | 5928 | 585.3 | - | - | 0 | - |
| - | - | 2192 | 585.8 | - | - | 0 | - |
| - | - | 1784 | 586.8 | - | - | 0 | - |
| - | - | 2340 | 587.3 | - | - | 0 | - |
| - | - | 842.1 | 587.8 | - | - | 0 | - |
| 5 | c | 1.003E+05 | 590.3 | 0.000211 | 0.3575 | +1 | 5 |
| - | - | 2.992E+04 | 591.3 | - | - | 0 | - |
| - | - | 743.1 | 591.8 | - | - | 0 | - |
| - | - | 6010 | 592.3 | - | - | 0 | - |
| - | - | 875.5 | 593.3 | - | - | 0 | - |
| 11 | c | 3986 | 600.8 | 0.0003157 | 0.5255 | +2 | 11 |
| - | - | 1587 | 601.3 | - | - | 0 | - |
| - | - | 1368 | 601.8 | - | - | 0 | - |
| - | - | 1020 | 607.3 | - | - | 0 | - |
| 8 | z | 5731 | 608.3 | 0.0005715 | 0.9395 | +1 | 5 |
| 11 | c | 4.049E+04 | 609.3 | 0.0007118 | 1.168 | +2 | 11 |
| - | - | 2.942E+04 | 609.8 | - | - | 0 | - |
| - | - | 1.592E+04 | 610.3 | - | - | 0 | - |
| - | - | 4776 | 610.8 | - | - | 0 | - |
| - | - | 988 | 611.3 | - | - | 0 | - |
| - | - | 1083 | 616.3 | - | - | 0 | - |
| 2 | w | 1906 | 618.3 | 0.0001207 | 0.1952 | +2 | 11 |
| - | - | 1978 | 618.8 | - | - | 0 | - |
| 8 | y | 8636 | 624.3 | 0.0006188 | 0.9912 | +1 | 5 |
| - | - | 2552 | 625.3 | - | - | 0 | - |
| - | - | 996.3 | 630.3 | - | - | 0 | - |
| - | - | 923.8 | 630.8 | - | - | 0 | - |
| 2 | z | 1279 | 632.8 | 0.002291 | 3.62 | +2 | 11 |
| - | - | 2924 | 633.3 | - | - | 0 | - |
| - | - | 713.3 | 633.8 | - | - | 0 | - |
| - | - | 961.4 | 636.3 | - | - | 0 | - |
| - | - | 836.2 | 638.3 | - | - | 0 | - |
| - | - | 1060 | 639.3 | - | - | 0 | - |
| 2 | y | 2689 | 640.8 | 0.00201 | 3.136 | +2 | 11 |
| - | - | 2603 | 641.3 | - | - | 0 | - |
| - | - | 1535 | 647.4 | - | - | 0 | - |
| - | - | 2310 | 647.9 | - | - | 0 | - |
| - | - | 602.4 | 648.3 | - | - | 0 | - |
| - | - | 795.9 | 652.8 | - | - | 0 | - |
| - | - | 945.7 | 653.8 | - | - | 0 | - |
| - | - | 662.4 | 654.3 | - | - | 0 | - |
| - | - | 650.6 | 654.8 | - | - | 0 | - |
| - | - | 3259 | 655.3 | - | - | 0 | - |
| - | - | 2635 | 655.8 | - | - | 0 | - |
| - | - | 936.4 | 656.3 | - | - | 0 | - |
| - | - | 3901 | 659.3 | - | - | 0 | - |
| - | - | 5566 | 660.3 | - | - | 0 | - |
| - | - | 4028 | 660.9 | - | - | 0 | - |
| 7 | z | 8762 | 661.3 | 0.00651 | 9.843 | +1 | 6 |
| - | - | 5231 | 661.8 | - | - | 0 | - |
| - | - | 6818 | 662.3 | - | - | 0 | - |
| - | - | 4796 | 662.8 | - | - | 0 | - |
| - | - | 2156 | 663.3 | - | - | 0 | - |
| - | - | 1062 | 663.8 | - | - | 0 | - |
| - | - | 3131 | 668.8 | - | - | 0 | - |
| - | - | 8772 | 669.3 | - | - | 0 | - |
| - | - | 7818 | 669.8 | - | - | 0 | - |
| - | - | 5688 | 670.3 | - | - | 0 | - |
| - | - | 3498 | 670.8 | - | - | 0 | - |
| - | - | 2585 | 671.3 | - | - | 0 | - |
| - | - | 778.9 | 673.9 | - | - | 0 | - |
| - | - | 2305 | 674.3 | - | - | 0 | - |
| - | - | 5238 | 674.8 | - | - | 0 | - |
| - | - | 4050 | 675.3 | - | - | 0 | - |
| - | - | 2991 | 675.8 | - | - | 0 | - |
| - | - | 2247 | 676.3 | - | - | 0 | - |
| - | - | 684.8 | 676.8 | - | - | 0 | - |
| 7 | z | 1.972E+04 | 679.4 | 0.0004539 | 0.6681 | +1 | 6 |
| - | - | 7415 | 680.4 | - | - | 0 | - |
| - | - | 2142 | 681.4 | - | - | 0 | - |
| - | - | 1211 | 682.9 | - | - | 0 | - |
| - | - | 2.427E+04 | 683.3 | - | - | 0 | - |
| - | - | 2.263E+04 | 683.8 | - | - | 0 | - |
| - | - | 1.269E+04 | 684.3 | - | - | 0 | - |
| - | - | 4621 | 684.8 | - | - | 0 | - |
| - | - | 987.1 | 685.3 | - | - | 0 | - |
| 6 | c | 854.5 | 687.3 | 0.01082 | 15.74 | +1 | 6 |
| - | - | 1345 | 689.2 | - | - | 0 | - |
| - | - | 2429 | 689.7 | - | - | 0 | - |
| - | - | 2060 | 690.2 | - | - | 0 | - |
| - | - | 2.289E+04 | 691.4 | - | - | 0 | - |
| - | - | 2.122E+04 | 691.9 | - | - | 0 | - |
| - | - | 1.239E+04 | 692.4 | - | - | 0 | - |
| - | - | 3532 | 692.9 | - | - | 0 | - |
| - | - | 1051 | 693.4 | - | - | 0 | - |
| 7 | y | 5408 | 695.4 | 0.001844 | 2.652 | +1 | 6 |
| - | - | 2215 | 696.4 | - | - | 0 | - |
| 6 | c | 2.615E+04 | 704.4 | 0.0003528 | 0.5009 | +1 | 6 |
| - | - | 9384 | 705.4 | - | - | 0 | - |
| - | - | 2376 | 706.4 | - | - | 0 | - |
| - | - | 631.5 | 707.4 | - | - | 0 | - |
| - | - | 936.2 | 722.4 | - | - | 0 | - |
| - | - | 4016 | 731.4 | - | - | 0 | - |
| - | - | 1997 | 732.4 | - | - | 0 | - |
| - | - | 1164 | 737.3 | - | - | 0 | - |
| - | - | 723.7 | 737.4 | - | - | 0 | - |
| 6 | w | 1.429E+04 | 749.4 | 0.0002867 | 0.3825 | +1 | 7 |
| - | - | 7603 | 750.4 | - | - | 0 | - |
| - | - | 2335 | 751.4 | - | - | 0 | - |
| 7 | c | 2280 | 758.4 | 0.001839 | 2.425 | +1 | 7 |
| - | - | 907.6 | 759.4 | - | - | 0 | - |
| - | - | 781.1 | 771.3 | - | - | 0 | - |
| 7 | c | 2.394E+04 | 775.4 | 0.001212 | 1.563 | +1 | 7 |
| - | - | 1.051E+04 | 776.4 | - | - | 0 | - |
| - | - | 3146 | 777.4 | - | - | 0 | - |
| - | - | 683.3 | 778.4 | - | - | 0 | - |
| - | - | 1324 | 787.4 | - | - | 0 | - |
| 6 | z | 5.584E+04 | 793.4 | 0.0008398 | 1.059 | +1 | 7 |
| - | - | 3.035E+04 | 794.4 | - | - | 0 | - |
| - | - | 8464 | 795.4 | - | - | 0 | - |
| - | - | 1639 | 796.4 | - | - | 0 | - |
| 6 | y | 796.4 | 809.4 | 0.004423 | 5.464 | +1 | 7 |
| - | - | 760.5 | 810.4 | - | - | 0 | - |
| - | - | 636.6 | 811.4 | - | - | 0 | - |
| - | - | 795.1 | 815.5 | - | - | 0 | - |
| - | - | 739.8 | 821.4 | - | - | 0 | - |
| - | - | 774.1 | 858.5 | - | - | 0 | - |
| - | - | 1923 | 859.5 | - | - | 0 | - |
| - | - | 2566 | 860.5 | - | - | 0 | - |
| - | - | 889.1 | 861.5 | - | - | 0 | - |
| - | - | 1.706E+04 | 864.4 | - | - | 0 | - |
| - | - | 1.069E+04 | 865.4 | - | - | 0 | - |
| - | - | 3272 | 866.4 | - | - | 0 | - |
| - | - | 821.5 | 884.5 | - | - | 0 | - |
| 8 | c | 782.1 | 885.5 | 0.004128 | 4.662 | +1 | 8 |
| 8 | c | 2509 | 886.5 | 0.001236 | 1.394 | +1 | 8 |
| - | - | 740.7 | 887.2 | - | - | 0 | - |
| - | - | 1065 | 887.5 | - | - | 0 | - |
| - | - | 675.7 | 895.4 | - | - | 0 | - |
| - | - | 2377 | 900.5 | - | - | 0 | - |
| - | - | 1654 | 901.5 | - | - | 0 | - |
| - | - | 9160 | 902.5 | - | - | 0 | - |
| 8 | c | 3.307E+04 | 903.5 | 0.002242 | 2.481 | +1 | 8 |
| - | - | 1.697E+04 | 904.5 | - | - | 0 | - |
| - | - | 5036 | 905.5 | - | - | 0 | - |
| - | - | 1287 | 906.5 | - | - | 0 | - |
| 5 | z | 1.822E+04 | 908.4 | 0.003125 | 3.44 | +1 | 8 |
| - | - | 1.123E+04 | 909.4 | - | - | 0 | - |
| - | - | 4441 | 910.4 | - | - | 0 | - |
| - | - | 1160 | 911.4 | - | - | 0 | - |
| - | - | 1189 | 919.5 | - | - | 0 | - |
| - | - | 751.3 | 921.4 | - | - | 0 | - |
| - | - | 816.4 | 921.5 | - | - | 0 | - |
| 5 | y | 1592 | 924.4 | 0.007505 | 8.119 | +1 | 8 |
| - | - | 1240 | 925.4 | - | - | 0 | - |
| - | - | 696.1 | 958.5 | - | - | 0 | - |
| - | - | 690 | 961.5 | - | - | 0 | - |
| - | - | 1255 | 986.5 | - | - | 0 | - |
| - | - | 3181 | 987.5 | - | - | 0 | - |
| - | - | 1491 | 988.5 | - | - | 0 | - |
| 9 | c | 1401 | 999.5 | 0.003787 | 3.788 | +1 | 9 |
| 9 | c | 2263 | 1001 | 0.002559 | 2.558 | +1 | 9 |
| - | - | 1615 | 1002 | - | - | 0 | - |
| - | - | 1497 | 1003 | - | - | 0 | - |
| - | - | 2126 | 1017 | - | - | 0 | - |
| 9 | c | 2.133E+04 | 1018 | 0.002139 | 2.102 | +1 | 9 |
| - | - | 1.425E+04 | 1019 | - | - | 0 | - |
| - | - | 5117 | 1020 | - | - | 0 | - |
| - | - | 1657 | 1021 | - | - | 0 | - |
| - | - | 810.4 | 1061 | - | - | 0 | - |
| 4 | y | 939.9 | 1063 | 0.02078 | 19.56 | +1 | 9 |
| 4 | z | 6547 | 1065 | 0.002673 | 2.511 | +1 | 9 |
| - | - | 1.109E+04 | 1066 | - | - | 0 | - |
| - | - | 4478 | 1067 | - | - | 0 | - |
| - | - | 2157 | 1068 | - | - | 0 | - |
| - | - | 2110 | 1089 | - | - | 0 | - |
| - | - | 1473 | 1090 | - | - | 0 | - |
| - | - | 763.2 | 1091 | - | - | 0 | - |
| - | - | 614.4 | 1096 | - | - | 0 | - |
| - | - | 758.9 | 1098 | - | - | 0 | - |
| 10 | c | 6296 | 1105 | 0.002246 | 2.034 | +1 | 10 |
| - | - | 5611 | 1106 | - | - | 0 | - |
| - | - | 2157 | 1107 | - | - | 0 | - |
| - | - | 792.2 | 1108 | - | - | 0 | - |
| - | - | 730.1 | 1137 | - | - | 0 | - |
| 3 | z | 3903 | 1152 | 0.002963 | 2.573 | +1 | 10 |
| - | - | 1.067E+04 | 1153 | - | - | 0 | - |
| - | - | 7094 | 1154 | - | - | 0 | - |
| - | - | 2571 | 1155 | - | - | 0 | - |
| - | - | 708.5 | 1156 | - | - | 0 | - |
| - | - | 844.6 | 1159 | - | - | 0 | - |
| - | - | 928.5 | 1160 | - | - | 0 | - |
| 3 | y | 792.6 | 1168 | 0.002238 | 1.917 | +1 | 10 |
| - | - | 1950 | 1174 | - | - | 0 | - |
| - | - | 3206 | 1175 | - | - | 0 | - |
| - | - | 3215 | 1176 | - | - | 0 | - |
| - | - | 1627 | 1177 | - | - | 0 | - |
| - | - | 5248 | 1202 | - | - | 0 | - |
| - | - | 3047 | 1203 | - | - | 0 | - |
| - | - | 1819 | 1204 | - | - | 0 | - |
| 11 | c | 3931 | 1218 | 0.002814 | 2.311 | +1 | 11 |
| - | - | 8848 | 1219 | - | - | 0 | - |
| - | - | 6182 | 1220 | - | - | 0 | - |
| - | - | 2256 | 1221 | - | - | 0 | - |
| - | - | 1637 | 1266 | - | - | 0 | - |
| - | - | 2045 | 1267 | - | - | 0 | - |
| - | - | 1226 | 1268 | - | - | 0 | - |
| - | - | 1553 | 1306 | - | - | 0 | - |
| - | - | 1919 | 1307 | - | - | 0 | - |
| - | - | 1654 | 1308 | - | - | 0 | - |
| - | - | 1408 | 1309 | - | - | 0 | - |
| - | - | 1383 | 1321 | - | - | 0 | - |
| - | - | 2496 | 1322 | - | - | 0 | - |
| - | - | 2223 | 1323 | - | - | 0 | - |
| - | - | 4906 | 1324 | - | - | 0 | - |
| - | - | 8575 | 1325 | - | - | 0 | - |
| - | - | 5789 | 1326 | - | - | 0 | - |
| - | - | 2374 | 1327 | - | - | 0 | - |
| - | - | 841.7 | 1328 | - | - | 0 | - |
| - | - | 2231 | 1338 | - | - | 0 | - |
| - | - | 1.148E+04 | 1339 | - | - | 0 | - |
| - | - | 9475 | 1340 | - | - | 0 | - |
| - | - | 4603 | 1341 | - | - | 0 | - |
| - | - | 2732 | 1342 | - | - | 0 | - |
| - | - | 3131 | 1349 | - | - | 0 | - |
| - | - | 4781 | 1350 | - | - | 0 | - |
| - | - | 3956 | 1351 | - | - | 0 | - |
| - | - | 1918 | 1352 | - | - | 0 | - |
| - | - | 978.8 | 1353 | - | - | 0 | - |
| - | - | 849.4 | 1356 | - | - | 0 | - |
| - | - | 770.6 | 1365 | - | - | 0 | - |
| - | - | 5636 | 1366 | - | - | 0 | - |
| - | - | 3.788E+04 | 1367 | - | - | 0 | - |
| - | - | 3.261E+04 | 1368 | - | - | 0 | - |
| - | - | 1.645E+04 | 1369 | - | - | 0 | - |
| - | - | 5825 | 1370 | - | - | 0 | - |
| - | - | 1437 | 1371 | - | - | 0 | - |
| - | - | 849.7 | 1378 | - | - | 0 | - |
| - | - | 3194 | 1379 | - | - | 0 | - |
| - | - | 1664 | 1380 | - | - | 0 | - |
| - | - | 843.8 | 1382 | - | - | 0 | - |
| - | - | 4305 | 1383 | - | - | 0 | - |
| - | - | 7786 | 1384 | - | - | 0 | - |
| - | - | 5627 | 1385 | - | - | 0 | - |
| - | - | 3327 | 1386 | - | - | 0 | - |

m/z Charge Intensity FragmentType MassShift Position
129.1024627685547 0 422.92093
133.08624267578125 0 512.35315
136.0758514404297 0 3457.1504
142.12254333496094 0 950.4897
143.11839294433594 0 654.5266
144.98500061035156 0 418.3118
148.92977905273438 0 502.5746
149.04498291015625 0 4698.465
160.7801513671875 0 446.8031
165.0546875 0 2831.1665
167.0553436279297 0 1792.8464
173.09231567382812 0 2870.586
173.1287078857422 0 531.56116
173.45278930664062 0 547.0715
182.08128356933594 0 11330.394 y 11
183.08480834960938 0 485.0945
183.16851806640625 0 475.88132
187.14427185058594 0 20544.209
188.1476287841797 0 1839.1086
189.96185302734375 0 574.3993
200.13973999023438 0 973.1955
201.12344360351562 0 678.5603
202.0827178955078 0 651.2816
207.71266174316406 0 546.94806
215.13919067382812 0 5183.809
216.14186096191406 0 858.6362
221.08473205566406 0 2773.9175
222.08485412597656 0 1398.0565
223.06385803222656 0 1312.7705
223.0814208984375 0 859.25366
225.04307556152344 0 2457.9526
226.04205322265625 0 659.70374
229.13002014160156 0 615.30615
235.11819458007812 0 555.08386
239.09510803222656 0 3322.359
239.10794067382812 0 731.4485
240.095947265625 0 2220.3462
241.09262084960938 0 708.8837
260.12420654296875 0 1712.1221
282.14422607421875 0 1212.9868
287.17144775390625 0 616.32214
295.1033935546875 0 2284.3674
295.1644287109375 0 760.89154 y 10
296.10406494140625 0 3000.1252
297.101806640625 0 1649.9592
299.06134033203125 0 1947.7654
300.0628662109375 0 1772.0247
300.15557861328125 0 3677.2463
301.05938720703125 0 1553.3164
301.1583557128906 0 671.6442
303.3233947753906 0 509.82047
313.1134033203125 0 745.2921
315.1665954589844 0 1045.7439
317.1820373535156 0 1750.1729
319.19842529296875 0 1442.8175 c 2
341.01812744140625 0 12300.057
355.0685119628906 0 803.4844
359.02825927734375 0 48420.254
369.12274169921875 0 790.1365
370.1224670410156 0 2680.037
371.1192932128906 0 1749.0494
372.11962890625 0 932.99335
375.86004638671875 0 845.4498
379.20709228515625 0 868.6314 c Water loss 6
382.19757080078125 0 903.3881 y 9
385.2193603515625 0 1901.1674
385.87933349609375 0 735.9023
386.2279052734375 0 1018.0405
391.5511169433594 0 1475.5511
391.88592529296875 0 1156.5037
415.26800537109375 0 605.9458
421.20379638671875 0 639.8768
427.2173156738281 0 1374.941
429.08880615234375 0 121952.21
429.755126953125 0 1534.5854
430.2455139160156 0 788.9154
430.279296875 0 735.17694
437.2120666503906 0 576.51544
442.74560546875 0 1518.9952
443.24298095703125 0 1470.8118 c Water loss 7
443.74127197265625 0 861.30444 c Ammonia loss 7
444.2437744140625 0 2004.3556
445.1202087402344 0 62859.625
445.2641906738281 0 4365.942
446.2126159667969 0 1171.4448
447.1006164550781 0 768.3983
451.7503356933594 0 5917.0264
452.2537536621094 0 18395.72 c 7
452.7520446777344 0 6677.1323
453.25360107421875 0 1856.3513
458.2720642089844 0 734.3283 c Ammonia loss 3
459.7003479003906 0 1234.1344
460.2010192871094 0 1161.8971
460.23785400390625 0 1084.3174
462.1162414550781 0 670.75366
472.2518615722656 0 700.05695
475.2989196777344 0 56556.906 c 3
476.3013000488281 0 11921.125
477.3054504394531 0 2063.2776
479.7574157714844 0 5649.1523
480.22216796875 0 717.6204 z 8
480.2582092285156 0 2564.2708
480.75299072265625 0 987.9416
481.229248046875 0 3324.5186
482.2323303222656 0 1008.9122
485.2409362792969 0 703.8787
486.76861572265625 0 1501.2767
487.268310546875 0 1759.9702
487.76739501953125 0 614.46906
491.7575378417969 0 695.75336
493.7544860839844 0 1407.1697
496.23968505859375 0 1938.4827 y 8
499.7664489746094 0 6304.2065
500.26220703125 0 3302.3591 c Water loss 8
500.7630920410156 0 3636.9968 c Ammonia loss 8
501.26397705078125 0 1537.12
501.7691955566406 0 1329.0054
502.2735595703125 0 5365.0664
503.2810363769531 0 949.747
508.251953125 0 1387.4716
508.7713928222656 0 12211.335
509.2734680175781 0 20401.254 c 8
509.77301025390625 0 13068.03
510.2724914550781 0 5191.9937
510.7719421386719 0 996.48584
526.2531127929688 0 845.0653
526.7568969726562 0 794.10986
530.2951049804688 0 879.7419
532.7665405273438 0 2128.1145 z 3
533.2588500976562 0 650.55035
535.27197265625 0 946.6709
535.7686767578125 0 1079.0212
541.2714233398438 0 854.589
544.2810668945312 0 1131.4982 c Ammonia loss 9
544.7774047851562 0 1661.963
545.3032836914062 0 2138.5908
546.3104858398438 0 1074.8959
548.2509155273438 0 2319.0986
548.748779296875 0 1620.3057
551.301025390625 0 611.0944
552.7913208007812 0 14693.241 c 9
553.2891845703125 0 13573.604
553.7861938476562 0 4385.6533
554.28662109375 0 3401.2205
554.7833862304688 0 899.48346
562.289794921875 0 546.9867
564.3042602539062 0 2471.0452
565.304443359375 0 790.6365
566.7783813476562 0 648.81915
568.773193359375 0 563.0472
573.3054809570312 0 790.33325 c Ammonia loss 4
573.8032836914062 0 595.99915
574.7962036132812 0 676.7342
575.7811889648438 0 627.18964 y Ammonia loss 2
576.2808227539062 0 6407.9326 z 2
576.7813110351562 0 4876.098
577.2833251953125 0 2486.7927
578.3154907226562 0 1663.1409
579.3129272460938 0 1155.6995
583.794677734375 0 1055.2821
584.290771484375 0 17033.37 y 2
584.788818359375 0 14167.958
585.2880859375 0 5928.076
585.7863159179688 0 2192.184
586.8206787109375 0 1783.741
587.3223876953125 0 2340.1338
587.8212890625 0 842.0553
590.325439453125 0 100269.36 c 4
591.3283081054688 0 29919.201
591.8121337890625 0 743.0865
592.3284912109375 0 6009.6523
593.333251953125 0 875.46136
600.8198852539062 0 3986.124 c Ammonia loss 10
601.3173828125 0 1586.7678
601.8271484375 0 1367.7255
607.3289794921875 0 1020.0762
608.3158569335938 0 5731.3647 z 7
609.332763671875 0 40488.047 c 10
609.8319091796875 0 29417.736
610.3303833007812 0 15923.573
610.8314208984375 0 4775.6113
611.3277587890625 0 988.0279
616.3449096679688 0 1082.9899
618.3045043945312 0 1905.5806 w 1
618.80126953125 0 1978.3174
624.3345336914062 0 8636.209 y 7
625.3330078125 0 2551.8335
630.3278198242188 0 996.31085
630.8243408203125 0 923.778
632.8216552734375 0 1278.994 z 1
633.3218383789062 0 2924.4177
633.8259887695312 0 713.3121
636.3471069335938 0 961.3784
638.3489379882812 0 836.1733
639.3433227539062 0 1059.6799
640.831298828125 0 2689.277 y 1
641.3317260742188 0 2603.3127
647.35205078125 0 1535.403
647.8504028320312 0 2310.3708
648.3416748046875 0 602.3821
652.8464965820312 0 795.9203
653.8331298828125 0 945.66223
654.329345703125 0 662.3576
654.8319702148438 0 650.5597
655.3176879882812 0 3258.882
655.8157348632812 0 2635.2378
656.3121948242188 0 936.4025
659.3466186523438 0 3901.3176
660.348876953125 0 5566.0127
660.8519287109375 0 4027.9922
661.3494873046875 0 8762.483 z Water loss 6
661.84912109375 0 5230.6265
662.3446655273438 0 6817.7476
662.8426513671875 0 4795.54
663.3435668945312 0 2155.9883
663.8466186523438 0 1061.5338
668.8279418945312 0 3130.911
669.34765625 0 8772.062
669.8480224609375 0 7817.7866
670.3473510742188 0 5687.882
670.8424682617188 0 3497.513
671.343994140625 0 2585.3342
673.8511352539062 0 778.9061
674.3450317382812 0 2305.0012
674.8435668945312 0 5237.8857
675.3441772460938 0 4049.8115
675.8373413085938 0 2990.5352
676.3364868164062 0 2247.377
676.8345947265625 0 684.7845
679.3530883789062 0 19724.512 z 6
680.35302734375 0 7414.559
681.357666015625 0 2141.867
682.8562622070312 0 1211.2609
683.3472900390625 0 24274.078
683.8463134765625 0 22629.967
684.345703125 0 12687.052
684.84619140625 0 4620.6914
685.3480834960938 0 987.0737
687.3528442382812 0 854.52356 c Ammonia loss 5
689.1546020507812 0 1344.5275
689.6566162109375 0 2428.9868
690.1580810546875 0 2060.4395
691.3565673828125 0 22892.768
691.8551025390625 0 21219.021
692.3549194335938 0 12386.752
692.85498046875 0 3532.3286
693.3612670898438 0 1050.5924
695.3704223632812 0 5407.752 y 6
696.3695678710938 0 2215.2751
704.3682250976562 0 26153.285 c 5
705.3659057617188 0 9383.781
706.3683471679688 0 2376.1902
707.3623657226562 0 631.4544
722.3500366210938 0 936.2272
731.390869140625 0 4015.5093
732.3917236328125 0 1997.4384
737.3291015625 0 1164.2262
737.3930053710938 0 723.69104
749.3831176757812 0 14293.315 w 5
750.3855590820312 0 7603.0864
751.3858642578125 0 2335.1462
758.3809814453125 0 2280.4639 c Ammonia loss 6
759.3787231445312 0 907.5746
771.3469848632812 0 781.1246
775.4044799804688 0 23939.684 c 6
776.402099609375 0 10507.162
777.4022827148438 0 3146.4832
778.4013061523438 0 683.2618
787.3905029296875 0 1323.8918
793.3956298828125 0 55840.652 z 5
794.3927612304688 0 30352.18
795.39306640625 0 8464.108
796.3960571289062 0 1639.4457
809.4196166992188 0 796.3671 y 5
810.4102172851562 0 760.52954
811.3975219726562 0 636.63947
815.4882202148438 0 795.09143
821.4264526367188 0 739.8103
858.4891967773438 0 774.0752
859.488037109375 0 1922.934
860.491455078125 0 2566.1091
861.4898071289062 0 889.13025
864.431884765625 0 17056.66
865.4275512695312 0 10690.093
866.427490234375 0 3271.9866
884.4807739257812 0 821.4872
885.4859619140625 0 782.12665 c Water loss 7
886.475341796875 0 2509.4358 c Ammonia loss 7
887.1654052734375 0 740.7416
887.4660034179688 0 1065.3279
895.4423217773438 0 675.66394
900.4764404296875 0 2377.0383
901.4716796875 0 1654.3584
902.4903564453125 0 9160.17
903.4984130859375 0 33069.57 c 7
904.4970703125 0 16973.537
905.49658203125 0 5035.7974
906.494873046875 0 1286.9967
908.4202880859375 0 18224.637 z 4
909.4188232421875 0 11230.513
910.4173583984375 0 4440.5034
911.4217529296875 0 1159.5496
919.4690551757812 0 1189.2211
921.3572387695312 0 751.2588
921.4617919921875 0 816.3586
924.4346313476562 0 1591.5022 y 4
925.4232788085938 0 1239.5042
958.502685546875 0 696.0671
961.5055541992188 0 689.99896
986.5015258789062 0 1255.4202
987.503662109375 0 3180.9304
988.5066528320312 0 1490.7494
999.5368041992188 0 1400.6086 c Water loss 8
1000.5195922851562 0 2262.6824 c Ammonia loss 8
1001.5192260742188 0 1615.4084
1002.525146484375 0 1497.0256
1016.5332641601562 0 2126.078
1017.5414428710938 0 21325.678 c 8
1018.5392456054688 0 14252.223
1019.538330078125 0 5117.2485
1020.5269775390625 0 1656.593
1060.5579833984375 0 810.41724
1062.553466796875 0 939.89655 y Water loss 3
1064.5218505859375 0 6546.918 z 3
1065.521484375 0 11093.558
1066.5198974609375 0 4477.9917
1067.52197265625 0 2157.2083
1088.555908203125 0 2109.6472
1089.5579833984375 0 1473.4613
1090.5430908203125 0 763.1789
1096.4971923828125 0 614.41223
1097.5045166015625 0 758.8618
1104.5733642578125 0 6295.5273 c 9
1105.571044921875 0 5610.962
1106.5679931640625 0 2156.786
1107.5657958984375 0 792.2259
1136.548095703125 0 730.1442
1151.5535888671875 0 3902.7104 z 2
1152.5574951171875 0 10670.762
1153.558837890625 0 7093.609
1154.559814453125 0 2570.5315
1155.5623779296875 0 708.5327
1158.62158203125 0 844.5628
1159.63232421875 0 928.50507
1167.5775146484375 0 792.5932 y 2
1173.6458740234375 0 1949.7273
1174.6485595703125 0 3206.4536
1175.653076171875 0 3215.205
1176.6495361328125 0 1627.1514
1201.6370849609375 0 5248.0137
1202.635009765625 0 3047.0596
1203.6357421875 0 1818.6705
1217.6568603515625 0 3931.2532 c 10
1218.6580810546875 0 8848.139
1219.659912109375 0 6181.747
1220.657958984375 0 2256.0828
1265.6484375 0 1637.1083
1266.6405029296875 0 2044.998
1267.63525390625 0 1225.6017
1305.6563720703125 0 1553.3674
1306.6546630859375 0 1919.1465
1307.6539306640625 0 1654.0778
1308.6595458984375 0 1407.8536
1320.6915283203125 0 1383.2081
1321.6868896484375 0 2496.1567
1322.6917724609375 0 2222.961
1323.6768798828125 0 4906.485
1324.673095703125 0 8574.802
1325.6756591796875 0 5788.7744
1326.6727294921875 0 2373.9836
1327.66943359375 0 841.7404
1337.6929931640625 0 2231.2026
1338.69677734375 0 11476.761
1339.6959228515625 0 9474.911
1340.695068359375 0 4603.18
1341.68603515625 0 2732.028
1348.679931640625 0 3130.7273
1349.6695556640625 0 4780.7017
1350.6697998046875 0 3955.6628
1351.667724609375 0 1918.4188
1352.6729736328125 0 978.8264
1355.7130126953125 0 849.4355
1364.7159423828125 0 770.5961
1365.6878662109375 0 5635.5225
1366.6912841796875 0 37875.94
1367.689208984375 0 32605.295
1368.68896484375 0 16447.67
1369.68994140625 0 5825.4824
1370.6971435546875 0 1437.2982
1378.3057861328125 0 849.67944
1379.311767578125 0 3193.7466
1380.313720703125 0 1663.656
1381.7135009765625 0 843.77435
1382.706298828125 0 4305.069
1383.712646484375 0 7785.8696
1384.7109375 0 5626.53
1385.7069091796875 0 3326.6238

Spectrum Details

|  |  |
| --- | --- |
| Matched peaks? Matched peaks The total absolute number of peaks matched. Additionally in brackets the total fraction of peaks matched and the total number of peaks is shown. | 54 (13.60% of 397) |
| FDR? FDR The false discovery rate estimated for this peptide. It is calculated by matching all theoretical fragments with a non-integer shift with the raw peaks for this spectrum. This is done with 40 different shifts. The resulting percentage is the average number of annotated peaks over the number of annotated peaks with the correct spectrum. | 1.32% |
| Satellite FDR? Satellite FDR See the FDR for details on its calculation. This satellite ion specific FDR only contains the satellite ions (d/w) for I/L/J positions. | 0.00% |
| PSM Score? PSM Score The PSM Score as given by Hecklib to this annotated spectrum. It is shown with three significant figures. | 465 |

#### Spectrum 4196? Spectrum 4196 The raw spectrum of this peptide as annotated by Hecklib. The fragments are coloured according to ion type (see legend). Any peaks with a star '\*' as text can be hovered over to see the full details, first the ion type second the mass shift type. By hovering over the amino acids in the peptide or ions in the legend the corresponding peaks are highlighted. By toggling the 'Unassigned' label you can turn the background (unassigned) peaks on or off in the plot. By updating the slider in the Ion legend you can update the spectrum to only show the top X% of the peaks with labels. The top X% means any peak that is within X% of the highest intensity. By dragging in the spectrum you can zoom in to a specific part of the spectrum and use 'Zoom Out' to get back to the original zoom level. The annotation of the spectrum is based on the given sequence in the peptides file and is done with different software so inconsistencies are likely. The peaks are annotated based on the given sequence, with 20 ppm tolerance.

Copy Data

##### Spectrum 4196 (TSV)

###### Preview

```
Loading example...
```

*Click on the button to copy the data to your clipboard.*

Mz MinMz MaxIntensity Max

WidthHeightPeptide font sizePeptide stroke widthSpectrum font sizeSpectrum stroke widthCompact peptide

Ion legend

wxyz

abcd

OtherUnassignedIon^Charge^_Position_Show for top:%

TISRDNAKNSJY

03.01e+46.02e+49.03e+41.20e+5

Zoom Out

y^+1^_1_y^+1^_2_c^+1^_3_c^+2^_7_y^+1^_3_c^+2^_8_c^+2^_8_c^+2^_8_c^+1^_4_c^+1^_4_y^+1^_4_c^+2^_9_c^+2^_9_c^+2^_9_z^+2^_9_c^+2^_10_c^+2^_10_z^+2^_10_w^+2^_10_c^+1^_5_y^+2^_10_z^+2^_10_y^+2^_10_c^+1^_5_c^+2^_11_z^+1^_5_c^+2^_11_w^+2^_11_y^+1^_5_z^+2^_11_y^+2^_11_z^+1^_6_z^+1^_6_c^+1^_6_y^+1^_6_c^+1^_6_w^+1^_7_c^+1^_7_c^+1^_7_z^+1^_7_c^+1^_8_c^+1^_8_c^+1^_8_z^+1^_8_y^+1^_8_c^+1^_9_c^+1^_9_c^+1^_9_z^+1^_9_c^+1^_10_w^+1^_10_z^+1^_10_c^+1^_11_

035070110511402

Fragment Matches Table

Show background peaks

| Position | Ion type | Intensity | mz Theoretical | mz Error (Th) | mz Error (ppm) | Charge | Series Number |
| --- | --- | --- | --- | --- | --- | --- | --- |
| - | - | 465.1 | 120.3 | - | - | 0 | - |
| - | - | 373.5 | 123.4 | - | - | 0 | - |
| - | - | 700.2 | 129.1 | - | - | 0 | - |
| - | - | 882.8 | 133.1 | - | - | 0 | - |
| - | - | 3545 | 136.1 | - | - | 0 | - |
| - | - | 444.2 | 142 | - | - | 0 | - |
| - | - | 544.7 | 142.1 | - | - | 0 | - |
| - | - | 3928 | 149 | - | - | 0 | - |
| - | - | 1412 | 150 | - | - | 0 | - |
| - | - | 501.1 | 151 | - | - | 0 | - |
| - | - | 504.8 | 157.1 | - | - | 0 | - |
| - | - | 407.6 | 157.4 | - | - | 0 | - |
| - | - | 3126 | 165.1 | - | - | 0 | - |
| - | - | 1470 | 167.1 | - | - | 0 | - |
| - | - | 521.4 | 169.1 | - | - | 0 | - |
| - | - | 2817 | 173.1 | - | - | 0 | - |
| - | - | 597.2 | 173.1 | - | - | 0 | - |
| - | - | 2823 | 173.4 | - | - | 0 | - |
| - | - | 500.7 | 176.6 | - | - | 0 | - |
| 12 | y | 1.221E+04 | 182.1 | 0.0002207 | 1.212 | +1 | 1 |
| - | - | 880.6 | 183.1 | - | - | 0 | - |
| - | - | 2.209E+04 | 187.1 | - | - | 0 | - |
| - | - | 2251 | 188.1 | - | - | 0 | - |
| - | - | 1224 | 201.1 | - | - | 0 | - |
| - | - | 483.3 | 209 | - | - | 0 | - |
| - | - | 6194 | 215.1 | - | - | 0 | - |
| - | - | 2030 | 221.1 | - | - | 0 | - |
| - | - | 1092 | 222.1 | - | - | 0 | - |
| - | - | 1407 | 223.1 | - | - | 0 | - |
| - | - | 654 | 223.1 | - | - | 0 | - |
| - | - | 2191 | 225 | - | - | 0 | - |
| - | - | 848.5 | 226 | - | - | 0 | - |
| - | - | 643.8 | 227 | - | - | 0 | - |
| - | - | 711.7 | 229.1 | - | - | 0 | - |
| - | - | 517.9 | 229.5 | - | - | 0 | - |
| - | - | 2800 | 239.1 | - | - | 0 | - |
| - | - | 1785 | 240.1 | - | - | 0 | - |
| - | - | 1051 | 241.1 | - | - | 0 | - |
| - | - | 1235 | 260.1 | - | - | 0 | - |
| - | - | 643.1 | 267.2 | - | - | 0 | - |
| - | - | 737.7 | 281.1 | - | - | 0 | - |
| - | - | 915 | 282.1 | - | - | 0 | - |
| - | - | 1358 | 282.1 | - | - | 0 | - |
| - | - | 1038 | 295.1 | - | - | 0 | - |
| 11 | y | 767.1 | 295.2 | 0.000904 | 3.063 | +1 | 2 |
| - | - | 2709 | 296.1 | - | - | 0 | - |
| - | - | 1647 | 297.1 | - | - | 0 | - |
| - | - | 502.4 | 297.5 | - | - | 0 | - |
| - | - | 1815 | 299.1 | - | - | 0 | - |
| - | - | 551.5 | 299.2 | - | - | 0 | - |
| - | - | 2337 | 300.1 | - | - | 0 | - |
| - | - | 4162 | 300.2 | - | - | 0 | - |
| - | - | 1448 | 301.1 | - | - | 0 | - |
| - | - | 541.9 | 301.2 | - | - | 0 | - |
| - | - | 639.2 | 314.1 | - | - | 0 | - |
| - | - | 585.6 | 315.2 | - | - | 0 | - |
| - | - | 2378 | 317.2 | - | - | 0 | - |
| 3 | c | 1927 | 319.2 | 0.001104 | 3.457 | +1 | 3 |
| - | - | 859.1 | 325 | - | - | 0 | - |
| - | - | 1.489E+04 | 341 | - | - | 0 | - |
| - | - | 662.3 | 343 | - | - | 0 | - |
| - | - | 519.8 | 351.4 | - | - | 0 | - |
| - | - | 1894 | 355.1 | - | - | 0 | - |
| - | - | 781.8 | 356.1 | - | - | 0 | - |
| - | - | 590.4 | 357.9 | - | - | 0 | - |
| - | - | 4.878E+04 | 359 | - | - | 0 | - |
| - | - | 904.1 | 369.1 | - | - | 0 | - |
| - | - | 2834 | 370.1 | - | - | 0 | - |
| - | - | 2065 | 371.1 | - | - | 0 | - |
| - | - | 618.6 | 373.1 | - | - | 0 | - |
| - | - | 552.5 | 374.2 | - | - | 0 | - |
| - | - | 495.7 | 375.3 | - | - | 0 | - |
| - | - | 847.5 | 376.9 | - | - | 0 | - |
| 7 | c | 796.1 | 379.2 | 0.006806 | 17.95 | +2 | 7 |
| - | - | 723.6 | 380.2 | - | - | 0 | - |
| 10 | y | 823.6 | 382.2 | 0.002323 | 6.078 | +1 | 3 |
| - | - | 3318 | 385.2 | - | - | 0 | - |
| - | - | 789.1 | 385.5 | - | - | 0 | - |
| - | - | 1036 | 385.9 | - | - | 0 | - |
| - | - | 674.9 | 386.2 | - | - | 0 | - |
| - | - | 1456 | 391.6 | - | - | 0 | - |
| - | - | 496.1 | 392.2 | - | - | 0 | - |
| - | - | 833.6 | 394.7 | - | - | 0 | - |
| - | - | 960.2 | 409.2 | - | - | 0 | - |
| - | - | 671.4 | 421.6 | - | - | 0 | - |
| - | - | 545.1 | 428.7 | - | - | 0 | - |
| - | - | 1.192E+05 | 429.1 | - | - | 0 | - |
| - | - | 771.1 | 429.8 | - | - | 0 | - |
| - | - | 1226 | 430.2 | - | - | 0 | - |
| - | - | 1046 | 433.2 | - | - | 0 | - |
| - | - | 531.2 | 442.2 | - | - | 0 | - |
| - | - | 2211 | 442.7 | - | - | 0 | - |
| 8 | c | 925.9 | 443.2 | 0.00677 | 15.27 | +2 | 8 |
| 8 | c | 1252 | 443.7 | 2.942E-05 | 0.0663 | +2 | 8 |
| - | - | 1750 | 444.2 | - | - | 0 | - |
| - | - | 6.564E+04 | 445.1 | - | - | 0 | - |
| - | - | 8274 | 445.3 | - | - | 0 | - |
| - | - | 1908 | 446.2 | - | - | 0 | - |
| - | - | 762.1 | 447.1 | - | - | 0 | - |
| - | - | 675.3 | 451.7 | - | - | 0 | - |
| - | - | 4464 | 451.8 | - | - | 0 | - |
| 8 | c | 1.806E+04 | 452.3 | 3.221E-05 | 0.07122 | +2 | 8 |
| - | - | 8455 | 452.8 | - | - | 0 | - |
| - | - | 3093 | 453.3 | - | - | 0 | - |
| 4 | c | 1011 | 458.3 | 0.0001805 | 0.3939 | +1 | 4 |
| - | - | 1213 | 459.7 | - | - | 0 | - |
| - | - | 4486 | 460.2 | - | - | 0 | - |
| - | - | 1777 | 460.7 | - | - | 0 | - |
| - | - | 982.4 | 461.1 | - | - | 0 | - |
| - | - | 782.4 | 461.2 | - | - | 0 | - |
| - | - | 643.4 | 461.2 | - | - | 0 | - |
| - | - | 642.1 | 462.1 | - | - | 0 | - |
| - | - | 815.4 | 470.7 | - | - | 0 | - |
| - | - | 1016 | 471.2 | - | - | 0 | - |
| 4 | c | 6.097E+04 | 475.3 | 0.00067 | 1.41 | +1 | 4 |
| - | - | 1.404E+04 | 476.3 | - | - | 0 | - |
| - | - | 2052 | 477.3 | - | - | 0 | - |
| - | - | 6167 | 479.8 | - | - | 0 | - |
| - | - | 2310 | 480.3 | - | - | 0 | - |
| - | - | 1742 | 480.8 | - | - | 0 | - |
| - | - | 3026 | 481.2 | - | - | 0 | - |
| - | - | 805.6 | 484.7 | - | - | 0 | - |
| - | - | 1384 | 485.2 | - | - | 0 | - |
| - | - | 737.3 | 486.8 | - | - | 0 | - |
| - | - | 1929 | 487.3 | - | - | 0 | - |
| - | - | 1041 | 493.8 | - | - | 0 | - |
| - | - | 778.3 | 494.3 | - | - | 0 | - |
| 9 | y | 2382 | 496.2 | 0.0005026 | 1.013 | +1 | 4 |
| - | - | 6076 | 499.8 | - | - | 0 | - |
| 9 | c | 3633 | 500.3 | 0.00498 | 9.954 | +2 | 9 |
| 9 | c | 3446 | 500.8 | 0.002066 | 4.127 | +2 | 9 |
| - | - | 1869 | 501.3 | - | - | 0 | - |
| - | - | 1175 | 501.8 | - | - | 0 | - |
| - | - | 6060 | 502.3 | - | - | 0 | - |
| - | - | 1696 | 503.3 | - | - | 0 | - |
| - | - | 639.5 | 504.3 | - | - | 0 | - |
| - | - | 2098 | 508.3 | - | - | 0 | - |
| - | - | 1.543E+04 | 508.8 | - | - | 0 | - |
| 9 | c | 2.491E+04 | 509.3 | 0.0008932 | 1.754 | +2 | 9 |
| - | - | 1.377E+04 | 509.8 | - | - | 0 | - |
| - | - | 5652 | 510.3 | - | - | 0 | - |
| - | - | 1298 | 510.8 | - | - | 0 | - |
| - | - | 587.4 | 521.3 | - | - | 0 | - |
| - | - | 568.1 | 523 | - | - | 0 | - |
| - | - | 595.8 | 526.3 | - | - | 0 | - |
| - | - | 717.1 | 526.8 | - | - | 0 | - |
| 4 | z | 2020 | 532.8 | 0.0005184 | 0.9729 | +2 | 9 |
| - | - | 1878 | 533.3 | - | - | 0 | - |
| - | - | 687.4 | 533.8 | - | - | 0 | - |
| - | - | 646.9 | 535.3 | - | - | 0 | - |
| - | - | 799.9 | 535.8 | - | - | 0 | - |
| - | - | 891.1 | 541.3 | - | - | 0 | - |
| 10 | c | 2881 | 544.3 | 0.0008227 | 1.512 | +2 | 10 |
| - | - | 1934 | 544.8 | - | - | 0 | - |
| - | - | 1792 | 545.3 | - | - | 0 | - |
| - | - | 1412 | 546.3 | - | - | 0 | - |
| - | - | 1909 | 548.3 | - | - | 0 | - |
| - | - | 1489 | 548.8 | - | - | 0 | - |
| 10 | c | 1.688E+04 | 552.8 | 0.0005487 | 0.9925 | +2 | 10 |
| - | - | 1.238E+04 | 553.3 | - | - | 0 | - |
| - | - | 5601 | 553.8 | - | - | 0 | - |
| - | - | 2099 | 554.3 | - | - | 0 | - |
| - | - | 1532 | 554.8 | - | - | 0 | - |
| - | - | 2357 | 564.3 | - | - | 0 | - |
| - | - | 882.6 | 565.3 | - | - | 0 | - |
| - | - | 1565 | 565.8 | - | - | 0 | - |
| - | - | 1230 | 566.3 | - | - | 0 | - |
| 3 | z | 852.8 | 567.3 | 0.002645 | 4.663 | +2 | 10 |
| 3 | w | 993.8 | 567.8 | 0.001316 | 2.318 | +2 | 10 |
| - | - | 972.9 | 568.3 | - | - | 0 | - |
| - | - | 659.9 | 569.3 | - | - | 0 | - |
| 5 | c | 914.9 | 573.3 | 0.005525 | 9.637 | +1 | 5 |
| 3 | y | 725.5 | 575.8 | 0.006056 | 10.52 | +2 | 10 |
| 3 | z | 8089 | 576.3 | 0.0003123 | 0.5418 | +2 | 10 |
| - | - | 5291 | 576.8 | - | - | 0 | - |
| - | - | 2605 | 577.3 | - | - | 0 | - |
| - | - | 1066 | 578.3 | - | - | 0 | - |
| - | - | 834.8 | 579.3 | - | - | 0 | - |
| - | - | 845.1 | 583.3 | - | - | 0 | - |
| 3 | y | 1.659E+04 | 584.3 | 0.0001055 | 0.1805 | +2 | 10 |
| - | - | 1.429E+04 | 584.8 | - | - | 0 | - |
| - | - | 8570 | 585.3 | - | - | 0 | - |
| - | - | 2233 | 585.8 | - | - | 0 | - |
| - | - | 1873 | 586.8 | - | - | 0 | - |
| - | - | 2259 | 587.3 | - | - | 0 | - |
| - | - | 734.1 | 587.8 | - | - | 0 | - |
| - | - | 1153 | 588.3 | - | - | 0 | - |
| - | - | 907.6 | 588.8 | - | - | 0 | - |
| - | - | 896.9 | 589.3 | - | - | 0 | - |
| 5 | c | 1.129E+05 | 590.3 | 0.0004604 | 0.7798 | +1 | 5 |
| - | - | 3.281E+04 | 591.3 | - | - | 0 | - |
| - | - | 1380 | 591.8 | - | - | 0 | - |
| - | - | 6493 | 592.3 | - | - | 0 | - |
| - | - | 960.1 | 592.8 | - | - | 0 | - |
| 11 | c | 2731 | 600.8 | 0.00121 | 2.014 | +2 | 11 |
| - | - | 2588 | 601.3 | - | - | 0 | - |
| - | - | 2789 | 601.8 | - | - | 0 | - |
| - | - | 1340 | 602.3 | - | - | 0 | - |
| - | - | 929.7 | 607.3 | - | - | 0 | - |
| 8 | z | 5912 | 608.3 | 0.0005271 | 0.8665 | +1 | 5 |
| 11 | c | 4.756E+04 | 609.3 | 2.058E-05 | 0.03378 | +2 | 11 |
| - | - | 3.154E+04 | 609.8 | - | - | 0 | - |
| - | - | 1.412E+04 | 610.3 | - | - | 0 | - |
| - | - | 5236 | 610.8 | - | - | 0 | - |
| - | - | 1068 | 611.3 | - | - | 0 | - |
| - | - | 1184 | 616.3 | - | - | 0 | - |
| 2 | w | 2367 | 618.3 | 0.0008531 | 1.38 | +2 | 11 |
| - | - | 2094 | 618.8 | - | - | 0 | - |
| - | - | 1377 | 619.3 | - | - | 0 | - |
| 8 | y | 6845 | 624.3 | 0.0005408 | 0.8662 | +1 | 5 |
| - | - | 2380 | 625.3 | - | - | 0 | - |
| - | - | 1378 | 630.3 | - | - | 0 | - |
| - | - | 1468 | 630.8 | - | - | 0 | - |
| - | - | 783.3 | 631.3 | - | - | 0 | - |
| 2 | z | 655.1 | 632.8 | 0.003207 | 5.067 | +2 | 11 |
| - | - | 1952 | 633.3 | - | - | 0 | - |
| - | - | 1247 | 633.8 | - | - | 0 | - |
| - | - | 833.3 | 636.3 | - | - | 0 | - |
| - | - | 1037 | 638.8 | - | - | 0 | - |
| - | - | 803.8 | 639.4 | - | - | 0 | - |
| - | - | 700.6 | 639.8 | - | - | 0 | - |
| - | - | 1154 | 640.3 | - | - | 0 | - |
| 2 | y | 3216 | 640.8 | 0.0001267 | 0.1977 | +2 | 11 |
| - | - | 1822 | 641.3 | - | - | 0 | - |
| - | - | 1378 | 641.8 | - | - | 0 | - |
| - | - | 970.2 | 642.3 | - | - | 0 | - |
| - | - | 1474 | 647.4 | - | - | 0 | - |
| - | - | 2504 | 647.9 | - | - | 0 | - |
| - | - | 848.8 | 653.3 | - | - | 0 | - |
| - | - | 1674 | 653.8 | - | - | 0 | - |
| - | - | 886.5 | 654.8 | - | - | 0 | - |
| - | - | 4044 | 655.3 | - | - | 0 | - |
| - | - | 3173 | 655.8 | - | - | 0 | - |
| - | - | 981 | 656.3 | - | - | 0 | - |
| - | - | 4941 | 659.3 | - | - | 0 | - |
| - | - | 6910 | 660.3 | - | - | 0 | - |
| - | - | 3373 | 660.9 | - | - | 0 | - |
| 7 | z | 7508 | 661.3 | 0.008524 | 12.89 | +1 | 6 |
| - | - | 7498 | 661.8 | - | - | 0 | - |
| - | - | 7543 | 662.3 | - | - | 0 | - |
| - | - | 5452 | 662.8 | - | - | 0 | - |
| - | - | 2788 | 663.3 | - | - | 0 | - |
| - | - | 711.9 | 663.8 | - | - | 0 | - |
| - | - | 3058 | 668.8 | - | - | 0 | - |
| - | - | 8092 | 669.3 | - | - | 0 | - |
| - | - | 5827 | 669.9 | - | - | 0 | - |
| - | - | 6809 | 670.3 | - | - | 0 | - |
| - | - | 6364 | 670.8 | - | - | 0 | - |
| - | - | 2833 | 671.3 | - | - | 0 | - |
| - | - | 1127 | 672.3 | - | - | 0 | - |
| - | - | 669.6 | 673.8 | - | - | 0 | - |
| - | - | 2243 | 674.3 | - | - | 0 | - |
| - | - | 4743 | 674.8 | - | - | 0 | - |
| - | - | 5392 | 675.3 | - | - | 0 | - |
| - | - | 4843 | 675.8 | - | - | 0 | - |
| - | - | 2804 | 676.3 | - | - | 0 | - |
| - | - | 886.2 | 676.8 | - | - | 0 | - |
| - | - | 1113 | 677.3 | - | - | 0 | - |
| 7 | z | 2.241E+04 | 679.4 | 0.0002785 | 0.41 | +1 | 6 |
| - | - | 8542 | 680.4 | - | - | 0 | - |
| - | - | 1972 | 681.4 | - | - | 0 | - |
| - | - | 1177 | 681.9 | - | - | 0 | - |
| - | - | 703.9 | 682.4 | - | - | 0 | - |
| - | - | 1010 | 682.9 | - | - | 0 | - |
| - | - | 2.732E+04 | 683.3 | - | - | 0 | - |
| - | - | 2.96E+04 | 683.8 | - | - | 0 | - |
| - | - | 1.364E+04 | 684.3 | - | - | 0 | - |
| - | - | 5165 | 684.8 | - | - | 0 | - |
| - | - | 815.7 | 685.3 | - | - | 0 | - |
| 6 | c | 1143 | 687.3 | 0.005444 | 7.921 | +1 | 6 |
| - | - | 1321 | 689.2 | - | - | 0 | - |
| - | - | 4467 | 689.7 | - | - | 0 | - |
| - | - | 2897 | 690.2 | - | - | 0 | - |
| - | - | 2.533E+04 | 691.4 | - | - | 0 | - |
| - | - | 2.199E+04 | 691.9 | - | - | 0 | - |
| - | - | 1.482E+04 | 692.4 | - | - | 0 | - |
| - | - | 5261 | 692.9 | - | - | 0 | - |
| - | - | 1837 | 693.4 | - | - | 0 | - |
| - | - | 2999 | 693.9 | - | - | 0 | - |
| - | - | 829 | 694.4 | - | - | 0 | - |
| 7 | y | 5668 | 695.4 | 0.000257 | 0.3697 | +1 | 6 |
| - | - | 1516 | 696.4 | - | - | 0 | - |
| 6 | c | 2.864E+04 | 704.4 | 0.0003796 | 0.5389 | +1 | 6 |
| - | - | 9668 | 705.4 | - | - | 0 | - |
| - | - | 2174 | 706.4 | - | - | 0 | - |
| - | - | 2206 | 722.3 | - | - | 0 | - |
| - | - | 4057 | 731.4 | - | - | 0 | - |
| - | - | 2593 | 732.4 | - | - | 0 | - |
| - | - | 1083 | 737.3 | - | - | 0 | - |
| - | - | 890.8 | 737.4 | - | - | 0 | - |
| 6 | w | 1.165E+04 | 749.4 | 0.001935 | 2.582 | +1 | 7 |
| - | - | 9269 | 750.4 | - | - | 0 | - |
| - | - | 2531 | 751.4 | - | - | 0 | - |
| 7 | c | 2151 | 758.4 | 0.002022 | 2.666 | +1 | 7 |
| - | - | 781.3 | 759.4 | - | - | 0 | - |
| - | - | 929.3 | 774.4 | - | - | 0 | - |
| 7 | c | 2.735E+04 | 775.4 | 0.0001741 | 0.2246 | +1 | 7 |
| - | - | 9816 | 776.4 | - | - | 0 | - |
| - | - | 3607 | 777.4 | - | - | 0 | - |
| - | - | 1196 | 787.4 | - | - | 0 | - |
| 6 | z | 6.366E+04 | 793.4 | 0.0003809 | 0.4801 | +1 | 7 |
| - | - | 3.3E+04 | 794.4 | - | - | 0 | - |
| - | - | 9715 | 795.4 | - | - | 0 | - |
| - | - | 1906 | 796.4 | - | - | 0 | - |
| - | - | 1180 | 810.4 | - | - | 0 | - |
| - | - | 711.1 | 843.2 | - | - | 0 | - |
| - | - | 685 | 858.5 | - | - | 0 | - |
| - | - | 1989 | 859.5 | - | - | 0 | - |
| - | - | 2522 | 860.5 | - | - | 0 | - |
| - | - | 2.15E+04 | 864.4 | - | - | 0 | - |
| - | - | 1.177E+04 | 865.4 | - | - | 0 | - |
| - | - | 3984 | 866.4 | - | - | 0 | - |
| - | - | 1529 | 867.4 | - | - | 0 | - |
| - | - | 1354 | 874.4 | - | - | 0 | - |
| - | - | 766.1 | 884.5 | - | - | 0 | - |
| 8 | c | 980 | 885.5 | 0.008828 | 9.969 | +1 | 8 |
| 8 | c | 3288 | 886.5 | 0.001297 | 1.463 | +1 | 8 |
| - | - | 1599 | 887.5 | - | - | 0 | - |
| - | - | 864.9 | 888.2 | - | - | 0 | - |
| - | - | 2500 | 900.5 | - | - | 0 | - |
| - | - | 1037 | 901.5 | - | - | 0 | - |
| - | - | 8770 | 902.5 | - | - | 0 | - |
| 8 | c | 3.439E+04 | 903.5 | 0.001204 | 1.333 | +1 | 8 |
| - | - | 1068 | 904.3 | - | - | 0 | - |
| - | - | 1.852E+04 | 904.5 | - | - | 0 | - |
| - | - | 6303 | 905.5 | - | - | 0 | - |
| - | - | 1618 | 906.5 | - | - | 0 | - |
| 5 | z | 1.865E+04 | 908.4 | 0.003613 | 3.977 | +1 | 8 |
| - | - | 1.296E+04 | 909.4 | - | - | 0 | - |
| - | - | 5232 | 910.4 | - | - | 0 | - |
| - | - | 1770 | 911.4 | - | - | 0 | - |
| - | - | 1498 | 919.5 | - | - | 0 | - |
| - | - | 1960 | 920.5 | - | - | 0 | - |
| 5 | y | 1894 | 924.4 | 0.004332 | 4.686 | +1 | 8 |
| - | - | 1441 | 925.4 | - | - | 0 | - |
| - | - | 1655 | 986.5 | - | - | 0 | - |
| - | - | 2080 | 987.5 | - | - | 0 | - |
| - | - | 973.1 | 988.5 | - | - | 0 | - |
| 9 | c | 1356 | 999.5 | 0.003359 | 3.361 | +1 | 9 |
| 9 | c | 1940 | 1001 | 0.004512 | 4.51 | +1 | 9 |
| - | - | 1230 | 1002 | - | - | 0 | - |
| - | - | 1079 | 1003 | - | - | 0 | - |
| - | - | 2295 | 1017 | - | - | 0 | - |
| 9 | c | 2.351E+04 | 1018 | 0.0009796 | 0.9628 | +1 | 9 |
| - | - | 1.262E+04 | 1019 | - | - | 0 | - |
| - | - | 4573 | 1020 | - | - | 0 | - |
| - | - | 2641 | 1021 | - | - | 0 | - |
| - | - | 1053 | 1023 | - | - | 0 | - |
| - | - | 842.9 | 1063 | - | - | 0 | - |
| 4 | z | 8544 | 1065 | 0.0006227 | 0.585 | +1 | 9 |
| - | - | 1.022E+04 | 1066 | - | - | 0 | - |
| - | - | 5961 | 1067 | - | - | 0 | - |
| - | - | 1836 | 1068 | - | - | 0 | - |
| - | - | 1918 | 1089 | - | - | 0 | - |
| - | - | 1841 | 1090 | - | - | 0 | - |
| - | - | 1209 | 1091 | - | - | 0 | - |
| - | - | 1002 | 1097 | - | - | 0 | - |
| 10 | c | 7089 | 1105 | 0.0004392 | 0.3976 | +1 | 10 |
| - | - | 7055 | 1106 | - | - | 0 | - |
| - | - | 2598 | 1107 | - | - | 0 | - |
| - | - | 1182 | 1108 | - | - | 0 | - |
| 3 | w | 1113 | 1135 | 0.000875 | 0.7712 | +1 | 10 |
| 3 | z | 4830 | 1152 | 0.001498 | 1.301 | +1 | 10 |
| - | - | 1.289E+04 | 1153 | - | - | 0 | - |
| - | - | 6175 | 1154 | - | - | 0 | - |
| - | - | 3244 | 1155 | - | - | 0 | - |
| - | - | 821 | 1159 | - | - | 0 | - |
| - | - | 805.7 | 1160 | - | - | 0 | - |
| - | - | 2270 | 1174 | - | - | 0 | - |
| - | - | 3201 | 1175 | - | - | 0 | - |
| - | - | 2464 | 1176 | - | - | 0 | - |
| - | - | 854.6 | 1177 | - | - | 0 | - |
| - | - | 5619 | 1202 | - | - | 0 | - |
| - | - | 3790 | 1203 | - | - | 0 | - |
| - | - | 1649 | 1204 | - | - | 0 | - |
| 11 | c | 5407 | 1218 | 0.0004949 | 0.4064 | +1 | 11 |
| - | - | 8359 | 1219 | - | - | 0 | - |
| - | - | 4996 | 1220 | - | - | 0 | - |
| - | - | 2498 | 1221 | - | - | 0 | - |
| - | - | 1184 | 1222 | - | - | 0 | - |
| - | - | 1315 | 1266 | - | - | 0 | - |
| - | - | 2168 | 1267 | - | - | 0 | - |
| - | - | 1541 | 1306 | - | - | 0 | - |
| - | - | 1748 | 1307 | - | - | 0 | - |
| - | - | 1422 | 1308 | - | - | 0 | - |
| - | - | 1128 | 1309 | - | - | 0 | - |
| - | - | 940 | 1310 | - | - | 0 | - |
| - | - | 833.7 | 1321 | - | - | 0 | - |
| - | - | 3113 | 1322 | - | - | 0 | - |
| - | - | 1648 | 1323 | - | - | 0 | - |
| - | - | 4150 | 1324 | - | - | 0 | - |
| - | - | 8451 | 1325 | - | - | 0 | - |
| - | - | 6736 | 1326 | - | - | 0 | - |
| - | - | 2290 | 1327 | - | - | 0 | - |
| - | - | 1956 | 1338 | - | - | 0 | - |
| - | - | 1.21E+04 | 1339 | - | - | 0 | - |
| - | - | 8347 | 1340 | - | - | 0 | - |
| - | - | 5066 | 1341 | - | - | 0 | - |
| - | - | 1922 | 1342 | - | - | 0 | - |
| - | - | 2354 | 1349 | - | - | 0 | - |
| - | - | 4529 | 1350 | - | - | 0 | - |
| - | - | 5110 | 1351 | - | - | 0 | - |
| - | - | 1941 | 1352 | - | - | 0 | - |
| - | - | 878.3 | 1365 | - | - | 0 | - |
| - | - | 7387 | 1366 | - | - | 0 | - |
| - | - | 3.996E+04 | 1367 | - | - | 0 | - |
| - | - | 3.303E+04 | 1368 | - | - | 0 | - |
| - | - | 1.755E+04 | 1369 | - | - | 0 | - |
| - | - | 6309 | 1370 | - | - | 0 | - |
| - | - | 1212 | 1371 | - | - | 0 | - |
| - | - | 1464 | 1378 | - | - | 0 | - |
| - | - | 2583 | 1379 | - | - | 0 | - |
| - | - | 2135 | 1380 | - | - | 0 | - |
| - | - | 2345 | 1382 | - | - | 0 | - |
| - | - | 5741 | 1383 | - | - | 0 | - |
| - | - | 7309 | 1384 | - | - | 0 | - |
| - | - | 6588 | 1385 | - | - | 0 | - |
| - | - | 3643 | 1386 | - | - | 0 | - |
| - | - | 964.8 | 1387 | - | - | 0 | - |
| - | - | 2248 | 1388 | - | - | 0 | - |

m/z Charge Intensity FragmentType MassShift Position
120.30294036865234 0 465.14633
123.43548583984375 0 373.45508
129.1023406982422 0 700.1869
133.0861053466797 0 882.8128
136.075927734375 0 3545.2107
142.03367614746094 0 444.20108
142.1233367919922 0 544.65137
149.04513549804688 0 3927.874
150.044677734375 0 1412.2996
151.04197692871094 0 501.0629
157.1343231201172 0 504.80777
157.4176483154297 0 407.64746
165.05477905273438 0 3125.522
167.0556182861328 0 1470.3878
169.1340789794922 0 521.43555
173.09230041503906 0 2817.3425
173.12945556640625 0 597.16943
173.44073486328125 0 2822.781
176.56056213378906 0 500.71866
182.08139038085938 0 12208.76 y 11
183.08438110351562 0 880.60724
187.1443634033203 0 22086.225
188.1476593017578 0 2250.6963
201.12362670898438 0 1223.7246
209.04127502441406 0 483.2515
215.13919067382812 0 6193.5474
221.08480834960938 0 2029.9227
222.08509826660156 0 1091.7163
223.0641326904297 0 1406.8182
223.08123779296875 0 654.008
225.0432586669922 0 2190.88
226.04293823242188 0 848.5141
227.03945922851562 0 643.81274
229.12957763671875 0 711.6746
229.5340118408203 0 517.90936
239.09506225585938 0 2799.8643
240.09608459472656 0 1784.5453
241.09259033203125 0 1051.366
260.124267578125 0 1235.1875
267.19683837890625 0 643.06696
281.0508728027344 0 737.73254
282.0506896972656 0 914.9574
282.1458740234375 0 1358.4275
295.103271484375 0 1038.4369
295.1661376953125 0 767.0653 y 10
296.1044921875 0 2708.641
297.10113525390625 0 1646.5566
297.53466796875 0 502.39557
299.061767578125 0 1814.6211
299.1711730957031 0 551.4679
300.0626525878906 0 2336.522
300.1556396484375 0 4161.5645
301.0594787597656 0 1447.5459
301.1595153808594 0 541.8987
314.1156311035156 0 639.19446
315.16583251953125 0 585.6439
317.1821594238281 0 2377.9814
319.1986999511719 0 1927.322 c 2
324.9871826171875 0 859.1097
341.0183410644531 0 14891.743
342.99639892578125 0 662.27374
351.4476623535156 0 519.81934
355.0700988769531 0 1894.2001
356.0710144042969 0 781.75946
357.8697204589844 0 590.43536
359.0286560058594 0 48781.82
369.1225891113281 0 904.1235
370.1230773925781 0 2834.3342
371.11962890625 0 2065.1367
373.1101989746094 0 618.6441
374.1800842285156 0 552.45044
375.25360107421875 0 495.6861
376.8973693847656 0 847.51953
379.2080078125 0 796.0762 c Water loss 6
380.2088317871094 0 723.5908
382.1995849609375 0 823.55347 y 9
385.2197570800781 0 3318.2983
385.5467529296875 0 789.0867
385.8798522949219 0 1035.8107
386.2275085449219 0 674.89136
391.55255126953125 0 1456.3373
392.2181396484375 0 496.06174
394.7034912109375 0 833.6261
409.2082214355469 0 960.1812
421.60980224609375 0 671.431
428.73370361328125 0 545.1317
429.0892333984375 0 119152.09
429.7549743652344 0 771.0771
430.24224853515625 0 1226.24
433.2204895019531 0 1045.9496
442.2215270996094 0 531.21356
442.7451171875 0 2210.879
443.2419128417969 0 925.9174 c Water loss 7
443.74066162109375 0 1252.0071 c Ammonia loss 7
444.24310302734375 0 1749.7942
445.1205749511719 0 65636.78
445.2648010253906 0 8274.188
446.2118835449219 0 1907.8054
447.09613037109375 0 762.0727
451.6850280761719 0 675.2615
451.7502746582031 0 4464.241
452.2539978027344 0 18063.889 c 7
452.7540588378906 0 8455.28
453.2541809082031 0 3093.4243
458.2723388671875 0 1010.9161 c Ammonia loss 3
459.6999206542969 0 1213.2008
460.23712158203125 0 4486.2905
460.7368469238281 0 1776.6677
461.1429443359375 0 982.3972
461.19598388671875 0 782.35876
461.2366943359375 0 643.4144
462.11260986328125 0 642.12976
470.74859619140625 0 815.4393
471.24896240234375 0 1016.2607
475.29937744140625 0 60967.26 c 3
476.3018798828125 0 14040.715
477.3046569824219 0 2051.7021
479.7591857910156 0 6166.685
480.2568664550781 0 2310.4766
480.7561340332031 0 1742.0084
481.22998046875 0 3025.8455
484.7486572265625 0 805.61725
485.2457275390625 0 1384.1809
486.77728271484375 0 737.3496
487.26898193359375 0 1929.2628
493.75421142578125 0 1041.2064
494.25439453125 0 778.2742
496.2406921386719 0 2381.674 y 8
499.76678466796875 0 6076.3325
500.2651672363281 0 3632.8555 c Water loss 8
500.76422119140625 0 3445.794 c Ammonia loss 8
501.2619323730469 0 1868.8927
501.76544189453125 0 1175.393
502.2746276855469 0 6060.06
503.2764892578125 0 1696.4956
504.27947998046875 0 639.528
508.2533874511719 0 2098.4368
508.7720642089844 0 15430.456
509.2745361328125 0 24910.342 c 8
509.77325439453125 0 13771.85
510.2740173339844 0 5652.285
510.7711486816406 0 1297.699
521.2734375 0 587.436
522.98388671875 0 568.11914
526.25439453125 0 595.7667
526.7568969726562 0 717.12305
532.7664184570312 0 2020.0083 z 3
533.2678833007812 0 1878.178
533.7666625976562 0 687.3654
535.2716674804688 0 646.946
535.7720947265625 0 799.938
541.2667846679688 0 891.0769
544.2789916992188 0 2881.0698 c Ammonia loss 9
544.7791137695312 0 1933.5406
545.3029174804688 0 1792.2874
546.3090209960938 0 1411.6757
548.2503662109375 0 1908.6338
548.7514038085938 0 1489.0175
552.7919921875 0 16884.852 c 9
553.289306640625 0 12380.343
553.7884521484375 0 5600.8687
554.2871704101562 0 2099.2578
554.7801513671875 0 1532.0203
564.3023681640625 0 2357.079
565.3092651367188 0 882.58685
565.8007202148438 0 1565.1578
566.3008422851562 0 1229.6531
567.2739868164062 0 852.78894 z Water loss 2
567.7818603515625 0 993.8482 w 2
568.279052734375 0 972.9215
569.324462890625 0 659.85034
573.3046264648438 0 914.8887 c Ammonia loss 4
575.7840576171875 0 725.4587 y Ammonia loss 2
576.2822265625 0 8088.539 z 2
576.7811279296875 0 5291.165
577.2799072265625 0 2605.4614
578.3147583007812 0 1066.4182
579.3125 0 834.8309
583.3080444335938 0 845.1034
584.2913818359375 0 16589.176 y 2
584.7903442382812 0 14292.828
585.2902221679688 0 8569.564
585.7890014648438 0 2233.4092
586.8228149414062 0 1872.8191
587.324951171875 0 2258.6985
587.8289794921875 0 734.10944
588.2754516601562 0 1153.2322
588.7758178710938 0 907.5635
589.32080078125 0 896.93274
590.3261108398438 0 112855.3 c 4
591.3287963867188 0 32805.742
591.8157958984375 0 1380.0837
592.3275756835938 0 6493.2305
592.8101806640625 0 960.0569
600.8214111328125 0 2731.3284 c Ammonia loss 10
601.3192138671875 0 2588.05
601.8256225585938 0 2788.801
602.32470703125 0 1340.447
607.3265991210938 0 929.7318
608.3169555664062 0 5912.128 z 7
609.33349609375 0 47555.797 c 10
609.8320922851562 0 31544.771
610.3321533203125 0 14119.332
610.8314208984375 0 5236.2104
611.3320922851562 0 1068.4344
616.3367919921875 0 1184.4551
618.3052368164062 0 2366.7947 w 1
618.8059692382812 0 2094.4387
619.306396484375 0 1377.4672
624.335693359375 0 6844.7646 y 7
625.3347778320312 0 2379.78
630.3291015625 0 1378.3435
630.8290405273438 0 1467.7957
631.327880859375 0 783.3206
632.8207397460938 0 655.08527 z 1
633.3204956054688 0 1951.9646
633.8212280273438 0 1247.2943
636.3468017578125 0 833.3334
638.84423828125 0 1036.987
639.3538208007812 0 803.8189
639.8212890625 0 700.6322
640.3427734375 0 1154.1807
640.8334350585938 0 3215.725 y 1
641.3275146484375 0 1821.6443
641.8260498046875 0 1378.0573
642.3244018554688 0 970.17554
647.3592529296875 0 1473.6881
647.8545532226562 0 2504.445
653.3424072265625 0 848.7512
653.8385620117188 0 1673.5453
654.8280639648438 0 886.5154
655.318115234375 0 4043.7441
655.8155517578125 0 3173.222
656.307861328125 0 981.01306
659.346923828125 0 4940.6553
660.3472900390625 0 6909.894
660.8526611328125 0 3372.5164
661.3515014648438 0 7507.7275 z Water loss 6
661.847900390625 0 7498.397
662.3460083007812 0 7542.9424
662.8433227539062 0 5451.9663
663.3442993164062 0 2787.7007
663.8495483398438 0 711.88245
668.8291625976562 0 3058.263
669.348876953125 0 8092.3877
669.8504028320312 0 5827.261
670.3472900390625 0 6809.4814
670.8456420898438 0 6363.9155
671.3462524414062 0 2832.8672
672.3421020507812 0 1127.3748
673.8473510742188 0 669.55286
674.3429565429688 0 2243.1545
674.8453369140625 0 4742.812
675.3438720703125 0 5392.248
675.8394775390625 0 4842.988
676.3375854492188 0 2804.4846
676.8350219726562 0 886.1749
677.338623046875 0 1113.4735
679.3538208007812 0 22413.893 z 6
680.3533325195312 0 8541.888
681.3607788085938 0 1971.6423
681.8735961914062 0 1177.4323
682.3733520507812 0 703.86115
682.8524169921875 0 1009.9798
683.3482666015625 0 27321.646
683.8464965820312 0 29598.816
684.346435546875 0 13637.367
684.846923828125 0 5164.881
685.3486938476562 0 815.664
687.3474731445312 0 1142.8643 c Ammonia loss 5
689.152587890625 0 1320.5348
689.6575317382812 0 4467.146
690.1580810546875 0 2896.8635
691.3572387695312 0 25332.805
691.856201171875 0 21989.568
692.3556518554688 0 14819.434
692.8572998046875 0 5261.1904
693.3604125976562 0 1837.2053
693.8741455078125 0 2999.46
694.3525390625 0 829.03033
695.3720092773438 0 5668.2515 y 6
696.3704833984375 0 1515.739
704.3689575195312 0 28644.162 c 5
705.3685302734375 0 9667.91
706.3695068359375 0 2173.5906
722.3492431640625 0 2205.5828
731.3919677734375 0 4057.059
732.39404296875 0 2592.6165
737.3264770507812 0 1082.608
737.3905029296875 0 890.7592
749.384765625 0 11646.015 w 5
750.387939453125 0 9268.959
751.388427734375 0 2530.672
758.3811645507812 0 2150.6677 c Ammonia loss 6
759.3788452148438 0 781.3016
774.3959350585938 0 929.27936
775.405517578125 0 27346.36 c 6
776.4027709960938 0 9816.487
777.4030151367188 0 3607.34
787.3917846679688 0 1196.3885
793.3968505859375 0 63658.57 z 5
794.3946533203125 0 32999.383
795.3948364257812 0 9714.938
796.40087890625 0 1906.0072
810.4065551757812 0 1179.607
843.2171020507812 0 711.0632
858.4951171875 0 684.98987
859.4857177734375 0 1989.2262
860.4857177734375 0 2522.1836
864.4339599609375 0 21497.818
865.4302978515625 0 11774.356
866.4296264648438 0 3984.1196
867.4274291992188 0 1528.6146
874.4227294921875 0 1353.5228
884.4806518554688 0 766.0825
885.4812622070312 0 979.9963 c Water loss 7
886.4754028320312 0 3287.7737 c Ammonia loss 7
887.470703125 0 1598.6692
888.1694946289062 0 864.86237
900.4764404296875 0 2499.6824
901.4761352539062 0 1036.9292
902.4916381835938 0 8770.272
903.4994506835938 0 34387.902 c 7
904.3447265625 0 1068.4115
904.4978637695312 0 18520.416
905.4962768554688 0 6303.353
906.49658203125 0 1618.3325
908.4197998046875 0 18647.945 z 4
909.4189453125 0 12962.522
910.421142578125 0 5231.689
911.4219970703125 0 1769.9832
919.4669189453125 0 1497.6332
920.4686279296875 0 1960.379
924.4378051757812 0 1893.6793 y 4
925.4282836914062 0 1441.4363
986.4976196289062 0 1654.9069
987.5057373046875 0 2080.2815
988.4955444335938 0 973.13007
999.536376953125 0 1356.1528 c Water loss 8
1000.5215454101562 0 1939.7256 c Ammonia loss 8
1001.5209350585938 0 1230.2322
1002.5270385742188 0 1078.7737
1016.5350341796875 0 2294.6975
1017.5426025390625 0 23505.156 c 8
1018.5396728515625 0 12620.767
1019.539794921875 0 4573.3955
1020.5364990234375 0 2640.696
1022.5173950195312 0 1053.2832
1062.5604248046875 0 842.88745
1064.525146484375 0 8543.702 z 3
1065.5240478515625 0 10221.525
1066.521728515625 0 5960.6733
1067.5262451171875 0 1836.4153
1088.556396484375 0 1917.5713
1089.552978515625 0 1840.5076
1090.5501708984375 0 1208.7289
1096.5003662109375 0 1001.6658
1104.5760498046875 0 7089.029 c 9
1105.5733642578125 0 7055.4165
1106.572509765625 0 2598.2227
1107.566162109375 0 1182.1337
1134.5546875 0 1113.4617 w 2
1151.5550537109375 0 4830.011 z 2
1152.5599365234375 0 12885.102
1153.55859375 0 6174.693
1154.5611572265625 0 3243.986
1158.616943359375 0 821.00946
1159.61083984375 0 805.6703
1173.6463623046875 0 2270.399
1174.651123046875 0 3201.4072
1175.649658203125 0 2463.5432
1176.6473388671875 0 854.63434
1201.640380859375 0 5619.123
1202.640625 0 3789.9182
1203.631591796875 0 1648.5703
1217.6591796875 0 5407.484 c 10
1218.662109375 0 8358.728
1219.6611328125 0 4995.658
1220.6629638671875 0 2497.625
1221.6654052734375 0 1183.9846
1265.6505126953125 0 1314.9636
1266.645263671875 0 2167.749
1305.6617431640625 0 1541.0164
1306.646484375 0 1748.3481
1307.6533203125 0 1421.5322
1308.6512451171875 0 1128.1565
1309.6561279296875 0 939.965
1320.6834716796875 0 833.65375
1321.6910400390625 0 3112.5762
1322.6925048828125 0 1647.5438
1323.676513671875 0 4150.4556
1324.677490234375 0 8450.679
1325.6796875 0 6735.928
1326.677734375 0 2289.8918
1337.697509765625 0 1955.6942
1338.69970703125 0 12102.202
1339.698486328125 0 8347.008
1340.701416015625 0 5065.542
1341.7010498046875 0 1922.169
1348.688232421875 0 2353.9653
1349.67578125 0 4528.8047
1350.6707763671875 0 5109.9863
1351.660888671875 0 1941.2885
1364.7113037109375 0 878.28876
1365.6912841796875 0 7386.766
1366.693359375 0 39957.21
1367.6927490234375 0 33025.34
1368.69189453125 0 17547.514
1369.6904296875 0 6309.3623
1370.69677734375 0 1211.7589
1378.3045654296875 0 1464.3722
1379.3138427734375 0 2583.0715
1380.319091796875 0 2135.2314
1381.7100830078125 0 2344.5583
1382.7122802734375 0 5741.415
1383.7156982421875 0 7309.3003
1384.712158203125 0 6587.6807
1385.7142333984375 0 3642.9988
1386.71484375 0 964.7904
1387.7520751953125 0 2247.6345

Spectrum Details

|  |  |
| --- | --- |
| Matched peaks? Matched peaks The total absolute number of peaks matched. Additionally in brackets the total fraction of peaks matched and the total number of peaks is shown. | 53 (12.62% of 420) |
| FDR? FDR The false discovery rate estimated for this peptide. It is calculated by matching all theoretical fragments with a non-integer shift with the raw peaks for this spectrum. This is done with 40 different shifts. The resulting percentage is the average number of annotated peaks over the number of annotated peaks with the correct spectrum. | 1.48% |
| Satellite FDR? Satellite FDR See the FDR for details on its calculation. This satellite ion specific FDR only contains the satellite ions (d/w) for I/L/J positions. | 0.00% |
| PSM Score? PSM Score The PSM Score as given by Hecklib to this annotated spectrum. It is shown with three significant figures. | 392 |

#### Spectrum 4924? Spectrum 4924 The raw spectrum of this peptide as annotated by Hecklib. The fragments are coloured according to ion type (see legend). Any peaks with a star '\*' as text can be hovered over to see the full details, first the ion type second the mass shift type. By hovering over the amino acids in the peptide or ions in the legend the corresponding peaks are highlighted. By toggling the 'Unassigned' label you can turn the background (unassigned) peaks on or off in the plot. By updating the slider in the Ion legend you can update the spectrum to only show the top X% of the peaks with labels. The top X% means any peak that is within X% of the highest intensity. By dragging in the spectrum you can zoom in to a specific part of the spectrum and use 'Zoom Out' to get back to the original zoom level. The annotation of the spectrum is based on the given sequence in the peptides file and is done with different software so inconsistencies are likely. The peaks are annotated based on the given sequence, with 20 ppm tolerance.

Copy Data

##### Spectrum 4924 (TSV)

###### Preview

```
Loading example...
```

*Click on the button to copy the data to your clipboard.*

Mz MinMz MaxIntensity Max

WidthHeightPeptide font sizePeptide stroke widthSpectrum font sizeSpectrum stroke widthCompact peptide

Ion legend

wxyz

abcd

OtherUnassignedIon^Charge^_Position_Show for top:%

TISRDNAKNSJY

04.54e+49.09e+41.36e+51.82e+5

Zoom Out

y^+1^_1_c^+1^_3_c^+2^_7_z^+2^_7_c^+2^_8_c^+1^_4_y^+1^_4_c^+2^_9_c^+2^_9_c^+2^_9_z^+2^_9_c^+2^_10_c^+2^_10_w^+2^_10_z^+2^_10_y^+2^_10_c^+1^_5_c^+2^_11_z^+1^_5_c^+2^_11_w^+2^_11_y^+1^_5_y^+2^_11_z^+1^_6_z^+1^_6_y^+1^_6_c^+1^_6_w^+1^_7_c^+1^_7_z^+1^_7_c^+1^_8_c^+1^_8_z^+1^_8_y^+1^_8_c^+1^_9_c^+1^_9_c^+1^_9_z^+1^_9_c^+1^_10_z^+1^_10_c^+1^_11_

0515103015452059

Fragment Matches Table

Show background peaks

| Position | Ion type | Intensity | mz Theoretical | mz Error (Th) | mz Error (ppm) | Charge | Series Number |
| --- | --- | --- | --- | --- | --- | --- | --- |
| - | - | 351.5 | 122.4 | - | - | 0 | - |
| - | - | 370.3 | 124.4 | - | - | 0 | - |
| - | - | 350.3 | 129.1 | - | - | 0 | - |
| - | - | 461.7 | 129.1 | - | - | 0 | - |
| - | - | 1689 | 136.1 | - | - | 0 | - |
| - | - | 436.4 | 136.7 | - | - | 0 | - |
| - | - | 382.1 | 143.1 | - | - | 0 | - |
| - | - | 424.5 | 143.2 | - | - | 0 | - |
| - | - | 420.6 | 146.5 | - | - | 0 | - |
| - | - | 546.6 | 147.1 | - | - | 0 | - |
| - | - | 1007 | 149 | - | - | 0 | - |
| - | - | 668.6 | 149 | - | - | 0 | - |
| - | - | 7154 | 149 | - | - | 0 | - |
| - | - | 602.3 | 150 | - | - | 0 | - |
| - | - | 715 | 151 | - | - | 0 | - |
| - | - | 482.8 | 162.9 | - | - | 0 | - |
| - | - | 2017 | 165.1 | - | - | 0 | - |
| - | - | 1874 | 167.1 | - | - | 0 | - |
| - | - | 3095 | 173.1 | - | - | 0 | - |
| - | - | 547.2 | 173.1 | - | - | 0 | - |
| - | - | 1043 | 173.4 | - | - | 0 | - |
| - | - | 522.4 | 174.1 | - | - | 0 | - |
| 12 | y | 4718 | 182.1 | 8.447E-05 | 0.4639 | +1 | 1 |
| - | - | 1.075E+04 | 187.1 | - | - | 0 | - |
| - | - | 646.8 | 188.1 | - | - | 0 | - |
| - | - | 1003 | 201.1 | - | - | 0 | - |
| - | - | 2046 | 215.1 | - | - | 0 | - |
| - | - | 3054 | 221.1 | - | - | 0 | - |
| - | - | 2178 | 222.1 | - | - | 0 | - |
| - | - | 1950 | 223.1 | - | - | 0 | - |
| - | - | 1205 | 223.1 | - | - | 0 | - |
| - | - | 3459 | 225 | - | - | 0 | - |
| - | - | 1005 | 226 | - | - | 0 | - |
| - | - | 621.6 | 227 | - | - | 0 | - |
| - | - | 884.2 | 227 | - | - | 0 | - |
| - | - | 1157 | 235.1 | - | - | 0 | - |
| - | - | 567.1 | 237.6 | - | - | 0 | - |
| - | - | 4410 | 239.1 | - | - | 0 | - |
| - | - | 3162 | 240.1 | - | - | 0 | - |
| - | - | 1216 | 241.1 | - | - | 0 | - |
| - | - | 1801 | 260.1 | - | - | 0 | - |
| - | - | 991.9 | 281.1 | - | - | 0 | - |
| - | - | 626.1 | 282.1 | - | - | 0 | - |
| - | - | 677.9 | 282.1 | - | - | 0 | - |
| - | - | 605.4 | 285 | - | - | 0 | - |
| - | - | 657.1 | 288.2 | - | - | 0 | - |
| - | - | 726.5 | 288.3 | - | - | 0 | - |
| - | - | 2891 | 295.1 | - | - | 0 | - |
| - | - | 4110 | 296.1 | - | - | 0 | - |
| - | - | 1809 | 297.1 | - | - | 0 | - |
| - | - | 757.3 | 298.1 | - | - | 0 | - |
| - | - | 2674 | 299.1 | - | - | 0 | - |
| - | - | 3065 | 300.1 | - | - | 0 | - |
| - | - | 1804 | 300.2 | - | - | 0 | - |
| - | - | 2135 | 301.1 | - | - | 0 | - |
| - | - | 508 | 313.1 | - | - | 0 | - |
| - | - | 904.4 | 314.1 | - | - | 0 | - |
| - | - | 731.7 | 317.2 | - | - | 0 | - |
| 3 | c | 1191 | 319.2 | 0.0004223 | 1.323 | +1 | 3 |
| - | - | 510.9 | 320.5 | - | - | 0 | - |
| - | - | 897.8 | 325 | - | - | 0 | - |
| - | - | 1.886E+04 | 341 | - | - | 0 | - |
| - | - | 947.4 | 343 | - | - | 0 | - |
| - | - | 565.5 | 351.6 | - | - | 0 | - |
| - | - | 2496 | 355.1 | - | - | 0 | - |
| - | - | 850 | 356.1 | - | - | 0 | - |
| - | - | 7.502E+04 | 359 | - | - | 0 | - |
| - | - | 1938 | 369.1 | - | - | 0 | - |
| - | - | 4088 | 370.1 | - | - | 0 | - |
| - | - | 3797 | 371.1 | - | - | 0 | - |
| - | - | 705.4 | 372.1 | - | - | 0 | - |
| - | - | 698.7 | 372.7 | - | - | 0 | - |
| - | - | 557.3 | 374.2 | - | - | 0 | - |
| - | - | 1386 | 375.9 | - | - | 0 | - |
| - | - | 614.2 | 376.9 | - | - | 0 | - |
| 7 | c | 972.4 | 379.2 | 0.007081 | 18.67 | +2 | 7 |
| - | - | 1554 | 385.2 | - | - | 0 | - |
| - | - | 1001 | 385.5 | - | - | 0 | - |
| 6 | z | 1461 | 397.2 | 0.006013 | 15.14 | +2 | 7 |
| - | - | 718.4 | 421.7 | - | - | 0 | - |
| - | - | 1.799E+05 | 429.1 | - | - | 0 | - |
| - | - | 630.6 | 430.2 | - | - | 0 | - |
| - | - | 650.8 | 431.1 | - | - | 0 | - |
| - | - | 2058 | 443.2 | - | - | 0 | - |
| - | - | 9.685E+04 | 445.1 | - | - | 0 | - |
| - | - | 1515 | 447.1 | - | - | 0 | - |
| - | - | 3222 | 451.7 | - | - | 0 | - |
| 8 | c | 7365 | 452.3 | 0.0008223 | 1.818 | +2 | 8 |
| - | - | 734 | 452.7 | - | - | 0 | - |
| - | - | 4184 | 452.8 | - | - | 0 | - |
| - | - | 1339 | 453.3 | - | - | 0 | - |
| - | - | 1919 | 460.2 | - | - | 0 | - |
| - | - | 665.6 | 461.2 | - | - | 0 | - |
| - | - | 719 | 461.7 | - | - | 0 | - |
| - | - | 896.7 | 472.3 | - | - | 0 | - |
| 4 | c | 2.895E+04 | 475.3 | 0.0001845 | 0.3882 | +1 | 4 |
| - | - | 6142 | 476.3 | - | - | 0 | - |
| - | - | 1216 | 477.3 | - | - | 0 | - |
| - | - | 663.7 | 479.3 | - | - | 0 | - |
| - | - | 3648 | 479.8 | - | - | 0 | - |
| - | - | 1396 | 480.3 | - | - | 0 | - |
| - | - | 738.8 | 480.8 | - | - | 0 | - |
| - | - | 1169 | 481.2 | - | - | 0 | - |
| - | - | 623.7 | 484.8 | - | - | 0 | - |
| - | - | 661.6 | 493.8 | - | - | 0 | - |
| 9 | y | 1196 | 496.2 | 0.001205 | 2.427 | +1 | 4 |
| - | - | 4650 | 499.8 | - | - | 0 | - |
| 9 | c | 2063 | 500.3 | 0.00675 | 13.49 | +2 | 9 |
| 9 | c | 1517 | 500.8 | 0.0007107 | 1.419 | +2 | 9 |
| - | - | 765.7 | 501.8 | - | - | 0 | - |
| - | - | 3856 | 502.3 | - | - | 0 | - |
| - | - | 887.5 | 503.3 | - | - | 0 | - |
| - | - | 1484 | 508.3 | - | - | 0 | - |
| - | - | 6152 | 508.8 | - | - | 0 | - |
| 9 | c | 1.107E+04 | 509.3 | 0.002297 | 4.51 | +2 | 9 |
| - | - | 5312 | 509.8 | - | - | 0 | - |
| - | - | 1803 | 510.3 | - | - | 0 | - |
| - | - | 644.4 | 510.8 | - | - | 0 | - |
| - | - | 498.3 | 524.5 | - | - | 0 | - |
| - | - | 1068 | 530.3 | - | - | 0 | - |
| 4 | z | 1156 | 532.8 | 0.002288 | 4.295 | +2 | 9 |
| - | - | 606.6 | 533.3 | - | - | 0 | - |
| 10 | c | 1477 | 544.3 | 0.0005786 | 1.063 | +2 | 10 |
| - | - | 599.8 | 544.8 | - | - | 0 | - |
| - | - | 616.2 | 548.2 | - | - | 0 | - |
| - | - | 577.3 | 550.8 | - | - | 0 | - |
| 10 | c | 7968 | 552.8 | 0.0001227 | 0.222 | +2 | 10 |
| - | - | 6610 | 553.3 | - | - | 0 | - |
| - | - | 1863 | 553.8 | - | - | 0 | - |
| - | - | 582.2 | 554.3 | - | - | 0 | - |
| - | - | 841.9 | 564.3 | - | - | 0 | - |
| - | - | 509 | 566 | - | - | 0 | - |
| - | - | 605.2 | 566.3 | - | - | 0 | - |
| 3 | w | 635.1 | 567.8 | 0.001926 | 3.393 | +2 | 10 |
| - | - | 700.6 | 568.3 | - | - | 0 | - |
| 3 | z | 2514 | 576.3 | 0.001885 | 3.271 | +2 | 10 |
| - | - | 2400 | 576.8 | - | - | 0 | - |
| - | - | 871.4 | 577.3 | - | - | 0 | - |
| 3 | y | 1.038E+04 | 584.3 | 0.000688 | 1.177 | +2 | 10 |
| - | - | 7224 | 584.8 | - | - | 0 | - |
| - | - | 2328 | 585.3 | - | - | 0 | - |
| - | - | 623.9 | 585.8 | - | - | 0 | - |
| - | - | 672.9 | 586.8 | - | - | 0 | - |
| - | - | 681 | 587.3 | - | - | 0 | - |
| - | - | 662.8 | 588.3 | - | - | 0 | - |
| - | - | 701.1 | 588.8 | - | - | 0 | - |
| 5 | c | 5.333E+04 | 590.3 | 0.000211 | 0.3575 | +1 | 5 |
| - | - | 1.717E+04 | 591.3 | - | - | 0 | - |
| - | - | 4140 | 592.3 | - | - | 0 | - |
| 11 | c | 1455 | 600.8 | 0.001231 | 2.049 | +2 | 11 |
| - | - | 863 | 601.3 | - | - | 0 | - |
| 8 | z | 3858 | 608.3 | 0.0006492 | 1.067 | +1 | 5 |
| 11 | c | 2.385E+04 | 609.3 | 0.001017 | 1.669 | +2 | 11 |
| - | - | 1.565E+04 | 609.8 | - | - | 0 | - |
| - | - | 6543 | 610.3 | - | - | 0 | - |
| - | - | 1636 | 610.8 | - | - | 0 | - |
| - | - | 673.5 | 614.3 | - | - | 0 | - |
| 2 | w | 1597 | 618.3 | 0.001039 | 1.68 | +2 | 11 |
| - | - | 980.7 | 618.8 | - | - | 0 | - |
| 8 | y | 3584 | 624.3 | 5.254E-05 | 0.08416 | +1 | 5 |
| - | - | 789.3 | 625.3 | - | - | 0 | - |
| - | - | 825.3 | 630.8 | - | - | 0 | - |
| - | - | 801.6 | 633.3 | - | - | 0 | - |
| - | - | 645.4 | 637.8 | - | - | 0 | - |
| - | - | 726.2 | 638.3 | - | - | 0 | - |
| - | - | 823.5 | 639.3 | - | - | 0 | - |
| - | - | 644 | 640.3 | - | - | 0 | - |
| 2 | y | 587.2 | 640.8 | 0.003169 | 4.945 | +2 | 11 |
| - | - | 808.2 | 641.3 | - | - | 0 | - |
| - | - | 1058 | 647.4 | - | - | 0 | - |
| - | - | 748.9 | 647.8 | - | - | 0 | - |
| - | - | 932.1 | 655.3 | - | - | 0 | - |
| - | - | 1272 | 655.8 | - | - | 0 | - |
| - | - | 2944 | 659.3 | - | - | 0 | - |
| - | - | 2986 | 660.3 | - | - | 0 | - |
| - | - | 1995 | 660.8 | - | - | 0 | - |
| 7 | z | 3582 | 661.3 | 0.009928 | 15.01 | +1 | 6 |
| - | - | 3326 | 661.8 | - | - | 0 | - |
| - | - | 4269 | 662.3 | - | - | 0 | - |
| - | - | 2116 | 662.8 | - | - | 0 | - |
| - | - | 1585 | 663.3 | - | - | 0 | - |
| - | - | 1137 | 668.8 | - | - | 0 | - |
| - | - | 4573 | 669.3 | - | - | 0 | - |
| - | - | 2636 | 669.8 | - | - | 0 | - |
| - | - | 3513 | 670.3 | - | - | 0 | - |
| - | - | 2281 | 670.8 | - | - | 0 | - |
| - | - | 797 | 674.3 | - | - | 0 | - |
| - | - | 2210 | 674.8 | - | - | 0 | - |
| - | - | 1932 | 675.3 | - | - | 0 | - |
| - | - | 1962 | 675.8 | - | - | 0 | - |
| - | - | 1741 | 676.3 | - | - | 0 | - |
| - | - | 620.4 | 678.3 | - | - | 0 | - |
| 7 | z | 9849 | 679.4 | 8.766E-05 | 0.129 | +1 | 6 |
| - | - | 4138 | 680.4 | - | - | 0 | - |
| - | - | 989 | 681.4 | - | - | 0 | - |
| - | - | 635.9 | 682.8 | - | - | 0 | - |
| - | - | 1.178E+04 | 683.3 | - | - | 0 | - |
| - | - | 1.061E+04 | 683.8 | - | - | 0 | - |
| - | - | 4552 | 684.3 | - | - | 0 | - |
| - | - | 1561 | 684.8 | - | - | 0 | - |
| - | - | 1146 | 689.7 | - | - | 0 | - |
| - | - | 1.253E+04 | 691.4 | - | - | 0 | - |
| - | - | 1.145E+04 | 691.9 | - | - | 0 | - |
| - | - | 4881 | 692.4 | - | - | 0 | - |
| - | - | 1638 | 692.9 | - | - | 0 | - |
| - | - | 1743 | 694.4 | - | - | 0 | - |
| 7 | y | 2825 | 695.4 | 0.004224 | 6.075 | +1 | 6 |
| - | - | 622.6 | 696.4 | - | - | 0 | - |
| 6 | c | 1.355E+04 | 704.4 | 0.0002308 | 0.3276 | +1 | 6 |
| - | - | 4530 | 705.4 | - | - | 0 | - |
| - | - | 2000 | 706.4 | - | - | 0 | - |
| - | - | 685 | 708.5 | - | - | 0 | - |
| - | - | 1762 | 731.4 | - | - | 0 | - |
| 6 | w | 7223 | 749.4 | 0.0001036 | 0.1382 | +1 | 7 |
| - | - | 4123 | 750.4 | - | - | 0 | - |
| - | - | 663.6 | 751.4 | - | - | 0 | - |
| - | - | 825.3 | 774.4 | - | - | 0 | - |
| 7 | c | 1.481E+04 | 775.4 | 0.001273 | 1.641 | +1 | 7 |
| - | - | 5483 | 776.4 | - | - | 0 | - |
| - | - | 2179 | 777.4 | - | - | 0 | - |
| 6 | z | 2.953E+04 | 793.4 | 0.0007788 | 0.9816 | +1 | 7 |
| - | - | 1.423E+04 | 794.4 | - | - | 0 | - |
| - | - | 4231 | 795.4 | - | - | 0 | - |
| - | - | 1008 | 796.4 | - | - | 0 | - |
| - | - | 628.7 | 841.1 | - | - | 0 | - |
| - | - | 1818 | 859.5 | - | - | 0 | - |
| - | - | 1449 | 860.5 | - | - | 0 | - |
| - | - | 8530 | 864.4 | - | - | 0 | - |
| - | - | 6047 | 865.4 | - | - | 0 | - |
| - | - | 1694 | 866.4 | - | - | 0 | - |
| - | - | 2329 | 880.4 | - | - | 0 | - |
| - | - | 855.1 | 881.4 | - | - | 0 | - |
| - | - | 911.7 | 884.5 | - | - | 0 | - |
| 8 | c | 1472 | 886.5 | 0.001388 | 1.566 | +1 | 8 |
| - | - | 722.8 | 888.4 | - | - | 0 | - |
| - | - | 923.8 | 900.5 | - | - | 0 | - |
| - | - | 3835 | 902.5 | - | - | 0 | - |
| 8 | c | 1.728E+04 | 903.5 | 0.002181 | 2.414 | +1 | 8 |
| - | - | 9238 | 904.5 | - | - | 0 | - |
| - | - | 2765 | 905.5 | - | - | 0 | - |
| - | - | 715.3 | 906.4 | - | - | 0 | - |
| 5 | z | 9262 | 908.4 | 0.002575 | 2.835 | +1 | 8 |
| - | - | 5505 | 909.4 | - | - | 0 | - |
| - | - | 1928 | 910.4 | - | - | 0 | - |
| - | - | 1028 | 919.3 | - | - | 0 | - |
| - | - | 950.3 | 919.5 | - | - | 0 | - |
| - | - | 2011 | 921 | - | - | 0 | - |
| - | - | 981.4 | 922 | - | - | 0 | - |
| - | - | 3725 | 924.3 | - | - | 0 | - |
| 5 | y | 1050 | 924.4 | 0.001894 | 2.049 | +1 | 8 |
| - | - | 839.4 | 925.3 | - | - | 0 | - |
| - | - | 652 | 974.5 | - | - | 0 | - |
| - | - | 714.3 | 987.5 | - | - | 0 | - |
| - | - | 948.9 | 988.5 | - | - | 0 | - |
| 9 | c | 861.9 | 999.5 | 0.003848 | 3.85 | +1 | 9 |
| 9 | c | 1321 | 1001 | 0.003902 | 3.9 | +1 | 9 |
| - | - | 663.7 | 1003 | - | - | 0 | - |
| - | - | 900.8 | 1017 | - | - | 0 | - |
| 9 | c | 1.285E+04 | 1018 | 0.001834 | 1.803 | +1 | 9 |
| - | - | 5809 | 1019 | - | - | 0 | - |
| - | - | 3143 | 1020 | - | - | 0 | - |
| 4 | z | 3505 | 1065 | 0.0008421 | 0.7911 | +1 | 9 |
| - | - | 5570 | 1066 | - | - | 0 | - |
| - | - | 2662 | 1067 | - | - | 0 | - |
| - | - | 736.2 | 1089 | - | - | 0 | - |
| 10 | c | 2813 | 1105 | 0.0007815 | 0.7075 | +1 | 10 |
| - | - | 2454 | 1106 | - | - | 0 | - |
| - | - | 1283 | 1107 | - | - | 0 | - |
| 3 | z | 2905 | 1152 | 0.001865 | 1.619 | +1 | 10 |
| - | - | 4909 | 1153 | - | - | 0 | - |
| - | - | 3210 | 1154 | - | - | 0 | - |
| - | - | 1238 | 1155 | - | - | 0 | - |
| - | - | 773.1 | 1174 | - | - | 0 | - |
| - | - | 2359 | 1175 | - | - | 0 | - |
| - | - | 3313 | 1202 | - | - | 0 | - |
| - | - | 1593 | 1203 | - | - | 0 | - |
| 11 | c | 2550 | 1218 | 0.004401 | 3.614 | +1 | 11 |
| - | - | 2949 | 1219 | - | - | 0 | - |
| - | - | 2410 | 1220 | - | - | 0 | - |
| - | - | 1093 | 1266 | - | - | 0 | - |
| - | - | 915.9 | 1267 | - | - | 0 | - |
| - | - | 655.8 | 1306 | - | - | 0 | - |
| - | - | 775.3 | 1309 | - | - | 0 | - |
| - | - | 981.5 | 1322 | - | - | 0 | - |
| - | - | 735.5 | 1323 | - | - | 0 | - |
| - | - | 2527 | 1324 | - | - | 0 | - |
| - | - | 3613 | 1325 | - | - | 0 | - |
| - | - | 3898 | 1326 | - | - | 0 | - |
| - | - | 1564 | 1327 | - | - | 0 | - |
| - | - | 1296 | 1338 | - | - | 0 | - |
| - | - | 4590 | 1339 | - | - | 0 | - |
| - | - | 4166 | 1340 | - | - | 0 | - |
| - | - | 1652 | 1341 | - | - | 0 | - |
| - | - | 891.3 | 1342 | - | - | 0 | - |
| - | - | 1666 | 1349 | - | - | 0 | - |
| - | - | 2324 | 1350 | - | - | 0 | - |
| - | - | 1579 | 1351 | - | - | 0 | - |
| - | - | 862.2 | 1352 | - | - | 0 | - |
| - | - | 4021 | 1366 | - | - | 0 | - |
| - | - | 1.9E+04 | 1367 | - | - | 0 | - |
| - | - | 1.593E+04 | 1368 | - | - | 0 | - |
| - | - | 7783 | 1369 | - | - | 0 | - |
| - | - | 3208 | 1370 | - | - | 0 | - |
| - | - | 1161 | 1371 | - | - | 0 | - |
| - | - | 1060 | 1379 | - | - | 0 | - |
| - | - | 880 | 1382 | - | - | 0 | - |
| - | - | 3018 | 1383 | - | - | 0 | - |
| - | - | 3910 | 1384 | - | - | 0 | - |
| - | - | 3357 | 1385 | - | - | 0 | - |
| - | - | 1882 | 1386 | - | - | 0 | - |
| - | - | 707.7 | 1796 | - | - | 0 | - |
| - | - | 801.2 | 2039 | - | - | 0 | - |

m/z Charge Intensity FragmentType MassShift Position
122.40363311767578 0 351.54962
124.43357849121094 0 370.34888
129.0542449951172 0 350.3402
129.1027374267578 0 461.67493
136.07582092285156 0 1689.3217
136.6658935546875 0 436.37796
143.1178436279297 0 382.13916
143.22927856445312 0 424.45587
146.4524688720703 0 420.61795
147.0655517578125 0 546.62885
148.9545440673828 0 1006.913
149.0393829345703 0 668.622
149.0448760986328 0 7153.7793
150.04449462890625 0 602.31055
151.0415496826172 0 715.0024
162.9347381591797 0 482.75995
165.05465698242188 0 2017.0441
167.05531311035156 0 1874.2985
173.092041015625 0 3095.1418
173.128173828125 0 547.2116
173.4387664794922 0 1042.7778
174.09503173828125 0 522.366
182.08108520507812 0 4718.278 y 11
187.1439208984375 0 10748.38
188.147705078125 0 646.7855
201.12344360351562 0 1003.3141
215.1388397216797 0 2046.4934
221.084228515625 0 3053.6538
222.0841827392578 0 2178.386
223.06338500976562 0 1949.538
223.08119201660156 0 1205.3656
225.0426788330078 0 3459.1997
226.04278564453125 0 1005.3134
227.0218505859375 0 621.63684
227.03977966308594 0 884.226
235.11767578125 0 1157.4232
237.6250457763672 0 567.0632
239.09469604492188 0 4409.7314
240.0955047607422 0 3162.463
241.09251403808594 0 1216.274
260.1239929199219 0 1800.8683
281.05108642578125 0 991.87494
282.052978515625 0 626.12036
282.1444396972656 0 677.87805
285.00787353515625 0 605.36115
288.1553649902344 0 657.1482
288.2846374511719 0 726.4871
295.1029968261719 0 2890.9678
296.1042175292969 0 4110.003
297.10113525390625 0 1808.5372
298.10198974609375 0 757.30664
299.0614929199219 0 2673.73
300.06243896484375 0 3064.9602
300.1556091308594 0 1804.4807
301.0589294433594 0 2135.2705
313.1125793457031 0 508.0257
314.1142272949219 0 904.3922
317.18145751953125 0 731.71173
319.1971740722656 0 1191.2346 c 2
320.46673583984375 0 510.89148
324.9845275878906 0 897.7605
341.01788330078125 0 18861.043
342.9962158203125 0 947.41504
351.5857238769531 0 565.4622
355.0693359375 0 2495.6458
356.0704345703125 0 849.96796
359.0280456542969 0 75021.64
369.1224670410156 0 1938.3795
370.1221008300781 0 4087.8967
371.1195983886719 0 3797.4324
372.1217346191406 0 705.38446
372.69451904296875 0 698.694
374.1786804199219 0 557.3371
375.8607177734375 0 1386.3448
376.8996887207031 0 614.1903
379.2082824707031 0 972.3553 c Water loss 6
385.22021484375 0 1553.5347
385.5464782714844 0 1001.06036
397.2078857421875 0 1461.272 z 5
421.65631103515625 0 718.42035
429.0885925292969 0 179901.25
430.24017333984375 0 630.5809
431.1397705078125 0 650.75464
443.2114562988281 0 2057.5427
445.1200256347656 0 96846.02
447.0990295410156 0 1515.2404
451.74969482421875 0 3221.7612
452.2531433105469 0 7364.594 c 7
452.7217102050781 0 733.9655
452.754150390625 0 4183.702
453.256103515625 0 1338.972
460.2402038574219 0 1918.5416
461.2403869628906 0 665.5889
461.71807861328125 0 719.01385
472.2519836425781 0 896.7139
475.29852294921875 0 28954.328 c 3
476.30169677734375 0 6141.7676
477.3022766113281 0 1215.5631
479.258056640625 0 663.6564
479.7570495605469 0 3647.992
480.2593994140625 0 1396.402
480.7523498535156 0 738.8
481.22906494140625 0 1169.1713
484.751220703125 0 623.65247
493.7542419433594 0 661.6097
496.24139404296875 0 1196.2919 y 8
499.7666015625 0 4650.396
500.2633972167969 0 2062.9204 c Water loss 8
500.7614440917969 0 1516.7281 c Ammonia loss 8
501.7660217285156 0 765.6532
502.273193359375 0 3856.2847
503.2784729003906 0 887.5138
508.2550048828125 0 1484.1045
508.77081298828125 0 6151.612
509.27313232421875 0 11069.735 c 8
509.7725524902344 0 5312.178
510.2735595703125 0 1803.1046
510.7696838378906 0 644.4187
524.5015258789062 0 498.30753
530.2916870117188 0 1067.8385
532.7681884765625 0 1155.501 z 3
533.2682495117188 0 606.62646
544.2787475585938 0 1477.2745 c Ammonia loss 9
544.7731323242188 0 599.7688
548.2481689453125 0 616.18353
550.8028564453125 0 577.3155
552.7913208007812 0 7967.509 c 9
553.2885131835938 0 6610.338
553.78857421875 0 1863.2751
554.2836303710938 0 582.19354
564.30224609375 0 841.8781
566.0166625976562 0 509.0371
566.3012084960938 0 605.2481
567.782470703125 0 635.09375 w 2
568.2791748046875 0 700.62177
576.280029296875 0 2513.8418 z 2
576.781982421875 0 2400.2122
577.2820434570312 0 871.41846
584.2905883789062 0 10384.01 y 2
584.7905883789062 0 7223.9917
585.2910766601562 0 2328.3057
585.7923583984375 0 623.91833
586.8232421875 0 672.9195
587.3277587890625 0 680.987
588.2772827148438 0 662.8164
588.7716674804688 0 701.0749
590.325439453125 0 53327.523 c 4
591.3279418945312 0 17166.498
592.3278198242188 0 4139.67
600.8189697265625 0 1455.4419 c Ammonia loss 10
601.3197631835938 0 862.97217
608.3170776367188 0 3857.8467 z 7
609.3324584960938 0 23849.701 c 10
609.8318481445312 0 15654.376
610.3317260742188 0 6543.234
610.8320922851562 0 1635.8143
614.34228515625 0 673.49146
618.3033447265625 0 1597.4203 w 1
618.8021240234375 0 980.7243
624.335205078125 0 3583.6548 y 7
625.3359985351562 0 789.3161
630.8230590820312 0 825.3298
633.3128662109375 0 801.59344
637.8369750976562 0 645.3586
638.3447265625 0 726.2473
639.3447875976562 0 823.49854
640.3264770507812 0 643.99005
640.8301391601562 0 587.1939 y 1
641.33203125 0 808.1718
647.3510131835938 0 1057.5363
647.8497314453125 0 748.9299
655.3201904296875 0 932.11237
655.8157958984375 0 1272.4976
659.3464965820312 0 2944.3074
660.347900390625 0 2986.1538
660.8487548828125 0 1994.5256
661.3529052734375 0 3582.2703 z Water loss 6
661.8477783203125 0 3326.2324
662.34375 0 4269.0728
662.8444213867188 0 2115.7834
663.34521484375 0 1585.3505
668.82470703125 0 1136.6742
669.347412109375 0 4573.225
669.8486328125 0 2636.2954
670.3463134765625 0 3512.6602
670.8460083007812 0 2280.734
674.3408813476562 0 796.9621
674.84423828125 0 2210.1375
675.3421020507812 0 1932.3201
675.8386840820312 0 1962.2349
676.3350830078125 0 1741.4148
678.3313598632812 0 620.36194
679.3534545898438 0 9849.335 z 6
680.3533935546875 0 4137.6743
681.3585815429688 0 989.0261
682.84912109375 0 635.8921
683.3474731445312 0 11777.447
683.8473510742188 0 10608.909
684.3429565429688 0 4552.398
684.8424072265625 0 1560.6653
689.6611938476562 0 1145.5642
691.356201171875 0 12525.327
691.8566284179688 0 11451.505
692.3554077148438 0 4880.8374
692.8560791015625 0 1638.4324
694.3521728515625 0 1743.4333
695.3680419921875 0 2824.5774 y 6
696.370361328125 0 622.62195
704.3683471679688 0 13553.824 c 5
705.3681640625 0 4529.913
706.3705444335938 0 2000.4966
708.4732055664062 0 685.021
731.3895263671875 0 1761.5111
749.3829345703125 0 7222.854 w 5
750.3856811523438 0 4123.055
751.3899536132812 0 663.63983
774.3742065429688 0 825.3406
775.4044189453125 0 14814.639 c 6
776.404541015625 0 5482.7207
777.40869140625 0 2179.471
793.3956909179688 0 29532.678 z 5
794.3934936523438 0 14225.055
795.3948974609375 0 4230.7466
796.3980712890625 0 1007.81213
841.1397094726562 0 628.74255
859.4883422851562 0 1818.2694
860.4883422851562 0 1448.6389
864.4317626953125 0 8529.887
865.43017578125 0 6047.023
866.429443359375 0 1694.4269
880.3929443359375 0 2328.8965
881.3914794921875 0 855.0797
884.4887084960938 0 911.73254
886.4727172851562 0 1471.8455 c Ammonia loss 7
888.3674926757812 0 722.80475
900.4742431640625 0 923.81067
902.4927978515625 0 3834.7317
903.4984741210938 0 17278.99 c 7
904.4989624023438 0 9237.523
905.5004272460938 0 2765.0967
906.4093627929688 0 715.2726
908.4208374023438 0 9262.461 z 4
909.4171752929688 0 5504.702
910.4205932617188 0 1928.0498
919.318603515625 0 1028.0682
919.5474243164062 0 950.27203
921.0123901367188 0 2011.4131
922.0179443359375 0 981.3908
924.3155517578125 0 3724.6697
924.4440307617188 0 1050.4576 y 4
925.3084716796875 0 839.37067
974.5241088867188 0 652.0335
987.5092163085938 0 714.26154
988.5025024414062 0 948.8592
999.536865234375 0 861.8683 c Water loss 8
1000.5209350585938 0 1321.2432 c Ammonia loss 8
1002.5135498046875 0 663.67676
1016.5315551757812 0 900.8103
1017.541748046875 0 12849.612 c 8
1018.5401000976562 0 5809.4717
1019.5400390625 0 3142.7937
1064.523681640625 0 3505.4604 z 3
1065.525146484375 0 5569.773
1066.5234375 0 2662.2393
1088.5614013671875 0 736.16016
1104.5748291015625 0 2812.669 c 9
1105.57470703125 0 2453.5503
1106.56787109375 0 1283.1061
1151.5546875 0 2904.9878 z 2
1152.5615234375 0 4908.682
1153.5601806640625 0 3209.6504
1154.5562744140625 0 1237.6201
1173.642333984375 0 773.1409
1174.6492919921875 0 2359.12
1201.6370849609375 0 3313.4812
1202.6400146484375 0 1592.9961
1217.6552734375 0 2550.3584 c 10
1218.6612548828125 0 2948.9783
1219.662841796875 0 2410.3955
1265.650390625 0 1092.9191
1266.6343994140625 0 915.86884
1306.394775390625 0 655.77313
1308.649169921875 0 775.2716
1321.70556640625 0 981.48
1322.691650390625 0 735.46735
1323.685302734375 0 2527.4673
1324.678466796875 0 3613.0134
1325.678466796875 0 3897.622
1326.677490234375 0 1563.8076
1337.68896484375 0 1296.3313
1338.699951171875 0 4589.933
1339.7010498046875 0 4165.611
1340.6995849609375 0 1651.8833
1341.7049560546875 0 891.28754
1348.6834716796875 0 1665.5344
1349.6690673828125 0 2324.063
1350.66796875 0 1579.2109
1351.6729736328125 0 862.2349
1365.69189453125 0 4021.3987
1366.693115234375 0 19000.006
1367.6920166015625 0 15931.7295
1368.692626953125 0 7782.766
1369.6898193359375 0 3208.0654
1370.70068359375 0 1160.7611
1379.3106689453125 0 1060.4312
1381.6953125 0 879.9909
1382.7069091796875 0 3018.2085
1383.71484375 0 3909.7454
1384.7152099609375 0 3356.7227
1385.7178955078125 0 1882.1147
1796.3057861328125 0 707.6748
2039.0460205078125 0 801.244

Spectrum Details

|  |  |
| --- | --- |
| Matched peaks? Matched peaks The total absolute number of peaks matched. Additionally in brackets the total fraction of peaks matched and the total number of peaks is shown. | 41 (13.14% of 312) |
| FDR? FDR The false discovery rate estimated for this peptide. It is calculated by matching all theoretical fragments with a non-integer shift with the raw peaks for this spectrum. This is done with 40 different shifts. The resulting percentage is the average number of annotated peaks over the number of annotated peaks with the correct spectrum. | 1.97% |
| Satellite FDR? Satellite FDR See the FDR for details on its calculation. This satellite ion specific FDR only contains the satellite ions (d/w) for I/L/J positions. | 0.00% |
| PSM Score? PSM Score The PSM Score as given by Hecklib to this annotated spectrum. It is shown with three significant figures. | 274 |

#### Spectrum 3892? Spectrum 3892 The raw spectrum of this peptide as annotated by Hecklib. The fragments are coloured according to ion type (see legend). Any peaks with a star '\*' as text can be hovered over to see the full details, first the ion type second the mass shift type. By hovering over the amino acids in the peptide or ions in the legend the corresponding peaks are highlighted. By toggling the 'Unassigned' label you can turn the background (unassigned) peaks on or off in the plot. By updating the slider in the Ion legend you can update the spectrum to only show the top X% of the peaks with labels. The top X% means any peak that is within X% of the highest intensity. By dragging in the spectrum you can zoom in to a specific part of the spectrum and use 'Zoom Out' to get back to the original zoom level. The annotation of the spectrum is based on the given sequence in the peptides file and is done with different software so inconsistencies are likely. The peaks are annotated based on the given sequence, with 20 ppm tolerance.

Copy Data

##### Spectrum 3892 (TSV)

###### Preview

```
Loading example...
```

*Click on the button to copy the data to your clipboard.*

Mz MinMz MaxIntensity Max

WidthHeightPeptide font sizePeptide stroke widthSpectrum font sizeSpectrum stroke widthCompact peptide

Ion legend

wxyz

abcd

OtherUnassignedIon^Charge^_Position_Show for top:%

TISRDNAKNSJY

04.64e+49.27e+41.39e+51.85e+5

Zoom Out

y^+1^_1_c^+1^_2_y^+1^_2_c^+1^_3_y^+1^_3_c^+2^_8_c^+2^_8_c^+2^_8_c^+1^_4_c^+1^_4_z^+1^_4_y^+1^_4_c^+2^_9_c^+2^_9_c^+2^_9_z^+2^_9_y^+2^_9_c^+2^_10_c^+2^_10_z^+2^_10_z^+2^_10_c^+1^_5_y^+2^_10_y^+2^_10_z^+2^_10_y^+2^_10_c^+1^_5_c^+2^_11_z^+1^_5_c^+2^_11_w^+2^_11_y^+1^_5_y^+2^_11_z^+2^_11_y^+2^_11_z^+1^_6_z^+1^_6_c^+1^_6_y^+1^_6_c^+1^_6_w^+1^_7_c^+1^_7_c^+1^_7_z^+1^_7_y^+1^_7_c^+1^_8_c^+1^_8_z^+1^_8_c^+1^_8_z^+1^_8_y^+1^_8_c^+1^_9_c^+1^_9_c^+1^_9_z^+1^_9_y^+1^_9_c^+1^_10_z^+1^_10_c^+1^_11_

035070010501401

Fragment Matches Table

Show background peaks

| Position | Ion type | Intensity | mz Theoretical | mz Error (Th) | mz Error (ppm) | Charge | Series Number |
| --- | --- | --- | --- | --- | --- | --- | --- |
| - | - | 420.6 | 120.1 | - | - | 0 | - |
| - | - | 394 | 121.1 | - | - | 0 | - |
| - | - | 586.8 | 123.1 | - | - | 0 | - |
| - | - | 396.7 | 128.6 | - | - | 0 | - |
| - | - | 3675 | 129.1 | - | - | 0 | - |
| - | - | 506.9 | 130.1 | - | - | 0 | - |
| - | - | 5613 | 136.1 | - | - | 0 | - |
| - | - | 913.2 | 137.1 | - | - | 0 | - |
| - | - | 491.8 | 142.1 | - | - | 0 | - |
| - | - | 493.5 | 143.1 | - | - | 0 | - |
| - | - | 390.5 | 143.9 | - | - | 0 | - |
| - | - | 409.6 | 144.2 | - | - | 0 | - |
| - | - | 393.4 | 144.8 | - | - | 0 | - |
| - | - | 4238 | 149 | - | - | 0 | - |
| - | - | 659.3 | 150 | - | - | 0 | - |
| - | - | 5246 | 165.1 | - | - | 0 | - |
| - | - | 623.6 | 166.1 | - | - | 0 | - |
| - | - | 1182 | 167.1 | - | - | 0 | - |
| - | - | 498 | 171.1 | - | - | 0 | - |
| - | - | 1957 | 173.1 | - | - | 0 | - |
| - | - | 557.4 | 173.1 | - | - | 0 | - |
| - | - | 679.5 | 174.1 | - | - | 0 | - |
| 12 | y | 1.824E+04 | 182.1 | 0.0002817 | 1.547 | +1 | 1 |
| - | - | 1774 | 183.1 | - | - | 0 | - |
| - | - | 568.8 | 186.1 | - | - | 0 | - |
| - | - | 3.865E+04 | 187.1 | - | - | 0 | - |
| - | - | 3249 | 188.1 | - | - | 0 | - |
| - | - | 438.1 | 193.5 | - | - | 0 | - |
| - | - | 682.8 | 200.1 | - | - | 0 | - |
| - | - | 1648 | 201.1 | - | - | 0 | - |
| - | - | 1.035E+04 | 215.1 | - | - | 0 | - |
| - | - | 1078 | 216.1 | - | - | 0 | - |
| - | - | 475.4 | 216.9 | - | - | 0 | - |
| - | - | 2364 | 221.1 | - | - | 0 | - |
| - | - | 1075 | 222.1 | - | - | 0 | - |
| - | - | 1289 | 223.1 | - | - | 0 | - |
| - | - | 1106 | 223.1 | - | - | 0 | - |
| - | - | 2574 | 225 | - | - | 0 | - |
| - | - | 545.5 | 226 | - | - | 0 | - |
| - | - | 1101 | 229.1 | - | - | 0 | - |
| 2 | c | 609.2 | 232.2 | 0.001211 | 5.214 | +1 | 2 |
| - | - | 629 | 235.1 | - | - | 0 | - |
| - | - | 3530 | 239.1 | - | - | 0 | - |
| - | - | 2022 | 240.1 | - | - | 0 | - |
| - | - | 1254 | 241.1 | - | - | 0 | - |
| - | - | 610.9 | 245 | - | - | 0 | - |
| - | - | 561.6 | 259.6 | - | - | 0 | - |
| - | - | 1211 | 260.1 | - | - | 0 | - |
| - | - | 772.4 | 270.1 | - | - | 0 | - |
| - | - | 777 | 279.1 | - | - | 0 | - |
| - | - | 2598 | 282.1 | - | - | 0 | - |
| - | - | 597.8 | 287.2 | - | - | 0 | - |
| - | - | 1972 | 295.1 | - | - | 0 | - |
| 11 | y | 1421 | 295.2 | 0.0004158 | 1.409 | +1 | 2 |
| - | - | 2960 | 296.1 | - | - | 0 | - |
| - | - | 1605 | 297.1 | - | - | 0 | - |
| - | - | 540.1 | 297.2 | - | - | 0 | - |
| - | - | 1478 | 299.1 | - | - | 0 | - |
| - | - | 862 | 299.2 | - | - | 0 | - |
| - | - | 2053 | 300.1 | - | - | 0 | - |
| - | - | 7054 | 300.2 | - | - | 0 | - |
| - | - | 845.2 | 301.1 | - | - | 0 | - |
| - | - | 647.4 | 301.2 | - | - | 0 | - |
| - | - | 1480 | 315.2 | - | - | 0 | - |
| - | - | 3698 | 317.2 | - | - | 0 | - |
| 3 | c | 2918 | 319.2 | 0.0005542 | 1.736 | +1 | 3 |
| - | - | 618.8 | 320.2 | - | - | 0 | - |
| - | - | 499.3 | 323.1 | - | - | 0 | - |
| - | - | 1.23E+04 | 341 | - | - | 0 | - |
| - | - | 1030 | 341 | - | - | 0 | - |
| - | - | 1785 | 350.7 | - | - | 0 | - |
| - | - | 1660 | 355.1 | - | - | 0 | - |
| - | - | 623.2 | 356.1 | - | - | 0 | - |
| - | - | 1784 | 358.7 | - | - | 0 | - |
| - | - | 4.55E+04 | 359 | - | - | 0 | - |
| - | - | 1934 | 359.7 | - | - | 0 | - |
| - | - | 938.7 | 360.2 | - | - | 0 | - |
| - | - | 763.2 | 364.2 | - | - | 0 | - |
| - | - | 1256 | 369.1 | - | - | 0 | - |
| - | - | 769.2 | 369.1 | - | - | 0 | - |
| - | - | 2876 | 370.1 | - | - | 0 | - |
| - | - | 3379 | 371.1 | - | - | 0 | - |
| - | - | 815.4 | 378.1 | - | - | 0 | - |
| - | - | 4765 | 379.2 | - | - | 0 | - |
| - | - | 1020 | 380.2 | - | - | 0 | - |
| 10 | y | 747.4 | 382.2 | 0.0002406 | 0.6295 | +1 | 3 |
| - | - | 697.1 | 385.2 | - | - | 0 | - |
| - | - | 3826 | 385.2 | - | - | 0 | - |
| - | - | 936.6 | 385.9 | - | - | 0 | - |
| - | - | 4204 | 386.2 | - | - | 0 | - |
| - | - | 1151 | 386.2 | - | - | 0 | - |
| - | - | 2856 | 386.7 | - | - | 0 | - |
| - | - | 1105 | 387.2 | - | - | 0 | - |
| - | - | 824.5 | 390.6 | - | - | 0 | - |
| - | - | 2457 | 391.6 | - | - | 0 | - |
| - | - | 2960 | 391.9 | - | - | 0 | - |
| - | - | 1247 | 399.7 | - | - | 0 | - |
| - | - | 763.6 | 408.2 | - | - | 0 | - |
| - | - | 652 | 412.7 | - | - | 0 | - |
| - | - | 909.5 | 415.3 | - | - | 0 | - |
| - | - | 691.9 | 418.7 | - | - | 0 | - |
| - | - | 2604 | 419.2 | - | - | 0 | - |
| - | - | 734 | 420.2 | - | - | 0 | - |
| - | - | 633.2 | 425.2 | - | - | 0 | - |
| - | - | 1480 | 427.7 | - | - | 0 | - |
| - | - | 1918 | 428.2 | - | - | 0 | - |
| - | - | 1.187E+05 | 429.1 | - | - | 0 | - |
| - | - | 1544 | 429.8 | - | - | 0 | - |
| - | - | 1880 | 430.2 | - | - | 0 | - |
| - | - | 1112 | 430.3 | - | - | 0 | - |
| - | - | 681.1 | 431.3 | - | - | 0 | - |
| - | - | 2059 | 433.7 | - | - | 0 | - |
| - | - | 1413 | 434.2 | - | - | 0 | - |
| - | - | 1460 | 435.2 | - | - | 0 | - |
| - | - | 3096 | 436.7 | - | - | 0 | - |
| - | - | 1291 | 437.2 | - | - | 0 | - |
| - | - | 920.8 | 437.2 | - | - | 0 | - |
| - | - | 5812 | 442.7 | - | - | 0 | - |
| - | - | 4154 | 442.7 | - | - | 0 | - |
| - | - | 4108 | 443.2 | - | - | 0 | - |
| 8 | c | 1579 | 443.2 | 0.008632 | 19.47 | +2 | 8 |
| - | - | 1314 | 443.7 | - | - | 0 | - |
| 8 | c | 1182 | 443.7 | 0.001893 | 4.266 | +2 | 8 |
| - | - | 664.1 | 444.7 | - | - | 0 | - |
| - | - | 5.961E+04 | 445.1 | - | - | 0 | - |
| - | - | 696.4 | 445.7 | - | - | 0 | - |
| - | - | 2095 | 446.2 | - | - | 0 | - |
| - | - | 1869 | 450.7 | - | - | 0 | - |
| - | - | 1219 | 450.7 | - | - | 0 | - |
| - | - | 855.7 | 451.2 | - | - | 0 | - |
| - | - | 1.841E+04 | 451.7 | - | - | 0 | - |
| - | - | 9725 | 451.8 | - | - | 0 | - |
| - | - | 1.067E+04 | 452.2 | - | - | 0 | - |
| 8 | c | 2.745E+04 | 452.3 | 0.0002764 | 0.6111 | +2 | 8 |
| - | - | 3714 | 452.7 | - | - | 0 | - |
| - | - | 1.405E+04 | 452.8 | - | - | 0 | - |
| - | - | 4407 | 453.3 | - | - | 0 | - |
| - | - | 1067 | 453.8 | - | - | 0 | - |
| - | - | 1108 | 455.2 | - | - | 0 | - |
| - | - | 608.7 | 455.2 | - | - | 0 | - |
| 4 | c | 1418 | 458.3 | 0.0007655 | 1.67 | +1 | 4 |
| - | - | 4.708E+04 | 459.7 | - | - | 0 | - |
| - | - | 2.053E+04 | 459.7 | - | - | 0 | - |
| - | - | 2.829E+04 | 460.2 | - | - | 0 | - |
| - | - | 1.516E+04 | 460.2 | - | - | 0 | - |
| - | - | 2.126E+04 | 460.7 | - | - | 0 | - |
| - | - | 5465 | 460.7 | - | - | 0 | - |
| - | - | 1.27E+04 | 461.2 | - | - | 0 | - |
| - | - | 1919 | 461.2 | - | - | 0 | - |
| - | - | 3542 | 461.7 | - | - | 0 | - |
| - | - | 672.3 | 462.2 | - | - | 0 | - |
| - | - | 772.9 | 462.2 | - | - | 0 | - |
| - | - | 658.7 | 465.3 | - | - | 0 | - |
| - | - | 826 | 470.8 | - | - | 0 | - |
| - | - | 1697 | 472.3 | - | - | 0 | - |
| 4 | c | 1.043E+05 | 475.3 | 0.0007921 | 1.666 | +1 | 4 |
| - | - | 2.398E+04 | 476.3 | - | - | 0 | - |
| - | - | 3514 | 477.3 | - | - | 0 | - |
| - | - | 645.8 | 479.3 | - | - | 0 | - |
| - | - | 7903 | 479.8 | - | - | 0 | - |
| 9 | z | 1514 | 480.2 | 6.042E-05 | 0.1258 | +1 | 4 |
| - | - | 2872 | 480.3 | - | - | 0 | - |
| - | - | 1835 | 480.8 | - | - | 0 | - |
| - | - | 7728 | 481.2 | - | - | 0 | - |
| - | - | 1020 | 482.2 | - | - | 0 | - |
| - | - | 1249 | 484.7 | - | - | 0 | - |
| - | - | 1173 | 485.2 | - | - | 0 | - |
| - | - | 763.8 | 485.7 | - | - | 0 | - |
| - | - | 1831 | 486.8 | - | - | 0 | - |
| - | - | 2182 | 487.3 | - | - | 0 | - |
| - | - | 1498 | 487.8 | - | - | 0 | - |
| - | - | 1067 | 488.3 | - | - | 0 | - |
| - | - | 2344 | 490.2 | - | - | 0 | - |
| - | - | 1507 | 493.8 | - | - | 0 | - |
| - | - | 1192 | 494.3 | - | - | 0 | - |
| 9 | y | 3728 | 496.2 | 0.0005026 | 1.013 | +1 | 4 |
| - | - | 1.032E+04 | 499.8 | - | - | 0 | - |
| 9 | c | 7675 | 500.3 | 0.004186 | 8.368 | +2 | 9 |
| 9 | c | 4879 | 500.8 | 0.001853 | 3.7 | +2 | 9 |
| - | - | 2792 | 501.3 | - | - | 0 | - |
| - | - | 842.2 | 501.8 | - | - | 0 | - |
| - | - | 1.04E+04 | 502.3 | - | - | 0 | - |
| - | - | 2319 | 503.3 | - | - | 0 | - |
| - | - | 826.5 | 506.1 | - | - | 0 | - |
| - | - | 776.5 | 508.3 | - | - | 0 | - |
| - | - | 2.219E+04 | 508.8 | - | - | 0 | - |
| 9 | c | 3.414E+04 | 509.3 | 0.001351 | 2.653 | +2 | 9 |
| - | - | 2.549E+04 | 509.8 | - | - | 0 | - |
| - | - | 8558 | 510.3 | - | - | 0 | - |
| - | - | 4138 | 510.8 | - | - | 0 | - |
| - | - | 3162 | 514.2 | - | - | 0 | - |
| - | - | 1628 | 515.2 | - | - | 0 | - |
| - | - | 604.3 | 522.3 | - | - | 0 | - |
| - | - | 691.6 | 523.3 | - | - | 0 | - |
| - | - | 1275 | 524.2 | - | - | 0 | - |
| - | - | 632.5 | 526.3 | - | - | 0 | - |
| - | - | 1108 | 526.8 | - | - | 0 | - |
| - | - | 989.2 | 530.2 | - | - | 0 | - |
| - | - | 1855 | 530.3 | - | - | 0 | - |
| 4 | z | 3923 | 532.8 | 0.0005794 | 1.088 | +2 | 9 |
| - | - | 1773 | 533.3 | - | - | 0 | - |
| - | - | 795 | 533.8 | - | - | 0 | - |
| - | - | 1326 | 534.3 | - | - | 0 | - |
| - | - | 1589 | 535.3 | - | - | 0 | - |
| - | - | 677.5 | 535.3 | - | - | 0 | - |
| - | - | 1283 | 535.8 | - | - | 0 | - |
| 4 | y | 705.8 | 540.8 | 0.0001767 | 0.3267 | +2 | 9 |
| - | - | 2234 | 542.2 | - | - | 0 | - |
| 10 | c | 3215 | 544.3 | 0.001006 | 1.848 | +2 | 10 |
| - | - | 715.4 | 544.3 | - | - | 0 | - |
| - | - | 2488 | 544.8 | - | - | 0 | - |
| - | - | 1129 | 545.3 | - | - | 0 | - |
| - | - | 3183 | 545.3 | - | - | 0 | - |
| - | - | 2273 | 546.3 | - | - | 0 | - |
| - | - | 5182 | 548.3 | - | - | 0 | - |
| - | - | 2388 | 548.7 | - | - | 0 | - |
| - | - | 1101 | 549.2 | - | - | 0 | - |
| 10 | c | 2.657E+04 | 552.8 | 0.0006097 | 1.103 | +2 | 10 |
| - | - | 2.374E+04 | 553.3 | - | - | 0 | - |
| - | - | 1.223E+04 | 553.8 | - | - | 0 | - |
| - | - | 4878 | 554.3 | - | - | 0 | - |
| - | - | 2533 | 554.8 | - | - | 0 | - |
| - | - | 649.2 | 555.3 | - | - | 0 | - |
| - | - | 982.6 | 556.2 | - | - | 0 | - |
| - | - | 3619 | 564.3 | - | - | 0 | - |
| - | - | 2190 | 565.3 | - | - | 0 | - |
| - | - | 2437 | 565.8 | - | - | 0 | - |
| - | - | 1591 | 566.3 | - | - | 0 | - |
| - | - | 653.8 | 566.8 | - | - | 0 | - |
| 3 | z | 685.4 | 567.3 | 0.002177 | 3.837 | +2 | 10 |
| 3 | z | 1565 | 567.8 | 0.009681 | 17.05 | +2 | 10 |
| - | - | 1725 | 568.3 | - | - | 0 | - |
| 5 | c | 1157 | 573.3 | 0.00815 | 14.22 | +1 | 5 |
| - | - | 1007 | 574.3 | - | - | 0 | - |
| 3 | y | 835.6 | 575.3 | 0.0001998 | 0.3474 | +2 | 10 |
| 3 | y | 1450 | 575.8 | 0.00392 | 6.807 | +2 | 10 |
| 3 | z | 1.01E+04 | 576.3 | 0.0006785 | 1.177 | +2 | 10 |
| - | - | 7994 | 576.8 | - | - | 0 | - |
| - | - | 5075 | 577.3 | - | - | 0 | - |
| - | - | 1837 | 577.8 | - | - | 0 | - |
| - | - | 1821 | 578.3 | - | - | 0 | - |
| - | - | 964.6 | 578.8 | - | - | 0 | - |
| - | - | 1101 | 579.3 | - | - | 0 | - |
| - | - | 780.5 | 579.8 | - | - | 0 | - |
| - | - | 1925 | 583.3 | - | - | 0 | - |
| 3 | y | 3.05E+04 | 584.3 | 0.0005327 | 0.9118 | +2 | 10 |
| - | - | 2.595E+04 | 584.8 | - | - | 0 | - |
| - | - | 1.162E+04 | 585.3 | - | - | 0 | - |
| - | - | 3548 | 585.8 | - | - | 0 | - |
| - | - | 3220 | 586.8 | - | - | 0 | - |
| - | - | 2970 | 587.3 | - | - | 0 | - |
| - | - | 2202 | 587.8 | - | - | 0 | - |
| - | - | 2000 | 588.3 | - | - | 0 | - |
| - | - | 1413 | 588.3 | - | - | 0 | - |
| - | - | 1307 | 588.8 | - | - | 0 | - |
| - | - | 907 | 589.3 | - | - | 0 | - |
| - | - | 2121 | 589.3 | - | - | 0 | - |
| 5 | c | 1.836E+05 | 590.3 | 0.0007045 | 1.193 | +1 | 5 |
| - | - | 5.017E+04 | 591.3 | - | - | 0 | - |
| - | - | 2352 | 591.8 | - | - | 0 | - |
| - | - | 1.032E+04 | 592.3 | - | - | 0 | - |
| - | - | 2440 | 592.8 | - | - | 0 | - |
| - | - | 1830 | 593.3 | - | - | 0 | - |
| - | - | 668.9 | 598.3 | - | - | 0 | - |
| 11 | c | 3666 | 600.8 | 0.001515 | 2.522 | +2 | 11 |
| - | - | 3198 | 601.3 | - | - | 0 | - |
| - | - | 2801 | 601.8 | - | - | 0 | - |
| - | - | 1230 | 607.3 | - | - | 0 | - |
| - | - | 858.7 | 607.8 | - | - | 0 | - |
| 8 | z | 1.18E+04 | 608.3 | 0.001443 | 2.372 | +1 | 5 |
| 11 | c | 7.367E+04 | 609.3 | 0.0002037 | 0.3343 | +2 | 11 |
| - | - | 5.745E+04 | 609.8 | - | - | 0 | - |
| - | - | 2.619E+04 | 610.3 | - | - | 0 | - |
| - | - | 8144 | 610.8 | - | - | 0 | - |
| - | - | 2218 | 611.3 | - | - | 0 | - |
| - | - | 694.3 | 613.2 | - | - | 0 | - |
| - | - | 1652 | 613.3 | - | - | 0 | - |
| - | - | 2333 | 614.2 | - | - | 0 | - |
| - | - | 1232 | 614.3 | - | - | 0 | - |
| - | - | 2149 | 616.3 | - | - | 0 | - |
| 2 | w | 4422 | 618.3 | 0.0001207 | 0.1952 | +2 | 11 |
| - | - | 4008 | 618.8 | - | - | 0 | - |
| - | - | 1085 | 619.3 | - | - | 0 | - |
| - | - | 1217 | 621.3 | - | - | 0 | - |
| 8 | y | 1.422E+04 | 624.3 | 0.0004798 | 0.7685 | +1 | 5 |
| - | - | 4190 | 625.3 | - | - | 0 | - |
| - | - | 964.9 | 626.3 | - | - | 0 | - |
| - | - | 2672 | 630.3 | - | - | 0 | - |
| - | - | 2193 | 630.8 | - | - | 0 | - |
| - | - | 1436 | 631.3 | - | - | 0 | - |
| 2 | y | 1026 | 631.8 | 0.002709 | 4.287 | +2 | 11 |
| 2 | z | 1708 | 632.8 | 0.00162 | 2.559 | +2 | 11 |
| - | - | 3763 | 633.3 | - | - | 0 | - |
| - | - | 1090 | 633.8 | - | - | 0 | - |
| - | - | 1569 | 634.3 | - | - | 0 | - |
| - | - | 635.9 | 635.4 | - | - | 0 | - |
| - | - | 2472 | 636.3 | - | - | 0 | - |
| - | - | 768.8 | 637.8 | - | - | 0 | - |
| - | - | 1712 | 638.3 | - | - | 0 | - |
| - | - | 1133 | 638.8 | - | - | 0 | - |
| - | - | 949.4 | 639.3 | - | - | 0 | - |
| - | - | 929.3 | 639.8 | - | - | 0 | - |
| - | - | 951.3 | 640.4 | - | - | 0 | - |
| - | - | 709.5 | 640.7 | - | - | 0 | - |
| 2 | y | 2916 | 640.8 | 0.001286 | 2.007 | +2 | 11 |
| - | - | 3405 | 641.3 | - | - | 0 | - |
| - | - | 1635 | 641.8 | - | - | 0 | - |
| - | - | 550.7 | 642.3 | - | - | 0 | - |
| - | - | 558.6 | 642.3 | - | - | 0 | - |
| - | - | 830.4 | 646.3 | - | - | 0 | - |
| - | - | 3144 | 647.4 | - | - | 0 | - |
| - | - | 2975 | 647.9 | - | - | 0 | - |
| - | - | 643.8 | 648.3 | - | - | 0 | - |
| - | - | 4438 | 649.3 | - | - | 0 | - |
| - | - | 2728 | 650.3 | - | - | 0 | - |
| - | - | 2626 | 653.3 | - | - | 0 | - |
| - | - | 1775 | 653.8 | - | - | 0 | - |
| - | - | 1702 | 654.3 | - | - | 0 | - |
| - | - | 6176 | 655.3 | - | - | 0 | - |
| - | - | 5204 | 655.8 | - | - | 0 | - |
| - | - | 2752 | 656.3 | - | - | 0 | - |
| - | - | 921.2 | 656.8 | - | - | 0 | - |
| - | - | 7060 | 659.3 | - | - | 0 | - |
| - | - | 1.108E+04 | 660.3 | - | - | 0 | - |
| - | - | 8128 | 660.9 | - | - | 0 | - |
| 7 | z | 1.506E+04 | 661.3 | 0.008341 | 12.61 | +1 | 6 |
| - | - | 1.184E+04 | 661.8 | - | - | 0 | - |
| - | - | 1.559E+04 | 662.3 | - | - | 0 | - |
| - | - | 9829 | 662.8 | - | - | 0 | - |
| - | - | 4596 | 663.3 | - | - | 0 | - |
| - | - | 1372 | 663.8 | - | - | 0 | - |
| - | - | 3640 | 668.8 | - | - | 0 | - |
| - | - | 1.259E+04 | 669.3 | - | - | 0 | - |
| - | - | 1.379E+04 | 669.8 | - | - | 0 | - |
| - | - | 1.384E+04 | 670.3 | - | - | 0 | - |
| - | - | 9008 | 670.8 | - | - | 0 | - |
| - | - | 4189 | 671.3 | - | - | 0 | - |
| - | - | 1350 | 671.8 | - | - | 0 | - |
| - | - | 1099 | 672.3 | - | - | 0 | - |
| - | - | 763.7 | 673.3 | - | - | 0 | - |
| - | - | 2844 | 674.3 | - | - | 0 | - |
| - | - | 8307 | 674.8 | - | - | 0 | - |
| - | - | 6791 | 675.3 | - | - | 0 | - |
| - | - | 7875 | 675.8 | - | - | 0 | - |
| - | - | 4875 | 676.3 | - | - | 0 | - |
| - | - | 2405 | 676.8 | - | - | 0 | - |
| 7 | z | 3.664E+04 | 679.4 | 0.0007058 | 1.039 | +1 | 6 |
| - | - | 1.436E+04 | 680.4 | - | - | 0 | - |
| - | - | 3894 | 681.4 | - | - | 0 | - |
| - | - | 3120 | 682.9 | - | - | 0 | - |
| - | - | 4.897E+04 | 683.3 | - | - | 0 | - |
| - | - | 4.34E+04 | 683.8 | - | - | 0 | - |
| - | - | 2.533E+04 | 684.3 | - | - | 0 | - |
| - | - | 7974 | 684.8 | - | - | 0 | - |
| - | - | 2282 | 685.3 | - | - | 0 | - |
| 6 | c | 1620 | 687.3 | 0.008984 | 13.07 | +1 | 6 |
| - | - | 1899 | 689.7 | - | - | 0 | - |
| - | - | 1283 | 690.2 | - | - | 0 | - |
| - | - | 4.456E+04 | 691.4 | - | - | 0 | - |
| - | - | 4.125E+04 | 691.9 | - | - | 0 | - |
| - | - | 2.495E+04 | 692.4 | - | - | 0 | - |
| - | - | 8890 | 692.9 | - | - | 0 | - |
| - | - | 2633 | 693.4 | - | - | 0 | - |
| - | - | 809.3 | 694.4 | - | - | 0 | - |
| 7 | y | 8343 | 695.4 | 0.0007195 | 1.035 | +1 | 6 |
| - | - | 3353 | 696.4 | - | - | 0 | - |
| - | - | 2795 | 700.3 | - | - | 0 | - |
| - | - | 3823 | 701.3 | - | - | 0 | - |
| - | - | 1.809E+04 | 702.3 | - | - | 0 | - |
| - | - | 1.156E+04 | 703.3 | - | - | 0 | - |
| - | - | 2902 | 704.3 | - | - | 0 | - |
| 6 | c | 4.148E+04 | 704.4 | 0.0008068 | 1.145 | +1 | 6 |
| - | - | 1.626E+04 | 705.4 | - | - | 0 | - |
| - | - | 3878 | 706.4 | - | - | 0 | - |
| - | - | 1019 | 707.4 | - | - | 0 | - |
| - | - | 917.2 | 711.3 | - | - | 0 | - |
| - | - | 4855 | 726.3 | - | - | 0 | - |
| - | - | 2482 | 727.3 | - | - | 0 | - |
| - | - | 5664 | 731.4 | - | - | 0 | - |
| - | - | 3420 | 732.4 | - | - | 0 | - |
| - | - | 1162 | 733.4 | - | - | 0 | - |
| - | - | 898.2 | 737.3 | - | - | 0 | - |
| - | - | 2557 | 737.4 | - | - | 0 | - |
| - | - | 789.7 | 738.3 | - | - | 0 | - |
| 6 | w | 2.152E+04 | 749.4 | 0.002362 | 3.152 | +1 | 7 |
| - | - | 1.464E+04 | 750.4 | - | - | 0 | - |
| - | - | 4999 | 751.4 | - | - | 0 | - |
| - | - | 1085 | 754.4 | - | - | 0 | - |
| - | - | 976.9 | 755.4 | - | - | 0 | - |
| 7 | c | 3062 | 758.4 | 0.001473 | 1.942 | +1 | 7 |
| - | - | 1085 | 759.4 | - | - | 0 | - |
| - | - | 8114 | 770.3 | - | - | 0 | - |
| - | - | 7948 | 771.3 | - | - | 0 | - |
| - | - | 1.951E+04 | 772.3 | - | - | 0 | - |
| - | - | 1.186E+04 | 773.3 | - | - | 0 | - |
| - | - | 2906 | 774.3 | - | - | 0 | - |
| 7 | c | 3.812E+04 | 775.4 | 0.0005583 | 0.72 | +1 | 7 |
| - | - | 1.959E+04 | 776.4 | - | - | 0 | - |
| - | - | 5802 | 777.4 | - | - | 0 | - |
| - | - | 796.7 | 778.4 | - | - | 0 | - |
| - | - | 897.4 | 781.3 | - | - | 0 | - |
| - | - | 1773 | 787.4 | - | - | 0 | - |
| - | - | 648.3 | 788.4 | - | - | 0 | - |
| 6 | z | 9.308E+04 | 793.4 | 0.0008692 | 1.096 | +1 | 7 |
| - | - | 5.348E+04 | 794.4 | - | - | 0 | - |
| - | - | 1.914E+04 | 795.4 | - | - | 0 | - |
| - | - | 4194 | 796.4 | - | - | 0 | - |
| - | - | 738.9 | 797.4 | - | - | 0 | - |
| - | - | 2887 | 799.3 | - | - | 0 | - |
| - | - | 1183 | 800.3 | - | - | 0 | - |
| - | - | 943.7 | 808.4 | - | - | 0 | - |
| 6 | y | 1560 | 809.4 | 0.000704 | 0.8698 | +1 | 7 |
| - | - | 1569 | 810.4 | - | - | 0 | - |
| - | - | 863.1 | 811.4 | - | - | 0 | - |
| - | - | 1621 | 815.3 | - | - | 0 | - |
| - | - | 1015 | 816.3 | - | - | 0 | - |
| - | - | 1.242E+04 | 817.3 | - | - | 0 | - |
| - | - | 4950 | 818.3 | - | - | 0 | - |
| - | - | 1213 | 819.3 | - | - | 0 | - |
| - | - | 834.5 | 820.4 | - | - | 0 | - |
| - | - | 1179 | 821.4 | - | - | 0 | - |
| - | - | 907.8 | 832.4 | - | - | 0 | - |
| - | - | 972.2 | 833.3 | - | - | 0 | - |
| - | - | 2018 | 848.5 | - | - | 0 | - |
| - | - | 1319 | 849.5 | - | - | 0 | - |
| - | - | 1173 | 852.4 | - | - | 0 | - |
| - | - | 1174 | 858.4 | - | - | 0 | - |
| - | - | 841.4 | 858.5 | - | - | 0 | - |
| - | - | 2044 | 859.4 | - | - | 0 | - |
| - | - | 4411 | 859.5 | - | - | 0 | - |
| - | - | 6563 | 860.4 | - | - | 0 | - |
| - | - | 3956 | 860.5 | - | - | 0 | - |
| - | - | 8477 | 861.4 | - | - | 0 | - |
| - | - | 1595 | 861.5 | - | - | 0 | - |
| - | - | 5740 | 862.4 | - | - | 0 | - |
| - | - | 2054 | 863.4 | - | - | 0 | - |
| - | - | 2.855E+04 | 864.4 | - | - | 0 | - |
| - | - | 1.998E+04 | 865.4 | - | - | 0 | - |
| - | - | 7401 | 866.4 | - | - | 0 | - |
| - | - | 1598 | 867.4 | - | - | 0 | - |
| - | - | 1401 | 874.4 | - | - | 0 | - |
| - | - | 2570 | 875.4 | - | - | 0 | - |
| - | - | 2202 | 876.4 | - | - | 0 | - |
| - | - | 1698 | 877.4 | - | - | 0 | - |
| - | - | 3363 | 878.3 | - | - | 0 | - |
| - | - | 1142 | 879.4 | - | - | 0 | - |
| - | - | 1899 | 884.5 | - | - | 0 | - |
| 8 | c | 1885 | 885.5 | 0.003152 | 3.559 | +1 | 8 |
| 8 | c | 4940 | 886.5 | 0.002762 | 3.116 | +1 | 8 |
| - | - | 1823 | 887.4 | - | - | 0 | - |
| - | - | 1021 | 887.5 | - | - | 0 | - |
| - | - | 1119 | 888.4 | - | - | 0 | - |
| - | - | 1634 | 888.5 | - | - | 0 | - |
| 5 | z | 811.9 | 891.4 | 0.01665 | 18.68 | +1 | 8 |
| - | - | 1975 | 896.4 | - | - | 0 | - |
| - | - | 4317 | 900.5 | - | - | 0 | - |
| - | - | 2562 | 901.5 | - | - | 0 | - |
| - | - | 1.264E+04 | 902.4 | - | - | 0 | - |
| - | - | 1.373E+04 | 902.5 | - | - | 0 | - |
| - | - | 9502 | 903.4 | - | - | 0 | - |
| 8 | c | 5.309E+04 | 903.5 | 0.0007158 | 0.7922 | +1 | 8 |
| - | - | 2.112E+04 | 904.4 | - | - | 0 | - |
| - | - | 3.088E+04 | 904.5 | - | - | 0 | - |
| - | - | 1.078E+04 | 905.4 | - | - | 0 | - |
| - | - | 1.204E+04 | 905.5 | - | - | 0 | - |
| - | - | 3097 | 906.4 | - | - | 0 | - |
| - | - | 2911 | 906.5 | - | - | 0 | - |
| 5 | z | 3.073E+04 | 908.4 | 0.0002933 | 0.3229 | +1 | 8 |
| - | - | 2.422E+04 | 909.4 | - | - | 0 | - |
| - | - | 8411 | 910.4 | - | - | 0 | - |
| - | - | 2590 | 911.4 | - | - | 0 | - |
| - | - | 4096 | 918.4 | - | - | 0 | - |
| - | - | 3359 | 918.5 | - | - | 0 | - |
| - | - | 9455 | 919.4 | - | - | 0 | - |
| - | - | 2.981E+04 | 919.5 | - | - | 0 | - |
| - | - | 8710 | 920.4 | - | - | 0 | - |
| - | - | 2.201E+04 | 920.5 | - | - | 0 | - |
| - | - | 2.209E+04 | 921.4 | - | - | 0 | - |
| - | - | 9040 | 921.5 | - | - | 0 | - |
| - | - | 9525 | 922.4 | - | - | 0 | - |
| - | - | 2486 | 922.5 | - | - | 0 | - |
| - | - | 3402 | 923.4 | - | - | 0 | - |
| 5 | y | 2353 | 924.4 | 0.004149 | 4.488 | +1 | 8 |
| - | - | 2697 | 925.4 | - | - | 0 | - |
| - | - | 1746 | 926.4 | - | - | 0 | - |
| - | - | 828.2 | 969.5 | - | - | 0 | - |
| - | - | 966.8 | 974.5 | - | - | 0 | - |
| - | - | 966.5 | 975.5 | - | - | 0 | - |
| - | - | 2596 | 986.5 | - | - | 0 | - |
| - | - | 3743 | 987.5 | - | - | 0 | - |
| - | - | 3035 | 988.5 | - | - | 0 | - |
| - | - | 868 | 989.5 | - | - | 0 | - |
| 9 | c | 1999 | 999.5 | 0.000857 | 0.8574 | +1 | 9 |
| 9 | c | 3364 | 1001 | 0.007137 | 7.133 | +1 | 9 |
| - | - | 2420 | 1002 | - | - | 0 | - |
| - | - | 1736 | 1003 | - | - | 0 | - |
| - | - | 1804 | 1004 | - | - | 0 | - |
| - | - | 3648 | 1017 | - | - | 0 | - |
| 9 | c | 3.159E+04 | 1018 | 0.0001252 | 0.123 | +1 | 9 |
| - | - | 2.156E+04 | 1019 | - | - | 0 | - |
| - | - | 9872 | 1020 | - | - | 0 | - |
| - | - | 3345 | 1021 | - | - | 0 | - |
| - | - | 1203 | 1022 | - | - | 0 | - |
| - | - | 799.2 | 1048 | - | - | 0 | - |
| - | - | 1270 | 1061 | - | - | 0 | - |
| - | - | 727.2 | 1062 | - | - | 0 | - |
| - | - | 680.6 | 1063 | - | - | 0 | - |
| - | - | 907.1 | 1064 | - | - | 0 | - |
| 4 | z | 1.047E+04 | 1065 | 0.0009889 | 0.929 | +1 | 9 |
| - | - | 1.782E+04 | 1066 | - | - | 0 | - |
| - | - | 9476 | 1067 | - | - | 0 | - |
| - | - | 3359 | 1068 | - | - | 0 | - |
| - | - | 809.7 | 1069 | - | - | 0 | - |
| 4 | y | 1749 | 1081 | 0.009068 | 8.392 | +1 | 9 |
| - | - | 2139 | 1089 | - | - | 0 | - |
| - | - | 2359 | 1090 | - | - | 0 | - |
| - | - | 1421 | 1091 | - | - | 0 | - |
| - | - | 873.1 | 1092 | - | - | 0 | - |
| - | - | 1302 | 1095 | - | - | 0 | - |
| - | - | 1233 | 1096 | - | - | 0 | - |
| - | - | 732.9 | 1102 | - | - | 0 | - |
| 10 | c | 8506 | 1105 | 0.0002932 | 0.2654 | +1 | 10 |
| - | - | 9148 | 1106 | - | - | 0 | - |
| - | - | 5059 | 1107 | - | - | 0 | - |
| - | - | 2171 | 1108 | - | - | 0 | - |
| - | - | 972.6 | 1109 | - | - | 0 | - |
| 3 | z | 6709 | 1152 | 0.0003326 | 0.2888 | +1 | 10 |
| - | - | 1.856E+04 | 1153 | - | - | 0 | - |
| - | - | 1.13E+04 | 1154 | - | - | 0 | - |
| - | - | 4430 | 1155 | - | - | 0 | - |
| - | - | 924.7 | 1156 | - | - | 0 | - |
| - | - | 1460 | 1159 | - | - | 0 | - |
| - | - | 959.4 | 1160 | - | - | 0 | - |
| - | - | 1343 | 1169 | - | - | 0 | - |
| - | - | 1091 | 1170 | - | - | 0 | - |
| - | - | 768.2 | 1171 | - | - | 0 | - |
| - | - | 2547 | 1174 | - | - | 0 | - |
| - | - | 4856 | 1175 | - | - | 0 | - |
| - | - | 5636 | 1176 | - | - | 0 | - |
| - | - | 1868 | 1177 | - | - | 0 | - |
| - | - | 1146 | 1185 | - | - | 0 | - |
| - | - | 7382 | 1202 | - | - | 0 | - |
| - | - | 4608 | 1203 | - | - | 0 | - |
| - | - | 3215 | 1204 | - | - | 0 | - |
| 11 | c | 6232 | 1218 | 0.003303 | 2.712 | +1 | 11 |
| - | - | 1.341E+04 | 1219 | - | - | 0 | - |
| - | - | 1.001E+04 | 1220 | - | - | 0 | - |
| - | - | 3336 | 1221 | - | - | 0 | - |
| - | - | 962.9 | 1222 | - | - | 0 | - |
| - | - | 2681 | 1266 | - | - | 0 | - |
| - | - | 3550 | 1267 | - | - | 0 | - |
| - | - | 1778 | 1268 | - | - | 0 | - |
| - | - | 990.9 | 1269 | - | - | 0 | - |
| - | - | 2289 | 1306 | - | - | 0 | - |
| - | - | 4612 | 1307 | - | - | 0 | - |
| - | - | 2118 | 1308 | - | - | 0 | - |
| - | - | 2143 | 1309 | - | - | 0 | - |
| - | - | 2153 | 1321 | - | - | 0 | - |
| - | - | 4275 | 1322 | - | - | 0 | - |
| - | - | 3996 | 1323 | - | - | 0 | - |
| - | - | 7773 | 1324 | - | - | 0 | - |
| - | - | 1.455E+04 | 1325 | - | - | 0 | - |
| - | - | 1.014E+04 | 1326 | - | - | 0 | - |
| - | - | 4963 | 1327 | - | - | 0 | - |
| - | - | 1507 | 1328 | - | - | 0 | - |
| - | - | 2926 | 1338 | - | - | 0 | - |
| - | - | 1.818E+04 | 1339 | - | - | 0 | - |
| - | - | 1.506E+04 | 1340 | - | - | 0 | - |
| - | - | 8021 | 1341 | - | - | 0 | - |
| - | - | 3573 | 1342 | - | - | 0 | - |
| - | - | 1066 | 1343 | - | - | 0 | - |
| - | - | 3827 | 1349 | - | - | 0 | - |
| - | - | 8293 | 1350 | - | - | 0 | - |
| - | - | 5198 | 1351 | - | - | 0 | - |
| - | - | 2930 | 1352 | - | - | 0 | - |
| - | - | 1211 | 1353 | - | - | 0 | - |
| - | - | 1442 | 1356 | - | - | 0 | - |
| - | - | 1477 | 1357 | - | - | 0 | - |
| - | - | 9540 | 1366 | - | - | 0 | - |
| - | - | 5.701E+04 | 1367 | - | - | 0 | - |
| - | - | 5.297E+04 | 1368 | - | - | 0 | - |
| - | - | 3.04E+04 | 1369 | - | - | 0 | - |
| - | - | 1.009E+04 | 1370 | - | - | 0 | - |
| - | - | 2296 | 1371 | - | - | 0 | - |
| - | - | 1134 | 1379 | - | - | 0 | - |
| - | - | 2123 | 1382 | - | - | 0 | - |
| - | - | 7105 | 1383 | - | - | 0 | - |
| - | - | 1.363E+04 | 1384 | - | - | 0 | - |
| - | - | 1.171E+04 | 1385 | - | - | 0 | - |
| - | - | 4690 | 1386 | - | - | 0 | - |
| - | - | 1741 | 1387 | - | - | 0 | - |

m/z Charge Intensity FragmentType MassShift Position
120.07127380371094 0 420.56067
121.13667297363281 0 394.0026
123.10444641113281 0 586.8235
128.5663299560547 0 396.6542
129.10252380371094 0 3675.234
130.105712890625 0 506.91302
136.0760040283203 0 5613.136
137.0793914794922 0 913.18646
142.1228485107422 0 491.76648
143.1181640625 0 493.5092
143.86541748046875 0 390.52197
144.16612243652344 0 409.55188
144.8082733154297 0 393.39523
149.0451202392578 0 4237.5063
150.0442657470703 0 659.2762
165.05490112304688 0 5245.7485
166.0581512451172 0 623.6248
167.05593872070312 0 1181.534
171.12501525878906 0 498.03366
173.09243774414062 0 1957.2543
173.1288299560547 0 557.37213
174.08787536621094 0 679.51794
182.08145141601562 0 18242.625 y 11
183.08497619628906 0 1774.4292
186.11231994628906 0 568.8148
187.14439392089844 0 38645.133
188.14781188964844 0 3248.8948
193.52252197265625 0 438.06528
200.13990783691406 0 682.79425
201.1233673095703 0 1647.9838
215.1392364501953 0 10345.386
216.14230346679688 0 1078.0643
216.85340881347656 0 475.41626
221.0849609375 0 2364.3843
222.08660888671875 0 1075.001
223.06405639648438 0 1288.8606
223.0816192626953 0 1106.2242
225.04339599609375 0 2573.581
226.04347229003906 0 545.4625
229.12942504882812 0 1100.5846
232.16677856445312 0 609.2487 c 1
235.14437866210938 0 629.03925
239.09536743164062 0 3529.736
240.09588623046875 0 2022.3805
241.09193420410156 0 1254.4349
245.0182342529297 0 610.90704
259.6243896484375 0 561.5982
260.1248474121094 0 1210.7234
270.1448974609375 0 772.4231
279.1333923339844 0 777.0389
282.1452331542969 0 2598.2266
287.1706848144531 0 597.84326
295.1034851074219 0 1971.95
295.1656494140625 0 1421.0953 y 10
296.10418701171875 0 2960.0825
297.1011657714844 0 1604.7072
297.1572570800781 0 540.1405
299.06219482421875 0 1477.636
299.1713562011719 0 861.9547
300.0627136230469 0 2053.4438
300.1558837890625 0 7054.0015
301.05877685546875 0 845.2299
301.1595764160156 0 647.3879
315.1669006347656 0 1479.6041
317.1823425292969 0 3697.5898
319.1981506347656 0 2917.626 c 2
320.2010498046875 0 618.813
323.1384582519531 0 499.29208
341.0183410644531 0 12302.078
341.0406494140625 0 1030.0142
350.6538391113281 0 1785.4917
355.0706787109375 0 1660.4617
356.0706787109375 0 623.15466
358.6698303222656 0 1783.7306
359.0286560058594 0 45497.184
359.6592102050781 0 1933.6842
360.16107177734375 0 938.6639
364.1974182128906 0 763.20544
369.1213684082031 0 1255.9395
369.1445007324219 0 769.2255
370.1227722167969 0 2875.8755
371.120361328125 0 3379.3623
378.1493835449219 0 815.35443
379.2090148925781 0 4764.767
380.2088623046875 0 1019.7039
382.197021484375 0 747.3555 y 9
385.1738586425781 0 697.1426
385.21942138671875 0 3826.3323
385.8817443847656 0 936.56445
386.16583251953125 0 4203.7153
386.22808837890625 0 1150.824
386.66741943359375 0 2856.0852
387.15203857421875 0 1104.9722
390.64923095703125 0 824.5287
391.55181884765625 0 2457.1526
391.8843078613281 0 2959.826
399.6551208496094 0 1247.4319
408.16668701171875 0 763.60077
412.66876220703125 0 651.9936
415.2657775878906 0 909.5159
418.6653747558594 0 691.86975
419.2173767089844 0 2604.0588
420.2232360839844 0 733.991
425.1599426269531 0 633.25
427.672119140625 0 1480.3867
428.1737060546875 0 1918.0797
429.0894470214844 0 118695.27
429.754638671875 0 1543.571
430.242431640625 0 1880.0922
430.27606201171875 0 1112.125
431.28265380859375 0 681.0621
433.6742858886719 0 2059.461
434.1730041503906 0 1413.1116
435.2362060546875 0 1459.598
436.6787109375 0 3096.3057
437.1800842285156 0 1290.849
437.21240234375 0 920.7735
442.6781921386719 0 5811.7646
442.7459716796875 0 4154.486
443.1795959472656 0 4108.37
443.24005126953125 0 1578.9961 c Water loss 7
443.6819152832031 0 1314.4785
443.7425842285156 0 1181.7843 c Ammonia loss 7
444.6914978027344 0 664.092
445.12066650390625 0 59606.535
445.682373046875 0 696.4122
446.21270751953125 0 2095.4219
450.6904602050781 0 1869.4375
450.7297058105469 0 1219.167
451.1910095214844 0 855.6551
451.6835021972656 0 18412.69
451.7506103515625 0 9724.996
452.1851501464844 0 10669.41
452.2542419433594 0 27445.125 c 7
452.68621826171875 0 3714.2039
452.752685546875 0 14051.956
453.25244140625 0 4406.8096
453.7537536621094 0 1067.4043
455.19976806640625 0 1107.9031
455.23907470703125 0 608.698
458.2713928222656 0 1418.477 c Ammonia loss 3
459.699951171875 0 47079.01
459.73822021484375 0 20530.729
460.2010498046875 0 28286.355
460.2400817871094 0 15155.414
460.69219970703125 0 21263.627
460.7408142089844 0 5465.3003
461.1913146972656 0 12700.887
461.2409362792969 0 1918.6069
461.6925964355469 0 3542.4922
462.194580078125 0 672.33
462.23931884765625 0 772.88806
465.2645568847656 0 658.7456
470.75 0 825.9577
472.2515869140625 0 1696.518
475.29949951171875 0 104288.08 c 3
476.3021240234375 0 23977.766
477.3048400878906 0 3514.0608
479.26348876953125 0 645.8147
479.75830078125 0 7903.3516
480.2214050292969 0 1514.4224 z 8
480.25921630859375 0 2872.1091
480.7587890625 0 1834.6539
481.22991943359375 0 7728.4365
482.2317199707031 0 1020.1805
484.74957275390625 0 1248.5802
485.24176025390625 0 1172.5056
485.74365234375 0 763.7824
486.7705993652344 0 1831.1289
487.2681884765625 0 2181.764
487.7662353515625 0 1498.3197
488.26385498046875 0 1066.66
490.2419128417969 0 2343.9648
493.7560119628906 0 1506.9203
494.25396728515625 0 1192.025
496.2406921386719 0 3727.532 y 8
499.76666259765625 0 10316.317
500.2659606933594 0 7675.0215 c Water loss 8
500.7640075683594 0 4879.093 c Ammonia loss 8
501.2610168457031 0 2792.118
501.7716369628906 0 842.1914
502.27423095703125 0 10397.862
503.2767028808594 0 2319.2058
506.14752197265625 0 826.5293
508.25836181640625 0 776.4669
508.7720031738281 0 22193.03
509.2740783691406 0 34137.977 c 8
509.7734680175781 0 25493.414
510.27142333984375 0 8557.662
510.7717590332031 0 4138.297
514.2152099609375 0 3161.8545
515.2193603515625 0 1627.8364
522.271728515625 0 604.3067
523.2669677734375 0 691.6104
524.1580810546875 0 1274.8138
526.2611083984375 0 632.4651
526.7547607421875 0 1107.6757
530.2323608398438 0 989.24066
530.2890625 0 1855.2202
532.7664794921875 0 3923.0308 z 3
533.2627563476562 0 1773.1716
533.7660522460938 0 795.003
534.30517578125 0 1326.1665
535.272705078125 0 1588.6418
535.3125610351562 0 677.4914
535.7691650390625 0 1282.9932
540.7750854492188 0 705.8212 y 3
542.1690063476562 0 2233.5225
544.2791748046875 0 3215.267 c Ammonia loss 9
544.3198852539062 0 715.4383
544.779052734375 0 2487.853
545.2659301757812 0 1128.8297
545.30517578125 0 3183.2285
546.3119506835938 0 2272.7024
548.250732421875 0 5181.5464
548.7496337890625 0 2387.6355
549.2484130859375 0 1100.8291
552.7920532226562 0 26571.996 c 9
553.288818359375 0 23736.781
553.7886962890625 0 12229.569
554.2870483398438 0 4878.116
554.7871704101562 0 2532.5342
555.2835693359375 0 649.2401
556.2085571289062 0 982.58594
564.3035278320312 0 3618.9497
565.3096923828125 0 2189.8403
565.8002319335938 0 2436.7014
566.2977905273438 0 1590.7854
566.7811889648438 0 653.78864
567.27880859375 0 685.40405 z Water loss 2
567.7783203125 0 1564.9417 z Ammonia loss 2
568.2783203125 0 1725.3718
573.3072509765625 0 1157.139 c Ammonia loss 4
574.3048095703125 0 1006.94867
575.2861938476562 0 835.58923 y Water loss 2
575.7819213867188 0 1449.5261 y Ammonia loss 2
576.2825927734375 0 10095.928 z 2
576.7807006835938 0 7993.6636
577.2791137695312 0 5074.9546
577.7802124023438 0 1836.9941
578.3137817382812 0 1821.0166
578.8128662109375 0 964.5966
579.31689453125 0 1100.9697
579.807373046875 0 780.48773
583.304931640625 0 1925.343
584.2918090820312 0 30501.879 y 2
584.7891235351562 0 25946.936
585.288330078125 0 11616.141
585.7879638671875 0 3547.6133
586.82373046875 0 3220.2905
587.3253173828125 0 2969.8147
587.8212280273438 0 2202.135
588.2759399414062 0 2000.2756
588.3226318359375 0 1412.7478
588.7741088867188 0 1307.0127
589.2736206054688 0 906.9882
589.3211059570312 0 2121.2393
590.3263549804688 0 183600.92 c 4
591.3290405273438 0 50173.58
591.8153686523438 0 2352.4539
592.3290405273438 0 10315.625
592.8109130859375 0 2439.6108
593.32958984375 0 1829.8527
598.2750244140625 0 668.907
600.8217163085938 0 3665.8152 c Ammonia loss 10
601.3197021484375 0 3198.381
601.8222045898438 0 2800.994
607.3250732421875 0 1230.0824
607.8257446289062 0 858.6745
608.31787109375 0 11800.321 z 7
609.3336791992188 0 73669.68 c 10
609.8326416015625 0 57454.18
610.3316040039062 0 26193.47
610.83056640625 0 8144.006
611.3311767578125 0 2217.6687
613.23193359375 0 694.2821
613.2827758789062 0 1652.4518
614.238525390625 0 2332.529
614.2888793945312 0 1231.8397
616.341796875 0 2148.6702
618.3045043945312 0 4422.113 w 1
618.80224609375 0 4008.0566
619.3045654296875 0 1084.8651
621.3242797851562 0 1216.5071
624.3356323242188 0 14215.139 y 7
625.3350219726562 0 4189.8105
626.3356323242188 0 964.8695
630.3278198242188 0 2672.0107
630.8270263671875 0 2192.6953
631.3253173828125 0 1435.6179
631.8253173828125 0 1025.8342 y Water loss 1
632.8223266601562 0 1708.3162 z 1
633.3228149414062 0 3762.7097
633.816650390625 0 1090.4128
634.3179931640625 0 1568.6984
635.3504638671875 0 635.8782
636.3475952148438 0 2471.5789
637.8311767578125 0 768.8172
638.3438720703125 0 1711.8948
638.845947265625 0 1133.2573
639.3463745117188 0 949.4478
639.8305053710938 0 929.26434
640.3553466796875 0 951.3484
640.6701049804688 0 709.47
640.8345947265625 0 2915.9426 y 1
641.3294067382812 0 3405.2075
641.826904296875 0 1634.8948
642.33251953125 0 550.6652
642.342041015625 0 558.6029
646.3217163085938 0 830.3549
647.3502807617188 0 3143.705
647.8522338867188 0 2974.8582
648.3392333984375 0 643.79065
649.3319702148438 0 4437.904
650.3333740234375 0 2727.51
653.3404541015625 0 2625.9897
653.8372192382812 0 1774.8551
654.3314819335938 0 1702.0316
655.3167724609375 0 6176.055
655.8151245117188 0 5204.3555
656.3116455078125 0 2752.3005
656.8221435546875 0 921.2422
659.3469848632812 0 7059.942
660.348388671875 0 11079.922
660.8513793945312 0 8127.7827
661.351318359375 0 15064.214 z Water loss 6
661.8491821289062 0 11838.7705
662.3450927734375 0 15590.147
662.8428344726562 0 9828.849
663.3434448242188 0 4595.6826
663.8427124023438 0 1371.7135
668.8302001953125 0 3640.0405
669.3484497070312 0 12591.614
669.8480224609375 0 13787.269
670.3475341796875 0 13844.35
670.8441772460938 0 9007.811
671.3425903320312 0 4189.0767
671.8417358398438 0 1349.8242
672.3426513671875 0 1099.1666
673.3494873046875 0 763.7397
674.34521484375 0 2843.6006
674.8448486328125 0 8307.49
675.345458984375 0 6791.187
675.8389892578125 0 7875.061
676.3368530273438 0 4875.108
676.8359985351562 0 2405.075
679.354248046875 0 36644.266 z 6
680.353271484375 0 14357.679
681.3545532226562 0 3893.8289
682.8529663085938 0 3119.9321
683.3486328125 0 48973
683.8465576171875 0 43398.016
684.34521484375 0 25331.49
684.845458984375 0 7974.4375
685.346923828125 0 2281.5427
687.3510131835938 0 1620.2837 c Ammonia loss 5
689.6597290039062 0 1898.8451
690.1611328125 0 1282.6439
691.3577270507812 0 44557.25
691.8555908203125 0 41253.008
692.3544311523438 0 24950.287
692.854736328125 0 8889.907
693.3546142578125 0 2633.4263
694.3534545898438 0 809.27924
695.3729858398438 0 8342.725 y 6
696.369384765625 0 3352.972
700.3126220703125 0 2794.8347
701.2841796875 0 3823.2974
702.291748046875 0 18091.352
703.2974243164062 0 11561.893
704.300048828125 0 2901.8674
704.369384765625 0 41476.164 c 5
705.3665771484375 0 16261.651
706.365478515625 0 3878.271
707.3671875 0 1018.75
711.3414306640625 0 917.16144
726.3272705078125 0 4855.4233
727.3287353515625 0 2481.7
731.3924560546875 0 5663.8193
732.38720703125 0 3419.957
733.388671875 0 1162.1239
737.3226928710938 0 898.1709
737.3848266601562 0 2556.9434
738.3369750976562 0 789.66455
749.3851928710938 0 21524.314 w 5
750.3876953125 0 14636.008
751.3871459960938 0 4999.0596
754.4204711914062 0 1085.0402
755.428955078125 0 976.9148
758.380615234375 0 3061.9043 c Ammonia loss 6
759.3860473632812 0 1085.2245
770.3388061523438 0 8114.415
771.3433227539062 0 7948.0425
772.3199462890625 0 19511.904
773.3241577148438 0 11860.269
774.3313598632812 0 2906.2803
775.40625 0 38123.742 c 6
776.4019165039062 0 19586.672
777.4034423828125 0 5802.189
778.4000244140625 0 796.7366
781.2960815429688 0 897.3937
787.3905029296875 0 1773.1285
788.3876953125 0 648.3286
793.3973388671875 0 93079.94 z 5
794.3943481445312 0 53475.293
795.3923950195312 0 19144.148
796.3934936523438 0 4194.3145
797.4024047851562 0 738.9317
799.3118286132812 0 2886.7463
800.311279296875 0 1182.6525
808.3736572265625 0 943.6798
809.4144897460938 0 1560.035 y 5
810.4089965820312 0 1568.73
811.4011840820312 0 863.11884
815.33740234375 0 1621.0894
816.3411254882812 0 1015.4882
817.3196411132812 0 12415.786
818.3207397460938 0 4950.3774
819.3272705078125 0 1212.5869
820.4292602539062 0 834.4739
821.4280395507812 0 1179.2775
832.4419555664062 0 907.83136
833.3408203125 0 972.24414
848.464111328125 0 2018.2487
849.4683837890625 0 1318.6185
852.3543090820312 0 1172.5798
858.3834838867188 0 1174.1229
858.49560546875 0 841.40814
859.3843994140625 0 2043.7501
859.4886474609375 0 4411.1904
860.3588256835938 0 6563.293
860.4879760742188 0 3955.5334
861.3539428710938 0 8477.493
861.4911499023438 0 1595.3466
862.35791015625 0 5739.5557
863.36181640625 0 2054.154
864.4336547851562 0 28547.795
865.4291381835938 0 19978.965
866.4296264648438 0 7401.183
867.4304809570312 0 1598.1616
874.4136352539062 0 1400.863
875.3668212890625 0 2570.298
876.3702392578125 0 2202.0645
877.3698120117188 0 1697.6698
878.3491821289062 0 3363.1956
879.355712890625 0 1142.0464
884.47998046875 0 1898.7166
885.4869384765625 0 1885.1869 c Water loss 7
886.4768676757812 0 4940.4136 c Ammonia loss 7
887.36865234375 0 1823.007
887.4677124023438 0 1021.15247
888.3770751953125 0 1119.267
888.4839477539062 0 1634.3585
891.4135131835938 0 811.9154 z Ammonia loss 4
896.4146728515625 0 1974.9071
900.4760131835938 0 4316.7085
901.4760131835938 0 2561.541
902.3753051757812 0 12641.348
902.4918212890625 0 13734.661
903.3737182617188 0 9502.037
903.4999389648438 0 53087.184 c 7
904.3543090820312 0 21116.982
904.4968872070312 0 30883.088
905.3558959960938 0 10782.785
905.4973754882812 0 12039.014
906.359130859375 0 3097.2527
906.4959716796875 0 2911.1013
908.4237060546875 0 30728.537 z 4
909.418212890625 0 24219.174
910.4166259765625 0 8410.831
911.4173583984375 0 2589.5095
918.3896484375 0 4095.8083
918.4727783203125 0 3359.4277
919.3916625976562 0 9454.847
919.4768676757812 0 29809.615
920.3818359375 0 8709.812
920.47998046875 0 22013.777
921.3779907226562 0 22090.47
921.4826049804688 0 9040.044
922.3804321289062 0 9524.904
922.4817504882812 0 2486.2993
923.38427734375 0 3401.6282
924.43798828125 0 2353.0762 y 4
925.4307250976562 0 2697.3726
926.4320678710938 0 1745.9141
969.479248046875 0 828.2336
974.5388793945312 0 966.8161
975.530029296875 0 966.5187
986.5014038085938 0 2596.4495
987.5034790039062 0 3743.058
988.5040893554688 0 3035.3096
989.5012817382812 0 867.9513
999.5338745117188 0 1999.3838 c Water loss 8
1000.524169921875 0 3364.4128 c Ammonia loss 8
1001.518310546875 0 2420.0571
1002.524169921875 0 1736.3138
1003.5283203125 0 1803.5919
1016.5377197265625 0 3647.823
1017.54345703125 0 31589.77 c 8
1018.5390625 0 21559.002
1019.5367431640625 0 9872.053
1020.5380859375 0 3345.3564
1021.5235595703125 0 1202.9851
1047.5491943359375 0 799.242
1060.56689453125 0 1269.5944
1061.5577392578125 0 727.19275
1062.567138671875 0 680.62616
1063.547607421875 0 907.06256
1064.5255126953125 0 10470.365 z 3
1065.5233154296875 0 17821.586
1066.5211181640625 0 9476.191
1067.5245361328125 0 3359.2761
1068.52978515625 0 809.6994
1080.5341796875 0 1749.1063 y 3
1088.5592041015625 0 2138.6096
1089.551025390625 0 2359.3477
1090.5479736328125 0 1421.3425
1091.5592041015625 0 873.091
1095.4964599609375 0 1301.872
1096.49560546875 0 1233.0402
1101.58447265625 0 732.88055
1104.5753173828125 0 8506.176 c 9
1105.5726318359375 0 9148.089
1106.5712890625 0 5058.5923
1107.56689453125 0 2171.4275
1108.5523681640625 0 972.6206
1151.556884765625 0 6708.706 z 2
1152.5604248046875 0 18555.715
1153.558837890625 0 11298.408
1154.5625 0 4430.29
1155.55126953125 0 924.71014
1158.62158203125 0 1459.6519
1159.63720703125 0 959.41565
1168.5794677734375 0 1342.7938
1169.56689453125 0 1091.467
1170.5675048828125 0 768.1924
1173.6488037109375 0 2546.5571
1174.6517333984375 0 4855.6323
1175.6453857421875 0 5635.9443
1176.6463623046875 0 1867.7972
1184.6177978515625 0 1146.2511
1201.6416015625 0 7381.7554
1202.6356201171875 0 4608.3193
1203.636962890625 0 3215.101
1217.6563720703125 0 6232.085 c 10
1218.6627197265625 0 13414.063
1219.6607666015625 0 10011.072
1220.660888671875 0 3335.559
1221.646240234375 0 962.8539
1265.6461181640625 0 2680.544
1266.6376953125 0 3550.2422
1267.635498046875 0 1777.7949
1268.63037109375 0 990.89276
1305.6585693359375 0 2288.7524
1306.6571044921875 0 4612.3774
1307.6495361328125 0 2118.4878
1308.6522216796875 0 2143.4653
1320.6922607421875 0 2153.499
1321.6929931640625 0 4274.609
1322.6915283203125 0 3996.2715
1323.6820068359375 0 7772.8936
1324.6773681640625 0 14551.378
1325.676513671875 0 10139.94
1326.6783447265625 0 4963.4
1327.6678466796875 0 1507.1814
1337.69482421875 0 2925.6372
1338.699951171875 0 18175.959
1339.6986083984375 0 15059.305
1340.69580078125 0 8021.113
1341.6986083984375 0 3573.435
1342.692626953125 0 1065.942
1348.6868896484375 0 3827.4282
1349.674560546875 0 8292.718
1350.6712646484375 0 5198.145
1351.667724609375 0 2929.9502
1352.66259765625 0 1210.9142
1355.7308349609375 0 1442.4395
1356.7108154296875 0 1476.8691
1365.691162109375 0 9540.089
1366.69384765625 0 57005.582
1367.691162109375 0 52968.613
1368.690673828125 0 30395.266
1369.687744140625 0 10087.133
1370.6871337890625 0 2295.6973
1379.3197021484375 0 1133.808
1381.7108154296875 0 2123.3994
1382.7086181640625 0 7104.6763
1383.7142333984375 0 13627.196
1384.71240234375 0 11710.063
1385.7103271484375 0 4689.816
1386.716552734375 0 1741.4272

Spectrum Details

|  |  |
| --- | --- |
| Matched peaks? Matched peaks The total absolute number of peaks matched. Additionally in brackets the total fraction of peaks matched and the total number of peaks is shown. | 59 (9.98% of 591) |
| FDR? FDR The false discovery rate estimated for this peptide. It is calculated by matching all theoretical fragments with a non-integer shift with the raw peaks for this spectrum. This is done with 40 different shifts. The resulting percentage is the average number of annotated peaks over the number of annotated peaks with the correct spectrum. | 2.46% |
| Satellite FDR? Satellite FDR See the FDR for details on its calculation. This satellite ion specific FDR only contains the satellite ions (d/w) for I/L/J positions. | 0.00% |
| PSM Score? PSM Score The PSM Score as given by Hecklib to this annotated spectrum. It is shown with three significant figures. | 483 |

#### Spectrum 5262? Spectrum 5262 The raw spectrum of this peptide as annotated by Hecklib. The fragments are coloured according to ion type (see legend). Any peaks with a star '\*' as text can be hovered over to see the full details, first the ion type second the mass shift type. By hovering over the amino acids in the peptide or ions in the legend the corresponding peaks are highlighted. By toggling the 'Unassigned' label you can turn the background (unassigned) peaks on or off in the plot. By updating the slider in the Ion legend you can update the spectrum to only show the top X% of the peaks with labels. The top X% means any peak that is within X% of the highest intensity. By dragging in the spectrum you can zoom in to a specific part of the spectrum and use 'Zoom Out' to get back to the original zoom level. The annotation of the spectrum is based on the given sequence in the peptides file and is done with different software so inconsistencies are likely. The peaks are annotated based on the given sequence, with 20 ppm tolerance.

Copy Data

##### Spectrum 5262 (TSV)

###### Preview

```
Loading example...
```

*Click on the button to copy the data to your clipboard.*

Mz MinMz MaxIntensity Max

WidthHeightPeptide font sizePeptide stroke widthSpectrum font sizeSpectrum stroke widthCompact peptide

Ion legend

wxyz

abcd

OtherUnassignedIon^Charge^_Position_Show for top:%

TISRDNAKNSJY

04.59e+49.18e+41.38e+51.84e+5

Zoom Out

y^+1^_1_c^+1^_2_y^+1^_2_c^+1^_3_c^+2^_7_c^+2^_8_c^+2^_8_c^+1^_4_y^+1^_4_c^+2^_9_c^+2^_9_c^+2^_9_c^+2^_10_c^+2^_10_z^+2^_10_y^+2^_10_c^+1^_5_c^+2^_11_z^+1^_5_c^+2^_11_w^+2^_11_y^+1^_5_y^+2^_11_z^+1^_6_y^+1^_6_z^+1^_6_y^+1^_6_c^+1^_6_w^+1^_7_c^+1^_7_c^+1^_7_z^+1^_7_c^+1^_8_c^+1^_8_z^+1^_8_y^+1^_8_c^+1^_9_c^+1^_9_c^+1^_9_z^+1^_9_c^+1^_10_z^+1^_10_c^+1^_11_

0679135820382717

Fragment Matches Table

Show background peaks

| Position | Ion type | Intensity | mz Theoretical | mz Error (Th) | mz Error (ppm) | Charge | Series Number |
| --- | --- | --- | --- | --- | --- | --- | --- |
| - | - | 373.2 | 120.1 | - | - | 0 | - |
| - | - | 405.6 | 124.1 | - | - | 0 | - |
| - | - | 337.7 | 125 | - | - | 0 | - |
| - | - | 378 | 127.1 | - | - | 0 | - |
| - | - | 329.2 | 128.8 | - | - | 0 | - |
| - | - | 1065 | 129.1 | - | - | 0 | - |
| - | - | 455 | 129.7 | - | - | 0 | - |
| - | - | 387.6 | 130.8 | - | - | 0 | - |
| - | - | 524.1 | 131.1 | - | - | 0 | - |
| - | - | 6903 | 133.1 | - | - | 0 | - |
| - | - | 2060 | 134.1 | - | - | 0 | - |
| - | - | 1745 | 136.1 | - | - | 0 | - |
| - | - | 475.8 | 143.2 | - | - | 0 | - |
| - | - | 391.7 | 146.1 | - | - | 0 | - |
| - | - | 498.7 | 148.6 | - | - | 0 | - |
| - | - | 1007 | 149 | - | - | 0 | - |
| - | - | 1.047E+04 | 149 | - | - | 0 | - |
| - | - | 843.8 | 150 | - | - | 0 | - |
| - | - | 1037 | 151 | - | - | 0 | - |
| - | - | 564.6 | 157.1 | - | - | 0 | - |
| - | - | 423.8 | 158.7 | - | - | 0 | - |
| - | - | 418.1 | 163.6 | - | - | 0 | - |
| - | - | 1044 | 165.1 | - | - | 0 | - |
| - | - | 395.9 | 165.8 | - | - | 0 | - |
| - | - | 2878 | 167.1 | - | - | 0 | - |
| - | - | 495.7 | 168.1 | - | - | 0 | - |
| - | - | 629.1 | 169.1 | - | - | 0 | - |
| - | - | 4634 | 173.1 | - | - | 0 | - |
| - | - | 671.4 | 173.1 | - | - | 0 | - |
| - | - | 2397 | 173.5 | - | - | 0 | - |
| - | - | 3106 | 177.1 | - | - | 0 | - |
| - | - | 1934 | 178.1 | - | - | 0 | - |
| 12 | y | 4227 | 182.1 | 0.0001597 | 0.8769 | +1 | 1 |
| - | - | 838.9 | 183.1 | - | - | 0 | - |
| - | - | 678.7 | 187.1 | - | - | 0 | - |
| - | - | 1.022E+04 | 187.1 | - | - | 0 | - |
| - | - | 812.4 | 188.1 | - | - | 0 | - |
| - | - | 727.4 | 201.1 | - | - | 0 | - |
| - | - | 1020 | 205.1 | - | - | 0 | - |
| - | - | 467.3 | 213.2 | - | - | 0 | - |
| - | - | 495.4 | 214.1 | - | - | 0 | - |
| - | - | 2753 | 215.1 | - | - | 0 | - |
| - | - | 6227 | 221.1 | - | - | 0 | - |
| - | - | 1216 | 221.1 | - | - | 0 | - |
| - | - | 2812 | 222.1 | - | - | 0 | - |
| - | - | 2195 | 223.1 | - | - | 0 | - |
| - | - | 1832 | 223.1 | - | - | 0 | - |
| - | - | 5757 | 225 | - | - | 0 | - |
| - | - | 1234 | 226 | - | - | 0 | - |
| - | - | 476.8 | 227 | - | - | 0 | - |
| - | - | 1023 | 227 | - | - | 0 | - |
| - | - | 529.7 | 229.1 | - | - | 0 | - |
| 2 | c | 480.6 | 232.2 | 0.0001272 | 0.5478 | +1 | 2 |
| - | - | 8441 | 239.1 | - | - | 0 | - |
| - | - | 4296 | 240.1 | - | - | 0 | - |
| - | - | 3070 | 241.1 | - | - | 0 | - |
| - | - | 614.7 | 242.1 | - | - | 0 | - |
| - | - | 2647 | 260.1 | - | - | 0 | - |
| - | - | 568.9 | 261.1 | - | - | 0 | - |
| - | - | 1214 | 281.1 | - | - | 0 | - |
| - | - | 753.7 | 282.1 | - | - | 0 | - |
| - | - | 567.5 | 282.1 | - | - | 0 | - |
| - | - | 610.8 | 283 | - | - | 0 | - |
| - | - | 524.6 | 292.6 | - | - | 0 | - |
| - | - | 5266 | 295.1 | - | - | 0 | - |
| 11 | y | 932.7 | 295.2 | 0.001057 | 3.58 | +1 | 2 |
| - | - | 7148 | 296.1 | - | - | 0 | - |
| - | - | 4057 | 297.1 | - | - | 0 | - |
| - | - | 716.4 | 298.1 | - | - | 0 | - |
| - | - | 5897 | 299.1 | - | - | 0 | - |
| - | - | 5451 | 300.1 | - | - | 0 | - |
| - | - | 2170 | 300.2 | - | - | 0 | - |
| - | - | 3593 | 301.1 | - | - | 0 | - |
| - | - | 593.1 | 301.2 | - | - | 0 | - |
| - | - | 830.7 | 306.2 | - | - | 0 | - |
| - | - | 1103 | 313.1 | - | - | 0 | - |
| - | - | 1371 | 314.1 | - | - | 0 | - |
| - | - | 490 | 314.9 | - | - | 0 | - |
| - | - | 1384 | 315.1 | - | - | 0 | - |
| - | - | 645.5 | 315.2 | - | - | 0 | - |
| - | - | 1149 | 317.2 | - | - | 0 | - |
| 3 | c | 713.9 | 319.2 | 0.001165 | 3.648 | +1 | 3 |
| - | - | 1147 | 341 | - | - | 0 | - |
| - | - | 2.1E+04 | 341 | - | - | 0 | - |
| - | - | 1315 | 343 | - | - | 0 | - |
| - | - | 2539 | 355.1 | - | - | 0 | - |
| - | - | 1247 | 356.1 | - | - | 0 | - |
| - | - | 1122 | 357.1 | - | - | 0 | - |
| - | - | 7.177E+04 | 359 | - | - | 0 | - |
| - | - | 564.4 | 365.9 | - | - | 0 | - |
| - | - | 3521 | 369.1 | - | - | 0 | - |
| - | - | 507.3 | 370 | - | - | 0 | - |
| - | - | 6480 | 370.1 | - | - | 0 | - |
| - | - | 5336 | 371.1 | - | - | 0 | - |
| - | - | 639.9 | 371.1 | - | - | 0 | - |
| - | - | 1721 | 372.1 | - | - | 0 | - |
| - | - | 638 | 375.9 | - | - | 0 | - |
| - | - | 2698 | 376.9 | - | - | 0 | - |
| 7 | c | 967.8 | 379.2 | 0.00644 | 16.98 | +2 | 7 |
| - | - | 1690 | 385.2 | - | - | 0 | - |
| - | - | 526 | 389.1 | - | - | 0 | - |
| - | - | 1485 | 391.6 | - | - | 0 | - |
| - | - | 1007 | 391.9 | - | - | 0 | - |
| - | - | 1.817E+05 | 429.1 | - | - | 0 | - |
| - | - | 655.7 | 429.8 | - | - | 0 | - |
| - | - | 962.2 | 433.2 | - | - | 0 | - |
| - | - | 617.9 | 442.7 | - | - | 0 | - |
| 8 | c | 2797 | 443.2 | 2.602E-05 | 0.05871 | +2 | 8 |
| - | - | 9.63E+04 | 445.1 | - | - | 0 | - |
| - | - | 902.2 | 446.2 | - | - | 0 | - |
| - | - | 1740 | 447.1 | - | - | 0 | - |
| - | - | 2808 | 451.8 | - | - | 0 | - |
| 8 | c | 9018 | 452.3 | 0.0004561 | 1.008 | +2 | 8 |
| - | - | 3745 | 452.8 | - | - | 0 | - |
| - | - | 1491 | 453.3 | - | - | 0 | - |
| - | - | 884.2 | 460.2 | - | - | 0 | - |
| - | - | 6621 | 460.3 | - | - | 0 | - |
| - | - | 3980 | 460.3 | - | - | 0 | - |
| - | - | 922.1 | 461.2 | - | - | 0 | - |
| - | - | 2242 | 461.3 | - | - | 0 | - |
| - | - | 1197 | 461.3 | - | - | 0 | - |
| - | - | 586.9 | 461.7 | - | - | 0 | - |
| 4 | c | 2.923E+04 | 475.3 | 0.0003648 | 0.7676 | +1 | 4 |
| - | - | 7032 | 476.3 | - | - | 0 | - |
| - | - | 1198 | 477.3 | - | - | 0 | - |
| - | - | 2186 | 479.8 | - | - | 0 | - |
| - | - | 1487 | 480.3 | - | - | 0 | - |
| - | - | 1194 | 481.2 | - | - | 0 | - |
| - | - | 640.8 | 482.2 | - | - | 0 | - |
| - | - | 704.3 | 487.8 | - | - | 0 | - |
| - | - | 641.3 | 493.2 | - | - | 0 | - |
| - | - | 766.9 | 494.2 | - | - | 0 | - |
| 9 | y | 810.2 | 496.2 | 0.0001974 | 0.3979 | +1 | 4 |
| - | - | 3121 | 499.8 | - | - | 0 | - |
| 9 | c | 2419 | 500.3 | 0.004247 | 8.49 | +2 | 9 |
| 9 | c | 1133 | 500.8 | 0.003196 | 6.381 | +2 | 9 |
| - | - | 859.9 | 501.8 | - | - | 0 | - |
| - | - | 3817 | 502.3 | - | - | 0 | - |
| - | - | 1142 | 503.3 | - | - | 0 | - |
| - | - | 2044 | 508.3 | - | - | 0 | - |
| - | - | 6801 | 508.8 | - | - | 0 | - |
| 9 | c | 1.345E+04 | 509.3 | 0.001443 | 2.832 | +2 | 9 |
| - | - | 6633 | 509.8 | - | - | 0 | - |
| - | - | 2711 | 510.3 | - | - | 0 | - |
| - | - | 1552 | 510.8 | - | - | 0 | - |
| - | - | 630.5 | 512.8 | - | - | 0 | - |
| 10 | c | 1178 | 544.3 | 0.002776 | 5.1 | +2 | 10 |
| - | - | 875 | 544.8 | - | - | 0 | - |
| - | - | 1210 | 545.3 | - | - | 0 | - |
| - | - | 645 | 546.3 | - | - | 0 | - |
| - | - | 1564 | 548.3 | - | - | 0 | - |
| 10 | c | 8266 | 552.8 | 6.169E-05 | 0.1116 | +2 | 10 |
| - | - | 7148 | 553.3 | - | - | 0 | - |
| - | - | 1937 | 553.8 | - | - | 0 | - |
| - | - | 540.4 | 558.3 | - | - | 0 | - |
| - | - | 1137 | 564.3 | - | - | 0 | - |
| - | - | 954.8 | 565.8 | - | - | 0 | - |
| - | - | 698.8 | 568.3 | - | - | 0 | - |
| - | - | 565.9 | 573.9 | - | - | 0 | - |
| 3 | z | 3067 | 576.3 | 0.0002981 | 0.5173 | +2 | 10 |
| - | - | 2374 | 576.8 | - | - | 0 | - |
| - | - | 1454 | 577.3 | - | - | 0 | - |
| 3 | y | 7264 | 584.3 | 0.0005327 | 0.9118 | +2 | 10 |
| - | - | 5664 | 584.8 | - | - | 0 | - |
| - | - | 2138 | 585.3 | - | - | 0 | - |
| - | - | 868.5 | 585.8 | - | - | 0 | - |
| - | - | 758.8 | 586.8 | - | - | 0 | - |
| - | - | 1502 | 587.3 | - | - | 0 | - |
| - | - | 668.1 | 587.8 | - | - | 0 | - |
| 5 | c | 5.524E+04 | 590.3 | 0.0002773 | 0.4697 | +1 | 5 |
| - | - | 1.596E+04 | 591.3 | - | - | 0 | - |
| - | - | 3157 | 592.3 | - | - | 0 | - |
| 11 | c | 1222 | 600.8 | 0.000804 | 1.338 | +2 | 11 |
| - | - | 769.2 | 601.3 | - | - | 0 | - |
| - | - | 708.5 | 601.8 | - | - | 0 | - |
| - | - | 801.3 | 607.3 | - | - | 0 | - |
| 8 | z | 2748 | 608.3 | 0.0006492 | 1.067 | +1 | 5 |
| 11 | c | 2.194E+04 | 609.3 | 0.0002236 | 0.3669 | +2 | 11 |
| - | - | 1.396E+04 | 609.8 | - | - | 0 | - |
| - | - | 7399 | 610.3 | - | - | 0 | - |
| - | - | 3082 | 610.8 | - | - | 0 | - |
| - | - | 722.5 | 613.3 | - | - | 0 | - |
| 2 | w | 913.8 | 618.3 | 0.0009169 | 1.483 | +2 | 11 |
| - | - | 932.2 | 618.8 | - | - | 0 | - |
| 8 | y | 3942 | 624.3 | 0.0002967 | 0.4752 | +1 | 5 |
| - | - | 950 | 625.3 | - | - | 0 | - |
| - | - | 674.3 | 625.8 | - | - | 0 | - |
| - | - | 823.7 | 630.3 | - | - | 0 | - |
| - | - | 980.4 | 633.3 | - | - | 0 | - |
| - | - | 1087 | 636.3 | - | - | 0 | - |
| 2 | y | 1505 | 640.8 | 0.001103 | 1.722 | +2 | 11 |
| - | - | 1530 | 641.3 | - | - | 0 | - |
| - | - | 913.5 | 647.3 | - | - | 0 | - |
| - | - | 638 | 653.4 | - | - | 0 | - |
| - | - | 1723 | 655.3 | - | - | 0 | - |
| - | - | 1447 | 655.8 | - | - | 0 | - |
| - | - | 916.4 | 656.3 | - | - | 0 | - |
| - | - | 2646 | 659.3 | - | - | 0 | - |
| - | - | 3761 | 660.3 | - | - | 0 | - |
| - | - | 1217 | 660.8 | - | - | 0 | - |
| 7 | z | 2232 | 661.3 | 0.009134 | 13.81 | +1 | 6 |
| - | - | 2787 | 661.9 | - | - | 0 | - |
| - | - | 4781 | 662.3 | - | - | 0 | - |
| - | - | 3385 | 662.8 | - | - | 0 | - |
| - | - | 917.3 | 663.3 | - | - | 0 | - |
| - | - | 3369 | 669.3 | - | - | 0 | - |
| - | - | 3279 | 669.8 | - | - | 0 | - |
| - | - | 3141 | 670.3 | - | - | 0 | - |
| - | - | 3232 | 670.8 | - | - | 0 | - |
| - | - | 770.9 | 671.3 | - | - | 0 | - |
| - | - | 928.4 | 672.4 | - | - | 0 | - |
| - | - | 1106 | 674.3 | - | - | 0 | - |
| - | - | 2917 | 674.8 | - | - | 0 | - |
| - | - | 2177 | 675.3 | - | - | 0 | - |
| - | - | 1757 | 675.8 | - | - | 0 | - |
| - | - | 1484 | 676.3 | - | - | 0 | - |
| - | - | 824 | 676.8 | - | - | 0 | - |
| 7 | y | 1001 | 678.3 | 0.01112 | 16.4 | +1 | 6 |
| 7 | z | 1.106E+04 | 679.4 | 0.0002785 | 0.41 | +1 | 6 |
| - | - | 5242 | 680.4 | - | - | 0 | - |
| - | - | 774.3 | 681.4 | - | - | 0 | - |
| - | - | 1197 | 682.9 | - | - | 0 | - |
| - | - | 1.575E+04 | 683.3 | - | - | 0 | - |
| - | - | 1.298E+04 | 683.8 | - | - | 0 | - |
| - | - | 5118 | 684.3 | - | - | 0 | - |
| - | - | 2767 | 684.8 | - | - | 0 | - |
| - | - | 2473 | 690.9 | - | - | 0 | - |
| - | - | 1.244E+04 | 691.4 | - | - | 0 | - |
| - | - | 1.131E+04 | 691.9 | - | - | 0 | - |
| - | - | 5846 | 692.4 | - | - | 0 | - |
| - | - | 2352 | 692.9 | - | - | 0 | - |
| - | - | 818.4 | 693.4 | - | - | 0 | - |
| - | - | 674.9 | 693.9 | - | - | 0 | - |
| - | - | 1764 | 694.4 | - | - | 0 | - |
| 7 | y | 2707 | 695.4 | 0.001417 | 2.037 | +1 | 6 |
| - | - | 714.9 | 696.4 | - | - | 0 | - |
| 6 | c | 1.354E+04 | 704.4 | 0.0005627 | 0.7989 | +1 | 6 |
| - | - | 5425 | 705.4 | - | - | 0 | - |
| - | - | 1655 | 709.4 | - | - | 0 | - |
| - | - | 1998 | 731.4 | - | - | 0 | - |
| - | - | 1208 | 732.4 | - | - | 0 | - |
| - | - | 915.2 | 733.4 | - | - | 0 | - |
| 6 | w | 7471 | 749.4 | 0.0007139 | 0.9527 | +1 | 7 |
| - | - | 2908 | 750.4 | - | - | 0 | - |
| - | - | 800.2 | 754.4 | - | - | 0 | - |
| 7 | c | 749 | 758.4 | 0.00428 | 5.644 | +1 | 7 |
| - | - | 1034 | 762.8 | - | - | 0 | - |
| - | - | 1140 | 774.4 | - | - | 0 | - |
| 7 | c | 1.33E+04 | 775.4 | 0.0005403 | 0.6969 | +1 | 7 |
| - | - | 4622 | 776.4 | - | - | 0 | - |
| - | - | 1763 | 777.4 | - | - | 0 | - |
| - | - | 732.1 | 787.4 | - | - | 0 | - |
| - | - | 897.3 | 788.4 | - | - | 0 | - |
| - | - | 893.2 | 790.1 | - | - | 0 | - |
| 6 | z | 2.841E+04 | 793.4 | 0.0001684 | 0.2123 | +1 | 7 |
| - | - | 1.627E+04 | 794.4 | - | - | 0 | - |
| - | - | 5810 | 795.4 | - | - | 0 | - |
| - | - | 1148 | 796.4 | - | - | 0 | - |
| - | - | 774.3 | 822.5 | - | - | 0 | - |
| - | - | 1287 | 837.5 | - | - | 0 | - |
| - | - | 1578 | 859.5 | - | - | 0 | - |
| - | - | 1713 | 860.5 | - | - | 0 | - |
| - | - | 9637 | 864.4 | - | - | 0 | - |
| - | - | 5615 | 865.4 | - | - | 0 | - |
| - | - | 1985 | 866.4 | - | - | 0 | - |
| - | - | 828.4 | 867.4 | - | - | 0 | - |
| - | - | 2842 | 880.4 | - | - | 0 | - |
| - | - | 695.6 | 881.4 | - | - | 0 | - |
| 8 | c | 1595 | 886.5 | 0.001663 | 1.876 | +1 | 8 |
| - | - | 1185 | 900.5 | - | - | 0 | - |
| - | - | 978.3 | 901.5 | - | - | 0 | - |
| - | - | 5347 | 902.5 | - | - | 0 | - |
| 8 | c | 1.785E+04 | 903.5 | 0.001143 | 1.265 | +1 | 8 |
| - | - | 9124 | 904.5 | - | - | 0 | - |
| - | - | 2081 | 905.5 | - | - | 0 | - |
| - | - | 779.1 | 906.4 | - | - | 0 | - |
| 5 | z | 9130 | 908.4 | 0.0006832 | 0.7521 | +1 | 8 |
| - | - | 5281 | 909.4 | - | - | 0 | - |
| - | - | 2008 | 910.4 | - | - | 0 | - |
| - | - | 775.4 | 914 | - | - | 0 | - |
| - | - | 1066 | 922 | - | - | 0 | - |
| - | - | 911.4 | 922.4 | - | - | 0 | - |
| - | - | 977.1 | 922.5 | - | - | 0 | - |
| - | - | 831.5 | 923.4 | - | - | 0 | - |
| 5 | y | 759.6 | 924.4 | 0.005003 | 5.412 | +1 | 8 |
| - | - | 814.1 | 925.4 | - | - | 0 | - |
| - | - | 825.2 | 938.6 | - | - | 0 | - |
| - | - | 816.6 | 986.5 | - | - | 0 | - |
| 9 | c | 983.8 | 999.5 | 0.006716 | 6.72 | +1 | 9 |
| 9 | c | 1075 | 1001 | 0.003056 | 3.054 | +1 | 9 |
| - | - | 960.1 | 1002 | - | - | 0 | - |
| - | - | 816.5 | 1003 | - | - | 0 | - |
| - | - | 1424 | 1017 | - | - | 0 | - |
| 9 | c | 1.075E+04 | 1018 | 0.001163 | 1.143 | +1 | 9 |
| - | - | 7805 | 1019 | - | - | 0 | - |
| - | - | 2605 | 1020 | - | - | 0 | - |
| 4 | z | 4799 | 1065 | 0.0002565 | 0.241 | +1 | 9 |
| - | - | 5230 | 1066 | - | - | 0 | - |
| - | - | 3435 | 1067 | - | - | 0 | - |
| - | - | 1026 | 1068 | - | - | 0 | - |
| - | - | 934.2 | 1089 | - | - | 0 | - |
| 10 | c | 3794 | 1105 | 0.001392 | 1.26 | +1 | 10 |
| - | - | 2801 | 1106 | - | - | 0 | - |
| - | - | 1362 | 1107 | - | - | 0 | - |
| - | - | 990.4 | 1114 | - | - | 0 | - |
| - | - | 876 | 1132 | - | - | 0 | - |
| - | - | 847.7 | 1144 | - | - | 0 | - |
| 3 | z | 1303 | 1152 | 0.0002778 | 0.2412 | +1 | 10 |
| - | - | 5235 | 1153 | - | - | 0 | - |
| - | - | 3446 | 1154 | - | - | 0 | - |
| - | - | 1071 | 1155 | - | - | 0 | - |
| - | - | 1566 | 1175 | - | - | 0 | - |
| - | - | 1325 | 1176 | - | - | 0 | - |
| - | - | 2398 | 1202 | - | - | 0 | - |
| - | - | 842 | 1203 | - | - | 0 | - |
| 11 | c | 1818 | 1218 | 0.003425 | 2.812 | +1 | 11 |
| - | - | 4327 | 1219 | - | - | 0 | - |
| - | - | 2709 | 1220 | - | - | 0 | - |
| - | - | 1217 | 1221 | - | - | 0 | - |
| - | - | 1017 | 1266 | - | - | 0 | - |
| - | - | 1200 | 1267 | - | - | 0 | - |
| - | - | 704.6 | 1305 | - | - | 0 | - |
| - | - | 1694 | 1306 | - | - | 0 | - |
| - | - | 1186 | 1322 | - | - | 0 | - |
| - | - | 952.4 | 1323 | - | - | 0 | - |
| - | - | 1846 | 1324 | - | - | 0 | - |
| - | - | 4089 | 1325 | - | - | 0 | - |
| - | - | 2873 | 1326 | - | - | 0 | - |
| - | - | 1452 | 1327 | - | - | 0 | - |
| - | - | 1614 | 1338 | - | - | 0 | - |
| - | - | 5916 | 1339 | - | - | 0 | - |
| - | - | 4887 | 1340 | - | - | 0 | - |
| - | - | 2741 | 1341 | - | - | 0 | - |
| - | - | 1200 | 1342 | - | - | 0 | - |
| - | - | 726.1 | 1343 | - | - | 0 | - |
| - | - | 1789 | 1349 | - | - | 0 | - |
| - | - | 2035 | 1350 | - | - | 0 | - |
| - | - | 1600 | 1351 | - | - | 0 | - |
| - | - | 1082 | 1352 | - | - | 0 | - |
| - | - | 785.7 | 1365 | - | - | 0 | - |
| - | - | 3795 | 1366 | - | - | 0 | - |
| - | - | 1.996E+04 | 1367 | - | - | 0 | - |
| - | - | 1.724E+04 | 1368 | - | - | 0 | - |
| - | - | 8283 | 1369 | - | - | 0 | - |
| - | - | 3301 | 1370 | - | - | 0 | - |
| - | - | 1368 | 1371 | - | - | 0 | - |
| - | - | 2473 | 1382 | - | - | 0 | - |
| - | - | 3334 | 1383 | - | - | 0 | - |
| - | - | 4592 | 1384 | - | - | 0 | - |
| - | - | 4005 | 1385 | - | - | 0 | - |
| - | - | 2995 | 1386 | - | - | 0 | - |
| - | - | 949.6 | 1387 | - | - | 0 | - |
| - | - | 984.7 | 1388 | - | - | 0 | - |
| - | - | 664.2 | 2690 | - | - | 0 | - |

m/z Charge Intensity FragmentType MassShift Position
120.08126831054688 0 373.22238
124.12974548339844 0 405.63184
124.95928955078125 0 337.74097
127.14202880859375 0 377.98944
128.75955200195312 0 329.20523
129.1022186279297 0 1065.3583
129.70144653320312 0 455.00815
130.7527618408203 0 387.56955
131.1177520751953 0 524.0748
133.0860137939453 0 6902.7217
134.08944702148438 0 2059.9849
136.0757293701172 0 1745.3374
143.23934936523438 0 475.8397
146.05857849121094 0 391.7142
148.5859375 0 498.7022
148.95448303222656 0 1006.74884
149.04495239257812 0 10466.904
150.0443115234375 0 843.7964
151.0417938232422 0 1036.8163
157.13372802734375 0 564.6081
158.65806579589844 0 423.75635
163.64111328125 0 418.063
165.05470275878906 0 1043.6156
165.82040405273438 0 395.93265
167.0554962158203 0 2877.8755
168.0553741455078 0 495.68964
169.05224609375 0 629.0555
173.0921630859375 0 4633.6455
173.1283416748047 0 671.3971
173.4509735107422 0 2396.5164
177.11228942871094 0 3105.753
178.11572265625 0 1933.6731
182.08132934570312 0 4226.7295 y 11
183.0843505859375 0 838.88135
187.1355438232422 0 678.6827
187.14419555664062 0 10221.335
188.14785766601562 0 812.3652
201.1230926513672 0 727.3528
205.10696411132812 0 1019.79297
213.23423767089844 0 467.3417
214.07545471191406 0 495.4108
215.13880920410156 0 2752.681
221.08453369140625 0 6226.689
221.13885498046875 0 1216.1133
222.08494567871094 0 2811.803
223.0638427734375 0 2195.2368
223.0811309814453 0 1832.185
225.0429229736328 0 5757.3496
226.04298400878906 0 1234.0264
227.0225067138672 0 476.80457
227.03976440429688 0 1023.2676
229.08273315429688 0 529.71173
232.1656951904297 0 480.63083 c 1
239.0950469970703 0 8440.794
240.095703125 0 4296.1367
241.09219360351562 0 3069.5808
242.09158325195312 0 614.73895
260.1237487792969 0 2647.1594
261.1280517578125 0 568.88525
281.05126953125 0 1213.6984
282.051513671875 0 753.67334
282.1453857421875 0 567.5416
283.04742431640625 0 610.82117
292.63958740234375 0 524.6111
295.103271484375 0 5265.511
295.1662902832031 0 932.711 y 10
296.1041564941406 0 7147.91
297.10113525390625 0 4057.0027
298.1019287109375 0 716.3714
299.0616760253906 0 5896.6816
300.06231689453125 0 5450.6675
300.15606689453125 0 2169.961
301.059326171875 0 3592.9175
301.1590270996094 0 593.0776
306.16839599609375 0 830.69977
313.1148376464844 0 1102.6357
314.1141357421875 0 1371.2495
314.8796081542969 0 489.9933
315.1105651855469 0 1383.7793
315.1665344238281 0 645.52515
317.1822204589844 0 1148.5475
319.1987609863281 0 713.8773 c 2
340.9943542480469 0 1146.6508
341.0180969238281 0 21004.225
342.9968566894531 0 1315.2484
355.0701904296875 0 2539.1567
356.07037353515625 0 1247.2081
357.0678405761719 0 1122.1012
359.0283203125 0 71772.664
365.9207458496094 0 564.407
369.1215515136719 0 3520.8267
370.0257263183594 0 507.25134
370.1225891113281 0 6480.002
371.1198425292969 0 5336.3237
371.14556884765625 0 639.903
372.1209411621094 0 1720.6107
375.86102294921875 0 637.9982
376.8988037109375 0 2698.115
379.2076416015625 0 967.7833 c Water loss 6
385.21954345703125 0 1690.1781
389.12969970703125 0 526.0314
391.55120849609375 0 1485.065
391.88397216796875 0 1007.0162
429.08905029296875 0 181736.97
429.7540588378906 0 655.6927
433.2215270996094 0 962.2128
442.7453308105469 0 617.917
443.2486572265625 0 2797.3604 c Water loss 7
445.12042236328125 0 96303.27
446.2123718261719 0 902.22266
447.1001281738281 0 1740.1859
451.7509765625 0 2808.0266
452.2535095214844 0 9017.5625 c 7
452.7543029785156 0 3744.7693
453.2557373046875 0 1491.2462
460.2458190917969 0 884.24457
460.28369140625 0 6620.694
460.3141784667969 0 3979.5957
461.2474670410156 0 922.0796
461.2864685058594 0 2241.8762
461.31842041015625 0 1197.3545
461.6830139160156 0 586.9499
475.299072265625 0 29225.861 c 3
476.3016662597656 0 7031.796
477.3028259277344 0 1197.8207
479.7569580078125 0 2186.492
480.2553405761719 0 1487.21
481.2276611328125 0 1193.584
482.2333679199219 0 640.79767
487.77215576171875 0 704.31616
493.2455749511719 0 641.32166
494.2491455078125 0 766.923
496.2403869628906 0 810.18665 y 8
499.76666259765625 0 3121.245
500.2658996582031 0 2419.4138 c Water loss 8
500.7653503417969 0 1132.9253 c Ammonia loss 8
501.7716369628906 0 859.91254
502.274169921875 0 3816.7886
503.2787780761719 0 1142.4955
508.2527160644531 0 2044.1724
508.77191162109375 0 6801.1094
509.27398681640625 0 13446.934 c 8
509.773681640625 0 6633.24
510.27447509765625 0 2711.0232
510.7732849121094 0 1551.678
512.772216796875 0 630.48706
544.2809448242188 0 1178.2108 c Ammonia loss 9
544.7822265625 0 875.04456
545.3037109375 0 1210.2085
546.3034057617188 0 645.039
548.2500610351562 0 1563.7433
552.7913818359375 0 8266.191 c 9
553.2897338867188 0 7148.3647
553.7869873046875 0 1936.9899
558.2882690429688 0 540.3636
564.30322265625 0 1137.4515
565.7998657226562 0 954.84955
568.2730102539062 0 698.799
573.876220703125 0 565.8829
576.2816162109375 0 3067.371 z 2
576.7799072265625 0 2373.9888
577.2817993164062 0 1453.8551
584.2918090820312 0 7263.75 y 2
584.79052734375 0 5663.724
585.2891845703125 0 2138.093
585.7894897460938 0 868.5151
586.8247680664062 0 758.78406
587.3237915039062 0 1501.7119
587.8250122070312 0 668.13367
590.325927734375 0 55244.83 c 4
591.3285522460938 0 15963.853
592.3323364257812 0 3156.9912
600.8193969726562 0 1221.7124 c Ammonia loss 10
601.314453125 0 769.1863
601.826416015625 0 708.5094
607.3248291015625 0 801.27136
608.3170776367188 0 2747.9438 z 7
609.333251953125 0 21940.799 c 10
609.8331909179688 0 13955.352
610.3324584960938 0 7398.531
610.8323364257812 0 3081.7534
613.3308715820312 0 722.4518
618.303466796875 0 913.8448 w 1
618.7990112304688 0 932.2219
624.33544921875 0 3942.067 y 7
625.3355712890625 0 950.0167
625.8220825195312 0 674.25665
630.3265991210938 0 823.74677
633.3187866210938 0 980.3851
636.3462524414062 0 1087.4156
640.8344116210938 0 1504.5315 y 1
641.3323364257812 0 1530.1144
647.3497314453125 0 913.4663
653.3504638671875 0 637.9795
655.3189697265625 0 1722.9148
655.8164672851562 0 1446.5717
656.3160400390625 0 916.40173
659.3467407226562 0 2646.3313
660.3466186523438 0 3760.7825
660.849609375 0 1217.3068
661.3521118164062 0 2231.6294 z Water loss 6
661.8516845703125 0 2786.6868
662.3448486328125 0 4780.5684
662.844970703125 0 3384.7893
663.3499145507812 0 917.299
669.3482666015625 0 3368.9019
669.8477172851562 0 3278.6929
670.3477783203125 0 3140.528
670.846435546875 0 3232.3616
671.3489990234375 0 770.8641
672.4388427734375 0 928.42737
674.3453369140625 0 1106.0751
674.8433837890625 0 2916.9397
675.343505859375 0 2177.0027
675.840087890625 0 1757.0616
676.3381958007812 0 1483.7378
676.8385620117188 0 823.95984
678.3345947265625 0 1001.2001 y Ammonia loss 6
679.3538208007812 0 11057.438 z 6
680.353759765625 0 5241.8325
681.3538208007812 0 774.2618
682.8590087890625 0 1196.8666
683.3480834960938 0 15752.11
683.8477783203125 0 12984.63
684.3482666015625 0 5117.5854
684.8494873046875 0 2767.3955
690.8753051757812 0 2472.9578
691.3584594726562 0 12440.547
691.8573608398438 0 11310.622
692.3568725585938 0 5846.2935
692.8591918945312 0 2351.8157
693.3583984375 0 818.4198
693.8812255859375 0 674.89685
694.3510131835938 0 1763.7712
695.370849609375 0 2707.079 y 6
696.3736572265625 0 714.9445
704.369140625 0 13543.911 c 5
705.368896484375 0 5424.962
709.4239501953125 0 1654.6237
731.3931884765625 0 1997.5778
732.3943481445312 0 1207.8778
733.3867797851562 0 915.1714
749.383544921875 0 7471.2295 w 5
750.385009765625 0 2908.2507
754.4376220703125 0 800.19305
758.3834228515625 0 748.9517 c Ammonia loss 6
762.753662109375 0 1033.53
774.3846435546875 0 1139.987
775.4051513671875 0 13304.249 c 6
776.4039306640625 0 4621.753
777.4039306640625 0 1763.026
787.3870849609375 0 732.1209
788.39208984375 0 897.3281
790.0662841796875 0 893.18494
793.3963012695312 0 28412.973 z 5
794.39453125 0 16267.319
795.3961181640625 0 5810.3193
796.3978881835938 0 1148.2565
822.5156860351562 0 774.2613
837.5180053710938 0 1286.7906
859.4854125976562 0 1577.9817
860.4947509765625 0 1712.5767
864.4327392578125 0 9637.015
865.4324340820312 0 5615.0938
866.427978515625 0 1984.9141
867.4201049804688 0 828.3952
880.3944091796875 0 2842.2925
881.4022216796875 0 695.55457
886.4757690429688 0 1595.3054 c Ammonia loss 7
900.4763793945312 0 1184.8788
901.47900390625 0 978.34863
902.4916381835938 0 5346.509
903.49951171875 0 17848.775 c 7
904.4994506835938 0 9124.372
905.5030517578125 0 2081.1328
906.4009399414062 0 779.13086
908.4227294921875 0 9130.452 z 4
909.421142578125 0 5281.0664
910.4185180664062 0 2008.2622
914.0162963867188 0 775.4203
922.0243530273438 0 1066.3635
922.4308471679688 0 911.36304
922.52978515625 0 977.0817
923.4337158203125 0 831.46515
924.4371337890625 0 759.62024 y 4
925.415771484375 0 814.06
938.5595703125 0 825.1817
986.50048828125 0 816.5604
999.5397338867188 0 983.83124 c Water loss 8
1000.5139770507812 0 1075.3811 c Ammonia loss 8
1001.5170288085938 0 960.06635
1002.5119018554688 0 816.529
1016.5305786132812 0 1424.3057
1017.5424194335938 0 10745.008 c 8
1018.5402221679688 0 7805.172
1019.5400390625 0 2605.1426
1064.5247802734375 0 4799.2153 z 3
1065.52685546875 0 5230.071
1066.52197265625 0 3435.0024
1067.524169921875 0 1026.0363
1088.5545654296875 0 934.2112
1104.57421875 0 3794.0217 c 9
1105.5767822265625 0 2800.997
1106.5770263671875 0 1362.0862
1113.580810546875 0 990.4359
1131.6553955078125 0 875.994
1143.62548828125 0 847.6929
1151.5562744140625 0 1303.489 z 2
1152.56103515625 0 5235.3286
1153.5595703125 0 3445.8909
1154.568603515625 0 1071.3994
1174.6517333984375 0 1566.3628
1175.639892578125 0 1324.659
1201.6383056640625 0 2398.4602
1202.6353759765625 0 841.96045
1217.65625 0 1817.515 c 10
1218.663330078125 0 4326.5356
1219.6624755859375 0 2709.419
1220.666015625 0 1217.4001
1265.64306640625 0 1017.4875
1266.636962890625 0 1199.6863
1304.7027587890625 0 704.6081
1305.6572265625 0 1694.2288
1321.6971435546875 0 1186.3595
1322.70166015625 0 952.388
1323.688720703125 0 1846.428
1324.679443359375 0 4088.9204
1325.6766357421875 0 2872.5034
1326.681396484375 0 1451.8306
1337.7008056640625 0 1614.2871
1338.701416015625 0 5916.0986
1339.69970703125 0 4887.4727
1340.697998046875 0 2740.913
1341.6998291015625 0 1200.193
1342.6768798828125 0 726.14685
1348.69091796875 0 1789.3606
1349.6700439453125 0 2035.1083
1350.6800537109375 0 1600.4897
1351.678955078125 0 1082.3367
1364.715087890625 0 785.73193
1365.69287109375 0 3795.2295
1366.6943359375 0 19960.197
1367.6937255859375 0 17244.137
1368.69287109375 0 8282.992
1369.7015380859375 0 3300.5894
1370.7216796875 0 1367.6378
1381.7423095703125 0 2473.043
1382.7197265625 0 3334.0544
1383.719970703125 0 4592.072
1384.7181396484375 0 4005.294
1385.7269287109375 0 2994.891
1386.746826171875 0 949.61835
1387.7689208984375 0 984.66187
2690.061279296875 0 664.2025

Spectrum Details

|  |  |
| --- | --- |
| Matched peaks? Matched peaks The total absolute number of peaks matched. Additionally in brackets the total fraction of peaks matched and the total number of peaks is shown. | 43 (12.15% of 354) |
| FDR? FDR The false discovery rate estimated for this peptide. It is calculated by matching all theoretical fragments with a non-integer shift with the raw peaks for this spectrum. This is done with 40 different shifts. The resulting percentage is the average number of annotated peaks over the number of annotated peaks with the correct spectrum. | 2.44% |
| Satellite FDR? Satellite FDR See the FDR for details on its calculation. This satellite ion specific FDR only contains the satellite ions (d/w) for I/L/J positions. | 0.00% |
| PSM Score? PSM Score The PSM Score as given by Hecklib to this annotated spectrum. It is shown with three significant figures. | 341 |

#### Spectrum 3238? Spectrum 3238 The raw spectrum of this peptide as annotated by Hecklib. The fragments are coloured according to ion type (see legend). Any peaks with a star '\*' as text can be hovered over to see the full details, first the ion type second the mass shift type. By hovering over the amino acids in the peptide or ions in the legend the corresponding peaks are highlighted. By toggling the 'Unassigned' label you can turn the background (unassigned) peaks on or off in the plot. By updating the slider in the Ion legend you can update the spectrum to only show the top X% of the peaks with labels. The top X% means any peak that is within X% of the highest intensity. By dragging in the spectrum you can zoom in to a specific part of the spectrum and use 'Zoom Out' to get back to the original zoom level. The annotation of the spectrum is based on the given sequence in the peptides file and is done with different software so inconsistencies are likely. The peaks are annotated based on the given sequence, with 20 ppm tolerance.

Copy Data

##### Spectrum 3238 (TSV)

###### Preview

```
Loading example...
```

*Click on the button to copy the data to your clipboard.*

Mz MinMz MaxIntensity Max

WidthHeightPeptide font sizePeptide stroke widthSpectrum font sizeSpectrum stroke widthCompact peptide

Ion legend

wxyz

abcd

OtherUnassignedIon^Charge^_Position_Show for top:%

TISRDNAKNSJY

04.52e+49.05e+41.36e+51.81e+5

Zoom Out

y^+1^_1_y^+1^_2_c^+1^_3_c^+2^_8_c^+2^_8_c^+2^_8_c^+1^_4_y^+1^_4_c^+2^_9_c^+2^_9_c^+2^_9_z^+2^_9_c^+2^_10_c^+2^_10_w^+2^_10_c^+1^_5_z^+2^_10_y^+2^_10_c^+1^_5_c^+2^_11_z^+1^_5_c^+2^_11_w^+2^_11_y^+1^_5_y^+2^_11_y^+2^_11_z^+1^_6_z^+1^_6_y^+1^_6_c^+1^_6_w^+1^_7_c^+1^_7_c^+1^_7_z^+1^_7_y^+1^_7_c^+1^_8_c^+1^_8_z^+1^_8_c^+1^_9_c^+1^_9_c^+1^_9_z^+1^_9_c^+1^_10_z^+1^_10_z^+1^_10_c^+1^_11_

0877175326303507

Fragment Matches Table

Show background peaks

| Position | Ion type | Intensity | mz Theoretical | mz Error (Th) | mz Error (ppm) | Charge | Series Number |
| --- | --- | --- | --- | --- | --- | --- | --- |
| - | - | 404.8 | 126.9 | - | - | 0 | - |
| - | - | 1529 | 129.1 | - | - | 0 | - |
| - | - | 2757 | 136.1 | - | - | 0 | - |
| - | - | 407.3 | 136.8 | - | - | 0 | - |
| - | - | 524.9 | 137.1 | - | - | 0 | - |
| - | - | 409.2 | 144.6 | - | - | 0 | - |
| - | - | 424.4 | 146.8 | - | - | 0 | - |
| - | - | 663.6 | 149 | - | - | 0 | - |
| - | - | 549.3 | 149 | - | - | 0 | - |
| - | - | 6660 | 149 | - | - | 0 | - |
| - | - | 529.9 | 150 | - | - | 0 | - |
| - | - | 407.7 | 157.1 | - | - | 0 | - |
| - | - | 1984 | 165.1 | - | - | 0 | - |
| - | - | 419.1 | 167 | - | - | 0 | - |
| - | - | 1947 | 167.1 | - | - | 0 | - |
| - | - | 4093 | 173.1 | - | - | 0 | - |
| - | - | 2633 | 173.4 | - | - | 0 | - |
| 12 | y | 7180 | 182.1 | 3.87E-05 | 0.2125 | +1 | 1 |
| - | - | 1.498E+04 | 187.1 | - | - | 0 | - |
| - | - | 1272 | 188.1 | - | - | 0 | - |
| - | - | 729.4 | 200.1 | - | - | 0 | - |
| - | - | 1617 | 201.1 | - | - | 0 | - |
| - | - | 529.8 | 203.7 | - | - | 0 | - |
| - | - | 492.9 | 209.6 | - | - | 0 | - |
| - | - | 4034 | 215.1 | - | - | 0 | - |
| - | - | 2737 | 221.1 | - | - | 0 | - |
| - | - | 1407 | 222.1 | - | - | 0 | - |
| - | - | 2329 | 223.1 | - | - | 0 | - |
| - | - | 865.6 | 223.1 | - | - | 0 | - |
| - | - | 564.3 | 224.1 | - | - | 0 | - |
| - | - | 3072 | 225 | - | - | 0 | - |
| - | - | 722 | 227 | - | - | 0 | - |
| - | - | 4738 | 239.1 | - | - | 0 | - |
| - | - | 3106 | 240.1 | - | - | 0 | - |
| - | - | 1800 | 241.1 | - | - | 0 | - |
| - | - | 999.8 | 259.1 | - | - | 0 | - |
| - | - | 2478 | 260.1 | - | - | 0 | - |
| - | - | 637.2 | 266.4 | - | - | 0 | - |
| - | - | 813.7 | 281.1 | - | - | 0 | - |
| - | - | 946.5 | 282.1 | - | - | 0 | - |
| - | - | 535.8 | 284.7 | - | - | 0 | - |
| - | - | 2914 | 295.1 | - | - | 0 | - |
| 11 | y | 1047 | 295.2 | 0.0002021 | 0.6848 | +1 | 2 |
| - | - | 3102 | 296.1 | - | - | 0 | - |
| - | - | 1977 | 297.1 | - | - | 0 | - |
| - | - | 2597 | 299.1 | - | - | 0 | - |
| - | - | 2365 | 300.1 | - | - | 0 | - |
| - | - | 2062 | 300.2 | - | - | 0 | - |
| - | - | 1326 | 301.1 | - | - | 0 | - |
| - | - | 706 | 313.1 | - | - | 0 | - |
| - | - | 836.4 | 315.2 | - | - | 0 | - |
| - | - | 1726 | 317.2 | - | - | 0 | - |
| 3 | c | 1374 | 319.2 | 6.593E-05 | 0.2066 | +1 | 3 |
| - | - | 1.952E+04 | 341 | - | - | 0 | - |
| - | - | 1308 | 343 | - | - | 0 | - |
| - | - | 590.8 | 344.2 | - | - | 0 | - |
| - | - | 1964 | 355.1 | - | - | 0 | - |
| - | - | 833.2 | 356.1 | - | - | 0 | - |
| - | - | 7.186E+04 | 359 | - | - | 0 | - |
| - | - | 1269 | 369.1 | - | - | 0 | - |
| - | - | 3952 | 370.1 | - | - | 0 | - |
| - | - | 3026 | 371.1 | - | - | 0 | - |
| - | - | 776.1 | 372.1 | - | - | 0 | - |
| - | - | 3395 | 375.9 | - | - | 0 | - |
| - | - | 552.3 | 376.9 | - | - | 0 | - |
| - | - | 1127 | 385.2 | - | - | 0 | - |
| - | - | 1180 | 386.2 | - | - | 0 | - |
| - | - | 659.5 | 391.5 | - | - | 0 | - |
| - | - | 1205 | 391.9 | - | - | 0 | - |
| - | - | 1021 | 393.9 | - | - | 0 | - |
| - | - | 1.791E+05 | 429.1 | - | - | 0 | - |
| - | - | 1574 | 429.8 | - | - | 0 | - |
| - | - | 860.1 | 430.2 | - | - | 0 | - |
| - | - | 534.6 | 441.5 | - | - | 0 | - |
| - | - | 1091 | 442.7 | - | - | 0 | - |
| 8 | c | 728.3 | 443.2 | 0.0006148 | 1.387 | +2 | 8 |
| 8 | c | 712.5 | 443.7 | 0.005189 | 11.69 | +2 | 8 |
| - | - | 9.547E+04 | 445.1 | - | - | 0 | - |
| - | - | 2105 | 446.2 | - | - | 0 | - |
| - | - | 1996 | 447.1 | - | - | 0 | - |
| - | - | 3891 | 451.8 | - | - | 0 | - |
| 8 | c | 1.432E+04 | 452.3 | 0.0001509 | 0.3337 | +2 | 8 |
| - | - | 4451 | 452.8 | - | - | 0 | - |
| - | - | 1884 | 453.3 | - | - | 0 | - |
| - | - | 584.3 | 459.9 | - | - | 0 | - |
| - | - | 1860 | 460.3 | - | - | 0 | - |
| - | - | 1461 | 461.2 | - | - | 0 | - |
| - | - | 865.6 | 461.8 | - | - | 0 | - |
| - | - | 1840 | 465.2 | - | - | 0 | - |
| - | - | 769.8 | 465.8 | - | - | 0 | - |
| 4 | c | 3.295E+04 | 475.3 | 0.0003648 | 0.7676 | +1 | 4 |
| - | - | 7353 | 476.3 | - | - | 0 | - |
| - | - | 840.4 | 477.3 | - | - | 0 | - |
| - | - | 2163 | 479.8 | - | - | 0 | - |
| - | - | 1426 | 480.3 | - | - | 0 | - |
| - | - | 1246 | 480.8 | - | - | 0 | - |
| - | - | 2733 | 481.2 | - | - | 0 | - |
| - | - | 694.3 | 482.2 | - | - | 0 | - |
| - | - | 642.7 | 487.8 | - | - | 0 | - |
| - | - | 748.1 | 493.8 | - | - | 0 | - |
| 9 | y | 1415 | 496.2 | 0.001601 | 3.227 | +1 | 4 |
| - | - | 595.9 | 497.2 | - | - | 0 | - |
| - | - | 2459 | 499.8 | - | - | 0 | - |
| 9 | c | 1791 | 500.3 | 0.003698 | 7.392 | +2 | 9 |
| 9 | c | 1849 | 500.8 | 0.001273 | 2.542 | +2 | 9 |
| - | - | 8886 | 502.3 | - | - | 0 | - |
| - | - | 1791 | 503.3 | - | - | 0 | - |
| - | - | 927.7 | 508.3 | - | - | 0 | - |
| - | - | 6745 | 508.8 | - | - | 0 | - |
| 9 | c | 1.119E+04 | 509.3 | 0.0003134 | 0.6153 | +2 | 9 |
| - | - | 4036 | 509.8 | - | - | 0 | - |
| - | - | 1658 | 510.3 | - | - | 0 | - |
| - | - | 624.1 | 521.3 | - | - | 0 | - |
| - | - | 772.5 | 522.8 | - | - | 0 | - |
| - | - | 657.6 | 530.3 | - | - | 0 | - |
| 4 | z | 1391 | 532.8 | 0.001617 | 3.035 | +2 | 9 |
| - | - | 886 | 533.8 | - | - | 0 | - |
| - | - | 697.4 | 535.8 | - | - | 0 | - |
| - | - | 782.1 | 540.3 | - | - | 0 | - |
| - | - | 959.2 | 541.3 | - | - | 0 | - |
| 10 | c | 1314 | 544.3 | 0.00302 | 5.549 | +2 | 10 |
| - | - | 1140 | 544.8 | - | - | 0 | - |
| - | - | 2137 | 545.3 | - | - | 0 | - |
| - | - | 647.5 | 546.3 | - | - | 0 | - |
| - | - | 1095 | 546.8 | - | - | 0 | - |
| - | - | 1586 | 547.3 | - | - | 0 | - |
| 10 | c | 2.021E+04 | 552.8 | 0.0001825 | 0.3301 | +2 | 10 |
| - | - | 1.156E+04 | 553.3 | - | - | 0 | - |
| - | - | 4296 | 553.8 | - | - | 0 | - |
| - | - | 830.6 | 554.3 | - | - | 0 | - |
| - | - | 595.9 | 554.8 | - | - | 0 | - |
| - | - | 1324 | 564.3 | - | - | 0 | - |
| - | - | 1120 | 565.8 | - | - | 0 | - |
| - | - | 694.7 | 566.3 | - | - | 0 | - |
| 3 | w | 897.1 | 567.8 | 0.003757 | 6.617 | +2 | 10 |
| - | - | 886.1 | 568.3 | - | - | 0 | - |
| 5 | c | 1083 | 573.3 | 0.008455 | 14.75 | +1 | 5 |
| 3 | z | 4170 | 576.3 | 0.0007864 | 1.365 | +2 | 10 |
| - | - | 4047 | 576.8 | - | - | 0 | - |
| - | - | 762.7 | 577.3 | - | - | 0 | - |
| - | - | 1327 | 578.3 | - | - | 0 | - |
| - | - | 810.1 | 578.8 | - | - | 0 | - |
| - | - | 711.8 | 579.3 | - | - | 0 | - |
| 3 | y | 1.852E+04 | 584.3 | 0.0002607 | 0.4462 | +2 | 10 |
| - | - | 8943 | 584.8 | - | - | 0 | - |
| - | - | 3195 | 585.3 | - | - | 0 | - |
| - | - | 1046 | 585.8 | - | - | 0 | - |
| - | - | 2135 | 586.8 | - | - | 0 | - |
| - | - | 2265 | 587.3 | - | - | 0 | - |
| - | - | 934.2 | 587.8 | - | - | 0 | - |
| - | - | 1335 | 588.3 | - | - | 0 | - |
| - | - | 988.6 | 589.3 | - | - | 0 | - |
| 5 | c | 6.339E+04 | 590.3 | 9.415E-05 | 0.1595 | +1 | 5 |
| - | - | 1.69E+04 | 591.3 | - | - | 0 | - |
| - | - | 1101 | 591.8 | - | - | 0 | - |
| - | - | 3334 | 592.3 | - | - | 0 | - |
| - | - | 1943 | 593.3 | - | - | 0 | - |
| - | - | 938.1 | 597.9 | - | - | 0 | - |
| - | - | 704 | 598.4 | - | - | 0 | - |
| 11 | c | 3758 | 600.8 | 0.0005388 | 0.8967 | +2 | 11 |
| - | - | 1576 | 601.3 | - | - | 0 | - |
| - | - | 1617 | 601.8 | - | - | 0 | - |
| 8 | z | 4303 | 608.3 | 0.001015 | 1.669 | +1 | 5 |
| 11 | c | 3.717E+04 | 609.3 | 0.0002236 | 0.3669 | +2 | 11 |
| - | - | 2.064E+04 | 609.8 | - | - | 0 | - |
| - | - | 7753 | 610.3 | - | - | 0 | - |
| - | - | 2144 | 610.8 | - | - | 0 | - |
| - | - | 1528 | 612.6 | - | - | 0 | - |
| - | - | 1772 | 612.9 | - | - | 0 | - |
| - | - | 1202 | 614.7 | - | - | 0 | - |
| - | - | 879 | 615 | - | - | 0 | - |
| - | - | 770.2 | 615.7 | - | - | 0 | - |
| - | - | 1521 | 616.3 | - | - | 0 | - |
| 2 | w | 1854 | 618.3 | 5.964E-05 | 0.09645 | +2 | 11 |
| - | - | 1577 | 618.8 | - | - | 0 | - |
| 8 | y | 3211 | 624.3 | 0.003304 | 5.293 | +1 | 5 |
| - | - | 1047 | 625.3 | - | - | 0 | - |
| - | - | 1173 | 630.3 | - | - | 0 | - |
| - | - | 1177 | 630.8 | - | - | 0 | - |
| 2 | y | 636.9 | 632.3 | 0.002102 | 3.324 | +2 | 11 |
| - | - | 799.1 | 633.3 | - | - | 0 | - |
| - | - | 1137 | 633.8 | - | - | 0 | - |
| - | - | 975.8 | 633.9 | - | - | 0 | - |
| - | - | 815.2 | 634.4 | - | - | 0 | - |
| - | - | 897.7 | 635.4 | - | - | 0 | - |
| - | - | 1263 | 638.3 | - | - | 0 | - |
| 2 | y | 1647 | 640.8 | 0.002812 | 4.388 | +2 | 11 |
| - | - | 811.8 | 641.3 | - | - | 0 | - |
| - | - | 869 | 644.3 | - | - | 0 | - |
| - | - | 1104 | 647.4 | - | - | 0 | - |
| - | - | 962.5 | 647.9 | - | - | 0 | - |
| - | - | 1166 | 651.9 | - | - | 0 | - |
| - | - | 969.6 | 652.8 | - | - | 0 | - |
| - | - | 792.9 | 653.3 | - | - | 0 | - |
| - | - | 722.1 | 653.8 | - | - | 0 | - |
| - | - | 974.1 | 654.8 | - | - | 0 | - |
| - | - | 2091 | 655.3 | - | - | 0 | - |
| - | - | 2556 | 659.3 | - | - | 0 | - |
| - | - | 5534 | 660.4 | - | - | 0 | - |
| - | - | 2179 | 660.9 | - | - | 0 | - |
| 7 | z | 6551 | 661.3 | 0.01023 | 15.47 | +1 | 6 |
| - | - | 5695 | 661.9 | - | - | 0 | - |
| - | - | 5656 | 662.3 | - | - | 0 | - |
| - | - | 3255 | 662.8 | - | - | 0 | - |
| - | - | 1851 | 663.3 | - | - | 0 | - |
| - | - | 1963 | 668.8 | - | - | 0 | - |
| - | - | 5414 | 669.3 | - | - | 0 | - |
| - | - | 3680 | 669.9 | - | - | 0 | - |
| - | - | 5691 | 670.3 | - | - | 0 | - |
| - | - | 2850 | 670.8 | - | - | 0 | - |
| - | - | 1581 | 671.4 | - | - | 0 | - |
| - | - | 880 | 674.3 | - | - | 0 | - |
| - | - | 3145 | 674.8 | - | - | 0 | - |
| - | - | 3697 | 675.3 | - | - | 0 | - |
| - | - | 3404 | 675.8 | - | - | 0 | - |
| - | - | 1993 | 676.3 | - | - | 0 | - |
| 7 | z | 2.149E+04 | 679.4 | 0.0001565 | 0.2303 | +1 | 6 |
| - | - | 6177 | 680.4 | - | - | 0 | - |
| - | - | 2033 | 681.4 | - | - | 0 | - |
| - | - | 1067 | 682.8 | - | - | 0 | - |
| - | - | 2.293E+04 | 683.3 | - | - | 0 | - |
| - | - | 1.641E+04 | 683.8 | - | - | 0 | - |
| - | - | 6943 | 684.4 | - | - | 0 | - |
| - | - | 2859 | 684.9 | - | - | 0 | - |
| - | - | 1032 | 689.9 | - | - | 0 | - |
| - | - | 843.3 | 690.9 | - | - | 0 | - |
| - | - | 2.392E+04 | 691.4 | - | - | 0 | - |
| - | - | 2.106E+04 | 691.9 | - | - | 0 | - |
| - | - | 1.274E+04 | 692.4 | - | - | 0 | - |
| - | - | 3534 | 692.9 | - | - | 0 | - |
| 7 | y | 8234 | 695.4 | 0.0002312 | 0.3325 | +1 | 6 |
| - | - | 2500 | 696.4 | - | - | 0 | - |
| - | - | 760.4 | 697.4 | - | - | 0 | - |
| 6 | c | 3.056E+04 | 704.4 | 0.0002575 | 0.3656 | +1 | 6 |
| - | - | 1.018E+04 | 705.4 | - | - | 0 | - |
| - | - | 3142 | 706.4 | - | - | 0 | - |
| - | - | 979.9 | 708.5 | - | - | 0 | - |
| - | - | 2670 | 731.4 | - | - | 0 | - |
| - | - | 1335 | 737.3 | - | - | 0 | - |
| 6 | w | 8525 | 749.4 | 0.001507 | 2.011 | +1 | 7 |
| - | - | 2274 | 750.4 | - | - | 0 | - |
| 7 | c | 1468 | 758.4 | 0.0005571 | 0.7346 | +1 | 7 |
| - | - | 691.9 | 766.4 | - | - | 0 | - |
| - | - | 1589 | 773.8 | - | - | 0 | - |
| - | - | 1478 | 774.3 | - | - | 0 | - |
| 7 | c | 2.774E+04 | 775.4 | 0.0003572 | 0.4607 | +1 | 7 |
| - | - | 9092 | 776.4 | - | - | 0 | - |
| - | - | 2028 | 777.4 | - | - | 0 | - |
| - | - | 893.7 | 778.4 | - | - | 0 | - |
| - | - | 1281 | 779.4 | - | - | 0 | - |
| - | - | 1105 | 787.4 | - | - | 0 | - |
| 6 | z | 4.206E+04 | 793.4 | 0.0001367 | 0.1724 | +1 | 7 |
| - | - | 1.957E+04 | 794.4 | - | - | 0 | - |
| - | - | 4122 | 795.4 | - | - | 0 | - |
| - | - | 904.4 | 796.4 | - | - | 0 | - |
| 6 | y | 1834 | 809.4 | 0.0005167 | 0.6383 | +1 | 7 |
| - | - | 749 | 822.5 | - | - | 0 | - |
| - | - | 1613 | 832.4 | - | - | 0 | - |
| - | - | 776.5 | 841.5 | - | - | 0 | - |
| - | - | 1382 | 857 | - | - | 0 | - |
| - | - | 2338 | 859.5 | - | - | 0 | - |
| - | - | 1537 | 860.5 | - | - | 0 | - |
| - | - | 1.333E+04 | 864.4 | - | - | 0 | - |
| - | - | 5399 | 865.4 | - | - | 0 | - |
| - | - | 1437 | 866.4 | - | - | 0 | - |
| - | - | 766.5 | 874.4 | - | - | 0 | - |
| - | - | 1293 | 884.5 | - | - | 0 | - |
| 8 | c | 2637 | 886.5 | 0.002762 | 3.116 | +1 | 8 |
| - | - | 880.2 | 887.5 | - | - | 0 | - |
| - | - | 819.8 | 892.5 | - | - | 0 | - |
| - | - | 1225 | 893 | - | - | 0 | - |
| - | - | 856.9 | 894 | - | - | 0 | - |
| - | - | 721.6 | 894.5 | - | - | 0 | - |
| - | - | 2629 | 900.5 | - | - | 0 | - |
| - | - | 1050 | 901.5 | - | - | 0 | - |
| - | - | 6084 | 902.5 | - | - | 0 | - |
| 8 | c | 2.585E+04 | 903.5 | 0.0005327 | 0.5896 | +1 | 8 |
| - | - | 9416 | 904.5 | - | - | 0 | - |
| - | - | 3514 | 905.5 | - | - | 0 | - |
| 5 | z | 1.121E+04 | 908.4 | 0.0007816 | 0.8604 | +1 | 8 |
| - | - | 4770 | 909.4 | - | - | 0 | - |
| - | - | 1392 | 910.4 | - | - | 0 | - |
| - | - | 866.2 | 910.9 | - | - | 0 | - |
| - | - | 838 | 914 | - | - | 0 | - |
| - | - | 1567 | 918.9 | - | - | 0 | - |
| - | - | 4285 | 919.4 | - | - | 0 | - |
| - | - | 1773 | 919.9 | - | - | 0 | - |
| - | - | 1577 | 921.5 | - | - | 0 | - |
| - | - | 2137 | 922.5 | - | - | 0 | - |
| - | - | 1750 | 923 | - | - | 0 | - |
| - | - | 1045 | 923.5 | - | - | 0 | - |
| - | - | 921 | 969.5 | - | - | 0 | - |
| - | - | 718.3 | 970.5 | - | - | 0 | - |
| - | - | 753.6 | 974.5 | - | - | 0 | - |
| - | - | 1177 | 986.5 | - | - | 0 | - |
| - | - | 924.7 | 988.5 | - | - | 0 | - |
| 9 | c | 758.7 | 999.5 | 0.005191 | 5.193 | +1 | 9 |
| 9 | c | 920.6 | 1001 | 0.0007367 | 0.7363 | +1 | 9 |
| - | - | 1014 | 1017 | - | - | 0 | - |
| 9 | c | 1.057E+04 | 1018 | 0.0007355 | 0.7228 | +1 | 9 |
| - | - | 5349 | 1019 | - | - | 0 | - |
| - | - | 2137 | 1020 | - | - | 0 | - |
| - | - | 839.6 | 1021 | - | - | 0 | - |
| - | - | 656.8 | 1063 | - | - | 0 | - |
| 4 | z | 7551 | 1065 | 0.0001344 | 0.1263 | +1 | 9 |
| - | - | 5611 | 1066 | - | - | 0 | - |
| - | - | 2956 | 1067 | - | - | 0 | - |
| - | - | 943.7 | 1067 | - | - | 0 | - |
| - | - | 753.9 | 1068 | - | - | 0 | - |
| - | - | 1887 | 1089 | - | - | 0 | - |
| - | - | 965.1 | 1090 | - | - | 0 | - |
| 10 | c | 5027 | 1105 | 4.907E-05 | 0.04442 | +1 | 10 |
| - | - | 5080 | 1106 | - | - | 0 | - |
| - | - | 1820 | 1107 | - | - | 0 | - |
| 3 | z | 834 | 1135 | 0.0176 | 15.52 | +1 | 10 |
| 3 | z | 5344 | 1152 | 0.000644 | 0.5592 | +1 | 10 |
| - | - | 9564 | 1153 | - | - | 0 | - |
| - | - | 4676 | 1154 | - | - | 0 | - |
| - | - | 1907 | 1155 | - | - | 0 | - |
| - | - | 844.7 | 1159 | - | - | 0 | - |
| - | - | 1012 | 1160 | - | - | 0 | - |
| - | - | 2814 | 1175 | - | - | 0 | - |
| - | - | 1383 | 1176 | - | - | 0 | - |
| - | - | 709.6 | 1177 | - | - | 0 | - |
| - | - | 4879 | 1202 | - | - | 0 | - |
| - | - | 2022 | 1203 | - | - | 0 | - |
| 11 | c | 3893 | 1218 | 0.001349 | 1.108 | +1 | 11 |
| - | - | 4896 | 1219 | - | - | 0 | - |
| - | - | 3026 | 1220 | - | - | 0 | - |
| - | - | 738.2 | 1221 | - | - | 0 | - |
| - | - | 774.2 | 1252 | - | - | 0 | - |
| - | - | 1914 | 1266 | - | - | 0 | - |
| - | - | 1194 | 1267 | - | - | 0 | - |
| - | - | 1089 | 1306 | - | - | 0 | - |
| - | - | 1086 | 1307 | - | - | 0 | - |
| - | - | 1175 | 1308 | - | - | 0 | - |
| - | - | 1893 | 1322 | - | - | 0 | - |
| - | - | 908.9 | 1323 | - | - | 0 | - |
| - | - | 5605 | 1324 | - | - | 0 | - |
| - | - | 7573 | 1325 | - | - | 0 | - |
| - | - | 3003 | 1326 | - | - | 0 | - |
| - | - | 1392 | 1327 | - | - | 0 | - |
| - | - | 1794 | 1338 | - | - | 0 | - |
| - | - | 9942 | 1339 | - | - | 0 | - |
| - | - | 5880 | 1340 | - | - | 0 | - |
| - | - | 2656 | 1341 | - | - | 0 | - |
| - | - | 2613 | 1349 | - | - | 0 | - |
| - | - | 3681 | 1350 | - | - | 0 | - |
| - | - | 1920 | 1351 | - | - | 0 | - |
| - | - | 1146 | 1352 | - | - | 0 | - |
| - | - | 5590 | 1366 | - | - | 0 | - |
| - | - | 3.246E+04 | 1367 | - | - | 0 | - |
| - | - | 1.922E+04 | 1368 | - | - | 0 | - |
| - | - | 1.02E+04 | 1369 | - | - | 0 | - |
| - | - | 2573 | 1370 | - | - | 0 | - |
| - | - | 793.3 | 1371 | - | - | 0 | - |
| - | - | 1222 | 1380 | - | - | 0 | - |
| - | - | 795.8 | 1381 | - | - | 0 | - |
| - | - | 1617 | 1382 | - | - | 0 | - |
| - | - | 4886 | 1383 | - | - | 0 | - |
| - | - | 9518 | 1384 | - | - | 0 | - |
| - | - | 5197 | 1385 | - | - | 0 | - |
| - | - | 3418 | 1386 | - | - | 0 | - |
| - | - | 955.8 | 1387 | - | - | 0 | - |
| - | - | 1369 | 1546 | - | - | 0 | - |
| - | - | 1039 | 1547 | - | - | 0 | - |
| - | - | 1050 | 1548 | - | - | 0 | - |
| - | - | 689.2 | 1803 | - | - | 0 | - |
| - | - | 890.6 | 1838 | - | - | 0 | - |
| - | - | 906 | 1844 | - | - | 0 | - |
| - | - | 839.3 | 1845 | - | - | 0 | - |
| - | - | 673 | 2105 | - | - | 0 | - |
| - | - | 663.6 | 2625 | - | - | 0 | - |
| - | - | 682 | 3081 | - | - | 0 | - |
| - | - | 715.4 | 3472 | - | - | 0 | - |

m/z Charge Intensity FragmentType MassShift Position
126.89169311523438 0 404.8257
129.1020965576172 0 1528.6046
136.07565307617188 0 2756.8423
136.7745819091797 0 407.25076
137.07870483398438 0 524.86945
144.5745391845703 0 409.1821
146.82801818847656 0 424.35983
148.9541015625 0 663.58307
149.03945922851562 0 549.2872
149.04489135742188 0 6660.2505
150.0442352294922 0 529.9441
157.13389587402344 0 407.65106
165.05459594726562 0 1983.8085
166.96351623535156 0 419.11636
167.05517578125 0 1947.1742
173.09201049804688 0 4093.4185
173.44046020507812 0 2632.5076
182.0811309814453 0 7180.184 y 11
187.14405822753906 0 14979.864
188.14755249023438 0 1272.1127
200.13893127441406 0 729.4168
201.1229248046875 0 1616.7886
203.65992736816406 0 529.76807
209.56423950195312 0 492.8972
215.13888549804688 0 4033.959
221.08428955078125 0 2737.3936
222.08445739746094 0 1406.8207
223.06324768066406 0 2328.9167
223.08114624023438 0 865.5725
224.0652618408203 0 564.3097
225.04278564453125 0 3072.0198
227.04052734375 0 722.04205
239.094970703125 0 4738.3364
240.0955047607422 0 3106.004
241.09173583984375 0 1799.6655
259.1170654296875 0 999.7977
260.1236267089844 0 2477.8035
266.3808288574219 0 637.1824
281.0509338378906 0 813.7193
282.05218505859375 0 946.454
284.7120666503906 0 535.8307
295.10302734375 0 2913.7056
295.1654357910156 0 1046.6866 y 10
296.10382080078125 0 3101.8533
297.100341796875 0 1976.6044
299.0616455078125 0 2597.4517
300.0627136230469 0 2365.0303
300.1551818847656 0 2061.93
301.0586853027344 0 1326.4148
313.114013671875 0 706.02625
315.16668701171875 0 836.4403
317.1820983886719 0 1726.3667
319.1976623535156 0 1373.8093 c 2
341.0180358886719 0 19522.842
342.997314453125 0 1308.2379
344.1832580566406 0 590.7575
355.0701599121094 0 1963.9354
356.070556640625 0 833.1639
359.0282287597656 0 71864.375
369.1226501464844 0 1268.8676
370.1223449707031 0 3951.549
371.12005615234375 0 3026.0125
372.1198425292969 0 776.14716
375.8599853515625 0 3394.5955
376.8987731933594 0 552.3003
385.2192077636719 0 1126.9508
386.2268981933594 0 1179.7821
391.5494689941406 0 659.5216
391.8541564941406 0 1205.1077
393.87200927734375 0 1021.0441
429.08892822265625 0 179121.06
429.7551574707031 0 1573.73
430.244140625 0 860.0734
441.5035400390625 0 534.63635
442.7444763183594 0 1091.3007
443.2492980957031 0 728.3326 c Water loss 7
443.7458801269531 0 712.5136 c Ammonia loss 7
445.1202697753906 0 95468.93
446.212646484375 0 2104.7817
447.099609375 0 1995.6935
451.7504577636719 0 3890.7097
452.2538146972656 0 14323.525 c 7
452.7554931640625 0 4451.103
453.25567626953125 0 1883.6637
459.8892822265625 0 584.26245
460.2884826660156 0 1859.8495
461.2474060058594 0 1460.6353
461.7501525878906 0 865.5564
465.2117919921875 0 1840.3877
465.7665710449219 0 769.8136
475.299072265625 0 32953.75 c 3
476.3017578125 0 7353.4067
477.3034973144531 0 840.37384
479.76080322265625 0 2162.6003
480.2587585449219 0 1426.1467
480.763427734375 0 1245.8226
481.2295837402344 0 2732.9915
482.2331237792969 0 694.2698
487.77313232421875 0 642.6957
493.7537536621094 0 748.0563
496.2417907714844 0 1415.2455 y 8
497.24432373046875 0 595.85724
499.7666320800781 0 2458.5295
500.2664489746094 0 1790.7222 c Water loss 8
500.763427734375 0 1848.6044 c Ammonia loss 8
502.27435302734375 0 8885.721
503.27685546875 0 1791.4105
508.2652587890625 0 927.7142
508.7713623046875 0 6744.915
509.2751159667969 0 11190.583 c 8
509.7757873535156 0 4036.1147
510.27655029296875 0 1658.0991
521.275146484375 0 624.09564
522.7723999023438 0 772.50037
530.294189453125 0 657.6409
532.7675170898438 0 1391.0981 z 3
533.8350219726562 0 886.0426
535.7710571289062 0 697.44495
540.2698974609375 0 782.14655
541.31396484375 0 959.22455
544.2811889648438 0 1314.0499 c Ammonia loss 9
544.779296875 0 1140.459
545.3037109375 0 2137.4265
546.3090209960938 0 647.4976
546.792236328125 0 1094.895
547.2966918945312 0 1586.0095
552.7916259765625 0 20213.611 c 9
553.29296875 0 11560.818
553.793701171875 0 4295.8823
554.2859497070312 0 830.6393
554.7857055664062 0 595.8835
564.2989501953125 0 1324.2355
565.798095703125 0 1120.1587
566.3020629882812 0 694.7317
567.7843017578125 0 897.10724 w 2
568.2760009765625 0 886.1008
573.3075561523438 0 1083.4159 c Ammonia loss 4
576.2811279296875 0 4170.109 z 2
576.7830200195312 0 4046.6636
577.2855834960938 0 762.71124
578.3131713867188 0 1327.4269
578.8152465820312 0 810.12103
579.3101196289062 0 711.7801
584.291015625 0 18521.65 y 2
584.7928466796875 0 8942.989
585.2933959960938 0 3194.6323
585.7977905273438 0 1045.7334
586.8228149414062 0 2134.7532
587.3265380859375 0 2265.421
587.8259887695312 0 934.2417
588.2735595703125 0 1334.8948
589.3206787109375 0 988.6292
590.3257446289062 0 63385.836 c 4
591.32861328125 0 16897.094
591.8128051757812 0 1100.8489
592.3261108398438 0 3333.689
593.3123779296875 0 1942.6843
597.86328125 0 938.12427
598.3654174804688 0 704.0246
600.8207397460938 0 3758.0325 c Ammonia loss 10
601.3218383789062 0 1575.6011
601.8261108398438 0 1616.7456
608.3174438476562 0 4302.5996 z 7
609.333251953125 0 37168.64 c 10
609.8348388671875 0 20637.611
610.3352661132812 0 7752.594
610.8391723632812 0 2143.878
612.5911254882812 0 1528.0713
612.9275512695312 0 1772.3821
614.677490234375 0 1201.7875
615.0138549804688 0 878.96466
615.6834106445312 0 770.24457
616.3375854492188 0 1520.8794
618.304443359375 0 1854.1477 w 1
618.8052368164062 0 1577.3103
624.3318481445312 0 3211.0657 y 7
625.3367919921875 0 1046.5405
630.3282470703125 0 1173.3518
630.828369140625 0 1177.2852
632.3179321289062 0 636.8513 y Ammonia loss 1
633.322265625 0 799.12787
633.819580078125 0 1137.2367
633.880859375 0 975.79974
634.3770751953125 0 815.2091
635.3629760742188 0 897.65875
638.3383178710938 0 1262.7699
640.8361206054688 0 1647.3049 y 1
641.32958984375 0 811.7516
644.3335571289062 0 869.0285
647.3564453125 0 1103.7834
647.85107421875 0 962.45056
651.8641357421875 0 1166.006
652.8472290039062 0 969.6005
653.3431396484375 0 792.8602
653.8263549804688 0 722.13837
654.8292236328125 0 974.09845
655.322998046875 0 2091.3274
659.3460693359375 0 2556.094
660.3511962890625 0 5533.899
660.851806640625 0 2179.1047
661.3532104492188 0 6551.371 z Water loss 6
661.8502807617188 0 5694.5254
662.3480224609375 0 5655.845
662.84521484375 0 3255.0242
663.3472290039062 0 1850.6665
668.8277587890625 0 1962.7881
669.3475952148438 0 5413.8623
669.8529052734375 0 3679.6003
670.3484497070312 0 5691.1377
670.8495483398438 0 2849.6157
671.3517456054688 0 1581.2831
674.3400268554688 0 880.02313
674.8457641601562 0 3144.6807
675.3475341796875 0 3697.0066
675.8394165039062 0 3404.348
676.338134765625 0 1992.5469
679.3536987304688 0 21490.014 z 6
680.3565673828125 0 6177.3823
681.3602905273438 0 2032.6572
682.8430786132812 0 1067.0692
683.3482055664062 0 22933.145
683.8494262695312 0 16407.223
684.3510131835938 0 6942.6826
684.8515014648438 0 2858.8843
689.9124145507812 0 1031.9333
690.9022827148438 0 843.28503
691.3572387695312 0 23916.725
691.860107421875 0 21057.672
692.362060546875 0 12742.753
692.8658447265625 0 3533.9473
695.3724975585938 0 8233.835 y 6
696.3751220703125 0 2499.887
697.3900756835938 0 760.3895
704.3688354492188 0 30557.895 c 5
705.3711547851562 0 10179.166
706.373779296875 0 3141.908
708.475341796875 0 979.8977
731.3904418945312 0 2670.3274
737.334228515625 0 1335.3868
749.3843383789062 0 8525.464 w 5
750.3877563476562 0 2274.293
758.3796997070312 0 1467.5961 c Ammonia loss 6
766.4208374023438 0 691.94415
773.8291015625 0 1589.2273
774.3269653320312 0 1478.1525
775.4053344726562 0 27735.766 c 6
776.4070434570312 0 9091.967
777.4103393554688 0 2027.6779
778.4114379882812 0 893.68774
779.4061889648438 0 1280.6849
787.393798828125 0 1104.6622
793.3966064453125 0 42059.523 z 5
794.3995971679688 0 19566.803
795.4017333984375 0 4122.0654
796.406494140625 0 904.36487
809.4157104492188 0 1833.6765 y 5
822.4879760742188 0 748.99176
832.4434204101562 0 1613.4332
841.4788208007812 0 776.5021
856.9730834960938 0 1382.3445
859.4852905273438 0 2338.2324
860.490966796875 0 1537.1594
864.4334716796875 0 13334.441
865.4359130859375 0 5398.5264
866.4408569335938 0 1437.2755
874.43017578125 0 766.5251
884.4871215820312 0 1293.337
886.4768676757812 0 2636.997 c Ammonia loss 7
887.4782104492188 0 880.19165
892.5009155273438 0 819.7674
893.0057983398438 0 1225.3597
894.015625 0 856.93915
894.501708984375 0 721.5965
900.4766235351562 0 2628.5347
901.4786376953125 0 1049.8546
902.4931030273438 0 6084.4883
903.5001220703125 0 25852.477 c 7
904.5026245117188 0 9415.895
905.5017700195312 0 3514.4924
908.4241943359375 0 11214.878 z 4
909.4273681640625 0 4769.8276
910.4285278320312 0 1391.7316
910.874267578125 0 866.1575
914.0092163085938 0 838.0108
918.8870849609375 0 1566.6083
919.3895874023438 0 4285.09
919.8871459960938 0 1772.92
921.460205078125 0 1576.6797
922.5098266601562 0 2136.8674
923.014404296875 0 1749.6854
923.5149536132812 0 1044.7413
969.477783203125 0 921.0365
970.48828125 0 718.26605
974.5422973632812 0 753.6156
986.5037841796875 0 1176.6855
988.517333984375 0 924.71344
999.5382080078125 0 758.7375 c Water loss 8
1000.5162963867188 0 920.5674 c Ammonia loss 8
1016.52734375 0 1014.19885
1017.5428466796875 0 10565.744 c 8
1018.546630859375 0 5348.8193
1019.5481567382812 0 2137.2454
1020.5377807617188 0 839.6129
1062.5650634765625 0 656.84314
1064.524658203125 0 7550.7627 z 3
1065.5291748046875 0 5611.336
1066.529541015625 0 2956.4312
1066.65283203125 0 943.71686
1067.531982421875 0 753.8759
1088.559326171875 0 1886.8464
1089.564208984375 0 965.05914
1104.5755615234375 0 5026.576 c 9
1105.5794677734375 0 5079.7627
1106.583251953125 0 1820.021
1134.547607421875 0 834.0122 z Ammonia loss 2
1151.555908203125 0 5344.4346 z 2
1152.5634765625 0 9563.803
1153.563720703125 0 4676.2026
1154.56396484375 0 1906.6711
1158.63525390625 0 844.69385
1159.6260986328125 0 1011.7896
1174.657470703125 0 2814.1658
1175.6529541015625 0 1382.7512
1176.6512451171875 0 709.5502
1201.6395263671875 0 4878.5586
1202.6424560546875 0 2022.0248
1217.6583251953125 0 3892.5198 c 10
1218.664794921875 0 4895.7446
1219.66552734375 0 3026.2605
1220.670166015625 0 738.1922
1251.718994140625 0 774.2076
1265.6497802734375 0 1914.1041
1266.6510009765625 0 1194.4271
1305.662109375 0 1089.485
1306.6617431640625 0 1085.6011
1307.648193359375 0 1174.5554
1321.692626953125 0 1892.595
1322.694580078125 0 908.91284
1323.6890869140625 0 5604.5996
1324.6865234375 0 7573.018
1325.6922607421875 0 3003.095
1326.6837158203125 0 1392.4299
1337.6949462890625 0 1793.7626
1338.702392578125 0 9942.323
1339.702880859375 0 5880.4434
1340.705078125 0 2656.172
1348.6805419921875 0 2613.1362
1349.6785888671875 0 3680.7915
1350.676513671875 0 1919.9008
1351.681640625 0 1145.6305
1365.6956787109375 0 5590.416
1366.6949462890625 0 32463.336
1367.6978759765625 0 19215.646
1368.7001953125 0 10203.786
1369.7032470703125 0 2573.4153
1370.7060546875 0 793.3479
1379.8292236328125 0 1221.6522
1380.848876953125 0 795.80597
1381.713623046875 0 1617.3553
1382.716796875 0 4886.1143
1383.7264404296875 0 9518.317
1384.7265625 0 5196.814
1385.7344970703125 0 3418.1
1386.7314453125 0 955.7731
1545.648681640625 0 1368.9268
1546.6497802734375 0 1038.578
1547.65234375 0 1049.5781
1802.7796630859375 0 689.1548
1837.7579345703125 0 890.60846
1843.982666015625 0 906.04834
1845.017578125 0 839.2748
2104.870849609375 0 672.9756
2625.24658203125 0 663.64825
3081.051025390625 0 682.03723
3471.990478515625 0 715.4447

Spectrum Details

|  |  |
| --- | --- |
| Matched peaks? Matched peaks The total absolute number of peaks matched. Additionally in brackets the total fraction of peaks matched and the total number of peaks is shown. | 46 (12.27% of 375) |
| FDR? FDR The false discovery rate estimated for this peptide. It is calculated by matching all theoretical fragments with a non-integer shift with the raw peaks for this spectrum. This is done with 40 different shifts. The resulting percentage is the average number of annotated peaks over the number of annotated peaks with the correct spectrum. | 2.33% |
| Satellite FDR? Satellite FDR See the FDR for details on its calculation. This satellite ion specific FDR only contains the satellite ions (d/w) for I/L/J positions. | 2.38% |
| PSM Score? PSM Score The PSM Score as given by Hecklib to this annotated spectrum. It is shown with three significant figures. | 341 |

#### Spectrum 4571? Spectrum 4571 The raw spectrum of this peptide as annotated by Hecklib. The fragments are coloured according to ion type (see legend). Any peaks with a star '\*' as text can be hovered over to see the full details, first the ion type second the mass shift type. By hovering over the amino acids in the peptide or ions in the legend the corresponding peaks are highlighted. By toggling the 'Unassigned' label you can turn the background (unassigned) peaks on or off in the plot. By updating the slider in the Ion legend you can update the spectrum to only show the top X% of the peaks with labels. The top X% means any peak that is within X% of the highest intensity. By dragging in the spectrum you can zoom in to a specific part of the spectrum and use 'Zoom Out' to get back to the original zoom level. The annotation of the spectrum is based on the given sequence in the peptides file and is done with different software so inconsistencies are likely. The peaks are annotated based on the given sequence, with 20 ppm tolerance.

Copy Data

##### Spectrum 4571 (TSV)

###### Preview

```
Loading example...
```

*Click on the button to copy the data to your clipboard.*

Mz MinMz MaxIntensity Max

WidthHeightPeptide font sizePeptide stroke widthSpectrum font sizeSpectrum stroke widthCompact peptide

Ion legend

wxyz

abcd

OtherUnassignedIon^Charge^_Position_Show for top:%

TISRDNAKNSJY

03.61e+47.21e+41.08e+51.44e+5

Zoom Out

y^+3^_4_y^+1^_1_z^+3^_7_y^+1^_2_c^+1^_3_c^+2^_7_c^+2^_8_y^+2^_8_z^+1^_4_c^+1^_4_y^+1^_4_c^+2^_9_c^+2^_9_c^+2^_9_z^+2^_9_c^+2^_10_c^+2^_10_c^+1^_5_y^+2^_10_z^+2^_10_y^+2^_10_c^+1^_5_c^+2^_11_z^+1^_5_c^+2^_11_w^+2^_11_y^+1^_5_z^+2^_11_y^+2^_11_z^+1^_6_y^+1^_6_z^+1^_6_y^+1^_6_c^+1^_6_w^+1^_7_c^+1^_7_c^+1^_7_z^+1^_7_c^+1^_8_c^+1^_8_y^+1^_8_z^+1^_8_y^+1^_8_c^+1^_9_c^+1^_9_c^+1^_9_z^+1^_9_c^+1^_10_z^+1^_10_c^+1^_11_

0830165924893318

Fragment Matches Table

Show background peaks

| Position | Ion type | Intensity | mz Theoretical | mz Error (Th) | mz Error (ppm) | Charge | Series Number |
| --- | --- | --- | --- | --- | --- | --- | --- |
| - | - | 638.8 | 120.1 | - | - | 0 | - |
| - | - | 321.2 | 121.4 | - | - | 0 | - |
| - | - | 498.8 | 125.5 | - | - | 0 | - |
| - | - | 567.5 | 129.1 | - | - | 0 | - |
| - | - | 1907 | 136.1 | - | - | 0 | - |
| - | - | 458.4 | 139.1 | - | - | 0 | - |
| - | - | 915.3 | 148.9 | - | - | 0 | - |
| - | - | 5667 | 149 | - | - | 0 | - |
| - | - | 807.5 | 151 | - | - | 0 | - |
| - | - | 444.4 | 157.3 | - | - | 0 | - |
| - | - | 1969 | 165.1 | - | - | 0 | - |
| 9 | y | 628.1 | 166.1 | 0.001359 | 8.183 | +3 | 4 |
| - | - | 1727 | 167.1 | - | - | 0 | - |
| - | - | 3476 | 173.1 | - | - | 0 | - |
| - | - | 625.9 | 173.1 | - | - | 0 | - |
| - | - | 859.3 | 173.5 | - | - | 0 | - |
| 12 | y | 7460 | 182.1 | 6.921E-05 | 0.3801 | +1 | 1 |
| - | - | 578 | 183.1 | - | - | 0 | - |
| - | - | 836.5 | 187.1 | - | - | 0 | - |
| - | - | 1.29E+04 | 187.1 | - | - | 0 | - |
| - | - | 1594 | 188.1 | - | - | 0 | - |
| - | - | 473.2 | 194.7 | - | - | 0 | - |
| - | - | 1480 | 201.1 | - | - | 0 | - |
| - | - | 1049 | 201.1 | - | - | 0 | - |
| - | - | 648.3 | 205.1 | - | - | 0 | - |
| - | - | 3366 | 215.1 | - | - | 0 | - |
| - | - | 633.3 | 221.1 | - | - | 0 | - |
| - | - | 2765 | 221.1 | - | - | 0 | - |
| - | - | 1828 | 222.1 | - | - | 0 | - |
| - | - | 1533 | 223.1 | - | - | 0 | - |
| - | - | 927.6 | 223.1 | - | - | 0 | - |
| - | - | 2628 | 225 | - | - | 0 | - |
| - | - | 1132 | 226 | - | - | 0 | - |
| - | - | 3233 | 229.1 | - | - | 0 | - |
| - | - | 625.4 | 230.7 | - | - | 0 | - |
| - | - | 596.8 | 232.3 | - | - | 0 | - |
| - | - | 588.4 | 236.9 | - | - | 0 | - |
| - | - | 3564 | 239.1 | - | - | 0 | - |
| - | - | 2428 | 240.1 | - | - | 0 | - |
| - | - | 804.3 | 241.1 | - | - | 0 | - |
| - | - | 504.5 | 249.2 | - | - | 0 | - |
| 6 | z | 474 | 259.1 | 0.0005167 | 1.994 | +3 | 7 |
| - | - | 1645 | 260.1 | - | - | 0 | - |
| - | - | 1055 | 281.1 | - | - | 0 | - |
| - | - | 721.4 | 282.1 | - | - | 0 | - |
| - | - | 637.4 | 282.1 | - | - | 0 | - |
| - | - | 2189 | 295.1 | - | - | 0 | - |
| 11 | y | 936.5 | 295.2 | 0.000927 | 3.141 | +1 | 2 |
| - | - | 2504 | 296.1 | - | - | 0 | - |
| - | - | 1576 | 297.1 | - | - | 0 | - |
| - | - | 662.5 | 298.1 | - | - | 0 | - |
| - | - | 2210 | 299.1 | - | - | 0 | - |
| - | - | 2460 | 300.1 | - | - | 0 | - |
| - | - | 2399 | 300.2 | - | - | 0 | - |
| - | - | 1411 | 301.1 | - | - | 0 | - |
| - | - | 511.6 | 310.9 | - | - | 0 | - |
| - | - | 638.5 | 315.2 | - | - | 0 | - |
| - | - | 1204 | 316.1 | - | - | 0 | - |
| 3 | c | 843.4 | 319.2 | 0.0001782 | 0.5583 | +1 | 3 |
| - | - | 1023 | 341 | - | - | 0 | - |
| - | - | 1.576E+04 | 341 | - | - | 0 | - |
| - | - | 1084 | 343 | - | - | 0 | - |
| - | - | 544.7 | 345.2 | - | - | 0 | - |
| - | - | 2049 | 355.1 | - | - | 0 | - |
| - | - | 713.9 | 356.1 | - | - | 0 | - |
| - | - | 5.81E+04 | 359 | - | - | 0 | - |
| - | - | 713.8 | 359.2 | - | - | 0 | - |
| - | - | 816.2 | 369.1 | - | - | 0 | - |
| - | - | 3499 | 370.1 | - | - | 0 | - |
| - | - | 3693 | 371.1 | - | - | 0 | - |
| - | - | 757.7 | 372.1 | - | - | 0 | - |
| - | - | 619.1 | 375.9 | - | - | 0 | - |
| 7 | c | 1489 | 379.2 | 0.007538 | 19.88 | +2 | 7 |
| - | - | 1429 | 385.2 | - | - | 0 | - |
| - | - | 674.9 | 386.2 | - | - | 0 | - |
| - | - | 569.4 | 391.6 | - | - | 0 | - |
| - | - | 775.7 | 391.9 | - | - | 0 | - |
| - | - | 678.6 | 397.2 | - | - | 0 | - |
| - | - | 601.9 | 409.5 | - | - | 0 | - |
| - | - | 599 | 415.2 | - | - | 0 | - |
| - | - | 1.428E+05 | 429.1 | - | - | 0 | - |
| - | - | 593.1 | 429.8 | - | - | 0 | - |
| - | - | 930.3 | 430.2 | - | - | 0 | - |
| - | - | 2894 | 433.2 | - | - | 0 | - |
| - | - | 818.5 | 442.7 | - | - | 0 | - |
| - | - | 7.281E+04 | 445.1 | - | - | 0 | - |
| - | - | 903.5 | 446.2 | - | - | 0 | - |
| - | - | 1118 | 447.1 | - | - | 0 | - |
| - | - | 2206 | 451.8 | - | - | 0 | - |
| 8 | c | 1.009E+04 | 452.3 | 0.0005476 | 1.211 | +2 | 8 |
| - | - | 953.6 | 452.7 | - | - | 0 | - |
| - | - | 7247 | 452.8 | - | - | 0 | - |
| - | - | 1529 | 453.3 | - | - | 0 | - |
| 5 | y | 781.9 | 453.7 | 0.00842 | 18.56 | +2 | 8 |
| - | - | 649.7 | 455.6 | - | - | 0 | - |
| - | - | 761.8 | 459.7 | - | - | 0 | - |
| - | - | 663.9 | 459.7 | - | - | 0 | - |
| - | - | 1253 | 460.2 | - | - | 0 | - |
| - | - | 3478 | 461.7 | - | - | 0 | - |
| 9 | z | 2318 | 462.2 | 0.007971 | 17.25 | +1 | 4 |
| - | - | 577.1 | 472.2 | - | - | 0 | - |
| 4 | c | 3.824E+04 | 475.3 | 0.0001512 | 0.3181 | +1 | 4 |
| - | - | 7429 | 476.3 | - | - | 0 | - |
| - | - | 1902 | 477.3 | - | - | 0 | - |
| - | - | 662.9 | 479.3 | - | - | 0 | - |
| - | - | 2423 | 479.8 | - | - | 0 | - |
| - | - | 2657 | 480.3 | - | - | 0 | - |
| - | - | 887.9 | 480.8 | - | - | 0 | - |
| - | - | 2021 | 481.2 | - | - | 0 | - |
| - | - | 2098 | 492.2 | - | - | 0 | - |
| - | - | 667.6 | 493.2 | - | - | 0 | - |
| 9 | y | 1237 | 496.2 | 0.00035 | 0.7054 | +1 | 4 |
| - | - | 3842 | 499.8 | - | - | 0 | - |
| 9 | c | 2106 | 500.3 | 0.004797 | 9.588 | +2 | 9 |
| 9 | c | 2653 | 500.8 | 0.0002834 | 0.566 | +2 | 9 |
| - | - | 912.4 | 501.3 | - | - | 0 | - |
| - | - | 3483 | 502.3 | - | - | 0 | - |
| - | - | 845.6 | 503.3 | - | - | 0 | - |
| - | - | 4715 | 508.3 | - | - | 0 | - |
| - | - | 9183 | 508.8 | - | - | 0 | - |
| 9 | c | 1.588E+04 | 509.3 | 0.001198 | 2.353 | +2 | 9 |
| - | - | 8382 | 509.8 | - | - | 0 | - |
| - | - | 3158 | 510.3 | - | - | 0 | - |
| 4 | z | 1691 | 532.8 | 3.097E-05 | 0.05812 | +2 | 9 |
| - | - | 1482 | 533.3 | - | - | 0 | - |
| - | - | 600 | 535.3 | - | - | 0 | - |
| - | - | 764 | 535.8 | - | - | 0 | - |
| 10 | c | 2118 | 544.3 | 3.176E-05 | 0.05836 | +2 | 10 |
| - | - | 1147 | 545.3 | - | - | 0 | - |
| - | - | 1612 | 548.3 | - | - | 0 | - |
| - | - | 1218 | 548.8 | - | - | 0 | - |
| 10 | c | 1.045E+04 | 552.8 | 0.0004279 | 0.7741 | +2 | 10 |
| - | - | 8915 | 553.3 | - | - | 0 | - |
| - | - | 3871 | 553.8 | - | - | 0 | - |
| - | - | 1770 | 554.3 | - | - | 0 | - |
| - | - | 624.2 | 554.8 | - | - | 0 | - |
| - | - | 879.7 | 564.3 | - | - | 0 | - |
| - | - | 812.5 | 565.8 | - | - | 0 | - |
| - | - | 653.7 | 566.3 | - | - | 0 | - |
| 5 | c | 557.6 | 573.3 | 0.009859 | 17.2 | +1 | 5 |
| 3 | y | 697.9 | 575.3 | 0.002425 | 4.215 | +2 | 10 |
| 3 | z | 4857 | 576.3 | 0.0003591 | 0.6232 | +2 | 10 |
| - | - | 2140 | 576.8 | - | - | 0 | - |
| - | - | 1871 | 577.3 | - | - | 0 | - |
| 3 | y | 1.124E+04 | 584.3 | 0.0002607 | 0.4462 | +2 | 10 |
| - | - | 7319 | 584.8 | - | - | 0 | - |
| - | - | 3237 | 585.3 | - | - | 0 | - |
| - | - | 710.8 | 585.8 | - | - | 0 | - |
| - | - | 1322 | 586.8 | - | - | 0 | - |
| - | - | 1943 | 587.3 | - | - | 0 | - |
| 5 | c | 6.884E+04 | 590.3 | 9.415E-05 | 0.1595 | +1 | 5 |
| - | - | 1.997E+04 | 591.3 | - | - | 0 | - |
| - | - | 931.7 | 591.8 | - | - | 0 | - |
| - | - | 3636 | 592.3 | - | - | 0 | - |
| - | - | 1045 | 592.8 | - | - | 0 | - |
| 11 | c | 1470 | 600.8 | 0.000743 | 1.237 | +2 | 11 |
| - | - | 1182 | 601.3 | - | - | 0 | - |
| - | - | 1394 | 601.8 | - | - | 0 | - |
| - | - | 2461 | 607.3 | - | - | 0 | - |
| 8 | z | 4876 | 608.3 | 0.001931 | 3.174 | +1 | 5 |
| 11 | c | 2.852E+04 | 609.3 | 0.0007729 | 1.268 | +2 | 11 |
| - | - | 2.191E+04 | 609.8 | - | - | 0 | - |
| - | - | 9000 | 610.3 | - | - | 0 | - |
| - | - | 2771 | 610.8 | - | - | 0 | - |
| - | - | 929.8 | 611.3 | - | - | 0 | - |
| 2 | w | 1794 | 618.3 | 0.0004897 | 0.792 | +2 | 11 |
| - | - | 1131 | 618.8 | - | - | 0 | - |
| 8 | y | 5381 | 624.3 | 0.0009851 | 1.578 | +1 | 5 |
| - | - | 1578 | 625.3 | - | - | 0 | - |
| - | - | 1387 | 630.3 | - | - | 0 | - |
| 2 | z | 701.5 | 632.8 | 0.003268 | 5.164 | +2 | 11 |
| - | - | 1144 | 633.3 | - | - | 0 | - |
| - | - | 1130 | 633.8 | - | - | 0 | - |
| - | - | 692.2 | 634.8 | - | - | 0 | - |
| - | - | 681.1 | 638.8 | - | - | 0 | - |
| - | - | 1714 | 640.7 | - | - | 0 | - |
| 2 | y | 1965 | 640.8 | 0.0009202 | 1.436 | +2 | 11 |
| - | - | 1489 | 641.3 | - | - | 0 | - |
| - | - | 767.2 | 642.8 | - | - | 0 | - |
| - | - | 837 | 647.4 | - | - | 0 | - |
| - | - | 712.8 | 648.3 | - | - | 0 | - |
| - | - | 3379 | 655.3 | - | - | 0 | - |
| - | - | 1714 | 655.8 | - | - | 0 | - |
| - | - | 3647 | 659.3 | - | - | 0 | - |
| - | - | 4446 | 660.3 | - | - | 0 | - |
| - | - | 2895 | 660.9 | - | - | 0 | - |
| 7 | z | 5314 | 661.3 | 0.007975 | 12.06 | +1 | 6 |
| - | - | 4325 | 661.9 | - | - | 0 | - |
| - | - | 4745 | 662.3 | - | - | 0 | - |
| - | - | 1970 | 662.8 | - | - | 0 | - |
| - | - | 983.9 | 663.3 | - | - | 0 | - |
| - | - | 1751 | 668.8 | - | - | 0 | - |
| - | - | 4915 | 669.3 | - | - | 0 | - |
| - | - | 3589 | 669.8 | - | - | 0 | - |
| - | - | 4732 | 670.3 | - | - | 0 | - |
| - | - | 1875 | 670.8 | - | - | 0 | - |
| - | - | 1216 | 671.3 | - | - | 0 | - |
| - | - | 840 | 672.3 | - | - | 0 | - |
| - | - | 1714 | 674.3 | - | - | 0 | - |
| - | - | 2457 | 674.8 | - | - | 0 | - |
| - | - | 2011 | 675.3 | - | - | 0 | - |
| - | - | 2636 | 675.8 | - | - | 0 | - |
| - | - | 2044 | 676.3 | - | - | 0 | - |
| - | - | 894.7 | 676.8 | - | - | 0 | - |
| 7 | y | 1300 | 678.3 | 0.01173 | 17.3 | +1 | 6 |
| 7 | z | 1.497E+04 | 679.4 | 3.441E-05 | 0.05065 | +1 | 6 |
| - | - | 5724 | 680.4 | - | - | 0 | - |
| - | - | 1904 | 681.4 | - | - | 0 | - |
| - | - | 1117 | 682.9 | - | - | 0 | - |
| - | - | 1.62E+04 | 683.3 | - | - | 0 | - |
| - | - | 1.613E+04 | 683.8 | - | - | 0 | - |
| - | - | 7508 | 684.3 | - | - | 0 | - |
| - | - | 2443 | 684.8 | - | - | 0 | - |
| - | - | 777.8 | 685.4 | - | - | 0 | - |
| - | - | 969 | 687.4 | - | - | 0 | - |
| - | - | 1368 | 689.2 | - | - | 0 | - |
| - | - | 5183 | 689.7 | - | - | 0 | - |
| - | - | 1935 | 690.2 | - | - | 0 | - |
| - | - | 1078 | 690.7 | - | - | 0 | - |
| - | - | 1.619E+04 | 691.4 | - | - | 0 | - |
| - | - | 1.459E+04 | 691.9 | - | - | 0 | - |
| - | - | 7691 | 692.4 | - | - | 0 | - |
| - | - | 2596 | 692.9 | - | - | 0 | - |
| - | - | 4960 | 694.4 | - | - | 0 | - |
| 7 | y | 4968 | 695.4 | 0.008863 | 12.75 | +1 | 6 |
| - | - | 997.4 | 696.4 | - | - | 0 | - |
| 6 | c | 1.722E+04 | 704.4 | 1.338E-05 | 0.019 | +1 | 6 |
| - | - | 6642 | 705.4 | - | - | 0 | - |
| - | - | 1093 | 706.4 | - | - | 0 | - |
| - | - | 799.6 | 707.8 | - | - | 0 | - |
| - | - | 873.8 | 719.3 | - | - | 0 | - |
| - | - | 2099 | 731.4 | - | - | 0 | - |
| - | - | 786.4 | 732.4 | - | - | 0 | - |
| 6 | w | 9132 | 749.4 | 0.0004698 | 0.6269 | +1 | 7 |
| - | - | 4414 | 750.4 | - | - | 0 | - |
| - | - | 1695 | 751.4 | - | - | 0 | - |
| 7 | c | 1447 | 758.4 | 0.001839 | 2.425 | +1 | 7 |
| - | - | 709.6 | 759.4 | - | - | 0 | - |
| - | - | 4935 | 774.4 | - | - | 0 | - |
| 7 | c | 1.6E+04 | 775.4 | 0.001151 | 1.484 | +1 | 7 |
| - | - | 6840 | 776.4 | - | - | 0 | - |
| - | - | 1783 | 777.4 | - | - | 0 | - |
| - | - | 679.8 | 788.4 | - | - | 0 | - |
| - | - | 803.9 | 788.8 | - | - | 0 | - |
| 6 | z | 4.015E+04 | 793.4 | 7.571E-05 | 0.09543 | +1 | 7 |
| - | - | 2.057E+04 | 794.4 | - | - | 0 | - |
| - | - | 7047 | 795.4 | - | - | 0 | - |
| - | - | 1589 | 796.4 | - | - | 0 | - |
| - | - | 859.5 | 810.4 | - | - | 0 | - |
| - | - | 973.5 | 859.5 | - | - | 0 | - |
| - | - | 937.3 | 860.5 | - | - | 0 | - |
| - | - | 1.207E+04 | 864.4 | - | - | 0 | - |
| - | - | 5484 | 865.4 | - | - | 0 | - |
| - | - | 2534 | 866.4 | - | - | 0 | - |
| - | - | 1.064E+04 | 880.4 | - | - | 0 | - |
| - | - | 5037 | 881.4 | - | - | 0 | - |
| 8 | c | 949.3 | 886.5 | 0.006485 | 7.316 | +1 | 8 |
| - | - | 1044 | 887.5 | - | - | 0 | - |
| - | - | 1085 | 888.4 | - | - | 0 | - |
| - | - | 1139 | 888.5 | - | - | 0 | - |
| - | - | 1323 | 889.4 | - | - | 0 | - |
| - | - | 2626 | 893.4 | - | - | 0 | - |
| - | - | 816.1 | 894.4 | - | - | 0 | - |
| - | - | 1801 | 900.5 | - | - | 0 | - |
| - | - | 1008 | 901.5 | - | - | 0 | - |
| - | - | 6229 | 902.5 | - | - | 0 | - |
| 8 | c | 2.458E+04 | 903.5 | 0.0008378 | 0.9273 | +1 | 8 |
| - | - | 1.245E+04 | 904.5 | - | - | 0 | - |
| - | - | 2947 | 905.5 | - | - | 0 | - |
| - | - | 3479 | 906.4 | - | - | 0 | - |
| 5 | y | 2042 | 907.4 | 0.006774 | 7.465 | +1 | 8 |
| 5 | z | 1.261E+04 | 908.4 | 0.0009884 | 1.088 | +1 | 8 |
| - | - | 8268 | 909.4 | - | - | 0 | - |
| - | - | 2605 | 910.4 | - | - | 0 | - |
| - | - | 919.4 | 914.9 | - | - | 0 | - |
| - | - | 1444 | 919.5 | - | - | 0 | - |
| - | - | 1178 | 920.5 | - | - | 0 | - |
| - | - | 1676 | 922.4 | - | - | 0 | - |
| - | - | 1031 | 922.9 | - | - | 0 | - |
| - | - | 2696 | 923.4 | - | - | 0 | - |
| 5 | y | 1153 | 924.4 | 0.01178 | 12.74 | +1 | 8 |
| - | - | 745.8 | 925.4 | - | - | 0 | - |
| - | - | 1718 | 938.3 | - | - | 0 | - |
| - | - | 3421 | 939.4 | - | - | 0 | - |
| - | - | 1670 | 940.4 | - | - | 0 | - |
| - | - | 806.9 | 986.5 | - | - | 0 | - |
| - | - | 1197 | 987.5 | - | - | 0 | - |
| - | - | 928.6 | 988.5 | - | - | 0 | - |
| 9 | c | 1091 | 999.5 | 0.004763 | 4.765 | +1 | 9 |
| 9 | c | 1187 | 1001 | 0.005681 | 5.678 | +1 | 9 |
| - | - | 855.2 | 1002 | - | - | 0 | - |
| - | - | 911.6 | 1003 | - | - | 0 | - |
| - | - | 2281 | 1017 | - | - | 0 | - |
| 9 | c | 1.453E+04 | 1018 | 0.001224 | 1.203 | +1 | 9 |
| - | - | 8918 | 1019 | - | - | 0 | - |
| - | - | 3108 | 1020 | - | - | 0 | - |
| - | - | 724.6 | 1021 | - | - | 0 | - |
| - | - | 846.1 | 1022 | - | - | 0 | - |
| - | - | 894.2 | 1061 | - | - | 0 | - |
| - | - | 957.7 | 1062 | - | - | 0 | - |
| 4 | z | 5304 | 1065 | 0.0008421 | 0.7911 | +1 | 9 |
| - | - | 6375 | 1066 | - | - | 0 | - |
| - | - | 4435 | 1067 | - | - | 0 | - |
| - | - | 1102 | 1068 | - | - | 0 | - |
| - | - | 1144 | 1089 | - | - | 0 | - |
| 10 | c | 4095 | 1105 | 0.0001711 | 0.1549 | +1 | 10 |
| - | - | 2870 | 1106 | - | - | 0 | - |
| - | - | 1767 | 1107 | - | - | 0 | - |
| - | - | 1117 | 1108 | - | - | 0 | - |
| - | - | 696.2 | 1136 | - | - | 0 | - |
| 3 | z | 3160 | 1152 | 0.001187 | 1.031 | +1 | 10 |
| - | - | 6592 | 1153 | - | - | 0 | - |
| - | - | 4966 | 1154 | - | - | 0 | - |
| - | - | 1393 | 1155 | - | - | 0 | - |
| - | - | 843.6 | 1156 | - | - | 0 | - |
| - | - | 1113 | 1174 | - | - | 0 | - |
| - | - | 2281 | 1175 | - | - | 0 | - |
| - | - | 2266 | 1176 | - | - | 0 | - |
| - | - | 3377 | 1202 | - | - | 0 | - |
| - | - | 2213 | 1203 | - | - | 0 | - |
| - | - | 823.4 | 1204 | - | - | 0 | - |
| 11 | c | 2942 | 1218 | 0.001336 | 1.097 | +1 | 11 |
| - | - | 6354 | 1219 | - | - | 0 | - |
| - | - | 4121 | 1220 | - | - | 0 | - |
| - | - | 1363 | 1221 | - | - | 0 | - |
| - | - | 1246 | 1266 | - | - | 0 | - |
| - | - | 927.5 | 1267 | - | - | 0 | - |
| - | - | 733.4 | 1268 | - | - | 0 | - |
| - | - | 948.3 | 1306 | - | - | 0 | - |
| - | - | 948.7 | 1321 | - | - | 0 | - |
| - | - | 2217 | 1322 | - | - | 0 | - |
| - | - | 889.8 | 1323 | - | - | 0 | - |
| - | - | 2479 | 1324 | - | - | 0 | - |
| - | - | 5078 | 1325 | - | - | 0 | - |
| - | - | 3873 | 1326 | - | - | 0 | - |
| - | - | 1792 | 1327 | - | - | 0 | - |
| - | - | 783.9 | 1328 | - | - | 0 | - |
| - | - | 1193 | 1338 | - | - | 0 | - |
| - | - | 6023 | 1339 | - | - | 0 | - |
| - | - | 5676 | 1340 | - | - | 0 | - |
| - | - | 3536 | 1341 | - | - | 0 | - |
| - | - | 1528 | 1342 | - | - | 0 | - |
| - | - | 2133 | 1349 | - | - | 0 | - |
| - | - | 3569 | 1350 | - | - | 0 | - |
| - | - | 2620 | 1351 | - | - | 0 | - |
| - | - | 1302 | 1352 | - | - | 0 | - |
| - | - | 907.4 | 1353 | - | - | 0 | - |
| - | - | 4837 | 1366 | - | - | 0 | - |
| - | - | 2.479E+04 | 1367 | - | - | 0 | - |
| - | - | 2.051E+04 | 1368 | - | - | 0 | - |
| - | - | 1.109E+04 | 1369 | - | - | 0 | - |
| - | - | 3859 | 1370 | - | - | 0 | - |
| - | - | 1104 | 1371 | - | - | 0 | - |
| - | - | 1686 | 1378 | - | - | 0 | - |
| - | - | 3325 | 1379 | - | - | 0 | - |
| - | - | 2582 | 1380 | - | - | 0 | - |
| - | - | 881.2 | 1382 | - | - | 0 | - |
| - | - | 3663 | 1383 | - | - | 0 | - |
| - | - | 6332 | 1384 | - | - | 0 | - |
| - | - | 4593 | 1385 | - | - | 0 | - |
| - | - | 2361 | 1386 | - | - | 0 | - |
| - | - | 909.7 | 1387 | - | - | 0 | - |
| - | - | 673.3 | 1767 | - | - | 0 | - |
| - | - | 727.5 | 2497 | - | - | 0 | - |
| - | - | 792.9 | 3084 | - | - | 0 | - |
| - | - | 765.8 | 3285 | - | - | 0 | - |

m/z Charge Intensity FragmentType MassShift Position
120.08096313476562 0 638.7553
121.380126953125 0 321.1968
125.50250244140625 0 498.7747
129.10256958007812 0 567.4659
136.07559204101562 0 1907.1296
139.07452392578125 0 458.3561
148.9471435546875 0 915.34216
149.0448455810547 0 5667.376
151.04176330566406 0 807.45844
157.33743286132812 0 444.42316
165.05433654785156 0 1968.8175
166.08627319335938 0 628.129 y 8
167.05523681640625 0 1727.2361
173.0921630859375 0 3476.356
173.1282958984375 0 625.94305
173.45204162597656 0 859.3402
182.0811004638672 0 7459.608 y 11
183.08505249023438 0 578.01404
187.1355438232422 0 836.5274
187.14401245117188 0 12899.438
188.14736938476562 0 1593.6133
194.72064208984375 0 473.15573
201.08673095703125 0 1479.6322
201.1232147216797 0 1048.6183
205.09716796875 0 648.343
215.1387939453125 0 3365.854
221.07391357421875 0 633.30536
221.08441162109375 0 2765.0098
222.0854949951172 0 1828.2257
223.0637664794922 0 1532.7952
223.08126831054688 0 927.57513
225.0428466796875 0 2628.4321
226.04269409179688 0 1131.5021
229.08187866210938 0 3232.5117
230.6939239501953 0 625.3997
232.3360595703125 0 596.78143
236.92160034179688 0 588.4222
239.09498596191406 0 3564.1648
240.09524536132812 0 2428.1597
241.09164428710938 0 804.3434
249.1990966796875 0 504.4558
259.1340026855469 0 474.0235 z Water loss 5
260.1236877441406 0 1645.3901
281.05126953125 0 1054.6112
282.0537414550781 0 721.4258
282.1436462402344 0 637.383
295.10308837890625 0 2189.1086
295.164306640625 0 936.4726 y 10
296.1044616699219 0 2503.8013
297.10107421875 0 1575.6787
298.10211181640625 0 662.5347
299.0614318847656 0 2209.6785
300.0614013671875 0 2460.4314
300.1555480957031 0 2399.0635
301.0586242675781 0 1410.7173
310.8862609863281 0 511.56995
315.16644287109375 0 638.46844
316.11334228515625 0 1204.0852
319.1974182128906 0 843.3846 c 2
340.9944763183594 0 1023.15686
341.01800537109375 0 15755.688
342.99652099609375 0 1084.2604
345.1801452636719 0 544.66187
355.0698547363281 0 2048.7815
356.06768798828125 0 713.899
359.0281677246094 0 58097.395
359.1542663574219 0 713.812
369.1203918457031 0 816.224
370.1221618652344 0 3499.3455
371.11993408203125 0 3693.2502
372.1194152832031 0 757.6608
375.8594665527344 0 619.1107
379.208740234375 0 1488.9683 c Water loss 6
385.2187805175781 0 1428.9075
386.2232971191406 0 674.85016
391.5521545410156 0 569.3609
391.8839111328125 0 775.74304
397.17230224609375 0 678.64685
409.4831237792969 0 601.8794
415.1813049316406 0 598.9961
429.08880615234375 0 142823.03
429.7548828125 0 593.0553
430.2421569824219 0 930.2788
433.2196044921875 0 2894.1897
442.74456787109375 0 818.5112
445.1202392578125 0 72807.27
446.2111511230469 0 903.4872
447.1007385253906 0 1117.6956
451.7507629394531 0 2205.7646
452.25341796875 0 10091.345 c 7
452.71405029296875 0 953.6229
452.7528076171875 0 7246.9375
453.25439453125 0 1529.2999
453.72784423828125 0 781.9069 y Water loss 4
455.5701904296875 0 649.6882
459.70068359375 0 761.795
459.7388000488281 0 663.90894
460.2397155761719 0 1252.6149
461.7167053222656 0 3478.037
462.2188720703125 0 2318.343 z Water loss 8
472.24713134765625 0 577.1358
475.2988586425781 0 38236.74 c 3
476.30145263671875 0 7428.944
477.30328369140625 0 1902.2572
479.2572326660156 0 662.89404
479.7581787109375 0 2423.1426
480.2563781738281 0 2656.8643
480.7536926269531 0 887.9484
481.2285461425781 0 2021.2585
492.23272705078125 0 2097.973
493.23577880859375 0 667.60345
496.24053955078125 0 1237.4817 y 8
499.76641845703125 0 3841.894
500.2653503417969 0 2106.4937 c Water loss 8
500.7618713378906 0 2653.237 c Ammonia loss 8
501.2641296386719 0 912.3691
502.27410888671875 0 3482.8538
503.2757873535156 0 845.6033
508.2514953613281 0 4715.053
508.7713928222656 0 9183.431
509.27423095703125 0 15884.66 c 8
509.77264404296875 0 8381.871
510.2721862792969 0 3158.4067
532.765869140625 0 1691.2428 z 3
533.263671875 0 1482.4001
535.2722778320312 0 600.04565
535.7675170898438 0 763.97156
544.2781372070312 0 2118.3867 c Ammonia loss 9
545.3055419921875 0 1147.3855
548.250244140625 0 1611.892
548.7510375976562 0 1217.9862
552.791015625 0 10445.035 c 9
553.2891235351562 0 8914.545
553.7883911132812 0 3871.0093
554.2869873046875 0 1770.1932
554.7841186523438 0 624.2
564.301025390625 0 879.6788
565.8013305664062 0 812.5216
566.3010864257812 0 653.6988
573.3089599609375 0 557.59406 c Ammonia loss 4
575.2835693359375 0 697.9432 y Water loss 2
576.2815551757812 0 4857.4956 z 2
576.7793579101562 0 2140.2158
577.2779541015625 0 1870.7313
584.291015625 0 11236.86 y 2
584.7904663085938 0 7319.237
585.2890014648438 0 3236.5388
585.79052734375 0 710.7745
586.823486328125 0 1321.7112
587.32275390625 0 1942.5332
590.3257446289062 0 68841.67 c 4
591.3285522460938 0 19969.29
591.8148803710938 0 931.67584
592.3281860351562 0 3635.5393
592.8087768554688 0 1044.662
600.8194580078125 0 1469.5682 c Ammonia loss 10
601.3203735351562 0 1181.7216
601.8223876953125 0 1394.0884
607.3219604492188 0 2461.0652
608.318359375 0 4875.9287 z 7
609.3327026367188 0 28522.332 c 10
609.8323364257812 0 21912.209
610.3330078125 0 8999.952
610.83251953125 0 2770.58
611.3253173828125 0 929.81573
618.3038940429688 0 1793.9595 w 1
618.8012084960938 0 1130.9858
624.3341674804688 0 5380.679 y 7
625.3389892578125 0 1578.379
630.3279418945312 0 1386.531
632.8206787109375 0 701.46295 z 1
633.3206787109375 0 1143.6749
633.8216552734375 0 1129.7373
634.8253784179688 0 692.24493
638.8462524414062 0 681.0521
640.6705932617188 0 1713.8848
640.834228515625 0 1964.506 y 1
641.332275390625 0 1488.7365
642.7754516601562 0 767.1678
647.3570556640625 0 836.95935
648.3482055664062 0 712.7569
655.31787109375 0 3378.8704
655.8131713867188 0 1713.6588
659.3455200195312 0 3647.0583
660.3475952148438 0 4446.274
660.8502197265625 0 2895.0278
661.3509521484375 0 5313.5337 z Water loss 6
661.8507690429688 0 4324.917
662.3435668945312 0 4744.578
662.8460083007812 0 1969.8053
663.3440551757812 0 983.8802
668.8330688476562 0 1751.0902
669.3455810546875 0 4914.836
669.8470458984375 0 3589.079
670.3460693359375 0 4732.0107
670.8458251953125 0 1875.3656
671.3464965820312 0 1216.3682
672.3469848632812 0 839.9532
674.34912109375 0 1714.0261
674.8427124023438 0 2456.5427
675.3448486328125 0 2010.7904
675.8383178710938 0 2636.0906
676.3323364257812 0 2043.9537
676.8377685546875 0 894.66986
678.333984375 0 1300.4984 y Ammonia loss 6
679.3535766601562 0 14974.532 z 6
680.353759765625 0 5724.3677
681.3567504882812 0 1903.5646
682.8555297851562 0 1116.9456
683.3482666015625 0 16199.575
683.8472290039062 0 16130.895
684.34619140625 0 7507.5703
684.8471069335938 0 2443.2974
685.3507690429688 0 777.7776
687.3564453125 0 968.9927
689.1571044921875 0 1367.6238
689.6580810546875 0 5183.4717
690.1571044921875 0 1934.6853
690.6597290039062 0 1078.4558
691.3569946289062 0 16191.777
691.8563232421875 0 14589.431
692.3560791015625 0 7691.4355
692.8551025390625 0 2595.5964
694.3521118164062 0 4959.847
695.3634033203125 0 4967.7827 y 6
696.3720092773438 0 997.4052
704.3685913085938 0 17223.941 c 5
705.36865234375 0 6641.778
706.3700561523438 0 1093.4624
707.8204956054688 0 799.6292
719.3424072265625 0 873.8417
731.3914794921875 0 2099.4646
732.397216796875 0 786.3865
749.38330078125 0 9131.683 w 5
750.384765625 0 4413.8286
751.3922729492188 0 1695.4772
758.3809814453125 0 1447.1465 c Ammonia loss 6
759.3870849609375 0 709.598
774.3753051757812 0 4934.818
775.404541015625 0 16004.847 c 6
776.4043579101562 0 6839.7656
777.403564453125 0 1783.4808
788.3967895507812 0 679.778
788.814697265625 0 803.9262
793.3965454101562 0 40152.258 z 5
794.3952026367188 0 20573.31
795.3929443359375 0 7046.796
796.3943481445312 0 1588.6663
810.406005859375 0 859.4812
859.4923095703125 0 973.49054
860.4872436523438 0 937.336
864.4323120117188 0 12071.185
865.4270629882812 0 5483.986
866.4314575195312 0 2533.5732
880.392822265625 0 10639.03
881.3975219726562 0 5036.7134
886.4805908203125 0 949.3477 c Ammonia loss 7
887.4714965820312 0 1044.2017
888.3787231445312 0 1085.4779
888.4859619140625 0 1138.504
889.3692626953125 0 1322.8752
893.4243774414062 0 2626.3328
894.4203491210938 0 816.0731
900.4757690429688 0 1800.6472
901.47314453125 0 1007.7012
902.4908447265625 0 6228.812
903.4998168945312 0 24580.828 c 7
904.4977416992188 0 12448.659
905.498779296875 0 2947.137
906.4094848632812 0 3479.1008
907.4088134765625 0 2042.4047 y Ammonia loss 4
908.4224243164062 0 12613.114 z 4
909.4193725585938 0 8268.034
910.4210815429688 0 2604.556
914.892333984375 0 919.37415
919.4735107421875 0 1443.9348
920.4690551757812 0 1177.5458
922.4249267578125 0 1675.9652
922.8921508789062 0 1030.7476
923.4165649414062 0 2695.6514
924.4303588867188 0 1152.63 y 4
925.4360961914062 0 745.79645
938.3497314453125 0 1718.2015
939.3548583984375 0 3421.1716
940.3558959960938 0 1669.5518
986.5012817382812 0 806.90985
987.512451171875 0 1196.8868
988.507080078125 0 928.6245
999.5377807617188 0 1091.4467 c Water loss 8
1000.5113525390625 0 1187.0233 c Ammonia loss 8
1001.5179443359375 0 855.16364
1002.512451171875 0 911.60614
1016.53369140625 0 2281.265
1017.5423583984375 0 14532.702 c 8
1018.5421142578125 0 8917.661
1019.5419921875 0 3107.5781
1020.551025390625 0 724.6062
1021.5314331054688 0 846.065
1060.5567626953125 0 894.23505
1061.5625 0 957.7434
1064.523681640625 0 5303.892 z 3
1065.5255126953125 0 6375.0093
1066.5257568359375 0 4434.977
1067.5224609375 0 1101.7338
1088.5599365234375 0 1143.9932
1104.575439453125 0 4094.8823 c 9
1105.5760498046875 0 2869.9172
1106.5745849609375 0 1767.4275
1107.568115234375 0 1117.491
1135.5504150390625 0 696.2297
1151.5577392578125 0 3160.2737 z 2
1152.5609130859375 0 6591.86
1153.56005859375 0 4965.7637
1154.560302734375 0 1393.2434
1155.5672607421875 0 843.576
1173.6534423828125 0 1113.4744
1174.64794921875 0 2281.064
1175.651123046875 0 2265.5374
1201.6435546875 0 3377.3613
1202.640625 0 2213.3418
1203.6500244140625 0 823.4138
1217.6610107421875 0 2941.5347 c 10
1218.663818359375 0 6354.114
1219.662109375 0 4120.547
1220.66162109375 0 1362.8358
1265.64794921875 0 1246.3029
1266.644287109375 0 927.5351
1267.6546630859375 0 733.39667
1305.668701171875 0 948.3293
1320.685546875 0 948.69147
1321.69677734375 0 2216.9531
1322.695068359375 0 889.8195
1323.67822265625 0 2478.6929
1324.6817626953125 0 5077.631
1325.68310546875 0 3873.0312
1326.6798095703125 0 1791.6105
1327.6953125 0 783.89136
1337.7044677734375 0 1192.6693
1338.7010498046875 0 6022.576
1339.6995849609375 0 5676.283
1340.699462890625 0 3536.2385
1341.694091796875 0 1528.164
1348.6885986328125 0 2133.3489
1349.6734619140625 0 3568.8147
1350.6766357421875 0 2619.9956
1351.6788330078125 0 1301.8192
1352.6846923828125 0 907.3782
1365.6888427734375 0 4837.3003
1366.6937255859375 0 24785.188
1367.6932373046875 0 20506.924
1368.6925048828125 0 11093.197
1369.693603515625 0 3858.9006
1370.69775390625 0 1103.518
1378.3096923828125 0 1685.9846
1379.318115234375 0 3325.2004
1380.316650390625 0 2582.0105
1381.707275390625 0 881.1983
1382.7086181640625 0 3662.7856
1383.7169189453125 0 6332.091
1384.7135009765625 0 4592.8125
1385.7225341796875 0 2361.005
1386.731201171875 0 909.6961
1766.7578125 0 673.26874
2497.347900390625 0 727.50665
3084.1533203125 0 792.8885
3285.318359375 0 765.8198

Spectrum Details

|  |  |
| --- | --- |
| Matched peaks? Matched peaks The total absolute number of peaks matched. Additionally in brackets the total fraction of peaks matched and the total number of peaks is shown. | 50 (13.66% of 366) |
| FDR? FDR The false discovery rate estimated for this peptide. It is calculated by matching all theoretical fragments with a non-integer shift with the raw peaks for this spectrum. This is done with 40 different shifts. The resulting percentage is the average number of annotated peaks over the number of annotated peaks with the correct spectrum. | 1.48% |
| Satellite FDR? Satellite FDR See the FDR for details on its calculation. This satellite ion specific FDR only contains the satellite ions (d/w) for I/L/J positions. | 2.38% |
| PSM Score? PSM Score The PSM Score as given by Hecklib to this annotated spectrum. It is shown with three significant figures. | 375 |

#### Spectrum 3162? Spectrum 3162 The raw spectrum of this peptide as annotated by Hecklib. The fragments are coloured according to ion type (see legend). Any peaks with a star '\*' as text can be hovered over to see the full details, first the ion type second the mass shift type. By hovering over the amino acids in the peptide or ions in the legend the corresponding peaks are highlighted. By toggling the 'Unassigned' label you can turn the background (unassigned) peaks on or off in the plot. By updating the slider in the Ion legend you can update the spectrum to only show the top X% of the peaks with labels. The top X% means any peak that is within X% of the highest intensity. By dragging in the spectrum you can zoom in to a specific part of the spectrum and use 'Zoom Out' to get back to the original zoom level. The annotation of the spectrum is based on the given sequence in the peptides file and is done with different software so inconsistencies are likely. The peaks are annotated based on the given sequence, with 20 ppm tolerance.

Copy Data

##### Spectrum 3162 (TSV)

###### Preview

```
Loading example...
```

*Click on the button to copy the data to your clipboard.*

Mz MinMz MaxIntensity Max

WidthHeightPeptide font sizePeptide stroke widthSpectrum font sizeSpectrum stroke widthCompact peptide

Ion legend

wxyz

abcd

OtherUnassignedIon^Charge^_Position_Show for top:%

TISRDNAKNSJY

03.93e+47.85e+41.18e+51.57e+5

Zoom Out

y^+1^_1_y^+1^_2_c^+1^_3_y^+1^_3_c^+2^_8_c^+1^_4_y^+1^_4_c^+2^_9_c^+2^_9_c^+2^_9_z^+2^_9_y^+2^_9_c^+2^_10_z^+2^_10_y^+2^_10_z^+2^_10_y^+2^_10_c^+1^_5_c^+2^_11_z^+1^_5_c^+2^_11_w^+2^_11_y^+1^_5_z^+2^_11_y^+2^_11_z^+1^_6_z^+1^_6_c^+1^_6_y^+1^_6_c^+1^_6_w^+1^_7_c^+1^_7_c^+1^_7_z^+1^_7_y^+1^_7_c^+1^_8_c^+1^_8_z^+1^_8_y^+1^_8_c^+1^_9_z^+1^_9_c^+1^_10_z^+1^_10_c^+1^_11_

039579011851580

Fragment Matches Table

Show background peaks

| Position | Ion type | Intensity | mz Theoretical | mz Error (Th) | mz Error (ppm) | Charge | Series Number |
| --- | --- | --- | --- | --- | --- | --- | --- |
| - | - | 454.9 | 129.1 | - | - | 0 | - |
| - | - | 438.1 | 131.7 | - | - | 0 | - |
| - | - | 612.6 | 133.1 | - | - | 0 | - |
| - | - | 1638 | 136.1 | - | - | 0 | - |
| - | - | 476.9 | 148.3 | - | - | 0 | - |
| - | - | 976 | 149 | - | - | 0 | - |
| - | - | 581.9 | 149 | - | - | 0 | - |
| - | - | 6468 | 149 | - | - | 0 | - |
| - | - | 662.1 | 150 | - | - | 0 | - |
| - | - | 438.2 | 157.4 | - | - | 0 | - |
| - | - | 384.8 | 159 | - | - | 0 | - |
| - | - | 383.8 | 163.1 | - | - | 0 | - |
| - | - | 1241 | 165.1 | - | - | 0 | - |
| - | - | 2691 | 167.1 | - | - | 0 | - |
| - | - | 421.4 | 171.8 | - | - | 0 | - |
| - | - | 4321 | 173.1 | - | - | 0 | - |
| - | - | 1821 | 173.4 | - | - | 0 | - |
| - | - | 460.4 | 181.9 | - | - | 0 | - |
| 12 | y | 4769 | 182.1 | 2.344E-05 | 0.1287 | +1 | 1 |
| - | - | 656.4 | 183.1 | - | - | 0 | - |
| - | - | 1.666E+04 | 187.1 | - | - | 0 | - |
| - | - | 1773 | 188.1 | - | - | 0 | - |
| - | - | 439.3 | 194.4 | - | - | 0 | - |
| - | - | 529.3 | 203.1 | - | - | 0 | - |
| - | - | 3626 | 215.1 | - | - | 0 | - |
| - | - | 3379 | 221.1 | - | - | 0 | - |
| - | - | 1570 | 222.1 | - | - | 0 | - |
| - | - | 1965 | 223.1 | - | - | 0 | - |
| - | - | 1053 | 223.1 | - | - | 0 | - |
| - | - | 2610 | 225 | - | - | 0 | - |
| - | - | 1168 | 226 | - | - | 0 | - |
| - | - | 4795 | 239.1 | - | - | 0 | - |
| - | - | 533.5 | 239.9 | - | - | 0 | - |
| - | - | 1512 | 240.1 | - | - | 0 | - |
| - | - | 780 | 241.1 | - | - | 0 | - |
| - | - | 687.5 | 244.1 | - | - | 0 | - |
| - | - | 2214 | 259.1 | - | - | 0 | - |
| - | - | 1829 | 260.1 | - | - | 0 | - |
| - | - | 1779 | 282.1 | - | - | 0 | - |
| - | - | 937.7 | 283 | - | - | 0 | - |
| - | - | 691.1 | 286 | - | - | 0 | - |
| - | - | 2930 | 295.1 | - | - | 0 | - |
| 11 | y | 648.8 | 295.2 | 0.0001336 | 0.4525 | +1 | 2 |
| - | - | 2686 | 296.1 | - | - | 0 | - |
| - | - | 1299 | 297.1 | - | - | 0 | - |
| - | - | 2323 | 299.1 | - | - | 0 | - |
| - | - | 2549 | 300.1 | - | - | 0 | - |
| - | - | 3281 | 300.2 | - | - | 0 | - |
| - | - | 1351 | 301.1 | - | - | 0 | - |
| - | - | 731 | 315.2 | - | - | 0 | - |
| - | - | 1536 | 317.2 | - | - | 0 | - |
| 3 | c | 853 | 319.2 | 0.001134 | 3.553 | +1 | 3 |
| - | - | 553.3 | 340.7 | - | - | 0 | - |
| - | - | 1.672E+04 | 341 | - | - | 0 | - |
| - | - | 598.2 | 343 | - | - | 0 | - |
| - | - | 1764 | 355.1 | - | - | 0 | - |
| - | - | 821 | 356.1 | - | - | 0 | - |
| - | - | 6.123E+04 | 359 | - | - | 0 | - |
| - | - | 483.9 | 360 | - | - | 0 | - |
| - | - | 1617 | 369.1 | - | - | 0 | - |
| - | - | 3164 | 370.1 | - | - | 0 | - |
| - | - | 592.3 | 370.8 | - | - | 0 | - |
| - | - | 3448 | 371.1 | - | - | 0 | - |
| - | - | 714.6 | 372.1 | - | - | 0 | - |
| - | - | 534.9 | 372.7 | - | - | 0 | - |
| - | - | 2545 | 375.9 | - | - | 0 | - |
| - | - | 1546 | 376.8 | - | - | 0 | - |
| - | - | 651.2 | 376.9 | - | - | 0 | - |
| 10 | y | 619.7 | 382.2 | 0.001919 | 5.021 | +1 | 3 |
| - | - | 3483 | 385.2 | - | - | 0 | - |
| - | - | 3868 | 386.2 | - | - | 0 | - |
| - | - | 1015 | 387.2 | - | - | 0 | - |
| - | - | 589 | 388.2 | - | - | 0 | - |
| - | - | 809.9 | 391.6 | - | - | 0 | - |
| - | - | 1273 | 393.9 | - | - | 0 | - |
| - | - | 671.3 | 400.3 | - | - | 0 | - |
| - | - | 1485 | 407.2 | - | - | 0 | - |
| - | - | 593.2 | 407.9 | - | - | 0 | - |
| - | - | 1600 | 425.2 | - | - | 0 | - |
| - | - | 566.9 | 426.2 | - | - | 0 | - |
| - | - | 1.555E+05 | 429.1 | - | - | 0 | - |
| - | - | 4644 | 430.2 | - | - | 0 | - |
| - | - | 773.9 | 442.7 | - | - | 0 | - |
| - | - | 617 | 442.8 | - | - | 0 | - |
| - | - | 8.361E+04 | 445.1 | - | - | 0 | - |
| - | - | 1028 | 446.2 | - | - | 0 | - |
| - | - | 1061 | 447.1 | - | - | 0 | - |
| - | - | 4723 | 451.7 | - | - | 0 | - |
| 8 | c | 1.24E+04 | 452.3 | 0.0005476 | 1.211 | +2 | 8 |
| - | - | 4533 | 452.8 | - | - | 0 | - |
| - | - | 779.1 | 453.3 | - | - | 0 | - |
| - | - | 1197 | 461.2 | - | - | 0 | - |
| - | - | 810 | 461.6 | - | - | 0 | - |
| - | - | 648.1 | 461.8 | - | - | 0 | - |
| - | - | 1430 | 462.2 | - | - | 0 | - |
| 4 | c | 3.597E+04 | 475.3 | 0.0001235 | 0.2598 | +1 | 4 |
| - | - | 8786 | 476.3 | - | - | 0 | - |
| - | - | 1234 | 477.3 | - | - | 0 | - |
| - | - | 3509 | 479.8 | - | - | 0 | - |
| - | - | 1267 | 480.3 | - | - | 0 | - |
| - | - | 2084 | 481.2 | - | - | 0 | - |
| - | - | 803.3 | 487.3 | - | - | 0 | - |
| - | - | 866.2 | 487.8 | - | - | 0 | - |
| - | - | 1066 | 493.8 | - | - | 0 | - |
| - | - | 652.5 | 494.3 | - | - | 0 | - |
| - | - | 565.9 | 495.8 | - | - | 0 | - |
| 9 | y | 1807 | 496.2 | 0.0001974 | 0.3979 | +1 | 4 |
| - | - | 559.9 | 497.4 | - | - | 0 | - |
| - | - | 2851 | 499.8 | - | - | 0 | - |
| 9 | c | 1127 | 500.3 | 0.003057 | 6.111 | +2 | 9 |
| 9 | c | 1049 | 500.8 | 0.0006016 | 1.201 | +2 | 9 |
| - | - | 668.3 | 501.8 | - | - | 0 | - |
| - | - | 6198 | 502.3 | - | - | 0 | - |
| - | - | 1642 | 503.3 | - | - | 0 | - |
| - | - | 5582 | 508.8 | - | - | 0 | - |
| 9 | c | 9060 | 509.3 | 0.001229 | 2.413 | +2 | 9 |
| - | - | 4176 | 509.8 | - | - | 0 | - |
| - | - | 1079 | 510.3 | - | - | 0 | - |
| - | - | 575.6 | 522.3 | - | - | 0 | - |
| 4 | z | 1536 | 532.8 | 0.002717 | 5.099 | +2 | 9 |
| - | - | 537.4 | 534.3 | - | - | 0 | - |
| 4 | y | 801.7 | 540.8 | 0.0009701 | 1.794 | +2 | 9 |
| - | - | 746.9 | 544.8 | - | - | 0 | - |
| - | - | 1176 | 545.3 | - | - | 0 | - |
| - | - | 630.6 | 550.8 | - | - | 0 | - |
| - | - | 692.7 | 551.3 | - | - | 0 | - |
| 10 | c | 2.318E+04 | 552.8 | 0.0004279 | 0.7741 | +2 | 10 |
| - | - | 1.095E+04 | 553.3 | - | - | 0 | - |
| - | - | 4257 | 553.8 | - | - | 0 | - |
| - | - | 2654 | 554.3 | - | - | 0 | - |
| - | - | 1433 | 554.8 | - | - | 0 | - |
| - | - | 671.6 | 559.8 | - | - | 0 | - |
| - | - | 562.8 | 564.3 | - | - | 0 | - |
| - | - | 1014 | 565.8 | - | - | 0 | - |
| - | - | 1182 | 566.8 | - | - | 0 | - |
| 3 | z | 1371 | 567.8 | 0.01017 | 17.91 | +2 | 10 |
| - | - | 582 | 568.3 | - | - | 0 | - |
| - | - | 605.5 | 568.8 | - | - | 0 | - |
| - | - | 844 | 570.3 | - | - | 0 | - |
| - | - | 823.4 | 574.8 | - | - | 0 | - |
| 3 | y | 1077 | 575.8 | 0.002794 | 4.853 | +2 | 10 |
| 3 | z | 6338 | 576.3 | 0.0007864 | 1.365 | +2 | 10 |
| - | - | 3664 | 576.8 | - | - | 0 | - |
| - | - | 1465 | 577.3 | - | - | 0 | - |
| - | - | 652.5 | 579 | - | - | 0 | - |
| 3 | y | 2.417E+04 | 584.3 | 0.0003218 | 0.5507 | +2 | 10 |
| - | - | 1.454E+04 | 584.8 | - | - | 0 | - |
| - | - | 4549 | 585.3 | - | - | 0 | - |
| - | - | 1278 | 585.8 | - | - | 0 | - |
| - | - | 1124 | 586.8 | - | - | 0 | - |
| - | - | 952.1 | 587.3 | - | - | 0 | - |
| - | - | 598.5 | 587.8 | - | - | 0 | - |
| - | - | 983.5 | 588.8 | - | - | 0 | - |
| - | - | 651.8 | 589.3 | - | - | 0 | - |
| 5 | c | 1.053E+05 | 590.3 | 0.000211 | 0.3575 | +1 | 5 |
| - | - | 3.142E+04 | 591.3 | - | - | 0 | - |
| - | - | 755 | 591.8 | - | - | 0 | - |
| - | - | 6307 | 592.3 | - | - | 0 | - |
| - | - | 772.8 | 593.3 | - | - | 0 | - |
| 11 | c | 2229 | 600.8 | 0.0001726 | 0.2872 | +2 | 11 |
| - | - | 796.5 | 601.3 | - | - | 0 | - |
| - | - | 807.3 | 607.3 | - | - | 0 | - |
| 8 | z | 3144 | 608.3 | 9.986E-05 | 0.1642 | +1 | 5 |
| 11 | c | 4.151E+04 | 609.3 | 0.0005898 | 0.9679 | +2 | 11 |
| - | - | 1.991E+04 | 609.8 | - | - | 0 | - |
| - | - | 802.1 | 609.9 | - | - | 0 | - |
| - | - | 7945 | 610.3 | - | - | 0 | - |
| - | - | 2406 | 610.8 | - | - | 0 | - |
| - | - | 815.8 | 616.3 | - | - | 0 | - |
| 2 | w | 2223 | 618.3 | 0.001158 | 1.873 | +2 | 11 |
| - | - | 1023 | 618.8 | - | - | 0 | - |
| - | - | 652.5 | 619.3 | - | - | 0 | - |
| 8 | y | 6323 | 624.3 | 0.00129 | 2.067 | +1 | 5 |
| - | - | 880 | 624.8 | - | - | 0 | - |
| - | - | 2286 | 625.3 | - | - | 0 | - |
| - | - | 592.8 | 626.3 | - | - | 0 | - |
| - | - | 941.6 | 630.3 | - | - | 0 | - |
| - | - | 561.9 | 631.3 | - | - | 0 | - |
| 2 | z | 774.4 | 632.8 | 0.004549 | 7.189 | +2 | 11 |
| - | - | 966.8 | 633.3 | - | - | 0 | - |
| - | - | 831.2 | 633.8 | - | - | 0 | - |
| - | - | 918 | 635.4 | - | - | 0 | - |
| - | - | 2243 | 636.3 | - | - | 0 | - |
| - | - | 603.6 | 639.3 | - | - | 0 | - |
| 2 | y | 2595 | 640.8 | 0.000554 | 0.8644 | +2 | 11 |
| - | - | 1691 | 641.3 | - | - | 0 | - |
| - | - | 737.7 | 641.8 | - | - | 0 | - |
| - | - | 870.7 | 642.3 | - | - | 0 | - |
| - | - | 1841 | 647.4 | - | - | 0 | - |
| - | - | 1651 | 653.3 | - | - | 0 | - |
| - | - | 937.1 | 654.3 | - | - | 0 | - |
| - | - | 1530 | 655.3 | - | - | 0 | - |
| - | - | 1365 | 655.8 | - | - | 0 | - |
| - | - | 720.5 | 656.3 | - | - | 0 | - |
| - | - | 5512 | 659.3 | - | - | 0 | - |
| - | - | 5272 | 660.3 | - | - | 0 | - |
| - | - | 3822 | 660.9 | - | - | 0 | - |
| 7 | z | 6055 | 661.3 | 0.009134 | 13.81 | +1 | 6 |
| - | - | 5216 | 661.9 | - | - | 0 | - |
| - | - | 5111 | 662.3 | - | - | 0 | - |
| - | - | 2349 | 662.8 | - | - | 0 | - |
| - | - | 933.5 | 663.3 | - | - | 0 | - |
| - | - | 2460 | 668.8 | - | - | 0 | - |
| - | - | 5756 | 669.3 | - | - | 0 | - |
| - | - | 4078 | 669.9 | - | - | 0 | - |
| - | - | 6214 | 670.3 | - | - | 0 | - |
| - | - | 1723 | 670.8 | - | - | 0 | - |
| - | - | 836.4 | 671.3 | - | - | 0 | - |
| - | - | 1215 | 672.3 | - | - | 0 | - |
| - | - | 724.8 | 673.3 | - | - | 0 | - |
| - | - | 1370 | 674.3 | - | - | 0 | - |
| - | - | 3475 | 674.8 | - | - | 0 | - |
| - | - | 3543 | 675.3 | - | - | 0 | - |
| - | - | 2467 | 675.8 | - | - | 0 | - |
| - | - | 1466 | 676.3 | - | - | 0 | - |
| 7 | z | 2.105E+04 | 679.4 | 0.0004539 | 0.6681 | +1 | 6 |
| - | - | 7132 | 680.4 | - | - | 0 | - |
| - | - | 2387 | 681.4 | - | - | 0 | - |
| - | - | 2325 | 682.9 | - | - | 0 | - |
| - | - | 2.459E+04 | 683.3 | - | - | 0 | - |
| - | - | 1.69E+04 | 683.8 | - | - | 0 | - |
| - | - | 7055 | 684.3 | - | - | 0 | - |
| - | - | 1759 | 684.9 | - | - | 0 | - |
| 6 | c | 1120 | 687.3 | 0.006604 | 9.608 | +1 | 6 |
| - | - | 2.413E+04 | 691.4 | - | - | 0 | - |
| - | - | 1.662E+04 | 691.9 | - | - | 0 | - |
| - | - | 414.5 | 691.9 | - | - | 0 | - |
| - | - | 1.647E+04 | 692.4 | - | - | 0 | - |
| - | - | 912.7 | 692.8 | - | - | 0 | - |
| - | - | 8788 | 692.9 | - | - | 0 | - |
| - | - | 3547 | 693.4 | - | - | 0 | - |
| - | - | 873.5 | 693.9 | - | - | 0 | - |
| 7 | y | 4027 | 695.4 | 0.001173 | 1.686 | +1 | 6 |
| - | - | 1939 | 696.4 | - | - | 0 | - |
| 6 | c | 2.65E+04 | 704.4 | 0.0003528 | 0.5009 | +1 | 6 |
| - | - | 7568 | 705.4 | - | - | 0 | - |
| - | - | 2345 | 706.4 | - | - | 0 | - |
| - | - | 880.1 | 708.3 | - | - | 0 | - |
| - | - | 3625 | 731.4 | - | - | 0 | - |
| - | - | 738 | 732.4 | - | - | 0 | - |
| - | - | 1182 | 737.3 | - | - | 0 | - |
| 6 | w | 1.412E+04 | 749.4 | 0.002911 | 3.885 | +1 | 7 |
| - | - | 6490 | 750.4 | - | - | 0 | - |
| - | - | 1444 | 751.4 | - | - | 0 | - |
| - | - | 675.5 | 754.4 | - | - | 0 | - |
| 7 | c | 878.5 | 758.4 | 0.007637 | 10.07 | +1 | 7 |
| 7 | c | 2.232E+04 | 775.4 | 0.0008455 | 1.09 | +1 | 7 |
| - | - | 8027 | 776.4 | - | - | 0 | - |
| - | - | 1994 | 777.4 | - | - | 0 | - |
| 6 | z | 6.242E+04 | 793.4 | 0.0005957 | 0.7508 | +1 | 7 |
| - | - | 2.603E+04 | 794.4 | - | - | 0 | - |
| - | - | 6172 | 795.4 | - | - | 0 | - |
| - | - | 937.9 | 796.4 | - | - | 0 | - |
| 6 | y | 1021 | 809.4 | 0.00162 | 2.001 | +1 | 7 |
| - | - | 692.8 | 832.4 | - | - | 0 | - |
| - | - | 2812 | 859.5 | - | - | 0 | - |
| - | - | 1771 | 860.5 | - | - | 0 | - |
| - | - | 612.9 | 862.5 | - | - | 0 | - |
| - | - | 1.667E+04 | 864.4 | - | - | 0 | - |
| - | - | 6479 | 865.4 | - | - | 0 | - |
| - | - | 2904 | 866.4 | - | - | 0 | - |
| - | - | 878.6 | 884.5 | - | - | 0 | - |
| 8 | c | 2237 | 886.5 | 0.000778 | 0.8776 | +1 | 8 |
| - | - | 1213 | 894.5 | - | - | 0 | - |
| - | - | 2254 | 900.5 | - | - | 0 | - |
| - | - | 1074 | 901.5 | - | - | 0 | - |
| - | - | 5903 | 902.5 | - | - | 0 | - |
| 8 | c | 2.493E+04 | 903.5 | 0.001021 | 1.13 | +1 | 8 |
| - | - | 1.027E+04 | 904.5 | - | - | 0 | - |
| - | - | 2762 | 905.5 | - | - | 0 | - |
| 5 | z | 1.108E+04 | 908.4 | 0.0009274 | 1.021 | +1 | 8 |
| - | - | 5495 | 909.4 | - | - | 0 | - |
| - | - | 1675 | 910.4 | - | - | 0 | - |
| - | - | 659 | 911.4 | - | - | 0 | - |
| - | - | 1467 | 922.5 | - | - | 0 | - |
| 5 | y | 1927 | 924.4 | 0.002073 | 2.243 | +1 | 8 |
| - | - | 790.9 | 973.5 | - | - | 0 | - |
| - | - | 1598 | 987.5 | - | - | 0 | - |
| - | - | 611.5 | 988.5 | - | - | 0 | - |
| 9 | c | 9668 | 1018 | 0.0006745 | 0.6628 | +1 | 9 |
| - | - | 3717 | 1019 | - | - | 0 | - |
| - | - | 1466 | 1020 | - | - | 0 | - |
| 4 | z | 5432 | 1065 | 0.001208 | 1.135 | +1 | 9 |
| - | - | 6034 | 1066 | - | - | 0 | - |
| - | - | 1602 | 1067 | - | - | 0 | - |
| - | - | 2151 | 1089 | - | - | 0 | - |
| - | - | 1329 | 1090 | - | - | 0 | - |
| 10 | c | 8951 | 1105 | 0.001392 | 1.26 | +1 | 10 |
| - | - | 5858 | 1106 | - | - | 0 | - |
| - | - | 2430 | 1107 | - | - | 0 | - |
| - | - | 710.1 | 1138 | - | - | 0 | - |
| 3 | z | 2522 | 1152 | 0.006625 | 5.753 | +1 | 10 |
| - | - | 9210 | 1153 | - | - | 0 | - |
| - | - | 3871 | 1154 | - | - | 0 | - |
| - | - | 1650 | 1155 | - | - | 0 | - |
| - | - | 759.6 | 1158 | - | - | 0 | - |
| - | - | 1837 | 1174 | - | - | 0 | - |
| - | - | 2216 | 1175 | - | - | 0 | - |
| - | - | 1408 | 1176 | - | - | 0 | - |
| - | - | 3948 | 1202 | - | - | 0 | - |
| - | - | 1953 | 1203 | - | - | 0 | - |
| - | - | 746.7 | 1204 | - | - | 0 | - |
| 11 | c | 4653 | 1218 | 0.002814 | 2.311 | +1 | 11 |
| - | - | 7616 | 1219 | - | - | 0 | - |
| - | - | 3698 | 1220 | - | - | 0 | - |
| - | - | 1525 | 1221 | - | - | 0 | - |
| - | - | 2173 | 1266 | - | - | 0 | - |
| - | - | 1031 | 1306 | - | - | 0 | - |
| - | - | 918.6 | 1307 | - | - | 0 | - |
| - | - | 1495 | 1321 | - | - | 0 | - |
| - | - | 1788 | 1322 | - | - | 0 | - |
| - | - | 1428 | 1323 | - | - | 0 | - |
| - | - | 3803 | 1324 | - | - | 0 | - |
| - | - | 4829 | 1325 | - | - | 0 | - |
| - | - | 3523 | 1326 | - | - | 0 | - |
| - | - | 1048 | 1327 | - | - | 0 | - |
| - | - | 1088 | 1338 | - | - | 0 | - |
| - | - | 9692 | 1339 | - | - | 0 | - |
| - | - | 6983 | 1340 | - | - | 0 | - |
| - | - | 2972 | 1341 | - | - | 0 | - |
| - | - | 1032 | 1342 | - | - | 0 | - |
| - | - | 875.2 | 1347 | - | - | 0 | - |
| - | - | 1967 | 1349 | - | - | 0 | - |
| - | - | 3428 | 1350 | - | - | 0 | - |
| - | - | 2801 | 1351 | - | - | 0 | - |
| - | - | 1111 | 1352 | - | - | 0 | - |
| - | - | 755.9 | 1357 | - | - | 0 | - |
| - | - | 660 | 1364 | - | - | 0 | - |
| - | - | 5893 | 1366 | - | - | 0 | - |
| - | - | 3.182E+04 | 1367 | - | - | 0 | - |
| - | - | 2.023E+04 | 1368 | - | - | 0 | - |
| - | - | 8307 | 1369 | - | - | 0 | - |
| - | - | 2055 | 1370 | - | - | 0 | - |
| - | - | 1471 | 1382 | - | - | 0 | - |
| - | - | 4870 | 1383 | - | - | 0 | - |
| - | - | 6289 | 1384 | - | - | 0 | - |
| - | - | 8544 | 1385 | - | - | 0 | - |
| - | - | 4555 | 1386 | - | - | 0 | - |
| - | - | 1752 | 1387 | - | - | 0 | - |
| - | - | 912.9 | 1388 | - | - | 0 | - |
| - | - | 763.2 | 1564 | - | - | 0 | - |

m/z Charge Intensity FragmentType MassShift Position
129.10218811035156 0 454.92654
131.67967224121094 0 438.13025
133.06089782714844 0 612.5828
136.07574462890625 0 1637.89
148.271728515625 0 476.91013
148.95460510253906 0 975.9797
149.03941345214844 0 581.864
149.04483032226562 0 6468.1216
150.04400634765625 0 662.1343
157.4095458984375 0 438.20322
158.9988250732422 0 384.83163
163.1098175048828 0 383.75098
165.05455017089844 0 1241.076
167.05535888671875 0 2690.5854
171.84585571289062 0 421.37628
173.09202575683594 0 4320.6904
173.43934631347656 0 1821.1348
181.92010498046875 0 460.44836
182.08114624023438 0 4768.784 y 11
183.0844268798828 0 656.3619
187.14401245117188 0 16664.803
188.147216796875 0 1772.5143
194.41455078125 0 439.27066
203.09188842773438 0 529.30914
215.13885498046875 0 3625.6975
221.084228515625 0 3378.925
222.08518981933594 0 1570.0087
223.06304931640625 0 1964.8872
223.081298828125 0 1053.0199
225.04283142089844 0 2610.3572
226.04286193847656 0 1168.3723
239.09481811523438 0 4794.7314
239.9251708984375 0 533.4719
240.09576416015625 0 1512.4059
241.09165954589844 0 780.0386
244.09422302246094 0 687.54
259.11724853515625 0 2213.8945
260.1234436035156 0 1828.8607
282.14471435546875 0 1778.8201
283.04779052734375 0 937.7255
286.0099182128906 0 691.13416
295.10308837890625 0 2929.9644
295.16510009765625 0 648.793 y 10
296.104248046875 0 2686.2666
297.09954833984375 0 1299.1427
299.0619201660156 0 2322.8506
300.06201171875 0 2548.867
300.15545654296875 0 3281.2002
301.0593566894531 0 1350.8282
315.1663513183594 0 731.00433
317.1817932128906 0 1535.5508
319.19873046875 0 853.0037 c 2
340.7305603027344 0 553.32025
341.0179138183594 0 16719.428
342.995849609375 0 598.1858
355.069091796875 0 1763.9622
356.07061767578125 0 821.01355
359.028076171875 0 61234.965
359.98193359375 0 483.87332
369.1219787597656 0 1616.742
370.12286376953125 0 3164.2031
370.80914306640625 0 592.3137
371.11932373046875 0 3447.594
372.1195983886719 0 714.56134
372.6959228515625 0 534.9288
375.8594055175781 0 2544.8687
376.8441162109375 0 1545.724
376.89971923828125 0 651.21313
382.1953430175781 0 619.70825 y 9
385.2194519042969 0 3483.0066
386.2259826660156 0 3868.156
387.2290954589844 0 1014.8586
388.2298889160156 0 588.97986
391.5505065917969 0 809.9285
393.86932373046875 0 1273.4794
400.2557678222656 0 671.3183
407.1914367675781 0 1485.4819
407.8541259765625 0 593.15204
425.20062255859375 0 1599.6702
426.20501708984375 0 566.8549
429.08868408203125 0 155512.4
430.2409973144531 0 4644.0264
442.74481201171875 0 773.93744
442.83270263671875 0 617.0139
445.12005615234375 0 83611.234
446.21246337890625 0 1027.7357
447.09912109375 0 1060.8696
451.7498779296875 0 4723.122
452.25341796875 0 12399.921 c 7
452.7555236816406 0 4533.4697
453.2580261230469 0 779.0598
461.2443542480469 0 1197.1501
461.5771484375 0 809.96387
461.7500305175781 0 648.0865
462.2301940917969 0 1429.6445
475.298583984375 0 35969.793 c 3
476.30126953125 0 8786.467
477.3021545410156 0 1234.15
479.7586975097656 0 3508.7732
480.2618103027344 0 1267.1848
481.2283935546875 0 2084.142
487.2651062011719 0 803.2734
487.7705383300781 0 866.1843
493.7545166015625 0 1066.1943
494.25384521484375 0 652.5408
495.8298034667969 0 565.91394
496.2403869628906 0 1807.0979 y 8
497.3872375488281 0 559.9202
499.7669372558594 0 2850.6506
500.26708984375 0 1127.3608 c Water loss 8
500.76275634765625 0 1049.3586 c Ammonia loss 8
501.7674865722656 0 668.3303
502.274658203125 0 6197.813
503.2749938964844 0 1642.1589
508.77166748046875 0 5582.3926
509.2742004394531 0 9060.253 c 8
509.7757263183594 0 4175.766
510.27899169921875 0 1079.0005
522.27880859375 0 575.6293
532.76318359375 0 1536.4797 z 3
534.3031616210938 0 537.4296
540.7742919921875 0 801.7199 y 3
544.7807006835938 0 746.87665
545.3042602539062 0 1175.8118
550.80126953125 0 630.585
551.3015747070312 0 692.6976
552.791015625 0 23177.98 c 9
553.2924194335938 0 10952.318
553.7933959960938 0 4256.873
554.2910766601562 0 2654.2595
554.7796020507812 0 1432.6162
559.8023071289062 0 671.5538
564.3204956054688 0 562.8395
565.7962036132812 0 1013.88995
566.7726440429688 0 1181.5162
567.77880859375 0 1371.2196 z Ammonia loss 2
568.276611328125 0 582.0404
568.7815551757812 0 605.4987
570.306396484375 0 843.9682
574.7952880859375 0 823.41907
575.7752075195312 0 1076.5815 y Ammonia loss 2
576.2811279296875 0 6337.952 z 2
576.7826538085938 0 3663.5298
577.2835693359375 0 1465.477
579.012451171875 0 652.5241
584.2909545898438 0 24173.76 y 2
584.7919921875 0 14537.669
585.29345703125 0 4549.067
585.7947998046875 0 1277.6028
586.8192138671875 0 1124.4354
587.327880859375 0 952.0968
587.82861328125 0 598.4987
588.777587890625 0 983.49744
589.3163452148438 0 651.84656
590.325439453125 0 105268.67 c 4
591.328125 0 31420.76
591.8123779296875 0 755.03046
592.328369140625 0 6306.6357
593.3316650390625 0 772.7932
600.8203735351562 0 2228.896 c Ammonia loss 10
601.3214721679688 0 796.4913
607.3320922851562 0 807.2879
608.3165283203125 0 3143.9924 z 7
609.3328857421875 0 41510.15 c 10
609.8345947265625 0 19907.25
609.8916015625 0 802.12866
610.3352661132812 0 7945.2563
610.8379516601562 0 2405.5896
616.3466186523438 0 815.7787
618.3055419921875 0 2223.2449 w 1
618.8051147460938 0 1023.4909
619.3057250976562 0 652.49005
624.3338623046875 0 6323.149 y 7
624.8124389648438 0 879.95056
625.3360595703125 0 2285.5337
626.3423461914062 0 592.8455
630.328125 0 941.6262
631.330078125 0 561.9042
632.8193969726562 0 774.39215 z 1
633.3214721679688 0 966.78503
633.825439453125 0 831.2298
635.3636474609375 0 918.0245
636.3485107421875 0 2242.6956
639.3489990234375 0 603.564
640.8338623046875 0 2594.5244 y 1
641.330810546875 0 1690.5337
641.8299560546875 0 737.7186
642.334228515625 0 870.68805
647.3505859375 0 1841.4944
653.3428344726562 0 1650.7886
654.3211669921875 0 937.1357
655.3179321289062 0 1529.9971
655.8189697265625 0 1364.571
656.3208618164062 0 720.5111
659.34619140625 0 5512.0967
660.34814453125 0 5271.688
660.8506469726562 0 3822.0054
661.3521118164062 0 6054.9756 z Water loss 6
661.8509521484375 0 5215.551
662.34814453125 0 5111.3066
662.8460083007812 0 2348.9824
663.3462524414062 0 933.4591
668.8309326171875 0 2459.9734
669.3484497070312 0 5755.717
669.8518676757812 0 4078.0398
670.3475952148438 0 6214.3438
670.84765625 0 1722.8741
671.3449096679688 0 836.3521
672.3441162109375 0 1215.2493
673.3436889648438 0 724.84204
674.3451538085938 0 1369.6903
674.8456420898438 0 3475.1655
675.34716796875 0 3542.7773
675.8402099609375 0 2467.4668
676.337646484375 0 1466.1129
679.3530883789062 0 21047.492 z 6
680.3561401367188 0 7132.297
681.35791015625 0 2386.8743
682.8532104492188 0 2325.179
683.3475952148438 0 24585.385
683.8486328125 0 16899.234
684.349365234375 0 7055.354
684.8515014648438 0 1759.4836
687.3486328125 0 1120.1881 c Ammonia loss 5
691.3565673828125 0 24126.004
691.85791015625 0 16620.672
691.9146118164062 0 414.46295
692.3558349609375 0 16471.98
692.7972412109375 0 912.6966
692.8558959960938 0 8788.464
693.3541870117188 0 3546.8958
693.8561401367188 0 873.46594
695.37109375 0 4026.9148 y 6
696.3742065429688 0 1938.9816
704.3682250976562 0 26501.213 c 5
705.3707885742188 0 7568.0034
706.3723754882812 0 2344.7493
708.28076171875 0 880.0571
731.3910522460938 0 3624.7812
732.3922119140625 0 737.98444
737.3359375 0 1182.2218
749.3857421875 0 14123.006 w 5
750.3892211914062 0 6489.7236
751.3903198242188 0 1443.7981
754.4196166992188 0 675.5294
758.3867797851562 0 878.50653 c Ammonia loss 6
775.4048461914062 0 22318.646 c 6
776.40673828125 0 8026.5146
777.40771484375 0 1994.0747
793.3958740234375 0 62419.863 z 5
794.3988647460938 0 26028.15
795.401611328125 0 6172.1035
796.4037475585938 0 937.9466
809.41357421875 0 1020.5532 y 5
832.4417114257812 0 692.75165
859.4874877929688 0 2811.9177
860.4923095703125 0 1770.814
862.4931030273438 0 612.88763
864.4328002929688 0 16667.535
865.4362182617188 0 6479.0444
866.4347534179688 0 2904.034
884.4803466796875 0 878.6259
886.4733276367188 0 2237.012 c Ammonia loss 7
894.5033569335938 0 1213.3545
900.47509765625 0 2254.1262
901.4864501953125 0 1074.3395
902.4913940429688 0 5903.21
903.4996337890625 0 24931.574 c 7
904.5020141601562 0 10269.591
905.5027465820312 0 2762.495
908.4224853515625 0 11081.388 z 4
909.4273681640625 0 5494.75
910.4259643554688 0 1675.3325
911.4361572265625 0 659.0019
922.4998168945312 0 1466.6182
924.4400634765625 0 1927.4833 y 4
973.5264282226562 0 790.91986
987.5079345703125 0 1597.7253
988.5227661132812 0 611.4797
1017.5429077148438 0 9668.413 c 8
1018.5440673828125 0 3716.5686
1019.5501098632812 0 1466.1272
1064.5233154296875 0 5431.5977 z 3
1065.5274658203125 0 6034.2827
1066.529052734375 0 1601.5037
1088.55615234375 0 2150.6948
1089.5604248046875 0 1328.8655
1104.57421875 0 8950.563 c 9
1105.578125 0 5858.4644
1106.580810546875 0 2429.8665
1137.5247802734375 0 710.06244
1151.5499267578125 0 2521.9966 z 2
1152.563232421875 0 9209.958
1153.56298828125 0 3870.6501
1154.566650390625 0 1650.4691
1157.642578125 0 759.62836
1173.64697265625 0 1836.9801
1174.653564453125 0 2216.1548
1175.65185546875 0 1408.2872
1201.638916015625 0 3947.8875
1202.639892578125 0 1952.5835
1203.651123046875 0 746.7336
1217.6568603515625 0 4652.788 c 10
1218.662353515625 0 7615.5107
1219.6649169921875 0 3697.6567
1220.6619873046875 0 1525.116
1265.6434326171875 0 2173.4646
1305.667236328125 0 1031.2632
1306.6600341796875 0 918.5599
1320.696044921875 0 1495.1047
1321.697998046875 0 1788.2584
1322.68212890625 0 1427.973
1323.682373046875 0 3803.4907
1324.682373046875 0 4829.4062
1325.6810302734375 0 3523.4312
1326.681640625 0 1047.5695
1337.6922607421875 0 1087.9407
1338.6978759765625 0 9692.223
1339.7005615234375 0 6982.8506
1340.697021484375 0 2971.9233
1341.70361328125 0 1032.499
1346.6763916015625 0 875.24713
1348.680419921875 0 1966.5015
1349.672119140625 0 3428.2927
1350.66845703125 0 2800.7612
1351.681396484375 0 1110.8444
1356.7130126953125 0 755.89246
1363.71044921875 0 660.0249
1365.6917724609375 0 5893.1016
1366.69287109375 0 31821.219
1367.6947021484375 0 20233.73
1368.6973876953125 0 8307.354
1369.701904296875 0 2054.5989
1381.6942138671875 0 1471.0182
1382.7122802734375 0 4870.2246
1383.716796875 0 6289.4688
1384.7130126953125 0 8543.555
1385.7113037109375 0 4555.206
1386.707763671875 0 1751.8685
1387.721923828125 0 912.93616
1564.260498046875 0 763.1688

Spectrum Details

|  |  |
| --- | --- |
| Matched peaks? Matched peaks The total absolute number of peaks matched. Additionally in brackets the total fraction of peaks matched and the total number of peaks is shown. | 44 (12.90% of 341) |
| FDR? FDR The false discovery rate estimated for this peptide. It is calculated by matching all theoretical fragments with a non-integer shift with the raw peaks for this spectrum. This is done with 40 different shifts. The resulting percentage is the average number of annotated peaks over the number of annotated peaks with the correct spectrum. | 1.79% |
| Satellite FDR? Satellite FDR See the FDR for details on its calculation. This satellite ion specific FDR only contains the satellite ions (d/w) for I/L/J positions. | 0.00% |
| PSM Score? PSM Score The PSM Score as given by Hecklib to this annotated spectrum. It is shown with three significant figures. | 324 |

#### Spectrum 4327? Spectrum 4327 The raw spectrum of this peptide as annotated by Hecklib. The fragments are coloured according to ion type (see legend). Any peaks with a star '\*' as text can be hovered over to see the full details, first the ion type second the mass shift type. By hovering over the amino acids in the peptide or ions in the legend the corresponding peaks are highlighted. By toggling the 'Unassigned' label you can turn the background (unassigned) peaks on or off in the plot. By updating the slider in the Ion legend you can update the spectrum to only show the top X% of the peaks with labels. The top X% means any peak that is within X% of the highest intensity. By dragging in the spectrum you can zoom in to a specific part of the spectrum and use 'Zoom Out' to get back to the original zoom level. The annotation of the spectrum is based on the given sequence in the peptides file and is done with different software so inconsistencies are likely. The peaks are annotated based on the given sequence, with 20 ppm tolerance.

Copy Data

##### Spectrum 4327 (TSV)

###### Preview

```
Loading example...
```

*Click on the button to copy the data to your clipboard.*

Mz MinMz MaxIntensity Max

WidthHeightPeptide font sizePeptide stroke widthSpectrum font sizeSpectrum stroke widthCompact peptide

Ion legend

wxyz

abcd

OtherUnassignedIon^Charge^_Position_Show for top:%

TISRDNAKNSJY

02.98e+45.96e+48.94e+41.19e+5

Zoom Out

y^+1^_1_y^+1^_2_c^+1^_3_c^+1^_3_y^+1^_3_c^+2^_8_c^+2^_8_c^+1^_4_y^+1^_4_c^+2^_9_c^+2^_9_c^+2^_9_z^+2^_9_c^+2^_10_c^+2^_10_y^+2^_10_z^+2^_10_y^+2^_10_c^+1^_5_c^+2^_11_z^+1^_5_c^+2^_11_w^+2^_11_y^+1^_5_z^+2^_11_y^+2^_11_z^+1^_6_z^+1^_6_c^+1^_6_y^+1^_6_c^+1^_6_w^+1^_7_c^+1^_7_c^+1^_7_z^+1^_7_c^+1^_8_c^+1^_8_c^+1^_8_z^+1^_8_y^+1^_8_c^+1^_9_c^+1^_9_c^+1^_9_z^+1^_9_c^+1^_10_z^+1^_10_y^+1^_10_c^+1^_11_

0778155623343112

Fragment Matches Table

Show background peaks

| Position | Ion type | Intensity | mz Theoretical | mz Error (Th) | mz Error (ppm) | Charge | Series Number |
| --- | --- | --- | --- | --- | --- | --- | --- |
| - | - | 407 | 124.5 | - | - | 0 | - |
| - | - | 440.4 | 127.6 | - | - | 0 | - |
| - | - | 1075 | 129.1 | - | - | 0 | - |
| - | - | 2827 | 136.1 | - | - | 0 | - |
| - | - | 395.2 | 140.9 | - | - | 0 | - |
| - | - | 402.2 | 144.7 | - | - | 0 | - |
| - | - | 4281 | 149 | - | - | 0 | - |
| - | - | 522.8 | 149.5 | - | - | 0 | - |
| - | - | 704.7 | 150 | - | - | 0 | - |
| - | - | 418 | 151 | - | - | 0 | - |
| - | - | 425.5 | 154 | - | - | 0 | - |
| - | - | 562.6 | 156.3 | - | - | 0 | - |
| - | - | 2649 | 165.1 | - | - | 0 | - |
| - | - | 1657 | 167.1 | - | - | 0 | - |
| - | - | 3202 | 173.1 | - | - | 0 | - |
| - | - | 1.041E+04 | 173.1 | - | - | 0 | - |
| - | - | 2613 | 173.5 | - | - | 0 | - |
| - | - | 437.7 | 173.6 | - | - | 0 | - |
| - | - | 849.7 | 174.1 | - | - | 0 | - |
| - | - | 757.2 | 182.1 | - | - | 0 | - |
| 12 | y | 1.093E+04 | 182.1 | 0.0002054 | 1.128 | +1 | 1 |
| - | - | 460.4 | 183.1 | - | - | 0 | - |
| - | - | 521.3 | 184.1 | - | - | 0 | - |
| - | - | 515.3 | 185.6 | - | - | 0 | - |
| - | - | 1.85E+04 | 187.1 | - | - | 0 | - |
| - | - | 2450 | 188.1 | - | - | 0 | - |
| - | - | 5330 | 201.1 | - | - | 0 | - |
| - | - | 543.9 | 202.1 | - | - | 0 | - |
| - | - | 518.2 | 204.7 | - | - | 0 | - |
| - | - | 467.7 | 205.9 | - | - | 0 | - |
| - | - | 474.7 | 210.2 | - | - | 0 | - |
| - | - | 5959 | 215.1 | - | - | 0 | - |
| - | - | 2992 | 221.1 | - | - | 0 | - |
| - | - | 1112 | 222.1 | - | - | 0 | - |
| - | - | 1412 | 223.1 | - | - | 0 | - |
| - | - | 504.8 | 223.4 | - | - | 0 | - |
| - | - | 1961 | 225 | - | - | 0 | - |
| - | - | 908 | 226 | - | - | 0 | - |
| - | - | 648.6 | 229.1 | - | - | 0 | - |
| - | - | 3878 | 239.1 | - | - | 0 | - |
| - | - | 1500 | 240.1 | - | - | 0 | - |
| - | - | 1118 | 241.1 | - | - | 0 | - |
| - | - | 1475 | 260.1 | - | - | 0 | - |
| - | - | 551.4 | 282.1 | - | - | 0 | - |
| - | - | 842.2 | 282.1 | - | - | 0 | - |
| - | - | 1764 | 295.1 | - | - | 0 | - |
| 11 | y | 1105 | 295.2 | 0.0005683 | 1.926 | +1 | 2 |
| - | - | 2703 | 296.1 | - | - | 0 | - |
| - | - | 1556 | 297.1 | - | - | 0 | - |
| - | - | 1770 | 299.1 | - | - | 0 | - |
| - | - | 976.8 | 299.2 | - | - | 0 | - |
| - | - | 2197 | 300.1 | - | - | 0 | - |
| - | - | 516.2 | 300.1 | - | - | 0 | - |
| - | - | 2787 | 300.2 | - | - | 0 | - |
| - | - | 1063 | 301.1 | - | - | 0 | - |
| 3 | c | 1846 | 301.2 | 0.003001 | 9.965 | +1 | 3 |
| - | - | 705.9 | 316.1 | - | - | 0 | - |
| - | - | 1316 | 317.2 | - | - | 0 | - |
| 3 | c | 1459 | 319.2 | 0.0004321 | 1.354 | +1 | 3 |
| - | - | 583.4 | 323.7 | - | - | 0 | - |
| - | - | 1.52E+04 | 341 | - | - | 0 | - |
| - | - | 850.9 | 343 | - | - | 0 | - |
| - | - | 2191 | 345.2 | - | - | 0 | - |
| - | - | 1052 | 346.2 | - | - | 0 | - |
| - | - | 543.3 | 353.9 | - | - | 0 | - |
| - | - | 1760 | 355.1 | - | - | 0 | - |
| - | - | 748.8 | 356.1 | - | - | 0 | - |
| - | - | 4.814E+04 | 359 | - | - | 0 | - |
| - | - | 933.6 | 369.1 | - | - | 0 | - |
| - | - | 2392 | 370.1 | - | - | 0 | - |
| - | - | 2462 | 371.1 | - | - | 0 | - |
| - | - | 727.5 | 376.9 | - | - | 0 | - |
| 10 | y | 588.2 | 382.2 | 0.0002477 | 0.6481 | +1 | 3 |
| - | - | 2166 | 385.2 | - | - | 0 | - |
| - | - | 1241 | 385.5 | - | - | 0 | - |
| - | - | 1202 | 385.9 | - | - | 0 | - |
| - | - | 882 | 386.2 | - | - | 0 | - |
| - | - | 943.2 | 391.6 | - | - | 0 | - |
| - | - | 543.1 | 425.2 | - | - | 0 | - |
| - | - | 806.8 | 427.2 | - | - | 0 | - |
| - | - | 1.18E+05 | 429.1 | - | - | 0 | - |
| - | - | 1037 | 429.8 | - | - | 0 | - |
| - | - | 2033 | 430.2 | - | - | 0 | - |
| - | - | 938 | 433.2 | - | - | 0 | - |
| - | - | 998.8 | 433.7 | - | - | 0 | - |
| - | - | 814.3 | 442.2 | - | - | 0 | - |
| - | - | 635.3 | 442.7 | - | - | 0 | - |
| - | - | 809.2 | 442.7 | - | - | 0 | - |
| 8 | c | 1236 | 443.7 | 0.003785 | 8.53 | +2 | 8 |
| - | - | 946.9 | 444.2 | - | - | 0 | - |
| - | - | 1739 | 444.2 | - | - | 0 | - |
| - | - | 6.475E+04 | 445.1 | - | - | 0 | - |
| - | - | 1048 | 445.3 | - | - | 0 | - |
| - | - | 1674 | 446.2 | - | - | 0 | - |
| - | - | 914.5 | 451.2 | - | - | 0 | - |
| - | - | 6021 | 451.8 | - | - | 0 | - |
| 8 | c | 1.641E+04 | 452.3 | 6.273E-05 | 0.1387 | +2 | 8 |
| - | - | 5690 | 452.8 | - | - | 0 | - |
| - | - | 2351 | 453.3 | - | - | 0 | - |
| - | - | 1326 | 459.7 | - | - | 0 | - |
| - | - | 3796 | 460.2 | - | - | 0 | - |
| - | - | 1498 | 460.7 | - | - | 0 | - |
| - | - | 604.2 | 461.2 | - | - | 0 | - |
| - | - | 1.049E+04 | 462.2 | - | - | 0 | - |
| - | - | 666.1 | 471.2 | - | - | 0 | - |
| - | - | 547.2 | 471.7 | - | - | 0 | - |
| 4 | c | 5.336E+04 | 475.3 | 0.0007005 | 1.474 | +1 | 4 |
| - | - | 1.367E+04 | 476.3 | - | - | 0 | - |
| - | - | 1898 | 477.3 | - | - | 0 | - |
| - | - | 4597 | 479.8 | - | - | 0 | - |
| - | - | 2944 | 480.3 | - | - | 0 | - |
| - | - | 3491 | 481.2 | - | - | 0 | - |
| - | - | 757.6 | 482.2 | - | - | 0 | - |
| - | - | 653.9 | 485.2 | - | - | 0 | - |
| - | - | 1406 | 487.3 | - | - | 0 | - |
| - | - | 1214 | 487.8 | - | - | 0 | - |
| - | - | 984 | 493.8 | - | - | 0 | - |
| - | - | 1092 | 494.3 | - | - | 0 | - |
| 9 | y | 1804 | 496.2 | 0.0002298 | 0.4631 | +1 | 4 |
| - | - | 6238 | 499.8 | - | - | 0 | - |
| 9 | c | 3259 | 500.3 | 0.002752 | 5.501 | +2 | 9 |
| 9 | c | 2249 | 500.8 | 0.002372 | 4.736 | +2 | 9 |
| - | - | 1557 | 501.3 | - | - | 0 | - |
| - | - | 785.6 | 501.8 | - | - | 0 | - |
| - | - | 6168 | 502.3 | - | - | 0 | - |
| - | - | 1229 | 503.3 | - | - | 0 | - |
| - | - | 1074 | 508.3 | - | - | 0 | - |
| - | - | 1.133E+04 | 508.8 | - | - | 0 | - |
| - | - | 419.3 | 508.8 | - | - | 0 | - |
| 9 | c | 2.296E+04 | 509.3 | 0.000649 | 1.274 | +2 | 9 |
| - | - | 1.128E+04 | 509.8 | - | - | 0 | - |
| - | - | 4122 | 510.3 | - | - | 0 | - |
| - | - | 1740 | 510.8 | - | - | 0 | - |
| - | - | 610.1 | 526.3 | - | - | 0 | - |
| - | - | 998.5 | 530.3 | - | - | 0 | - |
| - | - | 593.9 | 530.8 | - | - | 0 | - |
| 4 | z | 1977 | 532.8 | 0.001739 | 3.264 | +2 | 9 |
| - | - | 2587 | 533.3 | - | - | 0 | - |
| - | - | 804.1 | 535.3 | - | - | 0 | - |
| 10 | c | 2522 | 544.3 | 0.0006396 | 1.175 | +2 | 10 |
| - | - | 1406 | 544.8 | - | - | 0 | - |
| - | - | 781.5 | 545.3 | - | - | 0 | - |
| - | - | 1167 | 546.3 | - | - | 0 | - |
| - | - | 1080 | 548.3 | - | - | 0 | - |
| - | - | 958.5 | 548.8 | - | - | 0 | - |
| 10 | c | 1.243E+04 | 552.8 | 0.0004876 | 0.8821 | +2 | 10 |
| - | - | 1.041E+04 | 553.3 | - | - | 0 | - |
| - | - | 4236 | 553.8 | - | - | 0 | - |
| - | - | 1682 | 554.3 | - | - | 0 | - |
| - | - | 770.6 | 554.8 | - | - | 0 | - |
| - | - | 1686 | 564.3 | - | - | 0 | - |
| - | - | 759.8 | 565.3 | - | - | 0 | - |
| - | - | 875.1 | 565.8 | - | - | 0 | - |
| - | - | 918.6 | 574.3 | - | - | 0 | - |
| 3 | y | 2251 | 575.8 | 0.002272 | 3.945 | +2 | 10 |
| 3 | z | 6621 | 576.3 | 0.0002371 | 0.4114 | +2 | 10 |
| - | - | 5290 | 576.8 | - | - | 0 | - |
| - | - | 1407 | 577.3 | - | - | 0 | - |
| - | - | 968.5 | 578.3 | - | - | 0 | - |
| - | - | 1031 | 578.8 | - | - | 0 | - |
| 3 | y | 1.385E+04 | 584.3 | 0.0005327 | 0.9118 | +2 | 10 |
| - | - | 1.326E+04 | 584.8 | - | - | 0 | - |
| - | - | 4677 | 585.3 | - | - | 0 | - |
| - | - | 1800 | 585.8 | - | - | 0 | - |
| - | - | 2272 | 586.8 | - | - | 0 | - |
| - | - | 2473 | 587.3 | - | - | 0 | - |
| - | - | 1316 | 587.8 | - | - | 0 | - |
| - | - | 1153 | 588.3 | - | - | 0 | - |
| - | - | 1206 | 589.3 | - | - | 0 | - |
| 5 | c | 9.442E+04 | 590.3 | 0.0005214 | 0.8832 | +1 | 5 |
| - | - | 2.756E+04 | 591.3 | - | - | 0 | - |
| - | - | 1148 | 591.8 | - | - | 0 | - |
| - | - | 4948 | 592.3 | - | - | 0 | - |
| - | - | 939 | 592.8 | - | - | 0 | - |
| 11 | c | 2959 | 600.8 | 0.000804 | 1.338 | +2 | 11 |
| - | - | 2177 | 601.3 | - | - | 0 | - |
| - | - | 1170 | 601.8 | - | - | 0 | - |
| - | - | 911.3 | 602.3 | - | - | 0 | - |
| - | - | 860.7 | 604.3 | - | - | 0 | - |
| 8 | z | 4617 | 608.3 | 0.0006492 | 1.067 | +1 | 5 |
| 11 | c | 3.969E+04 | 609.3 | 2.058E-05 | 0.03378 | +2 | 11 |
| - | - | 2.764E+04 | 609.8 | - | - | 0 | - |
| - | - | 1.49E+04 | 610.3 | - | - | 0 | - |
| - | - | 5119 | 610.8 | - | - | 0 | - |
| - | - | 660.6 | 611.8 | - | - | 0 | - |
| - | - | 1232 | 616.3 | - | - | 0 | - |
| 2 | w | 1165 | 618.3 | 0.001586 | 2.564 | +2 | 11 |
| - | - | 1630 | 618.8 | - | - | 0 | - |
| - | - | 797.8 | 619.3 | - | - | 0 | - |
| 8 | y | 6617 | 624.3 | 5.254E-05 | 0.08416 | +1 | 5 |
| - | - | 3124 | 625.3 | - | - | 0 | - |
| - | - | 870.4 | 630.3 | - | - | 0 | - |
| - | - | 778.4 | 630.8 | - | - | 0 | - |
| 2 | z | 1075 | 632.8 | 0.002714 | 4.288 | +2 | 11 |
| - | - | 1723 | 633.3 | - | - | 0 | - |
| - | - | 1093 | 633.8 | - | - | 0 | - |
| - | - | 879.9 | 634.3 | - | - | 0 | - |
| - | - | 706.5 | 634.8 | - | - | 0 | - |
| - | - | 1040 | 636.4 | - | - | 0 | - |
| - | - | 650.6 | 637.8 | - | - | 0 | - |
| - | - | 913.2 | 638.3 | - | - | 0 | - |
| - | - | 696.2 | 638.8 | - | - | 0 | - |
| - | - | 1018 | 639.8 | - | - | 0 | - |
| - | - | 731 | 640.3 | - | - | 0 | - |
| - | - | 1049 | 640.7 | - | - | 0 | - |
| 2 | y | 1920 | 640.8 | 0.0007278 | 1.136 | +2 | 11 |
| - | - | 1920 | 641.3 | - | - | 0 | - |
| - | - | 611.2 | 641.8 | - | - | 0 | - |
| - | - | 2518 | 647.4 | - | - | 0 | - |
| - | - | 2737 | 647.9 | - | - | 0 | - |
| - | - | 918.6 | 653.3 | - | - | 0 | - |
| - | - | 768.8 | 653.8 | - | - | 0 | - |
| - | - | 2999 | 655.3 | - | - | 0 | - |
| - | - | 2105 | 655.8 | - | - | 0 | - |
| - | - | 1347 | 656.3 | - | - | 0 | - |
| - | - | 4603 | 659.3 | - | - | 0 | - |
| - | - | 6925 | 660.3 | - | - | 0 | - |
| - | - | 4328 | 660.9 | - | - | 0 | - |
| 7 | z | 7166 | 661.3 | 0.007853 | 11.87 | +1 | 6 |
| - | - | 5054 | 661.8 | - | - | 0 | - |
| - | - | 7145 | 662.3 | - | - | 0 | - |
| - | - | 4209 | 662.8 | - | - | 0 | - |
| - | - | 1795 | 663.3 | - | - | 0 | - |
| - | - | 1096 | 668.8 | - | - | 0 | - |
| - | - | 8611 | 669.3 | - | - | 0 | - |
| - | - | 5779 | 669.8 | - | - | 0 | - |
| - | - | 7596 | 670.3 | - | - | 0 | - |
| - | - | 3645 | 670.8 | - | - | 0 | - |
| - | - | 2744 | 671.3 | - | - | 0 | - |
| - | - | 1777 | 672.3 | - | - | 0 | - |
| - | - | 1353 | 674.4 | - | - | 0 | - |
| - | - | 4692 | 674.8 | - | - | 0 | - |
| - | - | 3357 | 675.3 | - | - | 0 | - |
| - | - | 4592 | 675.8 | - | - | 0 | - |
| - | - | 1301 | 676.3 | - | - | 0 | - |
| - | - | 965.8 | 676.8 | - | - | 0 | - |
| 7 | z | 1.928E+04 | 679.4 | 0.0005227 | 0.7694 | +1 | 6 |
| - | - | 7925 | 680.4 | - | - | 0 | - |
| - | - | 2065 | 681.4 | - | - | 0 | - |
| - | - | 766.4 | 682.4 | - | - | 0 | - |
| - | - | 1738 | 682.9 | - | - | 0 | - |
| - | - | 2.716E+04 | 683.3 | - | - | 0 | - |
| - | - | 2.134E+04 | 683.8 | - | - | 0 | - |
| - | - | 1.012E+04 | 684.3 | - | - | 0 | - |
| - | - | 3902 | 684.8 | - | - | 0 | - |
| - | - | 793.3 | 685.3 | - | - | 0 | - |
| 6 | c | 765.8 | 687.3 | 0.005749 | 8.365 | +1 | 6 |
| - | - | 1716 | 689.2 | - | - | 0 | - |
| - | - | 802.1 | 689.4 | - | - | 0 | - |
| - | - | 7505 | 689.7 | - | - | 0 | - |
| - | - | 2895 | 690.2 | - | - | 0 | - |
| - | - | 2.353E+04 | 691.4 | - | - | 0 | - |
| - | - | 1.936E+04 | 691.9 | - | - | 0 | - |
| - | - | 1.042E+04 | 692.4 | - | - | 0 | - |
| - | - | 5023 | 692.9 | - | - | 0 | - |
| - | - | 1798 | 693.4 | - | - | 0 | - |
| 7 | y | 4577 | 695.4 | 0.0008416 | 1.21 | +1 | 6 |
| - | - | 1538 | 696.4 | - | - | 0 | - |
| 6 | c | 2.418E+04 | 704.4 | 0.0003186 | 0.4523 | +1 | 6 |
| - | - | 9432 | 705.4 | - | - | 0 | - |
| - | - | 1793 | 706.4 | - | - | 0 | - |
| - | - | 1.405E+04 | 719.3 | - | - | 0 | - |
| - | - | 6251 | 720.3 | - | - | 0 | - |
| - | - | 1957 | 721.4 | - | - | 0 | - |
| - | - | 3125 | 731.4 | - | - | 0 | - |
| - | - | 5910 | 732.4 | - | - | 0 | - |
| - | - | 3219 | 733.4 | - | - | 0 | - |
| - | - | 1104 | 734.4 | - | - | 0 | - |
| - | - | 913.6 | 737.4 | - | - | 0 | - |
| 6 | w | 1.175E+04 | 749.4 | 0.001507 | 2.011 | +1 | 7 |
| - | - | 7116 | 750.4 | - | - | 0 | - |
| - | - | 2130 | 751.4 | - | - | 0 | - |
| 7 | c | 2392 | 758.4 | 0.002327 | 3.069 | +1 | 7 |
| - | - | 747.2 | 759.4 | - | - | 0 | - |
| 7 | c | 2.28E+04 | 775.4 | 0.0002531 | 0.3264 | +1 | 7 |
| - | - | 9557 | 776.4 | - | - | 0 | - |
| - | - | 2919 | 777.4 | - | - | 0 | - |
| - | - | 825 | 787.4 | - | - | 0 | - |
| 6 | z | 5.508E+04 | 793.4 | 0.000503 | 0.6339 | +1 | 7 |
| - | - | 2.614E+04 | 794.4 | - | - | 0 | - |
| - | - | 8564 | 795.4 | - | - | 0 | - |
| - | - | 1872 | 796.4 | - | - | 0 | - |
| - | - | 979.3 | 804.4 | - | - | 0 | - |
| - | - | 968.5 | 810.4 | - | - | 0 | - |
| - | - | 790.9 | 811.4 | - | - | 0 | - |
| - | - | 1145 | 821.4 | - | - | 0 | - |
| - | - | 656 | 853.4 | - | - | 0 | - |
| - | - | 843.3 | 858.4 | - | - | 0 | - |
| - | - | 732.8 | 858.5 | - | - | 0 | - |
| - | - | 2274 | 859.5 | - | - | 0 | - |
| - | - | 3130 | 860.5 | - | - | 0 | - |
| - | - | 2091 | 861.5 | - | - | 0 | - |
| - | - | 1195 | 862.5 | - | - | 0 | - |
| - | - | 1.749E+04 | 864.4 | - | - | 0 | - |
| - | - | 1.137E+04 | 865.4 | - | - | 0 | - |
| - | - | 3739 | 866.4 | - | - | 0 | - |
| - | - | 1346 | 867.4 | - | - | 0 | - |
| - | - | 902.8 | 884.5 | - | - | 0 | - |
| 8 | c | 862.7 | 885.5 | 0.003884 | 4.386 | +1 | 8 |
| 8 | c | 2858 | 886.5 | 0.0002287 | 0.258 | +1 | 8 |
| - | - | 941.3 | 887.5 | - | - | 0 | - |
| - | - | 1073 | 888.5 | - | - | 0 | - |
| - | - | 2118 | 891.5 | - | - | 0 | - |
| - | - | 2438 | 900.5 | - | - | 0 | - |
| - | - | 1639 | 901.5 | - | - | 0 | - |
| - | - | 7582 | 902.5 | - | - | 0 | - |
| - | - | 9895 | 903.4 | - | - | 0 | - |
| 8 | c | 2.381E+04 | 903.5 | 0.0006547 | 0.7247 | +1 | 8 |
| - | - | 4185 | 904.4 | - | - | 0 | - |
| - | - | 1.285E+04 | 904.5 | - | - | 0 | - |
| - | - | 3176 | 905.5 | - | - | 0 | - |
| - | - | 941.4 | 906.5 | - | - | 0 | - |
| 5 | z | 1.587E+04 | 908.4 | 0.001477 | 1.626 | +1 | 8 |
| - | - | 1.111E+04 | 909.4 | - | - | 0 | - |
| - | - | 2528 | 910.4 | - | - | 0 | - |
| - | - | 3216 | 919.5 | - | - | 0 | - |
| - | - | 5643 | 920.5 | - | - | 0 | - |
| - | - | 2908 | 921.5 | - | - | 0 | - |
| - | - | 1490 | 922.5 | - | - | 0 | - |
| - | - | 708.2 | 923.4 | - | - | 0 | - |
| 5 | y | 1827 | 924.4 | 0.005858 | 6.336 | +1 | 8 |
| - | - | 1788 | 925.4 | - | - | 0 | - |
| - | - | 1329 | 986.5 | - | - | 0 | - |
| - | - | 2536 | 987.5 | - | - | 0 | - |
| - | - | 1190 | 988.5 | - | - | 0 | - |
| 9 | c | 1124 | 999.5 | 0.001589 | 1.59 | +1 | 9 |
| 9 | c | 1903 | 1001 | 0.003658 | 3.656 | +1 | 9 |
| - | - | 1489 | 1002 | - | - | 0 | - |
| - | - | 953.8 | 1003 | - | - | 0 | - |
| - | - | 2868 | 1017 | - | - | 0 | - |
| 9 | c | 2.004E+04 | 1018 | 0.0006745 | 0.6628 | +1 | 9 |
| - | - | 1.349E+04 | 1019 | - | - | 0 | - |
| - | - | 4554 | 1020 | - | - | 0 | - |
| - | - | 1848 | 1021 | - | - | 0 | - |
| - | - | 1087 | 1049 | - | - | 0 | - |
| 4 | z | 6887 | 1065 | 0.0004759 | 0.4471 | +1 | 9 |
| - | - | 8077 | 1066 | - | - | 0 | - |
| - | - | 4757 | 1067 | - | - | 0 | - |
| - | - | 2324 | 1068 | - | - | 0 | - |
| - | - | 717.4 | 1069 | - | - | 0 | - |
| - | - | 805 | 1082 | - | - | 0 | - |
| - | - | 1508 | 1089 | - | - | 0 | - |
| - | - | 855 | 1090 | - | - | 0 | - |
| - | - | 707.2 | 1095 | - | - | 0 | - |
| 10 | c | 6458 | 1105 | 0.0007815 | 0.7075 | +1 | 10 |
| - | - | 5846 | 1106 | - | - | 0 | - |
| - | - | 2165 | 1107 | - | - | 0 | - |
| - | - | 979 | 1108 | - | - | 0 | - |
| - | - | 1194 | 1136 | - | - | 0 | - |
| 3 | z | 4104 | 1152 | 0.002841 | 2.467 | +1 | 10 |
| - | - | 9770 | 1153 | - | - | 0 | - |
| - | - | 7059 | 1154 | - | - | 0 | - |
| - | - | 1660 | 1155 | - | - | 0 | - |
| - | - | 683.6 | 1156 | - | - | 0 | - |
| - | - | 763.5 | 1161 | - | - | 0 | - |
| 3 | y | 948.5 | 1168 | 0.001546 | 1.324 | +1 | 10 |
| - | - | 1756 | 1174 | - | - | 0 | - |
| - | - | 2349 | 1175 | - | - | 0 | - |
| - | - | 1705 | 1176 | - | - | 0 | - |
| - | - | 1031 | 1177 | - | - | 0 | - |
| - | - | 5099 | 1202 | - | - | 0 | - |
| - | - | 3419 | 1203 | - | - | 0 | - |
| - | - | 1012 | 1204 | - | - | 0 | - |
| - | - | 890.3 | 1205 | - | - | 0 | - |
| 11 | c | 3194 | 1218 | 0.002692 | 2.211 | +1 | 11 |
| - | - | 7259 | 1219 | - | - | 0 | - |
| - | - | 4282 | 1220 | - | - | 0 | - |
| - | - | 1806 | 1221 | - | - | 0 | - |
| - | - | 1091 | 1222 | - | - | 0 | - |
| - | - | 1780 | 1266 | - | - | 0 | - |
| - | - | 1189 | 1267 | - | - | 0 | - |
| - | - | 968.7 | 1268 | - | - | 0 | - |
| - | - | 1562 | 1306 | - | - | 0 | - |
| - | - | 1416 | 1307 | - | - | 0 | - |
| - | - | 1893 | 1308 | - | - | 0 | - |
| - | - | 1339 | 1321 | - | - | 0 | - |
| - | - | 2190 | 1322 | - | - | 0 | - |
| - | - | 2090 | 1323 | - | - | 0 | - |
| - | - | 3400 | 1324 | - | - | 0 | - |
| - | - | 8592 | 1325 | - | - | 0 | - |
| - | - | 4558 | 1326 | - | - | 0 | - |
| - | - | 1871 | 1327 | - | - | 0 | - |
| - | - | 799.3 | 1328 | - | - | 0 | - |
| - | - | 2135 | 1338 | - | - | 0 | - |
| - | - | 1.029E+04 | 1339 | - | - | 0 | - |
| - | - | 8104 | 1340 | - | - | 0 | - |
| - | - | 3690 | 1341 | - | - | 0 | - |
| - | - | 2215 | 1342 | - | - | 0 | - |
| - | - | 3263 | 1349 | - | - | 0 | - |
| - | - | 4961 | 1350 | - | - | 0 | - |
| - | - | 2891 | 1351 | - | - | 0 | - |
| - | - | 2153 | 1352 | - | - | 0 | - |
| - | - | 5834 | 1366 | - | - | 0 | - |
| - | - | 3.497E+04 | 1367 | - | - | 0 | - |
| - | - | 2.812E+04 | 1368 | - | - | 0 | - |
| - | - | 1.398E+04 | 1369 | - | - | 0 | - |
| - | - | 5181 | 1370 | - | - | 0 | - |
| - | - | 805.8 | 1371 | - | - | 0 | - |
| - | - | 1575 | 1378 | - | - | 0 | - |
| - | - | 2535 | 1379 | - | - | 0 | - |
| - | - | 2158 | 1380 | - | - | 0 | - |
| - | - | 1828 | 1382 | - | - | 0 | - |
| - | - | 3782 | 1383 | - | - | 0 | - |
| - | - | 5698 | 1384 | - | - | 0 | - |
| - | - | 6279 | 1385 | - | - | 0 | - |
| - | - | 3374 | 1386 | - | - | 0 | - |
| - | - | 967.3 | 1387 | - | - | 0 | - |
| - | - | 698.4 | 3081 | - | - | 0 | - |

m/z Charge Intensity FragmentType MassShift Position
124.51460266113281 0 407.00027
127.6448974609375 0 440.415
129.1025848388672 0 1074.952
136.07591247558594 0 2827.0063
140.90402221679688 0 395.23477
144.66632080078125 0 402.19672
149.04510498046875 0 4281.0156
149.50430297851562 0 522.77344
150.0445098876953 0 704.711
151.0412139892578 0 417.9995
154.03912353515625 0 425.53278
156.3356170654297 0 562.57007
165.05484008789062 0 2649.4065
167.0556640625 0 1657.375
173.09231567382812 0 3202.2607
173.12872314453125 0 10411.7295
173.4505615234375 0 2612.9883
173.6227569580078 0 437.6555
174.1321563720703 0 849.6824
182.07308959960938 0 757.18823
182.0813751220703 0 10931.892 y 11
183.0847930908203 0 460.40842
184.07272338867188 0 521.29553
185.55307006835938 0 515.3492
187.1444091796875 0 18501.463
188.14779663085938 0 2449.66
201.1234893798828 0 5329.884
202.08253479003906 0 543.8504
204.67123413085938 0 518.21344
205.93191528320312 0 467.65485
210.2272186279297 0 474.65924
215.13938903808594 0 5958.905
221.08450317382812 0 2991.763
222.08526611328125 0 1112.0532
223.06356811523438 0 1411.8286
223.40359497070312 0 504.76385
225.04302978515625 0 1961.4082
226.0438690185547 0 907.9537
229.12950134277344 0 648.5919
239.09521484375 0 3878.2205
240.09654235839844 0 1500.3805
241.09182739257812 0 1118.0902
260.1241455078125 0 1474.6317
282.0501403808594 0 551.4012
282.1432189941406 0 842.2108
295.1034851074219 0 1764.3772
295.1658020019531 0 1104.8555 y 10
296.104248046875 0 2703.1697
297.1011657714844 0 1555.6438
299.0616455078125 0 1769.5735
299.17218017578125 0 976.7642
300.06298828125 0 2196.5173
300.1386413574219 0 516.2442
300.1557922363281 0 2787.2664
301.0596008300781 0 1063.4932
301.1900329589844 0 1846.212 c Water loss 2
316.14971923828125 0 705.9388
317.1823425292969 0 1316.0107
319.1980285644531 0 1458.5824 c 2
323.7435302734375 0 583.4254
341.0184631347656 0 15196.649
342.9971618652344 0 850.8574
345.1795349121094 0 2190.9192
346.1858825683594 0 1051.5253
353.93255615234375 0 543.3249
355.07049560546875 0 1760.2988
356.070068359375 0 748.77606
359.02862548828125 0 48137.34
369.12286376953125 0 933.5875
370.1230773925781 0 2392.3115
371.119873046875 0 2462.0898
376.8992004394531 0 727.5363
382.197509765625 0 588.1712 y 9
385.21954345703125 0 2165.9731
385.54766845703125 0 1241.3639
385.88201904296875 0 1201.9357
386.22772216796875 0 882.0247
391.55029296875 0 943.20905
425.2012939453125 0 543.0517
427.21734619140625 0 806.8263
429.08935546875 0 118045.06
429.757080078125 0 1036.5771
430.24383544921875 0 2033.2979
433.2209777832031 0 938.01465
433.7037658691406 0 998.77936
442.23095703125 0 814.3258
442.71173095703125 0 635.25214
442.7438049316406 0 809.16187
443.7444763183594 0 1235.6211 c Ammonia loss 7
444.1645202636719 0 946.9249
444.2438049316406 0 1738.7546
445.1206359863281 0 64750.508
445.264892578125 0 1048.0599
446.2114562988281 0 1674.0712
451.2320861816406 0 914.51
451.75018310546875 0 6021.0513
452.2540283203125 0 16414.594 c 7
452.7547912597656 0 5689.913
453.2540588378906 0 2350.7043
459.70245361328125 0 1326.2087
460.2369079589844 0 3796.2053
460.7379455566406 0 1497.8025
461.241455078125 0 604.1704
462.17669677734375 0 10493.807
471.24603271484375 0 666.05597
471.7438049316406 0 547.22925
475.2994079589844 0 53355.27 c 3
476.30224609375 0 13665.351
477.3033752441406 0 1898.3517
479.75799560546875 0 4596.604
480.25909423828125 0 2944.481
481.2296447753906 0 3490.808
482.2340087890625 0 757.60974
485.2453308105469 0 653.86633
487.2645568847656 0 1405.7123
487.7671813964844 0 1213.7463
493.75439453125 0 983.97485
494.25164794921875 0 1092.0496
496.2399597167969 0 1803.7211 y 8
499.766845703125 0 6237.5444
500.26739501953125 0 3258.8945 c Water loss 8
500.7645263671875 0 2249.4626 c Ammonia loss 8
501.2601318359375 0 1556.6938
501.7644958496094 0 785.62634
502.2746276855469 0 6167.815
503.28131103515625 0 1229.3364
508.2534484863281 0 1073.8484
508.77197265625 0 11328.699
508.8105163574219 0 419.2716
509.2747802734375 0 22958.877 c 8
509.7742004394531 0 11277.565
510.27313232421875 0 4122.0317
510.7709045410156 0 1740.1775
526.2607421875 0 610.1057
530.2835083007812 0 998.48724
530.779052734375 0 593.9365
532.7676391601562 0 1976.9395 z 3
533.2672119140625 0 2587.4683
535.2753295898438 0 804.12396
544.27880859375 0 2522.2156 c Ammonia loss 9
544.7772827148438 0 1406.4061
545.307861328125 0 781.4592
546.3078002929688 0 1167.2104
548.251953125 0 1079.6107
548.7505493164062 0 958.5498
552.7919311523438 0 12430.704 c 9
553.29052734375 0 10408.625
553.7885131835938 0 4235.9385
554.2872314453125 0 1682.2064
554.7861328125 0 770.60486
564.30419921875 0 1686.0485
565.310302734375 0 759.8139
565.80078125 0 875.1159
574.3033447265625 0 918.5879
575.7802734375 0 2251.1748 y Ammonia loss 2
576.2816772460938 0 6621.0474 z 2
576.7803955078125 0 5289.926
577.2813720703125 0 1406.6034
578.3123779296875 0 968.5193
578.8167114257812 0 1030.7389
584.2918090820312 0 13853.76 y 2
584.7904663085938 0 13259.646
585.291748046875 0 4676.8794
585.790283203125 0 1799.9106
586.8236083984375 0 2272.004
587.3222045898438 0 2472.618
587.8250122070312 0 1316.0515
588.306884765625 0 1153.3763
589.3130493164062 0 1206.1107
590.326171875 0 94419.31 c 4
591.3286743164062 0 27562.168
591.8154907226562 0 1148.0862
592.3287963867188 0 4947.542
592.8096923828125 0 938.99426
600.8193969726562 0 2958.8066 c Ammonia loss 10
601.3191528320312 0 2177.4968
601.8231201171875 0 1170.2056
602.3265991210938 0 911.2695
604.3175659179688 0 860.7399
608.3170776367188 0 4617.053 z 7
609.33349609375 0 39694.62 c 10
609.8333740234375 0 27639.777
610.3325805664062 0 14897.186
610.8324584960938 0 5118.8105
611.8192138671875 0 660.5565
616.3377685546875 0 1232.1643
618.3059692382812 0 1164.9178 w 1
618.806640625 0 1629.8314
619.3065185546875 0 797.83154
624.335205078125 0 6616.7495 y 7
625.3360595703125 0 3124.3
630.3276977539062 0 870.39905
630.824951171875 0 778.3796
632.82666015625 0 1074.7539 z 1
633.3190307617188 0 1723.0392
633.82080078125 0 1093.0935
634.3187866210938 0 879.9465
634.8231201171875 0 706.47394
636.3508911132812 0 1039.8246
637.8326416015625 0 650.62024
638.3411254882812 0 913.2085
638.8358764648438 0 696.2139
639.8355102539062 0 1018.1299
640.3185424804688 0 730.97424
640.6642456054688 0 1049.0135
640.8325805664062 0 1920.324 y 1
641.3303833007812 0 1919.7993
641.8309326171875 0 611.1932
647.3524169921875 0 2518.3423
647.8506469726562 0 2737.1174
653.3388671875 0 918.5504
653.83447265625 0 768.7899
655.3162231445312 0 2998.8965
655.8158569335938 0 2104.5813
656.3155517578125 0 1347.2041
659.3455200195312 0 4603.276
660.3479614257812 0 6925.263
660.8521728515625 0 4327.553
661.350830078125 0 7166.4766 z Water loss 6
661.8494262695312 0 5054.17
662.344970703125 0 7144.811
662.8446655273438 0 4208.84
663.3424682617188 0 1794.6912
668.8265991210938 0 1096.1682
669.347900390625 0 8611.349
669.8487548828125 0 5779.3125
670.346923828125 0 7595.5767
670.8455200195312 0 3645.1587
671.34521484375 0 2744.3914
672.3428955078125 0 1777.4597
674.350341796875 0 1352.5829
674.8448486328125 0 4691.6353
675.34521484375 0 3357.2192
675.839111328125 0 4591.6655
676.3428344726562 0 1300.5726
676.8367309570312 0 965.8301
679.3540649414062 0 19280.861 z 6
680.3541870117188 0 7925.456
681.3596801757812 0 2064.547
682.3648681640625 0 766.3601
682.8557739257812 0 1737.514
683.3486328125 0 27159.453
683.8475341796875 0 21341.758
684.3472900390625 0 10120.937
684.8473510742188 0 3902.145
685.3497314453125 0 793.272
687.3477783203125 0 765.8349 c Ammonia loss 5
689.1580200195312 0 1716.2319
689.3773803710938 0 802.0915
689.6586303710938 0 7504.8027
690.1598510742188 0 2895.35
691.357666015625 0 23533.3
691.8568115234375 0 19362.152
692.3550415039062 0 10416.2295
692.8599243164062 0 5022.8623
693.3589477539062 0 1798.365
695.3731079101562 0 4577.1943 y 6
696.3704833984375 0 1538.2001
704.368896484375 0 24182.996 c 5
705.3682861328125 0 9431.6875
706.3695678710938 0 1792.5375
719.3475952148438 0 14050.129
720.3494873046875 0 6251.414
721.3524780273438 0 1957.4523
731.3916625976562 0 3125.4004
732.3982543945312 0 5909.806
733.4005737304688 0 3219.2104
734.4041137695312 0 1104.1285
737.3897705078125 0 913.6345
749.3843383789062 0 11750.947 w 5
750.388916015625 0 7115.992
751.3936157226562 0 2129.6523
758.3814697265625 0 2392.401 c Ammonia loss 6
759.3869018554688 0 747.2403
775.4059448242188 0 22801.764 c 6
776.404052734375 0 9556.874
777.4041748046875 0 2919.0186
787.3910522460938 0 824.9673
793.39697265625 0 55077.254 z 5
794.3948974609375 0 26141.701
795.3946533203125 0 8564.419
796.3934326171875 0 1871.5807
804.3768920898438 0 979.28217
810.4116821289062 0 968.464
811.4014892578125 0 790.9364
821.42431640625 0 1144.6555
853.3601684570312 0 656.0462
858.3692626953125 0 843.2882
858.4869384765625 0 732.75446
859.478515625 0 2273.7422
860.4586791992188 0 3129.802
861.4566040039062 0 2091.4338
862.4555053710938 0 1195.2961
864.433837890625 0 17489.129
865.4299926757812 0 11367.579
866.4303588867188 0 3739.266
867.4247436523438 0 1345.7218
884.482421875 0 902.8211
885.4862060546875 0 862.74 c Water loss 7
886.473876953125 0 2858.185 c Ammonia loss 7
887.4794311523438 0 941.33765
888.48486328125 0 1073.2772
891.4662475585938 0 2118.0115
900.4760131835938 0 2437.6526
901.4788208007812 0 1639.4799
902.4915161132812 0 7582.309
903.4241333007812 0 9895.3
903.5 0 23810.428 c 7
904.4261474609375 0 4184.6675
904.4996948242188 0 12849.86
905.4989013671875 0 3175.9397
906.4916381835938 0 941.3777
908.4219360351562 0 15869.366 z 4
909.41943359375 0 11105.962
910.4144287109375 0 2527.6067
919.4660034179688 0 3215.835
920.4710083007812 0 5642.839
921.471923828125 0 2907.514
922.4785766601562 0 1490.3397
923.399169921875 0 708.19635
924.436279296875 0 1827.0087 y 4
925.42529296875 0 1788.1511
986.4981689453125 0 1329.211
987.504150390625 0 2536.0625
988.504150390625 0 1189.7534
999.5346069335938 0 1124.2628 c Water loss 8
1000.5206909179688 0 1902.5697 c Ammonia loss 8
1001.5167236328125 0 1489.0593
1002.5355224609375 0 953.82886
1016.5337524414062 0 2867.8667
1017.5429077148438 0 20036.068 c 8
1018.540283203125 0 13485.569
1019.5391845703125 0 4554.2437
1020.5369262695312 0 1847.8262
1048.54296875 0 1087.1691
1064.5240478515625 0 6886.8945 z 3
1065.525146484375 0 8077.2104
1066.5230712890625 0 4756.765
1067.52685546875 0 2323.6165
1068.5162353515625 0 717.39185
1081.547119140625 0 804.95966
1088.5548095703125 0 1507.753
1089.5562744140625 0 854.99786
1095.4937744140625 0 707.2385
1104.5748291015625 0 6458.063 c 9
1105.573974609375 0 5846.462
1106.5770263671875 0 2164.987
1107.571044921875 0 978.9948
1135.5572509765625 0 1193.6488
1151.5537109375 0 4104.3438 z 2
1152.55908203125 0 9769.814
1153.5606689453125 0 7059.0903
1154.5582275390625 0 1659.5559
1155.5604248046875 0 683.5717
1160.6290283203125 0 763.4572
1167.57373046875 0 948.5143 y 2
1173.6490478515625 0 1756.1411
1174.6517333984375 0 2348.9775
1175.6502685546875 0 1704.8981
1176.645751953125 0 1030.8278
1201.641845703125 0 5098.6704
1202.641845703125 0 3419.3525
1203.643798828125 0 1012.40607
1204.6361083984375 0 890.3088
1217.656982421875 0 3194.207 c 10
1218.661376953125 0 7259.1104
1219.66015625 0 4282.0117
1220.666748046875 0 1806.491
1221.66015625 0 1091.1936
1265.64306640625 0 1779.7557
1266.6436767578125 0 1188.547
1267.6552734375 0 968.6628
1305.6546630859375 0 1562.333
1306.6572265625 0 1416.0146
1307.6553955078125 0 1893.4253
1320.699462890625 0 1338.6042
1321.6959228515625 0 2189.5874
1322.68212890625 0 2089.934
1323.685302734375 0 3399.789
1324.6781005859375 0 8591.904
1325.6761474609375 0 4558.055
1326.6810302734375 0 1871.1079
1327.6834716796875 0 799.3004
1337.697021484375 0 2135.488
1338.69873046875 0 10292.57
1339.69921875 0 8103.598
1340.698486328125 0 3689.7727
1341.6983642578125 0 2215.0164
1348.684814453125 0 3262.9526
1349.6746826171875 0 4961.0464
1350.6739501953125 0 2891.1353
1351.6683349609375 0 2153.0706
1365.689697265625 0 5833.971
1366.693115234375 0 34972.12
1367.692138671875 0 28124.012
1368.6923828125 0 13983.177
1369.692626953125 0 5180.599
1370.6822509765625 0 805.8131
1378.3099365234375 0 1574.9335
1379.3133544921875 0 2534.7454
1380.3170166015625 0 2157.819
1381.706298828125 0 1828.3679
1382.7080078125 0 3781.9043
1383.717529296875 0 5697.523
1384.7171630859375 0 6279.4014
1385.71728515625 0 3373.507
1386.744140625 0 967.3022
3081.4736328125 0 698.35547

Spectrum Details

|  |  |
| --- | --- |
| Matched peaks? Matched peaks The total absolute number of peaks matched. Additionally in brackets the total fraction of peaks matched and the total number of peaks is shown. | 48 (11.76% of 408) |
| FDR? FDR The false discovery rate estimated for this peptide. It is calculated by matching all theoretical fragments with a non-integer shift with the raw peaks for this spectrum. This is done with 40 different shifts. The resulting percentage is the average number of annotated peaks over the number of annotated peaks with the correct spectrum. | 1.39% |
| Satellite FDR? Satellite FDR See the FDR for details on its calculation. This satellite ion specific FDR only contains the satellite ions (d/w) for I/L/J positions. | 0.00% |
| PSM Score? PSM Score The PSM Score as given by Hecklib to this annotated spectrum. It is shown with three significant figures. | 392 |

#### Spectrum 4073? Spectrum 4073 The raw spectrum of this peptide as annotated by Hecklib. The fragments are coloured according to ion type (see legend). Any peaks with a star '\*' as text can be hovered over to see the full details, first the ion type second the mass shift type. By hovering over the amino acids in the peptide or ions in the legend the corresponding peaks are highlighted. By toggling the 'Unassigned' label you can turn the background (unassigned) peaks on or off in the plot. By updating the slider in the Ion legend you can update the spectrum to only show the top X% of the peaks with labels. The top X% means any peak that is within X% of the highest intensity. By dragging in the spectrum you can zoom in to a specific part of the spectrum and use 'Zoom Out' to get back to the original zoom level. The annotation of the spectrum is based on the given sequence in the peptides file and is done with different software so inconsistencies are likely. The peaks are annotated based on the given sequence, with 20 ppm tolerance.

Copy Data

##### Spectrum 4073 (TSV)

###### Preview

```
Loading example...
```

*Click on the button to copy the data to your clipboard.*

Mz MinMz MaxIntensity Max

WidthHeightPeptide font sizePeptide stroke widthSpectrum font sizeSpectrum stroke widthCompact peptide

Ion legend

wxyz

abcd

OtherUnassignedIon^Charge^_Position_Show for top:%

TISRDNAKNSJY

03.38e+46.75e+41.01e+51.35e+5

Zoom Out

y^+1^_1_y^+1^_2_c^+1^_3_c^+3^_10_y^+1^_3_c^+2^_8_c^+2^_8_c^+2^_8_c^+1^_4_c^+1^_4_z^+1^_4_y^+1^_4_c^+2^_9_c^+2^_9_c^+2^_9_z^+2^_9_z^+2^_9_y^+2^_9_c^+2^_10_c^+2^_10_z^+2^_10_y^+2^_10_z^+2^_10_y^+2^_10_c^+1^_5_c^+2^_11_z^+1^_5_c^+2^_11_w^+2^_11_y^+1^_5_y^+2^_11_z^+2^_11_y^+2^_11_z^+1^_6_z^+1^_6_c^+1^_6_y^+1^_6_c^+1^_6_w^+1^_7_c^+1^_7_c^+1^_7_z^+1^_7_y^+1^_7_c^+1^_8_c^+1^_8_z^+1^_8_c^+1^_8_c^+1^_9_c^+1^_9_c^+1^_9_z^+1^_9_y^+1^_9_c^+1^_10_z^+1^_10_z^+1^_10_c^+1^_11_

0778155723353113

Fragment Matches Table

Show background peaks

| Position | Ion type | Intensity | mz Theoretical | mz Error (Th) | mz Error (ppm) | Charge | Series Number |
| --- | --- | --- | --- | --- | --- | --- | --- |
| - | - | 403.5 | 120.1 | - | - | 0 | - |
| - | - | 531.8 | 128.8 | - | - | 0 | - |
| - | - | 957.8 | 129.1 | - | - | 0 | - |
| - | - | 460 | 131 | - | - | 0 | - |
| - | - | 3914 | 136.1 | - | - | 0 | - |
| - | - | 444.4 | 142.1 | - | - | 0 | - |
| - | - | 470.9 | 144.7 | - | - | 0 | - |
| - | - | 418.8 | 145.1 | - | - | 0 | - |
| - | - | 664.8 | 148.9 | - | - | 0 | - |
| - | - | 632.8 | 148.9 | - | - | 0 | - |
| - | - | 898.1 | 148.9 | - | - | 0 | - |
| - | - | 867.9 | 148.9 | - | - | 0 | - |
| - | - | 1123 | 148.9 | - | - | 0 | - |
| - | - | 1543 | 148.9 | - | - | 0 | - |
| - | - | 3397 | 148.9 | - | - | 0 | - |
| - | - | 6115 | 149 | - | - | 0 | - |
| - | - | 3870 | 149 | - | - | 0 | - |
| - | - | 1384 | 149 | - | - | 0 | - |
| - | - | 1134 | 149 | - | - | 0 | - |
| - | - | 750.8 | 149 | - | - | 0 | - |
| - | - | 585.2 | 149 | - | - | 0 | - |
| - | - | 674.6 | 149 | - | - | 0 | - |
| - | - | 460.7 | 149 | - | - | 0 | - |
| - | - | 512.3 | 149 | - | - | 0 | - |
| - | - | 4263 | 149 | - | - | 0 | - |
| - | - | 445.9 | 151 | - | - | 0 | - |
| - | - | 1137 | 157.1 | - | - | 0 | - |
| - | - | 3530 | 165.1 | - | - | 0 | - |
| - | - | 508.6 | 166.1 | - | - | 0 | - |
| - | - | 1228 | 167.1 | - | - | 0 | - |
| - | - | 2227 | 173.1 | - | - | 0 | - |
| - | - | 2085 | 175.1 | - | - | 0 | - |
| 12 | y | 1.373E+04 | 182.1 | 0.0002207 | 1.212 | +1 | 1 |
| - | - | 1248 | 183.1 | - | - | 0 | - |
| - | - | 2.619E+04 | 187.1 | - | - | 0 | - |
| - | - | 1816 | 188.1 | - | - | 0 | - |
| - | - | 4001 | 199.1 | - | - | 0 | - |
| - | - | 1254 | 201.1 | - | - | 0 | - |
| - | - | 646.2 | 202.1 | - | - | 0 | - |
| - | - | 4578 | 203.1 | - | - | 0 | - |
| - | - | 7022 | 215.1 | - | - | 0 | - |
| - | - | 2312 | 221.1 | - | - | 0 | - |
| - | - | 804.6 | 222.1 | - | - | 0 | - |
| - | - | 1222 | 223.1 | - | - | 0 | - |
| - | - | 735.6 | 223.1 | - | - | 0 | - |
| - | - | 1578 | 225 | - | - | 0 | - |
| - | - | 8587 | 227.1 | - | - | 0 | - |
| - | - | 699.2 | 228.1 | - | - | 0 | - |
| - | - | 3388 | 239.1 | - | - | 0 | - |
| - | - | 1678 | 240.1 | - | - | 0 | - |
| - | - | 960.7 | 241.1 | - | - | 0 | - |
| - | - | 1102 | 260.1 | - | - | 0 | - |
| - | - | 515.1 | 266.6 | - | - | 0 | - |
| - | - | 592.1 | 271.8 | - | - | 0 | - |
| - | - | 1399 | 272.1 | - | - | 0 | - |
| - | - | 820.2 | 281.1 | - | - | 0 | - |
| - | - | 941.6 | 282.1 | - | - | 0 | - |
| - | - | 1757 | 282.1 | - | - | 0 | - |
| - | - | 2461 | 295.1 | - | - | 0 | - |
| 11 | y | 1637 | 295.2 | 4.954E-05 | 0.1679 | +1 | 2 |
| - | - | 3054 | 296.1 | - | - | 0 | - |
| - | - | 1535 | 297.1 | - | - | 0 | - |
| - | - | 885.5 | 297.2 | - | - | 0 | - |
| - | - | 556 | 298.1 | - | - | 0 | - |
| - | - | 1914 | 299.1 | - | - | 0 | - |
| - | - | 1535 | 300.1 | - | - | 0 | - |
| - | - | 5566 | 300.2 | - | - | 0 | - |
| - | - | 944.9 | 301.1 | - | - | 0 | - |
| - | - | 840.1 | 301.2 | - | - | 0 | - |
| - | - | 5400 | 306.2 | - | - | 0 | - |
| - | - | 2501 | 307.2 | - | - | 0 | - |
| - | - | 573 | 313.1 | - | - | 0 | - |
| - | - | 933 | 314.1 | - | - | 0 | - |
| - | - | 1113 | 315.2 | - | - | 0 | - |
| - | - | 3391 | 317.2 | - | - | 0 | - |
| - | - | 1641 | 318.1 | - | - | 0 | - |
| - | - | 563 | 318.8 | - | - | 0 | - |
| 3 | c | 1907 | 319.2 | 0.0004932 | 1.545 | +1 | 3 |
| - | - | 654.1 | 320.2 | - | - | 0 | - |
| - | - | 1.277E+04 | 341 | - | - | 0 | - |
| - | - | 2283 | 355.1 | - | - | 0 | - |
| - | - | 4.402E+04 | 359 | - | - | 0 | - |
| - | - | 573 | 359.2 | - | - | 0 | - |
| 10 | c | 867.5 | 363.2 | 0.002711 | 7.464 | +3 | 10 |
| - | - | 1811 | 364.2 | - | - | 0 | - |
| - | - | 1939 | 367.2 | - | - | 0 | - |
| - | - | 732.3 | 369.1 | - | - | 0 | - |
| - | - | 2413 | 370.1 | - | - | 0 | - |
| - | - | 1595 | 371.1 | - | - | 0 | - |
| - | - | 1809 | 371.2 | - | - | 0 | - |
| - | - | 848.7 | 372.1 | - | - | 0 | - |
| - | - | 561.7 | 373.2 | - | - | 0 | - |
| - | - | 741 | 375.9 | - | - | 0 | - |
| - | - | 5845 | 379.2 | - | - | 0 | - |
| 10 | y | 1232 | 382.2 | 0.001522 | 3.983 | +1 | 3 |
| - | - | 3422 | 385.2 | - | - | 0 | - |
| - | - | 899.7 | 385.5 | - | - | 0 | - |
| - | - | 728.7 | 385.9 | - | - | 0 | - |
| - | - | 596.2 | 391 | - | - | 0 | - |
| - | - | 1062 | 391.6 | - | - | 0 | - |
| - | - | 1804 | 391.9 | - | - | 0 | - |
| - | - | 972.6 | 392.2 | - | - | 0 | - |
| - | - | 647.6 | 395.2 | - | - | 0 | - |
| - | - | 1100 | 409.2 | - | - | 0 | - |
| - | - | 1.135E+05 | 429.1 | - | - | 0 | - |
| - | - | 1664 | 429.8 | - | - | 0 | - |
| - | - | 2288 | 430.2 | - | - | 0 | - |
| - | - | 638.8 | 431.2 | - | - | 0 | - |
| - | - | 726.6 | 431.3 | - | - | 0 | - |
| - | - | 5638 | 433.2 | - | - | 0 | - |
| - | - | 600.6 | 434.2 | - | - | 0 | - |
| - | - | 2943 | 442.7 | - | - | 0 | - |
| - | - | 580.7 | 442.8 | - | - | 0 | - |
| 8 | c | 1021 | 443.2 | 0.0079 | 17.82 | +2 | 8 |
| 8 | c | 1100 | 443.7 | 0.0003063 | 0.6902 | +2 | 8 |
| - | - | 622.8 | 444.7 | - | - | 0 | - |
| - | - | 6.253E+04 | 445.1 | - | - | 0 | - |
| - | - | 777.5 | 445.3 | - | - | 0 | - |
| - | - | 1819 | 446.2 | - | - | 0 | - |
| - | - | 683.7 | 447.1 | - | - | 0 | - |
| - | - | 1182 | 451.7 | - | - | 0 | - |
| - | - | 7005 | 451.8 | - | - | 0 | - |
| 8 | c | 2.113E+04 | 452.3 | 0.0002764 | 0.6111 | +2 | 8 |
| - | - | 1.044E+04 | 452.8 | - | - | 0 | - |
| - | - | 2697 | 453.3 | - | - | 0 | - |
| - | - | 1217 | 453.8 | - | - | 0 | - |
| 4 | c | 1299 | 458.3 | 2.585E-06 | 0.00564 | +1 | 4 |
| - | - | 668.3 | 459.2 | - | - | 0 | - |
| - | - | 2296 | 459.7 | - | - | 0 | - |
| - | - | 947.8 | 459.7 | - | - | 0 | - |
| - | - | 1002 | 460.2 | - | - | 0 | - |
| - | - | 1916 | 460.2 | - | - | 0 | - |
| - | - | 859.9 | 460.7 | - | - | 0 | - |
| - | - | 660 | 460.7 | - | - | 0 | - |
| - | - | 933.7 | 461.2 | - | - | 0 | - |
| - | - | 604.4 | 461.2 | - | - | 0 | - |
| - | - | 519.7 | 462.7 | - | - | 0 | - |
| - | - | 988.3 | 465.3 | - | - | 0 | - |
| - | - | 812.2 | 471.2 | - | - | 0 | - |
| 4 | c | 7.447E+04 | 475.3 | 0.0007005 | 1.474 | +1 | 4 |
| - | - | 602 | 476.2 | - | - | 0 | - |
| - | - | 1.578E+04 | 476.3 | - | - | 0 | - |
| - | - | 2499 | 477.3 | - | - | 0 | - |
| - | - | 6740 | 479.8 | - | - | 0 | - |
| 9 | z | 854.8 | 480.2 | 0.0004889 | 1.018 | +1 | 4 |
| - | - | 3324 | 480.3 | - | - | 0 | - |
| - | - | 1069 | 480.8 | - | - | 0 | - |
| - | - | 4928 | 481.2 | - | - | 0 | - |
| - | - | 1019 | 482.2 | - | - | 0 | - |
| - | - | 1419 | 484.8 | - | - | 0 | - |
| - | - | 668.9 | 485.2 | - | - | 0 | - |
| - | - | 569.5 | 485.7 | - | - | 0 | - |
| - | - | 988.8 | 486.8 | - | - | 0 | - |
| - | - | 2145 | 487.3 | - | - | 0 | - |
| - | - | 1093 | 487.8 | - | - | 0 | - |
| - | - | 713.2 | 490.2 | - | - | 0 | - |
| - | - | 607.6 | 491.8 | - | - | 0 | - |
| - | - | 3841 | 492.2 | - | - | 0 | - |
| - | - | 1616 | 493.2 | - | - | 0 | - |
| - | - | 1103 | 493.8 | - | - | 0 | - |
| - | - | 723.2 | 495.2 | - | - | 0 | - |
| 9 | y | 2050 | 496.2 | 0.0002298 | 0.4631 | +1 | 4 |
| - | - | 7847 | 499.8 | - | - | 0 | - |
| 9 | c | 5335 | 500.3 | 0.004217 | 8.429 | +2 | 9 |
| 9 | c | 5308 | 500.8 | 0.001303 | 2.603 | +2 | 9 |
| - | - | 2447 | 501.3 | - | - | 0 | - |
| - | - | 2052 | 501.8 | - | - | 0 | - |
| - | - | 7643 | 502.3 | - | - | 0 | - |
| - | - | 1680 | 503.3 | - | - | 0 | - |
| - | - | 1.447E+04 | 508.3 | - | - | 0 | - |
| - | - | 1.601E+04 | 508.8 | - | - | 0 | - |
| 9 | c | 2.805E+04 | 509.3 | 0.001259 | 2.473 | +2 | 9 |
| - | - | 1.775E+04 | 509.8 | - | - | 0 | - |
| - | - | 5852 | 510.3 | - | - | 0 | - |
| - | - | 2310 | 510.8 | - | - | 0 | - |
| - | - | 772.1 | 511.3 | - | - | 0 | - |
| - | - | 969.5 | 522.8 | - | - | 0 | - |
| 4 | z | 579.4 | 523.8 | 0.001162 | 2.219 | +2 | 9 |
| - | - | 1037 | 526.3 | - | - | 0 | - |
| - | - | 668.5 | 526.7 | - | - | 0 | - |
| - | - | 900.3 | 530.3 | - | - | 0 | - |
| - | - | 796.3 | 532.3 | - | - | 0 | - |
| 4 | z | 2778 | 532.8 | 0.0005192 | 0.9746 | +2 | 9 |
| - | - | 2632 | 533.3 | - | - | 0 | - |
| - | - | 1138 | 533.8 | - | - | 0 | - |
| - | - | 923.1 | 535.3 | - | - | 0 | - |
| - | - | 861.2 | 535.8 | - | - | 0 | - |
| 4 | y | 824.5 | 540.8 | 0.002435 | 4.503 | +2 | 9 |
| - | - | 785 | 541.3 | - | - | 0 | - |
| 10 | c | 2671 | 544.3 | 0.0001538 | 0.2826 | +2 | 10 |
| - | - | 683.2 | 544.3 | - | - | 0 | - |
| - | - | 2143 | 544.8 | - | - | 0 | - |
| - | - | 2073 | 545.3 | - | - | 0 | - |
| - | - | 1547 | 546.3 | - | - | 0 | - |
| - | - | 768.1 | 547.3 | - | - | 0 | - |
| - | - | 3412 | 548.3 | - | - | 0 | - |
| - | - | 2339 | 548.7 | - | - | 0 | - |
| - | - | 641.5 | 549.2 | - | - | 0 | - |
| - | - | 691.7 | 552.3 | - | - | 0 | - |
| - | - | 1226 | 552.7 | - | - | 0 | - |
| 10 | c | 2.214E+04 | 552.8 | 0.0006707 | 1.213 | +2 | 10 |
| - | - | 1.649E+04 | 553.3 | - | - | 0 | - |
| - | - | 7766 | 553.8 | - | - | 0 | - |
| - | - | 1862 | 554.3 | - | - | 0 | - |
| - | - | 1495 | 554.8 | - | - | 0 | - |
| - | - | 734.4 | 561.3 | - | - | 0 | - |
| - | - | 887.8 | 561.8 | - | - | 0 | - |
| - | - | 3613 | 564.3 | - | - | 0 | - |
| - | - | 1736 | 565.3 | - | - | 0 | - |
| - | - | 1524 | 565.8 | - | - | 0 | - |
| - | - | 1001 | 566.3 | - | - | 0 | - |
| - | - | 593.3 | 567.3 | - | - | 0 | - |
| 3 | z | 918.4 | 567.8 | 0.009131 | 16.08 | +2 | 10 |
| - | - | 1245 | 568.3 | - | - | 0 | - |
| - | - | 834.1 | 568.8 | - | - | 0 | - |
| - | - | 854.7 | 569.3 | - | - | 0 | - |
| - | - | 658 | 570.3 | - | - | 0 | - |
| - | - | 1155 | 573.3 | - | - | 0 | - |
| 3 | y | 2735 | 575.8 | 0.002577 | 4.475 | +2 | 10 |
| 3 | z | 7767 | 576.3 | 0.0003123 | 0.5418 | +2 | 10 |
| - | - | 5474 | 576.8 | - | - | 0 | - |
| - | - | 3259 | 577.3 | - | - | 0 | - |
| - | - | 1011 | 577.8 | - | - | 0 | - |
| - | - | 1437 | 578.3 | - | - | 0 | - |
| - | - | 1049 | 578.8 | - | - | 0 | - |
| - | - | 685.2 | 579.8 | - | - | 0 | - |
| - | - | 1346 | 583.3 | - | - | 0 | - |
| 3 | y | 2.077E+04 | 584.3 | 0.0005938 | 1.016 | +2 | 10 |
| - | - | 1.621E+04 | 584.8 | - | - | 0 | - |
| - | - | 8490 | 585.3 | - | - | 0 | - |
| - | - | 2561 | 585.8 | - | - | 0 | - |
| - | - | 2609 | 586.8 | - | - | 0 | - |
| - | - | 4086 | 587.3 | - | - | 0 | - |
| - | - | 1120 | 587.8 | - | - | 0 | - |
| - | - | 1095 | 588.3 | - | - | 0 | - |
| - | - | 987.3 | 588.8 | - | - | 0 | - |
| 5 | c | 1.337E+05 | 590.3 | 0.0004604 | 0.7798 | +1 | 5 |
| - | - | 3.549E+04 | 591.3 | - | - | 0 | - |
| - | - | 2133 | 591.8 | - | - | 0 | - |
| - | - | 6728 | 592.3 | - | - | 0 | - |
| - | - | 2013 | 592.8 | - | - | 0 | - |
| 11 | c | 3898 | 600.8 | 0.001027 | 1.709 | +2 | 11 |
| - | - | 2942 | 601.3 | - | - | 0 | - |
| - | - | 1508 | 601.8 | - | - | 0 | - |
| - | - | 5304 | 607.3 | - | - | 0 | - |
| 8 | z | 7829 | 608.3 | 0.001809 | 2.974 | +1 | 5 |
| 11 | c | 5.372E+04 | 609.3 | 4.045E-05 | 0.06639 | +2 | 11 |
| - | - | 4.06E+04 | 609.8 | - | - | 0 | - |
| - | - | 2.086E+04 | 610.3 | - | - | 0 | - |
| - | - | 5506 | 610.8 | - | - | 0 | - |
| - | - | 1185 | 611.3 | - | - | 0 | - |
| - | - | 1158 | 616.3 | - | - | 0 | - |
| 2 | w | 3294 | 618.3 | 0.0005479 | 0.8862 | +2 | 11 |
| - | - | 2941 | 618.8 | - | - | 0 | - |
| - | - | 1064 | 619.3 | - | - | 0 | - |
| 8 | y | 9742 | 624.3 | 5.254E-05 | 0.08416 | +1 | 5 |
| - | - | 3554 | 625.3 | - | - | 0 | - |
| - | - | 1202 | 630.3 | - | - | 0 | - |
| - | - | 1036 | 630.8 | - | - | 0 | - |
| - | - | 1665 | 631.3 | - | - | 0 | - |
| 2 | y | 667.5 | 632.3 | 0.0006368 | 1.007 | +2 | 11 |
| 2 | z | 2340 | 632.8 | 0.0005776 | 0.9128 | +2 | 11 |
| - | - | 2554 | 633.3 | - | - | 0 | - |
| - | - | 1562 | 633.8 | - | - | 0 | - |
| - | - | 1577 | 634.3 | - | - | 0 | - |
| - | - | 1828 | 636.3 | - | - | 0 | - |
| - | - | 895.3 | 637.8 | - | - | 0 | - |
| - | - | 540.4 | 638.3 | - | - | 0 | - |
| - | - | 542.7 | 638.4 | - | - | 0 | - |
| - | - | 709.7 | 639.8 | - | - | 0 | - |
| - | - | 594.6 | 640.3 | - | - | 0 | - |
| 2 | y | 2570 | 640.8 | 0.001216 | 1.898 | +2 | 11 |
| - | - | 2730 | 641.3 | - | - | 0 | - |
| - | - | 1712 | 641.8 | - | - | 0 | - |
| - | - | 705.4 | 642.3 | - | - | 0 | - |
| - | - | 2954 | 647.4 | - | - | 0 | - |
| - | - | 2658 | 647.9 | - | - | 0 | - |
| - | - | 1038 | 648.4 | - | - | 0 | - |
| - | - | 825.7 | 649.3 | - | - | 0 | - |
| - | - | 1229 | 652.8 | - | - | 0 | - |
| - | - | 1561 | 653.3 | - | - | 0 | - |
| - | - | 851.2 | 653.8 | - | - | 0 | - |
| - | - | 797.3 | 654.3 | - | - | 0 | - |
| - | - | 773 | 654.8 | - | - | 0 | - |
| - | - | 4526 | 655.3 | - | - | 0 | - |
| - | - | 4453 | 655.8 | - | - | 0 | - |
| - | - | 1645 | 656.3 | - | - | 0 | - |
| - | - | 5804 | 659.3 | - | - | 0 | - |
| - | - | 8164 | 660.3 | - | - | 0 | - |
| - | - | 5849 | 660.9 | - | - | 0 | - |
| 7 | z | 1.312E+04 | 661.3 | 0.00596 | 9.013 | +1 | 6 |
| - | - | 8246 | 661.9 | - | - | 0 | - |
| - | - | 1.032E+04 | 662.3 | - | - | 0 | - |
| - | - | 6173 | 662.8 | - | - | 0 | - |
| - | - | 2141 | 663.3 | - | - | 0 | - |
| - | - | 3503 | 668.8 | - | - | 0 | - |
| - | - | 1.076E+04 | 669.3 | - | - | 0 | - |
| - | - | 7902 | 669.8 | - | - | 0 | - |
| - | - | 1.002E+04 | 670.3 | - | - | 0 | - |
| - | - | 5580 | 670.8 | - | - | 0 | - |
| - | - | 3986 | 671.3 | - | - | 0 | - |
| - | - | 1760 | 672.3 | - | - | 0 | - |
| - | - | 2530 | 674.3 | - | - | 0 | - |
| - | - | 6054 | 674.8 | - | - | 0 | - |
| - | - | 4360 | 675.3 | - | - | 0 | - |
| - | - | 5670 | 675.8 | - | - | 0 | - |
| - | - | 4877 | 676.3 | - | - | 0 | - |
| - | - | 1947 | 676.8 | - | - | 0 | - |
| - | - | 1108 | 677.3 | - | - | 0 | - |
| - | - | 950.4 | 678.3 | - | - | 0 | - |
| 7 | z | 2.777E+04 | 679.4 | 0.0004617 | 0.6795 | +1 | 6 |
| - | - | 1.176E+04 | 680.4 | - | - | 0 | - |
| - | - | 3172 | 681.4 | - | - | 0 | - |
| - | - | 2559 | 682.9 | - | - | 0 | - |
| - | - | 3.461E+04 | 683.3 | - | - | 0 | - |
| - | - | 3.42E+04 | 683.8 | - | - | 0 | - |
| - | - | 1.469E+04 | 684.3 | - | - | 0 | - |
| - | - | 6009 | 684.8 | - | - | 0 | - |
| 6 | c | 1415 | 687.3 | 0.007092 | 10.32 | +1 | 6 |
| - | - | 778 | 689.2 | - | - | 0 | - |
| - | - | 4591 | 689.7 | - | - | 0 | - |
| - | - | 2129 | 690.2 | - | - | 0 | - |
| - | - | 3.205E+04 | 691.4 | - | - | 0 | - |
| - | - | 2.876E+04 | 691.9 | - | - | 0 | - |
| - | - | 1.568E+04 | 692.4 | - | - | 0 | - |
| - | - | 5942 | 692.9 | - | - | 0 | - |
| - | - | 1703 | 693.4 | - | - | 0 | - |
| - | - | 1.461E+04 | 694.4 | - | - | 0 | - |
| 7 | y | 1.041E+04 | 695.4 | 0.006849 | 9.849 | +1 | 6 |
| - | - | 3176 | 696.4 | - | - | 0 | - |
| - | - | 1334 | 697.4 | - | - | 0 | - |
| - | - | 1017 | 702.3 | - | - | 0 | - |
| 6 | c | 3.267E+04 | 704.4 | 0.0003186 | 0.4523 | +1 | 6 |
| - | - | 1.272E+04 | 705.4 | - | - | 0 | - |
| - | - | 2159 | 706.4 | - | - | 0 | - |
| - | - | 807.3 | 714.4 | - | - | 0 | - |
| - | - | 9898 | 722.3 | - | - | 0 | - |
| - | - | 2043 | 728.3 | - | - | 0 | - |
| - | - | 6138 | 731.4 | - | - | 0 | - |
| - | - | 1444 | 732.4 | - | - | 0 | - |
| - | - | 1181 | 737.4 | - | - | 0 | - |
| 6 | w | 1.622E+04 | 749.4 | 0.0004698 | 0.6269 | +1 | 7 |
| - | - | 1.035E+04 | 750.4 | - | - | 0 | - |
| - | - | 3748 | 751.4 | - | - | 0 | - |
| - | - | 928 | 752.4 | - | - | 0 | - |
| 7 | c | 2275 | 758.4 | 0.0009233 | 1.217 | +1 | 7 |
| - | - | 1072 | 759.4 | - | - | 0 | - |
| - | - | 7538 | 772.4 | - | - | 0 | - |
| - | - | 3164 | 773.4 | - | - | 0 | - |
| 7 | c | 3.271E+04 | 775.4 | 0.0004362 | 0.5626 | +1 | 7 |
| - | - | 8129 | 776.3 | - | - | 0 | - |
| - | - | 9304 | 776.4 | - | - | 0 | - |
| - | - | 3307 | 777.4 | - | - | 0 | - |
| - | - | 735.1 | 778.4 | - | - | 0 | - |
| - | - | 802.7 | 787.4 | - | - | 0 | - |
| - | - | 994.1 | 788.4 | - | - | 0 | - |
| 6 | z | 7.164E+04 | 793.4 | 0.0003199 | 0.4031 | +1 | 7 |
| - | - | 3.985E+04 | 794.4 | - | - | 0 | - |
| - | - | 1.07E+04 | 795.4 | - | - | 0 | - |
| - | - | 3129 | 796.4 | - | - | 0 | - |
| 6 | y | 1352 | 809.4 | 0.005465 | 6.751 | +1 | 7 |
| - | - | 934.1 | 810.4 | - | - | 0 | - |
| - | - | 1105 | 837.4 | - | - | 0 | - |
| - | - | 1099 | 858.5 | - | - | 0 | - |
| - | - | 2469 | 859.5 | - | - | 0 | - |
| - | - | 3040 | 860.5 | - | - | 0 | - |
| - | - | 805.1 | 861.4 | - | - | 0 | - |
| - | - | 1008 | 861.5 | - | - | 0 | - |
| - | - | 917.8 | 863.4 | - | - | 0 | - |
| - | - | 2.482E+04 | 864.4 | - | - | 0 | - |
| - | - | 1.504E+04 | 865.4 | - | - | 0 | - |
| - | - | 4116 | 866.4 | - | - | 0 | - |
| - | - | 965.8 | 867.4 | - | - | 0 | - |
| - | - | 754.4 | 873.4 | - | - | 0 | - |
| - | - | 1312 | 874.4 | - | - | 0 | - |
| - | - | 940.7 | 877.4 | - | - | 0 | - |
| - | - | 4169 | 879.4 | - | - | 0 | - |
| - | - | 1920 | 882.4 | - | - | 0 | - |
| - | - | 1002 | 884.5 | - | - | 0 | - |
| 8 | c | 1057 | 885.5 | 0.01609 | 18.17 | +1 | 8 |
| 8 | c | 2704 | 886.5 | 0.0009611 | 1.084 | +1 | 8 |
| - | - | 2242 | 887.5 | - | - | 0 | - |
| - | - | 977.8 | 888.5 | - | - | 0 | - |
| 5 | z | 1879 | 891.4 | 0.01238 | 13.89 | +1 | 8 |
| - | - | 927 | 892.4 | - | - | 0 | - |
| - | - | 3244 | 900.5 | - | - | 0 | - |
| - | - | 2433 | 901.5 | - | - | 0 | - |
| - | - | 1.072E+04 | 902.5 | - | - | 0 | - |
| 8 | c | 4.146E+04 | 903.5 | 0.0008989 | 0.9949 | +1 | 8 |
| - | - | 2075 | 904.4 | - | - | 0 | - |
| - | - | 2.319E+04 | 904.5 | - | - | 0 | - |
| - | - | 1129 | 905.4 | - | - | 0 | - |
| - | - | 7571 | 905.5 | - | - | 0 | - |
| - | - | 2050 | 906.5 | - | - | 0 | - |
| - | - | 3.189E+04 | 908.4 | - | - | 0 | - |
| - | - | 1.636E+04 | 909.4 | - | - | 0 | - |
| - | - | 5972 | 910.4 | - | - | 0 | - |
| - | - | 1013 | 911.4 | - | - | 0 | - |
| - | - | 737.8 | 916 | - | - | 0 | - |
| - | - | 2788 | 919.5 | - | - | 0 | - |
| - | - | 2285 | 920.4 | - | - | 0 | - |
| - | - | 1208 | 920.5 | - | - | 0 | - |
| - | - | 8565 | 921.4 | - | - | 0 | - |
| - | - | 5682 | 922.4 | - | - | 0 | - |
| - | - | 2238 | 923.4 | - | - | 0 | - |
| - | - | 5164 | 924.4 | - | - | 0 | - |
| - | - | 5482 | 925.4 | - | - | 0 | - |
| - | - | 795.3 | 961.5 | - | - | 0 | - |
| - | - | 710.9 | 969.5 | - | - | 0 | - |
| - | - | 1298 | 986.5 | - | - | 0 | - |
| - | - | 3742 | 987.5 | - | - | 0 | - |
| - | - | 1956 | 988.5 | - | - | 0 | - |
| - | - | 1168 | 989.5 | - | - | 0 | - |
| 9 | c | 2348 | 999.5 | 0.002017 | 2.018 | +1 | 9 |
| 9 | c | 3456 | 1001 | 0.006649 | 6.645 | +1 | 9 |
| - | - | 2446 | 1002 | - | - | 0 | - |
| - | - | 2105 | 1003 | - | - | 0 | - |
| - | - | 1559 | 1004 | - | - | 0 | - |
| - | - | 3916 | 1017 | - | - | 0 | - |
| 9 | c | 2.753E+04 | 1018 | 0.0005524 | 0.5429 | +1 | 9 |
| - | - | 1.709E+04 | 1019 | - | - | 0 | - |
| - | - | 7522 | 1020 | - | - | 0 | - |
| - | - | 2426 | 1021 | - | - | 0 | - |
| - | - | 1004 | 1062 | - | - | 0 | - |
| - | - | 786.5 | 1063 | - | - | 0 | - |
| 4 | z | 8310 | 1065 | 0.001355 | 1.273 | +1 | 9 |
| - | - | 1.218E+04 | 1066 | - | - | 0 | - |
| - | - | 8000 | 1067 | - | - | 0 | - |
| - | - | 2208 | 1068 | - | - | 0 | - |
| 4 | y | 1329 | 1081 | 0.004063 | 3.76 | +1 | 9 |
| - | - | 1054 | 1082 | - | - | 0 | - |
| - | - | 1539 | 1089 | - | - | 0 | - |
| - | - | 1954 | 1090 | - | - | 0 | - |
| - | - | 1112 | 1091 | - | - | 0 | - |
| 10 | c | 8258 | 1105 | 4.907E-05 | 0.04442 | +1 | 10 |
| - | - | 6746 | 1106 | - | - | 0 | - |
| - | - | 3869 | 1107 | - | - | 0 | - |
| - | - | 2001 | 1108 | - | - | 0 | - |
| 3 | z | 768.2 | 1135 | 0.02017 | 17.78 | +1 | 10 |
| - | - | 1449 | 1136 | - | - | 0 | - |
| 3 | z | 6021 | 1152 | 0.001187 | 1.031 | +1 | 10 |
| - | - | 1.337E+04 | 1153 | - | - | 0 | - |
| - | - | 7824 | 1154 | - | - | 0 | - |
| - | - | 4166 | 1155 | - | - | 0 | - |
| - | - | 819.6 | 1156 | - | - | 0 | - |
| - | - | 1203 | 1160 | - | - | 0 | - |
| - | - | 2829 | 1174 | - | - | 0 | - |
| - | - | 3325 | 1175 | - | - | 0 | - |
| - | - | 3610 | 1176 | - | - | 0 | - |
| - | - | 1034 | 1177 | - | - | 0 | - |
| - | - | 718.2 | 1178 | - | - | 0 | - |
| - | - | 5728 | 1202 | - | - | 0 | - |
| - | - | 4210 | 1203 | - | - | 0 | - |
| - | - | 1770 | 1204 | - | - | 0 | - |
| - | - | 1399 | 1205 | - | - | 0 | - |
| 11 | c | 4808 | 1218 | 0.000739 | 0.6069 | +1 | 11 |
| - | - | 1E+04 | 1219 | - | - | 0 | - |
| - | - | 6207 | 1220 | - | - | 0 | - |
| - | - | 2845 | 1221 | - | - | 0 | - |
| - | - | 2440 | 1266 | - | - | 0 | - |
| - | - | 2484 | 1267 | - | - | 0 | - |
| - | - | 962.7 | 1268 | - | - | 0 | - |
| - | - | 1315 | 1306 | - | - | 0 | - |
| - | - | 1575 | 1307 | - | - | 0 | - |
| - | - | 3323 | 1308 | - | - | 0 | - |
| - | - | 1186 | 1309 | - | - | 0 | - |
| - | - | 947.3 | 1310 | - | - | 0 | - |
| - | - | 1547 | 1321 | - | - | 0 | - |
| - | - | 3378 | 1322 | - | - | 0 | - |
| - | - | 2514 | 1323 | - | - | 0 | - |
| - | - | 5882 | 1324 | - | - | 0 | - |
| - | - | 9809 | 1325 | - | - | 0 | - |
| - | - | 7764 | 1326 | - | - | 0 | - |
| - | - | 3704 | 1327 | - | - | 0 | - |
| - | - | 2086 | 1338 | - | - | 0 | - |
| - | - | 1.377E+04 | 1339 | - | - | 0 | - |
| - | - | 1.082E+04 | 1340 | - | - | 0 | - |
| - | - | 7672 | 1341 | - | - | 0 | - |
| - | - | 2280 | 1342 | - | - | 0 | - |
| - | - | 2707 | 1349 | - | - | 0 | - |
| - | - | 5966 | 1350 | - | - | 0 | - |
| - | - | 5147 | 1351 | - | - | 0 | - |
| - | - | 1953 | 1352 | - | - | 0 | - |
| - | - | 959.5 | 1356 | - | - | 0 | - |
| - | - | 700.4 | 1357 | - | - | 0 | - |
| - | - | 8719 | 1366 | - | - | 0 | - |
| - | - | 4.412E+04 | 1367 | - | - | 0 | - |
| - | - | 4.037E+04 | 1368 | - | - | 0 | - |
| - | - | 2.13E+04 | 1369 | - | - | 0 | - |
| - | - | 7470 | 1370 | - | - | 0 | - |
| - | - | 1422 | 1371 | - | - | 0 | - |
| - | - | 1167 | 1378 | - | - | 0 | - |
| - | - | 2574 | 1379 | - | - | 0 | - |
| - | - | 1924 | 1380 | - | - | 0 | - |
| - | - | 2019 | 1382 | - | - | 0 | - |
| - | - | 5847 | 1383 | - | - | 0 | - |
| - | - | 1.087E+04 | 1384 | - | - | 0 | - |
| - | - | 9474 | 1385 | - | - | 0 | - |
| - | - | 4325 | 1386 | - | - | 0 | - |
| - | - | 1271 | 1387 | - | - | 0 | - |
| - | - | 766 | 3082 | - | - | 0 | - |
| - | - | 726.7 | 3083 | - | - | 0 | - |

m/z Charge Intensity FragmentType MassShift Position
120.08090209960938 0 403.5349
128.82406616210938 0 531.80853
129.10252380371094 0 957.77625
130.9553680419922 0 460.02142
136.07594299316406 0 3913.9495
142.12298583984375 0 444.369
144.7157745361328 0 470.93616
145.09207153320312 0 418.7609
148.89564514160156 0 664.8183
148.9026336669922 0 632.7684
148.9102783203125 0 898.06824
148.91749572753906 0 867.8682
148.92503356933594 0 1122.9015
148.93202209472656 0 1543.3232
148.9395751953125 0 3396.733
148.95614624023438 0 6115.088
148.9640655517578 0 3870.2825
148.97161865234375 0 1383.8481
148.97903442382812 0 1133.722
149.00083923339844 0 750.8317
149.0078887939453 0 585.18756
149.01519775390625 0 674.5746
149.02212524414062 0 460.6866
149.03024291992188 0 512.33795
149.0449981689453 0 4262.7627
151.0414276123047 0 445.9308
157.0610809326172 0 1136.9049
165.05479431152344 0 3530.3164
166.0577850341797 0 508.62305
167.05567932128906 0 1227.605
173.09226989746094 0 2226.9375
175.0716552734375 0 2084.7031
182.08139038085938 0 13732.665 y 11
183.08480834960938 0 1248.2152
187.14434814453125 0 26186.332
188.14779663085938 0 1815.994
199.0714874267578 0 4001.1882
201.1234588623047 0 1254.1567
202.0818328857422 0 646.216
203.0664520263672 0 4578.4897
215.13931274414062 0 7022.402
221.08482360839844 0 2311.7336
222.08541870117188 0 804.6001
223.0634002685547 0 1222.3859
223.0811004638672 0 735.6031
225.04306030273438 0 1578.2412
227.0665283203125 0 8587.045
228.06961059570312 0 699.16907
239.09556579589844 0 3387.5151
240.09671020507812 0 1678.368
241.0918731689453 0 960.72003
260.1243896484375 0 1102.4117
266.62689208984375 0 515.06744
271.7863464355469 0 592.1307
272.08782958984375 0 1399.488
281.0520935058594 0 820.2051
282.0520935058594 0 941.63745
282.14581298828125 0 1756.5577
295.10382080078125 0 2460.5034
295.165283203125 0 1637.0413 y 10
296.1044921875 0 3053.5
297.1011657714844 0 1534.7385
297.1567077636719 0 885.5273
298.1009826660156 0 555.9752
299.0619812011719 0 1913.9736
300.0624084472656 0 1534.5754
300.1558532714844 0 5566.2983
301.05828857421875 0 944.93036
301.1589660644531 0 840.0664
306.16888427734375 0 5399.8574
307.1755065917969 0 2501.172
313.1152038574219 0 572.9642
314.09832763671875 0 932.9937
315.1673583984375 0 1112.5007
317.1822814941406 0 3390.713
318.0937194824219 0 1640.7915
318.8009338378906 0 563.04083
319.1980895996094 0 1907.0994 c 2
320.2030029296875 0 654.1467
341.0184020996094 0 12766.729
355.0701904296875 0 2283.1143
359.0286560058594 0 44022.883
359.1634521484375 0 573.03955
363.1905822753906 0 867.5055 c Ammonia loss 9
364.1981201171875 0 1811.2899
367.16162109375 0 1939.2314
369.1212158203125 0 732.3497
370.1219482421875 0 2413.4019
371.1208190917969 0 1595.2191
371.1562194824219 0 1809.1965
372.1213684082031 0 848.66614
373.1974792480469 0 561.66656
375.8595886230469 0 741.048
379.2089538574219 0 5845.0986
382.19573974609375 0 1232.0817 y 9
385.2196960449219 0 3421.6633
385.5460510253906 0 899.70386
385.8833312988281 0 728.7388
390.9993591308594 0 596.219
391.551513671875 0 1061.8862
391.8841857910156 0 1804.0778
392.2195129394531 0 972.62274
395.1586608886719 0 647.6212
409.2079772949219 0 1099.7815
429.08935546875 0 113538.64
429.7550048828125 0 1663.9554
430.2418212890625 0 2288.3271
431.24639892578125 0 638.7684
431.2785339355469 0 726.556
433.2199401855469 0 5637.6895
434.2247314453125 0 600.5867
442.7457580566406 0 2942.9321
442.8352966308594 0 580.70496
443.24078369140625 0 1020.5712 c Water loss 7
443.7409973144531 0 1099.819 c Ammonia loss 7
444.69757080078125 0 622.7859
445.12060546875 0 62534.21
445.2666320800781 0 777.4536
446.21142578125 0 1818.5898
447.1003112792969 0 683.7333
451.6827697753906 0 1181.7133
451.7506408691406 0 7004.998
452.2542419433594 0 21125.838 c 7
452.7529296875 0 10443.786
453.2532043457031 0 2697.1348
453.7530517578125 0 1216.6819
458.27215576171875 0 1299.1317 c Ammonia loss 3
459.1849060058594 0 668.30164
459.7001037597656 0 2295.7576
459.73541259765625 0 947.844
460.2042541503906 0 1001.718
460.239501953125 0 1916.4812
460.6951904296875 0 859.9346
460.7348327636719 0 659.96063
461.1857604980469 0 933.66595
461.2390441894531 0 604.3542
462.7066650390625 0 519.6958
465.2623596191406 0 988.3156
471.2488098144531 0 812.15247
475.2994079589844 0 74470.63 c 3
476.2418212890625 0 601.97546
476.3020324707031 0 15777.223
477.3043518066406 0 2499.0205
479.75848388671875 0 6740.337
480.2219543457031 0 854.8291 z 8
480.2584533691406 0 3324.1025
480.7607727050781 0 1069.4181
481.23016357421875 0 4927.5605
482.2328796386719 0 1019.0408
484.7506103515625 0 1419.2194
485.24908447265625 0 668.86206
485.7402038574219 0 569.5418
486.7720031738281 0 988.7545
487.2686462402344 0 2144.919
487.77008056640625 0 1093.4458
490.24444580078125 0 713.21155
491.79791259765625 0 607.6418
492.233154296875 0 3841.4678
493.2398986816406 0 1616.0635
493.7541198730469 0 1103.2925
495.23223876953125 0 723.23157
496.2399597167969 0 2049.8557 y 8
499.7665100097656 0 7846.956
500.26593017578125 0 5334.888 c Water loss 8
500.7634582519531 0 5307.97 c Ammonia loss 8
501.2621154785156 0 2447.4626
501.76690673828125 0 2051.9958
502.2747802734375 0 7643.446
503.2743225097656 0 1680.1328
508.25238037109375 0 14473.449
508.7718811035156 0 16005.682
509.274169921875 0 28048.203 c 8
509.7731018066406 0 17748.13
510.2735290527344 0 5851.8955
510.7729797363281 0 2309.718
511.2661437988281 0 772.1055
522.7750854492188 0 969.4913
523.7617797851562 0 579.3542 z Water loss 3
526.2548828125 0 1036.7384
526.7471923828125 0 668.5213
530.293212890625 0 900.31433
532.2879028320312 0 796.26685
532.765380859375 0 2777.744 z 3
533.2645874023438 0 2631.532
533.7636108398438 0 1137.8339
535.2750854492188 0 923.0892
535.7692260742188 0 861.1666
540.7728271484375 0 824.53674 y 3
541.2752685546875 0 784.988
544.2780151367188 0 2671.1335 c Ammonia loss 9
544.3193359375 0 683.2269
544.7766723632812 0 2143.3396
545.3054809570312 0 2072.5205
546.309326171875 0 1547.425
547.3123779296875 0 768.0586
548.2509155273438 0 3412.4053
548.748291015625 0 2339.0286
549.2484741210938 0 641.4697
552.2518920898438 0 691.72144
552.74365234375 0 1225.7765
552.7921142578125 0 22137.178 c 9
553.288818359375 0 16488.64
553.7892456054688 0 7765.7476
554.2866821289062 0 1862.322
554.7809448242188 0 1495.4559
561.290283203125 0 734.3538
561.7864379882812 0 887.7554
564.3038330078125 0 3613.262
565.309326171875 0 1736.1666
565.7986450195312 0 1523.7568
566.2921142578125 0 1000.5091
567.3053588867188 0 593.3387
567.7777709960938 0 918.3727 z Ammonia loss 2
568.2736206054688 0 1244.7694
568.773193359375 0 834.0642
569.2801513671875 0 854.7289
570.3082275390625 0 657.9633
573.3113403320312 0 1155.1455
575.7805786132812 0 2734.7236 y Ammonia loss 2
576.2822265625 0 7767.1567 z 2
576.7821655273438 0 5473.7817
577.280029296875 0 3259.0393
577.8212280273438 0 1011.353
578.317138671875 0 1437.1978
578.8150634765625 0 1049.3191
579.8111572265625 0 685.1953
583.3043823242188 0 1345.8713
584.2918701171875 0 20769.41 y 2
584.7901611328125 0 16214.563
585.2894897460938 0 8490.17
585.7891845703125 0 2561.0752
586.8228149414062 0 2608.9878
587.3242797851562 0 4086.0232
587.8215942382812 0 1119.6145
588.276611328125 0 1095.1995
588.7693481445312 0 987.3284
590.3261108398438 0 133673.28 c 4
591.3287963867188 0 35486.52
591.8146362304688 0 2133.3135
592.3278198242188 0 6728.1187
592.8101806640625 0 2013.2255
600.8212280273438 0 3898.029 c Ammonia loss 10
601.3201904296875 0 2942.1802
601.820556640625 0 1508.4546
607.3212280273438 0 5304.2954
608.3182373046875 0 7828.548 z 7
609.3334350585938 0 53715.816 c 10
609.83251953125 0 40595.215
610.3317260742188 0 20855.283
610.8335571289062 0 5506.196
611.3319702148438 0 1185.2546
616.3364868164062 0 1158.4537
618.304931640625 0 3294.4973 w 1
618.806396484375 0 2941.4316
619.3030395507812 0 1063.7473
624.335205078125 0 9742.215 y 7
625.3363647460938 0 3553.9216
630.3301391601562 0 1201.9916
630.8240966796875 0 1036.1868
631.324951171875 0 1664.7708
632.3193969726562 0 667.5374 y Ammonia loss 1
632.8245239257812 0 2339.9053 z 1
633.3206787109375 0 2553.982
633.8172607421875 0 1561.649
634.3206176757812 0 1577.3413
636.3497314453125 0 1828.0121
637.828125 0 895.3121
638.325927734375 0 540.43555
638.3502807617188 0 542.739
639.8318481445312 0 709.7343
640.3402709960938 0 594.57684
640.8320922851562 0 2570.22 y 1
641.3323974609375 0 2729.635
641.828369140625 0 1712.4568
642.340087890625 0 705.40356
647.3501586914062 0 2954.1602
647.8515014648438 0 2658.2898
648.3536987304688 0 1038.3503
649.3291625976562 0 825.68335
652.8464965820312 0 1228.8749
653.3384399414062 0 1561.4531
653.8328857421875 0 851.1642
654.3333740234375 0 797.3135
654.8277587890625 0 772.957
655.3176879882812 0 4526.2744
655.8178100585938 0 4452.759
656.3141479492188 0 1644.8973
659.3480834960938 0 5803.6646
660.3484497070312 0 8164.1074
660.8525390625 0 5849.165
661.3489379882812 0 13124.057 z Water loss 6
661.851806640625 0 8245.766
662.3433837890625 0 10315.954
662.8434448242188 0 6173.1665
663.3414916992188 0 2141.2083
668.830810546875 0 3503.4631
669.3484497070312 0 10763.028
669.8490600585938 0 7901.82
670.3474731445312 0 10020.939
670.8455810546875 0 5579.5786
671.3419799804688 0 3985.8367
672.3458862304688 0 1760.0214
674.346923828125 0 2530.3418
674.844482421875 0 6054.031
675.3438110351562 0 4359.917
675.8411254882812 0 5669.5054
676.33837890625 0 4876.8027
676.83349609375 0 1947.0717
677.3345947265625 0 1107.6771
678.3307495117188 0 950.38367
679.35400390625 0 27769.033 z 6
680.3536376953125 0 11757.511
681.3544921875 0 3172.3977
682.8531494140625 0 2559.3726
683.348388671875 0 34606.76
683.8466796875 0 34196.28
684.3472900390625 0 14688.728
684.847900390625 0 6009.4595
687.34912109375 0 1415.0277 c Ammonia loss 5
689.159912109375 0 777.97095
689.6585083007812 0 4590.86
690.1613159179688 0 2129.238
691.3572387695312 0 32052.033
691.8560791015625 0 28761.693
692.3555908203125 0 15684.05
692.854736328125 0 5942.039
693.3567504882812 0 1703.3607
694.3522338867188 0 14609.29
695.3654174804688 0 10406.596 y 6
696.366943359375 0 3176.183
697.3634033203125 0 1334.3052
702.2886352539062 0 1016.9763
704.368896484375 0 32669.229 c 5
705.3670043945312 0 12715.702
706.3699340820312 0 2159.431
714.3723754882812 0 807.30994
722.347412109375 0 9898.378
728.3443603515625 0 2042.5779
731.3927612304688 0 6138.4067
732.3939819335938 0 1443.5836
737.387939453125 0 1180.5637
749.38330078125 0 16215.165 w 5
750.3865356445312 0 10350.048
751.3865356445312 0 3748.0679
752.3923950195312 0 928.01013
758.3800659179688 0 2274.695 c Ammonia loss 6
759.3844604492188 0 1071.8661
772.3585205078125 0 7538.2407
773.3607177734375 0 3164.2878
775.4061279296875 0 32708.668 c 6
776.34912109375 0 8129.186
776.40673828125 0 9303.865
777.4024658203125 0 3307.4954
778.40087890625 0 735.052
787.3937377929688 0 802.7047
788.3950805664062 0 994.1289
793.3967895507812 0 71642.4 z 5
794.3936767578125 0 39847.926
795.39404296875 0 10698.699
796.39501953125 0 3128.638
809.4097290039062 0 1352.386 y 5
810.4025268554688 0 934.08954
837.375 0 1105.4373
858.4882202148438 0 1099.2924
859.4859619140625 0 2469.0107
860.4893798828125 0 3039.865
861.3601684570312 0 805.0562
861.4817504882812 0 1008.4262
863.3902587890625 0 917.76886
864.4334106445312 0 24822.713
865.4290771484375 0 15041.819
866.4299926757812 0 4115.571
867.427490234375 0 965.8314
873.4212036132812 0 754.38403
874.4175415039062 0 1312.3635
877.4215087890625 0 940.65283
879.4075317382812 0 4169.012
882.3873901367188 0 1919.5055
884.4863891601562 0 1001.79425
885.4739990234375 0 1056.9293 c Water loss 7
886.47314453125 0 2704.0457 c Ammonia loss 7
887.4754028320312 0 2242.0986
888.4829711914062 0 977.8237
891.4092407226562 0 1879.1466 z Ammonia loss 4
892.4127197265625 0 927.0232
900.4755249023438 0 3244.2554
901.47509765625 0 2432.6775
902.4912719726562 0 10723.592
903.499755859375 0 41464.72 c 7
904.382568359375 0 2075.006
904.497802734375 0 23189.229
905.3984985351562 0 1129.1572
905.4959106445312 0 7570.6064
906.4920654296875 0 2049.7388
908.4019165039062 0 31885.484
909.4197998046875 0 16364.753
910.4193115234375 0 5972.214
911.418701171875 0 1012.7972
915.9605712890625 0 737.7656
919.4719848632812 0 2788.031
920.4057006835938 0 2285.0747
920.492919921875 0 1207.5453
921.4154052734375 0 8565.037
922.4181518554688 0 5681.8
923.4276123046875 0 2237.763
924.4137573242188 0 5163.8765
925.41650390625 0 5481.6543
961.4994506835938 0 795.33936
969.478271484375 0 710.899
986.5018310546875 0 1297.6903
987.5057373046875 0 3741.8713
988.5009155273438 0 1955.9841
989.500244140625 0 1168.1565
999.5350341796875 0 2348.153 c Water loss 8
1000.523681640625 0 3455.5647 c Ammonia loss 8
1001.5172119140625 0 2445.8015
1002.524658203125 0 2105.1677
1003.5223999023438 0 1559.206
1016.5348510742188 0 3916.4583
1017.5430297851562 0 27533.158 c 8
1018.5405883789062 0 17093.66
1019.5401611328125 0 7521.5083
1020.5367431640625 0 2426.258
1061.564453125 0 1003.8553
1062.565185546875 0 786.51544
1064.52587890625 0 8309.745 z 3
1065.524658203125 0 12179.549
1066.5224609375 0 7999.9375
1067.5264892578125 0 2208.285
1080.5391845703125 0 1328.7029 y 3
1081.5440673828125 0 1054.1461
1088.5509033203125 0 1539.2086
1089.552001953125 0 1954.478
1090.5570068359375 0 1111.741
1104.5755615234375 0 8258.495 c 9
1105.57470703125 0 6745.53
1106.571533203125 0 3868.5205
1107.5679931640625 0 2000.6396
1134.5501708984375 0 768.2017 z Ammonia loss 2
1135.5499267578125 0 1449.0759
1151.5577392578125 0 6020.89 z 2
1152.559326171875 0 13370.896
1153.55908203125 0 7824.039
1154.5596923828125 0 4166.1206
1155.5677490234375 0 819.6169
1159.624267578125 0 1202.9908
1173.6466064453125 0 2829.3457
1174.6546630859375 0 3324.5886
1175.648193359375 0 3609.5798
1176.6431884765625 0 1033.7799
1177.648193359375 0 718.1987
1201.638916015625 0 5728.092
1202.637939453125 0 4209.535
1203.64208984375 0 1769.7825
1204.6444091796875 0 1398.8699
1217.658935546875 0 4807.7437 c 10
1218.660888671875 0 10002.265
1219.662109375 0 6207.4517
1220.662841796875 0 2844.868
1265.6478271484375 0 2439.829
1266.6387939453125 0 2484.434
1267.64404296875 0 962.671
1305.6617431640625 0 1314.6189
1306.6492919921875 0 1574.7324
1307.6588134765625 0 3322.6082
1308.652099609375 0 1185.764
1309.6429443359375 0 947.2526
1320.7015380859375 0 1546.8508
1321.6851806640625 0 3377.5881
1322.694091796875 0 2514.016
1323.685546875 0 5881.8774
1324.6785888671875 0 9808.574
1325.676025390625 0 7764.3247
1326.6796875 0 3703.7217
1337.696044921875 0 2085.837
1338.7003173828125 0 13774.369
1339.69873046875 0 10821.009
1340.6971435546875 0 7671.5376
1341.6942138671875 0 2279.5256
1348.6851806640625 0 2706.5854
1349.6741943359375 0 5966.196
1350.66845703125 0 5147.4824
1351.6644287109375 0 1952.9272
1355.7132568359375 0 959.49054
1356.7357177734375 0 700.3709
1365.69140625 0 8719.433
1366.6934814453125 0 44115.887
1367.69189453125 0 40365.94
1368.689697265625 0 21304.93
1369.6922607421875 0 7469.7554
1370.6881103515625 0 1421.5083
1378.3067626953125 0 1166.5233
1379.3160400390625 0 2574.2134
1380.3193359375 0 1923.8948
1381.7076416015625 0 2018.7498
1382.7081298828125 0 5847.351
1383.7149658203125 0 10869.955
1384.714599609375 0 9474.397
1385.712646484375 0 4325.415
1386.71923828125 0 1270.9613
3081.830078125 0 765.99884
3082.523681640625 0 726.7383

Spectrum Details

|  |  |
| --- | --- |
| Matched peaks? Matched peaks The total absolute number of peaks matched. Additionally in brackets the total fraction of peaks matched and the total number of peaks is shown. | 56 (11.16% of 502) |
| FDR? FDR The false discovery rate estimated for this peptide. It is calculated by matching all theoretical fragments with a non-integer shift with the raw peaks for this spectrum. This is done with 40 different shifts. The resulting percentage is the average number of annotated peaks over the number of annotated peaks with the correct spectrum. | 1.74% |
| Satellite FDR? Satellite FDR See the FDR for details on its calculation. This satellite ion specific FDR only contains the satellite ions (d/w) for I/L/J positions. | 0.00% |
| PSM Score? PSM Score The PSM Score as given by Hecklib to this annotated spectrum. It is shown with three significant figures. | 428 |

#### Reverse Lookup? Reverse Lookup All places where this read could be placed.

| Group | Segment | Template | Template Part | Read Part | Score | Unique |
| --- | --- | --- | --- | --- | --- | --- |
| Homo sapiens Heavy Chain | IGHV | IGHV3-9 | [68..80] | [0..12] | 96 | False |
| Homo sapiens Heavy Chain | IGHV | IGHV3-48 | [68..80] | [0..12] | 96 | False |
| Homo sapiens Heavy Chain | IGHV | IGHV3-21 | [68..80] | [0..12] | 96 | False |
| Homo sapiens Heavy Chain | IGHV | IGHV3-7 | [68..80] | [0..12] | 96 | False |
| Homo sapiens Heavy Chain | IGHV | IGHV3-43 | [68..80] | [0..12] | 87 | False |
| Homo sapiens Heavy Chain | IGHV | IGHV3-13 | [67..79] | [0..12] | 87 | False |
| Homo sapiens Heavy Chain | IGHV | IGHV3-11 | [68..80] | [0..12] | 96 | False |
| Homo sapiens Heavy Chain | IGHV | IGHV3-74 | [68..80] | [0..12] | 87 | False |
| Homo sapiens Heavy Chain | IGHV | IGHV3-20 | [68..80] | [0..12] | 96 | False |

| Recombined | Template Part | Read Part | Score | Unique |
| --- | --- | --- | --- | --- |
| REC-0-1 | [68..80] | [0..12] | 96 | True |

#### Meta Information from Multiple reads

##### Number of combined reads

13

##### Intensity

0.9095

##### TotalArea

4.005E+09

##### Changes to the peptide sequence

TISRDNAKNSJY

L→JNo support for either Leucine or Isoleucine based on side chain ions (Position: 11)

L→ISupport for Isoleucine based on side chain ions (1 for I 0 for L) (Position: 2)

#### Positional Score

Copy Data

##### Positional Score (TSV)

###### Preview

```
Loading example...
```

*Click on the button to copy the data to your clipboard.*

1001234567891011

Label Value
"0" 0.808
"1" 0.815
"2" 0.818
"3" 0.824
"4" 0.835
"5" 0.825
"6" 0.842
"7" 0.839
"8" 0.825
"9" 0.825
"10" 0.836
"11" 0.836

#### Meta Information from PEAKS

##### Scan Identifier

F3:3437

##### Original sequence

T

L

S

R

D

N

A

K

N

S

L

Y

##### Posttranslational Modifications

##### Source File

D:\separate\_stitch\_analyses\xle-disambiguation\raw\20210323\_F1\_UM1\_Peng0013\_SA\_F59\_ingel\_3ug\_chymo.raw

##### Fraction

3

##### Scan Feature

F3:2673

##### De Novo Score

99

##### ConfidenceScore

99

### m/z

461.2412

##### Mass

1380.6997

##### Charge

3

##### Retention Time

17.97

##### Predicted Retention Time

-

##### Area

1.978E+09

##### Parts Per Million

1.6

##### Fragmentation mode

ETHCD

##### Originating file

01 D:\separate\_stitch\_analyses\xle-disambiguation\20210325\_F59\_3ug\_DENOVO\_12.csv

#### Meta Information from PEAKS

##### Scan Identifier

F3:3360

##### Original sequence

T

L

S

R

D

N

A

K

N

S

L

Y

##### Posttranslational Modifications

##### Source File

D:\separate\_stitch\_analyses\xle-disambiguation\raw\20210323\_F1\_UM1\_Peng0013\_SA\_F59\_ingel\_3ug\_chymo.raw

##### Fraction

3

##### Scan Feature

F3:2673

##### De Novo Score

99

##### ConfidenceScore

99

### m/z

461.2412

##### Mass

1380.6997

##### Charge

3

##### Retention Time

17.97

##### Predicted Retention Time

-

##### Area

1.978E+09

##### Parts Per Million

1.6

##### Fragmentation mode

ETHCD

##### Originating file

01 D:\separate\_stitch\_analyses\xle-disambiguation\20210325\_F59\_3ug\_DENOVO\_12.csv

#### Meta Information from PEAKS

##### Scan Identifier

F3:3614

##### Original sequence

T

L

S

R

D

N

A

K

N

S

L

Y

##### Posttranslational Modifications

##### Source File

D:\separate\_stitch\_analyses\xle-disambiguation\raw\20210323\_F1\_UM1\_Peng0013\_SA\_F59\_ingel\_3ug\_chymo.raw

##### Fraction

3

##### Scan Feature

-

##### De Novo Score

98

##### ConfidenceScore

98

### m/z

461.2418

##### Mass

1380.6997

##### Charge

3

##### Retention Time

19.78

##### Predicted Retention Time

-

##### Area

0

##### Parts Per Million

2.8

##### Fragmentation mode

ETHCD

##### Originating file

01 D:\separate\_stitch\_analyses\xle-disambiguation\20210325\_F59\_3ug\_DENOVO\_12.csv

#### Meta Information from PEAKS

##### Scan Identifier

F3:4261

##### Original sequence

T

L

S

R

D

N

A

K

N

S

L

Y

##### Posttranslational Modifications

##### Source File

D:\separate\_stitch\_analyses\xle-disambiguation\raw\20210323\_F1\_UM1\_Peng0013\_SA\_F59\_ingel\_3ug\_chymo.raw

##### Fraction

3

##### Scan Feature

-

##### De Novo Score

98

##### ConfidenceScore

98

### m/z

461.2411

##### Mass

1380.6997

##### Charge

3

##### Retention Time

23.43

##### Predicted Retention Time

-

##### Area

0

##### Parts Per Million

1.3

##### Fragmentation mode

ETHCD

##### Originating file

01 D:\separate\_stitch\_analyses\xle-disambiguation\20210325\_F59\_3ug\_DENOVO\_12.csv

#### Meta Information from PEAKS

##### Scan Identifier

F3:4196

##### Original sequence

T

L

S

R

D

N

A

K

N

S

L

Y

##### Posttranslational Modifications

##### Source File

D:\separate\_stitch\_analyses\xle-disambiguation\raw\20210323\_F1\_UM1\_Peng0013\_SA\_F59\_ingel\_3ug\_chymo.raw

##### Fraction

3

##### Scan Feature

-

##### De Novo Score

98

##### ConfidenceScore

98

### m/z

461.2417

##### Mass

1380.6997

##### Charge

3

##### Retention Time

23.06

##### Predicted Retention Time

-

##### Area

0

##### Parts Per Million

2.5

##### Fragmentation mode

ETHCD

##### Originating file

01 D:\separate\_stitch\_analyses\xle-disambiguation\20210325\_F59\_3ug\_DENOVO\_12.csv

#### Meta Information from PEAKS

##### Scan Identifier

F3:4924

##### Original sequence

T

L

S

R

D

N

A

K

N

S

L

Y

##### Posttranslational Modifications

##### Source File

D:\separate\_stitch\_analyses\xle-disambiguation\raw\20210323\_F1\_UM1\_Peng0013\_SA\_F59\_ingel\_3ug\_chymo.raw

##### Fraction

3

##### Scan Feature

F3:2674

##### De Novo Score

98

##### ConfidenceScore

98

### m/z

461.2412

##### Mass

1380.6997

##### Charge

3

##### Retention Time

26.97

##### Predicted Retention Time

-

##### Area

1.605E+07

##### Parts Per Million

1.6

##### Fragmentation mode

ETHCD

##### Originating file

01 D:\separate\_stitch\_analyses\xle-disambiguation\20210325\_F59\_3ug\_DENOVO\_12.csv

#### Meta Information from PEAKS

##### Scan Identifier

F3:3892

##### Original sequence

T

L

S

R

D

N

A

K

N

S

L

Y

##### Posttranslational Modifications

##### Source File

D:\separate\_stitch\_analyses\xle-disambiguation\raw\20210323\_F1\_UM1\_Peng0013\_SA\_F59\_ingel\_3ug\_chymo.raw

##### Fraction

3

##### Scan Feature

-

##### De Novo Score

97

##### ConfidenceScore

97

### m/z

461.2408

##### Mass

1380.6997

##### Charge

3

##### Retention Time

21.33

##### Predicted Retention Time

-

##### Area

0

##### Parts Per Million

0.7

##### Fragmentation mode

ETHCD

##### Originating file

01 D:\separate\_stitch\_analyses\xle-disambiguation\20210325\_F59\_3ug\_DENOVO\_12.csv

#### Meta Information from PEAKS

##### Scan Identifier

F3:5262

##### Original sequence

T

L

S

R

D

N

A

K

N

S

L

Y

##### Posttranslational Modifications

##### Source File

D:\separate\_stitch\_analyses\xle-disambiguation\raw\20210323\_F1\_UM1\_Peng0013\_SA\_F59\_ingel\_3ug\_chymo.raw

##### Fraction

3

##### Scan Feature

F3:2674

##### De Novo Score

97

##### ConfidenceScore

97

### m/z

461.2412

##### Mass

1380.6997

##### Charge

3

##### Retention Time

26.97

##### Predicted Retention Time

-

##### Area

1.605E+07

##### Parts Per Million

1.6

##### Fragmentation mode

ETHCD

##### Originating file

01 D:\separate\_stitch\_analyses\xle-disambiguation\20210325\_F59\_3ug\_DENOVO\_12.csv

#### Meta Information from PEAKS

##### Scan Identifier

F3:3238

##### Original sequence

T

L

S

R

D

N

A

K

N

S

L

Y

##### Posttranslational Modifications

##### Source File

D:\separate\_stitch\_analyses\xle-disambiguation\raw\20210323\_F1\_UM1\_Peng0013\_SA\_F59\_ingel\_3ug\_chymo.raw

##### Fraction

3

##### Scan Feature

-

##### De Novo Score

97

##### ConfidenceScore

97

### m/z

461.2415

##### Mass

1380.6997

##### Charge

3

##### Retention Time

17.7

##### Predicted Retention Time

-

##### Area

0

##### Parts Per Million

2.1

##### Fragmentation mode

ETHCD

##### Originating file

01 D:\separate\_stitch\_analyses\xle-disambiguation\20210325\_F59\_3ug\_DENOVO\_12.csv

#### Meta Information from PEAKS

##### Scan Identifier

F3:4571

##### Original sequence

T

L

S

R

D

N

A

K

N

S

L

Y

##### Posttranslational Modifications

##### Source File

D:\separate\_stitch\_analyses\xle-disambiguation\raw\20210323\_F1\_UM1\_Peng0013\_SA\_F59\_ingel\_3ug\_chymo.raw

##### Fraction

3

##### Scan Feature

F3:2674

##### De Novo Score

97

##### ConfidenceScore

97

### m/z

461.2412

##### Mass

1380.6997

##### Charge

3

##### Retention Time

26.97

##### Predicted Retention Time

-

##### Area

1.605E+07

##### Parts Per Million

1.6

##### Fragmentation mode

ETHCD

##### Originating file

01 D:\separate\_stitch\_analyses\xle-disambiguation\20210325\_F59\_3ug\_DENOVO\_12.csv

#### Meta Information from PEAKS

##### Scan Identifier

F3:3162

##### Original sequence

T

L

S

R

D

N

A

K

N

S

L

Y

##### Posttranslational Modifications

##### Source File

D:\separate\_stitch\_analyses\xle-disambiguation\raw\20210323\_F1\_UM1\_Peng0013\_SA\_F59\_ingel\_3ug\_chymo.raw

##### Fraction

3

##### Scan Feature

-

##### De Novo Score

97

##### ConfidenceScore

97

### m/z

461.2415

##### Mass

1380.6997

##### Charge

3

##### Retention Time

17.27

##### Predicted Retention Time

-

##### Area

0

##### Parts Per Million

2.1

##### Fragmentation mode

ETHCD

##### Originating file

01 D:\separate\_stitch\_analyses\xle-disambiguation\20210325\_F59\_3ug\_DENOVO\_12.csv

#### Meta Information from PEAKS

##### Scan Identifier

F3:4327

##### Original sequence

T

L

S

R

D

N

A

K

N

S

L

Y

##### Posttranslational Modifications

##### Source File

D:\separate\_stitch\_analyses\xle-disambiguation\raw\20210323\_F1\_UM1\_Peng0013\_SA\_F59\_ingel\_3ug\_chymo.raw

##### Fraction

3

##### Scan Feature

-

##### De Novo Score

97

##### ConfidenceScore

97

### m/z

461.2415

##### Mass

1380.6997

##### Charge

3

##### Retention Time

23.81

##### Predicted Retention Time

-

##### Area

0

##### Parts Per Million

2.1

##### Fragmentation mode

ETHCD

##### Originating file

01 D:\separate\_stitch\_analyses\xle-disambiguation\20210325\_F59\_3ug\_DENOVO\_12.csv

#### Meta Information from PEAKS

##### Scan Identifier

F3:4073

##### Original sequence

T

L

S

R

D

N

A

K

N

S

L

Y

##### Posttranslational Modifications

##### Source File

D:\separate\_stitch\_analyses\xle-disambiguation\raw\20210323\_F1\_UM1\_Peng0013\_SA\_F59\_ingel\_3ug\_chymo.raw

##### Fraction

3

##### Scan Feature

-

##### De Novo Score

97

##### ConfidenceScore

97

### m/z

461.2415

##### Mass

1380.6997

##### Charge

3

##### Retention Time

22.36

##### Predicted Retention Time

-

##### Area

0

##### Parts Per Million

2.1

##### Fragmentation mode

ETHCD

##### Originating file

01 D:\separate\_stitch\_analyses\xle-disambiguation\20210325\_F59\_3ug\_DENOVO\_12.csv
