## Supplementary material for "A handle on mass coincidence errors in *de novo* sequencing of antibodies by bottom-up proteomics": Combined_017.html

Details Combined\_017 | Stitch OverviewUndefined

### Read Combined\_017

#### Sequence (length=14)

JYLQMNSJRJDDTA

#### Spectrum 8058? Spectrum 8058 The raw spectrum of this peptide as annotated by Hecklib. The fragments are coloured according to ion type (see legend). Any peaks with a star '\*' as text can be hovered over to see the full details, first the ion type second the mass shift type. By hovering over the amino acids in the peptide or ions in the legend the corresponding peaks are highlighted. By toggling the 'Unassigned' label you can turn the background (unassigned) peaks on or off in the plot. By updating the slider in the Ion legend you can update the spectrum to only show the top X% of the peaks with labels. The top X% means any peak that is within X% of the highest intensity. By dragging in the spectrum you can zoom in to a specific part of the spectrum and use 'Zoom Out' to get back to the original zoom level. The annotation of the spectrum is based on the given sequence in the peptides file and is done with different software so inconsistencies are likely. The peaks are annotated based on the given sequence, with 20 ppm tolerance.

Copy Data

##### Spectrum 8058 (TSV)

###### Preview

```
Loading example...
```

*Click on the button to copy the data to your clipboard.*

Mz MinMz MaxIntensity Max

WidthHeightPeptide font sizePeptide stroke widthSpectrum font sizeSpectrum stroke widthCompact peptide

Ion legend

wxyz

abcd

OtherUnassignedIon^Charge^_Position_Show for top:%

JYLQMNSJRJDDTA

04.03e+48.06e+41.21e+51.61e+5

Zoom Out

y^+1^_2_y^+1^_2_a^+1^_2_b^+1^_2_y^+1^_3_y^+1^_3_b^+1^_3_y^+1^_4_y^+1^_4_y^+2^_8_y^+1^_5_b^+1^_4_y^+1^_5_b^+2^_9_y^+2^_10_b^+2^_9_y^+2^_10_b^+2^_10_y^+2^_11_y^+2^_11_y^+2^_11_b^+1^_5_y^+1^_6_y^+1^_6_b^+2^_11_b^+2^_11_y^+2^_12_y^+2^_12_y^+1^_6_y^+2^_12_b^+2^_12_b^+2^_12_b^+2^_12_b^+1^_6_y^+2^_13_y^+2^_13_y^+2^_13_b^+1^_6_b^+2^_13_b^+2^_13_y^+1^_7_y^+1^_7_b^+2^_13_y^+1^_7_\*\*\*b^+1^_7_b^+1^_7_b^+1^_7_y^+1^_8_y^+1^_8_y^+1^_8_b^+1^_8_b^+1^_8_y^+1^_9_y^+1^_9_y^+1^_9_b^+1^_9_y^+1^_10_y^+1^_10_b^+1^_9_y^+1^_10_b^+1^_10_y^+1^_11_y^+1^_11_y^+1^_11_b^+1^_11_b^+1^_11_y^+1^_12_y^+1^_12_y^+1^_12_b^+1^_12_b^+1^_12_y^+1^_13_

0823164624703293

Fragment Matches Table

Show background peaks

| Position | Ion type | Intensity | mz Theoretical | mz Error (Th) | mz Error (ppm) | Charge | Series Number |
| --- | --- | --- | --- | --- | --- | --- | --- |
| - | - | 6899 | 120.1 | - | - | 0 | - |
| - | - | 486.9 | 121.1 | - | - | 0 | - |
| - | - | 449.8 | 123.1 | - | - | 0 | - |
| - | - | 436.7 | 124.1 | - | - | 0 | - |
| - | - | 443.4 | 124.1 | - | - | 0 | - |
| - | - | 731.1 | 125.1 | - | - | 0 | - |
| - | - | 827 | 126.1 | - | - | 0 | - |
| - | - | 1248 | 127.1 | - | - | 0 | - |
| - | - | 2115 | 129.1 | - | - | 0 | - |
| - | - | 9327 | 129.1 | - | - | 0 | - |
| - | - | 410.1 | 130 | - | - | 0 | - |
| - | - | 560.6 | 130.1 | - | - | 0 | - |
| - | - | 446.9 | 130.5 | - | - | 0 | - |
| - | - | 1095 | 131.1 | - | - | 0 | - |
| - | - | 1.29E+04 | 131.1 | - | - | 0 | - |
| - | - | 442.1 | 132.1 | - | - | 0 | - |
| - | - | 914 | 133.1 | - | - | 0 | - |
| - | - | 909.2 | 134 | - | - | 0 | - |
| - | - | 1.338E+05 | 136.1 | - | - | 0 | - |
| - | - | 903.5 | 137.1 | - | - | 0 | - |
| - | - | 1.172E+04 | 137.1 | - | - | 0 | - |
| - | - | 543 | 138.1 | - | - | 0 | - |
| - | - | 390.9 | 138.1 | - | - | 0 | - |
| - | - | 767 | 139.1 | - | - | 0 | - |
| - | - | 851.9 | 139.1 | - | - | 0 | - |
| - | - | 916.5 | 140.1 | - | - | 0 | - |
| - | - | 1126 | 141.1 | - | - | 0 | - |
| - | - | 2296 | 141.1 | - | - | 0 | - |
| - | - | 1038 | 142.1 | - | - | 0 | - |
| - | - | 743.8 | 143 | - | - | 0 | - |
| - | - | 534 | 143.1 | - | - | 0 | - |
| - | - | 480 | 143.1 | - | - | 0 | - |
| - | - | 383.4 | 143.8 | - | - | 0 | - |
| - | - | 2166 | 144.1 | - | - | 0 | - |
| - | - | 459.1 | 144.6 | - | - | 0 | - |
| - | - | 779.7 | 145.1 | - | - | 0 | - |
| - | - | 825.5 | 145.1 | - | - | 0 | - |
| - | - | 3447 | 146.1 | - | - | 0 | - |
| - | - | 854.3 | 147 | - | - | 0 | - |
| - | - | 637.1 | 147.1 | - | - | 0 | - |
| - | - | 1407 | 147.1 | - | - | 0 | - |
| - | - | 539.2 | 148.9 | - | - | 0 | - |
| - | - | 531.7 | 148.9 | - | - | 0 | - |
| - | - | 536.6 | 148.9 | - | - | 0 | - |
| - | - | 616.9 | 148.9 | - | - | 0 | - |
| - | - | 574 | 148.9 | - | - | 0 | - |
| - | - | 887.6 | 148.9 | - | - | 0 | - |
| - | - | 1011 | 148.9 | - | - | 0 | - |
| - | - | 1180 | 148.9 | - | - | 0 | - |
| - | - | 1953 | 148.9 | - | - | 0 | - |
| - | - | 3586 | 148.9 | - | - | 0 | - |
| - | - | 3919 | 149 | - | - | 0 | - |
| - | - | 2310 | 149 | - | - | 0 | - |
| - | - | 1273 | 149 | - | - | 0 | - |
| - | - | 1168 | 149 | - | - | 0 | - |
| - | - | 861.1 | 149 | - | - | 0 | - |
| - | - | 736.2 | 149 | - | - | 0 | - |
| - | - | 487 | 149 | - | - | 0 | - |
| - | - | 526.4 | 149 | - | - | 0 | - |
| - | - | 624.7 | 149 | - | - | 0 | - |
| - | - | 582.4 | 149 | - | - | 0 | - |
| - | - | 577.1 | 149 | - | - | 0 | - |
| - | - | 401.9 | 149 | - | - | 0 | - |
| - | - | 493.9 | 149.2 | - | - | 0 | - |
| - | - | 475.8 | 150.1 | - | - | 0 | - |
| - | - | 2390 | 152.1 | - | - | 0 | - |
| - | - | 731.8 | 152.1 | - | - | 0 | - |
| - | - | 1505 | 153.1 | - | - | 0 | - |
| - | - | 506.9 | 153.1 | - | - | 0 | - |
| - | - | 1403 | 155.1 | - | - | 0 | - |
| - | - | 1388 | 155.1 | - | - | 0 | - |
| - | - | 2732 | 156.1 | - | - | 0 | - |
| - | - | 4107 | 157.1 | - | - | 0 | - |
| - | - | 1476 | 157.1 | - | - | 0 | - |
| - | - | 945.5 | 157.1 | - | - | 0 | - |
| - | - | 653.9 | 157.1 | - | - | 0 | - |
| - | - | 520.5 | 158.1 | - | - | 0 | - |
| - | - | 2149 | 159.1 | - | - | 0 | - |
| - | - | 744.3 | 159.1 | - | - | 0 | - |
| - | - | 507.5 | 159.1 | - | - | 0 | - |
| - | - | 445.3 | 165.1 | - | - | 0 | - |
| - | - | 540.2 | 165.1 | - | - | 0 | - |
| - | - | 842.2 | 166.1 | - | - | 0 | - |
| - | - | 3229 | 166.1 | - | - | 0 | - |
| - | - | 908.6 | 167 | - | - | 0 | - |
| - | - | 540.2 | 167.1 | - | - | 0 | - |
| - | - | 5330 | 167.1 | - | - | 0 | - |
| - | - | 962.7 | 167.1 | - | - | 0 | - |
| - | - | 806.7 | 168.1 | - | - | 0 | - |
| - | - | 682.4 | 168.1 | - | - | 0 | - |
| - | - | 974.8 | 168.1 | - | - | 0 | - |
| - | - | 501 | 168.1 | - | - | 0 | - |
| - | - | 583.3 | 169.1 | - | - | 0 | - |
| - | - | 2718 | 169.1 | - | - | 0 | - |
| - | - | 969.2 | 169.1 | - | - | 0 | - |
| - | - | 2621 | 171.1 | - | - | 0 | - |
| - | - | 685 | 171.1 | - | - | 0 | - |
| - | - | 614.2 | 173.1 | - | - | 0 | - |
| 13 | y | 5481 | 173.1 | 0.0003843 | 2.22 | +1 | 2 |
| - | - | 6737 | 173.1 | - | - | 0 | - |
| - | - | 719.8 | 173.4 | - | - | 0 | - |
| - | - | 9752 | 174.1 | - | - | 0 | - |
| - | - | 2174 | 174.1 | - | - | 0 | - |
| - | - | 505.2 | 174.1 | - | - | 0 | - |
| - | - | 3023 | 175.1 | - | - | 0 | - |
| - | - | 2162 | 175.1 | - | - | 0 | - |
| - | - | 801.7 | 177.1 | - | - | 0 | - |
| - | - | 459.3 | 177.7 | - | - | 0 | - |
| - | - | 1408 | 178 | - | - | 0 | - |
| - | - | 1019 | 180.1 | - | - | 0 | - |
| - | - | 1127 | 181.1 | - | - | 0 | - |
| - | - | 1459 | 181.1 | - | - | 0 | - |
| - | - | 544.6 | 182.1 | - | - | 0 | - |
| - | - | 1700 | 183.1 | - | - | 0 | - |
| - | - | 1825 | 183.1 | - | - | 0 | - |
| - | - | 4780 | 183.1 | - | - | 0 | - |
| - | - | 2067 | 184.1 | - | - | 0 | - |
| - | - | 2343 | 184.1 | - | - | 0 | - |
| - | - | 2746 | 185.1 | - | - | 0 | - |
| - | - | 1597 | 185.1 | - | - | 0 | - |
| - | - | 663.2 | 186.1 | - | - | 0 | - |
| - | - | 497.9 | 187.1 | - | - | 0 | - |
| - | - | 860.8 | 187.1 | - | - | 0 | - |
| - | - | 1.465E+04 | 187.1 | - | - | 0 | - |
| - | - | 1063 | 187.1 | - | - | 0 | - |
| - | - | 601 | 187.1 | - | - | 0 | - |
| - | - | 680.5 | 188.1 | - | - | 0 | - |
| - | - | 3714 | 188.1 | - | - | 0 | - |
| - | - | 5881 | 189.1 | - | - | 0 | - |
| - | - | 481.6 | 189.1 | - | - | 0 | - |
| - | - | 952.6 | 191.1 | - | - | 0 | - |
| 13 | y | 2.212E+04 | 191.1 | 0.0003176 | 1.662 | +1 | 2 |
| - | - | 1185 | 192.1 | - | - | 0 | - |
| - | - | 1117 | 192.2 | - | - | 0 | - |
| - | - | 524.5 | 193.1 | - | - | 0 | - |
| - | - | 1374 | 193.1 | - | - | 0 | - |
| - | - | 2337 | 194.1 | - | - | 0 | - |
| - | - | 1.226E+04 | 195.1 | - | - | 0 | - |
| - | - | 1929 | 195.1 | - | - | 0 | - |
| - | - | 778.9 | 196.1 | - | - | 0 | - |
| - | - | 916.6 | 196.1 | - | - | 0 | - |
| - | - | 572.6 | 196.1 | - | - | 0 | - |
| - | - | 1055 | 197.1 | - | - | 0 | - |
| - | - | 2054 | 197.1 | - | - | 0 | - |
| - | - | 4512 | 198.1 | - | - | 0 | - |
| - | - | 1769 | 199.1 | - | - | 0 | - |
| - | - | 637.7 | 199.1 | - | - | 0 | - |
| - | - | 1400 | 199.1 | - | - | 0 | - |
| - | - | 526.8 | 200.4 | - | - | 0 | - |
| - | - | 3941 | 201.1 | - | - | 0 | - |
| - | - | 5964 | 201.1 | - | - | 0 | - |
| - | - | 1260 | 202.1 | - | - | 0 | - |
| - | - | 6241 | 202.1 | - | - | 0 | - |
| - | - | 705.6 | 202.1 | - | - | 0 | - |
| - | - | 551.2 | 203.1 | - | - | 0 | - |
| - | - | 698.7 | 203.1 | - | - | 0 | - |
| - | - | 3.52E+04 | 203.1 | - | - | 0 | - |
| - | - | 2414 | 204.1 | - | - | 0 | - |
| - | - | 696 | 204.1 | - | - | 0 | - |
| - | - | 1008 | 207.1 | - | - | 0 | - |
| - | - | 1801 | 208.1 | - | - | 0 | - |
| - | - | 3182 | 208.1 | - | - | 0 | - |
| - | - | 760.8 | 208.1 | - | - | 0 | - |
| - | - | 1735 | 209.1 | - | - | 0 | - |
| - | - | 892.9 | 210.1 | - | - | 0 | - |
| - | - | 801.1 | 210.2 | - | - | 0 | - |
| - | - | 1079 | 211.1 | - | - | 0 | - |
| - | - | 1101 | 211.1 | - | - | 0 | - |
| - | - | 3687 | 211.1 | - | - | 0 | - |
| - | - | 3727 | 212.1 | - | - | 0 | - |
| - | - | 797.3 | 213.1 | - | - | 0 | - |
| - | - | 977.2 | 214.1 | - | - | 0 | - |
| - | - | 1651 | 214.2 | - | - | 0 | - |
| - | - | 1702 | 215.1 | - | - | 0 | - |
| - | - | 887.4 | 216.1 | - | - | 0 | - |
| - | - | 640 | 217.1 | - | - | 0 | - |
| - | - | 1.159E+04 | 217.1 | - | - | 0 | - |
| - | - | 894.7 | 217.1 | - | - | 0 | - |
| - | - | 536.8 | 218 | - | - | 0 | - |
| - | - | 821.8 | 218.1 | - | - | 0 | - |
| - | - | 528.8 | 219.1 | - | - | 0 | - |
| - | - | 1519 | 221.1 | - | - | 0 | - |
| - | - | 889.9 | 221.1 | - | - | 0 | - |
| - | - | 1747 | 223.1 | - | - | 0 | - |
| - | - | 951.3 | 224.1 | - | - | 0 | - |
| - | - | 1021 | 224.1 | - | - | 0 | - |
| - | - | 8255 | 225.1 | - | - | 0 | - |
| - | - | 6814 | 225.1 | - | - | 0 | - |
| - | - | 3959 | 225.2 | - | - | 0 | - |
| - | - | 1058 | 226.1 | - | - | 0 | - |
| - | - | 1089 | 226.1 | - | - | 0 | - |
| - | - | 618.7 | 226.1 | - | - | 0 | - |
| - | - | 1968 | 227.1 | - | - | 0 | - |
| - | - | 3658 | 228.1 | - | - | 0 | - |
| - | - | 717.9 | 228.1 | - | - | 0 | - |
| - | - | 1762 | 228.2 | - | - | 0 | - |
| - | - | 3779 | 229.1 | - | - | 0 | - |
| - | - | 734.4 | 229.2 | - | - | 0 | - |
| - | - | 1124 | 230.1 | - | - | 0 | - |
| - | - | 3733 | 231.1 | - | - | 0 | - |
| - | - | 605 | 231.2 | - | - | 0 | - |
| - | - | 957.2 | 233.1 | - | - | 0 | - |
| - | - | 660.9 | 233.1 | - | - | 0 | - |
| - | - | 795.1 | 235.1 | - | - | 0 | - |
| - | - | 1260 | 235.1 | - | - | 0 | - |
| - | - | 1027 | 236.1 | - | - | 0 | - |
| - | - | 969.7 | 237.1 | - | - | 0 | - |
| - | - | 776.9 | 238.2 | - | - | 0 | - |
| - | - | 2739 | 239.1 | - | - | 0 | - |
| - | - | 1366 | 240.1 | - | - | 0 | - |
| - | - | 522.2 | 240.1 | - | - | 0 | - |
| - | - | 770.4 | 241.1 | - | - | 0 | - |
| - | - | 2057 | 242.1 | - | - | 0 | - |
| - | - | 2.784E+04 | 242.2 | - | - | 0 | - |
| - | - | 1211 | 243.1 | - | - | 0 | - |
| - | - | 2155 | 243.2 | - | - | 0 | - |
| - | - | 853.3 | 244.1 | - | - | 0 | - |
| - | - | 735.6 | 244.1 | - | - | 0 | - |
| - | - | 1325 | 245.1 | - | - | 0 | - |
| - | - | 1424 | 245.1 | - | - | 0 | - |
| - | - | 633.2 | 246.1 | - | - | 0 | - |
| - | - | 1312 | 246.1 | - | - | 0 | - |
| 2 | a | 1.597E+05 | 249.2 | 0.0004629 | 1.858 | +1 | 2 |
| - | - | 2.07E+04 | 250.2 | - | - | 0 | - |
| - | - | 1827 | 251.2 | - | - | 0 | - |
| - | - | 857.2 | 252.1 | - | - | 0 | - |
| - | - | 1333 | 252.1 | - | - | 0 | - |
| - | - | 1304 | 253.1 | - | - | 0 | - |
| - | - | 7135 | 253.2 | - | - | 0 | - |
| - | - | 653.5 | 254.2 | - | - | 0 | - |
| - | - | 611.4 | 255.1 | - | - | 0 | - |
| - | - | 592.8 | 256.2 | - | - | 0 | - |
| - | - | 1733 | 258.1 | - | - | 0 | - |
| - | - | 4937 | 259.1 | - | - | 0 | - |
| - | - | 792.6 | 259.1 | - | - | 0 | - |
| - | - | 708.6 | 260.1 | - | - | 0 | - |
| - | - | 1250 | 261.1 | - | - | 0 | - |
| - | - | 719.3 | 261.1 | - | - | 0 | - |
| - | - | 824.3 | 261.2 | - | - | 0 | - |
| - | - | 2885 | 262.1 | - | - | 0 | - |
| - | - | 1457 | 262.2 | - | - | 0 | - |
| - | - | 1668 | 263.1 | - | - | 0 | - |
| - | - | 601.8 | 263.1 | - | - | 0 | - |
| - | - | 1154 | 264.1 | - | - | 0 | - |
| - | - | 807.8 | 264.1 | - | - | 0 | - |
| - | - | 2815 | 265.2 | - | - | 0 | - |
| - | - | 609 | 266.2 | - | - | 0 | - |
| - | - | 3530 | 267.1 | - | - | 0 | - |
| - | - | 2523 | 268.1 | - | - | 0 | - |
| - | - | 574.5 | 268.1 | - | - | 0 | - |
| - | - | 1026 | 269.2 | - | - | 0 | - |
| - | - | 915.6 | 270.1 | - | - | 0 | - |
| - | - | 917.2 | 270.1 | - | - | 0 | - |
| - | - | 1423 | 270.1 | - | - | 0 | - |
| - | - | 1739 | 270.2 | - | - | 0 | - |
| - | - | 728.1 | 274.1 | - | - | 0 | - |
| - | - | 4702 | 274.1 | - | - | 0 | - |
| - | - | 877.4 | 275.1 | - | - | 0 | - |
| - | - | 4917 | 276.1 | - | - | 0 | - |
| 2 | b | 1.234E+05 | 277.2 | 0.0003298 | 1.19 | +1 | 2 |
| - | - | 1325 | 278.1 | - | - | 0 | - |
| - | - | 1.885E+04 | 278.2 | - | - | 0 | - |
| - | - | 1116 | 279.2 | - | - | 0 | - |
| - | - | 1085 | 280.1 | - | - | 0 | - |
| - | - | 1912 | 280.2 | - | - | 0 | - |
| - | - | 565.3 | 282.1 | - | - | 0 | - |
| - | - | 675 | 282.1 | - | - | 0 | - |
| - | - | 3917 | 282.2 | - | - | 0 | - |
| - | - | 4422 | 285.1 | - | - | 0 | - |
| - | - | 1293 | 286.1 | - | - | 0 | - |
| - | - | 962 | 287.1 | - | - | 0 | - |
| - | - | 900.9 | 287.2 | - | - | 0 | - |
| - | - | 1120 | 288.1 | - | - | 0 | - |
| 12 | y | 8234 | 288.1 | 0.0004646 | 1.612 | +1 | 3 |
| - | - | 666.4 | 289.1 | - | - | 0 | - |
| - | - | 649.1 | 290.1 | - | - | 0 | - |
| - | - | 1558 | 291.1 | - | - | 0 | - |
| - | - | 1405 | 291.1 | - | - | 0 | - |
| - | - | 1319 | 293.1 | - | - | 0 | - |
| - | - | 609 | 294.1 | - | - | 0 | - |
| - | - | 1104 | 294.2 | - | - | 0 | - |
| - | - | 704.1 | 295.1 | - | - | 0 | - |
| - | - | 822.7 | 296.2 | - | - | 0 | - |
| - | - | 1954 | 297.2 | - | - | 0 | - |
| - | - | 592.8 | 297.2 | - | - | 0 | - |
| - | - | 700.8 | 298.1 | - | - | 0 | - |
| - | - | 1750 | 298.1 | - | - | 0 | - |
| - | - | 868 | 299.1 | - | - | 0 | - |
| - | - | 602.3 | 299.1 | - | - | 0 | - |
| - | - | 585.7 | 299.2 | - | - | 0 | - |
| - | - | 615.9 | 300.1 | - | - | 0 | - |
| - | - | 804.1 | 304.1 | - | - | 0 | - |
| - | - | 1125 | 304.1 | - | - | 0 | - |
| - | - | 3505 | 306.1 | - | - | 0 | - |
| 12 | y | 5467 | 306.1 | 0.0003369 | 1.101 | +1 | 3 |
| - | - | 1571 | 307.1 | - | - | 0 | - |
| - | - | 830.4 | 307.2 | - | - | 0 | - |
| - | - | 1226 | 308.1 | - | - | 0 | - |
| - | - | 923.4 | 308.2 | - | - | 0 | - |
| - | - | 3594 | 309.1 | - | - | 0 | - |
| - | - | 638.2 | 311.2 | - | - | 0 | - |
| - | - | 946.5 | 313.1 | - | - | 0 | - |
| - | - | 2418 | 314.1 | - | - | 0 | - |
| - | - | 3224 | 315.2 | - | - | 0 | - |
| - | - | 1116 | 317.2 | - | - | 0 | - |
| - | - | 1299 | 322.2 | - | - | 0 | - |
| - | - | 1900 | 323.2 | - | - | 0 | - |
| - | - | 666.2 | 324.1 | - | - | 0 | - |
| - | - | 2288 | 325.2 | - | - | 0 | - |
| - | - | 554.6 | 326.1 | - | - | 0 | - |
| - | - | 1229 | 326.2 | - | - | 0 | - |
| - | - | 686.3 | 328.1 | - | - | 0 | - |
| - | - | 666.1 | 328.1 | - | - | 0 | - |
| - | - | 1002 | 329.1 | - | - | 0 | - |
| - | - | 673.8 | 330.1 | - | - | 0 | - |
| - | - | 623.9 | 331.1 | - | - | 0 | - |
| - | - | 737.6 | 332.1 | - | - | 0 | - |
| - | - | 7355 | 332.1 | - | - | 0 | - |
| - | - | 1582 | 333.1 | - | - | 0 | - |
| - | - | 699 | 334.2 | - | - | 0 | - |
| - | - | 907.5 | 335.2 | - | - | 0 | - |
| - | - | 604.4 | 337.2 | - | - | 0 | - |
| - | - | 5578 | 340.2 | - | - | 0 | - |
| - | - | 2446 | 341.2 | - | - | 0 | - |
| - | - | 822.5 | 341.2 | - | - | 0 | - |
| - | - | 857 | 341.2 | - | - | 0 | - |
| - | - | 812.4 | 342.2 | - | - | 0 | - |
| - | - | 884.9 | 342.2 | - | - | 0 | - |
| - | - | 988.3 | 343.2 | - | - | 0 | - |
| - | - | 970.1 | 343.2 | - | - | 0 | - |
| - | - | 4507 | 344.1 | - | - | 0 | - |
| - | - | 775.7 | 345.2 | - | - | 0 | - |
| - | - | 2235 | 345.2 | - | - | 0 | - |
| - | - | 1446 | 349.1 | - | - | 0 | - |
| - | - | 633.5 | 350.2 | - | - | 0 | - |
| - | - | 1205 | 351.1 | - | - | 0 | - |
| - | - | 824.2 | 351.2 | - | - | 0 | - |
| - | - | 1506 | 352.2 | - | - | 0 | - |
| - | - | 3371 | 353.2 | - | - | 0 | - |
| - | - | 2564 | 354.1 | - | - | 0 | - |
| - | - | 778.3 | 354.2 | - | - | 0 | - |
| - | - | 1563 | 355.1 | - | - | 0 | - |
| - | - | 698.9 | 356.1 | - | - | 0 | - |
| - | - | 1759 | 357.2 | - | - | 0 | - |
| - | - | 1311 | 360.2 | - | - | 0 | - |
| - | - | 604.9 | 363.2 | - | - | 0 | - |
| - | - | 756.5 | 366.3 | - | - | 0 | - |
| - | - | 2097 | 368.2 | - | - | 0 | - |
| - | - | 5380 | 368.2 | - | - | 0 | - |
| - | - | 913 | 370.2 | - | - | 0 | - |
| - | - | 1304 | 370.2 | - | - | 0 | - |
| - | - | 1439 | 372.2 | - | - | 0 | - |
| - | - | 1624 | 372.2 | - | - | 0 | - |
| - | - | 2525 | 373.1 | - | - | 0 | - |
| - | - | 4325 | 378.1 | - | - | 0 | - |
| - | - | 1757 | 380.2 | - | - | 0 | - |
| - | - | 1180 | 381.2 | - | - | 0 | - |
| - | - | 1047 | 382.1 | - | - | 0 | - |
| - | - | 716.6 | 383.2 | - | - | 0 | - |
| - | - | 1447 | 385.1 | - | - | 0 | - |
| - | - | 5414 | 385.2 | - | - | 0 | - |
| - | - | 723.6 | 386.2 | - | - | 0 | - |
| - | - | 824.4 | 388.2 | - | - | 0 | - |
| - | - | 5047 | 389.2 | - | - | 0 | - |
| - | - | 1460 | 390.1 | - | - | 0 | - |
| 3 | b | 2.218E+04 | 390.2 | 0.0006469 | 1.658 | +1 | 3 |
| - | - | 782.8 | 391.2 | - | - | 0 | - |
| - | - | 4351 | 391.2 | - | - | 0 | - |
| - | - | 917.8 | 392.2 | - | - | 0 | - |
| - | - | 991.9 | 395.2 | - | - | 0 | - |
| - | - | 743.9 | 395.2 | - | - | 0 | - |
| - | - | 3710 | 396.2 | - | - | 0 | - |
| - | - | 921.2 | 397.2 | - | - | 0 | - |
| - | - | 963.2 | 398.2 | - | - | 0 | - |
| - | - | 661.8 | 402.2 | - | - | 0 | - |
| 11 | y | 7798 | 403.1 | 0.000438 | 1.087 | +1 | 4 |
| - | - | 892.6 | 404.2 | - | - | 0 | - |
| - | - | 6799 | 405.2 | - | - | 0 | - |
| - | - | 1715 | 406.2 | - | - | 0 | - |
| - | - | 5917 | 407.3 | - | - | 0 | - |
| - | - | 1045 | 408.3 | - | - | 0 | - |
| - | - | 717.3 | 409.2 | - | - | 0 | - |
| - | - | 650.9 | 410.2 | - | - | 0 | - |
| - | - | 1170 | 411.2 | - | - | 0 | - |
| - | - | 775.5 | 412.2 | - | - | 0 | - |
| - | - | 692.1 | 413.2 | - | - | 0 | - |
| - | - | 703.6 | 420.2 | - | - | 0 | - |
| - | - | 841.4 | 420.2 | - | - | 0 | - |
| 11 | y | 4413 | 421.2 | 0.0003409 | 0.8094 | +1 | 4 |
| - | - | 1503 | 424.2 | - | - | 0 | - |
| - | - | 1625 | 427.2 | - | - | 0 | - |
| - | - | 963.7 | 428.3 | - | - | 0 | - |
| - | - | 1711 | 430.2 | - | - | 0 | - |
| - | - | 920.5 | 430.7 | - | - | 0 | - |
| - | - | 2396 | 433.2 | - | - | 0 | - |
| 7 | y | 2332 | 437.2 | 0.004413 | 10.09 | +2 | 8 |
| - | - | 1157 | 440.2 | - | - | 0 | - |
| - | - | 3709 | 442.1 | - | - | 0 | - |
| - | - | 802.5 | 443.2 | - | - | 0 | - |
| - | - | 1139 | 444.2 | - | - | 0 | - |
| - | - | 6210 | 445.2 | - | - | 0 | - |
| - | - | 1647 | 446.2 | - | - | 0 | - |
| - | - | 734.2 | 453.3 | - | - | 0 | - |
| - | - | 3864 | 454.2 | - | - | 0 | - |
| - | - | 803.2 | 456.3 | - | - | 0 | - |
| - | - | 773.3 | 458.2 | - | - | 0 | - |
| - | - | 2038 | 460.2 | - | - | 0 | - |
| - | - | 831.6 | 462.2 | - | - | 0 | - |
| - | - | 1330 | 463.2 | - | - | 0 | - |
| - | - | 2387 | 464.2 | - | - | 0 | - |
| - | - | 1262 | 465.2 | - | - | 0 | - |
| - | - | 3505 | 469.2 | - | - | 0 | - |
| - | - | 637 | 469.3 | - | - | 0 | - |
| - | - | 1008 | 470.2 | - | - | 0 | - |
| - | - | 688.2 | 470.7 | - | - | 0 | - |
| - | - | 2197 | 471.3 | - | - | 0 | - |
| - | - | 806 | 472.3 | - | - | 0 | - |
| - | - | 888.7 | 473.2 | - | - | 0 | - |
| - | - | 1115 | 473.3 | - | - | 0 | - |
| - | - | 1029 | 477.2 | - | - | 0 | - |
| - | - | 834.8 | 479.2 | - | - | 0 | - |
| - | - | 1132 | 481.3 | - | - | 0 | - |
| - | - | 978.4 | 482.2 | - | - | 0 | - |
| - | - | 4549 | 483.2 | - | - | 0 | - |
| - | - | 2715 | 484.3 | - | - | 0 | - |
| - | - | 1067 | 485.3 | - | - | 0 | - |
| - | - | 1218 | 486.2 | - | - | 0 | - |
| - | - | 672 | 486.8 | - | - | 0 | - |
| - | - | 775.7 | 487.3 | - | - | 0 | - |
| - | - | 1369 | 490.3 | - | - | 0 | - |
| - | - | 2014 | 491.2 | - | - | 0 | - |
| - | - | 799.7 | 491.3 | - | - | 0 | - |
| - | - | 2575 | 498.3 | - | - | 0 | - |
| - | - | 6839 | 500.2 | - | - | 0 | - |
| - | - | 946.9 | 501.2 | - | - | 0 | - |
| - | - | 1459 | 503.2 | - | - | 0 | - |
| - | - | 1211 | 507.2 | - | - | 0 | - |
| - | - | 773.3 | 507.2 | - | - | 0 | - |
| - | - | 727 | 508.3 | - | - | 0 | - |
| - | - | 840.6 | 511.8 | - | - | 0 | - |
| - | - | 922.5 | 512.3 | - | - | 0 | - |
| - | - | 748.4 | 513.2 | - | - | 0 | - |
| 10 | y | 4030 | 516.2 | 0.00161 | 3.118 | +1 | 5 |
| - | - | 737.3 | 517.3 | - | - | 0 | - |
| 4 | b | 7802 | 518.3 | 0.0007242 | 1.397 | +1 | 4 |
| - | - | 3330 | 519.3 | - | - | 0 | - |
| - | - | 965.7 | 520.3 | - | - | 0 | - |
| - | - | 610.2 | 524.3 | - | - | 0 | - |
| - | - | 1727 | 526.3 | - | - | 0 | - |
| - | - | 605.5 | 526.8 | - | - | 0 | - |
| - | - | 916.4 | 528.2 | - | - | 0 | - |
| - | - | 631.8 | 528.7 | - | - | 0 | - |
| 10 | y | 1.079E+04 | 534.2 | 0.001055 | 1.974 | +1 | 5 |
| - | - | 3018 | 535.2 | - | - | 0 | - |
| - | - | 2411 | 536.2 | - | - | 0 | - |
| - | - | 691.1 | 536.7 | - | - | 0 | - |
| - | - | 1627 | 537.3 | - | - | 0 | - |
| - | - | 2813 | 544.3 | - | - | 0 | - |
| - | - | 979.7 | 544.8 | - | - | 0 | - |
| - | - | 1698 | 545.2 | - | - | 0 | - |
| - | - | 922.4 | 545.8 | - | - | 0 | - |
| - | - | 2186 | 552.2 | - | - | 0 | - |
| - | - | 986 | 554.3 | - | - | 0 | - |
| - | - | 2559 | 554.3 | - | - | 0 | - |
| - | - | 651.4 | 555.2 | - | - | 0 | - |
| - | - | 987.4 | 556.3 | - | - | 0 | - |
| - | - | 834.6 | 556.4 | - | - | 0 | - |
| 9 | b | 952.6 | 559.8 | 0.004064 | 7.261 | +2 | 9 |
| - | - | 1229 | 560.3 | - | - | 0 | - |
| - | - | 721.1 | 560.8 | - | - | 0 | - |
| - | - | 709.5 | 562.3 | - | - | 0 | - |
| - | - | 612.4 | 564.8 | - | - | 0 | - |
| - | - | 996.2 | 565.3 | - | - | 0 | - |
| 5 | y | 733.4 | 567.3 | 0.007956 | 14.03 | +2 | 10 |
| - | - | 1004 | 567.3 | - | - | 0 | - |
| 9 | b | 5599 | 568.3 | 0.004096 | 7.207 | +2 | 9 |
| - | - | 4350 | 568.8 | - | - | 0 | - |
| - | - | 2009 | 569.3 | - | - | 0 | - |
| - | - | 1057 | 569.8 | - | - | 0 | - |
| - | - | 1732 | 572.3 | - | - | 0 | - |
| - | - | 1166 | 573.2 | - | - | 0 | - |
| - | - | 2132 | 573.3 | - | - | 0 | - |
| - | - | 758.8 | 573.3 | - | - | 0 | - |
| - | - | 1541 | 574.3 | - | - | 0 | - |
| 5 | y | 783 | 576.3 | 0.003712 | 6.441 | +2 | 10 |
| - | - | 1088 | 576.8 | - | - | 0 | - |
| - | - | 756.4 | 582.3 | - | - | 0 | - |
| - | - | 1167 | 583.3 | - | - | 0 | - |
| - | - | 773.9 | 583.3 | - | - | 0 | - |
| - | - | 2533 | 584.3 | - | - | 0 | - |
| - | - | 1032 | 584.8 | - | - | 0 | - |
| - | - | 1086 | 585.3 | - | - | 0 | - |
| - | - | 3544 | 585.3 | - | - | 0 | - |
| - | - | 1504 | 586.8 | - | - | 0 | - |
| - | - | 2770 | 587.3 | - | - | 0 | - |
| - | - | 767.7 | 587.8 | - | - | 0 | - |
| - | - | 931.6 | 588.3 | - | - | 0 | - |
| - | - | 765 | 593.3 | - | - | 0 | - |
| - | - | 737.8 | 594.3 | - | - | 0 | - |
| - | - | 2165 | 595.8 | - | - | 0 | - |
| - | - | 835.7 | 597.3 | - | - | 0 | - |
| - | - | 3780 | 599.3 | - | - | 0 | - |
| - | - | 1914 | 599.8 | - | - | 0 | - |
| - | - | 956 | 600.3 | - | - | 0 | - |
| - | - | 3028 | 601.3 | - | - | 0 | - |
| - | - | 1796 | 601.3 | - | - | 0 | - |
| - | - | 6273 | 601.8 | - | - | 0 | - |
| - | - | 5443 | 602.3 | - | - | 0 | - |
| - | - | 2142 | 602.8 | - | - | 0 | - |
| - | - | 887.4 | 607.3 | - | - | 0 | - |
| - | - | 1731 | 608.3 | - | - | 0 | - |
| - | - | 1078 | 612.3 | - | - | 0 | - |
| - | - | 4813 | 613.3 | - | - | 0 | - |
| - | - | 1590 | 614.3 | - | - | 0 | - |
| - | - | 705.9 | 617.3 | - | - | 0 | - |
| - | - | 1138 | 618.3 | - | - | 0 | - |
| - | - | 830.8 | 619.2 | - | - | 0 | - |
| - | - | 4340 | 619.3 | - | - | 0 | - |
| - | - | 2688 | 620.3 | - | - | 0 | - |
| - | - | 1300 | 622.3 | - | - | 0 | - |
| - | - | 2771 | 622.8 | - | - | 0 | - |
| - | - | 2729 | 623.3 | - | - | 0 | - |
| 10 | b | 3235 | 624.8 | 0.003201 | 5.123 | +2 | 10 |
| - | - | 2487 | 625.3 | - | - | 0 | - |
| - | - | 893.7 | 625.8 | - | - | 0 | - |
| - | - | 746.1 | 626.3 | - | - | 0 | - |
| - | - | 788.6 | 626.8 | - | - | 0 | - |
| - | - | 640.3 | 627.3 | - | - | 0 | - |
| - | - | 1077 | 627.4 | - | - | 0 | - |
| - | - | 716.1 | 629.3 | - | - | 0 | - |
| - | - | 1763 | 630.3 | - | - | 0 | - |
| 4 | y | 1.356E+04 | 631.3 | 0.00296 | 4.688 | +2 | 11 |
| - | - | 909.9 | 631.4 | - | - | 0 | - |
| 4 | y | 7015 | 631.8 | 0.01077 | 17.04 | +2 | 11 |
| - | - | 5086 | 632.3 | - | - | 0 | - |
| - | - | 986.9 | 634.8 | - | - | 0 | - |
| - | - | 847.9 | 635.3 | - | - | 0 | - |
| - | - | 660 | 635.8 | - | - | 0 | - |
| - | - | 1024 | 636.2 | - | - | 0 | - |
| - | - | 1067 | 636.3 | - | - | 0 | - |
| - | - | 2321 | 637.3 | - | - | 0 | - |
| - | - | 1065 | 638.3 | - | - | 0 | - |
| - | - | 885.1 | 638.8 | - | - | 0 | - |
| - | - | 926.9 | 639.3 | - | - | 0 | - |
| 4 | y | 1.379E+04 | 640.3 | 0.003231 | 5.047 | +2 | 11 |
| - | - | 9578 | 640.8 | - | - | 0 | - |
| - | - | 3394 | 641.3 | - | - | 0 | - |
| - | - | 896.2 | 641.8 | - | - | 0 | - |
| - | - | 1865 | 643.3 | - | - | 0 | - |
| - | - | 1853 | 643.8 | - | - | 0 | - |
| - | - | 2178 | 644.3 | - | - | 0 | - |
| - | - | 4066 | 647.3 | - | - | 0 | - |
| - | - | 2074 | 647.8 | - | - | 0 | - |
| - | - | 2494 | 648.3 | - | - | 0 | - |
| - | - | 718 | 649.3 | - | - | 0 | - |
| - | - | 806.9 | 650.4 | - | - | 0 | - |
| - | - | 6133 | 652.3 | - | - | 0 | - |
| - | - | 2635 | 652.8 | - | - | 0 | - |
| - | - | 2029 | 653.3 | - | - | 0 | - |
| - | - | 1756 | 655.3 | - | - | 0 | - |
| - | - | 896.9 | 655.8 | - | - | 0 | - |
| - | - | 1701 | 656.3 | - | - | 0 | - |
| - | - | 1406 | 656.8 | - | - | 0 | - |
| - | - | 899.1 | 663.3 | - | - | 0 | - |
| - | - | 1284 | 663.8 | - | - | 0 | - |
| - | - | 3762 | 664.8 | - | - | 0 | - |
| 5 | b | 1.103E+04 | 665.3 | 0.008081 | 12.15 | +1 | 5 |
| - | - | 3745 | 666.3 | - | - | 0 | - |
| - | - | 1576 | 670.8 | - | - | 0 | - |
| 9 | y | 4017 | 672.3 | 0.0002301 | 0.3422 | +1 | 6 |
| 9 | y | 3595 | 673.3 | 0.005228 | 7.765 | +1 | 6 |
| 11 | b | 1569 | 673.8 | 0.004835 | 7.176 | +2 | 11 |
| - | - | 1036 | 674.3 | - | - | 0 | - |
| - | - | 682.1 | 675.3 | - | - | 0 | - |
| - | - | 5562 | 679.3 | - | - | 0 | - |
| - | - | 3805 | 679.8 | - | - | 0 | - |
| - | - | 1600 | 680.3 | - | - | 0 | - |
| - | - | 1519 | 680.8 | - | - | 0 | - |
| 11 | b | 1.739E+04 | 682.4 | 0.003402 | 4.985 | +2 | 11 |
| - | - | 1.112E+04 | 682.9 | - | - | 0 | - |
| - | - | 4796 | 683.4 | - | - | 0 | - |
| - | - | 1178 | 684.3 | - | - | 0 | - |
| - | - | 734.2 | 685.3 | - | - | 0 | - |
| - | - | 766.6 | 686.3 | - | - | 0 | - |
| 3 | y | 4438 | 687.8 | 0.003408 | 4.955 | +2 | 12 |
| 3 | y | 7092 | 688.3 | 0.007006 | 10.18 | +2 | 12 |
| - | - | 4825 | 688.8 | - | - | 0 | - |
| - | - | 1864 | 689.3 | - | - | 0 | - |
| 9 | y | 3.798E+04 | 690.3 | 0.0007738 | 1.121 | +1 | 6 |
| - | - | 1.283E+04 | 691.3 | - | - | 0 | - |
| - | - | 2321 | 692.3 | - | - | 0 | - |
| - | - | 1219 | 695.3 | - | - | 0 | - |
| - | - | 689.8 | 696.4 | - | - | 0 | - |
| 3 | y | 3.207E+04 | 696.8 | 0.003131 | 4.492 | +2 | 12 |
| - | - | 2.617E+04 | 697.3 | - | - | 0 | - |
| - | - | 9476 | 697.8 | - | - | 0 | - |
| - | - | 3301 | 698.3 | - | - | 0 | - |
| - | - | 6148 | 699.4 | - | - | 0 | - |
| - | - | 6669 | 700.4 | - | - | 0 | - |
| - | - | 2055 | 701.4 | - | - | 0 | - |
| - | - | 1124 | 703.3 | - | - | 0 | - |
| - | - | 1204 | 707.9 | - | - | 0 | - |
| - | - | 863.8 | 708.9 | - | - | 0 | - |
| - | - | 779.8 | 709.4 | - | - | 0 | - |
| - | - | 906.4 | 710.4 | - | - | 0 | - |
| - | - | 812.2 | 713.3 | - | - | 0 | - |
| - | - | 823.2 | 714.4 | - | - | 0 | - |
| - | - | 1093 | 715.4 | - | - | 0 | - |
| - | - | 686.6 | 716.4 | - | - | 0 | - |
| - | - | 725.9 | 717.4 | - | - | 0 | - |
| - | - | 765.6 | 717.9 | - | - | 0 | - |
| - | - | 685.6 | 721.3 | - | - | 0 | - |
| - | - | 914.2 | 722.8 | - | - | 0 | - |
| - | - | 1043 | 724.8 | - | - | 0 | - |
| - | - | 3052 | 725.4 | - | - | 0 | - |
| - | - | 737.7 | 725.8 | - | - | 0 | - |
| - | - | 961 | 726.3 | - | - | 0 | - |
| - | - | 668.3 | 728.4 | - | - | 0 | - |
| - | - | 1494 | 729.3 | - | - | 0 | - |
| 12 | b | 1608 | 730.9 | 0.002903 | 3.972 | +2 | 12 |
| 12 | b | 2571 | 731.4 | 0.007904 | 10.81 | +2 | 12 |
| - | - | 2605 | 731.9 | - | - | 0 | - |
| - | - | 1195 | 732.4 | - | - | 0 | - |
| - | - | 1252 | 733.4 | - | - | 0 | - |
| - | - | 3163 | 733.9 | - | - | 0 | - |
| - | - | 5626 | 734.4 | - | - | 0 | - |
| - | - | 2126 | 734.9 | - | - | 0 | - |
| 12 | b | 3.006E+04 | 739.9 | 0.003175 | 4.291 | +2 | 12 |
| - | - | 2.172E+04 | 740.4 | - | - | 0 | - |
| - | - | 9297 | 740.9 | - | - | 0 | - |
| - | - | 1318 | 741.4 | - | - | 0 | - |
| - | - | 812.4 | 742.4 | - | - | 0 | - |
| - | - | 2012 | 744.4 | - | - | 0 | - |
| - | - | 2776 | 746.4 | - | - | 0 | - |
| - | - | 1193 | 746.9 | - | - | 0 | - |
| - | - | 725.4 | 747.4 | - | - | 0 | - |
| - | - | 1566 | 756.9 | - | - | 0 | - |
| - | - | 1668 | 758.4 | - | - | 0 | - |
| - | - | 1634 | 758.9 | - | - | 0 | - |
| - | - | 1154 | 759.4 | - | - | 0 | - |
| - | - | 1551 | 760.9 | - | - | 0 | - |
| 6 | b | 2036 | 762.3 | 0.006413 | 8.412 | +1 | 6 |
| - | - | 889.9 | 763.4 | - | - | 0 | - |
| - | - | 874.8 | 765.4 | - | - | 0 | - |
| - | - | 1112 | 766.4 | - | - | 0 | - |
| - | - | 1309 | 767.3 | - | - | 0 | - |
| - | - | 1225 | 767.9 | - | - | 0 | - |
| - | - | 2997 | 768.4 | - | - | 0 | - |
| - | - | 1211 | 768.9 | - | - | 0 | - |
| 2 | y | 2353 | 769.4 | 0.0141 | 18.33 | +2 | 13 |
| 2 | y | 2966 | 769.9 | 0.007446 | 9.672 | +2 | 13 |
| - | - | 2769 | 770.4 | - | - | 0 | - |
| - | - | 1291 | 772.9 | - | - | 0 | - |
| - | - | 980.3 | 773.4 | - | - | 0 | - |
| - | - | 3774 | 776.4 | - | - | 0 | - |
| - | - | 3526 | 776.9 | - | - | 0 | - |
| - | - | 2187 | 777.4 | - | - | 0 | - |
| 2 | y | 6182 | 778.4 | 0.004059 | 5.215 | +2 | 13 |
| - | - | 5984 | 778.9 | - | - | 0 | - |
| 6 | b | 6678 | 779.4 | 0.006963 | 8.934 | +1 | 6 |
| - | - | 743.6 | 779.9 | - | - | 0 | - |
| - | - | 1792 | 780.4 | - | - | 0 | - |
| 13 | b | 8089 | 781.4 | 0.002745 | 3.513 | +2 | 13 |
| 13 | b | 8182 | 781.9 | 0.009944 | 12.72 | +2 | 13 |
| - | - | 5794 | 782.4 | - | - | 0 | - |
| - | - | 1771 | 782.9 | - | - | 0 | - |
| - | - | 2056 | 783.4 | - | - | 0 | - |
| - | - | 793.9 | 783.9 | - | - | 0 | - |
| 8 | y | 2555 | 785.4 | 0.007052 | 8.978 | +1 | 7 |
| 8 | y | 1645 | 786.4 | 0.01021 | 12.99 | +1 | 7 |
| 13 | b | 2.422E+04 | 790.4 | 0.0032 | 4.049 | +2 | 13 |
| - | - | 2.122E+04 | 790.9 | - | - | 0 | - |
| - | - | 9981 | 791.4 | - | - | 0 | - |
| - | - | 2402 | 791.9 | - | - | 0 | - |
| - | - | 2118 | 793.9 | - | - | 0 | - |
| - | - | 971.9 | 794.3 | - | - | 0 | - |
| - | - | 2620 | 794.4 | - | - | 0 | - |
| - | - | 2550 | 794.9 | - | - | 0 | - |
| - | - | 1988 | 795.4 | - | - | 0 | - |
| - | - | 918.5 | 796.5 | - | - | 0 | - |
| - | - | 2652 | 797.4 | - | - | 0 | - |
| - | - | 1626 | 799.4 | - | - | 0 | - |
| - | - | 984.3 | 800.4 | - | - | 0 | - |
| - | - | 1963 | 801.4 | - | - | 0 | - |
| - | - | 1119 | 802.4 | - | - | 0 | - |
| - | - | 8490 | 802.9 | - | - | 0 | - |
| 8 | y | 2.683E+04 | 803.4 | 0.001747 | 2.175 | +1 | 7 |
| - | - | 3765 | 803.9 | - | - | 0 | - |
| - | - | 8633 | 804.4 | - | - | 0 | - |
| - | - | 1036 | 805.4 | - | - | 0 | - |
| - | - | 861.1 | 808.4 | - | - | 0 | - |
| - | - | 1191 | 808.9 | - | - | 0 | - |
| - | - | 1657 | 809.4 | - | - | 0 | - |
| - | - | 2064 | 809.9 | - | - | 0 | - |
| - | - | 846.9 | 812.4 | - | - | 0 | - |
| - | - | 888.9 | 812.9 | - | - | 0 | - |
| - | - | 754.6 | 813.4 | - | - | 0 | - |
| - | - | 1.068E+04 | 814.4 | - | - | 0 | - |
| - | - | 5178 | 815.4 | - | - | 0 | - |
| - | - | 1068 | 815.9 | - | - | 0 | - |
| - | - | 3317 | 816.4 | - | - | 0 | - |
| - | - | 2746 | 816.9 | - | - | 0 | - |
| - | - | 6861 | 817.4 | - | - | 0 | - |
| - | - | 3061 | 817.9 | - | - | 0 | - |
| - | - | 2402 | 818.4 | - | - | 0 | - |
| - | - | 997.9 | 819.4 | - | - | 0 | - |
| - | - | 1124 | 824.4 | - | - | 0 | - |
| - | - | 1330 | 824.9 | - | - | 0 | - |
| - | - | 1095 | 825.4 | - | - | 0 | - |
| 0 | Precursor | 1.896E+04 | 825.9 | 0.003076 | 3.724 | +2 | -1 |
| 0 | Precursor | 2.645E+04 | 826.4 | 0.009847 | 11.92 | +2 | -1 |
| - | - | 1.586E+04 | 826.9 | - | - | 0 | - |
| - | - | 8832 | 827.4 | - | - | 0 | - |
| - | - | 1488 | 827.9 | - | - | 0 | - |
| - | - | 1180 | 828.4 | - | - | 0 | - |
| - | - | 717 | 829.4 | - | - | 0 | - |
| - | - | 1030 | 829.7 | - | - | 0 | - |
| - | - | 1603 | 831.4 | - | - | 0 | - |
| - | - | 904.4 | 832.4 | - | - | 0 | - |
| - | - | 3076 | 833.4 | - | - | 0 | - |
| - | - | 2910 | 833.9 | - | - | 0 | - |
| 0 | Precursor | 9.297E+04 | 834.9 | 0.003104 | 3.717 | +2 | -1 |
| - | - | 3214 | 835 | - | - | 0 | - |
| - | - | 8.726E+04 | 835.4 | - | - | 0 | - |
| - | - | 4104 | 835.7 | - | - | 0 | - |
| - | - | 4.839E+04 | 835.9 | - | - | 0 | - |
| - | - | 1338 | 836.1 | - | - | 0 | - |
| - | - | 1.02E+04 | 836.4 | - | - | 0 | - |
| - | - | 1797 | 840.4 | - | - | 0 | - |
| - | - | 2044 | 842.4 | - | - | 0 | - |
| - | - | 1058 | 844.4 | - | - | 0 | - |
| - | - | 3610 | 846.4 | - | - | 0 | - |
| - | - | 1084 | 847.4 | - | - | 0 | - |
| 7 | b | 1131 | 848.4 | 0.01107 | 13.05 | +1 | 7 |
| 7 | b | 2368 | 849.4 | 0.01381 | 16.26 | +1 | 7 |
| - | - | 1515 | 850.4 | - | - | 0 | - |
| - | - | 1072 | 857.4 | - | - | 0 | - |
| - | - | 2008 | 858.4 | - | - | 0 | - |
| - | - | 5173 | 859.4 | - | - | 0 | - |
| - | - | 2288 | 860.5 | - | - | 0 | - |
| - | - | 747.7 | 861.4 | - | - | 0 | - |
| 7 | b | 3650 | 866.4 | 0.007405 | 8.547 | +1 | 7 |
| - | - | 1824 | 867.4 | - | - | 0 | - |
| - | - | 1655 | 870.4 | - | - | 0 | - |
| - | - | 868.3 | 871.4 | - | - | 0 | - |
| 7 | y | 3990 | 872.4 | 0.001386 | 1.589 | +1 | 8 |
| 7 | y | 1080 | 873.4 | 0.01469 | 16.81 | +1 | 8 |
| - | - | 1417 | 874.5 | - | - | 0 | - |
| - | - | 1278 | 880.4 | - | - | 0 | - |
| 7 | y | 2.746E+04 | 890.5 | 0.0004651 | 0.5223 | +1 | 8 |
| - | - | 1.244E+04 | 891.5 | - | - | 0 | - |
| - | - | 2429 | 892.5 | - | - | 0 | - |
| - | - | 3854 | 893.4 | - | - | 0 | - |
| - | - | 993.8 | 894.4 | - | - | 0 | - |
| - | - | 8253 | 897.4 | - | - | 0 | - |
| - | - | 4866 | 898.4 | - | - | 0 | - |
| - | - | 2030 | 899.4 | - | - | 0 | - |
| - | - | 2032 | 900.4 | - | - | 0 | - |
| - | - | 1631 | 902.5 | - | - | 0 | - |
| - | - | 1037 | 903.4 | - | - | 0 | - |
| - | - | 4574 | 910.5 | - | - | 0 | - |
| - | - | 1801 | 911.5 | - | - | 0 | - |
| - | - | 2293 | 915.5 | - | - | 0 | - |
| - | - | 4283 | 916.5 | - | - | 0 | - |
| - | - | 2883 | 917.5 | - | - | 0 | - |
| - | - | 3491 | 918.5 | - | - | 0 | - |
| - | - | 727.6 | 919.5 | - | - | 0 | - |
| - | - | 1737 | 933.5 | - | - | 0 | - |
| - | - | 1301 | 934.5 | - | - | 0 | - |
| - | - | 1655 | 936.5 | - | - | 0 | - |
| - | - | 1534 | 939.5 | - | - | 0 | - |
| - | - | 2403 | 940.4 | - | - | 0 | - |
| - | - | 1026 | 944.5 | - | - | 0 | - |
| - | - | 1181 | 945.5 | - | - | 0 | - |
| - | - | 911.1 | 946.5 | - | - | 0 | - |
| - | - | 825.1 | 951.5 | - | - | 0 | - |
| - | - | 965.2 | 953.5 | - | - | 0 | - |
| - | - | 968.3 | 954.5 | - | - | 0 | - |
| - | - | 757.1 | 955.5 | - | - | 0 | - |
| - | - | 989.7 | 956.5 | - | - | 0 | - |
| - | - | 2443 | 957.4 | - | - | 0 | - |
| - | - | 1742 | 958.5 | - | - | 0 | - |
| - | - | 780.8 | 959.5 | - | - | 0 | - |
| - | - | 890.8 | 960.5 | - | - | 0 | - |
| - | - | 7514 | 961.4 | - | - | 0 | - |
| 8 | b | 5456 | 962.5 | 0.003296 | 3.424 | +1 | 8 |
| - | - | 2402 | 963.5 | - | - | 0 | - |
| - | - | 810.4 | 965.5 | - | - | 0 | - |
| - | - | 1118 | 970.5 | - | - | 0 | - |
| - | - | 1331 | 971.5 | - | - | 0 | - |
| - | - | 2296 | 972.5 | - | - | 0 | - |
| - | - | 1193 | 973.5 | - | - | 0 | - |
| - | - | 4318 | 974.5 | - | - | 0 | - |
| - | - | 2098 | 975.5 | - | - | 0 | - |
| 8 | b | 4573 | 979.5 | 0.004396 | 4.488 | +1 | 8 |
| - | - | 3208 | 980.5 | - | - | 0 | - |
| - | - | 1265 | 981.5 | - | - | 0 | - |
| 6 | y | 3203 | 986.5 | 0.001 | 1.014 | +1 | 9 |
| 6 | y | 6565 | 987.5 | 0.007097 | 7.187 | +1 | 9 |
| - | - | 3644 | 988.5 | - | - | 0 | - |
| - | - | 1204 | 989.5 | - | - | 0 | - |
| - | - | 1025 | 991.5 | - | - | 0 | - |
| - | - | 2031 | 998.5 | - | - | 0 | - |
| - | - | 966.4 | 1001 | - | - | 0 | - |
| 6 | y | 4.641E+04 | 1005 | 0.0006895 | 0.6864 | +1 | 9 |
| - | - | 2.161E+04 | 1006 | - | - | 0 | - |
| - | - | 6935 | 1007 | - | - | 0 | - |
| - | - | 1935 | 1007 | - | - | 0 | - |
| - | - | 5533 | 1008 | - | - | 0 | - |
| - | - | 3156 | 1009 | - | - | 0 | - |
| - | - | 1207 | 1010 | - | - | 0 | - |
| - | - | 2995 | 1017 | - | - | 0 | - |
| - | - | 2496 | 1017 | - | - | 0 | - |
| - | - | 2149 | 1018 | - | - | 0 | - |
| - | - | 3209 | 1018 | - | - | 0 | - |
| - | - | 801.4 | 1019 | - | - | 0 | - |
| - | - | 1450 | 1023 | - | - | 0 | - |
| - | - | 3266 | 1024 | - | - | 0 | - |
| - | - | 2109 | 1025 | - | - | 0 | - |
| - | - | 5462 | 1026 | - | - | 0 | - |
| - | - | 4310 | 1027 | - | - | 0 | - |
| - | - | 1521 | 1028 | - | - | 0 | - |
| - | - | 764.7 | 1030 | - | - | 0 | - |
| - | - | 2644 | 1031 | - | - | 0 | - |
| - | - | 2759 | 1032 | - | - | 0 | - |
| - | - | 1479 | 1033 | - | - | 0 | - |
| - | - | 1205 | 1034 | - | - | 0 | - |
| - | - | 721.3 | 1035 | - | - | 0 | - |
| - | - | 930.8 | 1042 | - | - | 0 | - |
| - | - | 809.2 | 1044 | - | - | 0 | - |
| - | - | 865.5 | 1050 | - | - | 0 | - |
| - | - | 837.5 | 1052 | - | - | 0 | - |
| - | - | 1271 | 1054 | - | - | 0 | - |
| - | - | 962 | 1060 | - | - | 0 | - |
| - | - | 3015 | 1061 | - | - | 0 | - |
| - | - | 2839 | 1062 | - | - | 0 | - |
| - | - | 3232 | 1062 | - | - | 0 | - |
| - | - | 1506 | 1063 | - | - | 0 | - |
| - | - | 1251 | 1066 | - | - | 0 | - |
| - | - | 3166 | 1070 | - | - | 0 | - |
| - | - | 3871 | 1071 | - | - | 0 | - |
| - | - | 3755 | 1072 | - | - | 0 | - |
| - | - | 4324 | 1072 | - | - | 0 | - |
| - | - | 2932 | 1073 | - | - | 0 | - |
| - | - | 3250 | 1073 | - | - | 0 | - |
| - | - | 1268 | 1074 | - | - | 0 | - |
| - | - | 1073 | 1075 | - | - | 0 | - |
| - | - | 1257 | 1077 | - | - | 0 | - |
| - | - | 1404 | 1080 | - | - | 0 | - |
| - | - | 982.1 | 1082 | - | - | 0 | - |
| - | - | 919.5 | 1083 | - | - | 0 | - |
| - | - | 1187 | 1085 | - | - | 0 | - |
| - | - | 3078 | 1086 | - | - | 0 | - |
| - | - | 3.805E+04 | 1088 | - | - | 0 | - |
| - | - | 1.902E+04 | 1089 | - | - | 0 | - |
| - | - | 8353 | 1090 | - | - | 0 | - |
| - | - | 4011 | 1091 | - | - | 0 | - |
| - | - | 991.6 | 1092 | - | - | 0 | - |
| - | - | 826.4 | 1096 | - | - | 0 | - |
| - | - | 1064 | 1099 | - | - | 0 | - |
| - | - | 1837 | 1102 | - | - | 0 | - |
| - | - | 1634 | 1104 | - | - | 0 | - |
| - | - | 1568 | 1110 | - | - | 0 | - |
| - | - | 703.5 | 1117 | - | - | 0 | - |
| 9 | b | 2281 | 1119 | 0.006616 | 5.915 | +1 | 9 |
| - | - | 1505 | 1120 | - | - | 0 | - |
| - | - | 1159 | 1122 | - | - | 0 | - |
| - | - | 1964 | 1127 | - | - | 0 | - |
| - | - | 1935 | 1128 | - | - | 0 | - |
| - | - | 1571 | 1129 | - | - | 0 | - |
| - | - | 3803 | 1131 | - | - | 0 | - |
| - | - | 2662 | 1132 | - | - | 0 | - |
| - | - | 871.6 | 1133 | - | - | 0 | - |
| 5 | y | 3577 | 1134 | 0.004634 | 4.088 | +1 | 10 |
| 5 | y | 3401 | 1135 | 0.01415 | 12.47 | +1 | 10 |
| 9 | b | 1.515E+04 | 1136 | 0.005214 | 4.591 | +1 | 9 |
| - | - | 1.066E+04 | 1137 | - | - | 0 | - |
| - | - | 3457 | 1138 | - | - | 0 | - |
| - | - | 5743 | 1139 | - | - | 0 | - |
| - | - | 3671 | 1140 | - | - | 0 | - |
| - | - | 831.6 | 1140 | - | - | 0 | - |
| - | - | 2107 | 1145 | - | - | 0 | - |
| - | - | 1220 | 1146 | - | - | 0 | - |
| - | - | 1507 | 1147 | - | - | 0 | - |
| - | - | 850 | 1150 | - | - | 0 | - |
| 5 | y | 4.593E+04 | 1152 | 0.005544 | 4.815 | +1 | 10 |
| - | - | 3.046E+04 | 1153 | - | - | 0 | - |
| - | - | 1.132E+04 | 1154 | - | - | 0 | - |
| - | - | 2367 | 1155 | - | - | 0 | - |
| - | - | 1087 | 1157 | - | - | 0 | - |
| - | - | 2099 | 1159 | - | - | 0 | - |
| - | - | 2484 | 1160 | - | - | 0 | - |
| - | - | 986.1 | 1161 | - | - | 0 | - |
| - | - | 941.9 | 1172 | - | - | 0 | - |
| - | - | 1032 | 1173 | - | - | 0 | - |
| - | - | 1142 | 1174 | - | - | 0 | - |
| - | - | 790.3 | 1176 | - | - | 0 | - |
| - | - | 1428 | 1181 | - | - | 0 | - |
| - | - | 2018 | 1182 | - | - | 0 | - |
| - | - | 1105 | 1188 | - | - | 0 | - |
| - | - | 2109 | 1191 | - | - | 0 | - |
| - | - | 1171 | 1192 | - | - | 0 | - |
| - | - | 3913 | 1198 | - | - | 0 | - |
| - | - | 1.513E+04 | 1199 | - | - | 0 | - |
| - | - | 8556 | 1200 | - | - | 0 | - |
| - | - | 2661 | 1201 | - | - | 0 | - |
| - | - | 6056 | 1203 | - | - | 0 | - |
| - | - | 4584 | 1204 | - | - | 0 | - |
| - | - | 989 | 1205 | - | - | 0 | - |
| - | - | 1872 | 1214 | - | - | 0 | - |
| - | - | 865.8 | 1215 | - | - | 0 | - |
| - | - | 2.456E+04 | 1216 | - | - | 0 | - |
| - | - | 1.525E+04 | 1217 | - | - | 0 | - |
| - | - | 4882 | 1218 | - | - | 0 | - |
| - | - | 979.5 | 1219 | - | - | 0 | - |
| - | - | 1210 | 1221 | - | - | 0 | - |
| - | - | 934.6 | 1222 | - | - | 0 | - |
| - | - | 1500 | 1227 | - | - | 0 | - |
| - | - | 805.1 | 1244 | - | - | 0 | - |
| - | - | 1005 | 1246 | - | - | 0 | - |
| - | - | 1189 | 1247 | - | - | 0 | - |
| 10 | b | 1.091E+04 | 1249 | 0.00428 | 3.427 | +1 | 10 |
| - | - | 7220 | 1250 | - | - | 0 | - |
| - | - | 2397 | 1251 | - | - | 0 | - |
| - | - | 1792 | 1258 | - | - | 0 | - |
| - | - | 1498 | 1259 | - | - | 0 | - |
| - | - | 1096 | 1260 | - | - | 0 | - |
| 4 | y | 3804 | 1262 | 0.005749 | 4.557 | +1 | 11 |
| 4 | y | 1.318E+04 | 1263 | 0.006963 | 5.515 | +1 | 11 |
| - | - | 8274 | 1264 | - | - | 0 | - |
| - | - | 4124 | 1265 | - | - | 0 | - |
| - | - | 890.8 | 1268 | - | - | 0 | - |
| - | - | 879.9 | 1271 | - | - | 0 | - |
| 4 | y | 3.46E+04 | 1280 | 0.004584 | 3.582 | +1 | 11 |
| - | - | 2.119E+04 | 1281 | - | - | 0 | - |
| - | - | 8947 | 1282 | - | - | 0 | - |
| - | - | 1004 | 1283 | - | - | 0 | - |
| - | - | 1068 | 1286 | - | - | 0 | - |
| - | - | 1183 | 1291 | - | - | 0 | - |
| - | - | 1084 | 1292 | - | - | 0 | - |
| - | - | 2113 | 1294 | - | - | 0 | - |
| - | - | 1893 | 1295 | - | - | 0 | - |
| - | - | 867.8 | 1296 | - | - | 0 | - |
| - | - | 2441 | 1300 | - | - | 0 | - |
| - | - | 2204 | 1301 | - | - | 0 | - |
| - | - | 1176 | 1302 | - | - | 0 | - |
| - | - | 1550 | 1303 | - | - | 0 | - |
| - | - | 1015 | 1304 | - | - | 0 | - |
| - | - | 1175 | 1305 | - | - | 0 | - |
| - | - | 960.6 | 1311 | - | - | 0 | - |
| - | - | 2097 | 1312 | - | - | 0 | - |
| - | - | 1195 | 1313 | - | - | 0 | - |
| - | - | 2883 | 1327 | - | - | 0 | - |
| - | - | 1142 | 1328 | - | - | 0 | - |
| - | - | 2.166E+04 | 1329 | - | - | 0 | - |
| - | - | 1.432E+04 | 1330 | - | - | 0 | - |
| - | - | 6749 | 1331 | - | - | 0 | - |
| - | - | 877.5 | 1332 | - | - | 0 | - |
| - | - | 1024 | 1340 | - | - | 0 | - |
| - | - | 1008 | 1341 | - | - | 0 | - |
| 11 | b | 1241 | 1347 | 0.003397 | 2.523 | +1 | 11 |
| - | - | 958.7 | 1349 | - | - | 0 | - |
| - | - | 2289 | 1358 | - | - | 0 | - |
| - | - | 1382 | 1359 | - | - | 0 | - |
| - | - | 1174 | 1360 | - | - | 0 | - |
| - | - | 2841 | 1361 | - | - | 0 | - |
| - | - | 1304 | 1362 | - | - | 0 | - |
| 11 | b | 7211 | 1364 | 0.004924 | 3.611 | +1 | 11 |
| - | - | 5367 | 1365 | - | - | 0 | - |
| - | - | 2888 | 1366 | - | - | 0 | - |
| - | - | 1578 | 1367 | - | - | 0 | - |
| 3 | y | 1970 | 1375 | 0.001031 | 0.7499 | +1 | 12 |
| 3 | y | 2782 | 1376 | 0.01665 | 12.1 | +1 | 12 |
| - | - | 1673 | 1377 | - | - | 0 | - |
| 3 | y | 2.532E+04 | 1393 | 0.004504 | 3.234 | +1 | 12 |
| - | - | 1.81E+04 | 1394 | - | - | 0 | - |
| - | - | 8601 | 1395 | - | - | 0 | - |
| - | - | 1913 | 1396 | - | - | 0 | - |
| - | - | 883.9 | 1404 | - | - | 0 | - |
| - | - | 2718 | 1415 | - | - | 0 | - |
| - | - | 2444 | 1416 | - | - | 0 | - |
| - | - | 870.5 | 1417 | - | - | 0 | - |
| - | - | 1158 | 1434 | - | - | 0 | - |
| 12 | b | 1927 | 1462 | 0.02843 | 19.45 | +1 | 12 |
| 12 | b | 5569 | 1479 | 0.004563 | 3.085 | +1 | 12 |
| - | - | 6558 | 1480 | - | - | 0 | - |
| - | - | 3743 | 1481 | - | - | 0 | - |
| - | - | 977.8 | 1482 | - | - | 0 | - |
| - | - | 3236 | 1492 | - | - | 0 | - |
| - | - | 2289 | 1493 | - | - | 0 | - |
| - | - | 1263 | 1534 | - | - | 0 | - |
| 2 | y | 1722 | 1556 | 0.002211 | 1.421 | +1 | 13 |
| - | - | 1794 | 1557 | - | - | 0 | - |
| - | - | 1152 | 1558 | - | - | 0 | - |
| - | - | 885.5 | 1567 | - | - | 0 | - |
| - | - | 687.7 | 2821 | - | - | 0 | - |
| - | - | 789.7 | 3038 | - | - | 0 | - |
| - | - | 689.7 | 3260 | - | - | 0 | - |

m/z Charge Intensity FragmentType MassShift Position
120.0810317993164 0 6899.098
121.08411407470703 0 486.85052
123.05565643310547 0 449.75656
124.0872573852539 0 436.69305
124.11251831054688 0 443.41367
125.0713119506836 0 731.14075
126.05538940429688 0 827.006
127.08683776855469 0 1247.8258
129.06607055664062 0 2114.8042
129.10252380371094 0 9327.088
130.04994201660156 0 410.1138
130.0654754638672 0 560.6244
130.482177734375 0 446.93
131.08187866210938 0 1095.0422
131.11814880371094 0 12898.492
132.1214599609375 0 442.0858
133.06103515625 0 914.0073
134.02731323242188 0 909.18616
136.0760040283203 0 133785.33
137.0733642578125 0 903.5052
137.079345703125 0 11720.573
138.05526733398438 0 542.97394
138.06666564941406 0 390.88516
139.05050659179688 0 767.0462
139.08689880371094 0 851.91504
140.0819549560547 0 916.5317
141.0662384033203 0 1126.4122
141.10255432128906 0 2296.0676
142.12315368652344 0 1038.3425
143.0450439453125 0 743.83405
143.08204650878906 0 533.95874
143.11810302734375 0 480.03687
143.77073669433594 0 383.40024
144.0657196044922 0 2166.009
144.5761260986328 0 459.11218
145.0609893798828 0 779.69244
145.0973358154297 0 825.48804
146.06031799316406 0 3447.374
147.04415893554688 0 854.2987
147.0640411376953 0 637.1206
147.0765838623047 0 1406.904
148.86216735839844 0 539.1652
148.88357543945312 0 531.72516
148.8911895751953 0 536.6404
148.89840698242188 0 616.8916
148.90577697753906 0 573.9786
148.912841796875 0 887.55945
148.92030334472656 0 1011.3282
148.92759704589844 0 1179.5825
148.9348907470703 0 1953.0652
148.9427947998047 0 3585.5486
148.95947265625 0 3918.7031
148.96730041503906 0 2309.5461
148.97462463378906 0 1272.6091
148.98191833496094 0 1167.6611
148.98899841308594 0 861.0696
148.9961700439453 0 736.2443
149.00352478027344 0 486.96683
149.0109405517578 0 526.3503
149.0182647705078 0 624.70703
149.0259246826172 0 582.4202
149.03309631347656 0 577.0611
149.0459442138672 0 401.90573
149.1636962890625 0 493.8668
150.0555877685547 0 475.75632
152.0708465576172 0 2390.2837
152.08216857910156 0 731.7701
153.06626892089844 0 1504.6763
153.10203552246094 0 506.8827
155.08189392089844 0 1403.2792
155.1179962158203 0 1388.0658
156.07713317871094 0 2731.7192
157.06103515625 0 4107.2197
157.09754943847656 0 1476.397
157.10882568359375 0 945.5418
157.13343811035156 0 653.92993
158.09228515625 0 520.4815
159.0765838623047 0 2148.7969
159.0920867919922 0 744.28314
159.11363220214844 0 507.5136
165.06619262695312 0 445.27383
165.10275268554688 0 540.22595
166.06137084960938 0 842.1876
166.09779357910156 0 3228.697
167.04562377929688 0 908.59155
167.05612182617188 0 540.1617
167.08184814453125 0 5330.0454
167.0934295654297 0 962.73413
168.06582641601562 0 806.7151
168.07717895507812 0 682.41583
168.08514404296875 0 974.76495
168.1137237548828 0 501.04337
169.06149291992188 0 583.3011
169.09754943847656 0 2717.5256
169.13389587402344 0 969.2354
171.07675170898438 0 2621.125
171.11312866210938 0 685.0255
173.07089233398438 0 614.1871
173.0924530029297 0 5481.3013 y Water loss 12
173.12876892089844 0 6736.8228
173.4407958984375 0 719.82935
174.0552520751953 0 9752.168
174.08761596679688 0 2174.3608
174.13287353515625 0 505.20898
175.05862426757812 0 3023.2505
175.07162475585938 0 2161.5493
177.0663604736328 0 801.74744
177.66238403320312 0 459.33722
178.04995727539062 0 1407.8682
180.0768280029297 0 1018.851
181.0609893798828 0 1127.2904
181.13388061523438 0 1458.6796
182.12925720214844 0 544.63556
183.0770721435547 0 1699.6177
183.11326599121094 0 1825.3665
183.14947509765625 0 4779.6436
184.0718536376953 0 2066.8958
184.1082000732422 0 2342.6038
185.0559844970703 0 2746.0698
185.09202575683594 0 1597.4956
186.12384033203125 0 663.23047
187.07215881347656 0 497.9083
187.09852600097656 0 860.76685
187.10801696777344 0 14647.203
187.11703491210938 0 1063.3708
187.14402770996094 0 600.9971
188.10304260253906 0 680.54144
188.11155700683594 0 3713.86
189.08726501464844 0 5881.3413
189.11312866210938 0 481.56033
191.08241271972656 0 952.6298
191.1029510498047 0 22119.842 y 12
192.1064453125 0 1184.5331
192.15020751953125 0 1116.938
193.0979461669922 0 524.5288
193.1082000732422 0 1373.7521
194.0927734375 0 2337.181
195.07672119140625 0 12264.696
195.08786010742188 0 1929.1428
196.07174682617188 0 778.8557
196.08058166503906 0 916.581
196.10885620117188 0 572.6355
197.1034698486328 0 1055.2527
197.12844848632812 0 2053.543
198.08755493164062 0 4512.162
199.0718536376953 0 1768.9777
199.09109497070312 0 637.72943
199.10816955566406 0 1399.8021
200.433837890625 0 526.81213
201.08726501464844 0 3941.2488
201.1236572265625 0 5964.3364
202.0504150390625 0 1260.407
202.08242797851562 0 6241.4443
202.0915069580078 0 705.6278
203.05447387695312 0 551.2261
203.06654357910156 0 698.7087
203.1029052734375 0 35203.445
204.1064453125 0 2414.0957
204.138916015625 0 696.01886
207.0878448486328 0 1008.1929
208.0718994140625 0 1801.3663
208.0970001220703 0 3182.3909
208.10726928710938 0 760.8205
209.1287078857422 0 1734.7719
210.08755493164062 0 892.8901
210.16049194335938 0 801.0515
211.1084747314453 0 1079.0665
211.11886596679688 0 1100.8354
211.14430236816406 0 3686.698
212.10321044921875 0 3727.416
213.08706665039062 0 797.2977
214.08251953125 0 977.19995
214.15492248535156 0 1650.8914
215.13926696777344 0 1701.8137
216.0980682373047 0 887.4202
217.06346130371094 0 639.99664
217.08218383789062 0 11591.666
217.0933074951172 0 894.7367
218.02769470214844 0 536.79285
218.08595275878906 0 821.7869
219.0974578857422 0 528.7969
221.103271484375 0 1519.096
221.12872314453125 0 889.8704
223.107666015625 0 1746.674
224.10284423828125 0 951.3298
224.13966369628906 0 1021.29645
225.09864807128906 0 8255.493
225.12367248535156 0 6813.5347
225.17138671875 0 3958.5188
226.10244750976562 0 1057.813
226.117919921875 0 1088.6821
226.12918090820312 0 618.7084
227.1027374267578 0 1967.8363
228.09815979003906 0 3658.3906
228.1342010498047 0 717.91003
228.1710205078125 0 1761.8538
229.11866760253906 0 3779.4534
229.15499877929688 0 734.3692
230.14988708496094 0 1124.0406
231.06129455566406 0 3732.8037
231.15013122558594 0 604.99194
233.0910186767578 0 957.1876
233.13084411621094 0 660.9492
235.09288024902344 0 795.12854
235.14443969726562 0 1260.175
236.13970947265625 0 1027.4733
237.13502502441406 0 969.7464
238.15525817871094 0 776.8755
239.114501953125 0 2738.939
240.09730529785156 0 1366.2704
240.13446044921875 0 522.17676
241.0936737060547 0 770.4064
242.1141815185547 0 2056.595
242.1503448486328 0 27840.236
243.13406372070312 0 1210.6469
243.15347290039062 0 2154.506
244.09364318847656 0 853.31506
244.12843322753906 0 735.59766
245.05978393554688 0 1325.2292
245.12852478027344 0 1424.3745
246.08763122558594 0 633.1906
246.10914611816406 0 1312.0671
249.16021728515625 0 159681.36 a 1
250.1634521484375 0 20698.172
251.1667022705078 0 1827.203
252.09765625 0 857.16797
252.13421630859375 0 1332.6608
253.11801147460938 0 1303.7153
253.1662139892578 0 7135.075
254.1691436767578 0 653.47375
255.11041259765625 0 611.3942
256.16558837890625 0 592.77136
258.09112548828125 0 1732.9923
259.0750732421875 0 4937.4116
259.1037902832031 0 792.63654
260.078857421875 0 708.5721
261.0903015136719 0 1250.3894
261.12664794921875 0 719.3018
261.1593017578125 0 824.3412
262.08563232421875 0 2885.3306
262.1554870605469 0 1456.693
263.0696716308594 0 1667.625
263.1387939453125 0 601.7773
264.09808349609375 0 1154.2468
264.1348571777344 0 807.7869
265.15478515625 0 2815.0293
266.1502685546875 0 609.03436
267.109130859375 0 3529.7976
268.0927734375 0 2523.197
268.1098937988281 0 574.4858
269.1608581542969 0 1026.0195
270.08953857421875 0 915.6222
270.10772705078125 0 917.2466
270.1449279785156 0 1423.155
270.1927490234375 0 1739.3586
274.1238708496094 0 728.0756
274.1400451660156 0 4701.58
275.1415100097656 0 877.3909
276.1016540527344 0 4916.829
277.1549987792969 0 123449.305 b 1
278.11376953125 0 1325.1799
278.158203125 0 18847.686
279.1603088378906 0 1115.5206
280.1287841796875 0 1084.6339
280.1659851074219 0 1912.0792
282.1087951660156 0 565.304
282.14520263671875 0 674.95294
282.1817626953125 0 3917.1746
285.1197204589844 0 4422.1157
286.1042175292969 0 1293.1033
287.1380920410156 0 961.9817
287.174560546875 0 900.92566
288.0999450683594 0 1120.1913
288.1194763183594 0 8234.331 y Water loss 11
289.1209411621094 0 666.4385
290.1493835449219 0 649.08057
291.06524658203125 0 1558.3186
291.1346130371094 0 1405.2549
293.1499328613281 0 1319.1002
294.1438293457031 0 608.9995
294.1822509765625 0 1104.3145
295.1036376953125 0 704.1432
296.2091979980469 0 822.7194
297.15679931640625 0 1953.7618
297.1924133300781 0 592.8151
298.0832824707031 0 700.8241
298.1404724121094 0 1750.3959
299.0629577636719 0 868.0476
299.1380920410156 0 602.3478
299.17242431640625 0 585.72363
300.0628356933594 0 615.8583
304.0956726074219 0 804.06665
304.1137390136719 0 1125.1317
306.1087341308594 0 3504.6055
306.1299133300781 0 5467.4062 y 11
307.11163330078125 0 1570.7366
307.17620849609375 0 830.4125
308.1347961425781 0 1225.6433
308.16015625 0 923.3898
309.12005615234375 0 3594.0098
311.17181396484375 0 638.19226
313.1154479980469 0 946.4506
314.09893798828125 0 2418.0005
315.1680603027344 0 3223.6467
317.2230224609375 0 1116.1733
322.1884460449219 0 1298.6987
323.17193603515625 0 1900.1753
324.1468811035156 0 666.1574
325.1874694824219 0 2287.5244
326.1487731933594 0 554.57513
326.17041015625 0 1229.19
328.097412109375 0 686.34735
328.1257629394531 0 666.06805
329.1490173339844 0 1001.85785
330.14227294921875 0 673.79944
331.10650634765625 0 623.902
332.08966064453125 0 737.6466
332.109375 0 7355.48
333.11651611328125 0 1582.0071
334.17724609375 0 698.9648
335.1734313964844 0 907.548
337.15203857421875 0 604.36816
340.19854736328125 0 5577.8936
341.1571350097656 0 2446.4285
341.179443359375 0 822.51184
341.202880859375 0 857.03656
342.1593933105469 0 812.4149
342.2131652832031 0 884.8908
343.1649475097656 0 988.3256
343.1985168457031 0 970.09705
344.1452941894531 0 4507.091
345.1504211425781 0 775.7281
345.2179260253906 0 2235.2288
349.11773681640625 0 1446.3604
350.18359375 0 633.464
351.13043212890625 0 1204.8303
351.1661376953125 0 824.1955
352.1981201171875 0 1506.0128
353.18218994140625 0 3371.3208
354.14190673828125 0 2564.1504
354.1815185546875 0 778.3218
355.0696716308594 0 1563.4915
356.0705261230469 0 698.9049
357.224853515625 0 1759.4648
360.1925964355469 0 1310.9401
363.1643371582031 0 604.89703
366.25030517578125 0 756.5286
368.1566162109375 0 2097.0466
368.1938171386719 0 5380.3926
370.1725158691406 0 913.0476
370.2098083496094 0 1304.1516
372.15850830078125 0 1439.0507
372.22930908203125 0 1624.3363
373.1168518066406 0 2525.3484
378.14117431640625 0 4325.3467
380.19287109375 0 1756.698
381.17694091796875 0 1180.1117
382.13592529296875 0 1047.2393
383.1920166015625 0 716.57056
385.1348876953125 0 1447.3492
385.2199401855469 0 5414.2437
386.2045593261719 0 723.56604
388.1851806640625 0 824.4181
389.18597412109375 0 5046.507
390.1435241699219 0 1459.7163
390.2393798828125 0 22180.484 b 2
391.2132263183594 0 782.81915
391.2423095703125 0 4350.8594
392.1910705566406 0 917.8069
395.1656188964844 0 991.9313
395.20465087890625 0 743.92706
396.15167236328125 0 3710.1885
397.15728759765625 0 921.1969
398.20440673828125 0 963.1864
402.174560546875 0 661.7513
403.1463928222656 0 7797.871 y Water loss 10
404.15057373046875 0 892.5642
405.2139587402344 0 6799.254
406.2158203125 0 1714.9297
407.2660827636719 0 5917.327
408.2680969238281 0 1045.006
409.2185363769531 0 717.2866
410.2043151855469 0 650.92755
411.23565673828125 0 1170.0623
412.219482421875 0 775.466
413.17681884765625 0 692.09296
420.1522521972656 0 703.61755
420.1878662109375 0 841.35956
421.1568603515625 0 4413.09 y 10
424.19134521484375 0 1502.6888
427.1825256347656 0 1625.1113
428.2583312988281 0 963.7171
430.22601318359375 0 1711.0975
430.7288513183594 0 920.53876
433.209228515625 0 2395.818
437.21484375 0 2331.83 y Ammonia loss 6
440.224853515625 0 1157.2794
442.1395568847656 0 3708.6558
443.2281799316406 0 802.5202
444.1887512207031 0 1138.8279
445.1940002441406 0 6210.315
446.1996154785156 0 1646.6244
453.2826232910156 0 734.2373
454.2413635253906 0 3863.727
456.2530517578125 0 803.24664
458.2083435058594 0 773.31433
460.1506042480469 0 2038.3906
462.2053527832031 0 831.6309
463.2293701171875 0 1329.6135
464.2166748046875 0 2386.596
465.21588134765625 0 1262.3589
469.1686706542969 0 3505.3296
469.2780456542969 0 636.9669
470.1699523925781 0 1007.94806
470.7159729003906 0 688.15015
471.26953125 0 2197.084
472.2736511230469 0 806.02686
473.2144470214844 0 888.6646
473.2755432128906 0 1114.768
477.1783752441406 0 1029.4935
479.2339172363281 0 834.75964
481.2793884277344 0 1131.7223
482.2350769042969 0 978.4431
483.22113037109375 0 4548.72
484.2762451171875 0 2715.482
485.2785339355469 0 1067.4247
486.2003479003906 0 1218.3445
486.7686767578125 0 672.0481
487.26910400390625 0 775.68506
490.3022155761719 0 1369.3851
491.224365234375 0 2014.4655
491.3057556152344 0 799.72723
498.3041687011719 0 2574.5125
500.24749755859375 0 6839.4785
501.2477722167969 0 946.8686
503.229248046875 0 1459.17
507.1842041015625 0 1211.2404
507.222900390625 0 773.3472
508.2522277832031 0 727.0317
511.7577209472656 0 840.6401
512.2625732421875 0 922.4842
513.2421264648438 0 748.3674
516.2316284179688 0 4030.041 y Water loss 9
517.27490234375 0 737.3185
518.2980346679688 0 7802.495 b 3
519.3014526367188 0 3330.3816
520.2515258789062 0 965.7334
524.2579345703125 0 610.19244
526.2639770507812 0 1727.1476
526.7594604492188 0 605.53357
528.210693359375 0 916.38495
528.737548828125 0 631.7983
534.2416381835938 0 10788.375 y 9
535.2440795898438 0 3017.5347
536.248046875 0 2411.1482
536.7457885742188 0 691.1361
537.2776489257812 0 1627.4309
544.2811279296875 0 2812.6157
544.7830200195312 0 979.7481
545.2482299804688 0 1697.8755
545.7564697265625 0 922.3661
552.2498168945312 0 2186.0894
554.257080078125 0 986.0301
554.304931640625 0 2559.204
555.22119140625 0 651.37067
556.27587890625 0 987.36566
556.3587036132812 0 834.62024
559.7883911132812 0 952.55084 b Ammonia loss 8
560.2889404296875 0 1229.405
560.7886962890625 0 721.11505
562.2864379882812 0 709.4838
564.760009765625 0 612.4337
565.2731323242188 0 996.17224
567.2719116210938 0 733.4053 y Water loss 4
567.3255615234375 0 1004.0025
568.3016967773438 0 5599.3623 b 8
568.8017578125 0 4349.543
569.3040161132812 0 2008.8157
569.7996826171875 0 1056.9834
572.2822875976562 0 1731.6934
573.2349243164062 0 1165.7911
573.2883911132812 0 2131.6658
573.3417358398438 0 758.818
574.2926025390625 0 1540.5347
576.27294921875 0 783.03986 y 4
576.7687377929688 0 1088.0675
582.2924194335938 0 756.432
583.2750244140625 0 1167.4191
583.3253784179688 0 773.9319
584.2797241210938 0 2533.0354
584.7783203125 0 1032.4332
585.2877197265625 0 1085.7871
585.3370361328125 0 3544.1035
586.7708129882812 0 1503.7762
587.2684326171875 0 2770.476
587.7654418945312 0 767.7348
588.2656860351562 0 931.64545
593.285400390625 0 765.04865
594.28564453125 0 737.8321
595.7770385742188 0 2164.794
597.3115844726562 0 835.6991
599.296875 0 3779.7446
599.7969360351562 0 1914.1438
600.296630859375 0 956.0164
601.2911987304688 0 3028.4001
601.336181640625 0 1796.0109
601.7953491210938 0 6272.827
602.2958984375 0 5443.401
602.7969360351562 0 2142.3152
607.2918701171875 0 887.4365
608.303955078125 0 1730.846
612.2987670898438 0 1077.8286
613.3313598632812 0 4813.0674
614.3346557617188 0 1590.443
617.2907104492188 0 705.8875
618.3037109375 0 1138.1277
619.22119140625 0 830.80475
619.3067626953125 0 4339.703
620.3095703125 0 2687.5015
622.2877197265625 0 1299.6742
622.7882080078125 0 2770.6753
623.2844848632812 0 2729.4966
624.8428344726562 0 3234.5232 b 9
625.3447875976562 0 2487.0789
625.8095092773438 0 893.65814
626.3242797851562 0 746.07825
626.8173217773438 0 788.6342
627.3298950195312 0 640.29224
627.3939819335938 0 1076.6094
629.31201171875 0 716.086
630.3110961914062 0 1763.276
631.2962036132812 0 13558.466 y Water loss 3
631.3518676757812 0 909.8677
631.7960205078125 0 7014.5293 y Ammonia loss 3
632.2965698242188 0 5085.5947
634.7976684570312 0 986.8887
635.3004150390625 0 847.8847
635.8057861328125 0 660.0419
636.24853515625 0 1024.2622
636.313720703125 0 1066.8152
637.3391723632812 0 2321.3835
638.3336181640625 0 1064.6182
638.8163452148438 0 885.06934
639.3413696289062 0 926.8727
640.3017578125 0 13790.834 y 3
640.8034057617188 0 9577.867
641.3023681640625 0 3393.6746
641.80078125 0 896.21027
643.3148193359375 0 1865.2247
643.8116455078125 0 1853.2188
644.3092041015625 0 2177.9368
647.3257446289062 0 4065.9922
647.8270263671875 0 2074.4692
648.3150024414062 0 2493.7063
649.3217163085938 0 718.0449
650.358642578125 0 806.87946
652.3201293945312 0 6132.6196
652.82080078125 0 2634.937
653.32080078125 0 2029.0653
655.3058471679688 0 1755.5586
655.8348388671875 0 896.9319
656.329833984375 0 1700.9746
656.832763671875 0 1406.1012
663.3308715820312 0 899.0564
663.8324584960938 0 1284.0286
664.84375 0 3761.5718
665.3358764648438 0 11027.487 b 4
666.3394165039062 0 3745.231
670.8191528320312 0 1575.7351
672.3313598632812 0 4017.1086 y Water loss 8
673.3203735351562 0 3595.2446 y Ammonia loss 8
673.8446655273438 0 1569.3156 b Ammonia loss 10
674.3193969726562 0 1036.1061
675.3213500976562 0 682.137
679.3258666992188 0 5561.6216
679.8260498046875 0 3804.7153
680.3283081054688 0 1599.5607
680.8341064453125 0 1518.6011
682.3565063476562 0 17385.375 b 10
682.8572387695312 0 11117.784
683.35546875 0 4795.6694
684.33642578125 0 1178.1548
685.3390502929688 0 734.15247
686.31689453125 0 766.6416
687.8386840820312 0 4437.658 y Water loss 2
688.3342895507812 0 7091.581 y Ammonia loss 2
688.8343505859375 0 4825.391
689.3297729492188 0 1863.6644
690.3424682617188 0 37983.098 y 8
691.3460083007812 0 12825.698
692.3460083007812 0 2320.5603
695.2681884765625 0 1219.4661
696.3690185546875 0 689.777
696.8436889648438 0 32065.25 y 2
697.3457641601562 0 26171.346
697.8456420898438 0 9476.139
698.347412109375 0 3301.3977
699.3780517578125 0 6148.432
700.3665161132812 0 6668.833
701.3677368164062 0 2054.6316
703.3404541015625 0 1124.0427
707.8741455078125 0 1203.8297
708.8570556640625 0 863.8192
709.3569946289062 0 779.83545
710.43408203125 0 906.3754
713.3424682617188 0 812.1598
714.3786010742188 0 823.2059
715.3746337890625 0 1092.788
716.3560180664062 0 686.5679
717.3954467773438 0 725.9404
717.8642578125 0 765.64246
721.3214111328125 0 685.5544
722.8400268554688 0 914.20245
724.84521484375 0 1043.4861
725.3519287109375 0 3051.7878
725.8433837890625 0 737.71625
726.3472900390625 0 960.9962
728.3806762695312 0 668.2943
729.3320922851562 0 1494.1567
730.8641967773438 0 1608.2894 b Water loss 11
731.3612060546875 0 2571.1501 b Ammonia loss 11
731.8562622070312 0 2604.783
732.3589477539062 0 1194.8191
733.3948974609375 0 1251.5989
733.8514404296875 0 3162.9858
734.3541259765625 0 5626.2344
734.854248046875 0 2125.9307
739.8697509765625 0 30057.102 b 11
740.37109375 0 21723.357
740.8712158203125 0 9297.21
741.374267578125 0 1318.3049
742.4254760742188 0 812.3808
744.3508911132812 0 2011.606
746.3642578125 0 2775.8315
746.876708984375 0 1192.5302
747.3632202148438 0 725.39655
756.865234375 0 1565.7051
758.39404296875 0 1667.6704
758.891845703125 0 1634.0206
759.3917846679688 0 1153.697
760.8580932617188 0 1551.2019
762.3505859375 0 2036.2234 b Ammonia loss 5
763.3583984375 0 889.9489
765.3888549804688 0 874.7986
766.3861694335938 0 1112.3165
767.3424072265625 0 1309.3959
767.889404296875 0 1225.0587
768.3909912109375 0 2997.0078
768.8884887695312 0 1211.4601
769.3810424804688 0 2353.2249 y Water loss 1
769.8663940429688 0 2966.373 y Ammonia loss 1
770.3682250976562 0 2768.9822
772.8859252929688 0 1291.373
773.3618774414062 0 980.31995
776.3973999023438 0 3774.2915
776.896728515625 0 3526.0837
777.398193359375 0 2186.9934
778.3762817382812 0 6181.5884 y 1
778.8785400390625 0 5983.944
779.377685546875 0 6677.8765 b 5
779.8778076171875 0 743.57385
780.3779907226562 0 1791.9803
781.3878784179688 0 8089.0483 b Water loss 12
781.8870849609375 0 8181.757 b Ammonia loss 12
782.3953857421875 0 5793.855
782.8837280273438 0 1771.1947
783.4136352539062 0 2056.139
783.86962890625 0 793.8758
785.4081420898438 0 2555.4504 y Water loss 7
786.409423828125 0 1645.4999 y Ammonia loss 7
790.3936157226562 0 24218.436 b 12
790.8956298828125 0 21222.123
791.396240234375 0 9980.994
791.8954467773438 0 2401.5242
793.9144897460938 0 2117.8228
794.3343505859375 0 971.8594
794.411865234375 0 2620.0747
794.9071655273438 0 2549.6501
795.4488525390625 0 1988.4749
796.4544067382812 0 918.498
797.3819580078125 0 2652.0762
799.3841552734375 0 1626.1858
800.4130249023438 0 984.2948
801.4083251953125 0 1963.4998
802.410888671875 0 1118.9326
802.9181518554688 0 8489.687
803.4240112304688 0 26829.617 y 7
803.9202270507812 0 3764.9517
804.4264526367188 0 8633.315
805.429443359375 0 1036.1016
808.4000854492188 0 861.0964
808.8981323242188 0 1191.1829
809.3984985351562 0 1657.059
809.902587890625 0 2064.2314
812.4039916992188 0 846.88184
812.9150390625 0 888.9338
813.413330078125 0 754.59534
814.4063720703125 0 10682.666
815.4071655273438 0 5177.8394
815.8779907226562 0 1067.5724
816.4117431640625 0 3316.9778
816.9053955078125 0 2746.0198
817.4046630859375 0 6860.578
817.9033203125 0 3060.5469
818.3944702148438 0 2402.0645
819.4203491210938 0 997.8506
824.3966064453125 0 1123.9221
824.8937377929688 0 1329.9152
825.3881225585938 0 1095.4469
825.9120483398438 0 18960.457 Precursor Water loss
826.4108276367188 0 26446.3 Precursor Ammonia loss
826.9112548828125 0 15863.732
827.41015625 0 8832.39
827.9120483398438 0 1487.5826
828.420654296875 0 1180.48
829.3763427734375 0 717.0353
829.708984375 0 1030.336
831.3806762695312 0 1603.0496
832.3688354492188 0 904.3795
833.389404296875 0 3076.0854
833.89990234375 0 2909.8247
834.9173583984375 0 92973.21 Precursor
835.03662109375 0 3214.102
835.4182739257812 0 87261.19
835.71484375 0 4104.222
835.9196166992188 0 48392.68
836.0530395507812 0 1337.5966
836.419677734375 0 10196.008
840.394287109375 0 1796.5206
842.4147338867188 0 2044.3145
844.4332275390625 0 1058.3562
846.4152221679688 0 3610.2378
847.4161987304688 0 1083.9531
848.4032592773438 0 1131.1212 b Water loss 6
849.3900146484375 0 2368.3284 b Ammonia loss 6
850.3844604492188 0 1515.0769
857.4398193359375 0 1071.8586
858.4332885742188 0 2007.9806
859.4439086914062 0 5172.941
860.4500122070312 0 2288.3333
861.4454345703125 0 747.7188
866.41015625 0 3649.8901 b 6
867.412109375 0 1823.5619
870.4237670898438 0 1655.3541
871.4332275390625 0 868.32245
872.4486083984375 0 3990.4114 y Water loss 6
873.4459228515625 0 1079.5641 y Ammonia loss 6
874.4583740234375 0 1416.7351
880.4216918945312 0 1277.7184
890.458251953125 0 27464.162 y 6
891.4612426757812 0 12442.399
892.4632568359375 0 2428.9746
893.4470825195312 0 3853.9468
894.4490356445312 0 993.7754
897.44384765625 0 8252.555
898.4456787109375 0 4865.707
899.442138671875 0 2029.966
900.4424438476562 0 2031.6422
902.455322265625 0 1630.7662
903.447265625 0 1036.6863
910.4750366210938 0 4573.661
911.4678955078125 0 1800.8458
915.4628295898438 0 2293.2703
916.4517822265625 0 4283.327
917.4558715820312 0 2882.9114
918.4585571289062 0 3491.369
919.4627075195312 0 727.5818
933.4664306640625 0 1736.6006
934.4713745117188 0 1300.8308
936.487548828125 0 1654.5956
939.4530029296875 0 1534.0858
940.4478149414062 0 2403.05
944.4660034179688 0 1025.666
945.4588012695312 0 1181.3674
946.4563598632812 0 911.1469
951.49169921875 0 825.13257
953.51220703125 0 965.19196
954.4813842773438 0 968.29803
955.5030517578125 0 757.09656
956.5064697265625 0 989.7482
957.4483642578125 0 2443.2676
958.452880859375 0 1741.9655
959.4827880859375 0 780.80963
960.4650268554688 0 890.8089
961.4469604492188 0 7513.6606
962.4569702148438 0 5456.0947 b Ammonia loss 7
963.4644165039062 0 2401.5986
965.4683837890625 0 810.3996
970.4661865234375 0 1117.646
971.458740234375 0 1331.4974
972.52587890625 0 2296.2424
973.5247192382812 0 1192.6135
974.4718627929688 0 4318.2373
975.47607421875 0 2097.8914
979.4912109375 0 4573.2954 b 7
980.4928588867188 0 3208.0723
981.4901733398438 0 1265.4529
986.4911499023438 0 3203.0125 y Water loss 5
987.4812622070312 0 6565.2563 y Ammonia loss 5
988.482177734375 0 3644.314
989.4808959960938 0 1203.8795
991.4524536132812 0 1025.4207
998.4872436523438 0 2031.1241
1001.4818115234375 0 966.35156
1004.5014038085938 0 46408.203 y 5
1005.5038452148438 0 21609.617
1006.50830078125 0 6935.2285
1007.4992065429688 0 1934.679
1008.4782104492188 0 5532.999
1009.4759521484375 0 3155.9714
1010.4813232421875 0 1207.0707
1016.5001831054688 0 2995.3264
1017.4985961914062 0 2495.9436
1017.9842529296875 0 2149.2898
1018.4888916015625 0 3208.9805
1018.9864501953125 0 801.39905
1022.5110473632812 0 1450.3347
1023.5560302734375 0 3265.6165
1024.55224609375 0 2109.21
1025.5028076171875 0 5461.5537
1026.501953125 0 4310.267
1027.504150390625 0 1520.7717
1029.52783203125 0 764.7293
1030.53662109375 0 2643.601
1031.5421142578125 0 2758.9216
1032.5396728515625 0 1479.4673
1034.4901123046875 0 1205.2546
1035.497802734375 0 721.3194
1041.5035400390625 0 930.7607
1044.482177734375 0 809.1721
1049.5255126953125 0 865.49286
1051.5164794921875 0 837.4823
1053.5191650390625 0 1270.5236
1059.517822265625 0 962.03766
1060.51611328125 0 3015.0278
1061.5186767578125 0 2839.0525
1062.4945068359375 0 3232.0212
1063.4893798828125 0 1505.6978
1065.5262451171875 0 1251.4498
1069.5235595703125 0 3166.2493
1070.5244140625 0 3870.7112
1071.595703125 0 3754.581
1072.4749755859375 0 4323.764
1072.597900390625 0 2931.982
1073.4803466796875 0 3250.1921
1074.4742431640625 0 1267.5333
1075.4857177734375 0 1072.6473
1076.5032958984375 0 1256.9486
1080.49267578125 0 1404.1141
1081.5115966796875 0 982.0941
1082.5484619140625 0 919.5348
1084.524169921875 0 1186.5608
1085.5211181640625 0 3078.3743
1087.5404052734375 0 38045.043
1088.54296875 0 19019.031
1089.52490234375 0 8353.195
1090.5030517578125 0 4010.9482
1091.5059814453125 0 991.6015
1096.49951171875 0 826.41473
1098.5123291015625 0 1064.186
1101.5673828125 0 1837.0348
1103.543701171875 0 1633.5836
1109.5228271484375 0 1568.1681
1116.5283203125 0 703.5154
1118.5679931640625 0 2281.2273 b Ammonia loss 8
1119.5643310546875 0 1504.844
1121.5572509765625 0 1158.7914
1126.546630859375 0 1964.4476
1127.5306396484375 0 1935.1122
1128.5313720703125 0 1571.1135
1130.5447998046875 0 3803.104
1131.54296875 0 2662.1265
1132.5426025390625 0 871.57245
1133.5252685546875 0 3576.9487 y Water loss 4
1134.518798828125 0 3401.3962 y Ammonia loss 4
1135.5931396484375 0 15148.28 b 8
1136.595458984375 0 10657.818
1137.5958251953125 0 3457.1077
1138.5855712890625 0 5743.256
1139.5897216796875 0 3671.4377
1140.0087890625 0 831.6435
1144.555419921875 0 2106.7898
1145.56103515625 0 1219.5739
1146.53955078125 0 1506.6508
1149.5274658203125 0 849.96497
1151.5367431640625 0 45926.023 y 4
1152.539794921875 0 30455.168
1153.538818359375 0 11318.171
1154.54638671875 0 2366.7974
1156.685791015625 0 1086.8491
1158.593505859375 0 2099.1936
1159.6009521484375 0 2484.3328
1160.6060791015625 0 986.11774
1171.5648193359375 0 941.87476
1172.5755615234375 0 1031.753
1173.5194091796875 0 1141.8627
1175.5689697265625 0 790.29047
1180.556396484375 0 1428.3185
1181.5509033203125 0 2018.4174
1187.5867919921875 0 1104.7849
1190.548095703125 0 2109.3264
1191.5445556640625 0 1170.501
1197.5833740234375 0 3912.795
1198.57373046875 0 15133.246
1199.5748291015625 0 8555.853
1200.578369140625 0 2661.1418
1202.5828857421875 0 6056.2134
1203.5867919921875 0 4583.884
1204.586181640625 0 988.9656
1213.5770263671875 0 1872.2059
1214.578125 0 865.8472
1215.596923828125 0 24563.738
1216.5994873046875 0 15254.547
1217.6026611328125 0 4881.71
1218.600830078125 0 979.47345
1220.67919921875 0 1209.9037
1221.685791015625 0 934.60284
1226.5653076171875 0 1500.0232
1243.6427001953125 0 805.1074
1245.5640869140625 0 1004.6587
1246.6470947265625 0 1189.042
1248.67626953125 0 10910.04 b 9
1249.6776123046875 0 7219.508
1250.6636962890625 0 2396.7583
1257.647216796875 0 1791.5916
1258.63818359375 0 1498.2988
1259.635498046875 0 1096.2838
1261.5849609375 0 3803.5923 y Water loss 3
1262.5701904296875 0 13183.837 y Ammonia loss 3
1263.57421875 0 8273.888
1264.5714111328125 0 4124.476
1267.5982666015625 0 890.81335
1270.6185302734375 0 879.90656
1279.5943603515625 0 34604.34 y 3
1280.59716796875 0 21192.459
1281.5982666015625 0 8947.332
1282.60009765625 0 1004.4501
1285.6383056640625 0 1068.1226
1290.5775146484375 0 1183.1434
1291.5625 0 1084.3175
1293.6478271484375 0 2112.646
1294.6461181640625 0 1893.2583
1295.6397705078125 0 867.8043
1299.703125 0 2441.3286
1300.7017822265625 0 2204.462
1301.6585693359375 0 1175.916
1302.649658203125 0 1550.3257
1303.6234130859375 0 1014.95636
1304.6346435546875 0 1175.1355
1310.6656494140625 0 960.6077
1311.6572265625 0 2097.2454
1312.66015625 0 1195.3291
1326.6636962890625 0 2883.01
1327.652587890625 0 1141.8849
1328.6800537109375 0 21657.807
1329.682373046875 0 14317.407
1330.6845703125 0 6748.8564
1331.6978759765625 0 877.4588
1339.6446533203125 0 1023.52606
1340.63525390625 0 1008.31464
1346.67578125 0 1240.5497 b Ammonia loss 10
1348.6829833984375 0 958.6809
1357.6480712890625 0 2289.3237
1358.6474609375 0 1381.7943
1359.665771484375 0 1174.1554
1360.670654296875 0 2840.7715
1361.6861572265625 0 1304.0101
1363.703857421875 0 7211.285 b 10
1364.7041015625 0 5367.1196
1365.694091796875 0 2887.993
1366.6517333984375 0 1578.0977
1374.664306640625 0 1970.361 y Water loss 2
1375.6639404296875 0 2781.5337 y Ammonia loss 2
1376.6427001953125 0 1672.7365
1392.6783447265625 0 25323.082 y 2
1393.681396484375 0 18101.074
1394.678955078125 0 8601.48
1395.677490234375 0 1913.4125
1403.6630859375 0 883.8513
1414.7276611328125 0 2718.3926
1415.7327880859375 0 2443.6123
1416.7362060546875 0 870.4875
1433.6700439453125 0 1157.7019
1461.6708984375 0 1927.4576 b Ammonia loss 11
1478.7213134765625 0 5569.259 b 11
1479.70947265625 0 6558.431
1480.7034912109375 0 3742.6309
1481.6868896484375 0 977.8452
1491.7423095703125 0 3235.8887
1492.742431640625 0 2289.3965
1533.8109130859375 0 1262.8894
1555.7393798828125 0 1722.1172 y 1
1556.74169921875 0 1793.9442
1557.7406005859375 0 1152.2249
1566.726318359375 0 885.4834
2821.138671875 0 687.70123
3038.22314453125 0 789.71295
3260.16796875 0 689.6752

Spectrum Details

|  |  |
| --- | --- |
| Matched peaks? Matched peaks The total absolute number of peaks matched. Additionally in brackets the total fraction of peaks matched and the total number of peaks is shown. | 75 (7.52% of 998) |
| FDR? FDR The false discovery rate estimated for this peptide. It is calculated by matching all theoretical fragments with a non-integer shift with the raw peaks for this spectrum. This is done with 40 different shifts. The resulting percentage is the average number of annotated peaks over the number of annotated peaks with the correct spectrum. | 0.76% |
| Satellite FDR? Satellite FDR See the FDR for details on its calculation. This satellite ion specific FDR only contains the satellite ions (d/w) for I/L/J positions. | ∞ |
| PSM Score? PSM Score The PSM Score as given by Hecklib to this annotated spectrum. It is shown with three significant figures. | 821 |

#### Spectrum 8370? Spectrum 8370 The raw spectrum of this peptide as annotated by Hecklib. The fragments are coloured according to ion type (see legend). Any peaks with a star '\*' as text can be hovered over to see the full details, first the ion type second the mass shift type. By hovering over the amino acids in the peptide or ions in the legend the corresponding peaks are highlighted. By toggling the 'Unassigned' label you can turn the background (unassigned) peaks on or off in the plot. By updating the slider in the Ion legend you can update the spectrum to only show the top X% of the peaks with labels. The top X% means any peak that is within X% of the highest intensity. By dragging in the spectrum you can zoom in to a specific part of the spectrum and use 'Zoom Out' to get back to the original zoom level. The annotation of the spectrum is based on the given sequence in the peptides file and is done with different software so inconsistencies are likely. The peaks are annotated based on the given sequence, with 20 ppm tolerance.

Copy Data

##### Spectrum 8370 (TSV)

###### Preview

```
Loading example...
```

*Click on the button to copy the data to your clipboard.*

Mz MinMz MaxIntensity Max

WidthHeightPeptide font sizePeptide stroke widthSpectrum font sizeSpectrum stroke widthCompact peptide

Ion legend

wxyz

abcd

OtherUnassignedIon^Charge^_Position_Show for top:%

JYLQMNSJRJDDTA

02.54e+45.08e+47.62e+41.02e+5

Zoom Out

y^+1^_2_y^+1^_2_a^+1^_2_b^+1^_2_y^+1^_3_y^+1^_3_b^+1^_3_y^+1^_4_y^+1^_4_y^+2^_8_y^+1^_5_b^+1^_4_y^+1^_5_b^+2^_9_b^+2^_10_y^+2^_11_y^+2^_11_y^+2^_11_b^+1^_5_y^+1^_6_y^+1^_6_b^+2^_11_y^+2^_12_y^+2^_12_y^+1^_6_y^+2^_12_b^+2^_12_b^+2^_12_b^+2^_12_b^+1^_6_y^+2^_13_y^+2^_13_y^+2^_13_b^+1^_6_b^+2^_13_b^+2^_13_b^+2^_13_y^+1^_7_\*\*\*b^+1^_7_b^+1^_7_b^+1^_7_y^+1^_8_y^+1^_8_y^+1^_8_b^+1^_8_b^+1^_8_y^+1^_9_y^+1^_9_y^+1^_9_b^+1^_9_y^+1^_10_y^+1^_10_b^+1^_9_y^+1^_10_b^+1^_10_y^+1^_11_y^+1^_11_y^+1^_11_b^+1^_11_y^+1^_12_y^+1^_12_y^+1^_12_b^+1^_12_b^+1^_12_y^+1^_13_

0767153423013068

Fragment Matches Table

Show background peaks

| Position | Ion type | Intensity | mz Theoretical | mz Error (Th) | mz Error (ppm) | Charge | Series Number |
| --- | --- | --- | --- | --- | --- | --- | --- |
| - | - | 607.7 | 120.1 | - | - | 0 | - |
| - | - | 3098 | 120.1 | - | - | 0 | - |
| - | - | 474 | 126.1 | - | - | 0 | - |
| - | - | 372.5 | 127.1 | - | - | 0 | - |
| - | - | 1018 | 129.1 | - | - | 0 | - |
| - | - | 3744 | 129.1 | - | - | 0 | - |
| - | - | 794.3 | 130.1 | - | - | 0 | - |
| - | - | 568.5 | 131.1 | - | - | 0 | - |
| - | - | 9154 | 131.1 | - | - | 0 | - |
| - | - | 410.9 | 132.1 | - | - | 0 | - |
| - | - | 610.2 | 133.1 | - | - | 0 | - |
| - | - | 8.84E+04 | 136.1 | - | - | 0 | - |
| - | - | 820.6 | 137.1 | - | - | 0 | - |
| - | - | 5738 | 137.1 | - | - | 0 | - |
| - | - | 485.6 | 139.1 | - | - | 0 | - |
| - | - | 343.3 | 139.7 | - | - | 0 | - |
| - | - | 916.2 | 140.1 | - | - | 0 | - |
| - | - | 615.5 | 141.1 | - | - | 0 | - |
| - | - | 647.6 | 142.1 | - | - | 0 | - |
| - | - | 1786 | 144.1 | - | - | 0 | - |
| - | - | 771.2 | 145.1 | - | - | 0 | - |
| - | - | 667.7 | 146.1 | - | - | 0 | - |
| - | - | 425.4 | 147.1 | - | - | 0 | - |
| - | - | 405.7 | 148.8 | - | - | 0 | - |
| - | - | 466.1 | 148.9 | - | - | 0 | - |
| - | - | 509.3 | 148.9 | - | - | 0 | - |
| - | - | 442.8 | 148.9 | - | - | 0 | - |
| - | - | 506.9 | 148.9 | - | - | 0 | - |
| - | - | 448.5 | 148.9 | - | - | 0 | - |
| - | - | 541.5 | 148.9 | - | - | 0 | - |
| - | - | 693.1 | 148.9 | - | - | 0 | - |
| - | - | 1095 | 148.9 | - | - | 0 | - |
| - | - | 1954 | 148.9 | - | - | 0 | - |
| - | - | 3552 | 148.9 | - | - | 0 | - |
| - | - | 4299 | 149 | - | - | 0 | - |
| - | - | 2613 | 149 | - | - | 0 | - |
| - | - | 1000 | 149 | - | - | 0 | - |
| - | - | 1032 | 149 | - | - | 0 | - |
| - | - | 707.1 | 149 | - | - | 0 | - |
| - | - | 573.2 | 149 | - | - | 0 | - |
| - | - | 724.5 | 149 | - | - | 0 | - |
| - | - | 517.9 | 149 | - | - | 0 | - |
| - | - | 862.3 | 149 | - | - | 0 | - |
| - | - | 1223 | 152.1 | - | - | 0 | - |
| - | - | 428.5 | 152.2 | - | - | 0 | - |
| - | - | 1244 | 153.1 | - | - | 0 | - |
| - | - | 440.4 | 153.1 | - | - | 0 | - |
| - | - | 810.9 | 155.1 | - | - | 0 | - |
| - | - | 1392 | 156.1 | - | - | 0 | - |
| - | - | 1465 | 157.1 | - | - | 0 | - |
| - | - | 781.1 | 157.1 | - | - | 0 | - |
| - | - | 654 | 159.1 | - | - | 0 | - |
| - | - | 881.3 | 159.1 | - | - | 0 | - |
| - | - | 1445 | 166.1 | - | - | 0 | - |
| - | - | 4217 | 167.1 | - | - | 0 | - |
| - | - | 738.7 | 169.1 | - | - | 0 | - |
| - | - | 710.2 | 169.1 | - | - | 0 | - |
| - | - | 1137 | 171.1 | - | - | 0 | - |
| - | - | 450.4 | 171.7 | - | - | 0 | - |
| 13 | y | 3230 | 173.1 | 0.0002927 | 1.691 | +1 | 2 |
| - | - | 4359 | 173.1 | - | - | 0 | - |
| - | - | 829.4 | 173.4 | - | - | 0 | - |
| - | - | 477.9 | 173.6 | - | - | 0 | - |
| - | - | 573.8 | 174 | - | - | 0 | - |
| - | - | 1862 | 174.1 | - | - | 0 | - |
| - | - | 1026 | 174.1 | - | - | 0 | - |
| - | - | 537.9 | 175.1 | - | - | 0 | - |
| - | - | 971.4 | 178.1 | - | - | 0 | - |
| - | - | 1098 | 181.1 | - | - | 0 | - |
| - | - | 1406 | 183.1 | - | - | 0 | - |
| - | - | 840.1 | 183.1 | - | - | 0 | - |
| - | - | 1644 | 184.1 | - | - | 0 | - |
| - | - | 1404 | 184.1 | - | - | 0 | - |
| - | - | 1226 | 185.1 | - | - | 0 | - |
| - | - | 515.1 | 186.1 | - | - | 0 | - |
| - | - | 3141 | 187.1 | - | - | 0 | - |
| - | - | 526.8 | 187.1 | - | - | 0 | - |
| - | - | 684.7 | 188.1 | - | - | 0 | - |
| - | - | 3815 | 189.1 | - | - | 0 | - |
| - | - | 775.3 | 191.1 | - | - | 0 | - |
| 13 | y | 1.326E+04 | 191.1 | 0.0002719 | 1.423 | +1 | 2 |
| - | - | 1063 | 192.1 | - | - | 0 | - |
| - | - | 1496 | 194.1 | - | - | 0 | - |
| - | - | 7416 | 195.1 | - | - | 0 | - |
| - | - | 696.6 | 196.1 | - | - | 0 | - |
| - | - | 1036 | 197.1 | - | - | 0 | - |
| - | - | 2569 | 198.1 | - | - | 0 | - |
| - | - | 1157 | 199.1 | - | - | 0 | - |
| - | - | 1376 | 199.1 | - | - | 0 | - |
| - | - | 828.9 | 201.1 | - | - | 0 | - |
| - | - | 3728 | 201.1 | - | - | 0 | - |
| - | - | 2264 | 202.1 | - | - | 0 | - |
| - | - | 1544 | 203.1 | - | - | 0 | - |
| - | - | 2528 | 203.1 | - | - | 0 | - |
| - | - | 1484 | 208.1 | - | - | 0 | - |
| - | - | 568 | 208.1 | - | - | 0 | - |
| - | - | 750.7 | 210.1 | - | - | 0 | - |
| - | - | 1054 | 211.1 | - | - | 0 | - |
| - | - | 3043 | 212.1 | - | - | 0 | - |
| - | - | 666.4 | 213.1 | - | - | 0 | - |
| - | - | 1350 | 214.2 | - | - | 0 | - |
| - | - | 3302 | 215.1 | - | - | 0 | - |
| - | - | 773.1 | 217.1 | - | - | 0 | - |
| - | - | 7361 | 217.1 | - | - | 0 | - |
| - | - | 959.8 | 223.1 | - | - | 0 | - |
| - | - | 738.2 | 224.1 | - | - | 0 | - |
| - | - | 693.7 | 225 | - | - | 0 | - |
| - | - | 1178 | 225.1 | - | - | 0 | - |
| - | - | 4742 | 225.1 | - | - | 0 | - |
| - | - | 2152 | 225.2 | - | - | 0 | - |
| - | - | 803.9 | 226.1 | - | - | 0 | - |
| - | - | 587.5 | 226.2 | - | - | 0 | - |
| - | - | 1726 | 227.1 | - | - | 0 | - |
| - | - | 675 | 228.2 | - | - | 0 | - |
| - | - | 2181 | 229.1 | - | - | 0 | - |
| - | - | 2304 | 231.1 | - | - | 0 | - |
| - | - | 554.2 | 236.1 | - | - | 0 | - |
| - | - | 943.9 | 239.1 | - | - | 0 | - |
| - | - | 768.3 | 239.1 | - | - | 0 | - |
| - | - | 989.4 | 240.1 | - | - | 0 | - |
| - | - | 913.5 | 240.1 | - | - | 0 | - |
| - | - | 621 | 242.1 | - | - | 0 | - |
| - | - | 675.3 | 242.1 | - | - | 0 | - |
| - | - | 1074 | 242.1 | - | - | 0 | - |
| - | - | 1.74E+04 | 242.2 | - | - | 0 | - |
| - | - | 1774 | 243.2 | - | - | 0 | - |
| - | - | 661.9 | 246.1 | - | - | 0 | - |
| 2 | a | 1.006E+05 | 249.2 | 0.0003866 | 1.552 | +1 | 2 |
| - | - | 1.366E+04 | 250.2 | - | - | 0 | - |
| - | - | 670.2 | 252.1 | - | - | 0 | - |
| - | - | 887 | 253.1 | - | - | 0 | - |
| - | - | 5576 | 253.2 | - | - | 0 | - |
| - | - | 745.4 | 258.1 | - | - | 0 | - |
| - | - | 3450 | 259.1 | - | - | 0 | - |
| - | - | 672.4 | 260.1 | - | - | 0 | - |
| - | - | 732.1 | 261.1 | - | - | 0 | - |
| - | - | 2481 | 262.1 | - | - | 0 | - |
| - | - | 844.5 | 262.2 | - | - | 0 | - |
| - | - | 618.1 | 264.1 | - | - | 0 | - |
| - | - | 1182 | 265.2 | - | - | 0 | - |
| - | - | 1612 | 267.1 | - | - | 0 | - |
| - | - | 1179 | 268.1 | - | - | 0 | - |
| - | - | 712.2 | 270.1 | - | - | 0 | - |
| - | - | 1436 | 270.1 | - | - | 0 | - |
| - | - | 779.4 | 270.2 | - | - | 0 | - |
| - | - | 917.4 | 274.1 | - | - | 0 | - |
| - | - | 3145 | 276.1 | - | - | 0 | - |
| 2 | b | 7.68E+04 | 277.2 | 0.0002688 | 0.9698 | +1 | 2 |
| - | - | 1.174E+04 | 278.2 | - | - | 0 | - |
| - | - | 695 | 279.1 | - | - | 0 | - |
| - | - | 1100 | 279.2 | - | - | 0 | - |
| - | - | 501.5 | 282.2 | - | - | 0 | - |
| - | - | 2183 | 285.1 | - | - | 0 | - |
| - | - | 1411 | 286.1 | - | - | 0 | - |
| - | - | 941.5 | 287.2 | - | - | 0 | - |
| 12 | y | 2898 | 288.1 | 0.0004035 | 1.401 | +1 | 3 |
| - | - | 677.3 | 290.1 | - | - | 0 | - |
| - | - | 763 | 294.2 | - | - | 0 | - |
| - | - | 1328 | 297.2 | - | - | 0 | - |
| - | - | 1438 | 298.1 | - | - | 0 | - |
| - | - | 1266 | 299.1 | - | - | 0 | - |
| - | - | 934.7 | 300.1 | - | - | 0 | - |
| 12 | y | 4161 | 306.1 | 0.0003064 | 1.001 | +1 | 3 |
| - | - | 671.1 | 307.2 | - | - | 0 | - |
| - | - | 755.2 | 308.1 | - | - | 0 | - |
| - | - | 1806 | 309.1 | - | - | 0 | - |
| - | - | 1328 | 314.1 | - | - | 0 | - |
| - | - | 2093 | 315.2 | - | - | 0 | - |
| - | - | 676 | 322.2 | - | - | 0 | - |
| - | - | 536.7 | 323.2 | - | - | 0 | - |
| - | - | 1321 | 325.2 | - | - | 0 | - |
| - | - | 1222 | 326.1 | - | - | 0 | - |
| - | - | 4909 | 332.1 | - | - | 0 | - |
| - | - | 2834 | 340.2 | - | - | 0 | - |
| - | - | 640.5 | 341.2 | - | - | 0 | - |
| - | - | 3249 | 344.1 | - | - | 0 | - |
| - | - | 798.7 | 345.2 | - | - | 0 | - |
| - | - | 814.1 | 349.1 | - | - | 0 | - |
| - | - | 664.2 | 350.2 | - | - | 0 | - |
| - | - | 1194 | 351.1 | - | - | 0 | - |
| - | - | 1628 | 353.2 | - | - | 0 | - |
| - | - | 2215 | 355.1 | - | - | 0 | - |
| - | - | 815.6 | 357.1 | - | - | 0 | - |
| - | - | 1833 | 357.2 | - | - | 0 | - |
| - | - | 683.5 | 359 | - | - | 0 | - |
| - | - | 1231 | 368.2 | - | - | 0 | - |
| - | - | 2827 | 368.2 | - | - | 0 | - |
| - | - | 627.2 | 369.2 | - | - | 0 | - |
| - | - | 1058 | 372.2 | - | - | 0 | - |
| - | - | 732.9 | 372.2 | - | - | 0 | - |
| - | - | 1744 | 373.1 | - | - | 0 | - |
| - | - | 2923 | 378.1 | - | - | 0 | - |
| - | - | 753 | 380.2 | - | - | 0 | - |
| - | - | 843.9 | 381.2 | - | - | 0 | - |
| - | - | 826.4 | 385.1 | - | - | 0 | - |
| - | - | 2727 | 385.2 | - | - | 0 | - |
| - | - | 919 | 386.2 | - | - | 0 | - |
| - | - | 3086 | 389.2 | - | - | 0 | - |
| - | - | 690.1 | 390.1 | - | - | 0 | - |
| 3 | b | 1.305E+04 | 390.2 | 0.0005859 | 1.501 | +1 | 3 |
| - | - | 3939 | 391.2 | - | - | 0 | - |
| - | - | 550.4 | 393.2 | - | - | 0 | - |
| - | - | 1004 | 395.2 | - | - | 0 | - |
| - | - | 1783 | 396.2 | - | - | 0 | - |
| 11 | y | 3947 | 403.1 | 0.0005296 | 1.314 | +1 | 4 |
| - | - | 796.3 | 404.1 | - | - | 0 | - |
| - | - | 646.2 | 404.2 | - | - | 0 | - |
| - | - | 4768 | 405.2 | - | - | 0 | - |
| - | - | 942.3 | 406.2 | - | - | 0 | - |
| - | - | 3935 | 407.3 | - | - | 0 | - |
| - | - | 693.1 | 417.2 | - | - | 0 | - |
| - | - | 641.5 | 420.2 | - | - | 0 | - |
| 11 | y | 2568 | 421.2 | 0.001287 | 3.056 | +1 | 4 |
| - | - | 1411 | 424.2 | - | - | 0 | - |
| - | - | 1865 | 427.2 | - | - | 0 | - |
| - | - | 866.9 | 428.3 | - | - | 0 | - |
| - | - | 1620 | 430.2 | - | - | 0 | - |
| - | - | 1226 | 430.7 | - | - | 0 | - |
| - | - | 553.1 | 431.9 | - | - | 0 | - |
| - | - | 1686 | 433.2 | - | - | 0 | - |
| 7 | y | 1940 | 437.2 | 0.004078 | 9.326 | +2 | 8 |
| - | - | 1542 | 442.1 | - | - | 0 | - |
| - | - | 4038 | 445.2 | - | - | 0 | - |
| - | - | 931.9 | 446.2 | - | - | 0 | - |
| - | - | 920.3 | 454.2 | - | - | 0 | - |
| - | - | 1126 | 460.1 | - | - | 0 | - |
| - | - | 984 | 464.2 | - | - | 0 | - |
| - | - | 649.8 | 466.9 | - | - | 0 | - |
| - | - | 535.4 | 468.6 | - | - | 0 | - |
| - | - | 1198 | 471.3 | - | - | 0 | - |
| - | - | 621.2 | 473.3 | - | - | 0 | - |
| - | - | 3455 | 483.2 | - | - | 0 | - |
| - | - | 568.9 | 486.2 | - | - | 0 | - |
| - | - | 1262 | 490.3 | - | - | 0 | - |
| - | - | 1016 | 491.2 | - | - | 0 | - |
| - | - | 2421 | 498.3 | - | - | 0 | - |
| - | - | 4519 | 500.2 | - | - | 0 | - |
| - | - | 972.2 | 501.3 | - | - | 0 | - |
| - | - | 1198 | 503.2 | - | - | 0 | - |
| - | - | 798 | 504.2 | - | - | 0 | - |
| - | - | 730.2 | 508.3 | - | - | 0 | - |
| - | - | 739.7 | 509.2 | - | - | 0 | - |
| 10 | y | 1892 | 516.2 | 0.0006941 | 1.345 | +1 | 5 |
| 4 | b | 6353 | 518.3 | 0.000419 | 0.8085 | +1 | 4 |
| - | - | 2510 | 519.3 | - | - | 0 | - |
| - | - | 757.1 | 520.3 | - | - | 0 | - |
| 10 | y | 5968 | 534.2 | 0.0009327 | 1.746 | +1 | 5 |
| - | - | 2092 | 535.2 | - | - | 0 | - |
| - | - | 1501 | 536.3 | - | - | 0 | - |
| - | - | 1058 | 536.8 | - | - | 0 | - |
| - | - | 571.2 | 538.3 | - | - | 0 | - |
| - | - | 1801 | 544.3 | - | - | 0 | - |
| - | - | 964.9 | 544.8 | - | - | 0 | - |
| - | - | 778.2 | 545.3 | - | - | 0 | - |
| - | - | 820.3 | 545.8 | - | - | 0 | - |
| - | - | 718.4 | 550.3 | - | - | 0 | - |
| - | - | 1079 | 552.2 | - | - | 0 | - |
| - | - | 744.9 | 552.8 | - | - | 0 | - |
| - | - | 644.9 | 554.3 | - | - | 0 | - |
| - | - | 1396 | 554.3 | - | - | 0 | - |
| - | - | 751.7 | 555.2 | - | - | 0 | - |
| 9 | b | 4394 | 568.3 | 0.00379 | 6.67 | +2 | 9 |
| - | - | 3110 | 568.8 | - | - | 0 | - |
| - | - | 592.4 | 569.3 | - | - | 0 | - |
| - | - | 817.4 | 573.3 | - | - | 0 | - |
| - | - | 564 | 573.8 | - | - | 0 | - |
| - | - | 856.1 | 584.3 | - | - | 0 | - |
| - | - | 2545 | 585.3 | - | - | 0 | - |
| - | - | 1565 | 587.3 | - | - | 0 | - |
| - | - | 974.1 | 587.8 | - | - | 0 | - |
| - | - | 810.3 | 590.8 | - | - | 0 | - |
| - | - | 1154 | 595.8 | - | - | 0 | - |
| - | - | 1971 | 599.3 | - | - | 0 | - |
| - | - | 1653 | 599.8 | - | - | 0 | - |
| - | - | 1768 | 601.3 | - | - | 0 | - |
| - | - | 1082 | 601.3 | - | - | 0 | - |
| - | - | 4375 | 601.8 | - | - | 0 | - |
| - | - | 2098 | 602.3 | - | - | 0 | - |
| - | - | 995.7 | 608.3 | - | - | 0 | - |
| - | - | 1193 | 610.8 | - | - | 0 | - |
| - | - | 2725 | 613.3 | - | - | 0 | - |
| - | - | 743.4 | 618.3 | - | - | 0 | - |
| - | - | 751.2 | 620.3 | - | - | 0 | - |
| - | - | 903.8 | 622.3 | - | - | 0 | - |
| - | - | 1373 | 622.8 | - | - | 0 | - |
| 10 | b | 2530 | 624.8 | 0.002835 | 4.537 | +2 | 10 |
| - | - | 1382 | 625.3 | - | - | 0 | - |
| - | - | 632.3 | 628.4 | - | - | 0 | - |
| - | - | 774.6 | 630.3 | - | - | 0 | - |
| 4 | y | 6078 | 631.3 | 0.002898 | 4.591 | +2 | 11 |
| 4 | y | 6847 | 631.8 | 0.01095 | 17.33 | +2 | 11 |
| - | - | 1867 | 632.3 | - | - | 0 | - |
| - | - | 930.8 | 634.8 | - | - | 0 | - |
| - | - | 683.2 | 635.8 | - | - | 0 | - |
| - | - | 1481 | 637.3 | - | - | 0 | - |
| - | - | 857.7 | 638.3 | - | - | 0 | - |
| - | - | 650.1 | 639.4 | - | - | 0 | - |
| 4 | y | 8323 | 640.3 | 0.003598 | 5.619 | +2 | 11 |
| - | - | 6316 | 640.8 | - | - | 0 | - |
| - | - | 1408 | 641.3 | - | - | 0 | - |
| - | - | 1028 | 643.3 | - | - | 0 | - |
| - | - | 2041 | 647.3 | - | - | 0 | - |
| - | - | 1514 | 647.8 | - | - | 0 | - |
| - | - | 1802 | 648.3 | - | - | 0 | - |
| - | - | 666.4 | 650.9 | - | - | 0 | - |
| - | - | 3585 | 652.3 | - | - | 0 | - |
| - | - | 1740 | 652.8 | - | - | 0 | - |
| - | - | 961.3 | 653.3 | - | - | 0 | - |
| - | - | 1144 | 655.3 | - | - | 0 | - |
| - | - | 1028 | 656.3 | - | - | 0 | - |
| - | - | 688.8 | 664.3 | - | - | 0 | - |
| - | - | 1849 | 664.8 | - | - | 0 | - |
| 5 | b | 6663 | 665.3 | 0.007715 | 11.6 | +1 | 5 |
| - | - | 1810 | 666.3 | - | - | 0 | - |
| - | - | 953 | 670.8 | - | - | 0 | - |
| - | - | 747.2 | 671.3 | - | - | 0 | - |
| 9 | y | 1716 | 672.3 | 0.001573 | 2.339 | +1 | 6 |
| 9 | y | 2786 | 673.3 | 0.001444 | 2.145 | +1 | 6 |
| - | - | 1010 | 674.3 | - | - | 0 | - |
| - | - | 4902 | 679.3 | - | - | 0 | - |
| - | - | 2876 | 679.8 | - | - | 0 | - |
| - | - | 1257 | 680.3 | - | - | 0 | - |
| 11 | b | 8066 | 682.4 | 0.00328 | 4.806 | +2 | 11 |
| - | - | 7278 | 682.9 | - | - | 0 | - |
| - | - | 3297 | 683.4 | - | - | 0 | - |
| - | - | 918.1 | 683.8 | - | - | 0 | - |
| 3 | y | 3090 | 687.8 | 0.003591 | 5.221 | +2 | 12 |
| 3 | y | 4584 | 688.3 | 0.005297 | 7.695 | +2 | 12 |
| - | - | 3353 | 688.8 | - | - | 0 | - |
| - | - | 1488 | 689.3 | - | - | 0 | - |
| 9 | y | 2.247E+04 | 690.3 | 0.0005907 | 0.8556 | +1 | 6 |
| - | - | 6992 | 691.3 | - | - | 0 | - |
| - | - | 1402 | 692.3 | - | - | 0 | - |
| 3 | y | 2.177E+04 | 696.8 | 0.002825 | 4.055 | +2 | 12 |
| - | - | 1.53E+04 | 697.3 | - | - | 0 | - |
| - | - | 6926 | 697.8 | - | - | 0 | - |
| - | - | 1696 | 698.3 | - | - | 0 | - |
| - | - | 3416 | 699.4 | - | - | 0 | - |
| - | - | 5051 | 700.4 | - | - | 0 | - |
| - | - | 1443 | 701.4 | - | - | 0 | - |
| - | - | 743.8 | 703.3 | - | - | 0 | - |
| - | - | 692.3 | 709.4 | - | - | 0 | - |
| - | - | 757.9 | 712.3 | - | - | 0 | - |
| - | - | 1073 | 714.4 | - | - | 0 | - |
| - | - | 1169 | 715.4 | - | - | 0 | - |
| - | - | 679 | 716.3 | - | - | 0 | - |
| - | - | 780 | 724.9 | - | - | 0 | - |
| - | - | 1542 | 725.3 | - | - | 0 | - |
| 12 | b | 1179 | 730.9 | 0.002598 | 3.554 | +2 | 12 |
| 12 | b | 1624 | 731.4 | 0.005463 | 7.47 | +2 | 12 |
| - | - | 1686 | 731.9 | - | - | 0 | - |
| - | - | 2807 | 733.9 | - | - | 0 | - |
| - | - | 2242 | 734.3 | - | - | 0 | - |
| - | - | 773.2 | 734.8 | - | - | 0 | - |
| - | - | 670.9 | 738.3 | - | - | 0 | - |
| 12 | b | 1.416E+04 | 739.9 | 0.002625 | 3.549 | +2 | 12 |
| - | - | 1.252E+04 | 740.4 | - | - | 0 | - |
| - | - | 5143 | 740.9 | - | - | 0 | - |
| - | - | 958.4 | 741.4 | - | - | 0 | - |
| - | - | 887.1 | 744.4 | - | - | 0 | - |
| - | - | 807.1 | 746.4 | - | - | 0 | - |
| 6 | b | 795.2 | 762.3 | 0.007755 | 10.17 | +1 | 6 |
| - | - | 909.6 | 766.4 | - | - | 0 | - |
| - | - | 1077 | 767.4 | - | - | 0 | - |
| - | - | 945 | 767.9 | - | - | 0 | - |
| - | - | 1793 | 768.4 | - | - | 0 | - |
| - | - | 1106 | 768.9 | - | - | 0 | - |
| 2 | y | 1546 | 769.4 | 0.006168 | 8.016 | +2 | 13 |
| 2 | y | 1747 | 769.9 | 0.008117 | 10.54 | +2 | 13 |
| - | - | 1260 | 770.4 | - | - | 0 | - |
| - | - | 890 | 776.4 | - | - | 0 | - |
| - | - | 1601 | 776.9 | - | - | 0 | - |
| - | - | 939.2 | 777.4 | - | - | 0 | - |
| 2 | y | 4460 | 778.4 | 0.005402 | 6.94 | +2 | 13 |
| - | - | 2550 | 778.9 | - | - | 0 | - |
| 6 | b | 3928 | 779.4 | 0.004461 | 5.723 | +1 | 6 |
| - | - | 1043 | 780.4 | - | - | 0 | - |
| 13 | b | 3488 | 781.4 | 0.002074 | 2.654 | +2 | 13 |
| 13 | b | 5249 | 781.9 | 0.007442 | 9.518 | +2 | 13 |
| - | - | 2703 | 782.4 | - | - | 0 | - |
| - | - | 1201 | 782.9 | - | - | 0 | - |
| - | - | 840.6 | 783.4 | - | - | 0 | - |
| 13 | b | 1.645E+04 | 790.4 | 0.002956 | 3.74 | +2 | 13 |
| - | - | 1.108E+04 | 790.9 | - | - | 0 | - |
| - | - | 7127 | 791.4 | - | - | 0 | - |
| - | - | 1702 | 791.9 | - | - | 0 | - |
| - | - | 1591 | 794.4 | - | - | 0 | - |
| - | - | 959.9 | 795.4 | - | - | 0 | - |
| - | - | 729.7 | 800.4 | - | - | 0 | - |
| - | - | 777 | 801.4 | - | - | 0 | - |
| - | - | 988.2 | 802.4 | - | - | 0 | - |
| - | - | 5147 | 802.9 | - | - | 0 | - |
| 8 | y | 1.683E+04 | 803.4 | 0.001442 | 1.795 | +1 | 7 |
| - | - | 2089 | 803.9 | - | - | 0 | - |
| - | - | 4666 | 804.4 | - | - | 0 | - |
| - | - | 955.6 | 805.4 | - | - | 0 | - |
| - | - | 1126 | 813.4 | - | - | 0 | - |
| - | - | 6089 | 814.4 | - | - | 0 | - |
| - | - | 3008 | 815.4 | - | - | 0 | - |
| - | - | 923.9 | 816.4 | - | - | 0 | - |
| - | - | 1869 | 817.4 | - | - | 0 | - |
| - | - | 1899 | 817.9 | - | - | 0 | - |
| - | - | 1256 | 818.4 | - | - | 0 | - |
| - | - | 1201 | 819.4 | - | - | 0 | - |
| 0 | Precursor | 1.098E+04 | 825.9 | 0.002588 | 3.133 | +2 | -1 |
| 0 | Precursor | 1.72E+04 | 826.4 | 0.009359 | 11.33 | +2 | -1 |
| - | - | 1.002E+04 | 826.9 | - | - | 0 | - |
| - | - | 4564 | 827.4 | - | - | 0 | - |
| - | - | 1061 | 833.4 | - | - | 0 | - |
| - | - | 819.3 | 834.4 | - | - | 0 | - |
| 0 | Precursor | 5.739E+04 | 834.9 | 0.002615 | 3.133 | +2 | -1 |
| - | - | 6.144E+04 | 835.4 | - | - | 0 | - |
| - | - | 3.321E+04 | 835.9 | - | - | 0 | - |
| - | - | 8283 | 836.4 | - | - | 0 | - |
| - | - | 661.8 | 842.4 | - | - | 0 | - |
| - | - | 720.5 | 843.4 | - | - | 0 | - |
| - | - | 1962 | 846.4 | - | - | 0 | - |
| - | - | 1216 | 847.4 | - | - | 0 | - |
| 7 | b | 940.6 | 848.4 | 0.01351 | 15.93 | +1 | 7 |
| 7 | b | 971.4 | 849.4 | 0.005512 | 6.49 | +1 | 7 |
| - | - | 1163 | 850.4 | - | - | 0 | - |
| - | - | 1192 | 858.4 | - | - | 0 | - |
| - | - | 2078 | 859.4 | - | - | 0 | - |
| - | - | 1453 | 860.4 | - | - | 0 | - |
| 7 | b | 1537 | 866.4 | 0.007527 | 8.688 | +1 | 7 |
| - | - | 1031 | 867.4 | - | - | 0 | - |
| 7 | y | 2284 | 872.4 | 0.0005669 | 0.6498 | +1 | 8 |
| 7 | y | 763.5 | 873.4 | 0.01243 | 14.23 | +1 | 8 |
| - | - | 1572 | 880.4 | - | - | 0 | - |
| 7 | y | 1.79E+04 | 890.5 | 0.0005872 | 0.6594 | +1 | 8 |
| - | - | 7192 | 891.5 | - | - | 0 | - |
| - | - | 2055 | 892.5 | - | - | 0 | - |
| - | - | 2219 | 893.5 | - | - | 0 | - |
| - | - | 1125 | 894.4 | - | - | 0 | - |
| - | - | 731 | 895.4 | - | - | 0 | - |
| - | - | 5406 | 897.4 | - | - | 0 | - |
| - | - | 3688 | 898.4 | - | - | 0 | - |
| - | - | 1265 | 899.4 | - | - | 0 | - |
| - | - | 815 | 908.4 | - | - | 0 | - |
| - | - | 2998 | 910.5 | - | - | 0 | - |
| - | - | 1459 | 911.5 | - | - | 0 | - |
| - | - | 697.3 | 915.5 | - | - | 0 | - |
| - | - | 1550 | 916.4 | - | - | 0 | - |
| - | - | 670.3 | 927.4 | - | - | 0 | - |
| - | - | 1167 | 933.5 | - | - | 0 | - |
| - | - | 1206 | 944.5 | - | - | 0 | - |
| - | - | 3801 | 961.4 | - | - | 0 | - |
| 8 | b | 3559 | 962.5 | 0.004699 | 4.883 | +1 | 8 |
| - | - | 946.1 | 963.5 | - | - | 0 | - |
| - | - | 1169 | 970.5 | - | - | 0 | - |
| - | - | 1201 | 972.5 | - | - | 0 | - |
| - | - | 1990 | 974.5 | - | - | 0 | - |
| - | - | 775.8 | 975.5 | - | - | 0 | - |
| 8 | b | 2665 | 979.5 | 0.004152 | 4.239 | +1 | 8 |
| - | - | 1193 | 980.5 | - | - | 0 | - |
| - | - | 1013 | 981.5 | - | - | 0 | - |
| 6 | y | 2223 | 986.5 | 0.003577 | 3.626 | +1 | 9 |
| 6 | y | 4198 | 987.5 | 0.00612 | 6.198 | +1 | 9 |
| - | - | 688.1 | 989.5 | - | - | 0 | - |
| - | - | 673.3 | 1002 | - | - | 0 | - |
| 6 | y | 2.945E+04 | 1005 | 0.0003843 | 0.3826 | +1 | 9 |
| - | - | 1.606E+04 | 1006 | - | - | 0 | - |
| - | - | 5584 | 1007 | - | - | 0 | - |
| - | - | 985.5 | 1008 | - | - | 0 | - |
| - | - | 2849 | 1008 | - | - | 0 | - |
| - | - | 1747 | 1009 | - | - | 0 | - |
| - | - | 1059 | 1010 | - | - | 0 | - |
| - | - | 2225 | 1017 | - | - | 0 | - |
| - | - | 1488 | 1018 | - | - | 0 | - |
| - | - | 1040 | 1018 | - | - | 0 | - |
| - | - | 776 | 1023 | - | - | 0 | - |
| - | - | 1133 | 1024 | - | - | 0 | - |
| - | - | 4139 | 1026 | - | - | 0 | - |
| - | - | 2346 | 1027 | - | - | 0 | - |
| - | - | 659.9 | 1056 | - | - | 0 | - |
| - | - | 2305 | 1061 | - | - | 0 | - |
| - | - | 1597 | 1062 | - | - | 0 | - |
| - | - | 1004 | 1062 | - | - | 0 | - |
| - | - | 1044 | 1063 | - | - | 0 | - |
| - | - | 986.7 | 1069 | - | - | 0 | - |
| - | - | 1775 | 1070 | - | - | 0 | - |
| - | - | 2570 | 1071 | - | - | 0 | - |
| - | - | 922.2 | 1072 | - | - | 0 | - |
| - | - | 2130 | 1072 | - | - | 0 | - |
| - | - | 989.8 | 1073 | - | - | 0 | - |
| - | - | 1729 | 1073 | - | - | 0 | - |
| - | - | 1.996E+04 | 1088 | - | - | 0 | - |
| - | - | 1.414E+04 | 1089 | - | - | 0 | - |
| - | - | 4456 | 1090 | - | - | 0 | - |
| - | - | 1911 | 1091 | - | - | 0 | - |
| - | - | 1000 | 1091 | - | - | 0 | - |
| - | - | 858.1 | 1111 | - | - | 0 | - |
| 9 | b | 1555 | 1119 | 0.003809 | 3.405 | +1 | 9 |
| - | - | 713.5 | 1121 | - | - | 0 | - |
| - | - | 872.7 | 1127 | - | - | 0 | - |
| - | - | 1541 | 1128 | - | - | 0 | - |
| - | - | 1260 | 1131 | - | - | 0 | - |
| - | - | 1477 | 1132 | - | - | 0 | - |
| 5 | y | 2214 | 1134 | 0.004878 | 4.304 | +1 | 10 |
| 5 | y | 3543 | 1135 | 0.009877 | 8.706 | +1 | 10 |
| 9 | b | 9134 | 1136 | 0.004603 | 4.054 | +1 | 9 |
| - | - | 6988 | 1137 | - | - | 0 | - |
| - | - | 1733 | 1138 | - | - | 0 | - |
| - | - | 2887 | 1139 | - | - | 0 | - |
| - | - | 1510 | 1140 | - | - | 0 | - |
| - | - | 711.5 | 1145 | - | - | 0 | - |
| - | - | 718 | 1147 | - | - | 0 | - |
| 5 | y | 2.769E+04 | 1152 | 0.004812 | 4.179 | +1 | 10 |
| - | - | 1.735E+04 | 1153 | - | - | 0 | - |
| - | - | 7530 | 1154 | - | - | 0 | - |
| - | - | 2328 | 1155 | - | - | 0 | - |
| - | - | 985.4 | 1175 | - | - | 0 | - |
| - | - | 830.9 | 1181 | - | - | 0 | - |
| - | - | 976.1 | 1191 | - | - | 0 | - |
| - | - | 2282 | 1198 | - | - | 0 | - |
| - | - | 8518 | 1199 | - | - | 0 | - |
| - | - | 4081 | 1200 | - | - | 0 | - |
| - | - | 1437 | 1201 | - | - | 0 | - |
| - | - | 3903 | 1203 | - | - | 0 | - |
| - | - | 3428 | 1204 | - | - | 0 | - |
| - | - | 1.28E+04 | 1216 | - | - | 0 | - |
| - | - | 8578 | 1217 | - | - | 0 | - |
| - | - | 2052 | 1218 | - | - | 0 | - |
| - | - | 1096 | 1221 | - | - | 0 | - |
| - | - | 1007 | 1227 | - | - | 0 | - |
| - | - | 1024 | 1246 | - | - | 0 | - |
| 10 | b | 6905 | 1249 | 0.005134 | 4.112 | +1 | 10 |
| - | - | 4230 | 1250 | - | - | 0 | - |
| - | - | 1501 | 1251 | - | - | 0 | - |
| - | - | 1132 | 1258 | - | - | 0 | - |
| 4 | y | 2230 | 1262 | 0.003674 | 2.912 | +1 | 11 |
| 4 | y | 7690 | 1263 | 0.008428 | 6.675 | +1 | 11 |
| - | - | 5940 | 1264 | - | - | 0 | - |
| - | - | 2383 | 1265 | - | - | 0 | - |
| 4 | y | 1.836E+04 | 1280 | 0.004828 | 3.773 | +1 | 11 |
| - | - | 1.485E+04 | 1281 | - | - | 0 | - |
| - | - | 6040 | 1282 | - | - | 0 | - |
| - | - | 928.8 | 1291 | - | - | 0 | - |
| - | - | 1328 | 1294 | - | - | 0 | - |
| - | - | 1472 | 1300 | - | - | 0 | - |
| - | - | 1018 | 1301 | - | - | 0 | - |
| - | - | 1134 | 1305 | - | - | 0 | - |
| - | - | 848 | 1312 | - | - | 0 | - |
| - | - | 1.159E+04 | 1329 | - | - | 0 | - |
| - | - | 9186 | 1330 | - | - | 0 | - |
| - | - | 3322 | 1331 | - | - | 0 | - |
| - | - | 875.5 | 1358 | - | - | 0 | - |
| - | - | 1042 | 1359 | - | - | 0 | - |
| 11 | b | 4136 | 1364 | 0.00346 | 2.537 | +1 | 11 |
| - | - | 3525 | 1365 | - | - | 0 | - |
| - | - | 2560 | 1366 | - | - | 0 | - |
| 3 | y | 1123 | 1375 | 0.006524 | 4.746 | +1 | 12 |
| 3 | y | 1084 | 1376 | 0.01531 | 11.13 | +1 | 12 |
| - | - | 1085 | 1377 | - | - | 0 | - |
| 3 | y | 1.455E+04 | 1393 | 0.003528 | 2.533 | +1 | 12 |
| - | - | 1.189E+04 | 1394 | - | - | 0 | - |
| - | - | 4983 | 1395 | - | - | 0 | - |
| - | - | 1259 | 1396 | - | - | 0 | - |
| - | - | 1629 | 1415 | - | - | 0 | - |
| - | - | 1478 | 1416 | - | - | 0 | - |
| 12 | b | 730.5 | 1462 | 0.008531 | 5.836 | +1 | 12 |
| 12 | b | 3578 | 1479 | 0.001663 | 1.125 | +1 | 12 |
| - | - | 2811 | 1480 | - | - | 0 | - |
| - | - | 1918 | 1481 | - | - | 0 | - |
| - | - | 1507 | 1492 | - | - | 0 | - |
| - | - | 952.8 | 1493 | - | - | 0 | - |
| - | - | 921.1 | 1494 | - | - | 0 | - |
| 2 | y | 1005 | 1556 | 0.006728 | 4.324 | +1 | 13 |
| - | - | 1105 | 1557 | - | - | 0 | - |
| - | - | 751.9 | 2958 | - | - | 0 | - |
| - | - | 1094 | 3037 | - | - | 0 | - |

m/z Charge Intensity FragmentType MassShift Position
120.06632995605469 0 607.7172
120.08099365234375 0 3097.6963
126.05497741699219 0 473.958
127.08650207519531 0 372.50497
129.06629943847656 0 1018.1017
129.10247802734375 0 3744.2454
130.06561279296875 0 794.2927
131.08172607421875 0 568.4983
131.1181182861328 0 9153.864
132.12164306640625 0 410.92896
133.06114196777344 0 610.21924
136.0759735107422 0 88402.234
137.07386779785156 0 820.6002
137.079345703125 0 5738.2773
139.0509796142578 0 485.55405
139.6966094970703 0 343.31717
140.08238220214844 0 916.1814
141.1026153564453 0 615.54706
142.12315368652344 0 647.6262
144.06578063964844 0 1786.0919
145.06143188476562 0 771.1757
146.0608673095703 0 667.73395
147.07701110839844 0 425.43362
148.84197998046875 0 405.74243
148.87686157226562 0 466.10138
148.88348388671875 0 509.32675
148.89108276367188 0 442.78625
148.8979034423828 0 506.93988
148.90567016601562 0 448.4849
148.91233825683594 0 541.5213
148.92039489746094 0 693.08057
148.92774963378906 0 1094.5159
148.93426513671875 0 1953.6741
148.94186401367188 0 3552
148.95848083496094 0 4298.763
148.96632385253906 0 2612.54
148.97296142578125 0 999.99146
148.98077392578125 0 1031.6201
148.98802185058594 0 707.058
148.99510192871094 0 573.20734
149.00221252441406 0 724.4857
149.00973510742188 0 517.85645
149.04530334472656 0 862.3386
152.0710906982422 0 1222.6818
152.17071533203125 0 428.45544
153.06629943847656 0 1244.1045
153.07699584960938 0 440.44873
155.08139038085938 0 810.91364
156.0772247314453 0 1391.8989
157.06105041503906 0 1464.5332
157.1086883544922 0 781.0847
159.07687377929688 0 653.9992
159.09164428710938 0 881.321
166.09774780273438 0 1445.3943
167.08177185058594 0 4216.726
169.09780883789062 0 738.67834
169.13412475585938 0 710.15186
171.0769500732422 0 1137.4619
171.7378387451172 0 450.43835
173.0923614501953 0 3230.088 y Water loss 12
173.12872314453125 0 4358.5938
173.4423370361328 0 829.413
173.645751953125 0 477.89355
174.04759216308594 0 573.82733
174.0552520751953 0 1862.2296
174.08782958984375 0 1026.0465
175.0869598388672 0 537.87415
178.05015563964844 0 971.42737
181.06149291992188 0 1098.3297
183.1131134033203 0 1406.0742
183.14898681640625 0 840.0864
184.0719451904297 0 1643.587
184.1080780029297 0 1403.925
185.05580139160156 0 1225.5656
186.12452697753906 0 515.0652
187.1079559326172 0 3141.3228
187.14463806152344 0 526.7659
188.1112060546875 0 684.74567
189.08709716796875 0 3815.2642
191.0817413330078 0 775.3263
191.1029052734375 0 13256.904 y 12
192.10641479492188 0 1062.6573
194.092529296875 0 1495.5521
195.07664489746094 0 7416.2
196.08001708984375 0 696.58167
197.12884521484375 0 1036.2526
198.087646484375 0 2568.7466
199.07150268554688 0 1156.5046
199.1077880859375 0 1376.4474
201.08755493164062 0 828.9403
201.12342834472656 0 3728.1372
202.08270263671875 0 2264.457
203.066650390625 0 1543.9794
203.1028594970703 0 2527.6467
208.09698486328125 0 1483.7649
208.14515686035156 0 568.0491
210.0886993408203 0 750.70764
211.1443328857422 0 1054.0482
212.10328674316406 0 3043.3174
213.0869598388672 0 666.37335
214.15548706054688 0 1350.3221
215.13926696777344 0 3301.9912
217.06417846679688 0 773.12964
217.08216857910156 0 7361.0107
223.1084442138672 0 959.82227
224.13955688476562 0 738.2464
225.04319763183594 0 693.7097
225.0986328125 0 1178.0311
225.12367248535156 0 4742.3247
225.17112731933594 0 2151.7793
226.11842346191406 0 803.9084
226.15628051757812 0 587.4515
227.1027374267578 0 1725.5447
228.1707000732422 0 675.0276
229.1185760498047 0 2181.2156
231.0616912841797 0 2303.691
236.1377716064453 0 554.23303
239.09422302246094 0 943.8583
239.11453247070312 0 768.268
240.09844970703125 0 989.4075
240.13479614257812 0 913.4932
242.0778350830078 0 621.02545
242.11325073242188 0 675.2622
242.13687133789062 0 1074.4875
242.15025329589844 0 17404.365
243.1536865234375 0 1773.7249
246.12425231933594 0 661.91956
249.16014099121094 0 100604.09 a 1
250.1634063720703 0 13664.389
252.13479614257812 0 670.20416
253.118408203125 0 887.00287
253.16619873046875 0 5576
258.0904846191406 0 745.39856
259.074951171875 0 3450.3303
260.1283264160156 0 672.3697
261.1268615722656 0 732.05316
262.08544921875 0 2480.6497
262.15447998046875 0 844.4651
264.1334228515625 0 618.13275
265.1546936035156 0 1181.7329
267.1092529296875 0 1611.8824
268.09307861328125 0 1178.72
270.1082763671875 0 712.19867
270.1443786621094 0 1436.1929
270.1932678222656 0 779.44904
274.1398620605469 0 917.39966
276.1015319824219 0 3144.7427
277.1549377441406 0 76801.62 b 1
278.1582336425781 0 11739.013
279.09912109375 0 694.97894
279.1611328125 0 1100.1592
282.1823425292969 0 501.51047
285.1195373535156 0 2182.6265
286.1037902832031 0 1410.8647
287.173828125 0 941.4913
288.1194152832031 0 2898.0325 y Water loss 11
290.14947509765625 0 677.2857
294.18096923828125 0 762.96924
297.15582275390625 0 1327.7477
298.1407470703125 0 1438.4238
299.06201171875 0 1265.6191
300.0624694824219 0 934.6756
306.1298828125 0 4161.0312 y 11
307.1759033203125 0 671.0905
308.1357727050781 0 755.1928
309.11981201171875 0 1806.2119
314.09844970703125 0 1328.0452
315.168212890625 0 2092.9834
322.1895751953125 0 676.04456
323.1685485839844 0 536.7356
325.1871643066406 0 1321.0068
326.1468200683594 0 1221.6561
332.10931396484375 0 4908.6016
340.19818115234375 0 2834.328
341.2022399902344 0 640.46564
344.1455078125 0 3249.169
345.2190246582031 0 798.6771
349.1158142089844 0 814.1126
350.1806640625 0 664.2184
351.1310119628906 0 1193.6525
353.1815490722656 0 1628.0879
355.0701904296875 0 2214.7507
357.06732177734375 0 815.5633
357.2250061035156 0 1832.5392
359.0284729003906 0 683.51306
368.1562194824219 0 1230.6753
368.19268798828125 0 2827.2288
369.1964416503906 0 627.2437
372.1579895019531 0 1058.2572
372.2271423339844 0 732.9129
373.1177978515625 0 1743.9188
378.1412353515625 0 2923.416
380.19378662109375 0 753.00916
381.1764831542969 0 843.8685
385.135986328125 0 826.374
385.21942138671875 0 2727.1968
386.2216796875 0 918.9733
389.18572998046875 0 3086.4373
390.1429443359375 0 690.065
390.23931884765625 0 13053.448 b 2
391.2418212890625 0 3939.4814
393.22021484375 0 550.389
395.1680908203125 0 1003.85144
396.1516418457031 0 1783.4677
403.146484375 0 3947.3965 y Water loss 10
404.1468505859375 0 796.2987
404.19573974609375 0 646.157
405.21368408203125 0 4767.7896
406.2174987792969 0 942.2941
407.26605224609375 0 3935.205
417.1800231933594 0 693.14166
420.1863098144531 0 641.4567
421.1578063964844 0 2568.4172 y 10
424.1908264160156 0 1411.3823
427.18450927734375 0 1864.6866
428.2587585449219 0 866.92365
430.2280578613281 0 1619.6476
430.72802734375 0 1225.5985
431.93902587890625 0 553.134
433.2102355957031 0 1685.637
437.2151794433594 0 1939.5334 y Ammonia loss 6
442.1399841308594 0 1542.1108
445.1933898925781 0 4038.026
446.1991882324219 0 931.909
454.2400207519531 0 920.2679
460.1476135253906 0 1126.1263
464.2139587402344 0 984.0316
466.85919189453125 0 649.8231
468.591796875 0 535.371
471.2724304199219 0 1198.2908
473.277587890625 0 621.1793
483.2208251953125 0 3455.4956
486.2042236328125 0 568.88824
490.3035888671875 0 1261.9047
491.2245178222656 0 1016.239
498.30389404296875 0 2421.2603
500.2467346191406 0 4518.867
501.25018310546875 0 972.1572
503.22772216796875 0 1198.1798
504.24774169921875 0 797.972
508.2517395019531 0 730.1671
509.2391357421875 0 739.6757
516.230712890625 0 1891.7074 y Water loss 9
518.2977294921875 0 6352.98 b 3
519.3011474609375 0 2510.1118
520.252685546875 0 757.13696
534.2415161132812 0 5967.509 y 9
535.2434692382812 0 2091.779
536.2501220703125 0 1501.176
536.7500610351562 0 1058.1805
538.2764892578125 0 571.2192
544.2822265625 0 1800.8845
544.7821044921875 0 964.9282
545.2532348632812 0 778.21594
545.754638671875 0 820.2986
550.3014526367188 0 718.4029
552.2481079101562 0 1079.0266
552.78369140625 0 744.94415
554.2604370117188 0 644.9044
554.305419921875 0 1396.0762
555.2203979492188 0 751.6693
568.3013916015625 0 4393.5254 b 8
568.8010864257812 0 3109.9639
569.2975463867188 0 592.4238
573.3411865234375 0 817.4074
573.7719116210938 0 564.0437
584.2753295898438 0 856.1452
585.3361206054688 0 2544.7283
587.2684936523438 0 1565.0715
587.7620849609375 0 974.0849
590.7853393554688 0 810.3163
595.7764282226562 0 1154.0225
599.297607421875 0 1971.2227
599.7938842773438 0 1652.5906
601.2914428710938 0 1767.53
601.338134765625 0 1081.7269
601.7954711914062 0 4375.115
602.2937622070312 0 2097.5413
608.3025512695312 0 995.6742
610.848388671875 0 1192.8394
613.331787109375 0 2724.6404
618.30712890625 0 743.3844
620.31494140625 0 751.1794
622.2893676757812 0 903.8151
622.7835693359375 0 1372.6228
624.8424682617188 0 2530.0688 b 9
625.3438110351562 0 1382.4741
628.3828125 0 632.3316
630.3184204101562 0 774.63416
631.296142578125 0 6077.924 y Water loss 3
631.7962036132812 0 6847.449 y Ammonia loss 3
632.2952880859375 0 1866.8772
634.8056030273438 0 930.75244
635.8134155273438 0 683.2351
637.3423461914062 0 1481.3687
638.325439453125 0 857.73785
639.3943481445312 0 650.1374
640.3021240234375 0 8322.783 y 3
640.802490234375 0 6316.2646
641.2972412109375 0 1408.3455
643.3102416992188 0 1028.3667
647.3282470703125 0 2041.4834
647.825927734375 0 1513.9292
648.3159790039062 0 1801.6243
650.8521118164062 0 666.36053
652.3197021484375 0 3584.8447
652.8196411132812 0 1739.6282
653.3201904296875 0 961.3007
655.3045043945312 0 1144.1272
656.336181640625 0 1027.618
664.349853515625 0 688.7813
664.8455200195312 0 1848.9962
665.3355102539062 0 6662.95 b 4
666.33447265625 0 1809.977
670.8193969726562 0 952.9985
671.3197021484375 0 747.21936
672.3327026367188 0 1715.6 y Water loss 8
673.3165893554688 0 2785.5276 y Ammonia loss 8
674.3395385742188 0 1010.0091
679.3257446289062 0 4902.182
679.8251953125 0 2876.2192
680.3255615234375 0 1256.6215
682.3563842773438 0 8065.6494 b 10
682.8570556640625 0 7278.271
683.3552856445312 0 3296.8296
683.831298828125 0 918.1401
687.8388671875 0 3089.5835 y Water loss 2
688.3325805664062 0 4583.941 y Ammonia loss 2
688.8308715820312 0 3353.0022
689.3312377929688 0 1488.4148
690.34228515625 0 22468.941 y 8
691.3450927734375 0 6992.442
692.3436889648438 0 1401.5204
696.8433837890625 0 21766.826 y 2
697.3441162109375 0 15303.695
697.8438110351562 0 6926.264
698.3455200195312 0 1695.9795
699.3782348632812 0 3415.5164
700.3641967773438 0 5051.198
701.365478515625 0 1443.272
703.3416137695312 0 743.8344
709.3715209960938 0 692.31445
712.313232421875 0 757.8738
714.3823852539062 0 1072.6156
715.3779907226562 0 1168.5023
716.3348388671875 0 679.0285
724.8532104492188 0 780.0373
725.3494873046875 0 1541.5015
730.8638916015625 0 1179.2261 b Water loss 11
731.3587646484375 0 1623.8508 b Ammonia loss 11
731.8578491210938 0 1685.7705
733.8534545898438 0 2806.8347
734.3497924804688 0 2242.4187
734.846923828125 0 773.16614
738.3489379882812 0 670.89325
739.8692016601562 0 14163.127 b 11
740.3700561523438 0 12524.589
740.8706665039062 0 5142.7812
741.3681030273438 0 958.3751
744.3533325195312 0 887.0714
746.3505859375 0 807.09717
762.3519287109375 0 795.1607 b Ammonia loss 5
766.381591796875 0 909.6495
767.38671875 0 1077.3188
767.8859252929688 0 945.0202
768.3905639648438 0 1793.2588
768.89306640625 0 1106.1779
769.3731079101562 0 1546.2971 y Water loss 1
769.8670654296875 0 1746.837 y Ammonia loss 1
770.367431640625 0 1260.4392
776.3921508789062 0 889.97253
776.8973999023438 0 1600.7476
777.3970336914062 0 939.21674
778.3776245117188 0 4459.9297 y 1
778.8753662109375 0 2550.2854
779.3751831054688 0 3927.6982 b 5
780.3750610351562 0 1042.9685
781.38720703125 0 3487.7615 b Water loss 12
781.8845825195312 0 5249.032 b Ammonia loss 12
782.4003295898438 0 2703.2498
782.8878173828125 0 1200.6929
783.4180297851562 0 840.6395
790.3933715820312 0 16451.232 b 12
790.89404296875 0 11077.927
791.394287109375 0 7126.907
791.8945922851562 0 1702.2457
794.4141235351562 0 1590.5679
795.4474487304688 0 959.8601
800.4013671875 0 729.69727
801.4149169921875 0 776.959
802.4112548828125 0 988.2101
802.91748046875 0 5147.284
803.42431640625 0 16829.785 y 7
803.9163818359375 0 2088.8616
804.4269409179688 0 4665.831
805.4224243164062 0 955.55194
813.4019165039062 0 1125.5457
814.4069213867188 0 6088.786
815.4046020507812 0 3007.727
816.405517578125 0 923.8892
817.401611328125 0 1868.8395
817.9027099609375 0 1899.4705
818.4027709960938 0 1255.625
819.4190063476562 0 1200.529
825.9115600585938 0 10975.359 Precursor Water loss
826.4103393554688 0 17196.527 Precursor Ammonia loss
826.9099731445312 0 10017.553
827.4052734375 0 4563.6875
833.3999633789062 0 1061.0902
834.417236328125 0 819.3474
834.9168701171875 0 57386.926 Precursor
835.4175415039062 0 61444.37
835.9175415039062 0 33207.266
836.4190063476562 0 8283.469
842.4262084960938 0 661.8064
843.4224243164062 0 720.46094
846.4138793945312 0 1962.3617
847.4259643554688 0 1215.834
848.4057006835938 0 940.56134 b Water loss 6
849.3817138671875 0 971.3828 b Ammonia loss 6
850.3895263671875 0 1163.4658
858.4317016601562 0 1192.2295
859.4428100585938 0 2078.2812
860.4470825195312 0 1453.1206
866.4102783203125 0 1537.3093 b 6
867.4132080078125 0 1031.3015
872.4466552734375 0 2284.427 y Water loss 6
873.4436645507812 0 763.5205 y Ammonia loss 6
880.4180908203125 0 1571.5991
890.4583740234375 0 17902.607 y 6
891.4610595703125 0 7192.338
892.4624633789062 0 2055.4006
893.450927734375 0 2219.3577
894.4491577148438 0 1125.3959
895.4259643554688 0 730.996
897.4430541992188 0 5405.573
898.4415283203125 0 3688.4702
899.4382934570312 0 1264.5691
908.4452514648438 0 815.03235
910.4749755859375 0 2997.9639
911.4728393554688 0 1458.6841
915.4500732421875 0 697.30237
916.4485473632812 0 1549.6976
927.4349975585938 0 670.3487
933.4723510742188 0 1167.2576
944.461669921875 0 1205.7229
961.4437255859375 0 3800.923
962.45556640625 0 3559.4521 b Ammonia loss 7
963.4578247070312 0 946.0794
970.4571533203125 0 1168.5228
972.5355224609375 0 1201.2401
974.4734497070312 0 1990.0051
975.4725341796875 0 775.8178
979.490966796875 0 2664.926 b 7
980.4833374023438 0 1192.7883
981.4766235351562 0 1012.5851
986.486572265625 0 2222.9502 y Water loss 5
987.4802856445312 0 4197.628 y Ammonia loss 5
989.4762573242188 0 688.1046
1002.487060546875 0 673.2942
1004.5010986328125 0 29450.057 y 5
1005.5010986328125 0 16058.555
1006.5029296875 0 5583.605
1007.5083618164062 0 985.51044
1008.4752807617188 0 2849.402
1009.4769287109375 0 1747.225
1010.4801635742188 0 1059.3198
1016.5013427734375 0 2225.1235
1017.5039672851562 0 1487.876
1018.49462890625 0 1040.1136
1022.5138549804688 0 776.02905
1023.555908203125 0 1132.8193
1025.5013427734375 0 4139.3164
1026.5067138671875 0 2346.4507
1056.455810546875 0 659.8809
1060.5126953125 0 2305.131
1061.5196533203125 0 1597.2501
1062.4930419921875 0 1004.20264
1063.4862060546875 0 1043.5117
1068.5069580078125 0 986.7015
1069.530517578125 0 1774.8685
1070.5184326171875 0 2570.3667
1071.60791015625 0 922.21466
1072.4757080078125 0 2130.4087
1072.599609375 0 989.83014
1073.4774169921875 0 1729.0969
1087.5394287109375 0 19957.969
1088.5400390625 0 14141.997
1089.5272216796875 0 4455.847
1090.5048828125 0 1911.0334
1091.494384765625 0 1000.4235
1110.5283203125 0 858.0688
1118.565185546875 0 1554.5983 b Ammonia loss 8
1120.5684814453125 0 713.5246
1126.55615234375 0 872.68805
1127.5438232421875 0 1540.9321
1130.5438232421875 0 1260.437
1131.5416259765625 0 1476.9633
1133.5255126953125 0 2213.743 y Water loss 4
1134.5145263671875 0 3543.0908 y Ammonia loss 4
1135.592529296875 0 9134.042 b 8
1136.5948486328125 0 6988.3613
1137.590576171875 0 1732.7133
1138.5836181640625 0 2887.0432
1139.58740234375 0 1509.8693
1144.5660400390625 0 711.4532
1146.5484619140625 0 718.02783
1151.5360107421875 0 27691.88 y 4
1152.5364990234375 0 17347.734
1153.535888671875 0 7529.998
1154.5445556640625 0 2327.6594
1174.5413818359375 0 985.38275
1180.56494140625 0 830.87787
1190.548583984375 0 976.05646
1197.5858154296875 0 2282.4482
1198.5733642578125 0 8517.747
1199.5697021484375 0 4080.605
1200.577392578125 0 1437.4673
1202.5828857421875 0 3903.0063
1203.5841064453125 0 3427.563
1215.5960693359375 0 12796.038
1216.596923828125 0 8577.532
1217.595458984375 0 2052.4353
1220.6837158203125 0 1096.3135
1226.566162109375 0 1007.3812
1245.5426025390625 0 1023.5272
1248.6771240234375 0 6904.673 b 9
1249.6771240234375 0 4229.7886
1250.68359375 0 1500.9702
1257.6414794921875 0 1131.773
1261.5828857421875 0 2230.017 y Water loss 3
1262.5716552734375 0 7690.06 y Ammonia loss 3
1263.570068359375 0 5939.685
1264.5694580078125 0 2382.7012
1279.5946044921875 0 18361.049 y 3
1280.5936279296875 0 14847.414
1281.59130859375 0 6040.38
1290.561279296875 0 928.8084
1293.633056640625 0 1328.2678
1299.70068359375 0 1471.9508
1300.702880859375 0 1018.3895
1304.6351318359375 0 1133.8545
1311.661865234375 0 848.02203
1328.678955078125 0 11594.078
1329.6805419921875 0 9185.739
1330.6844482421875 0 3322.2615
1357.6373291015625 0 875.4856
1358.6512451171875 0 1041.8379
1363.702392578125 0 4136.085 b 10
1364.7044677734375 0 3524.7236
1365.6976318359375 0 2559.699
1374.6697998046875 0 1123.2478 y Water loss 2
1375.66259765625 0 1083.508 y Ammonia loss 2
1376.667236328125 0 1085.1226
1392.6773681640625 0 14546.755 y 2
1393.6788330078125 0 11892.175
1394.6741943359375 0 4983.341
1395.68115234375 0 1259.1201
1414.7281494140625 0 1628.9739
1415.7271728515625 0 1477.7561
1461.6907958984375 0 730.45886 b Ammonia loss 11
1478.7275390625 0 3578.0037 b 11
1479.721435546875 0 2811.2786
1480.733154296875 0 1918.2483
1491.7393798828125 0 1507.1443
1492.7437744140625 0 952.8291
1493.74755859375 0 921.0787
1555.743896484375 0 1004.7965 y 1
1556.7401123046875 0 1105.0208
2957.8369140625 0 751.90393
3037.14013671875 0 1094.0383

Spectrum Details

|  |  |
| --- | --- |
| Matched peaks? Matched peaks The total absolute number of peaks matched. Additionally in brackets the total fraction of peaks matched and the total number of peaks is shown. | 68 (11.91% of 571) |
| FDR? FDR The false discovery rate estimated for this peptide. It is calculated by matching all theoretical fragments with a non-integer shift with the raw peaks for this spectrum. This is done with 40 different shifts. The resulting percentage is the average number of annotated peaks over the number of annotated peaks with the correct spectrum. | 0.39% |
| Satellite FDR? Satellite FDR See the FDR for details on its calculation. This satellite ion specific FDR only contains the satellite ions (d/w) for I/L/J positions. | ∞ |
| PSM Score? PSM Score The PSM Score as given by Hecklib to this annotated spectrum. It is shown with three significant figures. | 750 |

#### Spectrum 9788? Spectrum 9788 The raw spectrum of this peptide as annotated by Hecklib. The fragments are coloured according to ion type (see legend). Any peaks with a star '\*' as text can be hovered over to see the full details, first the ion type second the mass shift type. By hovering over the amino acids in the peptide or ions in the legend the corresponding peaks are highlighted. By toggling the 'Unassigned' label you can turn the background (unassigned) peaks on or off in the plot. By updating the slider in the Ion legend you can update the spectrum to only show the top X% of the peaks with labels. The top X% means any peak that is within X% of the highest intensity. By dragging in the spectrum you can zoom in to a specific part of the spectrum and use 'Zoom Out' to get back to the original zoom level. The annotation of the spectrum is based on the given sequence in the peptides file and is done with different software so inconsistencies are likely. The peaks are annotated based on the given sequence, with 20 ppm tolerance.

Copy Data

##### Spectrum 9788 (TSV)

###### Preview

```
Loading example...
```

*Click on the button to copy the data to your clipboard.*

Mz MinMz MaxIntensity Max

WidthHeightPeptide font sizePeptide stroke widthSpectrum font sizeSpectrum stroke widthCompact peptide

Ion legend

wxyz

abcd

OtherUnassignedIon^Charge^_Position_Show for top:%

JYLQMNSJRJDDTA

05.45e+41.09e+51.63e+52.18e+5

Zoom Out

y^+1^_2_y^+1^_3_y^+1^_3_y^+1^_4_c^+1^_3_y^+1^_4_y^+1^_5_c^+1^_4_y^+1^_5_c^+2^_9_y^+2^_11_y^+2^_11_z^+2^_11_y^+2^_11_c^+1^_5_y^+1^_6_z^+1^_6_c^+2^_11_y^+2^_12_y^+2^_12_z^+2^_12_y^+2^_12_y^+1^_6_c^+2^_12_w^+1^_7_y^+2^_13_z^+2^_13_c^+1^_6_y^+2^_13_c^+2^_13_z^+1^_7_y^+1^_7_c^+1^_7_y^+1^_8_z^+1^_8_y^+1^_8_c^+1^_8_y^+1^_9_y^+1^_9_z^+1^_9_y^+1^_9_w^+1^_10_y^+1^_10_y^+1^_10_z^+1^_10_y^+1^_10_c^+1^_9_w^+1^_11_c^+1^_10_y^+1^_11_y^+1^_11_z^+1^_11_c^+1^_10_y^+1^_11_w^+1^_12_c^+1^_11_y^+1^_12_y^+1^_12_z^+1^_12_c^+1^_11_y^+1^_12_c^+1^_12_c^+1^_12_y^+1^_13_z^+1^_13_y^+1^_13_c^+1^_13_c^+1^_13_c^+1^_13_

0745149122362982

Fragment Matches Table

Show background peaks

| Position | Ion type | Intensity | mz Theoretical | mz Error (Th) | mz Error (ppm) | Charge | Series Number |
| --- | --- | --- | --- | --- | --- | --- | --- |
| - | - | 353.1 | 121.1 | - | - | 0 | - |
| - | - | 411.9 | 125.7 | - | - | 0 | - |
| - | - | 555.6 | 131.1 | - | - | 0 | - |
| - | - | 400.7 | 134.3 | - | - | 0 | - |
| - | - | 3903 | 136.1 | - | - | 0 | - |
| - | - | 442 | 148.7 | - | - | 0 | - |
| - | - | 426.8 | 170.8 | - | - | 0 | - |
| - | - | 3429 | 173.4 | - | - | 0 | - |
| - | - | 599.5 | 189.7 | - | - | 0 | - |
| 13 | y | 6787 | 191.1 | 0.0001193 | 0.6242 | +1 | 2 |
| - | - | 457.7 | 200.9 | - | - | 0 | - |
| - | - | 4316 | 215.1 | - | - | 0 | - |
| - | - | 4055 | 217.1 | - | - | 0 | - |
| - | - | 8806 | 242.2 | - | - | 0 | - |
| - | - | 684.5 | 243.2 | - | - | 0 | - |
| - | - | 586 | 249.1 | - | - | 0 | - |
| - | - | 2.115E+04 | 249.2 | - | - | 0 | - |
| - | - | 2778 | 250.2 | - | - | 0 | - |
| - | - | 3.951E+04 | 277.2 | - | - | 0 | - |
| - | - | 7026 | 278.2 | - | - | 0 | - |
| 12 | y | 1131 | 288.1 | 0.000373 | 1.295 | +1 | 3 |
| 12 | y | 2475 | 306.1 | 6.224E-05 | 0.2033 | +1 | 3 |
| - | - | 588.4 | 325.1 | - | - | 0 | - |
| - | - | 2007 | 332.1 | - | - | 0 | - |
| - | - | 1334 | 373.2 | - | - | 0 | - |
| - | - | 628 | 374.2 | - | - | 0 | - |
| - | - | 722.6 | 385.1 | - | - | 0 | - |
| - | - | 1.161E+04 | 390.2 | - | - | 0 | - |
| - | - | 2304 | 391.2 | - | - | 0 | - |
| 11 | y | 2793 | 403.1 | 0.0007127 | 1.768 | +1 | 4 |
| - | - | 2557 | 405.2 | - | - | 0 | - |
| - | - | 820.5 | 406.2 | - | - | 0 | - |
| 3 | c | 3009 | 407.3 | 0.000465 | 1.142 | +1 | 3 |
| - | - | 594.1 | 415.1 | - | - | 0 | - |
| 11 | y | 1101 | 421.2 | 0.0001169 | 0.2775 | +1 | 4 |
| - | - | 1161 | 433.2 | - | - | 0 | - |
| - | - | 2320 | 445.2 | - | - | 0 | - |
| - | - | 652 | 479.1 | - | - | 0 | - |
| - | - | 632.8 | 487.2 | - | - | 0 | - |
| - | - | 1456 | 490.3 | - | - | 0 | - |
| - | - | 563.5 | 491.5 | - | - | 0 | - |
| - | - | 789.8 | 500.2 | - | - | 0 | - |
| - | - | 550 | 506.8 | - | - | 0 | - |
| 10 | y | 1622 | 516.2 | 0.0003279 | 0.6352 | +1 | 5 |
| 4 | c | 9042 | 518.3 | 0.0002359 | 0.4552 | +1 | 4 |
| - | - | 668.7 | 519.2 | - | - | 0 | - |
| - | - | 2197 | 519.3 | - | - | 0 | - |
| 10 | y | 4756 | 534.2 | 0.0004444 | 0.8318 | +1 | 5 |
| - | - | 638.1 | 535.2 | - | - | 0 | - |
| - | - | 1756 | 536.3 | - | - | 0 | - |
| 9 | c | 2110 | 560.3 | 0.0002553 | 0.4556 | +2 | 9 |
| - | - | 1251 | 560.8 | - | - | 0 | - |
| - | - | 2430 | 573.3 | - | - | 0 | - |
| - | - | 1118 | 593.8 | - | - | 0 | - |
| - | - | 844.6 | 594.3 | - | - | 0 | - |
| - | - | 867.7 | 601.3 | - | - | 0 | - |
| - | - | 746.8 | 614.8 | - | - | 0 | - |
| - | - | 1472 | 621.3 | - | - | 0 | - |
| 4 | y | 1937 | 623.3 | 0.0007061 | 1.133 | +2 | 11 |
| 4 | y | 2108 | 623.8 | 0.006867 | 11.01 | +2 | 11 |
| 4 | z | 644.3 | 624.3 | 0.009119 | 14.61 | +2 | 11 |
| - | - | 757.1 | 624.8 | - | - | 0 | - |
| 4 | y | 4638 | 632.3 | 0.001039 | 1.643 | +2 | 11 |
| - | - | 3304 | 632.8 | - | - | 0 | - |
| - | - | 1046 | 633.3 | - | - | 0 | - |
| - | - | 832.8 | 635.8 | - | - | 0 | - |
| - | - | 2852 | 644.3 | - | - | 0 | - |
| - | - | 1996 | 644.8 | - | - | 0 | - |
| - | - | 965.6 | 645.3 | - | - | 0 | - |
| 5 | c | 5481 | 649.3 | 0.0004007 | 0.6171 | +1 | 5 |
| - | - | 2124 | 650.3 | - | - | 0 | - |
| - | - | 845.7 | 665.8 | - | - | 0 | - |
| - | - | 633.9 | 666.4 | - | - | 0 | - |
| - | - | 891.1 | 671.3 | - | - | 0 | - |
| 9 | y | 1030 | 672.3 | 0.0007184 | 1.068 | +1 | 6 |
| 9 | z | 758.3 | 674.3 | 0.01181 | 17.52 | +1 | 6 |
| 11 | c | 2076 | 674.4 | 0.001209 | 1.793 | +2 | 11 |
| - | - | 1649 | 674.9 | - | - | 0 | - |
| - | - | 2261 | 675.3 | - | - | 0 | - |
| - | - | 944 | 676.3 | - | - | 0 | - |
| 3 | y | 2318 | 679.8 | 0.001155 | 1.698 | +2 | 12 |
| 3 | y | 4159 | 680.3 | 0.001884 | 2.769 | +2 | 12 |
| 3 | z | 2400 | 680.8 | 0.001907 | 2.801 | +2 | 12 |
| - | - | 1114 | 681.3 | - | - | 0 | - |
| 3 | y | 1.337E+04 | 688.8 | 0.000633 | 0.9189 | +2 | 12 |
| - | - | 7908 | 689.3 | - | - | 0 | - |
| - | - | 2924 | 689.8 | - | - | 0 | - |
| 9 | y | 1.295E+04 | 690.3 | 0.0005296 | 0.7672 | +1 | 6 |
| - | - | 3800 | 691.3 | - | - | 0 | - |
| - | - | 1082 | 692.3 | - | - | 0 | - |
| - | - | 782.1 | 699.4 | - | - | 0 | - |
| - | - | 617.2 | 700.4 | - | - | 0 | - |
| - | - | 619.2 | 701.4 | - | - | 0 | - |
| - | - | 749.1 | 717.3 | - | - | 0 | - |
| - | - | 610.9 | 718.4 | - | - | 0 | - |
| - | - | 1335 | 723.4 | - | - | 0 | - |
| - | - | 795.4 | 723.9 | - | - | 0 | - |
| - | - | 1237 | 725.9 | - | - | 0 | - |
| - | - | 1335 | 726.4 | - | - | 0 | - |
| 12 | c | 5935 | 731.9 | 5.813E-06 | 0.007943 | +2 | 12 |
| - | - | 4821 | 732.4 | - | - | 0 | - |
| - | - | 1978 | 732.9 | - | - | 0 | - |
| 8 | w | 1336 | 744.4 | 8.631E-05 | 0.116 | +1 | 7 |
| - | - | 1200 | 746.4 | - | - | 0 | - |
| - | - | 562.9 | 753.6 | - | - | 0 | - |
| - | - | 1199 | 759.9 | - | - | 0 | - |
| - | - | 1027 | 760.4 | - | - | 0 | - |
| - | - | 608.7 | 761.4 | - | - | 0 | - |
| 2 | y | 1209 | 761.9 | 0.00507 | 6.655 | +2 | 13 |
| 2 | z | 932.7 | 762.4 | 0.000429 | 0.5628 | +2 | 13 |
| 6 | c | 2350 | 763.4 | 0.0006251 | 0.8189 | +1 | 6 |
| - | - | 1466 | 764.4 | - | - | 0 | - |
| - | - | 857.5 | 765.4 | - | - | 0 | - |
| - | - | 903.7 | 768.4 | - | - | 0 | - |
| - | - | 988.6 | 768.9 | - | - | 0 | - |
| 2 | y | 2651 | 770.4 | 0.0001475 | 0.1915 | +2 | 13 |
| - | - | 2338 | 770.9 | - | - | 0 | - |
| - | - | 823.3 | 773.4 | - | - | 0 | - |
| - | - | 1540 | 773.9 | - | - | 0 | - |
| - | - | 1041 | 774.4 | - | - | 0 | - |
| 13 | c | 7354 | 782.4 | 0.0009468 | 1.21 | +2 | 13 |
| - | - | 8467 | 782.9 | - | - | 0 | - |
| - | - | 2966 | 783.4 | - | - | 0 | - |
| 8 | z | 1.266E+04 | 787.4 | 0.0005584 | 0.7091 | +1 | 7 |
| - | - | 6794 | 788.4 | - | - | 0 | - |
| - | - | 2145 | 789.4 | - | - | 0 | - |
| - | - | 979.4 | 802.4 | - | - | 0 | - |
| 8 | y | 6772 | 803.4 | 3.824E-05 | 0.0476 | +1 | 7 |
| - | - | 2689 | 804.4 | - | - | 0 | - |
| - | - | 1266 | 809.4 | - | - | 0 | - |
| - | - | 1484 | 809.9 | - | - | 0 | - |
| - | - | 1439 | 810.4 | - | - | 0 | - |
| - | - | 858 | 814.4 | - | - | 0 | - |
| - | - | 694.2 | 815.4 | - | - | 0 | - |
| - | - | 6945 | 817.9 | - | - | 0 | - |
| - | - | 8813 | 818.4 | - | - | 0 | - |
| - | - | 6160 | 818.9 | - | - | 0 | - |
| - | - | 2605 | 819.4 | - | - | 0 | - |
| - | - | 782.6 | 825.9 | - | - | 0 | - |
| - | - | 835.2 | 826.4 | - | - | 0 | - |
| - | - | 1.886E+04 | 826.9 | - | - | 0 | - |
| - | - | 1.998E+04 | 827.4 | - | - | 0 | - |
| - | - | 9240 | 827.9 | - | - | 0 | - |
| - | - | 1701 | 828.4 | - | - | 0 | - |
| - | - | 962.1 | 831.4 | - | - | 0 | - |
| - | - | 1033 | 833.4 | - | - | 0 | - |
| - | - | 737.4 | 843.5 | - | - | 0 | - |
| 7 | c | 2046 | 850.4 | 0.0007636 | 0.8979 | +1 | 7 |
| - | - | 838.4 | 854.4 | - | - | 0 | - |
| 7 | y | 1313 | 873.4 | 0.001685 | 1.929 | +1 | 8 |
| 7 | z | 2556 | 874.4 | 0.001611 | 1.842 | +1 | 8 |
| - | - | 2.319E+04 | 875.4 | - | - | 0 | - |
| - | - | 1.023E+04 | 876.5 | - | - | 0 | - |
| - | - | 2137 | 877.5 | - | - | 0 | - |
| 7 | y | 1.023E+04 | 890.5 | 0.000343 | 0.3852 | +1 | 8 |
| - | - | 4286 | 891.5 | - | - | 0 | - |
| - | - | 1471 | 945.4 | - | - | 0 | - |
| - | - | 2523 | 946.5 | - | - | 0 | - |
| - | - | 768.5 | 947.5 | - | - | 0 | - |
| - | - | 786.6 | 958.5 | - | - | 0 | - |
| 8 | c | 1553 | 963.5 | 0.0005381 | 0.5584 | +1 | 8 |
| 6 | y | 1388 | 986.5 | 0.002784 | 2.822 | +1 | 9 |
| 6 | y | 1708 | 987.5 | 0.001787 | 1.81 | +1 | 9 |
| 6 | z | 4476 | 988.5 | 0.001888 | 1.91 | +1 | 9 |
| - | - | 1.195E+05 | 989.5 | - | - | 0 | - |
| - | - | 5.864E+04 | 990.5 | - | - | 0 | - |
| - | - | 1.334E+04 | 991.5 | - | - | 0 | - |
| - | - | 1457 | 992.5 | - | - | 0 | - |
| - | - | 7706 | 1003 | - | - | 0 | - |
| 6 | y | 5.516E+04 | 1005 | 1.811E-05 | 0.01803 | +1 | 9 |
| - | - | 2.551E+04 | 1006 | - | - | 0 | - |
| - | - | 8759 | 1007 | - | - | 0 | - |
| - | - | 832.3 | 1046 | - | - | 0 | - |
| 5 | w | 1421 | 1059 | 0.003711 | 3.506 | +1 | 10 |
| - | - | 1022 | 1060 | - | - | 0 | - |
| - | - | 987.1 | 1072 | - | - | 0 | - |
| - | - | 1485 | 1074 | - | - | 0 | - |
| - | - | 938.9 | 1079 | - | - | 0 | - |
| - | - | 1.64E+04 | 1093 | - | - | 0 | - |
| - | - | 1.101E+04 | 1094 | - | - | 0 | - |
| - | - | 3662 | 1095 | - | - | 0 | - |
| - | - | 652 | 1101 | - | - | 0 | - |
| 5 | y | 2062 | 1118 | 0.001337 | 1.197 | +1 | 10 |
| 5 | y | 3303 | 1119 | 0.006468 | 5.783 | +1 | 10 |
| 5 | z | 1.578E+04 | 1120 | 0.001329 | 1.187 | +1 | 10 |
| - | - | 3.036E+04 | 1121 | - | - | 0 | - |
| - | - | 1.6E+04 | 1122 | - | - | 0 | - |
| - | - | 5218 | 1123 | - | - | 0 | - |
| - | - | 841.3 | 1135 | - | - | 0 | - |
| 5 | y | 2.897E+04 | 1136 | 0.0009153 | 0.8061 | +1 | 10 |
| 9 | c | 3.908E+04 | 1137 | 0.001306 | 1.149 | +1 | 9 |
| - | - | 2.369E+04 | 1138 | - | - | 0 | - |
| - | - | 8498 | 1139 | - | - | 0 | - |
| - | - | 891.8 | 1140 | - | - | 0 | - |
| - | - | 1098 | 1145 | - | - | 0 | - |
| - | - | 990.8 | 1146 | - | - | 0 | - |
| - | - | 1332 | 1175 | - | - | 0 | - |
| - | - | 2021 | 1175 | - | - | 0 | - |
| - | - | 1062 | 1176 | - | - | 0 | - |
| - | - | 1086 | 1187 | - | - | 0 | - |
| - | - | 940.1 | 1188 | - | - | 0 | - |
| - | - | 637 | 1189 | - | - | 0 | - |
| 4 | w | 778.4 | 1190 | 0.0004941 | 0.4153 | +1 | 11 |
| - | - | 1428 | 1206 | - | - | 0 | - |
| - | - | 1343 | 1211 | - | - | 0 | - |
| 10 | c | 4268 | 1233 | 0.0004714 | 0.3824 | +1 | 10 |
| - | - | 3069 | 1234 | - | - | 0 | - |
| - | - | 1201 | 1235 | - | - | 0 | - |
| 4 | y | 1516 | 1246 | 0.005027 | 4.035 | +1 | 11 |
| 4 | y | 5130 | 1247 | 0.0005955 | 0.4777 | +1 | 11 |
| 4 | z | 7799 | 1248 | 0.003416 | 2.738 | +1 | 11 |
| - | - | 3957 | 1249 | - | - | 0 | - |
| - | - | 1.461E+04 | 1249 | - | - | 0 | - |
| 10 | c | 2.703E+04 | 1250 | 0.002362 | 1.89 | +1 | 10 |
| - | - | 926.6 | 1251 | - | - | 0 | - |
| - | - | 1.489E+04 | 1251 | - | - | 0 | - |
| - | - | 5236 | 1252 | - | - | 0 | - |
| 4 | y | 2.236E+04 | 1264 | 0.001266 | 1.002 | +1 | 11 |
| - | - | 1.396E+04 | 1265 | - | - | 0 | - |
| - | - | 5270 | 1266 | - | - | 0 | - |
| - | - | 1114 | 1288 | - | - | 0 | - |
| - | - | 8125 | 1290 | - | - | 0 | - |
| - | - | 5997 | 1291 | - | - | 0 | - |
| - | - | 1397 | 1292 | - | - | 0 | - |
| - | - | 2312 | 1300 | - | - | 0 | - |
| - | - | 1309 | 1301 | - | - | 0 | - |
| - | - | 2255 | 1305 | - | - | 0 | - |
| - | - | 1015 | 1306 | - | - | 0 | - |
| 3 | w | 1.198E+04 | 1318 | 1.044E-05 | 0.007926 | +1 | 12 |
| - | - | 8036 | 1319 | - | - | 0 | - |
| - | - | 3254 | 1320 | - | - | 0 | - |
| - | - | 974.7 | 1325 | - | - | 0 | - |
| - | - | 653 | 1332 | - | - | 0 | - |
| - | - | 725.4 | 1342 | - | - | 0 | - |
| 11 | c | 4018 | 1348 | 0.002756 | 2.045 | +1 | 11 |
| - | - | 3448 | 1349 | - | - | 0 | - |
| - | - | 2080 | 1350 | - | - | 0 | - |
| - | - | 907.5 | 1351 | - | - | 0 | - |
| 3 | y | 898.3 | 1359 | 0.003232 | 2.379 | +1 | 12 |
| 3 | y | 1401 | 1360 | 0.004459 | 3.28 | +1 | 12 |
| 3 | z | 4.089E+04 | 1361 | 0.001542 | 1.133 | +1 | 12 |
| - | - | 3.567E+04 | 1362 | - | - | 0 | - |
| - | - | 1.489E+04 | 1363 | - | - | 0 | - |
| - | - | 2.331E+04 | 1364 | - | - | 0 | - |
| 11 | c | 3.56E+04 | 1365 | 0.003793 | 2.779 | +1 | 11 |
| - | - | 2.417E+04 | 1366 | - | - | 0 | - |
| - | - | 7906 | 1367 | - | - | 0 | - |
| - | - | 1070 | 1368 | - | - | 0 | - |
| 3 | y | 2.116E+04 | 1377 | 0.0009791 | 0.7112 | +1 | 12 |
| - | - | 1.583E+04 | 1378 | - | - | 0 | - |
| - | - | 7723 | 1379 | - | - | 0 | - |
| - | - | 1723 | 1380 | - | - | 0 | - |
| - | - | 931.5 | 1419 | - | - | 0 | - |
| - | - | 1451 | 1421 | - | - | 0 | - |
| - | - | 5038 | 1425 | - | - | 0 | - |
| - | - | 5129 | 1426 | - | - | 0 | - |
| - | - | 1920 | 1427 | - | - | 0 | - |
| - | - | 826.6 | 1433 | - | - | 0 | - |
| - | - | 926.9 | 1438 | - | - | 0 | - |
| - | - | 1066 | 1441 | - | - | 0 | - |
| - | - | 979.8 | 1447 | - | - | 0 | - |
| - | - | 1918 | 1450 | - | - | 0 | - |
| - | - | 1544 | 1451 | - | - | 0 | - |
| - | - | 1150 | 1452 | - | - | 0 | - |
| 12 | c | 3361 | 1463 | 0.003088 | 2.111 | +1 | 12 |
| - | - | 3421 | 1464 | - | - | 0 | - |
| - | - | 1810 | 1465 | - | - | 0 | - |
| - | - | 1812 | 1468 | - | - | 0 | - |
| - | - | 1295 | 1469 | - | - | 0 | - |
| - | - | 1063 | 1470 | - | - | 0 | - |
| 12 | c | 2.112E+04 | 1480 | 0.002049 | 1.385 | +1 | 12 |
| - | - | 1.615E+04 | 1481 | - | - | 0 | - |
| - | - | 8949 | 1482 | - | - | 0 | - |
| - | - | 1932 | 1483 | - | - | 0 | - |
| - | - | 1084 | 1519 | - | - | 0 | - |
| - | - | 793.6 | 1522 | - | - | 0 | - |
| 2 | y | 862.5 | 1523 | 0.01571 | 10.32 | +1 | 13 |
| 2 | z | 1.47E+04 | 1524 | 0.002004 | 1.315 | +1 | 13 |
| - | - | 1.326E+04 | 1525 | - | - | 0 | - |
| - | - | 4933 | 1526 | - | - | 0 | - |
| - | - | 937.8 | 1527 | - | - | 0 | - |
| - | - | 834 | 1535 | - | - | 0 | - |
| - | - | 913.6 | 1538 | - | - | 0 | - |
| 2 | y | 1068 | 1540 | 0.002906 | 1.887 | +1 | 13 |
| - | - | 1023 | 1541 | - | - | 0 | - |
| - | - | 897.9 | 1552 | - | - | 0 | - |
| 13 | c | 4636 | 1563 | 0.006082 | 3.892 | +1 | 13 |
| 13 | c | 4347 | 1564 | 0.01088 | 6.957 | +1 | 13 |
| - | - | 3958 | 1565 | - | - | 0 | - |
| - | - | 2200 | 1566 | - | - | 0 | - |
| - | - | 764.5 | 1567 | - | - | 0 | - |
| - | - | 1320 | 1576 | - | - | 0 | - |
| - | - | 959 | 1578 | - | - | 0 | - |
| - | - | 2563 | 1580 | - | - | 0 | - |
| 13 | c | 2.144E+04 | 1581 | 0.004195 | 2.654 | +1 | 13 |
| - | - | 2.173E+04 | 1582 | - | - | 0 | - |
| - | - | 1.178E+04 | 1583 | - | - | 0 | - |
| - | - | 2470 | 1584 | - | - | 0 | - |
| - | - | 1068 | 1590 | - | - | 0 | - |
| - | - | 1570 | 1591 | - | - | 0 | - |
| - | - | 1738 | 1592 | - | - | 0 | - |
| - | - | 2672 | 1593 | - | - | 0 | - |
| - | - | 1.792E+04 | 1594 | - | - | 0 | - |
| - | - | 2.093E+04 | 1595 | - | - | 0 | - |
| - | - | 1.346E+04 | 1596 | - | - | 0 | - |
| - | - | 4426 | 1597 | - | - | 0 | - |
| - | - | 3208 | 1598 | - | - | 0 | - |
| - | - | 2436 | 1599 | - | - | 0 | - |
| - | - | 766.8 | 1600 | - | - | 0 | - |
| - | - | 880.3 | 1607 | - | - | 0 | - |
| - | - | 3481 | 1608 | - | - | 0 | - |
| - | - | 4.815E+04 | 1609 | - | - | 0 | - |
| - | - | 4.494E+04 | 1610 | - | - | 0 | - |
| - | - | 2.531E+04 | 1611 | - | - | 0 | - |
| - | - | 7828 | 1612 | - | - | 0 | - |
| - | - | 1828 | 1613 | - | - | 0 | - |
| - | - | 1169 | 1620 | - | - | 0 | - |
| - | - | 1031 | 1622 | - | - | 0 | - |
| - | - | 985.5 | 1623 | - | - | 0 | - |
| - | - | 739.1 | 1624 | - | - | 0 | - |
| - | - | 3729 | 1626 | - | - | 0 | - |
| - | - | 3422 | 1627 | - | - | 0 | - |
| - | - | 1579 | 1628 | - | - | 0 | - |
| - | - | 856.5 | 1634 | - | - | 0 | - |
| - | - | 819.9 | 1635 | - | - | 0 | - |
| - | - | 8653 | 1636 | - | - | 0 | - |
| - | - | 1.132E+05 | 1637 | - | - | 0 | - |
| - | - | 1.098E+05 | 1638 | - | - | 0 | - |
| - | - | 1251 | 1638 | - | - | 0 | - |
| - | - | 5.592E+04 | 1639 | - | - | 0 | - |
| - | - | 1.384E+04 | 1640 | - | - | 0 | - |
| - | - | 1940 | 1651 | - | - | 0 | - |
| - | - | 4434 | 1652 | - | - | 0 | - |
| - | - | 1.293E+05 | 1653 | - | - | 0 | - |
| - | - | 2.158E+05 | 1654 | - | - | 0 | - |
| - | - | 1.518E+05 | 1655 | - | - | 0 | - |
| - | - | 5.955E+04 | 1656 | - | - | 0 | - |
| - | - | 1.083E+04 | 1657 | - | - | 0 | - |
| - | - | 709.3 | 2735 | - | - | 0 | - |
| - | - | 717 | 2952 | - | - | 0 | - |

m/z Charge Intensity FragmentType MassShift Position
121.1135025024414 0 353.10965
125.6550064086914 0 411.94827
131.11807250976562 0 555.56396
134.3105010986328 0 400.6626
136.0759735107422 0 3902.506
148.67945861816406 0 442.0351
170.76466369628906 0 426.8085
173.43954467773438 0 3429.2192
189.65235900878906 0 599.4757
191.10275268554688 0 6786.634 y 12
200.9423828125 0 457.71463
215.13916015625 0 4315.713
217.0820770263672 0 4055.2212
242.15020751953125 0 8805.606
243.1533966064453 0 684.4938
249.1449432373047 0 585.9593
249.15988159179688 0 21148.7
250.1631317138672 0 2778.0935
277.15472412109375 0 39507.496
278.1580810546875 0 7025.795
288.119384765625 0 1130.5844 y Water loss 11
306.129638671875 0 2474.6216 y 11
325.138916015625 0 588.3818
332.10870361328125 0 2007.0829
373.1900329589844 0 1334.3165
374.1953430175781 0 627.9853
385.1362609863281 0 722.614
390.2390441894531 0 11606.366
391.2427673339844 0 2304.4797
403.14666748046875 0 2793.4878 y Water loss 10
405.2137756347656 0 2557.4314
406.2167053222656 0 820.499
407.2657470703125 0 3008.5422 c 2
415.0577697753906 0 594.0517
421.1564025878906 0 1101.0986 y 10
433.2080383300781 0 1161.1171
445.19195556640625 0 2319.7764
479.1016540527344 0 651.9865
487.23382568359375 0 632.8324
490.30108642578125 0 1456.4414
491.537109375 0 563.54694
500.2475280761719 0 789.7943
506.7709045410156 0 549.98883
516.2303466796875 0 1621.8633 y Water loss 9
518.2975463867188 0 9042.153 c Ammonia loss 3
519.2291870117188 0 668.70715
519.3008422851562 0 2197.0386
534.2410278320312 0 4756.3574 y 9
535.2446899414062 0 638.0744
536.2540283203125 0 1755.7662
560.3028564453125 0 2110.2944 c Ammonia loss 8
560.80322265625 0 1250.9696
573.340576171875 0 2430.4246
593.8010864257812 0 1117.8972
594.2978515625 0 844.64154
601.292236328125 0 867.7208
614.7860717773438 0 746.7594
621.3436279296875 0 1472.0918
623.2989501953125 0 1936.6675 y Water loss 3
623.797119140625 0 2107.6982 y Ammonia loss 3
624.3032836914062 0 644.2524 z 3
624.8001098632812 0 757.08215
632.3045654296875 0 4638.0938 y 3
632.8053588867188 0 3304.0793
633.3095092773438 0 1045.825
635.8096313476562 0 832.83203
644.3200073242188 0 2851.652
644.8233032226562 0 1995.6077
645.326416015625 0 965.61847
649.3381958007812 0 5480.655 c Ammonia loss 4
650.3417358398438 0 2123.75
665.8477783203125 0 845.70483
666.3507080078125 0 633.9372
671.3289794921875 0 891.1351
672.3318481445312 0 1030.4784 y Water loss 8
674.3111572265625 0 758.2682 z 8
674.3593139648438 0 2076.1147 c Ammonia loss 10
674.8587646484375 0 1648.7211
675.3324584960938 0 2260.6016
676.3333129882812 0 944.01306
679.8414306640625 0 2317.7603 y Water loss 2
680.3341674804688 0 4159.1475 y Ammonia loss 2
680.8342895507812 0 2399.8748 z 2
681.336669921875 0 1114.1318
688.84619140625 0 13374.402 y 2
689.3473510742188 0 7907.83
689.8485107421875 0 2924.4058
690.3422241210938 0 12953.462 y 8
691.3448486328125 0 3799.6748
692.346435546875 0 1081.7345
699.3714599609375 0 782.0601
700.3663940429688 0 617.1755
701.3699951171875 0 619.2044
717.3455810546875 0 749.12103
718.3629760742188 0 610.8609
723.3590087890625 0 1335.479
723.859130859375 0 795.38074
725.8526000976562 0 1236.8228
726.3551025390625 0 1334.691
731.87158203125 0 5935.1313 c Ammonia loss 11
732.3734741210938 0 4821.1123
732.8709106445312 0 1977.6185
744.3521728515625 0 1335.6873 w 7
746.3539428710938 0 1199.5287
753.5846557617188 0 562.9128
759.8911743164062 0 1199.4973
760.3909912109375 0 1027.2294
761.3873291015625 0 608.69147
761.8690185546875 0 1209.0833 y Ammonia loss 1
762.367431640625 0 932.6878 z 1
763.38134765625 0 2349.8281 c Ammonia loss 5
764.385009765625 0 1465.8838
765.3768310546875 0 857.54614
768.4035034179688 0 903.6946
768.89990234375 0 988.58716
770.3770751953125 0 2650.6987 y 1
770.877685546875 0 2337.995
773.3904418945312 0 823.3269
773.8898315429688 0 1539.9891
774.3856811523438 0 1041.0398
782.3963623046875 0 7354.412 c Ammonia loss 12
782.8973999023438 0 8467.466
783.3984375 0 2966.3901
787.4075927734375 0 12660.918 z 7
788.4122924804688 0 6794.4766
789.4163208007812 0 2144.9048
802.4185791015625 0 979.423
803.4257202148438 0 6772.45 y 7
804.4285888671875 0 2689.3652
809.405029296875 0 1266.2628
809.9017944335938 0 1483.7568
810.4020385742188 0 1438.6996
814.4081420898438 0 858.00476
815.4111328125 0 694.1998
817.9138793945312 0 6945.0034
818.41259765625 0 8812.506
818.9113159179688 0 6160.1167
819.40966796875 0 2605.3965
825.9163208007812 0 782.6053
826.4260864257812 0 835.1921
826.9195556640625 0 18858.197
827.4213256835938 0 19978.947
827.922607421875 0 9240.077
828.422119140625 0 1700.555
831.4277954101562 0 962.05804
833.39306640625 0 1033.0781
843.4533081054688 0 737.3717
850.4119873046875 0 2046.0883 c Ammonia loss 6
854.4108276367188 0 838.4315
873.4329223632812 0 1312.972 y Ammonia loss 6
874.440673828125 0 2556.053 z 6
875.4470825195312 0 23193.688
876.4503784179688 0 10233.218
877.454345703125 0 2136.7878
890.4581298828125 0 10230.943 y 6
891.4620361328125 0 4285.801
945.4462890625 0 1470.6252
946.4667358398438 0 2522.7297
947.46826171875 0 768.5053
958.4735717773438 0 786.5699
963.4962768554688 0 1552.8997 c Ammonia loss 7
986.4873657226562 0 1388.311 y Water loss 5
987.4759521484375 0 1708.1493 y Ammonia loss 5
988.4801025390625 0 4475.518 z 5
989.4904174804688 0 119523.16
990.4931640625 0 58641.62
991.49560546875 0 13344.704
992.5010375976562 0 1456.7561
1003.494140625 0 7706.3286
1004.500732421875 0 55160.49 y 5
1005.5040893554688 0 25509.05
1006.5072631835938 0 8758.999
1046.497314453125 0 832.3468
1058.507568359375 0 1421.2052 w 4
1059.5120849609375 0 1022.0947
1071.5546875 0 987.0556
1073.514404296875 0 1484.5085
1078.5496826171875 0 938.9411
1092.6109619140625 0 16396.588
1093.613525390625 0 11008.251
1094.6160888671875 0 3662.4763
1100.5401611328125 0 652.04663
1117.529296875 0 2062.282 y Water loss 4
1118.5211181640625 0 3303.4878 y Ammonia loss 4
1119.5238037109375 0 15778.035 z 4
1120.529296875 0 30359.805
1121.5335693359375 0 16002.659
1122.5357666015625 0 5218.46
1134.52783203125 0 841.26
1135.5421142578125 0 28974.104 y 4
1136.6231689453125 0 39077.453 c 8
1137.627197265625 0 23688.514
1138.62744140625 0 8497.942
1139.6280517578125 0 891.8207
1144.585693359375 0 1098.1138
1145.57666015625 0 990.7903
1174.553466796875 0 1332.3026
1174.680908203125 0 2021.136
1175.689208984375 0 1061.6091
1186.5828857421875 0 1085.6178
1187.588134765625 0 940.12714
1188.5994873046875 0 636.9913
1189.55126953125 0 778.3707 w 3
1205.695556640625 0 1427.7509
1210.53857421875 0 1342.992
1232.6815185546875 0 4268.356 c Ammonia loss 9
1233.683349609375 0 3069.31
1234.680419921875 0 1200.8075
1245.59423828125 0 1515.5623 y Water loss 3
1246.5726318359375 0 5130.115 y Ammonia loss 3
1247.57763671875 0 7798.6255 z 3
1248.5751953125 0 3957.4727
1248.701904296875 0 14607.954
1249.7061767578125 0 27029.479 c 9
1250.5667724609375 0 926.6158
1250.70751953125 0 14886.774
1251.709716796875 0 5235.5303
1263.5985107421875 0 22361.39 y 3
1264.6019287109375 0 13960.661
1265.60205078125 0 5270.343
1287.632080078125 0 1113.61
1289.7066650390625 0 8125.3936
1290.7073974609375 0 5996.5557
1291.712890625 0 1396.7335
1299.65234375 0 2311.7178
1300.6575927734375 0 1309.3478
1304.603271484375 0 2255.3037
1305.5889892578125 0 1014.77344
1317.6103515625 0 11982.471 w 2
1318.61376953125 0 8035.829
1319.613037109375 0 3254.4866
1324.6234130859375 0 974.6645
1331.6845703125 0 653.0307
1341.648681640625 0 725.39984
1347.7061767578125 0 4017.5137 c Ammonia loss 10
1348.712890625 0 3448.3047
1349.705078125 0 2080.3672
1350.7015380859375 0 907.4518
1358.6700439453125 0 898.30853 y Water loss 2
1359.65283203125 0 1400.5836 y Ammonia loss 2
1360.66357421875 0 40887.14 z 2
1361.6669921875 0 35665.273
1362.67041015625 0 14885.85
1363.722412109375 0 23308.148
1364.731689453125 0 35604.742 c 10
1365.7342529296875 0 24170.832
1366.7294921875 0 7905.9155
1367.69677734375 0 1070.2675
1376.682861328125 0 21156.494 y 2
1377.685546875 0 15829.022
1378.685546875 0 7723.2974
1379.6871337890625 0 1723.1925
1418.7193603515625 0 931.48334
1420.7276611328125 0 1451.3606
1424.6751708984375 0 5037.951
1425.6790771484375 0 5129.063
1426.682373046875 0 1920.0513
1432.6756591796875 0 826.5892
1437.7618408203125 0 926.90125
1440.6534423828125 0 1066.1141
1446.7109375 0 979.84296
1449.6951904296875 0 1917.7129
1450.699951171875 0 1543.9072
1451.718994140625 0 1149.6421
1462.7327880859375 0 3361.3425 c Ammonia loss 11
1463.735107421875 0 3421.3909
1464.7445068359375 0 1810.0752
1467.6741943359375 0 1812.1433
1468.6697998046875 0 1294.5825
1469.6845703125 0 1062.5282
1479.7603759765625 0 21119.271 c 11
1480.7613525390625 0 16147.596
1481.7642822265625 0 8949.33
1482.770751953125 0 1931.6848
1518.7685546875 0 1083.8356
1521.7708740234375 0 793.6007
1522.736328125 0 862.5439 y Ammonia loss 1
1523.7264404296875 0 14698.987 z 1
1524.7293701171875 0 13262.253
1525.727783203125 0 4932.503
1526.7451171875 0 937.7755
1534.7957763671875 0 833.9653
1537.810546875 0 913.5708
1539.7442626953125 0 1068.2346 y 1
1540.7374267578125 0 1023.48706
1551.7288818359375 0 897.85516
1562.79345703125 0 4635.8647 c Water loss 12
1563.79443359375 0 4347.0396 c Ammonia loss 12
1564.80810546875 0 3957.777
1565.8048095703125 0 2199.9253
1566.8194580078125 0 764.46893
1575.8033447265625 0 1320.0676
1577.8212890625 0 959.0352
1579.80810546875 0 2562.8464
1580.805908203125 0 21441.256 c 12
1581.8099365234375 0 21728.324
1582.8114013671875 0 11782.788
1583.809814453125 0 2470.4612
1589.8048095703125 0 1067.8562
1590.8240966796875 0 1569.8425
1591.8193359375 0 1737.7617
1592.8204345703125 0 2672.4204
1593.798828125 0 17917.783
1594.7926025390625 0 20929.781
1595.7935791015625 0 13462.022
1596.7947998046875 0 4425.515
1597.775390625 0 3208.4019
1598.772705078125 0 2436.3691
1599.7728271484375 0 766.7523
1606.79833984375 0 880.32135
1607.824951171875 0 3481.133
1608.8154296875 0 48145.023
1609.81884765625 0 44935.81
1610.8154296875 0 25311.55
1611.810302734375 0 7827.93
1612.796630859375 0 1828.2305
1619.8092041015625 0 1169.1813
1621.7900390625 0 1030.971
1622.81689453125 0 985.4845
1623.765380859375 0 739.13196
1625.8389892578125 0 3728.5776
1626.841796875 0 3422.3354
1627.84326171875 0 1579.4573
1633.7662353515625 0 856.4969
1634.829345703125 0 819.91235
1635.821533203125 0 8653.305
1636.8115234375 0 113171.34
1637.8128662109375 0 109802.51
1638.080810546875 0 1250.631
1638.815185546875 0 55916.37
1639.81689453125 0 13835.331
1650.7974853515625 0 1939.5928
1651.8172607421875 0 4434.134
1652.829833984375 0 129279.34
1653.8355712890625 0 215751.16
1654.838134765625 0 151750.58
1655.8406982421875 0 59554.543
1656.8388671875 0 10828.231
2735.16455078125 0 709.3255
2952.269287109375 0 716.9906

Spectrum Details

|  |  |
| --- | --- |
| Matched peaks? Matched peaks The total absolute number of peaks matched. Additionally in brackets the total fraction of peaks matched and the total number of peaks is shown. | 69 (20.29% of 340) |
| FDR? FDR The false discovery rate estimated for this peptide. It is calculated by matching all theoretical fragments with a non-integer shift with the raw peaks for this spectrum. This is done with 40 different shifts. The resulting percentage is the average number of annotated peaks over the number of annotated peaks with the correct spectrum. | 1.52% |
| Satellite FDR? Satellite FDR See the FDR for details on its calculation. This satellite ion specific FDR only contains the satellite ions (d/w) for I/L/J positions. | 0.00% |
| PSM Score? PSM Score The PSM Score as given by Hecklib to this annotated spectrum. It is shown with three significant figures. | 748 |

#### Spectrum 9789? Spectrum 9789 The raw spectrum of this peptide as annotated by Hecklib. The fragments are coloured according to ion type (see legend). Any peaks with a star '\*' as text can be hovered over to see the full details, first the ion type second the mass shift type. By hovering over the amino acids in the peptide or ions in the legend the corresponding peaks are highlighted. By toggling the 'Unassigned' label you can turn the background (unassigned) peaks on or off in the plot. By updating the slider in the Ion legend you can update the spectrum to only show the top X% of the peaks with labels. The top X% means any peak that is within X% of the highest intensity. By dragging in the spectrum you can zoom in to a specific part of the spectrum and use 'Zoom Out' to get back to the original zoom level. The annotation of the spectrum is based on the given sequence in the peptides file and is done with different software so inconsistencies are likely. The peaks are annotated based on the given sequence, with 20 ppm tolerance.

Copy Data

##### Spectrum 9789 (TSV)

###### Preview

```
Loading example...
```

*Click on the button to copy the data to your clipboard.*

Mz MinMz MaxIntensity Max

WidthHeightPeptide font sizePeptide stroke widthSpectrum font sizeSpectrum stroke widthCompact peptide

Ion legend

wxyz

abcd

OtherUnassignedIon^Charge^_Position_Show for top:%

JYLQMNSJRJDDTA

09.41e+31.88e+42.82e+43.77e+4

Zoom Out

y^+1^_2_y^+1^_2_a^+1^_2_b^+1^_2_y^+1^_3_y^+1^_3_b^+1^_3_y^+1^_4_y^+1^_4_y^+2^_8_y^+1^_5_b^+1^_4_y^+1^_5_b^+2^_9_y^+2^_11_y^+2^_11_y^+2^_11_b^+1^_5_y^+1^_6_y^+1^_6_b^+2^_11_y^+2^_12_y^+1^_6_y^+2^_12_b^+2^_12_y^+2^_13_b^+1^_6_b^+2^_13_b^+2^_13_b^+2^_13_y^+1^_7_\*\*\*y^+1^_8_y^+1^_8_b^+1^_8_b^+1^_8_y^+1^_9_y^+1^_9_y^+1^_9_y^+1^_10_y^+1^_10_b^+1^_10_y^+1^_11_y^+1^_11_y^+1^_11_b^+1^_11_y^+1^_12_b^+1^_12_

0834166725013334

Fragment Matches Table

Show background peaks

| Position | Ion type | Intensity | mz Theoretical | mz Error (Th) | mz Error (ppm) | Charge | Series Number |
| --- | --- | --- | --- | --- | --- | --- | --- |
| - | - | 2991 | 120.1 | - | - | 0 | - |
| - | - | 423.2 | 120.1 | - | - | 0 | - |
| - | - | 469.6 | 122.1 | - | - | 0 | - |
| - | - | 325.7 | 124.6 | - | - | 0 | - |
| - | - | 677.1 | 127.1 | - | - | 0 | - |
| - | - | 618.4 | 127.1 | - | - | 0 | - |
| - | - | 328.8 | 127.7 | - | - | 0 | - |
| - | - | 493.3 | 129.1 | - | - | 0 | - |
| - | - | 3960 | 129.1 | - | - | 0 | - |
| - | - | 791.8 | 130.1 | - | - | 0 | - |
| - | - | 842.5 | 130.1 | - | - | 0 | - |
| - | - | 1057 | 131 | - | - | 0 | - |
| - | - | 6319 | 131.1 | - | - | 0 | - |
| - | - | 1091 | 133.1 | - | - | 0 | - |
| - | - | 456.7 | 135 | - | - | 0 | - |
| - | - | 3.515E+04 | 136.1 | - | - | 0 | - |
| - | - | 2517 | 137.1 | - | - | 0 | - |
| - | - | 1372 | 138.1 | - | - | 0 | - |
| - | - | 1722 | 141.1 | - | - | 0 | - |
| - | - | 406.5 | 141.9 | - | - | 0 | - |
| - | - | 450.1 | 142.1 | - | - | 0 | - |
| - | - | 1137 | 144.1 | - | - | 0 | - |
| - | - | 377.1 | 144.8 | - | - | 0 | - |
| - | - | 484 | 145.1 | - | - | 0 | - |
| - | - | 743.4 | 146.1 | - | - | 0 | - |
| - | - | 533.4 | 147.1 | - | - | 0 | - |
| - | - | 444 | 148.8 | - | - | 0 | - |
| - | - | 616.9 | 148.9 | - | - | 0 | - |
| - | - | 645.3 | 148.9 | - | - | 0 | - |
| - | - | 549.9 | 148.9 | - | - | 0 | - |
| - | - | 560.6 | 148.9 | - | - | 0 | - |
| - | - | 885.6 | 148.9 | - | - | 0 | - |
| - | - | 1040 | 148.9 | - | - | 0 | - |
| - | - | 2430 | 148.9 | - | - | 0 | - |
| - | - | 4133 | 148.9 | - | - | 0 | - |
| - | - | 3465 | 149 | - | - | 0 | - |
| - | - | 1873 | 149 | - | - | 0 | - |
| - | - | 1134 | 149 | - | - | 0 | - |
| - | - | 1091 | 149 | - | - | 0 | - |
| - | - | 1028 | 149 | - | - | 0 | - |
| - | - | 837.3 | 149 | - | - | 0 | - |
| - | - | 610.5 | 149 | - | - | 0 | - |
| - | - | 409.8 | 149.1 | - | - | 0 | - |
| - | - | 539.6 | 150.1 | - | - | 0 | - |
| - | - | 860.6 | 152.1 | - | - | 0 | - |
| - | - | 1.015E+04 | 152.1 | - | - | 0 | - |
| - | - | 635.4 | 155.1 | - | - | 0 | - |
| - | - | 593.8 | 155.1 | - | - | 0 | - |
| - | - | 1050 | 156.1 | - | - | 0 | - |
| - | - | 955.9 | 157.1 | - | - | 0 | - |
| - | - | 830.8 | 157.1 | - | - | 0 | - |
| - | - | 1424 | 159.1 | - | - | 0 | - |
| - | - | 534.8 | 162.1 | - | - | 0 | - |
| - | - | 553.8 | 165.1 | - | - | 0 | - |
| - | - | 652.1 | 166.1 | - | - | 0 | - |
| - | - | 562.7 | 167.1 | - | - | 0 | - |
| - | - | 2858 | 167.1 | - | - | 0 | - |
| - | - | 531.2 | 167.1 | - | - | 0 | - |
| - | - | 615.6 | 169.1 | - | - | 0 | - |
| - | - | 1073 | 171.1 | - | - | 0 | - |
| - | - | 557.4 | 171.1 | - | - | 0 | - |
| - | - | 728.7 | 172.1 | - | - | 0 | - |
| 13 | y | 1640 | 173.1 | 0.0002622 | 1.515 | +1 | 2 |
| - | - | 2234 | 173.1 | - | - | 0 | - |
| - | - | 729 | 174.1 | - | - | 0 | - |
| - | - | 1144 | 174.1 | - | - | 0 | - |
| - | - | 583.5 | 175.1 | - | - | 0 | - |
| - | - | 1507 | 178 | - | - | 0 | - |
| - | - | 599.7 | 180.1 | - | - | 0 | - |
| - | - | 1459 | 183.1 | - | - | 0 | - |
| - | - | 690.2 | 183.1 | - | - | 0 | - |
| - | - | 1213 | 184.1 | - | - | 0 | - |
| - | - | 597.2 | 185.1 | - | - | 0 | - |
| - | - | 664.3 | 186.1 | - | - | 0 | - |
| - | - | 1377 | 187.1 | - | - | 0 | - |
| - | - | 557 | 187.1 | - | - | 0 | - |
| - | - | 1497 | 189.1 | - | - | 0 | - |
| - | - | 735.5 | 191 | - | - | 0 | - |
| 13 | y | 6130 | 191.1 | 0.0002566 | 1.343 | +1 | 2 |
| - | - | 2343 | 195.1 | - | - | 0 | - |
| - | - | 609.6 | 195.1 | - | - | 0 | - |
| - | - | 889.4 | 195.1 | - | - | 0 | - |
| - | - | 513.1 | 197.1 | - | - | 0 | - |
| - | - | 696.9 | 197.1 | - | - | 0 | - |
| - | - | 1210 | 198.1 | - | - | 0 | - |
| - | - | 781.9 | 199.1 | - | - | 0 | - |
| - | - | 1404 | 199.1 | - | - | 0 | - |
| - | - | 1054 | 201.1 | - | - | 0 | - |
| - | - | 3285 | 201.1 | - | - | 0 | - |
| - | - | 1993 | 202.1 | - | - | 0 | - |
| - | - | 841.8 | 203.1 | - | - | 0 | - |
| - | - | 811.3 | 203.1 | - | - | 0 | - |
| - | - | 767.7 | 208.1 | - | - | 0 | - |
| - | - | 683.6 | 208.1 | - | - | 0 | - |
| - | - | 743.2 | 210.2 | - | - | 0 | - |
| - | - | 1092 | 212.1 | - | - | 0 | - |
| - | - | 1286 | 213.1 | - | - | 0 | - |
| - | - | 540.9 | 214.1 | - | - | 0 | - |
| - | - | 572 | 214.2 | - | - | 0 | - |
| - | - | 2.793E+04 | 215.1 | - | - | 0 | - |
| - | - | 2508 | 216.1 | - | - | 0 | - |
| - | - | 3562 | 217.1 | - | - | 0 | - |
| - | - | 599.9 | 218.1 | - | - | 0 | - |
| - | - | 2360 | 219 | - | - | 0 | - |
| - | - | 634.5 | 223.1 | - | - | 0 | - |
| - | - | 690.2 | 224.1 | - | - | 0 | - |
| - | - | 3012 | 225.1 | - | - | 0 | - |
| - | - | 1514 | 225.2 | - | - | 0 | - |
| - | - | 1054 | 226.1 | - | - | 0 | - |
| - | - | 2771 | 227.1 | - | - | 0 | - |
| - | - | 1116 | 229.1 | - | - | 0 | - |
| - | - | 1307 | 231.1 | - | - | 0 | - |
| - | - | 714.3 | 238.1 | - | - | 0 | - |
| - | - | 720.3 | 239.1 | - | - | 0 | - |
| - | - | 765.7 | 240.1 | - | - | 0 | - |
| - | - | 548.1 | 242.1 | - | - | 0 | - |
| - | - | 1.079E+04 | 242.2 | - | - | 0 | - |
| - | - | 915.5 | 243.1 | - | - | 0 | - |
| - | - | 879.4 | 243.2 | - | - | 0 | - |
| - | - | 947.6 | 244.1 | - | - | 0 | - |
| - | - | 732.1 | 245.1 | - | - | 0 | - |
| - | - | 1150 | 246.1 | - | - | 0 | - |
| - | - | 1994 | 246.1 | - | - | 0 | - |
| - | - | 661.8 | 247.1 | - | - | 0 | - |
| 2 | a | 3.728E+04 | 249.2 | 0.0003256 | 1.307 | +1 | 2 |
| - | - | 5228 | 250.2 | - | - | 0 | - |
| - | - | 527.3 | 252.1 | - | - | 0 | - |
| - | - | 3094 | 253.2 | - | - | 0 | - |
| - | - | 871 | 259.1 | - | - | 0 | - |
| - | - | 649.8 | 261.1 | - | - | 0 | - |
| - | - | 552 | 261.2 | - | - | 0 | - |
| - | - | 931.2 | 263.1 | - | - | 0 | - |
| - | - | 1.386E+04 | 265.2 | - | - | 0 | - |
| - | - | 2071 | 266.2 | - | - | 0 | - |
| - | - | 685.7 | 267.1 | - | - | 0 | - |
| - | - | 1127 | 268.1 | - | - | 0 | - |
| - | - | 722.8 | 270.2 | - | - | 0 | - |
| - | - | 721.9 | 273.1 | - | - | 0 | - |
| - | - | 1597 | 274.1 | - | - | 0 | - |
| - | - | 862.1 | 275.1 | - | - | 0 | - |
| - | - | 1207 | 276.1 | - | - | 0 | - |
| 2 | b | 2.658E+04 | 277.2 | 0.0001467 | 0.5293 | +1 | 2 |
| - | - | 4633 | 278.2 | - | - | 0 | - |
| - | - | 1036 | 280.1 | - | - | 0 | - |
| - | - | 881.9 | 285.1 | - | - | 0 | - |
| 12 | y | 1973 | 288.1 | 2.371E-05 | 0.08229 | +1 | 3 |
| - | - | 1149 | 290.2 | - | - | 0 | - |
| - | - | 9135 | 291.1 | - | - | 0 | - |
| - | - | 2032 | 292.1 | - | - | 0 | - |
| - | - | 1.058E+04 | 293.1 | - | - | 0 | - |
| - | - | 837.1 | 294.2 | - | - | 0 | - |
| - | - | 616.9 | 296.2 | - | - | 0 | - |
| - | - | 694 | 297.2 | - | - | 0 | - |
| - | - | 946 | 298.1 | - | - | 0 | - |
| - | - | 1031 | 299.1 | - | - | 0 | - |
| - | - | 766.9 | 300.1 | - | - | 0 | - |
| - | - | 861.1 | 300.1 | - | - | 0 | - |
| - | - | 625.9 | 301.1 | - | - | 0 | - |
| 12 | y | 1734 | 306.1 | 9.276E-05 | 0.303 | +1 | 3 |
| - | - | 698.5 | 309.1 | - | - | 0 | - |
| - | - | 1472 | 315.1 | - | - | 0 | - |
| - | - | 1101 | 315.2 | - | - | 0 | - |
| - | - | 574.2 | 318.1 | - | - | 0 | - |
| - | - | 535.4 | 323.2 | - | - | 0 | - |
| - | - | 514 | 328.2 | - | - | 0 | - |
| - | - | 1777 | 332.1 | - | - | 0 | - |
| - | - | 1648 | 332.1 | - | - | 0 | - |
| - | - | 878 | 333.1 | - | - | 0 | - |
| - | - | 2321 | 340.2 | - | - | 0 | - |
| - | - | 1346 | 344.1 | - | - | 0 | - |
| - | - | 581 | 345.1 | - | - | 0 | - |
| - | - | 536.4 | 351.1 | - | - | 0 | - |
| - | - | 1602 | 355.1 | - | - | 0 | - |
| - | - | 722.4 | 355.2 | - | - | 0 | - |
| - | - | 850.9 | 356.1 | - | - | 0 | - |
| - | - | 1007 | 357.1 | - | - | 0 | - |
| - | - | 726.5 | 357.2 | - | - | 0 | - |
| - | - | 543.5 | 360.6 | - | - | 0 | - |
| - | - | 1148 | 368.2 | - | - | 0 | - |
| - | - | 685 | 373.1 | - | - | 0 | - |
| - | - | 734.1 | 374.1 | - | - | 0 | - |
| - | - | 806.9 | 378.1 | - | - | 0 | - |
| - | - | 691.2 | 380.2 | - | - | 0 | - |
| - | - | 1745 | 385.2 | - | - | 0 | - |
| - | - | 684 | 386.2 | - | - | 0 | - |
| - | - | 870.1 | 389.2 | - | - | 0 | - |
| 3 | b | 4386 | 390.2 | 0.0005859 | 1.501 | +1 | 3 |
| - | - | 1075 | 391.2 | - | - | 0 | - |
| - | - | 685.2 | 396.2 | - | - | 0 | - |
| 11 | y | 1778 | 403.1 | 0.0004686 | 1.162 | +1 | 4 |
| - | - | 673.8 | 403.2 | - | - | 0 | - |
| - | - | 1687 | 404.2 | - | - | 0 | - |
| - | - | 2063 | 405.2 | - | - | 0 | - |
| - | - | 2042 | 406.2 | - | - | 0 | - |
| - | - | 914.5 | 407.2 | - | - | 0 | - |
| - | - | 1243 | 407.3 | - | - | 0 | - |
| 11 | y | 875.9 | 421.2 | 0.001134 | 2.693 | +1 | 4 |
| - | - | 692.5 | 423.3 | - | - | 0 | - |
| - | - | 755.9 | 427.2 | - | - | 0 | - |
| - | - | 859.5 | 431.2 | - | - | 0 | - |
| 7 | y | 717.3 | 437.2 | 0.004841 | 11.07 | +2 | 8 |
| - | - | 2979 | 441.2 | - | - | 0 | - |
| - | - | 1536 | 445.2 | - | - | 0 | - |
| - | - | 729.5 | 454.2 | - | - | 0 | - |
| - | - | 724.4 | 461.2 | - | - | 0 | - |
| - | - | 1965 | 483.2 | - | - | 0 | - |
| - | - | 1502 | 500.2 | - | - | 0 | - |
| 10 | y | 1182 | 516.2 | 0.002037 | 3.946 | +1 | 5 |
| 4 | b | 2105 | 518.3 | 0.0006796 | 1.311 | +1 | 4 |
| - | - | 914.5 | 519.3 | - | - | 0 | - |
| 10 | y | 3529 | 534.2 | 0.0003223 | 0.6033 | +1 | 5 |
| - | - | 956.5 | 534.3 | - | - | 0 | - |
| - | - | 702.1 | 537.3 | - | - | 0 | - |
| - | - | 1301 | 554.3 | - | - | 0 | - |
| - | - | 746.8 | 554.3 | - | - | 0 | - |
| 9 | b | 1125 | 568.3 | 0.005988 | 10.54 | +2 | 9 |
| - | - | 1151 | 568.8 | - | - | 0 | - |
| - | - | 625.2 | 584.3 | - | - | 0 | - |
| - | - | 1229 | 585.3 | - | - | 0 | - |
| - | - | 1317 | 601.8 | - | - | 0 | - |
| - | - | 1039 | 602.3 | - | - | 0 | - |
| - | - | 965.7 | 613.3 | - | - | 0 | - |
| - | - | 1143 | 623.3 | - | - | 0 | - |
| - | - | 601.1 | 627.3 | - | - | 0 | - |
| 4 | y | 2657 | 631.3 | 0.0004571 | 0.724 | +2 | 11 |
| 4 | y | 1614 | 631.8 | 0.01034 | 16.37 | +2 | 11 |
| - | - | 1206 | 632.3 | - | - | 0 | - |
| 4 | y | 1964 | 640.3 | 0.0003627 | 0.5665 | +2 | 11 |
| - | - | 1953 | 640.8 | - | - | 0 | - |
| - | - | 1040 | 641.3 | - | - | 0 | - |
| - | - | 1183 | 652.8 | - | - | 0 | - |
| - | - | 613.3 | 656.3 | - | - | 0 | - |
| 5 | b | 2265 | 665.3 | 0.007654 | 11.5 | +1 | 5 |
| - | - | 1035 | 666.3 | - | - | 0 | - |
| 9 | y | 981.8 | 672.3 | 0.004775 | 7.102 | +1 | 6 |
| 9 | y | 953.8 | 673.3 | 0.005167 | 7.674 | +1 | 6 |
| - | - | 1136 | 674.3 | - | - | 0 | - |
| - | - | 1264 | 679.3 | - | - | 0 | - |
| - | - | 939.7 | 679.8 | - | - | 0 | - |
| - | - | 561 | 680.3 | - | - | 0 | - |
| 11 | b | 3471 | 682.4 | 0.002242 | 3.286 | +2 | 11 |
| - | - | 2545 | 682.9 | - | - | 0 | - |
| - | - | 1498 | 683.4 | - | - | 0 | - |
| 3 | y | 1470 | 688.3 | 0.009508 | 13.81 | +2 | 12 |
| - | - | 1924 | 688.8 | - | - | 0 | - |
| - | - | 1790 | 689.3 | - | - | 0 | - |
| - | - | 1041 | 689.8 | - | - | 0 | - |
| 9 | y | 9826 | 690.3 | 0.0002855 | 0.4136 | +1 | 6 |
| - | - | 3777 | 691.3 | - | - | 0 | - |
| - | - | 807 | 692.3 | - | - | 0 | - |
| 3 | y | 5503 | 696.8 | 0.001605 | 2.303 | +2 | 12 |
| - | - | 4051 | 697.3 | - | - | 0 | - |
| - | - | 3007 | 697.8 | - | - | 0 | - |
| - | - | 1002 | 698.3 | - | - | 0 | - |
| - | - | 1977 | 699.4 | - | - | 0 | - |
| - | - | 2678 | 700.4 | - | - | 0 | - |
| - | - | 715.5 | 701.4 | - | - | 0 | - |
| - | - | 733 | 713.3 | - | - | 0 | - |
| - | - | 845.8 | 721.8 | - | - | 0 | - |
| - | - | 896 | 724.8 | - | - | 0 | - |
| - | - | 1128 | 725.3 | - | - | 0 | - |
| - | - | 830.3 | 734.4 | - | - | 0 | - |
| 12 | b | 3729 | 739.9 | 6.197E-05 | 0.08375 | +2 | 12 |
| - | - | 5523 | 740.4 | - | - | 0 | - |
| - | - | 1618 | 740.9 | - | - | 0 | - |
| - | - | 1043 | 741.4 | - | - | 0 | - |
| 2 | y | 1168 | 778.4 | 0.0003965 | 0.5094 | +2 | 13 |
| - | - | 1801 | 778.9 | - | - | 0 | - |
| 6 | b | 1070 | 779.4 | 0.005803 | 7.446 | +1 | 6 |
| - | - | 1894 | 780.4 | - | - | 0 | - |
| 13 | b | 1675 | 781.4 | 0.005189 | 6.641 | +2 | 13 |
| 13 | b | 1665 | 781.9 | 0.008845 | 11.31 | +2 | 13 |
| - | - | 1199 | 782.4 | - | - | 0 | - |
| 13 | b | 3522 | 790.4 | 0.0003316 | 0.4196 | +2 | 13 |
| - | - | 4663 | 790.9 | - | - | 0 | - |
| - | - | 3086 | 791.4 | - | - | 0 | - |
| - | - | 928.3 | 791.9 | - | - | 0 | - |
| - | - | 596 | 793.9 | - | - | 0 | - |
| - | - | 795.9 | 799.9 | - | - | 0 | - |
| - | - | 938.6 | 801.4 | - | - | 0 | - |
| - | - | 1708 | 802.9 | - | - | 0 | - |
| 8 | y | 7111 | 803.4 | 0.003273 | 4.074 | +1 | 7 |
| - | - | 820.8 | 803.9 | - | - | 0 | - |
| - | - | 2635 | 804.4 | - | - | 0 | - |
| - | - | 599.4 | 804.9 | - | - | 0 | - |
| - | - | 914.5 | 805.4 | - | - | 0 | - |
| - | - | 3173 | 814.4 | - | - | 0 | - |
| - | - | 1621 | 815.4 | - | - | 0 | - |
| - | - | 809.9 | 816.9 | - | - | 0 | - |
| - | - | 1153 | 817.4 | - | - | 0 | - |
| - | - | 758.5 | 818.4 | - | - | 0 | - |
| 0 | Precursor | 4469 | 825.9 | 0.0003904 | 0.4727 | +2 | -1 |
| 0 | Precursor | 6817 | 826.4 | 0.007528 | 9.11 | +2 | -1 |
| - | - | 4234 | 826.9 | - | - | 0 | - |
| - | - | 2008 | 827.4 | - | - | 0 | - |
| - | - | 817.4 | 827.9 | - | - | 0 | - |
| - | - | 1030 | 830.4 | - | - | 0 | - |
| - | - | 872.6 | 833.9 | - | - | 0 | - |
| - | - | 1130 | 834.4 | - | - | 0 | - |
| 0 | Precursor | 1.812E+04 | 834.9 | 0.001334 | 1.597 | +2 | -1 |
| - | - | 1.901E+04 | 835.4 | - | - | 0 | - |
| - | - | 895.9 | 835.7 | - | - | 0 | - |
| - | - | 9401 | 835.9 | - | - | 0 | - |
| - | - | 5049 | 836.4 | - | - | 0 | - |
| - | - | 1526 | 837.4 | - | - | 0 | - |
| - | - | 1014 | 846.4 | - | - | 0 | - |
| - | - | 893.9 | 858.4 | - | - | 0 | - |
| - | - | 1045 | 859.4 | - | - | 0 | - |
| 7 | y | 774.3 | 872.4 | 0.007795 | 8.935 | +1 | 8 |
| 7 | y | 8082 | 890.5 | 0.001061 | 1.191 | +1 | 8 |
| - | - | 4424 | 891.5 | - | - | 0 | - |
| - | - | 1626 | 897.4 | - | - | 0 | - |
| - | - | 796.3 | 898.4 | - | - | 0 | - |
| - | - | 1023 | 915.5 | - | - | 0 | - |
| - | - | 911.4 | 933.5 | - | - | 0 | - |
| - | - | 716.9 | 940.4 | - | - | 0 | - |
| - | - | 812.5 | 941.4 | - | - | 0 | - |
| - | - | 2034 | 945.4 | - | - | 0 | - |
| - | - | 948.2 | 946.5 | - | - | 0 | - |
| - | - | 1074 | 957.4 | - | - | 0 | - |
| - | - | 961 | 961.5 | - | - | 0 | - |
| 8 | b | 878.3 | 962.5 | 0.01666 | 17.31 | +1 | 8 |
| - | - | 636.1 | 963.5 | - | - | 0 | - |
| - | - | 1335 | 974.5 | - | - | 0 | - |
| 8 | b | 1144 | 979.5 | 0.004396 | 4.488 | +1 | 8 |
| - | - | 2485 | 984.4 | - | - | 0 | - |
| - | - | 1854 | 985.5 | - | - | 0 | - |
| 6 | y | 1510 | 986.5 | 0.01237 | 12.54 | +1 | 9 |
| 6 | y | 2213 | 987.5 | 0.001482 | 1.501 | +1 | 9 |
| - | - | 1134 | 988.5 | - | - | 0 | - |
| - | - | 946.8 | 995.5 | - | - | 0 | - |
| 6 | y | 1.495E+04 | 1005 | 0.001691 | 1.683 | +1 | 9 |
| - | - | 6666 | 1006 | - | - | 0 | - |
| - | - | 2288 | 1007 | - | - | 0 | - |
| - | - | 1072 | 1008 | - | - | 0 | - |
| - | - | 900.8 | 1009 | - | - | 0 | - |
| - | - | 740.1 | 1017 | - | - | 0 | - |
| - | - | 883.1 | 1018 | - | - | 0 | - |
| - | - | 1464 | 1026 | - | - | 0 | - |
| - | - | 1060 | 1026 | - | - | 0 | - |
| - | - | 717.2 | 1048 | - | - | 0 | - |
| - | - | 1263 | 1056 | - | - | 0 | - |
| - | - | 983.8 | 1072 | - | - | 0 | - |
| - | - | 1132 | 1073 | - | - | 0 | - |
| - | - | 6015 | 1088 | - | - | 0 | - |
| - | - | 3505 | 1089 | - | - | 0 | - |
| - | - | 1780 | 1090 | - | - | 0 | - |
| - | - | 839.1 | 1107 | - | - | 0 | - |
| - | - | 839.8 | 1120 | - | - | 0 | - |
| - | - | 884 | 1130 | - | - | 0 | - |
| - | - | 769.1 | 1131 | - | - | 0 | - |
| 5 | y | 1099 | 1134 | 0.01062 | 9.365 | +1 | 10 |
| - | - | 8257 | 1136 | - | - | 0 | - |
| - | - | 5237 | 1137 | - | - | 0 | - |
| - | - | 2395 | 1138 | - | - | 0 | - |
| - | - | 1475 | 1139 | - | - | 0 | - |
| 5 | y | 7637 | 1152 | 0.001638 | 1.422 | +1 | 10 |
| - | - | 4823 | 1153 | - | - | 0 | - |
| - | - | 2654 | 1154 | - | - | 0 | - |
| - | - | 915.3 | 1155 | - | - | 0 | - |
| - | - | 923.4 | 1173 | - | - | 0 | - |
| - | - | 772.1 | 1175 | - | - | 0 | - |
| - | - | 994.4 | 1188 | - | - | 0 | - |
| - | - | 1137 | 1189 | - | - | 0 | - |
| - | - | 1724 | 1199 | - | - | 0 | - |
| - | - | 1316 | 1200 | - | - | 0 | - |
| - | - | 1305 | 1203 | - | - | 0 | - |
| - | - | 1293 | 1204 | - | - | 0 | - |
| - | - | 5029 | 1211 | - | - | 0 | - |
| - | - | 3588 | 1212 | - | - | 0 | - |
| - | - | 1988 | 1213 | - | - | 0 | - |
| - | - | 3530 | 1216 | - | - | 0 | - |
| - | - | 2545 | 1217 | - | - | 0 | - |
| - | - | 2599 | 1247 | - | - | 0 | - |
| - | - | 2318 | 1248 | - | - | 0 | - |
| 10 | b | 2536 | 1249 | 0.003181 | 2.547 | +1 | 10 |
| - | - | 1646 | 1250 | - | - | 0 | - |
| 4 | y | 682.6 | 1262 | 0.02015 | 15.97 | +1 | 11 |
| 4 | y | 2354 | 1263 | 0.00501 | 3.968 | +1 | 11 |
| - | - | 6927 | 1264 | - | - | 0 | - |
| - | - | 4563 | 1265 | - | - | 0 | - |
| - | - | 2370 | 1266 | - | - | 0 | - |
| 4 | y | 5670 | 1280 | 0.0005556 | 0.4342 | +1 | 11 |
| - | - | 4084 | 1281 | - | - | 0 | - |
| - | - | 2135 | 1282 | - | - | 0 | - |
| - | - | 3241 | 1329 | - | - | 0 | - |
| - | - | 2448 | 1330 | - | - | 0 | - |
| - | - | 1330 | 1331 | - | - | 0 | - |
| - | - | 866.6 | 1360 | - | - | 0 | - |
| - | - | 817.1 | 1363 | - | - | 0 | - |
| 11 | b | 1654 | 1364 | 0.001057 | 0.7751 | +1 | 11 |
| - | - | 1462 | 1365 | - | - | 0 | - |
| - | - | 839.8 | 1366 | - | - | 0 | - |
| - | - | 6395 | 1377 | - | - | 0 | - |
| - | - | 6242 | 1378 | - | - | 0 | - |
| - | - | 2669 | 1379 | - | - | 0 | - |
| - | - | 1117 | 1380 | - | - | 0 | - |
| 3 | y | 4689 | 1393 | 0.00221 | 1.587 | +1 | 12 |
| - | - | 4175 | 1394 | - | - | 0 | - |
| - | - | 2379 | 1395 | - | - | 0 | - |
| - | - | 1686 | 1403 | - | - | 0 | - |
| - | - | 1735 | 1404 | - | - | 0 | - |
| - | - | 1058 | 1405 | - | - | 0 | - |
| 12 | b | 1939 | 1479 | 0.001999 | 1.352 | +1 | 12 |
| - | - | 2268 | 1480 | - | - | 0 | - |
| - | - | 1080 | 1481 | - | - | 0 | - |
| - | - | 742.9 | 3301 | - | - | 0 | - |

m/z Charge Intensity FragmentType MassShift Position
120.0810546875 0 2990.7441
120.08502960205078 0 423.1579
122.06024169921875 0 469.55463
124.55310821533203 0 325.69598
127.05059814453125 0 677.1423
127.08697509765625 0 618.4202
127.7225341796875 0 328.82755
129.06626892089844 0 493.31973
129.10250854492188 0 3959.6262
130.05006408691406 0 791.8172
130.06568908691406 0 842.5081
131.04551696777344 0 1056.9152
131.11814880371094 0 6319.49
133.06117248535156 0 1090.8734
135.04457092285156 0 456.72034
136.07598876953125 0 35145.117
137.07936096191406 0 2517.1064
138.05545043945312 0 1372.4045
141.10244750976562 0 1721.5262
141.8792266845703 0 406.47214
142.09754943847656 0 450.10257
144.0658721923828 0 1137.3546
144.75674438476562 0 377.1316
145.097412109375 0 483.987
146.0602264404297 0 743.4351
147.07681274414062 0 533.44824
148.8417205810547 0 443.98294
148.89254760742188 0 616.9081
148.900390625 0 645.29596
148.9072265625 0 549.8891
148.91407775878906 0 560.58246
148.92138671875 0 885.61224
148.928955078125 0 1040.3497
148.93609619140625 0 2430.254
148.94400024414062 0 4132.6387
148.9606170654297 0 3464.5588
148.96839904785156 0 1872.55
148.97557067871094 0 1134.4534
148.98306274414062 0 1090.9215
148.9902801513672 0 1028.4805
148.9976348876953 0 837.29736
149.00477600097656 0 610.52576
149.08322143554688 0 409.78333
150.0549774169922 0 539.6034
152.06468200683594 0 860.597
152.07090759277344 0 10146.114
155.08216857910156 0 635.3818
155.1180877685547 0 593.81055
156.07705688476562 0 1049.7007
157.0609588623047 0 955.93317
157.10914611816406 0 830.82
159.0919952392578 0 1423.5767
162.0555877685547 0 534.7851
165.1025390625 0 553.82007
166.0980987548828 0 652.1371
167.05596923828125 0 562.693
167.0816650390625 0 2857.728
167.1178741455078 0 531.23773
169.1336669921875 0 615.64404
171.0769500732422 0 1073.0054
171.11251831054688 0 557.35944
172.0720672607422 0 728.7427
173.0923309326172 0 1639.5951 y Water loss 12
173.12890625 0 2234.4592
174.0548553466797 0 728.9697
174.08721923828125 0 1144.3385
175.08705139160156 0 583.4655
178.04995727539062 0 1507.4921
180.0660858154297 0 599.6606
183.11312866210938 0 1458.794
183.14926147460938 0 690.15796
184.072021484375 0 1212.9669
185.0558319091797 0 597.17255
186.08590698242188 0 664.3189
187.10816955566406 0 1377.0969
187.14483642578125 0 556.95386
189.08750915527344 0 1496.9806
191.04867553710938 0 735.47015
191.10289001464844 0 6129.6978 y 12
195.07662963867188 0 2343.2693
195.08644104003906 0 609.56757
195.11338806152344 0 889.4497
197.09286499023438 0 513.1325
197.1286163330078 0 696.8789
198.08758544921875 0 1209.889
199.0721893310547 0 781.9373
199.10821533203125 0 1404.1869
201.0869598388672 0 1054.3441
201.12351989746094 0 3285.4424
202.08241271972656 0 1992.6577
203.06678771972656 0 841.82623
203.1029052734375 0 811.325
208.0978546142578 0 767.6959
208.14389038085938 0 683.63196
210.1598358154297 0 743.20905
212.10337829589844 0 1092.0328
213.12353515625 0 1285.7988
214.10098266601562 0 540.87427
214.155029296875 0 572.0338
215.1392822265625 0 27925.574
216.1426544189453 0 2507.8835
217.08197021484375 0 3561.5952
218.11695861816406 0 599.88007
219.04379272460938 0 2359.7227
223.1080780029297 0 634.5068
224.1392822265625 0 690.1578
225.1235809326172 0 3011.6248
225.1712188720703 0 1513.9849
226.11875915527344 0 1054.145
227.1029052734375 0 2770.9883
229.11827087402344 0 1116.3564
231.06137084960938 0 1306.7649
238.11904907226562 0 714.3087
239.09500122070312 0 720.3244
240.13380432128906 0 765.72174
242.09576416015625 0 548.09894
242.15020751953125 0 10792.299
243.1337127685547 0 915.4975
243.1536407470703 0 879.4238
244.12876892089844 0 947.55145
245.12832641601562 0 732.1412
246.09075927734375 0 1149.7888
246.11288452148438 0 1993.6841
247.1447296142578 0 661.7635
249.1600799560547 0 37283.348 a 1
250.1633758544922 0 5227.686
252.1090850830078 0 527.2829
253.166259765625 0 3094.0889
259.075927734375 0 871.0254
261.12640380859375 0 649.84296
261.1590881347656 0 551.9763
263.139404296875 0 931.15826
265.15484619140625 0 13863.657
266.157958984375 0 2071.0928
267.1087951660156 0 685.7463
268.1289978027344 0 1127.3636
270.1930847167969 0 722.77893
273.12298583984375 0 721.8566
274.107666015625 0 1596.653
275.1394348144531 0 862.05194
276.1004943847656 0 1207.3497
277.1548156738281 0 26576.545 b 1
278.15802001953125 0 4633.0815
280.1284484863281 0 1036.3098
285.11962890625 0 881.87177
288.1189880371094 0 1972.726 y Water loss 11
290.1502380371094 0 1149.0876
291.13433837890625 0 9134.586
292.1375427246094 0 2032.1559
293.14990234375 0 10582.366
294.15301513671875 0 837.08923
296.1964111328125 0 616.8634
297.1573791503906 0 693.99646
298.1396789550781 0 945.9892
299.0615539550781 0 1031.497
300.0622253417969 0 766.8729
300.1343994140625 0 861.06854
301.0583190917969 0 625.9001
306.1296691894531 0 1734.1343 y 11
309.12005615234375 0 698.54926
315.11187744140625 0 1471.6663
315.16900634765625 0 1101.0668
318.1448059082031 0 574.2462
323.1719665527344 0 535.39496
328.169921875 0 514.02716
332.10919189453125 0 1776.748
332.1286315917969 0 1648.2363
333.1248474121094 0 877.9874
340.1985168457031 0 2321.0266
344.14471435546875 0 1345.5651
345.1492919921875 0 581.02026
351.1322326660156 0 536.4376
355.07080078125 0 1601.5945
355.16259765625 0 722.40546
356.0718078613281 0 850.8802
357.06781005859375 0 1006.8312
357.2231750488281 0 726.50543
360.59515380859375 0 543.5444
368.19256591796875 0 1148.2465
373.1174621582031 0 685.0207
374.1490478515625 0 734.0533
378.141357421875 0 806.9091
380.19268798828125 0 691.2026
385.2191467285156 0 1744.5415
386.2044982910156 0 684.0409
389.18621826171875 0 870.08716
390.23931884765625 0 4386.49 b 2
391.2418212890625 0 1075.4799
396.1523742675781 0 685.2155
403.14642333984375 0 1778.2068 y Water loss 10
403.2301025390625 0 673.8184
404.21807861328125 0 1686.5402
405.2145690917969 0 2063.4878
406.2340393066406 0 2041.7408
407.2373046875 0 914.5143
407.26513671875 0 1243.2198
421.15765380859375 0 875.92615 y 10
423.2591857910156 0 692.4981
427.18212890625 0 755.92236
431.1963806152344 0 859.5285
437.21441650390625 0 717.3141 y Ammonia loss 6
441.2347412109375 0 2979.39
445.192626953125 0 1536.0405
454.2417297363281 0 729.54803
461.2052001953125 0 724.3581
483.2208251953125 0 1965.4602
500.2478332519531 0 1502.145
516.2320556640625 0 1182.4243 y Water loss 9
518.296630859375 0 2104.8923 b 3
519.298583984375 0 914.49866
534.2409057617188 0 3529.3345 y 9
534.2882690429688 0 956.4851
537.2734375 0 702.121
554.2600708007812 0 1300.84
554.3040161132812 0 746.8151
568.3035888671875 0 1125.1077 b 8
568.80126953125 0 1150.9315
584.27978515625 0 625.2198
585.337646484375 0 1228.5247
601.7954711914062 0 1316.8849
602.302001953125 0 1038.805
613.3296508789062 0 965.74615
623.2947998046875 0 1143.2133
627.260009765625 0 601.1409
631.293701171875 0 2657.0542 y Water loss 3
631.7955932617188 0 1613.9192 y Ammonia loss 3
632.2943115234375 0 1206.1888
640.2988891601562 0 1964.007 y 3
640.8034057617188 0 1953.4331
641.3031005859375 0 1039.588
652.8212890625 0 1183.1011
656.3287963867188 0 613.2518
665.33544921875 0 2265.2515 b 4
666.337646484375 0 1035.439
672.3263549804688 0 981.77875 y Water loss 8
673.3203125 0 953.75824 y Ammonia loss 8
674.3154907226562 0 1136.3828
679.3276977539062 0 1264.3744
679.8283081054688 0 939.6513
680.3367309570312 0 561.01794
682.3553466796875 0 3470.598 b 10
682.8563232421875 0 2545.239
683.358154296875 0 1498.4227
688.3367919921875 0 1470.041 y Ammonia loss 2
688.8384399414062 0 1924.1384
689.34228515625 0 1789.6896
689.8435668945312 0 1040.8333
690.3419799804688 0 9825.775 y 8
691.344970703125 0 3777.4136
692.3460083007812 0 806.963
696.8421630859375 0 5503.043 y 2
697.3432006835938 0 4051.1555
697.843505859375 0 3007.0122
698.3405151367188 0 1001.88525
699.3792724609375 0 1976.7191
700.3656616210938 0 2678.4417
701.3675537109375 0 715.4683
713.3474731445312 0 733.03204
721.8419189453125 0 845.8455
724.8236694335938 0 895.99567
725.3319091796875 0 1127.9918
734.3510131835938 0 830.2639
739.8666381835938 0 3728.5933 b 11
740.3690185546875 0 5522.5444
740.8738403320312 0 1618.0309
741.3705444335938 0 1042.7103
778.371826171875 0 1167.6171 y 1
778.87548828125 0 1801.2678
779.3765258789062 0 1069.9495 b 5
780.3611450195312 0 1893.7742
781.3799438476562 0 1675.2788 b Water loss 12
781.885986328125 0 1665.3198 b Ammonia loss 12
782.3935546875 0 1199.486
790.3907470703125 0 3521.7373 b 12
790.8926391601562 0 4663.0234
791.3930053710938 0 3085.5244
791.8939819335938 0 928.26294
793.9069213867188 0 595.9895
799.8974609375 0 795.86096
801.4059448242188 0 938.60516
802.9168701171875 0 1707.6915
803.4224853515625 0 7110.9614 y 7
803.91748046875 0 820.7854
804.4273071289062 0 2634.933
804.911376953125 0 599.4123
805.4324951171875 0 914.5003
814.4028930664062 0 3173.0247
815.4104614257812 0 1620.5326
816.896240234375 0 809.9061
817.4085693359375 0 1153.4907
818.3995361328125 0 758.4852
825.9093627929688 0 4469.327 Precursor Water loss
826.4085083007812 0 6817.083 Precursor Ammonia loss
826.9073486328125 0 4233.8647
827.4100341796875 0 2008.4324
827.9102783203125 0 817.37787
830.4208374023438 0 1029.9401
833.9039916992188 0 872.59503
834.404541015625 0 1130.053
834.9155883789062 0 18118.887 Precursor
835.416748046875 0 19014.71
835.71875 0 895.8904
835.9164428710938 0 9401.172
836.4185791015625 0 5049.408
837.3839111328125 0 1525.9423
846.416015625 0 1013.6436
858.4323120117188 0 893.8859
859.4476318359375 0 1044.749
872.4550170898438 0 774.3262 y Water loss 6
890.4567260742188 0 8082.1665 y 6
891.4609375 0 4424.262
897.4442749023438 0 1625.7101
898.433837890625 0 796.2819
915.4523315429688 0 1023.3753
933.4627075195312 0 911.43414
940.4384155273438 0 716.8831
941.4456787109375 0 812.5137
945.4472045898438 0 2034.2933
946.4511108398438 0 948.2309
957.4473266601562 0 1073.5088
961.4525756835938 0 961.047
962.443603515625 0 878.2806 b Ammonia loss 7
963.4635620117188 0 636.1071
974.470458984375 0 1334.6803
979.4912109375 0 1143.512 b 7
984.4478759765625 0 2485.2466
985.453857421875 0 1854.4126
986.477783203125 0 1509.9384 y Water loss 5
987.4756469726562 0 2212.8423 y Ammonia loss 5
988.4800415039062 0 1133.9434
995.4855346679688 0 946.78485
1004.4990234375 0 14950.227 y 5
1005.5012817382812 0 6665.889
1006.50390625 0 2287.5767
1007.5040283203125 0 1072.2543
1009.4773559570312 0 900.7883
1017.4894409179688 0 740.06805
1018.4838256835938 0 883.1176
1025.501708984375 0 1463.7238
1026.4832763671875 0 1060.2286
1048.0054931640625 0 717.1586
1056.4736328125 0 1263.3997
1071.569091796875 0 983.8101
1073.498779296875 0 1131.7664
1087.536865234375 0 6014.6807
1088.5404052734375 0 3504.5562
1089.5234375 0 1779.6666
1106.527099609375 0 839.05066
1119.5911865234375 0 839.7858
1129.5523681640625 0 883.98157
1130.55322265625 0 769.11395
1133.53125 0 1099.0964 y Water loss 4
1135.5469970703125 0 8257.381
1136.552001953125 0 5236.9727
1137.56103515625 0 2394.7524
1138.5732421875 0 1475.0051
1151.5328369140625 0 7636.6636 y 4
1152.5369873046875 0 4822.818
1153.537353515625 0 2653.5508
1154.54541015625 0 915.2578
1172.5592041015625 0 923.38116
1174.5203857421875 0 772.1369
1187.58544921875 0 994.4499
1188.5960693359375 0 1136.6924
1198.5665283203125 0 1723.9867
1199.567138671875 0 1315.7585
1202.5830078125 0 1304.8195
1203.5860595703125 0 1292.7025
1210.5418701171875 0 5028.718
1211.545654296875 0 3588.249
1212.550537109375 0 1987.7434
1215.5911865234375 0 3530.239
1216.5946044921875 0 2544.9834
1246.5721435546875 0 2598.5273
1247.5692138671875 0 2318.4277
1248.6751708984375 0 2536.165 b 9
1249.676513671875 0 1646.0564
1261.599365234375 0 682.5835 y Water loss 3
1262.5682373046875 0 2354.3152 y Ammonia loss 3
1263.5897216796875 0 6926.693
1264.592529296875 0 4563.4316
1265.5948486328125 0 2369.6633
1279.59033203125 0 5669.555 y 3
1280.5921630859375 0 4083.5273
1281.593994140625 0 2134.6536
1328.6737060546875 0 3241.4958
1329.67431640625 0 2447.5315
1330.6859130859375 0 1330.2611
1359.6612548828125 0 866.59174
1362.6717529296875 0 817.1454
1363.6978759765625 0 1654.2275 b 10
1364.6988525390625 0 1462.1283
1365.69775390625 0 839.7785
1376.677734375 0 6395.3843
1377.6781005859375 0 6242.3457
1378.6829833984375 0 2668.8994
1379.6761474609375 0 1116.6974
1392.671630859375 0 4688.933 y 2
1393.6739501953125 0 4175.002
1394.671630859375 0 2378.9941
1402.6580810546875 0 1686.1797
1403.66162109375 0 1734.6731
1404.6612548828125 0 1057.5891
1478.723876953125 0 1938.6144 b 11
1479.73095703125 0 2268.0015
1480.7266845703125 0 1079.7817
3301.04638671875 0 742.8937

Spectrum Details

|  |  |
| --- | --- |
| Matched peaks? Matched peaks The total absolute number of peaks matched. Additionally in brackets the total fraction of peaks matched and the total number of peaks is shown. | 50 (12.29% of 407) |
| FDR? FDR The false discovery rate estimated for this peptide. It is calculated by matching all theoretical fragments with a non-integer shift with the raw peaks for this spectrum. This is done with 40 different shifts. The resulting percentage is the average number of annotated peaks over the number of annotated peaks with the correct spectrum. | 0.43% |
| Satellite FDR? Satellite FDR See the FDR for details on its calculation. This satellite ion specific FDR only contains the satellite ions (d/w) for I/L/J positions. | - |
| PSM Score? PSM Score The PSM Score as given by Hecklib to this annotated spectrum. It is shown with three significant figures. | 490 |

#### Spectrum 9846? Spectrum 9846 The raw spectrum of this peptide as annotated by Hecklib. The fragments are coloured according to ion type (see legend). Any peaks with a star '\*' as text can be hovered over to see the full details, first the ion type second the mass shift type. By hovering over the amino acids in the peptide or ions in the legend the corresponding peaks are highlighted. By toggling the 'Unassigned' label you can turn the background (unassigned) peaks on or off in the plot. By updating the slider in the Ion legend you can update the spectrum to only show the top X% of the peaks with labels. The top X% means any peak that is within X% of the highest intensity. By dragging in the spectrum you can zoom in to a specific part of the spectrum and use 'Zoom Out' to get back to the original zoom level. The annotation of the spectrum is based on the given sequence in the peptides file and is done with different software so inconsistencies are likely. The peaks are annotated based on the given sequence, with 20 ppm tolerance.

Copy Data

##### Spectrum 9846 (TSV)

###### Preview

```
Loading example...
```

*Click on the button to copy the data to your clipboard.*

Mz MinMz MaxIntensity Max

WidthHeightPeptide font sizePeptide stroke widthSpectrum font sizeSpectrum stroke widthCompact peptide

Ion legend

wxyz

abcd

OtherUnassignedIon^Charge^_Position_Show for top:%

JYLQMNSJRJDDTA

01.47e+42.93e+44.40e+45.87e+4

Zoom Out

y^+1^_2_y^+1^_2_a^+1^_2_b^+1^_2_y^+1^_3_y^+1^_3_b^+1^_3_y^+1^_4_b^+1^_4_y^+1^_5_y^+2^_11_y^+2^_11_b^+1^_5_b^+1^_5_b^+2^_11_y^+2^_12_y^+2^_12_y^+2^_12_y^+1^_6_b^+2^_12_y^+2^_13_b^+2^_13_b^+2^_13_y^+1^_7_\*\*\*b^+1^_7_y^+1^_8_b^+1^_8_y^+1^_9_y^+1^_9_y^+1^_9_y^+1^_10_y^+1^_10_b^+1^_9_y^+1^_10_b^+1^_10_y^+1^_11_y^+1^_11_y^+1^_11_b^+1^_11_y^+1^_12_b^+1^_12_y^+1^_13_

0507101515222029

Fragment Matches Table

Show background peaks

| Position | Ion type | Intensity | mz Theoretical | mz Error (Th) | mz Error (ppm) | Charge | Series Number |
| --- | --- | --- | --- | --- | --- | --- | --- |
| - | - | 6618 | 120.1 | - | - | 0 | - |
| - | - | 339.2 | 123.1 | - | - | 0 | - |
| - | - | 398.1 | 126.1 | - | - | 0 | - |
| - | - | 429.3 | 126.1 | - | - | 0 | - |
| - | - | 476.3 | 127.1 | - | - | 0 | - |
| - | - | 543.3 | 127.1 | - | - | 0 | - |
| - | - | 450.9 | 128.1 | - | - | 0 | - |
| - | - | 984.1 | 129.1 | - | - | 0 | - |
| - | - | 7640 | 129.1 | - | - | 0 | - |
| - | - | 891.3 | 130 | - | - | 0 | - |
| - | - | 589.3 | 130.1 | - | - | 0 | - |
| - | - | 657 | 131 | - | - | 0 | - |
| - | - | 2853 | 131.1 | - | - | 0 | - |
| - | - | 655.9 | 133.1 | - | - | 0 | - |
| - | - | 859 | 134 | - | - | 0 | - |
| - | - | 2.551E+04 | 136.1 | - | - | 0 | - |
| - | - | 2207 | 137.1 | - | - | 0 | - |
| - | - | 777.3 | 138.1 | - | - | 0 | - |
| - | - | 587.3 | 139.1 | - | - | 0 | - |
| - | - | 1089 | 140.1 | - | - | 0 | - |
| - | - | 529.2 | 142.1 | - | - | 0 | - |
| - | - | 746.8 | 143.1 | - | - | 0 | - |
| - | - | 526 | 143.1 | - | - | 0 | - |
| - | - | 1029 | 145.1 | - | - | 0 | - |
| - | - | 555.1 | 147.1 | - | - | 0 | - |
| - | - | 520.6 | 148.9 | - | - | 0 | - |
| - | - | 763.7 | 148.9 | - | - | 0 | - |
| - | - | 975 | 148.9 | - | - | 0 | - |
| - | - | 1245 | 148.9 | - | - | 0 | - |
| - | - | 2672 | 148.9 | - | - | 0 | - |
| - | - | 4291 | 148.9 | - | - | 0 | - |
| - | - | 3492 | 149 | - | - | 0 | - |
| - | - | 1543 | 149 | - | - | 0 | - |
| - | - | 822.7 | 149 | - | - | 0 | - |
| - | - | 863.8 | 149 | - | - | 0 | - |
| - | - | 522.3 | 149 | - | - | 0 | - |
| - | - | 447.4 | 149 | - | - | 0 | - |
| - | - | 844.8 | 152.1 | - | - | 0 | - |
| - | - | 845.4 | 155.1 | - | - | 0 | - |
| - | - | 943.9 | 156.1 | - | - | 0 | - |
| - | - | 496.7 | 157.1 | - | - | 0 | - |
| - | - | 502.3 | 159 | - | - | 0 | - |
| - | - | 1422 | 159.1 | - | - | 0 | - |
| - | - | 1584 | 167.1 | - | - | 0 | - |
| - | - | 1205 | 167.1 | - | - | 0 | - |
| - | - | 1701 | 169.1 | - | - | 0 | - |
| - | - | 771.5 | 171.1 | - | - | 0 | - |
| - | - | 573.6 | 172.1 | - | - | 0 | - |
| - | - | 494.3 | 172.1 | - | - | 0 | - |
| 13 | y | 855.7 | 173.1 | 0.0005064 | 2.925 | +1 | 2 |
| - | - | 1590 | 173.1 | - | - | 0 | - |
| - | - | 766.8 | 174.1 | - | - | 0 | - |
| - | - | 498 | 174.3 | - | - | 0 | - |
| - | - | 653.4 | 181.1 | - | - | 0 | - |
| - | - | 844.4 | 183.1 | - | - | 0 | - |
| - | - | 658.3 | 183.1 | - | - | 0 | - |
| - | - | 564.7 | 184.1 | - | - | 0 | - |
| - | - | 1468 | 184.1 | - | - | 0 | - |
| - | - | 667.8 | 185.1 | - | - | 0 | - |
| - | - | 517.4 | 187.1 | - | - | 0 | - |
| - | - | 1551 | 187.1 | - | - | 0 | - |
| - | - | 1123 | 189.1 | - | - | 0 | - |
| 13 | y | 4356 | 191.1 | 0.0003939 | 2.061 | +1 | 2 |
| - | - | 1088 | 195.1 | - | - | 0 | - |
| - | - | 625.7 | 197.1 | - | - | 0 | - |
| - | - | 697.2 | 197.1 | - | - | 0 | - |
| - | - | 538.5 | 198.1 | - | - | 0 | - |
| - | - | 2323 | 199.1 | - | - | 0 | - |
| - | - | 564.1 | 201.1 | - | - | 0 | - |
| - | - | 1780 | 201.1 | - | - | 0 | - |
| - | - | 1678 | 202.1 | - | - | 0 | - |
| - | - | 1415 | 203.1 | - | - | 0 | - |
| - | - | 524 | 204.8 | - | - | 0 | - |
| - | - | 754.9 | 208.1 | - | - | 0 | - |
| - | - | 1165 | 209.1 | - | - | 0 | - |
| - | - | 589 | 211.1 | - | - | 0 | - |
| - | - | 601 | 213.1 | - | - | 0 | - |
| - | - | 2838 | 213.1 | - | - | 0 | - |
| - | - | 5.807E+04 | 215.1 | - | - | 0 | - |
| - | - | 5495 | 216.1 | - | - | 0 | - |
| - | - | 1519 | 217.1 | - | - | 0 | - |
| - | - | 790 | 225.1 | - | - | 0 | - |
| - | - | 1542 | 225.1 | - | - | 0 | - |
| - | - | 1064 | 225.2 | - | - | 0 | - |
| - | - | 1215 | 226.1 | - | - | 0 | - |
| - | - | 531.9 | 226.2 | - | - | 0 | - |
| - | - | 4466 | 227.1 | - | - | 0 | - |
| - | - | 658.7 | 228.1 | - | - | 0 | - |
| - | - | 1248 | 228.2 | - | - | 0 | - |
| - | - | 1141 | 229.1 | - | - | 0 | - |
| - | - | 520.2 | 231.1 | - | - | 0 | - |
| - | - | 822.3 | 231.1 | - | - | 0 | - |
| - | - | 634.8 | 232.1 | - | - | 0 | - |
| - | - | 1854 | 240.1 | - | - | 0 | - |
| - | - | 7390 | 242.2 | - | - | 0 | - |
| - | - | 1051 | 243.1 | - | - | 0 | - |
| - | - | 1098 | 243.2 | - | - | 0 | - |
| - | - | 1543 | 244.1 | - | - | 0 | - |
| - | - | 512.3 | 245.1 | - | - | 0 | - |
| - | - | 1424 | 246.1 | - | - | 0 | - |
| - | - | 4228 | 247.1 | - | - | 0 | - |
| 2 | a | 2.812E+04 | 249.2 | 0.0003561 | 1.429 | +1 | 2 |
| - | - | 4795 | 250.2 | - | - | 0 | - |
| - | - | 628.4 | 251.1 | - | - | 0 | - |
| - | - | 1509 | 253.2 | - | - | 0 | - |
| - | - | 580.6 | 257.9 | - | - | 0 | - |
| - | - | 791.6 | 260.1 | - | - | 0 | - |
| - | - | 712.5 | 270.1 | - | - | 0 | - |
| - | - | 3980 | 275.1 | - | - | 0 | - |
| 2 | b | 2.408E+04 | 277.2 | 0.0001162 | 0.4192 | +1 | 2 |
| - | - | 3027 | 278.2 | - | - | 0 | - |
| - | - | 698.3 | 283.1 | - | - | 0 | - |
| - | - | 751.3 | 287.2 | - | - | 0 | - |
| 12 | y | 865.2 | 288.1 | 0.0003425 | 1.189 | +1 | 3 |
| - | - | 506.4 | 292.1 | - | - | 0 | - |
| - | - | 640.5 | 297.2 | - | - | 0 | - |
| 12 | y | 1274 | 306.1 | 0.0002148 | 0.7018 | +1 | 3 |
| - | - | 795 | 315.1 | - | - | 0 | - |
| - | - | 946.4 | 315.2 | - | - | 0 | - |
| - | - | 548.9 | 324.2 | - | - | 0 | - |
| - | - | 979.9 | 326.2 | - | - | 0 | - |
| - | - | 944.7 | 333.1 | - | - | 0 | - |
| - | - | 1373 | 340.2 | - | - | 0 | - |
| - | - | 649.1 | 343.2 | - | - | 0 | - |
| - | - | 667 | 344.1 | - | - | 0 | - |
| - | - | 646.6 | 344.2 | - | - | 0 | - |
| - | - | 1098 | 357.1 | - | - | 0 | - |
| - | - | 1211 | 373.2 | - | - | 0 | - |
| - | - | 1341 | 374.2 | - | - | 0 | - |
| 3 | b | 4187 | 390.2 | 0.0002076 | 0.5319 | +1 | 3 |
| - | - | 630.9 | 391.2 | - | - | 0 | - |
| - | - | 683.7 | 391.2 | - | - | 0 | - |
| - | - | 2545 | 405.2 | - | - | 0 | - |
| - | - | 625.9 | 407.2 | - | - | 0 | - |
| - | - | 1042 | 407.3 | - | - | 0 | - |
| - | - | 595.4 | 408.2 | - | - | 0 | - |
| - | - | 572.5 | 408.3 | - | - | 0 | - |
| 11 | y | 736.3 | 421.2 | 0.004186 | 9.94 | +1 | 4 |
| - | - | 1061 | 423.2 | - | - | 0 | - |
| - | - | 1054 | 426.1 | - | - | 0 | - |
| - | - | 5222 | 441.2 | - | - | 0 | - |
| - | - | 1508 | 442.2 | - | - | 0 | - |
| - | - | 1022 | 444.2 | - | - | 0 | - |
| - | - | 1008 | 445.2 | - | - | 0 | - |
| - | - | 696.3 | 452.2 | - | - | 0 | - |
| - | - | 583.2 | 453.3 | - | - | 0 | - |
| - | - | 603.3 | 454.2 | - | - | 0 | - |
| - | - | 589.2 | 481.3 | - | - | 0 | - |
| - | - | 792.9 | 483.2 | - | - | 0 | - |
| 4 | b | 2652 | 518.3 | 0.001595 | 3.078 | +1 | 4 |
| - | - | 904.9 | 519.3 | - | - | 0 | - |
| 10 | y | 1148 | 534.2 | 0.0004711 | 0.8819 | +1 | 5 |
| - | - | 651.9 | 536.2 | - | - | 0 | - |
| - | - | 2030 | 554.3 | - | - | 0 | - |
| - | - | 680.7 | 557.2 | - | - | 0 | - |
| - | - | 765.1 | 573.3 | - | - | 0 | - |
| - | - | 603.1 | 585.3 | - | - | 0 | - |
| - | - | 1299 | 588.3 | - | - | 0 | - |
| - | - | 664.7 | 599.3 | - | - | 0 | - |
| - | - | 921.4 | 602.3 | - | - | 0 | - |
| - | - | 1012 | 617.3 | - | - | 0 | - |
| 4 | y | 986.7 | 623.3 | 0.003078 | 4.938 | +2 | 11 |
| 4 | y | 870.8 | 623.8 | 0.003388 | 5.432 | +2 | 11 |
| 5 | b | 2238 | 632.3 | 0.00784 | 12.4 | +1 | 5 |
| - | - | 680.9 | 633.3 | - | - | 0 | - |
| - | - | 612.9 | 635.3 | - | - | 0 | - |
| - | - | 773.9 | 644.3 | - | - | 0 | - |
| 5 | b | 971.3 | 649.3 | 0.0002707 | 0.4168 | +1 | 5 |
| - | - | 888.5 | 650.3 | - | - | 0 | - |
| 11 | b | 648.1 | 674.4 | 0.0008049 | 1.194 | +2 | 11 |
| - | - | 843.9 | 674.9 | - | - | 0 | - |
| 3 | y | 1699 | 679.8 | 0.001775 | 2.611 | +2 | 12 |
| 3 | y | 754 | 680.3 | 0.006522 | 9.587 | +2 | 12 |
| 3 | y | 3746 | 688.8 | 0.001686 | 2.448 | +2 | 12 |
| - | - | 2521 | 689.3 | - | - | 0 | - |
| - | - | 1816 | 689.8 | - | - | 0 | - |
| 9 | y | 5397 | 690.3 | 0.001301 | 1.885 | +1 | 6 |
| - | - | 1788 | 691.3 | - | - | 0 | - |
| - | - | 789.1 | 692.3 | - | - | 0 | - |
| - | - | 1042 | 699.4 | - | - | 0 | - |
| - | - | 932.4 | 700.4 | - | - | 0 | - |
| - | - | 1508 | 707.2 | - | - | 0 | - |
| - | - | 653.6 | 712.4 | - | - | 0 | - |
| 12 | b | 1996 | 731.9 | 0.001825 | 2.494 | +2 | 12 |
| - | - | 1232 | 732.4 | - | - | 0 | - |
| 2 | y | 669.9 | 770.4 | 0.000941 | 1.221 | +2 | 13 |
| 13 | b | 593.3 | 773.9 | 0.007385 | 9.543 | +2 | 13 |
| - | - | 2273 | 780.4 | - | - | 0 | - |
| - | - | 1130 | 781.4 | - | - | 0 | - |
| 13 | b | 2260 | 782.4 | 0.003081 | 3.939 | +2 | 13 |
| - | - | 1495 | 782.9 | - | - | 0 | - |
| - | - | 1113 | 783.4 | - | - | 0 | - |
| 8 | y | 4019 | 803.4 | 0.0002824 | 0.3515 | +1 | 7 |
| - | - | 1455 | 804.4 | - | - | 0 | - |
| - | - | 610.8 | 807.4 | - | - | 0 | - |
| - | - | 609.2 | 808.4 | - | - | 0 | - |
| - | - | 2300 | 814.4 | - | - | 0 | - |
| 0 | Precursor | 1073 | 817.9 | 0.001863 | 2.278 | +2 | -1 |
| 0 | Precursor | 2759 | 818.4 | 0.003077 | 3.76 | +2 | -1 |
| - | - | 1601 | 818.9 | - | - | 0 | - |
| - | - | 802.6 | 819.4 | - | - | 0 | - |
| - | - | 963.8 | 825.4 | - | - | 0 | - |
| - | - | 891.7 | 826.4 | - | - | 0 | - |
| 0 | Precursor | 5595 | 826.9 | 0.002324 | 2.81 | +2 | -1 |
| - | - | 5326 | 827.4 | - | - | 0 | - |
| - | - | 2834 | 827.9 | - | - | 0 | - |
| - | - | 1580 | 828.4 | - | - | 0 | - |
| - | - | 1568 | 830.4 | - | - | 0 | - |
| 7 | b | 750.3 | 850.4 | 0.0002753 | 0.3238 | +1 | 7 |
| 7 | y | 4599 | 890.5 | 0.00222 | 2.494 | +1 | 8 |
| - | - | 2154 | 891.5 | - | - | 0 | - |
| - | - | 860.5 | 892.5 | - | - | 0 | - |
| - | - | 3415 | 927.4 | - | - | 0 | - |
| - | - | 1898 | 928.4 | - | - | 0 | - |
| - | - | 804.3 | 929.4 | - | - | 0 | - |
| - | - | 2668 | 945.4 | - | - | 0 | - |
| 8 | b | 1782 | 946.5 | 0.01647 | 17.4 | +1 | 8 |
| - | - | 606.8 | 952.4 | - | - | 0 | - |
| - | - | 1006 | 957.4 | - | - | 0 | - |
| - | - | 1428 | 958.5 | - | - | 0 | - |
| - | - | 698.7 | 960.5 | - | - | 0 | - |
| 6 | y | 792.2 | 986.5 | 0.007484 | 7.586 | +1 | 9 |
| 6 | y | 1171 | 987.5 | 0.003279 | 3.321 | +1 | 9 |
| - | - | 867.7 | 994.5 | - | - | 0 | - |
| - | - | 683.7 | 1000 | - | - | 0 | - |
| 6 | y | 1.033E+04 | 1005 | 0.002973 | 2.959 | +1 | 9 |
| - | - | 4576 | 1005 | - | - | 0 | - |
| - | - | 1604 | 1007 | - | - | 0 | - |
| - | - | 766.7 | 1046 | - | - | 0 | - |
| - | - | 759 | 1072 | - | - | 0 | - |
| - | - | 1999 | 1073 | - | - | 0 | - |
| - | - | 1053 | 1074 | - | - | 0 | - |
| - | - | 753.9 | 1090 | - | - | 0 | - |
| 5 | y | 1687 | 1118 | 0.007563 | 6.768 | +1 | 10 |
| 5 | y | 2361 | 1119 | 0.01538 | 13.75 | +1 | 10 |
| 9 | b | 2386 | 1120 | 0.006495 | 5.801 | +1 | 9 |
| - | - | 2070 | 1121 | - | - | 0 | - |
| 5 | y | 1.612E+04 | 1136 | 0.005554 | 4.891 | +1 | 10 |
| - | - | 9401 | 1137 | - | - | 0 | - |
| - | - | 3955 | 1138 | - | - | 0 | - |
| - | - | 6860 | 1154 | - | - | 0 | - |
| - | - | 2776 | 1155 | - | - | 0 | - |
| - | - | 1363 | 1156 | - | - | 0 | - |
| - | - | 707.9 | 1165 | - | - | 0 | - |
| - | - | 2005 | 1187 | - | - | 0 | - |
| - | - | 1506 | 1188 | - | - | 0 | - |
| - | - | 1129 | 1211 | - | - | 0 | - |
| - | - | 814.1 | 1214 | - | - | 0 | - |
| - | - | 813.2 | 1215 | - | - | 0 | - |
| - | - | 965.2 | 1227 | - | - | 0 | - |
| 10 | b | 1505 | 1233 | 0.0008714 | 0.7069 | +1 | 10 |
| 4 | y | 1602 | 1246 | 0.00242 | 1.943 | +1 | 11 |
| 4 | y | 3284 | 1247 | 0.001816 | 1.457 | +1 | 11 |
| - | - | 2480 | 1248 | - | - | 0 | - |
| - | - | 819.7 | 1249 | - | - | 0 | - |
| 4 | y | 1.025E+04 | 1264 | 0.006515 | 5.156 | +1 | 11 |
| - | - | 6625 | 1265 | - | - | 0 | - |
| - | - | 2023 | 1266 | - | - | 0 | - |
| 11 | b | 1540 | 1348 | 0.005686 | 4.219 | +1 | 11 |
| - | - | 871 | 1349 | - | - | 0 | - |
| - | - | 815.9 | 1368 | - | - | 0 | - |
| 3 | y | 7432 | 1377 | 0.00867 | 6.297 | +1 | 12 |
| - | - | 4626 | 1378 | - | - | 0 | - |
| - | - | 2577 | 1379 | - | - | 0 | - |
| - | - | 1254 | 1440 | - | - | 0 | - |
| 12 | b | 1044 | 1463 | 0.01127 | 7.702 | +1 | 12 |
| - | - | 1403 | 1464 | - | - | 0 | - |
| 2 | y | 1199 | 1540 | 0.001686 | 1.095 | +1 | 13 |
| - | - | 813.1 | 2009 | - | - | 0 | - |

m/z Charge Intensity FragmentType MassShift Position
120.08116149902344 0 6618.08
123.0753402709961 0 339.2032
126.05506134033203 0 398.084
126.06649780273438 0 429.3048
127.05049133300781 0 476.2795
127.08683013916016 0 543.3081
128.071044921875 0 450.94608
129.06637573242188 0 984.1309
129.10260009765625 0 7640.289
130.04995727539062 0 891.29034
130.061279296875 0 589.27
131.04531860351562 0 656.9798
131.11825561523438 0 2853.097
133.06109619140625 0 655.9215
134.0272674560547 0 858.9856
136.0760498046875 0 25510.35
137.07945251464844 0 2206.9114
138.0552978515625 0 777.3393
139.06100463867188 0 587.34375
140.08241271972656 0 1089.0967
142.06150817871094 0 529.2317
143.08226013183594 0 746.81714
143.1190643310547 0 525.98395
145.06105041503906 0 1029.0574
147.07666015625 0 555.0624
148.9086151123047 0 520.5848
148.915283203125 0 763.74915
148.92291259765625 0 975.042
148.93016052246094 0 1244.8839
148.9374237060547 0 2671.8586
148.9452362060547 0 4290.5566
148.96177673339844 0 3492.0237
148.96957397460938 0 1542.5426
148.97654724121094 0 822.6777
148.9835662841797 0 863.78156
148.9982147216797 0 522.3158
149.02732849121094 0 447.40604
152.07095336914062 0 844.7932
155.08160400390625 0 845.43097
156.07650756835938 0 943.8779
157.09768676757812 0 496.73062
159.03936767578125 0 502.30786
159.09217834472656 0 1422.2877
167.08192443847656 0 1584.0796
167.11813354492188 0 1204.9525
169.13381958007812 0 1700.6768
171.0776824951172 0 771.53467
172.07171630859375 0 573.5622
172.1120147705078 0 494.27188
173.0925750732422 0 855.6802 y Water loss 12
173.12869262695312 0 1589.6067
174.08750915527344 0 766.8009
174.3274383544922 0 498.00934
181.09764099121094 0 653.41187
183.11370849609375 0 844.363
183.14939880371094 0 658.27527
184.07180786132812 0 564.7485
184.10842895507812 0 1468.3549
185.05535888671875 0 667.80865
187.07211303710938 0 517.4186
187.1081085205078 0 1551.3241
189.08778381347656 0 1123.2686
191.10302734375 0 4355.68 y 12
195.11297607421875 0 1088.3282
197.09237670898438 0 625.6952
197.129150390625 0 697.1976
198.12411499023438 0 538.5354
199.10787963867188 0 2323.412
201.08721923828125 0 564.13617
201.12351989746094 0 1779.8282
202.08248901367188 0 1677.8652
203.10317993164062 0 1415.1741
204.8045654296875 0 524.01953
208.09588623046875 0 754.9437
209.09300231933594 0 1165.3544
211.144287109375 0 588.996
213.09814453125 0 601.0118
213.1234893798828 0 2838.2517
215.1393280029297 0 58074.66
216.14276123046875 0 5495.3477
217.08204650878906 0 1518.7052
225.0867156982422 0 789.9926
225.12342834472656 0 1542.0408
225.1717071533203 0 1063.6368
226.11851501464844 0 1215.2874
226.15591430664062 0 531.8735
227.10302734375 0 4466.1562
228.1067352294922 0 658.65936
228.1712646484375 0 1247.5796
229.11911010742188 0 1140.881
231.13441467285156 0 520.2237
231.14987182617188 0 822.31494
232.1111297607422 0 634.7547
240.13479614257812 0 1854.3678
242.1503143310547 0 7390.3047
243.08001708984375 0 1051.2603
243.1539764404297 0 1097.6575
244.12948608398438 0 1543.1066
245.1131134033203 0 512.2813
246.09104919433594 0 1424.2197
247.1113739013672 0 4227.6123
249.1601104736328 0 28120.908 a 1
250.16336059570312 0 4795.246
251.1031036376953 0 628.36847
253.16647338867188 0 1508.8625
257.8699951171875 0 580.5713
260.1056213378906 0 791.59924
270.1441955566406 0 712.53296
275.1060485839844 0 3979.8574
277.15478515625 0 24078.629 b 1
278.1580810546875 0 3026.5332
283.14227294921875 0 698.2861
287.17535400390625 0 751.3251
288.1193542480469 0 865.1892 y Water loss 11
292.13153076171875 0 506.39505
297.15704345703125 0 640.5107
306.1297912597656 0 1273.7749 y 11
315.1125793457031 0 795.01154
315.1677551269531 0 946.43976
324.157470703125 0 548.94086
326.17144775390625 0 979.89764
333.1220703125 0 944.7219
340.19775390625 0 1373.2621
343.16375732421875 0 649.1144
344.1446533203125 0 666.9564
344.18017578125 0 646.56445
357.122314453125 0 1097.7279
373.19073486328125 0 1211.0017
374.1731262207031 0 1340.5117
390.238525390625 0 4186.9976 b 2
391.19683837890625 0 630.9004
391.2391052246094 0 683.70264
405.2134094238281 0 2544.8484
407.1916809082031 0 625.92474
407.26312255859375 0 1042.1101
408.1995544433594 0 595.3875
408.26654052734375 0 572.51306
421.16070556640625 0 736.2965 y 10
423.2239074707031 0 1060.5814
426.1429443359375 0 1054.356
441.2344970703125 0 5222.291
442.2377014160156 0 1508.1919
444.1551818847656 0 1022.4856
445.193603515625 0 1007.63354
452.2156066894531 0 696.28986
453.27716064453125 0 583.2153
454.23797607421875 0 603.29095
481.2688903808594 0 589.1942
483.22247314453125 0 792.90826
518.2957153320312 0 2652.289 b 3
519.2990112304688 0 904.9353
534.2401123046875 0 1148.1434 y 9
536.2491455078125 0 651.93774
554.2586059570312 0 2030.3334
557.2391967773438 0 680.7088
573.3377075195312 0 765.0947
585.283447265625 0 603.0794
588.3046875 0 1299.3295
599.2804565429688 0 664.7319
602.3049926757812 0 921.3543
617.2940673828125 0 1011.84076
623.295166015625 0 986.7429 y Water loss 3
623.7936401367188 0 870.80695 y Ammonia loss 3
632.3034057617188 0 2237.7625 b Ammonia loss 4
633.3013305664062 0 680.9342
635.3237915039062 0 612.92267
644.3226928710938 0 773.93207
649.3375244140625 0 971.3046 b 4
650.3370361328125 0 888.49963
674.3572998046875 0 648.0616 b 10
674.8583374023438 0 843.8533
679.8385009765625 0 1699.0795 y Water loss 2
680.3388061523438 0 754.01105 y Ammonia loss 2
688.8438720703125 0 3746.0137 y 2
689.3456420898438 0 2520.6987
689.8446044921875 0 1815.5787
690.3403930664062 0 5396.9487 y 8
691.344482421875 0 1787.6332
692.3480224609375 0 789.07983
699.3771362304688 0 1042.4515
700.3615112304688 0 932.3628
707.20703125 0 1507.8773
712.3554077148438 0 653.5996
731.8697509765625 0 1995.9846 b 11
732.3685302734375 0 1231.983
770.3762817382812 0 669.85736 y 1
773.8895263671875 0 593.2636 b Ammonia loss 12
780.355224609375 0 2272.5537
781.3618774414062 0 1129.56
782.392333984375 0 2260.3164 b 12
782.8930053710938 0 1494.8541
783.3923950195312 0 1112.8813
803.4254760742188 0 4018.626 y 7
804.4270629882812 0 1455.4419
807.3786010742188 0 610.821
808.4283447265625 0 609.1867
814.4046630859375 0 2299.6382
817.912109375 0 1073.3656 Precursor Water loss
818.4090576171875 0 2759.1025 Precursor Ammonia loss
818.9092407226562 0 1601.4059
819.40771484375 0 802.6047
825.3844604492188 0 963.8369
826.3782958984375 0 891.68646
826.9169311523438 0 5595.2324 Precursor
827.416748046875 0 5326.3696
827.9183959960938 0 2833.6133
828.4153442382812 0 1579.7194
830.4161987304688 0 1568.079
850.4124755859375 0 750.3211 b 6
890.45556640625 0 4599.01 y 6
891.458740234375 0 2153.6665
892.4518432617188 0 860.5462
927.4249267578125 0 3415.022
928.4292602539062 0 1898.4482
929.4356079101562 0 804.258
945.4464721679688 0 2668.4253
946.4537963867188 0 1781.8688 b Ammonia loss 7
952.4306030273438 0 606.765
957.4492797851562 0 1006.09125
958.4652099609375 0 1427.646
960.4800415039062 0 698.6961
986.482666015625 0 792.18097 y Water loss 5
987.4708862304688 0 1170.5857 y Ammonia loss 5
994.4838256835938 0 867.6865
1000.4536743164062 0 683.7073
1004.4977416992188 0 10326.007 y 5
1005.4988403320312 0 4575.735
1006.5011596679688 0 1604.0201
1046.4876708984375 0 766.7184
1071.5594482421875 0 758.9839
1073.4990234375 0 1999.4769
1074.49658203125 0 1053.4609
1089.5579833984375 0 753.90344
1117.5230712890625 0 1686.6644 y Water loss 4
1118.530029296875 0 2361.087 y Ammonia loss 4
1119.5914306640625 0 2386.096 b 8
1120.5899658203125 0 2070.1711
1135.53564453125 0 16122.1045 y 4
1136.5380859375 0 9401.051
1137.53857421875 0 3955.0461
1153.51953125 0 6859.9355
1154.521240234375 0 2776.4175
1155.511962890625 0 1363.1418
1164.529541015625 0 707.943
1186.5867919921875 0 2004.9318
1187.5894775390625 0 1505.6827
1210.5367431640625 0 1128.8918
1213.5545654296875 0 814.06274
1214.5474853515625 0 813.24536
1226.536865234375 0 965.20557
1232.682861328125 0 1505.2345 b 9
1245.5867919921875 0 1601.9349 y Water loss 3
1246.5714111328125 0 3284.321 y Ammonia loss 3
1247.5712890625 0 2480.1975
1248.5731201171875 0 819.68384
1263.59326171875 0 10250.035 y 3
1264.59521484375 0 6624.7466
1265.5968017578125 0 2023.0175
1347.7032470703125 0 1539.6227 b 10
1348.6964111328125 0 871.0205
1367.6419677734375 0 815.8828
1376.6751708984375 0 7431.6416 y 2
1377.6795654296875 0 4626.366
1378.6787109375 0 2577.4597
1439.6522216796875 0 1253.7821
1462.724609375 0 1043.9304 b 11
1463.7279052734375 0 1402.9087
1539.7454833984375 0 1199.0631 y 1
2009.357177734375 0 813.09155

Spectrum Details

|  |  |
| --- | --- |
| Matched peaks? Matched peaks The total absolute number of peaks matched. Additionally in brackets the total fraction of peaks matched and the total number of peaks is shown. | 45 (16.73% of 269) |
| FDR? FDR The false discovery rate estimated for this peptide. It is calculated by matching all theoretical fragments with a non-integer shift with the raw peaks for this spectrum. This is done with 40 different shifts. The resulting percentage is the average number of annotated peaks over the number of annotated peaks with the correct spectrum. | 0.21% |
| Satellite FDR? Satellite FDR See the FDR for details on its calculation. This satellite ion specific FDR only contains the satellite ions (d/w) for I/L/J positions. | - |
| PSM Score? PSM Score The PSM Score as given by Hecklib to this annotated spectrum. It is shown with three significant figures. | 450 |

#### Spectrum 9797? Spectrum 9797 The raw spectrum of this peptide as annotated by Hecklib. The fragments are coloured according to ion type (see legend). Any peaks with a star '\*' as text can be hovered over to see the full details, first the ion type second the mass shift type. By hovering over the amino acids in the peptide or ions in the legend the corresponding peaks are highlighted. By toggling the 'Unassigned' label you can turn the background (unassigned) peaks on or off in the plot. By updating the slider in the Ion legend you can update the spectrum to only show the top X% of the peaks with labels. The top X% means any peak that is within X% of the highest intensity. By dragging in the spectrum you can zoom in to a specific part of the spectrum and use 'Zoom Out' to get back to the original zoom level. The annotation of the spectrum is based on the given sequence in the peptides file and is done with different software so inconsistencies are likely. The peaks are annotated based on the given sequence, with 20 ppm tolerance.

Copy Data

##### Spectrum 9797 (TSV)

###### Preview

```
Loading example...
```

*Click on the button to copy the data to your clipboard.*

Mz MinMz MaxIntensity Max

WidthHeightPeptide font sizePeptide stroke widthSpectrum font sizeSpectrum stroke widthCompact peptide

Ion legend

wxyz

abcd

OtherUnassignedIon^Charge^_Position_Show for top:%

JYLQMNSJRJDDTA

02.47e+44.94e+47.40e+49.87e+4

Zoom Out

y^+1^_2_y^+1^_3_y^+1^_4_c^+1^_4_y^+1^_5_c^+2^_9_y^+2^_11_c^+1^_5_z^+2^_12_y^+1^_6_y^+2^_12_c^+2^_12_y^+2^_13_c^+1^_6_z^+1^_7_c^+2^_13_y^+1^_7_z^+1^_8_y^+1^_8_c^+1^_8_z^+1^_9_y^+1^_9_z^+1^_10_z^+1^_10_y^+1^_10_c^+1^_9_c^+1^_10_y^+1^_11_z^+1^_11_c^+1^_10_y^+1^_11_w^+1^_12_z^+1^_12_c^+1^_11_y^+1^_12_z^+1^_12_c^+1^_11_y^+1^_12_c^+1^_12_c^+1^_12_z^+1^_13_c^+1^_13_

042384512681691

Fragment Matches Table

Show background peaks

| Position | Ion type | Intensity | mz Theoretical | mz Error (Th) | mz Error (ppm) | Charge | Series Number |
| --- | --- | --- | --- | --- | --- | --- | --- |
| - | - | 336.9 | 120.1 | - | - | 0 | - |
| - | - | 413 | 120.4 | - | - | 0 | - |
| - | - | 352.3 | 121.2 | - | - | 0 | - |
| - | - | 351.9 | 128.1 | - | - | 0 | - |
| - | - | 376.5 | 128.1 | - | - | 0 | - |
| - | - | 472.1 | 130.4 | - | - | 0 | - |
| - | - | 687.6 | 136.1 | - | - | 0 | - |
| - | - | 758 | 141.1 | - | - | 0 | - |
| - | - | 464.5 | 144.3 | - | - | 0 | - |
| - | - | 432.4 | 147.8 | - | - | 0 | - |
| - | - | 576.3 | 148.9 | - | - | 0 | - |
| - | - | 474.4 | 148.9 | - | - | 0 | - |
| - | - | 585.3 | 148.9 | - | - | 0 | - |
| - | - | 1047 | 148.9 | - | - | 0 | - |
| - | - | 1146 | 148.9 | - | - | 0 | - |
| - | - | 1605 | 148.9 | - | - | 0 | - |
| - | - | 3208 | 148.9 | - | - | 0 | - |
| - | - | 5115 | 149 | - | - | 0 | - |
| - | - | 3314 | 149 | - | - | 0 | - |
| - | - | 1635 | 149 | - | - | 0 | - |
| - | - | 1128 | 149 | - | - | 0 | - |
| - | - | 860.4 | 149 | - | - | 0 | - |
| - | - | 905 | 149 | - | - | 0 | - |
| - | - | 588.3 | 149 | - | - | 0 | - |
| - | - | 508.8 | 149 | - | - | 0 | - |
| - | - | 520.5 | 149 | - | - | 0 | - |
| - | - | 417.9 | 149.1 | - | - | 0 | - |
| - | - | 496 | 149.4 | - | - | 0 | - |
| - | - | 496.9 | 152.1 | - | - | 0 | - |
| - | - | 458.3 | 169.7 | - | - | 0 | - |
| - | - | 936.2 | 173.4 | - | - | 0 | - |
| 13 | y | 2865 | 191.1 | 0.0004702 | 2.461 | +1 | 2 |
| - | - | 3931 | 215.1 | - | - | 0 | - |
| - | - | 1220 | 217.1 | - | - | 0 | - |
| - | - | 2931 | 242.2 | - | - | 0 | - |
| - | - | 4506 | 249.2 | - | - | 0 | - |
| - | - | 669.4 | 250.2 | - | - | 0 | - |
| - | - | 2173 | 265.2 | - | - | 0 | - |
| - | - | 7255 | 277.2 | - | - | 0 | - |
| - | - | 995.3 | 278.2 | - | - | 0 | - |
| 12 | y | 682.6 | 288.1 | 2.371E-05 | 0.08229 | +1 | 3 |
| - | - | 2713 | 291.1 | - | - | 0 | - |
| - | - | 4097 | 293.1 | - | - | 0 | - |
| - | - | 596.4 | 294.2 | - | - | 0 | - |
| - | - | 1113 | 299.1 | - | - | 0 | - |
| - | - | 606.2 | 310.4 | - | - | 0 | - |
| - | - | 588.9 | 316.9 | - | - | 0 | - |
| - | - | 720.4 | 332.1 | - | - | 0 | - |
| - | - | 814.2 | 355.1 | - | - | 0 | - |
| - | - | 1166 | 356.1 | - | - | 0 | - |
| - | - | 1041 | 357.1 | - | - | 0 | - |
| - | - | 2492 | 390.2 | - | - | 0 | - |
| 11 | y | 812.1 | 403.1 | 0.0008348 | 2.071 | +1 | 4 |
| - | - | 755.6 | 404.2 | - | - | 0 | - |
| - | - | 560.5 | 405.2 | - | - | 0 | - |
| - | - | 1017 | 406.2 | - | - | 0 | - |
| - | - | 1181 | 445.2 | - | - | 0 | - |
| - | - | 693.4 | 461.2 | - | - | 0 | - |
| - | - | 601.6 | 493.1 | - | - | 0 | - |
| 4 | c | 1005 | 518.3 | 0.001168 | 2.253 | +1 | 4 |
| 10 | y | 2168 | 534.2 | 0.0002613 | 0.4891 | +1 | 5 |
| - | - | 551.2 | 537 | - | - | 0 | - |
| 9 | c | 519.7 | 568.3 | 0.001654 | 2.911 | +2 | 9 |
| - | - | 1301 | 593.3 | - | - | 0 | - |
| 4 | y | 905.2 | 631.3 | 0.00296 | 4.688 | +2 | 11 |
| - | - | 990 | 640.8 | - | - | 0 | - |
| - | - | 720.5 | 652.8 | - | - | 0 | - |
| - | - | 842.5 | 665.3 | - | - | 0 | - |
| - | - | 977.4 | 666.3 | - | - | 0 | - |
| - | - | 610.9 | 673.4 | - | - | 0 | - |
| - | - | 642.4 | 675.3 | - | - | 0 | - |
| 5 | c | 1472 | 682.4 | 0.0002182 | 0.3198 | +1 | 5 |
| - | - | 1510 | 682.9 | - | - | 0 | - |
| 3 | z | 1592 | 688.8 | 0.01164 | 16.9 | +2 | 12 |
| - | - | 1353 | 689.3 | - | - | 0 | - |
| - | - | 1021 | 689.8 | - | - | 0 | - |
| 9 | y | 4361 | 690.3 | 0.0003859 | 0.559 | +1 | 6 |
| - | - | 2030 | 691.3 | - | - | 0 | - |
| 3 | y | 3228 | 696.8 | 0.004962 | 7.12 | +2 | 12 |
| - | - | 4109 | 697.3 | - | - | 0 | - |
| - | - | 1231 | 697.8 | - | - | 0 | - |
| - | - | 771.1 | 733.8 | - | - | 0 | - |
| 12 | c | 3240 | 739.9 | 0.002381 | 3.219 | +2 | 12 |
| - | - | 2960 | 740.4 | - | - | 0 | - |
| - | - | 1405 | 740.9 | - | - | 0 | - |
| - | - | 600.5 | 749.3 | - | - | 0 | - |
| 2 | y | 723.4 | 778.4 | 0.005463 | 7.018 | +2 | 13 |
| - | - | 1362 | 778.9 | - | - | 0 | - |
| 6 | c | 801.1 | 779.4 | 0.004339 | 5.567 | +1 | 6 |
| - | - | 927.8 | 781.4 | - | - | 0 | - |
| - | - | 1143 | 781.9 | - | - | 0 | - |
| - | - | 644.7 | 783.4 | - | - | 0 | - |
| 8 | z | 1880 | 787.4 | 0.001657 | 2.104 | +1 | 7 |
| - | - | 2651 | 788.4 | - | - | 0 | - |
| - | - | 806.3 | 789.4 | - | - | 0 | - |
| 13 | c | 2555 | 790.4 | 0.00375 | 4.744 | +2 | 13 |
| - | - | 2653 | 790.9 | - | - | 0 | - |
| - | - | 1872 | 791.4 | - | - | 0 | - |
| - | - | 778.7 | 794.4 | - | - | 0 | - |
| - | - | 784.6 | 802.9 | - | - | 0 | - |
| 8 | y | 2889 | 803.4 | 0.001442 | 1.795 | +1 | 7 |
| - | - | 2790 | 825.9 | - | - | 0 | - |
| - | - | 3739 | 826.4 | - | - | 0 | - |
| - | - | 2364 | 826.9 | - | - | 0 | - |
| - | - | 899.3 | 827.4 | - | - | 0 | - |
| - | - | 786.7 | 834.4 | - | - | 0 | - |
| - | - | 1.16E+04 | 834.9 | - | - | 0 | - |
| - | - | 1.068E+04 | 835.4 | - | - | 0 | - |
| - | - | 6899 | 835.9 | - | - | 0 | - |
| - | - | 2977 | 836.4 | - | - | 0 | - |
| 7 | z | 638.7 | 874.4 | 0.002099 | 2.401 | +1 | 8 |
| - | - | 3301 | 875.4 | - | - | 0 | - |
| - | - | 1505 | 876.5 | - | - | 0 | - |
| 7 | y | 1868 | 890.5 | 0.001915 | 2.151 | +1 | 8 |
| - | - | 1268 | 891.5 | - | - | 0 | - |
| - | - | 612.8 | 975.1 | - | - | 0 | - |
| 8 | c | 999.8 | 979.5 | 0.0007309 | 0.7462 | +1 | 8 |
| - | - | 1052 | 984.5 | - | - | 0 | - |
| 6 | z | 1453 | 988.5 | 0.0005537 | 0.5602 | +1 | 9 |
| - | - | 2.552E+04 | 989.5 | - | - | 0 | - |
| - | - | 1.349E+04 | 990.5 | - | - | 0 | - |
| - | - | 3082 | 991.5 | - | - | 0 | - |
| - | - | 2402 | 1003 | - | - | 0 | - |
| 6 | y | 1.134E+04 | 1005 | 0.0005922 | 0.5896 | +1 | 9 |
| - | - | 7125 | 1006 | - | - | 0 | - |
| - | - | 2180 | 1006 | - | - | 0 | - |
| - | - | 2577 | 1074 | - | - | 0 | - |
| - | - | 2070 | 1075 | - | - | 0 | - |
| - | - | 746.1 | 1089 | - | - | 0 | - |
| - | - | 3083 | 1109 | - | - | 0 | - |
| - | - | 1738 | 1110 | - | - | 0 | - |
| 5 | z | 815.3 | 1118 | 0.02018 | 18.06 | +1 | 10 |
| - | - | 787 | 1119 | - | - | 0 | - |
| - | - | 654.3 | 1120 | - | - | 0 | - |
| - | - | 2601 | 1121 | - | - | 0 | - |
| - | - | 1189 | 1122 | - | - | 0 | - |
| - | - | 663.9 | 1123 | - | - | 0 | - |
| 5 | z | 7076 | 1136 | 0.01499 | 13.2 | +1 | 10 |
| - | - | 7374 | 1137 | - | - | 0 | - |
| - | - | 3632 | 1138 | - | - | 0 | - |
| - | - | 1177 | 1139 | - | - | 0 | - |
| - | - | 621.4 | 1151 | - | - | 0 | - |
| 5 | y | 2487 | 1152 | 0.003713 | 3.225 | +1 | 10 |
| - | - | 2124 | 1152 | - | - | 0 | - |
| 9 | c | 9335 | 1153 | 0.002468 | 2.141 | +1 | 9 |
| - | - | 5018 | 1154 | - | - | 0 | - |
| - | - | 2758 | 1155 | - | - | 0 | - |
| - | - | 1019 | 1156 | - | - | 0 | - |
| - | - | 833.8 | 1212 | - | - | 0 | - |
| 10 | c | 876.3 | 1249 | 0.01453 | 11.64 | +1 | 10 |
| - | - | 1466 | 1250 | - | - | 0 | - |
| - | - | 772 | 1251 | - | - | 0 | - |
| 4 | y | 1379 | 1263 | 0.01197 | 9.479 | +1 | 11 |
| 4 | z | 4930 | 1264 | 0.01293 | 10.23 | +1 | 11 |
| - | - | 2567 | 1265 | - | - | 0 | - |
| - | - | 2346 | 1265 | - | - | 0 | - |
| - | - | 1057 | 1266 | - | - | 0 | - |
| 10 | c | 5391 | 1266 | 0.001029 | 0.8132 | +1 | 10 |
| - | - | 3334 | 1267 | - | - | 0 | - |
| - | - | 1176 | 1268 | - | - | 0 | - |
| 4 | y | 2783 | 1280 | 0.007269 | 5.681 | +1 | 11 |
| - | - | 2358 | 1281 | - | - | 0 | - |
| - | - | 896.9 | 1282 | - | - | 0 | - |
| - | - | 689.6 | 1318 | - | - | 0 | - |
| - | - | 726.2 | 1319 | - | - | 0 | - |
| - | - | 756.9 | 1320 | - | - | 0 | - |
| - | - | 748.2 | 1326 | - | - | 0 | - |
| 3 | w | 2178 | 1334 | 0.01062 | 7.964 | +1 | 12 |
| - | - | 2107 | 1335 | - | - | 0 | - |
| - | - | 853.2 | 1336 | - | - | 0 | - |
| 3 | z | 762.1 | 1359 | 0.01878 | 13.82 | +1 | 12 |
| - | - | 3303 | 1361 | - | - | 0 | - |
| - | - | 4145 | 1362 | - | - | 0 | - |
| - | - | 3163 | 1363 | - | - | 0 | - |
| 11 | c | 2015 | 1364 | 0.01156 | 8.473 | +1 | 11 |
| - | - | 1039 | 1365 | - | - | 0 | - |
| - | - | 1799 | 1367 | - | - | 0 | - |
| - | - | 1032 | 1368 | - | - | 0 | - |
| 3 | y | 901.3 | 1375 | 0.004584 | 3.335 | +1 | 12 |
| 3 | z | 1.407E+04 | 1377 | 0.01114 | 8.095 | +1 | 12 |
| - | - | 1.061E+04 | 1378 | - | - | 0 | - |
| - | - | 7258 | 1379 | - | - | 0 | - |
| - | - | 6513 | 1380 | - | - | 0 | - |
| 11 | c | 1.294E+04 | 1381 | 0.0008363 | 0.6057 | +1 | 11 |
| - | - | 9080 | 1382 | - | - | 0 | - |
| - | - | 4833 | 1383 | - | - | 0 | - |
| 3 | y | 3065 | 1393 | 0.005481 | 3.936 | +1 | 12 |
| - | - | 3010 | 1394 | - | - | 0 | - |
| - | - | 1416 | 1395 | - | - | 0 | - |
| - | - | 977.2 | 1404 | - | - | 0 | - |
| - | - | 3500 | 1425 | - | - | 0 | - |
| - | - | 4476 | 1426 | - | - | 0 | - |
| - | - | 1939 | 1427 | - | - | 0 | - |
| - | - | 827.2 | 1428 | - | - | 0 | - |
| - | - | 1019 | 1441 | - | - | 0 | - |
| 12 | c | 818.7 | 1479 | 0.001297 | 0.877 | +1 | 12 |
| - | - | 1599 | 1480 | - | - | 0 | - |
| - | - | 1657 | 1481 | - | - | 0 | - |
| - | - | 1084 | 1485 | - | - | 0 | - |
| - | - | 1141 | 1495 | - | - | 0 | - |
| 12 | c | 5443 | 1496 | 0.0002605 | 0.1741 | +1 | 12 |
| - | - | 6925 | 1497 | - | - | 0 | - |
| - | - | 3929 | 1498 | - | - | 0 | - |
| - | - | 1935 | 1499 | - | - | 0 | - |
| 2 | z | 4843 | 1540 | 0.001526 | 0.991 | +1 | 13 |
| - | - | 3991 | 1541 | - | - | 0 | - |
| - | - | 2648 | 1542 | - | - | 0 | - |
| - | - | 721.5 | 1543 | - | - | 0 | - |
| - | - | 1332 | 1553 | - | - | 0 | - |
| - | - | 1024 | 1578 | - | - | 0 | - |
| - | - | 1331 | 1581 | - | - | 0 | - |
| - | - | 934 | 1582 | - | - | 0 | - |
| - | - | 937.5 | 1583 | - | - | 0 | - |
| - | - | 719.3 | 1590 | - | - | 0 | - |
| - | - | 1613 | 1596 | - | - | 0 | - |
| 13 | c | 7016 | 1597 | 0.00152 | 0.9516 | +1 | 13 |
| - | - | 5952 | 1598 | - | - | 0 | - |
| - | - | 3355 | 1599 | - | - | 0 | - |
| - | - | 1825 | 1600 | - | - | 0 | - |
| - | - | 727.5 | 1606 | - | - | 0 | - |
| - | - | 808 | 1607 | - | - | 0 | - |
| - | - | 2111 | 1608 | - | - | 0 | - |
| - | - | 2260 | 1609 | - | - | 0 | - |
| - | - | 6465 | 1610 | - | - | 0 | - |
| - | - | 8029 | 1611 | - | - | 0 | - |
| - | - | 7019 | 1612 | - | - | 0 | - |
| - | - | 2556 | 1613 | - | - | 0 | - |
| - | - | 1660 | 1614 | - | - | 0 | - |
| - | - | 1378 | 1615 | - | - | 0 | - |
| - | - | 720.8 | 1622 | - | - | 0 | - |
| - | - | 1155 | 1623 | - | - | 0 | - |
| - | - | 1375 | 1624 | - | - | 0 | - |
| - | - | 1.411E+04 | 1625 | - | - | 0 | - |
| - | - | 1.322E+04 | 1626 | - | - | 0 | - |
| - | - | 8374 | 1627 | - | - | 0 | - |
| - | - | 3673 | 1628 | - | - | 0 | - |
| - | - | 1369 | 1629 | - | - | 0 | - |
| - | - | 936.3 | 1635 | - | - | 0 | - |
| - | - | 2019 | 1636 | - | - | 0 | - |
| - | - | 1722 | 1637 | - | - | 0 | - |
| - | - | 2646 | 1638 | - | - | 0 | - |
| - | - | 1448 | 1639 | - | - | 0 | - |
| - | - | 1156 | 1640 | - | - | 0 | - |
| - | - | 1586 | 1642 | - | - | 0 | - |
| - | - | 1197 | 1643 | - | - | 0 | - |
| - | - | 1140 | 1650 | - | - | 0 | - |
| - | - | 3410 | 1651 | - | - | 0 | - |
| - | - | 2.045E+04 | 1652 | - | - | 0 | - |
| - | - | 5.047E+04 | 1653 | - | - | 0 | - |
| - | - | 4.703E+04 | 1654 | - | - | 0 | - |
| - | - | 2.472E+04 | 1655 | - | - | 0 | - |
| - | - | 1.05E+04 | 1656 | - | - | 0 | - |
| - | - | 1019 | 1657 | - | - | 0 | - |
| - | - | 2912 | 1667 | - | - | 0 | - |
| - | - | 3.726E+04 | 1668 | - | - | 0 | - |
| - | - | 8.672E+04 | 1669 | - | - | 0 | - |
| - | - | 9.775E+04 | 1670 | - | - | 0 | - |
| - | - | 6.794E+04 | 1671 | - | - | 0 | - |
| - | - | 3.356E+04 | 1672 | - | - | 0 | - |
| - | - | 1.333E+04 | 1673 | - | - | 0 | - |
| - | - | 1975 | 1674 | - | - | 0 | - |

m/z Charge Intensity FragmentType MassShift Position
120.1077880859375 0 336.899
120.39189910888672 0 412.9885
121.1854019165039 0 352.27466
128.09544372558594 0 351.87
128.11390686035156 0 376.52612
130.3920440673828 0 472.14996
136.07623291015625 0 687.55194
141.1024169921875 0 757.9741
144.29527282714844 0 464.54834
147.78038024902344 0 432.3507
148.88168334960938 0 576.3116
148.90306091308594 0 474.43295
148.91073608398438 0 585.27094
148.9180908203125 0 1047.3555
148.9253692626953 0 1146.264
148.93231201171875 0 1604.592
148.94009399414062 0 3208.401
148.9567108154297 0 5115.09
148.96458435058594 0 3314.0974
148.97216796875 0 1635.3668
148.9791259765625 0 1128.239
148.9862823486328 0 860.4166
148.9936065673828 0 904.9559
149.0011749267578 0 588.2749
149.0232391357422 0 508.83643
149.04490661621094 0 520.4782
149.09483337402344 0 417.94394
149.3869171142578 0 495.95865
152.07144165039062 0 496.90854
169.73048400878906 0 458.26556
173.44985961914062 0 936.24664
191.1031036376953 0 2864.9387 y 12
215.13929748535156 0 3930.5222
217.08233642578125 0 1219.9683
242.1505126953125 0 2931.2283
249.1602020263672 0 4505.867
250.1638946533203 0 669.35645
265.1549377441406 0 2172.5303
277.1549072265625 0 7255.1636
278.1582946777344 0 995.2519
288.1189880371094 0 682.56226 y Water loss 11
291.1343994140625 0 2712.6265
293.1498718261719 0 4096.88
294.1523742675781 0 596.4282
299.06146240234375 0 1113.396
310.3763122558594 0 606.15485
316.8711853027344 0 588.8768
332.1077880859375 0 720.4351
355.0697326660156 0 814.24854
356.0699768066406 0 1166.3136
357.06842041015625 0 1040.8005
390.2392272949219 0 2491.753
403.14678955078125 0 812.0716 y Water loss 10
404.21759033203125 0 755.63074
405.2149963378906 0 560.4666
406.235107421875 0 1017.42664
445.1932373046875 0 1181.2086
461.20355224609375 0 693.4325
493.0553894042969 0 601.59534
518.296142578125 0 1004.7282 c Ammonia loss 3
534.2408447265625 0 2167.9495 y 9
536.9913330078125 0 551.1657
568.2992553710938 0 519.73596 c Ammonia loss 8
593.2915649414062 0 1300.5859
631.2962036132812 0 905.1959 y Water loss 3
640.8060913085938 0 989.95734
652.819091796875 0 720.5082
665.3445434570312 0 842.45374
666.3346557617188 0 977.3837
673.352294921875 0 610.9088
675.3307495117188 0 642.44037
682.3541259765625 0 1472.4237 c 4
682.8547973632812 0 1509.6589
688.8428344726562 0 1592.115 z 2
689.3463134765625 0 1353.2886
689.8467407226562 0 1020.93835
690.34130859375 0 4360.5923 y 8
691.345947265625 0 2030.3296
696.8455200195312 0 3227.883 y 2
697.3453979492188 0 4108.921
697.8438720703125 0 1230.8677
733.8463745117188 0 771.0927
739.8689575195312 0 3240.282 c Ammonia loss 11
740.3696899414062 0 2959.95
740.873046875 0 1405.1882
749.3150024414062 0 600.4913
778.377685546875 0 723.43207 y 1
778.8735961914062 0 1361.919
779.3750610351562 0 801.0838 c Ammonia loss 5
781.38330078125 0 927.81055
781.8793334960938 0 1143.0088
783.36865234375 0 644.7024
787.40869140625 0 1880.1497 z 7
788.4163208007812 0 2651.354
789.4152221679688 0 806.3468
790.3941650390625 0 2554.6387 c Ammonia loss 12
790.892822265625 0 2652.847
791.3948364257812 0 1871.6692
794.4057006835938 0 778.7112
802.9205932617188 0 784.593
803.42431640625 0 2888.7446 y 7
825.9107055664062 0 2790.1213
826.4122314453125 0 3738.965
826.9083251953125 0 2363.7031
827.4112548828125 0 899.2555
834.4107055664062 0 786.74414
834.9171752929688 0 11601.05
835.4183349609375 0 10678.079
835.9175415039062 0 6899.2163
836.4161987304688 0 2977.0032
874.441162109375 0 638.6533 z 6
875.4473266601562 0 3300.8508
876.4507446289062 0 1505.2573
890.4558715820312 0 1868.3049 y 6
891.459228515625 0 1267.9766
975.1422119140625 0 612.8225
979.486083984375 0 999.7889 c Ammonia loss 7
984.450927734375 0 1052.2471
988.4825439453125 0 1453.0153 z 5
989.4901733398438 0 25522.582
990.4931640625 0 13487.82
991.4945678710938 0 3082.294
1003.4849243164062 0 2401.838
1004.5001220703125 0 11341.017 y 5
1005.50048828125 0 7124.6177
1006.4990234375 0 2180.416
1073.53515625 0 2577.4976
1074.537353515625 0 2070.2031
1088.5194091796875 0 746.05115
1108.6043701171875 0 3082.7021
1109.6094970703125 0 1738.3964
1117.5220947265625 0 815.3383 z Water loss 4
1118.5633544921875 0 786.9933
1119.522216796875 0 654.3105
1120.5302734375 0 2601.2275
1121.530029296875 0 1188.6743
1122.5426025390625 0 663.8957
1135.5274658203125 0 7076.3784 z 4
1136.528076171875 0 7374.4546
1137.5311279296875 0 3631.5698
1138.532470703125 0 1177.1509
1150.616455078125 0 621.3517
1151.534912109375 0 2486.9062 y 4
1151.6199951171875 0 2123.9077
1152.616943359375 0 9334.745 c 8
1153.619384765625 0 5017.8247
1154.6226806640625 0 2757.831
1155.6268310546875 0 1019.33075
1211.535400390625 0 833.80975
1248.6865234375 0 876.2698 c Ammonia loss 9
1249.6719970703125 0 1466.4106
1250.6905517578125 0 772.0019
1262.5751953125 0 1379.2458 y Ammonia loss 3
1263.583984375 0 4929.805 z 3
1264.58203125 0 2566.7805
1264.702880859375 0 2346.1902
1265.5572509765625 0 1057.1372
1265.697509765625 0 5390.5967 c 9
1266.7027587890625 0 3333.7573
1267.7052001953125 0 1176.3256
1279.5970458984375 0 2782.8767 y 3
1280.5948486328125 0 2357.9878
1281.59765625 0 896.8596
1317.6126708984375 0 689.6428
1318.6158447265625 0 726.1849
1319.6357421875 0 756.8931
1325.644775390625 0 748.22656
1333.6109619140625 0 2177.7065 w 2
1334.609375 0 2106.6584
1335.5960693359375 0 853.188
1358.663330078125 0 762.1005 z Water loss 2
1360.6641845703125 0 3303.0298
1361.6690673828125 0 4145.2935
1362.67431640625 0 3163.319
1363.6873779296875 0 2015.0208 c Ammonia loss 10
1364.7012939453125 0 1039.3182
1366.6710205078125 0 1799.2161
1367.6739501953125 0 1032.3044
1374.65869140625 0 901.2575 y Water loss 2
1376.666259765625 0 14072.32 z 2
1377.6688232421875 0 10608.058
1378.6728515625 0 7258.2266
1379.7120361328125 0 6513.014
1380.726318359375 0 12937.348 c 10
1381.725830078125 0 9079.867
1382.72509765625 0 4832.679
1392.6793212890625 0 3064.873 y 2
1393.679931640625 0 3009.9788
1394.6795654296875 0 1415.9766
1403.6661376953125 0 977.2095
1424.6729736328125 0 3499.9453
1425.6768798828125 0 4476.1973
1426.683837890625 0 1938.8495
1427.6864013671875 0 827.2393
1440.66748046875 0 1019.4569
1478.7271728515625 0 818.68335 c Ammonia loss 11
1479.7308349609375 0 1599.1692
1480.728271484375 0 1656.6852
1484.650634765625 0 1083.7599
1494.733154296875 0 1141.0524
1495.752685546875 0 5443.0864 c 11
1496.7574462890625 0 6924.7427
1497.754638671875 0 3928.8813
1498.75048828125 0 1934.8911
1539.719970703125 0 4842.967 z 1
1540.7239990234375 0 3991.2942
1541.724365234375 0 2648.2456
1542.726806640625 0 721.5155
1552.779541015625 0 1331.791
1577.7994384765625 0 1023.51056
1580.7913818359375 0 1331.2476
1581.7781982421875 0 933.95776
1582.7701416015625 0 937.4721
1589.81591796875 0 719.27637
1595.7762451171875 0 1612.8735
1596.798583984375 0 7015.7466 c 12
1597.7999267578125 0 5951.867
1598.7957763671875 0 3355.3215
1599.8013916015625 0 1824.5735
1605.826416015625 0 727.456
1606.8284912109375 0 808.03723
1607.8272705078125 0 2110.711
1608.8038330078125 0 2260.1116
1609.8023681640625 0 6465.285
1610.7919921875 0 8028.5366
1611.794921875 0 7018.5894
1612.7879638671875 0 2556.3071
1613.7857666015625 0 1660.1051
1614.7825927734375 0 1378.4276
1621.7769775390625 0 720.7771
1622.809814453125 0 1154.9539
1623.8072509765625 0 1374.89
1624.8089599609375 0 14112.223
1625.81396484375 0 13217.554
1626.8079833984375 0 8373.575
1627.80908203125 0 3672.5955
1628.8077392578125 0 1368.7487
1634.794189453125 0 936.2742
1635.7874755859375 0 2019.0966
1636.788818359375 0 1722.2843
1637.8016357421875 0 2645.8652
1638.80029296875 0 1448.0123
1639.832763671875 0 1155.5027
1641.83447265625 0 1586.3253
1642.8348388671875 0 1196.6608
1649.81005859375 0 1140.2075
1650.7958984375 0 3409.9253
1651.8145751953125 0 20454.188
1652.8134765625 0 50474.168
1653.8138427734375 0 47029.953
1654.8145751953125 0 24719.914
1655.8138427734375 0 10497.321
1656.815185546875 0 1019.00464
1666.815673828125 0 2912.376
1667.8155517578125 0 37260
1668.821044921875 0 86718.15
1669.8275146484375 0 97745.01
1670.830078125 0 67942.664
1671.83056640625 0 33559.164
1672.831787109375 0 13329.594
1673.8253173828125 0 1975.0387

Spectrum Details

|  |  |
| --- | --- |
| Matched peaks? Matched peaks The total absolute number of peaks matched. Additionally in brackets the total fraction of peaks matched and the total number of peaks is shown. | 42 (16.09% of 261) |
| FDR? FDR The false discovery rate estimated for this peptide. It is calculated by matching all theoretical fragments with a non-integer shift with the raw peaks for this spectrum. This is done with 40 different shifts. The resulting percentage is the average number of annotated peaks over the number of annotated peaks with the correct spectrum. | 2.32% |
| Satellite FDR? Satellite FDR See the FDR for details on its calculation. This satellite ion specific FDR only contains the satellite ions (d/w) for I/L/J positions. | 0.00% |
| PSM Score? PSM Score The PSM Score as given by Hecklib to this annotated spectrum. It is shown with three significant figures. | 424 |

#### Spectrum 8157? Spectrum 8157 The raw spectrum of this peptide as annotated by Hecklib. The fragments are coloured according to ion type (see legend). Any peaks with a star '\*' as text can be hovered over to see the full details, first the ion type second the mass shift type. By hovering over the amino acids in the peptide or ions in the legend the corresponding peaks are highlighted. By toggling the 'Unassigned' label you can turn the background (unassigned) peaks on or off in the plot. By updating the slider in the Ion legend you can update the spectrum to only show the top X% of the peaks with labels. The top X% means any peak that is within X% of the highest intensity. By dragging in the spectrum you can zoom in to a specific part of the spectrum and use 'Zoom Out' to get back to the original zoom level. The annotation of the spectrum is based on the given sequence in the peptides file and is done with different software so inconsistencies are likely. The peaks are annotated based on the given sequence, with 20 ppm tolerance.

Copy Data

##### Spectrum 8157 (TSV)

###### Preview

```
Loading example...
```

*Click on the button to copy the data to your clipboard.*

Mz MinMz MaxIntensity Max

WidthHeightPeptide font sizePeptide stroke widthSpectrum font sizeSpectrum stroke widthCompact peptide

Ion legend

wxyz

abcd

OtherUnassignedIon^Charge^_Position_Show for top:%

JYLQMNSJRJDDTA

04.19e+48.38e+41.26e+51.68e+5

Zoom Out

y^+1^_2_y^+1^_3_y^+1^_4_c^+1^_3_y^+1^_4_y^+1^_5_c^+1^_4_y^+1^_5_c^+2^_9_c^+2^_10_y^+2^_11_y^+2^_11_z^+2^_11_y^+2^_11_c^+1^_5_z^+1^_6_z^+2^_12_c^+1^_5_y^+2^_12_y^+2^_12_z^+2^_12_y^+1^_6_y^+2^_12_c^+2^_12_y^+2^_13_y^+2^_13_c^+1^_6_z^+1^_7_c^+2^_13_y^+1^_7_y^+1^_8_z^+1^_8_y^+1^_8_c^+1^_8_y^+1^_9_z^+1^_9_c^+1^_8_y^+1^_9_y^+1^_10_z^+1^_10_y^+1^_10_c^+1^_9_c^+1^_10_y^+1^_11_y^+1^_11_z^+1^_11_c^+1^_10_y^+1^_11_w^+1^_12_c^+1^_11_y^+1^_12_z^+1^_12_c^+1^_11_y^+1^_12_c^+1^_12_c^+1^_12_c^+1^_12_z^+1^_13_y^+1^_13_c^+1^_13_c^+1^_13_

0739147822182957

Fragment Matches Table

Show background peaks

| Position | Ion type | Intensity | mz Theoretical | mz Error (Th) | mz Error (ppm) | Charge | Series Number |
| --- | --- | --- | --- | --- | --- | --- | --- |
| - | - | 426.5 | 125.7 | - | - | 0 | - |
| - | - | 335.9 | 128.7 | - | - | 0 | - |
| - | - | 463.5 | 129.9 | - | - | 0 | - |
| - | - | 388.1 | 130.4 | - | - | 0 | - |
| - | - | 393.2 | 134.8 | - | - | 0 | - |
| - | - | 2585 | 136.1 | - | - | 0 | - |
| - | - | 336.7 | 136.1 | - | - | 0 | - |
| - | - | 475.8 | 148.8 | - | - | 0 | - |
| - | - | 502.2 | 148.9 | - | - | 0 | - |
| - | - | 664.6 | 148.9 | - | - | 0 | - |
| - | - | 522.4 | 148.9 | - | - | 0 | - |
| - | - | 654.2 | 148.9 | - | - | 0 | - |
| - | - | 735.9 | 148.9 | - | - | 0 | - |
| - | - | 1053 | 148.9 | - | - | 0 | - |
| - | - | 1193 | 148.9 | - | - | 0 | - |
| - | - | 2297 | 148.9 | - | - | 0 | - |
| - | - | 3783 | 148.9 | - | - | 0 | - |
| - | - | 3624 | 149 | - | - | 0 | - |
| - | - | 1999 | 149 | - | - | 0 | - |
| - | - | 1156 | 149 | - | - | 0 | - |
| - | - | 993.9 | 149 | - | - | 0 | - |
| - | - | 536 | 149 | - | - | 0 | - |
| - | - | 565.6 | 149 | - | - | 0 | - |
| - | - | 460.4 | 149 | - | - | 0 | - |
| - | - | 457 | 170.6 | - | - | 0 | - |
| - | - | 450.8 | 170.9 | - | - | 0 | - |
| - | - | 785.3 | 173.4 | - | - | 0 | - |
| 13 | y | 4110 | 191.1 | 4.299E-05 | 0.225 | +1 | 2 |
| - | - | 3730 | 203.1 | - | - | 0 | - |
| - | - | 484.5 | 215.3 | - | - | 0 | - |
| - | - | 1813 | 217.1 | - | - | 0 | - |
| - | - | 818.8 | 227.1 | - | - | 0 | - |
| - | - | 3227 | 242.2 | - | - | 0 | - |
| - | - | 1091 | 245.2 | - | - | 0 | - |
| - | - | 1.053E+04 | 249.2 | - | - | 0 | - |
| - | - | 1036 | 250.2 | - | - | 0 | - |
| - | - | 634.2 | 265.2 | - | - | 0 | - |
| - | - | 743.4 | 274.1 | - | - | 0 | - |
| - | - | 1.945E+04 | 277.2 | - | - | 0 | - |
| - | - | 2430 | 278.2 | - | - | 0 | - |
| - | - | 592 | 293.1 | - | - | 0 | - |
| 12 | y | 1840 | 306.1 | 0.0004895 | 1.599 | +1 | 3 |
| - | - | 1569 | 332.1 | - | - | 0 | - |
| - | - | 643.6 | 340.2 | - | - | 0 | - |
| - | - | 846.6 | 355.1 | - | - | 0 | - |
| - | - | 1018 | 357.1 | - | - | 0 | - |
| - | - | 610.9 | 384.1 | - | - | 0 | - |
| - | - | 6036 | 390.2 | - | - | 0 | - |
| 11 | y | 1698 | 403.1 | 5.024E-05 | 0.1246 | +1 | 4 |
| - | - | 1048 | 405.2 | - | - | 0 | - |
| 3 | c | 1003 | 407.3 | 0.0002064 | 0.5067 | +1 | 3 |
| - | - | 639.6 | 409.1 | - | - | 0 | - |
| 11 | y | 1011 | 421.2 | 3.571E-05 | 0.08479 | +1 | 4 |
| - | - | 1131 | 445.2 | - | - | 0 | - |
| - | - | 2188 | 459.2 | - | - | 0 | - |
| - | - | 2230 | 471.3 | - | - | 0 | - |
| - | - | 530.9 | 472.3 | - | - | 0 | - |
| - | - | 563.4 | 484.2 | - | - | 0 | - |
| - | - | 894.3 | 500.2 | - | - | 0 | - |
| 10 | y | 1187 | 516.2 | 2.274E-05 | 0.04406 | +1 | 5 |
| 4 | c | 3243 | 518.3 | 0.0001303 | 0.2514 | +1 | 4 |
| - | - | 966.9 | 519.3 | - | - | 0 | - |
| - | - | 702.9 | 524.2 | - | - | 0 | - |
| 10 | y | 3744 | 534.2 | 0.0006885 | 1.289 | +1 | 5 |
| - | - | 1427 | 535.2 | - | - | 0 | - |
| - | - | 644.9 | 540.8 | - | - | 0 | - |
| 9 | c | 1633 | 568.3 | 0.001532 | 2.696 | +2 | 9 |
| - | - | 1375 | 568.8 | - | - | 0 | - |
| - | - | 711 | 575.2 | - | - | 0 | - |
| - | - | 633.5 | 587.8 | - | - | 0 | - |
| - | - | 677.1 | 595.8 | - | - | 0 | - |
| - | - | 783.7 | 601.3 | - | - | 0 | - |
| - | - | 1118 | 601.8 | - | - | 0 | - |
| 10 | c | 1019 | 624.8 | 3.358E-05 | 0.05374 | +2 | 10 |
| - | - | 661 | 629.3 | - | - | 0 | - |
| 4 | y | 1888 | 631.3 | 0.0018 | 2.851 | +2 | 11 |
| 4 | y | 1029 | 631.8 | 0.007961 | 12.6 | +2 | 11 |
| 4 | z | 633 | 632.3 | 0.006185 | 9.781 | +2 | 11 |
| 4 | y | 4823 | 640.3 | 0.00256 | 3.998 | +2 | 11 |
| - | - | 4482 | 640.8 | - | - | 0 | - |
| - | - | 2115 | 641.3 | - | - | 0 | - |
| - | - | 718.9 | 643.3 | - | - | 0 | - |
| - | - | 657.6 | 643.8 | - | - | 0 | - |
| - | - | 1890 | 652.3 | - | - | 0 | - |
| - | - | 1568 | 652.8 | - | - | 0 | - |
| - | - | 663.6 | 653.3 | - | - | 0 | - |
| 5 | c | 3053 | 665.3 | 0.01223 | 18.38 | +1 | 5 |
| - | - | 1530 | 666.3 | - | - | 0 | - |
| 9 | z | 1767 | 674.3 | 3.339E-05 | 0.04952 | +1 | 6 |
| - | - | 1380 | 675.3 | - | - | 0 | - |
| - | - | 1538 | 679.3 | - | - | 0 | - |
| 3 | z | 674.7 | 679.8 | 0.003648 | 5.367 | +2 | 12 |
| 5 | c | 5541 | 682.4 | 0.001308 | 1.916 | +1 | 5 |
| - | - | 2068 | 682.9 | - | - | 0 | - |
| - | - | 1628 | 683.4 | - | - | 0 | - |
| - | - | 857.2 | 683.9 | - | - | 0 | - |
| - | - | 605.4 | 684.3 | - | - | 0 | - |
| 3 | y | 2004 | 687.8 | 0.003286 | 4.777 | +2 | 12 |
| 3 | y | 2986 | 688.3 | 0.005663 | 8.227 | +2 | 12 |
| 3 | z | 1645 | 688.8 | 0.0008959 | 1.301 | +2 | 12 |
| - | - | 1403 | 689.3 | - | - | 0 | - |
| 9 | y | 8904 | 690.3 | 0.0004076 | 0.5904 | +1 | 6 |
| - | - | 2789 | 691.3 | - | - | 0 | - |
| 3 | y | 1.081E+04 | 696.8 | 0.001849 | 2.653 | +2 | 12 |
| - | - | 1.206E+04 | 697.3 | - | - | 0 | - |
| - | - | 6538 | 697.8 | - | - | 0 | - |
| - | - | 2633 | 698.3 | - | - | 0 | - |
| - | - | 693.2 | 709.4 | - | - | 0 | - |
| - | - | 905.2 | 724.9 | - | - | 0 | - |
| - | - | 1533 | 731.4 | - | - | 0 | - |
| - | - | 771.8 | 731.9 | - | - | 0 | - |
| 12 | c | 8112 | 739.9 | 0.002015 | 2.724 | +2 | 12 |
| - | - | 5924 | 740.4 | - | - | 0 | - |
| - | - | 3762 | 740.9 | - | - | 0 | - |
| - | - | 1887 | 741.4 | - | - | 0 | - |
| - | - | 894.6 | 762.3 | - | - | 0 | - |
| - | - | 676.7 | 767.9 | - | - | 0 | - |
| - | - | 1011 | 768.4 | - | - | 0 | - |
| 2 | y | 932.7 | 769.9 | 0.006713 | 8.72 | +2 | 13 |
| - | - | 937 | 772.9 | - | - | 0 | - |
| - | - | 802.8 | 776.4 | - | - | 0 | - |
| - | - | 673.6 | 776.9 | - | - | 0 | - |
| - | - | 843.3 | 777.4 | - | - | 0 | - |
| - | - | 682.9 | 777.5 | - | - | 0 | - |
| 2 | y | 1845 | 778.4 | 3.032E-05 | 0.03895 | +2 | 13 |
| - | - | 2477 | 778.9 | - | - | 0 | - |
| 6 | c | 2695 | 779.4 | 0.005559 | 7.133 | +1 | 6 |
| - | - | 1663 | 781.4 | - | - | 0 | - |
| - | - | 2451 | 781.9 | - | - | 0 | - |
| - | - | 2100 | 782.4 | - | - | 0 | - |
| - | - | 654.9 | 782.9 | - | - | 0 | - |
| 8 | z | 3576 | 787.4 | 0.0002532 | 0.3216 | +1 | 7 |
| - | - | 2836 | 788.4 | - | - | 0 | - |
| 13 | c | 7565 | 790.4 | 0.002651 | 3.354 | +2 | 13 |
| - | - | 8319 | 790.9 | - | - | 0 | - |
| - | - | 5294 | 791.4 | - | - | 0 | - |
| - | - | 1397 | 791.9 | - | - | 0 | - |
| - | - | 1436 | 794.4 | - | - | 0 | - |
| - | - | 699 | 799.4 | - | - | 0 | - |
| - | - | 890.7 | 802.4 | - | - | 0 | - |
| - | - | 2927 | 802.9 | - | - | 0 | - |
| 8 | y | 7372 | 803.4 | 0.005043 | 6.277 | +1 | 7 |
| - | - | 2032 | 803.9 | - | - | 0 | - |
| - | - | 2852 | 804.4 | - | - | 0 | - |
| - | - | 664.2 | 808.9 | - | - | 0 | - |
| - | - | 947.4 | 810.4 | - | - | 0 | - |
| - | - | 711.6 | 810.9 | - | - | 0 | - |
| - | - | 889.7 | 816.9 | - | - | 0 | - |
| - | - | 1655 | 817.4 | - | - | 0 | - |
| - | - | 1612 | 817.9 | - | - | 0 | - |
| - | - | 1130 | 818.4 | - | - | 0 | - |
| - | - | 5105 | 825.9 | - | - | 0 | - |
| - | - | 7751 | 826.4 | - | - | 0 | - |
| - | - | 6252 | 826.9 | - | - | 0 | - |
| - | - | 4938 | 827.4 | - | - | 0 | - |
| - | - | 1411 | 827.9 | - | - | 0 | - |
| - | - | 2.632E+04 | 834.9 | - | - | 0 | - |
| - | - | 3.094E+04 | 835.4 | - | - | 0 | - |
| - | - | 1.834E+04 | 835.9 | - | - | 0 | - |
| - | - | 1.126E+04 | 836.4 | - | - | 0 | - |
| - | - | 2125 | 836.9 | - | - | 0 | - |
| - | - | 1057 | 859.4 | - | - | 0 | - |
| 7 | y | 1441 | 873.4 | 0.005701 | 6.527 | +1 | 8 |
| 7 | z | 858.6 | 874.4 | 0.007361 | 8.418 | +1 | 8 |
| - | - | 5005 | 875.4 | - | - | 0 | - |
| - | - | 2385 | 876.4 | - | - | 0 | - |
| - | - | 1062 | 877.4 | - | - | 0 | - |
| 7 | y | 6533 | 890.5 | 0.0002673 | 0.3002 | +1 | 8 |
| - | - | 2223 | 891.5 | - | - | 0 | - |
| - | - | 856.5 | 892.5 | - | - | 0 | - |
| - | - | 2542 | 923.3 | - | - | 0 | - |
| - | - | 696.2 | 924.3 | - | - | 0 | - |
| - | - | 886.3 | 962.5 | - | - | 0 | - |
| - | - | 780.1 | 963.5 | - | - | 0 | - |
| - | - | 848.1 | 974.5 | - | - | 0 | - |
| 8 | c | 1256 | 979.5 | 0.00464 | 4.737 | +1 | 8 |
| - | - | 722.6 | 980.5 | - | - | 0 | - |
| - | - | 1037 | 981.5 | - | - | 0 | - |
| 6 | y | 1045 | 987.5 | 0.002702 | 2.737 | +1 | 9 |
| 6 | z | 2281 | 988.5 | 0.005794 | 5.861 | +1 | 9 |
| - | - | 4.812E+04 | 989.5 | - | - | 0 | - |
| - | - | 2.855E+04 | 990.5 | - | - | 0 | - |
| - | - | 9428 | 991.5 | - | - | 0 | - |
| - | - | 1405 | 992.5 | - | - | 0 | - |
| - | - | 971.2 | 994.5 | - | - | 0 | - |
| 8 | c | 671.9 | 996.5 | 0.01629 | 16.35 | +1 | 8 |
| - | - | 3312 | 1003 | - | - | 0 | - |
| 6 | y | 1.387E+04 | 1005 | 0.001996 | 1.987 | +1 | 9 |
| - | - | 8291 | 1005 | - | - | 0 | - |
| - | - | 3399 | 1007 | - | - | 0 | - |
| - | - | 745.3 | 1007 | - | - | 0 | - |
| - | - | 691.3 | 1037 | - | - | 0 | - |
| - | - | 764.5 | 1038 | - | - | 0 | - |
| - | - | 6456 | 1074 | - | - | 0 | - |
| - | - | 5929 | 1075 | - | - | 0 | - |
| - | - | 2442 | 1076 | - | - | 0 | - |
| - | - | 1148 | 1088 | - | - | 0 | - |
| - | - | 1855 | 1089 | - | - | 0 | - |
| - | - | 621.4 | 1090 | - | - | 0 | - |
| - | - | 695 | 1094 | - | - | 0 | - |
| - | - | 5074 | 1109 | - | - | 0 | - |
| - | - | 4715 | 1110 | - | - | 0 | - |
| - | - | 1579 | 1111 | - | - | 0 | - |
| - | - | 777.9 | 1121 | - | - | 0 | - |
| 5 | y | 667.1 | 1134 | 0.0003619 | 0.3192 | +1 | 10 |
| 5 | z | 1.576E+04 | 1136 | 0.005347 | 4.709 | +1 | 10 |
| - | - | 1.858E+04 | 1137 | - | - | 0 | - |
| - | - | 9577 | 1138 | - | - | 0 | - |
| - | - | 3754 | 1139 | - | - | 0 | - |
| - | - | 685.8 | 1140 | - | - | 0 | - |
| - | - | 910.8 | 1149 | - | - | 0 | - |
| 5 | y | 9548 | 1152 | 0.006277 | 5.451 | +1 | 10 |
| 9 | c | 2.135E+04 | 1153 | 0.0008814 | 0.7647 | +1 | 9 |
| - | - | 1.671E+04 | 1154 | - | - | 0 | - |
| - | - | 7986 | 1155 | - | - | 0 | - |
| - | - | 1967 | 1156 | - | - | 0 | - |
| - | - | 746.5 | 1162 | - | - | 0 | - |
| - | - | 1226 | 1166 | - | - | 0 | - |
| - | - | 1408 | 1167 | - | - | 0 | - |
| - | - | 751.9 | 1168 | - | - | 0 | - |
| - | - | 788.5 | 1187 | - | - | 0 | - |
| - | - | 842.6 | 1203 | - | - | 0 | - |
| - | - | 1004 | 1205 | - | - | 0 | - |
| - | - | 1417 | 1216 | - | - | 0 | - |
| - | - | 1102 | 1217 | - | - | 0 | - |
| - | - | 2611 | 1218 | - | - | 0 | - |
| - | - | 1397 | 1219 | - | - | 0 | - |
| - | - | 800.7 | 1220 | - | - | 0 | - |
| - | - | 685.4 | 1233 | - | - | 0 | - |
| 10 | c | 2853 | 1249 | 0.003059 | 2.45 | +1 | 10 |
| - | - | 2715 | 1250 | - | - | 0 | - |
| - | - | 1511 | 1251 | - | - | 0 | - |
| - | - | 1214 | 1254 | - | - | 0 | - |
| 4 | y | 825.5 | 1262 | 0.01283 | 10.17 | +1 | 11 |
| 4 | y | 1785 | 1263 | 0.005132 | 4.065 | +1 | 11 |
| 4 | z | 8124 | 1264 | 0.0006029 | 0.4771 | +1 | 11 |
| - | - | 9954 | 1265 | - | - | 0 | - |
| - | - | 6883 | 1265 | - | - | 0 | - |
| - | - | 5082 | 1266 | - | - | 0 | - |
| 10 | c | 1.195E+04 | 1266 | 0.0004189 | 0.331 | +1 | 10 |
| - | - | 1263 | 1267 | - | - | 0 | - |
| - | - | 8886 | 1267 | - | - | 0 | - |
| - | - | 4252 | 1268 | - | - | 0 | - |
| - | - | 1132 | 1269 | - | - | 0 | - |
| - | - | 785.3 | 1272 | - | - | 0 | - |
| 4 | y | 9601 | 1280 | 0.0002989 | 0.2336 | +1 | 11 |
| - | - | 7302 | 1281 | - | - | 0 | - |
| - | - | 4648 | 1282 | - | - | 0 | - |
| - | - | 2122 | 1283 | - | - | 0 | - |
| - | - | 4658 | 1288 | - | - | 0 | - |
| - | - | 5369 | 1289 | - | - | 0 | - |
| - | - | 3109 | 1290 | - | - | 0 | - |
| - | - | 1207 | 1291 | - | - | 0 | - |
| - | - | 957.1 | 1304 | - | - | 0 | - |
| - | - | 831.8 | 1306 | - | - | 0 | - |
| - | - | 938.3 | 1312 | - | - | 0 | - |
| - | - | 647.9 | 1318 | - | - | 0 | - |
| - | - | 816.5 | 1319 | - | - | 0 | - |
| - | - | 1110 | 1320 | - | - | 0 | - |
| - | - | 809.8 | 1321 | - | - | 0 | - |
| - | - | 746.4 | 1322 | - | - | 0 | - |
| - | - | 871.8 | 1325 | - | - | 0 | - |
| - | - | 1397 | 1329 | - | - | 0 | - |
| 3 | w | 8213 | 1334 | 0.004639 | 3.479 | +1 | 12 |
| - | - | 6670 | 1335 | - | - | 0 | - |
| - | - | 7216 | 1336 | - | - | 0 | - |
| - | - | 2296 | 1337 | - | - | 0 | - |
| - | - | 863.6 | 1338 | - | - | 0 | - |
| 11 | c | 1060 | 1364 | 0.01347 | 9.877 | +1 | 11 |
| - | - | 2233 | 1365 | - | - | 0 | - |
| - | - | 1416 | 1366 | - | - | 0 | - |
| 3 | y | 961.7 | 1376 | 0.005663 | 4.116 | +1 | 12 |
| 3 | z | 2.437E+04 | 1377 | 0.00211 | 1.533 | +1 | 12 |
| - | - | 2.174E+04 | 1378 | - | - | 0 | - |
| - | - | 1.081E+04 | 1379 | - | - | 0 | - |
| - | - | 1.358E+04 | 1380 | - | - | 0 | - |
| 11 | c | 3.203E+04 | 1381 | 0.0008727 | 0.632 | +1 | 11 |
| - | - | 2.679E+04 | 1382 | - | - | 0 | - |
| - | - | 1.38E+04 | 1383 | - | - | 0 | - |
| - | - | 4041 | 1384 | - | - | 0 | - |
| - | - | 814.2 | 1385 | - | - | 0 | - |
| 3 | y | 9589 | 1393 | 0.0002319 | 0.1665 | +1 | 12 |
| - | - | 9131 | 1394 | - | - | 0 | - |
| - | - | 9578 | 1395 | - | - | 0 | - |
| - | - | 5261 | 1396 | - | - | 0 | - |
| - | - | 2441 | 1397 | - | - | 0 | - |
| - | - | 906.8 | 1454 | - | - | 0 | - |
| - | - | 734.6 | 1463 | - | - | 0 | - |
| 12 | c | 912.4 | 1478 | 0.009072 | 6.139 | +1 | 12 |
| 12 | c | 1846 | 1479 | 0.0004424 | 0.2992 | +1 | 12 |
| - | - | 1755 | 1480 | - | - | 0 | - |
| - | - | 1868 | 1481 | - | - | 0 | - |
| - | - | 1095 | 1482 | - | - | 0 | - |
| - | - | 1715 | 1484 | - | - | 0 | - |
| - | - | 1492 | 1485 | - | - | 0 | - |
| 12 | c | 1.223E+04 | 1496 | 0.000594 | 0.3971 | +1 | 12 |
| - | - | 1.595E+04 | 1497 | - | - | 0 | - |
| - | - | 9163 | 1498 | - | - | 0 | - |
| - | - | 3495 | 1499 | - | - | 0 | - |
| - | - | 1050 | 1504 | - | - | 0 | - |
| - | - | 2613 | 1505 | - | - | 0 | - |
| - | - | 851.5 | 1506 | - | - | 0 | - |
| - | - | 891.7 | 1515 | - | - | 0 | - |
| - | - | 1464 | 1535 | - | - | 0 | - |
| 2 | z | 8310 | 1540 | 0.0007934 | 0.5153 | +1 | 13 |
| - | - | 1.199E+04 | 1541 | - | - | 0 | - |
| - | - | 6647 | 1542 | - | - | 0 | - |
| - | - | 2623 | 1543 | - | - | 0 | - |
| - | - | 1460 | 1553 | - | - | 0 | - |
| - | - | 1100 | 1554 | - | - | 0 | - |
| - | - | 741.4 | 1555 | - | - | 0 | - |
| 2 | y | 827.7 | 1556 | 0.003648 | 2.345 | +1 | 13 |
| - | - | 1278 | 1569 | - | - | 0 | - |
| - | - | 1090 | 1570 | - | - | 0 | - |
| - | - | 942.5 | 1578 | - | - | 0 | - |
| 13 | c | 944.2 | 1579 | 0.0132 | 8.358 | +1 | 13 |
| - | - | 1488 | 1581 | - | - | 0 | - |
| - | - | 1449 | 1582 | - | - | 0 | - |
| - | - | 1221 | 1583 | - | - | 0 | - |
| - | - | 1190 | 1590 | - | - | 0 | - |
| - | - | 1828 | 1591 | - | - | 0 | - |
| - | - | 760.3 | 1592 | - | - | 0 | - |
| - | - | 1098 | 1596 | - | - | 0 | - |
| 13 | c | 1.373E+04 | 1597 | 0.005182 | 3.245 | +1 | 13 |
| - | - | 1.645E+04 | 1598 | - | - | 0 | - |
| - | - | 9261 | 1599 | - | - | 0 | - |
| - | - | 3562 | 1600 | - | - | 0 | - |
| - | - | 1291 | 1601 | - | - | 0 | - |
| - | - | 1643 | 1606 | - | - | 0 | - |
| - | - | 2305 | 1607 | - | - | 0 | - |
| - | - | 3595 | 1608 | - | - | 0 | - |
| - | - | 3187 | 1609 | - | - | 0 | - |
| - | - | 1.292E+04 | 1610 | - | - | 0 | - |
| - | - | 2.446E+04 | 1611 | - | - | 0 | - |
| - | - | 1.722E+04 | 1612 | - | - | 0 | - |
| - | - | 9483 | 1613 | - | - | 0 | - |
| - | - | 5776 | 1614 | - | - | 0 | - |
| - | - | 3701 | 1615 | - | - | 0 | - |
| - | - | 2191 | 1616 | - | - | 0 | - |
| - | - | 1584 | 1621 | - | - | 0 | - |
| - | - | 1208 | 1622 | - | - | 0 | - |
| - | - | 824.7 | 1623 | - | - | 0 | - |
| - | - | 1023 | 1624 | - | - | 0 | - |
| - | - | 2.842E+04 | 1625 | - | - | 0 | - |
| - | - | 2.597E+04 | 1626 | - | - | 0 | - |
| - | - | 1.895E+04 | 1627 | - | - | 0 | - |
| - | - | 1.007E+04 | 1628 | - | - | 0 | - |
| - | - | 2758 | 1629 | - | - | 0 | - |
| - | - | 970.5 | 1630 | - | - | 0 | - |
| - | - | 752.9 | 1633 | - | - | 0 | - |
| - | - | 1710 | 1635 | - | - | 0 | - |
| - | - | 4636 | 1636 | - | - | 0 | - |
| - | - | 5833 | 1637 | - | - | 0 | - |
| - | - | 4517 | 1638 | - | - | 0 | - |
| - | - | 2589 | 1639 | - | - | 0 | - |
| - | - | 1541 | 1640 | - | - | 0 | - |
| - | - | 6057 | 1642 | - | - | 0 | - |
| - | - | 5396 | 1643 | - | - | 0 | - |
| - | - | 3353 | 1644 | - | - | 0 | - |
| - | - | 1545 | 1645 | - | - | 0 | - |
| - | - | 2969 | 1651 | - | - | 0 | - |
| - | - | 1.061E+04 | 1652 | - | - | 0 | - |
| - | - | 7.276E+04 | 1653 | - | - | 0 | - |
| - | - | 8.098E+04 | 1654 | - | - | 0 | - |
| - | - | 5.141E+04 | 1655 | - | - | 0 | - |
| - | - | 2.248E+04 | 1656 | - | - | 0 | - |
| - | - | 5604 | 1657 | - | - | 0 | - |
| - | - | 883.5 | 1667 | - | - | 0 | - |
| - | - | 5810 | 1668 | - | - | 0 | - |
| - | - | 8.677E+04 | 1669 | - | - | 0 | - |
| - | - | 1.66E+05 | 1670 | - | - | 0 | - |
| - | - | 1.44E+05 | 1671 | - | - | 0 | - |
| - | - | 8.625E+04 | 1672 | - | - | 0 | - |
| - | - | 3.118E+04 | 1673 | - | - | 0 | - |
| - | - | 4832 | 1674 | - | - | 0 | - |
| - | - | 706.7 | 2491 | - | - | 0 | - |
| - | - | 1331 | 2505 | - | - | 0 | - |
| - | - | 1040 | 2506 | - | - | 0 | - |
| - | - | 1393 | 2507 | - | - | 0 | - |
| - | - | 806.7 | 2510 | - | - | 0 | - |
| - | - | 727.6 | 2927 | - | - | 0 | - |

m/z Charge Intensity FragmentType MassShift Position
125.71434783935547 0 426.47607
128.74618530273438 0 335.87698
129.8897247314453 0 463.47174
130.41773986816406 0 388.0849
134.7777862548828 0 393.24564
136.07591247558594 0 2585.3389
136.08108520507812 0 336.6756
148.8481903076172 0 475.82516
148.8843994140625 0 502.22787
148.8919677734375 0 664.5811
148.8992919921875 0 522.4412
148.9069061279297 0 654.22437
148.913818359375 0 735.90076
148.92112731933594 0 1052.6171
148.9285430908203 0 1192.606
148.93572998046875 0 2296.5024
148.9436492919922 0 3782.7102
148.96029663085938 0 3624.3677
148.9681396484375 0 1998.5564
148.97543334960938 0 1155.9783
148.982666015625 0 993.9405
148.989990234375 0 535.9661
148.9968719482422 0 565.6084
149.01170349121094 0 460.36057
170.5755615234375 0 456.97772
170.9119415283203 0 450.81775
173.44009399414062 0 785.32166
191.10267639160156 0 4109.818 y 12
203.10264587402344 0 3730.4082
215.29840087890625 0 484.4988
217.0819854736328 0 1813.2183
227.102783203125 0 818.82806
242.1500701904297 0 3227.3313
245.18577575683594 0 1091.0001
249.15989685058594 0 10525.717
250.16371154785156 0 1035.7883
265.1540832519531 0 634.21295
274.13909912109375 0 743.4078
277.1546936035156 0 19450.72
278.15814208984375 0 2430.0908
293.14813232421875 0 592.0054
306.13006591796875 0 1839.8037 y 11
332.1092529296875 0 1569.0924
340.18585205078125 0 643.6491
355.06964111328125 0 846.61774
357.0683288574219 0 1017.7475
384.05438232421875 0 610.86414
390.23876953125 0 6036.327
403.1459045410156 0 1697.5956 y Water loss 10
405.2131042480469 0 1047.686
407.26507568359375 0 1003.0045 c 2
409.1310119628906 0 639.5629
421.15655517578125 0 1010.7999 y 10
445.1934814453125 0 1131.2783
459.2086181640625 0 2187.8562
471.2812194824219 0 2229.8987
472.34942626953125 0 530.9456
484.2132873535156 0 563.3727
500.2490539550781 0 894.27826
516.2300415039062 0 1187.2057 y Water loss 9
518.2971801757812 0 3242.6165 c Ammonia loss 3
519.3008422851562 0 966.9385
524.2094116210938 0 702.8586
534.2412719726562 0 3744.1 y 9
535.2445068359375 0 1427.0038
540.7546997070312 0 644.9237
568.2991333007812 0 1632.9385 c Ammonia loss 8
568.8026733398438 0 1374.7076
575.2182006835938 0 711.0106
587.7734375 0 633.4533
595.7807006835938 0 677.0718
601.3319702148438 0 783.7147
601.7966918945312 0 1117.6825
624.839599609375 0 1019.3657 c Ammonia loss 9
629.32470703125 0 660.97144
631.2950439453125 0 1887.504 y Water loss 3
631.793212890625 0 1029.4321 y Ammonia loss 3
632.2953491210938 0 632.95703 z 3
640.3010864257812 0 4823.1157 y 3
640.8012084960938 0 4481.772
641.301513671875 0 2115.383
643.3139038085938 0 718.8535
643.8092651367188 0 657.559
652.3189697265625 0 1890.363
652.8184204101562 0 1567.5359
653.3189697265625 0 663.57886
665.3400268554688 0 3052.5964 c Ammonia loss 4
666.3377685546875 0 1529.6744
674.3229370117188 0 1766.7352 z 8
675.3299560546875 0 1379.6675
679.3259887695312 0 1538.0629
679.822265625 0 674.65466 z Water loss 2
682.3556518554688 0 5540.8994 c 4
682.8565673828125 0 2068.2532
683.35107421875 0 1628.441
683.8508911132812 0 857.1978
684.327392578125 0 605.40375
687.8385620117188 0 2004.1145 y Water loss 2
688.3329467773438 0 2985.551 y Ammonia loss 2
688.8320922851562 0 1644.5911 z 2
689.3347778320312 0 1402.7888
690.3421020507812 0 8904.114 y 8
691.3431396484375 0 2789.483
696.8424072265625 0 10810.265 y 2
697.3421630859375 0 12058.983
697.84228515625 0 6537.9536
698.3366088867188 0 2632.8564
709.3690795898438 0 693.2254
724.8529052734375 0 905.1729
731.3607788085938 0 1533.0742
731.85546875 0 771.8044
739.8685913085938 0 8112.044 c Ammonia loss 11
740.3695068359375 0 5924.313
740.868896484375 0 3762.3416
741.3695678710938 0 1886.6797
762.3465576171875 0 894.5867
767.8864135742188 0 676.7365
768.3911743164062 0 1011.49384
769.8656616210938 0 932.69965 y Ammonia loss 1
772.872802734375 0 937.019
776.3972778320312 0 802.7771
776.8893432617188 0 673.5678
777.396240234375 0 843.3086
777.5327758789062 0 682.88226
778.3721923828125 0 1844.5807 y 1
778.8726196289062 0 2476.7515
779.3762817382812 0 2694.848 c Ammonia loss 5
781.3839721679688 0 1662.7355
781.8844604492188 0 2450.5276
782.3833618164062 0 2100.1492
782.8802490234375 0 654.9085
787.4072875976562 0 3576.108 z 7
788.4127197265625 0 2836.3115
790.39306640625 0 7564.653 c Ammonia loss 12
790.8924560546875 0 8318.663
791.39306640625 0 5294.0835
791.8941650390625 0 1397.3191
794.4072875976562 0 1435.6261
799.400390625 0 699.019
802.4108276367188 0 890.6591
802.916748046875 0 2926.774
803.4207153320312 0 7371.94 y 7
803.917724609375 0 2032.4216
804.424072265625 0 2851.5613
808.8939208984375 0 664.2079
810.4059448242188 0 947.35065
810.9070434570312 0 711.60876
816.9069213867188 0 889.682
817.4013671875 0 1655.3091
817.9022827148438 0 1612.365
818.396240234375 0 1129.6467
825.910888671875 0 5104.9473
826.4080200195312 0 7751.464
826.9083862304688 0 6252.139
827.4082641601562 0 4938.151
827.9082641601562 0 1411.1655
834.9163208007812 0 26316.938
835.4156494140625 0 30944.318
835.9163818359375 0 18340.098
836.417724609375 0 11262.974
836.917724609375 0 2124.5679
859.4420776367188 0 1057.4355
873.425537109375 0 1440.8225 y Ammonia loss 6
874.4317016601562 0 858.6253 z 6
875.4470825195312 0 5004.612
876.4473266601562 0 2384.7683
877.43798828125 0 1061.7369
890.45751953125 0 6532.5474 y 6
891.4601440429688 0 2222.8252
892.4682006835938 0 856.4783
923.3400268554688 0 2541.7798
924.3380737304688 0 696.17065
962.4700927734375 0 886.3422
963.4623413085938 0 780.0829
974.466796875 0 848.14307
979.491455078125 0 1256.1257 c Ammonia loss 7
980.4862060546875 0 722.64594
981.4769897460938 0 1036.9429
987.4768676757812 0 1044.791 y Ammonia loss 5
988.4761962890625 0 2281.1248 z 5
989.4867553710938 0 48122.074
990.488037109375 0 28553.717
991.4904174804688 0 9428.113
992.4921264648438 0 1405.1432
994.496826171875 0 971.1539
996.4970703125 0 671.91095 c 7
1003.4905395507812 0 3311.8606
1004.4987182617188 0 13872.411 y 5
1005.4988403320312 0 8290.575
1006.5012817382812 0 3398.8225
1007.4794311523438 0 745.287
1037.4200439453125 0 691.27356
1038.4169921875 0 764.51117
1073.53271484375 0 6456.228
1074.5335693359375 0 5929.394
1075.5340576171875 0 2441.7478
1087.529541015625 0 1147.5985
1088.532958984375 0 1855.0193
1089.5257568359375 0 621.4369
1093.5164794921875 0 694.969
1108.6036376953125 0 5073.913
1109.6065673828125 0 4714.804
1110.607666015625 0 1578.8787
1120.6072998046875 0 777.9382
1133.52099609375 0 667.0822 y Water loss 4
1135.517822265625 0 15758.765 z 4
1136.5185546875 0 18582.25
1137.519775390625 0 9576.633
1138.517822265625 0 3753.8499
1139.5145263671875 0 685.75854
1149.42041015625 0 910.80865
1151.5374755859375 0 9548.165 y 4
1152.6153564453125 0 21351.057 c 8
1153.615966796875 0 16713.297
1154.6146240234375 0 7985.703
1155.6156005859375 0 1967.4594
1161.5908203125 0 746.516
1166.4619140625 0 1225.6475
1167.4599609375 0 1408.1324
1168.4600830078125 0 751.9104
1186.5628662109375 0 788.5206
1202.5758056640625 0 842.61816
1204.5909423828125 0 1004.2264
1215.5997314453125 0 1416.9219
1216.588134765625 0 1101.9103
1217.6029052734375 0 2610.5798
1218.604248046875 0 1397.2847
1219.605712890625 0 800.7375
1232.609130859375 0 685.3981
1248.675048828125 0 2853.2935 c Ammonia loss 9
1249.67578125 0 2715.2285
1250.6640625 0 1510.7224
1253.6063232421875 0 1214.279
1261.592041015625 0 825.5496 y Water loss 3
1262.568359375 0 1785.4568 y Ammonia loss 3
1263.5716552734375 0 8124.4053 z 3
1264.5640869140625 0 9953.955
1264.6998291015625 0 6883.282
1265.5654296875 0 5082.4033
1265.6981201171875 0 11951.099 c 9
1266.570068359375 0 1263.079
1266.6964111328125 0 8886.487
1267.697265625 0 4251.83
1268.6912841796875 0 1131.6732
1271.6763916015625 0 785.3002
1279.5894775390625 0 9600.778 y 3
1280.58740234375 0 7302.454
1281.5892333984375 0 4647.9263
1282.593017578125 0 2122.2664
1287.68994140625 0 4658.3384
1288.693603515625 0 5368.6675
1289.698486328125 0 3108.6138
1290.7025146484375 0 1206.8058
1303.6368408203125 0 957.08624
1305.63232421875 0 831.7782
1311.658203125 0 938.2821
1317.6805419921875 0 647.9467
1318.687255859375 0 816.4737
1319.671142578125 0 1110.2906
1320.6968994140625 0 809.76465
1321.5823974609375 0 746.394
1324.6534423828125 0 871.8186
1328.6806640625 0 1396.7063
1333.60498046875 0 8212.519 w 2
1334.6072998046875 0 6670.0596
1335.6048583984375 0 7215.99
1336.603271484375 0 2295.8447
1337.6141357421875 0 863.6049
1363.71240234375 0 1060.2163 c Ammonia loss 10
1364.69921875 0 2233.1184
1365.7030029296875 0 1415.8273
1375.6529541015625 0 961.7289 y Ammonia loss 2
1376.6572265625 0 24370.209 z 2
1377.65673828125 0 21739.305
1378.6568603515625 0 10805.341
1379.70947265625 0 13581.914
1380.724609375 0 32027.197 c 10
1381.7249755859375 0 26790.63
1382.7254638671875 0 13798.433
1383.7218017578125 0 4040.9912
1384.7186279296875 0 814.2277
1392.674072265625 0 9589.191 y 2
1393.6500244140625 0 9131.369
1394.6309814453125 0 9577.852
1395.6287841796875 0 5261.235
1396.6180419921875 0 2440.7954
1453.755615234375 0 906.75464
1462.7137451171875 0 734.5964
1477.7327880859375 0 912.3923 c Water loss 11
1478.726318359375 0 1846.4274 c Ammonia loss 11
1479.7308349609375 0 1755.3877
1480.7340087890625 0 1867.5688
1481.734130859375 0 1095.2399
1483.6588134765625 0 1714.8821
1484.6651611328125 0 1492.1637
1495.7518310546875 0 12231.725 c 11
1496.7506103515625 0 15952.62
1497.7506103515625 0 9163.302
1498.74951171875 0 3495.486
1503.7049560546875 0 1050.264
1504.7149658203125 0 2612.8423
1505.7119140625 0 851.4683
1514.7666015625 0 891.6649
1534.7781982421875 0 1463.9127
1539.71923828125 0 8310.279 z 1
1540.7188720703125 0 11985.147
1541.718994140625 0 6647.216
1542.724853515625 0 2623.0847
1552.7664794921875 0 1459.5464
1553.779052734375 0 1099.5173
1554.784423828125 0 741.3898
1555.7335205078125 0 827.6568 y 1
1568.7578125 0 1278.3165
1569.7584228515625 0 1090.0732
1577.8028564453125 0 942.4577
1578.802734375 0 944.1756 c Water loss 12
1580.802490234375 0 1488.0846
1581.7950439453125 0 1448.9746
1582.798828125 0 1221.0144
1589.8101806640625 0 1190.0315
1590.8232421875 0 1827.9333
1591.816162109375 0 760.33
1595.78857421875 0 1098.05
1596.794921875 0 13732.675 c 12
1597.797119140625 0 16447.13
1598.7928466796875 0 9260.838
1599.797119140625 0 3561.6921
1600.7962646484375 0 1291.1792
1605.8204345703125 0 1642.5352
1606.8284912109375 0 2304.5085
1607.82421875 0 3595.4824
1608.824951171875 0 3186.9248
1609.7984619140625 0 12919.819
1610.7923583984375 0 24459.953
1611.7919921875 0 17221.258
1612.7955322265625 0 9482.936
1613.780517578125 0 5775.923
1614.7667236328125 0 3700.9246
1615.7657470703125 0 2191.1345
1620.802490234375 0 1584.2012
1621.8118896484375 0 1208.4424
1622.799072265625 0 824.73
1623.785400390625 0 1022.8548
1624.8076171875 0 28415.77
1625.8089599609375 0 25973.195
1626.8056640625 0 18950.559
1627.8033447265625 0 10068.049
1628.798095703125 0 2758.2627
1629.7845458984375 0 970.53735
1632.7664794921875 0 752.9244
1634.788818359375 0 1710.1393
1635.8018798828125 0 4636.231
1636.802001953125 0 5833.3813
1637.80224609375 0 4517.3584
1638.8006591796875 0 2588.8896
1639.8125 0 1541.1744
1641.833251953125 0 6057.376
1642.835693359375 0 5395.992
1643.8336181640625 0 3352.6558
1644.8392333984375 0 1544.8141
1650.812255859375 0 2968.8247
1651.8167724609375 0 10605.686
1652.808349609375 0 72757.125
1653.8070068359375 0 80978.43
1654.8096923828125 0 51410.668
1655.8084716796875 0 22483.611
1656.8077392578125 0 5603.6387
1666.80419921875 0 883.4617
1667.7921142578125 0 5809.6885
1668.8204345703125 0 86773.58
1669.8255615234375 0 166012.28
1670.8267822265625 0 143959.17
1671.8277587890625 0 86251.41
1672.83203125 0 31177.207
1673.8275146484375 0 4832.087
2491.172607421875 0 706.6523
2505.107666015625 0 1330.6416
2506.248046875 0 1039.882
2507.17529296875 0 1392.98
2510.225830078125 0 806.6893
2927.43994140625 0 727.5869

Spectrum Details

|  |  |
| --- | --- |
| Matched peaks? Matched peaks The total absolute number of peaks matched. Additionally in brackets the total fraction of peaks matched and the total number of peaks is shown. | 61 (16.01% of 381) |
| FDR? FDR The false discovery rate estimated for this peptide. It is calculated by matching all theoretical fragments with a non-integer shift with the raw peaks for this spectrum. This is done with 40 different shifts. The resulting percentage is the average number of annotated peaks over the number of annotated peaks with the correct spectrum. | 2.26% |
| Satellite FDR? Satellite FDR See the FDR for details on its calculation. This satellite ion specific FDR only contains the satellite ions (d/w) for I/L/J positions. | 0.00% |
| PSM Score? PSM Score The PSM Score as given by Hecklib to this annotated spectrum. It is shown with three significant figures. | 628 |

#### Reverse Lookup? Reverse Lookup All places where this read could be placed.

| Group | Segment | Template | Template Part | Read Part | Score | Unique |
| --- | --- | --- | --- | --- | --- | --- |
| Homo sapiens Heavy Chain | IGHV | IGHV3-9 | [78..92] | [0..14] | 94 | False |
| Homo sapiens Heavy Chain | IGHV | IGHV3-48 | [78..92] | [0..14] | 94 | False |
| Homo sapiens Heavy Chain | IGHV | IGHV3-21 | [78..92] | [0..14] | 94 | False |
| Homo sapiens Heavy Chain | IGHV | IGHV3-7 | [78..92] | [0..14] | 94 | False |
| Homo sapiens Heavy Chain | IGHV | IGHV3-43 | [78..92] | [0..14] | 94 | False |
| Homo sapiens Heavy Chain | IGHV | IGHV3-13 | [77..91] | [0..14] | 94 | False |
| Homo sapiens Heavy Chain | IGHV | IGHV3-11 | [78..92] | [0..14] | 94 | False |
| Homo sapiens Heavy Chain | IGHV | IGHV3-74 | [78..92] | [0..14] | 94 | False |
| Homo sapiens Heavy Chain | IGHV | IGHV3-20 | [78..92] | [0..14] | 94 | False |
| Homo sapiens Heavy Chain | IGHV | IGHV3-23 | [78..92] | [0..14] | 94 | False |
| Homo sapiens Heavy Chain | IGHV | IGHV3-53 | [77..91] | [0..14] | 94 | False |
| Homo sapiens Heavy Chain | IGHV | IGHV3-66 | [77..91] | [0..14] | 94 | False |
| Homo sapiens Heavy Chain | IGHV | IGHV3-NL1 | [78..92] | [0..14] | 94 | False |
| Homo sapiens Heavy Chain | IGHV | IGHV3-30-5 | [78..92] | [0..14] | 94 | False |
| Homo sapiens Heavy Chain | IGHV | IGHV3-72 | [80..94] | [0..14] | 85 | False |
| Homo sapiens Heavy Chain | IGHV | IGHV3-30 | [78..92] | [0..14] | 94 | False |
| Homo sapiens Heavy Chain | IGHV | IGHV3-33 | [78..92] | [0..14] | 94 | False |
| Homo sapiens Heavy Chain | IGHV | IGHV3-15 | [80..94] | [0..14] | 85 | False |

| Recombined | Template Part | Read Part | Score | Unique |
| --- | --- | --- | --- | --- |
| REC-0-1 | [78..92] | [0..14] | 112 | True |

#### Meta Information from Multiple reads

##### Number of combined reads

7

##### Intensity

0.6219

##### TotalArea

3.635E+08

##### Changes to the peptide sequence

JYLQMNSJRJDDTA

J→LSupport for Leucine based on side chain ions (1 for L 0 for I) (Position: 3)

L→JNo support for either Leucine or Isoleucine based on side chain ions (Position: 8)

L→JNo support for either Leucine or Isoleucine based on side chain ions (Position: 3)

J→LSupport for Leucine based on side chain ions (1 for L 0 for I) (Position: 8)

J→LSupport for Leucine based on side chain ions (1 for L 0 for I) (Position: 3)

L→JNo support for either Leucine or Isoleucine based on side chain ions (Position: 10)

L→JNo support for either Leucine or Isoleucine based on side chain ions (Position: 8)

L→JNo support for either Leucine or Isoleucine based on side chain ions (Position: 3)

L→JNo support for either Leucine or Isoleucine based on side chain ions (Position: 1)

#### Positional Score

Copy Data

##### Positional Score (TSV)

###### Preview

```
Loading example...
```

*Click on the button to copy the data to your clipboard.*

10012345678910111213

Label Value
"0" 0.611
"1" 0.714
"2" 0.709
"3" 0.687
"4" 0.701
"5" 0.686
"6" 0.681
"7" 0.69
"8" 0.69
"9" 0.7
"10" 0.707
"11" 0.714
"12" 0.704
"13" 0.704

#### Meta Information from PEAKS

##### Scan Identifier

F1:8058

##### Original sequence

L

Y

L

Q

M

+15.99

N

S

L

R

L

D

D

T

A

##### Posttranslational Modifications

Oxidation (M)

##### Source File

D:\separate\_stitch\_analyses\xle-disambiguation\raw\20210323\_F1\_UM1\_Peng0013\_SA\_F59\_ingel\_3ug\_ELA.raw

##### Fraction

1

##### Scan Feature

F1:17847

##### De Novo Score

99

##### ConfidenceScore

99

### m/z

834.9172

##### Mass

1667.8188

##### Charge

2

##### Retention Time

43.61

##### Predicted Retention Time

-

##### Area

1.16E+08

##### Parts Per Million

0.6

##### Fragmentation mode

HCD

##### Originating file

01 D:\separate\_stitch\_analyses\xle-disambiguation\20210325\_F59\_3ug\_DENOVO\_12.csv

#### Meta Information from PEAKS

##### Scan Identifier

F1:8370

##### Original sequence

L

Y

L

Q

M

+15.99

N

S

L

R

L

D

D

T

A

##### Posttranslational Modifications

Oxidation (M)

##### Source File

D:\separate\_stitch\_analyses\xle-disambiguation\raw\20210323\_F1\_UM1\_Peng0013\_SA\_F59\_ingel\_3ug\_ELA.raw

##### Fraction

1

##### Scan Feature

F1:17847

##### De Novo Score

99

##### ConfidenceScore

99

### m/z

834.9172

##### Mass

1667.8188

##### Charge

2

##### Retention Time

43.61

##### Predicted Retention Time

-

##### Area

1.16E+08

##### Parts Per Million

0.6

##### Fragmentation mode

HCD

##### Originating file

01 D:\separate\_stitch\_analyses\xle-disambiguation\20210325\_F59\_3ug\_DENOVO\_12.csv

#### Meta Information from PEAKS

##### Scan Identifier

F1:9788

##### Original sequence

L

Y

L

Q

M

N

S

L

R

L

D

D

T

A

##### Posttranslational Modifications

##### Source File

D:\separate\_stitch\_analyses\xle-disambiguation\raw\20210323\_F1\_UM1\_Peng0013\_SA\_F59\_ingel\_3ug\_ELA.raw

##### Fraction

1

##### Scan Feature

F1:17548

##### De Novo Score

98

##### ConfidenceScore

98

### m/z

826.9215

##### Mass

1651.824

##### Charge

2

##### Retention Time

54.35

##### Predicted Retention Time

-

##### Area

6.834E+06

##### Parts Per Million

2.7

##### Fragmentation mode

ETHCD

##### Originating file

01 D:\separate\_stitch\_analyses\xle-disambiguation\20210325\_F59\_3ug\_DENOVO\_12.csv

#### Meta Information from PEAKS

##### Scan Identifier

F1:9789

##### Original sequence

L

Y

L

Q

M

+15.99

N

S

L

R

L

D

D

T

A

##### Posttranslational Modifications

Oxidation (M)

##### Source File

D:\separate\_stitch\_analyses\xle-disambiguation\raw\20210323\_F1\_UM1\_Peng0013\_SA\_F59\_ingel\_3ug\_ELA.raw

##### Fraction

1

##### Scan Feature

F1:17851

##### De Novo Score

98

##### ConfidenceScore

98

### m/z

834.9178

##### Mass

1667.8188

##### Charge

2

##### Retention Time

54.34

##### Predicted Retention Time

-

##### Area

9.395E+05

##### Parts Per Million

1.4

##### Fragmentation mode

HCD

##### Originating file

01 D:\separate\_stitch\_analyses\xle-disambiguation\20210325\_F59\_3ug\_DENOVO\_12.csv

#### Meta Information from PEAKS

##### Scan Identifier

F1:9846

##### Original sequence

L

Y

L

Q

M

N

S

L

R

L

D

D

T

A

##### Posttranslational Modifications

##### Source File

D:\separate\_stitch\_analyses\xle-disambiguation\raw\20210323\_F1\_UM1\_Peng0013\_SA\_F59\_ingel\_3ug\_ELA.raw

##### Fraction

1

##### Scan Feature

F1:17548

##### De Novo Score

97

##### ConfidenceScore

97

### m/z

826.9215

##### Mass

1651.824

##### Charge

2

##### Retention Time

54.35

##### Predicted Retention Time

-

##### Area

6.834E+06

##### Parts Per Million

2.7

##### Fragmentation mode

HCD

##### Originating file

01 D:\separate\_stitch\_analyses\xle-disambiguation\20210325\_F59\_3ug\_DENOVO\_12.csv

#### Meta Information from PEAKS

##### Scan Identifier

F1:9797

##### Original sequence

L

Y

L

Q

M

+15.99

N

S

L

R

L

D

D

T

A

##### Posttranslational Modifications

Oxidation (M)

##### Source File

D:\separate\_stitch\_analyses\xle-disambiguation\raw\20210323\_F1\_UM1\_Peng0013\_SA\_F59\_ingel\_3ug\_ELA.raw

##### Fraction

1

##### Scan Feature

F1:17851

##### De Novo Score

95

##### ConfidenceScore

95

### m/z

834.9178

##### Mass

1667.8188

##### Charge

2

##### Retention Time

54.34

##### Predicted Retention Time

-

##### Area

9.395E+05

##### Parts Per Million

1.4

##### Fragmentation mode

ETHCD

##### Originating file

01 D:\separate\_stitch\_analyses\xle-disambiguation\20210325\_F59\_3ug\_DENOVO\_12.csv

#### Meta Information from PEAKS

##### Scan Identifier

F1:8157

##### Original sequence

L

Y

L

Q

M

+15.99

N

S

L

R

L

D

D

T

A

##### Posttranslational Modifications

Oxidation (M)

##### Source File

D:\separate\_stitch\_analyses\xle-disambiguation\raw\20210323\_F1\_UM1\_Peng0013\_SA\_F59\_ingel\_3ug\_ELA.raw

##### Fraction

1

##### Scan Feature

F1:17847

##### De Novo Score

95

##### ConfidenceScore

95

### m/z

834.9172

##### Mass

1667.8188

##### Charge

2

##### Retention Time

43.61

##### Predicted Retention Time

-

##### Area

1.16E+08

##### Parts Per Million

0.6

##### Fragmentation mode

ETHCD

##### Originating file

01 D:\separate\_stitch\_analyses\xle-disambiguation\20210325\_F59\_3ug\_DENOVO\_12.csv
