## Supplementary material for "A handle on mass coincidence errors in *de novo* sequencing of antibodies by bottom-up proteomics": Combined_018.html

Details Combined\_018 | Stitch OverviewUndefined

### Read Combined\_018

#### Sequence (length=11)

JSGGIDVVAHE

#### Spectrum 5525? Spectrum 5525 The raw spectrum of this peptide as annotated by Hecklib. The fragments are coloured according to ion type (see legend). Any peaks with a star '\*' as text can be hovered over to see the full details, first the ion type second the mass shift type. By hovering over the amino acids in the peptide or ions in the legend the corresponding peaks are highlighted. By toggling the 'Unassigned' label you can turn the background (unassigned) peaks on or off in the plot. By updating the slider in the Ion legend you can update the spectrum to only show the top X% of the peaks with labels. The top X% means any peak that is within X% of the highest intensity. By dragging in the spectrum you can zoom in to a specific part of the spectrum and use 'Zoom Out' to get back to the original zoom level. The annotation of the spectrum is based on the given sequence in the peptides file and is done with different software so inconsistencies are likely. The peaks are annotated based on the given sequence, with 20 ppm tolerance.

Copy Data

##### Spectrum 5525 (TSV)

###### Preview

```
Loading example...
```

*Click on the button to copy the data to your clipboard.*

Mz MinMz MaxIntensity Max

WidthHeightPeptide font sizePeptide stroke widthSpectrum font sizeSpectrum stroke widthCompact peptide

Ion legend

wxyz

abcd

OtherUnassignedIon^Charge^_Position_Show for top:%

JSGGIDVVAHE

02.04e+54.08e+56.12e+58.15e+5

Zoom Out

y^+1^_1_y^+1^_1_a^+1^_2_a^+1^_2_d^+1^_2_a^+1^_2_b^+1^_2_b^+1^_2_b^+1^_2_a^+1^_3_a^+1^_3_y^+2^_4_b^+1^_3_b^+1^_3_b^+1^_3_y^+1^_2_y^+2^_5_y^+1^_2_b^+1^_4_b^+1^_4_b^+1^_4_y^+2^_6_y^+1^_3_y^+1^_3_y^+2^_7_y^+2^_7_b^+2^_9_b^+1^_5_y^+2^_8_b^+1^_5_y^+1^_4_y^+2^_9_y^+2^_9_y^+1^_4_b^+2^_10_b^+2^_10_y^+2^_10_b^+1^_6_y^+1^_5_\*b^+1^_6_\*y^+1^_5_b^+1^_7_b^+1^_7_y^+1^_6_y^+1^_6_b^+1^_8_b^+1^_8_y^+1^_7_y^+1^_7_b^+1^_9_b^+1^_9_y^+1^_8_y^+1^_8_y^+1^_9_y^+1^_9_b^+1^_10_y^+1^_10_y^+1^_10_

0776155223283103

Fragment Matches Table

Show background peaks

| Position | Ion type | Intensity | mz Theoretical | mz Error (Th) | mz Error (ppm) | Charge | Series Number |
| --- | --- | --- | --- | --- | --- | --- | --- |
| - | - | 1455 | 120.1 | - | - | 0 | - |
| - | - | 615.2 | 121 | - | - | 0 | - |
| - | - | 2509 | 122.1 | - | - | 0 | - |
| - | - | 2051 | 125 | - | - | 0 | - |
| - | - | 6.575E+04 | 125.1 | - | - | 0 | - |
| - | - | 787.8 | 126.1 | - | - | 0 | - |
| - | - | 4639 | 126.1 | - | - | 0 | - |
| - | - | 1108 | 126.1 | - | - | 0 | - |
| - | - | 2394 | 127.1 | - | - | 0 | - |
| - | - | 1461 | 127.1 | - | - | 0 | - |
| - | - | 823.5 | 127.1 | - | - | 0 | - |
| - | - | 2.939E+04 | 128.1 | - | - | 0 | - |
| - | - | 4028 | 129.1 | - | - | 0 | - |
| - | - | 3118 | 129.1 | - | - | 0 | - |
| - | - | 2660 | 129.1 | - | - | 0 | - |
| 11 | y | 1896 | 130 | 0.0004997 | 3.842 | +1 | 1 |
| - | - | 937.7 | 130.1 | - | - | 0 | - |
| - | - | 997.4 | 131.1 | - | - | 0 | - |
| - | - | 430.4 | 132.1 | - | - | 0 | - |
| - | - | 2327 | 132.1 | - | - | 0 | - |
| - | - | 887.8 | 133.1 | - | - | 0 | - |
| - | - | 517.6 | 133.1 | - | - | 0 | - |
| - | - | 3366 | 136.1 | - | - | 0 | - |
| - | - | 1.176E+04 | 136.1 | - | - | 0 | - |
| - | - | 640 | 137.1 | - | - | 0 | - |
| - | - | 713.8 | 137.1 | - | - | 0 | - |
| - | - | 2.24E+04 | 138.1 | - | - | 0 | - |
| - | - | 1.224E+04 | 138.1 | - | - | 0 | - |
| - | - | 2304 | 139.1 | - | - | 0 | - |
| - | - | 1404 | 139.1 | - | - | 0 | - |
| - | - | 800.6 | 139.1 | - | - | 0 | - |
| - | - | 628.2 | 139.1 | - | - | 0 | - |
| - | - | 5422 | 140.1 | - | - | 0 | - |
| - | - | 1473 | 141.1 | - | - | 0 | - |
| - | - | 2825 | 141.1 | - | - | 0 | - |
| - | - | 1539 | 142.1 | - | - | 0 | - |
| - | - | 586.6 | 142.1 | - | - | 0 | - |
| - | - | 9.788E+04 | 143.1 | - | - | 0 | - |
| - | - | 961.4 | 144.1 | - | - | 0 | - |
| - | - | 6911 | 144.1 | - | - | 0 | - |
| - | - | 5.266E+04 | 145.1 | - | - | 0 | - |
| - | - | 3178 | 146.1 | - | - | 0 | - |
| - | - | 919.4 | 148.1 | - | - | 0 | - |
| 11 | y | 1.274E+04 | 148.1 | 0.0006467 | 4.368 | +1 | 1 |
| - | - | 2140 | 148.1 | - | - | 0 | - |
| - | - | 1007 | 149 | - | - | 0 | - |
| - | - | 1230 | 149 | - | - | 0 | - |
| - | - | 3335 | 150.1 | - | - | 0 | - |
| - | - | 491.4 | 151.1 | - | - | 0 | - |
| - | - | 1369 | 152.1 | - | - | 0 | - |
| - | - | 1241 | 152.1 | - | - | 0 | - |
| - | - | 803.7 | 153.1 | - | - | 0 | - |
| - | - | 806 | 154.1 | - | - | 0 | - |
| - | - | 558 | 154.1 | - | - | 0 | - |
| - | - | 1.337E+04 | 154.2 | - | - | 0 | - |
| - | - | 4142 | 155 | - | - | 0 | - |
| 2 | a | 7.971E+04 | 155.1 | 0.0006407 | 4.131 | +1 | 2 |
| - | - | 1485 | 155.2 | - | - | 0 | - |
| - | - | 3.975E+04 | 156.1 | - | - | 0 | - |
| 2 | a | 9520 | 156.1 | 0.0005882 | 3.768 | +1 | 2 |
| - | - | 1039 | 156.1 | - | - | 0 | - |
| - | - | 6797 | 156.1 | - | - | 0 | - |
| - | - | 1.654E+04 | 157.1 | - | - | 0 | - |
| - | - | 2480 | 157.1 | - | - | 0 | - |
| - | - | 2325 | 157.1 | - | - | 0 | - |
| 2 | d | 2745 | 157.1 | 0.0005852 | 3.724 | +1 | 2 |
| - | - | 991.2 | 158.1 | - | - | 0 | - |
| - | - | 819.7 | 158.1 | - | - | 0 | - |
| - | - | 1234 | 159.1 | - | - | 0 | - |
| - | - | 1144 | 159.1 | - | - | 0 | - |
| - | - | 1057 | 163.1 | - | - | 0 | - |
| - | - | 1915 | 163.1 | - | - | 0 | - |
| - | - | 4415 | 164.1 | - | - | 0 | - |
| - | - | 3847 | 164.1 | - | - | 0 | - |
| - | - | 624.8 | 165.1 | - | - | 0 | - |
| - | - | 3235 | 165.1 | - | - | 0 | - |
| - | - | 583.3 | 165.1 | - | - | 0 | - |
| - | - | 2.609E+04 | 166.1 | - | - | 0 | - |
| - | - | 850.9 | 166.1 | - | - | 0 | - |
| - | - | 3583 | 167 | - | - | 0 | - |
| - | - | 966.3 | 167.1 | - | - | 0 | - |
| - | - | 1697 | 167.1 | - | - | 0 | - |
| - | - | 644.3 | 168 | - | - | 0 | - |
| - | - | 1.979E+04 | 169.1 | - | - | 0 | - |
| - | - | 733.8 | 170.1 | - | - | 0 | - |
| - | - | 1653 | 170.1 | - | - | 0 | - |
| - | - | 678.8 | 171.1 | - | - | 0 | - |
| - | - | 4.354E+04 | 171.1 | - | - | 0 | - |
| - | - | 1.147E+04 | 171.1 | - | - | 0 | - |
| - | - | 848.6 | 172.1 | - | - | 0 | - |
| - | - | 3080 | 172.1 | - | - | 0 | - |
| - | - | 1132 | 172.2 | - | - | 0 | - |
| - | - | 1206 | 173.1 | - | - | 0 | - |
| 2 | a | 5.26E+05 | 173.1 | 0.0007114 | 4.109 | +1 | 2 |
| - | - | 6996 | 174.1 | - | - | 0 | - |
| - | - | 4.295E+04 | 174.1 | - | - | 0 | - |
| - | - | 862 | 175.1 | - | - | 0 | - |
| - | - | 643.1 | 175.1 | - | - | 0 | - |
| - | - | 1808 | 175.1 | - | - | 0 | - |
| - | - | 1587 | 176.1 | - | - | 0 | - |
| - | - | 1566 | 178.1 | - | - | 0 | - |
| - | - | 781.8 | 179.1 | - | - | 0 | - |
| - | - | 457.5 | 179.1 | - | - | 0 | - |
| - | - | 4448 | 180.1 | - | - | 0 | - |
| - | - | 2932 | 181.1 | - | - | 0 | - |
| - | - | 6.181E+04 | 181.1 | - | - | 0 | - |
| - | - | 1074 | 182.1 | - | - | 0 | - |
| - | - | 885 | 182.1 | - | - | 0 | - |
| - | - | 502.7 | 182.1 | - | - | 0 | - |
| - | - | 4887 | 182.1 | - | - | 0 | - |
| - | - | 857.6 | 182.1 | - | - | 0 | - |
| 2 | b | 8.175E+04 | 183.1 | 0.0007518 | 4.105 | +1 | 2 |
| - | - | 1.125E+04 | 184.1 | - | - | 0 | - |
| 2 | b | 930.9 | 184.1 | 0.0008212 | 4.461 | +1 | 2 |
| - | - | 6954 | 184.1 | - | - | 0 | - |
| - | - | 698.5 | 185.1 | - | - | 0 | - |
| - | - | 1033 | 185.1 | - | - | 0 | - |
| - | - | 746.7 | 185.1 | - | - | 0 | - |
| - | - | 5831 | 185.1 | - | - | 0 | - |
| - | - | 673.4 | 185.2 | - | - | 0 | - |
| - | - | 767.1 | 186.1 | - | - | 0 | - |
| - | - | 596 | 186.1 | - | - | 0 | - |
| - | - | 7.24E+04 | 187.1 | - | - | 0 | - |
| - | - | 1911 | 187.1 | - | - | 0 | - |
| - | - | 5764 | 188.1 | - | - | 0 | - |
| - | - | 602.1 | 189.1 | - | - | 0 | - |
| - | - | 545.5 | 191.1 | - | - | 0 | - |
| - | - | 3246 | 191.1 | - | - | 0 | - |
| - | - | 5.453E+04 | 192.1 | - | - | 0 | - |
| - | - | 1113 | 192.1 | - | - | 0 | - |
| - | - | 4488 | 193.1 | - | - | 0 | - |
| - | - | 2311 | 193.1 | - | - | 0 | - |
| - | - | 561.7 | 193.1 | - | - | 0 | - |
| - | - | 734.1 | 194.1 | - | - | 0 | - |
| - | - | 2982 | 194.1 | - | - | 0 | - |
| - | - | 509 | 194.1 | - | - | 0 | - |
| - | - | 936.3 | 195.1 | - | - | 0 | - |
| - | - | 7695 | 195.1 | - | - | 0 | - |
| - | - | 723.2 | 195.2 | - | - | 0 | - |
| - | - | 732.4 | 196.1 | - | - | 0 | - |
| - | - | 1329 | 197.1 | - | - | 0 | - |
| - | - | 1643 | 197.1 | - | - | 0 | - |
| - | - | 2632 | 197.2 | - | - | 0 | - |
| - | - | 1195 | 198.1 | - | - | 0 | - |
| - | - | 707.9 | 198.1 | - | - | 0 | - |
| - | - | 7169 | 199.1 | - | - | 0 | - |
| - | - | 1.277E+04 | 199.1 | - | - | 0 | - |
| - | - | 1860 | 199.2 | - | - | 0 | - |
| - | - | 1604 | 200.1 | - | - | 0 | - |
| - | - | 953.6 | 200.1 | - | - | 0 | - |
| - | - | 2.812E+04 | 200.1 | - | - | 0 | - |
| 2 | b | 4.699E+05 | 201.1 | 0.0007461 | 3.71 | +1 | 2 |
| - | - | 1.023E+05 | 202.1 | - | - | 0 | - |
| - | - | 4.366E+04 | 202.1 | - | - | 0 | - |
| - | - | 1140 | 203.1 | - | - | 0 | - |
| - | - | 6629 | 203.1 | - | - | 0 | - |
| - | - | 2552 | 203.1 | - | - | 0 | - |
| - | - | 1296 | 204.1 | - | - | 0 | - |
| - | - | 533.2 | 205.1 | - | - | 0 | - |
| - | - | 4321 | 207.1 | - | - | 0 | - |
| - | - | 583.4 | 207.1 | - | - | 0 | - |
| - | - | 723.6 | 207.1 | - | - | 0 | - |
| - | - | 4050 | 208.1 | - | - | 0 | - |
| - | - | 2.405E+05 | 209.1 | - | - | 0 | - |
| - | - | 2.102E+04 | 210.1 | - | - | 0 | - |
| - | - | 5321 | 210.1 | - | - | 0 | - |
| - | - | 1604 | 211.1 | - | - | 0 | - |
| - | - | 904.8 | 211.1 | - | - | 0 | - |
| - | - | 1549 | 211.1 | - | - | 0 | - |
| - | - | 795.1 | 211.2 | - | - | 0 | - |
| - | - | 6919 | 212.1 | - | - | 0 | - |
| 3 | a | 5016 | 212.1 | 0.0008598 | 4.053 | +1 | 3 |
| 3 | a | 1.383E+04 | 213.1 | 0.0007614 | 3.573 | +1 | 3 |
| - | - | 745.7 | 213.1 | - | - | 0 | - |
| - | - | 1002 | 213.2 | - | - | 0 | - |
| - | - | 3480 | 214.1 | - | - | 0 | - |
| - | - | 955 | 214.1 | - | - | 0 | - |
| - | - | 2050 | 214.2 | - | - | 0 | - |
| - | - | 8.077E+04 | 215.1 | - | - | 0 | - |
| - | - | 8630 | 216.1 | - | - | 0 | - |
| - | - | 1058 | 217.1 | - | - | 0 | - |
| - | - | 2167 | 219.1 | - | - | 0 | - |
| 8 | y | 1261 | 219.1 | 0.00154 | 7.028 | +2 | 4 |
| - | - | 4314 | 221.1 | - | - | 0 | - |
| - | - | 9027 | 221.1 | - | - | 0 | - |
| - | - | 1110 | 222.1 | - | - | 0 | - |
| - | - | 1249 | 222.1 | - | - | 0 | - |
| - | - | 3474 | 222.1 | - | - | 0 | - |
| - | - | 2107 | 223.1 | - | - | 0 | - |
| - | - | 1444 | 223.2 | - | - | 0 | - |
| - | - | 1907 | 224.1 | - | - | 0 | - |
| - | - | 3652 | 225 | - | - | 0 | - |
| - | - | 1150 | 225.1 | - | - | 0 | - |
| - | - | 705.8 | 225.1 | - | - | 0 | - |
| - | - | 3249 | 225.2 | - | - | 0 | - |
| - | - | 686.1 | 226.1 | - | - | 0 | - |
| - | - | 1735 | 226.1 | - | - | 0 | - |
| - | - | 913.4 | 226.1 | - | - | 0 | - |
| - | - | 1341 | 226.2 | - | - | 0 | - |
| - | - | 1584 | 227.1 | - | - | 0 | - |
| - | - | 1.707E+04 | 227.1 | - | - | 0 | - |
| - | - | 711.4 | 227.1 | - | - | 0 | - |
| - | - | 3239 | 227.2 | - | - | 0 | - |
| - | - | 2232 | 228.1 | - | - | 0 | - |
| - | - | 6.382E+04 | 228.1 | - | - | 0 | - |
| - | - | 5.213E+04 | 229.1 | - | - | 0 | - |
| - | - | 5283 | 229.1 | - | - | 0 | - |
| - | - | 9879 | 230.1 | - | - | 0 | - |
| - | - | 4709 | 230.1 | - | - | 0 | - |
| - | - | 589.7 | 231.1 | - | - | 0 | - |
| - | - | 731.3 | 233.1 | - | - | 0 | - |
| - | - | 740.8 | 235.1 | - | - | 0 | - |
| - | - | 931.6 | 235.1 | - | - | 0 | - |
| - | - | 1045 | 236.1 | - | - | 0 | - |
| - | - | 5084 | 237.1 | - | - | 0 | - |
| - | - | 945.9 | 237.2 | - | - | 0 | - |
| - | - | 3161 | 238.1 | - | - | 0 | - |
| - | - | 6998 | 239.1 | - | - | 0 | - |
| - | - | 2.227E+04 | 239.1 | - | - | 0 | - |
| - | - | 1193 | 239.2 | - | - | 0 | - |
| 3 | b | 1.163E+05 | 240.1 | 0.0009097 | 3.788 | +1 | 3 |
| 3 | b | 1354 | 241.1 | 0.0005215 | 2.163 | +1 | 3 |
| - | - | 1.452E+04 | 241.1 | - | - | 0 | - |
| - | - | 975.5 | 241.2 | - | - | 0 | - |
| - | - | 958.2 | 241.2 | - | - | 0 | - |
| - | - | 1.441E+04 | 241.2 | - | - | 0 | - |
| - | - | 1687 | 242.1 | - | - | 0 | - |
| - | - | 3492 | 242.1 | - | - | 0 | - |
| - | - | 607.2 | 242.1 | - | - | 0 | - |
| - | - | 2198 | 242.2 | - | - | 0 | - |
| - | - | 1090 | 242.2 | - | - | 0 | - |
| - | - | 1246 | 244.1 | - | - | 0 | - |
| - | - | 956.4 | 244.1 | - | - | 0 | - |
| - | - | 617.1 | 245.1 | - | - | 0 | - |
| - | - | 2385 | 245.1 | - | - | 0 | - |
| - | - | 550.8 | 245.2 | - | - | 0 | - |
| - | - | 3.287E+04 | 249.1 | - | - | 0 | - |
| - | - | 2221 | 250.1 | - | - | 0 | - |
| - | - | 3320 | 250.1 | - | - | 0 | - |
| - | - | 851.5 | 251.1 | - | - | 0 | - |
| - | - | 1314 | 251.2 | - | - | 0 | - |
| - | - | 1638 | 251.2 | - | - | 0 | - |
| - | - | 9465 | 252.1 | - | - | 0 | - |
| - | - | 1253 | 253.1 | - | - | 0 | - |
| - | - | 1268 | 253.1 | - | - | 0 | - |
| - | - | 919.6 | 253.2 | - | - | 0 | - |
| - | - | 627.9 | 254.1 | - | - | 0 | - |
| - | - | 1258 | 254.1 | - | - | 0 | - |
| - | - | 2203 | 254.2 | - | - | 0 | - |
| - | - | 2404 | 254.2 | - | - | 0 | - |
| - | - | 1402 | 255.1 | - | - | 0 | - |
| - | - | 812 | 255.1 | - | - | 0 | - |
| - | - | 6133 | 255.2 | - | - | 0 | - |
| - | - | 1.983E+04 | 256.1 | - | - | 0 | - |
| - | - | 838 | 256.2 | - | - | 0 | - |
| - | - | 1125 | 257.1 | - | - | 0 | - |
| - | - | 1669 | 257.1 | - | - | 0 | - |
| 3 | b | 1.748E+04 | 258.1 | 0.0008889 | 3.443 | +1 | 3 |
| - | - | 1766 | 259.1 | - | - | 0 | - |
| - | - | 722.1 | 262.1 | - | - | 0 | - |
| - | - | 864.4 | 263.2 | - | - | 0 | - |
| - | - | 1657 | 263.2 | - | - | 0 | - |
| - | - | 2774 | 264.2 | - | - | 0 | - |
| - | - | 1029 | 265.2 | - | - | 0 | - |
| - | - | 1132 | 266.1 | - | - | 0 | - |
| 10 | y | 8.514E+04 | 267.1 | 0.0008072 | 3.022 | +1 | 2 |
| - | - | 2968 | 267.1 | - | - | 0 | - |
| - | - | 955 | 268.1 | - | - | 0 | - |
| - | - | 9895 | 268.1 | - | - | 0 | - |
| - | - | 1513 | 268.1 | - | - | 0 | - |
| - | - | 3359 | 268.2 | - | - | 0 | - |
| - | - | 1628 | 269.1 | - | - | 0 | - |
| - | - | 4945 | 269.1 | - | - | 0 | - |
| - | - | 1.1E+04 | 269.2 | - | - | 0 | - |
| - | - | 1.382E+04 | 269.2 | - | - | 0 | - |
| - | - | 1.648E+04 | 270.1 | - | - | 0 | - |
| - | - | 1514 | 270.2 | - | - | 0 | - |
| - | - | 4130 | 270.2 | - | - | 0 | - |
| - | - | 6832 | 271.1 | - | - | 0 | - |
| - | - | 1964 | 271.1 | - | - | 0 | - |
| - | - | 4187 | 271.2 | - | - | 0 | - |
| - | - | 4172 | 272.1 | - | - | 0 | - |
| - | - | 627.7 | 272.2 | - | - | 0 | - |
| - | - | 2039 | 273.1 | - | - | 0 | - |
| - | - | 1452 | 273.1 | - | - | 0 | - |
| 7 | y | 622 | 277.7 | 0.0004748 | 1.71 | +2 | 5 |
| - | - | 1.433E+04 | 279.1 | - | - | 0 | - |
| - | - | 757.6 | 280.1 | - | - | 0 | - |
| - | - | 7762 | 280.1 | - | - | 0 | - |
| - | - | 2088 | 280.1 | - | - | 0 | - |
| - | - | 5757 | 280.2 | - | - | 0 | - |
| - | - | 877 | 281.1 | - | - | 0 | - |
| - | - | 1120 | 281.2 | - | - | 0 | - |
| - | - | 1.309E+04 | 282.2 | - | - | 0 | - |
| - | - | 1373 | 283.1 | - | - | 0 | - |
| - | - | 2366 | 283.1 | - | - | 0 | - |
| - | - | 9358 | 283.2 | - | - | 0 | - |
| - | - | 1583 | 283.2 | - | - | 0 | - |
| - | - | 2078 | 284.1 | - | - | 0 | - |
| - | - | 1216 | 284.2 | - | - | 0 | - |
| - | - | 1943 | 284.2 | - | - | 0 | - |
| 10 | y | 6.945E+05 | 285.1 | 0.001076 | 3.775 | +1 | 2 |
| - | - | 7.952E+04 | 286.1 | - | - | 0 | - |
| - | - | 6.955E+04 | 286.1 | - | - | 0 | - |
| - | - | 2533 | 286.2 | - | - | 0 | - |
| - | - | 9727 | 287.1 | - | - | 0 | - |
| - | - | 8315 | 287.1 | - | - | 0 | - |
| - | - | 1.542E+05 | 287.2 | - | - | 0 | - |
| - | - | 1759 | 288.1 | - | - | 0 | - |
| - | - | 2.127E+04 | 288.2 | - | - | 0 | - |
| - | - | 1518 | 289.2 | - | - | 0 | - |
| - | - | 1.024E+04 | 290.2 | - | - | 0 | - |
| - | - | 1067 | 291.1 | - | - | 0 | - |
| - | - | 1303 | 291.2 | - | - | 0 | - |
| - | - | 2665 | 291.2 | - | - | 0 | - |
| - | - | 740.2 | 294.1 | - | - | 0 | - |
| - | - | 1.133E+04 | 295.1 | - | - | 0 | - |
| - | - | 1708 | 295.1 | - | - | 0 | - |
| - | - | 912.5 | 296.1 | - | - | 0 | - |
| - | - | 3265 | 296.2 | - | - | 0 | - |
| - | - | 1374 | 296.2 | - | - | 0 | - |
| 4 | b | 5.755E+04 | 297.2 | 0.001159 | 3.901 | +1 | 4 |
| 4 | b | 2978 | 298.1 | 0.001244 | 4.173 | +1 | 4 |
| - | - | 6869 | 298.2 | - | - | 0 | - |
| - | - | 3955 | 298.2 | - | - | 0 | - |
| - | - | 3118 | 299.1 | - | - | 0 | - |
| - | - | 9662 | 299.1 | - | - | 0 | - |
| - | - | 2493 | 299.2 | - | - | 0 | - |
| - | - | 5414 | 299.2 | - | - | 0 | - |
| - | - | 1230 | 300.1 | - | - | 0 | - |
| - | - | 1484 | 300.2 | - | - | 0 | - |
| - | - | 3326 | 300.2 | - | - | 0 | - |
| - | - | 2541 | 301.1 | - | - | 0 | - |
| - | - | 5722 | 301.2 | - | - | 0 | - |
| - | - | 1841 | 302.1 | - | - | 0 | - |
| - | - | 1148 | 302.2 | - | - | 0 | - |
| - | - | 817.7 | 303.1 | - | - | 0 | - |
| - | - | 786.8 | 303.2 | - | - | 0 | - |
| - | - | 9885 | 307.1 | - | - | 0 | - |
| - | - | 6.037E+04 | 308.2 | - | - | 0 | - |
| - | - | 942.6 | 309.1 | - | - | 0 | - |
| - | - | 8087 | 309.2 | - | - | 0 | - |
| - | - | 626.9 | 310.2 | - | - | 0 | - |
| - | - | 2314 | 310.2 | - | - | 0 | - |
| - | - | 3733 | 311.1 | - | - | 0 | - |
| - | - | 1112 | 311.2 | - | - | 0 | - |
| - | - | 993.8 | 312.1 | - | - | 0 | - |
| - | - | 1391 | 312.2 | - | - | 0 | - |
| - | - | 1362 | 312.2 | - | - | 0 | - |
| - | - | 1092 | 313.2 | - | - | 0 | - |
| - | - | 2886 | 313.2 | - | - | 0 | - |
| - | - | 1592 | 313.2 | - | - | 0 | - |
| - | - | 1325 | 314.1 | - | - | 0 | - |
| - | - | 5.441E+04 | 314.2 | - | - | 0 | - |
| 4 | b | 2.242E+05 | 315.2 | 0.001398 | 4.435 | +1 | 4 |
| - | - | 3.277E+04 | 316.2 | - | - | 0 | - |
| - | - | 2.085E+04 | 317.1 | - | - | 0 | - |
| - | - | 3210 | 317.2 | - | - | 0 | - |
| - | - | 2469 | 318.1 | - | - | 0 | - |
| - | - | 889 | 318.2 | - | - | 0 | - |
| - | - | 3744 | 320.1 | - | - | 0 | - |
| - | - | 2488 | 320.2 | - | - | 0 | - |
| - | - | 3.332E+04 | 321.1 | - | - | 0 | - |
| - | - | 4443 | 322.1 | - | - | 0 | - |
| - | - | 1038 | 323.2 | - | - | 0 | - |
| - | - | 913.6 | 323.2 | - | - | 0 | - |
| - | - | 961.3 | 324.1 | - | - | 0 | - |
| - | - | 1.793E+04 | 325.2 | - | - | 0 | - |
| - | - | 923.8 | 325.2 | - | - | 0 | - |
| 6 | y | 2193 | 326.2 | 0.004327 | 13.27 | +2 | 6 |
| - | - | 6242 | 326.2 | - | - | 0 | - |
| - | - | 3313 | 326.2 | - | - | 0 | - |
| - | - | 1409 | 327.2 | - | - | 0 | - |
| - | - | 3385 | 327.2 | - | - | 0 | - |
| - | - | 1.487E+04 | 328.2 | - | - | 0 | - |
| - | - | 3.106E+04 | 329.1 | - | - | 0 | - |
| - | - | 2586 | 329.2 | - | - | 0 | - |
| - | - | 5048 | 330.1 | - | - | 0 | - |
| - | - | 973.2 | 335.2 | - | - | 0 | - |
| - | - | 4497 | 336.2 | - | - | 0 | - |
| - | - | 680.3 | 337.2 | - | - | 0 | - |
| - | - | 956.7 | 337.2 | - | - | 0 | - |
| - | - | 888.1 | 337.2 | - | - | 0 | - |
| 9 | y | 3.413E+04 | 338.1 | 0.001383 | 4.089 | +1 | 3 |
| - | - | 5262 | 339.2 | - | - | 0 | - |
| - | - | 3975 | 339.2 | - | - | 0 | - |
| - | - | 5074 | 340.2 | - | - | 0 | - |
| - | - | 2195 | 341.1 | - | - | 0 | - |
| - | - | 3084 | 341.2 | - | - | 0 | - |
| - | - | 1.194E+04 | 341.2 | - | - | 0 | - |
| - | - | 2724 | 342.2 | - | - | 0 | - |
| - | - | 1.195E+05 | 343.2 | - | - | 0 | - |
| - | - | 1.859E+04 | 344.2 | - | - | 0 | - |
| - | - | 3009 | 345.2 | - | - | 0 | - |
| - | - | 952.9 | 348.1 | - | - | 0 | - |
| - | - | 1275 | 349.2 | - | - | 0 | - |
| - | - | 913.1 | 351.2 | - | - | 0 | - |
| - | - | 1017 | 352.2 | - | - | 0 | - |
| - | - | 5149 | 353.1 | - | - | 0 | - |
| - | - | 2377 | 353.2 | - | - | 0 | - |
| - | - | 963.7 | 354.1 | - | - | 0 | - |
| - | - | 1318 | 354.2 | - | - | 0 | - |
| - | - | 1037 | 354.2 | - | - | 0 | - |
| - | - | 697.6 | 355.1 | - | - | 0 | - |
| - | - | 1292 | 355.2 | - | - | 0 | - |
| - | - | 1452 | 355.2 | - | - | 0 | - |
| - | - | 3668 | 355.2 | - | - | 0 | - |
| 9 | y | 4.046E+05 | 356.2 | 0.001285 | 3.609 | +1 | 3 |
| - | - | 6.875E+04 | 357.2 | - | - | 0 | - |
| - | - | 1.292E+04 | 357.2 | - | - | 0 | - |
| - | - | 7004 | 358.2 | - | - | 0 | - |
| - | - | 3151 | 358.2 | - | - | 0 | - |
| - | - | 734.4 | 359.2 | - | - | 0 | - |
| - | - | 794.7 | 362.2 | - | - | 0 | - |
| - | - | 910.7 | 365.2 | - | - | 0 | - |
| - | - | 3499 | 365.2 | - | - | 0 | - |
| - | - | 1.07E+04 | 366.1 | - | - | 0 | - |
| - | - | 1781 | 366.2 | - | - | 0 | - |
| - | - | 2515 | 367.2 | - | - | 0 | - |
| - | - | 1361 | 367.2 | - | - | 0 | - |
| - | - | 3656 | 368.2 | - | - | 0 | - |
| - | - | 1495 | 368.2 | - | - | 0 | - |
| - | - | 3064 | 369.1 | - | - | 0 | - |
| - | - | 6731 | 369.2 | - | - | 0 | - |
| - | - | 1.186E+04 | 369.2 | - | - | 0 | - |
| - | - | 1726 | 370.2 | - | - | 0 | - |
| - | - | 962.5 | 370.2 | - | - | 0 | - |
| - | - | 2037 | 370.2 | - | - | 0 | - |
| - | - | 2504 | 370.2 | - | - | 0 | - |
| - | - | 1717 | 371.2 | - | - | 0 | - |
| - | - | 3700 | 371.2 | - | - | 0 | - |
| - | - | 740.5 | 371.2 | - | - | 0 | - |
| - | - | 1641 | 372.2 | - | - | 0 | - |
| - | - | 790.2 | 377.1 | - | - | 0 | - |
| - | - | 1696 | 378.2 | - | - | 0 | - |
| - | - | 587.7 | 378.2 | - | - | 0 | - |
| - | - | 1.198E+04 | 379.2 | - | - | 0 | - |
| - | - | 2261 | 380.2 | - | - | 0 | - |
| - | - | 2162 | 380.2 | - | - | 0 | - |
| - | - | 3055 | 381.3 | - | - | 0 | - |
| - | - | 2110 | 382.2 | - | - | 0 | - |
| - | - | 1143 | 382.2 | - | - | 0 | - |
| - | - | 1.222E+04 | 382.2 | - | - | 0 | - |
| 5 | y | 1087 | 382.7 | 0.001767 | 4.618 | +2 | 7 |
| - | - | 1.647E+04 | 383.2 | - | - | 0 | - |
| - | - | 1.859E+04 | 384.2 | - | - | 0 | - |
| - | - | 3780 | 384.2 | - | - | 0 | - |
| - | - | 2661 | 385.2 | - | - | 0 | - |
| - | - | 3.376E+04 | 385.2 | - | - | 0 | - |
| - | - | 7871 | 386.2 | - | - | 0 | - |
| - | - | 1515 | 386.2 | - | - | 0 | - |
| - | - | 875.6 | 387.2 | - | - | 0 | - |
| - | - | 3877 | 388.2 | - | - | 0 | - |
| 5 | y | 1.536E+04 | 391.7 | 0.001337 | 3.414 | +2 | 7 |
| - | - | 1077 | 391.7 | - | - | 0 | - |
| - | - | 3952 | 392.2 | - | - | 0 | - |
| - | - | 813.9 | 392.2 | - | - | 0 | - |
| - | - | 2098 | 392.7 | - | - | 0 | - |
| - | - | 819.9 | 393.2 | - | - | 0 | - |
| - | - | 5433 | 394.2 | - | - | 0 | - |
| - | - | 1.947E+04 | 395.2 | - | - | 0 | - |
| - | - | 3725 | 396.2 | - | - | 0 | - |
| - | - | 4694 | 396.2 | - | - | 0 | - |
| - | - | 1.254E+04 | 396.2 | - | - | 0 | - |
| - | - | 1128 | 396.3 | - | - | 0 | - |
| - | - | 3.166E+04 | 397.2 | - | - | 0 | - |
| - | - | 8272 | 398.2 | - | - | 0 | - |
| 9 | b | 6820 | 398.2 | 0.00334 | 8.387 | +2 | 9 |
| - | - | 1801 | 398.2 | - | - | 0 | - |
| - | - | 874.1 | 399.2 | - | - | 0 | - |
| - | - | 3169 | 399.2 | - | - | 0 | - |
| - | - | 1405 | 399.3 | - | - | 0 | - |
| - | - | 3870 | 400.2 | - | - | 0 | - |
| - | - | 1.125E+05 | 400.3 | - | - | 0 | - |
| - | - | 2.411E+04 | 401.3 | - | - | 0 | - |
| - | - | 9824 | 402.2 | - | - | 0 | - |
| - | - | 2980 | 402.3 | - | - | 0 | - |
| - | - | 2407 | 403.2 | - | - | 0 | - |
| - | - | 5828 | 406.2 | - | - | 0 | - |
| - | - | 1960 | 407.2 | - | - | 0 | - |
| - | - | 1.628E+04 | 407.2 | - | - | 0 | - |
| - | - | 3586 | 408.2 | - | - | 0 | - |
| - | - | 1840 | 409.2 | - | - | 0 | - |
| - | - | 882.4 | 409.2 | - | - | 0 | - |
| - | - | 2858 | 410.2 | - | - | 0 | - |
| - | - | 2449 | 410.2 | - | - | 0 | - |
| 5 | b | 2.312E+04 | 410.2 | 0.001751 | 4.268 | +1 | 5 |
| - | - | 726.1 | 411.2 | - | - | 0 | - |
| - | - | 4152 | 411.2 | - | - | 0 | - |
| - | - | 736.3 | 411.7 | - | - | 0 | - |
| - | - | 5.164E+04 | 412.2 | - | - | 0 | - |
| - | - | 9836 | 413.2 | - | - | 0 | - |
| - | - | 8517 | 414.2 | - | - | 0 | - |
| - | - | 4.759E+04 | 414.2 | - | - | 0 | - |
| - | - | 1074 | 415.2 | - | - | 0 | - |
| - | - | 1.008E+04 | 415.2 | - | - | 0 | - |
| - | - | 1.625E+04 | 416.2 | - | - | 0 | - |
| - | - | 3144 | 417.2 | - | - | 0 | - |
| - | - | 887.6 | 418.7 | - | - | 0 | - |
| - | - | 4547 | 420.2 | - | - | 0 | - |
| 4 | y | 3564 | 420.2 | 0.003697 | 8.799 | +2 | 8 |
| - | - | 1757 | 420.7 | - | - | 0 | - |
| - | - | 728.8 | 421.2 | - | - | 0 | - |
| - | - | 1.303E+04 | 422.2 | - | - | 0 | - |
| - | - | 2088 | 423.2 | - | - | 0 | - |
| - | - | 2590 | 423.2 | - | - | 0 | - |
| - | - | 825.8 | 423.2 | - | - | 0 | - |
| - | - | 831.1 | 424.2 | - | - | 0 | - |
| - | - | 1.338E+04 | 424.2 | - | - | 0 | - |
| - | - | 3025 | 425.2 | - | - | 0 | - |
| - | - | 2732 | 425.3 | - | - | 0 | - |
| - | - | 770.8 | 425.7 | - | - | 0 | - |
| - | - | 1476 | 426.2 | - | - | 0 | - |
| - | - | 1048 | 427.2 | - | - | 0 | - |
| - | - | 3262 | 427.3 | - | - | 0 | - |
| - | - | 8972 | 428.2 | - | - | 0 | - |
| 5 | b | 6.814E+04 | 428.3 | 0.001806 | 4.218 | +1 | 5 |
| - | - | 1668 | 429.2 | - | - | 0 | - |
| - | - | 1.697E+04 | 429.3 | - | - | 0 | - |
| - | - | 7.152E+04 | 430.2 | - | - | 0 | - |
| - | - | 2157 | 430.3 | - | - | 0 | - |
| - | - | 1.372E+04 | 431.2 | - | - | 0 | - |
| - | - | 1838 | 432.2 | - | - | 0 | - |
| - | - | 9009 | 435.2 | - | - | 0 | - |
| - | - | 2130 | 435.2 | - | - | 0 | - |
| - | - | 1935 | 436.2 | - | - | 0 | - |
| - | - | 680.5 | 436.2 | - | - | 0 | - |
| 8 | y | 1.331E+04 | 437.2 | 0.001664 | 3.805 | +1 | 4 |
| - | - | 1685 | 438.2 | - | - | 0 | - |
| - | - | 1074 | 438.3 | - | - | 0 | - |
| - | - | 2318 | 439.3 | - | - | 0 | - |
| 3 | y | 5311 | 439.7 | 0.001818 | 4.135 | +2 | 9 |
| - | - | 2.298E+04 | 440.2 | - | - | 0 | - |
| - | - | 1847 | 440.2 | - | - | 0 | - |
| - | - | 2751 | 440.3 | - | - | 0 | - |
| - | - | 5029 | 441.2 | - | - | 0 | - |
| - | - | 6.035E+04 | 442.2 | - | - | 0 | - |
| - | - | 1.365E+04 | 443.2 | - | - | 0 | - |
| - | - | 1484 | 444.2 | - | - | 0 | - |
| 3 | y | 2.438E+04 | 448.7 | 0.001694 | 3.774 | +2 | 9 |
| - | - | 1.038E+04 | 449.2 | - | - | 0 | - |
| - | - | 2153 | 449.7 | - | - | 0 | - |
| - | - | 809.6 | 452.2 | - | - | 0 | - |
| - | - | 1392 | 452.3 | - | - | 0 | - |
| - | - | 1534 | 453.3 | - | - | 0 | - |
| - | - | 1145 | 453.7 | - | - | 0 | - |
| - | - | 2051 | 454.2 | - | - | 0 | - |
| - | - | 986.7 | 454.3 | - | - | 0 | - |
| - | - | 1048 | 454.3 | - | - | 0 | - |
| 8 | y | 2.861E+05 | 455.2 | 0.001841 | 4.045 | +1 | 4 |
| - | - | 5.861E+04 | 456.2 | - | - | 0 | - |
| - | - | 1657 | 456.3 | - | - | 0 | - |
| - | - | 8812 | 457.2 | - | - | 0 | - |
| - | - | 630 | 457.3 | - | - | 0 | - |
| - | - | 961.6 | 457.3 | - | - | 0 | - |
| - | - | 2244 | 461.3 | - | - | 0 | - |
| - | - | 1145 | 461.8 | - | - | 0 | - |
| - | - | 8154 | 465.2 | - | - | 0 | - |
| - | - | 1981 | 465.2 | - | - | 0 | - |
| - | - | 835.9 | 465.7 | - | - | 0 | - |
| - | - | 2493 | 466.2 | - | - | 0 | - |
| 10 | b | 3312 | 466.3 | 0.004744 | 10.18 | +2 | 10 |
| - | - | 1828 | 466.8 | - | - | 0 | - |
| - | - | 632.2 | 467.2 | - | - | 0 | - |
| - | - | 4131 | 467.3 | - | - | 0 | - |
| - | - | 8747 | 468.2 | - | - | 0 | - |
| - | - | 2079 | 468.3 | - | - | 0 | - |
| - | - | 1065 | 469.2 | - | - | 0 | - |
| - | - | 2283 | 469.2 | - | - | 0 | - |
| - | - | 979.6 | 470.2 | - | - | 0 | - |
| - | - | 1451 | 470.3 | - | - | 0 | - |
| - | - | 1283 | 471.3 | - | - | 0 | - |
| - | - | 1264 | 472.2 | - | - | 0 | - |
| 10 | b | 1.214E+04 | 475.3 | 0.002026 | 4.262 | +2 | 10 |
| - | - | 8251 | 475.8 | - | - | 0 | - |
| - | - | 2296 | 476.3 | - | - | 0 | - |
| - | - | 7584 | 477.2 | - | - | 0 | - |
| - | - | 1583 | 478.3 | - | - | 0 | - |
| - | - | 1518 | 479.3 | - | - | 0 | - |
| - | - | 1683 | 480.2 | - | - | 0 | - |
| - | - | 1955 | 481.2 | - | - | 0 | - |
| - | - | 2231 | 482.2 | - | - | 0 | - |
| - | - | 1671 | 482.3 | - | - | 0 | - |
| - | - | 1718 | 482.7 | - | - | 0 | - |
| - | - | 5797 | 483.3 | - | - | 0 | - |
| - | - | 5239 | 483.7 | - | - | 0 | - |
| - | - | 2.108E+04 | 484.2 | - | - | 0 | - |
| - | - | 4118 | 484.3 | - | - | 0 | - |
| - | - | 2751 | 485.2 | - | - | 0 | - |
| - | - | 6862 | 485.3 | - | - | 0 | - |
| - | - | 1239 | 486.3 | - | - | 0 | - |
| - | - | 1826 | 487.2 | - | - | 0 | - |
| - | - | 979.9 | 487.3 | - | - | 0 | - |
| - | - | 7842 | 488.2 | - | - | 0 | - |
| - | - | 3099 | 488.7 | - | - | 0 | - |
| - | - | 1431 | 489.2 | - | - | 0 | - |
| 2 | y | 4.88E+04 | 492.2 | 0.001915 | 3.89 | +2 | 10 |
| - | - | 3.23E+04 | 492.7 | - | - | 0 | - |
| - | - | 1.287E+04 | 493.2 | - | - | 0 | - |
| - | - | 2235 | 494.2 | - | - | 0 | - |
| - | - | 2.815E+04 | 494.3 | - | - | 0 | - |
| - | - | 2838 | 495.3 | - | - | 0 | - |
| - | - | 3906 | 495.3 | - | - | 0 | - |
| - | - | 7177 | 496.3 | - | - | 0 | - |
| - | - | 1.274E+04 | 497.2 | - | - | 0 | - |
| - | - | 4461 | 497.3 | - | - | 0 | - |
| - | - | 5844 | 497.7 | - | - | 0 | - |
| - | - | 1714 | 498.3 | - | - | 0 | - |
| - | - | 2309 | 498.3 | - | - | 0 | - |
| - | - | 9186 | 499.3 | - | - | 0 | - |
| - | - | 1343 | 499.3 | - | - | 0 | - |
| - | - | 1904 | 500.3 | - | - | 0 | - |
| - | - | 2514 | 501.2 | - | - | 0 | - |
| - | - | 2.558E+04 | 501.3 | - | - | 0 | - |
| - | - | 6634 | 502.3 | - | - | 0 | - |
| - | - | 941.1 | 504.3 | - | - | 0 | - |
| - | - | 1209 | 505.2 | - | - | 0 | - |
| - | - | 3373 | 506.3 | - | - | 0 | - |
| - | - | 3543 | 507.3 | - | - | 0 | - |
| - | - | 1153 | 508.2 | - | - | 0 | - |
| - | - | 970.3 | 509.2 | - | - | 0 | - |
| - | - | 856.7 | 509.3 | - | - | 0 | - |
| - | - | 2384 | 510.3 | - | - | 0 | - |
| - | - | 3.824E+04 | 511.3 | - | - | 0 | - |
| - | - | 9182 | 512.3 | - | - | 0 | - |
| - | - | 1.73E+04 | 513.3 | - | - | 0 | - |
| - | - | 1.098E+04 | 513.3 | - | - | 0 | - |
| - | - | 5004 | 514.3 | - | - | 0 | - |
| - | - | 3379 | 514.3 | - | - | 0 | - |
| - | - | 3631 | 515.2 | - | - | 0 | - |
| - | - | 4199 | 515.3 | - | - | 0 | - |
| - | - | 1683 | 516.3 | - | - | 0 | - |
| - | - | 1174 | 516.3 | - | - | 0 | - |
| - | - | 843 | 519.3 | - | - | 0 | - |
| - | - | 5253 | 521.2 | - | - | 0 | - |
| - | - | 3861 | 522.2 | - | - | 0 | - |
| - | - | 4.84E+04 | 522.3 | - | - | 0 | - |
| - | - | 1231 | 522.8 | - | - | 0 | - |
| - | - | 1.121E+04 | 523.3 | - | - | 0 | - |
| - | - | 2169 | 524.3 | - | - | 0 | - |
| 6 | b | 1.186E+04 | 525.3 | 0.002671 | 5.084 | +1 | 6 |
| - | - | 2856 | 525.8 | - | - | 0 | - |
| - | - | 3153 | 526.3 | - | - | 0 | - |
| - | - | 2560 | 527.3 | - | - | 0 | - |
| - | - | 5.231E+04 | 529.3 | - | - | 0 | - |
| - | - | 1.408E+04 | 530.3 | - | - | 0 | - |
| - | - | 7593 | 530.8 | - | - | 0 | - |
| - | - | 1.081E+04 | 531.3 | - | - | 0 | - |
| - | - | 3912 | 531.8 | - | - | 0 | - |
| - | - | 787.8 | 532.2 | - | - | 0 | - |
| - | - | 7478 | 534.3 | - | - | 0 | - |
| - | - | 1736 | 535.3 | - | - | 0 | - |
| 7 | y | 7128 | 536.3 | 0.0009684 | 1.806 | +1 | 5 |
| - | - | 1636 | 537.3 | - | - | 0 | - |
| - | - | 681.8 | 538.3 | - | - | 0 | - |
| - | - | 7178 | 539.2 | - | - | 0 | - |
| - | - | 990.7 | 539.3 | - | - | 0 | - |
| 0 | Precursor | 9.567E+04 | 539.8 | 0.001786 | 3.309 | +2 | -1 |
| - | - | 6.219E+04 | 540.3 | - | - | 0 | - |
| - | - | 523.4 | 540.7 | - | - | 0 | - |
| - | - | 1.913E+04 | 540.8 | - | - | 0 | - |
| - | - | 2.077E+04 | 541.3 | - | - | 0 | - |
| - | - | 6120 | 542.3 | - | - | 0 | - |
| 6 | b | 2.664E+04 | 543.3 | 0.00181 | 3.332 | +1 | 6 |
| - | - | 7911 | 544.3 | - | - | 0 | - |
| - | - | 1505 | 545.3 | - | - | 0 | - |
| - | - | 1141 | 547.8 | - | - | 0 | - |
| - | - | 1488 | 548.3 | - | - | 0 | - |
| 0 | Precursor | 2.3E+05 | 548.8 | 0.001936 | 3.528 | +2 | -1 |
| - | - | 1.436E+05 | 549.3 | - | - | 0 | - |
| - | - | 4.837E+04 | 549.8 | - | - | 0 | - |
| - | - | 1133 | 550 | - | - | 0 | - |
| - | - | 6801 | 550.3 | - | - | 0 | - |
| - | - | 1407 | 551.3 | - | - | 0 | - |
| - | - | 1074 | 552.3 | - | - | 0 | - |
| 7 | y | 1.654E+05 | 554.3 | 0.001878 | 3.389 | +1 | 5 |
| - | - | 4.668E+04 | 555.3 | - | - | 0 | - |
| - | - | 8482 | 556.3 | - | - | 0 | - |
| - | - | 3230 | 564.3 | - | - | 0 | - |
| - | - | 796.6 | 565.3 | - | - | 0 | - |
| - | - | 1044 | 566.3 | - | - | 0 | - |
| - | - | 4752 | 567.3 | - | - | 0 | - |
| - | - | 1442 | 568.3 | - | - | 0 | - |
| - | - | 1577 | 568.3 | - | - | 0 | - |
| - | - | 2436 | 572.3 | - | - | 0 | - |
| - | - | 1056 | 573.3 | - | - | 0 | - |
| - | - | 2111 | 576.3 | - | - | 0 | - |
| - | - | 1297 | 579.3 | - | - | 0 | - |
| - | - | 3670 | 582.3 | - | - | 0 | - |
| - | - | 1652 | 582.3 | - | - | 0 | - |
| - | - | 5094 | 583.3 | - | - | 0 | - |
| - | - | 8975 | 584.3 | - | - | 0 | - |
| - | - | 2618 | 585.3 | - | - | 0 | - |
| - | - | 6022 | 586.3 | - | - | 0 | - |
| - | - | 1547 | 587.3 | - | - | 0 | - |
| - | - | 773.7 | 590.3 | - | - | 0 | - |
| - | - | 1899 | 592.3 | - | - | 0 | - |
| - | - | 4152 | 593.3 | - | - | 0 | - |
| - | - | 3735 | 594.3 | - | - | 0 | - |
| - | - | 1375 | 595.3 | - | - | 0 | - |
| - | - | 2463 | 596.3 | - | - | 0 | - |
| - | - | 4367 | 597.3 | - | - | 0 | - |
| - | - | 954.8 | 598.3 | - | - | 0 | - |
| - | - | 8777 | 600.3 | - | - | 0 | - |
| - | - | 5429 | 600.3 | - | - | 0 | - |
| - | - | 2752 | 601.3 | - | - | 0 | - |
| - | - | 2271 | 601.3 | - | - | 0 | - |
| - | - | 1027 | 605.3 | - | - | 0 | - |
| - | - | 1673 | 606.3 | - | - | 0 | - |
| - | - | 1047 | 607.3 | - | - | 0 | - |
| - | - | 2532 | 607.4 | - | - | 0 | - |
| - | - | 1114 | 608.4 | - | - | 0 | - |
| - | - | 922.4 | 609.3 | - | - | 0 | - |
| - | - | 1.45E+04 | 610.3 | - | - | 0 | - |
| - | - | 3970 | 611.3 | - | - | 0 | - |
| - | - | 2.719E+04 | 612.3 | - | - | 0 | - |
| - | - | 9162 | 613.3 | - | - | 0 | - |
| - | - | 1.08E+04 | 614.4 | - | - | 0 | - |
| - | - | 2993 | 615.4 | - | - | 0 | - |
| - | - | 2832 | 620.3 | - | - | 0 | - |
| - | - | 4528 | 621.3 | - | - | 0 | - |
| - | - | 921.1 | 622.3 | - | - | 0 | - |
| - | - | 2514 | 623.3 | - | - | 0 | - |
| 7 | b | 1.147E+04 | 624.3 | 0.001456 | 2.333 | +1 | 7 |
| - | - | 3811 | 625.3 | - | - | 0 | - |
| - | - | 764.8 | 626.3 | - | - | 0 | - |
| - | - | 2.405E+04 | 628.3 | - | - | 0 | - |
| - | - | 8047 | 629.3 | - | - | 0 | - |
| - | - | 1172 | 630.3 | - | - | 0 | - |
| - | - | 7368 | 633.3 | - | - | 0 | - |
| - | - | 1538 | 634.3 | - | - | 0 | - |
| - | - | 1967 | 634.3 | - | - | 0 | - |
| - | - | 8627 | 635.4 | - | - | 0 | - |
| - | - | 1129 | 636.3 | - | - | 0 | - |
| - | - | 3300 | 636.4 | - | - | 0 | - |
| - | - | 4224 | 638.3 | - | - | 0 | - |
| - | - | 1769 | 639.3 | - | - | 0 | - |
| 7 | b | 1.999E+04 | 642.3 | 0.002061 | 3.209 | +1 | 7 |
| - | - | 7898 | 643.4 | - | - | 0 | - |
| - | - | 984.2 | 644.3 | - | - | 0 | - |
| - | - | 1725 | 647.4 | - | - | 0 | - |
| - | - | 7514 | 650.3 | - | - | 0 | - |
| 6 | y | 2.121E+04 | 651.3 | 0.002956 | 4.539 | +1 | 6 |
| - | - | 9323 | 652.3 | - | - | 0 | - |
| - | - | 2259 | 653.3 | - | - | 0 | - |
| - | - | 2936 | 653.4 | - | - | 0 | - |
| - | - | 3567 | 654.4 | - | - | 0 | - |
| - | - | 950.1 | 655.4 | - | - | 0 | - |
| - | - | 1386 | 663.3 | - | - | 0 | - |
| - | - | 4231 | 664.4 | - | - | 0 | - |
| - | - | 1268 | 665.4 | - | - | 0 | - |
| - | - | 1222 | 667.3 | - | - | 0 | - |
| 6 | y | 5.87E+05 | 669.3 | 0.002584 | 3.861 | +1 | 6 |
| - | - | 1.989E+05 | 670.3 | - | - | 0 | - |
| - | - | 1441 | 670.5 | - | - | 0 | - |
| - | - | 4.279E+04 | 671.3 | - | - | 0 | - |
| - | - | 2964 | 672.3 | - | - | 0 | - |
| - | - | 1747 | 672.4 | - | - | 0 | - |
| - | - | 3511 | 679.3 | - | - | 0 | - |
| - | - | 1174 | 680.3 | - | - | 0 | - |
| - | - | 1.093E+04 | 681.4 | - | - | 0 | - |
| - | - | 2429 | 681.8 | - | - | 0 | - |
| - | - | 3571 | 682.4 | - | - | 0 | - |
| - | - | 1263 | 683.4 | - | - | 0 | - |
| - | - | 1854 | 688.8 | - | - | 0 | - |
| - | - | 1198 | 689.3 | - | - | 0 | - |
| - | - | 1629 | 689.8 | - | - | 0 | - |
| - | - | 937.9 | 691.3 | - | - | 0 | - |
| - | - | 1.215E+04 | 692.4 | - | - | 0 | - |
| - | - | 4209 | 693.4 | - | - | 0 | - |
| - | - | 1028 | 695.4 | - | - | 0 | - |
| - | - | 1648 | 696.4 | - | - | 0 | - |
| - | - | 2.384E+04 | 699.4 | - | - | 0 | - |
| - | - | 7842 | 700.4 | - | - | 0 | - |
| - | - | 2237 | 701.4 | - | - | 0 | - |
| - | - | 6427 | 704.4 | - | - | 0 | - |
| - | - | 2637 | 705.4 | - | - | 0 | - |
| - | - | 1069 | 709.4 | - | - | 0 | - |
| - | - | 1012 | 710.4 | - | - | 0 | - |
| - | - | 974.1 | 713.4 | - | - | 0 | - |
| - | - | 609.7 | 713.4 | - | - | 0 | - |
| - | - | 793.8 | 714.4 | - | - | 0 | - |
| - | - | 4043 | 720.4 | - | - | 0 | - |
| - | - | 2.553E+04 | 721.4 | - | - | 0 | - |
| - | - | 9775 | 722.4 | - | - | 0 | - |
| 8 | b | 5928 | 723.4 | 0.002867 | 3.963 | +1 | 8 |
| - | - | 1734 | 724.4 | - | - | 0 | - |
| - | - | 3252 | 731.4 | - | - | 0 | - |
| - | - | 1957 | 732.4 | - | - | 0 | - |
| - | - | 3053 | 737.4 | - | - | 0 | - |
| - | - | 1068 | 738.4 | - | - | 0 | - |
| 8 | b | 7309 | 741.4 | 0.0028 | 3.777 | +1 | 8 |
| - | - | 2302 | 742.4 | - | - | 0 | - |
| - | - | 6.342E+04 | 749.4 | - | - | 0 | - |
| - | - | 2.905E+04 | 750.4 | - | - | 0 | - |
| - | - | 6218 | 751.4 | - | - | 0 | - |
| 5 | y | 3697 | 764.4 | 0.001778 | 2.326 | +1 | 7 |
| - | - | 1822 | 765.4 | - | - | 0 | - |
| - | - | 6345 | 767.4 | - | - | 0 | - |
| - | - | 3058 | 768.4 | - | - | 0 | - |
| - | - | 889.1 | 774.9 | - | - | 0 | - |
| - | - | 1047 | 776.4 | - | - | 0 | - |
| - | - | 1158 | 779.4 | - | - | 0 | - |
| 5 | y | 1.03E+05 | 782.4 | 0.00226 | 2.889 | +1 | 7 |
| - | - | 4.493E+04 | 783.4 | - | - | 0 | - |
| - | - | 1.215E+04 | 784.4 | - | - | 0 | - |
| - | - | 1155 | 790.4 | - | - | 0 | - |
| - | - | 3565 | 791.4 | - | - | 0 | - |
| - | - | 1681 | 792.4 | - | - | 0 | - |
| - | - | 1158 | 793.4 | - | - | 0 | - |
| 9 | b | 4685 | 794.4 | 0.001275 | 1.605 | +1 | 9 |
| - | - | 1867 | 795.4 | - | - | 0 | - |
| - | - | 805.1 | 801.4 | - | - | 0 | - |
| - | - | 1.652E+04 | 808.4 | - | - | 0 | - |
| - | - | 7438 | 809.4 | - | - | 0 | - |
| - | - | 2153 | 810.4 | - | - | 0 | - |
| 9 | b | 5633 | 812.5 | 0.002551 | 3.14 | +1 | 9 |
| - | - | 2362 | 813.5 | - | - | 0 | - |
| - | - | 6227 | 818.4 | - | - | 0 | - |
| - | - | 2631 | 819.4 | - | - | 0 | - |
| 4 | y | 4051 | 821.4 | 0.00192 | 2.338 | +1 | 8 |
| - | - | 1777 | 822.4 | - | - | 0 | - |
| - | - | 808.5 | 823.4 | - | - | 0 | - |
| - | - | 1112 | 828.4 | - | - | 0 | - |
| - | - | 4.423E+04 | 836.4 | - | - | 0 | - |
| - | - | 1.905E+04 | 837.4 | - | - | 0 | - |
| - | - | 4283 | 838.4 | - | - | 0 | - |
| 4 | y | 1.409E+05 | 839.4 | 0.002464 | 2.936 | +1 | 8 |
| - | - | 6.254E+04 | 840.4 | - | - | 0 | - |
| - | - | 1.725E+04 | 841.4 | - | - | 0 | - |
| - | - | 1362 | 842.4 | - | - | 0 | - |
| - | - | 3556 | 846.4 | - | - | 0 | - |
| - | - | 929.7 | 847.4 | - | - | 0 | - |
| - | - | 2894 | 850.4 | - | - | 0 | - |
| - | - | 1035 | 852.5 | - | - | 0 | - |
| - | - | 5136 | 854.4 | - | - | 0 | - |
| - | - | 2281 | 855.4 | - | - | 0 | - |
| 3 | y | 2.679E+04 | 878.4 | 0.00249 | 2.835 | +1 | 9 |
| - | - | 1.287E+04 | 879.4 | - | - | 0 | - |
| - | - | 2874 | 880.4 | - | - | 0 | - |
| - | - | 860.3 | 881.4 | - | - | 0 | - |
| - | - | 3106 | 894.4 | - | - | 0 | - |
| - | - | 2155 | 895.4 | - | - | 0 | - |
| 3 | y | 8.074E+05 | 896.4 | 0.00279 | 3.112 | +1 | 9 |
| - | - | 3.986E+05 | 897.5 | - | - | 0 | - |
| - | - | 1.077E+05 | 898.5 | - | - | 0 | - |
| - | - | 1.158E+04 | 899.5 | - | - | 0 | - |
| - | - | 1525 | 906.4 | - | - | 0 | - |
| - | - | 2834 | 937.5 | - | - | 0 | - |
| - | - | 753.6 | 938.5 | - | - | 0 | - |
| - | - | 2260 | 947.5 | - | - | 0 | - |
| - | - | 2074 | 948.4 | - | - | 0 | - |
| 10 | b | 2694 | 949.5 | 0.003454 | 3.638 | +1 | 10 |
| - | - | 1156 | 950.5 | - | - | 0 | - |
| - | - | 1922 | 963.5 | - | - | 0 | - |
| 2 | y | 6.023E+04 | 965.5 | 0.002444 | 2.532 | +1 | 10 |
| - | - | 3.342E+04 | 966.5 | - | - | 0 | - |
| - | - | 9986 | 967.5 | - | - | 0 | - |
| - | - | 982.3 | 968.5 | - | - | 0 | - |
| - | - | 2.559E+04 | 975.5 | - | - | 0 | - |
| - | - | 1.475E+04 | 976.5 | - | - | 0 | - |
| - | - | 4334 | 977.5 | - | - | 0 | - |
| - | - | 3030 | 981.5 | - | - | 0 | - |
| - | - | 964.1 | 982.5 | - | - | 0 | - |
| 2 | y | 6.036E+05 | 983.5 | 0.002927 | 2.976 | +1 | 10 |
| - | - | 3.189E+05 | 984.5 | - | - | 0 | - |
| - | - | 9.425E+04 | 985.5 | - | - | 0 | - |
| - | - | 1.017E+04 | 986.5 | - | - | 0 | - |
| - | - | 4.472E+04 | 993.5 | - | - | 0 | - |
| - | - | 2.485E+04 | 994.5 | - | - | 0 | - |
| - | - | 8834 | 995.5 | - | - | 0 | - |
| - | - | 695.7 | 3073 | - | - | 0 | - |

m/z Charge Intensity FragmentType MassShift Position
120.08136749267578 0 1455.2085
121.0402603149414 0 615.23663
122.07180786132812 0 2509.2327
125.03519439697266 0 2050.7207
125.10791015625 0 65751.45
126.09195709228516 0 787.8091
126.11128997802734 0 4638.9937
126.12828063964844 0 1108.0251
127.05074310302734 0 2393.8486
127.08727264404297 0 1460.6068
127.12368774414062 0 823.4832
128.10755920410156 0 29386.447
129.06649780273438 0 4027.9797
129.1029052734375 0 3118.3289
129.11090087890625 0 2660.2354
130.0503692626953 0 1896.3992 y Water loss 10
130.0869903564453 0 937.6785
131.11846923828125 0 997.4068
132.0768585205078 0 430.40363
132.1024627685547 0 2327.4307
133.0613555908203 0 887.8261
133.0863494873047 0 517.6482
136.07620239257812 0 3366.116
136.08755493164062 0 11761.543
137.09054565429688 0 640.03143
137.10800170898438 0 713.80096
138.0667724609375 0 22401.07
138.0919647216797 0 12243.171
139.05087280273438 0 2304.011
139.07008361816406 0 1404.065
139.0953826904297 0 800.6463
139.12327575683594 0 628.2033
140.1439971923828 0 5422.057
141.06668090820312 0 1472.7533
141.10279846191406 0 2824.7622
142.08689880371094 0 1538.8013
142.1234588623047 0 586.63416
143.1185302734375 0 97879.24
144.11582946777344 0 961.39874
144.12193298339844 0 6910.753
145.06138610839844 0 52662.223
146.06483459472656 0 3177.6401
148.0548095703125 0 919.35803
148.0610809326172 0 12741.898 y 10
148.0875244140625 0 2140.1655
148.95436096191406 0 1007.359
149.04562377929688 0 1229.542
150.06683349609375 0 3334.755
151.08726501464844 0 491.41565
152.07122802734375 0 1369.0724
152.08248901367188 0 1240.949
153.10279846191406 0 803.6569
154.0620574951172 0 806.0216
154.08734130859375 0 558.01
154.15968322753906 0 13368.843
155.0457000732422 0 4142.154
155.1185302734375 0 79711.125 a Water loss 1
155.1628875732422 0 1484.8041
156.077392578125 0 39752.695
156.1024932861328 0 9519.923 a Ammonia loss 1
156.115234375 0 1039.3495
156.1218719482422 0 6796.7095
157.06137084960938 0 16538.615
157.0806121826172 0 2479.7988
157.09764099121094 0 2324.5142
157.13412475585938 0 2744.6897 d 1
158.06494140625 0 991.2381
158.09271240234375 0 819.70386
159.09210205078125 0 1234.2527
159.113525390625 0 1143.9326
163.07176208496094 0 1056.6207
163.0985107421875 0 1914.6761
164.08248901367188 0 4415.1
164.1187286376953 0 3847.4539
165.10255432128906 0 624.77954
165.11410522460938 0 3234.8425
165.1208953857422 0 583.3396
166.06175231933594 0 26085.266
166.1346893310547 0 850.8682
167.0458221435547 0 3583.2349
167.05673217773438 0 966.27844
167.0651092529297 0 1696.9751
168.04920959472656 0 644.3275
169.09783935546875 0 19792.277
170.081787109375 0 733.7709
170.101318359375 0 1653.2308
171.07765197753906 0 678.7863
171.11354064941406 0 43538.383
171.14990234375 0 11466.253
172.07264709472656 0 848.5688
172.11695861816406 0 3079.908
172.15338134765625 0 1132.2781
173.05636596679688 0 1205.8716
173.12916564941406 0 526011.94 a 1
174.08798217773438 0 6996.3965
174.1325225830078 0 42945.977
175.07200622558594 0 861.9823
175.1190643310547 0 643.12646
175.1342315673828 0 1807.9579
176.0827178955078 0 1587.4136
178.1343536376953 0 1565.89
179.0821075439453 0 781.7975
179.0924530029297 0 457.54254
180.07745361328125 0 4447.891
181.0985870361328 0 2931.6865
181.10910034179688 0 61808.523
182.05670166015625 0 1074.3206
182.08209228515625 0 885.024
182.10630798339844 0 502.6988
182.1126251220703 0 4886.7607
182.12985229492188 0 857.6052
183.11355590820312 0 81747.84 b Water loss 1
184.0724334716797 0 11248.024
184.09764099121094 0 930.9013 b Ammonia loss 1
184.11695861816406 0 6953.679
185.05718994140625 0 698.4903
185.0758819580078 0 1033.1652
185.0935516357422 0 746.6734
185.12913513183594 0 5830.908
185.1654815673828 0 673.4399
186.0882110595703 0 767.059
186.13302612304688 0 596.0072
187.10848999023438 0 72395.76
187.14508056640625 0 1911.1711
188.1118927001953 0 5764.035
189.12411499023438 0 602.0863
191.0846710205078 0 545.5443
191.09347534179688 0 3246.1033
192.07749938964844 0 54528.332
192.11410522460938 0 1113.3639
193.0809326171875 0 4487.8276
193.0980224609375 0 2310.5308
193.10801696777344 0 561.6999
194.05645751953125 0 734.064
194.0931396484375 0 2981.7876
194.101806640625 0 508.99252
195.08811950683594 0 936.2513
195.1135711669922 0 7695.351
195.18618774414062 0 723.17786
196.1168212890625 0 732.43726
197.09259033203125 0 1329.15
197.12911987304688 0 1642.9923
197.16571044921875 0 2631.5186
198.07725524902344 0 1195.2056
198.0869598388672 0 707.92413
199.10841369628906 0 7168.7754
199.144775390625 0 12772.465
199.18141174316406 0 1860.1727
200.06735229492188 0 1604.0581
200.11244201660156 0 953.5703
200.14007568359375 0 28118.785
201.12411499023438 0 469863.75 b 1
202.08290100097656 0 102344.664
202.12745666503906 0 43659.58
203.06698608398438 0 1139.9437
203.08633422851562 0 6629.424
203.1293182373047 0 2552.375
204.0777130126953 0 1296.0432
205.06187438964844 0 533.1727
207.08824157714844 0 4321.2866
207.114013671875 0 583.3759
207.1496124267578 0 723.64044
208.07241821289062 0 4049.9128
209.10406494140625 0 240548.22
210.107421875 0 21015.959
210.12448120117188 0 5321.197
211.10862731933594 0 1603.8256
211.1280059814453 0 904.7592
211.14451599121094 0 1549.2637
211.18072509765625 0 795.0964
212.06739807128906 0 6918.926
212.1402130126953 0 5015.891 a Water loss 2
213.12413024902344 0 13826.205 a Ammonia loss 2
213.1427764892578 0 745.7033
213.1598358154297 0 1001.74976
214.119384765625 0 3479.992
214.12948608398438 0 954.99756
214.1556854248047 0 2049.9602
215.10340881347656 0 80766.09
216.10679626464844 0 8629.681
217.1085205078125 0 1057.8901
219.08804321289062 0 2166.5269
219.1092529296875 0 1261.043 y Water loss 7
221.08518981933594 0 4313.8896
221.104248046875 0 9026.759
222.08795166015625 0 1109.7019
222.107666015625 0 1249.4104
222.12457275390625 0 3474.168
223.10841369628906 0 2107.4387
223.1815185546875 0 1444.0227
224.14019775390625 0 1907.1826
225.04367065429688 0 3652.108
225.098876953125 0 1149.6155
225.12359619140625 0 705.76556
225.160400390625 0 3249.0642
226.08314514160156 0 686.0525
226.1195068359375 0 1735.4885
226.1309814453125 0 913.3644
226.15472412109375 0 1340.617
227.103515625 0 1583.8484
227.1147918701172 0 17069.648
227.1400909423828 0 711.3696
227.17637634277344 0 3238.9114
228.09881591796875 0 2232.1128
228.13516235351562 0 63815.395
229.11917114257812 0 52126.9
229.13882446289062 0 5282.715
230.07803344726562 0 9879.381
230.12249755859375 0 4709.3535
231.08111572265625 0 589.66644
233.11329650878906 0 731.305
235.10841369628906 0 740.8368
235.14491271972656 0 931.6377
236.1034393310547 0 1045.2542
237.0992889404297 0 5084.389
237.1606903076172 0 945.862
238.11959838867188 0 3161.4072
239.09588623046875 0 6997.759
239.11483764648438 0 22268.09
239.1757354736328 0 1192.9983
240.1351776123047 0 116301.27 b Water loss 2
241.11880493164062 0 1353.5432 b Ammonia loss 2
241.13853454589844 0 14516.696
241.15452575683594 0 975.49207
241.16830444335938 0 958.17413
241.1919708251953 0 14408.269
242.07766723632812 0 1687.1323
242.1143798828125 0 3491.7024
242.13864135742188 0 607.16016
242.15101623535156 0 2197.5981
242.19558715820312 0 1090.1775
244.09336853027344 0 1245.868
244.12954711914062 0 956.39
245.12789916992188 0 617.06415
245.14073181152344 0 2384.5725
245.1874237060547 0 550.7536
249.0991973876953 0 32870
250.08277893066406 0 2220.975
250.1020965576172 0 3319.6934
251.13885498046875 0 851.5303
251.15182495117188 0 1313.6447
251.17637634277344 0 1638.2903
252.13511657714844 0 9465.383
253.09414672851562 0 1253.0411
253.13894653320312 0 1267.931
253.15509033203125 0 919.615
254.0781707763672 0 627.94525
254.11427307128906 0 1258.0541
254.15084838867188 0 2202.5771
254.18714904785156 0 2404.2378
255.10960388183594 0 1402.4238
255.1339111328125 0 812.0164
255.17124938964844 0 6133.454
256.1300354003906 0 19830.959
256.1754150390625 0 838.0367
257.114013671875 0 1124.8438
257.1332092285156 0 1669.1073
258.1457214355469 0 17478.87 b 2
259.14898681640625 0 1765.8304
262.1195983886719 0 722.1439
263.1511535644531 0 864.35394
263.18743896484375 0 1657.0994
264.1827087402344 0 2774.4897
265.1544189453125 0 1028.9454
266.1261901855469 0 1132.3422
267.1095886230469 0 85143.05 y Water loss 9
267.1453552246094 0 2967.5225
268.09521484375 0 955.0039
268.1128845214844 0 9894.783
268.1305847167969 0 1513.1552
268.1665344238281 0 3358.6216
269.1148681640625 0 1627.537
269.14971923828125 0 4944.776
269.1619567871094 0 11004.907
269.1868896484375 0 13824.055
270.145751953125 0 16481.637
270.1647033691406 0 1514.0085
270.1827392578125 0 4129.7534
271.1045837402344 0 6832.376
271.149169921875 0 1963.7473
271.1773376464844 0 4187.3975
272.1250305175781 0 4171.642
272.1793212890625 0 627.7273
273.1202392578125 0 2039.041
273.13525390625 0 1452.4031
277.6507568359375 0 622.0383 y 6
279.14617919921875 0 14332.011
280.0929260253906 0 757.6203
280.13018798828125 0 7761.617
280.14898681640625 0 2088.1326
280.1778564453125 0 5757.2476
281.1336364746094 0 877.0442
281.1813659667969 0 1119.5593
282.18231201171875 0 13085.572
283.10491943359375 0 1372.7778
283.1413879394531 0 2365.8777
283.166259765625 0 9358.35
283.184814453125 0 1583.1526
284.125 0 2077.6792
284.17041015625 0 1215.5083
284.1990051269531 0 1943.4604
285.12042236328125 0 694461.56 y 9
286.1233215332031 0 79517.555
286.1409606933594 0 69551.02
286.1773986816406 0 2532.6155
287.1251220703125 0 9726.588
287.14349365234375 0 8315.34
287.17242431640625 0 154188.12
288.145751953125 0 1759.4695
288.1755676269531 0 21268.064
289.17755126953125 0 1518.273
290.162109375 0 10243.392
291.14666748046875 0 1066.9323
291.1648254394531 0 1303.4705
291.1827087402344 0 2665.1555
294.1200866699219 0 740.2365
295.104736328125 0 11333.713
295.14166259765625 0 1708.3455
296.10797119140625 0 912.4682
296.16162109375 0 3264.7632
296.1978454589844 0 1373.6582
297.1568908691406 0 57553.05 b Water loss 3
298.1409912109375 0 2978.0845 b Ammonia loss 3
298.1600341796875 0 6868.55
298.17755126953125 0 3955.197
299.0628662109375 0 3117.7969
299.09967041015625 0 9662.035
299.17242431640625 0 2492.5464
299.2088623046875 0 5413.613
300.1025695800781 0 1230.4742
300.1566162109375 0 1484.0095
300.19287109375 0 3326.099
301.1155090332031 0 2541.0103
301.151611328125 0 5721.7188
302.1357421875 0 1840.9264
302.1540832519531 0 1147.9457
303.1116943359375 0 817.6754
303.1676940917969 0 786.769
307.14129638671875 0 9884.774
308.1730041503906 0 60368.047
309.1194152832031 0 942.63184
309.1758117675781 0 8086.7896
310.1565246582031 0 626.8511
310.1783447265625 0 2313.5352
311.13592529296875 0 3732.731
311.158447265625 0 1111.5408
312.1392517089844 0 993.7515
312.1584777832031 0 1391.0309
312.192626953125 0 1361.9554
313.1521301269531 0 1091.5751
313.18841552734375 0 2886.1306
313.2256164550781 0 1591.7595
314.1377258300781 0 1325.0302
314.17236328125 0 54414.727
315.1676940917969 0 224197.8 b 3
316.17071533203125 0 32769.086
317.1104431152344 0 20852.281
317.17254638671875 0 3209.8481
318.11328125 0 2468.9119
318.1572265625 0 888.9682
320.1368713378906 0 3744.4363
320.1728210449219 0 2488.471
321.12066650390625 0 33322.785
322.12420654296875 0 4442.678
323.15020751953125 0 1038.2035
323.2091369628906 0 913.5671
324.133056640625 0 961.2749
325.1519775390625 0 17931.906
325.2232360839844 0 923.81256
326.1541442871094 0 2192.9148 y Water loss 5
326.1835021972656 0 6241.5684
326.2085266113281 0 3313.0566
327.1672058105469 0 1408.8834
327.2044372558594 0 3385.1267
328.1880798339844 0 14866.532
329.1468200683594 0 31061.629
329.19140625 0 2586.1443
330.1496276855469 0 5047.8013
335.2091369628906 0 973.1604
336.16790771484375 0 4496.5244
337.17010498046875 0 680.2675
337.2020263671875 0 956.7222
337.2244567871094 0 888.1208
338.14727783203125 0 34134.297 y Water loss 8
339.150146484375 0 5261.8975
339.20391845703125 0 3974.7
340.1882629394531 0 5073.8687
341.1468811035156 0 2194.507
341.1846008300781 0 3083.7778
341.21966552734375 0 11938.779
342.2228698730469 0 2724.0745
343.16259765625 0 119547.2
344.1656188964844 0 18585.605
345.168212890625 0 3009.4253
348.12908935546875 0 952.9494
349.150634765625 0 1274.6305
351.20306396484375 0 913.12744
352.1635437011719 0 1016.6351
353.1466064453125 0 5148.8125
353.2196960449219 0 2376.8828
354.14715576171875 0 963.6935
354.1788330078125 0 1317.7255
354.20196533203125 0 1037.0387
355.0706481933594 0 697.55536
355.1630554199219 0 1292.2229
355.1993408203125 0 1451.6022
355.23529052734375 0 3668.3625
356.1577453613281 0 404597.4 y 8
357.16058349609375 0 68750.11
357.2147216796875 0 12917.675
358.1623229980469 0 7004.053
358.2130126953125 0 3151.342
359.1587219238281 0 734.4148
362.2225341796875 0 794.7039
365.19549560546875 0 910.67944
365.2200012207031 0 3499.1475
366.1420593261719 0 10702.313
366.1785888671875 0 1781.352
367.1625671386719 0 2514.7031
367.1996154785156 0 1360.9742
368.1938781738281 0 3655.595
368.2308044433594 0 1494.757
369.1232604980469 0 3064.4116
369.17803955078125 0 6731.1245
369.2145080566406 0 11864.73
370.1755676269531 0 1725.663
370.19793701171875 0 962.45184
370.21826171875 0 2037.4944
370.24652099609375 0 2503.693
371.1575927734375 0 1716.6117
371.1940612792969 0 3700.3584
371.2283630371094 0 740.4767
372.1910095214844 0 1640.9908
377.1496276855469 0 790.2473
378.1778869628906 0 1695.7722
378.21917724609375 0 587.6972
379.2465515136719 0 11984.139
380.15777587890625 0 2260.6125
380.2498779296875 0 2162.4204
381.2512512207031 0 3055.2104
382.1733703613281 0 2110.4302
382.2086486816406 0 1142.8752
382.2452392578125 0 12223.543
382.7022705078125 0 1087.2632 y Water loss 4
383.2304382324219 0 16471.918
384.18902587890625 0 18594.215
384.2331237792969 0 3780.156
385.17279052734375 0 2660.5576
385.2095031738281 0 33755.797
386.2108154296875 0 7870.959
386.2411193847656 0 1514.5189
387.2107238769531 0 875.5942
388.1840515136719 0 3877.2026
391.7071228027344 0 15358.107 y 4
391.7342529296875 0 1077.3517
392.208740234375 0 3952.4836
392.2335510253906 0 813.8922
392.71087646484375 0 2097.6216
393.2145080566406 0 819.85986
394.173583984375 0 5432.6006
395.15753173828125 0 19465.125
396.16070556640625 0 3724.873
396.1895446777344 0 4693.57
396.22540283203125 0 12540.501
396.2511901855469 0 1128.2709
397.209716796875 0 31658.207
398.16864013671875 0 8271.728
398.212646484375 0 6820.2974 b Ammonia loss 8
398.2411193847656 0 1801.1787
399.17169189453125 0 874.1354
399.22369384765625 0 3169.0674
399.2625732421875 0 1405.166
400.1845397949219 0 3869.527
400.25701904296875 0 112478.53
401.26025390625 0 24110.912
402.19970703125 0 9823.992
402.2622985839844 0 2979.8267
403.2021179199219 0 2406.8376
406.2098693847656 0 5827.9746
407.2103576660156 0 1960.3325
407.2417297363281 0 16282.362
408.24371337890625 0 3585.8533
409.21624755859375 0 1839.7223
409.24822998046875 0 882.4395
410.1692199707031 0 2857.758
410.2081298828125 0 2449.2712
410.2415466308594 0 23116.523 b Water loss 4
411.1752624511719 0 726.09845
411.24468994140625 0 4152.482
411.7122802734375 0 736.3149
412.1842956542969 0 51636.68
413.18731689453125 0 9836.485
414.20037841796875 0 8517.095
414.23638916015625 0 47594.508
415.2062072753906 0 1074.0848
415.23944091796875 0 10084.611
416.1791076660156 0 16250.984
417.182861328125 0 3144.1858
418.71966552734375 0 887.6274
420.1890869140625 0 4547.034
420.22021484375 0 3564.4727 y 3
420.71917724609375 0 1756.9236
421.189453125 0 728.7931
422.1685485839844 0 13033.986
423.1707458496094 0 2087.728
423.2004699707031 0 2589.8108
423.2325744628906 0 825.76324
424.18896484375 0 831.12585
424.2209167480469 0 13377.73
425.2230529785156 0 3025.0327
425.2531433105469 0 2732.2983
425.7279968261719 0 770.84875
426.23553466796875 0 1476.1257
427.2251281738281 0 1048.2747
427.2572937011719 0 3261.8162
428.2164611816406 0 8972.342
428.2521667480469 0 68140.13 b 4
429.2201843261719 0 1667.5115
429.2551574707031 0 16970.389
430.19500732421875 0 71519.17
430.2585754394531 0 2156.6128
431.19805908203125 0 13724.723
432.19989013671875 0 1838.4299
435.20062255859375 0 9009.223
435.2358093261719 0 2129.8538
436.20318603515625 0 1934.8984
436.2329406738281 0 680.4846
437.2159729003906 0 13309.69 y Water loss 7
438.2200927734375 0 1685.3562
438.269775390625 0 1073.924
439.25701904296875 0 2317.959
439.7237854003906 0 5310.708 y Water loss 2
440.1795349121094 0 22976.736
440.2200012207031 0 1846.9419
440.2540588378906 0 2751.3384
441.18194580078125 0 5028.6655
442.2314453125 0 60346.99
443.23431396484375 0 13649.588
444.2376403808594 0 1484.3752
448.72894287109375 0 24383.129 y 2
449.23004150390625 0 10380.042
449.7318115234375 0 2152.7817
452.21954345703125 0 809.63153
452.2887268066406 0 1391.929
453.2755126953125 0 1534.1533
453.7196044921875 0 1144.5469
454.2308044433594 0 2051.3306
454.27294921875 0 986.7426
454.3040466308594 0 1048.1893
455.2267150878906 0 286112.03 y 7
456.22967529296875 0 58613.95
456.28619384765625 0 1657.373
457.2314453125 0 8812.184
457.2701721191406 0 630.0137
457.2838134765625 0 961.60547
461.2629089355469 0 2244.3257
461.76397705078125 0 1145.3229
465.2110290527344 0 8154.083
465.246826171875 0 1980.7454
465.7217102050781 0 835.9375
466.22857666015625 0 2493.268
466.2581787109375 0 3311.9924 b Water loss 9
466.7578125 0 1827.507
467.2314453125 0 632.23987
467.2624206542969 0 4131.3306
468.2467956542969 0 8747.156
468.2822265625 0 2078.7026
469.2069396972656 0 1065.0089
469.2498779296875 0 2282.6912
470.22393798828125 0 979.6327
470.3002624511719 0 1451.0089
471.2579650878906 0 1283.0453
472.239990234375 0 1264.1267
475.2607421875 0 12144.985 b 9
475.7624206542969 0 8250.762
476.2636413574219 0 2295.5378
477.2477722167969 0 7584.2065
478.2503356933594 0 1582.9513
479.2622985839844 0 1517.7593
480.22210693359375 0 1682.564
481.2442321777344 0 1955.3473
482.2316589355469 0 2231.2505
482.297607421875 0 1671.1921
482.7315979003906 0 1718.104
483.2532043457031 0 5796.9233
483.7414245605469 0 5238.9834
484.2420349121094 0 21078.688
484.276123046875 0 4117.6
485.24383544921875 0 2751.1372
485.2742614746094 0 6861.604
486.2779541015625 0 1239.022
487.2170715332031 0 1825.8995
487.252197265625 0 979.9325
488.23150634765625 0 7842.375
488.73358154296875 0 3098.7573
489.2354736328125 0 1431.1884
492.24517822265625 0 48800.246 y 1
492.74658203125 0 32301.262
493.2456970214844 0 12869.257
494.2303161621094 0 2235.0315
494.2740478515625 0 28150.564
495.25885009765625 0 2838.261
495.2872314453125 0 3906.4426
496.2779235839844 0 7176.5723
497.2370300292969 0 12739.319
497.2763977050781 0 4461.4995
497.7384948730469 0 5843.8403
498.25579833984375 0 1713.6002
498.29180908203125 0 2308.7397
499.2529602050781 0 9185.918
499.2934875488281 0 1342.7797
500.2550964355469 0 1903.6827
501.2314147949219 0 2514.1946
501.2684631347656 0 25580.793
502.2717590332031 0 6634.3276
504.25787353515625 0 941.1417
505.241943359375 0 1208.7444
506.2734069824219 0 3373.4321
507.2591552734375 0 3542.8054
508.2467956542969 0 1152.6632
509.2427978515625 0 970.3203
509.3088684082031 0 856.7244
510.2949523925781 0 2383.957
511.25299072265625 0 38241.137
512.2564697265625 0 9182.232
513.26708984375 0 17302.969
513.3059692382812 0 10983.388
514.271240234375 0 5004.0454
514.30859375 0 3379.1116
515.2469482421875 0 3630.7617
515.2835693359375 0 4198.907
516.2508544921875 0 1682.5846
516.289306640625 0 1174.1615
519.2537841796875 0 843.02466
521.23681640625 0 5253.3296
522.2344970703125 0 3861.3252
522.26904296875 0 48403.902
522.7648315429688 0 1231.0234
523.2733154296875 0 11206.992
524.2734375 0 2169.2053
525.2694091796875 0 11857.299 b Water loss 5
525.7841796875 0 2856.3704
526.2752075195312 0 3152.859
527.3213500976562 0 2560.063
529.2634887695312 0 52313.51
530.2666015625 0 14076.223
530.7760009765625 0 7592.84
531.2730102539062 0 10805.955
531.7744140625 0 3911.5442
532.246826171875 0 787.7521
534.2691650390625 0 7478.3594
535.2693481445312 0 1736.0891
536.28369140625 0 7127.643 y Water loss 6
537.2909545898438 0 1635.6909
538.2877197265625 0 681.84814
539.2476196289062 0 7177.8496
539.3215942382812 0 990.7165
539.7817993164062 0 95672.9 Precursor Water loss
540.2828979492188 0 62191.63
540.7391357421875 0 523.36646
540.784423828125 0 19129.342
541.2987060546875 0 20774.252
542.3027954101562 0 6119.51
543.2791137695312 0 26635.16 b 5
544.2822265625 0 7911.1597
545.2853393554688 0 1505.4138
547.7794799804688 0 1140.9374
548.2872314453125 0 1487.5118
548.7872314453125 0 230017.42 Precursor
549.2886352539062 0 143619.47
549.7899780273438 0 48367.266
549.9664916992188 0 1133.3466
550.2907104492188 0 6800.9443
551.254638671875 0 1406.8308
552.2825927734375 0 1074.408
554.295166015625 0 165403.97 y 6
555.2982788085938 0 46684.23
556.3007202148438 0 8481.93
564.2796630859375 0 3230.144
565.2767944335938 0 796.5778
566.3323364257812 0 1043.5571
567.3164672851562 0 4752.3574
568.2723999023438 0 1441.6688
568.3192138671875 0 1576.798
572.3050537109375 0 2435.6213
573.3109130859375 0 1056.2356
576.3141479492188 0 2111.3938
579.2901000976562 0 1296.6398
582.2900390625 0 3670.396
582.329345703125 0 1652.0079
583.3098754882812 0 5094.057
584.3414306640625 0 8974.86
585.3451538085938 0 2618.422
586.2847900390625 0 6022.001
587.2855834960938 0 1547.2295
590.3349609375 0 773.6621
592.3099365234375 0 1898.8557
593.3042602539062 0 4151.7725
594.3236694335938 0 3734.726
595.3318481445312 0 1375.4005
596.3394775390625 0 2463.499
597.3291015625 0 4366.59
598.3271484375 0 954.7739
600.2999267578125 0 8777.492
600.339599609375 0 5428.6353
601.3029174804688 0 2752.4673
601.343017578125 0 2270.7224
605.3478393554688 0 1026.9242
606.328125 0 1673.4298
607.3102416992188 0 1046.6959
607.3582763671875 0 2531.5342
608.3629760742188 0 1113.726
609.3076782226562 0 922.3557
610.3215942382812 0 14495.371
611.324462890625 0 3970.0312
612.3369140625 0 27192.328
613.33984375 0 9162.332
614.3511352539062 0 10796.896
615.3563842773438 0 2992.5034
620.305419921875 0 2832.023
621.3017578125 0 4527.7935
622.3046264648438 0 921.06476
623.317626953125 0 2514.0159
624.3366088867188 0 11471.222 b Water loss 6
625.3389892578125 0 3811.2725
626.3463134765625 0 764.7906
628.3320922851562 0 24046.535
629.3350830078125 0 8047.2603
630.3388671875 0 1172.317
633.3374633789062 0 7367.8027
634.2852172851562 0 1538.407
634.3418579101562 0 1966.5138
635.353271484375 0 8626.579
636.3035888671875 0 1128.7827
636.3564453125 0 3300.3625
638.3158569335938 0 4224.0044
639.3173217773438 0 1768.5812
642.3477783203125 0 19993.5 b 6
643.3512573242188 0 7897.5195
644.3499145507812 0 984.17163
647.3500366210938 0 1724.5721
650.3278198242188 0 7514.4707
651.3126220703125 0 21208.533 y Water loss 5
652.3087768554688 0 9322.853
653.3062133789062 0 2259.2444
653.3646850585938 0 2935.7783
654.3524169921875 0 3566.6338
655.3502807617188 0 950.09576
663.3466796875 0 1385.9048
664.3790283203125 0 4230.6846
665.3909301757812 0 1268.4395
667.31103515625 0 1222.0167
669.3228149414062 0 587042.25 y 5
670.3252563476562 0 198949.53
670.4607543945312 0 1441.2836
671.3277587890625 0 42787.715
672.3313598632812 0 2963.705
672.3799438476562 0 1747.3013
679.3075561523438 0 3510.974
680.3088989257812 0 1173.604
681.3585205078125 0 10927.941
681.8359375 0 2428.9592
682.3600463867188 0 3571.0354
683.36669921875 0 1263.3534
688.8260498046875 0 1854.4366
689.325927734375 0 1197.6824
689.8292236328125 0 1629.088
691.3419189453125 0 937.8748
692.374755859375 0 12153.816
693.376708984375 0 4209.4844
695.374755859375 0 1028.1676
696.3958740234375 0 1648.3853
699.3694458007812 0 23841.96
700.37255859375 0 7842.211
701.3736572265625 0 2237.2114
704.375 0 6426.538
705.3812255859375 0 2636.6619
709.35888671875 0 1069.1852
710.3804931640625 0 1011.9868
713.3798828125 0 974.08545
713.4291381835938 0 609.7042
714.38134765625 0 793.754
720.369384765625 0 4043.4622
721.4012451171875 0 25525.502
722.4047241210938 0 9775.426
723.4064331054688 0 5928.079 b Water loss 7
724.407958984375 0 1733.745
731.3880004882812 0 3251.6445
732.387451171875 0 1957.2585
737.35888671875 0 3053.0183
738.3649291992188 0 1067.6896
741.4169311523438 0 7309.243 b 7
742.4188842773438 0 2301.6592
749.3965454101562 0 63416.066
750.3992309570312 0 29048.77
751.4012451171875 0 6217.9087
764.3955078125 0 3697.18 y Water loss 4
765.39453125 0 1821.8141
767.4067993164062 0 6345.0815
768.4107055664062 0 3057.8567
774.9058227539062 0 889.1111
776.4305419921875 0 1047.0698
779.3665161132812 0 1158.0984
782.4065551757812 0 102953.4 y 4
783.409423828125 0 44927.113
784.4119873046875 0 12147.309
790.4166259765625 0 1155.0791
791.4088134765625 0 3565.2856
792.410400390625 0 1680.6571
793.4221801757812 0 1157.7279
794.4419555664062 0 4685.4893 b Water loss 8
795.4443969726562 0 1867.3735
801.4030151367188 0 805.07385
808.4330444335938 0 16520.621
809.4373779296875 0 7438.406
810.4373168945312 0 2152.8696
812.4537963867188 0 5633.138 b 8
813.4584350585938 0 2362.0217
818.4183349609375 0 6227.4863
819.4214477539062 0 2630.7766
821.4171142578125 0 4050.7188 y Water loss 3
822.4195556640625 0 1777.3057
823.4299926757812 0 808.5451
828.40380859375 0 1111.9263
836.4287109375 0 44233.242
837.4307861328125 0 19046.049
838.4364013671875 0 4282.729
839.42822265625 0 140879.1 y 3
840.4309692382812 0 62535.883
841.433837890625 0 17245.66
842.4342651367188 0 1361.5266
846.4140625 0 3555.6204
847.423583984375 0 929.7168
850.4443359375 0 2894.45
852.4508056640625 0 1035.3578
854.4382934570312 0 5135.533
855.4415893554688 0 2280.91
878.4391479492188 0 26792.906 y Water loss 2
879.4420776367188 0 12871.151
880.4425048828125 0 2873.871
881.4337768554688 0 860.26465
894.4336547851562 0 3106.1196
895.438232421875 0 2154.6829
896.4500122070312 0 807402.75 y 2
897.4522705078125 0 398555.47
898.4552001953125 0 107735.39
899.4568481445312 0 11582.367
906.4309692382812 0 1524.7693
937.4763793945312 0 2833.9539
938.4675903320312 0 753.6083
947.45703125 0 2260.0889
948.4452514648438 0 2073.83
949.5136108398438 0 2694.2302 b 9
950.516845703125 0 1156.3387
963.4561767578125 0 1921.6501
965.4711303710938 0 60233.625 y Water loss 1
966.4743041992188 0 33416.99
967.4768676757812 0 9985.585
968.4840698242188 0 982.3004
975.455322265625 0 25589.482
976.4583129882812 0 14751.996
977.4617919921875 0 4333.9243
981.4652099609375 0 3030.1873
982.4697265625 0 964.0651
983.482177734375 0 603585.5 y 1
984.4845581054688 0 318929.56
985.4873046875 0 94251.01
986.4889526367188 0 10171.209
993.466064453125 0 44721.39
994.46875 0 24852.016
995.47021484375 0 8833.712
3072.617919921875 0 695.66547

Spectrum Details

|  |  |
| --- | --- |
| Matched peaks? Matched peaks The total absolute number of peaks matched. Additionally in brackets the total fraction of peaks matched and the total number of peaks is shown. | 60 (6.88% of 872) |
| FDR? FDR The false discovery rate estimated for this peptide. It is calculated by matching all theoretical fragments with a non-integer shift with the raw peaks for this spectrum. This is done with 40 different shifts. The resulting percentage is the average number of annotated peaks over the number of annotated peaks with the correct spectrum. | 0.24% |
| Satellite FDR? Satellite FDR See the FDR for details on its calculation. This satellite ion specific FDR only contains the satellite ions (d/w) for I/L/J positions. | - |
| PSM Score? PSM Score The PSM Score as given by Hecklib to this annotated spectrum. It is shown with three significant figures. | 796 |

#### Spectrum 5530? Spectrum 5530 The raw spectrum of this peptide as annotated by Hecklib. The fragments are coloured according to ion type (see legend). Any peaks with a star '\*' as text can be hovered over to see the full details, first the ion type second the mass shift type. By hovering over the amino acids in the peptide or ions in the legend the corresponding peaks are highlighted. By toggling the 'Unassigned' label you can turn the background (unassigned) peaks on or off in the plot. By updating the slider in the Ion legend you can update the spectrum to only show the top X% of the peaks with labels. The top X% means any peak that is within X% of the highest intensity. By dragging in the spectrum you can zoom in to a specific part of the spectrum and use 'Zoom Out' to get back to the original zoom level. The annotation of the spectrum is based on the given sequence in the peptides file and is done with different software so inconsistencies are likely. The peaks are annotated based on the given sequence, with 20 ppm tolerance.

Copy Data

##### Spectrum 5530 (TSV)

###### Preview

```
Loading example...
```

*Click on the button to copy the data to your clipboard.*

Mz MinMz MaxIntensity Max

WidthHeightPeptide font sizePeptide stroke widthSpectrum font sizeSpectrum stroke widthCompact peptide

Ion legend

wxyz

abcd

OtherUnassignedIon^Charge^_Position_Show for top:%

JSGGIDVVAHE

01.99e+43.98e+45.97e+47.96e+4

Zoom Out

y^+1^_1_y^+1^_1_a^+1^_2_a^+1^_2_a^+1^_2_b^+1^_2_b^+1^_2_a^+1^_3_a^+1^_3_y^+2^_4_b^+1^_3_b^+1^_3_y^+1^_2_b^+1^_4_b^+1^_4_y^+1^_2_b^+1^_4_y^+2^_6_y^+1^_3_y^+1^_3_y^+2^_7_y^+2^_7_b^+1^_5_y^+2^_8_b^+1^_5_y^+2^_9_y^+1^_4_y^+1^_4_b^+2^_10_b^+2^_10_y^+2^_10_y^+2^_10_b^+1^_6_b^+1^_6_\*y^+1^_5_\*y^+1^_5_b^+1^_7_b^+1^_7_y^+1^_6_y^+1^_6_b^+1^_8_b^+1^_8_y^+1^_7_b^+1^_9_y^+1^_7_b^+1^_9_y^+1^_8_y^+1^_8_y^+1^_9_y^+1^_9_y^+1^_10_y^+1^_10_

02555107651021

Fragment Matches Table

Show background peaks

| Position | Ion type | Intensity | mz Theoretical | mz Error (Th) | mz Error (ppm) | Charge | Series Number |
| --- | --- | --- | --- | --- | --- | --- | --- |
| - | - | 4379 | 120.1 | - | - | 0 | - |
| - | - | 456 | 121.1 | - | - | 0 | - |
| - | - | 457.6 | 122 | - | - | 0 | - |
| - | - | 617.6 | 122.1 | - | - | 0 | - |
| - | - | 619.5 | 124.1 | - | - | 0 | - |
| - | - | 394.7 | 125 | - | - | 0 | - |
| - | - | 8089 | 125.1 | - | - | 0 | - |
| - | - | 1715 | 126.1 | - | - | 0 | - |
| - | - | 692.1 | 126.1 | - | - | 0 | - |
| - | - | 994.6 | 127.1 | - | - | 0 | - |
| - | - | 604.8 | 127.1 | - | - | 0 | - |
| - | - | 916.7 | 128.1 | - | - | 0 | - |
| - | - | 438.3 | 128.1 | - | - | 0 | - |
| - | - | 4590 | 128.1 | - | - | 0 | - |
| - | - | 367.3 | 129 | - | - | 0 | - |
| - | - | 1521 | 129.1 | - | - | 0 | - |
| - | - | 4555 | 129.1 | - | - | 0 | - |
| 11 | y | 4362 | 130 | 0.0003624 | 2.787 | +1 | 1 |
| - | - | 1615 | 130.1 | - | - | 0 | - |
| - | - | 4436 | 130.1 | - | - | 0 | - |
| - | - | 893.9 | 131.1 | - | - | 0 | - |
| - | - | 419.5 | 131.3 | - | - | 0 | - |
| - | - | 422 | 132 | - | - | 0 | - |
| - | - | 580.4 | 132.1 | - | - | 0 | - |
| - | - | 836.9 | 132.1 | - | - | 0 | - |
| - | - | 984 | 132.1 | - | - | 0 | - |
| - | - | 529.9 | 133.1 | - | - | 0 | - |
| - | - | 344.6 | 133.4 | - | - | 0 | - |
| - | - | 384.5 | 134.9 | - | - | 0 | - |
| - | - | 1993 | 136 | - | - | 0 | - |
| - | - | 800 | 136.1 | - | - | 0 | - |
| - | - | 6073 | 136.1 | - | - | 0 | - |
| - | - | 1506 | 136.1 | - | - | 0 | - |
| - | - | 1283 | 137 | - | - | 0 | - |
| - | - | 534.8 | 137.1 | - | - | 0 | - |
| - | - | 2402 | 138 | - | - | 0 | - |
| - | - | 3890 | 138.1 | - | - | 0 | - |
| - | - | 1371 | 138.1 | - | - | 0 | - |
| - | - | 2901 | 139.1 | - | - | 0 | - |
| - | - | 663.6 | 140.1 | - | - | 0 | - |
| - | - | 706.8 | 140.1 | - | - | 0 | - |
| - | - | 732.1 | 141.1 | - | - | 0 | - |
| - | - | 1687 | 141.1 | - | - | 0 | - |
| - | - | 685.8 | 142.1 | - | - | 0 | - |
| - | - | 598.3 | 142.1 | - | - | 0 | - |
| - | - | 1.527E+04 | 143.1 | - | - | 0 | - |
| - | - | 426.3 | 143.8 | - | - | 0 | - |
| - | - | 547.7 | 144.1 | - | - | 0 | - |
| - | - | 1032 | 144.1 | - | - | 0 | - |
| - | - | 1.008E+04 | 145.1 | - | - | 0 | - |
| 11 | y | 1.242E+04 | 148.1 | 0.0003415 | 2.307 | +1 | 1 |
| - | - | 688 | 150.1 | - | - | 0 | - |
| - | - | 1845 | 151.1 | - | - | 0 | - |
| - | - | 1272 | 152 | - | - | 0 | - |
| - | - | 1232 | 152.1 | - | - | 0 | - |
| - | - | 523.8 | 152.1 | - | - | 0 | - |
| - | - | 757.1 | 154.1 | - | - | 0 | - |
| - | - | 1176 | 154.1 | - | - | 0 | - |
| - | - | 1372 | 154.2 | - | - | 0 | - |
| - | - | 1075 | 155 | - | - | 0 | - |
| - | - | 723.7 | 155.1 | - | - | 0 | - |
| - | - | 480.9 | 155.1 | - | - | 0 | - |
| 2 | a | 8580 | 155.1 | 0.0003508 | 2.262 | +1 | 2 |
| - | - | 6048 | 156.1 | - | - | 0 | - |
| 2 | a | 1350 | 156.1 | 0.0004814 | 3.084 | +1 | 2 |
| - | - | 1358 | 156.1 | - | - | 0 | - |
| - | - | 3569 | 157.1 | - | - | 0 | - |
| - | - | 1486 | 157.1 | - | - | 0 | - |
| - | - | 3790 | 159.1 | - | - | 0 | - |
| - | - | 1522 | 159.1 | - | - | 0 | - |
| - | - | 1199 | 160.1 | - | - | 0 | - |
| - | - | 679.5 | 162.1 | - | - | 0 | - |
| - | - | 405.5 | 163 | - | - | 0 | - |
| - | - | 842.9 | 163.1 | - | - | 0 | - |
| - | - | 920.7 | 164.1 | - | - | 0 | - |
| - | - | 885.5 | 165.1 | - | - | 0 | - |
| - | - | 3078 | 166.1 | - | - | 0 | - |
| - | - | 888.4 | 167 | - | - | 0 | - |
| - | - | 2822 | 167.1 | - | - | 0 | - |
| - | - | 1788 | 167.1 | - | - | 0 | - |
| - | - | 542.8 | 169.1 | - | - | 0 | - |
| - | - | 8742 | 169.1 | - | - | 0 | - |
| - | - | 484.2 | 169.1 | - | - | 0 | - |
| - | - | 1016 | 170.1 | - | - | 0 | - |
| - | - | 8883 | 171.1 | - | - | 0 | - |
| - | - | 2396 | 171.1 | - | - | 0 | - |
| - | - | 944.7 | 172.1 | - | - | 0 | - |
| - | - | 554 | 172.1 | - | - | 0 | - |
| - | - | 526.5 | 172.1 | - | - | 0 | - |
| - | - | 1321 | 173.1 | - | - | 0 | - |
| 2 | a | 5.942E+04 | 173.1 | 0.000391 | 2.258 | +1 | 2 |
| - | - | 799.9 | 173.5 | - | - | 0 | - |
| - | - | 1135 | 174.1 | - | - | 0 | - |
| - | - | 4694 | 174.1 | - | - | 0 | - |
| - | - | 534.9 | 175.1 | - | - | 0 | - |
| - | - | 2386 | 179.1 | - | - | 0 | - |
| - | - | 2423 | 180.1 | - | - | 0 | - |
| - | - | 605.3 | 181.1 | - | - | 0 | - |
| - | - | 6214 | 181.1 | - | - | 0 | - |
| - | - | 8278 | 181.1 | - | - | 0 | - |
| - | - | 700.6 | 181.1 | - | - | 0 | - |
| - | - | 1710 | 182.1 | - | - | 0 | - |
| - | - | 820.2 | 182.1 | - | - | 0 | - |
| - | - | 535.8 | 182.1 | - | - | 0 | - |
| 2 | b | 9705 | 183.1 | 0.000355 | 1.939 | +1 | 2 |
| - | - | 2829 | 184.1 | - | - | 0 | - |
| - | - | 1168 | 184.1 | - | - | 0 | - |
| - | - | 525.2 | 185.1 | - | - | 0 | - |
| - | - | 1847 | 185.1 | - | - | 0 | - |
| - | - | 1507 | 185.1 | - | - | 0 | - |
| - | - | 1075 | 185.2 | - | - | 0 | - |
| - | - | 575.4 | 186.1 | - | - | 0 | - |
| - | - | 1.739E+04 | 187.1 | - | - | 0 | - |
| - | - | 947.8 | 187.1 | - | - | 0 | - |
| - | - | 1371 | 187.1 | - | - | 0 | - |
| - | - | 1789 | 188.1 | - | - | 0 | - |
| - | - | 560.2 | 189.1 | - | - | 0 | - |
| - | - | 1790 | 189.1 | - | - | 0 | - |
| - | - | 6978 | 192.1 | - | - | 0 | - |
| - | - | 526.3 | 193.1 | - | - | 0 | - |
| - | - | 807.7 | 193.1 | - | - | 0 | - |
| - | - | 873.6 | 194.1 | - | - | 0 | - |
| - | - | 730.8 | 195.1 | - | - | 0 | - |
| - | - | 484.5 | 195.1 | - | - | 0 | - |
| - | - | 1908 | 195.1 | - | - | 0 | - |
| - | - | 566.9 | 196.1 | - | - | 0 | - |
| - | - | 5115 | 197.1 | - | - | 0 | - |
| - | - | 593.6 | 197.1 | - | - | 0 | - |
| - | - | 703.2 | 198.1 | - | - | 0 | - |
| - | - | 604.7 | 199.1 | - | - | 0 | - |
| - | - | 5533 | 199.1 | - | - | 0 | - |
| - | - | 2663 | 199.1 | - | - | 0 | - |
| - | - | 553.5 | 199.2 | - | - | 0 | - |
| - | - | 3916 | 200.1 | - | - | 0 | - |
| 2 | b | 5.436E+04 | 201.1 | 0.0003342 | 1.661 | +1 | 2 |
| - | - | 1.597E+04 | 202.1 | - | - | 0 | - |
| - | - | 4415 | 202.1 | - | - | 0 | - |
| - | - | 1088 | 203.1 | - | - | 0 | - |
| - | - | 1736 | 203.1 | - | - | 0 | - |
| - | - | 678.3 | 204.1 | - | - | 0 | - |
| - | - | 990.7 | 206.1 | - | - | 0 | - |
| - | - | 1630 | 207.1 | - | - | 0 | - |
| - | - | 2194 | 208.1 | - | - | 0 | - |
| - | - | 555.2 | 208.1 | - | - | 0 | - |
| - | - | 1349 | 209.1 | - | - | 0 | - |
| - | - | 2.848E+04 | 209.1 | - | - | 0 | - |
| - | - | 686.5 | 210.1 | - | - | 0 | - |
| - | - | 2329 | 210.1 | - | - | 0 | - |
| - | - | 1652 | 210.1 | - | - | 0 | - |
| - | - | 528.9 | 211.1 | - | - | 0 | - |
| - | - | 1837 | 212.1 | - | - | 0 | - |
| 3 | a | 767.8 | 212.1 | 0.0005546 | 2.614 | +1 | 3 |
| - | - | 793 | 213.1 | - | - | 0 | - |
| 3 | a | 2456 | 213.1 | 0.0002731 | 1.282 | +1 | 3 |
| - | - | 767.1 | 213.2 | - | - | 0 | - |
| - | - | 760.3 | 214.1 | - | - | 0 | - |
| - | - | 545.9 | 215.1 | - | - | 0 | - |
| - | - | 1.234E+04 | 215.1 | - | - | 0 | - |
| - | - | 1072 | 215.1 | - | - | 0 | - |
| - | - | 821 | 216.1 | - | - | 0 | - |
| - | - | 1322 | 217.1 | - | - | 0 | - |
| - | - | 617.7 | 219.1 | - | - | 0 | - |
| - | - | 1161 | 221.1 | - | - | 0 | - |
| - | - | 660.1 | 222.1 | - | - | 0 | - |
| - | - | 1155 | 224.1 | - | - | 0 | - |
| - | - | 4862 | 225.1 | - | - | 0 | - |
| - | - | 1063 | 225.1 | - | - | 0 | - |
| - | - | 759 | 225.2 | - | - | 0 | - |
| - | - | 667.6 | 226.1 | - | - | 0 | - |
| - | - | 1859 | 226.1 | - | - | 0 | - |
| - | - | 2550 | 226.2 | - | - | 0 | - |
| 8 | y | 2277 | 227.1 | 0.002964 | 13.05 | +2 | 4 |
| - | - | 1742 | 227.1 | - | - | 0 | - |
| - | - | 911.3 | 227.1 | - | - | 0 | - |
| - | - | 660.3 | 228.1 | - | - | 0 | - |
| - | - | 9049 | 228.1 | - | - | 0 | - |
| - | - | 5973 | 229.1 | - | - | 0 | - |
| - | - | 626.1 | 229.1 | - | - | 0 | - |
| - | - | 1829 | 230.1 | - | - | 0 | - |
| - | - | 606.7 | 230.1 | - | - | 0 | - |
| - | - | 543.2 | 233.1 | - | - | 0 | - |
| - | - | 572 | 233.5 | - | - | 0 | - |
| - | - | 2844 | 236.1 | - | - | 0 | - |
| - | - | 695.3 | 238.1 | - | - | 0 | - |
| - | - | 5399 | 238.1 | - | - | 0 | - |
| - | - | 1074 | 238.2 | - | - | 0 | - |
| - | - | 797.7 | 239.1 | - | - | 0 | - |
| - | - | 670.8 | 239.1 | - | - | 0 | - |
| - | - | 2351 | 239.1 | - | - | 0 | - |
| 3 | b | 1.393E+04 | 240.1 | 0.000452 | 1.882 | +1 | 3 |
| - | - | 1684 | 241.1 | - | - | 0 | - |
| - | - | 1805 | 241.2 | - | - | 0 | - |
| - | - | 1429 | 242.1 | - | - | 0 | - |
| - | - | 775.9 | 243.1 | - | - | 0 | - |
| - | - | 2711 | 244.1 | - | - | 0 | - |
| - | - | 627.7 | 245.1 | - | - | 0 | - |
| - | - | 815.3 | 246.1 | - | - | 0 | - |
| - | - | 1428 | 246.1 | - | - | 0 | - |
| - | - | 4223 | 249.1 | - | - | 0 | - |
| - | - | 2092 | 252.1 | - | - | 0 | - |
| - | - | 1386 | 254.1 | - | - | 0 | - |
| - | - | 1231 | 255.2 | - | - | 0 | - |
| - | - | 2474 | 256.1 | - | - | 0 | - |
| - | - | 646.1 | 257.1 | - | - | 0 | - |
| - | - | 1481 | 257.2 | - | - | 0 | - |
| 3 | b | 1737 | 258.1 | 2.666E-05 | 0.1033 | +1 | 3 |
| - | - | 1277 | 265.1 | - | - | 0 | - |
| - | - | 1.064E+04 | 267.1 | - | - | 0 | - |
| - | - | 622.7 | 268.1 | - | - | 0 | - |
| - | - | 1045 | 269.2 | - | - | 0 | - |
| - | - | 1175 | 269.2 | - | - | 0 | - |
| - | - | 1614 | 270.1 | - | - | 0 | - |
| - | - | 904.2 | 271.1 | - | - | 0 | - |
| - | - | 888.3 | 272.1 | - | - | 0 | - |
| - | - | 916.3 | 274.1 | - | - | 0 | - |
| - | - | 664 | 275.1 | - | - | 0 | - |
| - | - | 705.7 | 275.2 | - | - | 0 | - |
| - | - | 1484 | 276.1 | - | - | 0 | - |
| - | - | 683.5 | 278.2 | - | - | 0 | - |
| - | - | 1068 | 279.1 | - | - | 0 | - |
| - | - | 736.1 | 280.1 | - | - | 0 | - |
| - | - | 976.2 | 280.1 | - | - | 0 | - |
| - | - | 638.7 | 280.2 | - | - | 0 | - |
| - | - | 885.9 | 281.1 | - | - | 0 | - |
| - | - | 1931 | 282.1 | - | - | 0 | - |
| - | - | 1322 | 282.1 | - | - | 0 | - |
| - | - | 2199 | 282.2 | - | - | 0 | - |
| - | - | 1.236E+04 | 283.1 | - | - | 0 | - |
| 10 | y | 2860 | 283.1 | 0.005345 | 18.88 | +1 | 2 |
| - | - | 1200 | 283.2 | - | - | 0 | - |
| - | - | 3127 | 284.1 | - | - | 0 | - |
| - | - | 572.8 | 284.1 | - | - | 0 | - |
| - | - | 1198 | 284.1 | - | - | 0 | - |
| - | - | 7.886E+04 | 285.1 | - | - | 0 | - |
| - | - | 2138 | 285.2 | - | - | 0 | - |
| - | - | 9485 | 286.1 | - | - | 0 | - |
| - | - | 8852 | 286.1 | - | - | 0 | - |
| - | - | 830.1 | 286.2 | - | - | 0 | - |
| - | - | 753.8 | 287.1 | - | - | 0 | - |
| - | - | 721.5 | 287.1 | - | - | 0 | - |
| - | - | 1.89E+04 | 287.2 | - | - | 0 | - |
| - | - | 921.8 | 287.2 | - | - | 0 | - |
| - | - | 3083 | 288.2 | - | - | 0 | - |
| - | - | 1061 | 290.2 | - | - | 0 | - |
| - | - | 674.4 | 293.1 | - | - | 0 | - |
| - | - | 1035 | 295.1 | - | - | 0 | - |
| - | - | 1116 | 295.1 | - | - | 0 | - |
| - | - | 1162 | 296.1 | - | - | 0 | - |
| - | - | 1668 | 296.2 | - | - | 0 | - |
| 4 | b | 6983 | 297.2 | 0.000732 | 2.463 | +1 | 4 |
| 4 | b | 679.1 | 298.1 | 0.001976 | 6.629 | +1 | 4 |
| - | - | 1607 | 298.2 | - | - | 0 | - |
| - | - | 3082 | 299.1 | - | - | 0 | - |
| 10 | y | 7533 | 301.1 | 0.005369 | 17.83 | +1 | 2 |
| - | - | 764.8 | 302.1 | - | - | 0 | - |
| - | - | 882.6 | 302.1 | - | - | 0 | - |
| - | - | 546 | 302.2 | - | - | 0 | - |
| - | - | 1881 | 304.1 | - | - | 0 | - |
| - | - | 848.7 | 307.1 | - | - | 0 | - |
| - | - | 6281 | 308.2 | - | - | 0 | - |
| - | - | 1395 | 309.1 | - | - | 0 | - |
| - | - | 908.7 | 309.2 | - | - | 0 | - |
| - | - | 913 | 310.1 | - | - | 0 | - |
| - | - | 894 | 311.1 | - | - | 0 | - |
| - | - | 3987 | 311.1 | - | - | 0 | - |
| - | - | 656.1 | 312.1 | - | - | 0 | - |
| - | - | 697.7 | 312.2 | - | - | 0 | - |
| - | - | 1844 | 313.2 | - | - | 0 | - |
| - | - | 6633 | 314.2 | - | - | 0 | - |
| 4 | b | 2.349E+04 | 315.2 | 0.0007874 | 2.499 | +1 | 4 |
| - | - | 3428 | 316.2 | - | - | 0 | - |
| - | - | 2313 | 317.1 | - | - | 0 | - |
| - | - | 720.8 | 317.2 | - | - | 0 | - |
| - | - | 3973 | 321.1 | - | - | 0 | - |
| - | - | 1364 | 322.2 | - | - | 0 | - |
| - | - | 3353 | 324.2 | - | - | 0 | - |
| - | - | 6896 | 325.2 | - | - | 0 | - |
| - | - | 2046 | 328.2 | - | - | 0 | - |
| - | - | 4907 | 329.1 | - | - | 0 | - |
| - | - | 2107 | 331.2 | - | - | 0 | - |
| - | - | 750.9 | 332.2 | - | - | 0 | - |
| - | - | 626.2 | 336.2 | - | - | 0 | - |
| - | - | 3395 | 338.1 | - | - | 0 | - |
| - | - | 1223 | 339.1 | - | - | 0 | - |
| - | - | 2148 | 340.1 | - | - | 0 | - |
| - | - | 1001 | 340.2 | - | - | 0 | - |
| - | - | 1678 | 341.1 | - | - | 0 | - |
| - | - | 1003 | 341.2 | - | - | 0 | - |
| 6 | y | 1.103E+04 | 343.2 | 0.00302 | 8.801 | +2 | 6 |
| - | - | 2113 | 344.2 | - | - | 0 | - |
| - | - | 2632 | 352.1 | - | - | 0 | - |
| - | - | 2483 | 353.1 | - | - | 0 | - |
| - | - | 946.6 | 353.2 | - | - | 0 | - |
| 9 | y | 2925 | 354.1 | 0.005951 | 16.8 | +1 | 3 |
| - | - | 3627 | 355.1 | - | - | 0 | - |
| - | - | 4.569E+04 | 356.2 | - | - | 0 | - |
| - | - | 6946 | 357.2 | - | - | 0 | - |
| - | - | 825.9 | 357.2 | - | - | 0 | - |
| - | - | 764.6 | 357.2 | - | - | 0 | - |
| - | - | 916.8 | 358.2 | - | - | 0 | - |
| - | - | 1360 | 359.2 | - | - | 0 | - |
| - | - | 666 | 365.2 | - | - | 0 | - |
| - | - | 1415 | 366.1 | - | - | 0 | - |
| - | - | 569.6 | 367.2 | - | - | 0 | - |
| - | - | 750.4 | 367.2 | - | - | 0 | - |
| - | - | 1399 | 369.2 | - | - | 0 | - |
| - | - | 2100 | 369.2 | - | - | 0 | - |
| - | - | 902.9 | 370.1 | - | - | 0 | - |
| - | - | 1057 | 370.2 | - | - | 0 | - |
| - | - | 615.1 | 371.2 | - | - | 0 | - |
| 9 | y | 4656 | 372.1 | 0.005396 | 14.5 | +1 | 3 |
| - | - | 806.8 | 373.2 | - | - | 0 | - |
| - | - | 887.3 | 374.2 | - | - | 0 | - |
| - | - | 691.3 | 377.2 | - | - | 0 | - |
| - | - | 627 | 379.2 | - | - | 0 | - |
| - | - | 665.3 | 380.2 | - | - | 0 | - |
| - | - | 748.6 | 383.2 | - | - | 0 | - |
| - | - | 1966 | 384.2 | - | - | 0 | - |
| - | - | 3830 | 385.2 | - | - | 0 | - |
| 5 | y | 705.6 | 390.7 | 0.001671 | 4.277 | +2 | 7 |
| - | - | 505.4 | 394.1 | - | - | 0 | - |
| - | - | 1068 | 394.2 | - | - | 0 | - |
| - | - | 1672 | 395.2 | - | - | 0 | - |
| - | - | 499.2 | 395.2 | - | - | 0 | - |
| - | - | 831.4 | 395.2 | - | - | 0 | - |
| - | - | 714.2 | 396.2 | - | - | 0 | - |
| - | - | 2327 | 396.2 | - | - | 0 | - |
| - | - | 2568 | 397.2 | - | - | 0 | - |
| - | - | 1379 | 398.2 | - | - | 0 | - |
| - | - | 1073 | 398.2 | - | - | 0 | - |
| 5 | y | 584.5 | 399.7 | 0.003316 | 8.296 | +2 | 7 |
| - | - | 785.7 | 400.2 | - | - | 0 | - |
| - | - | 539.9 | 400.2 | - | - | 0 | - |
| - | - | 7756 | 400.3 | - | - | 0 | - |
| - | - | 1432 | 401.3 | - | - | 0 | - |
| - | - | 1213 | 402.2 | - | - | 0 | - |
| - | - | 720.3 | 405.2 | - | - | 0 | - |
| - | - | 1263 | 407.2 | - | - | 0 | - |
| - | - | 1024 | 408.2 | - | - | 0 | - |
| - | - | 966.3 | 410.2 | - | - | 0 | - |
| - | - | 1197 | 410.2 | - | - | 0 | - |
| 5 | b | 1846 | 410.2 | 0.0006219 | 1.516 | +1 | 5 |
| - | - | 7168 | 412.2 | - | - | 0 | - |
| - | - | 1056 | 413.2 | - | - | 0 | - |
| - | - | 943.5 | 414.2 | - | - | 0 | - |
| - | - | 4046 | 414.2 | - | - | 0 | - |
| - | - | 1950 | 416.2 | - | - | 0 | - |
| - | - | 972.2 | 418.2 | - | - | 0 | - |
| - | - | 891.6 | 420.2 | - | - | 0 | - |
| - | - | 705.4 | 421.2 | - | - | 0 | - |
| - | - | 1010 | 422.2 | - | - | 0 | - |
| - | - | 2203 | 423.2 | - | - | 0 | - |
| - | - | 1143 | 424.2 | - | - | 0 | - |
| - | - | 2711 | 424.2 | - | - | 0 | - |
| - | - | 750.3 | 425.2 | - | - | 0 | - |
| - | - | 1691 | 426.2 | - | - | 0 | - |
| - | - | 1459 | 426.2 | - | - | 0 | - |
| - | - | 2248 | 428.2 | - | - | 0 | - |
| 4 | y | 1846 | 428.2 | 0.003204 | 7.483 | +2 | 8 |
| 5 | b | 5437 | 428.3 | 0.001074 | 2.508 | +1 | 5 |
| - | - | 703.6 | 429.3 | - | - | 0 | - |
| - | - | 7257 | 430.2 | - | - | 0 | - |
| - | - | 1577 | 431.2 | - | - | 0 | - |
| - | - | 1202 | 434.2 | - | - | 0 | - |
| - | - | 765.9 | 435.2 | - | - | 0 | - |
| - | - | 1467 | 437.2 | - | - | 0 | - |
| - | - | 1386 | 438.2 | - | - | 0 | - |
| - | - | 1537 | 439.2 | - | - | 0 | - |
| - | - | 1447 | 440.2 | - | - | 0 | - |
| - | - | 1297 | 440.2 | - | - | 0 | - |
| - | - | 583.9 | 440.3 | - | - | 0 | - |
| - | - | 6273 | 442.2 | - | - | 0 | - |
| - | - | 1739 | 443.2 | - | - | 0 | - |
| - | - | 900 | 444.2 | - | - | 0 | - |
| - | - | 2785 | 444.2 | - | - | 0 | - |
| - | - | 757.3 | 446.2 | - | - | 0 | - |
| - | - | 1782 | 446.7 | - | - | 0 | - |
| 3 | y | 2624 | 447.7 | 0.002546 | 5.687 | +2 | 9 |
| - | - | 1169 | 448.2 | - | - | 0 | - |
| - | - | 996.3 | 448.7 | - | - | 0 | - |
| - | - | 7180 | 451.2 | - | - | 0 | - |
| - | - | 1762 | 452.2 | - | - | 0 | - |
| 8 | y | 1216 | 453.2 | 0.005865 | 12.94 | +1 | 4 |
| - | - | 728.1 | 454.2 | - | - | 0 | - |
| - | - | 2.856E+04 | 455.2 | - | - | 0 | - |
| - | - | 7552 | 456.2 | - | - | 0 | - |
| - | - | 1690 | 457.2 | - | - | 0 | - |
| - | - | 653.1 | 458.2 | - | - | 0 | - |
| - | - | 780.2 | 461.3 | - | - | 0 | - |
| - | - | 704.7 | 464.2 | - | - | 0 | - |
| - | - | 1604 | 468.2 | - | - | 0 | - |
| - | - | 641.6 | 468.2 | - | - | 0 | - |
| - | - | 764.5 | 469.2 | - | - | 0 | - |
| 8 | y | 6444 | 471.2 | 0.00586 | 12.44 | +1 | 4 |
| - | - | 1029 | 472.2 | - | - | 0 | - |
| 10 | b | 851.8 | 474.2 | 0.001322 | 2.787 | +2 | 10 |
| - | - | 740.3 | 482.2 | - | - | 0 | - |
| 10 | b | 1673 | 483.3 | 0.0005733 | 1.186 | +2 | 10 |
| - | - | 1873 | 484.2 | - | - | 0 | - |
| - | - | 629.7 | 485.2 | - | - | 0 | - |
| - | - | 1013 | 487.6 | - | - | 0 | - |
| - | - | 761.6 | 487.9 | - | - | 0 | - |
| - | - | 2833 | 490.2 | - | - | 0 | - |
| - | - | 1602 | 490.7 | - | - | 0 | - |
| 2 | y | 2065 | 491.2 | 0.002798 | 5.695 | +2 | 10 |
| - | - | 1261 | 491.7 | - | - | 0 | - |
| - | - | 4047 | 492.2 | - | - | 0 | - |
| - | - | 1915 | 492.7 | - | - | 0 | - |
| - | - | 1663 | 493.2 | - | - | 0 | - |
| - | - | 3320 | 494.3 | - | - | 0 | - |
| - | - | 712.3 | 495.3 | - | - | 0 | - |
| - | - | 784.1 | 496.2 | - | - | 0 | - |
| - | - | 1220 | 497.2 | - | - | 0 | - |
| - | - | 1043 | 497.3 | - | - | 0 | - |
| - | - | 847.7 | 498.3 | - | - | 0 | - |
| - | - | 1634 | 499.2 | - | - | 0 | - |
| - | - | 691.5 | 499.3 | - | - | 0 | - |
| - | - | 1523 | 499.7 | - | - | 0 | - |
| 2 | y | 3229 | 500.2 | 0.001879 | 3.757 | +2 | 10 |
| - | - | 1009 | 500.7 | - | - | 0 | - |
| - | - | 2322 | 501.3 | - | - | 0 | - |
| - | - | 818.8 | 502.3 | - | - | 0 | - |
| - | - | 850.3 | 507.2 | - | - | 0 | - |
| - | - | 2112 | 509.2 | - | - | 0 | - |
| - | - | 654.2 | 510.2 | - | - | 0 | - |
| - | - | 5300 | 511.3 | - | - | 0 | - |
| - | - | 921.2 | 512.3 | - | - | 0 | - |
| - | - | 1418 | 513.3 | - | - | 0 | - |
| - | - | 1024 | 513.3 | - | - | 0 | - |
| - | - | 1291 | 515.2 | - | - | 0 | - |
| - | - | 1038 | 515.3 | - | - | 0 | - |
| - | - | 981.2 | 518.2 | - | - | 0 | - |
| - | - | 970.7 | 521.2 | - | - | 0 | - |
| - | - | 4140 | 522.3 | - | - | 0 | - |
| - | - | 1019 | 523.3 | - | - | 0 | - |
| - | - | 1188 | 525.2 | - | - | 0 | - |
| 6 | b | 2209 | 525.3 | 0.001755 | 3.341 | +1 | 6 |
| - | - | 1363 | 527.2 | - | - | 0 | - |
| - | - | 8253 | 529.3 | - | - | 0 | - |
| - | - | 2529 | 530.3 | - | - | 0 | - |
| - | - | 732.8 | 533.2 | - | - | 0 | - |
| - | - | 1405 | 533.8 | - | - | 0 | - |
| - | - | 2895 | 537.8 | - | - | 0 | - |
| - | - | 3549 | 538.3 | - | - | 0 | - |
| - | - | 947.2 | 538.8 | - | - | 0 | - |
| - | - | 2052 | 539.2 | - | - | 0 | - |
| - | - | 911.6 | 539.3 | - | - | 0 | - |
| - | - | 643.9 | 539.8 | - | - | 0 | - |
| - | - | 1320 | 540.3 | - | - | 0 | - |
| - | - | 1968 | 541.3 | - | - | 0 | - |
| 6 | b | 5071 | 543.3 | 0.000956 | 1.76 | +1 | 6 |
| - | - | 1016 | 544.3 | - | - | 0 | - |
| - | - | 1119 | 545.3 | - | - | 0 | - |
| - | - | 6364 | 546.8 | - | - | 0 | - |
| - | - | 6044 | 547.3 | - | - | 0 | - |
| 0 | Precursor | 1.393E+04 | 547.8 | 0.002147 | 3.92 | +2 | -1 |
| - | - | 7265 | 548.3 | - | - | 0 | - |
| - | - | 3656 | 548.8 | - | - | 0 | - |
| - | - | 933 | 549.3 | - | - | 0 | - |
| - | - | 5230 | 550.3 | - | - | 0 | - |
| - | - | 1482 | 551.3 | - | - | 0 | - |
| 7 | y | 778.6 | 552.3 | 0.005658 | 10.25 | +1 | 5 |
| - | - | 1704 | 553.3 | - | - | 0 | - |
| - | - | 1.454E+04 | 554.3 | - | - | 0 | - |
| - | - | 5622 | 555.3 | - | - | 0 | - |
| - | - | 7168 | 555.8 | - | - | 0 | - |
| - | - | 5288 | 556.3 | - | - | 0 | - |
| 0 | Precursor | 1.3E+04 | 556.8 | 0.002236 | 4.016 | +2 | -1 |
| - | - | 9505 | 557.3 | - | - | 0 | - |
| - | - | 2530 | 557.8 | - | - | 0 | - |
| - | - | 1107 | 559.3 | - | - | 0 | - |
| - | - | 759 | 567.2 | - | - | 0 | - |
| 7 | y | 4602 | 570.3 | 0.005714 | 10.02 | +1 | 5 |
| - | - | 1037 | 571.3 | - | - | 0 | - |
| - | - | 811.1 | 595.3 | - | - | 0 | - |
| - | - | 1598 | 597.3 | - | - | 0 | - |
| - | - | 1110 | 608.3 | - | - | 0 | - |
| - | - | 863.4 | 609.3 | - | - | 0 | - |
| - | - | 3193 | 610.3 | - | - | 0 | - |
| - | - | 953.1 | 611.3 | - | - | 0 | - |
| - | - | 2978 | 612.3 | - | - | 0 | - |
| - | - | 950.3 | 613.3 | - | - | 0 | - |
| - | - | 1804 | 614.4 | - | - | 0 | - |
| - | - | 1263 | 615.4 | - | - | 0 | - |
| 7 | b | 2615 | 624.3 | 0.0001746 | 0.2797 | +1 | 7 |
| - | - | 922.9 | 625.3 | - | - | 0 | - |
| - | - | 1125 | 626.3 | - | - | 0 | - |
| - | - | 4910 | 628.3 | - | - | 0 | - |
| - | - | 1775 | 629.3 | - | - | 0 | - |
| - | - | 1087 | 631.3 | - | - | 0 | - |
| - | - | 760 | 632.3 | - | - | 0 | - |
| - | - | 1102 | 638.3 | - | - | 0 | - |
| - | - | 1195 | 640.3 | - | - | 0 | - |
| 7 | b | 7635 | 642.3 | 0.0007794 | 1.213 | +1 | 7 |
| - | - | 2762 | 643.3 | - | - | 0 | - |
| - | - | 665.8 | 644.3 | - | - | 0 | - |
| - | - | 1362 | 645.8 | - | - | 0 | - |
| - | - | 1432 | 646.3 | - | - | 0 | - |
| - | - | 1048 | 651.3 | - | - | 0 | - |
| - | - | 580.9 | 652.3 | - | - | 0 | - |
| - | - | 1366 | 658.3 | - | - | 0 | - |
| - | - | 1.093E+04 | 665.3 | - | - | 0 | - |
| - | - | 3832 | 666.3 | - | - | 0 | - |
| 6 | y | 3073 | 667.3 | 0.0001997 | 0.2992 | +1 | 6 |
| - | - | 1039 | 668.3 | - | - | 0 | - |
| - | - | 4.876E+04 | 669.3 | - | - | 0 | - |
| - | - | 1.706E+04 | 670.3 | - | - | 0 | - |
| - | - | 4481 | 671.3 | - | - | 0 | - |
| - | - | 1303 | 681.4 | - | - | 0 | - |
| - | - | 979.4 | 681.8 | - | - | 0 | - |
| - | - | 2543 | 683.3 | - | - | 0 | - |
| - | - | 929.5 | 684.3 | - | - | 0 | - |
| 6 | y | 1.388E+04 | 685.3 | 0.00526 | 7.675 | +1 | 6 |
| - | - | 5263 | 686.3 | - | - | 0 | - |
| - | - | 715 | 687.3 | - | - | 0 | - |
| - | - | 683.2 | 690.4 | - | - | 0 | - |
| - | - | 742.6 | 692.4 | - | - | 0 | - |
| - | - | 729.8 | 694.4 | - | - | 0 | - |
| - | - | 877 | 697.4 | - | - | 0 | - |
| - | - | 2252 | 699.4 | - | - | 0 | - |
| - | - | 1929 | 700.4 | - | - | 0 | - |
| - | - | 738.4 | 708.4 | - | - | 0 | - |
| - | - | 1168 | 710.5 | - | - | 0 | - |
| - | - | 770.8 | 713.4 | - | - | 0 | - |
| - | - | 777.8 | 715.4 | - | - | 0 | - |
| - | - | 763.1 | 717.4 | - | - | 0 | - |
| - | - | 1007 | 721.4 | - | - | 0 | - |
| 8 | b | 1142 | 723.4 | 0.002565 | 3.546 | +1 | 8 |
| - | - | 672.1 | 735.4 | - | - | 0 | - |
| - | - | 770.8 | 738.4 | - | - | 0 | - |
| - | - | 1218 | 738.9 | - | - | 0 | - |
| - | - | 2061 | 739.4 | - | - | 0 | - |
| 8 | b | 3397 | 741.4 | 0.0008621 | 1.163 | +1 | 8 |
| - | - | 1094 | 742.4 | - | - | 0 | - |
| - | - | 1390 | 745.4 | - | - | 0 | - |
| - | - | 1121 | 746.4 | - | - | 0 | - |
| - | - | 965.5 | 747.4 | - | - | 0 | - |
| - | - | 1627 | 749.4 | - | - | 0 | - |
| - | - | 807.2 | 750.4 | - | - | 0 | - |
| - | - | 1594 | 763.4 | - | - | 0 | - |
| - | - | 2507 | 765.4 | - | - | 0 | - |
| - | - | 1208 | 766.4 | - | - | 0 | - |
| - | - | 805.2 | 767.4 | - | - | 0 | - |
| - | - | 1.647E+04 | 778.4 | - | - | 0 | - |
| - | - | 6403 | 779.4 | - | - | 0 | - |
| 5 | y | 2196 | 780.4 | 0.001345 | 1.723 | +1 | 7 |
| - | - | 1183 | 781.4 | - | - | 0 | - |
| - | - | 3765 | 782.4 | - | - | 0 | - |
| - | - | 1034 | 782.9 | - | - | 0 | - |
| - | - | 2965 | 783.4 | - | - | 0 | - |
| - | - | 679.5 | 783.9 | - | - | 0 | - |
| 9 | b | 1087 | 794.4 | 0.003717 | 4.678 | +1 | 9 |
| - | - | 1770 | 796.4 | - | - | 0 | - |
| - | - | 925.2 | 797.4 | - | - | 0 | - |
| 5 | y | 5281 | 798.4 | 0.004448 | 5.571 | +1 | 7 |
| - | - | 1874 | 799.4 | - | - | 0 | - |
| - | - | 852.9 | 808.4 | - | - | 0 | - |
| 9 | b | 2503 | 812.5 | 0.001209 | 1.488 | +1 | 9 |
| - | - | 1269 | 813.5 | - | - | 0 | - |
| - | - | 604.2 | 822.4 | - | - | 0 | - |
| - | - | 2472 | 829.5 | - | - | 0 | - |
| - | - | 762.1 | 832.4 | - | - | 0 | - |
| - | - | 743.8 | 833.4 | - | - | 0 | - |
| - | - | 997.3 | 834.4 | - | - | 0 | - |
| - | - | 2805 | 835.4 | - | - | 0 | - |
| - | - | 2481 | 836.4 | - | - | 0 | - |
| 4 | y | 1797 | 837.4 | 0.004108 | 4.906 | +1 | 8 |
| - | - | 2089 | 838.4 | - | - | 0 | - |
| - | - | 7629 | 839.4 | - | - | 0 | - |
| - | - | 3068 | 840.4 | - | - | 0 | - |
| - | - | 1001 | 841.4 | - | - | 0 | - |
| - | - | 1178 | 850.4 | - | - | 0 | - |
| - | - | 744.5 | 851.4 | - | - | 0 | - |
| - | - | 989.5 | 852.4 | - | - | 0 | - |
| - | - | 3973 | 853.4 | - | - | 0 | - |
| - | - | 2251 | 854.4 | - | - | 0 | - |
| 4 | y | 5739 | 855.4 | 0.004041 | 4.724 | +1 | 8 |
| - | - | 1683 | 856.4 | - | - | 0 | - |
| - | - | 633.9 | 866.4 | - | - | 0 | - |
| - | - | 1002 | 874.4 | - | - | 0 | - |
| - | - | 1.83E+04 | 892.4 | - | - | 0 | - |
| - | - | 8248 | 893.4 | - | - | 0 | - |
| 3 | y | 6951 | 894.4 | 0.001748 | 1.955 | +1 | 9 |
| - | - | 2999 | 895.4 | - | - | 0 | - |
| - | - | 1.89E+04 | 896.4 | - | - | 0 | - |
| - | - | 8881 | 897.4 | - | - | 0 | - |
| - | - | 2497 | 898.5 | - | - | 0 | - |
| - | - | 1.906E+04 | 910.4 | - | - | 0 | - |
| - | - | 1.228E+04 | 911.4 | - | - | 0 | - |
| 3 | y | 3.538E+04 | 912.4 | 0.003513 | 3.85 | +1 | 9 |
| - | - | 1.409E+04 | 913.4 | - | - | 0 | - |
| - | - | 4552 | 914.4 | - | - | 0 | - |
| - | - | 4182 | 953.5 | - | - | 0 | - |
| - | - | 1723 | 954.5 | - | - | 0 | - |
| - | - | 1932 | 961.4 | - | - | 0 | - |
| - | - | 906.4 | 962.4 | - | - | 0 | - |
| - | - | 1067 | 963.4 | - | - | 0 | - |
| - | - | 697.9 | 971.4 | - | - | 0 | - |
| - | - | 1.313E+04 | 979.4 | - | - | 0 | - |
| - | - | 8095 | 980.5 | - | - | 0 | - |
| 2 | y | 8065 | 981.5 | 0.0006036 | 0.6151 | +1 | 10 |
| - | - | 4922 | 982.5 | - | - | 0 | - |
| - | - | 6571 | 983.5 | - | - | 0 | - |
| - | - | 3314 | 984.5 | - | - | 0 | - |
| - | - | 784 | 985.5 | - | - | 0 | - |
| - | - | 1620 | 989.4 | - | - | 0 | - |
| - | - | 1651 | 991.4 | - | - | 0 | - |
| - | - | 942.2 | 992.5 | - | - | 0 | - |
| - | - | 985.6 | 993.5 | - | - | 0 | - |
| - | - | 1.657E+04 | 997.5 | - | - | 0 | - |
| - | - | 9904 | 998.5 | - | - | 0 | - |
| 2 | y | 2.419E+04 | 999.5 | 0.003284 | 3.285 | +1 | 10 |
| - | - | 1.152E+04 | 1000 | - | - | 0 | - |
| - | - | 4010 | 1001 | - | - | 0 | - |
| - | - | 1196 | 1007 | - | - | 0 | - |
| - | - | 2236 | 1009 | - | - | 0 | - |
| - | - | 839.7 | 1010 | - | - | 0 | - |

m/z Charge Intensity FragmentType MassShift Position
120.08109283447266 0 4378.629
121.08465576171875 0 455.99133
122.0478286743164 0 457.64056
122.07144165039062 0 617.5924
124.05059814453125 0 619.493
125.03492736816406 0 394.70282
125.1076889038086 0 8088.568
126.06660461425781 0 1715.4797
126.11125183105469 0 692.1273
127.05052185058594 0 994.56885
127.0869369506836 0 604.7643
128.07110595703125 0 916.6891
128.10325622558594 0 438.2636
128.10728454589844 0 4589.7393
129.0422821044922 0 367.29874
129.0662078857422 0 1520.9288
129.10260009765625 0 4555.097
130.05023193359375 0 4361.605 y Water loss 10
130.06558227539062 0 1614.7733
130.08660888671875 0 4435.932
131.1182403564453 0 893.8982
131.3487548828125 0 419.49802
132.0449981689453 0 422.04047
132.07638549804688 0 580.38416
132.0813446044922 0 836.94275
132.1022491455078 0 983.9815
133.0612030029297 0 529.9158
133.3936004638672 0 344.55188
134.85386657714844 0 384.52512
136.02720642089844 0 1993.431
136.05091857910156 0 800.02734
136.07606506347656 0 6073.393
136.0872344970703 0 1505.9357
137.03492736816406 0 1283.3676
137.07948303222656 0 534.7784
138.04275512695312 0 2401.863
138.06658935546875 0 3889.8025
138.0917510986328 0 1370.6937
139.0869903564453 0 2901.3528
140.07095336914062 0 663.5999
140.14358520507812 0 706.82837
141.0662078857422 0 732.06616
141.1025848388672 0 1686.5444
142.08645629882812 0 685.8252
142.1228485107422 0 598.34174
143.1182403564453 0 15270.439
143.81993103027344 0 426.32547
144.05398559570312 0 547.6775
144.12176513671875 0 1032.4368
145.06112670898438 0 10078.112
148.06077575683594 0 12420.859 y 10
150.0662078857422 0 688.03436
151.08682250976562 0 1845.2395
152.04600524902344 0 1271.6603
152.07107543945312 0 1231.6863
152.10777282714844 0 523.7697
154.06166076660156 0 757.08405
154.08663940429688 0 1176.1641
154.1593475341797 0 1371.5107
155.0453338623047 0 1074.7017
155.08197021484375 0 723.6597
155.09298706054688 0 480.93942
155.1182403564453 0 8579.647 a Water loss 1
156.0770263671875 0 6048.176
156.10238647460938 0 1349.5164 a Ammonia loss 1
156.12179565429688 0 1357.5275
157.0611114501953 0 3568.6177
157.09732055664062 0 1486.1456
159.0919952392578 0 3789.5518
159.11294555664062 0 1522.4336
160.09536743164062 0 1198.5526
162.05520629882812 0 679.49945
163.0037384033203 0 405.50787
163.07205200195312 0 842.9353
164.08238220214844 0 920.6915
165.10240173339844 0 885.5143
166.06143188476562 0 3077.7268
167.04566955566406 0 888.4096
167.081787109375 0 2822.3374
167.11849975585938 0 1788.0586
169.06072998046875 0 542.7915
169.09750366210938 0 8741.758
169.1342010498047 0 484.2146
170.1009979248047 0 1015.6041
171.1131591796875 0 8882.982
171.14955139160156 0 2396.2014
172.072265625 0 944.65094
172.09718322753906 0 554.01953
172.11656188964844 0 526.5413
173.09246826171875 0 1321.0352
173.12884521484375 0 59417.46 a 1
173.45217895507812 0 799.9424
174.08758544921875 0 1135.2573
174.13232421875 0 4693.645
175.0873260498047 0 534.85583
179.0820770263672 0 2386.0942
180.0771484375 0 2423.0728
181.07278442382812 0 605.2817
181.0976104736328 0 6214.3643
181.10874938964844 0 8278.048
181.13441467285156 0 700.58905
182.08135986328125 0 1709.6055
182.1009063720703 0 820.2184
182.11122131347656 0 535.7545
183.1131591796875 0 9704.621 b Water loss 1
184.0718994140625 0 2828.6785
184.11643981933594 0 1168.487
185.0556640625 0 525.17255
185.0923614501953 0 1847.2444
185.1288299560547 0 1506.9009
185.16546630859375 0 1075.2206
186.1239013671875 0 575.39056
187.10809326171875 0 17385.635
187.11708068847656 0 947.7655
187.1444854736328 0 1371.0461
188.1113739013672 0 1789.2787
189.08761596679688 0 560.15735
189.12380981445312 0 1789.9763
192.0771026611328 0 6977.6353
193.08116149902344 0 526.2767
193.10870361328125 0 807.72015
194.09237670898438 0 873.63367
195.07675170898438 0 730.7514
195.08694458007812 0 484.53452
195.11318969726562 0 1907.7411
196.1075897216797 0 566.8818
197.09234619140625 0 5114.858
197.12985229492188 0 593.6277
198.09666442871094 0 703.2154
199.07275390625 0 604.70764
199.10804748535156 0 5532.653
199.14439392089844 0 2663.106
199.1808624267578 0 553.47437
200.13966369628906 0 3916.3438
201.1237030029297 0 54364.168 b 1
202.0825653076172 0 15968.394
202.12722778320312 0 4415.439
203.067138671875 0 1088.05
203.08596801757812 0 1736.4508
204.0753173828125 0 678.2887
206.05621337890625 0 990.70404
207.08799743652344 0 1629.8224
208.0721435546875 0 2193.79
208.10772705078125 0 555.18365
209.0922088623047 0 1348.6987
209.10369873046875 0 28481.717
210.08782958984375 0 686.4602
210.10665893554688 0 2328.8906
210.12393188476562 0 1651.8774
211.10897827148438 0 528.8651
212.06678771972656 0 1836.7124
212.13990783691406 0 767.7997 a Water loss 2
213.08714294433594 0 792.9909
213.12364196777344 0 2455.7107 a Ammonia loss 2
213.16030883789062 0 767.0668
214.11810302734375 0 760.32733
215.09178161621094 0 545.8732
215.10305786132812 0 12339.633
215.13995361328125 0 1072.456
216.10623168945312 0 821.0334
217.11875915527344 0 1322.2495
219.10911560058594 0 617.7146
221.1040496826172 0 1161.2303
222.1240234375 0 660.0746
224.1033935546875 0 1154.7869
225.08738708496094 0 4861.775
225.09857177734375 0 1063.162
225.16030883789062 0 758.95404
226.08189392089844 0 667.63403
226.1190643310547 0 1858.505
226.15525817871094 0 2549.927
227.1028289794922 0 2276.662 y Water loss 7
227.11456298828125 0 1742.2426
227.12615966796875 0 911.3474
228.09783935546875 0 660.27313
228.1346893310547 0 9049.33
229.11866760253906 0 5972.7656
229.138427734375 0 626.13513
230.07760620117188 0 1828.67
230.12203979492188 0 606.68365
233.09324645996094 0 543.16614
233.45704650878906 0 571.95935
236.1033935546875 0 2844.139
238.0828094482422 0 695.2519
238.11920166015625 0 5398.5464
238.1540069580078 0 1073.8844
239.09078979492188 0 797.67505
239.1029510498047 0 670.8394
239.11448669433594 0 2350.8215
240.1347198486328 0 13932.739 b Water loss 2
241.1383819580078 0 1684.3259
241.19142150878906 0 1805.1985
242.11404418945312 0 1428.5132
243.0971221923828 0 775.9459
244.12945556640625 0 2710.8928
245.1136474609375 0 627.74927
246.09832763671875 0 815.33826
246.1238555908203 0 1427.6392
249.0985870361328 0 4223.1616
252.134765625 0 2091.91
254.11378479003906 0 1385.973
255.17027282714844 0 1230.6938
256.1297302246094 0 2474.1921
257.1139221191406 0 646.09607
257.1615295410156 0 1480.7339
258.1448059082031 0 1737.3787 b 2
265.06951904296875 0 1276.8539
267.1091003417969 0 10642.837
268.1128234863281 0 622.6943
269.16204833984375 0 1044.9779
269.1856689453125 0 1175.1866
270.14544677734375 0 1613.7244
271.10382080078125 0 904.21344
272.12451171875 0 888.26764
274.11761474609375 0 916.3199
275.12408447265625 0 664.0163
275.17059326171875 0 705.7057
276.1456604003906 0 1484.1143
278.1504821777344 0 683.5331
279.146484375 0 1067.5671
280.09356689453125 0 736.1307
280.12933349609375 0 976.2445
280.1777038574219 0 638.73865
281.12481689453125 0 885.9221
282.10894775390625 0 1930.666
282.14447021484375 0 1322.285
282.1821594238281 0 2198.9958
283.08038330078125 0 12355.104
283.1041259765625 0 2860.1973 y Water loss 9
283.1642761230469 0 1199.5757
284.0860290527344 0 3127.4106
284.1068420410156 0 572.815
284.1244201660156 0 1197.9656
285.119873046875 0 78860.53
285.15692138671875 0 2137.966
286.12274169921875 0 9485.029
286.14031982421875 0 8852.115
286.17669677734375 0 830.0604
287.1246337890625 0 753.8361
287.1439208984375 0 721.5184
287.171875 0 18904.229
287.1901550292969 0 921.77045
288.1760559082031 0 3082.8708
290.1614074707031 0 1061.4539
293.1248474121094 0 674.43384
295.1051940917969 0 1035.1554
295.140380859375 0 1115.5446
296.125 0 1161.9679
296.1606140136719 0 1668.332
297.1564636230469 0 6983.013 b Water loss 3
298.1417236328125 0 679.11993 b Ammonia loss 3
298.15985107421875 0 1606.7205
299.0994567871094 0 3081.762
301.1147155761719 0 7533.4277 y 9
302.1178283691406 0 764.7734
302.1355895996094 0 882.639
302.155517578125 0 546.00104
304.1409912109375 0 1881.4768
307.14031982421875 0 848.713
308.17236328125 0 6281.115
309.0840148925781 0 1395.3638
309.1743469238281 0 908.6715
310.1400451660156 0 912.954
311.1008605957031 0 893.9747
311.1355285644531 0 3987.0908
312.13848876953125 0 656.0859
312.1922607421875 0 697.7003
313.151611328125 0 1844.0161
314.1716613769531 0 6632.5312
315.1670837402344 0 23486.479 b 3
316.1703796386719 0 3428.127
317.10968017578125 0 2313.2498
317.17193603515625 0 720.83954
321.1200256347656 0 3973.0334
322.152099609375 0 1364.3586
324.1563415527344 0 3353.2168
325.15179443359375 0 6895.641
328.1878662109375 0 2045.8387
329.14599609375 0 4906.6123
331.1621398925781 0 2107.371
332.1928405761719 0 750.8767
336.1678771972656 0 626.24976
338.1468505859375 0 3394.8606
339.1304931640625 0 1222.9381
340.1367492675781 0 2147.9824
340.185302734375 0 1000.87146
341.1452331542969 0 1677.87
341.2195739746094 0 1003.2138
343.1617736816406 0 11034.396 y 5
344.1647644042969 0 2113.2932
352.125732421875 0 2632.4912
353.14556884765625 0 2483.3047
353.18194580078125 0 946.6004
354.141845703125 0 2925.3848 y Water loss 8
355.125244140625 0 3626.7358
356.1568603515625 0 45693.027
357.15997314453125 0 6946.3804
357.2134094238281 0 825.8732
357.2494812011719 0 764.6061
358.1597900390625 0 916.79974
359.15643310546875 0 1360.2212
365.1844787597656 0 665.98505
366.14111328125 0 1415.0902
367.1611633300781 0 569.56104
367.1971740722656 0 750.3788
369.1769714355469 0 1398.7023
369.2136535644531 0 2100
370.1374816894531 0 902.9043
370.24481201171875 0 1056.9642
371.1905822753906 0 615.06256
372.15185546875 0 4656.1763 y 8
373.1546630859375 0 806.7856
374.18023681640625 0 887.3207
377.2202453613281 0 691.317
379.2448425292969 0 627.0272
380.22637939453125 0 665.2513
383.2296142578125 0 748.56384
384.1890869140625 0 1965.9768
385.20880126953125 0 3830.4404
390.6971740722656 0 705.5945 y Water loss 4
394.13909912109375 0 505.39178
394.171142578125 0 1067.6891
395.15716552734375 0 1672.2972
395.1958923339844 0 499.16724
395.2301940917969 0 831.42664
396.19232177734375 0 714.2227
396.2252502441406 0 2327.2952
397.20806884765625 0 2567.6465
398.1669921875 0 1378.5518
398.2400817871094 0 1073.2432
399.7041015625 0 584.4606 y 4
400.18359375 0 785.73663
400.2269287109375 0 539.89484
400.2562561035156 0 7756.0703
401.2596740722656 0 1431.6044
402.20001220703125 0 1213.018
405.19061279296875 0 720.2746
407.2403259277344 0 1263.0571
408.1529846191406 0 1024.0454
410.1693115234375 0 966.26385
410.2044372558594 0 1197.3468
410.24041748046875 0 1846.1891 b Water loss 4
412.1835632324219 0 7167.9365
413.185791015625 0 1055.6063
414.20123291015625 0 943.4889
414.2353515625 0 4046.236
416.1788024902344 0 1950.3134
418.20172119140625 0 972.1719
420.18951416015625 0 891.5914
421.22088623046875 0 705.413
422.1668395996094 0 1010.3543
423.2273864746094 0 2203.4412
424.1846923828125 0 1142.5289
424.2196350097656 0 2710.74
425.2205810546875 0 750.3031
426.1631164550781 0 1690.5814
426.2341613769531 0 1459.1034
428.17822265625 0 2248.338
428.2147216796875 0 1845.8604 y 3
428.2514343261719 0 5437.2246 b 4
429.25616455078125 0 703.64075
430.1942138671875 0 7256.62
431.198486328125 0 1576.6345
434.1656494140625 0 1201.6453
435.1995544433594 0 765.9183
437.2149963378906 0 1466.7153
438.1996154785156 0 1385.6458
439.1989440917969 0 1537.088
440.1785888671875 0 1447.0137
440.2142333984375 0 1297.1804
440.2508850097656 0 583.8884
442.2301940917969 0 6273.177
443.2352600097656 0 1739.0763
444.17315673828125 0 899.99854
444.24664306640625 0 2785.4216
446.1888122558594 0 757.2586
446.7121887207031 0 1782.157
447.7195129394531 0 2623.7039 y Water loss 2
448.2214660644531 0 1169.1057
448.7254638671875 0 996.2726
451.19439697265625 0 7179.966
452.1978759765625 0 1761.998
453.2101745605469 0 1215.5801 y Water loss 7
454.191162109375 0 728.08673
455.22576904296875 0 28560.848
456.22821044921875 0 7552.092
457.2281494140625 0 1689.9088
458.2250061035156 0 653.1405
461.2612609863281 0 780.21375
464.2155456542969 0 704.689
468.17230224609375 0 1603.8356
468.24981689453125 0 641.5954
469.20556640625 0 764.49445
471.2207336425781 0 6443.7827 y 7
472.222412109375 0 1029.0378
474.249755859375 0 851.84344 b Water loss 9
482.239990234375 0 740.301
483.2531433105469 0 1672.6018 b 9
484.2416076660156 0 1873.3153
485.2389221191406 0 629.7131
487.5827331542969 0 1013.155
487.91827392578125 0 761.6463
490.2318420410156 0 2833.4497
490.73016357421875 0 1602.2168
491.23577880859375 0 2064.844 y Water loss 1
491.73834228515625 0 1261.3588
492.24365234375 0 4046.9028
492.7456359863281 0 1914.8196
493.2448425292969 0 1662.7792
494.2701110839844 0 3319.8008
495.2558288574219 0 712.34705
496.2292175292969 0 784.11194
497.23431396484375 0 1220.0969
497.273193359375 0 1043.0057
498.25714111328125 0 847.7276
499.2315979003906 0 1633.5697
499.2556457519531 0 691.4799
499.7352600097656 0 1523.074
500.2401428222656 0 3228.8542 y 1
500.7431640625 0 1008.8664
501.266845703125 0 2321.682
502.2715148925781 0 818.82623
507.2182922363281 0 850.3392
509.23626708984375 0 2112.4963
510.22418212890625 0 654.1685
511.25213623046875 0 5300.4697
512.2568359375 0 921.1858
513.263671875 0 1417.95
513.3035888671875 0 1024.0216
515.24609375 0 1290.9832
515.2842407226562 0 1038.1886
518.2400512695312 0 981.16565
521.2349853515625 0 970.68616
522.267578125 0 4139.9395
523.2532348632812 0 1018.9947
525.2303466796875 0 1188.0845
525.2684936523438 0 2209.4365 b Water loss 5
527.244873046875 0 1363.1464
529.2617797851562 0 8253.215
530.2645874023438 0 2529.0334
533.2393798828125 0 732.80194
533.7791748046875 0 1405.3763
537.7640991210938 0 2895.1943
538.2639770507812 0 3549.1301
538.7659301757812 0 947.2395
539.2482299804688 0 2051.9683
539.2779541015625 0 911.6414
539.7747802734375 0 643.9108
540.2811279296875 0 1319.9521
541.297607421875 0 1967.8174
543.2782592773438 0 5071.0674 b 5
544.28173828125 0 1015.50836
545.2568969726562 0 1119.3779
546.7697143554688 0 6363.947
547.271728515625 0 6044.0415
547.7771606445312 0 13929.32 Precursor Water loss
548.2793579101562 0 7265.1704
548.7810668945312 0 3656.2693
549.2858276367188 0 932.959
550.2622680664062 0 5230.22
551.2643432617188 0 1482.4137
552.2783813476562 0 778.5544 y Water loss 6
553.2628173828125 0 1703.89
554.2936401367188 0 14540.811
555.294921875 0 5621.8
555.7758178710938 0 7167.6177
556.279541015625 0 5288.4707
556.7825317382812 0 13001.064 Precursor
557.2859497070312 0 9504.964
557.783935546875 0 2530.104
559.27490234375 0 1107.1895
567.2431640625 0 759.0252
570.2890014648438 0 4601.954 y 6
571.2921142578125 0 1036.889
595.3275756835938 0 811.06433
597.32421875 0 1597.772
608.3035278320312 0 1110.4552
609.3046875 0 863.42706
610.3202514648438 0 3192.8787
611.3202514648438 0 953.0502
612.3348999023438 0 2977.7537
613.3406982421875 0 950.2743
614.350341796875 0 1803.8147
615.3556518554688 0 1263.0048
624.3353271484375 0 2615.3943 b Water loss 6
625.3414916992188 0 922.94025
626.3172607421875 0 1125.4438
628.3303833007812 0 4909.993
629.3348388671875 0 1774.7823
631.3209838867188 0 1087.4458
632.3218994140625 0 760.0186
638.314208984375 0 1102.1278
640.3306274414062 0 1194.6775
642.3464965820312 0 7635.3403 b 6
643.3497924804688 0 2761.681
644.3256225585938 0 665.7704
645.8424072265625 0 1361.9514
646.345947265625 0 1431.9438
651.3027954101562 0 1047.7965
652.29248046875 0 580.86444
658.335693359375 0 1366.0166
665.2901611328125 0 10928.091
666.2929077148438 0 3832.4487
667.2998657226562 0 3072.5251 y Water loss 5
668.2913208007812 0 1039.3843
669.3207397460938 0 48755.445
670.3238525390625 0 17059.777
671.3260498046875 0 4481.255
681.3572998046875 0 1302.7997
681.8330688476562 0 979.4132
683.3006591796875 0 2542.6133
684.3016357421875 0 929.5472
685.3154907226562 0 13881.042 y 5
686.318359375 0 5262.685
687.3223876953125 0 714.9895
690.3645629882812 0 683.1919
692.3705444335938 0 742.56464
694.3661499023438 0 729.77875
697.3516845703125 0 876.97577
699.3670043945312 0 2251.6978
700.3674926757812 0 1929.4084
708.3634643554688 0 738.4228
710.4566650390625 0 1167.717
713.4237060546875 0 770.8362
715.3653564453125 0 777.81354
717.3667602539062 0 763.125
721.3990478515625 0 1006.79144
723.4010009765625 0 1142.1605 b Water loss 7
735.3742065429688 0 672.1236
738.3795166015625 0 770.7811
738.8834838867188 0 1217.6608
739.3895263671875 0 2061.4727
741.4132690429688 0 3397.27 b 7
742.419189453125 0 1094.1724
745.3628540039062 0 1390.3652
746.3654174804688 0 1121.2356
747.3775634765625 0 965.52936
749.3955688476562 0 1627.2655
750.397216796875 0 807.1836
763.3746948242188 0 1593.5001
765.3890991210938 0 2507.4648
766.390869140625 0 1208.3225
767.4376831054688 0 805.188
778.3733520507812 0 16466.621
779.3760375976562 0 6402.6963
780.3823852539062 0 2195.8313 y Water loss 4
781.3826904296875 0 1182.5295
782.4038696289062 0 3764.5552
782.8997802734375 0 1033.8657
783.4056396484375 0 2964.792
783.8989868164062 0 679.5169
794.4443969726562 0 1086.6323 b Water loss 8
796.3856201171875 0 1770.466
797.3880615234375 0 925.2113
798.3987426757812 0 5281.3613 y 4
799.4027709960938 0 1874.2041
808.4296264648438 0 852.8858
812.4524536132812 0 2502.5994 b 8
813.4547119140625 0 1269.1432
822.4061279296875 0 604.2155
829.477783203125 0 2472.3726
832.3955078125 0 762.14935
833.3956298828125 0 743.8486
834.4075317382812 0 997.29913
835.3934936523438 0 2804.9387
836.40625 0 2481.2065
837.4093017578125 0 1797.1626 y Water loss 3
838.3974609375 0 2088.516
839.423828125 0 7629.165
840.4281005859375 0 3068.2107
841.428955078125 0 1001.30316
850.4053955078125 0 1178.2501
851.4071044921875 0 744.4924
852.4198608398438 0 989.49274
853.407470703125 0 3973
854.4080200195312 0 2251.4346
855.4197998046875 0 5738.841 y 3
856.4240112304688 0 1682.6913
866.4329223632812 0 633.91693
874.406494140625 0 1001.77405
892.416015625 0 18300.727
893.4188842773438 0 8248.088
894.4284057617188 0 6950.645 y Water loss 2
895.428955078125 0 2999.1328
896.4462280273438 0 18898.111
897.4496459960938 0 8881.151
898.4533081054688 0 2496.5054
910.42626953125 0 19064.873
911.4287109375 0 12280.474
912.4407348632812 0 35376.164 y 2
913.4441528320312 0 14091.172
914.44677734375 0 4552.4165
953.4660034179688 0 4182.135
954.4699096679688 0 1722.756
961.4398803710938 0 1932.1632
962.4401245117188 0 906.3777
963.442138671875 0 1067.3956
971.4151611328125 0 697.8523
979.4473266601562 0 13129.493
980.4507446289062 0 8094.673
981.4592895507812 0 8064.5693 y Water loss 1
982.4642944335938 0 4921.661
983.4761962890625 0 6571.244
984.4778442382812 0 3313.7422
985.4861450195312 0 784.0255
989.4351806640625 0 1620.0731
991.4445190429688 0 1650.6753
992.4515380859375 0 942.1674
993.4619140625 0 985.5807
997.4586791992188 0 16569.527
998.4605712890625 0 9903.721
999.4725341796875 0 24192.229 y 1
1000.4766235351562 0 11516.698
1001.4785766601562 0 4010.02
1007.4428100585938 0 1195.57
1009.4579467773438 0 2235.8135
1010.4530639648438 0 839.67456

Spectrum Details

|  |  |
| --- | --- |
| Matched peaks? Matched peaks The total absolute number of peaks matched. Additionally in brackets the total fraction of peaks matched and the total number of peaks is shown. | 54 (8.75% of 617) |
| FDR? FDR The false discovery rate estimated for this peptide. It is calculated by matching all theoretical fragments with a non-integer shift with the raw peaks for this spectrum. This is done with 40 different shifts. The resulting percentage is the average number of annotated peaks over the number of annotated peaks with the correct spectrum. | 0.31% |
| Satellite FDR? Satellite FDR See the FDR for details on its calculation. This satellite ion specific FDR only contains the satellite ions (d/w) for I/L/J positions. | - |
| PSM Score? PSM Score The PSM Score as given by Hecklib to this annotated spectrum. It is shown with three significant figures. | 719 |

#### Spectrum 5586? Spectrum 5586 The raw spectrum of this peptide as annotated by Hecklib. The fragments are coloured according to ion type (see legend). Any peaks with a star '\*' as text can be hovered over to see the full details, first the ion type second the mass shift type. By hovering over the amino acids in the peptide or ions in the legend the corresponding peaks are highlighted. By toggling the 'Unassigned' label you can turn the background (unassigned) peaks on or off in the plot. By updating the slider in the Ion legend you can update the spectrum to only show the top X% of the peaks with labels. The top X% means any peak that is within X% of the highest intensity. By dragging in the spectrum you can zoom in to a specific part of the spectrum and use 'Zoom Out' to get back to the original zoom level. The annotation of the spectrum is based on the given sequence in the peptides file and is done with different software so inconsistencies are likely. The peaks are annotated based on the given sequence, with 20 ppm tolerance.

Copy Data

##### Spectrum 5586 (TSV)

###### Preview

```
Loading example...
```

*Click on the button to copy the data to your clipboard.*

Mz MinMz MaxIntensity Max

WidthHeightPeptide font sizePeptide stroke widthSpectrum font sizeSpectrum stroke widthCompact peptide

Ion legend

wxyz

abcd

OtherUnassignedIon^Charge^_Position_Show for top:%

JSGGIDVVAHE

02.55e+45.10e+47.65e+41.02e+5

Zoom Out

c^+1^_2_y^+1^_2_c^+1^_4_z^+1^_3_y^+1^_3_c^+1^_5_z^+1^_4_y^+1^_4_y^+2^_10_w^+1^_5_z^+1^_5_c^+1^_6_y^+1^_5_z^+1^_6_c^+1^_7_y^+1^_6_w^+1^_7_c^+1^_8_c^+1^_8_z^+1^_7_y^+1^_7_z^+1^_8_c^+1^_9_y^+1^_8_z^+1^_9_y^+1^_9_y^+1^_10_c^+1^_10_y^+1^_10_

041783412511667

Fragment Matches Table

Show background peaks

| Position | Ion type | Intensity | mz Theoretical | mz Error (Th) | mz Error (ppm) | Charge | Series Number |
| --- | --- | --- | --- | --- | --- | --- | --- |
| - | - | 384.1 | 125.1 | - | - | 0 | - |
| - | - | 528.3 | 135.2 | - | - | 0 | - |
| - | - | 454.9 | 143.1 | - | - | 0 | - |
| - | - | 462 | 146.5 | - | - | 0 | - |
| - | - | 483.8 | 148.8 | - | - | 0 | - |
| - | - | 454.2 | 148.8 | - | - | 0 | - |
| - | - | 412.5 | 148.8 | - | - | 0 | - |
| - | - | 435.8 | 148.9 | - | - | 0 | - |
| - | - | 707.6 | 148.9 | - | - | 0 | - |
| - | - | 535.5 | 148.9 | - | - | 0 | - |
| - | - | 649.8 | 148.9 | - | - | 0 | - |
| - | - | 1352 | 148.9 | - | - | 0 | - |
| - | - | 1256 | 148.9 | - | - | 0 | - |
| - | - | 2322 | 148.9 | - | - | 0 | - |
| - | - | 3830 | 148.9 | - | - | 0 | - |
| - | - | 4435 | 149 | - | - | 0 | - |
| - | - | 2679 | 149 | - | - | 0 | - |
| - | - | 1499 | 149 | - | - | 0 | - |
| - | - | 1293 | 149 | - | - | 0 | - |
| - | - | 906.4 | 149 | - | - | 0 | - |
| - | - | 1004 | 149 | - | - | 0 | - |
| - | - | 662.1 | 149 | - | - | 0 | - |
| - | - | 663.2 | 149 | - | - | 0 | - |
| - | - | 567.1 | 149 | - | - | 0 | - |
| - | - | 688.3 | 149 | - | - | 0 | - |
| - | - | 1158 | 155.1 | - | - | 0 | - |
| - | - | 437.4 | 172.1 | - | - | 0 | - |
| - | - | 648.8 | 173.1 | - | - | 0 | - |
| - | - | 6736 | 173.1 | - | - | 0 | - |
| - | - | 605.5 | 187.1 | - | - | 0 | - |
| - | - | 488.5 | 187.1 | - | - | 0 | - |
| 2 | c | 9309 | 201.1 | 0.0003525 | 1.753 | +1 | 2 |
| - | - | 849.2 | 202.1 | - | - | 0 | - |
| - | - | 4884 | 221.1 | - | - | 0 | - |
| - | - | 1081 | 225 | - | - | 0 | - |
| - | - | 8636 | 239.1 | - | - | 0 | - |
| - | - | 1492 | 240.1 | - | - | 0 | - |
| - | - | 623.5 | 252.4 | - | - | 0 | - |
| - | - | 571.7 | 265.9 | - | - | 0 | - |
| - | - | 532.8 | 272.7 | - | - | 0 | - |
| 10 | y | 3010 | 285.1 | 0.000297 | 1.042 | +1 | 2 |
| - | - | 1300 | 287.2 | - | - | 0 | - |
| - | - | 6849 | 295.1 | - | - | 0 | - |
| - | - | 674.1 | 299.1 | - | - | 0 | - |
| - | - | 488.1 | 309.6 | - | - | 0 | - |
| - | - | 819.5 | 313.1 | - | - | 0 | - |
| 4 | c | 3053 | 315.2 | 0.0005248 | 1.665 | +1 | 4 |
| - | - | 529.3 | 316.2 | - | - | 0 | - |
| 9 | z | 2031 | 340.1 | 9.986E-06 | 0.02936 | +1 | 3 |
| - | - | 1017 | 355.1 | - | - | 0 | - |
| 9 | y | 2432 | 356.2 | 0.0007897 | 2.217 | +1 | 3 |
| - | - | 8345 | 369.1 | - | - | 0 | - |
| - | - | 1529 | 400.3 | - | - | 0 | - |
| - | - | 578.8 | 409 | - | - | 0 | - |
| 5 | c | 1835 | 428.3 | 0.0009401 | 2.195 | +1 | 5 |
| 8 | z | 1.025E+04 | 439.2 | 0.0001561 | 0.3554 | +1 | 4 |
| - | - | 3808 | 440.2 | - | - | 0 | - |
| - | - | 598.9 | 443.1 | - | - | 0 | - |
| 8 | y | 4705 | 455.2 | 8.132E-05 | 0.1786 | +1 | 4 |
| - | - | 1293 | 456.2 | - | - | 0 | - |
| - | - | 725 | 466.8 | - | - | 0 | - |
| - | - | 617.9 | 487.2 | - | - | 0 | - |
| 2 | y | 1084 | 492.2 | 0.0005572 | 1.132 | +2 | 10 |
| - | - | 1017 | 492.7 | - | - | 0 | - |
| 7 | w | 780 | 523.3 | 0.0004984 | 0.9524 | +1 | 5 |
| - | - | 628.4 | 529.3 | - | - | 0 | - |
| - | - | 733.2 | 530.3 | - | - | 0 | - |
| 7 | z | 5574 | 538.3 | 0.0002717 | 0.5047 | +1 | 5 |
| - | - | 1.801E+04 | 539.3 | - | - | 0 | - |
| - | - | 1693 | 539.8 | - | - | 0 | - |
| - | - | 6026 | 540.3 | - | - | 0 | - |
| - | - | 1121 | 541.3 | - | - | 0 | - |
| 6 | c | 624.7 | 543.3 | 0.001852 | 3.408 | +1 | 6 |
| - | - | 2068 | 547.4 | - | - | 0 | - |
| - | - | 5361 | 548.8 | - | - | 0 | - |
| - | - | 529.5 | 549 | - | - | 0 | - |
| - | - | 3596 | 549.3 | - | - | 0 | - |
| - | - | 2242 | 549.8 | - | - | 0 | - |
| 7 | y | 4092 | 554.3 | 0.000441 | 0.7957 | +1 | 5 |
| - | - | 1504 | 555.3 | - | - | 0 | - |
| - | - | 537.4 | 586.6 | - | - | 0 | - |
| - | - | 3539 | 609.3 | - | - | 0 | - |
| - | - | 2244 | 610.3 | - | - | 0 | - |
| - | - | 530.6 | 613.7 | - | - | 0 | - |
| 6 | z | 4842 | 653.3 | 0.0006644 | 1.017 | +1 | 6 |
| - | - | 1.644E+04 | 654.3 | - | - | 0 | - |
| - | - | 5735 | 655.3 | - | - | 0 | - |
| - | - | 1064 | 656.3 | - | - | 0 | - |
| - | - | 1917 | 658.4 | - | - | 0 | - |
| 7 | c | 3278 | 659.4 | 0.002149 | 3.259 | +1 | 7 |
| - | - | 596.2 | 662.2 | - | - | 0 | - |
| 6 | y | 1.169E+04 | 669.3 | 0.0008948 | 1.337 | +1 | 6 |
| - | - | 4740 | 670.3 | - | - | 0 | - |
| - | - | 938.8 | 671.3 | - | - | 0 | - |
| - | - | 5011 | 714.4 | - | - | 0 | - |
| - | - | 3537 | 715.4 | - | - | 0 | - |
| - | - | 1377 | 716.4 | - | - | 0 | - |
| - | - | 1200 | 722.4 | - | - | 0 | - |
| 5 | w | 5630 | 737.3 | 0.0005593 | 0.7585 | +1 | 7 |
| - | - | 2480 | 738.3 | - | - | 0 | - |
| 8 | c | 673 | 741.4 | 0.001335 | 1.801 | +1 | 8 |
| - | - | 553 | 742.4 | - | - | 0 | - |
| - | - | 1113 | 749.4 | - | - | 0 | - |
| - | - | 3926 | 757.4 | - | - | 0 | - |
| 8 | c | 9916 | 758.4 | 0.001166 | 1.537 | +1 | 8 |
| - | - | 3356 | 759.4 | - | - | 0 | - |
| - | - | 675.6 | 760.4 | - | - | 0 | - |
| 5 | z | 5027 | 766.4 | 0.0005377 | 0.7017 | +1 | 7 |
| - | - | 2234 | 767.4 | - | - | 0 | - |
| - | - | 1029 | 779.4 | - | - | 0 | - |
| - | - | 1137 | 781.4 | - | - | 0 | - |
| 5 | y | 2585 | 782.4 | 0.0004861 | 0.6213 | +1 | 7 |
| - | - | 1361 | 783.4 | - | - | 0 | - |
| - | - | 5394 | 785.5 | - | - | 0 | - |
| - | - | 613.4 | 786.4 | - | - | 0 | - |
| - | - | 3010 | 786.5 | - | - | 0 | - |
| - | - | 752.3 | 794.3 | - | - | 0 | - |
| - | - | 550 | 803 | - | - | 0 | - |
| - | - | 626.9 | 808.4 | - | - | 0 | - |
| - | - | 795.1 | 814.8 | - | - | 0 | - |
| - | - | 660.3 | 815.3 | - | - | 0 | - |
| - | - | 1275 | 818.4 | - | - | 0 | - |
| - | - | 672.5 | 823.3 | - | - | 0 | - |
| 4 | z | 6442 | 823.4 | 9.071E-06 | 0.01102 | +1 | 8 |
| - | - | 723.4 | 823.9 | - | - | 0 | - |
| - | - | 5757 | 824.4 | - | - | 0 | - |
| - | - | 2035 | 824.9 | - | - | 0 | - |
| - | - | 2484 | 825.4 | - | - | 0 | - |
| - | - | 7706 | 828.5 | - | - | 0 | - |
| 9 | c | 1.262E+04 | 829.5 | 0.002025 | 2.441 | +1 | 9 |
| - | - | 3745 | 830.5 | - | - | 0 | - |
| - | - | 1642 | 831.5 | - | - | 0 | - |
| 4 | y | 1.579E+04 | 839.4 | 0.0002824 | 0.3364 | +1 | 8 |
| - | - | 7268 | 840.4 | - | - | 0 | - |
| - | - | 2293 | 841.4 | - | - | 0 | - |
| - | - | 894.6 | 867.4 | - | - | 0 | - |
| - | - | 624.5 | 879.1 | - | - | 0 | - |
| 3 | z | 5125 | 880.4 | 0.0002144 | 0.2435 | +1 | 9 |
| - | - | 2902 | 881.4 | - | - | 0 | - |
| - | - | 757.5 | 882.4 | - | - | 0 | - |
| 3 | y | 2.31E+04 | 896.4 | 0.000689 | 0.7686 | +1 | 9 |
| - | - | 1.219E+04 | 897.4 | - | - | 0 | - |
| - | - | 2500 | 898.5 | - | - | 0 | - |
| - | - | 944.5 | 912.4 | - | - | 0 | - |
| - | - | 1412 | 922.5 | - | - | 0 | - |
| - | - | 623.6 | 960.5 | - | - | 0 | - |
| 2 | y | 1072 | 965.5 | 0.008115 | 8.405 | +1 | 10 |
| 10 | c | 4.466E+04 | 966.5 | 0.0006341 | 0.6561 | +1 | 10 |
| - | - | 2.211E+04 | 967.5 | - | - | 0 | - |
| - | - | 1622 | 968.5 | - | - | 0 | - |
| - | - | 5163 | 968.5 | - | - | 0 | - |
| - | - | 929.3 | 969.6 | - | - | 0 | - |
| - | - | 745.9 | 975.5 | - | - | 0 | - |
| 2 | y | 9881 | 983.5 | 0.0001246 | 0.1267 | +1 | 10 |
| - | - | 3948 | 984.5 | - | - | 0 | - |
| - | - | 1580 | 985.5 | - | - | 0 | - |
| - | - | 1343 | 993.5 | - | - | 0 | - |
| - | - | 633.6 | 1024 | - | - | 0 | - |
| - | - | 676.4 | 1034 | - | - | 0 | - |
| - | - | 6889 | 1037 | - | - | 0 | - |
| - | - | 3194 | 1038 | - | - | 0 | - |
| - | - | 1906 | 1039 | - | - | 0 | - |
| - | - | 892.2 | 1041 | - | - | 0 | - |
| - | - | 666.7 | 1048 | - | - | 0 | - |
| - | - | 1373 | 1052 | - | - | 0 | - |
| - | - | 969.9 | 1053 | - | - | 0 | - |
| - | - | 851.3 | 1054 | - | - | 0 | - |
| - | - | 812.7 | 1061 | - | - | 0 | - |
| - | - | 899.4 | 1070 | - | - | 0 | - |
| - | - | 781.2 | 1071 | - | - | 0 | - |
| - | - | 1024 | 1079 | - | - | 0 | - |
| - | - | 7187 | 1080 | - | - | 0 | - |
| - | - | 2.525E+04 | 1081 | - | - | 0 | - |
| - | - | 1.512E+04 | 1082 | - | - | 0 | - |
| - | - | 5515 | 1083 | - | - | 0 | - |
| - | - | 663.7 | 1084 | - | - | 0 | - |
| - | - | 958.1 | 1095 | - | - | 0 | - |
| - | - | 1521 | 1096 | - | - | 0 | - |
| - | - | 3.209E+04 | 1097 | - | - | 0 | - |
| - | - | 1.01E+05 | 1098 | - | - | 0 | - |
| - | - | 5.319E+04 | 1099 | - | - | 0 | - |
| - | - | 1.797E+04 | 1100 | - | - | 0 | - |
| - | - | 2567 | 1101 | - | - | 0 | - |
| - | - | 759 | 1122 | - | - | 0 | - |
| - | - | 607.9 | 1187 | - | - | 0 | - |
| - | - | 627.4 | 1219 | - | - | 0 | - |
| - | - | 660.4 | 1396 | - | - | 0 | - |
| - | - | 616.6 | 1527 | - | - | 0 | - |
| - | - | 803.3 | 1589 | - | - | 0 | - |
| - | - | 941.7 | 1629 | - | - | 0 | - |
| - | - | 1625 | 1648 | - | - | 0 | - |
| - | - | 1976 | 1649 | - | - | 0 | - |
| - | - | 3840 | 1650 | - | - | 0 | - |
| - | - | 1645 | 1651 | - | - | 0 | - |

m/z Charge Intensity FragmentType MassShift Position
125.1072998046875 0 384.12625
135.177734375 0 528.34937
143.117431640625 0 454.86945
146.48870849609375 0 462.04062
148.80767822265625 0 483.83777
148.82835388183594 0 454.1732
148.84324645996094 0 412.54913
148.89291381835938 0 435.82025
148.89990234375 0 707.6451
148.90695190429688 0 535.48846
148.91390991210938 0 649.8442
148.9210662841797 0 1352.087
148.92823791503906 0 1255.762
148.93528747558594 0 2321.9954
148.94287109375 0 3829.7483
148.9593048095703 0 4435.468
148.96693420410156 0 2679.3682
148.97421264648438 0 1499.2334
148.98129272460938 0 1292.7075
148.98831176757812 0 906.4172
148.99530029296875 0 1004.0647
149.00242614746094 0 662.11536
149.00990295410156 0 663.15155
149.0170440673828 0 567.0995
149.0454864501953 0 688.31934
155.11758422851562 0 1158.1538
172.09719848632812 0 437.40112
173.1214141845703 0 648.7757
173.1282501220703 0 6736.4546
187.10708618164062 0 605.4502
187.14369201660156 0 488.54214
201.12301635742188 0 9309.479 c Ammonia loss 1
202.12646484375 0 849.1862
221.08407592773438 0 4883.57
225.0424041748047 0 1080.5463
239.09461975097656 0 8636.102
240.13385009765625 0 1491.8865
252.3813934326172 0 623.52655
265.85980224609375 0 571.6885
272.73114013671875 0 532.8166
285.1190490722656 0 3009.5706 y 9
287.1713562011719 0 1299.6732
295.10296630859375 0 6848.736
299.0625915527344 0 674.12714
309.59344482421875 0 488.0754
313.1139221191406 0 819.51447
315.165771484375 0 3053.4143 c Ammonia loss 3
316.1705322265625 0 529.25244
340.1377258300781 0 2030.9087 z 8
355.0711364746094 0 1016.7188
356.1556701660156 0 2431.6174 y 8
369.1212463378906 0 8344.921
400.2543640136719 0 1529.3042
408.9764404296875 0 578.80005
428.2494201660156 0 1835.1133 c Ammonia loss 4
439.20599365234375 0 10247.055 z 7
440.2106628417969 0 3807.787
443.1412048339844 0 598.91046
455.22479248046875 0 4705.3438 y 7
456.22772216796875 0 1293.0355
466.7633361816406 0 725.0375
487.2386474609375 0 617.9189
492.2427062988281 0 1084.2518 y 1
492.7453918457031 0 1017.13684
523.2515869140625 0 779.9589 w 6
529.2620849609375 0 628.38556
530.3163452148438 0 733.21063
538.2742919921875 0 5573.776 z 6
539.2818603515625 0 18012.553
539.7796020507812 0 1693.3875
540.2842407226562 0 6025.6934
541.2880859375 0 1120.753
543.2754516601562 0 624.7386 c Ammonia loss 5
547.352294921875 0 2067.8691
548.7847290039062 0 5361.08
549.0353393554688 0 529.5294
549.2866821289062 0 3596.1492
549.787841796875 0 2241.791
554.2928466796875 0 4092.3755 y 6
555.29736328125 0 1503.5614
586.59375 0 537.4228
609.3114624023438 0 3538.7656
610.3167724609375 0 2244.393
613.6629638671875 0 530.61304
653.3008422851562 0 4842.4497 z 5
654.3084106445312 0 16444.797
655.3114013671875 0 5735.093
656.3157958984375 0 1063.7578
658.3630981445312 0 1916.8611
659.3701171875 0 3277.6477 c 6
662.240478515625 0 596.1804
669.3193359375 0 11691.069 y 5
670.3222045898438 0 4739.845
671.3240356445312 0 938.76074
714.4260864257812 0 5010.8003
715.4320068359375 0 3537.2014
716.4346313476562 0 1376.853
722.3938598632812 0 1199.7095
737.3458862304688 0 5630.104 w 4
738.3483276367188 0 2480.1794
741.4154663085938 0 672.95874 c Ammonia loss 7
742.3973388671875 0 553.0465
749.39794921875 0 1113.4141
757.4317016601562 0 3926.4805
758.4395141601562 0 9915.718 c 7
759.4432983398438 0 3356.1606
760.4390869140625 0 675.59863
766.3861083984375 0 5027.0176 z 4
767.39013671875 0 2233.9795
779.412353515625 0 1029.405
781.359375 0 1137.0599
782.40380859375 0 2585.4902 y 4
783.4085693359375 0 1360.9055
785.4627685546875 0 5394.0005
786.39599609375 0 613.41254
786.4672241210938 0 3009.889
794.3440551757812 0 752.2755
803.0391845703125 0 549.9799
808.3785400390625 0 626.9284
814.83544921875 0 795.1037
815.334228515625 0 660.3169
818.3914794921875 0 1275.045
823.3346557617188 0 672.4637
823.4070434570312 0 6442.0415 z 3
823.9203491210938 0 723.43945
824.4127197265625 0 5756.9985
824.9359741210938 0 2035.4723
825.4237060546875 0 2484.0625
828.469482421875 0 7705.878
829.4757690429688 0 12624.476 c 8
830.4786987304688 0 3745.3152
831.4884643554688 0 1641.8225
839.4254760742188 0 15787.771 y 3
840.4273681640625 0 7267.778
841.43212890625 0 2292.9097
867.4136962890625 0 894.57275
879.0712280273438 0 624.5132
880.4282836914062 0 5125.266 z 2
881.4307861328125 0 2902.033
882.4340209960938 0 757.51056
896.446533203125 0 23101.367 y 2
897.44921875 0 12190.366
898.4520874023438 0 2500.0623
912.3944091796875 0 944.5183
922.5219116210938 0 1412.1276
960.4817504882812 0 623.60406
965.4605712890625 0 1071.7086 y Water loss 1
966.5360717773438 0 44664.105 c 9
967.5394897460938 0 22113.28
968.4586181640625 0 1622.3105
968.5429077148438 0 5162.607
969.5570678710938 0 929.2521
975.4522094726562 0 745.8692
983.4791259765625 0 9881.487 y 1
984.4840698242188 0 3947.723
985.4869995117188 0 1580.0486
993.4617919921875 0 1342.5719
1024.46240234375 0 633.5693
1034.2327880859375 0 676.44946
1037.493896484375 0 6889.2534
1038.4969482421875 0 3193.5476
1039.4954833984375 0 1906.2502
1041.495849609375 0 892.23096
1047.5013427734375 0 666.6773
1051.5726318359375 0 1372.9985
1052.5767822265625 0 969.8612
1053.5621337890625 0 851.25604
1060.5086669921875 0 812.66235
1069.5716552734375 0 899.4052
1070.57177734375 0 781.2093
1078.5113525390625 0 1023.5066
1079.553955078125 0 7187.115
1080.545166015625 0 25250.652
1081.5465087890625 0 15120.2295
1082.54736328125 0 5515.0186
1083.565673828125 0 663.657
1094.50537109375 0 958.05347
1095.5279541015625 0 1520.814
1096.5625 0 32091.137
1097.5689697265625 0 101004.914
1098.5726318359375 0 53189.5
1099.5753173828125 0 17974.783
1100.572021484375 0 2567.432
1122.3963623046875 0 759.0194
1187.48193359375 0 607.92377
1218.60986328125 0 627.37146
1396.043701171875 0 660.42426
1527.1007080078125 0 616.59467
1588.675537109375 0 803.28357
1628.817626953125 0 941.692
1647.843994140625 0 1624.677
1648.8614501953125 0 1976.2836
1649.87548828125 0 3840.1128
1650.892578125 0 1645.0367

Spectrum Details

|  |  |
| --- | --- |
| Matched peaks? Matched peaks The total absolute number of peaks matched. Additionally in brackets the total fraction of peaks matched and the total number of peaks is shown. | 29 (14.95% of 194) |
| FDR? FDR The false discovery rate estimated for this peptide. It is calculated by matching all theoretical fragments with a non-integer shift with the raw peaks for this spectrum. This is done with 40 different shifts. The resulting percentage is the average number of annotated peaks over the number of annotated peaks with the correct spectrum. | 1.64% |
| Satellite FDR? Satellite FDR See the FDR for details on its calculation. This satellite ion specific FDR only contains the satellite ions (d/w) for I/L/J positions. | 9.52% |
| PSM Score? PSM Score The PSM Score as given by Hecklib to this annotated spectrum. It is shown with three significant figures. | 337 |

#### Reverse Lookup? Reverse Lookup All places where this read could be placed.

| Group | Segment | Template | Template Part | Read Part | Score | Unique |
| --- | --- | --- | --- | --- | --- | --- |
| Decoy | Decoy | THER | [364..375] | [0..11] | 88 | True |

| Recombined | Template Part | Read Part | Score | Unique |
| --- | --- | --- | --- | --- |
| THER | [364..375] | [0..11] | 88 | True |

#### Meta Information from Multiple reads

##### Number of combined reads

3

##### Intensity

0.6858

##### TotalArea

2.099E+07

##### Changes to the peptide sequence

JSGGIDVVAHE

J→ISupport for Isoleucine based on side chain ions (1 for I 0 for L) (Position: 5)

L→JNo support for either Leucine or Isoleucine based on side chain ions (Position: 5)

L→JNo support for either Leucine or Isoleucine based on side chain ions (Position: 1)

#### Positional Score

Copy Data

##### Positional Score (TSV)

###### Preview

```
Loading example...
```

*Click on the button to copy the data to your clipboard.*

10012345678910

Label Value
"0" 0.327
"1" 0.323
"2" 0.327
"3" 0.327
"4" 0.33
"5" 0.327
"6" 0.327
"7" 0.333
"8" 0.333
"9" 0.333
"10" 0.333

#### Meta Information from PEAKS

##### Scan Identifier

F2:5525

##### Original sequence

L

S

G

G

L

D

V

V

A

H

E

##### Posttranslational Modifications

##### Source File

D:\separate\_stitch\_analyses\xle-disambiguation\raw\20210323\_F1\_UM1\_Peng0013\_SA\_F59\_ingel\_3ug\_TL.raw

##### Fraction

2

##### Scan Feature

F2:6190

##### De Novo Score

99

##### ConfidenceScore

99

### m/z

548.7865

##### Mass

1095.5559

##### Charge

2

##### Retention Time

30.05

##### Predicted Retention Time

-

##### Area

2.048E+07

##### Parts Per Million

2.3

##### Fragmentation mode

HCD

##### Originating file

01 D:\separate\_stitch\_analyses\xle-disambiguation\20210325\_F59\_3ug\_DENOVO\_12.csv

#### Meta Information from PEAKS

##### Scan Identifier

F2:5530

##### Original sequence

L

S

G

G

L

D

V

V

A

H

+15.99

E

##### Posttranslational Modifications

Oxidation (HW)

##### Source File

D:\separate\_stitch\_analyses\xle-disambiguation\raw\20210323\_F1\_UM1\_Peng0013\_SA\_F59\_ingel\_3ug\_TL.raw

##### Fraction

2

##### Scan Feature

F2:6565

##### De Novo Score

99

##### ConfidenceScore

99

### m/z

556.783

##### Mass

1111.551

##### Charge

2

##### Retention Time

30.05

##### Predicted Retention Time

-

##### Area

5.125E+05

##### Parts Per Million

0.4

##### Fragmentation mode

HCD

##### Originating file

01 D:\separate\_stitch\_analyses\xle-disambiguation\20210325\_F59\_3ug\_DENOVO\_12.csv

#### Meta Information from PEAKS

##### Scan Identifier

F2:5586

##### Original sequence

L

S

G

G

L

D

V

V

A

H

E

##### Posttranslational Modifications

##### Source File

D:\separate\_stitch\_analyses\xle-disambiguation\raw\20210323\_F1\_UM1\_Peng0013\_SA\_F59\_ingel\_3ug\_TL.raw

##### Fraction

2

##### Scan Feature

-

##### De Novo Score

98

##### ConfidenceScore

98

### m/z

548.7865

##### Mass

1095.5559

##### Charge

2

##### Retention Time

30.4

##### Predicted Retention Time

-

##### Area

0

##### Parts Per Million

2.3

##### Fragmentation mode

ETHCD

##### Originating file

01 D:\separate\_stitch\_analyses\xle-disambiguation\20210325\_F59\_3ug\_DENOVO\_12.csv
