## Supplementary material for "A handle on mass coincidence errors in *de novo* sequencing of antibodies by bottom-up proteomics": Combined_019.html

Details Combined\_019 | Stitch OverviewUndefined

### Read Combined\_019

#### Sequence (length=9)

FTFDDYAMH

#### Spectrum 6605? Spectrum 6605 The raw spectrum of this peptide as annotated by Hecklib. The fragments are coloured according to ion type (see legend). Any peaks with a star '\*' as text can be hovered over to see the full details, first the ion type second the mass shift type. By hovering over the amino acids in the peptide or ions in the legend the corresponding peaks are highlighted. By toggling the 'Unassigned' label you can turn the background (unassigned) peaks on or off in the plot. By updating the slider in the Ion legend you can update the spectrum to only show the top X% of the peaks with labels. The top X% means any peak that is within X% of the highest intensity. By dragging in the spectrum you can zoom in to a specific part of the spectrum and use 'Zoom Out' to get back to the original zoom level. The annotation of the spectrum is based on the given sequence in the peptides file and is done with different software so inconsistencies are likely. The peaks are annotated based on the given sequence, with 20 ppm tolerance.

Copy Data

##### Spectrum 6605 (TSV)

###### Preview

```
Loading example...
```

*Click on the button to copy the data to your clipboard.*

Mz MinMz MaxIntensity Max

WidthHeightPeptide font sizePeptide stroke widthSpectrum font sizeSpectrum stroke widthCompact peptide

Ion legend

wxyz

abcd

OtherUnassignedIon^Charge^_Position_Show for top:%

FTFDDYAMH

01.46e+52.93e+54.39e+55.85e+5

Zoom Out

y^+1^_1_b^+2^_3_a^+1^_2_a^+1^_2_d^+1^_2_d^+1^_2_a^+1^_2_b^+1^_2_b^+1^_2_b^+2^_4_b^+1^_2_y^+1^_2_b^+2^_5_a^+1^_3_a^+1^_3_y^+1^_3_b^+1^_3_y^+2^_6_b^+2^_6_b^+1^_3_y^+2^_7_y^+2^_7_b^+1^_4_b^+2^_8_y^+2^_8_b^+2^_8_y^+2^_8_b^+1^_4_y^+1^_4_\*\*b^+1^_5_b^+1^_5_y^+1^_5_y^+1^_5_y^+1^_6_y^+1^_6_b^+1^_6_b^+1^_6_b^+1^_7_b^+1^_7_y^+1^_7_y^+1^_7_b^+1^_8_y^+1^_8_b^+1^_8_y^+1^_8_

0734146922032937

Fragment Matches Table

Show background peaks

| Position | Ion type | Intensity | mz Theoretical | mz Error (Th) | mz Error (ppm) | Charge | Series Number |
| --- | --- | --- | --- | --- | --- | --- | --- |
| - | - | 5.794E+05 | 120.1 | - | - | 0 | - |
| - | - | 4.735E+04 | 121.1 | - | - | 0 | - |
| - | - | 2007 | 122.1 | - | - | 0 | - |
| - | - | 1384 | 122.1 | - | - | 0 | - |
| - | - | 524.5 | 123 | - | - | 0 | - |
| - | - | 1236 | 124.1 | - | - | 0 | - |
| - | - | 1075 | 127.1 | - | - | 0 | - |
| - | - | 473 | 129.1 | - | - | 0 | - |
| - | - | 2391 | 129.1 | - | - | 0 | - |
| - | - | 554.5 | 130.1 | - | - | 0 | - |
| - | - | 2566 | 130.1 | - | - | 0 | - |
| - | - | 1362 | 131 | - | - | 0 | - |
| - | - | 1445 | 132.1 | - | - | 0 | - |
| - | - | 720.5 | 133.1 | - | - | 0 | - |
| - | - | 605.6 | 133.1 | - | - | 0 | - |
| - | - | 454.1 | 135 | - | - | 0 | - |
| - | - | 5.162E+04 | 136.1 | - | - | 0 | - |
| - | - | 3458 | 137.1 | - | - | 0 | - |
| - | - | 9117 | 138.1 | - | - | 0 | - |
| - | - | 928.7 | 138.1 | - | - | 0 | - |
| - | - | 459.8 | 139.1 | - | - | 0 | - |
| - | - | 389.2 | 139.1 | - | - | 0 | - |
| - | - | 405.2 | 139.1 | - | - | 0 | - |
| - | - | 859.4 | 140.1 | - | - | 0 | - |
| - | - | 615.3 | 141.1 | - | - | 0 | - |
| - | - | 446.7 | 142.1 | - | - | 0 | - |
| - | - | 1977 | 146.1 | - | - | 0 | - |
| - | - | 2046 | 146.1 | - | - | 0 | - |
| - | - | 636.6 | 148.1 | - | - | 0 | - |
| - | - | 1442 | 148.1 | - | - | 0 | - |
| - | - | 479.1 | 152.1 | - | - | 0 | - |
| - | - | 427.3 | 152.2 | - | - | 0 | - |
| - | - | 502.1 | 154 | - | - | 0 | - |
| - | - | 2144 | 155.1 | - | - | 0 | - |
| 9 | y | 1.85E+05 | 156.1 | 0.0003497 | 2.24 | +1 | 1 |
| - | - | 1418 | 157.1 | - | - | 0 | - |
| - | - | 1.053E+04 | 157.1 | - | - | 0 | - |
| - | - | 1.205E+04 | 158.1 | - | - | 0 | - |
| - | - | 1.811E+04 | 159.1 | - | - | 0 | - |
| - | - | 1432 | 159.1 | - | - | 0 | - |
| - | - | 1113 | 160.1 | - | - | 0 | - |
| - | - | 467.7 | 160.1 | - | - | 0 | - |
| - | - | 1546 | 160.1 | - | - | 0 | - |
| - | - | 1693 | 160.1 | - | - | 0 | - |
| - | - | 473.6 | 161.3 | - | - | 0 | - |
| - | - | 650 | 162.1 | - | - | 0 | - |
| - | - | 956.9 | 162.1 | - | - | 0 | - |
| - | - | 590.5 | 165.1 | - | - | 0 | - |
| - | - | 610.6 | 165.1 | - | - | 0 | - |
| - | - | 2889 | 166.1 | - | - | 0 | - |
| - | - | 3052 | 166.1 | - | - | 0 | - |
| - | - | 1559 | 166.1 | - | - | 0 | - |
| - | - | 573.9 | 167.1 | - | - | 0 | - |
| - | - | 1735 | 167.1 | - | - | 0 | - |
| - | - | 579.6 | 168.1 | - | - | 0 | - |
| - | - | 1359 | 169.1 | - | - | 0 | - |
| - | - | 1054 | 171.1 | - | - | 0 | - |
| - | - | 604.7 | 173.1 | - | - | 0 | - |
| - | - | 2384 | 173.1 | - | - | 0 | - |
| - | - | 3660 | 173.5 | - | - | 0 | - |
| - | - | 753 | 174.1 | - | - | 0 | - |
| - | - | 3639 | 175.1 | - | - | 0 | - |
| - | - | 1224 | 175.1 | - | - | 0 | - |
| - | - | 4295 | 176.1 | - | - | 0 | - |
| - | - | 2.792E+05 | 176.1 | - | - | 0 | - |
| - | - | 3568 | 177.1 | - | - | 0 | - |
| - | - | 3.136E+04 | 177.1 | - | - | 0 | - |
| - | - | 1116 | 178.1 | - | - | 0 | - |
| - | - | 481.3 | 180.1 | - | - | 0 | - |
| - | - | 2095 | 185.1 | - | - | 0 | - |
| - | - | 605.6 | 185.1 | - | - | 0 | - |
| - | - | 3.72E+04 | 186.1 | - | - | 0 | - |
| - | - | 1006 | 187.1 | - | - | 0 | - |
| - | - | 4701 | 187.1 | - | - | 0 | - |
| - | - | 628.4 | 187.1 | - | - | 0 | - |
| - | - | 587 | 187.1 | - | - | 0 | - |
| - | - | 791.1 | 188.1 | - | - | 0 | - |
| - | - | 654.2 | 189.1 | - | - | 0 | - |
| - | - | 848 | 189.1 | - | - | 0 | - |
| 3 | b | 964.6 | 190.1 | 0.0002623 | 1.38 | +2 | 3 |
| - | - | 860.6 | 191.1 | - | - | 0 | - |
| - | - | 1116 | 191.1 | - | - | 0 | - |
| - | - | 5070 | 191.1 | - | - | 0 | - |
| - | - | 643.3 | 192.1 | - | - | 0 | - |
| - | - | 928.9 | 192.1 | - | - | 0 | - |
| - | - | 771.4 | 193.1 | - | - | 0 | - |
| - | - | 6906 | 193.1 | - | - | 0 | - |
| - | - | 471.2 | 194.1 | - | - | 0 | - |
| - | - | 724.8 | 194.1 | - | - | 0 | - |
| - | - | 611.9 | 194.1 | - | - | 0 | - |
| - | - | 1161 | 195.1 | - | - | 0 | - |
| - | - | 478.2 | 199.1 | - | - | 0 | - |
| - | - | 1180 | 199.2 | - | - | 0 | - |
| - | - | 4519 | 202.1 | - | - | 0 | - |
| - | - | 3004 | 203.1 | - | - | 0 | - |
| 2 | a | 1.967E+04 | 203.1 | 0.0002898 | 1.427 | +1 | 2 |
| - | - | 3224 | 204.1 | - | - | 0 | - |
| 2 | a | 1033 | 204.1 | 0.0005576 | 2.732 | +1 | 2 |
| - | - | 2091 | 204.1 | - | - | 0 | - |
| - | - | 2048 | 205.1 | - | - | 0 | - |
| - | - | 933.1 | 205.1 | - | - | 0 | - |
| 2 | d | 1073 | 205.1 | 9.688E-05 | 0.4723 | +1 | 2 |
| 2 | d | 4313 | 207.1 | 0.0003703 | 1.788 | +1 | 2 |
| - | - | 5258 | 211.1 | - | - | 0 | - |
| - | - | 571 | 211.1 | - | - | 0 | - |
| - | - | 889.7 | 212.1 | - | - | 0 | - |
| - | - | 2082 | 212.1 | - | - | 0 | - |
| - | - | 2383 | 213.1 | - | - | 0 | - |
| - | - | 1436 | 214.1 | - | - | 0 | - |
| - | - | 1568 | 215.1 | - | - | 0 | - |
| - | - | 601.8 | 215.1 | - | - | 0 | - |
| - | - | 696.9 | 217.1 | - | - | 0 | - |
| - | - | 1.122E+04 | 217.1 | - | - | 0 | - |
| - | - | 1919 | 218.1 | - | - | 0 | - |
| - | - | 4529 | 219.1 | - | - | 0 | - |
| - | - | 1473 | 219.1 | - | - | 0 | - |
| - | - | 1578 | 220.1 | - | - | 0 | - |
| - | - | 2.89E+04 | 221.1 | - | - | 0 | - |
| 2 | a | 3.329E+05 | 221.1 | 0.0004673 | 2.113 | +1 | 2 |
| - | - | 1589 | 222.1 | - | - | 0 | - |
| - | - | 3259 | 222.1 | - | - | 0 | - |
| - | - | 4.065E+04 | 222.1 | - | - | 0 | - |
| - | - | 551.5 | 223.1 | - | - | 0 | - |
| - | - | 2372 | 223.1 | - | - | 0 | - |
| - | - | 1669 | 224.1 | - | - | 0 | - |
| - | - | 675.3 | 228.1 | - | - | 0 | - |
| - | - | 8122 | 231.1 | - | - | 0 | - |
| 2 | b | 7.265E+04 | 231.1 | 0.0004008 | 1.734 | +1 | 2 |
| 2 | b | 692.4 | 232.1 | 0.0005619 | 2.421 | +1 | 2 |
| - | - | 9977 | 232.1 | - | - | 0 | - |
| - | - | 819 | 233.1 | - | - | 0 | - |
| - | - | 926.8 | 233.1 | - | - | 0 | - |
| - | - | 2.415E+04 | 235.1 | - | - | 0 | - |
| - | - | 3526 | 236.1 | - | - | 0 | - |
| - | - | 1672 | 237.1 | - | - | 0 | - |
| - | - | 739.5 | 238.1 | - | - | 0 | - |
| - | - | 7.253E+04 | 239.1 | - | - | 0 | - |
| - | - | 771.3 | 240.1 | - | - | 0 | - |
| - | - | 7613 | 240.1 | - | - | 0 | - |
| - | - | 1471 | 245.1 | - | - | 0 | - |
| - | - | 740.7 | 245.1 | - | - | 0 | - |
| 4 | b | 545.6 | 247.1 | 0.0006326 | 2.56 | +2 | 4 |
| - | - | 1284 | 247.1 | - | - | 0 | - |
| - | - | 889 | 248.1 | - | - | 0 | - |
| 2 | b | 1.667E+05 | 249.1 | 0.000441 | 1.77 | +1 | 2 |
| - | - | 2.272E+04 | 250.1 | - | - | 0 | - |
| - | - | 2.195E+04 | 251.1 | - | - | 0 | - |
| - | - | 1518 | 251.1 | - | - | 0 | - |
| - | - | 2091 | 252.1 | - | - | 0 | - |
| - | - | 2665 | 257.1 | - | - | 0 | - |
| - | - | 4686 | 259.1 | - | - | 0 | - |
| - | - | 1724 | 261.1 | - | - | 0 | - |
| - | - | 1133 | 261.1 | - | - | 0 | - |
| - | - | 3.305E+04 | 263.1 | - | - | 0 | - |
| - | - | 3888 | 264.1 | - | - | 0 | - |
| - | - | 1041 | 267.1 | - | - | 0 | - |
| - | - | 712.6 | 267.1 | - | - | 0 | - |
| - | - | 1686 | 268.1 | - | - | 0 | - |
| - | - | 4121 | 268.1 | - | - | 0 | - |
| - | - | 697.5 | 269.1 | - | - | 0 | - |
| - | - | 568.5 | 271.1 | - | - | 0 | - |
| - | - | 806.6 | 273.1 | - | - | 0 | - |
| - | - | 1270 | 274.1 | - | - | 0 | - |
| - | - | 3062 | 275.1 | - | - | 0 | - |
| - | - | 1.643E+04 | 279.1 | - | - | 0 | - |
| - | - | 2634 | 280.1 | - | - | 0 | - |
| - | - | 726.4 | 281.1 | - | - | 0 | - |
| - | - | 1474 | 283.1 | - | - | 0 | - |
| - | - | 674.3 | 284.1 | - | - | 0 | - |
| - | - | 2.319E+04 | 285.1 | - | - | 0 | - |
| - | - | 2001 | 286.1 | - | - | 0 | - |
| - | - | 1043 | 287.1 | - | - | 0 | - |
| - | - | 806.7 | 289.1 | - | - | 0 | - |
| - | - | 1874 | 289.1 | - | - | 0 | - |
| - | - | 1823 | 289.1 | - | - | 0 | - |
| - | - | 2102 | 292.1 | - | - | 0 | - |
| - | - | 1662 | 293.1 | - | - | 0 | - |
| - | - | 1267 | 295.1 | - | - | 0 | - |
| - | - | 1157 | 299.1 | - | - | 0 | - |
| - | - | 814.2 | 300.1 | - | - | 0 | - |
| - | - | 1125 | 301.1 | - | - | 0 | - |
| - | - | 619 | 302.1 | - | - | 0 | - |
| - | - | 701.8 | 302.2 | - | - | 0 | - |
| 8 | y | 5.897E+04 | 303.1 | 0.005464 | 18.03 | +1 | 2 |
| - | - | 6714 | 304.1 | - | - | 0 | - |
| 5 | b | 1614 | 305.1 | 0.002785 | 9.129 | +2 | 5 |
| - | - | 712.4 | 305.2 | - | - | 0 | - |
| - | - | 511.1 | 306.1 | - | - | 0 | - |
| - | - | 1274 | 306.1 | - | - | 0 | - |
| - | - | 1675 | 306.2 | - | - | 0 | - |
| - | - | 592.7 | 307.1 | - | - | 0 | - |
| - | - | 1315 | 309.2 | - | - | 0 | - |
| - | - | 2.796E+04 | 310.2 | - | - | 0 | - |
| - | - | 5120 | 311.2 | - | - | 0 | - |
| - | - | 929.1 | 314.1 | - | - | 0 | - |
| - | - | 828.7 | 315.1 | - | - | 0 | - |
| - | - | 875.3 | 316.1 | - | - | 0 | - |
| - | - | 1131 | 317.1 | - | - | 0 | - |
| - | - | 5115 | 318.1 | - | - | 0 | - |
| - | - | 750.3 | 319.1 | - | - | 0 | - |
| - | - | 2028 | 320.1 | - | - | 0 | - |
| - | - | 830.2 | 323.2 | - | - | 0 | - |
| - | - | 1606 | 330.1 | - | - | 0 | - |
| - | - | 2202 | 332.1 | - | - | 0 | - |
| - | - | 873.1 | 333.1 | - | - | 0 | - |
| - | - | 5365 | 333.2 | - | - | 0 | - |
| - | - | 1113 | 334.1 | - | - | 0 | - |
| - | - | 2632 | 334.2 | - | - | 0 | - |
| - | - | 1021 | 336.2 | - | - | 0 | - |
| - | - | 699.9 | 337.2 | - | - | 0 | - |
| - | - | 691.2 | 338.1 | - | - | 0 | - |
| - | - | 878.9 | 339.1 | - | - | 0 | - |
| - | - | 2073 | 339.2 | - | - | 0 | - |
| - | - | 3857 | 346.1 | - | - | 0 | - |
| - | - | 616.9 | 347.1 | - | - | 0 | - |
| - | - | 4560 | 348.1 | - | - | 0 | - |
| - | - | 1.48E+04 | 350.1 | - | - | 0 | - |
| 3 | a | 2197 | 350.2 | 0.000159 | 0.4539 | +1 | 3 |
| - | - | 2429 | 351.1 | - | - | 0 | - |
| 3 | a | 1879 | 351.2 | 9.112E-05 | 0.2595 | +1 | 3 |
| - | - | 2030 | 356.1 | - | - | 0 | - |
| - | - | 1133 | 358.1 | - | - | 0 | - |
| - | - | 2648 | 360.1 | - | - | 0 | - |
| - | - | 677.5 | 361.1 | - | - | 0 | - |
| - | - | 9604 | 364.2 | - | - | 0 | - |
| - | - | 2420 | 365.2 | - | - | 0 | - |
| - | - | 3596 | 366.1 | - | - | 0 | - |
| - | - | 567.9 | 367.1 | - | - | 0 | - |
| - | - | 681.2 | 367.2 | - | - | 0 | - |
| 7 | y | 4.667E+04 | 374.1 | 0.005185 | 13.86 | +1 | 3 |
| - | - | 8901 | 375.2 | - | - | 0 | - |
| - | - | 4122 | 376.1 | - | - | 0 | - |
| - | - | 1162 | 376.1 | - | - | 0 | - |
| - | - | 1134 | 377.1 | - | - | 0 | - |
| - | - | 4.29E+04 | 378.1 | - | - | 0 | - |
| 3 | b | 2.683E+04 | 378.2 | 0.0003615 | 0.9559 | +1 | 3 |
| - | - | 7138 | 379.1 | - | - | 0 | - |
| - | - | 4987 | 379.2 | - | - | 0 | - |
| - | - | 1464 | 380.1 | - | - | 0 | - |
| - | - | 1158 | 380.2 | - | - | 0 | - |
| - | - | 869.4 | 381.1 | - | - | 0 | - |
| - | - | 7702 | 382.1 | - | - | 0 | - |
| - | - | 1081 | 383.1 | - | - | 0 | - |
| 4 | y | 1609 | 384.1 | 0.001413 | 3.677 | +2 | 6 |
| - | - | 2.289E+04 | 394.1 | - | - | 0 | - |
| - | - | 4355 | 395.1 | - | - | 0 | - |
| 6 | b | 903.2 | 395.2 | 0.003901 | 9.872 | +2 | 6 |
| - | - | 744.1 | 396.1 | - | - | 0 | - |
| 3 | b | 1236 | 396.2 | 0.0009663 | 2.439 | +1 | 3 |
| - | - | 1081 | 404.1 | - | - | 0 | - |
| - | - | 4070 | 404.1 | - | - | 0 | - |
| - | - | 1040 | 405.1 | - | - | 0 | - |
| - | - | 2457 | 410.1 | - | - | 0 | - |
| - | - | 859.1 | 415.2 | - | - | 0 | - |
| - | - | 2228 | 416.1 | - | - | 0 | - |
| - | - | 849 | 417.2 | - | - | 0 | - |
| - | - | 1425 | 419.2 | - | - | 0 | - |
| - | - | 817.4 | 420.2 | - | - | 0 | - |
| - | - | 954.1 | 423.2 | - | - | 0 | - |
| - | - | 2786 | 425.7 | - | - | 0 | - |
| - | - | 1461 | 426.2 | - | - | 0 | - |
| - | - | 1068 | 427.2 | - | - | 0 | - |
| - | - | 707.8 | 429.2 | - | - | 0 | - |
| - | - | 669.4 | 430.1 | - | - | 0 | - |
| - | - | 733.2 | 432.2 | - | - | 0 | - |
| - | - | 2279 | 433.2 | - | - | 0 | - |
| - | - | 753.1 | 439.7 | - | - | 0 | - |
| - | - | 954.9 | 443.2 | - | - | 0 | - |
| - | - | 733.5 | 445.1 | - | - | 0 | - |
| - | - | 3158 | 447.2 | - | - | 0 | - |
| - | - | 961.5 | 447.2 | - | - | 0 | - |
| - | - | 996.3 | 448.2 | - | - | 0 | - |
| 3 | y | 3053 | 448.7 | 0.00267 | 5.951 | +2 | 7 |
| - | - | 881 | 449.1 | - | - | 0 | - |
| - | - | 3649 | 449.2 | - | - | 0 | - |
| - | - | 1078 | 450.2 | - | - | 0 | - |
| - | - | 1171 | 455.2 | - | - | 0 | - |
| 3 | y | 3.223E+04 | 457.7 | 0.003125 | 6.828 | +2 | 7 |
| - | - | 1.955E+04 | 458.2 | - | - | 0 | - |
| - | - | 4969 | 458.7 | - | - | 0 | - |
| - | - | 578.7 | 459.2 | - | - | 0 | - |
| - | - | 5225 | 461.2 | - | - | 0 | - |
| - | - | 1122 | 462.2 | - | - | 0 | - |
| - | - | 2.137E+04 | 465.2 | - | - | 0 | - |
| - | - | 4344 | 466.2 | - | - | 0 | - |
| - | - | 1322 | 467.2 | - | - | 0 | - |
| - | - | 1.749E+04 | 473.2 | - | - | 0 | - |
| - | - | 4215 | 474.2 | - | - | 0 | - |
| - | - | 755.9 | 475.2 | - | - | 0 | - |
| - | - | 750.7 | 476.7 | - | - | 0 | - |
| - | - | 1133 | 477.2 | - | - | 0 | - |
| - | - | 5349 | 479.2 | - | - | 0 | - |
| - | - | 842.1 | 480.1 | - | - | 0 | - |
| - | - | 1301 | 480.2 | - | - | 0 | - |
| - | - | 1410 | 481.2 | - | - | 0 | - |
| - | - | 756.7 | 485.2 | - | - | 0 | - |
| - | - | 674.4 | 487.2 | - | - | 0 | - |
| - | - | 3640 | 489.2 | - | - | 0 | - |
| - | - | 738 | 490.2 | - | - | 0 | - |
| - | - | 1759 | 491.2 | - | - | 0 | - |
| - | - | 1084 | 491.7 | - | - | 0 | - |
| - | - | 621.4 | 493.2 | - | - | 0 | - |
| 4 | b | 2505 | 493.2 | 0.0008233 | 1.669 | +1 | 4 |
| 8 | b | 3575 | 495.2 | 0.0002969 | 0.5996 | +2 | 8 |
| - | - | 923.9 | 496.2 | - | - | 0 | - |
| - | - | 5355 | 497.2 | - | - | 0 | - |
| - | - | 1481 | 498.2 | - | - | 0 | - |
| 2 | y | 2213 | 499.2 | 0.002451 | 4.91 | +2 | 8 |
| - | - | 1832 | 499.7 | - | - | 0 | - |
| 8 | b | 985.3 | 504.2 | 0.009838 | 19.51 | +2 | 8 |
| - | - | 1124 | 505.2 | - | - | 0 | - |
| 2 | y | 1.522E+04 | 508.2 | 0.002845 | 5.598 | +2 | 8 |
| - | - | 1.016E+04 | 508.7 | - | - | 0 | - |
| - | - | 3449 | 509.2 | - | - | 0 | - |
| - | - | 646.2 | 509.7 | - | - | 0 | - |
| 4 | b | 9904 | 511.2 | 0.000543 | 1.062 | +1 | 4 |
| - | - | 2634 | 512.2 | - | - | 0 | - |
| - | - | 8555 | 513.2 | - | - | 0 | - |
| - | - | 695.5 | 513.2 | - | - | 0 | - |
| - | - | 691.9 | 513.7 | - | - | 0 | - |
| - | - | 2408 | 514.2 | - | - | 0 | - |
| - | - | 1363 | 519.2 | - | - | 0 | - |
| - | - | 1588 | 520.2 | - | - | 0 | - |
| - | - | 618.4 | 521.2 | - | - | 0 | - |
| - | - | 1028 | 523.2 | - | - | 0 | - |
| - | - | 2101 | 525.2 | - | - | 0 | - |
| - | - | 1474 | 530.2 | - | - | 0 | - |
| - | - | 1718 | 532.2 | - | - | 0 | - |
| - | - | 953.3 | 533.2 | - | - | 0 | - |
| 6 | y | 4.366E+04 | 537.2 | 0.005577 | 10.38 | +1 | 4 |
| - | - | 1.237E+04 | 538.2 | - | - | 0 | - |
| - | - | 4863 | 539.2 | - | - | 0 | - |
| - | - | 686 | 540.2 | - | - | 0 | - |
| - | - | 5074 | 540.7 | - | - | 0 | - |
| - | - | 3671 | 541.2 | - | - | 0 | - |
| - | - | 1960 | 541.2 | - | - | 0 | - |
| - | - | 1596 | 541.7 | - | - | 0 | - |
| - | - | 2179 | 542.2 | - | - | 0 | - |
| - | - | 1036 | 542.2 | - | - | 0 | - |
| - | - | 1234 | 544.3 | - | - | 0 | - |
| - | - | 826.5 | 547.2 | - | - | 0 | - |
| - | - | 4993 | 548.2 | - | - | 0 | - |
| - | - | 1522 | 549.2 | - | - | 0 | - |
| - | - | 3366 | 549.7 | - | - | 0 | - |
| - | - | 2476 | 550.2 | - | - | 0 | - |
| - | - | 897.3 | 550.7 | - | - | 0 | - |
| - | - | 822.2 | 551.2 | - | - | 0 | - |
| - | - | 657.9 | 551.7 | - | - | 0 | - |
| - | - | 610.5 | 557.2 | - | - | 0 | - |
| - | - | 1117 | 558.7 | - | - | 0 | - |
| - | - | 856.2 | 559.7 | - | - | 0 | - |
| - | - | 609 | 561.2 | - | - | 0 | - |
| - | - | 1443 | 562.2 | - | - | 0 | - |
| - | - | 1720 | 563.7 | - | - | 0 | - |
| - | - | 2693 | 564.2 | - | - | 0 | - |
| - | - | 1115 | 564.7 | - | - | 0 | - |
| - | - | 964.8 | 566.2 | - | - | 0 | - |
| - | - | 2711 | 567.2 | - | - | 0 | - |
| - | - | 722 | 568.2 | - | - | 0 | - |
| - | - | 1585 | 570.2 | - | - | 0 | - |
| 0 | Precursor | 2.471E+04 | 572.7 | 0.002912 | 5.085 | +2 | -1 |
| - | - | 1.891E+04 | 573.2 | - | - | 0 | - |
| - | - | 8185 | 573.7 | - | - | 0 | - |
| - | - | 2165 | 574.2 | - | - | 0 | - |
| - | - | 1902 | 579.2 | - | - | 0 | - |
| - | - | 1753 | 580.2 | - | - | 0 | - |
| 0 | Precursor | 3.394E+04 | 581.7 | 0.002818 | 4.844 | +2 | -1 |
| - | - | 2.269E+04 | 582.2 | - | - | 0 | - |
| - | - | 8403 | 582.7 | - | - | 0 | - |
| - | - | 1890 | 583.2 | - | - | 0 | - |
| - | - | 2891 | 584.2 | - | - | 0 | - |
| - | - | 929.7 | 585.2 | - | - | 0 | - |
| - | - | 2.085E+04 | 588.2 | - | - | 0 | - |
| - | - | 5484 | 589.2 | - | - | 0 | - |
| - | - | 935.7 | 590.2 | - | - | 0 | - |
| - | - | 1135 | 594.2 | - | - | 0 | - |
| - | - | 1163 | 596.2 | - | - | 0 | - |
| - | - | 758.8 | 598.3 | - | - | 0 | - |
| - | - | 1722 | 606.2 | - | - | 0 | - |
| - | - | 730.6 | 607.2 | - | - | 0 | - |
| 5 | b | 3712 | 608.2 | 0.001468 | 2.414 | +1 | 5 |
| - | - | 1396 | 609.2 | - | - | 0 | - |
| - | - | 3971 | 612.2 | - | - | 0 | - |
| - | - | 7364 | 612.2 | - | - | 0 | - |
| - | - | 1087 | 613.2 | - | - | 0 | - |
| - | - | 2373 | 613.2 | - | - | 0 | - |
| - | - | 2142 | 614.2 | - | - | 0 | - |
| - | - | 1008 | 615.2 | - | - | 0 | - |
| - | - | 1080 | 617.2 | - | - | 0 | - |
| - | - | 4776 | 624.2 | - | - | 0 | - |
| - | - | 1480 | 625.2 | - | - | 0 | - |
| 5 | b | 6163 | 626.2 | 0.0004249 | 0.6785 | +1 | 5 |
| - | - | 2115 | 627.2 | - | - | 0 | - |
| 5 | y | 2819 | 634.2 | 0.005251 | 8.279 | +1 | 5 |
| - | - | 1660 | 635.2 | - | - | 0 | - |
| - | - | 1842 | 642.2 | - | - | 0 | - |
| - | - | 783.5 | 643.2 | - | - | 0 | - |
| - | - | 1037 | 644.2 | - | - | 0 | - |
| - | - | 800 | 649.3 | - | - | 0 | - |
| - | - | 1722 | 650.3 | - | - | 0 | - |
| 5 | y | 6.98E+04 | 652.2 | 0.005428 | 8.323 | +1 | 5 |
| - | - | 2.227E+04 | 653.2 | - | - | 0 | - |
| - | - | 6354 | 654.2 | - | - | 0 | - |
| - | - | 1211 | 655.2 | - | - | 0 | - |
| - | - | 1019 | 657.3 | - | - | 0 | - |
| - | - | 845.4 | 662.2 | - | - | 0 | - |
| - | - | 2666 | 667.3 | - | - | 0 | - |
| - | - | 887.8 | 668.3 | - | - | 0 | - |
| - | - | 921.5 | 676.2 | - | - | 0 | - |
| - | - | 3463 | 677.3 | - | - | 0 | - |
| - | - | 1717 | 678.3 | - | - | 0 | - |
| - | - | 3459 | 685.3 | - | - | 0 | - |
| - | - | 1877 | 686.3 | - | - | 0 | - |
| - | - | 1036 | 688.2 | - | - | 0 | - |
| - | - | 1.214E+04 | 695.3 | - | - | 0 | - |
| - | - | 3747 | 696.3 | - | - | 0 | - |
| - | - | 1237 | 697.3 | - | - | 0 | - |
| - | - | 3.951E+04 | 703.3 | - | - | 0 | - |
| - | - | 1.552E+04 | 704.3 | - | - | 0 | - |
| - | - | 1143 | 704.3 | - | - | 0 | - |
| - | - | 3665 | 705.3 | - | - | 0 | - |
| - | - | 3406 | 713.3 | - | - | 0 | - |
| - | - | 832.6 | 717.3 | - | - | 0 | - |
| - | - | 1257 | 721.3 | - | - | 0 | - |
| - | - | 992.1 | 723.3 | - | - | 0 | - |
| - | - | 2000 | 731.3 | - | - | 0 | - |
| - | - | 1071 | 732.2 | - | - | 0 | - |
| - | - | 2083 | 741.3 | - | - | 0 | - |
| - | - | 1160 | 742.3 | - | - | 0 | - |
| - | - | 1772 | 743.3 | - | - | 0 | - |
| - | - | 718.2 | 744.3 | - | - | 0 | - |
| 4 | y | 7247 | 749.3 | 0.005529 | 7.38 | +1 | 6 |
| - | - | 3816 | 750.3 | - | - | 0 | - |
| - | - | 1135 | 751.3 | - | - | 0 | - |
| - | - | 1038 | 752.2 | - | - | 0 | - |
| - | - | 9190 | 759.3 | - | - | 0 | - |
| - | - | 4121 | 760.3 | - | - | 0 | - |
| 4 | y | 1.218E+05 | 767.3 | 0.00528 | 6.881 | +1 | 6 |
| - | - | 5.107E+04 | 768.3 | - | - | 0 | - |
| - | - | 1.45E+04 | 769.3 | - | - | 0 | - |
| - | - | 1641 | 770.3 | - | - | 0 | - |
| 6 | b | 2121 | 771.3 | 0.001433 | 1.858 | +1 | 6 |
| - | - | 1485 | 772.3 | - | - | 0 | - |
| - | - | 2753 | 777.3 | - | - | 0 | - |
| - | - | 978.9 | 778.3 | - | - | 0 | - |
| - | - | 1294 | 779.3 | - | - | 0 | - |
| 6 | b | 1484 | 789.3 | 0.0009529 | 1.207 | +1 | 6 |
| - | - | 1477 | 796.3 | - | - | 0 | - |
| - | - | 1966 | 804.3 | - | - | 0 | - |
| - | - | 768.5 | 805.3 | - | - | 0 | - |
| - | - | 1572 | 806.3 | - | - | 0 | - |
| - | - | 682.3 | 817.2 | - | - | 0 | - |
| - | - | 3315 | 823.3 | - | - | 0 | - |
| - | - | 2574 | 824.3 | - | - | 0 | - |
| - | - | 5786 | 832.3 | - | - | 0 | - |
| - | - | 2402 | 833.3 | - | - | 0 | - |
| - | - | 1035 | 834.3 | - | - | 0 | - |
| - | - | 1055 | 835.3 | - | - | 0 | - |
| 7 | b | 2066 | 842.3 | 0.01639 | 19.46 | +1 | 7 |
| 7 | b | 1519 | 843.3 | 0.004168 | 4.943 | +1 | 7 |
| - | - | 7.063E+04 | 850.3 | - | - | 0 | - |
| - | - | 3.236E+04 | 851.3 | - | - | 0 | - |
| - | - | 8382 | 852.3 | - | - | 0 | - |
| - | - | 1057 | 853.3 | - | - | 0 | - |
| - | - | 2204 | 860.3 | - | - | 0 | - |
| - | - | 1133 | 861.3 | - | - | 0 | - |
| - | - | 2906 | 868.3 | - | - | 0 | - |
| - | - | 1351 | 869.3 | - | - | 0 | - |
| 3 | y | 1.065E+04 | 896.3 | 0.005292 | 5.904 | +1 | 7 |
| - | - | 4638 | 897.3 | - | - | 0 | - |
| - | - | 1678 | 898.3 | - | - | 0 | - |
| - | - | 1201 | 899.3 | - | - | 0 | - |
| 3 | y | 2.307E+05 | 914.3 | 0.005347 | 5.848 | +1 | 7 |
| - | - | 1.239E+05 | 915.3 | - | - | 0 | - |
| - | - | 1227 | 915.4 | - | - | 0 | - |
| - | - | 4.41E+04 | 916.3 | - | - | 0 | - |
| - | - | 6854 | 917.3 | - | - | 0 | - |
| - | - | 919.8 | 924.3 | - | - | 0 | - |
| - | - | 2419 | 933.4 | - | - | 0 | - |
| - | - | 1165 | 934.4 | - | - | 0 | - |
| - | - | 718.8 | 935.4 | - | - | 0 | - |
| - | - | 864.8 | 943.4 | - | - | 0 | - |
| - | - | 1.582E+04 | 951.4 | - | - | 0 | - |
| - | - | 8065 | 952.4 | - | - | 0 | - |
| - | - | 2479 | 953.4 | - | - | 0 | - |
| - | - | 3264 | 961.4 | - | - | 0 | - |
| - | - | 2595 | 962.4 | - | - | 0 | - |
| - | - | 827.3 | 971.4 | - | - | 0 | - |
| - | - | 753.4 | 972.4 | - | - | 0 | - |
| - | - | 1189 | 979.4 | - | - | 0 | - |
| - | - | 764.3 | 980.4 | - | - | 0 | - |
| 8 | b | 1332 | 989.4 | 0.002743 | 2.773 | +1 | 8 |
| - | - | 1917 | 990.4 | - | - | 0 | - |
| 2 | y | 6829 | 997.4 | 0.005343 | 5.357 | +1 | 8 |
| - | - | 3788 | 998.4 | - | - | 0 | - |
| - | - | 1273 | 999.4 | - | - | 0 | - |
| 8 | b | 1162 | 1007 | 0.01106 | 10.98 | +1 | 8 |
| - | - | 1443 | 1008 | - | - | 0 | - |
| 2 | y | 5.152E+04 | 1015 | 0.005215 | 5.136 | +1 | 8 |
| - | - | 3.008E+04 | 1016 | - | - | 0 | - |
| - | - | 1.279E+04 | 1017 | - | - | 0 | - |
| - | - | 1620 | 1018 | - | - | 0 | - |
| - | - | 8243 | 1025 | - | - | 0 | - |
| - | - | 4259 | 1026 | - | - | 0 | - |
| - | - | 1457 | 1027 | - | - | 0 | - |
| - | - | 664.6 | 1666 | - | - | 0 | - |
| - | - | 671.6 | 1975 | - | - | 0 | - |
| - | - | 753.9 | 2908 | - | - | 0 | - |

m/z Charge Intensity FragmentType MassShift Position
120.08118438720703 0 579371.3
121.08442687988281 0 47347.273
122.07156372070312 0 2007.4412
122.08766174316406 0 1384.2568
123.00196838378906 0 524.5416
124.0871810913086 0 1235.549
127.0869369506836 0 1074.645
129.06626892089844 0 473.001
129.1025390625 0 2391.275
130.05006408691406 0 554.5346
130.06546020507812 0 2566.102
131.0493621826172 0 1361.6875
132.081298828125 0 1444.6152
133.0608673095703 0 720.47766
133.08580017089844 0 605.5665
135.0445098876953 0 454.14447
136.07601928710938 0 51621.64
137.07943725585938 0 3458.2373
138.05526733398438 0 9116.643
138.06622314453125 0 928.6596
139.05117797851562 0 459.78705
139.05813598632812 0 389.16867
139.09767150878906 0 405.21356
140.0824432373047 0 859.44806
141.06602478027344 0 615.34875
142.06544494628906 0 446.68893
146.06033325195312 0 1976.5182
146.09686279296875 0 2045.86
148.0760498046875 0 636.5967
148.0872802734375 0 1442.178
152.10714721679688 0 479.09116
152.15554809570312 0 427.28964
154.0498504638672 0 502.05685
155.0819549560547 0 2144.3315
156.0771026611328 0 184964.4 y 8
157.0738983154297 0 1418.229
157.08042907714844 0 10533.116
158.09669494628906 0 12051.154
159.0919952392578 0 18106.834
159.09957885742188 0 1431.5051
160.0760498046875 0 1112.6526
160.08837890625 0 467.74954
160.09542846679688 0 1545.7847
160.11233520507812 0 1692.9432
161.30999755859375 0 473.6382
162.05503845214844 0 650.0363
162.09201049804688 0 956.8729
165.07748413085938 0 590.5334
165.1027069091797 0 610.5665
166.05348205566406 0 2888.562
166.0615692138672 0 3051.8372
166.08645629882812 0 1558.6713
167.05596923828125 0 573.8897
167.09310913085938 0 1734.8666
168.0764617919922 0 579.5722
169.10887145996094 0 1358.6298
171.0768585205078 0 1053.5795
173.07083129882812 0 604.73096
173.09249877929688 0 2384.2373
173.45062255859375 0 3660.0562
174.06674194335938 0 753.03845
175.0869140625 0 3639.1372
175.12315368652344 0 1223.6322
176.0823974609375 0 4294.7544
176.10740661621094 0 279193.8
177.10252380371094 0 3568.129
177.11083984375 0 31355.69
178.11402893066406 0 1116.1445
180.0768585205078 0 481.26108
185.0559539794922 0 2095.0107
185.1076202392578 0 605.6438
186.09169006347656 0 37197.117
187.08677673339844 0 1006.2232
187.09512329101562 0 4701.4062
187.10853576660156 0 628.4335
187.14425659179688 0 587.02246
188.07095336914062 0 791.12476
189.08685302734375 0 654.20953
189.10220336914062 0 847.9524
190.08651733398438 0 964.6035 b Ammonia loss 2
191.08204650878906 0 860.5713
191.0933380126953 0 1116.048
191.11822509765625 0 5069.7593
192.10147094726562 0 643.31244
192.12139892578125 0 928.9176
193.07223510742188 0 771.43854
193.10867309570312 0 6905.6816
194.0801544189453 0 471.17184
194.09190368652344 0 724.7756
194.11172485351562 0 611.86835
195.1243896484375 0 1161.445
199.10964965820312 0 478.2142
199.18060302734375 0 1180.2385
202.053466796875 0 4518.9287
203.066650390625 0 3004.368
203.11817932128906 0 19667.877 a Water loss 1
204.0770721435547 0 3223.9983
204.1024627685547 0 1032.6416 a Ammonia loss 1
204.12135314941406 0 2091.143
205.09744262695312 0 2047.7534
205.1078338623047 0 933.14435
205.13363647460938 0 1072.9803 d 1
207.11317443847656 0 4313.0234 d 1
211.0828094482422 0 5257.623
211.11920166015625 0 571.0175
212.0912628173828 0 889.7348
212.1185302734375 0 2082.2954
213.0986785888672 0 2383.1577
214.0865478515625 0 1436.0851
215.08189392089844 0 1568.2721
215.1389617919922 0 601.7948
217.08421325683594 0 696.8602
217.0974578857422 0 11221.191
218.1012420654297 0 1918.7607
219.08013916015625 0 4529.258
219.11285400390625 0 1473.2006
220.07212829589844 0 1578.3923
221.10365295410156 0 28895.307
221.12892150878906 0 332926.75 a 1
222.0876922607422 0 1588.5927
222.10702514648438 0 3259.0303
222.13214111328125 0 40652.246
223.09539794921875 0 551.504
223.13473510742188 0 2372.243
224.12693786621094 0 1669.1392
228.11325073242188 0 675.325
231.06146240234375 0 8121.629
231.1132049560547 0 72653.37 b Water loss 1
232.09738159179688 0 692.3624 b Ammonia loss 1
232.1165008544922 0 9976.549
233.09226989746094 0 818.99146
233.1187286376953 0 926.8019
235.10813903808594 0 24151.31
236.1112060546875 0 3525.62
237.09849548339844 0 1671.7012
238.12208557128906 0 739.5215
239.11431884765625 0 72526.31
240.08148193359375 0 771.3205
240.11749267578125 0 7613.183
245.09280395507812 0 1471.242
245.129150390625 0 740.6599
247.10708618164062 0 545.6324 b Water loss 3
247.14398193359375 0 1284.4299
248.14990234375 0 888.9821
249.12380981445312 0 166716.78 b 1
250.1271209716797 0 22722.684
251.10299682617188 0 21950.13
251.129150390625 0 1517.7322
252.1063232421875 0 2091.2222
257.1065673828125 0 2665.075
259.10797119140625 0 4685.752
261.0872802734375 0 1723.5957
261.1233825683594 0 1132.8483
263.1029052734375 0 33050.59
264.1061706542969 0 3888.2031
267.0915222167969 0 1041.1589
267.11578369140625 0 712.60266
268.0750732421875 0 1686.2659
268.11688232421875 0 4121.073
269.0773620605469 0 697.54486
271.111083984375 0 568.4903
273.1246032714844 0 806.6086
274.13031005859375 0 1269.8535
275.114501953125 0 3061.969
279.09796142578125 0 16434.234
280.1012268066406 0 2634.3713
281.0795593261719 0 726.4088
283.12725830078125 0 1474.3102
284.13641357421875 0 674.33984
285.10198974609375 0 23191.113
286.10491943359375 0 2000.6133
287.101318359375 0 1043.1345
289.0824890136719 0 806.728
289.1184997558594 0 1874.4021
289.1339111328125 0 1822.7661
292.1409912109375 0 2101.5923
293.12530517578125 0 1662.4092
295.11102294921875 0 1266.5791
299.06146240234375 0 1156.9803
300.13470458984375 0 814.1955
301.1189880371094 0 1124.7985
302.1159973144531 0 619.0426
302.15087890625 0 701.80176
303.1127014160156 0 58968.9 y 7
304.1155700683594 0 6713.5127
305.11041259765625 0 1613.5675 b Ammonia loss 4
305.16363525390625 0 712.4032
306.1109924316406 0 511.11252
306.1444396972656 0 1273.5793
306.16156005859375 0 1674.8419
307.0930480957031 0 592.70294
309.16082763671875 0 1314.5012
310.15155029296875 0 27958.39
311.1548767089844 0 5119.864
314.1000061035156 0 929.05865
315.0962829589844 0 828.74615
316.130126953125 0 875.33014
317.12939453125 0 1131.3491
318.1453857421875 0 5115.139
319.1470947265625 0 750.3148
320.13568115234375 0 2027.8365
323.1767883300781 0 830.1926
330.1092834472656 0 1605.7034
332.1246643066406 0 2202.2097
333.1081848144531 0 873.1069
333.1601867675781 0 5365.38
334.1094970703125 0 1113.0443
334.1571960449219 0 2632.0244
336.1557922363281 0 1020.69135
337.1561279296875 0 699.8846
338.12664794921875 0 691.1896
339.1134948730469 0 878.85547
339.1712951660156 0 2073.1082
346.1401672363281 0 3856.6873
347.1424560546875 0 616.94666
348.11932373046875 0 4559.6196
350.13507080078125 0 14797.504
350.18646240234375 0 2196.83 a Water loss 2
351.1387634277344 0 2428.6875
351.17041015625 0 1879.4009 a Ammonia loss 2
356.13922119140625 0 2030.4845
358.10406494140625 0 1132.5098
360.119140625 0 2648.042
361.1214599609375 0 677.5291
364.15087890625 0 9604.083
365.1539001464844 0 2420.0708
366.1298522949219 0 3595.801
367.1305847167969 0 567.85944
367.1659851074219 0 681.2141
374.1495361328125 0 46667.184 y 6
375.15228271484375 0 8900.55
376.114013671875 0 4121.6904
376.14508056640625 0 1161.6827
377.1189270019531 0 1133.5332
378.1300048828125 0 42904.34
378.18157958984375 0 26827.104 b Water loss 2
379.1329650878906 0 7137.5957
379.184814453125 0 4987.2803
380.1374816894531 0 1464.335
380.1877746582031 0 1157.7637
381.1459655761719 0 869.36615
382.1435241699219 0 7702.4443
383.1462097167969 0 1080.8363
384.1358337402344 0 1608.6863 y 3
394.1251220703125 0 22885.041
395.1282958984375 0 4355.1675
395.15423583984375 0 903.22363 b 5
396.12957763671875 0 744.13275
396.1927490234375 0 1235.5446 b 2
404.1142578125 0 1081.4061
404.1463317871094 0 4069.9656
405.14727783203125 0 1039.9971
410.1388244628906 0 2457.4226
415.16363525390625 0 859.0692
416.1460266113281 0 2228.4338
417.1781005859375 0 849.0095
419.15704345703125 0 1424.7906
420.16094970703125 0 817.3545
423.20147705078125 0 954.10724
425.6718444824219 0 2785.6252
426.1737976074219 0 1461.0669
427.2093505859375 0 1067.5454
429.2269592285156 0 707.8099
430.1233215332031 0 669.432
432.1540222167969 0 733.24396
433.1723327636719 0 2278.539
439.6610107421875 0 753.08386
443.15484619140625 0 954.88586
445.1378479003906 0 733.4836
447.1520080566406 0 3157.691
447.20001220703125 0 961.4981
448.1568298339844 0 996.28296
448.666015625 0 3053.2363 y Water loss 2
449.1349792480469 0 881.04474
449.16705322265625 0 3649.1963
450.1668701171875 0 1077.5784
455.2037048339844 0 1171.47
457.6717529296875 0 32229.63 y 2
458.1733093261719 0 19547.002
458.67376708984375 0 4969.39
459.1728210449219 0 578.7446
461.16729736328125 0 5224.9556
462.1689758300781 0 1121.7687
465.16259765625 0 21374.12
466.165771484375 0 4344.227
467.1666564941406 0 1322.1763
473.2149658203125 0 17488.621
474.2181701660156 0 4215.301
475.2210693359375 0 755.8873
476.6967468261719 0 750.7155
477.16790771484375 0 1133.3389
479.1784362792969 0 5348.63
480.1430969238281 0 842.11224
480.1811828613281 0 1300.5935
481.1763916015625 0 1410.0438
485.1785888671875 0 756.7039
487.1822204589844 0 674.37897
489.161865234375 0 3640.1914
490.1845397949219 0 737.9726
491.17547607421875 0 1759.2495
491.67767333984375 0 1083.8965
493.16375732421875 0 621.38495
493.208984375 0 2504.6917 b Water loss 3
495.18695068359375 0 3574.5007 b Water loss 7
496.1699523925781 0 923.8622
497.1710205078125 0 5354.7246
498.1739196777344 0 1481.3485
499.18963623046875 0 2212.6697 y Water loss 1
499.6920166015625 0 1831.5045
504.1820983886719 0 985.315 b 7
505.1943054199219 0 1123.5775
508.1953125 0 15219.633 y 1
508.69683837890625 0 10160.576
509.19781494140625 0 3448.7612
509.69891357421875 0 646.15985
511.2192687988281 0 9904.267 b 3
512.2220458984375 0 2633.9424
513.1978149414062 0 8554.798
513.2308349609375 0 695.5315
513.689453125 0 691.8888
514.1997680664062 0 2408.269
519.2015991210938 0 1362.5226
520.204345703125 0 1587.7118
521.2081298828125 0 618.38135
523.1842041015625 0 1028.0198
525.1657104492188 0 2100.6548
530.1875610351562 0 1474.399
532.2080688476562 0 1717.5099
533.2062377929688 0 953.33636
537.2132568359375 0 43664.402 y 5
538.2162475585938 0 12373.045
539.2139282226562 0 4862.878
540.215576171875 0 686.0392
540.7252197265625 0 5073.776
541.1924438476562 0 3671.3916
541.2278442382812 0 1960.0474
541.7286376953125 0 1595.8644
542.1976318359375 0 2179.4626
542.2378540039062 0 1036.1471
544.2522583007812 0 1234.0084
547.196044921875 0 826.5401
548.2000732421875 0 4993.3223
549.2024536132812 0 1521.856
549.72900390625 0 3366.2673
550.2283935546875 0 2475.7935
550.734130859375 0 897.2801
551.2171630859375 0 822.20844
551.7048950195312 0 657.88
557.1971435546875 0 610.5267
558.728515625 0 1116.9393
559.7173461914062 0 856.2407
561.2100219726562 0 609.0442
562.2263793945312 0 1442.6924
563.7181396484375 0 1719.7944
564.2183837890625 0 2693.4255
564.7182006835938 0 1115.4037
566.2162475585938 0 964.7626
567.208984375 0 2711.0588
568.2186889648438 0 722.0179
570.2325439453125 0 1585.1976
572.7243041992188 0 24708.863 Precursor Water loss
573.225830078125 0 18914.854
573.7260131835938 0 8184.6816
574.22705078125 0 2165.2258
579.2107543945312 0 1902.053
580.2427368164062 0 1753.4017
581.7294921875 0 33940.484 Precursor
582.2308959960938 0 22687.89
582.7313232421875 0 8403.053
583.2306518554688 0 1890.0786
584.2360229492188 0 2890.902
585.2408447265625 0 929.6915
588.2418212890625 0 20848.861
589.2453002929688 0 5483.835
590.2470092773438 0 935.6858
594.2242431640625 0 1134.737
596.2393798828125 0 1163.4218
598.251708984375 0 758.8203
606.2281494140625 0 1722.17
607.2340087890625 0 730.605
608.236572265625 0 3712.224 b Water loss 4
609.2377319335938 0 1396.3934
612.1917114257812 0 3970.799
612.23193359375 0 7364.386
613.1917114257812 0 1086.7993
613.2340698242188 0 2373.013
614.2437744140625 0 2142.4968
615.2461547851562 0 1008.1439
617.2032470703125 0 1080.399
624.2302856445312 0 4775.866
625.232666015625 0 1479.9513
626.24609375 0 6162.836 b 4
627.249755859375 0 2115.391
634.2293090820312 0 2818.723 y Water loss 4
635.225341796875 0 1660.2196
642.2404174804688 0 1841.9089
643.244140625 0 783.54785
644.2412719726562 0 1036.6603
649.2635498046875 0 800.00507
650.2507934570312 0 1722.2919
652.2400512695312 0 69804.95 y 4
653.242919921875 0 22273.953
654.2421875 0 6353.6665
655.2412109375 0 1211.2649
657.265869140625 0 1018.93036
662.2227783203125 0 845.40906
667.2706298828125 0 2666.108
668.2769775390625 0 887.8454
676.2462158203125 0 921.489
677.2557983398438 0 3463.3933
678.2579345703125 0 1716.8954
685.2589721679688 0 3458.6453
686.2564086914062 0 1876.9333
688.2310180664062 0 1036.1807
695.2669677734375 0 12143.344
696.2694091796875 0 3747.3784
697.26953125 0 1236.6138
703.2684936523438 0 39507.605
704.2717895507812 0 15521.078
704.339599609375 0 1142.5331
705.2731323242188 0 3665.271
713.2706909179688 0 3406.185
717.2762451171875 0 832.5743
721.2611694335938 0 1257.202
723.2609252929688 0 992.142
731.268310546875 0 1999.6572
732.22509765625 0 1070.5848
741.2561645507812 0 2082.7437
742.2578125 0 1159.6034
743.3021240234375 0 1771.64
744.3003540039062 0 718.1609
749.2565307617188 0 7246.781 y Water loss 3
750.2533569335938 0 3815.6265
751.2523803710938 0 1134.9131
752.2431640625 0 1037.8273
759.2659301757812 0 9189.825
760.2697143554688 0 4120.806
767.266845703125 0 121823.73 y 3
768.2700805664062 0 51071.574
769.2684326171875 0 14502.838
770.2683715820312 0 1640.7924
771.2998657226562 0 2121.2366 b Water loss 5
772.301025390625 0 1484.61
777.2514038085938 0 2753.2126
778.3153076171875 0 978.9292
779.3211669921875 0 1293.9298
789.3080444335938 0 1483.6521 b 5
796.314453125 0 1477.2217
804.3320922851562 0 1965.841
805.3157958984375 0 768.478
806.347412109375 0 1571.9272
817.1873168945312 0 682.34186
823.3143920898438 0 3314.7183
824.315185546875 0 2574.461
832.326904296875 0 5786.0596
833.3281860351562 0 2402.2957
834.3263549804688 0 1035.1515
835.3081665039062 0 1054.8329
842.3191528320312 0 2066.4236 b Water loss 6
843.32373046875 0 1518.8983 b Ammonia loss 6
850.3369140625 0 70631.17
851.340087890625 0 32355.969
852.3422241210938 0 8381.54
853.3436889648438 0 1057.3453
860.3220825195312 0 2204.033
861.3389892578125 0 1132.9696
868.328125 0 2905.971
869.3323364257812 0 1351.196
896.32470703125 0 10648.114 y Water loss 2
897.326904296875 0 4637.604
898.3253784179688 0 1677.9125
899.3116455078125 0 1200.5249
914.3353271484375 0 230714.86 y 2
915.337890625 0 123918.14
915.4481201171875 0 1227.1559
916.337646484375 0 44099.67
917.3372192382812 0 6853.7144
924.3218383789062 0 919.7802
933.3765258789062 0 2419.24
934.375 0 1165.0057
935.3687133789062 0 718.7501
943.3684692382812 0 864.7763
951.3839111328125 0 15820.277
952.3872680664062 0 8065.375
953.3893432617188 0 2478.7925
961.3682250976562 0 3264.3901
962.3712158203125 0 2595.481
971.362548828125 0 827.3061
972.367431640625 0 753.3937
979.3612670898438 0 1188.893
980.3580322265625 0 764.3269
989.3687744140625 0 1332.4885 b Water loss 7
990.3740844726562 0 1916.6848
997.3724365234375 0 6828.852 y Water loss 1
998.3761596679688 0 3787.7603
999.3707885742188 0 1272.5223
1007.3655395507812 0 1161.6442 b 7
1008.3760986328125 0 1442.9084
1015.3828735351562 0 51517.45 y 1
1016.3853149414062 0 30083.697
1017.3857421875 0 12792.06
1018.3876953125 0 1619.9669
1025.3680419921875 0 8242.956
1026.36962890625 0 4259.278
1027.3720703125 0 1456.7566
1666.143798828125 0 664.6335
1974.631103515625 0 671.6293
2908.06396484375 0 753.93115

Spectrum Details

|  |  |
| --- | --- |
| Matched peaks? Matched peaks The total absolute number of peaks matched. Additionally in brackets the total fraction of peaks matched and the total number of peaks is shown. | 47 (9.25% of 508) |
| FDR? FDR The false discovery rate estimated for this peptide. It is calculated by matching all theoretical fragments with a non-integer shift with the raw peaks for this spectrum. This is done with 40 different shifts. The resulting percentage is the average number of annotated peaks over the number of annotated peaks with the correct spectrum. | 0.00% |
| Satellite FDR? Satellite FDR See the FDR for details on its calculation. This satellite ion specific FDR only contains the satellite ions (d/w) for I/L/J positions. | - |
| PSM Score? PSM Score The PSM Score as given by Hecklib to this annotated spectrum. It is shown with three significant figures. | 555 |

#### Spectrum 6682? Spectrum 6682 The raw spectrum of this peptide as annotated by Hecklib. The fragments are coloured according to ion type (see legend). Any peaks with a star '\*' as text can be hovered over to see the full details, first the ion type second the mass shift type. By hovering over the amino acids in the peptide or ions in the legend the corresponding peaks are highlighted. By toggling the 'Unassigned' label you can turn the background (unassigned) peaks on or off in the plot. By updating the slider in the Ion legend you can update the spectrum to only show the top X% of the peaks with labels. The top X% means any peak that is within X% of the highest intensity. By dragging in the spectrum you can zoom in to a specific part of the spectrum and use 'Zoom Out' to get back to the original zoom level. The annotation of the spectrum is based on the given sequence in the peptides file and is done with different software so inconsistencies are likely. The peaks are annotated based on the given sequence, with 20 ppm tolerance.

Copy Data

##### Spectrum 6682 (TSV)

###### Preview

```
Loading example...
```

*Click on the button to copy the data to your clipboard.*

Mz MinMz MaxIntensity Max

WidthHeightPeptide font sizePeptide stroke widthSpectrum font sizeSpectrum stroke widthCompact peptide

Ion legend

wxyz

abcd

OtherUnassignedIon^Charge^_Position_Show for top:%

FTFDDYAMH

02.61e+45.21e+47.82e+41.04e+5

Zoom Out

y^+1^_1_a^+1^_2_d^+1^_2_a^+1^_2_b^+1^_2_b^+1^_2_y^+1^_2_y^+1^_3_b^+1^_3_y^+2^_7_y^+2^_7_b^+2^_8_y^+2^_8_b^+1^_4_y^+1^_4_\*\*b^+1^_5_y^+1^_5_y^+1^_6_y^+1^_6_y^+1^_7_y^+1^_7_y^+1^_8_y^+1^_8_

0808161524233231

Fragment Matches Table

Show background peaks

| Position | Ion type | Intensity | mz Theoretical | mz Error (Th) | mz Error (ppm) | Charge | Series Number |
| --- | --- | --- | --- | --- | --- | --- | --- |
| - | - | 1.032E+05 | 120.1 | - | - | 0 | - |
| - | - | 7980 | 121.1 | - | - | 0 | - |
| - | - | 679.9 | 122.1 | - | - | 0 | - |
| - | - | 611.3 | 127.1 | - | - | 0 | - |
| - | - | 2837 | 129.1 | - | - | 0 | - |
| - | - | 1048 | 130.1 | - | - | 0 | - |
| - | - | 626.4 | 132.1 | - | - | 0 | - |
| - | - | 848 | 133.1 | - | - | 0 | - |
| - | - | 410 | 135.7 | - | - | 0 | - |
| - | - | 1.206E+04 | 136.1 | - | - | 0 | - |
| - | - | 924.6 | 137.1 | - | - | 0 | - |
| - | - | 1834 | 138.1 | - | - | 0 | - |
| - | - | 541.8 | 140.1 | - | - | 0 | - |
| - | - | 533.3 | 141.1 | - | - | 0 | - |
| - | - | 536.6 | 143.1 | - | - | 0 | - |
| - | - | 760.4 | 146.1 | - | - | 0 | - |
| - | - | 367.1 | 147.4 | - | - | 0 | - |
| - | - | 1076 | 149 | - | - | 0 | - |
| - | - | 649.7 | 152.1 | - | - | 0 | - |
| - | - | 1095 | 155.1 | - | - | 0 | - |
| 9 | y | 2.893E+04 | 156.1 | 0.0002276 | 1.458 | +1 | 1 |
| - | - | 585.3 | 157.1 | - | - | 0 | - |
| - | - | 1864 | 157.1 | - | - | 0 | - |
| - | - | 474.4 | 158 | - | - | 0 | - |
| - | - | 1397 | 158.1 | - | - | 0 | - |
| - | - | 3267 | 159.1 | - | - | 0 | - |
| - | - | 458.6 | 161.4 | - | - | 0 | - |
| - | - | 484.7 | 165.1 | - | - | 0 | - |
| - | - | 684.7 | 166.1 | - | - | 0 | - |
| - | - | 1055 | 167.1 | - | - | 0 | - |
| - | - | 539.4 | 167.3 | - | - | 0 | - |
| - | - | 483.6 | 171.1 | - | - | 0 | - |
| - | - | 656.1 | 173.1 | - | - | 0 | - |
| - | - | 785.6 | 173.1 | - | - | 0 | - |
| - | - | 2551 | 173.5 | - | - | 0 | - |
| - | - | 559 | 174.9 | - | - | 0 | - |
| - | - | 612.1 | 175.1 | - | - | 0 | - |
| - | - | 1140 | 175.1 | - | - | 0 | - |
| - | - | 4.429E+04 | 176.1 | - | - | 0 | - |
| - | - | 1019 | 177.1 | - | - | 0 | - |
| - | - | 5105 | 177.1 | - | - | 0 | - |
| - | - | 567.7 | 185.1 | - | - | 0 | - |
| - | - | 696 | 185.1 | - | - | 0 | - |
| - | - | 674.4 | 185.2 | - | - | 0 | - |
| - | - | 5872 | 186.1 | - | - | 0 | - |
| - | - | 556.8 | 187.1 | - | - | 0 | - |
| - | - | 572 | 187.1 | - | - | 0 | - |
| - | - | 1896 | 191.1 | - | - | 0 | - |
| - | - | 1008 | 193.1 | - | - | 0 | - |
| - | - | 1195 | 199.2 | - | - | 0 | - |
| - | - | 907.6 | 203.1 | - | - | 0 | - |
| 2 | a | 2410 | 203.1 | 0.0003203 | 1.577 | +1 | 2 |
| - | - | 512.5 | 204.1 | - | - | 0 | - |
| - | - | 644.8 | 205.1 | - | - | 0 | - |
| 2 | d | 1264 | 207.1 | 0.0009806 | 4.735 | +1 | 2 |
| - | - | 525.6 | 209.1 | - | - | 0 | - |
| - | - | 801.8 | 211.1 | - | - | 0 | - |
| - | - | 628.3 | 213.1 | - | - | 0 | - |
| - | - | 699.1 | 215.1 | - | - | 0 | - |
| - | - | 518.2 | 217.1 | - | - | 0 | - |
| - | - | 2798 | 217.1 | - | - | 0 | - |
| - | - | 555.6 | 218.1 | - | - | 0 | - |
| - | - | 641.6 | 219.1 | - | - | 0 | - |
| - | - | 744.8 | 219.1 | - | - | 0 | - |
| - | - | 5159 | 221.1 | - | - | 0 | - |
| 2 | a | 5.628E+04 | 221.1 | 0.0002384 | 1.078 | +1 | 2 |
| - | - | 818.9 | 222.1 | - | - | 0 | - |
| - | - | 7207 | 222.1 | - | - | 0 | - |
| - | - | 578.4 | 226.2 | - | - | 0 | - |
| - | - | 956.9 | 227.1 | - | - | 0 | - |
| - | - | 1998 | 231.1 | - | - | 0 | - |
| 2 | b | 1.12E+04 | 231.1 | 0.000294 | 1.272 | +1 | 2 |
| - | - | 1994 | 232.1 | - | - | 0 | - |
| - | - | 6129 | 235.1 | - | - | 0 | - |
| - | - | 952.2 | 237.1 | - | - | 0 | - |
| - | - | 1.022E+04 | 239.1 | - | - | 0 | - |
| - | - | 1070 | 240.1 | - | - | 0 | - |
| - | - | 555 | 245.1 | - | - | 0 | - |
| 2 | b | 2.527E+04 | 249.1 | 0.0003189 | 1.28 | +1 | 2 |
| - | - | 3212 | 250.1 | - | - | 0 | - |
| - | - | 3808 | 251.1 | - | - | 0 | - |
| - | - | 569.4 | 259.1 | - | - | 0 | - |
| - | - | 5644 | 263.1 | - | - | 0 | - |
| - | - | 902 | 264.1 | - | - | 0 | - |
| - | - | 772.2 | 275.1 | - | - | 0 | - |
| - | - | 530.2 | 275.5 | - | - | 0 | - |
| - | - | 3260 | 279.1 | - | - | 0 | - |
| - | - | 716 | 281.1 | - | - | 0 | - |
| - | - | 667.9 | 282.2 | - | - | 0 | - |
| - | - | 3905 | 285.1 | - | - | 0 | - |
| - | - | 664.5 | 296.4 | - | - | 0 | - |
| - | - | 2238 | 299.1 | - | - | 0 | - |
| 8 | y | 8056 | 303.1 | 0.005281 | 17.42 | +1 | 2 |
| - | - | 956.1 | 304.1 | - | - | 0 | - |
| - | - | 3862 | 310.2 | - | - | 0 | - |
| - | - | 545.6 | 318.1 | - | - | 0 | - |
| - | - | 597.7 | 320.5 | - | - | 0 | - |
| - | - | 640.4 | 333.2 | - | - | 0 | - |
| - | - | 610.3 | 336.9 | - | - | 0 | - |
| - | - | 932.5 | 346.1 | - | - | 0 | - |
| - | - | 2543 | 350.1 | - | - | 0 | - |
| - | - | 1321 | 364.2 | - | - | 0 | - |
| - | - | 915.3 | 366.1 | - | - | 0 | - |
| 7 | y | 6739 | 374.1 | 0.004757 | 12.72 | +1 | 3 |
| - | - | 994.6 | 375.2 | - | - | 0 | - |
| - | - | 7808 | 378.1 | - | - | 0 | - |
| 3 | b | 2919 | 378.2 | 0.000392 | 1.037 | +1 | 3 |
| - | - | 805.4 | 379.1 | - | - | 0 | - |
| - | - | 1222 | 379.2 | - | - | 0 | - |
| - | - | 1215 | 382.1 | - | - | 0 | - |
| - | - | 694.8 | 389.2 | - | - | 0 | - |
| - | - | 3772 | 394.1 | - | - | 0 | - |
| - | - | 873.6 | 404.1 | - | - | 0 | - |
| 3 | y | 966.1 | 448.7 | 0.001602 | 3.57 | +2 | 7 |
| - | - | 680.8 | 453.3 | - | - | 0 | - |
| 3 | y | 4809 | 457.7 | 0.002881 | 6.294 | +2 | 7 |
| - | - | 3251 | 458.2 | - | - | 0 | - |
| - | - | 615.9 | 458.2 | - | - | 0 | - |
| - | - | 1712 | 458.7 | - | - | 0 | - |
| - | - | 2946 | 465.2 | - | - | 0 | - |
| - | - | 2235 | 473.2 | - | - | 0 | - |
| - | - | 748.4 | 474.2 | - | - | 0 | - |
| - | - | 849.8 | 479.2 | - | - | 0 | - |
| - | - | 703.5 | 489.2 | - | - | 0 | - |
| 8 | b | 645.4 | 495.2 | 0.003532 | 7.132 | +2 | 8 |
| - | - | 920.1 | 497.2 | - | - | 0 | - |
| 2 | y | 2171 | 508.2 | 0.003822 | 7.52 | +2 | 8 |
| - | - | 812.9 | 508.7 | - | - | 0 | - |
| - | - | 1493 | 509.2 | - | - | 0 | - |
| 4 | b | 714 | 511.2 | 0.00271 | 5.301 | +1 | 4 |
| - | - | 1628 | 513.2 | - | - | 0 | - |
| 6 | y | 6962 | 537.2 | 0.0046 | 8.563 | +1 | 4 |
| - | - | 1665 | 538.2 | - | - | 0 | - |
| - | - | 853.3 | 540.7 | - | - | 0 | - |
| - | - | 746.3 | 548.2 | - | - | 0 | - |
| - | - | 751.4 | 549.7 | - | - | 0 | - |
| 0 | Precursor | 3698 | 572.7 | 0.002668 | 4.658 | +2 | -1 |
| - | - | 2215 | 573.2 | - | - | 0 | - |
| - | - | 964 | 573.7 | - | - | 0 | - |
| 0 | Precursor | 4524 | 581.7 | 0.002513 | 4.319 | +2 | -1 |
| - | - | 2608 | 582.2 | - | - | 0 | - |
| - | - | 1968 | 582.7 | - | - | 0 | - |
| - | - | 4158 | 588.2 | - | - | 0 | - |
| 5 | b | 745.8 | 626.2 | 5.871E-05 | 0.09376 | +1 | 5 |
| 5 | y | 9399 | 652.2 | 0.004879 | 7.48 | +1 | 5 |
| - | - | 2781 | 653.2 | - | - | 0 | - |
| - | - | 1066 | 654.2 | - | - | 0 | - |
| - | - | 611.3 | 656.9 | - | - | 0 | - |
| - | - | 1820 | 695.3 | - | - | 0 | - |
| - | - | 5374 | 703.3 | - | - | 0 | - |
| - | - | 2878 | 704.3 | - | - | 0 | - |
| 4 | y | 904.8 | 749.3 | 0.005102 | 6.81 | +1 | 6 |
| - | - | 1123 | 759.3 | - | - | 0 | - |
| - | - | 844.1 | 760.3 | - | - | 0 | - |
| 4 | y | 1.805E+04 | 767.3 | 0.004425 | 5.768 | +1 | 6 |
| - | - | 5975 | 768.3 | - | - | 0 | - |
| - | - | 2665 | 769.3 | - | - | 0 | - |
| - | - | 629.6 | 815.4 | - | - | 0 | - |
| - | - | 906 | 832.3 | - | - | 0 | - |
| - | - | 1.035E+04 | 850.3 | - | - | 0 | - |
| - | - | 4174 | 851.3 | - | - | 0 | - |
| - | - | 1369 | 852.3 | - | - | 0 | - |
| - | - | 630.1 | 855.8 | - | - | 0 | - |
| - | - | 754 | 868.3 | - | - | 0 | - |
| 3 | y | 1181 | 896.3 | 0.00163 | 1.818 | +1 | 7 |
| 3 | y | 3.51E+04 | 914.3 | 0.004676 | 5.114 | +1 | 7 |
| - | - | 1.606E+04 | 915.3 | - | - | 0 | - |
| - | - | 5929 | 916.3 | - | - | 0 | - |
| - | - | 906.6 | 934.4 | - | - | 0 | - |
| - | - | 2759 | 951.4 | - | - | 0 | - |
| - | - | 811.1 | 952.4 | - | - | 0 | - |
| 2 | y | 922.7 | 997.4 | 0.0006431 | 0.6448 | +1 | 8 |
| 2 | y | 8692 | 1015 | 0.003323 | 3.273 | +1 | 8 |
| - | - | 3205 | 1016 | - | - | 0 | - |
| - | - | 1259 | 1017 | - | - | 0 | - |
| - | - | 836.4 | 1025 | - | - | 0 | - |
| - | - | 674.8 | 1998 | - | - | 0 | - |
| - | - | 766.7 | 3199 | - | - | 0 | - |

m/z Charge Intensity FragmentType MassShift Position
120.08106994628906 0 103193.58
121.08438873291016 0 7980.28
122.07161712646484 0 679.9064
127.08695220947266 0 611.3302
129.1024932861328 0 2836.5537
130.06556701660156 0 1048.2698
132.08135986328125 0 626.39825
133.08619689941406 0 848.0223
135.7388916015625 0 410.03134
136.07595825195312 0 12055.274
137.0794219970703 0 924.60095
138.05523681640625 0 1833.8069
140.0821533203125 0 541.7863
141.06642150878906 0 533.3206
143.11807250976562 0 536.60547
146.0601806640625 0 760.3974
147.42529296875 0 367.09814
149.04510498046875 0 1075.9222
152.07057189941406 0 649.6658
155.0819854736328 0 1095.325
156.0769805908203 0 28934.076 y 8
157.0740509033203 0 585.33466
157.08050537109375 0 1864.2819
158.0494842529297 0 474.41452
158.09629821777344 0 1396.882
159.09197998046875 0 3266.6763
161.40093994140625 0 458.57138
165.07713317871094 0 484.6533
166.06130981445312 0 684.6915
167.0554656982422 0 1055.1343
167.34542846679688 0 539.3985
171.07676696777344 0 483.58667
173.0925750732422 0 656.1391
173.12896728515625 0 785.5689
173.45143127441406 0 2550.5115
174.9288330078125 0 558.9973
175.07177734375 0 612.12115
175.08677673339844 0 1140.4441
176.1072540283203 0 44293.523
177.102294921875 0 1019.3653
177.1107635498047 0 5105.386
185.05569458007812 0 567.69965
185.09234619140625 0 695.9566
185.1656036376953 0 674.3523
186.091552734375 0 5871.889
187.09536743164062 0 556.79987
187.1080322265625 0 572.00726
191.11788940429688 0 1895.5004
193.10826110839844 0 1007.8491
199.1807861328125 0 1194.9945
203.0664520263672 0 907.588
203.1182098388672 0 2409.882 a Water loss 1
204.0770263671875 0 512.53735
205.09657287597656 0 644.76416
207.11378479003906 0 1263.7214 d 1
209.0931854248047 0 525.56616
211.0826416015625 0 801.77734
213.09825134277344 0 628.28186
215.13938903808594 0 699.1319
217.08148193359375 0 518.16284
217.0973358154297 0 2798.243
218.10140991210938 0 555.61774
219.07958984375 0 641.6281
219.1133270263672 0 744.76807
221.10353088378906 0 5158.737
221.12869262695312 0 56277.44 a 1
222.10716247558594 0 818.8906
222.13201904296875 0 7206.7134
226.154296875 0 578.3958
227.10350036621094 0 956.9155
231.0615234375 0 1998.2627
231.11309814453125 0 11195.357 b Water loss 1
232.11666870117188 0 1994.4231
235.1080322265625 0 6129.4297
237.0979766845703 0 952.1809
239.1141815185547 0 10222.686
240.1168212890625 0 1070.2677
245.09197998046875 0 554.9629
249.12368774414062 0 25265.531 b 1
250.12710571289062 0 3211.6245
251.10292053222656 0 3808.4954
259.1068420410156 0 569.3685
263.1027526855469 0 5643.5947
264.10760498046875 0 902.0114
275.114501953125 0 772.22565
275.5014953613281 0 530.1659
279.0974426269531 0 3260.4292
281.0805969238281 0 716.0194
282.17852783203125 0 667.9
285.1019592285156 0 3905.2976
296.4035949707031 0 664.511
299.0617980957031 0 2237.7695
303.1125183105469 0 8056.3975 y 7
304.11541748046875 0 956.08
310.1515808105469 0 3862.0151
318.14495849609375 0 545.6248
320.4683837890625 0 597.72687
333.1597595214844 0 640.43774
336.86492919921875 0 610.31287
346.1392822265625 0 932.51086
350.1346435546875 0 2542.5713
364.1510925292969 0 1320.629
366.12994384765625 0 915.32043
374.14910888671875 0 6739.14 y 6
375.15252685546875 0 994.6118
378.12982177734375 0 7808.47
378.1816101074219 0 2919.394 b Water loss 2
379.13348388671875 0 805.4174
379.18365478515625 0 1221.6411
382.14373779296875 0 1214.9724
389.1664123535156 0 694.7534
394.12493896484375 0 3772.3037
404.146240234375 0 873.5662
448.6649475097656 0 966.11206 y Water loss 2
453.3440246582031 0 680.76544
457.6715087890625 0 4808.935 y 2
458.1729736328125 0 3251.28
458.20654296875 0 615.868
458.6730651855469 0 1711.7106
465.1625061035156 0 2946.4915
473.21527099609375 0 2235.25
474.2175598144531 0 748.40607
479.17523193359375 0 849.8381
489.1587219238281 0 703.5226
495.190185546875 0 645.4391 b Water loss 7
497.1717834472656 0 920.1308
508.1962890625 0 2170.815 y 1
508.6976318359375 0 812.9161
509.19403076171875 0 1492.8622
511.221435546875 0 713.98267 b 3
513.197265625 0 1627.668
537.2122802734375 0 6961.6753 y 5
538.2150268554688 0 1664.8735
540.7242431640625 0 853.273
548.1994018554688 0 746.33545
549.7286987304688 0 751.43317
572.7240600585938 0 3697.6614 Precursor Water loss
573.22705078125 0 2214.6707
573.7213134765625 0 964.0412
581.7291870117188 0 4524.4766 Precursor
582.23095703125 0 2608.448
582.73046875 0 1968.24
588.2415771484375 0 4158.1943
626.2457275390625 0 745.8314 b 4
652.239501953125 0 9398.686 y 4
653.2404174804688 0 2780.8057
654.2418212890625 0 1066.0741
656.8521728515625 0 611.3285
695.2674560546875 0 1820.0332
703.2682495117188 0 5374.134
704.2719116210938 0 2878.2437
749.256103515625 0 904.82654 y Water loss 3
759.2611694335938 0 1122.7709
760.26416015625 0 844.07336
767.2659912109375 0 18053.475 y 3
768.2691040039062 0 5975.483
769.2678833007812 0 2665.0764
815.3590087890625 0 629.5554
832.3272094726562 0 906.04456
850.3358154296875 0 10346.931
851.3400268554688 0 4174.136
852.3408203125 0 1368.9213
855.7626953125 0 630.083
868.3341674804688 0 753.982
896.321044921875 0 1180.5485 y Water loss 2
914.3346557617188 0 35098.133 y 2
915.3372802734375 0 16056.259
916.3378295898438 0 5928.9414
934.4334106445312 0 906.6401
951.3811645507812 0 2759.139
952.3914794921875 0 811.0903
997.3677368164062 0 922.6823 y Water loss 1
1015.3809814453125 0 8691.665 y 1
1016.3880004882812 0 3204.7793
1017.3899536132812 0 1258.8839
1025.3668212890625 0 836.40125
1998.4527587890625 0 674.7791
3198.60009765625 0 766.6529

Spectrum Details

|  |  |
| --- | --- |
| Matched peaks? Matched peaks The total absolute number of peaks matched. Additionally in brackets the total fraction of peaks matched and the total number of peaks is shown. | 25 (14.04% of 178) |
| FDR? FDR The false discovery rate estimated for this peptide. It is calculated by matching all theoretical fragments with a non-integer shift with the raw peaks for this spectrum. This is done with 40 different shifts. The resulting percentage is the average number of annotated peaks over the number of annotated peaks with the correct spectrum. | 0.10% |
| Satellite FDR? Satellite FDR See the FDR for details on its calculation. This satellite ion specific FDR only contains the satellite ions (d/w) for I/L/J positions. | - |
| PSM Score? PSM Score The PSM Score as given by Hecklib to this annotated spectrum. It is shown with three significant figures. | 248 |

#### Spectrum 6739? Spectrum 6739 The raw spectrum of this peptide as annotated by Hecklib. The fragments are coloured according to ion type (see legend). Any peaks with a star '\*' as text can be hovered over to see the full details, first the ion type second the mass shift type. By hovering over the amino acids in the peptide or ions in the legend the corresponding peaks are highlighted. By toggling the 'Unassigned' label you can turn the background (unassigned) peaks on or off in the plot. By updating the slider in the Ion legend you can update the spectrum to only show the top X% of the peaks with labels. The top X% means any peak that is within X% of the highest intensity. By dragging in the spectrum you can zoom in to a specific part of the spectrum and use 'Zoom Out' to get back to the original zoom level. The annotation of the spectrum is based on the given sequence in the peptides file and is done with different software so inconsistencies are likely. The peaks are annotated based on the given sequence, with 20 ppm tolerance.

Copy Data

##### Spectrum 6739 (TSV)

###### Preview

```
Loading example...
```

*Click on the button to copy the data to your clipboard.*

Mz MinMz MaxIntensity Max

WidthHeightPeptide font sizePeptide stroke widthSpectrum font sizeSpectrum stroke widthCompact peptide

Ion legend

wxyz

abcd

OtherUnassignedIon^Charge^_Position_Show for top:%

FTFDDYAMH

01.39e+42.78e+44.17e+45.56e+4

Zoom Out

y^+1^_1_a^+1^_2_d^+1^_2_a^+1^_2_b^+1^_2_b^+1^_2_y^+1^_2_y^+1^_3_b^+1^_3_y^+2^_7_y^+2^_8_b^+1^_4_y^+1^_4_\*\*y^+1^_5_y^+1^_6_y^+1^_7_y^+1^_8_y^+1^_8_

0695139020842779

Fragment Matches Table

Show background peaks

| Position | Ion type | Intensity | mz Theoretical | mz Error (Th) | mz Error (ppm) | Charge | Series Number |
| --- | --- | --- | --- | --- | --- | --- | --- |
| - | - | 5.504E+04 | 120.1 | - | - | 0 | - |
| - | - | 4031 | 121.1 | - | - | 0 | - |
| - | - | 427.3 | 125.1 | - | - | 0 | - |
| - | - | 438.8 | 127.1 | - | - | 0 | - |
| - | - | 574.3 | 128.1 | - | - | 0 | - |
| - | - | 502.6 | 129.1 | - | - | 0 | - |
| - | - | 3370 | 129.1 | - | - | 0 | - |
| - | - | 367.8 | 129.5 | - | - | 0 | - |
| - | - | 611.7 | 130.1 | - | - | 0 | - |
| - | - | 836 | 131.1 | - | - | 0 | - |
| - | - | 453.8 | 131.1 | - | - | 0 | - |
| - | - | 440.2 | 133.1 | - | - | 0 | - |
| - | - | 1748 | 133.1 | - | - | 0 | - |
| - | - | 481.6 | 134.3 | - | - | 0 | - |
| - | - | 8211 | 136.1 | - | - | 0 | - |
| - | - | 452.2 | 137.5 | - | - | 0 | - |
| - | - | 496.8 | 138.1 | - | - | 0 | - |
| - | - | 447.4 | 140.5 | - | - | 0 | - |
| - | - | 431.9 | 141.1 | - | - | 0 | - |
| - | - | 421.5 | 147.2 | - | - | 0 | - |
| - | - | 994.2 | 149 | - | - | 0 | - |
| - | - | 523.5 | 149 | - | - | 0 | - |
| - | - | 479.3 | 155.1 | - | - | 0 | - |
| - | - | 465 | 155.1 | - | - | 0 | - |
| 9 | y | 1.715E+04 | 156.1 | 0.0001818 | 1.165 | +1 | 1 |
| - | - | 761.2 | 157.1 | - | - | 0 | - |
| - | - | 544.2 | 158.1 | - | - | 0 | - |
| - | - | 2056 | 159.1 | - | - | 0 | - |
| - | - | 1040 | 167.1 | - | - | 0 | - |
| - | - | 484.9 | 171.2 | - | - | 0 | - |
| - | - | 807.4 | 173.1 | - | - | 0 | - |
| - | - | 2736 | 173.4 | - | - | 0 | - |
| - | - | 656.9 | 174.1 | - | - | 0 | - |
| - | - | 895.7 | 175.1 | - | - | 0 | - |
| - | - | 762 | 175.1 | - | - | 0 | - |
| - | - | 2.246E+04 | 176.1 | - | - | 0 | - |
| - | - | 2970 | 177.1 | - | - | 0 | - |
| - | - | 447.5 | 181.9 | - | - | 0 | - |
| - | - | 512.6 | 185.2 | - | - | 0 | - |
| - | - | 3120 | 186.1 | - | - | 0 | - |
| - | - | 668.4 | 191.1 | - | - | 0 | - |
| - | - | 502.9 | 193.1 | - | - | 0 | - |
| - | - | 531.5 | 199.1 | - | - | 0 | - |
| - | - | 1394 | 199.2 | - | - | 0 | - |
| - | - | 846.9 | 201.1 | - | - | 0 | - |
| - | - | 666.3 | 203.1 | - | - | 0 | - |
| 2 | a | 1609 | 203.1 | 6.09E-05 | 0.2998 | +1 | 2 |
| 2 | d | 965.2 | 207.1 | 0.000233 | 1.125 | +1 | 2 |
| - | - | 1385 | 217.1 | - | - | 0 | - |
| - | - | 2228 | 221.1 | - | - | 0 | - |
| 2 | a | 2.867E+04 | 221.1 | 7.056E-05 | 0.3191 | +1 | 2 |
| - | - | 2526 | 222.1 | - | - | 0 | - |
| - | - | 681 | 228.1 | - | - | 0 | - |
| - | - | 868.1 | 231.1 | - | - | 0 | - |
| 2 | b | 6934 | 231.1 | 0.0001414 | 0.6118 | +1 | 2 |
| - | - | 751.4 | 232.1 | - | - | 0 | - |
| - | - | 2875 | 235.1 | - | - | 0 | - |
| - | - | 6398 | 239.1 | - | - | 0 | - |
| - | - | 748.8 | 242.1 | - | - | 0 | - |
| 2 | b | 1.409E+04 | 249.1 | 0.0001816 | 0.7288 | +1 | 2 |
| - | - | 1943 | 250.1 | - | - | 0 | - |
| - | - | 1392 | 251.1 | - | - | 0 | - |
| - | - | 3159 | 263.1 | - | - | 0 | - |
| - | - | 1522 | 279.1 | - | - | 0 | - |
| - | - | 1205 | 281.1 | - | - | 0 | - |
| - | - | 2971 | 285.1 | - | - | 0 | - |
| - | - | 2057 | 299.1 | - | - | 0 | - |
| - | - | 533.3 | 300.1 | - | - | 0 | - |
| 8 | y | 4476 | 303.1 | 0.005098 | 16.82 | +1 | 2 |
| - | - | 886.6 | 304.1 | - | - | 0 | - |
| - | - | 2208 | 310.2 | - | - | 0 | - |
| - | - | 726.1 | 313.2 | - | - | 0 | - |
| - | - | 942.3 | 350.1 | - | - | 0 | - |
| - | - | 659.9 | 351.2 | - | - | 0 | - |
| - | - | 700.8 | 355.1 | - | - | 0 | - |
| - | - | 681.6 | 366.1 | - | - | 0 | - |
| 7 | y | 3892 | 374.1 | 0.004361 | 11.66 | +1 | 3 |
| - | - | 701.6 | 375.2 | - | - | 0 | - |
| - | - | 4599 | 378.1 | - | - | 0 | - |
| 3 | b | 2495 | 378.2 | 0.0008498 | 2.247 | +1 | 3 |
| - | - | 798.8 | 379.1 | - | - | 0 | - |
| - | - | 845 | 379.2 | - | - | 0 | - |
| - | - | 632.4 | 382.1 | - | - | 0 | - |
| - | - | 1338 | 394.1 | - | - | 0 | - |
| - | - | 611.9 | 416.1 | - | - | 0 | - |
| - | - | 597.6 | 449.2 | - | - | 0 | - |
| - | - | 1121 | 453.3 | - | - | 0 | - |
| - | - | 489.7 | 454.9 | - | - | 0 | - |
| 3 | y | 1973 | 457.7 | 0.002484 | 5.427 | +2 | 7 |
| - | - | 1910 | 458.2 | - | - | 0 | - |
| - | - | 595 | 458.7 | - | - | 0 | - |
| - | - | 2114 | 465.2 | - | - | 0 | - |
| - | - | 1426 | 473.2 | - | - | 0 | - |
| - | - | 643.4 | 477 | - | - | 0 | - |
| - | - | 583.9 | 479.2 | - | - | 0 | - |
| 2 | y | 709.6 | 508.2 | 0.00161 | 3.169 | +2 | 8 |
| 4 | b | 917.6 | 511.2 | 0.0004336 | 0.8481 | +1 | 4 |
| - | - | 876.7 | 513.2 | - | - | 0 | - |
| 6 | y | 3391 | 537.2 | 0.004356 | 8.109 | +1 | 4 |
| - | - | 939.1 | 538.2 | - | - | 0 | - |
| - | - | 728.6 | 539.2 | - | - | 0 | - |
| - | - | 599.2 | 546.8 | - | - | 0 | - |
| 0 | Precursor | 2295 | 572.7 | 0.001814 | 3.167 | +2 | -1 |
| - | - | 940.2 | 573.2 | - | - | 0 | - |
| 0 | Precursor | 2129 | 581.7 | 0.001292 | 2.221 | +2 | -1 |
| - | - | 1606 | 582.2 | - | - | 0 | - |
| - | - | 1284 | 588.2 | - | - | 0 | - |
| - | - | 681.2 | 612.2 | - | - | 0 | - |
| 5 | y | 5277 | 652.2 | 0.004086 | 6.264 | +1 | 5 |
| - | - | 2630 | 653.2 | - | - | 0 | - |
| - | - | 1357 | 695.3 | - | - | 0 | - |
| - | - | 842.7 | 696.3 | - | - | 0 | - |
| - | - | 2709 | 703.3 | - | - | 0 | - |
| - | - | 832.4 | 704.3 | - | - | 0 | - |
| 4 | y | 9980 | 767.3 | 0.003388 | 4.415 | +1 | 6 |
| - | - | 3220 | 768.3 | - | - | 0 | - |
| - | - | 1435 | 769.3 | - | - | 0 | - |
| - | - | 5158 | 850.3 | - | - | 0 | - |
| - | - | 2517 | 851.3 | - | - | 0 | - |
| - | - | 915.7 | 852.3 | - | - | 0 | - |
| - | - | 800.7 | 897.3 | - | - | 0 | - |
| 3 | y | 1.856E+04 | 914.3 | 0.00315 | 3.445 | +1 | 7 |
| - | - | 9530 | 915.3 | - | - | 0 | - |
| - | - | 3006 | 916.3 | - | - | 0 | - |
| - | - | 790.2 | 917.3 | - | - | 0 | - |
| - | - | 1111 | 933.4 | - | - | 0 | - |
| - | - | 1105 | 934.4 | - | - | 0 | - |
| - | - | 1243 | 951.4 | - | - | 0 | - |
| - | - | 785.9 | 961.5 | - | - | 0 | - |
| 2 | y | 878.6 | 997.4 | 0.007723 | 7.744 | +1 | 8 |
| 2 | y | 3710 | 1015 | 0.002041 | 2.01 | +1 | 8 |
| - | - | 1982 | 1016 | - | - | 0 | - |
| - | - | 909 | 1025 | - | - | 0 | - |
| - | - | 618.7 | 1326 | - | - | 0 | - |
| - | - | 673.6 | 1422 | - | - | 0 | - |
| - | - | 780.3 | 1606 | - | - | 0 | - |
| - | - | 660.2 | 2752 | - | - | 0 | - |

m/z Charge Intensity FragmentType MassShift Position
120.08102416992188 0 55044.39
121.08435821533203 0 4031.2654
125.0714111328125 0 427.3012
127.08707427978516 0 438.83282
128.10728454589844 0 574.3213
129.06622314453125 0 502.64093
129.1024932861328 0 3370.4922
129.5025634765625 0 367.7849
130.06533813476562 0 611.718
131.07044982910156 0 836.03845
131.10614013671875 0 453.7836
133.06141662597656 0 440.22586
133.08604431152344 0 1747.6536
134.28204345703125 0 481.59897
136.07589721679688 0 8210.874
137.54750061035156 0 452.19232
138.0553741455078 0 496.7805
140.53448486328125 0 447.3687
141.1025390625 0 431.91516
147.20614624023438 0 421.5392
148.95480346679688 0 994.18823
149.0447540283203 0 523.4603
155.08193969726562 0 479.2866
155.1181182861328 0 465.03625
156.07693481445312 0 17154.469 y 8
157.0801544189453 0 761.1891
158.0968475341797 0 544.1846
159.09202575683594 0 2055.6338
167.0554962158203 0 1040.0267
171.1500701904297 0 484.93515
173.1284637451172 0 807.4171
173.43984985351562 0 2735.651
174.0880889892578 0 656.8536
175.0869903564453 0 895.6936
175.0967254638672 0 762.03406
176.10720825195312 0 22460.906
177.11087036132812 0 2969.8027
181.87486267089844 0 447.5103
185.164794921875 0 512.6191
186.09164428710938 0 3119.8398
191.1178436279297 0 668.4462
193.1083221435547 0 502.87543
199.1075897216797 0 531.486
199.1807403564453 0 1393.88
201.12281799316406 0 846.8789
203.06674194335938 0 666.3397
203.11795043945312 0 1609.0426 a Water loss 1
207.113037109375 0 965.2407 d 1
217.0975341796875 0 1385.1245
221.10336303710938 0 2227.9424
221.12852478027344 0 28665.855 a 1
222.13182067871094 0 2525.5774
228.0979461669922 0 680.98376
231.0613555908203 0 868.0832
231.11294555664062 0 6934.0806 b Water loss 1
232.1168975830078 0 751.35895
235.10801696777344 0 2874.7185
239.11392211914062 0 6398.4033
242.11331176757812 0 748.805
249.12355041503906 0 14086.347 b 1
250.1265869140625 0 1943.1682
251.10250854492188 0 1391.6304
263.1021423339844 0 3159.06
279.0972900390625 0 1521.7576
281.0802917480469 0 1204.6101
285.1017150878906 0 2971.156
299.0617980957031 0 2056.821
300.0629577636719 0 533.2535
303.1123352050781 0 4475.794 y 7
304.11578369140625 0 886.57574
310.1510925292969 0 2207.6758
313.1867980957031 0 726.10474
350.1343688964844 0 942.27106
351.2000427246094 0 659.90845
355.0682373046875 0 700.8265
366.126953125 0 681.6368
374.1487121582031 0 3891.777 y 6
375.151611328125 0 701.57074
378.1295471191406 0 4599.335
378.18206787109375 0 2495.0083 b Water loss 2
379.1322326660156 0 798.8089
379.1839599609375 0 844.9743
382.1421203613281 0 632.4252
394.1232604980469 0 1338.228
416.12786865234375 0 611.9465
449.16583251953125 0 597.56586
453.3445129394531 0 1120.5385
454.8885192871094 0 489.72226
457.6711120605469 0 1972.7893 y 2
458.1713562011719 0 1910.4299
458.6715393066406 0 594.96954
465.1612548828125 0 2113.8547
473.21453857421875 0 1426.2299
477.001953125 0 643.3779
479.17877197265625 0 583.9424
508.19085693359375 0 709.63965 y 1
511.2182922363281 0 917.5965 b 3
513.1986694335938 0 876.6655
537.2120361328125 0 3390.6538 y 5
538.2120971679688 0 939.1042
539.2132568359375 0 728.58856
546.78466796875 0 599.2498
572.7232055664062 0 2295.0398 Precursor Water loss
573.2244873046875 0 940.16754
581.7279663085938 0 2129.21 Precursor
582.2299194335938 0 1605.9559
588.239990234375 0 1284.3334
612.2385864257812 0 681.1872
652.2387084960938 0 5277.4414 y 4
653.2409057617188 0 2630.3694
695.2693481445312 0 1357.2736
696.2767333984375 0 842.66693
703.2666625976562 0 2709.2942
704.2747192382812 0 832.3752
767.2649536132812 0 9979.581 y 3
768.2671508789062 0 3220.1057
769.2673950195312 0 1435.1293
850.3352661132812 0 5157.7207
851.3340454101562 0 2516.5251
852.3475952148438 0 915.7088
897.3272094726562 0 800.70905
914.3331298828125 0 18563.479 y 2
915.3356323242188 0 9530.291
916.3331909179688 0 3006.3062
917.3401489257812 0 790.24695
933.428955078125 0 1110.7528
934.4329833984375 0 1104.7604
951.3779907226562 0 1242.9332
961.4649658203125 0 785.8706
997.3748168945312 0 878.6464 y Water loss 1
1015.3796997070312 0 3709.8445 y 1
1016.382568359375 0 1981.8888
1025.361083984375 0 908.97345
1325.6263427734375 0 618.67804
1421.558349609375 0 673.6171
1605.9884033203125 0 780.287
2751.68310546875 0 660.2053

Spectrum Details

|  |  |
| --- | --- |
| Matched peaks? Matched peaks The total absolute number of peaks matched. Additionally in brackets the total fraction of peaks matched and the total number of peaks is shown. | 20 (14.60% of 137) |
| FDR? FDR The false discovery rate estimated for this peptide. It is calculated by matching all theoretical fragments with a non-integer shift with the raw peaks for this spectrum. This is done with 40 different shifts. The resulting percentage is the average number of annotated peaks over the number of annotated peaks with the correct spectrum. | 0.24% |
| Satellite FDR? Satellite FDR See the FDR for details on its calculation. This satellite ion specific FDR only contains the satellite ions (d/w) for I/L/J positions. | - |
| PSM Score? PSM Score The PSM Score as given by Hecklib to this annotated spectrum. It is shown with three significant figures. | 193 |

#### Spectrum 6798? Spectrum 6798 The raw spectrum of this peptide as annotated by Hecklib. The fragments are coloured according to ion type (see legend). Any peaks with a star '\*' as text can be hovered over to see the full details, first the ion type second the mass shift type. By hovering over the amino acids in the peptide or ions in the legend the corresponding peaks are highlighted. By toggling the 'Unassigned' label you can turn the background (unassigned) peaks on or off in the plot. By updating the slider in the Ion legend you can update the spectrum to only show the top X% of the peaks with labels. The top X% means any peak that is within X% of the highest intensity. By dragging in the spectrum you can zoom in to a specific part of the spectrum and use 'Zoom Out' to get back to the original zoom level. The annotation of the spectrum is based on the given sequence in the peptides file and is done with different software so inconsistencies are likely. The peaks are annotated based on the given sequence, with 20 ppm tolerance.

Copy Data

##### Spectrum 6798 (TSV)

###### Preview

```
Loading example...
```

*Click on the button to copy the data to your clipboard.*

Mz MinMz MaxIntensity Max

WidthHeightPeptide font sizePeptide stroke widthSpectrum font sizeSpectrum stroke widthCompact peptide

Ion legend

wxyz

abcd

OtherUnassignedIon^Charge^_Position_Show for top:%

FTFDDYAMH

09.33e+31.87e+42.80e+43.73e+4

Zoom Out

y^+1^_1_a^+1^_2_d^+1^_2_a^+1^_2_b^+1^_2_b^+1^_2_y^+1^_2_y^+1^_3_b^+1^_3_y^+2^_7_y^+2^_8_y^+1^_4_\*y^+1^_5_y^+1^_6_y^+1^_7_y^+1^_7_y^+1^_8_

0659131819772636

Fragment Matches Table

Show background peaks

| Position | Ion type | Intensity | mz Theoretical | mz Error (Th) | mz Error (ppm) | Charge | Series Number |
| --- | --- | --- | --- | --- | --- | --- | --- |
| - | - | 3.697E+04 | 120.1 | - | - | 0 | - |
| - | - | 2936 | 121.1 | - | - | 0 | - |
| - | - | 383 | 121.8 | - | - | 0 | - |
| - | - | 359.7 | 122.1 | - | - | 0 | - |
| - | - | 402 | 127.1 | - | - | 0 | - |
| - | - | 391.3 | 128.1 | - | - | 0 | - |
| - | - | 3185 | 129.1 | - | - | 0 | - |
| - | - | 496.7 | 130 | - | - | 0 | - |
| - | - | 525.5 | 133.1 | - | - | 0 | - |
| - | - | 505.6 | 133.1 | - | - | 0 | - |
| - | - | 4926 | 136.1 | - | - | 0 | - |
| - | - | 601.2 | 138.1 | - | - | 0 | - |
| - | - | 487.3 | 143.1 | - | - | 0 | - |
| - | - | 519.7 | 149 | - | - | 0 | - |
| - | - | 684.9 | 149 | - | - | 0 | - |
| - | - | 809.3 | 156.1 | - | - | 0 | - |
| 9 | y | 1.033E+04 | 156.1 | 0.0001055 | 0.6761 | +1 | 1 |
| - | - | 465.3 | 156.7 | - | - | 0 | - |
| - | - | 562.9 | 157.1 | - | - | 0 | - |
| - | - | 536.2 | 158.1 | - | - | 0 | - |
| - | - | 1585 | 159.1 | - | - | 0 | - |
| - | - | 836.8 | 167.1 | - | - | 0 | - |
| - | - | 522.9 | 173.1 | - | - | 0 | - |
| - | - | 4337 | 173.4 | - | - | 0 | - |
| - | - | 1.464E+04 | 176.1 | - | - | 0 | - |
| - | - | 1993 | 177.1 | - | - | 0 | - |
| - | - | 589.2 | 185.1 | - | - | 0 | - |
| - | - | 713.6 | 185.1 | - | - | 0 | - |
| - | - | 633.1 | 185.2 | - | - | 0 | - |
| - | - | 1804 | 186.1 | - | - | 0 | - |
| - | - | 926.5 | 187.1 | - | - | 0 | - |
| - | - | 574.2 | 193.1 | - | - | 0 | - |
| - | - | 1357 | 199.2 | - | - | 0 | - |
| 2 | a | 1048 | 203.1 | 0.0002745 | 1.352 | +1 | 2 |
| 2 | d | 649.1 | 207.1 | 0.0005076 | 2.451 | +1 | 2 |
| - | - | 488.4 | 210.4 | - | - | 0 | - |
| - | - | 673.6 | 215.1 | - | - | 0 | - |
| - | - | 783.5 | 217.1 | - | - | 0 | - |
| - | - | 486.5 | 218.9 | - | - | 0 | - |
| - | - | 1212 | 221.1 | - | - | 0 | - |
| 2 | a | 1.816E+04 | 221.1 | 0.0001316 | 0.5951 | +1 | 2 |
| - | - | 1522 | 222.1 | - | - | 0 | - |
| 2 | b | 3224 | 231.1 | 0.0002024 | 0.8759 | +1 | 2 |
| - | - | 561.1 | 233.8 | - | - | 0 | - |
| - | - | 1572 | 235.1 | - | - | 0 | - |
| - | - | 4310 | 239.1 | - | - | 0 | - |
| - | - | 525.1 | 249.1 | - | - | 0 | - |
| 2 | b | 8779 | 249.1 | 0.0001511 | 0.6063 | +1 | 2 |
| - | - | 1420 | 250.1 | - | - | 0 | - |
| - | - | 1250 | 251.1 | - | - | 0 | - |
| - | - | 1819 | 263.1 | - | - | 0 | - |
| - | - | 556.3 | 279.4 | - | - | 0 | - |
| - | - | 946.5 | 281.1 | - | - | 0 | - |
| - | - | 1150 | 285.1 | - | - | 0 | - |
| - | - | 582.7 | 292.6 | - | - | 0 | - |
| - | - | 674.2 | 295.3 | - | - | 0 | - |
| - | - | 1721 | 299.1 | - | - | 0 | - |
| - | - | 660.9 | 300.1 | - | - | 0 | - |
| 8 | y | 2494 | 303.1 | 0.005037 | 16.62 | +1 | 2 |
| - | - | 1640 | 310.2 | - | - | 0 | - |
| - | - | 719.8 | 331.5 | - | - | 0 | - |
| - | - | 521.5 | 350.1 | - | - | 0 | - |
| - | - | 638.4 | 356.1 | - | - | 0 | - |
| - | - | 581.3 | 364.2 | - | - | 0 | - |
| 7 | y | 1604 | 374.1 | 0.004757 | 12.72 | +1 | 3 |
| - | - | 2165 | 378.1 | - | - | 0 | - |
| 3 | b | 1312 | 378.2 | 0.0001573 | 0.4159 | +1 | 3 |
| - | - | 589.7 | 379.1 | - | - | 0 | - |
| - | - | 543.5 | 386 | - | - | 0 | - |
| - | - | 832.3 | 394.1 | - | - | 0 | - |
| - | - | 591.1 | 410.1 | - | - | 0 | - |
| - | - | 779.6 | 415 | - | - | 0 | - |
| - | - | 671.9 | 431.9 | - | - | 0 | - |
| 3 | y | 1824 | 457.7 | 0.003155 | 6.894 | +2 | 7 |
| - | - | 1447 | 458.2 | - | - | 0 | - |
| - | - | 903.4 | 465.2 | - | - | 0 | - |
| - | - | 565.4 | 475.7 | - | - | 0 | - |
| 2 | y | 881.3 | 508.2 | 0.001807 | 3.557 | +2 | 8 |
| 6 | y | 1895 | 537.2 | 0.0046 | 8.563 | +1 | 4 |
| - | - | 934.1 | 538.2 | - | - | 0 | - |
| - | - | 596 | 573.7 | - | - | 0 | - |
| 0 | Precursor | 846 | 581.7 | 0.006236 | 10.72 | +2 | -1 |
| - | - | 1387 | 582.2 | - | - | 0 | - |
| - | - | 913.1 | 588.2 | - | - | 0 | - |
| 5 | y | 3806 | 652.2 | 0.006039 | 9.258 | +1 | 5 |
| - | - | 863.7 | 653.2 | - | - | 0 | - |
| - | - | 624.4 | 695.3 | - | - | 0 | - |
| - | - | 1792 | 703.3 | - | - | 0 | - |
| - | - | 781.8 | 704.3 | - | - | 0 | - |
| - | - | 1181 | 738.4 | - | - | 0 | - |
| 4 | y | 6627 | 767.3 | 0.004975 | 6.483 | +1 | 6 |
| - | - | 2092 | 768.3 | - | - | 0 | - |
| - | - | 849.8 | 769.3 | - | - | 0 | - |
| - | - | 3487 | 850.3 | - | - | 0 | - |
| - | - | 1582 | 851.3 | - | - | 0 | - |
| 3 | y | 689.5 | 896.3 | 0.005292 | 5.904 | +1 | 7 |
| 3 | y | 1.106E+04 | 914.3 | 0.006141 | 6.716 | +1 | 7 |
| - | - | 4524 | 915.3 | - | - | 0 | - |
| - | - | 1769 | 916.3 | - | - | 0 | - |
| - | - | 1758 | 933.4 | - | - | 0 | - |
| - | - | 1126 | 934.4 | - | - | 0 | - |
| - | - | 708.8 | 951.4 | - | - | 0 | - |
| 2 | y | 2529 | 1015 | 0.005642 | 5.557 | +1 | 8 |
| - | - | 1153 | 1016 | - | - | 0 | - |
| - | - | 927.9 | 1017 | - | - | 0 | - |
| - | - | 649.3 | 2077 | - | - | 0 | - |
| - | - | 641.7 | 2436 | - | - | 0 | - |
| - | - | 711.3 | 2447 | - | - | 0 | - |
| - | - | 647.8 | 2610 | - | - | 0 | - |

m/z Charge Intensity FragmentType MassShift Position
120.0809326171875 0 36968.277
121.08425903320312 0 2935.8567
121.75404357910156 0 382.98663
122.07154846191406 0 359.66885
127.08728790283203 0 401.97745
128.10707092285156 0 391.31763
129.1023712158203 0 3185.4568
130.04981994628906 0 496.71423
133.06121826171875 0 525.4912
133.0860595703125 0 505.5754
136.0757598876953 0 4925.995
138.05514526367188 0 601.177
143.11785888671875 0 487.2708
149.02288818359375 0 519.6953
149.04486083984375 0 684.86224
156.0705108642578 0 809.3417
156.0768585205078 0 10326.476 y 8
156.700927734375 0 465.2962
157.0800323486328 0 562.91077
158.09640502929688 0 536.1793
159.09165954589844 0 1584.616
167.05572509765625 0 836.8122
173.12893676757812 0 522.9283
173.4406280517578 0 4337.2144
176.10708618164062 0 14637.8125
177.1107177734375 0 1992.7235
185.05596923828125 0 589.2123
185.0920867919922 0 713.631
185.1651611328125 0 633.05347
186.09146118164062 0 1803.6295
187.10829162597656 0 926.4534
193.1080780029297 0 574.1552
199.1802978515625 0 1357.3252
203.1181640625 0 1048.4795 a Water loss 1
207.11331176757812 0 649.1185 d 1
210.4039764404297 0 488.37
215.13946533203125 0 673.6461
217.09764099121094 0 783.4712
218.9051971435547 0 486.52246
221.10350036621094 0 1212.0835
221.1285858154297 0 18163.914 a 1
222.13182067871094 0 1521.7638
231.11300659179688 0 3223.6167 b Water loss 1
233.82249450683594 0 561.08417
235.1072540283203 0 1571.6218
239.11399841308594 0 4309.582
249.0996856689453 0 525.1201
249.12351989746094 0 8779.148 b 1
250.12660217285156 0 1420.3164
251.10301208496094 0 1250.0533
263.1029357910156 0 1819.4286
279.4374084472656 0 556.30853
281.0793762207031 0 946.4972
285.1024475097656 0 1149.735
292.6497802734375 0 582.7115
295.3057861328125 0 674.2435
299.0619201660156 0 1721.2448
300.0643310546875 0 660.8654
303.1122741699219 0 2494.2107 y 7
310.1513977050781 0 1639.8544
331.49114990234375 0 719.77484
350.13494873046875 0 521.45386
356.0693664550781 0 638.3728
364.15118408203125 0 581.30695
374.14910888671875 0 1604.1117 y 6
378.13006591796875 0 2165.2175
378.1810607910156 0 1312.4637 b Water loss 2
379.1314392089844 0 589.6526
385.9896240234375 0 543.46783
394.12640380859375 0 832.28937
410.1404724121094 0 591.0706
415.0390625 0 779.5586
431.8575134277344 0 671.91486
457.6717834472656 0 1824.123 y 2
458.1719665527344 0 1447.3939
465.164306640625 0 903.4216
475.67449951171875 0 565.39496
508.19427490234375 0 881.3367 y 1
537.2122802734375 0 1895.3838 y 5
538.2105102539062 0 934.0928
573.7192993164062 0 595.96454
581.73291015625 0 846.0337 Precursor
582.2286376953125 0 1387.4716
588.239501953125 0 913.14087
652.2406616210938 0 3806.0227 y 4
653.2427978515625 0 863.73175
695.2679443359375 0 624.42
703.2703247070312 0 1791.9917
704.2733764648438 0 781.7721
738.376708984375 0 1180.8514
767.2665405273438 0 6627.1484 y 3
768.2682495117188 0 2092.1108
769.2716064453125 0 849.8167
850.3375244140625 0 3487.2505
851.3407592773438 0 1582.1992
896.32470703125 0 689.5134 y Water loss 2
914.3361206054688 0 11059.022 y 2
915.3383178710938 0 4523.7217
916.3377075195312 0 1769.1204
933.427734375 0 1758.4933
934.423828125 0 1126.4307
951.3792724609375 0 708.7536
1015.38330078125 0 2529.247 y 1
1016.3833618164062 0 1152.6406
1017.3786010742188 0 927.9101
2077.2109375 0 649.3029
2435.7177734375 0 641.6512
2446.729248046875 0 711.30634
2610.29736328125 0 647.7889

Spectrum Details

|  |  |
| --- | --- |
| Matched peaks? Matched peaks The total absolute number of peaks matched. Additionally in brackets the total fraction of peaks matched and the total number of peaks is shown. | 18 (16.51% of 109) |
| FDR? FDR The false discovery rate estimated for this peptide. It is calculated by matching all theoretical fragments with a non-integer shift with the raw peaks for this spectrum. This is done with 40 different shifts. The resulting percentage is the average number of annotated peaks over the number of annotated peaks with the correct spectrum. | 0.26% |
| Satellite FDR? Satellite FDR See the FDR for details on its calculation. This satellite ion specific FDR only contains the satellite ions (d/w) for I/L/J positions. | - |
| PSM Score? PSM Score The PSM Score as given by Hecklib to this annotated spectrum. It is shown with three significant figures. | 176 |

#### Reverse Lookup? Reverse Lookup All places where this read could be placed.

| Group | Segment | Template | Template Part | Read Part | Score | Unique |
| --- | --- | --- | --- | --- | --- | --- |
| Homo sapiens Heavy Chain | IGHV | IGHV3-9 | [26..35] | [0..9] | 72 | True |

| Recombined | Template Part | Read Part | Score | Unique |
| --- | --- | --- | --- | --- |
| REC-0-1 | [26..35] | [0..9] | 72 | True |

#### Meta Information from Multiple reads

##### Number of combined reads

4

##### Intensity

0.6124

##### TotalArea

4.529E+07

#### Positional Score

Copy Data

##### Positional Score (TSV)

###### Preview

```
Loading example...
```

*Click on the button to copy the data to your clipboard.*

10012345678

Label Value
"0" 0.5
"1" 0.5
"2" 0.497
"3" 0.497
"4" 0.497
"5" 0.482
"6" 0.48
"7" 0.485
"8" 0.477

#### Meta Information from PEAKS

##### Scan Identifier

F2:6605

##### Original sequence

F

T

F

D

D

Y

A

M

+15.99

H

##### Posttranslational Modifications

Oxidation (M)

##### Source File

D:\separate\_stitch\_analyses\xle-disambiguation\raw\20210323\_F1\_UM1\_Peng0013\_SA\_F59\_ingel\_3ug\_TL.raw

##### Fraction

2

##### Scan Feature

F2:7517

##### De Novo Score

99

##### ConfidenceScore

99

### m/z

581.7293

##### Mass

1161.4436

##### Charge

2

##### Retention Time

36.56

##### Predicted Retention Time

-

##### Area

1.132E+07

##### Parts Per Million

0.4

##### Fragmentation mode

HCD

##### Originating file

01 D:\separate\_stitch\_analyses\xle-disambiguation\20210325\_F59\_3ug\_DENOVO\_12.csv

#### Meta Information from PEAKS

##### Scan Identifier

F2:6682

##### Original sequence

F

T

F

D

D

Y

A

M

+15.99

H

##### Posttranslational Modifications

Oxidation (M)

##### Source File

D:\separate\_stitch\_analyses\xle-disambiguation\raw\20210323\_F1\_UM1\_Peng0013\_SA\_F59\_ingel\_3ug\_TL.raw

##### Fraction

2

##### Scan Feature

F2:7517

##### De Novo Score

99

##### ConfidenceScore

99

### m/z

581.7293

##### Mass

1161.4436

##### Charge

2

##### Retention Time

36.56

##### Predicted Retention Time

-

##### Area

1.132E+07

##### Parts Per Million

0.4

##### Fragmentation mode

HCD

##### Originating file

01 D:\separate\_stitch\_analyses\xle-disambiguation\20210325\_F59\_3ug\_DENOVO\_12.csv

#### Meta Information from PEAKS

##### Scan Identifier

F2:6739

##### Original sequence

F

T

F

D

D

Y

A

M

+15.99

H

##### Posttranslational Modifications

Oxidation (M)

##### Source File

D:\separate\_stitch\_analyses\xle-disambiguation\raw\20210323\_F1\_UM1\_Peng0013\_SA\_F59\_ingel\_3ug\_TL.raw

##### Fraction

2

##### Scan Feature

F2:7517

##### De Novo Score

98

##### ConfidenceScore

98

### m/z

581.7293

##### Mass

1161.4436

##### Charge

2

##### Retention Time

36.56

##### Predicted Retention Time

-

##### Area

1.132E+07

##### Parts Per Million

0.4

##### Fragmentation mode

HCD

##### Originating file

01 D:\separate\_stitch\_analyses\xle-disambiguation\20210325\_F59\_3ug\_DENOVO\_12.csv

#### Meta Information from PEAKS

##### Scan Identifier

F2:6798

##### Original sequence

F

T

F

D

D

Y

A

M

+15.99

H

##### Posttranslational Modifications

Oxidation (M)

##### Source File

D:\separate\_stitch\_analyses\xle-disambiguation\raw\20210323\_F1\_UM1\_Peng0013\_SA\_F59\_ingel\_3ug\_TL.raw

##### Fraction

2

##### Scan Feature

F2:7517

##### De Novo Score

97

##### ConfidenceScore

97

### m/z

581.7293

##### Mass

1161.4436

##### Charge

2

##### Retention Time

36.56

##### Predicted Retention Time

-

##### Area

1.132E+07

##### Parts Per Million

0.4

##### Fragmentation mode

HCD

##### Originating file

01 D:\separate\_stitch\_analyses\xle-disambiguation\20210325\_F59\_3ug\_DENOVO\_12.csv
