## Supplementary material for "A handle on mass coincidence errors in *de novo* sequencing of antibodies by bottom-up proteomics": Combined_020.html

Details Combined\_020 | Stitch OverviewUndefined

### Read Combined\_020

#### Sequence (length=11)

JFPPKPKDTJM

#### Spectrum 5122? Spectrum 5122 The raw spectrum of this peptide as annotated by Hecklib. The fragments are coloured according to ion type (see legend). Any peaks with a star '\*' as text can be hovered over to see the full details, first the ion type second the mass shift type. By hovering over the amino acids in the peptide or ions in the legend the corresponding peaks are highlighted. By toggling the 'Unassigned' label you can turn the background (unassigned) peaks on or off in the plot. By updating the slider in the Ion legend you can update the spectrum to only show the top X% of the peaks with labels. The top X% means any peak that is within X% of the highest intensity. By dragging in the spectrum you can zoom in to a specific part of the spectrum and use 'Zoom Out' to get back to the original zoom level. The annotation of the spectrum is based on the given sequence in the peptides file and is done with different software so inconsistencies are likely. The peaks are annotated based on the given sequence, with 20 ppm tolerance.

Copy Data

##### Spectrum 5122 (TSV)

###### Preview

```
Loading example...
```

*Click on the button to copy the data to your clipboard.*

Mz MinMz MaxIntensity Max

WidthHeightPeptide font sizePeptide stroke widthSpectrum font sizeSpectrum stroke widthCompact peptide

Ion legend

wxyz

abcd

OtherUnassignedIon^Charge^_Position_Show for top:%

JFPPKPKDTJM

06.25e+51.25e+61.88e+62.50e+6

Zoom Out

y^+1^_2_y^+1^_3_y^+2^_7_y^+2^_8_y^+1^_4_y^+2^_9_z^+2^_9_y^+2^_9_y^+2^_10_z^+1^_5_y^+1^_5_c^+1^_6_c^+1^_6_y^+1^_6_w^+1^_7_c^+1^_7_c^+1^_7_z^+1^_7_y^+1^_7_c^+1^_8_c^+1^_8_y^+1^_8_c^+1^_9_c^+1^_9_y^+1^_9_c^+1^_10_c^+1^_10_z^+1^_10_y^+1^_10_

0803160724103213

Fragment Matches Table

Show background peaks

| Position | Ion type | Intensity | mz Theoretical | mz Error (Th) | mz Error (ppm) | Charge | Series Number |
| --- | --- | --- | --- | --- | --- | --- | --- |
| - | - | 4827 | 120.1 | - | - | 0 | - |
| - | - | 1.928E+04 | 120.1 | - | - | 0 | - |
| - | - | 1983 | 127.1 | - | - | 0 | - |
| - | - | 1855 | 127.9 | - | - | 0 | - |
| - | - | 1869 | 134.2 | - | - | 0 | - |
| - | - | 2429 | 135 | - | - | 0 | - |
| - | - | 7677 | 136.1 | - | - | 0 | - |
| - | - | 1931 | 137.2 | - | - | 0 | - |
| - | - | 4437 | 149 | - | - | 0 | - |
| - | - | 2134 | 150.8 | - | - | 0 | - |
| - | - | 6.306E+04 | 166.1 | - | - | 0 | - |
| - | - | 2265 | 167 | - | - | 0 | - |
| - | - | 3027 | 167.1 | - | - | 0 | - |
| - | - | 1.387E+04 | 173.4 | - | - | 0 | - |
| - | - | 2010 | 174.9 | - | - | 0 | - |
| - | - | 2300 | 191.6 | - | - | 0 | - |
| - | - | 4713 | 195.1 | - | - | 0 | - |
| - | - | 4749 | 197.1 | - | - | 0 | - |
| - | - | 2438 | 226.1 | - | - | 0 | - |
| - | - | 3171 | 226.2 | - | - | 0 | - |
| - | - | 1.599E+05 | 233.2 | - | - | 0 | - |
| - | - | 2.56E+04 | 234.2 | - | - | 0 | - |
| - | - | 1.489E+05 | 261.2 | - | - | 0 | - |
| - | - | 2.526E+04 | 262.2 | - | - | 0 | - |
| 10 | y | 1.093E+04 | 279.1 | 0.005458 | 19.55 | +1 | 2 |
| - | - | 2735 | 305.4 | - | - | 0 | - |
| - | - | 7524 | 316.2 | - | - | 0 | - |
| - | - | 3.331E+04 | 323.2 | - | - | 0 | - |
| - | - | 4146 | 324.2 | - | - | 0 | - |
| - | - | 2.006E+04 | 358.2 | - | - | 0 | - |
| - | - | 3952 | 359.2 | - | - | 0 | - |
| 9 | y | 7237 | 380.2 | 0.003709 | 9.755 | +1 | 3 |
| - | - | 2710 | 415.2 | - | - | 0 | - |
| 5 | y | 8457 | 424.7 | 0.002621 | 6.171 | +2 | 7 |
| - | - | 4600 | 425.2 | - | - | 0 | - |
| - | - | 2883 | 427.3 | - | - | 0 | - |
| - | - | 3159 | 430.3 | - | - | 0 | - |
| - | - | 4.129E+04 | 439.3 | - | - | 0 | - |
| - | - | 2.695E+04 | 439.8 | - | - | 0 | - |
| - | - | 5008 | 440.3 | - | - | 0 | - |
| - | - | 2452 | 450.5 | - | - | 0 | - |
| - | - | 2983 | 472.8 | - | - | 0 | - |
| 4 | y | 2.085E+04 | 473.3 | 0.003155 | 6.667 | +2 | 8 |
| - | - | 1.245E+04 | 473.8 | - | - | 0 | - |
| - | - | 9215 | 488.8 | - | - | 0 | - |
| - | - | 7.245E+04 | 489.8 | - | - | 0 | - |
| - | - | 3.369E+04 | 490.3 | - | - | 0 | - |
| - | - | 9784 | 490.8 | - | - | 0 | - |
| 8 | y | 3802 | 495.2 | 0.005208 | 10.52 | +1 | 4 |
| - | - | 3235 | 512.2 | - | - | 0 | - |
| 3 | y | 4.799E+04 | 512.8 | 0.003205 | 6.249 | +2 | 9 |
| - | - | 2.942E+04 | 513.3 | - | - | 0 | - |
| 3 | z | 1.006E+04 | 513.8 | 0.004355 | 8.476 | +2 | 9 |
| - | - | 9666 | 514.3 | - | - | 0 | - |
| - | - | 5190 | 514.8 | - | - | 0 | - |
| - | - | 2.162E+04 | 520.8 | - | - | 0 | - |
| - | - | 1.068E+04 | 521.3 | - | - | 0 | - |
| 3 | y | 2.047E+06 | 521.8 | 0.003659 | 7.013 | +2 | 9 |
| - | - | 1.181E+06 | 522.3 | - | - | 0 | - |
| - | - | 4.135E+05 | 522.8 | - | - | 0 | - |
| - | - | 5.114E+04 | 523.3 | - | - | 0 | - |
| - | - | 4652 | 565.4 | - | - | 0 | - |
| - | - | 3094 | 566.4 | - | - | 0 | - |
| 2 | y | 6.035E+04 | 595.3 | 0.003449 | 5.794 | +2 | 10 |
| - | - | 3.791E+04 | 595.8 | - | - | 0 | - |
| - | - | 1.535E+04 | 596.3 | - | - | 0 | - |
| 7 | z | 3.779E+04 | 607.3 | 0.006301 | 10.38 | +1 | 5 |
| - | - | 1.696E+05 | 608.3 | - | - | 0 | - |
| - | - | 4.661E+04 | 609.3 | - | - | 0 | - |
| - | - | 1.062E+04 | 610.3 | - | - | 0 | - |
| - | - | 3380 | 612.8 | - | - | 0 | - |
| - | - | 7441 | 615.3 | - | - | 0 | - |
| - | - | 4.496E+04 | 619.9 | - | - | 0 | - |
| - | - | 3.039E+04 | 620.4 | - | - | 0 | - |
| - | - | 1.237E+04 | 620.9 | - | - | 0 | - |
| - | - | 2839 | 620.9 | - | - | 0 | - |
| 7 | y | 7782 | 623.3 | 0.006681 | 10.72 | +1 | 5 |
| - | - | 6982 | 630.4 | - | - | 0 | - |
| - | - | 3123 | 634.8 | - | - | 0 | - |
| - | - | 1.251E+04 | 642.9 | - | - | 0 | - |
| - | - | 1.225E+04 | 643.4 | - | - | 0 | - |
| - | - | 6336 | 643.9 | - | - | 0 | - |
| - | - | 7684 | 644.4 | - | - | 0 | - |
| - | - | 5367 | 644.8 | - | - | 0 | - |
| - | - | 8875 | 650.9 | - | - | 0 | - |
| - | - | 2.339E+04 | 651.4 | - | - | 0 | - |
| - | - | 1.06E+06 | 651.9 | - | - | 0 | - |
| - | - | 8.047E+05 | 652.4 | - | - | 0 | - |
| - | - | 3.992E+05 | 652.9 | - | - | 0 | - |
| - | - | 7.007E+04 | 653.4 | - | - | 0 | - |
| - | - | 9214 | 653.4 | - | - | 0 | - |
| - | - | 4776 | 654.4 | - | - | 0 | - |
| - | - | 7080 | 663.4 | - | - | 0 | - |
| - | - | 3681 | 665.2 | - | - | 0 | - |
| 6 | c | 8035 | 680.4 | 0.002547 | 3.744 | +1 | 6 |
| - | - | 6264 | 687.4 | - | - | 0 | - |
| - | - | 8183 | 695.4 | - | - | 0 | - |
| - | - | 7.332E+05 | 696.4 | - | - | 0 | - |
| 6 | c | 1.037E+06 | 697.4 | 0.0001659 | 0.2379 | +1 | 6 |
| - | - | 3.672E+05 | 698.4 | - | - | 0 | - |
| - | - | 5.491E+04 | 699.4 | - | - | 0 | - |
| - | - | 8230 | 719.4 | - | - | 0 | - |
| 6 | y | 4.53E+04 | 720.4 | 0.005186 | 7.2 | +1 | 6 |
| - | - | 1.545E+04 | 721.4 | - | - | 0 | - |
| - | - | 5553 | 722.4 | - | - | 0 | - |
| - | - | 3242 | 728.4 | - | - | 0 | - |
| - | - | 1.367E+04 | 748.4 | - | - | 0 | - |
| - | - | 4611 | 749.4 | - | - | 0 | - |
| - | - | 5435 | 764.4 | - | - | 0 | - |
| - | - | 1.343E+04 | 768.5 | - | - | 0 | - |
| - | - | 5874 | 769.5 | - | - | 0 | - |
| 5 | w | 8086 | 774.4 | 0.006706 | 8.661 | +1 | 7 |
| - | - | 3375 | 780.5 | - | - | 0 | - |
| - | - | 4715 | 781.4 | - | - | 0 | - |
| - | - | 8.272E+04 | 781.5 | - | - | 0 | - |
| - | - | 4.439E+04 | 782.5 | - | - | 0 | - |
| - | - | 8583 | 783.5 | - | - | 0 | - |
| - | - | 1.469E+04 | 788.4 | - | - | 0 | - |
| - | - | 5564 | 789.4 | - | - | 0 | - |
| 7 | c | 5770 | 808.5 | 0.001807 | 2.235 | +1 | 7 |
| - | - | 1.066E+04 | 812.5 | - | - | 0 | - |
| - | - | 1.746E+04 | 823.5 | - | - | 0 | - |
| - | - | 6.256E+05 | 824.5 | - | - | 0 | - |
| 7 | c | 5.344E+05 | 825.5 | 0.001867 | 2.262 | +1 | 7 |
| - | - | 1.648E+05 | 826.5 | - | - | 0 | - |
| - | - | 2.927E+04 | 827.5 | - | - | 0 | - |
| - | - | 3118 | 830.4 | - | - | 0 | - |
| 5 | z | 1.911E+05 | 832.4 | 0.005913 | 7.103 | +1 | 7 |
| - | - | 9.948E+04 | 833.4 | - | - | 0 | - |
| - | - | 3.261E+04 | 834.4 | - | - | 0 | - |
| - | - | 6294 | 835.4 | - | - | 0 | - |
| - | - | 3907 | 847.3 | - | - | 0 | - |
| 5 | y | 1.796E+04 | 848.4 | 0.005621 | 6.625 | +1 | 7 |
| - | - | 7965 | 849.5 | - | - | 0 | - |
| - | - | 4148 | 868.5 | - | - | 0 | - |
| - | - | 6143 | 871.4 | - | - | 0 | - |
| - | - | 3.63E+04 | 877.5 | - | - | 0 | - |
| - | - | 1.924E+04 | 878.5 | - | - | 0 | - |
| - | - | 6806 | 879.5 | - | - | 0 | - |
| - | - | 9484 | 881.5 | - | - | 0 | - |
| - | - | 4219 | 882.5 | - | - | 0 | - |
| - | - | 8729 | 883.5 | - | - | 0 | - |
| - | - | 3373 | 888.4 | - | - | 0 | - |
| - | - | 1.852E+04 | 895.6 | - | - | 0 | - |
| - | - | 2.329E+04 | 896.6 | - | - | 0 | - |
| - | - | 1.379E+04 | 897.6 | - | - | 0 | - |
| - | - | 8528 | 898.6 | - | - | 0 | - |
| 8 | c | 5642 | 923.5 | 0.0009794 | 1.061 | +1 | 8 |
| - | - | 8129 | 924.5 | - | - | 0 | - |
| - | - | 1.687E+04 | 925.5 | - | - | 0 | - |
| - | - | 5276 | 926.6 | - | - | 0 | - |
| - | - | 9017 | 938.5 | - | - | 0 | - |
| - | - | 6.215E+05 | 939.6 | - | - | 0 | - |
| 8 | c | 6.498E+05 | 940.6 | 0.001589 | 1.689 | +1 | 8 |
| - | - | 2.602E+05 | 941.6 | - | - | 0 | - |
| - | - | 5.96E+04 | 942.6 | - | - | 0 | - |
| - | - | 1.095E+04 | 943.6 | - | - | 0 | - |
| - | - | 7298 | 944.5 | - | - | 0 | - |
| 4 | y | 1.878E+05 | 945.5 | 0.006202 | 6.559 | +1 | 8 |
| - | - | 9.571E+04 | 946.5 | - | - | 0 | - |
| - | - | 3.377E+04 | 947.5 | - | - | 0 | - |
| - | - | 3755 | 948.5 | - | - | 0 | - |
| - | - | 4703 | 972.4 | - | - | 0 | - |
| - | - | 4960 | 973.5 | - | - | 0 | - |
| - | - | 4232 | 974.5 | - | - | 0 | - |
| - | - | 5339 | 976.6 | - | - | 0 | - |
| - | - | 1.297E+04 | 978.6 | - | - | 0 | - |
| - | - | 1.183E+04 | 979.6 | - | - | 0 | - |
| - | - | 5307 | 980.6 | - | - | 0 | - |
| - | - | 1.028E+04 | 997.6 | - | - | 0 | - |
| - | - | 8558 | 998.6 | - | - | 0 | - |
| 9 | c | 8567 | 1025 | 0.01777 | 17.35 | +1 | 9 |
| - | - | 4504 | 1026 | - | - | 0 | - |
| - | - | 4647 | 1040 | - | - | 0 | - |
| - | - | 6.831E+04 | 1041 | - | - | 0 | - |
| 9 | c | 5.176E+05 | 1042 | 0.001209 | 1.161 | +1 | 9 |
| 3 | y | 3.56E+05 | 1043 | 0.01862 | 17.86 | +1 | 9 |
| - | - | 1.952E+05 | 1044 | - | - | 0 | - |
| - | - | 8.107E+04 | 1045 | - | - | 0 | - |
| - | - | 1.363E+04 | 1046 | - | - | 0 | - |
| - | - | 7240 | 1070 | - | - | 0 | - |
| - | - | 3.773E+04 | 1071 | - | - | 0 | - |
| - | - | 2.414E+04 | 1072 | - | - | 0 | - |
| - | - | 7825 | 1073 | - | - | 0 | - |
| - | - | 8600 | 1103 | - | - | 0 | - |
| - | - | 7275 | 1104 | - | - | 0 | - |
| - | - | 4.02E+04 | 1111 | - | - | 0 | - |
| - | - | 2.927E+04 | 1112 | - | - | 0 | - |
| - | - | 8032 | 1113 | - | - | 0 | - |
| - | - | 1.876E+04 | 1130 | - | - | 0 | - |
| - | - | 9700 | 1131 | - | - | 0 | - |
| - | - | 6462 | 1132 | - | - | 0 | - |
| 10 | c | 3.248E+04 | 1138 | 0.001678 | 1.475 | +1 | 10 |
| - | - | 1.781E+04 | 1139 | - | - | 0 | - |
| - | - | 1.038E+04 | 1140 | - | - | 0 | - |
| - | - | 6722 | 1153 | - | - | 0 | - |
| - | - | 4711 | 1154 | - | - | 0 | - |
| 10 | c | 9.183E+05 | 1155 | 0.001251 | 1.084 | +1 | 10 |
| - | - | 6.275E+05 | 1156 | - | - | 0 | - |
| - | - | 2.115E+05 | 1157 | - | - | 0 | - |
| - | - | 2.383E+04 | 1158 | - | - | 0 | - |
| - | - | 8507 | 1164 | - | - | 0 | - |
| 2 | z | 2.401E+05 | 1174 | 0.005921 | 5.045 | +1 | 10 |
| - | - | 1.672E+05 | 1175 | - | - | 0 | - |
| - | - | 7.226E+04 | 1176 | - | - | 0 | - |
| - | - | 8042 | 1177 | - | - | 0 | - |
| - | - | 9068 | 1184 | - | - | 0 | - |
| - | - | 6802 | 1185 | - | - | 0 | - |
| - | - | 7998 | 1186 | - | - | 0 | - |
| - | - | 4552 | 1187 | - | - | 0 | - |
| 2 | y | 3833 | 1190 | 0.005601 | 4.708 | +1 | 10 |
| - | - | 5720 | 1196 | - | - | 0 | - |
| - | - | 7792 | 1197 | - | - | 0 | - |
| - | - | 7731 | 1201 | - | - | 0 | - |
| - | - | 5060 | 1202 | - | - | 0 | - |
| - | - | 4300 | 1203 | - | - | 0 | - |
| - | - | 3845 | 1204 | - | - | 0 | - |
| - | - | 5.789E+04 | 1212 | - | - | 0 | - |
| - | - | 4.658E+04 | 1213 | - | - | 0 | - |
| - | - | 1.404E+04 | 1214 | - | - | 0 | - |
| - | - | 5632 | 1215 | - | - | 0 | - |
| - | - | 1.357E+04 | 1216 | - | - | 0 | - |
| - | - | 7520 | 1217 | - | - | 0 | - |
| - | - | 4959 | 1218 | - | - | 0 | - |
| - | - | 5047 | 1225 | - | - | 0 | - |
| - | - | 4624 | 1230 | - | - | 0 | - |
| - | - | 1.279E+04 | 1231 | - | - | 0 | - |
| - | - | 1.014E+04 | 1232 | - | - | 0 | - |
| - | - | 1.451E+04 | 1233 | - | - | 0 | - |
| - | - | 7558 | 1234 | - | - | 0 | - |
| - | - | 2.433E+04 | 1240 | - | - | 0 | - |
| - | - | 2.423E+04 | 1241 | - | - | 0 | - |
| - | - | 2257 | 1242 | - | - | 0 | - |
| - | - | 6.22E+04 | 1242 | - | - | 0 | - |
| - | - | 6.265E+04 | 1243 | - | - | 0 | - |
| - | - | 8.531E+04 | 1244 | - | - | 0 | - |
| - | - | 5.392E+04 | 1245 | - | - | 0 | - |
| - | - | 1.768E+04 | 1246 | - | - | 0 | - |
| - | - | 3666 | 1247 | - | - | 0 | - |
| - | - | 1.462E+04 | 1248 | - | - | 0 | - |
| - | - | 1.022E+04 | 1249 | - | - | 0 | - |
| - | - | 6093 | 1250 | - | - | 0 | - |
| - | - | 4255 | 1254 | - | - | 0 | - |
| - | - | 1.49E+04 | 1255 | - | - | 0 | - |
| - | - | 1.238E+04 | 1256 | - | - | 0 | - |
| - | - | 3.567E+04 | 1258 | - | - | 0 | - |
| - | - | 3.04E+04 | 1259 | - | - | 0 | - |
| - | - | 3.886E+04 | 1260 | - | - | 0 | - |
| - | - | 2.097E+04 | 1261 | - | - | 0 | - |
| - | - | 1.252E+04 | 1262 | - | - | 0 | - |
| - | - | 6546 | 1268 | - | - | 0 | - |
| - | - | 5619 | 1269 | - | - | 0 | - |
| - | - | 1.029E+04 | 1269 | - | - | 0 | - |
| - | - | 5.994E+04 | 1270 | - | - | 0 | - |
| - | - | 9146 | 1270 | - | - | 0 | - |
| - | - | 3.991E+04 | 1271 | - | - | 0 | - |
| - | - | 1.64E+04 | 1271 | - | - | 0 | - |
| - | - | 1.771E+04 | 1272 | - | - | 0 | - |
| - | - | 8922 | 1272 | - | - | 0 | - |
| - | - | 6914 | 1273 | - | - | 0 | - |
| - | - | 3910 | 1276 | - | - | 0 | - |
| - | - | 2.065E+04 | 1285 | - | - | 0 | - |
| - | - | 7912 | 1286 | - | - | 0 | - |
| - | - | 1.651E+05 | 1286 | - | - | 0 | - |
| - | - | 9.791E+05 | 1287 | - | - | 0 | - |
| - | - | 6.987E+05 | 1288 | - | - | 0 | - |
| - | - | 3.374E+05 | 1289 | - | - | 0 | - |
| - | - | 1.91E+04 | 1290 | - | - | 0 | - |
| - | - | 7.13E+04 | 1290 | - | - | 0 | - |
| - | - | 9823 | 1291 | - | - | 0 | - |
| - | - | 2.144E+04 | 1301 | - | - | 0 | - |
| - | - | 3.92E+04 | 1302 | - | - | 0 | - |
| - | - | 1.007E+06 | 1303 | - | - | 0 | - |
| - | - | 2.477E+06 | 1304 | - | - | 0 | - |
| - | - | 1.709E+06 | 1305 | - | - | 0 | - |
| - | - | 8854 | 1305 | - | - | 0 | - |
| - | - | 6.816E+05 | 1306 | - | - | 0 | - |
| - | - | 1.127E+05 | 1307 | - | - | 0 | - |
| - | - | 5393 | 1336 | - | - | 0 | - |
| - | - | 3959 | 3075 | - | - | 0 | - |
| - | - | 3574 | 3181 | - | - | 0 | - |

m/z Charge Intensity FragmentType MassShift Position
120.0657958984375 0 4826.661
120.08116149902344 0 19277.482
127.13186645507812 0 1982.5397
127.92845153808594 0 1855.3319
134.23460388183594 0 1868.9913
135.03265380859375 0 2428.5107
136.07614135742188 0 7676.835
137.16387939453125 0 1931.0896
148.95388793945312 0 4437.2104
150.84613037109375 0 2133.87
166.05360412597656 0 63062.875
167.03421020507812 0 2264.571
167.05670166015625 0 3026.6123
173.43968200683594 0 13872.144
174.9005889892578 0 2009.8663
191.5946807861328 0 2299.9163
195.1128692626953 0 4713.239
197.12835693359375 0 4748.5776
226.0706024169922 0 2437.6638
226.15457153320312 0 3171.0186
233.16526794433594 0 159854.36
234.168701171875 0 25600.684
261.16009521484375 0 148912.78
262.16351318359375 0 25257.965
279.1378479003906 0 10932.106 y 9
305.3903503417969 0 2734.6792
316.1869812011719 0 7524.206
323.2084655761719 0 33307.04
324.21221923828125 0 4145.7046
358.2129211425781 0 20064.094
359.2170104980469 0 3952.3572
380.18377685546875 0 7237.296 y 8
415.1632080078125 0 2710.0308
424.7311096191406 0 8456.638 y 4
425.23382568359375 0 4600.1377
427.2789306640625 0 2883.2844
430.25738525390625 0 3158.9707
439.2611389160156 0 41291.62
439.7635803222656 0 26954.51
440.264892578125 0 5008.378
450.5346374511719 0 2452.416
472.7589111328125 0 2983.2385
473.2580261230469 0 20846.244 y 3
473.7593078613281 0 12454.294
488.7785339355469 0 9215.398
489.7855224609375 0 72449.266
490.287353515625 0 33685.08
490.7886047363281 0 9784.382
495.21221923828125 0 3802.0178 y 7
512.2176513671875 0 3234.536
512.7791748046875 0 47987.484 y Water loss 2
513.2808227539062 0 29421.053
513.7762451171875 0 10058.966 z 2
514.2723388671875 0 9666.35
514.7759399414062 0 5189.77
520.7760620117188 0 21622.902
521.278076171875 0 10676.075
521.784912109375 0 2047190.2 y 2
522.2860717773438 0 1181218.1
522.7858276367188 0 413475.9
523.28564453125 0 51142.527
565.3817138671875 0 4651.5864
566.383544921875 0 3093.5315
595.3189086914062 0 60350.777 y 1
595.8203125 0 37914.812
596.3211059570312 0 15346.07
607.28955078125 0 37793.504 z 6
608.296875 0 169560.84
609.2990112304688 0 46612.203
610.2994384765625 0 10623.204
612.8238525390625 0 3380.2102
615.3472290039062 0 7441.089
619.861572265625 0 44960.96
620.362548828125 0 30391.912
620.8632202148438 0 12369.649
620.916015625 0 2838.7314
623.3086547851562 0 7781.7837 y 6
630.3600463867188 0 6982.2007
634.8462524414062 0 3122.8967
642.8543090820312 0 12510.035
643.35595703125 0 12246.729
643.8511352539062 0 6335.878
644.3514404296875 0 7683.6543
644.849365234375 0 5367.337
650.8513793945312 0 8874.683
651.3638916015625 0 23393.717
651.8610229492188 0 1060182
652.3624877929688 0 804714.5
652.86279296875 0 399213.78
653.361328125 0 70069.586
653.4251708984375 0 9213.605
654.4312133789062 0 4775.749
663.3817138671875 0 7080.204
665.1793212890625 0 3681.499
680.4104614257812 0 8035.241 c Ammonia loss 5
687.3807373046875 0 6264.212
695.4232788085938 0 8182.7944
696.4327392578125 0 733237.94
697.4393920898438 0 1036619.6 c 5
698.4426879882812 0 367223.62
699.4462280273438 0 54906.06
719.3541870117188 0 8230.046
720.3599243164062 0 45297.64 y 5
721.36279296875 0 15448.587
722.3630981445312 0 5553.1274
728.4253540039062 0 3241.9058
748.35546875 0 13668.191
749.3557739257812 0 4611.078
764.4404296875 0 5435.3716
768.4588623046875 0 13433.033
769.463134765625 0 5874.118
774.3720092773438 0 8086.1675 w 4
780.4608764648438 0 3374.5889
781.4476318359375 0 4715.448
781.5225219726562 0 82718.66
782.5260009765625 0 44388
783.5303344726562 0 8582.606
788.44775390625 0 14686.939
789.4493408203125 0 5564.319
808.5061645507812 0 5770.0464 c Ammonia loss 6
812.4668579101562 0 10659.258
823.51806640625 0 17460.523
824.5277099609375 0 625648.56
825.5326538085938 0 534444.8 c 6
826.5361938476562 0 164839.31
827.5404663085938 0 29268.455
830.420166015625 0 3117.6348
832.4368896484375 0 191148.53 z 4
833.4406127929688 0 99483.09
834.4409790039062 0 32605.268
835.4488525390625 0 6293.769
847.3301391601562 0 3906.9587
848.455322265625 0 17959.8 y 4
849.45654296875 0 7965.434
868.480224609375 0 4147.853
871.3719482421875 0 6143.0474
877.5145263671875 0 36301.547
878.5170288085938 0 19242.447
879.5147094726562 0 6805.884
881.5292358398438 0 9483.577
882.5386352539062 0 4218.7637
883.4951782226562 0 8728.922
888.367431640625 0 3373.436
895.5599975585938 0 18522.371
896.5516357421875 0 23288.152
897.55419921875 0 13787.542
898.5586547851562 0 8527.911
923.533935546875 0 5642.3247 c Ammonia loss 7
924.5419921875 0 8128.793
925.5484008789062 0 16869.021
926.5570068359375 0 5275.6094
938.5469360351562 0 9017.136
939.5545654296875 0 621543.25
940.5598754882812 0 649814.94 c 7
941.5635375976562 0 260204.3
942.5686645507812 0 59595.527
943.5761108398438 0 10950.169
944.5015258789062 0 7297.5566
945.5086669921875 0 187810.33 y 3
946.5115356445312 0 95710.2
947.510986328125 0 33768.938
948.5117797851562 0 3755.342
972.3870849609375 0 4703.3647
973.5010375976562 0 4959.925
974.508544921875 0 4232.1167
976.5540771484375 0 5339.3794
978.5632934570312 0 12966.696
979.5708618164062 0 11831.984
980.5912475585938 0 5307.3657
997.594970703125 0 10279.657
998.6044921875 0 8558.278
1024.5648193359375 0 8566.757 c Ammonia loss 8
1025.5635986328125 0 4504.161
1039.5933837890625 0 4647.279
1040.602783203125 0 68310.516
1041.6103515625 0 517609.3 c 8
1042.5738525390625 0 356009.28 y 2
1043.572265625 0 195184.95
1044.564208984375 0 81070.71
1045.5638427734375 0 13627.132
1069.5457763671875 0 7240.2837
1070.5555419921875 0 37728.43
1071.5587158203125 0 24138.742
1072.557373046875 0 7825.443
1102.53466796875 0 8600.325
1103.537109375 0 7275.318
1110.6796875 0 40204.434
1111.681396484375 0 29274.166
1112.6865234375 0 8031.5327
1129.6190185546875 0 18762.764
1130.6253662109375 0 9699.733
1131.612060546875 0 6461.635
1137.6683349609375 0 32481.893 c Ammonia loss 9
1138.6693115234375 0 17805.441
1139.674072265625 0 10384.2705
1152.6671142578125 0 6721.932
1153.6810302734375 0 4710.887
1154.6944580078125 0 918327.9 c 9
1155.6973876953125 0 627540.6
1156.6998291015625 0 211546.58
1157.70263671875 0 23828.857
1163.5218505859375 0 8507.245
1173.61083984375 0 240107.89 z 1
1174.6138916015625 0 167161.77
1175.61376953125 0 72261.05
1176.614013671875 0 8042.1006
1183.6624755859375 0 9068.2705
1184.664794921875 0 6802.268
1185.6553955078125 0 7997.7812
1186.6689453125 0 4552.3096
1189.6180419921875 0 3833.496 y 1
1195.729736328125 0 5719.7495
1196.7015380859375 0 7792.2534
1200.619873046875 0 7730.5347
1201.63671875 0 5060.031
1202.5377197265625 0 4299.9175
1203.5433349609375 0 3844.948
1211.6904296875 0 57890.688
1212.693603515625 0 46579.832
1213.6942138671875 0 14039.019
1214.63623046875 0 5632.3228
1215.623046875 0 13565.73
1216.61767578125 0 7519.5054
1217.6402587890625 0 4959.245
1224.7021484375 0 5046.812
1229.65283203125 0 4623.967
1230.635498046875 0 12791.18
1231.6341552734375 0 10141.384
1232.6490478515625 0 14505.598
1233.6536865234375 0 7557.5576
1239.720458984375 0 24328.873
1240.7232666015625 0 24230.29
1241.5693359375 0 2256.6035
1241.7205810546875 0 62195.246
1242.7144775390625 0 62647.562
1243.7012939453125 0 85313.34
1244.701171875 0 53924.96
1245.699462890625 0 17677.389
1246.68212890625 0 3665.8545
1247.6561279296875 0 14621.344
1248.6600341796875 0 10217.91
1249.660400390625 0 6093.182
1253.5517578125 0 4254.9253
1254.5712890625 0 14904.315
1255.568115234375 0 12382.343
1257.71484375 0 35668.72
1258.7161865234375 0 30404.451
1259.7247314453125 0 38860.766
1260.7303466796875 0 20971.744
1261.7376708984375 0 12521.358
1267.6865234375 0 6545.911
1268.540283203125 0 5619.129
1268.6812744140625 0 10292.953
1269.5611572265625 0 59942.906
1269.6959228515625 0 9145.803
1270.5628662109375 0 39914.543
1270.70068359375 0 16398.76
1271.559814453125 0 17705.223
1271.7008056640625 0 8922.341
1272.56201171875 0 6914.479
1275.7471923828125 0 3909.9556
1284.6817626953125 0 20650.652
1285.5594482421875 0 7912.102
1285.7066650390625 0 165143.83
1286.7021484375 0 979063.25
1287.7039794921875 0 698724.9
1288.7047119140625 0 337405.12
1289.5736083984375 0 19103.338
1289.704833984375 0 71298.7
1290.6983642578125 0 9822.581
1300.69091796875 0 21436.008
1301.7008056640625 0 39200.848
1302.71240234375 0 1007482.2
1303.719482421875 0 2477040
1304.72216796875 0 1709112.9
1305.082763671875 0 8853.731
1305.72265625 0 681609
1306.724609375 0 112742.01
1335.7099609375 0 5392.907
3075.175048828125 0 3959.3623
3181.2841796875 0 3573.6375

Spectrum Details

|  |  |
| --- | --- |
| Matched peaks? Matched peaks The total absolute number of peaks matched. Additionally in brackets the total fraction of peaks matched and the total number of peaks is shown. | 29 (10.32% of 281) |
| FDR? FDR The false discovery rate estimated for this peptide. It is calculated by matching all theoretical fragments with a non-integer shift with the raw peaks for this spectrum. This is done with 40 different shifts. The resulting percentage is the average number of annotated peaks over the number of annotated peaks with the correct spectrum. | 2.96% |
| Satellite FDR? Satellite FDR See the FDR for details on its calculation. This satellite ion specific FDR only contains the satellite ions (d/w) for I/L/J positions. | ∞ |
| PSM Score? PSM Score The PSM Score as given by Hecklib to this annotated spectrum. It is shown with three significant figures. | 269 |

#### Spectrum 5055? Spectrum 5055 The raw spectrum of this peptide as annotated by Hecklib. The fragments are coloured according to ion type (see legend). Any peaks with a star '\*' as text can be hovered over to see the full details, first the ion type second the mass shift type. By hovering over the amino acids in the peptide or ions in the legend the corresponding peaks are highlighted. By toggling the 'Unassigned' label you can turn the background (unassigned) peaks on or off in the plot. By updating the slider in the Ion legend you can update the spectrum to only show the top X% of the peaks with labels. The top X% means any peak that is within X% of the highest intensity. By dragging in the spectrum you can zoom in to a specific part of the spectrum and use 'Zoom Out' to get back to the original zoom level. The annotation of the spectrum is based on the given sequence in the peptides file and is done with different software so inconsistencies are likely. The peaks are annotated based on the given sequence, with 20 ppm tolerance.

Copy Data

##### Spectrum 5055 (TSV)

###### Preview

```
Loading example...
```

*Click on the button to copy the data to your clipboard.*

Mz MinMz MaxIntensity Max

WidthHeightPeptide font sizePeptide stroke widthSpectrum font sizeSpectrum stroke widthCompact peptide

Ion legend

wxyz

abcd

OtherUnassignedIon^Charge^_Position_Show for top:%

JFPPKPKDTJM

04.52e+59.04e+51.36e+61.81e+6

Zoom Out

y^+1^_2_y^+1^_3_y^+2^_7_y^+2^_8_y^+1^_4_y^+2^_9_z^+2^_9_y^+2^_9_c^+2^_10_y^+2^_10_y^+2^_10_z^+1^_5_y^+1^_5_c^+1^_6_c^+1^_6_y^+1^_6_w^+1^_7_c^+1^_7_c^+1^_7_z^+1^_7_y^+1^_7_c^+1^_8_c^+1^_8_y^+1^_8_c^+1^_9_c^+1^_9_y^+1^_9_c^+1^_10_c^+1^_10_y^+1^_10_z^+1^_10_

0691138220732765

Fragment Matches Table

Show background peaks

| Position | Ion type | Intensity | mz Theoretical | mz Error (Th) | mz Error (ppm) | Charge | Series Number |
| --- | --- | --- | --- | --- | --- | --- | --- |
| - | - | 1.145E+04 | 120.1 | - | - | 0 | - |
| - | - | 1349 | 121.1 | - | - | 0 | - |
| - | - | 1340 | 121.2 | - | - | 0 | - |
| - | - | 1139 | 121.8 | - | - | 0 | - |
| - | - | 1855 | 129.1 | - | - | 0 | - |
| - | - | 1245 | 136.6 | - | - | 0 | - |
| - | - | 1519 | 148.9 | - | - | 0 | - |
| - | - | 2136 | 148.9 | - | - | 0 | - |
| - | - | 2672 | 148.9 | - | - | 0 | - |
| - | - | 2590 | 148.9 | - | - | 0 | - |
| - | - | 3944 | 148.9 | - | - | 0 | - |
| - | - | 3789 | 148.9 | - | - | 0 | - |
| - | - | 4481 | 148.9 | - | - | 0 | - |
| - | - | 1.104E+04 | 148.9 | - | - | 0 | - |
| - | - | 2.129E+04 | 149 | - | - | 0 | - |
| - | - | 1.24E+04 | 149 | - | - | 0 | - |
| - | - | 5417 | 149 | - | - | 0 | - |
| - | - | 3643 | 149 | - | - | 0 | - |
| - | - | 3400 | 149 | - | - | 0 | - |
| - | - | 2469 | 149 | - | - | 0 | - |
| - | - | 1769 | 149 | - | - | 0 | - |
| - | - | 1611 | 149 | - | - | 0 | - |
| - | - | 1725 | 149 | - | - | 0 | - |
| - | - | 2510 | 149 | - | - | 0 | - |
| - | - | 1649 | 149 | - | - | 0 | - |
| - | - | 1397 | 151.2 | - | - | 0 | - |
| - | - | 4.923E+04 | 166.1 | - | - | 0 | - |
| - | - | 2874 | 167.1 | - | - | 0 | - |
| - | - | 1612 | 168.8 | - | - | 0 | - |
| - | - | 1378 | 170.8 | - | - | 0 | - |
| - | - | 3230 | 171.1 | - | - | 0 | - |
| - | - | 1.098E+04 | 173.1 | - | - | 0 | - |
| - | - | 2054 | 199.1 | - | - | 0 | - |
| - | - | 4269 | 201.1 | - | - | 0 | - |
| - | - | 1963 | 205.1 | - | - | 0 | - |
| - | - | 1832 | 207.6 | - | - | 0 | - |
| - | - | 1566 | 207.8 | - | - | 0 | - |
| - | - | 2280 | 217.1 | - | - | 0 | - |
| - | - | 3085 | 226.2 | - | - | 0 | - |
| - | - | 1464 | 232.1 | - | - | 0 | - |
| - | - | 1.286E+05 | 233.2 | - | - | 0 | - |
| - | - | 1.639E+04 | 234.2 | - | - | 0 | - |
| - | - | 3238 | 244.1 | - | - | 0 | - |
| - | - | 2051 | 245.1 | - | - | 0 | - |
| - | - | 8984 | 260.1 | - | - | 0 | - |
| - | - | 1.172E+05 | 261.2 | - | - | 0 | - |
| - | - | 1.783E+04 | 262.2 | - | - | 0 | - |
| - | - | 3130 | 274.1 | - | - | 0 | - |
| - | - | 1650 | 278.3 | - | - | 0 | - |
| 10 | y | 6918 | 279.1 | 0.004787 | 17.15 | +1 | 2 |
| - | - | 2.486E+04 | 323.2 | - | - | 0 | - |
| - | - | 5673 | 324.2 | - | - | 0 | - |
| - | - | 1.858E+04 | 358.2 | - | - | 0 | - |
| - | - | 4465 | 359.2 | - | - | 0 | - |
| 9 | y | 4393 | 380.2 | 0.004441 | 11.68 | +1 | 3 |
| - | - | 2349 | 399.2 | - | - | 0 | - |
| 5 | y | 6060 | 424.7 | 0.002712 | 6.386 | +2 | 7 |
| - | - | 2987 | 425.2 | - | - | 0 | - |
| - | - | 2645 | 425.3 | - | - | 0 | - |
| - | - | 2040 | 427.3 | - | - | 0 | - |
| - | - | 2508 | 433.2 | - | - | 0 | - |
| - | - | 1945 | 434.2 | - | - | 0 | - |
| - | - | 3.935E+04 | 439.3 | - | - | 0 | - |
| - | - | 1.67E+04 | 439.8 | - | - | 0 | - |
| 4 | y | 1.986E+04 | 473.3 | 0.003125 | 6.603 | +2 | 8 |
| - | - | 6025 | 473.8 | - | - | 0 | - |
| - | - | 4834 | 488.8 | - | - | 0 | - |
| - | - | 5.855E+04 | 489.8 | - | - | 0 | - |
| - | - | 2.743E+04 | 490.3 | - | - | 0 | - |
| - | - | 5574 | 490.8 | - | - | 0 | - |
| - | - | 1897 | 494.1 | - | - | 0 | - |
| 8 | y | 3157 | 495.2 | 0.004262 | 8.606 | +1 | 4 |
| - | - | 1852 | 499.3 | - | - | 0 | - |
| 3 | y | 3.779E+04 | 512.8 | 0.002777 | 5.416 | +2 | 9 |
| - | - | 1.818E+04 | 513.3 | - | - | 0 | - |
| 3 | z | 6283 | 513.8 | 0.006674 | 12.99 | +2 | 9 |
| - | - | 3997 | 514.3 | - | - | 0 | - |
| - | - | 2211 | 514.8 | - | - | 0 | - |
| - | - | 1.597E+04 | 520.8 | - | - | 0 | - |
| - | - | 4480 | 521.3 | - | - | 0 | - |
| 3 | y | 1.639E+06 | 521.8 | 0.002683 | 5.142 | +2 | 9 |
| - | - | 9.728E+05 | 522.3 | - | - | 0 | - |
| - | - | 3.287E+05 | 522.8 | - | - | 0 | - |
| - | - | 4.372E+04 | 523.3 | - | - | 0 | - |
| - | - | 1941 | 555.8 | - | - | 0 | - |
| - | - | 5193 | 565.4 | - | - | 0 | - |
| 10 | c | 2630 | 569.3 | 0.002616 | 4.595 | +2 | 10 |
| 2 | y | 2603 | 586.3 | 0.003543 | 6.044 | +2 | 10 |
| - | - | 2025 | 595 | - | - | 0 | - |
| - | - | 2323 | 595.3 | - | - | 0 | - |
| 2 | y | 4.229E+04 | 595.3 | 0.002167 | 3.641 | +2 | 10 |
| - | - | 2.868E+04 | 595.8 | - | - | 0 | - |
| - | - | 1.052E+04 | 596.3 | - | - | 0 | - |
| - | - | 1.341E+04 | 599.2 | - | - | 0 | - |
| - | - | 3339 | 600.2 | - | - | 0 | - |
| - | - | 1993 | 606.3 | - | - | 0 | - |
| 7 | z | 2.409E+04 | 607.3 | 0.004897 | 8.063 | +1 | 5 |
| - | - | 1.244E+05 | 608.3 | - | - | 0 | - |
| - | - | 3.344E+04 | 609.3 | - | - | 0 | - |
| - | - | 7302 | 610.3 | - | - | 0 | - |
| - | - | 2257 | 619.3 | - | - | 0 | - |
| - | - | 3.269E+04 | 619.9 | - | - | 0 | - |
| - | - | 2.005E+04 | 620.4 | - | - | 0 | - |
| - | - | 7100 | 620.9 | - | - | 0 | - |
| 7 | y | 3856 | 623.3 | 0.006986 | 11.21 | +1 | 5 |
| - | - | 2283 | 629.8 | - | - | 0 | - |
| - | - | 2019 | 635.3 | - | - | 0 | - |
| - | - | 3622 | 640.4 | - | - | 0 | - |
| - | - | 1.314E+04 | 642.9 | - | - | 0 | - |
| - | - | 1.138E+04 | 643.4 | - | - | 0 | - |
| - | - | 2556 | 643.9 | - | - | 0 | - |
| - | - | 3622 | 644.4 | - | - | 0 | - |
| - | - | 2805 | 644.9 | - | - | 0 | - |
| - | - | 5452 | 650.3 | - | - | 0 | - |
| - | - | 1.023E+04 | 650.8 | - | - | 0 | - |
| - | - | 2975 | 651.3 | - | - | 0 | - |
| - | - | 8082 | 651.4 | - | - | 0 | - |
| - | - | 8.358E+05 | 651.9 | - | - | 0 | - |
| - | - | 6.495E+05 | 652.4 | - | - | 0 | - |
| - | - | 3924 | 652.5 | - | - | 0 | - |
| - | - | 3.113E+05 | 652.9 | - | - | 0 | - |
| - | - | 2386 | 653.3 | - | - | 0 | - |
| - | - | 5.861E+04 | 653.4 | - | - | 0 | - |
| - | - | 7132 | 653.4 | - | - | 0 | - |
| - | - | 3851 | 654.4 | - | - | 0 | - |
| - | - | 2105 | 655.4 | - | - | 0 | - |
| - | - | 4173 | 663.4 | - | - | 0 | - |
| 6 | c | 6461 | 680.4 | 0.003829 | 5.628 | +1 | 6 |
| - | - | 2386 | 681.4 | - | - | 0 | - |
| - | - | 8809 | 695.4 | - | - | 0 | - |
| - | - | 5.465E+05 | 696.4 | - | - | 0 | - |
| 6 | c | 7.652E+05 | 697.4 | 0.001142 | 1.638 | +1 | 6 |
| - | - | 2.6E+05 | 698.4 | - | - | 0 | - |
| - | - | 4.794E+04 | 699.4 | - | - | 0 | - |
| - | - | 3580 | 700.4 | - | - | 0 | - |
| - | - | 6895 | 719.4 | - | - | 0 | - |
| 6 | y | 3.852E+04 | 720.4 | 0.004149 | 5.759 | +1 | 6 |
| - | - | 1.082E+04 | 721.4 | - | - | 0 | - |
| - | - | 4199 | 722.4 | - | - | 0 | - |
| - | - | 1.201E+04 | 748.4 | - | - | 0 | - |
| - | - | 4370 | 749.4 | - | - | 0 | - |
| - | - | 5684 | 764.4 | - | - | 0 | - |
| - | - | 8378 | 768.5 | - | - | 0 | - |
| - | - | 6795 | 769.5 | - | - | 0 | - |
| 5 | w | 2712 | 774.4 | 0.00335 | 4.326 | +1 | 7 |
| - | - | 2850 | 780.5 | - | - | 0 | - |
| - | - | 5.742E+04 | 781.5 | - | - | 0 | - |
| - | - | 3.624E+04 | 782.5 | - | - | 0 | - |
| - | - | 8775 | 783.5 | - | - | 0 | - |
| - | - | 1.129E+04 | 788.4 | - | - | 0 | - |
| - | - | 3282 | 789.5 | - | - | 0 | - |
| 7 | c | 2150 | 808.5 | 0.007056 | 8.728 | +1 | 7 |
| - | - | 6423 | 812.5 | - | - | 0 | - |
| - | - | 8184 | 823.5 | - | - | 0 | - |
| - | - | 4.784E+05 | 824.5 | - | - | 0 | - |
| 7 | c | 3.775E+05 | 825.5 | 0.002905 | 3.519 | +1 | 7 |
| - | - | 1.308E+05 | 826.5 | - | - | 0 | - |
| - | - | 2.466E+04 | 827.5 | - | - | 0 | - |
| 5 | z | 1.321E+05 | 832.4 | 0.004814 | 5.783 | +1 | 7 |
| - | - | 6.828E+04 | 833.4 | - | - | 0 | - |
| - | - | 2.135E+04 | 834.4 | - | - | 0 | - |
| - | - | 2970 | 835.4 | - | - | 0 | - |
| 5 | y | 1.177E+04 | 848.4 | 0.005255 | 6.194 | +1 | 7 |
| - | - | 6425 | 849.5 | - | - | 0 | - |
| - | - | 3231 | 868.5 | - | - | 0 | - |
| - | - | 2.759E+04 | 877.5 | - | - | 0 | - |
| - | - | 1.509E+04 | 878.5 | - | - | 0 | - |
| - | - | 8591 | 881.5 | - | - | 0 | - |
| - | - | 4426 | 882.5 | - | - | 0 | - |
| - | - | 6994 | 883.5 | - | - | 0 | - |
| - | - | 3780 | 884.5 | - | - | 0 | - |
| - | - | 8547 | 895.6 | - | - | 0 | - |
| - | - | 1.669E+04 | 896.5 | - | - | 0 | - |
| - | - | 9743 | 897.6 | - | - | 0 | - |
| - | - | 6476 | 898.6 | - | - | 0 | - |
| 8 | c | 3258 | 923.5 | 0.00488 | 5.284 | +1 | 8 |
| - | - | 5747 | 924.5 | - | - | 0 | - |
| - | - | 1.161E+04 | 925.5 | - | - | 0 | - |
| - | - | 2382 | 926.6 | - | - | 0 | - |
| - | - | 7092 | 938.5 | - | - | 0 | - |
| - | - | 4.606E+05 | 939.6 | - | - | 0 | - |
| 8 | c | 4.696E+05 | 940.6 | 0.002626 | 2.792 | +1 | 8 |
| - | - | 1.859E+05 | 941.6 | - | - | 0 | - |
| - | - | 4.969E+04 | 942.6 | - | - | 0 | - |
| - | - | 5064 | 943.6 | - | - | 0 | - |
| - | - | 2772 | 944.5 | - | - | 0 | - |
| 4 | y | 1.452E+05 | 945.5 | 0.005103 | 5.398 | +1 | 8 |
| - | - | 7.222E+04 | 946.5 | - | - | 0 | - |
| - | - | 3.015E+04 | 947.5 | - | - | 0 | - |
| - | - | 3066 | 948.5 | - | - | 0 | - |
| - | - | 2851 | 973.5 | - | - | 0 | - |
| - | - | 1.065E+04 | 978.6 | - | - | 0 | - |
| - | - | 7456 | 979.6 | - | - | 0 | - |
| - | - | 3400 | 980.6 | - | - | 0 | - |
| - | - | 2404 | 996.6 | - | - | 0 | - |
| - | - | 6210 | 997.6 | - | - | 0 | - |
| - | - | 8227 | 998.6 | - | - | 0 | - |
| 9 | c | 8150 | 1025 | 0.01191 | 11.63 | +1 | 9 |
| - | - | 4980 | 1026 | - | - | 0 | - |
| - | - | 2159 | 1027 | - | - | 0 | - |
| - | - | 2863 | 1040 | - | - | 0 | - |
| - | - | 4.663E+04 | 1041 | - | - | 0 | - |
| 9 | c | 3.901E+05 | 1042 | 0.0001338 | 0.1284 | +1 | 9 |
| 3 | y | 2.662E+05 | 1043 | 0.01728 | 16.58 | +1 | 9 |
| - | - | 1.506E+05 | 1044 | - | - | 0 | - |
| - | - | 5.797E+04 | 1045 | - | - | 0 | - |
| - | - | 8750 | 1046 | - | - | 0 | - |
| - | - | 4594 | 1070 | - | - | 0 | - |
| - | - | 2.707E+04 | 1071 | - | - | 0 | - |
| - | - | 1.472E+04 | 1072 | - | - | 0 | - |
| - | - | 5961 | 1073 | - | - | 0 | - |
| - | - | 2887 | 1084 | - | - | 0 | - |
| - | - | 4763 | 1103 | - | - | 0 | - |
| - | - | 7520 | 1104 | - | - | 0 | - |
| - | - | 2.779E+04 | 1111 | - | - | 0 | - |
| - | - | 2.112E+04 | 1112 | - | - | 0 | - |
| - | - | 6343 | 1113 | - | - | 0 | - |
| - | - | 1.38E+04 | 1130 | - | - | 0 | - |
| - | - | 6851 | 1131 | - | - | 0 | - |
| - | - | 2975 | 1132 | - | - | 0 | - |
| 10 | c | 2.273E+04 | 1138 | 0.000701 | 0.6161 | +1 | 10 |
| - | - | 1.649E+04 | 1139 | - | - | 0 | - |
| - | - | 6544 | 1140 | - | - | 0 | - |
| - | - | 3842 | 1153 | - | - | 0 | - |
| - | - | 6241 | 1154 | - | - | 0 | - |
| 10 | c | 6.68E+05 | 1155 | 3.077E-05 | 0.02665 | +1 | 10 |
| - | - | 4.245E+05 | 1156 | - | - | 0 | - |
| - | - | 1.523E+05 | 1157 | - | - | 0 | - |
| - | - | 1.835E+04 | 1158 | - | - | 0 | - |
| 2 | y | 2547 | 1173 | 0.0001698 | 0.1448 | +1 | 10 |
| 2 | z | 1.81E+05 | 1174 | 0.005067 | 4.317 | +1 | 10 |
| - | - | 1.221E+05 | 1175 | - | - | 0 | - |
| - | - | 5.049E+04 | 1176 | - | - | 0 | - |
| - | - | 6269 | 1177 | - | - | 0 | - |
| - | - | 8042 | 1184 | - | - | 0 | - |
| - | - | 7546 | 1185 | - | - | 0 | - |
| - | - | 6522 | 1186 | - | - | 0 | - |
| - | - | 3196 | 1187 | - | - | 0 | - |
| - | - | 3104 | 1196 | - | - | 0 | - |
| - | - | 5185 | 1197 | - | - | 0 | - |
| - | - | 3026 | 1200 | - | - | 0 | - |
| - | - | 7012 | 1201 | - | - | 0 | - |
| - | - | 2628 | 1202 | - | - | 0 | - |
| - | - | 4.278E+04 | 1212 | - | - | 0 | - |
| - | - | 3.167E+04 | 1213 | - | - | 0 | - |
| - | - | 1.412E+04 | 1214 | - | - | 0 | - |
| - | - | 4811 | 1215 | - | - | 0 | - |
| - | - | 8049 | 1216 | - | - | 0 | - |
| - | - | 6738 | 1217 | - | - | 0 | - |
| - | - | 3800 | 1218 | - | - | 0 | - |
| - | - | 3540 | 1219 | - | - | 0 | - |
| - | - | 4688 | 1224 | - | - | 0 | - |
| - | - | 5018 | 1225 | - | - | 0 | - |
| - | - | 2717 | 1230 | - | - | 0 | - |
| - | - | 7741 | 1231 | - | - | 0 | - |
| - | - | 7203 | 1232 | - | - | 0 | - |
| - | - | 6730 | 1233 | - | - | 0 | - |
| - | - | 3305 | 1234 | - | - | 0 | - |
| - | - | 2921 | 1239 | - | - | 0 | - |
| - | - | 1.959E+04 | 1240 | - | - | 0 | - |
| - | - | 1.561E+04 | 1241 | - | - | 0 | - |
| - | - | 4.043E+04 | 1242 | - | - | 0 | - |
| - | - | 3.928E+04 | 1243 | - | - | 0 | - |
| - | - | 6.291E+04 | 1244 | - | - | 0 | - |
| - | - | 4.356E+04 | 1245 | - | - | 0 | - |
| - | - | 1.578E+04 | 1246 | - | - | 0 | - |
| - | - | 2187 | 1247 | - | - | 0 | - |
| - | - | 1.212E+04 | 1248 | - | - | 0 | - |
| - | - | 1.204E+04 | 1249 | - | - | 0 | - |
| - | - | 5012 | 1250 | - | - | 0 | - |
| - | - | 2817 | 1252 | - | - | 0 | - |
| - | - | 2869 | 1256 | - | - | 0 | - |
| - | - | 2.769E+04 | 1258 | - | - | 0 | - |
| - | - | 1.859E+04 | 1259 | - | - | 0 | - |
| - | - | 3.009E+04 | 1260 | - | - | 0 | - |
| - | - | 1.843E+04 | 1261 | - | - | 0 | - |
| - | - | 7581 | 1262 | - | - | 0 | - |
| - | - | 3567 | 1268 | - | - | 0 | - |
| - | - | 3932 | 1269 | - | - | 0 | - |
| - | - | 7212 | 1270 | - | - | 0 | - |
| - | - | 1.277E+04 | 1271 | - | - | 0 | - |
| - | - | 5687 | 1272 | - | - | 0 | - |
| - | - | 4880 | 1273 | - | - | 0 | - |
| - | - | 3126 | 1276 | - | - | 0 | - |
| - | - | 2734 | 1284 | - | - | 0 | - |
| - | - | 1.203E+04 | 1285 | - | - | 0 | - |
| - | - | 1.201E+05 | 1286 | - | - | 0 | - |
| - | - | 7.025E+05 | 1287 | - | - | 0 | - |
| - | - | 5.277E+05 | 1288 | - | - | 0 | - |
| - | - | 3587 | 1288 | - | - | 0 | - |
| - | - | 2.458E+05 | 1289 | - | - | 0 | - |
| - | - | 5.42E+04 | 1290 | - | - | 0 | - |
| - | - | 4988 | 1291 | - | - | 0 | - |
| - | - | 1.046E+04 | 1301 | - | - | 0 | - |
| - | - | 2.675E+04 | 1302 | - | - | 0 | - |
| - | - | 7.081E+05 | 1303 | - | - | 0 | - |
| - | - | 1.791E+06 | 1304 | - | - | 0 | - |
| - | - | 1.214E+06 | 1305 | - | - | 0 | - |
| - | - | 6929 | 1305 | - | - | 0 | - |
| - | - | 5.069E+05 | 1306 | - | - | 0 | - |
| - | - | 7.853E+04 | 1307 | - | - | 0 | - |
| - | - | 3247 | 1337 | - | - | 0 | - |
| - | - | 2654 | 1740 | - | - | 0 | - |
| - | - | 2213 | 2396 | - | - | 0 | - |
| - | - | 2297 | 2737 | - | - | 0 | - |

m/z Charge Intensity FragmentType MassShift Position
120.08084106445312 0 11449.441
121.08451843261719 0 1349.2292
121.16686248779297 0 1340.1211
121.81196594238281 0 1139.3276
129.1022491455078 0 1855.1267
136.63946533203125 0 1245.4573
148.8899688720703 0 1519.115
148.8971710205078 0 2135.5967
148.90431213378906 0 2672.4192
148.91152954101562 0 2590.0537
148.91839599609375 0 3944.1855
148.92555236816406 0 3788.9146
148.93223571777344 0 4480.829
148.9397430419922 0 11037.724
148.95578002929688 0 21292.566
148.96334838867188 0 12400.1875
148.9707794189453 0 5417.1504
148.97763061523438 0 3642.588
148.98458862304688 0 3399.9863
148.99171447753906 0 2469.152
148.9990234375 0 1768.9303
149.00572204589844 0 1611.333
149.0130615234375 0 1724.9949
149.0200958251953 0 2510.4043
149.02691650390625 0 1648.8636
151.2084197998047 0 1397.1649
166.05316162109375 0 49231.996
167.05662536621094 0 2873.7358
168.78407287597656 0 1611.7793
170.80316162109375 0 1377.7778
171.0762939453125 0 3230.3718
173.09190368652344 0 10983.762
199.10789489746094 0 2053.59
201.12332153320312 0 4268.9653
205.08204650878906 0 1963.4812
207.5861358642578 0 1831.9249
207.80337524414062 0 1565.5674
217.13308715820312 0 2279.7378
226.15496826171875 0 3084.6707
232.12606811523438 0 1463.9596
233.1647491455078 0 128577.61
234.16807556152344 0 16390.65
244.13339233398438 0 3237.9165
245.1292266845703 0 2051.0786
260.1240539550781 0 8984.128
261.15948486328125 0 117215.09
262.162841796875 0 17830.92
274.1036376953125 0 3130.3118
278.2604064941406 0 1649.9795
279.1371765136719 0 6918.419 y 9
323.20782470703125 0 24864.967
324.2105712890625 0 5672.9653
358.2121276855469 0 18581.607
359.2164611816406 0 4465.179
380.18450927734375 0 4393.3945 y 8
399.2241516113281 0 2348.7932
424.731201171875 0 6059.5425 y 4
425.2320251464844 0 2987.4468
425.26385498046875 0 2644.9055
427.27349853515625 0 2040.4249
433.2286682128906 0 2508.0051
434.2322998046875 0 1945.0388
439.2609558105469 0 39347.836
439.7622375488281 0 16703.18
473.25799560546875 0 19861.482 y 3
473.760009765625 0 6024.7783
488.7782897949219 0 4833.6406
489.7846984863281 0 58554.504
490.2863464355469 0 27433.553
490.7877197265625 0 5574.141
494.126953125 0 1896.6864
495.2112731933594 0 3156.615 y 7
499.271240234375 0 1852.1165
512.7787475585938 0 37793.258 y Water loss 2
513.2802124023438 0 18175.258
513.778564453125 0 6283.3853 z 2
514.2739868164062 0 3996.7598
514.7685546875 0 2210.654
520.7761840820312 0 15973.067
521.2772216796875 0 4479.6367
521.783935546875 0 1639317.2 y 2
522.2852783203125 0 972833.5
522.784912109375 0 328725.47
523.2850341796875 0 43720.312
555.8414306640625 0 1940.5648
565.3829345703125 0 5192.6934
569.3343505859375 0 2629.823 c Ammonia loss 9
586.313720703125 0 2602.76 y Water loss 1
595.0433959960938 0 2025.4204
595.2634887695312 0 2323.1265
595.317626953125 0 42291.926 y 1
595.8197021484375 0 28677.594
596.31982421875 0 10517.931
599.2307739257812 0 13405.083
600.230224609375 0 3338.921
606.3181762695312 0 1993.4044
607.2881469726562 0 24089.96 z 6
608.2957153320312 0 124432.125
609.29833984375 0 33438.51
610.2969360351562 0 7301.643
619.2944946289062 0 2257.3909
619.8612670898438 0 32686.64
620.3621215820312 0 20047.295
620.8639526367188 0 7099.9585
623.3089599609375 0 3856.4324 y 6
629.843994140625 0 2283.2898
635.2842407226562 0 2019.2537
640.3698120117188 0 3621.8074
642.85302734375 0 13139.622
643.3533935546875 0 11382.011
643.8550415039062 0 2555.813
644.3528442382812 0 3622.1284
644.850830078125 0 2804.5767
650.3480834960938 0 5452.466
650.8493041992188 0 10229.658
651.283935546875 0 2975.4521
651.353515625 0 8081.577
651.8601684570312 0 835771.94
652.361572265625 0 649472.6
652.4892578125 0 3924.0837
652.861572265625 0 311267.47
653.298095703125 0 2386.1738
653.3617553710938 0 58612.89
653.4246826171875 0 7132.3438
654.4275512695312 0 3850.9077
655.4465942382812 0 2105.2566
663.385009765625 0 4172.7017
680.4091796875 0 6460.71 c Ammonia loss 5
681.4166259765625 0 2386.394
695.4230346679688 0 8808.676
696.431884765625 0 546513.8
697.4384155273438 0 765193.6 c 5
698.4417114257812 0 259995.12
699.4447631835938 0 47941.63
700.4496459960938 0 3579.9329
719.3511962890625 0 6895.146
720.35888671875 0 38517.918 y 5
721.3634643554688 0 10821.678
722.3594360351562 0 4198.5386
748.3528442382812 0 12014.674
749.3577880859375 0 4370.033
764.4346923828125 0 5683.6978
768.45361328125 0 8377.618
769.4577026367188 0 6795.037
774.36865234375 0 2712.0015 w 4
780.4586791992188 0 2849.928
781.5213012695312 0 57416.914
782.5252075195312 0 36242.17
783.5264282226562 0 8775.196
788.4458618164062 0 11289.887
789.4528198242188 0 3281.9275
808.5009155273438 0 2150.4927 c Ammonia loss 6
812.4642944335938 0 6422.7114
823.5183715820312 0 8183.997
824.526611328125 0 478390.06
825.5316162109375 0 377499.03 c 6
826.5354614257812 0 130828.48
827.5400390625 0 24663.398
832.435791015625 0 132073.72 z 4
833.4395751953125 0 68276.23
834.43994140625 0 21346.473
835.4387817382812 0 2970.1997
848.4549560546875 0 11772.906 y 4
849.4589233398438 0 6425.3213
868.4812622070312 0 3231.2405
877.5121459960938 0 27586.738
878.516845703125 0 15088.658
881.5304565429688 0 8591.464
882.53466796875 0 4425.5347
883.49267578125 0 6994.297
884.4979248046875 0 3779.6611
895.5558471679688 0 8547.211
896.5494995117188 0 16687.795
897.5528564453125 0 9742.742
898.5624389648438 0 6475.543
923.539794921875 0 3258.4036 c Ammonia loss 7
924.5377807617188 0 5746.5264
925.5476684570312 0 11612.644
926.5599975585938 0 2382.4124
938.5473022460938 0 7092.263
939.5533447265625 0 460561.7
940.558837890625 0 469590.88 c 7
941.5628662109375 0 185910.14
942.567626953125 0 49691.33
943.5779418945312 0 5063.7603
944.4987182617188 0 2772.204
945.507568359375 0 145231.03 y 3
946.5104370117188 0 72222.19
947.5103759765625 0 30150.562
948.5097045898438 0 3065.7603
973.5018310546875 0 2850.979
978.5647583007812 0 10653.812
979.5696411132812 0 7456.3813
980.5805053710938 0 3400.161
996.61083984375 0 2403.6592
997.5967407226562 0 6209.5386
998.6015014648438 0 8227.487
1024.5706787109375 0 8149.837 c Ammonia loss 8
1025.5673828125 0 4980.2373
1026.5594482421875 0 2159.3823
1039.60107421875 0 2863.3684
1040.6015625 0 46632.645
1041.6090087890625 0 390129.9 c 8
1042.572509765625 0 266207.28 y 2
1043.57080078125 0 150551.98
1044.564208984375 0 57967.477
1045.56103515625 0 8749.511
1069.5401611328125 0 4594.144
1070.5548095703125 0 27067.146
1071.55810546875 0 14720.174
1072.5592041015625 0 5961.2573
1083.59521484375 0 2887.1948
1102.5406494140625 0 4762.5884
1103.5419921875 0 7519.9844
1110.678466796875 0 27785.264
1111.6798095703125 0 21121.648
1112.6883544921875 0 6342.8267
1129.619140625 0 13795.791
1130.6236572265625 0 6851.4634
1131.61767578125 0 2974.8154
1137.6673583984375 0 22725.254 c Ammonia loss 9
1138.6697998046875 0 16491.414
1139.6702880859375 0 6543.5146
1152.6661376953125 0 3841.6277
1153.6768798828125 0 6240.7334
1154.6932373046875 0 668039 c 9
1155.6964111328125 0 424525.2
1156.6988525390625 0 152254.1
1157.701416015625 0 18350.822
1172.596923828125 0 2547.0017 y Ammonia loss 1
1173.6099853515625 0 180982.48 z 1
1174.6129150390625 0 122118.97
1175.6142578125 0 50487.434
1176.616455078125 0 6269.332
1183.6585693359375 0 8042.3726
1184.6591796875 0 7545.9814
1185.659423828125 0 6522.461
1186.664794921875 0 3195.6077
1195.7523193359375 0 3103.6448
1196.6942138671875 0 5185.1167
1199.6102294921875 0 3025.5994
1200.6214599609375 0 7012.222
1201.6273193359375 0 2628.2397
1211.6898193359375 0 42779.31
1212.6920166015625 0 31668.238
1213.7003173828125 0 14116.704
1214.639892578125 0 4811.0786
1215.629150390625 0 8048.8184
1216.626220703125 0 6737.9424
1217.6290283203125 0 3799.854
1218.6409912109375 0 3539.971
1223.7010498046875 0 4688.072
1224.7032470703125 0 5017.954
1229.67578125 0 2717.4143
1230.6383056640625 0 7740.7207
1231.636474609375 0 7202.6245
1232.64404296875 0 6730.4854
1233.6453857421875 0 3305.2358
1238.7073974609375 0 2920.6587
1239.718017578125 0 19588.064
1240.7232666015625 0 15605.877
1241.7208251953125 0 40427.746
1242.714599609375 0 39277.46
1243.6995849609375 0 62911.195
1244.6988525390625 0 43559.76
1245.69580078125 0 15777.141
1246.6883544921875 0 2187.3582
1247.65771484375 0 12121.261
1248.6585693359375 0 12041.286
1249.6593017578125 0 5011.7925
1251.676025390625 0 2817.0413
1255.7025146484375 0 2868.5117
1257.7147216796875 0 27687.988
1258.71630859375 0 18590.562
1259.725830078125 0 30092.19
1260.730712890625 0 18427.244
1261.7261962890625 0 7581.1616
1267.6864013671875 0 3567.1746
1268.6756591796875 0 3932.4202
1269.68798828125 0 7212.339
1270.6912841796875 0 12773.274
1271.689208984375 0 5686.5884
1272.6971435546875 0 4880.172
1275.7293701171875 0 3125.977
1283.68701171875 0 2733.503
1284.685546875 0 12026.17
1285.706298828125 0 120067.87
1286.7010498046875 0 702466.3
1287.7027587890625 0 527697.3
1288.0626220703125 0 3587.1218
1288.7025146484375 0 245755.6
1289.7005615234375 0 54198.95
1290.69775390625 0 4987.594
1300.6917724609375 0 10456.215
1301.7005615234375 0 26753.734
1302.711669921875 0 708097.8
1303.7188720703125 0 1790698.4
1304.7216796875 0 1213621.1
1305.087158203125 0 6929.1685
1305.7225341796875 0 506850.44
1306.723876953125 0 78529.3
1336.70947265625 0 3246.7488
1739.9178466796875 0 2654.1743
2396.1259765625 0 2212.9785
2737.16748046875 0 2297.431

Spectrum Details

|  |  |
| --- | --- |
| Matched peaks? Matched peaks The total absolute number of peaks matched. Additionally in brackets the total fraction of peaks matched and the total number of peaks is shown. | 31 (10.16% of 305) |
| FDR? FDR The false discovery rate estimated for this peptide. It is calculated by matching all theoretical fragments with a non-integer shift with the raw peaks for this spectrum. This is done with 40 different shifts. The resulting percentage is the average number of annotated peaks over the number of annotated peaks with the correct spectrum. | 2.23% |
| Satellite FDR? Satellite FDR See the FDR for details on its calculation. This satellite ion specific FDR only contains the satellite ions (d/w) for I/L/J positions. | ∞ |
| PSM Score? PSM Score The PSM Score as given by Hecklib to this annotated spectrum. It is shown with three significant figures. | 269 |

#### Spectrum 5239? Spectrum 5239 The raw spectrum of this peptide as annotated by Hecklib. The fragments are coloured according to ion type (see legend). Any peaks with a star '\*' as text can be hovered over to see the full details, first the ion type second the mass shift type. By hovering over the amino acids in the peptide or ions in the legend the corresponding peaks are highlighted. By toggling the 'Unassigned' label you can turn the background (unassigned) peaks on or off in the plot. By updating the slider in the Ion legend you can update the spectrum to only show the top X% of the peaks with labels. The top X% means any peak that is within X% of the highest intensity. By dragging in the spectrum you can zoom in to a specific part of the spectrum and use 'Zoom Out' to get back to the original zoom level. The annotation of the spectrum is based on the given sequence in the peptides file and is done with different software so inconsistencies are likely. The peaks are annotated based on the given sequence, with 20 ppm tolerance.

Copy Data

##### Spectrum 5239 (TSV)

###### Preview

```
Loading example...
```

*Click on the button to copy the data to your clipboard.*

Mz MinMz MaxIntensity Max

WidthHeightPeptide font sizePeptide stroke widthSpectrum font sizeSpectrum stroke widthCompact peptide

Ion legend

wxyz

abcd

OtherUnassignedIon^Charge^_Position_Show for top:%

JFPPKPKDTJM

05.43e+51.09e+61.63e+62.17e+6

Zoom Out

y^+1^_2_y^+1^_3_y^+2^_7_y^+2^_8_y^+2^_9_z^+2^_9_y^+2^_9_y^+2^_10_z^+1^_5_y^+1^_5_c^+1^_6_c^+1^_6_y^+1^_6_w^+1^_7_c^+1^_7_y^+1^_7_z^+1^_7_y^+1^_7_c^+1^_8_c^+1^_8_y^+1^_8_c^+1^_9_c^+1^_9_y^+1^_9_c^+1^_10_c^+1^_10_y^+1^_10_z^+1^_10_y^+1^_10_

0849169725463395

Fragment Matches Table

Show background peaks

| Position | Ion type | Intensity | mz Theoretical | mz Error (Th) | mz Error (ppm) | Charge | Series Number |
| --- | --- | --- | --- | --- | --- | --- | --- |
| - | - | 1321 | 120.1 | - | - | 0 | - |
| - | - | 1.595E+04 | 120.1 | - | - | 0 | - |
| - | - | 1619 | 121.1 | - | - | 0 | - |
| - | - | 4629 | 149 | - | - | 0 | - |
| - | - | 1779 | 150.8 | - | - | 0 | - |
| - | - | 5.065E+04 | 166.1 | - | - | 0 | - |
| - | - | 3153 | 167.1 | - | - | 0 | - |
| - | - | 2426 | 168 | - | - | 0 | - |
| - | - | 2249 | 169.1 | - | - | 0 | - |
| - | - | 1948 | 170.7 | - | - | 0 | - |
| - | - | 1.29E+04 | 173.5 | - | - | 0 | - |
| - | - | 2015 | 190.2 | - | - | 0 | - |
| - | - | 1982 | 194.5 | - | - | 0 | - |
| - | - | 2236 | 207.6 | - | - | 0 | - |
| - | - | 2045 | 209.4 | - | - | 0 | - |
| - | - | 3203 | 215.1 | - | - | 0 | - |
| - | - | 2827 | 226.2 | - | - | 0 | - |
| - | - | 1.292E+05 | 233.2 | - | - | 0 | - |
| - | - | 2.007E+04 | 234.2 | - | - | 0 | - |
| - | - | 2137 | 244.1 | - | - | 0 | - |
| - | - | 2150 | 245.1 | - | - | 0 | - |
| - | - | 1.251E+05 | 261.2 | - | - | 0 | - |
| - | - | 1.91E+04 | 262.2 | - | - | 0 | - |
| - | - | 2268 | 276.1 | - | - | 0 | - |
| 10 | y | 1.282E+04 | 279.1 | 0.005489 | 19.66 | +1 | 2 |
| - | - | 2.957E+04 | 323.2 | - | - | 0 | - |
| - | - | 3967 | 324.2 | - | - | 0 | - |
| - | - | 2392 | 339.5 | - | - | 0 | - |
| - | - | 1.53E+04 | 358.2 | - | - | 0 | - |
| - | - | 3184 | 359.2 | - | - | 0 | - |
| - | - | 2065 | 376.2 | - | - | 0 | - |
| 9 | y | 5554 | 380.2 | 0.006059 | 15.94 | +1 | 3 |
| - | - | 2054 | 393.5 | - | - | 0 | - |
| 5 | y | 1.048E+04 | 424.7 | 0.003658 | 8.614 | +2 | 7 |
| - | - | 4822 | 425.2 | - | - | 0 | - |
| - | - | 3630 | 427.3 | - | - | 0 | - |
| - | - | 5645 | 430.3 | - | - | 0 | - |
| - | - | 4.466E+04 | 439.3 | - | - | 0 | - |
| - | - | 2.343E+04 | 439.8 | - | - | 0 | - |
| - | - | 4095 | 440.3 | - | - | 0 | - |
| 4 | y | 1.833E+04 | 473.3 | 0.003583 | 7.57 | +2 | 8 |
| - | - | 9789 | 473.8 | - | - | 0 | - |
| - | - | 4103 | 474.3 | - | - | 0 | - |
| - | - | 2085 | 479 | - | - | 0 | - |
| - | - | 5747 | 488.8 | - | - | 0 | - |
| - | - | 5.752E+04 | 489.8 | - | - | 0 | - |
| - | - | 2.836E+04 | 490.3 | - | - | 0 | - |
| - | - | 7354 | 490.8 | - | - | 0 | - |
| 3 | y | 4.535E+04 | 512.8 | 0.003449 | 6.725 | +2 | 9 |
| - | - | 2.3E+04 | 513.3 | - | - | 0 | - |
| 3 | z | 1.055E+04 | 513.8 | 0.008932 | 17.39 | +2 | 9 |
| - | - | 9477 | 514.3 | - | - | 0 | - |
| - | - | 5841 | 514.8 | - | - | 0 | - |
| - | - | 1.626E+04 | 520.8 | - | - | 0 | - |
| - | - | 1.073E+04 | 521.3 | - | - | 0 | - |
| 3 | y | 1.795E+06 | 521.8 | 0.003537 | 6.779 | +2 | 9 |
| - | - | 9.827E+05 | 522.3 | - | - | 0 | - |
| - | - | 3.492E+05 | 522.8 | - | - | 0 | - |
| - | - | 4.621E+04 | 523.3 | - | - | 0 | - |
| - | - | 4328 | 548.4 | - | - | 0 | - |
| - | - | 5514 | 565.4 | - | - | 0 | - |
| - | - | 2807 | 566.4 | - | - | 0 | - |
| - | - | 3141 | 569.8 | - | - | 0 | - |
| 2 | y | 5.073E+04 | 595.3 | 0.003571 | 5.999 | +2 | 10 |
| - | - | 3.305E+04 | 595.8 | - | - | 0 | - |
| - | - | 1.264E+04 | 596.3 | - | - | 0 | - |
| 7 | z | 2.921E+04 | 607.3 | 0.005812 | 9.571 | +1 | 5 |
| - | - | 1.474E+05 | 608.3 | - | - | 0 | - |
| - | - | 4.101E+04 | 609.3 | - | - | 0 | - |
| - | - | 1.176E+04 | 610.3 | - | - | 0 | - |
| - | - | 3135 | 615.3 | - | - | 0 | - |
| - | - | 3.565E+04 | 619.9 | - | - | 0 | - |
| - | - | 1.709E+04 | 620.4 | - | - | 0 | - |
| - | - | 9621 | 620.9 | - | - | 0 | - |
| 7 | y | 5754 | 623.3 | 0.007291 | 11.7 | +1 | 5 |
| - | - | 3551 | 630.4 | - | - | 0 | - |
| - | - | 2556 | 640.4 | - | - | 0 | - |
| - | - | 1.455E+04 | 642.9 | - | - | 0 | - |
| - | - | 5989 | 643.4 | - | - | 0 | - |
| - | - | 7578 | 643.9 | - | - | 0 | - |
| - | - | 3852 | 644.3 | - | - | 0 | - |
| - | - | 3103 | 644.9 | - | - | 0 | - |
| - | - | 4642 | 650.3 | - | - | 0 | - |
| - | - | 1.357E+04 | 650.9 | - | - | 0 | - |
| - | - | 5684 | 651.4 | - | - | 0 | - |
| - | - | 9.272E+05 | 651.9 | - | - | 0 | - |
| - | - | 6.906E+05 | 652.4 | - | - | 0 | - |
| - | - | 3.408E+05 | 652.9 | - | - | 0 | - |
| - | - | 6.195E+04 | 653.4 | - | - | 0 | - |
| - | - | 1.04E+04 | 653.4 | - | - | 0 | - |
| - | - | 3821 | 654.4 | - | - | 0 | - |
| - | - | 6512 | 663.4 | - | - | 0 | - |
| 6 | c | 5490 | 680.4 | 0.002975 | 4.372 | +1 | 6 |
| - | - | 3398 | 681.4 | - | - | 0 | - |
| - | - | 6801 | 695.4 | - | - | 0 | - |
| - | - | 6.204E+05 | 696.4 | - | - | 0 | - |
| 6 | c | 9.128E+05 | 697.4 | 0.000227 | 0.3254 | +1 | 6 |
| - | - | 3.073E+05 | 698.4 | - | - | 0 | - |
| - | - | 5.687E+04 | 699.4 | - | - | 0 | - |
| - | - | 3344 | 700.4 | - | - | 0 | - |
| - | - | 9339 | 719.4 | - | - | 0 | - |
| 6 | y | 4.652E+04 | 720.4 | 0.004942 | 6.861 | +1 | 6 |
| - | - | 1.68E+04 | 721.4 | - | - | 0 | - |
| - | - | 3549 | 722.4 | - | - | 0 | - |
| - | - | 1.266E+04 | 748.4 | - | - | 0 | - |
| - | - | 6156 | 749.4 | - | - | 0 | - |
| - | - | 3044 | 761.4 | - | - | 0 | - |
| - | - | 1.013E+04 | 768.5 | - | - | 0 | - |
| - | - | 6088 | 769.5 | - | - | 0 | - |
| 5 | w | 4421 | 774.4 | 0.005242 | 6.769 | +1 | 7 |
| - | - | 8410 | 780.5 | - | - | 0 | - |
| - | - | 3996 | 781.4 | - | - | 0 | - |
| - | - | 7.365E+04 | 781.5 | - | - | 0 | - |
| - | - | 4.132E+04 | 782.5 | - | - | 0 | - |
| - | - | 1.003E+04 | 783.5 | - | - | 0 | - |
| - | - | 1.558E+04 | 788.4 | - | - | 0 | - |
| - | - | 7426 | 812.5 | - | - | 0 | - |
| - | - | 2788 | 822.5 | - | - | 0 | - |
| - | - | 1.008E+04 | 823.5 | - | - | 0 | - |
| - | - | 5.331E+05 | 824.5 | - | - | 0 | - |
| 7 | c | 4.355E+05 | 825.5 | 0.002111 | 2.558 | +1 | 7 |
| - | - | 1.35E+05 | 826.5 | - | - | 0 | - |
| - | - | 2.428E+04 | 827.5 | - | - | 0 | - |
| 5 | y | 3092 | 830.4 | 0.01641 | 19.76 | +1 | 7 |
| 5 | z | 1.466E+05 | 832.4 | 0.005607 | 6.736 | +1 | 7 |
| - | - | 8.164E+04 | 833.4 | - | - | 0 | - |
| - | - | 2.174E+04 | 834.4 | - | - | 0 | - |
| 5 | y | 1.189E+04 | 848.4 | 0.00733 | 8.639 | +1 | 7 |
| - | - | 6252 | 849.5 | - | - | 0 | - |
| - | - | 2693 | 868.5 | - | - | 0 | - |
| - | - | 2.353E+04 | 877.5 | - | - | 0 | - |
| - | - | 1.145E+04 | 878.5 | - | - | 0 | - |
| - | - | 8363 | 881.5 | - | - | 0 | - |
| - | - | 4929 | 882.5 | - | - | 0 | - |
| - | - | 5581 | 883.5 | - | - | 0 | - |
| - | - | 1.099E+04 | 895.6 | - | - | 0 | - |
| - | - | 1.755E+04 | 896.6 | - | - | 0 | - |
| - | - | 1.07E+04 | 897.6 | - | - | 0 | - |
| - | - | 6587 | 898.6 | - | - | 0 | - |
| - | - | 3449 | 899.6 | - | - | 0 | - |
| 8 | c | 3265 | 923.5 | 0.007382 | 7.994 | +1 | 8 |
| - | - | 4715 | 924.5 | - | - | 0 | - |
| - | - | 1.064E+04 | 925.6 | - | - | 0 | - |
| - | - | 7241 | 926.6 | - | - | 0 | - |
| - | - | 5774 | 938.5 | - | - | 0 | - |
| - | - | 5.425E+05 | 939.6 | - | - | 0 | - |
| 8 | c | 5.613E+05 | 940.6 | 0.001711 | 1.819 | +1 | 8 |
| - | - | 2.216E+05 | 941.6 | - | - | 0 | - |
| - | - | 4.734E+04 | 942.6 | - | - | 0 | - |
| - | - | 4106 | 943.6 | - | - | 0 | - |
| - | - | 4780 | 944.5 | - | - | 0 | - |
| 4 | y | 1.5E+05 | 945.5 | 0.006202 | 6.559 | +1 | 8 |
| - | - | 8.307E+04 | 946.5 | - | - | 0 | - |
| - | - | 2.939E+04 | 947.5 | - | - | 0 | - |
| - | - | 4132 | 973.5 | - | - | 0 | - |
| - | - | 1.17E+04 | 978.6 | - | - | 0 | - |
| - | - | 1.024E+04 | 979.6 | - | - | 0 | - |
| - | - | 1.141E+04 | 997.6 | - | - | 0 | - |
| - | - | 9538 | 998.6 | - | - | 0 | - |
| - | - | 3159 | 999.6 | - | - | 0 | - |
| 9 | c | 7383 | 1025 | 0.01399 | 13.65 | +1 | 9 |
| - | - | 5407 | 1026 | - | - | 0 | - |
| - | - | 4161 | 1040 | - | - | 0 | - |
| - | - | 5.716E+04 | 1041 | - | - | 0 | - |
| 9 | c | 4.629E+05 | 1042 | 0.0008428 | 0.8091 | +1 | 9 |
| 3 | y | 2.995E+05 | 1043 | 0.01875 | 17.98 | +1 | 9 |
| - | - | 1.717E+05 | 1044 | - | - | 0 | - |
| - | - | 7.229E+04 | 1045 | - | - | 0 | - |
| - | - | 1.367E+04 | 1046 | - | - | 0 | - |
| - | - | 6278 | 1070 | - | - | 0 | - |
| - | - | 3.032E+04 | 1071 | - | - | 0 | - |
| - | - | 1.982E+04 | 1072 | - | - | 0 | - |
| - | - | 7614 | 1073 | - | - | 0 | - |
| - | - | 9095 | 1103 | - | - | 0 | - |
| - | - | 3.096E+04 | 1111 | - | - | 0 | - |
| - | - | 2.297E+04 | 1112 | - | - | 0 | - |
| - | - | 7343 | 1113 | - | - | 0 | - |
| - | - | 1.639E+04 | 1130 | - | - | 0 | - |
| - | - | 9849 | 1131 | - | - | 0 | - |
| - | - | 7843 | 1132 | - | - | 0 | - |
| 10 | c | 2.76E+04 | 1138 | 9.062E-05 | 0.07965 | +1 | 10 |
| - | - | 1.55E+04 | 1139 | - | - | 0 | - |
| - | - | 4470 | 1140 | - | - | 0 | - |
| - | - | 4256 | 1153 | - | - | 0 | - |
| - | - | 7491 | 1154 | - | - | 0 | - |
| 10 | c | 8.093E+05 | 1155 | 0.001129 | 0.9781 | +1 | 10 |
| - | - | 5.614E+05 | 1156 | - | - | 0 | - |
| - | - | 1.886E+05 | 1157 | - | - | 0 | - |
| - | - | 2.159E+04 | 1158 | - | - | 0 | - |
| 2 | y | 3618 | 1173 | 0.001024 | 0.8735 | +1 | 10 |
| 2 | z | 2.209E+05 | 1174 | 0.005677 | 4.837 | +1 | 10 |
| - | - | 1.526E+05 | 1175 | - | - | 0 | - |
| - | - | 5.97E+04 | 1176 | - | - | 0 | - |
| - | - | 1.256E+04 | 1177 | - | - | 0 | - |
| - | - | 1.069E+04 | 1184 | - | - | 0 | - |
| - | - | 5857 | 1185 | - | - | 0 | - |
| - | - | 7948 | 1186 | - | - | 0 | - |
| 2 | y | 3694 | 1190 | 0.007217 | 6.066 | +1 | 10 |
| - | - | 3917 | 1196 | - | - | 0 | - |
| - | - | 8037 | 1197 | - | - | 0 | - |
| - | - | 5726 | 1201 | - | - | 0 | - |
| - | - | 4894 | 1202 | - | - | 0 | - |
| - | - | 4.498E+04 | 1212 | - | - | 0 | - |
| - | - | 3.913E+04 | 1213 | - | - | 0 | - |
| - | - | 1.504E+04 | 1214 | - | - | 0 | - |
| - | - | 6950 | 1215 | - | - | 0 | - |
| - | - | 1.18E+04 | 1216 | - | - | 0 | - |
| - | - | 7611 | 1217 | - | - | 0 | - |
| - | - | 4571 | 1218 | - | - | 0 | - |
| - | - | 3723 | 1219 | - | - | 0 | - |
| - | - | 4530 | 1224 | - | - | 0 | - |
| - | - | 7193 | 1225 | - | - | 0 | - |
| - | - | 3740 | 1230 | - | - | 0 | - |
| - | - | 9900 | 1231 | - | - | 0 | - |
| - | - | 9728 | 1232 | - | - | 0 | - |
| - | - | 8845 | 1233 | - | - | 0 | - |
| - | - | 8575 | 1234 | - | - | 0 | - |
| - | - | 3080 | 1235 | - | - | 0 | - |
| - | - | 2.538E+04 | 1240 | - | - | 0 | - |
| - | - | 1.844E+04 | 1241 | - | - | 0 | - |
| - | - | 5.298E+04 | 1242 | - | - | 0 | - |
| - | - | 4.961E+04 | 1243 | - | - | 0 | - |
| - | - | 6.902E+04 | 1244 | - | - | 0 | - |
| - | - | 4.302E+04 | 1245 | - | - | 0 | - |
| - | - | 1.829E+04 | 1246 | - | - | 0 | - |
| - | - | 9436 | 1248 | - | - | 0 | - |
| - | - | 9472 | 1249 | - | - | 0 | - |
| - | - | 3643 | 1250 | - | - | 0 | - |
| - | - | 4672 | 1255 | - | - | 0 | - |
| - | - | 3302 | 1256 | - | - | 0 | - |
| - | - | 2.908E+04 | 1258 | - | - | 0 | - |
| - | - | 2.386E+04 | 1259 | - | - | 0 | - |
| - | - | 3.197E+04 | 1260 | - | - | 0 | - |
| - | - | 1.795E+04 | 1261 | - | - | 0 | - |
| - | - | 6709 | 1262 | - | - | 0 | - |
| - | - | 3316 | 1268 | - | - | 0 | - |
| - | - | 3570 | 1269 | - | - | 0 | - |
| - | - | 4740 | 1269 | - | - | 0 | - |
| - | - | 1.496E+04 | 1270 | - | - | 0 | - |
| - | - | 7628 | 1270 | - | - | 0 | - |
| - | - | 1.339E+04 | 1271 | - | - | 0 | - |
| - | - | 9227 | 1271 | - | - | 0 | - |
| - | - | 4608 | 1272 | - | - | 0 | - |
| - | - | 7082 | 1272 | - | - | 0 | - |
| - | - | 4760 | 1277 | - | - | 0 | - |
| - | - | 3831 | 1278 | - | - | 0 | - |
| - | - | 1.58E+04 | 1285 | - | - | 0 | - |
| - | - | 1.393E+05 | 1286 | - | - | 0 | - |
| - | - | 8.095E+05 | 1287 | - | - | 0 | - |
| - | - | 5.982E+05 | 1288 | - | - | 0 | - |
| - | - | 2.886E+05 | 1289 | - | - | 0 | - |
| - | - | 5.895E+04 | 1290 | - | - | 0 | - |
| - | - | 6318 | 1291 | - | - | 0 | - |
| - | - | 2.339E+04 | 1301 | - | - | 0 | - |
| - | - | 3.539E+04 | 1302 | - | - | 0 | - |
| - | - | 8.418E+05 | 1303 | - | - | 0 | - |
| - | - | 2.152E+06 | 1304 | - | - | 0 | - |
| - | - | 1.464E+06 | 1305 | - | - | 0 | - |
| - | - | 5.876E+05 | 1306 | - | - | 0 | - |
| - | - | 9.29E+04 | 1307 | - | - | 0 | - |
| - | - | 3595 | 1336 | - | - | 0 | - |
| - | - | 3033 | 1828 | - | - | 0 | - |
| - | - | 3544 | 3361 | - | - | 0 | - |

m/z Charge Intensity FragmentType MassShift Position
120.07769012451172 0 1320.6727
120.08106231689453 0 15948.378
121.08460998535156 0 1618.6335
148.95370483398438 0 4629.13
150.7542266845703 0 1778.5695
166.05352783203125 0 50649.19
167.056640625 0 3153.0288
168.04905700683594 0 2425.779
169.1336212158203 0 2249.258
170.7303466796875 0 1947.7133
173.45106506347656 0 12900.621
190.18875122070312 0 2015.3574
194.53956604003906 0 1981.6077
207.56393432617188 0 2235.9119
209.4246368408203 0 2044.7394
215.13877868652344 0 3203.041
226.15469360351562 0 2826.8638
233.165283203125 0 129184.484
234.16859436035156 0 20067.084
244.13307189941406 0 2136.8945
245.12940979003906 0 2150.4214
261.1600036621094 0 125103.24
262.163330078125 0 19102.398
276.12841796875 0 2268.4153
279.13787841796875 0 12824.362 y 9
323.20843505859375 0 29567.955
324.2105407714844 0 3966.9153
339.5367431640625 0 2391.6304
358.21295166015625 0 15301.246
359.2160949707031 0 3184.4358
376.17333984375 0 2065.4216
380.1861267089844 0 5554.4014 y 8
393.5392150878906 0 2054.4917
424.7321472167969 0 10481.479 y 4
425.2331237792969 0 4822.313
427.27850341796875 0 3629.932
430.255859375 0 5645.4067
439.26165771484375 0 44662.195
439.7631530761719 0 23431.21
440.26422119140625 0 4094.5989
473.2584533691406 0 18326.158 y 3
473.75872802734375 0 9788.546
474.25946044921875 0 4102.8833
478.96014404296875 0 2085.0894
488.775390625 0 5746.619
489.7854309082031 0 57515.66
490.28662109375 0 28358.676
490.787841796875 0 7354.128
512.7794189453125 0 45345.973 y Water loss 2
513.2803955078125 0 23002.678
513.7808227539062 0 10545.669 z 2
514.2740478515625 0 9477.172
514.773193359375 0 5840.859
520.7767333984375 0 16262.318
521.2784423828125 0 10729.244
521.7847900390625 0 1794500.4 y 2
522.2860107421875 0 982677.1
522.7857666015625 0 349162.16
523.2852172851562 0 46207.074
548.3563232421875 0 4327.623
565.3816528320312 0 5513.975
566.3858032226562 0 2807.1643
569.8419189453125 0 3141.2993
595.3190307617188 0 50732.918 y 1
595.8201904296875 0 33048.445
596.3201904296875 0 12644.618
607.2890625 0 29208.525 z 6
608.2968139648438 0 147422.14
609.2991943359375 0 41012.188
610.2977294921875 0 11761.805
615.346435546875 0 3134.954
619.86181640625 0 35654.53
620.363037109375 0 17088.54
620.8645629882812 0 9620.739
623.3092651367188 0 5753.556 y 6
630.3577270507812 0 3551.0554
640.3630981445312 0 2555.51
642.8554077148438 0 14553.416
643.35693359375 0 5988.9834
643.8566284179688 0 7577.6665
644.3489990234375 0 3852.007
644.8543701171875 0 3103.4866
650.3462524414062 0 4642.346
650.8511352539062 0 13573.404
651.3543701171875 0 5683.687
651.8610229492188 0 927167.2
652.3623657226562 0 690580.7
652.8626708984375 0 340827.66
653.3627319335938 0 61950.473
653.4263916015625 0 10400.383
654.4306030273438 0 3820.627
663.3854370117188 0 6512.4746
680.4100341796875 0 5490.411 c Ammonia loss 5
681.4122314453125 0 3397.8604
695.4259033203125 0 6800.6685
696.4326171875 0 620363.4
697.4393310546875 0 912781.56 c 5
698.4426879882812 0 307347.6
699.4456787109375 0 56867.34
700.4429321289062 0 3343.7083
719.3524780273438 0 9338.524
720.3596801757812 0 46520.137 y 5
721.36328125 0 16802.92
722.3590087890625 0 3549.1401
748.3551635742188 0 12660.862
749.3561401367188 0 6155.8364
761.3649291992188 0 3044.1548
768.4584350585938 0 10125.2295
769.4547729492188 0 6087.6284
774.3705444335938 0 4420.9873 w 4
780.46044921875 0 8410.028
781.4481201171875 0 3995.56
781.522216796875 0 73647.375
782.5254516601562 0 41322.98
783.5289306640625 0 10027.07
788.44580078125 0 15579.792
812.4673461914062 0 7426.2656
822.5059204101562 0 2788.0254
823.5178833007812 0 10078.442
824.5275268554688 0 533110.06
825.5324096679688 0 435507.62 c 6
826.5364379882812 0 135005.64
827.5399169921875 0 24276.717
830.4227294921875 0 3092.1917 y Water loss 4
832.4365844726562 0 146641.66 z 4
833.4405517578125 0 81642.49
834.4404296875 0 21738.312
848.45703125 0 11886.4 y 4
849.4615478515625 0 6252.0137
868.4771728515625 0 2692.7861
877.513916015625 0 23531.275
878.5172119140625 0 11446.367
881.525146484375 0 8363.241
882.5363159179688 0 4929.39
883.49365234375 0 5580.772
895.5601196289062 0 10994.163
896.5513916015625 0 17549.326
897.5521850585938 0 10696.644
898.5613403320312 0 6586.997
899.5681762695312 0 3449.142
923.5422973632812 0 3264.6206 c Ammonia loss 7
924.541748046875 0 4714.514
925.550048828125 0 10643.669
926.5563354492188 0 7240.783
938.5464477539062 0 5774.192
939.554443359375 0 542474.56
940.5597534179688 0 561304.06 c 7
941.5635375976562 0 221596.02
942.5684204101562 0 47335.52
943.5806884765625 0 4106.108
944.50146484375 0 4780.1304
945.5086669921875 0 149991.3 y 3
946.5110473632812 0 83065.734
947.5109252929688 0 29388.79
973.5022583007812 0 4131.622
978.5648803710938 0 11704.191
979.5728759765625 0 10238.505
997.5971069335938 0 11413.45
998.6016845703125 0 9538.177
999.6146240234375 0 3158.9087
1024.568603515625 0 7383.016 c Ammonia loss 8
1025.5712890625 0 5406.507
1039.5987548828125 0 4161.246
1040.602294921875 0 57162.746
1041.6099853515625 0 462943.56 c 8
1042.573974609375 0 299474.3 y 2
1043.5711669921875 0 171702.84
1044.5634765625 0 72293.89
1045.56103515625 0 13668.332
1069.5462646484375 0 6278.4355
1070.555419921875 0 30320.482
1071.560302734375 0 19823.219
1072.559814453125 0 7613.9224
1102.5367431640625 0 9094.554
1110.678466796875 0 30961.135
1111.6829833984375 0 22974.434
1112.6849365234375 0 7343.0547
1129.617431640625 0 16390.484
1130.6243896484375 0 9849.368
1131.623779296875 0 7842.5103
1137.666748046875 0 27602.451 c Ammonia loss 9
1138.6693115234375 0 15503.304
1139.6593017578125 0 4470.27
1152.6793212890625 0 4256.128
1153.6788330078125 0 7490.5205
1154.6943359375 0 809340.1 c 9
1155.6971435546875 0 561367.75
1156.6998291015625 0 188577.34
1157.7020263671875 0 21586.307
1172.5960693359375 0 3618.1152 y Ammonia loss 1
1173.610595703125 0 220911.92 z 1
1174.6142578125 0 152564.78
1175.614013671875 0 59697.97
1176.61767578125 0 12563.596
1183.6600341796875 0 10690.219
1184.656982421875 0 5857.454
1185.663818359375 0 7948.1646
1189.630859375 0 3694.3367 y 1
1195.7415771484375 0 3917.4658
1196.692626953125 0 8036.895
1200.6278076171875 0 5726.439
1201.6397705078125 0 4893.782
1211.691162109375 0 44982.996
1212.6934814453125 0 39127.26
1213.7001953125 0 15036.547
1214.6431884765625 0 6949.883
1215.6268310546875 0 11804.979
1216.6282958984375 0 7611.211
1217.628173828125 0 4570.93
1218.6390380859375 0 3723.038
1223.6954345703125 0 4529.562
1224.7059326171875 0 7192.608
1229.65380859375 0 3740.2273
1230.638427734375 0 9900.4
1231.634765625 0 9727.76
1232.6448974609375 0 8845.247
1233.65478515625 0 8574.946
1234.6658935546875 0 3079.656
1239.7218017578125 0 25381.582
1240.723388671875 0 18438.863
1241.7220458984375 0 52976.914
1242.7135009765625 0 49611.53
1243.701171875 0 69018.516
1244.6988525390625 0 43021.254
1245.69580078125 0 18285.668
1247.6572265625 0 9435.805
1248.6629638671875 0 9472.103
1249.654541015625 0 3643.188
1254.5709228515625 0 4671.671
1255.5595703125 0 3302.3208
1257.7119140625 0 29082.674
1258.71435546875 0 23856.31
1259.7276611328125 0 31971.605
1260.728271484375 0 17948.963
1261.7318115234375 0 6709.2573
1267.6878662109375 0 3315.7346
1268.5379638671875 0 3570.0576
1268.6785888671875 0 4739.6533
1269.5577392578125 0 14964.691
1269.6971435546875 0 7627.728
1270.563720703125 0 13392.538
1270.6939697265625 0 9226.788
1271.552001953125 0 4608.2183
1271.687744140625 0 7081.9077
1276.7315673828125 0 4760.417
1277.7247314453125 0 3830.8364
1284.6903076171875 0 15796.653
1285.70703125 0 139318.36
1286.70166015625 0 809465.94
1287.70361328125 0 598168.2
1288.70263671875 0 288600.53
1289.702880859375 0 58945.24
1290.6976318359375 0 6318.289
1300.6937255859375 0 23385.182
1301.7008056640625 0 35388.05
1302.712158203125 0 841756.4
1303.719482421875 0 2152247.8
1304.72216796875 0 1463749.8
1305.722900390625 0 587620.2
1306.724609375 0 92901.63
1335.71044921875 0 3595.3064
1828.2760009765625 0 3033.412
3361.24072265625 0 3544.0461

Spectrum Details

|  |  |
| --- | --- |
| Matched peaks? Matched peaks The total absolute number of peaks matched. Additionally in brackets the total fraction of peaks matched and the total number of peaks is shown. | 29 (11.03% of 263) |
| FDR? FDR The false discovery rate estimated for this peptide. It is calculated by matching all theoretical fragments with a non-integer shift with the raw peaks for this spectrum. This is done with 40 different shifts. The resulting percentage is the average number of annotated peaks over the number of annotated peaks with the correct spectrum. | 2.55% |
| Satellite FDR? Satellite FDR See the FDR for details on its calculation. This satellite ion specific FDR only contains the satellite ions (d/w) for I/L/J positions. | ∞ |
| PSM Score? PSM Score The PSM Score as given by Hecklib to this annotated spectrum. It is shown with three significant figures. | 269 |

#### Spectrum 5301? Spectrum 5301 The raw spectrum of this peptide as annotated by Hecklib. The fragments are coloured according to ion type (see legend). Any peaks with a star '\*' as text can be hovered over to see the full details, first the ion type second the mass shift type. By hovering over the amino acids in the peptide or ions in the legend the corresponding peaks are highlighted. By toggling the 'Unassigned' label you can turn the background (unassigned) peaks on or off in the plot. By updating the slider in the Ion legend you can update the spectrum to only show the top X% of the peaks with labels. The top X% means any peak that is within X% of the highest intensity. By dragging in the spectrum you can zoom in to a specific part of the spectrum and use 'Zoom Out' to get back to the original zoom level. The annotation of the spectrum is based on the given sequence in the peptides file and is done with different software so inconsistencies are likely. The peaks are annotated based on the given sequence, with 20 ppm tolerance.

Copy Data

##### Spectrum 5301 (TSV)

###### Preview

```
Loading example...
```

*Click on the button to copy the data to your clipboard.*

Mz MinMz MaxIntensity Max

WidthHeightPeptide font sizePeptide stroke widthSpectrum font sizeSpectrum stroke widthCompact peptide

Ion legend

wxyz

abcd

OtherUnassignedIon^Charge^_Position_Show for top:%

JFPPKPKDTJM

03.56e+57.12e+51.07e+61.42e+6

Zoom Out

y^+1^_2_y^+1^_3_y^+2^_7_y^+2^_8_y^+1^_4_y^+2^_9_z^+2^_9_y^+2^_9_y^+2^_10_z^+1^_5_y^+1^_5_c^+1^_6_c^+1^_6_y^+1^_6_c^+1^_7_z^+1^_7_y^+1^_7_c^+1^_8_c^+1^_8_y^+1^_8_y^+1^_9_c^+1^_9_y^+1^_9_c^+1^_10_c^+1^_10_y^+1^_10_z^+1^_10_

0882176326453526

Fragment Matches Table

Show background peaks

| Position | Ion type | Intensity | mz Theoretical | mz Error (Th) | mz Error (ppm) | Charge | Series Number |
| --- | --- | --- | --- | --- | --- | --- | --- |
| - | - | 7762 | 120.1 | - | - | 0 | - |
| - | - | 1158 | 134.5 | - | - | 0 | - |
| - | - | 1239 | 136.1 | - | - | 0 | - |
| - | - | 1265 | 141.6 | - | - | 0 | - |
| - | - | 3.196E+04 | 166.1 | - | - | 0 | - |
| - | - | 1232 | 166.1 | - | - | 0 | - |
| - | - | 1544 | 167.1 | - | - | 0 | - |
| - | - | 1196 | 174.3 | - | - | 0 | - |
| - | - | 1335 | 180.7 | - | - | 0 | - |
| - | - | 1453 | 184.4 | - | - | 0 | - |
| - | - | 1569 | 189.3 | - | - | 0 | - |
| - | - | 1848 | 226.2 | - | - | 0 | - |
| - | - | 8.72E+04 | 233.2 | - | - | 0 | - |
| - | - | 1.113E+04 | 234.2 | - | - | 0 | - |
| - | - | 2189 | 244.1 | - | - | 0 | - |
| - | - | 1408 | 245.8 | - | - | 0 | - |
| - | - | 7.562E+04 | 261.2 | - | - | 0 | - |
| - | - | 1.057E+04 | 262.2 | - | - | 0 | - |
| 10 | y | 4794 | 279.1 | 0.005184 | 18.57 | +1 | 2 |
| - | - | 2.014E+04 | 323.2 | - | - | 0 | - |
| - | - | 4789 | 324.2 | - | - | 0 | - |
| - | - | 3905 | 355.1 | - | - | 0 | - |
| - | - | 1.106E+04 | 358.2 | - | - | 0 | - |
| - | - | 2725 | 359.2 | - | - | 0 | - |
| - | - | 1574 | 365.4 | - | - | 0 | - |
| 9 | y | 2774 | 380.2 | 0.005906 | 15.53 | +1 | 3 |
| - | - | 1511 | 410.1 | - | - | 0 | - |
| 5 | y | 4490 | 424.7 | 0.003628 | 8.542 | +2 | 7 |
| - | - | 3947 | 425.2 | - | - | 0 | - |
| - | - | 1992 | 437.3 | - | - | 0 | - |
| - | - | 2.397E+04 | 439.3 | - | - | 0 | - |
| - | - | 1.46E+04 | 439.8 | - | - | 0 | - |
| - | - | 2788 | 440.3 | - | - | 0 | - |
| 4 | y | 1.406E+04 | 473.3 | 0.003247 | 6.861 | +2 | 8 |
| - | - | 2868 | 473.8 | - | - | 0 | - |
| - | - | 4433 | 488.8 | - | - | 0 | - |
| - | - | 3.662E+04 | 489.8 | - | - | 0 | - |
| - | - | 2.185E+04 | 490.3 | - | - | 0 | - |
| - | - | 5027 | 490.8 | - | - | 0 | - |
| 8 | y | 2191 | 495.2 | 0.002492 | 5.032 | +1 | 4 |
| - | - | 1779 | 496.2 | - | - | 0 | - |
| - | - | 1503 | 499.8 | - | - | 0 | - |
| - | - | 1819 | 505.8 | - | - | 0 | - |
| 3 | y | 2.891E+04 | 512.8 | 0.002655 | 5.178 | +2 | 9 |
| - | - | 1.322E+04 | 513.3 | - | - | 0 | - |
| 3 | z | 7023 | 513.8 | 0.006186 | 12.04 | +2 | 9 |
| - | - | 6803 | 514.3 | - | - | 0 | - |
| - | - | 9610 | 520.8 | - | - | 0 | - |
| - | - | 4990 | 521.3 | - | - | 0 | - |
| - | - | 2104 | 521.4 | - | - | 0 | - |
| 3 | y | 1.158E+06 | 521.8 | 0.003415 | 6.546 | +2 | 9 |
| - | - | 6.764E+05 | 522.3 | - | - | 0 | - |
| - | - | 2.19E+05 | 522.8 | - | - | 0 | - |
| - | - | 2.925E+04 | 523.3 | - | - | 0 | - |
| - | - | 2565 | 548.4 | - | - | 0 | - |
| - | - | 1782 | 550.6 | - | - | 0 | - |
| - | - | 3743 | 565.4 | - | - | 0 | - |
| 2 | y | 2.952E+04 | 595.3 | 0.003449 | 5.794 | +2 | 10 |
| - | - | 2.042E+04 | 595.8 | - | - | 0 | - |
| - | - | 9695 | 596.3 | - | - | 0 | - |
| 7 | z | 1.714E+04 | 607.3 | 0.005934 | 9.772 | +1 | 5 |
| - | - | 1.008E+05 | 608.3 | - | - | 0 | - |
| - | - | 2.905E+04 | 609.3 | - | - | 0 | - |
| - | - | 6992 | 610.3 | - | - | 0 | - |
| - | - | 2.126E+04 | 619.9 | - | - | 0 | - |
| - | - | 1.144E+04 | 620.4 | - | - | 0 | - |
| - | - | 3991 | 620.9 | - | - | 0 | - |
| 7 | y | 4836 | 623.3 | 0.00546 | 8.76 | +1 | 5 |
| - | - | 2585 | 640.4 | - | - | 0 | - |
| - | - | 7923 | 642.9 | - | - | 0 | - |
| - | - | 6992 | 643.4 | - | - | 0 | - |
| - | - | 4193 | 643.9 | - | - | 0 | - |
| - | - | 3287 | 644.4 | - | - | 0 | - |
| - | - | 6415 | 650.9 | - | - | 0 | - |
| - | - | 3866 | 651.4 | - | - | 0 | - |
| - | - | 5.592E+05 | 651.9 | - | - | 0 | - |
| - | - | 4.36E+05 | 652.4 | - | - | 0 | - |
| - | - | 2.147E+05 | 652.9 | - | - | 0 | - |
| - | - | 3.588E+04 | 653.4 | - | - | 0 | - |
| - | - | 5053 | 653.4 | - | - | 0 | - |
| - | - | 2566 | 654.4 | - | - | 0 | - |
| - | - | 3495 | 663.4 | - | - | 0 | - |
| 6 | c | 3256 | 680.4 | 0.001876 | 2.757 | +1 | 6 |
| - | - | 2861 | 681.4 | - | - | 0 | - |
| - | - | 1806 | 694.4 | - | - | 0 | - |
| - | - | 6134 | 695.4 | - | - | 0 | - |
| - | - | 4.196E+05 | 696.4 | - | - | 0 | - |
| 6 | c | 5.963E+05 | 697.4 | 0.0004711 | 0.6755 | +1 | 6 |
| - | - | 2.053E+05 | 698.4 | - | - | 0 | - |
| - | - | 3.776E+04 | 699.4 | - | - | 0 | - |
| - | - | 2572 | 700.5 | - | - | 0 | - |
| - | - | 6990 | 719.4 | - | - | 0 | - |
| 6 | y | 2.674E+04 | 720.4 | 0.005613 | 7.793 | +1 | 6 |
| - | - | 8084 | 721.4 | - | - | 0 | - |
| - | - | 3174 | 722.4 | - | - | 0 | - |
| - | - | 1958 | 747.3 | - | - | 0 | - |
| - | - | 9837 | 748.4 | - | - | 0 | - |
| - | - | 3130 | 749.4 | - | - | 0 | - |
| - | - | 2866 | 764.4 | - | - | 0 | - |
| - | - | 8622 | 768.5 | - | - | 0 | - |
| - | - | 4330 | 769.5 | - | - | 0 | - |
| - | - | 2233 | 780.5 | - | - | 0 | - |
| - | - | 4.62E+04 | 781.5 | - | - | 0 | - |
| - | - | 2.74E+04 | 782.5 | - | - | 0 | - |
| - | - | 6621 | 783.5 | - | - | 0 | - |
| - | - | 9631 | 788.4 | - | - | 0 | - |
| - | - | 4770 | 789.4 | - | - | 0 | - |
| - | - | 6366 | 812.5 | - | - | 0 | - |
| - | - | 8068 | 823.5 | - | - | 0 | - |
| - | - | 3.643E+05 | 824.5 | - | - | 0 | - |
| 7 | c | 2.889E+05 | 825.5 | 0.002294 | 2.779 | +1 | 7 |
| - | - | 9.833E+04 | 826.5 | - | - | 0 | - |
| - | - | 1.74E+04 | 827.5 | - | - | 0 | - |
| 5 | z | 9.996E+04 | 832.4 | 0.005424 | 6.516 | +1 | 7 |
| - | - | 5.573E+04 | 833.4 | - | - | 0 | - |
| - | - | 1.706E+04 | 834.4 | - | - | 0 | - |
| 5 | y | 8477 | 848.4 | 0.005438 | 6.409 | +1 | 7 |
| - | - | 4227 | 849.5 | - | - | 0 | - |
| - | - | 2821 | 868.5 | - | - | 0 | - |
| - | - | 2.076E+04 | 877.5 | - | - | 0 | - |
| - | - | 1.121E+04 | 878.5 | - | - | 0 | - |
| - | - | 5694 | 881.5 | - | - | 0 | - |
| - | - | 3123 | 882.5 | - | - | 0 | - |
| - | - | 4776 | 883.5 | - | - | 0 | - |
| - | - | 2324 | 884.5 | - | - | 0 | - |
| - | - | 8632 | 895.6 | - | - | 0 | - |
| - | - | 1.291E+04 | 896.6 | - | - | 0 | - |
| - | - | 5228 | 897.6 | - | - | 0 | - |
| - | - | 4331 | 898.6 | - | - | 0 | - |
| 8 | c | 2385 | 923.5 | 0.004819 | 5.218 | +1 | 8 |
| - | - | 2729 | 924.5 | - | - | 0 | - |
| - | - | 7558 | 925.6 | - | - | 0 | - |
| - | - | 3052 | 926.6 | - | - | 0 | - |
| - | - | 5144 | 938.5 | - | - | 0 | - |
| - | - | 3.531E+05 | 939.6 | - | - | 0 | - |
| 8 | c | 3.661E+05 | 940.6 | 0.001894 | 2.013 | +1 | 8 |
| - | - | 1.525E+05 | 941.6 | - | - | 0 | - |
| - | - | 3.329E+04 | 942.6 | - | - | 0 | - |
| - | - | 3067 | 944.5 | - | - | 0 | - |
| 4 | y | 1.073E+05 | 945.5 | 0.005714 | 6.043 | +1 | 8 |
| - | - | 5.482E+04 | 946.5 | - | - | 0 | - |
| - | - | 1.513E+04 | 947.5 | - | - | 0 | - |
| - | - | 2983 | 948.5 | - | - | 0 | - |
| - | - | 1828 | 973.5 | - | - | 0 | - |
| - | - | 8156 | 978.6 | - | - | 0 | - |
| - | - | 7670 | 979.6 | - | - | 0 | - |
| - | - | 2219 | 980.6 | - | - | 0 | - |
| - | - | 7563 | 997.6 | - | - | 0 | - |
| - | - | 6470 | 998.6 | - | - | 0 | - |
| - | - | 3064 | 999.6 | - | - | 0 | - |
| 3 | y | 6077 | 1025 | 0.01454 | 14.19 | +1 | 9 |
| - | - | 2595 | 1026 | - | - | 0 | - |
| - | - | 3285 | 1040 | - | - | 0 | - |
| - | - | 3.469E+04 | 1041 | - | - | 0 | - |
| 9 | c | 2.977E+05 | 1042 | 0.0007207 | 0.6919 | +1 | 9 |
| 3 | y | 1.849E+05 | 1043 | 0.01899 | 18.21 | +1 | 9 |
| - | - | 1.069E+05 | 1044 | - | - | 0 | - |
| - | - | 4.642E+04 | 1045 | - | - | 0 | - |
| - | - | 7151 | 1046 | - | - | 0 | - |
| - | - | 1986 | 1068 | - | - | 0 | - |
| - | - | 4349 | 1070 | - | - | 0 | - |
| - | - | 2.249E+04 | 1071 | - | - | 0 | - |
| - | - | 1.277E+04 | 1072 | - | - | 0 | - |
| - | - | 6299 | 1073 | - | - | 0 | - |
| - | - | 2366 | 1084 | - | - | 0 | - |
| - | - | 3977 | 1103 | - | - | 0 | - |
| - | - | 4511 | 1104 | - | - | 0 | - |
| - | - | 2599 | 1105 | - | - | 0 | - |
| - | - | 2.075E+04 | 1111 | - | - | 0 | - |
| - | - | 1.479E+04 | 1112 | - | - | 0 | - |
| - | - | 2609 | 1113 | - | - | 0 | - |
| - | - | 6084 | 1130 | - | - | 0 | - |
| - | - | 7833 | 1131 | - | - | 0 | - |
| - | - | 1940 | 1132 | - | - | 0 | - |
| 10 | c | 1.552E+04 | 1138 | 0.0003977 | 0.3495 | +1 | 10 |
| - | - | 1.013E+04 | 1139 | - | - | 0 | - |
| - | - | 3857 | 1140 | - | - | 0 | - |
| - | - | 4237 | 1153 | - | - | 0 | - |
| - | - | 4786 | 1154 | - | - | 0 | - |
| 10 | c | 5.199E+05 | 1155 | 0.0005191 | 0.4495 | +1 | 10 |
| - | - | 3.615E+05 | 1156 | - | - | 0 | - |
| - | - | 1.293E+05 | 1157 | - | - | 0 | - |
| - | - | 1.426E+04 | 1158 | - | - | 0 | - |
| 2 | y | 1943 | 1172 | 0.01322 | 11.29 | +1 | 10 |
| 2 | z | 1.423E+05 | 1174 | 0.005433 | 4.629 | +1 | 10 |
| - | - | 9.414E+04 | 1175 | - | - | 0 | - |
| - | - | 2990 | 1175 | - | - | 0 | - |
| - | - | 4.073E+04 | 1176 | - | - | 0 | - |
| - | - | 6836 | 1177 | - | - | 0 | - |
| - | - | 3318 | 1184 | - | - | 0 | - |
| - | - | 4750 | 1185 | - | - | 0 | - |
| - | - | 2022 | 1186 | - | - | 0 | - |
| - | - | 2763 | 1187 | - | - | 0 | - |
| - | - | 3040 | 1196 | - | - | 0 | - |
| - | - | 3097 | 1197 | - | - | 0 | - |
| - | - | 2430 | 1198 | - | - | 0 | - |
| - | - | 5792 | 1201 | - | - | 0 | - |
| - | - | 3.144E+04 | 1212 | - | - | 0 | - |
| - | - | 1.872E+04 | 1213 | - | - | 0 | - |
| - | - | 9183 | 1214 | - | - | 0 | - |
| - | - | 2516 | 1215 | - | - | 0 | - |
| - | - | 7963 | 1216 | - | - | 0 | - |
| - | - | 6451 | 1217 | - | - | 0 | - |
| - | - | 3018 | 1218 | - | - | 0 | - |
| - | - | 2150 | 1219 | - | - | 0 | - |
| - | - | 3996 | 1224 | - | - | 0 | - |
| - | - | 6394 | 1225 | - | - | 0 | - |
| - | - | 2361 | 1226 | - | - | 0 | - |
| - | - | 2686 | 1230 | - | - | 0 | - |
| - | - | 5472 | 1231 | - | - | 0 | - |
| - | - | 4509 | 1232 | - | - | 0 | - |
| - | - | 8235 | 1233 | - | - | 0 | - |
| - | - | 3964 | 1234 | - | - | 0 | - |
| - | - | 2096 | 1238 | - | - | 0 | - |
| - | - | 1.86E+04 | 1240 | - | - | 0 | - |
| - | - | 1.094E+04 | 1241 | - | - | 0 | - |
| - | - | 3.293E+04 | 1242 | - | - | 0 | - |
| - | - | 3.642E+04 | 1243 | - | - | 0 | - |
| - | - | 5.13E+04 | 1244 | - | - | 0 | - |
| - | - | 3.097E+04 | 1245 | - | - | 0 | - |
| - | - | 1.053E+04 | 1246 | - | - | 0 | - |
| - | - | 2842 | 1247 | - | - | 0 | - |
| - | - | 1.036E+04 | 1248 | - | - | 0 | - |
| - | - | 7570 | 1249 | - | - | 0 | - |
| - | - | 2543 | 1250 | - | - | 0 | - |
| - | - | 3392 | 1255 | - | - | 0 | - |
| - | - | 1.97E+04 | 1258 | - | - | 0 | - |
| - | - | 1.597E+04 | 1259 | - | - | 0 | - |
| - | - | 1.951E+04 | 1260 | - | - | 0 | - |
| - | - | 1.454E+04 | 1261 | - | - | 0 | - |
| - | - | 5868 | 1262 | - | - | 0 | - |
| - | - | 2566 | 1268 | - | - | 0 | - |
| - | - | 4703 | 1269 | - | - | 0 | - |
| - | - | 3879 | 1270 | - | - | 0 | - |
| - | - | 6415 | 1270 | - | - | 0 | - |
| - | - | 2616 | 1271 | - | - | 0 | - |
| - | - | 1.102E+04 | 1271 | - | - | 0 | - |
| - | - | 5544 | 1272 | - | - | 0 | - |
| - | - | 2223 | 1276 | - | - | 0 | - |
| - | - | 3931 | 1284 | - | - | 0 | - |
| - | - | 8151 | 1285 | - | - | 0 | - |
| - | - | 9.309E+04 | 1286 | - | - | 0 | - |
| - | - | 5.461E+05 | 1287 | - | - | 0 | - |
| - | - | 4.122E+05 | 1288 | - | - | 0 | - |
| - | - | 1.983E+05 | 1289 | - | - | 0 | - |
| - | - | 3.942E+04 | 1290 | - | - | 0 | - |
| - | - | 6478 | 1291 | - | - | 0 | - |
| - | - | 1.198E+04 | 1301 | - | - | 0 | - |
| - | - | 2.008E+04 | 1302 | - | - | 0 | - |
| - | - | 5.564E+05 | 1303 | - | - | 0 | - |
| - | - | 1.41E+06 | 1304 | - | - | 0 | - |
| - | - | 9.551E+05 | 1305 | - | - | 0 | - |
| - | - | 3.725E+05 | 1306 | - | - | 0 | - |
| - | - | 6.087E+04 | 1307 | - | - | 0 | - |
| - | - | 2065 | 2238 | - | - | 0 | - |
| - | - | 2934 | 3075 | - | - | 0 | - |
| - | - | 1898 | 3309 | - | - | 0 | - |
| - | - | 2345 | 3491 | - | - | 0 | - |

m/z Charge Intensity FragmentType MassShift Position
120.0810317993164 0 7762.379
134.52401733398438 0 1158.4436
136.07586669921875 0 1239.1934
141.5788116455078 0 1264.57
166.05352783203125 0 31960.17
166.1212615966797 0 1232.0924
167.056640625 0 1544.4587
174.28379821777344 0 1196.0894
180.7252197265625 0 1335.0197
184.38731384277344 0 1452.6472
189.3246612548828 0 1569.4197
226.15576171875 0 1848.194
233.1651611328125 0 87200.22
234.16851806640625 0 11134.622
244.13352966308594 0 2189.0264
245.78138732910156 0 1407.5098
261.1600036621094 0 75621.98
262.16314697265625 0 10572.141
279.1375732421875 0 4794.044 y 9
323.2083435058594 0 20140.668
324.2110290527344 0 4789.4736
355.07000732421875 0 3904.7686
358.2125244140625 0 11058.736
359.2155456542969 0 2724.911
365.42633056640625 0 1573.6481
380.18597412109375 0 2774.4436 y 8
410.08294677734375 0 1510.7222
424.73211669921875 0 4490.013 y 4
425.2333679199219 0 3946.64
437.2872314453125 0 1992.143
439.26165771484375 0 23966.537
439.76251220703125 0 14603.404
440.26617431640625 0 2788.4731
473.25811767578125 0 14063.262 y 3
473.7611999511719 0 2868.037
488.7778625488281 0 4433.0625
489.7852783203125 0 36617.105
490.2864685058594 0 21846.37
490.7881164550781 0 5027.075
495.2095031738281 0 2190.5217 y 7
496.2115173339844 0 1779.393
499.79498291015625 0 1503.3596
505.77972412109375 0 1818.7128
512.7786254882812 0 28912.885 y Water loss 2
513.2799682617188 0 13218.095
513.778076171875 0 7023.1396 z 2
514.2725219726562 0 6803.0005
520.7770385742188 0 9609.844
521.2786865234375 0 4990.305
521.3750610351562 0 2103.7056
521.78466796875 0 1157502.8 y 2
522.2858276367188 0 676379.1
522.78564453125 0 218996.16
523.28466796875 0 29249.13
548.355712890625 0 2564.9163
550.5948486328125 0 1781.7982
565.3832397460938 0 3743.1377
595.3189086914062 0 29515.35 y 1
595.8204956054688 0 20420.336
596.3197631835938 0 9694.586
607.2891845703125 0 17137.621 z 6
608.2964477539062 0 100829.38
609.2987670898438 0 29047.527
610.2988891601562 0 6991.7
619.8621215820312 0 21263.973
620.3626098632812 0 11441.72
620.8622436523438 0 3990.5405
623.3074340820312 0 4836.1616 y 6
640.3719482421875 0 2585.3281
642.8546142578125 0 7923.2285
643.3554077148438 0 6992.164
643.85546875 0 4193.2637
644.3528442382812 0 3287.2632
650.8534545898438 0 6414.5527
651.3547973632812 0 3865.692
651.8607788085938 0 559241.9
652.3622436523438 0 436005
652.862548828125 0 214675.69
653.3618774414062 0 35883.938
653.4268188476562 0 5053.325
654.4335327148438 0 2566.3723
663.3840942382812 0 3495.204
680.4111328125 0 3256.364 c Ammonia loss 5
681.4170532226562 0 2861.3389
694.4121704101562 0 1805.6073
695.4244384765625 0 6134.4604
696.4324951171875 0 419555.84
697.4390869140625 0 596253.06 c 5
698.4423217773438 0 205282.19
699.446044921875 0 37762.547
700.452392578125 0 2572.302
719.3510131835938 0 6990.045
720.3603515625 0 26744.094 y 5
721.3623657226562 0 8084.276
722.357177734375 0 3173.614
747.33642578125 0 1958.3486
748.3558349609375 0 9837.173
749.3525390625 0 3130.1526
764.4379272460938 0 2866.2578
768.45849609375 0 8622.095
769.4569702148438 0 4330.1035
780.4612426757812 0 2233.0815
781.5219116210938 0 46199.457
782.5250854492188 0 27399.992
783.52978515625 0 6621.282
788.4464721679688 0 9631.333
789.445068359375 0 4770.159
812.4655151367188 0 6365.853
823.5176391601562 0 8067.757
824.5272827148438 0 364274.56
825.5322265625 0 288889.56 c 6
826.5363159179688 0 98328.19
827.5390014648438 0 17399.217
832.4364013671875 0 99963.76 z 4
833.4398193359375 0 55729.52
834.43994140625 0 17063.463
848.4551391601562 0 8477.046 y 4
849.4580688476562 0 4227.3804
868.4791259765625 0 2821.2898
877.5139770507812 0 20764.441
878.5172119140625 0 11209.038
881.5303344726562 0 5693.9106
882.5328369140625 0 3123.0388
883.4939575195312 0 4776.351
884.498291015625 0 2324.0662
895.5545043945312 0 8631.585
896.5509643554688 0 12913.054
897.5504760742188 0 5227.854
898.5586547851562 0 4331.074
923.5397338867188 0 2384.6128 c Ammonia loss 7
924.5325317382812 0 2728.5508
925.5509033203125 0 7557.726
926.557861328125 0 3052.1436
938.548583984375 0 5143.954
939.5540771484375 0 353090.56
940.5595703125 0 366065.84 c 7
941.5632934570312 0 152507.2
942.5684204101562 0 33286.348
944.5010986328125 0 3066.5747
945.5081787109375 0 107255.86 y 3
946.5108642578125 0 54821.125
947.509033203125 0 15126.766
948.5117797851562 0 2983.4453
973.5098266601562 0 1828.4774
978.5641479492188 0 8156.137
979.5777587890625 0 7669.7754
980.5735473632812 0 2218.6755
997.5922241210938 0 7563.0127
998.6012573242188 0 6469.902
999.611328125 0 3064.4187
1024.5592041015625 0 6076.6333 y Water loss 2
1025.5748291015625 0 2594.8792
1039.6007080078125 0 3284.813
1040.6016845703125 0 34691.414
1041.60986328125 0 297749.75 c 8
1042.57421875 0 184891.8 y 2
1043.5718994140625 0 106894.71
1044.5638427734375 0 46423.23
1045.5643310546875 0 7151.3745
1067.629150390625 0 1986.2083
1069.544921875 0 4348.9404
1070.5550537109375 0 22491.01
1071.560546875 0 12772.042
1072.5599365234375 0 6298.666
1083.5943603515625 0 2366.3313
1102.536376953125 0 3977.3567
1103.540283203125 0 4511.403
1104.5384521484375 0 2598.505
1110.6781005859375 0 20747.99
1111.6800537109375 0 14786.173
1112.6806640625 0 2608.7607
1129.6231689453125 0 6083.8193
1130.62255859375 0 7833.499
1131.61669921875 0 1940.0795
1137.666259765625 0 15520.324 c Ammonia loss 9
1138.67333984375 0 10125.427
1139.671875 0 3857.0762
1152.6771240234375 0 4236.65
1153.6800537109375 0 4785.666
1154.6937255859375 0 519877.5 c 9
1155.697265625 0 361489.34
1156.6998291015625 0 129293.81
1157.702392578125 0 14264.163
1171.599853515625 0 1942.6372 y Water loss 1
1173.6103515625 0 142291.97 z 1
1174.6136474609375 0 94144.31
1174.767333984375 0 2990.0356
1175.613525390625 0 40731.727
1176.617431640625 0 6836.469
1183.6636962890625 0 3317.8547
1184.65869140625 0 4749.5596
1185.6729736328125 0 2022.085
1186.6624755859375 0 2763.085
1195.73046875 0 3039.6042
1196.7021484375 0 3096.9543
1197.72314453125 0 2430.027
1200.6192626953125 0 5792.399
1211.6898193359375 0 31440.105
1212.6917724609375 0 18722.395
1213.69970703125 0 9183.452
1214.6356201171875 0 2516.1052
1215.62744140625 0 7963.115
1216.6273193359375 0 6451.3647
1217.625732421875 0 3018.1772
1218.6290283203125 0 2150.2078
1223.6961669921875 0 3996.317
1224.69921875 0 6393.7603
1225.708251953125 0 2360.6443
1229.6451416015625 0 2686.1665
1230.6363525390625 0 5471.5986
1231.6395263671875 0 4509.1787
1232.640380859375 0 8235.115
1233.65478515625 0 3963.751
1237.69677734375 0 2096.1355
1239.7208251953125 0 18601.924
1240.721923828125 0 10938.898
1241.7218017578125 0 32929.105
1242.7147216796875 0 36424.918
1243.7000732421875 0 51298.527
1244.7000732421875 0 30967.645
1245.6929931640625 0 10532.528
1246.695556640625 0 2842.3728
1247.659423828125 0 10364.557
1248.6622314453125 0 7570.443
1249.6605224609375 0 2542.945
1254.5694580078125 0 3391.7646
1257.7122802734375 0 19697.416
1258.7139892578125 0 15966.754
1259.72265625 0 19508.797
1260.7322998046875 0 14544.628
1261.728515625 0 5867.5605
1267.6763916015625 0 2565.5254
1268.682373046875 0 4703.3745
1269.5469970703125 0 3878.7422
1269.692138671875 0 6415.2456
1270.544189453125 0 2615.795
1270.690185546875 0 11019.345
1271.6822509765625 0 5543.881
1275.7095947265625 0 2223.0496
1283.682373046875 0 3930.7603
1284.6868896484375 0 8150.845
1285.7061767578125 0 93085.98
1286.7012939453125 0 546086.25
1287.70263671875 0 412205.1
1288.70361328125 0 198338.16
1289.7015380859375 0 39417.22
1290.701171875 0 6477.987
1300.689453125 0 11977.808
1301.7015380859375 0 20084.469
1302.7117919921875 0 556415.06
1303.718994140625 0 1409910.5
1304.7218017578125 0 955112.6
1305.722412109375 0 372528.66
1306.7232666015625 0 60870.797
2238.14013671875 0 2064.6404
3074.83740234375 0 2934.022
3308.667236328125 0 1898.008
3491.297119140625 0 2344.709

Spectrum Details

|  |  |
| --- | --- |
| Matched peaks? Matched peaks The total absolute number of peaks matched. Additionally in brackets the total fraction of peaks matched and the total number of peaks is shown. | 27 (10.47% of 258) |
| FDR? FDR The false discovery rate estimated for this peptide. It is calculated by matching all theoretical fragments with a non-integer shift with the raw peaks for this spectrum. This is done with 40 different shifts. The resulting percentage is the average number of annotated peaks over the number of annotated peaks with the correct spectrum. | 3.26% |
| Satellite FDR? Satellite FDR See the FDR for details on its calculation. This satellite ion specific FDR only contains the satellite ions (d/w) for I/L/J positions. | ∞ |
| PSM Score? PSM Score The PSM Score as given by Hecklib to this annotated spectrum. It is shown with three significant figures. | 252 |

#### Spectrum 4947? Spectrum 4947 The raw spectrum of this peptide as annotated by Hecklib. The fragments are coloured according to ion type (see legend). Any peaks with a star '\*' as text can be hovered over to see the full details, first the ion type second the mass shift type. By hovering over the amino acids in the peptide or ions in the legend the corresponding peaks are highlighted. By toggling the 'Unassigned' label you can turn the background (unassigned) peaks on or off in the plot. By updating the slider in the Ion legend you can update the spectrum to only show the top X% of the peaks with labels. The top X% means any peak that is within X% of the highest intensity. By dragging in the spectrum you can zoom in to a specific part of the spectrum and use 'Zoom Out' to get back to the original zoom level. The annotation of the spectrum is based on the given sequence in the peptides file and is done with different software so inconsistencies are likely. The peaks are annotated based on the given sequence, with 20 ppm tolerance.

Copy Data

##### Spectrum 4947 (TSV)

###### Preview

```
Loading example...
```

*Click on the button to copy the data to your clipboard.*

Mz MinMz MaxIntensity Max

WidthHeightPeptide font sizePeptide stroke widthSpectrum font sizeSpectrum stroke widthCompact peptide

Ion legend

wxyz

abcd

OtherUnassignedIon^Charge^_Position_Show for top:%

JFPPKPKDTJM

09.56e+41.91e+52.87e+53.82e+5

Zoom Out

y^+1^_2_y^+1^_3_y^+2^_7_y^+2^_8_y^+1^_4_y^+2^_9_z^+2^_9_y^+2^_9_y^+2^_10_z^+1^_5_y^+1^_5_c^+1^_6_c^+1^_6_y^+1^_6_c^+1^_7_z^+1^_7_y^+1^_7_c^+1^_8_y^+1^_8_c^+1^_9_c^+1^_9_y^+1^_9_c^+1^_10_c^+1^_10_z^+1^_10_y^+1^_10_

0776155223273103

Fragment Matches Table

Show background peaks

| Position | Ion type | Intensity | mz Theoretical | mz Error (Th) | mz Error (ppm) | Charge | Series Number |
| --- | --- | --- | --- | --- | --- | --- | --- |
| - | - | 2090 | 120.1 | - | - | 0 | - |
| - | - | 372 | 127.5 | - | - | 0 | - |
| - | - | 400.1 | 129.3 | - | - | 0 | - |
| - | - | 357.8 | 131.6 | - | - | 0 | - |
| - | - | 503.4 | 132.8 | - | - | 0 | - |
| - | - | 539 | 148.8 | - | - | 0 | - |
| - | - | 507.6 | 148.9 | - | - | 0 | - |
| - | - | 617.5 | 148.9 | - | - | 0 | - |
| - | - | 704.4 | 148.9 | - | - | 0 | - |
| - | - | 692.6 | 148.9 | - | - | 0 | - |
| - | - | 834.3 | 148.9 | - | - | 0 | - |
| - | - | 1086 | 148.9 | - | - | 0 | - |
| - | - | 1220 | 148.9 | - | - | 0 | - |
| - | - | 1365 | 148.9 | - | - | 0 | - |
| - | - | 2590 | 148.9 | - | - | 0 | - |
| - | - | 3991 | 148.9 | - | - | 0 | - |
| - | - | 4095 | 149 | - | - | 0 | - |
| - | - | 2366 | 149 | - | - | 0 | - |
| - | - | 1294 | 149 | - | - | 0 | - |
| - | - | 1229 | 149 | - | - | 0 | - |
| - | - | 907.9 | 149 | - | - | 0 | - |
| - | - | 476.2 | 149 | - | - | 0 | - |
| - | - | 529.7 | 149 | - | - | 0 | - |
| - | - | 643 | 149 | - | - | 0 | - |
| - | - | 648.1 | 149 | - | - | 0 | - |
| - | - | 516.7 | 149 | - | - | 0 | - |
| - | - | 8622 | 166.1 | - | - | 0 | - |
| - | - | 512.7 | 181 | - | - | 0 | - |
| - | - | 464 | 191.2 | - | - | 0 | - |
| - | - | 533.7 | 200.6 | - | - | 0 | - |
| - | - | 1.973E+04 | 233.2 | - | - | 0 | - |
| - | - | 3117 | 234.2 | - | - | 0 | - |
| - | - | 2.178E+04 | 261.2 | - | - | 0 | - |
| - | - | 2760 | 262.2 | - | - | 0 | - |
| - | - | 639 | 278.2 | - | - | 0 | - |
| 10 | y | 1127 | 279.1 | 0.005397 | 19.34 | +1 | 2 |
| - | - | 597.6 | 285.3 | - | - | 0 | - |
| - | - | 4166 | 323.2 | - | - | 0 | - |
| - | - | 621.6 | 324.2 | - | - | 0 | - |
| - | - | 535.8 | 332.3 | - | - | 0 | - |
| - | - | 749.7 | 344.2 | - | - | 0 | - |
| - | - | 1688 | 355.1 | - | - | 0 | - |
| - | - | 2779 | 358.2 | - | - | 0 | - |
| - | - | 692.6 | 359.2 | - | - | 0 | - |
| 9 | y | 771.8 | 380.2 | 0.003495 | 9.193 | +1 | 3 |
| 5 | y | 1383 | 424.7 | 0.002865 | 6.745 | +2 | 7 |
| - | - | 1026 | 429.1 | - | - | 0 | - |
| - | - | 637.9 | 430.1 | - | - | 0 | - |
| - | - | 640.8 | 431.1 | - | - | 0 | - |
| - | - | 6382 | 439.3 | - | - | 0 | - |
| - | - | 4113 | 439.8 | - | - | 0 | - |
| - | - | 579.6 | 455.3 | - | - | 0 | - |
| 4 | y | 2844 | 473.3 | 0.002972 | 6.281 | +2 | 8 |
| - | - | 1553 | 473.8 | - | - | 0 | - |
| 8 | y | 585.5 | 477.2 | 0.0005139 | 1.077 | +1 | 4 |
| - | - | 1355 | 488.8 | - | - | 0 | - |
| - | - | 9154 | 489.8 | - | - | 0 | - |
| - | - | 4329 | 490.3 | - | - | 0 | - |
| - | - | 1248 | 490.8 | - | - | 0 | - |
| 3 | y | 6614 | 512.8 | 0.002594 | 5.059 | +2 | 9 |
| - | - | 3808 | 513.3 | - | - | 0 | - |
| 3 | z | 2038 | 513.8 | 0.00765 | 14.89 | +2 | 9 |
| - | - | 1916 | 514.3 | - | - | 0 | - |
| - | - | 2151 | 520.8 | - | - | 0 | - |
| - | - | 822.4 | 521.3 | - | - | 0 | - |
| 3 | y | 2.789E+05 | 521.8 | 0.002988 | 5.727 | +2 | 9 |
| - | - | 1.669E+05 | 522.3 | - | - | 0 | - |
| - | - | 5.717E+04 | 522.8 | - | - | 0 | - |
| - | - | 8088 | 523.3 | - | - | 0 | - |
| - | - | 1119 | 565.4 | - | - | 0 | - |
| 2 | y | 7591 | 595.3 | 0.003083 | 5.179 | +2 | 10 |
| - | - | 5440 | 595.8 | - | - | 0 | - |
| - | - | 2679 | 596.3 | - | - | 0 | - |
| 7 | z | 5756 | 607.3 | 0.005934 | 9.772 | +1 | 5 |
| - | - | 2.633E+04 | 608.3 | - | - | 0 | - |
| - | - | 7736 | 609.3 | - | - | 0 | - |
| - | - | 2080 | 610.3 | - | - | 0 | - |
| - | - | 4636 | 619.9 | - | - | 0 | - |
| - | - | 4664 | 620.4 | - | - | 0 | - |
| - | - | 1526 | 620.9 | - | - | 0 | - |
| 7 | y | 1083 | 623.3 | 0.002774 | 4.451 | +1 | 5 |
| - | - | 668 | 624.3 | - | - | 0 | - |
| - | - | 2433 | 642.9 | - | - | 0 | - |
| - | - | 2251 | 643.4 | - | - | 0 | - |
| - | - | 767.8 | 643.9 | - | - | 0 | - |
| - | - | 2275 | 650.4 | - | - | 0 | - |
| - | - | 2968 | 650.9 | - | - | 0 | - |
| - | - | 1573 | 651.4 | - | - | 0 | - |
| - | - | 1.501E+05 | 651.9 | - | - | 0 | - |
| - | - | 1.124E+05 | 652.4 | - | - | 0 | - |
| - | - | 5.42E+04 | 652.9 | - | - | 0 | - |
| - | - | 9964 | 653.4 | - | - | 0 | - |
| - | - | 1474 | 653.4 | - | - | 0 | - |
| - | - | 1556 | 654.4 | - | - | 0 | - |
| - | - | 1638 | 663.4 | - | - | 0 | - |
| 6 | c | 1057 | 680.4 | 0.003402 | 5 | +1 | 6 |
| - | - | 679.9 | 681.4 | - | - | 0 | - |
| - | - | 1158 | 695.4 | - | - | 0 | - |
| - | - | 1.192E+05 | 696.4 | - | - | 0 | - |
| 6 | c | 1.63E+05 | 697.4 | 0.0008984 | 1.288 | +1 | 6 |
| - | - | 5.702E+04 | 698.4 | - | - | 0 | - |
| - | - | 9685 | 699.4 | - | - | 0 | - |
| - | - | 904.7 | 700.5 | - | - | 0 | - |
| 6 | y | 7636 | 720.4 | 0.004271 | 5.929 | +1 | 6 |
| - | - | 2474 | 721.4 | - | - | 0 | - |
| - | - | 2147 | 748.4 | - | - | 0 | - |
| - | - | 2120 | 768.5 | - | - | 0 | - |
| - | - | 787.9 | 780.5 | - | - | 0 | - |
| - | - | 987.3 | 781.4 | - | - | 0 | - |
| - | - | 1.286E+04 | 781.5 | - | - | 0 | - |
| - | - | 7341 | 782.5 | - | - | 0 | - |
| - | - | 1215 | 783.5 | - | - | 0 | - |
| - | - | 2015 | 788.4 | - | - | 0 | - |
| - | - | 737.7 | 789.4 | - | - | 0 | - |
| - | - | 1431 | 812.5 | - | - | 0 | - |
| - | - | 2909 | 823.5 | - | - | 0 | - |
| - | - | 9.603E+04 | 824.5 | - | - | 0 | - |
| 7 | c | 8.383E+04 | 825.5 | 0.002722 | 3.297 | +1 | 7 |
| - | - | 2.741E+04 | 826.5 | - | - | 0 | - |
| - | - | 4988 | 827.5 | - | - | 0 | - |
| 5 | z | 2.749E+04 | 832.4 | 0.005302 | 6.37 | +1 | 7 |
| - | - | 1.297E+04 | 833.4 | - | - | 0 | - |
| - | - | 4697 | 834.4 | - | - | 0 | - |
| 5 | y | 2805 | 848.4 | 0.0044 | 5.186 | +1 | 7 |
| - | - | 1009 | 849.5 | - | - | 0 | - |
| - | - | 4956 | 877.5 | - | - | 0 | - |
| - | - | 2147 | 878.5 | - | - | 0 | - |
| - | - | 734.8 | 879.5 | - | - | 0 | - |
| - | - | 1650 | 881.5 | - | - | 0 | - |
| - | - | 707.6 | 882.5 | - | - | 0 | - |
| - | - | 1080 | 883.5 | - | - | 0 | - |
| - | - | 909.2 | 884.5 | - | - | 0 | - |
| - | - | 2197 | 895.6 | - | - | 0 | - |
| - | - | 3263 | 896.6 | - | - | 0 | - |
| - | - | 2175 | 897.6 | - | - | 0 | - |
| - | - | 1260 | 898.6 | - | - | 0 | - |
| - | - | 1076 | 924.5 | - | - | 0 | - |
| - | - | 2637 | 925.6 | - | - | 0 | - |
| - | - | 785.7 | 926.6 | - | - | 0 | - |
| - | - | 2074 | 938.5 | - | - | 0 | - |
| - | - | 9.768E+04 | 939.6 | - | - | 0 | - |
| 8 | c | 1.01E+05 | 940.6 | 0.002626 | 2.792 | +1 | 8 |
| - | - | 4.207E+04 | 941.6 | - | - | 0 | - |
| - | - | 9286 | 942.6 | - | - | 0 | - |
| - | - | 1012 | 943.6 | - | - | 0 | - |
| 4 | y | 2.583E+04 | 945.5 | 0.00492 | 5.204 | +1 | 8 |
| - | - | 1.333E+04 | 946.5 | - | - | 0 | - |
| - | - | 4351 | 947.5 | - | - | 0 | - |
| - | - | 2108 | 978.6 | - | - | 0 | - |
| - | - | 1589 | 979.6 | - | - | 0 | - |
| - | - | 1929 | 997.6 | - | - | 0 | - |
| - | - | 1456 | 998.6 | - | - | 0 | - |
| 9 | c | 1434 | 1025 | 0.01021 | 9.961 | +1 | 9 |
| - | - | 1348 | 1026 | - | - | 0 | - |
| - | - | 9757 | 1041 | - | - | 0 | - |
| 9 | c | 7.967E+04 | 1042 | 0.0002558 | 0.2456 | +1 | 9 |
| 3 | y | 4.706E+04 | 1043 | 0.01923 | 18.45 | +1 | 9 |
| - | - | 2.727E+04 | 1044 | - | - | 0 | - |
| - | - | 1.056E+04 | 1045 | - | - | 0 | - |
| - | - | 1068 | 1046 | - | - | 0 | - |
| - | - | 700.9 | 1061 | - | - | 0 | - |
| - | - | 1606 | 1070 | - | - | 0 | - |
| - | - | 4821 | 1071 | - | - | 0 | - |
| - | - | 3504 | 1072 | - | - | 0 | - |
| - | - | 913.5 | 1097 | - | - | 0 | - |
| - | - | 1625 | 1103 | - | - | 0 | - |
| - | - | 692.7 | 1104 | - | - | 0 | - |
| - | - | 5566 | 1111 | - | - | 0 | - |
| - | - | 4590 | 1112 | - | - | 0 | - |
| - | - | 1397 | 1113 | - | - | 0 | - |
| - | - | 2564 | 1130 | - | - | 0 | - |
| - | - | 2368 | 1131 | - | - | 0 | - |
| - | - | 727.2 | 1132 | - | - | 0 | - |
| 10 | c | 3541 | 1138 | 0.00113 | 0.9933 | +1 | 10 |
| - | - | 2352 | 1139 | - | - | 0 | - |
| - | - | 836.8 | 1140 | - | - | 0 | - |
| - | - | 876.4 | 1153 | - | - | 0 | - |
| - | - | 1233 | 1154 | - | - | 0 | - |
| 10 | c | 1.347E+05 | 1155 | 0.0002134 | 0.1848 | +1 | 10 |
| - | - | 9.37E+04 | 1156 | - | - | 0 | - |
| - | - | 3.189E+04 | 1157 | - | - | 0 | - |
| - | - | 3720 | 1158 | - | - | 0 | - |
| 2 | z | 4.062E+04 | 1174 | 0.004578 | 3.901 | +1 | 10 |
| - | - | 2.638E+04 | 1175 | - | - | 0 | - |
| - | - | 1.128E+04 | 1176 | - | - | 0 | - |
| - | - | 1815 | 1177 | - | - | 0 | - |
| - | - | 1595 | 1184 | - | - | 0 | - |
| - | - | 1009 | 1185 | - | - | 0 | - |
| - | - | 1606 | 1186 | - | - | 0 | - |
| - | - | 1088 | 1187 | - | - | 0 | - |
| 2 | y | 1053 | 1190 | 0.007827 | 6.579 | +1 | 10 |
| - | - | 1649 | 1197 | - | - | 0 | - |
| - | - | 781.1 | 1200 | - | - | 0 | - |
| - | - | 915.1 | 1201 | - | - | 0 | - |
| - | - | 8998 | 1212 | - | - | 0 | - |
| - | - | 7498 | 1213 | - | - | 0 | - |
| - | - | 2837 | 1214 | - | - | 0 | - |
| - | - | 2076 | 1216 | - | - | 0 | - |
| - | - | 884.5 | 1217 | - | - | 0 | - |
| - | - | 890 | 1218 | - | - | 0 | - |
| - | - | 965.3 | 1219 | - | - | 0 | - |
| - | - | 1049 | 1224 | - | - | 0 | - |
| - | - | 1020 | 1225 | - | - | 0 | - |
| - | - | 813.1 | 1226 | - | - | 0 | - |
| - | - | 747.4 | 1228 | - | - | 0 | - |
| - | - | 874.9 | 1230 | - | - | 0 | - |
| - | - | 1722 | 1231 | - | - | 0 | - |
| - | - | 2199 | 1233 | - | - | 0 | - |
| - | - | 1238 | 1234 | - | - | 0 | - |
| - | - | 4366 | 1240 | - | - | 0 | - |
| - | - | 3771 | 1241 | - | - | 0 | - |
| - | - | 9378 | 1242 | - | - | 0 | - |
| - | - | 8927 | 1243 | - | - | 0 | - |
| - | - | 1.22E+04 | 1244 | - | - | 0 | - |
| - | - | 6886 | 1245 | - | - | 0 | - |
| - | - | 3668 | 1246 | - | - | 0 | - |
| - | - | 3550 | 1248 | - | - | 0 | - |
| - | - | 1586 | 1249 | - | - | 0 | - |
| - | - | 4550 | 1258 | - | - | 0 | - |
| - | - | 3617 | 1259 | - | - | 0 | - |
| - | - | 7174 | 1260 | - | - | 0 | - |
| - | - | 2754 | 1261 | - | - | 0 | - |
| - | - | 1667 | 1262 | - | - | 0 | - |
| - | - | 1030 | 1269 | - | - | 0 | - |
| - | - | 1935 | 1270 | - | - | 0 | - |
| - | - | 796.1 | 1271 | - | - | 0 | - |
| - | - | 3034 | 1271 | - | - | 0 | - |
| - | - | 2329 | 1272 | - | - | 0 | - |
| - | - | 924.5 | 1273 | - | - | 0 | - |
| - | - | 683.8 | 1277 | - | - | 0 | - |
| - | - | 1171 | 1284 | - | - | 0 | - |
| - | - | 3158 | 1285 | - | - | 0 | - |
| - | - | 2.444E+04 | 1286 | - | - | 0 | - |
| - | - | 1.512E+05 | 1287 | - | - | 0 | - |
| - | - | 1.071E+05 | 1288 | - | - | 0 | - |
| - | - | 5.351E+04 | 1289 | - | - | 0 | - |
| - | - | 883 | 1290 | - | - | 0 | - |
| - | - | 1.112E+04 | 1290 | - | - | 0 | - |
| - | - | 831.5 | 1291 | - | - | 0 | - |
| - | - | 881.4 | 1300 | - | - | 0 | - |
| - | - | 5148 | 1301 | - | - | 0 | - |
| - | - | 7013 | 1302 | - | - | 0 | - |
| - | - | 1.581E+05 | 1303 | - | - | 0 | - |
| - | - | 3.787E+05 | 1304 | - | - | 0 | - |
| - | - | 2.614E+05 | 1305 | - | - | 0 | - |
| - | - | 1643 | 1305 | - | - | 0 | - |
| - | - | 1.028E+05 | 1306 | - | - | 0 | - |
| - | - | 1.636E+04 | 1307 | - | - | 0 | - |
| - | - | 1087 | 1336 | - | - | 0 | - |
| - | - | 961.6 | 1952 | - | - | 0 | - |
| - | - | 931 | 3072 | - | - | 0 | - |

m/z Charge Intensity FragmentType MassShift Position
120.0810775756836 0 2090.4797
127.49662017822266 0 371.97604
129.34405517578125 0 400.07425
131.56492614746094 0 357.82275
132.7520751953125 0 503.4483
148.83407592773438 0 538.9989
148.86380004882812 0 507.61172
148.88540649414062 0 617.5258
148.89242553710938 0 704.41425
148.89889526367188 0 692.64526
148.90679931640625 0 834.2757
148.91395568847656 0 1085.832
148.9211883544922 0 1219.7465
148.9283905029297 0 1364.7028
148.9354248046875 0 2590.2537
148.9432373046875 0 3990.9177
148.9598846435547 0 4094.746
148.9676513671875 0 2366.3408
148.97470092773438 0 1293.5902
148.98199462890625 0 1228.7817
148.98907470703125 0 907.8862
148.9963836669922 0 476.20544
149.01124572753906 0 529.7077
149.01844787597656 0 643.00934
149.02598571777344 0 648.0713
149.03260803222656 0 516.7103
166.05345153808594 0 8621.651
181.0043487548828 0 512.7253
191.15843200683594 0 463.9602
200.59762573242188 0 533.7218
233.1651153564453 0 19728.838
234.1685333251953 0 3116.73
261.15985107421875 0 21783.324
262.1629638671875 0 2760.0686
278.2075500488281 0 638.9706
279.1377868652344 0 1127.2256 y 9
285.34197998046875 0 597.5818
323.2086181640625 0 4165.8164
324.20965576171875 0 621.555
332.2908020019531 0 535.7568
344.2472229003906 0 749.7217
355.0700378417969 0 1688.4583
358.2124938964844 0 2778.7283
359.2157287597656 0 692.6258
380.1835632324219 0 771.8381 y 8
424.7313537597656 0 1382.5659 y 4
429.0877685546875 0 1025.7208
430.0893249511719 0 637.9209
431.0867614746094 0 640.81213
439.26116943359375 0 6381.7637
439.7625427246094 0 4112.884
455.262451171875 0 579.60254
473.2578430175781 0 2844.151 y 3
473.7596740722656 0 1552.7244
477.19696044921875 0 585.48926 y Water loss 7
488.7783203125 0 1355.4224
489.7852783203125 0 9153.841
490.2869873046875 0 4328.736
490.7879943847656 0 1247.9819
512.778564453125 0 6613.678 y Water loss 2
513.280517578125 0 3807.8347
513.779541015625 0 2037.7329 z 2
514.2728271484375 0 1916.3954
520.7769775390625 0 2151.059
521.2763061523438 0 822.4091
521.7842407226562 0 278916.72 y 2
522.2855224609375 0 166921.33
522.7854614257812 0 57173.207
523.28564453125 0 8088.051
565.3802490234375 0 1118.6075
595.3185424804688 0 7591.4116 y 1
595.8203125 0 5439.8447
596.3204956054688 0 2679.1145
607.2891845703125 0 5755.753 z 6
608.2962646484375 0 26333.037
609.2985229492188 0 7736.4106
610.2979125976562 0 2080.4187
619.8612670898438 0 4636.489
620.3627319335938 0 4663.876
620.865234375 0 1525.943
623.3047485351562 0 1082.6578 y 6
624.3108520507812 0 667.97626
642.8550415039062 0 2432.7378
643.3545532226562 0 2251.4243
643.859375 0 767.8425
650.351318359375 0 2274.5469
650.8534545898438 0 2967.6328
651.3541870117188 0 1572.6484
651.8604736328125 0 150114.02
652.3619995117188 0 112428.336
652.8621826171875 0 54197.465
653.361572265625 0 9964.044
653.4263305664062 0 1474.2908
654.430419921875 0 1555.6895
663.3832397460938 0 1637.8209
680.4096069335938 0 1057.0917 c Ammonia loss 5
681.416748046875 0 679.8976
695.4226684570312 0 1157.9213
696.4321899414062 0 119207.914
697.4386596679688 0 163003.34 c 5
698.4420166015625 0 57019.434
699.4453125 0 9684.795
700.4503784179688 0 904.7149
720.3590087890625 0 7635.564 y 5
721.3609008789062 0 2473.7175
748.3530883789062 0 2146.709
768.4571533203125 0 2120.1692
780.4595336914062 0 787.9366
781.4479370117188 0 987.2739
781.5210571289062 0 12862.386
782.524169921875 0 7341.3633
783.5277099609375 0 1214.5262
788.4442138671875 0 2014.8577
789.448974609375 0 737.68726
812.4673461914062 0 1431.2576
823.5198364257812 0 2909.0488
824.5269165039062 0 96027.3
825.5317993164062 0 83832.945 c 6
826.5358276367188 0 27408.86
827.5400390625 0 4987.9126
832.436279296875 0 27491.537 z 4
833.4396362304688 0 12974.291
834.4397583007812 0 4697.254
848.4541015625 0 2804.8745 y 4
849.4613647460938 0 1008.88324
877.5139770507812 0 4956.2764
878.5162353515625 0 2146.5366
879.5076293945312 0 734.78546
881.529052734375 0 1649.9435
882.5411987304688 0 707.61395
883.4864501953125 0 1079.8369
884.5012817382812 0 909.21136
895.5639038085938 0 2197.1926
896.5510864257812 0 3263.0623
897.5518188476562 0 2174.8486
898.5606079101562 0 1259.5398
924.538330078125 0 1076.457
925.5504150390625 0 2636.5664
926.5518188476562 0 785.6871
938.5454711914062 0 2073.5542
939.5535888671875 0 97677.875
940.558837890625 0 101037.78 c 7
941.56298828125 0 42068.227
942.5669555664062 0 9285.523
943.57421875 0 1012.17645
945.5073852539062 0 25825.564 y 3
946.5103149414062 0 13330.15
947.5079956054688 0 4350.875
978.5623779296875 0 2107.9004
979.5756225585938 0 1589.2007
997.5960693359375 0 1928.5233
998.601318359375 0 1455.9469
1024.5723876953125 0 1434.056 c Ammonia loss 8
1025.5799560546875 0 1347.7816
1040.60205078125 0 9756.529
1041.60888671875 0 79674.8 c 8
1042.574462890625 0 47061.89 y 2
1043.5712890625 0 27270.018
1044.5633544921875 0 10555.942
1045.566650390625 0 1067.9554
1060.52783203125 0 700.8632
1069.5433349609375 0 1606.3597
1070.55322265625 0 4820.813
1071.557861328125 0 3503.5908
1096.631591796875 0 913.54706
1102.5341796875 0 1625.2065
1103.54248046875 0 692.7275
1110.677734375 0 5566.179
1111.6806640625 0 4590.006
1112.69580078125 0 1397.0481
1129.6240234375 0 2563.6204
1130.618896484375 0 2367.6458
1131.62548828125 0 727.191
1137.66552734375 0 3541.2673 c Ammonia loss 9
1138.669677734375 0 2351.7458
1139.6727294921875 0 836.7666
1152.6923828125 0 876.40765
1153.6826171875 0 1233.3699
1154.6929931640625 0 134691.19 c 9
1155.6964111328125 0 93697.78
1156.6988525390625 0 31892.723
1157.701416015625 0 3720.067
1173.6094970703125 0 40615.77 z 1
1174.613525390625 0 26375.238
1175.6124267578125 0 11276.305
1176.6165771484375 0 1814.5687
1183.653076171875 0 1594.94
1184.6517333984375 0 1009.4185
1185.6585693359375 0 1606.3507
1186.66845703125 0 1087.9849
1189.6314697265625 0 1052.9163 y 1
1196.6937255859375 0 1648.6393
1199.6114501953125 0 781.09595
1200.6287841796875 0 915.08826
1211.688232421875 0 8997.978
1212.691650390625 0 7497.534
1213.6954345703125 0 2836.588
1215.6226806640625 0 2076.1663
1216.6383056640625 0 884.5211
1217.6356201171875 0 889.97943
1218.6253662109375 0 965.2617
1223.6859130859375 0 1048.6786
1224.7059326171875 0 1019.5978
1225.703369140625 0 813.0936
1227.6634521484375 0 747.4099
1229.654541015625 0 874.91925
1230.63720703125 0 1722.0454
1232.6419677734375 0 2198.7783
1233.664306640625 0 1238.0215
1239.7186279296875 0 4366.362
1240.7200927734375 0 3771.239
1241.7200927734375 0 9378.473
1242.7158203125 0 8926.522
1243.7020263671875 0 12201.302
1244.697265625 0 6886.1133
1245.6910400390625 0 3667.6409
1247.6593017578125 0 3550.4802
1248.66259765625 0 1586.0299
1257.708740234375 0 4549.7173
1258.7122802734375 0 3617.0503
1259.7249755859375 0 7174.007
1260.7291259765625 0 2753.9165
1261.7208251953125 0 1667.2079
1268.697021484375 0 1029.6605
1269.6944580078125 0 1934.6798
1270.5487060546875 0 796.08453
1270.69287109375 0 3034.2498
1271.6846923828125 0 2328.544
1272.7081298828125 0 924.53143
1276.6993408203125 0 683.77325
1283.69189453125 0 1170.9261
1284.6900634765625 0 3157.9573
1285.7059326171875 0 24439.66
1286.699951171875 0 151190.02
1287.7021484375 0 107107.52
1288.702392578125 0 53511.082
1289.5633544921875 0 883.0497
1289.7017822265625 0 11117.452
1290.67724609375 0 831.45703
1299.6900634765625 0 881.43555
1300.6959228515625 0 5147.707
1301.6995849609375 0 7012.8047
1302.7110595703125 0 158060.67
1303.7181396484375 0 378695.2
1304.7210693359375 0 261407.88
1305.08056640625 0 1642.7645
1305.721435546875 0 102816.99
1306.7225341796875 0 16362.049
1335.7178955078125 0 1087.3296
1951.8870849609375 0 961.5731
3072.321533203125 0 930.97437

Spectrum Details

|  |  |
| --- | --- |
| Matched peaks? Matched peaks The total absolute number of peaks matched. Additionally in brackets the total fraction of peaks matched and the total number of peaks is shown. | 26 (10.36% of 251) |
| FDR? FDR The false discovery rate estimated for this peptide. It is calculated by matching all theoretical fragments with a non-integer shift with the raw peaks for this spectrum. This is done with 40 different shifts. The resulting percentage is the average number of annotated peaks over the number of annotated peaks with the correct spectrum. | 2.75% |
| Satellite FDR? Satellite FDR See the FDR for details on its calculation. This satellite ion specific FDR only contains the satellite ions (d/w) for I/L/J positions. | ∞ |
| PSM Score? PSM Score The PSM Score as given by Hecklib to this annotated spectrum. It is shown with three significant figures. | 236 |

#### Spectrum 5492? Spectrum 5492 The raw spectrum of this peptide as annotated by Hecklib. The fragments are coloured according to ion type (see legend). Any peaks with a star '\*' as text can be hovered over to see the full details, first the ion type second the mass shift type. By hovering over the amino acids in the peptide or ions in the legend the corresponding peaks are highlighted. By toggling the 'Unassigned' label you can turn the background (unassigned) peaks on or off in the plot. By updating the slider in the Ion legend you can update the spectrum to only show the top X% of the peaks with labels. The top X% means any peak that is within X% of the highest intensity. By dragging in the spectrum you can zoom in to a specific part of the spectrum and use 'Zoom Out' to get back to the original zoom level. The annotation of the spectrum is based on the given sequence in the peptides file and is done with different software so inconsistencies are likely. The peaks are annotated based on the given sequence, with 20 ppm tolerance.

Copy Data

##### Spectrum 5492 (TSV)

###### Preview

```
Loading example...
```

*Click on the button to copy the data to your clipboard.*

Mz MinMz MaxIntensity Max

WidthHeightPeptide font sizePeptide stroke widthSpectrum font sizeSpectrum stroke widthCompact peptide

Ion legend

wxyz

abcd

OtherUnassignedIon^Charge^_Position_Show for top:%

JFPPKPKDTJM

05.92e+41.18e+51.77e+52.37e+5

Zoom Out

y^+1^_2_y^+1^_3_y^+2^_7_y^+2^_8_y^+2^_9_z^+2^_9_y^+2^_9_y^+2^_10_z^+1^_5_y^+1^_5_c^+1^_6_c^+1^_6_y^+1^_6_c^+1^_7_z^+1^_7_y^+1^_7_c^+1^_8_y^+1^_8_c^+1^_9_c^+1^_9_c^+1^_10_c^+1^_10_z^+1^_10_

0842168425263368

Fragment Matches Table

Show background peaks

| Position | Ion type | Intensity | mz Theoretical | mz Error (Th) | mz Error (ppm) | Charge | Series Number |
| --- | --- | --- | --- | --- | --- | --- | --- |
| - | - | 1329 | 120.1 | - | - | 0 | - |
| - | - | 394.7 | 131.9 | - | - | 0 | - |
| - | - | 415.3 | 133.7 | - | - | 0 | - |
| - | - | 474.6 | 134.8 | - | - | 0 | - |
| - | - | 695.9 | 148.9 | - | - | 0 | - |
| - | - | 626.7 | 148.9 | - | - | 0 | - |
| - | - | 580.5 | 148.9 | - | - | 0 | - |
| - | - | 1006 | 148.9 | - | - | 0 | - |
| - | - | 1054 | 148.9 | - | - | 0 | - |
| - | - | 1390 | 148.9 | - | - | 0 | - |
| - | - | 2610 | 148.9 | - | - | 0 | - |
| - | - | 3823 | 148.9 | - | - | 0 | - |
| - | - | 4001 | 149 | - | - | 0 | - |
| - | - | 2611 | 149 | - | - | 0 | - |
| - | - | 1378 | 149 | - | - | 0 | - |
| - | - | 1228 | 149 | - | - | 0 | - |
| - | - | 729.3 | 149 | - | - | 0 | - |
| - | - | 567 | 149 | - | - | 0 | - |
| - | - | 579.3 | 149 | - | - | 0 | - |
| - | - | 598.6 | 149 | - | - | 0 | - |
| - | - | 528.7 | 149 | - | - | 0 | - |
| - | - | 612.8 | 149 | - | - | 0 | - |
| - | - | 417.9 | 149.4 | - | - | 0 | - |
| - | - | 431.7 | 159.6 | - | - | 0 | - |
| - | - | 4117 | 166.1 | - | - | 0 | - |
| - | - | 511.2 | 180.5 | - | - | 0 | - |
| - | - | 428.1 | 181.5 | - | - | 0 | - |
| - | - | 482.4 | 223.4 | - | - | 0 | - |
| - | - | 1.15E+04 | 233.2 | - | - | 0 | - |
| - | - | 1453 | 234.2 | - | - | 0 | - |
| - | - | 486.1 | 259.5 | - | - | 0 | - |
| - | - | 1.185E+04 | 261.2 | - | - | 0 | - |
| - | - | 1060 | 262.2 | - | - | 0 | - |
| - | - | 578.1 | 275.7 | - | - | 0 | - |
| 10 | y | 816.2 | 279.1 | 0.003475 | 12.45 | +1 | 2 |
| - | - | 2640 | 323.2 | - | - | 0 | - |
| - | - | 2932 | 355.1 | - | - | 0 | - |
| - | - | 1048 | 356.1 | - | - | 0 | - |
| - | - | 1598 | 358.2 | - | - | 0 | - |
| 9 | y | 858 | 380.2 | 0.004563 | 12 | +1 | 3 |
| - | - | 613.1 | 381.6 | - | - | 0 | - |
| 5 | y | 820.2 | 424.7 | 0.00314 | 7.392 | +2 | 7 |
| - | - | 1161 | 429.1 | - | - | 0 | - |
| - | - | 844.9 | 430.1 | - | - | 0 | - |
| - | - | 823.7 | 430.3 | - | - | 0 | - |
| - | - | 4781 | 439.3 | - | - | 0 | - |
| - | - | 620.8 | 440.3 | - | - | 0 | - |
| 4 | y | 1853 | 473.3 | 0.002698 | 5.7 | +2 | 8 |
| - | - | 715.5 | 473.8 | - | - | 0 | - |
| - | - | 1021 | 488.8 | - | - | 0 | - |
| - | - | 4417 | 489.8 | - | - | 0 | - |
| - | - | 2573 | 490.3 | - | - | 0 | - |
| - | - | 707.7 | 490.8 | - | - | 0 | - |
| 3 | y | 3618 | 512.8 | 0.001862 | 3.631 | +2 | 9 |
| - | - | 1889 | 513.3 | - | - | 0 | - |
| 3 | z | 644.8 | 513.8 | 0.001974 | 3.842 | +2 | 9 |
| - | - | 1147 | 520.8 | - | - | 0 | - |
| - | - | 1199 | 521.3 | - | - | 0 | - |
| 3 | y | 1.511E+05 | 521.8 | 0.0025 | 4.791 | +2 | 9 |
| - | - | 8.882E+04 | 522.3 | - | - | 0 | - |
| - | - | 3.356E+04 | 522.8 | - | - | 0 | - |
| - | - | 4808 | 523.3 | - | - | 0 | - |
| 2 | y | 4015 | 595.3 | 0.002289 | 3.846 | +2 | 10 |
| - | - | 2373 | 595.8 | - | - | 0 | - |
| - | - | 938.4 | 596.3 | - | - | 0 | - |
| 7 | z | 2858 | 607.3 | 0.005629 | 9.27 | +1 | 5 |
| - | - | 1.708E+04 | 608.3 | - | - | 0 | - |
| - | - | 4493 | 609.3 | - | - | 0 | - |
| - | - | 917.6 | 610.3 | - | - | 0 | - |
| - | - | 639.9 | 615.4 | - | - | 0 | - |
| - | - | 3293 | 619.9 | - | - | 0 | - |
| - | - | 2288 | 620.4 | - | - | 0 | - |
| 7 | y | 780 | 623.3 | 0.006009 | 9.641 | +1 | 5 |
| - | - | 1271 | 642.9 | - | - | 0 | - |
| - | - | 669.4 | 643.4 | - | - | 0 | - |
| - | - | 532.1 | 644.4 | - | - | 0 | - |
| - | - | 581.3 | 650.3 | - | - | 0 | - |
| - | - | 1181 | 650.9 | - | - | 0 | - |
| - | - | 1768 | 651.4 | - | - | 0 | - |
| - | - | 7.938E+04 | 651.9 | - | - | 0 | - |
| - | - | 6.227E+04 | 652.4 | - | - | 0 | - |
| - | - | 2.904E+04 | 652.9 | - | - | 0 | - |
| - | - | 4539 | 653.4 | - | - | 0 | - |
| - | - | 895.6 | 654.4 | - | - | 0 | - |
| 6 | c | 762.7 | 680.4 | 0.00212 | 3.116 | +1 | 6 |
| - | - | 774.8 | 695.4 | - | - | 0 | - |
| - | - | 7.075E+04 | 696.4 | - | - | 0 | - |
| 6 | c | 9.876E+04 | 697.4 | 0.001387 | 1.988 | +1 | 6 |
| - | - | 3.5E+04 | 698.4 | - | - | 0 | - |
| - | - | 6060 | 699.4 | - | - | 0 | - |
| 6 | y | 3598 | 720.4 | 0.003843 | 5.335 | +1 | 6 |
| - | - | 1114 | 721.4 | - | - | 0 | - |
| - | - | 1369 | 748.4 | - | - | 0 | - |
| - | - | 1651 | 768.5 | - | - | 0 | - |
| - | - | 927.3 | 769.5 | - | - | 0 | - |
| - | - | 7137 | 781.5 | - | - | 0 | - |
| - | - | 4692 | 782.5 | - | - | 0 | - |
| - | - | 957.8 | 783.5 | - | - | 0 | - |
| - | - | 844.9 | 788.4 | - | - | 0 | - |
| - | - | 1023 | 812.5 | - | - | 0 | - |
| - | - | 5.555E+04 | 824.5 | - | - | 0 | - |
| 7 | c | 4.65E+04 | 825.5 | 0.002783 | 3.371 | +1 | 7 |
| - | - | 1.388E+04 | 826.5 | - | - | 0 | - |
| - | - | 2025 | 827.5 | - | - | 0 | - |
| 5 | z | 1.732E+04 | 832.4 | 0.005363 | 6.443 | +1 | 7 |
| - | - | 8182 | 833.4 | - | - | 0 | - |
| - | - | 2552 | 834.4 | - | - | 0 | - |
| 5 | y | 1558 | 848.4 | 0.00617 | 7.273 | +1 | 7 |
| - | - | 2858 | 877.5 | - | - | 0 | - |
| - | - | 906.2 | 878.5 | - | - | 0 | - |
| - | - | 1402 | 881.5 | - | - | 0 | - |
| - | - | 1066 | 882.5 | - | - | 0 | - |
| - | - | 913.5 | 883.5 | - | - | 0 | - |
| - | - | 889.5 | 895.6 | - | - | 0 | - |
| - | - | 2462 | 896.6 | - | - | 0 | - |
| - | - | 1821 | 897.6 | - | - | 0 | - |
| - | - | 1471 | 898.6 | - | - | 0 | - |
| - | - | 2387 | 925.5 | - | - | 0 | - |
| - | - | 663.1 | 926.6 | - | - | 0 | - |
| - | - | 1040 | 938.5 | - | - | 0 | - |
| - | - | 5.916E+04 | 939.6 | - | - | 0 | - |
| 8 | c | 6.189E+04 | 940.6 | 0.002565 | 2.727 | +1 | 8 |
| - | - | 2.373E+04 | 941.6 | - | - | 0 | - |
| - | - | 5631 | 942.6 | - | - | 0 | - |
| - | - | 889 | 943.6 | - | - | 0 | - |
| 4 | y | 1.434E+04 | 945.5 | 0.005103 | 5.398 | +1 | 8 |
| - | - | 6902 | 946.5 | - | - | 0 | - |
| - | - | 3311 | 947.5 | - | - | 0 | - |
| - | - | 658.1 | 948.5 | - | - | 0 | - |
| - | - | 762.2 | 973.5 | - | - | 0 | - |
| - | - | 672.2 | 978.6 | - | - | 0 | - |
| - | - | 858.7 | 997.6 | - | - | 0 | - |
| 9 | c | 653.5 | 1025 | 0.01594 | 15.56 | +1 | 9 |
| - | - | 838 | 1026 | - | - | 0 | - |
| - | - | 6014 | 1041 | - | - | 0 | - |
| 9 | c | 5.086E+04 | 1042 | 0.0001338 | 0.1284 | +1 | 9 |
| - | - | 2.758E+04 | 1043 | - | - | 0 | - |
| - | - | 1.511E+04 | 1044 | - | - | 0 | - |
| - | - | 7773 | 1045 | - | - | 0 | - |
| - | - | 874.2 | 1046 | - | - | 0 | - |
| - | - | 858.9 | 1066 | - | - | 0 | - |
| - | - | 694.1 | 1070 | - | - | 0 | - |
| - | - | 2981 | 1071 | - | - | 0 | - |
| - | - | 2376 | 1072 | - | - | 0 | - |
| - | - | 771.1 | 1103 | - | - | 0 | - |
| - | - | 3290 | 1111 | - | - | 0 | - |
| - | - | 2297 | 1112 | - | - | 0 | - |
| - | - | 1890 | 1130 | - | - | 0 | - |
| - | - | 1466 | 1131 | - | - | 0 | - |
| 10 | c | 1781 | 1138 | 0.001374 | 1.208 | +1 | 10 |
| - | - | 1758 | 1139 | - | - | 0 | - |
| 10 | c | 8.427E+04 | 1155 | 0.0001528 | 0.1324 | +1 | 10 |
| - | - | 5.708E+04 | 1156 | - | - | 0 | - |
| - | - | 1.899E+04 | 1157 | - | - | 0 | - |
| - | - | 2333 | 1158 | - | - | 0 | - |
| 2 | z | 2.466E+04 | 1174 | 0.005189 | 4.421 | +1 | 10 |
| - | - | 1.675E+04 | 1175 | - | - | 0 | - |
| - | - | 6614 | 1176 | - | - | 0 | - |
| - | - | 1408 | 1177 | - | - | 0 | - |
| - | - | 918.6 | 1184 | - | - | 0 | - |
| - | - | 693.5 | 1186 | - | - | 0 | - |
| - | - | 1192 | 1197 | - | - | 0 | - |
| - | - | 724.1 | 1201 | - | - | 0 | - |
| - | - | 6113 | 1212 | - | - | 0 | - |
| - | - | 3232 | 1213 | - | - | 0 | - |
| - | - | 1941 | 1214 | - | - | 0 | - |
| - | - | 1431 | 1216 | - | - | 0 | - |
| - | - | 897 | 1218 | - | - | 0 | - |
| - | - | 688.4 | 1224 | - | - | 0 | - |
| - | - | 1069 | 1225 | - | - | 0 | - |
| - | - | 1392 | 1231 | - | - | 0 | - |
| - | - | 1162 | 1233 | - | - | 0 | - |
| - | - | 768.1 | 1234 | - | - | 0 | - |
| - | - | 2590 | 1240 | - | - | 0 | - |
| - | - | 2599 | 1241 | - | - | 0 | - |
| - | - | 5285 | 1242 | - | - | 0 | - |
| - | - | 679.7 | 1243 | - | - | 0 | - |
| - | - | 5923 | 1243 | - | - | 0 | - |
| - | - | 9092 | 1244 | - | - | 0 | - |
| - | - | 4719 | 1245 | - | - | 0 | - |
| - | - | 1752 | 1246 | - | - | 0 | - |
| - | - | 2203 | 1248 | - | - | 0 | - |
| - | - | 1416 | 1249 | - | - | 0 | - |
| - | - | 881.6 | 1255 | - | - | 0 | - |
| - | - | 3031 | 1258 | - | - | 0 | - |
| - | - | 2926 | 1259 | - | - | 0 | - |
| - | - | 3697 | 1260 | - | - | 0 | - |
| - | - | 2823 | 1261 | - | - | 0 | - |
| - | - | 1164 | 1262 | - | - | 0 | - |
| - | - | 765.2 | 1268 | - | - | 0 | - |
| - | - | 1622 | 1270 | - | - | 0 | - |
| - | - | 1565 | 1270 | - | - | 0 | - |
| - | - | 1756 | 1271 | - | - | 0 | - |
| - | - | 2663 | 1271 | - | - | 0 | - |
| - | - | 891.6 | 1272 | - | - | 0 | - |
| - | - | 2657 | 1285 | - | - | 0 | - |
| - | - | 1.615E+04 | 1286 | - | - | 0 | - |
| - | - | 8.855E+04 | 1287 | - | - | 0 | - |
| - | - | 7.029E+04 | 1288 | - | - | 0 | - |
| - | - | 3.029E+04 | 1289 | - | - | 0 | - |
| - | - | 6855 | 1290 | - | - | 0 | - |
| - | - | 851.2 | 1291 | - | - | 0 | - |
| - | - | 3023 | 1301 | - | - | 0 | - |
| - | - | 5093 | 1302 | - | - | 0 | - |
| - | - | 8.917E+04 | 1303 | - | - | 0 | - |
| - | - | 2.343E+05 | 1304 | - | - | 0 | - |
| - | - | 1.514E+05 | 1305 | - | - | 0 | - |
| - | - | 6.71E+04 | 1306 | - | - | 0 | - |
| - | - | 1.096E+04 | 1307 | - | - | 0 | - |
| - | - | 857.3 | 1320 | - | - | 0 | - |
| - | - | 691.7 | 1484 | - | - | 0 | - |
| - | - | 1119 | 1598 | - | - | 0 | - |
| - | - | 1507 | 1599 | - | - | 0 | - |
| - | - | 1476 | 1599 | - | - | 0 | - |
| - | - | 722.3 | 1628 | - | - | 0 | - |
| - | - | 1182 | 1629 | - | - | 0 | - |
| - | - | 766.1 | 1954 | - | - | 0 | - |
| - | - | 704.3 | 3335 | - | - | 0 | - |

m/z Charge Intensity FragmentType MassShift Position
120.08092498779297 0 1328.8225
131.9311981201172 0 394.71683
133.65713500976562 0 415.29178
134.81704711914062 0 474.56442
148.892333984375 0 695.9251
148.89974975585938 0 626.71063
148.9070587158203 0 580.51483
148.91400146484375 0 1006.1858
148.92129516601562 0 1053.8052
148.92831420898438 0 1389.6528
148.93548583984375 0 2609.921
148.943359375 0 3823.4045
148.95986938476562 0 4000.8945
148.96763610839844 0 2610.5078
148.9748077392578 0 1378.1207
148.98216247558594 0 1227.9426
148.9891357421875 0 729.26776
148.99607849121094 0 566.95404
149.00364685058594 0 579.29474
149.0111083984375 0 598.6185
149.01815795898438 0 528.7478
149.0250701904297 0 612.81525
149.44363403320312 0 417.94595
159.56216430664062 0 431.73816
166.05316162109375 0 4116.941
180.45053100585938 0 511.23203
181.49134826660156 0 428.05927
223.35630798339844 0 482.42447
233.1646728515625 0 11495.682
234.16778564453125 0 1452.8494
259.4545593261719 0 486.07312
261.159423828125 0 11846.132
262.1636962890625 0 1059.5469
275.7414855957031 0 578.0732
279.1358642578125 0 816.2459 y 9
323.20782470703125 0 2639.7515
355.0692138671875 0 2931.9858
356.07098388671875 0 1047.5889
358.21197509765625 0 1598.0189
380.18463134765625 0 858.01965 y 8
381.6330871582031 0 613.14294
424.73162841796875 0 820.18854 y 4
429.08819580078125 0 1161.3146
430.0891418457031 0 844.9374
430.255615234375 0 823.6965
439.2608337402344 0 4780.8716
440.2607727050781 0 620.78314
473.257568359375 0 1852.8492 y 3
473.7601318359375 0 715.47736
488.7778625488281 0 1021.27106
489.7840576171875 0 4417.253
490.2869873046875 0 2572.8384
490.7887268066406 0 707.66736
512.77783203125 0 3617.6494 y Water loss 2
513.2796020507812 0 1888.9271
513.7738647460938 0 644.8338 z 2
520.7785034179688 0 1146.7754
521.2771606445312 0 1199.2832
521.7837524414062 0 151134.72 y 2
522.28515625 0 88819.35
522.784912109375 0 33555.723
523.2855834960938 0 4807.9067
595.3177490234375 0 4015.21 y 1
595.8186645507812 0 2372.6772
596.3198852539062 0 938.37115
607.2888793945312 0 2858.4119 z 6
608.295654296875 0 17075.2
609.2992553710938 0 4492.679
610.2963256835938 0 917.60236
615.39599609375 0 639.8729
619.8594360351562 0 3293.1218
620.3616333007812 0 2288.33
623.3079833984375 0 780.04004 y 6
642.8518676757812 0 1270.9684
643.3563232421875 0 669.43774
644.3511962890625 0 532.1116
650.3458862304688 0 581.2999
650.8505249023438 0 1181.1692
651.3549194335938 0 1767.8457
651.8600463867188 0 79379.31
652.3615112304688 0 62273.22
652.8614501953125 0 29041.367
653.362060546875 0 4539.145
654.4287719726562 0 895.5868
680.410888671875 0 762.6929 c Ammonia loss 5
695.4251098632812 0 774.8397
696.4319458007812 0 70753.336
697.4381713867188 0 98761.125 c 5
698.441650390625 0 34996.16
699.444580078125 0 6060.415
720.3585815429688 0 3598.106 y 5
721.3595581054688 0 1114.048
748.3551635742188 0 1368.6394
768.455078125 0 1650.5593
769.45458984375 0 927.30774
781.5208129882812 0 7137.048
782.5238037109375 0 4691.7744
783.528564453125 0 957.81683
788.44970703125 0 844.8858
812.466552734375 0 1022.9234
824.526611328125 0 55546.285
825.53173828125 0 46503.23 c 6
826.5355224609375 0 13881.463
827.5387573242188 0 2025.0017
832.4363403320312 0 17319.475 z 4
833.4395141601562 0 8182.354
834.4411010742188 0 2551.802
848.4558715820312 0 1558.0692 y 4
877.5121459960938 0 2857.8464
878.5226440429688 0 906.2064
881.5346069335938 0 1402.263
882.5374755859375 0 1065.8138
883.4909057617188 0 913.5224
895.552734375 0 889.53033
896.5535278320312 0 2461.8564
897.5529174804688 0 1821.0549
898.5665893554688 0 1470.5978
925.5492553710938 0 2387.188
926.5569458007812 0 663.14484
938.5465087890625 0 1039.9863
939.5535888671875 0 59155.34
940.5588989257812 0 61892.36 c 7
941.5628662109375 0 23725.748
942.5680541992188 0 5631.0356
943.5789794921875 0 889.0212
945.507568359375 0 14338.401 y 3
946.5099487304688 0 6902.16
947.5125732421875 0 3311.0334
948.5182495117188 0 658.12384
973.5032348632812 0 762.17865
978.5687255859375 0 672.1523
997.6023559570312 0 858.6604
1024.566650390625 0 653.4638 c Ammonia loss 8
1025.56201171875 0 837.9841
1040.6005859375 0 6014.3247
1041.6090087890625 0 50864.883 c 8
1042.5914306640625 0 27575.234
1043.5721435546875 0 15106.068
1044.56201171875 0 7773.2188
1045.5513916015625 0 874.22546
1065.505615234375 0 858.9158
1069.5538330078125 0 694.0997
1070.553955078125 0 2981.1572
1071.5594482421875 0 2376.2212
1102.542724609375 0 771.0543
1110.677001953125 0 3289.6885
1111.679443359375 0 2297.107
1129.62060546875 0 1889.5767
1130.6192626953125 0 1465.5105
1137.665283203125 0 1781.4469 c Ammonia loss 9
1138.6666259765625 0 1757.8883
1154.693359375 0 84265.445 c 9
1155.69677734375 0 57076.664
1156.6995849609375 0 18985.729
1157.705810546875 0 2332.9526
1173.610107421875 0 24655.092 z 1
1174.61328125 0 16745.34
1175.613037109375 0 6613.782
1176.620849609375 0 1407.9733
1183.6591796875 0 918.5604
1185.6580810546875 0 693.5165
1196.6702880859375 0 1192.3076
1200.618408203125 0 724.10986
1211.6912841796875 0 6113.2075
1212.6920166015625 0 3231.6467
1213.697509765625 0 1941.4086
1215.62158203125 0 1431.3718
1217.623046875 0 896.99426
1223.6973876953125 0 688.35486
1224.70068359375 0 1069.2343
1230.63818359375 0 1392.4823
1232.6492919921875 0 1162.059
1233.6444091796875 0 768.1373
1239.7177734375 0 2590.2065
1240.7154541015625 0 2599.3193
1241.7191162109375 0 5285.22
1242.5762939453125 0 679.685
1242.711181640625 0 5922.5215
1243.6973876953125 0 9091.797
1244.6986083984375 0 4719.1978
1245.6944580078125 0 1752.1522
1247.651123046875 0 2202.7563
1248.6600341796875 0 1416.4321
1254.566162109375 0 881.59216
1257.7108154296875 0 3030.5034
1258.7176513671875 0 2926.4824
1259.720947265625 0 3697.1636
1260.7257080078125 0 2823.2275
1261.7301025390625 0 1163.9006
1267.682373046875 0 765.17163
1269.5567626953125 0 1621.6007
1269.696533203125 0 1564.5715
1270.5579833984375 0 1756.159
1270.6998291015625 0 2662.974
1271.709228515625 0 891.58734
1284.68017578125 0 2656.6921
1285.7039794921875 0 16151.951
1286.7005615234375 0 88548.3
1287.7027587890625 0 70285.11
1288.7034912109375 0 30288.725
1289.7025146484375 0 6855.29
1290.697265625 0 851.24945
1300.6942138671875 0 3022.5288
1301.6959228515625 0 5092.845
1302.711669921875 0 89165.83
1303.71875 0 234319.58
1304.7215576171875 0 151401.45
1305.721923828125 0 67104.11
1306.7222900390625 0 10956.708
1320.0684814453125 0 857.27527
1483.7149658203125 0 691.7445
1598.2303466796875 0 1118.634
1598.7613525390625 0 1507.4749
1599.26025390625 0 1475.7131
1628.268798828125 0 722.33636
1628.7747802734375 0 1182.1526
1953.919677734375 0 766.14136
3334.9912109375 0 704.30615

Spectrum Details

|  |  |
| --- | --- |
| Matched peaks? Matched peaks The total absolute number of peaks matched. Additionally in brackets the total fraction of peaks matched and the total number of peaks is shown. | 23 (10.55% of 218) |
| FDR? FDR The false discovery rate estimated for this peptide. It is calculated by matching all theoretical fragments with a non-integer shift with the raw peaks for this spectrum. This is done with 40 different shifts. The resulting percentage is the average number of annotated peaks over the number of annotated peaks with the correct spectrum. | 3.11% |
| Satellite FDR? Satellite FDR See the FDR for details on its calculation. This satellite ion specific FDR only contains the satellite ions (d/w) for I/L/J positions. | ∞ |
| PSM Score? PSM Score The PSM Score as given by Hecklib to this annotated spectrum. It is shown with three significant figures. | 188 |

#### Spectrum 5872? Spectrum 5872 The raw spectrum of this peptide as annotated by Hecklib. The fragments are coloured according to ion type (see legend). Any peaks with a star '\*' as text can be hovered over to see the full details, first the ion type second the mass shift type. By hovering over the amino acids in the peptide or ions in the legend the corresponding peaks are highlighted. By toggling the 'Unassigned' label you can turn the background (unassigned) peaks on or off in the plot. By updating the slider in the Ion legend you can update the spectrum to only show the top X% of the peaks with labels. The top X% means any peak that is within X% of the highest intensity. By dragging in the spectrum you can zoom in to a specific part of the spectrum and use 'Zoom Out' to get back to the original zoom level. The annotation of the spectrum is based on the given sequence in the peptides file and is done with different software so inconsistencies are likely. The peaks are annotated based on the given sequence, with 20 ppm tolerance.

Copy Data

##### Spectrum 5872 (TSV)

###### Preview

```
Loading example...
```

*Click on the button to copy the data to your clipboard.*

Mz MinMz MaxIntensity Max

WidthHeightPeptide font sizePeptide stroke widthSpectrum font sizeSpectrum stroke widthCompact peptide

Ion legend

wxyz

abcd

OtherUnassignedIon^Charge^_Position_Show for top:%

JFPPKPKDTJM

02.33e+44.66e+46.98e+49.31e+4

Zoom Out

y^+1^_2_y^+3^_9_z^+1^_3_w^+1^_4_z^+2^_8_z^+1^_4_c^+2^_8_c^+1^_4_y^+2^_8_z^+1^_4_y^+1^_4_w^+2^_9_y^+2^_9_c^+2^_9_y^+2^_9_c^+2^_10_z^+2^_10_y^+2^_10_z^+2^_10_y^+2^_10_z^+1^_5_w^+1^_6_c^+1^_6_c^+1^_6_y^+1^_6_c^+1^_7_c^+1^_7_z^+1^_7_y^+1^_7_c^+1^_8_c^+1^_8_y^+1^_8_c^+1^_9_c^+1^_9_c^+1^_10_c^+1^_10_z^+1^_10_y^+1^_10_

0776155123273102

Fragment Matches Table

Show background peaks

| Position | Ion type | Intensity | mz Theoretical | mz Error (Th) | mz Error (ppm) | Charge | Series Number |
| --- | --- | --- | --- | --- | --- | --- | --- |
| - | - | 5878 | 120.1 | - | - | 0 | - |
| - | - | 380.1 | 121.1 | - | - | 0 | - |
| - | - | 382.4 | 126.6 | - | - | 0 | - |
| - | - | 372.6 | 127.3 | - | - | 0 | - |
| - | - | 393.2 | 128.1 | - | - | 0 | - |
| - | - | 706.9 | 129.1 | - | - | 0 | - |
| - | - | 558.6 | 131 | - | - | 0 | - |
| - | - | 437.3 | 135.9 | - | - | 0 | - |
| - | - | 481 | 136.1 | - | - | 0 | - |
| - | - | 540.2 | 140.1 | - | - | 0 | - |
| - | - | 833.7 | 144.1 | - | - | 0 | - |
| - | - | 985.1 | 149 | - | - | 0 | - |
| - | - | 3993 | 149 | - | - | 0 | - |
| - | - | 567.8 | 149.1 | - | - | 0 | - |
| - | - | 820.2 | 156.1 | - | - | 0 | - |
| - | - | 675.5 | 156.1 | - | - | 0 | - |
| - | - | 646.3 | 158.1 | - | - | 0 | - |
| - | - | 454.1 | 161.6 | - | - | 0 | - |
| - | - | 2.378E+04 | 166.1 | - | - | 0 | - |
| - | - | 1314 | 167 | - | - | 0 | - |
| - | - | 937.3 | 167.1 | - | - | 0 | - |
| - | - | 1227 | 168 | - | - | 0 | - |
| - | - | 1433 | 172.1 | - | - | 0 | - |
| - | - | 1700 | 173.5 | - | - | 0 | - |
| - | - | 702.2 | 182.1 | - | - | 0 | - |
| - | - | 814.2 | 200.1 | - | - | 0 | - |
| - | - | 660.5 | 202.1 | - | - | 0 | - |
| - | - | 3875 | 209 | - | - | 0 | - |
| - | - | 1.199E+04 | 211 | - | - | 0 | - |
| - | - | 936.3 | 212 | - | - | 0 | - |
| - | - | 3.262E+04 | 212.1 | - | - | 0 | - |
| - | - | 2631 | 212.9 | - | - | 0 | - |
| - | - | 3925 | 213.1 | - | - | 0 | - |
| - | - | 677.1 | 215.1 | - | - | 0 | - |
| - | - | 1073 | 220.1 | - | - | 0 | - |
| - | - | 1319 | 226.1 | - | - | 0 | - |
| - | - | 804.8 | 226.2 | - | - | 0 | - |
| - | - | 1175 | 227.2 | - | - | 0 | - |
| - | - | 976.2 | 229.1 | - | - | 0 | - |
| - | - | 1.559E+04 | 233.2 | - | - | 0 | - |
| - | - | 1908 | 234.2 | - | - | 0 | - |
| - | - | 585 | 235.1 | - | - | 0 | - |
| - | - | 726.4 | 244.1 | - | - | 0 | - |
| - | - | 520.5 | 258.6 | - | - | 0 | - |
| - | - | 449.7 | 259.2 | - | - | 0 | - |
| - | - | 7890 | 260.2 | - | - | 0 | - |
| - | - | 6981 | 261.2 | - | - | 0 | - |
| - | - | 1207 | 262.2 | - | - | 0 | - |
| 10 | y | 7471 | 279.1 | 0.004909 | 17.59 | +1 | 2 |
| - | - | 634.5 | 291.2 | - | - | 0 | - |
| - | - | 1102 | 296.1 | - | - | 0 | - |
| - | - | 1020 | 297.1 | - | - | 0 | - |
| - | - | 868.8 | 297.2 | - | - | 0 | - |
| - | - | 4122 | 306.1 | - | - | 0 | - |
| - | - | 3329 | 308.1 | - | - | 0 | - |
| - | - | 1142 | 315.2 | - | - | 0 | - |
| - | - | 499.5 | 315.2 | - | - | 0 | - |
| - | - | 1814 | 320.9 | - | - | 0 | - |
| - | - | 7132 | 322.9 | - | - | 0 | - |
| - | - | 1182 | 323.2 | - | - | 0 | - |
| - | - | 675.6 | 323.9 | - | - | 0 | - |
| - | - | 1489 | 324.2 | - | - | 0 | - |
| - | - | 1875 | 324.9 | - | - | 0 | - |
| - | - | 925 | 326.9 | - | - | 0 | - |
| - | - | 807.3 | 327.2 | - | - | 0 | - |
| 3 | y | 1110 | 348.2 | 0.002547 | 7.315 | +3 | 9 |
| - | - | 787 | 358.2 | - | - | 0 | - |
| - | - | 1411 | 362.2 | - | - | 0 | - |
| - | - | 2718 | 363.2 | - | - | 0 | - |
| 9 | z | 1056 | 364.2 | 0.003512 | 9.644 | +1 | 3 |
| - | - | 7958 | 365.2 | - | - | 0 | - |
| - | - | 1640 | 366.2 | - | - | 0 | - |
| - | - | 1739 | 391.2 | - | - | 0 | - |
| - | - | 1126 | 391.7 | - | - | 0 | - |
| - | - | 693.1 | 401.3 | - | - | 0 | - |
| - | - | 870.9 | 416.2 | - | - | 0 | - |
| - | - | 678.5 | 425.3 | - | - | 0 | - |
| - | - | 749.6 | 426.3 | - | - | 0 | - |
| - | - | 914.9 | 428.3 | - | - | 0 | - |
| 8 | w | 1360 | 434.2 | 0.003611 | 8.318 | +1 | 4 |
| - | - | 668.9 | 434.2 | - | - | 0 | - |
| - | - | 1554 | 434.8 | - | - | 0 | - |
| - | - | 1396 | 434.9 | - | - | 0 | - |
| - | - | 1049 | 435.2 | - | - | 0 | - |
| - | - | 1063 | 435.3 | - | - | 0 | - |
| - | - | 2776 | 439.3 | - | - | 0 | - |
| - | - | 884.7 | 439.8 | - | - | 0 | - |
| - | - | 5579 | 447.8 | - | - | 0 | - |
| - | - | 1621 | 448.3 | - | - | 0 | - |
| - | - | 3338 | 448.8 | - | - | 0 | - |
| - | - | 2193 | 449.3 | - | - | 0 | - |
| - | - | 894.4 | 449.8 | - | - | 0 | - |
| 4 | z | 920.4 | 456.7 | 0.008946 | 19.59 | +2 | 8 |
| 8 | z | 1324 | 461.2 | 0.005322 | 11.54 | +1 | 4 |
| - | - | 6667 | 462.2 | - | - | 0 | - |
| - | - | 970.7 | 463.2 | - | - | 0 | - |
| - | - | 920.5 | 470.3 | - | - | 0 | - |
| 8 | c | 2837 | 470.8 | 0.0003737 | 0.7937 | +2 | 8 |
| - | - | 728.7 | 472.2 | - | - | 0 | - |
| 4 | c | 1303 | 472.3 | 0.0007103 | 1.504 | +1 | 4 |
| - | - | 743.7 | 472.8 | - | - | 0 | - |
| 4 | y | 2510 | 473.3 | 0.002576 | 5.442 | +2 | 8 |
| - | - | 677.4 | 473.3 | - | - | 0 | - |
| - | - | 1050 | 473.8 | - | - | 0 | - |
| 8 | z | 1683 | 479.2 | 0.006018 | 12.56 | +1 | 4 |
| - | - | 9443 | 480.2 | - | - | 0 | - |
| - | - | 2144 | 481.2 | - | - | 0 | - |
| - | - | 803.9 | 482.2 | - | - | 0 | - |
| - | - | 925.3 | 486.8 | - | - | 0 | - |
| - | - | 1011 | 487.3 | - | - | 0 | - |
| - | - | 3710 | 489.8 | - | - | 0 | - |
| - | - | 2257 | 490.3 | - | - | 0 | - |
| - | - | 872.5 | 490.8 | - | - | 0 | - |
| - | - | 917.3 | 492.3 | - | - | 0 | - |
| - | - | 654.8 | 492.8 | - | - | 0 | - |
| 8 | y | 874.9 | 495.2 | 0.004018 | 8.113 | +1 | 4 |
| 3 | w | 963.1 | 500.3 | 0.001669 | 3.337 | +2 | 9 |
| - | - | 1573 | 500.8 | - | - | 0 | - |
| - | - | 2399 | 507.8 | - | - | 0 | - |
| - | - | 738.5 | 508.3 | - | - | 0 | - |
| - | - | 883.3 | 508.8 | - | - | 0 | - |
| 3 | y | 1465 | 512.8 | 0.0001524 | 0.2973 | +2 | 9 |
| - | - | 3054 | 513.8 | - | - | 0 | - |
| - | - | 1194 | 514.3 | - | - | 0 | - |
| - | - | 1005 | 515.3 | - | - | 0 | - |
| - | - | 1461 | 520.8 | - | - | 0 | - |
| - | - | 1074 | 521.3 | - | - | 0 | - |
| 9 | c | 2821 | 521.3 | 0.0008725 | 1.674 | +2 | 9 |
| 3 | y | 3.279E+04 | 521.8 | 0.002744 | 5.259 | +2 | 9 |
| - | - | 1.862E+04 | 522.3 | - | - | 0 | - |
| - | - | 2.446E+04 | 522.8 | - | - | 0 | - |
| - | - | 9872 | 523.3 | - | - | 0 | - |
| - | - | 3182 | 523.8 | - | - | 0 | - |
| - | - | 946.5 | 524.3 | - | - | 0 | - |
| - | - | 1043 | 534.3 | - | - | 0 | - |
| - | - | 3456 | 535.3 | - | - | 0 | - |
| - | - | 1939 | 535.8 | - | - | 0 | - |
| - | - | 5337 | 536.3 | - | - | 0 | - |
| - | - | 3014 | 536.8 | - | - | 0 | - |
| - | - | 1045 | 537.3 | - | - | 0 | - |
| - | - | 1334 | 540.3 | - | - | 0 | - |
| - | - | 982.9 | 548.3 | - | - | 0 | - |
| - | - | 1729 | 548.8 | - | - | 0 | - |
| - | - | 908 | 555.8 | - | - | 0 | - |
| - | - | 1714 | 556.4 | - | - | 0 | - |
| - | - | 1069 | 557.4 | - | - | 0 | - |
| - | - | 1675 | 558.8 | - | - | 0 | - |
| - | - | 759.5 | 559.8 | - | - | 0 | - |
| - | - | 7345 | 563.3 | - | - | 0 | - |
| - | - | 1386 | 564.3 | - | - | 0 | - |
| - | - | 732.8 | 565.3 | - | - | 0 | - |
| - | - | 2165 | 565.4 | - | - | 0 | - |
| - | - | 915 | 569.8 | - | - | 0 | - |
| 10 | c | 2.109E+04 | 577.9 | 0.0001613 | 0.2792 | +2 | 10 |
| 2 | z | 879.6 | 578.3 | 0.009854 | 17.04 | +2 | 10 |
| - | - | 1.502E+04 | 578.4 | - | - | 0 | - |
| - | - | 5057 | 578.9 | - | - | 0 | - |
| - | - | 1518 | 579.4 | - | - | 0 | - |
| - | - | 791.8 | 585.4 | - | - | 0 | - |
| 2 | y | 1962 | 586.8 | 0.007996 | 13.63 | +2 | 10 |
| 2 | z | 1.146E+04 | 587.3 | 0.003351 | 5.705 | +2 | 10 |
| - | - | 7069 | 587.8 | - | - | 0 | - |
| - | - | 2457 | 588.3 | - | - | 0 | - |
| - | - | 1431 | 591.4 | - | - | 0 | - |
| - | - | 628.8 | 593.2 | - | - | 0 | - |
| - | - | 616.9 | 593.3 | - | - | 0 | - |
| 2 | y | 1485 | 595.3 | 0.002778 | 4.666 | +2 | 10 |
| - | - | 1143 | 595.8 | - | - | 0 | - |
| - | - | 751.7 | 600.3 | - | - | 0 | - |
| - | - | 1670 | 606.3 | - | - | 0 | - |
| 7 | z | 3856 | 607.3 | 0.004714 | 7.762 | +1 | 5 |
| - | - | 1117 | 608.3 | - | - | 0 | - |
| - | - | 2007 | 612.9 | - | - | 0 | - |
| - | - | 1054 | 613.4 | - | - | 0 | - |
| - | - | 859.9 | 613.9 | - | - | 0 | - |
| - | - | 910.6 | 614.4 | - | - | 0 | - |
| - | - | 698.4 | 615.3 | - | - | 0 | - |
| - | - | 799 | 615.8 | - | - | 0 | - |
| - | - | 722.1 | 616.3 | - | - | 0 | - |
| - | - | 1467 | 616.8 | - | - | 0 | - |
| - | - | 737.5 | 617.3 | - | - | 0 | - |
| - | - | 868.2 | 620.4 | - | - | 0 | - |
| - | - | 797.9 | 621.3 | - | - | 0 | - |
| - | - | 3.391E+04 | 622.3 | - | - | 0 | - |
| - | - | 2.195E+04 | 622.8 | - | - | 0 | - |
| - | - | 9584 | 623.3 | - | - | 0 | - |
| - | - | 4826 | 623.8 | - | - | 0 | - |
| - | - | 1664 | 624.3 | - | - | 0 | - |
| - | - | 830.7 | 624.8 | - | - | 0 | - |
| - | - | 899 | 626.4 | - | - | 0 | - |
| - | - | 1064 | 629.4 | - | - | 0 | - |
| - | - | 1641 | 630.4 | - | - | 0 | - |
| - | - | 1757 | 635.8 | - | - | 0 | - |
| - | - | 712.7 | 636.3 | - | - | 0 | - |
| - | - | 1360 | 641.8 | - | - | 0 | - |
| - | - | 1806 | 642.3 | - | - | 0 | - |
| - | - | 753.3 | 642.8 | - | - | 0 | - |
| - | - | 2964 | 643.4 | - | - | 0 | - |
| - | - | 3.588E+04 | 643.9 | - | - | 0 | - |
| - | - | 2.347E+04 | 644.4 | - | - | 0 | - |
| - | - | 1.159E+04 | 644.9 | - | - | 0 | - |
| - | - | 3461 | 645.4 | - | - | 0 | - |
| - | - | 778.5 | 645.9 | - | - | 0 | - |
| - | - | 1203 | 649.9 | - | - | 0 | - |
| - | - | 2135 | 650.4 | - | - | 0 | - |
| - | - | 2799 | 650.9 | - | - | 0 | - |
| - | - | 1287 | 651.4 | - | - | 0 | - |
| - | - | 6.828E+04 | 651.9 | - | - | 0 | - |
| - | - | 9.22E+04 | 652.4 | - | - | 0 | - |
| - | - | 5.278E+04 | 652.9 | - | - | 0 | - |
| - | - | 1.987E+04 | 653.4 | - | - | 0 | - |
| - | - | 7628 | 653.9 | - | - | 0 | - |
| - | - | 1370 | 654.4 | - | - | 0 | - |
| - | - | 683.7 | 674.3 | - | - | 0 | - |
| 6 | w | 1043 | 677.3 | 0.004356 | 6.431 | +1 | 6 |
| 6 | c | 2055 | 680.4 | 0.003341 | 4.91 | +1 | 6 |
| - | - | 1355 | 681.4 | - | - | 0 | - |
| - | - | 766.8 | 695.4 | - | - | 0 | - |
| - | - | 1957 | 696.4 | - | - | 0 | - |
| 6 | c | 2.946E+04 | 697.4 | 1.717E-05 | 0.02462 | +1 | 6 |
| - | - | 9456 | 698.4 | - | - | 0 | - |
| - | - | 2830 | 699.4 | - | - | 0 | - |
| 6 | y | 9719 | 720.4 | 0.004027 | 5.59 | +1 | 6 |
| - | - | 3545 | 721.4 | - | - | 0 | - |
| - | - | 842.6 | 722.4 | - | - | 0 | - |
| - | - | 1976 | 748.4 | - | - | 0 | - |
| - | - | 824.6 | 749.4 | - | - | 0 | - |
| - | - | 682.7 | 753.5 | - | - | 0 | - |
| - | - | 1028 | 780.5 | - | - | 0 | - |
| - | - | 1153 | 781.5 | - | - | 0 | - |
| - | - | 1190 | 781.5 | - | - | 0 | - |
| - | - | 4675 | 788.4 | - | - | 0 | - |
| - | - | 1610 | 789.4 | - | - | 0 | - |
| - | - | 1003 | 790.5 | - | - | 0 | - |
| 7 | c | 1270 | 808.5 | 0.005164 | 6.387 | +1 | 7 |
| - | - | 1179 | 809.5 | - | - | 0 | - |
| - | - | 1237 | 823.5 | - | - | 0 | - |
| - | - | 2.622E+04 | 824.5 | - | - | 0 | - |
| 7 | c | 6.747E+04 | 825.5 | 0.001135 | 1.375 | +1 | 7 |
| - | - | 3.186E+04 | 826.5 | - | - | 0 | - |
| - | - | 8748 | 827.5 | - | - | 0 | - |
| - | - | 1025 | 828.5 | - | - | 0 | - |
| 5 | z | 3.08E+04 | 832.4 | 0.004692 | 5.636 | +1 | 7 |
| - | - | 1.429E+04 | 833.4 | - | - | 0 | - |
| - | - | 4828 | 834.4 | - | - | 0 | - |
| - | - | 1218 | 835.4 | - | - | 0 | - |
| - | - | 629.9 | 838.7 | - | - | 0 | - |
| 5 | y | 1299 | 848.4 | 0.00556 | 6.553 | +1 | 7 |
| - | - | 945.6 | 868.6 | - | - | 0 | - |
| - | - | 2619 | 877.5 | - | - | 0 | - |
| - | - | 719 | 882.6 | - | - | 0 | - |
| - | - | 1732 | 894.5 | - | - | 0 | - |
| - | - | 903.4 | 895.5 | - | - | 0 | - |
| - | - | 1169 | 896.5 | - | - | 0 | - |
| - | - | 1086 | 897.6 | - | - | 0 | - |
| 8 | c | 779.3 | 923.5 | 0.002316 | 2.508 | +1 | 8 |
| - | - | 1173 | 925.6 | - | - | 0 | - |
| - | - | 678.2 | 926.5 | - | - | 0 | - |
| - | - | 1.993E+04 | 939.6 | - | - | 0 | - |
| 8 | c | 4.426E+04 | 940.6 | 0.001955 | 2.078 | +1 | 8 |
| - | - | 2.309E+04 | 941.6 | - | - | 0 | - |
| - | - | 5856 | 942.6 | - | - | 0 | - |
| - | - | 1488 | 943.6 | - | - | 0 | - |
| 4 | y | 4092 | 945.5 | 0.005348 | 5.656 | +1 | 8 |
| - | - | 2204 | 946.5 | - | - | 0 | - |
| - | - | 1192 | 947.5 | - | - | 0 | - |
| - | - | 721.1 | 981.6 | - | - | 0 | - |
| 9 | c | 728.4 | 1025 | 0.005811 | 5.672 | +1 | 9 |
| - | - | 881.1 | 1026 | - | - | 0 | - |
| - | - | 1672 | 1027 | - | - | 0 | - |
| - | - | 1069 | 1041 | - | - | 0 | - |
| 9 | c | 8253 | 1042 | 0.0009883 | 0.9488 | +1 | 9 |
| - | - | 6020 | 1043 | - | - | 0 | - |
| - | - | 2289 | 1044 | - | - | 0 | - |
| - | - | 5983 | 1045 | - | - | 0 | - |
| - | - | 2581 | 1046 | - | - | 0 | - |
| - | - | 1404 | 1047 | - | - | 0 | - |
| - | - | 643.9 | 1050 | - | - | 0 | - |
| - | - | 709.1 | 1095 | - | - | 0 | - |
| - | - | 1708 | 1111 | - | - | 0 | - |
| - | - | 5564 | 1112 | - | - | 0 | - |
| - | - | 2835 | 1113 | - | - | 0 | - |
| - | - | 1017 | 1114 | - | - | 0 | - |
| 10 | c | 735.6 | 1138 | 0.001374 | 1.208 | +1 | 10 |
| - | - | 1613 | 1139 | - | - | 0 | - |
| - | - | 1395 | 1140 | - | - | 0 | - |
| 10 | c | 7886 | 1155 | 0.001678 | 1.453 | +1 | 10 |
| - | - | 1.105E+04 | 1156 | - | - | 0 | - |
| - | - | 5706 | 1157 | - | - | 0 | - |
| - | - | 1863 | 1158 | - | - | 0 | - |
| 2 | z | 3018 | 1174 | 0.00116 | 0.9888 | +1 | 10 |
| - | - | 1.46E+04 | 1175 | - | - | 0 | - |
| - | - | 9021 | 1176 | - | - | 0 | - |
| - | - | 4252 | 1177 | - | - | 0 | - |
| - | - | 1493 | 1178 | - | - | 0 | - |
| 2 | y | 899.3 | 1190 | 0.0204 | 17.15 | +1 | 10 |
| - | - | 1024 | 1213 | - | - | 0 | - |
| - | - | 880.7 | 1214 | - | - | 0 | - |
| - | - | 985 | 1228 | - | - | 0 | - |
| - | - | 902.2 | 1229 | - | - | 0 | - |
| - | - | 867.4 | 1241 | - | - | 0 | - |
| - | - | 1919 | 1242 | - | - | 0 | - |
| - | - | 3956 | 1243 | - | - | 0 | - |
| - | - | 3402 | 1244 | - | - | 0 | - |
| - | - | 1.099E+04 | 1245 | - | - | 0 | - |
| - | - | 7331 | 1246 | - | - | 0 | - |
| - | - | 2496 | 1247 | - | - | 0 | - |
| - | - | 1406 | 1248 | - | - | 0 | - |
| - | - | 1399 | 1259 | - | - | 0 | - |
| - | - | 805 | 1260 | - | - | 0 | - |
| - | - | 904.2 | 1261 | - | - | 0 | - |
| - | - | 685.5 | 1269 | - | - | 0 | - |
| - | - | 2087 | 1270 | - | - | 0 | - |
| - | - | 3169 | 1271 | - | - | 0 | - |
| - | - | 2102 | 1272 | - | - | 0 | - |
| - | - | 668.2 | 1277 | - | - | 0 | - |
| - | - | 685.1 | 1278 | - | - | 0 | - |
| - | - | 1405 | 1284 | - | - | 0 | - |
| - | - | 1533 | 1285 | - | - | 0 | - |
| - | - | 2213 | 1286 | - | - | 0 | - |
| - | - | 1.532E+04 | 1287 | - | - | 0 | - |
| - | - | 4.7E+04 | 1288 | - | - | 0 | - |
| - | - | 3.174E+04 | 1289 | - | - | 0 | - |
| - | - | 1.391E+04 | 1290 | - | - | 0 | - |
| - | - | 6258 | 1291 | - | - | 0 | - |
| - | - | 1784 | 1301 | - | - | 0 | - |
| - | - | 1774 | 1302 | - | - | 0 | - |
| - | - | 7235 | 1303 | - | - | 0 | - |
| - | - | 2.355E+04 | 1304 | - | - | 0 | - |
| - | - | 5.561E+04 | 1305 | - | - | 0 | - |
| - | - | 3.951E+04 | 1306 | - | - | 0 | - |
| - | - | 1.817E+04 | 1307 | - | - | 0 | - |
| - | - | 5876 | 1308 | - | - | 0 | - |
| - | - | 1566 | 1309 | - | - | 0 | - |
| - | - | 701 | 1695 | - | - | 0 | - |
| - | - | 640.6 | 2158 | - | - | 0 | - |
| - | - | 937.9 | 3071 | - | - | 0 | - |

m/z Charge Intensity FragmentType MassShift Position
120.08099365234375 0 5878.4736
121.08422088623047 0 380.0725
126.6236801147461 0 382.3915
127.274658203125 0 372.6436
128.10726928710938 0 393.24948
129.1024932861328 0 706.8831
131.04541015625 0 558.58685
135.85040283203125 0 437.26752
136.0754852294922 0 481.0258
140.0816650390625 0 540.23315
144.06588745117188 0 833.7453
148.95448303222656 0 985.13715
149.02357482910156 0 3993.2349
149.0559539794922 0 567.8474
156.06578063964844 0 820.2199
156.07696533203125 0 675.50366
158.14202880859375 0 646.30896
161.56524658203125 0 454.14276
166.0535125732422 0 23783.377
167.0341339111328 0 1313.5143
167.056640625 0 937.3047
168.04904174804688 0 1227.4767
172.13331604003906 0 1432.6578
173.4525604248047 0 1700.1744
182.08140563964844 0 702.20386
200.12786865234375 0 814.16846
202.13197326660156 0 660.521
208.95327758789062 0 3875.0923
210.95046997070312 0 11993.839
211.95370483398438 0 936.31384
212.13958740234375 0 32624.15
212.9473876953125 0 2631.2502
213.1430206298828 0 3925.391
215.13929748535156 0 677.07153
220.06431579589844 0 1073.053
226.09507751464844 0 1318.9098
226.15541076660156 0 804.7659
227.1758575439453 0 1175.402
229.10946655273438 0 976.21674
233.1650390625 0 15592.25
234.1687469482422 0 1907.5729
235.07489013671875 0 585.04974
244.1060028076172 0 726.3716
258.6290283203125 0 520.4875
259.156494140625 0 449.68558
260.152099609375 0 7889.83
261.15960693359375 0 6981.364
262.16302490234375 0 1207.0865
279.1372985839844 0 7471.284 y 9
291.1708984375 0 634.45886
296.13629150390625 0 1101.6188
297.1446838378906 0 1019.87714
297.2287902832031 0 868.80396
306.0882263183594 0 4121.935
308.1038513183594 0 3328.9595
315.1556091308594 0 1142.0845
315.216552734375 0 499.5464
320.920166015625 0 1814.0883
322.9166259765625 0 7132.488
323.20684814453125 0 1182.4038
323.91851806640625 0 675.61346
324.21490478515625 0 1488.6415
324.9137268066406 0 1875.4014
326.8582763671875 0 924.9971
327.1937561035156 0 807.2913
348.1924743652344 0 1110.4205 y 2
358.2135009765625 0 787.02295
362.15081787109375 0 1410.7804
363.15863037109375 0 2717.9639
364.16485595703125 0 1056.0453 z 8
365.1739501953125 0 7958.254
366.177490234375 0 1639.8447
391.232421875 0 1739.3693
391.7369689941406 0 1126.2479
401.26458740234375 0 693.06866
416.2272644042969 0 870.86505
425.2610168457031 0 678.4911
426.2718200683594 0 749.55786
428.27899169921875 0 914.85455
434.1942443847656 0 1359.761 w 7
434.2373046875 0 668.86694
434.78094482421875 0 1553.9785
434.8829650878906 0 1395.6923
435.20526123046875 0 1048.761
435.2808837890625 0 1063.4606
439.2613220214844 0 2775.66
439.76226806640625 0 884.6558
447.7742004394531 0 5578.7593
448.2717590332031 0 1621.0304
448.7781066894531 0 3337.5354
449.2779846191406 0 2192.7253
449.7818298339844 0 894.3888
456.7411804199219 0 920.43243 z Ammonia loss 3
461.18304443359375 0 1323.9515 z Water loss 7
462.1907043457031 0 6667.4326
463.1935729980469 0 970.6808
470.27935791015625 0 920.46674
470.78399658203125 0 2837.1494 c 7
472.2484130859375 0 728.74493
472.29254150390625 0 1303.149 c 3
472.75225830078125 0 743.67926
473.2574462890625 0 2510.1707 y 3
473.2945556640625 0 677.41437
473.7570495605469 0 1049.5125
479.1943054199219 0 1682.9408 z 7
480.2012939453125 0 9443.478
481.20452880859375 0 2143.6624
482.1979675292969 0 803.86444
486.7517395019531 0 925.3308
487.2522277832031 0 1010.53723
489.7849426269531 0 3709.93
490.28680419921875 0 2256.6143
490.78924560546875 0 872.54517
492.27801513671875 0 917.3073
492.78155517578125 0 654.7857
495.2110290527344 0 874.9103 y 7
500.25848388671875 0 963.10767 w 2
500.7649230957031 0 1573.1837
507.7734069824219 0 2399.4104
508.279541015625 0 738.5058
508.7796630859375 0 883.261
512.7758178710938 0 1464.6932 y Water loss 2
513.7843017578125 0 3054.0957
514.2865600585938 0 1193.6207
515.283447265625 0 1005.13043
520.775390625 0 1460.5736
521.2714233398438 0 1074.0151
521.30908203125 0 2821.3428 c 8
521.7839965820312 0 32789.12 y 2
522.2853393554688 0 18618.398
522.7901000976562 0 24457.219
523.2921142578125 0 9871.877
523.7916870117188 0 3181.8687
524.2940673828125 0 946.47064
534.3150024414062 0 1042.5706
535.2777099609375 0 3456.3408
535.77978515625 0 1938.857
536.2840576171875 0 5337.4185
536.7862548828125 0 3014.4697
537.284423828125 0 1045.3611
540.3089599609375 0 1333.6255
548.3236694335938 0 982.946
548.8232421875 0 1728.6423
555.8419799804688 0 908.0289
556.3726196289062 0 1713.8013
557.3765258789062 0 1069.4338
558.7904052734375 0 1674.9846
559.7955322265625 0 759.48334
563.2987060546875 0 7344.5474
564.3015747070312 0 1385.9924
565.30712890625 0 732.7702
565.3817138671875 0 2165.26
569.834228515625 0 914.99524
577.8504028320312 0 21094.1 c 9
578.3106689453125 0 879.58026 z Water loss 1
578.3519897460938 0 15024.37
578.85400390625 0 5056.9736
579.35595703125 0 1518.3368
585.3754272460938 0 791.75934
586.8101806640625 0 1961.8159 y Ammonia loss 1
587.3094482421875 0 11463.133 z 1
587.8109741210938 0 7068.5996
588.3132934570312 0 2457.0994
591.35986328125 0 1431.1193
593.2466430664062 0 628.848
593.3190307617188 0 616.8764
595.3182373046875 0 1484.511 y 1
595.819580078125 0 1142.6438
600.3351440429688 0 751.7393
606.3407592773438 0 1669.8386
607.2879638671875 0 3855.5837 z 6
608.2929077148438 0 1116.656
612.85205078125 0 2006.8225
613.3551025390625 0 1053.775
613.8521728515625 0 859.93164
614.3638916015625 0 910.6487
615.3248291015625 0 698.35736
615.818359375 0 799.0061
616.3240966796875 0 722.10144
616.8287353515625 0 1467.3878
617.3301391601562 0 737.5272
620.3549194335938 0 868.1586
621.31005859375 0 797.8842
622.323974609375 0 33908.594
622.8251953125 0 21950.94
623.3252563476562 0 9583.606
623.828125 0 4826.24
624.3306274414062 0 1664.2142
624.8314208984375 0 830.7279
626.3723754882812 0 898.9615
629.3628540039062 0 1064.3861
630.36669921875 0 1640.8068
635.8475952148438 0 1757.0424
636.349609375 0 712.7376
641.8423461914062 0 1360.318
642.3418579101562 0 1806.0924
642.8486938476562 0 753.3257
643.3557739257812 0 2964.0427
643.8600463867188 0 35876.375
644.3607177734375 0 23470.596
644.8612670898438 0 11586.195
645.3619384765625 0 3461.2837
645.8657836914062 0 778.4767
649.8507080078125 0 1203.3722
650.3518676757812 0 2134.6086
650.8555297851562 0 2799.4336
651.3555908203125 0 1286.8103
651.860107421875 0 68282.99
652.362548828125 0 92201.04
652.8638916015625 0 52783.586
653.3643798828125 0 19870.535
653.86474609375 0 7628.434
654.3656005859375 0 1369.5791
674.337890625 0 683.7073
677.31689453125 0 1043.2152 w 5
680.40966796875 0 2055.3396 c Ammonia loss 5
681.4111328125 0 1355.2466
695.4273681640625 0 766.76544
696.43115234375 0 1956.8312
697.4395751953125 0 29459.334 c 5
698.4423217773438 0 9456.377
699.4442749023438 0 2830.0823
720.3587646484375 0 9719.245 y 5
721.3621215820312 0 3544.6038
722.3690795898438 0 842.58875
748.3521728515625 0 1975.7716
749.3630981445312 0 824.6395
753.4542236328125 0 682.7056
780.4585571289062 0 1027.8384
781.4525756835938 0 1152.8938
781.5234985351562 0 1189.901
788.44580078125 0 4675.02
789.4474487304688 0 1610.2235
790.4515991210938 0 1003.1795
808.5028076171875 0 1269.5376 c Ammonia loss 6
809.5119018554688 0 1179.2145
823.5167846679688 0 1236.6852
824.5261840820312 0 26223.05
825.5333862304688 0 67474.99 c 6
826.5368041992188 0 31863.004
827.5398559570312 0 8748.196
828.545654296875 0 1025.0276
832.4356689453125 0 30798.586 z 4
833.438720703125 0 14289.88
834.4397583007812 0 4828.2695
835.4440307617188 0 1218.132
838.6683959960938 0 629.8987
848.4552612304688 0 1299.4955 y 4
868.5568237304688 0 945.5795
877.5148315429688 0 2618.5435
882.5509033203125 0 719.012
894.5400390625 0 1732.0795
895.5419311523438 0 903.44366
896.5452880859375 0 1168.9355
897.5543823242188 0 1086.057
923.5372314453125 0 779.32227 c Ammonia loss 7
925.5509643554688 0 1172.5907
926.5413208007812 0 678.2213
939.55224609375 0 19926.395
940.5595092773438 0 44257.85 c 7
941.5631713867188 0 23092.01
942.5667114257812 0 5855.8135
943.5729370117188 0 1487.5751
945.5078125 0 4091.517 y 3
946.509521484375 0 2203.8098
947.5081787109375 0 1191.706
981.5828247070312 0 721.1286
1024.5767822265625 0 728.4337 c Ammonia loss 8
1025.5885009765625 0 881.1169
1026.589599609375 0 1672.3842
1040.598388671875 0 1068.7642
1041.608154296875 0 8253.287 c 8
1042.60498046875 0 6019.5283
1043.607421875 0 2288.9868
1044.57470703125 0 5983.2524
1045.5755615234375 0 2580.9136
1046.5770263671875 0 1404.4746
1050.236328125 0 643.9466
1094.660400390625 0 709.14624
1110.6800537109375 0 1708.4856
1111.6837158203125 0 5564.048
1112.690185546875 0 2834.9697
1113.6932373046875 0 1016.7387
1137.665283203125 0 735.6497 c Ammonia loss 9
1138.673828125 0 1612.597
1139.6796875 0 1395.4943
1154.6915283203125 0 7886.334 c 9
1155.6982421875 0 11050.727
1156.6993408203125 0 5705.932
1157.7054443359375 0 1863.0887
1173.6060791015625 0 3018.459 z 1
1174.6158447265625 0 14598.347
1175.61865234375 0 9021.405
1176.6177978515625 0 4252.394
1177.618896484375 0 1492.5752
1189.64404296875 0 899.3357 y 1
1212.6971435546875 0 1024.1006
1213.6971435546875 0 880.65265
1227.6837158203125 0 984.9577
1228.694091796875 0 902.19775
1240.68994140625 0 867.40875
1241.70947265625 0 1919.4974
1242.7216796875 0 3956.26
1243.7177734375 0 3402.2454
1244.706298828125 0 10989.949
1245.7083740234375 0 7330.597
1246.7054443359375 0 2496.0127
1247.7078857421875 0 1406.2622
1258.7105712890625 0 1399.124
1259.7169189453125 0 805.0015
1260.7310791015625 0 904.2413
1268.6986083984375 0 685.54895
1269.6893310546875 0 2086.8645
1270.6978759765625 0 3168.771
1271.697265625 0 2101.6665
1276.7242431640625 0 668.22955
1277.732666015625 0 685.12195
1283.6851806640625 0 1404.6761
1284.6834716796875 0 1532.8508
1285.700439453125 0 2212.7654
1286.705810546875 0 15324.847
1287.70458984375 0 47001.996
1288.70556640625 0 31736.584
1289.7066650390625 0 13912.051
1290.706298828125 0 6257.6704
1300.7061767578125 0 1783.6138
1301.7076416015625 0 1773.8365
1302.707763671875 0 7234.611
1303.7161865234375 0 23545.828
1304.72412109375 0 55610.41
1305.72705078125 0 39509.84
1306.7281494140625 0 18170.889
1307.7291259765625 0 5876.0894
1308.732666015625 0 1566.3364
1694.5440673828125 0 700.98895
2157.84619140625 0 640.5536
3071.340087890625 0 937.8993

Spectrum Details

|  |  |
| --- | --- |
| Matched peaks? Matched peaks The total absolute number of peaks matched. Additionally in brackets the total fraction of peaks matched and the total number of peaks is shown. | 38 (11.28% of 337) |
| FDR? FDR The false discovery rate estimated for this peptide. It is calculated by matching all theoretical fragments with a non-integer shift with the raw peaks for this spectrum. This is done with 40 different shifts. The resulting percentage is the average number of annotated peaks over the number of annotated peaks with the correct spectrum. | 2.13% |
| Satellite FDR? Satellite FDR See the FDR for details on its calculation. This satellite ion specific FDR only contains the satellite ions (d/w) for I/L/J positions. | ∞ |
| PSM Score? PSM Score The PSM Score as given by Hecklib to this annotated spectrum. It is shown with three significant figures. | 286 |

#### Spectrum 6145? Spectrum 6145 The raw spectrum of this peptide as annotated by Hecklib. The fragments are coloured according to ion type (see legend). Any peaks with a star '\*' as text can be hovered over to see the full details, first the ion type second the mass shift type. By hovering over the amino acids in the peptide or ions in the legend the corresponding peaks are highlighted. By toggling the 'Unassigned' label you can turn the background (unassigned) peaks on or off in the plot. By updating the slider in the Ion legend you can update the spectrum to only show the top X% of the peaks with labels. The top X% means any peak that is within X% of the highest intensity. By dragging in the spectrum you can zoom in to a specific part of the spectrum and use 'Zoom Out' to get back to the original zoom level. The annotation of the spectrum is based on the given sequence in the peptides file and is done with different software so inconsistencies are likely. The peaks are annotated based on the given sequence, with 20 ppm tolerance.

Copy Data

##### Spectrum 6145 (TSV)

###### Preview

```
Loading example...
```

*Click on the button to copy the data to your clipboard.*

Mz MinMz MaxIntensity Max

WidthHeightPeptide font sizePeptide stroke widthSpectrum font sizeSpectrum stroke widthCompact peptide

Ion legend

wxyz

abcd

OtherUnassignedIon^Charge^_Position_Show for top:%

JFPPKPKDTJM

01.66e+43.32e+44.98e+46.65e+4

Zoom Out

y^+1^_2_z^+1^_3_y^+1^_3_w^+1^_4_c^+2^_8_c^+1^_4_y^+2^_8_w^+2^_9_y^+2^_9_y^+2^_9_c^+2^_9_y^+2^_9_c^+2^_10_c^+2^_10_y^+2^_10_z^+2^_10_y^+2^_10_z^+1^_5_c^+1^_6_c^+1^_6_y^+1^_6_c^+1^_7_c^+1^_7_z^+1^_7_y^+1^_7_c^+1^_8_c^+1^_8_y^+1^_8_c^+1^_9_c^+1^_10_z^+1^_10_

0775155123263101

Fragment Matches Table

Show background peaks

| Position | Ion type | Intensity | mz Theoretical | mz Error (Th) | mz Error (ppm) | Charge | Series Number |
| --- | --- | --- | --- | --- | --- | --- | --- |
| - | - | 4810 | 120.1 | - | - | 0 | - |
| - | - | 511.2 | 121.1 | - | - | 0 | - |
| - | - | 414.8 | 124.7 | - | - | 0 | - |
| - | - | 385 | 126.7 | - | - | 0 | - |
| - | - | 512.8 | 128.1 | - | - | 0 | - |
| - | - | 686.6 | 129.1 | - | - | 0 | - |
| - | - | 684.7 | 133.1 | - | - | 0 | - |
| - | - | 422.9 | 136.1 | - | - | 0 | - |
| - | - | 469.9 | 139.3 | - | - | 0 | - |
| - | - | 451.2 | 143.5 | - | - | 0 | - |
| - | - | 832.3 | 144.1 | - | - | 0 | - |
| - | - | 1018 | 148.9 | - | - | 0 | - |
| - | - | 4376 | 149 | - | - | 0 | - |
| - | - | 695 | 149.1 | - | - | 0 | - |
| - | - | 724.8 | 156.1 | - | - | 0 | - |
| - | - | 811.5 | 158.1 | - | - | 0 | - |
| - | - | 1.772E+04 | 166.1 | - | - | 0 | - |
| - | - | 1727 | 167 | - | - | 0 | - |
| - | - | 927.8 | 168 | - | - | 0 | - |
| - | - | 528.2 | 169.4 | - | - | 0 | - |
| - | - | 448.1 | 170.1 | - | - | 0 | - |
| - | - | 446.4 | 170.2 | - | - | 0 | - |
| - | - | 1332 | 172.1 | - | - | 0 | - |
| - | - | 1515 | 173.4 | - | - | 0 | - |
| - | - | 1667 | 173.5 | - | - | 0 | - |
| - | - | 521 | 177 | - | - | 0 | - |
| - | - | 484.3 | 183.1 | - | - | 0 | - |
| - | - | 565.3 | 190.2 | - | - | 0 | - |
| - | - | 453.5 | 195.1 | - | - | 0 | - |
| - | - | 631.6 | 195.1 | - | - | 0 | - |
| - | - | 793.3 | 199.2 | - | - | 0 | - |
| - | - | 642.1 | 202.1 | - | - | 0 | - |
| - | - | 4891 | 209 | - | - | 0 | - |
| - | - | 1.163E+04 | 211 | - | - | 0 | - |
| - | - | 2.626E+04 | 212.1 | - | - | 0 | - |
| - | - | 3784 | 212.9 | - | - | 0 | - |
| - | - | 2489 | 213.1 | - | - | 0 | - |
| - | - | 820.3 | 220.1 | - | - | 0 | - |
| - | - | 575.7 | 226.1 | - | - | 0 | - |
| - | - | 883.4 | 226.2 | - | - | 0 | - |
| - | - | 532.6 | 227.2 | - | - | 0 | - |
| - | - | 479.4 | 227.2 | - | - | 0 | - |
| - | - | 1.234E+04 | 233.2 | - | - | 0 | - |
| - | - | 1477 | 234.2 | - | - | 0 | - |
| - | - | 771.7 | 246.1 | - | - | 0 | - |
| - | - | 772.1 | 255.2 | - | - | 0 | - |
| - | - | 7412 | 260.2 | - | - | 0 | - |
| - | - | 4558 | 261.2 | - | - | 0 | - |
| - | - | 1089 | 262.2 | - | - | 0 | - |
| 10 | y | 4566 | 279.1 | 0.00497 | 17.8 | +1 | 2 |
| - | - | 1002 | 296.1 | - | - | 0 | - |
| - | - | 693.6 | 297.1 | - | - | 0 | - |
| - | - | 733.1 | 302.1 | - | - | 0 | - |
| - | - | 900.5 | 303.1 | - | - | 0 | - |
| - | - | 2780 | 306.1 | - | - | 0 | - |
| - | - | 2637 | 308.1 | - | - | 0 | - |
| - | - | 2564 | 320.9 | - | - | 0 | - |
| - | - | 7480 | 322.9 | - | - | 0 | - |
| - | - | 556.1 | 323 | - | - | 0 | - |
| - | - | 591.1 | 323 | - | - | 0 | - |
| - | - | 899.7 | 323.2 | - | - | 0 | - |
| - | - | 726.4 | 323.9 | - | - | 0 | - |
| - | - | 906.3 | 324.2 | - | - | 0 | - |
| - | - | 1519 | 324.9 | - | - | 0 | - |
| - | - | 654.7 | 326.9 | - | - | 0 | - |
| - | - | 806.3 | 327.2 | - | - | 0 | - |
| - | - | 695.3 | 348.9 | - | - | 0 | - |
| - | - | 1304 | 355.1 | - | - | 0 | - |
| - | - | 1935 | 358.2 | - | - | 0 | - |
| - | - | 838 | 362.2 | - | - | 0 | - |
| - | - | 1576 | 363.2 | - | - | 0 | - |
| 9 | z | 792.7 | 364.2 | 0.002932 | 8.051 | +1 | 3 |
| - | - | 5699 | 365.2 | - | - | 0 | - |
| - | - | 1181 | 366.2 | - | - | 0 | - |
| 9 | y | 892 | 380.2 | 0.003465 | 9.113 | +1 | 3 |
| - | - | 1180 | 391.2 | - | - | 0 | - |
| - | - | 610 | 401.2 | - | - | 0 | - |
| - | - | 1027 | 425.3 | - | - | 0 | - |
| - | - | 694.7 | 425.8 | - | - | 0 | - |
| 8 | w | 1019 | 434.2 | 0.005443 | 12.53 | +1 | 4 |
| - | - | 655 | 434.2 | - | - | 0 | - |
| - | - | 2006 | 434.8 | - | - | 0 | - |
| - | - | 1783 | 434.9 | - | - | 0 | - |
| - | - | 741.2 | 435.2 | - | - | 0 | - |
| - | - | 825.3 | 435.2 | - | - | 0 | - |
| - | - | 779.2 | 435.3 | - | - | 0 | - |
| - | - | 3318 | 439.3 | - | - | 0 | - |
| - | - | 1960 | 439.8 | - | - | 0 | - |
| - | - | 2824 | 447.8 | - | - | 0 | - |
| - | - | 2117 | 448.3 | - | - | 0 | - |
| - | - | 3366 | 448.8 | - | - | 0 | - |
| - | - | 1773 | 449.3 | - | - | 0 | - |
| - | - | 4169 | 462.2 | - | - | 0 | - |
| - | - | 1151 | 463.2 | - | - | 0 | - |
| - | - | 1089 | 470.3 | - | - | 0 | - |
| 8 | c | 2833 | 470.8 | 0.0002211 | 0.4696 | +2 | 8 |
| - | - | 1463 | 471.3 | - | - | 0 | - |
| 4 | c | 859.3 | 472.3 | 0.0003272 | 0.6929 | +1 | 4 |
| 4 | y | 2085 | 473.3 | 0.00227 | 4.797 | +2 | 8 |
| - | - | 710.2 | 473.3 | - | - | 0 | - |
| - | - | 1220 | 473.8 | - | - | 0 | - |
| - | - | 7871 | 480.2 | - | - | 0 | - |
| - | - | 583.8 | 482.2 | - | - | 0 | - |
| - | - | 761.8 | 486.7 | - | - | 0 | - |
| - | - | 2572 | 489.8 | - | - | 0 | - |
| - | - | 2718 | 490.3 | - | - | 0 | - |
| - | - | 928.4 | 490.8 | - | - | 0 | - |
| - | - | 1118 | 492.8 | - | - | 0 | - |
| 3 | w | 1026 | 500.3 | 0.003092 | 6.18 | +2 | 9 |
| - | - | 1373 | 500.8 | - | - | 0 | - |
| - | - | 808.7 | 501.3 | - | - | 0 | - |
| - | - | 734.8 | 506.3 | - | - | 0 | - |
| - | - | 2543 | 507.8 | - | - | 0 | - |
| - | - | 704.3 | 508.3 | - | - | 0 | - |
| 3 | y | 805.8 | 512.8 | 0.002106 | 4.106 | +2 | 9 |
| 3 | y | 917.2 | 513.3 | 0.00845 | 16.46 | +2 | 9 |
| - | - | 1846 | 513.8 | - | - | 0 | - |
| - | - | 723.9 | 514.3 | - | - | 0 | - |
| - | - | 709.5 | 514.8 | - | - | 0 | - |
| - | - | 1455 | 520.8 | - | - | 0 | - |
| 9 | c | 3223 | 521.3 | 7.907E-05 | 0.1517 | +2 | 9 |
| 3 | y | 2.458E+04 | 521.8 | 0.002012 | 3.855 | +2 | 9 |
| - | - | 1.466E+04 | 522.3 | - | - | 0 | - |
| - | - | 1.65E+04 | 522.8 | - | - | 0 | - |
| - | - | 5896 | 523.3 | - | - | 0 | - |
| - | - | 3466 | 523.8 | - | - | 0 | - |
| - | - | 803 | 524.3 | - | - | 0 | - |
| - | - | 946.9 | 534.3 | - | - | 0 | - |
| - | - | 2528 | 535.3 | - | - | 0 | - |
| - | - | 2030 | 535.8 | - | - | 0 | - |
| - | - | 3566 | 536.3 | - | - | 0 | - |
| - | - | 1897 | 536.8 | - | - | 0 | - |
| - | - | 725.8 | 555.3 | - | - | 0 | - |
| - | - | 753.5 | 555.8 | - | - | 0 | - |
| - | - | 1408 | 558.8 | - | - | 0 | - |
| - | - | 4591 | 563.3 | - | - | 0 | - |
| - | - | 1161 | 564.3 | - | - | 0 | - |
| - | - | 986.8 | 565.3 | - | - | 0 | - |
| - | - | 1210 | 565.4 | - | - | 0 | - |
| 10 | c | 1012 | 569.3 | 0.0003133 | 0.5503 | +2 | 10 |
| 10 | c | 1.674E+04 | 577.9 | 0.000388 | 0.6714 | +2 | 10 |
| - | - | 1.185E+04 | 578.4 | - | - | 0 | - |
| - | - | 696.4 | 578.8 | - | - | 0 | - |
| - | - | 3948 | 578.9 | - | - | 0 | - |
| - | - | 625.2 | 585.4 | - | - | 0 | - |
| 2 | y | 2111 | 586.8 | 0.006592 | 11.23 | +2 | 10 |
| 2 | z | 8774 | 587.3 | 0.002618 | 4.458 | +2 | 10 |
| - | - | 4615 | 587.8 | - | - | 0 | - |
| - | - | 1698 | 588.3 | - | - | 0 | - |
| - | - | 1045 | 590.9 | - | - | 0 | - |
| - | - | 659.8 | 594.8 | - | - | 0 | - |
| 2 | y | 867.2 | 595.3 | 2.991E-05 | 0.05024 | +2 | 10 |
| - | - | 1086 | 595.8 | - | - | 0 | - |
| - | - | 678.2 | 596.3 | - | - | 0 | - |
| - | - | 720.1 | 599.8 | - | - | 0 | - |
| - | - | 1191 | 606.3 | - | - | 0 | - |
| 7 | z | 2883 | 607.3 | 0.003798 | 6.254 | +1 | 5 |
| - | - | 953 | 608.3 | - | - | 0 | - |
| - | - | 1826 | 612.9 | - | - | 0 | - |
| - | - | 647.5 | 613.4 | - | - | 0 | - |
| - | - | 1312 | 613.9 | - | - | 0 | - |
| - | - | 781.2 | 616.8 | - | - | 0 | - |
| - | - | 1064 | 621.4 | - | - | 0 | - |
| - | - | 2.687E+04 | 622.3 | - | - | 0 | - |
| - | - | 1.353E+04 | 622.8 | - | - | 0 | - |
| - | - | 6859 | 623.3 | - | - | 0 | - |
| - | - | 3047 | 623.8 | - | - | 0 | - |
| - | - | 869.4 | 624.3 | - | - | 0 | - |
| - | - | 903.7 | 629.4 | - | - | 0 | - |
| - | - | 1414 | 630.4 | - | - | 0 | - |
| - | - | 769.5 | 635.8 | - | - | 0 | - |
| - | - | 942.2 | 636.3 | - | - | 0 | - |
| - | - | 675.4 | 636.8 | - | - | 0 | - |
| - | - | 2030 | 643.4 | - | - | 0 | - |
| - | - | 2.413E+04 | 643.9 | - | - | 0 | - |
| - | - | 1.88E+04 | 644.4 | - | - | 0 | - |
| - | - | 7855 | 644.9 | - | - | 0 | - |
| - | - | 668.2 | 645.3 | - | - | 0 | - |
| - | - | 3318 | 645.4 | - | - | 0 | - |
| - | - | 1294 | 645.9 | - | - | 0 | - |
| - | - | 885.2 | 650.4 | - | - | 0 | - |
| - | - | 1441 | 650.9 | - | - | 0 | - |
| - | - | 1693 | 651.4 | - | - | 0 | - |
| - | - | 5.449E+04 | 651.9 | - | - | 0 | - |
| - | - | 6.579E+04 | 652.4 | - | - | 0 | - |
| - | - | 4.109E+04 | 652.9 | - | - | 0 | - |
| - | - | 1.735E+04 | 653.4 | - | - | 0 | - |
| - | - | 616 | 653.4 | - | - | 0 | - |
| - | - | 5207 | 653.9 | - | - | 0 | - |
| - | - | 858.2 | 654.4 | - | - | 0 | - |
| - | - | 753.3 | 663.4 | - | - | 0 | - |
| 6 | c | 1834 | 680.4 | 0.003158 | 4.641 | +1 | 6 |
| - | - | 795.9 | 684.4 | - | - | 0 | - |
| - | - | 1332 | 696.4 | - | - | 0 | - |
| 6 | c | 2.051E+04 | 697.4 | 0.0006542 | 0.938 | +1 | 6 |
| - | - | 9619 | 698.4 | - | - | 0 | - |
| - | - | 1842 | 699.4 | - | - | 0 | - |
| 6 | y | 6187 | 720.4 | 0.00366 | 5.081 | +1 | 6 |
| - | - | 2594 | 721.4 | - | - | 0 | - |
| - | - | 1195 | 722.4 | - | - | 0 | - |
| - | - | 937.9 | 748.4 | - | - | 0 | - |
| - | - | 984 | 749.4 | - | - | 0 | - |
| - | - | 692.9 | 753.5 | - | - | 0 | - |
| - | - | 896.9 | 768.5 | - | - | 0 | - |
| - | - | 688.3 | 781.4 | - | - | 0 | - |
| - | - | 991.2 | 781.5 | - | - | 0 | - |
| - | - | 2220 | 788.4 | - | - | 0 | - |
| - | - | 1549 | 789.4 | - | - | 0 | - |
| 7 | c | 1813 | 808.5 | 0.003699 | 4.576 | +1 | 7 |
| - | - | 681.5 | 817.5 | - | - | 0 | - |
| - | - | 745.5 | 823.5 | - | - | 0 | - |
| - | - | 2.079E+04 | 824.5 | - | - | 0 | - |
| 7 | c | 5.004E+04 | 825.5 | 0.002172 | 2.632 | +1 | 7 |
| - | - | 2.218E+04 | 826.5 | - | - | 0 | - |
| - | - | 5228 | 827.5 | - | - | 0 | - |
| - | - | 1155 | 828.5 | - | - | 0 | - |
| 5 | z | 2.395E+04 | 832.4 | 0.003654 | 4.39 | +1 | 7 |
| - | - | 1.187E+04 | 833.4 | - | - | 0 | - |
| - | - | 4226 | 834.4 | - | - | 0 | - |
| - | - | 1083 | 835.4 | - | - | 0 | - |
| 5 | y | 935.7 | 848.4 | 0.001276 | 1.504 | +1 | 7 |
| - | - | 1965 | 868.6 | - | - | 0 | - |
| - | - | 885.9 | 869.5 | - | - | 0 | - |
| - | - | 793.6 | 871.4 | - | - | 0 | - |
| - | - | 1650 | 877.5 | - | - | 0 | - |
| - | - | 839.8 | 895.5 | - | - | 0 | - |
| - | - | 1103 | 896.5 | - | - | 0 | - |
| 8 | c | 1011 | 923.5 | 0.005551 | 6.011 | +1 | 8 |
| - | - | 1.412E+04 | 939.6 | - | - | 0 | - |
| 8 | c | 3.404E+04 | 940.6 | 0.003359 | 3.571 | +1 | 8 |
| - | - | 1.792E+04 | 941.6 | - | - | 0 | - |
| - | - | 4793 | 942.6 | - | - | 0 | - |
| - | - | 964 | 943.6 | - | - | 0 | - |
| 4 | y | 3580 | 945.5 | 0.005225 | 5.527 | +1 | 8 |
| - | - | 1600 | 946.5 | - | - | 0 | - |
| - | - | 864.7 | 947.5 | - | - | 0 | - |
| - | - | 893.3 | 1026 | - | - | 0 | - |
| - | - | 1338 | 1027 | - | - | 0 | - |
| - | - | 734.5 | 1028 | - | - | 0 | - |
| 9 | c | 7168 | 1042 | 0.002331 | 2.238 | +1 | 9 |
| - | - | 4192 | 1043 | - | - | 0 | - |
| - | - | 2725 | 1044 | - | - | 0 | - |
| - | - | 4871 | 1045 | - | - | 0 | - |
| - | - | 2923 | 1046 | - | - | 0 | - |
| - | - | 840.7 | 1047 | - | - | 0 | - |
| - | - | 646.9 | 1095 | - | - | 0 | - |
| - | - | 1898 | 1111 | - | - | 0 | - |
| - | - | 4361 | 1112 | - | - | 0 | - |
| - | - | 2834 | 1113 | - | - | 0 | - |
| - | - | 766.9 | 1114 | - | - | 0 | - |
| - | - | 1516 | 1139 | - | - | 0 | - |
| - | - | 721.3 | 1140 | - | - | 0 | - |
| 10 | c | 6557 | 1155 | 0.003753 | 3.251 | +1 | 10 |
| - | - | 9085 | 1156 | - | - | 0 | - |
| - | - | 5024 | 1157 | - | - | 0 | - |
| - | - | 901.5 | 1158 | - | - | 0 | - |
| 2 | z | 1917 | 1174 | 0.002747 | 2.341 | +1 | 10 |
| - | - | 1.19E+04 | 1175 | - | - | 0 | - |
| - | - | 7103 | 1176 | - | - | 0 | - |
| - | - | 2928 | 1177 | - | - | 0 | - |
| - | - | 1061 | 1178 | - | - | 0 | - |
| - | - | 795 | 1214 | - | - | 0 | - |
| - | - | 1054 | 1241 | - | - | 0 | - |
| - | - | 941.5 | 1242 | - | - | 0 | - |
| - | - | 3395 | 1243 | - | - | 0 | - |
| - | - | 2123 | 1244 | - | - | 0 | - |
| - | - | 7756 | 1245 | - | - | 0 | - |
| - | - | 4815 | 1246 | - | - | 0 | - |
| - | - | 2061 | 1247 | - | - | 0 | - |
| - | - | 1101 | 1260 | - | - | 0 | - |
| - | - | 1090 | 1270 | - | - | 0 | - |
| - | - | 1339 | 1271 | - | - | 0 | - |
| - | - | 839.7 | 1272 | - | - | 0 | - |
| - | - | 941.1 | 1273 | - | - | 0 | - |
| - | - | 1128 | 1286 | - | - | 0 | - |
| - | - | 1.051E+04 | 1287 | - | - | 0 | - |
| - | - | 3.424E+04 | 1288 | - | - | 0 | - |
| - | - | 2.373E+04 | 1289 | - | - | 0 | - |
| - | - | 9433 | 1290 | - | - | 0 | - |
| - | - | 3597 | 1291 | - | - | 0 | - |
| - | - | 1179 | 1292 | - | - | 0 | - |
| - | - | 994.2 | 1300 | - | - | 0 | - |
| - | - | 722.3 | 1301 | - | - | 0 | - |
| - | - | 999.1 | 1302 | - | - | 0 | - |
| - | - | 4303 | 1303 | - | - | 0 | - |
| - | - | 1.833E+04 | 1304 | - | - | 0 | - |
| - | - | 4.558E+04 | 1305 | - | - | 0 | - |
| - | - | 2.892E+04 | 1306 | - | - | 0 | - |
| - | - | 1.417E+04 | 1307 | - | - | 0 | - |
| - | - | 5067 | 1308 | - | - | 0 | - |
| - | - | 831.8 | 1408 | - | - | 0 | - |
| - | - | 853.5 | 3070 | - | - | 0 | - |

m/z Charge Intensity FragmentType MassShift Position
120.08096313476562 0 4809.7124
121.08403778076172 0 511.20718
124.69488525390625 0 414.82925
126.67154693603516 0 385.02026
128.10733032226562 0 512.8014
129.1023406982422 0 686.59814
133.06069946289062 0 684.65607
136.07579040527344 0 422.86322
139.33953857421875 0 469.9192
143.5074920654297 0 451.20633
144.06570434570312 0 832.27606
148.94802856445312 0 1017.67896
149.02345275878906 0 4376.233
149.0554656982422 0 694.95
156.06549072265625 0 724.8244
158.1416015625 0 811.4554
166.05328369140625 0 17721.203
167.03372192382812 0 1726.7062
168.0488739013672 0 927.83154
169.38616943359375 0 528.2169
170.1183624267578 0 448.0791
170.15399169921875 0 446.42157
172.13314819335938 0 1331.9348
173.43772888183594 0 1515.0697
173.4514923095703 0 1667.2151
177.02511596679688 0 520.9538
183.12362670898438 0 484.3434
190.2366485595703 0 565.30396
195.08697509765625 0 453.55
195.1127166748047 0 631.627
199.1685791015625 0 793.28735
202.12232971191406 0 642.08704
208.95326232910156 0 4891.445
210.95022583007812 0 11631.668
212.13938903808594 0 26258.586
212.94732666015625 0 3784.378
213.1425018310547 0 2488.6323
220.06451416015625 0 820.27527
226.0959014892578 0 575.66895
226.15460205078125 0 883.41565
227.16224670410156 0 532.58624
227.17510986328125 0 479.4194
233.16488647460938 0 12343.917
234.16793823242188 0 1476.6176
246.1451416015625 0 771.7319
255.17037963867188 0 772.08594
260.1517639160156 0 7412.0728
261.1592712402344 0 4558.406
262.1625061035156 0 1088.844
279.1373596191406 0 4565.847 y 9
296.1366271972656 0 1002.1546
297.14459228515625 0 693.61914
302.1241149902344 0 733.14606
303.1065979003906 0 900.4845
306.08807373046875 0 2779.6357
308.1034240722656 0 2637.355
320.91888427734375 0 2563.7383
322.916259765625 0 7480.136
322.95025634765625 0 556.07214
323.0155944824219 0 591.12836
323.20721435546875 0 899.7233
323.91973876953125 0 726.4168
324.2138671875 0 906.2511
324.9129333496094 0 1518.9067
326.8594665527344 0 654.7015
327.1929016113281 0 806.27246
348.8601379394531 0 695.33636
355.0694580078125 0 1304.4167
358.21160888671875 0 1935.0951
362.1505432128906 0 837.9854
363.1593933105469 0 1576.3278
364.1642761230469 0 792.7155 z 8
365.1737365722656 0 5698.5835
366.17626953125 0 1180.8262
380.18353271484375 0 891.97186 y 8
391.2326354980469 0 1179.6946
401.21917724609375 0 609.9927
425.2630615234375 0 1026.6228
425.7633361816406 0 694.66736
434.1960754394531 0 1019.2213 w 7
434.23773193359375 0 655.0384
434.7799987792969 0 2005.9806
434.88250732421875 0 1782.6415
435.169677734375 0 741.1967
435.2220764160156 0 825.2759
435.27484130859375 0 779.2471
439.26116943359375 0 3317.9673
439.76318359375 0 1960.0641
447.7735595703125 0 2824.0437
448.2760009765625 0 2116.9326
448.7773132324219 0 3365.5364
449.27862548828125 0 1773.3276
462.1901550292969 0 4168.8584
463.1944274902344 0 1150.6913
470.2789611816406 0 1089.4602
470.7841491699219 0 2833.3457 c 7
471.2855224609375 0 1463.296
472.29150390625 0 859.2931 c 3
473.25714111328125 0 2084.9143 y 3
473.2944641113281 0 710.2468
473.75775146484375 0 1220.314
480.200927734375 0 7870.6025
482.20098876953125 0 583.7793
486.7489013671875 0 761.8067
489.783203125 0 2572.4568
490.2877502441406 0 2718.2212
490.79119873046875 0 928.404
492.78033447265625 0 1118.487
500.26324462890625 0 1026.2639 w 2
500.7620544433594 0 1372.9147
501.2667236328125 0 808.72485
506.2723083496094 0 734.8384
507.77313232421875 0 2543.3953
508.2748718261719 0 704.26825
512.7738647460938 0 805.8437 y Water loss 2
513.2764282226562 0 917.15356 y Ammonia loss 2
513.784912109375 0 1845.604
514.2864990234375 0 723.8629
514.784423828125 0 709.4741
520.7741088867188 0 1454.7075
521.3082885742188 0 3222.8271 c 8
521.7832641601562 0 24578.186 y 2
522.2848510742188 0 14657.082
522.789794921875 0 16500.582
523.2918701171875 0 5895.9106
523.792724609375 0 3465.8633
524.2967529296875 0 803.00574
534.3140258789062 0 946.9229
535.276611328125 0 2528.314
535.7793579101562 0 2030.0518
536.28369140625 0 3565.9587
536.7850952148438 0 1896.6655
555.339599609375 0 725.82605
555.8402709960938 0 753.5111
558.7914428710938 0 1408.232
563.2976684570312 0 4590.595
564.3008422851562 0 1161.1193
565.3093872070312 0 986.76373
565.3811645507812 0 1209.6973
569.3372802734375 0 1012.3718 c Ammonia loss 9
577.849853515625 0 16737.912 c 9
578.3512573242188 0 11849.95
578.8070678710938 0 696.394
578.8532104492188 0 3948.0242
585.3731689453125 0 625.2223
586.8087768554688 0 2110.8865 y Ammonia loss 1
587.3087158203125 0 8773.661 z 1
587.8095092773438 0 4614.6836
588.3116455078125 0 1698.0282
590.8565673828125 0 1044.8606
594.8251342773438 0 659.84235
595.3154296875 0 867.185 y 1
595.8196411132812 0 1086.2473
596.322265625 0 678.16174
599.8359375 0 720.06213
606.3411254882812 0 1190.6633
607.2870483398438 0 2882.6958 z 6
608.291748046875 0 953.02484
612.8534545898438 0 1825.7874
613.3549194335938 0 647.51483
613.8540649414062 0 1311.6699
616.828369140625 0 781.1731
621.3612670898438 0 1063.6299
622.3233032226562 0 26868.676
622.8253173828125 0 13532.662
623.323974609375 0 6859.0596
623.8262329101562 0 3046.543
624.329345703125 0 869.3954
629.3602905273438 0 903.73724
630.3659057617188 0 1413.5796
635.8452758789062 0 769.47186
636.3493041992188 0 942.1504
636.8493041992188 0 675.36414
643.356689453125 0 2030.2258
643.8590087890625 0 24129.87
644.3612670898438 0 18804.496
644.8610229492188 0 7854.703
645.3035888671875 0 668.2025
645.3621826171875 0 3317.9746
645.8651123046875 0 1294.4136
650.3541870117188 0 885.16296
650.8522338867188 0 1440.9138
651.3580932617188 0 1693.3215
651.859130859375 0 54492.23
652.36181640625 0 65792.21
652.86328125 0 41091.266
653.3634643554688 0 17348.08
653.42431640625 0 615.9693
653.865478515625 0 5206.593
654.3690185546875 0 858.2476
663.3775024414062 0 753.2597
680.4098510742188 0 1833.9712 c Ammonia loss 5
684.4154663085938 0 795.94714
696.4297485351562 0 1332.0195
697.4389038085938 0 20506.04 c 5
698.44189453125 0 9618.943
699.4464111328125 0 1842.1061
720.3583984375 0 6186.984 y 5
721.360595703125 0 2593.6626
722.36328125 0 1195.4375
748.3514404296875 0 937.8723
749.359130859375 0 983.97485
753.4535522460938 0 692.92755
768.4638671875 0 896.85767
781.4462890625 0 688.2893
781.5195922851562 0 991.1703
788.4459228515625 0 2220.4539
789.44775390625 0 1548.6731
808.5042724609375 0 1813.3125 c Ammonia loss 6
817.451171875 0 681.4509
823.515869140625 0 745.4528
824.5253295898438 0 20785.97
825.5323486328125 0 50036.344 c 6
826.5355224609375 0 22181.154
827.5390014648438 0 5227.7695
828.5454711914062 0 1155.3389
832.4346313476562 0 23945.57 z 4
833.4378051757812 0 11873.506
834.4374389648438 0 4226.146
835.44677734375 0 1082.8468
848.4484252929688 0 935.7061 y 4
868.5528564453125 0 1965.4537
869.5498657226562 0 885.92487
871.4202270507812 0 793.575
877.5149536132812 0 1650.188
895.5408325195312 0 839.7965
896.548095703125 0 1103.0135
923.5404663085938 0 1010.51276 c Ammonia loss 7
939.5512084960938 0 14124.623
940.55810546875 0 34037.812 c 7
941.5618896484375 0 17922.871
942.5661010742188 0 4793.2905
943.5728759765625 0 964.0036
945.5076904296875 0 3579.7017 y 3
946.5119018554688 0 1599.541
947.51123046875 0 864.67175
1025.581787109375 0 893.2832
1026.5872802734375 0 1337.899
1027.59375 0 734.53723
1041.6068115234375 0 7167.85 c 8
1042.602294921875 0 4191.6733
1043.5718994140625 0 2725.043
1044.574951171875 0 4871.466
1045.5760498046875 0 2922.7664
1046.5777587890625 0 840.6814
1094.6646728515625 0 646.8619
1110.676513671875 0 1897.6091
1111.6839599609375 0 4360.747
1112.6861572265625 0 2833.923
1113.6903076171875 0 766.94006
1138.6666259765625 0 1516.1805
1139.673583984375 0 721.27985
1154.689453125 0 6556.7896 c 9
1155.6954345703125 0 9085.241
1156.7008056640625 0 5023.775
1157.7142333984375 0 901.47345
1173.607666015625 0 1917.2136 z 1
1174.61474609375 0 11903.009
1175.6187744140625 0 7102.7065
1176.6181640625 0 2928.0396
1177.61962890625 0 1060.5621
1213.7010498046875 0 794.99915
1240.72119140625 0 1054.2051
1241.721435546875 0 941.49365
1242.72265625 0 3395.1064
1243.717529296875 0 2123.0896
1244.7042236328125 0 7755.513
1245.7049560546875 0 4815.3647
1246.70654296875 0 2061.1821
1259.72314453125 0 1101.1676
1269.689453125 0 1090.4023
1270.6959228515625 0 1339.0746
1271.7066650390625 0 839.7181
1272.7060546875 0 941.0878
1285.6959228515625 0 1127.7632
1286.7027587890625 0 10506.683
1287.702392578125 0 34235.207
1288.70556640625 0 23734.98
1289.705078125 0 9433.048
1290.7020263671875 0 3597.4138
1291.71728515625 0 1178.7535
1299.6947021484375 0 994.1768
1300.7003173828125 0 722.2918
1301.712890625 0 999.1175
1302.7081298828125 0 4303.3804
1303.71484375 0 18333.773
1304.7225341796875 0 45577.707
1305.724853515625 0 28916.457
1306.727294921875 0 14171.783
1307.7264404296875 0 5066.714
1407.906494140625 0 831.791
3070.328369140625 0 853.49335

Spectrum Details

|  |  |
| --- | --- |
| Matched peaks? Matched peaks The total absolute number of peaks matched. Additionally in brackets the total fraction of peaks matched and the total number of peaks is shown. | 31 (10.62% of 292) |
| FDR? FDR The false discovery rate estimated for this peptide. It is calculated by matching all theoretical fragments with a non-integer shift with the raw peaks for this spectrum. This is done with 40 different shifts. The resulting percentage is the average number of annotated peaks over the number of annotated peaks with the correct spectrum. | 2.38% |
| Satellite FDR? Satellite FDR See the FDR for details on its calculation. This satellite ion specific FDR only contains the satellite ions (d/w) for I/L/J positions. | ∞ |
| PSM Score? PSM Score The PSM Score as given by Hecklib to this annotated spectrum. It is shown with three significant figures. | 204 |

#### Spectrum 6095? Spectrum 6095 The raw spectrum of this peptide as annotated by Hecklib. The fragments are coloured according to ion type (see legend). Any peaks with a star '\*' as text can be hovered over to see the full details, first the ion type second the mass shift type. By hovering over the amino acids in the peptide or ions in the legend the corresponding peaks are highlighted. By toggling the 'Unassigned' label you can turn the background (unassigned) peaks on or off in the plot. By updating the slider in the Ion legend you can update the spectrum to only show the top X% of the peaks with labels. The top X% means any peak that is within X% of the highest intensity. By dragging in the spectrum you can zoom in to a specific part of the spectrum and use 'Zoom Out' to get back to the original zoom level. The annotation of the spectrum is based on the given sequence in the peptides file and is done with different software so inconsistencies are likely. The peaks are annotated based on the given sequence, with 20 ppm tolerance.

Copy Data

##### Spectrum 6095 (TSV)

###### Preview

```
Loading example...
```

*Click on the button to copy the data to your clipboard.*

Mz MinMz MaxIntensity Max

WidthHeightPeptide font sizePeptide stroke widthSpectrum font sizeSpectrum stroke widthCompact peptide

Ion legend

wxyz

abcd

OtherUnassignedIon^Charge^_Position_Show for top:%

JFPPKPKDTJM

01.60e+43.20e+44.79e+46.39e+4

Zoom Out

y^+1^_2_y^+3^_9_z^+1^_3_c^+2^_7_w^+1^_4_z^+2^_8_z^+2^_8_c^+2^_8_c^+1^_4_y^+2^_8_z^+1^_4_w^+2^_9_y^+2^_9_c^+2^_9_y^+2^_9_c^+2^_10_c^+2^_10_z^+2^_10_y^+2^_10_z^+2^_10_y^+2^_10_z^+1^_5_w^+1^_6_c^+1^_6_c^+1^_6_y^+1^_6_c^+1^_7_c^+1^_7_z^+1^_7_c^+1^_8_c^+1^_8_y^+1^_8_c^+1^_9_c^+1^_10_z^+1^_10_

0827165424803307

Fragment Matches Table

Show background peaks

| Position | Ion type | Intensity | mz Theoretical | mz Error (Th) | mz Error (ppm) | Charge | Series Number |
| --- | --- | --- | --- | --- | --- | --- | --- |
| - | - | 5409 | 120.1 | - | - | 0 | - |
| - | - | 489.9 | 121.1 | - | - | 0 | - |
| - | - | 360.5 | 121.9 | - | - | 0 | - |
| - | - | 457.3 | 124.1 | - | - | 0 | - |
| - | - | 386.2 | 125.3 | - | - | 0 | - |
| - | - | 482.7 | 129.1 | - | - | 0 | - |
| - | - | 545.1 | 131 | - | - | 0 | - |
| - | - | 489.4 | 136.1 | - | - | 0 | - |
| - | - | 4408 | 149 | - | - | 0 | - |
| - | - | 397.9 | 153.3 | - | - | 0 | - |
| - | - | 898.9 | 156.1 | - | - | 0 | - |
| - | - | 468.9 | 156.1 | - | - | 0 | - |
| - | - | 450.9 | 157.1 | - | - | 0 | - |
| - | - | 841.9 | 158.1 | - | - | 0 | - |
| - | - | 1.704E+04 | 166.1 | - | - | 0 | - |
| - | - | 1356 | 167 | - | - | 0 | - |
| - | - | 561 | 167.1 | - | - | 0 | - |
| - | - | 728 | 168 | - | - | 0 | - |
| - | - | 1147 | 172.1 | - | - | 0 | - |
| - | - | 1477 | 173.4 | - | - | 0 | - |
| - | - | 2058 | 173.5 | - | - | 0 | - |
| - | - | 584.9 | 173.5 | - | - | 0 | - |
| - | - | 586.6 | 173.5 | - | - | 0 | - |
| - | - | 489.4 | 184.4 | - | - | 0 | - |
| - | - | 548.7 | 185.2 | - | - | 0 | - |
| - | - | 527.2 | 189.6 | - | - | 0 | - |
| - | - | 581.7 | 199.2 | - | - | 0 | - |
| - | - | 4006 | 209 | - | - | 0 | - |
| - | - | 1.043E+04 | 211 | - | - | 0 | - |
| - | - | 2.455E+04 | 212.1 | - | - | 0 | - |
| - | - | 3123 | 212.9 | - | - | 0 | - |
| - | - | 2722 | 213.1 | - | - | 0 | - |
| - | - | 498.5 | 213.2 | - | - | 0 | - |
| - | - | 592.3 | 220.1 | - | - | 0 | - |
| - | - | 854.2 | 226.1 | - | - | 0 | - |
| - | - | 720.3 | 226.2 | - | - | 0 | - |
| - | - | 1025 | 227.2 | - | - | 0 | - |
| - | - | 933.4 | 229.1 | - | - | 0 | - |
| - | - | 1.039E+04 | 233.2 | - | - | 0 | - |
| - | - | 1071 | 234.2 | - | - | 0 | - |
| - | - | 743.6 | 255.2 | - | - | 0 | - |
| - | - | 5364 | 260.2 | - | - | 0 | - |
| - | - | 4897 | 261.2 | - | - | 0 | - |
| - | - | 788.9 | 262.2 | - | - | 0 | - |
| 10 | y | 5579 | 279.1 | 0.004817 | 17.26 | +1 | 2 |
| - | - | 752.4 | 279.2 | - | - | 0 | - |
| - | - | 596.8 | 280.1 | - | - | 0 | - |
| - | - | 574.3 | 287.2 | - | - | 0 | - |
| - | - | 550.3 | 291.4 | - | - | 0 | - |
| - | - | 716.7 | 296.1 | - | - | 0 | - |
| - | - | 568.4 | 297 | - | - | 0 | - |
| - | - | 858.9 | 297.1 | - | - | 0 | - |
| - | - | 2995 | 306.1 | - | - | 0 | - |
| - | - | 2731 | 308.1 | - | - | 0 | - |
| - | - | 992.2 | 315.2 | - | - | 0 | - |
| - | - | 528.8 | 319.1 | - | - | 0 | - |
| - | - | 2626 | 320.9 | - | - | 0 | - |
| - | - | 6411 | 322.9 | - | - | 0 | - |
| - | - | 540.5 | 323.1 | - | - | 0 | - |
| - | - | 819.8 | 323.9 | - | - | 0 | - |
| - | - | 619.6 | 324.2 | - | - | 0 | - |
| - | - | 2101 | 324.9 | - | - | 0 | - |
| - | - | 1734 | 326.9 | - | - | 0 | - |
| - | - | 531 | 327.8 | - | - | 0 | - |
| 3 | y | 1097 | 348.2 | 0.0003215 | 0.9234 | +3 | 9 |
| - | - | 1437 | 355.1 | - | - | 0 | - |
| - | - | 772.2 | 356.2 | - | - | 0 | - |
| - | - | 963.5 | 358.2 | - | - | 0 | - |
| - | - | 873.6 | 362.2 | - | - | 0 | - |
| - | - | 685.7 | 363.2 | - | - | 0 | - |
| 9 | z | 711.2 | 364.2 | 0.003756 | 10.31 | +1 | 3 |
| - | - | 5838 | 365.2 | - | - | 0 | - |
| - | - | 1225 | 391.2 | - | - | 0 | - |
| - | - | 862.7 | 391.7 | - | - | 0 | - |
| - | - | 1007 | 401.3 | - | - | 0 | - |
| 7 | c | 815.6 | 413.3 | 0.001734 | 4.195 | +2 | 7 |
| - | - | 1174 | 417.2 | - | - | 0 | - |
| - | - | 848.3 | 418 | - | - | 0 | - |
| - | - | 755.9 | 425.3 | - | - | 0 | - |
| - | - | 742.6 | 428.3 | - | - | 0 | - |
| 8 | w | 641.4 | 434.2 | 0.005961 | 13.73 | +1 | 4 |
| - | - | 2301 | 434.8 | - | - | 0 | - |
| - | - | 1273 | 434.9 | - | - | 0 | - |
| - | - | 996.4 | 435.2 | - | - | 0 | - |
| - | - | 1050 | 435.3 | - | - | 0 | - |
| - | - | 1865 | 439.3 | - | - | 0 | - |
| - | - | 1336 | 439.8 | - | - | 0 | - |
| - | - | 582.9 | 440 | - | - | 0 | - |
| - | - | 4116 | 447.8 | - | - | 0 | - |
| - | - | 2407 | 448.3 | - | - | 0 | - |
| - | - | 2656 | 448.8 | - | - | 0 | - |
| - | - | 993.9 | 449.3 | - | - | 0 | - |
| 4 | z | 711.1 | 456.2 | 0.006386 | 14 | +2 | 8 |
| 4 | z | 718.8 | 456.7 | 0.008458 | 18.52 | +2 | 8 |
| - | - | 4343 | 462.2 | - | - | 0 | - |
| - | - | 686.4 | 463.2 | - | - | 0 | - |
| - | - | 840.4 | 470.3 | - | - | 0 | - |
| 8 | c | 2330 | 470.8 | 0.001259 | 2.674 | +2 | 8 |
| - | - | 937.4 | 471.3 | - | - | 0 | - |
| 4 | c | 751.3 | 472.3 | 0.001962 | 4.153 | +1 | 4 |
| 4 | y | 1916 | 473.3 | 0.001721 | 3.637 | +2 | 8 |
| - | - | 836.4 | 473.8 | - | - | 0 | - |
| 8 | z | 1097 | 479.2 | 0.0026 | 5.426 | +1 | 4 |
| - | - | 7155 | 480.2 | - | - | 0 | - |
| - | - | 1061 | 481.2 | - | - | 0 | - |
| - | - | 1605 | 486.8 | - | - | 0 | - |
| - | - | 2789 | 489.8 | - | - | 0 | - |
| - | - | 1396 | 490.3 | - | - | 0 | - |
| - | - | 1227 | 490.8 | - | - | 0 | - |
| 3 | w | 1411 | 500.3 | 0.002084 | 4.167 | +2 | 9 |
| - | - | 1026 | 500.8 | - | - | 0 | - |
| - | - | 1591 | 507.8 | - | - | 0 | - |
| - | - | 733.1 | 508.8 | - | - | 0 | - |
| 3 | y | 941.3 | 513.3 | 0.007352 | 14.32 | +2 | 9 |
| - | - | 1707 | 513.8 | - | - | 0 | - |
| - | - | 682.7 | 514.3 | - | - | 0 | - |
| - | - | 1302 | 520.8 | - | - | 0 | - |
| - | - | 1042 | 521.3 | - | - | 0 | - |
| 9 | c | 2444 | 521.3 | 0.0001401 | 0.2688 | +2 | 9 |
| 3 | y | 2.299E+04 | 521.8 | 0.002195 | 4.206 | +2 | 9 |
| - | - | 1.186E+04 | 522.3 | - | - | 0 | - |
| - | - | 1.906E+04 | 522.8 | - | - | 0 | - |
| - | - | 8614 | 523.3 | - | - | 0 | - |
| - | - | 2020 | 523.8 | - | - | 0 | - |
| - | - | 686.2 | 524.3 | - | - | 0 | - |
| - | - | 798.3 | 534.3 | - | - | 0 | - |
| - | - | 2683 | 535.3 | - | - | 0 | - |
| - | - | 1314 | 535.8 | - | - | 0 | - |
| - | - | 2406 | 536.3 | - | - | 0 | - |
| - | - | 2189 | 536.8 | - | - | 0 | - |
| - | - | 799 | 539.8 | - | - | 0 | - |
| - | - | 523.8 | 545.2 | - | - | 0 | - |
| - | - | 611 | 550.6 | - | - | 0 | - |
| - | - | 652 | 555.8 | - | - | 0 | - |
| - | - | 1145 | 556.4 | - | - | 0 | - |
| - | - | 578.9 | 559.8 | - | - | 0 | - |
| - | - | 581.7 | 560.3 | - | - | 0 | - |
| - | - | 4015 | 563.3 | - | - | 0 | - |
| - | - | 1235 | 564.3 | - | - | 0 | - |
| - | - | 862.1 | 565.3 | - | - | 0 | - |
| - | - | 1263 | 565.4 | - | - | 0 | - |
| 10 | c | 654.6 | 569.3 | 0.008415 | 14.78 | +2 | 10 |
| - | - | 771.8 | 570.3 | - | - | 0 | - |
| 10 | c | 1.829E+04 | 577.9 | 0.0002659 | 0.4602 | +2 | 10 |
| 2 | z | 806.1 | 578.3 | 0.009121 | 15.77 | +2 | 10 |
| - | - | 8326 | 578.4 | - | - | 0 | - |
| - | - | 4050 | 578.9 | - | - | 0 | - |
| 2 | y | 1995 | 586.8 | 0.008667 | 14.77 | +2 | 10 |
| 2 | z | 7565 | 587.3 | 0.00213 | 3.627 | +2 | 10 |
| - | - | 4887 | 587.8 | - | - | 0 | - |
| - | - | 1487 | 588.3 | - | - | 0 | - |
| - | - | 967.7 | 590.9 | - | - | 0 | - |
| - | - | 793.9 | 591.4 | - | - | 0 | - |
| 2 | y | 697.1 | 595.3 | 0.001496 | 2.513 | +2 | 10 |
| - | - | 1455 | 606.3 | - | - | 0 | - |
| - | - | 1139 | 606.8 | - | - | 0 | - |
| 7 | z | 2627 | 607.3 | 0.00508 | 8.365 | +1 | 5 |
| - | - | 874.8 | 607.3 | - | - | 0 | - |
| - | - | 1069 | 608.3 | - | - | 0 | - |
| - | - | 758.4 | 612.8 | - | - | 0 | - |
| - | - | 1011 | 613.4 | - | - | 0 | - |
| - | - | 906 | 613.9 | - | - | 0 | - |
| - | - | 731.3 | 616.8 | - | - | 0 | - |
| - | - | 694.3 | 617.3 | - | - | 0 | - |
| - | - | 2.32E+04 | 622.3 | - | - | 0 | - |
| - | - | 1.376E+04 | 622.8 | - | - | 0 | - |
| - | - | 7969 | 623.3 | - | - | 0 | - |
| - | - | 3191 | 623.8 | - | - | 0 | - |
| - | - | 1474 | 624.3 | - | - | 0 | - |
| - | - | 1087 | 624.8 | - | - | 0 | - |
| - | - | 679.4 | 625.3 | - | - | 0 | - |
| - | - | 687.7 | 629.4 | - | - | 0 | - |
| - | - | 828.2 | 630.4 | - | - | 0 | - |
| - | - | 775.8 | 635.3 | - | - | 0 | - |
| - | - | 2416 | 643.4 | - | - | 0 | - |
| - | - | 2.254E+04 | 643.9 | - | - | 0 | - |
| - | - | 1.932E+04 | 644.4 | - | - | 0 | - |
| - | - | 8516 | 644.9 | - | - | 0 | - |
| - | - | 3394 | 645.4 | - | - | 0 | - |
| - | - | 744.5 | 649.9 | - | - | 0 | - |
| - | - | 611.8 | 650.4 | - | - | 0 | - |
| - | - | 1359 | 650.9 | - | - | 0 | - |
| - | - | 955.8 | 651.4 | - | - | 0 | - |
| - | - | 5.14E+04 | 651.9 | - | - | 0 | - |
| - | - | 6.329E+04 | 652.4 | - | - | 0 | - |
| - | - | 3.537E+04 | 652.9 | - | - | 0 | - |
| - | - | 1.419E+04 | 653.4 | - | - | 0 | - |
| - | - | 651.4 | 653.4 | - | - | 0 | - |
| - | - | 5211 | 653.9 | - | - | 0 | - |
| 6 | w | 1089 | 677.3 | 0.002464 | 3.637 | +1 | 6 |
| 6 | c | 2231 | 680.4 | 0.003219 | 4.731 | +1 | 6 |
| - | - | 1133 | 684.4 | - | - | 0 | - |
| - | - | 1889 | 696.4 | - | - | 0 | - |
| 6 | c | 1.968E+04 | 697.4 | 0.0007763 | 1.113 | +1 | 6 |
| - | - | 6750 | 698.4 | - | - | 0 | - |
| - | - | 1360 | 699.4 | - | - | 0 | - |
| 6 | y | 6483 | 720.4 | 0.003355 | 4.658 | +1 | 6 |
| - | - | 2858 | 721.4 | - | - | 0 | - |
| - | - | 1316 | 722.4 | - | - | 0 | - |
| - | - | 1112 | 748.4 | - | - | 0 | - |
| - | - | 746.6 | 749.3 | - | - | 0 | - |
| - | - | 917.8 | 768.5 | - | - | 0 | - |
| - | - | 775.4 | 780.5 | - | - | 0 | - |
| - | - | 843.2 | 781.5 | - | - | 0 | - |
| - | - | 959.4 | 781.5 | - | - | 0 | - |
| - | - | 3238 | 788.4 | - | - | 0 | - |
| - | - | 1270 | 789.4 | - | - | 0 | - |
| 7 | c | 919.1 | 808.5 | 0.003028 | 3.745 | +1 | 7 |
| - | - | 820.1 | 816.4 | - | - | 0 | - |
| - | - | 1051 | 823.5 | - | - | 0 | - |
| - | - | 2.055E+04 | 824.5 | - | - | 0 | - |
| 7 | c | 5.005E+04 | 825.5 | 0.001989 | 2.41 | +1 | 7 |
| - | - | 2.026E+04 | 826.5 | - | - | 0 | - |
| - | - | 4606 | 827.5 | - | - | 0 | - |
| - | - | 967.7 | 828.5 | - | - | 0 | - |
| 5 | z | 2.196E+04 | 832.4 | 0.004204 | 5.05 | +1 | 7 |
| - | - | 1.077E+04 | 833.4 | - | - | 0 | - |
| - | - | 4055 | 834.4 | - | - | 0 | - |
| - | - | 1050 | 835.4 | - | - | 0 | - |
| - | - | 1063 | 854.5 | - | - | 0 | - |
| - | - | 2192 | 868.6 | - | - | 0 | - |
| - | - | 1124 | 869.6 | - | - | 0 | - |
| - | - | 1350 | 877.5 | - | - | 0 | - |
| - | - | 773.7 | 894.5 | - | - | 0 | - |
| - | - | 2166 | 896.5 | - | - | 0 | - |
| - | - | 586.4 | 897.5 | - | - | 0 | - |
| 8 | c | 666.6 | 923.5 | 0.002378 | 2.574 | +1 | 8 |
| - | - | 960.2 | 924.5 | - | - | 0 | - |
| - | - | 1.337E+04 | 939.6 | - | - | 0 | - |
| 8 | c | 3.128E+04 | 940.6 | 0.002809 | 2.987 | +1 | 8 |
| - | - | 1.756E+04 | 941.6 | - | - | 0 | - |
| - | - | 5135 | 942.6 | - | - | 0 | - |
| - | - | 1158 | 943.6 | - | - | 0 | - |
| 4 | y | 2911 | 945.5 | 0.006019 | 6.366 | +1 | 8 |
| - | - | 1775 | 946.5 | - | - | 0 | - |
| - | - | 1161 | 1026 | - | - | 0 | - |
| - | - | 1776 | 1027 | - | - | 0 | - |
| 9 | c | 7713 | 1042 | 0.002087 | 2.004 | +1 | 9 |
| - | - | 4273 | 1043 | - | - | 0 | - |
| - | - | 1829 | 1044 | - | - | 0 | - |
| - | - | 4376 | 1045 | - | - | 0 | - |
| - | - | 2630 | 1046 | - | - | 0 | - |
| - | - | 828.6 | 1047 | - | - | 0 | - |
| - | - | 851.3 | 1096 | - | - | 0 | - |
| - | - | 1413 | 1111 | - | - | 0 | - |
| - | - | 4146 | 1112 | - | - | 0 | - |
| - | - | 2589 | 1113 | - | - | 0 | - |
| - | - | 897.6 | 1114 | - | - | 0 | - |
| - | - | 1778 | 1139 | - | - | 0 | - |
| - | - | 1292 | 1140 | - | - | 0 | - |
| 10 | c | 5480 | 1155 | 0.003509 | 3.039 | +1 | 10 |
| - | - | 8833 | 1156 | - | - | 0 | - |
| - | - | 3127 | 1157 | - | - | 0 | - |
| - | - | 1264 | 1158 | - | - | 0 | - |
| 2 | z | 1949 | 1174 | 0.0001839 | 0.1567 | +1 | 10 |
| - | - | 1.039E+04 | 1175 | - | - | 0 | - |
| - | - | 6516 | 1176 | - | - | 0 | - |
| - | - | 2596 | 1177 | - | - | 0 | - |
| - | - | 895.6 | 1178 | - | - | 0 | - |
| - | - | 1370 | 1213 | - | - | 0 | - |
| - | - | 715 | 1214 | - | - | 0 | - |
| - | - | 725.6 | 1224 | - | - | 0 | - |
| - | - | 666.8 | 1228 | - | - | 0 | - |
| - | - | 1394 | 1241 | - | - | 0 | - |
| - | - | 726.1 | 1242 | - | - | 0 | - |
| - | - | 2539 | 1243 | - | - | 0 | - |
| - | - | 2928 | 1244 | - | - | 0 | - |
| - | - | 8146 | 1245 | - | - | 0 | - |
| - | - | 5295 | 1246 | - | - | 0 | - |
| - | - | 1654 | 1247 | - | - | 0 | - |
| - | - | 709.7 | 1260 | - | - | 0 | - |
| - | - | 845.7 | 1261 | - | - | 0 | - |
| - | - | 1413 | 1270 | - | - | 0 | - |
| - | - | 2374 | 1271 | - | - | 0 | - |
| - | - | 1601 | 1272 | - | - | 0 | - |
| - | - | 882.3 | 1273 | - | - | 0 | - |
| - | - | 834.3 | 1284 | - | - | 0 | - |
| - | - | 1871 | 1286 | - | - | 0 | - |
| - | - | 1.095E+04 | 1287 | - | - | 0 | - |
| - | - | 3.473E+04 | 1288 | - | - | 0 | - |
| - | - | 2.228E+04 | 1289 | - | - | 0 | - |
| - | - | 1.121E+04 | 1290 | - | - | 0 | - |
| - | - | 4377 | 1291 | - | - | 0 | - |
| - | - | 1061 | 1301 | - | - | 0 | - |
| - | - | 813.4 | 1302 | - | - | 0 | - |
| - | - | 4898 | 1303 | - | - | 0 | - |
| - | - | 1.713E+04 | 1304 | - | - | 0 | - |
| - | - | 4.293E+04 | 1305 | - | - | 0 | - |
| - | - | 2.914E+04 | 1306 | - | - | 0 | - |
| - | - | 1.156E+04 | 1307 | - | - | 0 | - |
| - | - | 4695 | 1308 | - | - | 0 | - |
| - | - | 1729 | 1309 | - | - | 0 | - |
| - | - | 778.2 | 1313 | - | - | 0 | - |
| - | - | 655.1 | 1582 | - | - | 0 | - |
| - | - | 694.9 | 2768 | - | - | 0 | - |
| - | - | 678.2 | 3079 | - | - | 0 | - |
| - | - | 706 | 3274 | - | - | 0 | - |

m/z Charge Intensity FragmentType MassShift Position
120.08097839355469 0 5408.528
121.08454895019531 0 489.89822
121.9086685180664 0 360.54153
124.0870590209961 0 457.32022
125.33827209472656 0 386.18286
129.10218811035156 0 482.6717
131.04518127441406 0 545.0672
136.075927734375 0 489.43652
149.02346801757812 0 4408.088
153.3014373779297 0 397.94376
156.06582641601562 0 898.9093
156.0768280029297 0 468.90182
157.07681274414062 0 450.9243
158.14157104492188 0 841.9224
166.05335998535156 0 17040.863
167.03407287597656 0 1356.4559
167.0570526123047 0 560.984
168.04908752441406 0 728.00696
172.1333465576172 0 1146.904
173.43771362304688 0 1477.0217
173.45220947265625 0 2057.9915
173.46261596679688 0 584.86993
173.4720458984375 0 586.5519
184.37283325195312 0 489.38736
185.1647186279297 0 548.67346
189.6134033203125 0 527.19635
199.181396484375 0 581.74524
208.95321655273438 0 4005.8286
210.9503173828125 0 10430.797
212.13943481445312 0 24548.645
212.94735717773438 0 3123.024
213.14285278320312 0 2722.0947
213.1591033935547 0 498.49094
220.06333923339844 0 592.32367
226.09500122070312 0 854.23236
226.156005859375 0 720.298
227.17514038085938 0 1025.2496
229.11004638671875 0 933.42786
233.1648712158203 0 10386.203
234.16885375976562 0 1071.068
255.17010498046875 0 743.6426
260.1519470214844 0 5363.9023
261.15948486328125 0 4897.116
262.1600036621094 0 788.90283
279.13720703125 0 5578.697 y 9
279.1546325683594 0 752.3841
280.1410827636719 0 596.80383
287.1624755859375 0 574.34827
291.35626220703125 0 550.28314
296.13751220703125 0 716.6596
297.0384216308594 0 568.4468
297.1440734863281 0 858.9003
306.0880432128906 0 2995.173
308.1036376953125 0 2730.6982
315.1555480957031 0 992.233
319.0793762207031 0 528.79694
320.91937255859375 0 2626.3486
322.9163818359375 0 6410.7656
323.10882568359375 0 540.4668
323.9192199707031 0 819.75665
324.2141418457031 0 619.5553
324.9138488769531 0 2101.3872
326.858642578125 0 1734.3799
327.80902099609375 0 531.0084
348.1896057128906 0 1096.9316 y 2
355.0699462890625 0 1437.4109
356.2171936035156 0 772.16455
358.2115783691406 0 963.5398
362.1513977050781 0 873.6415
363.15985107421875 0 685.7146
364.16510009765625 0 711.1781 z 8
365.1741027832031 0 5838.3906
391.2321472167969 0 1225.3506
391.73443603515625 0 862.69653
401.2632141113281 0 1006.9604
413.2691650390625 0 815.6465 c 6
417.1920471191406 0 1173.5396
418.03521728515625 0 848.25006
425.2624816894531 0 755.92664
428.2793884277344 0 742.60596
434.19659423828125 0 641.4016 w 7
434.7800598144531 0 2300.7625
434.88079833984375 0 1273.0249
435.2203674316406 0 996.41174
435.2748718261719 0 1049.779
439.2617492675781 0 1865.2184
439.7614440917969 0 1336.0076
439.999755859375 0 582.91895
447.7740783691406 0 4116.4106
448.2743835449219 0 2407.165
448.7780456542969 0 2655.6646
449.2803955078125 0 993.8633
456.2466125488281 0 711.07336 z Water loss 3
456.7406921386719 0 718.7899 z Ammonia loss 3
462.1900939941406 0 4342.7065
463.192626953125 0 686.4312
470.2820129394531 0 840.4387
470.7831115722656 0 2329.99 c 7
471.28814697265625 0 937.39075
472.2937927246094 0 751.2847 c 3
473.256591796875 0 1916.3215 y 3
473.7578430175781 0 836.44086
479.1908874511719 0 1097.0756 z 7
480.2011413574219 0 7155.29
481.20184326171875 0 1061.0973
486.7513122558594 0 1605.1991
489.7848205566406 0 2788.7085
490.285400390625 0 1395.6324
490.79022216796875 0 1226.6532
500.2622375488281 0 1410.695 w 2
500.76361083984375 0 1026.4502
507.7754211425781 0 1591.229
508.7821044921875 0 733.0652
513.2753295898438 0 941.32367 y Ammonia loss 2
513.7839965820312 0 1706.9272
514.2871704101562 0 682.6785
520.7764892578125 0 1301.676
521.2716674804688 0 1041.811
521.308349609375 0 2444.117 c 8
521.783447265625 0 22988.854 y 2
522.2849731445312 0 11861.206
522.78955078125 0 19062.213
523.2918090820312 0 8613.646
523.7910766601562 0 2020.042
524.2966918945312 0 686.2108
534.3143920898438 0 798.2679
535.27734375 0 2682.5967
535.7783813476562 0 1313.5767
536.283203125 0 2406.3765
536.7853393554688 0 2189.3923
539.8067016601562 0 798.9528
545.2235107421875 0 523.8012
550.5935668945312 0 610.975
555.8407592773438 0 651.98755
556.3731079101562 0 1144.5404
559.7921752929688 0 578.854
560.2980346679688 0 581.7349
563.2986450195312 0 4015.0828
564.2984619140625 0 1234.6677
565.2971801757812 0 862.0531
565.3789672851562 0 1262.7532
569.3285522460938 0 654.5724 c Ammonia loss 9
570.339111328125 0 771.7845
577.8499755859375 0 18286.273 c 9
578.3099365234375 0 806.0526 z Water loss 1
578.3516845703125 0 8326.043
578.8526611328125 0 4050.4792
586.8108520507812 0 1994.8517 y Ammonia loss 1
587.3082275390625 0 7564.9946 z 1
587.8089599609375 0 4886.6626
588.3104248046875 0 1486.7737
590.859130859375 0 967.6524
591.3563232421875 0 793.90094
595.3169555664062 0 697.05743 y 1
606.3370361328125 0 1455.399
606.8353881835938 0 1138.631
607.288330078125 0 2626.54 z 6
607.342529296875 0 874.8263
608.2900390625 0 1069.156
612.8485107421875 0 758.432
613.3527221679688 0 1011.32086
613.8514404296875 0 906.0213
616.8224487304688 0 731.283
617.3292846679688 0 694.27783
622.323486328125 0 23200.453
622.8250732421875 0 13755.237
623.3239135742188 0 7969.051
623.8263549804688 0 3190.6343
624.3299560546875 0 1473.6898
624.8331909179688 0 1087.2545
625.3358154296875 0 679.433
629.3594360351562 0 687.68494
630.3694458007812 0 828.2211
635.2846069335938 0 775.82526
643.3574829101562 0 2415.7075
643.8594360351562 0 22539.66
644.3607177734375 0 19320.156
644.8607177734375 0 8515.919
645.3615112304688 0 3394.2969
649.8513793945312 0 744.5191
650.3563842773438 0 611.8139
650.8507690429688 0 1358.5818
651.3551025390625 0 955.7825
651.8594970703125 0 51397.7
652.3619995117188 0 63293.098
652.8630981445312 0 35367.62
653.3639526367188 0 14187.551
653.4230346679688 0 651.3502
653.8646240234375 0 5210.918
677.3150024414062 0 1088.5535 w 5
680.4097900390625 0 2230.5798 c Ammonia loss 5
684.4144287109375 0 1133.0848
696.429931640625 0 1888.956
697.4387817382812 0 19680.012 c 5
698.4418334960938 0 6750.374
699.4466552734375 0 1359.8353
720.3580932617188 0 6482.717 y 5
721.3616943359375 0 2857.5083
722.3601684570312 0 1315.9156
748.353271484375 0 1111.5018
749.3468627929688 0 746.60864
768.4656982421875 0 917.7974
780.4586791992188 0 775.3663
781.4526977539062 0 843.1624
781.5266723632812 0 959.36
788.4447631835938 0 3238.1758
789.4487915039062 0 1269.6874
808.5049438476562 0 919.1142 c Ammonia loss 6
816.4397583007812 0 820.082
823.5193481445312 0 1051.1052
824.5254516601562 0 20546.19
825.5325317382812 0 50052.633 c 6
826.5363159179688 0 20255.732
827.5392456054688 0 4605.8867
828.5445556640625 0 967.7064
832.4351806640625 0 21959.771 z 4
833.4379272460938 0 10769.006
834.4393310546875 0 4054.5059
835.4440307617188 0 1050.3398
854.50732421875 0 1062.6768
868.5540161132812 0 2191.833
869.5536499023438 0 1124.3198
877.5130004882812 0 1349.6171
894.5375366210938 0 773.6914
896.54833984375 0 2165.8303
897.5496215820312 0 586.4149
923.5372924804688 0 666.6444 c Ammonia loss 7
924.5363159179688 0 960.22296
939.551513671875 0 13371.8955
940.5586547851562 0 31281.2 c 7
941.5626220703125 0 17556.58
942.566650390625 0 5134.9478
943.5772094726562 0 1158.2626
945.5084838867188 0 2910.8054 y 3
946.5101318359375 0 1775.4817
1025.5841064453125 0 1160.5186
1026.5841064453125 0 1776.0077
1041.6070556640625 0 7713.193 c 8
1042.6055908203125 0 4272.783
1043.5892333984375 0 1829.4432
1044.573974609375 0 4376.299
1045.577392578125 0 2629.6763
1046.5809326171875 0 828.5907
1095.6727294921875 0 851.3361
1110.6710205078125 0 1413.1079
1111.683349609375 0 4146.3965
1112.6885986328125 0 2589.3909
1113.685302734375 0 897.5598
1138.66943359375 0 1777.9827
1139.669921875 0 1292.0314
1154.689697265625 0 5480.3423 c 9
1155.697021484375 0 8832.871
1156.69775390625 0 3127.4192
1157.7117919921875 0 1263.6533
1173.6051025390625 0 1949.4075 z 1
1174.61474609375 0 10387.37
1175.6177978515625 0 6516.496
1176.621337890625 0 2595.667
1177.6181640625 0 895.62366
1212.693359375 0 1369.7924
1213.71826171875 0 714.99884
1223.6922607421875 0 725.6379
1227.6883544921875 0 666.84894
1240.7125244140625 0 1394.2941
1241.7113037109375 0 726.0673
1242.7242431640625 0 2539.354
1243.712158203125 0 2927.6184
1244.70361328125 0 8146.1084
1245.7088623046875 0 5295.288
1246.707275390625 0 1654.2172
1259.728271484375 0 709.652
1260.7034912109375 0 845.7459
1269.686279296875 0 1412.6039
1270.6944580078125 0 2374.0408
1271.6993408203125 0 1600.7966
1272.698974609375 0 882.33813
1283.678466796875 0 834.25806
1285.695556640625 0 1871.4303
1286.703369140625 0 10953.555
1287.703369140625 0 34727.22
1288.7056884765625 0 22277.885
1289.7041015625 0 11212.792
1290.703857421875 0 4377.0303
1300.694580078125 0 1061.3119
1301.7144775390625 0 813.43665
1302.7100830078125 0 4898.034
1303.71533203125 0 17134.072
1304.7227783203125 0 42926.895
1305.725830078125 0 29142.35
1306.7264404296875 0 11555.49
1307.7274169921875 0 4694.922
1308.7222900390625 0 1728.5453
1313.2010498046875 0 778.2465
1581.64599609375 0 655.11475
2768.235107421875 0 694.86035
3078.821533203125 0 678.2243
3274.427490234375 0 705.9843

Spectrum Details

|  |  |
| --- | --- |
| Matched peaks? Matched peaks The total absolute number of peaks matched. Additionally in brackets the total fraction of peaks matched and the total number of peaks is shown. | 35 (11.78% of 297) |
| FDR? FDR The false discovery rate estimated for this peptide. It is calculated by matching all theoretical fragments with a non-integer shift with the raw peaks for this spectrum. This is done with 40 different shifts. The resulting percentage is the average number of annotated peaks over the number of annotated peaks with the correct spectrum. | 2.59% |
| Satellite FDR? Satellite FDR See the FDR for details on its calculation. This satellite ion specific FDR only contains the satellite ions (d/w) for I/L/J positions. | ∞ |
| PSM Score? PSM Score The PSM Score as given by Hecklib to this annotated spectrum. It is shown with three significant figures. | 188 |

#### Spectrum 6196? Spectrum 6196 The raw spectrum of this peptide as annotated by Hecklib. The fragments are coloured according to ion type (see legend). Any peaks with a star '\*' as text can be hovered over to see the full details, first the ion type second the mass shift type. By hovering over the amino acids in the peptide or ions in the legend the corresponding peaks are highlighted. By toggling the 'Unassigned' label you can turn the background (unassigned) peaks on or off in the plot. By updating the slider in the Ion legend you can update the spectrum to only show the top X% of the peaks with labels. The top X% means any peak that is within X% of the highest intensity. By dragging in the spectrum you can zoom in to a specific part of the spectrum and use 'Zoom Out' to get back to the original zoom level. The annotation of the spectrum is based on the given sequence in the peptides file and is done with different software so inconsistencies are likely. The peaks are annotated based on the given sequence, with 20 ppm tolerance.

Copy Data

##### Spectrum 6196 (TSV)

###### Preview

```
Loading example...
```

*Click on the button to copy the data to your clipboard.*

Mz MinMz MaxIntensity Max

WidthHeightPeptide font sizePeptide stroke widthSpectrum font sizeSpectrum stroke widthCompact peptide

Ion legend

wxyz

abcd

OtherUnassignedIon^Charge^_Position_Show for top:%

JFPPKPKDTJM

01.69e+43.39e+45.08e+46.78e+4

Zoom Out

z^+1^_1_y^+1^_2_y^+3^_9_z^+1^_3_y^+1^_3_z^+1^_4_c^+2^_8_c^+1^_4_y^+2^_8_z^+1^_4_y^+1^_4_w^+2^_9_y^+2^_9_c^+2^_9_y^+2^_9_c^+2^_10_y^+2^_10_z^+2^_10_z^+1^_5_c^+1^_6_c^+1^_6_y^+1^_6_c^+1^_7_c^+1^_7_z^+1^_7_c^+1^_8_y^+1^_8_c^+1^_9_c^+1^_10_z^+1^_10_

0586117317592345

Fragment Matches Table

Show background peaks

| Position | Ion type | Intensity | mz Theoretical | mz Error (Th) | mz Error (ppm) | Charge | Series Number |
| --- | --- | --- | --- | --- | --- | --- | --- |
| - | - | 5838 | 120.1 | - | - | 0 | - |
| - | - | 441.5 | 121.4 | - | - | 0 | - |
| - | - | 495.6 | 128.1 | - | - | 0 | - |
| - | - | 699 | 129.1 | - | - | 0 | - |
| - | - | 529.4 | 131 | - | - | 0 | - |
| - | - | 381.3 | 133.1 | - | - | 0 | - |
| - | - | 578.2 | 144.1 | - | - | 0 | - |
| - | - | 477.9 | 147.1 | - | - | 0 | - |
| - | - | 4744 | 149 | - | - | 0 | - |
| - | - | 502.1 | 149.1 | - | - | 0 | - |
| - | - | 460.7 | 149.5 | - | - | 0 | - |
| 11 | z | 491 | 150 | 0.002639 | 17.59 | +1 | 1 |
| - | - | 781 | 156.1 | - | - | 0 | - |
| - | - | 440.8 | 165.1 | - | - | 0 | - |
| - | - | 1.784E+04 | 166.1 | - | - | 0 | - |
| - | - | 1344 | 167 | - | - | 0 | - |
| - | - | 1321 | 167.1 | - | - | 0 | - |
| - | - | 559.8 | 172.1 | - | - | 0 | - |
| - | - | 2431 | 173.4 | - | - | 0 | - |
| - | - | 903.5 | 182.1 | - | - | 0 | - |
| - | - | 429 | 194.5 | - | - | 0 | - |
| - | - | 1031 | 199.2 | - | - | 0 | - |
| - | - | 4481 | 209 | - | - | 0 | - |
| - | - | 635 | 210 | - | - | 0 | - |
| - | - | 1.073E+04 | 211 | - | - | 0 | - |
| - | - | 537.8 | 211 | - | - | 0 | - |
| - | - | 1191 | 212 | - | - | 0 | - |
| - | - | 2.624E+04 | 212.1 | - | - | 0 | - |
| - | - | 3549 | 212.9 | - | - | 0 | - |
| - | - | 2596 | 213.1 | - | - | 0 | - |
| - | - | 1309 | 220.1 | - | - | 0 | - |
| - | - | 437.9 | 223 | - | - | 0 | - |
| - | - | 478 | 224.4 | - | - | 0 | - |
| - | - | 775.2 | 226.1 | - | - | 0 | - |
| - | - | 1009 | 226.2 | - | - | 0 | - |
| - | - | 686.2 | 227.2 | - | - | 0 | - |
| - | - | 691.6 | 229.1 | - | - | 0 | - |
| - | - | 549.1 | 231.7 | - | - | 0 | - |
| - | - | 1.003E+04 | 233.2 | - | - | 0 | - |
| - | - | 1975 | 234.2 | - | - | 0 | - |
| - | - | 470.3 | 244.1 | - | - | 0 | - |
| - | - | 874.6 | 255.2 | - | - | 0 | - |
| - | - | 5851 | 260.2 | - | - | 0 | - |
| - | - | 5496 | 261.2 | - | - | 0 | - |
| - | - | 1198 | 262.2 | - | - | 0 | - |
| - | - | 517 | 270 | - | - | 0 | - |
| - | - | 945 | 270.1 | - | - | 0 | - |
| 10 | y | 5053 | 279.1 | 0.005214 | 18.68 | +1 | 2 |
| - | - | 846 | 280.1 | - | - | 0 | - |
| - | - | 1039 | 296.1 | - | - | 0 | - |
| - | - | 963.2 | 297.1 | - | - | 0 | - |
| - | - | 3073 | 306.1 | - | - | 0 | - |
| - | - | 2597 | 308.1 | - | - | 0 | - |
| - | - | 629.5 | 309.1 | - | - | 0 | - |
| - | - | 1742 | 320.9 | - | - | 0 | - |
| - | - | 6005 | 322.9 | - | - | 0 | - |
| - | - | 1203 | 323.2 | - | - | 0 | - |
| - | - | 1078 | 323.9 | - | - | 0 | - |
| - | - | 867.4 | 324.2 | - | - | 0 | - |
| - | - | 2061 | 324.9 | - | - | 0 | - |
| - | - | 1292 | 326.9 | - | - | 0 | - |
| - | - | 753.9 | 336.2 | - | - | 0 | - |
| 3 | y | 653.6 | 348.2 | 0.003249 | 9.331 | +3 | 9 |
| - | - | 969.5 | 358.2 | - | - | 0 | - |
| - | - | 883.4 | 362.2 | - | - | 0 | - |
| - | - | 492.9 | 362.2 | - | - | 0 | - |
| - | - | 1704 | 363.2 | - | - | 0 | - |
| 9 | z | 574.7 | 364.2 | 0.003786 | 10.4 | +1 | 3 |
| - | - | 4715 | 365.2 | - | - | 0 | - |
| - | - | 855.7 | 366.2 | - | - | 0 | - |
| 9 | y | 603.4 | 380.2 | 0.003922 | 10.32 | +1 | 3 |
| - | - | 764.7 | 391.2 | - | - | 0 | - |
| - | - | 2032 | 391.2 | - | - | 0 | - |
| - | - | 753.2 | 391.7 | - | - | 0 | - |
| - | - | 609.7 | 392.2 | - | - | 0 | - |
| - | - | 640.3 | 401.3 | - | - | 0 | - |
| - | - | 661.5 | 416 | - | - | 0 | - |
| - | - | 930.1 | 417 | - | - | 0 | - |
| - | - | 882.6 | 418.2 | - | - | 0 | - |
| - | - | 801.9 | 434.8 | - | - | 0 | - |
| - | - | 953.6 | 434.9 | - | - | 0 | - |
| - | - | 963.5 | 435.2 | - | - | 0 | - |
| - | - | 784.7 | 435.2 | - | - | 0 | - |
| - | - | 1177 | 436.2 | - | - | 0 | - |
| - | - | 1079 | 436.2 | - | - | 0 | - |
| - | - | 4558 | 439.3 | - | - | 0 | - |
| - | - | 1948 | 439.8 | - | - | 0 | - |
| - | - | 843.7 | 440.3 | - | - | 0 | - |
| - | - | 3308 | 447.8 | - | - | 0 | - |
| - | - | 1939 | 448.3 | - | - | 0 | - |
| - | - | 3218 | 448.8 | - | - | 0 | - |
| - | - | 1173 | 449.3 | - | - | 0 | - |
| 8 | z | 1002 | 461.2 | 0.006482 | 14.05 | +1 | 4 |
| - | - | 6185 | 462.2 | - | - | 0 | - |
| - | - | 851.5 | 463.2 | - | - | 0 | - |
| - | - | 579.9 | 470.3 | - | - | 0 | - |
| 8 | c | 2558 | 470.8 | 9.902E-05 | 0.2103 | +2 | 8 |
| - | - | 2121 | 471.3 | - | - | 0 | - |
| - | - | 738.5 | 472.3 | - | - | 0 | - |
| 4 | c | 1169 | 472.3 | 0.0005104 | 1.081 | +1 | 4 |
| 4 | y | 1164 | 473.3 | 0.002118 | 4.475 | +2 | 8 |
| - | - | 960.1 | 473.8 | - | - | 0 | - |
| - | - | 642.1 | 474.3 | - | - | 0 | - |
| 8 | z | 955.3 | 479.2 | 0.004553 | 9.502 | +1 | 4 |
| - | - | 8415 | 480.2 | - | - | 0 | - |
| - | - | 2327 | 481.2 | - | - | 0 | - |
| - | - | 1114 | 486.8 | - | - | 0 | - |
| - | - | 762.3 | 487.8 | - | - | 0 | - |
| - | - | 2706 | 489.8 | - | - | 0 | - |
| - | - | 1852 | 490.3 | - | - | 0 | - |
| - | - | 1096 | 490.8 | - | - | 0 | - |
| - | - | 988.3 | 492.8 | - | - | 0 | - |
| 8 | y | 813.1 | 495.2 | 0.004506 | 9.099 | +1 | 4 |
| 3 | w | 1600 | 500.3 | 0.003488 | 6.973 | +2 | 9 |
| - | - | 952.3 | 500.8 | - | - | 0 | - |
| - | - | 1069 | 501.3 | - | - | 0 | - |
| - | - | 603.6 | 507.3 | - | - | 0 | - |
| - | - | 1678 | 507.8 | - | - | 0 | - |
| - | - | 1168 | 508.3 | - | - | 0 | - |
| 3 | y | 1575 | 512.8 | 3.068E-05 | 0.05983 | +2 | 9 |
| - | - | 1012 | 513.8 | - | - | 0 | - |
| - | - | 615.8 | 514.3 | - | - | 0 | - |
| - | - | 1539 | 520.8 | - | - | 0 | - |
| 9 | c | 2615 | 521.3 | 0.0008115 | 1.557 | +2 | 9 |
| 3 | y | 2.643E+04 | 521.8 | 0.002805 | 5.376 | +2 | 9 |
| - | - | 1.396E+04 | 522.3 | - | - | 0 | - |
| - | - | 1.715E+04 | 522.8 | - | - | 0 | - |
| - | - | 9327 | 523.3 | - | - | 0 | - |
| - | - | 3131 | 523.8 | - | - | 0 | - |
| - | - | 1059 | 524.3 | - | - | 0 | - |
| - | - | 1055 | 534.3 | - | - | 0 | - |
| - | - | 2364 | 535.3 | - | - | 0 | - |
| - | - | 1735 | 535.8 | - | - | 0 | - |
| - | - | 5380 | 536.3 | - | - | 0 | - |
| - | - | 2347 | 536.8 | - | - | 0 | - |
| - | - | 735.1 | 555.4 | - | - | 0 | - |
| - | - | 1036 | 555.8 | - | - | 0 | - |
| - | - | 1555 | 556.4 | - | - | 0 | - |
| - | - | 948.4 | 558.8 | - | - | 0 | - |
| - | - | 767.4 | 559.3 | - | - | 0 | - |
| - | - | 5484 | 563.3 | - | - | 0 | - |
| - | - | 1558 | 564.3 | - | - | 0 | - |
| - | - | 773.9 | 565.4 | - | - | 0 | - |
| 10 | c | 1.8E+04 | 577.9 | 0.0005276 | 0.913 | +2 | 10 |
| - | - | 1.184E+04 | 578.4 | - | - | 0 | - |
| - | - | 3331 | 578.9 | - | - | 0 | - |
| - | - | 705.3 | 579.4 | - | - | 0 | - |
| - | - | 1057 | 585.4 | - | - | 0 | - |
| 2 | y | 1017 | 586.8 | 0.0047 | 8.009 | +2 | 10 |
| 2 | z | 8735 | 587.3 | 0.002984 | 5.082 | +2 | 10 |
| - | - | 4142 | 587.8 | - | - | 0 | - |
| - | - | 1946 | 588.3 | - | - | 0 | - |
| - | - | 1161 | 588.8 | - | - | 0 | - |
| - | - | 1167 | 590.9 | - | - | 0 | - |
| - | - | 1112 | 606.3 | - | - | 0 | - |
| - | - | 837.3 | 606.8 | - | - | 0 | - |
| 7 | z | 3240 | 607.3 | 0.00508 | 8.365 | +1 | 5 |
| - | - | 869 | 607.3 | - | - | 0 | - |
| - | - | 962.6 | 608.3 | - | - | 0 | - |
| - | - | 2275 | 612.9 | - | - | 0 | - |
| - | - | 932.7 | 613.4 | - | - | 0 | - |
| - | - | 714.5 | 615.8 | - | - | 0 | - |
| - | - | 1343 | 616.8 | - | - | 0 | - |
| - | - | 676.8 | 621.9 | - | - | 0 | - |
| - | - | 2.648E+04 | 622.3 | - | - | 0 | - |
| - | - | 1.65E+04 | 622.8 | - | - | 0 | - |
| - | - | 8576 | 623.3 | - | - | 0 | - |
| - | - | 3409 | 623.8 | - | - | 0 | - |
| - | - | 818.8 | 624.8 | - | - | 0 | - |
| - | - | 769.9 | 626.4 | - | - | 0 | - |
| - | - | 1413 | 630.4 | - | - | 0 | - |
| - | - | 832.8 | 642.9 | - | - | 0 | - |
| - | - | 2307 | 643.4 | - | - | 0 | - |
| - | - | 2.599E+04 | 643.9 | - | - | 0 | - |
| - | - | 1.944E+04 | 644.4 | - | - | 0 | - |
| - | - | 8527 | 644.9 | - | - | 0 | - |
| - | - | 3585 | 645.4 | - | - | 0 | - |
| - | - | 1150 | 650.4 | - | - | 0 | - |
| - | - | 1595 | 650.9 | - | - | 0 | - |
| - | - | 798 | 651.4 | - | - | 0 | - |
| - | - | 781.3 | 651.8 | - | - | 0 | - |
| - | - | 5.57E+04 | 651.9 | - | - | 0 | - |
| - | - | 6.711E+04 | 652.4 | - | - | 0 | - |
| - | - | 4.084E+04 | 652.9 | - | - | 0 | - |
| - | - | 1.657E+04 | 653.4 | - | - | 0 | - |
| - | - | 5822 | 653.9 | - | - | 0 | - |
| - | - | 1131 | 654.4 | - | - | 0 | - |
| 6 | c | 1367 | 680.4 | 0.004989 | 7.332 | +1 | 6 |
| - | - | 1019 | 681.4 | - | - | 0 | - |
| - | - | 947.8 | 684.4 | - | - | 0 | - |
| - | - | 2516 | 696.4 | - | - | 0 | - |
| 6 | c | 2.354E+04 | 697.4 | 0.0003223 | 0.4622 | +1 | 6 |
| - | - | 8085 | 698.4 | - | - | 0 | - |
| - | - | 1696 | 699.4 | - | - | 0 | - |
| 6 | y | 7988 | 720.4 | 0.006346 | 8.809 | +1 | 6 |
| - | - | 2294 | 721.4 | - | - | 0 | - |
| - | - | 1045 | 722.4 | - | - | 0 | - |
| - | - | 1547 | 748.4 | - | - | 0 | - |
| - | - | 618.4 | 750.4 | - | - | 0 | - |
| - | - | 613.5 | 764.4 | - | - | 0 | - |
| - | - | 898.1 | 781.5 | - | - | 0 | - |
| - | - | 749.3 | 781.5 | - | - | 0 | - |
| - | - | 3162 | 788.4 | - | - | 0 | - |
| - | - | 975.4 | 789.5 | - | - | 0 | - |
| - | - | 750.3 | 790.5 | - | - | 0 | - |
| 7 | c | 1433 | 808.5 | 0.001441 | 1.783 | +1 | 7 |
| - | - | 849.1 | 809.5 | - | - | 0 | - |
| - | - | 1225 | 823.5 | - | - | 0 | - |
| - | - | 1006 | 824.4 | - | - | 0 | - |
| - | - | 2.281E+04 | 824.5 | - | - | 0 | - |
| 7 | c | 5.132E+04 | 825.5 | 0.0004024 | 0.4874 | +1 | 7 |
| - | - | 2.416E+04 | 826.5 | - | - | 0 | - |
| - | - | 6241 | 827.5 | - | - | 0 | - |
| - | - | 1357 | 828.5 | - | - | 0 | - |
| 5 | z | 2.469E+04 | 832.4 | 0.005119 | 6.15 | +1 | 7 |
| - | - | 1.2E+04 | 833.4 | - | - | 0 | - |
| - | - | 3362 | 834.4 | - | - | 0 | - |
| - | - | 983.8 | 835.4 | - | - | 0 | - |
| - | - | 1541 | 868.6 | - | - | 0 | - |
| - | - | 1339 | 877.5 | - | - | 0 | - |
| - | - | 1136 | 895.6 | - | - | 0 | - |
| - | - | 787 | 896.5 | - | - | 0 | - |
| - | - | 862.4 | 897.6 | - | - | 0 | - |
| - | - | 1.452E+04 | 939.6 | - | - | 0 | - |
| 8 | c | 3.415E+04 | 940.6 | 0.001222 | 1.3 | +1 | 8 |
| - | - | 1.654E+04 | 941.6 | - | - | 0 | - |
| - | - | 4389 | 942.6 | - | - | 0 | - |
| - | - | 986.4 | 943.6 | - | - | 0 | - |
| 4 | y | 3238 | 945.5 | 0.004066 | 4.3 | +1 | 8 |
| - | - | 1628 | 946.5 | - | - | 0 | - |
| - | - | 923.1 | 981.6 | - | - | 0 | - |
| - | - | 1192 | 1026 | - | - | 0 | - |
| - | - | 1453 | 1027 | - | - | 0 | - |
| 9 | c | 7112 | 1042 | 0.0005 | 0.48 | +1 | 9 |
| - | - | 5332 | 1043 | - | - | 0 | - |
| - | - | 1506 | 1044 | - | - | 0 | - |
| - | - | 4014 | 1045 | - | - | 0 | - |
| - | - | 3277 | 1046 | - | - | 0 | - |
| - | - | 955.5 | 1047 | - | - | 0 | - |
| - | - | 827.4 | 1095 | - | - | 0 | - |
| - | - | 2277 | 1111 | - | - | 0 | - |
| - | - | 5386 | 1112 | - | - | 0 | - |
| - | - | 2980 | 1113 | - | - | 0 | - |
| - | - | 1899 | 1139 | - | - | 0 | - |
| - | - | 1408 | 1140 | - | - | 0 | - |
| 10 | c | 4752 | 1155 | 0.000397 | 0.3438 | +1 | 10 |
| - | - | 8455 | 1156 | - | - | 0 | - |
| - | - | 4409 | 1157 | - | - | 0 | - |
| - | - | 1390 | 1158 | - | - | 0 | - |
| 2 | z | 2278 | 1174 | 0.004578 | 3.901 | +1 | 10 |
| - | - | 1.151E+04 | 1175 | - | - | 0 | - |
| - | - | 7444 | 1176 | - | - | 0 | - |
| - | - | 3040 | 1177 | - | - | 0 | - |
| - | - | 1161 | 1213 | - | - | 0 | - |
| - | - | 680 | 1214 | - | - | 0 | - |
| - | - | 762.9 | 1228 | - | - | 0 | - |
| - | - | 646.2 | 1230 | - | - | 0 | - |
| - | - | 661.8 | 1241 | - | - | 0 | - |
| - | - | 1471 | 1242 | - | - | 0 | - |
| - | - | 2376 | 1243 | - | - | 0 | - |
| - | - | 2352 | 1244 | - | - | 0 | - |
| - | - | 8631 | 1245 | - | - | 0 | - |
| - | - | 4581 | 1246 | - | - | 0 | - |
| - | - | 2203 | 1247 | - | - | 0 | - |
| - | - | 968.2 | 1248 | - | - | 0 | - |
| - | - | 635.4 | 1259 | - | - | 0 | - |
| - | - | 705.5 | 1260 | - | - | 0 | - |
| - | - | 1088 | 1261 | - | - | 0 | - |
| - | - | 1236 | 1270 | - | - | 0 | - |
| - | - | 2444 | 1271 | - | - | 0 | - |
| - | - | 1657 | 1272 | - | - | 0 | - |
| - | - | 770.3 | 1273 | - | - | 0 | - |
| - | - | 1057 | 1285 | - | - | 0 | - |
| - | - | 1428 | 1286 | - | - | 0 | - |
| - | - | 1.189E+04 | 1287 | - | - | 0 | - |
| - | - | 3.651E+04 | 1288 | - | - | 0 | - |
| - | - | 2.555E+04 | 1289 | - | - | 0 | - |
| - | - | 1.079E+04 | 1290 | - | - | 0 | - |
| - | - | 3851 | 1291 | - | - | 0 | - |
| - | - | 833.9 | 1292 | - | - | 0 | - |
| - | - | 3815 | 1303 | - | - | 0 | - |
| - | - | 1.715E+04 | 1304 | - | - | 0 | - |
| - | - | 4.481E+04 | 1305 | - | - | 0 | - |
| - | - | 3.083E+04 | 1306 | - | - | 0 | - |
| - | - | 1.387E+04 | 1307 | - | - | 0 | - |
| - | - | 4254 | 1308 | - | - | 0 | - |
| - | - | 852.9 | 1309 | - | - | 0 | - |
| - | - | 748.8 | 1970 | - | - | 0 | - |
| - | - | 763.2 | 2322 | - | - | 0 | - |

m/z Charge Intensity FragmentType MassShift Position
120.0811538696289 0 5837.874
121.402099609375 0 441.51294
128.10731506347656 0 495.58374
129.1026153564453 0 698.98083
131.0454559326172 0 529.4109
133.0857391357422 0 381.29727
144.0659942626953 0 578.19696
147.07760620117188 0 477.8984
149.02369689941406 0 4743.971
149.05613708496094 0 502.09036
149.4806671142578 0 460.66022
150.02696228027344 0 491.01456 z 10
156.06578063964844 0 781.0315
165.090576171875 0 440.81476
166.0535888671875 0 17839.795
167.0343017578125 0 1344.101
167.057373046875 0 1321.4092
172.13287353515625 0 559.7575
173.4403839111328 0 2430.8289
182.08103942871094 0 903.5139
194.46090698242188 0 429.00815
199.16941833496094 0 1030.6815
208.95352172851562 0 4481.1587
209.95681762695312 0 635.0024
210.9506072998047 0 10732.78
210.9617156982422 0 537.7517
211.95382690429688 0 1191.3057
212.13973999023438 0 26236.639
212.94760131835938 0 3549.1768
213.14349365234375 0 2596.4617
220.0637664794922 0 1308.6117
222.99517822265625 0 437.86533
224.4227752685547 0 477.95328
226.09495544433594 0 775.19824
226.15545654296875 0 1009.4418
227.1748809814453 0 686.1556
229.10968017578125 0 691.55896
231.7122039794922 0 549.10864
233.1652374267578 0 10033.403
234.16867065429688 0 1975.3931
244.10653686523438 0 470.3469
255.17144775390625 0 874.56635
260.1522216796875 0 5850.859
261.1595458984375 0 5495.7793
262.1634826660156 0 1198.2136
269.9535217285156 0 517.02747
270.12158203125 0 945.035
279.1376037597656 0 5052.6514 y 9
280.1422119140625 0 846.0417
296.1371765136719 0 1039.4866
297.14599609375 0 963.21655
306.08807373046875 0 3072.6982
308.1042785644531 0 2596.9758
309.1068115234375 0 629.53595
320.9197998046875 0 1741.893
322.9170837402344 0 6004.7275
323.2088317871094 0 1202.7988
323.9206237792969 0 1077.843
324.2156677246094 0 867.3926
324.9140625 0 2060.6882
326.8591003417969 0 1291.8066
336.1573181152344 0 753.8924
348.19317626953125 0 653.5728 y 2
358.2125244140625 0 969.46564
362.1512145996094 0 883.38586
362.21820068359375 0 492.8789
363.1601867675781 0 1704.336
364.1651306152344 0 574.6754 z 8
365.1739501953125 0 4714.771
366.175537109375 0 855.656
380.1839904785156 0 603.36035 y 8
391.177978515625 0 764.6931
391.2322692871094 0 2031.5135
391.7337951660156 0 753.1581
392.2346496582031 0 609.7156
401.2635192871094 0 640.3385
416.0351867675781 0 661.494
417.03448486328125 0 930.06006
418.1557312011719 0 882.63763
434.7815856933594 0 801.8698
434.8843994140625 0 953.6262
435.20269775390625 0 963.49457
435.2228698730469 0 784.71716
436.16839599609375 0 1177.2249
436.2250671386719 0 1079.074
439.26129150390625 0 4557.7715
439.76385498046875 0 1947.5966
440.2651672363281 0 843.7108
447.7743225097656 0 3308.0837
448.275390625 0 1938.5107
448.778076171875 0 3218.1938
449.2796630859375 0 1172.9829
461.1842041015625 0 1002.2572 z Water loss 7
462.1911315917969 0 6185.4834
463.1947937011719 0 851.46906
470.2780456542969 0 579.88763
470.7842712402344 0 2558.4456 c 7
471.2862243652344 0 2121.142
472.2536926269531 0 738.47986
472.29132080078125 0 1169.4321 c 3
473.2569885253906 0 1164.1648 y 3
473.75994873046875 0 960.1368
474.26129150390625 0 642.0814
479.1928405761719 0 955.34863 z 7
480.20166015625 0 8415.015
481.2043151855469 0 2327.386
486.7512512207031 0 1114.3951
487.755859375 0 762.2588
489.785400390625 0 2706.2593
490.28704833984375 0 1851.9775
490.78997802734375 0 1095.6454
492.7812194824219 0 988.31647
495.2115173339844 0 813.12494 y 7
500.2636413574219 0 1600.3485 w 2
500.76507568359375 0 952.2525
501.2690124511719 0 1069.3298
507.2730712890625 0 603.55963
507.7757568359375 0 1677.8441
508.2762451171875 0 1168.4717
512.7760009765625 0 1575.1854 y Water loss 2
513.7842407226562 0 1011.77814
514.285400390625 0 615.757
520.7767333984375 0 1538.6724
521.3090209960938 0 2614.6724 c 8
521.7840576171875 0 26428.62 y 2
522.2859497070312 0 13955.909
522.79052734375 0 17147.846
523.2925415039062 0 9326.589
523.793212890625 0 3131.106
524.2914428710938 0 1059.0343
534.3151245117188 0 1054.5575
535.27783203125 0 2364.1973
535.7810668945312 0 1735.016
536.2852172851562 0 5379.571
536.7874755859375 0 2346.663
555.36572265625 0 735.05695
555.8439331054688 0 1035.5156
556.3724365234375 0 1554.9706
558.7932739257812 0 948.3953
559.2972412109375 0 767.3671
563.2991943359375 0 5483.5537
564.3046875 0 1557.57
565.3822631835938 0 773.8669
577.8507690429688 0 18002.426 c 9
578.3523559570312 0 11837.707
578.8540649414062 0 3330.7156
579.3554077148438 0 705.2681
585.374267578125 0 1056.9056
586.806884765625 0 1016.9555 y Ammonia loss 1
587.30908203125 0 8735.038 z 1
587.8107299804688 0 4142.3213
588.3118896484375 0 1946.1089
588.8120727539062 0 1160.8568
590.8571166992188 0 1167.3303
606.3421020507812 0 1111.8962
606.8438110351562 0 837.28064
607.288330078125 0 3239.7776 z 6
607.3386840820312 0 869.0273
608.29541015625 0 962.6115
612.8536376953125 0 2275.3372
613.35302734375 0 932.6851
615.8248291015625 0 714.5222
616.8284912109375 0 1343.3157
621.8665161132812 0 676.84247
622.3240966796875 0 26478.943
622.8253173828125 0 16498.182
623.325927734375 0 8576.035
623.8289794921875 0 3408.7407
624.8282470703125 0 818.8044
626.3712158203125 0 769.94244
630.3692626953125 0 1412.8328
642.8519287109375 0 832.83966
643.3574829101562 0 2307.4485
643.8600463867188 0 25986.855
644.3612670898438 0 19443.58
644.8610229492188 0 8526.704
645.3621215820312 0 3585.2473
650.3505859375 0 1149.8698
650.8534545898438 0 1595.4355
651.3535766601562 0 798.02704
651.8133544921875 0 781.3373
651.8606567382812 0 55701.805
652.36279296875 0 67110.375
652.8641357421875 0 40842.66
653.3646850585938 0 16566.37
653.8657836914062 0 5821.5884
654.368896484375 0 1131.0305
680.4080200195312 0 1367.0582 c Ammonia loss 5
681.4143676757812 0 1019.0564
684.4171142578125 0 947.7608
696.4309692382812 0 2515.5576
697.4398803710938 0 23540.35 c 5
698.4429321289062 0 8084.7104
699.4429931640625 0 1695.8582
720.361083984375 0 7988.497 y 5
721.3621215820312 0 2293.551
722.3657836914062 0 1044.7407
748.3564453125 0 1547.1658
750.3555908203125 0 618.3684
764.4306640625 0 613.53345
781.454345703125 0 898.1092
781.52978515625 0 749.2645
788.446533203125 0 3161.8716
789.4541015625 0 975.4356
790.4500732421875 0 750.31604
808.5065307617188 0 1432.6534 c Ammonia loss 6
809.51171875 0 849.0884
823.5159912109375 0 1224.8685
824.438232421875 0 1005.68274
824.5267944335938 0 22813.625
825.5341186523438 0 51316.74 c 6
826.5369262695312 0 24163.195
827.5413208007812 0 6240.7188
828.5422973632812 0 1356.693
832.4360961914062 0 24687.086 z 4
833.439453125 0 11995.182
834.4390258789062 0 3362.26
835.445068359375 0 983.79834
868.5556030273438 0 1541.2473
877.5123901367188 0 1338.9045
895.5545043945312 0 1136.4167
896.5474243164062 0 787.0277
897.55322265625 0 862.4323
939.5526733398438 0 14520.176
940.5602416992188 0 34152.875 c 7
941.5634155273438 0 16539.672
942.56884765625 0 4389.4077
943.57568359375 0 986.38763
945.5065307617188 0 3238.379 y 3
946.505859375 0 1627.8351
981.5777587890625 0 923.0989
1025.58349609375 0 1191.9805
1026.5794677734375 0 1453.152
1041.608642578125 0 7111.9263 c 8
1042.607421875 0 5331.825
1043.617919921875 0 1506.0135
1044.5758056640625 0 4014.1565
1045.578857421875 0 3277.3704
1046.5692138671875 0 955.5413
1094.6619873046875 0 827.40594
1110.6785888671875 0 2276.9495
1111.6856689453125 0 5385.6836
1112.6888427734375 0 2979.9797
1138.67431640625 0 1898.7037
1139.681640625 0 1407.9432
1154.693603515625 0 4751.9854 c 9
1155.69873046875 0 8455.062
1156.7017822265625 0 4409.307
1157.7059326171875 0 1390.2263
1173.6094970703125 0 2277.6587 z 1
1174.6156005859375 0 11510.465
1175.6187744140625 0 7444.085
1176.62109375 0 3040.2205
1212.70703125 0 1160.7201
1213.6986083984375 0 679.9726
1227.7003173828125 0 762.921
1229.6749267578125 0 646.2029
1240.719970703125 0 661.79297
1241.7203369140625 0 1470.9646
1242.7279052734375 0 2375.758
1243.71435546875 0 2351.7683
1244.7078857421875 0 8631.474
1245.7078857421875 0 4581.3887
1246.7073974609375 0 2202.8872
1247.700927734375 0 968.1636
1258.720458984375 0 635.35834
1259.734375 0 705.4511
1260.722412109375 0 1088.37
1269.6988525390625 0 1236.389
1270.6927490234375 0 2444.147
1271.6973876953125 0 1656.8533
1272.6978759765625 0 770.2992
1284.6964111328125 0 1057.0878
1285.7034912109375 0 1427.9677
1286.7059326171875 0 11885.683
1287.7041015625 0 36512.984
1288.7061767578125 0 25553.188
1289.7059326171875 0 10789.021
1290.703369140625 0 3851.057
1291.7178955078125 0 833.862
1302.7088623046875 0 3815.41
1303.7156982421875 0 17152.564
1304.7242431640625 0 44812.52
1305.7274169921875 0 30826.668
1306.727783203125 0 13866.165
1307.7275390625 0 4254.03
1308.71337890625 0 852.8535
1970.453857421875 0 748.7682
2322.263671875 0 763.1896

Spectrum Details

|  |  |
| --- | --- |
| Matched peaks? Matched peaks The total absolute number of peaks matched. Additionally in brackets the total fraction of peaks matched and the total number of peaks is shown. | 30 (10.38% of 289) |
| FDR? FDR The false discovery rate estimated for this peptide. It is calculated by matching all theoretical fragments with a non-integer shift with the raw peaks for this spectrum. This is done with 40 different shifts. The resulting percentage is the average number of annotated peaks over the number of annotated peaks with the correct spectrum. | 2.94% |
| Satellite FDR? Satellite FDR See the FDR for details on its calculation. This satellite ion specific FDR only contains the satellite ions (d/w) for I/L/J positions. | ∞ |
| PSM Score? PSM Score The PSM Score as given by Hecklib to this annotated spectrum. It is shown with three significant figures. | 236 |

#### Reverse Lookup? Reverse Lookup All places where this read could be placed.

| Group | Segment | Template | Template Part | Read Part | Score | Unique |
| --- | --- | --- | --- | --- | --- | --- |
| Homo sapiens Heavy Chain | IGHC | IGHG1 | [124..135] | [0..11] | 88 | False |
| Homo sapiens Heavy Chain | IGHC | IGHG3 | [171..182] | [0..11] | 88 | False |
| Homo sapiens Heavy Chain | IGHC | IGHG2 | [120..131] | [0..11] | 88 | False |
| Homo sapiens Heavy Chain | IGHC | IGHG4 | [121..132] | [0..11] | 88 | False |

| Recombined | Template Part | Read Part | Score | Unique |
| --- | --- | --- | --- | --- |
| REC-0-1 | [247..258] | [0..11] | 88 | True |

#### Meta Information from Multiple reads

##### Number of combined reads

10

##### Intensity

0.9119

##### TotalArea

3.975E+09

##### Changes to the peptide sequence

JFPPKPKDTJM

L→JNo support for either Leucine or Isoleucine based on side chain ions (Position: 10)

L→JNo support for either Leucine or Isoleucine based on side chain ions (Position: 1)

#### Positional Score

Copy Data

##### Positional Score (TSV)

###### Preview

```
Loading example...
```

*Click on the button to copy the data to your clipboard.*

10012345678910

Label Value
"0" 0.786
"1" 0.743
"2" 0.778
"3" 0.776
"4" 0.752
"5" 0.789
"6" 0.79
"7" 0.794
"8" 0.792
"9" 0.795
"10" 0.782

#### Meta Information from PEAKS

##### Scan Identifier

F2:5122

##### Original sequence

L

F

P

P

K

P

K

D

T

L

M

+15.99

##### Posttranslational Modifications

Oxidation (M)

##### Source File

D:\separate\_stitch\_analyses\xle-disambiguation\raw\20210323\_F1\_UM1\_Peng0013\_SA\_F59\_ingel\_3ug\_TL.raw

##### Fraction

2

##### Scan Feature

F2:10471

##### De Novo Score

99

##### ConfidenceScore

99

### m/z

651.8606

##### Mass

1301.7053

##### Charge

2

##### Retention Time

27.68

##### Predicted Retention Time

-

##### Area

5.383E+08

##### Parts Per Million

1

##### Fragmentation mode

ETHCD

##### Originating file

01 D:\separate\_stitch\_analyses\xle-disambiguation\20210325\_F59\_3ug\_DENOVO\_12.csv

#### Meta Information from PEAKS

##### Scan Identifier

F2:5055

##### Original sequence

L

F

P

P

K

P

K

D

T

L

M

+15.99

##### Posttranslational Modifications

Oxidation (M)

##### Source File

D:\separate\_stitch\_analyses\xle-disambiguation\raw\20210323\_F1\_UM1\_Peng0013\_SA\_F59\_ingel\_3ug\_TL.raw

##### Fraction

2

##### Scan Feature

F2:10471

##### De Novo Score

99

##### ConfidenceScore

99

### m/z

651.8606

##### Mass

1301.7053

##### Charge

2

##### Retention Time

27.68

##### Predicted Retention Time

-

##### Area

5.383E+08

##### Parts Per Million

1

##### Fragmentation mode

ETHCD

##### Originating file

01 D:\separate\_stitch\_analyses\xle-disambiguation\20210325\_F59\_3ug\_DENOVO\_12.csv

#### Meta Information from PEAKS

##### Scan Identifier

F2:5239

##### Original sequence

L

F

P

P

K

P

K

D

T

L

M

+15.99

##### Posttranslational Modifications

Oxidation (M)

##### Source File

D:\separate\_stitch\_analyses\xle-disambiguation\raw\20210323\_F1\_UM1\_Peng0013\_SA\_F59\_ingel\_3ug\_TL.raw

##### Fraction

2

##### Scan Feature

F2:10471

##### De Novo Score

98

##### ConfidenceScore

98

### m/z

651.8606

##### Mass

1301.7053

##### Charge

2

##### Retention Time

27.68

##### Predicted Retention Time

-

##### Area

5.383E+08

##### Parts Per Million

1

##### Fragmentation mode

ETHCD

##### Originating file

01 D:\separate\_stitch\_analyses\xle-disambiguation\20210325\_F59\_3ug\_DENOVO\_12.csv

#### Meta Information from PEAKS

##### Scan Identifier

F2:5301

##### Original sequence

L

F

P

P

K

P

K

D

T

L

M

+15.99

##### Posttranslational Modifications

Oxidation (M)

##### Source File

D:\separate\_stitch\_analyses\xle-disambiguation\raw\20210323\_F1\_UM1\_Peng0013\_SA\_F59\_ingel\_3ug\_TL.raw

##### Fraction

2

##### Scan Feature

F2:10471

##### De Novo Score

98

##### ConfidenceScore

98

### m/z

651.8606

##### Mass

1301.7053

##### Charge

2

##### Retention Time

27.68

##### Predicted Retention Time

-

##### Area

5.383E+08

##### Parts Per Million

1

##### Fragmentation mode

ETHCD

##### Originating file

01 D:\separate\_stitch\_analyses\xle-disambiguation\20210325\_F59\_3ug\_DENOVO\_12.csv

#### Meta Information from PEAKS

##### Scan Identifier

F2:4947

##### Original sequence

L

F

P

P

K

P

K

D

T

L

M

+15.99

##### Posttranslational Modifications

Oxidation (M)

##### Source File

D:\separate\_stitch\_analyses\xle-disambiguation\raw\20210323\_F1\_UM1\_Peng0013\_SA\_F59\_ingel\_3ug\_TL.raw

##### Fraction

2

##### Scan Feature

F2:10471

##### De Novo Score

98

##### ConfidenceScore

98

### m/z

651.8606

##### Mass

1301.7053

##### Charge

2

##### Retention Time

27.68

##### Predicted Retention Time

-

##### Area

5.383E+08

##### Parts Per Million

1

##### Fragmentation mode

ETHCD

##### Originating file

01 D:\separate\_stitch\_analyses\xle-disambiguation\20210325\_F59\_3ug\_DENOVO\_12.csv

#### Meta Information from PEAKS

##### Scan Identifier

F2:5492

##### Original sequence

L

F

P

P

K

P

K

D

T

L

M

+15.99

##### Posttranslational Modifications

Oxidation (M)

##### Source File

D:\separate\_stitch\_analyses\xle-disambiguation\raw\20210323\_F1\_UM1\_Peng0013\_SA\_F59\_ingel\_3ug\_TL.raw

##### Fraction

2

##### Scan Feature

F2:10471

##### De Novo Score

98

##### ConfidenceScore

98

### m/z

651.8606

##### Mass

1301.7053

##### Charge

2

##### Retention Time

27.68

##### Predicted Retention Time

-

##### Area

5.383E+08

##### Parts Per Million

1

##### Fragmentation mode

ETHCD

##### Originating file

01 D:\separate\_stitch\_analyses\xle-disambiguation\20210325\_F59\_3ug\_DENOVO\_12.csv

#### Meta Information from PEAKS

##### Scan Identifier

F2:5872

##### Original sequence

L

F

P

P

K

P

K

D

T

L

M

+15.99

##### Posttranslational Modifications

Oxidation (M)

##### Source File

D:\separate\_stitch\_analyses\xle-disambiguation\raw\20210323\_F1\_UM1\_Peng0013\_SA\_F59\_ingel\_3ug\_TL.raw

##### Fraction

2

##### Scan Feature

F2:1738

##### De Novo Score

97

##### ConfidenceScore

97

### m/z

434.9093

##### Mass

1301.7053

##### Charge

3

##### Retention Time

27.95

##### Predicted Retention Time

-

##### Area

7.447E+08

##### Parts Per Million

0.6

##### Fragmentation mode

ETHCD

##### Originating file

01 D:\separate\_stitch\_analyses\xle-disambiguation\20210325\_F59\_3ug\_DENOVO\_12.csv

#### Meta Information from PEAKS

##### Scan Identifier

F2:6145

##### Original sequence

L

F

P

P

K

P

K

D

T

L

M

+15.99

##### Posttranslational Modifications

Oxidation (M)

##### Source File

D:\separate\_stitch\_analyses\xle-disambiguation\raw\20210323\_F1\_UM1\_Peng0013\_SA\_F59\_ingel\_3ug\_TL.raw

##### Fraction

2

##### Scan Feature

-

##### De Novo Score

96

##### ConfidenceScore

96

### m/z

434.9099

##### Mass

1301.7053

##### Charge

3

##### Retention Time

33.72

##### Predicted Retention Time

-

##### Area

0

##### Parts Per Million

1.8

##### Fragmentation mode

ETHCD

##### Originating file

01 D:\separate\_stitch\_analyses\xle-disambiguation\20210325\_F59\_3ug\_DENOVO\_12.csv

#### Meta Information from PEAKS

##### Scan Identifier

F2:6095

##### Original sequence

L

F

P

P

K

P

K

D

T

L

M

+15.99

##### Posttranslational Modifications

Oxidation (M)

##### Source File

D:\separate\_stitch\_analyses\xle-disambiguation\raw\20210323\_F1\_UM1\_Peng0013\_SA\_F59\_ingel\_3ug\_TL.raw

##### Fraction

2

##### Scan Feature

-

##### De Novo Score

96

##### ConfidenceScore

96

### m/z

434.9093

##### Mass

1301.7053

##### Charge

3

##### Retention Time

33.41

##### Predicted Retention Time

-

##### Area

0

##### Parts Per Million

0.5

##### Fragmentation mode

ETHCD

##### Originating file

01 D:\separate\_stitch\_analyses\xle-disambiguation\20210325\_F59\_3ug\_DENOVO\_12.csv

#### Meta Information from PEAKS

##### Scan Identifier

F2:6196

##### Original sequence

L

F

P

P

K

P

K

D

T

L

M

+15.99

##### Posttranslational Modifications

Oxidation (M)

##### Source File

D:\separate\_stitch\_analyses\xle-disambiguation\raw\20210323\_F1\_UM1\_Peng0013\_SA\_F59\_ingel\_3ug\_TL.raw

##### Fraction

2

##### Scan Feature

-

##### De Novo Score

96

##### ConfidenceScore

96

### m/z

434.91

##### Mass

1301.7053

##### Charge

3

##### Retention Time

34.03

##### Predicted Retention Time

-

##### Area

0

##### Parts Per Million

2.1

##### Fragmentation mode

ETHCD

##### Originating file

01 D:\separate\_stitch\_analyses\xle-disambiguation\20210325\_F59\_3ug\_DENOVO\_12.csv
