## Supplementary material for "A handle on mass coincidence errors in *de novo* sequencing of antibodies by bottom-up proteomics": Combined_021.html

Details Combined\_021 | Stitch OverviewUndefined

### Read Combined\_021

#### Sequence (length=8)

TLVGJVNY

#### Spectrum 10732? Spectrum 10732 The raw spectrum of this peptide as annotated by Hecklib. The fragments are coloured according to ion type (see legend). Any peaks with a star '\*' as text can be hovered over to see the full details, first the ion type second the mass shift type. By hovering over the amino acids in the peptide or ions in the legend the corresponding peaks are highlighted. By toggling the 'Unassigned' label you can turn the background (unassigned) peaks on or off in the plot. By updating the slider in the Ion legend you can update the spectrum to only show the top X% of the peaks with labels. The top X% means any peak that is within X% of the highest intensity. By dragging in the spectrum you can zoom in to a specific part of the spectrum and use 'Zoom Out' to get back to the original zoom level. The annotation of the spectrum is based on the given sequence in the peptides file and is done with different software so inconsistencies are likely. The peaks are annotated based on the given sequence, with 20 ppm tolerance.

Copy Data

##### Spectrum 10732 (TSV)

###### Preview

```
Loading example...
```

*Click on the button to copy the data to your clipboard.*

Mz MinMz MaxIntensity Max

WidthHeightPeptide font sizePeptide stroke widthSpectrum font sizeSpectrum stroke widthCompact peptide

Ion legend

wxyz

abcd

OtherUnassignedIon^Charge^_Position_Show for top:%

TLVGJVNY

01.52e+53.03e+54.55e+56.07e+5

Zoom Out

y^+1^_2_y^+1^_2_c^+1^_4_y^+2^_7_y^+1^_3_z^+1^_4_c^+1^_5_y^+1^_4_z^+1^_5_y^+1^_5_c^+1^_6_z^+1^_6_y^+1^_6_w^+1^_7_z^+1^_7_

0778155623343112

Fragment Matches Table

Show background peaks

| Position | Ion type | Intensity | mz Theoretical | mz Error (Th) | mz Error (ppm) | Charge | Series Number |
| --- | --- | --- | --- | --- | --- | --- | --- |
| - | - | 515.6 | 120 | - | - | 0 | - |
| - | - | 351.8 | 124.6 | - | - | 0 | - |
| - | - | 347.7 | 124.6 | - | - | 0 | - |
| - | - | 362.7 | 129.1 | - | - | 0 | - |
| - | - | 909.6 | 136.1 | - | - | 0 | - |
| - | - | 1139 | 143.1 | - | - | 0 | - |
| - | - | 4538 | 146.1 | - | - | 0 | - |
| - | - | 709.2 | 148.9 | - | - | 0 | - |
| - | - | 419.8 | 153.1 | - | - | 0 | - |
| - | - | 597.1 | 157.1 | - | - | 0 | - |
| - | - | 1980 | 163.1 | - | - | 0 | - |
| - | - | 442.2 | 166.3 | - | - | 0 | - |
| - | - | 669.3 | 169.1 | - | - | 0 | - |
| - | - | 2083 | 171.1 | - | - | 0 | - |
| - | - | 1697 | 171.1 | - | - | 0 | - |
| - | - | 3360 | 173.5 | - | - | 0 | - |
| - | - | 3387 | 174.1 | - | - | 0 | - |
| - | - | 1573 | 174.1 | - | - | 0 | - |
| - | - | 656.8 | 175.1 | - | - | 0 | - |
| - | - | 457.2 | 177.1 | - | - | 0 | - |
| - | - | 1735 | 185.1 | - | - | 0 | - |
| - | - | 795.9 | 185.2 | - | - | 0 | - |
| - | - | 2133 | 187.1 | - | - | 0 | - |
| - | - | 4.837E+04 | 187.1 | - | - | 0 | - |
| - | - | 3928 | 188.1 | - | - | 0 | - |
| - | - | 2868 | 191.1 | - | - | 0 | - |
| - | - | 1.323E+04 | 192.1 | - | - | 0 | - |
| - | - | 478 | 192.1 | - | - | 0 | - |
| - | - | 1142 | 193.1 | - | - | 0 | - |
| - | - | 546.2 | 197.8 | - | - | 0 | - |
| - | - | 728.8 | 199.1 | - | - | 0 | - |
| - | - | 6637 | 203.1 | - | - | 0 | - |
| - | - | 2.554E+04 | 209.1 | - | - | 0 | - |
| - | - | 2318 | 210.1 | - | - | 0 | - |
| - | - | 1578 | 213.2 | - | - | 0 | - |
| - | - | 3.749E+04 | 215.1 | - | - | 0 | - |
| - | - | 4071 | 216.1 | - | - | 0 | - |
| - | - | 729 | 221.1 | - | - | 0 | - |
| - | - | 1662 | 225.2 | - | - | 0 | - |
| - | - | 1498 | 242.2 | - | - | 0 | - |
| - | - | 2085 | 255.2 | - | - | 0 | - |
| - | - | 1590 | 260.1 | - | - | 0 | - |
| - | - | 505.7 | 261.1 | - | - | 0 | - |
| - | - | 1168 | 269.2 | - | - | 0 | - |
| - | - | 1.881E+04 | 270.2 | - | - | 0 | - |
| - | - | 1988 | 271.2 | - | - | 0 | - |
| - | - | 1613 | 278.1 | - | - | 0 | - |
| - | - | 1.097E+04 | 278.1 | - | - | 0 | - |
| 7 | y | 477.9 | 279.1 | 0.001418 | 5.079 | +1 | 2 |
| - | - | 2124 | 279.1 | - | - | 0 | - |
| - | - | 525.3 | 282.7 | - | - | 0 | - |
| - | - | 643.4 | 284.1 | - | - | 0 | - |
| - | - | 1310 | 286.2 | - | - | 0 | - |
| - | - | 2037 | 287.2 | - | - | 0 | - |
| - | - | 807.3 | 292.2 | - | - | 0 | - |
| - | - | 1029 | 293.1 | - | - | 0 | - |
| 7 | y | 2.777E+04 | 296.1 | 0.000323 | 1.091 | +1 | 2 |
| - | - | 896.2 | 296.2 | - | - | 0 | - |
| - | - | 3529 | 297.1 | - | - | 0 | - |
| - | - | 760.4 | 302.1 | - | - | 0 | - |
| - | - | 533.7 | 302.5 | - | - | 0 | - |
| - | - | 861.8 | 312.2 | - | - | 0 | - |
| - | - | 8306 | 314.2 | - | - | 0 | - |
| - | - | 568 | 315.2 | - | - | 0 | - |
| - | - | 834 | 326.7 | - | - | 0 | - |
| - | - | 537 | 327.9 | - | - | 0 | - |
| - | - | 779.9 | 339.2 | - | - | 0 | - |
| - | - | 3618 | 339.2 | - | - | 0 | - |
| - | - | 6196 | 341.3 | - | - | 0 | - |
| - | - | 1046 | 342.2 | - | - | 0 | - |
| - | - | 1494 | 342.3 | - | - | 0 | - |
| - | - | 1240 | 343.2 | - | - | 0 | - |
| - | - | 1207 | 345.3 | - | - | 0 | - |
| - | - | 4983 | 353.2 | - | - | 0 | - |
| - | - | 2847 | 354.2 | - | - | 0 | - |
| - | - | 1078 | 355.3 | - | - | 0 | - |
| - | - | 6343 | 357.2 | - | - | 0 | - |
| - | - | 1165 | 358.2 | - | - | 0 | - |
| - | - | 1162 | 367.2 | - | - | 0 | - |
| - | - | 1840 | 368.3 | - | - | 0 | - |
| - | - | 9431 | 369.3 | - | - | 0 | - |
| - | - | 2102 | 370.3 | - | - | 0 | - |
| - | - | 1.453E+04 | 371.2 | - | - | 0 | - |
| - | - | 1903 | 371.3 | - | - | 0 | - |
| - | - | 2302 | 372.2 | - | - | 0 | - |
| - | - | 568.5 | 377.2 | - | - | 0 | - |
| - | - | 2988 | 377.2 | - | - | 0 | - |
| - | - | 1031 | 382.3 | - | - | 0 | - |
| - | - | 1427 | 383.3 | - | - | 0 | - |
| - | - | 3102 | 385.2 | - | - | 0 | - |
| - | - | 785.1 | 386.2 | - | - | 0 | - |
| - | - | 1480 | 386.3 | - | - | 0 | - |
| - | - | 803.2 | 387.2 | - | - | 0 | - |
| - | - | 2664 | 387.2 | - | - | 0 | - |
| - | - | 687.3 | 387.3 | - | - | 0 | - |
| 4 | c | 1.85E+04 | 388.3 | 0.0001085 | 0.2796 | +1 | 4 |
| 2 | y | 589 | 389.2 | 0.004441 | 11.41 | +2 | 7 |
| - | - | 3229 | 389.3 | - | - | 0 | - |
| 6 | y | 1.181E+04 | 395.2 | 0.0003906 | 0.9883 | +1 | 3 |
| - | - | 2139 | 396.2 | - | - | 0 | - |
| - | - | 550.4 | 397.2 | - | - | 0 | - |
| - | - | 607.1 | 398.3 | - | - | 0 | - |
| - | - | 661.6 | 399.2 | - | - | 0 | - |
| - | - | 548.7 | 403.2 | - | - | 0 | - |
| - | - | 1794 | 403.3 | - | - | 0 | - |
| - | - | 1026 | 403.8 | - | - | 0 | - |
| - | - | 2172 | 406.2 | - | - | 0 | - |
| - | - | 2036 | 406.7 | - | - | 0 | - |
| - | - | 1246 | 421.8 | - | - | 0 | - |
| - | - | 743.2 | 422.8 | - | - | 0 | - |
| - | - | 2332 | 423.3 | - | - | 0 | - |
| - | - | 836.9 | 425.3 | - | - | 0 | - |
| - | - | 909.3 | 428.3 | - | - | 0 | - |
| - | - | 1649 | 431.2 | - | - | 0 | - |
| - | - | 1525 | 432.2 | - | - | 0 | - |
| - | - | 810.1 | 438.2 | - | - | 0 | - |
| - | - | 1996 | 438.3 | - | - | 0 | - |
| - | - | 2015 | 438.3 | - | - | 0 | - |
| - | - | 1295 | 439.1 | - | - | 0 | - |
| - | - | 1019 | 439.2 | - | - | 0 | - |
| - | - | 8910 | 439.2 | - | - | 0 | - |
| - | - | 1455 | 439.3 | - | - | 0 | - |
| - | - | 3252 | 439.8 | - | - | 0 | - |
| - | - | 777.4 | 440.2 | - | - | 0 | - |
| - | - | 1758 | 440.2 | - | - | 0 | - |
| - | - | 1297 | 440.3 | - | - | 0 | - |
| - | - | 728.3 | 440.3 | - | - | 0 | - |
| - | - | 848.4 | 440.3 | - | - | 0 | - |
| - | - | 1090 | 440.7 | - | - | 0 | - |
| - | - | 1036 | 440.8 | - | - | 0 | - |
| - | - | 2224 | 441.2 | - | - | 0 | - |
| - | - | 2013 | 441.2 | - | - | 0 | - |
| - | - | 1452 | 441.3 | - | - | 0 | - |
| - | - | 700.1 | 442.3 | - | - | 0 | - |
| - | - | 3905 | 449.7 | - | - | 0 | - |
| - | - | 2206 | 450.2 | - | - | 0 | - |
| - | - | 1643 | 452.3 | - | - | 0 | - |
| - | - | 757.3 | 453.3 | - | - | 0 | - |
| - | - | 4116 | 456.3 | - | - | 0 | - |
| - | - | 1.431E+04 | 456.3 | - | - | 0 | - |
| - | - | 848.8 | 457.3 | - | - | 0 | - |
| - | - | 4715 | 457.3 | - | - | 0 | - |
| - | - | 762.1 | 458.3 | - | - | 0 | - |
| - | - | 865.9 | 458.3 | - | - | 0 | - |
| - | - | 1631 | 460.3 | - | - | 0 | - |
| - | - | 1.196E+04 | 466.3 | - | - | 0 | - |
| - | - | 3199 | 467.3 | - | - | 0 | - |
| - | - | 2943 | 468.3 | - | - | 0 | - |
| - | - | 932.7 | 468.3 | - | - | 0 | - |
| - | - | 6487 | 469.3 | - | - | 0 | - |
| - | - | 2979 | 470.3 | - | - | 0 | - |
| - | - | 991.6 | 471.3 | - | - | 0 | - |
| - | - | 3060 | 483.3 | - | - | 0 | - |
| - | - | 4.471E+04 | 484.3 | - | - | 0 | - |
| - | - | 1727 | 484.3 | - | - | 0 | - |
| - | - | 1.078E+04 | 485.3 | - | - | 0 | - |
| - | - | 1370 | 486.3 | - | - | 0 | - |
| - | - | 797 | 487.3 | - | - | 0 | - |
| 5 | z | 5647 | 492.3 | 0.0002216 | 0.4501 | +1 | 4 |
| - | - | 2029 | 493.2 | - | - | 0 | - |
| - | - | 2487 | 493.3 | - | - | 0 | - |
| - | - | 1477 | 493.7 | - | - | 0 | - |
| - | - | 827.8 | 494.3 | - | - | 0 | - |
| - | - | 2311 | 500.3 | - | - | 0 | - |
| 5 | c | 1.031E+04 | 501.3 | 0.0001815 | 0.3621 | +1 | 5 |
| - | - | 1959 | 502.3 | - | - | 0 | - |
| - | - | 1634 | 505.3 | - | - | 0 | - |
| - | - | 8207 | 507.3 | - | - | 0 | - |
| 5 | y | 3459 | 508.3 | 0.001764 | 3.471 | +1 | 4 |
| - | - | 628.7 | 509.3 | - | - | 0 | - |
| - | - | 1335 | 510.3 | - | - | 0 | - |
| - | - | 2439 | 511.3 | - | - | 0 | - |
| - | - | 1350 | 512.3 | - | - | 0 | - |
| - | - | 771.1 | 514.3 | - | - | 0 | - |
| - | - | 692.2 | 516.3 | - | - | 0 | - |
| - | - | 1385 | 517.3 | - | - | 0 | - |
| - | - | 586.4 | 520.3 | - | - | 0 | - |
| - | - | 630.3 | 521.3 | - | - | 0 | - |
| - | - | 1996 | 522.2 | - | - | 0 | - |
| - | - | 1104 | 522.7 | - | - | 0 | - |
| - | - | 2659 | 530.3 | - | - | 0 | - |
| - | - | 1580 | 531.3 | - | - | 0 | - |
| - | - | 1978 | 532.3 | - | - | 0 | - |
| - | - | 928.5 | 533.3 | - | - | 0 | - |
| - | - | 1179 | 535.3 | - | - | 0 | - |
| - | - | 1499 | 536.8 | - | - | 0 | - |
| - | - | 722 | 538.4 | - | - | 0 | - |
| - | - | 1228 | 539.3 | - | - | 0 | - |
| - | - | 2066 | 539.4 | - | - | 0 | - |
| - | - | 890.7 | 540.4 | - | - | 0 | - |
| - | - | 2264 | 541.4 | - | - | 0 | - |
| - | - | 669.8 | 542.4 | - | - | 0 | - |
| - | - | 2240 | 546.3 | - | - | 0 | - |
| - | - | 2393 | 547.3 | - | - | 0 | - |
| 4 | z | 4372 | 549.3 | 0.0002619 | 0.4769 | +1 | 5 |
| - | - | 5191 | 550.3 | - | - | 0 | - |
| - | - | 850.6 | 551.3 | - | - | 0 | - |
| - | - | 820 | 552.3 | - | - | 0 | - |
| - | - | 2087 | 553.3 | - | - | 0 | - |
| - | - | 1698 | 554.3 | - | - | 0 | - |
| - | - | 4312 | 555.4 | - | - | 0 | - |
| - | - | 1.318E+04 | 556.4 | - | - | 0 | - |
| - | - | 3185 | 557.4 | - | - | 0 | - |
| - | - | 1.072E+04 | 564.3 | - | - | 0 | - |
| 4 | y | 1.815E+04 | 565.3 | 0.0009806 | 1.735 | +1 | 5 |
| - | - | 5887 | 565.4 | - | - | 0 | - |
| - | - | 5538 | 566.3 | - | - | 0 | - |
| - | - | 3542 | 566.4 | - | - | 0 | - |
| - | - | 852.1 | 567.3 | - | - | 0 | - |
| - | - | 1.419E+04 | 567.4 | - | - | 0 | - |
| - | - | 1.316E+04 | 568.4 | - | - | 0 | - |
| - | - | 4042 | 569.4 | - | - | 0 | - |
| - | - | 744.6 | 571.3 | - | - | 0 | - |
| - | - | 643.2 | 573.8 | - | - | 0 | - |
| - | - | 7584 | 578.8 | - | - | 0 | - |
| - | - | 3402 | 579.3 | - | - | 0 | - |
| - | - | 1530 | 580.8 | - | - | 0 | - |
| - | - | 1790 | 582.4 | - | - | 0 | - |
| - | - | 3.355E+04 | 583.4 | - | - | 0 | - |
| - | - | 1.055E+04 | 584.4 | - | - | 0 | - |
| - | - | 6666 | 585.4 | - | - | 0 | - |
| - | - | 1407 | 586.4 | - | - | 0 | - |
| - | - | 724.4 | 592.3 | - | - | 0 | - |
| 6 | c | 4.163E+04 | 600.4 | 0.0002185 | 0.364 | +1 | 6 |
| - | - | 1.473E+04 | 601.4 | - | - | 0 | - |
| - | - | 941.1 | 602.3 | - | - | 0 | - |
| - | - | 2683 | 602.4 | - | - | 0 | - |
| - | - | 810.4 | 607.4 | - | - | 0 | - |
| - | - | 1804 | 618.2 | - | - | 0 | - |
| - | - | 1093 | 624.3 | - | - | 0 | - |
| - | - | 1215 | 624.8 | - | - | 0 | - |
| - | - | 673.3 | 629.4 | - | - | 0 | - |
| - | - | 1252 | 631.3 | - | - | 0 | - |
| - | - | 851.5 | 631.8 | - | - | 0 | - |
| - | - | 729.9 | 632.3 | - | - | 0 | - |
| - | - | 1337 | 634.4 | - | - | 0 | - |
| - | - | 1698 | 635.4 | - | - | 0 | - |
| - | - | 1658 | 636.4 | - | - | 0 | - |
| - | - | 1212 | 637.4 | - | - | 0 | - |
| - | - | 1612 | 638.4 | - | - | 0 | - |
| - | - | 946 | 639.8 | - | - | 0 | - |
| - | - | 1713 | 642.4 | - | - | 0 | - |
| - | - | 1114 | 643.4 | - | - | 0 | - |
| - | - | 705.3 | 644.3 | - | - | 0 | - |
| - | - | 1679 | 646.4 | - | - | 0 | - |
| - | - | 1487 | 647.4 | - | - | 0 | - |
| 3 | z | 6362 | 648.3 | 4.973E-05 | 0.0767 | +1 | 6 |
| - | - | 2200 | 649.4 | - | - | 0 | - |
| - | - | 1453 | 649.9 | - | - | 0 | - |
| - | - | 903.3 | 650.4 | - | - | 0 | - |
| - | - | 1092 | 652.3 | - | - | 0 | - |
| - | - | 1151 | 652.4 | - | - | 0 | - |
| - | - | 2457 | 652.8 | - | - | 0 | - |
| - | - | 1106 | 653.4 | - | - | 0 | - |
| - | - | 3005 | 654.4 | - | - | 0 | - |
| - | - | 2689 | 655.4 | - | - | 0 | - |
| - | - | 1007 | 658.4 | - | - | 0 | - |
| - | - | 957 | 658.9 | - | - | 0 | - |
| - | - | 7425 | 660.8 | - | - | 0 | - |
| - | - | 4032 | 661.3 | - | - | 0 | - |
| - | - | 1167 | 663.4 | - | - | 0 | - |
| 3 | y | 1.697E+04 | 664.4 | 0.0003076 | 0.463 | +1 | 6 |
| - | - | 6012 | 665.4 | - | - | 0 | - |
| - | - | 906.3 | 666.4 | - | - | 0 | - |
| - | - | 3457 | 670.4 | - | - | 0 | - |
| - | - | 1601 | 671.4 | - | - | 0 | - |
| - | - | 1159 | 671.4 | - | - | 0 | - |
| - | - | 1264 | 672.4 | - | - | 0 | - |
| - | - | 926.9 | 672.4 | - | - | 0 | - |
| - | - | 2334 | 679.4 | - | - | 0 | - |
| - | - | 855.9 | 682.4 | - | - | 0 | - |
| - | - | 4368 | 688.4 | - | - | 0 | - |
| - | - | 1493 | 689.4 | - | - | 0 | - |
| - | - | 795.5 | 699.4 | - | - | 0 | - |
| - | - | 1202 | 708.4 | - | - | 0 | - |
| - | - | 904.9 | 709.4 | - | - | 0 | - |
| 2 | w | 2.507E+04 | 718.4 | 0.0001189 | 0.1655 | +1 | 7 |
| - | - | 1.235E+04 | 719.4 | - | - | 0 | - |
| - | - | 2726 | 720.4 | - | - | 0 | - |
| - | - | 9435 | 726.4 | - | - | 0 | - |
| - | - | 5816 | 727.4 | - | - | 0 | - |
| - | - | 2522 | 728.4 | - | - | 0 | - |
| - | - | 1018 | 729.4 | - | - | 0 | - |
| - | - | 4597 | 743.4 | - | - | 0 | - |
| - | - | 6.008E+05 | 744.4 | - | - | 0 | - |
| - | - | 2.392E+05 | 745.4 | - | - | 0 | - |
| - | - | 6.364E+04 | 746.4 | - | - | 0 | - |
| - | - | 3239 | 747.4 | - | - | 0 | - |
| 2 | z | 1.719E+04 | 761.4 | 0.0002753 | 0.3616 | +1 | 7 |
| - | - | 8792 | 762.4 | - | - | 0 | - |
| - | - | 8011 | 763.4 | - | - | 0 | - |
| - | - | 1947 | 764.4 | - | - | 0 | - |
| - | - | 1291 | 788.4 | - | - | 0 | - |
| - | - | 1756 | 790.3 | - | - | 0 | - |
| - | - | 1061 | 795.4 | - | - | 0 | - |
| - | - | 1866 | 800.5 | - | - | 0 | - |
| - | - | 2.058E+04 | 806.4 | - | - | 0 | - |
| - | - | 8159 | 807.4 | - | - | 0 | - |
| - | - | 2571 | 808.4 | - | - | 0 | - |
| - | - | 983.5 | 811.4 | - | - | 0 | - |
| - | - | 1220 | 819.4 | - | - | 0 | - |
| - | - | 1450 | 820.4 | - | - | 0 | - |
| - | - | 2.515E+04 | 844.5 | - | - | 0 | - |
| - | - | 1.467E+04 | 845.5 | - | - | 0 | - |
| - | - | 4143 | 846.5 | - | - | 0 | - |
| - | - | 2716 | 847.5 | - | - | 0 | - |
| - | - | 1439 | 848.5 | - | - | 0 | - |
| - | - | 3862 | 861.5 | - | - | 0 | - |
| - | - | 4.692E+04 | 862.5 | - | - | 0 | - |
| - | - | 2.059E+04 | 863.5 | - | - | 0 | - |
| - | - | 6633 | 864.5 | - | - | 0 | - |
| - | - | 2007 | 877.5 | - | - | 0 | - |
| - | - | 8.909E+04 | 878.5 | - | - | 0 | - |
| - | - | 4.603E+04 | 879.5 | - | - | 0 | - |
| - | - | 1.625E+04 | 880.5 | - | - | 0 | - |
| - | - | 2417 | 881.5 | - | - | 0 | - |
| - | - | 990 | 888.4 | - | - | 0 | - |
| - | - | 777.8 | 889.4 | - | - | 0 | - |
| - | - | 933 | 926.4 | - | - | 0 | - |
| - | - | 979.4 | 958.4 | - | - | 0 | - |
| - | - | 2419 | 959.4 | - | - | 0 | - |
| - | - | 4609 | 960.4 | - | - | 0 | - |
| - | - | 1241 | 961.4 | - | - | 0 | - |
| - | - | 1320 | 986.5 | - | - | 0 | - |
| - | - | 2216 | 1043 | - | - | 0 | - |
| - | - | 2897 | 1142 | - | - | 0 | - |
| - | - | 758.1 | 1143 | - | - | 0 | - |
| - | - | 924.3 | 1147 | - | - | 0 | - |
| - | - | 1745 | 1157 | - | - | 0 | - |
| - | - | 1203 | 1158 | - | - | 0 | - |
| - | - | 1211 | 1204 | - | - | 0 | - |
| - | - | 1.017E+04 | 1205 | - | - | 0 | - |
| - | - | 3409 | 1206 | - | - | 0 | - |
| - | - | 1486 | 1234 | - | - | 0 | - |
| - | - | 1829 | 1235 | - | - | 0 | - |
| - | - | 906.7 | 1264 | - | - | 0 | - |
| - | - | 1025 | 1265 | - | - | 0 | - |
| - | - | 854.7 | 1277 | - | - | 0 | - |
| - | - | 1611 | 1278 | - | - | 0 | - |
| - | - | 2213 | 1282 | - | - | 0 | - |
| - | - | 1206 | 1283 | - | - | 0 | - |
| - | - | 1040 | 1288 | - | - | 0 | - |
| - | - | 1873 | 1295 | - | - | 0 | - |
| - | - | 926.6 | 1296 | - | - | 0 | - |
| - | - | 1.993E+04 | 1299 | - | - | 0 | - |
| - | - | 1.236E+04 | 1300 | - | - | 0 | - |
| - | - | 5594 | 1301 | - | - | 0 | - |
| - | - | 936.3 | 1302 | - | - | 0 | - |
| - | - | 948.9 | 1304 | - | - | 0 | - |
| - | - | 3233 | 1305 | - | - | 0 | - |
| - | - | 2.067E+04 | 1306 | - | - | 0 | - |
| - | - | 9365 | 1307 | - | - | 0 | - |
| - | - | 9298 | 1322 | - | - | 0 | - |
| - | - | 3.974E+04 | 1323 | - | - | 0 | - |
| - | - | 1.597E+04 | 1324 | - | - | 0 | - |
| - | - | 693.4 | 2183 | - | - | 0 | - |
| - | - | 793.5 | 3081 | - | - | 0 | - |

m/z Charge Intensity FragmentType MassShift Position
120.04470825195312 0 515.56903
124.61660766601562 0 351.84357
124.64262390136719 0 347.74658
129.10218811035156 0 362.7497
136.0760955810547 0 909.5944
143.11817932128906 0 1139.3188
146.0603485107422 0 4537.7334
148.94776916503906 0 709.2294
153.0547332763672 0 419.81433
157.097900390625 0 597.1446
163.08709716796875 0 1980.3187
166.2740478515625 0 442.24933
169.13375854492188 0 669.27057
171.11306762695312 0 2082.7285
171.14944458007812 0 1696.7003
173.45025634765625 0 3360.1892
174.05511474609375 0 3387.2358
174.1238555908203 0 1573.2902
175.0587921142578 0 656.8287
177.1120147705078 0 457.17316
185.05564880371094 0 1734.7299
185.16473388671875 0 795.8943
187.10809326171875 0 2133.3774
187.14442443847656 0 48370.957
188.14776611328125 0 3927.8416
191.08181762695312 0 2868.2786
192.06578063964844 0 13225.646
192.08258056640625 0 477.96088
193.06935119628906 0 1142.3967
197.76473999023438 0 546.2408
199.1439666748047 0 728.7686
203.06640625 0 6636.8667
209.0923614501953 0 25536.994
210.09596252441406 0 2317.5212
213.15980529785156 0 1578.488
215.13931274414062 0 37491.863
216.14276123046875 0 4070.6196
221.1285400390625 0 729.02014
225.1600341796875 0 1661.7761
242.18609619140625 0 1498.3022
255.17066955566406 0 2085.1406
260.103515625 0 1589.9296
261.0870666503906 0 505.691
269.18603515625 0 1167.542
270.18145751953125 0 18806.75
271.1841125488281 0 1987.9581
278.0897216796875 0 1613.4447
278.11376953125 0 10972.628
279.09613037109375 0 477.93738 y Ammonia loss 6
279.1183776855469 0 2124.0742
282.7152404785156 0 525.2658
284.1253356933594 0 643.42316
286.21337890625 0 1309.6744
287.208251953125 0 2037.4574
292.195556640625 0 807.309
293.13616943359375 0 1028.8197
296.1244201660156 0 27773.842 y 6
296.1976318359375 0 896.2403
297.12744140625 0 3529.329
302.1349182128906 0 760.37634
302.5145568847656 0 533.7253
312.19293212890625 0 861.8442
314.20770263671875 0 8305.616
315.2105712890625 0 568.0246
326.70513916015625 0 833.99146
327.9349365234375 0 536.9771
339.16143798828125 0 779.91003
339.203125 0 3618.3953
341.2550048828125 0 6195.94
342.2026672363281 0 1045.7887
342.2585754394531 0 1494.3562
343.2342224121094 0 1240.1427
345.2503967285156 0 1206.5612
353.2190856933594 0 4982.6616
354.2388916015625 0 2846.9548
355.2701110839844 0 1078.2593
357.21343994140625 0 6343.228
358.21612548828125 0 1165.3138
367.2326354980469 0 1162.467
368.25311279296875 0 1840.1068
369.25018310546875 0 9430.503
370.2525634765625 0 2102.3882
371.22906494140625 0 14531.524
371.2641296386719 0 1903.4036
372.2325439453125 0 2302.0825
377.1585388183594 0 568.4514
377.1822509765625 0 2988.018
382.27032470703125 0 1030.8711
383.26593017578125 0 1426.882
385.2450256347656 0 3101.6707
386.2474365234375 0 785.06885
386.27667236328125 0 1479.9991
387.2228698730469 0 803.20874
387.2485046386719 0 2664.1682
387.27374267578125 0 687.277
388.25555419921875 0 18504.988 c 3
389.23333740234375 0 589.0468 y 1
389.2587890625 0 3228.6538
395.1929016113281 0 11812.919 y 5
396.1951904296875 0 2139.4026
397.1949462890625 0 550.4113
398.27679443359375 0 607.1341
399.1979675292969 0 661.56525
403.2061462402344 0 548.6907
403.2549743652344 0 1794.1593
403.8224182128906 0 1026.2148
406.198974609375 0 2171.7266
406.701904296875 0 2035.8887
421.8337097167969 0 1245.8899
422.8345642089844 0 743.1576
423.2604064941406 0 2332.3088
425.2754211425781 0 836.91833
428.2867126464844 0 909.2724
431.237548828125 0 1648.9436
432.24505615234375 0 1525.471
438.199462890625 0 810.1341
438.2715759277344 0 1995.7943
438.3089294433594 0 2014.9525
439.1480712890625 0 1295.4243
439.178955078125 0 1019.2228
439.21258544921875 0 8909.85
439.2914733886719 0 1455.2374
439.8442077636719 0 3252.4998
440.1839599609375 0 777.39557
440.2158508300781 0 1757.519
440.2500915527344 0 1297.1696
440.2960205078125 0 728.28937
440.3321838378906 0 848.3759
440.7085266113281 0 1089.8403
440.8457946777344 0 1036.2985
441.19195556640625 0 2224.1135
441.2279357910156 0 2012.924
441.30328369140625 0 1452.2474
442.3048400878906 0 700.08093
449.7151794433594 0 3904.846
450.21759033203125 0 2205.8503
452.288330078125 0 1642.7574
453.29144287109375 0 757.34406
456.2822265625 0 4116.448
456.31866455078125 0 14313.44
457.2872314453125 0 848.80566
457.3232421875 0 4715.0156
458.2602844238281 0 762.05493
458.33135986328125 0 865.85504
460.2770080566406 0 1631.3472
466.3058166503906 0 11957.633
467.3180847167969 0 3198.8567
468.2941589355469 0 2942.8342
468.32830810546875 0 932.6884
469.30145263671875 0 6487.4517
470.3006286621094 0 2979.1672
471.3027038574219 0 991.5902
483.30572509765625 0 3060.1675
484.3133544921875 0 44710.36
484.34368896484375 0 1727.2218
485.3167419433594 0 10777.564
486.3209533691406 0 1370.4875
487.2787170410156 0 797.0056
492.25762939453125 0 5647.2 z 4
493.22979736328125 0 2028.7168
493.26385498046875 0 2487.469
493.7353515625 0 1477.3604
494.2677917480469 0 827.82117
500.33282470703125 0 2310.9397
501.3396911621094 0 10305.675 c 4
502.3438415527344 0 1958.7932
505.28997802734375 0 1633.9694
507.26953125 0 8207.439
508.2748107910156 0 3458.9487 y 4
509.27569580078125 0 628.6581
510.2915344238281 0 1335.4578
511.299072265625 0 2439.3154
512.302978515625 0 1350.4469
514.2830810546875 0 771.0778
516.2854614257812 0 692.1543
517.2986450195312 0 1384.854
520.3025512695312 0 586.3874
521.3040771484375 0 630.3396
522.24658203125 0 1996.0853
522.7476196289062 0 1104.3668
530.306396484375 0 2658.923
531.31005859375 0 1579.7356
532.3072509765625 0 1977.6364
533.3018188476562 0 928.5015
535.33154296875 0 1179.0259
536.7522583007812 0 1499.0371
538.3510131835938 0 722.044
539.3186645507812 0 1227.5857
539.3676147460938 0 2066.334
540.3754272460938 0 890.69946
541.383056640625 0 2264.4243
542.3875122070312 0 669.7763
546.2799682617188 0 2240.0054
547.2848510742188 0 2393.4478
549.279052734375 0 4372.189 z 3
550.2862548828125 0 5190.676
551.2908325195312 0 850.6009
552.2918701171875 0 819.9847
553.33935546875 0 2087.3503
554.3491821289062 0 1698.3148
555.3867797851562 0 4311.979
556.3939819335938 0 13175.621
557.3978271484375 0 3185.2046
564.2906494140625 0 10721.55
565.2970581054688 0 18148.39 y 3
565.3701171875 0 5886.7505
566.3007202148438 0 5538.4536
566.3585205078125 0 3541.5308
567.3097534179688 0 852.11475
567.3623657226562 0 14193.059
568.3688354492188 0 13164.934
569.3726196289062 0 4042.4607
571.34326171875 0 744.62396
573.758056640625 0 643.1927
578.78759765625 0 7584.13
579.2882690429688 0 3402.1453
580.7515258789062 0 1530.2264
582.3742065429688 0 1790.1472
583.3814697265625 0 33554.35
584.3844604492188 0 10547.853
585.394287109375 0 6665.6836
586.4005126953125 0 1407.0968
592.2828979492188 0 724.4301
600.4081420898438 0 41629.65 c 5
601.4113159179688 0 14726.141
602.2811279296875 0 941.1401
602.4132690429688 0 2683.1685
607.3638916015625 0 810.37836
618.2078857421875 0 1803.8373
624.2859497070312 0 1092.8116
624.77197265625 0 1215.3678
629.3639526367188 0 673.3362
631.2774047851562 0 1251.7808
631.7777709960938 0 851.48425
632.3040771484375 0 729.94305
634.3922729492188 0 1337.269
635.350341796875 0 1698.1284
636.38427734375 0 1657.8784
637.3912353515625 0 1211.9734
638.3955688476562 0 1612.213
639.78857421875 0 945.9953
642.4192504882812 0 1713.3997
643.4213256835938 0 1113.9647
644.3004760742188 0 705.28815
646.3554077148438 0 1678.668
647.3556518554688 0 1487.4642
648.3477783203125 0 6361.6724 z 2
649.3504028320312 0 2200.1318
649.8973388671875 0 1453.2506
650.4037475585938 0 903.2958
652.3110961914062 0 1092.399
652.3969116210938 0 1150.6443
652.8023071289062 0 2456.7708
653.3751831054688 0 1106.4349
654.3948364257812 0 3004.5671
655.4014282226562 0 2689.3071
658.4102783203125 0 1007.048
658.91162109375 0 957
660.8113403320312 0 7425.076
661.31201171875 0 4031.6975
663.357177734375 0 1166.829
664.3667602539062 0 16968.248 y 2
665.3700561523438 0 6012.005
666.372314453125 0 906.2947
670.414794921875 0 3456.707
671.3506469726562 0 1601.0642
671.4141235351562 0 1159.1973
672.3511962890625 0 1263.7136
672.4205322265625 0 926.87823
679.3784790039062 0 2333.7883
682.3988037109375 0 855.92413
688.3750610351562 0 4367.8438
689.3779907226562 0 1493.4572
699.4331665039062 0 795.4775
708.416259765625 0 1201.6764
709.4082641601562 0 904.9135
718.3771362304688 0 25066.002 w 1
719.3799438476562 0 12353.575
720.3829956054688 0 2725.8071
726.4265747070312 0 9435.242
727.4207153320312 0 5815.908
728.4192504882812 0 2521.5178
729.4144897460938 0 1018.21234
743.4209594726562 0 4597.472
744.4381713867188 0 600797.6
745.440673828125 0 239244.11
746.4432983398438 0 63638.58
747.4462890625 0 3238.672
761.4320678710938 0 17192.2 z 1
762.434326171875 0 8791.768
763.4430541992188 0 8011.2695
764.4462280273438 0 1947.1597
788.4083251953125 0 1290.5027
790.3298950195312 0 1755.6755
795.3729248046875 0 1061.003
800.4803466796875 0 1866.1376
806.4171752929688 0 20577.871
807.4199829101562 0 8159.1074
808.421142578125 0 2570.512
811.3880004882812 0 983.54517
819.4220581054688 0 1220.4562
820.4290161132812 0 1449.9774
844.4681396484375 0 25146.918
845.4725952148438 0 14668.528
846.4744873046875 0 4142.7124
847.4625244140625 0 2715.8987
848.4605712890625 0 1439.4246
861.4951782226562 0 3862.2744
862.4801025390625 0 46917.375
863.4826049804688 0 20586.836
864.4844970703125 0 6632.5747
877.5111694335938 0 2006.5396
878.4981689453125 0 89089
879.501708984375 0 46026.918
880.5045776367188 0 16247.155
881.5079956054688 0 2417.2637
888.3809204101562 0 989.9944
889.3925170898438 0 777.8306
926.4164428710938 0 933.0456
958.4220581054688 0 979.4489
959.4257202148438 0 2418.6555
960.4327392578125 0 4609.388
961.439697265625 0 1240.9109
986.4597778320312 0 1320.0986
1043.4830322265625 0 2215.8145
1141.553955078125 0 2897.314
1142.54931640625 0 758.07916
1146.5462646484375 0 924.3201
1156.565185546875 0 1744.7865
1157.5703125 0 1202.5946
1203.5380859375 0 1210.5157
1204.5545654296875 0 10171.837
1205.5557861328125 0 3409.016
1233.58251953125 0 1485.8335
1234.587890625 0 1829.42
1263.5966796875 0 906.68317
1264.593505859375 0 1024.7621
1276.6298828125 0 854.665
1277.60498046875 0 1610.6984
1281.757080078125 0 2212.9597
1282.7547607421875 0 1205.6184
1287.59375 0 1039.9009
1294.6397705078125 0 1872.8176
1295.6419677734375 0 926.6301
1298.785888671875 0 19927.914
1299.7890625 0 12362.6455
1300.791259765625 0 5594.3516
1301.7847900390625 0 936.2761
1303.7274169921875 0 948.9231
1304.6060791015625 0 3232.8481
1305.6015625 0 20667.797
1306.6048583984375 0 9365.162
1321.6195068359375 0 9297.552
1322.627197265625 0 39735.91
1323.630859375 0 15971.987
2182.86376953125 0 693.42694
3081.471435546875 0 793.469

Spectrum Details

|  |  |
| --- | --- |
| Matched peaks? Matched peaks The total absolute number of peaks matched. Additionally in brackets the total fraction of peaks matched and the total number of peaks is shown. | 15 (4.20% of 357) |
| FDR? FDR The false discovery rate estimated for this peptide. It is calculated by matching all theoretical fragments with a non-integer shift with the raw peaks for this spectrum. This is done with 40 different shifts. The resulting percentage is the average number of annotated peaks over the number of annotated peaks with the correct spectrum. | 8.89% |
| Satellite FDR? Satellite FDR See the FDR for details on its calculation. This satellite ion specific FDR only contains the satellite ions (d/w) for I/L/J positions. | 9.52% |
| PSM Score? PSM Score The PSM Score as given by Hecklib to this annotated spectrum. It is shown with three significant figures. | 162 |

#### Spectrum 10783? Spectrum 10783 The raw spectrum of this peptide as annotated by Hecklib. The fragments are coloured according to ion type (see legend). Any peaks with a star '\*' as text can be hovered over to see the full details, first the ion type second the mass shift type. By hovering over the amino acids in the peptide or ions in the legend the corresponding peaks are highlighted. By toggling the 'Unassigned' label you can turn the background (unassigned) peaks on or off in the plot. By updating the slider in the Ion legend you can update the spectrum to only show the top X% of the peaks with labels. The top X% means any peak that is within X% of the highest intensity. By dragging in the spectrum you can zoom in to a specific part of the spectrum and use 'Zoom Out' to get back to the original zoom level. The annotation of the spectrum is based on the given sequence in the peptides file and is done with different software so inconsistencies are likely. The peaks are annotated based on the given sequence, with 20 ppm tolerance.

Copy Data

##### Spectrum 10783 (TSV)

###### Preview

```
Loading example...
```

*Click on the button to copy the data to your clipboard.*

Mz MinMz MaxIntensity Max

WidthHeightPeptide font sizePeptide stroke widthSpectrum font sizeSpectrum stroke widthCompact peptide

Ion legend

wxyz

abcd

OtherUnassignedIon^Charge^_Position_Show for top:%

TLVGJVNY

06.38e+51.28e+61.91e+62.55e+6

Zoom Out

c^+1^_2_y^+1^_2_y^+1^_2_c^+1^_3_y^+2^_6_z^+1^_3_c^+1^_4_y^+1^_3_z^+1^_4_c^+1^_5_y^+1^_4_z^+1^_5_y^+1^_5_c^+1^_6_w^+1^_6_z^+1^_6_y^+1^_6_w^+1^_7_z^+1^_7_y^+1^_7_

0778155623343113

Fragment Matches Table

Show background peaks

| Position | Ion type | Intensity | mz Theoretical | mz Error (Th) | mz Error (ppm) | Charge | Series Number |
| --- | --- | --- | --- | --- | --- | --- | --- |
| - | - | 2219 | 120 | - | - | 0 | - |
| - | - | 552.5 | 126.4 | - | - | 0 | - |
| - | - | 525 | 132.1 | - | - | 0 | - |
| - | - | 475.3 | 135.7 | - | - | 0 | - |
| - | - | 4795 | 136.1 | - | - | 0 | - |
| - | - | 633.4 | 137.1 | - | - | 0 | - |
| - | - | 1561 | 142.1 | - | - | 0 | - |
| - | - | 6725 | 143.1 | - | - | 0 | - |
| - | - | 2.849E+04 | 146.1 | - | - | 0 | - |
| - | - | 2082 | 147.1 | - | - | 0 | - |
| - | - | 1391 | 150.1 | - | - | 0 | - |
| - | - | 1785 | 157.1 | - | - | 0 | - |
| - | - | 3268 | 159.1 | - | - | 0 | - |
| - | - | 1.266E+04 | 163.1 | - | - | 0 | - |
| - | - | 1326 | 164.1 | - | - | 0 | - |
| - | - | 1356 | 169.1 | - | - | 0 | - |
| - | - | 689.2 | 169.1 | - | - | 0 | - |
| - | - | 1.059E+04 | 171.1 | - | - | 0 | - |
| - | - | 1339 | 171.1 | - | - | 0 | - |
| - | - | 4650 | 173.5 | - | - | 0 | - |
| - | - | 2.05E+04 | 174.1 | - | - | 0 | - |
| - | - | 4827 | 174.1 | - | - | 0 | - |
| - | - | 1425 | 175.1 | - | - | 0 | - |
| - | - | 721.8 | 176.1 | - | - | 0 | - |
| - | - | 1053 | 183.1 | - | - | 0 | - |
| - | - | 1.072E+04 | 185.1 | - | - | 0 | - |
| - | - | 4569 | 185.2 | - | - | 0 | - |
| - | - | 1032 | 186.1 | - | - | 0 | - |
| - | - | 1.344E+04 | 187.1 | - | - | 0 | - |
| - | - | 2.962E+05 | 187.1 | - | - | 0 | - |
| - | - | 2.471E+04 | 188.1 | - | - | 0 | - |
| - | - | 1399 | 189.2 | - | - | 0 | - |
| - | - | 843.5 | 191.1 | - | - | 0 | - |
| - | - | 2.134E+04 | 191.1 | - | - | 0 | - |
| - | - | 7.642E+04 | 192.1 | - | - | 0 | - |
| - | - | 6821 | 193.1 | - | - | 0 | - |
| - | - | 2707 | 197.1 | - | - | 0 | - |
| - | - | 1034 | 199.1 | - | - | 0 | - |
| - | - | 729.7 | 201.1 | - | - | 0 | - |
| - | - | 4.11E+04 | 203.1 | - | - | 0 | - |
| - | - | 3159 | 204.1 | - | - | 0 | - |
| - | - | 2564 | 204.1 | - | - | 0 | - |
| - | - | 692.1 | 208.1 | - | - | 0 | - |
| - | - | 1.464E+05 | 209.1 | - | - | 0 | - |
| - | - | 1.563E+04 | 210.1 | - | - | 0 | - |
| - | - | 8042 | 213.2 | - | - | 0 | - |
| - | - | 1072 | 214.2 | - | - | 0 | - |
| - | - | 2.034E+05 | 215.1 | - | - | 0 | - |
| - | - | 1.844E+04 | 216.1 | - | - | 0 | - |
| - | - | 1020 | 217.1 | - | - | 0 | - |
| - | - | 9921 | 225.2 | - | - | 0 | - |
| 2 | c | 836.7 | 232.2 | 0.0004781 | 2.059 | +1 | 2 |
| - | - | 782.9 | 234.1 | - | - | 0 | - |
| - | - | 791.6 | 238.9 | - | - | 0 | - |
| - | - | 1415 | 240.1 | - | - | 0 | - |
| - | - | 7462 | 242.2 | - | - | 0 | - |
| - | - | 857.9 | 243.2 | - | - | 0 | - |
| - | - | 3884 | 254.2 | - | - | 0 | - |
| - | - | 1832 | 254.2 | - | - | 0 | - |
| - | - | 9887 | 255.2 | - | - | 0 | - |
| - | - | 867.8 | 255.2 | - | - | 0 | - |
| - | - | 1771 | 256.2 | - | - | 0 | - |
| - | - | 3357 | 258.1 | - | - | 0 | - |
| - | - | 880.5 | 259.1 | - | - | 0 | - |
| - | - | 6826 | 260.1 | - | - | 0 | - |
| - | - | 3773 | 261.1 | - | - | 0 | - |
| - | - | 1131 | 268.2 | - | - | 0 | - |
| - | - | 1230 | 269.2 | - | - | 0 | - |
| - | - | 6373 | 269.2 | - | - | 0 | - |
| - | - | 9.696E+04 | 270.2 | - | - | 0 | - |
| - | - | 1.363E+04 | 271.2 | - | - | 0 | - |
| - | - | 5540 | 272.2 | - | - | 0 | - |
| - | - | 5273 | 278.1 | - | - | 0 | - |
| - | - | 6.229E+04 | 278.1 | - | - | 0 | - |
| - | - | 1053 | 278.1 | - | - | 0 | - |
| 7 | y | 1545 | 279.1 | 0.002669 | 9.563 | +1 | 2 |
| - | - | 8395 | 279.1 | - | - | 0 | - |
| - | - | 4452 | 283.2 | - | - | 0 | - |
| - | - | 5336 | 284.1 | - | - | 0 | - |
| - | - | 1555 | 284.2 | - | - | 0 | - |
| - | - | 1155 | 285.1 | - | - | 0 | - |
| - | - | 925.9 | 285.2 | - | - | 0 | - |
| - | - | 4803 | 286.2 | - | - | 0 | - |
| - | - | 9502 | 287.2 | - | - | 0 | - |
| - | - | 1815 | 288.2 | - | - | 0 | - |
| - | - | 2268 | 292.2 | - | - | 0 | - |
| - | - | 1183 | 292.7 | - | - | 0 | - |
| - | - | 1023 | 295.1 | - | - | 0 | - |
| 7 | y | 1.548E+05 | 296.1 | 0.0006893 | 2.328 | +1 | 2 |
| - | - | 5708 | 296.2 | - | - | 0 | - |
| - | - | 2.036E+04 | 297.1 | - | - | 0 | - |
| - | - | 2158 | 298.1 | - | - | 0 | - |
| - | - | 2294 | 298.2 | - | - | 0 | - |
| - | - | 3400 | 300.2 | - | - | 0 | - |
| - | - | 5064 | 302.1 | - | - | 0 | - |
| - | - | 3363 | 312.2 | - | - | 0 | - |
| - | - | 4.138E+04 | 314.2 | - | - | 0 | - |
| - | - | 7317 | 315.2 | - | - | 0 | - |
| - | - | 795.8 | 317.2 | - | - | 0 | - |
| - | - | 847.9 | 324.2 | - | - | 0 | - |
| - | - | 1797 | 324.2 | - | - | 0 | - |
| - | - | 1607 | 325.2 | - | - | 0 | - |
| - | - | 2239 | 326.2 | - | - | 0 | - |
| - | - | 880.2 | 326.2 | - | - | 0 | - |
| - | - | 4197 | 326.7 | - | - | 0 | - |
| - | - | 2708 | 327.2 | - | - | 0 | - |
| - | - | 1467 | 327.3 | - | - | 0 | - |
| - | - | 3185 | 329.2 | - | - | 0 | - |
| - | - | 1460 | 330.2 | - | - | 0 | - |
| 3 | c | 3874 | 331.2 | 0.0004541 | 1.371 | +1 | 3 |
| 3 | y | 2640 | 332.7 | 0.0003608 | 1.084 | +2 | 6 |
| - | - | 1619 | 333.2 | - | - | 0 | - |
| - | - | 3227 | 335.7 | - | - | 0 | - |
| - | - | 1330 | 336.2 | - | - | 0 | - |
| - | - | 3489 | 338.2 | - | - | 0 | - |
| - | - | 2.04E+04 | 339.2 | - | - | 0 | - |
| - | - | 2945 | 340.2 | - | - | 0 | - |
| - | - | 1770 | 340.3 | - | - | 0 | - |
| - | - | 3.923E+04 | 341.3 | - | - | 0 | - |
| - | - | 6559 | 342.2 | - | - | 0 | - |
| - | - | 6288 | 342.3 | - | - | 0 | - |
| - | - | 4372 | 343.2 | - | - | 0 | - |
| - | - | 5029 | 343.2 | - | - | 0 | - |
| - | - | 764 | 343.3 | - | - | 0 | - |
| - | - | 3512 | 345.3 | - | - | 0 | - |
| - | - | 2.571E+04 | 353.2 | - | - | 0 | - |
| - | - | 1096 | 354.2 | - | - | 0 | - |
| - | - | 2057 | 354.2 | - | - | 0 | - |
| - | - | 1.257E+04 | 354.2 | - | - | 0 | - |
| - | - | 1253 | 355.2 | - | - | 0 | - |
| - | - | 3026 | 355.2 | - | - | 0 | - |
| - | - | 4549 | 355.3 | - | - | 0 | - |
| - | - | 2783 | 356.2 | - | - | 0 | - |
| - | - | 3.419E+04 | 357.2 | - | - | 0 | - |
| - | - | 1359 | 357.3 | - | - | 0 | - |
| - | - | 5256 | 358.2 | - | - | 0 | - |
| - | - | 905.4 | 365.3 | - | - | 0 | - |
| - | - | 6327 | 367.2 | - | - | 0 | - |
| - | - | 9222 | 368.3 | - | - | 0 | - |
| - | - | 5.779E+04 | 369.3 | - | - | 0 | - |
| - | - | 1.213E+04 | 370.3 | - | - | 0 | - |
| - | - | 7.668E+04 | 371.2 | - | - | 0 | - |
| - | - | 8310 | 371.3 | - | - | 0 | - |
| - | - | 1.392E+04 | 372.2 | - | - | 0 | - |
| - | - | 1202 | 372.3 | - | - | 0 | - |
| - | - | 1158 | 372.3 | - | - | 0 | - |
| - | - | 1607 | 373.2 | - | - | 0 | - |
| - | - | 1021 | 374.2 | - | - | 0 | - |
| - | - | 1514 | 374.2 | - | - | 0 | - |
| - | - | 1309 | 377.2 | - | - | 0 | - |
| - | - | 1.239E+04 | 377.2 | - | - | 0 | - |
| - | - | 2561 | 378.2 | - | - | 0 | - |
| 6 | z | 1039 | 379.2 | 0.0008651 | 2.282 | +1 | 3 |
| - | - | 4214 | 382.3 | - | - | 0 | - |
| - | - | 2264 | 383.2 | - | - | 0 | - |
| - | - | 9706 | 383.3 | - | - | 0 | - |
| - | - | 2017 | 384.3 | - | - | 0 | - |
| - | - | 1.737E+04 | 385.2 | - | - | 0 | - |
| - | - | 3018 | 386.2 | - | - | 0 | - |
| - | - | 4314 | 386.3 | - | - | 0 | - |
| - | - | 1614 | 387.2 | - | - | 0 | - |
| - | - | 9945 | 387.2 | - | - | 0 | - |
| 4 | c | 6.885E+04 | 388.3 | 0.0006273 | 1.616 | +1 | 4 |
| - | - | 1.22E+04 | 389.3 | - | - | 0 | - |
| - | - | 1601 | 390.3 | - | - | 0 | - |
| - | - | 1194 | 393.2 | - | - | 0 | - |
| - | - | 1414 | 394.2 | - | - | 0 | - |
| 6 | y | 5.903E+04 | 395.2 | 0.0006347 | 1.606 | +1 | 3 |
| - | - | 1.004E+04 | 396.2 | - | - | 0 | - |
| - | - | 799.7 | 397.2 | - | - | 0 | - |
| - | - | 2782 | 401.2 | - | - | 0 | - |
| - | - | 1336 | 403.3 | - | - | 0 | - |
| - | - | 1136 | 406.2 | - | - | 0 | - |
| - | - | 1168 | 407.3 | - | - | 0 | - |
| - | - | 1101 | 410.2 | - | - | 0 | - |
| - | - | 1053 | 410.3 | - | - | 0 | - |
| - | - | 1593 | 410.3 | - | - | 0 | - |
| - | - | 3183 | 411.3 | - | - | 0 | - |
| - | - | 1158 | 412.3 | - | - | 0 | - |
| - | - | 1178 | 412.3 | - | - | 0 | - |
| - | - | 1664 | 413.2 | - | - | 0 | - |
| - | - | 1978 | 413.3 | - | - | 0 | - |
| - | - | 2025 | 416.2 | - | - | 0 | - |
| - | - | 1804 | 417.2 | - | - | 0 | - |
| - | - | 1979 | 421.3 | - | - | 0 | - |
| - | - | 2251 | 421.8 | - | - | 0 | - |
| - | - | 2302 | 422.2 | - | - | 0 | - |
| - | - | 9631 | 423.3 | - | - | 0 | - |
| - | - | 1638 | 424.3 | - | - | 0 | - |
| - | - | 971.2 | 424.3 | - | - | 0 | - |
| - | - | 4297 | 425.3 | - | - | 0 | - |
| - | - | 1360 | 426.3 | - | - | 0 | - |
| - | - | 1.033E+04 | 428.3 | - | - | 0 | - |
| - | - | 2171 | 429.3 | - | - | 0 | - |
| - | - | 8184 | 431.2 | - | - | 0 | - |
| - | - | 6975 | 432.2 | - | - | 0 | - |
| - | - | 1040 | 433.2 | - | - | 0 | - |
| - | - | 3629 | 437.3 | - | - | 0 | - |
| - | - | 1096 | 438.2 | - | - | 0 | - |
| - | - | 1.553E+04 | 438.3 | - | - | 0 | - |
| - | - | 8200 | 438.3 | - | - | 0 | - |
| - | - | 8547 | 439.2 | - | - | 0 | - |
| - | - | 3372 | 439.3 | - | - | 0 | - |
| - | - | 3003 | 439.3 | - | - | 0 | - |
| - | - | 1290 | 439.8 | - | - | 0 | - |
| - | - | 3391 | 439.8 | - | - | 0 | - |
| - | - | 2154 | 440.2 | - | - | 0 | - |
| - | - | 6742 | 440.3 | - | - | 0 | - |
| - | - | 3172 | 440.3 | - | - | 0 | - |
| - | - | 2879 | 440.3 | - | - | 0 | - |
| - | - | 2679 | 441.2 | - | - | 0 | - |
| - | - | 1600 | 441.2 | - | - | 0 | - |
| - | - | 2169 | 441.3 | - | - | 0 | - |
| - | - | 6068 | 441.3 | - | - | 0 | - |
| - | - | 1524 | 441.3 | - | - | 0 | - |
| - | - | 3564 | 442.3 | - | - | 0 | - |
| - | - | 937.1 | 444.5 | - | - | 0 | - |
| - | - | 2616 | 449.7 | - | - | 0 | - |
| - | - | 1281 | 450.2 | - | - | 0 | - |
| - | - | 1.217E+04 | 452.3 | - | - | 0 | - |
| - | - | 2476 | 453.3 | - | - | 0 | - |
| - | - | 1456 | 454.2 | - | - | 0 | - |
| - | - | 1466 | 454.3 | - | - | 0 | - |
| - | - | 1939 | 454.3 | - | - | 0 | - |
| - | - | 2.34E+04 | 456.3 | - | - | 0 | - |
| - | - | 6.935E+04 | 456.3 | - | - | 0 | - |
| - | - | 5179 | 457.3 | - | - | 0 | - |
| - | - | 1.983E+04 | 457.3 | - | - | 0 | - |
| - | - | 3115 | 458.3 | - | - | 0 | - |
| - | - | 4035 | 458.3 | - | - | 0 | - |
| - | - | 1474 | 460.3 | - | - | 0 | - |
| - | - | 1620 | 465.3 | - | - | 0 | - |
| - | - | 5.933E+04 | 466.3 | - | - | 0 | - |
| - | - | 4043 | 467.3 | - | - | 0 | - |
| - | - | 1.489E+04 | 467.3 | - | - | 0 | - |
| - | - | 1.26E+04 | 468.3 | - | - | 0 | - |
| - | - | 2330 | 468.3 | - | - | 0 | - |
| - | - | 2.467E+04 | 469.3 | - | - | 0 | - |
| - | - | 1.874E+04 | 470.3 | - | - | 0 | - |
| - | - | 3114 | 471.3 | - | - | 0 | - |
| - | - | 2681 | 473.3 | - | - | 0 | - |
| - | - | 1051 | 481.3 | - | - | 0 | - |
| - | - | 6332 | 482.3 | - | - | 0 | - |
| - | - | 1.03E+04 | 483.3 | - | - | 0 | - |
| - | - | 2.03E+05 | 484.3 | - | - | 0 | - |
| - | - | 4.736E+04 | 485.3 | - | - | 0 | - |
| - | - | 8110 | 486.3 | - | - | 0 | - |
| - | - | 3741 | 487.3 | - | - | 0 | - |
| - | - | 1149 | 490.3 | - | - | 0 | - |
| 5 | z | 2.201E+04 | 492.3 | 0.0009991 | 2.03 | +1 | 4 |
| - | - | 1004 | 493.2 | - | - | 0 | - |
| - | - | 1.237E+04 | 493.3 | - | - | 0 | - |
| - | - | 2245 | 494.3 | - | - | 0 | - |
| - | - | 1.024E+04 | 500.3 | - | - | 0 | - |
| 5 | c | 3.813E+04 | 501.3 | 0.0007308 | 1.458 | +1 | 5 |
| - | - | 9376 | 502.3 | - | - | 0 | - |
| - | - | 5246 | 505.3 | - | - | 0 | - |
| - | - | 975.9 | 506.3 | - | - | 0 | - |
| - | - | 3.551E+04 | 507.3 | - | - | 0 | - |
| 5 | y | 1.545E+04 | 508.3 | 0.001581 | 3.111 | +1 | 4 |
| - | - | 847.7 | 508.3 | - | - | 0 | - |
| - | - | 5055 | 509.3 | - | - | 0 | - |
| - | - | 5258 | 510.3 | - | - | 0 | - |
| - | - | 7997 | 511.3 | - | - | 0 | - |
| - | - | 2971 | 512.3 | - | - | 0 | - |
| - | - | 1089 | 513.3 | - | - | 0 | - |
| - | - | 1236 | 513.3 | - | - | 0 | - |
| - | - | 2380 | 515.3 | - | - | 0 | - |
| - | - | 1802 | 517.3 | - | - | 0 | - |
| - | - | 1972 | 520.4 | - | - | 0 | - |
| - | - | 1821 | 521.3 | - | - | 0 | - |
| - | - | 1738 | 521.4 | - | - | 0 | - |
| - | - | 1409 | 522.2 | - | - | 0 | - |
| - | - | 3135 | 526.3 | - | - | 0 | - |
| - | - | 1029 | 527.3 | - | - | 0 | - |
| - | - | 860.8 | 529.3 | - | - | 0 | - |
| - | - | 1778 | 529.3 | - | - | 0 | - |
| - | - | 1.35E+04 | 530.3 | - | - | 0 | - |
| - | - | 2845 | 531.3 | - | - | 0 | - |
| - | - | 4917 | 531.3 | - | - | 0 | - |
| - | - | 2733 | 532.3 | - | - | 0 | - |
| - | - | 2354 | 535.3 | - | - | 0 | - |
| - | - | 7805 | 535.3 | - | - | 0 | - |
| - | - | 2183 | 536.3 | - | - | 0 | - |
| - | - | 1977 | 537.4 | - | - | 0 | - |
| - | - | 7840 | 538.4 | - | - | 0 | - |
| - | - | 3795 | 539.3 | - | - | 0 | - |
| - | - | 1.096E+04 | 539.4 | - | - | 0 | - |
| - | - | 6974 | 540.4 | - | - | 0 | - |
| - | - | 6911 | 541.4 | - | - | 0 | - |
| - | - | 1836 | 542.4 | - | - | 0 | - |
| - | - | 2712 | 543.3 | - | - | 0 | - |
| - | - | 1166 | 544.4 | - | - | 0 | - |
| - | - | 9346 | 546.3 | - | - | 0 | - |
| - | - | 1.009E+04 | 547.3 | - | - | 0 | - |
| - | - | 1773 | 548.3 | - | - | 0 | - |
| 4 | z | 1.986E+04 | 549.3 | 0.0007757 | 1.412 | +1 | 5 |
| - | - | 2.257E+04 | 550.3 | - | - | 0 | - |
| - | - | 1075 | 550.4 | - | - | 0 | - |
| - | - | 5539 | 551.3 | - | - | 0 | - |
| - | - | 1392 | 551.4 | - | - | 0 | - |
| - | - | 1.2E+04 | 553.3 | - | - | 0 | - |
| - | - | 6083 | 554.4 | - | - | 0 | - |
| - | - | 1.927E+04 | 555.4 | - | - | 0 | - |
| - | - | 4.811E+04 | 556.4 | - | - | 0 | - |
| - | - | 3127 | 557.3 | - | - | 0 | - |
| - | - | 1.697E+04 | 557.4 | - | - | 0 | - |
| - | - | 2766 | 558.4 | - | - | 0 | - |
| - | - | 4.6E+04 | 564.3 | - | - | 0 | - |
| 4 | y | 7.903E+04 | 565.3 | 4.078E-06 | 0.007214 | +1 | 5 |
| - | - | 2.877E+04 | 565.4 | - | - | 0 | - |
| - | - | 2.203E+04 | 566.3 | - | - | 0 | - |
| - | - | 1.196E+04 | 566.4 | - | - | 0 | - |
| - | - | 2763 | 567.3 | - | - | 0 | - |
| - | - | 5.525E+04 | 567.4 | - | - | 0 | - |
| - | - | 5.457E+04 | 568.4 | - | - | 0 | - |
| - | - | 1545 | 568.4 | - | - | 0 | - |
| - | - | 1.456E+04 | 569.4 | - | - | 0 | - |
| - | - | 2249 | 570.4 | - | - | 0 | - |
| - | - | 867.8 | 571.3 | - | - | 0 | - |
| - | - | 6332 | 571.3 | - | - | 0 | - |
| - | - | 1107 | 571.4 | - | - | 0 | - |
| - | - | 1325 | 572.3 | - | - | 0 | - |
| - | - | 2937 | 578.8 | - | - | 0 | - |
| - | - | 1241 | 579.3 | - | - | 0 | - |
| - | - | 4750 | 582.4 | - | - | 0 | - |
| - | - | 1.456E+05 | 583.4 | - | - | 0 | - |
| - | - | 4.545E+04 | 584.4 | - | - | 0 | - |
| - | - | 2.247E+04 | 585.4 | - | - | 0 | - |
| - | - | 2927 | 586.3 | - | - | 0 | - |
| - | - | 6362 | 586.4 | - | - | 0 | - |
| - | - | 1524 | 587.4 | - | - | 0 | - |
| - | - | 2602 | 592.3 | - | - | 0 | - |
| - | - | 1639 | 596.3 | - | - | 0 | - |
| - | - | 1762 | 598.3 | - | - | 0 | - |
| - | - | 1418 | 599.3 | - | - | 0 | - |
| 6 | c | 1.756E+05 | 600.4 | 0.000951 | 1.584 | +1 | 6 |
| - | - | 5.469E+04 | 601.4 | - | - | 0 | - |
| - | - | 1.17E+04 | 602.4 | - | - | 0 | - |
| - | - | 1284 | 607.4 | - | - | 0 | - |
| - | - | 2649 | 624.4 | - | - | 0 | - |
| - | - | 1809 | 625.4 | - | - | 0 | - |
| - | - | 1666 | 628.3 | - | - | 0 | - |
| - | - | 2606 | 630.3 | - | - | 0 | - |
| 3 | w | 1275 | 633.3 | 0.005153 | 8.137 | +1 | 6 |
| - | - | 5291 | 634.4 | - | - | 0 | - |
| - | - | 1904 | 635.4 | - | - | 0 | - |
| - | - | 1423 | 635.4 | - | - | 0 | - |
| - | - | 9402 | 636.4 | - | - | 0 | - |
| - | - | 7527 | 637.4 | - | - | 0 | - |
| - | - | 4730 | 638.4 | - | - | 0 | - |
| - | - | 1458 | 639.4 | - | - | 0 | - |
| - | - | 1.047E+04 | 642.4 | - | - | 0 | - |
| - | - | 2578 | 643.4 | - | - | 0 | - |
| - | - | 1.028E+04 | 646.4 | - | - | 0 | - |
| - | - | 2981 | 647.4 | - | - | 0 | - |
| 3 | z | 2.883E+04 | 648.3 | 0.001148 | 1.771 | +1 | 6 |
| - | - | 8820 | 649.4 | - | - | 0 | - |
| - | - | 1674 | 650.4 | - | - | 0 | - |
| - | - | 1567 | 651.4 | - | - | 0 | - |
| - | - | 8377 | 652.4 | - | - | 0 | - |
| - | - | 4334 | 653.4 | - | - | 0 | - |
| - | - | 897.7 | 653.4 | - | - | 0 | - |
| - | - | 1.407E+04 | 654.4 | - | - | 0 | - |
| - | - | 1.279E+04 | 655.4 | - | - | 0 | - |
| - | - | 4684 | 656.4 | - | - | 0 | - |
| - | - | 2151 | 660.8 | - | - | 0 | - |
| - | - | 1309 | 661.3 | - | - | 0 | - |
| - | - | 1904 | 663.4 | - | - | 0 | - |
| 3 | y | 8.185E+04 | 664.4 | 0.00104 | 1.565 | +1 | 6 |
| - | - | 2.879E+04 | 665.4 | - | - | 0 | - |
| - | - | 4951 | 666.4 | - | - | 0 | - |
| - | - | 1333 | 669.4 | - | - | 0 | - |
| - | - | 1.376E+04 | 670.4 | - | - | 0 | - |
| - | - | 1.104E+04 | 671.3 | - | - | 0 | - |
| - | - | 5378 | 671.4 | - | - | 0 | - |
| - | - | 2865 | 672.4 | - | - | 0 | - |
| - | - | 3235 | 672.4 | - | - | 0 | - |
| - | - | 1252 | 673.3 | - | - | 0 | - |
| - | - | 3010 | 679.4 | - | - | 0 | - |
| - | - | 3970 | 682.4 | - | - | 0 | - |
| - | - | 2399 | 687.4 | - | - | 0 | - |
| - | - | 3460 | 687.4 | - | - | 0 | - |
| - | - | 1.66E+04 | 688.4 | - | - | 0 | - |
| - | - | 956.2 | 688.4 | - | - | 0 | - |
| - | - | 4759 | 689.4 | - | - | 0 | - |
| - | - | 2676 | 691.4 | - | - | 0 | - |
| - | - | 1424 | 692.3 | - | - | 0 | - |
| - | - | 5483 | 699.4 | - | - | 0 | - |
| - | - | 2007 | 700.4 | - | - | 0 | - |
| - | - | 3456 | 701.4 | - | - | 0 | - |
| - | - | 1657 | 702.4 | - | - | 0 | - |
| - | - | 4399 | 705.4 | - | - | 0 | - |
| - | - | 1691 | 706.4 | - | - | 0 | - |
| - | - | 5291 | 708.4 | - | - | 0 | - |
| - | - | 5028 | 709.4 | - | - | 0 | - |
| - | - | 1457 | 710.4 | - | - | 0 | - |
| - | - | 1413 | 714.4 | - | - | 0 | - |
| - | - | 2047 | 717.4 | - | - | 0 | - |
| 2 | w | 1.16E+05 | 718.4 | 0.0008513 | 1.185 | +1 | 7 |
| - | - | 4.262E+04 | 719.4 | - | - | 0 | - |
| - | - | 9386 | 720.4 | - | - | 0 | - |
| - | - | 3.787E+04 | 726.4 | - | - | 0 | - |
| - | - | 2.27E+04 | 727.4 | - | - | 0 | - |
| - | - | 7724 | 728.4 | - | - | 0 | - |
| - | - | 2848 | 729.4 | - | - | 0 | - |
| - | - | 2.297E+04 | 743.4 | - | - | 0 | - |
| - | - | 2.528E+06 | 744.4 | - | - | 0 | - |
| - | - | 9.206E+05 | 745.4 | - | - | 0 | - |
| - | - | 2.24E+05 | 746.4 | - | - | 0 | - |
| - | - | 1.578E+04 | 747.4 | - | - | 0 | - |
| - | - | 1571 | 757.4 | - | - | 0 | - |
| - | - | 1455 | 758.5 | - | - | 0 | - |
| 2 | z | 7.861E+04 | 761.4 | 0.0008246 | 1.083 | +1 | 7 |
| - | - | 3.27E+04 | 762.4 | - | - | 0 | - |
| - | - | 1.074E+04 | 763.4 | - | - | 0 | - |
| - | - | 2894 | 764.4 | - | - | 0 | - |
| 2 | y | 3057 | 777.5 | 0.001082 | 1.392 | +1 | 7 |
| - | - | 1760 | 778.5 | - | - | 0 | - |
| - | - | 1299 | 782.5 | - | - | 0 | - |
| - | - | 6285 | 788.4 | - | - | 0 | - |
| - | - | 2212 | 789.4 | - | - | 0 | - |
| - | - | 4625 | 800.5 | - | - | 0 | - |
| - | - | 2411 | 801.5 | - | - | 0 | - |
| - | - | 9.413E+04 | 806.4 | - | - | 0 | - |
| - | - | 3.521E+04 | 807.4 | - | - | 0 | - |
| - | - | 1.112E+04 | 808.4 | - | - | 0 | - |
| - | - | 2162 | 818.5 | - | - | 0 | - |
| - | - | 5287 | 819.4 | - | - | 0 | - |
| - | - | 5407 | 820.4 | - | - | 0 | - |
| - | - | 2013 | 821.4 | - | - | 0 | - |
| - | - | 2070 | 826.5 | - | - | 0 | - |
| - | - | 1479 | 832.5 | - | - | 0 | - |
| - | - | 1.077E+05 | 844.5 | - | - | 0 | - |
| - | - | 5.404E+04 | 845.5 | - | - | 0 | - |
| - | - | 1643 | 845.6 | - | - | 0 | - |
| - | - | 1.68E+04 | 846.5 | - | - | 0 | - |
| - | - | 1.303E+04 | 847.5 | - | - | 0 | - |
| - | - | 7641 | 848.5 | - | - | 0 | - |
| - | - | 2060 | 849.5 | - | - | 0 | - |
| - | - | 4302 | 861.5 | - | - | 0 | - |
| - | - | 1.951E+05 | 862.5 | - | - | 0 | - |
| - | - | 8.783E+04 | 863.5 | - | - | 0 | - |
| - | - | 2.496E+04 | 864.5 | - | - | 0 | - |
| - | - | 1506 | 865.5 | - | - | 0 | - |
| - | - | 3.691E+05 | 878.5 | - | - | 0 | - |
| - | - | 1.849E+05 | 879.5 | - | - | 0 | - |
| - | - | 5.818E+04 | 880.5 | - | - | 0 | - |
| - | - | 6824 | 881.5 | - | - | 0 | - |
| - | - | 2317 | 1205 | - | - | 0 | - |
| - | - | 4172 | 1299 | - | - | 0 | - |
| - | - | 1932 | 1300 | - | - | 0 | - |
| - | - | 1400 | 1301 | - | - | 0 | - |
| - | - | 4226 | 1306 | - | - | 0 | - |
| - | - | 2109 | 1307 | - | - | 0 | - |
| - | - | 1350 | 1322 | - | - | 0 | - |
| - | - | 8410 | 1323 | - | - | 0 | - |
| - | - | 2921 | 1324 | - | - | 0 | - |
| - | - | 1390 | 3082 | - | - | 0 | - |

m/z Charge Intensity FragmentType MassShift Position
120.04493713378906 0 2218.9646
126.37467956542969 0 552.455
132.12074279785156 0 525.01605
135.67539978027344 0 475.27148
136.0760498046875 0 4795.3584
137.07945251464844 0 633.36365
142.12294006347656 0 1561.0566
143.11834716796875 0 6725.1636
146.06045532226562 0 28492.83
147.0638427734375 0 2082.2478
150.05517578125 0 1390.6383
157.0976104736328 0 1785.4045
159.1131134033203 0 3267.511
163.08706665039062 0 12664.565
164.0711669921875 0 1326.3362
169.09750366210938 0 1355.5984
169.13417053222656 0 689.2094
171.11318969726562 0 10585.945
171.1493377685547 0 1338.9648
173.45059204101562 0 4649.6274
174.05540466308594 0 20501.271
174.12408447265625 0 4826.939
175.05877685546875 0 1425.3407
176.06097412109375 0 721.82935
183.11306762695312 0 1053.161
185.05609130859375 0 10724.433
185.1651611328125 0 4569.2285
186.05953979492188 0 1032.1031
187.10812377929688 0 13436.273
187.1446075439453 0 296224.22
188.14791870117188 0 24714.992
189.15049743652344 0 1398.5995
191.0586395263672 0 843.4994
191.08193969726562 0 21340.33
192.06593322753906 0 76422.27
193.0693817138672 0 6821.209
197.1287078857422 0 2706.8005
199.14439392089844 0 1033.6737
201.12362670898438 0 729.7444
203.06663513183594 0 41102.438
204.07025146484375 0 3159.2258
204.1346435546875 0 2564.115
208.08494567871094 0 692.12134
209.09254455566406 0 146403.62
210.0958709716797 0 15634.63
213.16012573242188 0 8041.6597
214.16375732421875 0 1071.846
215.13949584960938 0 203437.14
216.14283752441406 0 18435.67
217.14488220214844 0 1020.259
225.1602783203125 0 9920.692
232.16604614257812 0 836.72144 c 1
234.1000518798828 0 782.8667
238.85369873046875 0 791.591
240.135009765625 0 1415.1
242.18690490722656 0 7461.7847
243.1917266845703 0 857.852
254.15025329589844 0 3883.6443
254.16307067871094 0 1832.2577
255.1707305908203 0 9887.497
255.2071533203125 0 867.766
256.1661682128906 0 1771.0812
258.1451110839844 0 3357.4219
259.0944519042969 0 880.4822
260.1033630371094 0 6826.292
261.0874328613281 0 3773.077
268.1663513183594 0 1130.6445
269.1724853515625 0 1229.5216
269.1866149902344 0 6372.583
270.181640625 0 96962.87
271.18499755859375 0 13631.406
272.16119384765625 0 5540.0894
278.09002685546875 0 5273.339
278.1140441894531 0 62292.957
278.132080078125 0 1052.7787
279.0948791503906 0 1544.6062 y Ammonia loss 6
279.1172790527344 0 8394.578
283.2021789550781 0 4451.5615
284.1249084472656 0 5335.705
284.1600646972656 0 1554.6772
285.1277770996094 0 1154.6572
285.2065734863281 0 925.92566
286.2128601074219 0 4803.1304
287.2082824707031 0 9501.649
288.2121887207031 0 1814.8108
292.1944885253906 0 2268.2815
292.69622802734375 0 1182.6952
295.11669921875 0 1023.3453
296.1247863769531 0 154751.34 y 6
296.19757080078125 0 5707.677
297.12799072265625 0 20357.586
298.12957763671875 0 2157.8677
298.1766357421875 0 2293.839
300.19195556640625 0 3399.7498
302.13519287109375 0 5063.7075
312.1922302246094 0 3362.5654
314.2081604003906 0 41384.957
315.2113037109375 0 7316.794
317.2179870605469 0 795.8041
324.1922607421875 0 847.8876
324.229248046875 0 1797.196
325.1881103515625 0 1607.4706
326.20867919921875 0 2239.4744
326.24530029296875 0 880.16614
326.7059020996094 0 4197.0825
327.2061767578125 0 2708.0828
327.2508239746094 0 1467.2041
329.2190856933594 0 3185.1157
330.22454833984375 0 1459.5753
331.23443603515625 0 3874.0422 c 2
332.6872253417969 0 2639.9402 y 2
333.190185546875 0 1619.2064
335.71087646484375 0 3227.3828
336.21258544921875 0 1329.6467
338.24462890625 0 3488.819
339.2032775878906 0 20396.957
340.2080078125 0 2944.554
340.2598876953125 0 1770.4557
341.2554626464844 0 39232.02
342.203125 0 6558.8213
342.258056640625 0 6287.5767
343.1988525390625 0 4371.951
343.2347412109375 0 5028.543
343.2554626464844 0 763.9864
345.2502746582031 0 3511.6326
353.2191162109375 0 25705.535
354.20135498046875 0 1095.942
354.2221374511719 0 2056.766
354.2394104003906 0 12574.526
355.1989440917969 0 1253.473
355.24237060546875 0 3025.8801
355.2708435058594 0 4548.8193
356.21966552734375 0 2782.9246
357.2138366699219 0 34187.996
357.251220703125 0 1359.1304
358.216796875 0 5256.0137
365.259521484375 0 905.3771
367.2353515625 0 6327.0957
368.2540588378906 0 9222.369
369.25030517578125 0 57785.01
370.2535400390625 0 12131.93
371.2294006347656 0 76676.25
371.2652282714844 0 8310.07
372.23248291015625 0 13923.858
372.2611389160156 0 1202.1228
372.2686767578125 0 1157.6055
373.2403869628906 0 1606.6073
374.2178649902344 0 1020.6932
374.2409973144531 0 1514.426
377.158203125 0 1308.961
377.1825866699219 0 12387.085
378.18572998046875 0 2561.4941
379.1746520996094 0 1039.4376 z 5
382.2711181640625 0 4214.0522
383.2296447753906 0 2263.92
383.26641845703125 0 9705.912
384.26995849609375 0 2017.0674
385.2450866699219 0 17371.719
386.247802734375 0 3018.1514
386.27679443359375 0 4314.405
387.2226257324219 0 1613.642
387.2487487792969 0 9945.094
388.2560729980469 0 68849.34 c 3
389.2588195800781 0 12201.64
390.2608947753906 0 1601.3539
393.2487487792969 0 1194.059
394.18524169921875 0 1414.0659
395.1931457519531 0 59025.746 y 5
396.19622802734375 0 10041.025
397.1990966796875 0 799.67535
401.2396545410156 0 2782.1887
403.2569580078125 0 1335.5979
406.19842529296875 0 1136.0765
407.26666259765625 0 1168.387
410.2154846191406 0 1100.8181
410.25164794921875 0 1052.8529
410.277099609375 0 1592.9454
411.26080322265625 0 3183.3538
412.26287841796875 0 1157.8103
412.2920837402344 0 1177.5695
413.24102783203125 0 1663.5667
413.2761535644531 0 1977.521
416.2274169921875 0 2025.3304
417.2336730957031 0 1803.596
421.2841491699219 0 1978.6534
421.8334655761719 0 2250.5864
422.2419738769531 0 2302.0881
423.2608947753906 0 9631.497
424.26141357421875 0 1638.4246
424.2940368652344 0 971.1866
425.27667236328125 0 4296.701
426.2742919921875 0 1360.1506
428.2874755859375 0 10325.005
429.29229736328125 0 2170.982
431.23846435546875 0 8183.9395
432.24566650390625 0 6974.8374
433.24578857421875 0 1039.5941
437.31353759765625 0 3629.2183
438.200927734375 0 1096.1124
438.2723693847656 0 15533.013
438.3090515136719 0 8200.33
439.21282958984375 0 8546.96
439.28961181640625 0 3372.1553
439.3271484375 0 3003.4502
439.75445556640625 0 1289.5452
439.8443603515625 0 3390.6985
440.21539306640625 0 2153.9424
440.2530517578125 0 6742.0146
440.29534912109375 0 3172.3757
440.33544921875 0 2878.712
441.1903991699219 0 2678.815
441.22479248046875 0 1600.373
441.2712097167969 0 2169.162
441.30670166015625 0 6067.5312
441.34027099609375 0 1523.8485
442.3057556152344 0 3564.487
444.5133972167969 0 937.12714
449.7153015136719 0 2615.5076
450.21905517578125 0 1281.409
452.2878723144531 0 12170.865
453.29052734375 0 2476.1606
454.23004150390625 0 1455.8081
454.2660217285156 0 1465.5028
454.3386535644531 0 1938.7035
456.2825622558594 0 23400.861
456.3190002441406 0 69350.06
457.2856750488281 0 5179.4507
457.3230285644531 0 19834.81
458.2627868652344 0 3114.7864
458.330322265625 0 4035.1694
460.2739562988281 0 1473.7892
465.2968444824219 0 1620.101
466.30596923828125 0 59326.754
467.28546142578125 0 4043.0876
467.3189392089844 0 14890.751
468.2947692871094 0 12603.679
468.32489013671875 0 2330.123
469.3026123046875 0 24671.617
470.2995300292969 0 18743.584
471.30303955078125 0 3114.261
473.3092346191406 0 2681.4062
481.33984375 0 1050.7163
482.3348693847656 0 6332.4873
483.3060302734375 0 10296.544
484.31378173828125 0 202959.78
485.3169860839844 0 47362.965
486.3201599121094 0 8110.145
487.2796630859375 0 3740.6135
490.2703552246094 0 1148.6418
492.25885009765625 0 22012.188 z 4
493.2266845703125 0 1004.0469
493.2644958496094 0 12372.763
494.2658386230469 0 2245.4685
500.3322448730469 0 10237.984
501.3402404785156 0 38131.395 c 4
502.3434143066406 0 9375.705
505.2907409667969 0 5246.075
506.2955627441406 0 975.9189
507.269775390625 0 35508.39
508.2749938964844 0 15451.659 y 4
508.3110046386719 0 847.67255
509.27978515625 0 5054.982
510.2916259765625 0 5258.3345
511.3004455566406 0 7997.3813
512.3032836914062 0 2971.0027
513.2728881835938 0 1088.7405
513.311767578125 0 1235.7479
515.2947998046875 0 2380.38
517.2960205078125 0 1802.0747
520.35205078125 0 1972.3115
521.3069458007812 0 1820.5543
521.353271484375 0 1738.1819
522.2465209960938 0 1408.6229
526.323486328125 0 3134.7512
527.3284912109375 0 1029.2295
529.2854614257812 0 860.75525
529.3367919921875 0 1777.7972
530.3067626953125 0 13499.361
531.2682495117188 0 2844.908
531.3109741210938 0 4917.4062
532.3110961914062 0 2733.2668
535.2606811523438 0 2353.937
535.3320922851562 0 7805.393
536.340576171875 0 2183.086
537.38037109375 0 1976.919
538.3584594726562 0 7840.0806
539.31884765625 0 3795.126
539.3670043945312 0 10964.743
540.3738403320312 0 6974.22
541.3823852539062 0 6910.869
542.3911743164062 0 1836.0322
543.349365234375 0 2711.8225
544.3544921875 0 1165.9159
546.2803344726562 0 9346.484
547.2874145507812 0 10086.329
548.2926635742188 0 1773.024
549.2800903320312 0 19862.914 z 3
550.287109375 0 22570.092
550.3582153320312 0 1075.2019
551.2920532226562 0 5538.641
551.3536987304688 0 1392.1808
553.3395385742188 0 11997.729
554.3504028320312 0 6083.231
555.3871459960938 0 19273.328
556.3943481445312 0 48108.81
557.3294067382812 0 3127.4587
557.3980102539062 0 16966.965
558.4013671875 0 2766.4773
564.291015625 0 45996.195
565.2980346679688 0 79031.32 y 3
565.3709716796875 0 28773.463
566.3009643554688 0 22030.102
566.36083984375 0 11962.435
567.30712890625 0 2763.4304
567.3633422851562 0 55248.547
568.36962890625 0 54570.184
568.4219970703125 0 1544.7881
569.3727416992188 0 14557.161
570.3762817382812 0 2248.5034
571.3067016601562 0 867.7505
571.345458984375 0 6332.455
571.3912963867188 0 1106.857
572.3475341796875 0 1325.1896
578.7872924804688 0 2937.0867
579.2866821289062 0 1240.9304
582.3751220703125 0 4750.011
583.3820190429688 0 145564.39
584.3851928710938 0 45452.242
585.3953247070312 0 22467.867
586.3480834960938 0 2926.6367
586.4009399414062 0 6361.6665
587.39892578125 0 1524.236
592.2841186523438 0 2602.2603
596.3225708007812 0 1638.5358
598.3316040039062 0 1762.3997
599.33935546875 0 1418.0364
600.4088745117188 0 175589.69 c 5
601.4118041992188 0 54688.05
602.4143676757812 0 11696.154
607.3756713867188 0 1283.9061
624.4068603515625 0 2649.2693
625.3961181640625 0 1809.113
628.3400268554688 0 1665.6482
630.3363647460938 0 2606.3967
633.3294067382812 0 1275.2522 w 2
634.39306640625 0 5291.108
635.3502197265625 0 1903.6068
635.3984985351562 0 1423.455
636.3858032226562 0 9401.565
637.3914184570312 0 7526.6714
638.3975219726562 0 4730.16
639.4027099609375 0 1458.0216
642.419189453125 0 10473.012
643.4210815429688 0 2577.806
646.3565063476562 0 10281.68
647.3566284179688 0 2980.7568
648.348876953125 0 28825.172 z 2
649.3521118164062 0 8820.211
650.3523559570312 0 1673.6434
651.401123046875 0 1566.9977
652.4000854492188 0 8376.667
653.3782348632812 0 4333.739
653.4166870117188 0 897.6905
654.3956909179688 0 14066.3955
655.4021606445312 0 12789.721
656.40625 0 4683.825
660.8107299804688 0 2150.5796
661.3107299804688 0 1309.2615
663.3527221679688 0 1903.5514
664.3674926757812 0 81845.375 y 2
665.3706665039062 0 28794.639
666.3741455078125 0 4950.8896
669.4083251953125 0 1333.3368
670.4142456054688 0 13758.883
671.3493041992188 0 11040.894
671.4186401367188 0 5378.318
672.3547973632812 0 2865.2659
672.427734375 0 3234.8198
673.3488159179688 0 1251.9208
679.3778686523438 0 3010.133
682.4000244140625 0 3969.739
687.3603515625 0 2399.228
687.4376220703125 0 3459.6638
688.3751220703125 0 16604.734
688.437744140625 0 956.19214
689.3782958984375 0 4759.0884
691.3567504882812 0 2676.342
692.3497924804688 0 1424.2119
699.4317626953125 0 5483.118
700.43212890625 0 2006.9149
701.432861328125 0 3456.2861
702.3837280273438 0 1656.9757
705.3690795898438 0 4398.518
706.3671875 0 1690.5018
708.4174194335938 0 5290.9785
709.4031982421875 0 5028.144
710.4076538085938 0 1457.152
714.4186401367188 0 1412.6904
717.4487915039062 0 2047.2747
718.3778686523438 0 116036.85 w 1
719.381103515625 0 42618.76
720.3841552734375 0 9385.842
726.427490234375 0 37866.125
727.4210205078125 0 22702.086
728.4215698242188 0 7724.413
729.4205932617188 0 2847.6768
743.4224853515625 0 22969.959
744.4391479492188 0 2527779.2
745.4419555664062 0 920621.1
746.44384765625 0 223962.23
747.4470825195312 0 15776.488
757.4401245117188 0 1570.5378
758.455810546875 0 1454.7296
761.4326171875 0 78609.586 z 1
762.4356689453125 0 32703.512
763.4400024414062 0 10740.55
764.4456787109375 0 2894.1294
777.4515991210938 0 3057.47 y 1
778.4515380859375 0 1760.4835
782.4713745117188 0 1299.2628
788.4071655273438 0 6284.652
789.41162109375 0 2212.2156
800.48046875 0 4624.6616
801.4842529296875 0 2410.5513
806.4178466796875 0 94133.875
807.4203491210938 0 35205.254
808.4237060546875 0 11120.322
818.4905395507812 0 2161.509
819.42333984375 0 5286.9175
820.4298095703125 0 5406.9146
821.4328002929688 0 2012.5511
826.4613647460938 0 2070.3416
832.4688720703125 0 1478.7717
844.4696044921875 0 107666.47
845.4734497070312 0 54038.746
845.5672607421875 0 1642.8143
846.4761352539062 0 16799.96
847.45947265625 0 13029.714
848.460205078125 0 7640.884
849.465087890625 0 2060.082
861.4923095703125 0 4301.9985
862.4805297851562 0 195073.97
863.4833374023438 0 87833.13
864.4862060546875 0 24956.158
865.4928588867188 0 1505.8022
878.4989013671875 0 369125.66
879.502685546875 0 184885.8
880.5052490234375 0 58181.867
881.510009765625 0 6824.236
1204.556884765625 0 2316.6357
1298.78515625 0 4171.87
1299.787841796875 0 1932.4581
1300.7763671875 0 1399.5591
1305.606201171875 0 4226.4185
1306.6080322265625 0 2109.202
1321.635009765625 0 1350.4788
1322.62646484375 0 8410.288
1323.634521484375 0 2920.9746
3081.758056640625 0 1390.0924

Spectrum Details

|  |  |
| --- | --- |
| Matched peaks? Matched peaks The total absolute number of peaks matched. Additionally in brackets the total fraction of peaks matched and the total number of peaks is shown. | 20 (4.36% of 459) |
| FDR? FDR The false discovery rate estimated for this peptide. It is calculated by matching all theoretical fragments with a non-integer shift with the raw peaks for this spectrum. This is done with 40 different shifts. The resulting percentage is the average number of annotated peaks over the number of annotated peaks with the correct spectrum. | 8.81% |
| Satellite FDR? Satellite FDR See the FDR for details on its calculation. This satellite ion specific FDR only contains the satellite ions (d/w) for I/L/J positions. | 11.90% |
| PSM Score? PSM Score The PSM Score as given by Hecklib to this annotated spectrum. It is shown with three significant figures. | 234 |

#### Reverse Lookup? Reverse Lookup All places where this read could be placed.

| Group | Segment | Template | Template Part | Read Part | Score | Unique |
| --- | --- | --- | --- | --- | --- | --- |
| Homo sapiens Light Chain | IGLC | IGLC7 | [24..32] | [0..8] | 37 | True |

| Recombined | Template Part | Read Part | Score | Unique |
| --- | --- | --- | --- | --- |
| REC-0-1\_002 | [26..34] | [0..8] | 32 | True |

#### Meta Information from Multiple reads

##### Number of combined reads

2

##### Intensity

0.7078

##### TotalArea

9.415E+07

##### Changes to the peptide sequence

TLVGJVNY

L→JNo support for either Leucine or Isoleucine based on side chain ions (Position: 5)

#### Positional Score

Copy Data

##### Positional Score (TSV)

###### Preview

```
Loading example...
```

*Click on the button to copy the data to your clipboard.*

0001234567

Label Value
"0" 0
"1" 0
"2" 0
"3" 0
"4" 0
"5" 0
"6" 0
"7" 0

#### Meta Information from PEAKS

##### Scan Identifier

F3:10732

##### Original sequence

T

L

V

G

L

V

N

Y

##### Posttranslational Modifications

##### Source File

D:\separate\_stitch\_analyses\xle-disambiguation\raw\20210323\_F1\_UM1\_Peng0013\_SA\_F59\_ingel\_3ug\_chymo.raw

##### Fraction

3

##### Scan Feature

F3:1947

##### De Novo Score

99

##### ConfidenceScore

99

### m/z

439.7533

##### Mass

877.4909

##### Charge

2

##### Retention Time

60.78

##### Predicted Retention Time

-

##### Area

4.707E+07

##### Parts Per Million

1.3

##### Fragmentation mode

ETHCD

##### Originating file

01 D:\separate\_stitch\_analyses\xle-disambiguation\20210325\_F59\_3ug\_DENOVO\_12.csv

#### Meta Information from PEAKS

##### Scan Identifier

F3:10783

##### Original sequence

T

L

V

G

L

V

N

Y

##### Posttranslational Modifications

##### Source File

D:\separate\_stitch\_analyses\xle-disambiguation\raw\20210323\_F1\_UM1\_Peng0013\_SA\_F59\_ingel\_3ug\_chymo.raw

##### Fraction

3

##### Scan Feature

F3:1947

##### De Novo Score

99

##### ConfidenceScore

99

### m/z

439.7533

##### Mass

877.4909

##### Charge

2

##### Retention Time

60.78

##### Predicted Retention Time

-

##### Area

4.707E+07

##### Parts Per Million

1.3

##### Fragmentation mode

ETHCD

##### Originating file

01 D:\separate\_stitch\_analyses\xle-disambiguation\20210325\_F59\_3ug\_DENOVO\_12.csv
