## Supplementary material for "A handle on mass coincidence errors in *de novo* sequencing of antibodies by bottom-up proteomics": Combined_022.html

Details Combined\_022 | Stitch OverviewUndefined

### Read Combined\_022

#### Sequence (length=9)

TJJEGEESR

#### Spectrum 3697? Spectrum 3697 The raw spectrum of this peptide as annotated by Hecklib. The fragments are coloured according to ion type (see legend). Any peaks with a star '\*' as text can be hovered over to see the full details, first the ion type second the mass shift type. By hovering over the amino acids in the peptide or ions in the legend the corresponding peaks are highlighted. By toggling the 'Unassigned' label you can turn the background (unassigned) peaks on or off in the plot. By updating the slider in the Ion legend you can update the spectrum to only show the top X% of the peaks with labels. The top X% means any peak that is within X% of the highest intensity. By dragging in the spectrum you can zoom in to a specific part of the spectrum and use 'Zoom Out' to get back to the original zoom level. The annotation of the spectrum is based on the given sequence in the peptides file and is done with different software so inconsistencies are likely. The peaks are annotated based on the given sequence, with 20 ppm tolerance.

Copy Data

##### Spectrum 3697 (TSV)

###### Preview

```
Loading example...
```

*Click on the button to copy the data to your clipboard.*

Mz MinMz MaxIntensity Max

WidthHeightPeptide font sizePeptide stroke widthSpectrum font sizeSpectrum stroke widthCompact peptide

Ion legend

wxyz

abcd

OtherUnassignedIon^Charge^_Position_Show for top:%

TJJEGEESR

07.94e+41.59e+52.38e+53.18e+5

Zoom Out

y^+1^_1_a^+1^_2_d^+1^_2_y^+1^_1_a^+1^_2_b^+1^_2_b^+1^_2_b^+2^_4_y^+1^_2_y^+1^_2_y^+1^_2_b^+1^_3_b^+1^_3_y^+2^_6_y^+2^_6_y^+1^_3_y^+1^_3_y^+1^_3_y^+2^_7_y^+2^_7_b^+1^_4_b^+1^_4_y^+2^_8_y^+1^_4_\*\*\*y^+1^_4_y^+1^_5_y^+1^_5_y^+1^_5_b^+1^_6_y^+1^_6_y^+1^_6_y^+1^_6_y^+1^_7_y^+1^_7_y^+1^_7_y^+1^_8_

0778155623343112

Fragment Matches Table

Show background peaks

| Position | Ion type | Intensity | mz Theoretical | mz Error (Th) | mz Error (ppm) | Charge | Series Number |
| --- | --- | --- | --- | --- | --- | --- | --- |
| - | - | 590.4 | 120.1 | - | - | 0 | - |
| - | - | 338.9 | 120.9 | - | - | 0 | - |
| - | - | 422.5 | 124 | - | - | 0 | - |
| - | - | 565.1 | 124.1 | - | - | 0 | - |
| - | - | 396.7 | 125.6 | - | - | 0 | - |
| - | - | 1368 | 126.1 | - | - | 0 | - |
| - | - | 1483 | 126.1 | - | - | 0 | - |
| - | - | 399.7 | 127.1 | - | - | 0 | - |
| - | - | 627.9 | 129.1 | - | - | 0 | - |
| - | - | 362.9 | 129.1 | - | - | 0 | - |
| - | - | 2819 | 130.1 | - | - | 0 | - |
| - | - | 4330 | 130.1 | - | - | 0 | - |
| - | - | 804.9 | 130.1 | - | - | 0 | - |
| - | - | 426.7 | 134.1 | - | - | 0 | - |
| - | - | 400.2 | 135.1 | - | - | 0 | - |
| - | - | 867.3 | 136.1 | - | - | 0 | - |
| - | - | 430.3 | 141 | - | - | 0 | - |
| - | - | 3828 | 141.1 | - | - | 0 | - |
| - | - | 482.7 | 141.1 | - | - | 0 | - |
| - | - | 9065 | 142.1 | - | - | 0 | - |
| - | - | 1.179E+04 | 143.1 | - | - | 0 | - |
| - | - | 677.8 | 143.1 | - | - | 0 | - |
| - | - | 427.1 | 143.4 | - | - | 0 | - |
| - | - | 986.6 | 144.1 | - | - | 0 | - |
| - | - | 751.4 | 148.1 | - | - | 0 | - |
| - | - | 578.2 | 148.8 | - | - | 0 | - |
| - | - | 452.1 | 148.9 | - | - | 0 | - |
| - | - | 611.6 | 148.9 | - | - | 0 | - |
| - | - | 777.3 | 148.9 | - | - | 0 | - |
| - | - | 591.6 | 148.9 | - | - | 0 | - |
| - | - | 926 | 148.9 | - | - | 0 | - |
| - | - | 1084 | 148.9 | - | - | 0 | - |
| - | - | 1826 | 148.9 | - | - | 0 | - |
| - | - | 3733 | 148.9 | - | - | 0 | - |
| - | - | 5076 | 149 | - | - | 0 | - |
| - | - | 3299 | 149 | - | - | 0 | - |
| - | - | 1625 | 149 | - | - | 0 | - |
| - | - | 1086 | 149 | - | - | 0 | - |
| - | - | 841.7 | 149 | - | - | 0 | - |
| - | - | 807.8 | 149 | - | - | 0 | - |
| - | - | 639 | 149 | - | - | 0 | - |
| - | - | 547.7 | 149 | - | - | 0 | - |
| - | - | 450.6 | 149 | - | - | 0 | - |
| - | - | 470.9 | 149 | - | - | 0 | - |
| - | - | 589.2 | 149 | - | - | 0 | - |
| - | - | 1193 | 152.1 | - | - | 0 | - |
| - | - | 431.3 | 153.1 | - | - | 0 | - |
| - | - | 455.3 | 154 | - | - | 0 | - |
| - | - | 963.4 | 157.1 | - | - | 0 | - |
| 9 | y | 1.474E+04 | 158.1 | 0.0001873 | 1.185 | +1 | 1 |
| - | - | 5837 | 159.1 | - | - | 0 | - |
| - | - | 978.6 | 159.1 | - | - | 0 | - |
| - | - | 493.4 | 163 | - | - | 0 | - |
| - | - | 434.5 | 165.9 | - | - | 0 | - |
| - | - | 5258 | 169.1 | - | - | 0 | - |
| 2 | a | 3650 | 169.1 | 0.0001579 | 0.9336 | +1 | 2 |
| - | - | 928.9 | 170.1 | - | - | 0 | - |
| - | - | 2116 | 171.1 | - | - | 0 | - |
| - | - | 1116 | 171.1 | - | - | 0 | - |
| 2 | d | 623.7 | 173.1 | 0.0004825 | 2.787 | +1 | 2 |
| - | - | 861.4 | 173.4 | - | - | 0 | - |
| - | - | 466.8 | 174 | - | - | 0 | - |
| 9 | y | 2.066E+04 | 175.1 | 0.000219 | 1.25 | +1 | 1 |
| - | - | 1230 | 176.1 | - | - | 0 | - |
| - | - | 643.7 | 181.1 | - | - | 0 | - |
| - | - | 721.1 | 184.1 | - | - | 0 | - |
| - | - | 639.3 | 185.1 | - | - | 0 | - |
| - | - | 470.9 | 185.1 | - | - | 0 | - |
| - | - | 343.5 | 187.1 | - | - | 0 | - |
| - | - | 1.435E+04 | 187.1 | - | - | 0 | - |
| 2 | a | 3.145E+05 | 187.1 | 0.0003354 | 1.792 | +1 | 2 |
| - | - | 549.9 | 188.1 | - | - | 0 | - |
| - | - | 3.066E+04 | 188.1 | - | - | 0 | - |
| - | - | 1221 | 189.1 | - | - | 0 | - |
| - | - | 1403 | 189.2 | - | - | 0 | - |
| - | - | 472.7 | 193.1 | - | - | 0 | - |
| - | - | 1443 | 195.1 | - | - | 0 | - |
| 2 | b | 2790 | 197.1 | 0.0002384 | 1.209 | +1 | 2 |
| - | - | 690.8 | 198.1 | - | - | 0 | - |
| - | - | 1661 | 199.1 | - | - | 0 | - |
| - | - | 754.3 | 199.1 | - | - | 0 | - |
| - | - | 8945 | 199.2 | - | - | 0 | - |
| - | - | 634.2 | 200.2 | - | - | 0 | - |
| - | - | 2502 | 202.1 | - | - | 0 | - |
| - | - | 1430 | 203.1 | - | - | 0 | - |
| - | - | 553.2 | 209.2 | - | - | 0 | - |
| - | - | 1026 | 210.1 | - | - | 0 | - |
| - | - | 2252 | 213.1 | - | - | 0 | - |
| 2 | b | 1.591E+05 | 215.1 | 0.0002328 | 1.082 | +1 | 2 |
| - | - | 1.882E+04 | 216.1 | - | - | 0 | - |
| - | - | 1704 | 217.1 | - | - | 0 | - |
| - | - | 777.8 | 217.1 | - | - | 0 | - |
| 4 | b | 914.8 | 220.1 | 0.002349 | 10.67 | +2 | 4 |
| - | - | 773.9 | 221.1 | - | - | 0 | - |
| - | - | 735.6 | 225 | - | - | 0 | - |
| - | - | 2222 | 227.1 | - | - | 0 | - |
| - | - | 4081 | 227.2 | - | - | 0 | - |
| - | - | 805 | 228.1 | - | - | 0 | - |
| - | - | 774.1 | 228.2 | - | - | 0 | - |
| - | - | 1975 | 231.1 | - | - | 0 | - |
| - | - | 1130 | 239.1 | - | - | 0 | - |
| - | - | 4093 | 241.1 | - | - | 0 | - |
| - | - | 1015 | 242.1 | - | - | 0 | - |
| - | - | 2.294E+04 | 243.1 | - | - | 0 | - |
| 8 | y | 4433 | 244.1 | 0.002248 | 9.206 | +1 | 2 |
| 8 | y | 1.327E+04 | 245.1 | 0.0002023 | 0.8253 | +1 | 2 |
| - | - | 1082 | 245.1 | - | - | 0 | - |
| - | - | 594.7 | 246.1 | - | - | 0 | - |
| - | - | 1077 | 246.1 | - | - | 0 | - |
| - | - | 681.3 | 252.1 | - | - | 0 | - |
| - | - | 440 | 252.9 | - | - | 0 | - |
| - | - | 629.2 | 253.2 | - | - | 0 | - |
| - | - | 611.8 | 256.1 | - | - | 0 | - |
| - | - | 2836 | 259.1 | - | - | 0 | - |
| - | - | 595 | 259.4 | - | - | 0 | - |
| 8 | y | 4.235E+04 | 262.2 | 5.092E-05 | 0.1942 | +1 | 2 |
| - | - | 4877 | 263.2 | - | - | 0 | - |
| - | - | 685.2 | 266.2 | - | - | 0 | - |
| - | - | 3031 | 270.1 | - | - | 0 | - |
| - | - | 619.8 | 272.2 | - | - | 0 | - |
| - | - | 1034 | 282.1 | - | - | 0 | - |
| - | - | 3772 | 283.2 | - | - | 0 | - |
| - | - | 721.1 | 284.2 | - | - | 0 | - |
| - | - | 1120 | 288.1 | - | - | 0 | - |
| - | - | 1187 | 295.1 | - | - | 0 | - |
| - | - | 6636 | 298.1 | - | - | 0 | - |
| - | - | 900.2 | 299.1 | - | - | 0 | - |
| - | - | 4820 | 300.2 | - | - | 0 | - |
| - | - | 910.7 | 310.1 | - | - | 0 | - |
| 3 | b | 1696 | 310.2 | 0.000464 | 1.496 | +1 | 3 |
| - | - | 1552 | 313.2 | - | - | 0 | - |
| - | - | 6439 | 316.1 | - | - | 0 | - |
| - | - | 673.8 | 317.1 | - | - | 0 | - |
| - | - | 1170 | 326.2 | - | - | 0 | - |
| - | - | 586.3 | 327 | - | - | 0 | - |
| - | - | 619 | 327.2 | - | - | 0 | - |
| - | - | 1058 | 328.1 | - | - | 0 | - |
| 3 | b | 2.005E+04 | 328.2 | 0.0001532 | 0.4667 | +1 | 3 |
| - | - | 2813 | 329.2 | - | - | 0 | - |
| - | - | 662.6 | 331.2 | - | - | 0 | - |
| - | - | 1026 | 338.1 | - | - | 0 | - |
| - | - | 608.7 | 339.1 | - | - | 0 | - |
| - | - | 1207 | 344.2 | - | - | 0 | - |
| 4 | y | 638 | 344.6 | 3.018E-05 | 0.08756 | +2 | 6 |
| - | - | 688.6 | 346.1 | - | - | 0 | - |
| 4 | y | 584.1 | 353.7 | 0.0003992 | 1.129 | +2 | 6 |
| - | - | 617 | 355.4 | - | - | 0 | - |
| - | - | 5243 | 356.2 | - | - | 0 | - |
| - | - | 905.7 | 356.2 | - | - | 0 | - |
| - | - | 682.9 | 359 | - | - | 0 | - |
| - | - | 603.2 | 366.2 | - | - | 0 | - |
| - | - | 569.5 | 368.1 | - | - | 0 | - |
| 7 | y | 2708 | 373.2 | 0.0009716 | 2.604 | +1 | 3 |
| 7 | y | 3179 | 374.2 | 0.0003376 | 0.9022 | +1 | 3 |
| - | - | 821.4 | 375.2 | - | - | 0 | - |
| - | - | 927.4 | 381.1 | - | - | 0 | - |
| - | - | 776.1 | 385.1 | - | - | 0 | - |
| 7 | y | 1.768E+04 | 391.2 | 9.088E-05 | 0.2323 | +1 | 3 |
| - | - | 2631 | 392.2 | - | - | 0 | - |
| - | - | 578.8 | 393.3 | - | - | 0 | - |
| 3 | y | 2540 | 401.2 | 0.001029 | 2.564 | +2 | 7 |
| - | - | 637.1 | 401.7 | - | - | 0 | - |
| - | - | 980 | 403.1 | - | - | 0 | - |
| - | - | 527.5 | 407 | - | - | 0 | - |
| 3 | y | 2.249E+04 | 410.2 | 1.175E-05 | 0.02864 | +2 | 7 |
| - | - | 8901 | 410.7 | - | - | 0 | - |
| - | - | 3118 | 411.2 | - | - | 0 | - |
| - | - | 617.9 | 417.2 | - | - | 0 | - |
| - | - | 1661 | 427.1 | - | - | 0 | - |
| - | - | 1738 | 429.2 | - | - | 0 | - |
| - | - | 1745 | 429.3 | - | - | 0 | - |
| - | - | 709 | 430.3 | - | - | 0 | - |
| - | - | 661.3 | 432.2 | - | - | 0 | - |
| 4 | b | 907.2 | 439.3 | 0.001267 | 2.884 | +1 | 4 |
| - | - | 905.8 | 445.2 | - | - | 0 | - |
| - | - | 542.2 | 445.9 | - | - | 0 | - |
| - | - | 538.1 | 449.3 | - | - | 0 | - |
| 4 | b | 5324 | 457.3 | 0.000112 | 0.245 | +1 | 4 |
| - | - | 2343 | 458.3 | - | - | 0 | - |
| - | - | 666.8 | 463.2 | - | - | 0 | - |
| 2 | y | 3090 | 466.7 | 0.0005398 | 1.157 | +2 | 8 |
| - | - | 1611 | 467.2 | - | - | 0 | - |
| - | - | 762.5 | 469.8 | - | - | 0 | - |
| - | - | 697.5 | 479.2 | - | - | 0 | - |
| - | - | 660.6 | 483.3 | - | - | 0 | - |
| - | - | 1088 | 485.2 | - | - | 0 | - |
| - | - | 879.8 | 496.2 | - | - | 0 | - |
| 6 | y | 1057 | 502.2 | 0.0006247 | 1.244 | +1 | 4 |
| 0 | Precursor | 4038 | 508.3 | 0.0003284 | 0.6462 | +2 | -1 |
| 0 | Precursor | 1341 | 508.7 | 0.008183 | 16.08 | +2 | -1 |
| - | - | 1277 | 509.3 | - | - | 0 | - |
| - | - | 619.2 | 514.4 | - | - | 0 | - |
| 0 | Precursor | 6928 | 517.3 | 0.0004317 | 0.8346 | +2 | -1 |
| - | - | 5640 | 517.8 | - | - | 0 | - |
| - | - | 2228 | 518.3 | - | - | 0 | - |
| 6 | y | 7899 | 520.2 | 0.0001003 | 0.1928 | +1 | 4 |
| - | - | 1933 | 521.2 | - | - | 0 | - |
| - | - | 1791 | 542.2 | - | - | 0 | - |
| 5 | y | 3455 | 559.2 | 0.001033 | 1.847 | +1 | 5 |
| 5 | y | 4006 | 560.2 | 2.291E-06 | 0.00409 | +1 | 5 |
| 5 | y | 1.198E+05 | 577.3 | 0.0004894 | 0.8478 | +1 | 5 |
| - | - | 3.317E+04 | 578.3 | - | - | 0 | - |
| - | - | 5933 | 579.3 | - | - | 0 | - |
| - | - | 711.6 | 628.3 | - | - | 0 | - |
| 6 | b | 631.5 | 643.3 | 0.006449 | 10.02 | +1 | 6 |
| 4 | y | 1.837E+04 | 688.3 | 0.0009016 | 1.31 | +1 | 6 |
| 4 | y | 6320 | 689.3 | 0.01362 | 19.76 | +1 | 6 |
| - | - | 1582 | 690.3 | - | - | 0 | - |
| 4 | y | 1.283E+05 | 706.3 | 0.0007241 | 1.025 | +1 | 6 |
| - | - | 4.308E+04 | 707.3 | - | - | 0 | - |
| - | - | 9793 | 708.3 | - | - | 0 | - |
| - | - | 1023 | 716.3 | - | - | 0 | - |
| - | - | 838 | 717.3 | - | - | 0 | - |
| - | - | 683.1 | 741.3 | - | - | 0 | - |
| - | - | 675.5 | 766.3 | - | - | 0 | - |
| - | - | 2925 | 783.4 | - | - | 0 | - |
| - | - | 2824 | 784.4 | - | - | 0 | - |
| - | - | 1311 | 785.4 | - | - | 0 | - |
| 3 | y | 3081 | 801.4 | 0.001164 | 1.453 | +1 | 7 |
| 3 | y | 3479 | 802.4 | 0.005421 | 6.756 | +1 | 7 |
| - | - | 1735 | 803.4 | - | - | 0 | - |
| 3 | y | 2.218E+05 | 819.4 | 0.001231 | 1.502 | +1 | 7 |
| - | - | 9.56E+04 | 820.4 | - | - | 0 | - |
| - | - | 2.317E+04 | 821.4 | - | - | 0 | - |
| - | - | 1725 | 822.4 | - | - | 0 | - |
| - | - | 846.6 | 829.4 | - | - | 0 | - |
| 2 | y | 4277 | 932.5 | 0.002531 | 2.715 | +1 | 8 |
| - | - | 2734 | 933.5 | - | - | 0 | - |
| - | - | 762.8 | 934.5 | - | - | 0 | - |
| - | - | 994.3 | 3081 | - | - | 0 | - |

m/z Charge Intensity FragmentType MassShift Position
120.08106231689453 0 590.4454
120.86834716796875 0 338.9357
124.0395736694336 0 422.54218
124.1124267578125 0 565.08923
125.5521469116211 0 396.67346
126.09153747558594 0 1368.3694
126.12803649902344 0 1482.9655
127.13167572021484 0 399.65762
129.10227966308594 0 627.92114
129.1140594482422 0 362.91846
130.05007934570312 0 2819.0046
130.0613555908203 0 4330.0205
130.09774780273438 0 804.8634
134.1242218017578 0 426.68924
135.11607360839844 0 400.24332
136.0761260986328 0 867.26886
141.02513122558594 0 430.25302
141.0660858154297 0 3828.0674
141.1024169921875 0 482.69357
142.12286376953125 0 9064.883
143.11813354492188 0 11790.759
143.1253204345703 0 677.77435
143.3817138671875 0 427.08987
144.12152099609375 0 986.6453
148.0603485107422 0 751.361
148.8416748046875 0 578.2064
148.8763885498047 0 452.1192
148.89797973632812 0 611.59
148.90518188476562 0 777.3055
148.91229248046875 0 591.6119
148.9195098876953 0 925.962
148.92649841308594 0 1084.062
148.93341064453125 0 1826.4845
148.94100952148438 0 3733.0625
148.95733642578125 0 5075.613
148.96490478515625 0 3298.7847
148.97227478027344 0 1624.8955
148.9795684814453 0 1086.2876
148.98648071289062 0 841.71814
148.99359130859375 0 807.7773
149.0001983642578 0 638.96204
149.00804138183594 0 547.70184
149.01473999023438 0 450.55576
149.03663635253906 0 470.92978
149.04345703125 0 589.16974
152.1072998046875 0 1193.44
153.0904541015625 0 431.28906
153.95492553710938 0 455.3315
157.10877990722656 0 963.3725
158.09259033203125 0 14737.143 y Ammonia loss 8
159.07659912109375 0 5837.4097
159.09654235839844 0 978.5675
163.0299835205078 0 493.38788
165.94671630859375 0 434.535
169.0609588623047 0 5258.1313
169.13369750976562 0 3649.7715 a Water loss 1
170.11798095703125 0 928.92096
171.07667541503906 0 2116.1604
171.11279296875 0 1115.5242
173.12893676757812 0 623.6734 d 1
173.44105529785156 0 861.3747
173.9544677734375 0 466.75327
175.11917114257812 0 20661.762 y 8
176.1224365234375 0 1229.8995
181.06036376953125 0 643.704
184.10833740234375 0 721.13306
185.09243774414062 0 639.308
185.10255432128906 0 470.9308
187.06402587890625 0 343.46545
187.07156372070312 0 14350.877
187.14443969726562 0 314514.47 a 1
188.0746612548828 0 549.8854
188.14773559570312 0 30664.15
189.0873260498047 0 1220.875
189.151123046875 0 1403.1487
193.06056213378906 0 472.6618
195.0762939453125 0 1442.9758
197.12869262695312 0 2790.1494 b Water loss 1
198.0883331298828 0 690.7592
199.0716552734375 0 1660.7826
199.1078643798828 0 754.3023
199.18060302734375 0 8944.747
200.18385314941406 0 634.2429
202.1189422607422 0 2501.8984
203.10279846191406 0 1429.7085
209.1649932861328 0 553.1683
210.0871124267578 0 1025.5958
213.08700561523438 0 2251.5066
215.13925170898438 0 159132.94 b 1
216.14251708984375 0 18821.98
217.08193969726562 0 1703.8813
217.14442443847656 0 777.7782
220.12884521484375 0 914.768 b Water loss 3
221.08409118652344 0 773.9091
225.04356384277344 0 735.5545
227.114013671875 0 2222.4744
227.1756134033203 0 4081.2344
228.09771728515625 0 805.01044
228.17967224121094 0 774.13336
231.0972900390625 0 1975.4042
239.09500122070312 0 1129.828
241.08224487304688 0 4093.1575
242.0851593017578 0 1015.0085
243.13409423828125 0 22938.035
244.13816833496094 0 4432.751 y Water loss 7
245.1246337890625 0 13265.983 y Ammonia loss 7
245.13894653320312 0 1081.6562
246.108642578125 0 594.6829
246.1280059814453 0 1077.2427
252.0977783203125 0 681.29175
252.86532592773438 0 440.00327
253.15541076660156 0 629.1649
256.0939025878906 0 611.77106
259.09222412109375 0 2836.2163
259.35516357421875 0 595.0255
262.1510314941406 0 42349.883 y 7
263.1542053222656 0 4876.7505
266.1874694824219 0 685.1504
270.1084289550781 0 3030.5222
272.1609191894531 0 619.7975
282.1078796386719 0 1034.1135
283.20172119140625 0 3772.2808
284.2051086425781 0 721.12024
288.11907958984375 0 1120.0157
295.140380859375 0 1186.7891
298.1034851074219 0 6635.6206
299.1065979003906 0 900.2071
300.1556396484375 0 4820.275
310.1033630371094 0 910.6949
310.2129821777344 0 1695.739 b Water loss 2
313.1510009765625 0 1551.9092
316.114013671875 0 6439.1235
317.1182861328125 0 673.83167
326.17169189453125 0 1169.841
326.9976806640625 0 586.315
327.1542663574219 0 618.9527
328.1150817871094 0 1058.4421
328.2232360839844 0 20046.885 b 2
329.2263488769531 0 2812.6355
331.16217041015625 0 662.58496
338.1460876464844 0 1026.0774
339.12884521484375 0 608.69635
344.1817932128906 0 1207.4629
344.6484375 0 638.01624 y Water loss 3
346.125 0 688.5791
353.6533508300781 0 584.1382 y 3
355.3569030761719 0 617.03766
356.1561584472656 0 5243.324
356.21795654296875 0 905.70337
359.0279235839844 0 682.8993
366.1650695800781 0 603.163
368.1072082519531 0 569.51587
373.1820373535156 0 2707.9614 y Water loss 6
374.16668701171875 0 3179.0823 y Ammonia loss 6
375.1723937988281 0 821.3682
381.1418151855469 0 927.3779
385.13525390625 0 776.0903
391.19366455078125 0 17678.889 y 6
392.1954040527344 0 2630.924
393.2507629394531 0 578.79504
401.1915283203125 0 2540.426 y Water loss 2
401.69207763671875 0 637.0548
403.14642333984375 0 980.0171
407.0462951660156 0 527.46405
410.1957702636719 0 22489.81 y 2
410.69732666015625 0 8900.987
411.1978759765625 0 3118.2007
417.1598815917969 0 617.947
427.1466979980469 0 1661.215
429.1986389160156 0 1737.9697
429.2697448730469 0 1744.8689
430.27349853515625 0 709.032
432.15386962890625 0 661.28955
439.2563781738281 0 907.2024 b Water loss 3
445.15576171875 0 905.81775
445.9129333496094 0 542.1848
449.3047180175781 0 538.0976
457.26556396484375 0 5324.347 b 3
458.26702880859375 0 2343.056
463.1908874511719 0 666.7647
466.7372741699219 0 3089.516 y 1
467.2395935058594 0 1610.8196
469.7837219238281 0 762.45294
479.1861267089844 0 697.52637
483.3346252441406 0 660.6172
485.1971435546875 0 1088.4885
496.1677551269531 0 879.773
502.2262268066406 0 1057.0997 y Water loss 5
508.25604248046875 0 4038.2139 Precursor Water loss
508.7565612792969 0 1341.4341 Precursor Ammonia loss
509.25836181640625 0 1277.0673
514.447265625 0 619.19476
517.2620849609375 0 6927.9365 Precursor
517.7627563476562 0 5640.4336
518.264404296875 0 2228.4792
520.2362670898438 0 7898.5474 y 5
521.2387084960938 0 1933.2253
542.2202758789062 0 1791.0974
559.2460327148438 0 3454.9448 y Water loss 4
560.2310791015625 0 4006.4805 y Ammonia loss 4
577.2571411132812 0 119793.68 y 4
578.260009765625 0 33165.484
579.2622680664062 0 5932.84
628.2672119140625 0 711.63226
643.336181640625 0 631.49695 b 5
688.2887573242188 0 18373.64 y Water loss 3
689.2872924804688 0 6320.2397 y Ammonia loss 3
690.2896728515625 0 1582.1996
706.2994995117188 0 128323.81 y 3
707.3021240234375 0 43081.676
708.3040771484375 0 9793.306
716.2822875976562 0 1023.0168
717.2822265625 0 838.048
741.3490600585938 0 683.1169
766.3375854492188 0 675.5196
783.3628540039062 0 2924.639
784.355712890625 0 2823.6748
785.3560791015625 0 1310.6458
801.37255859375 0 3080.9663 y Water loss 2
802.3631591796875 0 3478.679 y Ammonia loss 2
803.3619384765625 0 1734.6481
819.383056640625 0 221783.53 y 2
820.3855590820312 0 95600.984
821.3883056640625 0 23173.84
822.3895263671875 0 1724.719
829.3600463867188 0 846.5871
932.4658203125 0 4277.3306 y 1
933.4677734375 0 2733.7307
934.4719848632812 0 762.81476
3081.2509765625 0 994.30396

Spectrum Details

|  |  |
| --- | --- |
| Matched peaks? Matched peaks The total absolute number of peaks matched. Additionally in brackets the total fraction of peaks matched and the total number of peaks is shown. | 39 (16.96% of 230) |
| FDR? FDR The false discovery rate estimated for this peptide. It is calculated by matching all theoretical fragments with a non-integer shift with the raw peaks for this spectrum. This is done with 40 different shifts. The resulting percentage is the average number of annotated peaks over the number of annotated peaks with the correct spectrum. | 0.92% |
| Satellite FDR? Satellite FDR See the FDR for details on its calculation. This satellite ion specific FDR only contains the satellite ions (d/w) for I/L/J positions. | 2.38% |
| PSM Score? PSM Score The PSM Score as given by Hecklib to this annotated spectrum. It is shown with three significant figures. | 467 |

#### Spectrum 3752? Spectrum 3752 The raw spectrum of this peptide as annotated by Hecklib. The fragments are coloured according to ion type (see legend). Any peaks with a star '\*' as text can be hovered over to see the full details, first the ion type second the mass shift type. By hovering over the amino acids in the peptide or ions in the legend the corresponding peaks are highlighted. By toggling the 'Unassigned' label you can turn the background (unassigned) peaks on or off in the plot. By updating the slider in the Ion legend you can update the spectrum to only show the top X% of the peaks with labels. The top X% means any peak that is within X% of the highest intensity. By dragging in the spectrum you can zoom in to a specific part of the spectrum and use 'Zoom Out' to get back to the original zoom level. The annotation of the spectrum is based on the given sequence in the peptides file and is done with different software so inconsistencies are likely. The peaks are annotated based on the given sequence, with 20 ppm tolerance.

Copy Data

##### Spectrum 3752 (TSV)

###### Preview

```
Loading example...
```

*Click on the button to copy the data to your clipboard.*

Mz MinMz MaxIntensity Max

WidthHeightPeptide font sizePeptide stroke widthSpectrum font sizeSpectrum stroke widthCompact peptide

Ion legend

wxyz

abcd

OtherUnassignedIon^Charge^_Position_Show for top:%

TJJEGEESR

03.65e+47.29e+41.09e+51.46e+5

Zoom Out

y^+1^_1_a^+1^_2_y^+1^_1_a^+1^_2_b^+1^_2_b^+1^_2_b^+2^_4_y^+1^_2_y^+1^_2_y^+1^_2_b^+1^_3_b^+1^_3_y^+1^_3_y^+1^_3_y^+1^_3_y^+2^_7_y^+2^_7_b^+1^_4_b^+1^_4_y^+2^_8_b^+1^_5_\*\*\*y^+1^_4_y^+1^_5_y^+1^_5_y^+1^_5_y^+1^_6_y^+1^_6_y^+1^_7_y^+1^_7_y^+1^_7_y^+1^_8_

0668133720052674

Fragment Matches Table

Show background peaks

| Position | Ion type | Intensity | mz Theoretical | mz Error (Th) | mz Error (ppm) | Charge | Series Number |
| --- | --- | --- | --- | --- | --- | --- | --- |
| - | - | 1806 | 120.1 | - | - | 0 | - |
| - | - | 529.4 | 126.1 | - | - | 0 | - |
| - | - | 644.7 | 126.1 | - | - | 0 | - |
| - | - | 1169 | 130.1 | - | - | 0 | - |
| - | - | 1891 | 130.1 | - | - | 0 | - |
| - | - | 374.3 | 140.4 | - | - | 0 | - |
| - | - | 1389 | 141.1 | - | - | 0 | - |
| - | - | 4078 | 142.1 | - | - | 0 | - |
| - | - | 4716 | 143.1 | - | - | 0 | - |
| - | - | 655.2 | 143.1 | - | - | 0 | - |
| - | - | 514 | 148.9 | - | - | 0 | - |
| - | - | 744.2 | 148.9 | - | - | 0 | - |
| - | - | 768.9 | 148.9 | - | - | 0 | - |
| - | - | 934.3 | 148.9 | - | - | 0 | - |
| - | - | 767.7 | 148.9 | - | - | 0 | - |
| - | - | 1065 | 148.9 | - | - | 0 | - |
| - | - | 1382 | 148.9 | - | - | 0 | - |
| - | - | 3268 | 148.9 | - | - | 0 | - |
| - | - | 5734 | 148.9 | - | - | 0 | - |
| - | - | 3733 | 149 | - | - | 0 | - |
| - | - | 1581 | 149 | - | - | 0 | - |
| - | - | 1167 | 149 | - | - | 0 | - |
| - | - | 1166 | 149 | - | - | 0 | - |
| - | - | 752.1 | 149 | - | - | 0 | - |
| - | - | 663.8 | 149 | - | - | 0 | - |
| - | - | 420.7 | 149 | - | - | 0 | - |
| - | - | 523 | 149 | - | - | 0 | - |
| - | - | 513.6 | 149 | - | - | 0 | - |
| - | - | 526 | 149 | - | - | 0 | - |
| - | - | 549.1 | 149.1 | - | - | 0 | - |
| - | - | 448.5 | 149.2 | - | - | 0 | - |
| - | - | 447.1 | 149.6 | - | - | 0 | - |
| - | - | 417.2 | 150.2 | - | - | 0 | - |
| - | - | 394.2 | 152.1 | - | - | 0 | - |
| - | - | 500 | 156.4 | - | - | 0 | - |
| - | - | 423.8 | 156.5 | - | - | 0 | - |
| - | - | 642.7 | 157.1 | - | - | 0 | - |
| 9 | y | 6773 | 158.1 | 0.000111 | 0.7019 | +1 | 1 |
| - | - | 1739 | 159.1 | - | - | 0 | - |
| - | - | 712.4 | 167.1 | - | - | 0 | - |
| - | - | 3048 | 169.1 | - | - | 0 | - |
| 2 | a | 1960 | 169.1 | 0.0001732 | 1.024 | +1 | 2 |
| - | - | 565.6 | 170.1 | - | - | 0 | - |
| - | - | 508.5 | 171.1 | - | - | 0 | - |
| - | - | 552.6 | 173.5 | - | - | 0 | - |
| 9 | y | 9931 | 175.1 | 0.0001122 | 0.6405 | +1 | 1 |
| - | - | 616.1 | 184.1 | - | - | 0 | - |
| - | - | 697.1 | 185.1 | - | - | 0 | - |
| - | - | 7153 | 187.1 | - | - | 0 | - |
| 2 | a | 1.444E+05 | 187.1 | 0.0001828 | 0.9769 | +1 | 2 |
| - | - | 1.253E+04 | 188.1 | - | - | 0 | - |
| - | - | 694.5 | 189.1 | - | - | 0 | - |
| - | - | 841.6 | 195.1 | - | - | 0 | - |
| 2 | b | 2127 | 197.1 | 0.0001163 | 0.5902 | +1 | 2 |
| - | - | 1361 | 199.1 | - | - | 0 | - |
| - | - | 4441 | 199.2 | - | - | 0 | - |
| - | - | 1067 | 202.1 | - | - | 0 | - |
| - | - | 1074 | 203.1 | - | - | 0 | - |
| - | - | 512.5 | 213.1 | - | - | 0 | - |
| 2 | b | 7.372E+04 | 215.1 | 8.021E-05 | 0.3728 | +1 | 2 |
| - | - | 7864 | 216.1 | - | - | 0 | - |
| - | - | 708.8 | 217.1 | - | - | 0 | - |
| 4 | b | 569 | 220.1 | 0.001754 | 7.966 | +2 | 4 |
| - | - | 496.6 | 221.2 | - | - | 0 | - |
| - | - | 928.1 | 225 | - | - | 0 | - |
| - | - | 1038 | 227.1 | - | - | 0 | - |
| - | - | 2200 | 227.2 | - | - | 0 | - |
| - | - | 598.4 | 231.5 | - | - | 0 | - |
| - | - | 647.5 | 232.1 | - | - | 0 | - |
| - | - | 2446 | 239.1 | - | - | 0 | - |
| - | - | 1866 | 241.1 | - | - | 0 | - |
| - | - | 1.069E+04 | 243.1 | - | - | 0 | - |
| 8 | y | 2128 | 244.1 | 0.001988 | 8.143 | +1 | 2 |
| 8 | y | 5388 | 245.1 | 9.55E-05 | 0.3896 | +1 | 2 |
| - | - | 673.4 | 246.1 | - | - | 0 | - |
| - | - | 1221 | 259.1 | - | - | 0 | - |
| 8 | y | 1.958E+04 | 262.2 | 4.064E-05 | 0.155 | +1 | 2 |
| - | - | 2202 | 263.2 | - | - | 0 | - |
| - | - | 473.1 | 269.3 | - | - | 0 | - |
| - | - | 1085 | 270.1 | - | - | 0 | - |
| - | - | 1301 | 283.2 | - | - | 0 | - |
| - | - | 608.1 | 295.1 | - | - | 0 | - |
| - | - | 2925 | 298.1 | - | - | 0 | - |
| - | - | 849.4 | 299.1 | - | - | 0 | - |
| - | - | 2223 | 300.2 | - | - | 0 | - |
| - | - | 743.4 | 302.1 | - | - | 0 | - |
| 3 | b | 1057 | 310.2 | 0.0003114 | 1.004 | +1 | 3 |
| - | - | 644.1 | 313.2 | - | - | 0 | - |
| - | - | 3109 | 316.1 | - | - | 0 | - |
| 3 | b | 8485 | 328.2 | 9.216E-05 | 0.2808 | +1 | 3 |
| - | - | 1921 | 329.2 | - | - | 0 | - |
| - | - | 658.8 | 331.2 | - | - | 0 | - |
| - | - | 549.3 | 335.3 | - | - | 0 | - |
| - | - | 1001 | 338.1 | - | - | 0 | - |
| - | - | 1964 | 356.2 | - | - | 0 | - |
| - | - | 1118 | 372.7 | - | - | 0 | - |
| 7 | y | 754.4 | 373.2 | 0.0006359 | 1.704 | +1 | 3 |
| 7 | y | 1316 | 374.2 | 8.969E-05 | 0.2397 | +1 | 3 |
| - | - | 651.9 | 380.2 | - | - | 0 | - |
| - | - | 921.1 | 381.1 | - | - | 0 | - |
| - | - | 1213 | 386.2 | - | - | 0 | - |
| 7 | y | 7403 | 391.2 | 3.119E-05 | 0.07974 | +1 | 3 |
| - | - | 1037 | 392.2 | - | - | 0 | - |
| - | - | 3134 | 395.2 | - | - | 0 | - |
| 3 | y | 1078 | 401.2 | 0.0002352 | 0.5862 | +2 | 7 |
| - | - | 745.3 | 409.1 | - | - | 0 | - |
| 3 | y | 1.078E+04 | 410.2 | 0.0001033 | 0.2518 | +2 | 7 |
| - | - | 4592 | 410.7 | - | - | 0 | - |
| - | - | 935.7 | 411.2 | - | - | 0 | - |
| - | - | 681.8 | 416.1 | - | - | 0 | - |
| - | - | 1986 | 417.2 | - | - | 0 | - |
| - | - | 836 | 419.2 | - | - | 0 | - |
| - | - | 1190 | 427.1 | - | - | 0 | - |
| - | - | 848.1 | 429.3 | - | - | 0 | - |
| 4 | b | 803.8 | 439.3 | 0.0004116 | 0.937 | +1 | 4 |
| - | - | 905.3 | 445.2 | - | - | 0 | - |
| 4 | b | 3339 | 457.3 | 0.0002951 | 0.6454 | +1 | 4 |
| 2 | y | 1101 | 466.7 | 0.0004178 | 0.8951 | +2 | 8 |
| 5 | b | 683.2 | 496.3 | 0.0009518 | 1.918 | +1 | 5 |
| 0 | Precursor | 1920 | 508.3 | 0.000343 | 0.6748 | +2 | -1 |
| 0 | Precursor | 981.3 | 508.7 | 0.01004 | 19.74 | +2 | -1 |
| - | - | 577.9 | 514.2 | - | - | 0 | - |
| 0 | Precursor | 4119 | 517.3 | 0.00085 | 1.643 | +2 | -1 |
| - | - | 922 | 517.3 | - | - | 0 | - |
| - | - | 2891 | 517.8 | - | - | 0 | - |
| - | - | 759 | 518.3 | - | - | 0 | - |
| 6 | y | 3293 | 520.2 | 0.0006321 | 1.215 | +1 | 4 |
| - | - | 1385 | 521.2 | - | - | 0 | - |
| - | - | 1616 | 542.2 | - | - | 0 | - |
| 5 | y | 1131 | 559.2 | 0.0009811 | 1.754 | +1 | 5 |
| 5 | y | 2049 | 560.2 | 0.0006081 | 1.085 | +1 | 5 |
| 5 | y | 5.69E+04 | 577.3 | 0.0007946 | 1.376 | +1 | 5 |
| - | - | 1.499E+04 | 578.3 | - | - | 0 | - |
| - | - | 2568 | 579.3 | - | - | 0 | - |
| - | - | 661.1 | 668 | - | - | 0 | - |
| 4 | y | 8061 | 688.3 | 0.001695 | 2.463 | +1 | 6 |
| - | - | 2626 | 689.3 | - | - | 0 | - |
| - | - | 659.8 | 701.1 | - | - | 0 | - |
| 4 | y | 6.25E+04 | 706.3 | 0.001029 | 1.457 | +1 | 6 |
| - | - | 2.041E+04 | 707.3 | - | - | 0 | - |
| - | - | 4126 | 708.3 | - | - | 0 | - |
| - | - | 620.9 | 744.3 | - | - | 0 | - |
| - | - | 1435 | 783.4 | - | - | 0 | - |
| - | - | 1677 | 784.3 | - | - | 0 | - |
| 3 | y | 2278 | 801.4 | 0.002568 | 3.205 | +1 | 7 |
| 3 | y | 2090 | 802.4 | 0.007374 | 9.19 | +1 | 7 |
| 3 | y | 1.075E+05 | 819.4 | 0.001597 | 1.949 | +1 | 7 |
| - | - | 4.219E+04 | 820.4 | - | - | 0 | - |
| - | - | 1.017E+04 | 821.4 | - | - | 0 | - |
| - | - | 1287 | 822.4 | - | - | 0 | - |
| 2 | y | 2767 | 932.5 | 0.004667 | 5.006 | +1 | 8 |
| - | - | 958.3 | 933.5 | - | - | 0 | - |
| - | - | 608.5 | 2647 | - | - | 0 | - |

m/z Charge Intensity FragmentType MassShift Position
120.0810775756836 0 1805.6283
126.09182739257812 0 529.4362
126.1279296875 0 644.6743
130.05001831054688 0 1168.7231
130.0613555908203 0 1891.2072
140.4221649169922 0 374.2708
141.06607055664062 0 1389.2212
142.12277221679688 0 4077.754
143.1179656982422 0 4715.9976
143.12445068359375 0 655.1708
148.88792419433594 0 513.995
148.8953094482422 0 744.1725
148.90206909179688 0 768.92303
148.9092254638672 0 934.2826
148.91651916503906 0 767.671
148.9237060546875 0 1065.1383
148.93093872070312 0 1381.618
148.93836975097656 0 3268.095
148.94606018066406 0 5734.202
148.96240234375 0 3733.4578
148.97012329101562 0 1580.9338
148.9771270751953 0 1167.4055
148.984130859375 0 1165.763
148.99169921875 0 752.1186
148.99893188476562 0 663.82605
149.00595092773438 0 420.73843
149.01316833496094 0 522.9786
149.02032470703125 0 513.5727
149.02774047851562 0 525.9723
149.0849151611328 0 549.11786
149.1642608642578 0 448.4534
149.64700317382812 0 447.07855
150.24180603027344 0 417.18854
152.10635375976562 0 394.15198
156.4262237548828 0 499.97388
156.5026397705078 0 423.8174
157.1087188720703 0 642.7155
158.09251403808594 0 6773.3105 y Ammonia loss 8
159.07650756835938 0 1739.4003
167.05528259277344 0 712.41547
169.06089782714844 0 3048.1682
169.1337127685547 0 1959.5428 a Water loss 1
170.1178741455078 0 565.6459
171.07691955566406 0 508.45996
173.4503631591797 0 552.61017
175.1190643310547 0 9930.568 y 8
184.1075897216797 0 616.1362
185.09181213378906 0 697.07947
187.07144165039062 0 7153.4487
187.144287109375 0 144428.31 a 1
188.1476287841797 0 12528.225
189.14894104003906 0 694.4667
195.07669067382812 0 841.63434
197.12857055664062 0 2127.0537 b Water loss 1
199.07135009765625 0 1360.9521
199.18067932128906 0 4440.945
202.11839294433594 0 1066.9061
203.1023712158203 0 1074.1924
213.08670043945312 0 512.45917
215.13909912109375 0 73715.13 b 1
216.1424102783203 0 7863.9614
217.14390563964844 0 708.77686
220.1294403076172 0 569.03204 b Water loss 3
221.2287139892578 0 496.61417
225.0430450439453 0 928.1067
227.1141815185547 0 1038.272
227.17532348632812 0 2199.7666
231.4586181640625 0 598.3696
232.139892578125 0 647.4752
239.09449768066406 0 2445.645
241.08184814453125 0 1865.5873
243.13406372070312 0 10691.308
244.138427734375 0 2128.067 y Water loss 7
245.12452697753906 0 5388.061 y Ammonia loss 7
246.12692260742188 0 673.3948
259.0922546386719 0 1221.2429
262.15093994140625 0 19581.666 y 7
263.1542053222656 0 2201.832
269.3113098144531 0 473.10303
270.1083068847656 0 1084.6323
283.2015075683594 0 1300.7162
295.1395263671875 0 608.1305
298.10321044921875 0 2925.095
299.0620422363281 0 849.38226
300.1555480957031 0 2223.052
302.1468505859375 0 743.42053
310.21282958984375 0 1056.7646 b Water loss 2
313.1501770019531 0 644.13855
316.1138000488281 0 3109.0442
328.2231750488281 0 8484.606 b 2
329.226806640625 0 1920.7173
331.1604309082031 0 658.8184
335.27191162109375 0 549.2588
338.1469421386719 0 1001.06384
356.15655517578125 0 1964.0432
372.6539306640625 0 1117.7601
373.182373046875 0 754.4163 y Water loss 6
374.1671142578125 0 1316.2238 y Ammonia loss 6
380.16943359375 0 651.9278
381.13995361328125 0 921.0769
386.16705322265625 0 1212.6217
391.19354248046875 0 7402.6685 y 6
392.1950378417969 0 1037.0262
395.1755676269531 0 3133.7473
401.19073486328125 0 1078.4208 y Water loss 2
409.1343994140625 0 745.2666
410.1956787109375 0 10784.1 y 2
410.69732666015625 0 4592.4546
411.197021484375 0 935.7366
416.1446838378906 0 681.8296
417.1881408691406 0 1986.1448
419.17828369140625 0 836.0205
427.1458435058594 0 1189.5023
429.2707824707031 0 848.1056
439.25469970703125 0 803.803 b Water loss 3
445.15753173828125 0 905.25073
457.265380859375 0 3338.8223 b 3
466.7373962402344 0 1100.8451 y 1
496.27752685546875 0 683.2061 b Water loss 4
508.2567138671875 0 1920.0028 Precursor Water loss
508.7584228515625 0 981.31885 Precursor Ammonia loss
514.1765747070312 0 577.9432
517.2608032226562 0 4118.5693 Precursor
517.3018188476562 0 922.0073
517.7620239257812 0 2891.23
518.2603149414062 0 758.9671
520.2355346679688 0 3292.94 y 5
521.23779296875 0 1384.5693
542.2222290039062 0 1616.475
559.248046875 0 1130.8768 y Water loss 4
560.231689453125 0 2049.3477 y Ammonia loss 4
577.2568359375 0 56896.344 y 4
578.259521484375 0 14988.151
579.2611694335938 0 2567.8904
667.9561157226562 0 661.0586
688.2879638671875 0 8061.1304 y Water loss 3
689.289306640625 0 2625.6807
701.06884765625 0 659.75653
706.2991943359375 0 62503.652 y 3
707.3018798828125 0 20409.572
708.3037719726562 0 4125.531
744.2646484375 0 620.92444
783.3623046875 0 1434.908
784.349609375 0 1676.6586
801.3711547851562 0 2278.4482 y Water loss 2
802.3651123046875 0 2089.8438 y Ammonia loss 2
819.3826904296875 0 107511.92 y 2
820.38525390625 0 42186.582
821.387939453125 0 10166.348
822.3869018554688 0 1286.7211
932.4636840820312 0 2767.3748 y 1
933.4635620117188 0 958.2768
2647.048095703125 0 608.4794

Spectrum Details

|  |  |
| --- | --- |
| Matched peaks? Matched peaks The total absolute number of peaks matched. Additionally in brackets the total fraction of peaks matched and the total number of peaks is shown. | 34 (22.22% of 153) |
| FDR? FDR The false discovery rate estimated for this peptide. It is calculated by matching all theoretical fragments with a non-integer shift with the raw peaks for this spectrum. This is done with 40 different shifts. The resulting percentage is the average number of annotated peaks over the number of annotated peaks with the correct spectrum. | 0.98% |
| Satellite FDR? Satellite FDR See the FDR for details on its calculation. This satellite ion specific FDR only contains the satellite ions (d/w) for I/L/J positions. | ∞ |
| PSM Score? PSM Score The PSM Score as given by Hecklib to this annotated spectrum. It is shown with three significant figures. | 422 |

#### Reverse Lookup? Reverse Lookup All places where this read could be placed.

| Group | Segment | Template | Template Part | Read Part | Score | Unique |
| --- | --- | --- | --- | --- | --- | --- |
| Homo sapiens Light Chain | IGLV | IGKV2D-26 | [49..58] | [0..9] | 36 | False |
| Homo sapiens Light Chain | IGLV | IGKV1-9 | [52..61] | [0..9] | 40 | False |
| Homo sapiens Light Chain | IGLV | IGKV1-8 | [52..61] | [0..9] | 40 | False |
| Homo sapiens Light Chain | IGLV | IGKV3-20 | [47..55] | [0..9] | 40 | False |
| Homo sapiens Light Chain | IGLV | IGKV1D-8 | [52..61] | [0..9] | 40 | False |
| Homo sapiens Light Chain | IGLV | IGKV1-27 | [52..61] | [0..9] | 40 | False |
| Decoy | Decoy | K1C20 | [372..380] | [0..9] | 40 | False |

| Recombined | Template Part | Read Part | Score | Unique |
| --- | --- | --- | --- | --- |
| K1C20 | [372..380] | [0..9] | 40 | True |

#### Meta Information from Multiple reads

##### Number of combined reads

2

##### Intensity

0.5691

##### TotalArea

1.185E+07

##### Changes to the peptide sequence

TJJEGEESR

I→JNo support for either Leucine or Isoleucine based on side chain ions (Position: 2)

L→JNo support for either Leucine or Isoleucine based on side chain ions (Position: 3)

L→ISupport for Isoleucine based on side chain ions (1 for I 0 for L) (Position: 2)

#### Positional Score

Copy Data

##### Positional Score (TSV)

###### Preview

```
Loading example...
```

*Click on the button to copy the data to your clipboard.*

00012345678

Label Value
"0" 0
"1" 0
"2" 0
"3" 0
"4" 0
"5" 0
"6" 0
"7" 0
"8" 0

#### Meta Information from PEAKS

##### Scan Identifier

F4:3697

##### Original sequence

T

L

L

E

G

E

E

S

R

##### Posttranslational Modifications

##### Source File

D:\separate\_stitch\_analyses\xle-disambiguation\raw\20210323\_F1\_UM1\_Peng0013\_SA\_F59\_ingel\_3ug\_tryp.raw

##### Fraction

4

##### Scan Feature

F4:3612

##### De Novo Score

99

##### ConfidenceScore

99

### m/z

517.2621

##### Mass

1032.5088

##### Charge

2

##### Retention Time

19.87

##### Predicted Retention Time

18.54

##### Area

5.926E+06

##### Parts Per Million

0.8

##### Fragmentation mode

HCD

##### Originating file

01 D:\separate\_stitch\_analyses\xle-disambiguation\20210325\_F59\_3ug\_DENOVO\_12.csv

#### Meta Information from PEAKS

##### Scan Identifier

F4:3752

##### Original sequence

T

L

L

E

G

E

E

S

R

##### Posttranslational Modifications

##### Source File

D:\separate\_stitch\_analyses\xle-disambiguation\raw\20210323\_F1\_UM1\_Peng0013\_SA\_F59\_ingel\_3ug\_tryp.raw

##### Fraction

4

##### Scan Feature

F4:3612

##### De Novo Score

99

##### ConfidenceScore

99

### m/z

517.2621

##### Mass

1032.5088

##### Charge

2

##### Retention Time

19.87

##### Predicted Retention Time

18.54

##### Area

5.926E+06

##### Parts Per Million

0.8

##### Fragmentation mode

HCD

##### Originating file

01 D:\separate\_stitch\_analyses\xle-disambiguation\20210325\_F59\_3ug\_DENOVO\_12.csv
