## Supplementary material for "A handle on mass coincidence errors in *de novo* sequencing of antibodies by bottom-up proteomics": Combined_023.html

Details Combined\_023 | Stitch OverviewUndefined

### Read Combined\_023

#### Sequence (length=10)

VVFGGGTKJT

#### Spectrum 5680? Spectrum 5680 The raw spectrum of this peptide as annotated by Hecklib. The fragments are coloured according to ion type (see legend). Any peaks with a star '\*' as text can be hovered over to see the full details, first the ion type second the mass shift type. By hovering over the amino acids in the peptide or ions in the legend the corresponding peaks are highlighted. By toggling the 'Unassigned' label you can turn the background (unassigned) peaks on or off in the plot. By updating the slider in the Ion legend you can update the spectrum to only show the top X% of the peaks with labels. The top X% means any peak that is within X% of the highest intensity. By dragging in the spectrum you can zoom in to a specific part of the spectrum and use 'Zoom Out' to get back to the original zoom level. The annotation of the spectrum is based on the given sequence in the peptides file and is done with different software so inconsistencies are likely. The peaks are annotated based on the given sequence, with 20 ppm tolerance.

Copy Data

##### Spectrum 5680 (TSV)

###### Preview

```
Loading example...
```

*Click on the button to copy the data to your clipboard.*

Mz MinMz MaxIntensity Max

WidthHeightPeptide font sizePeptide stroke widthSpectrum font sizeSpectrum stroke widthCompact peptide

Ion legend

wxyz

abcd

OtherUnassignedIon^Charge^_Position_Show for top:%

VVFGGGTKJT

01.40e+62.81e+64.21e+65.62e+6

Zoom Out

y^+1^_1_a^+1^_2_b^+1^_2_y^+1^_2_y^+1^_2_y^+2^_5_b^+2^_6_y^+2^_7_y^+1^_3_y^+1^_3_b^+1^_3_y^+1^_3_b^+2^_8_b^+2^_8_b^+2^_8_y^+2^_8_y^+2^_8_b^+1^_4_b^+2^_9_b^+2^_9_y^+2^_9_y^+2^_9_y^+1^_4_b^+1^_5_y^+1^_4_\*\*y^+1^_5_b^+1^_6_y^+1^_5_y^+1^_6_y^+1^_6_b^+1^_7_y^+1^_7_b^+1^_7_y^+1^_7_b^+1^_8_b^+1^_8_y^+1^_8_y^+1^_8_b^+1^_9_b^+1^_9_y^+1^_9_

0598119517932390

Fragment Matches Table

Show background peaks

| Position | Ion type | Intensity | mz Theoretical | mz Error (Th) | mz Error (ppm) | Charge | Series Number |
| --- | --- | --- | --- | --- | --- | --- | --- |
| 10 | y | 4.24E+05 | 120.1 | 0.0003831 | 3.191 | +1 | 1 |
| - | - | 2.857E+06 | 120.1 | - | - | 0 | - |
| - | - | 2877 | 121 | - | - | 0 | - |
| - | - | 2837 | 121.1 | - | - | 0 | - |
| - | - | 1.349E+04 | 121.1 | - | - | 0 | - |
| - | - | 2.204E+05 | 121.1 | - | - | 0 | - |
| - | - | 3994 | 122.1 | - | - | 0 | - |
| - | - | 1.991E+04 | 127.1 | - | - | 0 | - |
| - | - | 4438 | 128.1 | - | - | 0 | - |
| - | - | 2.24E+06 | 129.1 | - | - | 0 | - |
| - | - | 7616 | 130.1 | - | - | 0 | - |
| - | - | 1.614E+04 | 130.1 | - | - | 0 | - |
| - | - | 1.344E+05 | 130.1 | - | - | 0 | - |
| - | - | 9133 | 131 | - | - | 0 | - |
| - | - | 8.108E+04 | 131.1 | - | - | 0 | - |
| - | - | 3.638E+04 | 132.1 | - | - | 0 | - |
| - | - | 4587 | 136.1 | - | - | 0 | - |
| - | - | 1.238E+04 | 139.1 | - | - | 0 | - |
| - | - | 4.766E+04 | 141.1 | - | - | 0 | - |
| - | - | 8721 | 141.1 | - | - | 0 | - |
| - | - | 3648 | 142.1 | - | - | 0 | - |
| - | - | 4960 | 142.1 | - | - | 0 | - |
| - | - | 4334 | 143.1 | - | - | 0 | - |
| - | - | 8804 | 143.2 | - | - | 0 | - |
| - | - | 4741 | 146.1 | - | - | 0 | - |
| - | - | 9749 | 149 | - | - | 0 | - |
| - | - | 8160 | 151.1 | - | - | 0 | - |
| - | - | 2.018E+04 | 152.1 | - | - | 0 | - |
| - | - | 9851 | 153.1 | - | - | 0 | - |
| - | - | 6014 | 154.1 | - | - | 0 | - |
| - | - | 2.044E+04 | 155.1 | - | - | 0 | - |
| - | - | 6.129E+04 | 155.1 | - | - | 0 | - |
| - | - | 2.217E+04 | 158.1 | - | - | 0 | - |
| - | - | 7.638E+04 | 159.1 | - | - | 0 | - |
| - | - | 4484 | 160.1 | - | - | 0 | - |
| - | - | 7448 | 165.1 | - | - | 0 | - |
| - | - | 1.409E+04 | 167.1 | - | - | 0 | - |
| - | - | 6867 | 168.1 | - | - | 0 | - |
| - | - | 8861 | 169.1 | - | - | 0 | - |
| - | - | 4.424E+04 | 170.1 | - | - | 0 | - |
| - | - | 1.357E+04 | 171.1 | - | - | 0 | - |
| 2 | a | 5.56E+06 | 171.1 | 0.0004533 | 2.648 | +1 | 2 |
| - | - | 7.816E+04 | 172.1 | - | - | 0 | - |
| - | - | 4.903E+05 | 172.2 | - | - | 0 | - |
| - | - | 2.232E+04 | 173.2 | - | - | 0 | - |
| - | - | 1.027E+04 | 173.5 | - | - | 0 | - |
| - | - | 1.481E+04 | 174.1 | - | - | 0 | - |
| - | - | 1.109E+05 | 176.1 | - | - | 0 | - |
| - | - | 2.62E+05 | 177.1 | - | - | 0 | - |
| - | - | 1.151E+04 | 177.1 | - | - | 0 | - |
| - | - | 3.265E+04 | 178.1 | - | - | 0 | - |
| - | - | 9620 | 183.1 | - | - | 0 | - |
| - | - | 9617 | 183.1 | - | - | 0 | - |
| - | - | 3.311E+04 | 185.1 | - | - | 0 | - |
| - | - | 2.643E+04 | 186.1 | - | - | 0 | - |
| - | - | 7138 | 187.1 | - | - | 0 | - |
| - | - | 4462 | 188.1 | - | - | 0 | - |
| - | - | 2.961E+04 | 188.1 | - | - | 0 | - |
| - | - | 4677 | 188.1 | - | - | 0 | - |
| - | - | 7430 | 189.1 | - | - | 0 | - |
| - | - | 1.578E+04 | 194.1 | - | - | 0 | - |
| - | - | 9902 | 195.1 | - | - | 0 | - |
| - | - | 8094 | 196.1 | - | - | 0 | - |
| - | - | 5520 | 197.1 | - | - | 0 | - |
| - | - | 1.005E+05 | 197.2 | - | - | 0 | - |
| - | - | 8.881E+04 | 198.1 | - | - | 0 | - |
| - | - | 7326 | 198.2 | - | - | 0 | - |
| 2 | b | 2.772E+06 | 199.1 | 0.0003964 | 1.991 | +1 | 2 |
| - | - | 2.669E+05 | 200.1 | - | - | 0 | - |
| - | - | 1.052E+04 | 201.1 | - | - | 0 | - |
| - | - | 1.802E+04 | 201.1 | - | - | 0 | - |
| - | - | 2.736E+04 | 203.1 | - | - | 0 | - |
| - | - | 5.38E+05 | 205.1 | - | - | 0 | - |
| - | - | 6.013E+04 | 206.1 | - | - | 0 | - |
| - | - | 5470 | 206.1 | - | - | 0 | - |
| - | - | 3.87E+04 | 207.1 | - | - | 0 | - |
| - | - | 4.75E+04 | 208.1 | - | - | 0 | - |
| - | - | 1.555E+04 | 209.1 | - | - | 0 | - |
| - | - | 2.476E+04 | 210.1 | - | - | 0 | - |
| - | - | 1.755E+04 | 211.1 | - | - | 0 | - |
| - | - | 2.778E+05 | 212.1 | - | - | 0 | - |
| - | - | 6410 | 213.1 | - | - | 0 | - |
| - | - | 3.023E+04 | 213.1 | - | - | 0 | - |
| - | - | 1.02E+04 | 214.1 | - | - | 0 | - |
| - | - | 1.983E+04 | 214.2 | - | - | 0 | - |
| - | - | 4.76E+04 | 215.1 | - | - | 0 | - |
| 9 | y | 5.146E+04 | 215.1 | 0.0002481 | 1.153 | +1 | 2 |
| - | - | 5.792E+04 | 216.1 | - | - | 0 | - |
| - | - | 5.531E+04 | 217.1 | - | - | 0 | - |
| - | - | 4222 | 217.1 | - | - | 0 | - |
| - | - | 7557 | 218.1 | - | - | 0 | - |
| - | - | 2.538E+05 | 219.1 | - | - | 0 | - |
| - | - | 3.174E+04 | 220.2 | - | - | 0 | - |
| - | - | 2.277E+04 | 221.1 | - | - | 0 | - |
| - | - | 6427 | 221.2 | - | - | 0 | - |
| - | - | 4982 | 222.1 | - | - | 0 | - |
| - | - | 9.514E+04 | 224.2 | - | - | 0 | - |
| - | - | 2.757E+04 | 225.1 | - | - | 0 | - |
| - | - | 2.564E+04 | 225.2 | - | - | 0 | - |
| - | - | 6691 | 225.2 | - | - | 0 | - |
| - | - | 1.118E+05 | 227.1 | - | - | 0 | - |
| - | - | 5.579E+04 | 228.1 | - | - | 0 | - |
| - | - | 1.22E+04 | 228.1 | - | - | 0 | - |
| - | - | 7174 | 228.1 | - | - | 0 | - |
| - | - | 9677 | 228.1 | - | - | 0 | - |
| - | - | 1.829E+04 | 229.1 | - | - | 0 | - |
| - | - | 6126 | 229.1 | - | - | 0 | - |
| - | - | 4.884E+05 | 230.2 | - | - | 0 | - |
| - | - | 3.599E+04 | 231.1 | - | - | 0 | - |
| - | - | 5.171E+04 | 231.2 | - | - | 0 | - |
| - | - | 2.646E+04 | 233.1 | - | - | 0 | - |
| 9 | y | 1.548E+05 | 233.1 | 0.0003645 | 1.563 | +1 | 2 |
| - | - | 8948 | 233.2 | - | - | 0 | - |
| - | - | 1.537E+04 | 234.2 | - | - | 0 | - |
| - | - | 3.274E+04 | 237.1 | - | - | 0 | - |
| - | - | 1.628E+04 | 240.1 | - | - | 0 | - |
| - | - | 1.382E+04 | 241.2 | - | - | 0 | - |
| - | - | 1.001E+05 | 242.2 | - | - | 0 | - |
| - | - | 5.457E+04 | 243.1 | - | - | 0 | - |
| - | - | 1.149E+04 | 243.1 | - | - | 0 | - |
| - | - | 7353 | 243.2 | - | - | 0 | - |
| - | - | 7556 | 243.2 | - | - | 0 | - |
| - | - | 9059 | 244.1 | - | - | 0 | - |
| - | - | 5603 | 244.1 | - | - | 0 | - |
| - | - | 8.862E+04 | 245.1 | - | - | 0 | - |
| - | - | 1.06E+04 | 245.2 | - | - | 0 | - |
| - | - | 6012 | 246.1 | - | - | 0 | - |
| - | - | 1.174E+05 | 247.1 | - | - | 0 | - |
| - | - | 1.823E+04 | 248.1 | - | - | 0 | - |
| - | - | 1.065E+04 | 249.1 | - | - | 0 | - |
| 6 | y | 2.59E+04 | 251.2 | 0.004123 | 16.42 | +2 | 5 |
| - | - | 7251 | 252.2 | - | - | 0 | - |
| - | - | 8113 | 254.1 | - | - | 0 | - |
| - | - | 2.703E+05 | 255.1 | - | - | 0 | - |
| - | - | 2.461E+04 | 256.1 | - | - | 0 | - |
| - | - | 1.496E+04 | 256.2 | - | - | 0 | - |
| 6 | b | 1.439E+04 | 259.1 | 0.001889 | 7.29 | +2 | 6 |
| - | - | 1.336E+04 | 260.2 | - | - | 0 | - |
| - | - | 4.934E+04 | 261.1 | - | - | 0 | - |
| - | - | 7335 | 261.2 | - | - | 0 | - |
| - | - | 1.594E+05 | 262.1 | - | - | 0 | - |
| - | - | 2.269E+04 | 263.1 | - | - | 0 | - |
| - | - | 5113 | 263.2 | - | - | 0 | - |
| - | - | 5357 | 265.1 | - | - | 0 | - |
| - | - | 5586 | 268.2 | - | - | 0 | - |
| - | - | 2.738E+05 | 269.2 | - | - | 0 | - |
| - | - | 3.83E+04 | 270.2 | - | - | 0 | - |
| - | - | 3.121E+04 | 270.2 | - | - | 0 | - |
| - | - | 7723 | 271.1 | - | - | 0 | - |
| - | - | 7846 | 271.1 | - | - | 0 | - |
| - | - | 7155 | 271.2 | - | - | 0 | - |
| - | - | 6948 | 271.2 | - | - | 0 | - |
| - | - | 4.551E+04 | 272.1 | - | - | 0 | - |
| - | - | 3.031E+05 | 273.1 | - | - | 0 | - |
| - | - | 3.489E+04 | 273.2 | - | - | 0 | - |
| - | - | 7.12E+04 | 274.1 | - | - | 0 | - |
| - | - | 5880 | 274.2 | - | - | 0 | - |
| - | - | 1.4E+04 | 275.1 | - | - | 0 | - |
| - | - | 5844 | 276.2 | - | - | 0 | - |
| - | - | 8817 | 281.2 | - | - | 0 | - |
| - | - | 3.54E+04 | 282.2 | - | - | 0 | - |
| - | - | 6036 | 285.2 | - | - | 0 | - |
| - | - | 9.708E+04 | 287.2 | - | - | 0 | - |
| - | - | 1.166E+04 | 288.1 | - | - | 0 | - |
| - | - | 7278 | 288.2 | - | - | 0 | - |
| - | - | 5428 | 290.1 | - | - | 0 | - |
| - | - | 1.664E+04 | 290.1 | - | - | 0 | - |
| - | - | 6.481E+04 | 291.1 | - | - | 0 | - |
| - | - | 5868 | 292.1 | - | - | 0 | - |
| - | - | 6362 | 299.2 | - | - | 0 | - |
| - | - | 5486 | 299.2 | - | - | 0 | - |
| - | - | 6979 | 299.2 | - | - | 0 | - |
| - | - | 2.591E+04 | 300.1 | - | - | 0 | - |
| - | - | 8098 | 301.1 | - | - | 0 | - |
| - | - | 1.509E+05 | 301.2 | - | - | 0 | - |
| - | - | 3.773E+04 | 302.2 | - | - | 0 | - |
| - | - | 7134 | 303.2 | - | - | 0 | - |
| - | - | 6.262E+04 | 304.2 | - | - | 0 | - |
| - | - | 9170 | 305.2 | - | - | 0 | - |
| - | - | 1.253E+04 | 307.2 | - | - | 0 | - |
| 4 | y | 1.932E+04 | 308.2 | 0.002988 | 9.697 | +2 | 7 |
| - | - | 1.975E+04 | 308.2 | - | - | 0 | - |
| - | - | 5935 | 309.2 | - | - | 0 | - |
| - | - | 6111 | 310.2 | - | - | 0 | - |
| - | - | 1.987E+04 | 313.2 | - | - | 0 | - |
| - | - | 7910 | 315.2 | - | - | 0 | - |
| - | - | 9844 | 316.2 | - | - | 0 | - |
| - | - | 9627 | 317.2 | - | - | 0 | - |
| - | - | 3.899E+04 | 317.2 | - | - | 0 | - |
| - | - | 1.615E+04 | 317.7 | - | - | 0 | - |
| - | - | 3.041E+04 | 318.1 | - | - | 0 | - |
| - | - | 1.137E+05 | 319.1 | - | - | 0 | - |
| - | - | 1.687E+04 | 320.1 | - | - | 0 | - |
| - | - | 6639 | 321.2 | - | - | 0 | - |
| - | - | 1.396E+04 | 322.2 | - | - | 0 | - |
| - | - | 1.056E+04 | 323.2 | - | - | 0 | - |
| - | - | 5.458E+04 | 325.2 | - | - | 0 | - |
| - | - | 2.985E+05 | 326.2 | - | - | 0 | - |
| - | - | 9155 | 326.2 | - | - | 0 | - |
| - | - | 4.04E+04 | 327.2 | - | - | 0 | - |
| - | - | 1.229E+04 | 328.1 | - | - | 0 | - |
| - | - | 2.017E+04 | 328.2 | - | - | 0 | - |
| - | - | 1.377E+04 | 329.2 | - | - | 0 | - |
| - | - | 6066 | 330.2 | - | - | 0 | - |
| - | - | 7528 | 331.1 | - | - | 0 | - |
| - | - | 2.78E+04 | 331.2 | - | - | 0 | - |
| - | - | 2.186E+04 | 332.2 | - | - | 0 | - |
| - | - | 4755 | 333.2 | - | - | 0 | - |
| - | - | 2.613E+04 | 333.2 | - | - | 0 | - |
| - | - | 8231 | 336.2 | - | - | 0 | - |
| - | - | 6982 | 338.2 | - | - | 0 | - |
| - | - | 6501 | 338.2 | - | - | 0 | - |
| - | - | 2.734E+04 | 339.2 | - | - | 0 | - |
| - | - | 7371 | 340.2 | - | - | 0 | - |
| - | - | 1.214E+04 | 341.2 | - | - | 0 | - |
| - | - | 4045 | 343.2 | - | - | 0 | - |
| 8 | y | 9.755E+04 | 343.2 | 0.0005762 | 1.679 | +1 | 3 |
| - | - | 8.573E+04 | 344.2 | - | - | 0 | - |
| 8 | y | 1.059E+04 | 344.2 | 0.001048 | 3.045 | +1 | 3 |
| - | - | 1.181E+04 | 344.2 | - | - | 0 | - |
| - | - | 1.23E+04 | 345.2 | - | - | 0 | - |
| - | - | 1.762E+04 | 345.2 | - | - | 0 | - |
| 3 | b | 1.564E+05 | 346.2 | 0.0003114 | 0.8994 | +1 | 3 |
| - | - | 8913 | 347.2 | - | - | 0 | - |
| - | - | 2.682E+04 | 347.2 | - | - | 0 | - |
| - | - | 5.4E+04 | 348.2 | - | - | 0 | - |
| - | - | 8249 | 349.2 | - | - | 0 | - |
| - | - | 1.311E+04 | 354.2 | - | - | 0 | - |
| - | - | 9814 | 356.2 | - | - | 0 | - |
| - | - | 7767 | 356.2 | - | - | 0 | - |
| - | - | 8.322E+04 | 357.2 | - | - | 0 | - |
| - | - | 1.772E+04 | 358.2 | - | - | 0 | - |
| - | - | 1.833E+04 | 358.2 | - | - | 0 | - |
| - | - | 2.403E+04 | 361.2 | - | - | 0 | - |
| 8 | y | 3.6E+05 | 361.2 | 0.0002349 | 0.6502 | +1 | 3 |
| - | - | 6.162E+04 | 362.2 | - | - | 0 | - |
| - | - | 6846 | 363.3 | - | - | 0 | - |
| - | - | 7095 | 364.2 | - | - | 0 | - |
| 8 | b | 9604 | 364.7 | 0.001036 | 2.841 | +2 | 8 |
| 8 | b | 4.674E+04 | 365.2 | 0.006719 | 18.4 | +2 | 8 |
| - | - | 8112 | 366.2 | - | - | 0 | - |
| - | - | 1.219E+04 | 367.7 | - | - | 0 | - |
| - | - | 2.105E+04 | 368.2 | - | - | 0 | - |
| - | - | 7252 | 369.2 | - | - | 0 | - |
| - | - | 1.028E+04 | 372.3 | - | - | 0 | - |
| - | - | 5442 | 372.7 | - | - | 0 | - |
| - | - | 8294 | 373.2 | - | - | 0 | - |
| 8 | b | 3.201E+04 | 373.7 | 0.0004315 | 1.155 | +2 | 8 |
| - | - | 1.147E+05 | 374.2 | - | - | 0 | - |
| - | - | 1.156E+04 | 374.2 | - | - | 0 | - |
| - | - | 6.242E+04 | 375.2 | - | - | 0 | - |
| - | - | 3.103E+04 | 375.2 | - | - | 0 | - |
| - | - | 1.774E+04 | 376.2 | - | - | 0 | - |
| - | - | 5537 | 376.2 | - | - | 0 | - |
| - | - | 5779 | 377.2 | - | - | 0 | - |
| 3 | y | 1.38E+05 | 381.7 | 0.0004022 | 1.054 | +2 | 8 |
| - | - | 5.205E+04 | 382.2 | - | - | 0 | - |
| - | - | 6.893E+04 | 382.2 | - | - | 0 | - |
| - | - | 8470 | 382.7 | - | - | 0 | - |
| - | - | 2.809E+05 | 383.2 | - | - | 0 | - |
| - | - | 7201 | 383.2 | - | - | 0 | - |
| - | - | 1.27E+04 | 384.2 | - | - | 0 | - |
| - | - | 4.956E+04 | 384.2 | - | - | 0 | - |
| - | - | 1.453E+04 | 385.1 | - | - | 0 | - |
| - | - | 1.868E+04 | 386.2 | - | - | 0 | - |
| - | - | 1.95E+04 | 387.2 | - | - | 0 | - |
| - | - | 5610 | 388.2 | - | - | 0 | - |
| - | - | 6804 | 389.7 | - | - | 0 | - |
| - | - | 1.513E+04 | 390.2 | - | - | 0 | - |
| 3 | y | 9.801E+05 | 390.7 | 0.0003994 | 1.022 | +2 | 8 |
| - | - | 4.293E+05 | 391.2 | - | - | 0 | - |
| - | - | 1.014E+05 | 391.7 | - | - | 0 | - |
| - | - | 1.479E+05 | 392.2 | - | - | 0 | - |
| - | - | 9251 | 392.2 | - | - | 0 | - |
| - | - | 3.923E+04 | 393.2 | - | - | 0 | - |
| - | - | 8278 | 394.2 | - | - | 0 | - |
| - | - | 1.106E+04 | 395.2 | - | - | 0 | - |
| - | - | 9.714E+04 | 400.3 | - | - | 0 | - |
| - | - | 2.384E+05 | 401.2 | - | - | 0 | - |
| - | - | 1.847E+04 | 401.3 | - | - | 0 | - |
| - | - | 4.097E+05 | 402.2 | - | - | 0 | - |
| - | - | 4.674E+04 | 402.2 | - | - | 0 | - |
| - | - | 8.39E+04 | 403.2 | - | - | 0 | - |
| 4 | b | 7.156E+04 | 403.2 | 0.0001489 | 0.3693 | +1 | 4 |
| - | - | 8906 | 404.2 | - | - | 0 | - |
| - | - | 1.768E+04 | 404.2 | - | - | 0 | - |
| - | - | 1.146E+04 | 411.3 | - | - | 0 | - |
| - | - | 1.35E+04 | 413.3 | - | - | 0 | - |
| - | - | 1.014E+04 | 415.2 | - | - | 0 | - |
| - | - | 2.392E+04 | 416.3 | - | - | 0 | - |
| - | - | 1.613E+04 | 416.8 | - | - | 0 | - |
| - | - | 1.926E+04 | 418.2 | - | - | 0 | - |
| - | - | 2.633E+04 | 419.2 | - | - | 0 | - |
| - | - | 2.402E+05 | 420.2 | - | - | 0 | - |
| - | - | 5.097E+04 | 421.2 | - | - | 0 | - |
| 9 | b | 2.157E+04 | 421.3 | 0.004872 | 11.57 | +2 | 9 |
| - | - | 7029 | 422.2 | - | - | 0 | - |
| - | - | 5.258E+04 | 429.3 | - | - | 0 | - |
| 9 | b | 1.159E+04 | 430.3 | 0.0002917 | 0.6779 | +2 | 9 |
| 2 | y | 5491 | 431.2 | 0.0001003 | 0.2326 | +2 | 9 |
| - | - | 8003 | 431.7 | - | - | 0 | - |
| - | - | 5856 | 432.2 | - | - | 0 | - |
| - | - | 2.258E+04 | 432.3 | - | - | 0 | - |
| - | - | 2.51E+04 | 437.2 | - | - | 0 | - |
| - | - | 9542 | 438.2 | - | - | 0 | - |
| - | - | 1.538E+05 | 439.3 | - | - | 0 | - |
| 2 | y | 4.345E+04 | 440.3 | 0.001074 | 2.44 | +2 | 9 |
| - | - | 1.298E+04 | 440.3 | - | - | 0 | - |
| - | - | 2.801E+04 | 440.8 | - | - | 0 | - |
| - | - | 1.035E+04 | 442.2 | - | - | 0 | - |
| 7 | y | 2.127E+04 | 444.3 | 0.0005967 | 1.343 | +1 | 4 |
| - | - | 1.309E+04 | 445.2 | - | - | 0 | - |
| - | - | 1.009E+04 | 451.3 | - | - | 0 | - |
| - | - | 1.95E+04 | 452.3 | - | - | 0 | - |
| - | - | 2.799E+05 | 457.3 | - | - | 0 | - |
| - | - | 6.384E+04 | 458.3 | - | - | 0 | - |
| - | - | 1.049E+04 | 459.3 | - | - | 0 | - |
| 5 | b | 2.453E+04 | 460.3 | 0.0003222 | 0.7 | +1 | 5 |
| - | - | 1.213E+04 | 461.3 | - | - | 0 | - |
| 7 | y | 8.186E+04 | 462.3 | 0.0005606 | 1.213 | +1 | 4 |
| - | - | 2.044E+04 | 463.3 | - | - | 0 | - |
| - | - | 5140 | 464.3 | - | - | 0 | - |
| - | - | 5635 | 468.2 | - | - | 0 | - |
| - | - | 7304 | 468.3 | - | - | 0 | - |
| - | - | 6857 | 469.3 | - | - | 0 | - |
| - | - | 1.243E+04 | 470.2 | - | - | 0 | - |
| - | - | 1.174E+04 | 470.3 | - | - | 0 | - |
| - | - | 7266 | 472.3 | - | - | 0 | - |
| - | - | 1.445E+04 | 473.3 | - | - | 0 | - |
| - | - | 2.957E+04 | 478.3 | - | - | 0 | - |
| - | - | 6333 | 479.3 | - | - | 0 | - |
| 0 | Precursor | 4.223E+04 | 480.8 | 0.0005613 | 1.167 | +2 | -1 |
| - | - | 2.178E+04 | 481.3 | - | - | 0 | - |
| - | - | 8198 | 483.3 | - | - | 0 | - |
| - | - | 2.253E+04 | 486.2 | - | - | 0 | - |
| - | - | 1.523E+05 | 486.3 | - | - | 0 | - |
| - | - | 7670 | 487.3 | - | - | 0 | - |
| - | - | 2.796E+04 | 487.3 | - | - | 0 | - |
| - | - | 1.005E+04 | 488.3 | - | - | 0 | - |
| - | - | 5410 | 488.3 | - | - | 0 | - |
| - | - | 7385 | 489.3 | - | - | 0 | - |
| 0 | Precursor | 5.37E+04 | 489.8 | 0.0004975 | 1.016 | +2 | -1 |
| - | - | 3.634E+04 | 490.3 | - | - | 0 | - |
| - | - | 8414 | 491.3 | - | - | 0 | - |
| - | - | 1.411E+04 | 495.2 | - | - | 0 | - |
| - | - | 2.93E+05 | 496.3 | - | - | 0 | - |
| - | - | 6.894E+04 | 497.3 | - | - | 0 | - |
| - | - | 5961 | 498.3 | - | - | 0 | - |
| - | - | 4.014E+04 | 501.2 | - | - | 0 | - |
| 6 | y | 2.099E+04 | 501.3 | 0.0003732 | 0.7444 | +1 | 5 |
| - | - | 7915 | 502.2 | - | - | 0 | - |
| - | - | 7.481E+04 | 512.3 | - | - | 0 | - |
| - | - | 1.905E+04 | 513.3 | - | - | 0 | - |
| - | - | 7.247E+05 | 514.3 | - | - | 0 | - |
| - | - | 1.682E+04 | 515.3 | - | - | 0 | - |
| - | - | 1.755E+05 | 515.3 | - | - | 0 | - |
| - | - | 3.187E+04 | 516.3 | - | - | 0 | - |
| 6 | b | 1.693E+04 | 517.3 | 0.001472 | 2.846 | +1 | 6 |
| - | - | 7613 | 518.3 | - | - | 0 | - |
| - | - | 1.35E+04 | 519.3 | - | - | 0 | - |
| 6 | y | 1.609E+05 | 519.3 | 0.0005202 | 1.002 | +1 | 5 |
| - | - | 3.807E+04 | 520.3 | - | - | 0 | - |
| - | - | 2.687E+05 | 530.3 | - | - | 0 | - |
| - | - | 6.669E+04 | 531.3 | - | - | 0 | - |
| - | - | 1.46E+04 | 532.3 | - | - | 0 | - |
| - | - | 1.208E+04 | 533.3 | - | - | 0 | - |
| - | - | 1.04E+04 | 540.3 | - | - | 0 | - |
| - | - | 5.441E+05 | 548.3 | - | - | 0 | - |
| - | - | 1.539E+05 | 549.3 | - | - | 0 | - |
| - | - | 2.601E+04 | 550.3 | - | - | 0 | - |
| 5 | y | 6.043E+04 | 558.3 | 0.0002413 | 0.4321 | +1 | 6 |
| - | - | 1.702E+04 | 559.3 | - | - | 0 | - |
| - | - | 5999 | 560.3 | - | - | 0 | - |
| - | - | 1.741E+04 | 571.4 | - | - | 0 | - |
| - | - | 5520 | 573.3 | - | - | 0 | - |
| 5 | y | 7.156E+05 | 576.3 | 0.0001136 | 0.1971 | +1 | 6 |
| - | - | 1.944E+05 | 577.3 | - | - | 0 | - |
| - | - | 4.189E+04 | 578.3 | - | - | 0 | - |
| - | - | 8399 | 587.4 | - | - | 0 | - |
| - | - | 9869 | 588.4 | - | - | 0 | - |
| - | - | 1.552E+04 | 590.3 | - | - | 0 | - |
| - | - | 2.527E+04 | 597.3 | - | - | 0 | - |
| - | - | 2.087E+04 | 598.3 | - | - | 0 | - |
| - | - | 3.025E+04 | 599.3 | - | - | 0 | - |
| 7 | b | 3.732E+04 | 600.3 | 0.001834 | 3.055 | +1 | 7 |
| - | - | 9553 | 608.3 | - | - | 0 | - |
| 4 | y | 1.301E+05 | 615.3 | 0.001788 | 2.905 | +1 | 7 |
| - | - | 5.972E+04 | 616.3 | - | - | 0 | - |
| - | - | 2.109E+04 | 617.3 | - | - | 0 | - |
| 7 | b | 6.577E+04 | 618.3 | 0.000125 | 0.2021 | +1 | 7 |
| - | - | 1.963E+04 | 619.3 | - | - | 0 | - |
| - | - | 5.585E+04 | 625.3 | - | - | 0 | - |
| - | - | 2.064E+04 | 626.3 | - | - | 0 | - |
| - | - | 6186 | 627.3 | - | - | 0 | - |
| - | - | 1.282E+04 | 629.3 | - | - | 0 | - |
| - | - | 6437 | 630.3 | - | - | 0 | - |
| - | - | 1.189E+04 | 631.3 | - | - | 0 | - |
| - | - | 7519 | 632.3 | - | - | 0 | - |
| 4 | y | 2.106E+06 | 633.4 | 0.001355 | 2.139 | +1 | 7 |
| - | - | 6.44E+05 | 634.4 | - | - | 0 | - |
| - | - | 1.157E+05 | 635.4 | - | - | 0 | - |
| - | - | 1.193E+04 | 636.4 | - | - | 0 | - |
| - | - | 5.275E+05 | 643.4 | - | - | 0 | - |
| - | - | 1.781E+05 | 644.4 | - | - | 0 | - |
| - | - | 4.062E+04 | 645.4 | - | - | 0 | - |
| - | - | 2.674E+04 | 647.4 | - | - | 0 | - |
| - | - | 9829 | 648.4 | - | - | 0 | - |
| - | - | 7990 | 649.3 | - | - | 0 | - |
| - | - | 1.171E+06 | 661.4 | - | - | 0 | - |
| - | - | 4.266E+05 | 662.4 | - | - | 0 | - |
| - | - | 8.694E+04 | 663.4 | - | - | 0 | - |
| - | - | 3.087E+04 | 679.4 | - | - | 0 | - |
| - | - | 7728 | 680.4 | - | - | 0 | - |
| - | - | 6201 | 710.4 | - | - | 0 | - |
| - | - | 7192 | 712.4 | - | - | 0 | - |
| - | - | 4.124E+04 | 718.4 | - | - | 0 | - |
| - | - | 1.905E+04 | 719.4 | - | - | 0 | - |
| - | - | 5494 | 723.4 | - | - | 0 | - |
| - | - | 6370 | 726.4 | - | - | 0 | - |
| 8 | b | 5.194E+04 | 728.4 | 0.001699 | 2.332 | +1 | 8 |
| - | - | 1.804E+04 | 729.4 | - | - | 0 | - |
| - | - | 9160 | 732.4 | - | - | 0 | - |
| - | - | 1.21E+04 | 733.4 | - | - | 0 | - |
| - | - | 2.648E+04 | 734.4 | - | - | 0 | - |
| - | - | 2.267E+04 | 735.4 | - | - | 0 | - |
| - | - | 6872 | 736.4 | - | - | 0 | - |
| - | - | 2.675E+04 | 742.4 | - | - | 0 | - |
| - | - | 9949 | 743.4 | - | - | 0 | - |
| - | - | 4.416E+04 | 744.4 | - | - | 0 | - |
| - | - | 2.512E+04 | 745.4 | - | - | 0 | - |
| 8 | b | 1.71E+05 | 746.4 | 0.0008497 | 1.138 | +1 | 8 |
| - | - | 7.704E+04 | 747.4 | - | - | 0 | - |
| - | - | 1.423E+04 | 748.4 | - | - | 0 | - |
| - | - | 7.469E+04 | 760.4 | - | - | 0 | - |
| - | - | 3.706E+04 | 761.4 | - | - | 0 | - |
| 3 | y | 2.053E+05 | 762.4 | 3.678E-05 | 0.04824 | +1 | 8 |
| - | - | 9.289E+04 | 763.4 | - | - | 0 | - |
| - | - | 1.707E+04 | 764.4 | - | - | 0 | - |
| - | - | 2.66E+04 | 778.4 | - | - | 0 | - |
| - | - | 1.048E+04 | 779.4 | - | - | 0 | - |
| 3 | y | 4.08E+06 | 780.4 | 0.0001645 | 0.2107 | +1 | 8 |
| - | - | 1.724E+06 | 781.4 | - | - | 0 | - |
| - | - | 4.363E+05 | 782.4 | - | - | 0 | - |
| - | - | 3.58E+04 | 783.4 | - | - | 0 | - |
| - | - | 1.494E+04 | 831.5 | - | - | 0 | - |
| - | - | 6790 | 832.5 | - | - | 0 | - |
| 9 | b | 4.236E+04 | 841.5 | 0.0008627 | 1.025 | +1 | 9 |
| - | - | 3.26E+04 | 842.5 | - | - | 0 | - |
| 9 | b | 2.604E+05 | 859.5 | 0.00038 | 0.4421 | +1 | 9 |
| - | - | 1.303E+05 | 860.5 | - | - | 0 | - |
| - | - | 3.803E+04 | 861.5 | - | - | 0 | - |
| - | - | 7439 | 862.5 | - | - | 0 | - |
| 2 | y | 2.816E+05 | 879.5 | 0.0009514 | 1.082 | +1 | 9 |
| - | - | 8194 | 880.4 | - | - | 0 | - |
| - | - | 1.292E+05 | 880.5 | - | - | 0 | - |
| - | - | 4.158E+04 | 881.5 | - | - | 0 | - |
| - | - | 6341 | 1207 | - | - | 0 | - |
| - | - | 5773 | 1237 | - | - | 0 | - |
| - | - | 5627 | 1247 | - | - | 0 | - |
| - | - | 5918 | 1436 | - | - | 0 | - |
| - | - | 5385 | 1902 | - | - | 0 | - |
| - | - | 5207 | 2168 | - | - | 0 | - |
| - | - | 5753 | 2367 | - | - | 0 | - |

m/z Charge Intensity FragmentType MassShift Position
120.06590270996094 0 423982.66 y 9
120.08120727539062 0 2857335.5
121.0225830078125 0 2876.7695
121.06272888183594 0 2837.0366
121.06912994384766 0 13493.763
121.08448028564453 0 220413.27
122.08771514892578 0 3994.063
127.05062103271484 0 19906.752
128.1072998046875 0 4437.9497
129.10264587402344 0 2240369
130.05027770996094 0 7615.725
130.0997772216797 0 16141.431
130.10595703125 0 134392.31
131.04946899414062 0 9132.974
131.08187866210938 0 81076.21
132.08106994628906 0 36384.97
136.07623291015625 0 4586.9385
139.08694458007812 0 12377.401
141.0662841796875 0 47657.895
141.1029052734375 0 8720.808
142.06944274902344 0 3648.032
142.12261962890625 0 4959.6016
143.08209228515625 0 4334.4175
143.15438842773438 0 8803.56
146.1289520263672 0 4740.667
148.95382690429688 0 9749.447
151.08702087402344 0 8159.77
152.1437530517578 0 20175.947
153.06626892089844 0 9850.788
154.06134033203125 0 6014.426
155.0818328857422 0 20437.904
155.11825561523438 0 61288.824
158.0927734375 0 22172.535
159.07675170898438 0 76376.15
160.11160278320312 0 4483.9272
165.10195922851562 0 7447.9326
167.11842346191406 0 14093.048
168.11337280273438 0 6866.634
169.13412475585938 0 8860.694
170.0927734375 0 44240.223
171.07711791992188 0 13566.494
171.14964294433594 0 5559517 a 1
172.0719451904297 0 78159.28
172.15292358398438 0 490261.97
173.15586853027344 0 22316.115
173.45216369628906 0 10273.017
174.12777709960938 0 14814.129
176.10728454589844 0 110868.08
177.10260009765625 0 261952.72
177.1102752685547 0 11506.521
178.10601806640625 0 32648.494
183.11314392089844 0 9620.342
183.14952087402344 0 9616.965
185.12876892089844 0 33107.457
186.12413024902344 0 26430.615
187.1446075439453 0 7137.656
188.07196044921875 0 4461.973
188.10342407226562 0 29606.121
188.14364624023438 0 4677.3486
189.10040283203125 0 7429.99
194.12928771972656 0 15783.702
195.14959716796875 0 9901.766
196.10818481445312 0 8094.031
197.1289825439453 0 5519.9277
197.16514587402344 0 100495.45
198.08763122558594 0 88805.89
198.16831970214844 0 7326.3545
199.14450073242188 0 2772460.5 b 1
200.14779663085938 0 266882.88
201.09825134277344 0 10523.267
201.14988708496094 0 18019.607
203.1181640625 0 27361.54
205.0974884033203 0 537977.06
206.100830078125 0 60132.57
206.129638671875 0 5470.121
207.14959716796875 0 38700.504
208.10842895507812 0 47498.11
209.1033172607422 0 15548.536
210.0876922607422 0 24759.037
211.14451599121094 0 17551.387
212.13970947265625 0 277842.1
213.1242218017578 0 6410.3203
213.14317321777344 0 30229.996
214.11854553222656 0 10198.336
214.1915740966797 0 19826.688
215.11483764648438 0 47595.66
215.13926696777344 0 51455.14 y Water loss 8
216.09817504882812 0 57918.65
217.09756469726562 0 55306.56
217.1331024169922 0 4222.4497
218.1011962890625 0 7557.212
219.14956665039062 0 253790.31
220.15283203125 0 31743.785
221.12876892089844 0 22769.254
221.15606689453125 0 6426.909
222.1248779296875 0 4981.801
224.17611694335938 0 95140.75
225.13511657714844 0 27574.035
225.16004943847656 0 25640.05
225.17913818359375 0 6691.226
227.11422729492188 0 111847.29
228.0982208251953 0 55787.656
228.10951232910156 0 12201.052
228.11973571777344 0 7174.2114
228.1343231201172 0 9676.67
229.0935821533203 0 18286.404
229.1322784423828 0 6126.3877
230.15032958984375 0 488423.94
231.11322021484375 0 35991.773
231.15333557128906 0 51709.766
233.12828063964844 0 26463.35
233.1499481201172 0 154840.97 y 8
233.16351318359375 0 8948.389
234.15350341796875 0 15369.312
237.09854125976562 0 32736.285
240.13467407226562 0 16279.041
241.1549530029297 0 13820.356
242.18670654296875 0 100076.24
243.11334228515625 0 54571.32
243.14578247070312 0 11491.817
243.17111206054688 0 7353.1147
243.1890106201172 0 7555.5327
244.1071014404297 0 9059.382
244.11915588378906 0 5602.8105
245.1249237060547 0 88619.15
245.16500854492188 0 10598.112
246.1273956298828 0 6012.2534
247.14443969726562 0 117398.85
248.1475372314453 0 18230.326
249.1244354248047 0 10653.308
251.1510772705078 0 25898.697 y Water loss 5
252.17105102539062 0 7250.7305
254.1249542236328 0 8113.1245
255.109130859375 0 270317.38
256.1122131347656 0 24612.836
256.1663513183594 0 14955.437
259.14398193359375 0 14388.273 b 5
260.19683837890625 0 13356.932
261.12347412109375 0 49335.082
261.1595153808594 0 7334.967
262.1189270019531 0 159356.12
263.121826171875 0 22688.07
263.1764221191406 0 5112.617
265.1291809082031 0 5357.4453
268.16412353515625 0 5586.087
269.1611633300781 0 273752.16
270.1642761230469 0 38303.418
270.18170166015625 0 31211.43
271.1078186035156 0 7722.788
271.140380859375 0 7846.0415
271.1668701171875 0 7155.0083
271.18450927734375 0 6947.739
272.1360778808594 0 45513.69
273.11962890625 0 303063.62
273.1961364746094 0 34893.902
274.1199951171875 0 71198.664
274.16455078125 0 5880.322
275.1232604980469 0 14002.117
276.15557861328125 0 5844.4834
281.19781494140625 0 8816.828
282.1564025878906 0 35401.37
285.1566467285156 0 6036.353
287.1718444824219 0 97084.97
288.1347351074219 0 11660.833
288.1741638183594 0 7277.8457
290.1134948730469 0 5427.945
290.1467590332031 0 16644.71
291.1456604003906 0 64808.14
292.1498718261719 0 5868.2153
299.1732482910156 0 6361.9077
299.2090148925781 0 5486.07
299.2441711425781 0 6979.1235
300.1343078613281 0 25914.326
301.13067626953125 0 8098.315
301.19146728515625 0 150876.98
302.1947937011719 0 37734.742
303.18243408203125 0 7133.831
304.1658935546875 0 62616.016
305.1676025390625 0 9169.705
307.21331787109375 0 12526.685
308.1736755371094 0 19320.4 y Water loss 3
308.1969909667969 0 19753.047
309.1571350097656 0 5935.3003
310.15142822265625 0 6110.66
313.1878967285156 0 19865.13
315.23968505859375 0 7909.721
316.1663818359375 0 9843.595
317.16156005859375 0 9626.79
317.18951416015625 0 38991.348
317.6906433105469 0 16146.698
318.1451416015625 0 30406.244
319.14068603515625 0 113686.24
320.1429138183594 0 16865.113
321.1911315917969 0 6639.2197
322.18212890625 0 13959.666
323.171875 0 10559.622
325.22406005859375 0 54579.215
326.1827697753906 0 298507.47
326.22784423828125 0 9155.382
327.1856689453125 0 40399.645
328.1293640136719 0 12293.619
328.2017822265625 0 20165.75
329.1617431640625 0 13769.119
330.1819152832031 0 6066.0415
331.140625 0 7528.1294
331.18719482421875 0 27798.65
332.1609802246094 0 21861.516
333.1715087890625 0 4755.439
333.1927185058594 0 26132.963
336.1663818359375 0 8230.733
338.18255615234375 0 6982.4995
338.2196960449219 0 6501.111
339.1781921386719 0 27335.139
340.1986083984375 0 7370.721
341.18328857421875 0 12137.535
343.2120056152344 0 4044.9211
343.23455810546875 0 97545.87 y Water loss 7
344.193359375 0 85725.93
344.2169494628906 0 10588.174 y Ammonia loss 7
344.2377624511719 0 11809.828
345.1551818847656 0 12297.196
345.19561767578125 0 17620.07
346.21282958984375 0 156409.05 b 2
347.1711730957031 0 8912.851
347.21588134765625 0 26824.791
348.1671142578125 0 53998.883
349.1690368652344 0 8248.694
354.177978515625 0 13112.72
356.1732177734375 0 9814.203
356.2300720214844 0 7766.868
357.156005859375 0 83220.5
358.1560974121094 0 17717.914
358.210693359375 0 18328.969
361.1878356933594 0 24033.193
361.2447814941406 0 359979.88 y 7
362.2477722167969 0 61620.97
363.2500305175781 0 6846.3857
364.2348937988281 0 7095.479
364.70916748046875 0 9603.84 b Water loss 7
365.19342041015625 0 46740.145 b Ammonia loss 7
366.1976013183594 0 8112.157
367.71295166015625 0 12191.263
368.1924743652344 0 21047.346
369.1952819824219 0 7251.73
372.2613525390625 0 10284.38
372.70684814453125 0 5442.4824
373.1873779296875 0 8294.495
373.7129821777344 0 32008.412 b 7
374.1826171875 0 114708.21
374.2131042480469 0 11564.65
375.1671447753906 0 62420.23
375.23956298828125 0 31026.826
376.1640625 0 17738.174
376.24261474609375 0 5537.1035
377.1641845703125 0 5778.6636
381.7112731933594 0 138040.66 y Water loss 2
382.2121276855469 0 52046.195
382.24505615234375 0 68931.195
382.7146301269531 0 8469.923
383.2040710449219 0 280929.75
383.2479553222656 0 7200.66
384.1660461425781 0 12697.875
384.20721435546875 0 49563.71
385.14971923828125 0 14531.74
386.2040710449219 0 18684.174
387.2042541503906 0 19498.152
388.2039489746094 0 5609.647
389.7101745605469 0 6803.6494
390.2142639160156 0 15126.042
390.716552734375 0 980142.75 y 2
391.2179870605469 0 429308.9
391.71923828125 0 101428.5
392.1931457519531 0 147939.84
392.218994140625 0 9251.338
393.1949462890625 0 39232.12
394.244873046875 0 8277.918
395.2401428222656 0 11060.894
400.2558898925781 0 97142.36
401.2147216796875 0 238357.39
401.2611083984375 0 18472.678
402.17767333984375 0 409696.6
402.21697998046875 0 46741.07
403.1806335449219 0 83902.4
403.234130859375 0 71561.125 b 3
404.1845397949219 0 8906.233
404.2353210449219 0 17683.861
411.2716369628906 0 11459.568
413.250732421875 0 13499.203
415.2322998046875 0 10137.741
416.25885009765625 0 23919.668
416.75946044921875 0 16126.672
418.2091369628906 0 19255.74
419.2043762207031 0 26326.055
420.1883544921875 0 240232.22
421.19158935546875 0 50967.01
421.2550354003906 0 21569.893 b Water loss 8
422.1938781738281 0 7028.7505
429.28240966796875 0 52579.03
430.2557373046875 0 11585.099 b 8
431.24517822265625 0 5490.693 y Water loss 1
431.7467041015625 0 8002.613
432.22796630859375 0 5856.229
432.2611389160156 0 22577.436
437.2147521972656 0 25097.023
438.2183532714844 0 9541.875
439.2667541503906 0 153764.67
440.2514343261719 0 43453.195 y 1
440.275634765625 0 12984.932
440.7528381347656 0 28007.004
442.24737548828125 0 10353.828
444.2822570800781 0 21265.506 y Water loss 6
445.2417907714844 0 13090.204
451.2667236328125 0 10093.732
452.2620849609375 0 19504.709
457.2774353027344 0 279927.6
458.28045654296875 0 63844.152
459.2831726074219 0 10492.278
460.2557678222656 0 24525.006 b 4
461.2540588378906 0 12132.412
462.29278564453125 0 81859.914 y 6
463.2956237792969 0 20442.527
464.3025207519531 0 5139.55
468.2331848144531 0 5634.5293
468.29364013671875 0 7304.067
469.2767028808594 0 6856.828
470.2403564453125 0 12426.855
470.27386474609375 0 11744.23
472.25604248046875 0 7266.38
473.2539367675781 0 14454.85
478.2779235839844 0 29566.062
479.26544189453125 0 6333.1777
480.77984619140625 0 42227.996 Precursor Water loss
481.28204345703125 0 21783.766
483.2935791015625 0 8197.74
486.24615478515625 0 22530.277
486.3038330078125 0 152282.86
487.2696533203125 0 7669.801
487.3067932128906 0 27955.65
488.2508544921875 0 10047.051
488.3119812011719 0 5410.3467
489.28179931640625 0 7385.301
489.7850646972656 0 53697.992 Precursor
490.2861328125 0 36343.46
491.2611389160156 0 8414.134
495.2358703613281 0 14108.114
496.2883605957031 0 293026.8
497.2910461425781 0 68938.9
498.2935791015625 0 5960.51
501.2457580566406 0 40138.98
501.3034973144531 0 20988.588 y Water loss 5
502.2477722167969 0 7914.8154
512.2619018554688 0 74810.6
513.2627563476562 0 19045.309
514.298583984375 0 724681.2
515.261474609375 0 16819.408
515.3014526367188 0 175503.95
516.3040771484375 0 31866.06
517.2783813476562 0 16932.371 b 5
518.2793579101562 0 7612.562
519.256591796875 0 13501.737
519.314208984375 0 160859.53 y 5
520.3170166015625 0 38072.715
530.2724609375 0 268691.34
531.2752685546875 0 66685.86
532.3099365234375 0 14597.614
533.2721557617188 0 12084.682
540.3158569335938 0 10395.231
548.2830200195312 0 544114
549.2855834960938 0 153886.25
550.2891845703125 0 26013.338
558.3248291015625 0 60434.812 y Water loss 4
559.3275146484375 0 17015.549
560.3261108398438 0 5998.7686
571.3555908203125 0 17411.07
573.301025390625 0 5520.3193
576.3352661132812 0 715637.9 y 4
577.3385009765625 0 194376.33
578.3399658203125 0 41892.504
587.3500366210938 0 8398.664
588.3557739257812 0 9868.881
590.3311767578125 0 15524.817
597.3360595703125 0 25265.625
598.3303833007812 0 20872.82
599.33251953125 0 30245.39
600.3158569335938 0 37319.97 b Water loss 6
608.3187255859375 0 9552.64
615.3478393554688 0 130078.53 y Water loss 3
616.347900390625 0 59718.92
617.3433227539062 0 21090.889
618.324462890625 0 65770.15 b 6
619.3291015625 0 19634.172
625.3457641601562 0 55848.457
626.3436889648438 0 20636.541
627.3358764648438 0 6186.288
629.341796875 0 12822.755
630.3406982421875 0 6436.624
631.3362426757812 0 11886.168
632.3468627929688 0 7519.161
633.3579711914062 0 2105947.8 y 3
634.3607788085938 0 644030.7
635.3630981445312 0 115672.97
636.3639526367188 0 11932.225
643.35595703125 0 527523.5
644.3590087890625 0 178057.38
645.361572265625 0 40618.883
647.3519287109375 0 26736.812
648.3572387695312 0 9829.08
649.34033203125 0 7989.701
661.366943359375 0 1171225.4
662.369873046875 0 426641.22
663.372314453125 0 86941.85
679.376953125 0 30872.875
680.3814086914062 0 7727.7134
710.3980102539062 0 6200.548
712.4371948242188 0 7192.265
718.4232177734375 0 41239.293
719.4273681640625 0 19050.193
723.362548828125 0 5493.5464
726.3925170898438 0 6370.144
728.4072875976562 0 51944.902 b Water loss 7
729.4122314453125 0 18041.004
732.4417114257812 0 9159.824
733.44140625 0 12098.269
734.4203491210938 0 26475.535
735.4194946289062 0 22673.504
736.4017944335938 0 6871.9688
742.4236450195312 0 26752.352
743.4307250976562 0 9949.284
744.4055786132812 0 44159.633
745.400390625 0 25117.828
746.418701171875 0 170986.19 b 7
747.421630859375 0 77039.88
748.4252319335938 0 14228.027
760.4349975585938 0 74687.91
761.4368896484375 0 37062.996
762.4144287109375 0 205252.95 y Water loss 2
763.4156494140625 0 92885.7
764.4195556640625 0 17070.045
778.4094848632812 0 26600.453
779.4147338867188 0 10483.798
780.4248657226562 0 4080250.5 y 2
781.427734375 0 1723802
782.4301147460938 0 436330.6
783.431884765625 0 35802.914
831.5104370117188 0 14944.164
832.50830078125 0 6790.014
841.4921875 0 42359.81 b Water loss 8
842.4956665039062 0 32596.674
859.5032348632812 0 260371.25 b 8
860.505615234375 0 130345.984
861.5028076171875 0 38027.062
862.488525390625 0 7438.7764
879.4924926757812 0 281636.2 y 1
880.4072265625 0 8193.634
880.4949340820312 0 129237.79
881.4984130859375 0 41578.06
1207.4229736328125 0 6340.8784
1236.8106689453125 0 5773.26
1246.908447265625 0 5627.002
1435.970703125 0 5917.7554
1902.4832763671875 0 5384.525
2167.99755859375 0 5207.333
2366.6806640625 0 5752.8887

Spectrum Details

|  |  |
| --- | --- |
| Matched peaks? Matched peaks The total absolute number of peaks matched. Additionally in brackets the total fraction of peaks matched and the total number of peaks is shown. | 43 (9.29% of 463) |
| FDR? FDR The false discovery rate estimated for this peptide. It is calculated by matching all theoretical fragments with a non-integer shift with the raw peaks for this spectrum. This is done with 40 different shifts. The resulting percentage is the average number of annotated peaks over the number of annotated peaks with the correct spectrum. | 0.06% |
| Satellite FDR? Satellite FDR See the FDR for details on its calculation. This satellite ion specific FDR only contains the satellite ions (d/w) for I/L/J positions. | - |
| PSM Score? PSM Score The PSM Score as given by Hecklib to this annotated spectrum. It is shown with three significant figures. | 472 |

#### Spectrum 5744? Spectrum 5744 The raw spectrum of this peptide as annotated by Hecklib. The fragments are coloured according to ion type (see legend). Any peaks with a star '\*' as text can be hovered over to see the full details, first the ion type second the mass shift type. By hovering over the amino acids in the peptide or ions in the legend the corresponding peaks are highlighted. By toggling the 'Unassigned' label you can turn the background (unassigned) peaks on or off in the plot. By updating the slider in the Ion legend you can update the spectrum to only show the top X% of the peaks with labels. The top X% means any peak that is within X% of the highest intensity. By dragging in the spectrum you can zoom in to a specific part of the spectrum and use 'Zoom Out' to get back to the original zoom level. The annotation of the spectrum is based on the given sequence in the peptides file and is done with different software so inconsistencies are likely. The peaks are annotated based on the given sequence, with 20 ppm tolerance.

Copy Data

##### Spectrum 5744 (TSV)

###### Preview

```
Loading example...
```

*Click on the button to copy the data to your clipboard.*

Mz MinMz MaxIntensity Max

WidthHeightPeptide font sizePeptide stroke widthSpectrum font sizeSpectrum stroke widthCompact peptide

Ion legend

wxyz

abcd

OtherUnassignedIon^Charge^_Position_Show for top:%

VVFGGGTKJT

01.02e+52.04e+53.06e+54.07e+5

Zoom Out

y^+1^_1_y^+1^_2_y^+1^_2_z^+1^_3_y^+1^_3_c^+2^_8_y^+2^_8_z^+2^_8_y^+2^_8_w^+1^_4_c^+2^_9_y^+2^_9_y^+2^_9_y^+1^_4_z^+1^_4_y^+1^_4_c^+1^_5_y^+1^_5_z^+1^_5_y^+1^_5_c^+1^_6_z^+1^_6_y^+1^_6_y^+1^_6_z^+1^_6_y^+1^_6_z^+1^_7_z^+1^_7_y^+1^_7_y^+1^_7_c^+1^_7_y^+1^_7_c^+1^_7_c^+1^_8_y^+1^_8_c^+1^_8_z^+1^_8_y^+1^_8_z^+1^_9_w^+1^_9_c^+1^_9_z^+1^_9_c^+1^_9_y^+1^_9_

0775155123263101

Fragment Matches Table

Show background peaks

| Position | Ion type | Intensity | mz Theoretical | mz Error (Th) | mz Error (ppm) | Charge | Series Number |
| --- | --- | --- | --- | --- | --- | --- | --- |
| 10 | y | 1.948E+04 | 120.1 | 0.0003831 | 3.191 | +1 | 1 |
| - | - | 2.796E+04 | 120.1 | - | - | 0 | - |
| - | - | 624.6 | 121.1 | - | - | 0 | - |
| - | - | 2647 | 121.1 | - | - | 0 | - |
| - | - | 5893 | 129.1 | - | - | 0 | - |
| - | - | 729.8 | 133.1 | - | - | 0 | - |
| - | - | 1792 | 136.1 | - | - | 0 | - |
| - | - | 1042 | 149 | - | - | 0 | - |
| - | - | 413.9 | 149.4 | - | - | 0 | - |
| - | - | 1.659E+05 | 171.1 | - | - | 0 | - |
| - | - | 1151 | 172.1 | - | - | 0 | - |
| - | - | 1.604E+04 | 172.2 | - | - | 0 | - |
| - | - | 558.1 | 173.2 | - | - | 0 | - |
| - | - | 2188 | 177.1 | - | - | 0 | - |
| - | - | 816.3 | 177.1 | - | - | 0 | - |
| - | - | 478 | 177.1 | - | - | 0 | - |
| - | - | 556.6 | 179.2 | - | - | 0 | - |
| - | - | 654.5 | 197.1 | - | - | 0 | - |
| - | - | 480.4 | 198.8 | - | - | 0 | - |
| - | - | 1.526E+05 | 199.1 | - | - | 0 | - |
| - | - | 885.3 | 200.1 | - | - | 0 | - |
| - | - | 1.559E+04 | 200.1 | - | - | 0 | - |
| - | - | 1.404E+04 | 205.1 | - | - | 0 | - |
| - | - | 1522 | 206.1 | - | - | 0 | - |
| - | - | 1035 | 211.1 | - | - | 0 | - |
| 9 | y | 2607 | 215.1 | 0.0005227 | 2.43 | +1 | 2 |
| - | - | 2275 | 230.2 | - | - | 0 | - |
| 9 | y | 8933 | 233.1 | 0.0005171 | 2.218 | +1 | 2 |
| - | - | 1124 | 234.2 | - | - | 0 | - |
| - | - | 581.4 | 235.4 | - | - | 0 | - |
| - | - | 737.6 | 243.1 | - | - | 0 | - |
| - | - | 622.8 | 245 | - | - | 0 | - |
| - | - | 4864 | 247.1 | - | - | 0 | - |
| - | - | 660.4 | 256.2 | - | - | 0 | - |
| - | - | 754 | 258.1 | - | - | 0 | - |
| - | - | 2016 | 262.1 | - | - | 0 | - |
| - | - | 562.3 | 262.2 | - | - | 0 | - |
| - | - | 1700 | 273.1 | - | - | 0 | - |
| - | - | 2690 | 276.2 | - | - | 0 | - |
| - | - | 1169 | 287.2 | - | - | 0 | - |
| - | - | 1889 | 292.2 | - | - | 0 | - |
| - | - | 594.7 | 293.1 | - | - | 0 | - |
| - | - | 543 | 297.1 | - | - | 0 | - |
| - | - | 7823 | 301.2 | - | - | 0 | - |
| - | - | 1586 | 302.2 | - | - | 0 | - |
| - | - | 2102 | 304.2 | - | - | 0 | - |
| - | - | 737.5 | 308.2 | - | - | 0 | - |
| - | - | 3478 | 310.2 | - | - | 0 | - |
| - | - | 592.6 | 311.2 | - | - | 0 | - |
| - | - | 879.5 | 313.2 | - | - | 0 | - |
| - | - | 1763 | 317.2 | - | - | 0 | - |
| - | - | 1514 | 319.1 | - | - | 0 | - |
| - | - | 769.2 | 322.2 | - | - | 0 | - |
| - | - | 541.7 | 325.2 | - | - | 0 | - |
| - | - | 1184 | 326.2 | - | - | 0 | - |
| - | - | 1613 | 331.2 | - | - | 0 | - |
| - | - | 1399 | 332.2 | - | - | 0 | - |
| - | - | 1971 | 344.2 | - | - | 0 | - |
| 8 | z | 8258 | 345.2 | 0.0006179 | 1.79 | +1 | 3 |
| - | - | 9758 | 346.2 | - | - | 0 | - |
| - | - | 1.872E+04 | 346.2 | - | - | 0 | - |
| - | - | 2076 | 347.2 | - | - | 0 | - |
| - | - | 3745 | 347.2 | - | - | 0 | - |
| - | - | 1377 | 358.2 | - | - | 0 | - |
| - | - | 654.1 | 358.2 | - | - | 0 | - |
| - | - | 1107 | 359.2 | - | - | 0 | - |
| - | - | 560.5 | 359.3 | - | - | 0 | - |
| - | - | 1213 | 361.2 | - | - | 0 | - |
| 8 | y | 6484 | 361.2 | 0.0006316 | 1.748 | +1 | 3 |
| - | - | 1228 | 362.2 | - | - | 0 | - |
| - | - | 769.4 | 364.7 | - | - | 0 | - |
| - | - | 709.4 | 367.7 | - | - | 0 | - |
| - | - | 932.4 | 368.2 | - | - | 0 | - |
| 8 | c | 2263 | 373.7 | 8.732E-05 | 0.2336 | +2 | 8 |
| - | - | 1482 | 374.2 | - | - | 0 | - |
| - | - | 1793 | 375.2 | - | - | 0 | - |
| - | - | 659.6 | 380.7 | - | - | 0 | - |
| 3 | y | 9037 | 381.7 | 0.0009515 | 2.493 | +2 | 8 |
| - | - | 2698 | 382.2 | - | - | 0 | - |
| 3 | z | 1006 | 382.7 | 0.004818 | 12.59 | +2 | 8 |
| - | - | 1031 | 383.2 | - | - | 0 | - |
| - | - | 661.7 | 385.2 | - | - | 0 | - |
| - | - | 559.2 | 386.2 | - | - | 0 | - |
| 3 | y | 8.644E+04 | 390.7 | 0.0006436 | 1.647 | +2 | 8 |
| - | - | 3.652E+04 | 391.2 | - | - | 0 | - |
| - | - | 7263 | 391.7 | - | - | 0 | - |
| - | - | 770.4 | 392.2 | - | - | 0 | - |
| - | - | 1116 | 400.3 | - | - | 0 | - |
| - | - | 5560 | 401.2 | - | - | 0 | - |
| - | - | 3423 | 402.2 | - | - | 0 | - |
| - | - | 1097 | 402.2 | - | - | 0 | - |
| - | - | 6052 | 402.3 | - | - | 0 | - |
| - | - | 6292 | 403.2 | - | - | 0 | - |
| - | - | 1386 | 403.3 | - | - | 0 | - |
| - | - | 1085 | 404.2 | - | - | 0 | - |
| - | - | 1164 | 415.3 | - | - | 0 | - |
| - | - | 1356 | 415.3 | - | - | 0 | - |
| - | - | 1346 | 416.3 | - | - | 0 | - |
| - | - | 689.1 | 416.8 | - | - | 0 | - |
| - | - | 4079 | 420.2 | - | - | 0 | - |
| - | - | 883.9 | 421.2 | - | - | 0 | - |
| 7 | w | 651.4 | 429.3 | 0.001455 | 3.39 | +1 | 4 |
| 9 | c | 969.3 | 430.3 | 0.000841 | 1.955 | +2 | 9 |
| 2 | y | 721.4 | 431.2 | 0.0003575 | 0.8289 | +2 | 9 |
| - | - | 2791 | 437.2 | - | - | 0 | - |
| - | - | 1139 | 439.3 | - | - | 0 | - |
| 2 | y | 5191 | 440.3 | 0.001196 | 2.717 | +2 | 9 |
| - | - | 1494 | 440.8 | - | - | 0 | - |
| - | - | 1588 | 441.3 | - | - | 0 | - |
| - | - | 2664 | 442.3 | - | - | 0 | - |
| - | - | 1.198E+04 | 444.3 | - | - | 0 | - |
| 7 | y | 2.424E+04 | 445.3 | 0.0006204 | 1.393 | +1 | 4 |
| 7 | z | 5569 | 446.3 | 0.003604 | 8.075 | +1 | 4 |
| - | - | 3011 | 447.2 | - | - | 0 | - |
| - | - | 2.267E+04 | 447.3 | - | - | 0 | - |
| - | - | 1846 | 448.2 | - | - | 0 | - |
| - | - | 4279 | 448.3 | - | - | 0 | - |
| - | - | 787.5 | 449.3 | - | - | 0 | - |
| - | - | 1600 | 454.2 | - | - | 0 | - |
| - | - | 920.5 | 455.2 | - | - | 0 | - |
| - | - | 4119 | 457.3 | - | - | 0 | - |
| - | - | 768.8 | 458.3 | - | - | 0 | - |
| - | - | 4641 | 459.3 | - | - | 0 | - |
| - | - | 2932 | 460.3 | - | - | 0 | - |
| - | - | 1438 | 460.3 | - | - | 0 | - |
| 7 | y | 5138 | 462.3 | 0.001079 | 2.335 | +1 | 4 |
| - | - | 1011 | 463.3 | - | - | 0 | - |
| - | - | 4310 | 470.3 | - | - | 0 | - |
| - | - | 1303 | 471.3 | - | - | 0 | - |
| - | - | 2473 | 472.3 | - | - | 0 | - |
| - | - | 1693 | 473.3 | - | - | 0 | - |
| - | - | 922.2 | 473.3 | - | - | 0 | - |
| - | - | 5150 | 476.3 | - | - | 0 | - |
| 5 | c | 1.269E+04 | 477.3 | 0.0007201 | 1.509 | +1 | 5 |
| - | - | 2718 | 478.3 | - | - | 0 | - |
| - | - | 3647 | 480.8 | - | - | 0 | - |
| - | - | 3293 | 481.3 | - | - | 0 | - |
| - | - | 923.7 | 481.8 | - | - | 0 | - |
| - | - | 1363 | 486.2 | - | - | 0 | - |
| - | - | 729.7 | 486.3 | - | - | 0 | - |
| - | - | 798.8 | 487.3 | - | - | 0 | - |
| - | - | 766.6 | 489.3 | - | - | 0 | - |
| - | - | 8235 | 489.8 | - | - | 0 | - |
| - | - | 6987 | 490.3 | - | - | 0 | - |
| - | - | 1411 | 490.8 | - | - | 0 | - |
| - | - | 2266 | 491.3 | - | - | 0 | - |
| - | - | 2739 | 496.3 | - | - | 0 | - |
| - | - | 887.7 | 498.3 | - | - | 0 | - |
| - | - | 1161 | 498.3 | - | - | 0 | - |
| - | - | 5376 | 499.3 | - | - | 0 | - |
| - | - | 1090 | 500.3 | - | - | 0 | - |
| 6 | y | 590.4 | 501.3 | 0.002815 | 5.615 | +1 | 5 |
| 6 | z | 1.011E+04 | 503.3 | 0.0009337 | 1.855 | +1 | 5 |
| - | - | 1.176E+05 | 504.3 | - | - | 0 | - |
| - | - | 1.615E+05 | 504.3 | - | - | 0 | - |
| - | - | 3.032E+04 | 505.3 | - | - | 0 | - |
| - | - | 4.164E+04 | 505.3 | - | - | 0 | - |
| - | - | 6137 | 506.3 | - | - | 0 | - |
| - | - | 5518 | 506.3 | - | - | 0 | - |
| - | - | 806.5 | 512.3 | - | - | 0 | - |
| - | - | 1.037E+04 | 514.3 | - | - | 0 | - |
| - | - | 2445 | 515.3 | - | - | 0 | - |
| - | - | 6545 | 516.3 | - | - | 0 | - |
| - | - | 3025 | 517.3 | - | - | 0 | - |
| - | - | 1569 | 517.3 | - | - | 0 | - |
| - | - | 5716 | 518.3 | - | - | 0 | - |
| 6 | y | 1.41E+04 | 519.3 | 0.0005812 | 1.119 | +1 | 5 |
| - | - | 3522 | 520.3 | - | - | 0 | - |
| - | - | 679.5 | 521.3 | - | - | 0 | - |
| - | - | 570.4 | 523.1 | - | - | 0 | - |
| - | - | 2744 | 530.3 | - | - | 0 | - |
| - | - | 611.8 | 532.3 | - | - | 0 | - |
| - | - | 7855 | 533.3 | - | - | 0 | - |
| 6 | c | 6455 | 534.3 | 0.000602 | 1.127 | +1 | 6 |
| - | - | 1758 | 535.3 | - | - | 0 | - |
| 5 | z | 2490 | 542.3 | 0.0005659 | 1.044 | +1 | 6 |
| - | - | 1793 | 543.3 | - | - | 0 | - |
| - | - | 1231 | 543.3 | - | - | 0 | - |
| - | - | 630.9 | 544.3 | - | - | 0 | - |
| - | - | 1.493E+04 | 548.3 | - | - | 0 | - |
| - | - | 4303 | 549.3 | - | - | 0 | - |
| - | - | 812.5 | 550.3 | - | - | 0 | - |
| - | - | 1571 | 555.3 | - | - | 0 | - |
| - | - | 678 | 556.3 | - | - | 0 | - |
| 5 | y | 2140 | 558.3 | 0.0009794 | 1.754 | +1 | 6 |
| 5 | y | 831.5 | 559.3 | 0.00469 | 8.385 | +1 | 6 |
| - | - | 750.2 | 560.3 | - | - | 0 | - |
| - | - | 674.6 | 560.3 | - | - | 0 | - |
| 5 | z | 4.3E+04 | 560.3 | 0.0009543 | 1.703 | +1 | 6 |
| - | - | 4718 | 561.3 | - | - | 0 | - |
| - | - | 1.253E+05 | 561.3 | - | - | 0 | - |
| - | - | 3.391E+04 | 562.3 | - | - | 0 | - |
| - | - | 5300 | 563.3 | - | - | 0 | - |
| - | - | 893.3 | 573.3 | - | - | 0 | - |
| - | - | 5378 | 574.3 | - | - | 0 | - |
| - | - | 926.7 | 575.3 | - | - | 0 | - |
| - | - | 1.893E+04 | 575.3 | - | - | 0 | - |
| 5 | y | 4.583E+04 | 576.3 | 0.0001746 | 0.303 | +1 | 6 |
| - | - | 1.243E+04 | 577.3 | - | - | 0 | - |
| - | - | 1527 | 578.3 | - | - | 0 | - |
| - | - | 776.8 | 590.3 | - | - | 0 | - |
| - | - | 1664 | 592.3 | - | - | 0 | - |
| 4 | z | 1588 | 599.3 | 0.001591 | 2.655 | +1 | 7 |
| 4 | z | 2973 | 600.3 | 0.008054 | 13.42 | +1 | 7 |
| - | - | 762.5 | 606.3 | - | - | 0 | - |
| - | - | 1086 | 614.3 | - | - | 0 | - |
| 4 | y | 3557 | 615.3 | 0.0002264 | 0.3679 | +1 | 7 |
| 4 | y | 2511 | 616.3 | 0.005931 | 9.624 | +1 | 7 |
| 7 | c | 2.717E+04 | 617.3 | 0.001522 | 2.465 | +1 | 7 |
| - | - | 7.455E+04 | 618.3 | - | - | 0 | - |
| - | - | 2.404E+04 | 619.4 | - | - | 0 | - |
| - | - | 4200 | 620.4 | - | - | 0 | - |
| - | - | 677.8 | 630.3 | - | - | 0 | - |
| - | - | 1186 | 631.3 | - | - | 0 | - |
| - | - | 854.1 | 632.3 | - | - | 0 | - |
| - | - | 4.488E+04 | 632.3 | - | - | 0 | - |
| 4 | y | 1.946E+05 | 633.4 | 0.0008667 | 1.368 | +1 | 7 |
| - | - | 7.375E+04 | 634.4 | - | - | 0 | - |
| 7 | c | 2.47E+05 | 635.4 | 0.001158 | 1.823 | +1 | 7 |
| - | - | 8.621E+04 | 636.4 | - | - | 0 | - |
| - | - | 1.598E+04 | 637.4 | - | - | 0 | - |
| - | - | 1499 | 638.4 | - | - | 0 | - |
| - | - | 8435 | 643.4 | - | - | 0 | - |
| - | - | 2853 | 644.4 | - | - | 0 | - |
| - | - | 798.5 | 645.4 | - | - | 0 | - |
| - | - | 3301 | 646.3 | - | - | 0 | - |
| - | - | 2012 | 647.4 | - | - | 0 | - |
| - | - | 1075 | 659.3 | - | - | 0 | - |
| - | - | 1964 | 660.3 | - | - | 0 | - |
| - | - | 4.026E+04 | 661.4 | - | - | 0 | - |
| - | - | 1.384E+04 | 662.4 | - | - | 0 | - |
| - | - | 1.633E+04 | 663.4 | - | - | 0 | - |
| - | - | 5574 | 664.4 | - | - | 0 | - |
| - | - | 1022 | 685.4 | - | - | 0 | - |
| - | - | 946.9 | 690.3 | - | - | 0 | - |
| - | - | 1915 | 693.3 | - | - | 0 | - |
| - | - | 1354 | 694.3 | - | - | 0 | - |
| - | - | 6164 | 702.4 | - | - | 0 | - |
| - | - | 2957 | 703.4 | - | - | 0 | - |
| - | - | 3723 | 708.3 | - | - | 0 | - |
| - | - | 1311 | 709.3 | - | - | 0 | - |
| - | - | 3489 | 718.4 | - | - | 0 | - |
| - | - | 5518 | 719.4 | - | - | 0 | - |
| - | - | 1209 | 720.4 | - | - | 0 | - |
| - | - | 997.5 | 720.4 | - | - | 0 | - |
| - | - | 2385 | 721.4 | - | - | 0 | - |
| - | - | 2493 | 728.4 | - | - | 0 | - |
| - | - | 1.365E+04 | 729.4 | - | - | 0 | - |
| - | - | 5220 | 730.4 | - | - | 0 | - |
| - | - | 1652 | 731.4 | - | - | 0 | - |
| - | - | 1901 | 734.4 | - | - | 0 | - |
| - | - | 760.8 | 735.4 | - | - | 0 | - |
| - | - | 1685 | 745.4 | - | - | 0 | - |
| 8 | c | 1.706E+04 | 746.4 | 0.004878 | 6.535 | +1 | 8 |
| - | - | 9172 | 747.4 | - | - | 0 | - |
| - | - | 2011 | 748.4 | - | - | 0 | - |
| - | - | 1777 | 760.4 | - | - | 0 | - |
| - | - | 1873 | 761.4 | - | - | 0 | - |
| 3 | y | 9387 | 762.4 | 0.003381 | 4.435 | +1 | 8 |
| 8 | c | 2.97E+05 | 763.4 | 0.0008605 | 1.127 | +1 | 8 |
| 3 | z | 3.447E+04 | 764.4 | 0.00854 | 11.17 | +1 | 8 |
| - | - | 9.573E+04 | 764.5 | - | - | 0 | - |
| - | - | 2.393E+04 | 765.4 | - | - | 0 | - |
| - | - | 1.639E+04 | 765.5 | - | - | 0 | - |
| - | - | 6896 | 766.4 | - | - | 0 | - |
| - | - | 917.3 | 767.4 | - | - | 0 | - |
| - | - | 1924 | 778.4 | - | - | 0 | - |
| - | - | 1420 | 779.4 | - | - | 0 | - |
| 3 | y | 2.896E+05 | 780.4 | 0.001117 | 1.432 | +1 | 8 |
| - | - | 1.236E+05 | 781.4 | - | - | 0 | - |
| - | - | 2.936E+04 | 782.4 | - | - | 0 | - |
| - | - | 2075 | 783.4 | - | - | 0 | - |
| - | - | 3805 | 789.5 | - | - | 0 | - |
| - | - | 2110 | 790.5 | - | - | 0 | - |
| - | - | 4609 | 807.4 | - | - | 0 | - |
| - | - | 1919 | 808.4 | - | - | 0 | - |
| - | - | 3170 | 817.5 | - | - | 0 | - |
| - | - | 1605 | 818.5 | - | - | 0 | - |
| - | - | 1084 | 831.5 | - | - | 0 | - |
| - | - | 3.194E+04 | 832.5 | - | - | 0 | - |
| - | - | 1.675E+04 | 833.5 | - | - | 0 | - |
| - | - | 4178 | 834.5 | - | - | 0 | - |
| - | - | 1021 | 835.5 | - | - | 0 | - |
| - | - | 2320 | 841.5 | - | - | 0 | - |
| 2 | z | 872.6 | 845.5 | 0.003313 | 3.918 | +1 | 9 |
| 2 | w | 1143 | 848.5 | 0.002735 | 3.223 | +1 | 9 |
| 9 | c | 2.152E+04 | 859.5 | 0.0009018 | 1.049 | +1 | 9 |
| - | - | 1.08E+04 | 860.5 | - | - | 0 | - |
| - | - | 1.133E+04 | 861.5 | - | - | 0 | - |
| - | - | 4225 | 862.5 | - | - | 0 | - |
| 2 | z | 3.658E+04 | 863.5 | 0.0007438 | 0.8615 | +1 | 9 |
| - | - | 2.193E+04 | 864.5 | - | - | 0 | - |
| - | - | 7861 | 865.5 | - | - | 0 | - |
| - | - | 1929 | 874.5 | - | - | 0 | - |
| - | - | 1410 | 875.5 | - | - | 0 | - |
| 9 | c | 3.156E+05 | 876.5 | 0.0007809 | 0.8909 | +1 | 9 |
| - | - | 1.602E+05 | 877.5 | - | - | 0 | - |
| - | - | 4.442E+04 | 878.5 | - | - | 0 | - |
| 2 | y | 1.172E+04 | 879.5 | 0.004053 | 4.609 | +1 | 9 |
| - | - | 6398 | 880.5 | - | - | 0 | - |
| - | - | 1631 | 881.5 | - | - | 0 | - |
| - | - | 870.6 | 889.4 | - | - | 0 | - |
| - | - | 1287 | 900.5 | - | - | 0 | - |
| - | - | 5466 | 906.5 | - | - | 0 | - |
| - | - | 2794 | 907.5 | - | - | 0 | - |
| - | - | 998.8 | 908.5 | - | - | 0 | - |
| - | - | 1697 | 916.5 | - | - | 0 | - |
| - | - | 1147 | 917.5 | - | - | 0 | - |
| - | - | 1407 | 918.5 | - | - | 0 | - |
| - | - | 3603 | 919.5 | - | - | 0 | - |
| - | - | 1661 | 920.5 | - | - | 0 | - |
| - | - | 1.615E+04 | 923.5 | - | - | 0 | - |
| - | - | 7530 | 924.5 | - | - | 0 | - |
| - | - | 1713 | 925.5 | - | - | 0 | - |
| - | - | 799.5 | 932.6 | - | - | 0 | - |
| - | - | 4573 | 933.6 | - | - | 0 | - |
| - | - | 1601 | 934.6 | - | - | 0 | - |
| - | - | 800.5 | 935.6 | - | - | 0 | - |
| - | - | 847.7 | 936.5 | - | - | 0 | - |
| - | - | 1195 | 944.5 | - | - | 0 | - |
| - | - | 688.8 | 945.5 | - | - | 0 | - |
| - | - | 5488 | 947.5 | - | - | 0 | - |
| - | - | 1734 | 948.5 | - | - | 0 | - |
| - | - | 2224 | 951.6 | - | - | 0 | - |
| - | - | 1202 | 952.6 | - | - | 0 | - |
| - | - | 2661 | 960.5 | - | - | 0 | - |
| - | - | 6450 | 961.5 | - | - | 0 | - |
| - | - | 4.033E+05 | 962.5 | - | - | 0 | - |
| - | - | 2.144E+05 | 963.5 | - | - | 0 | - |
| - | - | 6.731E+04 | 964.5 | - | - | 0 | - |
| - | - | 8159 | 965.6 | - | - | 0 | - |
| - | - | 1597 | 976.5 | - | - | 0 | - |
| - | - | 733.5 | 977.5 | - | - | 0 | - |
| - | - | 5638 | 977.6 | - | - | 0 | - |
| - | - | 2.103E+05 | 978.6 | - | - | 0 | - |
| - | - | 2.715E+05 | 979.6 | - | - | 0 | - |
| - | - | 1.211E+05 | 980.6 | - | - | 0 | - |
| - | - | 3.33E+04 | 981.6 | - | - | 0 | - |
| - | - | 1035 | 982.5 | - | - | 0 | - |
| - | - | 712.9 | 982.5 | - | - | 0 | - |
| - | - | 3389 | 982.6 | - | - | 0 | - |
| - | - | 902.5 | 994.5 | - | - | 0 | - |
| - | - | 820.5 | 995.5 | - | - | 0 | - |
| - | - | 671.1 | 1453 | - | - | 0 | - |
| - | - | 868.3 | 3070 | - | - | 0 | - |

m/z Charge Intensity FragmentType MassShift Position
120.06590270996094 0 19479.293 y 9
120.08118438720703 0 27960.06
121.06893920898438 0 624.6173
121.08462524414062 0 2647.0166
129.1026153564453 0 5892.967
133.08621215820312 0 729.8323
136.07615661621094 0 1792.0742
148.9542999267578 0 1042.4749
149.4427032470703 0 413.8793
171.14967346191406 0 165869.61
172.1451873779297 0 1150.7023
172.15298461914062 0 16044.504
173.1564483642578 0 558.05237
177.1027069091797 0 2187.9343
177.1120147705078 0 816.2574
177.12075805664062 0 477.9821
179.1608123779297 0 556.6048
197.12802124023438 0 654.5181
198.78404235839844 0 480.3886
199.14456176757812 0 152624
200.1380615234375 0 885.3131
200.14788818359375 0 15590.027
205.0976104736328 0 14042.424
206.10096740722656 0 1521.8215
211.14439392089844 0 1035.3008
215.13954162597656 0 2606.8235 y Water loss 8
230.15036010742188 0 2275.154
233.1501007080078 0 8932.957 y 8
234.15394592285156 0 1124.2048
235.36451721191406 0 581.4488
243.13389587402344 0 737.60406
245.0293731689453 0 622.7771
247.1443634033203 0 4863.633
256.1659240722656 0 660.39905
258.1084289550781 0 753.95996
262.1191711425781 0 2015.9148
262.16278076171875 0 562.2789
273.1199951171875 0 1699.7476
276.1557312011719 0 2690.2925
287.1715087890625 0 1169.1467
292.202880859375 0 1888.8467
293.07244873046875 0 594.684
297.1111755371094 0 543.04614
301.191650390625 0 7822.596
302.1947021484375 0 1586.0425
304.16680908203125 0 2101.773
308.1959228515625 0 737.52783
310.2131652832031 0 3477.8416
311.1705627441406 0 592.56946
313.1880798339844 0 879.49963
317.18951416015625 0 1763.4678
319.1408386230469 0 1513.9371
322.18157958984375 0 769.19476
325.2251281738281 0 541.6674
326.1835632324219 0 1184.4583
331.18798828125 0 1613.283
332.1614685058594 0 1398.9077
344.1939392089844 0 1971.1443
345.2264404296875 0 8257.903 z 7
346.2126159667969 0 9757.73
346.2343444824219 0 18724.664
347.21588134765625 0 2075.5725
347.2375793457031 0 3744.626
358.2092590332031 0 1376.9012
358.2462158203125 0 654.12714
359.2165832519531 0 1106.519
359.2655029296875 0 560.5326
361.1878967285156 0 1213.3728
361.24517822265625 0 6483.6514 y 7
362.2496643066406 0 1228.1848
364.7091064453125 0 769.39355
367.7145690917969 0 709.41833
368.1916198730469 0 932.43414
373.7135009765625 0 2262.9795 c Ammonia loss 7
374.2148742675781 0 1481.7244
375.2391052246094 0 1792.8136
380.720947265625 0 659.596
381.7118225097656 0 9037.341 y Water loss 2
382.2130126953125 0 2698.2603
382.71160888671875 0 1005.8988 z 2
383.2040100097656 0 1030.7433
385.2437438964844 0 661.69617
386.2044677734375 0 559.2295
390.716796875 0 86438.81 y 2
391.2183837890625 0 36522.547
391.719970703125 0 7263.4834
392.2203063964844 0 770.4131
400.2563171386719 0 1115.5679
401.2151794433594 0 5559.539
402.17791748046875 0 3422.5508
402.2198791503906 0 1096.6663
402.2716979980469 0 6052.1123
403.2342224121094 0 6291.9575
403.2749328613281 0 1385.6389
404.2360534667969 0 1084.6661
415.2547302246094 0 1164.0156
415.2816467285156 0 1355.5635
416.2605895996094 0 1346.1517
416.7613525390625 0 689.14294
420.18902587890625 0 4079.4456
421.1913757324219 0 883.9029
429.272216796875 0 651.42554 w 6
430.25628662109375 0 969.28345 c Ammonia loss 8
431.2447204589844 0 721.4319 y Water loss 1
437.21588134765625 0 2790.7913
439.267333984375 0 1139.3363
440.2515563964844 0 5191.4165 y 1
440.753173828125 0 1494.3418
441.2563171386719 0 1588.036
442.2601623535156 0 2663.5723
444.259033203125 0 11980.479
445.26629638671875 0 24240.58 y Ammonia loss 6
446.2698974609375 0 5568.7437 z 6
447.23248291015625 0 3011.344
447.2821350097656 0 22674.15
448.2397766113281 0 1845.7981
448.2850646972656 0 4278.569
449.28887939453125 0 787.5289
454.234375 0 1600.281
455.2392883300781 0 920.54626
457.2780456542969 0 4119.0723
458.28125 0 768.8378
459.29351806640625 0 4641.012
460.2553405761719 0 2932.4
460.2958984375 0 1438.3864
462.2933044433594 0 5137.7676 y 6
463.2965393066406 0 1010.8162
470.2620849609375 0 4310.2153
471.2643737792969 0 1302.7037
472.302001953125 0 2472.805
473.2603454589844 0 1692.6512
473.3026428222656 0 922.2454
476.2749938964844 0 5149.7886
477.28271484375 0 12688.596 c 4
478.28643798828125 0 2717.6213
480.77984619140625 0 3646.9834
481.2812194824219 0 3293.1448
481.7828063964844 0 923.6585
486.2433166503906 0 1363.3563
486.296142578125 0 729.74286
487.3092041015625 0 798.75903
489.2772216796875 0 766.615
489.7857360839844 0 8234.644
490.2873229980469 0 6987.1187
490.7870788574219 0 1410.5502
491.2946472167969 0 2265.7444
496.2889404296875 0 2739.134
498.2779846191406 0 887.6556
498.31903076171875 0 1161.2944
499.2791442871094 0 5375.565
500.2820739746094 0 1090.4916
501.3059387207031 0 590.35175 y Water loss 5
503.2958984375 0 10106.288 z 5
504.2549743652344 0 117606.01
504.3035888671875 0 161494.6
505.258544921875 0 30315.725
505.3067321777344 0 41641.164
506.2610168457031 0 6136.6514
506.3090515136719 0 5517.683
512.283447265625 0 806.47546
514.2992553710938 0 10371.369
515.3026733398438 0 2444.879
516.3145141601562 0 6544.8164
517.2705078125 0 3024.9448
517.3189697265625 0 1568.9102
518.3074951171875 0 5715.718
519.3142700195312 0 14097.442 y 5
520.3167114257812 0 3521.9678
521.3194580078125 0 679.4652
523.0609130859375 0 570.38824
530.2733154296875 0 2743.6982
532.3155517578125 0 611.78595
533.2964477539062 0 7854.5454
534.3028564453125 0 6455.1147 c 5
535.30615234375 0 1757.5759
542.3052978515625 0 2490.1938 z Water loss 4
543.26416015625 0 1793.3119
543.3126220703125 0 1231.3954
544.2730712890625 0 630.86755
548.2833862304688 0 14933.792
549.2872314453125 0 4302.797
550.2893676757812 0 812.4893
555.339111328125 0 1571.0481
556.3411865234375 0 677.98486
558.3236083984375 0 2140.4048 y Water loss 4
559.3132934570312 0 831.5035 y Ammonia loss 4
560.265625 0 750.1608
560.2789306640625 0 674.5958
560.3173828125 0 42999.504 z 4
561.2787475585938 0 4717.5728
561.3247680664062 0 125310.25
562.3277587890625 0 33912.668
563.3305053710938 0 5299.7114
573.3231201171875 0 893.30914
574.283935546875 0 5378.0425
575.2882080078125 0 926.73724
575.3283081054688 0 18934.186
576.3353271484375 0 45833.035 y 4
577.3385009765625 0 12428.337
578.3402709960938 0 1527.1002
590.3329467773438 0 776.8196
592.3458251953125 0 1664.431
599.3289184570312 0 1588.178 z Water loss 3
600.3193969726562 0 2973.213 z Ammonia loss 3
606.3169555664062 0 762.4933
614.3380737304688 0 1085.6592
615.3458251953125 0 3557.315 y Water loss 3
616.3359985351562 0 2511.146 y Ammonia loss 3
617.3390502929688 0 27165.842 c Water loss 6
618.3453369140625 0 74546.92
619.3502807617188 0 24037.053
620.3573608398438 0 4200.1855
630.332275390625 0 677.84094
631.339111328125 0 1186.479
632.2868041992188 0 854.149
632.349853515625 0 44877.06
633.3574829101562 0 194559.03 y 3
634.3555908203125 0 73754.28
635.352294921875 0 246981.62 c 6
636.3551635742188 0 86207.9
637.3574829101562 0 15983.56
638.3609619140625 0 1499.4359
643.3566284179688 0 8435.406
644.3601684570312 0 2853.4683
645.3572998046875 0 798.50385
646.3441162109375 0 3301.1055
647.3502807617188 0 2011.5688
659.3482666015625 0 1074.5791
660.3474731445312 0 1964.1919
661.3675537109375 0 40255.145
662.3704833984375 0 13842.557
663.3817138671875 0 16329.601
664.3862915039062 0 5574.3887
685.3876342773438 0 1022.344
690.3316650390625 0 946.87524
693.3326416015625 0 1915.4484
694.336181640625 0 1353.6577
702.4086303710938 0 6163.8643
703.4100341796875 0 2956.598
708.3441772460938 0 3722.9612
709.3471069335938 0 1311.2141
718.4238891601562 0 3488.5054
719.4315795898438 0 5517.6426
720.3760375976562 0 1209.2153
720.4403686523438 0 997.50385
721.353515625 0 2385.1462
728.4102783203125 0 2493.299
729.4176635742188 0 13645.966
730.420166015625 0 5220.4478
731.4248046875 0 1651.7762
734.420166015625 0 1900.9508
735.4196166992188 0 760.83344
745.3970947265625 0 1685.4912
746.4146728515625 0 17056.096 c Ammonia loss 7
747.4202880859375 0 9171.626
748.424072265625 0 2011.2017
760.4354858398438 0 1776.8688
761.4345092773438 0 1872.8423
762.4178466796875 0 9387.073 y Water loss 2
763.4469604492188 0 296976.1 c 7
764.3977661132812 0 34470.746 z 2
764.450439453125 0 95729.93
765.4078369140625 0 23933.066
765.4590454101562 0 16393.94
766.41357421875 0 6896.0815
767.4136962890625 0 917.31085
778.4099731445312 0 1923.8788
779.4127807617188 0 1419.7301
780.4261474609375 0 289626.1 y 2
781.4290771484375 0 123570.78
782.4315795898438 0 29357.531
783.4339599609375 0 2074.5576
789.4620361328125 0 3805.0466
790.4657592773438 0 2110.2708
807.413818359375 0 4608.8853
808.4179077148438 0 1919.2826
817.5057373046875 0 3169.643
818.5082397460938 0 1605.2526
831.5115356445312 0 1084.352
832.5170288085938 0 31941.44
833.5203247070312 0 16748.178
834.5230712890625 0 4177.518
835.51220703125 0 1020.8113
841.4952392578125 0 2319.5205
845.4674682617188 0 872.59424 z Water loss 1
848.4539794921875 0 1142.6527 w 1
859.5045166015625 0 21522.803 c Ammonia loss 8
860.5074462890625 0 10800.674
861.51708984375 0 11326.89
862.5217895507812 0 4224.7847
863.4754638671875 0 36575.258 z 1
864.4786376953125 0 21934.043
865.4810180664062 0 7860.934
874.5173950195312 0 1929.0793
875.5232543945312 0 1410.3046
876.5309448242188 0 315611.28 c 8
877.5339965820312 0 160249.52
878.5369262695312 0 44417.883
879.4974975585938 0 11719.969 y 1
880.4985961914062 0 6397.8
881.5018920898438 0 1631.3616
889.43115234375 0 870.57806
900.5433959960938 0 1286.5532
906.4818115234375 0 5465.542
907.484619140625 0 2794.3394
908.496337890625 0 998.7725
916.538818359375 0 1696.6287
917.545654296875 0 1147.0481
918.51904296875 0 1406.5431
919.536376953125 0 3602.8826
920.5352172851562 0 1661.0032
923.507568359375 0 16145.276
924.510009765625 0 7530.227
925.5112915039062 0 1712.7218
932.5503540039062 0 799.5484
933.5634155273438 0 4573.456
934.5634155273438 0 1600.9169
935.5535278320312 0 800.502
936.5053100585938 0 847.74634
944.5394897460938 0 1194.8474
945.5330810546875 0 688.79596
947.5202026367188 0 5488.4785
948.5294799804688 0 1733.8213
951.5774536132812 0 2224.4392
952.5772094726562 0 1201.7019
960.5284423828125 0 2661.0054
961.548095703125 0 6449.978
962.54443359375 0 403311.8
963.5469970703125 0 214360.08
964.5496215820312 0 67312.62
965.5523071289062 0 8159.213
976.533935546875 0 1597.3978
977.4586791992188 0 733.5169
977.5551147460938 0 5637.6953
978.562744140625 0 210343.53
979.5686645507812 0 271515
980.572021484375 0 121109.78
981.5750732421875 0 33303.617
982.4793701171875 0 1035.1333
982.5131225585938 0 712.9306
982.5774536132812 0 3389.205
994.5316772460938 0 902.5026
995.5433349609375 0 820.49225
1452.7684326171875 0 671.0975
3070.46875 0 868.2661

Spectrum Details

|  |  |
| --- | --- |
| Matched peaks? Matched peaks The total absolute number of peaks matched. Additionally in brackets the total fraction of peaks matched and the total number of peaks is shown. | 44 (12.75% of 345) |
| FDR? FDR The false discovery rate estimated for this peptide. It is calculated by matching all theoretical fragments with a non-integer shift with the raw peaks for this spectrum. This is done with 40 different shifts. The resulting percentage is the average number of annotated peaks over the number of annotated peaks with the correct spectrum. | 0.22% |
| Satellite FDR? Satellite FDR See the FDR for details on its calculation. This satellite ion specific FDR only contains the satellite ions (d/w) for I/L/J positions. | - |
| PSM Score? PSM Score The PSM Score as given by Hecklib to this annotated spectrum. It is shown with three significant figures. | 523 |

#### Spectrum 6036? Spectrum 6036 The raw spectrum of this peptide as annotated by Hecklib. The fragments are coloured according to ion type (see legend). Any peaks with a star '\*' as text can be hovered over to see the full details, first the ion type second the mass shift type. By hovering over the amino acids in the peptide or ions in the legend the corresponding peaks are highlighted. By toggling the 'Unassigned' label you can turn the background (unassigned) peaks on or off in the plot. By updating the slider in the Ion legend you can update the spectrum to only show the top X% of the peaks with labels. The top X% means any peak that is within X% of the highest intensity. By dragging in the spectrum you can zoom in to a specific part of the spectrum and use 'Zoom Out' to get back to the original zoom level. The annotation of the spectrum is based on the given sequence in the peptides file and is done with different software so inconsistencies are likely. The peaks are annotated based on the given sequence, with 20 ppm tolerance.

Copy Data

##### Spectrum 6036 (TSV)

###### Preview

```
Loading example...
```

*Click on the button to copy the data to your clipboard.*

Mz MinMz MaxIntensity Max

WidthHeightPeptide font sizePeptide stroke widthSpectrum font sizeSpectrum stroke widthCompact peptide

Ion legend

wxyz

abcd

OtherUnassignedIon^Charge^_Position_Show for top:%

VVFGGGTKJT

04.00e+48.01e+41.20e+51.60e+5

Zoom Out

y^+1^_1_a^+1^_2_b^+1^_2_y^+1^_2_y^+1^_2_y^+2^_5_y^+1^_3_b^+1^_3_y^+1^_3_b^+2^_8_b^+2^_8_y^+2^_8_y^+2^_8_b^+1^_4_b^+2^_9_y^+2^_9_b^+1^_5_y^+1^_4_\*\*y^+1^_5_y^+1^_6_y^+1^_6_b^+1^_7_y^+1^_7_b^+1^_7_y^+1^_7_b^+1^_8_b^+1^_8_y^+1^_8_y^+1^_8_b^+1^_9_b^+1^_9_y^+1^_9_

0601120218032404

Fragment Matches Table

Show background peaks

| Position | Ion type | Intensity | mz Theoretical | mz Error (Th) | mz Error (ppm) | Charge | Series Number |
| --- | --- | --- | --- | --- | --- | --- | --- |
| 10 | y | 1.237E+04 | 120.1 | 0.0004441 | 3.699 | +1 | 1 |
| - | - | 8.635E+04 | 120.1 | - | - | 0 | - |
| - | - | 518.9 | 121.1 | - | - | 0 | - |
| - | - | 556.4 | 121.1 | - | - | 0 | - |
| - | - | 6734 | 121.1 | - | - | 0 | - |
| - | - | 1271 | 127.1 | - | - | 0 | - |
| - | - | 407.6 | 127.1 | - | - | 0 | - |
| - | - | 374.3 | 127.3 | - | - | 0 | - |
| - | - | 610.9 | 128.1 | - | - | 0 | - |
| - | - | 861.3 | 128.1 | - | - | 0 | - |
| - | - | 7.041E+04 | 129.1 | - | - | 0 | - |
| - | - | 727.7 | 130.1 | - | - | 0 | - |
| - | - | 3298 | 130.1 | - | - | 0 | - |
| - | - | 2133 | 131.1 | - | - | 0 | - |
| - | - | 1476 | 132.1 | - | - | 0 | - |
| - | - | 513.8 | 133.1 | - | - | 0 | - |
| - | - | 2792 | 133.1 | - | - | 0 | - |
| - | - | 1.335E+04 | 136.1 | - | - | 0 | - |
| - | - | 714.2 | 139.1 | - | - | 0 | - |
| - | - | 595.5 | 140.1 | - | - | 0 | - |
| - | - | 1510 | 141.1 | - | - | 0 | - |
| - | - | 500.5 | 141.1 | - | - | 0 | - |
| - | - | 1002 | 142.1 | - | - | 0 | - |
| - | - | 475.1 | 143.1 | - | - | 0 | - |
| - | - | 2498 | 145.1 | - | - | 0 | - |
| - | - | 1045 | 146.1 | - | - | 0 | - |
| - | - | 815.2 | 149 | - | - | 0 | - |
| - | - | 835.5 | 152.1 | - | - | 0 | - |
| - | - | 849.2 | 152.1 | - | - | 0 | - |
| - | - | 452.6 | 154.1 | - | - | 0 | - |
| - | - | 496.9 | 155.1 | - | - | 0 | - |
| - | - | 1973 | 155.1 | - | - | 0 | - |
| - | - | 1387 | 158.1 | - | - | 0 | - |
| - | - | 1655 | 159.1 | - | - | 0 | - |
| - | - | 503.4 | 160.9 | - | - | 0 | - |
| - | - | 1218 | 163.1 | - | - | 0 | - |
| - | - | 466.3 | 163.9 | - | - | 0 | - |
| - | - | 1848 | 165.1 | - | - | 0 | - |
| - | - | 763.8 | 166.1 | - | - | 0 | - |
| - | - | 1003 | 167.1 | - | - | 0 | - |
| - | - | 743.4 | 169.1 | - | - | 0 | - |
| - | - | 1167 | 170.1 | - | - | 0 | - |
| - | - | 659.5 | 171.1 | - | - | 0 | - |
| 2 | a | 1.586E+05 | 171.1 | 0.0005143 | 3.005 | +1 | 2 |
| - | - | 3094 | 172.1 | - | - | 0 | - |
| - | - | 693.3 | 172.1 | - | - | 0 | - |
| - | - | 1.578E+04 | 172.2 | - | - | 0 | - |
| - | - | 1114 | 173.1 | - | - | 0 | - |
| - | - | 515.8 | 173.2 | - | - | 0 | - |
| - | - | 1655 | 173.4 | - | - | 0 | - |
| - | - | 502.9 | 174.1 | - | - | 0 | - |
| - | - | 3357 | 176.1 | - | - | 0 | - |
| - | - | 7661 | 177.1 | - | - | 0 | - |
| - | - | 1312 | 177.1 | - | - | 0 | - |
| - | - | 867.9 | 178.1 | - | - | 0 | - |
| - | - | 1164 | 181.2 | - | - | 0 | - |
| - | - | 1037 | 183.1 | - | - | 0 | - |
| - | - | 673.1 | 183.1 | - | - | 0 | - |
| - | - | 873.5 | 185.1 | - | - | 0 | - |
| - | - | 1742 | 185.2 | - | - | 0 | - |
| - | - | 638.8 | 186.1 | - | - | 0 | - |
| - | - | 661.8 | 187.1 | - | - | 0 | - |
| - | - | 2282 | 187.1 | - | - | 0 | - |
| - | - | 787.1 | 188.1 | - | - | 0 | - |
| - | - | 559 | 189.1 | - | - | 0 | - |
| - | - | 823 | 195.1 | - | - | 0 | - |
| - | - | 526.9 | 197.1 | - | - | 0 | - |
| - | - | 1516 | 197.1 | - | - | 0 | - |
| - | - | 2907 | 197.2 | - | - | 0 | - |
| - | - | 2376 | 198.1 | - | - | 0 | - |
| - | - | 639.2 | 198.2 | - | - | 0 | - |
| 2 | b | 7.817E+04 | 199.1 | 0.0004575 | 2.297 | +1 | 2 |
| - | - | 7839 | 200.1 | - | - | 0 | - |
| - | - | 498.7 | 201.1 | - | - | 0 | - |
| - | - | 1440 | 201.1 | - | - | 0 | - |
| - | - | 828.1 | 201.1 | - | - | 0 | - |
| - | - | 1.571E+04 | 205.1 | - | - | 0 | - |
| - | - | 1207 | 206.1 | - | - | 0 | - |
| - | - | 851.6 | 207.1 | - | - | 0 | - |
| - | - | 880.9 | 208.1 | - | - | 0 | - |
| - | - | 672.3 | 209.1 | - | - | 0 | - |
| - | - | 1013 | 209.2 | - | - | 0 | - |
| - | - | 603.8 | 209.9 | - | - | 0 | - |
| - | - | 701.2 | 210.1 | - | - | 0 | - |
| - | - | 1082 | 211.1 | - | - | 0 | - |
| - | - | 6777 | 212.1 | - | - | 0 | - |
| - | - | 545.7 | 213.1 | - | - | 0 | - |
| - | - | 1014 | 213.1 | - | - | 0 | - |
| - | - | 1031 | 213.2 | - | - | 0 | - |
| - | - | 697.8 | 215.1 | - | - | 0 | - |
| 9 | y | 4150 | 215.1 | 0.0004617 | 2.146 | +1 | 2 |
| - | - | 1726 | 216.1 | - | - | 0 | - |
| - | - | 634.8 | 216.1 | - | - | 0 | - |
| - | - | 1431 | 217.1 | - | - | 0 | - |
| - | - | 7369 | 219.1 | - | - | 0 | - |
| - | - | 916.1 | 220.2 | - | - | 0 | - |
| - | - | 615.8 | 221.1 | - | - | 0 | - |
| - | - | 2361 | 224.2 | - | - | 0 | - |
| - | - | 639 | 225.1 | - | - | 0 | - |
| - | - | 885.9 | 225.1 | - | - | 0 | - |
| - | - | 3124 | 227.1 | - | - | 0 | - |
| - | - | 1642 | 228.1 | - | - | 0 | - |
| - | - | 2479 | 229.2 | - | - | 0 | - |
| - | - | 1.438E+04 | 230.2 | - | - | 0 | - |
| - | - | 722.8 | 230.2 | - | - | 0 | - |
| - | - | 1163 | 231.1 | - | - | 0 | - |
| - | - | 1523 | 231.2 | - | - | 0 | - |
| - | - | 901.4 | 233.1 | - | - | 0 | - |
| - | - | 882.6 | 233.1 | - | - | 0 | - |
| 9 | y | 5288 | 233.1 | 0.0005781 | 2.48 | +1 | 2 |
| - | - | 1412 | 233.2 | - | - | 0 | - |
| - | - | 574.9 | 234.1 | - | - | 0 | - |
| - | - | 657.7 | 240.1 | - | - | 0 | - |
| - | - | 828.6 | 241.2 | - | - | 0 | - |
| - | - | 2240 | 242.2 | - | - | 0 | - |
| - | - | 1549 | 243.1 | - | - | 0 | - |
| - | - | 549.1 | 243.2 | - | - | 0 | - |
| - | - | 768 | 244.1 | - | - | 0 | - |
| - | - | 2016 | 245.1 | - | - | 0 | - |
| - | - | 3522 | 247.1 | - | - | 0 | - |
| 6 | y | 967.9 | 251.2 | 0.004093 | 16.29 | +2 | 5 |
| - | - | 569.3 | 253.9 | - | - | 0 | - |
| - | - | 7549 | 255.1 | - | - | 0 | - |
| - | - | 622.3 | 256.1 | - | - | 0 | - |
| - | - | 1346 | 261.1 | - | - | 0 | - |
| - | - | 5219 | 262.1 | - | - | 0 | - |
| - | - | 1692 | 269.1 | - | - | 0 | - |
| - | - | 7066 | 269.2 | - | - | 0 | - |
| - | - | 788.6 | 270.2 | - | - | 0 | - |
| - | - | 978.5 | 270.2 | - | - | 0 | - |
| - | - | 611.4 | 271.1 | - | - | 0 | - |
| - | - | 1052 | 272.1 | - | - | 0 | - |
| - | - | 8946 | 273.1 | - | - | 0 | - |
| - | - | 1041 | 273.2 | - | - | 0 | - |
| - | - | 1711 | 274.1 | - | - | 0 | - |
| - | - | 2835 | 276.2 | - | - | 0 | - |
| - | - | 924 | 282.2 | - | - | 0 | - |
| - | - | 1039 | 283.1 | - | - | 0 | - |
| - | - | 532.2 | 287.1 | - | - | 0 | - |
| - | - | 2747 | 287.2 | - | - | 0 | - |
| - | - | 1412 | 291.1 | - | - | 0 | - |
| - | - | 648.1 | 298.1 | - | - | 0 | - |
| - | - | 3562 | 301.2 | - | - | 0 | - |
| - | - | 1618 | 304.2 | - | - | 0 | - |
| - | - | 3474 | 310.2 | - | - | 0 | - |
| - | - | 540.5 | 317.2 | - | - | 0 | - |
| - | - | 667.3 | 318.1 | - | - | 0 | - |
| - | - | 2159 | 319.1 | - | - | 0 | - |
| - | - | 1081 | 320.1 | - | - | 0 | - |
| - | - | 861.7 | 322.2 | - | - | 0 | - |
| - | - | 1064 | 325.2 | - | - | 0 | - |
| - | - | 6647 | 326.2 | - | - | 0 | - |
| - | - | 1164 | 327.2 | - | - | 0 | - |
| - | - | 692.3 | 339.2 | - | - | 0 | - |
| - | - | 899.7 | 341.2 | - | - | 0 | - |
| 8 | y | 2315 | 343.2 | 0.0004846 | 1.412 | +1 | 3 |
| - | - | 1340 | 344.2 | - | - | 0 | - |
| - | - | 634.9 | 344.2 | - | - | 0 | - |
| - | - | 625.4 | 345.2 | - | - | 0 | - |
| 3 | b | 2948 | 346.2 | 0.000586 | 1.693 | +1 | 3 |
| - | - | 783.6 | 347.2 | - | - | 0 | - |
| - | - | 899.9 | 348.2 | - | - | 0 | - |
| - | - | 2708 | 357.2 | - | - | 0 | - |
| - | - | 597.4 | 360 | - | - | 0 | - |
| - | - | 1211 | 361.2 | - | - | 0 | - |
| 8 | y | 9147 | 361.2 | 0.0003875 | 1.073 | +1 | 3 |
| - | - | 1598 | 362.2 | - | - | 0 | - |
| 8 | b | 1034 | 365.2 | 0.006047 | 16.56 | +2 | 8 |
| 8 | b | 594.1 | 373.7 | 0.0003315 | 0.8869 | +2 | 8 |
| - | - | 3957 | 374.2 | - | - | 0 | - |
| - | - | 694.9 | 375.2 | - | - | 0 | - |
| 3 | y | 3221 | 381.7 | 0.0004327 | 1.134 | +2 | 8 |
| - | - | 1065 | 382.2 | - | - | 0 | - |
| - | - | 2075 | 382.2 | - | - | 0 | - |
| - | - | 7487 | 383.2 | - | - | 0 | - |
| - | - | 871 | 384.2 | - | - | 0 | - |
| - | - | 1048 | 387.2 | - | - | 0 | - |
| 3 | y | 2.624E+04 | 390.7 | 0.0004299 | 1.1 | +2 | 8 |
| - | - | 1.031E+04 | 391.2 | - | - | 0 | - |
| - | - | 2471 | 391.7 | - | - | 0 | - |
| - | - | 3981 | 392.2 | - | - | 0 | - |
| - | - | 911.1 | 393.2 | - | - | 0 | - |
| - | - | 2027 | 400.3 | - | - | 0 | - |
| - | - | 5509 | 401.2 | - | - | 0 | - |
| - | - | 732.8 | 401.2 | - | - | 0 | - |
| - | - | 1.106E+04 | 402.2 | - | - | 0 | - |
| - | - | 896.6 | 402.2 | - | - | 0 | - |
| - | - | 2307 | 403.2 | - | - | 0 | - |
| 4 | b | 2524 | 403.2 | 0.0007593 | 1.883 | +1 | 4 |
| - | - | 631.8 | 404.2 | - | - | 0 | - |
| - | - | 1153 | 415.3 | - | - | 0 | - |
| - | - | 1124 | 416.3 | - | - | 0 | - |
| - | - | 906.8 | 419.2 | - | - | 0 | - |
| - | - | 7304 | 420.2 | - | - | 0 | - |
| - | - | 1631 | 421.2 | - | - | 0 | - |
| 9 | b | 1138 | 421.3 | 0.004994 | 11.86 | +2 | 9 |
| - | - | 1300 | 429.3 | - | - | 0 | - |
| - | - | 3505 | 439.3 | - | - | 0 | - |
| 2 | y | 1474 | 440.3 | 0.0003603 | 0.8183 | +2 | 9 |
| - | - | 735.9 | 452.3 | - | - | 0 | - |
| - | - | 8063 | 457.3 | - | - | 0 | - |
| - | - | 1247 | 458.3 | - | - | 0 | - |
| 5 | b | 820.7 | 460.3 | 0.00136 | 2.954 | +1 | 5 |
| 7 | y | 3116 | 462.3 | 0.0001719 | 0.3718 | +1 | 4 |
| - | - | 941.9 | 463.3 | - | - | 0 | - |
| - | - | 574.6 | 472.2 | - | - | 0 | - |
| - | - | 759.5 | 478.3 | - | - | 0 | - |
| 0 | Precursor | 1367 | 480.8 | 0.0001646 | 0.3423 | +2 | -1 |
| - | - | 641.5 | 486.2 | - | - | 0 | - |
| - | - | 3933 | 486.3 | - | - | 0 | - |
| - | - | 1040 | 487.3 | - | - | 0 | - |
| - | - | 1479 | 489.3 | - | - | 0 | - |
| 0 | Precursor | 2251 | 489.8 | 0.0008026 | 1.639 | +2 | -1 |
| - | - | 1440 | 490.3 | - | - | 0 | - |
| - | - | 8131 | 496.3 | - | - | 0 | - |
| - | - | 2554 | 497.3 | - | - | 0 | - |
| - | - | 667.1 | 501.2 | - | - | 0 | - |
| - | - | 1590 | 512.3 | - | - | 0 | - |
| - | - | 1.909E+04 | 514.3 | - | - | 0 | - |
| - | - | 768.5 | 515.3 | - | - | 0 | - |
| - | - | 5120 | 515.3 | - | - | 0 | - |
| 6 | y | 3913 | 519.3 | 0.000215 | 0.414 | +1 | 5 |
| - | - | 872.3 | 520.3 | - | - | 0 | - |
| - | - | 7275 | 530.3 | - | - | 0 | - |
| - | - | 2054 | 531.3 | - | - | 0 | - |
| - | - | 1.248E+04 | 548.3 | - | - | 0 | - |
| - | - | 4452 | 549.3 | - | - | 0 | - |
| 5 | y | 1705 | 558.3 | 0.0004244 | 0.7601 | +1 | 6 |
| 5 | y | 1.96E+04 | 576.3 | 0.0002356 | 0.4089 | +1 | 6 |
| - | - | 6317 | 577.3 | - | - | 0 | - |
| - | - | 746.6 | 597.3 | - | - | 0 | - |
| - | - | 769.8 | 599.3 | - | - | 0 | - |
| 7 | b | 1526 | 600.3 | 0.001706 | 2.842 | +1 | 7 |
| - | - | 911.5 | 601.3 | - | - | 0 | - |
| 4 | y | 2649 | 615.3 | 0.002764 | 4.492 | +1 | 7 |
| - | - | 877.4 | 616.3 | - | - | 0 | - |
| - | - | 587.5 | 616.8 | - | - | 0 | - |
| 7 | b | 1185 | 618.3 | 0.001645 | 2.661 | +1 | 7 |
| - | - | 1474 | 625.3 | - | - | 0 | - |
| 4 | y | 5.375E+04 | 633.4 | 0.001416 | 2.236 | +1 | 7 |
| - | - | 1.713E+04 | 634.4 | - | - | 0 | - |
| - | - | 2915 | 635.4 | - | - | 0 | - |
| - | - | 1.278E+04 | 643.4 | - | - | 0 | - |
| - | - | 5115 | 644.4 | - | - | 0 | - |
| - | - | 809.3 | 645.4 | - | - | 0 | - |
| - | - | 741.2 | 647.4 | - | - | 0 | - |
| - | - | 3.083E+04 | 661.4 | - | - | 0 | - |
| - | - | 1.191E+04 | 662.4 | - | - | 0 | - |
| - | - | 3002 | 663.4 | - | - | 0 | - |
| - | - | 1081 | 679.4 | - | - | 0 | - |
| - | - | 906.3 | 685.4 | - | - | 0 | - |
| - | - | 923.7 | 718.4 | - | - | 0 | - |
| 8 | b | 746.1 | 728.4 | 0.002574 | 3.534 | +1 | 8 |
| - | - | 926.5 | 729.4 | - | - | 0 | - |
| - | - | 936.6 | 734.4 | - | - | 0 | - |
| - | - | 622.8 | 742.4 | - | - | 0 | - |
| - | - | 1371 | 744.4 | - | - | 0 | - |
| - | - | 638.3 | 745.4 | - | - | 0 | - |
| 8 | b | 4427 | 746.4 | 0.001103 | 1.478 | +1 | 8 |
| - | - | 1374 | 747.4 | - | - | 0 | - |
| - | - | 1945 | 760.4 | - | - | 0 | - |
| 3 | y | 6267 | 762.4 | 0.0003294 | 0.4321 | +1 | 8 |
| - | - | 1917 | 763.4 | - | - | 0 | - |
| - | - | 707.9 | 778.4 | - | - | 0 | - |
| 3 | y | 1.066E+05 | 780.4 | 0.0002255 | 0.2889 | +1 | 8 |
| - | - | 4.614E+04 | 781.4 | - | - | 0 | - |
| - | - | 1.094E+04 | 782.4 | - | - | 0 | - |
| - | - | 1351 | 783.4 | - | - | 0 | - |
| 9 | b | 1355 | 841.5 | 0.001351 | 1.605 | +1 | 9 |
| 9 | b | 5699 | 859.5 | 0.0007797 | 0.9071 | +1 | 9 |
| - | - | 2746 | 860.5 | - | - | 0 | - |
| - | - | 1028 | 861.5 | - | - | 0 | - |
| 2 | y | 5790 | 879.5 | 3.589E-05 | 0.04081 | +1 | 9 |
| - | - | 3658 | 880.5 | - | - | 0 | - |
| - | - | 1023 | 881.5 | - | - | 0 | - |
| - | - | 619.6 | 1099 | - | - | 0 | - |
| - | - | 617.1 | 1315 | - | - | 0 | - |
| - | - | 650.4 | 1747 | - | - | 0 | - |
| - | - | 609.9 | 2380 | - | - | 0 | - |

m/z Charge Intensity FragmentType MassShift Position
120.06596374511719 0 12365.76 y 9
120.08123779296875 0 86346.87
121.07785034179688 0 518.8994
121.07954406738281 0 556.4312
121.08458709716797 0 6734.093
127.05075073242188 0 1270.9606
127.08718872070312 0 407.56436
127.3436508178711 0 374.25128
128.07101440429688 0 610.885
128.10748291015625 0 861.30396
129.10269165039062 0 70410.734
130.0503692626953 0 727.72205
130.10604858398438 0 3297.8372
131.08197021484375 0 2132.922
132.08140563964844 0 1476.3916
133.0612030029297 0 513.848
133.0863037109375 0 2791.8105
136.0761260986328 0 13352.339
139.0867462158203 0 714.244
140.0823211669922 0 595.52905
141.06639099121094 0 1509.6284
141.10284423828125 0 500.49716
142.12315368652344 0 1001.7796
143.1181182861328 0 475.13602
145.0501251220703 0 2497.9216
146.0602569580078 0 1045.0753
149.0233917236328 0 815.18054
152.07113647460938 0 835.4645
152.1439666748047 0 849.2234
154.06216430664062 0 452.6469
155.08193969726562 0 496.93988
155.11827087402344 0 1973.2236
158.09291076660156 0 1387.1895
159.07655334472656 0 1655.0225
160.9039306640625 0 503.35745
163.07159423828125 0 1218.4635
163.913330078125 0 466.31702
165.10272216796875 0 1848.0077
166.05380249023438 0 763.8448
167.1182403564453 0 1003.33417
169.13404846191406 0 743.3592
170.09280395507812 0 1166.6373
171.0771942138672 0 659.4578
171.1497039794922 0 158570.53 a 1
172.0720977783203 0 3093.8682
172.14596557617188 0 693.265
172.15306091308594 0 15777.2295
173.128662109375 0 1114.2742
173.1563720703125 0 515.84015
173.43894958496094 0 1655.2881
174.05540466308594 0 502.87875
176.10748291015625 0 3356.7295
177.10267639160156 0 7660.617
177.11154174804688 0 1312.1909
178.10621643066406 0 867.8624
181.17041015625 0 1164.2188
183.11317443847656 0 1036.8728
183.14981079101562 0 673.08185
185.12872314453125 0 873.4656
185.16554260253906 0 1741.7777
186.12350463867188 0 638.8336
187.10687255859375 0 661.75574
187.14453125 0 2281.8186
188.10279846191406 0 787.14166
189.0758819580078 0 558.9809
195.14993286132812 0 823.01465
197.09320068359375 0 526.886
197.12864685058594 0 1515.6272
197.1651611328125 0 2906.5046
198.0876922607422 0 2376.0972
198.1685333251953 0 639.1891
199.14456176757812 0 78169.47 b 1
200.14788818359375 0 7838.926
201.11392211914062 0 498.7124
201.12399291992188 0 1440.48
201.14938354492188 0 828.1315
205.0976104736328 0 15706.052
206.1010284423828 0 1206.6023
207.14955139160156 0 851.61035
208.10894775390625 0 880.9128
209.10316467285156 0 672.3025
209.16543579101562 0 1013.1
209.88717651367188 0 603.7958
210.0879669189453 0 701.2482
211.14466857910156 0 1081.7844
212.13973999023438 0 6777.378
213.12376403808594 0 545.7283
213.1436767578125 0 1014.4443
213.15982055664062 0 1031.3297
215.11497497558594 0 697.844
215.1394805908203 0 4149.6323 y Water loss 8
216.09841918945312 0 1725.9694
216.14434814453125 0 634.81647
217.0975341796875 0 1430.9434
219.14967346191406 0 7368.951
220.15286254882812 0 916.10986
221.1279296875 0 615.764
224.17623901367188 0 2361.2866
225.1238250732422 0 639.015
225.13528442382812 0 885.9364
227.11441040039062 0 3124.4534
228.09841918945312 0 1641.8656
229.15513610839844 0 2479.3242
230.15040588378906 0 14379.289
230.16329956054688 0 722.8336
231.11383056640625 0 1162.8301
231.15272521972656 0 1523.1438
233.1025848388672 0 901.42255
233.12838745117188 0 882.6376
233.15016174316406 0 5287.751 y 8
233.16522216796875 0 1412.3333
234.125 0 574.8838
240.13455200195312 0 657.66034
241.1553955078125 0 828.5726
242.18670654296875 0 2240.2463
243.11318969726562 0 1549.3019
243.19024658203125 0 549.1016
244.1079864501953 0 768.00287
245.12484741210938 0 2016.3381
247.14474487304688 0 3521.5916
251.15110778808594 0 967.8636 y Water loss 5
253.94097900390625 0 569.33453
255.10934448242188 0 7549.3643
256.1133117675781 0 622.3459
261.12420654296875 0 1345.5692
262.1191101074219 0 5218.9385
269.113525390625 0 1692.4045
269.16131591796875 0 7065.5903
270.1644592285156 0 788.6207
270.1812438964844 0 978.4663
271.1075744628906 0 611.3607
272.1361999511719 0 1052.0846
273.1196594238281 0 8946.032
273.1971435546875 0 1040.666
274.1206359863281 0 1711.1626
276.1562194824219 0 2835.2378
282.1563720703125 0 923.9855
283.14434814453125 0 1039.0913
287.07061767578125 0 532.1694
287.1716613769531 0 2747.3333
291.1456604003906 0 1412.2026
298.1398620605469 0 648.0563
301.1918640136719 0 3561.65
304.16558837890625 0 1617.9694
310.2129211425781 0 3474.0042
317.1904602050781 0 540.49426
318.1457214355469 0 667.3322
319.1405334472656 0 2159.1106
320.1435852050781 0 1081.2048
322.1816711425781 0 861.66626
325.2230529785156 0 1064.0116
326.1827697753906 0 6646.759
327.1872253417969 0 1164.1777
339.1792907714844 0 692.29266
341.1812438964844 0 899.70593
343.2344665527344 0 2315.2148 y Water loss 7
344.1932373046875 0 1340.2023
344.2392883300781 0 634.8987
345.1551208496094 0 625.3716
346.2131042480469 0 2948.358 b 2
347.2166748046875 0 783.61365
348.1669921875 0 899.8731
357.15594482421875 0 2708.1416
360.02972412109375 0 597.4159
361.18902587890625 0 1210.847
361.24493408203125 0 9146.701 y 7
362.2477111816406 0 1598.2574
365.194091796875 0 1034.2313 b Ammonia loss 7
373.7137451171875 0 594.0821 b 7
374.1825256347656 0 3957.1885
375.1656799316406 0 694.86633
381.7113037109375 0 3220.9463 y Water loss 2
382.2135009765625 0 1065.2837
382.24566650390625 0 2075.4998
383.20428466796875 0 7486.526
384.2062072753906 0 870.97473
387.2027587890625 0 1047.5577
390.7165832519531 0 26236.627 y 2
391.2182922363281 0 10309.294
391.7189636230469 0 2470.656
392.1932373046875 0 3980.9387
393.19378662109375 0 911.12164
400.2556457519531 0 2026.9354
401.215087890625 0 5509.0405
401.24676513671875 0 732.77826
402.1778869628906 0 11057.592
402.21014404296875 0 896.5797
403.18060302734375 0 2307.0503
403.2347412109375 0 2523.6672 b 3
404.2342224121094 0 631.80505
415.2551574707031 0 1152.838
416.2575378417969 0 1123.546
419.2049255371094 0 906.76575
420.1883544921875 0 7303.72
421.1906433105469 0 1631.1211
421.2551574707031 0 1138.4517 b Water loss 8
429.28314208984375 0 1299.9514
439.2677307128906 0 3505.391
440.25 0 1474.3938 y 1
452.26220703125 0 735.9216
457.2774658203125 0 8063.008
458.2809143066406 0 1247.2867
460.2568054199219 0 820.69354 b 4
462.29205322265625 0 3115.7983 y 6
463.2988586425781 0 941.9431
472.2454833984375 0 574.6467
478.27801513671875 0 759.4964
480.7794494628906 0 1367.199 Precursor Water loss
486.2460632324219 0 641.5224
486.3048095703125 0 3932.695
487.3051452636719 0 1039.8417
489.3085021972656 0 1479.1694
489.7853698730469 0 2250.7205 Precursor
490.28912353515625 0 1439.8972
496.2883605957031 0 8131.226
497.2912292480469 0 2554.0864
501.24627685546875 0 667.09344
512.261962890625 0 1589.6102
514.298583984375 0 19085.66
515.2619018554688 0 768.5359
515.3016357421875 0 5119.6245
519.3139038085938 0 3912.6777 y 5
520.3162841796875 0 872.3184
530.2725830078125 0 7275.3877
531.2766723632812 0 2054.4197
548.283203125 0 12475.086
549.2862548828125 0 4451.6753
558.3250122070312 0 1704.7919 y Water loss 4
576.3353881835938 0 19599.898 y 4
577.337890625 0 6316.6377
597.3355102539062 0 746.6363
599.3298950195312 0 769.7899
600.3123168945312 0 1525.9973 b Water loss 6
601.3153076171875 0 911.543
615.3488159179688 0 2649.129 y Water loss 3
616.3468627929688 0 877.4403
616.8321533203125 0 587.4633
618.3262329101562 0 1184.8198 b 6
625.3477172851562 0 1474.2263
633.3580322265625 0 53749.055 y 3
634.3612060546875 0 17133.52
635.3639526367188 0 2915.0168
643.3561401367188 0 12776.28
644.359130859375 0 5114.8325
645.3571166992188 0 809.25354
647.3515014648438 0 741.18494
661.3670654296875 0 30832.812
662.369873046875 0 11913.957
663.3738403320312 0 3002.1406
679.3810424804688 0 1080.6238
685.3898315429688 0 906.28357
718.4259033203125 0 923.7408
728.4115600585938 0 746.059 b Water loss 7
729.4155883789062 0 926.46106
734.4197998046875 0 936.63617
742.4267578125 0 622.8489
744.4039306640625 0 1371.4401
745.40087890625 0 638.25684
746.420654296875 0 4426.5103 b 7
747.4231567382812 0 1374.0616
760.4373168945312 0 1944.8915
762.414794921875 0 6266.6587 y Water loss 2
763.4149780273438 0 1916.8683
778.4083251953125 0 707.8723
780.4248046875 0 106634.734 y 2
781.4278564453125 0 46136.92
782.4303588867188 0 10937.398
783.4323120117188 0 1350.9248
841.49169921875 0 1355.2087 b Water loss 8
859.50439453125 0 5698.5254 b 8
860.50732421875 0 2745.594
861.5103759765625 0 1028.2424
879.493408203125 0 5790.234 y 1
880.4969482421875 0 3657.9106
881.4972534179688 0 1023.2498
1098.566162109375 0 619.58276
1314.5968017578125 0 617.0945
1746.606689453125 0 650.4332
2380.157958984375 0 609.9459

Spectrum Details

|  |  |
| --- | --- |
| Matched peaks? Matched peaks The total absolute number of peaks matched. Additionally in brackets the total fraction of peaks matched and the total number of peaks is shown. | 34 (12.19% of 279) |
| FDR? FDR The false discovery rate estimated for this peptide. It is calculated by matching all theoretical fragments with a non-integer shift with the raw peaks for this spectrum. This is done with 40 different shifts. The resulting percentage is the average number of annotated peaks over the number of annotated peaks with the correct spectrum. | 0.07% |
| Satellite FDR? Satellite FDR See the FDR for details on its calculation. This satellite ion specific FDR only contains the satellite ions (d/w) for I/L/J positions. | - |
| PSM Score? PSM Score The PSM Score as given by Hecklib to this annotated spectrum. It is shown with three significant figures. | 383 |

#### Spectrum 6156? Spectrum 6156 The raw spectrum of this peptide as annotated by Hecklib. The fragments are coloured according to ion type (see legend). Any peaks with a star '\*' as text can be hovered over to see the full details, first the ion type second the mass shift type. By hovering over the amino acids in the peptide or ions in the legend the corresponding peaks are highlighted. By toggling the 'Unassigned' label you can turn the background (unassigned) peaks on or off in the plot. By updating the slider in the Ion legend you can update the spectrum to only show the top X% of the peaks with labels. The top X% means any peak that is within X% of the highest intensity. By dragging in the spectrum you can zoom in to a specific part of the spectrum and use 'Zoom Out' to get back to the original zoom level. The annotation of the spectrum is based on the given sequence in the peptides file and is done with different software so inconsistencies are likely. The peaks are annotated based on the given sequence, with 20 ppm tolerance.

Copy Data

##### Spectrum 6156 (TSV)

###### Preview

```
Loading example...
```

*Click on the button to copy the data to your clipboard.*

Mz MinMz MaxIntensity Max

WidthHeightPeptide font sizePeptide stroke widthSpectrum font sizeSpectrum stroke widthCompact peptide

Ion legend

wxyz

abcd

OtherUnassignedIon^Charge^_Position_Show for top:%

VVFGGGTKJT

02.89e+45.77e+48.66e+41.15e+5

Zoom Out

y^+1^_1_a^+1^_2_b^+1^_2_y^+1^_2_y^+1^_2_y^+2^_5_y^+2^_7_y^+1^_3_b^+1^_3_y^+1^_3_b^+2^_8_y^+2^_8_y^+2^_8_b^+1^_4_b^+2^_9_y^+2^_9_y^+1^_4_\*\*y^+1^_5_y^+1^_6_y^+1^_6_y^+1^_7_y^+1^_7_b^+1^_8_b^+1^_8_y^+1^_8_y^+1^_8_b^+1^_9_b^+1^_9_b^+1^_9_y^+1^_9_y^+1^_9_

0818163724553273

Fragment Matches Table

Show background peaks

| Position | Ion type | Intensity | mz Theoretical | mz Error (Th) | mz Error (ppm) | Charge | Series Number |
| --- | --- | --- | --- | --- | --- | --- | --- |
| 10 | y | 7812 | 120.1 | 0.0001466 | 1.221 | +1 | 1 |
| - | - | 6.585E+04 | 120.1 | - | - | 0 | - |
| - | - | 400.7 | 121.1 | - | - | 0 | - |
| - | - | 447.5 | 121.1 | - | - | 0 | - |
| - | - | 640.8 | 121.1 | - | - | 0 | - |
| - | - | 5611 | 121.1 | - | - | 0 | - |
| - | - | 453.1 | 122.1 | - | - | 0 | - |
| - | - | 711.3 | 127.1 | - | - | 0 | - |
| - | - | 500.8 | 127.1 | - | - | 0 | - |
| - | - | 524.8 | 127.1 | - | - | 0 | - |
| - | - | 459.8 | 128.1 | - | - | 0 | - |
| - | - | 5.265E+04 | 129.1 | - | - | 0 | - |
| - | - | 475.7 | 130 | - | - | 0 | - |
| - | - | 500.1 | 130.1 | - | - | 0 | - |
| - | - | 3279 | 130.1 | - | - | 0 | - |
| - | - | 1116 | 131.1 | - | - | 0 | - |
| - | - | 2226 | 131.1 | - | - | 0 | - |
| - | - | 1071 | 132.1 | - | - | 0 | - |
| - | - | 571 | 133.1 | - | - | 0 | - |
| - | - | 1.562E+04 | 133.1 | - | - | 0 | - |
| - | - | 7480 | 136.1 | - | - | 0 | - |
| - | - | 417.3 | 141.1 | - | - | 0 | - |
| - | - | 1203 | 141.1 | - | - | 0 | - |
| - | - | 541 | 143.1 | - | - | 0 | - |
| - | - | 621.2 | 146.1 | - | - | 0 | - |
| - | - | 446.3 | 146.3 | - | - | 0 | - |
| - | - | 529 | 147.1 | - | - | 0 | - |
| - | - | 669.8 | 147.1 | - | - | 0 | - |
| - | - | 416.7 | 148.7 | - | - | 0 | - |
| - | - | 636.4 | 149 | - | - | 0 | - |
| - | - | 497 | 152.1 | - | - | 0 | - |
| - | - | 495.6 | 152.1 | - | - | 0 | - |
| - | - | 494.6 | 152.7 | - | - | 0 | - |
| - | - | 502.1 | 153.1 | - | - | 0 | - |
| - | - | 535.8 | 153.1 | - | - | 0 | - |
| - | - | 488.8 | 153.1 | - | - | 0 | - |
| - | - | 479.7 | 155.1 | - | - | 0 | - |
| - | - | 1351 | 155.1 | - | - | 0 | - |
| - | - | 488.8 | 156.1 | - | - | 0 | - |
| - | - | 450.3 | 157.1 | - | - | 0 | - |
| - | - | 706.1 | 158.1 | - | - | 0 | - |
| - | - | 2019 | 159.1 | - | - | 0 | - |
| - | - | 1499 | 163.1 | - | - | 0 | - |
| - | - | 496.9 | 169.1 | - | - | 0 | - |
| - | - | 861.8 | 170.1 | - | - | 0 | - |
| - | - | 489.1 | 171.1 | - | - | 0 | - |
| 2 | a | 1.143E+05 | 171.1 | 0.0001176 | 0.687 | +1 | 2 |
| - | - | 2018 | 172.1 | - | - | 0 | - |
| - | - | 937.3 | 172.1 | - | - | 0 | - |
| - | - | 1.023E+04 | 172.2 | - | - | 0 | - |
| - | - | 517.6 | 173.1 | - | - | 0 | - |
| - | - | 994.8 | 173.1 | - | - | 0 | - |
| - | - | 471.5 | 173.2 | - | - | 0 | - |
| - | - | 875 | 173.4 | - | - | 0 | - |
| - | - | 776 | 175.1 | - | - | 0 | - |
| - | - | 1156 | 175.1 | - | - | 0 | - |
| - | - | 3146 | 176.1 | - | - | 0 | - |
| - | - | 6189 | 177.1 | - | - | 0 | - |
| - | - | 5637 | 177.1 | - | - | 0 | - |
| - | - | 683.3 | 178.1 | - | - | 0 | - |
| - | - | 1151 | 181.2 | - | - | 0 | - |
| - | - | 592.5 | 183.1 | - | - | 0 | - |
| - | - | 849.8 | 183.1 | - | - | 0 | - |
| - | - | 1201 | 185.1 | - | - | 0 | - |
| - | - | 882.9 | 185.2 | - | - | 0 | - |
| - | - | 524.8 | 186.1 | - | - | 0 | - |
| - | - | 565.6 | 187.1 | - | - | 0 | - |
| - | - | 1708 | 187.1 | - | - | 0 | - |
| - | - | 1889 | 197.1 | - | - | 0 | - |
| - | - | 1852 | 197.2 | - | - | 0 | - |
| - | - | 537.7 | 197.3 | - | - | 0 | - |
| - | - | 1804 | 198.1 | - | - | 0 | - |
| 2 | b | 5.4E+04 | 199.1 | 1.554E-05 | 0.07804 | +1 | 2 |
| - | - | 487 | 200.1 | - | - | 0 | - |
| - | - | 5843 | 200.1 | - | - | 0 | - |
| - | - | 504.8 | 200.4 | - | - | 0 | - |
| - | - | 590.1 | 200.7 | - | - | 0 | - |
| - | - | 1810 | 201.1 | - | - | 0 | - |
| - | - | 721.4 | 203.1 | - | - | 0 | - |
| - | - | 1.016E+04 | 205.1 | - | - | 0 | - |
| - | - | 1356 | 206.1 | - | - | 0 | - |
| - | - | 1095 | 207.1 | - | - | 0 | - |
| - | - | 727.8 | 208.1 | - | - | 0 | - |
| - | - | 1317 | 209.2 | - | - | 0 | - |
| - | - | 5044 | 212.1 | - | - | 0 | - |
| - | - | 611.2 | 213.2 | - | - | 0 | - |
| - | - | 598.5 | 214.2 | - | - | 0 | - |
| - | - | 612.7 | 215.1 | - | - | 0 | - |
| 9 | y | 4124 | 215.1 | 4.186E-05 | 0.1946 | +1 | 2 |
| - | - | 1885 | 216.1 | - | - | 0 | - |
| - | - | 1078 | 217.1 | - | - | 0 | - |
| - | - | 680.8 | 219.1 | - | - | 0 | - |
| - | - | 4188 | 219.1 | - | - | 0 | - |
| - | - | 1169 | 221.1 | - | - | 0 | - |
| - | - | 1797 | 224.2 | - | - | 0 | - |
| - | - | 810.3 | 225.1 | - | - | 0 | - |
| - | - | 889.7 | 226.2 | - | - | 0 | - |
| - | - | 485.9 | 226.7 | - | - | 0 | - |
| - | - | 527.4 | 227.1 | - | - | 0 | - |
| - | - | 2346 | 227.1 | - | - | 0 | - |
| - | - | 1092 | 228.1 | - | - | 0 | - |
| - | - | 597.9 | 229.1 | - | - | 0 | - |
| - | - | 9449 | 230.1 | - | - | 0 | - |
| - | - | 685.2 | 231.1 | - | - | 0 | - |
| - | - | 1607 | 231.2 | - | - | 0 | - |
| 9 | y | 3204 | 233.1 | 0.0001085 | 0.4654 | +1 | 2 |
| - | - | 954.8 | 233.2 | - | - | 0 | - |
| - | - | 1201 | 237.2 | - | - | 0 | - |
| - | - | 787.1 | 239.1 | - | - | 0 | - |
| - | - | 492.6 | 240.8 | - | - | 0 | - |
| - | - | 2633 | 242.2 | - | - | 0 | - |
| - | - | 1214 | 243.1 | - | - | 0 | - |
| - | - | 1724 | 245.1 | - | - | 0 | - |
| - | - | 2182 | 247.1 | - | - | 0 | - |
| - | - | 571.2 | 249.1 | - | - | 0 | - |
| 6 | y | 646.5 | 251.2 | 0.004825 | 19.21 | +2 | 5 |
| - | - | 5494 | 255.1 | - | - | 0 | - |
| - | - | 623.6 | 256.1 | - | - | 0 | - |
| - | - | 1226 | 261.1 | - | - | 0 | - |
| - | - | 3483 | 262.1 | - | - | 0 | - |
| - | - | 630.6 | 263.1 | - | - | 0 | - |
| - | - | 5618 | 269.2 | - | - | 0 | - |
| - | - | 1163 | 270.2 | - | - | 0 | - |
| - | - | 738 | 272.1 | - | - | 0 | - |
| - | - | 5626 | 273.1 | - | - | 0 | - |
| - | - | 1162 | 274.1 | - | - | 0 | - |
| - | - | 2056 | 276.2 | - | - | 0 | - |
| - | - | 741.4 | 282.2 | - | - | 0 | - |
| - | - | 1474 | 283.2 | - | - | 0 | - |
| - | - | 2082 | 287.2 | - | - | 0 | - |
| - | - | 1439 | 291.1 | - | - | 0 | - |
| - | - | 679 | 300.1 | - | - | 0 | - |
| - | - | 564.7 | 301.1 | - | - | 0 | - |
| - | - | 3342 | 301.2 | - | - | 0 | - |
| - | - | 907.8 | 302.2 | - | - | 0 | - |
| - | - | 630.6 | 304.2 | - | - | 0 | - |
| - | - | 3239 | 310.2 | - | - | 0 | - |
| - | - | 536.2 | 311.2 | - | - | 0 | - |
| - | - | 594.9 | 313.2 | - | - | 0 | - |
| 4 | y | 1348 | 317.2 | 0.006317 | 19.91 | +2 | 7 |
| - | - | 612.1 | 318.1 | - | - | 0 | - |
| - | - | 2766 | 319.1 | - | - | 0 | - |
| - | - | 1227 | 325.2 | - | - | 0 | - |
| - | - | 6189 | 326.2 | - | - | 0 | - |
| - | - | 543 | 328.2 | - | - | 0 | - |
| - | - | 721.7 | 333.2 | - | - | 0 | - |
| - | - | 666.3 | 339.2 | - | - | 0 | - |
| 8 | y | 1004 | 343.2 | 3.417E-05 | 0.09956 | +1 | 3 |
| - | - | 1165 | 344.2 | - | - | 0 | - |
| 3 | b | 3572 | 346.2 | 0.000299 | 0.8635 | +1 | 3 |
| - | - | 670.9 | 347.2 | - | - | 0 | - |
| - | - | 728.4 | 353.2 | - | - | 0 | - |
| - | - | 1610 | 357.2 | - | - | 0 | - |
| - | - | 563.4 | 361.2 | - | - | 0 | - |
| 8 | y | 7377 | 361.2 | 0.0003755 | 1.039 | +1 | 3 |
| - | - | 1462 | 362.2 | - | - | 0 | - |
| 8 | b | 806.5 | 365.2 | 0.006322 | 17.31 | +2 | 8 |
| - | - | 705.4 | 368.2 | - | - | 0 | - |
| - | - | 1316 | 371.2 | - | - | 0 | - |
| - | - | 2133 | 374.2 | - | - | 0 | - |
| - | - | 1007 | 375.2 | - | - | 0 | - |
| 3 | y | 1694 | 381.7 | 0.0003912 | 1.025 | +2 | 8 |
| - | - | 549 | 382.2 | - | - | 0 | - |
| - | - | 684 | 382.2 | - | - | 0 | - |
| - | - | 767 | 382.2 | - | - | 0 | - |
| - | - | 5198 | 383.2 | - | - | 0 | - |
| - | - | 678.5 | 384.2 | - | - | 0 | - |
| - | - | 937.4 | 384.2 | - | - | 0 | - |
| - | - | 740.9 | 385.1 | - | - | 0 | - |
| 3 | y | 2.062E+04 | 390.7 | 0.0004246 | 1.087 | +2 | 8 |
| - | - | 7658 | 391.2 | - | - | 0 | - |
| - | - | 2625 | 391.7 | - | - | 0 | - |
| - | - | 2917 | 392.2 | - | - | 0 | - |
| - | - | 774.5 | 397.2 | - | - | 0 | - |
| - | - | 1802 | 400.3 | - | - | 0 | - |
| - | - | 4246 | 401.2 | - | - | 0 | - |
| - | - | 8697 | 402.2 | - | - | 0 | - |
| - | - | 622.3 | 402.2 | - | - | 0 | - |
| - | - | 579.8 | 402.2 | - | - | 0 | - |
| - | - | 1475 | 403.2 | - | - | 0 | - |
| 4 | b | 1602 | 403.2 | 9.521E-05 | 0.2361 | +1 | 4 |
| - | - | 3120 | 415.3 | - | - | 0 | - |
| - | - | 808.3 | 416.3 | - | - | 0 | - |
| - | - | 1366 | 419 | - | - | 0 | - |
| - | - | 4509 | 420.2 | - | - | 0 | - |
| - | - | 1124 | 421.2 | - | - | 0 | - |
| 9 | b | 613.3 | 421.3 | 0.00237 | 5.625 | +2 | 9 |
| - | - | 965 | 429.3 | - | - | 0 | - |
| - | - | 3251 | 439.3 | - | - | 0 | - |
| 2 | y | 1244 | 440.3 | 0.004144 | 9.414 | +2 | 9 |
| - | - | 767.8 | 440.8 | - | - | 0 | - |
| - | - | 846.7 | 443.2 | - | - | 0 | - |
| - | - | 2226 | 455.3 | - | - | 0 | - |
| - | - | 5546 | 457.3 | - | - | 0 | - |
| - | - | 1031 | 458.3 | - | - | 0 | - |
| 7 | y | 2576 | 462.3 | 7.228E-05 | 0.1563 | +1 | 4 |
| - | - | 8174 | 473.3 | - | - | 0 | - |
| 0 | Precursor | 595.6 | 481.3 | 0.001914 | 3.977 | +2 | -1 |
| - | - | 2466 | 486.3 | - | - | 0 | - |
| - | - | 805.9 | 487.3 | - | - | 0 | - |
| - | - | 940.3 | 489.3 | - | - | 0 | - |
| 0 | Precursor | 996 | 489.8 | 0.001181 | 2.411 | +2 | -1 |
| - | - | 1560 | 490.3 | - | - | 0 | - |
| - | - | 3041 | 490.3 | - | - | 0 | - |
| - | - | 5273 | 496.3 | - | - | 0 | - |
| - | - | 1629 | 497.3 | - | - | 0 | - |
| - | - | 995.4 | 512.3 | - | - | 0 | - |
| - | - | 1.133E+04 | 514.3 | - | - | 0 | - |
| - | - | 3667 | 515.3 | - | - | 0 | - |
| 6 | y | 2747 | 519.3 | 0.0008226 | 1.584 | +1 | 5 |
| - | - | 5318 | 530.3 | - | - | 0 | - |
| - | - | 1547 | 531.3 | - | - | 0 | - |
| - | - | 9120 | 548.3 | - | - | 0 | - |
| - | - | 2704 | 549.3 | - | - | 0 | - |
| 5 | y | 1511 | 558.3 | 0.0007295 | 1.307 | +1 | 6 |
| 5 | y | 1.283E+04 | 576.3 | 0.0009851 | 1.709 | +1 | 6 |
| - | - | 3814 | 577.3 | - | - | 0 | - |
| - | - | 618.3 | 578.3 | - | - | 0 | - |
| - | - | 773.5 | 603.3 | - | - | 0 | - |
| 4 | y | 3001 | 615.3 | 0.0002009 | 0.3264 | +1 | 7 |
| - | - | 1275 | 616.3 | - | - | 0 | - |
| - | - | 780.9 | 617.3 | - | - | 0 | - |
| - | - | 607.9 | 625.2 | - | - | 0 | - |
| - | - | 955.1 | 625.3 | - | - | 0 | - |
| 4 | y | 3.935E+04 | 633.4 | 0.0001709 | 0.2699 | +1 | 7 |
| - | - | 1.133E+04 | 634.4 | - | - | 0 | - |
| - | - | 2985 | 635.4 | - | - | 0 | - |
| - | - | 1.021E+04 | 643.4 | - | - | 0 | - |
| - | - | 3868 | 644.4 | - | - | 0 | - |
| - | - | 664.1 | 645.4 | - | - | 0 | - |
| - | - | 2.038E+04 | 661.4 | - | - | 0 | - |
| - | - | 8668 | 662.4 | - | - | 0 | - |
| - | - | 1571 | 663.4 | - | - | 0 | - |
| - | - | 1008 | 718.4 | - | - | 0 | - |
| 8 | b | 1100 | 728.4 | 0.001088 | 1.494 | +1 | 8 |
| - | - | 588.1 | 735.4 | - | - | 0 | - |
| - | - | 584.8 | 744.4 | - | - | 0 | - |
| - | - | 571.3 | 745.4 | - | - | 0 | - |
| 8 | b | 3382 | 746.4 | 0.001887 | 2.528 | +1 | 8 |
| - | - | 1356 | 747.4 | - | - | 0 | - |
| - | - | 1294 | 760.4 | - | - | 0 | - |
| 3 | y | 2975 | 762.4 | 0.00138 | 1.809 | +1 | 8 |
| - | - | 1933 | 763.4 | - | - | 0 | - |
| 3 | y | 7.912E+04 | 780.4 | 0.001812 | 2.322 | +1 | 8 |
| - | - | 3.001E+04 | 781.4 | - | - | 0 | - |
| - | - | 7978 | 782.4 | - | - | 0 | - |
| 9 | b | 947.7 | 841.5 | 0.003426 | 4.071 | +1 | 9 |
| 9 | b | 629.5 | 842.5 | 0.01579 | 18.75 | +1 | 9 |
| 9 | b | 3960 | 859.5 | 0.002821 | 3.283 | +1 | 9 |
| - | - | 1936 | 860.5 | - | - | 0 | - |
| 2 | y | 683.9 | 861.5 | 0.01663 | 19.31 | +1 | 9 |
| 2 | y | 5010 | 879.5 | 0.003271 | 3.719 | +1 | 9 |
| - | - | 1923 | 880.5 | - | - | 0 | - |
| - | - | 731.8 | 881.5 | - | - | 0 | - |
| - | - | 664.4 | 1570 | - | - | 0 | - |
| - | - | 616.1 | 1897 | - | - | 0 | - |
| - | - | 672.4 | 1982 | - | - | 0 | - |
| - | - | 753.3 | 2065 | - | - | 0 | - |
| - | - | 703.3 | 2842 | - | - | 0 | - |
| - | - | 786.4 | 3241 | - | - | 0 | - |

m/z Charge Intensity FragmentType MassShift Position
120.06566619873047 0 7812.3916 y 9
120.08096313476562 0 65848.664
121.06465148925781 0 400.68484
121.06916809082031 0 447.49313
121.07890319824219 0 640.77423
121.08428192138672 0 5611.372
122.0875244140625 0 453.08246
127.05049896240234 0 711.2634
127.0755386352539 0 500.8041
127.08696746826172 0 524.8285
128.10743713378906 0 459.84888
129.10238647460938 0 52649.2
130.0496368408203 0 475.74704
130.0608367919922 0 500.0602
130.10572814941406 0 3278.7224
131.07041931152344 0 1116.42
131.0816192626953 0 2226.1455
132.0810089111328 0 1071.3
133.06076049804688 0 570.976
133.08607482910156 0 15623.298
136.07579040527344 0 7480.101
141.05496215820312 0 417.2869
141.0660858154297 0 1202.7609
143.08145141601562 0 540.95526
146.0602569580078 0 621.19165
146.28477478027344 0 446.258
147.06463623046875 0 529.0318
147.1016845703125 0 669.81335
148.73611450195312 0 416.66522
148.9544219970703 0 636.4054
152.07093811035156 0 497.03772
152.14315795898438 0 495.56113
152.74266052246094 0 494.56815
153.0554656982422 0 502.05655
153.06593322753906 0 535.7575
153.0770721435547 0 488.78207
155.0818634033203 0 479.7165
155.1181640625 0 1351.1699
156.07730102539062 0 488.80515
157.085205078125 0 450.29013
158.0924530029297 0 706.06714
159.07659912109375 0 2018.8259
163.07159423828125 0 1499.0632
169.0974578857422 0 496.93088
170.0927276611328 0 861.7689
171.07614135742188 0 489.0807
171.14930725097656 0 114281.53 a 1
172.07179260253906 0 2017.6111
172.1451416015625 0 937.30695
172.1526641845703 0 10234.841
173.11746215820312 0 517.6031
173.1283416748047 0 994.8376
173.15570068359375 0 471.52176
173.43862915039062 0 874.9629
175.08770751953125 0 776.0133
175.09661865234375 0 1155.8839
176.1070098876953 0 3145.8882
177.1022186279297 0 6189.326
177.1121826171875 0 5636.9585
178.1057891845703 0 683.2553
181.169921875 0 1151.4916
183.08688354492188 0 592.5238
183.11293029785156 0 849.7637
185.12863159179688 0 1201.2587
185.1648406982422 0 882.88837
186.1237030029297 0 524.7928
187.1073455810547 0 565.6093
187.1441650390625 0 1708.4948
197.12843322753906 0 1889.3855
197.16464233398438 0 1852.3472
197.29843139648438 0 537.69183
198.08718872070312 0 1803.8441
199.1440887451172 0 54002.723 b 1
200.13973999023438 0 487.0454
200.14756774902344 0 5842.6533
200.40098571777344 0 504.8212
200.74465942382812 0 590.06146
201.123291015625 0 1809.6758
203.11721801757812 0 721.4299
205.0970916748047 0 10155.024
206.1005859375 0 1355.6387
207.1494903564453 0 1094.6349
208.10752868652344 0 727.81946
209.16452026367188 0 1316.7843
212.13929748535156 0 5044.252
213.15953063964844 0 611.2126
214.19142150878906 0 598.45636
215.1138916015625 0 612.7226
215.13897705078125 0 4124.3804 y Water loss 8
216.0980682373047 0 1884.8381
217.0972900390625 0 1077.5001
219.12265014648438 0 680.8233
219.14903259277344 0 4187.5835
221.13876342773438 0 1168.7703
224.1758270263672 0 1796.6577
225.1342010498047 0 810.2736
226.1553192138672 0 889.65533
226.67308044433594 0 485.91122
227.10215759277344 0 527.3629
227.11424255371094 0 2346.4316
228.0973663330078 0 1091.5073
229.0931396484375 0 597.85724
230.14990234375 0 9449.153
231.11326599121094 0 685.187
231.15252685546875 0 1606.7595
233.14947509765625 0 3203.5964 y 8
233.16314697265625 0 954.7684
237.1598663330078 0 1200.5621
239.14950561523438 0 787.1041
240.7532958984375 0 492.59537
242.18630981445312 0 2633.1062
243.11268615722656 0 1213.8092
245.1244659423828 0 1724.3523
247.14413452148438 0 2181.617
249.12403869628906 0 571.1792
251.15037536621094 0 646.53174 y Water loss 5
255.1087188720703 0 5494.1177
256.1116943359375 0 623.6035
261.122802734375 0 1226.4127
262.1182861328125 0 3482.73
263.12164306640625 0 630.59827
269.1605529785156 0 5617.68
270.1805419921875 0 1162.6511
272.13482666015625 0 737.9787
273.119140625 0 5625.9966
274.12030029296875 0 1162.2695
276.1549987792969 0 2055.7388
282.1556091308594 0 741.4304
283.1748962402344 0 1474.0009
287.17156982421875 0 2082.455
291.1455383300781 0 1439.0657
300.1347961425781 0 678.98865
301.1293029785156 0 564.66064
301.191162109375 0 3341.5317
302.1944274902344 0 907.8002
304.1665344238281 0 630.59875
310.2126770019531 0 3238.652
311.21722412109375 0 536.18604
313.185546875 0 594.8624
317.1882629394531 0 1348.2772 y 3
318.14410400390625 0 612.09174
319.14031982421875 0 2765.7844
325.22271728515625 0 1227.0422
326.1820983886719 0 6189.4688
328.2021179199219 0 542.9977
333.1908874511719 0 721.71564
339.1790466308594 0 666.3185
343.23394775390625 0 1004.2705 y Water loss 7
344.1928405761719 0 1164.5283
346.21221923828125 0 3572.1465 b 2
347.2152099609375 0 670.87787
353.2184753417969 0 728.40106
357.1559143066406 0 1609.6063
361.1880187988281 0 563.3545
361.2441711425781 0 7377.1055 y 7
362.2476501464844 0 1461.937
365.1938171386719 0 806.52527 b Ammonia loss 7
368.1930236816406 0 705.41205
371.22735595703125 0 1316.0475
374.1809997558594 0 2132.6052
375.16461181640625 0 1006.64746
381.7104797363281 0 1694.1271 y Water loss 2
382.166015625 0 549.01373
382.2110290527344 0 684.00586
382.24530029296875 0 767.01105
383.20281982421875 0 5197.5625
384.1674499511719 0 678.5222
384.2072448730469 0 937.41705
385.14971923828125 0 740.8619
390.7157287597656 0 20623.902 y 2
391.2173156738281 0 7657.949
391.7191467285156 0 2624.8665
392.19293212890625 0 2916.5608
397.2435302734375 0 774.5428
400.25445556640625 0 1802.188
401.2142333984375 0 4245.7744
402.17706298828125 0 8696.969
402.211181640625 0 622.2537
402.2160339355469 0 579.7624
403.1799621582031 0 1474.7601
403.23388671875 0 1602.2543 b 3
415.25341796875 0 3120.079
416.25732421875 0 808.2604
418.99468994140625 0 1365.6792
420.1872253417969 0 4508.8203
421.18951416015625 0 1124.4214
421.2525329589844 0 613.32245 b Water loss 8
429.2793884277344 0 964.99664
439.2662353515625 0 3251.0522
440.2462158203125 0 1244.3253 y 1
440.75103759765625 0 767.78485
443.2491760253906 0 846.67
455.2843017578125 0 2226.4353
457.2765808105469 0 5545.9365
458.279052734375 0 1030.6841
462.29229736328125 0 2576.0374 y 6
473.2951354980469 0 8174.129
481.2693786621094 0 595.5681 Precursor Ammonia loss
486.30218505859375 0 2466.0903
487.30450439453125 0 805.85205
489.2740478515625 0 940.27563
489.78338623046875 0 996.0499 Precursor
490.2840881347656 0 1559.8044
490.3218688964844 0 3041.0767
496.2870178222656 0 5272.5234
497.2901611328125 0 1628.6606
512.2612915039062 0 995.39685
514.2977294921875 0 11328.7
515.3003540039062 0 3667.3306
519.3128662109375 0 2746.8496 y 5
530.2716674804688 0 5317.571
531.2762451171875 0 1546.6428
548.2817993164062 0 9119.558
549.2844848632812 0 2704.1755
558.3253173828125 0 1510.5216 y Water loss 4
576.3341674804688 0 12831.864 y 4
577.33740234375 0 3813.7075
578.3373413085938 0 618.252
603.3081665039062 0 773.54443
615.3462524414062 0 3000.6294 y Water loss 3
616.3472900390625 0 1274.8779
617.3450317382812 0 780.90173
625.2266845703125 0 607.886
625.3447265625 0 955.09863
633.3564453125 0 39351.87 y 3
634.3589477539062 0 11334.753
635.3613891601562 0 2985.1921
643.3549194335938 0 10206.64
644.3582153320312 0 3868.035
645.3615112304688 0 664.08954
661.3654174804688 0 20384.268
662.368408203125 0 8667.907
663.3701171875 0 1570.5472
718.4235229492188 0 1008.1647
728.4078979492188 0 1099.8927 b Water loss 7
735.41748046875 0 588.05695
744.4072265625 0 584.795
745.4027099609375 0 571.2533
746.4176635742188 0 3381.5315 b 7
747.4198608398438 0 1356.3375
760.4364013671875 0 1293.9988
762.4130859375 0 2975.24 y Water loss 2
763.415283203125 0 1933.1635
780.4232177734375 0 79125 y 2
781.4259033203125 0 30009.088
782.428955078125 0 7978.069
841.4896240234375 0 947.71655 b Water loss 8
842.4928588867188 0 629.542 b Ammonia loss 8
859.5007934570312 0 3959.5845 b 8
860.5064697265625 0 1935.5408
861.49951171875 0 683.8862 y Water loss 1
879.4901733398438 0 5009.9497 y 1
880.4937133789062 0 1923.3438
881.489013671875 0 731.8364
1569.8533935546875 0 664.3772
1897.4599609375 0 616.14514
1981.6649169921875 0 672.36163
2064.89306640625 0 753.2543
2842.27099609375 0 703.2555
3240.7666015625 0 786.4481

Spectrum Details

|  |  |
| --- | --- |
| Matched peaks? Matched peaks The total absolute number of peaks matched. Additionally in brackets the total fraction of peaks matched and the total number of peaks is shown. | 33 (12.69% of 260) |
| FDR? FDR The false discovery rate estimated for this peptide. It is calculated by matching all theoretical fragments with a non-integer shift with the raw peaks for this spectrum. This is done with 40 different shifts. The resulting percentage is the average number of annotated peaks over the number of annotated peaks with the correct spectrum. | 0.14% |
| Satellite FDR? Satellite FDR See the FDR for details on its calculation. This satellite ion specific FDR only contains the satellite ions (d/w) for I/L/J positions. | - |
| PSM Score? PSM Score The PSM Score as given by Hecklib to this annotated spectrum. It is shown with three significant figures. | 362 |

#### Spectrum 5917? Spectrum 5917 The raw spectrum of this peptide as annotated by Hecklib. The fragments are coloured according to ion type (see legend). Any peaks with a star '\*' as text can be hovered over to see the full details, first the ion type second the mass shift type. By hovering over the amino acids in the peptide or ions in the legend the corresponding peaks are highlighted. By toggling the 'Unassigned' label you can turn the background (unassigned) peaks on or off in the plot. By updating the slider in the Ion legend you can update the spectrum to only show the top X% of the peaks with labels. The top X% means any peak that is within X% of the highest intensity. By dragging in the spectrum you can zoom in to a specific part of the spectrum and use 'Zoom Out' to get back to the original zoom level. The annotation of the spectrum is based on the given sequence in the peptides file and is done with different software so inconsistencies are likely. The peaks are annotated based on the given sequence, with 20 ppm tolerance.

Copy Data

##### Spectrum 5917 (TSV)

###### Preview

```
Loading example...
```

*Click on the button to copy the data to your clipboard.*

Mz MinMz MaxIntensity Max

WidthHeightPeptide font sizePeptide stroke widthSpectrum font sizeSpectrum stroke widthCompact peptide

Ion legend

wxyz

abcd

OtherUnassignedIon^Charge^_Position_Show for top:%

VVFGGGTKJT

02.24e+44.48e+46.71e+48.95e+4

Zoom Out

y^+1^_1_y^+1^_2_y^+1^_2_z^+1^_3_y^+1^_3_c^+2^_8_y^+2^_8_y^+2^_8_y^+2^_9_y^+1^_4_z^+1^_4_y^+1^_4_c^+1^_5_y^+1^_5_z^+1^_5_y^+1^_5_c^+1^_6_z^+1^_6_y^+1^_6_y^+1^_7_c^+1^_7_y^+1^_7_c^+1^_7_c^+1^_8_y^+1^_8_c^+1^_8_z^+1^_8_y^+1^_8_c^+1^_9_z^+1^_9_c^+1^_9_y^+1^_9_

03336659981331

Fragment Matches Table

Show background peaks

| Position | Ion type | Intensity | mz Theoretical | mz Error (Th) | mz Error (ppm) | Charge | Series Number |
| --- | --- | --- | --- | --- | --- | --- | --- |
| 10 | y | 3737 | 120.1 | 0.0002381 | 1.983 | +1 | 1 |
| - | - | 7504 | 120.1 | - | - | 0 | - |
| - | - | 675.9 | 121.1 | - | - | 0 | - |
| - | - | 521.4 | 126 | - | - | 0 | - |
| - | - | 426.3 | 127.7 | - | - | 0 | - |
| - | - | 1765 | 129.1 | - | - | 0 | - |
| - | - | 1858 | 136.1 | - | - | 0 | - |
| - | - | 406.7 | 138.8 | - | - | 0 | - |
| - | - | 370.5 | 141.8 | - | - | 0 | - |
| - | - | 516.2 | 148.9 | - | - | 0 | - |
| - | - | 475.3 | 153.1 | - | - | 0 | - |
| - | - | 436.9 | 161.6 | - | - | 0 | - |
| - | - | 489.2 | 164.7 | - | - | 0 | - |
| - | - | 3.451E+04 | 171.1 | - | - | 0 | - |
| - | - | 3469 | 172.2 | - | - | 0 | - |
| - | - | 1518 | 173.5 | - | - | 0 | - |
| - | - | 787.6 | 187.1 | - | - | 0 | - |
| - | - | 434.6 | 196.9 | - | - | 0 | - |
| - | - | 555.4 | 197.1 | - | - | 0 | - |
| - | - | 2.905E+04 | 199.1 | - | - | 0 | - |
| - | - | 3160 | 200.1 | - | - | 0 | - |
| - | - | 534.2 | 200.5 | - | - | 0 | - |
| - | - | 547.9 | 201.1 | - | - | 0 | - |
| - | - | 2208 | 205.1 | - | - | 0 | - |
| - | - | 591.3 | 214 | - | - | 0 | - |
| 9 | y | 2017 | 215.1 | 0.0002633 | 1.224 | +1 | 2 |
| - | - | 512.9 | 215.3 | - | - | 0 | - |
| - | - | 596.4 | 230.2 | - | - | 0 | - |
| 9 | y | 1718 | 233.1 | 0.0001051 | 0.4509 | +1 | 2 |
| - | - | 656.8 | 247.1 | - | - | 0 | - |
| - | - | 626.6 | 262.1 | - | - | 0 | - |
| - | - | 736.3 | 263.1 | - | - | 0 | - |
| - | - | 607.6 | 267.3 | - | - | 0 | - |
| - | - | 3435 | 276.2 | - | - | 0 | - |
| - | - | 765.3 | 277.1 | - | - | 0 | - |
| - | - | 1418 | 301.2 | - | - | 0 | - |
| - | - | 3926 | 310.2 | - | - | 0 | - |
| - | - | 892.1 | 311.2 | - | - | 0 | - |
| - | - | 788.8 | 334.2 | - | - | 0 | - |
| 8 | z | 1886 | 345.2 | 0.00132 | 3.823 | +1 | 3 |
| - | - | 1761 | 346.2 | - | - | 0 | - |
| - | - | 3837 | 346.2 | - | - | 0 | - |
| - | - | 708.6 | 347.2 | - | - | 0 | - |
| - | - | 1542 | 358.2 | - | - | 0 | - |
| 8 | y | 1120 | 361.2 | 0.0005095 | 1.411 | +1 | 3 |
| 8 | c | 1074 | 373.7 | 0.0005451 | 1.459 | +2 | 8 |
| 3 | y | 972 | 381.7 | 0.0009821 | 2.573 | +2 | 8 |
| - | - | 675.6 | 383.2 | - | - | 0 | - |
| - | - | 764.9 | 385.2 | - | - | 0 | - |
| 3 | y | 1.459E+04 | 390.7 | 0.0003689 | 0.9442 | +2 | 8 |
| - | - | 7795 | 391.2 | - | - | 0 | - |
| - | - | 2348 | 391.7 | - | - | 0 | - |
| - | - | 1185 | 401.2 | - | - | 0 | - |
| - | - | 1942 | 402.3 | - | - | 0 | - |
| - | - | 1074 | 403.2 | - | - | 0 | - |
| - | - | 1153 | 415.3 | - | - | 0 | - |
| - | - | 604.4 | 416.3 | - | - | 0 | - |
| - | - | 633.3 | 419 | - | - | 0 | - |
| - | - | 823.2 | 420.2 | - | - | 0 | - |
| 2 | y | 812.4 | 440.3 | 0.000189 | 0.4294 | +2 | 9 |
| - | - | 2833 | 444.3 | - | - | 0 | - |
| 7 | y | 5020 | 445.3 | 0.0009561 | 2.147 | +1 | 4 |
| 7 | z | 1475 | 446.3 | 0.00397 | 8.895 | +1 | 4 |
| - | - | 1152 | 447.2 | - | - | 0 | - |
| - | - | 5408 | 447.3 | - | - | 0 | - |
| - | - | 1516 | 448.3 | - | - | 0 | - |
| - | - | 835.7 | 457.3 | - | - | 0 | - |
| - | - | 735.1 | 459.3 | - | - | 0 | - |
| 7 | y | 1110 | 462.3 | 0.001476 | 3.193 | +1 | 4 |
| - | - | 1112 | 470.3 | - | - | 0 | - |
| - | - | 685.8 | 473.2 | - | - | 0 | - |
| - | - | 629.2 | 474.3 | - | - | 0 | - |
| - | - | 1034 | 476.3 | - | - | 0 | - |
| 5 | c | 2887 | 477.3 | 0.0004149 | 0.8693 | +1 | 5 |
| - | - | 939.3 | 478.3 | - | - | 0 | - |
| - | - | 1447 | 480.8 | - | - | 0 | - |
| - | - | 1497 | 489.2 | - | - | 0 | - |
| - | - | 1221 | 489.8 | - | - | 0 | - |
| - | - | 2893 | 490.3 | - | - | 0 | - |
| - | - | 769.2 | 491.3 | - | - | 0 | - |
| - | - | 1419 | 499.3 | - | - | 0 | - |
| 6 | y | 809.2 | 502.3 | 0.006622 | 13.18 | +1 | 5 |
| 6 | z | 2185 | 503.3 | 0.0001708 | 0.3393 | +1 | 5 |
| - | - | 2.87E+04 | 504.3 | - | - | 0 | - |
| - | - | 4.034E+04 | 504.3 | - | - | 0 | - |
| - | - | 7871 | 505.3 | - | - | 0 | - |
| - | - | 1.032E+04 | 505.3 | - | - | 0 | - |
| - | - | 1080 | 506.3 | - | - | 0 | - |
| - | - | 2282 | 506.3 | - | - | 0 | - |
| - | - | 2014 | 514.3 | - | - | 0 | - |
| - | - | 1166 | 516.3 | - | - | 0 | - |
| - | - | 992.9 | 517.3 | - | - | 0 | - |
| - | - | 1025 | 518.3 | - | - | 0 | - |
| 6 | y | 3359 | 519.3 | 0.0005785 | 1.114 | +1 | 5 |
| - | - | 1063 | 520.3 | - | - | 0 | - |
| - | - | 704.4 | 530.3 | - | - | 0 | - |
| - | - | 1671 | 533.3 | - | - | 0 | - |
| 6 | c | 1289 | 534.3 | 0.0004356 | 0.8152 | +1 | 6 |
| - | - | 2458 | 548.3 | - | - | 0 | - |
| - | - | 771.9 | 550.3 | - | - | 0 | - |
| 5 | z | 1.042E+04 | 560.3 | 0.000283 | 0.505 | +1 | 6 |
| - | - | 1234 | 561.3 | - | - | 0 | - |
| - | - | 2.933E+04 | 561.3 | - | - | 0 | - |
| - | - | 1.032E+04 | 562.3 | - | - | 0 | - |
| - | - | 1360 | 563.3 | - | - | 0 | - |
| - | - | 1506 | 574.3 | - | - | 0 | - |
| - | - | 5266 | 575.3 | - | - | 0 | - |
| 5 | y | 9932 | 576.3 | 0.0006188 | 1.074 | +1 | 6 |
| - | - | 3029 | 577.3 | - | - | 0 | - |
| - | - | 1299 | 589.3 | - | - | 0 | - |
| - | - | 686.3 | 607.3 | - | - | 0 | - |
| - | - | 550.2 | 613.8 | - | - | 0 | - |
| 4 | y | 761.3 | 615.3 | 0.001177 | 1.913 | +1 | 7 |
| - | - | 1015 | 616.3 | - | - | 0 | - |
| 7 | c | 8314 | 617.3 | 0.002682 | 4.344 | +1 | 7 |
| - | - | 1.991E+04 | 618.3 | - | - | 0 | - |
| - | - | 5362 | 619.4 | - | - | 0 | - |
| - | - | 1668 | 620.4 | - | - | 0 | - |
| - | - | 1.116E+04 | 632.3 | - | - | 0 | - |
| 4 | y | 4.097E+04 | 633.4 | 0.0001709 | 0.2699 | +1 | 7 |
| - | - | 1.846E+04 | 634.4 | - | - | 0 | - |
| 7 | c | 5.842E+04 | 635.4 | 0.0003645 | 0.5737 | +1 | 7 |
| - | - | 2.158E+04 | 636.4 | - | - | 0 | - |
| - | - | 4192 | 637.4 | - | - | 0 | - |
| - | - | 1684 | 643.4 | - | - | 0 | - |
| - | - | 1026 | 646.3 | - | - | 0 | - |
| - | - | 645.3 | 656.8 | - | - | 0 | - |
| - | - | 1143 | 659.3 | - | - | 0 | - |
| - | - | 1099 | 661.3 | - | - | 0 | - |
| - | - | 6870 | 661.4 | - | - | 0 | - |
| - | - | 2423 | 662.4 | - | - | 0 | - |
| - | - | 3617 | 663.4 | - | - | 0 | - |
| - | - | 1065 | 664.4 | - | - | 0 | - |
| - | - | 948.9 | 685.4 | - | - | 0 | - |
| - | - | 1771 | 702.4 | - | - | 0 | - |
| - | - | 869.8 | 708.3 | - | - | 0 | - |
| - | - | 624.7 | 708.5 | - | - | 0 | - |
| - | - | 622.6 | 709.3 | - | - | 0 | - |
| - | - | 870.6 | 715.4 | - | - | 0 | - |
| - | - | 762.5 | 716.4 | - | - | 0 | - |
| - | - | 832.8 | 718.4 | - | - | 0 | - |
| - | - | 824 | 719.4 | - | - | 0 | - |
| - | - | 983.9 | 721.3 | - | - | 0 | - |
| - | - | 3014 | 729.4 | - | - | 0 | - |
| - | - | 2049 | 730.4 | - | - | 0 | - |
| 8 | c | 3306 | 746.4 | 0.008906 | 11.93 | +1 | 8 |
| - | - | 1646 | 747.4 | - | - | 0 | - |
| - | - | 708.8 | 748.4 | - | - | 0 | - |
| 3 | y | 1814 | 762.4 | 0.002405 | 3.154 | +1 | 8 |
| 8 | c | 7.384E+04 | 763.4 | 0.0001161 | 0.152 | +1 | 8 |
| 3 | z | 6841 | 764.4 | 0.009028 | 11.81 | +1 | 8 |
| - | - | 2.751E+04 | 764.4 | - | - | 0 | - |
| - | - | 5148 | 765.4 | - | - | 0 | - |
| - | - | 5251 | 765.5 | - | - | 0 | - |
| - | - | 1812 | 766.4 | - | - | 0 | - |
| - | - | 691.3 | 773.5 | - | - | 0 | - |
| 3 | y | 6.01E+04 | 780.4 | 0.0002255 | 0.2889 | +1 | 8 |
| - | - | 2.234E+04 | 781.4 | - | - | 0 | - |
| - | - | 7754 | 782.4 | - | - | 0 | - |
| - | - | 1682 | 783.4 | - | - | 0 | - |
| - | - | 600.8 | 789 | - | - | 0 | - |
| - | - | 640.8 | 789.5 | - | - | 0 | - |
| - | - | 808 | 807.4 | - | - | 0 | - |
| - | - | 722.9 | 808.4 | - | - | 0 | - |
| - | - | 745.9 | 832.4 | - | - | 0 | - |
| - | - | 8714 | 832.5 | - | - | 0 | - |
| - | - | 3633 | 833.5 | - | - | 0 | - |
| - | - | 647.6 | 834.5 | - | - | 0 | - |
| 9 | c | 4957 | 859.5 | 0.001296 | 1.507 | +1 | 9 |
| - | - | 1737 | 860.5 | - | - | 0 | - |
| - | - | 2112 | 861.5 | - | - | 0 | - |
| - | - | 1144 | 862.5 | - | - | 0 | - |
| 2 | z | 8402 | 863.5 | 0.0004158 | 0.4816 | +1 | 9 |
| - | - | 5136 | 864.5 | - | - | 0 | - |
| - | - | 2440 | 865.5 | - | - | 0 | - |
| 9 | c | 7.294E+04 | 876.5 | 0.0004398 | 0.5018 | +1 | 9 |
| - | - | 4.171E+04 | 877.5 | - | - | 0 | - |
| - | - | 1.122E+04 | 878.5 | - | - | 0 | - |
| 2 | y | 2193 | 879.5 | 0.002039 | 2.319 | +1 | 9 |
| - | - | 1973 | 879.5 | - | - | 0 | - |
| - | - | 1018 | 880.5 | - | - | 0 | - |
| - | - | 849.3 | 888.5 | - | - | 0 | - |
| - | - | 778.2 | 889.5 | - | - | 0 | - |
| - | - | 914.8 | 893.5 | - | - | 0 | - |
| - | - | 1065 | 906.5 | - | - | 0 | - |
| - | - | 1202 | 917.5 | - | - | 0 | - |
| - | - | 3208 | 919.5 | - | - | 0 | - |
| - | - | 1577 | 920.5 | - | - | 0 | - |
| - | - | 3488 | 923.5 | - | - | 0 | - |
| - | - | 1880 | 924.5 | - | - | 0 | - |
| - | - | 1449 | 933.6 | - | - | 0 | - |
| - | - | 1672 | 934.5 | - | - | 0 | - |
| - | - | 2886 | 935.5 | - | - | 0 | - |
| - | - | 3347 | 936.5 | - | - | 0 | - |
| - | - | 1370 | 937.5 | - | - | 0 | - |
| - | - | 845.5 | 947.5 | - | - | 0 | - |
| - | - | 701.8 | 951.6 | - | - | 0 | - |
| - | - | 1698 | 961.6 | - | - | 0 | - |
| - | - | 8.862E+04 | 962.5 | - | - | 0 | - |
| - | - | 5.333E+04 | 963.5 | - | - | 0 | - |
| - | - | 1.575E+04 | 964.5 | - | - | 0 | - |
| - | - | 3095 | 965.6 | - | - | 0 | - |
| - | - | 844.7 | 977.6 | - | - | 0 | - |
| - | - | 4.783E+04 | 978.6 | - | - | 0 | - |
| - | - | 2994 | 979.5 | - | - | 0 | - |
| - | - | 6.257E+04 | 979.6 | - | - | 0 | - |
| - | - | 1635 | 980.5 | - | - | 0 | - |
| - | - | 3.038E+04 | 980.6 | - | - | 0 | - |
| - | - | 1648 | 981.5 | - | - | 0 | - |
| - | - | 1.021E+04 | 981.6 | - | - | 0 | - |
| - | - | 803.4 | 982.5 | - | - | 0 | - |
| - | - | 891.2 | 982.6 | - | - | 0 | - |
| - | - | 716.8 | 1288 | - | - | 0 | - |
| - | - | 1098 | 1289 | - | - | 0 | - |
| - | - | 759.7 | 1291 | - | - | 0 | - |
| - | - | 803.2 | 1318 | - | - | 0 | - |

m/z Charge Intensity FragmentType MassShift Position
120.06575775146484 0 3737.3677 y 9
120.0810546875 0 7504.0713
121.08441162109375 0 675.8651
125.97908020019531 0 521.3656
127.7051010131836 0 426.3066
129.10244750976562 0 1765.067
136.07598876953125 0 1858.4567
138.82870483398438 0 406.74847
141.78890991210938 0 370.49982
148.94778442382812 0 516.2002
153.06700134277344 0 475.2633
161.63055419921875 0 436.8516
164.6995391845703 0 489.18033
171.14950561523438 0 34514.12
172.15283203125 0 3469.2961
173.45176696777344 0 1517.8813
187.1078643798828 0 787.593
196.902099609375 0 434.58084
197.12884521484375 0 555.36487
199.1443634033203 0 29053.812
200.147705078125 0 3160.3174
200.5338592529297 0 534.15155
201.123291015625 0 547.9032
205.09744262695312 0 2208.3335
214.04061889648438 0 591.28674
215.1392822265625 0 2016.9354 y Water loss 8
215.26991271972656 0 512.898
230.15000915527344 0 596.35504
233.14968872070312 0 1718.287 y 8
247.1444854736328 0 656.7777
262.1193542480469 0 626.64966
263.1386413574219 0 736.296
267.2649230957031 0 607.5856
276.155517578125 0 3434.6162
277.1180114746094 0 765.34503
301.1914367675781 0 1418.0526
310.2126159667969 0 3925.9316
311.2151184082031 0 892.092
334.2095642089844 0 788.7951
345.2271423339844 0 1886.0591 z 7
346.2118225097656 0 1760.5319
346.23394775390625 0 3837.1165
347.23687744140625 0 708.61237
358.24468994140625 0 1541.753
361.24505615234375 0 1119.6552 y 7
373.7139587402344 0 1073.7673 c Ammonia loss 7
381.71185302734375 0 972.0082 y Water loss 2
383.2032775878906 0 675.61255
385.19610595703125 0 764.91125
390.7165222167969 0 14590.455 y 2
391.21820068359375 0 7795.343
391.7202453613281 0 2348.1924
401.2161865234375 0 1184.549
402.2705383300781 0 1941.538
403.2343444824219 0 1073.8855
415.25634765625 0 1152.7982
416.2601318359375 0 604.3912
419.00323486328125 0 633.28894
420.1893005371094 0 823.18365
440.25054931640625 0 812.4492 y 1
444.2585754394531 0 2833.294
445.2666320800781 0 5019.7817 y Ammonia loss 6
446.26953125 0 1474.7257 z 6
447.2322998046875 0 1151.8767
447.2817077636719 0 5408.4727
448.2853088378906 0 1516.2362
457.2758483886719 0 835.6617
459.29229736328125 0 735.0949
462.293701171875 0 1109.9473 y 6
470.2581481933594 0 1111.9171
473.2007751464844 0 685.7577
474.2662353515625 0 629.2444
476.27435302734375 0 1033.9656
477.28240966796875 0 2886.705 c 4
478.2818908691406 0 939.3196
480.7783508300781 0 1447.3915
489.2323303222656 0 1497.022
489.78424072265625 0 1221.2816
490.28680419921875 0 2893.3438
491.29095458984375 0 769.232
499.2760009765625 0 1418.8605
502.29376220703125 0 809.233 y Ammonia loss 5
503.2951354980469 0 2185.395 z 5
504.2544250488281 0 28700.053
504.3030090332031 0 40340.117
505.25823974609375 0 7870.7954
505.3065185546875 0 10321.923
506.26470947265625 0 1079.7227
506.3081970214844 0 2281.6414
514.298828125 0 2014.4594
516.31396484375 0 1166.0752
517.3165283203125 0 992.85535
518.3064575195312 0 1024.8264
519.3131103515625 0 3358.6272 y 5
520.3175659179688 0 1062.7725
530.2703247070312 0 704.4009
533.2958984375 0 1670.8177
534.3038940429688 0 1288.8097 c 5
548.28271484375 0 2458.4812
550.2824096679688 0 771.88513
560.3167114257812 0 10418.142 z 4
561.2777099609375 0 1234.4015
561.323974609375 0 29332.13
562.3270263671875 0 10316.409
563.3291015625 0 1359.9872
574.28271484375 0 1506.3181
575.32763671875 0 5265.9097
576.3345336914062 0 9931.773 y 4
577.3381958007812 0 3028.578
589.3308715820312 0 1298.9808
607.3040771484375 0 686.3347
613.834716796875 0 550.2379
615.3472290039062 0 761.2559 y Water loss 3
616.3433837890625 0 1014.8196
617.337890625 0 8313.696 c Water loss 6
618.3446655273438 0 19905.299
619.3515014648438 0 5362.1406
620.351806640625 0 1667.5077
632.3489379882812 0 11158.882
633.3564453125 0 40967.84 y 3
634.3538818359375 0 18455.098
635.3515014648438 0 58420.53 c 6
636.3545532226562 0 21575.596
637.3567504882812 0 4192.3135
643.3532104492188 0 1683.8325
646.3453369140625 0 1025.8331
656.7861328125 0 645.3005
659.2904052734375 0 1143.064
661.3035888671875 0 1099.1407
661.3671875 0 6870.1606
662.3689575195312 0 2422.7495
663.3806762695312 0 3616.941
664.38525390625 0 1065.3428
685.3894653320312 0 948.94135
702.4120483398438 0 1771.0065
708.3411865234375 0 869.7705
708.46630859375 0 624.73816
709.3475341796875 0 622.6164
715.388427734375 0 870.5963
716.3939819335938 0 762.47974
718.423095703125 0 832.76874
719.4295043945312 0 824.0131
721.3482055664062 0 983.91187
729.4169921875 0 3014.3337
730.4142456054688 0 2048.7812
746.41064453125 0 3305.5652 c Ammonia loss 7
747.4168090820312 0 1646.4978
748.4197387695312 0 708.8096
762.4168701171875 0 1813.6134 y Water loss 2
763.4459838867188 0 73838.04 c 7
764.3972778320312 0 6840.7627 z 2
764.4487915039062 0 27507.574
765.4046020507812 0 5148.054
765.455810546875 0 5250.641
766.4122924804688 0 1812.3293
773.5360107421875 0 691.2593
780.4248046875 0 60104.445 y 2
781.4281616210938 0 22342.62
782.4295654296875 0 7754.4688
783.4328002929688 0 1682.4066
788.9808959960938 0 600.80145
789.45654296875 0 640.8337
807.4105834960938 0 807.9744
808.41455078125 0 722.9063
832.4369506835938 0 745.94666
832.5167846679688 0 8714.071
833.520263671875 0 3632.9355
834.5255737304688 0 647.5607
859.5023193359375 0 4957.2305 c Ammonia loss 8
860.5068359375 0 1737.107
861.5152587890625 0 2112.0193
862.5235595703125 0 1143.9471
863.4743041992188 0 8402.397 z 1
864.47802734375 0 5135.95
865.477783203125 0 2440.2341
876.5297241210938 0 72943.35 c 8
877.5325927734375 0 41710.812
878.5357055664062 0 11222.768
879.4954833984375 0 2193.282 y 1
879.5313110351562 0 1973.443
880.5051879882812 0 1017.8252
888.4613037109375 0 849.33466
889.4539184570312 0 778.2067
893.4734497070312 0 914.8245
906.4798583984375 0 1064.8096
917.4561157226562 0 1201.6222
919.4850463867188 0 3207.5718
920.4903564453125 0 1577.4535
923.5087280273438 0 3487.6206
924.5122680664062 0 1880.3875
933.5633544921875 0 1448.5292
934.4694213867188 0 1672.3009
935.4800415039062 0 2885.5024
936.4878540039062 0 3347.2007
937.4855346679688 0 1370.4166
947.5120239257812 0 845.49677
951.5818481445312 0 701.824
961.5518798828125 0 1698.2008
962.5427856445312 0 88623.48
963.5458374023438 0 53328.117
964.5482788085938 0 15750.935
965.5506591796875 0 3094.8801
977.5574951171875 0 844.7487
978.5610961914062 0 47831.305
979.4760131835938 0 2994.2346
979.5671997070312 0 62571.77
980.48193359375 0 1634.544
980.570556640625 0 30375.58
981.4823608398438 0 1647.5968
981.5731811523438 0 10208.285
982.4834594726562 0 803.4348
982.5875854492188 0 891.2393
1287.6358642578125 0 716.81085
1288.617919921875 0 1098.2749
1290.6336669921875 0 759.7346
1317.5653076171875 0 803.2275

Spectrum Details

|  |  |
| --- | --- |
| Matched peaks? Matched peaks The total absolute number of peaks matched. Additionally in brackets the total fraction of peaks matched and the total number of peaks is shown. | 32 (14.81% of 216) |
| FDR? FDR The false discovery rate estimated for this peptide. It is calculated by matching all theoretical fragments with a non-integer shift with the raw peaks for this spectrum. This is done with 40 different shifts. The resulting percentage is the average number of annotated peaks over the number of annotated peaks with the correct spectrum. | 0.45% |
| Satellite FDR? Satellite FDR See the FDR for details on its calculation. This satellite ion specific FDR only contains the satellite ions (d/w) for I/L/J positions. | - |
| PSM Score? PSM Score The PSM Score as given by Hecklib to this annotated spectrum. It is shown with three significant figures. | 383 |

#### Spectrum 5975? Spectrum 5975 The raw spectrum of this peptide as annotated by Hecklib. The fragments are coloured according to ion type (see legend). Any peaks with a star '\*' as text can be hovered over to see the full details, first the ion type second the mass shift type. By hovering over the amino acids in the peptide or ions in the legend the corresponding peaks are highlighted. By toggling the 'Unassigned' label you can turn the background (unassigned) peaks on or off in the plot. By updating the slider in the Ion legend you can update the spectrum to only show the top X% of the peaks with labels. The top X% means any peak that is within X% of the highest intensity. By dragging in the spectrum you can zoom in to a specific part of the spectrum and use 'Zoom Out' to get back to the original zoom level. The annotation of the spectrum is based on the given sequence in the peptides file and is done with different software so inconsistencies are likely. The peaks are annotated based on the given sequence, with 20 ppm tolerance.

Copy Data

##### Spectrum 5975 (TSV)

###### Preview

```
Loading example...
```

*Click on the button to copy the data to your clipboard.*

Mz MinMz MaxIntensity Max

WidthHeightPeptide font sizePeptide stroke widthSpectrum font sizeSpectrum stroke widthCompact peptide

Ion legend

wxyz

abcd

OtherUnassignedIon^Charge^_Position_Show for top:%

VVFGGGTKJT

04.77e+49.53e+41.43e+51.91e+5

Zoom Out

y^+1^_1_a^+1^_2_b^+1^_2_y^+1^_2_y^+1^_2_y^+1^_3_b^+1^_3_y^+1^_3_b^+2^_8_y^+2^_8_y^+2^_8_b^+1^_4_y^+2^_9_y^+1^_4_\*\*\*y^+1^_5_y^+1^_5_y^+1^_6_y^+1^_6_y^+1^_7_b^+1^_7_y^+1^_7_b^+1^_8_b^+1^_8_y^+1^_8_y^+1^_8_b^+1^_9_b^+1^_9_y^+1^_9_

048997914681958

Fragment Matches Table

Show background peaks

| Position | Ion type | Intensity | mz Theoretical | mz Error (Th) | mz Error (ppm) | Charge | Series Number |
| --- | --- | --- | --- | --- | --- | --- | --- |
| 10 | y | 1.513E+04 | 120.1 | 0.0003068 | 2.555 | +1 | 1 |
| - | - | 1.063E+05 | 120.1 | - | - | 0 | - |
| - | - | 463.1 | 121.1 | - | - | 0 | - |
| - | - | 7973 | 121.1 | - | - | 0 | - |
| - | - | 359.4 | 121.5 | - | - | 0 | - |
| - | - | 334.9 | 122.4 | - | - | 0 | - |
| - | - | 442.6 | 125.1 | - | - | 0 | - |
| - | - | 402 | 125.1 | - | - | 0 | - |
| - | - | 416.9 | 126.1 | - | - | 0 | - |
| - | - | 1190 | 127.1 | - | - | 0 | - |
| - | - | 556.4 | 127.1 | - | - | 0 | - |
| - | - | 397.2 | 127.6 | - | - | 0 | - |
| - | - | 517.7 | 128.1 | - | - | 0 | - |
| - | - | 735.5 | 128.1 | - | - | 0 | - |
| - | - | 8.442E+04 | 129.1 | - | - | 0 | - |
| - | - | 620.6 | 130 | - | - | 0 | - |
| - | - | 2566 | 130.1 | - | - | 0 | - |
| - | - | 794.6 | 130.1 | - | - | 0 | - |
| - | - | 5070 | 130.1 | - | - | 0 | - |
| - | - | 3584 | 131.1 | - | - | 0 | - |
| - | - | 1174 | 131.1 | - | - | 0 | - |
| - | - | 1489 | 132.1 | - | - | 0 | - |
| - | - | 1180 | 133.1 | - | - | 0 | - |
| - | - | 478.3 | 134.1 | - | - | 0 | - |
| - | - | 1.004E+05 | 136.1 | - | - | 0 | - |
| - | - | 381.5 | 137.2 | - | - | 0 | - |
| - | - | 592.1 | 139.1 | - | - | 0 | - |
| - | - | 412 | 139.1 | - | - | 0 | - |
| - | - | 378.1 | 139.2 | - | - | 0 | - |
| - | - | 1202 | 140.1 | - | - | 0 | - |
| - | - | 1496 | 141.1 | - | - | 0 | - |
| - | - | 905.2 | 141.1 | - | - | 0 | - |
| - | - | 451.6 | 143.1 | - | - | 0 | - |
| - | - | 714.8 | 143.1 | - | - | 0 | - |
| - | - | 717.4 | 145.1 | - | - | 0 | - |
| - | - | 522.7 | 145.1 | - | - | 0 | - |
| - | - | 410 | 145.7 | - | - | 0 | - |
| - | - | 1543 | 146.1 | - | - | 0 | - |
| - | - | 938.7 | 147 | - | - | 0 | - |
| - | - | 1042 | 147.1 | - | - | 0 | - |
| - | - | 865.8 | 149 | - | - | 0 | - |
| - | - | 564.8 | 149 | - | - | 0 | - |
| - | - | 879.3 | 152.1 | - | - | 0 | - |
| - | - | 622.9 | 152.1 | - | - | 0 | - |
| - | - | 696.9 | 153.1 | - | - | 0 | - |
| - | - | 386.4 | 153.1 | - | - | 0 | - |
| - | - | 1073 | 154.2 | - | - | 0 | - |
| - | - | 1450 | 155.1 | - | - | 0 | - |
| - | - | 2830 | 155.1 | - | - | 0 | - |
| - | - | 427.4 | 155.6 | - | - | 0 | - |
| - | - | 427.7 | 155.7 | - | - | 0 | - |
| - | - | 687 | 157.1 | - | - | 0 | - |
| - | - | 428.6 | 157.1 | - | - | 0 | - |
| - | - | 1026 | 158.1 | - | - | 0 | - |
| - | - | 2588 | 159.1 | - | - | 0 | - |
| - | - | 406 | 159.1 | - | - | 0 | - |
| - | - | 1268 | 159.1 | - | - | 0 | - |
| - | - | 907.3 | 159.1 | - | - | 0 | - |
| - | - | 513.6 | 162.1 | - | - | 0 | - |
| - | - | 1209 | 163.1 | - | - | 0 | - |
| - | - | 751.2 | 166.1 | - | - | 0 | - |
| - | - | 552.6 | 166.1 | - | - | 0 | - |
| - | - | 497.3 | 167 | - | - | 0 | - |
| - | - | 1244 | 167.1 | - | - | 0 | - |
| - | - | 1448 | 169.1 | - | - | 0 | - |
| - | - | 2092 | 170.1 | - | - | 0 | - |
| 2 | a | 1.888E+05 | 171.1 | 0.0003922 | 2.292 | +1 | 2 |
| - | - | 3440 | 172.1 | - | - | 0 | - |
| - | - | 988.8 | 172.1 | - | - | 0 | - |
| - | - | 1.757E+04 | 172.2 | - | - | 0 | - |
| - | - | 1487 | 173.1 | - | - | 0 | - |
| - | - | 849.6 | 173.2 | - | - | 0 | - |
| - | - | 1555 | 173.5 | - | - | 0 | - |
| - | - | 568.3 | 174.1 | - | - | 0 | - |
| - | - | 603.7 | 174.1 | - | - | 0 | - |
| - | - | 531.8 | 175.1 | - | - | 0 | - |
| - | - | 3500 | 176.1 | - | - | 0 | - |
| - | - | 9069 | 177.1 | - | - | 0 | - |
| - | - | 699.3 | 177.1 | - | - | 0 | - |
| - | - | 897.5 | 178.1 | - | - | 0 | - |
| - | - | 555.2 | 180.1 | - | - | 0 | - |
| - | - | 1743 | 181.1 | - | - | 0 | - |
| - | - | 992.8 | 181.2 | - | - | 0 | - |
| - | - | 1165 | 183.1 | - | - | 0 | - |
| - | - | 863.9 | 183.1 | - | - | 0 | - |
| - | - | 509.6 | 184.1 | - | - | 0 | - |
| - | - | 1135 | 185.1 | - | - | 0 | - |
| - | - | 1.51E+04 | 185.2 | - | - | 0 | - |
| - | - | 501.5 | 187.1 | - | - | 0 | - |
| - | - | 2877 | 187.1 | - | - | 0 | - |
| - | - | 964.7 | 188.1 | - | - | 0 | - |
| - | - | 471.8 | 190.5 | - | - | 0 | - |
| - | - | 642.6 | 191.1 | - | - | 0 | - |
| - | - | 663.9 | 194.1 | - | - | 0 | - |
| - | - | 566.1 | 196.1 | - | - | 0 | - |
| - | - | 539.2 | 197.1 | - | - | 0 | - |
| - | - | 554.9 | 197.1 | - | - | 0 | - |
| - | - | 2557 | 197.1 | - | - | 0 | - |
| - | - | 3592 | 197.2 | - | - | 0 | - |
| - | - | 4811 | 198.1 | - | - | 0 | - |
| - | - | 768.6 | 199.1 | - | - | 0 | - |
| 2 | b | 9.778E+04 | 199.1 | 0.0002896 | 1.454 | +1 | 2 |
| - | - | 9533 | 200.1 | - | - | 0 | - |
| - | - | 3112 | 201.1 | - | - | 0 | - |
| - | - | 505.1 | 202.1 | - | - | 0 | - |
| - | - | 576.1 | 203.1 | - | - | 0 | - |
| - | - | 1.806E+04 | 205.1 | - | - | 0 | - |
| - | - | 1913 | 206.1 | - | - | 0 | - |
| - | - | 1660 | 207.1 | - | - | 0 | - |
| - | - | 1028 | 208.1 | - | - | 0 | - |
| - | - | 540.1 | 208.9 | - | - | 0 | - |
| - | - | 738.3 | 209.1 | - | - | 0 | - |
| - | - | 733.3 | 209.2 | - | - | 0 | - |
| - | - | 989.3 | 210.1 | - | - | 0 | - |
| - | - | 1012 | 211.1 | - | - | 0 | - |
| - | - | 8682 | 212.1 | - | - | 0 | - |
| - | - | 846.2 | 212.1 | - | - | 0 | - |
| - | - | 1053 | 213.1 | - | - | 0 | - |
| - | - | 6895 | 213.2 | - | - | 0 | - |
| - | - | 711.5 | 214.1 | - | - | 0 | - |
| - | - | 1642 | 215.1 | - | - | 0 | - |
| 9 | y | 2931 | 215.1 | 0.0002023 | 0.9402 | +1 | 2 |
| - | - | 2706 | 216.1 | - | - | 0 | - |
| - | - | 2687 | 217.1 | - | - | 0 | - |
| - | - | 7939 | 219.1 | - | - | 0 | - |
| - | - | 1231 | 220.2 | - | - | 0 | - |
| - | - | 936.5 | 221.1 | - | - | 0 | - |
| - | - | 527.2 | 222.1 | - | - | 0 | - |
| - | - | 610 | 223.1 | - | - | 0 | - |
| - | - | 2918 | 224.1 | - | - | 0 | - |
| - | - | 1539 | 224.1 | - | - | 0 | - |
| - | - | 2605 | 224.2 | - | - | 0 | - |
| - | - | 1185 | 225.1 | - | - | 0 | - |
| - | - | 1039 | 225.1 | - | - | 0 | - |
| - | - | 820.8 | 225.2 | - | - | 0 | - |
| - | - | 649.7 | 225.2 | - | - | 0 | - |
| - | - | 738 | 226.1 | - | - | 0 | - |
| - | - | 1331 | 226.1 | - | - | 0 | - |
| - | - | 935.8 | 226.2 | - | - | 0 | - |
| - | - | 3390 | 227.1 | - | - | 0 | - |
| - | - | 1358 | 228.1 | - | - | 0 | - |
| - | - | 799.1 | 229.1 | - | - | 0 | - |
| - | - | 1.727E+04 | 230.2 | - | - | 0 | - |
| - | - | 989.3 | 231.1 | - | - | 0 | - |
| - | - | 2208 | 231.2 | - | - | 0 | - |
| 9 | y | 5370 | 233.1 | 0.0004103 | 1.76 | +1 | 2 |
| - | - | 1057 | 233.2 | - | - | 0 | - |
| - | - | 684.3 | 234.1 | - | - | 0 | - |
| - | - | 929.3 | 237.1 | - | - | 0 | - |
| - | - | 700.6 | 237.2 | - | - | 0 | - |
| - | - | 1578 | 240.1 | - | - | 0 | - |
| - | - | 577.8 | 241.2 | - | - | 0 | - |
| - | - | 2044 | 242.1 | - | - | 0 | - |
| - | - | 4348 | 242.2 | - | - | 0 | - |
| - | - | 3860 | 242.2 | - | - | 0 | - |
| - | - | 570.3 | 242.4 | - | - | 0 | - |
| - | - | 2121 | 243.1 | - | - | 0 | - |
| - | - | 1486 | 244.1 | - | - | 0 | - |
| - | - | 691.4 | 244.1 | - | - | 0 | - |
| - | - | 2242 | 245.1 | - | - | 0 | - |
| - | - | 4117 | 247.1 | - | - | 0 | - |
| - | - | 681.1 | 249.1 | - | - | 0 | - |
| - | - | 1547 | 249.2 | - | - | 0 | - |
| - | - | 1619 | 250.1 | - | - | 0 | - |
| - | - | 4821 | 251.1 | - | - | 0 | - |
| - | - | 1112 | 251.1 | - | - | 0 | - |
| - | - | 1762 | 252.1 | - | - | 0 | - |
| - | - | 535.4 | 252.1 | - | - | 0 | - |
| - | - | 2649 | 253.2 | - | - | 0 | - |
| - | - | 9250 | 255.1 | - | - | 0 | - |
| - | - | 1087 | 256.1 | - | - | 0 | - |
| - | - | 2008 | 258.1 | - | - | 0 | - |
| - | - | 1196 | 261.1 | - | - | 0 | - |
| - | - | 6180 | 262.1 | - | - | 0 | - |
| - | - | 547.5 | 263 | - | - | 0 | - |
| - | - | 584.5 | 263.1 | - | - | 0 | - |
| - | - | 1455 | 263.1 | - | - | 0 | - |
| - | - | 1058 | 263.1 | - | - | 0 | - |
| - | - | 789.1 | 268.1 | - | - | 0 | - |
| - | - | 652 | 268.2 | - | - | 0 | - |
| - | - | 1.613E+04 | 269.1 | - | - | 0 | - |
| - | - | 9875 | 269.2 | - | - | 0 | - |
| - | - | 1438 | 270.2 | - | - | 0 | - |
| - | - | 1683 | 270.2 | - | - | 0 | - |
| - | - | 585 | 271.1 | - | - | 0 | - |
| - | - | 996.4 | 272.1 | - | - | 0 | - |
| - | - | 1.037E+04 | 273.1 | - | - | 0 | - |
| - | - | 1261 | 273.2 | - | - | 0 | - |
| - | - | 2510 | 274.1 | - | - | 0 | - |
| - | - | 2365 | 276.2 | - | - | 0 | - |
| - | - | 540.6 | 277.2 | - | - | 0 | - |
| - | - | 579.1 | 277.9 | - | - | 0 | - |
| - | - | 584.7 | 281.2 | - | - | 0 | - |
| - | - | 1654 | 281.2 | - | - | 0 | - |
| - | - | 1005 | 282.2 | - | - | 0 | - |
| - | - | 642.3 | 282.2 | - | - | 0 | - |
| - | - | 535 | 285.2 | - | - | 0 | - |
| - | - | 2624 | 287.2 | - | - | 0 | - |
| - | - | 949.5 | 288.1 | - | - | 0 | - |
| - | - | 658.9 | 288.2 | - | - | 0 | - |
| - | - | 2573 | 291.1 | - | - | 0 | - |
| - | - | 605.3 | 292.8 | - | - | 0 | - |
| - | - | 636.9 | 296.1 | - | - | 0 | - |
| - | - | 853.7 | 296.2 | - | - | 0 | - |
| - | - | 851.6 | 297.1 | - | - | 0 | - |
| - | - | 656.8 | 299.2 | - | - | 0 | - |
| - | - | 773.2 | 300.1 | - | - | 0 | - |
| - | - | 4027 | 301.2 | - | - | 0 | - |
| - | - | 1211 | 302.2 | - | - | 0 | - |
| - | - | 2343 | 304.2 | - | - | 0 | - |
| - | - | 993.7 | 310.1 | - | - | 0 | - |
| - | - | 3952 | 310.2 | - | - | 0 | - |
| - | - | 917.2 | 311.2 | - | - | 0 | - |
| - | - | 1912 | 313.2 | - | - | 0 | - |
| - | - | 1114 | 313.2 | - | - | 0 | - |
| - | - | 659.4 | 315.2 | - | - | 0 | - |
| - | - | 1524 | 317.2 | - | - | 0 | - |
| - | - | 1319 | 318.1 | - | - | 0 | - |
| - | - | 3371 | 319.1 | - | - | 0 | - |
| - | - | 911.8 | 322.2 | - | - | 0 | - |
| - | - | 762 | 322.7 | - | - | 0 | - |
| - | - | 881.5 | 325.1 | - | - | 0 | - |
| - | - | 1617 | 325.2 | - | - | 0 | - |
| - | - | 9617 | 326.2 | - | - | 0 | - |
| - | - | 6171 | 327.2 | - | - | 0 | - |
| - | - | 1042 | 327.2 | - | - | 0 | - |
| - | - | 685.7 | 328.1 | - | - | 0 | - |
| - | - | 753.5 | 331.2 | - | - | 0 | - |
| - | - | 855.4 | 332.2 | - | - | 0 | - |
| - | - | 1057 | 333.2 | - | - | 0 | - |
| - | - | 1097 | 339.2 | - | - | 0 | - |
| - | - | 778 | 339.2 | - | - | 0 | - |
| - | - | 2280 | 341.1 | - | - | 0 | - |
| - | - | 896.2 | 341.2 | - | - | 0 | - |
| - | - | 3035 | 341.2 | - | - | 0 | - |
| - | - | 645.8 | 342.1 | - | - | 0 | - |
| 8 | y | 3504 | 343.2 | 0.000271 | 0.7896 | +1 | 3 |
| - | - | 2259 | 344.2 | - | - | 0 | - |
| - | - | 828.5 | 345.2 | - | - | 0 | - |
| 3 | b | 5766 | 346.2 | 0.0004029 | 1.164 | +1 | 3 |
| - | - | 665.3 | 347.1 | - | - | 0 | - |
| - | - | 998.8 | 347.2 | - | - | 0 | - |
| - | - | 1704 | 348.2 | - | - | 0 | - |
| - | - | 1998 | 353.1 | - | - | 0 | - |
| - | - | 750.2 | 354.2 | - | - | 0 | - |
| - | - | 5416 | 355.2 | - | - | 0 | - |
| - | - | 2163 | 357.2 | - | - | 0 | - |
| - | - | 1252 | 359.2 | - | - | 0 | - |
| - | - | 1460 | 361.2 | - | - | 0 | - |
| 8 | y | 1.285E+04 | 361.2 | 0.0001433 | 0.3968 | +1 | 3 |
| - | - | 2805 | 362.2 | - | - | 0 | - |
| - | - | 745 | 364.2 | - | - | 0 | - |
| - | - | 792.7 | 365.2 | - | - | 0 | - |
| - | - | 699.6 | 367.2 | - | - | 0 | - |
| - | - | 841.1 | 368.2 | - | - | 0 | - |
| - | - | 1795 | 370.1 | - | - | 0 | - |
| - | - | 2930 | 371.1 | - | - | 0 | - |
| - | - | 628.9 | 372.3 | - | - | 0 | - |
| - | - | 950.9 | 372.6 | - | - | 0 | - |
| - | - | 4401 | 373.2 | - | - | 0 | - |
| 8 | b | 729.8 | 373.7 | 0.0003704 | 0.9913 | +2 | 8 |
| - | - | 3859 | 374.2 | - | - | 0 | - |
| - | - | 1577 | 375.2 | - | - | 0 | - |
| - | - | 731.2 | 375.2 | - | - | 0 | - |
| - | - | 1791 | 376.2 | - | - | 0 | - |
| 3 | y | 3895 | 381.7 | 8.606E-05 | 0.2255 | +2 | 8 |
| - | - | 878.8 | 382.2 | - | - | 0 | - |
| - | - | 1767 | 382.2 | - | - | 0 | - |
| - | - | 8561 | 383.2 | - | - | 0 | - |
| - | - | 1164 | 383.2 | - | - | 0 | - |
| - | - | 1534 | 384.2 | - | - | 0 | - |
| - | - | 2018 | 388.1 | - | - | 0 | - |
| 3 | y | 3.017E+04 | 390.7 | 0.0002163 | 0.5536 | +2 | 8 |
| - | - | 1.368E+04 | 391.2 | - | - | 0 | - |
| - | - | 3971 | 391.7 | - | - | 0 | - |
| - | - | 4120 | 392.2 | - | - | 0 | - |
| - | - | 944.8 | 393.2 | - | - | 0 | - |
| - | - | 3305 | 400.3 | - | - | 0 | - |
| - | - | 7228 | 401.2 | - | - | 0 | - |
| - | - | 1027 | 401.3 | - | - | 0 | - |
| - | - | 1.365E+04 | 402.2 | - | - | 0 | - |
| - | - | 946.3 | 402.2 | - | - | 0 | - |
| - | - | 2935 | 403.2 | - | - | 0 | - |
| 4 | b | 2389 | 403.2 | 0.0003015 | 0.7478 | +1 | 4 |
| - | - | 714.1 | 405.2 | - | - | 0 | - |
| - | - | 828.8 | 408.2 | - | - | 0 | - |
| - | - | 909.1 | 409.2 | - | - | 0 | - |
| - | - | 798.2 | 409.3 | - | - | 0 | - |
| - | - | 635.5 | 410.2 | - | - | 0 | - |
| - | - | 1004 | 415.3 | - | - | 0 | - |
| - | - | 651 | 416.1 | - | - | 0 | - |
| - | - | 661.8 | 416.3 | - | - | 0 | - |
| - | - | 938 | 416.8 | - | - | 0 | - |
| - | - | 1638 | 419 | - | - | 0 | - |
| - | - | 863.5 | 419.2 | - | - | 0 | - |
| - | - | 7949 | 420.2 | - | - | 0 | - |
| - | - | 1548 | 421.2 | - | - | 0 | - |
| - | - | 3491 | 426.2 | - | - | 0 | - |
| - | - | 1381 | 429.3 | - | - | 0 | - |
| - | - | 1237 | 432.7 | - | - | 0 | - |
| - | - | 1506 | 437.2 | - | - | 0 | - |
| - | - | 1244 | 438.2 | - | - | 0 | - |
| - | - | 7043 | 439.3 | - | - | 0 | - |
| 2 | y | 2300 | 440.3 | 0.001227 | 2.786 | +2 | 9 |
| - | - | 695.4 | 440.8 | - | - | 0 | - |
| - | - | 712.5 | 441.3 | - | - | 0 | - |
| - | - | 933.3 | 442.2 | - | - | 0 | - |
| - | - | 1194 | 444.2 | - | - | 0 | - |
| - | - | 5649 | 454.2 | - | - | 0 | - |
| - | - | 2019 | 455.2 | - | - | 0 | - |
| - | - | 3731 | 456.2 | - | - | 0 | - |
| - | - | 9396 | 457.3 | - | - | 0 | - |
| - | - | 1754 | 458.3 | - | - | 0 | - |
| 7 | y | 3137 | 462.3 | 0.0004995 | 1.081 | +1 | 4 |
| - | - | 996.8 | 463.3 | - | - | 0 | - |
| - | - | 875.7 | 468.3 | - | - | 0 | - |
| - | - | 1321 | 470.2 | - | - | 0 | - |
| - | - | 3789 | 472.2 | - | - | 0 | - |
| - | - | 1449 | 473.2 | - | - | 0 | - |
| - | - | 685.9 | 478.3 | - | - | 0 | - |
| 0 | Precursor | 1971 | 480.8 | 0.001813 | 3.77 | +2 | -1 |
| 0 | Precursor | 1137 | 481.3 | 0.009469 | 19.68 | +2 | -1 |
| - | - | 630.8 | 482.2 | - | - | 0 | - |
| - | - | 1384 | 483.2 | - | - | 0 | - |
| - | - | 5078 | 484.2 | - | - | 0 | - |
| - | - | 1319 | 486.3 | - | - | 0 | - |
| - | - | 5125 | 486.3 | - | - | 0 | - |
| - | - | 1649 | 487.3 | - | - | 0 | - |
| - | - | 1079 | 489.2 | - | - | 0 | - |
| 0 | Precursor | 3237 | 489.8 | 0.0002533 | 0.5172 | +2 | -1 |
| - | - | 1378 | 490.2 | - | - | 0 | - |
| - | - | 1831 | 490.3 | - | - | 0 | - |
| - | - | 793.2 | 491.3 | - | - | 0 | - |
| - | - | 9259 | 496.3 | - | - | 0 | - |
| - | - | 2847 | 497.3 | - | - | 0 | - |
| - | - | 2054 | 501.2 | - | - | 0 | - |
| 6 | y | 780.8 | 501.3 | 0.00135 | 2.692 | +1 | 5 |
| - | - | 684.8 | 504.3 | - | - | 0 | - |
| - | - | 1869 | 512.3 | - | - | 0 | - |
| - | - | 2.275E+04 | 514.3 | - | - | 0 | - |
| - | - | 1006 | 515.3 | - | - | 0 | - |
| - | - | 6163 | 515.3 | - | - | 0 | - |
| - | - | 810.3 | 516.3 | - | - | 0 | - |
| - | - | 1737 | 518.2 | - | - | 0 | - |
| - | - | 1180 | 519.3 | - | - | 0 | - |
| 6 | y | 4974 | 519.3 | 0.0003371 | 0.6491 | +1 | 5 |
| - | - | 1211 | 520.3 | - | - | 0 | - |
| - | - | 1182 | 522.3 | - | - | 0 | - |
| - | - | 7904 | 530.3 | - | - | 0 | - |
| - | - | 2334 | 531.3 | - | - | 0 | - |
| - | - | 560 | 536.2 | - | - | 0 | - |
| - | - | 796.9 | 537.3 | - | - | 0 | - |
| - | - | 1420 | 539.3 | - | - | 0 | - |
| - | - | 1.737E+04 | 548.3 | - | - | 0 | - |
| - | - | 4718 | 549.3 | - | - | 0 | - |
| - | - | 858.9 | 550.3 | - | - | 0 | - |
| - | - | 2709 | 554.3 | - | - | 0 | - |
| - | - | 1614 | 555.3 | - | - | 0 | - |
| 5 | y | 1266 | 558.3 | 0.001035 | 1.853 | +1 | 6 |
| - | - | 1850 | 567.3 | - | - | 0 | - |
| - | - | 1183 | 569.3 | - | - | 0 | - |
| - | - | 1592 | 572.3 | - | - | 0 | - |
| 5 | y | 2.316E+04 | 576.3 | 0.0001306 | 0.2265 | +1 | 6 |
| - | - | 6944 | 577.3 | - | - | 0 | - |
| - | - | 1630 | 578.3 | - | - | 0 | - |
| - | - | 2254 | 582.3 | - | - | 0 | - |
| - | - | 3003 | 583.3 | - | - | 0 | - |
| - | - | 1680 | 585.3 | - | - | 0 | - |
| - | - | 1453 | 586.3 | - | - | 0 | - |
| - | - | 616.1 | 589.3 | - | - | 0 | - |
| - | - | 975.6 | 590.3 | - | - | 0 | - |
| - | - | 893.9 | 596.3 | - | - | 0 | - |
| - | - | 890.3 | 597.3 | - | - | 0 | - |
| - | - | 5758 | 600.3 | - | - | 0 | - |
| - | - | 1368 | 605.3 | - | - | 0 | - |
| - | - | 1633 | 614.3 | - | - | 0 | - |
| 4 | y | 3883 | 615.3 | 7.88E-05 | 0.1281 | +1 | 7 |
| - | - | 2186 | 616.3 | - | - | 0 | - |
| - | - | 965.6 | 617.3 | - | - | 0 | - |
| 7 | b | 1507 | 618.3 | 0.000125 | 0.2021 | +1 | 7 |
| - | - | 1356 | 625.3 | - | - | 0 | - |
| 4 | y | 6.886E+04 | 633.4 | 0.00105 | 1.657 | +1 | 7 |
| - | - | 1.998E+04 | 634.4 | - | - | 0 | - |
| - | - | 3101 | 635.4 | - | - | 0 | - |
| - | - | 1.766E+04 | 643.4 | - | - | 0 | - |
| - | - | 7300 | 644.4 | - | - | 0 | - |
| - | - | 2124 | 645.4 | - | - | 0 | - |
| - | - | 1631 | 647.3 | - | - | 0 | - |
| - | - | 1225 | 649.4 | - | - | 0 | - |
| - | - | 1484 | 650.4 | - | - | 0 | - |
| - | - | 3.855E+04 | 661.4 | - | - | 0 | - |
| - | - | 1.464E+04 | 662.4 | - | - | 0 | - |
| - | - | 2583 | 663.4 | - | - | 0 | - |
| - | - | 2187 | 667.4 | - | - | 0 | - |
| - | - | 2069 | 668.4 | - | - | 0 | - |
| - | - | 994.7 | 679.4 | - | - | 0 | - |
| - | - | 4829 | 685.4 | - | - | 0 | - |
| - | - | 1900 | 695.4 | - | - | 0 | - |
| - | - | 2085 | 696.4 | - | - | 0 | - |
| - | - | 6153 | 713.4 | - | - | 0 | - |
| - | - | 1421 | 718.4 | - | - | 0 | - |
| 8 | b | 1430 | 728.4 | 0.0008441 | 1.159 | +1 | 8 |
| - | - | 1045 | 733.3 | - | - | 0 | - |
| - | - | 2301 | 734.3 | - | - | 0 | - |
| - | - | 836.3 | 742.4 | - | - | 0 | - |
| - | - | 865.8 | 744.4 | - | - | 0 | - |
| 8 | b | 5202 | 746.4 | 0.001399 | 1.874 | +1 | 8 |
| - | - | 2964 | 747.4 | - | - | 0 | - |
| - | - | 2621 | 751.3 | - | - | 0 | - |
| - | - | 2555 | 760.4 | - | - | 0 | - |
| 3 | y | 6466 | 762.4 | 0.0005251 | 0.6887 | +1 | 8 |
| - | - | 2577 | 763.4 | - | - | 0 | - |
| - | - | 836.5 | 764.4 | - | - | 0 | - |
| - | - | 868.2 | 769.3 | - | - | 0 | - |
| - | - | 869.3 | 779.4 | - | - | 0 | - |
| 3 | y | 1.335E+05 | 780.4 | 0.0007138 | 0.9146 | +1 | 8 |
| - | - | 6.169E+04 | 781.4 | - | - | 0 | - |
| - | - | 1.433E+04 | 782.4 | - | - | 0 | - |
| - | - | 1177 | 783.4 | - | - | 0 | - |
| 9 | b | 2449 | 841.5 | 0.0005575 | 0.6625 | +1 | 9 |
| - | - | 753 | 842.5 | - | - | 0 | - |
| 9 | b | 7101 | 859.5 | 0.001723 | 2.004 | +1 | 9 |
| - | - | 4406 | 860.5 | - | - | 0 | - |
| - | - | 930.1 | 861.5 | - | - | 0 | - |
| - | - | 1066 | 876.4 | - | - | 0 | - |
| 2 | y | 7798 | 879.5 | 0.0009514 | 1.082 | +1 | 9 |
| - | - | 4084 | 880.5 | - | - | 0 | - |
| - | - | 1128 | 881.5 | - | - | 0 | - |
| - | - | 778.9 | 1045 | - | - | 0 | - |
| - | - | 709.4 | 1938 | - | - | 0 | - |

m/z Charge Intensity FragmentType MassShift Position
120.06582641601562 0 15133.282 y 9
120.08114624023438 0 106255.95
121.0692367553711 0 463.103
121.08448028564453 0 7972.7393
121.5196304321289 0 359.4432
122.36387634277344 0 334.859
125.07147979736328 0 442.62064
125.10783386230469 0 402.0187
126.0551986694336 0 416.89124
127.05046081542969 0 1190.4209
127.08685302734375 0 556.4114
127.62518310546875 0 397.23157
128.0712127685547 0 517.6804
128.1070556640625 0 735.48145
129.1025848388672 0 84418
130.04995727539062 0 620.5821
130.06552124023438 0 2565.9617
130.10015869140625 0 794.594
130.10594177246094 0 5070.4473
131.0818328857422 0 3583.7688
131.1182861328125 0 1174.0203
132.08102416992188 0 1488.8461
133.0862274169922 0 1180.4683
134.06080627441406 0 478.29196
136.0760498046875 0 100423.1
137.21206665039062 0 381.49268
139.05079650878906 0 592.0949
139.08717346191406 0 412.0063
139.15565490722656 0 378.11807
140.14337158203125 0 1202.2892
141.06619262695312 0 1496.0588
141.1026611328125 0 905.157
143.08169555664062 0 451.5695
143.1181182861328 0 714.75183
145.06109619140625 0 717.44116
145.0677032470703 0 522.6568
145.66604614257812 0 409.97015
146.0605010986328 0 1542.5164
147.04747009277344 0 938.6891
147.0767059326172 0 1042.0134
148.9544219970703 0 865.80554
149.0238800048828 0 564.8324
152.0711212158203 0 879.3157
152.1436767578125 0 622.88086
153.06643676757812 0 696.86115
153.07742309570312 0 386.40097
154.15940856933594 0 1072.8315
155.0818634033203 0 1449.8328
155.11807250976562 0 2829.6284
155.6244659423828 0 427.4267
155.7075653076172 0 427.7129
157.0612030029297 0 687.0285
157.0979766845703 0 428.5587
158.0926513671875 0 1025.7222
159.0764617919922 0 2587.9368
159.08277893066406 0 405.99548
159.09193420410156 0 1267.9916
159.11289978027344 0 907.34515
162.08709716796875 0 513.55927
163.0718536376953 0 1209.2422
166.05307006835938 0 751.15936
166.08656311035156 0 552.599
167.04507446289062 0 497.31787
167.11827087402344 0 1244.0175
169.13389587402344 0 1448.4541
170.0928955078125 0 2091.8198
171.1495819091797 0 188788.86 a 1
172.0718231201172 0 3440.3171
172.14630126953125 0 988.8266
172.15292358398438 0 17566.152
173.12884521484375 0 1487.0333
173.15586853027344 0 849.6367
173.45162963867188 0 1554.6666
174.05517578125 0 568.3163
174.12782287597656 0 603.7469
175.0868377685547 0 531.80133
176.10733032226562 0 3499.8699
177.10255432128906 0 9068.588
177.11102294921875 0 699.3302
178.10589599609375 0 897.4652
180.07708740234375 0 555.17664
181.0612335205078 0 1742.9727
181.17030334472656 0 992.8173
183.1127166748047 0 1164.6178
183.14952087402344 0 863.9017
184.071533203125 0 509.57034
185.12879943847656 0 1135.4216
185.16514587402344 0 15101.134
187.10800170898438 0 501.52887
187.14459228515625 0 2876.9885
188.1033172607422 0 964.7311
190.47528076171875 0 471.75854
191.0736083984375 0 642.59155
194.129638671875 0 663.8712
196.07212829589844 0 566.14703
197.09214782714844 0 539.16
197.10342407226562 0 554.8904
197.12879943847656 0 2557.4548
197.16510009765625 0 3592.4707
198.08750915527344 0 4810.856
199.07135009765625 0 768.5979
199.14439392089844 0 97775.87 b 1
200.1477813720703 0 9533.329
201.1234893798828 0 3111.8843
202.0831756591797 0 505.1147
203.1186981201172 0 576.12177
205.09742736816406 0 18057.25
206.10121154785156 0 1912.895
207.14944458007812 0 1660.1167
208.10861206054688 0 1027.6489
208.8999786376953 0 540.11273
209.1031036376953 0 738.27673
209.16561889648438 0 733.29913
210.0878448486328 0 989.29663
211.14454650878906 0 1012.1283
212.13963317871094 0 8681.692
212.1499481201172 0 846.2227
213.12408447265625 0 1052.8555
213.16012573242188 0 6894.8965
214.08226013183594 0 711.5058
215.1140594482422 0 1641.8379
215.13922119140625 0 2931.2732 y Water loss 8
216.09829711914062 0 2706.4272
217.09739685058594 0 2686.5403
219.14952087402344 0 7939.4697
220.15321350097656 0 1230.8787
221.12860107421875 0 936.5261
222.1246337890625 0 527.19653
223.1074676513672 0 609.9811
224.06680297851562 0 2917.94
224.09205627441406 0 1538.794
224.17623901367188 0 2604.825
225.12379455566406 0 1184.901
225.1353302001953 0 1038.6901
225.1603240966797 0 820.7774
225.1796875 0 649.65704
226.08343505859375 0 737.97076
226.11920166015625 0 1331.0244
226.15513610839844 0 935.79236
227.1143798828125 0 3390.1157
228.0983123779297 0 1358.4072
229.0929412841797 0 799.05554
230.15028381347656 0 17273.404
231.11314392089844 0 989.26196
231.15301513671875 0 2208.123
233.14999389648438 0 5370.1895 y 8
233.1639862060547 0 1056.8125
234.10873413085938 0 684.34674
237.0986328125 0 929.26117
237.15933227539062 0 700.56665
240.1346893310547 0 1577.9836
241.15403747558594 0 577.7664
242.077392578125 0 2043.8289
242.15011596679688 0 4347.9863
242.18679809570312 0 3859.9888
242.36085510253906 0 570.2718
243.11326599121094 0 2120.7366
244.09312438964844 0 1485.84
244.1074676513672 0 691.41254
245.12440490722656 0 2242.4695
247.1444091796875 0 4116.923
249.12399291992188 0 681.10736
249.1599578857422 0 1546.8506
250.0825653076172 0 1619.0728
251.10299682617188 0 4820.715
251.14987182617188 0 1112.2609
252.0872344970703 0 1762.207
252.13458251953125 0 535.3651
253.22793579101562 0 2648.7917
255.1089630126953 0 9250.436
256.11138916015625 0 1087.1735
258.1452941894531 0 2008.264
261.1236572265625 0 1196.4926
262.11883544921875 0 6180.474
262.9559326171875 0 547.4611
263.1032409667969 0 584.53357
263.1230163574219 0 1455.4075
263.13861083984375 0 1057.6019
268.0929870605469 0 789.05035
268.2029113769531 0 652.0103
269.1133728027344 0 16130.655
269.16094970703125 0 9875.494
270.1644287109375 0 1437.8572
270.1814270019531 0 1682.9569
271.10748291015625 0 584.9909
272.1356201171875 0 996.4336
273.1195373535156 0 10374.641
273.19598388671875 0 1260.54
274.11968994140625 0 2509.8203
276.15582275390625 0 2365.085
277.1539306640625 0 540.59033
277.9388427734375 0 579.14386
281.1606140136719 0 584.7222
281.22216796875 0 1654.2928
282.1573791503906 0 1004.67194
282.18145751953125 0 642.26904
285.18975830078125 0 535.0266
287.17156982421875 0 2624.2747
288.13531494140625 0 949.514
288.1753234863281 0 658.9388
291.1457214355469 0 2572.9338
292.8443298339844 0 605.33795
296.0876159667969 0 636.86694
296.19622802734375 0 853.6981
297.10797119140625 0 851.55286
299.17169189453125 0 656.792
300.1351318359375 0 773.1594
301.1911926269531 0 4026.8164
302.1946716308594 0 1211.1672
304.16595458984375 0 2343.3342
310.1400146484375 0 993.7218
310.2129211425781 0 3951.7573
311.171142578125 0 917.1874
313.15057373046875 0 1912.1499
313.1863708496094 0 1114.0223
315.2395324707031 0 659.41724
317.1883239746094 0 1523.8916
318.14483642578125 0 1318.9272
319.1408386230469 0 3371.4685
322.18212890625 0 911.83044
322.6838684082031 0 761.98114
325.1129455566406 0 881.46124
325.2240295410156 0 1616.843
326.1826477050781 0 9617.17
327.1665344238281 0 6171.008
327.18603515625 0 1042.2605
328.1282958984375 0 685.72095
331.1858825683594 0 753.4869
332.16131591796875 0 855.4137
333.19354248046875 0 1056.6671
339.1786804199219 0 1097.2726
339.2019958496094 0 777.9778
341.1465148925781 0 2280.3794
341.18206787109375 0 896.2047
341.2186584472656 0 3034.8823
342.1433410644531 0 645.82733
343.2342529296875 0 3503.8196 y Water loss 7
344.1934814453125 0 2258.757
345.1763916015625 0 828.4762
346.2129211425781 0 5766.4824 b 2
347.1360778808594 0 665.2802
347.2166748046875 0 998.8036
348.1669616699219 0 1703.751
353.10943603515625 0 1997.7305
354.17706298828125 0 750.217
355.16143798828125 0 5415.9883
357.1552429199219 0 2162.8362
359.1553039550781 0 1251.6664
361.1882629394531 0 1460.2842
361.24468994140625 0 12846.841 y 7
362.248291015625 0 2804.8057
364.1866760253906 0 745.0219
365.19244384765625 0 792.7431
367.1618347167969 0 699.56714
368.1923828125 0 841.1126
370.1356506347656 0 1795.0262
371.1202697753906 0 2930.4019
372.25921630859375 0 628.9121
372.6363525390625 0 950.9287
373.1718444824219 0 4400.7256
373.7130432128906 0 729.77844 b 7
374.18231201171875 0 3858.9465
375.1658935546875 0 1576.9999
375.2384948730469 0 731.2452
376.1663513183594 0 1790.8079
381.7107849121094 0 3894.9316 y Water loss 2
382.2132568359375 0 878.82477
382.2452392578125 0 1767.0859
383.20367431640625 0 8561.0625
383.24871826171875 0 1164.0194
384.2066955566406 0 1533.7195
388.1463928222656 0 2017.878
390.71636962890625 0 30173.117 y 2
391.2178649902344 0 13684.313
391.71893310546875 0 3970.5051
392.19281005859375 0 4119.8945
393.1945495605469 0 944.8018
400.25634765625 0 3305.1475
401.21453857421875 0 7228.2153
401.258544921875 0 1026.7896
402.1775207519531 0 13647.804
402.2148132324219 0 946.2815
403.18072509765625 0 2935.3877
403.2342834472656 0 2388.511 b 3
405.17486572265625 0 714.14655
408.2224426269531 0 828.7787
409.2070617675781 0 909.12115
409.28216552734375 0 798.18335
410.2396240234375 0 635.50073
415.257080078125 0 1004.19037
416.14581298828125 0 650.9937
416.2585144042969 0 661.8111
416.7595520019531 0 937.9666
418.996337890625 0 1638.4617
419.2063293457031 0 863.53986
420.1883850097656 0 7949.4126
421.1913757324219 0 1547.9514
426.2345886230469 0 3490.6716
429.2821350097656 0 1380.7397
432.7090759277344 0 1236.7115
437.21331787109375 0 1505.5449
438.19793701171875 0 1243.8894
439.26654052734375 0 7042.5884
440.2515869140625 0 2299.6562 y 1
440.7532653808594 0 695.3521
441.25189208984375 0 712.50385
442.1949768066406 0 933.3161
444.2453308105469 0 1193.987
454.2303161621094 0 5648.559
455.2245788574219 0 2018.8044
456.2088317871094 0 3730.8406
457.2772521972656 0 9395.558
458.2794189453125 0 1753.5522
462.292724609375 0 3137.0964 y 6
463.2958068847656 0 996.818
468.2937927246094 0 875.6765
470.1888427734375 0 1320.6212
472.2413024902344 0 3788.9788
473.23577880859375 0 1449.4509
478.27581787109375 0 685.94965
480.7810974121094 0 1970.5244 Precursor Water loss
481.28076171875 0 1136.9652 Precursor Ammonia loss
482.2065124511719 0 630.75854
483.2198486328125 0 1384.2013
484.203857421875 0 5078.4346
486.25079345703125 0 1319.2957
486.3037109375 0 5124.53
487.306396484375 0 1648.7616
489.1870422363281 0 1078.6804
489.7848205566406 0 3236.5623 Precursor
490.2287902832031 0 1378.1051
490.28533935546875 0 1831.4235
491.26092529296875 0 793.2469
496.28802490234375 0 9259.148
497.29156494140625 0 2847.297
501.2335510253906 0 2054.074
501.3044738769531 0 780.83636 y Water loss 5
504.2523498535156 0 684.8095
512.26318359375 0 1869.3445
514.2982788085938 0 22749.701
515.26123046875 0 1006.1849
515.30126953125 0 6163.169
516.3049926757812 0 810.2853
518.2264404296875 0 1736.5343
519.2562255859375 0 1179.9646
519.3140258789062 0 4974.331 y 5
520.316650390625 0 1211.0685
522.29052734375 0 1182.2062
530.2724609375 0 7904.109
531.2750244140625 0 2334.297
536.2319946289062 0 560.01715
537.26416015625 0 796.9473
539.3189697265625 0 1420.3721
548.2828369140625 0 17367.682
549.2852172851562 0 4717.9355
550.2852783203125 0 858.9037
554.29296875 0 2709.2148
555.2763671875 0 1614.3656
558.3256225585938 0 1266.428 y Water loss 4
567.3134765625 0 1850.0574
569.296630859375 0 1183.3871
572.3038940429688 0 1591.8147
576.3350219726562 0 23163.352 y 4
577.33837890625 0 6943.991
578.3416748046875 0 1629.9268
582.2883911132812 0 2254.1428
583.270751953125 0 3003.4033
585.323974609375 0 1680.4062
586.31982421875 0 1452.9998
589.2940673828125 0 616.0951
590.3311157226562 0 975.6
596.3049926757812 0 893.94434
597.337646484375 0 890.28217
600.3013916015625 0 5757.9204
605.256103515625 0 1368.3418
614.3111572265625 0 1632.9669
615.3461303710938 0 3882.608 y Water loss 3
616.3474731445312 0 2185.7246
617.3439331054688 0 965.58215
618.324462890625 0 1507.4529 b 6
625.3449096679688 0 1356.3832
633.357666015625 0 68863.06 y 3
634.360595703125 0 19975.057
635.3622436523438 0 3101.2014
643.3554077148438 0 17657.37
644.3585205078125 0 7300.2285
645.3604125976562 0 2124.41
647.26953125 0 1631.2983
649.3688354492188 0 1224.7771
650.3524780273438 0 1484.4756
661.3663940429688 0 38545.008
662.3692626953125 0 14639.82
663.3713989257812 0 2582.831
667.3765869140625 0 2187.2273
668.3612670898438 0 2069.138
679.38037109375 0 994.69025
685.38720703125 0 4829.1694
695.3726806640625 0 1899.9305
696.3557739257812 0 2085.0425
713.3815307617188 0 6153.1953
718.4244995117188 0 1421.2668
728.4081420898438 0 1430.0739 b Water loss 7
733.3134155273438 0 1044.8867
734.2992553710938 0 2301.056
742.4242553710938 0 836.2743
744.4046020507812 0 865.8239
746.4181518554688 0 5202.1074 b 7
747.421630859375 0 2963.7952
751.3250122070312 0 2621.2139
760.4336547851562 0 2555.2502
762.4139404296875 0 6466.156 y Water loss 2
763.4166259765625 0 2577.1714
764.4130249023438 0 836.49347
769.3342895507812 0 868.177
779.4111328125 0 869.2863
780.42431640625 0 133463.84 y 2
781.427001953125 0 61687.215
782.4302978515625 0 14334.992
783.4329223632812 0 1176.5977
841.4924926757812 0 2448.6343 b Water loss 8
842.4969482421875 0 752.974
859.5018920898438 0 7100.834 b 8
860.505859375 0 4406.0146
861.5020751953125 0 930.144
876.446533203125 0 1066.3663
879.4924926757812 0 7797.7036 y 1
880.494140625 0 4084.251
881.4978637695312 0 1127.5146
1045.3974609375 0 778.939
1938.295166015625 0 709.39014

Spectrum Details

|  |  |
| --- | --- |
| Matched peaks? Matched peaks The total absolute number of peaks matched. Additionally in brackets the total fraction of peaks matched and the total number of peaks is shown. | 31 (7.21% of 430) |
| FDR? FDR The false discovery rate estimated for this peptide. It is calculated by matching all theoretical fragments with a non-integer shift with the raw peaks for this spectrum. This is done with 40 different shifts. The resulting percentage is the average number of annotated peaks over the number of annotated peaks with the correct spectrum. | 1.46% |
| Satellite FDR? Satellite FDR See the FDR for details on its calculation. This satellite ion specific FDR only contains the satellite ions (d/w) for I/L/J positions. | - |
| PSM Score? PSM Score The PSM Score as given by Hecklib to this annotated spectrum. It is shown with three significant figures. | 362 |

#### Spectrum 5854? Spectrum 5854 The raw spectrum of this peptide as annotated by Hecklib. The fragments are coloured according to ion type (see legend). Any peaks with a star '\*' as text can be hovered over to see the full details, first the ion type second the mass shift type. By hovering over the amino acids in the peptide or ions in the legend the corresponding peaks are highlighted. By toggling the 'Unassigned' label you can turn the background (unassigned) peaks on or off in the plot. By updating the slider in the Ion legend you can update the spectrum to only show the top X% of the peaks with labels. The top X% means any peak that is within X% of the highest intensity. By dragging in the spectrum you can zoom in to a specific part of the spectrum and use 'Zoom Out' to get back to the original zoom level. The annotation of the spectrum is based on the given sequence in the peptides file and is done with different software so inconsistencies are likely. The peaks are annotated based on the given sequence, with 20 ppm tolerance.

Copy Data

##### Spectrum 5854 (TSV)

###### Preview

```
Loading example...
```

*Click on the button to copy the data to your clipboard.*

Mz MinMz MaxIntensity Max

WidthHeightPeptide font sizePeptide stroke widthSpectrum font sizeSpectrum stroke widthCompact peptide

Ion legend

wxyz

abcd

OtherUnassignedIon^Charge^_Position_Show for top:%

VVFGGGTKJT

03.00e+46.00e+49.01e+41.20e+5

Zoom Out

y^+1^_1_y^+1^_2_y^+1^_2_z^+1^_3_y^+1^_3_c^+2^_8_y^+2^_8_y^+2^_8_y^+2^_9_y^+1^_4_z^+1^_4_y^+1^_4_c^+1^_5_z^+1^_5_y^+1^_5_c^+1^_6_z^+1^_6_y^+1^_6_z^+1^_7_y^+1^_7_y^+1^_7_c^+1^_7_y^+1^_7_c^+1^_7_c^+1^_8_y^+1^_8_c^+1^_8_z^+1^_8_y^+1^_8_c^+1^_9_z^+1^_9_c^+1^_9_y^+1^_9_

0870174026103480

Fragment Matches Table

Show background peaks

| Position | Ion type | Intensity | mz Theoretical | mz Error (Th) | mz Error (ppm) | Charge | Series Number |
| --- | --- | --- | --- | --- | --- | --- | --- |
| 10 | y | 4371 | 120.1 | 0.0003068 | 2.555 | +1 | 1 |
| - | - | 8414 | 120.1 | - | - | 0 | - |
| - | - | 550.8 | 121.1 | - | - | 0 | - |
| - | - | 547.3 | 122.1 | - | - | 0 | - |
| - | - | 1698 | 129.1 | - | - | 0 | - |
| - | - | 2314 | 136.1 | - | - | 0 | - |
| - | - | 448.5 | 169.1 | - | - | 0 | - |
| - | - | 4.348E+04 | 171.1 | - | - | 0 | - |
| - | - | 3789 | 172.2 | - | - | 0 | - |
| - | - | 702.2 | 173.4 | - | - | 0 | - |
| - | - | 897.4 | 177.1 | - | - | 0 | - |
| - | - | 510.5 | 195.5 | - | - | 0 | - |
| - | - | 3.703E+04 | 199.1 | - | - | 0 | - |
| - | - | 4035 | 200.1 | - | - | 0 | - |
| - | - | 1377 | 201.1 | - | - | 0 | - |
| - | - | 3525 | 205.1 | - | - | 0 | - |
| - | - | 636.5 | 211.1 | - | - | 0 | - |
| - | - | 529.4 | 213.4 | - | - | 0 | - |
| 9 | y | 2026 | 215.1 | 0.0002175 | 1.011 | +1 | 2 |
| - | - | 435.2 | 220 | - | - | 0 | - |
| 9 | y | 1370 | 233.1 | 0.0003187 | 1.367 | +1 | 2 |
| - | - | 540.5 | 233.2 | - | - | 0 | - |
| - | - | 793.5 | 247.1 | - | - | 0 | - |
| - | - | 550.3 | 254.4 | - | - | 0 | - |
| - | - | 3303 | 276.2 | - | - | 0 | - |
| - | - | 1508 | 290.1 | - | - | 0 | - |
| - | - | 2379 | 301.2 | - | - | 0 | - |
| - | - | 746 | 302.2 | - | - | 0 | - |
| - | - | 794.1 | 304.2 | - | - | 0 | - |
| - | - | 3614 | 310.2 | - | - | 0 | - |
| - | - | 761.5 | 331.2 | - | - | 0 | - |
| - | - | 541.6 | 332.5 | - | - | 0 | - |
| 8 | z | 3380 | 345.2 | 0.0003432 | 0.9942 | +1 | 3 |
| - | - | 1641 | 346.2 | - | - | 0 | - |
| - | - | 5515 | 346.2 | - | - | 0 | - |
| - | - | 694.9 | 347.2 | - | - | 0 | - |
| - | - | 813.9 | 349.1 | - | - | 0 | - |
| - | - | 1792 | 358.2 | - | - | 0 | - |
| 8 | y | 2545 | 361.2 | 0.0004485 | 1.242 | +1 | 3 |
| - | - | 553.9 | 362.2 | - | - | 0 | - |
| - | - | 527.1 | 366.1 | - | - | 0 | - |
| 8 | c | 961 | 373.7 | 0.001805 | 4.829 | +2 | 8 |
| 3 | y | 1885 | 381.7 | 0.000799 | 2.093 | +2 | 8 |
| - | - | 624.6 | 389.2 | - | - | 0 | - |
| 3 | y | 2.142E+04 | 390.7 | 0.0002774 | 0.7099 | +2 | 8 |
| - | - | 1.016E+04 | 391.2 | - | - | 0 | - |
| - | - | 1458 | 391.7 | - | - | 0 | - |
| - | - | 744.2 | 397.2 | - | - | 0 | - |
| - | - | 1242 | 401.2 | - | - | 0 | - |
| - | - | 603.5 | 402.2 | - | - | 0 | - |
| - | - | 2256 | 402.3 | - | - | 0 | - |
| - | - | 1527 | 403.2 | - | - | 0 | - |
| - | - | 766.7 | 403.3 | - | - | 0 | - |
| - | - | 1185 | 415.3 | - | - | 0 | - |
| - | - | 801 | 416.3 | - | - | 0 | - |
| - | - | 1418 | 420.2 | - | - | 0 | - |
| - | - | 1093 | 437.2 | - | - | 0 | - |
| 2 | y | 1409 | 440.3 | 0.001062 | 2.413 | +2 | 9 |
| - | - | 1043 | 442.3 | - | - | 0 | - |
| - | - | 3137 | 444.3 | - | - | 0 | - |
| 7 | y | 7578 | 445.3 | 0.0004068 | 0.9136 | +1 | 4 |
| 7 | z | 1490 | 446.3 | 0.004336 | 9.716 | +1 | 4 |
| - | - | 863.8 | 447.2 | - | - | 0 | - |
| - | - | 687.7 | 447.2 | - | - | 0 | - |
| - | - | 7093 | 447.3 | - | - | 0 | - |
| - | - | 1376 | 448.3 | - | - | 0 | - |
| - | - | 1568 | 457.3 | - | - | 0 | - |
| - | - | 1157 | 459.3 | - | - | 0 | - |
| 7 | y | 1033 | 462.3 | 0.0006601 | 1.428 | +1 | 4 |
| - | - | 1090 | 470.3 | - | - | 0 | - |
| - | - | 1658 | 476.3 | - | - | 0 | - |
| 5 | c | 3684 | 477.3 | 0.0003539 | 0.7415 | +1 | 5 |
| - | - | 896.5 | 478.3 | - | - | 0 | - |
| - | - | 1134 | 480.8 | - | - | 0 | - |
| - | - | 899.8 | 481.3 | - | - | 0 | - |
| - | - | 745.7 | 486.2 | - | - | 0 | - |
| - | - | 2396 | 489.8 | - | - | 0 | - |
| - | - | 4176 | 490.3 | - | - | 0 | - |
| - | - | 1182 | 499.3 | - | - | 0 | - |
| - | - | 603.4 | 500.3 | - | - | 0 | - |
| 6 | z | 2237 | 503.3 | 0.001208 | 2.401 | +1 | 5 |
| - | - | 3.855E+04 | 504.3 | - | - | 0 | - |
| - | - | 5.056E+04 | 504.3 | - | - | 0 | - |
| - | - | 1.013E+04 | 505.3 | - | - | 0 | - |
| - | - | 1.259E+04 | 505.3 | - | - | 0 | - |
| - | - | 1283 | 506.3 | - | - | 0 | - |
| - | - | 2382 | 506.3 | - | - | 0 | - |
| - | - | 2615 | 514.3 | - | - | 0 | - |
| - | - | 1016 | 515.3 | - | - | 0 | - |
| - | - | 1885 | 516.3 | - | - | 0 | - |
| - | - | 635.4 | 517.3 | - | - | 0 | - |
| - | - | 1642 | 518.3 | - | - | 0 | - |
| 6 | y | 4986 | 519.3 | 0.0005174 | 0.9964 | +1 | 5 |
| - | - | 1361 | 520.3 | - | - | 0 | - |
| - | - | 2337 | 533.3 | - | - | 0 | - |
| 6 | c | 2441 | 534.3 | 0.001151 | 2.155 | +1 | 6 |
| - | - | 4039 | 548.3 | - | - | 0 | - |
| - | - | 1315 | 549.3 | - | - | 0 | - |
| 5 | z | 1.414E+04 | 560.3 | 0.0005881 | 1.05 | +1 | 6 |
| - | - | 3.826E+04 | 561.3 | - | - | 0 | - |
| - | - | 1.087E+04 | 562.3 | - | - | 0 | - |
| - | - | 1163 | 563.3 | - | - | 0 | - |
| - | - | 2033 | 574.3 | - | - | 0 | - |
| - | - | 6219 | 575.3 | - | - | 0 | - |
| 5 | y | 1.343E+04 | 576.3 | 5.254E-05 | 0.09117 | +1 | 6 |
| - | - | 3383 | 577.3 | - | - | 0 | - |
| - | - | 823.2 | 589.3 | - | - | 0 | - |
| 4 | z | 1379 | 600.3 | 0.006528 | 10.87 | +1 | 7 |
| - | - | 1825 | 604.4 | - | - | 0 | - |
| - | - | 793.1 | 605.4 | - | - | 0 | - |
| 4 | y | 970.8 | 615.3 | 0.005389 | 8.757 | +1 | 7 |
| 4 | y | 631.2 | 616.3 | 0.005687 | 9.228 | +1 | 7 |
| 7 | c | 6734 | 617.3 | 0.00317 | 5.135 | +1 | 7 |
| - | - | 2.305E+04 | 618.3 | - | - | 0 | - |
| - | - | 7827 | 619.3 | - | - | 0 | - |
| - | - | 1122 | 620.4 | - | - | 0 | - |
| - | - | 1.253E+04 | 632.3 | - | - | 0 | - |
| 4 | y | 5.584E+04 | 633.4 | 0.0001342 | 0.2119 | +1 | 7 |
| - | - | 2.103E+04 | 634.4 | - | - | 0 | - |
| 7 | c | 7.586E+04 | 635.4 | 0.0008528 | 1.342 | +1 | 7 |
| - | - | 2.765E+04 | 636.4 | - | - | 0 | - |
| - | - | 5267 | 637.4 | - | - | 0 | - |
| - | - | 681.5 | 638.4 | - | - | 0 | - |
| - | - | 848.9 | 641.3 | - | - | 0 | - |
| - | - | 2089 | 643.4 | - | - | 0 | - |
| - | - | 1143 | 646.3 | - | - | 0 | - |
| - | - | 723.6 | 647.4 | - | - | 0 | - |
| - | - | 872.4 | 660.3 | - | - | 0 | - |
| - | - | 8624 | 661.4 | - | - | 0 | - |
| - | - | 3162 | 662.4 | - | - | 0 | - |
| - | - | 678.6 | 662.6 | - | - | 0 | - |
| - | - | 4642 | 663.4 | - | - | 0 | - |
| - | - | 1352 | 664.4 | - | - | 0 | - |
| - | - | 698 | 670.4 | - | - | 0 | - |
| - | - | 759.8 | 671.4 | - | - | 0 | - |
| - | - | 657.3 | 685.4 | - | - | 0 | - |
| - | - | 643.2 | 687.4 | - | - | 0 | - |
| - | - | 697.1 | 690.3 | - | - | 0 | - |
| - | - | 2076 | 702.4 | - | - | 0 | - |
| - | - | 780.9 | 708.3 | - | - | 0 | - |
| - | - | 1115 | 718.4 | - | - | 0 | - |
| - | - | 1184 | 719.4 | - | - | 0 | - |
| - | - | 843.4 | 721.3 | - | - | 0 | - |
| - | - | 761.7 | 722.4 | - | - | 0 | - |
| - | - | 747.4 | 728.4 | - | - | 0 | - |
| - | - | 4443 | 729.4 | - | - | 0 | - |
| - | - | 2654 | 730.4 | - | - | 0 | - |
| - | - | 789.9 | 734.4 | - | - | 0 | - |
| 8 | c | 4082 | 746.4 | 0.009028 | 12.1 | +1 | 8 |
| - | - | 2133 | 747.4 | - | - | 0 | - |
| - | - | 1024 | 761.4 | - | - | 0 | - |
| 3 | y | 1905 | 762.4 | 0.004907 | 6.436 | +1 | 8 |
| 8 | c | 1.007E+05 | 763.4 | 0.0001891 | 0.2477 | +1 | 8 |
| 3 | z | 1.132E+04 | 764.4 | 0.008479 | 11.09 | +1 | 8 |
| - | - | 3.003E+04 | 764.5 | - | - | 0 | - |
| - | - | 9155 | 765.4 | - | - | 0 | - |
| - | - | 5074 | 765.5 | - | - | 0 | - |
| - | - | 1621 | 766.4 | - | - | 0 | - |
| - | - | 814.5 | 778.4 | - | - | 0 | - |
| 3 | y | 7.331E+04 | 780.4 | 0.0001407 | 0.1803 | +1 | 8 |
| - | - | 3.306E+04 | 781.4 | - | - | 0 | - |
| - | - | 8070 | 782.4 | - | - | 0 | - |
| - | - | 752.7 | 789.5 | - | - | 0 | - |
| - | - | 1359 | 807.4 | - | - | 0 | - |
| - | - | 1.035E+04 | 832.5 | - | - | 0 | - |
| - | - | 5298 | 833.5 | - | - | 0 | - |
| - | - | 1083 | 834.5 | - | - | 0 | - |
| - | - | 757.4 | 841.5 | - | - | 0 | - |
| 9 | c | 5476 | 859.5 | 0.0002579 | 0.3001 | +1 | 9 |
| - | - | 2504 | 860.5 | - | - | 0 | - |
| - | - | 3474 | 861.5 | - | - | 0 | - |
| - | - | 989.9 | 862.5 | - | - | 0 | - |
| 2 | z | 1.216E+04 | 863.5 | 0.0003776 | 0.4373 | +1 | 9 |
| - | - | 5455 | 864.5 | - | - | 0 | - |
| - | - | 3016 | 865.5 | - | - | 0 | - |
| - | - | 835.5 | 866.4 | - | - | 0 | - |
| - | - | 1124 | 874.5 | - | - | 0 | - |
| - | - | 721.9 | 875.5 | - | - | 0 | - |
| 9 | c | 9.676E+04 | 876.5 | 1.258E-05 | 0.01435 | +1 | 9 |
| - | - | 4.837E+04 | 877.5 | - | - | 0 | - |
| - | - | 1.225E+04 | 878.5 | - | - | 0 | - |
| 2 | y | 2907 | 879.5 | 0.003321 | 3.776 | +1 | 9 |
| - | - | 1161 | 880.5 | - | - | 0 | - |
| - | - | 1528 | 887.5 | - | - | 0 | - |
| - | - | 2110 | 893.5 | - | - | 0 | - |
| - | - | 1975 | 906.5 | - | - | 0 | - |
| - | - | 812.1 | 909.4 | - | - | 0 | - |
| - | - | 859.7 | 916.5 | - | - | 0 | - |
| - | - | 829.9 | 919.5 | - | - | 0 | - |
| - | - | 4219 | 923.5 | - | - | 0 | - |
| - | - | 2919 | 924.5 | - | - | 0 | - |
| - | - | 697.6 | 925.5 | - | - | 0 | - |
| - | - | 1399 | 933.6 | - | - | 0 | - |
| - | - | 847.5 | 936.5 | - | - | 0 | - |
| - | - | 787.6 | 937.5 | - | - | 0 | - |
| - | - | 1651 | 947.5 | - | - | 0 | - |
| - | - | 682.4 | 948.5 | - | - | 0 | - |
| - | - | 855.1 | 951.6 | - | - | 0 | - |
| - | - | 1753 | 960.5 | - | - | 0 | - |
| - | - | 2011 | 961.6 | - | - | 0 | - |
| - | - | 1.189E+05 | 962.5 | - | - | 0 | - |
| - | - | 6.214E+04 | 963.5 | - | - | 0 | - |
| - | - | 1.965E+04 | 964.5 | - | - | 0 | - |
| - | - | 1376 | 965.6 | - | - | 0 | - |
| - | - | 2888 | 976.5 | - | - | 0 | - |
| - | - | 1180 | 977.5 | - | - | 0 | - |
| - | - | 4275 | 977.5 | - | - | 0 | - |
| - | - | 6.452E+04 | 978.6 | - | - | 0 | - |
| - | - | 7.489E+04 | 979.6 | - | - | 0 | - |
| - | - | 3.672E+04 | 980.6 | - | - | 0 | - |
| - | - | 768 | 981.5 | - | - | 0 | - |
| - | - | 8993 | 981.6 | - | - | 0 | - |
| - | - | 868.7 | 982.5 | - | - | 0 | - |
| - | - | 653.8 | 1360 | - | - | 0 | - |
| - | - | 708.1 | 1449 | - | - | 0 | - |
| - | - | 661.4 | 1829 | - | - | 0 | - |
| - | - | 871.7 | 3071 | - | - | 0 | - |
| - | - | 771.5 | 3446 | - | - | 0 | - |

m/z Charge Intensity FragmentType MassShift Position
120.06582641601562 0 4371.225 y 9
120.08110809326172 0 8414.104
121.08460235595703 0 550.8088
122.06005859375 0 547.2897
129.1026153564453 0 1697.8344
136.07591247558594 0 2314.2637
169.13409423828125 0 448.48605
171.14955139160156 0 43475.9
172.1529998779297 0 3789.3289
173.43821716308594 0 702.24084
177.10264587402344 0 897.4217
195.48403930664062 0 510.48218
199.14443969726562 0 37027.285
200.147705078125 0 4035.004
201.12355041503906 0 1376.931
205.09765625 0 3525.1187
211.1439208984375 0 636.48254
213.4066619873047 0 529.39685
215.1392364501953 0 2025.9315 y Water loss 8
220.0250701904297 0 435.24677
233.14990234375 0 1369.5399 y 8
233.21560668945312 0 540.487
247.14468383789062 0 793.5182
254.3744354248047 0 550.3361
276.15576171875 0 3302.949
290.13525390625 0 1507.6487
301.1917419433594 0 2379.2476
302.19525146484375 0 746.02655
304.1658020019531 0 794.126
310.2130432128906 0 3614.3674
331.1886291503906 0 761.5254
332.5350036621094 0 541.64136
345.2261657714844 0 3379.7378 z 7
346.21234130859375 0 1640.8776
346.2336730957031 0 5515.149
347.23699951171875 0 694.946
349.1189270019531 0 813.9064
358.24517822265625 0 1791.9456
361.2449951171875 0 2544.6838 y 7
362.24774169921875 0 553.8866
366.146728515625 0 527.0793
373.71160888671875 0 961.02325 c Ammonia loss 7
381.711669921875 0 1884.5312 y Water loss 2
389.20306396484375 0 624.5962
390.7164306640625 0 21415.184 y 2
391.2180480957031 0 10156.991
391.71942138671875 0 1457.8813
397.2447509765625 0 744.2353
401.21490478515625 0 1241.6357
402.1784362792969 0 603.51483
402.2718200683594 0 2255.8245
403.2335510253906 0 1526.7872
403.275390625 0 766.702
415.2564697265625 0 1185.347
416.25799560546875 0 800.97046
420.189453125 0 1418.0237
437.21575927734375 0 1093.1855
440.2492980957031 0 1408.8885 y 1
442.2567138671875 0 1043.0812
444.25897216796875 0 3136.6265
445.2660827636719 0 7578.443 y Ammonia loss 6
446.2691650390625 0 1489.7451 z 6
447.231201171875 0 863.82385
447.2441101074219 0 687.728
447.28179931640625 0 7092.5522
448.2847595214844 0 1376.1823
457.2777099609375 0 1568.4366
459.2938232421875 0 1156.8005
462.29156494140625 0 1033.2585 y 6
470.2596740722656 0 1090.4825
476.27447509765625 0 1658.138
477.2823486328125 0 3683.7349 c 4
478.2862854003906 0 896.4561
480.7790222167969 0 1134.0225
481.27899169921875 0 899.822
486.24542236328125 0 745.69135
489.784912109375 0 2396.188
490.2871398925781 0 4175.83
499.2777404785156 0 1182.3408
500.281494140625 0 603.41846
503.2961730957031 0 2236.6501 z 5
504.2545166015625 0 38548.406
504.30316162109375 0 50563.086
505.25823974609375 0 10125.465
505.3064880371094 0 12594.58
506.2610168457031 0 1283.2125
506.309326171875 0 2381.8718
514.298583984375 0 2614.9487
515.3042602539062 0 1015.50555
516.3125 0 1884.87
517.2769165039062 0 635.3624
518.3072509765625 0 1641.7467
519.3131713867188 0 4986.303 y 5
520.3173828125 0 1361.066
533.2962036132812 0 2337.1812
534.3023071289062 0 2441.254 c 5
548.283203125 0 4038.9678
549.2870483398438 0 1314.6288
560.3170166015625 0 14139.153 z 4
561.324462890625 0 38255.617
562.3272094726562 0 10869.688
563.328369140625 0 1162.5415
574.2822265625 0 2032.9583
575.328369140625 0 6218.518
576.335205078125 0 13426.029 y 4
577.3394775390625 0 3382.8047
589.3297729492188 0 823.15314
600.31787109375 0 1379.4045 z Ammonia loss 3
604.3673706054688 0 1824.779
605.371826171875 0 793.14386
615.3514404296875 0 970.81506 y Water loss 3
616.3357543945312 0 631.17126 y Ammonia loss 3
617.33740234375 0 6734.1074 c Water loss 6
618.3447875976562 0 23047.451
619.3497314453125 0 7826.999
620.3500366210938 0 1121.9498
632.3491821289062 0 12526.983
633.3567504882812 0 55843.797 y 3
634.3539428710938 0 21033.031
635.3519897460938 0 75864.1 c 6
636.3546752929688 0 27645.484
637.3571166992188 0 5266.7295
638.3599243164062 0 681.45135
641.2783813476562 0 848.8721
643.354736328125 0 2089.382
646.3447265625 0 1143.3049
647.3517456054688 0 723.62256
660.348388671875 0 872.38776
661.3670043945312 0 8624.224
662.3692016601562 0 3162.4006
662.610107421875 0 678.5822
663.382080078125 0 4642.27
664.38623046875 0 1352.1545
670.3858032226562 0 697.9982
671.3947143554688 0 759.7884
685.386474609375 0 657.3347
687.4144897460938 0 643.1587
690.34033203125 0 697.07684
702.4097290039062 0 2076.298
708.34228515625 0 780.88806
718.4215087890625 0 1114.6671
719.428466796875 0 1183.7628
721.3490600585938 0 843.4458
722.3549194335938 0 761.7128
728.40771484375 0 747.42365
729.4170532226562 0 4442.8604
730.4168701171875 0 2653.573
734.4234008789062 0 789.87775
746.4105224609375 0 4081.8206 c Ammonia loss 7
747.4148559570312 0 2132.8105
761.431396484375 0 1024.4872
762.4193725585938 0 1904.5009 y Water loss 2
763.4462890625 0 100717.3 c 7
764.3978271484375 0 11322.1455 z 2
764.4500732421875 0 30031.9
765.407958984375 0 9155.397
765.4591674804688 0 5073.678
766.412353515625 0 1620.8805
778.4099731445312 0 814.5094
780.4251708984375 0 73305.55 y 2
781.4282836914062 0 33059.03
782.4307250976562 0 8070.409
789.4618530273438 0 752.69836
807.4115600585938 0 1359.101
832.5167846679688 0 10351.481
833.5192260742188 0 5298.176
834.5210571289062 0 1082.8016
841.4933471679688 0 757.38293
859.5033569335938 0 5475.802 c Ammonia loss 8
860.50390625 0 2503.5928
861.5148315429688 0 3474.28
862.5264282226562 0 989.94696
863.47509765625 0 12155.253 z 1
864.478515625 0 5455.381
865.4808959960938 0 3015.5461
866.4019775390625 0 835.52313
874.514404296875 0 1123.9214
875.5174560546875 0 721.90063
876.5301513671875 0 96761.64 c 8
877.5330810546875 0 48367.1
878.5365600585938 0 12254.273
879.4967651367188 0 2906.8066 y 1
880.494384765625 0 1160.9554
887.5191040039062 0 1527.6332
893.4734497070312 0 2109.6567
906.4793701171875 0 1975.055
909.4080200195312 0 812.1401
916.5397338867188 0 859.66174
919.5380249023438 0 829.8972
923.5050659179688 0 4219.2505
924.5086669921875 0 2918.726
925.5148315429688 0 697.56335
933.5612182617188 0 1398.734
936.49072265625 0 847.5118
937.4879760742188 0 787.60986
947.52001953125 0 1650.7812
948.5291748046875 0 682.36035
951.5791625976562 0 855.06836
960.5255737304688 0 1752.7578
961.5521240234375 0 2011.2479
962.5433349609375 0 118907.15
963.5459594726562 0 62141.21
964.5488891601562 0 19649.793
965.5527954101562 0 1376.2129
976.5421752929688 0 2887.9731
977.4525146484375 0 1180.0604
977.5497436523438 0 4275.1646
978.5615234375 0 64516.06
979.5675048828125 0 74891.664
980.5712890625 0 36724.176
981.4929809570312 0 768.049
981.574951171875 0 8993.204
982.4894409179688 0 868.6786
1359.9801025390625 0 653.7652
1448.715576171875 0 708.0959
1828.6090087890625 0 661.36707
3070.998046875 0 871.6905
3445.8896484375 0 771.5108

Spectrum Details

|  |  |
| --- | --- |
| Matched peaks? Matched peaks The total absolute number of peaks matched. Additionally in brackets the total fraction of peaks matched and the total number of peaks is shown. | 33 (15.14% of 218) |
| FDR? FDR The false discovery rate estimated for this peptide. It is calculated by matching all theoretical fragments with a non-integer shift with the raw peaks for this spectrum. This is done with 40 different shifts. The resulting percentage is the average number of annotated peaks over the number of annotated peaks with the correct spectrum. | 0.14% |
| Satellite FDR? Satellite FDR See the FDR for details on its calculation. This satellite ion specific FDR only contains the satellite ions (d/w) for I/L/J positions. | - |
| PSM Score? PSM Score The PSM Score as given by Hecklib to this annotated spectrum. It is shown with three significant figures. | 403 |

#### Spectrum 6339? Spectrum 6339 The raw spectrum of this peptide as annotated by Hecklib. The fragments are coloured according to ion type (see legend). Any peaks with a star '\*' as text can be hovered over to see the full details, first the ion type second the mass shift type. By hovering over the amino acids in the peptide or ions in the legend the corresponding peaks are highlighted. By toggling the 'Unassigned' label you can turn the background (unassigned) peaks on or off in the plot. By updating the slider in the Ion legend you can update the spectrum to only show the top X% of the peaks with labels. The top X% means any peak that is within X% of the highest intensity. By dragging in the spectrum you can zoom in to a specific part of the spectrum and use 'Zoom Out' to get back to the original zoom level. The annotation of the spectrum is based on the given sequence in the peptides file and is done with different software so inconsistencies are likely. The peaks are annotated based on the given sequence, with 20 ppm tolerance.

Copy Data

##### Spectrum 6339 (TSV)

###### Preview

```
Loading example...
```

*Click on the button to copy the data to your clipboard.*

Mz MinMz MaxIntensity Max

WidthHeightPeptide font sizePeptide stroke widthSpectrum font sizeSpectrum stroke widthCompact peptide

Ion legend

wxyz

abcd

OtherUnassignedIon^Charge^_Position_Show for top:%

VVFGGGTKJT

01.96e+43.92e+45.87e+47.83e+4

Zoom Out

y^+1^_1_d^+1^_2_a^+1^_2_b^+1^_2_y^+1^_2_y^+1^_2_y^+2^_5_y^+1^_3_b^+1^_3_y^+1^_3_y^+2^_8_y^+2^_8_y^+1^_4_\*\*y^+1^_5_y^+1^_6_b^+1^_7_y^+1^_7_y^+1^_7_b^+1^_8_y^+1^_8_y^+1^_8_b^+1^_9_b^+1^_9_y^+1^_9_

0754150722613014

Fragment Matches Table

Show background peaks

| Position | Ion type | Intensity | mz Theoretical | mz Error (Th) | mz Error (ppm) | Charge | Series Number |
| --- | --- | --- | --- | --- | --- | --- | --- |
| 10 | y | 6973 | 120.1 | 0.0003297 | 2.746 | +1 | 1 |
| - | - | 4.748E+04 | 120.1 | - | - | 0 | - |
| - | - | 3671 | 121.1 | - | - | 0 | - |
| - | - | 406.9 | 121.2 | - | - | 0 | - |
| - | - | 355.3 | 121.3 | - | - | 0 | - |
| - | - | 329.4 | 122.9 | - | - | 0 | - |
| - | - | 395.5 | 125.7 | - | - | 0 | - |
| - | - | 439.1 | 126.1 | - | - | 0 | - |
| - | - | 1090 | 127.1 | - | - | 0 | - |
| - | - | 645 | 129.1 | - | - | 0 | - |
| - | - | 3.646E+04 | 129.1 | - | - | 0 | - |
| - | - | 1647 | 130.1 | - | - | 0 | - |
| - | - | 2381 | 130.1 | - | - | 0 | - |
| - | - | 492.6 | 131.1 | - | - | 0 | - |
| - | - | 1762 | 131.1 | - | - | 0 | - |
| - | - | 518.6 | 132.1 | - | - | 0 | - |
| - | - | 2038 | 133.1 | - | - | 0 | - |
| - | - | 5667 | 136.1 | - | - | 0 | - |
| - | - | 712.2 | 140.1 | - | - | 0 | - |
| - | - | 968.4 | 141.1 | - | - | 0 | - |
| - | - | 492.1 | 141.1 | - | - | 0 | - |
| - | - | 559.8 | 142.1 | - | - | 0 | - |
| - | - | 599.3 | 143.1 | - | - | 0 | - |
| - | - | 703.2 | 145.1 | - | - | 0 | - |
| - | - | 1670 | 146.1 | - | - | 0 | - |
| - | - | 4025 | 147 | - | - | 0 | - |
| - | - | 471.2 | 147.1 | - | - | 0 | - |
| - | - | 438.9 | 150.5 | - | - | 0 | - |
| - | - | 592.7 | 152.1 | - | - | 0 | - |
| - | - | 460.8 | 153.1 | - | - | 0 | - |
| - | - | 453.6 | 153.1 | - | - | 0 | - |
| - | - | 468.7 | 154.1 | - | - | 0 | - |
| - | - | 1511 | 155.1 | - | - | 0 | - |
| - | - | 1650 | 155.1 | - | - | 0 | - |
| - | - | 979.2 | 156.1 | - | - | 0 | - |
| 2 | d | 499.7 | 157.1 | 5.11E-05 | 0.3252 | +1 | 2 |
| - | - | 515.4 | 158.1 | - | - | 0 | - |
| - | - | 2095 | 159.1 | - | - | 0 | - |
| - | - | 1302 | 159.1 | - | - | 0 | - |
| - | - | 449.1 | 159.1 | - | - | 0 | - |
| - | - | 508.3 | 160.9 | - | - | 0 | - |
| - | - | 5206 | 163.1 | - | - | 0 | - |
| - | - | 575.3 | 169.1 | - | - | 0 | - |
| - | - | 929.7 | 170.1 | - | - | 0 | - |
| - | - | 743 | 171.1 | - | - | 0 | - |
| 2 | a | 7.755E+04 | 171.1 | 0.0004075 | 2.381 | +1 | 2 |
| - | - | 1460 | 172.1 | - | - | 0 | - |
| - | - | 7072 | 172.2 | - | - | 0 | - |
| - | - | 594.1 | 173.1 | - | - | 0 | - |
| - | - | 2076 | 173.5 | - | - | 0 | - |
| - | - | 1560 | 175.1 | - | - | 0 | - |
| - | - | 1604 | 176.1 | - | - | 0 | - |
| - | - | 502.2 | 177 | - | - | 0 | - |
| - | - | 4099 | 177.1 | - | - | 0 | - |
| - | - | 1282 | 177.1 | - | - | 0 | - |
| - | - | 818.4 | 181.1 | - | - | 0 | - |
| - | - | 2568 | 181.2 | - | - | 0 | - |
| - | - | 1036 | 185.1 | - | - | 0 | - |
| - | - | 1012 | 185.2 | - | - | 0 | - |
| - | - | 874.9 | 186.1 | - | - | 0 | - |
| - | - | 2303 | 187.1 | - | - | 0 | - |
| - | - | 651.1 | 188.1 | - | - | 0 | - |
| - | - | 508.6 | 195.5 | - | - | 0 | - |
| - | - | 734.2 | 197.1 | - | - | 0 | - |
| - | - | 475 | 197.1 | - | - | 0 | - |
| - | - | 2617 | 197.1 | - | - | 0 | - |
| - | - | 1090 | 197.2 | - | - | 0 | - |
| - | - | 446.8 | 198.1 | - | - | 0 | - |
| - | - | 1771 | 198.1 | - | - | 0 | - |
| 2 | b | 3.893E+04 | 199.1 | 0.0003354 | 1.684 | +1 | 2 |
| - | - | 640.2 | 200.1 | - | - | 0 | - |
| - | - | 4082 | 200.1 | - | - | 0 | - |
| - | - | 1608 | 201.1 | - | - | 0 | - |
| - | - | 3313 | 204.1 | - | - | 0 | - |
| - | - | 7410 | 205.1 | - | - | 0 | - |
| - | - | 701.9 | 205.1 | - | - | 0 | - |
| - | - | 1014 | 206.1 | - | - | 0 | - |
| - | - | 738.6 | 207.1 | - | - | 0 | - |
| - | - | 1834 | 208.1 | - | - | 0 | - |
| - | - | 554.2 | 208.9 | - | - | 0 | - |
| - | - | 1387 | 209.1 | - | - | 0 | - |
| - | - | 1349 | 209.2 | - | - | 0 | - |
| - | - | 499.1 | 211.1 | - | - | 0 | - |
| - | - | 3987 | 212.1 | - | - | 0 | - |
| - | - | 462.7 | 213.2 | - | - | 0 | - |
| - | - | 783.6 | 215.1 | - | - | 0 | - |
| 9 | y | 3779 | 215.1 | 0.0003854 | 1.791 | +1 | 2 |
| - | - | 1904 | 216.1 | - | - | 0 | - |
| - | - | 652 | 217.1 | - | - | 0 | - |
| - | - | 3246 | 219.1 | - | - | 0 | - |
| - | - | 1209 | 220.1 | - | - | 0 | - |
| - | - | 790.4 | 220.2 | - | - | 0 | - |
| - | - | 912.1 | 221.1 | - | - | 0 | - |
| - | - | 613.1 | 224.1 | - | - | 0 | - |
| - | - | 1161 | 224.2 | - | - | 0 | - |
| - | - | 567.9 | 226.1 | - | - | 0 | - |
| - | - | 544.5 | 226.5 | - | - | 0 | - |
| - | - | 1687 | 227.1 | - | - | 0 | - |
| - | - | 959.8 | 228.1 | - | - | 0 | - |
| - | - | 717.5 | 228.2 | - | - | 0 | - |
| - | - | 6636 | 230.2 | - | - | 0 | - |
| - | - | 610 | 231.2 | - | - | 0 | - |
| - | - | 605.1 | 233.1 | - | - | 0 | - |
| 9 | y | 1780 | 233.1 | 0.0003187 | 1.367 | +1 | 2 |
| - | - | 1018 | 237.2 | - | - | 0 | - |
| - | - | 2030 | 239.2 | - | - | 0 | - |
| - | - | 1822 | 242.2 | - | - | 0 | - |
| - | - | 1502 | 245.1 | - | - | 0 | - |
| - | - | 2322 | 247.1 | - | - | 0 | - |
| - | - | 637.5 | 250.1 | - | - | 0 | - |
| 6 | y | 708 | 251.2 | 0.004733 | 18.85 | +2 | 5 |
| - | - | 4317 | 255.1 | - | - | 0 | - |
| - | - | 674.4 | 256.1 | - | - | 0 | - |
| - | - | 694.4 | 256.2 | - | - | 0 | - |
| - | - | 699.3 | 261.1 | - | - | 0 | - |
| - | - | 2504 | 262.1 | - | - | 0 | - |
| - | - | 4146 | 269.2 | - | - | 0 | - |
| - | - | 989.1 | 272.1 | - | - | 0 | - |
| - | - | 4038 | 273.1 | - | - | 0 | - |
| - | - | 1089 | 274.1 | - | - | 0 | - |
| - | - | 723.2 | 275.2 | - | - | 0 | - |
| - | - | 1856 | 276.2 | - | - | 0 | - |
| - | - | 559.3 | 282.4 | - | - | 0 | - |
| - | - | 994.3 | 287.2 | - | - | 0 | - |
| - | - | 1056 | 291.1 | - | - | 0 | - |
| - | - | 2703 | 292.2 | - | - | 0 | - |
| - | - | 539.1 | 292.6 | - | - | 0 | - |
| - | - | 1561 | 301.2 | - | - | 0 | - |
| - | - | 658 | 304.2 | - | - | 0 | - |
| - | - | 1430 | 308.2 | - | - | 0 | - |
| - | - | 725.3 | 309.2 | - | - | 0 | - |
| - | - | 3126 | 310.2 | - | - | 0 | - |
| - | - | 808.1 | 317.2 | - | - | 0 | - |
| - | - | 1426 | 319.1 | - | - | 0 | - |
| - | - | 629.7 | 323.2 | - | - | 0 | - |
| - | - | 646.9 | 324.2 | - | - | 0 | - |
| - | - | 805 | 325.2 | - | - | 0 | - |
| - | - | 3474 | 326.2 | - | - | 0 | - |
| - | - | 719.7 | 328.1 | - | - | 0 | - |
| - | - | 691.8 | 342.2 | - | - | 0 | - |
| - | - | 1075 | 343.2 | - | - | 0 | - |
| 8 | y | 1142 | 343.2 | 0.0001184 | 0.345 | +1 | 3 |
| - | - | 1345 | 344.2 | - | - | 0 | - |
| 3 | b | 1905 | 346.2 | 0.0003905 | 1.128 | +1 | 3 |
| - | - | 594.5 | 346.2 | - | - | 0 | - |
| - | - | 1412 | 357.2 | - | - | 0 | - |
| - | - | 919.3 | 357.2 | - | - | 0 | - |
| 8 | y | 4887 | 361.2 | 0.0002959 | 0.8192 | +1 | 3 |
| - | - | 672.9 | 362.2 | - | - | 0 | - |
| - | - | 900 | 365.2 | - | - | 0 | - |
| - | - | 803.1 | 367.2 | - | - | 0 | - |
| - | - | 1785 | 374.2 | - | - | 0 | - |
| - | - | 718.9 | 375.2 | - | - | 0 | - |
| - | - | 625.3 | 376.2 | - | - | 0 | - |
| 3 | y | 1035 | 381.7 | 0.0006964 | 1.824 | +2 | 8 |
| - | - | 1065 | 382.2 | - | - | 0 | - |
| - | - | 4066 | 383.2 | - | - | 0 | - |
| - | - | 669 | 384.2 | - | - | 0 | - |
| - | - | 811.6 | 384.2 | - | - | 0 | - |
| 3 | y | 1.278E+04 | 390.7 | 0.0003079 | 0.788 | +2 | 8 |
| - | - | 4983 | 391.2 | - | - | 0 | - |
| - | - | 2111 | 391.7 | - | - | 0 | - |
| - | - | 1275 | 392.2 | - | - | 0 | - |
| - | - | 1390 | 400.3 | - | - | 0 | - |
| - | - | 3499 | 401.2 | - | - | 0 | - |
| - | - | 5978 | 402.2 | - | - | 0 | - |
| - | - | 1170 | 403.2 | - | - | 0 | - |
| - | - | 1057 | 415.3 | - | - | 0 | - |
| - | - | 858.9 | 419 | - | - | 0 | - |
| - | - | 2810 | 420.2 | - | - | 0 | - |
| - | - | 614.9 | 422.7 | - | - | 0 | - |
| - | - | 820.9 | 429.3 | - | - | 0 | - |
| - | - | 1567 | 439.3 | - | - | 0 | - |
| - | - | 592.2 | 447.4 | - | - | 0 | - |
| - | - | 1015 | 454.2 | - | - | 0 | - |
| - | - | 650.4 | 455.2 | - | - | 0 | - |
| - | - | 3399 | 457.3 | - | - | 0 | - |
| - | - | 1011 | 458.3 | - | - | 0 | - |
| 7 | y | 1018 | 462.3 | 0.001301 | 2.814 | +1 | 4 |
| - | - | 833.3 | 470.3 | - | - | 0 | - |
| - | - | 3323 | 471.2 | - | - | 0 | - |
| - | - | 876.9 | 478.3 | - | - | 0 | - |
| 0 | Precursor | 863.8 | 480.8 | 0.001483 | 3.085 | +2 | -1 |
| - | - | 628.5 | 484 | - | - | 0 | - |
| - | - | 1869 | 486.3 | - | - | 0 | - |
| - | - | 879.2 | 487.3 | - | - | 0 | - |
| - | - | 804.6 | 488.2 | - | - | 0 | - |
| - | - | 1122 | 489.2 | - | - | 0 | - |
| 0 | Precursor | 987.3 | 489.8 | 0.001199 | 2.449 | +2 | -1 |
| - | - | 1891 | 490.3 | - | - | 0 | - |
| - | - | 4070 | 496.3 | - | - | 0 | - |
| - | - | 1356 | 497.3 | - | - | 0 | - |
| - | - | 757 | 512.3 | - | - | 0 | - |
| - | - | 8239 | 514.3 | - | - | 0 | - |
| - | - | 2002 | 515.3 | - | - | 0 | - |
| 6 | y | 1870 | 519.3 | 0.0003343 | 0.6438 | +1 | 5 |
| - | - | 1321 | 525.8 | - | - | 0 | - |
| - | - | 3746 | 530.3 | - | - | 0 | - |
| - | - | 1109 | 531.3 | - | - | 0 | - |
| - | - | 2396 | 539.8 | - | - | 0 | - |
| - | - | 2840 | 540.3 | - | - | 0 | - |
| - | - | 6841 | 548.3 | - | - | 0 | - |
| - | - | 566 | 548.8 | - | - | 0 | - |
| - | - | 1225 | 549.3 | - | - | 0 | - |
| - | - | 646 | 554.3 | - | - | 0 | - |
| - | - | 635.3 | 559.3 | - | - | 0 | - |
| - | - | 660.1 | 572.4 | - | - | 0 | - |
| 5 | y | 9113 | 576.3 | 0.0001306 | 0.2265 | +1 | 6 |
| - | - | 2592 | 577.3 | - | - | 0 | - |
| - | - | 694.8 | 589.3 | - | - | 0 | - |
| 7 | b | 722.4 | 600.3 | 0.001162 | 1.936 | +1 | 7 |
| 4 | y | 2157 | 615.3 | 0.003192 | 5.187 | +1 | 7 |
| 4 | y | 2.793E+04 | 633.4 | 0.0009277 | 1.465 | +1 | 7 |
| - | - | 8899 | 634.4 | - | - | 0 | - |
| - | - | 1584 | 635.4 | - | - | 0 | - |
| - | - | 6962 | 643.4 | - | - | 0 | - |
| - | - | 614.1 | 643.9 | - | - | 0 | - |
| - | - | 2221 | 644.4 | - | - | 0 | - |
| - | - | 1.491E+04 | 661.4 | - | - | 0 | - |
| - | - | 5087 | 662.4 | - | - | 0 | - |
| - | - | 866.3 | 663.4 | - | - | 0 | - |
| - | - | 655.5 | 690.4 | - | - | 0 | - |
| - | - | 700 | 706.8 | - | - | 0 | - |
| - | - | 837.4 | 718.4 | - | - | 0 | - |
| 8 | b | 2421 | 746.4 | 0.002131 | 2.856 | +1 | 8 |
| - | - | 765.2 | 747.4 | - | - | 0 | - |
| - | - | 650.5 | 760.4 | - | - | 0 | - |
| 3 | y | 2568 | 762.4 | 0.0009398 | 1.233 | +1 | 8 |
| - | - | 1402 | 763.4 | - | - | 0 | - |
| 3 | y | 5.494E+04 | 780.4 | 0.0005917 | 0.7582 | +1 | 8 |
| - | - | 2.476E+04 | 781.4 | - | - | 0 | - |
| - | - | 5514 | 782.4 | - | - | 0 | - |
| 9 | b | 795.2 | 841.5 | 0.006661 | 7.916 | +1 | 9 |
| 9 | b | 3700 | 859.5 | 0.001723 | 2.004 | +1 | 9 |
| - | - | 1932 | 860.5 | - | - | 0 | - |
| 2 | y | 3924 | 879.5 | 0.002294 | 2.609 | +1 | 9 |
| - | - | 614.6 | 880.5 | - | - | 0 | - |
| - | - | 603.7 | 1073 | - | - | 0 | - |
| - | - | 743.1 | 1628 | - | - | 0 | - |
| - | - | 595.4 | 2434 | - | - | 0 | - |
| - | - | 592.2 | 2443 | - | - | 0 | - |
| - | - | 648 | 2582 | - | - | 0 | - |
| - | - | 604.7 | 2984 | - | - | 0 | - |

m/z Charge Intensity FragmentType MassShift Position
120.06584930419922 0 6973.053 y 9
120.08116149902344 0 47479.715
121.08453369140625 0 3670.6294
121.16876983642578 0 406.86975
121.28961944580078 0 355.33685
122.91858673095703 0 329.36676
125.68621063232422 0 395.50677
126.0666275024414 0 439.05588
127.05072021484375 0 1090.4191
129.06602478027344 0 645.0214
129.1025848388672 0 36458.746
130.0655059814453 0 1647.3429
130.1060028076172 0 2380.8635
131.0699920654297 0 492.62546
131.08169555664062 0 1762.3055
132.08139038085938 0 518.6462
133.08628845214844 0 2038.0095
136.07603454589844 0 5666.7676
140.08206176757812 0 712.19116
141.0660400390625 0 968.42786
141.10243225097656 0 492.11142
142.0986328125 0 559.8411
143.0817413330078 0 599.26843
145.0608367919922 0 703.21796
146.0604248046875 0 1669.8799
147.0444793701172 0 4025.365
147.07675170898438 0 471.17148
150.50462341308594 0 438.93427
152.0704345703125 0 592.6591
153.06646728515625 0 460.78217
153.0772705078125 0 453.57806
154.06146240234375 0 468.72223
155.08189392089844 0 1511.256
155.11819458007812 0 1650.4062
156.07705688476562 0 979.2211
157.1335906982422 0 499.66406 d 1
158.09300231933594 0 515.4058
159.07688903808594 0 2094.6294
159.091796875 0 1302.2345
159.11300659179688 0 449.10608
160.88592529296875 0 508.27695
163.0717315673828 0 5205.941
169.13369750976562 0 575.33466
170.09283447265625 0 929.679
171.07638549804688 0 743.003
171.14959716796875 0 77551.21 a 1
172.07205200195312 0 1459.796
172.15293884277344 0 7072.2007
173.12901306152344 0 594.14825
173.45150756835938 0 2075.849
175.0868682861328 0 1560.4288
176.10726928710938 0 1603.5885
176.96363830566406 0 502.19113
177.10264587402344 0 4098.614
177.11163330078125 0 1282.4615
181.0975799560547 0 818.4096
181.1704559326172 0 2567.783
185.12924194335938 0 1035.729
185.1649169921875 0 1011.6733
186.12350463867188 0 874.8724
187.14419555664062 0 2303.026
188.1038055419922 0 651.1421
195.487060546875 0 508.55414
197.0924072265625 0 734.2223
197.10238647460938 0 474.96823
197.12884521484375 0 2616.9385
197.1652069091797 0 1090.4988
198.0771942138672 0 446.84387
198.087646484375 0 1771.4011
199.14443969726562 0 38933.055 b 1
200.10238647460938 0 640.2037
200.14785766601562 0 4081.8752
201.123779296875 0 1607.6016
204.10226440429688 0 3313.128
205.0974884033203 0 7409.6006
205.1064453125 0 701.9286
206.10089111328125 0 1013.62274
207.14984130859375 0 738.6138
208.1084442138672 0 1833.9558
208.86631774902344 0 554.19885
209.09239196777344 0 1387.1255
209.16517639160156 0 1349.1902
211.14523315429688 0 499.13092
212.13966369628906 0 3987.0693
213.16067504882812 0 462.7433
215.11520385742188 0 783.5915
215.139404296875 0 3778.5984 y Water loss 8
216.09800720214844 0 1904.1075
217.09820556640625 0 651.974
219.14962768554688 0 3245.732
220.09703063964844 0 1208.7897
220.15269470214844 0 790.4275
221.1289520263672 0 912.114
224.1024627685547 0 613.1035
224.17579650878906 0 1160.7097
226.1188507080078 0 567.8858
226.4524383544922 0 544.454
227.1142578125 0 1687.254
228.09817504882812 0 959.79596
228.1825408935547 0 717.5452
230.1503143310547 0 6636.1475
231.15243530273438 0 610.04565
233.12840270996094 0 605.1092
233.14990234375 0 1780.3704 y 8
237.1602020263672 0 1018.0726
239.15029907226562 0 2030.1676
242.18678283691406 0 1822.1862
245.12522888183594 0 1502.1378
247.14407348632812 0 2321.571
250.08360290527344 0 637.5444
251.1504669189453 0 707.9574 y Water loss 5
255.10910034179688 0 4317.0786
256.1302490234375 0 674.3931
256.1773681640625 0 694.3851
261.1246337890625 0 699.31
262.1190490722656 0 2503.5535
269.1611633300781 0 4146.254
272.1365051269531 0 989.066
273.1195373535156 0 4037.988
274.1198425292969 0 1089.4766
275.1746520996094 0 723.2115
276.156005859375 0 1856.0795
282.3924560546875 0 559.2517
287.1715393066406 0 994.26025
291.1449890136719 0 1055.7496
292.202392578125 0 2702.7375
292.644775390625 0 539.1249
301.191650390625 0 1560.793
304.1661071777344 0 658.045
308.1974182128906 0 1430.4031
309.2016296386719 0 725.3341
310.2122802734375 0 3126.1338
317.18988037109375 0 808.0587
319.1407775878906 0 1425.59
323.1720886230469 0 629.74146
324.1922912597656 0 646.86676
325.22308349609375 0 805.0241
326.1831359863281 0 3473.9626
328.13037109375 0 719.6993
342.201171875 0 691.8354
343.1627197265625 0 1074.6631
343.2341003417969 0 1142.0222 y Water loss 7
344.19384765625 0 1344.7555
346.2121276855469 0 1905.2012 b 2
346.2358093261719 0 594.50494
357.15582275390625 0 1411.6521
357.2460632324219 0 919.25507
361.2448425292969 0 4887.452 y 7
362.24737548828125 0 672.8546
365.1924743652344 0 899.9778
367.1613464355469 0 803.0797
374.18243408203125 0 1784.8743
375.16741943359375 0 718.8837
376.1640625 0 625.26013
381.7101745605469 0 1035.086 y Water loss 2
382.24493408203125 0 1064.7485
383.20391845703125 0 4065.8816
384.20660400390625 0 668.9519
384.23431396484375 0 811.58765
390.7164611816406 0 12778.933 y 2
391.2179260253906 0 4982.528
391.7195129394531 0 2111.3132
392.1927185058594 0 1275.0254
400.2555847167969 0 1390.4974
401.2144775390625 0 3499.4814
402.17767333984375 0 5977.714
403.180419921875 0 1169.5343
415.25604248046875 0 1057.1584
418.9960632324219 0 858.93634
420.1887512207031 0 2810.189
422.7127990722656 0 614.8726
429.2838439941406 0 820.87946
439.266357421875 0 1566.8752
447.40863037109375 0 592.2043
454.2339172363281 0 1015.27515
455.233154296875 0 650.36163
457.2774658203125 0 3399.1655
458.2810974121094 0 1011.43365
462.2909240722656 0 1018.44293 y 6
470.2743225097656 0 833.27026
471.2200927734375 0 3323.204
478.2786560058594 0 876.9026
480.7778015136719 0 863.8479 Precursor Water loss
483.9583435058594 0 628.4798
486.3042297363281 0 1868.5577
487.30377197265625 0 879.24194
488.2420654296875 0 804.6286
489.2331237792969 0 1122.2523
489.7857666015625 0 987.32806 Precursor
490.2879638671875 0 1890.5157
496.2886962890625 0 4069.6648
497.2903747558594 0 1355.7548
512.2620239257812 0 757.0226
514.2986450195312 0 8238.85
515.3013916015625 0 2002.4819
519.3133544921875 0 1870.4867 y 5
525.8109741210938 0 1320.8478
530.2732543945312 0 3746.1223
531.2752685546875 0 1109.3408
539.8106079101562 0 2396.4893
540.3109130859375 0 2840.1235
548.2825317382812 0 6841.1294
548.8220825195312 0 566.0294
549.2861938476562 0 1225.4838
554.252197265625 0 645.98413
559.3247680664062 0 635.26245
572.4217529296875 0 660.05634
576.3350219726562 0 9113.12 y 4
577.338134765625 0 2591.9539
589.3192749023438 0 694.783
600.315185546875 0 722.3685 b Water loss 6
615.3492431640625 0 2157.1694 y Water loss 3
633.3575439453125 0 27925.205 y 3
634.3600463867188 0 8898.938
635.3623046875 0 1584.4344
643.3554077148438 0 6961.68
643.8522338867188 0 614.1057
644.3590087890625 0 2220.504
661.36669921875 0 14907.267
662.3689575195312 0 5087.4917
663.3743286132812 0 866.3436
690.3812866210938 0 655.4824
706.804931640625 0 699.9647
718.4241943359375 0 837.43353
746.4174194335938 0 2421.3108 b 7
747.4232177734375 0 765.17535
760.4368286132812 0 650.54474
762.4154052734375 0 2567.7625 y Water loss 2
763.4146118164062 0 1402.3741
780.4244384765625 0 54944.832 y 2
781.4273071289062 0 24762.459
782.4297485351562 0 5513.5825
841.4863891601562 0 795.24084 b Water loss 8
859.5018920898438 0 3699.6475 b 8
860.5073852539062 0 1931.9714
879.4911499023438 0 3924.1313 y 1
880.4879760742188 0 614.63586
1072.7750244140625 0 603.7266
1627.74365234375 0 743.0939
2433.64404296875 0 595.39764
2442.69580078125 0 592.15295
2582.04150390625 0 647.98566
2984.391845703125 0 604.7111

Spectrum Details

|  |  |
| --- | --- |
| Matched peaks? Matched peaks The total absolute number of peaks matched. Additionally in brackets the total fraction of peaks matched and the total number of peaks is shown. | 26 (10.70% of 243) |
| FDR? FDR The false discovery rate estimated for this peptide. It is calculated by matching all theoretical fragments with a non-integer shift with the raw peaks for this spectrum. This is done with 40 different shifts. The resulting percentage is the average number of annotated peaks over the number of annotated peaks with the correct spectrum. | 0.46% |
| Satellite FDR? Satellite FDR See the FDR for details on its calculation. This satellite ion specific FDR only contains the satellite ions (d/w) for I/L/J positions. | - |
| PSM Score? PSM Score The PSM Score as given by Hecklib to this annotated spectrum. It is shown with three significant figures. | 281 |

#### Spectrum 6400? Spectrum 6400 The raw spectrum of this peptide as annotated by Hecklib. The fragments are coloured according to ion type (see legend). Any peaks with a star '\*' as text can be hovered over to see the full details, first the ion type second the mass shift type. By hovering over the amino acids in the peptide or ions in the legend the corresponding peaks are highlighted. By toggling the 'Unassigned' label you can turn the background (unassigned) peaks on or off in the plot. By updating the slider in the Ion legend you can update the spectrum to only show the top X% of the peaks with labels. The top X% means any peak that is within X% of the highest intensity. By dragging in the spectrum you can zoom in to a specific part of the spectrum and use 'Zoom Out' to get back to the original zoom level. The annotation of the spectrum is based on the given sequence in the peptides file and is done with different software so inconsistencies are likely. The peaks are annotated based on the given sequence, with 20 ppm tolerance.

Copy Data

##### Spectrum 6400 (TSV)

###### Preview

```
Loading example...
```

*Click on the button to copy the data to your clipboard.*

Mz MinMz MaxIntensity Max

WidthHeightPeptide font sizePeptide stroke widthSpectrum font sizeSpectrum stroke widthCompact peptide

Ion legend

wxyz

abcd

OtherUnassignedIon^Charge^_Position_Show for top:%

VVFGGGTKJT

01.92e+43.85e+45.77e+47.69e+4

Zoom Out

y^+1^_1_a^+1^_2_b^+1^_2_y^+1^_2_y^+1^_2_y^+1^_3_b^+1^_3_y^+1^_3_y^+2^_8_y^+2^_8_b^+1^_4_y^+1^_4_\*\*y^+1^_5_y^+1^_6_y^+1^_6_y^+1^_7_b^+1^_7_y^+1^_7_b^+1^_8_y^+1^_8_y^+1^_8_y^+1^_8_b^+1^_9_y^+1^_9_

0776155123273102

Fragment Matches Table

Show background peaks

| Position | Ion type | Intensity | mz Theoretical | mz Error (Th) | mz Error (ppm) | Charge | Series Number |
| --- | --- | --- | --- | --- | --- | --- | --- |
| 10 | y | 5859 | 120.1 | 0.0002153 | 1.793 | +1 | 1 |
| - | - | 4.081E+04 | 120.1 | - | - | 0 | - |
| - | - | 3673 | 121.1 | - | - | 0 | - |
| - | - | 375.5 | 126.1 | - | - | 0 | - |
| - | - | 411.4 | 127.1 | - | - | 0 | - |
| - | - | 477.7 | 127.1 | - | - | 0 | - |
| - | - | 458.6 | 128.1 | - | - | 0 | - |
| - | - | 548.3 | 129.1 | - | - | 0 | - |
| - | - | 3.455E+04 | 129.1 | - | - | 0 | - |
| - | - | 662.9 | 130 | - | - | 0 | - |
| - | - | 767.7 | 130.1 | - | - | 0 | - |
| - | - | 1819 | 130.1 | - | - | 0 | - |
| - | - | 1682 | 131.1 | - | - | 0 | - |
| - | - | 588.5 | 132.1 | - | - | 0 | - |
| - | - | 460.4 | 133.1 | - | - | 0 | - |
| - | - | 1326 | 133.1 | - | - | 0 | - |
| - | - | 403.8 | 135.9 | - | - | 0 | - |
| - | - | 6069 | 136.1 | - | - | 0 | - |
| - | - | 505.2 | 138.1 | - | - | 0 | - |
| - | - | 475.1 | 140 | - | - | 0 | - |
| - | - | 694 | 140.1 | - | - | 0 | - |
| - | - | 1065 | 141.1 | - | - | 0 | - |
| - | - | 713.8 | 142.1 | - | - | 0 | - |
| - | - | 496.8 | 143.2 | - | - | 0 | - |
| - | - | 576 | 144.1 | - | - | 0 | - |
| - | - | 518.2 | 144.1 | - | - | 0 | - |
| - | - | 1043 | 145.1 | - | - | 0 | - |
| - | - | 752.2 | 146.1 | - | - | 0 | - |
| - | - | 721.3 | 147 | - | - | 0 | - |
| - | - | 411 | 148.9 | - | - | 0 | - |
| - | - | 902.4 | 149 | - | - | 0 | - |
| - | - | 505.8 | 152.1 | - | - | 0 | - |
| - | - | 888.6 | 155.1 | - | - | 0 | - |
| - | - | 847.1 | 155.1 | - | - | 0 | - |
| - | - | 608 | 156.1 | - | - | 0 | - |
| - | - | 432.2 | 156.6 | - | - | 0 | - |
| - | - | 736.4 | 158.1 | - | - | 0 | - |
| - | - | 994 | 159.1 | - | - | 0 | - |
| - | - | 2274 | 163.1 | - | - | 0 | - |
| - | - | 471.7 | 165.1 | - | - | 0 | - |
| - | - | 551.5 | 165.1 | - | - | 0 | - |
| - | - | 483.7 | 166.1 | - | - | 0 | - |
| - | - | 494.7 | 167.1 | - | - | 0 | - |
| - | - | 578.6 | 169.1 | - | - | 0 | - |
| - | - | 748.8 | 170.1 | - | - | 0 | - |
| 2 | a | 7.615E+04 | 171.1 | 0.0002244 | 1.311 | +1 | 2 |
| - | - | 1608 | 172.1 | - | - | 0 | - |
| - | - | 588.1 | 172.1 | - | - | 0 | - |
| - | - | 7087 | 172.2 | - | - | 0 | - |
| - | - | 470.3 | 173.1 | - | - | 0 | - |
| - | - | 625.3 | 173.1 | - | - | 0 | - |
| - | - | 3089 | 173.4 | - | - | 0 | - |
| - | - | 638.5 | 174.1 | - | - | 0 | - |
| - | - | 565.9 | 175.1 | - | - | 0 | - |
| - | - | 1015 | 176.1 | - | - | 0 | - |
| - | - | 4150 | 177.1 | - | - | 0 | - |
| - | - | 878.2 | 177.1 | - | - | 0 | - |
| - | - | 2357 | 181.2 | - | - | 0 | - |
| - | - | 509.7 | 185.1 | - | - | 0 | - |
| - | - | 1247 | 185.2 | - | - | 0 | - |
| - | - | 653.8 | 187.1 | - | - | 0 | - |
| - | - | 2556 | 187.1 | - | - | 0 | - |
| - | - | 493.8 | 188.1 | - | - | 0 | - |
| - | - | 481.2 | 188.3 | - | - | 0 | - |
| - | - | 672.6 | 191.1 | - | - | 0 | - |
| - | - | 548.1 | 197.1 | - | - | 0 | - |
| - | - | 1699 | 197.1 | - | - | 0 | - |
| - | - | 685.5 | 197.2 | - | - | 0 | - |
| - | - | 1741 | 198.1 | - | - | 0 | - |
| 2 | b | 3.673E+04 | 199.1 | 9.127E-05 | 0.4583 | +1 | 2 |
| - | - | 3947 | 200.1 | - | - | 0 | - |
| - | - | 1948 | 201.1 | - | - | 0 | - |
| - | - | 561.6 | 203.1 | - | - | 0 | - |
| - | - | 6865 | 205.1 | - | - | 0 | - |
| - | - | 1057 | 206.1 | - | - | 0 | - |
| - | - | 628.7 | 207.1 | - | - | 0 | - |
| - | - | 1011 | 208.1 | - | - | 0 | - |
| - | - | 1370 | 209.2 | - | - | 0 | - |
| - | - | 928.3 | 211.1 | - | - | 0 | - |
| - | - | 4279 | 212.1 | - | - | 0 | - |
| - | - | 518.2 | 213.2 | - | - | 0 | - |
| 9 | y | 5173 | 215.1 | 0.0001718 | 0.7984 | +1 | 2 |
| - | - | 1211 | 216.1 | - | - | 0 | - |
| - | - | 1036 | 217.1 | - | - | 0 | - |
| - | - | 2343 | 219.1 | - | - | 0 | - |
| - | - | 742.4 | 221.1 | - | - | 0 | - |
| - | - | 1308 | 224.2 | - | - | 0 | - |
| - | - | 1663 | 227.1 | - | - | 0 | - |
| - | - | 1326 | 228.1 | - | - | 0 | - |
| - | - | 6587 | 230.2 | - | - | 0 | - |
| - | - | 519 | 231.2 | - | - | 0 | - |
| 9 | y | 2357 | 233.1 | 0.0001509 | 0.6472 | +1 | 2 |
| - | - | 709.2 | 233.2 | - | - | 0 | - |
| - | - | 1550 | 237.2 | - | - | 0 | - |
| - | - | 780.2 | 239.1 | - | - | 0 | - |
| - | - | 505.7 | 239.2 | - | - | 0 | - |
| - | - | 725.6 | 240.1 | - | - | 0 | - |
| - | - | 1964 | 242.2 | - | - | 0 | - |
| - | - | 618.2 | 243.1 | - | - | 0 | - |
| - | - | 636 | 244.1 | - | - | 0 | - |
| - | - | 711.3 | 245.1 | - | - | 0 | - |
| - | - | 1388 | 247.1 | - | - | 0 | - |
| - | - | 571.2 | 251.2 | - | - | 0 | - |
| - | - | 573.8 | 253.2 | - | - | 0 | - |
| - | - | 1230 | 254 | - | - | 0 | - |
| - | - | 504.4 | 254.2 | - | - | 0 | - |
| - | - | 3785 | 255.1 | - | - | 0 | - |
| - | - | 568.8 | 261.1 | - | - | 0 | - |
| - | - | 2263 | 262.1 | - | - | 0 | - |
| - | - | 4047 | 269.2 | - | - | 0 | - |
| - | - | 589.8 | 270.2 | - | - | 0 | - |
| - | - | 3085 | 273.1 | - | - | 0 | - |
| - | - | 873.6 | 274.1 | - | - | 0 | - |
| - | - | 2696 | 276.2 | - | - | 0 | - |
| - | - | 1052 | 287.2 | - | - | 0 | - |
| - | - | 599.9 | 291.1 | - | - | 0 | - |
| - | - | 595.1 | 292.2 | - | - | 0 | - |
| - | - | 1686 | 301.2 | - | - | 0 | - |
| - | - | 663.4 | 302.2 | - | - | 0 | - |
| - | - | 1141 | 304.2 | - | - | 0 | - |
| - | - | 3151 | 310.2 | - | - | 0 | - |
| - | - | 1349 | 319.1 | - | - | 0 | - |
| - | - | 685.9 | 325.2 | - | - | 0 | - |
| - | - | 3384 | 326.2 | - | - | 0 | - |
| - | - | 532.6 | 337.6 | - | - | 0 | - |
| 8 | y | 634.1 | 343.2 | 0.001713 | 4.99 | +1 | 3 |
| - | - | 1069 | 344.2 | - | - | 0 | - |
| 3 | b | 1508 | 346.2 | 0.0008607 | 2.486 | +1 | 3 |
| - | - | 725.8 | 347.2 | - | - | 0 | - |
| - | - | 631.9 | 347.4 | - | - | 0 | - |
| - | - | 831.3 | 348.2 | - | - | 0 | - |
| - | - | 1032 | 349.1 | - | - | 0 | - |
| - | - | 1039 | 357.2 | - | - | 0 | - |
| 8 | y | 3723 | 361.2 | 0.0003144 | 0.8704 | +1 | 3 |
| - | - | 1184 | 362.2 | - | - | 0 | - |
| - | - | 789.3 | 365.2 | - | - | 0 | - |
| - | - | 844.5 | 367.1 | - | - | 0 | - |
| - | - | 770.6 | 373.2 | - | - | 0 | - |
| - | - | 1520 | 374.2 | - | - | 0 | - |
| - | - | 1018 | 375.2 | - | - | 0 | - |
| 3 | y | 1454 | 381.7 | 0.0003302 | 0.865 | +2 | 8 |
| - | - | 1061 | 382.2 | - | - | 0 | - |
| - | - | 848.6 | 382.2 | - | - | 0 | - |
| - | - | 3186 | 383.2 | - | - | 0 | - |
| - | - | 586.8 | 383.2 | - | - | 0 | - |
| - | - | 838.6 | 384.2 | - | - | 0 | - |
| 3 | y | 1.4E+04 | 390.7 | 0.0001194 | 0.3055 | +2 | 8 |
| - | - | 4844 | 391.2 | - | - | 0 | - |
| - | - | 799.2 | 391.7 | - | - | 0 | - |
| - | - | 1686 | 392.2 | - | - | 0 | - |
| - | - | 1480 | 400.3 | - | - | 0 | - |
| - | - | 3180 | 401.2 | - | - | 0 | - |
| - | - | 4872 | 402.2 | - | - | 0 | - |
| - | - | 580.3 | 402.7 | - | - | 0 | - |
| - | - | 1523 | 403.2 | - | - | 0 | - |
| 4 | b | 584.8 | 403.2 | 0.001621 | 4.02 | +1 | 4 |
| - | - | 1212 | 415.3 | - | - | 0 | - |
| - | - | 595.3 | 416.3 | - | - | 0 | - |
| - | - | 804.9 | 419 | - | - | 0 | - |
| - | - | 3648 | 420.2 | - | - | 0 | - |
| - | - | 975.2 | 421.2 | - | - | 0 | - |
| - | - | 669 | 429.3 | - | - | 0 | - |
| - | - | 1449 | 439.3 | - | - | 0 | - |
| - | - | 1057 | 455.2 | - | - | 0 | - |
| - | - | 3378 | 457.3 | - | - | 0 | - |
| - | - | 709 | 458.3 | - | - | 0 | - |
| 7 | y | 703.6 | 462.3 | 0.0008657 | 1.873 | +1 | 4 |
| 0 | Precursor | 1392 | 480.8 | 0.0006899 | 1.435 | +2 | -1 |
| - | - | 2249 | 486.3 | - | - | 0 | - |
| 0 | Precursor | 1533 | 489.8 | 0.001913 | 3.907 | +2 | -1 |
| - | - | 628.2 | 490.1 | - | - | 0 | - |
| - | - | 1494 | 490.2 | - | - | 0 | - |
| - | - | 1614 | 490.3 | - | - | 0 | - |
| - | - | 3866 | 496.3 | - | - | 0 | - |
| - | - | 958.1 | 497.3 | - | - | 0 | - |
| - | - | 9134 | 514.3 | - | - | 0 | - |
| - | - | 2030 | 515.3 | - | - | 0 | - |
| 6 | y | 1403 | 519.3 | 0.000215 | 0.414 | +1 | 5 |
| - | - | 830.2 | 528.3 | - | - | 0 | - |
| - | - | 2558 | 530.3 | - | - | 0 | - |
| - | - | 1325 | 531.3 | - | - | 0 | - |
| - | - | 7155 | 548.3 | - | - | 0 | - |
| - | - | 1255 | 549.3 | - | - | 0 | - |
| 5 | y | 755.6 | 558.3 | 0.001767 | 3.165 | +1 | 6 |
| 5 | y | 8057 | 576.3 | 0.000924 | 1.603 | +1 | 6 |
| - | - | 2560 | 577.3 | - | - | 0 | - |
| - | - | 604.4 | 578.3 | - | - | 0 | - |
| 4 | y | 2263 | 615.3 | 0.0001043 | 0.1695 | +1 | 7 |
| - | - | 809.2 | 616.3 | - | - | 0 | - |
| - | - | 715.7 | 617.3 | - | - | 0 | - |
| 7 | b | 648.8 | 618.3 | 0.0006132 | 0.9918 | +1 | 7 |
| 4 | y | 2.316E+04 | 633.4 | 0.0002563 | 0.4047 | +1 | 7 |
| - | - | 8645 | 634.4 | - | - | 0 | - |
| - | - | 1419 | 635.4 | - | - | 0 | - |
| - | - | 6578 | 643.4 | - | - | 0 | - |
| - | - | 2047 | 644.4 | - | - | 0 | - |
| - | - | 648.6 | 645.4 | - | - | 0 | - |
| - | - | 1.41E+04 | 661.4 | - | - | 0 | - |
| - | - | 5870 | 662.4 | - | - | 0 | - |
| - | - | 679.9 | 663.9 | - | - | 0 | - |
| - | - | 554.2 | 693.1 | - | - | 0 | - |
| - | - | 746.4 | 714.4 | - | - | 0 | - |
| - | - | 706.4 | 718.4 | - | - | 0 | - |
| - | - | 1086 | 744.4 | - | - | 0 | - |
| 8 | b | 1877 | 746.4 | 0.001704 | 2.283 | +1 | 8 |
| - | - | 719.3 | 761.4 | - | - | 0 | - |
| 3 | y | 2238 | 762.4 | 0.0026 | 3.411 | +1 | 8 |
| 3 | y | 777.9 | 763.4 | 0.01399 | 18.33 | +1 | 8 |
| 3 | y | 5.055E+04 | 780.4 | 0.001385 | 1.775 | +1 | 8 |
| - | - | 1.992E+04 | 781.4 | - | - | 0 | - |
| - | - | 4081 | 782.4 | - | - | 0 | - |
| - | - | 1091 | 785.4 | - | - | 0 | - |
| - | - | 1417 | 813.4 | - | - | 0 | - |
| 9 | b | 1865 | 859.5 | 0.001234 | 1.436 | +1 | 9 |
| - | - | 1982 | 860.5 | - | - | 0 | - |
| - | - | 589.2 | 864.4 | - | - | 0 | - |
| 2 | y | 2573 | 879.5 | 0.004186 | 4.76 | +1 | 9 |
| - | - | 1447 | 880.5 | - | - | 0 | - |
| - | - | 660.8 | 1591 | - | - | 0 | - |
| - | - | 635.4 | 1645 | - | - | 0 | - |
| - | - | 602.3 | 1659 | - | - | 0 | - |
| - | - | 663.6 | 2030 | - | - | 0 | - |
| - | - | 637.1 | 2085 | - | - | 0 | - |
| - | - | 731.4 | 3071 | - | - | 0 | - |

m/z Charge Intensity FragmentType MassShift Position
120.06573486328125 0 5858.802 y 9
120.08102416992188 0 40809.137
121.08436584472656 0 3673.1475
126.103271484375 0 375.52063
127.075927734375 0 411.3504
127.08683013916016 0 477.72018
128.0707550048828 0 458.63223
129.06591796875 0 548.2848
129.10247802734375 0 34552.375
130.04966735839844 0 662.933
130.06568908691406 0 767.6641
130.10581970214844 0 1819.201
131.0819091796875 0 1682.3208
132.0811004638672 0 588.5493
133.0609893798828 0 460.3856
133.08609008789062 0 1325.8231
135.89952087402344 0 403.8002
136.07591247558594 0 6069.027
138.06651306152344 0 505.17755
140.04583740234375 0 475.08588
140.0821075439453 0 693.9735
141.0659942626953 0 1065.0125
142.1228485107422 0 713.8262
143.24642944335938 0 496.80347
144.0658721923828 0 575.9788
144.07716369628906 0 518.22864
145.0611114501953 0 1043.0603
146.060302734375 0 752.16364
147.04478454589844 0 721.3017
148.946533203125 0 410.98242
149.02365112304688 0 902.4376
152.10736083984375 0 505.76492
155.08168029785156 0 888.61365
155.11813354492188 0 847.0568
156.07644653320312 0 607.98724
156.57723999023438 0 432.23785
158.0927276611328 0 736.4239
159.0767364501953 0 994.04346
163.071533203125 0 2274.461
165.10316467285156 0 471.69412
165.11338806152344 0 551.45013
166.0531463623047 0 483.69507
167.1185302734375 0 494.652
169.1338653564453 0 578.5896
170.09190368652344 0 748.77026
171.1494140625 0 76153.2 a 1
172.07192993164062 0 1607.5352
172.14508056640625 0 588.1027
172.15283203125 0 7086.58
173.09213256835938 0 470.31238
173.1283416748047 0 625.25714
173.4404754638672 0 3089.1997
174.05543518066406 0 638.5018
175.08670043945312 0 565.85175
176.1072998046875 0 1015.03705
177.10235595703125 0 4150.279
177.11094665527344 0 878.1681
181.1702117919922 0 2356.9937
185.1280517578125 0 509.68182
185.1646270751953 0 1247.4637
187.1079864501953 0 653.7504
187.1444549560547 0 2556.021
188.1022491455078 0 493.84344
188.3090057373047 0 481.2129
191.0814971923828 0 672.55194
197.091552734375 0 548.13446
197.1287384033203 0 1699.1036
197.16461181640625 0 685.4905
198.08767700195312 0 1740.522
199.14419555664062 0 36731.605 b 1
200.14747619628906 0 3946.6387
201.12350463867188 0 1947.717
203.1180419921875 0 561.5614
205.09722900390625 0 6865.0776
206.1005401611328 0 1056.9495
207.1490936279297 0 628.741
208.10841369628906 0 1011.39453
209.16510009765625 0 1370.2964
211.1443328857422 0 928.27704
212.1394805908203 0 4279.1772
213.16079711914062 0 518.2394
215.13919067382812 0 5172.702 y Water loss 8
216.09808349609375 0 1211.0161
217.0972442626953 0 1036.1095
219.14930725097656 0 2342.7583
221.1289520263672 0 742.4412
224.17599487304688 0 1307.6448
227.1135711669922 0 1663.4539
228.0985107421875 0 1325.8206
230.1500701904297 0 6586.5356
231.15292358398438 0 518.985
233.1497344970703 0 2356.9966 y 8
233.16357421875 0 709.184
237.15957641601562 0 1549.678
239.13873291015625 0 780.2079
239.15081787109375 0 505.71246
240.13417053222656 0 725.60596
242.18630981445312 0 1963.7181
243.11341857910156 0 618.1654
244.1078643798828 0 635.95404
245.12460327148438 0 711.30865
247.14410400390625 0 1388.0388
251.198974609375 0 571.24304
253.22781372070312 0 573.788
254.03895568847656 0 1229.5808
254.2090301513672 0 504.38745
255.10873413085938 0 3785.4902
261.122314453125 0 568.8417
262.11871337890625 0 2262.7026
269.1609191894531 0 4047.4888
270.1796875 0 589.8121
273.119384765625 0 3085.1074
274.1186218261719 0 873.62054
276.1553955078125 0 2696.0852
287.1717529296875 0 1052.0466
291.1446838378906 0 599.8836
292.2015075683594 0 595.11523
301.1910705566406 0 1685.541
302.17071533203125 0 663.42725
304.1651916503906 0 1141.1342
310.21282958984375 0 3150.6865
319.1402282714844 0 1348.6733
325.2239074707031 0 685.9103
326.1822509765625 0 3383.946
337.6119079589844 0 532.6332
343.2322692871094 0 634.05255 y Water loss 7
344.1929626464844 0 1068.87
346.21337890625 0 1507.6664 b 2
347.2167663574219 0 725.7696
347.4217834472656 0 631.941
348.1663818359375 0 831.31335
349.1136474609375 0 1031.6606
357.1559143066406 0 1039.1796
361.2442321777344 0 3722.8164 y 7
362.2483215332031 0 1184.0919
365.1919860839844 0 789.30206
367.12286376953125 0 844.5295
373.16558837890625 0 770.62146
374.1826477050781 0 1520.1292
375.1661376953125 0 1017.65424
381.7105407714844 0 1453.8348 y Water loss 2
382.21209716796875 0 1061.3319
382.24261474609375 0 848.60406
383.20361328125 0 3185.9683
383.24688720703125 0 586.7629
384.2067565917969 0 838.5777
390.7160339355469 0 14002.041 y 2
391.2176818847656 0 4843.784
391.7194519042969 0 799.2098
392.1927795410156 0 1685.7826
400.2561340332031 0 1479.9042
401.2144775390625 0 3179.869
402.17724609375 0 4871.992
402.67755126953125 0 580.2946
403.17974853515625 0 1522.5068
403.23236083984375 0 584.8239 b 3
415.2547912597656 0 1211.9066
416.2598876953125 0 595.25977
418.996826171875 0 804.86993
420.187744140625 0 3648.4265
421.1919860839844 0 975.1582
429.28167724609375 0 668.9743
439.2664489746094 0 1448.7776
455.23040771484375 0 1057.2844
457.27703857421875 0 3378.0515
458.2793884277344 0 708.9632
462.2930908203125 0 703.6155 y 6
480.7785949707031 0 1392.4382 Precursor Water loss
486.30364990234375 0 2249.1287
489.78265380859375 0 1532.7493 Precursor
490.0687255859375 0 628.24255
490.2080993652344 0 1493.9609
490.2840270996094 0 1614.355
496.287353515625 0 3865.826
497.2916564941406 0 958.10394
514.2977905273438 0 9133.502
515.300048828125 0 2030.2205
519.3139038085938 0 1403.2529 y 5
528.2971801757812 0 830.23926
530.2720336914062 0 2557.9026
531.275146484375 0 1324.6451
548.2822875976562 0 7155.3564
549.2847290039062 0 1255.2521
558.3263549804688 0 755.583 y Water loss 4
576.334228515625 0 8056.785 y 4
577.3381958007812 0 2560.3818
578.3362426757812 0 604.38666
615.345947265625 0 2263.1738 y Water loss 3
616.3461303710938 0 809.171
617.3492431640625 0 715.73944
618.323974609375 0 648.7728 b 6
633.3568725585938 0 23158.393 y 3
634.3598022460938 0 8645.164
635.3638305664062 0 1418.9702
643.3544921875 0 6577.8096
644.3580322265625 0 2047.2167
645.36376953125 0 648.6247
661.365966796875 0 14099.464
662.3685913085938 0 5870.363
663.9440307617188 0 679.9339
693.0884399414062 0 554.1798
714.3709716796875 0 746.39307
718.4208984375 0 706.4245
744.4080810546875 0 1085.9738
746.4178466796875 0 1877.183 b 7
761.4364624023438 0 719.2698
762.411865234375 0 2238.3474 y Water loss 2
763.4124755859375 0 777.9394 y Ammonia loss 2
780.4236450195312 0 50554.203 y 2
781.4262084960938 0 19922.336
782.4292602539062 0 4081.1814
785.448486328125 0 1090.8755
813.4440307617188 0 1416.5719
859.5023803710938 0 1865.4285 b 8
860.5051879882812 0 1981.7001
864.4431762695312 0 589.21173
879.4892578125 0 2573.269 y 1
880.4899291992188 0 1447.1566
1591.01904296875 0 660.78064
1644.618896484375 0 635.4362
1659.3343505859375 0 602.33844
2029.508056640625 0 663.6413
2084.946044921875 0 637.1261
3071.49609375 0 731.437

Spectrum Details

|  |  |
| --- | --- |
| Matched peaks? Matched peaks The total absolute number of peaks matched. Additionally in brackets the total fraction of peaks matched and the total number of peaks is shown. | 26 (11.61% of 224) |
| FDR? FDR The false discovery rate estimated for this peptide. It is calculated by matching all theoretical fragments with a non-integer shift with the raw peaks for this spectrum. This is done with 40 different shifts. The resulting percentage is the average number of annotated peaks over the number of annotated peaks with the correct spectrum. | 0.27% |
| Satellite FDR? Satellite FDR See the FDR for details on its calculation. This satellite ion specific FDR only contains the satellite ions (d/w) for I/L/J positions. | - |
| PSM Score? PSM Score The PSM Score as given by Hecklib to this annotated spectrum. It is shown with three significant figures. | 321 |

#### Spectrum 6461? Spectrum 6461 The raw spectrum of this peptide as annotated by Hecklib. The fragments are coloured according to ion type (see legend). Any peaks with a star '\*' as text can be hovered over to see the full details, first the ion type second the mass shift type. By hovering over the amino acids in the peptide or ions in the legend the corresponding peaks are highlighted. By toggling the 'Unassigned' label you can turn the background (unassigned) peaks on or off in the plot. By updating the slider in the Ion legend you can update the spectrum to only show the top X% of the peaks with labels. The top X% means any peak that is within X% of the highest intensity. By dragging in the spectrum you can zoom in to a specific part of the spectrum and use 'Zoom Out' to get back to the original zoom level. The annotation of the spectrum is based on the given sequence in the peptides file and is done with different software so inconsistencies are likely. The peaks are annotated based on the given sequence, with 20 ppm tolerance.

Copy Data

##### Spectrum 6461 (TSV)

###### Preview

```
Loading example...
```

*Click on the button to copy the data to your clipboard.*

Mz MinMz MaxIntensity Max

WidthHeightPeptide font sizePeptide stroke widthSpectrum font sizeSpectrum stroke widthCompact peptide

Ion legend

wxyz

abcd

OtherUnassignedIon^Charge^_Position_Show for top:%

VVFGGGTKJT

01.75e+43.51e+45.26e+47.01e+4

Zoom Out

y^+1^_1_a^+1^_2_b^+1^_2_y^+1^_2_y^+1^_2_y^+2^_7_y^+1^_3_b^+1^_3_y^+1^_3_b^+2^_8_y^+2^_8_y^+2^_8_b^+1^_4_y^+1^_4_\*y^+1^_5_y^+1^_6_y^+1^_7_y^+1^_7_b^+1^_8_y^+1^_8_y^+1^_8_b^+1^_9_y^+1^_9_

0883176626493531

Fragment Matches Table

Show background peaks

| Position | Ion type | Intensity | mz Theoretical | mz Error (Th) | mz Error (ppm) | Charge | Series Number |
| --- | --- | --- | --- | --- | --- | --- | --- |
| 10 | y | 5688 | 120.1 | 0.000261 | 2.174 | +1 | 1 |
| - | - | 4.621E+04 | 120.1 | - | - | 0 | - |
| - | - | 374.3 | 120.4 | - | - | 0 | - |
| - | - | 3245 | 121.1 | - | - | 0 | - |
| - | - | 649.5 | 127.1 | - | - | 0 | - |
| - | - | 817.4 | 128.1 | - | - | 0 | - |
| - | - | 930.7 | 129.1 | - | - | 0 | - |
| - | - | 3.293E+04 | 129.1 | - | - | 0 | - |
| - | - | 440.5 | 130.1 | - | - | 0 | - |
| - | - | 2346 | 130.1 | - | - | 0 | - |
| - | - | 1396 | 131.1 | - | - | 0 | - |
| - | - | 690 | 132.1 | - | - | 0 | - |
| - | - | 671.4 | 133.1 | - | - | 0 | - |
| - | - | 1060 | 133.1 | - | - | 0 | - |
| - | - | 6736 | 136.1 | - | - | 0 | - |
| - | - | 411.9 | 137.3 | - | - | 0 | - |
| - | - | 507 | 139.1 | - | - | 0 | - |
| - | - | 550.4 | 140.1 | - | - | 0 | - |
| - | - | 696.2 | 141.1 | - | - | 0 | - |
| - | - | 467.8 | 141.1 | - | - | 0 | - |
| - | - | 535.6 | 142.1 | - | - | 0 | - |
| - | - | 907.8 | 142.1 | - | - | 0 | - |
| - | - | 583.8 | 146.1 | - | - | 0 | - |
| - | - | 558 | 152.1 | - | - | 0 | - |
| - | - | 419.9 | 153.1 | - | - | 0 | - |
| - | - | 650 | 154.1 | - | - | 0 | - |
| - | - | 1296 | 155.1 | - | - | 0 | - |
| - | - | 560.8 | 156.1 | - | - | 0 | - |
| - | - | 488.5 | 157.1 | - | - | 0 | - |
| - | - | 826.7 | 158.1 | - | - | 0 | - |
| - | - | 698.3 | 159.1 | - | - | 0 | - |
| - | - | 1524 | 163.1 | - | - | 0 | - |
| - | - | 586.1 | 165.1 | - | - | 0 | - |
| - | - | 752.8 | 169.1 | - | - | 0 | - |
| - | - | 542.6 | 170 | - | - | 0 | - |
| - | - | 1294 | 170.1 | - | - | 0 | - |
| 2 | a | 6.943E+04 | 171.1 | 0.0002549 | 1.489 | +1 | 2 |
| - | - | 1107 | 172.1 | - | - | 0 | - |
| - | - | 6619 | 172.2 | - | - | 0 | - |
| - | - | 1531 | 173.1 | - | - | 0 | - |
| - | - | 1465 | 176.1 | - | - | 0 | - |
| - | - | 455.7 | 176.7 | - | - | 0 | - |
| - | - | 3067 | 177.1 | - | - | 0 | - |
| - | - | 2789 | 181.2 | - | - | 0 | - |
| - | - | 538.5 | 182.2 | - | - | 0 | - |
| - | - | 495.5 | 183.4 | - | - | 0 | - |
| - | - | 657 | 185.1 | - | - | 0 | - |
| - | - | 1204 | 185.2 | - | - | 0 | - |
| - | - | 697.3 | 186.1 | - | - | 0 | - |
| - | - | 1730 | 187.1 | - | - | 0 | - |
| - | - | 941.7 | 191.1 | - | - | 0 | - |
| - | - | 635.1 | 197.1 | - | - | 0 | - |
| - | - | 1958 | 197.1 | - | - | 0 | - |
| - | - | 1374 | 197.2 | - | - | 0 | - |
| - | - | 1428 | 198.1 | - | - | 0 | - |
| - | - | 556.9 | 199.1 | - | - | 0 | - |
| 2 | b | 3.527E+04 | 199.1 | 0.0001523 | 0.7648 | +1 | 2 |
| - | - | 3167 | 200.1 | - | - | 0 | - |
| - | - | 1342 | 201.1 | - | - | 0 | - |
| - | - | 479.1 | 202.4 | - | - | 0 | - |
| - | - | 8014 | 205.1 | - | - | 0 | - |
| - | - | 660.5 | 206.1 | - | - | 0 | - |
| - | - | 557.8 | 207.1 | - | - | 0 | - |
| - | - | 607.3 | 208.1 | - | - | 0 | - |
| - | - | 606.1 | 209.1 | - | - | 0 | - |
| - | - | 1576 | 209.2 | - | - | 0 | - |
| - | - | 984.1 | 211.1 | - | - | 0 | - |
| - | - | 3632 | 212.1 | - | - | 0 | - |
| - | - | 611 | 213.1 | - | - | 0 | - |
| - | - | 547.2 | 213.2 | - | - | 0 | - |
| 9 | y | 3634 | 215.1 | 0.0002481 | 1.153 | +1 | 2 |
| - | - | 1370 | 216.1 | - | - | 0 | - |
| - | - | 535.1 | 217.1 | - | - | 0 | - |
| - | - | 3444 | 219.1 | - | - | 0 | - |
| - | - | 600.7 | 223.1 | - | - | 0 | - |
| - | - | 815.6 | 224.2 | - | - | 0 | - |
| - | - | 1361 | 227.1 | - | - | 0 | - |
| - | - | 6655 | 230.2 | - | - | 0 | - |
| 9 | y | 2268 | 233.1 | 0.0001051 | 0.4509 | +1 | 2 |
| - | - | 1130 | 233.2 | - | - | 0 | - |
| - | - | 1096 | 235.1 | - | - | 0 | - |
| - | - | 701.1 | 235.1 | - | - | 0 | - |
| - | - | 504.5 | 237.1 | - | - | 0 | - |
| - | - | 1708 | 237.2 | - | - | 0 | - |
| - | - | 1485 | 242.2 | - | - | 0 | - |
| - | - | 679.9 | 243.1 | - | - | 0 | - |
| - | - | 1238 | 245.1 | - | - | 0 | - |
| - | - | 1238 | 247.1 | - | - | 0 | - |
| - | - | 495.9 | 251.8 | - | - | 0 | - |
| - | - | 4492 | 255.1 | - | - | 0 | - |
| - | - | 798 | 261.1 | - | - | 0 | - |
| - | - | 2227 | 262.1 | - | - | 0 | - |
| - | - | 629.7 | 269.1 | - | - | 0 | - |
| - | - | 3615 | 269.2 | - | - | 0 | - |
| - | - | 4311 | 273.1 | - | - | 0 | - |
| - | - | 1354 | 274.1 | - | - | 0 | - |
| - | - | 2364 | 276.2 | - | - | 0 | - |
| - | - | 852.2 | 287.2 | - | - | 0 | - |
| - | - | 791.6 | 291.1 | - | - | 0 | - |
| - | - | 890.2 | 301.1 | - | - | 0 | - |
| - | - | 1314 | 301.2 | - | - | 0 | - |
| - | - | 597.4 | 302.2 | - | - | 0 | - |
| - | - | 872.9 | 304.2 | - | - | 0 | - |
| - | - | 2775 | 310.2 | - | - | 0 | - |
| - | - | 584.7 | 312.1 | - | - | 0 | - |
| - | - | 640.8 | 313.2 | - | - | 0 | - |
| 4 | y | 670.1 | 317.2 | 0.005828 | 18.38 | +2 | 7 |
| - | - | 1136 | 319.1 | - | - | 0 | - |
| - | - | 583.4 | 320.1 | - | - | 0 | - |
| - | - | 637.6 | 325.2 | - | - | 0 | - |
| - | - | 3189 | 326.2 | - | - | 0 | - |
| - | - | 618.2 | 328.1 | - | - | 0 | - |
| - | - | 614.1 | 339.2 | - | - | 0 | - |
| - | - | 697.3 | 341.2 | - | - | 0 | - |
| 8 | y | 661.5 | 343.2 | 0.001919 | 5.591 | +1 | 3 |
| - | - | 1566 | 344.2 | - | - | 0 | - |
| 3 | b | 2348 | 346.2 | 0.0001893 | 0.5468 | +1 | 3 |
| - | - | 1371 | 357.2 | - | - | 0 | - |
| 8 | y | 4266 | 361.2 | 0.0001618 | 0.448 | +1 | 3 |
| - | - | 753.1 | 362.2 | - | - | 0 | - |
| 8 | b | 895.7 | 365.2 | 0.006871 | 18.82 | +2 | 8 |
| - | - | 856.4 | 372.2 | - | - | 0 | - |
| - | - | 1884 | 374.2 | - | - | 0 | - |
| - | - | 633.5 | 375.2 | - | - | 0 | - |
| 3 | y | 1307 | 381.7 | 0.0002081 | 0.5453 | +2 | 8 |
| - | - | 1047 | 382.2 | - | - | 0 | - |
| - | - | 677.6 | 382.2 | - | - | 0 | - |
| - | - | 2400 | 383.2 | - | - | 0 | - |
| 3 | y | 9767 | 390.7 | 2.783E-05 | 0.07122 | +2 | 8 |
| - | - | 4737 | 391.2 | - | - | 0 | - |
| - | - | 1778 | 392.2 | - | - | 0 | - |
| - | - | 1423 | 400.3 | - | - | 0 | - |
| - | - | 2897 | 401.2 | - | - | 0 | - |
| - | - | 5624 | 402.2 | - | - | 0 | - |
| - | - | 764.1 | 403.2 | - | - | 0 | - |
| 4 | b | 768.7 | 403.2 | 0.000271 | 0.6721 | +1 | 4 |
| - | - | 1513 | 415.3 | - | - | 0 | - |
| - | - | 564.8 | 416.3 | - | - | 0 | - |
| - | - | 1343 | 419 | - | - | 0 | - |
| - | - | 2434 | 420.2 | - | - | 0 | - |
| - | - | 1438 | 439.3 | - | - | 0 | - |
| - | - | 3261 | 457.3 | - | - | 0 | - |
| 7 | y | 1399 | 462.3 | 0.0002554 | 0.5524 | +1 | 4 |
| - | - | 659 | 476.9 | - | - | 0 | - |
| - | - | 1628 | 486.3 | - | - | 0 | - |
| 0 | Precursor | 619 | 489.8 | 0.003165 | 6.461 | +2 | -1 |
| - | - | 1580 | 490.3 | - | - | 0 | - |
| - | - | 3019 | 496.3 | - | - | 0 | - |
| - | - | 1277 | 497.3 | - | - | 0 | - |
| - | - | 624.9 | 513.3 | - | - | 0 | - |
| - | - | 6457 | 514.3 | - | - | 0 | - |
| - | - | 1614 | 515.3 | - | - | 0 | - |
| 6 | y | 781.8 | 519.3 | 0.001314 | 2.53 | +1 | 5 |
| - | - | 794 | 521.8 | - | - | 0 | - |
| - | - | 2952 | 530.3 | - | - | 0 | - |
| - | - | 774.7 | 531.3 | - | - | 0 | - |
| - | - | 6309 | 548.3 | - | - | 0 | - |
| - | - | 1657 | 549.3 | - | - | 0 | - |
| - | - | 1060 | 566.3 | - | - | 0 | - |
| 5 | y | 8101 | 576.3 | 0.0004357 | 0.756 | +1 | 6 |
| - | - | 2410 | 577.3 | - | - | 0 | - |
| 4 | y | 1698 | 615.3 | 0.001422 | 2.31 | +1 | 7 |
| - | - | 924.7 | 616.3 | - | - | 0 | - |
| 4 | y | 2.37E+04 | 633.4 | 0.0002563 | 0.4047 | +1 | 7 |
| - | - | 8182 | 634.4 | - | - | 0 | - |
| - | - | 1387 | 635.4 | - | - | 0 | - |
| - | - | 5681 | 643.4 | - | - | 0 | - |
| - | - | 2965 | 644.4 | - | - | 0 | - |
| - | - | 1.337E+04 | 661.4 | - | - | 0 | - |
| - | - | 4177 | 662.4 | - | - | 0 | - |
| - | - | 1174 | 663.4 | - | - | 0 | - |
| - | - | 665.1 | 695.4 | - | - | 0 | - |
| 8 | b | 2925 | 746.4 | 0.002192 | 2.937 | +1 | 8 |
| - | - | 1156 | 747.4 | - | - | 0 | - |
| - | - | 988 | 760.4 | - | - | 0 | - |
| 3 | y | 1994 | 762.4 | 0.001807 | 2.37 | +1 | 8 |
| - | - | 764.4 | 763.4 | - | - | 0 | - |
| - | - | 800.9 | 778.4 | - | - | 0 | - |
| 3 | y | 4.82E+04 | 780.4 | 0.001263 | 1.618 | +1 | 8 |
| - | - | 1.915E+04 | 781.4 | - | - | 0 | - |
| - | - | 5190 | 782.4 | - | - | 0 | - |
| - | - | 697 | 796.3 | - | - | 0 | - |
| 9 | b | 2568 | 859.5 | 0.002211 | 2.572 | +1 | 9 |
| - | - | 1564 | 860.5 | - | - | 0 | - |
| 2 | y | 2983 | 879.5 | 0.002599 | 2.956 | +1 | 9 |
| - | - | 1246 | 880.5 | - | - | 0 | - |
| - | - | 646.8 | 1228 | - | - | 0 | - |
| - | - | 704.3 | 2025 | - | - | 0 | - |
| - | - | 661.1 | 2137 | - | - | 0 | - |
| - | - | 658.4 | 2525 | - | - | 0 | - |
| - | - | 691.3 | 3424 | - | - | 0 | - |
| - | - | 822.3 | 3496 | - | - | 0 | - |

m/z Charge Intensity FragmentType MassShift Position
120.06578063964844 0 5688.1733 y 9
120.0810546875 0 46208.418
120.38567352294922 0 374.26358
121.08440399169922 0 3244.7769
127.05062866210938 0 649.4713
128.10714721679688 0 817.3699
129.06617736816406 0 930.6936
129.10250854492188 0 32925.754
130.0501251220703 0 440.54456
130.10580444335938 0 2346.4675
131.0817108154297 0 1396.016
132.0813446044922 0 689.98346
133.06126403808594 0 671.4209
133.08604431152344 0 1059.9276
136.07594299316406 0 6736.275
137.32965087890625 0 411.9074
139.0869140625 0 507.00598
140.0816192626953 0 550.437
141.06614685058594 0 696.2009
141.1025390625 0 467.83167
142.0609130859375 0 535.60345
142.09793090820312 0 907.7681
146.06028747558594 0 583.8188
152.07086181640625 0 558.0238
153.06625366210938 0 419.88962
154.0605010986328 0 650.044
155.11805725097656 0 1296.1174
156.07696533203125 0 560.84033
157.07598876953125 0 488.45303
158.09262084960938 0 826.69385
159.07627868652344 0 698.25885
163.07174682617188 0 1523.9865
165.10263061523438 0 586.11926
169.13368225097656 0 752.7964
170.0237579345703 0 542.5829
170.0926971435547 0 1294.2485
171.14944458007812 0 69432.04 a 1
172.0720977783203 0 1107.3374
172.15280151367188 0 6619.2227
173.12887573242188 0 1531.3326
176.1073455810547 0 1465.19
176.7014923095703 0 455.73914
177.10244750976562 0 3067.3071
181.17022705078125 0 2789.2075
182.172607421875 0 538.53534
183.35069274902344 0 495.46585
185.081298828125 0 657.0298
185.16505432128906 0 1203.6625
186.1243133544922 0 697.252
187.1439971923828 0 1729.6442
191.1180877685547 0 941.7398
197.10415649414062 0 635.12317
197.12869262695312 0 1958.2733
197.1651153564453 0 1374.276
198.08726501464844 0 1428.2211
199.07188415527344 0 556.85785
199.14425659179688 0 35267.4 b 1
200.14764404296875 0 3166.6062
201.1236114501953 0 1342.1993
202.43141174316406 0 479.05084
205.09730529785156 0 8013.5693
206.10079956054688 0 660.51337
207.149658203125 0 557.7837
208.10842895507812 0 607.28015
209.1038818359375 0 606.10065
209.1648712158203 0 1576.0582
211.14447021484375 0 984.0699
212.1395263671875 0 3632.0872
213.14353942871094 0 610.9693
213.16033935546875 0 547.19855
215.13926696777344 0 3634.4502 y Water loss 8
216.09805297851562 0 1369.9229
217.0991973876953 0 535.1097
219.14939880371094 0 3444.1448
223.10745239257812 0 600.6829
224.17498779296875 0 815.60815
227.11415100097656 0 1360.7804
230.1501007080078 0 6655.27
233.14968872070312 0 2268.4233 y 8
233.16412353515625 0 1130.0677
235.09242248535156 0 1095.6396
235.12979125976562 0 701.0718
237.09913635253906 0 504.45053
237.16001892089844 0 1707.9462
242.1864471435547 0 1485.2445
243.11305236816406 0 679.8509
245.12503051757812 0 1238.0709
247.1444854736328 0 1237.6028
251.78504943847656 0 495.90753
255.1088409423828 0 4491.828
261.12298583984375 0 798.0351
262.11871337890625 0 2226.63
269.1145935058594 0 629.67255
269.1610107421875 0 3615.3994
273.1193542480469 0 4311.0273
274.11932373046875 0 1354.1176
276.1554260253906 0 2363.9941
287.17138671875 0 852.1521
291.144775390625 0 791.62585
301.12982177734375 0 890.24115
301.1912841796875 0 1313.616
302.1949157714844 0 597.4115
304.1665344238281 0 872.89844
310.2126159667969 0 2774.897
312.0976867675781 0 584.71735
313.1864318847656 0 640.8132
317.1877746582031 0 670.1234 y 3
319.1391296386719 0 1136.1337
320.1424865722656 0 583.3541
325.2242736816406 0 637.64453
326.1822509765625 0 3188.7349
328.12921142578125 0 618.2312
339.177734375 0 614.0788
341.1810607910156 0 697.3233
343.23590087890625 0 661.4781 y Water loss 7
344.1933288574219 0 1566.0864
346.21270751953125 0 2347.6099 b 2
357.1561279296875 0 1371.031
361.244384765625 0 4266.1665 y 7
362.2471618652344 0 753.1288
365.1932678222656 0 895.7127 b Ammonia loss 7
372.1969909667969 0 856.4086
374.1824645996094 0 1883.8109
375.1663513183594 0 633.4975
381.7106628417969 0 1306.6354 y Water loss 2
382.2135925292969 0 1046.6986
382.2413330078125 0 677.59436
383.2032775878906 0 2399.8918
390.71612548828125 0 9767.316 y 2
391.2178649902344 0 4737.1787
392.1922912597656 0 1778.0321
400.25567626953125 0 1422.7456
401.21453857421875 0 2897.4702
402.1772155761719 0 5623.9126
403.1808166503906 0 764.05206
403.2342529296875 0 768.65045 b 3
415.2543640136719 0 1512.586
416.26043701171875 0 564.75073
418.9952087402344 0 1343.0687
420.18817138671875 0 2434.182
439.26605224609375 0 1437.7229
457.2768249511719 0 3260.5676
462.29248046875 0 1399.3896 y 6
476.9361877441406 0 659.0306
486.3012390136719 0 1628.0525
489.7814025878906 0 619.00793 Precursor
490.2861633300781 0 1579.6013
496.28814697265625 0 3018.6794
497.2901611328125 0 1277.136
513.2655029296875 0 624.88257
514.2982788085938 0 6456.9175
515.2999877929688 0 1614.2615
519.3150024414062 0 781.78534 y 5
521.7810668945312 0 794.03705
530.2724609375 0 2951.7542
531.27685546875 0 774.7137
548.282470703125 0 6308.595
549.2838745117188 0 1656.7798
566.3087158203125 0 1060.0771
576.334716796875 0 8101.3267 y 4
577.3375854492188 0 2409.613
615.3474731445312 0 1698.1289 y Water loss 3
616.345947265625 0 924.699
633.3568725585938 0 23699.129 y 3
634.3598022460938 0 8181.5166
635.3611450195312 0 1386.6931
643.354736328125 0 5681.005
644.3580932617188 0 2964.819
661.3660888671875 0 13373.225
662.3692016601562 0 4177.2417
663.3721923828125 0 1173.6992
695.3750610351562 0 665.1222
746.4173583984375 0 2925.2275 b 7
747.4207763671875 0 1156.0645
760.434814453125 0 987.973
762.4126586914062 0 1994.1414 y Water loss 2
763.4192504882812 0 764.44824
778.4083862304688 0 800.85315
780.4237670898438 0 48196.68 y 2
781.4263305664062 0 19146.486
782.4304809570312 0 5190.27
796.3203125 0 696.9803
859.5014038085938 0 2568.3672 b 8
860.50341796875 0 1564.043
879.4908447265625 0 2983.1255 y 1
880.4951782226562 0 1245.8922
1227.93017578125 0 646.8348
2024.9432373046875 0 704.27167
2136.6923828125 0 661.0837
2524.904541015625 0 658.40845
3423.8837890625 0 691.28735
3496.47265625 0 822.335

Spectrum Details

|  |  |
| --- | --- |
| Matched peaks? Matched peaks The total absolute number of peaks matched. Additionally in brackets the total fraction of peaks matched and the total number of peaks is shown. | 24 (12.50% of 192) |
| FDR? FDR The false discovery rate estimated for this peptide. It is calculated by matching all theoretical fragments with a non-integer shift with the raw peaks for this spectrum. This is done with 40 different shifts. The resulting percentage is the average number of annotated peaks over the number of annotated peaks with the correct spectrum. | 0.20% |
| Satellite FDR? Satellite FDR See the FDR for details on its calculation. This satellite ion specific FDR only contains the satellite ions (d/w) for I/L/J positions. | - |
| PSM Score? PSM Score The PSM Score as given by Hecklib to this annotated spectrum. It is shown with three significant figures. | 262 |

#### Spectrum 6214? Spectrum 6214 The raw spectrum of this peptide as annotated by Hecklib. The fragments are coloured according to ion type (see legend). Any peaks with a star '\*' as text can be hovered over to see the full details, first the ion type second the mass shift type. By hovering over the amino acids in the peptide or ions in the legend the corresponding peaks are highlighted. By toggling the 'Unassigned' label you can turn the background (unassigned) peaks on or off in the plot. By updating the slider in the Ion legend you can update the spectrum to only show the top X% of the peaks with labels. The top X% means any peak that is within X% of the highest intensity. By dragging in the spectrum you can zoom in to a specific part of the spectrum and use 'Zoom Out' to get back to the original zoom level. The annotation of the spectrum is based on the given sequence in the peptides file and is done with different software so inconsistencies are likely. The peaks are annotated based on the given sequence, with 20 ppm tolerance.

Copy Data

##### Spectrum 6214 (TSV)

###### Preview

```
Loading example...
```

*Click on the button to copy the data to your clipboard.*

Mz MinMz MaxIntensity Max

WidthHeightPeptide font sizePeptide stroke widthSpectrum font sizeSpectrum stroke widthCompact peptide

Ion legend

wxyz

abcd

OtherUnassignedIon^Charge^_Position_Show for top:%

VVFGGGTKJT

02.49e+44.99e+47.48e+49.97e+4

Zoom Out

y^+1^_1_a^+1^_2_b^+1^_2_y^+1^_2_y^+1^_2_y^+2^_5_y^+2^_7_y^+1^_3_b^+1^_3_y^+1^_3_b^+2^_8_b^+2^_8_y^+2^_8_y^+2^_8_b^+1^_4_y^+2^_9_y^+1^_4_\*\*y^+1^_5_y^+1^_6_y^+1^_6_b^+1^_7_y^+1^_7_b^+1^_7_y^+1^_7_b^+1^_8_y^+1^_8_y^+1^_8_b^+1^_9_y^+1^_9_

0816163224473263

Fragment Matches Table

Show background peaks

| Position | Ion type | Intensity | mz Theoretical | mz Error (Th) | mz Error (ppm) | Charge | Series Number |
| --- | --- | --- | --- | --- | --- | --- | --- |
| 10 | y | 7672 | 120.1 | 0.0002229 | 1.856 | +1 | 1 |
| - | - | 5.395E+04 | 120.1 | - | - | 0 | - |
| - | - | 429.2 | 120.5 | - | - | 0 | - |
| - | - | 4451 | 121.1 | - | - | 0 | - |
| - | - | 367.9 | 125.1 | - | - | 0 | - |
| - | - | 386 | 125.4 | - | - | 0 | - |
| - | - | 526.3 | 126.1 | - | - | 0 | - |
| - | - | 842.9 | 127.1 | - | - | 0 | - |
| - | - | 401.4 | 127.1 | - | - | 0 | - |
| - | - | 1362 | 128.1 | - | - | 0 | - |
| - | - | 659.7 | 129.1 | - | - | 0 | - |
| - | - | 4.257E+04 | 129.1 | - | - | 0 | - |
| - | - | 542.6 | 129.1 | - | - | 0 | - |
| - | - | 346 | 129.3 | - | - | 0 | - |
| - | - | 2321 | 130.1 | - | - | 0 | - |
| - | - | 602 | 130.1 | - | - | 0 | - |
| - | - | 2844 | 130.1 | - | - | 0 | - |
| - | - | 411.1 | 130.2 | - | - | 0 | - |
| - | - | 1728 | 131.1 | - | - | 0 | - |
| - | - | 1214 | 132.1 | - | - | 0 | - |
| - | - | 879.6 | 133.1 | - | - | 0 | - |
| - | - | 4274 | 133.1 | - | - | 0 | - |
| - | - | 367 | 133.1 | - | - | 0 | - |
| - | - | 7581 | 136.1 | - | - | 0 | - |
| - | - | 454 | 137.1 | - | - | 0 | - |
| - | - | 460 | 140.1 | - | - | 0 | - |
| - | - | 692.6 | 141.1 | - | - | 0 | - |
| - | - | 544.7 | 141.1 | - | - | 0 | - |
| - | - | 655.2 | 142.1 | - | - | 0 | - |
| - | - | 441.5 | 142.4 | - | - | 0 | - |
| - | - | 493.5 | 143.1 | - | - | 0 | - |
| - | - | 521.8 | 143.1 | - | - | 0 | - |
| - | - | 377.1 | 144 | - | - | 0 | - |
| - | - | 643.4 | 146.2 | - | - | 0 | - |
| - | - | 3828 | 147 | - | - | 0 | - |
| - | - | 696.8 | 148 | - | - | 0 | - |
| - | - | 436.5 | 150.1 | - | - | 0 | - |
| - | - | 733.1 | 152.1 | - | - | 0 | - |
| - | - | 448.4 | 154.1 | - | - | 0 | - |
| - | - | 660.2 | 155.1 | - | - | 0 | - |
| - | - | 1200 | 155.1 | - | - | 0 | - |
| - | - | 694.9 | 156.1 | - | - | 0 | - |
| - | - | 840.1 | 158.1 | - | - | 0 | - |
| - | - | 1898 | 159.1 | - | - | 0 | - |
| - | - | 4549 | 159.1 | - | - | 0 | - |
| - | - | 725.3 | 159.1 | - | - | 0 | - |
| - | - | 529.9 | 160.1 | - | - | 0 | - |
| - | - | 397.6 | 161.8 | - | - | 0 | - |
| - | - | 561.9 | 163 | - | - | 0 | - |
| - | - | 1539 | 163.1 | - | - | 0 | - |
| - | - | 491.6 | 167.1 | - | - | 0 | - |
| - | - | 720.5 | 169.1 | - | - | 0 | - |
| - | - | 954.8 | 169.1 | - | - | 0 | - |
| - | - | 1121 | 170.1 | - | - | 0 | - |
| - | - | 961.9 | 171.1 | - | - | 0 | - |
| 2 | a | 9.874E+04 | 171.1 | 0.0002549 | 1.489 | +1 | 2 |
| - | - | 1968 | 172.1 | - | - | 0 | - |
| - | - | 1072 | 172.1 | - | - | 0 | - |
| - | - | 9393 | 172.2 | - | - | 0 | - |
| - | - | 716.7 | 173.1 | - | - | 0 | - |
| - | - | 678.2 | 175 | - | - | 0 | - |
| - | - | 489.1 | 175.1 | - | - | 0 | - |
| - | - | 1933 | 176.1 | - | - | 0 | - |
| - | - | 4806 | 177.1 | - | - | 0 | - |
| - | - | 1888 | 177.1 | - | - | 0 | - |
| - | - | 469.2 | 181.1 | - | - | 0 | - |
| - | - | 899.9 | 181.2 | - | - | 0 | - |
| - | - | 767.3 | 182.1 | - | - | 0 | - |
| - | - | 570 | 185.1 | - | - | 0 | - |
| - | - | 542.1 | 185.1 | - | - | 0 | - |
| - | - | 1480 | 185.2 | - | - | 0 | - |
| - | - | 1039 | 186.1 | - | - | 0 | - |
| - | - | 1977 | 187.1 | - | - | 0 | - |
| - | - | 1774 | 197.1 | - | - | 0 | - |
| - | - | 1961 | 197.2 | - | - | 0 | - |
| - | - | 1431 | 198.1 | - | - | 0 | - |
| 2 | b | 4.739E+04 | 199.1 | 0.0001218 | 0.6116 | +1 | 2 |
| - | - | 5297 | 200.1 | - | - | 0 | - |
| - | - | 2188 | 201.1 | - | - | 0 | - |
| - | - | 2062 | 204.1 | - | - | 0 | - |
| - | - | 9097 | 205.1 | - | - | 0 | - |
| - | - | 587.2 | 205.1 | - | - | 0 | - |
| - | - | 1180 | 206.1 | - | - | 0 | - |
| - | - | 1020 | 207.1 | - | - | 0 | - |
| - | - | 1476 | 209.2 | - | - | 0 | - |
| - | - | 634.3 | 211.1 | - | - | 0 | - |
| - | - | 5398 | 212.1 | - | - | 0 | - |
| - | - | 639.9 | 215.1 | - | - | 0 | - |
| 9 | y | 3270 | 215.1 | 0.0002023 | 0.9402 | +1 | 2 |
| - | - | 2008 | 216.1 | - | - | 0 | - |
| - | - | 1361 | 217.1 | - | - | 0 | - |
| - | - | 3804 | 219.1 | - | - | 0 | - |
| - | - | 882.8 | 220.1 | - | - | 0 | - |
| - | - | 678.1 | 220.2 | - | - | 0 | - |
| - | - | 604.3 | 221.1 | - | - | 0 | - |
| - | - | 953.2 | 224.2 | - | - | 0 | - |
| - | - | 593.2 | 225.1 | - | - | 0 | - |
| - | - | 968.9 | 226.2 | - | - | 0 | - |
| - | - | 2201 | 227.1 | - | - | 0 | - |
| - | - | 1054 | 228.1 | - | - | 0 | - |
| - | - | 9073 | 230.2 | - | - | 0 | - |
| - | - | 745.5 | 231.1 | - | - | 0 | - |
| - | - | 962.4 | 231.2 | - | - | 0 | - |
| 9 | y | 2691 | 233.1 | 0.000334 | 1.433 | +1 | 2 |
| - | - | 679.6 | 233.2 | - | - | 0 | - |
| - | - | 1019 | 237.2 | - | - | 0 | - |
| - | - | 1208 | 239.2 | - | - | 0 | - |
| - | - | 2027 | 242.2 | - | - | 0 | - |
| - | - | 922.5 | 243.1 | - | - | 0 | - |
| - | - | 1161 | 245.1 | - | - | 0 | - |
| - | - | 1754 | 247.1 | - | - | 0 | - |
| 6 | y | 543.7 | 251.2 | 0.004138 | 16.48 | +2 | 5 |
| - | - | 5063 | 255.1 | - | - | 0 | - |
| - | - | 514.7 | 258.1 | - | - | 0 | - |
| - | - | 580.5 | 260.2 | - | - | 0 | - |
| - | - | 864.9 | 261.1 | - | - | 0 | - |
| - | - | 942.2 | 261.2 | - | - | 0 | - |
| - | - | 2665 | 262.1 | - | - | 0 | - |
| - | - | 553 | 269.1 | - | - | 0 | - |
| - | - | 5925 | 269.2 | - | - | 0 | - |
| - | - | 890.4 | 270.2 | - | - | 0 | - |
| - | - | 4250 | 273.1 | - | - | 0 | - |
| - | - | 1317 | 274.1 | - | - | 0 | - |
| - | - | 925.1 | 275.2 | - | - | 0 | - |
| - | - | 1916 | 276.2 | - | - | 0 | - |
| - | - | 1716 | 287.2 | - | - | 0 | - |
| - | - | 660.6 | 288.8 | - | - | 0 | - |
| - | - | 885.4 | 291.1 | - | - | 0 | - |
| - | - | 2352 | 292.2 | - | - | 0 | - |
| - | - | 839.6 | 293.2 | - | - | 0 | - |
| - | - | 954.3 | 294.2 | - | - | 0 | - |
| - | - | 2576 | 301.2 | - | - | 0 | - |
| - | - | 1216 | 304.2 | - | - | 0 | - |
| - | - | 832.2 | 306.2 | - | - | 0 | - |
| 4 | y | 599.9 | 308.2 | 0.002073 | 6.726 | +2 | 7 |
| - | - | 1198 | 308.2 | - | - | 0 | - |
| - | - | 2696 | 310.2 | - | - | 0 | - |
| - | - | 908.9 | 317.2 | - | - | 0 | - |
| - | - | 579.1 | 318.1 | - | - | 0 | - |
| - | - | 1420 | 319.1 | - | - | 0 | - |
| - | - | 5149 | 326.2 | - | - | 0 | - |
| - | - | 799.5 | 330.2 | - | - | 0 | - |
| - | - | 816.4 | 331.2 | - | - | 0 | - |
| - | - | 701.3 | 339.2 | - | - | 0 | - |
| 8 | y | 2075 | 343.2 | 0.0003393 | 0.9887 | +1 | 3 |
| - | - | 1167 | 344.2 | - | - | 0 | - |
| 3 | b | 2582 | 346.2 | 0.00036 | 1.04 | +1 | 3 |
| - | - | 1054 | 347.2 | - | - | 0 | - |
| - | - | 1302 | 357.2 | - | - | 0 | - |
| - | - | 599 | 358.2 | - | - | 0 | - |
| - | - | 785.5 | 361.2 | - | - | 0 | - |
| 8 | y | 5007 | 361.2 | 0.0001924 | 0.5325 | +1 | 3 |
| - | - | 771.2 | 362.2 | - | - | 0 | - |
| 8 | b | 594.2 | 365.2 | 0.006963 | 19.07 | +2 | 8 |
| - | - | 680.7 | 367.2 | - | - | 0 | - |
| - | - | 1884 | 371.2 | - | - | 0 | - |
| 8 | b | 1088 | 373.7 | 0.001225 | 3.278 | +2 | 8 |
| - | - | 1876 | 374.2 | - | - | 0 | - |
| - | - | 941.8 | 375.2 | - | - | 0 | - |
| 3 | y | 3300 | 381.7 | 0.0002081 | 0.5453 | +2 | 8 |
| - | - | 1007 | 382.2 | - | - | 0 | - |
| - | - | 845.7 | 382.2 | - | - | 0 | - |
| - | - | 4093 | 383.2 | - | - | 0 | - |
| - | - | 1189 | 383.2 | - | - | 0 | - |
| 3 | y | 1.607E+04 | 390.7 | 2.693E-06 | 0.006891 | +2 | 8 |
| - | - | 7805 | 391.2 | - | - | 0 | - |
| - | - | 2326 | 391.7 | - | - | 0 | - |
| - | - | 2153 | 392.2 | - | - | 0 | - |
| - | - | 755 | 397.2 | - | - | 0 | - |
| - | - | 1081 | 400.3 | - | - | 0 | - |
| - | - | 4341 | 401.2 | - | - | 0 | - |
| - | - | 7041 | 402.2 | - | - | 0 | - |
| - | - | 1198 | 403.2 | - | - | 0 | - |
| 4 | b | 1117 | 403.2 | 0.00021 | 0.5207 | +1 | 4 |
| - | - | 849.6 | 404.7 | - | - | 0 | - |
| - | - | 1356 | 415.3 | - | - | 0 | - |
| - | - | 810 | 419 | - | - | 0 | - |
| - | - | 3659 | 420.2 | - | - | 0 | - |
| - | - | 624.4 | 421.2 | - | - | 0 | - |
| - | - | 643.5 | 422.2 | - | - | 0 | - |
| - | - | 2351 | 439.3 | - | - | 0 | - |
| 2 | y | 1613 | 440.3 | 0.0002077 | 0.4717 | +2 | 9 |
| - | - | 572.1 | 444 | - | - | 0 | - |
| - | - | 629.4 | 451.3 | - | - | 0 | - |
| - | - | 2252 | 454.2 | - | - | 0 | - |
| - | - | 1449 | 455.2 | - | - | 0 | - |
| - | - | 4767 | 457.3 | - | - | 0 | - |
| - | - | 819.7 | 458.3 | - | - | 0 | - |
| 7 | y | 1677 | 462.3 | 0.001293 | 2.797 | +1 | 4 |
| - | - | 567.8 | 473.3 | - | - | 0 | - |
| - | - | 2690 | 473.3 | - | - | 0 | - |
| - | - | 933.4 | 475.3 | - | - | 0 | - |
| 0 | Precursor | 1152 | 480.8 | 0.0005003 | 1.041 | +2 | -1 |
| - | - | 755.6 | 484.6 | - | - | 0 | - |
| - | - | 2746 | 486.3 | - | - | 0 | - |
| - | - | 716 | 487.3 | - | - | 0 | - |
| - | - | 622.3 | 489.2 | - | - | 0 | - |
| 0 | Precursor | 819.4 | 489.8 | 0.0008332 | 1.701 | +2 | -1 |
| - | - | 2564 | 490.3 | - | - | 0 | - |
| - | - | 819 | 490.3 | - | - | 0 | - |
| - | - | 814 | 490.8 | - | - | 0 | - |
| - | - | 757.7 | 491.2 | - | - | 0 | - |
| - | - | 4329 | 496.3 | - | - | 0 | - |
| - | - | 1265 | 497.3 | - | - | 0 | - |
| - | - | 912.5 | 512.3 | - | - | 0 | - |
| - | - | 1.073E+04 | 514.3 | - | - | 0 | - |
| - | - | 2468 | 515.3 | - | - | 0 | - |
| 6 | y | 2377 | 519.3 | 9.293E-05 | 0.1789 | +1 | 5 |
| - | - | 848.5 | 521.8 | - | - | 0 | - |
| - | - | 2389 | 525.8 | - | - | 0 | - |
| - | - | 1648 | 526.3 | - | - | 0 | - |
| - | - | 4711 | 530.3 | - | - | 0 | - |
| - | - | 1530 | 531.3 | - | - | 0 | - |
| - | - | 666.1 | 532.3 | - | - | 0 | - |
| - | - | 8367 | 539.8 | - | - | 0 | - |
| - | - | 4634 | 540.3 | - | - | 0 | - |
| - | - | 2387 | 540.8 | - | - | 0 | - |
| - | - | 7494 | 548.3 | - | - | 0 | - |
| - | - | 2377 | 549.3 | - | - | 0 | - |
| 5 | y | 809.9 | 558.3 | 0.00159 | 2.847 | +1 | 6 |
| 5 | y | 1.241E+04 | 576.3 | 0.0004968 | 0.862 | +1 | 6 |
| - | - | 3648 | 577.3 | - | - | 0 | - |
| 7 | b | 655.6 | 600.3 | 0.006961 | 11.6 | +1 | 7 |
| 4 | y | 1707 | 615.3 | 0.002337 | 3.798 | +1 | 7 |
| - | - | 1111 | 616.4 | - | - | 0 | - |
| 7 | b | 1130 | 618.3 | 0.002811 | 4.545 | +1 | 7 |
| - | - | 4168 | 618.9 | - | - | 0 | - |
| - | - | 1743 | 619.4 | - | - | 0 | - |
| - | - | 1410 | 619.9 | - | - | 0 | - |
| - | - | 798.7 | 625.3 | - | - | 0 | - |
| - | - | 7478 | 632.8 | - | - | 0 | - |
| 4 | y | 3.762E+04 | 633.4 | 0.0006592 | 1.041 | +1 | 7 |
| - | - | 2679 | 633.9 | - | - | 0 | - |
| - | - | 1.095E+04 | 634.4 | - | - | 0 | - |
| - | - | 1928 | 635.4 | - | - | 0 | - |
| - | - | 7210 | 643.4 | - | - | 0 | - |
| - | - | 2976 | 644.4 | - | - | 0 | - |
| - | - | 1041 | 645.4 | - | - | 0 | - |
| - | - | 1.938E+04 | 661.4 | - | - | 0 | - |
| - | - | 6219 | 662.4 | - | - | 0 | - |
| - | - | 1142 | 663.4 | - | - | 0 | - |
| - | - | 3709 | 668.4 | - | - | 0 | - |
| - | - | 3382 | 668.9 | - | - | 0 | - |
| - | - | 2087 | 669.4 | - | - | 0 | - |
| - | - | 909.4 | 679.4 | - | - | 0 | - |
| - | - | 6789 | 682.4 | - | - | 0 | - |
| - | - | 3726 | 682.9 | - | - | 0 | - |
| - | - | 1842 | 683.4 | - | - | 0 | - |
| - | - | 654 | 692.1 | - | - | 0 | - |
| - | - | 642 | 718.4 | - | - | 0 | - |
| 8 | b | 2458 | 746.4 | 0.001826 | 2.447 | +1 | 8 |
| - | - | 1804 | 747.4 | - | - | 0 | - |
| - | - | 1411 | 760.4 | - | - | 0 | - |
| 3 | y | 2140 | 762.4 | 0.00138 | 1.809 | +1 | 8 |
| - | - | 1515 | 763.4 | - | - | 0 | - |
| - | - | 670.6 | 766.1 | - | - | 0 | - |
| 3 | y | 6.187E+04 | 780.4 | 0.001324 | 1.697 | +1 | 8 |
| - | - | 3.049E+04 | 781.4 | - | - | 0 | - |
| - | - | 7409 | 782.4 | - | - | 0 | - |
| - | - | 799.3 | 783.4 | - | - | 0 | - |
| 9 | b | 3970 | 859.5 | 0.001967 | 2.288 | +1 | 9 |
| - | - | 1836 | 860.5 | - | - | 0 | - |
| - | - | 745.6 | 861.5 | - | - | 0 | - |
| - | - | 731.1 | 876.4 | - | - | 0 | - |
| 2 | y | 4226 | 879.5 | 0.00144 | 1.637 | +1 | 9 |
| - | - | 1366 | 880.5 | - | - | 0 | - |
| - | - | 608.8 | 1008 | - | - | 0 | - |
| - | - | 640.4 | 1235 | - | - | 0 | - |
| - | - | 636.6 | 2508 | - | - | 0 | - |
| - | - | 700.1 | 2534 | - | - | 0 | - |
| - | - | 739.8 | 3231 | - | - | 0 | - |

m/z Charge Intensity FragmentType MassShift Position
120.06574249267578 0 7671.89 y 9
120.08103942871094 0 53946.258
120.53569793701172 0 429.20486
121.08436584472656 0 4450.7173
125.07105255126953 0 367.87167
125.43244934082031 0 386.0118
126.06655883789062 0 526.28204
127.0504379272461 0 842.9084
127.0866470336914 0 401.36047
128.10726928710938 0 1361.8518
129.0659942626953 0 659.73364
129.10247802734375 0 42567.92
129.13894653320312 0 542.5533
129.3145751953125 0 345.99078
130.0652618408203 0 2320.733
130.10072326660156 0 602.0432
130.1058349609375 0 2844.4775
130.1593780517578 0 411.075
131.0817108154297 0 1727.8835
132.0812225341797 0 1214.0975
133.06094360351562 0 879.6271
133.08612060546875 0 4274.2725
133.1020965576172 0 366.9515
136.07591247558594 0 7581.484
137.0791778564453 0 454.03723
140.0823211669922 0 460.04578
141.06613159179688 0 692.57935
141.10226440429688 0 544.6662
142.12306213378906 0 655.1806
142.3741912841797 0 441.50082
143.0814971923828 0 493.5038
143.11795043945312 0 521.75476
144.0227508544922 0 377.12158
146.1649627685547 0 643.4278
147.04429626464844 0 3828.1458
148.04763793945312 0 696.81335
150.0553741455078 0 436.45828
152.07077026367188 0 733.149
154.0502166748047 0 448.38632
155.0814666748047 0 660.1814
155.11819458007812 0 1199.5664
156.0767822265625 0 694.86536
158.09275817871094 0 840.12665
159.07669067382812 0 1897.7109
159.0919189453125 0 4549.46
159.11309814453125 0 725.2557
160.09507751464844 0 529.9292
161.8136749267578 0 397.59094
163.03884887695312 0 561.918
163.07147216796875 0 1539.2206
167.11781311035156 0 491.62592
169.09762573242188 0 720.5366
169.1339569091797 0 954.7707
170.09243774414062 0 1121.2263
171.07630920410156 0 961.899
171.14944458007812 0 98743.11 a 1
172.07192993164062 0 1967.7811
172.14491271972656 0 1072.0721
172.15280151367188 0 9393.212
173.12811279296875 0 716.65674
175.02952575683594 0 678.187
175.06982421875 0 489.11276
176.10679626464844 0 1933.1235
177.10252380371094 0 4805.92
177.11207580566406 0 1888.1298
181.09669494628906 0 469.16086
181.17018127441406 0 899.94104
182.08108520507812 0 767.27936
185.10333251953125 0 570.01556
185.12884521484375 0 542.1392
185.16481018066406 0 1480.0673
186.12359619140625 0 1039.1511
187.1441650390625 0 1977.279
197.1287841796875 0 1773.8118
197.1649169921875 0 1960.7085
198.08737182617188 0 1430.8257
199.14422607421875 0 47387.38 b 1
200.1476287841797 0 5297.2793
201.1235809326172 0 2188.3525
204.10191345214844 0 2061.5996
205.0972137451172 0 9096.558
205.106689453125 0 587.20856
206.1001739501953 0 1180.4047
207.1493682861328 0 1020.30133
209.1649169921875 0 1475.878
211.14419555664062 0 634.3003
212.13943481445312 0 5397.6904
215.11460876464844 0 639.90137
215.13922119140625 0 3270.1123 y Water loss 8
216.09805297851562 0 2007.5168
217.0967559814453 0 1360.8984
219.14944458007812 0 3804.131
220.09739685058594 0 882.81384
220.15267944335938 0 678.14154
221.12820434570312 0 604.3274
224.17587280273438 0 953.1679
225.1346435546875 0 593.1929
226.15493774414062 0 968.85345
227.11407470703125 0 2200.9277
228.09817504882812 0 1054.0948
230.15011596679688 0 9073.177
231.1130828857422 0 745.45123
231.15309143066406 0 962.4213
233.14991760253906 0 2690.675 y 8
233.1633758544922 0 679.6263
237.1600799560547 0 1019.06726
239.1505889892578 0 1207.6969
242.1865234375 0 2026.8425
243.1129150390625 0 922.47437
245.12574768066406 0 1160.9541
247.14430236816406 0 1754.1062
251.15106201171875 0 543.74414 y Water loss 5
255.10877990722656 0 5063.2935
258.14495849609375 0 514.74774
260.19677734375 0 580.5004
261.1225280761719 0 864.85175
261.158935546875 0 942.225
262.11865234375 0 2664.792
269.11236572265625 0 553.00397
269.160888671875 0 5924.784
270.16375732421875 0 890.36475
273.1194763183594 0 4250.4336
274.1178283691406 0 1316.5287
275.1751708984375 0 925.0852
276.15582275390625 0 1915.9196
287.17144775390625 0 1715.748
288.7704772949219 0 660.62616
291.1451416015625 0 885.4224
292.20208740234375 0 2352.4438
293.20538330078125 0 839.63617
294.1557312011719 0 954.33295
301.1911926269531 0 2575.9285
304.1673889160156 0 1215.6578
306.1805419921875 0 832.20233
308.1745910644531 0 599.8584 y Water loss 3
308.1964111328125 0 1197.807
310.2126770019531 0 2696.0916
317.1897277832031 0 908.8527
318.1442565917969 0 579.14825
319.14068603515625 0 1419.5436
326.1824035644531 0 5148.9116
330.18096923828125 0 799.49054
331.18804931640625 0 816.37915
339.17626953125 0 701.3461
343.233642578125 0 2075.412 y Water loss 7
344.1911315917969 0 1167.3804
346.212158203125 0 2581.6511 b 2
347.2152404785156 0 1053.634
357.1550598144531 0 1301.9094
358.2097473144531 0 598.97504
361.186767578125 0 785.50494
361.2443542480469 0 5006.797 y 7
362.24908447265625 0 771.1594
365.19317626953125 0 594.16797 b Ammonia loss 7
367.20721435546875 0 680.7185
371.2270202636719 0 1884.3713
373.7121887207031 0 1087.8568 b 7
374.1824645996094 0 1876.0211
375.1686706542969 0 941.75574
381.7106628417969 0 3300.3325 y Water loss 2
382.2144775390625 0 1006.8109
382.244384765625 0 845.719
383.20318603515625 0 4092.8953
383.2285461425781 0 1188.6213
390.7161560058594 0 16074.865 y 2
391.2176208496094 0 7804.947
391.7182312011719 0 2326.1882
392.1927795410156 0 2152.6904
397.2449035644531 0 755.0035
400.256591796875 0 1080.599
401.2143249511719 0 4341.3794
402.177001953125 0 7041.4185
403.1805725097656 0 1198.1091
403.23419189453125 0 1116.614 b 3
404.7463073730469 0 849.573
415.255126953125 0 1355.6722
418.9938659667969 0 810.0438
420.1884765625 0 3659.113
421.1911926269531 0 624.419
422.2371826171875 0 643.5411
439.26629638671875 0 2351.4463
440.2501525878906 0 1613.3763 y 1
443.97332763671875 0 572.0575
451.26568603515625 0 629.4195
454.2334289550781 0 2251.7822
455.2360534667969 0 1449.3469
457.2769775390625 0 4766.753
458.2806396484375 0 819.6535
462.29351806640625 0 1676.9904 y 6
473.2586669921875 0 567.81
473.29571533203125 0 2690.2686
475.2872619628906 0 933.4372
480.77978515625 0 1151.6274 Precursor Water loss
484.601318359375 0 755.5586
486.3031311035156 0 2746.235
487.30657958984375 0 716.02313
489.2408142089844 0 622.3306
489.785400390625 0 819.39545 Precursor
490.28607177734375 0 2564.4822
490.3214416503906 0 819.00757
490.78741455078125 0 813.95447
491.2192687988281 0 757.71436
496.2870788574219 0 4328.8696
497.29254150390625 0 1265.0577
512.2628173828125 0 912.5129
514.2981567382812 0 10728.374
515.3009033203125 0 2467.51
519.3137817382812 0 2377.1777 y 5
521.780517578125 0 848.48444
525.8114624023438 0 2388.9243
526.3131713867188 0 1648.482
530.2714233398438 0 4711.41
531.2745971679688 0 1529.8138
532.3082275390625 0 666.0787
539.8094482421875 0 8367.242
540.3099975585938 0 4634.116
540.811279296875 0 2387.003
548.28173828125 0 7494.0015
549.2836303710938 0 2377.2407
558.322998046875 0 809.86176 y Water loss 4
576.3346557617188 0 12406.756 y 4
577.3372192382812 0 3648.2166
600.3209838867188 0 655.63385 b Water loss 6
615.348388671875 0 1706.8212 y Water loss 3
616.3508911132812 0 1111.1091
618.32177734375 0 1129.7778 b 6
618.8515014648438 0 4168.186
619.350341796875 0 1743.1696
619.854248046875 0 1410.1005
625.34814453125 0 798.74414
632.8484497070312 0 7477.646
633.35595703125 0 37624.383 y 3
633.8517456054688 0 2678.8228
634.359375 0 10949.52
635.361572265625 0 1928.345
643.3553466796875 0 7210.372
644.358154296875 0 2976.4487
645.3583984375 0 1040.7736
661.3662109375 0 19378.785
662.3694458007812 0 6219.093
663.3743286132812 0 1142.38
668.3829345703125 0 3708.6963
668.8872680664062 0 3382.449
669.3903198242188 0 2087.309
679.3771362304688 0 909.40204
682.38232421875 0 6788.908
682.8834838867188 0 3725.8467
683.387451171875 0 1842.2751
692.09033203125 0 653.9929
718.4239501953125 0 641.974
746.417724609375 0 2457.9082 b 7
747.4215698242188 0 1804.1313
760.434814453125 0 1410.6584
762.4130859375 0 2140.2192 y Water loss 2
763.4151611328125 0 1514.6963
766.1044311523438 0 670.5923
780.4237060546875 0 61865.438 y 2
781.426513671875 0 30490.213
782.4297485351562 0 7409.2266
783.4325561523438 0 799.32294
859.5016479492188 0 3969.657 b 8
860.501708984375 0 1836.2124
861.502197265625 0 745.63885
876.4014892578125 0 731.1462
879.4920043945312 0 4225.8516 y 1
880.5003662109375 0 1366.033
1008.10888671875 0 608.8236
1235.3211669921875 0 640.39215
2508.2255859375 0 636.6002
2533.780029296875 0 700.1198
3230.733154296875 0 739.80383

Spectrum Details

|  |  |
| --- | --- |
| Matched peaks? Matched peaks The total absolute number of peaks matched. Additionally in brackets the total fraction of peaks matched and the total number of peaks is shown. | 31 (11.44% of 271) |
| FDR? FDR The false discovery rate estimated for this peptide. It is calculated by matching all theoretical fragments with a non-integer shift with the raw peaks for this spectrum. This is done with 40 different shifts. The resulting percentage is the average number of annotated peaks over the number of annotated peaks with the correct spectrum. | 0.23% |
| Satellite FDR? Satellite FDR See the FDR for details on its calculation. This satellite ion specific FDR only contains the satellite ions (d/w) for I/L/J positions. | - |
| PSM Score? PSM Score The PSM Score as given by Hecklib to this annotated spectrum. It is shown with three significant figures. | 321 |

#### Spectrum 7091? Spectrum 7091 The raw spectrum of this peptide as annotated by Hecklib. The fragments are coloured according to ion type (see legend). Any peaks with a star '\*' as text can be hovered over to see the full details, first the ion type second the mass shift type. By hovering over the amino acids in the peptide or ions in the legend the corresponding peaks are highlighted. By toggling the 'Unassigned' label you can turn the background (unassigned) peaks on or off in the plot. By updating the slider in the Ion legend you can update the spectrum to only show the top X% of the peaks with labels. The top X% means any peak that is within X% of the highest intensity. By dragging in the spectrum you can zoom in to a specific part of the spectrum and use 'Zoom Out' to get back to the original zoom level. The annotation of the spectrum is based on the given sequence in the peptides file and is done with different software so inconsistencies are likely. The peaks are annotated based on the given sequence, with 20 ppm tolerance.

Copy Data

##### Spectrum 7091 (TSV)

###### Preview

```
Loading example...
```

*Click on the button to copy the data to your clipboard.*

Mz MinMz MaxIntensity Max

WidthHeightPeptide font sizePeptide stroke widthSpectrum font sizeSpectrum stroke widthCompact peptide

Ion legend

wxyz

abcd

OtherUnassignedIon^Charge^_Position_Show for top:%

VVFGGGTKJT

09.59e+31.92e+42.88e+43.84e+4

Zoom Out

y^+1^_1_a^+1^_2_b^+1^_2_y^+1^_2_y^+1^_2_b^+1^_3_y^+1^_3_y^+2^_8_y^+2^_8_y^+1^_4_\*y^+1^_5_y^+1^_6_y^+1^_7_b^+1^_8_y^+1^_8_y^+1^_8_b^+1^_9_y^+1^_9_

0801160224033204

Fragment Matches Table

Show background peaks

| Position | Ion type | Intensity | mz Theoretical | mz Error (Th) | mz Error (ppm) | Charge | Series Number |
| --- | --- | --- | --- | --- | --- | --- | --- |
| 10 | y | 3077 | 120.1 | 0.0002 | 1.666 | +1 | 1 |
| - | - | 2.272E+04 | 120.1 | - | - | 0 | - |
| - | - | 2246 | 121.1 | - | - | 0 | - |
| - | - | 583.4 | 123.1 | - | - | 0 | - |
| - | - | 369.7 | 123.3 | - | - | 0 | - |
| - | - | 411.4 | 127.1 | - | - | 0 | - |
| - | - | 808.6 | 127.1 | - | - | 0 | - |
| - | - | 1105 | 128.1 | - | - | 0 | - |
| - | - | 423.4 | 129.1 | - | - | 0 | - |
| - | - | 1.715E+04 | 129.1 | - | - | 0 | - |
| - | - | 596.8 | 130.1 | - | - | 0 | - |
| - | - | 931 | 130.1 | - | - | 0 | - |
| - | - | 347.2 | 131.1 | - | - | 0 | - |
| - | - | 468.3 | 131.1 | - | - | 0 | - |
| - | - | 911.6 | 133.1 | - | - | 0 | - |
| - | - | 1855 | 133.1 | - | - | 0 | - |
| - | - | 416.7 | 134.6 | - | - | 0 | - |
| - | - | 4574 | 136.1 | - | - | 0 | - |
| - | - | 2730 | 137.1 | - | - | 0 | - |
| - | - | 514.2 | 138.1 | - | - | 0 | - |
| - | - | 495 | 139.1 | - | - | 0 | - |
| - | - | 838.3 | 140.1 | - | - | 0 | - |
| - | - | 481.6 | 141.1 | - | - | 0 | - |
| - | - | 639.6 | 144.1 | - | - | 0 | - |
| - | - | 959.5 | 146.1 | - | - | 0 | - |
| - | - | 623.7 | 147.1 | - | - | 0 | - |
| - | - | 980.7 | 149 | - | - | 0 | - |
| - | - | 397.3 | 151.1 | - | - | 0 | - |
| - | - | 635.6 | 152.1 | - | - | 0 | - |
| - | - | 634.9 | 153.1 | - | - | 0 | - |
| - | - | 601.5 | 153.1 | - | - | 0 | - |
| - | - | 664.9 | 155.1 | - | - | 0 | - |
| - | - | 650.3 | 155.1 | - | - | 0 | - |
| - | - | 630.8 | 156.1 | - | - | 0 | - |
| - | - | 724.4 | 157.1 | - | - | 0 | - |
| - | - | 974.9 | 162.1 | - | - | 0 | - |
| - | - | 1206 | 163.1 | - | - | 0 | - |
| - | - | 1510 | 165.1 | - | - | 0 | - |
| - | - | 622.2 | 167.1 | - | - | 0 | - |
| - | - | 517.6 | 169.1 | - | - | 0 | - |
| - | - | 661.8 | 170.1 | - | - | 0 | - |
| 2 | a | 3.799E+04 | 171.1 | 0.0002702 | 1.579 | +1 | 2 |
| - | - | 1297 | 172.1 | - | - | 0 | - |
| - | - | 2454 | 172.2 | - | - | 0 | - |
| - | - | 626.8 | 173.5 | - | - | 0 | - |
| - | - | 840.6 | 175.1 | - | - | 0 | - |
| - | - | 1005 | 176.1 | - | - | 0 | - |
| - | - | 1521 | 177.1 | - | - | 0 | - |
| - | - | 819.4 | 177.1 | - | - | 0 | - |
| - | - | 1285 | 181.2 | - | - | 0 | - |
| - | - | 737.1 | 185.2 | - | - | 0 | - |
| - | - | 2049 | 187.1 | - | - | 0 | - |
| - | - | 561.2 | 193.1 | - | - | 0 | - |
| - | - | 1679 | 193.1 | - | - | 0 | - |
| - | - | 1912 | 197.1 | - | - | 0 | - |
| - | - | 877.2 | 197.2 | - | - | 0 | - |
| - | - | 1046 | 198.1 | - | - | 0 | - |
| - | - | 743.1 | 199.1 | - | - | 0 | - |
| 2 | b | 1.932E+04 | 199.1 | 0.0002133 | 1.071 | +1 | 2 |
| - | - | 1759 | 200.1 | - | - | 0 | - |
| - | - | 537.5 | 201.1 | - | - | 0 | - |
| - | - | 1467 | 201.1 | - | - | 0 | - |
| - | - | 501.3 | 203.1 | - | - | 0 | - |
| - | - | 729.4 | 203.1 | - | - | 0 | - |
| - | - | 3397 | 205.1 | - | - | 0 | - |
| - | - | 528.6 | 205.4 | - | - | 0 | - |
| - | - | 898 | 209.2 | - | - | 0 | - |
| - | - | 2405 | 212.1 | - | - | 0 | - |
| 9 | y | 3193 | 215.1 | 9.547E-05 | 0.4438 | +1 | 2 |
| - | - | 682.6 | 216.1 | - | - | 0 | - |
| - | - | 583.9 | 217.1 | - | - | 0 | - |
| - | - | 1615 | 219.1 | - | - | 0 | - |
| - | - | 911.1 | 227.1 | - | - | 0 | - |
| - | - | 892.2 | 227.1 | - | - | 0 | - |
| - | - | 583.8 | 228.1 | - | - | 0 | - |
| - | - | 559.5 | 229.1 | - | - | 0 | - |
| - | - | 3769 | 230.2 | - | - | 0 | - |
| - | - | 481.2 | 232.1 | - | - | 0 | - |
| 9 | y | 1131 | 233.1 | 0.0005019 | 2.153 | +1 | 2 |
| - | - | 1053 | 237.2 | - | - | 0 | - |
| - | - | 711.3 | 242.2 | - | - | 0 | - |
| - | - | 541.4 | 245.1 | - | - | 0 | - |
| - | - | 681.3 | 247.1 | - | - | 0 | - |
| - | - | 578.5 | 251.1 | - | - | 0 | - |
| - | - | 2177 | 255.1 | - | - | 0 | - |
| - | - | 777.1 | 262.1 | - | - | 0 | - |
| - | - | 546.7 | 265.2 | - | - | 0 | - |
| - | - | 594.5 | 267.1 | - | - | 0 | - |
| - | - | 1898 | 269.2 | - | - | 0 | - |
| - | - | 1280 | 273.1 | - | - | 0 | - |
| - | - | 555.3 | 274.1 | - | - | 0 | - |
| - | - | 1920 | 276.2 | - | - | 0 | - |
| - | - | 517.5 | 288.1 | - | - | 0 | - |
| - | - | 932.6 | 301.2 | - | - | 0 | - |
| - | - | 2651 | 310.2 | - | - | 0 | - |
| - | - | 526.7 | 311.2 | - | - | 0 | - |
| - | - | 1447 | 326.2 | - | - | 0 | - |
| - | - | 838 | 341.2 | - | - | 0 | - |
| - | - | 636.5 | 344.2 | - | - | 0 | - |
| 3 | b | 1151 | 346.2 | 0.000421 | 1.216 | +1 | 3 |
| - | - | 616.1 | 357.2 | - | - | 0 | - |
| - | - | 633.3 | 361.2 | - | - | 0 | - |
| 8 | y | 2392 | 361.2 | 0.000345 | 0.9549 | +1 | 3 |
| - | - | 620.1 | 362.2 | - | - | 0 | - |
| - | - | 522.1 | 370.4 | - | - | 0 | - |
| 3 | y | 705.6 | 381.7 | 0.0003107 | 0.8139 | +2 | 8 |
| - | - | 982 | 382.2 | - | - | 0 | - |
| - | - | 1044 | 382.2 | - | - | 0 | - |
| - | - | 1801 | 383.2 | - | - | 0 | - |
| 3 | y | 6488 | 390.7 | 0.0001858 | 0.4755 | +2 | 8 |
| - | - | 3228 | 391.2 | - | - | 0 | - |
| - | - | 731.4 | 391.7 | - | - | 0 | - |
| - | - | 1248 | 392.2 | - | - | 0 | - |
| - | - | 735.5 | 400.3 | - | - | 0 | - |
| - | - | 1487 | 401.2 | - | - | 0 | - |
| - | - | 2598 | 402.2 | - | - | 0 | - |
| - | - | 509.2 | 402.2 | - | - | 0 | - |
| - | - | 577.3 | 415.2 | - | - | 0 | - |
| - | - | 1244 | 415.3 | - | - | 0 | - |
| - | - | 1039 | 419 | - | - | 0 | - |
| - | - | 1476 | 420.2 | - | - | 0 | - |
| - | - | 504.5 | 436.3 | - | - | 0 | - |
| - | - | 592.8 | 438.2 | - | - | 0 | - |
| - | - | 862.4 | 439.3 | - | - | 0 | - |
| - | - | 1920 | 457.3 | - | - | 0 | - |
| 7 | y | 633.2 | 462.3 | 0.001087 | 2.352 | +1 | 4 |
| - | - | 774.3 | 486.3 | - | - | 0 | - |
| - | - | 874.7 | 488.8 | - | - | 0 | - |
| - | - | 689.8 | 489.2 | - | - | 0 | - |
| 0 | Precursor | 975.9 | 489.8 | 0.003183 | 6.499 | +2 | -1 |
| - | - | 1356 | 496.3 | - | - | 0 | - |
| - | - | 3458 | 514.3 | - | - | 0 | - |
| 6 | y | 914.9 | 519.3 | 0.0004591 | 0.8841 | +1 | 5 |
| - | - | 1362 | 530.3 | - | - | 0 | - |
| - | - | 2871 | 548.3 | - | - | 0 | - |
| - | - | 1087 | 549.3 | - | - | 0 | - |
| 5 | y | 4786 | 576.3 | 0.0006629 | 1.15 | +1 | 6 |
| - | - | 968.7 | 577.3 | - | - | 0 | - |
| 4 | y | 1.161E+04 | 633.4 | 0.001294 | 2.043 | +1 | 7 |
| - | - | 4161 | 634.4 | - | - | 0 | - |
| - | - | 744.5 | 635.4 | - | - | 0 | - |
| - | - | 3584 | 643.4 | - | - | 0 | - |
| - | - | 6746 | 661.4 | - | - | 0 | - |
| - | - | 2552 | 662.4 | - | - | 0 | - |
| 8 | b | 811.3 | 746.4 | 0.006465 | 8.661 | +1 | 8 |
| - | - | 734.9 | 760.4 | - | - | 0 | - |
| 3 | y | 1222 | 762.4 | 0.001074 | 1.409 | +1 | 8 |
| - | - | 668.2 | 763.4 | - | - | 0 | - |
| - | - | 1555 | 769.4 | - | - | 0 | - |
| 3 | y | 2.357E+04 | 780.4 | 0.0004086 | 0.5236 | +1 | 8 |
| - | - | 9902 | 781.4 | - | - | 0 | - |
| - | - | 1811 | 782.4 | - | - | 0 | - |
| 9 | b | 1840 | 859.5 | 0.001451 | 1.688 | +1 | 9 |
| - | - | 690 | 860.5 | - | - | 0 | - |
| 2 | y | 1162 | 879.5 | 2.515E-05 | 0.02859 | +1 | 9 |
| - | - | 708.7 | 881.5 | - | - | 0 | - |
| - | - | 598.4 | 1038 | - | - | 0 | - |
| - | - | 677.3 | 1271 | - | - | 0 | - |
| - | - | 612.9 | 1571 | - | - | 0 | - |
| - | - | 623.1 | 1771 | - | - | 0 | - |
| - | - | 639 | 2456 | - | - | 0 | - |
| - | - | 625.4 | 3173 | - | - | 0 | - |

m/z Charge Intensity FragmentType MassShift Position
120.06571960449219 0 3077.147 y 9
120.0810546875 0 22722.605
121.08440399169922 0 2246.1184
123.11734008789062 0 583.40985
123.3079605102539 0 369.6923
127.05030059814453 0 411.37433
127.07568359375 0 808.56573
128.10736083984375 0 1105.2638
129.0662078857422 0 423.38904
129.1024932861328 0 17146.45
130.06112670898438 0 596.8351
130.10617065429688 0 931.04144
131.07130432128906 0 347.1994
131.08157348632812 0 468.3143
133.06094360351562 0 911.55524
133.08612060546875 0 1854.7799
134.6354522705078 0 416.72324
136.07598876953125 0 4573.829
137.10757446289062 0 2729.8013
138.06654357910156 0 514.22705
139.0750732421875 0 494.95956
140.08203125 0 838.314
141.10215759277344 0 481.60812
144.07644653320312 0 639.60614
146.06060791015625 0 959.52203
147.06491088867188 0 623.67456
148.95465087890625 0 980.743
151.08895874023438 0 397.2598
152.07061767578125 0 635.6208
153.05499267578125 0 634.93585
153.07752990722656 0 601.5189
155.08126831054688 0 664.9219
155.11822509765625 0 650.2981
156.07675170898438 0 630.82825
157.08599853515625 0 724.4189
162.05487060546875 0 974.9342
163.07147216796875 0 1205.5701
165.10263061523438 0 1510.0057
167.0558624267578 0 622.1548
169.13380432128906 0 517.57336
170.0928955078125 0 661.7796
171.1494598388672 0 37993.11 a 1
172.07215881347656 0 1296.9777
172.1529541015625 0 2453.8499
173.45285034179688 0 626.7994
175.08668518066406 0 840.6109
176.10696411132812 0 1004.86
177.10275268554688 0 1521.2983
177.11215209960938 0 819.41327
181.17022705078125 0 1284.5498
185.1651153564453 0 737.0868
187.14447021484375 0 2049.225
193.0876007080078 0 561.22253
193.0974884033203 0 1679.2637
197.128662109375 0 1912.3755
197.16510009765625 0 877.16077
198.08741760253906 0 1045.681
199.10791015625 0 743.07904
199.14431762695312 0 19319.914 b 1
200.14808654785156 0 1759.4813
201.11370849609375 0 537.46545
201.12362670898438 0 1467.3062
203.09278869628906 0 501.31573
203.1029510498047 0 729.4221
205.0975341796875 0 3397.1453
205.3946075439453 0 528.5589
209.16554260253906 0 897.95105
212.13943481445312 0 2405.0244
215.1391143798828 0 3192.8257 y Water loss 8
216.0977783203125 0 682.58655
217.09841918945312 0 583.8501
219.14947509765625 0 1614.7184
227.10302734375 0 911.1208
227.1146697998047 0 892.1967
228.09817504882812 0 583.7874
229.0930633544922 0 559.5217
230.1502685546875 0 3769.43
232.055908203125 0 481.19147
233.15008544921875 0 1130.6554 y 8
237.160400390625 0 1053.4044
242.18719482421875 0 711.2618
245.1250457763672 0 541.4134
247.14413452148438 0 681.30914
251.1491241455078 0 578.46155
255.10865783691406 0 2176.5276
262.1184387207031 0 777.06055
265.19024658203125 0 546.66565
267.1331787109375 0 594.46674
269.1612548828125 0 1898.1956
273.1197204589844 0 1280.1061
274.1180114746094 0 555.3133
276.15545654296875 0 1920.1361
288.1358947753906 0 517.5067
301.1910400390625 0 932.6045
310.2127685546875 0 2650.758
311.2198791503906 0 526.68567
326.1827697753906 0 1446.8738
341.18170166015625 0 838.0074
344.1929016113281 0 636.5143
346.21209716796875 0 1150.9408 b 2
357.15325927734375 0 616.0604
361.1849670410156 0 633.2584
361.24420166015625 0 2392.0305 y 7
362.2492980957031 0 620.1019
370.4265441894531 0 522.0943
381.711181640625 0 705.6125 y Water loss 2
382.2124328613281 0 981.9906
382.244873046875 0 1043.8846
383.20306396484375 0 1800.9519
390.7163391113281 0 6488.112 y 2
391.21820068359375 0 3227.9744
391.7186279296875 0 731.4048
392.1939392089844 0 1247.6593
400.25665283203125 0 735.48474
401.21392822265625 0 1487.3433
402.177490234375 0 2598.058
402.2047119140625 0 509.1779
415.221435546875 0 577.333
415.252685546875 0 1244.2245
418.9945983886719 0 1038.7026
420.1872253417969 0 1476.2793
436.2761535644531 0 504.47543
438.1552734375 0 592.8376
439.26641845703125 0 862.39374
457.27850341796875 0 1920.4823
462.2911376953125 0 633.23645 y 6
486.3053894042969 0 774.28613
488.7843322753906 0 874.65857
489.22894287109375 0 689.83594
489.7877502441406 0 975.89185 Precursor
496.28955078125 0 1355.9384
514.2987670898438 0 3458.407
519.3141479492188 0 914.9494 y 5
530.2737426757812 0 1361.6881
548.283203125 0 2870.7915
549.2841186523438 0 1087.4011
576.3358154296875 0 4785.546 y 4
577.3411254882812 0 968.6525
633.35791015625 0 11607.773 y 3
634.3604125976562 0 4160.6606
635.3681030273438 0 744.46765
643.3565063476562 0 3583.8145
661.3663330078125 0 6745.599
662.3699951171875 0 2552.106
746.4130859375 0 811.33954 b 7
760.4336547851562 0 734.919
762.4133911132812 0 1222.1782 y Water loss 2
763.41455078125 0 668.24536
769.4105834960938 0 1555.0555
780.4246215820312 0 23572.662 y 2
781.4279174804688 0 9901.799
782.4276733398438 0 1811.2587
859.5050659179688 0 1840.4305 b 8
860.507568359375 0 689.9534
879.4934692382812 0 1162.4467 y 1
881.4906616210938 0 708.7494
1037.8983154296875 0 598.37146
1270.702880859375 0 677.3211
1570.5159912109375 0 612.8799
1771.4395751953125 0 623.1058
2456.406005859375 0 639.01624
3172.52783203125 0 625.4101

Spectrum Details

|  |  |
| --- | --- |
| Matched peaks? Matched peaks The total absolute number of peaks matched. Additionally in brackets the total fraction of peaks matched and the total number of peaks is shown. | 19 (11.73% of 162) |
| FDR? FDR The false discovery rate estimated for this peptide. It is calculated by matching all theoretical fragments with a non-integer shift with the raw peaks for this spectrum. This is done with 40 different shifts. The resulting percentage is the average number of annotated peaks over the number of annotated peaks with the correct spectrum. | 0.25% |
| Satellite FDR? Satellite FDR See the FDR for details on its calculation. This satellite ion specific FDR only contains the satellite ions (d/w) for I/L/J positions. | - |
| PSM Score? PSM Score The PSM Score as given by Hecklib to this annotated spectrum. It is shown with three significant figures. | 207 |

#### Spectrum 5801? Spectrum 5801 The raw spectrum of this peptide as annotated by Hecklib. The fragments are coloured according to ion type (see legend). Any peaks with a star '\*' as text can be hovered over to see the full details, first the ion type second the mass shift type. By hovering over the amino acids in the peptide or ions in the legend the corresponding peaks are highlighted. By toggling the 'Unassigned' label you can turn the background (unassigned) peaks on or off in the plot. By updating the slider in the Ion legend you can update the spectrum to only show the top X% of the peaks with labels. The top X% means any peak that is within X% of the highest intensity. By dragging in the spectrum you can zoom in to a specific part of the spectrum and use 'Zoom Out' to get back to the original zoom level. The annotation of the spectrum is based on the given sequence in the peptides file and is done with different software so inconsistencies are likely. The peaks are annotated based on the given sequence, with 20 ppm tolerance.

Copy Data

##### Spectrum 5801 (TSV)

###### Preview

```
Loading example...
```

*Click on the button to copy the data to your clipboard.*

Mz MinMz MaxIntensity Max

WidthHeightPeptide font sizePeptide stroke widthSpectrum font sizeSpectrum stroke widthCompact peptide

Ion legend

wxyz

abcd

OtherUnassignedIon^Charge^_Position_Show for top:%

VVFGGGTKJT

05.05e+41.01e+51.52e+52.02e+5

Zoom Out

y^+1^_1_y^+1^_2_y^+1^_2_z^+1^_3_y^+1^_3_c^+2^_8_y^+2^_8_y^+2^_8_z^+1^_4_y^+2^_9_y^+1^_4_z^+1^_4_y^+1^_4_c^+1^_5_z^+1^_5_y^+1^_5_c^+1^_6_y^+1^_6_z^+1^_6_y^+1^_6_z^+1^_7_y^+1^_7_y^+1^_7_c^+1^_7_y^+1^_7_c^+1^_7_c^+1^_8_c^+1^_8_y^+1^_8_c^+1^_8_z^+1^_8_y^+1^_8_z^+1^_9_w^+1^_9_c^+1^_9_z^+1^_9_c^+1^_9_y^+1^_9_

037174211131484

Fragment Matches Table

Show background peaks

| Position | Ion type | Intensity | mz Theoretical | mz Error (Th) | mz Error (ppm) | Charge | Series Number |
| --- | --- | --- | --- | --- | --- | --- | --- |
| 10 | y | 9080 | 120.1 | 0.0002381 | 1.983 | +1 | 1 |
| - | - | 1.485E+04 | 120.1 | - | - | 0 | - |
| - | - | 900.6 | 121.1 | - | - | 0 | - |
| - | - | 553.4 | 128.1 | - | - | 0 | - |
| - | - | 5354 | 129.1 | - | - | 0 | - |
| - | - | 384 | 129.5 | - | - | 0 | - |
| - | - | 2672 | 133.1 | - | - | 0 | - |
| - | - | 534.2 | 134 | - | - | 0 | - |
| - | - | 2.02E+04 | 136.1 | - | - | 0 | - |
| - | - | 508.1 | 136.7 | - | - | 0 | - |
| - | - | 410.6 | 149.3 | - | - | 0 | - |
| - | - | 486.8 | 153.9 | - | - | 0 | - |
| - | - | 891.4 | 166.1 | - | - | 0 | - |
| - | - | 7.253E+04 | 171.1 | - | - | 0 | - |
| - | - | 7385 | 172.2 | - | - | 0 | - |
| - | - | 2988 | 173.4 | - | - | 0 | - |
| - | - | 648.4 | 175.1 | - | - | 0 | - |
| - | - | 1157 | 177.1 | - | - | 0 | - |
| - | - | 1170 | 177.1 | - | - | 0 | - |
| - | - | 492.6 | 181.8 | - | - | 0 | - |
| - | - | 488.5 | 184.5 | - | - | 0 | - |
| - | - | 3912 | 185.2 | - | - | 0 | - |
| - | - | 7.227E+04 | 199.1 | - | - | 0 | - |
| - | - | 5822 | 200.1 | - | - | 0 | - |
| - | - | 6259 | 205.1 | - | - | 0 | - |
| - | - | 1095 | 205.1 | - | - | 0 | - |
| - | - | 2755 | 213.2 | - | - | 0 | - |
| 9 | y | 1804 | 215.1 | 0.0002481 | 1.153 | +1 | 2 |
| - | - | 527.8 | 227.2 | - | - | 0 | - |
| - | - | 952.2 | 230.1 | - | - | 0 | - |
| 9 | y | 3177 | 233.1 | 0.0001051 | 0.4509 | +1 | 2 |
| - | - | 957.3 | 242.2 | - | - | 0 | - |
| - | - | 959.7 | 244.1 | - | - | 0 | - |
| - | - | 2865 | 247.1 | - | - | 0 | - |
| - | - | 589 | 251.1 | - | - | 0 | - |
| - | - | 785.5 | 262.1 | - | - | 0 | - |
| - | - | 4060 | 269.1 | - | - | 0 | - |
| - | - | 860.9 | 271.1 | - | - | 0 | - |
| - | - | 537.6 | 273.1 | - | - | 0 | - |
| - | - | 2395 | 276.2 | - | - | 0 | - |
| - | - | 3047 | 297.1 | - | - | 0 | - |
| - | - | 2837 | 301.2 | - | - | 0 | - |
| - | - | 1150 | 302.2 | - | - | 0 | - |
| - | - | 641.8 | 304.2 | - | - | 0 | - |
| - | - | 592.3 | 310 | - | - | 0 | - |
| - | - | 3820 | 310.2 | - | - | 0 | - |
| - | - | 669.9 | 311.2 | - | - | 0 | - |
| - | - | 1561 | 315.2 | - | - | 0 | - |
| - | - | 1096 | 317.2 | - | - | 0 | - |
| - | - | 815.8 | 325.2 | - | - | 0 | - |
| - | - | 1055 | 326.2 | - | - | 0 | - |
| - | - | 513.3 | 327.8 | - | - | 0 | - |
| - | - | 739.5 | 329.2 | - | - | 0 | - |
| - | - | 785.6 | 331.2 | - | - | 0 | - |
| - | - | 551.3 | 334.2 | - | - | 0 | - |
| - | - | 1062 | 340.2 | - | - | 0 | - |
| - | - | 5306 | 341.2 | - | - | 0 | - |
| - | - | 593 | 343.2 | - | - | 0 | - |
| - | - | 512.5 | 343.8 | - | - | 0 | - |
| - | - | 903.8 | 344.2 | - | - | 0 | - |
| - | - | 870.3 | 345.2 | - | - | 0 | - |
| 8 | z | 5259 | 345.2 | 0.0002212 | 0.6406 | +1 | 3 |
| - | - | 4664 | 346.2 | - | - | 0 | - |
| - | - | 9421 | 346.2 | - | - | 0 | - |
| - | - | 837.5 | 347.2 | - | - | 0 | - |
| - | - | 1872 | 347.2 | - | - | 0 | - |
| - | - | 4002 | 357.2 | - | - | 0 | - |
| - | - | 1096 | 358.2 | - | - | 0 | - |
| - | - | 2.422E+04 | 358.2 | - | - | 0 | - |
| - | - | 776.2 | 359.2 | - | - | 0 | - |
| - | - | 684.5 | 360.2 | - | - | 0 | - |
| - | - | 1171 | 361.2 | - | - | 0 | - |
| 8 | y | 3234 | 361.2 | 8.229E-05 | 0.2278 | +1 | 3 |
| - | - | 1616 | 373.2 | - | - | 0 | - |
| 8 | c | 1635 | 373.7 | 0.0007672 | 2.053 | +2 | 8 |
| - | - | 625.7 | 375.2 | - | - | 0 | - |
| 3 | y | 4337 | 381.7 | 0.0004218 | 1.105 | +2 | 8 |
| - | - | 577.9 | 382.2 | - | - | 0 | - |
| - | - | 939.7 | 382.2 | - | - | 0 | - |
| - | - | 835.2 | 383.2 | - | - | 0 | - |
| - | - | 918.9 | 383.2 | - | - | 0 | - |
| - | - | 670.4 | 384.2 | - | - | 0 | - |
| - | - | 1404 | 384.3 | - | - | 0 | - |
| 3 | y | 3.777E+04 | 390.7 | 0.0001553 | 0.3974 | +2 | 8 |
| - | - | 1.421E+04 | 391.2 | - | - | 0 | - |
| - | - | 937.7 | 391.2 | - | - | 0 | - |
| - | - | 4695 | 391.7 | - | - | 0 | - |
| - | - | 642.4 | 392.2 | - | - | 0 | - |
| - | - | 817.8 | 397.2 | - | - | 0 | - |
| - | - | 734.9 | 400.3 | - | - | 0 | - |
| - | - | 1975 | 401.2 | - | - | 0 | - |
| - | - | 1436 | 402.2 | - | - | 0 | - |
| - | - | 5503 | 402.3 | - | - | 0 | - |
| - | - | 2681 | 403.2 | - | - | 0 | - |
| - | - | 1076 | 403.3 | - | - | 0 | - |
| - | - | 1850 | 405.2 | - | - | 0 | - |
| - | - | 2490 | 415.3 | - | - | 0 | - |
| - | - | 1331 | 416.3 | - | - | 0 | - |
| - | - | 2383 | 420.2 | - | - | 0 | - |
| - | - | 693.7 | 426.3 | - | - | 0 | - |
| - | - | 2139 | 428.3 | - | - | 0 | - |
| 7 | z | 776.5 | 429.2 | 0.001926 | 4.488 | +1 | 4 |
| - | - | 884.3 | 432.7 | - | - | 0 | - |
| - | - | 565.6 | 435.2 | - | - | 0 | - |
| - | - | 1454 | 436.2 | - | - | 0 | - |
| - | - | 1214 | 437.2 | - | - | 0 | - |
| - | - | 696.8 | 439.3 | - | - | 0 | - |
| 2 | y | 2236 | 440.3 | 0.0003722 | 0.8453 | +2 | 9 |
| - | - | 798.2 | 440.8 | - | - | 0 | - |
| - | - | 1112 | 442.3 | - | - | 0 | - |
| - | - | 1544 | 443.3 | - | - | 0 | - |
| - | - | 7751 | 444.3 | - | - | 0 | - |
| 7 | y | 2.578E+04 | 445.3 | 0.008433 | 18.94 | +1 | 4 |
| 7 | z | 3083 | 446.3 | 0.003481 | 7.801 | +1 | 4 |
| - | - | 1.07E+04 | 447.3 | - | - | 0 | - |
| - | - | 2433 | 448.3 | - | - | 0 | - |
| - | - | 1735 | 457.3 | - | - | 0 | - |
| - | - | 689.8 | 458.2 | - | - | 0 | - |
| - | - | 1960 | 459.3 | - | - | 0 | - |
| - | - | 817.6 | 460.3 | - | - | 0 | - |
| - | - | 679.8 | 460.3 | - | - | 0 | - |
| 7 | y | 1986 | 462.3 | 0.0004995 | 1.081 | +1 | 4 |
| - | - | 2253 | 470.3 | - | - | 0 | - |
| - | - | 2889 | 471.3 | - | - | 0 | - |
| - | - | 1207 | 472.2 | - | - | 0 | - |
| - | - | 1023 | 472.3 | - | - | 0 | - |
| - | - | 2770 | 476.3 | - | - | 0 | - |
| 5 | c | 4785 | 477.3 | 7.336E-05 | 0.1537 | +1 | 5 |
| - | - | 1524 | 478.3 | - | - | 0 | - |
| - | - | 707.5 | 479.3 | - | - | 0 | - |
| - | - | 1042 | 480.8 | - | - | 0 | - |
| - | - | 1905 | 481.3 | - | - | 0 | - |
| - | - | 918.3 | 484.2 | - | - | 0 | - |
| - | - | 1465 | 486.2 | - | - | 0 | - |
| - | - | 1456 | 488.3 | - | - | 0 | - |
| - | - | 1630 | 488.3 | - | - | 0 | - |
| - | - | 1003 | 489.3 | - | - | 0 | - |
| - | - | 1222 | 489.3 | - | - | 0 | - |
| - | - | 4073 | 489.8 | - | - | 0 | - |
| - | - | 746.9 | 490.2 | - | - | 0 | - |
| - | - | 4758 | 490.3 | - | - | 0 | - |
| - | - | 891.1 | 490.3 | - | - | 0 | - |
| - | - | 692.3 | 496.3 | - | - | 0 | - |
| - | - | 784 | 497.3 | - | - | 0 | - |
| - | - | 2524 | 499.3 | - | - | 0 | - |
| - | - | 809.9 | 501.2 | - | - | 0 | - |
| - | - | 1663 | 503.2 | - | - | 0 | - |
| 6 | z | 5305 | 503.3 | 7.338E-05 | 0.1458 | +1 | 5 |
| - | - | 6.018E+04 | 504.3 | - | - | 0 | - |
| - | - | 7.974E+04 | 504.3 | - | - | 0 | - |
| - | - | 1.533E+04 | 505.3 | - | - | 0 | - |
| - | - | 2.058E+04 | 505.3 | - | - | 0 | - |
| - | - | 2416 | 506.3 | - | - | 0 | - |
| - | - | 2806 | 506.3 | - | - | 0 | - |
| - | - | 737.9 | 508.2 | - | - | 0 | - |
| - | - | 875.2 | 513.3 | - | - | 0 | - |
| - | - | 6143 | 514.3 | - | - | 0 | - |
| - | - | 782.1 | 515.3 | - | - | 0 | - |
| - | - | 3435 | 516.3 | - | - | 0 | - |
| - | - | 1572 | 517.3 | - | - | 0 | - |
| - | - | 1052 | 517.3 | - | - | 0 | - |
| - | - | 3261 | 518.3 | - | - | 0 | - |
| 6 | y | 6501 | 519.3 | 0.0002733 | 0.5262 | +1 | 5 |
| - | - | 1682 | 520.3 | - | - | 0 | - |
| - | - | 1043 | 522.8 | - | - | 0 | - |
| - | - | 1137 | 523.3 | - | - | 0 | - |
| - | - | 1672 | 530.3 | - | - | 0 | - |
| - | - | 2659 | 533.3 | - | - | 0 | - |
| 6 | c | 1945 | 534.3 | 0.001029 | 1.926 | +1 | 6 |
| - | - | 807.6 | 535.3 | - | - | 0 | - |
| - | - | 842.8 | 536.2 | - | - | 0 | - |
| - | - | 764.2 | 543.3 | - | - | 0 | - |
| - | - | 7008 | 548.3 | - | - | 0 | - |
| - | - | 1708 | 549.3 | - | - | 0 | - |
| - | - | 2264 | 551.3 | - | - | 0 | - |
| - | - | 5324 | 551.8 | - | - | 0 | - |
| - | - | 3460 | 552.3 | - | - | 0 | - |
| - | - | 1572 | 553.2 | - | - | 0 | - |
| - | - | 876.1 | 554.2 | - | - | 0 | - |
| - | - | 1874 | 554.3 | - | - | 0 | - |
| 5 | y | 1092 | 558.3 | 0.000125 | 0.2238 | +1 | 6 |
| - | - | 2655 | 559.3 | - | - | 0 | - |
| - | - | 1168 | 559.3 | - | - | 0 | - |
| 5 | z | 2.219E+04 | 560.3 | 0.0003274 | 0.5843 | +1 | 6 |
| - | - | 2851 | 561.3 | - | - | 0 | - |
| - | - | 6.402E+04 | 561.3 | - | - | 0 | - |
| - | - | 989.8 | 562.3 | - | - | 0 | - |
| - | - | 1.767E+04 | 562.3 | - | - | 0 | - |
| - | - | 2347 | 563.3 | - | - | 0 | - |
| - | - | 968.3 | 564.3 | - | - | 0 | - |
| - | - | 950.6 | 571.3 | - | - | 0 | - |
| - | - | 1827 | 572.3 | - | - | 0 | - |
| - | - | 2756 | 574.3 | - | - | 0 | - |
| - | - | 955.5 | 575.3 | - | - | 0 | - |
| - | - | 9337 | 575.3 | - | - | 0 | - |
| 5 | y | 2.108E+04 | 576.3 | 0.0002526 | 0.4383 | +1 | 6 |
| - | - | 6008 | 577.3 | - | - | 0 | - |
| - | - | 1153 | 578.3 | - | - | 0 | - |
| - | - | 1205 | 583.3 | - | - | 0 | - |
| - | - | 883.3 | 585.3 | - | - | 0 | - |
| - | - | 773.5 | 588.3 | - | - | 0 | - |
| - | - | 5187 | 589.3 | - | - | 0 | - |
| - | - | 1376 | 598.3 | - | - | 0 | - |
| - | - | 2267 | 599.3 | - | - | 0 | - |
| 4 | z | 3370 | 600.3 | 0.009158 | 15.26 | +1 | 7 |
| - | - | 2647 | 603.4 | - | - | 0 | - |
| - | - | 1112 | 604.4 | - | - | 0 | - |
| - | - | 1213 | 608.8 | - | - | 0 | - |
| - | - | 974.3 | 609.3 | - | - | 0 | - |
| - | - | 1097 | 614.3 | - | - | 0 | - |
| 4 | y | 1587 | 615.3 | 0.003217 | 5.228 | +1 | 7 |
| 4 | y | 4621 | 616.3 | 0.001171 | 1.899 | +1 | 7 |
| 7 | c | 2.729E+04 | 617.3 | 0.0108 | 17.49 | +1 | 7 |
| - | - | 899.7 | 617.8 | - | - | 0 | - |
| - | - | 3.808E+04 | 618.3 | - | - | 0 | - |
| - | - | 1.11E+04 | 619.4 | - | - | 0 | - |
| - | - | 2346 | 620.4 | - | - | 0 | - |
| - | - | 2.08E+04 | 632.3 | - | - | 0 | - |
| - | - | 814.4 | 632.4 | - | - | 0 | - |
| 4 | y | 9.124E+04 | 633.4 | 0.0001342 | 0.2119 | +1 | 7 |
| - | - | 3.36E+04 | 634.4 | - | - | 0 | - |
| 7 | c | 1.265E+05 | 635.4 | 0.0005476 | 0.8619 | +1 | 7 |
| - | - | 4.484E+04 | 636.4 | - | - | 0 | - |
| - | - | 7377 | 637.4 | - | - | 0 | - |
| - | - | 767.6 | 638.4 | - | - | 0 | - |
| - | - | 1609 | 640.3 | - | - | 0 | - |
| - | - | 1.228E+04 | 641.3 | - | - | 0 | - |
| - | - | 3748 | 643.4 | - | - | 0 | - |
| - | - | 1788 | 644.4 | - | - | 0 | - |
| - | - | 1749 | 646.3 | - | - | 0 | - |
| - | - | 707.7 | 647.3 | - | - | 0 | - |
| - | - | 857.3 | 647.4 | - | - | 0 | - |
| - | - | 867.6 | 648.3 | - | - | 0 | - |
| - | - | 1928 | 649.8 | - | - | 0 | - |
| - | - | 973.7 | 650.3 | - | - | 0 | - |
| - | - | 5458 | 658.3 | - | - | 0 | - |
| - | - | 5684 | 658.8 | - | - | 0 | - |
| - | - | 2896 | 659.3 | - | - | 0 | - |
| - | - | 844.1 | 660.3 | - | - | 0 | - |
| - | - | 1.791E+04 | 661.4 | - | - | 0 | - |
| - | - | 6642 | 662.4 | - | - | 0 | - |
| - | - | 6465 | 663.4 | - | - | 0 | - |
| - | - | 2024 | 664.4 | - | - | 0 | - |
| - | - | 1711 | 667.4 | - | - | 0 | - |
| - | - | 672 | 668.4 | - | - | 0 | - |
| - | - | 828 | 669.4 | - | - | 0 | - |
| - | - | 7872 | 685.4 | - | - | 0 | - |
| - | - | 643.4 | 686.4 | - | - | 0 | - |
| - | - | 692.2 | 687.4 | - | - | 0 | - |
| - | - | 2568 | 692.3 | - | - | 0 | - |
| - | - | 2582 | 693.3 | - | - | 0 | - |
| - | - | 925.1 | 694.3 | - | - | 0 | - |
| - | - | 2475 | 695.4 | - | - | 0 | - |
| - | - | 2429 | 696.4 | - | - | 0 | - |
| - | - | 3700 | 697.4 | - | - | 0 | - |
| - | - | 1272 | 698.4 | - | - | 0 | - |
| - | - | 1044 | 702.4 | - | - | 0 | - |
| - | - | 7838 | 702.4 | - | - | 0 | - |
| - | - | 2174 | 702.9 | - | - | 0 | - |
| - | - | 1099 | 703.4 | - | - | 0 | - |
| - | - | 1130 | 703.4 | - | - | 0 | - |
| - | - | 1254 | 703.9 | - | - | 0 | - |
| - | - | 841.5 | 707.3 | - | - | 0 | - |
| - | - | 1817 | 708.3 | - | - | 0 | - |
| - | - | 798.7 | 709.3 | - | - | 0 | - |
| - | - | 961 | 710.4 | - | - | 0 | - |
| - | - | 1414 | 711.4 | - | - | 0 | - |
| - | - | 1187 | 711.9 | - | - | 0 | - |
| - | - | 1109 | 712.4 | - | - | 0 | - |
| - | - | 1.211E+04 | 713.4 | - | - | 0 | - |
| - | - | 887.4 | 715.4 | - | - | 0 | - |
| - | - | 705.8 | 717.4 | - | - | 0 | - |
| - | - | 945.9 | 718.3 | - | - | 0 | - |
| - | - | 1502 | 718.4 | - | - | 0 | - |
| - | - | 1974 | 719.4 | - | - | 0 | - |
| - | - | 774.1 | 720.4 | - | - | 0 | - |
| - | - | 863.9 | 721.4 | - | - | 0 | - |
| - | - | 4003 | 724.4 | - | - | 0 | - |
| - | - | 4324 | 724.9 | - | - | 0 | - |
| - | - | 1751 | 725.4 | - | - | 0 | - |
| - | - | 1359 | 728.4 | - | - | 0 | - |
| - | - | 6912 | 729.4 | - | - | 0 | - |
| - | - | 2.824E+04 | 730.4 | - | - | 0 | - |
| - | - | 4464 | 732.4 | - | - | 0 | - |
| - | - | 5130 | 732.9 | - | - | 0 | - |
| - | - | 2392 | 733.4 | - | - | 0 | - |
| - | - | 899.5 | 733.9 | - | - | 0 | - |
| - | - | 2963 | 735.3 | - | - | 0 | - |
| 8 | c | 2190 | 745.4 | 0.01378 | 18.49 | +1 | 8 |
| 8 | c | 8658 | 746.4 | 0.003352 | 4.491 | +1 | 8 |
| - | - | 4019 | 747.4 | - | - | 0 | - |
| - | - | 708.6 | 748.4 | - | - | 0 | - |
| - | - | 1342 | 751.3 | - | - | 0 | - |
| - | - | 6200 | 753.3 | - | - | 0 | - |
| - | - | 4687 | 761.4 | - | - | 0 | - |
| - | - | 1430 | 761.4 | - | - | 0 | - |
| 3 | y | 3993 | 762.4 | 0.001672 | 2.193 | +1 | 8 |
| 8 | c | 1.576E+05 | 763.4 | 5.989E-06 | 0.007845 | +1 | 8 |
| 3 | z | 1.735E+04 | 764.4 | 0.009211 | 12.05 | +1 | 8 |
| - | - | 5.209E+04 | 764.4 | - | - | 0 | - |
| - | - | 1.298E+04 | 765.4 | - | - | 0 | - |
| - | - | 7445 | 765.5 | - | - | 0 | - |
| - | - | 3862 | 766.4 | - | - | 0 | - |
| - | - | 3133 | 769.3 | - | - | 0 | - |
| - | - | 1043 | 779.4 | - | - | 0 | - |
| 3 | y | 1.338E+05 | 780.4 | 4.238E-05 | 0.05431 | +1 | 8 |
| - | - | 5.626E+04 | 781.4 | - | - | 0 | - |
| - | - | 1.525E+04 | 782.4 | - | - | 0 | - |
| - | - | 833.2 | 788.4 | - | - | 0 | - |
| - | - | 4677 | 789.4 | - | - | 0 | - |
| - | - | 1187 | 790.4 | - | - | 0 | - |
| - | - | 1115 | 793.4 | - | - | 0 | - |
| - | - | 1406 | 804.4 | - | - | 0 | - |
| - | - | 1725 | 807.4 | - | - | 0 | - |
| - | - | 1427 | 808.4 | - | - | 0 | - |
| - | - | 755.7 | 818.5 | - | - | 0 | - |
| - | - | 6439 | 823.3 | - | - | 0 | - |
| - | - | 905.8 | 830.4 | - | - | 0 | - |
| - | - | 1417 | 831.4 | - | - | 0 | - |
| - | - | 1056 | 832.4 | - | - | 0 | - |
| - | - | 1.714E+04 | 832.5 | - | - | 0 | - |
| - | - | 7441 | 833.5 | - | - | 0 | - |
| - | - | 2288 | 834.5 | - | - | 0 | - |
| - | - | 960.5 | 841.5 | - | - | 0 | - |
| 2 | z | 772.9 | 845.5 | 0.008284 | 9.798 | +1 | 9 |
| - | - | 945.9 | 848.4 | - | - | 0 | - |
| 2 | w | 931.7 | 848.5 | 0.01287 | 15.16 | +1 | 9 |
| - | - | 3156 | 849.5 | - | - | 0 | - |
| - | - | 1025 | 858.4 | - | - | 0 | - |
| - | - | 925.3 | 859.4 | - | - | 0 | - |
| 9 | c | 8580 | 859.5 | 1.377E-05 | 0.01602 | +1 | 9 |
| - | - | 3894 | 860.4 | - | - | 0 | - |
| - | - | 4390 | 860.5 | - | - | 0 | - |
| - | - | 4953 | 861.5 | - | - | 0 | - |
| - | - | 1873 | 862.5 | - | - | 0 | - |
| 2 | z | 1.761E+04 | 863.5 | 0.0001335 | 0.1546 | +1 | 9 |
| - | - | 9067 | 864.5 | - | - | 0 | - |
| - | - | 1.494E+04 | 865.5 | - | - | 0 | - |
| - | - | 1.501E+04 | 866.4 | - | - | 0 | - |
| - | - | 959.3 | 873.5 | - | - | 0 | - |
| - | - | 870.8 | 874.5 | - | - | 0 | - |
| 9 | c | 1.6E+05 | 876.5 | 4.845E-05 | 0.05528 | +1 | 9 |
| - | - | 8.155E+04 | 877.5 | - | - | 0 | - |
| - | - | 758.1 | 878.4 | - | - | 0 | - |
| - | - | 2.403E+04 | 878.5 | - | - | 0 | - |
| - | - | 1040 | 879.4 | - | - | 0 | - |
| 2 | y | 4457 | 879.5 | 0.005518 | 6.274 | +1 | 9 |
| - | - | 1959 | 880.5 | - | - | 0 | - |
| - | - | 2604 | 881.4 | - | - | 0 | - |
| - | - | 1113 | 889.4 | - | - | 0 | - |
| - | - | 1459 | 891.5 | - | - | 0 | - |
| - | - | 5.087E+04 | 893.5 | - | - | 0 | - |
| - | - | 2492 | 906.5 | - | - | 0 | - |
| - | - | 1661 | 907.5 | - | - | 0 | - |
| - | - | 942.6 | 908.5 | - | - | 0 | - |
| - | - | 6596 | 909.4 | - | - | 0 | - |
| - | - | 640 | 916.5 | - | - | 0 | - |
| - | - | 7089 | 917.5 | - | - | 0 | - |
| - | - | 7713 | 918.5 | - | - | 0 | - |
| - | - | 1892 | 919.5 | - | - | 0 | - |
| - | - | 7943 | 923.5 | - | - | 0 | - |
| - | - | 3151 | 924.5 | - | - | 0 | - |
| - | - | 832.5 | 925.5 | - | - | 0 | - |
| - | - | 1488 | 933.6 | - | - | 0 | - |
| - | - | 1035 | 934.6 | - | - | 0 | - |
| - | - | 9890 | 936.5 | - | - | 0 | - |
| - | - | 5456 | 937.5 | - | - | 0 | - |
| - | - | 1805 | 943.5 | - | - | 0 | - |
| - | - | 1969 | 944.5 | - | - | 0 | - |
| - | - | 1020 | 945.5 | - | - | 0 | - |
| - | - | 1830 | 947.5 | - | - | 0 | - |
| - | - | 1109 | 948.5 | - | - | 0 | - |
| - | - | 1759 | 951.6 | - | - | 0 | - |
| - | - | 1202 | 952.6 | - | - | 0 | - |
| - | - | 798.5 | 953.5 | - | - | 0 | - |
| - | - | 1004 | 954.5 | - | - | 0 | - |
| - | - | 1954 | 960.5 | - | - | 0 | - |
| - | - | 3484 | 961.5 | - | - | 0 | - |
| - | - | 2.002E+05 | 962.5 | - | - | 0 | - |
| - | - | 1.062E+05 | 963.5 | - | - | 0 | - |
| - | - | 3.582E+04 | 964.5 | - | - | 0 | - |
| - | - | 8406 | 965.5 | - | - | 0 | - |
| - | - | 2842 | 965.6 | - | - | 0 | - |
| - | - | 846.4 | 977.5 | - | - | 0 | - |
| - | - | 3724 | 977.6 | - | - | 0 | - |
| - | - | 1.031E+05 | 978.6 | - | - | 0 | - |
| - | - | 1.364E+05 | 979.6 | - | - | 0 | - |
| - | - | 6.11E+04 | 980.6 | - | - | 0 | - |
| - | - | 1.826E+04 | 981.5 | - | - | 0 | - |
| - | - | 1.262E+04 | 981.6 | - | - | 0 | - |
| - | - | 1.655E+04 | 982.5 | - | - | 0 | - |
| - | - | 724.2 | 985.5 | - | - | 0 | - |
| - | - | 1237 | 1002 | - | - | 0 | - |
| - | - | 3856 | 1003 | - | - | 0 | - |
| - | - | 1771 | 1004 | - | - | 0 | - |
| - | - | 7679 | 1018 | - | - | 0 | - |
| - | - | 4640 | 1020 | - | - | 0 | - |
| - | - | 1899 | 1021 | - | - | 0 | - |
| - | - | 996.5 | 1045 | - | - | 0 | - |
| - | - | 1971 | 1046 | - | - | 0 | - |
| - | - | 1144 | 1047 | - | - | 0 | - |
| - | - | 1044 | 1086 | - | - | 0 | - |
| - | - | 853.8 | 1087 | - | - | 0 | - |
| - | - | 1046 | 1102 | - | - | 0 | - |
| - | - | 7461 | 1103 | - | - | 0 | - |
| - | - | 6144 | 1104 | - | - | 0 | - |
| - | - | 1493 | 1105 | - | - | 0 | - |
| - | - | 4646 | 1106 | - | - | 0 | - |
| - | - | 4086 | 1107 | - | - | 0 | - |
| - | - | 2050 | 1108 | - | - | 0 | - |
| - | - | 1100 | 1218 | - | - | 0 | - |
| - | - | 781.6 | 1219 | - | - | 0 | - |
| - | - | 894.4 | 1234 | - | - | 0 | - |
| - | - | 5490 | 1235 | - | - | 0 | - |
| - | - | 4769 | 1236 | - | - | 0 | - |
| - | - | 1628 | 1237 | - | - | 0 | - |
| - | - | 923.4 | 1272 | - | - | 0 | - |
| - | - | 1388 | 1273 | - | - | 0 | - |
| - | - | 850.2 | 1274 | - | - | 0 | - |
| - | - | 2111 | 1300 | - | - | 0 | - |
| - | - | 849.4 | 1301 | - | - | 0 | - |
| - | - | 1211 | 1316 | - | - | 0 | - |
| - | - | 2917 | 1317 | - | - | 0 | - |
| - | - | 1176 | 1318 | - | - | 0 | - |
| - | - | 2498 | 1350 | - | - | 0 | - |
| - | - | 2174 | 1351 | - | - | 0 | - |
| - | - | 923.9 | 1352 | - | - | 0 | - |
| - | - | 2284 | 1406 | - | - | 0 | - |
| - | - | 3271 | 1407 | - | - | 0 | - |
| - | - | 2402 | 1408 | - | - | 0 | - |
| - | - | 980.1 | 1409 | - | - | 0 | - |
| - | - | 3852 | 1421 | - | - | 0 | - |
| - | - | 2523 | 1422 | - | - | 0 | - |
| - | - | 1899 | 1423 | - | - | 0 | - |
| - | - | 1158 | 1433 | - | - | 0 | - |
| - | - | 2415 | 1448 | - | - | 0 | - |
| - | - | 1.322E+04 | 1449 | - | - | 0 | - |
| - | - | 1.107E+04 | 1450 | - | - | 0 | - |
| - | - | 5656 | 1451 | - | - | 0 | - |
| - | - | 2691 | 1452 | - | - | 0 | - |
| - | - | 3190 | 1465 | - | - | 0 | - |
| - | - | 1.513E+04 | 1466 | - | - | 0 | - |
| - | - | 1.456E+04 | 1467 | - | - | 0 | - |
| - | - | 6610 | 1468 | - | - | 0 | - |
| - | - | 2207 | 1469 | - | - | 0 | - |
| - | - | 1205 | 1470 | - | - | 0 | - |

m/z Charge Intensity FragmentType MassShift Position
120.06575775146484 0 9080.106 y 9
120.0810317993164 0 14849.098
121.08451080322266 0 900.604
128.0821533203125 0 553.38605
129.10256958007812 0 5353.609
129.53421020507812 0 384.0204
133.086181640625 0 2672.475
134.0450439453125 0 534.2445
136.0759735107422 0 20201.559
136.7425537109375 0 508.05814
149.32876586914062 0 410.63797
153.90077209472656 0 486.82776
166.0865020751953 0 891.4305
171.1494903564453 0 72527.95
172.15289306640625 0 7385.2515
173.4404296875 0 2987.5806
175.09646606445312 0 648.4135
177.10279846191406 0 1156.9637
177.11224365234375 0 1169.5748
181.8274688720703 0 492.644
184.52569580078125 0 488.53003
185.16505432128906 0 3912.285
199.14434814453125 0 72273.85
200.1478271484375 0 5822.1147
205.097412109375 0 6259.0713
205.10723876953125 0 1095.1503
213.16015625 0 2754.822
215.13926696777344 0 1804.3832 y Water loss 8
227.1751708984375 0 527.7658
230.1497344970703 0 952.18274
233.14968872070312 0 3177.4019 y 8
242.1501007080078 0 957.27466
244.12864685058594 0 959.71783
247.14434814453125 0 2865.381
251.10301208496094 0 589.03906
262.11895751953125 0 785.48804
269.11334228515625 0 4059.9392
271.1397705078125 0 860.8767
273.11895751953125 0 537.64465
276.1551513671875 0 2395.3184
297.10833740234375 0 3046.604
301.1913146972656 0 2837.062
302.1947326660156 0 1149.9922
304.165771484375 0 641.8144
309.9974365234375 0 592.2581
310.2128601074219 0 3819.8574
311.17169189453125 0 669.85205
315.2399597167969 0 1561.1921
317.1899108886719 0 1096.3746
325.15191650390625 0 815.7616
326.1833190917969 0 1055.3798
327.76544189453125 0 513.26416
329.1817626953125 0 739.4595
331.1867980957031 0 785.58466
334.1748352050781 0 551.3433
340.2105407714844 0 1061.5159
341.2189025878906 0 5306.3467
343.1983642578125 0 592.9702
343.7999572753906 0 512.4637
344.19329833984375 0 903.7686
345.1770935058594 0 870.31384
345.2260437011719 0 5258.708 z 7
346.2122497558594 0 4663.9004
346.2339172363281 0 9420.732
347.2144775390625 0 837.48083
347.2361755371094 0 1871.6959
357.2371826171875 0 4001.7776
358.2083740234375 0 1096.4453
358.2450866699219 0 24216.38
359.2153625488281 0 776.2369
360.2251281738281 0 684.46094
361.188720703125 0 1170.5542
361.24462890625 0 3234.1482 y 7
373.17181396484375 0 1616.3243
373.712646484375 0 1635.095 c Ammonia loss 7
375.18896484375 0 625.7235
381.71044921875 0 4337.31 y Water loss 2
382.1737060546875 0 577.9373
382.2115783691406 0 939.7177
383.17877197265625 0 835.2154
383.2288513183594 0 918.8876
384.2337951660156 0 670.4083
384.2608947753906 0 1403.5253
390.71630859375 0 37766.805 y 2
391.2181701660156 0 14214.247
391.2434997558594 0 937.70435
391.7191162109375 0 4694.792
392.1929626464844 0 642.4128
397.2444152832031 0 817.84686
400.2542419433594 0 734.87286
401.214111328125 0 1975.1913
402.177490234375 0 1436.2935
402.2716064453125 0 5502.5815
403.2337951660156 0 2680.6448
403.2740783691406 0 1076.0973
405.17437744140625 0 1849.8878
415.25390625 0 2489.5415
416.2588806152344 0 1330.82
420.18878173828125 0 2383.201
426.2596435546875 0 693.7387
428.2513122558594 0 2139.3284
429.2450256347656 0 776.5415 z Ammonia loss 6
432.7083435058594 0 884.32074
435.2099304199219 0 565.6386
436.218017578125 0 1453.833
437.21575927734375 0 1213.6095
439.26751708984375 0 696.79144
440.250732421875 0 2235.637 y 1
440.7528076171875 0 798.16534
442.2598876953125 0 1112.0892
443.2501525878906 0 1544.0823
444.2649230957031 0 7751.452
445.27410888671875 0 25782.953 y Ammonia loss 6
446.27001953125 0 3082.8188 z 6
447.2816467285156 0 10696.578
448.2850036621094 0 2433.0623
457.27825927734375 0 1735.0986
458.2240905761719 0 689.783
459.2937316894531 0 1960.4934
460.25653076171875 0 817.5687
460.29327392578125 0 679.8229
462.292724609375 0 1986.1677 y 6
470.26165771484375 0 2253.0442
471.29290771484375 0 2888.5273
472.2405700683594 0 1206.9081
472.3023376464844 0 1022.9167
476.2743835449219 0 2769.5154
477.28192138671875 0 4784.9297 c 4
478.2851867675781 0 1523.5957
479.2568054199219 0 707.51324
480.7789001464844 0 1042.3778
481.28082275390625 0 1904.9619
484.20343017578125 0 918.3156
486.2187805175781 0 1465.1512
488.2689514160156 0 1456.0819
488.3082580566406 0 1630.105
489.27374267578125 0 1002.8717
489.3103332519531 0 1222.1714
489.7851867675781 0 4073.3838
490.2298889160156 0 746.8662
490.28729248046875 0 4758.2656
490.3213195800781 0 891.1209
496.2871398925781 0 692.3014
497.27239990234375 0 784.0328
499.2793884277344 0 2523.7708
501.2315368652344 0 809.9076
503.2461853027344 0 1663.1965
503.2948913574219 0 5305.0713 z 5
504.2543640136719 0 60176.69
504.3030700683594 0 79736.11
505.2579345703125 0 15331.552
505.3061218261719 0 20577.77
506.259521484375 0 2416.331
506.30780029296875 0 2805.5393
508.2391052246094 0 737.85876
513.2902221679688 0 875.16315
514.2982177734375 0 6143.0063
515.3024291992188 0 782.1139
516.3142700195312 0 3434.8074
517.2665405273438 0 1572.28
517.3163452148438 0 1051.8038
518.3072509765625 0 3260.8887
519.3134155273438 0 6501.133 y 5
520.3153076171875 0 1682.4515
522.7761840820312 0 1043.0178
523.2807006835938 0 1136.7424
530.2724609375 0 1672.3236
533.296875 0 2658.5686
534.3024291992188 0 1944.8695 c 5
535.3085327148438 0 807.5958
536.2333984375 0 842.7791
543.2744140625 0 764.25
548.282958984375 0 7008.356
549.2859497070312 0 1707.6973
551.2872924804688 0 2263.7793
551.7903442382812 0 5323.8623
552.2916870117188 0 3460.1777
553.238525390625 0 1572.0433
554.2459106445312 0 876.10834
554.2940673828125 0 1874.4152
558.324462890625 0 1091.712 y Water loss 4
559.2973022460938 0 2655.4453
559.3414916992188 0 1168.4574
560.3161010742188 0 22185.164 z 4
561.2769775390625 0 2851.2188
561.3240966796875 0 64019.004
562.2776489257812 0 989.81366
562.3272094726562 0 17668.246
563.3302612304688 0 2346.7893
564.2805786132812 0 968.2501
571.3131103515625 0 950.5544
572.3038330078125 0 1827.0472
574.28173828125 0 2755.6543
575.2817993164062 0 955.5045
575.3275756835938 0 9337.198
576.3348999023438 0 21076.518 y 4
577.3379516601562 0 6008.0884
578.3417358398438 0 1153.2869
583.2703857421875 0 1205.3325
585.3216552734375 0 883.2681
588.3386840820312 0 773.49365
589.3308715820312 0 5186.599
598.2958984375 0 1375.9996
599.305908203125 0 2267.1394
600.3021850585938 0 3370.006 z Ammonia loss 3
603.3573608398438 0 2647.01
604.3599853515625 0 1111.5516
608.8126831054688 0 1213.2587
609.3151245117188 0 974.26117
614.3292846679688 0 1096.545
615.3428344726562 0 1586.8973 y Water loss 3
616.3312377929688 0 4620.543 y Ammonia loss 3
617.3297729492188 0 27287.43 c Water loss 6
617.8151245117188 0 899.6627
618.3445434570312 0 38081.086
619.3501586914062 0 11101.098
620.3534545898438 0 2346.359
632.34912109375 0 20801.148
632.4092407226562 0 814.3886
633.3567504882812 0 91240.125 y 3
634.3541259765625 0 33596.59
635.3516845703125 0 126493.65 c 6
636.3541259765625 0 44839.47
637.356689453125 0 7377.135
638.3604736328125 0 767.5969
640.2691650390625 0 1609.4415
641.2783813476562 0 12277.463
643.3551635742188 0 3747.6711
644.3580322265625 0 1788.2335
646.345947265625 0 1749.1218
647.270751953125 0 707.7457
647.3508911132812 0 857.3427
648.2713012695312 0 867.63824
649.8338012695312 0 1927.6611
650.33935546875 0 973.7246
658.3466186523438 0 5457.6333
658.8489379882812 0 5683.9365
659.3497924804688 0 2896.0547
660.3411254882812 0 844.06946
661.3668212890625 0 17913.367
662.3696899414062 0 6641.614
663.3805541992188 0 6464.619
664.3861083984375 0 2024.102
667.375732421875 0 1710.8428
668.3611450195312 0 672.0062
669.3740234375 0 828.0167
685.387939453125 0 7872.2446
686.408935546875 0 643.37354
687.40478515625 0 692.16473
692.335693359375 0 2567.6267
693.3360595703125 0 2582.3054
694.3374633789062 0 925.0827
695.3693237304688 0 2474.6458
696.35498046875 0 2428.8704
697.3649291992188 0 3700.35
698.37060546875 0 1271.9961
702.3564453125 0 1043.6445
702.4132080078125 0 7838.4424
702.8672485351562 0 2173.9695
703.3590087890625 0 1099.4774
703.4185180664062 0 1130.4397
703.8648071289062 0 1253.5841
707.316650390625 0 841.52435
708.3464965820312 0 1816.5952
709.3426513671875 0 798.6695
710.3681030273438 0 960.98376
711.3665771484375 0 1414.4841
711.8662719726562 0 1187.4983
712.3717651367188 0 1109.4813
713.3821411132812 0 12111.041
715.3994750976562 0 887.40393
717.3515625 0 705.7533
718.3043823242188 0 945.8789
718.4234008789062 0 1502.1667
719.4309692382812 0 1973.9441
720.438720703125 0 774.1336
721.350341796875 0 863.86707
724.364013671875 0 4002.7285
724.8656616210938 0 4324.073
725.3666381835938 0 1750.8438
728.407470703125 0 1358.9153
729.4159545898438 0 6912.0527
730.4102783203125 0 28241.219
732.3726806640625 0 4463.7524
732.8746948242188 0 5130.0825
733.3737182617188 0 2391.989
733.87744140625 0 899.4582
735.3062744140625 0 2963.2952
745.4217529296875 0 2190.476 c Water loss 7
746.4161987304688 0 8658.22 c Ammonia loss 7
747.4210815429688 0 4019.1973
748.4076538085938 0 708.6084
751.3245239257812 0 1342.0536
753.3174438476562 0 6199.5684
761.3588256835938 0 4686.6567
761.4361572265625 0 1430.2933
762.4161376953125 0 3993.0632 y Water loss 2
763.4461059570312 0 157570.23 c 7
764.3970947265625 0 17346.459 z 2
764.4495849609375 0 52086.312
765.40771484375 0 12977.516
765.45947265625 0 7445.139
766.4131469726562 0 3861.6755
769.3353271484375 0 3132.8381
779.4019165039062 0 1042.9524
780.4249877929688 0 133809.34 y 2
781.4280395507812 0 56263.246
782.4307861328125 0 15249.164
788.4292602539062 0 833.23425
789.44384765625 0 4676.8613
790.4478149414062 0 1186.8641
793.3949584960938 0 1114.9001
804.3690795898438 0 1406.4539
807.4099731445312 0 1724.5293
808.4111938476562 0 1426.8918
818.5072021484375 0 755.6961
823.3475341796875 0 6439.315
830.44140625 0 905.77637
831.4389038085938 0 1416.5063
832.4342041015625 0 1055.888
832.516357421875 0 17135.57
833.5195922851562 0 7440.735
834.5230102539062 0 2287.8816
841.4852294921875 0 960.4671
845.4558715820312 0 772.9339 z Water loss 1
848.381591796875 0 945.8905
848.464111328125 0 931.73975 w 1
849.4631958007812 0 3156.1575
858.4314575195312 0 1024.7053
859.4155883789062 0 925.3012
859.5036010742188 0 8579.786 c Ammonia loss 8
860.4246215820312 0 3894.054
860.5082397460938 0 4389.678
861.516357421875 0 4953.4404
862.5194702148438 0 1872.9773
863.474853515625 0 17607.928 z 1
864.4769287109375 0 9066.781
865.478515625 0 14935.191
866.4009399414062 0 15013.518
873.4862060546875 0 959.33307
874.5098266601562 0 870.81903
876.5302124023438 0 160022.31 c 8
877.532958984375 0 81551.9
878.431396484375 0 758.06055
878.5363159179688 0 24032.236
879.4103393554688 0 1040.0786
879.4989624023438 0 4457.1055 y 1
880.497314453125 0 1958.6061
881.4125366210938 0 2604.3638
889.437255859375 0 1112.8185
891.4644165039062 0 1458.5548
893.4728393554688 0 50867.312
906.4820556640625 0 2492.204
907.4835815429688 0 1661.1941
908.4830932617188 0 942.62335
909.4066772460938 0 6596.1226
916.48046875 0 639.9682
917.494873046875 0 7089.3735
918.4873657226562 0 7712.521
919.5245971679688 0 1891.8627
923.5071411132812 0 7942.6743
924.5112915039062 0 3151.2651
925.5062255859375 0 832.4603
933.5612182617188 0 1488.0073
934.5664672851562 0 1035.2754
936.4905395507812 0 9889.934
937.4766845703125 0 5456.106
943.5083618164062 0 1804.8076
944.5236206054688 0 1969.4164
945.5186767578125 0 1019.681
947.5184326171875 0 1830.4343
948.5234375 0 1109.3214
951.5751953125 0 1758.8759
952.5757446289062 0 1202.1115
953.4942016601562 0 798.48486
954.4971313476562 0 1004.2082
960.5297241210938 0 1954.1769
961.5485229492188 0 3484.1677
962.5431518554688 0 200164.22
963.5459594726562 0 106215.35
964.5485229492188 0 35819.84
965.4685668945312 0 8405.939
965.5564575195312 0 2842.3772
977.4542846679688 0 846.42773
977.552734375 0 3724.346
978.561279296875 0 103105.234
979.5675659179688 0 136399.53
980.5706787109375 0 61099.062
981.4857788085938 0 18264.95
981.5787963867188 0 12619.332
982.494873046875 0 16553.14
985.5043334960938 0 724.23145
1001.5372314453125 0 1236.6571
1002.5470581054688 0 3855.583
1003.552001953125 0 1770.8975
1018.497314453125 0 7678.6606
1019.5004272460938 0 4640.281
1020.50048828125 0 1899.3297
1044.546875 0 996.45935
1045.5516357421875 0 1970.5938
1046.5489501953125 0 1143.9333
1085.5582275390625 0 1043.9597
1086.5679931640625 0 853.76135
1101.5616455078125 0 1046.2921
1102.5726318359375 0 7461.2134
1103.5772705078125 0 6144.333
1104.5802001953125 0 1493.1544
1105.5313720703125 0 4646.4175
1106.5328369140625 0 4086.1743
1107.5343017578125 0 2049.5242
1217.62890625 0 1100.4252
1218.60888671875 0 781.5592
1233.6171875 0 894.3788
1234.6297607421875 0 5490.47
1235.6337890625 0 4769.1816
1236.6348876953125 0 1628.1774
1271.6683349609375 0 923.4492
1272.670166015625 0 1388.0752
1273.6746826171875 0 850.17847
1299.667724609375 0 2111.0947
1300.66650390625 0 849.4116
1315.673828125 0 1211.4203
1316.68896484375 0 2917.4578
1317.6845703125 0 1176.1388
1349.654296875 0 2497.7285
1350.6580810546875 0 2174.3523
1351.662353515625 0 923.9304
1405.717041015625 0 2284.4827
1406.716064453125 0 3271.1052
1407.7216796875 0 2402.4119
1408.7186279296875 0 980.1265
1420.7265625 0 3851.9568
1421.7344970703125 0 2522.6306
1422.732421875 0 1898.7045
1432.68701171875 0 1158.1857
1447.7305908203125 0 2415.342
1448.72607421875 0 13216.535
1449.7294921875 0 11072.405
1450.72998046875 0 5656.231
1451.7327880859375 0 2690.769
1464.7403564453125 0 3189.831
1465.75 0 15128.996
1466.75341796875 0 14560.854
1467.755615234375 0 6609.98
1468.7584228515625 0 2207.0173
1469.760986328125 0 1204.5457

Spectrum Details

|  |  |
| --- | --- |
| Matched peaks? Matched peaks The total absolute number of peaks matched. Additionally in brackets the total fraction of peaks matched and the total number of peaks is shown. | 38 (8.52% of 446) |
| FDR? FDR The false discovery rate estimated for this peptide. It is calculated by matching all theoretical fragments with a non-integer shift with the raw peaks for this spectrum. This is done with 40 different shifts. The resulting percentage is the average number of annotated peaks over the number of annotated peaks with the correct spectrum. | 1.00% |
| Satellite FDR? Satellite FDR See the FDR for details on its calculation. This satellite ion specific FDR only contains the satellite ions (d/w) for I/L/J positions. | - |
| PSM Score? PSM Score The PSM Score as given by Hecklib to this annotated spectrum. It is shown with three significant figures. | 482 |

#### Spectrum 6098? Spectrum 6098 The raw spectrum of this peptide as annotated by Hecklib. The fragments are coloured according to ion type (see legend). Any peaks with a star '\*' as text can be hovered over to see the full details, first the ion type second the mass shift type. By hovering over the amino acids in the peptide or ions in the legend the corresponding peaks are highlighted. By toggling the 'Unassigned' label you can turn the background (unassigned) peaks on or off in the plot. By updating the slider in the Ion legend you can update the spectrum to only show the top X% of the peaks with labels. The top X% means any peak that is within X% of the highest intensity. By dragging in the spectrum you can zoom in to a specific part of the spectrum and use 'Zoom Out' to get back to the original zoom level. The annotation of the spectrum is based on the given sequence in the peptides file and is done with different software so inconsistencies are likely. The peaks are annotated based on the given sequence, with 20 ppm tolerance.

Copy Data

##### Spectrum 6098 (TSV)

###### Preview

```
Loading example...
```

*Click on the button to copy the data to your clipboard.*

Mz MinMz MaxIntensity Max

WidthHeightPeptide font sizePeptide stroke widthSpectrum font sizeSpectrum stroke widthCompact peptide

Ion legend

wxyz

abcd

OtherUnassignedIon^Charge^_Position_Show for top:%

VVFGGGTKJT

01.31e+42.61e+43.92e+45.22e+4

Zoom Out

y^+1^_1_y^+1^_2_y^+1^_2_c^+2^_5_z^+1^_3_y^+1^_3_y^+2^_8_w^+1^_4_y^+1^_4_z^+1^_4_c^+1^_5_z^+1^_5_y^+1^_5_c^+1^_6_z^+1^_6_y^+1^_6_y^+1^_7_c^+1^_7_y^+1^_7_c^+1^_7_c^+1^_8_y^+1^_8_c^+1^_8_z^+1^_8_y^+1^_8_c^+1^_9_z^+1^_9_c^+1^_9_

0695138920842779

Fragment Matches Table

Show background peaks

| Position | Ion type | Intensity | mz Theoretical | mz Error (Th) | mz Error (ppm) | Charge | Series Number |
| --- | --- | --- | --- | --- | --- | --- | --- |
| 10 | y | 1882 | 120.1 | 0.0001466 | 1.221 | +1 | 1 |
| - | - | 3433 | 120.1 | - | - | 0 | - |
| - | - | 398.1 | 123.9 | - | - | 0 | - |
| - | - | 1018 | 129.1 | - | - | 0 | - |
| - | - | 470.9 | 130.1 | - | - | 0 | - |
| - | - | 492.5 | 131 | - | - | 0 | - |
| - | - | 707.3 | 131.1 | - | - | 0 | - |
| - | - | 6426 | 133.1 | - | - | 0 | - |
| - | - | 433.6 | 134.4 | - | - | 0 | - |
| - | - | 424.9 | 134.6 | - | - | 0 | - |
| - | - | 417.1 | 135.5 | - | - | 0 | - |
| - | - | 843.6 | 136.1 | - | - | 0 | - |
| - | - | 871.8 | 147.1 | - | - | 0 | - |
| - | - | 559.2 | 157.4 | - | - | 0 | - |
| - | - | 470 | 158.3 | - | - | 0 | - |
| - | - | 603.8 | 161.1 | - | - | 0 | - |
| - | - | 470.3 | 162.8 | - | - | 0 | - |
| - | - | 1.722E+04 | 171.1 | - | - | 0 | - |
| - | - | 1444 | 172.2 | - | - | 0 | - |
| - | - | 1294 | 175.1 | - | - | 0 | - |
| - | - | 899.5 | 177.1 | - | - | 0 | - |
| - | - | 4554 | 177.1 | - | - | 0 | - |
| - | - | 625.8 | 179.1 | - | - | 0 | - |
| - | - | 508.4 | 194.2 | - | - | 0 | - |
| - | - | 1.702E+04 | 199.1 | - | - | 0 | - |
| - | - | 1659 | 200.1 | - | - | 0 | - |
| - | - | 643 | 201.1 | - | - | 0 | - |
| - | - | 1216 | 205.1 | - | - | 0 | - |
| - | - | 2963 | 205.1 | - | - | 0 | - |
| 9 | y | 965.3 | 215.1 | 0.0004159 | 1.933 | +1 | 2 |
| - | - | 780.5 | 221.1 | - | - | 0 | - |
| - | - | 1372 | 223.1 | - | - | 0 | - |
| - | - | 515.1 | 230.2 | - | - | 0 | - |
| 9 | y | 805.8 | 233.1 | 6.272E-05 | 0.269 | +1 | 2 |
| 5 | c | 1419 | 239.1 | 0.00458 | 19.15 | +2 | 5 |
| - | - | 773 | 249.1 | - | - | 0 | - |
| - | - | 583.9 | 265.2 | - | - | 0 | - |
| - | - | 1066 | 267.1 | - | - | 0 | - |
| - | - | 1183 | 268.1 | - | - | 0 | - |
| - | - | 548.9 | 275.9 | - | - | 0 | - |
| - | - | 3238 | 276.2 | - | - | 0 | - |
| - | - | 876.8 | 283.2 | - | - | 0 | - |
| - | - | 544.2 | 293.2 | - | - | 0 | - |
| - | - | 678 | 301.2 | - | - | 0 | - |
| - | - | 4350 | 310.2 | - | - | 0 | - |
| - | - | 654.3 | 311.2 | - | - | 0 | - |
| - | - | 895.4 | 312.2 | - | - | 0 | - |
| - | - | 725.5 | 326.2 | - | - | 0 | - |
| - | - | 1260 | 327.2 | - | - | 0 | - |
| - | - | 871.9 | 328.2 | - | - | 0 | - |
| 8 | z | 957.3 | 345.2 | 0.0005723 | 1.658 | +1 | 3 |
| - | - | 701.8 | 346.2 | - | - | 0 | - |
| - | - | 1723 | 346.2 | - | - | 0 | - |
| - | - | 722.2 | 351.2 | - | - | 0 | - |
| - | - | 794.8 | 358.2 | - | - | 0 | - |
| 8 | y | 691.1 | 361.2 | 0.0003264 | 0.9036 | +1 | 3 |
| - | - | 1113 | 371.2 | - | - | 0 | - |
| - | - | 568.2 | 373.2 | - | - | 0 | - |
| - | - | 576.8 | 389.6 | - | - | 0 | - |
| 3 | y | 9363 | 390.7 | 8.886E-05 | 0.2274 | +2 | 8 |
| - | - | 4151 | 391.2 | - | - | 0 | - |
| - | - | 798.9 | 391.7 | - | - | 0 | - |
| - | - | 648.4 | 399.2 | - | - | 0 | - |
| - | - | 573 | 401.2 | - | - | 0 | - |
| - | - | 859.6 | 402.3 | - | - | 0 | - |
| - | - | 9869 | 415.3 | - | - | 0 | - |
| - | - | 1490 | 416.3 | - | - | 0 | - |
| - | - | 734.3 | 420.2 | - | - | 0 | - |
| - | - | 547.3 | 429.2 | - | - | 0 | - |
| 7 | w | 1591 | 429.3 | 0.0012 | 2.794 | +1 | 4 |
| - | - | 9673 | 443.2 | - | - | 0 | - |
| - | - | 2469 | 444.3 | - | - | 0 | - |
| 7 | y | 2712 | 445.3 | 0.0006308 | 1.417 | +1 | 4 |
| 7 | z | 1162 | 446.3 | 0.005831 | 13.07 | +1 | 4 |
| - | - | 3431 | 447.3 | - | - | 0 | - |
| - | - | 680.2 | 448.3 | - | - | 0 | - |
| - | - | 1411 | 457.2 | - | - | 0 | - |
| - | - | 790 | 459.3 | - | - | 0 | - |
| - | - | 627.4 | 476.3 | - | - | 0 | - |
| 5 | c | 1437 | 477.3 | 0.0002623 | 0.5496 | +1 | 5 |
| - | - | 8350 | 488.3 | - | - | 0 | - |
| - | - | 3470 | 489.3 | - | - | 0 | - |
| - | - | 995.3 | 490.2 | - | - | 0 | - |
| - | - | 3674 | 490.3 | - | - | 0 | - |
| - | - | 617.9 | 491.3 | - | - | 0 | - |
| - | - | 632.1 | 499.3 | - | - | 0 | - |
| 6 | z | 897.7 | 503.3 | 0.0005922 | 1.177 | +1 | 5 |
| - | - | 1.576E+04 | 504.3 | - | - | 0 | - |
| - | - | 2.328E+04 | 504.3 | - | - | 0 | - |
| - | - | 4485 | 505.3 | - | - | 0 | - |
| - | - | 5459 | 505.3 | - | - | 0 | - |
| - | - | 682.5 | 506.3 | - | - | 0 | - |
| - | - | 893.9 | 506.3 | - | - | 0 | - |
| - | - | 807.3 | 514.3 | - | - | 0 | - |
| - | - | 636.6 | 516.3 | - | - | 0 | - |
| 6 | y | 1751 | 519.3 | 0.0003981 | 0.7666 | +1 | 5 |
| - | - | 987.6 | 533.3 | - | - | 0 | - |
| 6 | c | 1152 | 534.3 | 0.002311 | 4.325 | +1 | 6 |
| - | - | 1472 | 548.3 | - | - | 0 | - |
| - | - | 674.6 | 553.3 | - | - | 0 | - |
| 5 | z | 4948 | 560.3 | 0.0002219 | 0.3961 | +1 | 6 |
| - | - | 1058 | 561.3 | - | - | 0 | - |
| - | - | 1.572E+04 | 561.3 | - | - | 0 | - |
| - | - | 825.2 | 561.4 | - | - | 0 | - |
| - | - | 4593 | 562.3 | - | - | 0 | - |
| - | - | 999.9 | 563.3 | - | - | 0 | - |
| - | - | 933.7 | 574.3 | - | - | 0 | - |
| - | - | 1643 | 575.3 | - | - | 0 | - |
| 5 | y | 4719 | 576.3 | 0.001168 | 2.027 | +1 | 6 |
| - | - | 1407 | 577.3 | - | - | 0 | - |
| 4 | y | 873.9 | 616.3 | 0.004833 | 7.841 | +1 | 7 |
| 7 | c | 3057 | 617.3 | 0.004085 | 6.618 | +1 | 7 |
| - | - | 9121 | 618.3 | - | - | 0 | - |
| - | - | 3330 | 619.4 | - | - | 0 | - |
| - | - | 5672 | 632.3 | - | - | 0 | - |
| 4 | y | 2.332E+04 | 633.4 | 0.0009644 | 1.523 | +1 | 7 |
| - | - | 9227 | 634.4 | - | - | 0 | - |
| 7 | c | 3.384E+04 | 635.4 | 0.0001848 | 0.2909 | +1 | 7 |
| - | - | 9145 | 636.4 | - | - | 0 | - |
| - | - | 2523 | 637.4 | - | - | 0 | - |
| - | - | 851.5 | 643.4 | - | - | 0 | - |
| - | - | 3442 | 661.4 | - | - | 0 | - |
| - | - | 1466 | 662.4 | - | - | 0 | - |
| - | - | 1964 | 663.4 | - | - | 0 | - |
| - | - | 828.2 | 702.4 | - | - | 0 | - |
| - | - | 951.8 | 719.4 | - | - | 0 | - |
| - | - | 1207 | 729.4 | - | - | 0 | - |
| - | - | 961.2 | 730.4 | - | - | 0 | - |
| 8 | c | 1860 | 746.4 | 0.008357 | 11.2 | +1 | 8 |
| - | - | 689.4 | 747.4 | - | - | 0 | - |
| 3 | y | 1179 | 762.4 | 0.0009398 | 1.233 | +1 | 8 |
| 8 | c | 3.919E+04 | 763.4 | 0.0009095 | 1.191 | +1 | 8 |
| 3 | z | 4616 | 764.4 | 0.009456 | 12.37 | +1 | 8 |
| - | - | 1.133E+04 | 764.4 | - | - | 0 | - |
| - | - | 2829 | 765.4 | - | - | 0 | - |
| - | - | 1938 | 765.5 | - | - | 0 | - |
| - | - | 1284 | 766.4 | - | - | 0 | - |
| 3 | y | 2.631E+04 | 780.4 | 0.0008969 | 1.149 | +1 | 8 |
| - | - | 1.397E+04 | 781.4 | - | - | 0 | - |
| - | - | 3713 | 782.4 | - | - | 0 | - |
| - | - | 3618 | 832.5 | - | - | 0 | - |
| - | - | 1399 | 833.5 | - | - | 0 | - |
| 9 | c | 1967 | 859.5 | 0.001173 | 1.365 | +1 | 9 |
| - | - | 771.3 | 860.5 | - | - | 0 | - |
| - | - | 807.1 | 861.5 | - | - | 0 | - |
| 2 | z | 5268 | 863.5 | 0.0007438 | 0.8615 | +1 | 9 |
| - | - | 2236 | 864.5 | - | - | 0 | - |
| - | - | 633.5 | 865.5 | - | - | 0 | - |
| 9 | c | 4.159E+04 | 876.5 | 0.0009281 | 1.059 | +1 | 9 |
| - | - | 1.892E+04 | 877.5 | - | - | 0 | - |
| - | - | 6268 | 878.5 | - | - | 0 | - |
| - | - | 1696 | 893.5 | - | - | 0 | - |
| - | - | 1141 | 923.5 | - | - | 0 | - |
| - | - | 664.1 | 924.5 | - | - | 0 | - |
| - | - | 644.8 | 933.6 | - | - | 0 | - |
| - | - | 964.8 | 947.5 | - | - | 0 | - |
| - | - | 962.2 | 961.6 | - | - | 0 | - |
| - | - | 5.17E+04 | 962.5 | - | - | 0 | - |
| - | - | 2.742E+04 | 963.5 | - | - | 0 | - |
| - | - | 7497 | 964.5 | - | - | 0 | - |
| - | - | 751 | 965.5 | - | - | 0 | - |
| - | - | 835.2 | 977.6 | - | - | 0 | - |
| - | - | 2.433E+04 | 978.6 | - | - | 0 | - |
| - | - | 3.229E+04 | 979.6 | - | - | 0 | - |
| - | - | 1.347E+04 | 980.6 | - | - | 0 | - |
| - | - | 3712 | 981.6 | - | - | 0 | - |
| - | - | 723.6 | 1449 | - | - | 0 | - |
| - | - | 643.2 | 1465 | - | - | 0 | - |
| - | - | 713.5 | 2734 | - | - | 0 | - |
| - | - | 725.9 | 2751 | - | - | 0 | - |

m/z Charge Intensity FragmentType MassShift Position
120.06566619873047 0 1881.5671 y 9
120.0809097290039 0 3433.14
123.85073852539062 0 398.09702
129.1024169921875 0 1017.8876
130.14080810546875 0 470.87317
131.04515075683594 0 492.48004
131.07061767578125 0 707.32916
133.0860137939453 0 6426.4116
134.42396545410156 0 433.558
134.5571746826172 0 424.85385
135.46534729003906 0 417.13907
136.0755615234375 0 843.6326
147.10177612304688 0 871.8253
157.40853881835938 0 559.15497
158.30435180664062 0 469.9799
161.08099365234375 0 603.8087
162.78292846679688 0 470.3106
171.14932250976562 0 17216.85
172.1524658203125 0 1443.6716
175.0963592529297 0 1293.9373
177.10391235351562 0 899.4859
177.11224365234375 0 4554.0034
179.09149169921875 0 625.8225
194.19064331054688 0 508.40207
199.1440887451172 0 17015.953
200.14772033691406 0 1659.4855
201.12286376953125 0 643.04205
205.09710693359375 0 1216.4049
205.10720825195312 0 2962.5156
215.13943481445312 0 965.3008 y Water loss 8
221.1384735107422 0 780.4521
223.11758422851562 0 1371.808
230.1510467529297 0 515.13995
233.14952087402344 0 805.75555 y 8
239.1492156982422 0 1419.0879 c 4
249.13311767578125 0 772.9603
265.1655578613281 0 583.8836
267.1431884765625 0 1066.0054
268.1383056640625 0 1182.684
275.9171447753906 0 548.9352
276.1552734375 0 3237.6675
283.17425537109375 0 876.7548
293.16168212890625 0 544.2142
301.19140625 0 677.9585
310.21240234375 0 4349.866
311.1702880859375 0 654.2979
312.16522216796875 0 895.369
326.17852783203125 0 725.53613
327.2018737792969 0 1260.3611
328.2235107421875 0 871.93097
345.2252502441406 0 957.25055 z 7
346.2108459472656 0 701.7505
346.2324523925781 0 1723.3818
351.20318603515625 0 722.1754
358.2447814941406 0 794.76373
361.244873046875 0 691.12915 y 7
371.227294921875 0 1113.1241
373.23046875 0 568.2323
389.5953674316406 0 576.7642
390.716064453125 0 9362.794 y 2
391.2179870605469 0 4151.2227
391.7198181152344 0 798.8915
399.22503662109375 0 648.4446
401.214599609375 0 572.97327
402.2718505859375 0 859.5518
415.25408935546875 0 9868.655
416.2557678222656 0 1490.4874
420.1872253417969 0 734.28595
429.23516845703125 0 547.302
429.2695617675781 0 1590.6769 w 6
443.2486877441406 0 9672.727
444.2548522949219 0 2468.5464
445.2650451660156 0 2712.2163 y Ammonia loss 6
446.2676696777344 0 1162.186 z 6
447.2809143066406 0 3430.5474
448.2857971191406 0 680.24365
457.2268371582031 0 1411.4922
459.2911376953125 0 790.0358
476.2722473144531 0 627.3851
477.2822570800781 0 1437.0192 c 4
488.3066711425781 0 8349.686
489.3097839355469 0 3469.5215
490.24932861328125 0 995.3397
490.286865234375 0 3673.9272
491.2871398925781 0 617.9305
499.2796630859375 0 632.12427
503.29437255859375 0 897.6719 z 5
504.2540283203125 0 15761.606
504.3025207519531 0 23284.13
505.25732421875 0 4485.363
505.3060302734375 0 5458.6177
506.2593688964844 0 682.52386
506.30950927734375 0 893.908
514.2977905273438 0 807.2929
516.310791015625 0 636.597
519.3140869140625 0 1750.7878 y 5
533.2911987304688 0 987.60236
534.3011474609375 0 1151.7587 c 5
548.282470703125 0 1471.9568
553.2817993164062 0 674.6028
560.316650390625 0 4948.395 z 4
561.2767333984375 0 1058.4077
561.32373046875 0 15724.267
561.3727416992188 0 825.1518
562.3261108398438 0 4593.4272
563.3298950195312 0 999.8839
574.2813720703125 0 933.67957
575.3256225585938 0 1643.0371
576.333984375 0 4719.318 y 4
577.3350830078125 0 1406.7319
616.3348999023438 0 873.85754 y Ammonia loss 3
617.3364868164062 0 3057.1675 c Water loss 6
618.3438720703125 0 9120.582
619.3508911132812 0 3330.256
632.3485107421875 0 5672.067
633.3556518554688 0 23318.264 y 3
634.3526000976562 0 9227.112
635.3509521484375 0 33841.555 c 6
636.3530883789062 0 9144.87
637.35595703125 0 2522.9858
643.3522338867188 0 851.5285
661.3650512695312 0 3441.9724
662.3677368164062 0 1466.3706
663.379638671875 0 1964.3435
702.405517578125 0 828.15656
719.4296875 0 951.7594
729.4168090820312 0 1207.221
730.4185791015625 0 961.16095
746.4111938476562 0 1859.7947 c Ammonia loss 7
747.4189453125 0 689.4225
762.4154052734375 0 1179.4115 y Water loss 2
763.4451904296875 0 39185.895 c 7
764.3968505859375 0 4616.219 z 2
764.449462890625 0 11331.524
765.405517578125 0 2829.4727
765.4567260742188 0 1938.4338
766.4129028320312 0 1283.5192
780.4241333007812 0 26306.584 y 2
781.4274291992188 0 13971.69
782.4295654296875 0 3712.9185
832.515380859375 0 3617.57
833.51953125 0 1399.0415
859.50244140625 0 1967.072 c Ammonia loss 8
860.5055541992188 0 771.31696
861.5166015625 0 807.1157
863.4754638671875 0 5268.197 z 1
864.4765625 0 2235.559
865.4781494140625 0 633.5317
876.5292358398438 0 41585.863 c 8
877.5319213867188 0 18923.707
878.5342407226562 0 6268.3496
893.4714965820312 0 1696.2721
923.504638671875 0 1140.5049
924.5042114257812 0 664.0752
933.5657348632812 0 644.79944
947.5253295898438 0 964.79443
961.55029296875 0 962.20685
962.5418090820312 0 51696.066
963.54443359375 0 27418.904
964.5474243164062 0 7496.5874
965.5460815429688 0 751.03436
977.5545654296875 0 835.1697
978.559814453125 0 24326.705
979.566162109375 0 32292.584
980.5695190429688 0 13472.483
981.5733642578125 0 3712.0488
1448.7490234375 0 723.594
1464.7880859375 0 643.20807
2733.854736328125 0 713.51904
2751.30615234375 0 725.8977

Spectrum Details

|  |  |
| --- | --- |
| Matched peaks? Matched peaks The total absolute number of peaks matched. Additionally in brackets the total fraction of peaks matched and the total number of peaks is shown. | 28 (16.47% of 170) |
| FDR? FDR The false discovery rate estimated for this peptide. It is calculated by matching all theoretical fragments with a non-integer shift with the raw peaks for this spectrum. This is done with 40 different shifts. The resulting percentage is the average number of annotated peaks over the number of annotated peaks with the correct spectrum. | 0.00% |
| Satellite FDR? Satellite FDR See the FDR for details on its calculation. This satellite ion specific FDR only contains the satellite ions (d/w) for I/L/J positions. | - |
| PSM Score? PSM Score The PSM Score as given by Hecklib to this annotated spectrum. It is shown with three significant figures. | 328 |

#### Spectrum 7033? Spectrum 7033 The raw spectrum of this peptide as annotated by Hecklib. The fragments are coloured according to ion type (see legend). Any peaks with a star '\*' as text can be hovered over to see the full details, first the ion type second the mass shift type. By hovering over the amino acids in the peptide or ions in the legend the corresponding peaks are highlighted. By toggling the 'Unassigned' label you can turn the background (unassigned) peaks on or off in the plot. By updating the slider in the Ion legend you can update the spectrum to only show the top X% of the peaks with labels. The top X% means any peak that is within X% of the highest intensity. By dragging in the spectrum you can zoom in to a specific part of the spectrum and use 'Zoom Out' to get back to the original zoom level. The annotation of the spectrum is based on the given sequence in the peptides file and is done with different software so inconsistencies are likely. The peaks are annotated based on the given sequence, with 20 ppm tolerance.

Copy Data

##### Spectrum 7033 (TSV)

###### Preview

```
Loading example...
```

*Click on the button to copy the data to your clipboard.*

Mz MinMz MaxIntensity Max

WidthHeightPeptide font sizePeptide stroke widthSpectrum font sizeSpectrum stroke widthCompact peptide

Ion legend

wxyz

abcd

OtherUnassignedIon^Charge^_Position_Show for top:%

VVFGGGTKJT

09.89e+31.98e+42.97e+43.95e+4

Zoom Out

y^+1^_1_a^+1^_2_b^+1^_2_y^+1^_2_y^+1^_2_y^+1^_3_y^+1^_3_y^+2^_8_y^+1^_4_y^+1^_5_y^+1^_6_y^+1^_6_y^+1^_7_y^+1^_7_b^+1^_8_y^+1^_8_y^+1^_8_b^+1^_9_y^+1^_9_

0566113116972263

Fragment Matches Table

Show background peaks

| Position | Ion type | Intensity | mz Theoretical | mz Error (Th) | mz Error (ppm) | Charge | Series Number |
| --- | --- | --- | --- | --- | --- | --- | --- |
| 10 | y | 2850 | 120.1 | 0.0002305 | 1.92 | +1 | 1 |
| - | - | 2.484E+04 | 120.1 | - | - | 0 | - |
| - | - | 1768 | 121.1 | - | - | 0 | - |
| - | - | 394.4 | 121.1 | - | - | 0 | - |
| - | - | 379.7 | 121.4 | - | - | 0 | - |
| - | - | 387.4 | 124.1 | - | - | 0 | - |
| - | - | 353 | 126.7 | - | - | 0 | - |
| - | - | 544 | 127.1 | - | - | 0 | - |
| - | - | 1445 | 127.1 | - | - | 0 | - |
| - | - | 363.5 | 128.9 | - | - | 0 | - |
| - | - | 1.748E+04 | 129.1 | - | - | 0 | - |
| - | - | 523.3 | 130.1 | - | - | 0 | - |
| - | - | 1245 | 130.1 | - | - | 0 | - |
| - | - | 979.3 | 131 | - | - | 0 | - |
| - | - | 653.1 | 131.1 | - | - | 0 | - |
| - | - | 471.4 | 131.1 | - | - | 0 | - |
| - | - | 970.9 | 133.1 | - | - | 0 | - |
| - | - | 2057 | 133.1 | - | - | 0 | - |
| - | - | 418.3 | 134.1 | - | - | 0 | - |
| - | - | 6835 | 136.1 | - | - | 0 | - |
| - | - | 483.6 | 139.1 | - | - | 0 | - |
| - | - | 460.2 | 140.1 | - | - | 0 | - |
| - | - | 537.8 | 142.1 | - | - | 0 | - |
| - | - | 667.9 | 142.1 | - | - | 0 | - |
| - | - | 1153 | 146.1 | - | - | 0 | - |
| - | - | 640.7 | 149 | - | - | 0 | - |
| - | - | 646.1 | 152.1 | - | - | 0 | - |
| - | - | 422.1 | 152.1 | - | - | 0 | - |
| - | - | 483.2 | 153.1 | - | - | 0 | - |
| - | - | 477.3 | 153.1 | - | - | 0 | - |
| - | - | 672.3 | 155.1 | - | - | 0 | - |
| - | - | 802.7 | 156.1 | - | - | 0 | - |
| - | - | 775.9 | 157.1 | - | - | 0 | - |
| - | - | 474.6 | 157.1 | - | - | 0 | - |
| - | - | 712 | 158.1 | - | - | 0 | - |
| - | - | 720.4 | 159.1 | - | - | 0 | - |
| - | - | 852.4 | 162.1 | - | - | 0 | - |
| - | - | 1306 | 163.1 | - | - | 0 | - |
| - | - | 603.8 | 166.1 | - | - | 0 | - |
| - | - | 578.9 | 166.1 | - | - | 0 | - |
| - | - | 606.6 | 167.1 | - | - | 0 | - |
| - | - | 954.2 | 170.1 | - | - | 0 | - |
| 2 | a | 3.915E+04 | 171.1 | 0.0002549 | 1.489 | +1 | 2 |
| - | - | 775.8 | 172.1 | - | - | 0 | - |
| - | - | 3211 | 172.2 | - | - | 0 | - |
| - | - | 549.2 | 173.1 | - | - | 0 | - |
| - | - | 1566 | 173.1 | - | - | 0 | - |
| - | - | 861.7 | 175.1 | - | - | 0 | - |
| - | - | 551.8 | 175.1 | - | - | 0 | - |
| - | - | 957.4 | 176.1 | - | - | 0 | - |
| - | - | 2040 | 177.1 | - | - | 0 | - |
| - | - | 770.4 | 177.1 | - | - | 0 | - |
| - | - | 503.8 | 177.4 | - | - | 0 | - |
| - | - | 478.9 | 181 | - | - | 0 | - |
| - | - | 1708 | 181.2 | - | - | 0 | - |
| - | - | 522.7 | 183.1 | - | - | 0 | - |
| - | - | 498.5 | 185.1 | - | - | 0 | - |
| - | - | 521.1 | 186 | - | - | 0 | - |
| - | - | 1912 | 187.1 | - | - | 0 | - |
| - | - | 682.9 | 190.1 | - | - | 0 | - |
| - | - | 601.5 | 191.1 | - | - | 0 | - |
| - | - | 1812 | 197.1 | - | - | 0 | - |
| - | - | 977.3 | 197.2 | - | - | 0 | - |
| - | - | 1098 | 198.1 | - | - | 0 | - |
| 2 | b | 1.666E+04 | 199.1 | 0.0001523 | 0.7648 | +1 | 2 |
| - | - | 2014 | 200.1 | - | - | 0 | - |
| - | - | 1826 | 201.1 | - | - | 0 | - |
| - | - | 3662 | 205.1 | - | - | 0 | - |
| - | - | 972.2 | 207.1 | - | - | 0 | - |
| - | - | 627.5 | 208.1 | - | - | 0 | - |
| - | - | 1086 | 209.2 | - | - | 0 | - |
| - | - | 1495 | 212.1 | - | - | 0 | - |
| - | - | 782.4 | 213.1 | - | - | 0 | - |
| 9 | y | 2570 | 215.1 | 0.0002938 | 1.366 | +1 | 2 |
| - | - | 469.6 | 215.5 | - | - | 0 | - |
| - | - | 961.6 | 216.1 | - | - | 0 | - |
| - | - | 618.2 | 217.1 | - | - | 0 | - |
| - | - | 1391 | 219.1 | - | - | 0 | - |
| - | - | 593.1 | 221.1 | - | - | 0 | - |
| - | - | 705.4 | 223.6 | - | - | 0 | - |
| - | - | 818.5 | 227.1 | - | - | 0 | - |
| - | - | 2923 | 230.2 | - | - | 0 | - |
| 9 | y | 953.7 | 233.1 | 0.0001085 | 0.4654 | +1 | 2 |
| - | - | 1414 | 237.2 | - | - | 0 | - |
| - | - | 693.7 | 239.2 | - | - | 0 | - |
| - | - | 825.2 | 245.1 | - | - | 0 | - |
| - | - | 1744 | 255.1 | - | - | 0 | - |
| - | - | 2257 | 269.2 | - | - | 0 | - |
| - | - | 1375 | 273.1 | - | - | 0 | - |
| - | - | 647.6 | 274.1 | - | - | 0 | - |
| - | - | 2367 | 276.2 | - | - | 0 | - |
| - | - | 1093 | 301.2 | - | - | 0 | - |
| - | - | 639.6 | 304.2 | - | - | 0 | - |
| - | - | 2437 | 310.2 | - | - | 0 | - |
| - | - | 939.4 | 319.1 | - | - | 0 | - |
| - | - | 671.7 | 324.1 | - | - | 0 | - |
| - | - | 1810 | 326.2 | - | - | 0 | - |
| 8 | y | 711.1 | 343.2 | 0.0008508 | 2.479 | +1 | 3 |
| - | - | 835.9 | 357.2 | - | - | 0 | - |
| 8 | y | 2368 | 361.2 | 3.978E-05 | 0.1101 | +1 | 3 |
| - | - | 540.7 | 371.2 | - | - | 0 | - |
| - | - | 577.9 | 374.2 | - | - | 0 | - |
| - | - | 824.4 | 382.2 | - | - | 0 | - |
| - | - | 1938 | 383.2 | - | - | 0 | - |
| 3 | y | 5038 | 390.7 | 0.0004299 | 1.1 | +2 | 8 |
| - | - | 2099 | 391.2 | - | - | 0 | - |
| - | - | 1128 | 391.7 | - | - | 0 | - |
| - | - | 634.8 | 392.2 | - | - | 0 | - |
| - | - | 1133 | 401.2 | - | - | 0 | - |
| - | - | 1756 | 402.2 | - | - | 0 | - |
| - | - | 614.3 | 415.3 | - | - | 0 | - |
| - | - | 964.7 | 419 | - | - | 0 | - |
| - | - | 688.1 | 420 | - | - | 0 | - |
| - | - | 1094 | 420.2 | - | - | 0 | - |
| - | - | 720.7 | 421.2 | - | - | 0 | - |
| - | - | 1080 | 439.3 | - | - | 0 | - |
| - | - | 1061 | 453.2 | - | - | 0 | - |
| - | - | 1428 | 457.3 | - | - | 0 | - |
| - | - | 574.4 | 462.1 | - | - | 0 | - |
| 7 | y | 587.6 | 462.3 | 0.001545 | 3.342 | +1 | 4 |
| - | - | 747 | 473.3 | - | - | 0 | - |
| - | - | 837.7 | 486.3 | - | - | 0 | - |
| - | - | 576.6 | 489.3 | - | - | 0 | - |
| - | - | 758.1 | 490.3 | - | - | 0 | - |
| - | - | 2057 | 496.3 | - | - | 0 | - |
| - | - | 734.4 | 497.3 | - | - | 0 | - |
| - | - | 3264 | 514.3 | - | - | 0 | - |
| - | - | 626.2 | 515.3 | - | - | 0 | - |
| 6 | y | 1136 | 519.3 | 0.0004591 | 0.8841 | +1 | 5 |
| - | - | 993.3 | 530.3 | - | - | 0 | - |
| - | - | 1872 | 548.3 | - | - | 0 | - |
| - | - | 1232 | 549.3 | - | - | 0 | - |
| - | - | 584.5 | 558.3 | - | - | 0 | - |
| 5 | y | 895.4 | 558.3 | 0.001584 | 2.837 | +1 | 6 |
| 5 | y | 4235 | 576.3 | 0.0007409 | 1.286 | +1 | 6 |
| - | - | 1006 | 577.3 | - | - | 0 | - |
| 4 | y | 614.6 | 615.3 | 0.001727 | 2.806 | +1 | 7 |
| 4 | y | 1.131E+04 | 633.4 | 0.0008056 | 1.272 | +1 | 7 |
| - | - | 2718 | 634.4 | - | - | 0 | - |
| - | - | 768.9 | 635.4 | - | - | 0 | - |
| - | - | 3333 | 643.4 | - | - | 0 | - |
| - | - | 1300 | 644.4 | - | - | 0 | - |
| - | - | 667.7 | 651.9 | - | - | 0 | - |
| - | - | 6969 | 661.4 | - | - | 0 | - |
| - | - | 2455 | 662.4 | - | - | 0 | - |
| 8 | b | 904.5 | 746.4 | 0.007441 | 9.97 | +1 | 8 |
| 3 | y | 795.8 | 762.4 | 0.002222 | 2.914 | +1 | 8 |
| 3 | y | 2.345E+04 | 780.4 | 0.0005917 | 0.7582 | +1 | 8 |
| - | - | 1.005E+04 | 781.4 | - | - | 0 | - |
| - | - | 2451 | 782.4 | - | - | 0 | - |
| 9 | b | 755 | 859.5 | 0.002977 | 3.464 | +1 | 9 |
| - | - | 1070 | 860.5 | - | - | 0 | - |
| 2 | y | 1064 | 879.5 | 0.004247 | 4.829 | +1 | 9 |
| - | - | 566.5 | 1326 | - | - | 0 | - |
| - | - | 636.2 | 1347 | - | - | 0 | - |
| - | - | 612.6 | 1529 | - | - | 0 | - |
| - | - | 578.6 | 1807 | - | - | 0 | - |
| - | - | 632.2 | 2149 | - | - | 0 | - |
| - | - | 629.5 | 2240 | - | - | 0 | - |

m/z Charge Intensity FragmentType MassShift Position
120.06575012207031 0 2850.4937 y 9
120.0810317993164 0 24843.29
121.08454132080078 0 1768.262
121.10989379882812 0 394.3656
121.39811706542969 0 379.74988
124.08724212646484 0 387.44324
126.70362091064453 0 352.9714
127.05065155029297 0 543.97296
127.07574462890625 0 1444.6123
128.88128662109375 0 363.52335
129.1024932861328 0 17479.79
130.05043029785156 0 523.34564
130.10572814941406 0 1245.2063
131.04513549804688 0 979.32367
131.0702362060547 0 653.0547
131.08155822753906 0 471.406
133.0607452392578 0 970.89874
133.08619689941406 0 2056.764
134.06007385253906 0 418.2815
136.07594299316406 0 6835.258
139.0504913330078 0 483.64105
140.0823516845703 0 460.18222
142.09811401367188 0 537.8216
142.122802734375 0 667.8821
146.0601043701172 0 1152.98
149.02285766601562 0 640.68085
152.07106018066406 0 646.059
152.10787963867188 0 422.14728
153.05474853515625 0 483.2272
153.07748413085938 0 477.34875
155.11793518066406 0 672.34906
156.07708740234375 0 802.6822
157.08657836914062 0 775.8549
157.09719848632812 0 474.56033
158.09231567382812 0 712.0245
159.0766143798828 0 720.40625
162.0550537109375 0 852.446
163.0715789794922 0 1306.2258
166.05348205566406 0 603.75183
166.0863037109375 0 578.85236
167.08180236816406 0 606.64
170.09275817871094 0 954.2375
171.14944458007812 0 39152.945 a 1
172.07211303710938 0 775.7998
172.1528778076172 0 3211.329
173.0917205810547 0 549.1669
173.1283416748047 0 1565.5425
175.0876007080078 0 861.6613
175.0966033935547 0 551.8082
176.10714721679688 0 957.41235
177.10238647460938 0 2039.5464
177.111083984375 0 770.41003
177.3594512939453 0 503.80066
180.95579528808594 0 478.90714
181.1703643798828 0 1707.9376
183.11314392089844 0 522.727
185.0950927734375 0 498.49762
185.99551391601562 0 521.11145
187.14434814453125 0 1911.6384
190.05072021484375 0 682.8725
191.08233642578125 0 601.5436
197.1287078857422 0 1811.7262
197.16525268554688 0 977.2508
198.08729553222656 0 1097.6387
199.14425659179688 0 16655.717 b 1
200.14756774902344 0 2013.75
201.1236114501953 0 1826.148
205.097412109375 0 3662.4614
207.07655334472656 0 972.15643
208.10816955566406 0 627.45447
209.16510009765625 0 1085.5724
212.1397705078125 0 1494.6511
213.08750915527344 0 782.3556
215.13931274414062 0 2570.2651 y Water loss 8
215.45626831054688 0 469.6472
216.0976104736328 0 961.63214
217.0975341796875 0 618.2278
219.1490478515625 0 1391.1262
221.1271514892578 0 593.10944
223.5814666748047 0 705.4199
227.11456298828125 0 818.4741
230.15005493164062 0 2922.6025
233.14947509765625 0 953.71625 y 8
237.1604461669922 0 1413.9093
239.15098571777344 0 693.6939
245.12477111816406 0 825.2203
255.10928344726562 0 1743.9756
269.16107177734375 0 2257.3972
273.1191711425781 0 1375.433
274.1183776855469 0 647.593
276.1556701660156 0 2367.3083
301.1915283203125 0 1093.3391
304.16510009765625 0 639.56647
310.2128601074219 0 2436.7886
319.14007568359375 0 939.3982
324.11822509765625 0 671.69037
326.18304443359375 0 1809.5803
343.2348327636719 0 711.0584 y Water loss 7
357.1554870605469 0 835.9466
361.2445068359375 0 2367.8167 y 7
371.23162841796875 0 540.7273
374.178955078125 0 577.90137
382.2115478515625 0 824.418
383.2037658691406 0 1938.0732
390.7165832519531 0 5037.693 y 2
391.2174987792969 0 2098.651
391.7193908691406 0 1128.1953
392.1922302246094 0 634.76636
401.2146911621094 0 1132.8815
402.1770324707031 0 1756.4495
415.2588195800781 0 614.29504
418.996337890625 0 964.6844
419.9939270019531 0 688.11035
420.188720703125 0 1094.251
421.19696044921875 0 720.7475
439.2657165527344 0 1079.8373
453.232421875 0 1060.6929
457.2764892578125 0 1428.345
462.14093017578125 0 574.39777
462.2906799316406 0 587.64307 y 6
473.2526550292969 0 746.97894
486.3044738769531 0 837.70886
489.28277587890625 0 576.57227
490.28851318359375 0 758.1222
496.2889709472656 0 2056.7876
497.2879638671875 0 734.3605
514.2988891601562 0 3264.0198
515.3009643554688 0 626.2324
519.3141479492188 0 1136.0552 y 5
530.2728881835938 0 993.3351
548.2826538085938 0 1871.8367
549.28564453125 0 1231.8851
558.268310546875 0 584.47864
558.326171875 0 895.42255 y Water loss 4
576.3344116210938 0 4235.372 y 4
577.3377075195312 0 1005.59875
615.3477783203125 0 614.599 y Water loss 3
633.357421875 0 11308.524 y 3
634.362060546875 0 2717.586
635.362548828125 0 768.86163
643.3563232421875 0 3332.5984
644.3583374023438 0 1300.3259
651.8742065429688 0 667.69653
661.3668823242188 0 6969.401
662.3697509765625 0 2455.0134
746.412109375 0 904.54254 b 7
762.4166870117188 0 795.7654 y Water loss 2
780.4244384765625 0 23451.967 y 2
781.4282836914062 0 10051.743
782.4302368164062 0 2450.7117
859.506591796875 0 755.04285 b 8
860.5059814453125 0 1069.8368
879.4891967773438 0 1063.641 y 1
1326.2442626953125 0 566.48346
1347.3359375 0 636.1686
1529.454833984375 0 612.5792
1807.0721435546875 0 578.57086
2148.682373046875 0 632.1973
2240.4853515625 0 629.49255

Spectrum Details

|  |  |
| --- | --- |
| Matched peaks? Matched peaks The total absolute number of peaks matched. Additionally in brackets the total fraction of peaks matched and the total number of peaks is shown. | 19 (11.95% of 159) |
| FDR? FDR The false discovery rate estimated for this peptide. It is calculated by matching all theoretical fragments with a non-integer shift with the raw peaks for this spectrum. This is done with 40 different shifts. The resulting percentage is the average number of annotated peaks over the number of annotated peaks with the correct spectrum. | 0.13% |
| Satellite FDR? Satellite FDR See the FDR for details on its calculation. This satellite ion specific FDR only contains the satellite ions (d/w) for I/L/J positions. | - |
| PSM Score? PSM Score The PSM Score as given by Hecklib to this annotated spectrum. It is shown with three significant figures. | 243 |

#### Spectrum 6276? Spectrum 6276 The raw spectrum of this peptide as annotated by Hecklib. The fragments are coloured according to ion type (see legend). Any peaks with a star '\*' as text can be hovered over to see the full details, first the ion type second the mass shift type. By hovering over the amino acids in the peptide or ions in the legend the corresponding peaks are highlighted. By toggling the 'Unassigned' label you can turn the background (unassigned) peaks on or off in the plot. By updating the slider in the Ion legend you can update the spectrum to only show the top X% of the peaks with labels. The top X% means any peak that is within X% of the highest intensity. By dragging in the spectrum you can zoom in to a specific part of the spectrum and use 'Zoom Out' to get back to the original zoom level. The annotation of the spectrum is based on the given sequence in the peptides file and is done with different software so inconsistencies are likely. The peaks are annotated based on the given sequence, with 20 ppm tolerance.

Copy Data

##### Spectrum 6276 (TSV)

###### Preview

```
Loading example...
```

*Click on the button to copy the data to your clipboard.*

Mz MinMz MaxIntensity Max

WidthHeightPeptide font sizePeptide stroke widthSpectrum font sizeSpectrum stroke widthCompact peptide

Ion legend

wxyz

abcd

OtherUnassignedIon^Charge^_Position_Show for top:%

VVFGGGTKJT

01.74e+43.48e+45.23e+46.97e+4

Zoom Out

y^+1^_1_a^+1^_2_b^+1^_2_y^+1^_2_y^+1^_2_y^+1^_3_b^+1^_3_y^+1^_3_b^+2^_8_y^+2^_8_y^+2^_8_b^+1^_4_y^+2^_9_y^+1^_4_\*y^+1^_5_y^+1^_6_y^+1^_6_y^+1^_7_b^+1^_7_y^+1^_7_b^+1^_8_y^+1^_8_y^+1^_8_y^+1^_8_b^+1^_9_y^+1^_9_

0658131719752633

Fragment Matches Table

Show background peaks

| Position | Ion type | Intensity | mz Theoretical | mz Error (Th) | mz Error (ppm) | Charge | Series Number |
| --- | --- | --- | --- | --- | --- | --- | --- |
| 10 | y | 5263 | 120.1 | 0.0003373 | 2.809 | +1 | 1 |
| - | - | 4.113E+04 | 120.1 | - | - | 0 | - |
| - | - | 343.7 | 121.1 | - | - | 0 | - |
| - | - | 2723 | 121.1 | - | - | 0 | - |
| - | - | 427.9 | 122.6 | - | - | 0 | - |
| - | - | 394.5 | 124.1 | - | - | 0 | - |
| - | - | 405.6 | 126.1 | - | - | 0 | - |
| - | - | 949 | 127.1 | - | - | 0 | - |
| - | - | 620.8 | 127.1 | - | - | 0 | - |
| - | - | 1380 | 127.1 | - | - | 0 | - |
| - | - | 1343 | 128.1 | - | - | 0 | - |
| - | - | 2523 | 129.1 | - | - | 0 | - |
| - | - | 3.206E+04 | 129.1 | - | - | 0 | - |
| - | - | 775.8 | 130.1 | - | - | 0 | - |
| - | - | 485.9 | 130.1 | - | - | 0 | - |
| - | - | 1702 | 130.1 | - | - | 0 | - |
| - | - | 679.5 | 131 | - | - | 0 | - |
| - | - | 2179 | 131.1 | - | - | 0 | - |
| - | - | 1.321E+04 | 131.1 | - | - | 0 | - |
| - | - | 882.6 | 131.1 | - | - | 0 | - |
| - | - | 461.6 | 132.1 | - | - | 0 | - |
| - | - | 763.1 | 132.1 | - | - | 0 | - |
| - | - | 684 | 133.1 | - | - | 0 | - |
| - | - | 1.904E+04 | 133.1 | - | - | 0 | - |
| - | - | 769.2 | 134.1 | - | - | 0 | - |
| - | - | 5378 | 136.1 | - | - | 0 | - |
| - | - | 457.2 | 137.7 | - | - | 0 | - |
| - | - | 2004 | 141.1 | - | - | 0 | - |
| - | - | 788.1 | 141.1 | - | - | 0 | - |
| - | - | 389 | 142.8 | - | - | 0 | - |
| - | - | 458.2 | 143 | - | - | 0 | - |
| - | - | 714.2 | 143.1 | - | - | 0 | - |
| - | - | 933 | 143.1 | - | - | 0 | - |
| - | - | 1026 | 144.1 | - | - | 0 | - |
| - | - | 3402 | 145.1 | - | - | 0 | - |
| - | - | 946.6 | 145.1 | - | - | 0 | - |
| - | - | 735.3 | 146.1 | - | - | 0 | - |
| - | - | 6769 | 147 | - | - | 0 | - |
| - | - | 1013 | 147.1 | - | - | 0 | - |
| - | - | 1419 | 147.1 | - | - | 0 | - |
| - | - | 790.1 | 147.1 | - | - | 0 | - |
| - | - | 784.8 | 148 | - | - | 0 | - |
| - | - | 1017 | 149 | - | - | 0 | - |
| - | - | 478.6 | 149 | - | - | 0 | - |
| - | - | 546.1 | 152.1 | - | - | 0 | - |
| - | - | 813.2 | 153.1 | - | - | 0 | - |
| - | - | 554.6 | 154.1 | - | - | 0 | - |
| - | - | 3077 | 155.1 | - | - | 0 | - |
| - | - | 525.7 | 155.1 | - | - | 0 | - |
| - | - | 1975 | 155.1 | - | - | 0 | - |
| - | - | 569.5 | 157.1 | - | - | 0 | - |
| - | - | 442.7 | 157.9 | - | - | 0 | - |
| - | - | 818.8 | 158.1 | - | - | 0 | - |
| - | - | 9574 | 159.1 | - | - | 0 | - |
| - | - | 453.6 | 160.1 | - | - | 0 | - |
| - | - | 558.4 | 161.1 | - | - | 0 | - |
| - | - | 939.1 | 163 | - | - | 0 | - |
| - | - | 1997 | 163.1 | - | - | 0 | - |
| - | - | 458.6 | 163.6 | - | - | 0 | - |
| - | - | 468.3 | 164.2 | - | - | 0 | - |
| - | - | 591.9 | 167.1 | - | - | 0 | - |
| - | - | 661.2 | 169.1 | - | - | 0 | - |
| - | - | 630.7 | 170.1 | - | - | 0 | - |
| - | - | 2270 | 171.1 | - | - | 0 | - |
| 2 | a | 6.899E+04 | 171.1 | 0.0003922 | 2.292 | +1 | 2 |
| - | - | 2007 | 172.1 | - | - | 0 | - |
| - | - | 6937 | 172.2 | - | - | 0 | - |
| - | - | 485.4 | 173.1 | - | - | 0 | - |
| - | - | 463.1 | 173.1 | - | - | 0 | - |
| - | - | 1417 | 173.1 | - | - | 0 | - |
| - | - | 3065 | 173.4 | - | - | 0 | - |
| - | - | 448.7 | 175.1 | - | - | 0 | - |
| - | - | 683.4 | 175.1 | - | - | 0 | - |
| - | - | 2191 | 175.1 | - | - | 0 | - |
| - | - | 1430 | 176.1 | - | - | 0 | - |
| - | - | 3471 | 177.1 | - | - | 0 | - |
| - | - | 7925 | 177.1 | - | - | 0 | - |
| - | - | 738.1 | 178.1 | - | - | 0 | - |
| - | - | 530.1 | 178.1 | - | - | 0 | - |
| - | - | 567.2 | 179.1 | - | - | 0 | - |
| - | - | 6269 | 180.1 | - | - | 0 | - |
| - | - | 826.2 | 181.1 | - | - | 0 | - |
| - | - | 1.005E+04 | 181.1 | - | - | 0 | - |
| - | - | 734.4 | 181.2 | - | - | 0 | - |
| - | - | 661.7 | 182.1 | - | - | 0 | - |
| - | - | 654.8 | 183.1 | - | - | 0 | - |
| - | - | 810.3 | 184.1 | - | - | 0 | - |
| - | - | 1378 | 185.1 | - | - | 0 | - |
| - | - | 701.2 | 185.2 | - | - | 0 | - |
| - | - | 1923 | 187.1 | - | - | 0 | - |
| - | - | 649.6 | 188.1 | - | - | 0 | - |
| - | - | 3238 | 189.1 | - | - | 0 | - |
| - | - | 1993 | 191.1 | - | - | 0 | - |
| - | - | 806.4 | 191.1 | - | - | 0 | - |
| - | - | 4193 | 197.1 | - | - | 0 | - |
| - | - | 1647 | 197.1 | - | - | 0 | - |
| - | - | 1251 | 197.2 | - | - | 0 | - |
| - | - | 1605 | 198.1 | - | - | 0 | - |
| - | - | 541 | 198.1 | - | - | 0 | - |
| - | - | 514.3 | 198.1 | - | - | 0 | - |
| - | - | 504.7 | 199.1 | - | - | 0 | - |
| - | - | 609.3 | 199.1 | - | - | 0 | - |
| 2 | b | 3.577E+04 | 199.1 | 0.0003202 | 1.608 | +1 | 2 |
| - | - | 3750 | 200.1 | - | - | 0 | - |
| - | - | 5051 | 200.1 | - | - | 0 | - |
| - | - | 2173 | 201.1 | - | - | 0 | - |
| - | - | 1691 | 202.1 | - | - | 0 | - |
| - | - | 4545 | 204.1 | - | - | 0 | - |
| - | - | 6400 | 205.1 | - | - | 0 | - |
| - | - | 1953 | 205.1 | - | - | 0 | - |
| - | - | 623.2 | 206.1 | - | - | 0 | - |
| - | - | 2786 | 207.1 | - | - | 0 | - |
| - | - | 675.1 | 207.1 | - | - | 0 | - |
| - | - | 593.3 | 207.1 | - | - | 0 | - |
| - | - | 1.604E+04 | 208.1 | - | - | 0 | - |
| - | - | 1.333E+04 | 209.1 | - | - | 0 | - |
| - | - | 1909 | 209.1 | - | - | 0 | - |
| - | - | 764.5 | 209.2 | - | - | 0 | - |
| - | - | 1041 | 210.1 | - | - | 0 | - |
| - | - | 1454 | 210.1 | - | - | 0 | - |
| - | - | 683.1 | 212.1 | - | - | 0 | - |
| - | - | 3673 | 212.1 | - | - | 0 | - |
| - | - | 627.7 | 213.1 | - | - | 0 | - |
| - | - | 751.8 | 213.1 | - | - | 0 | - |
| - | - | 565 | 214.1 | - | - | 0 | - |
| - | - | 1131 | 215.1 | - | - | 0 | - |
| - | - | 588.2 | 215.1 | - | - | 0 | - |
| 9 | y | 3953 | 215.1 | 0.0002023 | 0.9402 | +1 | 2 |
| - | - | 1360 | 216.1 | - | - | 0 | - |
| - | - | 681.6 | 217.1 | - | - | 0 | - |
| - | - | 600.9 | 218.1 | - | - | 0 | - |
| - | - | 836 | 219.1 | - | - | 0 | - |
| - | - | 3599 | 219.1 | - | - | 0 | - |
| - | - | 967.3 | 220.1 | - | - | 0 | - |
| - | - | 1031 | 221.1 | - | - | 0 | - |
| - | - | 1939 | 221.1 | - | - | 0 | - |
| - | - | 1065 | 224.1 | - | - | 0 | - |
| - | - | 1028 | 224.2 | - | - | 0 | - |
| - | - | 3456 | 225.1 | - | - | 0 | - |
| - | - | 1082 | 226.1 | - | - | 0 | - |
| - | - | 1132 | 227.1 | - | - | 0 | - |
| - | - | 897.8 | 227.1 | - | - | 0 | - |
| - | - | 4155 | 228.1 | - | - | 0 | - |
| - | - | 1852 | 228.1 | - | - | 0 | - |
| - | - | 869.9 | 229.1 | - | - | 0 | - |
| - | - | 5617 | 230.2 | - | - | 0 | - |
| - | - | 789.2 | 233.1 | - | - | 0 | - |
| 9 | y | 1985 | 233.1 | 0.0004103 | 1.76 | +1 | 2 |
| - | - | 772.7 | 233.2 | - | - | 0 | - |
| - | - | 792.4 | 237.2 | - | - | 0 | - |
| - | - | 588.2 | 238.1 | - | - | 0 | - |
| - | - | 2395 | 239.1 | - | - | 0 | - |
| - | - | 631.2 | 241.1 | - | - | 0 | - |
| - | - | 768.2 | 241.1 | - | - | 0 | - |
| - | - | 1535 | 242.1 | - | - | 0 | - |
| - | - | 1683 | 242.2 | - | - | 0 | - |
| - | - | 645.4 | 243.1 | - | - | 0 | - |
| - | - | 525.8 | 243.2 | - | - | 0 | - |
| - | - | 719.1 | 245.1 | - | - | 0 | - |
| - | - | 534.3 | 245.2 | - | - | 0 | - |
| - | - | 6640 | 246.1 | - | - | 0 | - |
| - | - | 1617 | 247.1 | - | - | 0 | - |
| - | - | 805.6 | 255.1 | - | - | 0 | - |
| - | - | 3895 | 255.1 | - | - | 0 | - |
| - | - | 994.4 | 256.1 | - | - | 0 | - |
| - | - | 3726 | 256.1 | - | - | 0 | - |
| - | - | 889.8 | 257.2 | - | - | 0 | - |
| - | - | 2672 | 259.1 | - | - | 0 | - |
| - | - | 652.3 | 261.2 | - | - | 0 | - |
| - | - | 2411 | 262.1 | - | - | 0 | - |
| - | - | 2425 | 264.1 | - | - | 0 | - |
| - | - | 994.3 | 265.1 | - | - | 0 | - |
| - | - | 726.6 | 266.1 | - | - | 0 | - |
| - | - | 674.8 | 269.1 | - | - | 0 | - |
| - | - | 3211 | 269.2 | - | - | 0 | - |
| - | - | 787.4 | 270.1 | - | - | 0 | - |
| - | - | 640.3 | 271.2 | - | - | 0 | - |
| - | - | 3201 | 273.1 | - | - | 0 | - |
| - | - | 1020 | 274.1 | - | - | 0 | - |
| - | - | 1135 | 275.2 | - | - | 0 | - |
| - | - | 1942 | 276.2 | - | - | 0 | - |
| - | - | 758.1 | 279.2 | - | - | 0 | - |
| - | - | 955.7 | 283.1 | - | - | 0 | - |
| - | - | 1740 | 283.2 | - | - | 0 | - |
| - | - | 608.9 | 284.1 | - | - | 0 | - |
| - | - | 627.8 | 284.2 | - | - | 0 | - |
| - | - | 811.7 | 285.1 | - | - | 0 | - |
| - | - | 848.6 | 287.2 | - | - | 0 | - |
| - | - | 4633 | 292.2 | - | - | 0 | - |
| - | - | 1621 | 293.2 | - | - | 0 | - |
| - | - | 652.1 | 299.1 | - | - | 0 | - |
| - | - | 1881 | 301.2 | - | - | 0 | - |
| - | - | 840.4 | 304.2 | - | - | 0 | - |
| - | - | 663.1 | 306.2 | - | - | 0 | - |
| - | - | 2031 | 308.2 | - | - | 0 | - |
| - | - | 631.2 | 309.2 | - | - | 0 | - |
| - | - | 689.9 | 309.2 | - | - | 0 | - |
| - | - | 2820 | 310.2 | - | - | 0 | - |
| - | - | 844.8 | 313.2 | - | - | 0 | - |
| - | - | 638.4 | 324.2 | - | - | 0 | - |
| - | - | 514.2 | 324.6 | - | - | 0 | - |
| - | - | 926.7 | 325.2 | - | - | 0 | - |
| - | - | 636 | 325.2 | - | - | 0 | - |
| - | - | 3914 | 326.2 | - | - | 0 | - |
| - | - | 819.5 | 327.2 | - | - | 0 | - |
| - | - | 990.7 | 330.2 | - | - | 0 | - |
| - | - | 1222 | 331.2 | - | - | 0 | - |
| - | - | 2711 | 338.1 | - | - | 0 | - |
| - | - | 2184 | 339.2 | - | - | 0 | - |
| - | - | 577.8 | 341.2 | - | - | 0 | - |
| - | - | 6702 | 343.2 | - | - | 0 | - |
| 8 | y | 1415 | 343.2 | 0.0004919 | 1.433 | +1 | 3 |
| - | - | 1133 | 344.2 | - | - | 0 | - |
| - | - | 3142 | 344.2 | - | - | 0 | - |
| 3 | b | 1592 | 346.2 | 0.0002504 | 0.7231 | +1 | 3 |
| - | - | 1039 | 348.2 | - | - | 0 | - |
| - | - | 1051 | 348.2 | - | - | 0 | - |
| - | - | 1830 | 349.2 | - | - | 0 | - |
| - | - | 967.5 | 354.2 | - | - | 0 | - |
| - | - | 640.7 | 355.2 | - | - | 0 | - |
| - | - | 1039 | 357.2 | - | - | 0 | - |
| - | - | 564.8 | 358.2 | - | - | 0 | - |
| - | - | 6417 | 361.2 | - | - | 0 | - |
| 8 | y | 4128 | 361.2 | 0.0001128 | 0.3123 | +1 | 3 |
| - | - | 1924 | 362.2 | - | - | 0 | - |
| - | - | 658.4 | 362.2 | - | - | 0 | - |
| 8 | b | 694.6 | 365.2 | 0.005986 | 16.39 | +2 | 8 |
| - | - | 1535 | 366.2 | - | - | 0 | - |
| - | - | 1.16E+04 | 367.2 | - | - | 0 | - |
| - | - | 2492 | 368.2 | - | - | 0 | - |
| - | - | 496.4 | 368.2 | - | - | 0 | - |
| - | - | 524 | 370.1 | - | - | 0 | - |
| - | - | 1989 | 372.2 | - | - | 0 | - |
| - | - | 1432 | 374.2 | - | - | 0 | - |
| - | - | 1118 | 375.2 | - | - | 0 | - |
| - | - | 870.7 | 375.2 | - | - | 0 | - |
| 3 | y | 943 | 381.7 | 0.001714 | 4.492 | +2 | 8 |
| - | - | 2282 | 383.2 | - | - | 0 | - |
| - | - | 1595 | 384.2 | - | - | 0 | - |
| 3 | y | 1.086E+04 | 390.7 | 9.425E-05 | 0.2412 | +2 | 8 |
| - | - | 4103 | 391.2 | - | - | 0 | - |
| - | - | 1364 | 391.7 | - | - | 0 | - |
| - | - | 1600 | 392.2 | - | - | 0 | - |
| - | - | 1397 | 400.3 | - | - | 0 | - |
| - | - | 2862 | 401.2 | - | - | 0 | - |
| - | - | 4741 | 402.2 | - | - | 0 | - |
| 4 | b | 810.4 | 403.2 | 0.001156 | 2.867 | +1 | 4 |
| - | - | 590 | 415.2 | - | - | 0 | - |
| - | - | 3345 | 415.3 | - | - | 0 | - |
| - | - | 845.2 | 416.3 | - | - | 0 | - |
| - | - | 1038 | 418.2 | - | - | 0 | - |
| - | - | 867.1 | 419 | - | - | 0 | - |
| - | - | 2405 | 420.2 | - | - | 0 | - |
| - | - | 732.7 | 423.2 | - | - | 0 | - |
| - | - | 1149 | 425.2 | - | - | 0 | - |
| - | - | 973.6 | 429.2 | - | - | 0 | - |
| - | - | 620.6 | 429.3 | - | - | 0 | - |
| - | - | 4575 | 436.2 | - | - | 0 | - |
| - | - | 1837 | 439.3 | - | - | 0 | - |
| 2 | y | 861.8 | 440.3 | 0.0005129 | 1.165 | +2 | 9 |
| - | - | 965.3 | 443.2 | - | - | 0 | - |
| - | - | 831.4 | 443.3 | - | - | 0 | - |
| - | - | 1840 | 453.2 | - | - | 0 | - |
| - | - | 2787 | 454.2 | - | - | 0 | - |
| - | - | 3060 | 454.2 | - | - | 0 | - |
| - | - | 1154 | 455.2 | - | - | 0 | - |
| - | - | 2040 | 455.2 | - | - | 0 | - |
| - | - | 3141 | 457.3 | - | - | 0 | - |
| - | - | 864.8 | 458.3 | - | - | 0 | - |
| 7 | y | 986.6 | 462.3 | 0.002209 | 4.777 | +1 | 4 |
| - | - | 3.463E+04 | 471.2 | - | - | 0 | - |
| - | - | 7273 | 472.2 | - | - | 0 | - |
| - | - | 639.5 | 473.2 | - | - | 0 | - |
| - | - | 618 | 473.2 | - | - | 0 | - |
| - | - | 5850 | 473.3 | - | - | 0 | - |
| - | - | 1162 | 482.2 | - | - | 0 | - |
| - | - | 1549 | 486.3 | - | - | 0 | - |
| - | - | 702.2 | 488.2 | - | - | 0 | - |
| - | - | 1203 | 488.3 | - | - | 0 | - |
| - | - | 1.395E+04 | 489.2 | - | - | 0 | - |
| - | - | 711.3 | 489.3 | - | - | 0 | - |
| 0 | Precursor | 1043 | 489.8 | 0.0005096 | 1.04 | +2 | -1 |
| - | - | 3414 | 490.2 | - | - | 0 | - |
| - | - | 1394 | 490.3 | - | - | 0 | - |
| - | - | 1189 | 490.3 | - | - | 0 | - |
| - | - | 3166 | 496.3 | - | - | 0 | - |
| - | - | 794.3 | 497.3 | - | - | 0 | - |
| - | - | 8467 | 514.3 | - | - | 0 | - |
| - | - | 1711 | 515.3 | - | - | 0 | - |
| 6 | y | 1592 | 519.3 | 0.0004591 | 0.8841 | +1 | 5 |
| - | - | 3284 | 530.3 | - | - | 0 | - |
| - | - | 1264 | 531.3 | - | - | 0 | - |
| - | - | 6004 | 548.3 | - | - | 0 | - |
| - | - | 1488 | 549.3 | - | - | 0 | - |
| - | - | 863.3 | 550.3 | - | - | 0 | - |
| 5 | y | 784 | 558.3 | 0.00134 | 2.4 | +1 | 6 |
| 5 | y | 8755 | 576.3 | 6.953E-05 | 0.1206 | +1 | 6 |
| - | - | 3039 | 577.3 | - | - | 0 | - |
| 4 | y | 991 | 615.3 | 0.0002874 | 0.4671 | +1 | 7 |
| 7 | b | 1238 | 618.3 | 0.003537 | 5.721 | +1 | 7 |
| - | - | 830.4 | 625.3 | - | - | 0 | - |
| - | - | 729.9 | 632.8 | - | - | 0 | - |
| 4 | y | 2.36E+04 | 633.4 | 0.0006835 | 1.079 | +1 | 7 |
| - | - | 7402 | 634.4 | - | - | 0 | - |
| - | - | 1039 | 635.4 | - | - | 0 | - |
| - | - | 6593 | 643.4 | - | - | 0 | - |
| - | - | 1778 | 644.4 | - | - | 0 | - |
| - | - | 1.364E+04 | 661.4 | - | - | 0 | - |
| - | - | 5798 | 662.4 | - | - | 0 | - |
| - | - | 1443 | 663.4 | - | - | 0 | - |
| - | - | 764.8 | 718.4 | - | - | 0 | - |
| 8 | b | 1779 | 746.4 | 0.0001783 | 0.2389 | +1 | 8 |
| - | - | 950.5 | 747.4 | - | - | 0 | - |
| - | - | 1157 | 760.4 | - | - | 0 | - |
| 3 | y | 2037 | 762.4 | 0.001319 | 1.729 | +1 | 8 |
| 3 | y | 775.5 | 763.4 | 0.01168 | 15.29 | +1 | 8 |
| 3 | y | 4.907E+04 | 780.4 | 0.0006527 | 0.8364 | +1 | 8 |
| - | - | 2E+04 | 781.4 | - | - | 0 | - |
| - | - | 4598 | 782.4 | - | - | 0 | - |
| 9 | b | 3378 | 859.5 | 0.001357 | 1.578 | +1 | 9 |
| - | - | 1472 | 860.5 | - | - | 0 | - |
| - | - | 1147 | 861.5 | - | - | 0 | - |
| 2 | y | 2774 | 879.5 | 0.0004524 | 0.5144 | +1 | 9 |
| - | - | 1303 | 880.5 | - | - | 0 | - |
| - | - | 742 | 1010 | - | - | 0 | - |
| - | - | 599.9 | 1830 | - | - | 0 | - |
| - | - | 565.3 | 1897 | - | - | 0 | - |
| - | - | 661.7 | 1911 | - | - | 0 | - |
| - | - | 629.6 | 2028 | - | - | 0 | - |
| - | - | 694.6 | 2607 | - | - | 0 | - |

m/z Charge Intensity FragmentType MassShift Position
120.06585693359375 0 5262.97 y 9
120.08116149902344 0 41132.887
121.07987213134766 0 343.73547
121.08454895019531 0 2723.423
122.6209487915039 0 427.8947
124.07620239257812 0 394.5352
126.10271453857422 0 405.59338
127.05057525634766 0 949.02136
127.07527923583984 0 620.8442
127.0869140625 0 1380.0385
128.10752868652344 0 1343.2046
129.06617736816406 0 2522.943
129.10260009765625 0 32059.21
130.05027770996094 0 775.816
130.098876953125 0 485.924
130.10581970214844 0 1702.4008
131.03402709960938 0 679.4807
131.07052612304688 0 2179.4597
131.08187866210938 0 13212.714
131.08724975585938 0 882.6025
132.06541442871094 0 461.62714
132.08558654785156 0 763.1324
133.06117248535156 0 683.9545
133.0862579345703 0 19038.625
134.089599609375 0 769.20123
136.07603454589844 0 5378.1943
137.66082763671875 0 457.15173
141.066162109375 0 2003.5226
141.10308837890625 0 788.0898
142.77557373046875 0 389.01236
143.04576110839844 0 458.23312
143.08155822753906 0 714.17896
143.11827087402344 0 932.99
144.07723999023438 0 1025.7269
145.06114196777344 0 3401.5586
145.08644104003906 0 946.61304
146.0605010986328 0 735.3005
147.04443359375 0 6769.0977
147.06558227539062 0 1012.9857
147.07662963867188 0 1419.2614
147.1017608642578 0 790.0774
148.04776000976562 0 784.8126
148.95480346679688 0 1017.4845
149.02340698242188 0 478.57303
152.143798828125 0 546.1136
153.06605529785156 0 813.2269
154.0612335205078 0 554.55035
155.0818634033203 0 3077.47
155.09292602539062 0 525.73553
155.11822509765625 0 1974.6125
157.0615234375 0 569.50183
157.9469757080078 0 442.72018
158.0923309326172 0 818.8163
159.07676696777344 0 9574.011
160.07200622558594 0 453.61737
161.08155822753906 0 558.41614
163.0390167236328 0 939.1376
163.07159423828125 0 1997.3279
163.55128479003906 0 458.57983
164.21148681640625 0 468.30075
167.1186981201172 0 591.8877
169.134033203125 0 661.2499
170.09255981445312 0 630.74347
171.07684326171875 0 2270.0886
171.1495819091797 0 68988.54 a 1
172.0718994140625 0 2007.004
172.15292358398438 0 6937.228
173.05645751953125 0 485.3613
173.11868286132812 0 463.1459
173.12875366210938 0 1416.5497
173.4404296875 0 3064.999
175.0598907470703 0 448.7023
175.0880126953125 0 683.3924
175.09678649902344 0 2191.4482
176.1072998046875 0 1429.8334
177.10260009765625 0 3471.3147
177.11239624023438 0 7925
178.10606384277344 0 738.1103
178.11537170410156 0 530.0558
179.08164978027344 0 567.1664
180.1134033203125 0 6269.294
181.06100463867188 0 826.2256
181.09754943847656 0 10049.085
181.1700439453125 0 734.4007
182.1013641357422 0 661.68835
183.07728576660156 0 654.7544
184.07174682617188 0 810.28284
185.1289825439453 0 1378.4521
185.16531372070312 0 701.20557
187.1444549560547 0 1922.5719
188.103759765625 0 649.56085
189.08741760253906 0 3238.4368
191.08184814453125 0 1992.7852
191.09144592285156 0 806.41327
197.0924072265625 0 4193.041
197.1284942626953 0 1646.9156
197.1652069091797 0 1250.5146
198.08746337890625 0 1604.6316
198.09671020507812 0 540.97955
198.123779296875 0 514.34814
199.07183837890625 0 504.70398
199.10855102539062 0 609.25507
199.14442443847656 0 35768.73 b 1
200.10321044921875 0 3750.4424
200.14788818359375 0 5050.9453
201.12387084960938 0 2172.9348
202.08273315429688 0 1690.6896
204.10206604003906 0 4545.462
205.09738159179688 0 6399.7847
205.10714721679688 0 1952.5027
206.10098266601562 0 623.1733
207.09811401367188 0 2786.2588
207.12359619140625 0 675.0716
207.14987182617188 0 593.2803
208.10838317871094 0 16043.28
209.09243774414062 0 13330.4
209.11148071289062 0 1909.3104
209.16514587402344 0 764.5117
210.0867156982422 0 1040.7816
210.0962371826172 0 1453.6447
212.1027069091797 0 683.1485
212.1397247314453 0 3672.8438
213.12364196777344 0 627.67285
213.14341735839844 0 751.8204
214.08229064941406 0 565.0279
215.10333251953125 0 1130.8401
215.11434936523438 0 588.22516
215.13922119140625 0 3953.2883 y Water loss 8
216.0978546142578 0 1360.437
217.09823608398438 0 681.64777
218.0812530517578 0 600.9159
219.12277221679688 0 836.0366
219.14952087402344 0 3599.0918
220.09788513183594 0 967.3307
221.12802124023438 0 1031.313
221.13882446289062 0 1939.2241
224.103271484375 0 1064.5802
224.17544555664062 0 1028.035
225.0872344970703 0 3455.9133
226.1189727783203 0 1081.9355
227.1029052734375 0 1132.031
227.1142578125 0 897.78326
228.09820556640625 0 4154.7085
228.13450622558594 0 1852.1887
229.11849975585938 0 869.9401
230.1502227783203 0 5617.072
233.12844848632812 0 789.17633
233.14999389648438 0 1984.8212 y 8
233.1644744873047 0 772.71936
237.1597137451172 0 792.3893
238.0817108154297 0 588.18964
239.14947509765625 0 2395.1177
241.08177185058594 0 631.1581
241.09439086914062 0 768.2477
242.11375427246094 0 1535.4215
242.18687438964844 0 1682.7765
243.11251831054688 0 645.423
243.18994140625 0 525.75354
245.1250457763672 0 719.06366
245.166015625 0 534.28815
246.10897827148438 0 6640.1846
247.144775390625 0 1617.4965
255.0940704345703 0 805.63617
255.10922241210938 0 3895.0945
256.11431884765625 0 994.43805
256.12933349609375 0 3725.8955
257.16058349609375 0 889.8147
259.1042785644531 0 2672.2908
261.15911865234375 0 652.2705
262.11883544921875 0 2410.585
264.1192932128906 0 2424.6667
265.1291809082031 0 994.2528
266.1141357421875 0 726.628
269.1134338378906 0 674.77405
269.16119384765625 0 3211.0615
270.14447021484375 0 787.4462
271.2083740234375 0 640.33276
273.1196594238281 0 3200.626
274.1194763183594 0 1020.2845
275.1759338378906 0 1135.119
276.1559143066406 0 1942.3567
279.18231201171875 0 758.1178
283.1408386230469 0 955.73083
283.1753234863281 0 1739.9512
284.1227111816406 0 608.8616
284.1598815917969 0 627.8011
285.1197814941406 0 811.74854
287.17138671875 0 848.6324
292.202392578125 0 4633.2866
293.2059020996094 0 1620.9286
299.1332092285156 0 652.1003
301.1913146972656 0 1881.2828
304.16650390625 0 840.3894
306.1817321777344 0 663.0844
308.1977233886719 0 2030.7874
309.1540832519531 0 631.23206
309.2001037597656 0 689.9386
310.21295166015625 0 2820.2014
313.18670654296875 0 844.8291
324.1918029785156 0 638.43097
324.63092041015625 0 514.2161
325.150146484375 0 926.6827
325.2244567871094 0 635.9793
326.18212890625 0 3914.194
327.2023010253906 0 819.5171
330.1820068359375 0 990.6808
331.1867370605469 0 1221.7593
338.13543701171875 0 2711.4685
339.16778564453125 0 2184.2805
341.180419921875 0 577.7562
343.1615905761719 0 6702.485
343.2334899902344 0 1415.1572 y Water loss 7
344.16754150390625 0 1132.9752
344.1932678222656 0 3141.525
346.2127685546875 0 1592.1157 b 2
348.16766357421875 0 1039.2062
348.19189453125 0 1051.2803
349.1514892578125 0 1829.8336
354.1764831542969 0 967.54456
355.16168212890625 0 640.7087
357.1555480957031 0 1038.8237
358.20672607421875 0 564.7768
361.1718444824219 0 6417.213
361.2446594238281 0 4128.015 y 7
362.1744384765625 0 1923.8735
362.2446594238281 0 658.4145
365.19415283203125 0 694.63495 b Ammonia loss 7
366.1777038574219 0 1534.5364
367.1612854003906 0 11604.898
368.16412353515625 0 2491.7085
368.19036865234375 0 496.42322
370.1371154785156 0 523.9765
372.18804931640625 0 1988.6183
374.1839294433594 0 1432.3241
375.1663818359375 0 1117.7021
375.2412414550781 0 870.7472
381.71258544921875 0 943.0365 y Water loss 2
383.2039794921875 0 2281.6357
384.18792724609375 0 1595.2279
390.71624755859375 0 10859.145 y 2
391.2174987792969 0 4103.0933
391.7195739746094 0 1364.4615
392.1918640136719 0 1600.261
400.2551574707031 0 1397.2501
401.2150573730469 0 2861.584
402.1773376464844 0 4740.9463
403.2351379394531 0 810.4276 b 3
415.2277526855469 0 589.9569
415.25439453125 0 3344.6257
416.25738525390625 0 845.21295
418.1713562011719 0 1038.3132
418.99468994140625 0 867.1396
420.1881103515625 0 2405.1423
423.2231140136719 0 732.6563
425.2143249511719 0 1148.8701
429.2329406738281 0 973.5764
429.2817687988281 0 620.5851
436.182861328125 0 4575.3516
439.26678466796875 0 1837.093
440.2498474121094 0 861.79193 y 1
443.2212829589844 0 965.2733
443.25299072265625 0 831.3942
453.20947265625 0 1840.3534
454.1947326660156 0 2786.5107
454.233154296875 0 3059.63
455.1956481933594 0 1153.798
455.2364807128906 0 2039.5071
457.2781677246094 0 3140.8525
458.2825927734375 0 864.7811
462.29443359375 0 986.62286 y 6
471.2203369140625 0 34633.57
472.22216796875 0 7273.0137
473.2056579589844 0 639.4508
473.2273254394531 0 618.00275
473.2962951660156 0 5850.281
482.19073486328125 0 1161.7584
486.3039245605469 0 1549.0946
488.23712158203125 0 702.18726
488.2721252441406 0 1203.338
489.2307434082031 0 13951.556
489.2720642089844 0 711.32446
489.7840576171875 0 1043.4403 Precursor
490.2340393066406 0 3413.6565
490.28399658203125 0 1393.5864
490.32135009765625 0 1188.6099
496.2884216308594 0 3166.124
497.2920227050781 0 794.25854
514.298583984375 0 8467.423
515.301025390625 0 1711.4702
519.3141479492188 0 1591.6633 y 5
530.2721557617188 0 3284.3708
531.2725830078125 0 1263.7566
548.2824096679688 0 6004.1416
549.2864379882812 0 1488.2126
550.28759765625 0 863.32025
558.325927734375 0 783.99554 y Water loss 4
576.3350830078125 0 8754.883 y 4
577.3385620117188 0 3038.8894
615.3457641601562 0 991.0414 y Water loss 3
618.328125 0 1238.0144 b 6
625.3450927734375 0 830.4123
632.844482421875 0 729.94244
633.3572998046875 0 23595.78 y 3
634.3603515625 0 7402.2734
635.361083984375 0 1039.2571
643.3565673828125 0 6593.3203
644.3585205078125 0 1778.4441
661.3667602539062 0 13639.845
662.3698120117188 0 5798.1533
663.3707275390625 0 1443.2295
718.4251708984375 0 764.8396
746.4193725585938 0 1778.5547 b 7
747.4215698242188 0 950.4962
760.433837890625 0 1157.0846
762.4131469726562 0 2037.3413 y Water loss 2
763.41015625 0 775.4995 y Ammonia loss 2
780.4243774414062 0 49069.742 y 2
781.4273071289062 0 20000.293
782.4306640625 0 4597.907
859.5022583007812 0 3377.5708 b 8
860.5040283203125 0 1471.8486
861.5046997070312 0 1146.7672
879.493896484375 0 2773.8372 y 1
880.49560546875 0 1302.8984
1010.4130859375 0 741.9635
1829.66357421875 0 599.88995
1897.3707275390625 0 565.2602
1910.93701171875 0 661.7273
2027.5819091796875 0 629.5511
2607.359375 0 694.6452

Spectrum Details

|  |  |
| --- | --- |
| Matched peaks? Matched peaks The total absolute number of peaks matched. Additionally in brackets the total fraction of peaks matched and the total number of peaks is shown. | 27 (8.18% of 330) |
| FDR? FDR The false discovery rate estimated for this peptide. It is calculated by matching all theoretical fragments with a non-integer shift with the raw peaks for this spectrum. This is done with 40 different shifts. The resulting percentage is the average number of annotated peaks over the number of annotated peaks with the correct spectrum. | 0.79% |
| Satellite FDR? Satellite FDR See the FDR for details on its calculation. This satellite ion specific FDR only contains the satellite ions (d/w) for I/L/J positions. | - |
| PSM Score? PSM Score The PSM Score as given by Hecklib to this annotated spectrum. It is shown with three significant figures. | 321 |

#### Reverse Lookup? Reverse Lookup All places where this read could be placed.

| Group | Segment | Template | Template Part | Read Part | Score | Unique |
| --- | --- | --- | --- | --- | --- | --- |
| Homo sapiens Light Chain | IGLJ | IGLJ2 | [0..10] | [0..10] | 80 | True |

| Recombined | Template Part | Read Part | Score | Unique |
| --- | --- | --- | --- | --- |
| REC-0-1\_002 | [99..109] | [0..10] | 80 | True |

#### Meta Information from Multiple reads

##### Number of combined reads

16

##### Intensity

1

##### TotalArea

0

##### Changes to the peptide sequence

VVFGGGTKJT

L→JNo support for either Leucine or Isoleucine based on side chain ions (Position: 9)

#### Positional Score

Copy Data

##### Positional Score (TSV)

###### Preview

```
Loading example...
```

*Click on the button to copy the data to your clipboard.*

100123456789

Label Value
"0" 0.875
"1" 0.875
"2" 0.875
"3" 0.87
"4" 0.835
"5" 0.782
"6" 0.821
"7" 0.863
"8" 0.859
"9" 0.866

#### Meta Information from PEAKS

##### Scan Identifier

F2:5680

##### Original sequence

V

V

F

G

G

G

T

K

L

T

##### Posttranslational Modifications

##### Source File

D:\separate\_stitch\_analyses\xle-disambiguation\raw\20210323\_F1\_UM1\_Peng0013\_SA\_F59\_ingel\_3ug\_TL.raw

##### Fraction

2

##### Scan Feature

-

##### De Novo Score

99

##### ConfidenceScore

99

### m/z

489.7861

##### Mass

977.5546

##### Charge

2

##### Retention Time

30.95

##### Predicted Retention Time

-

##### Area

0

##### Parts Per Million

3.2

##### Fragmentation mode

HCD

##### Originating file

01 D:\separate\_stitch\_analyses\xle-disambiguation\20210325\_F59\_3ug\_DENOVO\_12.csv

#### Meta Information from PEAKS

##### Scan Identifier

F2:5744

##### Original sequence

V

V

F

G

G

G

T

K

L

T

##### Posttranslational Modifications

##### Source File

D:\separate\_stitch\_analyses\xle-disambiguation\raw\20210323\_F1\_UM1\_Peng0013\_SA\_F59\_ingel\_3ug\_TL.raw

##### Fraction

2

##### Scan Feature

-

##### De Novo Score

99

##### ConfidenceScore

99

### m/z

489.7856

##### Mass

977.5546

##### Charge

2

##### Retention Time

31.32

##### Predicted Retention Time

-

##### Area

0

##### Parts Per Million

2.2

##### Fragmentation mode

ETHCD

##### Originating file

01 D:\separate\_stitch\_analyses\xle-disambiguation\20210325\_F59\_3ug\_DENOVO\_12.csv

#### Meta Information from PEAKS

##### Scan Identifier

F2:6036

##### Original sequence

V

V

F

G

G

G

T

K

L

T

##### Posttranslational Modifications

##### Source File

D:\separate\_stitch\_analyses\xle-disambiguation\raw\20210323\_F1\_UM1\_Peng0013\_SA\_F59\_ingel\_3ug\_TL.raw

##### Fraction

2

##### Scan Feature

-

##### De Novo Score

98

##### ConfidenceScore

98

### m/z

489.7855

##### Mass

977.5546

##### Charge

2

##### Retention Time

33.05

##### Predicted Retention Time

-

##### Area

0

##### Parts Per Million

1.8

##### Fragmentation mode

HCD

##### Originating file

01 D:\separate\_stitch\_analyses\xle-disambiguation\20210325\_F59\_3ug\_DENOVO\_12.csv

#### Meta Information from PEAKS

##### Scan Identifier

F2:6156

##### Original sequence

V

V

F

G

G

G

T

K

L

T

##### Posttranslational Modifications

##### Source File

D:\separate\_stitch\_analyses\xle-disambiguation\raw\20210323\_F1\_UM1\_Peng0013\_SA\_F59\_ingel\_3ug\_TL.raw

##### Fraction

2

##### Scan Feature

-

##### De Novo Score

98

##### ConfidenceScore

98

### m/z

489.7853

##### Mass

977.5546

##### Charge

2

##### Retention Time

33.79

##### Predicted Retention Time

-

##### Area

0

##### Parts Per Million

1.5

##### Fragmentation mode

HCD

##### Originating file

01 D:\separate\_stitch\_analyses\xle-disambiguation\20210325\_F59\_3ug\_DENOVO\_12.csv

#### Meta Information from PEAKS

##### Scan Identifier

F2:5917

##### Original sequence

V

V

F

G

G

G

T

K

L

T

##### Posttranslational Modifications

##### Source File

D:\separate\_stitch\_analyses\xle-disambiguation\raw\20210323\_F1\_UM1\_Peng0013\_SA\_F59\_ingel\_3ug\_TL.raw

##### Fraction

2

##### Scan Feature

-

##### De Novo Score

98

##### ConfidenceScore

98

### m/z

489.7851

##### Mass

977.5546

##### Charge

2

##### Retention Time

32.35

##### Predicted Retention Time

-

##### Area

0

##### Parts Per Million

1

##### Fragmentation mode

ETHCD

##### Originating file

01 D:\separate\_stitch\_analyses\xle-disambiguation\20210325\_F59\_3ug\_DENOVO\_12.csv

#### Meta Information from PEAKS

##### Scan Identifier

F2:5975

##### Original sequence

V

V

F

G

G

G

T

K

L

T

##### Posttranslational Modifications

##### Source File

D:\separate\_stitch\_analyses\xle-disambiguation\raw\20210323\_F1\_UM1\_Peng0013\_SA\_F59\_ingel\_3ug\_TL.raw

##### Fraction

2

##### Scan Feature

-

##### De Novo Score

98

##### ConfidenceScore

98

### m/z

489.7849

##### Mass

977.5546

##### Charge

2

##### Retention Time

32.68

##### Predicted Retention Time

-

##### Area

0

##### Parts Per Million

0.8

##### Fragmentation mode

HCD

##### Originating file

01 D:\separate\_stitch\_analyses\xle-disambiguation\20210325\_F59\_3ug\_DENOVO\_12.csv

#### Meta Information from PEAKS

##### Scan Identifier

F2:5854

##### Original sequence

V

V

F

G

G

G

T

K

L

T

##### Posttranslational Modifications

##### Source File

D:\separate\_stitch\_analyses\xle-disambiguation\raw\20210323\_F1\_UM1\_Peng0013\_SA\_F59\_ingel\_3ug\_TL.raw

##### Fraction

2

##### Scan Feature

-

##### De Novo Score

98

##### ConfidenceScore

98

### m/z

489.7856

##### Mass

977.5546

##### Charge

2

##### Retention Time

31.98

##### Predicted Retention Time

-

##### Area

0

##### Parts Per Million

2

##### Fragmentation mode

ETHCD

##### Originating file

01 D:\separate\_stitch\_analyses\xle-disambiguation\20210325\_F59\_3ug\_DENOVO\_12.csv

#### Meta Information from PEAKS

##### Scan Identifier

F2:6339

##### Original sequence

V

V

F

G

G

G

T

K

L

T

##### Posttranslational Modifications

##### Source File

D:\separate\_stitch\_analyses\xle-disambiguation\raw\20210323\_F1\_UM1\_Peng0013\_SA\_F59\_ingel\_3ug\_TL.raw

##### Fraction

2

##### Scan Feature

-

##### De Novo Score

97

##### ConfidenceScore

97

### m/z

489.7853

##### Mass

977.5546

##### Charge

2

##### Retention Time

34.88

##### Predicted Retention Time

-

##### Area

0

##### Parts Per Million

1.5

##### Fragmentation mode

HCD

##### Originating file

01 D:\separate\_stitch\_analyses\xle-disambiguation\20210325\_F59\_3ug\_DENOVO\_12.csv

#### Meta Information from PEAKS

##### Scan Identifier

F2:6400

##### Original sequence

V

V

F

G

G

G

T

K

L

T

##### Posttranslational Modifications

##### Source File

D:\separate\_stitch\_analyses\xle-disambiguation\raw\20210323\_F1\_UM1\_Peng0013\_SA\_F59\_ingel\_3ug\_TL.raw

##### Fraction

2

##### Scan Feature

-

##### De Novo Score

97

##### ConfidenceScore

97

### m/z

489.7853

##### Mass

977.5546

##### Charge

2

##### Retention Time

35.24

##### Predicted Retention Time

-

##### Area

0

##### Parts Per Million

1.6

##### Fragmentation mode

HCD

##### Originating file

01 D:\separate\_stitch\_analyses\xle-disambiguation\20210325\_F59\_3ug\_DENOVO\_12.csv

#### Meta Information from PEAKS

##### Scan Identifier

F2:6461

##### Original sequence

V

V

F

G

G

G

T

K

L

T

##### Posttranslational Modifications

##### Source File

D:\separate\_stitch\_analyses\xle-disambiguation\raw\20210323\_F1\_UM1\_Peng0013\_SA\_F59\_ingel\_3ug\_TL.raw

##### Fraction

2

##### Scan Feature

-

##### De Novo Score

97

##### ConfidenceScore

97

### m/z

489.7852

##### Mass

977.5546

##### Charge

2

##### Retention Time

35.6

##### Predicted Retention Time

-

##### Area

0

##### Parts Per Million

1.3

##### Fragmentation mode

HCD

##### Originating file

01 D:\separate\_stitch\_analyses\xle-disambiguation\20210325\_F59\_3ug\_DENOVO\_12.csv

#### Meta Information from PEAKS

##### Scan Identifier

F2:6214

##### Original sequence

V

V

F

G

G

G

T

K

L

T

##### Posttranslational Modifications

##### Source File

D:\separate\_stitch\_analyses\xle-disambiguation\raw\20210323\_F1\_UM1\_Peng0013\_SA\_F59\_ingel\_3ug\_TL.raw

##### Fraction

2

##### Scan Feature

-

##### De Novo Score

96

##### ConfidenceScore

96

### m/z

489.7851

##### Mass

977.5546

##### Charge

2

##### Retention Time

34.15

##### Predicted Retention Time

-

##### Area

0

##### Parts Per Million

1.2

##### Fragmentation mode

HCD

##### Originating file

01 D:\separate\_stitch\_analyses\xle-disambiguation\20210325\_F59\_3ug\_DENOVO\_12.csv

#### Meta Information from PEAKS

##### Scan Identifier

F2:7091

##### Original sequence

V

V

F

G

G

G

T

K

L

T

##### Posttranslational Modifications

##### Source File

D:\separate\_stitch\_analyses\xle-disambiguation\raw\20210323\_F1\_UM1\_Peng0013\_SA\_F59\_ingel\_3ug\_TL.raw

##### Fraction

2

##### Scan Feature

-

##### De Novo Score

96

##### ConfidenceScore

96

### m/z

489.7854

##### Mass

977.5546

##### Charge

2

##### Retention Time

39.48

##### Predicted Retention Time

-

##### Area

0

##### Parts Per Million

1.7

##### Fragmentation mode

HCD

##### Originating file

01 D:\separate\_stitch\_analyses\xle-disambiguation\20210325\_F59\_3ug\_DENOVO\_12.csv

#### Meta Information from PEAKS

##### Scan Identifier

F2:5801

##### Original sequence

V

V

F

G

G

G

T

K

L

T

##### Posttranslational Modifications

##### Source File

D:\separate\_stitch\_analyses\xle-disambiguation\raw\20210323\_F1\_UM1\_Peng0013\_SA\_F59\_ingel\_3ug\_TL.raw

##### Fraction

2

##### Scan Feature

-

##### De Novo Score

96

##### ConfidenceScore

96

### m/z

489.7851

##### Mass

977.5546

##### Charge

2

##### Retention Time

31.66

##### Predicted Retention Time

-

##### Area

0

##### Parts Per Million

1.2

##### Fragmentation mode

ETHCD

##### Originating file

01 D:\separate\_stitch\_analyses\xle-disambiguation\20210325\_F59\_3ug\_DENOVO\_12.csv

#### Meta Information from PEAKS

##### Scan Identifier

F2:6098

##### Original sequence

V

V

F

G

G

G

T

K

L

T

##### Posttranslational Modifications

##### Source File

D:\separate\_stitch\_analyses\xle-disambiguation\raw\20210323\_F1\_UM1\_Peng0013\_SA\_F59\_ingel\_3ug\_TL.raw

##### Fraction

2

##### Scan Feature

-

##### De Novo Score

96

##### ConfidenceScore

96

### m/z

489.7849

##### Mass

977.5546

##### Charge

2

##### Retention Time

33.43

##### Predicted Retention Time

-

##### Area

0

##### Parts Per Million

0.7

##### Fragmentation mode

ETHCD

##### Originating file

01 D:\separate\_stitch\_analyses\xle-disambiguation\20210325\_F59\_3ug\_DENOVO\_12.csv

#### Meta Information from PEAKS

##### Scan Identifier

F2:7033

##### Original sequence

V

V

F

G

G

G

T

K

L

T

##### Posttranslational Modifications

##### Source File

D:\separate\_stitch\_analyses\xle-disambiguation\raw\20210323\_F1\_UM1\_Peng0013\_SA\_F59\_ingel\_3ug\_TL.raw

##### Fraction

2

##### Scan Feature

-

##### De Novo Score

95

##### ConfidenceScore

95

### m/z

489.7857

##### Mass

977.5546

##### Charge

2

##### Retention Time

39.1

##### Predicted Retention Time

-

##### Area

0

##### Parts Per Million

2.3

##### Fragmentation mode

HCD

##### Originating file

01 D:\separate\_stitch\_analyses\xle-disambiguation\20210325\_F59\_3ug\_DENOVO\_12.csv

#### Meta Information from PEAKS

##### Scan Identifier

F2:6276

##### Original sequence

V

V

F

G

G

G

T

K

L

T

##### Posttranslational Modifications

##### Source File

D:\separate\_stitch\_analyses\xle-disambiguation\raw\20210323\_F1\_UM1\_Peng0013\_SA\_F59\_ingel\_3ug\_TL.raw

##### Fraction

2

##### Scan Feature

-

##### De Novo Score

95

##### ConfidenceScore

95

### m/z

489.7854

##### Mass

977.5546

##### Charge

2

##### Retention Time

34.53

##### Predicted Retention Time

-

##### Area

0

##### Parts Per Million

1.8

##### Fragmentation mode

HCD

##### Originating file

01 D:\separate\_stitch\_analyses\xle-disambiguation\20210325\_F59\_3ug\_DENOVO\_12.csv
