## Supplementary material for "A handle on mass coincidence errors in *de novo* sequencing of antibodies by bottom-up proteomics": Combined_024.html

Details Combined\_024 | Stitch OverviewUndefined

### Read Combined\_024

#### Sequence (length=18)

SWYQHHPGKAPKJJJSEV

#### Spectrum 7437? Spectrum 7437 The raw spectrum of this peptide as annotated by Hecklib. The fragments are coloured according to ion type (see legend). Any peaks with a star '\*' as text can be hovered over to see the full details, first the ion type second the mass shift type. By hovering over the amino acids in the peptide or ions in the legend the corresponding peaks are highlighted. By toggling the 'Unassigned' label you can turn the background (unassigned) peaks on or off in the plot. By updating the slider in the Ion legend you can update the spectrum to only show the top X% of the peaks with labels. The top X% means any peak that is within X% of the highest intensity. By dragging in the spectrum you can zoom in to a specific part of the spectrum and use 'Zoom Out' to get back to the original zoom level. The annotation of the spectrum is based on the given sequence in the peptides file and is done with different software so inconsistencies are likely. The peaks are annotated based on the given sequence, with 20 ppm tolerance.

Copy Data

##### Spectrum 7437 (TSV)

###### Preview

```
Loading example...
```

*Click on the button to copy the data to your clipboard.*

Mz MinMz MaxIntensity Max

WidthHeightPeptide font sizePeptide stroke widthSpectrum font sizeSpectrum stroke widthCompact peptide

Ion legend

wxyz

abcd

OtherUnassignedIon^Charge^_Position_Show for top:%

SWYQHHPGKAPKJJJSEV

04.29e+58.59e+51.29e+61.72e+6

Zoom Out

c^+3^_4_y^+1^_2_y^+1^_2_y^+1^_3_y^+1^_3_y^+1^_4_y^+1^_5_c^+2^_9_z^+2^_11_c^+1^_4_c^+1^_4_c^+3^_15_c^+2^_10_y^+3^_16_w^+1^_6_c^+3^_16_y^+2^_12_z^+1^_6_c^+3^_17_y^+3^_17_y^+1^_6_y^+2^_13_z^+2^_13_y^+2^_13_c^+1^_5_c^+2^_12_c^+2^_12_c^+1^_5_z^+2^_14_y^+2^_14_c^+2^_13_c^+2^_13_z^+1^_7_w^+2^_15_y^+1^_7_y^+2^_15_z^+2^_15_c^+2^_14_y^+2^_15_c^+2^_14_c^+1^_6_c^+2^_15_c^+2^_15_c^+2^_15_z^+2^_16_y^+1^_8_y^+2^_16_y^+2^_16_z^+2^_16_y^+2^_16_c^+2^_16_c^+2^_16_c^+2^_16_c^+1^_7_z^+1^_9_y^+1^_9_c^+2^_17_y^+2^_17_y^+2^_17_z^+2^_17_c^+2^_17_y^+2^_17_c^+1^_8_z^+1^_10_y^+1^_10_c^+1^_9_z^+1^_11_y^+1^_11_y^+1^_12_c^+1^_11_z^+1^_13_y^+1^_13_c^+1^_12_z^+1^_14_y^+1^_14_c^+1^_13_z^+1^_15_y^+1^_15_c^+1^_14_c^+1^_15_c^+1^_15_y^+1^_16_z^+1^_16_y^+1^_16_c^+1^_16_z^+1^_17_c^+1^_17_

0580116017392319

Fragment Matches Table

Show background peaks

| Position | Ion type | Intensity | mz Theoretical | mz Error (Th) | mz Error (ppm) | Charge | Series Number |
| --- | --- | --- | --- | --- | --- | --- | --- |
| - | - | 2153 | 122.3 | - | - | 0 | - |
| - | - | 2524 | 130.1 | - | - | 0 | - |
| - | - | 2181 | 136.1 | - | - | 0 | - |
| - | - | 2383 | 138.6 | - | - | 0 | - |
| - | - | 2443 | 149 | - | - | 0 | - |
| - | - | 2702 | 184.2 | - | - | 0 | - |
| - | - | 2957 | 186.1 | - | - | 0 | - |
| - | - | 5678 | 187.1 | - | - | 0 | - |
| 4 | c | 4652 | 189.1 | 0.001644 | 8.696 | +3 | 4 |
| - | - | 3126 | 199.4 | - | - | 0 | - |
| - | - | 2716 | 203.9 | - | - | 0 | - |
| - | - | 2636 | 212.6 | - | - | 0 | - |
| - | - | 2980 | 216.7 | - | - | 0 | - |
| - | - | 2.792E+04 | 217.1 | - | - | 0 | - |
| - | - | 2606 | 218.6 | - | - | 0 | - |
| 17 | y | 2.374E+04 | 229.1 | 0.0002567 | 1.121 | +1 | 2 |
| - | - | 2.73E+04 | 246.1 | - | - | 0 | - |
| 17 | y | 1.662E+04 | 247.1 | 0.0007964 | 3.223 | +1 | 2 |
| - | - | 3176 | 273.1 | - | - | 0 | - |
| - | - | 4.546E+04 | 274.1 | - | - | 0 | - |
| - | - | 8537 | 275.1 | - | - | 0 | - |
| - | - | 5677 | 292.1 | - | - | 0 | - |
| - | - | 4614 | 309.2 | - | - | 0 | - |
| - | - | 3167 | 314 | - | - | 0 | - |
| 16 | y | 4076 | 316.2 | 1.28E-05 | 0.0405 | +1 | 3 |
| - | - | 2702 | 328.5 | - | - | 0 | - |
| 16 | y | 2.306E+04 | 334.2 | 0.0004151 | 1.242 | +1 | 3 |
| - | - | 3299 | 369.1 | - | - | 0 | - |
| - | - | 3444 | 434.2 | - | - | 0 | - |
| - | - | 7297 | 437.2 | - | - | 0 | - |
| 15 | y | 1.121E+04 | 447.2 | 0.0005558 | 1.243 | +1 | 4 |
| - | - | 1.264E+04 | 465.2 | - | - | 0 | - |
| - | - | 9493 | 477.2 | - | - | 0 | - |
| - | - | 5921 | 478.2 | - | - | 0 | - |
| - | - | 6707 | 480.2 | - | - | 0 | - |
| - | - | 3272 | 490.2 | - | - | 0 | - |
| - | - | 5890 | 528.2 | - | - | 0 | - |
| - | - | 7665 | 538.3 | - | - | 0 | - |
| - | - | 1.552E+04 | 539.3 | - | - | 0 | - |
| - | - | 2972 | 541.3 | - | - | 0 | - |
| 14 | y | 1.153E+04 | 560.3 | 0.0005133 | 0.9161 | +1 | 5 |
| 9 | c | 1.771E+04 | 561.3 | 0.001233 | 2.197 | +2 | 9 |
| 8 | z | 2984 | 561.3 | 0.005711 | 10.17 | +2 | 11 |
| - | - | 1.076E+04 | 561.8 | - | - | 0 | - |
| - | - | 4315 | 562.3 | - | - | 0 | - |
| - | - | 2.491E+04 | 563.2 | - | - | 0 | - |
| - | - | 7954 | 564.2 | - | - | 0 | - |
| 4 | c | 1.31E+04 | 565.2 | 0.0001988 | 0.3517 | +1 | 4 |
| - | - | 3863 | 566.2 | - | - | 0 | - |
| - | - | 9204 | 567.8 | - | - | 0 | - |
| - | - | 5013 | 568.3 | - | - | 0 | - |
| - | - | 3569 | 577.7 | - | - | 0 | - |
| - | - | 2.082E+04 | 581.3 | - | - | 0 | - |
| 4 | c | 1.105E+05 | 582.3 | 0.0008376 | 1.439 | +1 | 4 |
| - | - | 3.79E+04 | 583.3 | - | - | 0 | - |
| - | - | 6544 | 584.3 | - | - | 0 | - |
| 15 | c | 2.504E+04 | 586.3 | 0.001545 | 2.636 | +3 | 15 |
| - | - | 2.574E+04 | 586.7 | - | - | 0 | - |
| - | - | 1.822E+04 | 587 | - | - | 0 | - |
| - | - | 3742 | 587.3 | - | - | 0 | - |
| 10 | c | 1.686E+04 | 596.8 | 0.0005348 | 0.8962 | +2 | 10 |
| - | - | 8901 | 597.3 | - | - | 0 | - |
| - | - | 3995 | 597.8 | - | - | 0 | - |
| - | - | 4804 | 601.3 | - | - | 0 | - |
| 3 | y | 7571 | 606.3 | 0.001349 | 2.226 | +3 | 16 |
| - | - | 5998 | 606.7 | - | - | 0 | - |
| - | - | 3339 | 609.7 | - | - | 0 | - |
| 13 | w | 1.02E+04 | 614.3 | 0.003327 | 5.415 | +1 | 6 |
| 16 | c | 2.056E+04 | 615.3 | 0.000686 | 1.115 | +3 | 16 |
| - | - | 2.515E+04 | 615.7 | - | - | 0 | - |
| - | - | 1.079E+04 | 616 | - | - | 0 | - |
| 7 | y | 2.827E+04 | 626.4 | 0.001141 | 1.821 | +2 | 12 |
| - | - | 2.241E+04 | 626.9 | - | - | 0 | - |
| - | - | 6522 | 627.4 | - | - | 0 | - |
| - | - | 1.286E+04 | 649 | - | - | 0 | - |
| - | - | 1.164E+04 | 649.4 | - | - | 0 | - |
| - | - | 6292 | 649.7 | - | - | 0 | - |
| - | - | 1.076E+04 | 652.3 | - | - | 0 | - |
| - | - | 1.928E+04 | 652.7 | - | - | 0 | - |
| - | - | 1.345E+04 | 653 | - | - | 0 | - |
| - | - | 1.049E+04 | 653.3 | - | - | 0 | - |
| - | - | 9045 | 657.3 | - | - | 0 | - |
| 13 | z | 1.398E+04 | 657.4 | 0.0005456 | 0.83 | +1 | 6 |
| 17 | c | 1.156E+05 | 658.4 | 0.0007235 | 1.099 | +3 | 17 |
| - | - | 1.362E+05 | 658.7 | - | - | 0 | - |
| - | - | 6.858E+04 | 659 | - | - | 0 | - |
| - | - | 1.135E+04 | 659.3 | - | - | 0 | - |
| - | - | 3.504E+04 | 659.4 | - | - | 0 | - |
| - | - | 1.028E+04 | 659.7 | - | - | 0 | - |
| - | - | 5356 | 660.3 | - | - | 0 | - |
| - | - | 7189 | 664.4 | - | - | 0 | - |
| - | - | 6365 | 664.7 | - | - | 0 | - |
| 2 | y | 5480 | 668.4 | 0.001115 | 1.668 | +3 | 17 |
| - | - | 3975 | 668.7 | - | - | 0 | - |
| 13 | y | 1.076E+04 | 673.4 | 0.0007499 | 1.114 | +1 | 6 |
| - | - | 5613 | 674.3 | - | - | 0 | - |
| - | - | 3.457E+04 | 675.3 | - | - | 0 | - |
| - | - | 2.067E+04 | 676.3 | - | - | 0 | - |
| - | - | 5553 | 677.3 | - | - | 0 | - |
| - | - | 6443 | 682 | - | - | 0 | - |
| - | - | 6675 | 682.4 | - | - | 0 | - |
| 6 | y | 3343 | 685.9 | 0.007553 | 11.01 | +2 | 13 |
| - | - | 3641 | 686 | - | - | 0 | - |
| 6 | z | 3490 | 686.9 | 0.000677 | 0.9856 | +2 | 13 |
| - | - | 4436 | 687.7 | - | - | 0 | - |
| - | - | 3.935E+04 | 691.4 | - | - | 0 | - |
| - | - | 5.433E+04 | 691.7 | - | - | 0 | - |
| - | - | 3.18E+04 | 692 | - | - | 0 | - |
| - | - | 1.488E+04 | 692.4 | - | - | 0 | - |
| - | - | 6121 | 692.7 | - | - | 0 | - |
| 6 | y | 4170 | 694.9 | 0.003888 | 5.595 | +2 | 13 |
| - | - | 7447 | 695.4 | - | - | 0 | - |
| - | - | 1.348E+04 | 696.7 | - | - | 0 | - |
| - | - | 1.234E+04 | 697 | - | - | 0 | - |
| - | - | 7.196E+05 | 697.4 | - | - | 0 | - |
| - | - | 8.802E+05 | 697.7 | - | - | 0 | - |
| - | - | 5.502E+05 | 698 | - | - | 0 | - |
| - | - | 2.713E+05 | 698.4 | - | - | 0 | - |
| - | - | 8.964E+04 | 698.7 | - | - | 0 | - |
| - | - | 1.836E+04 | 699 | - | - | 0 | - |
| 5 | c | 1.445E+04 | 702.3 | 0.0001859 | 0.2646 | +1 | 5 |
| - | - | 4617 | 703.3 | - | - | 0 | - |
| 12 | c | 4138 | 708.9 | 0.008689 | 12.26 | +2 | 12 |
| - | - | 4958 | 709.9 | - | - | 0 | - |
| - | - | 3.47E+04 | 717.4 | - | - | 0 | - |
| 12 | c | 3.597E+05 | 717.9 | 0.001093 | 1.522 | +2 | 12 |
| - | - | 3.079E+05 | 718.4 | - | - | 0 | - |
| - | - | 1.491E+05 | 718.9 | - | - | 0 | - |
| 5 | c | 1.18E+05 | 719.3 | 0.001217 | 1.692 | +1 | 5 |
| - | - | 2.088E+04 | 719.4 | - | - | 0 | - |
| - | - | 8380 | 719.9 | - | - | 0 | - |
| - | - | 4.587E+04 | 720.3 | - | - | 0 | - |
| - | - | 3657 | 720.4 | - | - | 0 | - |
| - | - | 1.309E+04 | 721.3 | - | - | 0 | - |
| - | - | 5179 | 742.4 | - | - | 0 | - |
| - | - | 4391 | 742.9 | - | - | 0 | - |
| - | - | 5425 | 743.4 | - | - | 0 | - |
| - | - | 1.523E+04 | 752.4 | - | - | 0 | - |
| - | - | 1.546E+04 | 752.9 | - | - | 0 | - |
| - | - | 4440 | 753.4 | - | - | 0 | - |
| 5 | z | 4.271E+04 | 755.4 | 0.001496 | 1.981 | +2 | 14 |
| - | - | 6.701E+04 | 755.9 | - | - | 0 | - |
| - | - | 4.388E+04 | 756.4 | - | - | 0 | - |
| - | - | 8834 | 756.9 | - | - | 0 | - |
| - | - | 5586 | 757.4 | - | - | 0 | - |
| 5 | y | 4.608E+04 | 763.4 | 0.0004823 | 0.6317 | +2 | 14 |
| - | - | 3.925E+04 | 763.9 | - | - | 0 | - |
| - | - | 1.369E+04 | 764.4 | - | - | 0 | - |
| - | - | 4847 | 765 | - | - | 0 | - |
| 13 | c | 7511 | 765.9 | 0.001225 | 1.599 | +2 | 13 |
| - | - | 7414 | 766.4 | - | - | 0 | - |
| - | - | 3961 | 766.9 | - | - | 0 | - |
| - | - | 2.717E+04 | 773.9 | - | - | 0 | - |
| 13 | c | 2.149E+05 | 774.4 | 0.001194 | 1.541 | +2 | 13 |
| - | - | 2.027E+05 | 774.9 | - | - | 0 | - |
| - | - | 1.002E+05 | 775.4 | - | - | 0 | - |
| - | - | 2.476E+04 | 775.9 | - | - | 0 | - |
| - | - | 6436 | 776.4 | - | - | 0 | - |
| - | - | 3532 | 780.9 | - | - | 0 | - |
| - | - | 6200 | 783.5 | - | - | 0 | - |
| - | - | 5684 | 783.9 | - | - | 0 | - |
| 12 | z | 1.444E+04 | 785.5 | 0.0006828 | 0.8692 | +1 | 7 |
| - | - | 1.29E+04 | 786.5 | - | - | 0 | - |
| - | - | 6891 | 787.5 | - | - | 0 | - |
| 4 | w | 3.688E+04 | 790.5 | 0.0009429 | 1.193 | +2 | 15 |
| - | - | 3.021E+04 | 791 | - | - | 0 | - |
| - | - | 2.113E+04 | 791.5 | - | - | 0 | - |
| - | - | 9249 | 791.9 | - | - | 0 | - |
| - | - | 3356 | 792.4 | - | - | 0 | - |
| - | - | 1.017E+04 | 796.4 | - | - | 0 | - |
| - | - | 3657 | 797.4 | - | - | 0 | - |
| 12 | y | 7385 | 801.5 | 3.592E-05 | 0.04481 | +1 | 7 |
| - | - | 8443 | 809 | - | - | 0 | - |
| - | - | 8290 | 809.5 | - | - | 0 | - |
| - | - | 7706 | 810 | - | - | 0 | - |
| - | - | 3.075E+04 | 811.4 | - | - | 0 | - |
| - | - | 1.838E+04 | 812.4 | - | - | 0 | - |
| - | - | 5136 | 813.4 | - | - | 0 | - |
| 4 | y | 7452 | 819 | 0.004805 | 5.867 | +2 | 15 |
| 4 | z | 1.539E+05 | 819.5 | 0.001122 | 1.369 | +2 | 15 |
| - | - | 2.432E+05 | 820 | - | - | 0 | - |
| - | - | 1.658E+05 | 820.5 | - | - | 0 | - |
| - | - | 7.693E+04 | 821 | - | - | 0 | - |
| - | - | 2.629E+04 | 821.5 | - | - | 0 | - |
| - | - | 3755 | 822 | - | - | 0 | - |
| 14 | c | 1.484E+04 | 822.4 | 0.0006884 | 0.8371 | +2 | 14 |
| - | - | 2.26E+04 | 822.9 | - | - | 0 | - |
| - | - | 4338 | 823.4 | - | - | 0 | - |
| - | - | 5574 | 824 | - | - | 0 | - |
| - | - | 3641 | 827 | - | - | 0 | - |
| 4 | y | 3.616E+04 | 827.5 | 1.411E-05 | 0.01705 | +2 | 15 |
| - | - | 4.948E+04 | 828 | - | - | 0 | - |
| - | - | 2.076E+04 | 828.5 | - | - | 0 | - |
| - | - | 8771 | 829 | - | - | 0 | - |
| - | - | 3485 | 830.4 | - | - | 0 | - |
| 14 | c | 7.032E+04 | 831 | 0.001356 | 1.631 | +2 | 14 |
| - | - | 7.004E+04 | 831.5 | - | - | 0 | - |
| - | - | 2216 | 831.5 | - | - | 0 | - |
| - | - | 3.506E+04 | 832 | - | - | 0 | - |
| - | - | 1.848E+04 | 832.5 | - | - | 0 | - |
| - | - | 7192 | 833 | - | - | 0 | - |
| - | - | 3408 | 836 | - | - | 0 | - |
| 6 | c | 3.189E+04 | 839.4 | 0.0009868 | 1.176 | +1 | 6 |
| - | - | 1.63E+04 | 840.4 | - | - | 0 | - |
| - | - | 3747 | 841.4 | - | - | 0 | - |
| - | - | 3871 | 842.5 | - | - | 0 | - |
| - | - | 5261 | 850.5 | - | - | 0 | - |
| - | - | 8861 | 851 | - | - | 0 | - |
| - | - | 9133 | 851.5 | - | - | 0 | - |
| - | - | 9614 | 852 | - | - | 0 | - |
| - | - | 6538 | 858 | - | - | 0 | - |
| - | - | 5895 | 858.5 | - | - | 0 | - |
| - | - | 3712 | 859 | - | - | 0 | - |
| - | - | 4408 | 860 | - | - | 0 | - |
| - | - | 8436 | 865 | - | - | 0 | - |
| - | - | 1.91E+04 | 865.5 | - | - | 0 | - |
| - | - | 2.53E+04 | 866 | - | - | 0 | - |
| - | - | 1.153E+04 | 866.5 | - | - | 0 | - |
| - | - | 6085 | 869.5 | - | - | 0 | - |
| - | - | 7240 | 870 | - | - | 0 | - |
| - | - | 4025 | 870.5 | - | - | 0 | - |
| - | - | 1.029E+04 | 872 | - | - | 0 | - |
| - | - | 1.063E+04 | 872.5 | - | - | 0 | - |
| - | - | 8010 | 873 | - | - | 0 | - |
| - | - | 3669 | 873.5 | - | - | 0 | - |
| - | - | 3298 | 874.4 | - | - | 0 | - |
| - | - | 1.166E+04 | 878 | - | - | 0 | - |
| 15 | c | 1.573E+04 | 878.5 | 0.007832 | 8.915 | +2 | 15 |
| 15 | c | 6.665E+04 | 879 | 8.378E-05 | 0.09531 | +2 | 15 |
| - | - | 7.387E+04 | 879.5 | - | - | 0 | - |
| - | - | 4.138E+04 | 880 | - | - | 0 | - |
| - | - | 2.276E+04 | 880.5 | - | - | 0 | - |
| - | - | 9863 | 881 | - | - | 0 | - |
| - | - | 3626 | 884.8 | - | - | 0 | - |
| - | - | 9.718E+04 | 887 | - | - | 0 | - |
| 15 | c | 1.955E+05 | 887.5 | 0.002372 | 2.673 | +2 | 15 |
| - | - | 1.584E+05 | 888 | - | - | 0 | - |
| - | - | 8.812E+04 | 888.5 | - | - | 0 | - |
| - | - | 3.687E+04 | 889 | - | - | 0 | - |
| - | - | 7272 | 889.5 | - | - | 0 | - |
| 3 | z | 4984 | 892.5 | 0.006795 | 7.613 | +2 | 16 |
| - | - | 1.737E+04 | 897.5 | - | - | 0 | - |
| 11 | y | 3.591E+04 | 898.6 | 0.002873 | 3.197 | +1 | 8 |
| - | - | 1.494E+04 | 899.6 | - | - | 0 | - |
| 3 | y | 1.214E+04 | 900 | 0.00171 | 1.9 | +2 | 16 |
| 3 | y | 1.778E+04 | 900.5 | 0.009823 | 10.91 | +2 | 16 |
| 3 | z | 1.116E+05 | 901 | 0.0004986 | 0.5534 | +2 | 16 |
| - | - | 1.439E+05 | 901.5 | - | - | 0 | - |
| - | - | 7.723E+04 | 902 | - | - | 0 | - |
| - | - | 4.073E+04 | 902.5 | - | - | 0 | - |
| - | - | 8185 | 903 | - | - | 0 | - |
| - | - | 4388 | 908.5 | - | - | 0 | - |
| 3 | y | 1.161E+05 | 909 | 0.001011 | 1.112 | +2 | 16 |
| - | - | 1.158E+05 | 909.5 | - | - | 0 | - |
| - | - | 6.281E+04 | 910 | - | - | 0 | - |
| - | - | 2.52E+04 | 910.5 | - | - | 0 | - |
| - | - | 9540 | 911 | - | - | 0 | - |
| - | - | 3936 | 913.5 | - | - | 0 | - |
| - | - | 9493 | 914 | - | - | 0 | - |
| 16 | c | 5572 | 922 | 0.006634 | 7.195 | +2 | 16 |
| 16 | c | 2.816E+04 | 922.5 | 0.0007171 | 0.7774 | +2 | 16 |
| - | - | 3.789E+04 | 923 | - | - | 0 | - |
| - | - | 2.921E+04 | 923.5 | - | - | 0 | - |
| - | - | 5420 | 924 | - | - | 0 | - |
| - | - | 4473 | 928.5 | - | - | 0 | - |
| - | - | 4513 | 929.1 | - | - | 0 | - |
| - | - | 4448 | 929.5 | - | - | 0 | - |
| - | - | 4943 | 930.2 | - | - | 0 | - |
| - | - | 2.008E+04 | 930.5 | - | - | 0 | - |
| - | - | 4008 | 930.8 | - | - | 0 | - |
| 16 | c | 2.21E+05 | 931 | 0.001602 | 1.72 | +2 | 16 |
| - | - | 2.564E+05 | 931.5 | - | - | 0 | - |
| - | - | 1.685E+05 | 932 | - | - | 0 | - |
| - | - | 5.493E+04 | 932.5 | - | - | 0 | - |
| - | - | 1.645E+04 | 933 | - | - | 0 | - |
| - | - | 4715 | 933.5 | - | - | 0 | - |
| - | - | 1.428E+04 | 944 | - | - | 0 | - |
| - | - | 1.203E+04 | 944.5 | - | - | 0 | - |
| - | - | 1.537E+04 | 945 | - | - | 0 | - |
| - | - | 1.079E+04 | 952.4 | - | - | 0 | - |
| 7 | c | 8.865E+04 | 953.4 | 0.0008319 | 0.8726 | +1 | 7 |
| 10 | z | 1.438E+05 | 953.6 | 0.000877 | 0.9197 | +1 | 9 |
| - | - | 4.272E+04 | 954.4 | - | - | 0 | - |
| - | - | 7.968E+04 | 954.6 | - | - | 0 | - |
| - | - | 1.502E+04 | 955.4 | - | - | 0 | - |
| - | - | 2.789E+04 | 955.6 | - | - | 0 | - |
| - | - | 4893 | 956.6 | - | - | 0 | - |
| - | - | 3576 | 958.5 | - | - | 0 | - |
| - | - | 7565 | 965 | - | - | 0 | - |
| - | - | 5704 | 965.5 | - | - | 0 | - |
| - | - | 3846 | 966 | - | - | 0 | - |
| - | - | 1.41E+04 | 966.4 | - | - | 0 | - |
| - | - | 9605 | 967.5 | - | - | 0 | - |
| - | - | 5806 | 968.4 | - | - | 0 | - |
| 10 | y | 3.632E+04 | 969.6 | 0.001474 | 1.52 | +1 | 9 |
| - | - | 2.032E+04 | 970.6 | - | - | 0 | - |
| - | - | 6057 | 971.6 | - | - | 0 | - |
| - | - | 3580 | 972 | - | - | 0 | - |
| - | - | 1.451E+04 | 973 | - | - | 0 | - |
| - | - | 4.37E+04 | 973.5 | - | - | 0 | - |
| - | - | 4.589E+04 | 974 | - | - | 0 | - |
| - | - | 2.661E+04 | 974.5 | - | - | 0 | - |
| - | - | 1.348E+04 | 975 | - | - | 0 | - |
| - | - | 3851 | 978 | - | - | 0 | - |
| - | - | 5516 | 978.5 | - | - | 0 | - |
| - | - | 6112 | 979 | - | - | 0 | - |
| - | - | 5771 | 980 | - | - | 0 | - |
| - | - | 1.042E+04 | 980.5 | - | - | 0 | - |
| - | - | 1.227E+04 | 981 | - | - | 0 | - |
| - | - | 1.21E+04 | 981.5 | - | - | 0 | - |
| - | - | 4959 | 984.5 | - | - | 0 | - |
| - | - | 5023 | 986.6 | - | - | 0 | - |
| 17 | c | 4.422E+04 | 987 | 0.001607 | 1.628 | +2 | 17 |
| - | - | 6.454E+04 | 987.5 | - | - | 0 | - |
| - | - | 4.101E+04 | 988 | - | - | 0 | - |
| - | - | 2.72E+04 | 988.5 | - | - | 0 | - |
| - | - | 5468 | 989 | - | - | 0 | - |
| 2 | y | 9294 | 993 | 0.002975 | 2.996 | +2 | 17 |
| 2 | y | 1.041E+04 | 993.5 | 0.007092 | 7.139 | +2 | 17 |
| 2 | z | 3.75E+04 | 994 | 0.002997 | 3.015 | +2 | 17 |
| - | - | 4.502E+04 | 994.5 | - | - | 0 | - |
| - | - | 2.565E+04 | 995 | - | - | 0 | - |
| 17 | c | 2.346E+05 | 995.5 | 0.0008644 | 0.8683 | +2 | 17 |
| - | - | 2.664E+05 | 996 | - | - | 0 | - |
| - | - | 1.618E+05 | 996.5 | - | - | 0 | - |
| - | - | 9.058E+04 | 997 | - | - | 0 | - |
| - | - | 2.821E+04 | 997.5 | - | - | 0 | - |
| - | - | 1.014E+04 | 998 | - | - | 0 | - |
| - | - | 6131 | 1002 | - | - | 0 | - |
| 2 | y | 1.982E+04 | 1002 | 0.01027 | 10.25 | +2 | 17 |
| - | - | 2.383E+04 | 1003 | - | - | 0 | - |
| - | - | 1.23E+04 | 1003 | - | - | 0 | - |
| - | - | 4538 | 1004 | - | - | 0 | - |
| - | - | 1.977E+04 | 1005 | - | - | 0 | - |
| - | - | 1.828E+04 | 1006 | - | - | 0 | - |
| - | - | 1.55E+04 | 1006 | - | - | 0 | - |
| - | - | 1.41E+04 | 1007 | - | - | 0 | - |
| - | - | 2.808E+04 | 1008 | - | - | 0 | - |
| - | - | 2.621E+04 | 1009 | - | - | 0 | - |
| - | - | 2.416E+04 | 1009 | - | - | 0 | - |
| - | - | 1.355E+04 | 1009 | - | - | 0 | - |
| - | - | 3.441E+04 | 1010 | - | - | 0 | - |
| - | - | 4.362E+04 | 1010 | - | - | 0 | - |
| 8 | c | 1.28E+05 | 1010 | 0.001483 | 1.467 | +1 | 8 |
| - | - | 2.385E+04 | 1011 | - | - | 0 | - |
| - | - | 8.257E+04 | 1011 | - | - | 0 | - |
| - | - | 4983 | 1012 | - | - | 0 | - |
| - | - | 2.481E+04 | 1012 | - | - | 0 | - |
| - | - | 6315 | 1013 | - | - | 0 | - |
| - | - | 2.45E+04 | 1015 | - | - | 0 | - |
| - | - | 3.637E+04 | 1015 | - | - | 0 | - |
| - | - | 4.298E+04 | 1016 | - | - | 0 | - |
| - | - | 3.582E+04 | 1016 | - | - | 0 | - |
| - | - | 3.946E+04 | 1017 | - | - | 0 | - |
| - | - | 2.244E+04 | 1017 | - | - | 0 | - |
| - | - | 9884 | 1018 | - | - | 0 | - |
| - | - | 1.574E+04 | 1018 | - | - | 0 | - |
| - | - | 1.802E+04 | 1019 | - | - | 0 | - |
| - | - | 1.323E+04 | 1019 | - | - | 0 | - |
| - | - | 7065 | 1020 | - | - | 0 | - |
| - | - | 1.078E+04 | 1023 | - | - | 0 | - |
| - | - | 7.466E+04 | 1024 | - | - | 0 | - |
| - | - | 8.037E+04 | 1024 | - | - | 0 | - |
| - | - | 1.029E+05 | 1025 | - | - | 0 | - |
| - | - | 9.737E+04 | 1025 | - | - | 0 | - |
| - | - | 4.657E+04 | 1026 | - | - | 0 | - |
| - | - | 1.819E+04 | 1026 | - | - | 0 | - |
| - | - | 5323 | 1028 | - | - | 0 | - |
| - | - | 1.339E+04 | 1029 | - | - | 0 | - |
| - | - | 1.517E+04 | 1029 | - | - | 0 | - |
| - | - | 9114 | 1030 | - | - | 0 | - |
| - | - | 4755 | 1030 | - | - | 0 | - |
| - | - | 7671 | 1031 | - | - | 0 | - |
| - | - | 5781 | 1036 | - | - | 0 | - |
| - | - | 9023 | 1037 | - | - | 0 | - |
| - | - | 3.745E+04 | 1037 | - | - | 0 | - |
| - | - | 3.105E+05 | 1038 | - | - | 0 | - |
| - | - | 3.73E+05 | 1038 | - | - | 0 | - |
| - | - | 2.268E+05 | 1039 | - | - | 0 | - |
| - | - | 1.012E+05 | 1039 | - | - | 0 | - |
| - | - | 5.313E+04 | 1040 | - | - | 0 | - |
| - | - | 8722 | 1040 | - | - | 0 | - |
| - | - | 1.86E+04 | 1045 | - | - | 0 | - |
| - | - | 4.063E+05 | 1046 | - | - | 0 | - |
| - | - | 1.616E+06 | 1046 | - | - | 0 | - |
| - | - | 1.701E+06 | 1047 | - | - | 0 | - |
| - | - | 9.993E+05 | 1047 | - | - | 0 | - |
| - | - | 4.876E+05 | 1048 | - | - | 0 | - |
| - | - | 1.493E+05 | 1048 | - | - | 0 | - |
| - | - | 3.356E+04 | 1049 | - | - | 0 | - |
| 9 | z | 8.488E+04 | 1082 | 0.001663 | 1.537 | +1 | 10 |
| - | - | 1.151E+05 | 1083 | - | - | 0 | - |
| - | - | 4.923E+04 | 1084 | - | - | 0 | - |
| - | - | 1.403E+04 | 1085 | - | - | 0 | - |
| - | - | 2.215E+04 | 1095 | - | - | 0 | - |
| - | - | 1.229E+04 | 1096 | - | - | 0 | - |
| 9 | y | 2.601E+04 | 1098 | 0.002198 | 2.003 | +1 | 10 |
| - | - | 1.513E+04 | 1099 | - | - | 0 | - |
| - | - | 4675 | 1100 | - | - | 0 | - |
| 9 | c | 1.903E+05 | 1139 | 0.001231 | 1.081 | +1 | 9 |
| 8 | z | 4.952E+04 | 1139 | 0.0006655 | 0.5845 | +1 | 11 |
| - | - | 1.225E+05 | 1140 | - | - | 0 | - |
| - | - | 5.319E+04 | 1140 | - | - | 0 | - |
| - | - | 4.849E+04 | 1141 | - | - | 0 | - |
| - | - | 3.129E+04 | 1141 | - | - | 0 | - |
| - | - | 1.276E+04 | 1142 | - | - | 0 | - |
| - | - | 8742 | 1142 | - | - | 0 | - |
| 8 | y | 8929 | 1155 | 0.005229 | 4.529 | +1 | 11 |
| - | - | 4192 | 1156 | - | - | 0 | - |
| - | - | 2.107E+04 | 1166 | - | - | 0 | - |
| - | - | 1.354E+04 | 1167 | - | - | 0 | - |
| - | - | 5245 | 1168 | - | - | 0 | - |
| - | - | 4501 | 1251 | - | - | 0 | - |
| 7 | y | 1.073E+04 | 1252 | 0.0008644 | 0.6905 | +1 | 12 |
| - | - | 7749 | 1253 | - | - | 0 | - |
| - | - | 4883 | 1263 | - | - | 0 | - |
| - | - | 1.204E+04 | 1279 | - | - | 0 | - |
| - | - | 1.248E+04 | 1280 | - | - | 0 | - |
| - | - | 9822 | 1281 | - | - | 0 | - |
| - | - | 4768 | 1282 | - | - | 0 | - |
| - | - | 5221 | 1306 | - | - | 0 | - |
| 11 | c | 2.298E+04 | 1307 | 0.0002001 | 0.1531 | +1 | 11 |
| - | - | 1.493E+04 | 1308 | - | - | 0 | - |
| - | - | 6402 | 1309 | - | - | 0 | - |
| 6 | z | 9.528E+04 | 1373 | 0.002112 | 1.538 | +1 | 13 |
| - | - | 9.8E+04 | 1374 | - | - | 0 | - |
| - | - | 5.212E+04 | 1375 | - | - | 0 | - |
| - | - | 1.911E+04 | 1376 | - | - | 0 | - |
| - | - | 6111 | 1378 | - | - | 0 | - |
| - | - | 6327 | 1385 | - | - | 0 | - |
| - | - | 7074 | 1386 | - | - | 0 | - |
| - | - | 7537 | 1386 | - | - | 0 | - |
| - | - | 5844 | 1388 | - | - | 0 | - |
| 6 | y | 6.284E+04 | 1389 | 0.002525 | 1.818 | +1 | 13 |
| - | - | 4.023E+04 | 1390 | - | - | 0 | - |
| - | - | 1.872E+04 | 1391 | - | - | 0 | - |
| - | - | 6372 | 1392 | - | - | 0 | - |
| - | - | 1.169E+04 | 1393 | - | - | 0 | - |
| - | - | 5723 | 1393 | - | - | 0 | - |
| - | - | 1.005E+04 | 1394 | - | - | 0 | - |
| - | - | 8716 | 1394 | - | - | 0 | - |
| - | - | 5723 | 1395 | - | - | 0 | - |
| - | - | 7656 | 1395 | - | - | 0 | - |
| - | - | 8317 | 1396 | - | - | 0 | - |
| - | - | 6379 | 1396 | - | - | 0 | - |
| - | - | 9335 | 1397 | - | - | 0 | - |
| - | - | 6924 | 1397 | - | - | 0 | - |
| - | - | 1.444E+04 | 1434 | - | - | 0 | - |
| 12 | c | 7.296E+04 | 1435 | 0.004187 | 2.918 | +1 | 12 |
| - | - | 6.143E+04 | 1436 | - | - | 0 | - |
| - | - | 2.676E+04 | 1437 | - | - | 0 | - |
| - | - | 4747 | 1438 | - | - | 0 | - |
| - | - | 5018 | 1504 | - | - | 0 | - |
| - | - | 5938 | 1505 | - | - | 0 | - |
| - | - | 5980 | 1506 | - | - | 0 | - |
| 5 | z | 7.027E+04 | 1510 | 0.00182 | 1.205 | +1 | 14 |
| - | - | 1.734E+05 | 1511 | - | - | 0 | - |
| - | - | 1.404E+05 | 1512 | - | - | 0 | - |
| - | - | 5.933E+04 | 1513 | - | - | 0 | - |
| - | - | 2.053E+04 | 1514 | - | - | 0 | - |
| - | - | 5822 | 1525 | - | - | 0 | - |
| 5 | y | 4.136E+04 | 1526 | 0.002965 | 1.943 | +1 | 14 |
| - | - | 3.474E+04 | 1527 | - | - | 0 | - |
| - | - | 1.764E+04 | 1528 | - | - | 0 | - |
| - | - | 6930 | 1529 | - | - | 0 | - |
| - | - | 4858 | 1547 | - | - | 0 | - |
| 13 | c | 2.57E+04 | 1548 | 0.004755 | 3.072 | +1 | 13 |
| - | - | 3.068E+04 | 1549 | - | - | 0 | - |
| - | - | 1.236E+04 | 1550 | - | - | 0 | - |
| - | - | 5200 | 1551 | - | - | 0 | - |
| - | - | 4088 | 1618 | - | - | 0 | - |
| 4 | z | 2.004E+04 | 1638 | 0.005343 | 3.262 | +1 | 15 |
| - | - | 1.264E+05 | 1639 | - | - | 0 | - |
| - | - | 1.22E+05 | 1640 | - | - | 0 | - |
| - | - | 6.899E+04 | 1641 | - | - | 0 | - |
| - | - | 1.871E+04 | 1642 | - | - | 0 | - |
| - | - | 6698 | 1643 | - | - | 0 | - |
| 4 | y | 1.223E+04 | 1654 | 0.000957 | 0.5786 | +1 | 15 |
| - | - | 1.009E+04 | 1655 | - | - | 0 | - |
| - | - | 6210 | 1656 | - | - | 0 | - |
| 14 | c | 1.769E+04 | 1661 | 0.007398 | 4.454 | +1 | 14 |
| - | - | 1.394E+04 | 1662 | - | - | 0 | - |
| - | - | 6675 | 1663 | - | - | 0 | - |
| - | - | 5014 | 1702 | - | - | 0 | - |
| 15 | c | 3940 | 1757 | 0.0112 | 6.376 | +1 | 15 |
| - | - | 2.031E+04 | 1773 | - | - | 0 | - |
| 15 | c | 6.895E+04 | 1774 | 0.003937 | 2.219 | +1 | 15 |
| - | - | 6.558E+04 | 1775 | - | - | 0 | - |
| - | - | 3.641E+04 | 1776 | - | - | 0 | - |
| - | - | 1.864E+04 | 1777 | - | - | 0 | - |
| - | - | 5858 | 1778 | - | - | 0 | - |
| 3 | y | 4247 | 1800 | 0.01252 | 6.954 | +1 | 16 |
| 3 | z | 2.288E+04 | 1801 | 0.0008008 | 0.4446 | +1 | 16 |
| - | - | 7.487E+04 | 1802 | - | - | 0 | - |
| - | - | 6.969E+04 | 1803 | - | - | 0 | - |
| - | - | 4.066E+04 | 1804 | - | - | 0 | - |
| - | - | 1.427E+04 | 1805 | - | - | 0 | - |
| 3 | y | 7517 | 1817 | 0.0002377 | 0.1308 | +1 | 16 |
| - | - | 1.613E+04 | 1818 | - | - | 0 | - |
| - | - | 1.297E+04 | 1819 | - | - | 0 | - |
| - | - | 5493 | 1820 | - | - | 0 | - |
| - | - | 4474 | 1846 | - | - | 0 | - |
| 16 | c | 6.905E+04 | 1861 | 0.00264 | 1.419 | +1 | 16 |
| - | - | 8.83E+04 | 1862 | - | - | 0 | - |
| - | - | 5.557E+04 | 1863 | - | - | 0 | - |
| - | - | 2.47E+04 | 1864 | - | - | 0 | - |
| - | - | 5311 | 1865 | - | - | 0 | - |
| - | - | 1.151E+04 | 1946 | - | - | 0 | - |
| - | - | 2.116E+04 | 1947 | - | - | 0 | - |
| - | - | 2.352E+04 | 1948 | - | - | 0 | - |
| - | - | 1.274E+04 | 1949 | - | - | 0 | - |
| - | - | 6370 | 1950 | - | - | 0 | - |
| - | - | 5256 | 1974 | - | - | 0 | - |
| - | - | 6868 | 1975 | - | - | 0 | - |
| - | - | 4758 | 1976 | - | - | 0 | - |
| - | - | 5035 | 1978 | - | - | 0 | - |
| 2 | z | 8730 | 1987 | 0.01322 | 6.653 | +1 | 17 |
| - | - | 2.548E+04 | 1988 | - | - | 0 | - |
| - | - | 2.668E+04 | 1989 | - | - | 0 | - |
| 17 | c | 3.558E+04 | 1990 | 0.001519 | 0.7635 | +1 | 17 |
| - | - | 6.564E+04 | 1991 | - | - | 0 | - |
| - | - | 5.743E+04 | 1992 | - | - | 0 | - |
| - | - | 2.514E+04 | 1993 | - | - | 0 | - |
| - | - | 1.484E+04 | 1994 | - | - | 0 | - |
| - | - | 5692 | 1995 | - | - | 0 | - |
| - | - | 8866 | 2030 | - | - | 0 | - |
| - | - | 9545 | 2031 | - | - | 0 | - |
| - | - | 6071 | 2032 | - | - | 0 | - |
| - | - | 1.426E+04 | 2046 | - | - | 0 | - |
| - | - | 5.088E+04 | 2047 | - | - | 0 | - |
| - | - | 5.838E+04 | 2048 | - | - | 0 | - |
| - | - | 2.954E+04 | 2049 | - | - | 0 | - |
| - | - | 1.233E+04 | 2050 | - | - | 0 | - |
| - | - | 4607 | 2051 | - | - | 0 | - |
| - | - | 2.046E+04 | 2057 | - | - | 0 | - |
| - | - | 2.508E+04 | 2058 | - | - | 0 | - |
| - | - | 1.625E+04 | 2059 | - | - | 0 | - |
| - | - | 5499 | 2060 | - | - | 0 | - |
| - | - | 1.855E+04 | 2063 | - | - | 0 | - |
| - | - | 6.219E+04 | 2064 | - | - | 0 | - |
| - | - | 7.81E+04 | 2065 | - | - | 0 | - |
| - | - | 4.331E+04 | 2066 | - | - | 0 | - |
| - | - | 1.882E+04 | 2067 | - | - | 0 | - |
| - | - | 7065 | 2068 | - | - | 0 | - |
| - | - | 6443 | 2073 | - | - | 0 | - |
| - | - | 4.426E+04 | 2074 | - | - | 0 | - |
| - | - | 1.964E+05 | 2075 | - | - | 0 | - |
| - | - | 2.134E+05 | 2076 | - | - | 0 | - |
| - | - | 1.223E+05 | 2077 | - | - | 0 | - |
| - | - | 5.657E+04 | 2078 | - | - | 0 | - |
| - | - | 2.267E+04 | 2079 | - | - | 0 | - |
| - | - | 7089 | 2080 | - | - | 0 | - |
| - | - | 4.566E+04 | 2090 | - | - | 0 | - |
| - | - | 2.425E+05 | 2091 | - | - | 0 | - |
| - | - | 9.038E+05 | 2092 | - | - | 0 | - |
| - | - | 9.753E+05 | 2093 | - | - | 0 | - |
| - | - | 5.933E+05 | 2094 | - | - | 0 | - |
| - | - | 2.744E+05 | 2095 | - | - | 0 | - |
| - | - | 8.727E+04 | 2096 | - | - | 0 | - |
| - | - | 2.471E+04 | 2097 | - | - | 0 | - |
| - | - | 8358 | 2295 | - | - | 0 | - |
| - | - | 5793 | 2296 | - | - | 0 | - |

m/z Charge Intensity FragmentType MassShift Position
122.33216857910156 0 2153.0276
130.06483459472656 0 2524.4446
136.07574462890625 0 2181.309
138.61068725585938 0 2383.2966
148.95452880859375 0 2443.2344
184.23052978515625 0 2702.062
186.07943725585938 0 2957.0486
187.0862579345703 0 5677.901
189.086669921875 0 4652.1006 c Ammonia loss 3
199.3961181640625 0 3126.2969
203.86627197265625 0 2716.3914
212.5716552734375 0 2636.0469
216.7153778076172 0 2979.8477
217.08160400390625 0 27923.398
218.5696563720703 0 2606.1506
229.11802673339844 0 23744.178 y Water loss 16
246.12332153320312 0 27302.145
247.1280517578125 0 16618.803 y 16
273.1073303222656 0 3175.8274
274.1180725097656 0 45457.8
275.12188720703125 0 8536.907
292.1296081542969 0 5677.3276
309.15545654296875 0 4614.051
313.9812316894531 0 3166.7026
316.1502990722656 0 4076.238 y Water loss 15
328.4894714355469 0 2701.543
334.16046142578125 0 23058.502 y 15
369.1227111816406 0 3298.5312
434.1933898925781 0 3443.5288
437.18304443359375 0 7296.5146
447.244384765625 0 11210.91 y 14
465.2084655761719 0 12638.843
477.1999206542969 0 9493.094
478.2041015625 0 5920.5166
480.22332763671875 0 6707.418
490.1791687011719 0 3271.7224
528.244873046875 0 5889.533
538.2513427734375 0 7664.8667
539.2598266601562 0 15521.292
541.319091796875 0 2972.335
560.3284912109375 0 11530.902 y 13
561.2661743164062 0 17706.254 c Ammonia loss 8
561.3324584960938 0 2983.7905 z Ammonia loss 7
561.7675170898438 0 10763.826
562.265380859375 0 4314.5293
563.2482299804688 0 24907.99
564.244873046875 0 7953.649
565.24072265625 0 13103.085 c Ammonia loss 3
566.2446899414062 0 3863.1143
567.8469848632812 0 9204.193
568.3494873046875 0 5012.8975
577.661376953125 0 3569.1616
581.2587280273438 0 20822.281
582.2662353515625 0 110547.805 c 3
583.268798828125 0 37899.957
584.2708129882812 0 6544.3315
586.3248291015625 0 25037.617 c Ammonia loss 14
586.6599731445312 0 25735.066
586.9942016601562 0 18216.408
587.3298950195312 0 3742.14
596.7864990234375 0 16855.35 c Ammonia loss 9
597.2871704101562 0 8900.981
597.7881469726562 0 3994.822
601.3338623046875 0 4803.98
606.339111328125 0 7571.3765 y 2
606.6730346679688 0 5998.2754
609.666259765625 0 3338.8167
614.3362426757812 0 10200.112 w 12
615.3363647460938 0 20560.004 c Ammonia loss 15
615.6704711914062 0 25149.074
616.0048828125 0 10787.149
626.3860473632812 0 28272.574 y 6
626.888671875 0 22408.29
627.3906860351562 0 6521.7
649.02099609375 0 12859.624
649.352783203125 0 11640.873
649.6843872070312 0 6292
652.345458984375 0 10756.674
652.6804809570312 0 19276.182
653.0166015625 0 13448.908
653.3499755859375 0 10485.375
657.2982177734375 0 9045.316
657.393798828125 0 13981.682 z 12
658.3505249023438 0 115640.72 c Ammonia loss 16
658.6851196289062 0 136216.19
659.018798828125 0 68581.39
659.2913818359375 0 11353.147
659.352294921875 0 35042.516
659.6874389648438 0 10277.45
660.2969970703125 0 5356.0957
664.354248046875 0 7188.9644
664.6868286132812 0 6364.6426
668.3657836914062 0 5480.3545 y 1
668.7026977539062 0 3975.408
673.413818359375 0 10760.517 y 12
674.3045043945312 0 5613.4727
675.311279296875 0 34569.414
676.3162841796875 0 20673.375
677.3201904296875 0 5552.9463
682.0399780273438 0 6442.991
682.37646484375 0 6675.485
685.90380859375 0 3342.6758 y Water loss 5
686.0366821289062 0 3641.0269
686.907958984375 0 3489.709 z 5
687.7097778320312 0 4436.483
691.3732299804688 0 39354.41
691.7072143554688 0 54325.99
692.0418701171875 0 31804.305
692.3741455078125 0 14882.121
692.7076416015625 0 6120.6685
694.9205322265625 0 4169.9927 y 5
695.4165649414062 0 7446.607
696.7068481445312 0 13480.439
697.0364379882812 0 12339.277
697.376708984375 0 719648.75
697.7113647460938 0 880244.94
698.0451049804688 0 550154.94
698.3795776367188 0 271298.25
698.713623046875 0 89639.34
699.0473022460938 0 18362.502
702.2996215820312 0 14454.445 c Ammonia loss 4
703.3023681640625 0 4617.3813
708.859130859375 0 4138.3257 c Water loss 11
709.8590698242188 0 4958.388
717.3678588867188 0 34695.39
717.8720092773438 0 359706.34 c 11
718.3737182617188 0 307942.28
718.8748168945312 0 149064.69
719.3247680664062 0 117959.61 c 4
719.3792114257812 0 20879.326
719.8763427734375 0 8380.205
720.3278198242188 0 45873.76
720.3930053710938 0 3656.7441
721.3285522460938 0 13090.985
742.4296264648438 0 5179.3716
742.9310913085938 0 4390.779
743.4361572265625 0 5425.3857
752.4082641601562 0 15230.9375
752.9076538085938 0 15457.869
753.4046630859375 0 4439.954
755.4352416992188 0 42709.52 z 4
755.9382934570312 0 67006.03
756.4400024414062 0 43878.3
756.94482421875 0 8834.341
757.4461059570312 0 5585.6274
763.4456176757812 0 46083.02 y 4
763.946533203125 0 39248.195
764.4481811523438 0 13689.933
764.951904296875 0 4846.709
765.900634765625 0 7511.442 c Ammonia loss 12
766.40283203125 0 7413.661
766.9019775390625 0 3961.1738
773.9108276367188 0 27169.893
774.4139404296875 0 214871.5 c 12
774.9158325195312 0 202680.73
775.4172973632812 0 100190.625
775.9185791015625 0 24760.104
776.4232177734375 0 6436.4224
780.9082641601562 0 3532.06
783.4529418945312 0 6200.3867
783.9403076171875 0 5683.5884
785.489990234375 0 14442.36 z 11
786.4960327148438 0 12901.924
787.4959716796875 0 6890.983
790.450439453125 0 36884.04 w 3
790.9509887695312 0 30206.45
791.450439453125 0 21127.459
791.942626953125 0 9249.339
792.4402465820312 0 3356.098
796.3536376953125 0 10172.976
797.355224609375 0 3657.2085
801.5079956054688 0 7384.7188 y 11
808.9508666992188 0 8443.452
809.4501953125 0 8290.32
809.953125 0 7705.9175
811.3639526367188 0 30747.201
812.365966796875 0 18375.734
813.369873046875 0 5135.5537
818.9669189453125 0 7452.388 y Ammonia loss 3
819.4649047851562 0 153867.19 z 3
819.9676513671875 0 243240.17
820.4696044921875 0 165807.55
820.9713745117188 0 76931.63
821.471435546875 0 26288.346
821.9694213867188 0 3754.6575
822.444580078125 0 14841.358 c Ammonia loss 13
822.94580078125 0 22595.268
823.4425659179688 0 4338.106
823.9512329101562 0 5574.2344
826.971923828125 0 3641.158
827.4754028320312 0 36160.965 y 3
827.975830078125 0 49482.992
828.4777221679688 0 20764.582
828.9793090820312 0 8770.902
830.44873046875 0 3484.868
830.955810546875 0 70321.086 c 13
831.4576416015625 0 70044.16
831.5389404296875 0 2216.3281
831.9594116210938 0 35058.035
832.4609985351562 0 18477.422
832.9592895507812 0 7192.022
835.9762573242188 0 3408.1865
839.3573608398438 0 31887.01 c Ammonia loss 5
840.360107421875 0 16304.283
841.3609008789062 0 3746.9324
842.4591064453125 0 3870.603
850.4666137695312 0 5261.4937
850.9696044921875 0 8861.355
851.4691772460938 0 9132.687
851.9688720703125 0 9614.062
857.9822998046875 0 6537.834
858.4833374023438 0 5894.629
858.962646484375 0 3712.1855
859.9767456054688 0 4407.8657
864.9881591796875 0 8435.652
865.48681640625 0 19100.475
865.9853515625 0 25297.695
866.4849853515625 0 11534.373
869.477294921875 0 6085.353
869.9781494140625 0 7240.037
870.4780883789062 0 4024.5947
871.9758911132812 0 10286.644
872.4819946289062 0 10630.578
872.9722900390625 0 8009.9214
873.4713745117188 0 3668.534
874.4439697265625 0 3297.7031
877.989990234375 0 11657.392
878.486083984375 0 15733.463 c Water loss 14
878.98583984375 0 66646.45 c Ammonia loss 14
879.4866333007812 0 73871.7
879.9865112304688 0 41376.617
880.4879150390625 0 22761.443
880.9929809570312 0 9862.547
884.791015625 0 3626.3223
886.994140625 0 97183.33
887.496826171875 0 195478.67 c 14
887.998779296875 0 158408.56
888.5005493164062 0 88119.57
889.002685546875 0 36867.836
889.5025024414062 0 7271.543
892.4912109375 0 4983.5474 z Ammonia loss 2
897.514404296875 0 17371.459
898.5579223632812 0 35911.086 y 10
899.5641479492188 0 14935.015
900.0000610351562 0 12139.263 y Water loss 2
900.5036010742188 0 17780.447 y Ammonia loss 2
900.9971923828125 0 111634.125 z 2
901.4988403320312 0 143903.3
902.0001831054688 0 77230.35
902.5015258789062 0 40728.465
902.9981689453125 0 8185.3564
908.5018920898438 0 4387.985
909.0060424804688 0 116084.586 y 2
909.5068359375 0 115805.1
910.0079345703125 0 62812.996
910.5101318359375 0 25197.21
911.0095825195312 0 9539.801
913.4823608398438 0 3936.068
913.996337890625 0 9493.483
922.0032958984375 0 5571.878 c Water loss 15
922.501220703125 0 28164.162 c Ammonia loss 15
923.0022583007812 0 37888.047
923.5030517578125 0 29206.848
924.0023803710938 0 5419.8013
928.494873046875 0 4473.398
929.1478271484375 0 4513.302
929.4762573242188 0 4447.6235
930.15234375 0 4942.605
930.4974365234375 0 20076.135
930.8345336914062 0 4007.9421
931.0136108398438 0 220953.44 c 15
931.5150756835938 0 256441.58
932.0169067382812 0 168475.67
932.5178833007812 0 54929.03
933.0188598632812 0 16451.611
933.5234375 0 4715.2925
944.0231323242188 0 14282.171
944.521484375 0 12030.049
945.0205078125 0 15365.048
952.4277954101562 0 10786.222
953.4368286132812 0 88651.47 c 6
953.5783081054688 0 143767.9 z 9
954.4402465820312 0 42722.406
954.5812377929688 0 79675.59
955.4434204101562 0 15017.737
955.585205078125 0 27886.246
956.5894775390625 0 4893.196
958.5196533203125 0 3576.1362
965.0191650390625 0 7565.0347
965.5175170898438 0 5704.301
966.0252685546875 0 3845.768
966.4442138671875 0 14098.789
967.452880859375 0 9605.131
968.4447021484375 0 5806.0454
969.596435546875 0 36322.652 y 9
970.5990600585938 0 20316.361
971.6083374023438 0 6056.6753
972.0181884765625 0 3580.1558
973.025146484375 0 14508.838
973.5267333984375 0 43701.89
974.0283813476562 0 45888.656
974.5326538085938 0 26612.354
975.0309448242188 0 13483.1875
978.0157470703125 0 3850.776
978.5234985351562 0 5516.0034
979.015869140625 0 6112.0312
980.0349731445312 0 5771.414
980.5386352539062 0 10415.101
981.033935546875 0 12272.996
981.5279541015625 0 12104.708
984.5398559570312 0 4958.708
986.5574951171875 0 5023.1387
987.0248413085938 0 44218.367 c Ammonia loss 16
987.525146484375 0 64535.492
988.0260009765625 0 41011.01
988.5310668945312 0 27197.547
989.0317993164062 0 5468.4463
993.0384521484375 0 9293.873 y Water loss 1
993.54052734375 0 10407.887 y Ammonia loss 1
994.0403442382812 0 37501.277 z 1
994.5380859375 0 45023.008
995.03857421875 0 25651.129
995.53564453125 0 234613.97 c 16
996.0370483398438 0 266360
996.5382690429688 0 161838.61
997.0392456054688 0 90583.25
997.5408325195312 0 28212.494
998.0443725585938 0 10136.354
1001.5526733398438 0 6131.0557
1002.0364379882812 0 19816.916 y 1
1002.5372314453125 0 23829.014
1003.0399169921875 0 12299.158
1003.529052734375 0 4537.949
1005.04150390625 0 19767.164
1005.5413208007812 0 18275.123
1006.0450439453125 0 15500.841
1006.5393676757812 0 14095.44
1008.0455932617188 0 28076.826
1008.5436401367188 0 26210.086
1009.044677734375 0 24160.375
1009.44140625 0 13549.875
1009.5347290039062 0 34405.336
1010.0335693359375 0 43620.312
1010.4576416015625 0 128017.875 c 7
1011.0400390625 0 23854.756
1011.460205078125 0 82573.04
1012.0513916015625 0 4982.557
1012.4644165039062 0 24809.768
1013.462646484375 0 6314.8306
1014.5530395507812 0 24503.943
1015.0543823242188 0 36372.04
1015.5572509765625 0 42981.496
1016.0576171875 0 35824.824
1016.5555419921875 0 39457.97
1017.0431518554688 0 22440.35
1017.5429077148438 0 9883.676
1018.0374755859375 0 15740.961
1018.5361938476562 0 18020.506
1019.0368041992188 0 13230.064
1019.5397338867188 0 7064.571
1023.0618286132812 0 10784.875
1023.5556640625 0 74664.12
1024.0574951171875 0 80371.34
1024.5472412109375 0 102895.37
1025.0443115234375 0 97372.5
1025.54443359375 0 46570.965
1026.0447998046875 0 18194.445
1028.06591796875 0 5323.4624
1028.5469970703125 0 13388.092
1029.0523681640625 0 15171.418
1029.5531005859375 0 9114.283
1030.05224609375 0 4755.4604
1030.541015625 0 7670.772
1036.05224609375 0 5781.034
1036.5615234375 0 9023.411
1037.0582275390625 0 37448.055
1037.5540771484375 0 310483.28
1038.054931640625 0 372984.44
1038.55615234375 0 226786.61
1039.0556640625 0 101191.01
1039.558837890625 0 53133.31
1040.055419921875 0 8721.789
1045.0589599609375 0 18599.459
1045.5618896484375 0 406298.38
1046.065185546875 0 1615780.9
1046.5670166015625 0 1700635.4
1047.068115234375 0 999284.5
1047.5694580078125 0 487556.4
1048.0699462890625 0 149339.3
1048.571044921875 0 33557.062
1081.6724853515625 0 84878.28 z 8
1082.6781005859375 0 115135.42
1083.6805419921875 0 49233.113
1084.683837890625 0 14026.282
1094.5384521484375 0 22147.965
1095.53955078125 0 12294.692
1097.690673828125 0 26013.967 y 8
1098.69287109375 0 15133.369
1099.7000732421875 0 4675.489
1138.5528564453125 0 190325.36 c 8
1138.6949462890625 0 49515.117 z 7
1139.555419921875 0 122451.19
1139.70068359375 0 53186.367
1140.5570068359375 0 48486.09
1140.70361328125 0 31290.63
1141.561767578125 0 12756.13
1141.710205078125 0 8741.698
1154.7091064453125 0 8928.854 y 7
1155.72265625 0 4192.4956
1165.5750732421875 0 21072.512
1166.579833984375 0 13543.914
1167.588134765625 0 5244.806
1250.7584228515625 0 4501.2695
1251.7662353515625 0 10729.676 y 6
1252.7689208984375 0 7749.38
1262.63134765625 0 4883.4634
1278.752685546875 0 12040.297
1279.7542724609375 0 12484.603
1280.765380859375 0 9821.664
1281.77197265625 0 4767.59
1305.64501953125 0 5221.2695
1306.6441650390625 0 22979.79 c 10
1307.6424560546875 0 14927.379
1308.6451416015625 0 6402.3984
1372.80517578125 0 95277.14 z 5
1373.809326171875 0 97995.54
1374.8118896484375 0 52120.074
1375.815185546875 0 19110.965
1377.703125 0 6110.819
1385.212890625 0 6326.7563
1385.7261962890625 0 7073.5996
1386.2186279296875 0 7537.2397
1387.8214111328125 0 5843.7817
1388.823486328125 0 62837.645 y 5
1389.8250732421875 0 40230.43
1390.8297119140625 0 18722.416
1391.7080078125 0 6371.816
1392.7056884765625 0 11690.663
1393.236572265625 0 5723.229
1393.7137451171875 0 10050.095
1394.2279052734375 0 8716.461
1394.7144775390625 0 5722.9463
1395.2281494140625 0 7655.8115
1395.71337890625 0 8316.826
1396.238525390625 0 6378.6167
1396.736328125 0 9335.276
1397.2430419921875 0 6924.173
1433.729736328125 0 14441.734
1434.7347412109375 0 72960.2 c 11
1435.73828125 0 61428.69
1436.7423095703125 0 26761.135
1437.723388671875 0 4747.013
1503.809814453125 0 5017.78
1504.818359375 0 5937.76
1505.81005859375 0 5979.894
1509.8643798828125 0 70267.18 z 4
1510.8695068359375 0 173386.8
1511.8729248046875 0 140401.31
1512.8768310546875 0 59327.48
1513.880615234375 0 20532.309
1524.8692626953125 0 5821.9463
1525.8819580078125 0 41360.516 y 4
1526.8848876953125 0 34743.168
1527.8883056640625 0 17637.498
1528.89599609375 0 6930.3813
1546.8099365234375 0 4858.2705
1547.8182373046875 0 25701.59 c 12
1548.8212890625 0 30676.053
1549.82666015625 0 12362.747
1550.83154296875 0 5199.889
1617.9217529296875 0 4087.6707
1637.91943359375 0 20040.053 z 3
1638.929931640625 0 126449.414
1639.933349609375 0 122014.33
1640.9366455078125 0 68994.44
1641.9454345703125 0 18707.969
1642.94873046875 0 6698.298
1653.9444580078125 0 12226.059 y 3
1654.9476318359375 0 10089.782
1655.9329833984375 0 6210.3955
1660.899658203125 0 17686.771 c 13
1661.9134521484375 0 13939.915
1662.9193115234375 0 6674.807
1701.9346923828125 0 5014.055
1756.953369140625 0 3940.386 c Ammonia loss 14
1772.9814453125 0 20308.768
1773.9871826171875 0 68945.51 c 14
1774.9912109375 0 65581.47
1775.9913330078125 0 36407.883
1776.9989013671875 0 18641.846
1778.0003662109375 0 5858.085
1799.9927978515625 0 4246.773 y Ammonia loss 2
1800.9873046875 0 22883.287 z 2
1801.991943359375 0 74865.32
1802.9952392578125 0 69691.15
1803.9974365234375 0 40656.42
1805.0003662109375 0 14269.932
1817.006591796875 0 7516.72 y 2
1818.0067138671875 0 16133.044
1819.012939453125 0 12968.208
1820.0262451171875 0 5493.1655
1846.007080078125 0 4473.943
1861.0205078125 0 69047.22 c 15
1862.0238037109375 0 88299.11
1863.0283203125 0 55568.58
1864.0328369140625 0 24702.672
1865.04052734375 0 5311.4824
1946.0435791015625 0 11511.748
1947.0531005859375 0 21161.932
1948.0577392578125 0 23524.725
1949.0614013671875 0 12742.124
1950.0570068359375 0 6370.4634
1974.040771484375 0 5256.027
1975.0418701171875 0 6868.396
1976.0556640625 0 4757.623
1978.0572509765625 0 5035.1895
1987.05419921875 0 8729.944 z 1
1988.066650390625 0 25483.81
1989.07080078125 0 26677.453
1990.0672607421875 0 35577.93 c 16
1991.068359375 0 65641.625
1992.07177734375 0 57431.49
1993.0732421875 0 25143.912
1994.08056640625 0 14835.646
1995.0399169921875 0 5692.0176
2030.1083984375 0 8866.111
2031.1114501953125 0 9545.114
2032.122802734375 0 6071.4946
2046.11181640625 0 14258.986
2047.1087646484375 0 50882.4
2048.11376953125 0 58380.35
2049.112060546875 0 29538.625
2050.109130859375 0 12328.465
2051.123046875 0 4606.7393
2057.09033203125 0 20457.557
2058.087646484375 0 25078.719
2059.0947265625 0 16248.488
2060.09521484375 0 5499.272
2063.132080078125 0 18550.432
2064.134765625 0 62185.293
2065.13720703125 0 78103.64
2066.13818359375 0 43305.164
2067.1435546875 0 18819.305
2068.159912109375 0 7064.852
2073.09765625 0 6442.8916
2074.098876953125 0 44264.508
2075.10400390625 0 196449.27
2076.10546875 0 213406.81
2077.10693359375 0 122327.695
2078.109130859375 0 56574.27
2079.111083984375 0 22665.521
2080.106689453125 0 7088.748
2090.1162109375 0 45657.195
2091.12060546875 0 242505.4
2092.127685546875 0 903773.8
2093.1318359375 0 975279.06
2094.133544921875 0 593346.4
2095.13671875 0 274363.47
2096.1396484375 0 87269.41
2097.136962890625 0 24708.697
2295.21630859375 0 8358.337
2296.22509765625 0 5792.9507

Spectrum Details

|  |  |
| --- | --- |
| Matched peaks? Matched peaks The total absolute number of peaks matched. Additionally in brackets the total fraction of peaks matched and the total number of peaks is shown. | 87 (15.48% of 562) |
| FDR? FDR The false discovery rate estimated for this peptide. It is calculated by matching all theoretical fragments with a non-integer shift with the raw peaks for this spectrum. This is done with 40 different shifts. The resulting percentage is the average number of annotated peaks over the number of annotated peaks with the correct spectrum. | 3.34% |
| Satellite FDR? Satellite FDR See the FDR for details on its calculation. This satellite ion specific FDR only contains the satellite ions (d/w) for I/L/J positions. | 0.00% |
| PSM Score? PSM Score The PSM Score as given by Hecklib to this annotated spectrum. It is shown with three significant figures. | 527 |

#### Spectrum 7499? Spectrum 7499 The raw spectrum of this peptide as annotated by Hecklib. The fragments are coloured according to ion type (see legend). Any peaks with a star '\*' as text can be hovered over to see the full details, first the ion type second the mass shift type. By hovering over the amino acids in the peptide or ions in the legend the corresponding peaks are highlighted. By toggling the 'Unassigned' label you can turn the background (unassigned) peaks on or off in the plot. By updating the slider in the Ion legend you can update the spectrum to only show the top X% of the peaks with labels. The top X% means any peak that is within X% of the highest intensity. By dragging in the spectrum you can zoom in to a specific part of the spectrum and use 'Zoom Out' to get back to the original zoom level. The annotation of the spectrum is based on the given sequence in the peptides file and is done with different software so inconsistencies are likely. The peaks are annotated based on the given sequence, with 20 ppm tolerance.

Copy Data

##### Spectrum 7499 (TSV)

###### Preview

```
Loading example...
```

*Click on the button to copy the data to your clipboard.*

Mz MinMz MaxIntensity Max

WidthHeightPeptide font sizePeptide stroke widthSpectrum font sizeSpectrum stroke widthCompact peptide

Ion legend

wxyz

abcd

OtherUnassignedIon^Charge^_Position_Show for top:%

SWYQHHPGKAPKJJJSEV

01.21e+52.42e+53.63e+54.84e+5

Zoom Out

c^+3^_4_y^+1^_2_y^+1^_2_y^+1^_3_y^+1^_3_y^+1^_4_y^+1^_5_c^+2^_9_z^+2^_11_c^+1^_4_c^+1^_4_c^+1^_4_c^+3^_15_c^+2^_10_y^+3^_16_y^+3^_16_w^+1^_6_c^+3^_16_y^+2^_12_z^+1^_6_c^+3^_17_y^+3^_17_y^+1^_6_y^+2^_13_c^+1^_5_c^+2^_12_c^+2^_12_c^+2^_12_c^+1^_5_z^+2^_14_y^+2^_14_c^+2^_13_c^+2^_13_z^+1^_7_w^+2^_15_y^+1^_7_z^+2^_15_y^+2^_15_z^+2^_15_c^+2^_14_y^+2^_15_c^+2^_14_c^+1^_6_c^+2^_15_c^+2^_15_c^+2^_15_z^+2^_16_z^+2^_16_y^+1^_8_y^+2^_16_y^+2^_16_z^+2^_16_y^+2^_16_c^+2^_16_c^+2^_16_c^+2^_16_c^+1^_7_z^+1^_9_y^+1^_9_c^+2^_17_c^+2^_17_y^+2^_17_z^+2^_17_c^+2^_17_y^+2^_17_c^+1^_8_z^+1^_10_y^+1^_10_c^+1^_9_z^+1^_11_y^+1^_11_w^+1^_12_y^+1^_12_c^+1^_11_z^+1^_13_y^+1^_13_c^+1^_12_z^+1^_14_y^+1^_14_c^+1^_13_z^+1^_15_y^+1^_15_c^+1^_14_c^+1^_15_z^+1^_16_y^+1^_16_c^+1^_16_z^+1^_17_c^+1^_17_

0580115917392318

Fragment Matches Table

Show background peaks

| Position | Ion type | Intensity | mz Theoretical | mz Error (Th) | mz Error (ppm) | Charge | Series Number |
| --- | --- | --- | --- | --- | --- | --- | --- |
| - | - | 3578 | 120.1 | - | - | 0 | - |
| - | - | 1095 | 120.1 | - | - | 0 | - |
| - | - | 509.1 | 125.8 | - | - | 0 | - |
| - | - | 447.5 | 129.1 | - | - | 0 | - |
| - | - | 436.7 | 129.2 | - | - | 0 | - |
| - | - | 467 | 130.6 | - | - | 0 | - |
| - | - | 511.1 | 132.1 | - | - | 0 | - |
| - | - | 801 | 132.1 | - | - | 0 | - |
| - | - | 1021 | 136.1 | - | - | 0 | - |
| - | - | 678.7 | 146.1 | - | - | 0 | - |
| - | - | 645.2 | 162.4 | - | - | 0 | - |
| - | - | 1052 | 173.4 | - | - | 0 | - |
| - | - | 584.7 | 183.1 | - | - | 0 | - |
| - | - | 766.5 | 187.1 | - | - | 0 | - |
| 4 | c | 840.5 | 189.1 | 0.002285 | 12.09 | +3 | 4 |
| - | - | 719 | 201.1 | - | - | 0 | - |
| - | - | 7880 | 217.1 | - | - | 0 | - |
| - | - | 978.9 | 219.1 | - | - | 0 | - |
| - | - | 1015 | 223.1 | - | - | 0 | - |
| 17 | y | 6401 | 229.1 | 1.259E-05 | 0.05496 | +1 | 2 |
| - | - | 699.9 | 230.1 | - | - | 0 | - |
| - | - | 1674 | 232.1 | - | - | 0 | - |
| - | - | 1447 | 239.1 | - | - | 0 | - |
| - | - | 9142 | 246.1 | - | - | 0 | - |
| 17 | y | 3734 | 247.1 | 0.0007811 | 3.161 | +1 | 2 |
| - | - | 1582 | 247.1 | - | - | 0 | - |
| - | - | 1533 | 249.1 | - | - | 0 | - |
| - | - | 695 | 255.1 | - | - | 0 | - |
| - | - | 604.7 | 260.5 | - | - | 0 | - |
| - | - | 1421 | 267.1 | - | - | 0 | - |
| - | - | 1.226E+04 | 274.1 | - | - | 0 | - |
| - | - | 1905 | 275.1 | - | - | 0 | - |
| - | - | 1101 | 292.1 | - | - | 0 | - |
| - | - | 3838 | 295.1 | - | - | 0 | - |
| - | - | 1613 | 309.2 | - | - | 0 | - |
| - | - | 719.8 | 310.2 | - | - | 0 | - |
| - | - | 724.2 | 312.6 | - | - | 0 | - |
| 16 | y | 855.7 | 316.2 | 0.00044 | 1.392 | +1 | 3 |
| - | - | 878.9 | 320.1 | - | - | 0 | - |
| 16 | y | 5584 | 334.2 | 0.0001037 | 0.3102 | +1 | 3 |
| - | - | 797.2 | 335.2 | - | - | 0 | - |
| - | - | 862.9 | 365.5 | - | - | 0 | - |
| - | - | 713.5 | 366.1 | - | - | 0 | - |
| - | - | 1939 | 369.1 | - | - | 0 | - |
| - | - | 829.6 | 396.8 | - | - | 0 | - |
| - | - | 1674 | 403.2 | - | - | 0 | - |
| - | - | 691.7 | 434.2 | - | - | 0 | - |
| - | - | 1185 | 435.2 | - | - | 0 | - |
| - | - | 2460 | 437.2 | - | - | 0 | - |
| - | - | 753.5 | 445 | - | - | 0 | - |
| 15 | y | 2098 | 447.2 | 6.461E-06 | 0.01445 | +1 | 4 |
| - | - | 1439 | 453.2 | - | - | 0 | - |
| - | - | 2101 | 465.2 | - | - | 0 | - |
| - | - | 788.5 | 474.3 | - | - | 0 | - |
| - | - | 2093 | 477.2 | - | - | 0 | - |
| - | - | 2188 | 480.2 | - | - | 0 | - |
| - | - | 880 | 481.2 | - | - | 0 | - |
| - | - | 798.7 | 524.3 | - | - | 0 | - |
| - | - | 1703 | 528.2 | - | - | 0 | - |
| - | - | 1060 | 529.2 | - | - | 0 | - |
| - | - | 788.6 | 536.3 | - | - | 0 | - |
| - | - | 2412 | 538.3 | - | - | 0 | - |
| - | - | 3893 | 539.3 | - | - | 0 | - |
| - | - | 859.6 | 540.3 | - | - | 0 | - |
| - | - | 916.5 | 541.3 | - | - | 0 | - |
| - | - | 835.2 | 546.3 | - | - | 0 | - |
| - | - | 1225 | 549 | - | - | 0 | - |
| 14 | y | 4005 | 560.3 | 0.0005243 | 0.9357 | +1 | 5 |
| 9 | c | 3813 | 561.3 | 0.001208 | 2.153 | +2 | 9 |
| 8 | z | 1050 | 561.3 | 0.003209 | 5.716 | +2 | 11 |
| - | - | 1425 | 561.8 | - | - | 0 | - |
| - | - | 6444 | 563.2 | - | - | 0 | - |
| 4 | c | 1314 | 564.3 | 0.00486 | 8.614 | +1 | 4 |
| 4 | c | 2701 | 565.2 | 0.0009922 | 1.755 | +1 | 4 |
| - | - | 2623 | 566.2 | - | - | 0 | - |
| - | - | 2173 | 567.8 | - | - | 0 | - |
| - | - | 832.7 | 577 | - | - | 0 | - |
| - | - | 937.6 | 578 | - | - | 0 | - |
| - | - | 1106 | 580.3 | - | - | 0 | - |
| - | - | 5387 | 581.3 | - | - | 0 | - |
| 4 | c | 2.806E+04 | 582.3 | 0.0002 | 0.3435 | +1 | 4 |
| - | - | 8857 | 583.3 | - | - | 0 | - |
| - | - | 2084 | 584.3 | - | - | 0 | - |
| 15 | c | 6979 | 586.3 | 0.0004077 | 0.6953 | +3 | 15 |
| - | - | 8286 | 586.7 | - | - | 0 | - |
| - | - | 3869 | 587 | - | - | 0 | - |
| - | - | 1220 | 587.8 | - | - | 0 | - |
| 10 | c | 3910 | 596.8 | 0.0001976 | 0.3311 | +2 | 10 |
| - | - | 2304 | 597.3 | - | - | 0 | - |
| 3 | y | 873.7 | 600.7 | 0.002536 | 4.222 | +3 | 16 |
| - | - | 887.9 | 601.3 | - | - | 0 | - |
| 3 | y | 1512 | 606.3 | 0.002313 | 3.814 | +3 | 16 |
| - | - | 1151 | 606.7 | - | - | 0 | - |
| - | - | 1969 | 607 | - | - | 0 | - |
| - | - | 1520 | 609.3 | - | - | 0 | - |
| 13 | w | 1746 | 614.3 | 0.0005187 | 0.8443 | +1 | 6 |
| 16 | c | 5938 | 615.3 | 0.001328 | 2.159 | +3 | 16 |
| - | - | 6900 | 615.7 | - | - | 0 | - |
| - | - | 4037 | 616 | - | - | 0 | - |
| - | - | 1581 | 616.3 | - | - | 0 | - |
| - | - | 1220 | 620.7 | - | - | 0 | - |
| - | - | 1609 | 621 | - | - | 0 | - |
| - | - | 1333 | 621.3 | - | - | 0 | - |
| - | - | 977.4 | 623.7 | - | - | 0 | - |
| 7 | y | 7835 | 626.4 | 0.0009955 | 1.589 | +2 | 12 |
| - | - | 5403 | 626.9 | - | - | 0 | - |
| - | - | 2422 | 627.4 | - | - | 0 | - |
| - | - | 785 | 627.9 | - | - | 0 | - |
| - | - | 1385 | 639.9 | - | - | 0 | - |
| - | - | 938.9 | 644.3 | - | - | 0 | - |
| - | - | 735 | 647.7 | - | - | 0 | - |
| - | - | 3105 | 649 | - | - | 0 | - |
| - | - | 3448 | 649.4 | - | - | 0 | - |
| - | - | 1124 | 649.7 | - | - | 0 | - |
| - | - | 888 | 650 | - | - | 0 | - |
| - | - | 4728 | 652.3 | - | - | 0 | - |
| - | - | 6157 | 652.7 | - | - | 0 | - |
| - | - | 4637 | 653 | - | - | 0 | - |
| - | - | 1955 | 653.4 | - | - | 0 | - |
| - | - | 1713 | 657.3 | - | - | 0 | - |
| 13 | z | 4103 | 657.4 | 0.0001868 | 0.2842 | +1 | 6 |
| - | - | 859.9 | 658.3 | - | - | 0 | - |
| 17 | c | 2.4E+04 | 658.4 | 0.0006193 | 0.9406 | +3 | 17 |
| - | - | 3.775E+04 | 658.7 | - | - | 0 | - |
| - | - | 2.265E+04 | 659 | - | - | 0 | - |
| - | - | 2828 | 659.3 | - | - | 0 | - |
| - | - | 1.199E+04 | 659.4 | - | - | 0 | - |
| - | - | 1046 | 659.4 | - | - | 0 | - |
| - | - | 3036 | 659.7 | - | - | 0 | - |
| - | - | 1168 | 660 | - | - | 0 | - |
| - | - | 1334 | 660.3 | - | - | 0 | - |
| 2 | y | 1020 | 662.7 | 0.00509 | 7.681 | +3 | 17 |
| - | - | 1223 | 664.4 | - | - | 0 | - |
| - | - | 1227 | 664.7 | - | - | 0 | - |
| - | - | 917.4 | 665 | - | - | 0 | - |
| - | - | 1159 | 665.4 | - | - | 0 | - |
| - | - | 846.4 | 669.4 | - | - | 0 | - |
| 13 | y | 2400 | 673.4 | 0.0002005 | 0.2978 | +1 | 6 |
| - | - | 2586 | 674.3 | - | - | 0 | - |
| - | - | 912.7 | 674.4 | - | - | 0 | - |
| - | - | 1.062E+04 | 675.3 | - | - | 0 | - |
| - | - | 5438 | 676.3 | - | - | 0 | - |
| - | - | 1346 | 677.3 | - | - | 0 | - |
| - | - | 938.9 | 678.4 | - | - | 0 | - |
| - | - | 886.6 | 680.4 | - | - | 0 | - |
| - | - | 3229 | 682 | - | - | 0 | - |
| - | - | 1875 | 682.4 | - | - | 0 | - |
| - | - | 1205 | 682.7 | - | - | 0 | - |
| - | - | 1617 | 685.7 | - | - | 0 | - |
| - | - | 1394 | 687.4 | - | - | 0 | - |
| - | - | 1.188E+04 | 691.4 | - | - | 0 | - |
| - | - | 1.401E+04 | 691.7 | - | - | 0 | - |
| - | - | 1.528E+04 | 692 | - | - | 0 | - |
| - | - | 4900 | 692.4 | - | - | 0 | - |
| - | - | 3402 | 692.7 | - | - | 0 | - |
| - | - | 1565 | 693 | - | - | 0 | - |
| 6 | y | 1781 | 694.9 | 0.0003232 | 0.4651 | +2 | 13 |
| - | - | 1984 | 695.4 | - | - | 0 | - |
| - | - | 937.2 | 695.9 | - | - | 0 | - |
| - | - | 1384 | 696.7 | - | - | 0 | - |
| - | - | 3553 | 697 | - | - | 0 | - |
| - | - | 2.066E+05 | 697.4 | - | - | 0 | - |
| - | - | 2.386E+05 | 697.7 | - | - | 0 | - |
| - | - | 1.682E+05 | 698 | - | - | 0 | - |
| - | - | 7.816E+04 | 698.4 | - | - | 0 | - |
| - | - | 2.864E+04 | 698.7 | - | - | 0 | - |
| - | - | 8484 | 699 | - | - | 0 | - |
| 5 | c | 3468 | 702.3 | 0.002139 | 3.046 | +1 | 5 |
| - | - | 973.5 | 703.3 | - | - | 0 | - |
| 12 | c | 818.7 | 708.9 | 0.008933 | 12.6 | +2 | 12 |
| 12 | c | 1238 | 709.4 | 0.005834 | 8.224 | +2 | 12 |
| - | - | 914.7 | 709.9 | - | - | 0 | - |
| - | - | 9586 | 717.4 | - | - | 0 | - |
| 12 | c | 9.928E+04 | 717.9 | 0.0002499 | 0.348 | +2 | 12 |
| - | - | 7.956E+04 | 718.4 | - | - | 0 | - |
| - | - | 3.871E+04 | 718.9 | - | - | 0 | - |
| 5 | c | 2.844E+04 | 719.3 | 0.0001792 | 0.2491 | +1 | 5 |
| - | - | 6150 | 719.4 | - | - | 0 | - |
| - | - | 2661 | 719.9 | - | - | 0 | - |
| - | - | 1.513E+04 | 720.3 | - | - | 0 | - |
| - | - | 3665 | 721.3 | - | - | 0 | - |
| - | - | 1104 | 730.9 | - | - | 0 | - |
| - | - | 1062 | 742.4 | - | - | 0 | - |
| - | - | 1667 | 742.9 | - | - | 0 | - |
| - | - | 1580 | 743.4 | - | - | 0 | - |
| - | - | 4800 | 752.4 | - | - | 0 | - |
| - | - | 3150 | 752.9 | - | - | 0 | - |
| - | - | 958.6 | 753.4 | - | - | 0 | - |
| 5 | z | 1.507E+04 | 755.4 | 0.0004569 | 0.6048 | +2 | 14 |
| - | - | 2.039E+04 | 755.9 | - | - | 0 | - |
| - | - | 1.273E+04 | 756.4 | - | - | 0 | - |
| - | - | 4325 | 756.9 | - | - | 0 | - |
| - | - | 1439 | 762.9 | - | - | 0 | - |
| 5 | y | 1.256E+04 | 763.4 | 0.0002501 | 0.3276 | +2 | 14 |
| - | - | 7931 | 763.9 | - | - | 0 | - |
| - | - | 5076 | 764.4 | - | - | 0 | - |
| 13 | c | 1104 | 765.9 | 0.004948 | 6.46 | +2 | 13 |
| - | - | 1551 | 766.4 | - | - | 0 | - |
| - | - | 1221 | 768.4 | - | - | 0 | - |
| - | - | 913.6 | 769.4 | - | - | 0 | - |
| - | - | 1234 | 770.4 | - | - | 0 | - |
| - | - | 6761 | 773.9 | - | - | 0 | - |
| 13 | c | 5.532E+04 | 774.4 | 0.0006983 | 0.9018 | +2 | 13 |
| - | - | 5.325E+04 | 774.9 | - | - | 0 | - |
| - | - | 2.49E+04 | 775.4 | - | - | 0 | - |
| - | - | 1.009E+04 | 775.9 | - | - | 0 | - |
| - | - | 1764 | 776.4 | - | - | 0 | - |
| - | - | 1164 | 777.5 | - | - | 0 | - |
| - | - | 1068 | 780.9 | - | - | 0 | - |
| - | - | 1034 | 781.4 | - | - | 0 | - |
| - | - | 998.8 | 783.5 | - | - | 0 | - |
| - | - | 2747 | 783.9 | - | - | 0 | - |
| - | - | 2503 | 784.4 | - | - | 0 | - |
| 12 | z | 3370 | 785.5 | 0.0002327 | 0.2963 | +1 | 7 |
| - | - | 4146 | 786.5 | - | - | 0 | - |
| - | - | 1537 | 787.5 | - | - | 0 | - |
| 4 | w | 8039 | 790.5 | 0.000522 | 0.6604 | +2 | 15 |
| - | - | 8054 | 791 | - | - | 0 | - |
| - | - | 6446 | 791.5 | - | - | 0 | - |
| - | - | 3189 | 791.9 | - | - | 0 | - |
| - | - | 1001 | 792.4 | - | - | 0 | - |
| - | - | 3343 | 796.4 | - | - | 0 | - |
| 12 | y | 1305 | 801.5 | 0.0021 | 2.62 | +1 | 7 |
| - | - | 1074 | 802.5 | - | - | 0 | - |
| - | - | 992.6 | 803 | - | - | 0 | - |
| - | - | 1118 | 805.5 | - | - | 0 | - |
| - | - | 1331 | 808.4 | - | - | 0 | - |
| - | - | 2122 | 809 | - | - | 0 | - |
| - | - | 2978 | 809.5 | - | - | 0 | - |
| - | - | 2794 | 810 | - | - | 0 | - |
| 4 | z | 2098 | 810.5 | 0.01574 | 19.42 | +2 | 15 |
| - | - | 9901 | 811.4 | - | - | 0 | - |
| - | - | 5011 | 812.4 | - | - | 0 | - |
| - | - | 2226 | 813.4 | - | - | 0 | - |
| 4 | y | 2407 | 819 | 0.002031 | 2.48 | +2 | 15 |
| 4 | z | 4.31E+04 | 819.5 | 0.0007702 | 0.9399 | +2 | 15 |
| - | - | 7.212E+04 | 820 | - | - | 0 | - |
| - | - | 4.672E+04 | 820.5 | - | - | 0 | - |
| - | - | 2.446E+04 | 821 | - | - | 0 | - |
| - | - | 5761 | 821.5 | - | - | 0 | - |
| - | - | 1493 | 822 | - | - | 0 | - |
| 14 | c | 5380 | 822.4 | 0.001421 | 1.728 | +2 | 14 |
| - | - | 4139 | 822.9 | - | - | 0 | - |
| - | - | 1826 | 823.4 | - | - | 0 | - |
| - | - | 1245 | 823.9 | - | - | 0 | - |
| - | - | 1094 | 824.5 | - | - | 0 | - |
| - | - | 1885 | 827 | - | - | 0 | - |
| 4 | y | 1.194E+04 | 827.5 | 0.0006245 | 0.7547 | +2 | 15 |
| - | - | 1.228E+04 | 828 | - | - | 0 | - |
| - | - | 6393 | 828.5 | - | - | 0 | - |
| - | - | 2207 | 829 | - | - | 0 | - |
| - | - | 1931 | 830.5 | - | - | 0 | - |
| 14 | c | 2.032E+04 | 831 | 0.001147 | 1.38 | +2 | 14 |
| - | - | 1.764E+04 | 831.5 | - | - | 0 | - |
| - | - | 1.034E+04 | 832 | - | - | 0 | - |
| - | - | 3990 | 832.5 | - | - | 0 | - |
| - | - | 957 | 833 | - | - | 0 | - |
| - | - | 1534 | 835.5 | - | - | 0 | - |
| - | - | 989.4 | 836.5 | - | - | 0 | - |
| - | - | 1221 | 837.5 | - | - | 0 | - |
| - | - | 1030 | 838.5 | - | - | 0 | - |
| 6 | c | 8830 | 839.4 | 0.000356 | 0.4242 | +1 | 6 |
| - | - | 4289 | 840.4 | - | - | 0 | - |
| - | - | 1304 | 841.4 | - | - | 0 | - |
| - | - | 2058 | 842 | - | - | 0 | - |
| - | - | 1284 | 843.5 | - | - | 0 | - |
| - | - | 824.1 | 844 | - | - | 0 | - |
| - | - | 1342 | 845.5 | - | - | 0 | - |
| - | - | 987 | 848 | - | - | 0 | - |
| - | - | 2494 | 850.5 | - | - | 0 | - |
| - | - | 2770 | 851 | - | - | 0 | - |
| - | - | 1522 | 851.5 | - | - | 0 | - |
| - | - | 2661 | 852 | - | - | 0 | - |
| - | - | 1746 | 853.4 | - | - | 0 | - |
| - | - | 1413 | 853.7 | - | - | 0 | - |
| - | - | 1295 | 854.1 | - | - | 0 | - |
| - | - | 1369 | 855.5 | - | - | 0 | - |
| - | - | 1883 | 858 | - | - | 0 | - |
| - | - | 2713 | 858.5 | - | - | 0 | - |
| - | - | 1976 | 859 | - | - | 0 | - |
| - | - | 2088 | 859.5 | - | - | 0 | - |
| - | - | 960.1 | 863.5 | - | - | 0 | - |
| - | - | 1926 | 865 | - | - | 0 | - |
| - | - | 5188 | 865.5 | - | - | 0 | - |
| - | - | 5374 | 866 | - | - | 0 | - |
| - | - | 2965 | 866.5 | - | - | 0 | - |
| - | - | 2152 | 867 | - | - | 0 | - |
| - | - | 1269 | 867.5 | - | - | 0 | - |
| - | - | 1069 | 869.5 | - | - | 0 | - |
| - | - | 2278 | 870 | - | - | 0 | - |
| - | - | 1762 | 870.5 | - | - | 0 | - |
| - | - | 1213 | 871 | - | - | 0 | - |
| - | - | 3028 | 872 | - | - | 0 | - |
| - | - | 3978 | 872.5 | - | - | 0 | - |
| - | - | 3228 | 873 | - | - | 0 | - |
| - | - | 2134 | 873.5 | - | - | 0 | - |
| - | - | 1032 | 874 | - | - | 0 | - |
| - | - | 3337 | 878 | - | - | 0 | - |
| 15 | c | 5730 | 878.5 | 0.00301 | 3.426 | +2 | 15 |
| 15 | c | 2.258E+04 | 879 | 0.002113 | 2.404 | +2 | 15 |
| - | - | 1.846E+04 | 879.5 | - | - | 0 | - |
| - | - | 1.496E+04 | 880 | - | - | 0 | - |
| - | - | 5623 | 880.5 | - | - | 0 | - |
| - | - | 2431 | 881 | - | - | 0 | - |
| - | - | 1059 | 882.5 | - | - | 0 | - |
| - | - | 1119 | 882.6 | - | - | 0 | - |
| - | - | 1402 | 886.5 | - | - | 0 | - |
| - | - | 2.604E+04 | 887 | - | - | 0 | - |
| 15 | c | 5.622E+04 | 887.5 | 0.0005409 | 0.6095 | +2 | 15 |
| - | - | 4.594E+04 | 888 | - | - | 0 | - |
| - | - | 2.769E+04 | 888.5 | - | - | 0 | - |
| - | - | 9106 | 889 | - | - | 0 | - |
| - | - | 3162 | 889.5 | - | - | 0 | - |
| - | - | 905.1 | 891 | - | - | 0 | - |
| - | - | 1220 | 891.5 | - | - | 0 | - |
| 3 | z | 1518 | 892 | 0.006859 | 7.689 | +2 | 16 |
| 3 | z | 3271 | 892.5 | 0.01577 | 17.67 | +2 | 16 |
| - | - | 1716 | 895.4 | - | - | 0 | - |
| - | - | 1180 | 896 | - | - | 0 | - |
| - | - | 1742 | 896.5 | - | - | 0 | - |
| - | - | 5191 | 897.5 | - | - | 0 | - |
| 11 | y | 9520 | 898.6 | 0.0033 | 3.673 | +1 | 8 |
| - | - | 3971 | 899.6 | - | - | 0 | - |
| 3 | y | 4702 | 900 | 0.001343 | 1.493 | +2 | 16 |
| 3 | y | 5311 | 900.5 | 0.008419 | 9.349 | +2 | 16 |
| 3 | z | 3.504E+04 | 901 | 0.00121 | 1.343 | +2 | 16 |
| - | - | 4.037E+04 | 901.5 | - | - | 0 | - |
| - | - | 2.122E+04 | 902 | - | - | 0 | - |
| - | - | 9540 | 902.5 | - | - | 0 | - |
| - | - | 3429 | 903 | - | - | 0 | - |
| - | - | 1137 | 908 | - | - | 0 | - |
| - | - | 1484 | 908.5 | - | - | 0 | - |
| 3 | y | 3.041E+04 | 909 | 0.0008816 | 0.9698 | +2 | 16 |
| - | - | 2.923E+04 | 909.5 | - | - | 0 | - |
| - | - | 1.835E+04 | 910 | - | - | 0 | - |
| - | - | 7628 | 910.5 | - | - | 0 | - |
| - | - | 1864 | 911 | - | - | 0 | - |
| - | - | 1243 | 912.5 | - | - | 0 | - |
| - | - | 1873 | 913.5 | - | - | 0 | - |
| - | - | 2047 | 914 | - | - | 0 | - |
| - | - | 1005 | 914.5 | - | - | 0 | - |
| - | - | 1302 | 915 | - | - | 0 | - |
| - | - | 985.8 | 918.6 | - | - | 0 | - |
| - | - | 985.4 | 919.6 | - | - | 0 | - |
| - | - | 1184 | 920.5 | - | - | 0 | - |
| - | - | 1066 | 921.5 | - | - | 0 | - |
| 16 | c | 1404 | 922 | 0.002056 | 2.23 | +2 | 16 |
| 16 | c | 1.01E+04 | 922.5 | 0.0009308 | 1.009 | +2 | 16 |
| - | - | 1.016E+04 | 923 | - | - | 0 | - |
| - | - | 5829 | 923.5 | - | - | 0 | - |
| - | - | 2606 | 924 | - | - | 0 | - |
| - | - | 1002 | 924.2 | - | - | 0 | - |
| - | - | 1474 | 924.5 | - | - | 0 | - |
| - | - | 1234 | 928.5 | - | - | 0 | - |
| - | - | 1091 | 929.2 | - | - | 0 | - |
| - | - | 3870 | 929.5 | - | - | 0 | - |
| - | - | 1252 | 930 | - | - | 0 | - |
| - | - | 6683 | 930.5 | - | - | 0 | - |
| - | - | 2862 | 930.8 | - | - | 0 | - |
| 16 | c | 6.346E+04 | 931 | 0.0005347 | 0.5743 | +2 | 16 |
| - | - | 6.621E+04 | 931.5 | - | - | 0 | - |
| - | - | 3.768E+04 | 932 | - | - | 0 | - |
| - | - | 1.708E+04 | 932.5 | - | - | 0 | - |
| - | - | 5595 | 933 | - | - | 0 | - |
| - | - | 1306 | 933.5 | - | - | 0 | - |
| - | - | 966.8 | 934.4 | - | - | 0 | - |
| - | - | 1208 | 937.5 | - | - | 0 | - |
| - | - | 1068 | 938 | - | - | 0 | - |
| - | - | 1023 | 939.5 | - | - | 0 | - |
| - | - | 2831 | 944 | - | - | 0 | - |
| - | - | 5467 | 944.5 | - | - | 0 | - |
| - | - | 3330 | 945 | - | - | 0 | - |
| - | - | 1347 | 945.5 | - | - | 0 | - |
| - | - | 1226 | 946 | - | - | 0 | - |
| - | - | 1464 | 952 | - | - | 0 | - |
| - | - | 3771 | 952.4 | - | - | 0 | - |
| 7 | c | 2.017E+04 | 953.4 | 0.002403 | 2.52 | +1 | 7 |
| 10 | z | 3.805E+04 | 953.6 | 0.001442 | 1.513 | +1 | 9 |
| - | - | 1.302E+04 | 954.4 | - | - | 0 | - |
| - | - | 2.159E+04 | 954.6 | - | - | 0 | - |
| - | - | 3269 | 955.4 | - | - | 0 | - |
| - | - | 7743 | 955.6 | - | - | 0 | - |
| - | - | 2055 | 956.6 | - | - | 0 | - |
| - | - | 1917 | 965 | - | - | 0 | - |
| - | - | 1514 | 966 | - | - | 0 | - |
| - | - | 3572 | 966.4 | - | - | 0 | - |
| - | - | 1189 | 967 | - | - | 0 | - |
| - | - | 3174 | 967.5 | - | - | 0 | - |
| 10 | y | 8213 | 969.6 | 0.0008022 | 0.8274 | +1 | 9 |
| - | - | 6149 | 970.6 | - | - | 0 | - |
| - | - | 1421 | 971.6 | - | - | 0 | - |
| - | - | 1442 | 972 | - | - | 0 | - |
| - | - | 2251 | 972.5 | - | - | 0 | - |
| - | - | 3060 | 973 | - | - | 0 | - |
| - | - | 1.271E+04 | 973.5 | - | - | 0 | - |
| - | - | 1.321E+04 | 974 | - | - | 0 | - |
| - | - | 8326 | 974.5 | - | - | 0 | - |
| - | - | 3698 | 975 | - | - | 0 | - |
| - | - | 2166 | 975.5 | - | - | 0 | - |
| - | - | 1624 | 977 | - | - | 0 | - |
| - | - | 1093 | 977.6 | - | - | 0 | - |
| - | - | 2513 | 978 | - | - | 0 | - |
| - | - | 2811 | 978.5 | - | - | 0 | - |
| - | - | 2204 | 979 | - | - | 0 | - |
| - | - | 2247 | 979.5 | - | - | 0 | - |
| - | - | 2546 | 980 | - | - | 0 | - |
| - | - | 5808 | 980.5 | - | - | 0 | - |
| - | - | 4349 | 981 | - | - | 0 | - |
| - | - | 2233 | 981.5 | - | - | 0 | - |
| - | - | 1394 | 982 | - | - | 0 | - |
| - | - | 1300 | 984 | - | - | 0 | - |
| - | - | 1381 | 985.6 | - | - | 0 | - |
| - | - | 1453 | 986.1 | - | - | 0 | - |
| 17 | c | 2409 | 986.5 | 0.01778 | 18.03 | +2 | 17 |
| 17 | c | 1.431E+04 | 987 | 0.003743 | 3.792 | +2 | 17 |
| - | - | 1.543E+04 | 987.5 | - | - | 0 | - |
| - | - | 1.146E+04 | 988 | - | - | 0 | - |
| - | - | 5424 | 988.5 | - | - | 0 | - |
| - | - | 2252 | 989 | - | - | 0 | - |
| - | - | 1212 | 989.5 | - | - | 0 | - |
| - | - | 2870 | 992.6 | - | - | 0 | - |
| 2 | y | 4097 | 993.5 | 0.007703 | 7.753 | +2 | 17 |
| 2 | z | 1.133E+04 | 994 | 0.002692 | 2.708 | +2 | 17 |
| - | - | 1.314E+04 | 994.5 | - | - | 0 | - |
| - | - | 8610 | 995 | - | - | 0 | - |
| 17 | c | 7.04E+04 | 995.5 | 0.001638 | 1.645 | +2 | 17 |
| - | - | 7.428E+04 | 996 | - | - | 0 | - |
| - | - | 5.233E+04 | 996.5 | - | - | 0 | - |
| - | - | 1.881E+04 | 997 | - | - | 0 | - |
| - | - | 8562 | 997.5 | - | - | 0 | - |
| - | - | 2269 | 998 | - | - | 0 | - |
| - | - | 1186 | 1000 | - | - | 0 | - |
| - | - | 2555 | 1001 | - | - | 0 | - |
| - | - | 2980 | 1002 | - | - | 0 | - |
| 2 | y | 6552 | 1002 | 0.004656 | 4.647 | +2 | 17 |
| - | - | 5758 | 1003 | - | - | 0 | - |
| - | - | 4053 | 1003 | - | - | 0 | - |
| - | - | 1392 | 1004 | - | - | 0 | - |
| - | - | 3845 | 1005 | - | - | 0 | - |
| - | - | 5631 | 1006 | - | - | 0 | - |
| - | - | 6041 | 1006 | - | - | 0 | - |
| - | - | 3522 | 1007 | - | - | 0 | - |
| - | - | 1294 | 1007 | - | - | 0 | - |
| - | - | 1331 | 1008 | - | - | 0 | - |
| - | - | 8120 | 1008 | - | - | 0 | - |
| - | - | 8609 | 1009 | - | - | 0 | - |
| - | - | 6653 | 1009 | - | - | 0 | - |
| - | - | 5028 | 1009 | - | - | 0 | - |
| - | - | 1.076E+04 | 1010 | - | - | 0 | - |
| - | - | 1.143E+04 | 1010 | - | - | 0 | - |
| 8 | c | 3.675E+04 | 1010 | 0.0002874 | 0.2844 | +1 | 8 |
| - | - | 5039 | 1011 | - | - | 0 | - |
| - | - | 1.836E+04 | 1011 | - | - | 0 | - |
| - | - | 1947 | 1012 | - | - | 0 | - |
| - | - | 6201 | 1012 | - | - | 0 | - |
| - | - | 8017 | 1015 | - | - | 0 | - |
| - | - | 1.326E+04 | 1015 | - | - | 0 | - |
| - | - | 1.541E+04 | 1016 | - | - | 0 | - |
| - | - | 1.384E+04 | 1016 | - | - | 0 | - |
| - | - | 8721 | 1017 | - | - | 0 | - |
| - | - | 6417 | 1017 | - | - | 0 | - |
| - | - | 3106 | 1018 | - | - | 0 | - |
| - | - | 5369 | 1018 | - | - | 0 | - |
| - | - | 4937 | 1019 | - | - | 0 | - |
| - | - | 1600 | 1019 | - | - | 0 | - |
| - | - | 1221 | 1022 | - | - | 0 | - |
| - | - | 3602 | 1022 | - | - | 0 | - |
| - | - | 3941 | 1023 | - | - | 0 | - |
| - | - | 6880 | 1023 | - | - | 0 | - |
| - | - | 2.252E+04 | 1024 | - | - | 0 | - |
| - | - | 2.395E+04 | 1024 | - | - | 0 | - |
| - | - | 2.703E+04 | 1025 | - | - | 0 | - |
| - | - | 2.55E+04 | 1025 | - | - | 0 | - |
| - | - | 1.382E+04 | 1026 | - | - | 0 | - |
| - | - | 5615 | 1026 | - | - | 0 | - |
| - | - | 3573 | 1027 | - | - | 0 | - |
| - | - | 2586 | 1028 | - | - | 0 | - |
| - | - | 4414 | 1029 | - | - | 0 | - |
| - | - | 6462 | 1029 | - | - | 0 | - |
| - | - | 7533 | 1030 | - | - | 0 | - |
| - | - | 8241 | 1030 | - | - | 0 | - |
| - | - | 7676 | 1031 | - | - | 0 | - |
| - | - | 5600 | 1031 | - | - | 0 | - |
| - | - | 3636 | 1032 | - | - | 0 | - |
| - | - | 2584 | 1032 | - | - | 0 | - |
| - | - | 2877 | 1037 | - | - | 0 | - |
| - | - | 1.233E+04 | 1037 | - | - | 0 | - |
| - | - | 8.417E+04 | 1038 | - | - | 0 | - |
| - | - | 9.962E+04 | 1038 | - | - | 0 | - |
| - | - | 7.713E+04 | 1039 | - | - | 0 | - |
| - | - | 4.284E+04 | 1039 | - | - | 0 | - |
| - | - | 1.761E+04 | 1040 | - | - | 0 | - |
| - | - | 5075 | 1040 | - | - | 0 | - |
| - | - | 2224 | 1043 | - | - | 0 | - |
| - | - | 1384 | 1044 | - | - | 0 | - |
| - | - | 3114 | 1045 | - | - | 0 | - |
| - | - | 6599 | 1045 | - | - | 0 | - |
| - | - | 1.16E+05 | 1046 | - | - | 0 | - |
| - | - | 4.54E+05 | 1046 | - | - | 0 | - |
| - | - | 4.792E+05 | 1047 | - | - | 0 | - |
| - | - | 2.976E+05 | 1047 | - | - | 0 | - |
| - | - | 1.357E+05 | 1048 | - | - | 0 | - |
| - | - | 4.795E+04 | 1048 | - | - | 0 | - |
| - | - | 1.116E+04 | 1049 | - | - | 0 | - |
| 9 | z | 2.432E+04 | 1082 | 0.001999 | 1.848 | +1 | 10 |
| - | - | 2.662E+04 | 1083 | - | - | 0 | - |
| - | - | 1.25E+04 | 1084 | - | - | 0 | - |
| - | - | 4212 | 1085 | - | - | 0 | - |
| - | - | 7157 | 1095 | - | - | 0 | - |
| - | - | 5272 | 1096 | - | - | 0 | - |
| - | - | 1724 | 1097 | - | - | 0 | - |
| 9 | y | 7895 | 1098 | 0.0002431 | 0.2214 | +1 | 10 |
| - | - | 4781 | 1099 | - | - | 0 | - |
| - | - | 1543 | 1100 | - | - | 0 | - |
| 9 | c | 5E+04 | 1139 | 0.001577 | 1.385 | +1 | 9 |
| 8 | z | 1.258E+04 | 1139 | 0.00202 | 1.774 | +1 | 11 |
| - | - | 3.22E+04 | 1140 | - | - | 0 | - |
| - | - | 1.764E+04 | 1140 | - | - | 0 | - |
| - | - | 1.312E+04 | 1141 | - | - | 0 | - |
| - | - | 7794 | 1141 | - | - | 0 | - |
| - | - | 2876 | 1142 | - | - | 0 | - |
| - | - | 2021 | 1142 | - | - | 0 | - |
| - | - | 2766 | 1147 | - | - | 0 | - |
| 8 | y | 2346 | 1155 | 0.002788 | 2.414 | +1 | 11 |
| - | - | 2222 | 1156 | - | - | 0 | - |
| - | - | 5930 | 1166 | - | - | 0 | - |
| - | - | 3528 | 1167 | - | - | 0 | - |
| - | - | 1101 | 1168 | - | - | 0 | - |
| 7 | w | 937 | 1209 | 0.009446 | 7.815 | +1 | 12 |
| - | - | 1002 | 1210 | - | - | 0 | - |
| 7 | y | 3676 | 1252 | 0.0006004 | 0.4797 | +1 | 12 |
| - | - | 3308 | 1253 | - | - | 0 | - |
| - | - | 1019 | 1254 | - | - | 0 | - |
| - | - | 2937 | 1279 | - | - | 0 | - |
| - | - | 2989 | 1280 | - | - | 0 | - |
| - | - | 2175 | 1280 | - | - | 0 | - |
| - | - | 1225 | 1281 | - | - | 0 | - |
| - | - | 3137 | 1281 | - | - | 0 | - |
| - | - | 2378 | 1281 | - | - | 0 | - |
| - | - | 2062 | 1282 | - | - | 0 | - |
| - | - | 959.4 | 1288 | - | - | 0 | - |
| - | - | 2037 | 1306 | - | - | 0 | - |
| 11 | c | 7540 | 1307 | 0.0003222 | 0.2466 | +1 | 11 |
| - | - | 4748 | 1308 | - | - | 0 | - |
| - | - | 1617 | 1309 | - | - | 0 | - |
| - | - | 1527 | 1338 | - | - | 0 | - |
| 6 | z | 2.574E+04 | 1373 | 0.0004517 | 0.329 | +1 | 13 |
| - | - | 2.762E+04 | 1374 | - | - | 0 | - |
| - | - | 1.386E+04 | 1375 | - | - | 0 | - |
| - | - | 4251 | 1376 | - | - | 0 | - |
| - | - | 1316 | 1386 | - | - | 0 | - |
| - | - | 1221 | 1388 | - | - | 0 | - |
| 6 | y | 1.521E+04 | 1389 | 0.001259 | 0.9064 | +1 | 13 |
| - | - | 1.242E+04 | 1390 | - | - | 0 | - |
| - | - | 6241 | 1391 | - | - | 0 | - |
| - | - | 1334 | 1392 | - | - | 0 | - |
| - | - | 2612 | 1393 | - | - | 0 | - |
| - | - | 2190 | 1394 | - | - | 0 | - |
| - | - | 1297 | 1394 | - | - | 0 | - |
| - | - | 1541 | 1395 | - | - | 0 | - |
| - | - | 3958 | 1396 | - | - | 0 | - |
| - | - | 5479 | 1396 | - | - | 0 | - |
| - | - | 3641 | 1397 | - | - | 0 | - |
| - | - | 1911 | 1397 | - | - | 0 | - |
| - | - | 2865 | 1434 | - | - | 0 | - |
| 12 | c | 1.894E+04 | 1435 | 0.0002078 | 0.1448 | +1 | 12 |
| - | - | 1.532E+04 | 1436 | - | - | 0 | - |
| - | - | 6981 | 1437 | - | - | 0 | - |
| - | - | 1850 | 1438 | - | - | 0 | - |
| - | - | 2698 | 1504 | - | - | 0 | - |
| - | - | 1949 | 1505 | - | - | 0 | - |
| 5 | z | 1.871E+04 | 1510 | 0.001476 | 0.9778 | +1 | 14 |
| - | - | 4.408E+04 | 1511 | - | - | 0 | - |
| - | - | 3.013E+04 | 1512 | - | - | 0 | - |
| - | - | 1.509E+04 | 1513 | - | - | 0 | - |
| - | - | 4329 | 1514 | - | - | 0 | - |
| - | - | 2362 | 1525 | - | - | 0 | - |
| 5 | y | 1.07E+04 | 1526 | 0.0001579 | 0.1035 | +1 | 14 |
| - | - | 8558 | 1527 | - | - | 0 | - |
| - | - | 4132 | 1528 | - | - | 0 | - |
| - | - | 2036 | 1529 | - | - | 0 | - |
| 13 | c | 6850 | 1548 | 0.003412 | 2.204 | +1 | 13 |
| - | - | 7376 | 1549 | - | - | 0 | - |
| - | - | 3723 | 1550 | - | - | 0 | - |
| - | - | 1362 | 1551 | - | - | 0 | - |
| 4 | z | 4287 | 1638 | 0.000394 | 0.2405 | +1 | 15 |
| - | - | 2.792E+04 | 1639 | - | - | 0 | - |
| - | - | 2.482E+04 | 1640 | - | - | 0 | - |
| - | - | 1.174E+04 | 1641 | - | - | 0 | - |
| - | - | 5432 | 1642 | - | - | 0 | - |
| - | - | 1376 | 1643 | - | - | 0 | - |
| 4 | y | 3337 | 1654 | 0.005962 | 3.605 | +1 | 15 |
| - | - | 3071 | 1655 | - | - | 0 | - |
| - | - | 1440 | 1656 | - | - | 0 | - |
| - | - | 1198 | 1657 | - | - | 0 | - |
| 14 | c | 5477 | 1661 | 0.003955 | 2.381 | +1 | 14 |
| - | - | 4109 | 1662 | - | - | 0 | - |
| - | - | 2085 | 1663 | - | - | 0 | - |
| - | - | 2621 | 1758 | - | - | 0 | - |
| - | - | 1339 | 1759 | - | - | 0 | - |
| - | - | 2683 | 1773 | - | - | 0 | - |
| 15 | c | 1.501E+04 | 1774 | 0.001678 | 0.9459 | +1 | 15 |
| - | - | 1.315E+04 | 1775 | - | - | 0 | - |
| - | - | 8755 | 1776 | - | - | 0 | - |
| - | - | 3395 | 1777 | - | - | 0 | - |
| 3 | z | 4072 | 1801 | 0.00396 | 2.199 | +1 | 16 |
| - | - | 1.588E+04 | 1802 | - | - | 0 | - |
| - | - | 1.709E+04 | 1803 | - | - | 0 | - |
| - | - | 8383 | 1804 | - | - | 0 | - |
| - | - | 3201 | 1805 | - | - | 0 | - |
| 3 | y | 2500 | 1817 | 0.0127 | 6.99 | +1 | 16 |
| - | - | 4688 | 1818 | - | - | 0 | - |
| - | - | 2904 | 1819 | - | - | 0 | - |
| - | - | 1377 | 1820 | - | - | 0 | - |
| - | - | 1386 | 1860 | - | - | 0 | - |
| 16 | c | 1.73E+04 | 1861 | 0.002975 | 1.598 | +1 | 16 |
| - | - | 1.794E+04 | 1862 | - | - | 0 | - |
| - | - | 1.224E+04 | 1863 | - | - | 0 | - |
| - | - | 5467 | 1864 | - | - | 0 | - |
| - | - | 1488 | 1865 | - | - | 0 | - |
| - | - | 2467 | 1946 | - | - | 0 | - |
| - | - | 4938 | 1947 | - | - | 0 | - |
| - | - | 5069 | 1948 | - | - | 0 | - |
| - | - | 2879 | 1949 | - | - | 0 | - |
| - | - | 1129 | 1965 | - | - | 0 | - |
| - | - | 1840 | 1975 | - | - | 0 | - |
| 2 | z | 1779 | 1987 | 0.00204 | 1.026 | +1 | 17 |
| - | - | 6178 | 1988 | - | - | 0 | - |
| - | - | 4774 | 1989 | - | - | 0 | - |
| 17 | c | 9929 | 1990 | 0.008355 | 4.198 | +1 | 17 |
| - | - | 1.358E+04 | 1991 | - | - | 0 | - |
| - | - | 1.123E+04 | 1992 | - | - | 0 | - |
| - | - | 5810 | 1993 | - | - | 0 | - |
| - | - | 2844 | 1994 | - | - | 0 | - |
| - | - | 1229 | 1995 | - | - | 0 | - |
| - | - | 1930 | 2030 | - | - | 0 | - |
| - | - | 2892 | 2031 | - | - | 0 | - |
| - | - | 1918 | 2045 | - | - | 0 | - |
| - | - | 3316 | 2046 | - | - | 0 | - |
| - | - | 1.082E+04 | 2047 | - | - | 0 | - |
| - | - | 1.182E+04 | 2048 | - | - | 0 | - |
| - | - | 7749 | 2049 | - | - | 0 | - |
| - | - | 2840 | 2050 | - | - | 0 | - |
| - | - | 1553 | 2051 | - | - | 0 | - |
| - | - | 3528 | 2057 | - | - | 0 | - |
| - | - | 6402 | 2058 | - | - | 0 | - |
| - | - | 5188 | 2059 | - | - | 0 | - |
| - | - | 3287 | 2060 | - | - | 0 | - |
| - | - | 2135 | 2061 | - | - | 0 | - |
| - | - | 1925 | 2062 | - | - | 0 | - |
| - | - | 3641 | 2063 | - | - | 0 | - |
| - | - | 1.669E+04 | 2064 | - | - | 0 | - |
| - | - | 1.51E+04 | 2065 | - | - | 0 | - |
| - | - | 9744 | 2066 | - | - | 0 | - |
| - | - | 4446 | 2067 | - | - | 0 | - |
| - | - | 1649 | 2068 | - | - | 0 | - |
| - | - | 1315 | 2073 | - | - | 0 | - |
| - | - | 8855 | 2074 | - | - | 0 | - |
| - | - | 4.257E+04 | 2075 | - | - | 0 | - |
| - | - | 4.952E+04 | 2076 | - | - | 0 | - |
| - | - | 3.241E+04 | 2077 | - | - | 0 | - |
| - | - | 1.798E+04 | 2078 | - | - | 0 | - |
| - | - | 7696 | 2079 | - | - | 0 | - |
| - | - | 1579 | 2089 | - | - | 0 | - |
| - | - | 1.029E+04 | 2090 | - | - | 0 | - |
| - | - | 5.319E+04 | 2091 | - | - | 0 | - |
| - | - | 1.963E+05 | 2092 | - | - | 0 | - |
| - | - | 2.16E+05 | 2093 | - | - | 0 | - |
| - | - | 1.336E+05 | 2094 | - | - | 0 | - |
| - | - | 6.312E+04 | 2095 | - | - | 0 | - |
| - | - | 2.305E+04 | 2096 | - | - | 0 | - |
| - | - | 6931 | 2097 | - | - | 0 | - |
| - | - | 1900 | 2294 | - | - | 0 | - |
| - | - | 2156 | 2295 | - | - | 0 | - |

m/z Charge Intensity FragmentType MassShift Position
120.06543731689453 0 3577.9106
120.08071899414062 0 1095.0812
125.80577087402344 0 509.11505
129.1022186279297 0 447.48383
129.1880340576172 0 436.74875
130.56668090820312 0 467.0091
132.07476806640625 0 511.11154
132.08056640625 0 801.00635
136.07577514648438 0 1021.4047
146.0920867919922 0 678.72766
162.3561553955078 0 645.2172
173.43798828125 0 1052.3127
183.0762939453125 0 584.6598
187.08644104003906 0 766.46356
189.08731079101562 0 840.47296 c Ammonia loss 3
201.12364196777344 0 718.99335
217.0817108154297 0 7879.6973
219.14944458007812 0 978.87616
223.11920166015625 0 1015.32764
229.11827087402344 0 6400.8184 y Water loss 16
230.12294006347656 0 699.93036
232.11952209472656 0 1674.4578
239.09532165527344 0 1447.4939
246.12359619140625 0 9142.34
247.12806701660156 0 3733.8074 y 16
247.1431121826172 0 1582.3436
249.05422973632812 0 1533.3331
255.1011505126953 0 694.9501
260.4669189453125 0 604.73114
267.06427001953125 0 1421.3252
274.11846923828125 0 12256.459
275.1225891113281 0 1904.6837
292.1287536621094 0 1101.2076
295.1032409667969 0 3838.1538
309.1558837890625 0 1613.2571
310.1597595214844 0 719.81726
312.56317138671875 0 724.1588
316.1498718261719 0 855.735 y Water loss 15
320.12451171875 0 878.8829
334.1609802246094 0 5584.293 y 15
335.1638488769531 0 797.2035
365.5375671386719 0 862.9326
366.13238525390625 0 713.4621
369.1204833984375 0 1938.6417
396.824951171875 0 829.6097
403.21893310546875 0 1674.499
434.19488525390625 0 691.66626
435.2352294921875 0 1184.6187
437.1815185546875 0 2459.7385
445.0373229980469 0 753.5343
447.24493408203125 0 2097.8826 y 14
453.2007751464844 0 1439.2776
465.2070617675781 0 2101.2102
474.25567626953125 0 788.49976
477.20037841796875 0 2092.7004
480.2247314453125 0 2187.553
481.22705078125 0 879.96063
524.2998046875 0 798.71985
528.24560546875 0 1703.1184
529.2473754882812 0 1059.8573
536.2831420898438 0 788.6462
538.2537841796875 0 2412.2646
539.2613525390625 0 3893.493
540.2659301757812 0 859.59894
541.3245239257812 0 916.5131
546.2564086914062 0 835.1703
548.9672241210938 0 1224.5847
560.3295288085938 0 4004.9573 y 13
561.2686157226562 0 3812.6033 c Ammonia loss 8
561.3349609375 0 1049.684 z Ammonia loss 7
561.7665405273438 0 1425.0626
563.2490844726562 0 6443.7646
564.2516479492188 0 1313.594 c Water loss 3
565.2415161132812 0 2700.5999 c Ammonia loss 3
566.2457275390625 0 2622.958
567.847900390625 0 2173.4487
576.995849609375 0 832.70575
577.9803466796875 0 937.63873
580.2568969726562 0 1105.7633
581.2592163085938 0 5386.7534
582.2672729492188 0 28055.48 c 3
583.2703247070312 0 8857.146
584.2733764648438 0 2083.6624
586.3267822265625 0 6979.339 c Ammonia loss 14
586.6614990234375 0 8285.65
586.9950561523438 0 3868.7634
587.7823486328125 0 1220.1656
596.7857666015625 0 3910.347 c Ammonia loss 9
597.28955078125 0 2303.7373
600.66748046875 0 873.72974 y Ammonia loss 2
601.3323364257812 0 887.8984
606.3427734375 0 1512.0795 y 2
606.6747436523438 0 1150.5939
607.0103149414062 0 1968.744
609.3341064453125 0 1520.4674
614.340087890625 0 1746.3632 w 12
615.33837890625 0 5937.8027 c Ammonia loss 15
615.671630859375 0 6900.317
616.0056762695312 0 4037.25
616.3405151367188 0 1580.961
620.6626586914062 0 1219.9722
620.9930419921875 0 1609.4952
621.3290405273438 0 1333.021
623.6672973632812 0 977.3721
626.38818359375 0 7834.85 y 6
626.8885498046875 0 5402.8657
627.3924560546875 0 2422.0913
627.892822265625 0 784.9748
639.896728515625 0 1384.598
644.34619140625 0 938.9299
647.6752319335938 0 735.0277
649.01953125 0 3104.9778
649.3533325195312 0 3447.778
649.6875610351562 0 1123.7885
650.0248413085938 0 887.9704
652.3473510742188 0 4728.49
652.6822509765625 0 6157.2275
653.0164794921875 0 4636.846
653.3544311523438 0 1954.9736
657.3046264648438 0 1712.9708
657.39453125 0 4102.543 z 12
658.3006591796875 0 859.8733
658.3518676757812 0 23995.346 c Ammonia loss 16
658.6859741210938 0 37745.547
659.0198974609375 0 22651.48
659.2920532226562 0 2828.1787
659.3541870117188 0 11989.451
659.4117431640625 0 1046.3322
659.6876220703125 0 3035.6426
660.0222778320312 0 1168.4766
660.2965698242188 0 1333.9264
662.6964721679688 0 1019.61237 y Ammonia loss 1
664.3600463867188 0 1222.5736
664.6883544921875 0 1227.0685
665.0260009765625 0 917.3647
665.36083984375 0 1158.502
669.370849609375 0 846.3991
673.4132690429688 0 2400.441 y 12
674.3027954101562 0 2585.8445
674.4178466796875 0 912.71625
675.3128051757812 0 10615.0625
676.317138671875 0 5438.4414
677.3204956054688 0 1346.1116
678.355224609375 0 938.85126
680.3722534179688 0 886.6252
682.0430297851562 0 3229.1333
682.374267578125 0 1875.4081
682.7093505859375 0 1205.2146
685.7045288085938 0 1617.488
687.4078979492188 0 1394.4607
691.3753051757812 0 11882.291
691.7081909179688 0 14007.183
692.0420532226562 0 15280.536
692.375 0 4900.221
692.7091064453125 0 3401.97
693.0368041992188 0 1565.069
694.9163208007812 0 1781.3605 y 5
695.4193725585938 0 1984.2451
695.9260864257812 0 937.2218
696.7077026367188 0 1383.5132
697.0409545898438 0 3552.9658
697.3780517578125 0 206612.9
697.7125244140625 0 238624.81
698.046630859375 0 168228.03
698.3803100585938 0 78159.164
698.7151489257812 0 28639.445
699.04931640625 0 8483.547
702.3015747070312 0 3468.1003 c Ammonia loss 4
703.3034057617188 0 973.4629
708.85888671875 0 818.70056 c Water loss 11
709.3656616210938 0 1238.0778 c Ammonia loss 11
709.8651733398438 0 914.68506
717.3700561523438 0 9586.435
717.8733520507812 0 99278.92 c 11
718.3748779296875 0 79558.57
718.8760986328125 0 38706.168
719.3258056640625 0 28439.02 c 4
719.3798217773438 0 6149.6255
719.8792724609375 0 2660.8472
720.32958984375 0 15126.535
721.33203125 0 3665.0415
730.8826293945312 0 1104.0983
742.4382934570312 0 1061.826
742.9325561523438 0 1666.5667
743.4368286132812 0 1579.9005
752.40869140625 0 4799.6216
752.9099731445312 0 3150.2095
753.4163208007812 0 958.5512
755.4371948242188 0 15074.738 z 4
755.9398193359375 0 20391.459
756.4413452148438 0 12727.148
756.9430541992188 0 4325.297
762.9423828125 0 1439.1777
763.4463500976562 0 12557.269 y 4
763.9471435546875 0 7931.4233
764.4495239257812 0 5076.455
765.8969116210938 0 1104.0521 c Ammonia loss 12
766.4005126953125 0 1551.0599
768.4287719726562 0 1221.2684
769.4243774414062 0 913.57513
770.4393920898438 0 1234.467
773.9124755859375 0 6760.7944
774.4158325195312 0 55315.73 c 12
774.9174194335938 0 53245.32
775.418701171875 0 24900.377
775.9200439453125 0 10090.501
776.4202270507812 0 1764.1848
777.45068359375 0 1163.9565
780.92333984375 0 1068.3763
781.4203491210938 0 1033.9764
783.4585571289062 0 998.7635
783.9417114257812 0 2747.1226
784.4329223632812 0 2503.3853
785.4890747070312 0 3369.8694 z 11
786.4949340820312 0 4146.0996
787.5006713867188 0 1536.8976
790.451904296875 0 8038.982 w 3
790.9542236328125 0 8054.1353
791.4515991210938 0 6445.8374
791.9490356445312 0 3189.1064
792.440185546875 0 1001.1965
796.3538818359375 0 3343.4446
801.5101318359375 0 1304.8322 y 11
802.5099487304688 0 1074.3075
802.9600219726562 0 992.58124
805.4595336914062 0 1117.9825
808.4478149414062 0 1330.9993
808.95263671875 0 2121.8594
809.4507446289062 0 2978.1726
809.95361328125 0 2794.4756
810.4450073242188 0 2097.7905 z Water loss 3
811.3641967773438 0 9900.91
812.3692016601562 0 5011.012
813.3768920898438 0 2226.3604
818.9600830078125 0 2407.219 y Ammonia loss 3
819.466796875 0 43103.52 z 3
819.96923828125 0 72118.27
820.4710083007812 0 46721.668
820.9724731445312 0 24463.752
821.4740600585938 0 5760.5293
821.9737548828125 0 1493.0232
822.4453125 0 5379.536 c Ammonia loss 13
822.945556640625 0 4139.417
823.445068359375 0 1826.4634
823.94921875 0 1244.962
824.4514770507812 0 1093.5039
826.9763793945312 0 1885.4723
827.4760131835938 0 11940.801 y 3
827.9777221679688 0 12275.601
828.4771728515625 0 6393.3604
828.9810180664062 0 2207.3442
830.4608764648438 0 1930.7534
830.9583129882812 0 20322.68 c 13
831.4592895507812 0 17635.436
831.961181640625 0 10340.116
832.4619750976562 0 3990.2092
832.9638061523438 0 956.9953
835.4716796875 0 1533.5323
836.4675903320312 0 989.4079
837.4633178710938 0 1221.1625
838.4592895507812 0 1029.9541
839.3587036132812 0 8829.619 c Ammonia loss 5
840.3644409179688 0 4289.4585
841.3696899414062 0 1303.9944
841.9713134765625 0 2057.7996
843.4741821289062 0 1283.5978
843.9786376953125 0 824.0763
845.4735717773438 0 1342.2755
847.962646484375 0 987.0173
850.4667358398438 0 2493.7417
850.9694213867188 0 2769.702
851.479248046875 0 1521.6993
851.97265625 0 2660.875
853.4051513671875 0 1745.5873
853.7308959960938 0 1413.108
854.0761108398438 0 1294.6503
855.4978637695312 0 1368.8871
857.9840698242188 0 1883.282
858.4901123046875 0 2713.3818
858.9730224609375 0 1976.3904
859.4719848632812 0 2087.8228
863.4842529296875 0 960.0835
864.9888305664062 0 1926.125
865.4807739257812 0 5188.4854
865.98486328125 0 5373.8076
866.489990234375 0 2965.3755
866.9930419921875 0 2151.9192
867.5010986328125 0 1268.5758
869.4794311523438 0 1069.0952
869.9808959960938 0 2278.2231
870.4838256835938 0 1761.7234
870.9931030273438 0 1212.5806
871.984619140625 0 3027.5835
872.4828491210938 0 3978.2139
872.97607421875 0 3227.9487
873.4785766601562 0 2134.4329
873.9534912109375 0 1031.5416
877.9910888671875 0 3337.0542
878.4909057617188 0 5729.775 c Water loss 14
878.988037109375 0 22577.582 c Ammonia loss 14
879.4891967773438 0 18458.186
879.9893798828125 0 14957.017
880.4921264648438 0 5622.9893
880.9890747070312 0 2430.7566
882.4844360351562 0 1058.7695
882.5656127929688 0 1119.2313
886.4844360351562 0 1402.326
886.9953002929688 0 26040.49
887.4986572265625 0 56222.27 c 14
888.0001831054688 0 45943.527
888.5029907226562 0 27686.848
889.0042114257812 0 9105.756
889.5031127929688 0 3162.4082
891.0012817382812 0 905.1094
891.496337890625 0 1220.3512
891.999267578125 0 1517.9691 z Water loss 2
892.5001831054688 0 3270.6707 z Ammonia loss 2
895.4449462890625 0 1715.6301
895.996826171875 0 1179.7712
896.4533081054688 0 1742.1278
897.5183715820312 0 5190.6685
898.5574951171875 0 9519.757 y 10
899.5624389648438 0 3971.1963
900.0004272460938 0 4701.8784 y Water loss 2
900.502197265625 0 5311.188 y Ammonia loss 2
900.9989013671875 0 35038.812 z 2
901.5003662109375 0 40366.125
902.00146484375 0 21223.936
902.5031127929688 0 9539.565
903.0054321289062 0 3429.2769
907.993408203125 0 1136.7601
908.5005493164062 0 1483.9491
909.0079345703125 0 30407.59 y 2
909.509033203125 0 29230.705
910.010986328125 0 18348.414
910.5125732421875 0 7627.7593
911.0054321289062 0 1863.9895
912.48095703125 0 1242.9395
913.489990234375 0 1873.4838
913.9962768554688 0 2046.9808
914.49755859375 0 1004.53503
915.0034790039062 0 1302.3865
918.61083984375 0 985.8464
919.614013671875 0 985.4185
920.5040283203125 0 1184.346
921.5023803710938 0 1065.6481
922.0078735351562 0 1404.2607 c Water loss 15
922.5028686523438 0 10097.341 c Ammonia loss 15
923.0029296875 0 10158.476
923.5027465820312 0 5829.366
924.0090942382812 0 2605.8167
924.1544189453125 0 1002.24457
924.500732421875 0 1474.4946
928.4913330078125 0 1233.6271
929.1585693359375 0 1091.1085
929.4972534179688 0 3869.8655
930.0133666992188 0 1252.3958
930.5015869140625 0 6683.2275
930.8319091796875 0 2861.942
931.0157470703125 0 63460.613 c 15
931.5170288085938 0 66214.24
932.0188598632812 0 37684.34
932.5203247070312 0 17076.44
933.0200805664062 0 5595.3306
933.5224609375 0 1306.4874
934.41748046875 0 966.8098
937.5018310546875 0 1207.7438
938.0017700195312 0 1067.6191
939.5067138671875 0 1023.4484
944.0230102539062 0 2831.0981
944.5230712890625 0 5467.101
945.0204467773438 0 3329.871
945.5188598632812 0 1347.2933
946.011962890625 0 1226.4067
952.0135498046875 0 1464.0825
952.4303588867188 0 3770.536
953.4400634765625 0 20165.082 c 6
953.5806274414062 0 38048.51 z 9
954.4428100585938 0 13015.234
954.5839233398438 0 21593.982
955.4453735351562 0 3269.2456
955.5861206054688 0 7742.9126
956.5891723632812 0 2054.975
965.0220947265625 0 1916.9
966.026123046875 0 1514.1205
966.4466552734375 0 3572.0515
967.0159301757812 0 1189.4781
967.4528198242188 0 3174.3828
969.5971069335938 0 8212.551 y 9
970.6031494140625 0 6149.321
971.6127319335938 0 1420.5686
972.0433349609375 0 1442.4412
972.52880859375 0 2250.8262
973.025146484375 0 3059.85
973.5299072265625 0 12713.997
974.03125 0 13207.112
974.5316162109375 0 8325.813
975.0313110351562 0 3697.5942
975.5321655273438 0 2166.4685
977.03662109375 0 1624.1981
977.5634155273438 0 1093.2279
978.0248413085938 0 2512.7214
978.5256958007812 0 2810.8464
979.0260009765625 0 2203.9272
979.5387573242188 0 2246.849
980.0416870117188 0 2545.9219
980.5321044921875 0 5808.0527
981.0316772460938 0 4349.238
981.5286865234375 0 2233.1533
982.0376586914062 0 1394.4681
984.021728515625 0 1299.7231
985.5501098632812 0 1380.6527
986.0508422851562 0 1452.5593
986.5490112304688 0 2408.9 c Water loss 16
987.0269775390625 0 14309.525 c Ammonia loss 16
987.5294189453125 0 15432.872
988.0308837890625 0 11464.823
988.53125 0 5424.468
989.03466796875 0 2252.1685
989.5322875976562 0 1211.85
992.5534057617188 0 2869.7634
993.5411376953125 0 4096.609 y Ammonia loss 1
994.0400390625 0 11330.736 z 1
994.5404663085938 0 13136.86
995.0407104492188 0 8610.311
995.5381469726562 0 70400.375 c 16
996.0396118164062 0 74279.76
996.5408935546875 0 52325.96
997.04150390625 0 18805.664
997.5436401367188 0 8561.689
998.0386352539062 0 2269.335
1000.0509033203125 0 1186.342
1001.048095703125 0 2554.6936
1001.547119140625 0 2979.6707
1002.0420532226562 0 6552.1074 y 1
1002.5394897460938 0 5757.9243
1003.0352783203125 0 4052.969
1003.5443725585938 0 1392.1274
1005.03955078125 0 3845.2854
1005.5438842773438 0 5630.587
1006.04248046875 0 6040.651
1006.5489501953125 0 3521.606
1007.05810546875 0 1294.4215
1007.547607421875 0 1330.7625
1008.0468139648438 0 8120.312
1008.5468139648438 0 8609.404
1009.0458984375 0 6652.873
1009.4453735351562 0 5027.6245
1009.536865234375 0 10756.275
1010.0357666015625 0 11430.881
1010.4594116210938 0 36750.93 c 7
1011.0440673828125 0 5039.4263
1011.4629516601562 0 18355.053
1012.0440063476562 0 1946.8044
1012.4683837890625 0 6201.404
1014.5559692382812 0 8017.0737
1015.0598754882812 0 13262.455
1015.5613403320312 0 15405.544
1016.0560913085938 0 13836.284
1016.5611572265625 0 8720.811
1017.0525512695312 0 6417.426
1017.5481567382812 0 3106.3845
1018.0415649414062 0 5368.916
1018.53857421875 0 4937.474
1019.0479736328125 0 1600.131
1021.5536499023438 0 1220.8065
1022.0634765625 0 3601.8562
1022.5680541992188 0 3941.4724
1023.0615844726562 0 6879.9756
1023.5607299804688 0 22524.89
1024.0616455078125 0 23954.588
1024.5516357421875 0 27030.104
1025.04833984375 0 25496.793
1025.549072265625 0 13818.689
1026.0457763671875 0 5615.0728
1026.545166015625 0 3573.1018
1028.05517578125 0 2586.4602
1028.5543212890625 0 4413.558
1029.0576171875 0 6461.673
1029.56201171875 0 7532.8506
1030.065673828125 0 8241.255
1030.5670166015625 0 7676.312
1031.0697021484375 0 5599.816
1031.5721435546875 0 3636.2427
1032.0682373046875 0 2584.2776
1036.560546875 0 2877.3503
1037.06005859375 0 12333.071
1037.55615234375 0 84166.34
1038.057861328125 0 99623.234
1038.560302734375 0 77133.25
1039.062255859375 0 42836.926
1039.566650390625 0 17614.547
1040.0704345703125 0 5075.3975
1042.5230712890625 0 2223.774
1044.0479736328125 0 1384.3381
1044.5570068359375 0 3114.3264
1045.0606689453125 0 6599.412
1045.5645751953125 0 116006.22
1046.0673828125 0 454005.5
1046.5693359375 0 479203.6
1047.070556640625 0 297550.16
1047.5718994140625 0 135650.8
1048.0726318359375 0 47952.184
1048.5699462890625 0 11159.229
1081.6761474609375 0 24324.38 z 8
1082.6800537109375 0 26617.34
1083.683349609375 0 12499.073
1084.6865234375 0 4212.191
1094.5423583984375 0 7157.0264
1095.5472412109375 0 5272.2363
1096.54736328125 0 1724.0764
1097.693115234375 0 7895.2217 y 8
1098.6953125 0 4780.7666
1099.6971435546875 0 1543.1014
1138.5556640625 0 49995.207 c 8
1138.6976318359375 0 12579.143 z 7
1139.5574951171875 0 32204.773
1139.7032470703125 0 17644.309
1140.561279296875 0 13118.083
1140.7060546875 0 7794.3003
1141.5662841796875 0 2875.7434
1141.7100830078125 0 2021.2695
1146.5877685546875 0 2765.8118
1154.7115478515625 0 2345.7644 y 7
1155.718994140625 0 2222.4744
1165.578369140625 0 5929.629
1166.5831298828125 0 3527.9949
1167.5806884765625 0 1101.4917
1208.7154541015625 0 936.9696 w 6
1209.7255859375 0 1002.0586
1251.7677001953125 0 3675.9097 y 6
1252.7708740234375 0 3307.8596
1253.775146484375 0 1018.96936
1278.7481689453125 0 2936.7559
1279.756591796875 0 2989.0237
1280.1080322265625 0 2174.7173
1280.6162109375 0 1225.0375
1280.7708740234375 0 3136.696
1281.1082763671875 0 2378.021
1281.770751953125 0 2062.2017
1288.1141357421875 0 959.41986
1305.6273193359375 0 2036.5209
1306.644287109375 0 7540.3228 c 10
1307.634765625 0 4747.8843
1308.629150390625 0 1617.0613
1337.5889892578125 0 1527.251
1372.8077392578125 0 25738.848 z 5
1373.8128662109375 0 27620.633
1374.8143310546875 0 13855.634
1375.8175048828125 0 4251.0586
1385.7088623046875 0 1316.1455
1387.7470703125 0 1220.9224
1388.8272705078125 0 15208.877 y 5
1389.8310546875 0 12423.753
1390.8330078125 0 6241.3794
1391.8443603515625 0 1334.4851
1392.7301025390625 0 2611.7068
1393.7059326171875 0 2189.9585
1394.2388916015625 0 1297.1959
1394.727783203125 0 1541.1709
1395.732177734375 0 3957.5728
1396.232177734375 0 5479.139
1396.74267578125 0 3641.3562
1397.2442626953125 0 1910.6301
1433.7275390625 0 2865.1328
1434.7391357421875 0 18940.219 c 11
1435.74267578125 0 15316.085
1436.74755859375 0 6980.8003
1437.7474365234375 0 1849.8611
1503.807861328125 0 2697.5217
1504.8179931640625 0 1948.7576
1509.86767578125 0 18708.742 z 4
1510.874267578125 0 44083
1511.8770751953125 0 30130.316
1512.8787841796875 0 15093.081
1513.88525390625 0 4329.1865
1524.8731689453125 0 2361.8623
1525.884765625 0 10696.278 y 4
1526.886962890625 0 8558.432
1527.89404296875 0 4131.8735
1528.8973388671875 0 2036.0001
1547.819580078125 0 6850.3374 c 12
1548.8275146484375 0 7375.7114
1549.8270263671875 0 3722.7715
1550.8408203125 0 1361.8497
1637.9251708984375 0 4286.7695 z 3
1638.935546875 0 27920.861
1639.9384765625 0 24822.512
1640.939453125 0 11739.536
1641.9415283203125 0 5432.232
1642.9447021484375 0 1375.8104
1653.949462890625 0 3336.6116 y 3
1654.9454345703125 0 3071.167
1655.9478759765625 0 1440.3856
1656.9368896484375 0 1197.5565
1660.9110107421875 0 5476.644 c 13
1661.915283203125 0 4109.4653
1662.906494140625 0 2085.0955
1757.9739990234375 0 2620.671
1758.9796142578125 0 1338.6646
1772.9830322265625 0 2683.4727
1773.9927978515625 0 15005.456 c 14
1774.9952392578125 0 13147.07
1775.997802734375 0 8755.19
1777.01416015625 0 3395.2603
1800.9920654296875 0 4071.773 z 2
1801.9967041015625 0 15880.31
1803.000732421875 0 17090.24
1804.00732421875 0 8383.09
1805.00634765625 0 3201.2424
1817.01953125 0 2500.0127 y 2
1818.0057373046875 0 4688.497
1819.0164794921875 0 2903.786
1820.02587890625 0 1377.4547
1860.00341796875 0 1385.6492
1861.026123046875 0 17303.686 c 15
1862.028564453125 0 17940.838
1863.0321044921875 0 12244.572
1864.034423828125 0 5466.5117
1865.0390625 0 1488.4033
1946.0487060546875 0 2467.358
1947.0556640625 0 4938.2554
1948.057861328125 0 5069.195
1949.0732421875 0 2878.6882
1965.0511474609375 0 1129.0619
1975.0394287109375 0 1840.1793
1987.0694580078125 0 1778.6313 z 1
1988.0728759765625 0 6178.1763
1989.080810546875 0 4773.6943
1990.0740966796875 0 9929.254 c 16
1991.0743408203125 0 13575.703
1992.0745849609375 0 11228.075
1993.0806884765625 0 5809.6523
1994.085693359375 0 2843.9626
1995.0731201171875 0 1228.8876
2030.1231689453125 0 1929.9036
2031.123779296875 0 2891.9487
2045.134033203125 0 1917.9181
2046.12060546875 0 3315.933
2047.1207275390625 0 10818.528
2048.121337890625 0 11824.099
2049.12109375 0 7749.385
2050.11328125 0 2840.456
2051.13671875 0 1553.184
2057.092529296875 0 3528.3987
2058.10693359375 0 6401.8794
2059.111083984375 0 5187.7495
2060.12646484375 0 3286.6077
2061.11962890625 0 2134.75
2062.113525390625 0 1924.987
2063.137939453125 0 3641.1545
2064.137939453125 0 16686.162
2065.14501953125 0 15097.086
2066.146240234375 0 9744.046
2067.145751953125 0 4446.3027
2068.15673828125 0 1648.5182
2073.141357421875 0 1314.8085
2074.1083984375 0 8854.691
2075.110107421875 0 42570.58
2076.113525390625 0 49521.56
2077.114013671875 0 32412.293
2078.119140625 0 17975.926
2079.131591796875 0 7696.244
2089.125732421875 0 1579.4366
2090.119873046875 0 10288.268
2091.1279296875 0 53187.375
2092.13330078125 0 196286.16
2093.137939453125 0 216035.19
2094.1396484375 0 133627.22
2095.143310546875 0 63123.89
2096.143310546875 0 23054.271
2097.1455078125 0 6931.032
2294.2236328125 0 1899.6287
2295.20751953125 0 2156.3833

Spectrum Details

|  |  |
| --- | --- |
| Matched peaks? Matched peaks The total absolute number of peaks matched. Additionally in brackets the total fraction of peaks matched and the total number of peaks is shown. | 89 (13.20% of 674) |
| FDR? FDR The false discovery rate estimated for this peptide. It is calculated by matching all theoretical fragments with a non-integer shift with the raw peaks for this spectrum. This is done with 40 different shifts. The resulting percentage is the average number of annotated peaks over the number of annotated peaks with the correct spectrum. | 3.77% |
| Satellite FDR? Satellite FDR See the FDR for details on its calculation. This satellite ion specific FDR only contains the satellite ions (d/w) for I/L/J positions. | 0.00% |
| PSM Score? PSM Score The PSM Score as given by Hecklib to this annotated spectrum. It is shown with three significant figures. | 510 |

#### Spectrum 7638? Spectrum 7638 The raw spectrum of this peptide as annotated by Hecklib. The fragments are coloured according to ion type (see legend). Any peaks with a star '\*' as text can be hovered over to see the full details, first the ion type second the mass shift type. By hovering over the amino acids in the peptide or ions in the legend the corresponding peaks are highlighted. By toggling the 'Unassigned' label you can turn the background (unassigned) peaks on or off in the plot. By updating the slider in the Ion legend you can update the spectrum to only show the top X% of the peaks with labels. The top X% means any peak that is within X% of the highest intensity. By dragging in the spectrum you can zoom in to a specific part of the spectrum and use 'Zoom Out' to get back to the original zoom level. The annotation of the spectrum is based on the given sequence in the peptides file and is done with different software so inconsistencies are likely. The peaks are annotated based on the given sequence, with 20 ppm tolerance.

Copy Data

##### Spectrum 7638 (TSV)

###### Preview

```
Loading example...
```

*Click on the button to copy the data to your clipboard.*

Mz MinMz MaxIntensity Max

WidthHeightPeptide font sizePeptide stroke widthSpectrum font sizeSpectrum stroke widthCompact peptide

Ion legend

wxyz

abcd

OtherUnassignedIon^Charge^_Position_Show for top:%

SWYQHHPGKAPKJJJSEV

04.77e+49.53e+41.43e+51.91e+5

Zoom Out

y^+1^_2_y^+1^_2_y^+1^_3_y^+1^_4_y^+1^_5_c^+2^_9_c^+1^_4_c^+1^_4_c^+1^_4_c^+3^_15_y^+3^_16_w^+1^_6_c^+3^_16_y^+2^_12_z^+1^_6_c^+3^_17_y^+3^_17_y^+1^_6_y^+2^_13_c^+1^_5_c^+2^_12_c^+2^_12_c^+1^_5_z^+2^_14_y^+2^_14_c^+2^_13_z^+1^_7_w^+2^_15_y^+1^_7_y^+2^_15_z^+2^_15_c^+2^_14_y^+2^_15_c^+2^_14_c^+1^_6_c^+2^_15_c^+2^_15_c^+2^_15_y^+1^_8_y^+2^_16_y^+2^_16_z^+2^_16_y^+2^_16_c^+2^_16_c^+2^_16_c^+1^_7_z^+1^_9_y^+1^_9_c^+2^_17_y^+2^_17_z^+2^_17_c^+2^_17_y^+2^_17_c^+1^_8_z^+1^_10_y^+1^_10_c^+1^_9_z^+1^_11_y^+1^_11_y^+1^_12_c^+1^_11_z^+1^_13_y^+1^_13_c^+1^_12_z^+1^_14_y^+1^_14_c^+1^_13_z^+1^_15_y^+1^_15_c^+1^_14_c^+1^_15_z^+1^_16_y^+1^_16_c^+1^_16_c^+1^_17_

0705141021152819

Fragment Matches Table

Show background peaks

| Position | Ion type | Intensity | mz Theoretical | mz Error (Th) | mz Error (ppm) | Charge | Series Number |
| --- | --- | --- | --- | --- | --- | --- | --- |
| - | - | 442.7 | 128.1 | - | - | 0 | - |
| - | - | 349 | 129.1 | - | - | 0 | - |
| - | - | 393.3 | 129.1 | - | - | 0 | - |
| - | - | 1181 | 136.1 | - | - | 0 | - |
| - | - | 447.4 | 155.6 | - | - | 0 | - |
| - | - | 2751 | 217.1 | - | - | 0 | - |
| - | - | 1616 | 221.1 | - | - | 0 | - |
| 17 | y | 2142 | 229.1 | 0.0003483 | 1.52 | +1 | 2 |
| - | - | 743.6 | 232.1 | - | - | 0 | - |
| - | - | 2215 | 239.1 | - | - | 0 | - |
| - | - | 484.6 | 239.1 | - | - | 0 | - |
| - | - | 3396 | 246.1 | - | - | 0 | - |
| 17 | y | 1516 | 247.1 | 0.0008879 | 3.593 | +1 | 2 |
| - | - | 4715 | 274.1 | - | - | 0 | - |
| - | - | 815.1 | 275.1 | - | - | 0 | - |
| - | - | 1890 | 295.1 | - | - | 0 | - |
| - | - | 4185 | 300.1 | - | - | 0 | - |
| 16 | y | 2104 | 334.2 | 1.841E-05 | 0.05509 | +1 | 3 |
| - | - | 599.1 | 335.2 | - | - | 0 | - |
| - | - | 651.9 | 360.2 | - | - | 0 | - |
| - | - | 1965 | 369.1 | - | - | 0 | - |
| - | - | 645.5 | 397.2 | - | - | 0 | - |
| - | - | 1157 | 398.2 | - | - | 0 | - |
| - | - | 1332 | 437.2 | - | - | 0 | - |
| 15 | y | 1325 | 447.2 | 2.406E-05 | 0.05379 | +1 | 4 |
| - | - | 2114 | 463.2 | - | - | 0 | - |
| - | - | 953.7 | 465.2 | - | - | 0 | - |
| - | - | 641 | 471.3 | - | - | 0 | - |
| - | - | 790.8 | 477.2 | - | - | 0 | - |
| - | - | 1008 | 480.2 | - | - | 0 | - |
| - | - | 570.8 | 483.2 | - | - | 0 | - |
| - | - | 895.3 | 528.2 | - | - | 0 | - |
| - | - | 1060 | 538.3 | - | - | 0 | - |
| - | - | 1613 | 539.3 | - | - | 0 | - |
| - | - | 611.2 | 541.3 | - | - | 0 | - |
| 14 | y | 1326 | 560.3 | 0.001002 | 1.788 | +1 | 5 |
| 9 | c | 1685 | 561.3 | 0.001453 | 2.588 | +2 | 9 |
| - | - | 2401 | 563.2 | - | - | 0 | - |
| 4 | c | 889 | 564.3 | 0.007363 | 13.05 | +1 | 4 |
| 4 | c | 913.7 | 565.2 | 0.0005336 | 0.9441 | +1 | 4 |
| - | - | 706.5 | 567.8 | - | - | 0 | - |
| - | - | 664.8 | 577 | - | - | 0 | - |
| - | - | 987.2 | 580.3 | - | - | 0 | - |
| - | - | 2165 | 581.3 | - | - | 0 | - |
| 4 | c | 1.242E+04 | 582.3 | 0.0005324 | 0.9144 | +1 | 4 |
| - | - | 3950 | 583.3 | - | - | 0 | - |
| 15 | c | 2223 | 586.3 | 0.0005079 | 0.8662 | +3 | 15 |
| - | - | 3108 | 586.7 | - | - | 0 | - |
| - | - | 720.7 | 587 | - | - | 0 | - |
| 3 | y | 840 | 606.3 | 0.004083 | 6.733 | +3 | 16 |
| - | - | 619.3 | 606.7 | - | - | 0 | - |
| 13 | w | 856.6 | 614.3 | 0.0018 | 2.931 | +1 | 6 |
| 16 | c | 2564 | 615.3 | 0.0002587 | 0.4204 | +3 | 16 |
| - | - | 2737 | 615.7 | - | - | 0 | - |
| - | - | 929.2 | 616 | - | - | 0 | - |
| - | - | 753.2 | 616.3 | - | - | 0 | - |
| - | - | 764.8 | 621 | - | - | 0 | - |
| 7 | y | 3470 | 626.4 | 0.0004694 | 0.7493 | +2 | 12 |
| - | - | 2972 | 626.9 | - | - | 0 | - |
| - | - | 1668 | 627.4 | - | - | 0 | - |
| - | - | 703.8 | 646.3 | - | - | 0 | - |
| - | - | 1053 | 649.4 | - | - | 0 | - |
| - | - | 1626 | 652.3 | - | - | 0 | - |
| - | - | 1731 | 652.7 | - | - | 0 | - |
| - | - | 1551 | 653 | - | - | 0 | - |
| - | - | 830.8 | 657.3 | - | - | 0 | - |
| 13 | z | 829.3 | 657.4 | 0.000431 | 0.6556 | +1 | 6 |
| - | - | 934.9 | 658.3 | - | - | 0 | - |
| 17 | c | 1.117E+04 | 658.4 | 0.0002352 | 0.3573 | +3 | 17 |
| - | - | 966.8 | 658.4 | - | - | 0 | - |
| - | - | 1.338E+04 | 658.7 | - | - | 0 | - |
| - | - | 5896 | 659 | - | - | 0 | - |
| - | - | 1125 | 659.3 | - | - | 0 | - |
| - | - | 2519 | 659.4 | - | - | 0 | - |
| - | - | 2579 | 659.7 | - | - | 0 | - |
| - | - | 1235 | 664.4 | - | - | 0 | - |
| 2 | y | 937.9 | 668.4 | 0.0003213 | 0.4807 | +3 | 17 |
| - | - | 710.5 | 669 | - | - | 0 | - |
| 13 | y | 1279 | 673.4 | 0.0008719 | 1.295 | +1 | 6 |
| - | - | 1545 | 674.3 | - | - | 0 | - |
| - | - | 3195 | 675.3 | - | - | 0 | - |
| - | - | 2212 | 676.3 | - | - | 0 | - |
| - | - | 4093 | 691.4 | - | - | 0 | - |
| - | - | 5204 | 691.7 | - | - | 0 | - |
| - | - | 3495 | 692 | - | - | 0 | - |
| - | - | 954.9 | 692.4 | - | - | 0 | - |
| 6 | y | 952.5 | 694.9 | 0.001447 | 2.082 | +2 | 13 |
| - | - | 1119 | 696.7 | - | - | 0 | - |
| - | - | 889.8 | 697 | - | - | 0 | - |
| - | - | 6.895E+04 | 697.4 | - | - | 0 | - |
| - | - | 8.402E+04 | 697.7 | - | - | 0 | - |
| - | - | 5.565E+04 | 698 | - | - | 0 | - |
| - | - | 1201 | 698.1 | - | - | 0 | - |
| - | - | 2.791E+04 | 698.4 | - | - | 0 | - |
| - | - | 8776 | 698.7 | - | - | 0 | - |
| - | - | 4382 | 699 | - | - | 0 | - |
| 5 | c | 1591 | 702.3 | 0.002627 | 3.741 | +1 | 5 |
| 12 | c | 844.7 | 709.4 | 2.54E-05 | 0.03581 | +2 | 12 |
| - | - | 3849 | 717.4 | - | - | 0 | - |
| 12 | c | 4.251E+04 | 717.9 | 0.0007877 | 1.097 | +2 | 12 |
| - | - | 3.147E+04 | 718.4 | - | - | 0 | - |
| - | - | 1.366E+04 | 718.9 | - | - | 0 | - |
| 5 | c | 1.304E+04 | 719.3 | 0.0009116 | 1.267 | +1 | 5 |
| - | - | 3269 | 719.4 | - | - | 0 | - |
| - | - | 1035 | 719.9 | - | - | 0 | - |
| - | - | 6224 | 720.3 | - | - | 0 | - |
| - | - | 1966 | 721.3 | - | - | 0 | - |
| - | - | 717.8 | 743.9 | - | - | 0 | - |
| - | - | 1640 | 752.4 | - | - | 0 | - |
| - | - | 1068 | 752.9 | - | - | 0 | - |
| - | - | 881 | 753.4 | - | - | 0 | - |
| 5 | z | 3941 | 755.4 | 0.0005807 | 0.7687 | +2 | 14 |
| - | - | 7314 | 755.9 | - | - | 0 | - |
| - | - | 4525 | 756.4 | - | - | 0 | - |
| - | - | 2539 | 756.9 | - | - | 0 | - |
| - | - | 869.2 | 757.4 | - | - | 0 | - |
| - | - | 905.5 | 762.9 | - | - | 0 | - |
| 5 | y | 4770 | 763.4 | 0.001337 | 1.751 | +2 | 14 |
| - | - | 3642 | 763.9 | - | - | 0 | - |
| - | - | 1577 | 764.5 | - | - | 0 | - |
| - | - | 1087 | 764.9 | - | - | 0 | - |
| - | - | 2891 | 773.9 | - | - | 0 | - |
| 13 | c | 2.545E+04 | 774.4 | 0.0009496 | 1.226 | +2 | 13 |
| - | - | 2.189E+04 | 774.9 | - | - | 0 | - |
| - | - | 9347 | 775.4 | - | - | 0 | - |
| - | - | 3362 | 775.9 | - | - | 0 | - |
| - | - | 724 | 776.4 | - | - | 0 | - |
| - | - | 703.7 | 783.9 | - | - | 0 | - |
| 12 | z | 1637 | 785.5 | 0.001881 | 2.394 | +1 | 7 |
| - | - | 1198 | 786.5 | - | - | 0 | - |
| - | - | 841.8 | 787.5 | - | - | 0 | - |
| 4 | w | 3665 | 790.5 | 0.0009429 | 1.193 | +2 | 15 |
| - | - | 2794 | 791 | - | - | 0 | - |
| - | - | 2033 | 791.5 | - | - | 0 | - |
| - | - | 1585 | 791.9 | - | - | 0 | - |
| - | - | 1215 | 796.4 | - | - | 0 | - |
| 12 | y | 836.8 | 801.5 | 0.004247 | 5.299 | +1 | 7 |
| - | - | 741.5 | 802.5 | - | - | 0 | - |
| - | - | 848.9 | 808.9 | - | - | 0 | - |
| - | - | 999.8 | 809.5 | - | - | 0 | - |
| - | - | 997.7 | 810 | - | - | 0 | - |
| - | - | 4125 | 811.4 | - | - | 0 | - |
| - | - | 1816 | 812.4 | - | - | 0 | - |
| 4 | y | 1190 | 818.5 | 0.005751 | 7.026 | +2 | 15 |
| 4 | z | 1.379E+04 | 819.5 | 0.0009388 | 1.146 | +2 | 15 |
| - | - | 2.824E+04 | 820 | - | - | 0 | - |
| - | - | 1.494E+04 | 820.5 | - | - | 0 | - |
| - | - | 7363 | 821 | - | - | 0 | - |
| - | - | 3280 | 821.5 | - | - | 0 | - |
| 14 | c | 1653 | 822.4 | 0.001997 | 2.428 | +2 | 14 |
| - | - | 2046 | 822.9 | - | - | 0 | - |
| - | - | 700.5 | 823.4 | - | - | 0 | - |
| 4 | y | 5075 | 827.5 | 0.001634 | 1.974 | +2 | 15 |
| - | - | 3489 | 828 | - | - | 0 | - |
| - | - | 2196 | 828.5 | - | - | 0 | - |
| 14 | c | 8514 | 831 | 0.0016 | 1.925 | +2 | 14 |
| - | - | 6105 | 831.5 | - | - | 0 | - |
| - | - | 3719 | 832 | - | - | 0 | - |
| - | - | 1252 | 832.5 | - | - | 0 | - |
| - | - | 709.6 | 839 | - | - | 0 | - |
| 6 | c | 3206 | 839.4 | 0.00178 | 2.121 | +1 | 6 |
| - | - | 1406 | 840.4 | - | - | 0 | - |
| - | - | 734.6 | 842.5 | - | - | 0 | - |
| - | - | 1098 | 850.5 | - | - | 0 | - |
| - | - | 1776 | 851 | - | - | 0 | - |
| - | - | 1032 | 851.5 | - | - | 0 | - |
| - | - | 716.8 | 852 | - | - | 0 | - |
| - | - | 659.4 | 858.5 | - | - | 0 | - |
| - | - | 2014 | 865.5 | - | - | 0 | - |
| - | - | 1475 | 866 | - | - | 0 | - |
| - | - | 987.6 | 866.5 | - | - | 0 | - |
| - | - | 847.3 | 867 | - | - | 0 | - |
| - | - | 943.9 | 869.5 | - | - | 0 | - |
| - | - | 1204 | 872 | - | - | 0 | - |
| - | - | 1042 | 872.5 | - | - | 0 | - |
| - | - | 1171 | 873 | - | - | 0 | - |
| - | - | 928.2 | 873.5 | - | - | 0 | - |
| - | - | 1140 | 878 | - | - | 0 | - |
| 15 | c | 2165 | 878.5 | 0.003376 | 3.843 | +2 | 15 |
| 15 | c | 5341 | 879 | 0.0008318 | 0.9463 | +2 | 15 |
| - | - | 7469 | 879.5 | - | - | 0 | - |
| - | - | 5427 | 880 | - | - | 0 | - |
| - | - | 1091 | 880.5 | - | - | 0 | - |
| - | - | 838.9 | 881 | - | - | 0 | - |
| - | - | 9763 | 887 | - | - | 0 | - |
| 15 | c | 2.15E+04 | 887.5 | 0.001884 | 2.123 | +2 | 15 |
| - | - | 1.826E+04 | 888 | - | - | 0 | - |
| - | - | 9056 | 888.5 | - | - | 0 | - |
| - | - | 2872 | 889 | - | - | 0 | - |
| - | - | 1066 | 889.5 | - | - | 0 | - |
| - | - | 626.5 | 890 | - | - | 0 | - |
| - | - | 1779 | 897.5 | - | - | 0 | - |
| 11 | y | 3702 | 898.6 | 0.003605 | 4.012 | +1 | 8 |
| - | - | 1008 | 899.6 | - | - | 0 | - |
| 3 | y | 2150 | 900 | 0.001588 | 1.764 | +2 | 16 |
| 3 | y | 2284 | 900.5 | 0.008602 | 9.552 | +2 | 16 |
| 3 | z | 1.123E+04 | 901 | 0.0007427 | 0.8243 | +2 | 16 |
| - | - | 1.433E+04 | 901.5 | - | - | 0 | - |
| - | - | 1.019E+04 | 902 | - | - | 0 | - |
| - | - | 3080 | 902.5 | - | - | 0 | - |
| - | - | 1223 | 903 | - | - | 0 | - |
| - | - | 754.3 | 908.5 | - | - | 0 | - |
| 3 | y | 1.17E+04 | 909 | 0.001072 | 1.179 | +2 | 16 |
| - | - | 1.164E+04 | 909.5 | - | - | 0 | - |
| - | - | 7413 | 910 | - | - | 0 | - |
| - | - | 3182 | 910.5 | - | - | 0 | - |
| - | - | 686.5 | 911.5 | - | - | 0 | - |
| - | - | 710.3 | 914.5 | - | - | 0 | - |
| 16 | c | 3280 | 922.5 | 0.001389 | 1.505 | +2 | 16 |
| - | - | 2430 | 923 | - | - | 0 | - |
| - | - | 2948 | 923.5 | - | - | 0 | - |
| - | - | 1499 | 924 | - | - | 0 | - |
| - | - | 798.5 | 928.5 | - | - | 0 | - |
| - | - | 3800 | 930.5 | - | - | 0 | - |
| - | - | 1907 | 930.8 | - | - | 0 | - |
| 16 | c | 2.328E+04 | 931 | 0.001602 | 1.72 | +2 | 16 |
| - | - | 3.221E+04 | 931.5 | - | - | 0 | - |
| - | - | 1.625E+04 | 932 | - | - | 0 | - |
| - | - | 6093 | 932.5 | - | - | 0 | - |
| - | - | 2379 | 933 | - | - | 0 | - |
| - | - | 680.9 | 944 | - | - | 0 | - |
| - | - | 1997 | 944.5 | - | - | 0 | - |
| - | - | 670.1 | 945 | - | - | 0 | - |
| - | - | 592.9 | 945.5 | - | - | 0 | - |
| - | - | 1215 | 952.4 | - | - | 0 | - |
| 7 | c | 8552 | 953.4 | 0.0002826 | 0.2964 | +1 | 7 |
| 10 | z | 1.823E+04 | 953.6 | 0.000816 | 0.8557 | +1 | 9 |
| - | - | 5603 | 954.4 | - | - | 0 | - |
| - | - | 8008 | 954.6 | - | - | 0 | - |
| - | - | 1579 | 955.4 | - | - | 0 | - |
| - | - | 2399 | 955.6 | - | - | 0 | - |
| - | - | 1017 | 966.4 | - | - | 0 | - |
| - | - | 1925 | 967.5 | - | - | 0 | - |
| - | - | 874.1 | 968.5 | - | - | 0 | - |
| 10 | y | 2715 | 969.6 | 0.0001743 | 0.1798 | +1 | 9 |
| - | - | 1913 | 970.6 | - | - | 0 | - |
| - | - | 1187 | 973 | - | - | 0 | - |
| - | - | 5370 | 973.5 | - | - | 0 | - |
| - | - | 3920 | 974 | - | - | 0 | - |
| - | - | 2594 | 974.5 | - | - | 0 | - |
| - | - | 1827 | 975 | - | - | 0 | - |
| - | - | 995.1 | 979 | - | - | 0 | - |
| - | - | 1570 | 979.5 | - | - | 0 | - |
| - | - | 716.4 | 980 | - | - | 0 | - |
| - | - | 2021 | 980.5 | - | - | 0 | - |
| - | - | 1506 | 981 | - | - | 0 | - |
| 17 | c | 5825 | 987 | 0.001119 | 1.133 | +2 | 17 |
| - | - | 4904 | 987.5 | - | - | 0 | - |
| - | - | 3056 | 988 | - | - | 0 | - |
| - | - | 2047 | 988.5 | - | - | 0 | - |
| - | - | 817.9 | 989 | - | - | 0 | - |
| 2 | y | 1144 | 993.5 | 0.002271 | 2.285 | +2 | 17 |
| 2 | z | 4030 | 994 | 0.0001892 | 0.1903 | +2 | 17 |
| - | - | 3833 | 994.5 | - | - | 0 | - |
| - | - | 3184 | 995 | - | - | 0 | - |
| 17 | c | 2.647E+04 | 995.5 | 0.0003151 | 0.3165 | +2 | 17 |
| - | - | 3.118E+04 | 996 | - | - | 0 | - |
| - | - | 1.629E+04 | 996.5 | - | - | 0 | - |
| - | - | 7904 | 997 | - | - | 0 | - |
| - | - | 2922 | 997.5 | - | - | 0 | - |
| - | - | 1052 | 998 | - | - | 0 | - |
| 2 | y | 2119 | 1002 | 0.01155 | 11.53 | +2 | 17 |
| - | - | 1423 | 1003 | - | - | 0 | - |
| - | - | 962.1 | 1003 | - | - | 0 | - |
| - | - | 2299 | 1005 | - | - | 0 | - |
| - | - | 1798 | 1006 | - | - | 0 | - |
| - | - | 2153 | 1006 | - | - | 0 | - |
| - | - | 1719 | 1007 | - | - | 0 | - |
| - | - | 1427 | 1007 | - | - | 0 | - |
| - | - | 1132 | 1008 | - | - | 0 | - |
| - | - | 2421 | 1009 | - | - | 0 | - |
| - | - | 2389 | 1009 | - | - | 0 | - |
| - | - | 1671 | 1009 | - | - | 0 | - |
| - | - | 4711 | 1010 | - | - | 0 | - |
| - | - | 4171 | 1010 | - | - | 0 | - |
| 8 | c | 1.313E+04 | 1010 | 0.001483 | 1.467 | +1 | 8 |
| - | - | 2126 | 1011 | - | - | 0 | - |
| - | - | 7915 | 1011 | - | - | 0 | - |
| - | - | 2310 | 1012 | - | - | 0 | - |
| - | - | 2761 | 1015 | - | - | 0 | - |
| - | - | 4286 | 1015 | - | - | 0 | - |
| - | - | 5417 | 1016 | - | - | 0 | - |
| - | - | 4952 | 1016 | - | - | 0 | - |
| - | - | 3057 | 1017 | - | - | 0 | - |
| - | - | 1923 | 1017 | - | - | 0 | - |
| - | - | 1037 | 1018 | - | - | 0 | - |
| - | - | 1600 | 1019 | - | - | 0 | - |
| - | - | 802.4 | 1019 | - | - | 0 | - |
| - | - | 1837 | 1023 | - | - | 0 | - |
| - | - | 7022 | 1024 | - | - | 0 | - |
| - | - | 8436 | 1024 | - | - | 0 | - |
| - | - | 9488 | 1025 | - | - | 0 | - |
| - | - | 7916 | 1025 | - | - | 0 | - |
| - | - | 5048 | 1026 | - | - | 0 | - |
| - | - | 2668 | 1026 | - | - | 0 | - |
| - | - | 801.2 | 1027 | - | - | 0 | - |
| - | - | 1247 | 1029 | - | - | 0 | - |
| - | - | 801 | 1030 | - | - | 0 | - |
| - | - | 909.2 | 1031 | - | - | 0 | - |
| - | - | 1082 | 1032 | - | - | 0 | - |
| - | - | 1776 | 1037 | - | - | 0 | - |
| - | - | 4319 | 1037 | - | - | 0 | - |
| - | - | 3.554E+04 | 1038 | - | - | 0 | - |
| - | - | 3.584E+04 | 1038 | - | - | 0 | - |
| - | - | 2.25E+04 | 1039 | - | - | 0 | - |
| - | - | 871.3 | 1039 | - | - | 0 | - |
| - | - | 1.245E+04 | 1039 | - | - | 0 | - |
| - | - | 3676 | 1040 | - | - | 0 | - |
| - | - | 949.9 | 1040 | - | - | 0 | - |
| - | - | 896.6 | 1045 | - | - | 0 | - |
| - | - | 3136 | 1045 | - | - | 0 | - |
| - | - | 5.002E+04 | 1046 | - | - | 0 | - |
| - | - | 1.735E+05 | 1046 | - | - | 0 | - |
| - | - | 1.887E+05 | 1047 | - | - | 0 | - |
| - | - | 1.079E+05 | 1047 | - | - | 0 | - |
| - | - | 5.187E+04 | 1048 | - | - | 0 | - |
| - | - | 1.898E+04 | 1048 | - | - | 0 | - |
| - | - | 4493 | 1049 | - | - | 0 | - |
| 9 | z | 1.038E+04 | 1082 | 0.00032 | 0.2958 | +1 | 10 |
| - | - | 1.114E+04 | 1083 | - | - | 0 | - |
| - | - | 5278 | 1084 | - | - | 0 | - |
| - | - | 1889 | 1085 | - | - | 0 | - |
| - | - | 2422 | 1095 | - | - | 0 | - |
| - | - | 1752 | 1096 | - | - | 0 | - |
| 9 | y | 2426 | 1098 | 0.003907 | 3.56 | +1 | 10 |
| - | - | 1756 | 1099 | - | - | 0 | - |
| - | - | 1123 | 1100 | - | - | 0 | - |
| 9 | c | 2.072E+04 | 1139 | 0.0008646 | 0.7594 | +1 | 9 |
| 8 | z | 7395 | 1139 | 0.0001889 | 0.1659 | +1 | 11 |
| - | - | 1.352E+04 | 1140 | - | - | 0 | - |
| - | - | 7216 | 1140 | - | - | 0 | - |
| - | - | 5314 | 1141 | - | - | 0 | - |
| - | - | 3575 | 1141 | - | - | 0 | - |
| - | - | 1580 | 1142 | - | - | 0 | - |
| - | - | 1191 | 1142 | - | - | 0 | - |
| 8 | y | 1007 | 1155 | 0.008564 | 7.417 | +1 | 11 |
| - | - | 2130 | 1166 | - | - | 0 | - |
| - | - | 1099 | 1167 | - | - | 0 | - |
| - | - | 1091 | 1218 | - | - | 0 | - |
| - | - | 1017 | 1219 | - | - | 0 | - |
| 7 | y | 1682 | 1252 | 0.006358 | 5.079 | +1 | 12 |
| - | - | 761.6 | 1253 | - | - | 0 | - |
| - | - | 2167 | 1279 | - | - | 0 | - |
| - | - | 1032 | 1280 | - | - | 0 | - |
| - | - | 1419 | 1281 | - | - | 0 | - |
| 11 | c | 2958 | 1307 | 0.001543 | 1.181 | +1 | 11 |
| - | - | 2058 | 1308 | - | - | 0 | - |
| - | - | 760.7 | 1337 | - | - | 0 | - |
| 6 | z | 1.024E+04 | 1373 | 0.003088 | 2.25 | +1 | 13 |
| - | - | 1.242E+04 | 1374 | - | - | 0 | - |
| - | - | 5827 | 1375 | - | - | 0 | - |
| - | - | 1035 | 1376 | - | - | 0 | - |
| - | - | 905.2 | 1379 | - | - | 0 | - |
| - | - | 1995 | 1386 | - | - | 0 | - |
| - | - | 1215 | 1386 | - | - | 0 | - |
| - | - | 1675 | 1387 | - | - | 0 | - |
| - | - | 864.6 | 1388 | - | - | 0 | - |
| 6 | y | 6244 | 1389 | 0.005577 | 4.016 | +1 | 13 |
| - | - | 4235 | 1390 | - | - | 0 | - |
| - | - | 1185 | 1391 | - | - | 0 | - |
| - | - | 1911 | 1393 | - | - | 0 | - |
| - | - | 2299 | 1394 | - | - | 0 | - |
| - | - | 3118 | 1395 | - | - | 0 | - |
| - | - | 1891 | 1395 | - | - | 0 | - |
| - | - | 4218 | 1396 | - | - | 0 | - |
| - | - | 4343 | 1396 | - | - | 0 | - |
| - | - | 5097 | 1397 | - | - | 0 | - |
| - | - | 2196 | 1397 | - | - | 0 | - |
| - | - | 1662 | 1398 | - | - | 0 | - |
| - | - | 928.5 | 1434 | - | - | 0 | - |
| 12 | c | 8585 | 1435 | 0.003576 | 2.493 | +1 | 12 |
| - | - | 7484 | 1436 | - | - | 0 | - |
| - | - | 3156 | 1437 | - | - | 0 | - |
| - | - | 1500 | 1505 | - | - | 0 | - |
| 5 | z | 7301 | 1510 | 0.002186 | 1.448 | +1 | 14 |
| - | - | 2.129E+04 | 1511 | - | - | 0 | - |
| - | - | 1.491E+04 | 1512 | - | - | 0 | - |
| - | - | 6986 | 1513 | - | - | 0 | - |
| - | - | 2143 | 1514 | - | - | 0 | - |
| - | - | 948.1 | 1525 | - | - | 0 | - |
| 5 | y | 5926 | 1526 | 0.003698 | 2.423 | +1 | 14 |
| - | - | 4187 | 1527 | - | - | 0 | - |
| - | - | 1854 | 1528 | - | - | 0 | - |
| 13 | c | 3041 | 1548 | 0.002679 | 1.731 | +1 | 13 |
| - | - | 3126 | 1549 | - | - | 0 | - |
| - | - | 2316 | 1550 | - | - | 0 | - |
| 4 | z | 2303 | 1638 | 0.003146 | 1.921 | +1 | 15 |
| - | - | 1.563E+04 | 1639 | - | - | 0 | - |
| - | - | 1.629E+04 | 1640 | - | - | 0 | - |
| - | - | 6843 | 1641 | - | - | 0 | - |
| - | - | 2804 | 1642 | - | - | 0 | - |
| - | - | 1058 | 1643 | - | - | 0 | - |
| 4 | y | 785.2 | 1654 | 0.002095 | 1.267 | +1 | 15 |
| - | - | 1415 | 1655 | - | - | 0 | - |
| 14 | c | 2033 | 1661 | 0.003247 | 1.955 | +1 | 14 |
| - | - | 2121 | 1662 | - | - | 0 | - |
| - | - | 1385 | 1663 | - | - | 0 | - |
| - | - | 919.8 | 1759 | - | - | 0 | - |
| - | - | 1770 | 1773 | - | - | 0 | - |
| 15 | c | 8257 | 1774 | 0.00235 | 1.325 | +1 | 15 |
| - | - | 7897 | 1775 | - | - | 0 | - |
| - | - | 4782 | 1776 | - | - | 0 | - |
| - | - | 2360 | 1777 | - | - | 0 | - |
| 3 | z | 3168 | 1801 | 0.0009082 | 0.5043 | +1 | 16 |
| - | - | 1.039E+04 | 1802 | - | - | 0 | - |
| - | - | 9312 | 1803 | - | - | 0 | - |
| - | - | 3804 | 1804 | - | - | 0 | - |
| - | - | 1873 | 1805 | - | - | 0 | - |
| 3 | y | 1553 | 1817 | 0.001092 | 0.6011 | +1 | 16 |
| - | - | 2365 | 1818 | - | - | 0 | - |
| - | - | 1685 | 1819 | - | - | 0 | - |
| - | - | 1099 | 1820 | - | - | 0 | - |
| 16 | c | 9347 | 1861 | 0.001542 | 0.8285 | +1 | 16 |
| - | - | 1.114E+04 | 1862 | - | - | 0 | - |
| - | - | 6724 | 1863 | - | - | 0 | - |
| - | - | 2917 | 1864 | - | - | 0 | - |
| - | - | 1513 | 1865 | - | - | 0 | - |
| - | - | 1252 | 1946 | - | - | 0 | - |
| - | - | 3839 | 1947 | - | - | 0 | - |
| - | - | 3580 | 1948 | - | - | 0 | - |
| - | - | 1689 | 1949 | - | - | 0 | - |
| - | - | 753.1 | 1976 | - | - | 0 | - |
| - | - | 2569 | 1988 | - | - | 0 | - |
| - | - | 3276 | 1989 | - | - | 0 | - |
| 17 | c | 4226 | 1990 | 0.003961 | 1.99 | +1 | 17 |
| - | - | 7646 | 1991 | - | - | 0 | - |
| - | - | 6775 | 1992 | - | - | 0 | - |
| - | - | 3355 | 1993 | - | - | 0 | - |
| - | - | 1129 | 1994 | - | - | 0 | - |
| - | - | 955.6 | 2011 | - | - | 0 | - |
| - | - | 904.4 | 2029 | - | - | 0 | - |
| - | - | 1539 | 2030 | - | - | 0 | - |
| - | - | 2402 | 2046 | - | - | 0 | - |
| - | - | 6862 | 2047 | - | - | 0 | - |
| - | - | 6370 | 2048 | - | - | 0 | - |
| - | - | 4238 | 2049 | - | - | 0 | - |
| - | - | 2562 | 2050 | - | - | 0 | - |
| - | - | 1884 | 2057 | - | - | 0 | - |
| - | - | 1908 | 2058 | - | - | 0 | - |
| - | - | 1862 | 2059 | - | - | 0 | - |
| - | - | 1137 | 2060 | - | - | 0 | - |
| - | - | 2284 | 2063 | - | - | 0 | - |
| - | - | 8643 | 2064 | - | - | 0 | - |
| - | - | 9930 | 2065 | - | - | 0 | - |
| - | - | 5411 | 2066 | - | - | 0 | - |
| - | - | 3303 | 2067 | - | - | 0 | - |
| - | - | 5270 | 2074 | - | - | 0 | - |
| - | - | 2.497E+04 | 2075 | - | - | 0 | - |
| - | - | 2.747E+04 | 2076 | - | - | 0 | - |
| - | - | 1.798E+04 | 2077 | - | - | 0 | - |
| - | - | 7083 | 2078 | - | - | 0 | - |
| - | - | 2538 | 2079 | - | - | 0 | - |
| - | - | 6398 | 2090 | - | - | 0 | - |
| - | - | 3.086E+04 | 2091 | - | - | 0 | - |
| - | - | 1.156E+05 | 2092 | - | - | 0 | - |
| - | - | 1.281E+05 | 2093 | - | - | 0 | - |
| - | - | 7.843E+04 | 2094 | - | - | 0 | - |
| - | - | 3.873E+04 | 2095 | - | - | 0 | - |
| - | - | 1.26E+04 | 2096 | - | - | 0 | - |
| - | - | 2973 | 2097 | - | - | 0 | - |
| - | - | 895.6 | 2295 | - | - | 0 | - |
| - | - | 1074 | 2296 | - | - | 0 | - |
| - | - | 909 | 2790 | - | - | 0 | - |
| - | - | 1213 | 2791 | - | - | 0 | - |

m/z Charge Intensity FragmentType MassShift Position
128.14625549316406 0 442.7388
129.05523681640625 0 349.03812
129.10232543945312 0 393.2813
136.07557678222656 0 1180.599
155.5974578857422 0 447.43616
217.08160400390625 0 2750.839
221.08433532714844 0 1616.2615
229.11793518066406 0 2142.351 y Water loss 16
232.11865234375 0 743.56256
239.09474182128906 0 2215.205
239.11056518554688 0 484.64236
246.12384033203125 0 3395.984
247.12796020507812 0 1516.3779 y 16
274.1183166503906 0 4714.769
275.122802734375 0 815.1379
295.10284423828125 0 1890.1278
300.1186828613281 0 4185.1997
334.1608581542969 0 2104.22 y 15
335.16168212890625 0 599.1071
360.17852783203125 0 651.90875
369.1217956542969 0 1965.0573
397.242431640625 0 645.5163
398.171630859375 0 1156.8427
437.18121337890625 0 1331.7845
447.2449645996094 0 1324.7798 y 14
463.1823425292969 0 2113.6812
465.207763671875 0 953.7275
471.2802734375 0 640.96814
477.2019958496094 0 790.8331
480.22210693359375 0 1008.3995
483.21978759765625 0 570.79315
528.2437744140625 0 895.3397
538.2530517578125 0 1060.3479
539.2593994140625 0 1612.692
541.3123168945312 0 611.17236
560.3280029296875 0 1326.4161 y 13
561.2688598632812 0 1685.2954 c Ammonia loss 8
563.2485961914062 0 2401.462
564.2491455078125 0 889.01105 c Water loss 3
565.239990234375 0 913.67944 c Ammonia loss 3
567.8462524414062 0 706.4975
576.9967041015625 0 664.83405
580.2612915039062 0 987.2467
581.258544921875 0 2164.5078
582.2665405273438 0 12420.761 c 3
583.270263671875 0 3950.4104
586.3258666992188 0 2223.3733 c Ammonia loss 14
586.6607055664062 0 3108.46
586.995849609375 0 720.7471
606.3445434570312 0 840.0331 y 2
606.6712646484375 0 619.3245
614.3413696289062 0 856.6079 w 12
615.3367919921875 0 2563.912 c Ammonia loss 15
615.6704711914062 0 2737.351
616.0062866210938 0 929.1837
616.3387451171875 0 753.22595
620.9900512695312 0 764.8442
626.38671875 0 3470.041 y 6
626.8869018554688 0 2972.266
627.3917236328125 0 1668.2186
646.33837890625 0 703.7529
649.3526611328125 0 1053.2057
652.34619140625 0 1626.4562
652.6824340820312 0 1730.9727
653.0144653320312 0 1551.3398
657.2978515625 0 830.8204
657.394775390625 0 829.2593 z 12
658.2919921875 0 934.9009
658.3510131835938 0 11169.417 c Ammonia loss 16
658.406005859375 0 966.8139
658.68505859375 0 13381.483
659.01953125 0 5896.182
659.2957153320312 0 1125.3145
659.3536376953125 0 2519.425
659.6889038085938 0 2579.0876
664.35546875 0 1235.3738
668.3665771484375 0 937.90155 y 1
669.0347900390625 0 710.49536
673.4139404296875 0 1278.9269 y 12
674.30908203125 0 1545.3799
675.3118896484375 0 3194.9878
676.3182983398438 0 2212.4583
691.3728637695312 0 4093.203
691.7073364257812 0 5203.771
692.0411987304688 0 3494.5767
692.3806762695312 0 954.90063
694.9180908203125 0 952.5168 y 5
696.7046508789062 0 1118.5615
697.039794921875 0 889.8273
697.376953125 0 68952.07
697.71142578125 0 84021.43
698.0453491210938 0 55654.86
698.1181640625 0 1201.0298
698.3798217773438 0 27913.596
698.7136840820312 0 8776.436
699.0468139648438 0 4381.6055
702.3020629882812 0 1591.1051 c Ammonia loss 4
709.3598022460938 0 844.70483 c Ammonia loss 11
717.369384765625 0 3848.6953
717.872314453125 0 42508.906 c 11
718.3739624023438 0 31473.258
718.8748168945312 0 13659.093
719.3250732421875 0 13038.911 c 4
719.3795776367188 0 3268.5466
719.8768310546875 0 1035.271
720.328369140625 0 6223.9707
721.3284912109375 0 1965.8077
743.931884765625 0 717.7877
752.4075317382812 0 1639.9565
752.9109497070312 0 1068.2625
753.406494140625 0 880.953
755.4361572265625 0 3941.171 z 4
755.9376220703125 0 7314.429
756.4400024414062 0 4524.877
756.9427490234375 0 2538.853
757.4461059570312 0 869.1705
762.9420776367188 0 905.5451
763.4447631835938 0 4769.974 y 4
763.9462890625 0 3641.6929
764.4503173828125 0 1576.5131
764.9464111328125 0 1087.4454
773.910400390625 0 2890.5503
774.4141845703125 0 25448.143 c 12
774.9158325195312 0 21889.123
775.4169311523438 0 9347.064
775.9189453125 0 3362.0986
776.4221801757812 0 723.9557
783.94287109375 0 703.7182
785.4874267578125 0 1636.8478 z 11
786.498291015625 0 1198.3154
787.4999389648438 0 841.79376
790.450439453125 0 3664.709 w 3
790.9528198242188 0 2794.2812
791.450439453125 0 2032.8217
791.9463500976562 0 1585.1912
796.3551025390625 0 1214.7533
801.5037841796875 0 836.83826 y 11
802.5119018554688 0 741.49677
808.9407958984375 0 848.85754
809.4517211914062 0 999.8172
809.9518432617188 0 997.7052
811.3623657226562 0 4125.381
812.3692626953125 0 1816.0491
818.46435546875 0 1190.2843 y Water loss 3
819.465087890625 0 13788.812 z 3
819.9678955078125 0 28242.297
820.4701538085938 0 14939
820.9710083007812 0 7363.1855
821.4750366210938 0 3279.9453
822.44189453125 0 1652.7168 c Ammonia loss 13
822.946044921875 0 2045.902
823.4459228515625 0 700.5064
827.4737548828125 0 5074.7134 y 3
827.9744262695312 0 3489.357
828.4760131835938 0 2195.9795
830.95556640625 0 8514.217 c 13
831.4584350585938 0 6105.031
831.9607543945312 0 3719.334
832.4594116210938 0 1251.7655
838.9522094726562 0 709.62994
839.3565673828125 0 3205.6152 c Ammonia loss 5
840.3590698242188 0 1406.3524
842.4653930664062 0 734.6092
850.4713745117188 0 1098.277
850.9677124023438 0 1775.5575
851.4690551757812 0 1031.9891
851.9666748046875 0 716.75415
858.4965209960938 0 659.36176
865.4837036132812 0 2013.8643
865.9867553710938 0 1475.3922
866.489013671875 0 987.5926
866.9921264648438 0 847.31165
869.4742431640625 0 943.91876
871.9786987304688 0 1204.2932
872.47412109375 0 1042.4296
872.9884033203125 0 1171.2921
873.4661865234375 0 928.19055
877.9808349609375 0 1140.1674
878.4905395507812 0 2164.7927 c Water loss 14
878.9867553710938 0 5341.102 c Ammonia loss 14
879.486572265625 0 7468.999
879.9871826171875 0 5427.12
880.4844970703125 0 1091.414
880.9808959960938 0 838.88837
886.993896484375 0 9763.31
887.497314453125 0 21498.547 c 14
887.9989624023438 0 18262.193
888.5005493164062 0 9055.832
889.003662109375 0 2871.77
889.501953125 0 1066.1104
890.0059814453125 0 626.5177
897.5173950195312 0 1779.396
898.5571899414062 0 3701.6785 y 10
899.565673828125 0 1007.6926
900.0001831054688 0 2150.041 y Water loss 2
900.5023803710938 0 2284.166 y Ammonia loss 2
900.9969482421875 0 11233.843 z 2
901.4987182617188 0 14330.602
902.0007934570312 0 10185.825
902.5010375976562 0 3080.1794
903.0042724609375 0 1222.722
908.5025634765625 0 754.2544
909.0059814453125 0 11700.891 y 2
909.5073852539062 0 11639.902
910.0089721679688 0 7413.047
910.5126342773438 0 3182.2268
911.5255126953125 0 686.4904
914.5044555664062 0 710.2777
922.5005493164062 0 3279.915 c Ammonia loss 15
923.0022583007812 0 2430.005
923.5025634765625 0 2947.808
924.008056640625 0 1498.6934
928.481689453125 0 798.5043
930.4960327148438 0 3800.3093
930.827392578125 0 1907.144
931.0136108398438 0 23280.965 c 15
931.5145874023438 0 32213.771
932.0167846679688 0 16249.574
932.5182495117188 0 6092.6025
933.0192260742188 0 2378.8787
944.0192260742188 0 680.89545
944.5225219726562 0 1996.6633
945.0248413085938 0 670.1351
945.4822998046875 0 592.85345
952.4286499023438 0 1215.0126
953.4373779296875 0 8551.662 c 6
953.578369140625 0 18225.26 z 9
954.4408569335938 0 5603.456
954.5809326171875 0 8008.1533
955.4464721679688 0 1579.3517
955.583740234375 0 2399.3997
966.4425659179688 0 1017.0471
967.454345703125 0 1924.7162
968.45263671875 0 874.0555
969.5980834960938 0 2715.3435 y 9
970.601318359375 0 1913.3558
973.0289916992188 0 1186.9258
973.5293579101562 0 5370.1914
974.0272216796875 0 3920.316
974.530517578125 0 2593.7815
975.03173828125 0 1827.1488
979.0391845703125 0 995.09485
979.5184326171875 0 1569.6204
980.0386352539062 0 716.35626
980.5272216796875 0 2020.8132
981.0280151367188 0 1506.1033
987.0243530273438 0 5825.183 c Ammonia loss 16
987.5244750976562 0 4903.583
988.0254516601562 0 3056.3647
988.5249633789062 0 2046.708
989.0330200195312 0 817.9344
993.5357055664062 0 1143.5653 y Ammonia loss 1
994.0375366210938 0 4030.0793 z 1
994.5381469726562 0 3833.1138
995.042236328125 0 3183.9602
995.5361938476562 0 26467.393 c 16
996.036865234375 0 31180.77
996.5386962890625 0 16290.932
997.03857421875 0 7903.549
997.5407104492188 0 2921.6887
998.0341796875 0 1052.4852
1002.03515625 0 2118.5117 y 1
1002.5402221679688 0 1422.7434
1003.0368041992188 0 962.0766
1005.0418701171875 0 2299.1777
1005.5390014648438 0 1797.9379
1006.041015625 0 2152.9023
1006.555419921875 0 1719.4805
1007.0543823242188 0 1426.8065
1008.0488891601562 0 1131.582
1008.5484008789062 0 2421.0527
1009.0436401367188 0 2389.3918
1009.43603515625 0 1671.1815
1009.5350952148438 0 4710.7197
1010.0341796875 0 4171.0073
1010.4576416015625 0 13134.845 c 7
1011.03564453125 0 2126.1978
1011.4608154296875 0 7914.6797
1012.4638061523438 0 2309.7085
1014.5506591796875 0 2760.9646
1015.0536499023438 0 4285.762
1015.5573120117188 0 5416.909
1016.0535888671875 0 4952.3433
1016.5506591796875 0 3057.256
1017.05029296875 0 1922.6202
1018.0411376953125 0 1037.2931
1018.5335083007812 0 1599.589
1019.0379638671875 0 802.39575
1023.0570678710938 0 1837.1619
1023.5557861328125 0 7021.9014
1024.0592041015625 0 8435.709
1024.5479736328125 0 9488.26
1025.047607421875 0 7915.6226
1025.5443115234375 0 5047.8975
1026.0389404296875 0 2667.5847
1026.5418701171875 0 801.2498
1028.548583984375 0 1247.1094
1029.555419921875 0 801.00256
1031.0491943359375 0 909.18304
1031.55029296875 0 1081.7345
1036.5579833984375 0 1775.7755
1037.0567626953125 0 4318.848
1037.553955078125 0 35540.656
1038.055419921875 0 35836.918
1038.55615234375 0 22496.635
1038.68212890625 0 871.2854
1039.056396484375 0 12449.999
1039.5572509765625 0 3675.7793
1040.0606689453125 0 949.908
1044.5477294921875 0 896.5869
1045.05615234375 0 3136.451
1045.5616455078125 0 50022.71
1046.0653076171875 0 173517.31
1046.5670166015625 0 188728.83
1047.06787109375 0 107908.64
1047.5692138671875 0 51866.67
1048.0701904296875 0 18982.791
1048.564697265625 0 4493.1646
1081.673828125 0 10381.82 z 8
1082.677490234375 0 11137.428
1083.680419921875 0 5278.327
1084.682861328125 0 1888.6501
1094.5411376953125 0 2422.2031
1095.5433349609375 0 1751.7366
1097.68896484375 0 2425.524 y 8
1098.694580078125 0 1756.294
1099.6904296875 0 1122.973
1138.55322265625 0 20720.871 c 8
1138.69580078125 0 7394.595 z 7
1139.5552978515625 0 13524.011
1139.70068359375 0 7215.6875
1140.5579833984375 0 5314.1094
1140.7037353515625 0 3575.3892
1141.5589599609375 0 1579.8125
1141.7027587890625 0 1191.355
1154.722900390625 0 1006.8451 y 7
1165.5765380859375 0 2130.4104
1166.5831298828125 0 1099.0052
1217.599853515625 0 1090.6013
1218.6065673828125 0 1016.625
1251.7607421875 0 1681.6364 y 6
1252.7764892578125 0 761.5802
1278.7501220703125 0 2166.609
1279.751953125 0 1031.7279
1280.7576904296875 0 1419.3248
1306.6455078125 0 2957.801 c 10
1307.6451416015625 0 2058.108
1336.615234375 0 760.6778
1372.80419921875 0 10244.367 z 5
1373.8101806640625 0 12422.301
1374.8135986328125 0 5826.9336
1375.826416015625 0 1035.2573
1378.6727294921875 0 905.19855
1385.7239990234375 0 1995.2009
1386.214111328125 0 1215.3656
1387.2193603515625 0 1674.7114
1387.69580078125 0 864.5513
1388.8204345703125 0 6244.485 y 5
1389.8277587890625 0 4235.2974
1390.84765625 0 1184.6654
1392.7032470703125 0 1910.5352
1393.6932373046875 0 2299.4849
1394.6875 0 3118.4165
1395.21435546875 0 1891.2666
1395.6953125 0 4218.29
1396.23291015625 0 4343.162
1396.7047119140625 0 5097.4883
1397.2381591796875 0 2196.0632
1397.692626953125 0 1661.9081
1433.7230224609375 0 928.48535
1434.7353515625 0 8584.835 c 11
1435.73779296875 0 7483.7437
1436.7349853515625 0 3156.1343
1504.80712890625 0 1499.7495
1509.864013671875 0 7301.064 z 4
1510.8687744140625 0 21294.51
1511.8729248046875 0 14911.028
1512.874755859375 0 6986.3315
1513.881103515625 0 2142.8418
1524.86083984375 0 948.1321
1525.8812255859375 0 5926.4253 y 4
1526.885498046875 0 4186.563
1527.8916015625 0 1854.1912
1547.8203125 0 3040.86 c 12
1548.8250732421875 0 3126.4185
1549.8162841796875 0 2316.3914
1637.921630859375 0 2302.642 z 3
1638.9300537109375 0 15632.94
1639.93408203125 0 16291.153
1640.936279296875 0 6843.213
1641.9385986328125 0 2803.7812
1642.9375 0 1057.6749
1653.94140625 0 785.2063 y 3
1654.9471435546875 0 1414.9683
1660.90380859375 0 2032.5173 c 13
1661.9083251953125 0 2121.4792
1662.9156494140625 0 1385.4585
1758.988525390625 0 919.83093
1772.9847412109375 0 1770.4431
1773.98876953125 0 8257.229 c 14
1774.9884033203125 0 7896.5566
1775.9923095703125 0 4782.0107
1777.0010986328125 0 2360.3015
1800.989013671875 0 3167.8013 z 2
1801.9925537109375 0 10392.847
1802.992919921875 0 9311.701
1804.0029296875 0 3804.457
1805.0013427734375 0 1872.9828
1817.0057373046875 0 1553.3573 y 2
1818.0029296875 0 2364.9907
1819.007568359375 0 1684.8618
1820.0208740234375 0 1099.2358
1861.0216064453125 0 9347.224 c 15
1862.0205078125 0 11141.604
1863.0262451171875 0 6723.913
1864.025390625 0 2916.6504
1865.03662109375 0 1513.1249
1946.04736328125 0 1252.0447
1947.0579833984375 0 3839.3665
1948.058837890625 0 3580.4304
1949.0711669921875 0 1689.1492
1976.040283203125 0 753.13464
1988.076171875 0 2569.0886
1989.080322265625 0 3275.6418
1990.0697021484375 0 4225.691 c 16
1991.0643310546875 0 7646.402
1992.0692138671875 0 6774.645
1993.0732421875 0 3355.0298
1994.0667724609375 0 1128.736
2011.0655517578125 0 955.58234
2029.112060546875 0 904.3709
2030.1112060546875 0 1539.1418
2046.1123046875 0 2402.0103
2047.1083984375 0 6862.0327
2048.111083984375 0 6370.4126
2049.1171875 0 4237.6973
2050.1171875 0 2561.8047
2057.083984375 0 1883.7068
2058.090576171875 0 1908.151
2059.095458984375 0 1861.943
2060.108642578125 0 1136.7332
2063.125732421875 0 2283.53
2064.131591796875 0 8642.531
2065.134521484375 0 9929.797
2066.1376953125 0 5410.8027
2067.137939453125 0 3302.629
2074.10009765625 0 5269.801
2075.10205078125 0 24970.012
2076.105224609375 0 27469.895
2077.106689453125 0 17983.096
2078.10888671875 0 7082.6616
2079.106689453125 0 2537.6582
2090.11474609375 0 6398.2617
2091.121337890625 0 30860.861
2092.126953125 0 115574.45
2093.131591796875 0 128093.91
2094.13330078125 0 78428.34
2095.137451171875 0 38733.543
2096.136962890625 0 12595.154
2097.1337890625 0 2973.0237
2295.223388671875 0 895.5543
2296.220703125 0 1074.3301
2790.449951171875 0 908.9613
2791.4453125 0 1213.4009

Spectrum Details

|  |  |
| --- | --- |
| Matched peaks? Matched peaks The total absolute number of peaks matched. Additionally in brackets the total fraction of peaks matched and the total number of peaks is shown. | 75 (16.16% of 464) |
| FDR? FDR The false discovery rate estimated for this peptide. It is calculated by matching all theoretical fragments with a non-integer shift with the raw peaks for this spectrum. This is done with 40 different shifts. The resulting percentage is the average number of annotated peaks over the number of annotated peaks with the correct spectrum. | 3.05% |
| Satellite FDR? Satellite FDR See the FDR for details on its calculation. This satellite ion specific FDR only contains the satellite ions (d/w) for I/L/J positions. | 0.00% |
| PSM Score? PSM Score The PSM Score as given by Hecklib to this annotated spectrum. It is shown with three significant figures. | 477 |

#### Spectrum 7373? Spectrum 7373 The raw spectrum of this peptide as annotated by Hecklib. The fragments are coloured according to ion type (see legend). Any peaks with a star '\*' as text can be hovered over to see the full details, first the ion type second the mass shift type. By hovering over the amino acids in the peptide or ions in the legend the corresponding peaks are highlighted. By toggling the 'Unassigned' label you can turn the background (unassigned) peaks on or off in the plot. By updating the slider in the Ion legend you can update the spectrum to only show the top X% of the peaks with labels. The top X% means any peak that is within X% of the highest intensity. By dragging in the spectrum you can zoom in to a specific part of the spectrum and use 'Zoom Out' to get back to the original zoom level. The annotation of the spectrum is based on the given sequence in the peptides file and is done with different software so inconsistencies are likely. The peaks are annotated based on the given sequence, with 20 ppm tolerance.

Copy Data

##### Spectrum 7373 (TSV)

###### Preview

```
Loading example...
```

*Click on the button to copy the data to your clipboard.*

Mz MinMz MaxIntensity Max

WidthHeightPeptide font sizePeptide stroke widthSpectrum font sizeSpectrum stroke widthCompact peptide

Ion legend

wxyz

abcd

OtherUnassignedIon^Charge^_Position_Show for top:%

SWYQHHPGKAPKJJJSEV

09.06e+51.81e+62.72e+63.62e+6

Zoom Out

b^+3^_4_a^+1^_2_b^+3^_5_y^+1^_2_a^+1^_2_y^+1^_2_b^+1^_2_b^+1^_2_y^+1^_3_b^+3^_8_y^+2^_6_y^+1^_3_y^+2^_6_a^+1^_3_b^+2^_6_b^+1^_3_b^+2^_6_y^+1^_4_b^+1^_3_y^+1^_4_y^+2^_8_b^+2^_8_b^+3^_13_b^+3^_13_b^+3^_13_y^+2^_10_y^+1^_5_b^+3^_14_b^+3^_14_b^+1^_4_b^+3^_14_y^+2^_10_y^+1^_5_b^+2^_9_b^+1^_4_y^+2^_11_b^+3^_15_b^+3^_15_b^+2^_10_b^+2^_10_b^+2^_10_y^+3^_16_y^+3^_16_y^+3^_16_b^+3^_16_b^+3^_16_b^+3^_16_y^+2^_12_y^+2^_12_y^+2^_12_b^+2^_11_b^+3^_17_b^+3^_17_b^+3^_17_y^+3^_17_y^+3^_17_y^+3^_17_y^+1^_6_b^+1^_5_b^+1^_5_y^+2^_13_\*\*y^+2^_13_\*b^+1^_5_b^+2^_12_y^+2^_14_b^+2^_13_b^+2^_13_y^+2^_14_b^+2^_13_y^+1^_7_y^+1^_7_b^+2^_14_b^+2^_14_y^+2^_15_y^+2^_15_b^+1^_6_b^+1^_6_b^+2^_14_y^+2^_15_b^+1^_6_b^+2^_15_b^+2^_15_b^+2^_15_y^+1^_8_y^+2^_16_y^+2^_16_y^+2^_16_b^+2^_16_b^+2^_16_b^+2^_16_b^+1^_7_y^+1^_9_b^+2^_17_b^+2^_17_b^+2^_17_b^+1^_8_y^+1^_10_y^+1^_10_b^+1^_9_y^+1^_11_y^+1^_11_y^+1^_11_b^+1^_10_b^+1^_10_b^+1^_10_y^+1^_12_y^+1^_12_y^+1^_12_y^+1^_13_y^+1^_13_b^+1^_12_

0548109616442192

Fragment Matches Table

Show background peaks

| Position | Ion type | Intensity | mz Theoretical | mz Error (Th) | mz Error (ppm) | Charge | Series Number |
| --- | --- | --- | --- | --- | --- | --- | --- |
| - | - | 2375 | 120.1 | - | - | 0 | - |
| - | - | 3831 | 120.1 | - | - | 0 | - |
| - | - | 2000 | 120.1 | - | - | 0 | - |
| - | - | 1928 | 120.2 | - | - | 0 | - |
| - | - | 2146 | 120.3 | - | - | 0 | - |
| - | - | 6203 | 122.1 | - | - | 0 | - |
| - | - | 8368 | 123.1 | - | - | 0 | - |
| - | - | 4.768E+04 | 126.1 | - | - | 0 | - |
| - | - | 4120 | 127.1 | - | - | 0 | - |
| - | - | 1.054E+04 | 127.1 | - | - | 0 | - |
| - | - | 2211 | 128 | - | - | 0 | - |
| - | - | 2563 | 128.1 | - | - | 0 | - |
| - | - | 2.025E+04 | 129.1 | - | - | 0 | - |
| - | - | 2.738E+05 | 129.1 | - | - | 0 | - |
| - | - | 4785 | 130.1 | - | - | 0 | - |
| - | - | 1.218E+05 | 130.1 | - | - | 0 | - |
| - | - | 3553 | 130.1 | - | - | 0 | - |
| - | - | 1.881E+04 | 130.1 | - | - | 0 | - |
| - | - | 1933 | 130.1 | - | - | 0 | - |
| - | - | 1.269E+04 | 131.1 | - | - | 0 | - |
| - | - | 3693 | 131.1 | - | - | 0 | - |
| - | - | 4.419E+05 | 132.1 | - | - | 0 | - |
| - | - | 4.726E+04 | 133.1 | - | - | 0 | - |
| - | - | 3916 | 134.1 | - | - | 0 | - |
| - | - | 2.821E+05 | 136.1 | - | - | 0 | - |
| - | - | 2.343E+04 | 137.1 | - | - | 0 | - |
| - | - | 1.375E+04 | 138.1 | - | - | 0 | - |
| - | - | 3675 | 138.1 | - | - | 0 | - |
| - | - | 2539 | 139.1 | - | - | 0 | - |
| - | - | 2.387E+04 | 141.1 | - | - | 0 | - |
| - | - | 2141 | 143.1 | - | - | 0 | - |
| - | - | 1.254E+04 | 144.1 | - | - | 0 | - |
| - | - | 5952 | 144.1 | - | - | 0 | - |
| - | - | 2484 | 146.1 | - | - | 0 | - |
| - | - | 3898 | 146.1 | - | - | 0 | - |
| - | - | 3254 | 146.1 | - | - | 0 | - |
| - | - | 4544 | 147 | - | - | 0 | - |
| - | - | 3779 | 148.1 | - | - | 0 | - |
| - | - | 2209 | 148.9 | - | - | 0 | - |
| - | - | 6434 | 151.1 | - | - | 0 | - |
| - | - | 2.364E+04 | 152.1 | - | - | 0 | - |
| - | - | 2.916E+04 | 152.1 | - | - | 0 | - |
| - | - | 9152 | 153.1 | - | - | 0 | - |
| - | - | 2460 | 153.1 | - | - | 0 | - |
| - | - | 1.47E+04 | 153.1 | - | - | 0 | - |
| - | - | 2595 | 153.1 | - | - | 0 | - |
| - | - | 2243 | 154 | - | - | 0 | - |
| - | - | 5.797E+04 | 154.1 | - | - | 0 | - |
| - | - | 1.159E+04 | 154.2 | - | - | 0 | - |
| - | - | 4806 | 155.1 | - | - | 0 | - |
| - | - | 2.062E+04 | 155.1 | - | - | 0 | - |
| - | - | 1.54E+04 | 155.1 | - | - | 0 | - |
| - | - | 1981 | 155.1 | - | - | 0 | - |
| - | - | 2.694E+04 | 155.1 | - | - | 0 | - |
| - | - | 1.885E+04 | 156.1 | - | - | 0 | - |
| - | - | 2380 | 156.1 | - | - | 0 | - |
| - | - | 3090 | 156.1 | - | - | 0 | - |
| - | - | 3309 | 157.1 | - | - | 0 | - |
| - | - | 3448 | 157.1 | - | - | 0 | - |
| - | - | 5932 | 158.1 | - | - | 0 | - |
| - | - | 3.578E+04 | 158.1 | - | - | 0 | - |
| - | - | 1.827E+05 | 159.1 | - | - | 0 | - |
| - | - | 6285 | 159.1 | - | - | 0 | - |
| - | - | 9134 | 160.1 | - | - | 0 | - |
| - | - | 1978 | 160.1 | - | - | 0 | - |
| - | - | 2.275E+04 | 160.1 | - | - | 0 | - |
| - | - | 3231 | 165.1 | - | - | 0 | - |
| - | - | 1.091E+04 | 165.1 | - | - | 0 | - |
| - | - | 6959 | 166.1 | - | - | 0 | - |
| - | - | 2549 | 166.1 | - | - | 0 | - |
| - | - | 1.02E+04 | 168.1 | - | - | 0 | - |
| - | - | 6281 | 168.1 | - | - | 0 | - |
| - | - | 2.836E+04 | 169.1 | - | - | 0 | - |
| - | - | 3.554E+04 | 169.1 | - | - | 0 | - |
| - | - | 4745 | 170 | - | - | 0 | - |
| - | - | 6.102E+04 | 170.1 | - | - | 0 | - |
| - | - | 5416 | 171.1 | - | - | 0 | - |
| - | - | 6.478E+04 | 171.1 | - | - | 0 | - |
| - | - | 2987 | 171.1 | - | - | 0 | - |
| - | - | 2.366E+04 | 171.1 | - | - | 0 | - |
| - | - | 6368 | 172.1 | - | - | 0 | - |
| - | - | 2861 | 172.1 | - | - | 0 | - |
| - | - | 3.222E+04 | 172.1 | - | - | 0 | - |
| - | - | 2.722E+04 | 173.1 | - | - | 0 | - |
| - | - | 1.267E+04 | 173.5 | - | - | 0 | - |
| - | - | 4213 | 174.1 | - | - | 0 | - |
| - | - | 5085 | 175.1 | - | - | 0 | - |
| - | - | 8834 | 176.1 | - | - | 0 | - |
| - | - | 3415 | 178.1 | - | - | 0 | - |
| - | - | 3085 | 179.1 | - | - | 0 | - |
| - | - | 2654 | 179.2 | - | - | 0 | - |
| - | - | 8292 | 180.1 | - | - | 0 | - |
| - | - | 7849 | 181.1 | - | - | 0 | - |
| - | - | 8.693E+04 | 181.1 | - | - | 0 | - |
| - | - | 4779 | 181.1 | - | - | 0 | - |
| - | - | 3393 | 181.2 | - | - | 0 | - |
| - | - | 9339 | 182.1 | - | - | 0 | - |
| - | - | 1.522E+04 | 182.1 | - | - | 0 | - |
| - | - | 2766 | 182.1 | - | - | 0 | - |
| - | - | 6.359E+04 | 183.1 | - | - | 0 | - |
| - | - | 2.436E+05 | 183.1 | - | - | 0 | - |
| - | - | 7215 | 184.1 | - | - | 0 | - |
| - | - | 9441 | 184.1 | - | - | 0 | - |
| - | - | 2.913E+04 | 184.1 | - | - | 0 | - |
| - | - | 1.731E+04 | 185.1 | - | - | 0 | - |
| - | - | 3272 | 185.1 | - | - | 0 | - |
| - | - | 3224 | 185.1 | - | - | 0 | - |
| - | - | 2958 | 186.1 | - | - | 0 | - |
| - | - | 5.75E+04 | 186.1 | - | - | 0 | - |
| - | - | 4.63E+05 | 187.1 | - | - | 0 | - |
| - | - | 5426 | 187.1 | - | - | 0 | - |
| - | - | 6.092E+04 | 188.1 | - | - | 0 | - |
| 4 | b | 3.277E+05 | 189.1 | 0.00259 | 13.7 | +3 | 4 |
| - | - | 2.845E+04 | 190.1 | - | - | 0 | - |
| - | - | 4499 | 191.1 | - | - | 0 | - |
| - | - | 1.363E+04 | 195.1 | - | - | 0 | - |
| - | - | 6585 | 195.1 | - | - | 0 | - |
| - | - | 4.378E+04 | 197.2 | - | - | 0 | - |
| - | - | 5465 | 198.2 | - | - | 0 | - |
| - | - | 3.543E+04 | 199.1 | - | - | 0 | - |
| - | - | 1.037E+04 | 199.1 | - | - | 0 | - |
| - | - | 6660 | 199.1 | - | - | 0 | - |
| - | - | 1.691E+05 | 199.2 | - | - | 0 | - |
| - | - | 3143 | 200.1 | - | - | 0 | - |
| - | - | 1.066E+04 | 200.1 | - | - | 0 | - |
| - | - | 2.113E+04 | 200.2 | - | - | 0 | - |
| - | - | 3.69E+04 | 201.1 | - | - | 0 | - |
| - | - | 4.351E+04 | 201.1 | - | - | 0 | - |
| - | - | 5137 | 202.1 | - | - | 0 | - |
| - | - | 5.536E+04 | 202.1 | - | - | 0 | - |
| - | - | 3400 | 202.1 | - | - | 0 | - |
| - | - | 6811 | 203.1 | - | - | 0 | - |
| - | - | 3073 | 204.1 | - | - | 0 | - |
| - | - | 5641 | 205.1 | - | - | 0 | - |
| - | - | 2911 | 207.1 | - | - | 0 | - |
| - | - | 1.781E+04 | 207.1 | - | - | 0 | - |
| - | - | 3912 | 208.1 | - | - | 0 | - |
| - | - | 3091 | 209.1 | - | - | 0 | - |
| - | - | 5604 | 209.1 | - | - | 0 | - |
| - | - | 5938 | 209.2 | - | - | 0 | - |
| - | - | 9658 | 211.1 | - | - | 0 | - |
| - | - | 2.731E+04 | 211.1 | - | - | 0 | - |
| - | - | 3748 | 211.1 | - | - | 0 | - |
| - | - | 3173 | 212.1 | - | - | 0 | - |
| - | - | 4800 | 213.1 | - | - | 0 | - |
| - | - | 2.394E+04 | 214.1 | - | - | 0 | - |
| - | - | 3333 | 214.1 | - | - | 0 | - |
| - | - | 2967 | 214.2 | - | - | 0 | - |
| - | - | 3666 | 216.1 | - | - | 0 | - |
| - | - | 5948 | 216.1 | - | - | 0 | - |
| - | - | 5.42E+05 | 217.1 | - | - | 0 | - |
| - | - | 1.084E+04 | 217.1 | - | - | 0 | - |
| - | - | 4.956E+04 | 218.1 | - | - | 0 | - |
| - | - | 3043 | 218.2 | - | - | 0 | - |
| - | - | 1.235E+04 | 219.1 | - | - | 0 | - |
| - | - | 3.727E+04 | 220.1 | - | - | 0 | - |
| - | - | 6.199E+04 | 221.1 | - | - | 0 | - |
| - | - | 4460 | 221.1 | - | - | 0 | - |
| - | - | 2852 | 221.2 | - | - | 0 | - |
| - | - | 4922 | 222.1 | - | - | 0 | - |
| - | - | 3280 | 222.1 | - | - | 0 | - |
| - | - | 3696 | 222.1 | - | - | 0 | - |
| - | - | 4343 | 223.1 | - | - | 0 | - |
| - | - | 4524 | 223.1 | - | - | 0 | - |
| - | - | 2460 | 223.2 | - | - | 0 | - |
| - | - | 1.344E+04 | 224.1 | - | - | 0 | - |
| - | - | 2.67E+04 | 224.2 | - | - | 0 | - |
| - | - | 8651 | 225.1 | - | - | 0 | - |
| - | - | 3840 | 225.1 | - | - | 0 | - |
| - | - | 5769 | 225.2 | - | - | 0 | - |
| - | - | 3.29E+05 | 226.2 | - | - | 0 | - |
| - | - | 2753 | 227 | - | - | 0 | - |
| - | - | 3010 | 227.1 | - | - | 0 | - |
| - | - | 2731 | 227.1 | - | - | 0 | - |
| - | - | 3.182E+04 | 227.2 | - | - | 0 | - |
| - | - | 1.084E+05 | 227.2 | - | - | 0 | - |
| 2 | a | 3.68E+04 | 228.1 | 0.0007073 | 3.101 | +1 | 2 |
| - | - | 3944 | 228.1 | - | - | 0 | - |
| - | - | 1.301E+04 | 228.2 | - | - | 0 | - |
| 5 | b | 4.779E+04 | 229.1 | 0.002087 | 9.111 | +3 | 5 |
| 17 | y | 4.927E+05 | 229.1 | 0.0006893 | 3.009 | +1 | 2 |
| - | - | 1.177E+04 | 230.1 | - | - | 0 | - |
| - | - | 5.641E+04 | 230.1 | - | - | 0 | - |
| - | - | 2.038E+04 | 231.1 | - | - | 0 | - |
| - | - | 3190 | 231.1 | - | - | 0 | - |
| - | - | 7830 | 231.1 | - | - | 0 | - |
| - | - | 2719 | 232.1 | - | - | 0 | - |
| - | - | 4628 | 232.1 | - | - | 0 | - |
| - | - | 9275 | 233.1 | - | - | 0 | - |
| - | - | 2605 | 233.2 | - | - | 0 | - |
| - | - | 5705 | 234.1 | - | - | 0 | - |
| - | - | 7098 | 235.1 | - | - | 0 | - |
| - | - | 1.18E+05 | 235.1 | - | - | 0 | - |
| - | - | 9986 | 236.1 | - | - | 0 | - |
| - | - | 4021 | 236.1 | - | - | 0 | - |
| - | - | 3202 | 237.1 | - | - | 0 | - |
| - | - | 1.065E+04 | 237.2 | - | - | 0 | - |
| - | - | 1.489E+04 | 238.1 | - | - | 0 | - |
| - | - | 2.897E+04 | 239.2 | - | - | 0 | - |
| - | - | 5184 | 239.2 | - | - | 0 | - |
| - | - | 4213 | 240.1 | - | - | 0 | - |
| - | - | 6375 | 241.2 | - | - | 0 | - |
| - | - | 3505 | 241.2 | - | - | 0 | - |
| - | - | 1.056E+04 | 242.2 | - | - | 0 | - |
| - | - | 1.115E+04 | 243.2 | - | - | 0 | - |
| - | - | 6594 | 244.1 | - | - | 0 | - |
| - | - | 2770 | 244.2 | - | - | 0 | - |
| - | - | 2667 | 245.2 | - | - | 0 | - |
| 2 | a | 1.249E+06 | 246.1 | 0.0008238 | 3.347 | +1 | 2 |
| 17 | y | 4.009E+05 | 247.1 | 0.0005159 | 2.087 | +1 | 2 |
| - | - | 7.216E+04 | 248.1 | - | - | 0 | - |
| - | - | 3.661E+04 | 248.1 | - | - | 0 | - |
| - | - | 8.365E+04 | 249.1 | - | - | 0 | - |
| - | - | 5474 | 249.1 | - | - | 0 | - |
| - | - | 6147 | 249.1 | - | - | 0 | - |
| - | - | 7540 | 249.1 | - | - | 0 | - |
| - | - | 1.003E+04 | 250.1 | - | - | 0 | - |
| - | - | 4365 | 250.1 | - | - | 0 | - |
| - | - | 3210 | 251.1 | - | - | 0 | - |
| - | - | 7314 | 251.1 | - | - | 0 | - |
| - | - | 2981 | 251.2 | - | - | 0 | - |
| - | - | 2718 | 251.2 | - | - | 0 | - |
| - | - | 2.549E+04 | 252.2 | - | - | 0 | - |
| - | - | 5640 | 254.2 | - | - | 0 | - |
| - | - | 5528 | 255.1 | - | - | 0 | - |
| - | - | 2850 | 255.2 | - | - | 0 | - |
| - | - | 2456 | 256.1 | - | - | 0 | - |
| 2 | b | 2.687E+04 | 256.1 | 0.0007726 | 3.017 | +1 | 2 |
| - | - | 5715 | 257.1 | - | - | 0 | - |
| - | - | 8698 | 257.1 | - | - | 0 | - |
| - | - | 3474 | 257.1 | - | - | 0 | - |
| - | - | 8550 | 257.2 | - | - | 0 | - |
| - | - | 1.656E+05 | 258.1 | - | - | 0 | - |
| - | - | 2.267E+04 | 259.1 | - | - | 0 | - |
| - | - | 8355 | 259.1 | - | - | 0 | - |
| - | - | 4345 | 259.2 | - | - | 0 | - |
| - | - | 6529 | 260.1 | - | - | 0 | - |
| - | - | 7132 | 263.1 | - | - | 0 | - |
| - | - | 6082 | 264.1 | - | - | 0 | - |
| - | - | 3565 | 265.1 | - | - | 0 | - |
| - | - | 8079 | 265.1 | - | - | 0 | - |
| - | - | 5270 | 265.2 | - | - | 0 | - |
| - | - | 6789 | 265.2 | - | - | 0 | - |
| - | - | 9.394E+04 | 266.1 | - | - | 0 | - |
| - | - | 1.312E+04 | 267.1 | - | - | 0 | - |
| - | - | 5055 | 267.1 | - | - | 0 | - |
| - | - | 7340 | 267.2 | - | - | 0 | - |
| - | - | 5329 | 268.2 | - | - | 0 | - |
| - | - | 2.118E+04 | 269.1 | - | - | 0 | - |
| - | - | 4188 | 270.1 | - | - | 0 | - |
| - | - | 3985 | 270.1 | - | - | 0 | - |
| - | - | 4.459E+04 | 270.2 | - | - | 0 | - |
| - | - | 5215 | 271.2 | - | - | 0 | - |
| - | - | 3101 | 273.1 | - | - | 0 | - |
| 2 | b | 1.005E+06 | 274.1 | 0.0007364 | 2.687 | +1 | 2 |
| - | - | 1.691E+04 | 275.1 | - | - | 0 | - |
| - | - | 4.167E+05 | 275.1 | - | - | 0 | - |
| - | - | 1.002E+04 | 276.1 | - | - | 0 | - |
| - | - | 4.978E+04 | 276.1 | - | - | 0 | - |
| - | - | 4074 | 277.1 | - | - | 0 | - |
| - | - | 1.081E+04 | 277.1 | - | - | 0 | - |
| - | - | 3603 | 277.2 | - | - | 0 | - |
| - | - | 3384 | 278.2 | - | - | 0 | - |
| - | - | 1.231E+04 | 279.2 | - | - | 0 | - |
| - | - | 3609 | 282.2 | - | - | 0 | - |
| - | - | 1.88E+04 | 283.2 | - | - | 0 | - |
| - | - | 1.308E+05 | 283.2 | - | - | 0 | - |
| - | - | 4445 | 284.2 | - | - | 0 | - |
| - | - | 1.786E+04 | 284.2 | - | - | 0 | - |
| - | - | 9306 | 285.1 | - | - | 0 | - |
| - | - | 2.806E+04 | 285.2 | - | - | 0 | - |
| - | - | 3253 | 286.1 | - | - | 0 | - |
| - | - | 5195 | 286.2 | - | - | 0 | - |
| - | - | 4877 | 290.1 | - | - | 0 | - |
| - | - | 7142 | 290.2 | - | - | 0 | - |
| - | - | 1.022E+04 | 291.2 | - | - | 0 | - |
| - | - | 9.647E+04 | 292.1 | - | - | 0 | - |
| - | - | 1.407E+04 | 293.1 | - | - | 0 | - |
| - | - | 3520 | 293.2 | - | - | 0 | - |
| - | - | 4.101E+04 | 294.1 | - | - | 0 | - |
| - | - | 1.234E+04 | 294.2 | - | - | 0 | - |
| - | - | 3664 | 295.1 | - | - | 0 | - |
| - | - | 6572 | 295.2 | - | - | 0 | - |
| - | - | 3695 | 295.2 | - | - | 0 | - |
| - | - | 1.464E+04 | 296.2 | - | - | 0 | - |
| - | - | 5.104E+04 | 297.2 | - | - | 0 | - |
| - | - | 1.447E+04 | 298.1 | - | - | 0 | - |
| - | - | 7876 | 298.2 | - | - | 0 | - |
| - | - | 2618 | 299.2 | - | - | 0 | - |
| - | - | 5928 | 300.2 | - | - | 0 | - |
| - | - | 2.065E+04 | 303.1 | - | - | 0 | - |
| - | - | 3889 | 303.2 | - | - | 0 | - |
| - | - | 2807 | 304.1 | - | - | 0 | - |
| - | - | 2.001E+04 | 305.1 | - | - | 0 | - |
| - | - | 4031 | 305.7 | - | - | 0 | - |
| - | - | 8927 | 308.1 | - | - | 0 | - |
| - | - | 3375 | 309.2 | - | - | 0 | - |
| - | - | 6682 | 309.2 | - | - | 0 | - |
| - | - | 4281 | 310.2 | - | - | 0 | - |
| - | - | 2.115E+04 | 311.2 | - | - | 0 | - |
| - | - | 4.05E+04 | 312.2 | - | - | 0 | - |
| - | - | 3567 | 312.2 | - | - | 0 | - |
| - | - | 7663 | 313.2 | - | - | 0 | - |
| - | - | 9558 | 313.2 | - | - | 0 | - |
| - | - | 3434 | 314.1 | - | - | 0 | - |
| - | - | 1.23E+04 | 314.2 | - | - | 0 | - |
| - | - | 4453 | 314.7 | - | - | 0 | - |
| - | - | 4205 | 315.2 | - | - | 0 | - |
| 16 | y | 6.387E+04 | 316.2 | 0.001086 | 3.435 | +1 | 3 |
| - | - | 8331 | 317.2 | - | - | 0 | - |
| - | - | 3.878E+04 | 320.1 | - | - | 0 | - |
| - | - | 8283 | 320.2 | - | - | 0 | - |
| - | - | 7346 | 321.1 | - | - | 0 | - |
| - | - | 1.065E+05 | 321.2 | - | - | 0 | - |
| - | - | 3.512E+04 | 322.2 | - | - | 0 | - |
| - | - | 2.306E+04 | 322.2 | - | - | 0 | - |
| - | - | 1.043E+04 | 323.2 | - | - | 0 | - |
| - | - | 2905 | 323.2 | - | - | 0 | - |
| - | - | 6154 | 325.2 | - | - | 0 | - |
| 8 | b | 3777 | 326.1 | 0.004024 | 12.34 | +3 | 8 |
| - | - | 2.575E+04 | 326.2 | - | - | 0 | - |
| - | - | 3589 | 327.2 | - | - | 0 | - |
| - | - | 5782 | 327.2 | - | - | 0 | - |
| - | - | 3156 | 327.3 | - | - | 0 | - |
| 13 | y | 2931 | 328.2 | 0.003392 | 10.34 | +2 | 6 |
| - | - | 3240 | 329.1 | - | - | 0 | - |
| - | - | 2911 | 329.2 | - | - | 0 | - |
| - | - | 2.255E+04 | 330.2 | - | - | 0 | - |
| - | - | 4231 | 330.2 | - | - | 0 | - |
| - | - | 3605 | 331.2 | - | - | 0 | - |
| - | - | 4290 | 332.7 | - | - | 0 | - |
| - | - | 6763 | 333.1 | - | - | 0 | - |
| 16 | y | 1.824E+05 | 334.2 | 0.00108 | 3.233 | +1 | 3 |
| - | - | 2.408E+04 | 335.2 | - | - | 0 | - |
| - | - | 9268 | 336.1 | - | - | 0 | - |
| - | - | 5608 | 336.2 | - | - | 0 | - |
| - | - | 1.257E+05 | 336.2 | - | - | 0 | - |
| - | - | 5742 | 337.2 | - | - | 0 | - |
| 13 | y | 1.806E+04 | 337.2 | 0.002622 | 7.777 | +2 | 6 |
| - | - | 6096 | 337.3 | - | - | 0 | - |
| - | - | 1.027E+04 | 337.7 | - | - | 0 | - |
| - | - | 9204 | 338.2 | - | - | 0 | - |
| - | - | 3750 | 338.2 | - | - | 0 | - |
| - | - | 9980 | 338.2 | - | - | 0 | - |
| - | - | 3554 | 339.2 | - | - | 0 | - |
| - | - | 5822 | 339.2 | - | - | 0 | - |
| - | - | 8.201E+04 | 339.2 | - | - | 0 | - |
| - | - | 1.063E+04 | 340.2 | - | - | 0 | - |
| - | - | 1.56E+04 | 340.2 | - | - | 0 | - |
| - | - | 9353 | 340.3 | - | - | 0 | - |
| - | - | 5792 | 341.2 | - | - | 0 | - |
| - | - | 3573 | 342.2 | - | - | 0 | - |
| - | - | 3328 | 344.1 | - | - | 0 | - |
| - | - | 6130 | 346.7 | - | - | 0 | - |
| - | - | 4564 | 347.2 | - | - | 0 | - |
| - | - | 4538 | 348.1 | - | - | 0 | - |
| - | - | 4948 | 348.2 | - | - | 0 | - |
| - | - | 1.66E+04 | 350.2 | - | - | 0 | - |
| - | - | 3785 | 351.2 | - | - | 0 | - |
| - | - | 2.499E+04 | 351.2 | - | - | 0 | - |
| - | - | 5761 | 352.2 | - | - | 0 | - |
| - | - | 1.11E+04 | 353.2 | - | - | 0 | - |
| - | - | 1.711E+05 | 354.2 | - | - | 0 | - |
| - | - | 2.9E+04 | 355.2 | - | - | 0 | - |
| - | - | 1.888E+04 | 355.3 | - | - | 0 | - |
| - | - | 4796 | 356.2 | - | - | 0 | - |
| - | - | 6829 | 356.3 | - | - | 0 | - |
| - | - | 7680 | 357.2 | - | - | 0 | - |
| - | - | 5362 | 357.2 | - | - | 0 | - |
| - | - | 1.95E+04 | 358.2 | - | - | 0 | - |
| - | - | 5623 | 359.2 | - | - | 0 | - |
| - | - | 4785 | 359.2 | - | - | 0 | - |
| - | - | 3932 | 360.7 | - | - | 0 | - |
| - | - | 1.311E+04 | 365.3 | - | - | 0 | - |
| - | - | 3.41E+04 | 366.2 | - | - | 0 | - |
| - | - | 3648 | 366.2 | - | - | 0 | - |
| - | - | 1.168E+04 | 367.2 | - | - | 0 | - |
| - | - | 1.859E+04 | 368.1 | - | - | 0 | - |
| - | - | 4869 | 368.2 | - | - | 0 | - |
| - | - | 4043 | 369.2 | - | - | 0 | - |
| - | - | 8948 | 370.2 | - | - | 0 | - |
| - | - | 3322 | 370.7 | - | - | 0 | - |
| - | - | 3914 | 371.2 | - | - | 0 | - |
| - | - | 3762 | 372.2 | - | - | 0 | - |
| - | - | 3285 | 374.3 | - | - | 0 | - |
| - | - | 7388 | 375.2 | - | - | 0 | - |
| - | - | 3278 | 378.3 | - | - | 0 | - |
| - | - | 3751 | 379.2 | - | - | 0 | - |
| - | - | 7733 | 380.2 | - | - | 0 | - |
| - | - | 4019 | 380.3 | - | - | 0 | - |
| - | - | 5269 | 382.2 | - | - | 0 | - |
| - | - | 9160 | 382.3 | - | - | 0 | - |
| - | - | 4.838E+04 | 383.3 | - | - | 0 | - |
| - | - | 4372 | 383.3 | - | - | 0 | - |
| - | - | 3498 | 384.1 | - | - | 0 | - |
| - | - | 3.239E+04 | 384.2 | - | - | 0 | - |
| - | - | 3938 | 384.2 | - | - | 0 | - |
| - | - | 9477 | 384.3 | - | - | 0 | - |
| - | - | 6.88E+04 | 385.2 | - | - | 0 | - |
| - | - | 2.462E+04 | 385.2 | - | - | 0 | - |
| - | - | 4.752E+04 | 386.2 | - | - | 0 | - |
| - | - | 4891 | 386.2 | - | - | 0 | - |
| - | - | 1.036E+04 | 387.2 | - | - | 0 | - |
| - | - | 8487 | 389.3 | - | - | 0 | - |
| - | - | 3218 | 389.8 | - | - | 0 | - |
| - | - | 3004 | 390.2 | - | - | 0 | - |
| - | - | 3895 | 390.3 | - | - | 0 | - |
| - | - | 1.16E+04 | 392.2 | - | - | 0 | - |
| - | - | 2.732E+04 | 392.3 | - | - | 0 | - |
| - | - | 8527 | 393.2 | - | - | 0 | - |
| - | - | 7631 | 393.3 | - | - | 0 | - |
| - | - | 4.611E+04 | 394.2 | - | - | 0 | - |
| - | - | 7425 | 394.3 | - | - | 0 | - |
| - | - | 1.105E+04 | 395.2 | - | - | 0 | - |
| - | - | 4685 | 396.2 | - | - | 0 | - |
| - | - | 2.314E+04 | 396.3 | - | - | 0 | - |
| - | - | 4419 | 396.3 | - | - | 0 | - |
| - | - | 5225 | 397.2 | - | - | 0 | - |
| - | - | 1.09E+04 | 398.2 | - | - | 0 | - |
| - | - | 4344 | 398.3 | - | - | 0 | - |
| - | - | 4460 | 400.2 | - | - | 0 | - |
| - | - | 1.005E+04 | 401.2 | - | - | 0 | - |
| - | - | 2.097E+04 | 402.2 | - | - | 0 | - |
| - | - | 9.24E+04 | 403.2 | - | - | 0 | - |
| - | - | 5335 | 403.2 | - | - | 0 | - |
| - | - | 2.085E+04 | 403.3 | - | - | 0 | - |
| - | - | 1.347E+04 | 403.8 | - | - | 0 | - |
| - | - | 1.776E+04 | 404.2 | - | - | 0 | - |
| - | - | 3809 | 404.3 | - | - | 0 | - |
| - | - | 2.21E+04 | 406.2 | - | - | 0 | - |
| - | - | 7175 | 406.3 | - | - | 0 | - |
| - | - | 8702 | 406.7 | - | - | 0 | - |
| - | - | 5785 | 407.2 | - | - | 0 | - |
| - | - | 9342 | 407.3 | - | - | 0 | - |
| - | - | 3133 | 408.2 | - | - | 0 | - |
| - | - | 8513 | 408.3 | - | - | 0 | - |
| 3 | a | 3496 | 409.2 | 0.002147 | 5.246 | +1 | 3 |
| - | - | 5367 | 409.3 | - | - | 0 | - |
| - | - | 5.121E+04 | 410.3 | - | - | 0 | - |
| 6 | b | 8.008E+04 | 411.2 | 0.00112 | 2.724 | +2 | 6 |
| - | - | 3545 | 411.2 | - | - | 0 | - |
| - | - | 9902 | 411.3 | - | - | 0 | - |
| - | - | 1.418E+04 | 412.2 | - | - | 0 | - |
| - | - | 1.19E+04 | 412.2 | - | - | 0 | - |
| - | - | 7780 | 413.2 | - | - | 0 | - |
| - | - | 6571 | 413.3 | - | - | 0 | - |
| - | - | 2962 | 414.2 | - | - | 0 | - |
| - | - | 5065 | 415.3 | - | - | 0 | - |
| 3 | b | 8621 | 419.2 | 0.001531 | 3.652 | +1 | 3 |
| 6 | b | 2.288E+04 | 420.2 | 0.0006291 | 1.497 | +2 | 6 |
| - | - | 1.504E+04 | 420.2 | - | - | 0 | - |
| - | - | 9846 | 420.7 | - | - | 0 | - |
| - | - | 5028 | 421.2 | - | - | 0 | - |
| - | - | 3.765E+04 | 422.3 | - | - | 0 | - |
| - | - | 9629 | 423.3 | - | - | 0 | - |
| - | - | 3743 | 423.3 | - | - | 0 | - |
| - | - | 7235 | 424.2 | - | - | 0 | - |
| - | - | 3.55E+04 | 424.3 | - | - | 0 | - |
| - | - | 8908 | 424.7 | - | - | 0 | - |
| - | - | 4488 | 425.2 | - | - | 0 | - |
| - | - | 1.259E+04 | 425.2 | - | - | 0 | - |
| - | - | 7714 | 425.3 | - | - | 0 | - |
| - | - | 7279 | 425.3 | - | - | 0 | - |
| - | - | 5099 | 426.2 | - | - | 0 | - |
| - | - | 9839 | 426.3 | - | - | 0 | - |
| - | - | 4922 | 426.3 | - | - | 0 | - |
| - | - | 5638 | 427.3 | - | - | 0 | - |
| - | - | 2.984E+05 | 429.2 | - | - | 0 | - |
| 15 | y | 8621 | 429.2 | 0.0036 | 8.388 | +1 | 4 |
| - | - | 7.541E+04 | 430.2 | - | - | 0 | - |
| - | - | 3.624E+04 | 431.2 | - | - | 0 | - |
| - | - | 9426 | 432.2 | - | - | 0 | - |
| - | - | 5405 | 433.3 | - | - | 0 | - |
| - | - | 3583 | 433.3 | - | - | 0 | - |
| - | - | 6619 | 434.3 | - | - | 0 | - |
| - | - | 2849 | 436.2 | - | - | 0 | - |
| 3 | b | 1.035E+05 | 437.2 | 0.001373 | 3.14 | +1 | 3 |
| - | - | 3.187E+04 | 438.2 | - | - | 0 | - |
| - | - | 5441 | 439.2 | - | - | 0 | - |
| - | - | 1.09E+04 | 439.3 | - | - | 0 | - |
| - | - | 1.038E+04 | 442.2 | - | - | 0 | - |
| - | - | 6250 | 442.3 | - | - | 0 | - |
| - | - | 1.575E+04 | 443.3 | - | - | 0 | - |
| - | - | 3332 | 444.3 | - | - | 0 | - |
| - | - | 3344 | 445.3 | - | - | 0 | - |
| - | - | 1.239E+04 | 445.8 | - | - | 0 | - |
| - | - | 1.253E+04 | 446.2 | - | - | 0 | - |
| - | - | 7096 | 446.3 | - | - | 0 | - |
| 15 | y | 3.759E+04 | 447.2 | 0.001062 | 2.374 | +1 | 4 |
| - | - | 6219 | 447.3 | - | - | 0 | - |
| - | - | 8816 | 448.3 | - | - | 0 | - |
| - | - | 5086 | 449.3 | - | - | 0 | - |
| 11 | y | 5017 | 449.8 | 0.00167 | 3.712 | +2 | 8 |
| - | - | 9636 | 450.3 | - | - | 0 | - |
| - | - | 3496 | 450.3 | - | - | 0 | - |
| - | - | 5342 | 451.2 | - | - | 0 | - |
| - | - | 1.89E+04 | 451.3 | - | - | 0 | - |
| - | - | 4940 | 451.3 | - | - | 0 | - |
| - | - | 4803 | 451.7 | - | - | 0 | - |
| - | - | 7026 | 452.3 | - | - | 0 | - |
| - | - | 7.341E+04 | 452.3 | - | - | 0 | - |
| - | - | 4888 | 453.3 | - | - | 0 | - |
| - | - | 2.172E+04 | 453.3 | - | - | 0 | - |
| - | - | 4.507E+04 | 454.2 | - | - | 0 | - |
| - | - | 4236 | 454.3 | - | - | 0 | - |
| - | - | 1.13E+04 | 455.2 | - | - | 0 | - |
| - | - | 2.354E+04 | 455.3 | - | - | 0 | - |
| - | - | 5699 | 456.3 | - | - | 0 | - |
| - | - | 1.382E+04 | 457.2 | - | - | 0 | - |
| - | - | 3214 | 457.6 | - | - | 0 | - |
| - | - | 2.376E+04 | 459.8 | - | - | 0 | - |
| - | - | 9955 | 460.2 | - | - | 0 | - |
| - | - | 1.378E+04 | 460.3 | - | - | 0 | - |
| - | - | 5511 | 460.8 | - | - | 0 | - |
| - | - | 8450 | 461.2 | - | - | 0 | - |
| - | - | 3483 | 462.8 | - | - | 0 | - |
| - | - | 6641 | 463.3 | - | - | 0 | - |
| - | - | 4.606E+04 | 467.3 | - | - | 0 | - |
| - | - | 3559 | 467.3 | - | - | 0 | - |
| - | - | 4236 | 468.2 | - | - | 0 | - |
| - | - | 2.441E+04 | 468.3 | - | - | 0 | - |
| - | - | 1.898E+04 | 468.4 | - | - | 0 | - |
| - | - | 2979 | 469.3 | - | - | 0 | - |
| - | - | 6134 | 469.4 | - | - | 0 | - |
| - | - | 8932 | 470.3 | - | - | 0 | - |
| - | - | 6311 | 471.8 | - | - | 0 | - |
| - | - | 3.436E+04 | 473.3 | - | - | 0 | - |
| - | - | 6013 | 473.6 | - | - | 0 | - |
| - | - | 1.134E+04 | 474.3 | - | - | 0 | - |
| - | - | 4123 | 474.8 | - | - | 0 | - |
| - | - | 3777 | 476.3 | - | - | 0 | - |
| - | - | 2.787E+04 | 478.2 | - | - | 0 | - |
| - | - | 7839 | 478.3 | - | - | 0 | - |
| - | - | 6757 | 479.2 | - | - | 0 | - |
| - | - | 3826 | 482.3 | - | - | 0 | - |
| - | - | 8027 | 482.3 | - | - | 0 | - |
| - | - | 4163 | 482.8 | - | - | 0 | - |
| - | - | 4055 | 483.2 | - | - | 0 | - |
| - | - | 3470 | 483.8 | - | - | 0 | - |
| - | - | 3731 | 484.3 | - | - | 0 | - |
| - | - | 3954 | 488.3 | - | - | 0 | - |
| - | - | 1.413E+04 | 491.3 | - | - | 0 | - |
| - | - | 6172 | 492.3 | - | - | 0 | - |
| - | - | 1.024E+04 | 495.2 | - | - | 0 | - |
| - | - | 1.073E+04 | 495.4 | - | - | 0 | - |
| - | - | 3990 | 496 | - | - | 0 | - |
| - | - | 1.707E+04 | 496.4 | - | - | 0 | - |
| 8 | b | 1.964E+04 | 497.2 | 0.001479 | 2.975 | +2 | 8 |
| - | - | 7156 | 497.3 | - | - | 0 | - |
| - | - | 4635 | 497.4 | - | - | 0 | - |
| - | - | 1.385E+04 | 497.7 | - | - | 0 | - |
| - | - | 6138 | 498.2 | - | - | 0 | - |
| - | - | 6500 | 501.6 | - | - | 0 | - |
| - | - | 8295 | 501.9 | - | - | 0 | - |
| - | - | 1.528E+04 | 503.2 | - | - | 0 | - |
| - | - | 1.57E+04 | 503.3 | - | - | 0 | - |
| - | - | 1.003E+04 | 503.8 | - | - | 0 | - |
| - | - | 5041 | 504.2 | - | - | 0 | - |
| - | - | 3301 | 504.3 | - | - | 0 | - |
| 13 | b | 9016 | 504.9 | 0.003443 | 6.819 | +3 | 13 |
| 13 | b | 4483 | 505.3 | 0.007805 | 15.45 | +3 | 13 |
| - | - | 1.244E+04 | 505.6 | - | - | 0 | - |
| - | - | 6343 | 509.3 | - | - | 0 | - |
| - | - | 3812 | 510.3 | - | - | 0 | - |
| - | - | 3306 | 510.4 | - | - | 0 | - |
| 13 | b | 2.213E+04 | 510.9 | 0.002363 | 4.625 | +3 | 13 |
| - | - | 1.81E+04 | 511.3 | - | - | 0 | - |
| - | - | 7225 | 511.6 | - | - | 0 | - |
| - | - | 3578 | 511.9 | - | - | 0 | - |
| - | - | 3989 | 514.2 | - | - | 0 | - |
| - | - | 4680 | 514.3 | - | - | 0 | - |
| - | - | 3828 | 517.7 | - | - | 0 | - |
| - | - | 4161 | 518.9 | - | - | 0 | - |
| - | - | 7414 | 519.3 | - | - | 0 | - |
| - | - | 4261 | 519.8 | - | - | 0 | - |
| - | - | 3940 | 520 | - | - | 0 | - |
| - | - | 4427 | 520.2 | - | - | 0 | - |
| - | - | 8710 | 520.3 | - | - | 0 | - |
| - | - | 9789 | 520.4 | - | - | 0 | - |
| - | - | 1.495E+04 | 521.2 | - | - | 0 | - |
| - | - | 4293 | 521.3 | - | - | 0 | - |
| - | - | 5957 | 522.2 | - | - | 0 | - |
| - | - | 4.078E+04 | 523.4 | - | - | 0 | - |
| - | - | 1.405E+04 | 524.4 | - | - | 0 | - |
| - | - | 5534 | 524.9 | - | - | 0 | - |
| - | - | 3778 | 525.4 | - | - | 0 | - |
| - | - | 5080 | 527.3 | - | - | 0 | - |
| - | - | 9242 | 528.2 | - | - | 0 | - |
| - | - | 8765 | 528.3 | - | - | 0 | - |
| - | - | 9907 | 528.8 | - | - | 0 | - |
| - | - | 1.709E+04 | 529.2 | - | - | 0 | - |
| - | - | 8983 | 530.2 | - | - | 0 | - |
| - | - | 4743 | 530.3 | - | - | 0 | - |
| - | - | 5581 | 530.9 | - | - | 0 | - |
| - | - | 1.511E+04 | 531.2 | - | - | 0 | - |
| - | - | 3432 | 532.2 | - | - | 0 | - |
| - | - | 5489 | 535.4 | - | - | 0 | - |
| - | - | 4517 | 535.8 | - | - | 0 | - |
| - | - | 4727 | 536.4 | - | - | 0 | - |
| - | - | 5570 | 537.2 | - | - | 0 | - |
| - | - | 5225 | 537.3 | - | - | 0 | - |
| - | - | 3.478E+04 | 537.4 | - | - | 0 | - |
| - | - | 1.807E+05 | 538.3 | - | - | 0 | - |
| - | - | 6984 | 538.4 | - | - | 0 | - |
| - | - | 8882 | 538.4 | - | - | 0 | - |
| - | - | 4763 | 538.6 | - | - | 0 | - |
| - | - | 6106 | 538.8 | - | - | 0 | - |
| - | - | 4586 | 538.9 | - | - | 0 | - |
| - | - | 5.632E+04 | 539.3 | - | - | 0 | - |
| - | - | 1.999E+04 | 539.3 | - | - | 0 | - |
| - | - | 9382 | 539.4 | - | - | 0 | - |
| - | - | 2.153E+04 | 539.6 | - | - | 0 | - |
| - | - | 7538 | 539.8 | - | - | 0 | - |
| - | - | 1.107E+04 | 540 | - | - | 0 | - |
| - | - | 5028 | 540.3 | - | - | 0 | - |
| - | - | 9074 | 540.3 | - | - | 0 | - |
| 9 | y | 8917 | 540.8 | 0.008736 | 16.15 | +2 | 10 |
| - | - | 6004 | 541.3 | - | - | 0 | - |
| 14 | y | 4343 | 542.3 | 0.002422 | 4.466 | +1 | 5 |
| 14 | b | 3846 | 542.6 | 0.0008023 | 1.479 | +3 | 14 |
| 14 | b | 4767 | 543 | 0.01057 | 19.46 | +3 | 14 |
| - | - | 4217 | 544.6 | - | - | 0 | - |
| - | - | 5731 | 544.8 | - | - | 0 | - |
| 4 | b | 5575 | 547.2 | 0.001242 | 2.27 | +1 | 4 |
| - | - | 5284 | 548 | - | - | 0 | - |
| - | - | 8456 | 548.2 | - | - | 0 | - |
| 14 | b | 4.749E+04 | 548.6 | 0.001126 | 2.052 | +3 | 14 |
| - | - | 3.982E+04 | 549 | - | - | 0 | - |
| - | - | 9960 | 549.2 | - | - | 0 | - |
| - | - | 2.48E+04 | 549.3 | - | - | 0 | - |
| 9 | y | 5416 | 549.4 | 0.002906 | 5.291 | +2 | 10 |
| - | - | 8946 | 549.6 | - | - | 0 | - |
| - | - | 4964 | 550.2 | - | - | 0 | - |
| - | - | 4463 | 550.4 | - | - | 0 | - |
| - | - | 6064 | 553.9 | - | - | 0 | - |
| - | - | 6860 | 555.3 | - | - | 0 | - |
| - | - | 3607 | 556.3 | - | - | 0 | - |
| - | - | 9630 | 557.3 | - | - | 0 | - |
| - | - | 7429 | 557.7 | - | - | 0 | - |
| - | - | 3916 | 558 | - | - | 0 | - |
| - | - | 5932 | 558.8 | - | - | 0 | - |
| - | - | 4023 | 559.3 | - | - | 0 | - |
| 14 | y | 1.713E+04 | 560.3 | 0.002477 | 4.421 | +1 | 5 |
| 9 | b | 8.733E+04 | 561.3 | 0.001086 | 1.935 | +2 | 9 |
| - | - | 3464 | 561.3 | - | - | 0 | - |
| - | - | 5088 | 561.3 | - | - | 0 | - |
| - | - | 9612 | 561.6 | - | - | 0 | - |
| - | - | 5.753E+04 | 561.8 | - | - | 0 | - |
| - | - | 3887 | 562 | - | - | 0 | - |
| - | - | 1.993E+04 | 562.3 | - | - | 0 | - |
| - | - | 1.067E+04 | 562.6 | - | - | 0 | - |
| - | - | 4804 | 562.8 | - | - | 0 | - |
| - | - | 4547 | 563 | - | - | 0 | - |
| - | - | 3219 | 563.4 | - | - | 0 | - |
| - | - | 4245 | 564.4 | - | - | 0 | - |
| 4 | b | 4.355E+04 | 565.2 | 0.001053 | 1.863 | +1 | 4 |
| - | - | 5421 | 565.3 | - | - | 0 | - |
| - | - | 5410 | 565.4 | - | - | 0 | - |
| - | - | 9.094E+04 | 565.4 | - | - | 0 | - |
| - | - | 6567 | 565.6 | - | - | 0 | - |
| - | - | 5924 | 565.9 | - | - | 0 | - |
| - | - | 2.198E+05 | 566.2 | - | - | 0 | - |
| - | - | 3.706E+04 | 566.4 | - | - | 0 | - |
| - | - | 7.357E+04 | 567.3 | - | - | 0 | - |
| - | - | 1.9E+04 | 567.3 | - | - | 0 | - |
| - | - | 9484 | 567.4 | - | - | 0 | - |
| - | - | 9550 | 567.6 | - | - | 0 | - |
| - | - | 4.377E+04 | 567.8 | - | - | 0 | - |
| - | - | 4781 | 568 | - | - | 0 | - |
| - | - | 1.241E+04 | 568.3 | - | - | 0 | - |
| - | - | 6589 | 568.3 | - | - | 0 | - |
| - | - | 2.271E+04 | 568.4 | - | - | 0 | - |
| - | - | 9602 | 568.6 | - | - | 0 | - |
| 8 | y | 7286 | 568.9 | 0.002924 | 5.14 | +2 | 11 |
| - | - | 3598 | 569 | - | - | 0 | - |
| - | - | 2.397E+04 | 570.2 | - | - | 0 | - |
| - | - | 6332 | 571.2 | - | - | 0 | - |
| - | - | 3358 | 571.3 | - | - | 0 | - |
| - | - | 3744 | 571.7 | - | - | 0 | - |
| - | - | 4612 | 571.9 | - | - | 0 | - |
| - | - | 3863 | 572 | - | - | 0 | - |
| - | - | 5609 | 572.8 | - | - | 0 | - |
| - | - | 8244 | 573.3 | - | - | 0 | - |
| - | - | 4175 | 573.8 | - | - | 0 | - |
| - | - | 3780 | 574 | - | - | 0 | - |
| - | - | 1.2E+04 | 576.6 | - | - | 0 | - |
| - | - | 3.533E+04 | 577 | - | - | 0 | - |
| - | - | 3.296E+04 | 577.3 | - | - | 0 | - |
| - | - | 1.845E+04 | 577.4 | - | - | 0 | - |
| - | - | 1.833E+04 | 577.7 | - | - | 0 | - |
| - | - | 1.868E+04 | 578 | - | - | 0 | - |
| - | - | 8157 | 578.3 | - | - | 0 | - |
| - | - | 6733 | 578.4 | - | - | 0 | - |
| - | - | 4.843E+04 | 579.4 | - | - | 0 | - |
| - | - | 2.13E+04 | 580.4 | - | - | 0 | - |
| 15 | b | 6753 | 580.7 | 0.003011 | 5.186 | +3 | 15 |
| - | - | 6623 | 581.4 | - | - | 0 | - |
| - | - | 1.015E+04 | 582.8 | - | - | 0 | - |
| - | - | 9545 | 583.3 | - | - | 0 | - |
| - | - | 6590 | 583.4 | - | - | 0 | - |
| - | - | 5448 | 585.7 | - | - | 0 | - |
| 15 | b | 1.387E+05 | 586.3 | 0.001384 | 2.361 | +3 | 15 |
| - | - | 1.713E+05 | 586.7 | - | - | 0 | - |
| - | - | 3622 | 586.8 | - | - | 0 | - |
| - | - | 1.048E+05 | 587 | - | - | 0 | - |
| - | - | 9911 | 587.3 | - | - | 0 | - |
| - | - | 3.356E+04 | 587.3 | - | - | 0 | - |
| - | - | 6726 | 587.7 | - | - | 0 | - |
| 10 | b | 4.439E+04 | 587.8 | 0.001606 | 2.732 | +2 | 10 |
| 10 | b | 4.11E+04 | 588.3 | 0.009659 | 16.42 | +2 | 10 |
| - | - | 1.177E+04 | 588.8 | - | - | 0 | - |
| - | - | 4245 | 591.4 | - | - | 0 | - |
| - | - | 5866 | 592 | - | - | 0 | - |
| - | - | 4189 | 592.3 | - | - | 0 | - |
| - | - | 5196 | 594.2 | - | - | 0 | - |
| - | - | 1.055E+04 | 595.4 | - | - | 0 | - |
| - | - | 1.707E+04 | 595.4 | - | - | 0 | - |
| - | - | 8231 | 595.9 | - | - | 0 | - |
| - | - | 6016 | 596.3 | - | - | 0 | - |
| - | - | 7757 | 596.4 | - | - | 0 | - |
| 10 | b | 8.548E+04 | 596.8 | 0.001572 | 2.635 | +2 | 10 |
| - | - | 1.773E+04 | 596.9 | - | - | 0 | - |
| - | - | 6.086E+04 | 597.3 | - | - | 0 | - |
| - | - | 6879 | 597.4 | - | - | 0 | - |
| - | - | 2.054E+04 | 597.8 | - | - | 0 | - |
| - | - | 8738 | 597.9 | - | - | 0 | - |
| - | - | 1.189E+04 | 598.2 | - | - | 0 | - |
| - | - | 7703 | 599.6 | - | - | 0 | - |
| 3 | y | 1.277E+04 | 600.3 | 0.003687 | 6.142 | +3 | 16 |
| 3 | y | 1.015E+04 | 600.7 | 0.004977 | 8.287 | +3 | 16 |
| - | - | 1.454E+04 | 604.4 | - | - | 0 | - |
| - | - | 7398 | 604.9 | - | - | 0 | - |
| - | - | 4807 | 605.3 | - | - | 0 | - |
| - | - | 6485 | 605.4 | - | - | 0 | - |
| - | - | 9302 | 605.7 | - | - | 0 | - |
| - | - | 1.109E+04 | 606 | - | - | 0 | - |
| 3 | y | 5.366E+04 | 606.3 | 0.0008478 | 1.398 | +3 | 16 |
| - | - | 4044 | 606.4 | - | - | 0 | - |
| - | - | 4.336E+04 | 606.7 | - | - | 0 | - |
| - | - | 2.594E+04 | 607 | - | - | 0 | - |
| - | - | 8207 | 607.3 | - | - | 0 | - |
| - | - | 5980 | 608.5 | - | - | 0 | - |
| - | - | 1.096E+04 | 609 | - | - | 0 | - |
| 16 | b | 1.285E+04 | 609.3 | 0.007479 | 12.27 | +3 | 16 |
| 16 | b | 1.603E+04 | 609.7 | 0.0002699 | 0.4426 | +3 | 16 |
| - | - | 5174 | 610 | - | - | 0 | - |
| - | - | 2.162E+04 | 610.3 | - | - | 0 | - |
| - | - | 4877 | 610.4 | - | - | 0 | - |
| - | - | 3750 | 610.7 | - | - | 0 | - |
| - | - | 1.994E+04 | 611.3 | - | - | 0 | - |
| - | - | 1.152E+04 | 611.7 | - | - | 0 | - |
| - | - | 1.146E+04 | 612 | - | - | 0 | - |
| - | - | 8085 | 612.3 | - | - | 0 | - |
| - | - | 5170 | 614.3 | - | - | 0 | - |
| - | - | 1.635E+04 | 614.7 | - | - | 0 | - |
| - | - | 1.655E+04 | 615 | - | - | 0 | - |
| - | - | 1.105E+04 | 615.3 | - | - | 0 | - |
| 16 | b | 9.935E+04 | 615.3 | 0.001816 | 2.952 | +3 | 16 |
| - | - | 1.11E+05 | 615.7 | - | - | 0 | - |
| - | - | 8138 | 615.9 | - | - | 0 | - |
| - | - | 7.149E+04 | 616 | - | - | 0 | - |
| - | - | 5779 | 616.3 | - | - | 0 | - |
| - | - | 2.327E+04 | 616.3 | - | - | 0 | - |
| - | - | 1.155E+04 | 616.7 | - | - | 0 | - |
| 7 | y | 1.153E+04 | 617.4 | 0.002188 | 3.545 | +2 | 12 |
| 7 | y | 1.294E+04 | 617.9 | 0.0111 | 17.96 | +2 | 12 |
| - | - | 3500 | 619 | - | - | 0 | - |
| - | - | 6209 | 619.3 | - | - | 0 | - |
| - | - | 2.205E+04 | 620.3 | - | - | 0 | - |
| - | - | 2.565E+04 | 620.7 | - | - | 0 | - |
| - | - | 1.782E+04 | 620.8 | - | - | 0 | - |
| - | - | 3.629E+04 | 621 | - | - | 0 | - |
| - | - | 1.662E+04 | 621.3 | - | - | 0 | - |
| - | - | 4507 | 621.4 | - | - | 0 | - |
| - | - | 7845 | 621.7 | - | - | 0 | - |
| - | - | 8964 | 623.4 | - | - | 0 | - |
| - | - | 4250 | 623.5 | - | - | 0 | - |
| - | - | 4132 | 623.7 | - | - | 0 | - |
| - | - | 4621 | 624.4 | - | - | 0 | - |
| - | - | 5910 | 624.5 | - | - | 0 | - |
| - | - | 4503 | 625.5 | - | - | 0 | - |
| 7 | y | 1.425E+05 | 626.4 | 0.001667 | 2.661 | +2 | 12 |
| - | - | 1.185E+05 | 626.9 | - | - | 0 | - |
| - | - | 4.567E+04 | 627.4 | - | - | 0 | - |
| - | - | 1.284E+04 | 627.9 | - | - | 0 | - |
| - | - | 8438 | 628.3 | - | - | 0 | - |
| - | - | 5441 | 628.4 | - | - | 0 | - |
| - | - | 5047 | 629.3 | - | - | 0 | - |
| - | - | 4.63E+04 | 629.3 | - | - | 0 | - |
| - | - | 1.451E+04 | 629.7 | - | - | 0 | - |
| - | - | 3.301E+04 | 629.8 | - | - | 0 | - |
| - | - | 5353 | 630 | - | - | 0 | - |
| - | - | 1.28E+04 | 630.3 | - | - | 0 | - |
| - | - | 8371 | 632.3 | - | - | 0 | - |
| - | - | 5818 | 633.3 | - | - | 0 | - |
| - | - | 4331 | 633.4 | - | - | 0 | - |
| - | - | 6486 | 634.4 | - | - | 0 | - |
| - | - | 9144 | 636.4 | - | - | 0 | - |
| - | - | 5.308E+04 | 636.4 | - | - | 0 | - |
| - | - | 4887 | 636.9 | - | - | 0 | - |
| - | - | 4033 | 637.3 | - | - | 0 | - |
| - | - | 1.59E+04 | 637.4 | - | - | 0 | - |
| - | - | 3911 | 637.7 | - | - | 0 | - |
| - | - | 4139 | 638.4 | - | - | 0 | - |
| - | - | 4158 | 638.4 | - | - | 0 | - |
| - | - | 7968 | 640.4 | - | - | 0 | - |
| - | - | 1.216E+04 | 643 | - | - | 0 | - |
| - | - | 8467 | 643.3 | - | - | 0 | - |
| - | - | 1.25E+04 | 643.3 | - | - | 0 | - |
| - | - | 1.487E+04 | 643.7 | - | - | 0 | - |
| 11 | b | 6354 | 645.3 | 0.001802 | 2.792 | +2 | 11 |
| - | - | 6128 | 645.8 | - | - | 0 | - |
| - | - | 1.747E+04 | 646.7 | - | - | 0 | - |
| - | - | 1.435E+04 | 647 | - | - | 0 | - |
| - | - | 5789 | 647.3 | - | - | 0 | - |
| - | - | 8677 | 647.4 | - | - | 0 | - |
| - | - | 5525 | 647.7 | - | - | 0 | - |
| - | - | 4735 | 648 | - | - | 0 | - |
| - | - | 8009 | 648.4 | - | - | 0 | - |
| - | - | 3918 | 648.4 | - | - | 0 | - |
| - | - | 6227 | 648.7 | - | - | 0 | - |
| - | - | 5.764E+04 | 649 | - | - | 0 | - |
| - | - | 6.019E+04 | 649.4 | - | - | 0 | - |
| - | - | 3.832E+04 | 649.7 | - | - | 0 | - |
| - | - | 1.737E+04 | 650 | - | - | 0 | - |
| - | - | 7928 | 650.4 | - | - | 0 | - |
| - | - | 1.002E+04 | 651.5 | - | - | 0 | - |
| - | - | 1.34E+04 | 651.9 | - | - | 0 | - |
| 17 | b | 8.701E+04 | 652.3 | 0.0007839 | 1.202 | +3 | 17 |
| - | - | 2.59E+04 | 652.4 | - | - | 0 | - |
| 17 | b | 1.242E+05 | 652.7 | 0.007679 | 11.76 | +3 | 17 |
| - | - | 1.017E+04 | 652.9 | - | - | 0 | - |
| - | - | 7.928E+04 | 653 | - | - | 0 | - |
| - | - | 3.08E+04 | 653.3 | - | - | 0 | - |
| - | - | 1.026E+04 | 653.4 | - | - | 0 | - |
| - | - | 2.232E+04 | 653.7 | - | - | 0 | - |
| - | - | 1.615E+04 | 653.8 | - | - | 0 | - |
| - | - | 5699 | 654 | - | - | 0 | - |
| - | - | 9810 | 654.3 | - | - | 0 | - |
| - | - | 1.224E+04 | 656.3 | - | - | 0 | - |
| - | - | 4.506E+04 | 657.3 | - | - | 0 | - |
| - | - | 6054 | 658 | - | - | 0 | - |
| 17 | b | 5.481E+05 | 658.4 | 0.001718 | 2.609 | +3 | 17 |
| - | - | 6.077E+05 | 658.7 | - | - | 0 | - |
| - | - | 4.082E+05 | 659 | - | - | 0 | - |
| - | - | 1.646E+05 | 659.4 | - | - | 0 | - |
| - | - | 4.672E+04 | 659.7 | - | - | 0 | - |
| - | - | 5278 | 660 | - | - | 0 | - |
| - | - | 9806 | 660.9 | - | - | 0 | - |
| - | - | 1.16E+04 | 661.4 | - | - | 0 | - |
| 2 | y | 5332 | 662.4 | 0.003575 | 5.397 | +3 | 17 |
| 2 | y | 4548 | 662.7 | 0.00155 | 2.339 | +3 | 17 |
| - | - | 5473 | 663.3 | - | - | 0 | - |
| - | - | 8344 | 663.4 | - | - | 0 | - |
| - | - | 2.938E+04 | 664.4 | - | - | 0 | - |
| - | - | 3.011E+04 | 664.5 | - | - | 0 | - |
| - | - | 3.184E+04 | 664.7 | - | - | 0 | - |
| - | - | 1.868E+04 | 665 | - | - | 0 | - |
| - | - | 6131 | 665.4 | - | - | 0 | - |
| - | - | 1.117E+04 | 665.5 | - | - | 0 | - |
| - | - | 6360 | 666.4 | - | - | 0 | - |
| - | - | 6122 | 667.3 | - | - | 0 | - |
| 2 | y | 2.05E+04 | 668.4 | 0.004623 | 6.916 | +3 | 17 |
| - | - | 3.207E+04 | 668.7 | - | - | 0 | - |
| - | - | 1.582E+04 | 669 | - | - | 0 | - |
| - | - | 3859 | 669.4 | - | - | 0 | - |
| - | - | 2.244E+04 | 671.9 | - | - | 0 | - |
| - | - | 1.42E+04 | 672.4 | - | - | 0 | - |
| - | - | 5779 | 672.9 | - | - | 0 | - |
| 13 | y | 1.074E+04 | 673.4 | 0.003034 | 4.506 | +1 | 6 |
| - | - | 1.291E+05 | 674.3 | - | - | 0 | - |
| - | - | 9.123E+04 | 674.4 | - | - | 0 | - |
| - | - | 5.044E+04 | 675.3 | - | - | 0 | - |
| - | - | 3.923E+04 | 675.4 | - | - | 0 | - |
| - | - | 1.198E+04 | 676.3 | - | - | 0 | - |
| - | - | 6238 | 676.4 | - | - | 0 | - |
| - | - | 1.025E+04 | 676.4 | - | - | 0 | - |
| - | - | 7433 | 676.7 | - | - | 0 | - |
| - | - | 6144 | 676.9 | - | - | 0 | - |
| - | - | 5991 | 677.4 | - | - | 0 | - |
| - | - | 4285 | 677.7 | - | - | 0 | - |
| - | - | 5685 | 678.4 | - | - | 0 | - |
| - | - | 5132 | 678.7 | - | - | 0 | - |
| - | - | 5606 | 679.4 | - | - | 0 | - |
| - | - | 6596 | 679.7 | - | - | 0 | - |
| - | - | 7334 | 680 | - | - | 0 | - |
| - | - | 2.094E+04 | 682 | - | - | 0 | - |
| - | - | 1.098E+04 | 682.4 | - | - | 0 | - |
| - | - | 2.063E+04 | 682.7 | - | - | 0 | - |
| - | - | 1.068E+04 | 683 | - | - | 0 | - |
| - | - | 7639 | 683.4 | - | - | 0 | - |
| - | - | 6188 | 683.7 | - | - | 0 | - |
| 5 | b | 2.812E+04 | 684.3 | 0.002023 | 2.956 | +1 | 5 |
| 5 | b | 1.858E+04 | 685.3 | 0.01001 | 14.61 | +1 | 5 |
| - | - | 1.572E+04 | 685.4 | - | - | 0 | - |
| - | - | 2.591E+04 | 685.7 | - | - | 0 | - |
| - | - | 7.794E+04 | 685.9 | - | - | 0 | - |
| - | - | 1.496E+04 | 686 | - | - | 0 | - |
| - | - | 5880 | 686.3 | - | - | 0 | - |
| 6 | y | 6.399E+04 | 686.4 | 0.01177 | 17.14 | +2 | 13 |
| - | - | 3.101E+04 | 686.9 | - | - | 0 | - |
| - | - | 1.227E+04 | 687.4 | - | - | 0 | - |
| - | - | 6176 | 687.7 | - | - | 0 | - |
| - | - | 5469 | 688 | - | - | 0 | - |
| - | - | 7958 | 690.5 | - | - | 0 | - |
| - | - | 5352 | 691 | - | - | 0 | - |
| 0 | Precursor | 1.874E+05 | 691.4 | 0.001679 | 2.429 | +3 | -1 |
| 0 | Precursor | 2.376E+05 | 691.7 | 0.007781 | 11.25 | +3 | -1 |
| - | - | 1.916E+05 | 692 | - | - | 0 | - |
| - | - | 8.724E+04 | 692.4 | - | - | 0 | - |
| - | - | 7.177E+04 | 692.4 | - | - | 0 | - |
| - | - | 3.308E+04 | 692.7 | - | - | 0 | - |
| - | - | 1.731E+04 | 693 | - | - | 0 | - |
| - | - | 3.48E+04 | 693.5 | - | - | 0 | - |
| - | - | 9928 | 694.5 | - | - | 0 | - |
| 6 | y | 5.082E+04 | 694.9 | 0.001386 | 1.994 | +2 | 13 |
| - | - | 4.343E+04 | 695.4 | - | - | 0 | - |
| - | - | 2.074E+04 | 695.9 | - | - | 0 | - |
| - | - | 1.188E+04 | 696.4 | - | - | 0 | - |
| - | - | 1.95E+04 | 696.7 | - | - | 0 | - |
| - | - | 5.064E+04 | 697 | - | - | 0 | - |
| 0 | Precursor | 2.838E+06 | 697.4 | 0.002064 | 2.96 | +3 | -1 |
| - | - | 3.589E+06 | 697.7 | - | - | 0 | - |
| - | - | 2.417E+06 | 698 | - | - | 0 | - |
| - | - | 1.103E+06 | 698.4 | - | - | 0 | - |
| - | - | 3.909E+05 | 698.7 | - | - | 0 | - |
| - | - | 1.105E+05 | 699 | - | - | 0 | - |
| 5 | b | 2.253E+05 | 702.3 | 0.00159 | 2.264 | +1 | 5 |
| - | - | 8.459E+04 | 703.3 | - | - | 0 | - |
| - | - | 2.476E+04 | 704.3 | - | - | 0 | - |
| - | - | 6560 | 704.4 | - | - | 0 | - |
| - | - | 1.238E+04 | 704.9 | - | - | 0 | - |
| - | - | 5260 | 705.3 | - | - | 0 | - |
| - | - | 1.11E+04 | 705.4 | - | - | 0 | - |
| - | - | 1.432E+04 | 707.3 | - | - | 0 | - |
| - | - | 6022 | 708.3 | - | - | 0 | - |
| - | - | 5768 | 708.4 | - | - | 0 | - |
| - | - | 2.588E+04 | 708.5 | - | - | 0 | - |
| 12 | b | 4.064E+04 | 709.4 | 0.0002798 | 0.3944 | +2 | 12 |
| - | - | 1.645E+04 | 709.5 | - | - | 0 | - |
| - | - | 4.009E+04 | 709.9 | - | - | 0 | - |
| - | - | 2.667E+04 | 710.4 | - | - | 0 | - |
| - | - | 1.37E+04 | 710.9 | - | - | 0 | - |
| - | - | 4995 | 711.4 | - | - | 0 | - |
| - | - | 9058 | 716.4 | - | - | 0 | - |
| - | - | 6691 | 719.9 | - | - | 0 | - |
| - | - | 1.097E+04 | 720.3 | - | - | 0 | - |
| - | - | 6178 | 720.4 | - | - | 0 | - |
| - | - | 9603 | 721.4 | - | - | 0 | - |
| - | - | 6112 | 723.4 | - | - | 0 | - |
| - | - | 1.567E+04 | 723.5 | - | - | 0 | - |
| - | - | 1.71E+04 | 724.3 | - | - | 0 | - |
| - | - | 7642 | 724.5 | - | - | 0 | - |
| - | - | 8021 | 725.3 | - | - | 0 | - |
| - | - | 6344 | 725.4 | - | - | 0 | - |
| - | - | 1.816E+04 | 728.4 | - | - | 0 | - |
| - | - | 1.638E+04 | 728.9 | - | - | 0 | - |
| - | - | 1.17E+04 | 729.4 | - | - | 0 | - |
| - | - | 1.151E+04 | 729.9 | - | - | 0 | - |
| - | - | 4396 | 732.5 | - | - | 0 | - |
| - | - | 5670 | 733.4 | - | - | 0 | - |
| - | - | 1.602E+04 | 735.3 | - | - | 0 | - |
| - | - | 5462 | 736.5 | - | - | 0 | - |
| - | - | 4103 | 737.4 | - | - | 0 | - |
| - | - | 1.121E+04 | 738.4 | - | - | 0 | - |
| - | - | 7991 | 739.4 | - | - | 0 | - |
| - | - | 4945 | 740.4 | - | - | 0 | - |
| - | - | 8244 | 741.4 | - | - | 0 | - |
| - | - | 1.044E+05 | 742.4 | - | - | 0 | - |
| - | - | 1.002E+05 | 742.9 | - | - | 0 | - |
| - | - | 3.894E+04 | 743.4 | - | - | 0 | - |
| - | - | 9823 | 743.9 | - | - | 0 | - |
| - | - | 7069 | 751.4 | - | - | 0 | - |
| - | - | 2.21E+04 | 751.9 | - | - | 0 | - |
| - | - | 2.021E+04 | 752.3 | - | - | 0 | - |
| - | - | 2.623E+04 | 752.4 | - | - | 0 | - |
| - | - | 7286 | 752.9 | - | - | 0 | - |
| - | - | 8859 | 753.3 | - | - | 0 | - |
| - | - | 4404 | 753.4 | - | - | 0 | - |
| - | - | 9875 | 753.5 | - | - | 0 | - |
| 5 | y | 8264 | 754.9 | 0.006872 | 9.103 | +2 | 14 |
| 13 | b | 4.608E+04 | 756.9 | 0.001616 | 2.135 | +2 | 13 |
| 13 | b | 4.825E+04 | 757.4 | 0.009974 | 13.17 | +2 | 13 |
| - | - | 2.247E+04 | 757.9 | - | - | 0 | - |
| - | - | 1.505E+04 | 758.4 | - | - | 0 | - |
| - | - | 5424 | 758.9 | - | - | 0 | - |
| - | - | 1.306E+04 | 759.9 | - | - | 0 | - |
| - | - | 1.07E+04 | 760.4 | - | - | 0 | - |
| - | - | 9311 | 760.5 | - | - | 0 | - |
| - | - | 1.094E+04 | 760.9 | - | - | 0 | - |
| - | - | 4485 | 762.9 | - | - | 0 | - |
| 5 | y | 8.682E+04 | 763.4 | 0.002081 | 2.726 | +2 | 14 |
| - | - | 5.91E+04 | 763.9 | - | - | 0 | - |
| - | - | 2.841E+04 | 764.5 | - | - | 0 | - |
| - | - | 1.661E+04 | 764.5 | - | - | 0 | - |
| - | - | 1.108E+04 | 764.9 | - | - | 0 | - |
| - | - | 7415 | 765.4 | - | - | 0 | - |
| 13 | b | 8.802E+04 | 765.9 | 0.001461 | 1.907 | +2 | 13 |
| - | - | 7.669E+04 | 766.4 | - | - | 0 | - |
| - | - | 3.498E+04 | 766.9 | - | - | 0 | - |
| - | - | 1.564E+04 | 767.4 | - | - | 0 | - |
| - | - | 5610 | 767.9 | - | - | 0 | - |
| - | - | 1.376E+04 | 768.9 | - | - | 0 | - |
| - | - | 4847 | 769.4 | - | - | 0 | - |
| - | - | 5036 | 769.7 | - | - | 0 | - |
| - | - | 5384 | 770.4 | - | - | 0 | - |
| - | - | 5297 | 776.9 | - | - | 0 | - |
| - | - | 1.135E+04 | 777.4 | - | - | 0 | - |
| - | - | 3.011E+04 | 777.5 | - | - | 0 | - |
| - | - | 1.218E+04 | 778.5 | - | - | 0 | - |
| - | - | 1.216E+04 | 778.9 | - | - | 0 | - |
| - | - | 8814 | 779.5 | - | - | 0 | - |
| - | - | 5247 | 779.9 | - | - | 0 | - |
| - | - | 9108 | 780.3 | - | - | 0 | - |
| - | - | 5153 | 780.9 | - | - | 0 | - |
| - | - | 4357 | 781.3 | - | - | 0 | - |
| - | - | 4.394E+04 | 781.5 | - | - | 0 | - |
| - | - | 2.153E+04 | 782.5 | - | - | 0 | - |
| 12 | y | 1.249E+04 | 783.5 | 0.005768 | 7.361 | +1 | 7 |
| - | - | 8090 | 785.4 | - | - | 0 | - |
| - | - | 4.76E+04 | 785.9 | - | - | 0 | - |
| - | - | 4.78E+04 | 786.4 | - | - | 0 | - |
| - | - | 2.017E+04 | 786.9 | - | - | 0 | - |
| - | - | 4300 | 787.5 | - | - | 0 | - |
| - | - | 1.829E+04 | 787.5 | - | - | 0 | - |
| - | - | 1.09E+04 | 788.5 | - | - | 0 | - |
| - | - | 6722 | 792.4 | - | - | 0 | - |
| - | - | 1.012E+04 | 793.4 | - | - | 0 | - |
| - | - | 1.526E+04 | 793.9 | - | - | 0 | - |
| - | - | 2.547E+04 | 794.3 | - | - | 0 | - |
| - | - | 1.295E+04 | 794.4 | - | - | 0 | - |
| - | - | 1.182E+04 | 794.9 | - | - | 0 | - |
| - | - | 1.507E+04 | 795.4 | - | - | 0 | - |
| - | - | 5975 | 795.4 | - | - | 0 | - |
| - | - | 9195 | 799.9 | - | - | 0 | - |
| - | - | 4676 | 800.9 | - | - | 0 | - |
| - | - | 6044 | 801.4 | - | - | 0 | - |
| 12 | y | 1.325E+04 | 801.5 | 0.002894 | 3.61 | +1 | 7 |
| - | - | 6173 | 802.5 | - | - | 0 | - |
| - | - | 6298 | 804.3 | - | - | 0 | - |
| - | - | 5582 | 805.4 | - | - | 0 | - |
| - | - | 1.39E+05 | 805.5 | - | - | 0 | - |
| - | - | 5.98E+04 | 806.5 | - | - | 0 | - |
| - | - | 1.783E+04 | 807.5 | - | - | 0 | - |
| - | - | 2.023E+04 | 808.4 | - | - | 0 | - |
| - | - | 5725 | 808.5 | - | - | 0 | - |
| - | - | 2.942E+04 | 808.9 | - | - | 0 | - |
| - | - | 1.711E+04 | 809.4 | - | - | 0 | - |
| - | - | 1.922E+04 | 809.9 | - | - | 0 | - |
| - | - | 5954 | 810.4 | - | - | 0 | - |
| - | - | 1.978E+05 | 811.4 | - | - | 0 | - |
| - | - | 1.976E+04 | 811.5 | - | - | 0 | - |
| - | - | 1.051E+05 | 812.4 | - | - | 0 | - |
| - | - | 8965 | 812.5 | - | - | 0 | - |
| - | - | 1.957E+04 | 813.4 | - | - | 0 | - |
| 14 | b | 1.268E+04 | 813.4 | 0.004689 | 5.764 | +2 | 14 |
| 14 | b | 2.624E+04 | 813.9 | 0.007127 | 8.756 | +2 | 14 |
| - | - | 1.149E+04 | 814.4 | - | - | 0 | - |
| - | - | 9094 | 814.9 | - | - | 0 | - |
| - | - | 5498 | 817.4 | - | - | 0 | - |
| - | - | 9231 | 817.9 | - | - | 0 | - |
| - | - | 3798 | 818.4 | - | - | 0 | - |
| 4 | y | 2.065E+04 | 818.5 | 0.003465 | 4.234 | +2 | 15 |
| 4 | y | 5.095E+04 | 819 | 0.005293 | 6.463 | +2 | 15 |
| - | - | 4.187E+04 | 819.5 | - | - | 0 | - |
| - | - | 2.467E+04 | 820 | - | - | 0 | - |
| - | - | 1.026E+04 | 820.5 | - | - | 0 | - |
| 6 | b | 2.64E+04 | 821.3 | 0.001216 | 1.481 | +1 | 6 |
| - | - | 3.204E+04 | 821.6 | - | - | 0 | - |
| - | - | 7910 | 821.9 | - | - | 0 | - |
| 6 | b | 2.348E+04 | 822.3 | 0.01415 | 17.21 | +1 | 6 |
| 14 | b | 1.702E+05 | 822.4 | 0.001177 | 1.431 | +2 | 14 |
| - | - | 1.452E+04 | 822.6 | - | - | 0 | - |
| - | - | 1.711E+05 | 822.9 | - | - | 0 | - |
| - | - | 5296 | 823.4 | - | - | 0 | - |
| - | - | 8.698E+04 | 823.4 | - | - | 0 | - |
| - | - | 3.805E+04 | 823.9 | - | - | 0 | - |
| - | - | 1.041E+04 | 824.5 | - | - | 0 | - |
| 4 | y | 6.482E+04 | 827.5 | 0.0003803 | 0.4596 | +2 | 15 |
| - | - | 5.84E+04 | 828 | - | - | 0 | - |
| - | - | 3.448E+04 | 828.5 | - | - | 0 | - |
| - | - | 1.386E+04 | 829 | - | - | 0 | - |
| - | - | 1.046E+04 | 829.5 | - | - | 0 | - |
| - | - | 4944 | 830.4 | - | - | 0 | - |
| - | - | 7428 | 830.5 | - | - | 0 | - |
| - | - | 5206 | 833 | - | - | 0 | - |
| - | - | 8793 | 833.5 | - | - | 0 | - |
| - | - | 6098 | 834.9 | - | - | 0 | - |
| - | - | 1.266E+04 | 835.5 | - | - | 0 | - |
| - | - | 7635 | 836 | - | - | 0 | - |
| - | - | 1.451E+04 | 836.5 | - | - | 0 | - |
| - | - | 1.106E+04 | 837 | - | - | 0 | - |
| 6 | b | 4.185E+05 | 839.4 | 0.001699 | 2.024 | +1 | 6 |
| - | - | 2.183E+05 | 840.4 | - | - | 0 | - |
| - | - | 4605 | 840.4 | - | - | 0 | - |
| - | - | 5.38E+04 | 841.4 | - | - | 0 | - |
| - | - | 2.233E+04 | 841.5 | - | - | 0 | - |
| - | - | 2.067E+04 | 842 | - | - | 0 | - |
| - | - | 8963 | 842.4 | - | - | 0 | - |
| - | - | 9973 | 842.5 | - | - | 0 | - |
| - | - | 6156 | 842.9 | - | - | 0 | - |
| - | - | 5277 | 843.4 | - | - | 0 | - |
| - | - | 2.847E+04 | 848.4 | - | - | 0 | - |
| - | - | 1.553E+04 | 849.4 | - | - | 0 | - |
| - | - | 8.514E+04 | 850.5 | - | - | 0 | - |
| - | - | 7.972E+04 | 851 | - | - | 0 | - |
| - | - | 4.86E+04 | 851.5 | - | - | 0 | - |
| - | - | 5537 | 851.6 | - | - | 0 | - |
| - | - | 1.612E+04 | 852 | - | - | 0 | - |
| - | - | 9288 | 852.5 | - | - | 0 | - |
| - | - | 1.009E+04 | 852.5 | - | - | 0 | - |
| - | - | 1.038E+04 | 856.4 | - | - | 0 | - |
| - | - | 1.687E+04 | 856.9 | - | - | 0 | - |
| - | - | 6076 | 857.4 | - | - | 0 | - |
| - | - | 6308 | 859.5 | - | - | 0 | - |
| - | - | 7902 | 860 | - | - | 0 | - |
| - | - | 4796 | 860.5 | - | - | 0 | - |
| - | - | 7954 | 864.5 | - | - | 0 | - |
| - | - | 1.656E+04 | 865 | - | - | 0 | - |
| - | - | 1.876E+04 | 865.5 | - | - | 0 | - |
| - | - | 2.23E+04 | 866 | - | - | 0 | - |
| - | - | 2.086E+04 | 866.5 | - | - | 0 | - |
| - | - | 1.147E+04 | 867 | - | - | 0 | - |
| 15 | b | 1.028E+04 | 870 | 0.0006608 | 0.7596 | +2 | 15 |
| 15 | b | 1.225E+04 | 870.5 | 0.007576 | 8.703 | +2 | 15 |
| - | - | 1.674E+04 | 871 | - | - | 0 | - |
| - | - | 5406 | 871.5 | - | - | 0 | - |
| - | - | 7389 | 872 | - | - | 0 | - |
| - | - | 5566 | 873.6 | - | - | 0 | - |
| - | - | 2.374E+04 | 873.9 | - | - | 0 | - |
| - | - | 1.962E+04 | 874.4 | - | - | 0 | - |
| - | - | 1.12E+04 | 874.9 | - | - | 0 | - |
| - | - | 5426 | 875.4 | - | - | 0 | - |
| - | - | 1.046E+04 | 878 | - | - | 0 | - |
| - | - | 1.256E+04 | 878.5 | - | - | 0 | - |
| 15 | b | 4.035E+05 | 879 | 0.001503 | 1.71 | +2 | 15 |
| - | - | 3.757E+05 | 879.5 | - | - | 0 | - |
| - | - | 2.065E+05 | 880 | - | - | 0 | - |
| - | - | 7.081E+04 | 880.5 | - | - | 0 | - |
| - | - | 2.129E+04 | 881 | - | - | 0 | - |
| - | - | 5301 | 887 | - | - | 0 | - |
| - | - | 5461 | 887.5 | - | - | 0 | - |
| - | - | 2.136E+04 | 890.6 | - | - | 0 | - |
| - | - | 1.669E+04 | 891.5 | - | - | 0 | - |
| - | - | 9692 | 891.6 | - | - | 0 | - |
| - | - | 8979 | 892 | - | - | 0 | - |
| - | - | 6533 | 892.5 | - | - | 0 | - |
| - | - | 1.268E+04 | 892.6 | - | - | 0 | - |
| - | - | 8995 | 893.6 | - | - | 0 | - |
| 11 | y | 1.164E+05 | 898.6 | 0.001705 | 1.897 | +1 | 8 |
| - | - | 3170 | 899.5 | - | - | 0 | - |
| - | - | 6.153E+04 | 899.6 | - | - | 0 | - |
| 3 | y | 3.387E+04 | 900 | 0.0009759 | 1.084 | +2 | 16 |
| 3 | y | 3.589E+04 | 900.5 | 0.007808 | 8.671 | +2 | 16 |
| - | - | 1.181E+04 | 900.6 | - | - | 0 | - |
| - | - | 2.021E+04 | 901 | - | - | 0 | - |
| - | - | 2.672E+04 | 901.4 | - | - | 0 | - |
| - | - | 8183 | 901.6 | - | - | 0 | - |
| - | - | 5925 | 902 | - | - | 0 | - |
| - | - | 2.114E+04 | 902.4 | - | - | 0 | - |
| - | - | 7658 | 903.5 | - | - | 0 | - |
| - | - | 1.47E+04 | 908.5 | - | - | 0 | - |
| - | - | 9585 | 908.6 | - | - | 0 | - |
| 3 | y | 3.99E+05 | 909 | 0.00137 | 1.507 | +2 | 16 |
| - | - | 4.175E+05 | 909.5 | - | - | 0 | - |
| - | - | 2.436E+05 | 910 | - | - | 0 | - |
| - | - | 9.589E+04 | 910.5 | - | - | 0 | - |
| - | - | 2.261E+04 | 911 | - | - | 0 | - |
| - | - | 6494 | 911.5 | - | - | 0 | - |
| - | - | 1.688E+04 | 913 | - | - | 0 | - |
| 16 | b | 3.802E+04 | 913.5 | 0.009656 | 10.57 | +2 | 16 |
| 16 | b | 3.129E+04 | 914 | 0.003463 | 3.789 | +2 | 16 |
| - | - | 2.002E+04 | 914.5 | - | - | 0 | - |
| - | - | 4497 | 914.6 | - | - | 0 | - |
| - | - | 9326 | 915 | - | - | 0 | - |
| - | - | 5480 | 915.5 | - | - | 0 | - |
| - | - | 8826 | 916.5 | - | - | 0 | - |
| - | - | 6305 | 917 | - | - | 0 | - |
| - | - | 2.074E+05 | 918.6 | - | - | 0 | - |
| - | - | 2.118E+04 | 919.5 | - | - | 0 | - |
| - | - | 1.264E+05 | 919.6 | - | - | 0 | - |
| - | - | 1.144E+04 | 920.5 | - | - | 0 | - |
| - | - | 3.61E+04 | 920.6 | - | - | 0 | - |
| - | - | 2.319E+04 | 921.5 | - | - | 0 | - |
| - | - | 7909 | 921.6 | - | - | 0 | - |
| - | - | 1.431E+04 | 922 | - | - | 0 | - |
| 16 | b | 1.515E+05 | 922.5 | 0.0001374 | 0.1489 | +2 | 16 |
| - | - | 2.023E+05 | 923 | - | - | 0 | - |
| - | - | 1.122E+05 | 923.5 | - | - | 0 | - |
| - | - | 4.159E+04 | 924 | - | - | 0 | - |
| - | - | 9335 | 924.6 | - | - | 0 | - |
| - | - | 6273 | 925.6 | - | - | 0 | - |
| - | - | 4.153E+04 | 930.5 | - | - | 0 | - |
| - | - | 3.43E+04 | 931 | - | - | 0 | - |
| - | - | 1.87E+04 | 931.5 | - | - | 0 | - |
| - | - | 1.047E+04 | 932 | - | - | 0 | - |
| 7 | b | 7950 | 936.4 | 0.001303 | 1.392 | +1 | 7 |
| - | - | 5829 | 937.4 | - | - | 0 | - |
| - | - | 2.12E+04 | 942.6 | - | - | 0 | - |
| - | - | 5811 | 943.5 | - | - | 0 | - |
| - | - | 7929 | 943.6 | - | - | 0 | - |
| - | - | 7478 | 944.5 | - | - | 0 | - |
| - | - | 6466 | 948.6 | - | - | 0 | - |
| - | - | 9001 | 966.6 | - | - | 0 | - |
| - | - | 7708 | 967.6 | - | - | 0 | - |
| - | - | 5221 | 969.5 | - | - | 0 | - |
| 10 | y | 5.326E+04 | 969.6 | 0.002066 | 2.131 | +1 | 9 |
| - | - | 3.71E+04 | 970.6 | - | - | 0 | - |
| - | - | 9001 | 971.6 | - | - | 0 | - |
| - | - | 7789 | 973 | - | - | 0 | - |
| - | - | 9037 | 973.5 | - | - | 0 | - |
| - | - | 8162 | 974 | - | - | 0 | - |
| 17 | b | 3.713E+04 | 978 | 0.00219 | 2.239 | +2 | 17 |
| 17 | b | 4.462E+04 | 978.5 | 0.009938 | 10.16 | +2 | 17 |
| - | - | 2.983E+04 | 979 | - | - | 0 | - |
| - | - | 8145 | 979.5 | - | - | 0 | - |
| - | - | 5522 | 980 | - | - | 0 | - |
| - | - | 5955 | 980.5 | - | - | 0 | - |
| - | - | 5158 | 981.5 | - | - | 0 | - |
| 17 | b | 2.338E+05 | 987 | 0.001668 | 1.69 | +2 | 17 |
| - | - | 2.914E+05 | 987.5 | - | - | 0 | - |
| - | - | 1.434E+04 | 987.6 | - | - | 0 | - |
| - | - | 1.619E+05 | 988 | - | - | 0 | - |
| - | - | 6.647E+04 | 988.5 | - | - | 0 | - |
| - | - | 7899 | 988.6 | - | - | 0 | - |
| - | - | 2.326E+04 | 989 | - | - | 0 | - |
| 8 | b | 1.785E+04 | 993.4 | 0.001751 | 1.763 | +1 | 8 |
| - | - | 1.118E+04 | 994.4 | - | - | 0 | - |
| - | - | 2.311E+04 | 996 | - | - | 0 | - |
| - | - | 3.003E+04 | 996.5 | - | - | 0 | - |
| - | - | 1.768E+04 | 997 | - | - | 0 | - |
| - | - | 6505 | 997.5 | - | - | 0 | - |
| - | - | 5574 | 1004 | - | - | 0 | - |
| - | - | 7.141E+04 | 1006 | - | - | 0 | - |
| - | - | 5.314E+04 | 1007 | - | - | 0 | - |
| - | - | 1.042E+04 | 1008 | - | - | 0 | - |
| - | - | 5943 | 1010 | - | - | 0 | - |
| - | - | 1.806E+04 | 1010 | - | - | 0 | - |
| - | - | 8015 | 1011 | - | - | 0 | - |
| - | - | 1.867E+04 | 1022 | - | - | 0 | - |
| - | - | 1.231E+04 | 1023 | - | - | 0 | - |
| - | - | 1.007E+04 | 1033 | - | - | 0 | - |
| - | - | 1.429E+04 | 1034 | - | - | 0 | - |
| - | - | 5109 | 1036 | - | - | 0 | - |
| - | - | 1.793E+04 | 1038 | - | - | 0 | - |
| - | - | 1.457E+04 | 1039 | - | - | 0 | - |
| - | - | 3.443E+04 | 1056 | - | - | 0 | - |
| - | - | 2.221E+04 | 1057 | - | - | 0 | - |
| - | - | 1.333E+04 | 1080 | - | - | 0 | - |
| 9 | y | 1.055E+04 | 1081 | 0.01203 | 11.13 | +1 | 10 |
| - | - | 5513 | 1089 | - | - | 0 | - |
| 9 | y | 1.927E+04 | 1098 | 0.0009777 | 0.8906 | +1 | 10 |
| - | - | 1.395E+04 | 1099 | - | - | 0 | - |
| - | - | 1.069E+04 | 1107 | - | - | 0 | - |
| - | - | 7788 | 1108 | - | - | 0 | - |
| - | - | 1.916E+04 | 1117 | - | - | 0 | - |
| - | - | 1.042E+04 | 1118 | - | - | 0 | - |
| - | - | 7182 | 1119 | - | - | 0 | - |
| 9 | b | 3.033E+04 | 1122 | 0.001881 | 1.677 | +1 | 9 |
| - | - | 1.639E+04 | 1123 | - | - | 0 | - |
| - | - | 8419 | 1124 | - | - | 0 | - |
| - | - | 5318 | 1126 | - | - | 0 | - |
| - | - | 8.464E+04 | 1135 | - | - | 0 | - |
| - | - | 5.022E+04 | 1136 | - | - | 0 | - |
| 8 | y | 2.316E+04 | 1137 | 0.004797 | 4.22 | +1 | 11 |
| 8 | y | 7477 | 1138 | 0.01082 | 9.512 | +1 | 11 |
| - | - | 1.244E+04 | 1143 | - | - | 0 | - |
| - | - | 8908 | 1145 | - | - | 0 | - |
| - | - | 1.32E+04 | 1146 | - | - | 0 | - |
| - | - | 1.017E+04 | 1147 | - | - | 0 | - |
| - | - | 5495 | 1153 | - | - | 0 | - |
| - | - | 6049 | 1154 | - | - | 0 | - |
| 8 | y | 2.711E+04 | 1155 | 0.001973 | 1.708 | +1 | 11 |
| - | - | 1.628E+04 | 1156 | - | - | 0 | - |
| - | - | 7524 | 1165 | - | - | 0 | - |
| 10 | b | 4.418E+04 | 1175 | 0.001211 | 1.031 | +1 | 10 |
| 10 | b | 3.362E+04 | 1176 | 0.01964 | 16.7 | +1 | 10 |
| - | - | 1.649E+04 | 1177 | - | - | 0 | - |
| - | - | 5165 | 1178 | - | - | 0 | - |
| 10 | b | 2.472E+04 | 1193 | 0.0005336 | 0.4474 | +1 | 10 |
| - | - | 1.899E+04 | 1193 | - | - | 0 | - |
| - | - | 1.959E+04 | 1194 | - | - | 0 | - |
| - | - | 1.179E+04 | 1194 | - | - | 0 | - |
| - | - | 5370 | 1208 | - | - | 0 | - |
| 7 | y | 1.58E+04 | 1234 | 0.002751 | 2.23 | +1 | 12 |
| - | - | 7154 | 1235 | - | - | 0 | - |
| 7 | y | 1.23E+04 | 1235 | 0.02202 | 17.84 | +1 | 12 |
| - | - | 6740 | 1236 | - | - | 0 | - |
| - | - | 4976 | 1240 | - | - | 0 | - |
| - | - | 6288 | 1241 | - | - | 0 | - |
| 7 | y | 1.255E+05 | 1252 | 0.0004784 | 0.3822 | +1 | 12 |
| - | - | 9.092E+04 | 1253 | - | - | 0 | - |
| - | - | 3.927E+04 | 1254 | - | - | 0 | - |
| - | - | 1.282E+04 | 1255 | - | - | 0 | - |
| - | - | 5667 | 1256 | - | - | 0 | - |
| - | - | 1.346E+04 | 1258 | - | - | 0 | - |
| - | - | 9334 | 1259 | - | - | 0 | - |
| - | - | 5501 | 1260 | - | - | 0 | - |
| - | - | 5871 | 1261 | - | - | 0 | - |
| - | - | 9589 | 1272 | - | - | 0 | - |
| - | - | 8706 | 1273 | - | - | 0 | - |
| - | - | 1.214E+04 | 1306 | - | - | 0 | - |
| - | - | 2.117E+04 | 1307 | - | - | 0 | - |
| - | - | 1.442E+04 | 1308 | - | - | 0 | - |
| - | - | 1.323E+04 | 1371 | - | - | 0 | - |
| 6 | y | 1.09E+04 | 1372 | 0.02187 | 15.95 | +1 | 13 |
| - | - | 4546 | 1373 | - | - | 0 | - |
| 6 | y | 7526 | 1389 | 0.001137 | 0.8185 | +1 | 13 |
| - | - | 9507 | 1390 | - | - | 0 | - |
| - | - | 4154 | 1391 | - | - | 0 | - |
| - | - | 6219 | 1409 | - | - | 0 | - |
| - | - | 5105 | 1410 | - | - | 0 | - |
| 12 | b | 6095 | 1418 | 0.005592 | 3.944 | +1 | 12 |
| - | - | 9528 | 1419 | - | - | 0 | - |
| - | - | 1.122E+04 | 1420 | - | - | 0 | - |
| - | - | 4798 | 1533 | - | - | 0 | - |
| - | - | 3627 | 2170 | - | - | 0 | - |

m/z Charge Intensity FragmentType MassShift Position
120.05614471435547 0 2374.6692
120.08135223388672 0 3831.4648
120.09076690673828 0 2000.436
120.15461730957031 0 1927.64
120.2545166015625 0 2145.951
122.07200622558594 0 6202.9766
123.05584716796875 0 8368.218
126.0554428100586 0 47677.934
127.05854797363281 0 4120.071
127.08712768554688 0 10540.783
128.03488159179688 0 2210.5518
128.10755920410156 0 2563.137
129.06637573242188 0 20253.324
129.10275268554688 0 273822.03
130.05055236816406 0 4785.1553
130.06564331054688 0 121795.29
130.10093688964844 0 3552.8499
130.1060791015625 0 18812.371
130.10997009277344 0 1933.0543
131.0689697265625 0 12690.692
131.11825561523438 0 3693.1733
132.081298828125 0 441858.66
133.08468627929688 0 47255.64
134.07183837890625 0 3916.254
136.07620239257812 0 282059.25
137.07952880859375 0 23431.896
138.06663513183594 0 13749.715
138.09190368652344 0 3675.4033
139.05052185058594 0 2538.637
141.10279846191406 0 23868.06
143.11871337890625 0 2141.1772
144.06625366210938 0 12544.492
144.0813446044922 0 5951.512
146.060302734375 0 2483.8843
146.0931854248047 0 3897.9343
146.12925720214844 0 3253.7344
147.04441833496094 0 4543.8506
148.06094360351562 0 3778.5469
148.94613647460938 0 2209.0488
151.08724975585938 0 6434.403
152.08245849609375 0 23639.334
152.14389038085938 0 29155.05
153.06640625 0 9152.047
153.09698486328125 0 2459.93
153.10287475585938 0 14697.266
153.14730834960938 0 2594.962
154.04421997070312 0 2243.4297
154.05039978027344 0 57968.54
154.1594696044922 0 11594.096
155.05392456054688 0 4806.494
155.0819854736328 0 20623.076
155.09329223632812 0 15401.032
155.09906005859375 0 1981.3433
155.1184844970703 0 26938.98
156.07748413085938 0 18849.281
156.0962677001953 0 2379.9224
156.12196350097656 0 3089.503
157.0760498046875 0 3308.9983
157.0977020263672 0 3447.8992
158.0606689453125 0 5932.08
158.08438110351562 0 35776.66
159.09219360351562 0 182720.7
159.11317443847656 0 6285.1895
160.07595825195312 0 9134.484
160.09054565429688 0 1977.5507
160.0955810546875 0 22749.951
165.06640625 0 3231.4546
165.10276794433594 0 10910.016
166.06161499023438 0 6958.513
166.09829711914062 0 2549.2573
168.1024932861328 0 10199.175
168.11375427246094 0 6280.5747
169.07656860351562 0 28363.053
169.09774780273438 0 35537.46
170.0460662841797 0 4745.104
170.06063842773438 0 61022.97
171.06442260742188 0 5416.1055
171.0769805908203 0 64781.98
171.09181213378906 0 2986.9946
171.11334228515625 0 23660.88
172.06092834472656 0 6367.506
172.0803680419922 0 2860.9128
172.1085968017578 0 32216.994
173.1290283203125 0 27217.898
173.4512176513672 0 12670.98
174.1325225830078 0 4212.7026
175.09835815429688 0 5085.102
176.0823211669922 0 8833.652
178.13465881347656 0 3414.5955
179.08241271972656 0 3084.6367
179.15476989746094 0 2654.4207
180.07737731933594 0 8291.933
181.06124877929688 0 7848.645
181.0977783203125 0 86931.37
181.13418579101562 0 4779.3516
181.17066955566406 0 3392.6663
182.10121154785156 0 9338.865
182.1293487548828 0 15221.205
182.1373291015625 0 2765.563
183.09231567382812 0 63589.04
183.11341857910156 0 243556.56
184.0762481689453 0 7215.268
184.09576416015625 0 9441.122
184.11679077148438 0 29126.367
185.07159423828125 0 17308.486
185.08070373535156 0 3271.9832
185.11911010742188 0 3224.0166
186.0794677734375 0 2958.1606
186.12428283691406 0 57495.445
187.0872039794922 0 463020.4
187.1280517578125 0 5426.2236
188.090576171875 0 60923.902
189.08761596679688 0 327690.78 b 3
190.09109497070312 0 28449.863
191.09255981445312 0 4498.897
195.11354064941406 0 13632.233
195.1497344970703 0 6585.1333
197.16539001464844 0 43777.617
198.16934204101562 0 5465.182
199.07196044921875 0 35430.19
199.08694458007812 0 10371.778
199.10816955566406 0 6659.5566
199.1810302734375 0 169089.9
200.07542419433594 0 3142.944
200.1400909423828 0 10657.9795
200.18446350097656 0 21128.455
201.1027374267578 0 36904.645
201.12396240234375 0 43511.625
202.0871124267578 0 5137.2144
202.10911560058594 0 55358.64
202.1265869140625 0 3400.1985
203.11293029785156 0 6810.814
204.0773468017578 0 3073.1724
205.10897827148438 0 5640.725
207.12448120117188 0 2910.506
207.14976501464844 0 17805.094
208.14471435546875 0 3912.2659
209.1038360595703 0 3091.214
209.12913513183594 0 5603.7114
209.16529846191406 0 5937.837
211.0870819091797 0 9657.546
211.10830688476562 0 27305.9
211.14476013183594 0 3748.1763
212.11221313476562 0 3172.5898
213.12408447265625 0 4799.748
214.1094207763672 0 23941.85
214.11932373046875 0 3333.4868
214.19236755371094 0 2967.409
216.098388671875 0 3666
216.1134033203125 0 5947.8604
217.08258056640625 0 542030.1
217.10804748535156 0 10844.928
218.08592224121094 0 49556.97
218.15072631835938 0 3043.1477
219.08824157714844 0 12353.38
220.11996459960938 0 37270.207
221.10398864746094 0 61994.832
221.1224365234375 0 4460.0635
221.16603088378906 0 2852.0957
222.1074676513672 0 4921.6416
222.1243438720703 0 3279.694
222.13546752929688 0 3695.6084
223.10824584960938 0 4343.141
223.1443328857422 0 4524.388
223.18218994140625 0 2460.393
224.11883544921875 0 13437.133
224.17630004882812 0 26696.793
225.0990753173828 0 8651.425
225.12261962890625 0 3840.4216
225.16098022460938 0 5768.6885
226.1557159423828 0 328950.94
226.96258544921875 0 2752.6958
227.0671844482422 0 3009.7026
227.0828094482422 0 2730.9722
227.15916442871094 0 31820.346
227.1761016845703 0 108417.5
228.1138458251953 0 36799.773 a Water loss 1
228.13429260253906 0 3943.535
228.17947387695312 0 13005.144
229.097900390625 0 47785.05 b Ammonia loss 4
229.1189727783203 0 492690.8 y Water loss 16
230.1028289794922 0 11770.301
230.12225341796875 0 56412.875
231.08819580078125 0 20381.195
231.1245880126953 0 3190.4382
231.13607788085938 0 7830.09
232.0922088623047 0 2719.154
232.11917114257812 0 4627.7085
233.10426330566406 0 9274.563
233.1640625 0 2604.8628
234.1357421875 0 5704.6963
235.0930633544922 0 7098.207
235.1197052001953 0 118040.29
236.12310791015625 0 9986.122
236.14048767089844 0 4020.8376
237.12423706054688 0 3202.3093
237.1602020263672 0 10654.889
238.13070678710938 0 14893.162
239.15101623535156 0 28969.928
239.17616271972656 0 5183.6084
240.13453674316406 0 4212.756
241.15567016601562 0 6375.1606
241.19212341308594 0 3505.3472
242.1868133544922 0 10561.314
243.18206787109375 0 11154.67
244.1309814453125 0 6593.723
244.1838836669922 0 2770.126
245.18544006347656 0 2667.1401
246.12452697753906 0 1248843 a 1
247.12936401367188 0 400876.6 y 16
248.11477661132812 0 72155.195
248.13217163085938 0 36611.04
249.09898376464844 0 83654.625
249.1156005859375 0 5473.512
249.11972045898438 0 6146.706
249.1353759765625 0 7539.696
250.1020965576172 0 10031.67
250.1221923828125 0 4364.654
251.10398864746094 0 3209.7344
251.12945556640625 0 7314.142
251.15142822265625 0 2981.0679
251.17518615722656 0 2717.7183
252.17140197753906 0 25489.197
254.18739318847656 0 5639.7817
255.0768280029297 0 5528.2407
255.1714324951172 0 2849.781
256.0799865722656 0 2456.457
256.10882568359375 0 26869.576 b Water loss 1
257.0925598144531 0 5715.0796
257.1148376464844 0 8697.631
257.14862060546875 0 3474.0752
257.16156005859375 0 8550.355
258.0992431640625 0 165622.03
259.102294921875 0 22673.203
259.1306457519531 0 8354.951
259.21295166015625 0 4344.7188
260.1076965332031 0 6528.848
263.11456298828125 0 7131.621
264.1352233886719 0 6081.6772
265.1204528808594 0 3565.054
265.14105224609375 0 8078.8916
265.16571044921875 0 5270.306
265.2287902832031 0 6789.1714
266.1253967285156 0 93938.82
267.12884521484375 0 13119.97
267.14501953125 0 5055.1177
267.2436828613281 0 7339.6826
268.1664123535156 0 5329.257
269.1139831542969 0 21182.441
270.11895751953125 0 4188.3057
270.1451416015625 0 3985.2637
270.1820983886719 0 44585.7
271.1851501464844 0 5215.2466
273.1097412109375 0 3101.1729
274.1193542480469 0 1005013.44 b 1
275.1039733886719 0 16909.402
275.12469482421875 0 416667.53
276.1084899902344 0 10024.403
276.1281433105469 0 49778.082
277.0935974121094 0 4074.1584
277.1335754394531 0 10808.613
277.2292175292969 0 3603.0322
278.1514587402344 0 3384.3408
279.1824951171875 0 12311.215
282.18194580078125 0 3608.7317
283.1515197753906 0 18796.703
283.17730712890625 0 130845.125
284.1642761230469 0 4444.532
284.1805725097656 0 17863.559
285.1103820800781 0 9306.363
285.1566467285156 0 28056.736
286.1165466308594 0 3253.4368
286.16009521484375 0 5194.881
290.1142578125 0 4877.0747
290.1853332519531 0 7141.9097
291.1576843261719 0 10223.455
292.13006591796875 0 96473.086
293.1337890625 0 14065.076
293.1614990234375 0 3519.5227
294.1205139160156 0 41012.395
294.2184143066406 0 12341.606
295.1246643066406 0 3664.0786
295.1763916015625 0 6572.2505
295.2395935058594 0 3694.67
296.1976623535156 0 14639.808
297.1932678222656 0 51040.26
298.140625 0 14469.06
298.19635009765625 0 7876.2124
299.2077331542969 0 2618.4568
300.2043151855469 0 5927.573
303.1208190917969 0 20646.488
303.21954345703125 0 3889.4937
304.1256408691406 0 2806.669
305.1293029785156 0 20007.514
305.6646423339844 0 4031.0254
308.125732421875 0 8927.18
309.16986083984375 0 3375.1265
309.1934814453125 0 6682.158
310.1767578125 0 4281.4053
311.24517822265625 0 21147.328
312.1564025878906 0 40497.12
312.248779296875 0 3567.105
313.1597900390625 0 7662.7666
313.1883544921875 0 9558.146
314.135986328125 0 3434.4753
314.20849609375 0 12304.055
314.66925048828125 0 4453.081
315.2109375 0 4205.3726
316.1513977050781 0 63865.348 y Water loss 15
317.15484619140625 0 8331.406
320.1250915527344 0 38782.09
320.1504821777344 0 8282.986
321.1297912597656 0 7346.475
321.2294616699219 0 106501.57
322.1560974121094 0 35115.613
322.2326354980469 0 23062.203
323.1595458984375 0 10425.827
323.2341613769531 0 2904.6975
325.224853515625 0 6154.477
326.13616943359375 0 3777.4487 b Ammonia loss 7
326.2198791503906 0 25747.736
327.2028503417969 0 3589.11
327.2231140136719 0 5782.271
327.2770080566406 0 3156.0908
328.2082824707031 0 2931.0547 y Water loss 12
329.13543701171875 0 3239.8965
329.2194519042969 0 2911.128
330.1669006347656 0 22554.217
330.21453857421875 0 4230.6855
331.1690368652344 0 3605.0095
332.7299499511719 0 4289.6836
333.1242980957031 0 6763.3086
334.1619567871094 0 182369.34 y 15
335.16510009765625 0 24080.514
336.14691162109375 0 9267.521
336.16693115234375 0 5607.8784
336.2041015625 0 125681.48
337.18792724609375 0 5742.475
337.2075500488281 0 18055.645 y 12
337.2605895996094 0 6095.9644
337.7221374511719 0 10269.486
338.1620788574219 0 9204.282
338.2253112792969 0 3749.5613
338.2458190917969 0 9979.53
339.1663818359375 0 3553.686
339.20428466796875 0 5821.692
339.2402038574219 0 82011.96
340.15240478515625 0 10628.137
340.24273681640625 0 15603.431
340.2610168457031 0 9353.428
341.2197570800781 0 5791.683
342.1680908203125 0 3572.997
344.1440734863281 0 3328.2185
346.72650146484375 0 6130.2837
347.2272033691406 0 4563.803
348.1470642089844 0 4538.462
348.1756591796875 0 4947.934
350.15106201171875 0 16603.549
351.1533508300781 0 3784.6013
351.2398986816406 0 24992.023
352.2424011230469 0 5761.0264
353.2193298339844 0 11104.68
354.2145690917969 0 171059.14
355.2174377441406 0 29003.953
355.27154541015625 0 18879.598
356.2218322753906 0 4796.4873
356.2690734863281 0 6828.852
357.1798400878906 0 7680.3257
357.2138671875 0 5361.9087
358.1630554199219 0 19495.234
359.16571044921875 0 5622.5947
359.19549560546875 0 4785.255
360.6912841796875 0 3931.5474
365.25567626953125 0 13107.858
366.1571044921875 0 34100.246
366.2152404785156 0 3647.698
367.16241455078125 0 11684.472
368.1474914550781 0 18588.32
368.1690368652344 0 4869.0327
369.1758117675781 0 4042.7046
370.2460021972656 0 8948.448
370.6869812011719 0 3321.8652
371.2427062988281 0 3914.1902
372.22515869140625 0 3761.687
374.255615234375 0 3285.175
375.1903381347656 0 7388.0303
378.2521667480469 0 3278.2898
379.19818115234375 0 3751.4526
380.2308654785156 0 7733.1855
380.26641845703125 0 4019.486
382.2087097167969 0 5269.2773
382.281982421875 0 9159.97
383.26629638671875 0 48378.332
383.2904357910156 0 4371.716
384.14117431640625 0 3498.203
384.1678161621094 0 32388.338
384.22601318359375 0 3937.575
384.26959228515625 0 9476.713
385.1737060546875 0 68803.68
385.1995544433594 0 24615.342
386.1581115722656 0 47517.7
386.2018127441406 0 4890.8696
387.1603698730469 0 10364.419
389.271484375 0 8486.956
389.7728576660156 0 3217.8323
390.2457275390625 0 3003.8894
390.274658203125 0 3894.6274
392.1620178222656 0 11601.612
392.2666931152344 0 27324.03
393.1677551269531 0 8526.642
393.2694091796875 0 7631.1514
394.152099609375 0 46114.73
394.2828369140625 0 7425.057
395.155517578125 0 11049.545
396.23321533203125 0 4685.4385
396.2618103027344 0 23143.43
396.2882080078125 0 4419.0703
397.2444763183594 0 5224.9707
398.2405090332031 0 10899.295
398.2764892578125 0 4344.1953
400.17218017578125 0 4459.561
401.1947021484375 0 10046.001
402.2260437011719 0 20967.262
403.18475341796875 0 92402.664
403.22998046875 0 5334.51
403.2692565917969 0 20848.555
403.7709655761719 0 13469.546
404.1871032714844 0 17764.273
404.26922607421875 0 3808.735
406.18634033203125 0 22099.559
406.2502136230469 0 7174.8223
406.68768310546875 0 8702.101
407.1930236816406 0 5785.0415
407.3026123046875 0 9341.789
408.2289733886719 0 3132.6409
408.26220703125 0 8512.874
409.1891784667969 0 3496.0684 a 2
409.2805480957031 0 5366.8467
410.2775573730469 0 51210.34
411.17864990234375 0 80075.266 b Water loss 5
411.22528076171875 0 3545.3752
411.27978515625 0 9901.733
412.16168212890625 0 14184.883
412.1840515136719 0 11900.657
413.168212890625 0 7780.2773
413.287353515625 0 6570.8237
414.2334289550781 0 2962.4224
415.25848388671875 0 5065.24
419.17291259765625 0 8620.552 b Water loss 2
420.1834411621094 0 22884.758 b 5
420.23663330078125 0 15039.618
420.6854248046875 0 9846.206
421.1867370605469 0 5027.8906
422.2774353027344 0 37654.77
423.2779541015625 0 9629.363
423.33514404296875 0 3742.6255
424.2196350097656 0 7235.2144
424.3298645019531 0 35504.29
424.712158203125 0 8908.142
425.2137145996094 0 4488.287
425.24176025390625 0 12591.084
425.2900390625 0 7714.423
425.3320007324219 0 7279.2524
426.2420654296875 0 5099.216
426.27313232421875 0 9839.491
426.3066101074219 0 4922.2954
427.295654296875 0 5638.018
429.1895446777344 0 298421.78
429.23797607421875 0 8621.193 y Water loss 14
430.1925964355469 0 75407.56
431.18048095703125 0 36235.438
432.1832580566406 0 9425.624
433.257080078125 0 5405.0845
433.2884521484375 0 3583.3218
434.3135986328125 0 6619.3164
436.1923828125 0 2848.5972
437.1833190917969 0 103507.33 b 2
438.1866455078125 0 31869.01
439.18914794921875 0 5440.5737
439.30364990234375 0 10904.516
442.2315368652344 0 10383.978
442.3039855957031 0 6250.477
443.2508544921875 0 15754.973
444.2536315917969 0 3331.98
445.2691955566406 0 3343.7761
445.8150329589844 0 12388.066
446.21746826171875 0 12533.893
446.3156433105469 0 7095.6543
447.2460021972656 0 37591.508 y 14
447.28192138671875 0 6218.907
448.250732421875 0 8816.129
449.29150390625 0 5085.927
449.78570556640625 0 5016.704 y 10
450.276611328125 0 9635.789
450.3443908691406 0 3496.1418
451.2300720214844 0 5342.2397
451.2681884765625 0 18898.889
451.3316955566406 0 4939.67
451.7272033691406 0 4802.6313
452.2908020019531 0 7026.4434
452.3249206542969 0 73413.81
453.29302978515625 0 4888.0864
453.3283996582031 0 21717.35
454.2098388671875 0 45071.22
454.33062744140625 0 4235.94
455.2128601074219 0 11300.457
455.25323486328125 0 23540.48
456.25360107421875 0 5699.359
457.18475341796875 0 13815.242
457.5970764160156 0 3214.1113
459.8116760253906 0 23761.154
460.2315979003906 0 9955.327
460.3133850097656 0 13781.932
460.8144836425781 0 5511.0234
461.1832580566406 0 8449.792
462.79229736328125 0 3482.7156
463.27960205078125 0 6641.017
467.2989196777344 0 46061.07
467.33416748046875 0 3558.964
468.2482604980469 0 4235.9
468.29644775390625 0 24411.018
468.3561096191406 0 18982.047
469.3020324707031 0 2978.839
469.356689453125 0 6134.429
470.2970886230469 0 8932.218
471.7984924316406 0 6311.382
473.2626953125 0 34357.617
473.57733154296875 0 6012.552
474.26513671875 0 11338.927
474.7801818847656 0 4122.972
476.2542724609375 0 3776.792
478.2104187011719 0 27867.857
478.3408203125 0 7838.895
479.2137756347656 0 6756.719
482.26422119140625 0 3825.895
482.30804443359375 0 8027.409
482.76214599609375 0 4162.917
483.21832275390625 0 4054.7605
483.78546142578125 0 3469.9783
484.2890319824219 0 3730.5266
488.30218505859375 0 3953.7085
491.2733459472656 0 14127.846
492.27764892578125 0 6171.565
495.2386169433594 0 10241.762
495.36700439453125 0 10732.024
495.9579772949219 0 3989.9216
496.3514099121094 0 17068.164
497.2214050292969 0 19642.846 b 7
497.3109130859375 0 7155.59
497.3519287109375 0 4635.1636
497.7226867675781 0 13852.495
498.21923828125 0 6138.1353
501.6050109863281 0 6499.541
501.9425048828125 0 8295.155
503.21661376953125 0 15278.578
503.3277587890625 0 15704.775
503.8296203613281 0 10030.657
504.221435546875 0 5041.1885
504.324951171875 0 3301.1218
504.9369201660156 0 9015.891 b Water loss 12
505.269287109375 0 4482.8066 b Ammonia loss 12
505.6018371582031 0 12439.024
509.3461608886719 0 6342.5874
510.2945556640625 0 3812.4382
510.3792419433594 0 3305.6665
510.9393615722656 0 22134.982 b 12
511.2721252441406 0 18097.11
511.6093444824219 0 7224.796
511.9432373046875 0 3578.4036
514.2086791992188 0 3988.7336
514.2804565429688 0 4680.301
517.7488403320312 0 3828.4915
518.9388427734375 0 4160.977
519.33349609375 0 7413.8325
519.8370361328125 0 4260.6914
519.955078125 0 3940.2476
520.2410888671875 0 4426.711
520.2827758789062 0 8710.178
520.3606567382812 0 9788.95
521.2273559570312 0 14951.68
521.3479614257812 0 4292.5396
522.226318359375 0 5956.5063
523.3613891601562 0 40775.457
524.3626098632812 0 14052.071
524.9459838867188 0 5534.285
525.3655395507812 0 3778.2712
527.3187866210938 0 5080.3965
528.2327270507812 0 9242.3545
528.3407592773438 0 8765.371
528.8438720703125 0 9907.39
529.2219848632812 0 17093.258
530.2227172851562 0 8982.963
530.3021850585938 0 4743.2314
530.9482421875 0 5580.9077
531.2112426757812 0 15111.488
532.210205078125 0 3431.835
535.3604125976562 0 5489.109
535.75341796875 0 4517.131
536.3555908203125 0 4726.552
537.2463989257812 0 5569.5396
537.30322265625 0 5224.6055
537.4138793945312 0 34784.348
538.2532958984375 0 180714.75
538.3746337890625 0 6984.2466
538.4168090820312 0 8881.844
538.6181030273438 0 4762.804
538.8106079101562 0 6105.7324
538.9479370117188 0 4585.825
539.2559814453125 0 56323.758
539.302978515625 0 19988.912
539.3577880859375 0 9381.705
539.634521484375 0 21532.844
539.8045043945312 0 7537.965
539.9676513671875 0 11072.518
540.2568359375 0 5028.0757
540.3281860351562 0 9073.961
540.8280639648438 0 8916.588 y Ammonia loss 8
541.3297729492188 0 6004.1025
542.3208618164062 0 4343.057 y Water loss 13
542.6289672851562 0 3846.2842 b Water loss 13
542.9667358398438 0 4767.1924 b Ammonia loss 13
544.6283569335938 0 4216.8394
544.7604370117188 0 5730.7026
547.231201171875 0 5574.5933 b Water loss 3
547.950439453125 0 5283.5415
548.232666015625 0 8456.234
548.6328125 0 47486.29 b 13
548.96728515625 0 39817.887
549.2235107421875 0 9959.801
549.3017578125 0 24801.883
549.34716796875 0 5416.056 y 8
549.6355590820312 0 8946.052
550.2230224609375 0 4963.507
550.3723754882812 0 4462.91
553.8517456054688 0 6064.3657
555.3158569335938 0 6859.9624
556.3363647460938 0 3606.8274
557.2967529296875 0 9630.49
557.65234375 0 7429.051
557.9845581054688 0 3916.496
558.84228515625 0 5932.203
559.3389892578125 0 4023.354
560.3314819335938 0 17132.691 y 13
561.2684936523438 0 87327.78 b 8
561.3150634765625 0 3464.358
561.346923828125 0 5088.384
561.6441040039062 0 9612.146
561.77001953125 0 57528.33
561.9557495117188 0 3886.8984
562.2724609375 0 19934.629
562.6366577148438 0 10673.259
562.7693481445312 0 4803.824
562.968505859375 0 4547.2227
563.3582763671875 0 3219.1348
564.3546752929688 0 4244.821
565.2415771484375 0 43548.76 b 3
565.28466796875 0 5420.5054
565.3591918945312 0 5409.7227
565.4083862304688 0 90938.414
565.6148071289062 0 6567.246
565.9468994140625 0 5924.1636
566.2482299804688 0 219818.97
566.4119262695312 0 37063.49
567.2511596679688 0 73565.26
567.3126220703125 0 19003.879
567.4086303710938 0 9484.088
567.647216796875 0 9549.8955
567.8487548828125 0 43773.184
567.982666015625 0 4780.6904
568.2531127929688 0 12406.817
568.305419921875 0 6589.318
568.3504028320312 0 22713.123
568.6473388671875 0 9602.446
568.8526000976562 0 7285.9204 y Water loss 7
568.9762573242188 0 3598.1416
570.2477416992188 0 23965.822
571.2481079101562 0 6331.521
571.3294067382812 0 3358.2031
571.650634765625 0 3744.4717
571.85498046875 0 4612.2544
571.9796142578125 0 3862.5703
572.8031005859375 0 5609.2544
573.3086547851562 0 8243.601
573.8099365234375 0 4174.95
573.9644775390625 0 3779.9082
576.6443481445312 0 12001.519
576.9932250976562 0 35333.312
577.330078125 0 32956.387
577.3836059570312 0 18445.709
577.6587524414062 0 18331.28
577.97802734375 0 18675.426
578.3129272460938 0 8157.103
578.3828125 0 6733.3594
579.3626708984375 0 48432.29
580.376708984375 0 21304.516
580.6538696289062 0 6752.61 b Ammonia loss 14
581.3778686523438 0 6622.7627
582.79052734375 0 10150.997
583.29248046875 0 9545.2705
583.37451171875 0 6590.3535
585.6517944335938 0 5447.745
586.3277587890625 0 138656.33 b 14
586.662109375 0 171316.64
586.84375 0 3622.3718
586.9959716796875 0 104776.99
587.274169921875 0 9911.074
587.3320922851562 0 33562.61
587.6655883789062 0 6725.6206
587.7822875976562 0 44385.902 b Water loss 9
588.2823486328125 0 41100.54 b Ammonia loss 9
588.7854614257812 0 11769.809
591.4200439453125 0 4244.937
591.9868774414062 0 5865.6016
592.3154907226562 0 4188.984
594.2427978515625 0 5196.1504
595.3507080078125 0 10548.662
595.3946533203125 0 17074.273
595.8515625 0 8231.499
596.348876953125 0 6016.437
596.3978271484375 0 7757.4604
596.7875366210938 0 85479.87 b 9
596.8709106445312 0 17731.51
597.2890625 0 60863.773
597.371337890625 0 6879.052
597.7889404296875 0 20544.057
597.874755859375 0 8738.126
598.2412719726562 0 11892.309
599.649658203125 0 7703.2812
600.333251953125 0 12768.718 y Water loss 2
600.669921875 0 10150.409 y Ammonia loss 2
604.3590698242188 0 14535.936
604.8592529296875 0 7398.076
605.3165283203125 0 4806.6777
605.3663330078125 0 6484.843
605.6565551757812 0 9302.411
606.0003662109375 0 11093.601
606.34130859375 0 53655.156 y 2
606.4314575195312 0 4044.2349
606.6756591796875 0 43363.965
607.0106201171875 0 25940.531
607.3444213867188 0 8207.469
608.4501953125 0 5980.2783
608.9849243164062 0 10959.896
609.3260498046875 0 12849.49 b Water loss 15
609.6618041992188 0 16028.046 b Ammonia loss 15
610.000732421875 0 5174.4136
610.3237915039062 0 21624.92
610.3938598632812 0 4876.7104
610.6723022460938 0 3749.5828
611.3265380859375 0 19939.605
611.6605224609375 0 11515.581
611.995361328125 0 11455.669
612.3286743164062 0 8084.842
614.3342895507812 0 5170.4316
614.6624145507812 0 16353.303
614.9942626953125 0 16554.332
615.2704467773438 0 11051.285
615.3388671875 0 99353.7 b 15
615.6723022460938 0 111024.39
615.850341796875 0 8138.2007
616.0065307617188 0 71492.44
616.2728271484375 0 5778.945
616.3412475585938 0 23273.766
616.6740112304688 0 11546.622
617.3840942382812 0 11531.284 y Water loss 6
617.885009765625 0 12940.095 y Ammonia loss 6
619.0023193359375 0 3499.5967
619.34814453125 0 6208.829
620.3432006835938 0 22049.518
620.6581420898438 0 25646.357
620.8433837890625 0 17822.98
620.9930419921875 0 36288.117
621.3292846679688 0 16624.977
621.4080200195312 0 4507.222
621.6612548828125 0 7845.466
623.38720703125 0 8963.729
623.4588012695312 0 4249.5596
623.674560546875 0 4131.999
624.3935546875 0 4621.486
624.4505615234375 0 5909.816
625.4542236328125 0 4502.804
626.3888549804688 0 142451.02 y 6
626.8902587890625 0 118472.85
627.3904418945312 0 45667.934
627.8889770507812 0 12836.269
628.33056640625 0 8437.838
628.388427734375 0 5440.7036
629.2882080078125 0 5047.077
629.3468017578125 0 46304.996
629.6751098632812 0 14508.703
629.8486938476562 0 33010.258
630.0153198242188 0 5353.2183
630.34912109375 0 12798.1455
632.2958984375 0 8371.425
633.2999877929688 0 5818.1387
633.3613891601562 0 4331.2153
634.4300537109375 0 6486.1436
636.3797607421875 0 9144.081
636.4454956054688 0 53080.332
636.8760986328125 0 4886.5254
637.3423461914062 0 4032.5955
637.4480590820312 0 15903.805
637.6806030273438 0 3910.525
638.396484375 0 4138.5234
638.4491577148438 0 4157.866
640.3924560546875 0 7968.3403
643.0180053710938 0 12161.664
643.265625 0 8467.399
643.3491821289062 0 12498.36
643.6827392578125 0 14869.69
645.3141479492188 0 6353.974 b 10
645.8168334960938 0 6128.0684
646.6764526367188 0 17470.35
647.012451171875 0 14348.566
647.343505859375 0 5788.7085
647.4251098632812 0 8677.333
647.6771240234375 0 5524.979
648.0148315429688 0 4734.634
648.3665161132812 0 8009.299
648.4304809570312 0 3918.186
648.6851806640625 0 6226.6787
649.02099609375 0 57638.316
649.3556518554688 0 60185.613
649.6893920898438 0 38316.465
650.0239868164062 0 17366.658
650.354736328125 0 7927.6694
651.4577026367188 0 10018.541
651.893798828125 0 13397.891
652.3485107421875 0 87011.2 b Water loss 16
652.4442749023438 0 25904.428
652.6834106445312 0 124155.78 b Ammonia loss 16
652.8888549804688 0 10167.666
653.0176391601562 0 79275.734
653.3486938476562 0 30796.36
653.4463500976562 0 10257.42
653.6863403320312 0 22324.404
653.82958984375 0 16150.203
654.0158081054688 0 5698.7485
654.333251953125 0 9810.301
656.294189453125 0 12239.314
657.2799682617188 0 45062.164
658.015380859375 0 6054.4897
658.3529663085938 0 548091.75 b 16
658.68701171875 0 607655.4
659.0216674804688 0 408171.94
659.3556518554688 0 164613.05
659.689697265625 0 46722.66
660.0225219726562 0 5277.742
660.8998413085938 0 9805.543
661.4032592773438 0 11603.273
662.3598022460938 0 5332.202 y Water loss 1
662.6929321289062 0 4547.5347 y Ammonia loss 1
663.3016357421875 0 5472.731
663.3757934570312 0 8344.069
664.3571166992188 0 29382.516
664.4524536132812 0 30109.188
664.6903076171875 0 31836.469
665.02587890625 0 18681.32
665.3627319335938 0 6130.793
665.4558715820312 0 11165.471
666.3986206054688 0 6359.987
667.3455810546875 0 6122.492
668.3715209960938 0 20497.027 y 1
668.7025756835938 0 32067.893
669.0364990234375 0 15819.677
669.3921508789062 0 3859.4321
671.8914794921875 0 22436.793
672.3936767578125 0 14203.839
672.8948364257812 0 5778.6323
673.4100341796875 0 10742.907 y 12
674.30615234375 0 129053.89
674.4364624023438 0 91226.94
675.3086547851562 0 50442.066
675.4388427734375 0 39232
676.31103515625 0 11976.47
676.375 0 6238.106
676.44482421875 0 10249.009
676.705810546875 0 7432.812
676.8869018554688 0 6143.8613
677.3811645507812 0 5990.702
677.7030639648438 0 4285.2456
678.3765869140625 0 5684.6763
678.6904907226562 0 5132.483
679.3626098632812 0 5606.088
679.6976928710938 0 6596.483
680.0296630859375 0 7334.4995
682.0430908203125 0 20943.438
682.3782958984375 0 10983.795
682.7122192382812 0 20630.629
683.04150390625 0 10677.126
683.3646850585938 0 7639.1206
683.7044677734375 0 6187.8423
684.2908935546875 0 28121.482 b Water loss 4
685.2828979492188 0 18576.58 b Ammonia loss 4
685.3678588867188 0 15719.86
685.7061157226562 0 25910.662
685.8922119140625 0 77940.11
686.0380249023438 0 14962.785
686.2926635742188 0 5879.569
686.3916015625 0 63985.004 y Ammonia loss 5
686.8954467773438 0 31005.484
687.3904418945312 0 12267.564
687.7062377929688 0 6175.775
688.043212890625 0 5468.983
690.46875 0 7958.419
691.0408935546875 0 5352.293
691.375732421875 0 187391.73 Precursor Water loss
691.7098388671875 0 237603.16 Precursor Ammonia loss
692.0443115234375 0 191553.9
692.37744140625 0 87243.734
692.447998046875 0 71766.94
692.7094116210938 0 33076.516
693.0408935546875 0 17313.23
693.4513549804688 0 34804.805
694.4552612304688 0 9928.396
694.9180297851562 0 50818.04 y 5
695.4186401367188 0 43427.906
695.9178466796875 0 20739.809
696.4121704101562 0 11882.741
696.70556640625 0 19495.734
697.0423583984375 0 50638.49
697.379638671875 0 2838229 Precursor
697.7137451171875 0 3588539.2
698.0480346679688 0 2416834.8
698.3817749023438 0 1103313.8
698.7161254882812 0 390900.25
699.049560546875 0 110481.7
702.301025390625 0 225297.31 b 4
703.3040771484375 0 84586.57
704.3071899414062 0 24761.236
704.407470703125 0 6560.132
704.91064453125 0 12381.824
705.3101806640625 0 5259.966
705.4093017578125 0 11104.274
707.306884765625 0 14318.884
708.3085327148438 0 6022.187
708.4077758789062 0 5768.1816
708.4783935546875 0 25882.213
709.360107421875 0 40636.598 b 11
709.479736328125 0 16451.824
709.8638916015625 0 40085.945
710.3662719726562 0 26672.797
710.8671264648438 0 13696.297
711.3621215820312 0 4995.0854
716.4227905273438 0 9058.145
719.8651123046875 0 6691.4785
720.32080078125 0 10968.747
720.4307250976562 0 6177.744
721.35546875 0 9603.175
723.3905029296875 0 6112.206
723.4776611328125 0 15666.588
724.3333129882812 0 17097.146
724.48046875 0 7642.0503
725.3345947265625 0 8020.9194
725.4051513671875 0 6343.6455
728.4310913085938 0 18163.533
728.9353637695312 0 16380.291
729.4091796875 0 11697.935
729.9068603515625 0 11511.52
732.5159301757812 0 4396.455
733.4164428710938 0 5669.863
735.3023071289062 0 16024.164
736.4759521484375 0 5461.6416
737.3580932617188 0 4102.702
738.3810424804688 0 11207.746
739.371826171875 0 7991.117
740.44140625 0 4944.7593
741.41259765625 0 8244.012
742.4310913085938 0 104418.914
742.9332885742188 0 100235
743.4347534179688 0 38944.027
743.9346923828125 0 9823.402
751.4219360351562 0 7068.5015
751.9077758789062 0 22099.68
752.328125 0 20206.566
752.4093017578125 0 26228.797
752.9096069335938 0 7286.411
753.3280639648438 0 8858.7295
753.3956909179688 0 4403.5513
753.4891357421875 0 9875.336
754.939697265625 0 8264.373 y Ammonia loss 4
756.898193359375 0 46075.305 b Water loss 12
757.3985595703125 0 48254.477 b Ammonia loss 12
757.8999633789062 0 22469.398
758.3997192382812 0 15045.752
758.9002075195312 0 5424.173
759.931396484375 0 13064.1875
760.4290771484375 0 10704.04
760.5103759765625 0 9311.046
760.9286499023438 0 10941.558
762.9140014648438 0 4484.601
763.4481811523438 0 86815.44 y 4
763.9495849609375 0 59098.754
764.4533081054688 0 28408.424
764.5429077148438 0 16608.432
764.9498901367188 0 11079.805
765.3800048828125 0 7414.7656
765.9033203125 0 88021.31 b 12
766.4053344726562 0 76692.79
766.90625 0 34983.047
767.4104614257812 0 15635.424
767.9110717773438 0 5609.6904
768.9348754882812 0 13762.656
769.4378051757812 0 4846.7104
769.6996459960938 0 5036.022
770.4274291992188 0 5383.8047
776.941162109375 0 5296.952
777.4375 0 11349.198
777.5369873046875 0 30111.139
778.5401000976562 0 12177.133
778.9248657226562 0 12160.688
779.4805908203125 0 8813.881
779.93310546875 0 5246.5527
780.3204956054688 0 9107.678
780.9277954101562 0 5152.6353
781.3472290039062 0 4356.881
781.4829711914062 0 43942.355
782.4864501953125 0 21534.611
783.49169921875 0 12490.168 y Water loss 11
785.4285278320312 0 8090.441
785.9473876953125 0 47597.965
786.4483642578125 0 47796.42
786.9490966796875 0 20168.512
787.455322265625 0 4300.4463
787.521728515625 0 18289.062
788.5238647460938 0 10896.901
792.3936767578125 0 6721.701
793.3543090820312 0 10115.351
793.9270629882812 0 15260.537
794.3429565429688 0 25470.28
794.427490234375 0 12954.442
794.9299926757812 0 11816.536
795.3529663085938 0 15067.202
795.43212890625 0 5975.064
799.879638671875 0 9194.98
800.8959350585938 0 4676.2485
801.4103393554688 0 6044.1504
801.5109252929688 0 13249.878 y 11
802.5167236328125 0 6172.9683
804.3256225585938 0 6298.104
805.4442138671875 0 5582.139
805.5315551757812 0 138998.05
806.5340576171875 0 59795.613
807.5361938476562 0 17827.475
808.44580078125 0 20229.182
808.53466796875 0 5725.367
808.9483032226562 0 29415.22
809.4324340820312 0 17105.637
809.92724609375 0 19219.41
810.4241943359375 0 5954.2305
811.3648681640625 0 197801.39
811.4967041015625 0 19762.557
812.3681640625 0 105125.85
812.5 0 8964.651
813.3724975585938 0 19565.998
813.4432983398438 0 12677.59 b Water loss 13
813.937744140625 0 26237.312 b Ammonia loss 13
814.4384765625 0 11485.39
814.9470825195312 0 9094.316
817.4010620117188 0 5498.495
817.9010620117188 0 9231.328
818.396728515625 0 3798.3806
818.4735717773438 0 20645.855 y Water loss 3
818.9674072265625 0 50948.066 y Ammonia loss 3
819.4672241210938 0 41870.69
819.9691772460938 0 24673.684
820.4652709960938 0 10258.282
821.3489990234375 0 26397.717 b Water loss 5
821.5626220703125 0 32040.02
821.920166015625 0 7910.2227
822.345947265625 0 23478.7 b Ammonia loss 5
822.445068359375 0 170222.03 b 13
822.56103515625 0 14520.402
822.946533203125 0 171103.88
823.35546875 0 5295.643
823.4486083984375 0 86977.15
823.9490356445312 0 38052.375
824.4506225585938 0 10410.788
827.4757690429688 0 64820.66 y 3
827.9763793945312 0 58404.504
828.4757080078125 0 34481.156
828.9788818359375 0 13858.922
829.5094604492188 0 10459.111
830.4227294921875 0 4944.27
830.5093383789062 0 7427.6846
832.957275390625 0 5205.515
833.4532470703125 0 8793.499
834.947998046875 0 6097.8633
835.468017578125 0 12661.978
835.9688720703125 0 7635.358
836.4725341796875 0 14505.583
836.97021484375 0 11061.059
839.3600463867188 0 418490.4 b 5
840.3631591796875 0 218253.77
840.4407348632812 0 4605.3296
841.365966796875 0 53802.88
841.4634399414062 0 22332.385
841.963134765625 0 20665.9
842.3685913085938 0 8962.83
842.4627075195312 0 9973.054
842.9459228515625 0 6155.7163
843.4495239257812 0 5276.852
848.4224853515625 0 28472.104
849.423828125 0 15527.842
850.4691772460938 0 85139.99
850.9703979492188 0 79719.55
851.471435546875 0 48604.223
851.5654296875 0 5536.865
851.97021484375 0 16116.994
852.4586181640625 0 9287.575
852.5265502929688 0 10085.547
856.4234619140625 0 10383.45
856.9277954101562 0 16867.955
857.442626953125 0 6076.4805
859.4728393554688 0 6307.7256
859.9714965820312 0 7902.422
860.4520263671875 0 4795.661
864.4595947265625 0 7953.6787
864.9824829101562 0 16563.658
865.4889526367188 0 18759.742
865.979248046875 0 22295.06
866.4623413085938 0 20860.744
866.9652099609375 0 11470.811
869.97998046875 0 10281.346 b Water loss 14
870.480224609375 0 12247.395 b Ammonia loss 14
870.9776611328125 0 16739.162
871.474609375 0 5406.1
871.9774780273438 0 7389.3696
873.5931396484375 0 5566.451
873.9422607421875 0 23736.568
874.4419555664062 0 19619.197
874.9439697265625 0 11204.466
875.4436645507812 0 5425.831
877.9631958007812 0 10460.931
878.4706420898438 0 12563.411
878.9874267578125 0 403524.8 b 14
879.4888916015625 0 375673.03
879.9903564453125 0 206503
880.491943359375 0 70811.77
880.9927368164062 0 21285.326
886.961181640625 0 5301.3823
887.4691162109375 0 5461.2925
890.617919921875 0 21355.467
891.4842529296875 0 16689.734
891.6199951171875 0 9691.839
891.9915771484375 0 8979.117
892.476318359375 0 6533.257
892.5645751953125 0 12675.005
893.5660400390625 0 8994.733
898.5625 0 116370.96 y 10
899.4907836914062 0 3169.6492
899.5651245117188 0 61525.645
900.0027465820312 0 33874.695 y Water loss 2
900.5015869140625 0 35885.133 y Ammonia loss 2
900.5830688476562 0 11811.1045
901.0045166015625 0 20212.564
901.4439086914062 0 26717.928
901.5952758789062 0 8182.831
901.995849609375 0 5924.83
902.4483642578125 0 21137.537
903.4526977539062 0 7658.4473
908.5095825195312 0 14699.756
908.5980834960938 0 9585.483
909.0084228515625 0 399003.22 y 2
909.5098266601562 0 417529.88
910.0111083984375 0 243554.7
910.5128784179688 0 95891.1
911.0148315429688 0 22606.994
911.5186157226562 0 6493.9097
912.9660034179688 0 16875.832
913.4869995117188 0 38015.645 b Water loss 15
913.9921264648438 0 31293.012 b Ammonia loss 15
914.4951171875 0 20021.305
914.5963134765625 0 4496.536
914.9979248046875 0 9325.724
915.4996948242188 0 5479.72
916.4821166992188 0 8825.997
916.9820556640625 0 6305.4526
918.615234375 0 207350.5
919.453369140625 0 21180.44
919.6181640625 0 126399.93
920.4575805664062 0 11441.352
920.621826171875 0 36099.188
921.4793090820312 0 23190.787
921.6265869140625 0 7908.6714
921.9822998046875 0 14305.278
922.5020751953125 0 151540.77 b 15
923.0048828125 0 202271.44
923.5060424804688 0 112241.13
924.0065307617188 0 41588.344
924.580078125 0 9334.675
925.575927734375 0 6273.2114
930.4828491210938 0 41528.613
930.9853515625 0 34295.074
931.4864501953125 0 18697.773
931.9844360351562 0 10471.179
936.4124145507812 0 7950.3057 b 6
937.418212890625 0 5828.5376
942.5899658203125 0 21197.14
943.4996948242188 0 5810.9634
943.5946044921875 0 7928.6553
944.5067138671875 0 7477.6855
948.5572509765625 0 6465.7456
966.5590209960938 0 9001.015
967.569091796875 0 7707.671
969.5091552734375 0 5221.4585
969.5999755859375 0 53262.76 y 9
970.6022338867188 0 37099.934
971.6085815429688 0 9001.327
973.0249633789062 0 7789.149
973.5299682617188 0 9037.334
974.0283813476562 0 8161.6123
978.0201416015625 0 37127.734 b Water loss 16
978.5198974609375 0 44616.16 b Ammonia loss 16
979.0218505859375 0 29828.54
979.5220947265625 0 8145.464
980.0202026367188 0 5521.8013
980.5177612304688 0 5955.068
981.5347900390625 0 5158.1714
987.02490234375 0 233779.75 b 16
987.5260009765625 0 291361.75
987.6337890625 0 14342.074
988.027587890625 0 161917.81
988.5286254882812 0 66470.97
988.6416625976562 0 7899.269
989.0296020507812 0 23264.045
993.434326171875 0 17846.662 b 7
994.4373168945312 0 11182.593
996.0294799804688 0 23105.576
996.5322875976562 0 30025.508
997.03173828125 0 17679.275
997.5352783203125 0 6505.117
1003.6038818359375 0 5574.497
1005.6469116210938 0 71409.77
1006.6499633789062 0 53136.805
1007.6522216796875 0 10420.778
1009.6634521484375 0 5942.5327
1010.4598999023438 0 18061.354
1011.461669921875 0 8015.136
1021.6094970703125 0 18668.785
1022.6084594726562 0 12305.846
1032.5272216796875 0 10071.386
1033.532470703125 0 14287.048
1035.5301513671875 0 5108.5933
1037.65380859375 0 17930.912
1038.6558837890625 0 14570.541
1055.673095703125 0 34430.76
1056.6773681640625 0 22212.988
1079.650390625 0 13327.012
1080.654296875 0 10551.316 y Ammonia loss 8
1088.52001953125 0 5513.072
1097.69189453125 0 19267.742 y 8
1098.69482421875 0 13946.37
1106.693115234375 0 10689.483
1107.6943359375 0 7788.4194
1116.6806640625 0 19157.96
1117.6805419921875 0 10422.222
1118.6815185546875 0 7182.4844
1121.5294189453125 0 30331.834 b 8
1122.532470703125 0 16391.85
1123.533935546875 0 8419.038
1125.6966552734375 0 5317.97
1134.690185546875 0 84638.31
1135.6927490234375 0 50221.31
1136.698974609375 0 23156.098 y Water loss 7
1137.6986083984375 0 7476.773 y Ammonia loss 7
1142.703857421875 0 12436.147
1144.6029052734375 0 8907.798
1145.606689453125 0 13204.933
1146.6182861328125 0 10174.671
1152.7078857421875 0 5495.005
1153.6990966796875 0 6049.2285
1154.71630859375 0 27111.635 y 7
1155.720703125 0 16281.462
1164.5760498046875 0 7523.571
1174.5552978515625 0 44183.734 b Water loss 9
1175.5577392578125 0 33617.566 b Ammonia loss 9
1176.561767578125 0 16490.766
1177.562744140625 0 5164.7773
1192.565185546875 0 24718.822 b 9
1192.7298583984375 0 18994.8
1193.5667724609375 0 19588.95
1193.735595703125 0 11785.845
1207.7137451171875 0 5370.188
1233.7537841796875 0 15801.228 y Water loss 6
1234.608154296875 0 7153.705
1234.7625732421875 0 12304.856 y Ammonia loss 6
1235.7615966796875 0 6740.0073
1239.66455078125 0 4976.107
1240.6734619140625 0 6288.434
1251.767578125 0 125458.94 y 6
1252.7713623046875 0 90924.375
1253.768310546875 0 39267.125
1254.7669677734375 0 12823.455
1255.76220703125 0 5666.8623
1257.6868896484375 0 13459.844
1258.6915283203125 0 9334.315
1259.712890625 0 5500.9575
1260.6326904296875 0 5870.6367
1271.7459716796875 0 9588.916
1272.746826171875 0 8706.293
1305.6488037109375 0 12143.118
1306.6480712890625 0 21169.629
1307.647216796875 0 14420.325
1370.7744140625 0 13233.425
1371.777587890625 0 10901.365 y Ammonia loss 5
1372.7921142578125 0 4546.4336
1388.8271484375 0 7526.2334 y 5
1389.83056640625 0 9506.706
1390.8387451171875 0 4154.251
1408.8067626953125 0 6218.5376
1409.812744140625 0 5105.0776
1417.706787109375 0 6095.222 b 11
1418.7305908203125 0 9527.974
1419.7266845703125 0 11224.231
1532.8150634765625 0 4798.1416
2170.064697265625 0 3626.7698

Spectrum Details

|  |  |
| --- | --- |
| Matched peaks? Matched peaks The total absolute number of peaks matched. Additionally in brackets the total fraction of peaks matched and the total number of peaks is shown. | 114 (8.66% of 1317) |
| FDR? FDR The false discovery rate estimated for this peptide. It is calculated by matching all theoretical fragments with a non-integer shift with the raw peaks for this spectrum. This is done with 40 different shifts. The resulting percentage is the average number of annotated peaks over the number of annotated peaks with the correct spectrum. | 6.16% |
| Satellite FDR? Satellite FDR See the FDR for details on its calculation. This satellite ion specific FDR only contains the satellite ions (d/w) for I/L/J positions. | - |
| PSM Score? PSM Score The PSM Score as given by Hecklib to this annotated spectrum. It is shown with three significant figures. | 622 |

#### Reverse Lookup? Reverse Lookup All places where this read could be placed.

| Group | Segment | Template | Template Part | Read Part | Score | Unique |
| --- | --- | --- | --- | --- | --- | --- |
| Homo sapiens Light Chain | IGLV | IGLV2-14 | [35..53] | [0..18] | 117 | False |
| Homo sapiens Light Chain | IGLV | IGLV2-23 | [35..53] | [0..18] | 108 | False |
| Homo sapiens Light Chain | IGLV | IGLV2-8 | [35..53] | [0..18] | 117 | False |
| Homo sapiens Light Chain | IGLV | IGLV2-11 | [35..53] | [0..18] | 112 | False |

| Recombined | Template Part | Read Part | Score | Unique |
| --- | --- | --- | --- | --- |
| REC-0-1\_002 | [35..53] | [0..18] | 144 | True |

#### Meta Information from Multiple reads

##### Number of combined reads

4

##### Intensity

0.809

##### TotalArea

8.536E+08

##### Changes to the peptide sequence

SWYQHHPGKAPKJJJSEV

L→JNo support for either Leucine or Isoleucine based on side chain ions (Position: 13)

L→JNo support for either Leucine or Isoleucine based on side chain ions (Position: 15)

L→JNo support for either Leucine or Isoleucine based on side chain ions (Position: 14)

#### Positional Score

Copy Data

##### Positional Score (TSV)

###### Preview

```
Loading example...
```

*Click on the button to copy the data to your clipboard.*

1001234567891011121314151617

Label Value
"0" 0.473
"1" 0.47
"2" 0.495
"3" 0.49
"4" 0.497
"5" 0.497
"6" 0.497
"7" 0.497
"8" 0.495
"9" 0.48
"10" 0.44
"11" 0.43
"12" 0.478
"13" 0.5
"14" 0.5
"15" 0.5
"16" 0.497
"17" 0.492

#### Meta Information from PEAKS

##### Scan Identifier

F1:7437

##### Original sequence

S

W

Y

Q

H

H

P

G

K

A

P

K

L

L

L

S

E

V

##### Posttranslational Modifications

##### Source File

D:\separate\_stitch\_analyses\xle-disambiguation\raw\20210323\_F1\_UM1\_Peng0013\_SA\_F59\_ingel\_3ug\_ELA.raw

##### Fraction

1

##### Scan Feature

F1:12656

##### De Novo Score

99

##### ConfidenceScore

99

### m/z

697.3787

##### Mass

2089.1108

##### Charge

3

##### Retention Time

40.03

##### Predicted Retention Time

-

##### Area

2.134E+08

##### Parts Per Million

1.6

##### Fragmentation mode

ETHCD

##### Originating file

01 D:\separate\_stitch\_analyses\xle-disambiguation\20210325\_F59\_3ug\_DENOVO\_12.csv

#### Meta Information from PEAKS

##### Scan Identifier

F1:7499

##### Original sequence

S

W

Y

Q

H

H

P

G

K

A

P

K

L

L

L

S

E

V

##### Posttranslational Modifications

##### Source File

D:\separate\_stitch\_analyses\xle-disambiguation\raw\20210323\_F1\_UM1\_Peng0013\_SA\_F59\_ingel\_3ug\_ELA.raw

##### Fraction

1

##### Scan Feature

F1:12656

##### De Novo Score

99

##### ConfidenceScore

99

### m/z

697.3787

##### Mass

2089.1108

##### Charge

3

##### Retention Time

40.03

##### Predicted Retention Time

-

##### Area

2.134E+08

##### Parts Per Million

1.6

##### Fragmentation mode

ETHCD

##### Originating file

01 D:\separate\_stitch\_analyses\xle-disambiguation\20210325\_F59\_3ug\_DENOVO\_12.csv

#### Meta Information from PEAKS

##### Scan Identifier

F1:7638

##### Original sequence

S

W

Y

Q

H

H

P

G

K

A

P

K

L

L

L

S

E

V

##### Posttranslational Modifications

##### Source File

D:\separate\_stitch\_analyses\xle-disambiguation\raw\20210323\_F1\_UM1\_Peng0013\_SA\_F59\_ingel\_3ug\_ELA.raw

##### Fraction

1

##### Scan Feature

F1:12656

##### De Novo Score

98

##### ConfidenceScore

98

### m/z

697.3787

##### Mass

2089.1108

##### Charge

3

##### Retention Time

40.03

##### Predicted Retention Time

-

##### Area

2.134E+08

##### Parts Per Million

1.6

##### Fragmentation mode

ETHCD

##### Originating file

01 D:\separate\_stitch\_analyses\xle-disambiguation\20210325\_F59\_3ug\_DENOVO\_12.csv

#### Meta Information from PEAKS

##### Scan Identifier

F1:7373

##### Original sequence

S

W

Y

Q

H

H

P

G

K

A

P

K

L

L

L

S

E

V

##### Posttranslational Modifications

##### Source File

D:\separate\_stitch\_analyses\xle-disambiguation\raw\20210323\_F1\_UM1\_Peng0013\_SA\_F59\_ingel\_3ug\_ELA.raw

##### Fraction

1

##### Scan Feature

F1:12656

##### De Novo Score

96

##### ConfidenceScore

96

### m/z

697.3787

##### Mass

2089.1108

##### Charge

3

##### Retention Time

40.03

##### Predicted Retention Time

-

##### Area

2.134E+08

##### Parts Per Million

1.6

##### Fragmentation mode

HCD

##### Originating file

01 D:\separate\_stitch\_analyses\xle-disambiguation\20210325\_F59\_3ug\_DENOVO\_12.csv
