## Supplementary material for "A handle on mass coincidence errors in *de novo* sequencing of antibodies by bottom-up proteomics": Combined_025.html

Details Combined\_025 | Stitch OverviewUndefined

### Read Combined\_025

#### Sequence (length=16)

JHQDWJDGKEYKCKVS

#### Spectrum 5024? Spectrum 5024 The raw spectrum of this peptide as annotated by Hecklib. The fragments are coloured according to ion type (see legend). Any peaks with a star '\*' as text can be hovered over to see the full details, first the ion type second the mass shift type. By hovering over the amino acids in the peptide or ions in the legend the corresponding peaks are highlighted. By toggling the 'Unassigned' label you can turn the background (unassigned) peaks on or off in the plot. By updating the slider in the Ion legend you can update the spectrum to only show the top X% of the peaks with labels. The top X% means any peak that is within X% of the highest intensity. By dragging in the spectrum you can zoom in to a specific part of the spectrum and use 'Zoom Out' to get back to the original zoom level. The annotation of the spectrum is based on the given sequence in the peptides file and is done with different software so inconsistencies are likely. The peaks are annotated based on the given sequence, with 20 ppm tolerance.

Copy Data

##### Spectrum 5024 (TSV)

###### Preview

```
Loading example...
```

*Click on the button to copy the data to your clipboard.*

Mz MinMz MaxIntensity Max

WidthHeightPeptide font sizePeptide stroke widthSpectrum font sizeSpectrum stroke widthCompact peptide

Ion legend

wxyz

abcd

OtherUnassignedIon^Charge^_Position_Show for top:%

JHQDWJDGKEYKCKVS

02.24e+54.49e+56.73e+58.98e+5

Zoom Out

y^+1^_2_c^+1^_2_z^+1^_3_y^+1^_3_c^+1^_3_c^+1^_3_c^+1^_4_c^+1^_4_w^+1^_5_y^+2^_9_w^+2^_10_y^+2^_10_c^+3^_14_z^+1^_5_c^+3^_14_y^+2^_10_c^+2^_10_y^+1^_5_y^+3^_15_y^+3^_15_c^+3^_15_y^+2^_11_c^+1^_5_c^+1^_5_c^+2^_12_y^+1^_6_z^+1^_6_y^+1^_6_c^+1^_6_c^+1^_6_y^+2^_13_w^+1^_7_w^+2^_14_c^+2^_13_z^+2^_14_y^+2^_14_y^+2^_14_z^+2^_14_y^+2^_14_z^+1^_7_y^+1^_7_z^+1^_7_c^+2^_14_c^+2^_14_c^+1^_7_c^+2^_14_y^+1^_7_c^+1^_7_z^+2^_15_y^+2^_15_c^+2^_15_c^+2^_15_c^+2^_15_c^+1^_8_c^+1^_8_z^+1^_8_y^+1^_8_z^+1^_9_y^+1^_9_c^+1^_9_z^+1^_10_y^+1^_10_c^+1^_10_w^+1^_11_y^+1^_11_z^+1^_11_y^+1^_11_c^+1^_11_y^+1^_12_y^+1^_12_z^+1^_12_c^+1^_12_c^+1^_12_c^+1^_12_w^+1^_13_y^+1^_13_z^+1^_13_y^+1^_13_w^+1^_14_c^+1^_13_z^+1^_14_y^+1^_14_c^+1^_14_c^+1^_15_c^+1^_15_

0769153823073076

Fragment Matches Table

Show background peaks

| Position | Ion type | Intensity | mz Theoretical | mz Error (Th) | mz Error (ppm) | Charge | Series Number |
| --- | --- | --- | --- | --- | --- | --- | --- |
| - | - | 1797 | 120.3 | - | - | 0 | - |
| - | - | 1489 | 121 | - | - | 0 | - |
| - | - | 2143 | 122.3 | - | - | 0 | - |
| - | - | 2362 | 128.1 | - | - | 0 | - |
| - | - | 3052 | 129.1 | - | - | 0 | - |
| - | - | 5144 | 129.1 | - | - | 0 | - |
| - | - | 1944 | 133.6 | - | - | 0 | - |
| - | - | 2222 | 136 | - | - | 0 | - |
| - | - | 3554 | 138.1 | - | - | 0 | - |
| - | - | 2616 | 146.1 | - | - | 0 | - |
| - | - | 1.086E+04 | 155.1 | - | - | 0 | - |
| - | - | 2967 | 164.1 | - | - | 0 | - |
| - | - | 2114 | 166.1 | - | - | 0 | - |
| - | - | 2.363E+04 | 166.1 | - | - | 0 | - |
| - | - | 5835 | 168.1 | - | - | 0 | - |
| - | - | 6558 | 178.1 | - | - | 0 | - |
| - | - | 1.388E+04 | 185.2 | - | - | 0 | - |
| 15 | y | 7645 | 205.1 | 0.0002468 | 1.203 | +1 | 2 |
| - | - | 5785 | 207.1 | - | - | 0 | - |
| - | - | 4265 | 207.2 | - | - | 0 | - |
| - | - | 2.764E+04 | 223.2 | - | - | 0 | - |
| - | - | 5861 | 224.2 | - | - | 0 | - |
| - | - | 2425 | 227.9 | - | - | 0 | - |
| - | - | 3191 | 230.2 | - | - | 0 | - |
| - | - | 4598 | 233.1 | - | - | 0 | - |
| - | - | 3.622E+04 | 234.1 | - | - | 0 | - |
| - | - | 2764 | 234.1 | - | - | 0 | - |
| - | - | 2780 | 234.1 | - | - | 0 | - |
| - | - | 5856 | 235.1 | - | - | 0 | - |
| - | - | 4260 | 248.1 | - | - | 0 | - |
| - | - | 5711 | 250.1 | - | - | 0 | - |
| - | - | 1.5E+05 | 251.2 | - | - | 0 | - |
| - | - | 1.721E+04 | 252.2 | - | - | 0 | - |
| - | - | 2902 | 262.9 | - | - | 0 | - |
| - | - | 1.043E+04 | 266.1 | - | - | 0 | - |
| 2 | c | 8.403E+04 | 268.2 | 0.0002311 | 0.8616 | +1 | 2 |
| - | - | 1.038E+04 | 269.2 | - | - | 0 | - |
| - | - | 9400 | 283.2 | - | - | 0 | - |
| - | - | 2677 | 293.1 | - | - | 0 | - |
| 14 | z | 1.119E+05 | 317.2 | 0.0006985 | 2.202 | +1 | 3 |
| - | - | 2.906E+04 | 318.2 | - | - | 0 | - |
| - | - | 3756 | 319.2 | - | - | 0 | - |
| 14 | y | 1.132E+04 | 333.2 | 0.0004071 | 1.222 | +1 | 3 |
| - | - | 3667 | 351.2 | - | - | 0 | - |
| - | - | 8106 | 352.2 | - | - | 0 | - |
| - | - | 3206 | 361.2 | - | - | 0 | - |
| - | - | 3002 | 361.8 | - | - | 0 | - |
| 3 | c | 2.849E+04 | 379.2 | 0.0003072 | 0.81 | +1 | 3 |
| - | - | 3640 | 380.2 | - | - | 0 | - |
| - | - | 1.442E+05 | 387.2 | - | - | 0 | - |
| - | - | 2.573E+04 | 388.2 | - | - | 0 | - |
| - | - | 3227 | 389.2 | - | - | 0 | - |
| - | - | 2796 | 390.2 | - | - | 0 | - |
| - | - | 1.668E+04 | 395.2 | - | - | 0 | - |
| 3 | c | 1.782E+05 | 396.2 | 0.0006135 | 1.548 | +1 | 3 |
| - | - | 3.808E+04 | 397.2 | - | - | 0 | - |
| - | - | 2919 | 398.2 | - | - | 0 | - |
| - | - | 3857 | 409.2 | - | - | 0 | - |
| - | - | 5467 | 429.1 | - | - | 0 | - |
| - | - | 4870 | 430.1 | - | - | 0 | - |
| - | - | 4387 | 431.1 | - | - | 0 | - |
| - | - | 4304 | 466.2 | - | - | 0 | - |
| - | - | 6.739E+04 | 467.2 | - | - | 0 | - |
| - | - | 2.151E+04 | 468.3 | - | - | 0 | - |
| - | - | 3544 | 469.3 | - | - | 0 | - |
| 4 | c | 3.969E+04 | 494.2 | 0.004266 | 8.633 | +1 | 4 |
| - | - | 8954 | 495.2 | - | - | 0 | - |
| - | - | 3117 | 496.3 | - | - | 0 | - |
| - | - | 6526 | 497.8 | - | - | 0 | - |
| - | - | 2625 | 505.7 | - | - | 0 | - |
| - | - | 3838 | 510.3 | - | - | 0 | - |
| 4 | c | 1.385E+05 | 511.3 | 0.0007701 | 1.506 | +1 | 4 |
| - | - | 3.181E+04 | 512.3 | - | - | 0 | - |
| - | - | 4503 | 513.3 | - | - | 0 | - |
| - | - | 1.875E+04 | 515.3 | - | - | 0 | - |
| - | - | 6496 | 516.3 | - | - | 0 | - |
| - | - | 3544 | 522.3 | - | - | 0 | - |
| - | - | 8836 | 547.3 | - | - | 0 | - |
| 12 | w | 5386 | 548.2 | 0.004165 | 7.597 | +1 | 5 |
| - | - | 2605 | 548.3 | - | - | 0 | - |
| 8 | y | 4.76E+04 | 550.3 | 0.0007384 | 1.342 | +2 | 9 |
| - | - | 2.967E+04 | 550.8 | - | - | 0 | - |
| - | - | 8299 | 551.3 | - | - | 0 | - |
| - | - | 3860 | 555.3 | - | - | 0 | - |
| - | - | 4974 | 567.3 | - | - | 0 | - |
| 7 | w | 2578 | 577.3 | 0.005471 | 9.478 | +2 | 10 |
| - | - | 6738 | 597.3 | - | - | 0 | - |
| 7 | y | 2844 | 598.8 | 0.0003827 | 0.6391 | +2 | 10 |
| 14 | c | 1.688E+04 | 601.6 | 0.0009041 | 1.503 | +3 | 14 |
| - | - | 1.528E+04 | 602 | - | - | 0 | - |
| - | - | 7720 | 602.3 | - | - | 0 | - |
| - | - | 3590 | 602.6 | - | - | 0 | - |
| 12 | z | 1.142E+05 | 606.3 | 0.003494 | 5.763 | +1 | 5 |
| 14 | c | 4.941E+04 | 607.3 | 0.009531 | 15.69 | +3 | 14 |
| 7 | y | 2.804E+04 | 607.8 | 0.001026 | 1.689 | +2 | 10 |
| - | - | 2.03E+04 | 608.3 | - | - | 0 | - |
| - | - | 8638 | 608.8 | - | - | 0 | - |
| - | - | 2937 | 609.3 | - | - | 0 | - |
| 10 | c | 4703 | 611.8 | 0.00184 | 3.008 | +2 | 10 |
| - | - | 3428 | 612.3 | - | - | 0 | - |
| 12 | y | 5.082E+04 | 622.3 | 0.003297 | 5.298 | +1 | 5 |
| - | - | 1.511E+04 | 623.3 | - | - | 0 | - |
| - | - | 3735 | 624.3 | - | - | 0 | - |
| - | - | 3391 | 625.3 | - | - | 0 | - |
| - | - | 7217 | 625.6 | - | - | 0 | - |
| 2 | y | 4930 | 626.3 | 0.007019 | 11.21 | +3 | 15 |
| - | - | 1.329E+04 | 628.6 | - | - | 0 | - |
| - | - | 1.497E+04 | 629 | - | - | 0 | - |
| - | - | 1.026E+04 | 629.3 | - | - | 0 | - |
| - | - | 2987 | 629.6 | - | - | 0 | - |
| - | - | 2595 | 631.6 | - | - | 0 | - |
| 2 | y | 4.798E+04 | 632 | 0.0001017 | 0.1609 | +3 | 15 |
| - | - | 4.408E+04 | 632.3 | - | - | 0 | - |
| - | - | 2.458E+04 | 632.6 | - | - | 0 | - |
| - | - | 8258 | 633 | - | - | 0 | - |
| 15 | c | 7.406E+04 | 634.6 | 0.000111 | 0.1749 | +3 | 15 |
| - | - | 7.826E+04 | 635 | - | - | 0 | - |
| - | - | 4.122E+04 | 635.3 | - | - | 0 | - |
| - | - | 2.099E+04 | 635.6 | - | - | 0 | - |
| - | - | 5780 | 636 | - | - | 0 | - |
| - | - | 4271 | 637.3 | - | - | 0 | - |
| - | - | 5548 | 637.4 | - | - | 0 | - |
| - | - | 2772 | 638.4 | - | - | 0 | - |
| - | - | 2857 | 641 | - | - | 0 | - |
| - | - | 5803 | 652.3 | - | - | 0 | - |
| - | - | 4335 | 653.3 | - | - | 0 | - |
| - | - | 4133 | 654.3 | - | - | 0 | - |
| - | - | 3931 | 655.6 | - | - | 0 | - |
| - | - | 4921 | 657.7 | - | - | 0 | - |
| - | - | 1.274E+04 | 658 | - | - | 0 | - |
| - | - | 1.839E+04 | 658.3 | - | - | 0 | - |
| - | - | 8739 | 658.7 | - | - | 0 | - |
| - | - | 6.027E+04 | 663.7 | - | - | 0 | - |
| - | - | 7.496E+04 | 664 | - | - | 0 | - |
| 6 | y | 5.79E+04 | 664.3 | 0.01083 | 16.3 | +2 | 11 |
| - | - | 2.982E+04 | 664.7 | - | - | 0 | - |
| - | - | 5234 | 664.8 | - | - | 0 | - |
| - | - | 1.105E+04 | 665 | - | - | 0 | - |
| - | - | 5025 | 665.3 | - | - | 0 | - |
| - | - | 3148 | 668.3 | - | - | 0 | - |
| - | - | 4105 | 668.7 | - | - | 0 | - |
| - | - | 8857 | 669 | - | - | 0 | - |
| - | - | 3.881E+04 | 669.3 | - | - | 0 | - |
| - | - | 8.106E+05 | 669.7 | - | - | 0 | - |
| - | - | 8.888E+05 | 670 | - | - | 0 | - |
| - | - | 5.309E+05 | 670.3 | - | - | 0 | - |
| - | - | 2.594E+05 | 670.7 | - | - | 0 | - |
| - | - | 9.105E+04 | 671 | - | - | 0 | - |
| - | - | 7601 | 678.4 | - | - | 0 | - |
| - | - | 7228 | 679.4 | - | - | 0 | - |
| 5 | c | 1.08E+04 | 680.3 | 0.001504 | 2.21 | +1 | 5 |
| - | - | 7633 | 681.3 | - | - | 0 | - |
| - | - | 6715 | 696.3 | - | - | 0 | - |
| 5 | c | 1.488E+05 | 697.3 | 0.0007723 | 1.108 | +1 | 5 |
| - | - | 5.777E+04 | 698.3 | - | - | 0 | - |
| - | - | 1.621E+04 | 699.3 | - | - | 0 | - |
| - | - | 4032 | 711.3 | - | - | 0 | - |
| 12 | c | 2.132E+04 | 757.4 | 0.002354 | 3.109 | +2 | 12 |
| - | - | 1.272E+04 | 757.9 | - | - | 0 | - |
| - | - | 7950 | 758.4 | - | - | 0 | - |
| - | - | 1.616E+04 | 766.4 | - | - | 0 | - |
| - | - | 1.623E+04 | 767.4 | - | - | 0 | - |
| 11 | y | 3683 | 768.4 | 0.007789 | 10.14 | +1 | 6 |
| - | - | 4267 | 768.4 | - | - | 0 | - |
| 11 | z | 1.494E+05 | 769.4 | 0.003468 | 4.508 | +1 | 6 |
| - | - | 9.139E+04 | 770.4 | - | - | 0 | - |
| - | - | 3.174E+04 | 771.4 | - | - | 0 | - |
| - | - | 9401 | 772.4 | - | - | 0 | - |
| - | - | 3537 | 777.4 | - | - | 0 | - |
| 11 | y | 5.048E+04 | 785.4 | 0.003088 | 3.932 | +1 | 6 |
| - | - | 2.114E+04 | 786.4 | - | - | 0 | - |
| - | - | 5062 | 787.4 | - | - | 0 | - |
| 6 | c | 1.232E+04 | 793.4 | 0.00179 | 2.256 | +1 | 6 |
| - | - | 4911 | 794.4 | - | - | 0 | - |
| - | - | 4557 | 807.4 | - | - | 0 | - |
| - | - | 1.152E+04 | 809.4 | - | - | 0 | - |
| 6 | c | 1.987E+05 | 810.4 | 0.001242 | 1.533 | +1 | 6 |
| - | - | 8.341E+04 | 811.4 | - | - | 0 | - |
| - | - | 2.464E+04 | 812.4 | - | - | 0 | - |
| - | - | 4550 | 813.4 | - | - | 0 | - |
| 4 | y | 1.359E+04 | 814.9 | 0.001155 | 1.417 | +2 | 13 |
| - | - | 1.815E+04 | 815.4 | - | - | 0 | - |
| - | - | 9659 | 815.9 | - | - | 0 | - |
| - | - | 6226 | 816.4 | - | - | 0 | - |
| - | - | 3363 | 817.9 | - | - | 0 | - |
| - | - | 6580 | 818.4 | - | - | 0 | - |
| - | - | 4782 | 824.4 | - | - | 0 | - |
| - | - | 9428 | 826.4 | - | - | 0 | - |
| - | - | 9065 | 826.9 | - | - | 0 | - |
| 10 | w | 2.852E+04 | 839.4 | 0.002911 | 3.468 | +1 | 7 |
| - | - | 1.131E+04 | 840.4 | - | - | 0 | - |
| - | - | 7318 | 841.4 | - | - | 0 | - |
| 3 | w | 3453 | 841.9 | 0.001376 | 1.634 | +2 | 14 |
| - | - | 1.34E+04 | 842.9 | - | - | 0 | - |
| - | - | 1.438E+04 | 843.4 | - | - | 0 | - |
| - | - | 8255 | 843.9 | - | - | 0 | - |
| 13 | c | 8249 | 846.4 | 0.001451 | 1.714 | +2 | 13 |
| - | - | 6797 | 846.9 | - | - | 0 | - |
| - | - | 3496 | 847.4 | - | - | 0 | - |
| - | - | 4548 | 849.4 | - | - | 0 | - |
| - | - | 4807 | 858.9 | - | - | 0 | - |
| - | - | 5610 | 859.4 | - | - | 0 | - |
| - | - | 4065 | 859.9 | - | - | 0 | - |
| - | - | 3789 | 861.4 | - | - | 0 | - |
| 3 | z | 3693 | 862.4 | 0.01081 | 12.53 | +2 | 14 |
| - | - | 4283 | 864.4 | - | - | 0 | - |
| - | - | 5643 | 866.4 | - | - | 0 | - |
| - | - | 4026 | 866.9 | - | - | 0 | - |
| - | - | 5.33E+04 | 867.4 | - | - | 0 | - |
| - | - | 2.757E+04 | 868.4 | - | - | 0 | - |
| - | - | 5390 | 869.4 | - | - | 0 | - |
| 3 | y | 1.471E+04 | 869.9 | 0.00331 | 3.805 | +2 | 14 |
| 3 | y | 3.858E+04 | 870.4 | 0.002301 | 2.644 | +2 | 14 |
| 3 | z | 3.269E+04 | 870.9 | 0.0008787 | 1.009 | +2 | 14 |
| - | - | 2.26E+04 | 871.4 | - | - | 0 | - |
| - | - | 1.04E+04 | 871.9 | - | - | 0 | - |
| - | - | 6408 | 878.4 | - | - | 0 | - |
| 3 | y | 8.384E+04 | 878.9 | 0.0007192 | 0.8183 | +2 | 14 |
| - | - | 8.143E+04 | 879.4 | - | - | 0 | - |
| - | - | 4.285E+04 | 879.9 | - | - | 0 | - |
| 10 | z | 1.746E+04 | 880.4 | 0.01382 | 15.7 | +1 | 7 |
| - | - | 5786 | 880.9 | - | - | 0 | - |
| - | - | 2.228E+04 | 881.4 | - | - | 0 | - |
| - | - | 1.34E+04 | 882.4 | - | - | 0 | - |
| - | - | 3848 | 883.5 | - | - | 0 | - |
| - | - | 4273 | 886.9 | - | - | 0 | - |
| - | - | 5593 | 887.9 | - | - | 0 | - |
| - | - | 1.951E+04 | 888.4 | - | - | 0 | - |
| - | - | 2.164E+04 | 888.9 | - | - | 0 | - |
| - | - | 1.307E+04 | 889.4 | - | - | 0 | - |
| - | - | 4945 | 889.9 | - | - | 0 | - |
| - | - | 4530 | 892.5 | - | - | 0 | - |
| - | - | 6295 | 893 | - | - | 0 | - |
| - | - | 5058 | 893.5 | - | - | 0 | - |
| - | - | 8455 | 893.9 | - | - | 0 | - |
| - | - | 5136 | 894.4 | - | - | 0 | - |
| - | - | 3298 | 894.9 | - | - | 0 | - |
| - | - | 4430 | 895.4 | - | - | 0 | - |
| - | - | 3502 | 895.9 | - | - | 0 | - |
| 10 | y | 4532 | 896.4 | 0.012 | 13.39 | +1 | 7 |
| 10 | z | 1.486E+05 | 898.4 | 0.002909 | 3.238 | +1 | 7 |
| - | - | 6.671E+04 | 899.4 | - | - | 0 | - |
| - | - | 2.969E+04 | 900.4 | - | - | 0 | - |
| 14 | c | 3483 | 901.4 | 0.01524 | 16.9 | +2 | 14 |
| 14 | c | 1.028E+04 | 901.9 | 0.0007735 | 0.8575 | +2 | 14 |
| - | - | 1.292E+04 | 902.4 | - | - | 0 | - |
| - | - | 6809 | 902.9 | - | - | 0 | - |
| - | - | 8262 | 903.9 | - | - | 0 | - |
| - | - | 3825 | 904.4 | - | - | 0 | - |
| - | - | 3702 | 905.4 | - | - | 0 | - |
| 7 | c | 3.525E+04 | 908.4 | 0.004205 | 4.629 | +1 | 7 |
| - | - | 7817 | 909 | - | - | 0 | - |
| - | - | 1.638E+04 | 909.4 | - | - | 0 | - |
| - | - | 1.46E+04 | 909.9 | - | - | 0 | - |
| 14 | c | 1.35E+05 | 910.4 | 0.001841 | 2.022 | +2 | 14 |
| - | - | 1.277E+05 | 910.9 | - | - | 0 | - |
| - | - | 8.079E+04 | 911.4 | - | - | 0 | - |
| - | - | 3.071E+04 | 911.9 | - | - | 0 | - |
| - | - | 1.093E+04 | 912.4 | - | - | 0 | - |
| 10 | y | 2.493E+04 | 914.4 | 0.00198 | 2.165 | +1 | 7 |
| - | - | 1.644E+04 | 915.4 | - | - | 0 | - |
| - | - | 3374 | 916 | - | - | 0 | - |
| - | - | 8526 | 916.5 | - | - | 0 | - |
| - | - | 4727 | 917 | - | - | 0 | - |
| - | - | 1.548E+04 | 917.4 | - | - | 0 | - |
| - | - | 7753 | 917.9 | - | - | 0 | - |
| - | - | 5374 | 918.4 | - | - | 0 | - |
| - | - | 3334 | 923.5 | - | - | 0 | - |
| - | - | 2.56E+04 | 924.4 | - | - | 0 | - |
| 7 | c | 5.232E+04 | 925.5 | 0.0009714 | 1.05 | +1 | 7 |
| - | - | 2.625E+04 | 926.5 | - | - | 0 | - |
| - | - | 5488 | 927.5 | - | - | 0 | - |
| - | - | 3236 | 928.5 | - | - | 0 | - |
| - | - | 6242 | 929 | - | - | 0 | - |
| - | - | 5571 | 929.5 | - | - | 0 | - |
| - | - | 3580 | 930 | - | - | 0 | - |
| - | - | 7636 | 930.5 | - | - | 0 | - |
| - | - | 7865 | 931.5 | - | - | 0 | - |
| - | - | 1.196E+04 | 935.5 | - | - | 0 | - |
| - | - | 9241 | 936.5 | - | - | 0 | - |
| - | - | 3498 | 937 | - | - | 0 | - |
| - | - | 5127 | 937.5 | - | - | 0 | - |
| - | - | 3.315E+04 | 938 | - | - | 0 | - |
| - | - | 4.373E+04 | 938.5 | - | - | 0 | - |
| - | - | 3.815E+04 | 939 | - | - | 0 | - |
| 2 | z | 8.264E+04 | 939.4 | 0.008789 | 9.356 | +2 | 15 |
| - | - | 8.013E+04 | 939.9 | - | - | 0 | - |
| - | - | 4.6E+04 | 940.4 | - | - | 0 | - |
| - | - | 1.564E+04 | 940.9 | - | - | 0 | - |
| - | - | 5767 | 941.4 | - | - | 0 | - |
| - | - | 5026 | 943 | - | - | 0 | - |
| - | - | 6872 | 943.5 | - | - | 0 | - |
| - | - | 4569 | 944 | - | - | 0 | - |
| - | - | 3224 | 945 | - | - | 0 | - |
| - | - | 7609 | 946 | - | - | 0 | - |
| - | - | 1.28E+04 | 946.5 | - | - | 0 | - |
| 2 | y | 5872 | 947.4 | 0.003516 | 3.711 | +2 | 15 |
| - | - | 6421 | 947.9 | - | - | 0 | - |
| - | - | 2827 | 948.4 | - | - | 0 | - |
| - | - | 2.266E+04 | 950.5 | - | - | 0 | - |
| 15 | c | 1.37E+04 | 951 | 0.004635 | 4.874 | +2 | 15 |
| 15 | c | 1.554E+04 | 951.5 | 0.01476 | 15.52 | +2 | 15 |
| - | - | 9261 | 952 | - | - | 0 | - |
| - | - | 1.465E+04 | 952.5 | - | - | 0 | - |
| - | - | 8209 | 953 | - | - | 0 | - |
| - | - | 9118 | 953.5 | - | - | 0 | - |
| - | - | 7616 | 954 | - | - | 0 | - |
| - | - | 3574 | 959 | - | - | 0 | - |
| - | - | 1.642E+04 | 959.5 | - | - | 0 | - |
| 15 | c | 2.299E+05 | 960 | 2.385E-05 | 0.02484 | +2 | 15 |
| - | - | 2.341E+05 | 960.5 | - | - | 0 | - |
| - | - | 1.455E+05 | 961 | - | - | 0 | - |
| - | - | 5.352E+04 | 961.5 | - | - | 0 | - |
| - | - | 1.515E+04 | 962 | - | - | 0 | - |
| - | - | 1.125E+04 | 964.5 | - | - | 0 | - |
| - | - | 1.161E+04 | 965 | - | - | 0 | - |
| 8 | c | 1.61E+04 | 965.4 | 0.01894 | 19.61 | +1 | 8 |
| - | - | 1.561E+04 | 966 | - | - | 0 | - |
| - | - | 1.607E+04 | 966.5 | - | - | 0 | - |
| - | - | 1.485E+04 | 967 | - | - | 0 | - |
| - | - | 1.532E+04 | 967.5 | - | - | 0 | - |
| - | - | 1.727E+04 | 968 | - | - | 0 | - |
| - | - | 2.402E+04 | 968.5 | - | - | 0 | - |
| - | - | 1.442E+04 | 969 | - | - | 0 | - |
| - | - | 4399 | 969.5 | - | - | 0 | - |
| - | - | 5056 | 972.5 | - | - | 0 | - |
| - | - | 1.054E+04 | 973 | - | - | 0 | - |
| - | - | 1.704E+04 | 973.5 | - | - | 0 | - |
| - | - | 3.295E+04 | 974 | - | - | 0 | - |
| - | - | 1.401E+05 | 974.5 | - | - | 0 | - |
| - | - | 4.054E+05 | 975 | - | - | 0 | - |
| - | - | 3.343E+05 | 975.5 | - | - | 0 | - |
| - | - | 2.122E+05 | 976 | - | - | 0 | - |
| - | - | 7.432E+04 | 976.5 | - | - | 0 | - |
| - | - | 2.901E+04 | 977 | - | - | 0 | - |
| - | - | 5338 | 978 | - | - | 0 | - |
| - | - | 4698 | 980.5 | - | - | 0 | - |
| - | - | 9.876E+04 | 981.5 | - | - | 0 | - |
| - | - | 6.444E+04 | 982 | - | - | 0 | - |
| 8 | c | 2.368E+05 | 982.5 | 0.004959 | 5.048 | +1 | 8 |
| - | - | 3.423E+04 | 983 | - | - | 0 | - |
| - | - | 1.044E+05 | 983.5 | - | - | 0 | - |
| - | - | 3945 | 984 | - | - | 0 | - |
| - | - | 2.888E+04 | 984.5 | - | - | 0 | - |
| - | - | 6657 | 985.5 | - | - | 0 | - |
| - | - | 1.165E+04 | 986.5 | - | - | 0 | - |
| - | - | 1.082E+04 | 987 | - | - | 0 | - |
| - | - | 9230 | 987.5 | - | - | 0 | - |
| - | - | 8817 | 988 | - | - | 0 | - |
| - | - | 5165 | 988.5 | - | - | 0 | - |
| - | - | 8843 | 995 | - | - | 0 | - |
| - | - | 4.136E+04 | 995.5 | - | - | 0 | - |
| - | - | 1.208E+05 | 996 | - | - | 0 | - |
| - | - | 1.039E+05 | 996.5 | - | - | 0 | - |
| - | - | 7.566E+04 | 997 | - | - | 0 | - |
| - | - | 4.692E+04 | 997.5 | - | - | 0 | - |
| - | - | 1.016E+04 | 998 | - | - | 0 | - |
| - | - | 6867 | 1002 | - | - | 0 | - |
| - | - | 5778 | 1003 | - | - | 0 | - |
| - | - | 2.262E+04 | 1003 | - | - | 0 | - |
| - | - | 4.364E+05 | 1004 | - | - | 0 | - |
| - | - | 7.188E+05 | 1004 | - | - | 0 | - |
| - | - | 6.097E+05 | 1005 | - | - | 0 | - |
| - | - | 3.591E+05 | 1005 | - | - | 0 | - |
| - | - | 1.316E+05 | 1006 | - | - | 0 | - |
| - | - | 3.555E+04 | 1006 | - | - | 0 | - |
| 9 | z | 1.259E+05 | 1027 | 0.003085 | 3.005 | +1 | 8 |
| - | - | 1.069E+05 | 1028 | - | - | 0 | - |
| - | - | 4.221E+04 | 1029 | - | - | 0 | - |
| - | - | 1.528E+04 | 1030 | - | - | 0 | - |
| - | - | 3587 | 1031 | - | - | 0 | - |
| - | - | 8802 | 1042 | - | - | 0 | - |
| 9 | y | 1.406E+04 | 1043 | 0.004963 | 4.761 | +1 | 8 |
| - | - | 8834 | 1044 | - | - | 0 | - |
| - | - | 1.13E+04 | 1067 | - | - | 0 | - |
| - | - | 9664 | 1068 | - | - | 0 | - |
| 8 | z | 3.393E+04 | 1084 | 0.002576 | 2.377 | +1 | 9 |
| - | - | 4.516E+04 | 1085 | - | - | 0 | - |
| - | - | 1.929E+04 | 1086 | - | - | 0 | - |
| - | - | 7014 | 1087 | - | - | 0 | - |
| 8 | y | 3.258E+04 | 1100 | 0.003356 | 3.052 | +1 | 9 |
| - | - | 1.783E+04 | 1101 | - | - | 0 | - |
| - | - | 7707 | 1102 | - | - | 0 | - |
| - | - | 4415 | 1108 | - | - | 0 | - |
| - | - | 6858 | 1109 | - | - | 0 | - |
| - | - | 9917 | 1110 | - | - | 0 | - |
| 9 | c | 1.856E+05 | 1111 | 0.001488 | 1.34 | +1 | 9 |
| - | - | 1.085E+05 | 1112 | - | - | 0 | - |
| - | - | 4.042E+04 | 1113 | - | - | 0 | - |
| - | - | 8126 | 1114 | - | - | 0 | - |
| - | - | 1.527E+04 | 1137 | - | - | 0 | - |
| - | - | 1.256E+04 | 1138 | - | - | 0 | - |
| - | - | 4230 | 1139 | - | - | 0 | - |
| - | - | 5.342E+04 | 1142 | - | - | 0 | - |
| - | - | 3.4E+04 | 1143 | - | - | 0 | - |
| - | - | 1.846E+04 | 1144 | - | - | 0 | - |
| - | - | 1.179E+04 | 1155 | - | - | 0 | - |
| - | - | 6116 | 1156 | - | - | 0 | - |
| - | - | 1.991E+04 | 1196 | - | - | 0 | - |
| - | - | 1.788E+04 | 1197 | - | - | 0 | - |
| - | - | 7435 | 1198 | - | - | 0 | - |
| 7 | z | 1.192E+05 | 1199 | 0.002663 | 2.222 | +1 | 10 |
| - | - | 9.061E+04 | 1200 | - | - | 0 | - |
| - | - | 4.582E+04 | 1201 | - | - | 0 | - |
| - | - | 1.069E+04 | 1202 | - | - | 0 | - |
| - | - | 5806 | 1214 | - | - | 0 | - |
| 7 | y | 4.167E+04 | 1215 | 0.002711 | 2.232 | +1 | 10 |
| - | - | 2.381E+04 | 1216 | - | - | 0 | - |
| - | - | 1.007E+04 | 1217 | - | - | 0 | - |
| - | - | 3.011E+04 | 1239 | - | - | 0 | - |
| 10 | c | 1.849E+05 | 1240 | 0.0006429 | 0.5187 | +1 | 10 |
| - | - | 1.194E+05 | 1241 | - | - | 0 | - |
| - | - | 4622 | 1241 | - | - | 0 | - |
| - | - | 3.835E+04 | 1242 | - | - | 0 | - |
| - | - | 1.028E+04 | 1243 | - | - | 0 | - |
| - | - | 1.725E+04 | 1268 | - | - | 0 | - |
| 6 | w | 1.038E+04 | 1269 | 0.0148 | 11.67 | +1 | 11 |
| - | - | 5531 | 1270 | - | - | 0 | - |
| 6 | y | 5126 | 1311 | 0.01729 | 13.19 | +1 | 11 |
| 6 | z | 7.732E+04 | 1312 | 0.002499 | 1.905 | +1 | 11 |
| - | - | 5.915E+04 | 1313 | - | - | 0 | - |
| - | - | 3.907E+04 | 1314 | - | - | 0 | - |
| - | - | 1.07E+04 | 1315 | - | - | 0 | - |
| 6 | y | 1.126E+04 | 1328 | 0.01011 | 7.618 | +1 | 11 |
| - | - | 3794 | 1329 | - | - | 0 | - |
| - | - | 4815 | 1330 | - | - | 0 | - |
| - | - | 3.104E+04 | 1359 | - | - | 0 | - |
| - | - | 2.057E+04 | 1360 | - | - | 0 | - |
| - | - | 7289 | 1361 | - | - | 0 | - |
| - | - | 1.138E+04 | 1402 | - | - | 0 | - |
| 11 | c | 1.34E+05 | 1403 | 0.001279 | 0.912 | +1 | 11 |
| - | - | 9.531E+04 | 1404 | - | - | 0 | - |
| - | - | 3.619E+04 | 1405 | - | - | 0 | - |
| - | - | 1.034E+04 | 1406 | - | - | 0 | - |
| - | - | 5711 | 1442 | - | - | 0 | - |
| - | - | 4559 | 1443 | - | - | 0 | - |
| - | - | 3300 | 1482 | - | - | 0 | - |
| - | - | 1.356E+04 | 1487 | - | - | 0 | - |
| - | - | 1.421E+04 | 1488 | - | - | 0 | - |
| - | - | 6198 | 1489 | - | - | 0 | - |
| 5 | y | 4738 | 1496 | 0.01038 | 6.941 | +1 | 12 |
| 5 | y | 5892 | 1497 | 0.02697 | 18.02 | +1 | 12 |
| 5 | z | 6.785E+04 | 1498 | 0.001612 | 1.076 | +1 | 12 |
| - | - | 7.159E+04 | 1499 | - | - | 0 | - |
| - | - | 3.049E+04 | 1500 | - | - | 0 | - |
| - | - | 1.834E+04 | 1501 | - | - | 0 | - |
| 12 | c | 7717 | 1513 | 0.0172 | 11.37 | +1 | 12 |
| 12 | c | 6.578E+04 | 1514 | 0.007564 | 4.997 | +1 | 12 |
| - | - | 4.858E+04 | 1515 | - | - | 0 | - |
| - | - | 2.887E+04 | 1516 | - | - | 0 | - |
| - | - | 7972 | 1517 | - | - | 0 | - |
| - | - | 3625 | 1522 | - | - | 0 | - |
| - | - | 4003 | 1523 | - | - | 0 | - |
| - | - | 5474 | 1526 | - | - | 0 | - |
| - | - | 3786 | 1527 | - | - | 0 | - |
| - | - | 1.035E+04 | 1530 | - | - | 0 | - |
| 12 | c | 1.276E+05 | 1531 | 0.001043 | 0.6812 | +1 | 12 |
| - | - | 1.11E+05 | 1532 | - | - | 0 | - |
| - | - | 6.03E+04 | 1533 | - | - | 0 | - |
| - | - | 1.607E+04 | 1534 | - | - | 0 | - |
| - | - | 8478 | 1551 | - | - | 0 | - |
| - | - | 6721 | 1552 | - | - | 0 | - |
| - | - | 4716 | 1555 | - | - | 0 | - |
| - | - | 6816 | 1556 | - | - | 0 | - |
| - | - | 1.041E+04 | 1557 | - | - | 0 | - |
| - | - | 1.037E+04 | 1558 | - | - | 0 | - |
| - | - | 4771 | 1559 | - | - | 0 | - |
| 4 | w | 4338 | 1568 | 0.007341 | 4.682 | +1 | 13 |
| - | - | 4.571E+04 | 1569 | - | - | 0 | - |
| - | - | 4.056E+04 | 1570 | - | - | 0 | - |
| - | - | 3.25E+04 | 1571 | - | - | 0 | - |
| - | - | 1.226E+04 | 1572 | - | - | 0 | - |
| - | - | 5607 | 1573 | - | - | 0 | - |
| 4 | y | 7240 | 1612 | 0.009422 | 5.846 | +1 | 13 |
| 4 | z | 1.213E+05 | 1613 | 0.004141 | 2.567 | +1 | 13 |
| - | - | 1.623E+05 | 1614 | - | - | 0 | - |
| - | - | 9.326E+04 | 1615 | - | - | 0 | - |
| - | - | 3.966E+04 | 1616 | - | - | 0 | - |
| - | - | 1.572E+04 | 1617 | - | - | 0 | - |
| - | - | 3925 | 1628 | - | - | 0 | - |
| 4 | y | 2.593E+04 | 1629 | 0.00431 | 2.646 | +1 | 13 |
| - | - | 2.22E+04 | 1630 | - | - | 0 | - |
| - | - | 1.154E+04 | 1631 | - | - | 0 | - |
| - | - | 4115 | 1633 | - | - | 0 | - |
| - | - | 5478 | 1634 | - | - | 0 | - |
| - | - | 3362 | 1647 | - | - | 0 | - |
| - | - | 3438 | 1653 | - | - | 0 | - |
| - | - | 3665 | 1655 | - | - | 0 | - |
| - | - | 4276 | 1678 | - | - | 0 | - |
| 3 | w | 5766 | 1683 | 0.02101 | 12.49 | +1 | 14 |
| - | - | 8670 | 1684 | - | - | 0 | - |
| - | - | 6831 | 1685 | - | - | 0 | - |
| - | - | 2.249E+04 | 1691 | - | - | 0 | - |
| 13 | c | 1.548E+05 | 1692 | 0.003127 | 1.848 | +1 | 13 |
| - | - | 1.534E+05 | 1693 | - | - | 0 | - |
| - | - | 8.706E+04 | 1694 | - | - | 0 | - |
| - | - | 3.381E+04 | 1695 | - | - | 0 | - |
| - | - | 8794 | 1696 | - | - | 0 | - |
| - | - | 6167 | 1697 | - | - | 0 | - |
| - | - | 5116 | 1698 | - | - | 0 | - |
| - | - | 3878 | 1699 | - | - | 0 | - |
| 3 | z | 9.889E+04 | 1741 | 0.001927 | 1.107 | +1 | 14 |
| - | - | 1.469E+05 | 1742 | - | - | 0 | - |
| - | - | 8.613E+04 | 1743 | - | - | 0 | - |
| - | - | 4.239E+04 | 1744 | - | - | 0 | - |
| - | - | 1.018E+04 | 1745 | - | - | 0 | - |
| 3 | y | 2.592E+04 | 1757 | 0.002219 | 1.263 | +1 | 14 |
| - | - | 2.394E+04 | 1758 | - | - | 0 | - |
| - | - | 1.706E+04 | 1759 | - | - | 0 | - |
| - | - | 8025 | 1760 | - | - | 0 | - |
| - | - | 6339 | 1761 | - | - | 0 | - |
| - | - | 6152 | 1762 | - | - | 0 | - |
| - | - | 6757 | 1763 | - | - | 0 | - |
| - | - | 5326 | 1764 | - | - | 0 | - |
| - | - | 2E+04 | 1777 | - | - | 0 | - |
| - | - | 1.835E+04 | 1778 | - | - | 0 | - |
| - | - | 7897 | 1779 | - | - | 0 | - |
| - | - | 5035 | 1804 | - | - | 0 | - |
| - | - | 5621 | 1805 | - | - | 0 | - |
| - | - | 3668 | 1819 | - | - | 0 | - |
| 14 | c | 2.096E+04 | 1820 | 0.009101 | 5.001 | +1 | 14 |
| - | - | 3.853E+04 | 1821 | - | - | 0 | - |
| - | - | 2.798E+04 | 1822 | - | - | 0 | - |
| - | - | 1.658E+04 | 1823 | - | - | 0 | - |
| - | - | 6819 | 1824 | - | - | 0 | - |
| - | - | 4074 | 1860 | - | - | 0 | - |
| - | - | 1.09E+04 | 1861 | - | - | 0 | - |
| - | - | 7598 | 1862 | - | - | 0 | - |
| - | - | 4017 | 1863 | - | - | 0 | - |
| - | - | 9342 | 1875 | - | - | 0 | - |
| - | - | 1.99E+04 | 1876 | - | - | 0 | - |
| - | - | 1.94E+04 | 1877 | - | - | 0 | - |
| - | - | 2.209E+04 | 1878 | - | - | 0 | - |
| - | - | 5.642E+04 | 1879 | - | - | 0 | - |
| - | - | 5.468E+04 | 1880 | - | - | 0 | - |
| - | - | 3.586E+04 | 1881 | - | - | 0 | - |
| - | - | 1.36E+04 | 1882 | - | - | 0 | - |
| - | - | 5921 | 1890 | - | - | 0 | - |
| - | - | 4614 | 1891 | - | - | 0 | - |
| - | - | 7682 | 1892 | - | - | 0 | - |
| - | - | 9253 | 1893 | - | - | 0 | - |
| - | - | 7424 | 1894 | - | - | 0 | - |
| 15 | c | 8851 | 1901 | 0.02924 | 15.38 | +1 | 15 |
| - | - | 5298 | 1902 | - | - | 0 | - |
| - | - | 5900 | 1903 | - | - | 0 | - |
| - | - | 8371 | 1904 | - | - | 0 | - |
| - | - | 5169 | 1905 | - | - | 0 | - |
| - | - | 5298 | 1906 | - | - | 0 | - |
| - | - | 1.622E+04 | 1917 | - | - | 0 | - |
| - | - | 3.392E+04 | 1918 | - | - | 0 | - |
| 15 | c | 5.426E+04 | 1919 | 0.02466 | 12.85 | +1 | 15 |
| - | - | 6.125E+04 | 1920 | - | - | 0 | - |
| - | - | 5.457E+04 | 1921 | - | - | 0 | - |
| - | - | 3.731E+04 | 1922 | - | - | 0 | - |
| - | - | 2.173E+04 | 1923 | - | - | 0 | - |
| - | - | 1.235E+04 | 1924 | - | - | 0 | - |
| - | - | 1.3E+04 | 1932 | - | - | 0 | - |
| - | - | 2.329E+04 | 1933 | - | - | 0 | - |
| - | - | 2.296E+04 | 1934 | - | - | 0 | - |
| - | - | 1.118E+04 | 1935 | - | - | 0 | - |
| - | - | 9073 | 1936 | - | - | 0 | - |
| - | - | 7825 | 1946 | - | - | 0 | - |
| - | - | 5839 | 1947 | - | - | 0 | - |
| - | - | 2.376E+04 | 1948 | - | - | 0 | - |
| - | - | 8.34E+04 | 1949 | - | - | 0 | - |
| - | - | 1.492E+05 | 1950 | - | - | 0 | - |
| - | - | 1.086E+05 | 1951 | - | - | 0 | - |
| - | - | 5.992E+04 | 1952 | - | - | 0 | - |
| - | - | 2.898E+04 | 1953 | - | - | 0 | - |
| - | - | 6770 | 1954 | - | - | 0 | - |
| - | - | 7713 | 1962 | - | - | 0 | - |
| - | - | 4.236E+04 | 1963 | - | - | 0 | - |
| - | - | 7.821E+04 | 1964 | - | - | 0 | - |
| - | - | 6.422E+04 | 1965 | - | - | 0 | - |
| - | - | 3.54E+04 | 1966 | - | - | 0 | - |
| - | - | 1.36E+04 | 1967 | - | - | 0 | - |
| - | - | 4640 | 1968 | - | - | 0 | - |
| - | - | 6905 | 1973 | - | - | 0 | - |
| - | - | 8787 | 1974 | - | - | 0 | - |
| - | - | 5887 | 1975 | - | - | 0 | - |
| - | - | 3.273E+04 | 1980 | - | - | 0 | - |
| - | - | 9.008E+04 | 1981 | - | - | 0 | - |
| - | - | 8.928E+04 | 1982 | - | - | 0 | - |
| - | - | 5.711E+04 | 1983 | - | - | 0 | - |
| - | - | 2.028E+04 | 1984 | - | - | 0 | - |
| - | - | 7685 | 1985 | - | - | 0 | - |
| - | - | 4728 | 1989 | - | - | 0 | - |
| - | - | 1.441E+04 | 1990 | - | - | 0 | - |
| - | - | 6.711E+04 | 1991 | - | - | 0 | - |
| - | - | 1.704E+05 | 1992 | - | - | 0 | - |
| - | - | 1.63E+05 | 1993 | - | - | 0 | - |
| - | - | 9.784E+04 | 1994 | - | - | 0 | - |
| - | - | 4.527E+04 | 1995 | - | - | 0 | - |
| - | - | 1.13E+04 | 1996 | - | - | 0 | - |
| - | - | 4303 | 2005 | - | - | 0 | - |
| - | - | 6614 | 2006 | - | - | 0 | - |
| - | - | 5.01E+04 | 2007 | - | - | 0 | - |
| - | - | 2.183E+05 | 2008 | - | - | 0 | - |
| - | - | 7.08E+05 | 2009 | - | - | 0 | - |
| - | - | 6.621E+05 | 2010 | - | - | 0 | - |
| - | - | 4.123E+05 | 2011 | - | - | 0 | - |
| - | - | 1.82E+05 | 2012 | - | - | 0 | - |
| - | - | 6.071E+04 | 2013 | - | - | 0 | - |
| - | - | 4702 | 2211 | - | - | 0 | - |
| - | - | 4412 | 2212 | - | - | 0 | - |
| - | - | 3628 | 3046 | - | - | 0 | - |

m/z Charge Intensity FragmentType MassShift Position
120.27899169921875 0 1797.1187
121.01175689697266 0 1489.3013
122.3408432006836 0 2142.6545
128.08238220214844 0 2362.201
129.06637573242188 0 3052.2378
129.10255432128906 0 5144.0903
133.62940979003906 0 1943.9524
136.00030517578125 0 2221.8662
138.06661987304688 0 3553.6016
146.09304809570312 0 2615.8118
155.09298706054688 0 10863.127
164.08233642578125 0 2967.3643
166.05593872070312 0 2114.0125
166.06138610839844 0 23628.047
168.13861083984375 0 5835.252
178.134033203125 0 6558.3306
185.1652069091797 0 13881.655
205.1185302734375 0 7644.7397 y 14
207.13665771484375 0 5784.7837
207.1605987548828 0 4264.9585
223.15570068359375 0 27641.166
224.16249084472656 0 5860.902
227.9366455078125 0 2424.5361
230.18594360351562 0 3190.7449
233.13978576660156 0 4598.4336
234.12408447265625 0 36220.797
234.1388702392578 0 2764.0176
234.14309692382812 0 2779.5244
235.12744140625 0 5855.589
248.11537170410156 0 4260.2397
250.14276123046875 0 5710.5054
251.15065002441406 0 150026.73
252.15396118164062 0 17205.578
262.88140869140625 0 2902.383
266.1249694824219 0 10430.946
268.1770324707031 0 84031.836 c 1
269.1803894042969 0 10380.701
283.15167236328125 0 9400.108
293.13555908203125 0 2676.579
317.1952209472656 0 111890.16 z 13
318.2005615234375 0 29062.574
319.20166015625 0 3756.3376
333.2136535644531 0 11324.22 y 13
351.2152404785156 0 3666.8909
352.22247314453125 0 8106.3125
361.2002868652344 0 3205.8823
361.75006103515625 0 3002.336
379.2091369628906 0 28486.643 c Ammonia loss 2
380.2113952636719 0 3640.0833
387.224365234375 0 144237.84
388.227294921875 0 25728.266
389.23089599609375 0 3227.3298
390.1946716308594 0 2796.4355
395.2281799316406 0 16677.72
396.2359924316406 0 178192.92 c 2
397.2392272949219 0 38083.215
398.2403259277344 0 2918.637
409.2208251953125 0 3856.603
429.09136962890625 0 5466.9673
430.0901184082031 0 4870.4155
431.0881652832031 0 4387.4995
466.2397766113281 0 4304.3584
467.24957275390625 0 67393.19
468.2536926269531 0 21508.158
469.2590026855469 0 3543.7493
494.23150634765625 0 39688.473 c Ammonia loss 3
495.2345275878906 0 8953.923
496.253173828125 0 3117.4497
497.75567626953125 0 6526.399
505.7311706542969 0 2625.1248
510.25347900390625 0 3837.5745
511.2630920410156 0 138463.75 c 3
512.26611328125 0 31807.549
513.2687377929688 0 4502.846
515.3198852539062 0 18749.229
516.3229370117188 0 6496.0684
522.2694702148438 0 3543.9883
547.2928466796875 0 8836.169
548.2388305664062 0 5386.3984 w 11
548.2918090820312 0 2605.117
550.2777709960938 0 47600.527 y 7
550.7786865234375 0 29674.607
551.278076171875 0 8298.995
555.26806640625 0 3859.5552
567.276611328125 0 4974.3276
577.2783203125 0 2578.458 w 6
597.28466796875 0 6737.5156
598.7863159179688 0 2844.0408 y Water loss 6
601.6249389648438 0 16883.572 c Ammonia loss 13
601.9579467773438 0 15279.824
602.2927856445312 0 7719.6924
602.6233520507812 0 3589.79
606.30517578125 0 114195.26 z 11
607.30908203125 0 49408.46 c 13
607.7909545898438 0 28044.504 y 6
608.301513671875 0 20304.303
608.79296875 0 8638.007
609.2985229492188 0 2937.1914
611.7980346679688 0 4702.97 c Ammonia loss 9
612.2974243164062 0 3427.8142
622.3240966796875 0 50815.64 y 11
623.325927734375 0 15111.716
624.3251342773438 0 3734.9402
625.310791015625 0 3390.6958
625.6480102539062 0 7217.2646
626.2978515625 0 4929.8574 y Ammonia loss 1
628.6435546875 0 13293.178
628.9778442382812 0 14973.32
629.3087768554688 0 10262.403
629.6444091796875 0 2987.0554
631.63427734375 0 2594.506
631.9662475585938 0 47983.02 y 1
632.2999877929688 0 44082.72
632.6346435546875 0 24584.59
632.972412109375 0 8257.993
634.646728515625 0 74059.68 c Ammonia loss 14
634.98046875 0 78257.9
635.3135986328125 0 41224.68
635.6473388671875 0 20988.807
635.9857177734375 0 5780.3384
637.3043823242188 0 4270.89
637.3568725585938 0 5547.697
638.3604736328125 0 2771.8826
640.9851684570312 0 2856.9602
652.3209838867188 0 5802.634
653.3295288085938 0 4334.591
654.331787109375 0 4132.687
655.6436767578125 0 3931.0815
657.65283203125 0 4921.2754
657.9808959960938 0 12738.881
658.3153076171875 0 18387.72
658.6538696289062 0 8739.335
663.656982421875 0 60265.67
663.989501953125 0 74955.125
664.3231811523438 0 57895.11 y 5
664.6558227539062 0 29815.027
664.8357543945312 0 5233.7617
664.9901123046875 0 11048.88
665.3305053710938 0 5024.792
668.3036499023438 0 3148.259
668.6539916992188 0 4105.47
668.98876953125 0 8856.653
669.3316650390625 0 38806.434
669.6609497070312 0 810588
669.9949951171875 0 888803.2
670.3291625976562 0 530911.5
670.662841796875 0 259402.6
670.996337890625 0 91045.18
678.3819580078125 0 7601.183
679.3861694335938 0 7227.882
680.3165893554688 0 10798.608 c Ammonia loss 4
681.3167114257812 0 7633.037
696.3351440429688 0 6714.9893
697.3424072265625 0 148775.12 c 4
698.345947265625 0 57767.23
699.349365234375 0 16212.173
711.2994384765625 0 4032.3845
757.3729858398438 0 21321.123 c Ammonia loss 11
757.8724365234375 0 12718.573
758.3792724609375 0 7950.2163
766.4140625 0 16156.421
767.4196166992188 0 16231.374
768.3563842773438 0 3682.8672 y Ammonia loss 10
768.4271850585938 0 4267.451
769.3685302734375 0 149429.34 z 10
770.3736572265625 0 91388.92
771.376220703125 0 31737.293
772.373291015625 0 9401.011
777.3644409179688 0 3536.8171
785.3876342773438 0 50482.996 y 10
786.389892578125 0 21144.537
787.3803100585938 0 5061.897
793.4009399414062 0 12319.603 c Ammonia loss 5
794.4027709960938 0 4910.5396
807.4221801757812 0 4556.719
809.4180297851562 0 11519.747
810.4269409179688 0 198657.36 c 5
811.4298095703125 0 83405.02
812.4315185546875 0 24639.084
813.4346923828125 0 4549.511
814.885986328125 0 13585.422 y 3
815.3862915039062 0 18148.979
815.8883666992188 0 9658.92
816.3825073242188 0 6226.485
817.8851928710938 0 3362.8887
818.3861694335938 0 6580.0654
824.4066162109375 0 4782.105
826.3966674804688 0 9428.302
826.8967895507812 0 9065.295
839.3983764648438 0 28524.639 w 9
840.4030151367188 0 11305.699
841.4025268554688 0 7318.3516
841.893798828125 0 3452.8132 w 2
842.94580078125 0 13400.996
843.4476318359375 0 14376.251
843.946044921875 0 8254.887
846.399658203125 0 8249.259 c 12
846.8973999023438 0 6796.6934
847.3969116210938 0 3496.3872
849.4122924804688 0 4547.854
858.9308471679688 0 4807.0903
859.4312744140625 0 5609.658
859.9271850585938 0 4064.8853
861.40087890625 0 3788.7139
862.4046020507812 0 3693.182 z Ammonia loss 2
864.4490356445312 0 4283.197
866.4389038085938 0 5643.0967
866.9431762695312 0 4025.914
867.4251098632812 0 53298.742
868.426513671875 0 27574.998
869.4307861328125 0 5390.332
869.9078369140625 0 14705.162 y Water loss 2
870.4054565429688 0 38583.3 y Ammonia loss 2
870.9061889648438 0 32687.406 z 2
871.4061279296875 0 22601.533
871.9100952148438 0 10404.718
878.4204711914062 0 6408.4536
878.9157104492188 0 83835.22 y 2
879.417724609375 0 81430.76
879.9176025390625 0 42852.156
880.4178466796875 0 17459.008 z Water loss 9
880.9185180664062 0 5786.116
881.4412841796875 0 22284.145
882.445068359375 0 13397.891
883.4532470703125 0 3848.3586
886.9420776367188 0 4272.564
887.9356689453125 0 5593.186
888.4371948242188 0 19505.5
888.93798828125 0 21635.77
889.4378051757812 0 13070.625
889.938720703125 0 4945.3623
892.4839477539062 0 4530.1304
892.9813842773438 0 6294.856
893.4888305664062 0 5058.0483
893.94677734375 0 8454.742
894.4447021484375 0 5136.1865
894.9457397460938 0 3298.3005
895.4446411132812 0 4429.977
895.9307861328125 0 3501.8896
896.4347534179688 0 4532.3604 y Water loss 9
898.4116821289062 0 148609.23 z 9
899.4146118164062 0 66711.19
900.41552734375 0 29694.986
901.4251708984375 0 3482.8752 c Water loss 13
901.931640625 0 10276.76 c Ammonia loss 13
902.4359741210938 0 12923.86
902.9332275390625 0 6809.2153
903.91796875 0 8262.279
904.415771484375 0 3825.05
905.4156494140625 0 3701.704
908.4302978515625 0 35251.375 c Ammonia loss 6
908.9527587890625 0 7817.1934
909.4371337890625 0 16381.82
909.9423217773438 0 14603.487
910.44384765625 0 134967.19 c 13
910.9456787109375 0 127682.78
911.4453125 0 80792.836
911.947998046875 0 30708.838
912.4494018554688 0 10934.767
914.4313354492188 0 24927.525 y 9
915.4328002929688 0 16440.418
915.9718017578125 0 3373.855
916.4603881835938 0 8525.67
916.95947265625 0 4727.0493
917.4426879882812 0 15481.859
917.940185546875 0 7753.421
918.4376220703125 0 5374.1816
923.4584350585938 0 3334.2532
924.4483642578125 0 25599.209
925.45361328125 0 52319.957 c 6
926.4566040039062 0 26245.916
927.4525756835938 0 5488.4907
928.4864501953125 0 3235.563
928.9852294921875 0 6242.1816
929.4820556640625 0 5571.415
929.9658813476562 0 3579.6313
930.464111328125 0 7636.4316
931.4572143554688 0 7865.025
935.5187377929688 0 11959.684
936.5051879882812 0 9241.294
936.9895629882812 0 3497.958
937.470458984375 0 5126.7993
937.9745483398438 0 33147.51
938.4702758789062 0 43733.805
938.9637451171875 0 38149.637
939.4453125 0 82636.7 z 1
939.9390258789062 0 80132.65
940.442626953125 0 45999
940.9393920898438 0 15639.498
941.4418334960938 0 5767.3613
942.9825439453125 0 5026.2915
943.47705078125 0 6871.7134
943.973876953125 0 4569.439
944.9520874023438 0 3223.6548
945.9619750976562 0 7608.6826
946.462646484375 0 12801.71
947.4494018554688 0 5871.822 y 1
947.9472045898438 0 6420.731
948.43798828125 0 2826.9976
950.4807739257812 0 22659.482
950.979248046875 0 13703.208 c Water loss 14
951.4813842773438 0 15540.551 c Ammonia loss 14
951.9813232421875 0 9261.18
952.4765625 0 14651.065
952.9761962890625 0 8209.485
953.4780883789062 0 9118.275
953.9623413085938 0 7615.7983
958.9822387695312 0 3573.7993
959.4837646484375 0 16418.924
959.9799194335938 0 229938.83 c 14
960.4810180664062 0 234073.4
960.9814453125 0 145548.86
961.481689453125 0 53524.35
961.982177734375 0 15145.281
964.4866943359375 0 11246.34
964.975830078125 0 11612.982
965.4664916992188 0 16099.399 c Ammonia loss 7
965.9782104492188 0 15608.743
966.4747924804688 0 16065.481
966.976806640625 0 14850.19
967.4813842773438 0 15319.598
967.9630737304688 0 17266.486
968.4660034179688 0 24023.576
968.9697875976562 0 14419.306
969.4644165039062 0 4398.9414
972.4803466796875 0 5056.064
972.983642578125 0 10537.485
973.4869995117188 0 17042.32
973.9805908203125 0 32948.5
974.4707641601562 0 140121.7
974.980712890625 0 405418.22
975.4830322265625 0 334257.7
975.9850463867188 0 212197.66
976.485595703125 0 74319.87
976.9893798828125 0 29013.348
977.97216796875 0 5338.4536
980.4757080078125 0 4698.3413
981.4800415039062 0 98758.84
981.9860229492188 0 64442.516
982.4790649414062 0 236777.22 c 7
982.9866943359375 0 34233.887
983.4798583984375 0 104371.53
983.98828125 0 3944.6726
984.4819946289062 0 28876.578
985.4850463867188 0 6656.636
986.4820556640625 0 11650.901
986.9769287109375 0 10819.148
987.4783935546875 0 9229.716
987.9837646484375 0 8816.886
988.4693603515625 0 5164.9434
994.976806640625 0 8843.285
995.4859619140625 0 41362.117
995.9796752929688 0 120826.58
996.48095703125 0 103909.67
996.9810791015625 0 75664.76
997.481201171875 0 46920.48
997.9819946289062 0 10160.197
1002.478515625 0 6867.283
1002.97802734375 0 5778.459
1003.4898681640625 0 22624.053
1003.9876708984375 0 436397.84
1004.490234375 0 718761.25
1004.9912719726562 0 609722.94
1005.492431640625 0 359079.2
1005.992431640625 0 131596.77
1006.49462890625 0 35552.32
1026.5064697265625 0 125929.984 z 8
1027.5115966796875 0 106949.78
1028.5115966796875 0 42211.67
1029.513427734375 0 15276.56
1030.5313720703125 0 3586.629
1041.52099609375 0 8801.807
1042.5233154296875 0 14063.119 y 8
1043.52685546875 0 8833.649
1066.552490234375 0 11298.295
1067.554931640625 0 9664.185
1083.5284423828125 0 33927.09 z 7
1084.5330810546875 0 45164.28
1085.5372314453125 0 19292.695
1086.5404052734375 0 7013.95
1099.54638671875 0 32579.557 y 7
1100.549560546875 0 17830.945
1101.5482177734375 0 7707.311
1107.5684814453125 0 4414.5464
1108.5716552734375 0 6857.816
1109.57958984375 0 9916.814
1110.570556640625 0 185585.62 c 8
1111.5728759765625 0 108501.57
1112.5750732421875 0 40422.992
1113.572998046875 0 8125.8755
1136.5848388671875 0 15266.298
1137.58837890625 0 12555.81
1138.5830078125 0 4230.423
1141.5574951171875 0 53419.227
1142.559814453125 0 33996.652
1143.559814453125 0 18455.709
1154.5670166015625 0 11791.567
1155.5631103515625 0 6115.6084
1195.60107421875 0 19911.16
1196.6011962890625 0 17880.043
1197.59521484375 0 7434.784
1198.5552978515625 0 119238.44 z 6
1199.5594482421875 0 90611.22
1200.5601806640625 0 45818.887
1201.56298828125 0 10692.02
1213.5726318359375 0 5805.9263
1214.573974609375 0 41672.19 y 6
1215.57861328125 0 23805.744
1216.5745849609375 0 10068.315
1238.606689453125 0 30108.105
1239.6123046875 0 184912.72 c 9
1240.6153564453125 0 119354.12
1240.7813720703125 0 4622.21
1241.6175537109375 0 38347.3
1242.6204833984375 0 10277.979
1267.6470947265625 0 17250.596
1268.60205078125 0 10382.379 w 5
1269.626953125 0 5530.742
1310.6514892578125 0 5125.8096 y Ammonia loss 5
1311.6395263671875 0 77324.414 z 5
1312.641357421875 0 59152.574
1313.646728515625 0 39072.844
1314.649658203125 0 10698.202
1327.650634765625 0 11255.933 y 5
1328.6646728515625 0 3793.5452
1329.6651611328125 0 4815.2266
1358.6634521484375 0 31040.592
1359.6666259765625 0 20565.12
1360.6641845703125 0 7288.905
1401.6788330078125 0 11379.9795
1402.67626953125 0 133960.69 c 10
1403.6790771484375 0 95306.98
1404.680908203125 0 36186.145
1405.6854248046875 0 10344.597
1441.654052734375 0 5710.641
1442.6715087890625 0 4558.6743
1481.732177734375 0 3300.3162
1486.75732421875 0 13556.549
1487.761962890625 0 14209.763
1488.7635498046875 0 6197.5215
1495.7191162109375 0 4737.839 y Water loss 4
1496.740478515625 0 5892.033 y Ammonia loss 4
1497.7197265625 0 67848.33 z 4
1498.7205810546875 0 71588.29
1499.7239990234375 0 30488.404
1500.728271484375 0 18344.967
1512.7421875 0 7717.159 c Water loss 11
1513.73583984375 0 65777.57 c Ammonia loss 11
1514.7403564453125 0 48583.12
1515.7430419921875 0 28865.848
1516.7401123046875 0 7971.7456
1521.778564453125 0 3625.4922
1522.751220703125 0 4003.237
1525.74365234375 0 5473.731
1526.7374267578125 0 3785.5613
1529.776611328125 0 10351.189
1530.77099609375 0 127611.09 c 11
1531.77392578125 0 110998.39
1532.77490234375 0 60302.45
1533.7784423828125 0 16071.575
1550.743896484375 0 8477.67
1551.7435302734375 0 6720.9546
1554.7264404296875 0 4715.7153
1555.7415771484375 0 6816.2896
1556.7742919921875 0 10412.448
1557.7750244140625 0 10366.617
1558.783935546875 0 4771.3657
1567.7432861328125 0 4337.8066 w 3
1568.7532958984375 0 45705.066
1569.756591796875 0 40564.344
1570.7591552734375 0 32504.246
1571.7618408203125 0 12264.462
1572.7657470703125 0 5607.3843
1611.7498779296875 0 7240.1147 y Ammonia loss 3
1612.744140625 0 121291.35 z 3
1613.747802734375 0 162279.17
1614.7509765625 0 93264.04
1615.7535400390625 0 39658.168
1616.757568359375 0 15721.625
1627.7578125 0 3924.504
1628.7626953125 0 25925.695 y 3
1629.7647705078125 0 22196.281
1630.7723388671875 0 11541.898
1632.76025390625 0 4114.9937
1633.7816162109375 0 5478.007
1646.73388671875 0 3362.076
1652.8184814453125 0 3438.0938
1654.8037109375 0 3665.1255
1677.788818359375 0 4276.387
1682.798583984375 0 5766.0264 w 2
1683.7974853515625 0 8669.97
1684.788330078125 0 6831.298
1690.782958984375 0 22494.875
1691.7860107421875 0 154796.12 c 12
1692.7884521484375 0 153407.84
1693.7906494140625 0 87059.8
1694.791748046875 0 33813.375
1695.7882080078125 0 8794.21
1696.807861328125 0 6166.8735
1697.80615234375 0 5116.1177
1698.822265625 0 3878.4246
1740.804931640625 0 98887.51 z 2
1741.80859375 0 146948.11
1742.8111572265625 0 86131.29
1743.8126220703125 0 42386.293
1744.812744140625 0 10179.094
1756.8233642578125 0 25920.963 y 2
1757.824951171875 0 23942.578
1758.8277587890625 0 17059.344
1759.828369140625 0 8025.178
1760.85888671875 0 6339.4746
1761.8760986328125 0 6151.622
1762.86865234375 0 6757.0054
1763.8780517578125 0 5325.9824
1776.87353515625 0 19999.24
1777.8685302734375 0 18352.064
1778.8658447265625 0 7896.9243
1803.8638916015625 0 5034.978
1804.86865234375 0 5621.0405
1818.8807373046875 0 3667.534
1819.875 0 20964.133 c 13
1820.8824462890625 0 38528.973
1821.885986328125 0 27982.729
1822.8895263671875 0 16580.848
1823.87646484375 0 6818.7656
1859.9493408203125 0 4074.041
1860.931396484375 0 10897.164
1861.931884765625 0 7598.2397
1862.926513671875 0 4017.1948
1874.9395751953125 0 9342.356
1875.9400634765625 0 19903.92
1876.944580078125 0 19400.615
1877.905029296875 0 22090.248
1878.8741455078125 0 56415.27
1879.8743896484375 0 54677.28
1880.874755859375 0 35857.887
1881.8782958984375 0 13599.847
1889.9810791015625 0 5921.471
1891.0101318359375 0 4614.4917
1891.9700927734375 0 7681.7393
1892.9471435546875 0 9253.237
1893.9483642578125 0 7424.225
1900.97119140625 0 8850.906 c Water loss 14
1901.98779296875 0 5298.119
1902.94775390625 0 5900.1304
1903.9378662109375 0 8370.724
1904.94677734375 0 5168.509
1905.9395751953125 0 5298.003
1916.9814453125 0 16224.9795
1917.9942626953125 0 33916.11
1918.9771728515625 0 54263.453 c 14
1919.96484375 0 61246.824
1920.96484375 0 54567.215
1921.9671630859375 0 37311
1922.969482421875 0 21726.271
1923.9669189453125 0 12352.989
1931.9515380859375 0 12999.791
1932.953857421875 0 23288.121
1933.95361328125 0 22961.014
1934.9500732421875 0 11179.233
1935.950927734375 0 9072.915
1945.9500732421875 0 7825.2896
1946.962890625 0 5838.738
1947.951171875 0 23757.426
1948.958251953125 0 83401.09
1949.9608154296875 0 149210.45
1950.9644775390625 0 108561.38
1951.9677734375 0 59923.65
1952.97021484375 0 28976.703
1953.9775390625 0 6770.4185
1961.9571533203125 0 7712.559
1962.96630859375 0 42358.297
1963.9664306640625 0 78214.9
1964.9637451171875 0 64216.496
1965.9664306640625 0 35396.55
1966.96240234375 0 13597.107
1967.9686279296875 0 4639.9307
1972.9681396484375 0 6904.8896
1973.9622802734375 0 8787.183
1974.976318359375 0 5887.1226
1979.9794921875 0 32732.408
1980.9818115234375 0 90083.625
1981.9859619140625 0 89283.85
1982.987548828125 0 57108.84
1983.986572265625 0 20277.404
1985.0009765625 0 7684.6157
1988.9119873046875 0 4727.686
1989.953369140625 0 14406.847
1990.9593505859375 0 67107.14
1991.954345703125 0 170432
1992.9573974609375 0 163002.11
1993.9581298828125 0 97839.99
1994.960205078125 0 45267.6
1995.9493408203125 0 11297.103
2004.9581298828125 0 4302.5405
2005.9580078125 0 6614.266
2006.964111328125 0 50102.25
2007.97265625 0 218307.44
2008.978515625 0 708045.3
2009.9805908203125 0 662149.4
2010.9832763671875 0 412288.75
2011.9852294921875 0 181966.17
2012.9830322265625 0 60710.973
2211.085205078125 0 4702.4883
2212.069580078125 0 4411.896
3045.9716796875 0 3628.2747

Spectrum Details

|  |  |
| --- | --- |
| Matched peaks? Matched peaks The total absolute number of peaks matched. Additionally in brackets the total fraction of peaks matched and the total number of peaks is shown. | 85 (14.03% of 606) |
| FDR? FDR The false discovery rate estimated for this peptide. It is calculated by matching all theoretical fragments with a non-integer shift with the raw peaks for this spectrum. This is done with 40 different shifts. The resulting percentage is the average number of annotated peaks over the number of annotated peaks with the correct spectrum. | 1.34% |
| Satellite FDR? Satellite FDR See the FDR for details on its calculation. This satellite ion specific FDR only contains the satellite ions (d/w) for I/L/J positions. | 0.00% |
| PSM Score? PSM Score The PSM Score as given by Hecklib to this annotated spectrum. It is shown with three significant figures. | 686 |

#### Spectrum 5098? Spectrum 5098 The raw spectrum of this peptide as annotated by Hecklib. The fragments are coloured according to ion type (see legend). Any peaks with a star '\*' as text can be hovered over to see the full details, first the ion type second the mass shift type. By hovering over the amino acids in the peptide or ions in the legend the corresponding peaks are highlighted. By toggling the 'Unassigned' label you can turn the background (unassigned) peaks on or off in the plot. By updating the slider in the Ion legend you can update the spectrum to only show the top X% of the peaks with labels. The top X% means any peak that is within X% of the highest intensity. By dragging in the spectrum you can zoom in to a specific part of the spectrum and use 'Zoom Out' to get back to the original zoom level. The annotation of the spectrum is based on the given sequence in the peptides file and is done with different software so inconsistencies are likely. The peaks are annotated based on the given sequence, with 20 ppm tolerance.

Copy Data

##### Spectrum 5098 (TSV)

###### Preview

```
Loading example...
```

*Click on the button to copy the data to your clipboard.*

Mz MinMz MaxIntensity Max

WidthHeightPeptide font sizePeptide stroke widthSpectrum font sizeSpectrum stroke widthCompact peptide

Ion legend

wxyz

abcd

OtherUnassignedIon^Charge^_Position_Show for top:%

JHQDWJDGKEYKCKVS

01.66e+43.32e+44.98e+46.64e+4

Zoom Out

y^+1^_2_c^+1^_2_z^+1^_3_y^+1^_3_c^+1^_3_c^+1^_3_c^+1^_4_c^+1^_4_w^+1^_5_y^+2^_9_c^+3^_14_z^+1^_5_c^+3^_14_y^+2^_10_y^+1^_5_y^+3^_15_c^+3^_15_c^+3^_15_y^+2^_11_y^+2^_11_c^+1^_5_c^+2^_12_z^+1^_6_y^+1^_6_c^+1^_6_c^+1^_6_y^+2^_13_w^+1^_7_c^+2^_13_y^+2^_14_y^+2^_14_z^+2^_14_y^+2^_14_z^+1^_7_c^+2^_14_c^+2^_14_c^+1^_7_c^+2^_14_y^+1^_7_c^+1^_7_z^+2^_15_c^+2^_15_c^+2^_15_c^+1^_8_z^+1^_8_z^+1^_9_y^+1^_9_c^+1^_9_z^+1^_10_y^+1^_10_c^+1^_10_y^+1^_11_z^+1^_11_y^+1^_11_c^+1^_11_y^+1^_12_z^+1^_12_c^+1^_12_c^+1^_12_c^+1^_12_y^+1^_13_z^+1^_13_y^+1^_13_c^+1^_13_z^+1^_14_y^+1^_14_c^+1^_14_z^+1^_15_c^+1^_15_

0843168725303374

Fragment Matches Table

Show background peaks

| Position | Ion type | Intensity | mz Theoretical | mz Error (Th) | mz Error (ppm) | Charge | Series Number |
| --- | --- | --- | --- | --- | --- | --- | --- |
| - | - | 1595 | 129.1 | - | - | 0 | - |
| - | - | 799.3 | 131.1 | - | - | 0 | - |
| - | - | 372.2 | 133.1 | - | - | 0 | - |
| - | - | 413.8 | 136.7 | - | - | 0 | - |
| - | - | 522.9 | 143.1 | - | - | 0 | - |
| - | - | 391.5 | 144.6 | - | - | 0 | - |
| - | - | 621.5 | 146.1 | - | - | 0 | - |
| - | - | 614 | 148.9 | - | - | 0 | - |
| - | - | 581.4 | 148.9 | - | - | 0 | - |
| - | - | 772.5 | 148.9 | - | - | 0 | - |
| - | - | 816 | 148.9 | - | - | 0 | - |
| - | - | 753 | 148.9 | - | - | 0 | - |
| - | - | 1049 | 148.9 | - | - | 0 | - |
| - | - | 1141 | 148.9 | - | - | 0 | - |
| - | - | 2706 | 148.9 | - | - | 0 | - |
| - | - | 4425 | 148.9 | - | - | 0 | - |
| - | - | 3784 | 149 | - | - | 0 | - |
| - | - | 1391 | 149 | - | - | 0 | - |
| - | - | 1261 | 149 | - | - | 0 | - |
| - | - | 1088 | 149 | - | - | 0 | - |
| - | - | 829.7 | 149 | - | - | 0 | - |
| - | - | 613.8 | 149 | - | - | 0 | - |
| - | - | 692.3 | 149 | - | - | 0 | - |
| - | - | 409.9 | 149.1 | - | - | 0 | - |
| - | - | 501.8 | 153.1 | - | - | 0 | - |
| - | - | 894.3 | 155.1 | - | - | 0 | - |
| - | - | 459.9 | 164.9 | - | - | 0 | - |
| - | - | 1887 | 166.1 | - | - | 0 | - |
| - | - | 518.5 | 168.1 | - | - | 0 | - |
| - | - | 836.7 | 173.1 | - | - | 0 | - |
| - | - | 582.1 | 178.1 | - | - | 0 | - |
| - | - | 1020 | 185.2 | - | - | 0 | - |
| 15 | y | 934 | 205.1 | 0.000201 | 0.9801 | +1 | 2 |
| - | - | 577 | 214.2 | - | - | 0 | - |
| - | - | 1920 | 223.2 | - | - | 0 | - |
| - | - | 1092 | 228.1 | - | - | 0 | - |
| - | - | 1203 | 229.1 | - | - | 0 | - |
| - | - | 573.2 | 229.2 | - | - | 0 | - |
| - | - | 2013 | 230.1 | - | - | 0 | - |
| - | - | 2173 | 234.1 | - | - | 0 | - |
| - | - | 572.4 | 237.2 | - | - | 0 | - |
| - | - | 1.08E+04 | 242.2 | - | - | 0 | - |
| - | - | 1364 | 243.2 | - | - | 0 | - |
| - | - | 7907 | 246.1 | - | - | 0 | - |
| - | - | 1569 | 247.1 | - | - | 0 | - |
| - | - | 829.4 | 247.1 | - | - | 0 | - |
| - | - | 3329 | 248.1 | - | - | 0 | - |
| - | - | 550.5 | 249.1 | - | - | 0 | - |
| - | - | 1184 | 249.2 | - | - | 0 | - |
| - | - | 649.2 | 250.1 | - | - | 0 | - |
| - | - | 9494 | 251.2 | - | - | 0 | - |
| - | - | 1711 | 252.2 | - | - | 0 | - |
| - | - | 9713 | 264.1 | - | - | 0 | - |
| - | - | 812.5 | 265.1 | - | - | 0 | - |
| - | - | 1641 | 265.2 | - | - | 0 | - |
| - | - | 618.3 | 266.1 | - | - | 0 | - |
| - | - | 618.8 | 268.1 | - | - | 0 | - |
| 2 | c | 6127 | 268.2 | 0.0001351 | 0.5039 | +1 | 2 |
| - | - | 683.6 | 269.2 | - | - | 0 | - |
| - | - | 5097 | 277.2 | - | - | 0 | - |
| - | - | 689.7 | 278.2 | - | - | 0 | - |
| - | - | 2666 | 291.1 | - | - | 0 | - |
| - | - | 7244 | 293.1 | - | - | 0 | - |
| - | - | 1190 | 294.2 | - | - | 0 | - |
| - | - | 487.2 | 298.8 | - | - | 0 | - |
| - | - | 585.4 | 306.1 | - | - | 0 | - |
| - | - | 961.4 | 309.1 | - | - | 0 | - |
| - | - | 927.3 | 316.2 | - | - | 0 | - |
| 14 | z | 7803 | 317.2 | 8.818E-05 | 0.278 | +1 | 3 |
| - | - | 3794 | 318.2 | - | - | 0 | - |
| 14 | y | 1189 | 333.2 | 0.0001629 | 0.489 | +1 | 3 |
| - | - | 762.2 | 341 | - | - | 0 | - |
| - | - | 1455 | 342.2 | - | - | 0 | - |
| - | - | 934.8 | 343 | - | - | 0 | - |
| - | - | 1298 | 343.2 | - | - | 0 | - |
| - | - | 762.5 | 344.2 | - | - | 0 | - |
| - | - | 597.4 | 353.2 | - | - | 0 | - |
| - | - | 1908 | 355.1 | - | - | 0 | - |
| - | - | 1176 | 356.1 | - | - | 0 | - |
| - | - | 2079 | 357.1 | - | - | 0 | - |
| - | - | 1825 | 359 | - | - | 0 | - |
| - | - | 6548 | 359.2 | - | - | 0 | - |
| - | - | 1629 | 360 | - | - | 0 | - |
| - | - | 867.2 | 360.2 | - | - | 0 | - |
| - | - | 1367 | 360.2 | - | - | 0 | - |
| - | - | 1789 | 361 | - | - | 0 | - |
| - | - | 2929 | 361.2 | - | - | 0 | - |
| - | - | 593.5 | 362.2 | - | - | 0 | - |
| - | - | 737.9 | 375.2 | - | - | 0 | - |
| - | - | 9504 | 377.2 | - | - | 0 | - |
| - | - | 1434 | 378.2 | - | - | 0 | - |
| 3 | c | 1624 | 379.2 | 0.0004292 | 1.132 | +1 | 3 |
| - | - | 1.08E+04 | 387.2 | - | - | 0 | - |
| - | - | 2947 | 388.2 | - | - | 0 | - |
| - | - | 1465 | 389.2 | - | - | 0 | - |
| - | - | 4083 | 390.2 | - | - | 0 | - |
| - | - | 1088 | 395.2 | - | - | 0 | - |
| 3 | c | 1.31E+04 | 396.2 | 2.735E-05 | 0.06902 | +1 | 3 |
| - | - | 2643 | 397.2 | - | - | 0 | - |
| - | - | 763.1 | 404.2 | - | - | 0 | - |
| - | - | 837.6 | 405.2 | - | - | 0 | - |
| - | - | 4653 | 406.2 | - | - | 0 | - |
| - | - | 1340 | 407.2 | - | - | 0 | - |
| - | - | 706.9 | 407.3 | - | - | 0 | - |
| - | - | 1720 | 411.2 | - | - | 0 | - |
| - | - | 982.5 | 412.2 | - | - | 0 | - |
| - | - | 629.8 | 419.2 | - | - | 0 | - |
| - | - | 881.8 | 421.2 | - | - | 0 | - |
| - | - | 673.1 | 422.2 | - | - | 0 | - |
| - | - | 763.7 | 423.3 | - | - | 0 | - |
| - | - | 2324 | 429.1 | - | - | 0 | - |
| - | - | 2019 | 430.1 | - | - | 0 | - |
| - | - | 5218 | 431.1 | - | - | 0 | - |
| - | - | 1316 | 432.1 | - | - | 0 | - |
| - | - | 609.5 | 439.3 | - | - | 0 | - |
| - | - | 616.7 | 461.2 | - | - | 0 | - |
| - | - | 5122 | 467.2 | - | - | 0 | - |
| - | - | 1354 | 468.3 | - | - | 0 | - |
| - | - | 566.8 | 490.3 | - | - | 0 | - |
| 4 | c | 2656 | 494.2 | 0.00445 | 9.003 | +1 | 4 |
| - | - | 1082 | 495.2 | - | - | 0 | - |
| - | - | 1334 | 506.3 | - | - | 0 | - |
| 4 | c | 1.505E+04 | 511.3 | 6.82E-05 | 0.1334 | +1 | 4 |
| - | - | 4000 | 512.3 | - | - | 0 | - |
| - | - | 2071 | 515.3 | - | - | 0 | - |
| - | - | 701.8 | 516.3 | - | - | 0 | - |
| - | - | 1217 | 517.3 | - | - | 0 | - |
| - | - | 3575 | 518.3 | - | - | 0 | - |
| - | - | 1543 | 519.3 | - | - | 0 | - |
| - | - | 839.8 | 519.3 | - | - | 0 | - |
| - | - | 673.1 | 519.8 | - | - | 0 | - |
| - | - | 8969 | 534.3 | - | - | 0 | - |
| - | - | 2513 | 535.3 | - | - | 0 | - |
| - | - | 757.6 | 547.3 | - | - | 0 | - |
| 12 | w | 931.6 | 548.2 | 0.007583 | 13.83 | +1 | 5 |
| - | - | 1586 | 548.3 | - | - | 0 | - |
| 8 | y | 3563 | 550.3 | 0.002069 | 3.76 | +2 | 9 |
| - | - | 993.3 | 550.8 | - | - | 0 | - |
| - | - | 1031 | 551.3 | - | - | 0 | - |
| - | - | 1136 | 556.3 | - | - | 0 | - |
| - | - | 573.5 | 563.2 | - | - | 0 | - |
| - | - | 833.3 | 564.2 | - | - | 0 | - |
| - | - | 2780 | 592.3 | - | - | 0 | - |
| - | - | 1238 | 593.3 | - | - | 0 | - |
| - | - | 1020 | 600.3 | - | - | 0 | - |
| 14 | c | 847.6 | 601.3 | 0.006827 | 11.35 | +3 | 14 |
| - | - | 845.1 | 602 | - | - | 0 | - |
| 12 | z | 8186 | 606.3 | 0.004898 | 8.078 | +1 | 5 |
| 14 | c | 3705 | 607.3 | 0.009653 | 15.89 | +3 | 14 |
| 7 | y | 881.2 | 607.8 | 0.002125 | 3.496 | +2 | 10 |
| - | - | 1412 | 608.3 | - | - | 0 | - |
| - | - | 1175 | 617.3 | - | - | 0 | - |
| - | - | 1152 | 618.3 | - | - | 0 | - |
| - | - | 2163 | 619.2 | - | - | 0 | - |
| 12 | y | 4170 | 622.3 | 0.004274 | 6.868 | +1 | 5 |
| - | - | 709.3 | 623.3 | - | - | 0 | - |
| - | - | 1052 | 629.3 | - | - | 0 | - |
| - | - | 1002 | 631.6 | - | - | 0 | - |
| 2 | y | 3345 | 632 | 2.042E-05 | 0.03231 | +3 | 15 |
| - | - | 2645 | 632.3 | - | - | 0 | - |
| - | - | 964.2 | 632.6 | - | - | 0 | - |
| - | - | 611.8 | 633 | - | - | 0 | - |
| 15 | c | 1584 | 634.3 | 0.001574 | 2.481 | +3 | 15 |
| 15 | c | 4699 | 634.6 | 0.0009266 | 1.46 | +3 | 15 |
| - | - | 5117 | 635 | - | - | 0 | - |
| - | - | 5056 | 635.3 | - | - | 0 | - |
| - | - | 1083 | 635.6 | - | - | 0 | - |
| - | - | 2509 | 636.3 | - | - | 0 | - |
| - | - | 1127 | 637.2 | - | - | 0 | - |
| - | - | 1767 | 637.4 | - | - | 0 | - |
| - | - | 1124 | 651.3 | - | - | 0 | - |
| - | - | 3628 | 651.3 | - | - | 0 | - |
| - | - | 2860 | 652.3 | - | - | 0 | - |
| - | - | 1024 | 653.3 | - | - | 0 | - |
| 6 | y | 646.9 | 655.3 | 0.01049 | 16.01 | +2 | 11 |
| - | - | 1361 | 658 | - | - | 0 | - |
| - | - | 1499 | 658.3 | - | - | 0 | - |
| - | - | 778 | 658.7 | - | - | 0 | - |
| - | - | 2391 | 663.7 | - | - | 0 | - |
| - | - | 3144 | 664 | - | - | 0 | - |
| 6 | y | 1804 | 664.3 | 0.01156 | 17.41 | +2 | 11 |
| - | - | 2829 | 664.7 | - | - | 0 | - |
| - | - | 3920 | 668.3 | - | - | 0 | - |
| - | - | 3.121E+04 | 669.3 | - | - | 0 | - |
| - | - | 5.672E+04 | 669.7 | - | - | 0 | - |
| - | - | 4.986E+04 | 670 | - | - | 0 | - |
| - | - | 3.825E+04 | 670.3 | - | - | 0 | - |
| - | - | 1.356E+04 | 670.7 | - | - | 0 | - |
| - | - | 5693 | 671 | - | - | 0 | - |
| - | - | 3952 | 671.3 | - | - | 0 | - |
| 5 | c | 1.126E+04 | 697.3 | 0.0002653 | 0.3804 | +1 | 5 |
| - | - | 4375 | 698.3 | - | - | 0 | - |
| - | - | 602 | 732.7 | - | - | 0 | - |
| 12 | c | 1149 | 757.4 | 0.002721 | 3.592 | +2 | 12 |
| - | - | 1298 | 757.9 | - | - | 0 | - |
| - | - | 1393 | 766.4 | - | - | 0 | - |
| - | - | 790.5 | 767.4 | - | - | 0 | - |
| 11 | z | 1.173E+04 | 769.4 | 0.004811 | 6.253 | +1 | 6 |
| - | - | 6163 | 770.4 | - | - | 0 | - |
| - | - | 3381 | 771.4 | - | - | 0 | - |
| - | - | 611.4 | 780.4 | - | - | 0 | - |
| 11 | y | 2997 | 785.4 | 0.003638 | 4.632 | +1 | 6 |
| - | - | 1491 | 786.4 | - | - | 0 | - |
| 6 | c | 1013 | 793.4 | 0.00468 | 5.898 | +1 | 6 |
| - | - | 640.8 | 795.4 | - | - | 0 | - |
| 6 | c | 1.112E+04 | 810.4 | 0.001138 | 1.405 | +1 | 6 |
| - | - | 6256 | 811.4 | - | - | 0 | - |
| - | - | 1563 | 812.4 | - | - | 0 | - |
| 4 | y | 1364 | 814.9 | 0.0001882 | 0.2309 | +2 | 13 |
| - | - | 639 | 815.4 | - | - | 0 | - |
| - | - | 759 | 824.4 | - | - | 0 | - |
| - | - | 696.8 | 826.4 | - | - | 0 | - |
| - | - | 783.3 | 826.9 | - | - | 0 | - |
| - | - | 632.2 | 836.4 | - | - | 0 | - |
| 10 | w | 1726 | 839.4 | 0.009808 | 11.68 | +1 | 7 |
| - | - | 1233 | 840.4 | - | - | 0 | - |
| - | - | 621.7 | 841.4 | - | - | 0 | - |
| 13 | c | 809.6 | 846.4 | 0.002638 | 3.117 | +2 | 13 |
| - | - | 760.6 | 847.4 | - | - | 0 | - |
| - | - | 3091 | 867.4 | - | - | 0 | - |
| - | - | 1974 | 868.4 | - | - | 0 | - |
| 3 | y | 973.5 | 869.9 | 0.001145 | 1.316 | +2 | 14 |
| 3 | y | 3909 | 870.4 | 0.001874 | 2.153 | +2 | 14 |
| 3 | z | 1605 | 870.9 | 0.003747 | 4.303 | +2 | 14 |
| - | - | 1504 | 871.4 | - | - | 0 | - |
| - | - | 1536 | 878.4 | - | - | 0 | - |
| 3 | y | 4966 | 878.9 | 0.0001699 | 0.1933 | +2 | 14 |
| - | - | 5704 | 879.4 | - | - | 0 | - |
| - | - | 3409 | 879.9 | - | - | 0 | - |
| - | - | 1090 | 880.4 | - | - | 0 | - |
| - | - | 1988 | 881.4 | - | - | 0 | - |
| - | - | 1699 | 882.4 | - | - | 0 | - |
| - | - | 664 | 887.9 | - | - | 0 | - |
| - | - | 1398 | 888.4 | - | - | 0 | - |
| - | - | 658.8 | 889.4 | - | - | 0 | - |
| - | - | 834.4 | 889.9 | - | - | 0 | - |
| 10 | z | 1.111E+04 | 898.4 | 0.004924 | 5.48 | +1 | 7 |
| - | - | 5957 | 899.4 | - | - | 0 | - |
| - | - | 1683 | 900.4 | - | - | 0 | - |
| 14 | c | 1124 | 901.4 | 0.0167 | 18.53 | +2 | 14 |
| 14 | c | 954.1 | 901.9 | 0.001119 | 1.24 | +2 | 14 |
| - | - | 1406 | 902.4 | - | - | 0 | - |
| 7 | c | 1835 | 908.4 | 0.006036 | 6.645 | +1 | 7 |
| - | - | 840.8 | 909.4 | - | - | 0 | - |
| - | - | 2345 | 909.9 | - | - | 0 | - |
| 14 | c | 8574 | 910.4 | 0.003794 | 4.167 | +2 | 14 |
| - | - | 7956 | 910.9 | - | - | 0 | - |
| - | - | 4794 | 911.4 | - | - | 0 | - |
| - | - | 1819 | 911.9 | - | - | 0 | - |
| 10 | y | 2713 | 914.4 | 0.00375 | 4.101 | +1 | 7 |
| - | - | 975 | 917.4 | - | - | 0 | - |
| - | - | 1151 | 917.9 | - | - | 0 | - |
| - | - | 1152 | 918.4 | - | - | 0 | - |
| - | - | 2429 | 924.5 | - | - | 0 | - |
| 7 | c | 3874 | 925.5 | 0.0001883 | 0.2035 | +1 | 7 |
| - | - | 1516 | 926.5 | - | - | 0 | - |
| - | - | 769.2 | 927.5 | - | - | 0 | - |
| - | - | 557.7 | 931.4 | - | - | 0 | - |
| - | - | 1224 | 935.5 | - | - | 0 | - |
| - | - | 873.1 | 936.5 | - | - | 0 | - |
| - | - | 2137 | 938 | - | - | 0 | - |
| - | - | 3632 | 938.5 | - | - | 0 | - |
| - | - | 3289 | 939 | - | - | 0 | - |
| 2 | z | 6191 | 939.4 | 0.007568 | 8.056 | +2 | 15 |
| - | - | 4943 | 939.9 | - | - | 0 | - |
| - | - | 3407 | 940.4 | - | - | 0 | - |
| - | - | 804 | 940.9 | - | - | 0 | - |
| - | - | 671 | 945.5 | - | - | 0 | - |
| - | - | 878.4 | 946 | - | - | 0 | - |
| 15 | c | 922.4 | 951 | 0.004818 | 5.066 | +2 | 15 |
| - | - | 1221 | 952 | - | - | 0 | - |
| - | - | 1632 | 952.5 | - | - | 0 | - |
| - | - | 1478 | 953 | - | - | 0 | - |
| - | - | 1035 | 953.5 | - | - | 0 | - |
| - | - | 596.6 | 955 | - | - | 0 | - |
| - | - | 4581 | 959.5 | - | - | 0 | - |
| 15 | c | 1.639E+04 | 960 | 0.0006342 | 0.6606 | +2 | 15 |
| - | - | 1.596E+04 | 960.5 | - | - | 0 | - |
| - | - | 9281 | 961 | - | - | 0 | - |
| - | - | 4319 | 961.5 | - | - | 0 | - |
| - | - | 2277 | 962 | - | - | 0 | - |
| - | - | 1111 | 962.5 | - | - | 0 | - |
| - | - | 760.8 | 965 | - | - | 0 | - |
| - | - | 1680 | 965.5 | - | - | 0 | - |
| - | - | 1183 | 966 | - | - | 0 | - |
| - | - | 1138 | 966.5 | - | - | 0 | - |
| - | - | 1187 | 967 | - | - | 0 | - |
| - | - | 1161 | 967.5 | - | - | 0 | - |
| - | - | 591.9 | 967.6 | - | - | 0 | - |
| - | - | 1633 | 968 | - | - | 0 | - |
| - | - | 1949 | 968.5 | - | - | 0 | - |
| - | - | 1708 | 969 | - | - | 0 | - |
| - | - | 893 | 969.5 | - | - | 0 | - |
| - | - | 2039 | 973.5 | - | - | 0 | - |
| - | - | 3241 | 974 | - | - | 0 | - |
| - | - | 1.268E+04 | 974.5 | - | - | 0 | - |
| - | - | 2.57E+04 | 975 | - | - | 0 | - |
| - | - | 2.522E+04 | 975.5 | - | - | 0 | - |
| - | - | 1.378E+04 | 976 | - | - | 0 | - |
| - | - | 8099 | 976.5 | - | - | 0 | - |
| - | - | 2900 | 977 | - | - | 0 | - |
| - | - | 967.4 | 977.5 | - | - | 0 | - |
| - | - | 664.6 | 979.5 | - | - | 0 | - |
| - | - | 1039 | 980.5 | - | - | 0 | - |
| - | - | 1.349E+04 | 981.5 | - | - | 0 | - |
| - | - | 9175 | 982 | - | - | 0 | - |
| 8 | c | 1.983E+04 | 982.5 | 0.00734 | 7.471 | +1 | 8 |
| - | - | 3648 | 983 | - | - | 0 | - |
| - | - | 7621 | 983.5 | - | - | 0 | - |
| - | - | 1949 | 984.5 | - | - | 0 | - |
| - | - | 1059 | 986.5 | - | - | 0 | - |
| - | - | 1164 | 987 | - | - | 0 | - |
| - | - | 6390 | 995.5 | - | - | 0 | - |
| - | - | 8897 | 996 | - | - | 0 | - |
| - | - | 9361 | 996.5 | - | - | 0 | - |
| - | - | 5872 | 997 | - | - | 0 | - |
| - | - | 2686 | 997.5 | - | - | 0 | - |
| - | - | 1517 | 998 | - | - | 0 | - |
| - | - | 745.8 | 1001 | - | - | 0 | - |
| - | - | 1532 | 1003 | - | - | 0 | - |
| - | - | 8237 | 1003 | - | - | 0 | - |
| - | - | 3.404E+04 | 1004 | - | - | 0 | - |
| - | - | 5.39E+04 | 1004 | - | - | 0 | - |
| - | - | 4.227E+04 | 1005 | - | - | 0 | - |
| - | - | 2.337E+04 | 1005 | - | - | 0 | - |
| - | - | 9967 | 1006 | - | - | 0 | - |
| - | - | 4131 | 1006 | - | - | 0 | - |
| - | - | 1733 | 1007 | - | - | 0 | - |
| 9 | z | 1.257E+04 | 1027 | 0.004794 | 4.67 | +1 | 8 |
| - | - | 1.056E+04 | 1028 | - | - | 0 | - |
| - | - | 4301 | 1029 | - | - | 0 | - |
| - | - | 1172 | 1030 | - | - | 0 | - |
| - | - | 2010 | 1037 | - | - | 0 | - |
| - | - | 1167 | 1039 | - | - | 0 | - |
| - | - | 1579 | 1067 | - | - | 0 | - |
| 8 | z | 2651 | 1084 | 0.004773 | 4.405 | +1 | 9 |
| - | - | 4225 | 1085 | - | - | 0 | - |
| - | - | 1746 | 1086 | - | - | 0 | - |
| - | - | 1058 | 1087 | - | - | 0 | - |
| - | - | 808.7 | 1097 | - | - | 0 | - |
| 8 | y | 3823 | 1100 | 0.004454 | 4.051 | +1 | 9 |
| - | - | 2459 | 1101 | - | - | 0 | - |
| - | - | 872.3 | 1102 | - | - | 0 | - |
| - | - | 4444 | 1110 | - | - | 0 | - |
| - | - | 1057 | 1110 | - | - | 0 | - |
| 9 | c | 1.326E+04 | 1111 | 0.0008776 | 0.7903 | +1 | 9 |
| - | - | 7361 | 1112 | - | - | 0 | - |
| - | - | 2732 | 1113 | - | - | 0 | - |
| - | - | 1120 | 1114 | - | - | 0 | - |
| - | - | 850.6 | 1116 | - | - | 0 | - |
| - | - | 1087 | 1117 | - | - | 0 | - |
| - | - | 770.7 | 1137 | - | - | 0 | - |
| - | - | 897.1 | 1138 | - | - | 0 | - |
| - | - | 3753 | 1142 | - | - | 0 | - |
| - | - | 1498 | 1143 | - | - | 0 | - |
| - | - | 1083 | 1155 | - | - | 0 | - |
| - | - | 1457 | 1196 | - | - | 0 | - |
| - | - | 1645 | 1198 | - | - | 0 | - |
| 7 | z | 7291 | 1199 | 0.00657 | 5.481 | +1 | 10 |
| - | - | 6231 | 1200 | - | - | 0 | - |
| - | - | 3865 | 1201 | - | - | 0 | - |
| - | - | 919.5 | 1202 | - | - | 0 | - |
| 7 | y | 3193 | 1215 | 0.003687 | 3.036 | +1 | 10 |
| - | - | 1732 | 1216 | - | - | 0 | - |
| - | - | 4391 | 1239 | - | - | 0 | - |
| 10 | c | 1.208E+04 | 1240 | 0.001554 | 1.254 | +1 | 10 |
| - | - | 8979 | 1241 | - | - | 0 | - |
| - | - | 3877 | 1242 | - | - | 0 | - |
| - | - | 902.6 | 1243 | - | - | 0 | - |
| - | - | 1158 | 1268 | - | - | 0 | - |
| - | - | 895.4 | 1269 | - | - | 0 | - |
| - | - | 753.8 | 1270 | - | - | 0 | - |
| 6 | y | 3182 | 1311 | 0.01582 | 12.07 | +1 | 11 |
| 6 | z | 5303 | 1312 | 0.001041 | 0.7937 | +1 | 11 |
| - | - | 4938 | 1313 | - | - | 0 | - |
| - | - | 2661 | 1314 | - | - | 0 | - |
| - | - | 1404 | 1315 | - | - | 0 | - |
| 6 | y | 983.1 | 1328 | 0.005144 | 3.875 | +1 | 11 |
| - | - | 1038 | 1329 | - | - | 0 | - |
| - | - | 813.6 | 1334 | - | - | 0 | - |
| - | - | 995.6 | 1342 | - | - | 0 | - |
| - | - | 2015 | 1359 | - | - | 0 | - |
| - | - | 902.3 | 1360 | - | - | 0 | - |
| - | - | 740.4 | 1361 | - | - | 0 | - |
| - | - | 842.8 | 1401 | - | - | 0 | - |
| - | - | 4100 | 1402 | - | - | 0 | - |
| 11 | c | 1.04E+04 | 1403 | 0.001035 | 0.738 | +1 | 11 |
| - | - | 7248 | 1404 | - | - | 0 | - |
| - | - | 2452 | 1405 | - | - | 0 | - |
| - | - | 801.3 | 1406 | - | - | 0 | - |
| - | - | 1220 | 1487 | - | - | 0 | - |
| - | - | 1140 | 1488 | - | - | 0 | - |
| - | - | 702.3 | 1489 | - | - | 0 | - |
| 5 | y | 2962 | 1497 | 0.01439 | 9.616 | +1 | 12 |
| 5 | z | 7196 | 1498 | 0.001368 | 0.9131 | +1 | 12 |
| - | - | 5844 | 1499 | - | - | 0 | - |
| - | - | 3002 | 1500 | - | - | 0 | - |
| - | - | 1485 | 1501 | - | - | 0 | - |
| 12 | c | 1366 | 1513 | 0.009144 | 6.045 | +1 | 12 |
| 12 | c | 5529 | 1514 | 0.005855 | 3.868 | +1 | 12 |
| - | - | 4718 | 1515 | - | - | 0 | - |
| - | - | 1991 | 1516 | - | - | 0 | - |
| - | - | 862.7 | 1517 | - | - | 0 | - |
| - | - | 3420 | 1530 | - | - | 0 | - |
| 12 | c | 1.082E+04 | 1531 | 0.0006662 | 0.4352 | +1 | 12 |
| - | - | 8331 | 1532 | - | - | 0 | - |
| - | - | 4027 | 1533 | - | - | 0 | - |
| - | - | 1321 | 1534 | - | - | 0 | - |
| - | - | 1061 | 1557 | - | - | 0 | - |
| - | - | 3300 | 1569 | - | - | 0 | - |
| - | - | 2480 | 1570 | - | - | 0 | - |
| - | - | 1953 | 1571 | - | - | 0 | - |
| - | - | 822.8 | 1572 | - | - | 0 | - |
| - | - | 991.5 | 1608 | - | - | 0 | - |
| 4 | y | 1841 | 1612 | 0.007835 | 4.861 | +1 | 13 |
| 4 | z | 1.02E+04 | 1613 | 0.003408 | 2.113 | +1 | 13 |
| - | - | 1.164E+04 | 1614 | - | - | 0 | - |
| - | - | 7897 | 1615 | - | - | 0 | - |
| - | - | 3603 | 1616 | - | - | 0 | - |
| - | - | 1199 | 1617 | - | - | 0 | - |
| 4 | y | 1936 | 1629 | 0.01066 | 6.543 | +1 | 13 |
| - | - | 1597 | 1630 | - | - | 0 | - |
| - | - | 803.8 | 1632 | - | - | 0 | - |
| - | - | 699.5 | 1633 | - | - | 0 | - |
| - | - | 979.3 | 1665 | - | - | 0 | - |
| - | - | 1158 | 1665 | - | - | 0 | - |
| - | - | 1207 | 1666 | - | - | 0 | - |
| - | - | 793 | 1668 | - | - | 0 | - |
| - | - | 846.8 | 1672 | - | - | 0 | - |
| - | - | 1005 | 1673 | - | - | 0 | - |
| - | - | 1502 | 1674 | - | - | 0 | - |
| - | - | 883.5 | 1675 | - | - | 0 | - |
| - | - | 859.6 | 1684 | - | - | 0 | - |
| - | - | 4615 | 1691 | - | - | 0 | - |
| 13 | c | 1.394E+04 | 1692 | 0.00508 | 3.003 | +1 | 13 |
| - | - | 1.264E+04 | 1693 | - | - | 0 | - |
| - | - | 5612 | 1694 | - | - | 0 | - |
| - | - | 2890 | 1695 | - | - | 0 | - |
| 3 | z | 8274 | 1741 | 0.005589 | 3.211 | +1 | 14 |
| - | - | 1.069E+04 | 1742 | - | - | 0 | - |
| - | - | 7426 | 1743 | - | - | 0 | - |
| - | - | 4545 | 1744 | - | - | 0 | - |
| - | - | 1589 | 1745 | - | - | 0 | - |
| 3 | y | 2078 | 1757 | 0.006491 | 3.695 | +1 | 14 |
| - | - | 1374 | 1758 | - | - | 0 | - |
| - | - | 1069 | 1759 | - | - | 0 | - |
| - | - | 1015 | 1762 | - | - | 0 | - |
| - | - | 1204 | 1777 | - | - | 0 | - |
| - | - | 1344 | 1778 | - | - | 0 | - |
| - | - | 982.1 | 1819 | - | - | 0 | - |
| 14 | c | 2298 | 1820 | 0.007514 | 4.129 | +1 | 14 |
| - | - | 2738 | 1821 | - | - | 0 | - |
| - | - | 1229 | 1822 | - | - | 0 | - |
| - | - | 1417 | 1823 | - | - | 0 | - |
| - | - | 1211 | 1860 | - | - | 0 | - |
| - | - | 1009 | 1875 | - | - | 0 | - |
| - | - | 2147 | 1876 | - | - | 0 | - |
| - | - | 1931 | 1877 | - | - | 0 | - |
| 2 | z | 2004 | 1878 | 0.03389 | 18.05 | +1 | 15 |
| - | - | 4816 | 1879 | - | - | 0 | - |
| - | - | 4531 | 1880 | - | - | 0 | - |
| - | - | 2401 | 1881 | - | - | 0 | - |
| - | - | 895.4 | 1882 | - | - | 0 | - |
| - | - | 813.6 | 1902 | - | - | 0 | - |
| - | - | 1085 | 1904 | - | - | 0 | - |
| - | - | 1531 | 1917 | - | - | 0 | - |
| - | - | 2926 | 1918 | - | - | 0 | - |
| 15 | c | 5038 | 1919 | 0.01709 | 8.906 | +1 | 15 |
| - | - | 6402 | 1920 | - | - | 0 | - |
| - | - | 4060 | 1921 | - | - | 0 | - |
| - | - | 3587 | 1922 | - | - | 0 | - |
| - | - | 1716 | 1923 | - | - | 0 | - |
| - | - | 983.5 | 1924 | - | - | 0 | - |
| - | - | 1736 | 1932 | - | - | 0 | - |
| - | - | 2067 | 1933 | - | - | 0 | - |
| - | - | 2201 | 1934 | - | - | 0 | - |
| - | - | 814.9 | 1947 | - | - | 0 | - |
| - | - | 3047 | 1948 | - | - | 0 | - |
| - | - | 7319 | 1949 | - | - | 0 | - |
| - | - | 1.259E+04 | 1950 | - | - | 0 | - |
| - | - | 9152 | 1951 | - | - | 0 | - |
| - | - | 4461 | 1952 | - | - | 0 | - |
| - | - | 2496 | 1953 | - | - | 0 | - |
| - | - | 1433 | 1954 | - | - | 0 | - |
| - | - | 1468 | 1962 | - | - | 0 | - |
| - | - | 4992 | 1963 | - | - | 0 | - |
| - | - | 7179 | 1964 | - | - | 0 | - |
| - | - | 5369 | 1965 | - | - | 0 | - |
| - | - | 2921 | 1966 | - | - | 0 | - |
| - | - | 1300 | 1967 | - | - | 0 | - |
| - | - | 840.9 | 1973 | - | - | 0 | - |
| - | - | 1056 | 1979 | - | - | 0 | - |
| - | - | 4317 | 1980 | - | - | 0 | - |
| - | - | 8500 | 1981 | - | - | 0 | - |
| - | - | 7173 | 1982 | - | - | 0 | - |
| - | - | 3689 | 1983 | - | - | 0 | - |
| - | - | 1874 | 1984 | - | - | 0 | - |
| - | - | 1919 | 1990 | - | - | 0 | - |
| - | - | 9072 | 1991 | - | - | 0 | - |
| - | - | 1.467E+04 | 1992 | - | - | 0 | - |
| - | - | 1.277E+04 | 1993 | - | - | 0 | - |
| - | - | 7942 | 1994 | - | - | 0 | - |
| - | - | 4311 | 1995 | - | - | 0 | - |
| - | - | 1947 | 1996 | - | - | 0 | - |
| - | - | 1931 | 2006 | - | - | 0 | - |
| - | - | 6832 | 2007 | - | - | 0 | - |
| - | - | 2.861E+04 | 2008 | - | - | 0 | - |
| - | - | 6.576E+04 | 2009 | - | - | 0 | - |
| - | - | 6.082E+04 | 2010 | - | - | 0 | - |
| - | - | 3.511E+04 | 2011 | - | - | 0 | - |
| - | - | 1.273E+04 | 2012 | - | - | 0 | - |
| - | - | 6707 | 2013 | - | - | 0 | - |
| - | - | 2372 | 2014 | - | - | 0 | - |
| - | - | 730.2 | 3136 | - | - | 0 | - |
| - | - | 814.2 | 3340 | - | - | 0 | - |

m/z Charge Intensity FragmentType MassShift Position
129.1023406982422 0 1595.4075
131.11819458007812 0 799.31775
133.06076049804688 0 372.18225
136.73361206054688 0 413.75217
143.1182861328125 0 522.8993
144.5810546875 0 391.4636
146.0923309326172 0 621.5145
148.8863525390625 0 614.04004
148.8936004638672 0 581.3791
148.9008026123047 0 772.4953
148.9080047607422 0 816.00854
148.91517639160156 0 752.96106
148.9228057861328 0 1048.6404
148.9298553466797 0 1141.0881
148.93731689453125 0 2705.52
148.9451141357422 0 4424.7056
148.9617462158203 0 3784.4458
148.96957397460938 0 1390.98
148.97683715820312 0 1260.7562
148.9840850830078 0 1088.1587
148.99105834960938 0 829.7018
148.9984893798828 0 613.8021
149.0056610107422 0 692.2753
149.12921142578125 0 409.85983
153.09036254882812 0 501.7764
155.09274291992188 0 894.26184
164.91709899902344 0 459.87146
166.06100463867188 0 1886.5402
168.1389923095703 0 518.4581
173.0919189453125 0 836.7313
178.1338653564453 0 582.05426
185.16500854492188 0 1020.4675
205.1184844970703 0 933.9519 y 14
214.1536102294922 0 576.9631
223.1554718017578 0 1920.1818
228.0981903076172 0 1092.0344
229.0818634033203 0 1202.5707
229.15496826171875 0 573.2088
230.11358642578125 0 2012.6818
234.12353515625 0 2172.8152
237.2140350341797 0 572.4098
242.1500244140625 0 10803.819
243.15382385253906 0 1364.3542
246.10848999023438 0 7907.2993
247.0920867919922 0 1569.2091
247.11080932617188 0 829.4429
248.12403869628906 0 3329.0508
249.0977783203125 0 550.5301
249.15919494628906 0 1184.381
250.1414794921875 0 649.2046
251.150146484375 0 9494.126
252.1534881591797 0 1710.8204
264.118896484375 0 9713.34
265.12152099609375 0 812.4687
265.1546630859375 0 1641.4579
266.1247253417969 0 618.28644
268.1290283203125 0 618.79614
268.1766662597656 0 6126.5034 c 1
269.1806640625 0 683.5553
277.15460205078125 0 5097.119
278.1591796875 0 689.72125
291.1339111328125 0 2666.459
293.1494445800781 0 7243.731
294.1529846191406 0 1190.0333
298.794677734375 0 487.1698
306.10772705078125 0 585.3715
309.1450500488281 0 961.368
316.1745300292969 0 927.26654
317.1946105957031 0 7803.0903 z 13
318.2012023925781 0 3794.2163
333.2134094238281 0 1188.9213 y 13
341.01885986328125 0 762.19885
342.1668395996094 0 1454.7233
343.0146484375 0 934.7709
343.19757080078125 0 1298.3398
344.1820068359375 0 762.53894
353.2289733886719 0 597.3909
355.0697937011719 0 1908.1282
356.0706787109375 0 1176.059
357.06805419921875 0 2079.254
359.0285339355469 0 1824.5017
359.19244384765625 0 6548.2744
360.028076171875 0 1628.6329
360.17474365234375 0 867.2238
360.19659423828125 0 1367.1229
361.0264892578125 0 1788.9178
361.20758056640625 0 2928.8347
362.21026611328125 0 593.5108
375.188720703125 0 737.8983
377.20257568359375 0 9504.415
378.2052001953125 0 1433.6903
379.2092590332031 0 1623.698 c Ammonia loss 2
387.2239074707031 0 10800.3
388.2265319824219 0 2946.858
389.2299499511719 0 1465.181
390.23883056640625 0 4082.7656
395.2281188964844 0 1087.7249
396.2353515625 0 13095.751 c 2
397.2375793457031 0 2642.874
404.2181091308594 0 763.0879
405.21466064453125 0 837.5644
406.2334899902344 0 4653.0894
407.23663330078125 0 1339.512
407.26568603515625 0 706.9203
411.1509704589844 0 1719.5303
412.1531982421875 0 982.5245
419.19305419921875 0 629.8327
421.2073059082031 0 881.7945
422.2291259765625 0 673.0938
423.2596740722656 0 763.655
429.0889892578125 0 2323.6187
430.0886535644531 0 2018.6757
431.0871276855469 0 5218.4414
432.08770751953125 0 1316.207
439.2566223144531 0 609.5212
461.2236633300781 0 616.66956
467.2489013671875 0 5122.3184
468.2519836425781 0 1354.2274
490.30108642578125 0 566.76013
494.2313232421875 0 2655.7915 c Ammonia loss 3
495.23272705078125 0 1081.843
506.2976989746094 0 1334.3662
511.26239013671875 0 15047.525 c 3
512.2651977539062 0 4000.3079
515.3186645507812 0 2071.0098
516.322265625 0 701.8253
517.2900390625 0 1217.2639
518.296630859375 0 3574.8833
519.2507934570312 0 1542.7308
519.296142578125 0 839.83264
519.7544555664062 0 673.12994
534.2918701171875 0 8969.432
535.2947387695312 0 2512.758
547.2907104492188 0 757.6091
548.2354125976562 0 931.6358 w 11
548.30517578125 0 1585.61
550.2805786132812 0 3563.3896 y 7
550.7752075195312 0 993.2636
551.28857421875 0 1031.1495
556.258544921875 0 1136.0203
563.2492065429688 0 573.47906
564.248779296875 0 833.34235
592.321533203125 0 2779.5637
593.3225708007812 0 1237.9519
600.3016967773438 0 1020.41656
601.3028564453125 0 847.5682 c Water loss 13
601.9561157226562 0 845.14276
606.3037719726562 0 8185.665 z 11
607.3092041015625 0 3704.8481 c 13
607.7898559570312 0 881.2051 y 6
608.2963256835938 0 1412.1199
617.32861328125 0 1174.6694
618.328857421875 0 1152.1467
619.2277221679688 0 2163.3162
622.3231201171875 0 4169.6387 y 11
623.3223266601562 0 709.33246
629.30810546875 0 1052.1815
631.6336669921875 0 1001.7882
631.9663696289062 0 3344.7432 y 1
632.3016967773438 0 2644.6868
632.6349487304688 0 964.16315
632.9679565429688 0 611.8406
634.3172607421875 0 1584.3071 c Water loss 14
634.6477661132812 0 4699.4736 c Ammonia loss 14
634.9794311523438 0 5117.3506
635.3358764648438 0 5055.97
635.6461791992188 0 1083.1542
636.3421020507812 0 2509.262
637.2380981445312 0 1127.0186
637.3556518554688 0 1766.5178
651.2835693359375 0 1123.9324
651.3349609375 0 3627.6204
652.3292846679688 0 2860.424
653.3218994140625 0 1024.152
655.3182373046875 0 646.88446 y Water loss 5
657.9818725585938 0 1361.0337
658.3171997070312 0 1499.4714
658.6510620117188 0 778.00073
663.6561889648438 0 2390.9026
663.987548828125 0 3143.5723
664.3224487304688 0 1804.3417 y 5
664.656982421875 0 2828.7227
668.288818359375 0 3919.8374
669.3394165039062 0 31209.998
669.6611328125 0 56718.918
669.9945068359375 0 49863.14
670.3318481445312 0 38249.66
670.6620483398438 0 13558.995
670.997314453125 0 5692.5654
671.3488159179688 0 3952.3584
697.3413696289062 0 11263.8545 c 4
698.3450317382812 0 4374.571
732.7398071289062 0 601.9711
757.3726196289062 0 1149.4097 c Ammonia loss 11
757.8724365234375 0 1298.2551
766.410400390625 0 1393.1251
767.4171752929688 0 790.4577
769.3671875 0 11729.803 z 10
770.3720092773438 0 6162.927
771.3718872070312 0 3380.5596
780.3684692382812 0 611.38446
785.3870849609375 0 2996.844 y 10
786.3892211914062 0 1490.6606
793.3944702148438 0 1012.6153 c Ammonia loss 5
795.4050903320312 0 640.7749
810.424560546875 0 11117.804 c 5
811.4274291992188 0 6255.5483
812.4304809570312 0 1563.0872
814.8873291015625 0 1363.8685 y 3
815.3858032226562 0 639.00134
824.391357421875 0 759.00323
826.3959350585938 0 696.7759
826.895263671875 0 783.33386
836.43017578125 0 632.1721
839.3914794921875 0 1726.2085 w 9
840.4002075195312 0 1232.8561
841.392578125 0 621.7214
846.3955688476562 0 809.5881 c 12
847.4022827148438 0 760.56616
867.4228515625 0 3090.9478
868.4287109375 0 1973.9955
869.9122924804688 0 973.52405 y Water loss 2
870.405029296875 0 3908.6921 y Ammonia loss 2
870.9033203125 0 1604.6293 z 2
871.40576171875 0 1503.8108
878.4271850585938 0 1536.4072
878.916259765625 0 4966.4326 y 2
879.416748046875 0 5703.8433
879.9186401367188 0 3409.1948
880.4432373046875 0 1089.8318
881.443359375 0 1987.799
882.4442138671875 0 1698.5895
887.9398803710938 0 664.01483
888.4297485351562 0 1398.0984
889.4204711914062 0 658.8245
889.9420776367188 0 834.4333
898.40966796875 0 11105.1875 z 9
899.4126586914062 0 5957.1523
900.413330078125 0 1683.0065
901.4237060546875 0 1123.5972 c Water loss 13
901.9335327148438 0 954.0646 c Ammonia loss 13
902.4386596679688 0 1405.633
908.43212890625 0 1834.9581 c Ammonia loss 6
909.43505859375 0 840.7753
909.9479370117188 0 2344.7195
910.44189453125 0 8574.021 c 13
910.9437255859375 0 7955.7056
911.4458618164062 0 4793.9746
911.9439697265625 0 1819.0106
914.4295654296875 0 2712.6458 y 9
917.44677734375 0 974.96545
917.9347534179688 0 1151.3486
918.4419555664062 0 1151.6863
924.4525146484375 0 2428.7776
925.4524536132812 0 3874.4521 c 6
926.4515380859375 0 1516.4386
927.451904296875 0 769.2006
931.4364013671875 0 557.7274
935.5181884765625 0 1223.5831
936.5009155273438 0 873.09784
937.9752197265625 0 2137.1567
938.4692993164062 0 3632.0735
938.9549560546875 0 3288.7976
939.444091796875 0 6190.6465 z 1
939.9380493164062 0 4942.821
940.44384765625 0 3407.1736
940.9325561523438 0 803.98364
945.4777221679688 0 670.96185
945.9616088867188 0 878.43744
950.9794311523438 0 922.38477 c Water loss 14
951.9619750976562 0 1221.0314
952.4764404296875 0 1632.4033
952.9713134765625 0 1478.1869
953.47216796875 0 1035.3207
954.9614868164062 0 596.55316
959.483642578125 0 4580.8574
959.9805297851562 0 16385.486 c 14
960.4802856445312 0 15964.853
960.9806518554688 0 9281.328
961.4788818359375 0 4319.2964
961.984130859375 0 2276.6523
962.476318359375 0 1110.852
964.9808959960938 0 760.76447
965.4733276367188 0 1679.8439
965.9761352539062 0 1182.8843
966.4793090820312 0 1138.0315
966.96728515625 0 1187.2058
967.477294921875 0 1160.503
967.6199951171875 0 591.94147
967.9719848632812 0 1632.9329
968.4695434570312 0 1948.9496
968.9646606445312 0 1708.1127
969.4688110351562 0 892.96606
973.4853515625 0 2039.067
973.9739379882812 0 3241.359
974.4765014648438 0 12678.118
974.980712890625 0 25699.834
975.482177734375 0 25216.703
975.9820556640625 0 13782.321
976.4838256835938 0 8098.647
976.9835205078125 0 2900.4324
977.486572265625 0 967.38196
979.4833984375 0 664.6251
980.4764404296875 0 1038.5417
981.4855346679688 0 13488.293
981.9851684570312 0 9175.29
982.4814453125 0 19828.04 c 7
982.9871826171875 0 3648.2942
983.480224609375 0 7620.5264
984.4745483398438 0 1948.9857
986.4857788085938 0 1058.9117
986.9769287109375 0 1164.1221
995.48046875 0 6390.44
995.9790649414062 0 8896.679
996.4799194335938 0 9361.223
996.9779052734375 0 5871.883
997.480224609375 0 2685.7227
997.977294921875 0 1516.7754
1001.4915771484375 0 745.7605
1002.977294921875 0 1531.6067
1003.4908447265625 0 8237.467
1003.9874877929688 0 34042.742
1004.4884033203125 0 53900.402
1004.989990234375 0 42268.566
1005.4906616210938 0 23370.893
1005.9905395507812 0 9967.264
1006.4884033203125 0 4130.831
1006.9778442382812 0 1733.3926
1026.5047607421875 0 12573.623 z 8
1027.50830078125 0 10558.272
1028.509521484375 0 4301.3667
1029.513916015625 0 1172.4938
1037.49169921875 0 2009.8502
1038.5037841796875 0 1166.9723
1066.55419921875 0 1578.5908
1083.5262451171875 0 2650.9355 z 7
1084.5333251953125 0 4224.528
1085.53759765625 0 1745.9757
1086.537353515625 0 1057.9103
1096.5224609375 0 808.705
1099.5452880859375 0 3823.0889 y 7
1100.541259765625 0 2458.876
1101.548583984375 0 872.32666
1109.5828857421875 0 4444.2964
1110.240966796875 0 1056.5433
1110.5699462890625 0 13257.448 c 8
1111.572998046875 0 7361.211
1112.572265625 0 2732.3892
1113.576416015625 0 1119.5698
1116.2529296875 0 850.5631
1116.91015625 0 1087.3335
1136.5965576171875 0 770.6953
1137.58056640625 0 897.14484
1141.552978515625 0 3753.3862
1142.55859375 0 1498.1161
1154.5496826171875 0 1083.3923
1195.5986328125 0 1456.6025
1197.5811767578125 0 1644.9358
1198.5513916015625 0 7291.3354 z 6
1199.5584716796875 0 6231.036
1200.5587158203125 0 3865.295
1201.550048828125 0 919.4671
1214.572998046875 0 3192.659 y 6
1215.575439453125 0 1732.0514
1238.6168212890625 0 4390.715
1239.610107421875 0 12084.594 c 9
1240.61279296875 0 8979.268
1241.6156005859375 0 3876.761
1242.6190185546875 0 902.6458
1267.636962890625 0 1157.9811
1268.6351318359375 0 895.36426
1269.568603515625 0 753.8149
1310.6500244140625 0 3182.2363 y Ammonia loss 5
1311.64306640625 0 5303.477 z 5
1312.6439208984375 0 4938.44
1313.642822265625 0 2661.1392
1314.6474609375 0 1404.3264
1327.6658935546875 0 983.09576 y 5
1328.652099609375 0 1038.0747
1333.605712890625 0 813.5611
1341.6400146484375 0 995.5707
1358.6622314453125 0 2015.3671
1359.662841796875 0 902.32184
1360.66455078125 0 740.36786
1400.6824951171875 0 842.8015
1401.6817626953125 0 4099.521
1402.676025390625 0 10399.516 c 10
1403.6767578125 0 7248.215
1404.6776123046875 0 2451.7654
1405.694091796875 0 801.34576
1486.746826171875 0 1219.7665
1487.771728515625 0 1139.7734
1488.76806640625 0 702.2932
1496.7279052734375 0 2961.5754 y Ammonia loss 4
1497.719970703125 0 7196.2974 z 4
1498.7200927734375 0 5843.657
1499.7230224609375 0 3002.1726
1500.7322998046875 0 1485.3486
1512.750244140625 0 1365.5378 c Water loss 11
1513.737548828125 0 5528.736 c Ammonia loss 11
1514.738037109375 0 4718.1357
1515.73486328125 0 1991.0117
1516.7520751953125 0 862.71173
1529.7779541015625 0 3420.0564
1530.769287109375 0 10818.247 c 11
1531.7723388671875 0 8330.95
1532.774658203125 0 4027.163
1533.775146484375 0 1321.2124
1556.7454833984375 0 1060.787
1568.751708984375 0 3300.3142
1569.7607421875 0 2480.4397
1570.75244140625 0 1952.6418
1571.77001953125 0 822.7757
1607.7913818359375 0 991.5013
1611.748291015625 0 1840.8657 y Ammonia loss 3
1612.744873046875 0 10197.3955 z 3
1613.7462158203125 0 11639.862
1614.7513427734375 0 7896.9873
1615.753173828125 0 3602.763
1616.750244140625 0 1198.884
1628.75634765625 0 1935.9888 y 3
1629.7498779296875 0 1596.5566
1631.7681884765625 0 803.8395
1632.763671875 0 699.49927
1664.8115234375 0 979.3155
1665.3563232421875 0 1157.9436
1666.3480224609375 0 1207.0793
1667.8243408203125 0 792.96375
1672.3233642578125 0 846.78204
1672.8656005859375 0 1004.7794
1673.819580078125 0 1501.9966
1674.894287109375 0 883.5308
1683.7977294921875 0 859.64746
1690.7935791015625 0 4615.28
1691.7840576171875 0 13938.74 c 12
1692.785400390625 0 12636.343
1693.78662109375 0 5612.1777
1694.7874755859375 0 2890.006
1740.80126953125 0 8274.368 z 2
1741.8057861328125 0 10693.155
1742.811767578125 0 7425.8657
1743.809814453125 0 4544.872
1744.8140869140625 0 1589.4873
1756.819091796875 0 2077.7285 y 2
1757.8209228515625 0 1374.2401
1758.840576171875 0 1068.7906
1761.8563232421875 0 1014.7475
1776.872802734375 0 1203.9055
1777.8673095703125 0 1343.7047
1818.89111328125 0 982.08234
1819.8765869140625 0 2297.8943 c 13
1820.880126953125 0 2737.6182
1821.879150390625 0 1229.283
1822.8909912109375 0 1417.0293
1859.932373046875 0 1211.2616
1874.9383544921875 0 1008.86304
1875.93994140625 0 2147.0122
1876.94140625 0 1931.4971
1877.899658203125 0 2004.1013 z 1
1878.8746337890625 0 4816.2803
1879.8677978515625 0 4530.7363
1880.876220703125 0 2401.4448
1881.8748779296875 0 895.44763
1901.9940185546875 0 813.6421
1903.9659423828125 0 1084.6539
1916.987060546875 0 1531.4512
1917.9842529296875 0 2926.1616
1918.9696044921875 0 5038.1006 c 14
1919.965576171875 0 6401.675
1920.9561767578125 0 4060.2312
1921.958984375 0 3587.2097
1922.9630126953125 0 1716.1556
1923.9501953125 0 983.4587
1931.9422607421875 0 1735.9248
1932.9449462890625 0 2067.462
1933.9461669921875 0 2201.1055
1946.9501953125 0 814.8916
1947.9486083984375 0 3047.3677
1948.9573974609375 0 7318.771
1949.9576416015625 0 12585.936
1950.9598388671875 0 9151.68
1951.96142578125 0 4461.2515
1952.9573974609375 0 2495.8005
1953.9571533203125 0 1432.6387
1961.94921875 0 1468.0159
1962.966064453125 0 4991.5605
1963.9671630859375 0 7178.5728
1964.9599609375 0 5369.468
1965.96337890625 0 2920.603
1966.97998046875 0 1299.7888
1972.9775390625 0 840.91986
1978.964599609375 0 1056.1927
1979.9810791015625 0 4317.2427
1980.975830078125 0 8500.354
1981.98291015625 0 7173.3213
1982.9869384765625 0 3688.6306
1983.98828125 0 1874.1843
1989.9598388671875 0 1918.8889
1990.9569091796875 0 9071.985
1991.9530029296875 0 14669.127
1992.954833984375 0 12767.934
1993.9510498046875 0 7941.9907
1994.949951171875 0 4311.4565
1995.9429931640625 0 1946.7386
2005.9622802734375 0 1930.5361
2006.9732666015625 0 6832.225
2007.9759521484375 0 28608.818
2008.977294921875 0 65761.26
2009.9765625 0 60816.77
2010.9786376953125 0 35105.934
2011.980224609375 0 12728.602
2012.9781494140625 0 6706.92
2013.953857421875 0 2371.649
3135.609130859375 0 730.17676
3340.49267578125 0 814.19086

Spectrum Details

|  |  |
| --- | --- |
| Matched peaks? Matched peaks The total absolute number of peaks matched. Additionally in brackets the total fraction of peaks matched and the total number of peaks is shown. | 69 (13.40% of 515) |
| FDR? FDR The false discovery rate estimated for this peptide. It is calculated by matching all theoretical fragments with a non-integer shift with the raw peaks for this spectrum. This is done with 40 different shifts. The resulting percentage is the average number of annotated peaks over the number of annotated peaks with the correct spectrum. | 1.17% |
| Satellite FDR? Satellite FDR See the FDR for details on its calculation. This satellite ion specific FDR only contains the satellite ions (d/w) for I/L/J positions. | - |
| PSM Score? PSM Score The PSM Score as given by Hecklib to this annotated spectrum. It is shown with three significant figures. | 560 |

#### Spectrum 5274? Spectrum 5274 The raw spectrum of this peptide as annotated by Hecklib. The fragments are coloured according to ion type (see legend). Any peaks with a star '\*' as text can be hovered over to see the full details, first the ion type second the mass shift type. By hovering over the amino acids in the peptide or ions in the legend the corresponding peaks are highlighted. By toggling the 'Unassigned' label you can turn the background (unassigned) peaks on or off in the plot. By updating the slider in the Ion legend you can update the spectrum to only show the top X% of the peaks with labels. The top X% means any peak that is within X% of the highest intensity. By dragging in the spectrum you can zoom in to a specific part of the spectrum and use 'Zoom Out' to get back to the original zoom level. The annotation of the spectrum is based on the given sequence in the peptides file and is done with different software so inconsistencies are likely. The peaks are annotated based on the given sequence, with 20 ppm tolerance.

Copy Data

##### Spectrum 5274 (TSV)

###### Preview

```
Loading example...
```

*Click on the button to copy the data to your clipboard.*

Mz MinMz MaxIntensity Max

WidthHeightPeptide font sizePeptide stroke widthSpectrum font sizeSpectrum stroke widthCompact peptide

Ion legend

wxyz

abcd

OtherUnassignedIon^Charge^_Position_Show for top:%

JHQDWJDGKEYKCKVS

02.27e+54.54e+56.81e+59.08e+5

Zoom Out

b^+4^_4_a^+2^_3_y^+1^_2_b^+4^_6_y^+1^_2_a^+1^_2_b^+2^_4_b^+2^_4_b^+1^_2_y^+4^_8_y^+2^_5_y^+2^_5_y^+1^_3_y^+4^_11_b^+2^_5_y^+1^_3_a^+1^_3_b^+2^_5_y^+3^_8_y^+3^_9_b^+1^_3_y^+3^_9_b^+4^_12_b^+1^_3_y^+2^_6_y^+2^_6_b^+2^_6_b^+2^_6_y^+2^_6_b^+2^_6_y^+3^_10_y^+3^_10_y^+3^_10_y^+3^_11_y^+3^_11_y^+3^_11_b^+2^_7_b^+2^_7_b^+4^_14_y^+2^_7_b^+2^_7_y^+2^_7_y^+4^_15_b^+4^_15_b^+4^_15_b^+2^_8_b^+1^_4_b^+1^_4_b^+1^_4_\*\*b^+3^_12_\*b^+3^_12_y^+2^_8_y^+2^_8_y^+2^_8_b^+2^_9_b^+2^_9_y^+2^_9_y^+2^_9_y^+3^_13_b^+2^_9_y^+2^_9_y^+3^_14_y^+3^_14_y^+3^_14_b^+3^_14_b^+3^_14_y^+2^_10_y^+2^_10_b^+3^_14_y^+1^_5_y^+2^_10_b^+2^_10_y^+1^_5_y^+3^_15_y^+3^_15_b^+3^_15_b^+3^_15_y^+3^_15_b^+3^_15_y^+2^_11_y^+2^_11_b^+1^_5_y^+2^_11_\*b^+1^_5_b^+2^_11_b^+2^_11_b^+2^_12_b^+2^_12_b^+2^_12_y^+1^_6_b^+1^_6_b^+1^_6_y^+1^_6_b^+1^_6_y^+2^_13_y^+2^_13_y^+2^_13_y^+2^_14_y^+2^_14_b^+1^_7_b^+1^_7_y^+1^_7_b^+1^_7_y^+1^_7_b^+1^_8_b^+1^_8_y^+1^_8_y^+1^_9_b^+1^_9_y^+1^_9_y^+1^_10_b^+1^_10_y^+1^_10_

0724144721712894

Fragment Matches Table

Show background peaks

| Position | Ion type | Intensity | mz Theoretical | mz Error (Th) | mz Error (ppm) | Charge | Series Number |
| --- | --- | --- | --- | --- | --- | --- | --- |
| 4 | b | 1661 | 120.1 | 0.001382 | 11.51 | +4 | 4 |
| - | - | 875.8 | 120.1 | - | - | 0 | - |
| - | - | 1751 | 121 | - | - | 0 | - |
| - | - | 3263 | 122.1 | - | - | 0 | - |
| - | - | 2661 | 123.1 | - | - | 0 | - |
| - | - | 668.4 | 124.1 | - | - | 0 | - |
| - | - | 524.1 | 124.7 | - | - | 0 | - |
| - | - | 3251 | 125.1 | - | - | 0 | - |
| - | - | 5829 | 127.1 | - | - | 0 | - |
| - | - | 4891 | 127.1 | - | - | 0 | - |
| - | - | 1290 | 128.1 | - | - | 0 | - |
| - | - | 1911 | 128.1 | - | - | 0 | - |
| - | - | 609.7 | 128.1 | - | - | 0 | - |
| - | - | 4807 | 129.1 | - | - | 0 | - |
| - | - | 8.99E+05 | 129.1 | - | - | 0 | - |
| - | - | 3.313E+05 | 130.1 | - | - | 0 | - |
| - | - | 6165 | 130.1 | - | - | 0 | - |
| - | - | 6.199E+04 | 130.1 | - | - | 0 | - |
| - | - | 1757 | 131.1 | - | - | 0 | - |
| - | - | 3.085E+04 | 131.1 | - | - | 0 | - |
| - | - | 1412 | 131.1 | - | - | 0 | - |
| - | - | 804.3 | 131.1 | - | - | 0 | - |
| - | - | 1408 | 132.1 | - | - | 0 | - |
| - | - | 4.548E+04 | 132.1 | - | - | 0 | - |
| - | - | 1320 | 132.1 | - | - | 0 | - |
| - | - | 535.7 | 132.9 | - | - | 0 | - |
| - | - | 2245 | 133.1 | - | - | 0 | - |
| - | - | 4556 | 133.1 | - | - | 0 | - |
| - | - | 9699 | 134 | - | - | 0 | - |
| - | - | 634.4 | 135.1 | - | - | 0 | - |
| - | - | 837.9 | 136 | - | - | 0 | - |
| - | - | 6.613E+04 | 136.1 | - | - | 0 | - |
| - | - | 6145 | 137.1 | - | - | 0 | - |
| - | - | 893.2 | 137.1 | - | - | 0 | - |
| - | - | 1.063E+04 | 138.1 | - | - | 0 | - |
| - | - | 1002 | 138.1 | - | - | 0 | - |
| - | - | 1.035E+04 | 138.1 | - | - | 0 | - |
| - | - | 2538 | 139.1 | - | - | 0 | - |
| - | - | 1.389E+04 | 139.1 | - | - | 0 | - |
| - | - | 954.5 | 139.1 | - | - | 0 | - |
| - | - | 1223 | 140.1 | - | - | 0 | - |
| - | - | 1.619E+04 | 141.1 | - | - | 0 | - |
| - | - | 6392 | 142.1 | - | - | 0 | - |
| - | - | 1172 | 142.1 | - | - | 0 | - |
| - | - | 6534 | 143 | - | - | 0 | - |
| - | - | 698.1 | 143.1 | - | - | 0 | - |
| - | - | 1352 | 143.1 | - | - | 0 | - |
| - | - | 3015 | 144.1 | - | - | 0 | - |
| - | - | 1.187E+04 | 145.1 | - | - | 0 | - |
| - | - | 3157 | 146.1 | - | - | 0 | - |
| - | - | 872.1 | 146.1 | - | - | 0 | - |
| - | - | 2271 | 146.1 | - | - | 0 | - |
| - | - | 3895 | 147 | - | - | 0 | - |
| - | - | 1549 | 147.1 | - | - | 0 | - |
| - | - | 1280 | 148.1 | - | - | 0 | - |
| - | - | 978.5 | 148.1 | - | - | 0 | - |
| - | - | 892 | 149 | - | - | 0 | - |
| - | - | 882 | 150.1 | - | - | 0 | - |
| - | - | 7923 | 151.1 | - | - | 0 | - |
| - | - | 1167 | 152.1 | - | - | 0 | - |
| - | - | 5974 | 153.1 | - | - | 0 | - |
| - | - | 670.7 | 154.1 | - | - | 0 | - |
| - | - | 2671 | 154.1 | - | - | 0 | - |
| - | - | 930.1 | 154.1 | - | - | 0 | - |
| - | - | 2440 | 154.2 | - | - | 0 | - |
| - | - | 1677 | 155 | - | - | 0 | - |
| - | - | 5967 | 155.1 | - | - | 0 | - |
| - | - | 3.752E+04 | 155.1 | - | - | 0 | - |
| - | - | 2227 | 155.1 | - | - | 0 | - |
| - | - | 2382 | 155.2 | - | - | 0 | - |
| - | - | 1419 | 156.1 | - | - | 0 | - |
| - | - | 2539 | 156.1 | - | - | 0 | - |
| - | - | 839 | 156.1 | - | - | 0 | - |
| - | - | 2268 | 157.1 | - | - | 0 | - |
| - | - | 1388 | 157.1 | - | - | 0 | - |
| - | - | 2921 | 157.1 | - | - | 0 | - |
| - | - | 6881 | 158.1 | - | - | 0 | - |
| - | - | 1.202E+04 | 158.1 | - | - | 0 | - |
| - | - | 5.554E+05 | 159.1 | - | - | 0 | - |
| - | - | 4121 | 160.1 | - | - | 0 | - |
| - | - | 3176 | 160.1 | - | - | 0 | - |
| - | - | 5.529E+04 | 160.1 | - | - | 0 | - |
| - | - | 1904 | 161 | - | - | 0 | - |
| - | - | 888.6 | 161.1 | - | - | 0 | - |
| - | - | 2318 | 161.1 | - | - | 0 | - |
| - | - | 678.8 | 161.1 | - | - | 0 | - |
| - | - | 1174 | 162.1 | - | - | 0 | - |
| - | - | 736.2 | 164.1 | - | - | 0 | - |
| - | - | 3676 | 165.1 | - | - | 0 | - |
| - | - | 1952 | 165.1 | - | - | 0 | - |
| - | - | 1196 | 165.1 | - | - | 0 | - |
| - | - | 1.539E+05 | 166.1 | - | - | 0 | - |
| - | - | 2099 | 167.1 | - | - | 0 | - |
| - | - | 1.117E+04 | 167.1 | - | - | 0 | - |
| - | - | 1356 | 167.1 | - | - | 0 | - |
| - | - | 1936 | 167.1 | - | - | 0 | - |
| - | - | 1.503E+04 | 168.1 | - | - | 0 | - |
| - | - | 2505 | 168.1 | - | - | 0 | - |
| - | - | 5954 | 168.1 | - | - | 0 | - |
| - | - | 1.189E+04 | 169.1 | - | - | 0 | - |
| - | - | 2196 | 169.1 | - | - | 0 | - |
| - | - | 2.088E+04 | 169.1 | - | - | 0 | - |
| - | - | 645.3 | 169.1 | - | - | 0 | - |
| - | - | 4.836E+04 | 170.1 | - | - | 0 | - |
| - | - | 2091 | 170.1 | - | - | 0 | - |
| - | - | 5633 | 171.1 | - | - | 0 | - |
| - | - | 1429 | 171.1 | - | - | 0 | - |
| - | - | 1659 | 171.1 | - | - | 0 | - |
| - | - | 2710 | 171.1 | - | - | 0 | - |
| - | - | 5038 | 172.1 | - | - | 0 | - |
| - | - | 1680 | 172.1 | - | - | 0 | - |
| - | - | 931 | 172.1 | - | - | 0 | - |
| - | - | 1.08E+04 | 173.1 | - | - | 0 | - |
| - | - | 614.1 | 173.1 | - | - | 0 | - |
| - | - | 1196 | 173.1 | - | - | 0 | - |
| - | - | 815.5 | 173.1 | - | - | 0 | - |
| - | - | 737.5 | 174.1 | - | - | 0 | - |
| - | - | 1072 | 174.1 | - | - | 0 | - |
| - | - | 2008 | 175.1 | - | - | 0 | - |
| - | - | 965.1 | 175.1 | - | - | 0 | - |
| - | - | 2111 | 176.1 | - | - | 0 | - |
| 3 | a | 789.4 | 176.1 | 0.0005645 | 3.205 | +2 | 3 |
| - | - | 1301 | 177.1 | - | - | 0 | - |
| - | - | 6.596E+04 | 178.1 | - | - | 0 | - |
| - | - | 2293 | 179 | - | - | 0 | - |
| - | - | 1767 | 179.1 | - | - | 0 | - |
| - | - | 1098 | 179.1 | - | - | 0 | - |
| - | - | 7136 | 179.1 | - | - | 0 | - |
| - | - | 993.1 | 180.1 | - | - | 0 | - |
| - | - | 1.807E+04 | 180.1 | - | - | 0 | - |
| - | - | 915.8 | 180.1 | - | - | 0 | - |
| - | - | 1.73E+04 | 181.1 | - | - | 0 | - |
| - | - | 7514 | 181.1 | - | - | 0 | - |
| - | - | 1836 | 181.1 | - | - | 0 | - |
| - | - | 3686 | 181.1 | - | - | 0 | - |
| - | - | 759.7 | 182 | - | - | 0 | - |
| - | - | 1428 | 182.1 | - | - | 0 | - |
| - | - | 652.1 | 182.1 | - | - | 0 | - |
| - | - | 2126 | 182.6 | - | - | 0 | - |
| - | - | 2156 | 183.1 | - | - | 0 | - |
| - | - | 1393 | 183.1 | - | - | 0 | - |
| - | - | 1.509E+04 | 183.1 | - | - | 0 | - |
| - | - | 3.112E+04 | 183.2 | - | - | 0 | - |
| - | - | 1502 | 184.1 | - | - | 0 | - |
| - | - | 3189 | 184.2 | - | - | 0 | - |
| - | - | 1592 | 185.1 | - | - | 0 | - |
| - | - | 991 | 185.2 | - | - | 0 | - |
| - | - | 1.47E+05 | 186.1 | - | - | 0 | - |
| - | - | 5.664E+04 | 187.1 | - | - | 0 | - |
| - | - | 1159 | 187.1 | - | - | 0 | - |
| 15 | y | 8911 | 187.1 | 0.0008933 | 4.774 | +1 | 2 |
| - | - | 904.4 | 187.1 | - | - | 0 | - |
| - | - | 968.9 | 187.1 | - | - | 0 | - |
| - | - | 1.466E+04 | 187.1 | - | - | 0 | - |
| - | - | 2047 | 188.1 | - | - | 0 | - |
| - | - | 6455 | 188.1 | - | - | 0 | - |
| - | - | 962.1 | 188.1 | - | - | 0 | - |
| - | - | 809.8 | 188.1 | - | - | 0 | - |
| - | - | 5481 | 190.1 | - | - | 0 | - |
| - | - | 1.45E+04 | 190.1 | - | - | 0 | - |
| - | - | 1135 | 191.1 | - | - | 0 | - |
| - | - | 1558 | 191.1 | - | - | 0 | - |
| - | - | 1482 | 191.1 | - | - | 0 | - |
| - | - | 1315 | 192.1 | - | - | 0 | - |
| - | - | 1161 | 193.1 | - | - | 0 | - |
| - | - | 1.014E+04 | 193.1 | - | - | 0 | - |
| - | - | 939.5 | 194.1 | - | - | 0 | - |
| - | - | 1470 | 194.1 | - | - | 0 | - |
| - | - | 1925 | 195.1 | - | - | 0 | - |
| - | - | 2.974E+04 | 195.1 | - | - | 0 | - |
| - | - | 634.1 | 195.1 | - | - | 0 | - |
| - | - | 2.564E+04 | 196.1 | - | - | 0 | - |
| - | - | 1261 | 196.6 | - | - | 0 | - |
| - | - | 1925 | 197.1 | - | - | 0 | - |
| - | - | 2480 | 197.1 | - | - | 0 | - |
| - | - | 807.1 | 197.1 | - | - | 0 | - |
| - | - | 1.284E+04 | 198.1 | - | - | 0 | - |
| - | - | 4.373E+04 | 198.1 | - | - | 0 | - |
| - | - | 8393 | 199.1 | - | - | 0 | - |
| - | - | 2052 | 199.1 | - | - | 0 | - |
| 6 | b | 1141 | 199.1 | 0.002696 | 13.54 | +4 | 6 |
| - | - | 6873 | 199.1 | - | - | 0 | - |
| - | - | 3035 | 200.1 | - | - | 0 | - |
| - | - | 3358 | 200.1 | - | - | 0 | - |
| - | - | 4036 | 200.2 | - | - | 0 | - |
| - | - | 1.833E+04 | 201.1 | - | - | 0 | - |
| - | - | 2314 | 202.1 | - | - | 0 | - |
| - | - | 2739 | 202.1 | - | - | 0 | - |
| - | - | 723.5 | 203.1 | - | - | 0 | - |
| - | - | 1800 | 203.1 | - | - | 0 | - |
| - | - | 2567 | 203.1 | - | - | 0 | - |
| - | - | 1020 | 204.1 | - | - | 0 | - |
| - | - | 1545 | 204.1 | - | - | 0 | - |
| - | - | 937.3 | 204.1 | - | - | 0 | - |
| - | - | 1.479E+04 | 205.1 | - | - | 0 | - |
| 15 | y | 6.904E+04 | 205.1 | 0.001147 | 5.592 | +1 | 2 |
| - | - | 1.119E+04 | 205.1 | - | - | 0 | - |
| - | - | 1213 | 206.1 | - | - | 0 | - |
| - | - | 3853 | 206.1 | - | - | 0 | - |
| - | - | 7197 | 206.1 | - | - | 0 | - |
| - | - | 1592 | 206.1 | - | - | 0 | - |
| - | - | 1926 | 207.1 | - | - | 0 | - |
| - | - | 739.6 | 207.1 | - | - | 0 | - |
| - | - | 998.8 | 207.1 | - | - | 0 | - |
| - | - | 1373 | 207.2 | - | - | 0 | - |
| - | - | 1.235E+04 | 207.2 | - | - | 0 | - |
| - | - | 861.8 | 208.1 | - | - | 0 | - |
| - | - | 3471 | 208.1 | - | - | 0 | - |
| - | - | 1251 | 208.2 | - | - | 0 | - |
| - | - | 4745 | 209.1 | - | - | 0 | - |
| - | - | 856.2 | 209.1 | - | - | 0 | - |
| - | - | 1070 | 209.1 | - | - | 0 | - |
| - | - | 919.3 | 209.1 | - | - | 0 | - |
| - | - | 1.904E+04 | 210.1 | - | - | 0 | - |
| - | - | 1241 | 210.2 | - | - | 0 | - |
| - | - | 1.708E+04 | 210.2 | - | - | 0 | - |
| - | - | 691 | 211.1 | - | - | 0 | - |
| - | - | 881.2 | 211.1 | - | - | 0 | - |
| - | - | 2898 | 211.1 | - | - | 0 | - |
| - | - | 3354 | 211.1 | - | - | 0 | - |
| - | - | 4436 | 211.1 | - | - | 0 | - |
| - | - | 1874 | 211.2 | - | - | 0 | - |
| - | - | 3997 | 211.6 | - | - | 0 | - |
| - | - | 1832 | 212.1 | - | - | 0 | - |
| - | - | 3246 | 212.1 | - | - | 0 | - |
| - | - | 1021 | 212.2 | - | - | 0 | - |
| - | - | 1.604E+04 | 213.1 | - | - | 0 | - |
| - | - | 1128 | 214.1 | - | - | 0 | - |
| - | - | 1131 | 214.1 | - | - | 0 | - |
| - | - | 1392 | 215.1 | - | - | 0 | - |
| - | - | 1.195E+04 | 216.1 | - | - | 0 | - |
| - | - | 637.3 | 217.1 | - | - | 0 | - |
| - | - | 1366 | 219.1 | - | - | 0 | - |
| - | - | 1629 | 220.1 | - | - | 0 | - |
| - | - | 1536 | 220.1 | - | - | 0 | - |
| - | - | 3337 | 220.1 | - | - | 0 | - |
| - | - | 3277 | 221.1 | - | - | 0 | - |
| - | - | 2026 | 221.1 | - | - | 0 | - |
| - | - | 8148 | 221.1 | - | - | 0 | - |
| - | - | 897.8 | 221.1 | - | - | 0 | - |
| - | - | 6496 | 222.1 | - | - | 0 | - |
| - | - | 1882 | 223.1 | - | - | 0 | - |
| - | - | 1.058E+04 | 223.1 | - | - | 0 | - |
| 2 | a | 1.263E+05 | 223.2 | 0.001156 | 5.182 | +1 | 2 |
| - | - | 4623 | 224.1 | - | - | 0 | - |
| - | - | 6856 | 224.1 | - | - | 0 | - |
| - | - | 934.4 | 224.1 | - | - | 0 | - |
| - | - | 1.508E+04 | 224.2 | - | - | 0 | - |
| - | - | 604.5 | 225.1 | - | - | 0 | - |
| - | - | 1861 | 225.1 | - | - | 0 | - |
| - | - | 3.086E+04 | 226.1 | - | - | 0 | - |
| - | - | 2590 | 226.1 | - | - | 0 | - |
| - | - | 2852 | 226.2 | - | - | 0 | - |
| - | - | 2.814E+04 | 227.1 | - | - | 0 | - |
| - | - | 2.735E+04 | 227.1 | - | - | 0 | - |
| - | - | 5906 | 227.1 | - | - | 0 | - |
| - | - | 1.014E+04 | 227.2 | - | - | 0 | - |
| - | - | 2656 | 228.1 | - | - | 0 | - |
| - | - | 2312 | 228.1 | - | - | 0 | - |
| - | - | 1328 | 228.2 | - | - | 0 | - |
| - | - | 9901 | 228.2 | - | - | 0 | - |
| - | - | 3.665E+04 | 229.1 | - | - | 0 | - |
| - | - | 1881 | 229.2 | - | - | 0 | - |
| - | - | 1050 | 229.2 | - | - | 0 | - |
| - | - | 1977 | 229.6 | - | - | 0 | - |
| - | - | 789.7 | 230.1 | - | - | 0 | - |
| - | - | 3665 | 230.1 | - | - | 0 | - |
| - | - | 1337 | 230.2 | - | - | 0 | - |
| - | - | 1857 | 231.1 | - | - | 0 | - |
| - | - | 970.4 | 233.1 | - | - | 0 | - |
| - | - | 1344 | 233.1 | - | - | 0 | - |
| - | - | 1.872E+04 | 233.1 | - | - | 0 | - |
| - | - | 2.021E+05 | 234.1 | - | - | 0 | - |
| - | - | 1100 | 235.1 | - | - | 0 | - |
| - | - | 2.377E+04 | 235.1 | - | - | 0 | - |
| - | - | 947.8 | 235.1 | - | - | 0 | - |
| - | - | 4086 | 235.2 | - | - | 0 | - |
| - | - | 1356 | 236.1 | - | - | 0 | - |
| - | - | 2231 | 237.1 | - | - | 0 | - |
| - | - | 881.8 | 238.1 | - | - | 0 | - |
| - | - | 909.2 | 238.1 | - | - | 0 | - |
| - | - | 1.826E+04 | 238.1 | - | - | 0 | - |
| - | - | 1603 | 238.1 | - | - | 0 | - |
| - | - | 3278 | 238.2 | - | - | 0 | - |
| - | - | 3303 | 238.6 | - | - | 0 | - |
| - | - | 5006 | 239.1 | - | - | 0 | - |
| 4 | b | 2572 | 239.1 | 0.003529 | 14.76 | +2 | 4 |
| - | - | 1887 | 239.1 | - | - | 0 | - |
| - | - | 1.971E+04 | 240.1 | - | - | 0 | - |
| - | - | 1504 | 241.1 | - | - | 0 | - |
| - | - | 5267 | 241.1 | - | - | 0 | - |
| - | - | 1538 | 241.1 | - | - | 0 | - |
| - | - | 1240 | 243.1 | - | - | 0 | - |
| - | - | 6588 | 243.1 | - | - | 0 | - |
| - | - | 2334 | 243.2 | - | - | 0 | - |
| - | - | 4998 | 243.6 | - | - | 0 | - |
| - | - | 6.201E+04 | 244.1 | - | - | 0 | - |
| - | - | 2.796E+04 | 245.1 | - | - | 0 | - |
| - | - | 3142 | 246.1 | - | - | 0 | - |
| - | - | 2143 | 246.1 | - | - | 0 | - |
| - | - | 6764 | 246.2 | - | - | 0 | - |
| - | - | 2152 | 246.6 | - | - | 0 | - |
| - | - | 1344 | 247.1 | - | - | 0 | - |
| - | - | 1.361E+04 | 247.1 | - | - | 0 | - |
| - | - | 1310 | 247.1 | - | - | 0 | - |
| - | - | 1.649E+04 | 247.1 | - | - | 0 | - |
| 4 | b | 1.076E+04 | 247.6 | 0.0003168 | 1.279 | +2 | 4 |
| - | - | 1434 | 248.1 | - | - | 0 | - |
| - | - | 2.172E+04 | 248.1 | - | - | 0 | - |
| - | - | 2552 | 248.1 | - | - | 0 | - |
| - | - | 3.573E+04 | 249.1 | - | - | 0 | - |
| - | - | 1943 | 249.1 | - | - | 0 | - |
| - | - | 1153 | 249.1 | - | - | 0 | - |
| - | - | 4253 | 250.1 | - | - | 0 | - |
| - | - | 1538 | 250.1 | - | - | 0 | - |
| - | - | 1265 | 250.2 | - | - | 0 | - |
| 2 | b | 5.724E+05 | 251.2 | 0.001344 | 5.35 | +1 | 2 |
| - | - | 1579 | 252.1 | - | - | 0 | - |
| - | - | 7.206E+04 | 252.2 | - | - | 0 | - |
| - | - | 1.93E+04 | 252.6 | - | - | 0 | - |
| - | - | 4981 | 253.1 | - | - | 0 | - |
| - | - | 1297 | 253.1 | - | - | 0 | - |
| - | - | 4101 | 253.2 | - | - | 0 | - |
| - | - | 1710 | 254.1 | - | - | 0 | - |
| - | - | 2421 | 254.1 | - | - | 0 | - |
| - | - | 4520 | 255.1 | - | - | 0 | - |
| - | - | 2.189E+04 | 255.1 | - | - | 0 | - |
| - | - | 1200 | 256.1 | - | - | 0 | - |
| - | - | 6656 | 256.1 | - | - | 0 | - |
| - | - | 1018 | 256.1 | - | - | 0 | - |
| - | - | 2160 | 256.2 | - | - | 0 | - |
| - | - | 1.675E+04 | 256.2 | - | - | 0 | - |
| - | - | 1652 | 257.1 | - | - | 0 | - |
| - | - | 7399 | 257.1 | - | - | 0 | - |
| 9 | y | 2678 | 257.1 | 0.001373 | 5.34 | +4 | 8 |
| - | - | 1574 | 257.2 | - | - | 0 | - |
| - | - | 2476 | 257.6 | - | - | 0 | - |
| - | - | 1480 | 258.1 | - | - | 0 | - |
| - | - | 3044 | 258.1 | - | - | 0 | - |
| - | - | 6632 | 258.1 | - | - | 0 | - |
| - | - | 1205 | 259.1 | - | - | 0 | - |
| - | - | 2948 | 260.6 | - | - | 0 | - |
| - | - | 843.5 | 261.1 | - | - | 0 | - |
| - | - | 2.049E+04 | 261.1 | - | - | 0 | - |
| - | - | 2742 | 261.1 | - | - | 0 | - |
| - | - | 6683 | 261.6 | - | - | 0 | - |
| - | - | 2039 | 261.6 | - | - | 0 | - |
| - | - | 5142 | 262.1 | - | - | 0 | - |
| - | - | 1034 | 262.1 | - | - | 0 | - |
| - | - | 1833 | 262.2 | - | - | 0 | - |
| - | - | 606.2 | 262.6 | - | - | 0 | - |
| - | - | 2084 | 263.1 | - | - | 0 | - |
| - | - | 1122 | 263.1 | - | - | 0 | - |
| - | - | 1287 | 264.2 | - | - | 0 | - |
| - | - | 2.15E+04 | 265.1 | - | - | 0 | - |
| - | - | 1.123E+04 | 265.1 | - | - | 0 | - |
| - | - | 1511 | 265.1 | - | - | 0 | - |
| - | - | 8.284E+04 | 266.1 | - | - | 0 | - |
| - | - | 1986 | 266.6 | - | - | 0 | - |
| - | - | 1511 | 267.1 | - | - | 0 | - |
| - | - | 1.058E+04 | 267.1 | - | - | 0 | - |
| - | - | 1487 | 267.1 | - | - | 0 | - |
| - | - | 5390 | 268.1 | - | - | 0 | - |
| - | - | 4016 | 268.2 | - | - | 0 | - |
| - | - | 2619 | 269.2 | - | - | 0 | - |
| - | - | 771.4 | 269.2 | - | - | 0 | - |
| - | - | 1666 | 269.2 | - | - | 0 | - |
| - | - | 2.02E+05 | 270.1 | - | - | 0 | - |
| - | - | 5.835E+04 | 270.6 | - | - | 0 | - |
| - | - | 1121 | 271.1 | - | - | 0 | - |
| - | - | 1.011E+04 | 271.1 | - | - | 0 | - |
| - | - | 1218 | 271.6 | - | - | 0 | - |
| - | - | 1.183E+05 | 272.1 | - | - | 0 | - |
| - | - | 1.684E+04 | 272.2 | - | - | 0 | - |
| - | - | 3.798E+04 | 273.1 | - | - | 0 | - |
| - | - | 1.346E+04 | 273.1 | - | - | 0 | - |
| - | - | 3574 | 273.2 | - | - | 0 | - |
| - | - | 4158 | 273.6 | - | - | 0 | - |
| - | - | 1354 | 274.1 | - | - | 0 | - |
| - | - | 3513 | 274.1 | - | - | 0 | - |
| - | - | 2.309E+04 | 274.1 | - | - | 0 | - |
| - | - | 5177 | 274.1 | - | - | 0 | - |
| - | - | 1471 | 274.2 | - | - | 0 | - |
| - | - | 1551 | 274.2 | - | - | 0 | - |
| - | - | 1314 | 275.1 | - | - | 0 | - |
| - | - | 3400 | 275.1 | - | - | 0 | - |
| - | - | 4932 | 275.1 | - | - | 0 | - |
| - | - | 5990 | 275.1 | - | - | 0 | - |
| - | - | 3328 | 275.6 | - | - | 0 | - |
| - | - | 9315 | 276.1 | - | - | 0 | - |
| - | - | 1274 | 277.1 | - | - | 0 | - |
| - | - | 795.1 | 279.1 | - | - | 0 | - |
| - | - | 2761 | 279.1 | - | - | 0 | - |
| - | - | 4076 | 279.2 | - | - | 0 | - |
| - | - | 5298 | 280.1 | - | - | 0 | - |
| - | - | 984.2 | 280.1 | - | - | 0 | - |
| - | - | 4878 | 280.2 | - | - | 0 | - |
| - | - | 798.6 | 280.6 | - | - | 0 | - |
| - | - | 1069 | 281.1 | - | - | 0 | - |
| - | - | 709.2 | 281.1 | - | - | 0 | - |
| - | - | 1330 | 281.7 | - | - | 0 | - |
| - | - | 2658 | 282.1 | - | - | 0 | - |
| - | - | 4349 | 282.2 | - | - | 0 | - |
| - | - | 935.9 | 282.2 | - | - | 0 | - |
| - | - | 969.3 | 282.6 | - | - | 0 | - |
| - | - | 4135 | 283.1 | - | - | 0 | - |
| - | - | 1.032E+05 | 283.1 | - | - | 0 | - |
| - | - | 2178 | 284.1 | - | - | 0 | - |
| - | - | 1.303E+04 | 284.1 | - | - | 0 | - |
| - | - | 1.228E+04 | 284.1 | - | - | 0 | - |
| - | - | 1.338E+04 | 284.1 | - | - | 0 | - |
| - | - | 5052 | 284.6 | - | - | 0 | - |
| - | - | 1336 | 285.1 | - | - | 0 | - |
| - | - | 1771 | 285.1 | - | - | 0 | - |
| - | - | 1315 | 285.1 | - | - | 0 | - |
| - | - | 2385 | 285.2 | - | - | 0 | - |
| - | - | 1.602E+04 | 286.1 | - | - | 0 | - |
| - | - | 654.4 | 286.2 | - | - | 0 | - |
| - | - | 2686 | 287.1 | - | - | 0 | - |
| - | - | 3253 | 287.2 | - | - | 0 | - |
| - | - | 2097 | 289.1 | - | - | 0 | - |
| - | - | 9.749E+04 | 290.1 | - | - | 0 | - |
| - | - | 1044 | 291.1 | - | - | 0 | - |
| - | - | 1.219E+04 | 291.1 | - | - | 0 | - |
| - | - | 863.7 | 291.2 | - | - | 0 | - |
| - | - | 6130 | 291.2 | - | - | 0 | - |
| - | - | 3671 | 292.1 | - | - | 0 | - |
| - | - | 4.767E+04 | 292.2 | - | - | 0 | - |
| - | - | 7293 | 293.1 | - | - | 0 | - |
| - | - | 8453 | 293.2 | - | - | 0 | - |
| - | - | 719.1 | 294.1 | - | - | 0 | - |
| - | - | 1.184E+04 | 294.1 | - | - | 0 | - |
| - | - | 1143 | 295.1 | - | - | 0 | - |
| - | - | 1123 | 295.1 | - | - | 0 | - |
| - | - | 3.21E+04 | 297.2 | - | - | 0 | - |
| - | - | 5774 | 297.2 | - | - | 0 | - |
| - | - | 3097 | 298.1 | - | - | 0 | - |
| - | - | 4081 | 298.2 | - | - | 0 | - |
| - | - | 908.6 | 298.2 | - | - | 0 | - |
| - | - | 1424 | 298.2 | - | - | 0 | - |
| - | - | 3552 | 299.1 | - | - | 0 | - |
| - | - | 1195 | 299.1 | - | - | 0 | - |
| - | - | 5045 | 300.1 | - | - | 0 | - |
| - | - | 2245 | 300.1 | - | - | 0 | - |
| - | - | 4950 | 300.2 | - | - | 0 | - |
| - | - | 1181 | 300.6 | - | - | 0 | - |
| - | - | 5974 | 301.1 | - | - | 0 | - |
| - | - | 4.211E+04 | 301.2 | - | - | 0 | - |
| - | - | 1.879E+04 | 302.1 | - | - | 0 | - |
| - | - | 7157 | 302.2 | - | - | 0 | - |
| - | - | 1257 | 303.1 | - | - | 0 | - |
| - | - | 2963 | 303.1 | - | - | 0 | - |
| 12 | y | 4025 | 303.2 | 0.001625 | 5.362 | +2 | 5 |
| - | - | 910.2 | 303.7 | - | - | 0 | - |
| - | - | 2084 | 304.2 | - | - | 0 | - |
| - | - | 2100 | 304.2 | - | - | 0 | - |
| - | - | 1.036E+04 | 307.1 | - | - | 0 | - |
| - | - | 1036 | 307.2 | - | - | 0 | - |
| - | - | 1497 | 308.1 | - | - | 0 | - |
| - | - | 1330 | 308.1 | - | - | 0 | - |
| - | - | 860.8 | 308.2 | - | - | 0 | - |
| - | - | 912.7 | 308.7 | - | - | 0 | - |
| - | - | 1.688E+04 | 309.1 | - | - | 0 | - |
| - | - | 1160 | 309.2 | - | - | 0 | - |
| - | - | 1354 | 309.2 | - | - | 0 | - |
| - | - | 3426 | 309.6 | - | - | 0 | - |
| - | - | 3631 | 309.6 | - | - | 0 | - |
| - | - | 1936 | 310.1 | - | - | 0 | - |
| - | - | 694.9 | 310.2 | - | - | 0 | - |
| - | - | 6868 | 311.1 | - | - | 0 | - |
| - | - | 2229 | 311.2 | - | - | 0 | - |
| 12 | y | 7501 | 311.7 | 0.0006787 | 2.178 | +2 | 5 |
| - | - | 1096 | 312.1 | - | - | 0 | - |
| - | - | 1926 | 312.2 | - | - | 0 | - |
| - | - | 844.6 | 312.7 | - | - | 0 | - |
| - | - | 1441 | 313.1 | - | - | 0 | - |
| - | - | 927.1 | 314.1 | - | - | 0 | - |
| - | - | 812.7 | 314.2 | - | - | 0 | - |
| - | - | 461.4 | 315.2 | - | - | 0 | - |
| - | - | 4.338E+04 | 315.2 | - | - | 0 | - |
| 14 | y | 3121 | 315.2 | 0.001694 | 5.376 | +1 | 3 |
| - | - | 6496 | 316.2 | - | - | 0 | - |
| - | - | 2.244E+04 | 317.7 | - | - | 0 | - |
| - | - | 1.042E+04 | 318.1 | - | - | 0 | - |
| - | - | 4.545E+04 | 318.2 | - | - | 0 | - |
| - | - | 1.565E+04 | 318.7 | - | - | 0 | - |
| - | - | 9540 | 319.2 | - | - | 0 | - |
| - | - | 6057 | 319.2 | - | - | 0 | - |
| - | - | 1029 | 320.2 | - | - | 0 | - |
| - | - | 882.8 | 321.2 | - | - | 0 | - |
| - | - | 860.7 | 322.7 | - | - | 0 | - |
| - | - | 1877 | 322.8 | - | - | 0 | - |
| - | - | 9893 | 323.1 | - | - | 0 | - |
| - | - | 1332 | 323.2 | - | - | 0 | - |
| - | - | 4172 | 323.6 | - | - | 0 | - |
| - | - | 1035 | 324.1 | - | - | 0 | - |
| - | - | 1041 | 324.6 | - | - | 0 | - |
| - | - | 2446 | 324.7 | - | - | 0 | - |
| - | - | 6941 | 325.2 | - | - | 0 | - |
| - | - | 2153 | 325.2 | - | - | 0 | - |
| - | - | 2507 | 326.2 | - | - | 0 | - |
| - | - | 984.4 | 326.2 | - | - | 0 | - |
| - | - | 4.16E+05 | 326.7 | - | - | 0 | - |
| - | - | 1.553E+05 | 327.2 | - | - | 0 | - |
| - | - | 3.631E+04 | 327.7 | - | - | 0 | - |
| 6 | y | 5372 | 328.2 | 0.001522 | 4.637 | +4 | 11 |
| - | - | 5795 | 329.2 | - | - | 0 | - |
| - | - | 1622 | 330.2 | - | - | 0 | - |
| 5 | b | 5.319E+04 | 331.7 | 0.001725 | 5.2 | +2 | 5 |
| - | - | 1.422E+04 | 332.2 | - | - | 0 | - |
| - | - | 775.6 | 332.2 | - | - | 0 | - |
| - | - | 3993 | 332.7 | - | - | 0 | - |
| - | - | 1343 | 333.2 | - | - | 0 | - |
| 14 | y | 3.732E+04 | 333.2 | 0.001719 | 5.16 | +1 | 3 |
| 3 | a | 2517 | 334.2 | 0.00108 | 3.232 | +1 | 3 |
| - | - | 5230 | 334.2 | - | - | 0 | - |
| - | - | 2288 | 335.2 | - | - | 0 | - |
| - | - | 903.2 | 336.1 | - | - | 0 | - |
| - | - | 1262 | 338.2 | - | - | 0 | - |
| - | - | 1556 | 338.7 | - | - | 0 | - |
| - | - | 760.8 | 339.7 | - | - | 0 | - |
| 5 | b | 1.202E+05 | 340.7 | 0.001813 | 5.323 | +2 | 5 |
| - | - | 4.805E+04 | 341.2 | - | - | 0 | - |
| - | - | 1.035E+04 | 341.7 | - | - | 0 | - |
| - | - | 2159 | 342.2 | - | - | 0 | - |
| 9 | y | 715.7 | 342.5 | 0.0007981 | 2.33 | +3 | 8 |
| - | - | 1890 | 343.2 | - | - | 0 | - |
| - | - | 4523 | 343.2 | - | - | 0 | - |
| - | - | 8540 | 344.2 | - | - | 0 | - |
| - | - | 9663 | 345.1 | - | - | 0 | - |
| - | - | 1027 | 345.2 | - | - | 0 | - |
| - | - | 3.668E+04 | 346.1 | - | - | 0 | - |
| - | - | 2849 | 346.2 | - | - | 0 | - |
| - | - | 1184 | 346.7 | - | - | 0 | - |
| - | - | 6629 | 347.1 | - | - | 0 | - |
| - | - | 1013 | 347.2 | - | - | 0 | - |
| - | - | 906.7 | 348.1 | - | - | 0 | - |
| - | - | 2187 | 349.1 | - | - | 0 | - |
| - | - | 1477 | 349.2 | - | - | 0 | - |
| - | - | 1127 | 349.2 | - | - | 0 | - |
| - | - | 1453 | 349.7 | - | - | 0 | - |
| - | - | 1561 | 350.2 | - | - | 0 | - |
| - | - | 1504 | 351.1 | - | - | 0 | - |
| - | - | 1939 | 351.2 | - | - | 0 | - |
| - | - | 1007 | 351.2 | - | - | 0 | - |
| - | - | 1007 | 351.9 | - | - | 0 | - |
| - | - | 1833 | 352.2 | - | - | 0 | - |
| - | - | 3374 | 353.2 | - | - | 0 | - |
| - | - | 1474 | 355.2 | - | - | 0 | - |
| - | - | 852.2 | 356.3 | - | - | 0 | - |
| - | - | 1112 | 357.2 | - | - | 0 | - |
| - | - | 1340 | 358.2 | - | - | 0 | - |
| - | - | 885.8 | 359.1 | - | - | 0 | - |
| - | - | 1047 | 359.2 | - | - | 0 | - |
| - | - | 1.427E+04 | 360.7 | - | - | 0 | - |
| - | - | 2.146E+04 | 361.2 | - | - | 0 | - |
| 8 | y | 4102 | 361.5 | 0.002219 | 6.138 | +3 | 9 |
| - | - | 1733 | 361.7 | - | - | 0 | - |
| - | - | 2076 | 361.8 | - | - | 0 | - |
| 3 | b | 8670 | 362.2 | 0.002869 | 7.923 | +1 | 3 |
| - | - | 5.538E+04 | 363.1 | - | - | 0 | - |
| - | - | 2517 | 363.2 | - | - | 0 | - |
| - | - | 8163 | 364.1 | - | - | 0 | - |
| - | - | 8026 | 364.1 | - | - | 0 | - |
| - | - | 3933 | 364.2 | - | - | 0 | - |
| - | - | 1095 | 364.2 | - | - | 0 | - |
| - | - | 1531 | 365.1 | - | - | 0 | - |
| - | - | 947.2 | 365.2 | - | - | 0 | - |
| - | - | 4990 | 365.7 | - | - | 0 | - |
| - | - | 2975 | 366.2 | - | - | 0 | - |
| - | - | 5077 | 366.2 | - | - | 0 | - |
| - | - | 1.439E+04 | 367.1 | - | - | 0 | - |
| 8 | y | 7.734E+04 | 367.2 | 0.000134 | 0.3649 | +3 | 9 |
| - | - | 4.851E+04 | 367.5 | - | - | 0 | - |
| - | - | 1.964E+04 | 367.9 | - | - | 0 | - |
| - | - | 3001 | 368.1 | - | - | 0 | - |
| - | - | 1168 | 368.2 | - | - | 0 | - |
| - | - | 6908 | 368.2 | - | - | 0 | - |
| - | - | 687.4 | 368.5 | - | - | 0 | - |
| - | - | 1041 | 369.2 | - | - | 0 | - |
| - | - | 2447 | 369.2 | - | - | 0 | - |
| - | - | 996.4 | 369.2 | - | - | 0 | - |
| - | - | 2249 | 370.2 | - | - | 0 | - |
| - | - | 1534 | 370.7 | - | - | 0 | - |
| - | - | 3.335E+04 | 371.2 | - | - | 0 | - |
| - | - | 4581 | 372.2 | - | - | 0 | - |
| - | - | 5273 | 372.2 | - | - | 0 | - |
| - | - | 1164 | 373.2 | - | - | 0 | - |
| - | - | 1501 | 373.2 | - | - | 0 | - |
| - | - | 805.4 | 373.2 | - | - | 0 | - |
| - | - | 1453 | 374.1 | - | - | 0 | - |
| - | - | 1.215E+04 | 374.2 | - | - | 0 | - |
| 12 | b | 2.937E+04 | 374.7 | 0.006371 | 17 | +4 | 12 |
| - | - | 1.266E+04 | 375.2 | - | - | 0 | - |
| - | - | 1031 | 375.7 | - | - | 0 | - |
| - | - | 1643 | 376.2 | - | - | 0 | - |
| - | - | 1881 | 377.1 | - | - | 0 | - |
| - | - | 1646 | 377.1 | - | - | 0 | - |
| - | - | 974.7 | 377.2 | - | - | 0 | - |
| - | - | 1078 | 378.2 | - | - | 0 | - |
| - | - | 3353 | 378.2 | - | - | 0 | - |
| 3 | b | 1.047E+05 | 379.2 | 0.001864 | 4.914 | +1 | 3 |
| - | - | 2605 | 380.2 | - | - | 0 | - |
| - | - | 1.932E+04 | 380.2 | - | - | 0 | - |
| - | - | 1.32E+05 | 381.2 | - | - | 0 | - |
| - | - | 3178 | 381.2 | - | - | 0 | - |
| - | - | 2.275E+04 | 382.2 | - | - | 0 | - |
| - | - | 2766 | 382.2 | - | - | 0 | - |
| - | - | 2548 | 382.7 | - | - | 0 | - |
| - | - | 3217 | 383.2 | - | - | 0 | - |
| - | - | 5.726E+04 | 383.2 | - | - | 0 | - |
| - | - | 2.987E+04 | 383.7 | - | - | 0 | - |
| - | - | 1.434E+04 | 384.2 | - | - | 0 | - |
| 11 | y | 3075 | 384.2 | 0.003074 | 8.001 | +2 | 6 |
| - | - | 3995 | 384.2 | - | - | 0 | - |
| - | - | 1478 | 384.3 | - | - | 0 | - |
| 11 | y | 1589 | 384.7 | 0.001246 | 3.239 | +2 | 6 |
| - | - | 931.1 | 384.7 | - | - | 0 | - |
| - | - | 2.588E+04 | 385.2 | - | - | 0 | - |
| - | - | 1572 | 385.2 | - | - | 0 | - |
| - | - | 3124 | 386.2 | - | - | 0 | - |
| - | - | 3534 | 386.2 | - | - | 0 | - |
| - | - | 1653 | 386.2 | - | - | 0 | - |
| - | - | 1208 | 386.7 | - | - | 0 | - |
| - | - | 1313 | 387.2 | - | - | 0 | - |
| - | - | 2097 | 387.2 | - | - | 0 | - |
| - | - | 1013 | 387.7 | - | - | 0 | - |
| 6 | b | 2738 | 388.2 | 0.001959 | 5.047 | +2 | 6 |
| 6 | b | 7436 | 388.7 | 0.00278 | 7.152 | +2 | 6 |
| - | - | 7.334E+04 | 389.2 | - | - | 0 | - |
| - | - | 1158 | 389.7 | - | - | 0 | - |
| - | - | 1.41E+04 | 390.2 | - | - | 0 | - |
| - | - | 1.454E+04 | 391.1 | - | - | 0 | - |
| - | - | 3671 | 391.2 | - | - | 0 | - |
| - | - | 1953 | 391.2 | - | - | 0 | - |
| - | - | 2960 | 392.1 | - | - | 0 | - |
| - | - | 1452 | 392.1 | - | - | 0 | - |
| - | - | 1148 | 392.2 | - | - | 0 | - |
| - | - | 1.925E+04 | 392.2 | - | - | 0 | - |
| 11 | y | 1.402E+05 | 393.2 | 0.0001164 | 0.2961 | +2 | 6 |
| - | - | 6.103E+04 | 393.7 | - | - | 0 | - |
| - | - | 1499 | 394.2 | - | - | 0 | - |
| - | - | 1.151E+04 | 394.2 | - | - | 0 | - |
| - | - | 1.758E+04 | 394.2 | - | - | 0 | - |
| - | - | 5690 | 394.7 | - | - | 0 | - |
| - | - | 1.329E+04 | 395.1 | - | - | 0 | - |
| - | - | 1968 | 395.2 | - | - | 0 | - |
| - | - | 1533 | 395.2 | - | - | 0 | - |
| - | - | 2899 | 396.1 | - | - | 0 | - |
| - | - | 1157 | 396.2 | - | - | 0 | - |
| - | - | 7476 | 396.2 | - | - | 0 | - |
| - | - | 9574 | 396.2 | - | - | 0 | - |
| - | - | 2018 | 396.7 | - | - | 0 | - |
| 6 | b | 2.529E+04 | 397.2 | 0.001804 | 4.542 | +2 | 6 |
| - | - | 947.5 | 397.2 | - | - | 0 | - |
| - | - | 1.005E+04 | 397.7 | - | - | 0 | - |
| - | - | 1.126E+04 | 398.2 | - | - | 0 | - |
| - | - | 2683 | 398.2 | - | - | 0 | - |
| - | - | 3652 | 399.2 | - | - | 0 | - |
| 7 | y | 3421 | 399.5 | 0.001002 | 2.508 | +3 | 10 |
| 7 | y | 3526 | 399.9 | 0.003685 | 9.216 | +3 | 10 |
| - | - | 3629 | 401.2 | - | - | 0 | - |
| - | - | 3.069E+04 | 402.2 | - | - | 0 | - |
| - | - | 1854 | 403.1 | - | - | 0 | - |
| - | - | 1.413E+04 | 403.2 | - | - | 0 | - |
| - | - | 3382 | 404.2 | - | - | 0 | - |
| - | - | 1186 | 405.2 | - | - | 0 | - |
| - | - | 1655 | 405.2 | - | - | 0 | - |
| 7 | y | 4.732E+04 | 405.5 | 0.0005931 | 1.463 | +3 | 10 |
| - | - | 2.966E+04 | 405.9 | - | - | 0 | - |
| - | - | 1.254E+04 | 406.2 | - | - | 0 | - |
| - | - | 4616 | 406.5 | - | - | 0 | - |
| - | - | 1242 | 406.9 | - | - | 0 | - |
| - | - | 3601 | 407.2 | - | - | 0 | - |
| - | - | 1037 | 407.3 | - | - | 0 | - |
| - | - | 2289 | 408.2 | - | - | 0 | - |
| - | - | 2.841E+04 | 409.1 | - | - | 0 | - |
| - | - | 1462 | 409.2 | - | - | 0 | - |
| - | - | 5063 | 410.2 | - | - | 0 | - |
| - | - | 3557 | 411.2 | - | - | 0 | - |
| - | - | 1819 | 411.7 | - | - | 0 | - |
| - | - | 8812 | 412.2 | - | - | 0 | - |
| - | - | 5.655E+04 | 412.2 | - | - | 0 | - |
| - | - | 1.639E+04 | 413.1 | - | - | 0 | - |
| - | - | 9904 | 413.2 | - | - | 0 | - |
| - | - | 3764 | 414.2 | - | - | 0 | - |
| - | - | 5078 | 414.2 | - | - | 0 | - |
| - | - | 7196 | 414.2 | - | - | 0 | - |
| - | - | 1288 | 415 | - | - | 0 | - |
| - | - | 809.5 | 415.1 | - | - | 0 | - |
| - | - | 9354 | 415.2 | - | - | 0 | - |
| - | - | 2135 | 415.2 | - | - | 0 | - |
| - | - | 2263 | 416.2 | - | - | 0 | - |
| - | - | 925.6 | 416.5 | - | - | 0 | - |
| - | - | 900.5 | 416.9 | - | - | 0 | - |
| - | - | 2890 | 417.2 | - | - | 0 | - |
| - | - | 1011 | 417.5 | - | - | 0 | - |
| - | - | 1513 | 418.1 | - | - | 0 | - |
| - | - | 4.333E+04 | 418.2 | - | - | 0 | - |
| - | - | 7734 | 419.2 | - | - | 0 | - |
| - | - | 7408 | 419.7 | - | - | 0 | - |
| - | - | 1795 | 420.2 | - | - | 0 | - |
| - | - | 5946 | 420.2 | - | - | 0 | - |
| - | - | 1189 | 420.7 | - | - | 0 | - |
| - | - | 1.878E+04 | 421.2 | - | - | 0 | - |
| - | - | 4452 | 422.2 | - | - | 0 | - |
| - | - | 1897 | 422.3 | - | - | 0 | - |
| - | - | 1053 | 423.2 | - | - | 0 | - |
| - | - | 2086 | 424.2 | - | - | 0 | - |
| - | - | 2007 | 425.3 | - | - | 0 | - |
| - | - | 1416 | 427.7 | - | - | 0 | - |
| - | - | 1586 | 428.2 | - | - | 0 | - |
| - | - | 898.7 | 429.2 | - | - | 0 | - |
| - | - | 9.407E+04 | 430.2 | - | - | 0 | - |
| - | - | 1.958E+04 | 431.2 | - | - | 0 | - |
| - | - | 4124 | 432.2 | - | - | 0 | - |
| - | - | 1588 | 432.2 | - | - | 0 | - |
| - | - | 2115 | 432.7 | - | - | 0 | - |
| - | - | 1294 | 433.2 | - | - | 0 | - |
| - | - | 1290 | 434.2 | - | - | 0 | - |
| - | - | 5644 | 435.2 | - | - | 0 | - |
| - | - | 1416 | 436.2 | - | - | 0 | - |
| - | - | 901.1 | 436.7 | - | - | 0 | - |
| - | - | 1441 | 437.2 | - | - | 0 | - |
| 6 | y | 6918 | 437.2 | 0.004831 | 11.05 | +3 | 11 |
| 6 | y | 4386 | 437.5 | 0.006751 | 15.43 | +3 | 11 |
| - | - | 1891 | 437.9 | - | - | 0 | - |
| - | - | 3797 | 438.2 | - | - | 0 | - |
| - | - | 2398 | 438.2 | - | - | 0 | - |
| - | - | 1152 | 439.2 | - | - | 0 | - |
| - | - | 2191 | 439.2 | - | - | 0 | - |
| - | - | 2576 | 439.7 | - | - | 0 | - |
| - | - | 2196 | 440.2 | - | - | 0 | - |
| - | - | 1431 | 440.2 | - | - | 0 | - |
| - | - | 2064 | 440.7 | - | - | 0 | - |
| - | - | 1400 | 441.2 | - | - | 0 | - |
| - | - | 1276 | 441.7 | - | - | 0 | - |
| 6 | y | 4.823E+04 | 443.2 | 0.0008514 | 1.921 | +3 | 11 |
| - | - | 3.616E+04 | 443.6 | - | - | 0 | - |
| - | - | 1.62E+04 | 443.9 | - | - | 0 | - |
| - | - | 1973 | 444.2 | - | - | 0 | - |
| - | - | 3506 | 444.2 | - | - | 0 | - |
| - | - | 1827 | 444.6 | - | - | 0 | - |
| 7 | b | 2406 | 445.7 | 0.0007254 | 1.628 | +2 | 7 |
| 7 | b | 5735 | 446.2 | 0.005025 | 11.26 | +2 | 7 |
| - | - | 1072 | 446.2 | - | - | 0 | - |
| - | - | 3664 | 446.7 | - | - | 0 | - |
| 14 | b | 1117 | 447.2 | 0.006268 | 14.02 | +4 | 14 |
| - | - | 2936 | 447.3 | - | - | 0 | - |
| - | - | 1685 | 448.2 | - | - | 0 | - |
| - | - | 3.321E+04 | 448.2 | - | - | 0 | - |
| 10 | y | 2.05E+04 | 448.7 | 0.006452 | 14.38 | +2 | 7 |
| - | - | 9808 | 449.2 | - | - | 0 | - |
| - | - | 3956 | 449.7 | - | - | 0 | - |
| - | - | 1904 | 450.2 | - | - | 0 | - |
| - | - | 1502 | 450.2 | - | - | 0 | - |
| - | - | 8938 | 450.2 | - | - | 0 | - |
| - | - | 1613 | 451.2 | - | - | 0 | - |
| - | - | 3497 | 451.7 | - | - | 0 | - |
| - | - | 1220 | 451.8 | - | - | 0 | - |
| - | - | 1055 | 451.9 | - | - | 0 | - |
| - | - | 1969 | 452.2 | - | - | 0 | - |
| - | - | 1269 | 452.2 | - | - | 0 | - |
| - | - | 2.376E+04 | 453.2 | - | - | 0 | - |
| - | - | 1114 | 453.2 | - | - | 0 | - |
| - | - | 1887 | 453.7 | - | - | 0 | - |
| - | - | 4421 | 454.2 | - | - | 0 | - |
| - | - | 3697 | 454.2 | - | - | 0 | - |
| - | - | 1211 | 454.2 | - | - | 0 | - |
| 7 | b | 7665 | 454.7 | 0.002004 | 4.408 | +2 | 7 |
| - | - | 3395 | 455.2 | - | - | 0 | - |
| - | - | 3654 | 455.2 | - | - | 0 | - |
| - | - | 1302 | 455.7 | - | - | 0 | - |
| - | - | 1487 | 455.8 | - | - | 0 | - |
| - | - | 1045 | 456.2 | - | - | 0 | - |
| - | - | 1505 | 457.3 | - | - | 0 | - |
| 10 | y | 6467 | 457.7 | 0.001567 | 3.423 | +2 | 7 |
| - | - | 1657 | 458.2 | - | - | 0 | - |
| - | - | 5277 | 458.2 | - | - | 0 | - |
| - | - | 1976 | 458.7 | - | - | 0 | - |
| - | - | 1.418E+04 | 459.2 | - | - | 0 | - |
| - | - | 984.1 | 459.7 | - | - | 0 | - |
| - | - | 1226 | 460.2 | - | - | 0 | - |
| - | - | 5611 | 460.2 | - | - | 0 | - |
| - | - | 6178 | 460.7 | - | - | 0 | - |
| - | - | 1887 | 461.2 | - | - | 0 | - |
| - | - | 2203 | 461.2 | - | - | 0 | - |
| - | - | 887 | 461.7 | - | - | 0 | - |
| - | - | 2417 | 462.2 | - | - | 0 | - |
| - | - | 1261 | 462.7 | - | - | 0 | - |
| - | - | 2619 | 463.2 | - | - | 0 | - |
| - | - | 875.7 | 463.2 | - | - | 0 | - |
| - | - | 4743 | 464.2 | - | - | 0 | - |
| - | - | 1165 | 464.2 | - | - | 0 | - |
| - | - | 2091 | 464.6 | - | - | 0 | - |
| - | - | 2712 | 465.2 | - | - | 0 | - |
| - | - | 2205 | 465.5 | - | - | 0 | - |
| - | - | 1372 | 465.7 | - | - | 0 | - |
| - | - | 6640 | 466.2 | - | - | 0 | - |
| - | - | 3819 | 466.2 | - | - | 0 | - |
| - | - | 1325 | 467.2 | - | - | 0 | - |
| - | - | 1098 | 467.2 | - | - | 0 | - |
| - | - | 1.606E+04 | 469.2 | - | - | 0 | - |
| - | - | 6769 | 469.5 | - | - | 0 | - |
| - | - | 8985 | 469.7 | - | - | 0 | - |
| 2 | y | 7165 | 470 | 0.007107 | 15.12 | +4 | 15 |
| - | - | 3558 | 470.2 | - | - | 0 | - |
| - | - | 1290 | 470.3 | - | - | 0 | - |
| - | - | 2309 | 470.5 | - | - | 0 | - |
| 15 | b | 4216 | 471.7 | 0.002313 | 4.903 | +4 | 15 |
| 15 | b | 2697 | 472 | 0.005528 | 11.71 | +4 | 15 |
| - | - | 7291 | 472.2 | - | - | 0 | - |
| - | - | 7222 | 472.3 | - | - | 0 | - |
| - | - | 1243 | 473.2 | - | - | 0 | - |
| - | - | 2155 | 473.3 | - | - | 0 | - |
| - | - | 1378 | 473.5 | - | - | 0 | - |
| - | - | 3206 | 474 | - | - | 0 | - |
| 8 | b | 8.006E+04 | 474.2 | 0.005832 | 12.3 | +2 | 8 |
| - | - | 8.507E+04 | 474.5 | - | - | 0 | - |
| - | - | 5.685E+04 | 474.7 | - | - | 0 | - |
| - | - | 2.65E+04 | 475 | - | - | 0 | - |
| - | - | 6437 | 475.2 | - | - | 0 | - |
| - | - | 5582 | 475.3 | - | - | 0 | - |
| - | - | 2432 | 475.5 | - | - | 0 | - |
| - | - | 3345 | 475.7 | - | - | 0 | - |
| 4 | b | 1.947E+04 | 476.2 | 0.001904 | 3.997 | +1 | 4 |
| - | - | 2203 | 476.5 | - | - | 0 | - |
| - | - | 3034 | 476.7 | - | - | 0 | - |
| - | - | 1206 | 477 | - | - | 0 | - |
| 4 | b | 2.098E+04 | 477.2 | 0.003728 | 7.812 | +1 | 4 |
| - | - | 2042 | 477.7 | - | - | 0 | - |
| - | - | 1.632E+04 | 478.2 | - | - | 0 | - |
| - | - | 941.2 | 478.6 | - | - | 0 | - |
| - | - | 4931 | 479.2 | - | - | 0 | - |
| - | - | 1230 | 479.5 | - | - | 0 | - |
| - | - | 3230 | 479.7 | - | - | 0 | - |
| - | - | 2973 | 480.2 | - | - | 0 | - |
| - | - | 6534 | 480.7 | - | - | 0 | - |
| - | - | 6912 | 481 | - | - | 0 | - |
| - | - | 4054 | 481.2 | - | - | 0 | - |
| - | - | 3764 | 481.2 | - | - | 0 | - |
| - | - | 1320 | 482.2 | - | - | 0 | - |
| - | - | 1293 | 483.5 | - | - | 0 | - |
| - | - | 2.716E+04 | 483.8 | - | - | 0 | - |
| - | - | 1.554E+04 | 484.3 | - | - | 0 | - |
| - | - | 1459 | 484.5 | - | - | 0 | - |
| - | - | 4919 | 484.8 | - | - | 0 | - |
| - | - | 1604 | 485 | - | - | 0 | - |
| - | - | 2423 | 485.3 | - | - | 0 | - |
| - | - | 1493 | 486.2 | - | - | 0 | - |
| - | - | 1451 | 487.2 | - | - | 0 | - |
| - | - | 912.9 | 487.5 | - | - | 0 | - |
| - | - | 1336 | 487.7 | - | - | 0 | - |
| - | - | 1721 | 488 | - | - | 0 | - |
| - | - | 4049 | 488.2 | - | - | 0 | - |
| - | - | 1612 | 488.5 | - | - | 0 | - |
| - | - | 3.081E+04 | 488.8 | - | - | 0 | - |
| - | - | 1.825E+04 | 489.3 | - | - | 0 | - |
| - | - | 8801 | 489.3 | - | - | 0 | - |
| - | - | 8029 | 489.8 | - | - | 0 | - |
| - | - | 1139 | 490 | - | - | 0 | - |
| - | - | 4698 | 490.2 | - | - | 0 | - |
| - | - | 2705 | 490.3 | - | - | 0 | - |
| - | - | 1601 | 490.6 | - | - | 0 | - |
| - | - | 1246 | 490.7 | - | - | 0 | - |
| - | - | 1816 | 491 | - | - | 0 | - |
| - | - | 4001 | 491.2 | - | - | 0 | - |
| - | - | 2997 | 491.5 | - | - | 0 | - |
| - | - | 3484 | 491.7 | - | - | 0 | - |
| - | - | 2078 | 492 | - | - | 0 | - |
| - | - | 2416 | 492.2 | - | - | 0 | - |
| - | - | 2118 | 493.3 | - | - | 0 | - |
| - | - | 1863 | 493.5 | - | - | 0 | - |
| - | - | 1.496E+04 | 493.7 | - | - | 0 | - |
| - | - | 1.281E+04 | 494 | - | - | 0 | - |
| 4 | b | 3.748E+05 | 494.2 | 0.00141 | 2.852 | +1 | 4 |
| - | - | 6010 | 494.5 | - | - | 0 | - |
| - | - | 3188 | 494.7 | - | - | 0 | - |
| - | - | 9.222E+04 | 495.2 | - | - | 0 | - |
| - | - | 1.617E+04 | 496.2 | - | - | 0 | - |
| - | - | 1.002E+04 | 496.7 | - | - | 0 | - |
| - | - | 7113 | 497.2 | - | - | 0 | - |
| - | - | 7.723E+04 | 497.8 | - | - | 0 | - |
| 0 | Precursor | 5.15E+04 | 498 | 0.001657 | 3.327 | +4 | -1 |
| 0 | Precursor | 8.885E+04 | 498.2 | 0.009193 | 18.45 | +4 | -1 |
| - | - | 4.891E+04 | 498.5 | - | - | 0 | - |
| - | - | 3.15E+04 | 498.7 | - | - | 0 | - |
| - | - | 1.293E+04 | 499 | - | - | 0 | - |
| - | - | 2.338E+04 | 499.3 | - | - | 0 | - |
| 12 | b | 1.573E+04 | 499.6 | 0.0007449 | 1.491 | +3 | 12 |
| - | - | 1.473E+04 | 499.9 | - | - | 0 | - |
| - | - | 5857 | 500.2 | - | - | 0 | - |
| - | - | 5885 | 500.3 | - | - | 0 | - |
| - | - | 2620 | 500.6 | - | - | 0 | - |
| - | - | 2732 | 501.3 | - | - | 0 | - |
| - | - | 974.8 | 501.8 | - | - | 0 | - |
| - | - | 4252 | 502 | - | - | 0 | - |
| - | - | 1.005E+04 | 502.3 | - | - | 0 | - |
| 0 | Precursor | 6.18E+05 | 502.5 | 0.00164 | 3.264 | +4 | -1 |
| - | - | 7.653E+05 | 502.7 | - | - | 0 | - |
| - | - | 4.425E+05 | 503 | - | - | 0 | - |
| - | - | 2.411E+05 | 503.3 | - | - | 0 | - |
| - | - | 7.752E+04 | 503.5 | - | - | 0 | - |
| - | - | 2.331E+04 | 503.8 | - | - | 0 | - |
| - | - | 2584 | 504 | - | - | 0 | - |
| - | - | 5988 | 504.2 | - | - | 0 | - |
| - | - | 1125 | 505.2 | - | - | 0 | - |
| 12 | b | 3.269E+04 | 505.3 | 0.0005772 | 1.142 | +3 | 12 |
| - | - | 2.956E+04 | 505.6 | - | - | 0 | - |
| - | - | 7.211E+04 | 505.7 | - | - | 0 | - |
| - | - | 1.245E+04 | 505.9 | - | - | 0 | - |
| - | - | 4.027E+04 | 506.2 | - | - | 0 | - |
| - | - | 1.508E+04 | 506.7 | - | - | 0 | - |
| - | - | 5940 | 507.2 | - | - | 0 | - |
| - | - | 2189 | 507.3 | - | - | 0 | - |
| - | - | 3249 | 509.3 | - | - | 0 | - |
| - | - | 1011 | 510.3 | - | - | 0 | - |
| - | - | 608.5 | 511.2 | - | - | 0 | - |
| - | - | 1.828E+04 | 511.3 | - | - | 0 | - |
| - | - | 3487 | 512.3 | - | - | 0 | - |
| - | - | 1516 | 512.6 | - | - | 0 | - |
| 9 | y | 3985 | 512.8 | 0.002608 | 5.085 | +2 | 8 |
| 9 | y | 4203 | 513.3 | 4.743E-05 | 0.0924 | +2 | 8 |
| - | - | 2745 | 514.2 | - | - | 0 | - |
| - | - | 2396 | 514.3 | - | - | 0 | - |
| - | - | 1464 | 514.7 | - | - | 0 | - |
| - | - | 1154 | 515.2 | - | - | 0 | - |
| - | - | 4.032E+04 | 517.3 | - | - | 0 | - |
| - | - | 1.107E+04 | 518.3 | - | - | 0 | - |
| - | - | 4712 | 519.3 | - | - | 0 | - |
| - | - | 1619 | 519.6 | - | - | 0 | - |
| - | - | 1757 | 519.9 | - | - | 0 | - |
| - | - | 9122 | 520.3 | - | - | 0 | - |
| - | - | 2414 | 521.2 | - | - | 0 | - |
| - | - | 2664 | 521.3 | - | - | 0 | - |
| 9 | y | 2.454E+04 | 521.8 | 0.0001668 | 0.3197 | +2 | 8 |
| - | - | 3616 | 522.2 | - | - | 0 | - |
| - | - | 1.258E+04 | 522.3 | - | - | 0 | - |
| - | - | 6884 | 522.8 | - | - | 0 | - |
| - | - | 1.033E+04 | 523.3 | - | - | 0 | - |
| - | - | 4011 | 523.8 | - | - | 0 | - |
| - | - | 6299 | 524.3 | - | - | 0 | - |
| - | - | 1852 | 524.8 | - | - | 0 | - |
| - | - | 1.162E+04 | 525.3 | - | - | 0 | - |
| - | - | 4199 | 526.2 | - | - | 0 | - |
| - | - | 4039 | 526.3 | - | - | 0 | - |
| - | - | 2453 | 527.3 | - | - | 0 | - |
| - | - | 1359 | 528.3 | - | - | 0 | - |
| - | - | 1942 | 528.7 | - | - | 0 | - |
| - | - | 1230 | 529.2 | - | - | 0 | - |
| - | - | 2471 | 530.2 | - | - | 0 | - |
| - | - | 1208 | 531.2 | - | - | 0 | - |
| - | - | 7907 | 532.2 | - | - | 0 | - |
| - | - | 1.022E+04 | 532.3 | - | - | 0 | - |
| - | - | 1.454E+04 | 532.8 | - | - | 0 | - |
| - | - | 1537 | 533.2 | - | - | 0 | - |
| - | - | 7907 | 533.3 | - | - | 0 | - |
| - | - | 3113 | 533.8 | - | - | 0 | - |
| - | - | 2515 | 534.2 | - | - | 0 | - |
| - | - | 1446 | 535.3 | - | - | 0 | - |
| - | - | 2332 | 535.9 | - | - | 0 | - |
| - | - | 3936 | 536.2 | - | - | 0 | - |
| - | - | 4153 | 536.3 | - | - | 0 | - |
| - | - | 2328 | 536.6 | - | - | 0 | - |
| - | - | 8165 | 537.3 | - | - | 0 | - |
| - | - | 9315 | 537.8 | - | - | 0 | - |
| 9 | b | 4850 | 538.3 | 0.00985 | 18.3 | +2 | 9 |
| 9 | b | 1522 | 538.8 | 0.004795 | 8.9 | +2 | 9 |
| - | - | 3.752E+04 | 539.2 | - | - | 0 | - |
| - | - | 2309 | 539.6 | - | - | 0 | - |
| - | - | 1513 | 539.9 | - | - | 0 | - |
| - | - | 1.162E+04 | 540.2 | - | - | 0 | - |
| - | - | 1019 | 540.8 | - | - | 0 | - |
| 8 | y | 6.804E+04 | 541.3 | 0.0003935 | 0.7271 | +2 | 9 |
| 8 | y | 4.276E+04 | 541.8 | 0.009301 | 17.17 | +2 | 9 |
| - | - | 1.955E+04 | 542.3 | - | - | 0 | - |
| - | - | 5591 | 542.8 | - | - | 0 | - |
| - | - | 1.652E+04 | 543.3 | - | - | 0 | - |
| 4 | y | 8678 | 543.6 | 0.001606 | 2.955 | +3 | 13 |
| - | - | 7128 | 543.9 | - | - | 0 | - |
| - | - | 7695 | 544.3 | - | - | 0 | - |
| - | - | 1807 | 544.6 | - | - | 0 | - |
| - | - | 1.229E+04 | 545.3 | - | - | 0 | - |
| - | - | 6807 | 545.6 | - | - | 0 | - |
| - | - | 5048 | 545.9 | - | - | 0 | - |
| - | - | 6.837E+04 | 546.3 | - | - | 0 | - |
| - | - | 5.156E+04 | 546.8 | - | - | 0 | - |
| 9 | b | 2.529E+04 | 547.3 | 0.009517 | 17.39 | +2 | 9 |
| - | - | 6994 | 547.8 | - | - | 0 | - |
| - | - | 1747 | 548.2 | - | - | 0 | - |
| - | - | 2760 | 548.3 | - | - | 0 | - |
| - | - | 1.148E+04 | 549.2 | - | - | 0 | - |
| - | - | 5837 | 549.3 | - | - | 0 | - |
| 8 | y | 2.022E+05 | 550.3 | 0.0004213 | 0.7655 | +2 | 9 |
| - | - | 1.154E+05 | 550.8 | - | - | 0 | - |
| - | - | 4.158E+04 | 551.3 | - | - | 0 | - |
| - | - | 2200 | 551.6 | - | - | 0 | - |
| - | - | 1.364E+04 | 551.8 | - | - | 0 | - |
| - | - | 1601 | 551.9 | - | - | 0 | - |
| - | - | 1.778E+04 | 552.3 | - | - | 0 | - |
| - | - | 7772 | 553.3 | - | - | 0 | - |
| - | - | 1218 | 553.3 | - | - | 0 | - |
| - | - | 2130 | 553.8 | - | - | 0 | - |
| - | - | 1325 | 554.2 | - | - | 0 | - |
| - | - | 2885 | 554.3 | - | - | 0 | - |
| - | - | 2171 | 554.6 | - | - | 0 | - |
| - | - | 2373 | 554.8 | - | - | 0 | - |
| - | - | 1464 | 555 | - | - | 0 | - |
| - | - | 1.917E+05 | 555.3 | - | - | 0 | - |
| - | - | 1.145E+05 | 555.8 | - | - | 0 | - |
| - | - | 4.963E+04 | 556.3 | - | - | 0 | - |
| - | - | 1.823E+04 | 556.8 | - | - | 0 | - |
| - | - | 3406 | 557.3 | - | - | 0 | - |
| - | - | 928.5 | 557.9 | - | - | 0 | - |
| - | - | 3162 | 560.2 | - | - | 0 | - |
| - | - | 1236 | 560.3 | - | - | 0 | - |
| - | - | 1002 | 561.2 | - | - | 0 | - |
| - | - | 1135 | 561.3 | - | - | 0 | - |
| - | - | 4631 | 561.8 | - | - | 0 | - |
| - | - | 1.635E+04 | 562.3 | - | - | 0 | - |
| - | - | 1.322E+04 | 562.8 | - | - | 0 | - |
| - | - | 1.249E+04 | 563.3 | - | - | 0 | - |
| - | - | 1855 | 563.8 | - | - | 0 | - |
| - | - | 6119 | 563.9 | - | - | 0 | - |
| - | - | 2.217E+04 | 564.2 | - | - | 0 | - |
| - | - | 1.605E+04 | 564.3 | - | - | 0 | - |
| - | - | 1369 | 564.6 | - | - | 0 | - |
| - | - | 7305 | 564.8 | - | - | 0 | - |
| - | - | 1811 | 564.9 | - | - | 0 | - |
| - | - | 5757 | 565.2 | - | - | 0 | - |
| - | - | 1.356E+04 | 565.3 | - | - | 0 | - |
| - | - | 1930 | 566.2 | - | - | 0 | - |
| - | - | 4802 | 566.3 | - | - | 0 | - |
| - | - | 5.739E+04 | 567.2 | - | - | 0 | - |
| - | - | 1.706E+04 | 568.2 | - | - | 0 | - |
| - | - | 6513 | 568.3 | - | - | 0 | - |
| - | - | 1364 | 568.8 | - | - | 0 | - |
| - | - | 1862 | 569.2 | - | - | 0 | - |
| - | - | 7425 | 569.3 | - | - | 0 | - |
| - | - | 2256 | 570.3 | - | - | 0 | - |
| - | - | 1832 | 570.3 | - | - | 0 | - |
| - | - | 1525 | 571.3 | - | - | 0 | - |
| - | - | 1241 | 572.3 | - | - | 0 | - |
| - | - | 1295 | 573.2 | - | - | 0 | - |
| - | - | 6059 | 574.3 | - | - | 0 | - |
| - | - | 7316 | 574.6 | - | - | 0 | - |
| - | - | 5158 | 574.9 | - | - | 0 | - |
| - | - | 1.039E+04 | 575.3 | - | - | 0 | - |
| - | - | 1186 | 575.8 | - | - | 0 | - |
| - | - | 951.6 | 575.9 | - | - | 0 | - |
| - | - | 1376 | 576.2 | - | - | 0 | - |
| - | - | 1343 | 576.3 | - | - | 0 | - |
| - | - | 3337 | 576.8 | - | - | 0 | - |
| - | - | 1639 | 577.2 | - | - | 0 | - |
| - | - | 2289 | 577.3 | - | - | 0 | - |
| - | - | 2411 | 578.2 | - | - | 0 | - |
| - | - | 2300 | 578.3 | - | - | 0 | - |
| - | - | 1271 | 578.8 | - | - | 0 | - |
| - | - | 1511 | 579.8 | - | - | 0 | - |
| 3 | y | 4.378E+04 | 580.3 | 0.001491 | 2.57 | +3 | 14 |
| 3 | y | 8.598E+04 | 580.6 | 0.004297 | 7.401 | +3 | 14 |
| - | - | 7.011E+04 | 580.9 | - | - | 0 | - |
| - | - | 6.45E+04 | 581.3 | - | - | 0 | - |
| - | - | 1.828E+04 | 581.6 | - | - | 0 | - |
| - | - | 4368 | 581.9 | - | - | 0 | - |
| - | - | 2.514E+04 | 582.2 | - | - | 0 | - |
| - | - | 1.165E+04 | 582.3 | - | - | 0 | - |
| - | - | 1359 | 582.6 | - | - | 0 | - |
| - | - | 7731 | 583.2 | - | - | 0 | - |
| - | - | 5319 | 583.3 | - | - | 0 | - |
| - | - | 2770 | 584.2 | - | - | 0 | - |
| - | - | 2005 | 584.8 | - | - | 0 | - |
| - | - | 3095 | 585.3 | - | - | 0 | - |
| - | - | 2464 | 585.6 | - | - | 0 | - |
| - | - | 1205 | 585.8 | - | - | 0 | - |
| 3 | y | 5.107E+04 | 586.3 | 0.00151 | 2.575 | +3 | 14 |
| - | - | 4.337E+04 | 586.6 | - | - | 0 | - |
| - | - | 2.518E+04 | 586.9 | - | - | 0 | - |
| - | - | 9370 | 587.3 | - | - | 0 | - |
| - | - | 5397 | 587.6 | - | - | 0 | - |
| - | - | 6547 | 588 | - | - | 0 | - |
| - | - | 2149 | 588.3 | - | - | 0 | - |
| - | - | 3798 | 588.8 | - | - | 0 | - |
| - | - | 2021 | 588.9 | - | - | 0 | - |
| - | - | 1549 | 589.3 | - | - | 0 | - |
| - | - | 2590 | 589.3 | - | - | 0 | - |
| - | - | 3722 | 589.8 | - | - | 0 | - |
| - | - | 1203 | 589.9 | - | - | 0 | - |
| - | - | 7624 | 590.3 | - | - | 0 | - |
| - | - | 2536 | 590.8 | - | - | 0 | - |
| - | - | 6593 | 591 | - | - | 0 | - |
| - | - | 1280 | 591.2 | - | - | 0 | - |
| - | - | 7027 | 591.3 | - | - | 0 | - |
| - | - | 4369 | 591.6 | - | - | 0 | - |
| - | - | 1624 | 592 | - | - | 0 | - |
| - | - | 1616 | 592.2 | - | - | 0 | - |
| - | - | 1971 | 592.3 | - | - | 0 | - |
| - | - | 4.389E+04 | 593.3 | - | - | 0 | - |
| - | - | 4747 | 593.8 | - | - | 0 | - |
| - | - | 1.325E+04 | 594.3 | - | - | 0 | - |
| - | - | 2247 | 594.8 | - | - | 0 | - |
| - | - | 1.727E+04 | 595.2 | - | - | 0 | - |
| 14 | b | 2656 | 595.6 | 0.001617 | 2.715 | +3 | 14 |
| 14 | b | 2506 | 595.9 | 0.005156 | 8.651 | +3 | 14 |
| - | - | 5482 | 596.2 | - | - | 0 | - |
| - | - | 1532 | 596.3 | - | - | 0 | - |
| - | - | 1.358E+04 | 597 | - | - | 0 | - |
| - | - | 933.9 | 597.2 | - | - | 0 | - |
| - | - | 1.765E+04 | 597.3 | - | - | 0 | - |
| - | - | 8484 | 597.6 | - | - | 0 | - |
| - | - | 1.548E+04 | 597.8 | - | - | 0 | - |
| - | - | 4131 | 598 | - | - | 0 | - |
| - | - | 9327 | 598.3 | - | - | 0 | - |
| 7 | y | 6.461E+04 | 598.8 | 0.000838 | 1.4 | +2 | 10 |
| 7 | y | 5.387E+04 | 599.3 | 0.007549 | 12.6 | +2 | 10 |
| - | - | 2.685E+04 | 599.8 | - | - | 0 | - |
| - | - | 8620 | 600.3 | - | - | 0 | - |
| - | - | 1139 | 600.6 | - | - | 0 | - |
| - | - | 3317 | 600.8 | - | - | 0 | - |
| - | - | 2664 | 601.3 | - | - | 0 | - |
| 14 | b | 1.091E+04 | 601.6 | 0.001514 | 2.517 | +3 | 14 |
| - | - | 1.089E+04 | 602 | - | - | 0 | - |
| - | - | 8195 | 602.3 | - | - | 0 | - |
| - | - | 3516 | 602.6 | - | - | 0 | - |
| - | - | 2.397E+04 | 602.8 | - | - | 0 | - |
| - | - | 3363 | 603 | - | - | 0 | - |
| - | - | 1.745E+04 | 603.3 | - | - | 0 | - |
| - | - | 2192 | 603.6 | - | - | 0 | - |
| - | - | 7006 | 603.8 | - | - | 0 | - |
| 12 | y | 4703 | 604.3 | 0.005367 | 8.881 | +1 | 5 |
| - | - | 1.409E+04 | 606.3 | - | - | 0 | - |
| - | - | 1527 | 606.8 | - | - | 0 | - |
| - | - | 7406 | 607.3 | - | - | 0 | - |
| 7 | y | 7.019E+05 | 607.8 | 0.0008658 | 1.424 | +2 | 10 |
| - | - | 4.461E+05 | 608.3 | - | - | 0 | - |
| - | - | 1.877E+05 | 608.8 | - | - | 0 | - |
| - | - | 6.694E+04 | 609.3 | - | - | 0 | - |
| - | - | 1.699E+04 | 609.8 | - | - | 0 | - |
| - | - | 4666 | 610.3 | - | - | 0 | - |
| - | - | 2293 | 611.3 | - | - | 0 | - |
| 10 | b | 7.81E+04 | 611.8 | 0.00947 | 15.48 | +2 | 10 |
| - | - | 2909 | 612 | - | - | 0 | - |
| - | - | 5.456E+04 | 612.3 | - | - | 0 | - |
| - | - | 1578 | 612.6 | - | - | 0 | - |
| - | - | 2.156E+04 | 612.8 | - | - | 0 | - |
| - | - | 5479 | 613.3 | - | - | 0 | - |
| - | - | 1531 | 613.8 | - | - | 0 | - |
| - | - | 1994 | 614.3 | - | - | 0 | - |
| - | - | 1534 | 616.3 | - | - | 0 | - |
| - | - | 4852 | 617.3 | - | - | 0 | - |
| - | - | 1852 | 618.3 | - | - | 0 | - |
| - | - | 5597 | 618.3 | - | - | 0 | - |
| - | - | 4950 | 619.3 | - | - | 0 | - |
| - | - | 1994 | 620 | - | - | 0 | - |
| - | - | 2451 | 620.3 | - | - | 0 | - |
| - | - | 1919 | 620.6 | - | - | 0 | - |
| - | - | 2736 | 620.8 | - | - | 0 | - |
| - | - | 2813 | 621 | - | - | 0 | - |
| - | - | 1349 | 621.3 | - | - | 0 | - |
| - | - | 1258 | 621.8 | - | - | 0 | - |
| 12 | y | 4.614E+04 | 622.3 | 0.001955 | 3.141 | +1 | 5 |
| - | - | 2194 | 623 | - | - | 0 | - |
| - | - | 1.377E+04 | 623.3 | - | - | 0 | - |
| - | - | 4706 | 624.3 | - | - | 0 | - |
| - | - | 2654 | 624.8 | - | - | 0 | - |
| - | - | 6556 | 625.3 | - | - | 0 | - |
| - | - | 5279 | 625.7 | - | - | 0 | - |
| 2 | y | 5651 | 626 | 0.004519 | 7.219 | +3 | 15 |
| 2 | y | 8134 | 626.3 | 0.01123 | 17.93 | +3 | 15 |
| - | - | 7942 | 626.6 | - | - | 0 | - |
| - | - | 5285 | 627 | - | - | 0 | - |
| - | - | 4602 | 627.3 | - | - | 0 | - |
| - | - | 2375 | 627.6 | - | - | 0 | - |
| 15 | b | 7098 | 628.6 | 0.001763 | 2.804 | +3 | 15 |
| 15 | b | 8775 | 629 | 0.005911 | 9.397 | +3 | 15 |
| - | - | 7045 | 629.3 | - | - | 0 | - |
| - | - | 2319 | 629.6 | - | - | 0 | - |
| - | - | 1418 | 630.3 | - | - | 0 | - |
| - | - | 1227 | 630.4 | - | - | 0 | - |
| - | - | 1963 | 631.6 | - | - | 0 | - |
| 2 | y | 5.994E+04 | 632 | 0.001546 | 2.447 | +3 | 15 |
| - | - | 6.469E+04 | 632.3 | - | - | 0 | - |
| - | - | 3.675E+04 | 632.6 | - | - | 0 | - |
| - | - | 1.939E+04 | 633 | - | - | 0 | - |
| - | - | 5693 | 633.3 | - | - | 0 | - |
| - | - | 1.014E+04 | 634.3 | - | - | 0 | - |
| 15 | b | 3.167E+04 | 634.6 | 0.001659 | 2.614 | +3 | 15 |
| - | - | 3.565E+04 | 635 | - | - | 0 | - |
| - | - | 3.312E+04 | 635.3 | - | - | 0 | - |
| - | - | 8740 | 635.6 | - | - | 0 | - |
| - | - | 1983 | 636 | - | - | 0 | - |
| - | - | 2.238E+04 | 636.3 | - | - | 0 | - |
| - | - | 1457 | 636.8 | - | - | 0 | - |
| - | - | 8363 | 637.3 | - | - | 0 | - |
| - | - | 1993 | 638.3 | - | - | 0 | - |
| - | - | 8930 | 640.3 | - | - | 0 | - |
| - | - | 6331 | 640.7 | - | - | 0 | - |
| - | - | 8996 | 641 | - | - | 0 | - |
| - | - | 2338 | 641.3 | - | - | 0 | - |
| - | - | 2481 | 641.3 | - | - | 0 | - |
| - | - | 2341 | 641.7 | - | - | 0 | - |
| - | - | 5964 | 645.3 | - | - | 0 | - |
| - | - | 4962 | 646.3 | - | - | 0 | - |
| - | - | 2881 | 646.8 | - | - | 0 | - |
| - | - | 7641 | 647.3 | - | - | 0 | - |
| - | - | 1529 | 648.3 | - | - | 0 | - |
| - | - | 1.218E+05 | 652.3 | - | - | 0 | - |
| - | - | 4.975E+04 | 653.3 | - | - | 0 | - |
| - | - | 1.055E+04 | 654.3 | - | - | 0 | - |
| 6 | y | 2.375E+04 | 655.3 | 0.0004835 | 0.7378 | +2 | 11 |
| 6 | y | 1.938E+04 | 655.8 | 0.008424 | 12.85 | +2 | 11 |
| - | - | 8217 | 656.3 | - | - | 0 | - |
| - | - | 3429 | 656.8 | - | - | 0 | - |
| - | - | 6041 | 657.3 | - | - | 0 | - |
| - | - | 3706 | 658.3 | - | - | 0 | - |
| - | - | 1960 | 658.3 | - | - | 0 | - |
| - | - | 1506 | 659.3 | - | - | 0 | - |
| - | - | 3.162E+04 | 662.3 | - | - | 0 | - |
| 5 | b | 2.095E+04 | 663.3 | 0.005531 | 8.338 | +1 | 5 |
| - | - | 7349 | 663.3 | - | - | 0 | - |
| - | - | 1335 | 663.8 | - | - | 0 | - |
| 6 | y | 2.336E+05 | 664.3 | 0.002718 | 4.091 | +2 | 11 |
| - | - | 1.35E+05 | 664.8 | - | - | 0 | - |
| - | - | 7.311E+04 | 665.3 | - | - | 0 | - |
| - | - | 2.163E+04 | 665.8 | - | - | 0 | - |
| - | - | 1.019E+04 | 666.3 | - | - | 0 | - |
| - | - | 4785 | 669.3 | - | - | 0 | - |
| 0 | Precursor | 1456 | 669.7 | 0.002049 | 3.059 | +3 | -1 |
| - | - | 2317 | 669.8 | - | - | 0 | - |
| - | - | 4873 | 670.3 | - | - | 0 | - |
| - | - | 2560 | 670.8 | - | - | 0 | - |
| - | - | 1612 | 671.3 | - | - | 0 | - |
| - | - | 2126 | 678.3 | - | - | 0 | - |
| - | - | 2214 | 679.3 | - | - | 0 | - |
| 5 | b | 3.895E+05 | 680.3 | 0.003823 | 5.619 | +1 | 5 |
| - | - | 1.537E+05 | 681.3 | - | - | 0 | - |
| - | - | 3.521E+04 | 682.3 | - | - | 0 | - |
| - | - | 8157 | 683.3 | - | - | 0 | - |
| - | - | 1141 | 683.8 | - | - | 0 | - |
| 11 | b | 3455 | 684.3 | 0.004938 | 7.216 | +2 | 11 |
| - | - | 1627 | 684.8 | - | - | 0 | - |
| - | - | 1587 | 685.3 | - | - | 0 | - |
| - | - | 3228 | 688.3 | - | - | 0 | - |
| - | - | 1165 | 689.3 | - | - | 0 | - |
| - | - | 2061 | 690.3 | - | - | 0 | - |
| - | - | 1810 | 691.4 | - | - | 0 | - |
| - | - | 1.248E+04 | 692.3 | - | - | 0 | - |
| 11 | b | 6120 | 693.3 | 0.01072 | 15.46 | +2 | 11 |
| - | - | 1987 | 693.8 | - | - | 0 | - |
| - | - | 2557 | 694.3 | - | - | 0 | - |
| - | - | 1326 | 695.9 | - | - | 0 | - |
| - | - | 2396 | 696.3 | - | - | 0 | - |
| - | - | 1939 | 698.4 | - | - | 0 | - |
| - | - | 2447 | 699.4 | - | - | 0 | - |
| - | - | 1341 | 700.4 | - | - | 0 | - |
| - | - | 4199 | 703.3 | - | - | 0 | - |
| - | - | 1503 | 704.3 | - | - | 0 | - |
| - | - | 4040 | 704.9 | - | - | 0 | - |
| - | - | 3354 | 705.4 | - | - | 0 | - |
| - | - | 2345 | 705.9 | - | - | 0 | - |
| - | - | 1.026E+04 | 706.3 | - | - | 0 | - |
| - | - | 3051 | 707.3 | - | - | 0 | - |
| - | - | 2388 | 708.3 | - | - | 0 | - |
| - | - | 1468 | 709.3 | - | - | 0 | - |
| - | - | 2.719E+04 | 710.3 | - | - | 0 | - |
| - | - | 9334 | 711.3 | - | - | 0 | - |
| - | - | 5870 | 712.3 | - | - | 0 | - |
| - | - | 2122 | 712.8 | - | - | 0 | - |
| - | - | 3335 | 713.3 | - | - | 0 | - |
| - | - | 1139 | 715.3 | - | - | 0 | - |
| - | - | 2219 | 717.4 | - | - | 0 | - |
| - | - | 2.841E+04 | 721.4 | - | - | 0 | - |
| - | - | 1.319E+04 | 722.4 | - | - | 0 | - |
| - | - | 3915 | 723.4 | - | - | 0 | - |
| - | - | 3074 | 729.4 | - | - | 0 | - |
| - | - | 1275 | 730.4 | - | - | 0 | - |
| - | - | 1229 | 739.4 | - | - | 0 | - |
| - | - | 1375 | 740.4 | - | - | 0 | - |
| - | - | 1435 | 745.4 | - | - | 0 | - |
| - | - | 2060 | 746.4 | - | - | 0 | - |
| - | - | 1381 | 747.4 | - | - | 0 | - |
| 12 | b | 3992 | 748.4 | 0.004271 | 5.707 | +2 | 12 |
| 12 | b | 2113 | 748.9 | 0.01135 | 15.15 | +2 | 12 |
| - | - | 1084 | 749.3 | - | - | 0 | - |
| - | - | 1182 | 749.4 | - | - | 0 | - |
| - | - | 1332 | 749.9 | - | - | 0 | - |
| - | - | 1639 | 750.3 | - | - | 0 | - |
| - | - | 1865 | 753.4 | - | - | 0 | - |
| - | - | 2237 | 753.9 | - | - | 0 | - |
| - | - | 1956 | 754.4 | - | - | 0 | - |
| 12 | b | 1.197E+04 | 757.4 | 0.0006455 | 0.8523 | +2 | 12 |
| - | - | 1.208E+04 | 757.9 | - | - | 0 | - |
| - | - | 6078 | 758.4 | - | - | 0 | - |
| - | - | 1699 | 758.9 | - | - | 0 | - |
| - | - | 1160 | 759.4 | - | - | 0 | - |
| - | - | 1597 | 760.3 | - | - | 0 | - |
| - | - | 2847 | 762.4 | - | - | 0 | - |
| - | - | 2322 | 762.9 | - | - | 0 | - |
| - | - | 5379 | 763.4 | - | - | 0 | - |
| - | - | 5210 | 764.4 | - | - | 0 | - |
| - | - | 4315 | 765.4 | - | - | 0 | - |
| - | - | 2454 | 766.4 | - | - | 0 | - |
| - | - | 5135 | 767.3 | - | - | 0 | - |
| - | - | 2055 | 767.9 | - | - | 0 | - |
| 11 | y | 8259 | 768.4 | 0.009498 | 12.36 | +1 | 6 |
| - | - | 2097 | 768.9 | - | - | 0 | - |
| - | - | 2270 | 769.4 | - | - | 0 | - |
| - | - | 1171 | 771.4 | - | - | 0 | - |
| - | - | 9706 | 772.4 | - | - | 0 | - |
| - | - | 1.663E+04 | 773.4 | - | - | 0 | - |
| - | - | 7227 | 774.4 | - | - | 0 | - |
| 6 | b | 4332 | 775.4 | 0.007665 | 9.885 | +1 | 6 |
| 6 | b | 3755 | 776.4 | 0.005512 | 7.1 | +1 | 6 |
| - | - | 2369 | 776.9 | - | - | 0 | - |
| - | - | 2443 | 777.4 | - | - | 0 | - |
| - | - | 1427 | 777.9 | - | - | 0 | - |
| - | - | 1477 | 778.3 | - | - | 0 | - |
| - | - | 1.162E+04 | 781.4 | - | - | 0 | - |
| - | - | 3989 | 782.4 | - | - | 0 | - |
| - | - | 1422 | 783.4 | - | - | 0 | - |
| - | - | 2766 | 784.4 | - | - | 0 | - |
| 11 | y | 4.832E+04 | 785.4 | 0.001623 | 2.067 | +1 | 6 |
| - | - | 1.999E+04 | 786.4 | - | - | 0 | - |
| - | - | 6973 | 787.4 | - | - | 0 | - |
| - | - | 1268 | 788.4 | - | - | 0 | - |
| - | - | 2436 | 790.4 | - | - | 0 | - |
| - | - | 5.815E+04 | 791.4 | - | - | 0 | - |
| - | - | 2.563E+04 | 792.4 | - | - | 0 | - |
| 6 | b | 2.793E+04 | 793.4 | 0.001872 | 2.359 | +1 | 6 |
| - | - | 1.188E+04 | 794.4 | - | - | 0 | - |
| - | - | 1.379E+04 | 795.3 | - | - | 0 | - |
| - | - | 7444 | 796.3 | - | - | 0 | - |
| - | - | 2723 | 797.4 | - | - | 0 | - |
| - | - | 2112 | 804.4 | - | - | 0 | - |
| 4 | y | 1564 | 805.9 | 0.007835 | 9.722 | +2 | 13 |
| 4 | y | 1547 | 806.4 | 0.006627 | 8.218 | +2 | 13 |
| - | - | 2841 | 808.4 | - | - | 0 | - |
| - | - | 2511 | 808.9 | - | - | 0 | - |
| - | - | 4.403E+04 | 809.4 | - | - | 0 | - |
| - | - | 1195 | 809.9 | - | - | 0 | - |
| - | - | 1.795E+04 | 810.4 | - | - | 0 | - |
| - | - | 5983 | 811.4 | - | - | 0 | - |
| - | - | 3215 | 812.4 | - | - | 0 | - |
| - | - | 1421 | 813.4 | - | - | 0 | - |
| - | - | 3790 | 814.4 | - | - | 0 | - |
| 4 | y | 6814 | 814.9 | 0.001043 | 1.28 | +2 | 13 |
| - | - | 6364 | 815.4 | - | - | 0 | - |
| - | - | 2802 | 815.9 | - | - | 0 | - |
| - | - | 4134 | 817.4 | - | - | 0 | - |
| - | - | 7296 | 817.9 | - | - | 0 | - |
| - | - | 6364 | 818.4 | - | - | 0 | - |
| - | - | 1841 | 818.9 | - | - | 0 | - |
| - | - | 2571 | 819.4 | - | - | 0 | - |
| - | - | 2167 | 820.4 | - | - | 0 | - |
| - | - | 1302 | 825.4 | - | - | 0 | - |
| - | - | 7924 | 826.4 | - | - | 0 | - |
| - | - | 5506 | 826.9 | - | - | 0 | - |
| - | - | 4658 | 827.4 | - | - | 0 | - |
| - | - | 1490 | 827.9 | - | - | 0 | - |
| - | - | 1566 | 828.4 | - | - | 0 | - |
| - | - | 4801 | 834.4 | - | - | 0 | - |
| - | - | 2027 | 835.4 | - | - | 0 | - |
| - | - | 5272 | 838.4 | - | - | 0 | - |
| - | - | 1346 | 839.4 | - | - | 0 | - |
| - | - | 1137 | 843.4 | - | - | 0 | - |
| - | - | 3957 | 844.4 | - | - | 0 | - |
| - | - | 2731 | 845.4 | - | - | 0 | - |
| - | - | 1567 | 845.9 | - | - | 0 | - |
| - | - | 1413 | 846.4 | - | - | 0 | - |
| - | - | 2369 | 852.4 | - | - | 0 | - |
| - | - | 1323 | 853.4 | - | - | 0 | - |
| - | - | 1426 | 854.4 | - | - | 0 | - |
| - | - | 4601 | 864.4 | - | - | 0 | - |
| - | - | 1680 | 865.4 | - | - | 0 | - |
| - | - | 1202 | 866.3 | - | - | 0 | - |
| - | - | 1224 | 869.4 | - | - | 0 | - |
| 3 | y | 3345 | 870.4 | 0.002154 | 2.475 | +2 | 14 |
| - | - | 1454 | 871.4 | - | - | 0 | - |
| - | - | 2779 | 878.4 | - | - | 0 | - |
| 3 | y | 1041 | 878.9 | 0.0006846 | 0.7789 | +2 | 14 |
| - | - | 2782 | 879.4 | - | - | 0 | - |
| - | - | 1339 | 880.4 | - | - | 0 | - |
| - | - | 1.254E+04 | 882.4 | - | - | 0 | - |
| - | - | 4620 | 883.4 | - | - | 0 | - |
| - | - | 1966 | 884.4 | - | - | 0 | - |
| - | - | 1968 | 885.4 | - | - | 0 | - |
| - | - | 1200 | 886.4 | - | - | 0 | - |
| 7 | b | 1104 | 890.4 | 0.002503 | 2.811 | +1 | 7 |
| 7 | b | 1786 | 891.4 | 0.005302 | 5.948 | +1 | 7 |
| - | - | 1515 | 892.4 | - | - | 0 | - |
| - | - | 969.5 | 894.4 | - | - | 0 | - |
| - | - | 1.362E+04 | 895.4 | - | - | 0 | - |
| 10 | y | 1.955E+04 | 896.4 | 0.003885 | 4.334 | +1 | 7 |
| - | - | 9607 | 897.4 | - | - | 0 | - |
| - | - | 4274 | 898.4 | - | - | 0 | - |
| - | - | 4498 | 899.4 | - | - | 0 | - |
| - | - | 2817 | 900.4 | - | - | 0 | - |
| - | - | 956.3 | 901.4 | - | - | 0 | - |
| - | - | 1097 | 902.4 | - | - | 0 | - |
| 7 | b | 2.499E+04 | 908.4 | 0.00274 | 3.016 | +1 | 7 |
| - | - | 1.114E+04 | 909.4 | - | - | 0 | - |
| - | - | 3301 | 910.4 | - | - | 0 | - |
| 10 | y | 5.419E+04 | 914.4 | 0.001858 | 2.032 | +1 | 7 |
| - | - | 2.54E+04 | 915.4 | - | - | 0 | - |
| - | - | 1.022E+04 | 916.4 | - | - | 0 | - |
| - | - | 2925 | 917.4 | - | - | 0 | - |
| - | - | 2175 | 919.5 | - | - | 0 | - |
| - | - | 1286 | 924.4 | - | - | 0 | - |
| - | - | 2172 | 936.4 | - | - | 0 | - |
| - | - | 1637 | 937.4 | - | - | 0 | - |
| - | - | 2363 | 937.5 | - | - | 0 | - |
| 8 | b | 1251 | 947.4 | 0.006978 | 7.365 | +1 | 8 |
| - | - | 3627 | 954.4 | - | - | 0 | - |
| - | - | 7244 | 955.4 | - | - | 0 | - |
| - | - | 2583 | 956.4 | - | - | 0 | - |
| - | - | 1096 | 963.4 | - | - | 0 | - |
| 8 | b | 5930 | 965.4 | 0.004409 | 4.567 | +1 | 8 |
| - | - | 2915 | 966.5 | - | - | 0 | - |
| - | - | 1175 | 967.5 | - | - | 0 | - |
| - | - | 3825 | 972.4 | - | - | 0 | - |
| - | - | 2492 | 973.4 | - | - | 0 | - |
| - | - | 4620 | 976.5 | - | - | 0 | - |
| - | - | 1326 | 977.5 | - | - | 0 | - |
| - | - | 1657 | 980.5 | - | - | 0 | - |
| - | - | 3134 | 981.4 | - | - | 0 | - |
| - | - | 3036 | 982.5 | - | - | 0 | - |
| - | - | 1090 | 983.5 | - | - | 0 | - |
| - | - | 3349 | 992.5 | - | - | 0 | - |
| - | - | 2767 | 993.4 | - | - | 0 | - |
| - | - | 8828 | 994.5 | - | - | 0 | - |
| - | - | 4176 | 995.5 | - | - | 0 | - |
| - | - | 1015 | 996.5 | - | - | 0 | - |
| - | - | 1091 | 997.5 | - | - | 0 | - |
| - | - | 2.425E+04 | 1010 | - | - | 0 | - |
| - | - | 1.112E+04 | 1011 | - | - | 0 | - |
| - | - | 7116 | 1012 | - | - | 0 | - |
| - | - | 2065 | 1013 | - | - | 0 | - |
| 9 | y | 2545 | 1043 | 0.00301 | 2.887 | +1 | 8 |
| - | - | 1644 | 1044 | - | - | 0 | - |
| 8 | y | 1531 | 1082 | 6.9E-06 | 0.006379 | +1 | 9 |
| - | - | 5188 | 1092 | - | - | 0 | - |
| - | - | 3536 | 1093 | - | - | 0 | - |
| 9 | b | 3272 | 1094 | 0.007607 | 6.957 | +1 | 9 |
| - | - | 2572 | 1095 | - | - | 0 | - |
| 8 | y | 5597 | 1100 | 0.003356 | 3.052 | +1 | 9 |
| - | - | 3240 | 1101 | - | - | 0 | - |
| - | - | 1984 | 1102 | - | - | 0 | - |
| - | - | 8957 | 1110 | - | - | 0 | - |
| - | - | 5444 | 1111 | - | - | 0 | - |
| - | - | 2403 | 1112 | - | - | 0 | - |
| - | - | 4161 | 1124 | - | - | 0 | - |
| - | - | 2587 | 1125 | - | - | 0 | - |
| - | - | 1413 | 1126 | - | - | 0 | - |
| - | - | 927.1 | 1137 | - | - | 0 | - |
| 7 | y | 1000 | 1198 | 0.007847 | 6.553 | +1 | 10 |
| 10 | b | 1167 | 1205 | 0.01835 | 15.23 | +1 | 10 |
| 7 | y | 3798 | 1215 | 0.004542 | 3.739 | +1 | 10 |
| - | - | 2248 | 1216 | - | - | 0 | - |
| - | - | 978 | 1217 | - | - | 0 | - |
| - | - | 806.4 | 1218 | - | - | 0 | - |
| - | - | 1438 | 1223 | - | - | 0 | - |
| - | - | 1175 | 1224 | - | - | 0 | - |
| - | - | 1048 | 1247 | - | - | 0 | - |
| - | - | 780.4 | 2866 | - | - | 0 | - |

m/z Charge Intensity FragmentType MassShift Position
120.05638122558594 0 1660.9021 b Ammonia loss 3
120.08167266845703 0 875.76575
121.04031372070312 0 1751.2965
122.07208251953125 0 3262.643
123.05603790283203 0 2661.2117
124.07624816894531 0 668.3914
124.66868591308594 0 524.1327
125.10807800292969 0 3250.6785
127.05108642578125 0 5829.383
127.0874252319336 0 4890.512
128.07162475585938 0 1289.6066
128.082763671875 0 1910.8245
128.11953735351562 0 609.68115
129.06689453125 0 4807.1816
129.10308837890625 0 898992.44
130.06588745117188 0 331302.38
130.10020446777344 0 6165.399
130.1063232421875 0 61993.195
131.06344604492188 0 1757.357
131.06922912597656 0 30852.695
131.1076202392578 0 1412.1545
131.1182861328125 0 804.31274
132.07266235351562 0 1408.3145
132.08152770996094 0 45481.992
132.10264587402344 0 1320.0609
132.94798278808594 0 535.65405
133.06170654296875 0 2245.3657
133.0850067138672 0 4555.969
134.02783203125 0 9698.972
135.0922088623047 0 634.3636
136.03981018066406 0 837.86926
136.07652282714844 0 66133.97
137.07989501953125 0 6145.465
137.10841369628906 0 893.17755
138.06700134277344 0 10633.049
138.09214782714844 0 1002.3672
138.12855529785156 0 10348.889
139.05093383789062 0 2537.9602
139.08741760253906 0 13885.787
139.13145446777344 0 954.5161
140.09085083007812 0 1223.4747
141.1030731201172 0 16192.184
142.0660400390625 0 6391.7954
142.10643005371094 0 1172.3142
143.0460205078125 0 6534.4917
143.06887817382812 0 698.0635
143.1190948486328 0 1352.1211
144.0813751220703 0 3015.0034
145.06167602539062 0 11867.728
146.06089782714844 0 3156.55
146.09303283691406 0 872.1013
146.12969970703125 0 2270.9812
147.0449676513672 0 3895.2139
147.11367797851562 0 1549.2924
148.05116271972656 0 1279.7926
148.07676696777344 0 978.474
149.04498291015625 0 891.9811
150.09251403808594 0 882.002
151.0874786376953 0 7922.99
152.1197509765625 0 1166.8854
153.1031036376953 0 5973.985
154.06149291992188 0 670.70624
154.09840393066406 0 2670.6282
154.1348114013672 0 930.09863
154.1597137451172 0 2439.668
155.04638671875 0 1677.3438
155.0823516845703 0 5967.4487
155.09364318847656 0 37517.04
155.11854553222656 0 2226.9492
155.15512084960938 0 2382.138
156.0771026611328 0 1419.3622
156.09693908691406 0 2539.1174
156.10316467285156 0 838.9701
157.06163024902344 0 2268.366
157.09820556640625 0 1387.7948
157.1344451904297 0 2920.9429
158.06092834472656 0 6881.0435
158.08477783203125 0 12017.055
159.09255981445312 0 555355.4
160.0763702392578 0 4120.6895
160.08953857421875 0 3176.2507
160.09591674804688 0 55293.082
161.03883361816406 0 1903.6935
161.0797882080078 0 888.6095
161.09939575195312 0 2317.5896
161.10665893554688 0 678.8441
162.05528259277344 0 1173.6677
164.08311462402344 0 736.16345
165.0779266357422 0 3676.0366
165.10311889648438 0 1952.3605
165.13995361328125 0 1195.6848
166.06204223632812 0 153859.88
167.05860900878906 0 2099.2937
167.0654754638672 0 11171.298
167.09323120117188 0 1356.1902
167.11865234375 0 1936.1934
168.0817108154297 0 15026.493
168.10275268554688 0 2505.0977
168.11399841308594 0 5953.595
169.076904296875 0 11892.474
169.0848846435547 0 2195.6199
169.09811401367188 0 20883.74
169.11749267578125 0 645.2865
170.0609893798828 0 48358.98
170.1297607421875 0 2090.687
171.0643310546875 0 5633.2925
171.0768585205078 0 1429.221
171.0919952392578 0 1659.3673
171.11378479003906 0 2710.4592
172.0725555419922 0 5037.976
172.109375 0 1680.078
172.14427185058594 0 930.97064
173.05665588378906 0 10796.975
173.07225036621094 0 614.1199
173.09280395507812 0 1195.714
173.12998962402344 0 815.4955
174.06008911132812 0 737.54254
174.08847045898438 0 1072.3705
175.08705139160156 0 2008.2284
175.0985107421875 0 965.1037
176.08279418945312 0 2110.7283
176.1111602783203 0 789.35114 a 2
177.10301208496094 0 1301.0802
178.1348419189453 0 65962.46
179.0493927001953 0 2292.6504
179.1188507080078 0 1766.6935
179.13046264648438 0 1097.6792
179.13827514648438 0 7136.197
180.0777130126953 0 993.0608
180.11407470703125 0 18070.402
180.1229705810547 0 915.80774
181.06173706054688 0 17295.537
181.09815979003906 0 7514.201
181.1173858642578 0 1836.0126
181.13442993164062 0 3685.6843
182.0466766357422 0 759.67426
182.06503295898438 0 1427.5239
182.1299591064453 0 652.06085
182.6113739013672 0 2126.4124
183.08876037597656 0 2155.6108
183.10475158691406 0 1393.3132
183.1138153076172 0 15091.75
183.15016174316406 0 31120.17
184.1174774169922 0 1502.4032
184.15371704101562 0 3189.0364
185.0723114013672 0 1591.6702
185.1665802001953 0 990.9703
186.12472534179688 0 147012.55
187.08758544921875 0 56640.695
187.0977783203125 0 1159.0989
187.10861206054688 0 8910.823 y Water loss 14
187.1177978515625 0 904.405
187.119873046875 0 968.87946
187.12814331054688 0 14662.833
188.07130432128906 0 2047.0332
188.09104919433594 0 6454.617
188.1126708984375 0 962.09753
188.13072204589844 0 809.7566
190.0833282470703 0 5480.545
190.1349334716797 0 14499.949
191.0836181640625 0 1134.5963
191.10324096679688 0 1558.2893
191.13800048828125 0 1481.6744
192.07742309570312 0 1315.2275
193.0979461669922 0 1161.0582
193.134521484375 0 10141.882
194.12913513183594 0 939.4582
194.13858032226562 0 1469.5117
195.0927276611328 0 1924.5692
195.11378479003906 0 29743.775
195.1498565673828 0 634.1319
196.1090545654297 0 25642.895
196.6086883544922 0 1260.791
197.07180786132812 0 1924.84
197.11160278320312 0 2480.419
197.12977600097656 0 807.1238
198.08834838867188 0 12844.52
198.12844848632812 0 43731.836
199.07223510742188 0 8392.713
199.09019470214844 0 2052.2524
199.10794067382812 0 1141.1581 b 5
199.1319122314453 0 6873.4736
200.07505798339844 0 3035.4978
200.1442108154297 0 3358.1055
200.1766815185547 0 4035.5918
201.12440490722656 0 18327.455
202.0868377685547 0 2313.7317
202.12417602539062 0 2739.081
203.08172607421875 0 723.52
203.10389709472656 0 1799.8545
203.11703491210938 0 2567.4795
204.0786590576172 0 1020.2084
204.11422729492188 0 1545.0747
204.1361541748047 0 937.3052
205.0981903076172 0 14790.393
205.1194305419922 0 69037.984 y 14
205.14573669433594 0 11193.204
206.1013641357422 0 1212.5192
206.12191772460938 0 3852.92
206.12985229492188 0 7197.39
206.14903259277344 0 1592.2319
207.11367797851562 0 1926.0829
207.12351989746094 0 739.56256
207.13368225097656 0 998.752
207.15074157714844 0 1373.1307
207.16152954101562 0 12353.764
208.0728759765625 0 861.7986
208.10903930664062 0 3471.4512
208.1653594970703 0 1250.8107
209.0568389892578 0 4744.943
209.07623291015625 0 856.1888
209.0933380126953 0 1069.5726
209.1295166015625 0 919.32855
210.12869262695312 0 19044.049
210.1505126953125 0 1240.9025
210.1612091064453 0 17084.523
211.0723876953125 0 691.0446
211.086669921875 0 881.1526
211.10888671875 0 2898.1116
211.1322021484375 0 3354.1865
211.1452178955078 0 4435.952
211.16419982910156 0 1873.9263
211.63026428222656 0 3996.7974
212.09300231933594 0 1831.9557
212.14073181152344 0 3246.1648
212.17672729492188 0 1021.2232
213.12437438964844 0 16038.36
214.1187744140625 0 1128.0647
214.12840270996094 0 1130.9784
215.14010620117188 0 1392.1165
216.0990447998047 0 11949.634
217.10191345214844 0 637.32263
219.12510681152344 0 1366.0953
220.09751892089844 0 1629.1101
220.10894775390625 0 1535.9591
220.120361328125 0 3337.2498
221.07200622558594 0 3277.191
221.09307861328125 0 2026.4775
221.10452270507812 0 8147.575
221.139404296875 0 897.8379
222.12477111816406 0 6495.6777
223.0880126953125 0 1882.001
223.10887145996094 0 10584.559
223.156494140625 0 126326.7 a 1
224.10438537597656 0 4623.445
224.1193389892578 0 6856.237
224.1409454345703 0 934.42413
224.15985107421875 0 15083.999
225.09889221191406 0 604.4541
225.12307739257812 0 1860.8816
226.0834197998047 0 30862.686
226.11997985839844 0 2589.6687
226.15623474121094 0 2852.4893
227.0673828125 0 28138.406
227.08616638183594 0 27346.188
227.14027404785156 0 5905.5825
227.1553955078125 0 10142.535
228.07034301757812 0 2656.2612
228.0896453857422 0 2311.733
228.15957641601562 0 1327.9788
228.17190551757812 0 9900.671
229.11952209472656 0 36646.54
229.15589904785156 0 1880.7587
229.1751708984375 0 1050.4425
229.60177612304688 0 1976.8984
230.10498046875 0 789.72107
230.12258911132812 0 3665.0408
230.15090942382812 0 1336.8884
231.08836364746094 0 1857.036
233.1055145263672 0 970.402
233.12799072265625 0 1344.2407
233.14088439941406 0 18716.117
234.12493896484375 0 202147.81
235.10888671875 0 1100.4169
235.12826538085938 0 23774.502
235.14404296875 0 947.82275
235.15676879882812 0 4085.7888
236.13108825683594 0 1356.3793
237.1361541748047 0 2230.7266
238.08412170410156 0 881.76764
238.0994415283203 0 909.213
238.11985778808594 0 18262.254
238.1309814453125 0 1602.8989
238.15628051757812 0 3277.8884
238.60708618164062 0 3302.8325
239.0826873779297 0 5006.205
239.10472106933594 0 2572.3909 b Ammonia loss 3
239.12286376953125 0 1886.8201
240.1354522705078 0 19714.975
241.09786987304688 0 1503.8094
241.11962890625 0 5267.42
241.13912963867188 0 1537.785
243.08029174804688 0 1240.4504
243.11004638671875 0 6588.26
243.15049743652344 0 2334.3083
243.5991668701172 0 4997.8994
244.09413146972656 0 62012.52
245.0968780517578 0 27959.52
246.0997314453125 0 3142.2742
246.12484741210938 0 2142.6914
246.18252563476562 0 6764.2705
246.6409912109375 0 2152.098
247.09420776367188 0 1344.252
247.10891723632812 0 13607.971
247.13172912597656 0 1310.455
247.14541625976562 0 16492.38
247.62184143066406 0 10759.137 b 3
248.1055145263672 0 1434.4625
248.115478515625 0 21723.605
248.14877319335938 0 2552.1948
249.09957885742188 0 35727.91
249.11837768554688 0 1942.749
249.13592529296875 0 1152.8411
250.10287475585938 0 4253.474
250.11929321289062 0 1537.9639
250.16709899902344 0 1265.1477
251.15159606933594 0 572444.2 b 1
252.1132049560547 0 1579.3186
252.1547088623047 0 72062.82
252.6043701171875 0 19298.004
253.105712890625 0 4981.4077
253.11965942382812 0 1296.5077
253.15650939941406 0 4101.0283
254.07872009277344 0 1710.1346
254.09693908691406 0 2420.9697
255.0809783935547 0 4520.4253
255.14915466308594 0 21889.922
256.09503173828125 0 1200.3652
256.1095275878906 0 6655.5747
256.133056640625 0 1017.80475
256.1512756347656 0 2159.5981
256.1668395996094 0 16753.703
257.0934143066406 0 1651.5215
257.11407470703125 0 7398.5396
257.1295166015625 0 2677.9934 y Ammonia loss 8
257.17169189453125 0 1573.5171
257.59698486328125 0 2476.1865
258.0987243652344 0 1479.5375
258.1246032714844 0 3043.6123
258.1462707519531 0 6632.3696
259.1483154296875 0 1204.5167
260.637939453125 0 2948.344
261.09283447265625 0 843.4625
261.1183166503906 0 20488.328
261.1368408203125 0 2741.8442
261.6097106933594 0 6682.5967
261.6218566894531 0 2038.9609
262.1214599609375 0 5141.9077
262.1369934082031 0 1033.771
262.1564025878906 0 1833.4683
262.6128845214844 0 606.22015
263.1036376953125 0 2084.081
263.12701416015625 0 1121.7083
264.17156982421875 0 1287.2947
265.1191711425781 0 21500.316
265.1312561035156 0 11230.448
265.1438903808594 0 1510.8394
266.1260986328125 0 82838.82
266.6016540527344 0 1985.894
267.07763671875 0 1510.7877
267.12896728515625 0 10579.558
267.14581298828125 0 1487.2849
268.13092041015625 0 5390.007
268.177978515625 0 4016.2107
269.1620178222656 0 2619.2493
269.1802978515625 0 771.43256
269.1983337402344 0 1666.2698
270.1228942871094 0 202007.92
270.6243896484375 0 58350.37
271.0749816894531 0 1120.6136
271.12591552734375 0 10109.364
271.6260986328125 0 1218.3567
272.107666015625 0 118305.914
272.1769714355469 0 16843.785
273.0916442871094 0 37979.168
273.1104736328125 0 13464.781
273.1800537109375 0 3574.3784
273.64593505859375 0 4157.707
274.0916442871094 0 1353.8092
274.10333251953125 0 3513.2786
274.11993408203125 0 23091.045
274.13134765625 0 5177.4907
274.1503601074219 0 1471.0613
274.1556396484375 0 1550.7015
275.087158203125 0 1314.1838
275.1029357910156 0 3399.9126
275.1166076660156 0 4932.2207
275.1399841308594 0 5989.931
275.6167297363281 0 3328.2234
276.11016845703125 0 9315.421
277.0941467285156 0 1273.7253
279.1249084472656 0 795.0924
279.1466979980469 0 2760.919
279.18304443359375 0 4075.6655
280.1305847167969 0 5297.5938
280.1466064453125 0 984.2084
280.1670227050781 0 4877.9404
280.60955810546875 0 798.6329
281.11480712890625 0 1069.4176
281.13311767578125 0 709.2157
281.6535949707031 0 1329.7
282.133544921875 0 2658.2566
282.1595764160156 0 4349.0005
282.18072509765625 0 935.866
282.63629150390625 0 969.32404
283.10504150390625 0 4135.3467
283.1416320800781 0 103160.7
284.08843994140625 0 2178.419
284.1043701171875 0 13034.916
284.1213073730469 0 12280.939
284.1446228027344 0 13375.48
284.6219177246094 0 5052.097
285.0891418457031 0 1336.1719
285.1067810058594 0 1771.4807
285.12530517578125 0 1314.7906
285.1568908691406 0 2385.2632
286.1412353515625 0 16015.613
286.1583251953125 0 654.35736
287.1423645019531 0 2685.9482
287.1729736328125 0 3252.8367
289.0859680175781 0 2097.0713
290.118408203125 0 97487.81
291.1017761230469 0 1043.6879
291.1215515136719 0 12185.095
291.183837890625 0 863.73944
291.2193298339844 0 6130.421
292.1147155761719 0 3671.233
292.1670837402344 0 47667.863
293.1145935058594 0 7293.3716
293.1701354980469 0 8453.439
294.0899963378906 0 719.0563
294.1208801269531 0 11836.751
295.10498046875 0 1142.9344
295.1222229003906 0 1122.9786
297.1573181152344 0 32100.623
297.1937255859375 0 5773.799
298.14093017578125 0 3096.7334
298.1601867675781 0 4080.7373
298.17864990234375 0 908.5611
298.1969299316406 0 1423.8795
299.0635681152344 0 3552.244
299.1444396972656 0 1194.6367
300.1031188964844 0 5044.59
300.13189697265625 0 2244.7021
300.17169189453125 0 4950.0254
300.6334228515625 0 1181.0903
301.1156311035156 0 5974.007
301.1521911621094 0 42112.6
302.11541748046875 0 18789.533
302.15509033203125 0 7157.2
303.0997009277344 0 1256.9159
303.1185302734375 0 2963.1777
303.1524353027344 0 4024.5762 y Ammonia loss 11
303.6528625488281 0 910.1555
304.15301513671875 0 2084.0447
304.1683044433594 0 2100.343
307.1446838378906 0 10362.66
307.1773681640625 0 1036.4805
308.127685546875 0 1496.8807
308.1459045410156 0 1329.5457
308.2104187011719 0 860.8434
308.6558532714844 0 912.6779
309.14666748046875 0 16876.32
309.16552734375 0 1160.0734
309.193115234375 0 1354.4902
309.6393737792969 0 3426.3704
309.64813232421875 0 3631.282
310.1402282714844 0 1935.6594
310.17559814453125 0 694.85736
311.1367492675781 0 6867.571
311.24566650390625 0 2228.6628
311.6666564941406 0 7500.768 y 11
312.1381530761719 0 1096.0298
312.1684875488281 0 1925.9335
312.6661071777344 0 844.61847
313.1153259277344 0 1440.6157
314.1196594238281 0 927.10516
314.1726379394531 0 812.7371
315.1512145996094 0 461.4041
315.16796875 0 43375.844
315.2043762207031 0 3120.6155 y Water loss 13
316.1717834472656 0 6495.6987
317.66015625 0 22438.127
318.11419677734375 0 10422.578
318.1524353027344 0 45450.07
318.65380859375 0 15648.838
319.15411376953125 0 9540.172
319.2141418457031 0 6057
320.21844482421875 0 1028.6509
321.1576232910156 0 882.7785
322.6526794433594 0 860.7377
322.8431701660156 0 1877.4982
323.1445007324219 0 9892.79
323.1768798828125 0 1332.2784
323.645751953125 0 4172.037
324.1447448730469 0 1035.458
324.64892578125 0 1041.2549
324.6783447265625 0 2445.836
325.1524353027344 0 6940.7637
325.1875 0 2153.4753
326.15447998046875 0 2507.0515
326.220703125 0 984.37915
326.6654968261719 0 416044.9
327.16680908203125 0 155333.23
327.66815185546875 0 36312.12
328.1695251464844 0 5372.164 y Water loss 5
329.187744140625 0 5794.694
330.1910095214844 0 1621.5077
331.6576232910156 0 53191.125 b Water loss 4
332.15716552734375 0 14220.817
332.1931457519531 0 775.6157
332.6600036621094 0 3992.9124
333.155517578125 0 1342.6158
333.2149658203125 0 37315.652 y 13
334.1884460449219 0 2516.5793 a Ammonia loss 2
334.21826171875 0 5229.537
335.21075439453125 0 2288.1914
336.1352233886719 0 903.22235
338.1776428222656 0 1262.1068
338.67584228515625 0 1555.8096
339.6553039550781 0 760.8137
340.6629943847656 0 120219.484 b 4
341.1644592285156 0 48045.3
341.66583251953125 0 10350.507
342.16748046875 0 2159.2542
342.5062255859375 0 715.7272 y Ammonia loss 8
343.160400390625 0 1890.0508
343.179931640625 0 4522.6387
344.17181396484375 0 8540.0205
345.13214111328125 0 9663.1
345.17413330078125 0 1027.318
346.1163635253906 0 36681.535
346.21392822265625 0 2849.038
346.6599426269531 0 1184.3365
347.1192626953125 0 6629.3193
347.2181701660156 0 1012.5611
348.0993347167969 0 906.6852
349.13043212890625 0 2187.0027
349.15252685546875 0 1477.1793
349.18939208984375 0 1127.4364
349.683349609375 0 1453.1567
350.1836853027344 0 1560.6151
351.14697265625 0 1504.4521
351.1676330566406 0 1938.6265
351.2040100097656 0 1006.70905
351.8529052734375 0 1006.9326
352.1657409667969 0 1833.2701
353.16522216796875 0 3374.4143
355.18035888671875 0 1473.9961
356.26702880859375 0 852.19806
357.1758728027344 0 1112.4318
358.1787109375 0 1340.2875
359.1395568847656 0 885.83624
359.17486572265625 0 1047.3358
360.69671630859375 0 14268.547
361.19622802734375 0 21462.807
361.5148010253906 0 4101.8335 y Ammonia loss 7
361.7023010253906 0 1733.2273
361.8489685058594 0 2075.5225
362.1851501464844 0 8670.421 b Ammonia loss 2
363.142822265625 0 55383.46
363.1864929199219 0 2517.015
364.12640380859375 0 8163.1895
364.146484375 0 8026.017
364.21478271484375 0 3932.9204
364.2369079589844 0 1095.2927
365.128173828125 0 1531.4009
365.2189025878906 0 947.2246
365.6893310546875 0 4990.2246
366.1565246582031 0 2974.8699
366.1826171875 0 5077.293
367.1421813964844 0 14385.357
367.188232421875 0 77343.625 y 7
367.5226745605469 0 48513.53
367.8564453125 0 19637.852
368.14495849609375 0 3000.5842
368.1681213378906 0 1168.2458
368.1911926269531 0 6907.808
368.52423095703125 0 687.4417
369.1559143066406 0 1040.556
369.1782531738281 0 2447.4873
369.2173767089844 0 996.3821
370.1772155761719 0 2248.7104
370.6873779296875 0 1533.6594
371.17669677734375 0 33345.543
372.1603088378906 0 4580.7437
372.1802673339844 0 5273.4404
373.16693115234375 0 1164.1582
373.193359375 0 1500.9854
373.2256164550781 0 805.37115
374.1112060546875 0 1453.4573
374.20245361328125 0 12146.547
374.6950378417969 0 29365.13 b Water loss 11
375.1963806152344 0 12658.572
375.69891357421875 0 1030.8014
376.163818359375 0 1642.5929
377.12567138671875 0 1880.6826
377.1481018066406 0 1645.8247
377.1886901855469 0 974.67725
378.1812744140625 0 1077.8918
378.21630859375 0 3353.1511
379.210693359375 0 104736.65 b 2
380.16461181640625 0 2605.076
380.2137451171875 0 19319.873
381.1535949707031 0 132032.2
381.2139587402344 0 3178.0503
382.156494140625 0 22750.453
382.1954650878906 0 2765.6426
382.6971435546875 0 2548.0947
383.159423828125 0 3216.9565
383.20751953125 0 57264.02
383.7091064453125 0 29873.375
384.1682434082031 0 14341.75
384.1906433105469 0 3074.9648 y Water loss 10
384.211669921875 0 3994.7273
384.26129150390625 0 1478.0778
384.6844787597656 0 1589.118 y Ammonia loss 10
384.71246337890625 0 931.0941
385.1522521972656 0 25879.166
385.1932067871094 0 1571.5864
386.15570068359375 0 3124.2336
386.1805725097656 0 3534.1938
386.2061462402344 0 1652.8569
386.6859130859375 0 1207.7007
387.1585998535156 0 1313.0514
387.1861877441406 0 2097.3928
387.6854553222656 0 1013.2393
388.19989013671875 0 2737.6123 b Water loss 5
388.6927185058594 0 7435.9287 b Ammonia loss 5
389.1875915527344 0 73340.33
389.6958923339844 0 1157.6484
390.19036865234375 0 14103.851
391.1378479003906 0 14535.059
391.18505859375 0 3671.081
391.2366638183594 0 1952.5356
392.12249755859375 0 2960.3909
392.144287109375 0 1452.3904
392.1825256347656 0 1147.94
392.21014404296875 0 19246.736
393.1988830566406 0 140194.12 y 10
393.7002258300781 0 61025.156
394.1515197753906 0 1499.4087
394.17352294921875 0 11510.505
394.2005615234375 0 17581.79
394.7012634277344 0 5689.9966
395.1366882324219 0 13288.751
395.156494140625 0 1967.9102
395.2039489746094 0 1533.456
396.1400146484375 0 2899.4902
396.1668395996094 0 1157.1549
396.1923828125 0 7475.567
396.2269287109375 0 9574.368
396.6942138671875 0 2018.2786
397.20501708984375 0 25293.207 b 5
397.22711181640625 0 947.45734
397.7065734863281 0 10046.662
398.17791748046875 0 11259.393
398.20831298828125 0 2683.1775
399.1769104003906 0 3651.5044
399.52789306640625 0 3421.0999 y Water loss 6
399.85858154296875 0 3525.8367 y Ammonia loss 6
401.1868591308594 0 3629.2666
402.1792297363281 0 30691.617
403.1369323730469 0 1853.9231
403.199951171875 0 14133.609
404.2032775878906 0 3382.025
405.1951904296875 0 1185.6868
405.2255554199219 0 1654.9779
405.531005859375 0 47320.324 y 6
405.8654479980469 0 29658.889
406.19915771484375 0 12537.826
406.5331726074219 0 4616.125
406.86688232421875 0 1242.1813
407.1986389160156 0 3601.3772
407.27783203125 0 1036.8416
408.1637268066406 0 2288.7605
409.1488037109375 0 28412.758
409.2364501953125 0 1462.2659
410.1513366699219 0 5062.9595
411.1997375488281 0 3556.667
411.7012634277344 0 1819.3917
412.1653747558594 0 8811.85
412.18505859375 0 56547.445
413.14727783203125 0 16387.574
413.18829345703125 0 9903.546
414.1506042480469 0 3764.1711
414.1982421875 0 5077.5635
414.2369384765625 0 7196.289
415.03900146484375 0 1288.091
415.14996337890625 0 809.45154
415.1996154785156 0 9353.863
415.2375793457031 0 2134.923
416.20269775390625 0 2262.6646
416.482421875 0 925.6459
416.88397216796875 0 900.47003
417.1826171875 0 2889.5747
417.54803466796875 0 1010.7693
418.1470031738281 0 1512.5272
418.2139892578125 0 43326.277
419.2181396484375 0 7734.1562
419.7121887207031 0 7407.9316
420.1656494140625 0 1794.7467
420.21392822265625 0 5945.902
420.71429443359375 0 1189.16
421.2103271484375 0 18779.361
422.2139892578125 0 4451.6875
422.25323486328125 0 1896.5256
423.2026062011719 0 1053.0907
424.2204895019531 0 2085.8594
425.2889709472656 0 2006.9594
427.6905822753906 0 1416.4071
428.197021484375 0 1585.6139
429.2151794433594 0 898.74304
430.1955261230469 0 94071.875
431.1987609863281 0 19577.25
432.20074462890625 0 4123.9185
432.2301940917969 0 1588.4546
432.6836853027344 0 2115.3618
433.2095031738281 0 1293.5614
434.2128601074219 0 1290.445
435.1697082519531 0 5644.0063
436.15576171875 0 1415.7513
436.70751953125 0 901.05835
437.19195556640625 0 1441.0214
437.2264099121094 0 6917.6084 y Water loss 5
437.55633544921875 0 4385.699 y Ammonia loss 5
437.89013671875 0 1891.2859
438.17547607421875 0 3797.155
438.2336730957031 0 2397.7078
439.1789245605469 0 1152.2799
439.219482421875 0 2191.4585
439.7118225097656 0 2576.2524
440.1794738769531 0 2195.6519
440.2117614746094 0 1431.1367
440.7186279296875 0 2064.3005
441.2226257324219 0 1399.7311
441.6888427734375 0 1276.2665
443.2259521484375 0 48228.36 y 5
443.5603332519531 0 36163.188
443.89385986328125 0 16197.06
444.15625 0 1973.0392
444.22900390625 0 3505.8452
444.5606994628906 0 1826.5419
445.7121276855469 0 2405.6626 b Water loss 6
446.20843505859375 0 5734.849 b Ammonia loss 6
446.2396240234375 0 1071.6261
446.73577880859375 0 3663.911
447.2069396972656 0 1116.6083 b Ammonia loss 13
447.27362060546875 0 2935.6792
448.161865234375 0 1684.9421
448.22344970703125 0 33205.27
448.7214660644531 0 20500.02 y Water loss 9
449.2191162109375 0 9808.293
449.7195129394531 0 3956.1836
450.1755676269531 0 1903.7681
450.20648193359375 0 1501.8983
450.2369689941406 0 8937.958
451.2381286621094 0 1612.928
451.72882080078125 0 3497.0483
451.76220703125 0 1220.4042
451.88507080078125 0 1055.3632
452.1911315917969 0 1969.0677
452.2271728515625 0 1269.4811
453.1827697753906 0 23761.955
453.21466064453125 0 1114.1044
453.71893310546875 0 1887.4291
454.1847229003906 0 4421.4165
454.21136474609375 0 3696.9187
454.2450256347656 0 1211.1755
454.71868896484375 0 7664.9653 b 6
455.2181091308594 0 3394.89
455.24945068359375 0 3653.9902
455.7207336425781 0 1302.3488
455.7520751953125 0 1486.5256
456.21112060546875 0 1045.4232
457.2508850097656 0 1505.0448
457.72186279296875 0 6467.3115 y 9
458.1716003417969 0 1657.2839
458.2234191894531 0 5277.3037
458.72979736328125 0 1975.5564
459.20074462890625 0 14175.638
459.7237854003906 0 984.0894
460.20147705078125 0 1226.4712
460.2403869628906 0 5610.8506
460.7378845214844 0 6177.8926
461.20654296875 0 1887.1162
461.2405700683594 0 2202.9702
461.73492431640625 0 887.0405
462.1663513183594 0 2416.873
462.7197265625 0 1260.6976
463.16717529296875 0 2619.2886
463.2200927734375 0 875.7076
464.2165832519531 0 4743.439
464.24969482421875 0 1165.1229
464.5638427734375 0 2091.3206
465.226318359375 0 2712.2075
465.4695739746094 0 2204.5598
465.72503662109375 0 1372.1648
466.1705322265625 0 6639.9736
466.24200439453125 0 3819.0828
467.1727600097656 0 1325.3596
467.2474365234375 0 1097.8187
469.2445983886719 0 16059.853
469.48968505859375 0 6769.1987
469.7440490722656 0 8984.8125
469.97705078125 0 7164.5234 y Ammonia loss 1
470.2210693359375 0 3557.6873
470.2508544921875 0 1290.488
470.4761657714844 0 2309.266
471.73199462890625 0 4215.976 b Water loss 14
471.98583984375 0 2696.762 b Ammonia loss 14
472.2223815917969 0 7290.9775
472.2614440917969 0 7222.0786
473.2308654785156 0 1242.8774
473.2642822265625 0 2155.1826
473.4850769042969 0 1378.0225
473.97943115234375 0 3205.8555
474.22796630859375 0 80056.71 b Water loss 7
474.4786376953125 0 85069.96
474.72918701171875 0 56851.438
474.9797668457031 0 26498.592
475.2305603027344 0 6437.113
475.2554016113281 0 5582.0376
475.48028564453125 0 2431.5527
475.74554443359375 0 3345.3254
476.22711181640625 0 19470.32 b Water loss 3
476.4892272949219 0 2203.3342
476.7395324707031 0 3033.8845
476.98394775390625 0 1206.1399
477.21295166015625 0 20975.898 b Ammonia loss 3
477.7237854003906 0 2042.0851
478.2301330566406 0 16317.066
478.57183837890625 0 941.1549
479.2362976074219 0 4931.35
479.502685546875 0 1230.2755
479.7483825683594 0 3230.1738
480.2312316894531 0 2972.8752
480.7406921386719 0 6534.3687
480.9915466308594 0 6911.633
481.17742919921875 0 4054.425
481.24127197265625 0 3764.3647
482.2367858886719 0 1319.8069
483.4903869628906 0 1292.8536
483.760009765625 0 27158.047
484.2607421875 0 15539.103
484.4883728027344 0 1458.5039
484.7611389160156 0 4918.8716
484.9923095703125 0 1604.1449
485.2613220214844 0 2422.5652
486.1914978027344 0 1493.092
487.2460021972656 0 1451.4873
487.495849609375 0 912.85034
487.7384033203125 0 1335.7494
487.99749755859375 0 1721.4053
488.22503662109375 0 4049.4534
488.4875183105469 0 1612.4443
488.7521667480469 0 30806.543
489.25201416015625 0 18247.83
489.28857421875 0 8800.732
489.7528381347656 0 8029.397
489.9893493652344 0 1138.8817
490.2448425292969 0 4697.5825
490.2912292480469 0 2705.4653
490.5804748535156 0 1601.2542
490.7380065917969 0 1246.3574
490.9963684082031 0 1816.3304
491.2472839355469 0 4000.571
491.49749755859375 0 2996.5134
491.7460632324219 0 3484.3455
491.9918518066406 0 2077.6077
492.2474060058594 0 2416.0342
493.2527160644531 0 2118.1218
493.4957580566406 0 1863.3958
493.7423095703125 0 14964.508
493.9910888671875 0 12811.855
494.2371826171875 0 374765.2 b 3
494.4910583496094 0 6009.874
494.74615478515625 0 3188.1174
495.24005126953125 0 92219.38
496.2408447265625 0 16170.479
496.7320556640625 0 10022.644
497.2284240722656 0 7112.6455
497.75762939453125 0 77226.75
497.9966125488281 0 51498.023 Precursor Water loss
498.2501525878906 0 88850.35 Precursor Ammonia loss
498.49591064453125 0 48909.793
498.7499694824219 0 31496.32
498.9961853027344 0 12928.365
499.2712097167969 0 23376.031
499.577880859375 0 15730.878 b Ammonia loss 11
499.91229248046875 0 14734.7295
500.2463073730469 0 5856.8364
500.27606201171875 0 5884.5244
500.5795593261719 0 2619.6821
501.2677917480469 0 2732.1362
501.7534484863281 0 974.77264
501.99432373046875 0 4252.042
502.25469970703125 0 10050.862
502.4992370605469 0 617969 Precursor
502.7497253417969 0 765311.94
503.0002136230469 0 442518.72
503.25054931640625 0 241130.83
503.5009460449219 0 77519.4
503.75115966796875 0 23310.729
504.0008544921875 0 2583.7688
504.2024841308594 0 5987.896
505.2071228027344 0 1124.7548
505.2520751953125 0 32691.494 b 11
505.58673095703125 0 29558.668
505.73663330078125 0 72111.03
505.9206237792969 0 12446.94
506.2384948730469 0 40270.69
506.7384033203125 0 15083.197
507.2364807128906 0 5940.2314
507.2686462402344 0 2188.653
509.2732238769531 0 3248.7239
510.2753601074219 0 1010.7013
511.2249450683594 0 608.4706
511.2646484375 0 18280.34
512.26611328125 0 3487.179
512.5747680664062 0 1515.8817
512.7598876953125 0 3985.3474 y Water loss 8
513.2544555664062 0 4203.0054 y Ammonia loss 8
514.1864624023438 0 2745.457
514.264892578125 0 2395.7585
514.7489624023438 0 1464.4287
515.2384643554688 0 1153.7948
517.2826538085938 0 40315.137
518.285400390625 0 11067.864
519.2810668945312 0 4711.8296
519.6043701171875 0 1618.7135
519.9360961914062 0 1757.3246
520.273681640625 0 9121.847
521.23095703125 0 2413.9604
521.2756958007812 0 2664.486
521.7679443359375 0 24536.299 y 8
522.2152709960938 0 3616.0474
522.2701416015625 0 12577.459
522.7691040039062 0 6884.1377
523.2642822265625 0 10329.313
523.75927734375 0 4011.182
524.259521484375 0 6298.9277
524.755126953125 0 1852.0468
525.267333984375 0 11617.77
526.2325439453125 0 4198.6465
526.2700805664062 0 4039.2668
527.27099609375 0 2453.4434
528.257568359375 0 1359.1938
528.7365112304688 0 1941.9009
529.232666015625 0 1230.2261
530.2274169921875 0 2471.3318
531.2164306640625 0 1208.1376
532.1968994140625 0 7906.6416
532.2689208984375 0 10218.468
532.7634887695312 0 14543.377
533.2024536132812 0 1536.7023
533.2566528320312 0 7906.517
533.76123046875 0 3113.438
534.2376098632812 0 2515.325
535.294677734375 0 1445.7227
535.9329223632812 0 2332.2703
536.2200927734375 0 3935.6265
536.2659912109375 0 4153.307
536.5948486328125 0 2328.0374
537.2601928710938 0 8164.763
537.757080078125 0 9314.547
538.259765625 0 4849.6074 b Water loss 8
538.7664184570312 0 1521.7767 b Ammonia loss 8
539.2387084960938 0 37521.91
539.5914916992188 0 2308.606
539.92626953125 0 1512.7307
540.2425537109375 0 11616.919
540.7794189453125 0 1018.87024
541.2736206054688 0 68039.06 y Water loss 7
541.7745361328125 0 42761.582 y Ammonia loss 7
542.2749633789062 0 19550.217
542.77587890625 0 5590.942
543.2771606445312 0 16523.418
543.595458984375 0 8677.821 y 3
543.9301147460938 0 7127.939
544.2675170898438 0 7695.0957
544.5985107421875 0 1806.852
545.2700805664062 0 12290.793
545.5951538085938 0 6807.483
545.9297485351562 0 5047.7812
546.265380859375 0 68370.945
546.76513671875 0 51563.586
547.265380859375 0 25290.943 b 8
547.7672729492188 0 6994.212
548.2348022460938 0 1747.187
548.27197265625 0 2759.5095
549.2236328125 0 11480.694
549.30615234375 0 5836.6274
550.2789306640625 0 202175.64 y 7
550.7802734375 0 115380.01
551.2776489257812 0 41577.11
551.5966796875 0 2200.344
551.7811279296875 0 13640.616
551.9331665039062 0 1601.2958
552.2608642578125 0 17783.314
553.2670288085938 0 7772.016
553.3134765625 0 1218.4233
553.774169921875 0 2129.7302
554.2275390625 0 1325.3982
554.2710571289062 0 2884.5898
554.6378173828125 0 2170.779
554.7741088867188 0 2372.9329
554.9725341796875 0 1464.2461
555.2706909179688 0 191687.31
555.7719116210938 0 114533.69
556.2725219726562 0 49629.496
556.7734985351562 0 18229.74
557.2747192382812 0 3406.0544
557.9266357421875 0 928.54706
560.1904907226562 0 3162.1875
560.2625122070312 0 1235.5713
561.1959228515625 0 1001.9973
561.2681274414062 0 1135.3492
561.7958374023438 0 4631.3647
562.2832641601562 0 16347.93
562.7800903320312 0 13221.382
563.2619018554688 0 12489.971
563.780517578125 0 1855.0873
563.930908203125 0 6118.654
564.2144165039062 0 22167.186
564.2701416015625 0 16054.806
564.599853515625 0 1369.1411
564.777587890625 0 7305.0864
564.9291381835938 0 1811.3833
565.2157592773438 0 5757.331
565.2666625976562 0 13556.962
566.2138671875 0 1929.9381
566.2669677734375 0 4802.0537
567.2334594726562 0 57385.395
568.2362060546875 0 17061.607
568.28515625 0 6512.687
568.7794799804688 0 1363.8713
569.2389526367188 0 1861.5701
569.284912109375 0 7424.7495
570.2869873046875 0 2255.694
570.3439331054688 0 1832.2954
571.287841796875 0 1525.2136
572.2979125976562 0 1240.6337
573.1983032226562 0 1295.0813
574.2743530273438 0 6058.6606
574.6065673828125 0 7316.418
574.93701171875 0 5157.73
575.250732421875 0 10386.561
575.7828979492188 0 1186.3495
575.9390869140625 0 951.58746
576.2440185546875 0 1376.0082
576.28759765625 0 1343.278
576.7866821289062 0 3337.3413
577.2161865234375 0 1638.6158
577.2880249023438 0 2289.3513
578.205322265625 0 2410.579
578.2816772460938 0 2299.6357
578.7922973632812 0 1270.9586
579.7904052734375 0 1510.7705
580.2780151367188 0 43779.47 y Water loss 2
580.6088256835938 0 85980.7 y Ammonia loss 2
580.9421997070312 0 70108.96
581.276611328125 0 64499.13
581.60986328125 0 18277.482
581.9461059570312 0 4367.947
582.2255249023438 0 25136.232
582.2813110351562 0 11649.592
582.6183471679688 0 1358.5222
583.2281494140625 0 7731.4365
583.2835083007812 0 5319.4453
584.227294921875 0 2770.2666
584.7880249023438 0 2004.6315
585.2845458984375 0 3094.825
585.6109619140625 0 2463.619
585.7836303710938 0 1204.5973
586.2815551757812 0 51071.316 y 2
586.6156616210938 0 43371.984
586.9497680664062 0 25179.27
587.2840576171875 0 9369.635
587.6203002929688 0 5397.1104
587.9556274414062 0 6546.7217
588.2901000976562 0 2148.9744
588.8084716796875 0 3798.256
588.949462890625 0 2021.1832
589.2791748046875 0 1548.5658
589.3077392578125 0 2589.9285
589.7827758789062 0 3722.4521
589.9486694335938 0 1203.142
590.2777099609375 0 7624.4517
590.7783813476562 0 2535.8499
590.9507446289062 0 6593.4727
591.2103881835938 0 1280.3041
591.2814331054688 0 7026.8896
591.6195678710938 0 4368.8354
591.9561767578125 0 1623.6083
592.2122192382812 0 1615.6678
592.2749633789062 0 1971.4967
593.259521484375 0 43886.633
593.801025390625 0 4746.9146
594.2640991210938 0 13249.335
594.791259765625 0 2247.482
595.228759765625 0 17273.693
595.618896484375 0 2656.0518 b Water loss 13
595.9536743164062 0 2505.5437 b Ammonia loss 13
596.2316284179688 0 5482.124
596.2808227539062 0 1532.2179
596.95361328125 0 13580.998
597.2366333007812 0 933.9188
597.2881469726562 0 17649.213
597.6229248046875 0 8484.135
597.8125610351562 0 15482.816
597.9558715820312 0 4131.4717
598.3121948242188 0 9327.319
598.7875366210938 0 64612.137 y Water loss 6
599.2862548828125 0 53872.64 y Ammonia loss 6
599.7864379882812 0 26849.268
600.284912109375 0 8619.941
600.6116943359375 0 1139.2866
600.7854614257812 0 3316.5796
601.3062744140625 0 2664.089
601.6255493164062 0 10911.521 b 13
601.9603271484375 0 10892.071
602.2955932617188 0 8195.244
602.62841796875 0 3516.0505
602.8034057617188 0 23969.207
602.9584350585938 0 3363.0938
603.3005981445312 0 17448.34
603.6256713867188 0 2191.501
603.8034057617188 0 7005.632
604.3114624023438 0 4703.1157 y Water loss 11
606.326904296875 0 14092.185
606.7816772460938 0 1527.0333
607.3006591796875 0 7405.5425
607.7928466796875 0 701885.06 y 6
608.2941284179688 0 446076.44
608.7946166992188 0 187674.03
609.2951049804688 0 66936.68
609.7954711914062 0 16986.361
610.2930297851562 0 4666.1055
611.2996826171875 0 2293.3108
611.8056640625 0 78100.875 b 9
611.958251953125 0 2908.6462
612.3067016601562 0 54561.92
612.624267578125 0 1578.375
612.8084716796875 0 21559.129
613.30859375 0 5479.2056
613.7985229492188 0 1531.1577
614.2927856445312 0 1994.1185
616.30517578125 0 1534.0145
617.2892456054688 0 4851.549
618.2672729492188 0 1852.4304
618.3273315429688 0 5597.024
619.3209228515625 0 4949.691
619.9779663085938 0 1993.5009
620.317138671875 0 2451.0627
620.6244506835938 0 1919.2404
620.8157958984375 0 2735.7776
620.9639282226562 0 2813.4385
621.3165283203125 0 1348.9147
621.8186645507812 0 1257.8942
622.325439453125 0 46143.125 y 11
622.9718017578125 0 2193.6912
623.3275146484375 0 13771.286
624.3264770507812 0 4705.8423
624.8207397460938 0 2653.6597
625.3190307617188 0 6555.7866
625.6513061523438 0 5279.405
625.9673461914062 0 5650.8247 y Water loss 1
626.3020629882812 0 8134.171 y Ammonia loss 1
626.6394653320312 0 7941.992
626.9752807617188 0 5285.1123
627.3090209960938 0 4602.3306
627.6431884765625 0 2375.2625
628.6450805664062 0 7097.823 b Water loss 14
628.9772338867188 0 8774.9375 b Ammonia loss 14
629.3103637695312 0 7044.8115
629.644287109375 0 2319.4734
630.3092651367188 0 1417.749
630.3656005859375 0 1226.9049
631.6366577148438 0 1963.2137
631.9678955078125 0 59943.93 y 1
632.3021850585938 0 64689.605
632.6358642578125 0 36754.81
632.9703369140625 0 19391.113
633.30419921875 0 5693.0063
634.31396484375 0 10142.969
634.6484985351562 0 31669.098 b 14
634.9826049804688 0 35648.516
635.3076171875 0 33122.387
635.6488037109375 0 8739.808
635.9724731445312 0 1982.9529
636.3341674804688 0 22384.537
636.7896728515625 0 1456.8794
637.3355712890625 0 8363.419
638.3387451171875 0 1992.6501
640.2752685546875 0 8929.887
640.6507568359375 0 6331.2046
640.98583984375 0 8996.159
641.2691040039062 0 2338.0645
641.3191528320312 0 2480.9019
641.6506958007812 0 2341.497
645.2814331054688 0 5963.7573
646.3306884765625 0 4961.776
646.8168334960938 0 2881.4023
647.3195190429688 0 7640.838
648.3172607421875 0 1528.6776
652.3236083984375 0 121770.336
653.3262939453125 0 49748.875
654.329833984375 0 10549.396
655.3282470703125 0 23747.146 y Water loss 5
655.8291625976562 0 19377.855 y Ammonia loss 5
656.3316040039062 0 8216.749
656.8270874023438 0 3428.9473
657.3386840820312 0 6040.738
658.285400390625 0 3706.4854
658.3412475585938 0 1960.1952
659.28759765625 0 1505.7528
662.3323364257812 0 31624.86
663.2940673828125 0 20954.918 b Ammonia loss 4
663.343017578125 0 7348.6416
663.8361206054688 0 1334.9678
664.3367309570312 0 233590.6 y 5
664.83642578125 0 135045.94
665.3391723632812 0 73114.836
665.8377685546875 0 21632.09
666.342529296875 0 10189.305
669.3248901367188 0 4784.6626
669.6630859375 0 1456.0627 Precursor
669.8273315429688 0 2317.1401
670.3317260742188 0 4873.2676
670.8335571289062 0 2560.4548
671.3271484375 0 1612.4363
678.3453979492188 0 2125.7163
679.3341674804688 0 2213.741
680.3189086914062 0 389451.94 b 4
681.3214721679688 0 153652.73
682.32470703125 0 35214.395
683.3271484375 0 8156.851
683.8255004882812 0 1141.158
684.3275146484375 0 3455.2825 b Water loss 10
684.8306884765625 0 1627.1809
685.3328857421875 0 1586.9954
688.3331298828125 0 3227.9119
689.3352661132812 0 1165.1193
690.3384399414062 0 2061.2764
691.3556518554688 0 1810.424
692.3110961914062 0 12480.506
693.317138671875 0 6120.38 b 10
693.8306884765625 0 1987.0094
694.318359375 0 2556.6067
695.8502807617188 0 1325.9896
696.3411865234375 0 2395.953
698.3554077148438 0 1938.9615
699.3807373046875 0 2447.3484
700.3828735351562 0 1341.2422
703.3458862304688 0 4198.5166
704.33837890625 0 1503.3622
704.8535766601562 0 4040.4797
705.35400390625 0 3353.5056
705.8561401367188 0 2345.4656
706.3440551757812 0 10257.953
707.345947265625 0 3051.21
708.3412475585938 0 2388.356
709.3471069335938 0 1467.8419
710.3211669921875 0 27189.977
711.3251953125 0 9334.012
712.324951171875 0 5870.086
712.8257446289062 0 2121.9644
713.3306274414062 0 3334.6387
715.3229370117188 0 1139.4844
717.39794921875 0 2218.9707
721.3544921875 0 28413.674
722.356689453125 0 13194.203
723.3599853515625 0 3915.1238
729.3564453125 0 3074.3884
730.3646240234375 0 1275.2764
739.3550415039062 0 1229.3351
740.3514404296875 0 1375.3567
745.3682250976562 0 1435.275
746.3614501953125 0 2060.2517
747.3915405273438 0 1381.3054
748.3743286132812 0 3992.2693 b Water loss 11
748.8734130859375 0 2113.3684 b Ammonia loss 11
749.3306274414062 0 1084.4974
749.3726806640625 0 1182.2153
749.8709106445312 0 1331.5833
750.341064453125 0 1639.0446
753.36279296875 0 1864.7122
753.8619995117188 0 2237.2732
754.3601684570312 0 1956.4283
757.3746948242188 0 11969.18 b 11
757.8768310546875 0 12078.611
758.3763427734375 0 6077.5366
758.8775024414062 0 1698.9779
759.3731079101562 0 1159.7164
760.3097534179688 0 1597.0767
762.3655395507812 0 2846.572
762.8684692382812 0 2321.9075
763.3809814453125 0 5378.6685
764.3788452148438 0 5210.343
765.4030151367188 0 4314.58
766.4059448242188 0 2454.2842
767.3461303710938 0 5135.4653
767.85693359375 0 2055.368
768.3546752929688 0 8258.727 y Ammonia loss 10
768.851806640625 0 2096.5886
769.3551635742188 0 2269.5645
771.378173828125 0 1170.7598
772.3651123046875 0 9706.218
773.368896484375 0 16631.297
774.3673095703125 0 7227.4175
775.3809204101562 0 4331.6943 b Water loss 5
776.3781127929688 0 3754.7515 b Ammonia loss 5
776.8605346679688 0 2369.4177
777.3598022460938 0 2442.9048
777.8652954101562 0 1426.6398
778.329833984375 0 1477.2258
781.3946533203125 0 11616.052
782.3969116210938 0 3988.6233
783.3983764648438 0 1422.4403
784.3697509765625 0 2766.0952
785.3890991210938 0 48321.934 y 10
786.3917846679688 0 19991.879
787.3907470703125 0 6973.3604
788.3920288085938 0 1268.3422
790.3744506835938 0 2435.75
791.3786010742188 0 58145.992
792.3804321289062 0 25629.908
793.3972778320312 0 27928.758 b 5
794.40185546875 0 11876.128
795.3457641601562 0 13785.801
796.3486328125 0 7444.033
797.3560791015625 0 2723.275
804.3868408203125 0 2112.037
805.8740234375 0 1563.6848 y Water loss 3
806.3804931640625 0 1546.8185 y Ammonia loss 3
808.3775634765625 0 2841.313
808.8825073242188 0 2510.5737
809.3887939453125 0 44025.3
809.8843383789062 0 1195.3811
810.3914184570312 0 17948.81
811.394287109375 0 5983.1567
812.3953857421875 0 3215.11
813.4015502929688 0 1421.4242
814.4048461914062 0 3790.4526
814.88818359375 0 6814.163 y 3
815.3939208984375 0 6364.385
815.8919067382812 0 2802.029
817.3942260742188 0 4133.841
817.8876953125 0 7296.1636
818.3880004882812 0 6363.6855
818.8870849609375 0 1841.0272
819.3801879882812 0 2571.0757
820.4234619140625 0 2166.9814
825.3930053710938 0 1301.69
826.3914794921875 0 7924.272
826.8965454101562 0 5506.0576
827.3971557617188 0 4657.79
827.8947143554688 0 1489.5863
828.390625 0 1565.5853
834.4365844726562 0 4801.3228
835.4397583007812 0 2026.7219
838.412109375 0 5271.993
839.4195556640625 0 1345.8129
843.3935546875 0 1136.6406
844.3850708007812 0 3957.2405
845.3909301757812 0 2730.9363
845.8945922851562 0 1567.1151
846.3987426757812 0 1412.9377
852.3671875 0 2368.5398
853.3644409179688 0 1322.7072
854.3797607421875 0 1426.2745
864.3593139648438 0 4601.206
865.3604736328125 0 1679.6686
866.3451538085938 0 1202.1223
869.3960571289062 0 1224.175
870.4010009765625 0 3345.3757 y Ammonia loss 2
871.4085083007812 0 1453.5977
878.41162109375 0 2778.908
878.9171142578125 0 1040.81 y 2
879.4148559570312 0 2782.255
880.417724609375 0 1339.4446
882.3692016601562 0 12543.053
883.3720092773438 0 4619.883
884.372802734375 0 1965.5212
885.4489135742188 0 1968.3131
886.4419555664062 0 1199.7775
890.4130249023438 0 1104.0204 b Water loss 6
891.4048461914062 0 1785.7428 b Ammonia loss 6
892.4212646484375 0 1515.4967
894.4247436523438 0 969.53015
895.4361572265625 0 13624.745
896.4266357421875 0 19547.686 y Water loss 9
897.4268798828125 0 9607.371
898.4234619140625 0 4274.1025
899.3990478515625 0 4498.2944
900.400146484375 0 2817.3132
901.3876342773438 0 956.2659
902.4290161132812 0 1096.6738
908.4288330078125 0 24992.15 b 6
909.432373046875 0 11142.506
910.4282836914062 0 3301.3267
914.4314575195312 0 54194.203 y 9
915.4349975585938 0 25401.75
916.4369506835938 0 10216.86
917.435791015625 0 2924.8037
919.47314453125 0 2174.6606
924.4215087890625 0 1285.8486
936.4251098632812 0 2171.9658
937.4012451171875 0 1636.709
937.4859008789062 0 2362.831
947.4439697265625 0 1251.4473 b Water loss 7
954.4344482421875 0 3627.0632
955.4221801757812 0 7243.7285
956.4301147460938 0 2582.8245
963.427978515625 0 1095.6066
965.4519653320312 0 5929.913 b 7
966.45703125 0 2915.119
967.4556884765625 0 1175.2937
972.44580078125 0 3824.6614
973.4481201171875 0 2492.4321
976.4954223632812 0 4620.368
977.4955444335938 0 1325.7896
980.4669189453125 0 1656.6523
981.4378051757812 0 3133.9724
982.4715576171875 0 3036.4639
983.4738159179688 0 1089.9059
992.4533081054688 0 3349.2
993.4457397460938 0 2766.6262
994.5029907226562 0 8827.524
995.5045776367188 0 4176.3257
996.460693359375 0 1015.45306
997.5104370117188 0 1091.4958
1010.46435546875 0 24250.588
1011.4683837890625 0 11118.293
1012.4712524414062 0 7115.9766
1013.4710693359375 0 2065.4495
1042.5252685546875 0 2544.6863 y 8
1043.5330810546875 0 1644.2207
1081.5391845703125 0 1531.2584 y Water loss 7
1091.5189208984375 0 5187.761
1092.511474609375 0 3536.3171
1093.534912109375 0 3272.0317 b 8
1094.542236328125 0 2571.633
1099.54638671875 0 5597.3916 y 7
1100.54931640625 0 3240.3687
1101.5531005859375 0 1984.2701
1109.530517578125 0 8956.626
1110.5345458984375 0 5444.378
1111.5438232421875 0 2403.1443
1123.5474853515625 0 4161.168
1124.5521240234375 0 2586.92
1125.5531005859375 0 1412.8214
1136.528076171875 0 927.11584
1197.5579833984375 0 999.9617 y Ammonia loss 6
1204.5928955078125 0 1166.7612 b Water loss 9
1214.5721435546875 0 3797.7808 y 6
1215.58154296875 0 2248.285
1216.5755615234375 0 978.03046
1217.585205078125 0 806.4009
1222.6151123046875 0 1437.6995
1223.617431640625 0 1175.3778
1246.581787109375 0 1047.7904
2865.501953125 0 780.3992

Spectrum Details

|  |  |
| --- | --- |
| Matched peaks? Matched peaks The total absolute number of peaks matched. Additionally in brackets the total fraction of peaks matched and the total number of peaks is shown. | 117 (7.99% of 1464) |
| FDR? FDR The false discovery rate estimated for this peptide. It is calculated by matching all theoretical fragments with a non-integer shift with the raw peaks for this spectrum. This is done with 40 different shifts. The resulting percentage is the average number of annotated peaks over the number of annotated peaks with the correct spectrum. | 0.69% |
| Satellite FDR? Satellite FDR See the FDR for details on its calculation. This satellite ion specific FDR only contains the satellite ions (d/w) for I/L/J positions. | - |
| PSM Score? PSM Score The PSM Score as given by Hecklib to this annotated spectrum. It is shown with three significant figures. | 456 |

#### Reverse Lookup? Reverse Lookup All places where this read could be placed.

| Group | Segment | Template | Template Part | Read Part | Score | Unique |
| --- | --- | --- | --- | --- | --- | --- |
| Homo sapiens Heavy Chain | IGHC | IGHG1 | [191..207] | [0..16] | 123 | False |
| Homo sapiens Heavy Chain | IGHC | IGHG3 | [238..254] | [0..16] | 123 | False |
| Homo sapiens Heavy Chain | IGHC | IGHG2 | [188..203] | [0..16] | 114 | False |
| Homo sapiens Heavy Chain | IGHC | IGHG4 | [188..204] | [0..16] | 123 | False |

| Recombined | Template Part | Read Part | Score | Unique |
| --- | --- | --- | --- | --- |
| REC-0-1 | [316..332] | [0..16] | 128 | True |

#### Meta Information from Multiple reads

##### Number of combined reads

3

##### Intensity

0.6926

##### TotalArea

1.137E+08

##### Changes to the peptide sequence

JHQDWJDGKEYKCKVS

L→JNo support for either Leucine or Isoleucine based on side chain ions (Position: 6)

L→JNo support for either Leucine or Isoleucine based on side chain ions (Position: 1)

#### Positional Score

Copy Data

##### Positional Score (TSV)

###### Preview

```
Loading example...
```

*Click on the button to copy the data to your clipboard.*

100123456789101112131415

Label Value
"0" 0.333
"1" 0.327
"2" 0.313
"3" 0.317
"4" 0.323
"5" 0.333
"6" 0.333
"7" 0.327
"8" 0.323
"9" 0.33
"10" 0.323
"11" 0.3
"12" 0.253
"13" 0.317
"14" 0.297
"15" 0.32

#### Meta Information from PEAKS

##### Scan Identifier

F1:5024

##### Original sequence

L

H

Q

D

W

L

D

G

K

E

Y

K

C

+58.01

K

V

S

##### Posttranslational Modifications

Carboxymethyl

##### Source File

D:\separate\_stitch\_analyses\xle-disambiguation\raw\20210323\_F1\_UM1\_Peng0013\_SA\_F59\_ingel\_3ug\_ELA.raw

##### Fraction

1

##### Scan Feature

F1:11461

##### De Novo Score

99

##### ConfidenceScore

99

### m/z

669.6603

##### Mass

2005.9568

##### Charge

3

##### Retention Time

26.79

##### Predicted Retention Time

-

##### Area

3.382E+07

##### Parts Per Million

1.1

##### Fragmentation mode

ETHCD

##### Originating file

01 D:\separate\_stitch\_analyses\xle-disambiguation\20210325\_F59\_3ug\_DENOVO\_12.csv

#### Meta Information from PEAKS

##### Scan Identifier

F1:5098

##### Original sequence

L

H

Q

D

W

L

D

G

K

E

Y

K

C

+58.01

K

V

S

##### Posttranslational Modifications

Carboxymethyl

##### Source File

D:\separate\_stitch\_analyses\xle-disambiguation\raw\20210323\_F1\_UM1\_Peng0013\_SA\_F59\_ingel\_3ug\_ELA.raw

##### Fraction

1

##### Scan Feature

F1:11461

##### De Novo Score

99

##### ConfidenceScore

99

### m/z

669.6603

##### Mass

2005.9568

##### Charge

3

##### Retention Time

26.79

##### Predicted Retention Time

-

##### Area

3.382E+07

##### Parts Per Million

1.1

##### Fragmentation mode

ETHCD

##### Originating file

01 D:\separate\_stitch\_analyses\xle-disambiguation\20210325\_F59\_3ug\_DENOVO\_12.csv

#### Meta Information from PEAKS

##### Scan Identifier

F1:5274

##### Original sequence

L

H

Q

D

W

L

D

G

K

E

Y

K

C

+58.01

K

V

S

##### Posttranslational Modifications

Carboxymethyl

##### Source File

D:\separate\_stitch\_analyses\xle-disambiguation\raw\20210323\_F1\_UM1\_Peng0013\_SA\_F59\_ingel\_3ug\_ELA.raw

##### Fraction

1

##### Scan Feature

F1:4129

##### De Novo Score

95

##### ConfidenceScore

95

### m/z

502.4968

##### Mass

2005.9568

##### Charge

4

##### Retention Time

27.6

##### Predicted Retention Time

-

##### Area

4.602E+07

##### Parts Per Million

0.6

##### Fragmentation mode

HCD

##### Originating file

01 D:\separate\_stitch\_analyses\xle-disambiguation\20210325\_F59\_3ug\_DENOVO\_12.csv
