## Supplementary material for "A handle on mass coincidence errors in *de novo* sequencing of antibodies by bottom-up proteomics": Combined_026.html

Details Combined\_026 | Stitch OverviewUndefined

### Read Combined\_026

#### Sequence (length=7)

JASYLDK

#### Spectrum 3928? Spectrum 3928 The raw spectrum of this peptide as annotated by Hecklib. The fragments are coloured according to ion type (see legend). Any peaks with a star '\*' as text can be hovered over to see the full details, first the ion type second the mass shift type. By hovering over the amino acids in the peptide or ions in the legend the corresponding peaks are highlighted. By toggling the 'Unassigned' label you can turn the background (unassigned) peaks on or off in the plot. By updating the slider in the Ion legend you can update the spectrum to only show the top X% of the peaks with labels. The top X% means any peak that is within X% of the highest intensity. By dragging in the spectrum you can zoom in to a specific part of the spectrum and use 'Zoom Out' to get back to the original zoom level. The annotation of the spectrum is based on the given sequence in the peptides file and is done with different software so inconsistencies are likely. The peaks are annotated based on the given sequence, with 20 ppm tolerance.

Copy Data

##### Spectrum 3928 (TSV)

###### Preview

```
Loading example...
```

*Click on the button to copy the data to your clipboard.*

Mz MinMz MaxIntensity Max

WidthHeightPeptide font sizePeptide stroke widthSpectrum font sizeSpectrum stroke widthCompact peptide

Ion legend

wxyz

abcd

OtherUnassignedIon^Charge^_Position_Show for top:%

JASYLDK

01.30e+42.60e+43.90e+45.19e+4

Zoom Out

y^+1^_1_y^+1^_1_a^+1^_2_b^+1^_2_y^+1^_2_y^+1^_2_y^+1^_2_b^+1^_3_y^+1^_3_y^+1^_3_\*\*b^+1^_4_y^+1^_4_y^+1^_5_y^+1^_5_y^+1^_6_y^+1^_6_

0537107416112148

Fragment Matches Table

Show background peaks

| Position | Ion type | Intensity | mz Theoretical | mz Error (Th) | mz Error (ppm) | Charge | Series Number |
| --- | --- | --- | --- | --- | --- | --- | --- |
| - | - | 559.5 | 120.1 | - | - | 0 | - |
| - | - | 355.2 | 124.1 | - | - | 0 | - |
| - | - | 454.8 | 129 | - | - | 0 | - |
| - | - | 9840 | 129 | - | - | 0 | - |
| - | - | 1.257E+04 | 129.1 | - | - | 0 | - |
| - | - | 3235 | 130 | - | - | 0 | - |
| 7 | y | 9871 | 130.1 | 0.0002012 | 1.547 | +1 | 1 |
| - | - | 613.5 | 130.1 | - | - | 0 | - |
| - | - | 565.8 | 131.1 | - | - | 0 | - |
| - | - | 549.7 | 133.1 | - | - | 0 | - |
| - | - | 4.167E+04 | 136.1 | - | - | 0 | - |
| - | - | 3305 | 137.1 | - | - | 0 | - |
| - | - | 372.2 | 138.5 | - | - | 0 | - |
| - | - | 1919 | 139 | - | - | 0 | - |
| - | - | 1231 | 140 | - | - | 0 | - |
| - | - | 517.2 | 140.1 | - | - | 0 | - |
| - | - | 755.6 | 141.1 | - | - | 0 | - |
| 7 | y | 1.58E+04 | 147.1 | 0.0001567 | 1.065 | +1 | 1 |
| - | - | 1053 | 148.1 | - | - | 0 | - |
| - | - | 3907 | 149 | - | - | 0 | - |
| - | - | 4594 | 157 | - | - | 0 | - |
| - | - | 535.7 | 157.1 | - | - | 0 | - |
| 2 | a | 3.768E+04 | 157.1 | 0.0001427 | 0.9078 | +1 | 2 |
| - | - | 1844 | 158 | - | - | 0 | - |
| - | - | 541.1 | 158.1 | - | - | 0 | - |
| - | - | 2649 | 158.1 | - | - | 0 | - |
| - | - | 1587 | 159.1 | - | - | 0 | - |
| - | - | 1608 | 160.1 | - | - | 0 | - |
| - | - | 485.8 | 167.1 | - | - | 0 | - |
| - | - | 470.6 | 171.1 | - | - | 0 | - |
| - | - | 523.3 | 171.2 | - | - | 0 | - |
| - | - | 2298 | 173.5 | - | - | 0 | - |
| - | - | 785.8 | 175.1 | - | - | 0 | - |
| - | - | 1962 | 178.1 | - | - | 0 | - |
| - | - | 852.2 | 179 | - | - | 0 | - |
| - | - | 718.9 | 183.1 | - | - | 0 | - |
| - | - | 863.1 | 184.1 | - | - | 0 | - |
| - | - | 5471 | 185.1 | - | - | 0 | - |
| - | - | 977.5 | 185.1 | - | - | 0 | - |
| 2 | b | 2.127E+04 | 185.1 | 0.0001163 | 0.6284 | +1 | 2 |
| - | - | 3507 | 186.1 | - | - | 0 | - |
| - | - | 2227 | 186.1 | - | - | 0 | - |
| - | - | 688.7 | 188.1 | - | - | 0 | - |
| - | - | 538.3 | 193.9 | - | - | 0 | - |
| - | - | 675.2 | 198.1 | - | - | 0 | - |
| - | - | 1181 | 201.1 | - | - | 0 | - |
| - | - | 1207 | 205.1 | - | - | 0 | - |
| - | - | 2547 | 209.1 | - | - | 0 | - |
| - | - | 823.1 | 217 | - | - | 0 | - |
| - | - | 2957 | 223.1 | - | - | 0 | - |
| - | - | 2.894E+04 | 223.1 | - | - | 0 | - |
| - | - | 594.8 | 224.1 | - | - | 0 | - |
| - | - | 3543 | 224.1 | - | - | 0 | - |
| - | - | 1581 | 225 | - | - | 0 | - |
| - | - | 3826 | 226.1 | - | - | 0 | - |
| - | - | 640.9 | 227 | - | - | 0 | - |
| - | - | 1018 | 227 | - | - | 0 | - |
| - | - | 1388 | 229.1 | - | - | 0 | - |
| - | - | 1092 | 233.1 | - | - | 0 | - |
| - | - | 533.2 | 240.1 | - | - | 0 | - |
| 6 | y | 4501 | 244.1 | 4.737E-05 | 0.194 | +1 | 2 |
| 6 | y | 3509 | 245.1 | 5.264E-05 | 0.2148 | +1 | 2 |
| - | - | 3429 | 248 | - | - | 0 | - |
| - | - | 3402 | 249.2 | - | - | 0 | - |
| - | - | 1.263E+04 | 251.1 | - | - | 0 | - |
| - | - | 1349 | 252.1 | - | - | 0 | - |
| - | - | 1205 | 259.2 | - | - | 0 | - |
| - | - | 1470 | 260.2 | - | - | 0 | - |
| 6 | y | 2.908E+04 | 262.1 | 9.875E-05 | 0.3767 | +1 | 2 |
| - | - | 3610 | 263.1 | - | - | 0 | - |
| - | - | 752.8 | 264.1 | - | - | 0 | - |
| - | - | 690.1 | 266 | - | - | 0 | - |
| 3 | b | 2026 | 272.2 | 5.172E-05 | 0.19 | +1 | 3 |
| - | - | 1165 | 276.1 | - | - | 0 | - |
| - | - | 1479 | 277.2 | - | - | 0 | - |
| - | - | 599.1 | 284 | - | - | 0 | - |
| - | - | 589.6 | 285.8 | - | - | 0 | - |
| - | - | 607 | 297.1 | - | - | 0 | - |
| - | - | 555.5 | 297.1 | - | - | 0 | - |
| - | - | 2839 | 299.1 | - | - | 0 | - |
| - | - | 1038 | 300.1 | - | - | 0 | - |
| - | - | 992.1 | 301.1 | - | - | 0 | - |
| - | - | 9583 | 304.1 | - | - | 0 | - |
| - | - | 1438 | 305.1 | - | - | 0 | - |
| - | - | 627.2 | 316.9 | - | - | 0 | - |
| - | - | 3404 | 322.1 | - | - | 0 | - |
| - | - | 820 | 323.1 | - | - | 0 | - |
| - | - | 864.7 | 346.2 | - | - | 0 | - |
| - | - | 658.1 | 351.2 | - | - | 0 | - |
| 5 | y | 1239 | 357.2 | 0.0004779 | 1.338 | +1 | 3 |
| - | - | 2081 | 364.2 | - | - | 0 | - |
| - | - | 585 | 374.3 | - | - | 0 | - |
| - | - | 859.6 | 374.7 | - | - | 0 | - |
| 5 | y | 1.58E+04 | 375.2 | 0.0003309 | 0.882 | +1 | 3 |
| - | - | 2984 | 376.2 | - | - | 0 | - |
| - | - | 903 | 389.2 | - | - | 0 | - |
| - | - | 2473 | 392.2 | - | - | 0 | - |
| 0 | Precursor | 3912 | 396.2 | 6.859E-05 | 0.1731 | +2 | -1 |
| 0 | Precursor | 939.1 | 405.2 | 0.0001784 | 0.4401 | +2 | -1 |
| - | - | 930.9 | 407.2 | - | - | 0 | - |
| - | - | 1164 | 412.2 | - | - | 0 | - |
| 4 | b | 1385 | 435.2 | 0.0006056 | 1.391 | +1 | 4 |
| - | - | 620.6 | 469.9 | - | - | 0 | - |
| - | - | 2169 | 479.2 | - | - | 0 | - |
| - | - | 743.6 | 480.2 | - | - | 0 | - |
| - | - | 658.6 | 532.2 | - | - | 0 | - |
| - | - | 574.2 | 536.4 | - | - | 0 | - |
| 4 | y | 6640 | 538.3 | 0.0005186 | 0.9635 | +1 | 4 |
| - | - | 1811 | 539.3 | - | - | 0 | - |
| - | - | 561.1 | 542.5 | - | - | 0 | - |
| - | - | 1586 | 550.2 | - | - | 0 | - |
| - | - | 563.9 | 570.4 | - | - | 0 | - |
| 3 | y | 1054 | 607.3 | 0.0006617 | 1.09 | +1 | 5 |
| 3 | y | 5.142E+04 | 625.3 | 0.0009308 | 1.489 | +1 | 5 |
| - | - | 1.89E+04 | 626.3 | - | - | 0 | - |
| - | - | 647.6 | 626.4 | - | - | 0 | - |
| - | - | 4230 | 627.3 | - | - | 0 | - |
| 2 | y | 6168 | 678.3 | 0.001357 | 2 | +1 | 6 |
| - | - | 1877 | 679.3 | - | - | 0 | - |
| - | - | 656.1 | 680.3 | - | - | 0 | - |
| 2 | y | 3.859E+04 | 696.4 | 0.001546 | 2.22 | +1 | 6 |
| - | - | 1.42E+04 | 697.4 | - | - | 0 | - |
| - | - | 4092 | 698.4 | - | - | 0 | - |
| - | - | 685.9 | 1711 | - | - | 0 | - |
| - | - | 665.3 | 1943 | - | - | 0 | - |
| - | - | 639.5 | 2127 | - | - | 0 | - |

m/z Charge Intensity FragmentType MassShift Position
120.0807876586914 0 559.5138
124.13679504394531 0 355.19254
129.01290893554688 0 454.83765
129.01841735839844 0 9840.478
129.1024169921875 0 12567.581
130.02191162109375 0 3234.8215
130.08645629882812 0 9871.079 y Ammonia loss 6
130.10552978515625 0 613.5336
131.0707550048828 0 565.79694
133.08584594726562 0 549.7343
136.0758514404297 0 41669.71
137.0791778564453 0 3305.446
138.5235137939453 0 372.24725
139.00279235839844 0 1918.9174
140.00607299804688 0 1230.5154
140.1075897216797 0 517.15094
141.06605529785156 0 755.5946
147.1129608154297 0 15796.565 y 6
148.11627197265625 0 1052.7246
149.02340698242188 0 3907.2234
157.0132293701172 0 4593.56
157.09768676757812 0 535.713
157.13368225097656 0 37676.742 a 1
158.01661682128906 0 1843.5094
158.1306610107422 0 541.1173
158.1370849609375 0 2649.3665
159.07652282714844 0 1587.0896
160.0759735107422 0 1608.206
167.05612182617188 0 485.7788
171.13771057128906 0 470.55676
171.16770935058594 0 523.33075
173.45053100585938 0 2297.5627
175.0972442626953 0 785.8262
178.08628845214844 0 1962.1626
179.00038146972656 0 852.2161
183.11228942871094 0 718.8764
184.1079559326172 0 863.086
185.08087158203125 0 5470.588
185.09017944335938 0 977.5431
185.12857055664062 0 21274.145 b 1
186.08421325683594 0 3506.5032
186.13211059570312 0 2226.6887
188.0701446533203 0 688.65356
193.93475341796875 0 538.3264
198.12338256835938 0 675.19977
201.12351989746094 0 1180.6456
205.09762573242188 0 1207.0486
209.09225463867188 0 2547.1294
217.03396606445312 0 823.1245
223.06370544433594 0 2956.5063
223.10772705078125 0 28940.945
224.06317138671875 0 594.79706
224.11106872558594 0 3543.157
225.0430450439453 0 1581.2361
226.1185760498047 0 3826.173
227.0218963623047 0 640.9384
227.03807067871094 0 1017.6736
229.11831665039062 0 1388.019
233.092041015625 0 1091.6484
240.080810546875 0 533.15985
244.12913513183594 0 4501.331 y Water loss 5
245.11325073242188 0 3508.8032 y Ammonia loss 5
248.0343017578125 0 3428.691
249.1597137451172 0 3401.7544
251.1027069091797 0 12629.317
252.1058349609375 0 1349.4203
259.1544189453125 0 1205.4846
260.1573791503906 0 1469.9017
262.1396484375 0 29077.13 y 5
263.1425476074219 0 3609.9705
264.1441955566406 0 752.8431
266.0433349609375 0 690.08307
272.1604309082031 0 2026.118 b 2
276.1343078613281 0 1164.9661
277.1543884277344 0 1479.4193
284.029052734375 0 599.0828
285.81292724609375 0 589.6471
297.0825500488281 0 606.992
297.0997619628906 0 555.4904
299.0617980957031 0 2838.571
300.0626220703125 0 1037.9607
301.05859375 0 992.10034
304.12908935546875 0 9582.578
305.132568359375 0 1438.0509
316.8788757324219 0 627.1934
322.1397399902344 0 3403.9343
323.14166259765625 0 819.9628
346.17486572265625 0 864.68756
351.201904296875 0 658.07477
357.2127685546875 0 1239.0344 y Water loss 4
364.18609619140625 0 2080.7532
374.31439208984375 0 585.0159
374.6972961425781 0 859.6221
375.2234802246094 0 15804.989 y 4
376.22686767578125 0 2984.0674
389.2184753417969 0 903.0282
392.181396484375 0 2472.8796
396.2185974121094 0 3911.592 Precursor Water loss
405.2236328125 0 939.0849 Precursor
407.2283630371094 0 930.94946
412.22015380859375 0 1163.8545
435.22320556640625 0 1384.5431 b 3
469.8988342285156 0 620.5808
479.21295166015625 0 2168.749
480.2149353027344 0 743.58105
532.2391357421875 0 658.5918
536.39794921875 0 574.20184
538.28662109375 0 6640.2744 y 3
539.2901611328125 0 1811.4309
542.4983520507812 0 561.1283
550.2499389648438 0 1585.7563
570.4082641601562 0 563.9074
607.3092651367188 0 1053.753 y Water loss 2
625.3182373046875 0 51422.188 y 2
626.3212280273438 0 18898.46
626.3814697265625 0 647.62823
627.3231811523438 0 4229.653
678.3443603515625 0 6168.039 y Water loss 1
679.3486328125 0 1876.901
680.3488159179688 0 656.1134
696.354736328125 0 38594.227 y 1
697.3578491210938 0 14195.15
698.3609619140625 0 4092.2297
1711.08154296875 0 685.94684
1943.1783447265625 0 665.2829
2126.744140625 0 639.51135

Spectrum Details

|  |  |
| --- | --- |
| Matched peaks? Matched peaks The total absolute number of peaks matched. Additionally in brackets the total fraction of peaks matched and the total number of peaks is shown. | 18 (14.29% of 126) |
| FDR? FDR The false discovery rate estimated for this peptide. It is calculated by matching all theoretical fragments with a non-integer shift with the raw peaks for this spectrum. This is done with 40 different shifts. The resulting percentage is the average number of annotated peaks over the number of annotated peaks with the correct spectrum. | 2.12% |
| Satellite FDR? Satellite FDR See the FDR for details on its calculation. This satellite ion specific FDR only contains the satellite ions (d/w) for I/L/J positions. | - |
| PSM Score? PSM Score The PSM Score as given by Hecklib to this annotated spectrum. It is shown with three significant figures. | 233 |

#### Spectrum 3979? Spectrum 3979 The raw spectrum of this peptide as annotated by Hecklib. The fragments are coloured according to ion type (see legend). Any peaks with a star '\*' as text can be hovered over to see the full details, first the ion type second the mass shift type. By hovering over the amino acids in the peptide or ions in the legend the corresponding peaks are highlighted. By toggling the 'Unassigned' label you can turn the background (unassigned) peaks on or off in the plot. By updating the slider in the Ion legend you can update the spectrum to only show the top X% of the peaks with labels. The top X% means any peak that is within X% of the highest intensity. By dragging in the spectrum you can zoom in to a specific part of the spectrum and use 'Zoom Out' to get back to the original zoom level. The annotation of the spectrum is based on the given sequence in the peptides file and is done with different software so inconsistencies are likely. The peaks are annotated based on the given sequence, with 20 ppm tolerance.

Copy Data

##### Spectrum 3979 (TSV)

###### Preview

```
Loading example...
```

*Click on the button to copy the data to your clipboard.*

Mz MinMz MaxIntensity Max

WidthHeightPeptide font sizePeptide stroke widthSpectrum font sizeSpectrum stroke widthCompact peptide

Ion legend

wxyz

abcd

OtherUnassignedIon^Charge^_Position_Show for top:%

JASYLDK

01.83e+43.67e+45.50e+47.33e+4

Zoom Out

y^+1^_1_z^+1^_1_y^+1^_1_z^+1^_2_y^+1^_2_c^+1^_3_y^+2^_5_y^+2^_5_w^+1^_3_z^+1^_3_y^+1^_3_c^+1^_4_z^+1^_4_y^+1^_4_c^+1^_5_w^+1^_5_y^+1^_5_z^+1^_5_y^+1^_5_y^+1^_6_c^+1^_6_y^+1^_6_

0640128019202560

Fragment Matches Table

Show background peaks

| Position | Ion type | Intensity | mz Theoretical | mz Error (Th) | mz Error (ppm) | Charge | Series Number |
| --- | --- | --- | --- | --- | --- | --- | --- |
| - | - | 343.2 | 128 | - | - | 0 | - |
| - | - | 463.1 | 128.6 | - | - | 0 | - |
| - | - | 2640 | 129 | - | - | 0 | - |
| - | - | 426.3 | 129 | - | - | 0 | - |
| - | - | 694.6 | 129.1 | - | - | 0 | - |
| - | - | 563.4 | 130 | - | - | 0 | - |
| 7 | y | 1024 | 130.1 | 0.0001707 | 1.312 | +1 | 1 |
| 7 | z | 519.8 | 131.1 | 3.612E-05 | 0.2756 | +1 | 1 |
| - | - | 401.9 | 133.6 | - | - | 0 | - |
| - | - | 1334 | 139 | - | - | 0 | - |
| - | - | 543.9 | 140 | - | - | 0 | - |
| 7 | y | 2721 | 147.1 | 0.000118 | 0.8021 | +1 | 1 |
| - | - | 475.5 | 148.9 | - | - | 0 | - |
| - | - | 550.7 | 148.9 | - | - | 0 | - |
| - | - | 785.8 | 148.9 | - | - | 0 | - |
| - | - | 628.7 | 148.9 | - | - | 0 | - |
| - | - | 671.8 | 148.9 | - | - | 0 | - |
| - | - | 1539 | 148.9 | - | - | 0 | - |
| - | - | 2208 | 148.9 | - | - | 0 | - |
| - | - | 4843 | 149 | - | - | 0 | - |
| - | - | 2472 | 149 | - | - | 0 | - |
| - | - | 1787 | 149 | - | - | 0 | - |
| - | - | 967.7 | 149 | - | - | 0 | - |
| - | - | 586.1 | 149 | - | - | 0 | - |
| - | - | 884.7 | 149 | - | - | 0 | - |
| - | - | 1420 | 149 | - | - | 0 | - |
| - | - | 401.1 | 149.1 | - | - | 0 | - |
| - | - | 2739 | 157 | - | - | 0 | - |
| - | - | 1.271E+04 | 157.1 | - | - | 0 | - |
| - | - | 1239 | 158 | - | - | 0 | - |
| - | - | 754.4 | 158.1 | - | - | 0 | - |
| - | - | 408.8 | 173.5 | - | - | 0 | - |
| - | - | 604.1 | 175.1 | - | - | 0 | - |
| - | - | 4377 | 185.1 | - | - | 0 | - |
| - | - | 8867 | 185.1 | - | - | 0 | - |
| - | - | 3246 | 186.1 | - | - | 0 | - |
| - | - | 468.1 | 201.1 | - | - | 0 | - |
| - | - | 1340 | 217 | - | - | 0 | - |
| - | - | 849.2 | 218 | - | - | 0 | - |
| - | - | 916.4 | 223.1 | - | - | 0 | - |
| - | - | 1128 | 223.1 | - | - | 0 | - |
| 6 | z | 3084 | 246.1 | 0.000204 | 0.829 | +1 | 2 |
| - | - | 1514 | 247.1 | - | - | 0 | - |
| - | - | 2841 | 248 | - | - | 0 | - |
| - | - | 586.7 | 249 | - | - | 0 | - |
| - | - | 2387 | 251.1 | - | - | 0 | - |
| - | - | 1473 | 259.2 | - | - | 0 | - |
| - | - | 3056 | 260.2 | - | - | 0 | - |
| 6 | y | 5999 | 262.1 | 0.0001903 | 0.726 | +1 | 2 |
| - | - | 639.7 | 266 | - | - | 0 | - |
| - | - | 571 | 270.1 | - | - | 0 | - |
| - | - | 991 | 272.2 | - | - | 0 | - |
| 3 | c | 519.5 | 289.2 | 0.0005693 | 1.969 | +1 | 3 |
| - | - | 561.8 | 293 | - | - | 0 | - |
| - | - | 1338 | 299.1 | - | - | 0 | - |
| - | - | 1084 | 304.1 | - | - | 0 | - |
| 3 | y | 548.6 | 304.2 | 0.000843 | 2.772 | +2 | 5 |
| 3 | y | 507.7 | 313.2 | 0.001664 | 5.314 | +2 | 5 |
| 5 | w | 1.598E+04 | 316.2 | 0.0001044 | 0.3301 | +1 | 3 |
| - | - | 2182 | 317.2 | - | - | 0 | - |
| - | - | 1019 | 322.1 | - | - | 0 | - |
| - | - | 526.9 | 343.2 | - | - | 0 | - |
| - | - | 615.7 | 353.7 | - | - | 0 | - |
| 5 | z | 2.906E+04 | 359.2 | 0.0005278 | 1.469 | +1 | 3 |
| - | - | 2.232E+04 | 360.2 | - | - | 0 | - |
| - | - | 4763 | 361.2 | - | - | 0 | - |
| - | - | 637.5 | 362.2 | - | - | 0 | - |
| - | - | 964.3 | 364.2 | - | - | 0 | - |
| 5 | y | 5013 | 375.2 | 0.000392 | 1.045 | +1 | 3 |
| - | - | 600.4 | 381.2 | - | - | 0 | - |
| - | - | 1322 | 396.2 | - | - | 0 | - |
| - | - | 577.7 | 404.8 | - | - | 0 | - |
| - | - | 680.8 | 405.2 | - | - | 0 | - |
| - | - | 839.2 | 407.2 | - | - | 0 | - |
| - | - | 799.9 | 422.2 | - | - | 0 | - |
| - | - | 2474 | 435.2 | - | - | 0 | - |
| - | - | 2430 | 451.2 | - | - | 0 | - |
| 4 | c | 1670 | 452.3 | 0.001337 | 2.956 | +1 | 4 |
| - | - | 3778 | 466.2 | - | - | 0 | - |
| - | - | 1282 | 467.2 | - | - | 0 | - |
| - | - | 664.7 | 478.3 | - | - | 0 | - |
| - | - | 1.194E+04 | 479.2 | - | - | 0 | - |
| - | - | 2534 | 480.2 | - | - | 0 | - |
| - | - | 609.7 | 480.3 | - | - | 0 | - |
| - | - | 700.8 | 507.2 | - | - | 0 | - |
| - | - | 1836 | 520.3 | - | - | 0 | - |
| - | - | 730.7 | 521.3 | - | - | 0 | - |
| 4 | z | 1.122E+04 | 522.3 | 0.0005323 | 1.019 | +1 | 4 |
| - | - | 4606 | 523.3 | - | - | 0 | - |
| - | - | 1483 | 530.3 | - | - | 0 | - |
| - | - | 860.1 | 531.3 | - | - | 0 | - |
| 4 | y | 4355 | 538.3 | 0.0005797 | 1.077 | +1 | 4 |
| - | - | 1403 | 539.3 | - | - | 0 | - |
| - | - | 1672 | 547.3 | - | - | 0 | - |
| - | - | 1.022E+04 | 548.3 | - | - | 0 | - |
| - | - | 3403 | 549.3 | - | - | 0 | - |
| - | - | 678 | 550.3 | - | - | 0 | - |
| - | - | 686.3 | 553.2 | - | - | 0 | - |
| - | - | 1.154E+04 | 564.3 | - | - | 0 | - |
| 5 | c | 2.722E+04 | 565.3 | 0.0009892 | 1.75 | +1 | 5 |
| - | - | 4430 | 566.2 | - | - | 0 | - |
| - | - | 1.037E+04 | 566.3 | - | - | 0 | - |
| - | - | 1347 | 567.2 | - | - | 0 | - |
| - | - | 2054 | 567.3 | - | - | 0 | - |
| - | - | 562.7 | 582.1 | - | - | 0 | - |
| 3 | w | 1849 | 592.3 | 0.0008294 | 1.4 | +1 | 5 |
| 3 | y | 780.6 | 607.3 | 0.002024 | 3.332 | +1 | 5 |
| 3 | z | 1.101E+04 | 609.3 | 0.0005173 | 0.849 | +1 | 5 |
| - | - | 3663 | 610.3 | - | - | 0 | - |
| - | - | 630.5 | 611.3 | - | - | 0 | - |
| - | - | 1518 | 624.3 | - | - | 0 | - |
| 3 | y | 4.189E+04 | 625.3 | 0.0004425 | 0.7077 | +1 | 5 |
| - | - | 1.168E+04 | 626.3 | - | - | 0 | - |
| - | - | 2684 | 627.3 | - | - | 0 | - |
| - | - | 856.2 | 636.3 | - | - | 0 | - |
| - | - | 670.2 | 637.3 | - | - | 0 | - |
| - | - | 575.7 | 638.3 | - | - | 0 | - |
| - | - | 6171 | 663.3 | - | - | 0 | - |
| - | - | 1834 | 664.3 | - | - | 0 | - |
| 2 | y | 3495 | 678.3 | 0.0008686 | 1.28 | +1 | 6 |
| - | - | 1950 | 679.3 | - | - | 0 | - |
| 6 | c | 7.259E+04 | 680.4 | 0.001443 | 2.121 | +1 | 6 |
| - | - | 2.438E+04 | 681.4 | - | - | 0 | - |
| - | - | 4624 | 682.4 | - | - | 0 | - |
| - | - | 2697 | 694.4 | - | - | 0 | - |
| - | - | 815.4 | 695.4 | - | - | 0 | - |
| 2 | y | 1.834E+04 | 696.4 | 0.0006911 | 0.9924 | +1 | 6 |
| - | - | 6003 | 697.4 | - | - | 0 | - |
| - | - | 983 | 698.4 | - | - | 0 | - |
| - | - | 2783 | 737.4 | - | - | 0 | - |
| - | - | 776.1 | 739.4 | - | - | 0 | - |
| - | - | 1.594E+04 | 750.4 | - | - | 0 | - |
| - | - | 7722 | 750.4 | - | - | 0 | - |
| - | - | 6331 | 751.4 | - | - | 0 | - |
| - | - | 2860 | 751.4 | - | - | 0 | - |
| - | - | 1497 | 752.4 | - | - | 0 | - |
| - | - | 693 | 764.4 | - | - | 0 | - |
| - | - | 1.664E+04 | 793.4 | - | - | 0 | - |
| - | - | 6561 | 794.4 | - | - | 0 | - |
| - | - | 1862 | 795.4 | - | - | 0 | - |
| - | - | 3.379E+04 | 809.4 | - | - | 0 | - |
| - | - | 2.706E+04 | 810.4 | - | - | 0 | - |
| - | - | 9115 | 811.4 | - | - | 0 | - |
| - | - | 1507 | 812.5 | - | - | 0 | - |
| - | - | 640.7 | 1002 | - | - | 0 | - |
| - | - | 676 | 1409 | - | - | 0 | - |
| - | - | 623.8 | 1500 | - | - | 0 | - |
| - | - | 749.1 | 2331 | - | - | 0 | - |
| - | - | 821.3 | 2366 | - | - | 0 | - |
| - | - | 628.9 | 2393 | - | - | 0 | - |
| - | - | 613.1 | 2535 | - | - | 0 | - |

m/z Charge Intensity FragmentType MassShift Position
127.99368286132812 0 343.1597
128.55947875976562 0 463.0643
129.01820373535156 0 2640.0762
129.0226593017578 0 426.33057
129.10247802734375 0 694.5839
130.02210998535156 0 563.4133
130.08642578125 0 1023.7832 y Ammonia loss 6
131.0941162109375 0 519.7643 z 6
133.5539093017578 0 401.93216
139.00247192382812 0 1333.8317
140.0060577392578 0 543.8547
147.11268615722656 0 2720.5444 y 6
148.86834716796875 0 475.52582
148.9104766845703 0 550.73914
148.9208984375 0 785.8319
148.92628479003906 0 628.6529
148.93133544921875 0 671.8146
148.9365997314453 0 1538.7993
148.94276428222656 0 2208.1567
148.95472717285156 0 4843.159
148.96006774902344 0 2471.75
148.96597290039062 0 1786.6161
148.97140502929688 0 967.732
148.9764404296875 0 586.12354
148.98202514648438 0 884.73724
149.02349853515625 0 1420.3026
149.0518035888672 0 401.1267
157.01304626464844 0 2738.8643
157.1334228515625 0 12707.413
158.01661682128906 0 1238.9998
158.13671875 0 754.35815
173.4566650390625 0 408.84406
175.09666442871094 0 604.12274
185.0806427001953 0 4376.6113
185.12832641601562 0 8866.574
186.0841064453125 0 3245.5142
201.05908203125 0 468.08105
217.03408813476562 0 1339.8645
218.0375213623047 0 849.1917
223.0640106201172 0 916.44257
223.10723876953125 0 1128.3965
246.12081909179688 0 3084.211 z 5
247.1284942626953 0 1514.3712
248.0338897705078 0 2840.9773
249.03611755371094 0 586.7479
251.10260009765625 0 2387.4202
259.1535949707031 0 1473.1685
260.15692138671875 0 3055.6604
262.1395568847656 0 5999.4326 y 5
266.0447692871094 0 639.7208
270.0913391113281 0 571.013
272.16033935546875 0 991.0416
289.18646240234375 0 519.5101 c 2
293.04815673828125 0 561.8415
299.0619201660156 0 1337.903
304.1289978027344 0 1083.844
304.1587829589844 0 548.6421 y Water loss 2
313.1648864746094 0 507.74075 y 2
316.15020751953125 0 15978.592 w 4
317.153564453125 0 2181.9072
322.1391296386719 0 1019.23175
343.1773681640625 0 526.9019
353.673828125 0 615.7071
359.2045593261719 0 29055.234 z 4
360.2118835449219 0 22323.463
361.2151794433594 0 4762.501
362.2193298339844 0 637.5188
364.1869201660156 0 964.3376
375.2234191894531 0 5012.832 y 4
381.2140808105469 0 600.38806
396.2186279296875 0 1322.1558
404.81585693359375 0 577.66296
405.22381591796875 0 680.82135
407.22845458984375 0 839.20337
422.2163391113281 0 799.85034
435.2231750488281 0 2474.0667
451.242431640625 0 2429.7454
452.2490234375 0 1669.9329 c 3
466.2054443359375 0 3778.377
467.20880126953125 0 1282.0773
478.2793884277344 0 664.7101
479.2135314941406 0 11941.215
480.21630859375 0 2533.802
480.25079345703125 0 609.6707
507.2446594238281 0 700.84344
520.3131103515625 0 1836.2043
521.3189086914062 0 730.71094
522.2678833007812 0 11216.905 z 3
523.2723999023438 0 4606.3354
530.297607421875 0 1483.1719
531.3013305664062 0 860.1487
538.2865600585938 0 4355.421 y 3
539.2902221679688 0 1403.3972
547.30126953125 0 1672.1998
548.3073120117188 0 10217.499
549.3103637695312 0 3403.2146
550.3099975585938 0 678.03516
553.2371826171875 0 686.26697
564.326171875 0 11535.976
565.3334350585938 0 27219.848 c 4
566.2457275390625 0 4430.489
566.3363037109375 0 10365.314
567.2476196289062 0 1347.1124
567.338623046875 0 2053.6326
582.1104125976562 0 562.7326
592.296875 0 1848.5398 w 2
607.3065795898438 0 780.5682 y Water loss 2
609.2999267578125 0 11005.656 z 2
610.3042602539062 0 3663.4666
611.308837890625 0 630.47125
624.2742919921875 0 1518.4293
625.3187255859375 0 41886.96 y 2
626.3215942382812 0 11680.252
627.3242797851562 0 2684.1309
636.34765625 0 856.1875
637.3456420898438 0 670.2316
638.34619140625 0 575.6537
663.3348999023438 0 6171.0547
664.3370361328125 0 1834.4288
678.3448486328125 0 3494.5654 y Water loss 1
679.349365234375 0 1949.6893
680.3599243164062 0 72588.125 c 5
681.3629150390625 0 24384.635
682.3646850585938 0 4623.504
694.3646240234375 0 2697.267
695.370849609375 0 815.4063
696.3555908203125 0 18337.172 y 1
697.3583984375 0 6003.0728
698.3601684570312 0 983.04663
737.3578491210938 0 2783.3447
739.3701171875 0 776.1319
750.3652954101562 0 15938.501
750.4293823242188 0 7721.5776
751.3684692382812 0 6331.2446
751.4317016601562 0 2859.7964
752.371826171875 0 1496.7067
764.440673828125 0 693.00964
793.4212036132812 0 16640.082
794.42431640625 0 6561.21
795.4275512695312 0 1862.4526
809.439697265625 0 33789.137
810.4451904296875 0 27057.984
811.4490966796875 0 9114.622
812.451904296875 0 1506.6163
1002.0197143554688 0 640.7243
1409.4969482421875 0 675.9762
1499.7510986328125 0 623.77875
2331.280517578125 0 749.09393
2366.405029296875 0 821.3057
2393.41796875 0 628.8602
2534.52685546875 0 613.09515

Spectrum Details

|  |  |
| --- | --- |
| Matched peaks? Matched peaks The total absolute number of peaks matched. Additionally in brackets the total fraction of peaks matched and the total number of peaks is shown. | 22 (14.57% of 151) |
| FDR? FDR The false discovery rate estimated for this peptide. It is calculated by matching all theoretical fragments with a non-integer shift with the raw peaks for this spectrum. This is done with 40 different shifts. The resulting percentage is the average number of annotated peaks over the number of annotated peaks with the correct spectrum. | 1.08% |
| Satellite FDR? Satellite FDR See the FDR for details on its calculation. This satellite ion specific FDR only contains the satellite ions (d/w) for I/L/J positions. | 2.38% |
| PSM Score? PSM Score The PSM Score as given by Hecklib to this annotated spectrum. It is shown with three significant figures. | 243 |

#### Reverse Lookup? Reverse Lookup All places where this read could be placed.

| Group | Segment | Template | Template Part | Read Part | Score | Unique |
| --- | --- | --- | --- | --- | --- | --- |
| Decoy | Decoy | K1C20 | [80..87] | [0..7] | 47 | True |

| Recombined | Template Part | Read Part | Score | Unique |
| --- | --- | --- | --- | --- |
| K1C20 | [80..87] | [0..7] | 47 | True |

#### Meta Information from Multiple reads

##### Number of combined reads

2

##### Intensity

0.5499

##### TotalArea

8.895E+06

##### Changes to the peptide sequence

JASYLDK

J→LSupport for Leucine based on side chain ions (1 for L 0 for I) (Position: 5)

L→JNo support for either Leucine or Isoleucine based on side chain ions (Position: 5)

L→JNo support for either Leucine or Isoleucine based on side chain ions (Position: 1)

#### Positional Score

Copy Data

##### Positional Score (TSV)

###### Preview

```
Loading example...
```

*Click on the button to copy the data to your clipboard.*

000123456

Label Value
"0" 0
"1" 0
"2" 0
"3" 0
"4" 0
"5" 0
"6" 0

#### Meta Information from PEAKS

##### Scan Identifier

F4:3928

##### Original sequence

L

A

S

Y

L

D

K

##### Posttranslational Modifications

##### Source File

D:\separate\_stitch\_analyses\xle-disambiguation\raw\20210323\_F1\_UM1\_Peng0013\_SA\_F59\_ingel\_3ug\_tryp.raw

##### Fraction

4

##### Scan Feature

F4:792

##### De Novo Score

99

##### ConfidenceScore

99

### m/z

405.2246

##### Mass

808.433

##### Charge

2

##### Retention Time

21.13

##### Predicted Retention Time

22.30

##### Area

4.448E+06

##### Parts Per Million

2

##### Fragmentation mode

HCD

##### Originating file

01 D:\separate\_stitch\_analyses\xle-disambiguation\20210325\_F59\_3ug\_DENOVO\_12.csv

#### Meta Information from PEAKS

##### Scan Identifier

F4:3979

##### Original sequence

L

A

S

Y

L

D

K

##### Posttranslational Modifications

##### Source File

D:\separate\_stitch\_analyses\xle-disambiguation\raw\20210323\_F1\_UM1\_Peng0013\_SA\_F59\_ingel\_3ug\_tryp.raw

##### Fraction

4

##### Scan Feature

F4:792

##### De Novo Score

99

##### ConfidenceScore

98

### m/z

405.2246

##### Mass

808.433

##### Charge

2

##### Retention Time

21.13

##### Predicted Retention Time

22.30

##### Area

4.448E+06

##### Parts Per Million

2

##### Fragmentation mode

ETHCD

##### Originating file

01 D:\separate\_stitch\_analyses\xle-disambiguation\20210325\_F59\_3ug\_DENOVO\_12.csv
